# The lipid phosphatase activity of PTEN dampens FRA1 expression via AKT/mTOR signaling to suppress melanoma

**DOI:** 10.1101/2023.06.01.543131

**Authors:** Xiaonan Xu, Ilah Bok, Neel Jasani, Kaizhen Wang, Manon Chadourne, Ou Deng, Eric A. Welsh, Fumi Kinose, Uwe Rix, Florian A. Karreth

## Abstract

PTEN, a phosphatase frequently inactivated in melanoma, opposes PI3K/AKT/mTOR pathway activation. However, AKT- and mTOR-targeted therapies have so far yielded insufficient results in preclinical models and clinical trials of melanoma. We therefore examined whether PTEN suppresses melanoma through lipid phosphatase-independent functions or by opposing lipid phosphatase-dependent, AKT-independent pathways. Restoring different PTEN functions in PTEN-deficient cells or mouse models revealed that PTEN lipid phosphatase activity predominantly suppresses melanoma with minimal contribution from its protein phosphatase and scaffold functions. A drug screen highlighted the dependence of PTEN-deficient melanoma cells on the AKT/mTOR pathway. Moreover, activation of AKT was sufficient to overcome several aspects of PTEN-mediated melanoma suppression. Phosphoproteomics analysis of the PTEN lipid phosphatase activity identified the AP-1 transcription factor FRA1 as a downstream effector. PTEN regulates FRA1 translation via AKT/mTOR and FRA1 overexpression overcomes PTEN-mediated melanoma suppression. Our study affirms AKT as the key mediator of PTEN inactivation in melanoma and identifies an AKT/mTOR/FRA1 axis as a driver of melanomagenesis.

## INTRODUCTION

Recurrent somatic mutation or copy number alteration of common driver genes promote the initiation and early stages of progression of cutaneous malignant melanoma. One of the earliest events in melanomagenesis is the hyperactivation of the MAPK pathway (Shain et al., 2018), which is most frequently caused by activating mutations in BRAF or NRAS or inactivating mutations in NF1 (Network, 2015). Melanoma progression requires the additional inactivation of tumor suppressor genes, and *CDKN2A* (encoding the cell cycle regulators p16INK4a and p14ARF) and *PTEN* are most commonly affected (Network, 2015; Shain et al., 2018). PTEN encodes a phosphatase that dephosphorylates the lipid second messenger Phosphatidylinositol-3,4,5-triphosphate (PIP_3_) to Phosphatidylinositol-4,5-bisphosphate (PIP_2_), thereby opposing the function of PI3K, which phosphorylates PIP_2_ at the 3 position. Anchored in the cell membrane, PIP_3_ recruits pleckstrin homology (PH) domain-containing proteins, enabling their activation and downstream signaling. AKT and its downstream effector mTOR (as part of the mTORC1 complex) constitute the most widely studied pathway that is activated by PI3K via PIP_3_. The importance of the PI3K/AKT/mTOR pathway in maintaining normal physiology is highlighted by its frequent alteration in disease, and activating mutations and copy number gains of PI3K or AKT as well as inactivating mutations and copy number losses of PTEN are frequently observed in several cancers (Manning and Toker, 2017; Fruman et al., 2017; Thorpe et al., 2015; Song et al., 2012; Lee et al., 2018), including melanoma. Notably, alterations of PTEN are far more frequently observed in melanoma than alterations of PI3K or AKT (Zhang et al., 2016), suggesting that PTEN inactivation more potently promotes melanomagenesis.

Given the frequent hyperactivation of the PI3K/AKT pathway, significant effort has been invested into gaining a better understanding of the role of this pathway in melanomagenesis and exploring its suitability as a therapeutic target. AKT is considered the key mediator of elevated PIP_3_ levels, regulating numerous effectors to promote melanomagenesis. Indeed, melanoma mouse models overexpressing myristoylated AKT1 or the activated AKT1^E17K^ mutant revealed a pro-metastatic effect of AKT (Cho et al., 2015; Kircher et al., 2019). Interestingly, while independent AKT1 activation or PTEN deletion resulted in primary melanomas with similar latency and penetrance, activated AKT1 synergized with PTEN deficiency to promote metastasis in this mouse model (Cho et al., 2015). This suggests that the activation of AKT and possibly other downstream signaling pathways in response to the loss of PTEN changes during melanoma progression and metastasis. Several mouse melanoma models driven by BRAF^V600E^ and melanocyte-specific deletion of PTEN demonstrated that pharmacological inhibition of PI3K (Deuker et al., 2015; Durban et al., 2013) or mTOR (Dankort et al., 2009) as well as genetic restoration of PTEN (Bok et al., 2019) had superior antitumor effects than AKT inhibition (Durban et al., 2013). Moreover, PI3K maintains melanoma cell proliferation by regulating protein synthesis via mTORC1 independently of AKT (Silva et al., 2014). Elevated PIP_3_ levels stimulate several kinases in addition to AKT (Lien et al., 2017) of which PDK1 and SGKs have been reported as drivers of a malignant phenotype in PTEN-deficient melanoma (Scortegagna et al., 2015, 2014). Thus, although AKT is widely considered a key signaling node in PTEN-deficient melanoma, it remains to be determined if AKT critically mediates the pro-tumorigenic effects of PTEN loss in melanoma.

PTEN possesses non-canonical functions that are independent of its lipid phosphatase activity. Indeed, PTEN was originally described as a protein phosphatase (Li et al., 1997; Myers et al., 1997). Several protein phosphatase targets for PTEN have been identified, including FAK (Tamura et al., 1998), SRC (Dey et al., 2008), IRS1 (Shi et al., 2014), PGK1 (Qian et al., 2019) RAB7 (Shinde and Maddika, 2016), and the glucocorticoid receptor (Yip et al., 2020). Interestingly, PTEN protein phosphatase activity has been associated with cancer cell migration (Tamura et al., 1998; Dey et al., 2008; Tibarewal et al., 2012; Gildea et al., 2004; Raftopoulou et al., 2004), and a recent study suggested a role for the protein phosphatase activity of PTEN in melanoma metastasis (Yu et al., 2023). Even phosphatase-independent functions for PTEN as a scaffold have been discovered (Kuchay et al., 2017; Zhao et al., 2017). These non-canonical functions raise the possibility that the loss or inactivation of PTEN promote melanoma, at least in part, through PIP_3_- and AKT-independent mechanisms. These findings highlight the critical need for a better understanding of the mechanisms whereby PTEN suppresses melanoma, which promises to reveal new avenues for therapeutic intervention.

Here, we report that the lipid phosphatase activity confers PTEN with the ability to suppress melanoma, with negligible contribution of the protein phosphatase activity. The PTEN scaffold function is dispensable for melanoma suppression. Moreover, AKT is sufficient to overcome melanoma suppression mediated by restoring PTEN expression, affirming AKT as the predominant downstream effector of PTEN loss-of-function. AKT controls the expression of the AP1 transcription factor FRA1 via mTOR-mediated translation, and FRA1 partially negates the effects of PTEN restoration. Thus, our results nominate a PTEN/AKT/mTOR/FRA1 axis as a key driver of melanoma.

## RESULTS

### The lipid phosphatase activity mediates the melanoma suppressor function of PTEN in vitro

Using an ESC-GEMM platform, we showed previously that restoring endogenous Pten expression in melanomas and melanoma-derived cell lines significantly diminished tumor growth and cell proliferation, respectively (Bok et al., 2019). Similarly, in murine Braf^V600E^; Pten^Δ/Δ^ (BPP) melanoma cell lines harboring the rtTA3 reverse transactivator to enable Dox-mediated transcriptional control of expression cassettes, we demonstrated that expression of ectopic Pten reduces proliferation in vitro and allograft tumor growth (Bok et al., 2019, 2021). This represents an ideal system to assess the melanoma suppressive potential of the different PTEN functions. We therefore generated Dox-inducible expression constructs harboring either wildtype Pten (Pten^WT^), one of three mutant Pten isoforms (Pten^C124S^, Pten^G129E^, or Pten^Y138L^), or a GFP control cDNA. Pten^C124S^ has no phosphatase activity, Pten^G129E^ is lipid phosphatase-deficient but retains protein phosphatase activity, and Pten^Y138L^ is protein phosphatase-deficient but retains lipid phosphatase activity. All Pten isoforms retain scaffolding activity.

To examine the contribution of PTEN functions to melanoma suppression, these constructs were expressed in BPP mouse melanoma cell lines M10M1 and M10M6 (**Figure 1A, Supplementary Figure 1**). Similar to our previous findings (Bok et al., 2019), the restoration of Pten^WT^ expression significantly attenuated proliferation of both cell lines (**Figure 1B**). Protein phosphatase-deficient Pten^Y138L^ moderately reduced proliferation of M10M1 cells, while it phenocopied the robust suppression of proliferation in M10M6 cells (**Figure 1B**). Neither phosphatase-dead Pten^C124S^ nor lipid phosphatase-deficient Pten^G129E^ affected melanoma cell proliferation (**Figure 1B**). Pten^WT^ robustly reduced low-density colony formation of M10M1 and M10M6 cells, which was partially recapitulated by Pten^Y138L^, while Pten^C124S^ and Pten^G129E^ had no effect (**Figure 1C**). Moreover, the colony size and number under anchorage-independent growth conditions in soft agar were markedly reduced in M10M1 and M10M6 cells expressing Pten^WT^ or Pten^Y138L^ (**Figure 1D**). Interestingly, Pten^G129E^ moderately reduced colony size and number in both cell lines, while Pten^C124S^ had again no effect (**Figure 1D**). Taken together, the PTEN lipid phosphatase function accounts for most of the suppressive activity in several cell proliferation assays, while the PTEN protein phosphatase function only contributes to suppression of anchorage-independent growth and putative PTEN scaffold functions play no role.

**Figure 1.**
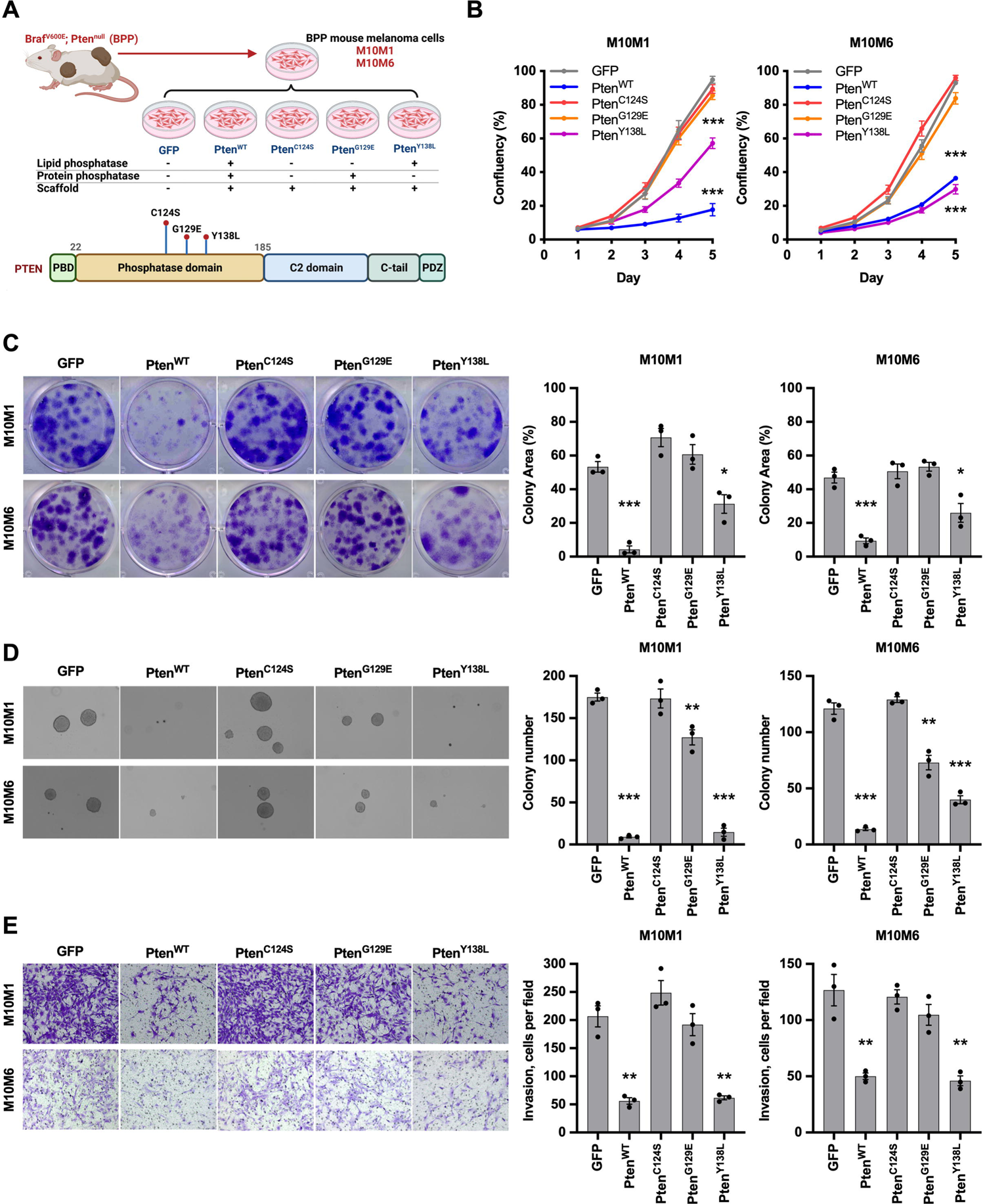
Comparison of different PTEN functions in melanoma suppression in vitro. **(A)** Schematic diagram depicting the generation of mouse melanoma cell lines harboring inducible expression constructs of different PTEN constructs (created with BioRender.com). **(B)** Proliferation as measured by relative confluence of M10M6 and M10M1 cells expressing PTEN constructs. **(C)** Representative images (left) and quantification of percentage occupied area (right) of low-density colony formation assays of M10M6 and M10M1 cells expressing PTEN constructs. **(D)** Representative images (left) and quantification of colony number (right) of anchorage-independent growth agar assays of M10M6 and M10M1 cells expressing PTEN constructs. **(E)** Representative images (left) and quantification of cell numbers (right) of transwell invasion assays of M10M6 and M10M1 cells expressing PTEN constructs. Mean ± SEM are shown in (B-E). Data are analyzed with Student’s unpaired *t* test, * P<0.05, ** P<0.01, *** P<0.001.

Interestingly, we observed cell morphological changes in M10M1 and M10M6 cells expressing the different PTEN mutants under low serum (M10M6) or normal (M10M1) growth conditions. Pten^WT^- and Pten^Y138L^-expressing cells appeared more epithelial, were smaller and exhibited contact inhibition, while cells expressing Pten^G129E^ and Pten^C124S^ are more elongated and spindle-like (**Supplementary Figure 1B**). This suggested that PTEN restoration may also affect the invasive capability of melanoma cells. Indeed, expression of Pten^WT^ markedly inhibited melanoma cell invasion in transwell assays, and this was completely phenocopied by Pten^Y138L^ (**Figure 1E**). Pten^G129E^ and Pten^C124S^ exhibited no effect on invasion of M10M1 and M10M6 cells (**Figure 1E**). Thus, PTEN suppresses melanoma cell invasion entirely through its lipid phosphatase function.

### The lipid phosphatase function of PTEN suppresses melanoma initiation and growth in vivo

We next determined which function of PTEN suppresses melanoma in vivo by performing allografts in immunocompromised Nu/Nu mice. We subcutaneously injected M10M6 cells harboring the Doxycycline (Dox)-inducible Pten expression constructs, placed the mice on a Dox diet immediately following the transplantation, and monitored tumor growth. Tumors expressing Pten^G129E^ or Pten^C124S^ grew rapidly, requiring euthanasia after 3 weeks (**Figure 2A**). By contrast, activating the expression of Pten^WT^ or Pten^Y138L^ markedly slowed tumor growth and prolonged survival to approximately 4.5 weeks (**Figure 2A**). At endpoint, we collected tumors to determine the expression of PTEN. Notably, Pten^WT^ or Pten^Y138L^ tumors had lost expression of PTEN, while Pten^C124S^ and Pten^G129E^ tumors maintained PTEN expression (**Figure 2B**). This finding suggests that there is strong selective pressure against PTEN lipid phosphatase activity to allow for melanoma growth. Accordingly, the levels of AKT phosphorylated at Ser473 was comparable between tumors expressing lipid phosphatase active (Pten^WT^ or Pten^Y138L^) and inactive (Pten^C124S^ and Pten^G129E^) PTEN (**Figure 2B**).

**Figure 2.**
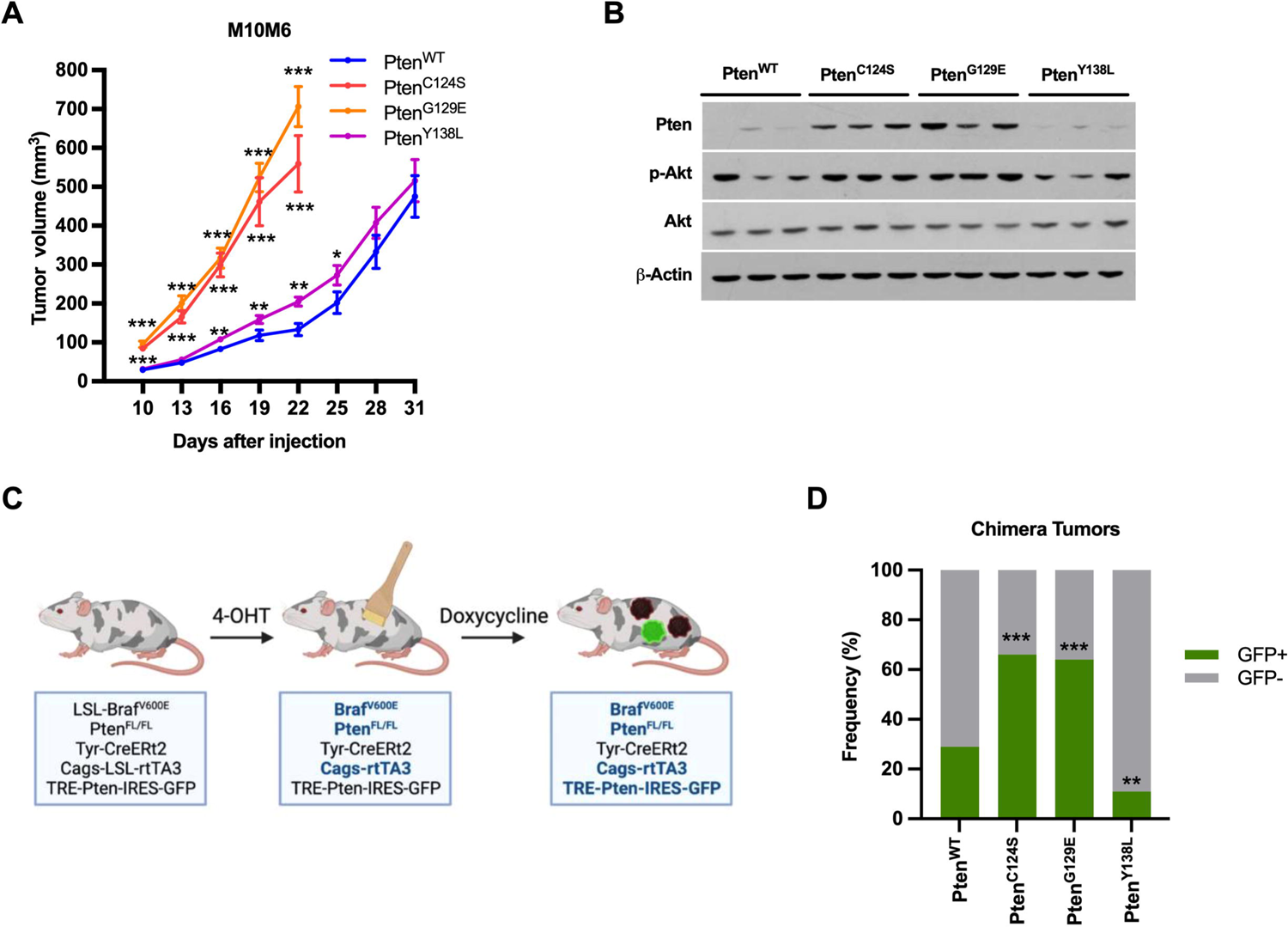
Lipid phosphatase activity is essential for PTEN to suppress melanoma in vivo. **(A)** M10M6 melanoma cells expressing PTEN constructs were subcutaneously injected into nude mice (n=10). Mice were fed chow containing 200 mg/kg Doxycycline to induce the expression of PTEN mutant constructs. Tumor volumes were measured every 3 days. Tumor volumes are shown as mean ± SEM. Data are analyzed with Student’s unpaired *t* test, * P<0.05, ** P<0.01, *** P<0.001. **(B)** Western blot analysis of tumors shown in (A). Pten, p-Akt (S473), pan-Akt, and β-Actin were detected. **(C)** Schematic diagram of the ESC-GEMMs where Pten-IRES-GFP constructs are expressed in the Braf^V600E^; Pten^null^ (BPP) mouse melanoma model. **(D)** Melanomas were isolated and GFP expression was detected as a surrogate for ectopic PTEN expression. The percentages of GFP+ and GFP-tumors in the different PTEN cohorts are shown. Data are analyzed with Chi-square test. * P<0.05, ** P<0.01, *** P<0.001.

To further determine the role of the PTEN lipid and protein phosphatase activities in the initiation and early stages of spontaneous melanomagenesis, we used our ESC-GEMM platform. We targeted BPP (LSL-Braf^V600E^; Pten^FL/FL^; Tyr-CreERt2; CAGs-LSL-rtTA3; CHC) ES cells (Bok et al., 2019) with Dox-inducible PTEN expression constructs (TRE-PTEN^WT^-IRES-GFP, TRE-PTEN^C124S^-IRES-GFP, TRE-PTEN^G129E^-IRES-GFP, and TRE-PTEN^Y138L^-IRES-GFP) (**Figure 2C**). Because we suspected that there could be negative selection against ectopic PTEN expression in tumors, we linked a GFP cDNA to the PTEN cDNAs via an internal ribosome entry site (IRES). This enables the use of GFP as in indicator of ectopic PTEN expression and circumvents the difficulties with distinguishing between endogenous wildtype and ectopic mutant PTEN expression. Targeted BPP ES cells were used to generate experimental chimeras and melanomagenesis was induced by topical administration of 4-Hydroxytamoxifen (4OHT) on shaved backs at 3 weeks of age. 4OHT activates the melanocyte-specific Cre, which in turn induces Braf^V600E^ expression, deletes both Pten alleles, and activates expression of the rtTA3 reverse transactivator. The mice were placed on a Dox-containing diet immediately following 4OHT administration to activate expression of the transgenic PTEN cDNA constructs (**Figure 2C**). Melanomas formed readily in each of the four PTEN cohorts, leading to indistinguishable tumor-free survival and overall survival as well as tumor numbers (**Supplementary Figure 2A-C**). To test if expression of the ectopic PTEN alleles was induced, we collected tumor tissue and determined GFP expression as a surrogate for transgenic activation by Western blot. Notably, the frequency of GFP-positive tumors in the PTEN^C124S^ and PTEN^G129E^ cohorts was significantly higher than in the PTEN^WT^ and PTEN^Y138L^ cohorts (66% in PTEN^C124S^ and 64% in PTEN^G129E^ vs. 29% in PTEN^WT^ and 11% in PTEN^Y138L^) (**Figure 2D**), again suggesting that there is strong selective pressure against the expression of PTEN having lipid phosphatase activity. These in vivo findings corroborate the in vitro data indicating that PTEN suppresses melanoma primarily through its lipid phosphatase function.

### A small molecule inhibitor screen highlights the importance of the AKT/mTOR axis in PTEN-deficient melanoma

After establishing that PTEN suppresses melanoma through its lipid phosphatase activity, we sought to determine key effectors that mediate the suppressive effects on PTEN. To this end, we performed a small molecule inhibitor screen in M10M6 cells in which PTEN is inactive (Pten^C124S^) or in which PTEN activity is restored (Pten^WT^). We reasoned that PTEN restoration would inactivate critical pathways, thereby reducing the sensitivity to drugs targeting those pathways. Thus, the differential drug sensitivity between Pten^C124S^ and Pten^WT^ melanoma cells may identify critical effectors. We used an expanded drug panel that is based on an earlier library (Sumi et al., 2019), which contains 500 small molecule inhibitors (of which approximately 20% are FDA-approved, 40% are currently in clinical trials, 20% were previously in clinical trials, and 20% are preclinical compounds/chemical probes for unique targets) at four concentrations (0.1 µM, 0.5 µM, 2.5 µM, and 10 µM). We induced Pten^C124S^ and Pten^WT^ expression by incubating M10M6 cells with Dox for 24 hours, followed by treating cells with the drug library for 5 days. Drug sensitivity was measured via cell viability and compared to DMSO treated cells. At the 2.5 µM concentration, 287 out of 500 inhibitors were more effective in Pten^C124S^ cells compared to Pten^WT^ cells (**Supplementary Table 1**), possibly reflecting the widespread signaling changes that occur upon PTEN inactivation. Notably, Pten^C124S^ cells were more sensitive to all AKT inhibitors (7/7), all mTORC1 inhibitors (4/4), and most mTOR inhibitors (8/10) compared to Pten^WT^ cells, suggesting robust addiction of PTEN-deficient cells to the AKT/mTOR pathway (**Figure 3A, Supplementary Table 1**).

**Figure 3.**
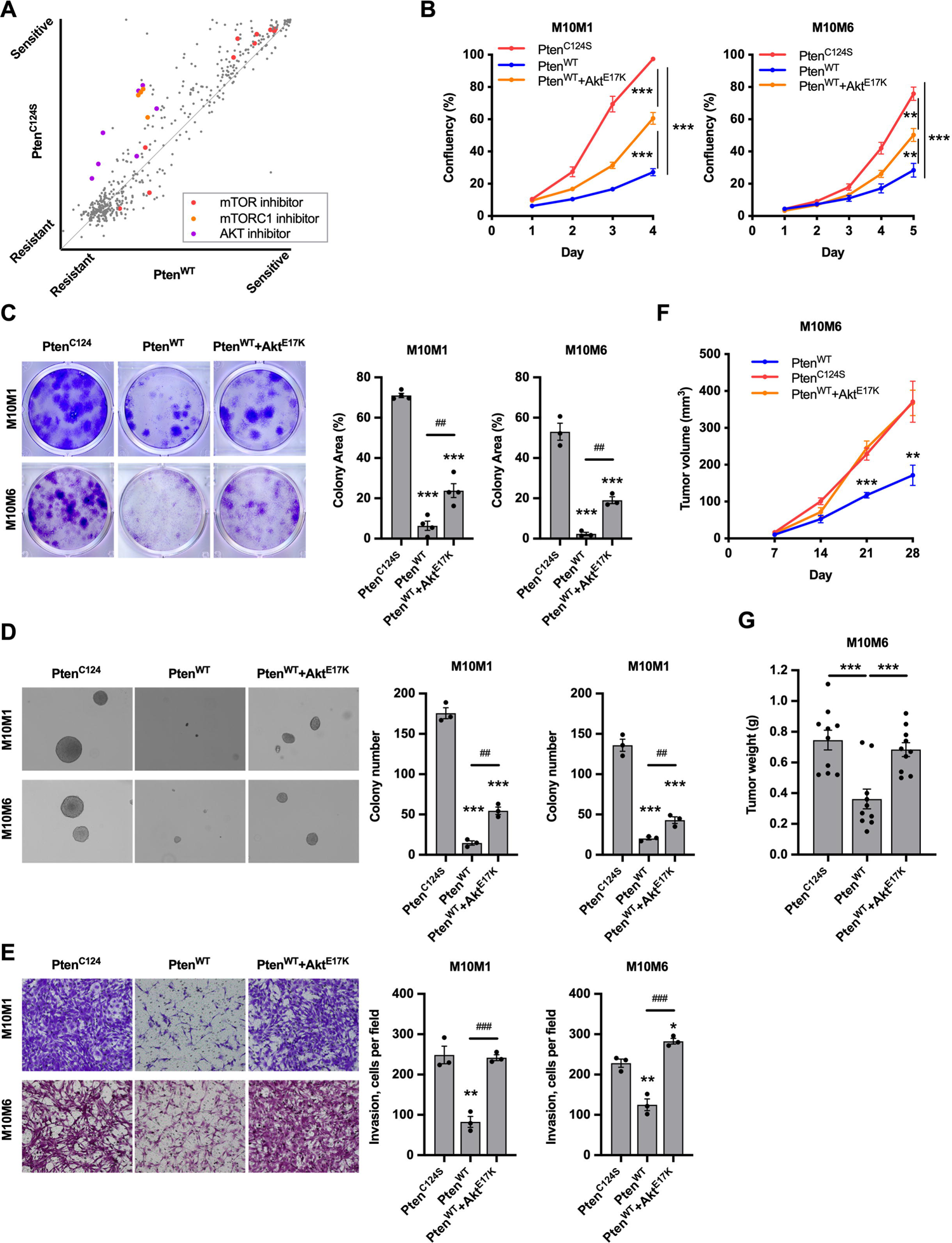
AKT reactivation partially abrogates melanoma suppression by PTEN. **(A)** M10M6 cells having Pten^WT^ or phosphatase deficient Pten^C124S^ were subjected to a small molecule inhibitor library screen containing 500 inhibitors. The sensitivity of each inhibitor towards Pten^WT^ and Pten^C124S^ cells is calculated by comparing cell viability of inhibitor treatment to DMSO control (see Materials and Methods) and plotted. **(B)** Proliferation as measured by relative confluence of M10M6 and M10M1 cells expressing Pten^WT^, Pten^C124S^, or wildtype Pten together with constitutively activated Akt (Pten^WT^+Akt^E17K^). **(C)** Representative images (left) and quantification of percentage occupied area (right) of low-density colony formation assays of M10M6 and M10M1 cells expressing Pten^WT^, Pten^C124S^, or Pten^WT^+Akt^E17K^. **(D)** Representative images (left) and quantification of colony number (right) of anchorage-independent growth assays of M10M6 and M10M1 cells expressing Pten^WT^, Pten^C124S^, or Pten^WT^+Akt^E17K^. **(E)** Representative images (left) and quantification of cell numbers (right) of transwell invasion assays of M10M6 and M10M1 cells expressing Pten^WT^, Pten^C124S^, or Pten^WT^+Akt^E17K^. **(F, G)** M10M6 cells expressing Pten^WT^, Pten^C124S^, or Pten^WT^+Akt^E17K^ were subcutaneously injected into NSG mice (n=10). Mice were fed chow containing 200 mg/kg Doxycycline to induce expression of PTEN constructs. Tumor volumes were measured every 3 days. The curves of tumor volume **(F)** and the tumor weight at the end point **(G)** are shown. Mean ± SEM are shown in (B-G). Data are analyzed with Student’s unpaired *t* test, * P<0.05, ** P<0.01, *** P<0.001, ^##^ P<0.01.

### Reactivating AKT partially rescues PTEN induced melanoma suppression

We next determined whether inhibiting the AKT/mTOR pathway is sufficient to mediate the tumor suppressive effects of PTEN. To this end, we expressed constitutively active AKT (Akt^E17K^) in Pten^WT^ M10M1 and M10M6 cells (**Supplementary Figure 3A**) and examined whether this negates the tumor suppressive effects of PTEN. Reactivation of AKT partially rescued the suppression of cell proliferation induced by Pten^WT^ in both melanoma cell lines (**Figure 3B**). Moreover, AKT reactivation was less potent in low density and anchorage-independent growth assays where it only moderately rescued the effects of Pten^WT^ restoration (**Figure 3C,D**). Conversely, reactivating AKT fully rescued the Pten^WT^-mediated suppression of cell invasion in transwell assays (**Figure 3E**). To determine whether active AKT can overcome Pten^WT^-mediated melanoma suppression, we performed subcutaneous allografts in NSG mice. Notably, active AKT completely rescued the repression of melanoma growth by Pten^WT^ (**Figure 3F,G, Supplementary Figure 3B**). These results, combined with the data obtained with the PTEN phosphatase mutants, suggest that PTEN suppresses melanoma invasion in vitro and tumor growth by inhibiting AKT, while melanoma cell proliferation under different in vitro conditions is suppressed through additional pathways.

A previous study reported minimal effects of the AKT inhibitor MK2206 on melanoma cell proliferation (Silva et al., 2014), while our results indicate that PTEN suppresses proliferation through its lipid phosphatase activity at least in part by inactivating AKT. We therefore evaluated this further by examining the effect of MK2206 in our murine melanoma cell lines. Even at low doses of the MK2206 (0.1-0.5 µM), AKT activation as measured by Ser473 phosphorylation was reduced to an extent similar to that elicited by Pten^WT^ restoration (**Figure 4A**). In agreement with the study by Silva et al. (Silva et al., 2014), even the highest dose (1 µM) of MK2206 failed to inhibit melanoma cell proliferation and low density growth, while Pten^WT^ restoration elicited marked suppression (**Figure 4B,C, Supplementary Figure 4A**). By contrast, MK2206 inhibited melanoma cell invasivon in a dose dependent manner, with the highest dose eliciting effects comparable to Pten^WT^ restoration (**Figure 4D, Supplementary Figure 4B**). Taken together, these findings further support the notion that the PTEN lipid phosphatase activity suppresses some aspects of melanomagenesis predominantly by inactivating AKT signaling.

**Figure 4.**
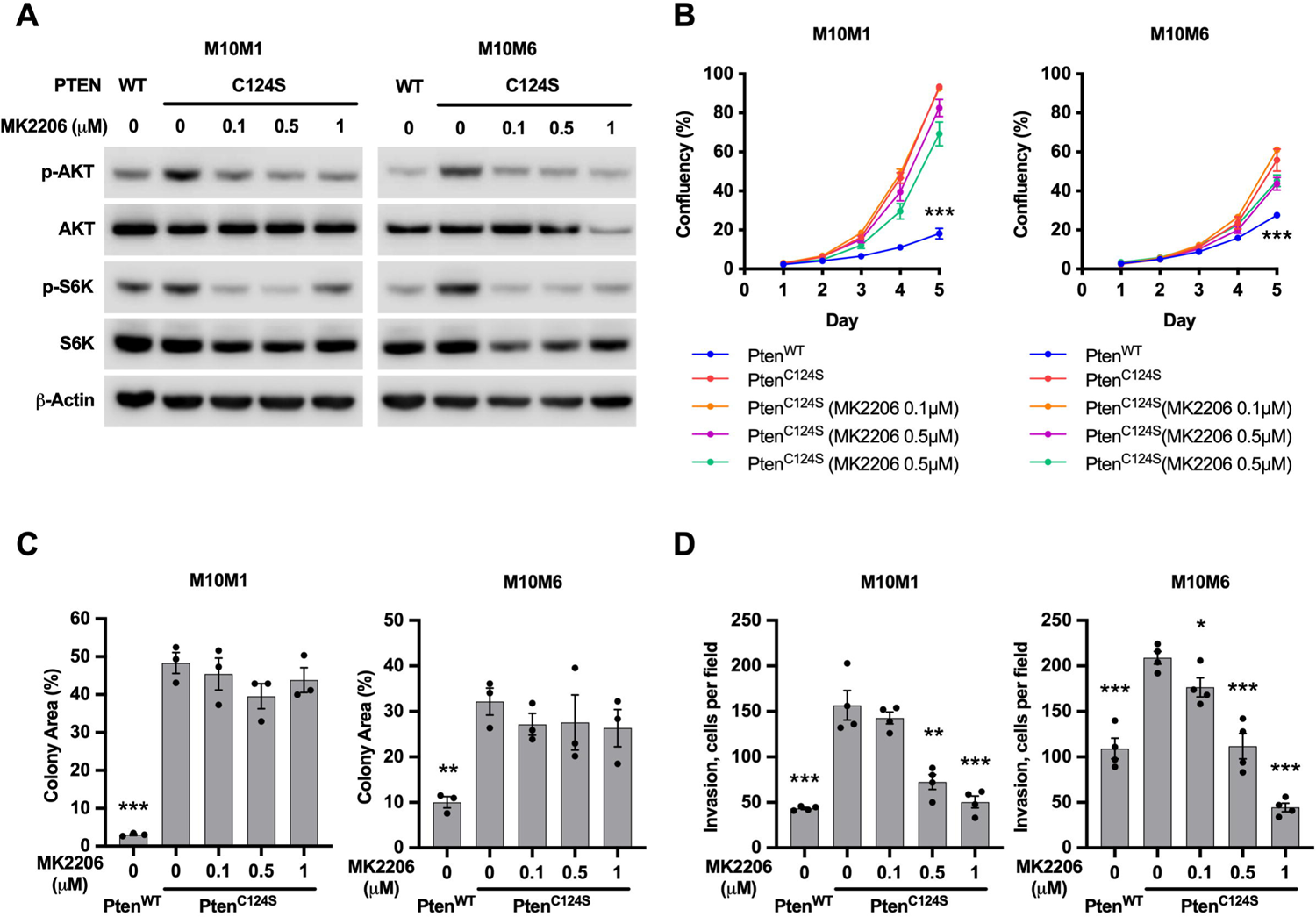
Comparison of tumor suppressive effects of AKT inhibitor vs. PTEN restoration. **(A)** Western blot showing the changes of AKT activity in M10M1 and M10M6 melanoma cells expressing Pten^WT^ or Pten^C124S^ and treated with the AKT inhibitor MK2206. p-Akt (Ser473), pan-Akt, p-S6K (Thr389), S6K, and β-Actin were detected. **(B)** Proliferation as measured by relative confluence of M10M6 and M10M1 cells expressing Pten^WT^ or Pten^C124S^ and treated with the AKT inhibitor MK2206. **(C)** Quantification of percentage occupied area of low-density colony formation assays of M10M6 and M10M1 cells expressing Pten^WT^ or Pten^C124S^ and treated with the AKT inhibitor MK2206. **(D)** Quantification of cell numbers of transwell invasion assays of M10M6 and M10M1 cells expressing Pten^WT^ or Pten^C124S^ and treated with the AKT inhibitor MK2206. Mean ± SEM are shown in (B-D). Data are analyzed with Student’s unpaired *t* test, * P<0.05, ** P<0.01, *** P<0.001

### Phosphoproteomics analysis reveals potential PTEN lipid phosphatase targets

To comprehensively identify the pathways downstream of PTEN lipid phosphatase activity, we performed phosphoproteomics in M10M6 melanoma cells expressing GFP, Pten^WT^, Pten^C124S^, Pten^G129E^, or Pten^Y138L^. We induced PTEN expression for 16 hours with Dox (**Supplementary Figure 5A**) and then denatured, reduced, alkylated, and digested the isolated proteins. Using LC-MS/MS, we performed 16-plexed global phosphoproteomics to detect phosphorylation changes at Serine, Threonine, and Tyrosine residues (pS/T/Y) and an additional 16-plexed phospho-Tyrosine proteomics analysis to enrich for phosphorylation changes at Tyrosine residues (pY) (**Figure 5A**). Principle component analysis (PCA) showed that the phosphorylation patterns in the GFP control and Pten^C124S^ cells were highly similar, as expected given the lack of a suppressive effect of the PTEN scaffold function. Phosphorylation in Pten^G129E^ cells was also similar to GFP/Pten^C124S^ cells, suggesting that the PTEN protein phosphatase function does not provoke pronounced phosphorylation changes of the proteome. By contrast, phosphorylation changes in Pten^WT^ and Pten^Y138L^ cells were remarkably similar to each other but distinct from GFP/Pten^C124S^/Pten^G129E^ cells (**Figure 5B**). Given the high similarity between GFP and Pten^C124S^ cells, we combined them to a single negative control and compared it to Pten^WT^, Pten^G129E^, and Pten^Y138L^ cells to identify proteins differentially phosphorylated in response to PTEN lipid phosphatase activity. We found that phosphorylation of 594 proteins (a total of 1160 sites) were decreased while 119 proteins (240 sites) were increased by Pten^WT^ (**Figure 5C, Supplementary Table 2**), and Pten^Y138L^ led to decreased phosphorylation of 264 proteins (451 sites) and increased phosphorylation of 80 proteins (161 sites) (**Figure 5D, Supplementary Table 2**). Phosphorylation of 105 proteins (138 sites) was decreased while phosphorylation of 30 proteins (40 sites) was increased by Pten^G129E^ (**Figure 5E, Supplementary Table 2**). Notably, the extent of phosphorylation changes and the affected proteins largely overlap in Pten^WT^ and Pten^Y138L^ cells (**Figure 5C,D, Supplementary Figure 5B, Supplementary Table 2**). Kyoto Encyclopedia of Genes and Genomes (KEGG) analysis revealed that ErbB signaling, insulin signaling, and autophagy pathways – pathways that have been associated with AKT signaling – are enriched in Pten^WT^ and Pten^Y138L^ cells (**Figure 5F,G**). Conversely, the pathways enriched in Pten^G129E^ cells are different from those identified in Pten^WT^ and Pten^Y138L^ cells (**Figure 5H**). Thus, the lipid phosphatase activity predominantly shapes the phospho-proteome in response to PTEN expression.

**Figure 5.**
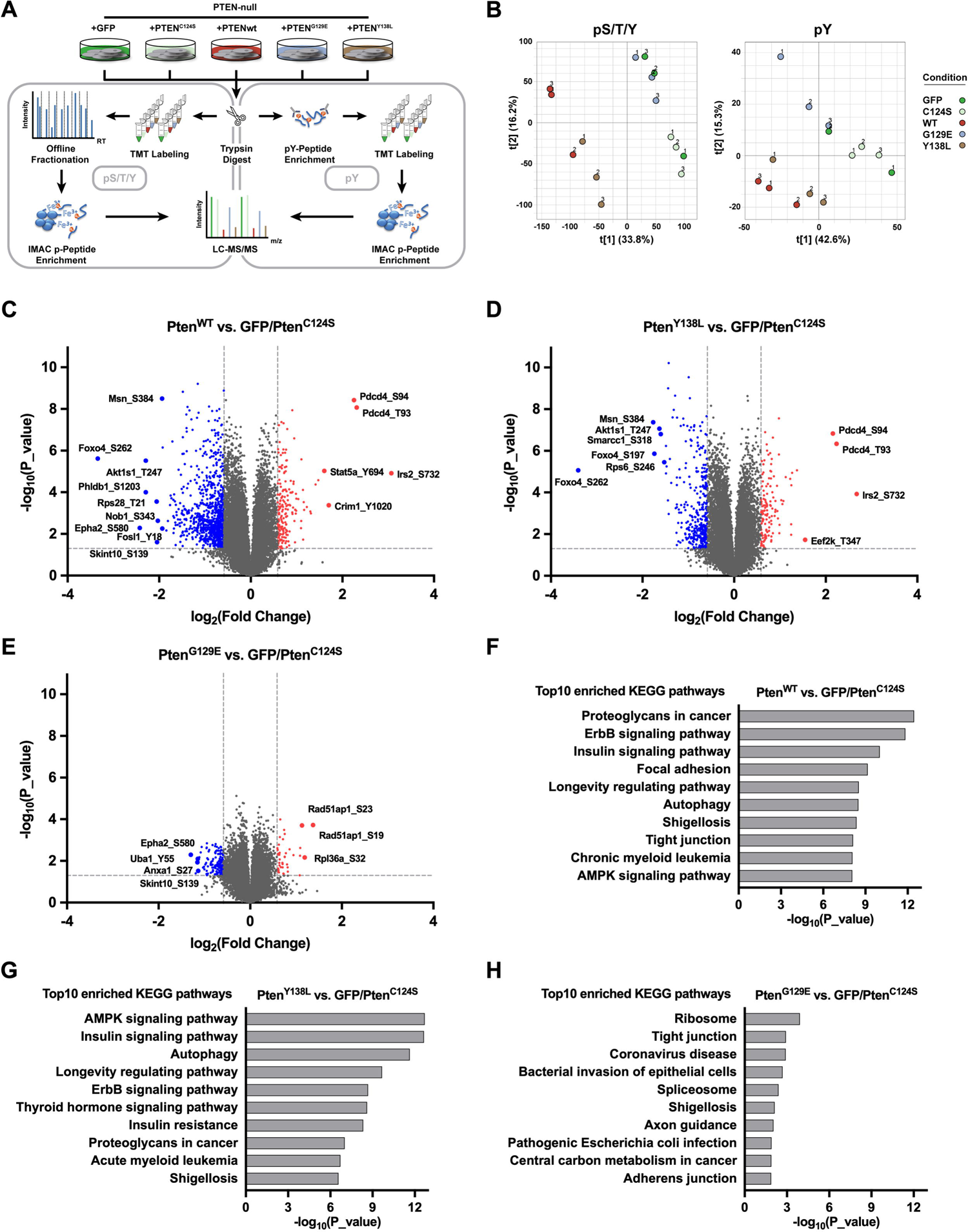
Quantitative global and tyrosine phosphoproteomics analysis of network-wide signaling effects of PTEN lipid phosphatase and protein phosphatase activities. **(A)** Schematic of quantitative global (pS/T/Y) and tyrosine (pY) phosphoproteomics. **(B)** Principal Component Analysis of pS/T/Y (left panel) and pY (right panel) phosphoproteomics data showing all biological replicates for each condition on the first two components. **(C-E)** Volcano plots indicating phosphorylation ratios of pS/T/Y and pY peptides between Pten^WT^ vs Pten^C124S^/GFP **(C)**, Pten^Y138L^ vs Pten^C124S^/GFP **(D)**, and Pten^G129E^ vs Pten^C124S^/GFP **(E)**. **(F-H)** KEGG pathway analysis of differentially phosphorylated proteins from Pten^WT^ vs Pten^C124S^/GFP **(F)**, Pten^Y138L^ vs Pten^C124S^/GFP **(G)**, and Pten^G129E^ vs Pten^C124S^/GFP **(H)** using Enrichr. Displayed are the top 10 enriched pathways ranked by P-value.

### PTEN inhibits FRA1 via AKT-mTOR pathway

We next sought to discover key proteins that mediate the effects of the PTEN lipid phosphatase activity, either downstream of AKT or through AKT-independent pathways. The phosphoproteomics analysis identified 143 proteins whose phosphorylation is regulated by the PTEN lipid phosphatase function in melanoma cells (**Supplementary Table 2**). To further prioritize these candidates, we reasoned that critical drivers of melanomagenesis may also be transcriptionally deregulated. We therefore analyzed the expression of the 143 candidates in a publicly available dataset (GSE112509) containing RNA expression data from 23 nevi and 57 melanomas. Seven genes exhibited significantly increased expression in melanoma compared to nevi, indicating they may be oncogenic in melanoma (**Figure 6A**). Interestingly, FOSL1, which encodes that AP-1 transcription factor FRA1, is the most upregulated gene in melanoma (fold change = 7.574) (**Figure 6A,B**). Moreover, an analysis of the TCGA-SKCM dataset revealed that high expression of FOSL1 negatively correlates with the overall survival of melanoma patients (**Figure 6C**).

**Figure 6.**
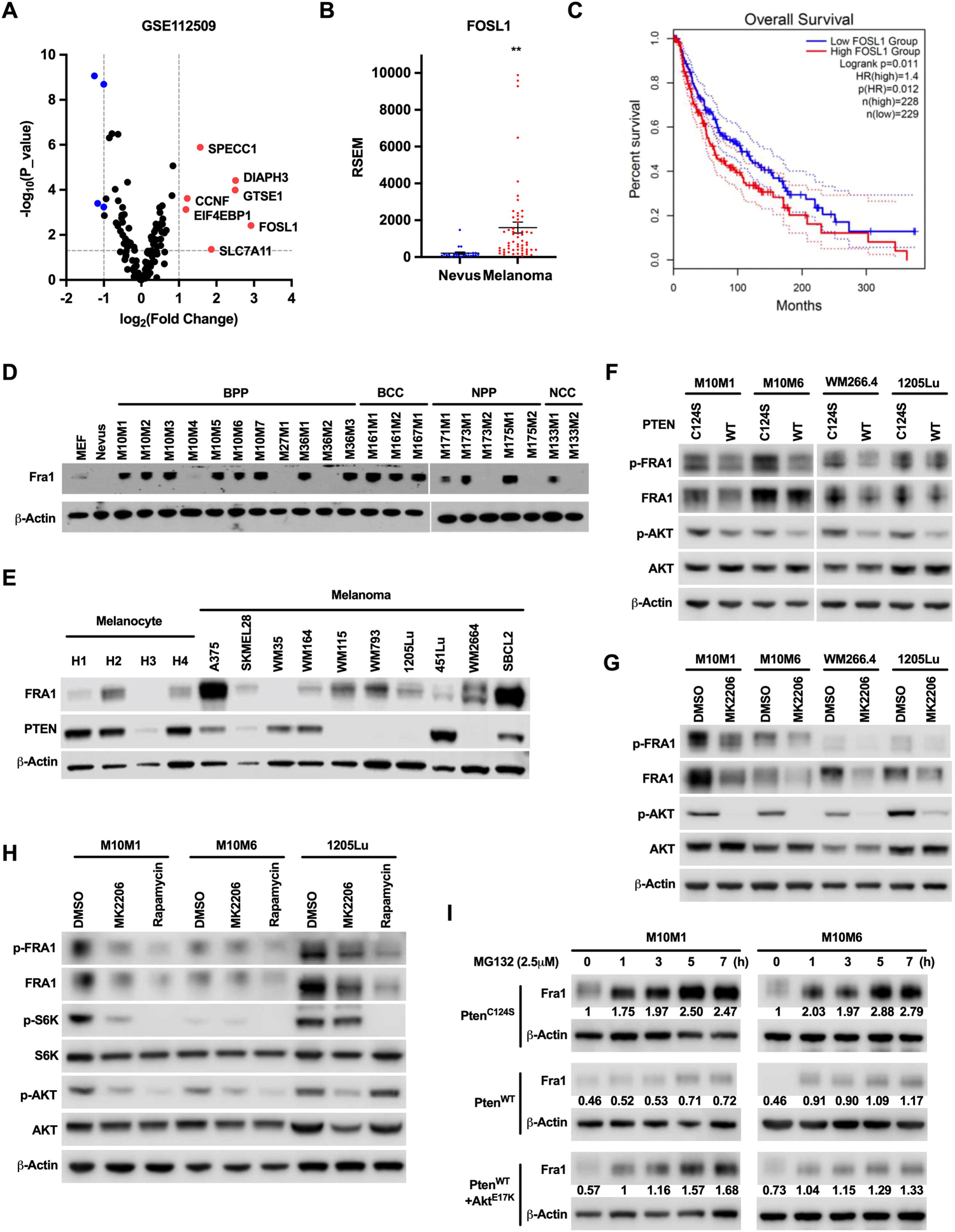
FRA1 is suppressed by PTEN through AKT-mTOR signaling. **(A)** 143 proteins were identified as potential targets of PTEN lipid phosphatase activity (see Table S3) and their differential mRNA expression between melanoma and nevi was analyzed in the GSE112509 dataset and shown as volcano plot. P-value <0.05 and fold-change >2 were used as thresholds. **(B)** Expression of FRA1 in nevi and melanoma samples in the GSE112509 dataset. **(C)** Overall survival curve of melanoma patients with high or low FRA1 expression from TCGA. **(D, E)** Western blot analysis of FRA1 protein level in mouse nevus and melanoma cells **(D)** and in human melanocytes and melanoma cells **(E)**. FRA1, PTEN, and β-Actin were detected. **(F, G)** Western blot analysis of FRA1 in mouse and human melanoma cells expressing PTEN **(F)** or treated with the AKT inhibitor MK2206 **(G)**. FRA1, p-FRA1 (Ser 265), pan-AKT, p-AKT (Ser473), and β-Actin were detected. **(H)** Western blot analysis of FRA1 expression in mouse and human melanoma cells treated with the AKT inhibitor MK2206 the mTOR inhibitor Rapamycin. FRA1, p-FRA1 (Ser 265), pan-AKT, p-AKT (Ser473), S6K, p-S6K (Thr389), and β-Actin were detected. **(I)** Western blot analysis of FRA1 protein levels in melanoma cells treated with the proteasome inhibitor MG132 for the indicated time points. FRA1 and β-Actin were detected.

The phosphoproteomic analysis identified five phosphosites within FRA1, two of which, Ser254 and Ser267 (corresponding to human Ser252 and Ser265), are associated with FRA1 protein stability (Basbous et al., 2007). It is therefore possible that the difference in phosphopeptide abundance reflects alterations in total FRA1 protein levels. Interestingly, Fra1 protein levels are increased in 15 out of 21 mouse melanoma cell lines compared to mouse nevus, while the remaining six cell lines exhibited very low Fra1 levels (**Figure 6D**). FRA1 protein levels are similarly increased in human melanoma cell lines compared to human melanocytes, and several of the cell lines exhibiting high FRA1 levels also have low or absent PTEN (**Figure 6E**). We next assessed if PTEN expression influences FRA1 abundance. Consistent with the phosphoproteomic analysis, Ser265-phosphorylated and total FRA1 levels decreased upon wildtype PTEN expression in human and mouse melanoma cell lines (**Figure 6F**). FOSL1 mRNA levels were unaffected by PTEN expression in both human and mouse melanoma cell lines, indicating post-translational regulation of FRA1 by PTEN (**Supplementary Figure 6A**). Additionally, co-immunoprecipitation assays failed to detect an interaction between PTEN and FRA1 (**Supplementary Figure 6B**), suggesting that FRA1 is not a direct substrate of PTEN. Given the role of Ser252 and Ser265 in FRA1 protein stability, we examined if PTEN expression promotes turnover of FRA1 by mediating the dephosphorylation of these sites. To this end, we treated melanoma cells with the translation inhibitor Cycloheximide and assessed FRA1 degradation in response to PTEN expression. FRA1 degradation was not accelerated by PTEN expression (**Supplementary Figure 6C**). Moreover, while active Akt^E17K^ rescued the reduction of FRA1 levels in PTEN expressing cells, it had no effect on FRA1 turnover (**Supplementary Figure 6C**). Thus, PTEN does not regulate the stability of FRA1.

To characterize the signaling pathway connecting PTEN to FRA1, we firstly treated human and mouse melanoma cell lines with MK2206 and found that Ser265-phosphorylated and total FRA1 levels were decreased by the AKT inhibitor (**Figure 6G**). Since mTOR is a major downstream effector of AKT, we assessed the effect of the mTORC1 inhibitor Rapamycin on FRA1 expression. mTOR inhibition reduced pSer265 and total FRA1 levels in human and mouse melanoma cell lines (**Figure 6H**). The mTORC1 complex is a critical regulator of protein translation and we next examined whether PTEN attenuates FRA1 protein levels by reducing its translation. Notably, following the treatment with the proteasome inhibitor MG132, FRA1 protein rapidly accumulated in Pten^C124S^ cells but not Pten^WT^ cells (**Figure 6I**). Moreover, restoring AKT signaling with Akt^E17K^ in Pten^WT^ cells promoted the accumulation of FRA1 protein in response to MG132 treatment (**Figure 6I**). These results suggest that PTEN diminishes FRA1 translation by suppressing the AKT-mTOR-axis.

### FRA1 repression mediates melanoma suppression by PTEN

We next examined whether PTEN provokes melanoma suppression by diminishing FRA1 protein levels. To this end, we overexpressed FRA1 in melanoma cells in which PTEN function was restored by expressing Pten^WT^ (**Supplementary Figure 7A)**. FRA1 overexpression was unable to rescue the suppressive effect of PTEN on cell proliferation and low-density growth (**Figure 7A,B, Supplementary Figure 7B**). By contrast, overexpression of FRA1 partially rescued the anchorage-independent growth ability of melanoma cells expressing Pten^WT^ (**Figure 7C**). Similar to Akt^E17K^, FRA1 fully restored cell invasion (**Figure 7D**) and subcutaneous tumor growth in allograft models of melanoma cells in which PTEN activity was restored (**Figure 7E,F, Supplementary Figure 3B**). Thus, FRA1 expression potently overcomes most aspects of PTEN-mediated melanoma suppression that are also overcome by restoring AKT signaling. Taken together, PTEN exerts its melanoma suppressive functions at least in part by diminishing AKT-mediated FRA1 expression.

**Figure 7.**
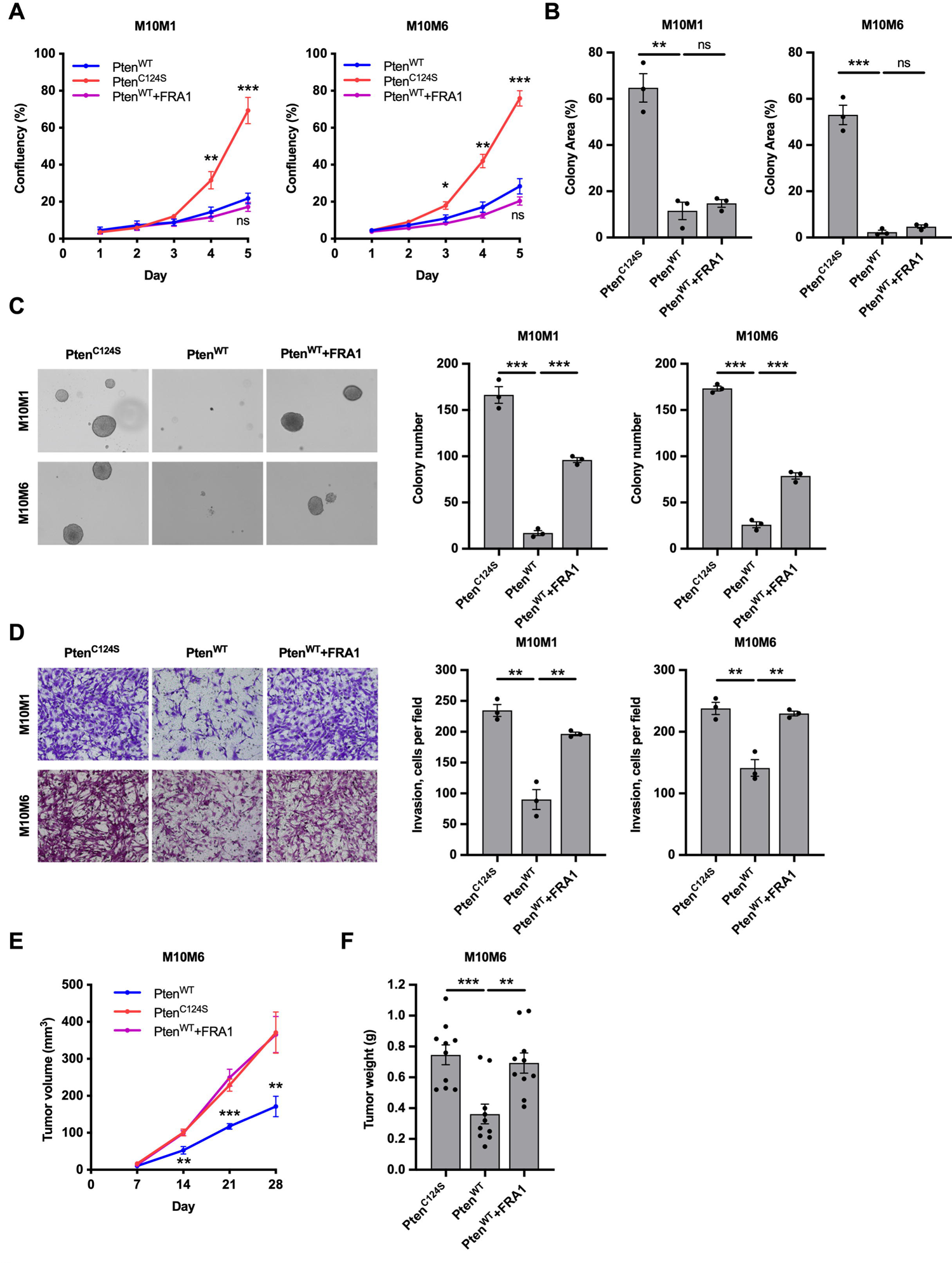
FRA1 overexpression partially abrogates melanoma suppression by PTEN. **(A)** Proliferation as measured by relative confluence of M10M6 and M10M1 cells expressing Pten^WT^ or Pten^C124S^ or Pten^WT^+FRA1. **(B)** Quantification of percentage occupied area of low-density colony formation assays of M10M6 and M10M1 cells expressing Pten^WT^ or Pten^C124S^ or Pten^WT^+FRA1. **(C)** Representative images (left) and quantification of colony number (right) of anchorage-independent growth of M10M6 and M10M1 cells expressing Pten^WT^ or Pten^C124S^ or Pten^WT^+FRA1. **(D)** Representative images (left) and quantification of cell numbers (right) of transwell invasion assays of M10M6 and M10M1 cells expressing Pten^WT^ or Pten^C124S^ or Pten^WT^+FRA1. **(E, F)** M10M6 cells expressing Pten^WT^ or Pten^C124S^ or Pten^WT^+FRA1 were subcutaneously injected into NSG mice (n=10). Mice were fed chow containing 200 mg/kg Doxycycline to induce expression of PTEN mutants. Tumor volumes were measured every 3 days. The curves of tumor volume **(E)** and the tumor weight at the end point **(F)** are shown. Mean ± SEM are shown in (A-F). Data are analyzed with Student’s unpaired *t* test, * P<0.05, ** P<0.01, *** P<0.001.

## DISCUSSION

PTEN loss-of-function occurs in up to half of all malignant melanomas, and the resulting disruptions in downstream pathways may offer exploitable vulnerabilities for targeted therapies. Although drugs targeting AKT or mTOR have been developed, their routine clinical use for treating melanoma has not been realized. In fact, the underwhelming preclinical and clinical performances of these inhibitors have prompted two fundamental questions regarding PTEN-mediated melanoma suppression. First, does PTEN suppress melanoma through its lipid phosphatase activity? And second, is AKT a crucial signaling node in PTEN-deficient melanoma? Our study provides affirmative answers to these questions, demonstrating that loss of PTEN’s lipid phosphatase activity promotes melanomagenesis predominantly through AKT activation. Moreover, we uncovered an AKT/mTOR/FRA1 axis that plays a key role in PTEN-mediated melanoma suppression.

It is widely assumed that the loss of PTEN promotes melanomagenesis through increased PIP_3_ levels and the consequential activation of downstream signaling pathways. However, this assumption discounts the contribution of the protein phosphatase and scaffold functions of PTEN, neither of which has previously been characterized in melanoma. Our collection of murine melanoma cell lines and ESC-GEMM platform (Bok et al., 2021, 2019) enabled the analysis of different PTEN functions in an otherwise PTEN deficient background, revealing lipid phosphatase activity as the predominant melanoma suppressive function. The scaffold function retained by the Pten^C124S^ mutant had no effect in vitro and in vivo, while the protein phosphatase active Pten^G129E^ mutant only moderately impacted anchorage-independent growth. Besides its cytoplasmic localization, PTEN also enters the nucleus where it controls DNA damage repair, genomic integrity, transcription, and chromatin structure (Langdon, 2023). The role of the nuclear functions of PTEN in melanoma suppression remains to be investigated. However, PTEN shifts from the nucleus to the cytoplasm in melanomagenesis (Tsao et al., 2003; Whiteman et al., 2002) and our findings highlight PTEN-mediated control of FRA1 translation via AKT/mTOR signaling, indicating that PTEN suppresses melanomagenesis primarily in the cytoplasm.

If the PTEN lipid phosphatase activity is lost in melanoma, the resulting increase in PIP_3_ levels will activate several independent signaling pathways (Lien et al., 2017). It is conceivable to these pathways work in concert to promote melanomagenesis, offering a potential explanation as to why AKT and mTOR inhibitors are not sufficient to target melanoma. However, our unbiased drug screening and phosphoproteomics approaches aimed at identifying pathways activated by the loss of PTEN lipid phosphatase activity suggested the critical importance of AKT signaling. Moreover, activated AKT was sufficient to rescue the suppression of PTEN-deficient melanoma by acute PTEN restoration. In a complementary study, Parkman et al. demonstrate that genetic silencing of all three AKT paralogs diminishes the proliferation and induces death of PTEN-deficient melanoma cells (Parkman et al., 2022). These findings support the notion that AKT is the predominant effector downstream of PIP_3_, promoting melanomagenesis upon the loss of the PTEN lipid phosphatase activity.

Interestingly, we observed different potencies of AKT in overcoming PTEN-mediated melanoma suppression. While AKT only moderately rescued the growth of melanoma cells in proliferation, low density, and anchorage-independence assays, it completely rescued cell invasion in vitro and tumor growth in allograft experiments. The effects of AKT on invasion are in line with previous reports on the pro-invasive and pro-metastatic role of AKT1 in melanoma (Cho et al., 2015; Kircher et al., 2019). We were unable to test the effect of PTEN and AKT on spontaneous metastasis in our study due to the rapid growth of the primary tumors in transplant models. Similarly, in our genetic BPP models, most tumors that developed were escapers in which ectopic PTEN was not expressed, precluding a meaningful analysis of the metastatic burden in these mice. While PTEN alterations occur late in melanomagenesis (Shain et al., 2018; Cabrita et al., 2020), the relative contribution of AKT to the metastatic spreading of PTEN-deficient melanoma remains to be investigated in more detail. While the effect of PTEN restoration is completely dependent on inactivation of AKT, other in vitro phenotypes are likely influenced by other effector pathways. Our results using our ESC-GEMM models to restore PTEN at tumor initiation further demonstrate that PTEN suppresses the early steps of melanocyte transformation and melanoma progression, which corroborates the drastically accelerated melanoma formation in Braf^V600E^; Pten^null^ mice (Dankort et al., 2009). The effects of PTEN in early melanomagenesis are also dependent on its lipid phosphatase activity, but whether AKT is required in these stages remains to be determined.

Our study and the study by Parkman et al. aimed to provide an answer to the question of whether AKT is an inadequate therapeutic target in PTEN-deficient melanoma or if AKT inhibitors lack efficacy to elicit anti-melanoma effects. Our results show AKT-independent pathway contribution to melanoma proliferation in vitro, while in vivo tumor growth in transplant models completely relies on AKT. Parkman et al. demonstrate that depletion of all three AKT paralogs kills melanoma cells and outperforms current AKT inhibitors. These findings suggest that AKT inhibitors having improved potency may perform better in the clinic. By contrast, we show that the AKT inhibitor MK2206 fully mimics the effect of PTEN restoration on cell invasion, an in vitro phenotype that is entirely dependent on AKT. Thus, whether current AKT inhibitors are simply not potent enough molecularly is difficult to reconcile, but regardless, their lack of efficacy in the clinic calls for better and improved treatment strategies. The efficacy of any future AKT inhibitors must be determined in assays that rely predominantly on AKT activity, while the effectiveness of these drugs is best evaluated in in vivo models that encompass the full spectrum of PTEN loss-induced downstream effectors. However, whether pan-AKT inhibitors that recapitulate the genetic silencing of all three AKT paralogs while eliciting low toxicity can be accomplished is questionable. Alternative treatment strategies for PTEN-deficient melanomas therefore need to be devised. One approach could be the combinatorial inhibition of AKT and parallel PIP_3_-dependent pathways that contribute to melanomagenesis. One such parallel pathway could be the SGK pathway, which has previously been implicated in melanoma (Scortegagna et al., 2015). Indeed, Parkman et al. identified a compensatory increase in SGK1 expression upon pan-AKT silencing and demonstrate superior melanoma cell killing by combined AKT and SGK inhibition (Parkman et al., 2022).

In addition to parallel pathways, downstream effectors of AKT may prove useful for combination therapy strategies. We aimed to identify such downstream effectors through phosphoproteomics, which revealed 143 candidate proteins that are potentially regulated by the PTEN lipid phosphatase activity. Only 18 of these proteins are considered to participate in canonical PI3K/AKT signaling, while 101 proteins have never or rarely (<10 studies) been associated with AKT signaling. Our strategy to prioritize candidates by analyzing their expression in melanoma revealed the AP-1 transcription factor FRA1 as the top candidate. We found remarkably elevated FRA1 protein abundance in murine and human melanoma cell lines. However, these increases also occurred in some cell lines having intact PTEN, suggesting that FRA1 is regulated by additional pathways. Indeed, ERK and PKC phosphorylate FRA1, preventing its proteasomal degradation (Belguise et al., 2012; Basbous et al., 2007). Our results instead point toward translational regulation of FRA1 via AKT and mTOR. Thus, multiple pathways converge on FRA1, enhancing FRA1 protein abundance by regulating translation and turnover. FRA1 is capable of transforming melanocytes in vitro leading to a reprogrammed cell state that even in the absence of FRA1 expression enables tumor formation in transplant models (Maurus et al., 2017). Whether FRA1 contributes to spontaneous melanoma formation in genetic mouse models and what aspects of melanomagenesis are affect by FRA1 remains to be determined. We showed that FRA1 overexpression recapitulates the effects of AKT in overcoming PTEN-mediated melanoma suppression, specifically invasion and tumor growth. FRA1 expression has also previously been associated with an invasive phenotype in melanoma cell lines (Maurus et al., 2017) and we also observed that FRA1 is associated with a melanoma metastasis signature in TCGA samples (**Supplementary Figure 7C**). These findings suggest that FRA1 is a key AKT effector expected to play important roles in the same steps of melanomagenesis as AKT. Interestingly, FRA1 upregulation is critical for the survival or MAPK inhibitor-addicted melanoma cells (Hong et al., 2017; Kong et al., 2017), indicating that FRA1 targeting may forestall or overcome acquired resistance to BRAF and MEK inhibitors. While therapeutic targeting of FRA1 will need innovative approaches (Casalino et al., 2022), early efforts with compounds that target the ERK-FRA1 interface have resulted in diminished FRA1 protein levels and enhanced apoptosis of melanoma cells (Samadani et al., 2015). Taken together, not only does the discovery of the AKT/mTOR/FRA1 axis add to our understanding of AKT-mediated melanomagenesis, but it may also offer new opportunities for targeting PTEN-deficient melanoma.

## MATERIALS AND METHODS

### Cell lines and culture condition

Mouse melanoma cell lines M10M1 and M10M6 were isolated from melanomas from Braf^V600E^; Pten^null^ (BPP) mice as described before (Bok et al., 2019). Human melanoma cell lines WM266-4 (RRID:CVCL_2765), and 1205Lu (RRID:CVCL_5239) were provided by M. Herlyn (Wistar Institute, Philadelphia, PA). A375 (RRID: CVCL_6233) and HEK293T Lenti-X were obtained from ATCC and Takara, respectively. All cell lines were cultured in a humidified atmosphere at 37°C and 5% CO2. Melanoma cell lines were cultured in RPMI1640 (Lonza) containing 5% FBS (Sigma). HEK293T Lenti-X were cultured in DMEM (VWR) containing 10% FBS. All cell lines were confirmed bimonthly to be Mycoplasma-free with the MycoAlert Mycoplasma Detection Kit (Lonza) and STR authenticated when they were initially obtained. Cell lines were used for experiments within 15 passages after thawing.

### Animal models

All animal experiments were conducted in accordance with an Institutional Animal Care and Use Committee protocol approved by the University of South Florida. The development of Braf^V600E^; Pten^null^ (BPP) ESC-GEMM approach was described previously (Bok et al., 2019). BPP ES cells were targeted with TRE-Pten^WT^ or mutant constructs (TRE-Pten^C124S^, TRE-Pten^G129E^, TRE-Pten^Y138L^) designed to co-express GFP via an IRES. To induce melanomagenesis, mice were anesthetized with Isoflurane and their back hair was shaved with clippers. 4OHT (Sigma Aldrich) dissolved in DMSO (2.5 mg/mL) was administered with a paintbrush on the back skin of 3-4 week old mice on two consecutive days. Nude mice (Foxn1^Nu/Nu^) were obtained from JAX (Stock #007850). NSG mice were obtained from JAX (Stock #005557) and bred in-house. 2.5×10^5^ M10M6 melanoma cells were subcutaneously injected into the flank of nude mice or NSG mice. Tumors were measured using calipers and volume was calculated using the formula (width^2^ x length)/2. Mice were fed chow containing 200 mg/kg Doxycycline (Envigo).

### TCGA data and GEO dataset

The GEO dataset GEO112509 containing RNA expression data from 23 nevi and 57 melanomas was used to analyze differential expression of potential targets of PTEN lipid phosphatase in nevi and melanomas using T-test. A P value <0.05 and a fold-change of Mean > 2 were used as the threshold. Gene expression data and patient follow-up data from 457 skin cutaneous melanoma (SKCM) cases in The Cancer Genome Atlas (TCGA) were analyzed for the correlation between FOSL1 expression and patient survival using Kaplan–Meier analysis.

### Gene set enrichment analysis

Global mRNA expression profiles of TCGA skin cutaneous melanoma dataset (TCGA-SKCM) were subjected to gene set enrichment analysis (GSEA; RRID:SCR_003199) to identify the association of FOSL1 with a melanoma metastasis gene signature (WINNEPENNINCKX_MELANOMA_METASTASIS_UP). For GSEA, expression of FOSL1 was used as phenotype, and “No_Collapse” was used for gene symbol. The metric for ranking genes in GSEA was set as “Pearson,” otherwise default parameters were used. GSEA was performed using GSEA 4.3.2 software.

### Plasmid and lentivirus production

pLenti-TRE-GFP-Blast was generated by replacing the CMV promoter in pLenti-GFP-Blast with the TRE3G promoter from pRRL-TRE3G-GFP-PGK-Puro-IRES-rtTA3 (from J. Zuber). pLenti-TRE-Pten^WT^-Blast and pLenti-TRE-Pten^C124S^-Blast were created by replacing GFP in pLenti-TRE-GFP-Blast with mouse Pten wildtype or mutant cDNA. pLenti-TRE-Pten^G129E^-Blast and pLenti-TRE-Pten^Y138L^-Blast were generated by site-directed mutagenesis using pLenti-TRE-Pten^WT^-Blast as a template. Q5 Site-Directed Mutagenesis Kit (New England BioLabs) was used following the manufacturer’s instruction. pLenti-CMV-Akt^E17K^-Hygro was generated by replacing GFP in pLenti-CMV-GFP-Hygro with AKT^E17K^ cDNA from pFBD-Akt1(E17K) (Addgene plasmid #86563). pLenti-CMV-Fosl1-Hygro was generated by replacing GFP in pLenti-CMV-GFP-Hygro with Fosl1 cDNA from mouse embryonic fibroblasts (MEF). To produce lentivirus supernatants, HEK293T cells were transfected with lentiviral vector and Δ8.2 and VSVG helper plasmids at a 9:8:1 ratio. Lentiviral supernatant was cleared using 0.45 µm syringe filters and used to infect melanoma cells in the presence of 8 µg/mL polybrene. Infected cells were selected with the appropriate antibiotics (1 µg/mL puromycin, 100 µg/mL hygromycin, or 10 µg/mL blasticidin).

### Proliferation assays

1×10^3^ M10M1 or M10M6 cells/well were seeded in 96 well plates in 200µL RPMI-1640 medium containing 5% FBS. After 24 hours, the plate was loaded into Cellcyte-X live cell analyzer (ECHO). Images of each well were taken daily for 4-5 days and the cell confluency of each image was quantified.

### Low density growth assays

0.5×10^3^ M10M1 or M10M6 cells were seeded in 6-well plates in triplicates and incubated for 8 days. Cells were fixed in cold 4% paraformaldehyde (VWR) and stained with 0.5% crystal violet (VWR). Percent area of crystal violet staining was quantified using ImageJ.

### Soft agar assays

1.6% SeaPlaque Agarose in PBS was mixed 1:1 with RPMI-1640 and 1mL of the 0.8% agarose was used to pour a bottom layer in 6-well plates. Melanoma cells were trypsinized, counted, and 5×10^3^ cells were resuspended in 1 mL of RPMI-1640 medium containing 20% FBS. Cells were then mixed with 1 mL of 0.8% SeaPlaque Agarose in PBS and plated atop the solidified agar. The top layer was allowed to solidify at room temperature, followed by culture at 37°C with 100 µL of fresh medium added every 3 days. Pictures were taken after 10 to 15 days and analyzed with ImageJ.

### Transwell invasion assays

To pre-coat transwells, Matrigel (Corning) was thawed on ice for at least 2 hours, diluted to 5% with ice-cold RPMI-1640 medium, and gently added to transwell inserts (50 µL/insert) and solidified at 37°C for 30 min. Melanoma cells were trypsinized and resuspend in RPMI-1640 medium. For transwell invasion assays, 2×10^4^ M10M1 or M10M6 cells were plated per Matrigel-coated insert in 200 µl of RPMI-1640 without FBS. 500 µl RPMI-1640 medium containing 15% FBS were added into the bottom well. Plates were incubated at 37°C in a humidified incubator for 48 hr. Media was discarded, inserts were gently washed once with PBS, and cells were fixed with 500 µl fixing solution (ethanol:acetic acid = 3:1) in the bottom well for 10 minutes. Inserts were washed once with PBS, and cells were stained in 500 µL staining solution (0.5% crystal violet) for 30 minutes. Inserts were washed with tap water twice and non-migrated cells or Matrigel on the top side of the insert were carefully removed with cotton swabs, and inserts were air dried overnight. Pictures were taken at 200x magnification and cell numbers quantified.

### Small molecule inhibitor library screening

250 M10M6-Pten^C124S^ cells and 750 M10M6-Pten^WT^ cells were seeded in 384 well plate with 40 µL RPMI-1640 medium containing 5% FBS using Bravo automated liquid handling platform (Agilent). After 24 hours, the drug library was added to 40 µL cells with final concentrations of 0.1 µM, 0.5 µM, 2.5 µM, and 10 µM drug using Echo650 (Labcyte/Beckman Coulter). The same volume of DMSO was used as negative control. After 5 days, 10 µL CellTiter Glo (Promega) were added to 40µL cells and gently shaken at room temperature for 10 minutes, luminescence was recorded using a PerkinElmer EnVision plate reader. Cell viability = [luminescence (drug) / luminescence (DMSO)] x 100%. Sensitivity = [100% - (cell viability)] / 100%.

### Phosphoproteomics

Cells were harvested, lysed with urea lysis buffer, sonicated using a microtip sonicator, and digested using the PTMScan Kit Protocol (Cell Signaling). Peptide purification was performed using Sep-Pak® C18 columns (Waters). Eluates were collected in the same tube and then aliquoted for pY enrichment (1 mg per sample) or global phosphoproteomics (200 μg per sample). Samples were frozen at −80°C overnight and lyophilized for two days to ensure that there was no TFA left in the samples. Phosphotyrosine-containing peptides were immunoprecipitated using PTMScan p-Tyr-1000 immunoaffinity beads (Cell Signaling #8803). Subsequently, samples were labeled using TMT reagents according to the manufacturer instructions (TMTpro™ 16plex Label Reagent Set, # A44521). Labeling efficiency was confirmed by MS. After sample combination and lyophilization, peptides were redissolved in 250 μl of 20 mM ammonium formate buffer (pH 10.0). Basic pH reversed-phase liquid chromatography separation was performed on an XBridge column (Waters). Fifteen concatenated peptide fractions were dried by vacuum. TMT channel 16 was used for boosting of pY signals with a 10 mg bulk GFP sample and channel 14 was left empty (Fang et al., 2020). Peptides for global phosphoproteomics (pSTY) were labeled using TMT 16-plex reagents as described above. Phosphopeptide enrichment was achieved using an IMAC enrichment kit (Sigma I1408). Samples were evaporated using a speed vacuum centrifuge and resuspended in 2% ACN / 0.1% FA, which contained 50 fmol/μl of PRTC (Thermo Scientific Pierce Retention Time Calibration Mixture, PIERCE) to confirm consistent performance of the LC-MS analyses. The acquired pY and IMAC data were searched with MaxQuant with the murine UniProt database using the embedded search engine, Andromeda. Carbamidomethylated cysteines were set as fixed modification and oxidation of methionine, N-terminal protein acetylation, and phosphorylation of serine, threonine, and tyrosine as variable modifications. Further, the MaxQuant initial search precursor and fragment ion tolerance were set to 20 ppm and m/z 0.05, respectively. The MaxQuant main search precursor ion mass tolerance was 4.5 ppm. Resolution was set to 70,000 at 200 m/z for MS1 and 17,500 for MS2. MaxQuant automatically filters out any TMT-labeled peptides that are isolated with less than 75% purity. The data were then filtered for 1% protein FDR, plus common contaminants. For the analysis and comparison of TMT 16-plex global pSTY data and pY data, the reporter ion intensity was used for the relative quantification of each peptide. IRON (iterative rank-order normalization) of MaxQuant data was performed as described before (Welsh et al., 2013). The experiment was performed with three biological replicates. Each treated sample was run as two technical replicates. Biological triplicates were then averaged, and the log2 ratio was determined between the samples. The phosphorylated peptides were filtered for further analysis with absolute value of log2 fold change (> 0.585) and p value (<0.05). KEGG pathway enrichment analysis of differentially phosphorylated proteins from phosphoproteomics data sets was performed with Enrichr (https://maayanlab.cloud/Enrichr/)(3, 4, 5). Volcano Plots were created by GraphPad Prism 9.

### qRT-PCR

RNA was extracted from cells using TRIzol (Invitrogen) following protocols supplied by the manufacturer. cDNA was generated with PrimeScript RT Master Mix (Takara). qPCR was performed on StepOnePlus Real-Time PCR System (Thermo Fisher Scientific) using PerfeCTa SYBR Green Fastmix (Quantabio). mRNA expression was normalized to 18S rRNA. Primer sequences were obtained from PrimerBank (https://pga.mgh.harvard.edu/primerbank/index.html; RRID:SCR_006898): Mouse Fosl1 forward: ATGTACCGAGACTACGGGGAA. Mouse Fosl1 reverse: CTGCTGCTGTCGATGCTTG. Human FOSL1 forward: CAGGCGGAGACTGACAAACTG. Human FOSL1 reverse: TCCTTCCGGGATTTTGCAGAT. RNA18S forward: CTAAATACCGGCACGAGACC. RNA18S reverse: TTCACGCCCTCTTGAACTCT.

### Nuclear/Cytoplasm Co-Immunoprecipitation

Nuclear and cytoplasmic fractions of A375 cells were lysed and isolated using NE-PER Nuclear and Cytoplasmic Extraction Kit (Thermo Fisher Scientific). Nuclear and cytoplasmic lysates were incubated with PTEN antibody (1:100) at 4°C overnight. 20 µL Pierce™ Protein A/G Magnetic Beads (Thermo Fisher Scientific) were washed with TBST (Tris-buffered saline containing 0.05% Tween-20) and incubated with lysate-antibody solution at 4°C for 1 hour. Supernatants were discarded and the beads were washed 3 times with TBST. Protein bound to the beads were eluted with SDS-PAGE sample buffer and subjected to Western Blot analysis.

### Western blot analysis

15µg of total protein were separated on NuPAGE 4-12% precast gels (Thermo Fisher Scientific) and transferred to nitrocellulose membranes. Membranes were blocked in 5% non-fat dry milk in TBST and incubated with one of the following primary antibodies overnight at 4°C: PTEN (1:1,000), phospho-AKT (Ser 473) (1:1,000), AKT (pan) (1:1,000), phospho-p70 S6 kinase (Thr389) (1:1,000), p70 S6 kinase (1:1,000), phospho-S6 ribosomal protein (Ser235/236) (1:1000), S6 ribosomal protein (1:1000), phospho-FAK (Try397) (1:1,000), phospho-PRAS40 (Thr246) (1:1000), PRASS40 (1:1000), FAK (1:1000), phospho-FRA1 (Ser 265) (1:1000), or FRA1 (1:1,000). β-Actin (1:3,000) was blotted as a loading control. Membranes were washed 3 times with TBST for 10 min, followed by incubation with HRP-conjugated secondary antibodies (1:3,000) for 1 hr at room temperature. After three washes in TBST, chemiluminescence substrate (1:1) was applied to the blots for 4 min and chemiluminescence signal was captured using a LI-COR imaging system.

### Quantification and statistical analysis

Proliferation assays, low density growth assays, and soft agar assays are presented as mean ± SEM. For transwell invasion assays, three or four random fields were quantified for each well and data are presented as mean ± SEM. All experiments were performed at least three times with three to four technical replicates. Statistical analyses were performed by Student’s unpaired t test. Data of subcutaneous tumor growth curve and tumor weight at endpoint are presented as mean ± SEM. Graphpad Prism 9 (RRID:SCR_002798) was used for statistical analyses and a P value of <0.05 was considered statistically significant.

### Data availability

The data generated in this study are available within the article and its supplementary data files.

## Supporting information

Supplementary Table 2 (pSTY)

## ACKNOWLEDGMENTS

We are grateful to Karreth lab members for helpful discussions. This work was supported by a Miles for Moffitt Award, a Harry J. Lloyd Charitable Trust Career Development Award, and a Bankhead-Coley Grant from the Florida Department of Health (F. A. Karreth). This work was also supported by the Gene Targeting Core, the Proteomics Core, and the Biostatistics and Bioinformatics Shared Resource, which are funded in part by Moffitt’s Cancer Center Support Grant (P30CA076292).

## Author Contributions

X. Xu, I. Bok. And F. A. Karreth conceived the study and designed all experiments. X. Xu and I. Bok performed all experiments with contributions from N. Jasani, K. Wang., and M. Chadourne. O. Deng performed the phosphoproteomics with I. Bok and the drug library screen with X. Xu. U. Rix helped design and supervised phosphoproteomics and the drug library screen and provided the COCTAIL 2.0 library. O. Deng and U. Rix analyzed the drug screen results. E. A. Welsh analyzed the phosphoproteomics results. X. Xu, I. Bok., and F. A. Karreth analyzed all data. X. Xu and F. A. Karreth wrote the manuscript with input from all authors. F. A. Karreth supervised the study and acquired funding.

## SUPPLEMENTARY FIGURE LEGENDS

**Supplementary Figure 1.**
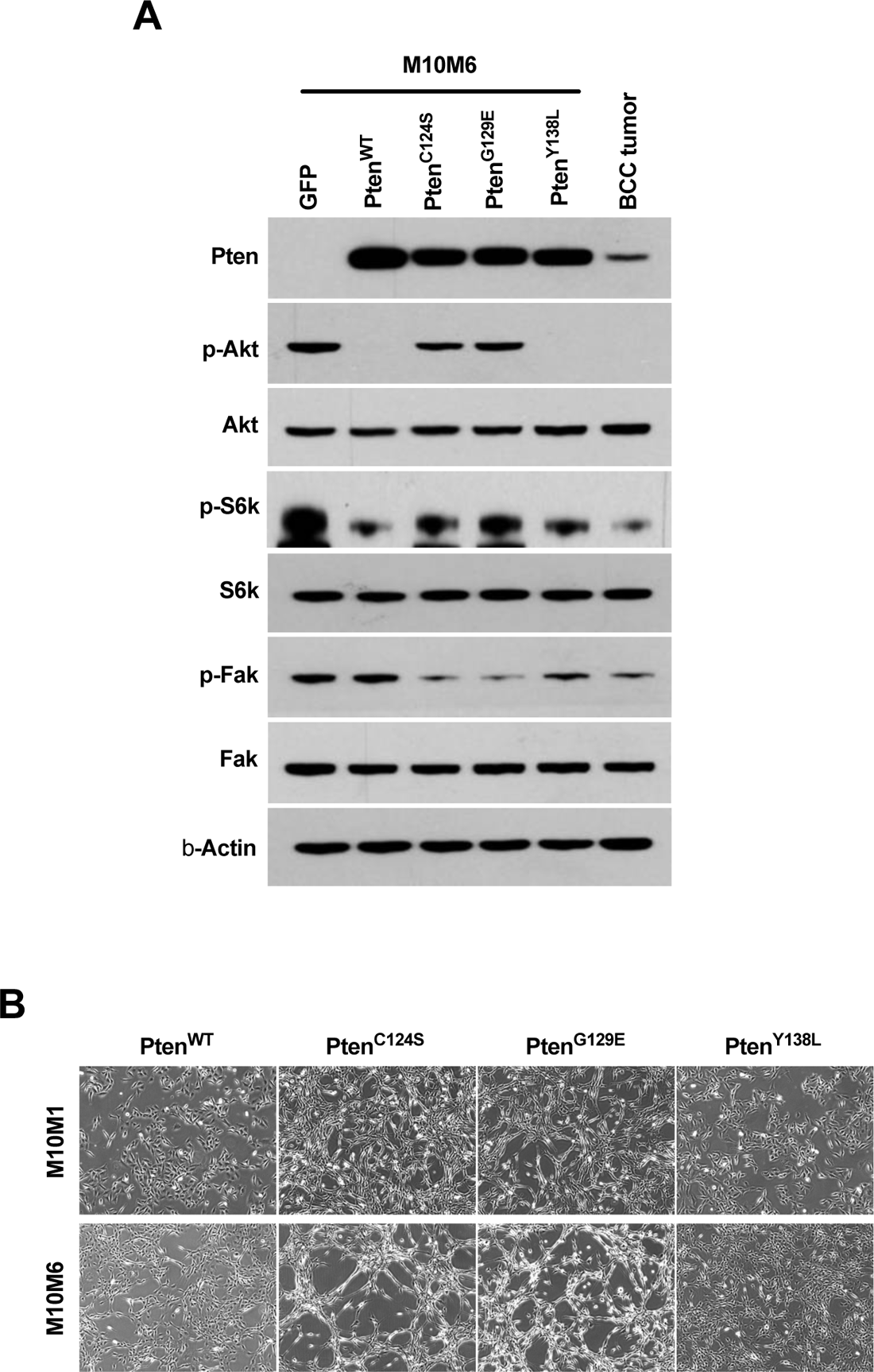
**(A)** Western blot confirming the expression and activity of different PTEN mutants expressed in M10M6 cells. Pten, p-AKT (Ser473), pan-Akt, p-S6k (Thr389), S6K, p-Fak (Tyr397), Fak and β-Actin were detected. Pten^WT^ and Pten^Y138L^ reduce p-Akt and p-S6k, indicative of lipid phosphatase activity. p-Fak was used as an indicator of protein phosphatase activity, but neither Pten^WT^ nor Pten^G138E^ reduce p-Fak levels suggesting that PTEN does not dephosphorylate the Tyr397 site of FAK in melanoma. **(B)** Representative images showing the morphology of M10M6 and M10M1 cells expressing PTEN constructs.

**Supplementary Figure 2.**
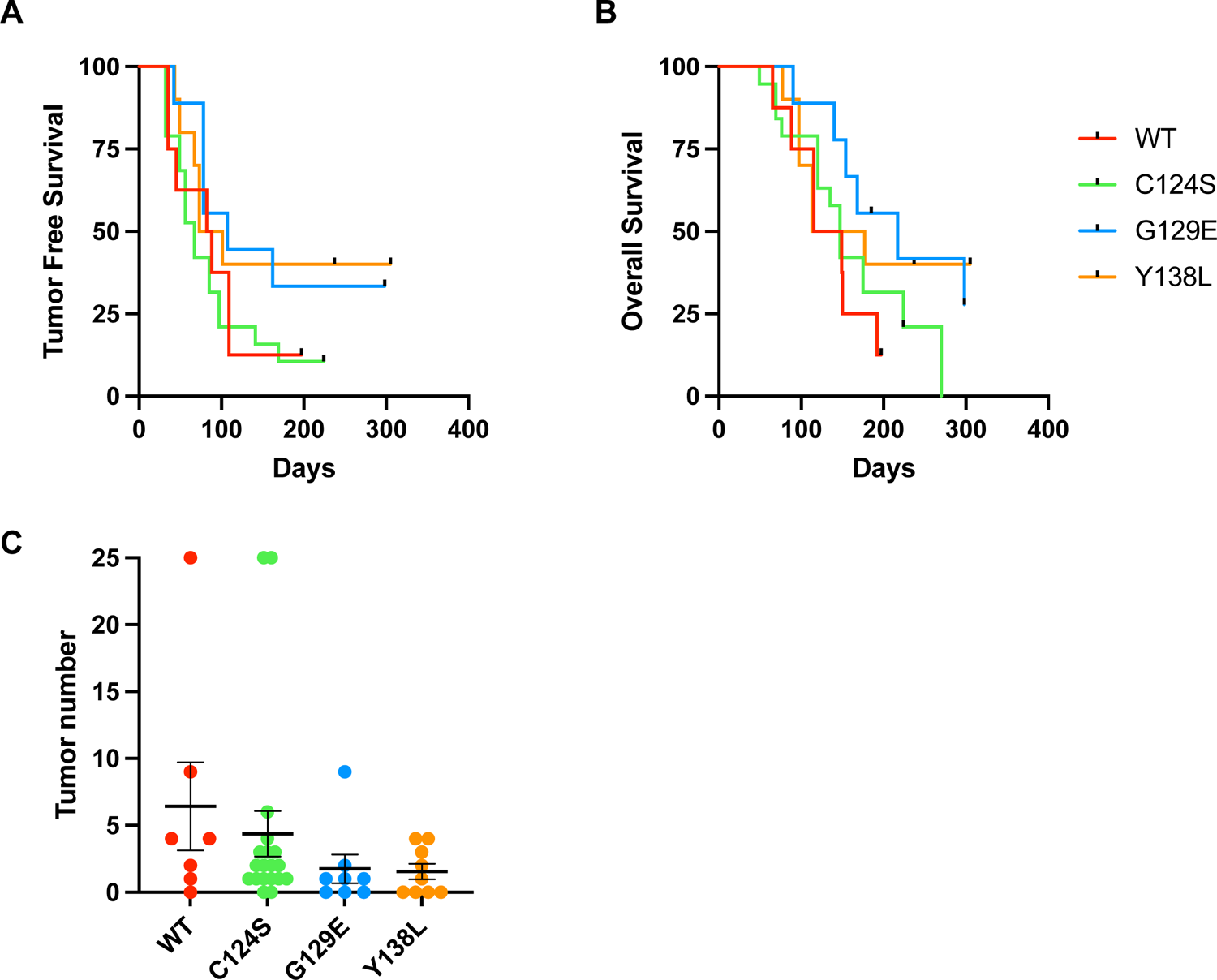
**(A, B)** Tumor-free survival curve **(A)** and overall survival **(B)** of Braf^V600E^; Pten^null^ (BPP) mice expressing PTEN mutants. **(C)** Tumor numbers of each cohort are shown.

**Supplementary Figure 3.**
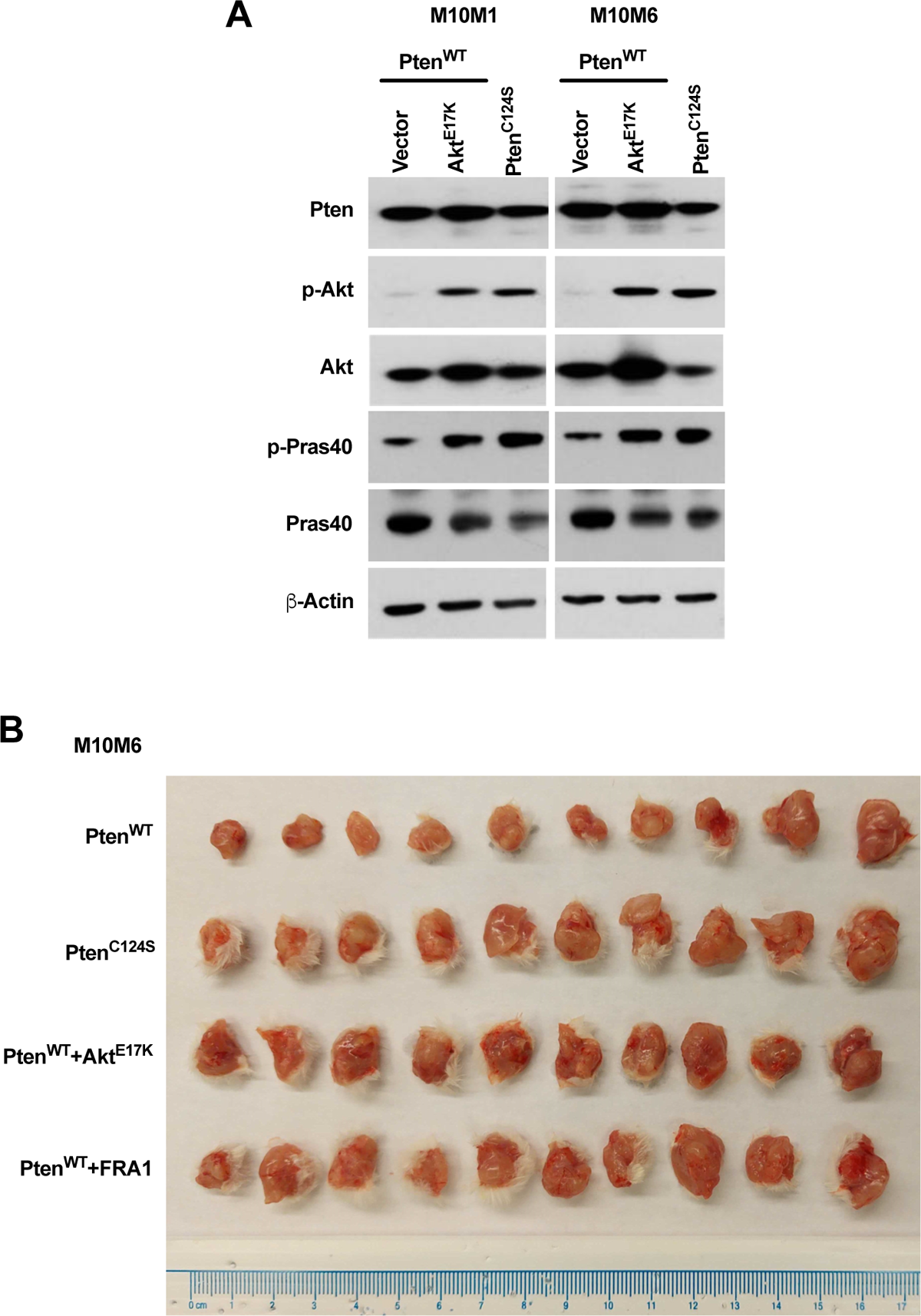
**(A)** Western blot showing AKT activation in M10M1 and M10M6 cells expressing Pten^WT^, Pten^C124S^, or Pten^WT^+Akt^E17K^. Pten, p-Akt (Ser473), pan-Akt, p-Pras40 (Thr246), Pras40, and β-Actin were detected. **(B)** Subcutaneous tumors from the indicated cell lines at end point are shown.

**Supplementary Figure 4.**
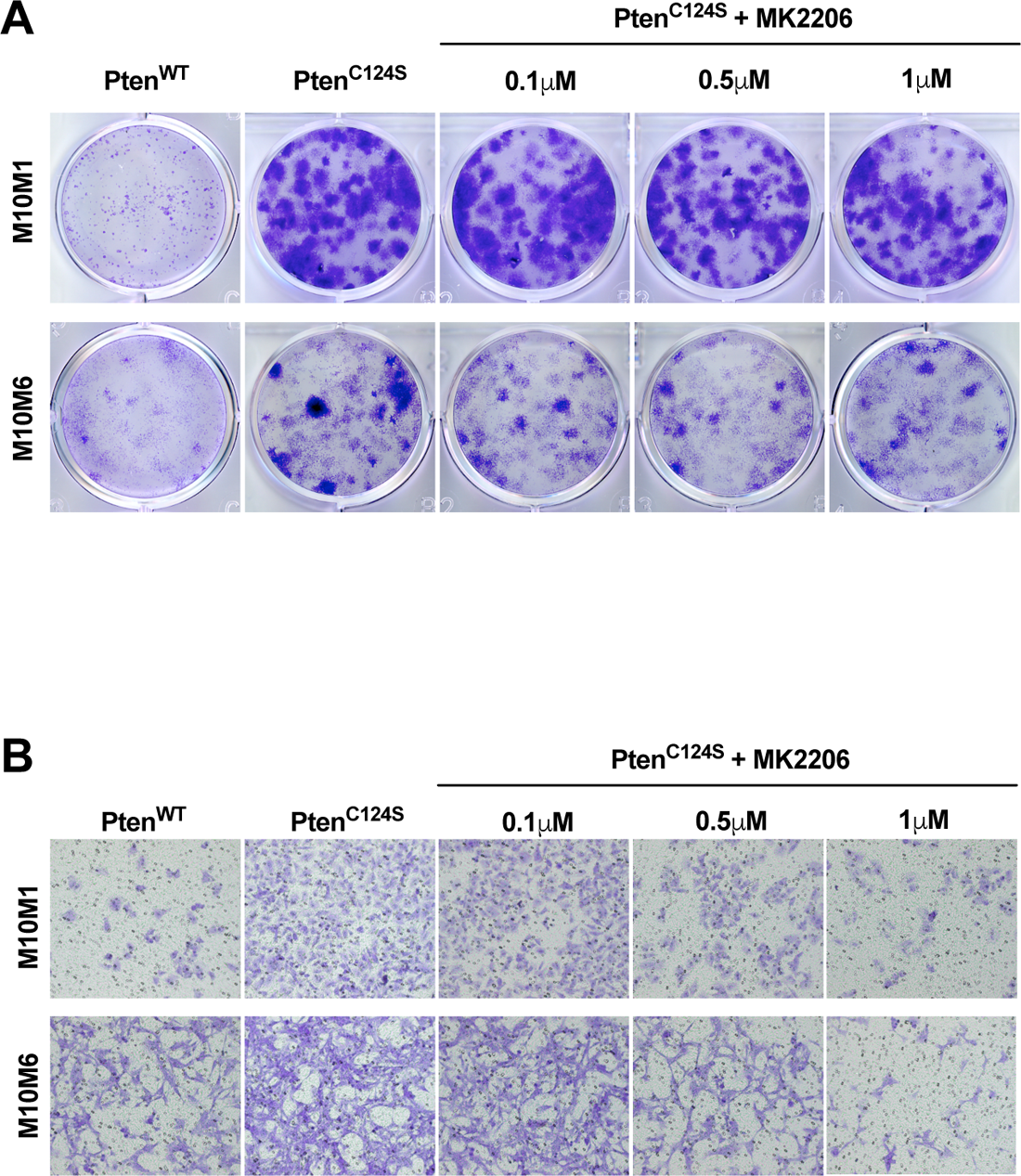
**(A)** Representative images of low-density colony formation assays of M10M1 and M10M6 melanoma cells expressing Pten^WT^, or Pten^C124S^ treated with the indicated doses of MK2206. **(B)** Representative images of cell numbers of transwell invasion assays of M10M1 and M10M6 melanoma cells expressing Pten^WT^, or Pten^C124S^ treated with the indicated doses of MK2206.

**Supplementary Figure 5.**
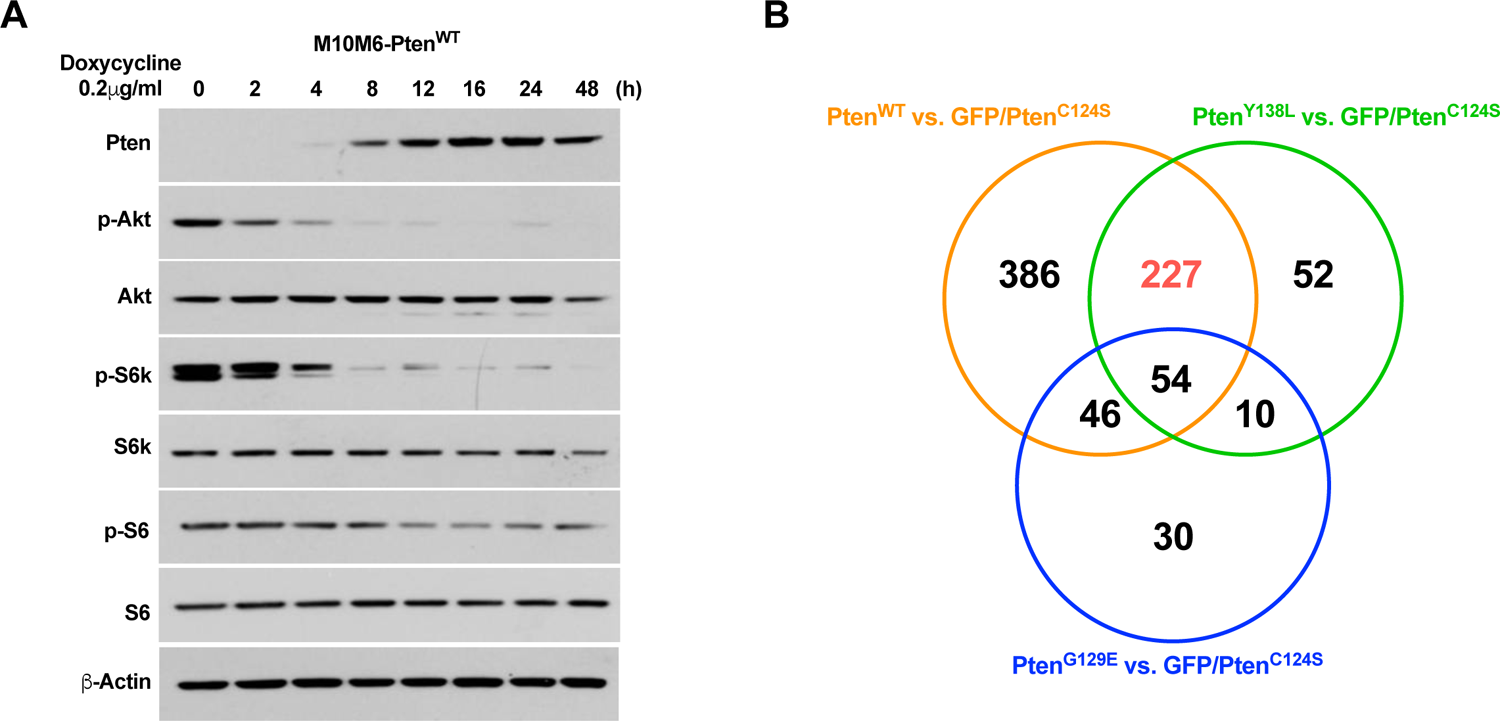
**(A)** Western blot showing AKT pathway inhibition in M10M6 melanoma cells treated with Doxycycline for the indicated time points to induce ectopic PTEN expression. Pten, p-Akt (Ser473), pan-Akt, p-S6k (Thr389), S6k, p-S6 (Ser235/236), S6, and β-Actin were detected. **(B)** Venn diagram showing the overlapping genes in Figure 5C-E.

**Supplementary Figure 6.**
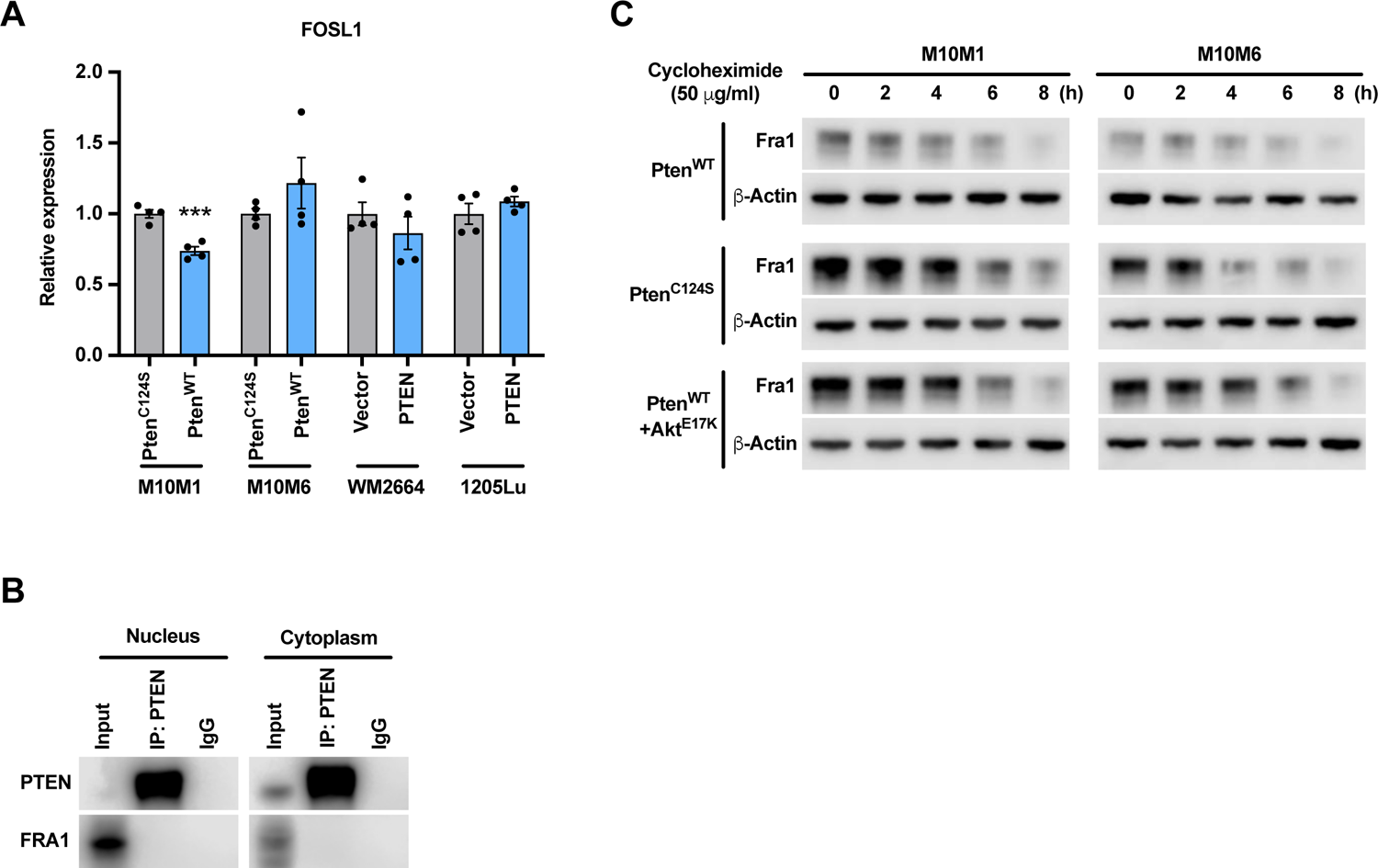
**(A)** qRT-PCR analysis of FOSL1 mRNA levels in mouse and human melanoma cell lines expressing Pten^WT^ or Pten^C124S^. **(B)** Co-IP analysis in nuclear and cytoplasmic fractions of A375 cells showed FRA1 does not directly bind to PTEN. **(C)** Western blot analysis of FRA1 protein level in M10M1 and M10M6 cells expressing Pten^WT^, Pten^C124S^, or Pten^WT^+Akt^E17K^ and treated with Cycloheximide for the indicated time points. Fra1 and β-Actin were detected.

**Supplementary Figure 7.**
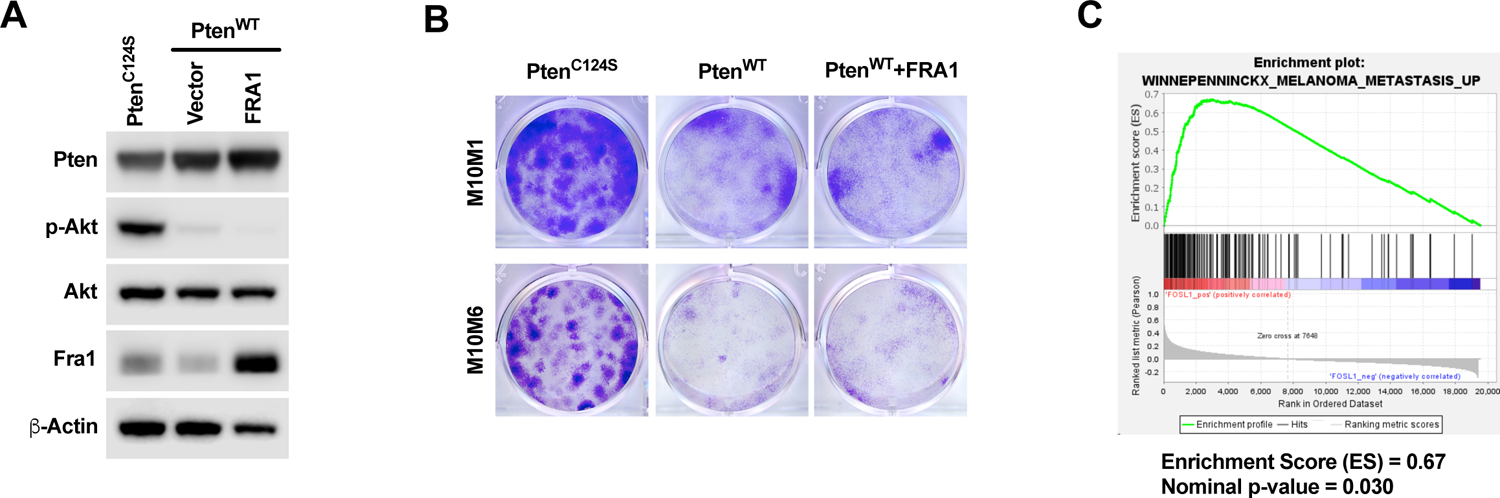
**(A)** Western blot confirming FRA1 overexpression in M10M1 cells expressing Pten^WT^, Pten^C124S^, or Pten^WT^+FRA1. **(B)** Representative images of low-density colony formation assays of M10M1 and M10M6 melanoma cells expressing Pten^WT^, Pten^C124S^, or Pten^WT^+FRA1. **(C)** GSEA of TCGA-SKCM samples associates FOSL1 expression with a melanoma metastasis gene signature.

## Notes

### Competing Interest Statement

The authors have declared no competing interest.

