## Supplementary Table 1 for "The lipid phosphatase activity of PTEN dampens FRA1 expression via AKT/mTOR signaling to suppress melanoma"

| Drug, DMSO or aq=H20 | Alt Name | Target 1 | Target | Target | Target | M10M6-WT 0.1uM | M10M6-WT 0.5uM | M10M6-WT 2.5uM | M10M6-WT 10uM | M10M6-C124S 0.1uM | M10M6-C124S 0.5uM | M10M6-C124S 2.5uM | M10M6-C124S 10uM | C124S/wt_0.1 | C124S/wt_0.5 | C124S/wt_2.5 | C124S/wt_10 |
|---|---|---|---|---|---|---|---|---|---|---|---|---|---|---|---|---|---|
| AZD4635 | HTL1071 | A2A Receptor | | | | 94.40 | 97.32 | 100.29 | 90.54 | 101.11 | 101.15 | 102.59 | 97.79 | 0.93 | 0.96 | 0.98 | 0.93 |
| XMD16-5 | | ACK1 | | | | 100.35 | 92.53 | 46.32 | 36.14 | 96.83 | 95.59 | 36.42 | 20.49 | 1.04 | 0.97 | 1.27 | 1.76 |
| K02288 | | ACVR1 (ALK2) | ACVRL | BMPR1B (ALK | | 104.38 | 100.86 | 86.43 | 44.65 | 100.44 | 96.01 | 92.77 | 50.45 | 1.04 | 1.05 | 0.93 | 0.88 |
| LDN-193189, aq | | ACVR1 (ALK2) | | | | 96.66 | 99.78 | 0.57 | 110.28 | 109.07 | 97.41 | 0.58 | 111.73 | 0.89 | 1.02 | 0.99 | 0.99 |
| Afuresertib | GSK2110183 | AKT | | | | 93.75 | 88.69 | 76.09 | 0.73 | 77.24 | 64.10 | 35.83 | 0.55 | 1.21 | 1.38 | 2.12 | 1.32 |
| Uprosertib | GSK2141795, GS | AKT | | | | 94.73 | 84.94 | 73.96 | 0.75 | 68.53 | 67.60 | 33.04 | 0.46 | 1.38 | 1.26 | 2.24 | 1.66 |
| Miransertib | ARQ092 | AKT | | | | 95.70 | 86.81 | 67.04 | 0.50 | 83.92 | 70.98 | 44.62 | 0.49 | 1.14 | 1.22 | 1.50 | 1.02 |
| Ipatasertib | GDC-0068 | AKT | | | | 98.43 | 97.22 | 93.78 | 83.13 | 93.85 | 86.07 | 56.69 | 46.37 | 1.05 | 1.13 | 1.65 | 1.79 |
| Capivasertib | AZD-5363, AZD5363 | AKT | | | | 98.09 | 101.90 | 99.39 | 75.09 | 91.66 | 91.03 | 79.54 | 40.96 | 1.07 | 1.12 | 1.25 | 1.83 |
| GSK690693 | | AKT | | | | 104.67 | 92.63 | 96.18 | 92.28 | 93.44 | 93.56 | 72.31 | 44.19 | 1.12 | 0.99 | 1.33 | 2.09 |
| MK-2206 | | AKT | | | | 94.76 | 82.73 | 77.01 | 0.55 | 96.70 | 87.62 | 68.37 | 0.48 | 0.98 | 0.94 | 1.13 | 1.16 |
| NCT-501 | | ALDH1A | | | | 101.40 | 103.42 | 94.29 | 77.71 | 103.09 | 103.11 | 92.75 | 94.07 | 0.98 | 1.00 | 1.02 | 0.83 |
| GSK1838705A | | ALK | IGF1R | IR | | 102.10 | 101.40 | 82.28 | 0.42 | 103.45 | 95.52 | 0.37 | 0.29 | 0.99 | 1.06 | 221.55 | 1.45 |
| Brigatinib | AP26113 | ALK | ROS1 | EGFR | | 104.42 | 106.18 | 94.93 | 85.41 | 92.06 | 100.10 | 88.61 | 52.65 | 1.13 | 1.06 | 1.07 | 1.62 |
| Ensartinib | X-396 | ALK | ROS1 | TRKA-C | | 95.25 | 85.71 | 28.85 | 6.08 | 98.95 | 81.66 | 24.12 | 2.25 | 0.96 | 1.05 | 1.20 | 2.70 |
| Lorlatinib | PF-06463922 | ALK | ROS1 | | | 87.70 | 83.29 | 70.53 | 34.65 | 80.11 | 80.34 | 65.31 | 31.06 | 1.09 | 1.04 | 1.08 | 1.12 |
| Alectinib | CH5424802 | ALK | | | | 102.35 | 95.54 | 68.74 | 42.00 | 92.10 | 92.18 | 35.23 | 2.64 | 1.11 | 1.04 | 1.95 | 15.89 |
| Belizatinib | TSR-011 | ALK | TRKA-C | | | 101.16 | 98.87 | 90.36 | 22.50 | 102.64 | 96.67 | 89.87 | 42.15 | 0.99 | 1.02 | 1.01 | 0.53 |
| Ceritinib | LDK378 | ALK | IGF1R | | | 92.71 | 88.45 | 0.51 | 0.53 | 94.66 | 92.37 | 0.57 | 0.51 | 0.98 | 0.96 | 0.89 | 1.03 |
| Crizotinib | PF-02341066 | ALK | MET | | | 74.50 | 76.03 | 40.23 | 0.36 | 99.19 | 90.67 | 38.05 | 0.44 | 0.75 | 0.84 | 1.06 | 0.83 |
| Levamisole, aq | | Alkaline phosphatase | | | | 91.35 | 91.46 | 98.99 | 97.58 | 102.38 | 89.83 | 97.51 | 104.18 | 0.89 | 1.02 | 1.02 | 0.94 |
| MK-3903 | | AMPK activator | | | | 98.28 | 91.50 | 71.41 | 42.84 | 100.34 | 96.67 | 77.94 | 43.14 | 0.98 | 0.95 | 0.92 | 0.99 |
| PF-06409577 | | AMPK activator | | | | 94.09 | 89.30 | 81.92 | 65.83 | 97.65 | 99.26 | 91.70 | 64.95 | 0.96 | 0.90 | 0.89 | 1.01 |
| Eprosartan Mesylate | | Angiotensin II R | | | | 107.88 | 100.06 | 102.73 | 101.98 | 103.41 | 100.06 | 96.12 | 95.15 | 1.04 | 1.00 | 1.07 | 1.07 |
| TAME | | APC | | | | 101.36 | 91.30 | 96.46 | 93.65 | 102.16 | 96.21 | 95.03 | 94.58 | 0.99 | 0.95 | 1.02 | 0.99 |
| Selonsertib | GS-4997 | ASK1 | | | | 98.09 | 93.13 | 88.32 | 77.42 | 97.21 | 99.91 | 99.30 | 87.91 | 1.01 | 0.93 | 0.89 | 0.88 |
| AZD1390 | | ATM | | | | 99.31 | 95.24 | 89.02 | 64.48 | 70.12 | 77.41 | 76.13 | 49.13 | 1.42 | 1.23 | 1.17 | 1.31 |
| AZD0156 | | ATM | | | | 107.21 | 79.83 | 49.82 | 34.65 | 99.98 | 68.93 | 35.08 | 24.21 | 1.07 | 1.16 | 1.42 | 1.43 |
| KU60019 | | ATM | | | | 100.31 | 96.79 | 86.48 | 53.95 | 103.34 | 97.03 | 70.73 | 15.38 | 0.97 | 1.00 | 1.22 | 3.51 |
| Berzosertib | VX-970, VE-822, I | ATR | ATM | | | 86.59 | 36.38 | 1.62 | 0.42 | 36.10 | 11.28 | 0.43 | 0.39 | 2.40 | 3.22 | 3.79 | 1.10 |
| BAY-1895344 | BAY 1895344 | ATR | | | | 19.50 | 12.40 | 7.48 | 3.74 | 15.95 | 8.89 | 4.66 | 2.48 | 1.22 | 1.39 | 1.60 | 1.51 |
| Ceralasertib | AZD6738 | ATR | | | | 94.57 | 89.04 | 42.03 | 11.13 | 95.54 | 84.87 | 33.91 | 11.77 | 0.99 | 1.05 | 1.24 | 0.95 |
| CYC-116 | | AURKA | AURKI | VEGFR | | 108.37 | 73.43 | 36.65 | 3.09 | 102.06 | 45.80 | 22.49 | 1.18 | 1.06 | 1.60 | 1.63 | 2.62 |
| AMG 900 | | AURKA | AURKI | AURKC | | 37.53 | 36.95 | 33.83 | 22.63 | 25.95 | 28.66 | 25.97 | 18.00 | 1.45 | 1.29 | 1.30 | 1.26 |
| Tozasertib | VX-680, MK-045 | AURKA | AURKI | BCR-ABL | | 78.09 | 48.88 | 30.62 | 24.28 | 75.12 | 47.56 | 29.21 | 16.35 | 1.04 | 1.03 | 1.05 | 1.49 |
| ENMD-2076 | | AURKA | SRC | | | 84.72 | 72.03 | 34.74 | 0.45 | 101.46 | 72.63 | 20.51 | 0.48 | 0.84 | 0.99 | 1.69 | 0.93 |
| MK-5108 | | AURKA | | | | 91.30 | 72.88 | 48.87 | 40.99 | 98.51 | 74.42 | 34.41 | 27.14 | 0.93 | 0.98 | 1.42 | 1.51 |
| Alisertib | MLN-8237 | AURKA | | | | 38.21 | 21.54 | 7.85 | 9.76 | 32.87 | 22.22 | 14.45 | 15.29 | 1.16 | 0.97 | 0.54 | 0.64 |
| LY-3295668 | AK-01 | AURKA | | | | 86.25 | 65.02 | 52.19 | 49.54 | 100.73 | 73.56 | 56.22 | 61.83 | 0.86 | 0.88 | 0.93 | 0.80 |
| Danusertib | | AURKA | BCR-ABL | | | 93.89 | 73.75 | 59.51 | 38.33 | 101.48 | 84.14 | 52.18 | 25.40 | 0.93 | 0.88 | 1.14 | 1.51 |
| Barasertib | AZD1152-HQPA( | AURKB | | | | 105.90 | 81.88 | 64.20 | 65.38 | 96.49 | 84.49 | 43.51 | 34.52 | 1.10 | 0.97 | 1.48 | 1.89 |
| TAK-901 | | AURKB | | | | 93.71 | 83.43 | 64.19 | 30.43 | 98.19 | 94.09 | 54.26 | 19.63 | 0.95 | 0.89 | 1.18 | 1.55 |
| Bemcentinib | BGB324, R428 | AXL | | | | 102.39 | 59.89 | 0.52 | 0.52 | 102.86 | 36.35 | 0.50 | 0.52 | 1.00 | 1.65 | 1.04 | 1.00 |
| BCL201 | | BCL2 | | | | 103.31 | 103.58 | 83.65 | 80.14 | 100.10 | 93.69 | 83.89 | 69.05 | 1.03 | 1.11 | 1.00 | 1.16 |
| S55746 | | BCL2 | | | | 99.29 | 100.19 | 74.31 | 62.92 | 105.35 | 93.20 | 80.45 | 68.68 | 0.94 | 1.08 | 0.92 | 0.92 |
| AZD4320 | | BCL2 | BCL-XL | | | 104.88 | 98.69 | 93.19 | 70.79 | 96.37 | 98.10 | 95.73 | 71.45 | 1.09 | 1.01 | 0.97 | 0.99 |
| Navitoclax | ABT-263 | BCL2 | | | | 98.09 | 79.83 | 63.09 | 0.50 | 103.77 | 94.63 | 77.04 | 0.72 | 0.95 | 0.84 | 0.82 | 0.70 |
| Obatoclax | GX15-070MS | BCL2 | | | | 6.97 | 1.83 | 0.68 | 0.47 | 9.91 | 3.84 | 1.71 | 0.94 | 0.70 | 0.48 | 0.40 | 0.50 |
| Venetoclax | ABT-199 | BCL2 (not Bcl-xl) | | | | 104.45 | 90.17 | 47.77 | 1.08 | 94.09 | 90.47 | 23.53 | 0.98 | 1.11 | 1.00 | 2.03 | 1.10 |
| BI-3812 | | BCL6 | | | | 97.55 | 97.78 | 74.42 | 27.10 | 96.61 | 92.23 | 67.22 | 45.87 | 1.01 | 1.06 | 1.11 | 0.59 |
| Nilotinib | | BCR-ABL | KIT | PDGFR | | 91.95 | 85.28 | 55.41 | 8.31 | 92.66 | 84.20 | 57.82 | 1.97 | 0.99 | 1.01 | 0.96 | 4.22 |
| Imatinib (free base or mesylate) | | BCR-ABL | PDGFR | | | 100.55 | 99.38 | 92.29 | 57.65 | 95.19 | 100.52 | 95.21 | 49.44 | 1.06 | 0.99 | 0.97 | 1.17 |
| Ponatinib | AP24534 | BCR-ABL | FGFR | PDGFF | VEGFR | 49.74 | 35.24 | 0.49 | 0.47 | 64.64 | 36.59 | 0.47 | 0.55 | 0.77 | 0.96 | 1.05 | 0.86 |
| Asciminib | ABL001 | BCR-ABL | | | | 96.10 | 93.41 | 92.76 | 64.81 | 101.14 | 101.31 | 93.53 | 76.77 | 0.95 | 0.92 | 0.99 | 0.84 |
| Bosutinib | SKI-606 | BCR-ABL | SRC | | | 84.34 | 80.00 | 66.91 | 0.40 | 105.92 | 92.28 | 89.05 | 0.52 | 0.80 | 0.87 | 0.75 | 0.78 |
| Dasatinib | | BCR-ABL | SRC | KIT | | 47.16 | 43.16 | 40.48 | 7.38 | 65.74 | 54.49 | 41.41 | 5.95 | 0.72 | 0.79 | 0.98 | 1.24 |
| Rebastinib | DCC-2036 | BCR-ABL | | | | 82.53 | 52.33 | 27.77 | 4.27 | 85.87 | 75.90 | 28.64 | 3.00 | 0.96 | 0.69 | 0.97 | 1.42 |
| TAK-580 | MLN2480 | BRAF | | | | 98.15 | 93.46 | 64.50 | 45.09 | 95.92 | 72.37 | 49.27 | 30.21 | 1.02 | 1.29 | 1.31 | 1.49 |
| RO5126766 | CH5126766 | BRAF | CRAF | MEK | | 42.53 | 36.55 | 31.28 | 25.48 | 30.55 | 30.07 | 26.33 | 24.04 | 1.39 | 1.22 | 1.19 | 1.06 |
| Belvarafenib | GDC-5573, HM9 | BRAF | CRAF | | | 38.74 | 35.19 | 29.87 | 11.91 | 34.31 | 31.44 | 27.28 | 8.17 | 1.13 | 1.12 | 1.10 | 1.46 |
| LY3009120 | | BRAF | CRAF | ARAF | | 34.28 | 31.63 | 26.88 | 12.21 | 28.49 | 28.37 | 20.78 | 11.41 | 1.20 | 1.12 | 1.29 | 1.07 |
| Encorafenib | LGX818 | BRAF | | | | 49.48 | 39.65 | 33.97 | 17.03 | 48.79 | 36.02 | 39.94 | 19.49 | 1.01 | 1.10 | 0.85 | 0.87 |
| Agerafenib | CEP-32496, RXD | BRAF | RET | | | 57.88 | 44.96 | 31.22 | 4.92 | 55.91 | 41.96 | 28.02 | 6.79 | 1.04 | 1.07 | 1.11 | 0.72 |
| PLX8394 | | BRAF | CRAF | | | 59.25 | 42.69 | 30.02 | 14.87 | 45.59 | 40.15 | 23.44 | 8.52 | 1.30 | 1.06 | 1.28 | 1.74 |
| Lifirafenib | BGB-283 | BRAF | EGFR | | | 47.68 | 37.65 | 29.17 | 19.94 | 50.88 | 38.34 | 31.87 | 18.30 | 0.94 | 0.98 | 0.92 | 1.09 |
| LXH254 | | BRAF | CRAF | | | 67.76 | 49.39 | 23.89 | 13.15 | 83.09 | 50.74 | 17.35 | 7.95 | 0.82 | 0.97 | 1.38 | 1.66 |
| Dabrafenib | GSK2118436A | BRAF | | | | 47.64 | 44.81 | 41.84 | 25.87 | 49.21 | 46.96 | 40.45 | 15.10 | 0.97 | 0.95 | 1.03 | 1.71 |
| TAK-632 | | BRAF | CRAF | VEGFR | | 52.17 | 41.04 | 28.70 | 5.24 | 60.98 | 47.39 | 31.30 | 10.25 | 0.86 | 0.87 | 0.92 | 0.51 |
| Regorafenib | BAY 73-4506 | BRAF | VEGFR | | | 83.89 | 73.41 | 43.67 | 7.59 | 96.66 | 92.52 | 43.48 | 4.24 | 0.87 | 0.79 | 1.00 | 1.79 |
| Vemurafenib | PLX4032 | BRAF | | | | 38.80 | 35.34 | 30.28 | 7.36 | 59.45 | 45.77 | 29.47 | 3.19 | 0.65 | 0.77 | 1.03 | 2.31 |
| Mivebresib | ABBV-075 | BRD2 | BRD4 | BRDT | | 27.73 | 24.79 | 9.08 | 3.09 | 17.17 | 11.37 | 4.99 | 2.07 | 1.62 | 2.18 | 1.82 | 1.49 |
| PLX51107 | | BRD2 | BRD3 | BRD4 | | 48.95 | 43.13 | 34.43 | 21.56 | 44.17 | 34.19 | 24.58 | 14.15 | 1.11 | 1.26 | 1.40 | 1.52 |
| Birabresib | OTX015, Y-803 | BRD2 | BRD3 | BRD4 | | 53.25 | 39.42 | 53.25 | 28.64 | 45.50 | 34.31 | 17.49 | 18.56 | 1.17 | 1.15 | 3.04 | 1.54 |
| ABBV-744 | | BRD2 | BRD3 | BRD4 | | 76.54 | 61.89 | 28.13 | 7.67 | 59.38 | 60.76 | 15.71 | 3.59 | 1.29 | 1.02 | 1.79 | 2.13 |
| INCB057643 | | BRD2 | BRD3 | BRD4 | | 66.10 | 48.89 | 31.93 | 10.67 | 52.40 | 54.72 | 25.73 | 6.26 | 1.26 | 0.89 | 1.24 | 1.70 |
| BMS-986158 | | BRD4 | | | | 33.38 | 22.61 | 11.42 | 2.39 | 20.22 | 18.24 | 9.00 | 4.81 | 1.65 | 1.24 | 1.27 | 0.50 |
| AZD5153 | | BRD4 | | | | 41.72 | 42.64 | 34.07 | 27.74 | 49.86 | 35.10 | 27.82 | 28.65 | 0.84 | 1.21 | 1.22 | 0.97 |
| Bromosporine | | BRD4 | BRD9 | BRD2 | | 90.05 | 65.84 | 34.96 | 14.46 | 88.99 | 59.95 | 30.86 | 10.09 | 1.01 | 1.10 | 1.13 | 1.43 |
| Molibresib | I-BET762, GSK52 | BRD4 | BRD3 | BRD2 | | 86.76 | 68.16 | 51.22 | 41.05 | 75.14 | 63.51 | 48.81 | 30.23 | 1.15 | 1.07 | 1.05 | 1.36 |
| Apabetalone | RVX-208, RVX00 | BRD4 | | | | 99.61 | 96.20 | 83.98 | 70.63 | 96.90 | 98.54 | 88.40 | 60.12 | 1.03 | 0.98 | 0.95 | 1.17 |
| BI-7273 | | BRD9 | | | | 101.45 | 98.21 | 98.67 | 96.93 | 99.28 | 102.50 | 95.28 | 100.18 | 1.02 | 0.96 | 1.04 | 0.97 |
| GSK6853 | | BRPF1 | | | | 102.44 | 98.73 | 93.94 | 94.27 | 100.22 | 99.26 | 94.02 | 90.57 | 1.02 | 0.99 | 1.00 | 1.04 |
| OF-1 | | BRPF1 | BRPF2 | | | 94.29 | 84.31 | 78.71 | 55.17 | 92.51 | 90.85 | 79.61 | 30.31 | 1.02 | 0.93 | 0.99 | 1.82 |
| ARQ531 | | BTK | | | | 39.33 | 33.95 | 15.53 | 0.53 | 34.60 | 24.45 | 8.25 | 0.80 | 1.14 | 1.39 | 1.88 | 0.66 |
| Ibrutinib | PCI-32765 | BTK | | | | 109.97 | 110.15 | 99.87 | 55.73 | 105.74 | 97.28 | 92.78 | 36.19 | 1.04 | 1.13 | 1.08 | 1.54 |
| Acalabrutinib | | BTK | TEC | | | 102.59 | 96.18 | 94.46 | 87.72 | 98.69 | 96.58 | 91.30 | 76.30 | 1.04 | 1.00 | 1.03 | 1.15 |
| Tirabrutinib | ONO-4059, GS-4 | BTK | | | | 99.58 | 99.14 | 92.97 | 81.50 | 99.26 | 100.16 | 84.37 | 65.79 | 1.00 | 0.99 | 1.10 | 1.24 |
| Zanubrutinib | BGB-3111 | BTK | | | | 102.65 | 97.69 | 91.98 | 73.87 | 102.62 | 102.07 | 100.55 | 86.28 | 1.00 | 0.96 | 0.91 | 0.86 |
| Fenebrutinib | GDC-0853 | BTK | | | | 98.02 | 93.02 | 91.48 | 98.25 | 106.12 | 100.55 | 95.93 | 94.87 | 0.92 | 0.93 | 0.95 | 1.04 |
| Pimecrolimus | | Calcineurin | | | | 101.38 | 96.98 | 77.23 | 23.20 | 95.86 | 90.54 | 80.64 | 24.39 | 1.06 | 1.07 | 0.96 | 0.95 |
| Gossypol | AT-101 | Calcineurin | MCL1 | BCL2 | BCL-X | 81.31 | 28.62 | 11.26 | 0.67 | 79.24 | 28.03 | 8.90 | 0.57 | 1.03 | 1.02 | 1.26 | 1.17 |
| Tacrolimus | FK-506 | Calcineurin | | | | 102.09 | 99.14 | 89.27 | 88.97 | 99.18 | 97.67 | 88.87 | 87.29 | 1.03 | 1.02 | 1.00 | 1.02 |
| Amcasertib | BBI503 | cancer stemness pathway (NANOG) | | | | 100.85 | 85.66 | 0.55 | 0.55 | 103.94 | 69.05 | 0.53 | 0.53 | 0.97 | 1.24 | 1.02 | 1.02 |
| Belnacasan | VX-765 | Caspase1 | | | | 96.71 | 105.11 | 96.00 | 94.44 | 94.68 | 101.17 | 95.66 | 100.15 | 1.02 | 1.04 | 1.00 | 0.94 |
| SGC-CBP30 | | CBP (CREBBP) | EP300 | | | 104.42 | 97.58 | 82.64 | 47.51 | 90.33 | 94.29 | 83.25 | 51.32 | 1.16 | 1.03 | 0.99 | 0.93 |
| CPI-637 | | CBP (CREBBP) | EP300 | | | 87.78 | 85.15 | 74.14 | 56.33 | 86.80 | 84.96 | 68.86 | 54.44 | 1.01 | 1.00 | 1.08 | 1.03 |
| E7386 | E-7386 | CBP/b-catenin PPI | | | | 16.22 | 6.41 | 3.99 | 2.39 | 6.46 | 5.38 | 4.62 | 3.45 | 2.51 | 1.19 | 0.86 | 0.69 |
| Alendronate, aq | | CD45 | | | | 95.10 | 98.52 | 96.31 | 23.14 | 93.60 | 99.19 | 91.20 | 4.94 | 1.02 | 0.99 | 1.06 | 4.68 |
| MBQ-167 | | CDC42 | RAC | | | 24.90 | 18.03 | 14.66 | 17.12 | 11.77 | 13.80 | 15.19 | 17.25 | 2.12 | 1.31 | 0.97 | 0.99 |
| ML141 | CID-2950007 | CDC42 | | | | 94.38 | 93.73 | 90.30 | 61.09 | 95.46 | 96.70 | 91.78 | 44.59 | 0.99 | 0.97 | 0.98 | 1.37 |
| Simurosertib | TAK-931 | CDC7 | | | | 93.24 | 93.32 | 93.40 | 72.34 | 97.35 | 96.89 | 85.31 | 67.91 | 0.96 | 0.96 | 1.09 | 1.07 |
| CYC065 | | CDK | | | | 113.84 | 93.77 | 1.23 | 1.65 | 102.70 | 48.44 | 1.51 | 1.44 | 1.11 | 1.94 | 0.82 | 1.14 |
| AT7519 | | CDK | | | | 97.03 | 94.54 | 104.58 | 86.72 | 102.71 | 102.11 | 97.93 | 54.88 | 0.94 | 0.93 | 1.07 | 1.58 |
| Roscovitine | Seliciclib | CDK | | | | 89.08 | 90.15 | 99.54 | 101.92 | 104.56 | 104.80 | 100.40 | 95.29 | 0.85 | 0.86 | 0.99 | 1.07 |
| Dinaciclib | SCH727965 | CDK | | | | 2.44 | 1.37 | 1.37 | 1.58 | 1.79 | 1.88 | 1.26 | 1.24 | 1.36 | 0.73 | 1.09 | 1.27 |
| Ribociclib | LEE-011 | CDK4/6 | | | | 97.10 | 95.30 | 92.91 | 83.67 | 102.64 | 90.00 | 87.54 | 80.58 | 0.95 | 1.06 | 1.06 | 1.04 |
| Palbociclib, aq | PD-0332991 | CDK4/6 | | | | 90.39 | 91.75 | 69.08 | 106.53 | 96.64 | 90.21 | 37.52 | 112.29 | 0.94 | 1.02 | 1.84 | 0.95 |
| Lerociclib | G1T38 | CDK4/6 | CDK9 | | | 98.15 | 97.01 | 73.91 | 77.81 | 92.30 | 98.92 | 81.48 | 86.06 | 1.06 | 0.98 | 0.91 | 0.90 |
| Abemaciclib | LY2835219 | CDK4/6 | | | | 98.07 | 86.47 | 70.07 | 69.16 | 97.96 | 92.12 | 33.11 | 18.52 | 1.00 | 0.94 | 2.12 | 3.73 |
| AZD4573 | | CDK9 | | | | 3.22 | 2.04 | 3.26 | 2.36 | 4.09 | 2.42 | 2.55 | 2.34 | 0.79 | 0.84 | 1.28 | 1.01 |
| BAY1251152 | BAY-1251152 | CDK9 | | | | 4.57 | 1.56 | 1.53 | 2.53 | 2.79 | 1.86 | 1.44 | 0.97 | 1.64 | 0.84 | 1.06 | 2.61 |
| GSK923295 | | CENPE | | | | 85.60 | 65.53 | 53.90 | 7.70 | 74.64 | 61.08 | 43.52 | 3.05 | 1.15 | 1.07 | 1.24 | 2.53 |
| Rabusertib | LY2603618, IC-8 | CHK1 | | | | 88.08 | 55.55 | 9.62 | 6.19 | 97.66 | 34.34 | 17.04 | 8.64 | 0.90 | 1.62 | 0.56 | 0.72 |
| SCH900776 | MK-8776 | CHK1 | | | | 99.97 | 98.22 | 44.78 | 35.93 | 93.26 | 91.63 | 33.66 | 21.74 | 1.07 | 1.07 | 1.33 | 1.65 |
| Prexasertib | LY2606368 | CHK1 | | | | 15.57 | 20.32 | 18.82 | 15.51 | 27.86 | 19.43 | 25.11 | 24.65 | 0.56 | 1.05 | 0.75 | 0.63 |
| AZD7762 | | CHK1 | | | | 46.51 | 24.72 | 11.91 | 6.78 | 36.83 | 29.81 | 19.79 | 5.94 | 1.26 | 0.83 | 0.60 | 1.14 |
| SR-3029 | | CK1d | CK1e | | | 31.79 | 4.26 | 0.49 | 0.38 | 12.03 | 1.20 | 0.37 | 0.45 | 2.64 | 3.56 | 1.33 | 0.86 |
| Silmitasertib | CX-4945 | CK2 | | | | 93.00 | 103.93 | 84.84 | 50.69 | 96.17 | 100.77 | 92.51 | 60.93 | 0.97 | 1.03 | 0.92 | 0.83 |
| Lorecivivint | Adavivint, SM046 | CLK2 | DYRK1A | | | 105.30 | 98.32 | 25.27 | 24.83 | 106.32 | 96.36 | 9.71 | 11.11 | 0.99 | 1.02 | 2.60 | 2.24 |

| | | | | | | | | | | | | | | | |
|---|---|---|---|---|---|---|---|---|---|---|---|---|---|---|---|
| Selinexor | KPT-330 | CRM1 (XPO1) | | 24.52 | 22.16 | 15.13 | 12.74 | 19.15 | 14.32 | 11.73 | 9.56 | 1.28 | 1.55 | 1.29 | 1.33 |
| Eltanexor | KPT-8602 | CRM1 (XPO1) | | 42.44 | 20.86 | 11.16 | 4.92 | 31.52 | 15.21 | 8.66 | 1.31 | 1.35 | 1.37 | 1.29 | 3.74 |
| Pexidartinib | PLX3397 | CSF1R | KIT FLT3 | 103.02 | 100.18 | 95.95 | 60.56 | 100.78 | 100.03 | 98.29 | 44.21 | 1.02 | 1.00 | 0.98 | 1.37 |
| SX-682 | | CXCR1 | CXCR2 | 96.66 | 98.45 | 79.91 | 32.35 | 97.94 | 101.21 | 83.89 | 16.49 | 0.99 | 0.97 | 0.95 | 1.96 |
| AZD5069 | | CXCR2 | | 109.34 | 112.87 | 104.72 | 105.00 | 96.35 | 105.14 | 97.63 | 102.11 | 1.13 | 1.07 | 1.07 | 1.03 |
| Brequinar | | Dihydroorotate dehydrogenase | | 99.96 | 79.52 | 26.97 | 12.93 | 97.15 | 60.39 | 26.99 | 19.17 | 1.03 | 1.32 | 1.00 | 0.67 |
| ML390 | | Dihydroorotate dehydrogenase | | 102.84 | 90.80 | 44.18 | 26.64 | 99.58 | 81.47 | 33.76 | 22.46 | 1.03 | 1.11 | 1.31 | 1.19 |
| AZD7648 | | DNAPK | | 106.90 | 106.23 | 89.38 | 82.25 | 101.78 | 104.58 | 96.85 | 83.81 | 1.05 | 1.02 | 0.92 | 0.98 |
| Nedisertib | M-3814 | DNAPK | | 94.86 | 94.64 | 73.44 | 61.75 | 99.48 | 97.46 | 87.80 | 64.53 | 0.95 | 0.97 | 0.84 | 0.96 |
| CC-115 | | DNAPK | mTOR | 95.41 | 83.95 | 22.63 | 4.27 | 103.24 | 87.05 | 15.56 | 2.97 | 0.92 | 0.96 | 1.45 | 1.44 |
| KU-0060648 | | DNAPK | PI3K | 95.80 | 83.15 | 38.29 | 0.46 | 102.88 | 95.47 | 41.63 | 0.50 | 0.93 | 0.87 | 0.92 | 0.92 |
| Decitabine | | DNMT | | 80.17 | 64.25 | 50.15 | 30.64 | 73.76 | 47.50 | 36.89 | 29.19 | 1.09 | 1.35 | 1.36 | 1.05 |
| RG108 | | DNMT | | 103.05 | 107.85 | 100.31 | 96.99 | 99.73 | 91.81 | 94.08 | 92.64 | 1.03 | 1.17 | 1.07 | 1.05 |
| Azacitidine | | DNMT | | 102.92 | 94.28 | 68.95 | 30.47 | 99.07 | 94.45 | 53.61 | 19.55 | 1.04 | 1.00 | 1.29 | 1.56 |
| SGC0946 | | DOT1L | | 94.51 | 95.26 | 91.96 | 88.93 | 94.87 | 95.60 | 89.20 | 77.63 | 1.00 | 1.00 | 1.03 | 1.15 |
| Pinometostat | EPZ-5676 | DOT1L | | 89.40 | 93.36 | 86.82 | 84.63 | 100.61 | 98.38 | 92.01 | 91.11 | 0.89 | 0.95 | 0.94 | 0.93 |
| Bioymifi | | DR5 | | 103.15 | 103.04 | 37.07 | 4.35 | 98.32 | 103.05 | 1.93 | 1.15 | 1.05 | 1.00 | 19.25 | 3.78 |
| AZ191 | | DYRK1B | DYRK1A | 103.90 | 102.20 | 93.75 | 0.86 | 100.40 | 98.58 | 90.57 | 0.49 | 1.03 | 1.04 | 1.04 | 1.73 |
| EED226 | | EED (PRC2 complex) | | 99.96 | 100.68 | 98.56 | 87.63 | 106.24 | 103.32 | 99.01 | 95.31 | 0.94 | 0.97 | 1.00 | 0.92 |
| MAK683 | | EED (PRC2 complex) | | 97.06 | 100.61 | 93.80 | 95.32 | 102.59 | 104.45 | 99.63 | 109.35 | 0.95 | 0.96 | 0.94 | 0.87 |
| ARQ621 | | Eg5 | | 83.12 | 41.53 | 26.15 | 0.48 | 32.54 | 26.93 | 12.56 | 0.47 | 2.55 | 1.54 | 2.08 | 1.01 |
| BMS-599626 | AC480 | EGFR | ERBB2 | 111.01 | 108.13 | 102.33 | 0.63 | 106.56 | 99.80 | 96.04 | 0.64 | 1.04 | 1.08 | 1.07 | 0.98 |
| Sapitinib | AZD8931 | EGFR | ERBB2 ERRB3 | 97.88 | 108.04 | 81.70 | 54.74 | 102.05 | 102.34 | 94.88 | 74.37 | 0.96 | 1.06 | 0.86 | 0.74 |
| Gefitinib | | EGFR | | 94.60 | 104.16 | 85.90 | 3.02 | 96.93 | 99.88 | 86.84 | 1.77 | 0.98 | 1.04 | 0.99 | 1.70 |
| Varlitinib | ARRY334543 | EGFR | ERBB2 | 97.38 | 96.21 | 86.69 | 31.37 | 97.84 | 93.36 | 94.12 | 23.25 | 1.00 | 1.03 | 0.92 | 1.35 |
| Vandetanib | | EGFR | VEGFR RET | 105.13 | 92.89 | 85.89 | 0.55 | 92.18 | 91.32 | 93.14 | 0.50 | 1.14 | 1.02 | 0.92 | 1.09 |
| Lapatinib (free base) | | EGFR | ERBB2 | 99.56 | 93.57 | 40.77 | 0.40 | 101.24 | 95.52 | 43.17 | 0.53 | 0.98 | 0.98 | 0.94 | 0.74 |
| Lazertinib | YH25448,GNS-14 | EGFR | EGFR T790M | 97.56 | 99.71 | 94.62 | 94.80 | 102.07 | 103.12 | 94.44 | 95.54 | 0.96 | 0.97 | 1.00 | 0.99 |
| Osimertinib | AZD9291, Merele | EGFR | EGFR T790M | 104.23 | 92.80 | 2.74 | 0.58 | 105.74 | 99.42 | 8.54 | 0.54 | 0.99 | 0.93 | 0.32 | 1.07 |
| Dacomitinib | PF-00299804 | EGFR | ERBB2 | 96.68 | 91.95 | 55.63 | 0.45 | 99.85 | 99.21 | 36.76 | 0.39 | 0.97 | 0.93 | 1.51 | 1.17 |
| Erlotinib | | EGFR | | 100.51 | 95.25 | 87.00 | 48.16 | 97.81 | 102.77 | 97.10 | 76.90 | 1.03 | 0.93 | 0.90 | 0.63 |
| Poziotinib | HM781-36B | EGFR | ERBB2 ERRB4 | 98.59 | 94.10 | 68.08 | 94.02 | 103.76 | 101.94 | 89.80 | 99.34 | 0.95 | 0.92 | 0.76 | 0.95 |
| Afatinib | BIBW-2992 | EGFR | ERBB2 | 66.74 | 71.03 | 35.74 | 0.47 | 101.46 | 102.06 | 39.22 | 0.46 | 0.66 | 0.70 | 0.91 | 1.02 |
| Icotinib | BPI-2009H | EGFR T790M | | 98.83 | 97.56 | 93.42 | 89.96 | 97.27 | 94.89 | 96.60 | 98.40 | 1.02 | 1.03 | 0.97 | 0.91 |
| HS-10296 | | EGFR T790M | | 101.41 | 98.53 | 0.38 | 0.38 | 102.70 | 101.49 | 0.47 | 0.39 | 0.99 | 0.97 | 0.80 | 0.97 |
| Nazartinib | EGF816 | EGFR T790M | | 95.12 | 95.94 | 86.58 | 2.40 | 102.67 | 100.27 | 90.42 | 0.58 | 0.93 | 0.96 | 0.96 | 4.10 |
| Avitinib | AC0010 | EGFR T790M | | 95.82 | 62.76 | 16.03 | 0.40 | 106.75 | 72.17 | 10.78 | 0.41 | 0.90 | 0.87 | 1.49 | 0.97 |
| NVP-BHG712 | | EPHB4 | | 96.20 | 56.26 | 3.08 | 0.53 | 98.58 | 48.16 | 1.76 | 0.47 | 0.98 | 1.17 | 1.75 | 1.14 |
| Neratinib | | ERBB2 | EGFR | 81.81 | 56.36 | 53.51 | 43.28 | 74.11 | 33.61 | 19.34 | 0.55 | 1.10 | 1.68 | 2.77 | 78.44 |
| Canertinib | CI-1033, PD1838 | ERBB2 | | 91.37 | 86.66 | 46.90 | 30.36 | 93.64 | 80.81 | 12.55 | 8.59 | 0.98 | 1.07 | 3.74 | 3.53 |
| Tucatinib | ARRY-380, ONT-3 | ERBB2 | | 100.40 | 99.56 | 95.45 | 75.63 | 97.92 | 93.18 | 92.89 | 62.48 | 1.03 | 1.07 | 1.03 | 1.21 |
| Ulixertinib | BVD-523 | ERK | | 53.33 | 37.96 | 29.07 | 6.51 | 46.82 | 31.13 | 27.13 | 4.60 | 1.14 | 1.22 | 1.07 | 1.41 |
| LY3214996 | | ERK1/2 | | 74.63 | 52.94 | 32.33 | 7.90 | 78.91 | 48.46 | 27.76 | 7.08 | 0.95 | 1.09 | 1.16 | 1.12 |
| MK-8353 | SCH900353 | ERK1/2 | | 43.32 | 31.35 | 21.32 | 1.76 | 33.32 | 30.47 | 16.88 | 1.25 | 1.30 | 1.03 | 1.26 | 1.41 |
| Ravoxertinib | GDC-0994 | ERK1/2 | | 82.77 | 53.67 | 39.47 | 35.31 | 97.69 | 63.44 | 46.31 | 37.98 | 0.85 | 0.85 | 0.85 | 0.93 |
| LSZ102 | | ESR1 degrader | | 106.44 | 107.23 | 101.66 | 53.08 | 99.65 | 102.21 | 97.71 | 38.48 | 1.07 | 1.05 | 1.04 | 1.38 |
| CPI-1205 | | EZH2 | EZH1 | 101.31 | 105.61 | 99.04 | 96.03 | 98.54 | 101.14 | 91.47 | 98.17 | 1.03 | 1.04 | 1.08 | 0.98 |
| EBI-2511 | | EZH2 | | 103.18 | 103.43 | 99.90 | 101.27 | 105.99 | 104.00 | 100.40 | 79.62 | 0.97 | 0.99 | 0.99 | 1.27 |
| Tazemetostat | EPZ6438, E7438 | EZH2 | | 103.53 | 99.91 | 92.62 | 89.68 | 106.62 | 104.51 | 86.76 | 92.73 | 0.97 | 0.96 | 1.07 | 0.97 |
| PF-06821497 | | EZH2 | | 102.17 | 99.71 | 100.15 | 96.72 | 108.56 | 104.85 | 99.09 | 102.05 | 0.94 | 0.95 | 1.01 | 0.95 |
| Defactinib | VS-6063, PF-045 | FAK | | 93.94 | 98.61 | 53.66 | 28.76 | 102.55 | 89.13 | 31.09 | 24.30 | 0.92 | 1.11 | 1.73 | 1.18 |
| GSK2256098 | | FAK | | 94.84 | 90.70 | 84.50 | 81.05 | 93.14 | 93.42 | 85.99 | 84.72 | 1.02 | 0.97 | 0.98 | 0.96 |
| PND-1186 | VS-4718 | FAK | | 81.15 | 74.67 | 48.68 | 31.93 | 83.19 | 78.62 | 42.24 | 17.57 | 0.98 | 0.95 | 1.15 | 1.82 |
| PF573228 | | FAK | | 99.20 | 90.65 | 47.92 | 39.50 | 94.02 | 96.17 | 51.47 | 30.63 | 1.06 | 0.94 | 0.93 | 1.29 |
| Tipifarnib | | Farnesyltransferase | | 55.65 | 57.05 | 50.18 | 16.82 | 51.08 | 45.50 | 37.61 | 8.64 | 1.09 | 1.25 | 1.33 | 1.95 |
| Derazantinib | ARQ-087 | FGFR | RET DDR2 PDGF | 105.74 | 68.91 | 0.65 | 0.59 | 106.13 | 4.64 | 0.55 | 0.62 | 1.00 | 14.85 | 1.17 | 0.96 |
| LY2874455 | | FGFR | VEGFR2 | 51.36 | 39.23 | 16.86 | 0.71 | 36.12 | 28.97 | 17.46 | 1.02 | 1.42 | 1.35 | 0.97 | 0.70 |
| ASP5878 | | FGFR | | 105.07 | 82.93 | 28.46 | 18.40 | 106.70 | 81.18 | 23.02 | 17.59 | 0.98 | 1.02 | 1.24 | 1.05 |
| AZD4547 | | FGFR | | 96.09 | 97.40 | 90.28 | 1.78 | 98.93 | 97.57 | 91.66 | 1.93 | 0.97 | 1.00 | 0.99 | 0.92 |
| Infigratinib | BGJ398, NVP-BG. | FGFR | | 102.33 | 101.14 | 83.67 | 0.53 | 101.32 | 106.38 | 67.03 | 0.53 | 1.01 | 0.95 | 1.25 | 1.00 |
| Rogaratinib | BAY1163877 | FGFR | | 96.30 | 93.06 | 96.30 | 85.51 | 103.41 | 99.40 | 102.61 | 104.20 | 0.93 | 0.94 | 0.94 | 0.82 |
| PRN1371 | | FGFR (pan) | | 104.29 | 102.15 | 53.05 | 20.00 | 103.24 | 101.80 | 46.86 | 16.22 | 1.01 | 1.00 | 1.13 | 1.23 |
| Erdafitinib | JNJ-42756493 | FGFR (pan) | | 99.00 | 103.18 | 82.71 | 8.40 | 95.46 | 103.83 | 99.34 | 15.56 | 1.04 | 0.99 | 0.83 | 0.54 |
| BLU-554 | | FGFR4 | | 106.49 | 108.83 | 103.56 | 93.00 | 99.73 | 102.73 | 97.20 | 95.09 | 1.07 | 1.06 | 1.07 | 0.98 |
| H3B-6527 | | FGFR4 | | 97.55 | 106.77 | 93.07 | 86.45 | 105.37 | 106.83 | 91.31 | 74.16 | 0.93 | 1.00 | 1.02 | 1.17 |
| Roblitinib | FGF401 | FGFR4 | | 96.90 | 90.63 | 93.61 | 91.22 | 106.77 | 93.00 | 99.62 | 100.51 | 0.91 | 0.97 | 0.94 | 0.91 |
| Quizartinib | AC220 | FLT3 | | 78.30 | 24.96 | 25.28 | 16.98 | 76.98 | 18.08 | 7.70 | 8.54 | 1.02 | 1.38 | 3.28 | 1.99 |
| Midostaurin | PKC412 | FLT3 | PKC KIT | 83.97 | 25.40 | 7.91 | 8.11 | 92.99 | 23.33 | 14.17 | 14.17 | 0.90 | 1.09 | 0.56 | 0.57 |
| Tandutinib | MLN-518 | FLT3 | PDGF KIT | 97.98 | 99.80 | 92.71 | 68.80 | 92.85 | 99.71 | 92.30 | 57.82 | 1.06 | 1.00 | 1.00 | 1.19 |
| Gilteritinib | ASP2215 | FLT3 | AXL multikinase | 92.66 | 51.92 | 36.47 | 1.76 | 101.14 | 57.92 | 28.22 | 7.97 | 0.92 | 0.90 | 1.29 | 0.22 |
| Linifanib | ABT-869 | FLT3 | VEGFR PDGFR | 68.35 | 66.57 | 88.44 | 36.06 | 100.02 | 96.48 | 80.85 | 13.16 | 0.68 | 0.69 | 1.09 | 2.74 |
| JK184 | | GLI1 | | 23.61 | 26.82 | 25.26 | 25.63 | 16.95 | 17.77 | 18.03 | 17.79 | 1.39 | 1.51 | 1.40 | 1.44 |
| GANT61 | | GLI1 | GLI2 | 101.71 | 109.53 | 91.38 | 57.91 | 103.05 | 103.80 | 94.83 | 69.79 | 0.99 | 1.06 | 0.96 | 0.83 |
| Telaglenastat | CB-839 | Glutaminase I | | 36.09 | 22.09 | 26.87 | 35.76 | 6.89 | 3.72 | 2.81 | 4.99 | 5.24 | 5.94 | 9.57 | 7.17 |
| DAPT (GSI-IX) | | g-Secretase | | 94.17 | 101.24 | 95.13 | 60.39 | 103.91 | 94.88 | 96.39 | 31.10 | 0.91 | 1.07 | 0.99 | 1.94 |
| Crenigacestat | LY3039478 | g-Secretase | | 98.29 | 96.32 | 94.50 | 90.40 | 97.34 | 94.96 | 94.33 | 90.83 | 1.01 | 1.01 | 1.00 | 1.00 |
| RO4929097 | | g-Secretase | | 100.99 | 95.50 | 92.42 | 86.37 | 104.55 | 99.75 | 97.14 | 93.03 | 0.97 | 0.96 | 0.95 | 0.93 |
| Nirogacestat | PF-03084014 | g-Secretase | | 96.30 | 91.04 | 85.48 | 71.09 | 94.21 | 101.89 | 91.16 | 72.99 | 1.02 | 0.89 | 0.94 | 0.97 |
| Avagacestat | BMS-708163 | g-Secretase | | 71.20 | 77.94 | 78.52 | 67.29 | 100.52 | 94.01 | 94.95 | 70.46 | 0.71 | 0.83 | 0.83 | 0.96 |
| LY2090314 | | GSK3 | | 78.28 | 79.39 | 74.50 | 74.68 | 33.56 | 48.89 | 30.43 | 33.03 | 2.33 | 1.62 | 2.45 | 2.26 |
| Tideglusib | NP031112 | GSK3 | | 104.07 | 98.63 | 97.06 | 92.93 | 94.99 | 95.56 | 90.70 | 81.19 | 1.10 | 1.03 | 1.07 | 1.14 |
| C646 | | HAT | | 95.40 | 99.30 | 92.83 | 81.11 | 95.42 | 92.85 | 89.11 | 56.55 | 1.00 | 1.07 | 1.04 | 1.43 |
| Remodelin | | HAT | | 98.21 | 99.40 | 92.68 | 60.51 | 101.25 | 100.48 | 96.11 | 52.15 | 0.97 | 0.99 | 0.96 | 1.16 |
| A-485 | | HAT | | 98.37 | 96.16 | 10.09 | 0.58 | 106.39 | 103.66 | 76.53 | 1.75 | 0.92 | 0.93 | 0.13 | 0.33 |
| Belinostat | PDX101 | HDAC | | 34.91 | 20.84 | 6.70 | 0.43 | 52.84 | 11.11 | 2.99 | 0.50 | 0.66 | 1.88 | 2.24 | 0.85 |
| Panobinostat | LBH-589 | HDAC | | 0.72 | 0.91 | 0.44 | 0.42 | 0.62 | 0.52 | 0.47 | 0.41 | 1.16 | 1.75 | 0.93 | 1.01 |
| Givinostat | ITF2357 | HDAC | | 56.82 | 23.64 | 1.54 | 0.51 | 37.45 | 14.51 | 1.19 | 0.53 | 1.52 | 1.63 | 1.30 | 0.97 |
| CUDC-907 | | HDAC | PI3K | 0.89 | 0.73 | 0.61 | 0.46 | 0.53 | 0.47 | 0.51 | 0.41 | 1.68 | 1.56 | 1.19 | 1.14 |
| Romidepsin | FK-228 | HDAC | | 0.82 | 0.80 | 0.70 | 0.46 | 0.48 | 0.58 | 0.63 | 0.40 | 1.71 | 1.37 | 1.12 | 1.16 |
| CUDC-101 | CUDC101 | HDAC | ERBB2 EGFR | 104.54 | 67.26 | 19.31 | 0.72 | 99.40 | 53.54 | 6.31 | 0.54 | 1.05 | 1.26 | 3.06 | 1.34 |
| Mocetinostat | MGCD0103 | HDAC | | 89.26 | 39.97 | 2.76 | 2.14 | 98.19 | 32.56 | 0.79 | 1.00 | 0.91 | 1.23 | 3.48 | 2.13 |
| Tucidinostat | Chidamide | HDAC | | 98.30 | 92.00 | 63.47 | 26.58 | 100.67 | 88.33 | 44.25 | 13.82 | 0.98 | 1.04 | 1.43 | 1.92 |
| PCI-34051 | | HDAC | | 103.77 | 96.00 | 71.30 | 15.47 | 103.38 | 96.62 | 52.94 | 14.51 | 1.00 | 0.99 | 1.35 | 1.07 |
| Vorinostat | SAHA | HDAC | | 94.29 | 73.14 | 36.59 | 4.70 | 96.90 | 79.64 | 38.41 | 1.69 | 0.97 | 0.92 | 0.95 | 2.78 |
| Entinostat | MS-275, SNDX-2 | HDAC | | 85.36 | 65.27 | 34.29 | 5.27 | 100.38 | 82.72 | 28.99 | 2.35 | 0.85 | 0.79 | 1.18 | 2.24 |
| Abexinostat | PCI-24781 | HDAC1 | | 36.57 | 22.47 | 1.96 | 0.47 | 40.75 | 18.99 | 1.21 | 0.54 | 0.90 | 1.18 | 1.61 | 0.87 |
| ACY-957 | CP-0006946-157 | HDAC1 | HDAC2 | 96.41 | 40.29 | 18.24 | 0.56 | 97.79 | 44.19 | 8.40 | 0.47 | 0.99 | 0.91 | 2.17 | 1.17 |
| Rocilinostat | Ricilinostat, ACY-1 | HDAC6 | | 101.24 | 97.92 | 85.98 | 29.15 | 99.57 | 90.95 | 82.18 | 20.43 | 1.02 | 1.08 | 1.05 | 1.43 |
| Citarinostat | ACY-241, CP-000 | HDAC6 | HDAC3 | 101.03 | 101.18 | 68.81 | 17.61 | 104.80 | 100.88 | 48.27 | 9.32 | 0.96 | 1.00 | 1.43 | 1.89 |
| TAS0728 | | HER2 | | 99.93 | 112.94 | 92.25 | 84.92 | 99.50 | 95.73 | 96.40 | 86.90 | 1.00 | 1.18 | 0.96 | 0.98 |
| Daprodustat | GSK1278863 | HIF Prolyl hydroxylase | | 91.56 | 90.15 | 70.24 | 48.41 | 94.57 | 82.80 | 62.59 | 45.42 | 0.97 | 1.09 | 1.12 | 1.07 |
| Roxadustat | FG-4592 | HIF Prolyl hydroxylase | | 98.50 | 99.52 | 105.96 | 67.31 | 99.97 | 99.29 | 92.88 | 69.60 | 0.99 | 1.00 | 1.14 | 0.97 |
| BAY-872243 | BAY 87-2243 | HIF1 | | 87.26 | 77.93 | 61.72 | 58.96 | 87.45 | 70.37 | 39.33 | 51.04 | 1.00 | 1.11 | 1.57 | 1.16 |
| PX-478 | | HIF1 | | 98.21 | 92.95 | 92.12 | 83.35 | 96.88 | 96.35 | 93.26 | 83.17 | 1.01 | 0.96 | 0.99 | 1.00 |
| PT2385 | | HIF2a | | 102.00 | 102.23 | 98.77 | 97.13 | 92.79 | 99.95 | 96.05 | 94.94 | 1.10 | 1.02 | 1.03 | 1.02 |
| A-366 | | HMT | | 108.06 | 108.06 | 103.47 | 91.79 | 99.24 | 87.89 | 96.18 | 92.20 | 1.09 | 1.23 | 1.08 | 1.00 |
| UNC0638 | | HMT | | 96.33 | 101.86 | 89.64 | 0.57 | 103.53 | 87.21 | 82.75 | 0.65 | 0.93 | 1.17 | 1.08 | 0.88 |
| UNC0642 | | HMT | | 103.86 | 108.23 | 84.16 | 5.06 | 105.24 | 105.51 | 81.04 | 0.64 | 0.99 | 1.03 | 1.04 | 7.91 |
| UNC0646 | | HMT | | 98.69 | 93.22 | 59.17 | 5.12 | 97.79 | 94.72 | 67.43 | 2.13 | 1.01 | 0.98 | 0.88 | 2.40 |
| BIX-01294 | | HMT (G9a) | | 100.89 | 101.88 | 1.47 | 0.48 | 102.63 | 92.50 | 0.48 | 0.48 | 0.98 | 1.10 | 3.06 | 0.99 |
| A-196 | | HMT (KMT5B HMT (KMT5C) | | 98.40 | 95.33 | 90.13 | 68.85 | 101.02 | 98.61 | 92.27 | 58.11 | 0.97 | 0.97 | 0.98 | 1.18 |
| PFI-2 | | HMT (SETD7; KMT7) | | 95.97 | 100.90 | 92.98 | 91.05 | 96.94 | 99.00 | 92.11 | 88.67 | 0.99 | 1.02 | 1.01 | 1.03 |
| Tanespimycin | Retaspimycin, IPI- | HSP90 | | 94.64 | 62.11 | 32.97 | 5.62 | 47.60 | 17.06 | 2.83 | 0.60 | 1.99 | 3.64 | 11.65 | 9.36 |
| Luminespib | AUY922, NVP-AU | HSP90 | | 0.68 | 0.74 | 0.60 | 0.38 | 0.50 | 0.38 | 0.56 | 0.46 | 1.37 | 1.97 | 1.07 | 0.84 |
| PU-H71 | | HSP90 | | 93.96 | 40.93 | 10.20 | 1.51 | 60.72 | 24.51 | 3.16 | 0.61 | 1.55 | 1.67 | 3.23 | 2.49 |
| GDC-0152 | | IAP | | 103.64 | 103.42 | 99.44 | 85.96 | 99.28 | 98.83 | 91.96 | 86.31 | 1.04 | 1.05 | 1.08 | 1.00 |
| LCL161 | LCL-161 | IAP | | 106.94 | 102.13 | 97.14 | 94.53 | 99.79 | 101.37 | 95.84 | 93.04 | 1.07 | 1.01 | 1.01 | 1.02 |
| Birinapant | TL32711 | IAP | | 103.93 | 98.55 | 99.85 | 103.16 | 96.60 | 98.53 | 97.49 | 100.58 | 1.08 | 1.00 | 1.02 | 1.03 |
| AT-406 | SM-406 | IAP | | 100.77 | 97.56 | 97.48 | 94.75 | 100.68 | 98.73 | 94.81 | 95.31 | 1.00 | 0.99 | 1.03 | 0.99 |
| **AZD5582, aq** | | IAP | | 113.14 | 104.10 | 113.66 | 97.14 | 109.19 | 108.81 | 105.74 | 104.75 | 1.04 | 0.96 | 1.07 | 0.93 |
| AGI-5198 | IDH-C35 | IDH1 | | 98.04 | 98.39 | 93.71 | 81.72 | 101.77 | 93.70 | 96.39 | 92.07 | 0.96 | 1.05 | 0.97 | 0.89 |
| Vorasidenib | AG-881 | IDH1 | IDH2 | 101.54 | 106.50 | 99.59 | 101.88 | 104.12 | 106.71 | 99.34 | 102.62 | 0.98 | 1.00 | 1.00 | 0.99 |
| BAY-1436032 | BAY 1436032, S8 | IDH1 | | 93.54 | 98.53 | 93.62 | 91.48 | 105.03 | 101.92 | 96.02 | 87.15 | 0.89 | 0.97 | 0.98 | 1.05 |

| | | | | | | | | | | | | | | | | | |
|---|---|---|---|---|---|---|---|---|---|---|---|---|---|---|---|---|---|
| Ivosidenib | AG-120 | IDH1 | | | | 96.76 | 96.33 | 89.92 | 70.93 | 103.11 | 99.76 | 92.91 | 78.37 | 0.94 | 0.97 | 0.97 | 0.91 |
| AGI-6780 | | IDH2 | | | | 104.56 | 106.02 | 94.32 | 58.51 | 105.55 | 102.89 | 92.86 | 35.40 | 0.99 | 1.03 | 1.02 | 1.65 |
| Enasidenib | AG-221, CC-9000 | IDH2 | | | | 99.64 | 100.70 | 97.11 | 72.19 | 101.14 | 99.09 | 96.70 | 90.09 | 0.99 | 1.02 | 1.00 | 0.80 |
| BMS-986205 | | IDO1 | | | | 103.83 | 103.39 | 86.94 | 53.83 | 103.07 | 97.10 | 87.02 | 42.21 | 1.01 | 1.06 | 1.00 | 1.28 |
| Epacadostat | INCB024360 | IDO1 | | | | 106.69 | 103.81 | 98.98 | 95.37 | 99.36 | 99.34 | 92.68 | 95.49 | 1.07 | 1.04 | 1.07 | 1.00 |
| BMS-754807 | | IGF1R | | | | 80.26 | 72.22 | 28.38 | 25.30 | 96.08 | 80.85 | 25.68 | 17.76 | 0.84 | 0.89 | 1.11 | 1.42 |
| Linsitinib | OSI-906 | IGF1R | IR | | | 80.64 | 89.55 | 82.51 | 69.65 | 104.84 | 100.50 | 97.44 | 84.69 | 0.77 | 0.89 | 0.85 | 0.82 |
| Mycophenolic acid | | IMPDH1 | IMPDH2 | | | 93.06 | 47.17 | 13.54 | 8.77 | 89.37 | 32.32 | 10.09 | 5.58 | 1.04 | 1.46 | 1.34 | 1.57 |
| Cilengitide | EMD 121974 | Integrin avb3 | Integrin avb5 | | | 110.31 | 101.61 | 86.83 | 50.67 | 102.85 | 91.85 | 70.09 | 43.64 | 1.07 | 1.11 | 1.24 | 1.16 |
| SB273005 | | Integrin avb3 | Integrin avb5 | | | 56.70 | 44.88 | 44.54 | 39.31 | 55.43 | 46.37 | 40.24 | 29.50 | 1.02 | 0.97 | 1.11 | 1.33 |
| MKC-3946 | | IRE1 (endonuclease activity) | | | | 98.38 | 100.10 | 92.43 | 84.90 | 103.72 | 95.47 | 96.10 | 18.67 | 0.95 | 1.05 | 0.96 | 4.55 |
| Cerdulatinib | PRT062070, PRT- | JAK (pan) | SYK | | | 87.83 | 64.73 | 33.81 | 24.21 | 99.12 | 68.18 | 37.10 | 35.31 | 0.89 | 0.95 | 0.91 | 0.69 |
| Itacitinib | INCB39110 | JAK1 | | | | 101.07 | 103.04 | 99.83 | 91.91 | 105.24 | 99.17 | 98.59 | 96.70 | 0.96 | 1.04 | 1.01 | 0.95 |
| AZD4205 | AZD-4205 | JAK1 | | | | 100.68 | 106.51 | 100.22 | 96.09 | 101.43 | 104.00 | 100.54 | 97.38 | 0.99 | 1.02 | 1.00 | 0.99 |
| Filgotinib | GLPG0634 | JAK1 | JAK2 | | | 96.07 | 96.16 | 95.30 | 93.92 | 97.75 | 95.14 | 90.69 | 87.07 | 0.98 | 1.01 | 1.05 | 1.08 |
| Peficitinib | ASP015K, JNJ-54 | JAK1 | JAK3 | | | 96.47 | 96.45 | 93.60 | 81.01 | 98.23 | 97.57 | 93.36 | 86.12 | 0.98 | 0.99 | 1.00 | 0.94 |
| Upadacitinib | ABT-494 | JAK1 | | | | 102.35 | 95.14 | 101.13 | 88.42 | 105.78 | 98.27 | 97.98 | 93.45 | 0.97 | 0.97 | 1.03 | 0.95 |
| Pacritinib | SB1518 | JAK2 | FLT3 | | | 99.54 | 86.17 | 27.60 | 0.68 | 106.48 | 73.11 | 5.40 | 0.69 | 0.93 | 1.18 | 5.11 | 0.99 |
| Fedratinib | TG101348 | JAK2 | BRD4 | | | 103.07 | 95.53 | 27.29 | 0.49 | 97.75 | 94.18 | 24.53 | 0.50 | 1.05 | 1.01 | 1.11 | 0.98 |
| Baricitinib | LY3009104, INCB | JAK2 | JAK1 | | | 102.67 | 98.13 | 95.44 | 91.68 | 94.68 | 97.02 | 93.34 | 88.48 | 1.08 | 1.01 | 1.02 | 1.04 |
| Momelotinib | CYT-387 | JAK2 | JAK1 | TBK1 | | 103.87 | 99.05 | 54.85 | 7.53 | 101.07 | 102.34 | 45.01 | 2.86 | 1.03 | 0.97 | 1.22 | 2.63 |
| Ruxolitinib | INCB018424 | JAK2 | JAK1 | | | 94.76 | 94.42 | 91.55 | 89.34 | 98.73 | 98.77 | 93.02 | 90.23 | 0.96 | 0.96 | 0.98 | 0.99 |
| Tofacitinib | CP-690550 | JAK3 | | | | 94.91 | 100.98 | 94.10 | 98.05 | 99.73 | 103.20 | 91.21 | 98.97 | 0.95 | 0.98 | 1.03 | 0.99 |
| JIB-04 | | JARID1A (KDM | JMJD2 | JMJD3 | JMJD2 | 8.45 | 9.96 | 9.73 | 2.93 | 0.49 | 0.42 | 0.51 | 0.42 | 17.09 | 23.97 | 18.91 | 7.05 |
| CPI-455 | | JARID1A (KDM5A) | | | | 95.83 | 101.09 | 96.39 | 91.16 | 100.41 | 97.54 | 92.29 | 100.43 | 0.95 | 1.04 | 1.04 | 0.91 |
| IOX1 | | JMJC | | | | 104.61 | 104.80 | 100.32 | 98.62 | 103.23 | 98.13 | 100.42 | 95.89 | 1.01 | 1.07 | 1.00 | 1.03 |
| ML324 | | JMJD2 | | | | 96.85 | 100.82 | 88.92 | 15.21 | 96.90 | 87.31 | 54.41 | 6.45 | 1.00 | 1.15 | 1.63 | 2.36 |
| GSK-J4 | | JMJD3 | UTX | | | 96.81 | 90.81 | 9.56 | 2.47 | 98.83 | 57.02 | 3.27 | 0.65 | 0.98 | 1.59 | 2.92 | 3.79 |
| Ispinesib | SB-715992 | Kinesin | | | | 41.91 | 36.20 | 26.16 | 0.36 | 28.85 | 26.19 | 21.13 | 0.41 | 1.45 | 1.38 | 1.24 | 0.87 |
| SB743921 | | Kinesin | | | | 33.67 | 35.65 | 19.62 | 0.49 | 30.87 | 26.74 | 4.49 | 0.49 | 1.09 | 1.33 | 4.37 | 1.01 |
| Dovitinib | TKI-258, CHIR-25 | KIT | VEGFR | | | 73.52 | 35.06 | 36.55 | 0.34 | 106.06 | 36.37 | 33.87 | 0.46 | 0.69 | 0.96 | 1.08 | 0.74 |
| Ripretinib | DCC-2618 | KIT | PDGFRA | | | 77.59 | 47.36 | 33.30 | 18.64 | 87.55 | 52.75 | 33.04 | 12.24 | 0.89 | 0.90 | 1.01 | 1.52 |
| Avapritinib | BLU-285 | KIT | PDGFRa | | | 94.90 | 89.90 | 65.46 | 0.54 | 107.12 | 101.61 | 74.94 | 0.54 | 0.89 | 0.88 | 0.87 | 0.99 |
| Masitinib | AB1010 | KIT | | | | 85.23 | 81.62 | 83.66 | 0.45 | 106.85 | 103.85 | 86.15 | 0.48 | 0.80 | 0.79 | 0.97 | 0.93 |
| MRTX1257 | | KRAS G12C | | | | 106.88 | 97.30 | 71.83 | 0.57 | 100.07 | 93.24 | 7.27 | 0.60 | 1.07 | 1.04 | 9.88 | 0.95 |
| Adagrasib | MRTX849 | KRAS G12C | | | | 98.23 | 99.79 | 15.63 | 0.56 | 101.78 | 100.21 | 8.03 | 0.69 | 0.97 | 1.00 | 1.94 | 0.81 |
| ARS-1620 | | KRAS G12C | | | | 98.79 | 106.17 | 104.53 | 83.11 | 106.49 | 107.41 | 108.73 | 90.07 | 0.93 | 0.99 | 0.96 | 0.92 |
| Sotorasib | AMG-510 | KRAS G12C | | | | 105.37 | 98.19 | 97.08 | 91.34 | 107.83 | 103.99 | 103.05 | 100.46 | 0.98 | 0.94 | 0.94 | 0.91 |
| Deltarasin | | KRAS-PDEd PPI | | | | 101.97 | 105.40 | 0.56 | 0.50 | 101.75 | 95.28 | 0.42 | 0.51 | 1.00 | 1.11 | 1.36 | 0.98 |
| Lifitegrast | | LFA1-ICAM-1 PPI | | | | 98.61 | 97.68 | 92.47 | 91.05 | 98.61 | 99.50 | 88.22 | 91.49 | 1.00 | 0.98 | 1.05 | 1.00 |
| GSK2578215A | | LRRK2 | | | | 95.11 | 96.95 | 90.97 | 51.28 | 95.56 | 95.77 | 94.39 | 36.88 | 1.00 | 1.01 | 0.96 | 1.39 |
| GNE-9605 | | LRRK2 | | | | 101.32 | 101.57 | 92.49 | 99.48 | 103.11 | 102.34 | 92.81 | 98.11 | 0.98 | 0.99 | 1.00 | 1.01 |
| CZC-54252 | | LRRK2 | | | | 90.96 | 91.14 | 50.78 | 23.02 | 98.63 | 94.93 | 59.73 | 13.84 | 0.92 | 0.96 | 0.85 | 1.66 |
| SP2509 | | LSD1 | | | | 26.19 | 13.47 | 11.80 | 9.60 | 20.24 | 13.04 | 7.48 | 6.90 | 1.29 | 1.03 | 1.58 | 1.39 |
| GSK-LSD1 | | LSD1 | | | | 88.64 | 93.27 | 90.26 | 91.51 | 87.21 | 95.06 | 91.86 | 91.40 | 1.02 | 0.98 | 0.98 | 1.00 |
| T-3775440 | | LSD1 | | | | 99.71 | 97.34 | 95.93 | 98.28 | 101.64 | 100.08 | 99.38 | 95.43 | 0.98 | 0.97 | 0.97 | 1.03 |
| OG-L002 | | LSD1 | | | | 100.90 | 96.36 | 93.69 | 94.71 | 101.14 | 99.75 | 94.41 | 98.98 | 1.00 | 0.97 | 0.99 | 0.96 |
| SR9243 | | LXR | | | | 81.96 | 75.26 | 67.84 | 66.43 | 65.41 | 57.10 | 60.76 | 53.94 | 1.25 | 1.32 | 1.12 | 1.23 |
| RGX-104 | | LXR | | | | 102.97 | 108.07 | 96.60 | 22.82 | 104.86 | 102.85 | 98.68 | 23.14 | 0.98 | 1.05 | 0.98 | 0.99 |
| Safinamide | | MAO B | | | | 96.04 | 99.80 | 89.30 | 93.73 | 104.89 | 102.97 | 94.37 | 97.29 | 0.92 | 0.97 | 0.95 | 0.96 |
| UNC669 | | MBT domain | | | | 100.44 | 98.73 | 100.40 | 92.54 | 99.28 | 101.35 | 99.69 | 95.91 | 1.01 | 0.97 | 1.01 | 0.96 |
| UNC1215 | | MBT domain | | | | 96.27 | 96.15 | 92.37 | 92.45 | 98.16 | 99.58 | 96.29 | 91.90 | 0.98 | 0.97 | 0.96 | 1.01 |
| AZD5991 | | MCL1 | | | | 92.56 | 102.83 | 88.20 | 15.59 | 104.68 | 104.20 | 100.25 | 13.96 | 0.88 | 0.99 | 0.88 | 1.12 |
| MIK665 | S64315 | MCL1 | | | | 103.79 | 98.70 | 97.93 | 50.46 | 102.98 | 100.70 | 91.96 | 35.73 | 1.01 | 0.98 | 1.06 | 1.41 |
| S63845 | | MCL1 | | | | 100.19 | 101.43 | 93.36 | 57.03 | 97.71 | 103.74 | 98.04 | 63.17 | 1.03 | 0.98 | 0.95 | 0.90 |
| BAY-8002 | BAY 8002 | MCT1 | | | | 96.41 | 66.47 | 8.85 | 2.27 | 91.84 | 35.87 | 8.30 | 6.59 | 1.05 | 1.85 | 1.07 | 0.34 |
| AZD3965 | | MCT1 | | | | 7.39 | 4.10 | 2.36 | 0.84 | 7.10 | 9.60 | 5.42 | 2.12 | 1.04 | 0.43 | 0.44 | 0.40 |
| DS-3032 | DS-3032b | MDM2 | | | | 57.28 | 30.64 | 7.01 | 0.37 | 35.67 | 14.50 | 3.96 | 0.29 | 1.61 | 2.11 | 1.77 | 1.29 |
| Siremadlin | HDM201 | MDM2 | | | | 23.88 | 7.43 | 0.54 | 0.42 | 22.62 | 3.80 | 0.63 | 0.50 | 1.06 | 1.96 | 0.86 | 0.84 |
| Idasanutlin | RG7388 | MDM2 | | | | 83.64 | 54.02 | 21.40 | 22.37 | 85.05 | 38.48 | 15.19 | 14.88 | 0.98 | 1.40 | 1.41 | 1.50 |
| NVP-CGM097 | CGM097 | MDM2 | | | | 97.61 | 88.20 | 64.77 | 0.68 | 97.79 | 75.77 | 45.30 | 0.48 | 1.00 | 1.16 | 1.43 | 1.42 |
| AMG-232 | | MDM2 | | | | 98.85 | 80.54 | 8.91 | 1.45 | 98.06 | 84.65 | 4.01 | 0.77 | 1.01 | 0.95 | 2.22 | 1.89 |
| Cobimetinib | GDC-0973 | MEK | | | | 26.75 | 21.38 | 14.66 | 0.50 | 22.25 | 16.20 | 10.21 | 0.44 | 1.20 | 1.32 | 1.44 | 1.13 |
| Trametinib | GSK1120212 | MEK | | | | 19.39 | 15.77 | 8.32 | 5.91 | 17.16 | 12.59 | 7.29 | 6.09 | 1.13 | 1.25 | 1.14 | 0.97 |
| TAK-733 | | MEK | | | | 27.23 | 24.74 | 12.13 | 9.03 | 23.25 | 20.35 | 12.97 | 8.68 | 1.17 | 1.22 | 0.94 | 1.04 |
| PD0325901 | | MEK | | | | 28.32 | 24.88 | 16.12 | 5.40 | 25.21 | 21.42 | 12.48 | 6.69 | 1.12 | 1.16 | 1.29 | 0.81 |
| Binimetinib | MEK162, Arry-16 | MEK | | | | 45.19 | 38.54 | 31.87 | 24.42 | 44.58 | 33.62 | 28.15 | 18.88 | 1.01 | 1.15 | 1.13 | 1.29 |
| GDC-0623 | | MEK | | | | 27.28 | 23.73 | 18.83 | 10.27 | 21.64 | 20.89 | 13.93 | 10.31 | 1.26 | 1.14 | 1.35 | 1.00 |
| Selumetinib | AZD6244 | MEK | | | | 49.47 | 38.74 | 33.25 | 25.43 | 45.21 | 36.45 | 32.85 | 23.55 | 1.09 | 1.06 | 1.01 | 1.08 |
| Pimasertib | AS-703026, MSC: | MEK | | | | 33.04 | 30.37 | 22.65 | 16.72 | 31.42 | 28.93 | 21.33 | 13.93 | 1.05 | 1.05 | 1.06 | 1.20 |
| BI-847325 | | MEK | AURKA | AURKB | | 18.78 | 12.13 | 0.58 | 0.37 | 16.77 | 13.00 | 1.30 | 0.36 | 1.12 | 0.93 | 0.45 | 1.05 |
| UNC2025 | | MERTK | FLT3 | | | 99.78 | 60.73 | 7.39 | 0.61 | 92.37 | 41.84 | 1.78 | 0.46 | 1.08 | 1.45 | 4.16 | 1.35 |
| Capmatinib | INC280 | MET | | | | 109.76 | 49.12 | 96.46 | 90.17 | 102.31 | 38.03 | 100.75 | 95.13 | 1.07 | 1.29 | 0.96 | 0.95 |
| Merestinib | LY2801653 | MET | TRKA | multikinase | | 103.65 | 93.46 | 44.86 | 27.41 | 90.72 | 75.68 | 56.12 | 39.84 | 1.14 | 1.24 | 0.80 | 0.69 |
| Foretinib | XL880 | MET | VEGFR | | | 98.90 | 59.76 | 1.35 | 0.53 | 88.18 | 53.17 | 0.54 | 0.47 | 1.12 | 1.12 | 2.51 | 1.12 |
| Amuvatinib | MP-470 | MET | FLT3 | PDGFR | | 73.41 | 69.37 | 50.41 | 32.78 | 89.03 | 69.03 | 47.34 | 16.05 | 0.82 | 1.01 | 1.06 | 2.04 |
| BMS-777607 | BMS777607 | MET | AXL | RON | TYRO3 | 106.93 | 94.88 | 86.68 | 41.62 | 98.15 | 95.47 | 89.71 | 32.20 | 1.09 | 0.99 | 0.97 | 1.29 |
| Savolitinib | Volitinib, AZD609 | MET | | | | 94.67 | 94.95 | 92.41 | 89.65 | 98.78 | 98.80 | 92.57 | 94.84 | 0.96 | 0.96 | 1.00 | 0.95 |
| Tepotinib | EMD1214063 | MET | | | | 110.15 | 97.34 | 104.40 | 19.92 | 110.65 | 99.04 | 99.88 | 0.72 | 1.00 | 0.98 | 1.05 | 27.67 |
| AMG 337 | | MET | | | | 95.35 | 93.30 | 92.20 | 86.02 | 100.51 | 95.87 | 96.68 | 94.09 | 0.95 | 0.97 | 0.95 | 0.91 |
| Glumetinib | SCC244 | MET | | | | 95.03 | 98.95 | 79.08 | 45.30 | 97.53 | 103.11 | 88.13 | 71.88 | 0.97 | 0.96 | 0.90 | 0.63 |
| Altiratinib | DCC-2701 | MET | TIE2 | VEGFR | TRKA-( | 104.35 | 92.08 | 70.86 | 13.52 | 98.71 | 97.49 | 71.82 | 5.85 | 1.06 | 0.94 | 0.99 | 2.31 |
| Glesatinib | MGCD265 | MET | VEGFR | AXL | | 99.71 | 95.10 | 93.06 | 76.77 | 104.31 | 101.20 | 100.54 | 71.14 | 0.96 | 0.94 | 0.93 | 1.08 |
| Sitravatinib | MGCD516 | MET | TRKA-( | AXL | PDGFF | 98.58 | 94.19 | 0.44 | 0.46 | 103.01 | 100.55 | 0.64 | 0.57 | 0.96 | 0.94 | 0.69 | 0.80 |
| JZL 184 | | MGLL | | | | 97.46 | 94.95 | 95.36 | 73.57 | 94.40 | 93.61 | 88.18 | 75.25 | 1.03 | 1.01 | 1.08 | 0.98 |
| ABX-1431 | | MGLL | | | | 96.64 | 95.68 | 95.76 | 94.48 | 95.89 | 101.92 | 99.46 | 95.41 | 1.01 | 0.94 | 0.96 | 0.99 |
| MM-102 | | MLL1 | | | | 103.52 | 101.28 | 95.86 | 94.50 | 94.35 | 96.57 | 93.78 | 86.73 | 1.10 | 1.05 | 1.02 | 1.09 |
| MI-503 | | MLL1 | | | | 96.00 | 96.86 | 0.62 | 0.44 | 96.87 | 97.37 | 0.44 | 0.46 | 0.99 | 0.99 | 1.41 | 0.96 |
| Ilomastat | Galardin, GM600 | MMP | | | | 96.05 | 100.34 | 92.85 | 91.39 | 104.40 | 100.29 | 98.33 | 96.55 | 0.92 | 1.00 | 0.94 | 0.95 |
| Batimastat | BB-94 | MMP | | | | 100.27 | 93.01 | 90.41 | 89.88 | 101.11 | 96.27 | 95.09 | 90.98 | 0.99 | 0.97 | 0.95 | 0.99 |
| Marimastat | BB-2516 | MMP | | | | 94.51 | 92.57 | 93.66 | 92.31 | 96.74 | 96.90 | 94.30 | 93.40 | 0.98 | 0.96 | 0.99 | 0.99 |
| Tomivosertib | eFT-508 | MNK2 | MNK1 | | | 99.56 | 99.77 | 94.71 | 87.22 | 97.94 | 97.04 | 95.32 | 97.52 | 1.02 | 1.03 | 0.99 | 0.89 |
| BAY-1217389 | BAY 1217389 | MPS1 | | | | 38.23 | 38.19 | 30.71 | 18.10 | 35.27 | 30.02 | 21.37 | 3.45 | 1.08 | 1.27 | 1.44 | 5.24 |
| MPI-0479605 | | MPS1 | | | | 36.80 | 43.27 | 39.92 | 36.15 | 42.04 | 36.36 | 35.63 | 31.87 | 0.88 | 1.19 | 1.12 | 1.13 |
| AZ3146 | | MPS1 | | | | 97.56 | 96.02 | 48.79 | 35.43 | 97.52 | 98.60 | 35.49 | 33.36 | 1.00 | 0.97 | 1.37 | 1.06 |
| Dactolisib | NVP-BEZ235, BEZ | mTOR | PI3K | | | 17.24 | 9.37 | 10.31 | 7.37 | 6.17 | 4.58 | 5.45 | 3.29 | 2.80 | 2.04 | 1.89 | 2.24 |
| Gedatolisib | PF-05212384, PK | mTOR | PI3K | | | 62.83 | 39.75 | 17.13 | 10.25 | 25.79 | 21.33 | 7.56 | 5.25 | 2.44 | 1.86 | 2.27 | 1.95 |
| AZD8055 | | mTOR | | | | 32.19 | 13.38 | 8.36 | 7.59 | 12.26 | 7.29 | 5.95 | 5.05 | 2.63 | 1.83 | 1.41 | 1.50 |
| Sapanisertib | INK128 | mTOR | | | | 14.29 | 9.20 | 7.76 | 7.24 | 10.51 | 6.42 | 4.96 | 5.61 | 1.36 | 1.43 | 1.56 | 1.29 |
| PF-04691502 | | mTOR | PI3K | | | 82.09 | 75.89 | 28.63 | 10.95 | 85.79 | 63.81 | 19.12 | 6.13 | 0.96 | 1.19 | 1.50 | 1.79 |
| Apitolisib | GDC-0980 | mTOR | PI3K | | | 88.87 | 71.93 | 26.66 | 10.18 | 95.47 | 69.67 | 9.77 | 4.90 | 0.93 | 1.03 | 2.73 | 2.08 |
| Vistusertib | AZD2014 | mTOR | | | | 87.60 | 58.55 | 18.13 | 7.52 | 92.51 | 60.88 | 12.09 | 3.21 | 0.95 | 0.96 | 1.50 | 2.34 |
| Bimiralisib | PQR309 | mTOR | PI3K | | | 96.48 | 91.58 | 72.69 | 22.86 | 101.66 | 95.58 | 64.16 | 12.36 | 0.95 | 0.96 | 1.13 | 1.85 |
| Pilaralisib | XL147, SAR-2454 | mTOR | PI3K | | | 96.34 | 93.77 | 85.44 | 58.27 | 99.67 | 100.92 | 94.51 | 70.47 | 0.97 | 0.93 | 0.90 | 0.83 |
| OSI-027 | | mTOR | | | | 92.80 | 85.71 | 70.61 | 13.04 | 98.83 | 97.44 | 86.57 | 17.08 | 0.94 | 0.88 | 0.82 | 0.76 |
| Everolimus | | mTORC1 | | | | 73.31 | 71.99 | 76.24 | 57.47 | 48.26 | 41.93 | 37.93 | 28.69 | 1.52 | 1.72 | 2.01 | 2.00 |
| Ridaforolimus | Deforolimus, MK- | mTORC1 | | | | 77.01 | 73.63 | 74.92 | 68.37 | 45.33 | 49.91 | 36.45 | 51.21 | 1.70 | 1.48 | 2.06 | 1.34 |
| Rapamycin | Sirolimus | mTORC1 | | | | 63.97 | 76.07 | 73.80 | 76.86 | 38.39 | 53.04 | 34.92 | 41.97 | 1.67 | 1.43 | 2.11 | 1.83 |
| Temsirolimus | | mTORC1 | | | | 74.92 | 68.63 | 71.56 | 32.47 | 54.33 | 49.42 | 49.08 | 20.98 | 1.38 | 1.39 | 1.46 | 1.55 |
| DMSO | NA | NA | NA | NA | NA | 98.52 | 104.97 | 92.73 | 91.20 | 92.32 | 92.95 | 97.53 | 94.41 | 1.07 | 1.13 | 0.95 | 0.97 |
| DMSO | NA | NA | NA | NA | NA | 94.47 | 107.48 | 93.26 | 95.37 | 102.31 | 99.50 | 94.60 | 99.26 | 0.92 | 1.08 | 0.99 | 0.96 |
| DMSO | NA | NA | NA | NA | NA | 104.32 | 112.39 | 97.54 | 102.28 | 99.64 | 104.90 | 90.43 | 95.73 | 1.05 | 1.07 | 1.08 | 1.07 |
| DMSO | NA | NA | NA | NA | NA | 113.61 | 108.85 | 102.61 | 103.03 | 97.44 | 101.63 | 97.26 | 101.30 | 1.17 | 1.07 | 1.06 | 1.02 |
| DMSO | NA | NA | NA | NA | NA | 103.99 | 102.08 | 95.41 | 95.28 | 102.91 | 95.37 | 98.92 | 90.07 | 1.01 | 1.07 | 0.96 | 1.06 |
| DMSO | NA | NA | NA | NA | NA | 99.80 | 96.59 | 91.37 | 92.80 | 99.64 | 90.36 | 100.84 | 98.78 | 1.00 | 1.07 | 0.91 | 0.94 |
| DMSO | NA | NA | NA | NA | NA | 102.56 | 104.12 | 98.90 | 97.01 | 104.58 | 97.46 | 94.46 | 93.93 | 0.98 | 1.07 | 1.05 | 1.03 |
| DMSO | NA | NA | NA | NA | NA | 96.74 | 106.27 | 90.06 | 94.96 | 103.59 | 101.15 | 95.16 | 98.04 | 0.93 | 1.05 | 0.95 | 0.97 |
| DMSO | NA | NA | NA | NA | NA | 97.73 | 101.95 | 94.49 | 101.44 | 95.04 | 97.24 | 90.26 | 97.24 | 1.03 | 1.05 | 1.05 | 1.04 |
| DMSO | NA | NA | NA | NA | NA | 110.18 | 107.27 | 106.18 | 106.07 | 99.73 | 102.39 | 93.36 | 99.01 | 1.10 | 1.05 | 1.14 | 1.07 |
| DMSO | NA | NA | NA | NA | NA | 98.90 | 103.57 | 96.90 | 95.58 | 90.93 | 98.87 | 102.19 | 94.89 | 1.09 | 1.05 | 0.95 | 1.01 |
| DMSO | NA | NA | NA | NA | NA | 103.97 | 103.09 | 98.88 | 100.35 | 102.43 | 98.59 | 99.04 | 85.87 | 1.02 | 1.05 | 1.00 | 1.17 |
| DMSO | NA | NA | NA | NA | NA | 100.50 | 100.76 | 94.25 | 102.19 | 110.39 | 96.76 | 102.09 | 99.83 | 0.91 | 1.04 | 0.92 | 1.02 |
| DMSO | NA | NA | NA | NA | NA | 103.32 | 99.90 | 101.81 | 93.85 | 103.36 | 96.58 | 96.48 | 100.34 | 1.00 | 1.03 | 1.06 | 0.94 |

| | | | | | | | | | | | | | | | | |
|---|---|---|---|---|---|---|---|---|---|---|---|---|---|---|---|---|
| DMSO | NA | NA | NA | NA | NA | 100.29 | 99.10 | 93.22 | 91.41 | 96.91 | 95.85 | 93.02 | 96.93 | 1.03 | 1.03 | 1.00 | 0.94 |
| DMSO | NA | NA | NA | NA | NA | 100.01 | 96.19 | 93.50 | 91.44 | 86.61 | 93.32 | 91.19 | 93.63 | 1.15 | 1.03 | 1.03 | 0.98 |
| DMSO | NA | NA | NA | NA | NA | 95.26 | 104.80 | 93.79 | 99.93 | 104.94 | 101.78 | 95.71 | 99.16 | 0.91 | 1.03 | 0.98 | 1.01 |
| DMSO | NA | NA | NA | NA | NA | 99.95 | 91.18 | 93.08 | 97.27 | 101.42 | 88.63 | 94.96 | 96.52 | 0.99 | 1.03 | 0.98 | 1.01 |
| DMSO | NA | NA | NA | NA | NA | 102.84 | 103.52 | 98.61 | 95.33 | 102.42 | 100.97 | 97.48 | 97.12 | 1.00 | 1.03 | 1.01 | 0.98 |
| DMSO | NA | NA | NA | NA | NA | 96.64 | 98.44 | 92.66 | 92.87 | 98.84 | 96.51 | 90.71 | 93.95 | 0.98 | 1.02 | 1.02 | 0.99 |
| DMSO | NA | NA | NA | NA | NA | 100.38 | 99.61 | 94.46 | 96.24 | 102.74 | 98.93 | 99.03 | 100.44 | 0.98 | 1.01 | 0.95 | 0.96 |
| DMSO | NA | NA | NA | NA | NA | 101.99 | 99.02 | 91.95 | 98.04 | 101.00 | 98.82 | 98.37 | 95.36 | 1.01 | 1.00 | 0.93 | 1.03 |
| DMSO | NA | NA | NA | NA | NA | 100.59 | 96.73 | 93.86 | 95.50 | 95.44 | 96.72 | 94.83 | 91.62 | 1.05 | 1.00 | 0.99 | 1.04 |
| DMSO | NA | NA | NA | NA | NA | 93.68 | 97.05 | 100.48 | 89.09 | 97.94 | 97.36 | 98.08 | 97.10 | 0.96 | 1.00 | 1.02 | 0.92 |
| DMSO | NA | NA | NA | NA | NA | 95.98 | 95.28 | 102.87 | 100.74 | 101.66 | 96.28 | 88.46 | 90.43 | 0.94 | 0.99 | 1.16 | 1.11 |
| DMSO | NA | NA | NA | NA | NA | 100.51 | 97.56 | 94.03 | 95.94 | 100.30 | 98.66 | 95.70 | 94.86 | 1.00 | 0.99 | 0.98 | 1.01 |
| DMSO | NA | NA | NA | NA | NA | 96.37 | 98.06 | 87.91 | 93.54 | 101.15 | 99.38 | 97.29 | 89.54 | 0.95 | 0.99 | 0.90 | 1.04 |
| DMSO | NA | NA | NA | NA | NA | 99.97 | 96.40 | 100.15 | 95.85 | 106.55 | 98.15 | 100.53 | 93.48 | 0.94 | 0.98 | 1.00 | 1.03 |
| DMSO | NA | NA | NA | NA | NA | 99.91 | 100.26 | 98.48 | 93.59 | 100.55 | 102.59 | 98.27 | 96.50 | 0.99 | 0.98 | 1.00 | 0.97 |
| DMSO | NA | NA | NA | NA | NA | 108.48 | 104.80 | 102.85 | 94.60 | 102.40 | 107.87 | 100.51 | 105.40 | 1.06 | 0.97 | 1.02 | 0.90 |
| DMSO | NA | NA | NA | NA | NA | 99.60 | 106.20 | 94.44 | 97.19 | 100.36 | 110.05 | 95.59 | 99.86 | 0.99 | 0.96 | 0.99 | 0.97 |
| DMSO | NA | NA | NA | NA | NA | 106.53 | 98.09 | 98.57 | 100.21 | 107.37 | 101.95 | 95.88 | 102.07 | 0.99 | 0.96 | 1.03 | 0.98 |
| DMSO | NA | NA | NA | NA | NA | 105.74 | 103.11 | 100.41 | 96.62 | 100.00 | 107.37 | 97.89 | 104.58 | 1.06 | 0.96 | 1.03 | 0.92 |
| DMSO | NA | NA | NA | NA | NA | 98.55 | 97.60 | 100.43 | 93.98 | 100.17 | 101.92 | 97.67 | 101.59 | 0.98 | 0.96 | 1.03 | 0.93 |
| DMSO | NA | NA | NA | NA | NA | 95.50 | 99.84 | 93.98 | 102.71 | 103.67 | 104.54 | 102.95 | 99.23 | 0.92 | 0.96 | 0.91 | 1.04 |
| DMSO | NA | NA | NA | NA | NA | 95.88 | 95.00 | 88.79 | 89.74 | 99.95 | 99.83 | 91.61 | 92.92 | 0.96 | 0.95 | 0.97 | 0.97 |
| DMSO | NA | NA | NA | NA | NA | 102.23 | 96.22 | 96.59 | 90.30 | 104.10 | 101.63 | 95.49 | 100.53 | 0.98 | 0.95 | 1.01 | 0.90 |
| DMSO | NA | NA | NA | NA | NA | 93.19 | 91.57 | 91.08 | 97.93 | 101.52 | 98.47 | 95.13 | 99.47 | 0.92 | 0.93 | 0.96 | 0.98 |
| DMSO | NA | NA | NA | NA | NA | 96.66 | 93.74 | 93.16 | 94.10 | 100.88 | 101.82 | 97.75 | 98.83 | 0.96 | 0.92 | 0.95 | 0.95 |
| DMSO | NA | NA | NA | NA | NA | 97.57 | 89.81 | 97.84 | 94.23 | 101.38 | 100.58 | 95.76 | 96.74 | 0.96 | 0.89 | 1.02 | 0.97 |
| Pevonedistat | MLN-4924 | NAE | | | | 94.22 | 83.71 | 49.62 | 16.57 | 100.21 | 68.99 | 22.35 | 5.76 | 0.94 | 1.21 | 2.22 | 2.88 |
| Nodinitib-1 | ML130 | NOD1 | | | | 97.24 | 100.61 | 93.72 | 87.07 | 98.71 | 99.24 | 97.14 | 90.64 | 0.99 | 1.01 | 0.96 | 0.96 |
| ML385 | | NRF2 | | | | 100.17 | 97.08 | 78.22 | 74.46 | 102.66 | 100.81 | 75.97 | 72.58 | 0.98 | 0.96 | 1.03 | 1.03 |
| MK-3697 | | Orexin receptor | | | | 96.86 | 100.35 | 90.00 | 94.34 | 103.36 | 102.17 | 97.48 | 95.54 | 0.94 | 0.98 | 0.92 | 0.99 |
| **Ralimetinib, aq** | LY2228820 | p38 | | | | 110.13 | 5.95 | 75.49 | 99.05 | 95.95 | 2.28 | 81.34 | 103.60 | 1.15 | 2.61 | 0.93 | 0.96 |
| Pexmetinib | ARRY-614 | p38 | TIE2 | | | 90.67 | 57.39 | 40.76 | 26.06 | 96.10 | 57.33 | 46.70 | 25.85 | 0.94 | 1.00 | 0.87 | 1.01 |
| VX-702 | | p38a | | | | 93.91 | 100.38 | 99.62 | 91.03 | 97.22 | 99.92 | 98.81 | 93.54 | 0.97 | 1.00 | 1.01 | 0.97 |
| PF-03758309 | PF-3758309 | PAK | | | | 28.56 | 30.46 | 31.55 | 17.99 | 18.01 | 24.78 | 23.88 | 8.68 | 1.59 | 1.23 | 1.32 | 2.07 |
| FRAX486 | | PAK | | | | 82.52 | 66.68 | 0.54 | 0.54 | 104.13 | 90.53 | 0.50 | 0.48 | 0.79 | 0.74 | 1.07 | 1.12 |
| KPT-9274 | | PAK4 | NAMPT | | | 81.54 | 24.52 | 5.06 | 0.99 | 46.49 | 23.80 | 5.91 | 0.64 | 1.75 | 1.03 | 0.86 | 1.55 |
| AZ3451 | | PAR2 | | | | 111.99 | 95.58 | 79.49 | 76.13 | 100.52 | 96.00 | 90.92 | 77.99 | 1.11 | 1.00 | 0.87 | 0.98 |
| Talazoparib | BMN673 | PARP1 | | | | 66.24 | 46.66 | 35.59 | 24.20 | 53.47 | 38.82 | 35.66 | 30.30 | 1.24 | 1.20 | 1.00 | 0.80 |
| Niraparib | MK-4827 | PARP1 | | | | 96.54 | 92.05 | 81.32 | 42.29 | 100.19 | 84.89 | 65.34 | 33.24 | 0.96 | 1.08 | 1.24 | 1.27 |
| Pamiparib | BGB-290 | PARP1 | | | | 98.16 | 96.24 | 83.72 | 51.62 | 99.75 | 92.46 | 70.30 | 29.18 | 0.98 | 1.04 | 1.19 | 1.77 |
| Veliparib | ABT-888 | PARP1 | | | | 99.86 | 92.91 | 96.44 | 90.12 | 96.10 | 96.90 | 95.11 | 95.05 | 1.04 | 0.96 | 1.01 | 0.95 |
| Olaparib | AZD2281 | PARP1 | | | | 100.14 | 83.70 | 86.81 | 57.81 | 102.08 | 100.32 | 98.01 | 42.90 | 0.98 | 0.83 | 0.89 | 1.35 |
| Rucaparib | AG-014699, PF-0 | PARP1 | | | | 106.77 | 78.73 | 90.32 | 77.90 | 103.15 | 98.03 | 95.05 | 60.41 | 1.04 | 0.80 | 0.95 | 1.29 |
| Crisaborole | AN2728 | PDE | | | | 105.52 | 96.83 | 95.70 | 89.64 | 103.75 | 92.16 | 92.22 | 88.06 | 1.02 | 1.05 | 1.04 | 1.02 |
| Cilomilast | Ariflo, SB-207499 | PDE | | | | 100.55 | 95.51 | 93.14 | 97.15 | 103.69 | 95.49 | 89.86 | 93.82 | 0.97 | 1.00 | 1.04 | 1.04 |
| Apremilast | CC-10004 | PDE | | | | 102.38 | 100.43 | 101.14 | 91.45 | 110.79 | 102.71 | 96.81 | 95.30 | 0.92 | 0.98 | 1.04 | 0.96 |
| Roflumilast | | PDE4 | | | | 109.46 | 98.97 | 92.29 | 88.04 | 101.36 | 99.85 | 99.25 | 98.42 | 1.08 | 0.99 | 0.93 | 0.89 |
| Sildenafil | | PDE5 | | | | 104.40 | 93.55 | 94.40 | 92.39 | 103.73 | 97.58 | 86.85 | 93.80 | 1.01 | 0.96 | 1.09 | 0.98 |
| Tadalafil | | PDE5 | PDE11 | | | 99.86 | 95.67 | 95.31 | 90.33 | 99.83 | 100.70 | 94.37 | 95.33 | 1.00 | 0.95 | 1.01 | 0.95 |
| Vardenafil | | PDE5 | | | | 98.10 | 94.22 | 88.97 | 87.15 | 98.40 | 102.34 | 95.94 | 93.29 | 1.00 | 0.92 | 0.93 | 0.93 |
| Orantinib | TSU-68 | PDGFR | FGFR | KDR | | 99.03 | 95.13 | 93.02 | 82.58 | 100.42 | 96.11 | 95.58 | 92.66 | 0.99 | 0.99 | 0.97 | 0.89 |
| Sunitinib (free base) | | PDGFR | KIT | VEGFR | | 100.78 | 99.81 | 74.06 | 0.60 | 99.51 | 103.17 | 75.71 | 0.54 | 1.01 | 0.97 | 0.98 | 1.10 |
| Sorafenib (free base) | | PDGFR | VEGFR | BRAF | | 102.24 | 95.93 | 70.21 | 1.15 | 98.80 | 99.57 | 80.62 | 0.81 | 1.03 | 0.96 | 0.87 | 1.42 |
| Crenolanib | CP-868569 | PDGFR | FLT3 | | | 100.96 | 93.10 | 84.26 | 42.82 | 102.32 | 98.17 | 91.86 | 23.13 | 0.99 | 0.95 | 0.92 | 1.85 |
| CCF642 | | PDI | | | | 92.76 | 67.61 | 2.07 | 0.35 | 97.92 | 15.01 | 2.55 | 0.48 | 0.95 | 4.50 | 0.81 | 0.73 |
| GSK2334470 | | PDK1 | | | | 94.87 | 90.55 | 82.46 | 27.61 | 99.85 | 93.87 | 89.05 | 27.38 | 0.95 | 0.96 | 0.93 | 1.01 |
| BX-912 | | PDK1 | | | | 95.82 | 89.23 | 84.79 | 37.59 | 105.58 | 101.02 | 96.52 | 22.12 | 0.91 | 0.88 | 0.88 | 1.70 |
| BMS202 | | PD-L1 | | | | 106.17 | 101.63 | 83.01 | 0.44 | 94.63 | 98.38 | 78.62 | 0.50 | 1.12 | 1.03 | 1.06 | 0.87 |
| BMS-1166 | | PD-L1 | | | | 98.32 | 95.09 | 90.24 | 91.39 | 95.19 | 96.44 | 91.53 | 89.75 | 1.03 | 0.99 | 0.99 | 1.02 |
| PF-04418948 | | PGE2 receptor | | | | 102.70 | 101.64 | 92.85 | 92.72 | 101.22 | 100.58 | 97.48 | 94.75 | 1.01 | 1.01 | 0.95 | 0.98 |
| ONO-AE3-208 | | PGE2 receptor | | | | 101.17 | 95.48 | 96.74 | 89.74 | 100.80 | 97.29 | 96.12 | 93.91 | 1.00 | 0.98 | 1.01 | 0.96 |
| Zosuquidar | LY335979 | PGP | | | | 103.65 | 96.30 | 90.51 | 0.43 | 100.01 | 101.11 | 93.33 | 0.53 | 1.04 | 0.95 | 0.97 | 0.82 |
| Buparlisib | BKM-120 | PI3K (pan) | | | | 97.96 | 79.88 | 31.72 | 4.60 | 98.76 | 88.92 | 16.10 | 4.54 | 0.99 | 0.90 | 1.97 | 1.02 |
| Alpelisib | NVP-BYL719, BYL | PI3Ka | | | | 98.17 | 94.79 | 87.27 | 75.41 | 97.00 | 97.04 | 90.66 | 80.01 | 1.01 | 0.98 | 0.96 | 0.94 |
| Taselisib | GDC-0032 | PI3Ka | | | | 87.82 | 86.38 | 77.65 | 72.90 | 93.73 | 91.70 | 77.30 | 65.60 | 0.94 | 0.94 | 1.00 | 1.11 |
| Serabelisib | TAK-117, MLN11 | PI3Ka | | | | 99.35 | 96.86 | 95.72 | 81.39 | 104.43 | 103.09 | 99.94 | 100.19 | 0.95 | 0.94 | 0.96 | 0.81 |
| Copanlisib | BAY80-6946 | PI3Ka/d | | | | 79.39 | 72.97 | 40.39 | 0.48 | 90.79 | 52.16 | 4.97 | 0.43 | 0.87 | 1.40 | 8.13 | 1.12 |
| AZD8835 | | PI3Ka/d | | | | 104.02 | 97.47 | 82.99 | 41.71 | 103.84 | 102.17 | 94.53 | 87.77 | 1.00 | 0.95 | 0.88 | 0.48 |
| Pictilisib | GDC-0941 (free b | PI3Ka/d | | | | 78.24 | 74.77 | 57.81 | 4.78 | 104.31 | 82.74 | 50.57 | 1.00 | 0.75 | 0.90 | 1.14 | 4.81 |
| GSK2636771 | | PI3Kb | | | | 94.92 | 105.25 | 90.08 | 90.28 | 98.86 | 98.63 | 95.52 | 91.89 | 0.96 | 1.07 | 0.94 | 0.98 |
| AZD6482 | | PI3Kb/d | | | | 97.35 | 98.63 | 94.82 | 21.16 | 98.95 | 93.39 | 89.67 | 51.16 | 0.98 | 1.06 | 1.06 | 0.41 |
| AZD8186 | | PI3Kb/d | | | | 99.21 | 93.69 | 84.36 | 58.86 | 97.88 | 100.28 | 84.65 | 67.06 | 1.01 | 0.93 | 1.00 | 0.88 |
| Acalisib | GS-9820 | PI3Kd | | | | 99.62 | 98.90 | 97.22 | 79.88 | 94.85 | 98.69 | 94.87 | 78.32 | 1.05 | 1.00 | 1.02 | 1.02 |
| Idelalisib | CAL-101, GS-110 | PI3Kd | | | | 107.37 | 93.91 | 93.26 | 91.66 | 96.09 | 97.44 | 97.68 | 89.45 | 1.12 | 0.96 | 0.95 | 1.02 |
| Umbralisib | TGR-1202 | PI3Kd | CSNK1E | | | 99.17 | 95.93 | 91.52 | 55.15 | 99.28 | 100.32 | 92.50 | 41.49 | 1.00 | 0.96 | 0.99 | 1.33 |
| IPI-549 | | PI3Kg | | | | 104.88 | 97.53 | 81.14 | 61.62 | 100.13 | 85.02 | 67.28 | 26.66 | 1.05 | 1.15 | 1.21 | 2.31 |
| Duvelisib | IPI-145 | PI3Kg/d | | | | 99.22 | 96.39 | 88.47 | 83.78 | 95.07 | 93.51 | 85.05 | 79.62 | 1.04 | 1.03 | 1.04 | 1.05 |
| Apilimod | LAM-002A | PIKfyve | | | | 14.67 | 12.21 | 7.30 | 1.21 | 3.01 | 2.64 | 1.18 | 0.45 | 4.88 | 4.62 | 6.19 | 2.71 |
| YM201636 | | PIKfyve | | | | 56.51 | 33.38 | 3.13 | 0.68 | 90.87 | 20.23 | 0.72 | 0.46 | 0.62 | 1.65 | 4.36 | 1.49 |
| SGI-1776 | | PIM | | | | 110.25 | 97.09 | 79.74 | 0.47 | 97.54 | 100.97 | 91.72 | 0.43 | 1.13 | 0.96 | 0.87 | 1.11 |
| INCB053914 | | PIM (pan) | | | | 102.21 | 100.73 | 92.42 | 19.66 | 102.73 | 98.66 | 96.45 | 3.92 | 1.00 | 1.02 | 0.96 | 5.01 |
| AZD1208 | | PIM (pan) | | | | 95.88 | 90.10 | 88.57 | 76.50 | 98.38 | 94.54 | 87.31 | 79.07 | 0.97 | 0.95 | 1.01 | 0.97 |
| LGH447 | PIM447 | PIM (pan) | | | | 91.62 | 89.46 | 84.53 | 68.04 | 103.70 | 100.88 | 97.19 | 86.31 | 0.88 | 0.89 | 0.87 | 0.79 |
| Sotrastaurin | AEB071 | PKC | | | | 100.49 | 100.69 | 92.79 | 87.25 | 100.13 | 100.17 | 95.78 | 95.68 | 1.00 | 1.01 | 0.97 | 0.91 |
| LXS196 | LXS-196, NVP-LX | PKC | | | | 102.82 | 98.20 | 95.58 | 39.78 | 98.66 | 101.40 | 100.91 | 11.10 | 1.04 | 0.97 | 0.95 | 3.58 |
| Ruboxistaurin | LY333531 | PKCb | | | | 99.52 | 89.67 | 72.03 | 23.10 | 97.34 | 97.85 | 88.04 | 1.23 | 1.02 | 0.92 | 0.82 | 18.83 |
| **Perifosine, aq** | | PLC | PKC | AKT | | 106.22 | 107.72 | 88.45 | 87.36 | 96.48 | 103.46 | 95.18 | 86.37 | 1.10 | 1.04 | 0.93 | 1.01 |
| Rigosertib | ON-01910 | PLK1 | | | | 22.35 | 17.60 | 14.46 | 16.29 | 13.44 | 13.62 | 14.63 | 13.20 | 1.66 | 1.29 | 0.99 | 1.23 |
| Volasertib | BI6727 | PLK1 | | | | 26.44 | 27.25 | 12.58 | 0.46 | 39.96 | 28.62 | 13.41 | 0.54 | 0.66 | 0.95 | 0.94 | 0.87 |
| BI2536 | | PLK1 | | | | 27.60 | 29.94 | 23.11 | 0.47 | 37.77 | 34.03 | 24.13 | 0.52 | 0.73 | 0.88 | 0.96 | 0.90 |
| CFI-400945 | | PLK4 | | | | 37.72 | 35.42 | 32.28 | 1.07 | 31.11 | 31.42 | 20.07 | 0.47 | 1.21 | 1.13 | 1.61 | 2.25 |
| CX-5461 | | POL1 | | | | 47.61 | 39.58 | 35.36 | 26.37 | 23.25 | 15.98 | 13.92 | 11.32 | 2.05 | 2.48 | 2.54 | 2.33 |
| ETC-159 | ETC-1922159 | PORCN | | | | 108.50 | 105.93 | 108.60 | 110.64 | 102.19 | 99.11 | 95.11 | 90.70 | 1.06 | 1.07 | 1.14 | 1.22 |
| GNF-6231 | | PORCN | | | | 106.64 | 100.70 | 103.47 | 98.24 | 97.02 | 99.03 | 96.89 | 92.87 | 1.10 | 1.02 | 1.07 | 1.06 |
| WNT974 | LGK974, LGK-97 | PORCN | | | | 99.70 | 97.26 | 94.91 | 80.28 | 97.29 | 98.49 | 90.42 | 84.58 | 1.02 | 0.99 | 1.05 | 0.95 |
| **LB-100, aq** | | PP2A | | | | 100.85 | 100.48 | 93.43 | 99.88 | 100.29 | 100.77 | 108.14 | 104.01 | 1.01 | 1.00 | 0.86 | 0.96 |
| FH535 | | PPARd (Wnt/ | PPARg | | | 100.67 | 93.98 | 75.21 | 45.94 | 96.09 | 90.29 | 81.31 | 33.81 | 1.05 | 1.04 | 0.92 | 1.36 |
| GSK0660 | | PPARd (Wnt/ | PPARg | | | 102.78 | 104.51 | 93.50 | 89.94 | 103.12 | 103.75 | 96.09 | 91.49 | 1.00 | 1.01 | 0.97 | 0.98 |
| GSK3787 | | PPARd (Wnt/ | PPARg | | | 105.63 | 101.65 | 94.73 | 95.39 | 103.46 | 103.55 | 96.37 | 98.57 | 1.02 | 0.98 | 0.98 | 0.97 |
| JNJ-64619178 | | PRMT5 | | | | 76.64 | 70.12 | 57.10 | 39.34 | 61.34 | 51.72 | 24.32 | 9.20 | 1.25 | 1.36 | 2.35 | 4.28 |
| GSK3326595 | EPZ015938 | PRMT5 | | | | 83.91 | 83.28 | 72.90 | 72.08 | 87.20 | 66.67 | 54.55 | 58.72 | 0.96 | 1.25 | 1.34 | 1.23 |
| GSK591 | | PRMT5 | | | | 84.74 | 70.64 | 63.67 | 60.58 | 64.30 | 61.16 | 63.02 | 58.38 | 1.32 | 1.15 | 1.01 | 1.04 |
| MS023 | | PRMT6 | PRMT | PRMT1 | | 91.65 | 87.55 | 75.45 | 46.05 | 89.54 | 80.44 | 61.63 | 21.85 | 1.02 | 1.09 | 1.22 | 2.11 |
| Carfilzomib | PX-171 | Proteasome | | | | 0.38 | 0.49 | 0.36 | 0.53 | 0.35 | 0.41 | 0.30 | 0.43 | 1.11 | 1.19 | 1.19 | 1.23 |
| Ixazomib | MLN9708 | Proteasome | | | | 27.13 | 3.34 | 0.65 | 0.41 | 23.50 | 3.11 | 0.49 | 0.47 | 1.15 | 1.08 | 1.33 | 0.88 |
| Bortezomib | | Proteasome | | | | 0.47 | 0.47 | 0.53 | 0.49 | 0.44 | 0.46 | 0.54 | 0.48 | 1.07 | 1.02 | 0.99 | 1.02 |
| EHT1864 | | RAC | | | | 101.18 | 90.26 | 86.30 | 64.27 | 99.14 | 90.49 | 81.39 | 69.11 | 1.02 | 1.00 | 1.06 | 0.93 |
| Selpercatinib | LOXO-292 | RET | | | | 100.79 | 105.79 | 89.96 | 54.93 | 102.78 | 104.14 | 97.92 | 38.84 | 0.98 | 1.02 | 0.92 | 1.41 |
| Pralsetinib | BLU-667 | RET | | | | 99.09 | 91.96 | 73.62 | 48.10 | 102.13 | 96.85 | 94.31 | 56.95 | 0.97 | 0.95 | 0.78 | 0.84 |
| GSK2982772 | | RIPK1 | | | | 97.69 | 101.10 | 97.95 | 94.73 | 100.75 | 101.25 | 97.94 | 96.36 | 0.97 | 1.00 | 1.00 | 0.98 |
| GSK2606414 | | RIPK1 | PERK | | | 100.61 | 98.25 | 84.40 | 22.97 | 97.54 | 99.06 | 86.18 | 27.52 | 1.03 | 0.99 | 0.98 | 0.83 |
| Ripasudil | K-115 | ROCK | | | | 106.63 | 101.80 | 99.39 | 93.51 | 102.75 | 98.86 | 97.08 | 87.24 | 1.04 | 1.03 | 1.02 | 1.07 |
| Netarsudil | AR-13324 | ROCK | | | | 96.14 | 102.51 | 82.11 | 0.52 | 103.13 | 100.15 | 70.91 | 0.42 | 0.93 | 1.02 | 1.16 | 1.24 |
| LJH685 | | RSK1 | RSK2 | RSK3 | | 98.29 | 94.42 | 87.08 | 81.83 | 102.25 | 103.02 | 96.37 | 100.95 | 0.96 | 0.92 | 0.90 | 0.81 |
| Adapalene | | RXR | | | | 83.87 | 92.70 | 14.05 | 8.78 | 97.95 | 81.81 | 8.58 | 6.67 | 0.86 | 1.13 | 1.64 | 1.32 |
| Bexarotene | | RXR | | | | 92.96 | 96.06 | 90.47 | 72.66 | 88.90 | 93.45 | 87.20 | 77.05 | 1.05 | 1.03 | 1.04 | 0.94 |
| Acitretin | | RXR | | | | 79.83 | 79.62 | 79.03 | 86.15 | 97.38 | 90.27 | 88.86 | 90.25 | 0.82 | 0.88 | 0.89 | 0.95 |
| AT13148 | | S6K | PKA | ROCK | | 97.80 | 99.29 | 63.06 | 32.96 | 94.08 | 84.65 | 49.04 | 11.22 | 1.04 | 1.17 | 1.29 | 2.94 |
| LY2584702 | | S6K | | | | 106.35 | 109.87 | 95.83 | 106.58 | 104.54 | 101.69 | 97.81 | 99.77 | 1.02 | 1.08 | 0.98 | 1.07 |
| SHP099 | | SHP2 | | | | 99.06 | 99.10 | 95.66 | 92.05 | 100.85 | 98.02 | 94.95 | 90.15 | 0.98 | 1.01 | 1.01 | 1.02 |
| RMC-4550 | | SHP2 | | | | 95.70 | 94.74 | 97.94 | 85.49 | 102.65 | 99.36 | 95.95 | 86.67 | 0.93 | 0.95 | 1.02 | 0.99 |
| SRT1720 | | SIRT1 activator | | | | 106.00 | 106.95 | 0.53 | 0.51 | 92.66 | 96.76 | 0.52 | 0.52 | 1.14 | 1.11 | 1.03 | 0.99 |

| | | | | | | | | | | | | | | | | |
|---|---|---|---|---|---|---|---|---|---|---|---|---|---|---|---|---|
| GSK2245840 | SRT2104 | SIRT1 activator | | | 97.34 | 99.35 | 92.39 | 93.77 | 97.12 | 98.06 | 99.55 | 97.69 | 1.00 | 1.01 | 0.93 | 0.96 |
| Selisistat | EX-527 | SIRT1 inhibitor | | | 99.42 | 110.15 | 95.43 | 96.38 | 99.54 | 99.05 | 96.09 | 95.21 | 1.00 | 1.11 | 0.99 | 1.01 |
| BMS-833923 | | SMO | | | 98.53 | 97.22 | 0.49 | 0.38 | 98.73 | 79.78 | 0.43 | 0.33 | 1.00 | 1.22 | 1.14 | 1.18 |
| Erismodegib | Sonidegib, LDE22 | SMO | | | 105.30 | 105.79 | 90.63 | 41.68 | 99.26 | 93.50 | 81.35 | 12.26 | 1.06 | 1.13 | 1.11 | 3.40 |
| Taladegib | LY2940680, LY-2! | SMO | | | 94.80 | 98.35 | 92.51 | 67.13 | 102.86 | 95.66 | 87.98 | 73.49 | 0.92 | 1.03 | 1.05 | 0.91 |
| Vismodegib | GDC-0449 | SMO | | | 91.27 | 102.09 | 94.40 | 95.57 | 100.98 | 100.48 | 98.09 | 100.22 | 0.90 | 1.02 | 0.96 | 0.95 |
| Glasdegib | PF-04449913 | SMO | | | 98.62 | 96.91 | 98.60 | 91.03 | 96.25 | 97.02 | 96.76 | 101.35 | 1.02 | 1.00 | 1.02 | 0.90 |
| BAY-293 | | SOS1-KRAS PPI | | | 102.09 | 92.61 | 25.00 | 97.93 | 98.00 | 94.86 | 2.40 | 101.31 | 1.04 | 0.98 | 10.41 | 0.97 |
| Fingolimod | FTY720 | Sphingosine 1-phosphate receptor | | | 104.33 | 99.46 | 39.19 | 0.59 | 96.25 | 87.81 | 0.43 | 0.41 | 1.08 | 1.13 | 91.81 | 1.44 |
| Siponimod | BAF312 | Sphingosine 1-phosphate receptor | | | 101.03 | 102.06 | 98.08 | 96.24 | 101.47 | 97.63 | 96.96 | 59.29 | 1.00 | 1.05 | 1.01 | 1.62 |
| Ozanimod | RPC1063 | Sphingosine 1-phosphate receptor | | | 100.91 | 100.74 | 81.73 | 0.58 | 100.99 | 102.90 | 71.13 | 0.67 | 1.00 | 0.98 | 1.15 | 0.87 |
| Ponesimod | ACT-128800 | Sphingosine 1-phosphate receptor | | | 91.39 | 87.66 | 65.50 | 2.11 | 101.14 | 99.63 | 83.50 | 8.67 | 0.90 | 0.88 | 0.78 | 0.24 |
| KX2-391 | | SRC | | | 21.98 | 22.75 | 19.90 | 20.79 | 13.57 | 16.04 | 13.20 | 17.00 | 1.62 | 1.42 | 1.51 | 1.22 |
| Saracatinib | AZD0530 | SRC | TGFBR | | 85.11 | 72.99 | 59.59 | 53.28 | 103.11 | 96.00 | 72.79 | 54.21 | 0.83 | 0.76 | 0.82 | 0.98 |
| SPHINX31 | | SRPK1 | | | 97.72 | 96.14 | 84.27 | 82.61 | 101.27 | 104.82 | 86.91 | 9.57 | 0.96 | 0.92 | 0.97 | 8.63 |
| Niclosamide | | STAT3 | | | 77.11 | 37.90 | 1.56 | 0.61 | 83.38 | 29.78 | 1.40 | 0.55 | 0.92 | 1.27 | 1.11 | 1.10 |
| Napabucasin | BBI608 | STAT3 | | | 91.76 | 0.46 | 0.56 | 0.48 | 8.06 | 0.59 | 0.50 | 0.54 | 11.39 | 0.78 | 1.11 | 0.88 |
| **ADU-S100, aq** | | STING | | | 98.22 | 95.56 | 93.15 | 99.67 | 97.46 | 92.30 | 85.05 | 107.08 | 1.01 | 1.04 | 1.10 | 0.93 |
| Entospletinib | GS-9973 | SYK | | | 105.32 | 104.58 | 93.60 | 52.15 | 97.84 | 102.21 | 90.53 | 33.84 | 1.08 | 1.02 | 1.03 | 1.54 |
| **TAK-659, aq** | | SYK | FLT3 | | 107.06 | 101.60 | 92.10 | 101.62 | 96.43 | 100.43 | 100.70 | 109.60 | 1.11 | 1.01 | 0.91 | 0.93 |
| Fostamatinib | R788 | SYK | | | 90.49 | 83.28 | 57.11 | 4.21 | 100.54 | 98.80 | 60.44 | 1.65 | 0.90 | 0.84 | 0.95 | 2.54 |
| Galunisertib | LY2157299 | TGFBR | | | 95.31 | 93.30 | 85.55 | 76.10 | 97.33 | 99.40 | 90.71 | 90.49 | 0.98 | 0.94 | 0.94 | 0.84 |
| LY3200882 | | TGFBR | | | 92.17 | 87.69 | 76.73 | 63.53 | 103.22 | 94.35 | 77.47 | 64.92 | 0.89 | 0.93 | 0.99 | 0.98 |
| Vactosertib | TEW-7197, EW-7: | TGFBR | | | 77.33 | 64.79 | 66.64 | 67.87 | 76.58 | 74.01 | 76.39 | 82.78 | 1.01 | 0.88 | 0.87 | 0.82 |
| Ramatroban | Bay u3405 | Thromboxane receptor | | | 94.99 | 97.18 | 94.65 | 89.03 | 95.40 | 95.03 | 91.03 | 92.31 | 1.00 | 1.02 | 1.04 | 0.96 |
| AZ6102 | | TNKS1/2 | | | 96.99 | 96.06 | 91.04 | 0.50 | 78.03 | 93.88 | 89.65 | 0.73 | 1.24 | 1.02 | 1.02 | 0.68 |
| E7449 | 2X-121 | TNKS1/2 | PARP1 | PARP2 | 100.68 | 94.96 | 79.75 | 52.61 | 102.05 | 94.70 | 71.39 | 42.24 | 0.99 | 1.00 | 1.12 | 1.25 |
| NVP-TNKS656 | TNKS656 | TNKS1/2 | | | 95.72 | 93.25 | 89.63 | 88.75 | 92.59 | 95.08 | 86.00 | 88.75 | 1.03 | 0.98 | 1.04 | 1.00 |
| XAV-939 | | TNKS1/2 | | | 67.65 | 64.42 | 63.70 | 80.77 | 96.88 | 93.12 | 88.20 | 76.89 | 0.70 | 0.69 | 0.72 | 1.05 |
| Entrectinib | RXDX-101 | TRKA-C | ROS1 | ALK | 106.97 | 102.98 | 19.91 | 0.42 | 98.87 | 96.04 | 0.99 | 0.52 | 1.08 | 1.07 | 20.11 | 0.81 |
| Larotrectinib | LOXO-101 | TRKA-C | | | 102.23 | 102.01 | 98.36 | 94.52 | 97.29 | 100.37 | 94.32 | 92.68 | 1.05 | 1.02 | 1.04 | 1.02 |
| Repotrectinib | Ropotrectinib, TP | TRKA-C | ROS1 | ALK | 97.68 | 91.44 | 35.81 | 7.25 | 99.48 | 91.45 | 16.36 | 5.09 | 0.98 | 1.00 | 2.19 | 1.42 |
| Selitrectinib | LOXO-195 | TRKA-C | | | 95.12 | 102.49 | 99.55 | 80.91 | 103.86 | 102.57 | 96.29 | 93.45 | 0.92 | 1.00 | 1.03 | 0.87 |
| Telotristat (ethyl) | LX1606 | Tryptophan hydroxylase | | | 103.79 | 96.34 | 93.18 | 0.41 | 102.20 | 99.32 | 88.57 | 0.39 | 1.02 | 0.97 | 1.05 | 1.07 |
| ABT-751 | E7010 | Tubulin | | | 54.43 | 21.71 | 22.43 | 27.04 | 101.92 | 26.70 | 15.19 | 19.11 | 0.53 | 0.81 | 1.48 | 1.41 |
| TAK-243 | MLN7243 | UAE | | | 95.00 | 72.78 | 1.14 | 0.48 | 103.68 | 91.18 | 3.36 | 0.54 | 0.92 | 0.80 | 0.34 | 0.88 |
| MRT68921 | | ULK1 | ULK2 | | 99.27 | 88.65 | 53.97 | 19.78 | 95.01 | 75.42 | 33.47 | 9.21 | 1.04 | 1.18 | 1.61 | 2.15 |
| Anlotinib | AL3818 | VEGFR | EGFR | multikinase | 108.76 | 84.48 | 18.96 | 0.60 | 98.45 | 78.98 | 11.26 | 0.50 | 1.10 | 1.07 | 1.68 | 1.22 |
| Axitinib | AG-013736 | VEGFR | PDGFI | KIT | 94.16 | 79.79 | 47.16 | 34.63 | 95.62 | 77.29 | 32.33 | 27.53 | 0.98 | 1.03 | 1.46 | 1.26 |
| Cabozantinib | XL184 | VEGFR | MET | FLT3 | 106.04 | 98.80 | 78.68 | 17.57 | 99.44 | 96.41 | 77.99 | 13.74 | 1.07 | 1.02 | 1.01 | 1.28 |
| Apatinib | YN968D1 | VEGFR | | | 100.71 | 100.27 | 98.90 | 73.99 | 94.73 | 99.99 | 93.89 | 76.41 | 1.06 | 1.00 | 1.05 | 0.97 |
| Brivanib (alaninat | BMS-582664 (pr | VEGFR | | | 78.05 | 105.26 | 94.19 | 80.03 | 103.75 | 106.85 | 103.17 | 75.54 | 0.75 | 0.99 | 0.91 | 1.06 |
| Telatinib | BAY 57-9352 | VEGFR | PDGFI | KIT | 98.83 | 97.19 | 86.66 | 80.15 | 97.50 | 100.38 | 94.82 | 79.62 | 1.01 | 0.97 | 0.91 | 1.01 |
| Motesanib | AMG-706 | VEGFR | PDGFI | KIT | 98.13 | 95.98 | 94.66 | 91.66 | 104.39 | 102.09 | 98.23 | 98.04 | 0.94 | 0.94 | 0.96 | 0.93 |
| Cediranib | AZD2171 | VEGFR | PDGFI | FGFR | 93.40 | 88.23 | 84.13 | 0.38 | 97.00 | 99.19 | 90.69 | 0.44 | 0.96 | 0.89 | 0.93 | 0.87 |
| Lenvatinib | E7080 | VEGFR | EGFR | | 78.39 | 85.96 | 92.59 | 87.66 | 101.12 | 100.00 | 98.57 | 94.55 | 0.78 | 0.86 | 0.94 | 0.93 |
| Pazopanib | GW-786034 | VEGFR | PDGFI | KIT | 88.76 | 82.94 | 95.35 | 49.92 | 98.87 | 97.02 | 92.92 | 51.98 | 0.90 | 0.85 | 1.03 | 0.96 |
| Tivozanib | AV-951 | VEGFR | | | 62.23 | 82.92 | 73.60 | 31.70 | 97.54 | 99.49 | 87.16 | 31.78 | 0.64 | 0.83 | 0.84 | 1.00 |
| Nintedanib | BIBF1120 | VEGFR | PDGFI | FGFR | 73.18 | 73.97 | 64.04 | 0.53 | 100.98 | 93.81 | 75.52 | 0.50 | 0.72 | 0.79 | 0.85 | 1.07 |
| Valbenazine | | VMAT2 | | | 97.32 | 97.90 | 94.94 | 88.85 | 99.48 | 93.67 | 93.79 | 91.88 | 0.98 | 1.05 | 1.01 | 0.97 |
| NMS-873 | | VPS/p97 | | | 68.89 | 41.65 | 1.97 | 0.48 | 55.38 | 29.40 | 2.69 | 0.38 | 1.24 | 1.42 | 0.73 | 1.28 |
| Adavosertib | AZD-1775, MK-1: | WEE1 | PLK1 | | 38.65 | 23.16 | 25.39 | 20.16 | 51.55 | 22.43 | 16.41 | 12.97 | 0.75 | 1.03 | 1.55 | 1.55 |
| GSK2830371 | | WIP1 Phosphatase | | | 90.65 | 90.70 | 82.41 | 73.73 | 96.72 | 98.48 | 89.72 | 79.51 | 0.94 | 0.92 | 0.92 | 0.93 |
