## Supplementary Table 2 (pSTY) for "The lipid phosphatase activity of PTEN dampens FRA1 expression via AKT/mTOR signaling to suppress melanoma"

| | | | *Log2 ratio vs. C124S+GFP* | | | WT vs. other | | *T-tests* | | | | |
|---|---|---|---|---|---|---|---|---|---|---|---|---|
| | | | G129E | Y138L | WT | WT | WT | G129E | Y138L | WT | WT | WT |
| | | | vs | vs | vs | vs | vs | vs | vs | vs | vs | vs |
| | | | C124S+GFP | C124S+GFP | C124S+GFP | G129E | Y138L | C124S+GFP | C124S+GFP | C124S+GFP | G129E | Y138L |
| **Symbol** | **Amino** | **Positions Within Proteins** | **G129EvsC124S+GFP _log2ratio** | **Y138LvsC124S+GFP _log2ratio** | **WTvsC124S+GFP _log2ratio** | **WTvsG129E _log2ratio** | **WTvsY138L _log2ratio** | **G129EvsC124S+GFP _pvalue** | **Y138LvsC124S+GFP _pvalue** | **WTvsC124S+GFP _pvalue** | **WTvsG129E _pvalue** | **WTvsY138L _pvalue** |
| Rps28 | T | 21 | -0.720881004 | -1.173620014 | -2.054796878 | -1.333915875 | -0.881176865 | 0.041173535 | 0.025019607 | 0.00027957 | 0.007077639 | 0.067787418 |
| Rps28 | S | 23 | -0.679719872 | -1.049154314 | -1.747158885 | -1.067439012 | -0.698004571 | 0.036401755 | 0.014741683 | 0.000144213 | 0.00823599 | 0.063886834 |
| Cdc42se1 | S | 77 | -0.710788957 | -1.002065351 | -1.621225669 | -0.910436712 | -0.619160318 | 0.04775846 | 0.015707274 | 0.000884171 | 0.009998699 | 0.056597528 |
| Nsd2 | S | 555 | -0.724414313 | -0.882512919 | -1.552270266 | -0.827855953 | -0.669757347 | 0.034125624 | 0.0419485 | 0.003075937 | 0.043997646 | 0.110989974 |
| Sema4b | S | 741 | -0.917144965 | -0.93830643 | -1.383160891 | -0.466015925 | -0.44485446 | 0.008553131 | 0.016011074 | 0.001064961 | 0.024713924 | 0.121020541 |
| Igf2r | S | 2344 | -0.823220461 | -0.927840809 | -1.277657311 | -0.454436849 | -0.349816502 | 0.014998507 | 0.013912507 | 0.001333249 | 0.121022621 | 0.226993104 |
| Epha2 | S | 578 | -0.819888908 | -0.696939701 | -1.036670566 | -0.216781658 | -0.339730866 | 0.001742089 | 0.035833828 | 0.002403725 | 0.274477299 | 0.22430899 |
| Snrnp27 | S | 132 | -0.764421166 | -0.716716519 | -1.408073237 | -0.64365207 | -0.691356718 | 0.020433506 | 0.045203481 | 0.002133596 | 0.054735876 | 0.060386979 |
| Slc2a1 | S | 473 | -0.7326462 | -0.719864011 | -1.085608192 | -0.352961992 | -0.365744181 | 0.017274349 | 0.023000601 | 0.002026475 | 0.10774526 | 0.123412329 |
| Epha2 | Y | 595 | -0.610987929 | -0.813416702 | -1.06977842 | -0.458790491 | -0.256361718 | 0.021453946 | 0.019679099 | 0.001467186 | 0.075298877 | 0.317497845 |
| Epha2 | T | 594 | -0.605498677 | -0.712976341 | -0.689243077 | -0.0837444 | 0.023733264 | 0.004423326 | 0.007924408 | 0.002331479 | 0.502229313 | 0.883567649 |
| Tom1l2 | S | 394 | -0.637928433 | -0.700574567 | -1.137026207 | -0.499097775 | -0.43645164 | 0.027381601 | 0.039347497 | 0.002067724 | 0.037338706 | 0.122865184 |
| Emd | S | 54 | -0.754355152 | -0.615623661 | -0.714881411 | 0.039473741 | -0.099257749 | 0.001557047 | 0.010761084 | 0.037008419 | 0.855644993 | 0.682862183 |
| Atp10a | S | 1433 | -0.639675153 | -0.626705127 | -1.459585945 | -0.819910792 | -0.832880817 | 0.047329669 | 0.035296884 | 0.011310727 | 0.072176883 | 0.068617207 |
| Slc39a6 | S | 481 | -0.717124012 | -0.606700223 | -0.976367143 | -0.259243131 | -0.36966692 | 0.02530359 | 0.042688735 | 0.003712182 | 0.315448788 | 0.174217258 |
| Slc22a23 | S | 12 | -0.686074244 | -0.612551695 | -0.779139431 | -0.093065187 | -0.166587736 | 0.002455563 | 0.045126674 | 0.014155407 | 0.624667778 | 0.521374566 |
| Arhgef2 | S | 718 | -0.645291821 | -0.597369165 | -1.012909842 | -0.36761802 | -0.415540676 | 0.022106038 | 0.03657287 | 0.016431318 | 0.22390921 | 0.186298752 |
| A430005L14Rik | S | 22 | -1.189044147 | -1.248180215 | -0.033941506 | 1.155102641 | 1.214238709 | 0.039881674 | 0.033611469 | 0.948360083 | 0.045540071 | 0.036612034 |
| Rbbp6 | T | 1469 | 0.743178754 | 1.126829486 | 0.34612451 | -0.397054244 | -0.780704975 | 0.000326079 | 1.82788E-05 | 0.014913169 | 0.010607079 | 0.000665451 |
| Epha2 | Y | 589 | -0.700769532 | -0.775712409 | -0.581194245 | 0.119575287 | 0.194518164 | 0.003212726 | 0.005373027 | 0.026011476 | 0.498754295 | 0.352092041 |
| Osbpl11 | S | 15 | -0.72647879 | -0.612031735 | -0.535008771 | 0.191470019 | 0.077022964 | 0.003188511 | 0.044194789 | 0.01355002 | 0.094655995 | 0.712009838 |
| Rpl10l Rpl10 Rpl10-p | T | 125;125 | 0.946347792 | 0.846136051 | 0.841583788 | -0.104764004 | -0.004552263 | 0.005331804 | 0.008997967 | 0.072358624 | 0.753846955 | 0.988958615 |
| Synpo | S | 805 | 0.754962877 | 0.78807893 | 0.247426121 | -0.507536756 | -0.540652809 | 0.000509845 | 0.000400543 | 0.503097815 | 0.229608007 | 0.210293915 |
| Rad51ap1 | S | 19 | 1.371997853 | 0.755367472 | -0.094025643 | -1.466023496 | -0.849393115 | 0.000190364 | 0.011590018 | 0.918121592 | 0.206795471 | 0.400281926 |
| Rad51ap1 | S | 23 | 1.128661441 | 0.626249846 | 0.082502352 | -1.04615909 | -0.543747494 | 0.00019694 | 0.015351516 | 0.903497015 | 0.218308263 | 0.458661788 |
| Sgo2a | S | 253 | 0.752089565 | 0.768119824 | 0.142083915 | -0.61000565 | -0.62603591 | 0.006499232 | 0.004765623 | 0.792043448 | 0.311637767 | 0.302842393 |
| Dnm1 | T | 776 | 0.752232201 | 0.67142709 | -0.054770886 | -0.807003087 | -0.726197975 | 0.006260947 | 0.016436336 | 0.903069728 | 0.163160335 | 0.185003369 |
| Ddx3x | S | 594 | 1.093308154 | 0.588516748 | -0.473668036 | -1.56697619 | -1.062184784 | 0.002375145 | 0.035967876 | 0.528567296 | 0.121961957 | 0.225183275 |
| Kif2c | S | 183 | -0.819765362 | -0.774652051 | -1.689968098 | -0.870202736 | -0.915316047 | 0.033174439 | 0.077466746 | 0.001389182 | 0.024029856 | 0.045319105 |
| Cd44 | T | 726 | -0.811933061 | -0.802995125 | -1.618720853 | -0.806787792 | -0.815725727 | 0.027773458 | 0.075363643 | 0.001245645 | 0.032923079 | 0.075521519 |
| Hnrnpab | S | 266 | -0.925110984 | -0.835513584 | -1.810276026 | -0.885165042 | -0.974762442 | 0.040505677 | 0.114382396 | 0.00241311 | 0.049610338 | 0.078168074 |
| Epha2 | S | 580 | -1.294383632 | -1.008599527 | -2.420613612 | -1.12622998 | -1.412014085 | 0.005060643 | 0.119492805 | 0.005144992 | 0.082731487 | 0.066955986 |
| Abcb1a Abcb1b | S | 16;16 | -1.111377441 | -0.945910348 | -1.638872716 | -0.527495275 | -0.692962368 | 0.006720244 | 0.056641888 | 0.003732177 | 0.153761732 | 0.143406216 |
| Rab21 | S | 194 | -0.949333448 | -0.806434601 | -1.67507526 | -0.725741813 | -0.868640659 | 0.030051793 | 0.076859446 | 0.005515929 | 0.107640039 | 0.082351454 |
| Epb41l2 | S | 68 | -1.027378818 | -0.762537155 | -1.842804438 | -0.81542562 | -1.080267283 | 0.020804742 | 0.19165237 | 0.006810982 | 0.106598563 | 0.100997442 |
| Anxa1 | S | 27 | -1.152712672 | -0.756726836 | -1.389077335 | -0.236364663 | -0.632350499 | 0.01177174 | 0.111515658 | 0.012075951 | 0.466959272 | 0.168057685 |
| Arpp19 | S | 104 | -0.961336436 | -0.768029776 | -1.512622597 | -0.551286161 | -0.744592821 | 0.012572796 | 0.122438142 | 0.011888358 | 0.196392788 | 0.161320245 |
| Skint10 | S | 139 | -1.133273802 | -0.821356458 | -2.047083162 | -0.91380936 | -1.225726705 | 0.029984976 | 0.231686504 | 0.024517902 | 0.186554797 | 0.144856572 |
| Tom1 | S | 182 | -0.860958923 | -0.765000556 | -1.491684001 | -0.630725078 | -0.726683445 | 0.037553578 | 0.052409734 | 0.008751988 | 0.14707181 | 0.105536456 |
| Slc5a3 | S | 557 | -0.907062805 | -0.622155091 | -1.08963916 | -0.182576356 | -0.467484069 | 0.003527987 | 0.120282186 | 0.00473556 | 0.423680745 | 0.217845399 |
| Grsf1 | S | 334 | -0.999854521 | -0.662980151 | -1.190553463 | -0.190698943 | -0.527573312 | 0.025328829 | 0.060565559 | 0.005321646 | 0.504810535 | 0.040309585 |
| Nptn | S | 382 | -0.923973831 | -0.75243483 | -1.395215162 | -0.471241331 | -0.642780332 | 0.018271962 | 0.130855964 | 0.007455679 | 0.196578316 | 0.192183908 |
| Ankrd11 | S | 1070 | -0.942782291 | -0.847370102 | -0.835276343 | 0.107505949 | 0.01209376 | 0.001436667 | 0.060718919 | 0.028869898 | 0.65790598 | 0.973221707 |
| Piwil4 | S | 499 | -0.848307387 | -0.503509287 | -1.241997706 | -0.393690319 | -0.738488419 | 0.010833137 | 0.15930625 | 0.001835832 | 0.095669335 | 0.058944372 |
| Ocrl | S | 170 | -0.945672557 | -0.911187997 | -1.641000618 | -0.695328061 | -0.729812621 | 0.048538884 | 0.055065953 | 0.034185224 | 0.247250273 | 0.229189385 |
| Ocrl | S | 172 | -0.945672557 | -0.911187997 | -1.641000618 | -0.695328061 | -0.729812621 | 0.048538884 | 0.055065953 | 0.034185224 | 0.247250273 | 0.229189385 |
| Tom1 | S | 180 | -0.796064824 | -0.681570154 | -1.483832193 | -0.687767369 | -0.802262039 | 0.020745527 | 0.112530285 | 0.007694508 | 0.100710969 | 0.099495557 |
| Ddx19a | S | 402 | -1.009870263 | -0.521799231 | -1.685712232 | -0.675842068 | -1.163913101 | 0.019205593 | 0.375173903 | 0.009090387 | 0.150846334 | 0.104162187 |
| Hells | S | 178 | -0.662097494 | -0.601615073 | -1.014852988 | -0.352755494 | -0.413237915 | 0.014169067 | 0.053700633 | 0.001974781 | 0.069860915 | 0.127019565 |
| Snx9 | S | 200 | -0.710037962 | -0.75079411 | -1.154825202 | -0.44478724 | -0.404031091 | 0.024569687 | 0.072590892 | 0.004691124 | 0.141857094 | 0.273458408 |
| Ifitm3 | S | 40 | -0.587725416 | -0.574237236 | -1.066854758 | -0.479129341 | -0.492617522 | 0.019178103 | 0.031659078 | 0.002174004 | 0.064057426 | 0.070769229 |
| Slc5a3 | S | 559 | -0.744611305 | -0.521432993 | -0.818547182 | -0.073935877 | -0.297114189 | 0.003398171 | 0.104908163 | 0.00251289 | 0.567904098 | 0.283532476 |
| Snap23 | S | 95 | -0.671689418 | -0.575667143 | -1.059071921 | -0.387382502 | -0.483404778 | 0.014628313 | 0.090186421 | 0.003170862 | 0.101477324 | 0.136237001 |
| Uhrf1bp1l | S | 952 | -0.761441001 | -0.586815056 | -0.96347139 | -0.202030389 | -0.376656334 | 0.004092317 | 0.101351491 | 0.012755054 | 0.407016657 | 0.276007484 |
| Eif4h | S | 21 | -0.747990384 | -0.620036469 | -1.223583708 | -0.475593323 | -0.603547238 | 0.04316211 | 0.091427647 | 0.006625666 | 0.140292308 | 0.091032013 |
| Sema6d | S | 744 | -0.609209347 | -0.638732286 | -1.274544213 | -0.665334867 | -0.635811928 | 0.015822485 | 0.087874223 | 0.014563157 | 0.093440878 | 0.132438577 |
| Ccdc43 | S | 186 | -0.710542931 | -0.550598573 | -0.951901542 | -0.241358611 | -0.401302969 | 0.016827161 | 0.06382429 | 0.006416513 | 0.225520404 | 0.118565258 |
| Atp1a1 | S | 16 | -0.613416385 | -0.473057076 | -0.718832183 | -0.105415797 | -0.245775107 | 0.008330461 | 0.026010658 | 0.003842271 | 0.183179528 | 0.057163339 |
| Sh3pxd2b | S | 528 | -0.617807567 | -0.476796485 | -0.864458047 | -0.24665048 | -0.387661563 | 0.004402992 | 0.112110269 | 0.008995868 | 0.253830285 | 0.197691577 |
| Uhrf1bp1l | S | 932 | -0.711559389 | -0.49813884 | -0.848779402 | -0.137220013 | -0.350640562 | 0.011834945 | 0.107936623 | 0.008643585 | 0.473849178 | 0.210891906 |
| Epb41l2 | T | 597 | -0.727481739 | -0.371435606 | -1.177706473 | -0.450224734 | -0.806270867 | 0.016612506 | 0.343367402 | 0.005595323 | 0.12871505 | 0.086023558 |
| Uhrf1bp1l | S | 934 | -0.702724064 | -0.507277689 | -0.848752385 | -0.146028321 | -0.341474695 | 0.013216752 | 0.102637794 | 0.008869923 | 0.450335404 | 0.21871191 |
| Cttn | S | 261 | -0.626042867 | -0.379624961 | -1.043515328 | -0.417472461 | -0.663890367 | 0.03571586 | 0.228621366 | 0.004968954 | 0.094287426 | 0.05473674 |
| Ppp1r12a | S | 911 | -0.677824312 | -0.417698421 | -0.853465993 | -0.175641681 | -0.435767572 | 0.014205372 | 0.132573549 | 0.017554513 | 0.454567745 | 0.146108011 |
| Ppp1r12a | S | 909 | -0.66018934 | -0.396667123 | -0.827698147 | -0.167508807 | -0.431031024 | 0.02174101 | 0.204535854 | 0.023754095 | 0.500620637 | 0.191226388 |
| Cdc42ep3 | S | 134 | -0.622794892 | -0.19678765 | -0.804565921 | -0.181771029 | -0.607778271 | 0.026261492 | 0.546855203 | 0.004517012 | 0.381072888 | 0.121753371 |
| Dbnl | T | 299 | -0.63842118 | -0.281048129 | -0.717638337 | -0.079217157 | -0.436590208 | 0.015943792 | 0.26885419 | 0.041351696 | 0.749559246 | 0.163824887 |
| Eef2 | T | 54 | -0.618845496 | -0.249448567 | -0.828045671 | -0.209200176 | -0.578597104 | 0.030205712 | 0.418201654 | 0.020534193 | 0.411692461 | 0.110429813 |
| Ppp3cb | S | 479 | -0.606437157 | -0.208421359 | -0.665875686 | -0.05943853 | -0.457454327 | 0.015965673 | 0.410606452 | 0.037940404 | 0.772492594 | 0.12768354 |
| Hnrnpm | S | 452 | -0.592449948 | -0.247317005 | -0.621938061 | -0.029488113 | -0.374621056 | 0.039317455 | 0.407871029 | 0.035111169 | 0.878441691 | 0.194178988 |
| Itprid2 | S | 151 | 0.702051895 | 0.520698926 | 0.970276917 | 0.268225022 | 0.449577991 | 0.012559186 | 0.117426671 | 0.000695359 | 0.197626869 | 0.161458123 |
| Itprid2 | S | 154 | 0.702051895 | 0.520698926 | 0.970276917 | 0.268225022 | 0.449577991 | 0.012559186 | 0.117426671 | 0.000695359 | 0.197626869 | 0.161458123 |
| Ugp2 | S | 13 | -0.953500369 | -0.109955161 | -0.276837899 | 0.67666247 | -0.166882738 | 0.0024619 | 0.755482535 | 0.240025765 | 0.010995812 | 0.617271042 |
| Eif6 | S | 175 | -0.633547189 | -0.134509645 | 0.072079933 | 0.705627122 | 0.206589577 | 0.012185879 | 0.27948516 | 0.693021217 | 0.024360221 | 0.326507301 |
| Rbm27 | T | 889 | 0.648425704 | 0.323321627 | -0.027117186 | -0.675542891 | -0.350438813 | 0.000723511 | 0.072238006 | 0.806997591 | 0.000466891 | 0.06478865 |
| Gmeb2 | S | 385 | 0.62765908 | -0.030780487 | -0.177877672 | -0.805536752 | -0.147097185 | 0.001646989 | 0.906405473 | 0.360022342 | 0.01466545 | 0.601329403 |
| Dmxl1 | S | 1903 | -0.635747654 | -0.349787286 | -0.606353339 | 0.029394314 | -0.256566053 | 0.002176707 | 0.287007157 | 0.074125191 | 0.902774937 | 0.466280852 |
| Abi1 | S | 292 | -0.626330119 | -0.119594987 | -0.97260789 | -0.346277771 | -0.853012904 | 0.041152043 | 0.69727546 | 0.066139201 | 0.399172202 | 0.098987276 |
| Usp47 | S | 849 | -0.589826935 | -0.235687245 | -0.679559294 | -0.089732358 | -0.443872048 | 0.010809506 | 0.286049732 | 0.24945184 | 0.856108579 | 0.414076672 |
| Tmsb10 | T | 23;23 | -0.645277391 | -0.115485339 | -0.367988787 | 0.277288604 | -0.252503448 | 0.00210741 | 0.665359209 | 0.297235483 | 0.407543394 | 0.503315682 |
| Pacs1 | S | 526 | -0.601840471 | -0.114871265 | -0.412527909 | 0.189312562 | -0.297656644 | 0.010495404 | 0.635168649 | 0.19998374 | 0.484544963 | 0.347029485 |
| Noc2l | T | 680 | -0.658555443 | -0.279297663 | -0.146130171 | 0.512425272 | 0.133167492 | 0.010601021 | 0.392730185 | 0.646633499 | 0.164120695 | 0.718170365 |
| Pip4k2b | S | 319 | -0.629331427 | 0.029845525 | -0.258332893 | 0.370998534 | -0.288178418 | 0.002940682 | 0.821074261 | 0.481889093 | 0.333991367 | 0.431155407 |
| Kif2a | S | 432 | 0.857676475 | 0.185130713 | 0.536253053 | -0.321423422 | 0.35112234 | 0.002749239 | 0.309403497 | 0.01898773 | 0.093390342 | 0.043715718 |

| | | | | | | | | | | | | |
|---|---|---|---|---|---|---|---|---|---|---|---|---|
| Tpx2 | T | 113 | 0.713963663 | 0.527886808 | 0.076683166 | -0.637280497 | -0.451203642 | 0.000914543 | 0.004741358 | 0.865177023 | 0.243096779 | 0.366656378 |
| Srek1 | S | 238 | 0.599506836 | 0.536756129 | 0.112145515 | -0.487361321 | -0.424610613 | 0.002944026 | 0.004034668 | 0.787438897 | 0.302558193 | 0.354707331 |
| Oxsr1 | S | 359 | 0.656141904 | 0.152463511 | 0.178304394 | -0.477837511 | 0.025840883 | 0.000102022 | 0.458840491 | 0.059666812 | 0.001322289 | 0.890045382 |
| Rpl23a | S | 43 | 0.636921316 | 0.46623873 | -0.17769932 | -0.814620637 | -0.64393805 | 0.005068075 | 0.016799369 | 0.69089861 | 0.156451591 | 0.226623651 |
| Rfc4 | S | 29 | 0.633811734 | 0.538265764 | -0.120113642 | -0.753925376 | -0.658379406 | 0.012459468 | 0.025012169 | 0.843658576 | 0.28453064 | 0.333478456 |
| Cenpa | S | 15 | 0.611024142 | 0.455752301 | -0.004186293 | -0.615210435 | -0.459938594 | 0.023115038 | 0.013398263 | 0.99231821 | 0.23606617 | 0.34311669 |
| Faf1 | T | 579 | 0.614788834 | 0.471985289 | -0.263685504 | -0.878474338 | -0.735670793 | 0.005031719 | 0.028991019 | 0.580236564 | 0.151177526 | 0.197777378 |
| Rbm27 | S | 128 | 0.638734292 | 0.439338033 | -0.191638551 | -0.830372843 | -0.630976584 | 0.016367858 | 0.065009484 | 0.696332214 | 0.175394131 | 0.256901133 |
| D1Pas1 Ddx3y | S | 592;592 | 0.643033156 | 0.198540717 | -0.580942612 | -1.223975768 | -0.779483329 | 0.004077745 | 0.21887849 | 0.248335844 | 0.072003926 | 0.162446289 |
| Akt1s1 | T | 247 | -0.122612614 | -1.629697936 | -2.289125992 | -2.166513378 | -0.659428056 | 0.265489441 | 8.62669E-08 | 3.05091E-06 | 6.70178E-05 | 0.001035012 |
| Phldb1 | S | 1203 | -0.782126554 | -1.426707218 | -2.291759576 | -1.509633022 | -0.865052358 | 0.092259774 | 0.001085975 | 9.93503E-05 | 0.017333021 | 0.022445174 |
| Acly | T | 453 | 0.279583827 | -0.71720894 | -1.387187841 | -1.666771668 | -0.669978901 | 0.127881177 | 0.027086267 | 0.000919038 | 0.00090924 | 0.040010463 |
| Foxo4 | S | 262 | -0.285308847 | -3.406516409 | -3.34147644 | -3.056167593 | 0.06503997 | 0.398394839 | 8.57309E-06 | 2.3756E-06 | 0.001631776 | 0.620713587 |
| Msn | S | 384 | -0.089763918 | -1.765311645 | -1.936788102 | -1.847024183 | -0.171476456 | 0.183477563 | 4.26028E-08 | 3.20226E-09 | 8.03028E-06 | 0.012272475 |
| Hnrnpa1 | S | 199 | -0.056216834 | -1.440155183 | -1.481251345 | -1.425034511 | -0.041096163 | 0.124798451 | 1.00362E-09 | 2.28249E-07 | 3.92787E-05 | 0.324862561 |
| Foxo4 | S | 197 | -0.104014914 | -1.741650122 | -1.924137589 | -1.820122675 | -0.182487467 | 0.321514911 | 1.38047E-06 | 7.01297E-07 | 6.71757E-05 | 0.096475456 |
| Myef2 Gm9833 | S | 511 | -0.031014499 | -1.425869763 | -1.398210289 | -1.36719579 | 0.027659474 | 0.68215244 | 6.07864E-11 | 1.07051E-06 | 0.000698421 | 0.390271581 |
| Arhgap22 | S | 397 | -0.036279052 | -1.109327432 | -1.482255733 | -1.445976681 | -0.372928301 | 0.620616859 | 2.83726E-08 | 1.4994E-09 | 0.000604963 | 0.00032972 |
| Rps27a | T | 87 | -0.035739265 | -0.990135273 | -1.155294498 | -1.119555232 | -0.165159224 | 0.227421802 | 2.9505E-10 | 6.25793E-10 | 0.000108074 | 0.000377799 |
| Dlgap4 | T | 714 | -0.120343248 | -1.318642193 | -1.466900132 | -1.346556884 | -0.148257939 | 0.061367974 | 1.0809E-08 | 5.4222E-05 | 0.000120218 | 0.107139863 |
| Eif4b | S | 425 | -0.061963605 | -1.342541469 | -1.525630987 | -1.463667382 | -0.183089518 | 0.223445559 | 2.49705E-08 | 1.44406E-05 | 0.000380408 | 0.067518757 |
| Akt1s1 | S | 184 | -0.146981121 | -1.182607956 | -1.382228544 | -1.235247423 | -0.199620588 | 0.092257388 | 1.94251E-05 | 7.9545E-09 | 0.001300416 | 0.037232817 |
| Stip1 | S | 16 | -0.202546254 | -1.250291699 | -1.336175117 | -1.133628863 | -0.085883418 | 0.002632332 | 0.000375947 | 1.17522E-08 | 7.84219E-06 | 0.18056343 |
| Hnrnpa1 | S | 197 | 0.020798609 | -1.475298042 | -1.404580826 | -1.425379435 | 0.070717216 | 0.672760046 | 5.62882E-06 | 1.85609E-08 | 8.01037E-06 | 0.250343832 |
| Prrc2c | S | 901 | -0.12093171 | -1.249932107 | -1.396290614 | -1.275358904 | -0.146358507 | 0.085077978 | 1.91351E-05 | 1.21253E-07 | 8.32556E-06 | 0.127628923 |
| Wnk1 | T | 58 | -0.164945685 | -0.926110227 | -1.272612425 | -1.10766674 | -0.346502198 | 0.057855975 | 3.23577E-07 | 1.0025E-07 | 0.000157986 | 0.001643379 |
| Rrn3 | S | 42 | 0.025241345 | -1.100655622 | -1.29568279 | -1.320924135 | -0.195027168 | 0.686466338 | 5.75164E-08 | 3.12271E-08 | 0.00012094 | 0.006514416 |
| Prrc2c | S | 904 | -0.118506485 | -1.248416012 | -1.400905561 | -1.282399076 | -0.152489549 | 0.068905937 | 4.42572E-05 | 8.29294E-08 | 0.000170738 | 0.130107686 |
| Usp36 | S | 841 | -0.060859916 | -1.486362255 | -1.507811803 | -1.446951887 | -0.021449548 | 0.367475145 | 5.80177E-07 | 6.91446E-05 | 0.000297544 | 0.79217269 |
| Lmna | S | 429 | -0.060823799 | -1.195015599 | -1.139802861 | -1.078979062 | 0.055212739 | 0.134091138 | 7.86108E-06 | 1.5066E-08 | 1.57838E-05 | 0.326533082 |
| Zfp36l2 | S | 147 | -0.301126602 | -0.948635936 | -1.024072111 | -0.722945509 | -0.075436175 | 0.059202895 | 2.94506E-07 | 3.51499E-07 | 0.008827362 | 0.129458504 |
| Cct2 | S | 260 | -0.104729981 | -1.370335116 | -1.57609279 | -1.471362809 | -0.205757674 | 0.080558505 | 3.47262E-05 | 8.30342E-06 | 0.000274518 | 0.068376985 |
| Gab2 | S | 220 | -0.196605278 | -1.206705522 | -1.440482204 | -1.243876925 | -0.233776682 | 0.000936461 | 0.000661827 | 4.45933E-06 | 4.73601E-05 | 0.043566561 |
| Smarcc1 | S | 318 | -0.189521733 | -1.599858045 | -1.528493674 | -1.33897194 | 0.071364371 | 0.071151152 | 1.58438E-07 | 0.012861903 | 0.015101832 | 0.759661862 |
| Rps6 Rps6-ps4 | S | 246 | 0.191596148 | -1.523647801 | -1.492327304 | -1.683923452 | 0.031320497 | 0.147628664 | 3.53754E-06 | 1.43014E-06 | 0.000292276 | 0.714576079 |
| Eif3a | S | 584 | -0.093542462 | -0.942204622 | -0.934318694 | -0.840776232 | 0.007885928 | 0.05282331 | 7.38741E-08 | 2.9673E-07 | 2.63663E-05 | 0.75300799 |
| Eif4b | S | 422 | -0.024998035 | -1.052808355 | -1.196172907 | -1.171174872 | -0.143364552 | 0.634482417 | 3.49076E-06 | 1.01032E-06 | 0.000586896 | 0.057673524 |
| Rasa3 | S | 207 | 0.098915548 | -1.021551887 | -1.173628222 | -1.27254377 | -0.152076334 | 0.033117786 | 6.35568E-08 | 2.6608E-08 | 2.76858E-05 | 0.01245766 |
| Afdn | S | 1795 | 0.06691214 | -1.287675844 | -1.310817762 | -1.377729902 | -0.023141918 | 0.070081235 | 2.2428E-09 | 0.00014458 | 0.000192016 | 0.678788515 |
| Rps27a | T | 86 | -0.031899438 | -0.966277223 | -1.132254989 | -1.10035555 | -0.165977766 | 0.49263116 | 2.95653E-09 | 1.86583E-05 | 2.43846E-05 | 0.030390735 |
| Larp1 | S | 743 | -0.037992661 | -0.974087288 | -0.978977927 | -0.940985266 | -0.00489064 | 0.368326692 | 1.02331E-07 | 3.41321E-07 | 0.000638061 | 0.633823571 |
| Phldb2 | S | 46 | -0.476900705 | -1.154908587 | -1.632618041 | -1.155717336 | -0.477709453 | 0.042306609 | 0.000406597 | 0.000400682 | 0.004661859 | 0.069188469 |
| Diaph3 | T | 675 | -0.078840999 | -0.948913977 | -0.990241394 | -0.911400395 | -0.041327416 | 0.283033524 | 2.86525E-07 | 1.15157E-06 | 0.000271388 | 0.418382057 |
| Rps6kb2 | S | 370 | -0.140747713 | -1.156323229 | -1.467253929 | -1.326506216 | -0.310930699 | 0.283462924 | 0.00093453 | 1.39049E-06 | 0.000689877 | 0.077778575 |
| Rps6kb2 | S | 379 | -0.140747713 | -1.156323229 | -1.467253929 | -1.326506216 | -0.310930699 | 0.283462924 | 0.00093453 | 1.39049E-06 | 0.000689877 | 0.077778575 |
| Pfkfb2 | S | 486 | 0.008860629 | -1.145574418 | -1.71068471 | -1.719545338 | -0.565110292 | 0.944447826 | 0.002866531 | 3.66103E-07 | 0.000790236 | 0.023796015 |
| Rps6 Rps6-ps4 | S | 244 | 0.242290779 | -1.40155202 | -1.300376195 | -1.542666974 | 0.101175825 | 0.18291522 | 2.42859E-06 | 5.88654E-06 | 0.001744278 | 0.305069406 |
| Eif4b | T | 420 | 0.05376665 | -1.244998251 | -1.481585372 | -1.535352022 | -0.236587121 | 0.299210545 | 2.55807E-08 | 0.001174787 | 0.002042821 | 0.120934223 |
| Sqstm1 | T | 269 | -0.127082516 | -0.742958486 | -0.805119696 | -0.67803718 | -0.062161209 | 0.036092712 | 6.49755E-08 | 1.31206E-06 | 0.000135327 | 0.136449584 |
| Pfkl | S | 775 | -0.51723566 | -1.217879523 | -1.493041446 | -0.975805786 | -0.275161923 | 0.011426752 | 0.002923715 | 0.000770592 | 0.009852569 | 0.28505225 |
| Cad | S | 1859 | -0.133187517 | -0.6976637 | -0.841676869 | -0.708489352 | -0.144013169 | 0.053988523 | 1.10386E-05 | 1.26031E-08 | 0.000922916 | 0.013012947 |
| Pds5a | S | 1274 | -0.128559243 | -1.080158848 | -1.033011297 | -0.904452054 | 0.047147551 | 0.084828052 | 0.000376343 | 6.00078E-07 | 7.64208E-05 | 0.613304108 |
| Eif4b | S | 424 | 0.075875717 | -1.16268228 | -1.418231845 | -1.494107562 | -0.255549565 | 0.259701789 | 6.40381E-08 | 0.000696345 | 0.000660719 | 0.090123075 |
| Riox1 | S | 107 | -0.160791495 | -0.879882438 | -0.885226543 | -0.724435048 | -0.005344105 | 0.030294227 | 1.50479E-07 | 0.000196529 | 0.000380736 | 0.922828866 |
| Prim1 | S | 59 | -0.118599619 | -0.940215134 | -0.76892142 | -0.6503218 | 0.171293714 | 0.37523155 | 5.68071E-07 | 1.17676E-06 | 0.024243592 | 0.000115875 |
| Bad | S | 170 | -0.250594061 | -1.250281846 | -1.542865207 | -1.292271147 | -0.292583361 | 0.221462322 | 0.000514784 | 0.000332504 | 3.14908E-06 | 0.097661967 |
| Numa1 | S | 1974 | -0.054296017 | -0.817032221 | -0.79013228 | -0.735836263 | 0.026899941 | 0.256174289 | 1.05675E-06 | 1.02213E-07 | 0.000323385 | 0.396526653 |
| Rps6 Rps6-ps4 | S | 247 | 0.278046797 | -1.487717255 | -1.314629402 | -1.592676199 | 0.173087853 | 0.219810277 | 1.2096E-05 | 2.91005E-05 | 0.003228967 | 0.20235402 |
| Rps6kb1 | S | 452 | -0.099736414 | -1.05034787 | -1.04772753 | -0.947991116 | 0.00262034 | 0.198712447 | 0.000289648 | 1.78593E-06 | 0.000111448 | 0.977389176 |
| Larp1 | S | 845 | -0.041437849 | -0.7845791 | -0.705259615 | -0.663821766 | 0.079319485 | 0.201279808 | 5.49269E-07 | 1.35852E-07 | 1.47322E-05 | 0.061022769 |
| Rps6 Rps6-ps4 | T | 241 | 0.001610146 | -1.276165253 | -1.731298934 | -1.73290908 | -0.455133681 | 0.954289243 | 0.000573502 | 0.000196176 | 0.000425774 | 0.004552445 |
| Kif13a | T | 1717 | -0.083484322 | -1.199789026 | -1.39099829 | -1.307513969 | -0.191209264 | 0.238966801 | 0.000923213 | 6.4077E-05 | 7.15983E-05 | 0.051417087 |
| Pfkfb2 | S | 469 | -0.069812075 | -0.822546758 | -1.171787422 | -1.101975347 | -0.349240664 | 0.536204624 | 9.67671E-06 | 1.49787E-05 | 0.000821531 | 0.011160811 |
| Eaf1 Eaf2 | S | 21;22 | -0.584620453 | -0.974440083 | -1.443370519 | -0.858750066 | -0.468930436 | 0.032454712 | 0.006501294 | 0.000653875 | 0.011607349 | 0.0946667 |
| Acly | S | 457 | -0.125885205 | -0.967154706 | -1.087406518 | -0.961521313 | -0.120251813 | 0.058702764 | 4.05383E-05 | 0.000115567 | 6.63164E-05 | 0.085399048 |
| Map2k4 | S | 78 | -0.350325454 | -0.84237999 | -1.177052027 | -0.826726573 | -0.334672037 | 0.014158375 | 0.003901042 | 3.67827E-05 | 0.001196926 | 0.071890805 |
| Plec | S | 4397 | -0.219975023 | -0.672000684 | -1.134115418 | -0.914140395 | -0.462114734 | 0.164527428 | 0.000115312 | 7.90959E-06 | 0.004420985 | 0.003810125 |
| Gsk3b | S | 9 | -0.269861479 | -0.817972732 | -1.217611181 | -0.947749703 | -0.39963845 | 0.147232936 | 0.000731729 | 2.53789E-05 | 0.005113937 | 0.024617836 |
| Fkbp15 | S | 1127 | 0.028769876 | -0.726209252 | -0.810679871 | -0.839449747 | -0.08447062 | 0.703219298 | 1.0834E-07 | 5.29382E-07 | 0.001926828 | 0.060817649 |
| Eef2 | S | 595 | -0.025828618 | -0.870515664 | -1.180022537 | -1.154193919 | -0.309506873 | 0.477714207 | 6.76401E-08 | 0.001625297 | 0.004195782 | 0.056098707 |
| Zc3hav1 | S | 347 | -0.209027358 | -1.04840389 | -1.346724127 | -1.137696769 | -0.298320237 | 0.252275598 | 3.95969E-06 | 0.00984878 | 0.005752294 | 0.193718219 |
| Ranbp3 | S | 58 | 0.020881396 | -0.795690794 | -1.105799308 | -1.126680704 | -0.310108514 | 0.709879038 | 3.99457E-07 | 5.40428E-05 | 5.46672E-05 | 0.007716828 |
| Rps6 Rps6-ps4 | S | 235 | -0.039803846 | -1.091740729 | -1.462644199 | -1.422840352 | -0.370903469 | 0.600893986 | 3.23676E-05 | 0.001230402 | 0.000596133 | 0.039078065 |
| Dab2 | S | 401 | -0.00651357 | -0.699023716 | -0.855337758 | -0.848824188 | -0.156314042 | 0.911161971 | 7.28849E-07 | 1.06129E-06 | 0.000164634 | 0.020537922 |
| Foxo3 | S | 252 | -0.149218462 | -0.695377547 | -0.760458433 | -0.611239972 | -0.065080887 | 0.167286081 | 6.63743E-06 | 2.4849E-06 | 0.007084216 | 0.146371517 |
| Eif4ebp1 | T | 69 | -0.331072717 | -0.959585817 | -1.073115348 | -0.742042631 | -0.113529531 | 0.164076625 | 0.001248698 | 7.92773E-05 | 0.029891864 | 0.350876274 |
| Thoc5 | T | 30 | -0.009333045 | -1.060708571 | -0.836902483 | -0.827569439 | 0.223806088 | 0.907486615 | 6.65884E-06 | 4.64408E-05 | 1.04495E-05 | 0.006441314 |
| Ltv1 | S | 17 | -0.500677511 | -1.000763983 | -1.655445942 | -1.154768431 | -0.654681959 | 0.127246313 | 0.047083328 | 0.000275564 | 0.013691046 | 0.135274832 |
| Sqstm1 | T | 272 | -0.139915004 | -0.75774079 | -0.795160488 | -0.655245484 | -0.037419698 | 0.126870164 | 8.11009E-07 | 0.000154311 | 0.001574647 | 0.435973226 |
| Shroom3 | S | 1671 | 0.02767249 | -0.866363348 | -0.832211366 | -0.859883856 | 0.034151982 | 0.779387789 | 5.85675E-08 | 0.00017387 | 0.002660234 | 0.543054363 |
| Irs1 | S | 444 | -0.017880358 | -0.665938357 | -0.869193252 | -0.851312894 | -0.203254894 | 0.788956788 | 4.74848E-06 | 9.72091E-07 | 0.000216645 | 0.007835396 |
| Irs1 | T | 441 | -0.017880358 | -0.665938357 | -0.869193252 | -0.851312894 | -0.203254894 | 0.788956788 | 4.74848E-06 | 9.72091E-07 | 0.000216645 | 0.007835396 |
| Irs1 | T | 448 | -0.017880358 | -0.665938357 | -0.869193252 | -0.851312894 | -0.203254894 | 0.788956788 | 4.74848E-06 | 9.72091E-07 | 0.000216645 | 0.007835396 |
| Bad | S | 155 | 0.076438478 | -1.015559148 | -1.369380805 | -1.445819284 | -0.353821658 | 0.616773772 | 0.000401662 | 3.91984E-05 | 0.00011212 | 0.013490571 |
| Usp36 | S | 578 | 0.054270862 | -0.723257169 | -0.710837436 | -0.765108298 | 0.012419733 | 0.536212059 | 6.86794E-07 | 1.80492E-07 | 0.005728123 | 0.590324479 |
| Phldb2 | S | 413 | -0.162740682 | -0.684524288 | -0.8487888 | -0.686048118 | -0.164264512 | 0.064050807 | 2.45285E-05 | 3.64154E-05 | 0.000701597 | 0.077721849 |
| Mdm2 | S | 183 | -0.021812669 | -0.643870418 | -0.910575816 | -0.888763147 | -0.266705398 | 0.605012879 | 0.001414051 | 7.6411E-09 | 0.00015609 | 0.024971852 |
| Plekhf2 | S | 226 | -0.243327953 | -0.969041836 | -1.221351556 | -0.978023603 | -0.25230972 | 0.010681766 | 0.01018215 | 0.000369868 | 0.000868541 | 0.177490291 |
| Cbx5 | S | 93 | -0.054938299 | -0.693941728 | -0.722163661 | -0.667225362 | -0.028221933 | 0.117929995 | 1.70671E-05 | 1.28767E-06 | 0.00033795 | 0.538340667 |
| Pum3 | S | 13 | -0.049975707 | -0.605635043 | -0.775335192 | -0.725359485 | -0.169700149 | 0.52698665 | 1.75384E-05 | 1.89714E-07 | 0.004023811 | 0.017451673 |
| Lrrc47 | S | 293 | -0.09303779 | -0.670021627 | -1.015884864 | -0.922847074 | -0.345863237 | 0.413699857 | 0.000660414 | 1.36852E-06 | 0.005941349 | 0.004490438 |
| Nsd3 | S | 585 | -0.120340754 | -1.06198959 | -1.116731827 | -0.996391073 | -0.054742237 | 0.025863522 | 0.00070881 | 0.001118237 | 0.003321449 | 0.62564004 |
| Aff4 | S | 831 | -0.037656168 | -0.587511737 | -0.724579711 | -0.686923543 | -0.137067974 | 0.184119294 | 4.94728E-05 | 3.6962E-08 | 3.79925E-05 | 0.028617557 |
| Sqstm1 | S | 368 | 0.033123251 | -0.696218438 | -0.849999308 | -0.883122559 | -0.15378087 | 0.489376233 | 4.20711E-08 | 6.26416E-05 | 6.80486E-05 | 0.038935266 |

| | | | | | | | | | | | | | |
|---|---|---|---|---|---|---|---|---|---|---|---|---|---|
| Phldb1 | S | | 535 | -0.05387948 | -0.700981495 | -1.100274497 | -1.046395017 | -0.399293002 | 0.687205874 | 1.63343E-05 | 0.000107113 | 0.002249538 | 0.019059808 |
| Fosl1 | S | | 256 | -0.093642261 | -0.993071372 | -1.185585166 | -1.091942904 | -0.192513794 | 0.266833145 | 0.006123425 | 3.50594E-05 | 0.0002257 | 0.190392231 |
| Fam13b | S | | 492 | -0.263262801 | -0.747358141 | -0.970442944 | -0.707180142 | -0.223084802 | 0.14710227 | 0.000672594 | 9.06611E-05 | 0.01123746 | 0.093605466 |
| Srpk2 | S | | 489 | -0.042859656 | -0.739880505 | -0.776041968 | -0.733182312 | -0.036161462 | 0.385070717 | 1.54255E-06 | 8.67721E-05 | 0.000164323 | 0.375898351 |
| Rrp15 | S | | 194 | -0.255242861 | -0.5868715 | -0.956915634 | -0.701672774 | -0.370044135 | 0.040987211 | 0.001479904 | 2.77987E-05 | 0.001449713 | 0.018250744 |
| Phldb2 | S | | 414 | -0.198833367 | -0.820013521 | -0.950609449 | -0.751776082 | -0.130595928 | 0.043521158 | 0.000889935 | 0.000211533 | 0.001419678 | 0.283695389 |
| Surf2 | S | | 156 | 0.061539441 | -0.979200248 | -0.858232809 | -0.91977225 | 0.120967439 | 0.546367804 | 1.74109E-05 | 4.55657E-05 | 0.000516738 | 0.206267242 |
| Surf2 | S | | 157 | 0.061539441 | -0.979200248 | -0.858232809 | -0.91977225 | 0.120967439 | 0.546367804 | 1.74109E-05 | 4.55657E-05 | 0.000516738 | 0.206267242 |
| Phldb1 | S | | 537 | 0.048669858 | -0.807881486 | -1.171752727 | -1.220422585 | -0.363871241 | 0.737164487 | 0.000980109 | 5.48712E-06 | 0.003396439 | 0.02505683 |
| Acly | S | | 455 | -0.03053823 | -0.617654563 | -0.855591354 | -0.825053124 | -0.237936791 | 0.647301228 | 3.42098E-05 | 1.88249E-06 | 0.000417214 | 0.002077301 |
| Sipa1l3 | T | | 1157 | 0.036304992 | -0.78872757 | -0.882021854 | -0.918326846 | -0.093294284 | 0.355796319 | 5.84899E-07 | 0.000130644 | 0.002830026 | 0.201318654 |
| Frmd4a | S | | 900 | -0.422718125 | -0.859194561 | -1.124453674 | -0.701735549 | -0.265259113 | 0.026754628 | 0.008331954 | 0.001601203 | 0.021433001 | 0.272357768 |
| Cbx4 | S | | 410 | -0.005639678 | -0.677486436 | -0.68544053 | -0.679800852 | -0.007954094 | 0.894281379 | 9.78786E-07 | 1.00951E-08 | 0.001017566 | 0.84112101 |
| Ccdc88a | S | | 1387 | 0.008317142 | -0.614541757 | -0.780985712 | -0.789302854 | -0.166443955 | 0.813125049 | 0.000332789 | 3.45073E-08 | 0.000833326 | 0.037037987 |
| Gtse1 | S | | 311 | -0.449338535 | -0.796819786 | -1.222863535 | -0.773524999 | -0.426043749 | 0.059809671 | 0.011513055 | 0.001658957 | 0.020074241 | 0.112542643 |
| Rps6 Rps6-ps4 | S | | 240 | 0.164749201 | -0.823091159 | -1.026482697 | -1.191231899 | -0.203391538 | 0.104978581 | 6.7656E-06 | 2.75003E-05 | 0.000181555 | 0.059007493 |
| Uck2 | S | | 256 | -0.042697819 | -0.962956022 | -1.12588835 | -1.083190531 | -0.162932329 | 0.438993439 | 0.003621643 | 6.94675E-05 | 7.99587E-05 | 0.188336239 |
| Pcyt1a Pcyt1b | S | 233;233 | | -0.171971917 | -0.602425004 | -0.957970866 | -0.785998948 | -0.355545862 | 0.110948878 | 0.005042034 | 6.52559E-06 | 0.001196628 | 0.033992137 |
| Mast4 | T | | 2551 | -0.296468643 | -0.612509054 | -0.885592441 | -0.589123798 | -0.273083386 | 0.017105478 | 0.017593747 | 2.08063E-05 | 0.001545301 | 0.150353336 |
| Rbbp6 | S | | 1699 | -0.04680118 | -0.814474115 | -0.824776278 | -0.777975098 | -0.010302163 | 0.631422564 | 1.03491E-05 | 0.000470645 | 0.001611839 | 0.910200077 |
| Rps6 Rps6-ps4 | S | | 236 | -0.106646618 | -1.022401848 | -1.596089405 | -1.489442787 | -0.573687557 | 0.229161317 | 0.005225921 | 0.003687578 | 0.002034879 | 0.026528979 |
| Gsk3a | S | | 21 | -0.320703084 | -0.660261923 | -1.061344938 | -0.740641853 | -0.401083015 | 0.149765115 | 0.003555852 | 0.000241054 | 0.016431378 | 0.025487531 |
| Gnl2 | S | | 515 | 0.044727439 | -0.627889824 | -0.652782444 | -0.697509883 | -0.02489352 | 0.232399301 | 9.78786E-07 | 1.20079E-06 | 6.94476E-05 | 0.428960313 |
| Foxo4 | T | | 32 | -0.082758704 | -0.870575928 | -0.918786947 | -0.836028243 | -0.048211019 | 0.230152346 | 2.01173E-06 | 0.018003269 | 0.023495084 | 0.777616246 |
| Sema4b | S | | 816 | 0.036816192 | -0.65118885 | -0.918531941 | -0.955348133 | -0.267343091 | 0.290978329 | 0.003901665 | 3.82551E-08 | 1.16368E-06 | 0.05268479 |
| Sdc2 | S | | 188 | -0.132091992 | -0.785866621 | -1.08048624 | -0.948394248 | -0.294619619 | 0.01354326 | 0.002420444 | 0.001386205 | 0.003580483 | 0.047171029 |
| Mast4 | S | | 2517 | -0.32415421 | -0.662887134 | -1.029760274 | -0.705606064 | -0.36687314 | 0.075816364 | 0.039016408 | 4.52648E-05 | 0.009412694 | 0.156070461 |
| Phldb1 | S | | 565 | -0.122370763 | -0.619560098 | -0.871320578 | -0.748949814 | -0.25176048 | 0.070222337 | 5.59733E-05 | 0.000552314 | 0.001393636 | 0.037363271 |
| Tbc1d4 | S | | 348 | -0.1699786 | -0.63081313 | -1.055258527 | -0.885279926 | -0.424445397 | 0.023386121 | 0.016071182 | 7.49925E-05 | 0.000294541 | 0.03524765 |
| Larp1 | T | | 842 | -0.027659808 | -0.652580519 | -0.625534425 | -0.597874617 | 0.027046094 | 0.279019625 | 1.74514E-05 | 9.71342E-06 | 0.000197983 | 0.489630485 |
| Phldb1 | S | | 431 | 0.10586816 | -0.60754908 | -0.812583952 | -0.918452112 | -0.205034872 | 0.31958113 | 1.53863E-05 | 1.86085E-06 | 0.004038524 | 0.008150934 |
| Ccnf | S | | 577 | -0.091128207 | -0.756497296 | -0.878110054 | -0.786981847 | -0.121612759 | 0.080648316 | 0.000388838 | 0.000502068 | 0.00066989 | 0.162280251 |
| Lims1 | S | | 319 | -0.046783822 | -0.705042393 | -0.909100924 | -0.862317102 | -0.204058531 | 0.477130707 | 1.66574E-05 | 0.000915895 | 0.001515118 | 0.109996817 |
| Npm1 | S | | 139 | -0.071250717 | -0.62726499 | -0.680886582 | -0.609635865 | -0.053621591 | 0.671562222 | 2.17364E-05 | 1.88879E-05 | 0.040890861 | 0.320182102 |
| Sorbs1 | S | | 62 | -0.167948447 | -0.720783788 | -0.818071566 | -0.650123119 | -0.097287778 | 0.11264086 | 0.004390994 | 3.76939E-05 | 0.000804469 | 0.458682486 |
| Eif4b | S | | 418 | 0.034861762 | -0.926416272 | -1.170460115 | -1.205321877 | -0.244043843 | 0.316608915 | 0.000467958 | 0.000474352 | 0.000474426 | 0.027579353 |
| Znrf2 | S | | 20 | -0.107893865 | -0.727606331 | -0.955643162 | -0.847749298 | -0.228036832 | 0.251035894 | 0.009376981 | 2.0054E-05 | 0.000423629 | 0.18019701 |
| Rpl14 | T | | 202 | 0.258833572 | -0.670910187 | -1.101548838 | -1.36038241 | -0.430638652 | 0.004344304 | 1.92986E-05 | 8.45159E-07 | 0.000203589 | 0.001990235 |
| Rps6 Rps6-ps4 | S | | 242 | -0.132107371 | -0.828126646 | -1.240913055 | -1.108805684 | -0.412786409 | 0.030898643 | 0.005192617 | 0.003254864 | 0.002530118 | 0.023122223 |
| Mepce | S | | 347 | -0.161570828 | -0.905854338 | -0.939701059 | -0.778130231 | -0.033846721 | 0.120285582 | 0.002138097 | 0.001428294 | 0.002754683 | 0.803380527 |
| Kdm2b | S | | 741 | -0.01203441 | -0.590465172 | -0.66462369 | -0.65258928 | -0.074158518 | 0.788370813 | 0.000138668 | 6.35627E-07 | 8.41243E-05 | 0.19997508 |
| Rhoj | S | | 9 | -0.093901886 | -0.609754514 | -1.054171923 | -0.960270037 | -0.444417409 | 0.045643613 | 6.90585E-05 | 0.003174233 | 0.007173408 | 0.025573452 |
| Emg1 | T | | 24 | -0.17015865 | -0.732808378 | -0.883771578 | -0.713612928 | -0.1509632 | 0.122862459 | 0.001855644 | 0.000318589 | 0.001987366 | 0.103497216 |
| Slc7a11 | S | | 21 | -0.443371958 | -0.630764546 | -1.076618994 | -0.633247037 | -0.445854448 | 0.011465077 | 0.01359224 | 0.005960976 | 0.037744252 | 0.095334217 |
| Cep170 Plin5 | S | 1152;220 | | -0.507865932 | -0.743436628 | -1.309908125 | -0.802042194 | -0.566471497 | 0.098516158 | 0.049277151 | 0.002456626 | 0.027190102 | 0.108807202 |
| Gtse1 | S | | 310 | -0.337675997 | -0.774131456 | -1.070504464 | -0.732828467 | -0.296373008 | 0.099349638 | 0.012519384 | 0.001479624 | 0.020052796 | 0.205181644 |
| Alkbh5 | S | | 362 | -0.188420898 | -0.600614168 | -0.830744964 | -0.642324066 | -0.230130795 | 0.035731891 | 0.001591942 | 0.000316479 | 0.001842397 | 0.06921159 |
| Gas7 | T | | 98 | -0.03574228 | -0.624556046 | -0.903902578 | -0.868160298 | -0.279346532 | 0.713945853 | 0.00097265 | 1.76048E-05 | 0.000339314 | 0.03883314 |
| Rangap1 | S | | 429 | -0.033456862 | -0.65190336 | -0.768436452 | -0.73497959 | -0.116533092 | 0.608868638 | 1.01086E-05 | 0.000381629 | 0.001098233 | 0.192739259 |
| Rtkn | S | | 520 | -0.036449852 | -0.728397638 | -0.84098816 | -0.804538308 | -0.112590522 | 0.794782477 | 8.96935E-05 | 0.000361321 | 0.004812131 | 0.301386503 |
| Akt1 | T | | 308 | 0.198672167 | -1.042023348 | -0.963617736 | -1.162289904 | 0.078405612 | 0.034248882 | 0.002865638 | 1.52922E-06 | 0.001650341 | 0.497209264 |
| Phf8-ps Phf8 | S | | 984 | -0.143170904 | -0.804265745 | -1.171796778 | -1.028625874 | -0.367531033 | 0.281368644 | 0.017072382 | 0.000470529 | 0.005637457 | 0.153283705 |
| Zfp36l1 Zfp36l2 | S | 480;334 | | -0.071306937 | -0.599016075 | -0.74109963 | -0.669792693 | -0.142083555 | 0.463205486 | 0.002282288 | 3.63627E-06 | 0.005797532 | 0.139646529 |
| Fosl1 | S | | 253 | -0.130908354 | -0.64496413 | -0.746318374 | -0.61541002 | -0.101354244 | 0.117886332 | 6.96018E-06 | 0.008264889 | 0.012983847 | 0.427423979 |
| Zfp609 | S | | 467 | 0.096261427 | -0.611223844 | -0.890662746 | -0.986924174 | -0.279438903 | 0.124004448 | 0.00035192 | 2.16081E-06 | 5.52149E-05 | 0.017302354 |
| Abi1 | S | | 243 | -0.19293204 | -0.688119475 | -0.78502546 | -0.592093421 | -0.096905986 | 0.087053932 | 5.19886E-05 | 0.01283801 | 0.02929425 | 0.548462972 |
| Sdc2 | S | | 189 | -0.147189655 | -0.84073122 | -0.873296339 | -0.726106685 | -0.032565119 | 0.017712927 | 0.001519155 | 0.009365329 | 0.012670998 | 0.799459749 |
| Gabarapl2 | S | | 10 | -0.013739252 | -0.678851935 | -0.760847049 | -0.747107798 | -0.081995114 | 0.711660125 | 0.00663437 | 1.52626E-06 | 0.00010698 | 0.418179761 |
| Prkaa1 | S | | 496 | -0.239591008 | -0.674734059 | -1.050482625 | -0.810891617 | -0.375748566 | 0.057939953 | 0.017617911 | 0.001786762 | 0.004679462 | 0.088733375 |
| Khdrbs1 | S | | 322 | -0.414317444 | -0.621832507 | -1.030629676 | -0.616312232 | -0.408797168 | 0.05170256 | 0.017136345 | 0.00460234 | 0.040276636 | 0.121690677 |
| Phldb2 | S | | 241 | -0.147256387 | -0.590113317 | -0.776674751 | -0.629418364 | -0.186561434 | 0.274284588 | 0.000300462 | 0.000665743 | 0.006978759 | 0.134255706 |
| Zc3hav1 | S | | 345 | 0.129951279 | -0.623636806 | -0.805219228 | -0.935170506 | -0.181582422 | 0.063525478 | 1.66186E-05 | 1.87365E-05 | 0.000720959 | 0.021444903 |
| Zc3hav1 | T | | 346 | 0.129951279 | -0.623636806 | -0.805219228 | -0.935170506 | -0.181582422 | 0.063525478 | 1.66186E-05 | 1.87365E-05 | 0.000720959 | 0.021444903 |
| Eif4b | S | | 131 | -0.015885877 | -0.909041167 | -0.952837037 | -0.93695116 | -0.04379587 | 0.837060105 | 0.001549919 | 0.001432838 | 0.000883643 | 0.707955558 |
| Ranbp3 | T | | 56 | 0.085921762 | -0.645498293 | -0.847970414 | -0.933892176 | -0.202472121 | 0.095949025 | 0.009961332 | 1.05063E-07 | 3.29805E-05 | 0.14170666 |
| Gnl2 | S | | 504 | -0.028252998 | -0.665905231 | -0.644736051 | -0.616483054 | 0.021169179 | 0.430747343 | 0.000117876 | 0.000179725 | 0.000395615 | 0.700173943 |
| Gtf2f1 | T | | 384 | -0.065187103 | -0.587634486 | -0.967248205 | -0.902061102 | -0.379613719 | 0.595902129 | 0.003180484 | 0.000133441 | 0.00240257 | 0.022204091 |
| Akt3 | S | | 472 | -0.050978114 | -0.61436441 | -0.699143537 | -0.648165422 | -0.084779127 | 0.669655098 | 0.000594804 | 3.71172E-05 | 0.016861051 | 0.183697823 |
| Akt3 | S | | 476 | -0.050978114 | -0.61436441 | -0.699143537 | -0.648165422 | -0.084779127 | 0.669655098 | 0.000594804 | 3.71172E-05 | 0.016861051 | 0.183697823 |
| Errfi1 | S | | 368 | -0.251174027 | -0.603909046 | -0.967345612 | -0.716171585 | -0.363436566 | 0.131693301 | 0.004722951 | 0.002518355 | 0.015117453 | 0.093356681 |
| Bad | S | | 136 | -0.004968495 | -0.625543674 | -0.800666201 | -0.795697707 | -0.175122527 | 0.952899147 | 0.005392894 | 1.52197E-05 | 0.005120242 | 0.07266875 |
| Dock7 | S | | 434 | -0.041638617 | -0.626868152 | -0.882292043 | -0.840653426 | -0.255423892 | 0.604382939 | 0.000144917 | 0.002525633 | 0.002395837 | 0.082272516 |
| Nav3 | S | | 1537 | 0.004584691 | -0.686317189 | -0.783775486 | -0.788360177 | -0.097458297 | 0.947320754 | 0.001391867 | 0.000111073 | 0.000550979 | 0.24869311 |
| Bnip3l | S | | 165 | -0.131604698 | -0.586822866 | -0.801315022 | -0.669710325 | -0.214492156 | 0.011983468 | 0.005718785 | 0.001832704 | 0.001392368 | 0.045787935 |
| Trim33 | S | | 1134 | -0.291880393 | -0.875408723 | -1.090440058 | -0.798559665 | -0.215031335 | 0.150570075 | 0.017702798 | 0.016729017 | 0.03574529 | 0.461343945 |
| Fosl1 | S | | 254 | -0.069772903 | -0.905627416 | -1.104176599 | -1.034403696 | -0.198549182 | 0.60727288 | 0.015477712 | 0.002407271 | 0.002434499 | 0.331832511 |
| Sqstm1 | S | | 152 | 0.022084077 | -0.604170934 | -0.688219241 | -0.710303317 | -0.084048307 | 0.588305299 | 9.24012E-06 | 0.001081686 | 0.00065567 | 0.222240419 |
| Specc1l | S | | 923 | 0.219623944 | -0.736756193 | -0.917097561 | -1.136721504 | -0.180341368 | 0.109916388 | 5.26679E-05 | 0.000442108 | 0.000693356 | 0.120574743 |
| Mettl1 | S | | 21 | 7.71755E-05 | -0.715036673 | -0.88742372 | -0.887500895 | -0.172387047 | 0.999347788 | 0.002946344 | 0.000561941 | 0.001280172 | 0.13131981 |
| Nfil3 | S | | 301 | -0.027040344 | -0.698397559 | -0.676218019 | -0.649177675 | 0.02217954 | 0.517249955 | 0.00053217 | 0.000833298 | 0.007625917 | 0.798670923 |
| Mycbp2 | S | | 2943 | -0.027269693 | -0.591426639 | -0.749851683 | -0.72258199 | -0.158425044 | 0.298080813 | 0.000711694 | 0.000720043 | 0.001113618 | 0.046306091 |
| Mepce | S | | 348 | 0.075431466 | -0.697883495 | -0.811508096 | -0.886939563 | -0.113624602 | 0.271781278 | 1.81886E-05 | 0.003681982 | 0.009822401 | 0.374991309 |
| Grb14 | S | | 364 | -0.022350273 | -0.742477858 | -0.954867944 | -0.932517672 | -0.212390086 | 0.743215663 | 0.000178146 | 0.018308021 | 0.019044244 | 0.300615318 |
| Pin4 | S | | 21 | -0.304515476 | -0.607273291 | -1.042080237 | -0.737564761 | -0.434806946 | 0.187975547 | 0.043130832 | 0.002836604 | 0.025441408 | 0.124238616 |
| Gab2 | S | | 211 | -0.103131917 | -0.666835474 | -0.824844581 | -0.721712664 | -0.158009107 | 0.217977725 | 0.010117617 | 0.00086795 | 0.001036597 | 0.152305727 |
| Larp4 | S | | 586 | 0.074184329 | -0.589750027 | -0.6115399 | -0.685724229 | -0.021789873 | 0.423661243 | 7.63684E-06 | 0.001002313 | 0.002104501 | 0.752430352 |
| Zfp507 | T | | 473 | 0.076442159 | -1.063044597 | -1.27643113 | -1.352873289 | -0.213386533 | 0.437107799 | 0.010261353 | 0.008659329 | 0.002058128 | 0.293618099 |
| Specc1 | S | | 37 | -0.229482131 | -0.603550228 | -1.023110941 | -0.79362881 | -0.419560714 | 0.12141542 | 0.04972477 | 0.006446822 | 0.019654268 | 0.147924152 |
| Zfp507 | S | | 475 | 0.067237675 | -0.913754354 | -1.037742391 | -1.104980066 | -0.123988037 | 0.332238947 | 0.00457707 | 0.005507893 | 0.006553006 | 0.479532656 |
| Akap10 | S | | 278 | 0.240050402 | -0.62706627 | -0.720628995 | -0.960679397 | -0.093562725 | 0.011692447 | 5.76482E-05 | 4.54115E-05 | 0.000149027 | 0.180333389 |
| Foxo4 | S | | 30 | 0.059798532 | -0.653901879 | -0.654278538 | -0.71407707 | -0.000376659 | 0.402364816 | 6.35385E-05 | 0.007072656 | 0.01366202 | 0.99735363 |
| Ehbp1l1 | S | | 1460 | -0.23046846 | -0.593728302 | -1.006815623 | -0.776347164 | -0.413087322 | 0.177073523 | 0.030759434 | 0.022350247 | 0.029387775 | 0.145800754 |
| Ei24 | S | | 47 | 0.002675149 | -0.69578143 | -0.862976368 | -0.865651517 | -0.167194938 | 0.974206399 | 0.004861608 | 0.005877281 | 0.001263775 | 0.172754938 |
| Psmc3 | S | | 12 | 0.068045834 | -0.697546189 | -0.639080709 | -0.707126543 | 0.058465481 | 0.465018054 | 0.000648875 | 0.002074801 | 0.00253833 | 0.422000925 |

| | | | | | | | | | | | | |
|---|---|---|---|---|---|---|---|---|---|---|---|---|
| Larp1 | T | 1047 | 0.048895541 | -0.673378603 | -0.860052401 | -0.908947942 | -0.186673797 | 0.545312612 | 0.003714858 | 0.006364389 | 0.00771817 | 0.274298219 |
| Wdhd1 | S | 1077 | -0.109666109 | -0.623037512 | -0.808812409 | -0.699146299 | -0.185774897 | 0.249763217 | 0.043836682 | 0.005298655 | 0.003884034 | 0.346774574 |
| Usp36 | S | 899 | 0.077594713 | -0.750758932 | -0.683871327 | -0.76146604 | 0.066887605 | 0.144229025 | 0.003611037 | 0.001005657 | 0.001455044 | 0.532935338 |
| Phldb1 | S | 976 | 0.122752019 | -0.610185646 | -0.647498016 | -0.770250035 | -0.03731237 | 0.176600884 | 0.00050198 | 0.001451308 | 0.001406491 | 0.711602775 |
| Slc38a2 | S | 19 | -0.008530057 | -0.636941517 | -0.71929656 | -0.710766503 | -0.082355043 | 0.909069393 | 0.015038084 | 0.004070888 | 0.006267842 | 0.601988737 |
| Fkbp15 | S | 340 | 0.059152283 | -0.684226398 | -0.86928414 | -0.928436423 | -0.185057742 | 0.448934784 | 0.008362093 | 0.008371573 | 0.004437316 | 0.26449498 |
| Akt1 | S | 473 | 0.070361776 | -0.760444895 | -0.717150982 | -0.787512758 | 0.043293913 | 0.483852091 | 0.013560597 | 0.009544233 | 0.002479783 | 0.754059953 |
| Akt1 | T | 479 | 0.088829303 | -0.748423944 | -0.690323167 | -0.779152471 | 0.058100777 | 0.268134986 | 0.014612329 | 0.011105625 | 0.003585698 | 0.700942246 |
| Plekha2 | S | 308 | 0.258925919 | -0.677586254 | -0.687652768 | -0.946578687 | -0.010066513 | 0.201086866 | 0.009744411 | 0.009145832 | 0.001116943 | 0.827702252 |
| Zc3hav1 | S | 344 | 0.556424913 | -0.675559507 | -1.408814942 | -1.965239855 | -0.733255435 | 0.010287259 | 0.004113745 | 0.01833467 | 0.017446068 | 0.10957856 |
| Pdcd4 | S | 94 | 0.064381512 | 2.154493627 | 2.251504349 | 2.187122836 | 0.097010721 | 0.405060737 | 1.46656E-07 | 3.78675E-09 | 0.000117491 | 0.175368189 |
| Pdcd4 | T | 93 | 0.064085926 | 2.234105477 | 2.314215538 | 2.250129612 | 0.080110061 | 0.60658333 | 4.54196E-07 | 8.51782E-09 | 0.000563314 | 0.30115015 |
| Nrcam | S | 1190 | 0.208168861 | 0.977454681 | 0.948081532 | 0.739912671 | -0.02937315 | 0.008382448 | 1.49755E-06 | 2.54699E-07 | 0.000203339 | 0.544694389 |
| Synpo | T | 769 | 0.187961908 | 0.78821738 | 0.931194687 | 0.743232779 | 0.142977307 | 0.007754799 | 1.28509E-06 | 3.99881E-05 | 0.004058833 | 0.092714596 |
| Eif4g1 | S | 1096 | -0.052369492 | 0.979321951 | 0.937009339 | 0.989378831 | -0.042312612 | 0.428643272 | 2.76192E-08 | 3.04868E-05 | 0.000161572 | 0.389603235 |
| Tmpo | S | 155;155 | 0.098165981 | 0.678419827 | 0.736548974 | 0.638382993 | 0.058129147 | 0.127461806 | 1.25583E-07 | 1.07277E-05 | 0.000504772 | 0.169687032 |
| Jup | S | 665 | 0.206121202 | 0.8234506 | 0.951398911 | 0.745277709 | 0.127948311 | 0.039895817 | 2.46643E-06 | 0.002396261 | 0.003678931 | 0.282781494 |
| Zfp280c | S | 528 | -0.039144571 | 0.800887778 | 0.904602685 | 0.943747256 | 0.103714907 | 0.60733706 | 0.000434659 | 1.14886E-08 | 0.002727559 | 0.146849738 |
| Foxk1 | T | 422 | 0.005737228 | 0.803237719 | 0.872364617 | 0.866627389 | 0.069126898 | 0.872578953 | 2.12305E-05 | 2.80459E-05 | 3.51776E-05 | 0.096885522 |
| Tmpo | S | 157;157 | 0.058024126 | 0.711730478 | 0.760843848 | 0.702819722 | 0.049113371 | 0.304139269 | 4.87546E-07 | 0.000635748 | 0.000785895 | 0.466220605 |
| Carhsp1 | S | 59 | 0.047706187 | 0.848268072 | 0.740835272 | 0.693129085 | -0.1074328 | 0.368244859 | 2.2922E-07 | 0.005162755 | 0.007471349 | 0.328851068 |
| Eif4g1 | S | 1211 | -0.080229412 | 0.791614631 | 0.750790533 | 0.831019945 | -0.040824099 | 0.063776461 | 9.2679E-06 | 1.2478E-06 | 0.000302355 | 0.352840474 |
| Plec | S | 4620 | 0.020286565 | 0.625103735 | 0.796705553 | 0.776418988 | 0.171601819 | 0.725756869 | 0.001016377 | 9.16499E-08 | 0.000908375 | 0.065522541 |
| Eif4g1 | T | 1213 | -0.052219042 | 0.762748068 | 0.712885572 | 0.765104614 | -0.049862496 | 0.245044721 | 2.09497E-05 | 8.76583E-07 | 3.2019E-05 | 0.358656808 |
| Eif4g1 | S | 708 | 0.024444956 | 0.958649807 | 0.912583338 | 0.888138382 | -0.046066469 | 0.65698465 | 0.001783476 | 9.70438E-05 | 0.000190181 | 0.528553808 |
| Irs2 | S | 980 | -0.062385548 | 0.644519891 | 0.764239359 | 0.826624907 | 0.119719468 | 0.43560983 | 8.27787E-05 | 9.8324E-06 | 0.000263022 | 0.116471601 |
| Eif4g3 | S | 240 | -0.052908151 | 0.662931105 | 0.667917616 | 0.720825768 | 0.004986511 | 0.090171868 | 0.000730399 | 4.24837E-08 | 3.3892E-06 | 0.922020811 |
| Tsc2 | S | 1412 | 0.101627419 | 0.674753102 | 0.802651662 | 0.701024243 | 0.12789856 | 0.587227672 | 0.00042278 | 0.00014493 | 0.031844405 | 0.135838895 |
| Filip1l | S | 789 | -0.043606321 | 0.672920424 | 0.656807605 | 0.700413925 | -0.016112819 | 0.420521657 | 0.000219769 | 1.63935E-06 | 0.000277009 | 0.769942203 |
| Pphln1 | S | 221 | 0.0768107 | 0.595131403 | 0.807778807 | 0.730968107 | 0.212647404 | 0.463967524 | 0.000351682 | 0.000407346 | 0.002313814 | 0.094454156 |
| Prex2 | S | 1065 | -0.035784125 | 0.661394846 | 0.592882998 | 0.628667124 | -0.068511847 | 0.661845306 | 2.86328E-06 | 0.004548521 | 0.002504969 | 0.393351646 |
| Pdcd4 | S | 457 | -0.014078395 | 0.690917307 | 0.690938721 | 0.705017116 | 2.14141E-05 | 0.766820961 | 0.001207256 | 0.000157202 | 0.001873586 | 0.999804223 |
| Foxo1 | S | 467 | -0.298443126 | 0.627722153 | 0.622782093 | 0.921225219 | -0.004940061 | 0.06441173 | 2.9192E-05 | 6.10543E-06 | 0.005776148 | 0.923977038 |
| Dpysl2 | T | 509 | -0.05193307 | 0.680430762 | 0.609571467 | 0.661504537 | -0.070859295 | 0.034524647 | 0.001178167 | 0.00250754 | 0.005103165 | 0.335530371 |
| Irs2 | S | 965 | -0.186608299 | 0.70343416 | 0.981497334 | 1.168105633 | 0.278063174 | 0.014674109 | 0.00611021 | 0.00105955 | 0.000384557 | 0.054834711 |
| Dpysl3 | T | 508 | -0.17116638 | 0.807198188 | 0.587636618 | 0.758802998 | -0.21956157 | 0.002762429 | 9.79366E-06 | 0.028582094 | 0.016730208 | 0.17721553 |
| Lrrfip1 | S | 614 | -0.166431437 | -0.600511103 | -0.719047003 | -0.552615566 | -0.1185359 | 0.000442474 | 2.19583E-08 | 1.42176E-07 | 0.00067459 | 0.006793114 |
| Tfrc | S | 19 | -0.49323722 | -0.6191931 | -1.016559945 | -0.523322725 | -0.397366846 | 0.003430827 | 0.001090774 | 0.003681375 | 0.039894338 | 0.077050772 |
| Eif4b | T | 412 | -0.144471984 | -0.620460056 | -0.71345009 | -0.568978106 | -0.092990034 | 0.104628981 | 1.55645E-07 | 0.000382422 | 0.00174686 | 0.105661545 |
| Hells | S | 498 | -0.294448352 | -0.640319987 | -0.867707361 | -0.57325901 | -0.227387374 | 0.029742171 | 0.004873401 | 5.08539E-05 | 0.003078793 | 0.136772164 |
| Slc2a1 | S | 226 | -0.5800578 | -0.777192243 | -1.152224165 | -0.572166365 | -0.375031922 | 0.004030083 | 0.047968047 | 0.005172582 | 0.076210608 | 0.253701383 |
| Rwdd1 | T | 144 | -0.325889785 | -0.618266912 | -0.735427752 | -0.409537967 | -0.11716084 | 0.005889041 | 4.31354E-05 | 0.015978327 | 0.06300451 | 0.448067789 |
| Abl2 | Y | 719 | -0.427483315 | -0.625317302 | -0.93255208 | -0.505068765 | -0.307234778 | 0.060234449 | 0.008998941 | 0.000533108 | 0.037035695 | 0.086439079 |
| Slc4a7 | S | 994 | -0.474916846 | -0.594553695 | -0.716204423 | -0.241287577 | -0.121650728 | 0.00926403 | 0.0063003 | 0.001445278 | 0.075773469 | 0.380712059 |
| Dennd1b | S | 633 | -0.134893825 | -0.705803103 | -0.702769352 | -0.567875528 | 0.00303375 | 0.142857012 | 0.016477037 | 9.17175E-07 | 0.006126991 | 0.981232186 |
| Tpr | T | 2184 | -0.308524534 | -0.585944119 | -0.718146863 | -0.409622329 | -0.132202744 | 0.025601067 | 0.00092113 | 0.001355397 | 0.022743406 | 0.292857078 |
| Dennd1b | S | 632 | -0.258423775 | -0.711192566 | -0.620671063 | -0.362247288 | 0.090521503 | 0.04900532 | 0.005297206 | 0.000117478 | 0.022042858 | 0.467301525 |
| Phldb2 | S | 527 | -0.45979184 | -0.590458561 | -0.993543766 | -0.533751926 | -0.403085206 | 0.015174887 | 0.012986756 | 0.015583727 | 0.096941005 | 0.167357708 |
| Map1b | S | 2001 | -0.194963168 | -0.608287811 | -0.760087366 | -0.565124198 | -0.151799555 | 0.026933969 | 0.001699438 | 0.002927734 | 0.004725047 | 0.180151566 |
| Tepsin | S | 520 | -0.392801954 | -0.857824284 | -0.984482777 | -0.591680824 | -0.126658493 | 0.397034888 | 0.01127382 | 0.004882671 | 0.243522097 | 0.55576608 |
| Nsd2 | S | 172 | -0.262018866 | -0.637993118 | -0.797862668 | -0.535843801 | -0.15986955 | 0.122871057 | 0.001424243 | 0.005331297 | 0.032321669 | 0.34215552 |
| Eloa | S | 243 | -0.433359616 | -0.60181638 | -1.028247422 | -0.594887806 | -0.426431043 | 0.2034468 | 0.039025531 | 0.003925563 | 0.103593129 | 0.040654512 |
| Npm1 | T | 235 | -0.070546598 | -0.644308129 | -0.653404009 | -0.582857411 | -0.00909588 | 0.622717943 | 0.000145454 | 0.002094074 | 0.015276682 | 0.929080958 |
| Npm1 | T | 232 | -0.078571883 | -0.59423019 | -0.623206366 | -0.544634484 | -0.028976176 | 0.458874694 | 0.000235383 | 0.000235383 | 0.009697137 | 0.724615557 |
| Itsn2 | Y | 922 | -0.420519422 | -0.611410305 | -0.696365962 | -0.27584654 | -0.084955657 | 0.03435361 | 0.025847168 | 0.010318217 | 0.181372101 | 0.688241524 |
| Aagab | S | 211 | -0.167855372 | -0.647565697 | -0.646213426 | -0.478358054 | 0.00135227 | 0.012160788 | 0.008211606 | 0.013118682 | 0.040448411 | 0.992715025 |
| Gys1 | S | 653 | -0.145851767 | -0.662282531 | -0.654710347 | -0.50885858 | 0.007572184 | 0.046813996 | 0.018610146 | 0.00286313 | 0.002695876 | 0.952009101 |
| Gys1 | S | 657 | -0.145851767 | -0.662282531 | -0.654710347 | -0.50885858 | 0.007572184 | 0.046813996 | 0.018610146 | 0.00286313 | 0.002695876 | 0.952009101 |
| Larp4 | S | 642 | -0.085324094 | -0.649722758 | -0.597993314 | -0.51266922 | 0.051729445 | 0.3876436 | 0.001276754 | 0.028497684 | 0.045431444 | 0.759944314 |
| Racgap1 | S | 226 | -0.145331159 | -0.651962448 | -0.665872173 | -0.520541014 | -0.013909725 | 0.077003403 | 0.011424913 | 0.043184933 | 0.068418118 | 0.944290648 |
| Eef2k | T | 347 | 0.573187171 | 1.549762458 | 1.474139804 | 0.900952633 | -0.075622654 | 0.132919341 | 0.01877945 | 2.61694E-05 | 0.068335836 | 0.807973696 |
| Synpo | T | 826 | 0.371258829 | 0.869240533 | 0.807861902 | 0.436603072 | -0.061378631 | 0.002865432 | 1.70259E-05 | 0.000238779 | 0.017333497 | 0.502765433 |
| Synpo | T | 822 | 0.338499406 | 0.809140145 | 0.779913021 | 0.441413615 | -0.029227125 | 0.001008161 | 4.37044E-06 | 0.000456203 | 0.016345113 | 0.722677836 |
| Eif4g3 | S | 1126 | 0.396235335 | 1.370238208 | 0.691137865 | 0.294902531 | -0.679100342 | 0.029326261 | 8.73815E-06 | 0.048652645 | 0.271204436 | 0.063052464 |
| Tsc22d4 | S | 123 | 0.494344796 | 0.806996197 | 0.621691063 | 0.127346268 | -0.185305134 | 6.4155E-05 | 0.022884573 | 2.51312E-05 | 0.068501615 | 0.327091668 |
| Synpo | S | 828 | 0.305739982 | 0.749039758 | 0.75196414 | 0.446224157 | 0.002924382 | 0.012069188 | 1.5271E-06 | 0.002748242 | 0.012007457 | 0.973722594 |
| Synpo | S | 740 | 0.226406828 | 0.73828942 | 0.780954003 | 0.554547175 | 0.042664583 | 0.065981175 | 3.91263E-05 | 8.89363E-05 | 0.004252136 | 0.570253556 |
| Synpo | S | 744 | 0.226406828 | 0.73828942 | 0.780954003 | 0.554547175 | 0.042664583 | 0.065981175 | 3.91263E-05 | 8.89363E-05 | 0.004252136 | 0.570253556 |
| Nrcam | S | 1194 | 0.121466917 | 0.640688422 | 0.621272365 | 0.499805448 | -0.019416057 | 0.067051542 | 4.46838E-06 | 9.9175E-06 | 0.000646189 | 0.681340519 |
| Nrcam | T | 1192 | 0.121466917 | 0.640688422 | 0.621272365 | 0.499805448 | -0.019416057 | 0.067051542 | 4.46838E-06 | 9.9175E-06 | 0.000646189 | 0.681340519 |
| Eif4g1 | S | 1187 | 0.039074657 | 0.607101874 | 0.598329719 | 0.559255062 | -0.000772155 | 0.308281916 | 3.09484E-06 | 8.61257E-07 | 0.000559308 | 0.687327596 |
| Fgfrl1 | S | 128 | 0.156124556 | 0.790718887 | 0.710546196 | 0.554421639 | -0.080172692 | 0.360908399 | 5.06965E-05 | 7.71134E-05 | 0.047029195 | 0.046772247 |
| Fgfrl1 | S | 129 | 0.156124556 | 0.790718887 | 0.710546196 | 0.554421639 | -0.080172692 | 0.360908399 | 5.06965E-05 | 7.71134E-05 | 0.047029195 | 0.046772247 |
| Arhgap22 | S | 482 | 0.273545469 | 0.71863904 | 0.619190846 | 0.345645377 | -0.099448194 | 0.00165294 | 0.00040166 | 0.000963543 | 0.027415301 | 0.316131436 |
| Dapk3 | S | 301 | 0.44836509 | 0.652189318 | 0.656261766 | 0.207896646 | 0.004072418 | 0.005458122 | 0.000761846 | 0.001707441 | 0.108003669 | 0.968558412 |
| Rab11fip5 | S | 209 | 0.580246365 | 0.808007732 | 0.687775706 | 0.10752934 | -0.120223027 | 0.006569627 | 0.002265101 | 0.014215341 | 0.564270365 | 0.53828967 |
| Mrnip | S | 140 | 0.324790443 | 0.701420066 | 0.601765194 | 0.276974751 | -0.099654872 | 0.02959541 | 5.01999E-05 | 0.00407489 | 0.081201821 | 0.383789849 |
| Synpo | S | 535 | 0.246956931 | 0.679974285 | 0.616978307 | 0.370021376 | -0.062995978 | 0.033788235 | 0.000232522 | 0.000665287 | 0.011959411 | 0.436146183 |
| Rb1cc1 | T | 630 | 0.35573906 | 0.769361877 | 0.806275497 | 0.450536438 | 0.03691362 | 0.012219111 | 0.001604602 | 0.01128184 | 0.029904146 | 0.779235733 |
| Arhgap22 | S | 483 | 0.143766547 | 0.815049605 | 0.698547318 | 0.554780771 | -0.116502287 | 0.258382661 | 0.003267348 | 8.69221E-05 | 0.011538958 | 0.362413531 |
| Sorbs3 | S | 412 | 0.295177666 | 0.643051449 | 0.611576586 | 0.316398919 | -0.031474863 | 0.012728774 | 0.023385755 | 0.000486255 | 0.017315214 | 0.832515812 |
| Il16 | S | 967 | 0.04776075 | 0.714288202 | 0.620977755 | 0.573217008 | -0.093310444 | 0.664546163 | 0.000471686 | 0.001134047 | 0.004098206 | 0.31495506 |
| Ctdspl2 | T | 89 | 0.161824157 | -0.67528415 | -0.479519179 | -0.641343336 | 0.195764971 | 0.00755541 | 5.97403E-07 | 0.009160606 | 0.004142998 | 0.082660026 |
| Washc2 | S | 1003 | 0.066118471 | -0.605707793 | -0.572617017 | -0.638735488 | 0.033090775 | 0.245208127 | 0.007098079 | 0.005515811 | 0.010188339 | 0.792403984 |
| Ccdc88a | S | 1680 | 0.318498387 | -0.848600227 | -1.089028388 | -1.407526775 | -0.240428161 | 0.11887921 | 0.007213312 | 0.074109233 | 0.037495604 | 0.556959363 |
| Plec | S | 4623 | -0.016239604 | 0.648621332 | 0.577191354 | 0.593430958 | -0.071429977 | 0.820189964 | 0.000424188 | 5.80907E-05 | 0.001168487 | 0.35630217 |
| Rhox5 | S | 22 | -0.209596018 | -1.444790197 | -0.058405262 | 0.151190755 | 1.386384935 | 0.284654566 | 0.000239146 | 0.750421759 | 0.247359185 | 0.001365773 |
| Rhox5 | S | 141 | -0.034441451 | -0.786247355 | 0.043481342 | 0.077922793 | 0.829728697 | 0.750704976 | 0.000301515 | 0.740997809 | 0.451186122 | 0.005639875 |
| Myo9a | S | 2502 | -0.373661764 | -0.600760716 | 0.134697764 | 0.508359528 | 0.73545848 | 0.105665726 | 0.006592291 | 0.377129078 | 0.0584941 | 0.00054722 |
| Reps1 | S | 145 | 0.429104411 | 1.224584857 | -0.135903468 | -0.565007879 | -1.360488325 | 0.12496958 | 2.22686E-05 | 0.227782714 | 0.088757862 | 1.03077E-05 |
| Akap12 | S | 598 | 0.094446894 | 0.69191513 | 0.090615997 | -0.003830896 | -0.601299133 | 0.345078902 | 0.000284246 | 0.392318378 | 0.946563665 | 0.001239559 |
| Ell2 | S | 314 | -0.206617299 | -0.641856453 | -0.529417998 | -0.322800699 | 0.112438456 | 0.008169049 | 5.36349E-05 | 5.21118E-05 | 0.000239786 | 0.110095769 |
| Prune2 | S | 1075 | -0.053264853 | -0.599918052 | -0.572533775 | -0.519268923 | 0.027384277 | 0.513743077 | 1.6961E-07 | 5.55673E-05 | 0.00675637 | 0.492456217 |
| Map3k4 | S | 1217 | -0.138524712 | -0.828847467 | -0.566722277 | -0.428197565 | 0.26212519 | 0.124929867 | 5.83735E-05 | 0.000148006 | 0.003512111 | 0.024304665 |

| | | | | | | | | | | | | |
|---|---|---|---|---|---|---|---|---|---|---|---|---|
| Tepsin | S | 519 | -0.281533253 | -0.859957684 | -1.352702182 | -1.071168928 | -0.492744498 | 0.091915301 | 7.98895E-05 | 0.075361742 | 0.113667557 | 0.353540068 |
| Disp2 | S | 387 | -0.013340847 | -0.620589608 | -0.575158819 | -0.561817972 | 0.045430789 | 0.744926394 | 4.70967E-06 | 3.21935E-06 | 0.000155991 | 0.035417959 |
| Disp2 | T | 380 | -0.013340847 | -0.620589608 | -0.575158819 | -0.561817972 | 0.045430789 | 0.744926394 | 4.70967E-06 | 3.21935E-06 | 0.000155991 | 0.035417959 |
| Gtf2f1 | S | 377 | -0.029989162 | -0.636676365 | -0.58001655 | -0.550027388 | 0.056659814 | 0.57458994 | 1.2164E-06 | 4.94076E-05 | 0.000535024 | 0.248835874 |
| Kdm4c | S | 105 | -0.328810012 | -0.806637354 | -0.476320422 | -0.14751041 | 0.330316932 | 0.000946923 | 0.000114518 | 0.049829497 | 0.372495287 | 0.109298379 |
| Sec61b | S | 49 | 0.004561035 | -0.597099623 | -0.510125039 | -0.514686075 | 0.086974584 | 0.925167544 | 2.39604E-06 | 5.88068E-06 | 0.000426777 | 0.046140041 |
| Bclaf1 | S | 882 | -0.1919495 | -0.628236283 | -0.4944374 | -0.3024879 | 0.133798883 | 0.047654374 | 0.000151969 | 0.000546434 | 0.003386791 | 0.04510221 |
| Psmf1 | T | 251 | -0.232088926 | -0.762171687 | -1.250585504 | -1.018496578 | -0.488413817 | 0.393119857 | 0.000303416 | 0.082081477 | 0.109169184 | 0.347084617 |
| Pdpk1 | S | 37 | -0.179179683 | -0.592122143 | -0.518658385 | -0.339478702 | 0.073463758 | 0.205024308 | 0.004487904 | 4.05986E-05 | 0.075013276 | 0.435478047 |
| Espl1 | S | 1922 | 0.005874213 | -0.609152172 | -0.498783992 | -0.504658205 | 0.11036818 | 0.895850434 | 1.01995E-05 | 0.000697329 | 0.007187062 | 0.147444073 |
| Espl1 | T | 1924 | 0.005874213 | -0.609152172 | -0.498783992 | -0.504658205 | 0.11036818 | 0.895850434 | 1.01995E-05 | 0.000697329 | 0.007187062 | 0.147444073 |
| Espl1 | Y | 1923 | 0.005874213 | -0.609152172 | -0.498783992 | -0.504658205 | 0.11036818 | 0.895850434 | 1.01995E-05 | 0.000697329 | 0.007187062 | 0.147444073 |
| 1810030O07Rik | S | 281 | -0.46461127 | -0.631864134 | -0.583077073 | -0.118465804 | 0.04878706 | 0.023148306 | 0.006948638 | 0.034262307 | 0.515205918 | 0.793656507 |
| Gtf2f1 | S | 380 | -0.008203333 | -0.611022488 | -0.549786471 | -0.541583138 | 0.061236017 | 0.890668168 | 3.88461E-05 | 0.002361763 | 0.005160491 | 0.480106692 |
| Mllt11 | S | 84 | -0.527088202 | -0.667951174 | -1.26945761 | -0.742369409 | -0.601506436 | 0.199797891 | 0.049746874 | 0.052917407 | 0.190909485 | 0.24366016 |
| Nfil3 | S | 286 | -0.058578795 | -0.600672445 | -0.570413406 | -0.511834611 | 0.030259039 | 0.247631619 | 3.93205E-05 | 0.019256722 | 0.031917154 | 0.802667191 |
| Slc20a1 | S | 273 | 0.048765872 | -0.611261761 | -0.526674229 | -0.575440101 | 0.084587531 | 0.070349364 | 4.17746E-05 | 0.000216629 | 0.000449553 | 0.094695054 |
| Uba1 | S | 820 | -0.094676589 | -0.591834496 | -0.584733365 | -0.490056776 | 0.007101131 | 0.24385535 | 0.000522004 | 0.006513122 | 0.0083256 | 0.942293362 |
| Iws1 | T | 668 | -0.248013065 | -0.628573433 | -0.435483285 | -0.18747022 | 0.193090148 | 0.107833812 | 0.002244101 | 0.013073383 | 0.081224055 | 0.050064257 |
| Zfp703 | S | 187 | -0.090353293 | -0.651738344 | -0.876263123 | -0.78590983 | -0.224524779 | 0.643046046 | 0.000302785 | 0.072667393 | 0.074563299 | 0.481682738 |
| Ranbp2 | S | 954 | 0.005761936 | -0.586798031 | -0.495599634 | -0.50136157 | 0.091198396 | 0.908101205 | 0.000246305 | 0.001682304 | 0.011145777 | 0.303737976 |
| Mapre2 | S | 227 | -0.312485274 | -0.588152318 | -0.574601302 | -0.262116028 | 0.013551016 | 0.124585924 | 0.045528556 | 0.039504353 | 0.223811666 | 0.954684445 |
| Abi1 | S | 242 | -0.175427592 | -0.610748297 | -0.801872231 | -0.626444639 | -0.191123934 | 0.451889258 | 0.025896764 | 0.051440034 | 0.08284527 | 0.457320616 |
| Adrm1 | T | 217 | -0.282915626 | -0.779660411 | -0.021305742 | 0.261609883 | 0.758354668 | 0.082134351 | 0.001225697 | 0.93328717 | 0.319870554 | 0.056101276 |
| Prrc2b | S | 1006 | 0.064003728 | -0.604882097 | -0.447639204 | -0.511642932 | 0.157242892 | 0.352086984 | 0.012145713 | 0.000810578 | 0.003775867 | 0.277358825 |
| Cavin4 | T | 334 | -0.190250862 | -0.619137519 | -0.297764302 | -0.10751344 | 0.321373217 | 0.298997267 | 0.012017423 | 0.125215954 | 0.115691016 | 0.007836044 |
| Abl2 | S | 936 | -0.022367265 | -0.590204237 | -0.387296025 | -0.36492876 | 0.202908212 | 0.868641141 | 0.037779633 | 0.048416317 | 0.078235745 | 0.290532832 |
| Agfg1 | T | 177 | 0.370801743 | 0.585491071 | 0.550545608 | 0.179743864 | -0.034945463 | 0.001172026 | 0.000222747 | 6.99678E-05 | 0.031375385 | 0.60450733 |
| Cdc42ep4 | S | 64 | 0.208220555 | 0.618646385 | 0.486221641 | 0.278001086 | -0.132424744 | 0.007260901 | 1.34282E-05 | 0.000219336 | 0.012987963 | 0.068491627 |
| Dpysl3 | T | 514 | 0.054070402 | 0.718921163 | 0.502311194 | 0.448240793 | -0.216609969 | 0.03999983 | 9.81349E-07 | 0.005202704 | 0.010453041 | 0.037986176 |
| Slain2 | S | 350 | 0.405379037 | 0.704350839 | 0.536992825 | 0.131613787 | -0.167358014 | 0.062097121 | 0.006816544 | 0.000377869 | 0.413307054 | 0.26652319 |
| Slain2 | S | 358 | 0.410416867 | 0.673360666 | 0.525029013 | 0.114612146 | -0.148331653 | 0.042474377 | 0.011474661 | 0.000392204 | 0.425256562 | 0.340118454 |
| Ezh2 | S | 363 | 0.197644414 | 0.638601157 | 0.554892038 | 0.357247624 | -0.083709119 | 0.005791692 | 1.0965E-05 | 0.12418267 | 0.247364377 | 0.743299823 |
| Numa1 | S | 1221 | 0.219462111 | 0.943616693 | 0.560289175 | 0.340827064 | -0.383327518 | 0.30061912 | 0.000788074 | 0.009781791 | 0.10924551 | 0.021200718 |
| Bcl2l13 | T | 389 | 0.256868106 | 0.632504405 | 0.437519254 | 0.180651148 | -0.194985151 | 0.063845196 | 0.001063649 | 0.007302163 | 0.069685048 | 0.056690138 |
| Rb1cc1 | S | 628 | 0.082764896 | 0.587351061 | 0.536317531 | 0.453552635 | -0.05103353 | 0.387805111 | 1.91843E-05 | 0.032337473 | 0.033137155 | 0.694155172 |
| Bcl2l13 | S | 387 | 0.182505462 | 0.596147953 | 0.326086392 | 0.14358093 | -0.270061562 | 0.095299185 | 0.000280413 | 0.007959838 | 0.128666422 | 0.015115464 |
| Carmil1 | S | 1153 | 0.340305687 | 0.730113801 | 0.27586891 | -0.064436777 | -0.45424489 | 0.033467062 | 0.001079841 | 0.190718516 | 0.711693687 | 0.085037084 |
| Bzw1 | S | 413 | 0.677394998 | 0.899823474 | 0.265653664 | -0.411741334 | -0.63416981 | 0.209313477 | 0.000860429 | 0.507307501 | 0.456961555 | 0.181617222 |
| Ikzf4 | S | 326 | 0.096721176 | 0.6513721 | 0.6179261 | 0.521204923 | -0.033446 | 0.614878444 | 0.000136555 | 0.078482729 | 0.116451775 | 0.88593674 |
| Ikzf4 | T | 327 | 0.096721176 | 0.6513721 | 0.6179261 | 0.521204923 | -0.033446 | 0.614878444 | 0.000136555 | 0.078482729 | 0.116451775 | 0.88593674 |
| Dnm1 | T | 780 | 0.509971564 | 0.62209431 | 0.147086095 | -0.362885469 | -0.475008215 | 0.01139614 | 0.002170113 | 0.687306655 | 0.359392962 | 0.258555726 |
| Tnrc18 | S | 385 | 0.318140415 | 0.617872106 | 0.264830539 | -0.053309876 | -0.353041567 | 0.027640527 | 0.000991987 | 0.348528223 | 0.834007296 | 0.240570304 |
| Uvssa | S | 403 | 0.50814882 | 0.763367843 | -0.011857888 | -0.520006708 | -0.775225732 | 0.003274505 | 0.006943853 | 0.978249471 | 0.30221649 | 0.16729016 |
| Plpp7 | S | 43 | 0.233505805 | 0.698275252 | 0.300222843 | 0.066717038 | -0.398052409 | 0.099287507 | 0.012335199 | 0.051126603 | 0.346706818 | 0.087116644 |
| Hdac4 | S | 338 | 0.233500012 | 0.604134114 | 0.702302649 | 0.468802638 | 0.098168535 | 0.52701592 | 0.008591307 | 0.097822514 | 0.287351305 | 0.666855723 |
| Dpysl3 | T | 519 | -0.063704503 | 0.608864744 | 0.245805698 | 0.309510201 | -0.363059046 | 0.510443383 | 4.98689E-05 | 0.0094396 | 0.033471894 | 0.004170497 |
| Bzw1 | S | 411 | 0.457091028 | 0.587962191 | 0.169800114 | -0.287290914 | -0.418162077 | 0.190927661 | 0.000891345 | 0.486801154 | 0.409956101 | 0.155413427 |
| Afap1l2 | S | 484 | 0.066328383 | 0.720048687 | 0.281508231 | 0.215179848 | -0.438540456 | 0.630730961 | 0.000726093 | 0.175627437 | 0.272232012 | 0.068870832 |
| Plec | S | 4641 | 0.339127173 | 0.622976007 | 0.076719299 | -0.262407874 | -0.546256709 | 0.035437575 | 0.002439835 | 0.753471734 | 0.304824717 | 0.09739375 |
| Fez1 | S | 298 | -0.096747328 | 0.602154616 | 0.376609724 | 0.473357052 | -0.225544892 | 0.07426055 | 0.00080783 | 0.001858089 | 0.00734684 | 0.041269603 |
| Pdcd4 | S | 76 | -0.050860692 | 0.811079363 | 0.450479263 | 0.501339956 | -0.360600099 | 0.680742367 | 0.012967952 | 0.021643429 | 0.020188564 | 0.130875075 |
| Cnst | S | 522 | 0.053916711 | 0.611627181 | 0.445652847 | 0.391736136 | -0.165974333 | 0.660373355 | 0.001136502 | 0.222535275 | 0.269552887 | 0.593338327 |
| Mycbp2 | S | 2910 | 0.419643939 | 0.622224733 | 0.147912468 | -0.271731471 | -0.474312264 | 0.222053528 | 0.035709035 | 0.089849189 | 0.377080225 | 0.054620572 |
| Snx1 | S | 188 | -0.026410873 | 0.639238359 | 0.317298235 | 0.343709108 | -0.321940124 | 0.829138003 | 0.001993544 | 0.1829654 | 0.165757455 | 0.174392301 |
| Cpsf7 | S | 427 | 0.056051114 | 0.585748234 | 0.300203466 | 0.244152352 | -0.285544768 | 0.729302066 | 0.009690827 | 0.156858293 | 0.159696945 | 0.119670688 |
| Arhgef6 | S | 679 | 0.053187445 | 0.623079088 | 0.061425823 | 0.008238378 | -0.561653265 | 0.617678749 | 0.001068164 | 0.592747156 | 0.872174857 | 0.00087537 |
| Hdac4 | Y | 345 | 0.216913957 | 0.595728324 | 1.053163148 | 0.836249191 | 0.457434824 | 0.549458922 | 0.022710244 | NA | NA | NA |
| Zfp609 | S | 604 | 0.146315099 | 0.639021141 | -0.084530807 | -0.230845906 | -0.723551948 | 0.619408526 | 0.035229976 | 0.833000024 | 0.56529577 | 0.139639968 |
| Zfp609 | S | 608 | 0.146315099 | 0.639021141 | -0.084530807 | -0.230845906 | -0.723551948 | 0.619408526 | 0.035229976 | 0.833000024 | 0.56529577 | 0.139639968 |
| Zfp609 | S | 613 | 0.146315099 | 0.639021141 | -0.084530807 | -0.230845906 | -0.723551948 | 0.619408526 | 0.035229976 | 0.833000024 | 0.56529577 | 0.139639968 |
| Zfp609 | S | 619 | 0.146315099 | 0.639021141 | -0.084530807 | -0.230845906 | -0.723551948 | 0.619408526 | 0.035229976 | 0.833000024 | 0.56529577 | 0.139639968 |
| Rpl14 | S | 139 | 0.032812135 | -0.464400444 | -1.06087384 | -1.093685975 | -0.596473395 | 0.824763034 | 0.000648226 | 1.1143E-05 | 0.011730672 | 1.29236E-06 |
| Specc1 | S | 126 | -0.396716273 | -0.422827074 | -1.175900019 | -0.779183746 | -0.753072945 | 0.155433798 | 0.112321889 | 0.000398226 | 0.037442669 | 0.0300917 |
| Sart1 | S | 744 | -0.414353318 | -0.505217849 | -1.116356059 | -0.70200274 | -0.611138209 | 0.108448578 | 0.073077977 | 0.001646475 | 0.019752059 | 0.039302432 |
| Ndrg1 | T | 366 | -0.137120638 | -0.274700114 | -1.001982287 | -0.864861649 | -0.727282173 | 0.138622071 | 0.022260136 | 5.60709E-05 | 0.000570902 | 0.001266318 |
| Dcun1d3 | S | 25 | -0.426522623 | -0.334692377 | -1.509520206 | -1.082997583 | -1.174827829 | 0.172309575 | 0.395330888 | 0.002811855 | 0.020234407 | 0.039344366 |
| Cul4a | S | 10 | -0.310462395 | -0.312700321 | -0.967009246 | -0.656546851 | -0.654308926 | 0.068663261 | 0.112813744 | 0.00073287 | 0.009351129 | 0.015484218 |
| Rps7 | S | 119 | -0.171420342 | -0.29493062 | -0.905198439 | -0.733778097 | -0.610267819 | 0.311647551 | 0.167066229 | 3.21668E-05 | 0.022885066 | 0.048433628 |
| Cmc2 | S | 59 | -0.213212187 | -0.446717528 | -1.166330805 | -0.953118618 | -0.719613277 | 0.281410635 | 0.061478306 | 0.007708852 | 0.009906562 | 0.021993284 |
| Ddx5 | S | 24 | -0.260921813 | -0.4493705 | -1.232834778 | -0.971912965 | -0.783464278 | 0.349539568 | 0.144895897 | 0.004952027 | 0.020072662 | 0.042412433 |
| Serpinb1a Serpinb1c | S | 83;83 | -0.237814227 | -0.374856607 | -0.983355521 | -0.745541294 | -0.608498914 | 0.162549245 | 0.120514747 | 0.002332231 | 0.011953322 | 0.041993561 |
| Ttpal | S | 16 | -0.076822201 | -0.159880741 | -0.98121411 | -0.904391909 | -0.821333369 | 0.524365656 | 0.17131794 | 0.012844523 | 0.01739941 | 0.025346576 |
| Ndrg1 | S | 362 | -0.109319704 | -0.083079497 | -0.725701145 | -0.616381441 | -0.642621648 | 0.300436368 | 0.439291039 | 0.005767449 | 0.011861583 | 0.010122661 |
| Pea15a | S | 104 | -0.035632472 | 0.039389464 | -1.485162773 | -1.449530301 | -1.524552237 | 0.935101995 | 0.93438908 | 0.038298152 | 0.047825321 | 0.043144996 |
| Mark3 | S | 638 | 0.195680656 | 0.144948043 | -0.712199678 | -0.907880334 | -0.857147721 | 0.315917921 | 0.369927734 | 0.021086079 | 0.013092488 | 0.021661167 |
| Nefl | S | 532 | -0.080017163 | -0.332867146 | 0.745151371 | 0.825168534 | 1.078018517 | 0.307465988 | 0.02420187 | 6.23916E-06 | 0.000649099 | 0.001083274 |
| Wdr20rt | S | 345 | -0.099465623 | -0.506796918 | -0.687660476 | -0.588194853 | -0.180863558 | 0.005985549 | 9.51479E-08 | 2.78886E-08 | 0.000123383 | 0.000978875 |
| Nob1 | S | 343 | -0.869374564 | -1.072711201 | -2.02944915 | -1.160074586 | -0.956737949 | 0.057657978 | 0.064328686 | 0.002312941 | 0.037163604 | 0.100626137 |
| Nolc1 | S | 585 | -0.09432676 | -0.549299395 | -0.742344349 | -0.648017589 | -0.193044954 | 0.122220365 | 0.000114506 | 1.21858E-08 | 0.002023969 | 0.014532318 |
| Jpt1 | S | 17 | -0.808005522 | -0.738612097 | -1.673807067 | -0.865801545 | -0.93519497 | 0.06867862 | 0.132157565 | 0.001670624 | 0.041297195 | 0.068629589 |
| Sh3pxd2a | S | 721 | -0.073078051 | -0.51384597 | -0.713085402 | -0.64000735 | -0.199239432 | 0.12860828 | 1.01459E-05 | 1.22917E-06 | 0.000123833 | 0.00490808 |
| Frmd4a | S | 342 | -0.101452644 | -0.542126126 | -0.763801954 | -0.66234931 | -0.221675828 | 0.068684132 | 1.37124E-05 | 1.23475E-05 | 0.000299343 | 0.012143482 |
| Pcnp | S | 87 | -0.797555009 | -0.697489833 | -1.589140839 | -0.791585831 | -0.891651007 | 0.058743301 | 0.117215518 | 0.002001144 | 0.039573348 | 0.050674849 |
| Zc3hav1 | T | 349 | -0.000759103 | -0.533100834 | -0.831871506 | -0.831112403 | -0.298770672 | 0.991094812 | 1.21805E-06 | 1.69352E-05 | 0.000729572 | 0.008996475 |
| Ddx5 | S | 30 | -0.523373885 | -0.69633023 | -1.441284054 | -0.917910169 | -0.744953824 | 0.086760769 | 0.079397504 | 0.00063526 | 0.012347917 | 0.06945799 |
| Ddx10 | S | 665 | -0.044169615 | -0.464026535 | -0.765293302 | -0.721123687 | -0.301266767 | 0.350975268 | 0.00016957 | 1.45769E-07 | 0.000127809 | 0.003656286 |
| Npm1 | S | 225 | -0.581545971 | -0.648546125 | -1.350691008 | -0.769145037 | -0.702144884 | 0.038164033 | 0.092140502 | 0.000996233 | 0.018087391 | 0.079488411 |
| Gtf2e2 | S | 265 | -0.623362123 | -0.841355812 | -1.546210628 | -0.922848505 | -0.704854816 | 0.119223615 | 0.063277444 | 0.004267226 | 0.046010525 | 0.111553322 |
| Serf2 | S | 21 | -0.641877568 | -0.609151278 | -1.507209588 | -0.86533202 | -0.89805831 | 0.071946072 | 0.156711896 | 0.001060186 | 0.030308097 | 0.065952617 |
| Katnb1 | S | 354 | 0.012447582 | -0.581548206 | -0.811788517 | -0.824236099 | -0.230240311 | 0.879053343 | 7.84974E-05 | 5.23121E-06 | 0.004301887 | 0.025372393 |
| Ndrg1 | S | 352 | -0.173208404 | -0.484756618 | -1.020566217 | -0.847357813 | -0.535809599 | 0.212158439 | 2.44549E-05 | 0.002452548 | 0.003229026 | 0.019392134 |
| Zc3hav1 | S | 350 | 0.044728185 | -0.521169594 | -0.660239607 | -0.704967792 | -0.139070013 | 0.434461391 | 7.71926E-06 | 1.68565E-06 | 0.000320175 | 0.022743888 |
| Srrm2 | S | 1475 | 0.053076656 | -0.560816746 | -0.65378485 | -0.706861506 | -0.092968104 | 0.101542094 | 2.92138E-05 | 2.32317E-07 | 0.000203183 | 0.069468049 |

| | | | | | | | | | | | | |
|---|---|---|---|---|---|---|---|---|---|---|---|---|
| H1f1 | S | 123 | -0.586866526 | -0.558343687 | -1.501459158 | -0.914592632 | -0.943115471 | 0.136155933 | 0.216321287 | 0.000997634 | 0.041285109 | 0.073527297 |
| Gtse1 | S | 434 | -0.475835922 | -0.555416055 | -1.111023203 | -0.635187282 | -0.555607148 | 0.050463144 | 0.046766006 | 0.001330426 | 0.016926406 | 0.043465804 |
| Zfand3 | S | 124 | -0.104856544 | -0.533308539 | -0.69700555 | -0.592149006 | -0.163697011 | 0.058224876 | 0.006198123 | 2.82819E-06 | 0.000188981 | 0.108098747 |
| Spred1 | S | 309 | -0.551442107 | -0.517745747 | -1.541795241 | -0.990353134 | -1.024049494 | 0.080785471 | 0.221284663 | 0.002055558 | 0.021170204 | 0.051643857 |
| Rangap1 | S | 430 | -0.118094906 | -0.575230123 | -0.7083573 | -0.590262394 | -0.133127177 | 0.029135828 | 0.000142584 | 0.00087071 | 0.00072106 | 0.073626917 |
| Gja1 | S | 297 | 0.015879491 | -0.573414151 | -0.939401818 | -0.955281308 | -0.365987667 | 0.873179854 | 0.000539201 | 4.8294E-05 | 0.000377287 | 0.011924257 |
| Gja1 | T | 326 | 0.00456864 | -0.504332332 | -0.778691651 | -0.783260292 | -0.274359319 | 0.922780662 | 1.75341E-05 | 6.12943E-05 | 0.001296431 | 0.013848318 |
| Rassf1 | S | 203 | -0.217639228 | -0.461987404 | -0.862607347 | -0.644968119 | -0.400619943 | 0.029481301 | 0.034152838 | 4.25051E-05 | 0.001057145 | 0.050509172 |
| Fhdc1 | S | 970 | -0.196589788 | -0.555297868 | -0.949190251 | -0.752600462 | -0.393892382 | 0.274482299 | 0.012846739 | 7.4002E-05 | 0.013256083 | 0.050259247 |
| H1f1 | S | 106 | -0.549836892 | -0.413870105 | -1.46736949 | -0.917532598 | -1.053499384 | 0.153306129 | 0.349337407 | 0.000468013 | 0.048561264 | 0.07050784 |
| Gja1 | S | 328 | -0.003335738 | -0.463124982 | -0.72618872 | -0.722852982 | -0.263063738 | 0.944686179 | 3.35181E-05 | 1.68965E-05 | 0.000461925 | 0.007671696 |
| Cep170b | S | 421 | -0.073060612 | -0.473758895 | -0.745252248 | -0.672191636 | -0.271493353 | 0.495311788 | 0.009501762 | 4.14216E-07 | 0.012342875 | 0.054400265 |
| Prkaa1 | T | 490 | -0.097394943 | -0.468784827 | -0.716087499 | -0.618692556 | -0.247302672 | 0.155678635 | 1.10232E-05 | 0.000939601 | 0.001653478 | 0.043738557 |
| Map4k4 | S | 701 | -0.151633946 | -0.52722412 | -0.770935961 | -0.619302015 | -0.243711842 | 0.045288947 | 0.0017554 | 0.000407191 | 0.000511443 | 0.017924114 |
| Hmgn2 Hmgn3 | S | 29;31 | -0.359655703 | -0.572774389 | -1.269081828 | -0.909426125 | -0.696307439 | 0.207911877 | 0.110526419 | 0.000597337 | 0.017251291 | 0.072610442 |
| Erc1 | S | 824 | 0.002259161 | -0.526795287 | -0.750421657 | -0.752680818 | -0.22362637 | 0.975314105 | 0.000511811 | 9.38188E-06 | 0.000532222 | 0.027602461 |
| Uvrag | S | 548 | -0.068096557 | -0.566684662 | -0.727704925 | -0.659608368 | -0.161020263 | 0.392785125 | 0.000209072 | 0.000203133 | 0.002197564 | 0.028885969 |
| 1110059G10Rik | S | 130 | -0.520651096 | -0.467913683 | -1.160777749 | -0.640126652 | -0.692864066 | 0.075231306 | 0.15726306 | 0.001039691 | 0.045581445 | 0.074117823 |
| 1110059G10Rik | S | 115 | -0.395367086 | -0.452464284 | -1.048632693 | -0.653265607 | -0.59616841 | 0.06015305 | 0.134897438 | 0.000325513 | 0.016704967 | 0.089678486 |
| Troap | S | 39 | -0.37004197 | -0.529288155 | -0.991554913 | -0.621512943 | -0.462266758 | 0.046751049 | 0.016957561 | 0.003629301 | 0.032449131 | 0.068558028 |
| Gja1 | S | 325 | -0.015547897 | -0.497991507 | -0.757466228 | -0.741918331 | -0.259474721 | 0.752904031 | 2.54685E-05 | 0.000222643 | 0.003245046 | 0.029548368 |
| Cep170 | T | 1133 | -0.601264262 | -0.648713581 | -1.403664348 | -0.802400086 | -0.754950767 | 0.11459302 | 0.20789401 | 0.005714697 | 0.048227251 | 0.15091997 |
| Irs1 | S | 265 | -0.03253973 | -0.486788196 | -0.919993997 | -0.887454267 | -0.433205801 | 0.779710444 | 0.002512075 | 4.20721E-05 | 0.001542702 | 0.006681686 |
| Tsc2 | S | 939 | -0.016278525 | -0.549340716 | -0.963655189 | -0.947376664 | -0.414314473 | 0.843069532 | 0.014841793 | 1.61061E-05 | 0.001407919 | 0.023366633 |
| Mybbp1a | S | 1219 | -0.067667167 | -0.426332648 | -0.781134796 | -0.71346763 | -0.354802148 | 0.472922618 | 0.002659136 | 1.11563E-05 | 0.002083744 | 0.012437463 |
| Tbc1d5 | S | 563 | -0.015539413 | -0.419435314 | -0.656868275 | -0.641328862 | -0.237432961 | 0.754451689 | 0.000168737 | 6.17801E-06 | 0.000137332 | 0.00718771 |
| Pxk | S | 448 | -0.469222121 | -0.477432448 | -1.313945776 | -0.844723655 | -0.836513328 | 0.062635307 | 0.163660955 | 0.004201339 | 0.031875975 | 0.051274009 |
| Fam98a | S | 283 | -0.095292981 | -0.519000972 | -0.683183811 | -0.58789083 | -0.164182839 | 0.279249131 | 0.000661917 | 0.000149836 | 0.005247445 | 0.063090952 |
| Rpl37 Rpl37rt | T | 83 | -0.259797651 | -0.457395565 | -0.870133595 | -0.610335945 | -0.41273803 | 0.062826748 | 0.057062935 | 0.000187764 | 0.005385768 | 0.075344878 |
| Cdkn1a | S | 78 | 0.020985759 | -0.358930807 | -0.605251137 | -0.626236895 | -0.24632033 | 0.522220767 | 1.66849E-05 | 1.63706E-06 | 1.0613E-06 | 0.000740023 |
| Snx16 | S | 29 | 0.01113861 | -0.49157943 | -0.708063014 | -0.719201624 | -0.216483584 | 0.824595182 | 0.011790758 | 2.73688E-07 | 0.000497455 | 0.095902195 |
| Tcof1 | S | 309 | -0.141991406 | -0.507240712 | -0.877485566 | -0.73549416 | -0.370244855 | 0.153659318 | 0.08703368 | 3.18782E-05 | 0.000557801 | 0.171582286 |
| Nolc1 | T | 411 | -0.014899765 | -0.429235178 | -0.799389266 | -0.784489501 | -0.370154088 | 0.719694282 | 0.003796002 | 1.53839E-05 | 0.001472756 | 0.008260326 |
| Gja1 | S | 296 | -0.01956078 | -0.502172316 | -0.953174381 | -0.9336136 | -0.451002064 | 0.852489339 | 0.00199137 | 0.000344022 | 0.002520921 | 0.018644494 |
| Gnl3l | S | 459 | 0.079801896 | -0.575404628 | -0.654568532 | -0.734370429 | -0.079163904 | 0.249781616 | 0.000177647 | 1.70119E-05 | 8.47621E-05 | 0.245273737 |
| Tmcc3 | T | 48 | -0.174082585 | -0.563320021 | -0.87976715 | -0.705684564 | -0.316447129 | 0.075080727 | 0.000682674 | 0.01849878 | 0.025112333 | 0.145407336 |
| Ahctf1 | S | 2144 | -0.139610552 | -0.445109115 | -0.859923158 | -0.720312606 | -0.414814044 | 0.226660202 | 0.014165838 | 0.00016953 | 0.002227187 | 0.025624855 |
| Serbp1 | S | 391 | 0.03092426 | -0.505478842 | -0.607011061 | -0.637935321 | -0.101532219 | 0.475657506 | 5.39731E-06 | 0.00038649 | 0.001900722 | 0.157946803 |
| Phldb1 | S | 566 | -0.059130427 | -0.379008673 | -0.673487652 | -0.614357225 | -0.294478979 | 0.522209787 | 0.000802479 | 2.71876E-05 | 0.002643975 | 0.004973663 |
| Hmmr | S | 50 | -0.168317391 | -0.52998045 | -1.031042186 | -0.862724796 | -0.501061737 | 0.330694225 | 0.018207023 | 0.001389064 | 0.008478602 | 0.036184161 |
| Mark3 | S | 601 | -0.171600103 | -0.566197293 | -1.368612509 | -1.197012406 | -0.802415216 | 0.589009287 | 0.132123648 | 0.000923713 | 0.01886844 | 0.062132422 |
| Wnk1 | S | 29 | 0.119372285 | -0.449371101 | -0.910482284 | -1.02985457 | -0.461111183 | 0.387079633 | 0.001304063 | 3.30122E-05 | 0.002207392 | 0.00413373 |
| Hmgn1 | S | 24 | -0.273651912 | -0.437143272 | -0.921039075 | -0.647387163 | -0.483895804 | 0.127289655 | 0.050481274 | 0.000841742 | 0.001544622 | 0.033607794 |
| Nifk | T | 299 | 0.032317166 | -0.574388527 | -0.682198256 | -0.714515422 | -0.107809729 | 0.620962135 | 6.25833E-05 | 0.000853418 | 0.001233692 | 0.233540137 |
| Mid1ip1 | S | 110 | 0.005890777 | -0.466900971 | -0.635528352 | -0.641419129 | -0.16862738 | 0.934394309 | 2.81871E-05 | 0.000434613 | 0.001076032 | 0.076432343 |
| Hdgfl2 | S | 450 | -0.024050853 | -0.57022234 | -0.649000248 | -0.624949395 | -0.078777907 | 0.28910235 | 0.007168541 | 4.70144E-05 | 1.85529E-05 | 0.279389834 |
| Ica1l | S | 316 | 0.018578653 | -0.416928769 | -0.750458746 | -0.769037399 | -0.333529976 | 0.484742408 | 0.008201599 | 4.09089E-06 | 0.00060641 | 0.013796652 |
| Dmtn | S | 156 | 0.016883819 | -0.425985363 | -0.631268881 | -0.6481527 | -0.205283518 | 0.725334319 | 0.001698685 | 5.69543E-06 | 0.000104709 | 0.034761454 |
| Tbc1d4 | S | 673 | -0.073403079 | -0.507069157 | -0.674309075 | -0.600905996 | -0.167239917 | 0.347071717 | 0.000307383 | 0.003127521 | 0.002465404 | 0.106496271 |
| Cep170 Plin5 | S | 1150;218 | -0.324765548 | -0.509547239 | -1.018442013 | -0.693676465 | -0.508894773 | 0.199998572 | 0.069295039 | 0.003733461 | 0.026427102 | 0.071393337 |
| 1810009A15Rik | S | 65 | -0.225072851 | -0.497056656 | -0.909810701 | -0.684737851 | -0.412754045 | 0.240701766 | 0.014642041 | 0.004820407 | 0.022725595 | 0.08145074 |
| Rps10 | T | 148 | 0.033076578 | -0.429609205 | -0.586432953 | -0.619509531 | -0.156823747 | 0.473437107 | 2.64904E-06 | 0.000910278 | 0.000511799 | 0.054605918 |
| Rps25 | S | 26 | 0.025392113 | -0.483011031 | -0.680283096 | -0.70567521 | -0.197272066 | 0.687079705 | 6.40293E-06 | 0.004685508 | 0.002261113 | 0.103090378 |
| Anln | S | 683 | -0.39929964 | -0.319500409 | -1.085040026 | -0.685740387 | -0.765539618 | 0.120247791 | 0.316578283 | 0.001525303 | 0.034561044 | 0.06605359 |
| Afdn | S | 216 | -0.35397937 | -0.423801709 | -1.000351255 | -0.646371885 | -0.576549546 | 0.076551692 | 0.127505372 | 0.005591829 | 0.035810534 | 0.076631841 |
| Mark3 | S | 456 | -0.276490516 | -0.438879591 | -0.863690989 | -0.587200473 | -0.424811399 | 0.097195711 | 0.095949157 | 0.004836126 | 0.028398902 | 0.121658372 |
| Rrm2 | S | 20 | -0.030877465 | -0.428364139 | -0.675804526 | -0.644927061 | -0.247440387 | 0.776526484 | 0.000233507 | 0.007799153 | 0.007155744 | 0.093179701 |
| Map4 | S | 840 | -0.314966257 | -0.314665762 | -1.250725081 | -0.935758824 | -0.936059319 | 0.351919572 | 0.434210287 | 0.003135751 | 0.023063192 | 0.058912612 |
| Nolc1 | S | 527 | -0.024181262 | -0.543577108 | -0.708737229 | -0.684555967 | -0.165160121 | 0.601298413 | 0.004289023 | 0.005300473 | 0.01142472 | 0.206955657 |
| Pes1 | S | 574 | 0.136536865 | -0.571952578 | -0.74381781 | -0.880354675 | -0.171865232 | 0.009985594 | 0.016096343 | 4.98406E-06 | 2.79112E-05 | 0.183752064 |
| Adam10 | S | 741 | -0.352343441 | -0.296033173 | -1.039620937 | -0.687277497 | -0.743587764 | 0.118697242 | 0.317771102 | 0.006674204 | 0.037987737 | 0.058382137 |
| Dock5 | S | 1656 | -0.058081987 | -0.280763328 | -0.832125402 | -0.774043415 | -0.551362074 | 0.762170757 | 0.094637821 | 0.000138218 | 0.025272604 | 0.016766385 |
| Dtl | S | 489 | -0.12329854 | -0.395254522 | -0.711681685 | -0.588383145 | -0.316427164 | 0.378524395 | 0.118878419 | 0.000322453 | 0.008851623 | 0.186237342 |
| Eef1a2 Eef1a1 Eef1a | S | 454;454 | -0.08045088 | -0.481305397 | -0.728023767 | -0.647572887 | -0.24671837 | 0.424588411 | 0.003268759 | 0.017006913 | 0.047294243 | 0.23170004 |
| Stk35 | T | 501 | 0.104529738 | -0.49846799 | -0.643323158 | -0.747852896 | -0.144855168 | 0.424532539 | 1.08289E-05 | 0.023251458 | 0.009303694 | 0.344568088 |
| Frmd4a | S | 699 | -0.079074854 | -0.462406561 | -0.86955642 | -0.790481566 | -0.407149859 | 0.351150191 | 0.052241677 | 0.004498484 | 0.01148384 | 0.082126914 |
| Rps5 | S | 142 | 0.09303545 | -0.507561799 | -0.587272479 | -0.68030793 | -0.079710681 | 0.260159623 | 0.000347104 | 0.001441608 | 0.001195579 | 0.240358128 |
| Irs1 | S | 263 | 0.019110771 | -0.421090229 | -0.835899154 | -0.855009924 | -0.414808924 | 0.896888422 | 0.003184054 | 0.005157726 | 0.00688375 | 0.062613001 |
| Spred1 | S | 234 | -0.225468411 | -0.289542438 | -1.139105391 | -0.91363698 | -0.849562953 | 0.315204286 | 0.399415935 | 0.003127833 | 0.013964176 | 0.061729405 |
| Nolc1 | T | 410 | 0.010407478 | -0.391383572 | -0.772646391 | -0.783053869 | -0.381262819 | 0.875213046 | 0.022835626 | 0.000613549 | 0.00127652 | 0.026462315 |
| Snx7 | S | 383 | -0.140493239 | -0.44608307 | -0.728533239 | -0.58804 | -0.282450169 | 0.148009762 | 0.05191698 | 0.006990756 | 0.016626313 | 0.169032415 |
| Clasp2 | S | 495 | 0.031365713 | -0.326079115 | -0.586229003 | -0.617594716 | -0.260149888 | 0.722699976 | 0.003677417 | 0.000137449 | 0.001202254 | 0.027783868 |
| Eml1 | S | 159 | 0.038484135 | -0.377489672 | -0.761533627 | -0.800017762 | -0.384043955 | 0.755109218 | 0.018864056 | 0.000498426 | 0.00535522 | 0.032142874 |
| Eif4b | S | 230 | 0.02910214 | -0.543262581 | -0.790113918 | -0.819216058 | -0.246851337 | 0.726240038 | 0.020848392 | 0.002589384 | 0.001436285 | 0.128715225 |
| Stk35 | S | 502 | 0.184347415 | -0.538448888 | -0.647746952 | -0.832094367 | -0.109298064 | 0.28303518 | 0.002558659 | 0.001140212 | 0.011112996 | 0.2809896 |
| Pald1 | S | 45 | -0.251543747 | -0.205984043 | -0.870973313 | -0.619429566 | -0.66498927 | 0.107835328 | 0.479769769 | 0.002991895 | 0.017404072 | 0.084384469 |
| Rpl18 | S | 130 | 0.116833289 | -0.457438834 | -0.682564116 | -0.799397405 | -0.225125282 | 0.621538247 | 0.022226659 | 0.001353534 | 0.047289101 | 0.149289252 |
| Nfatc1 | S | 271 | -0.044228784 | -0.216755823 | -0.743142726 | -0.698913942 | -0.526386903 | 0.759423669 | 0.087504362 | 0.000556693 | 0.006029039 | 0.006192932 |
| Marcks | S | 160 | -0.090276051 | -0.337617435 | -0.850645505 | -0.760369453 | -0.51302807 | 0.396976221 | 0.153636527 | 0.006383092 | 0.013565538 | 0.068791484 |
| Tsc2 | S | 1806 | 0.039844115 | -0.405395678 | -0.677690188 | -0.717534303 | -0.27229451 | 0.584978526 | 0.000868312 | 0.022797261 | 0.01793436 | 0.153071004 |
| Uckl1 | S | 63 | 0.105520795 | -0.317487971 | -0.641910942 | -0.747431738 | -0.324422971 | 0.294841448 | 0.004723421 | 0.000574401 | 0.001205556 | 0.030366412 |
| Csnk1g1 | S | 373 | -0.135158854 | -0.137751667 | -0.766364393 | -0.631205539 | -0.628612726 | 0.370503159 | 0.586161738 | 0.000310792 | 0.024756598 | 0.092192016 |
| Mark3 | T | 507 | -0.120160215 | -0.153892614 | -0.848560787 | -0.728400572 | -0.694668172 | 0.572533918 | 0.571252416 | 0.000481738 | 0.028058568 | 0.071288265 |
| Supt16 | S | 1023 | 0.068988599 | -0.221632089 | -0.692777558 | -0.761766157 | -0.471145469 | 0.54291945 | 0.059700021 | 0.000117983 | 0.006011768 | 0.009845343 |
| Mapre2 | S | 215 | 0.051626923 | -0.501491887 | -0.791306935 | -0.842933857 | -0.289815047 | 0.662657421 | 0.007842088 | 0.030935209 | 0.021670872 | 0.22795748 |
| Ncl | T | 129 | 0.284236829 | -0.477335762 | -0.639499663 | -0.923736492 | -0.162163901 | 0.022816615 | 0.002083072 | 0.000243359 | 0.00031131 | 0.112374165 |
| Marcks | S | 156 | -0.237094299 | -0.177346658 | -1.014395766 | -0.777301468 | -0.837049108 | 0.214935282 | 0.532815685 | 0.022515323 | 0.046877213 | 0.055106338 |
| Eif2ak3 | S | 1092 | 0.280775847 | -0.5309272 | -1.016895793 | -1.297671639 | -0.485968592 | 0.018563514 | 0.000858654 | 0.019846446 | 0.020246739 | 0.123597846 |
| Dab2 | S | 220 | -0.006771044 | -0.379533193 | -1.050751368 | -1.043980324 | -0.671218175 | 0.982421201 | 0.504207861 | 0.006579723 | 0.023520136 | 0.274082101 |
| Rpl18 | T | 158 | 0.167069291 | -0.350251212 | -0.610366945 | -0.777436236 | -0.260115733 | 0.378359814 | 0.01358575 | 0.00163824 | 0.01707432 | 0.066309436 |
| Phldb1 | S | 1022 | 0.305270022 | -0.543740929 | -0.619780992 | -0.925051013 | -0.076040063 | 0.007828342 | 0.000537869 | 0.001629757 | 0.000625549 | 0.439181873 |
| Jmjd1c | S | 1799 | 0.04163123 | -0.304509044 | -0.634185195 | -0.675816425 | -0.329676151 | 0.685597315 | 0.001454638 | 0.034170475 | 0.023656906 | 0.145138085 |
| Mvb12a | S | 193 | -0.202772418 | -0.122469173 | -0.842938834 | -0.640166415 | -0.72046966 | 0.280625375 | 0.66395934 | 0.008529021 | 0.031207921 | 0.063123991 |
| Dvl2 | S | 255 | -0.055555698 | -0.244779517 | -0.808296147 | -0.75274045 | -0.56351663 | 0.819562702 | 0.288988437 | 0.008252652 | 0.014886131 | 0.021395439 |

| | | | | | | | | | | | | |
|---|---|---|---|---|---|---|---|---|---|---|---|---|
| Phldb1 | S | 580 | 0.008697167 | -0.187866199 | -0.65890667 | -0.667603837 | -0.471040472 | 0.933997043 | 0.168505837 | 0.004087035 | 0.005828683 | 0.022686625 |
| Setmar | S | 282 | -0.023697721 | -0.191331054 | -0.66347751 | -0.639779789 | -0.472146456 | 0.722628937 | 0.082127133 | 0.016435765 | 0.014578391 | 0.027883245 |
| Phldb1 | S | 1021 | 0.289702854 | -0.433576836 | -0.597354506 | -0.88705736 | -0.16377767 | 0.009196683 | 0.00168641 | 0.002790751 | 0.006903267 | 0.184947642 |
| Marcks | S | 163 | -0.151860073 | -0.069353995 | -1.038067259 | -0.886207186 | -0.968713263 | 0.446246427 | 0.825614243 | 0.035574004 | 0.042076656 | 0.054936017 |
| Phldb1 | T | 550 | 0.205713449 | -0.462862969 | -0.619782446 | -0.825495895 | -0.156919477 | 0.005806553 | 0.00911636 | 0.006686917 | 0.002130072 | 0.1650956 |
| Htt | S | 1843 | 0.31362161 | -0.574108387 | -0.615518935 | -0.929140545 | -0.041410547 | 0.025436174 | 0.011684 | 0.012825953 | 0.002696707 | 0.783770594 |
| Phldb1 | S | 491 | 0.344487053 | -0.483390718 | -0.674234952 | -1.018722005 | -0.190844234 | 0.000855481 | 0.008019218 | 0.001272442 | 0.000707243 | 0.133699406 |
| Nectin1 | S | 389 | 0.049790953 | -0.05537798 | -0.637775748 | -0.687566701 | -0.582397768 | 0.632910588 | 0.765208934 | 0.004307162 | 0.004700923 | 0.043363307 |
| Rgs12 | S | 909 | 0.192646276 | -0.224091746 | -0.597424137 | -0.790070414 | -0.373332392 | 0.388934619 | 0.275361184 | 0.019005917 | 0.008085521 | 0.043623021 |
| Synpo | S | 773 | 0.115745359 | 0.535030531 | 0.769543343 | 0.653797984 | 0.234512813 | 0.208144517 | 1.05072E-05 | 8.89151E-07 | 0.007587453 | 0.000219845 |
| Atrx | S | 801 | -0.03693944 | 0.519150557 | 0.615678977 | 0.652618417 | 0.09652842 | 0.59299389 | 6.01934E-07 | 6.03233E-08 | 0.005339121 | 0.007939834 |
| Sap30l | S | 98 | 0.009972517 | 0.502348141 | 0.714649472 | 0.704676955 | 0.212301332 | 0.936098663 | 7.58616E-05 | 4.85989E-08 | 0.021561074 | 0.003509111 |
| Ncoa2 | S | 699 | 0.047100074 | 0.566486987 | 0.860063917 | 0.812963842 | 0.293576929 | 0.46384351 | 1.58781E-05 | 0.00042518 | 0.000544349 | 0.037099476 |
| Dcun1d4 | S | 103 | -0.022041292 | 0.503258724 | 0.679063247 | 0.701104539 | 0.175804523 | 0.806862192 | 8.44318E-06 | 7.13512E-06 | 0.004522321 | 0.015100275 |
| Actn1 Actn2 Actn3 Ac | S | 160;140;147;153 | 0.027879797 | 0.354197592 | 0.630346323 | 0.602466525 | 0.27614873 | 0.429109089 | 1.88664E-05 | 3.85975E-07 | 0.001171622 | 0.003687976 |
| Filip1l | T | 984 | 0.024329603 | 0.491270628 | 0.772909163 | 0.748579559 | 0.281638535 | 0.780567229 | 0.005771385 | 2.95144E-06 | 0.003101117 | 0.049566877 |
| Sorbs2 | S | 378 | 0.019469667 | 0.577288927 | 0.712721844 | 0.693252178 | 0.135432917 | 0.666751043 | 1.16036E-06 | 0.007512582 | 0.009135999 | 0.252842044 |
| Pphln1 | S | 215 | -0.075414853 | 0.482157182 | 0.593009393 | 0.668424246 | 0.110852211 | 0.148177662 | 3.96372E-07 | 3.79757E-05 | 0.000150985 | 0.047152099 |
| Epb41l3 | S | 456 | -0.020388047 | 0.382917626 | 0.651978615 | 0.672366662 | 0.269060989 | 0.719153879 | 0.000195855 | 6.13955E-06 | 6.54993E-05 | 0.001860561 |
| Thrap3 | S | 746 | 0.175124558 | 0.476005906 | 0.761124218 | 0.58587666 | 0.285118313 | 0.140236258 | 0.021887251 | 0.000354284 | 0.004792116 | 0.096722947 |
| Vcam1 | S | 166 | -0.170669059 | 0.387933481 | 0.772902309 | 0.943571369 | 0.384968828 | 0.037906829 | 0.000458797 | 5.95158E-06 | 6.14846E-05 | 0.001729148 |
| Zc3h6 | S | 1150 | -0.032675653 | 0.496106595 | 0.656453898 | 0.689129551 | 0.160347302 | 0.765899011 | 0.000825761 | 0.000478089 | 0.003634084 | 0.130584509 |
| Tent4b | S | 54 | -0.060402537 | 0.473884256 | 0.684845306 | 0.745247843 | 0.210961049 | 0.250261875 | 0.021342389 | 5.04512E-06 | 0.00013007 | 0.144158818 |
| Tent4b | S | 57 | -0.060402537 | 0.473884256 | 0.684845306 | 0.745247843 | 0.210961049 | 0.250261875 | 0.021342389 | 5.04512E-06 | 0.00013007 | 0.144158818 |
| Irs2 | S | 1089 | -0.191050163 | 0.453653576 | 0.72735481 | 0.918404973 | 0.273701234 | 0.063024916 | 0.000971836 | 1.19383E-05 | 0.000557332 | 0.015330413 |
| Filip1l | T | 992 | -0.090570511 | 0.418665997 | 0.635680226 | 0.726250737 | 0.217014229 | 0.17229504 | 0.00028168 | 0.000552324 | 0.001694237 | 0.063883408 |
| Foxo1 | S | 284 | -0.095713826 | 0.392247781 | 0.73184101 | 0.827554836 | 0.339593229 | 0.164708416 | 0.003209609 | 0.000282875 | 0.000209301 | 0.00506797 |
| Rps6ka5 | T | 839 | -0.068409478 | 0.58186769 | 0.601966286 | 0.670375765 | 0.020098596 | 0.531288102 | 0.001155622 | 0.008818561 | 0.009047926 | 0.88206823 |
| Zc3h13 | S | 1347 | 0.009405137 | 0.326637345 | 0.635146852 | 0.625741715 | 0.308509507 | 0.935161636 | 0.029641324 | 0.000359828 | 0.005860306 | 0.043487024 |
| Rtp4 | S | 181 | 0.012881133 | 0.22119483 | 0.612982824 | 0.600101691 | 0.391787994 | 0.912132899 | 0.094462534 | 0.002250963 | 0.002576035 | 0.002540705 |
| Plag1 | S | 389 | -0.01669743 | 0.485589026 | 0.590154085 | 0.606851516 | 0.104565059 | 0.828059198 | 0.103650312 | 0.008311665 | 0.006920818 | 0.635115968 |
| Zrsr1 | S | 417 | -0.107017527 | 0.336406085 | 0.622446088 | 0.729463616 | 0.286040004 | 0.139828319 | 0.061440837 | 0.003734227 | 0.010161694 | 0.105792538 |
| Mylk | T | 1031 | -0.283796311 | 0.258481214 | 0.587643285 | 0.871439596 | 0.329162071 | 0.054643311 | 0.07290132 | 0.003420344 | 0.000259672 | 0.003873461 |
| Mylk | T | 1033 | -0.283796311 | 0.258481214 | 0.587643285 | 0.871439596 | 0.329162071 | 0.054643311 | 0.07290132 | 0.003420344 | 0.000259672 | 0.003873461 |
| Mylk | T | 1040 | -0.283796311 | 0.258481214 | 0.587643285 | 0.871439596 | 0.329162071 | 0.054643311 | 0.07290132 | 0.003420344 | 0.000259672 | 0.003873461 |
| Gpatch11 | S | 50 | -0.560259184 | -0.508074625 | -1.142490988 | -0.582231804 | -0.634416364 | 0.068551791 | 0.12337007 | 0.004124241 | 0.020311691 | 0.037117791 |
| Cep170 | S | 122 | -0.533135509 | -0.335823742 | -0.996601865 | -0.463466356 | -0.660778123 | 0.028219103 | 0.164278244 | 0.005809094 | 0.08665264 | 0.036757419 |
| Rasa3 | S | 530 | -0.386159739 | -0.417231297 | -1.004696986 | -0.618537248 | -0.58746569 | 0.21841428 | 0.074780614 | 0.001999302 | 0.086896689 | 0.033926419 |
| Slc7a6 | T | 23 | -0.47138914 | -0.330271834 | -0.954231192 | -0.482842052 | -0.623959358 | 0.044400936 | 0.131437098 | 0.004820483 | 0.07171849 | 0.035766415 |
| Dcun1d3 | S | 30 | -0.499652139 | -0.22217485 | -1.647903033 | -1.148250894 | -1.425728183 | 0.297092807 | 0.570578607 | 0.002827498 | 0.062802629 | 0.02262722 |
| Plpp2 | S | 267 | -0.290688779 | -0.229668152 | -0.851789969 | -0.561101189 | -0.622121816 | 0.010059656 | 0.086841389 | 0.002874702 | 0.013705732 | 0.008848074 |
| Tsr3 | S | 24 | -0.399751483 | -0.295800337 | -1.037479917 | -0.637728434 | -0.741679581 | 0.401122784 | 0.116459288 | 0.005698508 | 0.226205392 | 0.023773212 |
| Map4 | S | 841 | -0.399681564 | -0.313496441 | -1.17090607 | -0.771224506 | -0.857409629 | 0.240028709 | 0.367736515 | 0.011177784 | 0.058766376 | 0.048972367 |
| Ccnb1 | S | 53 | -0.348272372 | -0.216554038 | -0.808879456 | -0.460607084 | -0.592325418 | 0.075620524 | 0.30037774 | 0.007451929 | 0.063214391 | 0.039200851 |
| Syn1 | S | 605 | -0.420049859 | 0.049808259 | -0.780808312 | -0.360758453 | -0.830616572 | 0.064344624 | 0.844681957 | 0.005359345 | 0.083609188 | 0.025085975 |
| Cdk18 | S | 109 | -0.318136508 | -0.130017104 | -0.937784434 | -0.619647926 | -0.80776733 | 0.222449074 | 0.656389898 | 0.018999378 | 0.077201148 | 0.048710716 |
| Marcks | S | 152 | -0.283115517 | -0.129965905 | -1.177515342 | -0.894399825 | -1.047549436 | 0.26634663 | 0.676028533 | 0.032284067 | 0.064718823 | 0.049667911 |
| Gli3 | S | 980 | -0.404495483 | 0.166537124 | -0.688453621 | -0.283958138 | -0.854990745 | 0.041846 | 0.521593617 | 0.034027744 | 0.25600744 | 0.032954771 |
| Cdk18 | S | 66 | -0.22947881 | 0.037438597 | -0.755659633 | -0.526180823 | -0.79309823 | 0.288081853 | 0.884656975 | 0.021043446 | 0.068701241 | 0.026531298 |
| Lima1 | S | 581 | -0.169076425 | -0.068627597 | -0.696325806 | -0.527249381 | -0.627698209 | 0.428419906 | 0.691859566 | 0.026941413 | 0.081230813 | 0.04418821 |
| Ppp1r12c | S | 431 | -0.331413908 | 0.167631224 | -0.658316145 | -0.326902237 | -0.825947369 | 0.156304747 | 0.598717553 | 0.035151011 | 0.17422241 | 0.049729689 |
| Nr2f6 | S | 143 | -0.091665758 | -0.029932414 | -0.676874536 | -0.585208778 | -0.646942122 | 0.55590067 | 0.857163463 | 0.039866512 | 0.05203921 | 0.041758875 |
| Bcl7c | S | 103 | 0.026579845 | 0.171017129 | 1.300355652 | 1.273775807 | 1.129338523 | 0.955425443 | 0.525684211 | 0.000831975 | 0.072432957 | 0.005712339 |
| Baz1a | S | 284 | -0.053033707 | -0.517992775 | -0.631749266 | -0.578715558 | -0.113756491 | 0.320771439 | 4.30328E-07 | 5.64379E-08 | 0.001909317 | 0.007372208 |
| Larp1 | S | 1032 | -0.177830936 | -0.57566109 | -0.610927134 | -0.433096198 | -0.035266044 | 0.023848801 | 1.6861E-05 | 2.04113E-06 | 0.002276016 | 0.434471232 |
| Eif4b | S | 495 | -0.091515852 | -0.567627436 | -0.64180368 | -0.550287828 | -0.074176244 | 0.012504299 | 0.000199676 | 6.13806E-08 | 1.46795E-05 | 0.153350342 |
| Parp3 | S | 14 | -0.757544551 | -0.542876886 | -1.76046465 | -1.002920099 | -1.217587764 | 0.102502967 | 0.25678449 | 0.000413874 | 0.055443609 | 0.051884019 |
| Fnip1 | S | 594 | -0.515961497 | -0.761149727 | -0.943820247 | -0.42785875 | -0.18267052 | 0.00076054 | 0.056088373 | 0.002295327 | 0.056322547 | 0.503737364 |
| Khdrbs1 | S | 18 | -0.709397835 | -0.723687572 | -1.307575632 | -0.598177797 | -0.583888061 | 0.052661443 | 0.061038896 | 0.001738362 | 0.078092616 | 0.101513519 |
| Pip5k1a | T | 34 | -0.654092268 | -0.718442239 | -1.380907512 | -0.726815245 | -0.662465273 | 0.083093198 | 0.088368124 | 0.000839578 | 0.062467858 | 0.10864313 |
| Utp18 | S | 27 | -0.136437011 | -0.506478278 | -0.619679204 | -0.483242193 | -0.113200925 | 0.039406326 | 1.57782E-05 | 9.16514E-06 | 0.001770488 | 0.018371247 |
| Ddx17 | S | 520 | -0.370976234 | -0.544940835 | -0.876909918 | -0.505933684 | -0.331969083 | 0.002203845 | 0.000268027 | 0.007219837 | 0.051391385 | 0.115261981 |
| Zfp622 | S | 145 | -0.099888625 | -0.384699955 | -0.590039727 | -0.490151102 | -0.205339772 | 0.027215105 | 0.001405452 | 1.32266E-09 | 0.00152119 | 0.01312276 |
| Nolc1 | S | 312 | -0.075650762 | -0.472286656 | -0.618179766 | -0.542529004 | -0.14589311 | 0.067712089 | 4.21494E-06 | 4.22337E-06 | 0.000200758 | 0.015428574 |
| Plec | S | 4396 | -0.089436722 | -0.337225223 | -0.618381142 | -0.52894442 | -0.281155919 | 0.035392178 | 9.41685E-06 | 1.68671E-07 | 0.000228355 | 0.000273347 |
| Zfp26 Zfp28 Zfp39 Zf | T | 381;460;297;625 | -0.379478006 | -0.499902298 | -0.660804163 | -0.281326157 | -0.160901866 | 0.029728438 | 0.000524911 | 7.92566E-05 | 0.087546235 | 0.052823707 |
| Hspbp1 | S | 349 | -0.155625271 | -0.540264843 | -0.607396862 | -0.451771592 | -0.06713202 | 0.008869163 | 6.12116E-06 | 0.000392687 | 0.00999452 | 0.312784775 |
| Sqstm1 | S | 334 | -0.054007284 | -0.544277365 | -0.638120325 | -0.58411304 | -0.093842959 | 0.114714544 | 3.28094E-07 | 0.000199766 | 0.000192409 | 0.090686282 |
| Dcaf10 | S | 356 | -0.362027825 | -0.55047982 | -0.617941598 | -0.255913772 | -0.067461777 | 0.005601357 | 0.00311623 | 0.000172135 | 0.031287478 | 0.511827472 |
| Rangap1 | S | 444 | -0.127838029 | -0.479979404 | -0.619892913 | -0.492054884 | -0.139913509 | 0.040951489 | 3.98828E-06 | 0.000134952 | 0.000486529 | 0.027642491 |
| Errfi1 | S | 97 | -0.074285986 | -0.526521103 | -0.606438447 | -0.532152462 | -0.079917344 | 0.310016378 | 2.22942E-05 | 8.01137E-06 | 0.002902747 | 0.093723568 |
| Cd2ap | S | 233 | -0.150146402 | -0.405145063 | -0.674086461 | -0.523940059 | -0.268941397 | 0.00819748 | 0.002904036 | 2.70471E-06 | 0.000367772 | 0.018408492 |
| Ddx3x | S | 90 | -0.490329025 | -0.55620529 | -1.033526202 | -0.543197177 | -0.477320911 | 0.034252704 | 0.030643332 | 0.001279127 | 0.019809444 | 0.043976697 |
| Lrrc41 | S | 326 | -0.688622465 | -0.527116118 | -1.285513137 | -0.596890672 | -0.75839702 | 0.054716542 | 0.160568918 | 0.00150055 | 0.079013973 | 0.06828205 |
| Lrrc41 | S | 327 | -0.688622465 | -0.527116118 | -1.285513137 | -0.596890672 | -0.75839702 | 0.054716542 | 0.160568918 | 0.00150055 | 0.079013973 | 0.06828205 |
| Vars | S | 1224 | -0.374038958 | -0.517848805 | -0.702556884 | -0.328517926 | -0.184708079 | 0.015072813 | 0.003273518 | 0.000682946 | 0.005414664 | 0.029388188 |
| Fnip1 | S | 593 | -0.375431489 | -0.565996191 | -0.729247504 | -0.353816015 | -0.163251313 | 0.006461248 | 0.032974575 | 0.000238645 | 0.01673369 | 0.35241632 |
| Ppfibp1 | S | 403 | -0.179447144 | -0.383581474 | -0.585042184 | -0.40559504 | -0.201460709 | 0.001237727 | 0.002418225 | 1.45465E-05 | 0.00117746 | 0.026252757 |
| Shc1 | S | 139 | -0.439730568 | -0.507454949 | -0.945797197 | -0.506066629 | -0.438342248 | 0.012737914 | 0.080575169 | 0.001119093 | 0.032578946 | 0.120256067 |
| Irs2 | S | 590 | -0.085350381 | -0.563099972 | -0.585644955 | -0.500294574 | -0.022544982 | 0.259969237 | 5.23728E-06 | 0.000607283 | 0.002706895 | 0.65541666 |
| Troap | S | 37 | -0.401409898 | -0.489524724 | -0.95410969 | -0.552699792 | -0.464584966 | 0.027396846 | 0.01820489 | 0.002056798 | 0.027811637 | 0.047269631 |
| Dnaja1 | S | 335 | -0.080811709 | -0.522449791 | -0.649790491 | -0.568978782 | -0.1273407 | 0.201405249 | 4.80797E-07 | 0.003678808 | 0.001583866 | 0.14731311 |
| Ddx50 | S | 39 | -0.397295065 | -0.532277738 | -0.682592407 | -0.285297342 | -0.150314669 | 0.203574547 | 0.000171612 | 0.001753913 | 0.321753881 | 0.198447248 |
| Tjp1 | S | 1411 | -0.377548308 | -0.437597689 | -0.742018844 | -0.364470537 | -0.304421155 | 0.00393827 | 0.050971763 | 0.000441679 | 0.020931216 | 0.127853119 |
| Rpl12 | S | 38 | -0.076552371 | -0.498234787 | -0.647036074 | -0.570483703 | -0.148801287 | 0.168334647 | 6.1894E-05 | 0.000122963 | 0.003527831 | 0.083347714 |
| Emg1 | S | 16 | -0.184221943 | -0.582314947 | -0.712910284 | -0.528688341 | -0.130595338 | 0.132302739 | 0.004335747 | 9.08229E-05 | 0.009365622 | 0.197054186 |
| Spry4 | S | 281 | -0.425041471 | -0.50907835 | -0.904400854 | -0.479359382 | -0.395322504 | 0.021904115 | 0.038069344 | 0.002170071 | 0.036071317 | 0.096648227 |
| Cep170 | S | 1135 | -0.773842079 | -0.607115708 | -1.536731508 | -0.762889429 | -0.9296158 | 0.059314426 | 0.185243453 | 0.024033853 | 0.15876125 | 0.116277536 |
| Arpp19 | S | 23 | -0.22497878 | -0.55534482 | -0.610411929 | -0.38543315 | -0.055067109 | 0.117606874 | 0.001260682 | 0.000308036 | 0.033493756 | 0.55457691 |
| Gsk3b | S | 25 | -0.431372619 | -0.49689926 | -0.894084553 | -0.462711934 | -0.397185293 | 0.028217558 | 0.034520498 | 0.002636902 | 0.046174801 | 0.09164403 |
| Eif4ebp2 | T | 37 | -0.261441948 | -0.490839224 | -0.717468591 | -0.456026644 | -0.226629368 | 0.050261924 | 0.001642641 | 0.001334704 | 0.011432119 | 0.065518434 |
| Tfrc | S | 24 | -0.40696125 | -0.530678151 | -0.97482601 | -0.567864759 | -0.444147859 | 0.016454224 | 0.008640405 | 0.020328043 | 0.091158305 | 0.143143053 |
| Trip12 | S | 1111 | -0.303071138 | -0.392200804 | -0.664602949 | -0.361531811 | -0.272402145 | 0.012754071 | 0.004378592 | 0.0007223 | 0.018325257 | 0.043158639 |
| Nolc1 | S | 554 | -0.021504952 | -0.394557424 | -0.60511372 | -0.583608768 | -0.210556296 | 0.520128998 | 1.90411E-06 | 8.00963E-05 | 0.000144923 | 0.016025325 |

| | | | | | | | | | | | | |
|---|---|---|---|---|---|---|---|---|---|---|---|---|
| Scrib | S | 1271 | -0.380756857 | -0.407291059 | -0.835501036 | -0.454744179 | -0.428209977 | 0.026395875 | 0.037308619 | 0.000781111 | 0.015024009 | 0.032007892 |
| Phldb2 | S | 411 | -0.189531642 | -0.54033668 | -0.647699019 | -0.458167377 | -0.107362339 | 0.002946467 | 0.002736261 | 0.003180562 | 0.017796111 | 0.318076152 |
| Nop2 | S | 59 | -0.252948802 | -0.548343611 | -0.728782766 | -0.475833964 | -0.180439156 | 0.056717994 | 0.003424379 | 0.001807462 | 0.031809768 | 0.207725443 |
| Ubap2 | S | 114 | -0.38618542 | -0.498943308 | -0.725692411 | -0.339506991 | -0.226749103 | 0.029881191 | 0.032341953 | 0.000770326 | 0.038203972 | 0.213827306 |
| Pus7 | S | 657 | -0.068395691 | -0.492190375 | -0.642305995 | -0.573910304 | -0.15011562 | 0.469645957 | 0.000281819 | 5.51031E-05 | 0.004295946 | 0.033375237 |
| Ankrd17 | S | 1453 | -0.542675335 | -0.622150274 | -0.629574041 | -0.086898706 | -0.007423768 | 0.032754771 | 0.115926654 | 0.000407227 | 0.592491381 | 0.978821723 |
| Cpd | S | 1352 | -0.508385718 | -0.504299904 | -0.939464856 | -0.431079138 | -0.435164952 | 0.024319826 | 0.077299195 | 0.005787126 | 0.0911434 | 0.134795358 |
| Ddx3x | S | 612 | -0.505653859 | -0.583369781 | -1.084920707 | -0.579266848 | -0.501550926 | 0.048222144 | 0.116953917 | 0.006196033 | 0.065159552 | 0.173730825 |
| Rrp12 | S | 1214 | -0.239442834 | -0.499037255 | -0.65747333 | -0.418030497 | -0.158436075 | 0.032485146 | 0.003626052 | 0.00109442 | 0.01541325 | 0.212198807 |
| Mtmr10 | S | 565 | -0.48512827 | -0.419177246 | -1.018095552 | -0.532967283 | -0.598918306 | 0.07804574 | 0.14766794 | 0.000819825 | 0.065536819 | 0.068411268 |
| Rab5a | S | 193 | -0.57393201 | -0.518820312 | -0.980496201 | -0.406564191 | -0.461675888 | 0.038464972 | 0.165189765 | 0.003616048 | 0.035367203 | 0.190500395 |
| Dap | T | 50 | -0.427602833 | -0.496522419 | -0.84954134 | -0.421938507 | -0.353018921 | 0.016097058 | 0.016639176 | 0.012876519 | 0.078161695 | 0.121883738 |
| Errfi1 | S | 327 | -0.109329374 | -0.489694116 | -0.636661802 | -0.527332428 | -0.146967686 | 0.207539358 | 0.000722228 | 0.000152095 | 0.002558186 | 0.052668235 |
| Pphln1 | S | 223 | -0.30832829 | -0.443496425 | -0.598469936 | -0.290141646 | -0.15497351 | 0.005519834 | 0.001048383 | 0.011261709 | 0.109030642 | 0.286266179 |
| Samd4 | S | 652 | -0.193943345 | -0.453108031 | -0.696997291 | -0.503053946 | -0.24388926 | 0.229298563 | 0.003837289 | 0.000106059 | 0.033419804 | 0.046505184 |
| Plec | S | 4275 | -0.392510659 | -0.381289411 | -0.915370302 | -0.522859643 | -0.534080892 | 0.077368828 | 0.06550957 | 0.000558956 | 0.038569885 | 0.023512636 |
| Knstrn | S | 159 | -0.501047578 | -0.550665536 | -1.035467658 | -0.53442008 | -0.484802121 | 0.105097633 | 0.096265933 | 0.003196916 | 0.064922694 | 0.106698375 |
| Iws1 | S | 667 | -0.074321742 | -0.446029334 | -0.615314921 | -0.540993179 | -0.169285587 | 0.295018561 | 0.000346643 | 5.88214E-05 | 9.56318E-05 | 0.011156608 |
| Parp3 | S | 8 | -0.513519026 | -0.57429983 | -1.025674707 | -0.512155681 | -0.451374877 | 0.103861876 | 0.095618021 | 0.003883347 | 0.073913067 | 0.13574181 |
| Kif2a | S | 156 | -0.283820213 | -0.475987753 | -0.774474529 | -0.490654316 | -0.298486776 | 0.059603879 | 0.02357432 | 0.000690318 | 0.011008836 | 0.090061976 |
| Dap | S | 51 | -0.442246215 | -0.504998493 | -0.907915359 | -0.465669144 | -0.402916866 | 0.021679849 | 0.030423803 | 0.013996832 | 0.073685332 | 0.112460913 |
| Slc16a1 | S | 491 | -0.074972383 | -0.343527641 | -0.586234784 | -0.511262401 | -0.242707144 | 0.604907345 | 0.000150979 | 5.70216E-06 | 0.050013612 | 0.00019045 |
| Jpt2 | S | 30 | -0.159518967 | -0.455987266 | -0.589222709 | -0.429703742 | -0.133235442 | 0.08639796 | 0.000994191 | 0.000291409 | 0.000240483 | 0.030347011 |
| Frmd4a | S | 882 | -0.370051084 | -0.526854242 | -0.923252857 | -0.553201774 | -0.396398615 | 0.053374926 | 0.04627998 | 0.00384391 | 0.038188188 | 0.124495091 |
| Zfp787 | T | 89 | -0.396481054 | -0.411625784 | -0.666451558 | -0.269970504 | -0.254825774 | 0.058200362 | 0.012707048 | 0.000836527 | 0.156381246 | 0.061361424 |
| Slc38a2 | S | 29 | -0.418148401 | -0.670556144 | -0.862027921 | -0.44387952 | -0.191471777 | 0.079706148 | 0.05200913 | 0.004994401 | 0.035801101 | 0.456182441 |
| Pak1ip1 | S | 374 | -0.365798649 | -0.621720512 | -1.240584831 | -0.874786182 | -0.618864319 | 0.323408872 | 0.117304744 | 0.002114894 | 0.060844656 | 0.115985401 |
| Tagln2 Tagln3 | S | 7;7 | -0.473024693 | -0.435789187 | -0.727334148 | -0.254309455 | -0.291544961 | 0.014652981 | 0.0275308 | 0.007387748 | 0.173210075 | 0.137496363 |
| Top2a | S | 1211 | -0.347964154 | -0.385751479 | -0.671700322 | -0.323736168 | -0.285948843 | 0.019671841 | 0.035707752 | 0.000536832 | 0.018406779 | 0.08560458 |
| Eef2 | T | 435 | -0.226750261 | -0.525376939 | -0.729858709 | -0.503108448 | -0.20448177 | 0.009416675 | 0.011597912 | 0.004826787 | 0.009960818 | 0.135186597 |
| Nolc1 | S | 424 | -0.149036205 | -0.548368118 | -0.654501514 | -0.505465309 | -0.106133396 | 0.111401191 | 0.007652762 | 0.00026764 | 0.003359145 | 0.271760229 |
| Unc5b | S | 528 | -0.101415653 | -0.561018001 | -0.611736546 | -0.510320894 | -0.050718545 | 0.141104135 | 0.000500141 | 0.001340522 | 0.013510923 | 0.603290924 |
| Unc5b | S | 531 | -0.101415653 | -0.561018001 | -0.611736546 | -0.510320894 | -0.050718545 | 0.141104135 | 0.000500141 | 0.001340522 | 0.013510923 | 0.603290924 |
| Kif2c | S | 188 | -0.434497428 | -0.508620336 | -1.042039333 | -0.607541905 | -0.533418997 | 0.154115705 | 0.076513055 | 0.00279028 | 0.053639419 | 0.036919592 |
| Slc4a7 | S | 60 | -0.31464724 | -0.460060166 | -0.784037723 | -0.469390483 | -0.323977557 | 0.012360072 | 0.022146764 | 0.00558294 | 0.024982566 | 0.084803143 |
| Twf1 | S | 322 | -0.55028843 | -0.548871616 | -1.042596084 | -0.492307654 | -0.493724468 | 0.056874382 | 0.147121427 | 0.008523086 | 0.11475737 | 0.194947712 |
| Cr1l | S | 454 | -0.572465922 | -0.446867498 | -0.688049532 | -0.115583609 | -0.241182034 | 0.011899679 | 0.093343978 | 0.014678119 | 0.501059641 | 0.226635112 |
| Prkd1 | S | 255 | -0.22529378 | -0.338544604 | -0.766414372 | -0.541120593 | -0.427869768 | 0.034648296 | 0.018823967 | 0.000452105 | 0.015130133 | 0.01508317 |
| Chrm1 | S | 356 | -0.532324598 | -0.419018249 | -0.608566686 | -0.076242088 | -0.189548438 | 0.001065528 | 0.17294483 | 0.009440141 | 0.558308613 | 0.481701725 |
| Chrm1 | T | 354 | -0.532324598 | -0.419018249 | -0.608566686 | -0.076242088 | -0.189548438 | 0.001065528 | 0.17294483 | 0.009440141 | 0.558308613 | 0.481701725 |
| Pld5 | S | 187 | -0.443735753 | -0.410723126 | -0.718269966 | -0.274534212 | -0.30754684 | 0.015993609 | 0.031054907 | 0.006848373 | 0.142228925 | 0.119929689 |
| Arhgdia | S | 148 | -0.159966884 | -0.47267654 | -0.648708898 | -0.488742014 | -0.176032358 | 0.216592605 | 0.003193418 | 0.000292558 | 0.014464556 | 0.124682167 |
| Il1rap | S | 566 | -0.407945478 | -0.378984287 | -0.706169176 | -0.298223698 | -0.327184889 | 0.004025434 | 0.094630861 | 0.003593833 | 0.070544205 | 0.137469606 |
| Pcnp | S | 147 | -0.569196978 | -0.443846504 | -1.085337037 | -0.516140059 | -0.641490533 | 0.072126425 | 0.22533825 | 0.005125472 | 0.064284545 | 0.092165914 |
| Pacsin2 | S | 313 | -0.409571684 | -0.353897153 | -0.612668961 | -0.203097276 | -0.258771808 | 0.009687202 | 0.041402482 | 0.001635809 | 0.099370027 | 0.098917219 |
| Pacsin2 | T | 311 | -0.409571684 | -0.353897153 | -0.612668961 | -0.203097276 | -0.258771808 | 0.009687202 | 0.041402482 | 0.001635809 | 0.099370027 | 0.098917219 |
| Hmgn1 | S | 20 | -0.281513185 | -0.421971311 | -0.83361854 | -0.552105355 | -0.41164723 | 0.094263484 | 0.032718804 | 0.001033128 | 0.005965002 | 0.018818486 |
| H1f2 H1f3 H1f4 H1f5 | S | 104;104;106;104; | -0.553896929 | -0.398305156 | -1.341543713 | -0.787646784 | -0.943238557 | 0.103925224 | 0.330149939 | 0.005885785 | 0.055082348 | 0.064088253 |
| Rps6kb1 | S | 403 | -0.190868228 | -0.57561486 | -0.742600393 | -0.551732165 | -0.166985533 | 0.068812805 | 0.02142586 | 0.001860874 | 0.007179629 | 0.311953288 |
| Mllt3 | S | 302 | -0.212858745 | -0.553263207 | -0.591735045 | -0.3788763 | -0.038471837 | 0.186147811 | 0.006703113 | 0.00060887 | 0.058566509 | 0.732725249 |
| Rassf8 | S | 129 | -0.265921522 | -0.382661378 | -0.780351803 | -0.514430281 | -0.397690424 | 0.104910172 | 0.027102381 | 0.000607312 | 0.010538229 | 0.015213134 |
| Ssrp1 | S | 120 | -0.408177414 | -0.407619473 | -0.883944976 | -0.475767562 | -0.476325503 | 0.114580319 | 0.231238634 | 0.000381627 | 0.086458735 | 0.183019013 |
| Ufd1 | S | 299 | -0.400934763 | -0.56600912 | -1.046803563 | -0.645868801 | -0.480794444 | 0.033193399 | 0.032351233 | 0.047073154 | 0.138283442 | 0.226041522 |
| Tnk2 | S | 951 | -0.491296822 | -0.404556587 | -0.78205647 | -0.290759648 | -0.377499883 | 0.017548732 | 0.139832981 | 0.004966169 | 0.146993395 | 0.171427703 |
| Myl12b Myl9 | S | 20;20 | -0.569296447 | -0.375934749 | -1.085320627 | -0.51602418 | -0.709385878 | 0.093971305 | 0.289746239 | 0.003237644 | 0.093820612 | 0.076785903 |
| Hecw2 | S | 1046 | -0.185782757 | -0.430835952 | -0.705357464 | -0.519574707 | -0.274521512 | 0.178903187 | 0.018215461 | 0.000194108 | 0.01566228 | 0.086288884 |
| Utp14a | S | 531 | -0.317074836 | -0.405375637 | -0.615189711 | -0.298114875 | -0.209814075 | 0.025474022 | 0.039261081 | 0.00075892 | 0.042010242 | 0.19410223 |
| Rin1 | Y | 35 | -0.424127852 | -0.472541442 | -0.746347515 | -0.322219663 | -0.273806073 | 0.019622021 | 0.035933778 | 0.01464605 | 0.154386347 | 0.236399895 |
| Fiz1 Zfp768 | T | 508;80 | -0.238341398 | -0.482509929 | -0.588484962 | -0.350143563 | -0.105975033 | 0.047982266 | 0.001405503 | 0.007849254 | 0.055677185 | 0.412568888 |
| Dnajb2 | S | 302 | -0.545120157 | -0.502992558 | -1.118135103 | -0.573014945 | -0.615142545 | 0.058998316 | 0.126744694 | 0.022615595 | 0.141921441 | 0.137978516 |
| Zfp507 | S | 185 | -0.269463958 | -0.25338263 | -0.707739343 | -0.438275385 | -0.454356713 | 0.016459417 | 0.135910007 | 9.89012E-05 | 0.005427085 | 0.052387117 |
| Klf10 Klf11 Klf13 Sp1 | T | 678;439;682;423; | -0.488347685 | -0.524932859 | -0.855790152 | -0.367442466 | -0.330857293 | 0.183918437 | 0.043355309 | 0.005184221 | 0.29185083 | 0.172788771 |
| Nolc1 | T | 594 | -0.150583038 | -0.467401465 | -0.599588288 | -0.449005249 | -0.132186823 | 0.012845335 | 0.002989411 | 0.003269732 | 0.006638758 | 0.163580336 |
| Vim | S | 10 | -0.385514096 | -0.289543786 | -0.594544211 | -0.209030115 | -0.305000424 | 0.005283564 | 0.111551522 | 0.00065368 | 0.043152966 | 0.103085451 |
| Errfi1 | S | 367 | -0.229808012 | -0.452415486 | -0.732253025 | -0.502445013 | -0.279837539 | 0.108438289 | 0.009125655 | 0.002169206 | 0.017407477 | 0.091521448 |
| Hivep3 | S | 669 | -0.536898848 | -0.272835175 | -0.791184405 | -0.254285557 | -0.518349231 | 0.009246876 | 0.170073889 | 0.00619842 | 0.197017327 | 0.046789511 |
| Cdk17 | S | 92 | -0.286144621 | -0.323266609 | -0.794521706 | -0.508377085 | -0.471255096 | 0.13088123 | 0.22418202 | 4.38318E-05 | 0.04607176 | 0.127024918 |
| Iqsec1 | S | 106 | -0.226497413 | -0.33741572 | -0.615834555 | -0.389337143 | -0.278418835 | 0.036856236 | 0.029750198 | 0.000157111 | 0.006072537 | 0.061033844 |
| Nifk | S | 249 | -0.183070111 | -0.418219624 | -0.682888565 | -0.499818454 | -0.264668942 | 0.171960685 | 0.007647478 | 0.000502609 | 0.006987279 | 0.022632461 |
| Rasa3 | S | 532 | -0.418409836 | -0.354373268 | -0.635962264 | -0.217552428 | -0.281588996 | 0.031238304 | 0.036468418 | 0.002118886 | 0.158713672 | 0.060754756 |
| 4930579G24Rik | S | 41 | -0.199350401 | -0.516193435 | -0.656959193 | -0.457608792 | -0.140765758 | 0.149160063 | 0.000104625 | 0.037445213 | 0.069116182 | 0.441954249 |
| Mapre1 | S | 155 | -0.358160808 | -0.412250654 | -0.668836113 | -0.310675305 | -0.256585459 | 0.062124757 | 0.032024473 | 0.001676227 | 0.08480647 | 0.102609951 |
| Rpl37 Rpl37rt | T | 82 | -0.223137153 | -0.379791092 | -0.788397162 | -0.565260008 | -0.40860607 | 0.019431896 | 0.066798835 | 0.0013448 | 0.005357668 | 0.052265228 |
| Cep170 | S | 353 | -0.236808463 | -0.381026113 | -0.590150276 | -0.353341814 | -0.209124164 | 0.17051971 | 0.004729654 | 0.000379316 | 0.077895067 | 0.046379644 |
| Eef1b2 | S | 128 | -0.255260768 | -0.487588049 | -0.746706429 | -0.491445661 | -0.259118379 | 0.152536753 | 0.020633793 | 0.001903238 | 0.010981205 | 0.085522956 |
| Nolc1 | S | 689 | -0.303428229 | -0.304441237 | -0.674322078 | -0.370893849 | -0.369880842 | 0.014559171 | 0.001066927 | 0.019829995 | 0.077236019 | 0.091712885 |
| Lrp1 | T | 4461 | -0.583044789 | -0.41348669 | -0.946551356 | -0.363506567 | -0.533064666 | 0.029697016 | 0.192276944 | 0.018655854 | 0.216528472 | 0.140019961 |
| Tnk2 | S | 74 | -0.488955258 | -0.366377203 | -0.807903408 | -0.31894815 | -0.441526205 | 0.020102525 | 0.127030573 | 0.009663216 | 0.157700642 | 0.104843848 |
| Cdc42bpb | S | 481 | -0.538512105 | -0.334522269 | -0.803666758 | -0.265154653 | -0.469144489 | 0.012988296 | 0.183498606 | 0.011195549 | 0.230807232 | 0.108981776 |
| Ddx17 | S | 492 | -0.138864348 | -0.577124296 | -0.668804493 | -0.529940146 | -0.091680197 | 0.034714051 | 0.001412229 | 0.027117547 | 0.045341489 | 0.563462167 |
| Irf3 | S | 123 | -0.192447176 | -0.505360175 | -0.738526911 | -0.546079736 | -0.233166737 | 0.06124229 | 0.012329167 | 0.006429175 | 0.021819567 | 0.184999345 |
| Ybx3 | S | 335 | -0.40378018 | -0.392622118 | -0.612850198 | -0.209070018 | -0.22022808 | 0.02481261 | 0.051033675 | 0.003945672 | 0.043041759 | 0.15925422 |
| Mtm1 | S | 18 | -0.581894985 | -0.28398099 | -1.066268915 | -0.48437393 | -0.782287926 | 0.041068879 | 0.378758176 | 0.008596561 | 0.108826311 | 0.069438464 |
| Nav1 | S | 93 | -0.458816692 | -0.495442977 | -0.986385693 | -0.527569001 | -0.490942716 | 0.065102387 | 0.114896943 | 0.019553719 | 0.116093146 | 0.168086719 |
| Abi1 | Y | 213 | -0.492255741 | -0.449055374 | -0.71244706 | -0.220191319 | -0.263391686 | 0.025263247 | 0.122952682 | 0.010159581 | 0.282915656 | 0.327998997 |
| Slc7a6os | S | 256 | -0.13485303 | -0.436734678 | -0.66337716 | -0.52852413 | -0.226642482 | 0.160503812 | 0.006206035 | 0.001026087 | 0.005089826 | 0.087302978 |
| Sfr1 | S | 139 | -0.430583893 | -0.419165501 | -0.791322988 | -0.360739095 | -0.372157487 | 0.063346176 | 0.112967386 | 0.004418204 | 0.05201768 | 0.126021641 |
| Zc3hav1 | Y | 508 | -0.411940398 | -0.44111938 | -0.679114953 | -0.267174555 | -0.237995573 | 0.04008742 | 0.054525318 | 0.007353608 | 0.137097129 | 0.229488611 |
| Slc4a2 | S | 494 | -0.370711126 | -0.369334004 | -0.668590661 | -0.297879534 | -0.299256657 | 0.000454086 | 0.144006015 | 0.024621079 | 0.142449677 | 0.23001828 |
| Slc4a2 | S | 496 | -0.330779131 | -0.456598308 | -0.682334308 | -0.351555177 | -0.225736 | 0.010692166 | 0.088122415 | 0.00741962 | 0.040079307 | 0.290599482 |
| Pdia6 | S | 148 | -0.50227038 | -0.374055471 | -0.77214594 | -0.26987556 | -0.398090469 | 0.038236124 | 0.179663295 | 0.004954319 | 0.114071846 | 0.148411257 |
| Trir | S | 21 | -0.294825149 | -0.409709409 | -0.698165346 | -0.403340198 | -0.288455938 | 0.066381404 | 0.06663243 | 0.001475276 | 0.03493377 | 0.162832815 |
| Orc1 | S | 376 | -0.313060538 | -0.353462931 | -0.6268105 | -0.313749962 | -0.273347569 | 0.0274878 | 0.041820726 | 0.002333738 | 0.042308311 | 0.09474633 |
| Notch2 | T | 1718 | -0.376844879 | -0.261703697 | -0.771449198 | -0.39460432 | -0.509745501 | 0.087986996 | 0.091276312 | 0.000986018 | 0.08472016 | 0.014686268 |

| | | | | | | | | | | | | |
|---|---|---|---|---|---|---|---|---|---|---|---|---|
| Septin9 | S | 60 | -0.399348012 | -0.280269693 | -0.901258815 | -0.501910803 | -0.620989122 | 0.015526494 | 0.241587564 | 0.006257889 | 0.047239134 | 0.050984609 |
| Anln | S | 131 | -0.146994964 | -0.402206487 | -0.704596784 | -0.55760182 | -0.302390297 | 0.199910194 | 0.03275822 | 0.000291082 | 0.002210864 | 0.081170298 |
| Epb41l2 | S | 630 | -0.544802794 | -0.234007419 | -1.064245775 | -0.519442981 | -0.830238356 | 0.014784615 | 0.521008855 | 0.013090526 | 0.107306077 | 0.087845724 |
| Rbm3 | S | 117 | -0.46455258 | -0.464898225 | -0.795921749 | -0.331369169 | -0.331023524 | 0.095690829 | 0.074651564 | 0.010310362 | 0.106401742 | 0.023760675 |
| Hmga1 Hmga1b | T | 21 | -0.570415084 | -0.462951066 | -0.767712042 | -0.197296958 | -0.304760977 | 0.038534394 | 0.211017538 | 0.012291316 | 0.215385728 | 0.347518201 |
| Hnrnpa0 | S | 88 | -0.529296012 | -0.392276484 | -0.755007899 | -0.225711886 | -0.362731415 | 0.033181546 | 0.145570045 | 0.011454027 | 0.239032235 | 0.152798765 |
| Arhgef2 | S | 781 | -0.228552499 | -0.550832959 | -0.74655922 | -0.518006721 | -0.195726261 | 0.196130652 | 0.018714548 | 0.005918744 | 0.028978256 | 0.289844912 |
| Pdlim5 | S | 356 | -0.450589877 | -0.431904435 | -0.731328989 | -0.280739112 | -0.299424553 | 0.051431097 | 0.094473615 | 0.008082918 | 0.177319728 | 0.208232034 |
| Hdgf | S | 103 | -0.416588932 | -0.334352771 | -0.77650293 | -0.359913998 | -0.442150159 | 0.054722406 | 0.15747857 | 0.003011686 | 0.040276036 | 0.069109133 |
| Gas7 | S | 101 | -0.389539112 | -0.505432674 | -0.596495435 | -0.206956323 | -0.091062761 | 0.066935344 | 0.062877409 | 0.004395588 | 0.225759043 | 0.644273573 |
| Plekha2 | S | 312 | -0.249787097 | -0.426832821 | -0.726063615 | -0.476276519 | -0.299230794 | 0.07778829 | 0.057754014 | 0.002224966 | 0.020256118 | 0.14947835 |
| Crk | S | 125 | -0.133484155 | -0.558694293 | -0.714963841 | -0.581479686 | -0.156269548 | 0.249205979 | 0.015923221 | 0.002718591 | 0.006543241 | 0.22508923 |
| Oxr1 | S | 503 | -0.528756576 | -0.331064695 | -0.736484536 | -0.20772796 | -0.405419841 | 0.045345304 | 0.140897192 | 0.006649375 | 0.272316034 | 0.043017434 |
| Rnf169 | S | 689 | -0.396667835 | -0.475461256 | -0.850355833 | -0.453687998 | -0.374894577 | 0.068729964 | 0.13242321 | 0.009291895 | 0.080801874 | 0.223271701 |
| Racgap1 | S | 611 | -0.466769872 | -0.260550603 | -0.708770767 | -0.242000895 | -0.448220164 | 0.021613805 | 0.27033041 | 0.002211186 | 0.089084859 | 0.09704492 |
| Sord | S | 169 | -0.200133547 | -0.313936959 | -0.668983758 | -0.468850211 | -0.355046799 | 0.042773012 | 0.04079673 | 0.000698974 | 0.011644776 | 0.036154307 |
| Nifk | S | 270 | -0.306621766 | -0.455382339 | -0.817066123 | -0.510444356 | -0.361683784 | 0.170200754 | 0.052769828 | 0.004704696 | 0.062082666 | 0.126065889 |
| Ddhd2 | S | 589 | -0.456517877 | -0.451978095 | -1.264775748 | -0.808257871 | -0.812797654 | 0.199584662 | 0.189460513 | 0.017602257 | 0.082571716 | 0.079066938 |
| Nme2 | S | 120 | -0.339633252 | -0.304311674 | -0.759280763 | -0.419647511 | -0.454969089 | 0.098090267 | 0.135580578 | 0.000877706 | 0.061728609 | 0.054299364 |
| Slc9a1 | S | 801 | -0.373842603 | -0.341325724 | -0.867950021 | -0.494107418 | -0.526624297 | 0.041923365 | 0.145482261 | 0.004702662 | 0.041282042 | 0.094369286 |
| Eif4ebp1 | T | 36 | -0.266863242 | -0.394773449 | -0.634953978 | -0.368090737 | -0.240180529 | 0.027302345 | 0.007165996 | 0.017859875 | 0.074577567 | 0.179782797 |
| Spred1 | S | 288 | -0.2900607 | -0.313333093 | -0.78021477 | -0.49015407 | -0.466881676 | 0.12306765 | 0.098327211 | 0.000951883 | 0.034655632 | 0.038079511 |
| Mlh1 | S | 388 | -0.550397456 | -0.33456855 | -0.810394456 | -0.2605469 | -0.476375805 | 0.024628703 | 0.255110693 | 0.014706285 | 0.274096151 | 0.14677899 |
| Stard7 | S | 356 | -0.262814868 | -0.408278994 | -0.670044913 | -0.407230045 | -0.261765919 | 0.016411858 | 0.030401808 | 0.012663211 | 0.055355033 | 0.165695732 |
| Drosha | S | 279 | -0.249155938 | -0.279542399 | -0.738742984 | -0.489587045 | -0.459200584 | 0.045892114 | 0.105523278 | 0.0009117 | 0.010469611 | 0.032592925 |
| Drosha | S | 283 | -0.249155938 | -0.279542399 | -0.738742984 | -0.489587045 | -0.459200584 | 0.045892114 | 0.105523278 | 0.0009117 | 0.010469611 | 0.032592925 |
| Crip2 | Y | 77 | -0.472535001 | -0.341645197 | -0.595613694 | -0.123078693 | -0.253968497 | 0.023890696 | 0.09409982 | 0.00728202 | 0.285341794 | 0.126587877 |
| Epb41l1 | T | 517 | -0.49909532 | -0.313132427 | -1.061426556 | -0.562331235 | -0.748294129 | 0.135450051 | 0.373165547 | 0.005142295 | 0.101541518 | 0.074525001 |
| Prrc2a | S | 18 | -0.384318452 | -0.278791761 | -0.699292227 | -0.314973775 | -0.420500466 | 0.044341212 | 0.16277812 | 0.001681084 | 0.061970991 | 0.062151566 |
| Jpt1 | S | 87 | -0.403684661 | -0.440959677 | -0.627005483 | -0.223320821 | -0.186045805 | 0.024948353 | 0.03398349 | 0.032561983 | 0.288090216 | 0.38148324 |
| Septin9 | S | 111 | -0.427443978 | -0.314395819 | -0.774799533 | -0.347355555 | -0.460403714 | 0.04288776 | 0.190792598 | 0.005188905 | 0.069891388 | 0.070082437 |
| Vipas39 | S | 119 | -0.494238216 | -0.372676313 | -0.79711252 | -0.302874304 | -0.424436208 | 0.024490894 | 0.24447631 | 0.015681248 | 0.213739587 | 0.216394717 |
| Nup133 | S | 55 | -0.508826174 | -0.371045771 | -0.673243573 | -0.164417398 | -0.302197802 | 0.042568218 | 0.178601435 | 0.006872144 | 0.328491029 | 0.226245861 |
| Plec | S | 4279 | -0.360530587 | -0.203588008 | -0.710364369 | -0.349833781 | -0.50677636 | 0.02336614 | 0.28723474 | 0.000953809 | 0.017548351 | 0.040180366 |
| Ubap2 | S | 254 | -0.515693598 | -0.345254082 | -0.681677039 | -0.165983441 | -0.336422957 | 0.031838386 | 0.087465076 | 0.019349998 | 0.447932354 | 0.15044757 |
| Sltm | S | 887 | -0.389905936 | -0.41312183 | -0.609362387 | -0.219456451 | -0.196240557 | 0.046114455 | 0.044380602 | 0.010972694 | 0.143383823 | 0.197425524 |
| Abi1 | S | 240 | -0.166458558 | -0.555533009 | -0.688589436 | -0.522130877 | -0.133056427 | 0.156080002 | 0.007442009 | 0.016935228 | 0.038912172 | 0.464096338 |
| Nhsl1 | S | 1364 | -0.391314572 | -0.314322897 | -0.742108208 | -0.350793636 | -0.427785311 | 0.056715835 | 0.171662344 | 0.003069392 | 0.046843266 | 0.072460085 |
| Pdia6 | S | 428 | -0.326536625 | -0.310142911 | -0.603631712 | -0.277095087 | -0.293488801 | 0.03314649 | 0.090682353 | 0.00183002 | 0.056537454 | 0.104116939 |
| Prkaa1 | S | 486 | -0.093720722 | -0.328030836 | -0.589302416 | -0.495581694 | -0.26127158 | 0.184803426 | 0.000690956 | 0.002006707 | 0.006406308 | 0.044753084 |
| Cd2ap | S | 234 | -0.16224321 | -0.277798814 | -0.589388258 | -0.427145048 | -0.311589444 | 0.04646578 | 0.015245711 | 0.000737659 | 0.006517184 | 0.020802471 |
| Spn | S | 339 | -0.502484451 | -0.280597463 | -0.64034164 | -0.137857189 | -0.359744177 | 0.021279596 | 0.261158581 | 0.005315439 | 0.420105893 | 0.180272827 |
| Lbh | S | 63 | -0.380973163 | -0.412796897 | -0.666738869 | -0.285765706 | -0.253941972 | 0.03489395 | 0.043132588 | 0.025899678 | 0.210873585 | 0.269384325 |
| Lima1 | S | 599 | -0.459731319 | -0.230897895 | -0.825005179 | -0.36527386 | -0.594107284 | 0.021989889 | 0.38716352 | 0.00679496 | 0.108620194 | 0.073884895 |
| Eif3g | S | 189 | -0.230310588 | -0.388803206 | -0.661465607 | -0.431155019 | -0.272662401 | 0.037642584 | 0.011967901 | 0.017512064 | 0.048800951 | 0.137679777 |
| Eif3g | T | 190 | -0.230310588 | -0.388803206 | -0.661465607 | -0.431155019 | -0.272662401 | 0.037642584 | 0.011967901 | 0.017512064 | 0.048800951 | 0.137679777 |
| Atp5e | S | 42 | -0.319372523 | -0.295624339 | -0.646421531 | -0.327040008 | -0.350788192 | 0.091029451 | 0.107810233 | 0.000932317 | 0.089196853 | 0.073686347 |
| Cep170 | S | 662 | -0.471345771 | -0.221039935 | -0.715629273 | -0.244283502 | -0.494589338 | 0.016773855 | 0.310739835 | 0.006108776 | 0.179597525 | 0.065208926 |
| Alms1 Alms1-ps2 | S | 2416 | -0.464382273 | -0.310017478 | -0.615294104 | -0.150911832 | -0.305276626 | 0.052909055 | 0.061976111 | 0.008935966 | 0.444250161 | 0.090240205 |
| Myl12b Myl9 | T | 19;19 | -0.454976684 | -0.28994054 | -0.700435542 | -0.245458858 | -0.410495002 | 0.035855751 | 0.242262275 | 0.004967342 | 0.079357063 | 0.109522765 |
| Morf4l2 | S | 71 | -0.327233627 | -0.401792188 | -0.75724419 | -0.430010562 | -0.355452002 | 0.172364123 | 0.085088373 | 0.004193587 | 0.071171173 | 0.076160938 |
| Pkp4 | S | 457 | -0.395353223 | -0.130554977 | -0.618948432 | -0.223595209 | -0.488393455 | 0.012689435 | 0.607731237 | 0.000391944 | 0.082731784 | 0.141310867 |
| Nufip2 | S | 160 | -0.447147987 | -0.343712421 | -0.704732929 | -0.257584942 | -0.361020508 | 0.050543907 | 0.164755497 | 0.008698236 | 0.063329704 | 0.098759395 |
| Alkbh5 | S | 376 | -0.324752599 | -0.380919433 | -0.626642737 | -0.301890138 | -0.245723304 | 0.031939994 | 0.084186069 | 0.008094138 | 0.077881284 | 0.213597559 |
| Zzef1 | S | 1535 | -0.301789091 | -0.332980536 | -0.668352722 | -0.366563632 | -0.335372186 | 0.070978655 | 0.097746067 | 0.002797636 | 0.046979406 | 0.101414888 |
| Ahctf1 | S | 2142 | -0.120953034 | -0.3502545 | -0.646881 | -0.525927966 | -0.2966265 | 0.266889559 | 0.01730702 | 0.00073165 | 0.000369229 | 0.022193467 |
| Myadm | T | 9 | -0.468059948 | -0.220721513 | -0.700634249 | -0.232574301 | -0.479912736 | 0.017360786 | 0.318594946 | 0.00674896 | 0.210286672 | 0.081207706 |
| Hadh | S | 14 | -0.601786927 | -0.328791549 | -0.823607367 | -0.22182044 | -0.494815818 | 0.060735737 | 0.216353773 | 0.025446395 | 0.46352546 | 0.133423038 |
| Scrib | T | 475 | -0.433382315 | -0.243111099 | -0.809515449 | -0.376133134 | -0.56640435 | 0.045401643 | 0.422133789 | 0.003663936 | 0.07053159 | 0.113583792 |
| Nav3 | S | 336 | -0.226784472 | -0.360606371 | -0.600561551 | -0.373777079 | -0.23995518 | 0.005038911 | 0.053726899 | 0.013242645 | 0.049109299 | 0.164435643 |
| Pin4 | S | 19 | -0.25393054 | -0.558449379 | -0.985141759 | -0.731211219 | -0.42669238 | 0.271174061 | 0.067281468 | 0.023954418 | 0.058394458 | 0.209228211 |
| Cnn3 | Y | 261 | -0.381133561 | -0.34458051 | -0.655024507 | -0.273890946 | -0.310443997 | 0.050044707 | 0.240982847 | 0.003066545 | 0.084754668 | 0.272859557 |
| Eef2 | T | 59 | -0.473102238 | -0.235316951 | -0.862112811 | -0.389010572 | -0.62679586 | 0.035624408 | 0.334127424 | 0.012059852 | 0.134002783 | 0.05231313 |
| Tnks1bp1 | S | 783 | -0.387260165 | -0.225216648 | -0.652996959 | -0.265736794 | -0.427780311 | 0.036562447 | 0.198047979 | 0.002515587 | 0.063893721 | 0.026794003 |
| Zwint | S | 216 | -0.390675658 | -0.277738534 | -0.666176533 | -0.275500874 | -0.388437998 | 0.071457816 | 0.128320968 | 0.003485966 | 0.173271082 | 0.062135436 |
| Cdv3 | S | 66 | -0.208561669 | -0.460105832 | -0.599779513 | -0.391217844 | -0.139673681 | 0.270765244 | 0.004961421 | 0.012453394 | 0.092752488 | 0.36561609 |
| Pimreg | S | 16 | -0.331980627 | -0.34323631 | -0.642484829 | -0.310504203 | -0.29924852 | 0.021198373 | 0.050811562 | 0.023558363 | 0.159718111 | 0.177185619 |
| Cavin1 | S | 302 | -0.524050723 | -0.321214926 | -0.700372714 | -0.176321992 | -0.379157789 | 0.02752668 | 0.306436067 | 0.015724266 | 0.354306992 | 0.234135059 |
| Apbb2 | S | 718 | -0.298999256 | -0.31665973 | -0.776702905 | -0.477703649 | -0.460043175 | 0.017846054 | 0.203614484 | 0.010251763 | 0.053577539 | 0.110109138 |
| Mapk14 | T | 180 | -0.532547455 | -0.335212886 | -0.603437766 | -0.070890311 | -0.26822488 | 0.01950979 | 0.178932171 | 0.026769358 | 0.667020332 | 0.26076887 |
| Notch1 | S | 1769 | -0.350943386 | -0.243185628 | -0.811711234 | -0.460767848 | -0.568525605 | 0.027126689 | 0.315375287 | 0.006275952 | 0.058612701 | 0.06749641 |
| Pa2g4 | S | 363 | -0.360740157 | -0.354559928 | -0.762149734 | -0.401409577 | -0.407589806 | 0.129300656 | 0.195858491 | 0.003906737 | 0.086154041 | 0.140605268 |
| Jpt1 | S | 14 | -0.502208039 | -0.320519764 | -1.051336499 | -0.54912846 | -0.730816735 | 0.151494871 | 0.390680548 | 0.012777506 | 0.091462509 | 0.070253571 |
| Papola | S | 732 | -0.286765166 | -0.300834316 | -0.679135297 | -0.39237013 | -0.378300981 | 0.077849649 | 0.100403894 | 0.003212355 | 0.027518147 | 0.043976125 |
| Vangl1 | S | 42 | -0.276751787 | -0.45686123 | -0.701535256 | -0.424783469 | -0.244674026 | 0.135786388 | 0.135031538 | 0.004280562 | 0.030453856 | 0.355491958 |
| Wwc2 | S | 1145 | -0.34772153 | -0.305286838 | -0.653673127 | -0.305951598 | -0.34838629 | 0.045161208 | 0.088302624 | 0.008976849 | 0.106615573 | 0.077527191 |
| Snap23 | S | 120 | -0.320703352 | -0.28048715 | -0.694188327 | -0.373484975 | -0.413701177 | 0.04544935 | 0.173868377 | 0.004003754 | 0.050625092 | 0.078599214 |
| Rock1 | S | 1336 | -0.468724179 | -0.318338131 | -1.04956457 | -0.580840391 | -0.731226438 | 0.091762648 | 0.390661389 | 0.018348781 | 0.091289846 | 0.115745565 |
| Mdm2 | S | 185 | -0.167095598 | -0.462567317 | -0.626719106 | -0.459623508 | -0.164151789 | 0.476495247 | 0.001248715 | 0.028201339 | 0.124432546 | 0.274834891 |
| Cttn | S | 150 | -0.425824228 | -0.309222605 | -0.788208377 | -0.36238415 | -0.478985773 | 0.108026133 | 0.247861846 | 0.007100706 | 0.100950942 | 0.072768588 |
| Tpx2 | S | 60 | -0.307881781 | -0.372032833 | -0.7191454 | -0.411263619 | -0.347112566 | 0.141857646 | 0.089858642 | 0.007347028 | 0.068162592 | 0.108119275 |
| Map4k4 | S | 649 | -0.436060411 | -0.235965216 | -0.613937877 | -0.177877466 | -0.377972661 | 0.022394338 | 0.32646727 | 0.005161844 | 0.109030641 | 0.152922548 |
| Lnpep | S | 80 | -0.344839434 | -0.321834396 | -0.799503712 | -0.454664279 | -0.477669316 | 0.034136143 | 0.208482899 | 0.017067264 | 0.098324386 | 0.122565682 |
| Eif3a | S | 978 | -0.192879854 | -0.325833835 | -0.608989049 | -0.416109195 | -0.283155213 | 0.105802662 | 0.009101489 | 0.009566684 | 0.023649888 | 0.062835154 |
| Clasp1 | S | 720 | -0.407630689 | -0.232425841 | -0.823419772 | -0.415789083 | -0.590993931 | 0.088763993 | 0.390555694 | 0.004322811 | 0.057829527 | 0.062757951 |
| Nf2 | S | 13 | -0.235752713 | -0.338392483 | -0.681755606 | -0.446002892 | -0.343363123 | 0.136131325 | 0.113037831 | 0.002435207 | 0.024480865 | 0.113178285 |
| Cep170 | S | 1195 | -0.230728096 | -0.390085465 | -0.679281439 | -0.448553343 | -0.289195974 | 0.059557519 | 0.058330282 | 0.01416485 | 0.038535818 | 0.151142476 |
| Cep170 | S | 1202 | -0.230728096 | -0.390085465 | -0.679281439 | -0.448553343 | -0.289195974 | 0.059557519 | 0.058330282 | 0.01416485 | 0.038535818 | 0.151142476 |
| Bcr | S | 697 | -0.19512398 | -0.409139689 | -0.729342492 | -0.534218512 | -0.320202803 | 0.082716037 | 0.09113396 | 0.00837148 | 0.034077603 | 0.177930843 |
| Fyn | S | 26 | -0.438725586 | -0.300083031 | -0.980775668 | -0.542050082 | -0.680692637 | 0.075531533 | 0.37763902 | 0.021339447 | 0.09717281 | 0.113931587 |
| Cbl | S | 838 | -0.025328722 | -0.513828294 | -1.32653128 | -1.301202558 | -0.812702986 | 0.959267973 | 0.228230312 | 0.005225389 | 0.05597041 | 0.073540388 |
| Itprid2 | S | 441 | -0.406049723 | -0.293871464 | -0.729046441 | -0.322996718 | -0.435174977 | 0.057143075 | 0.164741076 | 0.016300543 | 0.173528369 | 0.098189135 |
| Spats2 | S | 472 | -0.422028659 | -0.173830864 | -0.887290341 | -0.465261682 | -0.713459477 | 0.050991714 | 0.527970776 | 0.007691048 | 0.079383474 | 0.060637165 |
| Ppp1r14c | S | 33 | -0.281061966 | -0.278454506 | -0.681490814 | -0.400428848 | -0.403036308 | 0.008426244 | 0.09374399 | 0.028012748 | 0.09158872 | 0.0884694 |

| | | | | | | | | | | | | |
|---|---|---|---|---|---|---|---|---|---|---|---|---|
| Hnrnpm | S | 617 | -0.586873911 | -0.2810473 | -0.773505971 | -0.18663206 | -0.492458671 | 0.053920069 | 0.390757619 | 0.023974465 | 0.402992764 | 0.137376065 |
| Abi1 | S | 222 | -0.369179957 | -0.242522112 | -0.864679957 | -0.4955 | -0.622157845 | 0.052198064 | 0.385411464 | 0.010228284 | 0.066532342 | 0.084889046 |
| Snx17 | S | 437 | -0.273827985 | -0.34861127 | -0.607476876 | -0.333648891 | -0.258865605 | 0.040394997 | 0.112889678 | 0.008347563 | 0.046211463 | 0.197842826 |
| Eef2 | T | 355 | -1.053376427 | -0.31030598 | -0.971044504 | 0.082331923 | -0.660738524 | 0.417428038 | 0.524386559 | 0.010348696 | 0.937842714 | 0.214908216 |
| Jpt1 | S | 70 | -0.32545128 | -0.289101876 | -0.589776792 | -0.264325511 | -0.300674916 | 0.009380738 | 0.059286396 | 0.041259798 | 0.229807449 | 0.188459092 |
| Lrp1 | Y | 4474 | -0.37249637 | -0.412908623 | -0.635549597 | -0.263053227 | -0.222640973 | 0.137861977 | 0.123336427 | 0.009783716 | 0.226653578 | 0.320604726 |
| Gys1 | S | 698 | -0.164954862 | -0.294076858 | -0.629177048 | -0.464222186 | -0.33510019 | 0.079877726 | 0.035427391 | 0.004104411 | 0.03001607 | 0.051715348 |
| Dtl | S | 484 | -0.219554419 | -0.386211808 | -0.744608313 | -0.525053894 | -0.358396505 | 0.222066376 | 0.161667893 | 0.003030028 | 0.014949802 | 0.180167518 |
| Epha2 | Y | 773 | -0.102329481 | -0.344067334 | -0.645102664 | -0.542773182 | -0.301035329 | 0.430781565 | 0.099392413 | 0.000287511 | 0.018718075 | 0.125347043 |
| Rrn3 | T | 290 | -0.05670193 | -0.43975977 | -0.606640669 | -0.549938739 | -0.166880899 | 0.400729924 | 0.00256197 | 0.012491537 | 0.007017084 | 0.161402409 |
| Rps7 | T | 123 | -0.207786684 | -0.339266074 | -0.711711249 | -0.503924565 | -0.372445175 | 0.332082034 | 0.162410719 | 0.001024663 | 0.077941563 | 0.147374185 |
| Hnrnpa3 | S | 376 | -0.298908869 | -0.360116711 | -0.771617411 | -0.472708542 | -0.411500699 | 0.176586925 | 0.038249414 | 0.029582852 | 0.116948433 | 0.13523944 |
| Serpinh1 | S | 140 | -0.336682849 | -0.127370419 | -0.791352104 | -0.454669255 | -0.663981686 | 0.06336673 | 0.613545798 | 0.001851936 | 0.024894321 | 0.056208108 |
| Arfgap2 | S | 432 | -0.434334441 | -0.255565359 | -0.703786971 | -0.269452531 | -0.448221613 | 0.058605984 | 0.261972834 | 0.01312249 | 0.236947359 | 0.101435452 |
| Stk10 | S | 13 | -0.341383912 | -0.350916238 | -0.890068511 | -0.548684599 | -0.539152273 | 0.045827002 | 0.232760082 | 0.038903052 | 0.100423724 | 0.142251461 |
| Map1b | S | 2323 | -0.024250575 | -0.3705722 | -0.617825941 | -0.593575366 | -0.247253741 | 0.888406985 | 0.036732899 | 0.000221694 | 0.054063844 | 0.12833688 |
| Dok1 | Y | 340 | -0.424764302 | -0.343221362 | -0.659913697 | -0.235149395 | -0.316692335 | 0.026224521 | 0.203659979 | 0.041369212 | 0.332474315 | 0.292465967 |
| Specc1 | S | 119 | -0.260801765 | -0.242394562 | -0.745969245 | -0.485167479 | -0.503574683 | 0.127863023 | 0.263208791 | 0.002286962 | 0.022936127 | 0.056213612 |
| Jcad | S | 693 | -0.460350448 | -0.181074831 | -0.66719947 | -0.206849022 | -0.486124639 | 0.006324595 | 0.325042612 | 0.04320616 | 0.377908535 | 0.103386667 |
| Baiap2 | S | 455 | -0.299728891 | -0.282518173 | -0.647305742 | -0.347576851 | -0.364787568 | 0.127733928 | 0.168808063 | 0.00324875 | 0.094946905 | 0.104320113 |
| Kdm4a | S | 572 | -0.281782608 | -0.399001361 | -0.606973874 | -0.325191266 | -0.207972513 | 0.148762558 | 0.099920596 | 0.007522776 | 0.055197749 | 0.280397113 |
| Ap3d1 | S | 9 | -0.519555264 | -0.231797272 | -0.668881421 | -0.149326156 | -0.437084148 | 0.03145591 | 0.338679418 | 0.025773133 | 0.431672875 | 0.100404019 |
| Plpp1 | S | 278 | -0.275892966 | -0.248300916 | -0.6423722 | -0.366479234 | -0.394071284 | 0.088332728 | 0.230235811 | 0.002214966 | 0.047925926 | 0.096477923 |
| Mt2 | S | 45 | -0.236865744 | -0.290350817 | -0.597586532 | -0.360720788 | -0.307235715 | 0.097193146 | 0.080235447 | 0.004528495 | 0.044756639 | 0.087827482 |
| Nadk | S | 46 | -0.155987231 | -0.205235869 | -0.586811826 | -0.430824595 | -0.381575957 | 0.157722204 | 0.14611915 | 0.000220583 | 0.007994736 | 0.037863474 |
| Mxra7 | S | 125 | -0.317090106 | -0.249582465 | -0.74510568 | -0.428015574 | -0.495523215 | 0.062131551 | 0.286353934 | 0.008378437 | 0.071772663 | 0.079424916 |
| Arfgap2 | S | 431 | -0.422216152 | -0.244387466 | -0.694206226 | -0.271990074 | -0.44981876 | 0.062444455 | 0.277380118 | 0.013048387 | 0.228726969 | 0.097999102 |
| Eif4ebp1 | S | 64 | -0.159788976 | -0.37139938 | -0.679320071 | -0.519531095 | -0.30792069 | 0.308628682 | 0.035970209 | 0.007231454 | 0.03041131 | 0.105457847 |
| Cdv3 | S | 68 | -0.285223247 | -0.220759704 | -0.745169682 | -0.459946436 | -0.524409978 | 0.127655352 | 0.30898006 | 0.003024832 | 0.008930605 | 0.039681952 |
| Hectd1 | S | 1384 | -0.253111063 | -0.352931172 | -0.606392803 | -0.35328174 | -0.25346163 | 0.017309602 | 0.095495644 | 0.035132555 | 0.123790052 | 0.266192174 |
| Ccdc124 | S | 102 | -0.320085001 | -0.289389641 | -0.647065722 | -0.326980721 | -0.357676082 | 0.108953594 | 0.276335693 | 0.004513453 | 0.066464343 | 0.186283799 |
| Slc20a2 | S | 425 | -0.175203313 | -0.346184271 | -0.661582672 | -0.486379359 | -0.315398402 | 0.278151068 | 0.067053316 | 0.00494318 | 0.022345276 | 0.090922215 |
| Hnrnpm Plvap | S | 4;527 | -0.407633337 | -0.24692815 | -0.720405028 | -0.312771691 | -0.473476878 | 0.086597929 | 0.380739442 | 0.009994254 | 0.124854418 | 0.118427741 |
| Stom | S | 12 | -0.324383204 | -0.280066421 | -0.707463408 | -0.383080204 | -0.427396987 | 0.090014365 | 0.205222059 | 0.012733889 | 0.091475257 | 0.091443458 |
| Nedd1 | S | 379 | -0.515374033 | -0.206430734 | -0.655251245 | -0.139877212 | -0.44882051 | 0.031548823 | 0.44349965 | 0.024319517 | 0.449909057 | 0.126942617 |
| Map7d1 | S | 444 | -0.220743741 | -0.234181725 | -0.587556961 | -0.36681322 | -0.353375236 | 0.075092158 | 0.156800335 | 0.00239357 | 0.023505507 | 0.059578218 |
| Cep170 | S | 960 | -0.347021499 | -0.214604128 | -0.993259728 | -0.646238229 | -0.7786556 | 0.119321793 | 0.469327503 | 0.017378285 | 0.088083446 | 0.058871625 |
| Stx5a | S | 9 | -0.097894181 | -0.306886037 | -0.605765281 | -0.5078711 | -0.298879244 | 0.506139189 | 0.032774228 | 0.001526598 | 0.013829589 | 0.019170623 |
| Pard3 | S | 1255 | -0.378563614 | -0.176567707 | -0.687791463 | -0.309227849 | -0.511223756 | 0.069362637 | 0.464799871 | 0.005629312 | 0.10402014 | 0.077387358 |
| Map2k2 | T | 395 | -0.329733669 | -0.260190652 | -0.590799494 | -0.261065825 | -0.330608843 | 0.021603478 | 0.133473603 | 0.035707224 | 0.223733441 | 0.161764058 |
| Atp2b1 | S | 1140 | -0.315438844 | -0.286738364 | -0.591871096 | -0.276432252 | -0.305132732 | 0.069504119 | 0.156871997 | 0.012893109 | 0.116345058 | 0.135135537 |
| Ppp1r12b | S | 833 | -0.263237104 | -0.350826178 | -0.624865459 | -0.361628355 | -0.274039281 | 0.171862904 | 0.125928444 | 0.009135969 | 0.061106203 | 0.186408113 |
| Mast4 | S | 1281 | -0.297093825 | -0.324021841 | -0.66979766 | -0.372703835 | -0.345775819 | 0.058885975 | 0.126487141 | 0.033509915 | 0.143860893 | 0.190167767 |
| Fhod1 | S | 1177 | -0.429116107 | -0.178070999 | -0.790463991 | -0.361347884 | -0.612392992 | 0.083591016 | 0.630245058 | 0.009957902 | 0.143086014 | 0.162461651 |
| Afdn | S | 1119 | -0.342029602 | -0.199880437 | -0.661467199 | -0.319437597 | -0.461586762 | 0.01238199 | 0.391850118 | 0.027559877 | 0.136215663 | 0.12120116 |
| Sh3pxd2b | S | 672 | -0.463346329 | -0.208569456 | -0.594665326 | -0.131318997 | -0.38609587 | 0.032417577 | 0.39230504 | 0.023226569 | 0.435576725 | 0.139004726 |
| Eef2 | T | 57 | -0.423486199 | -0.192879373 | -0.683226782 | -0.259740583 | -0.49034741 | 0.060162052 | 0.406522207 | 0.015432713 | 0.223147255 | 0.077496952 |
| Smtn | S | 798 | -0.180556566 | -0.228535161 | -0.621251942 | -0.440695376 | -0.392716781 | 0.293909185 | 0.084020762 | 0.001936731 | 0.048226122 | 0.023015418 |
| Adam15 | T | 757 | -0.213026997 | -0.244488604 | -0.605101972 | -0.392074976 | -0.360613368 | 0.096353717 | 0.123884152 | 0.005722104 | 0.037115562 | 0.062374718 |
| Arfgap3 | S | 374 | -0.429255301 | -0.149686299 | -0.599870786 | -0.170615485 | -0.450184486 | 0.014182572 | 0.393682199 | 0.028328882 | 0.37446585 | 0.07812058 |
| Psmg1 | T | 55 | -0.372053696 | -0.279104305 | -0.586042417 | -0.213988721 | -0.306938112 | 0.140879099 | 0.15095581 | 0.01297181 | 0.34236551 | 0.112608693 |
| Rbm10 | S | 710 | -0.404858341 | -0.277531721 | -0.601460398 | -0.196602057 | -0.323928678 | 0.086183016 | 0.276391509 | 0.016288061 | 0.278983824 | 0.180732472 |
| Sp100 | S | 365 | -0.31437007 | -0.257644592 | -0.626515058 | -0.312144988 | -0.368870466 | 0.133122495 | 0.245580269 | 0.007346896 | 0.127085331 | 0.116262253 |
| Sphk1 | S | 373 | -0.430523479 | -0.20721419 | -0.71989869 | -0.289375211 | -0.5126845 | 0.039455445 | 0.499388445 | 0.030378417 | 0.256056896 | 0.170772917 |
| Cdc20 | S | 86 | -0.179251975 | -0.184778249 | -0.600070083 | -0.420818107 | -0.415291834 | 0.119868085 | 0.258382863 | 0.001146505 | 0.009349284 | 0.043284122 |
| Sh3pxd2a | S | 1007 | -0.194220354 | -0.326333429 | -0.704027449 | -0.509807096 | -0.37769402 | 0.297148551 | 0.140205647 | 0.008582762 | 0.041897537 | 0.11684925 |
| Cdv3 | S | 76 | -0.176659631 | -0.396420774 | -0.611859945 | -0.435200314 | -0.215439172 | 0.185926594 | 0.180161424 | 0.007527588 | 0.032628451 | 0.412938813 |
| Nme1 | S | 122 | -0.27598663 | -0.220907154 | -0.590384606 | -0.314397976 | -0.369477452 | 0.05900151 | 0.210682824 | 0.011463084 | 0.093057129 | 0.08699844 |
| Slc38a2 | S | 21 | -0.033581538 | -0.537188239 | -0.586057401 | -0.552475863 | -0.048869162 | 0.60594732 | 0.012078521 | 0.034111015 | 0.040723992 | 0.774559238 |
| Pald1 | T | 44 | -0.238645583 | -0.318768745 | -1.024986777 | -0.786341194 | -0.706218032 | 0.177014589 | 0.353373395 | 0.034659647 | 0.08177514 | 0.124990332 |
| Rbsn | S | 642 | 0.025788303 | -0.423462117 | -0.718216686 | -0.744004989 | -0.294754568 | 0.929540485 | 0.012782204 | 0.013847242 | 0.080465719 | 0.133968416 |
| Lnpep | S | 91 | -0.249498417 | -0.214742891 | -0.590778932 | -0.341280516 | -0.376036041 | 0.145907543 | 0.220562912 | 0.004811384 | 0.050049382 | 0.047415072 |
| Cdv3 | S | 75 | -0.143580735 | -0.418370703 | -0.600003387 | -0.456422652 | -0.181632684 | 0.296461047 | 0.136756892 | 0.007889702 | 0.031393675 | 0.450979193 |
| Epb41l2 | S | 86 | -0.367122406 | -0.102569447 | -0.701172535 | -0.334050129 | -0.598603088 | 0.08594683 | 0.668595528 | 0.00644637 | 0.132723205 | 0.067806455 |
| Cwc15 | T | 47 | -0.352211416 | -0.232549205 | -0.65140723 | -0.299195814 | -0.418858025 | 0.065167807 | 0.352033418 | 0.024018085 | 0.181590051 | 0.148352663 |
| Dock7 | S | 1445 | -0.458610366 | -0.084592785 | -0.601442151 | -0.142831785 | -0.516849366 | 0.025679566 | 0.691671105 | 0.017744317 | 0.408235309 | 0.057541625 |
| Limd2 | S | 30 | -0.433923966 | -0.254582181 | -0.757929276 | -0.324005309 | -0.503347095 | 0.117460264 | 0.416711999 | 0.035533263 | 0.279671844 | 0.171098901 |
| Nedd4l | S | 364 | -0.276415923 | -0.242770496 | -0.637482427 | -0.361066504 | -0.394711931 | 0.097833682 | 0.12003309 | 0.030479816 | 0.123043301 | 0.099187983 |
| Adgra2 | S | 1163 | -0.353146349 | -0.179473513 | -0.715309956 | -0.362163607 | -0.535836443 | 0.10662333 | 0.506492004 | 0.012645139 | 0.133131012 | 0.107113225 |
| Aprt | S | 15 | -0.227882795 | -0.34106935 | -0.639503259 | -0.411620464 | -0.29843391 | 0.195045491 | 0.119278638 | 0.024383518 | 0.097749206 | 0.219485911 |
| Ptpn12 | S | 448 | -0.43849623 | -0.221669687 | -0.641912479 | -0.203416249 | -0.420242793 | 0.056805268 | 0.363866103 | 0.049580678 | 0.3875186 | 0.150749673 |
| Ppp1r12c | S | 454 | -0.458754755 | -0.177977473 | -0.634718649 | -0.175963894 | -0.456741176 | 0.061807733 | 0.459554827 | 0.032575833 | 0.445991088 | 0.115256625 |
| Map4 | S | 975 | -0.176673237 | -0.267322059 | -0.697650005 | -0.520976768 | -0.430327945 | 0.478347611 | 0.172261 | 0.005329434 | 0.076369046 | 0.020771741 |
| Arpp19 Ensa | S | 62;67 | -0.403284815 | -0.187990203 | -0.680519101 | -0.277234285 | -0.492528897 | 0.09406808 | 0.449844993 | 0.028065863 | 0.284942072 | 0.122579496 |
| Dbnl | S | 237 | -0.18363841 | -0.19485902 | -0.585507905 | -0.401869495 | -0.390648885 | 0.044279864 | 0.134905069 | 0.018782836 | 0.025793616 | 0.030548887 |
| Lima1 | S | 720 | -0.405481028 | -0.136620765 | -0.592151758 | -0.18667073 | -0.455530994 | 0.044246845 | 0.470975359 | 0.029864262 | 0.324255794 | 0.066640966 |
| Cep170 | T | 1261 | -0.349020498 | -0.135686339 | -0.63158277 | -0.282562272 | -0.495896431 | 0.045815462 | 0.41803012 | 0.030277177 | 0.212006088 | 0.069565821 |
| Taok1 | S | 445 | -0.248224269 | -0.196391683 | -0.779249601 | -0.531025332 | -0.582857918 | 0.088425537 | 0.246342619 | 0.04881852 | 0.095443778 | 0.076419213 |
| Tiam2 | S | 29 | -0.204412688 | -0.20175095 | -0.710492551 | -0.506079863 | -0.508741601 | 0.103719041 | 0.235881559 | 0.026218488 | 0.072556861 | 0.068798552 |
| Arhgap22 | S | 22 | -0.046599674 | -0.322199881 | -0.61355341 | -0.566953736 | -0.291353528 | 0.667152803 | 0.097479782 | 0.006171753 | 0.009456448 | 0.121467376 |
| Mpp1 | S | 110 | -0.397321897 | -0.086769879 | -0.607971235 | -0.210649338 | -0.521201356 | 0.04490294 | 0.673521456 | 0.034847552 | 0.334772152 | 0.073253109 |
| Cdk11b | S | 212 | -0.14889308 | -0.171613669 | -0.597301007 | -0.448407927 | -0.425687338 | 0.257247476 | 0.220799982 | 0.005677809 | 0.026091134 | 0.026456128 |
| Rab13 | T | 180 | -0.327343984 | -0.113139601 | -0.686329109 | -0.358985125 | -0.573189508 | 0.059096705 | 0.610589807 | 0.035978958 | 0.183203245 | 0.078448739 |
| Arhgef19 | S | 127 | -0.250458588 | -0.183189402 | -0.864700011 | -0.614241423 | -0.68151061 | 0.371228215 | 0.525724024 | 0.018323716 | 0.093732077 | 0.084581531 |
| Neo1 | S | 1179 | -0.097371366 | -0.249591614 | -0.604009145 | -0.506637778 | -0.354417531 | 0.141537347 | 0.055021635 | 0.038520746 | 0.069912701 | 0.117199666 |
| Agap1 | T | 535 | -0.22816004 | -0.236860941 | -0.668801104 | -0.440641064 | -0.431940162 | 0.30076256 | 0.274017703 | 0.022502588 | 0.060716819 | 0.064751241 |
| Braf | S | 463 | -0.206079018 | -0.319453293 | -0.608884568 | -0.40280555 | -0.289431275 | 0.268763088 | 0.158006708 | 0.045112018 | 0.136979114 | 0.27422999 |
| Adgrl3 | S | 1238 | -0.229940331 | -0.208683377 | -0.660759324 | -0.430818993 | -0.452075947 | 0.089409997 | 0.332618971 | 0.045350717 | 0.112310896 | 0.129692901 |
| Rgl1 | S | 391 | -0.323493553 | -0.042854709 | -0.980475248 | -0.656981695 | -0.937620538 | 0.249469709 | 0.899044345 | 0.027101586 | 0.096735915 | 0.063159264 |
| Rgl1 | S | 392 | -0.323493553 | -0.042854709 | -0.980475248 | -0.656981695 | -0.937620538 | 0.249469709 | 0.899044345 | 0.027101586 | 0.096735915 | 0.063159264 |
| Eml3 | S | 157 | -0.148858969 | -0.152632024 | -0.664155581 | -0.515296612 | -0.511523557 | 0.275184858 | 0.182164079 | 0.017734099 | 0.041932435 | 0.047573328 |
| Cep55 | S | 20 | -0.256105721 | -0.055954849 | -0.60015107 | -0.344045349 | -0.544196221 | 0.116149218 | 0.793805018 | 0.007432812 | 0.049496808 | 0.053310089 |
| Zgpat | S | 353 | -0.096596652 | -0.265595082 | -0.661706221 | -0.565109569 | -0.396111139 | 0.64138707 | 0.205227205 | 0.012363563 | 0.02469874 | 0.059257208 |
| Tpd52 | S | 175 | -0.346334919 | -0.059648249 | -0.767053346 | -0.420718427 | -0.707405097 | 0.223439625 | 0.83730811 | 0.021614875 | 0.182266798 | 0.071552175 |

| | | | | | | | | | | | | |
|---|---|---|---|---|---|---|---|---|---|---|---|---|
| Cotl1 | S | 141 | -0.333841252 | -0.077411212 | -0.60397808 | -0.270136828 | -0.526566869 | 0.091815079 | 0.724183512 | 0.040174733 | 0.24807908 | 0.080169306 |
| Nolc1 | S | 691 | -0.107696712 | -0.274903883 | -0.595884642 | -0.48818793 | -0.320980759 | 0.570597949 | 0.404436272 | 0.008467984 | 0.064393268 | 0.346619182 |
| Abi1 | S | 231 | -0.228490261 | -0.091663077 | -0.607377167 | -0.378886906 | -0.51571409 | 0.146625731 | 0.699139495 | 0.016321987 | 0.081689571 | 0.100587652 |
| Sptbn1 | S | 2357 | -0.201688037 | -0.01899816 | -0.762644214 | -0.560956177 | -0.743646054 | 0.349444956 | 0.948692599 | 0.008980097 | 0.055788604 | 0.075789271 |
| Plcl1 | S | 78 | -0.343027306 | -0.006408506 | -0.708893844 | -0.365866538 | -0.702485338 | 0.13076512 | 0.982927322 | 0.041178789 | 0.195115234 | 0.078705371 |
| Epha2 | T | 772 | -0.089485073 | -0.247778375 | -0.602245787 | -0.512760714 | -0.354467412 | 0.644424743 | 0.241331561 | 0.017986 | 0.06575662 | 0.149507102 |
| Erbb2 | Y | 878 | -0.15796172 | -0.156483178 | -0.65410585 | -0.496144131 | -0.497622673 | 0.444290288 | 0.58116216 | 0.012902542 | 0.069796827 | 0.14991235 |
| Septin9 | S | 58 | -0.197503444 | 0.001242411 | -0.607574544 | -0.4100711 | -0.608816955 | 0.324288267 | 0.995441917 | 0.003560549 | 0.093212709 | 0.060132591 |
| Calu | S | 277 | -0.082510014 | -0.057593915 | -0.698951949 | -0.616441934 | -0.641358033 | 0.76855899 | 0.861477468 | 0.002972426 | 0.096225636 | 0.133618707 |
| Ensa | S | 109 | -0.223770761 | -0.125670427 | -0.812893001 | -0.58912224 | -0.687222574 | 0.488797576 | 0.690422453 | 0.037000506 | 0.124152728 | 0.084513112 |
| Nolc1 | S | 672 | 0.324882135 | -0.267773847 | -0.938606233 | -1.263488368 | -0.670832386 | 0.612308343 | 0.402514868 | 0.031545965 | 0.124617445 | 0.105368404 |
| Nbr1 | S | 891 | 0.036814826 | -0.093154373 | -0.610892678 | -0.647707504 | -0.517738305 | 0.937433918 | 0.648431951 | 0.011734296 | 0.241885233 | 0.023748309 |
| Epb41l1 | S | 510 | -0.130506607 | 0.018233566 | -0.627677592 | -0.497170985 | -0.645911157 | 0.389322189 | 0.935175078 | 0.034407112 | 0.068374182 | 0.058523893 |
| Cenpv | S | 6 | 0.081145827 | -0.07335848 | -0.595332236 | -0.676478063 | -0.521973756 | 0.754550472 | 0.768794698 | 0.023552823 | 0.051768558 | 0.088058289 |
| Hnrnpab | S | 272 | 0.074546826 | 0.120341675 | -0.593677609 | -0.668224435 | -0.714019284 | 0.901854678 | 0.873555281 | 0.032559942 | 0.321244931 | 0.387075098 |
| Tmpo | S | 158;158 | 0.071327484 | 0.576395004 | 0.652686303 | 0.581358818 | 0.076291298 | 0.012571147 | 9.30006E-07 | 6.10677E-06 | 0.000113247 | 0.055423029 |
| Specc1l | S | 393 | 0.140397178 | 0.500486939 | 0.697715492 | 0.557318314 | 0.197228553 | 0.179123144 | 3.27381E-05 | 1.29133E-06 | 0.012942482 | 0.011651815 |
| Epb41l3 | S | 99 | 0.342447506 | 0.584904734 | 0.744518298 | 0.402070792 | 0.159613563 | 0.016814576 | 0.011998838 | 7.63337E-05 | 0.01398856 | 0.268868543 |
| Pdlim2 | S | 127 | 0.072791008 | 0.409228684 | 0.606835755 | 0.534044748 | 0.197607071 | 0.06182365 | 2.1881E-05 | 1.24097E-06 | 3.02765E-05 | 0.00667435 |
| Ttc7 | T | 693 | 0.332614424 | 0.491033193 | 0.686439305 | 0.353824881 | 0.195406112 | 0.033500585 | 0.000269104 | 0.000828083 | 0.036442256 | 0.096513546 |
| Phf2 | S | 650 | 0.368929579 | 0.419775957 | 0.6442046 | 0.27527502 | 0.224428642 | 0.001984925 | 0.040356085 | 0.000118445 | 0.016149349 | 0.169796563 |
| Zranb2 | S | 280 | 0.168567085 | 0.576223584 | 0.692211111 | 0.523644026 | 0.115987527 | 0.084551885 | 1.92632E-05 | 0.006018752 | 0.012777879 | 0.335311426 |
| Pxn | S | 260 | 0.262047783 | 0.516241463 | 0.671073407 | 0.409025623 | 0.154831944 | 0.023474083 | 0.010595081 | 0.000185598 | 0.003671741 | 0.253161842 |
| Srsf4 | S | 361 | 0.439490037 | 0.471608979 | 0.907555626 | 0.468065588 | 0.435946647 | 0.026153898 | 0.109144737 | 0.000924417 | 0.03185369 | 0.131172081 |
| Srsf4 | S | 363 | 0.439490037 | 0.471608979 | 0.907555626 | 0.468065588 | 0.435946647 | 0.026153898 | 0.109144737 | 0.000924417 | 0.03185369 | 0.131172081 |
| Arhgap21 | S | 609 | 0.518747644 | 0.356751808 | 0.610955905 | 0.092208261 | 0.254204097 | 0.000235736 | 0.128049277 | 0.004443941 | 0.415234912 | 0.236762493 |
| Adnp | S | 737 | 0.515475196 | 0.497751794 | 0.711837065 | 0.196361869 | 0.214085271 | 0.01083332 | 0.03054616 | 0.009328431 | 0.296787119 | 0.302199982 |
| Thrap3 | S | 913 | 0.501043744 | 0.251496911 | 0.65659309 | 0.155549346 | 0.405096179 | 0.011791426 | 0.004289667 | 0.004500968 | 0.28220338 | 0.038390139 |
| Lamtor1 | S | 56 | 0.223920429 | 0.571923779 | 0.606975609 | 0.383055181 | 0.03505183 | 0.192943633 | 0.004161537 | 0.001101294 | 0.057236771 | 0.746589999 |
| Dapk3 | S | 307 | 0.18662665 | 0.35233999 | 0.587176388 | 0.400549738 | 0.234836399 | 0.037604931 | 0.006334665 | 0.000129193 | 0.006402464 | 0.034177841 |
| Cpsf7 | T | 203 | 0.387702446 | 0.451392229 | 0.608931393 | 0.221228947 | 0.157539165 | 0.013527449 | 0.051651343 | 0.002000761 | 0.069906375 | 0.377314996 |
| Synpo | S | 230 | 0.285757612 | 0.333340654 | 0.66820336 | 0.382445748 | 0.334862706 | 0.016731778 | 0.036001768 | 0.0014064 | 0.021450719 | 0.04951529 |
| Grk5 | T | 485 | 0.054115279 | 0.477698757 | 0.586969859 | 0.532854581 | 0.109271102 | 0.533796697 | 0.000732097 | 0.000305632 | 0.002467132 | 0.218604477 |
| Thrap3 | S | 744 | 0.17748376 | 0.553361511 | 0.699131661 | 0.521647901 | 0.14577015 | 0.048232571 | 0.005308527 | 0.0105237 | 0.022383392 | 0.330885541 |
| Tnrc6b | S | 1793 | 0.193754104 | 0.45698134 | 0.592230199 | 0.398476095 | 0.135248859 | 0.162896626 | 0.008878011 | 0.002720586 | 0.011024929 | 0.174496588 |
| Atf2 | S | 310 | 0.309702037 | 0.365593631 | 0.626986019 | 0.317283983 | 0.261392389 | 0.060758837 | 0.03385861 | 0.010243749 | 0.098945148 | 0.149793665 |
| Zranb2 | S | 279 | 0.066970382 | 0.523583703 | 0.632397347 | 0.565426965 | 0.108813644 | 0.628865621 | 0.002390389 | 0.01064356 | 0.022087081 | 0.456861486 |
| Elf1 | S | 334 | 0.0293069 | 0.343335172 | 0.605202584 | 0.575895684 | 0.261867412 | 0.587167523 | 0.003663733 | 0.000715315 | 0.000795061 | 0.014934496 |
| Phc3 | S | 759 | 0.246320985 | 0.535884614 | 0.66878151 | 0.422460525 | 0.132896896 | 0.621799495 | 0.065930237 | 0.005753196 | 0.415525423 | 0.530166654 |
| Phc3 | S | 760 | 0.246320985 | 0.535884614 | 0.66878151 | 0.422460525 | 0.132896896 | 0.621799495 | 0.065930237 | 0.005753196 | 0.415525423 | 0.530166654 |
| Phc3 | T | 762 | 0.204537992 | 0.486120135 | 0.61913495 | 0.414596958 | 0.133014815 | 0.665998847 | 0.076747803 | 0.006238532 | 0.405352716 | 0.510403553 |
| Adgra2 | S | 963 | 0.335437479 | 0.303861405 | 0.587671174 | 0.252233695 | 0.283809768 | 0.131029103 | 0.056352156 | 0.037441988 | 0.246092377 | 0.140487909 |
| Tjp1 | T | 840 | 0.057017327 | 0.457419094 | 0.639542313 | 0.582524986 | 0.182123219 | 0.790521281 | 0.018994239 | 0.018651834 | 0.067521105 | 0.280027927 |
| Cavin2 | S | 296 | 0.102471012 | 0.260127877 | 0.635087841 | 0.53261683 | 0.374959965 | 0.355253348 | 0.042969327 | 0.006972037 | 0.018717629 | 0.04949798 |
| Larp1 | T | 196 | 0.385425734 | 0.068248576 | 0.60605943 | 0.220633696 | 0.537810855 | 0.055200088 | 0.527196566 | 0.009213688 | 0.245337158 | 0.031879804 |
| Clasp2 | T | 546 | 0.335624544 | 0.177621459 | 0.60649231 | 0.270867766 | 0.428870851 | 0.149953737 | 0.451061788 | 0.005636327 | 0.212355688 | 0.11535597 |
| Zfp692 | S | 267 | 0.457245418 | 0.386642495 | 0.637339538 | 0.18009412 | 0.250697043 | 0.237316064 | 0.322377253 | 0.041640165 | 0.54760835 | 0.442915077 |
| Ankib1 | S | 908 | -0.105428223 | -0.043720541 | 0.605903311 | 0.711331534 | 0.649623851 | 0.815218014 | 0.894634251 | 0.008658834 | 0.199209962 | 0.127589528 |
| Apc | S | 2774 | -0.42434189 | -0.381327687 | 0.25303402 | 0.67737591 | 0.634361707 | 0.087807091 | 0.127116934 | 0.278053178 | 0.013261491 | 0.019712063 |
| Hsph1 | S | 852 | 0.233634729 | 0.28005271 | -0.394388456 | -0.628023185 | -0.674441166 | 0.27820341 | 0.034485991 | 0.017526051 | 0.038942579 | 0.004146346 |
| H2ax | S | 121 | 0.048095552 | -0.439640593 | -0.582563916 | -0.630659468 | -0.142923323 | 0.437780027 | 2.80578E-05 | 7.93993E-07 | 0.001838317 | 0.015114507 |
| Srpk2 | S | 490 | 0.059629625 | -0.580425932 | -0.583188004 | -0.642817629 | -0.002762072 | 0.333676177 | 4.05707E-06 | 8.21367E-05 | 0.00096408 | 0.939291385 |
| Adipor2 | T | 14 | 0.046248968 | -0.485967537 | -0.572793338 | -0.619042306 | -0.086825801 | 0.493593398 | 2.0357E-06 | 0.001058934 | 0.001021786 | 0.203234585 |
| Stx12 | S | 139 | 0.069687041 | -0.401089818 | -0.519301218 | -0.588988258 | -0.1182114 | 0.1101631 | 1.7924E-05 | 3.16911E-06 | 2.53185E-05 | 0.010255318 |
| Ddx21 | S | 192 | 0.085772183 | -0.386497777 | -0.562906153 | -0.648678337 | -0.176408376 | 0.026122719 | 9.90063E-06 | 5.26753E-06 | 0.00011231 | 0.0062875 |
| Ncl | T | 58 | 0.067420539 | -0.366257042 | -0.544172905 | -0.611593443 | -0.177915863 | 0.218735359 | 0.000144382 | 1.19483E-05 | 0.000136699 | 0.006988326 |
| Acaca | S | 23 | 0.064953725 | -0.543190831 | -0.576829114 | -0.641782839 | -0.033638283 | 0.322551342 | 0.000164177 | 0.000950832 | 0.005224759 | 0.650439127 |
| Ddx21 | S | 155 | 0.093983801 | -0.347832844 | -0.518559585 | -0.612543386 | -0.170726742 | 0.06780335 | 7.34725E-05 | 7.12255E-06 | 0.000105701 | 0.001709659 |
| Cd44 | S | 771 | 0.055030585 | -0.290487962 | -0.560022253 | -0.615052838 | -0.269534291 | 0.211398889 | 0.00037148 | 7.64694E-06 | 8.97857E-05 | 0.000125199 |
| Pi4k2a | S | 462 | 0.08621469 | -0.390472077 | -0.501205605 | -0.587420295 | -0.110733528 | 0.03477614 | 4.81352E-06 | 0.000539561 | 0.00068513 | 0.093524378 |
| Nop2 | S | 68 | 0.089530723 | -0.322183306 | -0.515092541 | -0.604623263 | -0.192909235 | 0.073763597 | 0.00025003 | 1.39366E-05 | 0.000391245 | 0.009423572 |
| Gtf2f1 | S | 391 | 0.221277674 | -0.568740891 | -0.48068037 | -0.701958043 | 0.088060521 | 0.343558492 | 0.000235211 | 0.001359109 | 0.04854102 | 0.275449684 |
| Irs2 | S | 573 | 0.040026221 | -0.380649112 | -0.567426671 | -0.607452892 | -0.186777559 | 0.576093598 | 0.0013118 | 0.002964049 | 0.006854453 | 0.118496793 |
| Cpeb4 | S | 332 | 0.193761881 | -0.46686542 | -0.519024068 | -0.712785949 | -0.052158648 | 0.086970236 | 0.003382207 | 0.000160223 | 0.003217058 | 0.539915724 |
| Rsl1d1 | T | 385 | 0.186798681 | -0.369033678 | -0.545978364 | -0.732777044 | -0.176944686 | 0.116620152 | 0.017607618 | 1.41886E-05 | 0.008341494 | 0.132634531 |
| Mybbp1a | S | 1187 | 0.017035242 | -0.399772873 | -0.576446349 | -0.593481591 | -0.176673476 | 0.813225156 | 0.005092945 | 0.026006678 | 0.027157848 | 0.285364452 |
| Mybbp1a | T | 1190 | 0.017035242 | -0.399772873 | -0.576446349 | -0.593481591 | -0.176673476 | 0.813225156 | 0.005092945 | 0.026006678 | 0.027157848 | 0.285364452 |
| Tmem74 | S | 124 | 0.078882488 | -0.437942293 | -0.571332502 | -0.65021489 | -0.133390209 | 0.225491314 | 0.001489804 | 0.020644543 | 0.016864095 | 0.347432878 |
| Rsl1d1 | S | 326 | 0.114013477 | -0.386435925 | -0.496839544 | -0.610853021 | -0.110403619 | 0.05968593 | 0.002190479 | 0.001274678 | 0.000529308 | 0.144593283 |
| Nolc1 | T | 551 | 0.211486207 | -0.278614394 | -0.513271416 | -0.724757624 | -0.234657022 | 0.022074622 | 0.001653144 | 7.59819E-05 | 0.00040281 | 0.017060427 |
| Cox8a | S | 15 | 0.055714198 | -0.334707084 | -0.560194933 | -0.615909132 | -0.225487849 | 0.428254076 | 0.01296856 | 0.009579453 | 0.005729915 | 0.09976824 |
| Gja1 | T | 251 | 0.068925058 | -0.431986345 | -0.517146349 | -0.586071407 | -0.085160004 | 0.680796867 | 0.00795501 | 0.041081376 | 0.035989521 | 0.565105166 |
| Bcl6 | S | 405 | 0.08434254 | -0.267946933 | -0.530650442 | -0.614992981 | -0.262703509 | 0.246908921 | 0.004024151 | 0.005496737 | 0.002959125 | 0.043625266 |
| Mybbp1a | S | 6 | 0.173015503 | -0.371357734 | -0.445042495 | -0.618057998 | -0.073684761 | 0.16203078 | 0.000506511 | 0.015975795 | 0.006672864 | 0.489514891 |
| Phldb1 | S | 445 | 0.208138366 | -0.331203393 | -0.422398058 | -0.630536423 | -0.091194664 | 0.00563053 | 0.003850151 | 3.86868E-05 | 0.000195906 | 0.193270193 |
| Npm1 | T | 198 | 0.254326546 | -0.350350008 | -0.575596317 | -0.829922863 | -0.225246309 | 0.004049034 | 0.028752298 | 3.33722E-05 | 7.25138E-05 | 0.099220043 |
| Utp18 | S | 29 | 0.149360835 | -0.429791113 | -0.529655649 | -0.679016484 | -0.099864536 | 0.215659491 | 0.093370661 | 0.001774477 | 0.003717039 | 0.598116336 |
| Rragc | T | 393 | 0.248661728 | -0.487238946 | -0.439727521 | -0.688389249 | 0.047511425 | 0.066877229 | 0.004238302 | 0.005004795 | 0.003572077 | 0.561364856 |
| Ncl | T | 106 | 0.213965767 | -0.305729978 | -0.462560896 | -0.676526663 | -0.156830918 | 0.021151021 | 0.001579882 | 0.000682839 | 0.000594182 | 0.067113157 |
| Map1b | T | 702 | 0.209119915 | -0.486629636 | -0.401846631 | -0.610966547 | 0.084783005 | 0.027851382 | 0.007174543 | 0.00374899 | 0.00283868 | 0.457817779 |
| Pnn | S | 288 | 0.07079648 | -0.25334474 | -0.546085049 | -0.616881529 | -0.292740309 | 0.451553596 | 0.025879618 | 0.030374745 | 0.037220402 | 0.157742262 |
| Phrf1 | S | 457 | 0.231396987 | -0.458436919 | -0.550287479 | -0.781684466 | -0.09185056 | 0.349836905 | 0.242759365 | 0.003561926 | 0.042310525 | 0.779381091 |
| Phldb1 | S | 463 | 0.196682941 | -0.317537721 | -0.390038112 | -0.586721053 | -0.072500391 | 0.138248755 | 0.014040665 | 0.005664418 | 0.007713721 | 0.389158648 |
| Ulk1 | S | 614 | 0.118093395 | -0.408988371 | -0.477895487 | -0.595988883 | -0.068907116 | 0.467707188 | 0.126127923 | 0.027204891 | 0.025108392 | 0.756306121 |
| Fnip1 | S | 284 | 0.206926314 | -0.233040456 | -0.571642833 | -0.778569147 | -0.338602377 | 0.079143178 | 0.065602242 | 0.002767503 | 0.009526252 | 0.02893298 |
| Jun | S | 63 | 0.254122454 | -0.289142465 | -0.408468415 | -0.662590869 | -0.11932595 | 0.008256655 | 0.004665567 | 0.001561166 | 0.001485493 | 0.123964458 |
| Mybbp1a | S | 1321 | 0.240281426 | -0.360535926 | -0.496281379 | -0.736562805 | -0.135745452 | 0.0057494 | 0.005718497 | 0.014125275 | 0.014989272 | 0.293025157 |
| Eml1 | S | 162 | 0.183666407 | -0.200200483 | -0.476024428 | -0.659690834 | -0.275823945 | 0.049467773 | 0.06127316 | 0.003281607 | 0.005726958 | 0.03814011 |
| Rsl1d1 | S | 384 | 0.241124979 | -0.295325363 | -0.541052164 | -0.782177143 | -0.245726801 | 0.038703107 | 0.05219518 | 0.011797603 | 0.002569356 | 0.111742662 |
| Layn | S | 299 | 0.066229705 | 0.00130506 | -0.525913279 | -0.592142984 | -0.52721834 | 0.550290447 | 0.990592584 | 0.002631168 | 0.002566086 | 0.002788226 |
| Phldb1 | S | 420 | 0.326051413 | -0.302720125 | -0.401133594 | -0.727185006 | -0.098413468 | 0.002358564 | 0.017702175 | 0.00065636 | 6.27504E-05 | 0.270028106 |
| Phldb1 | S | 387 | 0.327740523 | -0.350202643 | -0.400686613 | -0.728427136 | -0.05048397 | 0.005245959 | 0.074533329 | 0.001895407 | 0.000166419 | 0.720176203 |
| Eif3c | S | 207 | 0.323381989 | -0.285690222 | -0.294390258 | -0.617772248 | -0.008700036 | 0.0299292 | 0.025954775 | 0.007179812 | 0.005062873 | 0.924290598 |
| Rpl19 | S | 13 | 0.20304433 | -0.181664307 | -0.430349895 | -0.633393325 | -0.248685588 | 0.033484538 | 0.105982496 | 0.019790158 | 0.009972396 | 0.104973903 |

| | | | | | | | | | | | | |
|---|---|---|---|---|---|---|---|---|---|---|---|---|
| Jpt2 | S | 133 | 0.228209952 | -0.112126138 | -0.415241231 | -0.643451183 | -0.303115093 | 0.025447523 | 0.34701712 | 0.001220171 | 0.001100636 | 0.060001746 |
| Psrc1 | S | 107 | 0.208423327 | -0.115493526 | -0.38813154 | -0.596554867 | -0.272638013 | 0.021579917 | 0.186382382 | 0.003315295 | 0.001438526 | 0.026478133 |
| Rpl23a | S | 34 | 0.261444517 | -0.028289983 | -0.580904487 | -0.842349004 | -0.552614505 | 0.103433184 | 0.850199275 | 0.008821788 | 0.004111081 | 0.019620584 |
| Arhgap35 | S | 589 | 0.177232432 | -0.082766611 | -0.420128897 | -0.597361329 | -0.337362286 | 0.080817406 | 0.442262281 | 0.017033351 | 0.015630789 | 0.043521966 |
| Dop1b | S | 719 | 0.201757857 | -0.072924394 | -0.416402525 | -0.618160383 | -0.343478131 | 0.132734521 | 0.392875961 | 0.022537547 | 0.008890361 | 0.060293959 |
| Svil | S | 831 | 0.52910206 | -0.215843485 | -0.442632423 | -0.971734484 | -0.226788938 | 0.126796229 | 0.813316801 | 0.032783198 | 0.049838635 | 0.804381209 |
| Tpst2 | T | 38 | 0.383173594 | -0.045498412 | -0.401649418 | -0.784823012 | -0.356151006 | 0.238676912 | 0.85273059 | 0.160290184 | 0.031218536 | 0.06979426 |
| Syne2 | T | 6489 | -0.405622668 | 0.291709158 | 0.25137566 | 0.656998328 | -0.040333498 | 0.000992965 | 0.143031058 | 0.115846219 | 0.019049922 | 0.823306686 |
| Osbpl6 | S | 69 | 0.317283105 | -0.113092596 | -0.383904283 | -0.701187388 | -0.270811687 | 0.008353692 | 0.407715979 | 0.104443205 | 0.034879691 | 0.216256512 |
| Rsl1d1 | T | 429 | 0.416050657 | -0.141310322 | -0.202831498 | -0.618882155 | -0.061521175 | 0.017046124 | 0.311204009 | 0.018217618 | 0.007672172 | 0.627312672 |
| Fn1 | S | 2475 | -0.036136558 | 0.53376214 | 0.554406946 | 0.590543504 | 0.020644806 | 0.300141873 | 1.51159E-06 | 0.000730701 | 0.004978086 | 0.708079925 |
| Lmna | T | 416 | -0.152513157 | 0.403360177 | 0.437761129 | 0.590274286 | 0.034400952 | 0.368587956 | 0.0001141 | 9.97451E-06 | 0.043299315 | 0.411174092 |
| Pxn | S | 250 | -0.015370698 | 0.445273898 | 0.58442548 | 0.599796178 | 0.139151582 | 0.826806075 | 0.00024482 | 0.002612945 | 0.001934266 | 0.134069714 |
| Cep170 | S | 358 | -0.084584329 | 0.339791867 | 0.512741192 | 0.597325522 | 0.172949325 | 0.24839496 | 0.001032772 | 0.000106254 | 0.001575867 | 0.022342611 |
| Prr14 | S | 135 | -0.095777015 | 0.405931525 | 0.556434466 | 0.652211481 | 0.150502941 | 0.242345359 | 0.002041253 | 0.000966728 | 0.001290931 | 0.071656777 |
| Arid5b | S | 1032 | -0.080607977 | 0.264451154 | 0.571502931 | 0.652110908 | 0.307051776 | 0.137390937 | 0.000107907 | 0.005176738 | 0.003008378 | 0.038566647 |
| Pbrm1 | S | 378 | -0.038475909 | 0.31742617 | 0.556850443 | 0.595326352 | 0.239424273 | 0.739799315 | 0.059199563 | 0.000617126 | 0.004510444 | 0.121968537 |
| Prickle2 | S | 807 | -0.139918991 | 0.364997885 | 0.467570875 | 0.607489866 | 0.10257299 | 0.111393349 | 0.001795422 | 0.001065638 | 0.001125887 | 0.184481643 |
| Lrrk2 | S | 976 | -0.114771947 | 0.275052907 | 0.499567935 | 0.614339882 | 0.224515028 | 0.399501229 | 0.033317366 | 0.002508833 | 0.01214748 | 0.081437871 |
| Cdc42ep4 | S | 136 | -0.132903106 | 0.481641628 | 0.453355486 | 0.586258591 | -0.028286142 | 0.012881078 | 0.026186565 | 0.00102717 | 0.000498596 | 0.79276104 |
| Tcf20 | S | 1819 | -0.147831615 | 0.306433481 | 0.495993072 | 0.643824688 | 0.189559591 | 0.007466406 | 0.000151754 | 0.02018182 | 0.011484343 | 0.146740018 |
| Reps1 | S | 559 | 0.735143175 | 0.335714675 | -0.087876686 | -0.823019861 | -0.423591361 | 0.051701177 | 0.228738337 | 0.744018517 | 0.040178131 | 0.11657613 |
| Tcf20 | S | 669 | -0.172125353 | 0.189575745 | 0.461715656 | 0.633841009 | 0.272139911 | 0.098154211 | 0.001474758 | 0.001370538 | 0.025440054 |  |
| Mndal | S | 276 | -0.158683856 | 0.16949256 | 0.456389624 | 0.61507348 | 0.286897064 | 0.027494335 | 0.038224638 | 0.006362148 | 0.00435123 | 0.032477714 |
| Stard8 | S | 732 | -0.303863585 | 0.254969238 | 0.407018856 | 0.710882441 | 0.152049618 | 0.001344905 | 0.017629641 | 0.000346624 | 0.000626005 | 0.093227502 |
| D5Ertd579e | S | 226 | -0.241850824 | 0.211537155 | 0.349512362 | 0.591363187 | 0.137975207 | 0.050111682 | 0.001957696 | 0.04669587 | 0.006615526 | 0.24538559 |
| Plekha2 | S | 401 | -0.384280892 | 0.558858837 | 0.308126452 | 0.692407344 | -0.250732385 | 0.037737837 | 0.015840299 | 0.114466272 | 0.010919884 | 0.203147986 |
| Trpm4 | S | 538 | -0.292762314 | 0.012526286 | 0.407541292 | 0.700303606 | 0.395015005 | 0.130654827 | 0.762536251 | 0.000122335 | 0.023327059 | 0.000907842 |
| Ptpn13 | S | 1456 | 0.438232482 | 0.233696767 | -0.23604648 | -0.674278963 | -0.469743247 | 0.034399467 | 0.249164745 | 0.187870321 | 0.015335178 | 0.067743102 |
| Ttk | S | 820 | 0.41639777 | 0.111797867 | -0.249576278 | -0.665974047 | -0.361374145 | 0.002947098 | 0.227076732 | 0.042947919 | 0.001201351 | 0.025923395 |
| Kif23 | S | 715 | 0.414905107 | 0.146999697 | -0.260243417 | -0.675148524 | -0.407243114 | 0.026643655 | 0.129319225 | 0.06269138 | 0.006998519 | 0.020327253 |
| Srgap2 | S | 1053 | 0.531730725 | 0.172335182 | -0.302516072 | -0.834246797 | -0.474851254 | 0.130644506 | 0.2568655 | 0.147879307 | 0.047328098 | 0.052551822 |
| Stard3nl | S | 210 | -0.404236141 | 0.129708807 | 0.410751116 | 0.814987256 | 0.281042309 | 0.056682939 | 0.106421158 | 0.02063228 | 0.007188833 | 0.079801469 |
| Stard3nl | S | 214 | -0.404236141 | 0.129708807 | 0.410751116 | 0.814987256 | 0.281042309 | 0.056682939 | 0.106421158 | 0.02063228 | 0.007188833 | 0.079801469 |
| Stard3nl | S | 218 | -0.404236141 | 0.129708807 | 0.410751116 | 0.814987256 | 0.281042309 | 0.056682939 | 0.106421158 | 0.02063228 | 0.007188833 | 0.079801469 |
| Atxn2l | T | 684 | 0.501720269 | -0.123051869 | -0.158750949 | -0.660471217 | -0.035699079 | 0.029930204 | 0.425716337 | 0.089111763 | 0.022450242 | 0.796741648 |
| Phf11d | S | 17 | -0.362944069 | 0.023239372 | 0.486407278 | 0.849351346 | 0.463173305 | 0.058045035 | 0.88625276 | 0.024364369 | 0.004184891 | 0.014253817 |
| Top2b | T | 1458 | -0.272677018 | -0.051550664 | 0.463709285 | 0.736386303 | 0.515259948 | 0.123873427 | 0.737045728 | 0.04236269 | 0.012570058 | 0.042532848 |
| Cep170 | S | 355 | -0.251101227 | 0.006926401 | 0.393164034 | 0.644265261 | 0.386237633 | 0.107810359 | 0.968034513 | 0.103256698 | 0.029677998 | 0.138230865 |
| Phf11d | S | 240 | -0.289218879 | -0.01876017 | 0.302704917 | 0.591923796 | 0.321465087 | 0.139345491 | 0.914946797 | 0.122398663 | 0.003172511 | 0.012279744 |
| Tanc2 | S | 1726 | -0.318453193 | 0.042024464 | -0.551806189 | -0.233352996 | -0.593830653 | 0.083723982 | 0.784260441 | 0.022893664 | 0.240383574 | 0.024514208 |
| Akap12 | T | 272 | -0.30565473 | 0.131447918 | -0.513463373 | -0.207808643 | -0.644911291 | 0.029141535 | 0.486334649 | 0.043999033 | 0.286514825 | 0.033966001 |
| Bloc1s4 | S | 177 | -0.335134322 | 0.163876451 | -0.615240144 | -0.280105822 | -0.779116595 | 0.308713405 | 0.562767694 | 0.071919226 | 0.410629013 | 0.049056033 |
| Scarf1 | S | 590 | -0.165770023 | 0.24543045 | -0.579790056 | -0.414020033 | -0.825220506 | 0.092673373 | 0.221654657 | 0.039607061 | 0.074735901 | 0.013870139 |
| Eml3 | S | 177 | -0.161816497 | 0.224490514 | -0.378378242 | -0.216561745 | -0.602868756 | 0.24489799 | 0.26192598 | 0.045111112 | 0.206963756 | 0.033529405 |
| Dcaf5 | S | 821 | -0.162351617 | -0.37076942 | 0.240619672 | 0.402971288 | 0.611389092 | 0.35579516 | 0.036662625 | 0.139887964 | 0.033996311 | 0.003035343 |
| Srrm3 | S | 516 | -0.062740861 | 0.185915767 | -0.400094681 | -0.337353819 | -0.586010447 | 0.626248107 | 0.104164786 | 0.08181839 | 0.12454968 | 0.031148259 |
| Srrm3 | S | 518 | -0.062740861 | 0.185915767 | -0.400094681 | -0.337353819 | -0.586010447 | 0.626248107 | 0.104164786 | 0.08181839 | 0.12454968 | 0.031148259 |
| Srrm3 | S | 520 | -0.062740861 | 0.185915767 | -0.400094681 | -0.337353819 | -0.586010447 | 0.626248107 | 0.104164786 | 0.08181839 | 0.12454968 | 0.031148259 |
| Clec16a | S | 848 | 0.004047983 | 0.264950637 | -0.422104541 | -0.426152524 | -0.687055178 | 0.986283983 | 0.111946564 | 0.020691541 | 0.152883129 | 0.00882952 |
| Cep55 | S | 40 | -0.018945587 | 0.324227319 | -0.473403191 | -0.454457603 | -0.79763051 | 0.933280787 | 0.220724482 | 0.144518594 | 0.169855431 | 0.047777507 |
| Cep55 | T | 39 | -0.018945587 | 0.324227319 | -0.473403191 | -0.454457603 | -0.79763051 | 0.933280787 | 0.220724482 | 0.144518594 | 0.169855431 | 0.047777507 |
| Luzp1 | S | 261 | 0.127076161 | 0.514804085 | -0.080059557 | -0.207135717 | -0.594863642 | 0.177513703 | 0.000797913 | 0.480591417 | 0.090595136 | 0.005433365 |
| Plcl1 | S | 570 | 0.090056053 | 0.483420448 | -0.142010659 | -0.232066712 | -0.625431107 | 0.505056968 | 0.010695886 | 0.380953669 | 0.109793039 | 0.019812298 |
| Gpr149 | T | 239 | 0.169971192 | -0.200283274 | 0.55364272 | 0.383671528 | 0.753925994 | 0.063818829 | 0.080736664 | 0.053460005 | 0.119364239 | 0.024346381 |
| Ino80 | S | 85 | 0.186707183 | 0.59063094 | -0.2696779 | -0.456385084 | -0.86030884 | 0.751599424 | 0.155098286 | 0.45369584 | 0.469874678 | 0.002370524 |
| Ino80 | S | 88 | 0.186707183 | 0.59063094 | -0.2696779 | -0.456385084 | -0.86030884 | 0.751599424 | 0.155098286 | 0.45369584 | 0.469874678 | 0.002370524 |
| Ino80 | S | 89 | 0.186707183 | 0.59063094 | -0.2696779 | -0.456385084 | -0.86030884 | 0.751599424 | 0.155098286 | 0.45369584 | 0.469874678 | 0.002370524 |
| Ino80 | S | 90 | 0.186707183 | 0.59063094 | -0.2696779 | -0.456385084 | -0.86030884 | 0.751599424 | 0.155098286 | 0.45369584 | 0.469874678 | 0.002370524 |
| Ino80 | T | 84 | 0.186707183 | 0.59063094 | -0.2696779 | -0.456385084 | -0.86030884 | 0.751599424 | 0.155098286 | 0.45369584 | 0.469874678 | 0.002370524 |
| Dock11 | S | 797 | -0.072421217 | 0.481629912 | -0.278416945 | -0.205995728 | -0.760046857 | 0.592545214 | 0.020495133 | 0.124074288 | 0.214543638 | 0.00821619 |
| Gm13889 | T | 179 | 0.125035227 | 0.450191316 | -0.294551639 | -0.419586866 | -0.744742955 | 0.516500322 | 0.101226867 | 0.052426402 | 0.079044839 | 0.042315611 |
| Gpr149 | T | 242 | 0.138006145 | -0.333431358 | 0.368182368 | 0.230176223 | 0.701613726 | 0.131635191 | 0.007642351 | 0.108605488 | 0.252683736 | 0.029222552 |
| Gab2 | T | 388 | -0.120725712 | -0.5177795 | -0.546142601 | -0.425416889 | -0.028363101 | 0.022781453 | 1.02112E-07 | 1.67504E-07 | 0.003537893 | 0.157230229 |
| Pdcd11 | S | 1468 | -0.085263049 | -0.52548364 | -0.533450458 | -0.448187409 | -0.007966818 | 0.051642999 | 3.68049E-07 | 1.3306E-07 | 0.000308904 | 0.69974519 |
| Setd1a | T | 1174 | -0.03578665 | -0.516058731 | -0.528405651 | -0.492619001 | -0.01234692 | 0.261462636 | 1.3448E-07 | 3.57019E-07 | 6.19765E-05 | 0.564324814 |
| Rbm15 | S | 723 | -0.083925275 | -0.492133586 | -0.543716724 | -0.45979145 | -0.051583138 | 0.000265073 | 0.004561431 | 3.73158E-09 | 2.5266E-05 | 0.336274593 |
| Larp4 | S | 579 | -0.052628757 | -0.541525127 | -0.505638081 | -0.452811324 | 0.035887046 | 0.108744698 | 5.54042E-07 | 2.55364E-06 | 2.02328E-05 | 0.121878308 |
| Impdh2 | S | 416 | -0.161364063 | -0.502019727 | -0.511118757 | -0.349754694 | -0.009099031 | 0.014959317 | 7.31912E-06 | 1.25464E-05 | 0.001348866 | 0.791417633 |
| Serbp1 | S | 234 | -0.108647608 | -0.519669532 | -0.48896763 | -0.380320022 | 0.030701902 | 0.016152566 | 6.06923E-06 | 4.86677E-06 | 3.34624E-05 | 0.096618932 |
| Jmy | S | 58 | -0.068173056 | -0.489751126 | -0.582676656 | -0.5145036 | -0.09292553 | 0.108322289 | 6.68381E-06 | 1.81548E-06 | 0.000332321 | 0.023135611 |
| Larp4 | S | 717 | -0.036292211 | -0.442860075 | -0.54221125 | -0.505919039 | -0.099351175 | 0.333478263 | 3.62868E-06 | 1.05596E-07 | 0.000461621 | 0.01554697 |
| Atr | S | 438 | -0.08384177 | -0.538509681 | -0.570268023 | -0.486426253 | -0.031758342 | 0.11841295 | 2.12742E-05 | 3.78118E-06 | 0.000702371 | 0.164528257 |
| Pdcd11 | S | 1470 | -0.114617313 | -0.452032644 | -0.477154182 | -0.362536869 | -0.025121537 | 0.008303174 | 5.3376E-06 | 6.89695E-06 | 0.00202365 | 0.403156145 |
| Pcbp2 | S | 268 | -0.117998586 | -0.495202099 | -0.55344959 | -0.435451004 | -0.058247491 | 0.098597775 | 1.23944E-05 | 6.4638E-06 | 0.002549208 | 0.198279062 |
| Gtf2f1 | S | 385 | -0.013004745 | -0.527153354 | -0.562615885 | -0.549611139 | -0.035462531 | 0.86622888 | 2.06184E-06 | 1.26447E-06 | 0.009754426 | 0.265198342 |
| Pcbp1 | S | 264 | -0.051307411 | -0.465603225 | -0.545453116 | -0.494145706 | -0.079849892 | 0.36838733 | 9.33226E-07 | 2.28664E-06 | 0.002053556 | 0.048733007 |
| Lmna | S | 404 | -0.0545843 | -0.542561194 | -0.533066203 | -0.478481903 | 0.009494991 | 0.247660154 | 6.7826E-06 | 3.10078E-06 | 0.001452757 | 0.704055419 |
| Eif4b | S | 409 | -0.016774625 | -0.556773182 | -0.504994255 | -0.488219631 | 0.051778927 | 0.855154694 | 9.39478E-07 | 1.8001E-06 | 0.019540725 | 0.138238285 |
| Rpl31 | S | 98 | -0.173194799 | -0.387410643 | -0.365250723 | -0.192055924 | 0.022159919 | 0.005783814 | 9.11363E-07 | 5.38025E-06 | 0.017392 | 0.171966206 |
| Rtcb | S | 208 | -0.110889442 | -0.51636045 | -0.567966575 | -0.457077133 | -0.051606126 | 0.014971927 | 0.000351042 | 6.61395E-06 | 0.000262141 | 0.345007817 |
| Aif1l | S | 133 | -0.248362012 | -0.364494132 | -0.37373683 | -0.125374818 | -0.009242698 | 5.26958E-05 | 0.000174538 | 3.67647E-05 | 0.02017455 | 0.812025362 |
| Serbp1 | S | 197 | -0.020161622 | -0.439307084 | -0.468242638 | -0.448081016 | -0.028935554 | 0.7121909 | 4.80238E-06 | 2.52002E-08 | 0.009176855 | 0.125447025 |
| Ybx1 | S | 100 | -0.066713065 | -0.525864825 | -0.500105802 | -0.433392737 | 0.025759023 | 0.031813244 | 1.40573E-07 | 0.000940196 | 0.00193722 | 0.539052715 |
| Ybx2 Ybx3 | S | 126;139 | -0.059282773 | -0.503791669 | -0.480814031 | -0.421531258 | 0.022977638 | 0.041587303 | 6.82401E-07 | 0.002273986 | 0.002343658 | 0.591030169 |
| Hectd1 | S | 252 | -0.119870216 | -0.473014834 | -0.521235749 | -0.401365534 | -0.048220916 | 0.111031817 | 2.05307E-05 | 1.18788E-05 | 0.0065994 | 0.161997696 |
| Lrrfip1 | S | 16 | -0.139936055 | -0.373594409 | -0.429539812 | -0.289603757 | -0.055945403 | 0.004600483 | 7.30494E-07 | 0.000101205 | 0.001104721 | 0.146678308 |
| Cfdp1 | T | 202 | -0.151778856 | -0.430619978 | -0.538111057 | -0.3863322 | -0.107491079 | 0.008265741 | 0.001017866 | 4.88389E-06 | 0.000389306 | 0.129595741 |
| Ncl | T | 99 | -0.087204105 | -0.389148186 | -0.528337109 | -0.441133003 | -0.139188923 | 0.013494705 | 1.73052E-07 | 0.000335044 | 0.000158664 | 0.013684828 |
| Rps4x | S | 262 | -0.050991485 | -0.40575412 | -0.425543487 | -0.374552001 | -0.019789366 | 0.325660959 | 7.06891E-07 | 4.95141E-07 | 0.005785247 | 0.298604782 |
| Ccdc86 | S | 283 | -0.005425187 | -0.522371789 | -0.57179596 | -0.566370773 | -0.049424171 | 0.852561715 | 0.000101743 | 4.1187E-07 | 9.75785E-06 | 0.258134205 |
| Dab2 | S | 227 | -0.150377361 | -0.537220921 | -0.556986923 | -0.406609562 | -0.019766002 | 0.027082864 | 2.68401E-09 | 0.004056946 | 0.008048386 | 0.775834991 |
| Ccz1 | S | 74 | -0.160723785 | -0.456593975 | -0.582004176 | -0.421280391 | -0.125410201 | 0.001277007 | 0.000375109 | 0.000287103 | 0.002287513 | 0.069310273 |
| Usp33 | S | 407 | -0.045499829 | -0.533392283 | -0.568280252 | -0.522780422 | -0.034887969 | 0.377407765 | 1.81569E-05 | 2.16066E-05 | 0.000773632 | 0.366200031 |

| | | | | | | | | | | | | |
|---|---|---|---|---|---|---|---|---|---|---|---|---|
| Tpcn1 | S | 767 | -0.02315114 | -0.395525435 | -0.47348161 | -0.45033047 | -0.077956175 | 0.516640997 | 2.34948E-06 | 3.18215E-07 | 0.000714864 | 0.022164603 |
| Pls3 | S | 339 | -0.256894517 | -0.509711979 | -0.421369686 | -0.164475169 | 0.088342293 | 0.015666598 | 0.00220952 | 6.13998E-06 | 0.063126162 | 0.217731671 |
| Cdc6 | S | 108 | -0.073018053 | -0.498045218 | -0.434157831 | -0.361139778 | 0.063887387 | 0.29669138 | 4.35825E-07 | 6.45902E-05 | 0.008623746 | 0.128303869 |
| Prrc2a | S | 759 | 0.014008052 | -0.465991646 | -0.498639553 | -0.512647605 | -0.032647907 | 0.776837325 | 1.40871E-06 | 1.40554E-06 | 0.00316636 | 0.161448645 |
| Gtpbp4 | S | 558 | -0.006898928 | -0.410371466 | -0.582055728 | -0.5751568 | -0.171684261 | 0.918075685 | 1.53162E-05 | 9.19107E-07 | 0.004326522 | 0.00490326 |
| Nifk | S | 304 | -0.061413597 | -0.45502424 | -0.520028322 | -0.458614725 | -0.065004082 | 0.186085289 | 0.001694949 | 4.04705E-08 | 0.002199638 | 0.211858507 |
| Prrc2a | S | 761 | 0.018736414 | -0.463561638 | -0.50219572 | -0.520932134 | -0.038634081 | 0.572425497 | 2.78829E-06 | 7.08709E-07 | 0.000431563 | 0.025210637 |
| Tmem87b | S | 497 | -0.301788031 | -0.519801291 | -0.430451956 | -0.128663925 | 0.089349336 | 0.007949102 | 0.00020302 | 0.001586041 | 0.161275408 | 0.266330202 |
| Rnf219 | S | 210 | -0.146589926 | -0.491577215 | -0.475522799 | -0.328932873 | 0.016054415 | 0.00016731 | 0.000304848 | 0.000762352 | 0.003220422 | 0.704834555 |
| Mybbp1a | T | 1220 | -0.146229587 | -0.500223541 | -0.490216358 | -0.343986771 | 0.010007184 | 0.147011275 | 0.000103743 | 2.17653E-05 | 0.031146191 | 0.825289703 |
| Mybbp1a | T | 1191 | -0.0120787 | -0.409531491 | -0.509498331 | -0.497419631 | -0.09996684 | 0.773882084 | 3.70058E-05 | 1.88856E-07 | 0.002632625 | 0.035855642 |
| Sqstm1 | T | 271 | -0.15928055 | -0.544438642 | -0.524086198 | -0.364805648 | 0.020352443 | 0.075044081 | 2.33249E-06 | 0.007950346 | 0.011838116 | 0.782141177 |
| Zfp106 | S | 878 | 0.008847348 | -0.435764968 | -0.468894108 | -0.477741456 | -0.03312914 | 0.783738985 | 1.65776E-07 | 1.01553E-05 | 0.000140737 | 0.239671711 |
| Ddx24 | S | 170 | -0.098050524 | -0.465552705 | -0.51854252 | -0.420491995 | -0.052989815 | 0.158649813 | 1.25403E-06 | 0.00080106 | 0.002619938 | 0.274142693 |
| Dnttip2 | S | 353 | -0.397978776 | -0.393719355 | -0.348632404 | 0.049346372 | 0.045086951 | 0.00076334 | 4.29806E-05 | 0.019124071 | 0.584960231 | 0.59301531 |
| Abl2 | S | 822 | -0.212334315 | -0.569252104 | -0.551397436 | -0.339063121 | 0.017854668 | 0.09244789 | 0.000266885 | 0.000836406 | 0.026461161 | 0.822226183 |
| Ap3d1 | S | 825 | -0.08922788 | -0.321453273 | -0.523901165 | -0.434673285 | -0.202447891 | 0.028522625 | 3.89344E-05 | 8.04816E-06 | 0.000854865 | 0.004420861 |
| Scaf8 | S | 617 | -0.069996074 | -0.493417129 | -0.393069987 | -0.323073913 | 0.100347143 | 0.033975368 | 1.85857E-06 | 0.000263792 | 0.002107281 | 0.071687955 |
| Jmy | S | 56 | -0.067292835 | -0.386236629 | -0.496398213 | -0.429105378 | -0.110161584 | 0.189956856 | 9.58383E-06 | 1.06143E-05 | 0.001039105 | 0.025173075 |
| Gpatch4 | T | 339 | -0.131995743 | -0.405524326 | -0.571192971 | -0.439197227 | -0.165668644 | 0.029646615 | 0.000127343 | 0.00015858 | 0.00393911 | 0.057743951 |
| Gnl2 | S | 647 | -0.169743556 | -0.464428814 | -0.517422573 | -0.347679018 | -0.052993759 | 0.023172883 | 0.002897698 | 1.88589E-05 | 0.002649642 | 0.477520261 |
| Bmi1 | S | 314 | -0.07233583 | -0.48224955 | -0.541192853 | -0.468857023 | -0.058943302 | 0.218759717 | 5.65242E-06 | 0.000371314 | 0.00130988 | 0.305898161 |
| Ddx21 | S | 152 | -0.001798483 | -0.480127945 | -0.568334073 | -0.56653559 | -0.088206128 | 0.980810628 | 0.00034943 | 6.6762E-07 | 0.009949555 | 0.132067215 |
| Nup50 1700123L14Ri | S | 322 | -0.248578638 | -0.507702761 | -0.500855124 | -0.252276486 | 0.006847637 | 0.109827505 | 0.000262365 | 0.000310765 | 0.120006722 | 0.876486468 |
| Prune2 | T | 2576 | -0.022927807 | -0.476823973 | -0.500150334 | -0.477222527 | -0.023326361 | 0.576604688 | 2.64674E-05 | 7.12414E-06 | 0.002859849 | 0.487435303 |
| Prrc2c | S | 899 | -0.025199624 | -0.457124811 | -0.579920299 | -0.554720675 | -0.122795488 | 0.5062848 | 9.02376E-05 | 9.42352E-06 | 0.000115503 | 0.038964129 |
| Stx12 | S | 142 | 0.021961398 | -0.510880647 | -0.513753501 | -0.535714899 | -0.002872854 | 0.499571399 | 5.64103E-06 | 1.11827E-05 | 0.00010475 | 0.729493304 |
| Map1b | S | 2267 | -0.117781124 | -0.470959481 | -0.461763471 | -0.343982347 | 0.00919601 | 0.086178655 | 9.36899E-06 | 0.000391659 | 0.004295286 | 0.854581169 |
| H2ax | S | 122 | -0.019624171 | -0.451323258 | -0.436922778 | -0.417298607 | 0.014400479 | 0.717855951 | 2.55526E-08 | 0.000350226 | 0.003717598 | 0.630948804 |
| Gja1 | S | 314 | -0.030691555 | -0.426923431 | -0.422199959 | -0.391508404 | 0.004723473 | 0.540525281 | 0.000136487 | 5.61885E-08 | 0.007998715 | 0.860088382 |
| Srrt | S | 21 | -0.2811012 | -0.43726875 | -0.41389614 | -0.13279494 | 0.02337261 | 0.001800768 | 0.001980734 | 0.000586886 | 0.074710497 | 0.759132928 |
| Nup98 | S | 839 | -0.082606607 | -0.281104808 | -0.335423622 | -0.252817015 | -0.054318814 | 0.053398652 | 2.49467E-07 | 7.53298E-07 | 0.008701348 | 0.020256378 |
| Rrp8 | S | 105 | -0.154311978 | -0.340773435 | -0.394069648 | -0.239757671 | -0.053296213 | 0.024755634 | 3.81353E-05 | 1.19302E-05 | 0.014741262 | 0.03396329 |
| Gphn | S | 338 | -0.571817433 | -0.282432888 | -0.560753874 | 0.011063558 | -0.278320986 | 0.00027244 | 0.116714394 | 0.004021029 | 0.914541311 | 0.132827943 |
| Palld | S | 988 | -0.122455108 | -0.421598142 | -0.418804353 | -0.296349245 | 0.00279379 | 0.352962741 | 1.53539E-05 | 1.16734E-05 | 0.094772717 | 0.914464592 |
| Rab3ip | S | 111 | -0.35861486 | -0.438191324 | -0.582386163 | -0.223771303 | -0.144194839 | 0.009054567 | 0.003994735 | 0.002909419 | 0.095356361 | 0.227671891 |
| Dlg5 | S | 1669 | -0.098304537 | -0.541418115 | -0.567570901 | -0.469266364 | -0.026152785 | 0.055145771 | 2.144E-06 | 0.018547524 | 0.031518466 | 0.811743 |
| Mybbp1a | S | 1303 | -0.011529163 | -0.359583293 | -0.481080879 | -0.469551715 | -0.121497586 | 0.66051986 | 1.39355E-05 | 7.09459E-07 | 5.93745E-05 | 0.015992283 |
| Sbno1 | S | 695 | -0.032338488 | -0.481626054 | -0.515537941 | -0.483199454 | -0.033911887 | 0.555027461 | 1.46587E-06 | 0.00042451 | 0.001129385 | 0.474042497 |
| Nolc1 | T | 610 | -0.092374154 | -0.459936523 | -0.519466358 | -0.427092204 | -0.059529835 | 0.030721587 | 0.000109293 | 0.000336853 | 0.000680035 | 0.229638982 |
| Nolc1 | T | 613 | -0.092374154 | -0.459936523 | -0.519466358 | -0.427092204 | -0.059529835 | 0.030721587 | 0.000109293 | 0.000336853 | 0.000680035 | 0.229638982 |
| Sh3pxd2b | S | 717 | -0.203932646 | -0.353963455 | -0.399739122 | -0.195806476 | -0.045775668 | 0.002402528 | 0.001123926 | 5.08847E-05 | 0.002527913 | 0.38419101 |
| Nolc1 | T | 600 | 0.001406976 | -0.412706934 | -0.523053751 | -0.524460727 | -0.110346817 | 0.974740637 | 5.03171E-06 | 1.60711E-05 | 0.00029174 | 0.042553893 |
| Trip12 | S | 340 | -0.124493344 | -0.397308659 | -0.520528008 | -0.396034664 | -0.12321935 | 0.030070544 | 0.000123773 | 0.000247022 | 0.001973592 | 0.07748914 |
| Rrp1b | S | 528 | -0.052160503 | -0.415690769 | -0.407020881 | -0.354860378 | 0.008669888 | 0.292977986 | 8.11891E-06 | 1.11369E-05 | 0.001756433 | 0.743844076 |
| Tacc3 | S | 71 | -0.006823742 | -0.347596154 | -0.460760564 | -0.453936823 | -0.113164411 | 0.875114983 | 5.35528E-06 | 9.74161E-08 | 0.003282761 | 0.014058568 |
| Trip12 | T | 618 | -0.300650073 | -0.395419875 | -0.443779145 | -0.143129072 | -0.04835927 | 0.006288627 | 0.007029758 | 0.000210951 | 0.105186539 | 0.533738836 |
| Rpl6 | S | 21 | -0.07384605 | -0.376823397 | -0.352060859 | -0.278214809 | 0.024762538 | 0.070444839 | 2.24581E-05 | 2.82666E-06 | 0.001639179 | 0.368782396 |
| Clip2 | S | 43 | -0.017983384 | -0.395879199 | -0.511651192 | -0.493667808 | -0.115771993 | 0.798620937 | 2.05547E-05 | 8.85245E-06 | 0.005114111 | 0.030053484 |
| Lmna | S | 424 | -0.043890441 | -0.429394528 | -0.386830792 | -0.342940351 | 0.042563736 | 0.144605072 | 6.60567E-05 | 2.18071E-06 | 0.000122338 | 0.239175083 |
| Trip12 | S | 1024 | -0.045814687 | -0.309069403 | -0.405715961 | -0.359901275 | -0.096646558 | 0.202398071 | 5.83237E-06 | 9.99331E-07 | 0.000650518 | 0.010126509 |
| Tnks1bp1 | S | 1643 | -0.019834299 | -0.437161682 | -0.494512361 | -0.474678062 | -0.057350679 | 0.647915896 | 3.30185E-05 | 1.4803E-05 | 0.000893185 | 0.134057364 |
| Polm | S | 12 | -0.013218615 | -0.493040963 | -0.485409444 | -0.472190829 | 0.00763152 | 0.806487187 | 1.58598E-05 | 6.54435E-05 | 0.00139555 | 0.85663535 |
| Polr3g | S | 158 | -0.030032538 | -0.437089473 | -0.379159012 | -0.349126474 | 0.057930461 | 0.441285057 | 0.000113622 | 2.53134E-07 | 0.004219637 | 0.125807806 |
| Rpl19 | S | 189 | -0.185896696 | -0.496445671 | -0.442176942 | -0.256280246 | 0.054268729 | 0.006757668 | 0.001978242 | 0.000565092 | 0.006826789 | 0.425828787 |
| Ddx50 | S | 41 | -0.449925864 | -0.506340458 | -0.554449248 | -0.104523384 | -0.04810879 | 0.261889456 | 0.001796755 | 0.000954523 | 0.758231571 | 0.469509111 |
| Eaf1 | S | 158 | -0.025378752 | -0.449937976 | -0.439970248 | -0.414591497 | 0.009967727 | 0.631037559 | 3.23614E-06 | 8.23649E-05 | 0.001546579 | 0.785225944 |
| Sipa1l3 | S | 1169 | -0.027293149 | -0.382246645 | -0.417689764 | -0.390396615 | -0.035443119 | 0.539557247 | 1.26098E-05 | 2.52506E-06 | 0.001923074 | 0.249316945 |
| Tpr | S | 2204 | -0.201140552 | -0.528493251 | -0.56918496 | -0.368044408 | -0.040691709 | 0.024942895 | 0.000125616 | 0.021099733 | 0.068619823 | 0.755276011 |
| Arfgap1 | S | 342 | -0.156750046 | -0.424211949 | -0.503376934 | -0.346626887 | -0.079164985 | 0.058759855 | 0.001952865 | 5.42709E-05 | 0.005621211 | 0.291703714 |
| Itprid2 | S | 299 | -0.38908318 | -0.389737903 | -0.548379834 | -0.159296653 | -0.158641931 | 0.002755799 | 0.015953283 | 0.004318689 | 0.191901665 | 0.249625127 |
| Ddx24 | T | 169 | -0.03785798 | -0.481025366 | -0.563241648 | -0.525383668 | -0.082216283 | 0.593042559 | 0.000217853 | 5.5442E-05 | 0.000572549 | 0.07227631 |
| Uck2 | S | 254 | -0.089291772 | -0.424537891 | -0.516572568 | -0.427280795 | -0.092034677 | 0.177101396 | 9.28378E-06 | 0.000992028 | 0.002413275 | 0.161459929 |
| Pdpk1 | S | 38 | -0.081312371 | -0.523728879 | -0.541271376 | -0.459959005 | -0.017542496 | 0.000706891 | 0.000862937 | 0.002347252 | 0.003445845 | 0.700690387 |
| Eif4g3 | S | 230 | -0.204102843 | -0.399876011 | -0.461855362 | -0.257752519 | -0.06197935 | 0.014794987 | 0.002339444 | 0.000151356 | 0.004770046 | 0.202056443 |
| Ube2o | S | 82 | -0.068200868 | -0.337680325 | -0.419103251 | -0.350902382 | -0.081422926 | 0.277521594 | 2.06169E-05 | 3.61508E-06 | 0.008853834 | 0.03867405 |
| Ube2o | S | 84 | -0.068200868 | -0.337680325 | -0.419103251 | -0.350902382 | -0.081422926 | 0.277521594 | 2.06169E-05 | 3.61508E-06 | 0.008853834 | 0.03867405 |
| Fgd6 | S | 1262 | -0.114543062 | -0.46828419 | -0.478278288 | -0.363735226 | -0.009994098 | 0.183489375 | 0.000319919 | 9.02514E-05 | 0.009681875 | 0.867541625 |
| Map2k3 | T | 39 | -0.123615505 | -0.362713019 | -0.45932806 | -0.335712555 | -0.096615041 | 0.000683811 | 1.73482E-05 | 0.005500253 | 0.018480149 | 0.180543486 |
| Bsg | S | 372 | -0.274240245 | -0.526725553 | -0.453860638 | -0.179620394 | 0.072864914 | 0.034085699 | 0.001317967 | 0.002414661 | 0.074829364 | 0.116704597 |
| Rrn3 | S | 168 | 0.015464735 | -0.464262617 | -0.515188996 | -0.53065373 | -0.050926379 | 0.733970381 | 5.49665E-05 | 1.08058E-05 | 0.000106928 | 0.283065201 |
| Uvrag | S | 549 | -0.117265292 | -0.574090159 | -0.542609975 | -0.425344683 | 0.031480184 | 0.005359361 | 0.000626084 | 0.006996073 | 0.015681378 | 0.699310449 |
| Ngef | S | 696 | -0.444638208 | -0.304246505 | -0.397243856 | 0.047394351 | -0.092997351 | 0.000198099 | 0.015342952 | 0.007761206 | 0.571718759 | 0.377807539 |
| Ranbp2 | S | 1600 | -0.373989107 | -0.509629215 | -0.493012432 | -0.119023325 | 0.016616783 | 0.000969512 | 0.01530392 | 0.020238086 | 0.340775278 | 0.904848324 |
| Larp4 | S | 69 | -0.143760963 | -0.329572922 | -0.458127286 | -0.314366323 | -0.128554364 | 0.012579829 | 0.002915043 | 1.31982E-05 | 0.001013646 | 0.041141263 |
| Bag3 | S | 360 | -0.256419782 | -0.285736042 | -0.313955065 | -0.057535282 | -0.028219023 | 0.000270582 | 0.001966953 | 0.000149871 | 0.137678297 | 0.56535648 |
| Nsrp1 | S | 442 | 0.089640105 | -0.454703619 | -0.46534115 | -0.554981255 | -0.010637531 | 0.207370781 | 4.74533E-06 | 2.2149E-06 | 0.004338498 | 0.711950599 |
| Nsrp1 | S | 443 | 0.089640105 | -0.454703619 | -0.46534115 | -0.554981255 | -0.010637531 | 0.207370781 | 4.74533E-06 | 2.2149E-06 | 0.004338498 | 0.711950599 |
| Riok2 | S | 478 | -0.142545213 | -0.359778609 | -0.425255325 | -0.282710112 | -0.065476716 | 0.023311856 | 0.000510899 | 4.87524E-05 | 0.011738452 | 0.206667239 |
| Nav3 | S | 1193 | -0.212156019 | -0.346865982 | -0.246439134 | -0.034283115 | 0.100426848 | 0.000806233 | 6.65849E-06 | 0.003812026 | 0.462962806 | 0.085241633 |
| Tut1 | S | 655 | 0.044087622 | -0.434666335 | -0.54027801 | -0.584365633 | -0.105611675 | 0.263392471 | 2.27444E-05 | 8.79303E-06 | 0.000145659 | 0.04930439 |
| Luzp1 | S | 866 | -0.152115027 | -0.392980006 | -0.376276556 | -0.22416153 | 0.016703449 | 0.009833018 | 0.000996617 | 4.97404E-05 | 0.003560694 | 0.751839818 |
| Gja1 | S | 372 | -0.113540795 | -0.400106091 | -0.538494841 | -0.424954046 | -0.13838875 | 0.162734411 | 0.005028085 | 8.7581E-06 | 0.006635594 | 0.114955751 |
| Pin4 | S | 23 | -0.214996601 | -0.405098617 | -0.497524472 | -0.28252787 | -0.092425854 | 0.008261643 | 0.003781395 | 0.000767189 | 0.014542859 | 0.303802022 |
| Cdca3 | S | 220 | -0.327465075 | -0.37406908 | -0.504804028 | -0.177338953 | -0.130734948 | 0.005083384 | 0.024931367 | 0.000687318 | 0.056515963 | 0.287745315 |
| Nmt1 | S | 44 | -0.138027816 | -0.317406323 | -0.515695535 | -0.377667719 | -0.198289212 | 0.117231771 | 0.000276508 | 5.40879E-05 | 0.021781468 | 0.006717198 |
| Polq | S | 1572 | 0.00865385 | -0.367591221 | -0.49377111 | -0.502424961 | -0.126179889 | 0.803119011 | 1.15042E-05 | 8.67033E-06 | 0.000121292 | 0.024498603 |
| Ranbp3 | S | 283 | -0.180930166 | -0.264824306 | -0.32865007 | -0.147719904 | -0.063825763 | 0.00076257 | 0.000135481 | 0.000135722 | 0.014782526 | 0.131798113 |
| Fasn | S | 207 | -0.092932135 | -0.433911953 | -0.404743229 | -0.311811094 | 0.029168724 | 0.036006734 | 6.53313E-06 | 0.002629835 | 0.007296355 | 0.590172463 |
| Lrrfip1 | S | 547 | -0.083616848 | -0.28899244 | -0.368392822 | -0.284775974 | -0.079400381 | 0.003933798 | 4.71767E-07 | 0.000622282 | 0.002014965 | 0.070077092 |
| Rrn3 | S | 170 | 0.013781765 | -0.43110912 | -0.489669776 | -0.503451541 | -0.058560656 | 0.740705393 | 5.07938E-05 | 9.76255E-06 | 0.000131141 | 0.205466768 |
| Psma5 | S | 16 | -0.140604853 | -0.475973514 | -0.551832383 | -0.411227529 | -0.075858869 | 0.025286304 | 0.001136457 | 0.001910875 | 0.00584841 | 0.354479097 |
| Eif3c | S | 204 | -0.104952513 | -0.474443282 | -0.486469939 | -0.381517426 | -0.012026657 | 0.127253038 | 0.003496819 | 2.57982E-05 | 0.010994259 | 0.858348371 |
| Sipa1l3 | S | 1154 | 0.054271845 | -0.336624194 | -0.462911321 | -0.517183166 | -0.126287127 | 0.106385674 | 3.86649E-06 | 2.13182E-07 | 0.000427997 | 0.005001891 |

| | | | | | | | | | | | | |
|---|---|---|---|---|---|---|---|---|---|---|---|---|
| Ranbp3 | S | 148 | -0.082197137 | -0.348757396 | -0.386042716 | -0.303845578 | -0.037285319 | 0.020947691 | 1.37865E-05 | 0.000184815 | 0.000470065 | 0.237542472 |
| Dync1h1 | T | 4367 | -0.041613852 | -0.419530986 | -0.535317175 | -0.493703323 | -0.115786188 | 0.327380037 | 1.23659E-06 | 0.003203278 | 0.002741091 | 0.148280691 |
| Larp1 | T | 822 | -0.054948641 | -0.43892344 | -0.407663233 | -0.352714592 | 0.031260207 | 0.037813493 | 1.83106E-05 | 0.000575513 | 0.000215445 | 0.25860119 |
| Jpt2 | S | 97 | 0.001596639 | -0.454359756 | -0.55090945 | -0.55250609 | -0.096549694 | 0.976381227 | 1.88962E-05 | 0.000251611 | 0.000771367 | 0.095176922 |
| Eif4g1 | S | 1189 | -0.109376845 | -0.333087578 | -0.397659728 | -0.288282883 | -0.06457215 | 0.079631146 | 5.69359E-05 | 3.5082E-05 | 0.016697227 | 0.103624697 |
| Glyr1 | S | 130 | -0.081529672 | -0.327735772 | -0.461024364 | -0.379494692 | -0.133288593 | 0.007158573 | 0.000355441 | 7.52196E-05 | 0.000623563 | 0.016946004 |
| Eif5b | S | 571 | 0.008393862 | -0.497200863 | -0.545621262 | -0.554015124 | -0.048420399 | 0.859625533 | 2.10769E-05 | 0.000430158 | 0.00116759 | 0.432541528 |
| Bod1l | S | 864 | -0.165672006 | -0.53637057 | -0.439737711 | -0.274065705 | 0.096632859 | 0.057873285 | 4.24024E-05 | 0.019816656 | 0.048667895 | 0.336871783 |
| Tnks1bp1 | S | 214 | -0.083802896 | -0.348146295 | -0.537103308 | -0.453300411 | -0.188957013 | 0.013717316 | 0.00146747 | 0.000112891 | 0.003955554 | 0.016431417 |
| Fkbp15 | S | 344 | -0.037477679 | -0.422427248 | -0.504306917 | -0.466829238 | -0.081879669 | 0.781426665 | 0.000119741 | 4.10518E-05 | 0.049580594 | 0.05148961 |
| Zfp36l2 | S | 127 | -0.047276958 | -0.478924194 | -0.584733572 | -0.537456615 | -0.105809379 | 0.448660607 | 0.00051274 | 0.000241982 | 0.001050735 | 0.157194153 |
| Tpr | S | 2223 | -0.174634309 | -0.450157742 | -0.482765797 | -0.308131488 | -0.032608056 | 0.034734551 | 0.008459109 | 0.000138784 | 0.017197678 | 0.705300169 |
| Cenpe | S | 2383 | 0.022249546 | -0.351496565 | -0.446741787 | -0.468991333 | -0.095245222 | 0.371423254 | 1.44168E-05 | 1.17215E-06 | 0.000502787 | 0.021416115 |
| Ap2m1 | T | 229 | 0.023702096 | -0.441750159 | -0.515979206 | -0.539681301 | -0.074229047 | 0.787709193 | 4.43492E-05 | 4.94153E-05 | 0.007389897 | 0.1580167 |
| Eif4g1 | S | 15 | -0.169177829 | -0.317979553 | -0.434286897 | -0.265109069 | -0.116307345 | 0.005979307 | 0.000223112 | 0.001128629 | 0.014942055 | 0.113353603 |
| Sipa1l3 | S | 1129 | -0.100449348 | -0.344608444 | -0.393921156 | -0.293471808 | -0.049312712 | 0.210793216 | 9.18135E-06 | 0.000118816 | 0.020000006 | 0.224347418 |
| Mff | S | 146 | -0.246755386 | -0.324106201 | -0.467726174 | -0.220970787 | -0.143619973 | 0.016363141 | 0.006453042 | 0.000207536 | 0.045817631 | 0.115314194 |
| Hsph1 | T | 816 | -0.09799677 | -0.43460798 | -0.406309436 | -0.308312666 | 0.028298544 | 0.297616807 | 0.002486853 | 1.85642E-06 | 0.042999695 | 0.593411224 |
| Larp4 | S | 626 | -0.076899161 | -0.452301575 | -0.405015758 | -0.328116597 | 0.047285817 | 0.296895793 | 9.98023E-06 | 0.000903107 | 0.01062607 | 0.341739758 |
| Tanc1 | S | 462 | -0.044386904 | -0.429607793 | -0.400226013 | -0.355839108 | 0.02938178 | 0.328609998 | 4.02289E-05 | 6.44332E-05 | 0.0024994 | 0.373533052 |
| Gpatch4 | S | 130 | -0.093818629 | -0.470771513 | -0.416717325 | -0.322898695 | 0.054054188 | 0.182815131 | 2.49019E-05 | 0.001623482 | 0.0086999 | 0.401166405 |
| Spata5 | S | 244 | -0.150121425 | -0.38753304 | -0.351433244 | -0.201311819 | 0.036099796 | 0.034324688 | 7.34595E-06 | 0.003361135 | 0.027692449 | 0.504587631 |
| Errfi1 | S | 301 | -0.1910463 | -0.423891562 | -0.440948677 | -0.249902377 | -0.017057115 | 0.048574714 | 1.7681E-05 | 0.015997714 | 0.066217422 | 0.849798933 |
| Ercc6l | S | 942 | -0.111193983 | -0.376521265 | -0.392073464 | -0.280879481 | -0.015552198 | 0.037066414 | 0.000591508 | 4.79604E-05 | 0.000702201 | 0.748955786 |
| Nmd3 | T | 433 | -0.215941486 | -0.510202634 | -0.555646841 | -0.339705355 | -0.045444207 | 0.256258328 | 0.000755254 | 0.003054885 | 0.118666053 | 0.612628056 |
| Errfi1 | S | 329 | -0.109693904 | -0.498009552 | -0.489187383 | -0.379493479 | 0.008822169 | 0.18541783 | 0.000701557 | 0.000553138 | 0.007017087 | 0.894306289 |
| Gja1 | S | 262 | -0.009805728 | -0.400028721 | -0.56446766 | -0.554661932 | -0.164438939 | 0.88301084 | 6.45096E-05 | 0.000130621 | 0.001422002 | 0.03338289 |
| Larp1 | S | 751 | 0.002829538 | -0.42138088 | -0.397300622 | -0.40013016 | 0.024080257 | 0.948557499 | 1.51253E-05 | 1.58802E-05 | 0.003377009 | 0.096207326 |
| Errfi1 | S | 333 | -0.089100546 | -0.499891253 | -0.477967042 | -0.388866496 | 0.021924211 | 0.240718048 | 0.0004373 | 0.000449559 | 0.004602423 | 0.70006522 |
| Eml4 | S | 891 | -0.031138809 | -0.42665237 | -0.354375585 | -0.323236776 | 0.072276785 | 0.517727691 | 6.86363E-06 | 6.69874E-05 | 0.001898166 | 0.088331199 |
| Eml4 | T | 888 | -0.031138809 | -0.42665237 | -0.354375585 | -0.323236776 | 0.072276785 | 0.517727691 | 6.86363E-06 | 6.69874E-05 | 0.001898166 | 0.088331199 |
| Usp15 | T | 226 | -0.109898251 | -0.359529213 | -0.387562035 | -0.277663784 | -0.028032821 | 0.071684065 | 0.000281276 | 3.74294E-05 | 0.005471846 | 0.49253224 |
| Map1b | S | 1391 | -0.146982212 | -0.334195463 | -0.314039424 | -0.167057212 | 0.020156039 | 0.006984911 | 0.000200452 | 0.00010347 | 0.002009985 | 0.364945843 |
| Mapre2 | S | 207 | -0.05410176 | -0.333842499 | -0.501374806 | -0.447273046 | -0.167532107 | 0.133514077 | 2.71681E-06 | 0.001180501 | 0.001345205 | 0.043690151 |
| Ankrd17 Muc20 | S | 1705;618 | -0.049640031 | -0.293217459 | -0.349003759 | -0.299363728 | -0.0557863 | 0.060239024 | 1.79391E-05 | 4.58474E-06 | 3.53941E-06 | 0.002358649 |
| Sgip1 | S | 558 | -0.444227597 | -0.446949015 | -0.561606554 | -0.117378957 | -0.114657539 | 0.001362235 | 0.105341102 | 0.01760694 | 0.406809866 | 0.603657465 |
| Ctdspl2 | S | 28 | -0.133677937 | -0.441441609 | -0.421684979 | -0.288007042 | 0.019756631 | 0.18872975 | 3.6371E-06 | 0.009737973 | 0.038861299 | 0.786171648 |
| Rps27 Rps27rt | S | 11 | -0.026253958 | -0.404875906 | -0.35853581 | -0.332281852 | 0.046340096 | 0.601234581 | 6.07302E-05 | 3.60529E-06 | 0.007591183 | 0.187082026 |
| Tnks1bp1 | S | 178 | -0.062924022 | -0.398587271 | -0.518811935 | -0.455887912 | -0.120224664 | 0.171416006 | 0.000140115 | 0.000434989 | 0.000655574 | 0.055670619 |
| Dsn1 | S | 79 | -0.001314478 | -0.422651831 | -0.530478445 | -0.529163967 | -0.107826615 | 0.946517377 | 0.001335341 | 5.43491E-06 | 3.46358E-05 | 0.050430317 |
| Kpna2 | S | 62 | -0.01924763 | -0.254090347 | -0.465323985 | -0.446076355 | -0.211233638 | 0.552190012 | 3.15076E-05 | 1.69191E-06 | 0.000781679 | 0.000210132 |
| Pdlim2 | S | 117 | 0.031511766 | -0.44823704 | -0.507737794 | -0.53924956 | -0.059500754 | 0.814676976 | 0.000134712 | 3.09997E-05 | 0.035745055 | 0.266623283 |
| Tax1bp1 | S | 632 | -0.062510888 | -0.423779102 | -0.389907779 | -0.32739689 | 0.033871323 | 0.086420226 | 0.00079227 | 1.49802E-05 | 0.000369635 | 0.476539518 |
| Plec | S | 1055 | -0.06631009 | -0.578136699 | -0.527651604 | -0.461341514 | 0.050485095 | 0.554124466 | 0.000543943 | 0.000991395 | 0.009152465 | 0.47821119 |
| Nolc1 | S | 394 | -0.019045617 | -0.434943494 | -0.465511543 | -0.446465926 | -0.030568049 | 0.71483084 | 5.74688E-06 | 0.000868801 | 0.001606694 | 0.559640476 |
| Pnn | S | 702 | 0.005125977 | -0.361247714 | -0.423871937 | -0.428997914 | -0.062624224 | 0.845984422 | 4.87532E-06 | 2.77856E-05 | 0.002300042 | 0.111182751 |
| Bclaf1 | T | 724 | -0.06233164 | -0.462099322 | -0.502381689 | -0.440050049 | -0.040282367 | 0.436900442 | 0.011033822 | 3.57298E-06 | 0.016933784 | 0.625085521 |
| Plec | S | 4393 | -0.04965624 | -0.276945983 | -0.495482685 | -0.445826445 | -0.218536702 | 0.122803672 | 0.000126885 | 2.81531E-05 | 0.003798225 | 0.01518552 |
| 2310022B05Rik | S | 278 | -0.258186599 | -0.343879872 | -0.328310748 | -0.070124149 | 0.015569124 | 0.050324819 | 0.000364243 | 0.000255265 | 0.447958248 | 0.691486427 |
| Map1b | S | 1395 | -0.14245935 | -0.31467343 | -0.298570339 | -0.15611099 | 0.016103091 | 0.006094006 | 0.000167442 | 8.94757E-05 | 0.003011307 | 0.45476899 |
| Phrf1 | S | 622 | -0.047088932 | -0.333083926 | -0.353716398 | -0.306627466 | -0.020632472 | 0.460831679 | 7.00068E-06 | 1.56754E-05 | 0.015539661 | 0.4431194 |
| Cep170 | S | 702 | -0.07273384 | -0.416968324 | -0.535869366 | -0.463135525 | -0.118901042 | 0.186322167 | 0.00176808 | 0.000139817 | 0.00104433 | 0.135570756 |
| Gja1 | S | 330 | -0.027722228 | -0.392588042 | -0.563218886 | -0.535496658 | -0.170630845 | 0.510636389 | 7.28794E-05 | 0.000448925 | 0.004299579 | 0.0658155 |
| Ajuba | S | 147 | -0.082840559 | -0.310103747 | -0.433804206 | -0.350963646 | -0.123700458 | 0.097667222 | 9.16252E-05 | 8.46586E-05 | 0.001213243 | 0.042116532 |
| Phldb2 | T | 546 | -0.011003129 | -0.557492412 | -0.535837165 | -0.524834036 | 0.021655247 | 0.890611067 | 0.00052975 | 0.000347692 | 0.000197928 | 0.509125039 |
| Csde1 | S | 445 | -0.279546619 | -0.391332608 | -0.536538662 | -0.256992043 | -0.145206054 | 0.02942823 | 0.000596429 | 0.019568001 | 0.116879666 | 0.293020801 |
| Usp10 | S | 359 | -0.059087989 | -0.330988053 | -0.314587865 | -0.255499876 | 0.016400188 | 0.120392657 | 6.74668E-06 | 3.03652E-05 | 0.007851599 | 0.410605473 |
| Errfi1 | T | 331 | -0.097994039 | -0.533757815 | -0.434703648 | -0.336709609 | 0.099054166 | 0.18503366 | 0.000910481 | 0.000559586 | 0.007162705 | 0.241022063 |
| Tnks1bp1 | S | 1437 | -0.159128933 | -0.441431256 | -0.539071158 | -0.379942226 | -0.097639903 | 0.022547789 | 0.000107705 | 0.032974033 | 0.089417622 | 0.510844671 |
| Prrc2b | S | 479 | -0.135632742 | -0.319867 | -0.28807362 | -0.152440878 | 0.031793381 | 0.012258167 | 4.87085E-05 | 0.000181155 | 0.01982885 | 0.214797268 |
| Ubap2 | S | 113 | -0.222021119 | -0.389825422 | -0.573810973 | -0.351789853 | -0.18398555 | 0.035190475 | 0.019695594 | 0.000774681 | 0.012856858 | 0.168830862 |
| Knop1 | T | 141 | -0.020429411 | -0.347070118 | -0.387587906 | -0.367158495 | -0.040517789 | 0.345362677 | 4.88825E-07 | 0.000432982 | 0.001211631 | 0.254897064 |
| Nup35 | S | 260 | -0.330973858 | -0.370654677 | -0.474023949 | -0.143050091 | -0.103369271 | 0.209315889 | 0.0067425 | 0.000153967 | 0.522013665 | 0.239845727 |
| Cpeb4 | S | 97 | -0.044529006 | -0.359522548 | -0.29971124 | -0.255182234 | 0.059811307 | 0.139224627 | 1.49018E-05 | 1.55717E-05 | 0.000293458 | 0.079191126 |
| Tmem200a | S | 350 | -0.170532839 | -0.381683998 | -0.483262643 | -0.312729804 | -0.101578645 | 0.083739791 | 0.016472375 | 4.04119E-05 | 0.028077472 | 0.309384879 |
| Nvl | S | 214 | -0.105327665 | -0.434573025 | -0.563383122 | -0.458055457 | -0.128810097 | 0.022130413 | 0.01446523 | 0.000339414 | 0.000367449 | 0.157546899 |
| Atp7a | S | 1454 | -0.435650147 | -0.363395762 | -0.543193501 | -0.107543354 | -0.179797739 | 0.011267962 | 0.076344414 | 0.003730126 | 0.379448076 | 0.299449096 |
| Atp7a | S | 1457 | -0.435650147 | -0.363395762 | -0.543193501 | -0.107543354 | -0.179797739 | 0.011267962 | 0.076344414 | 0.003730126 | 0.379448076 | 0.299449096 |
| Prrc2a | S | 764 | 0.028339683 | -0.425589487 | -0.463548381 | -0.491888065 | -0.037958894 | 0.579388497 | 1.71111E-05 | 9.64007E-05 | 0.000552757 | 0.413581055 |
| Ddx10 | T | 831 | -0.024451764 | -0.384108858 | -0.446584083 | -0.422132318 | -0.062475225 | 0.590477056 | 0.000282955 | 1.58007E-05 | 0.000232164 | 0.219007635 |
| Phax | S | 155 | -0.441897239 | -0.36397626 | -0.494649578 | -0.052752339 | -0.130673318 | 0.002047423 | 0.038664978 | 0.023296524 | 0.697310737 | 0.437915921 |
| Agap3 | S | 478 | -0.082440106 | -0.394923848 | -0.468229527 | -0.385789421 | -0.07330568 | 0.046037726 | 0.012851984 | 1.41791E-05 | 0.000328332 | 0.270698967 |
| Mybbp1a | T | 1277 | -0.002306741 | -0.371845818 | -0.457989492 | -0.455682751 | -0.086143674 | 0.952763968 | 1.00916E-05 | 0.000136636 | 0.000498705 | 0.10482756 |
| Mybbp1a | S | 1323 | 0.022800926 | -0.340301464 | -0.432439872 | -0.455240798 | -0.092138407 | 0.575950931 | 2.35301E-05 | 4.61314E-06 | 0.000310615 | 0.017918807 |
| Sde2 | S | 204 | -0.119389683 | -0.369420753 | -0.496127091 | -0.376737408 | -0.126706338 | 0.198070149 | 0.007218284 | 2.19359E-05 | 0.022774071 | 0.161929051 |
| Smtn | S | 304 | 0.042637525 | -0.398528829 | -0.454001162 | -0.496638687 | -0.055472332 | 0.131632405 | 1.70325E-06 | 6.72979E-05 | 0.001509769 | 0.171257403 |
| Ddx21 | S | 144 | -0.003809982 | -0.327954102 | -0.547527935 | -0.543717954 | -0.219573833 | 0.949566668 | 0.000132097 | 3.54922E-05 | 0.000896517 | 0.008271276 |
| Dync1li1 | S | 421 | -0.076815215 | -0.318795592 | -0.476149111 | -0.399333897 | -0.157353519 | 0.04362177 | 7.09387E-06 | 0.003174238 | 0.004651402 | 0.07076198 |
| Tacc2 | S | 763 | -0.059806748 | -0.370340414 | -0.356329704 | -0.296522956 | 0.01401071 | 0.157111241 | 6.62688E-06 | 0.000461119 | 0.001835365 | 0.707050344 |
| Baz1a | S | 1302 | -0.085881815 | -0.414453781 | -0.581128162 | -0.495246347 | -0.166674381 | 0.305971767 | 8.66144E-05 | 0.003832459 | 0.006708755 | 0.141939587 |
| Thoc2 | S | 1222 | -0.259116771 | -0.203009599 | -0.371299312 | -0.112182541 | -0.168289713 | 0.000265141 | 0.146602468 | 2.08673E-05 | 0.019924444 | 0.204361898 |
| 8030462N17Rik | T | 149 | 0.017173646 | -0.539171569 | -0.515247311 | -0.532420957 | 0.023924259 | 0.711647512 | 8.9267E-06 | 0.005076235 | 0.009487707 | 0.760286486 |
| Mindy1 | S | 382 | -0.205126765 | -0.425837448 | -0.508570308 | -0.303443543 | -0.08273286 | 0.063484272 | 0.000450723 | 0.01148501 | 0.025345525 | 0.328949209 |
| Atxn2l | S | 597 | -0.032819036 | -0.221417625 | -0.225341516 | -0.19252248 | -0.003923891 | 0.025742794 | 1.44386E-07 | 2.36454E-07 | 0.000188316 | 0.59239247 |
| Lrrfip1 | S | 302 | -0.055844846 | -0.300365061 | -0.299003756 | -0.243158909 | 0.001361306 | 0.156145244 | 0.000232666 | 1.65449E-07 | 0.007173434 | 0.944192014 |
| Tsc2 | S | 960 | -0.149895671 | -0.349727743 | -0.424034649 | -0.274138978 | -0.074306906 | 0.018715755 | 0.000489404 | 0.001488688 | 0.008456126 | 0.244894639 |
| Slc20a1 | S | 269 | 0.035008117 | -0.554125216 | -0.463645219 | -0.498653336 | 0.090479997 | 0.21153218 | 3.31078E-05 | 0.00041312 | 0.000632787 | 0.087161272 |
| Fbxw11 | S | 65 | -0.106348356 | -0.317751889 | -0.408928193 | -0.302579837 | -0.091176304 | 0.003554379 | 0.003785632 | 8.93339E-05 | 0.006499425 | 0.126186886 |
| Dis3l2 | S | 31 | 0.060346101 | -0.273540182 | -0.325228205 | -0.385574306 | -0.051688023 | 0.215986467 | 2.22482E-07 | 7.37375E-08 | 0.006951867 | 0.0021114 |
| Zeb1 Zeb2 | S | 933;1050 | -0.105632514 | -0.315229839 | -0.4375573 | -0.331924786 | -0.12232746 | 0.284309057 | 0.006264602 | 1.47459E-06 | 0.041772823 | 0.100958044 |
| Wapl | T | 286 | -0.44252788 | -0.420615627 | -0.546929049 | -0.104401169 | -0.126313422 | 0.016756282 | 0.02811139 | 0.018199494 | 0.524819035 | 0.46482152 |
| Map7d1 | S | 460 | -0.060110068 | -0.379483394 | -0.450549393 | -0.390439326 | -0.071066089 | 0.154518325 | 0.000600817 | 0.000100422 | 0.000638727 | 0.188807426 |
| Cenpe | T | 422 | -0.128396062 | -0.440479693 | -0.541484788 | -0.413088725 | -0.101005094 | 0.118685599 | 0.002928736 | 0.001081876 | 0.005179246 | 0.24782926 |
| Rpl18a | S | 123 | -0.062439751 | -0.395008951 | -0.503169629 | -0.440729878 | -0.108160678 | 0.352087623 | 0.003362095 | 2.96131E-05 | 0.005637254 | 0.096882037 |

| | | | | | | | | | | | | |
|---|---|---|---|---|---|---|---|---|---|---|---|---|
| Gemin5 | S | 757 | -0.300805763 | -0.257872881 | -0.376612449 | -0.075806686 | -0.118739568 | 0.001548245 | 0.073957654 | 0.000136607 | 0.175571907 | 0.300581873 |
| Ddx21 | S | 218 | 0.039000482 | -0.337777873 | -0.444383161 | -0.483383642 | -0.106605287 | 0.608688152 | 3.89958E-05 | 4.86394E-06 | 0.012706269 | 0.018553811 |
| Tbc1d1 | S | 565 | -0.116794562 | -0.359371853 | -0.401833279 | -0.285038717 | -0.042461426 | 0.039652482 | 0.000153431 | 0.001132439 | 0.013615435 | 0.478167089 |
| Tbc1d1 | S | 567 | -0.116794562 | -0.359371853 | -0.401833279 | -0.285038717 | -0.042461426 | 0.039652482 | 0.000153431 | 0.001132439 | 0.013615435 | 0.478167089 |
| Aif1l | S | 134 | -0.204811804 | -0.286974111 | -0.302264764 | -0.09745296 | -0.015290653 | 0.001865343 | 0.001934487 | 0.000304459 | 0.049219721 | 0.757620049 |
| Rprd2 | T | 844 | -0.116340443 | -0.475781405 | -0.496566429 | -0.380225986 | -0.020785024 | 0.172096735 | 0.009706445 | 0.000139019 | 0.006764596 | 0.831319328 |
| Wfs1 | S | 32 | -0.477887357 | -0.336253782 | -0.302438908 | 0.175448449 | 0.033814874 | 0.000193532 | 0.007781331 | 0.165331322 | 0.356929891 | 0.84727563 |
| Chmp7 | S | 442 | -0.144147056 | -0.44397734 | -0.570512707 | -0.426365651 | -0.126535366 | 0.185113065 | 0.00328553 | 0.001317658 | 0.008226642 | 0.179979288 |
| Tnks1bp1 | S | 1440 | -0.175222513 | -0.483998839 | -0.362687988 | -0.187465475 | 0.121310851 | 0.04554891 | 0.000440667 | 0.006123367 | 0.074192986 | 0.178624609 |
| Zfand3 | S | 122 | -0.047673332 | -0.277328514 | -0.434547031 | -0.386873699 | -0.157218517 | 0.391955993 | 0.000556017 | 2.29613E-06 | 0.006043186 | 0.008348124 |
| Sipa1l1 | T | 1562 | -0.086538522 | -0.254151529 | -0.389368958 | -0.302830437 | -0.135217429 | 0.262073349 | 0.000112689 | 5.80387E-06 | 0.027361617 | 0.002012072 |
| Cep170 | T | 1376 | -0.038197089 | -0.360085983 | -0.45149108 | -0.413293991 | -0.091405097 | 0.551724503 | 0.000281815 | 3.34896E-05 | 0.004272461 | 0.10162922 |
| Espl1 | S | 1391 | -0.219531343 | -0.346067022 | -0.528809548 | -0.309278205 | -0.182742525 | 0.002797904 | 0.001252536 | 0.029410107 | 0.089941924 | 0.220277241 |
| Tnks1bp1 | S | 692 | -0.061637586 | -0.270663971 | -0.365590431 | -0.303952845 | -0.09492646 | 0.123285795 | 7.23342E-06 | 6.79967E-05 | 0.00097435 | 0.039277497 |
| Ncl | T | 59 | 0.048080254 | -0.35638324 | -0.528750484 | -0.576830739 | -0.172367244 | 0.319286764 | 0.000180096 | 7.17607E-06 | 0.000244295 | 0.011406837 |
| Macf1 | S | 1500 | -0.466660888 | -0.399303429 | -0.400690977 | 0.065969911 | -0.001387549 | 0.114322868 | 0.017715262 | 0.000498719 | 0.75128118 | 0.98844312 |
| Nav1 | S | 148 | -0.396279414 | -0.38546956 | -0.577608003 | -0.181328589 | -0.192138443 | 0.068031723 | 0.055284368 | 0.001576726 | 0.306173739 | 0.246533924 |
| Tbc1d5 | S | 582 | -0.04425195 | -0.412579966 | -0.514636783 | -0.470384833 | -0.102056817 | 0.472757857 | 0.00074866 | 0.000172089 | 0.001241244 | 0.145419678 |
| Jmjd6 | S | 381 | -0.082908779 | -0.370892279 | -0.325053112 | -0.242144333 | 0.045839167 | 0.016135148 | 0.001592548 | 1.86425E-05 | 0.000595328 | 0.293285339 |
| Cep170b | S | 1166 | -0.208897044 | -0.500380664 | -0.571748028 | -0.362850984 | -0.071367364 | 0.30082478 | 0.005788584 | 0.001891918 | 0.113882954 | 0.516631584 |
| Abi1 | S | 216 | 0.052680342 | -0.397246762 | -0.494845482 | -0.547525824 | -0.09759872 | 0.121420169 | 0.000162544 | 4.46874E-06 | 0.000596541 | 0.068318216 |
| Mindy1 | S | 103 | 0.043639127 | -0.384601092 | -0.390811929 | -0.434451055 | -0.006210836 | 0.477461072 | 5.75414E-06 | 2.84504E-05 | 0.004055046 | 0.834826186 |
| Picalm | S | 16 | -0.05832551 | -0.319233963 | -0.484149936 | -0.425824425 | -0.164915973 | 0.377336735 | 2.68969E-05 | 0.00047271 | 0.003274926 | 0.031053502 |
| Dnttip2 | S | 352 | -0.344531599 | -0.361192817 | -0.299854547 | 0.044677052 | 0.061338269 | 0.010573426 | 0.000372248 | 0.024294622 | 0.658316373 | 0.469083158 |
| Mtrex | S | 56 | -0.094961012 | -0.321592825 | -0.420545695 | -0.325584683 | -0.098952871 | 0.036349832 | 0.001100746 | 0.000114709 | 0.000862405 | 0.069649496 |
| Utp11 | S | 241 | 0.005816904 | -0.322874242 | -0.418632044 | -0.424448948 | -0.095757801 | 0.891307938 | 7.73332E-05 | 4.52412E-06 | 0.000724396 | 0.031305979 |
| C2cd5 | S | 321 | 0.003442542 | -0.334486647 | -0.495083424 | -0.498525967 | -0.160596778 | 0.925394327 | 2.77036E-06 | 0.000107746 | 0.00034607 | 0.018234829 |
| Atxn2l | S | 304 | -0.122045961 | -0.350527009 | -0.387331231 | -0.26528527 | -0.036804222 | 0.049115863 | 4.30428E-05 | 0.003167319 | 0.005628844 | 0.425482966 |
| Ddx24 | S | 288 | 0.021376297 | -0.322218204 | -0.392846381 | -0.414222679 | -0.070628178 | 0.57416333 | 4.40026E-05 | 1.72032E-06 | 0.00167669 | 0.059751176 |
| Larp4 | S | 578 | -0.038274143 | -0.564489545 | -0.569624158 | -0.531350016 | -0.005134613 | 0.438244799 | 0.000268824 | 0.009133486 | 0.017736091 | 0.96050879 |
| Csdc2 | S | 48 | -0.25774541 | -0.347873046 | -0.540811005 | -0.283065594 | -0.192937959 | 0.056456634 | 0.001183663 | 0.010264413 | 0.084378267 | 0.169115428 |
| Uba1 | T | 800 | -0.094104186 | -0.365052055 | -0.370437957 | -0.276333772 | -0.005385902 | 0.096217787 | 0.000311337 | 0.000196852 | 0.000400381 | 0.809635788 |
| Ddx21 | S | 243 | -0.033674887 | -0.312036855 | -0.389862257 | -0.35618737 | -0.077825401 | 0.408758525 | 7.18591E-05 | 1.79219E-05 | 0.000897781 | 0.046779034 |
| Taf3 | S | 199 | -0.49930984 | -0.36051576 | -0.58042894 | -0.081119101 | -0.21991318 | 0.017637476 | 0.133873518 | 0.007475873 | 0.488452899 | 0.279081444 |
| Taf3 | S | 204 | -0.49930984 | -0.36051576 | -0.58042894 | -0.081119101 | -0.21991318 | 0.017637476 | 0.133873518 | 0.007475873 | 0.488452899 | 0.279081444 |
| Slc7a1 | S | 616 | -0.020441743 | -0.348328678 | -0.426239386 | -0.405797644 | -0.077910709 | 0.330686373 | 0.002210915 | 2.43701E-06 | 0.000459253 | 0.134541945 |
| Larp1 | S | 838 | -0.033016698 | -0.362490193 | -0.391008681 | -0.357991983 | -0.028518488 | 0.461074798 | 0.000722196 | 5.68621E-06 | 0.008865017 | 0.449998756 |
| Eif5 | S | 417 | -0.116121618 | -0.402630804 | -0.390720837 | -0.27459922 | 0.011909967 | 0.168655244 | 0.00098983 | 0.000201252 | 0.022936497 | 0.835369847 |
| Rabl6 | S | 414 | -0.152711556 | -0.353178686 | -0.454728715 | -0.302017159 | -0.101550029 | 0.031761361 | 0.000811213 | 0.002789425 | 0.024602347 | 0.240152094 |
| Ranbp2 | T | 799 | 0.013798587 | -0.325550524 | -0.326985614 | -0.340784201 | -0.00143509 | 0.542469208 | 6.17836E-06 | 2.25427E-06 | 6.04529E-05 | 0.92447082 |
| Rbm3 | S | 135 | -0.201653824 | -0.391921458 | -0.402398535 | -0.20074471 | -0.010477076 | 0.056736879 | 0.000124771 | 0.014377753 | 0.089587805 | 0.895909578 |
| Nab1 | S | 456 | 0.022737792 | -0.354627125 | -0.47757221 | -0.500310001 | -0.122945085 | 0.784387226 | 0.002193316 | 1.3724E-06 | 0.014938053 | 0.077419672 |
| Prune2 | S | 2648 | -0.006624958 | -0.475182709 | -0.372248212 | -0.365623253 | 0.102934497 | 0.946535093 | 9.88161E-05 | 0.000133699 | 0.035898403 | 0.091323583 |
| Bysl | S | 97 | -0.004861566 | -0.283820818 | -0.267864518 | -0.263002951 | 0.015956301 | 0.889248323 | 8.85758E-07 | 1.02701E-06 | 0.008059207 | 0.193867706 |
| Zfp703 | S | 231 | -0.063834933 | -0.339061856 | -0.48099878 | -0.417163846 | -0.141936924 | 0.425989552 | 0.011450713 | 3.15509E-06 | 0.014631859 | 0.109982329 |
| Dnaja1 | S | 83 | 0.018461655 | -0.534126979 | -0.555146283 | -0.573607938 | -0.021019305 | 0.739556867 | 0.000192402 | 0.001842774 | 0.00551253 | 0.802793782 |
| Irs2 | T | 347 | -0.032524448 | -0.356017057 | -0.393393742 | -0.360869294 | -0.037376685 | 0.393345927 | 0.000245725 | 2.66614E-05 | 0.000410941 | 0.387473843 |
| Map1b | T | 1908 | -0.1826461 | -0.353375578 | -0.495848976 | -0.313202877 | -0.142473399 | 0.040135337 | 0.00015296 | 0.019299505 | 0.058590065 | 0.265033817 |
| Tor1aip1 | S | 153 | -0.293262553 | -0.362250355 | -0.38882312 | -0.095560568 | -0.026572765 | 0.034064368 | 0.00026332 | 0.02468199 | 0.430596067 | 0.78478071 |
| Eif4ebp2 | S | 65 | -0.082490877 | -0.511748789 | -0.37369365 | -0.291202773 | 0.138055139 | 0.296683838 | 0.000893659 | 0.00040371 | 0.020116996 | 0.078307434 |
| Ssrp1 | S | 659 | -0.143002492 | -0.348630673 | -0.353182203 | -0.210179711 | -0.00455153 | 0.024608675 | 0.001833849 | 0.000270821 | 0.010418847 | 0.93587219 |
| Slc2a1 | T | 234 | -0.008212204 | -0.303253687 | -0.407560922 | -0.399348718 | -0.104307235 | 0.839481702 | 0.001145591 | 4.48834E-07 | 0.003957794 | 0.058460427 |
| Pld1 | S | 629 | -0.214694806 | -0.25385353 | -0.307975264 | -0.093280459 | -0.054121734 | 6.34076E-05 | 0.00039223 | 0.027366182 | 0.263040071 | 0.47671426 |
| Mybbp1a | S | 1164 | -0.042288552 | -0.275107209 | -0.318344665 | -0.276056113 | -0.043237456 | 0.054960883 | 4.79469E-07 | 0.000512875 | 0.001504495 | 0.183849778 |
| Flcn | S | 72 | -0.138921756 | -0.45363736 | -0.443814278 | -0.304892523 | 0.009823082 | 0.016883501 | 0.002522274 | 0.006204786 | 0.041404233 | 0.914859961 |
| Dnmt1 | S | 140 | -0.037848994 | -0.21304343 | -0.348496703 | -0.310647708 | -0.135453273 | 0.223101636 | 0.000305735 | 1.40425E-07 | 0.001656496 | 0.00578498 |
| Sipa1 | S | 72 | -0.071099681 | -0.443062731 | -0.440912401 | -0.36981272 | 0.002150329 | 0.137326913 | 0.001033511 | 0.000789089 | 0.013650701 | 0.975832102 |
| Slc9a1 | S | 707 | 0.063709949 | -0.336564295 | -0.483537759 | -0.547247708 | -0.146973465 | 0.20811894 | 0.000355601 | 1.11275E-06 | 0.000854867 | 0.016594669 |
| Sgta | T | 305 | -0.052757968 | -0.235915365 | -0.266895526 | -0.214137558 | -0.030980161 | 0.02485733 | 2.34004E-05 | 2.29566E-06 | 5.71889E-05 | 0.14770549 |
| Rpl17 | S | 142 | -0.284816944 | -0.345651796 | -0.445723415 | -0.160906471 | -0.100071619 | 0.10195226 | 0.013986866 | 0.000417651 | 0.28080906 | 0.293514275 |
| Sall1 Sall3 Zfp143 | T | 322;715;728 | -0.302212912 | -0.459607655 | -0.561173284 | -0.258960371 | -0.101565629 | 0.160828179 | 0.01416958 | 0.004585274 | 0.200980625 | 0.363497595 |
| Nolc1 | S | 418 | -0.09720373 | -0.335721817 | -0.412992035 | -0.315788305 | -0.077270218 | 0.250537895 | 0.000147476 | 0.000423966 | 0.019774477 | 0.137123416 |
| Errfi1 | S | 249 | -0.103600152 | -0.324942454 | -0.553774881 | -0.450174729 | -0.228832427 | 0.105766369 | 0.010742973 | 0.000189101 | 0.002835577 | 0.04500303 |
| Zcchc2 | S | 482 | -0.244075442 | -0.13340003 | -0.524934737 | -0.280859295 | -0.391534707 | 0.021596305 | 0.043322034 | 6.28846E-05 | 0.018771114 | 0.002600424 |
| Zcchc2 | S | 483 | -0.244075442 | -0.13340003 | -0.524934737 | -0.280859295 | -0.391534707 | 0.021596305 | 0.043322034 | 6.28846E-05 | 0.018771114 | 0.002600424 |
| Zcchc2 | S | 486 | -0.244075442 | -0.13340003 | -0.524934737 | -0.280859295 | -0.391534707 | 0.021596305 | 0.043322034 | 6.28846E-05 | 0.018771114 | 0.002600424 |
| Zcchc2 | S | 487 | -0.244075442 | -0.13340003 | -0.524934737 | -0.280859295 | -0.391534707 | 0.021596305 | 0.043322034 | 6.28846E-05 | 0.018771114 | 0.002600424 |
| Ptk2 | Y | 861 | -0.583751158 | -0.430535977 | -0.412978222 | 0.170772937 | 0.017557755 | 0.003053873 | 0.028801291 | 0.2192885 | 0.55077399 | 0.950945649 |
| Ptpn12 | S | 396 | -0.079585173 | -0.354994552 | -0.51189841 | -0.432313237 | -0.156903858 | 0.286099005 | 0.000473371 | 0.000874123 | 0.00437674 | 0.088461319 |
| Grk2 | S | 670 | -0.350400481 | -0.480582278 | -0.415952965 | -0.065552484 | 0.064629313 | 0.020638152 | 0.004386146 | 0.060275489 | 0.680602666 | 0.679915222 |
| Riok2 | S | 484 | -0.096074995 | -0.291821214 | -0.325026512 | -0.228951518 | -0.033205298 | 0.009485799 | 4.91613E-05 | 0.001067393 | 0.009677374 | 0.412747386 |
| Ncl | T | 84 | 0.059932049 | -0.382324465 | -0.459446555 | -0.519378604 | -0.07712209 | 0.460852993 | 9.49087E-05 | 2.8224E-05 | 0.010359953 | 0.018079989 |
| Bmp2k | S | 807 | -0.291451283 | -0.316247706 | -0.526415365 | -0.234964083 | -0.210167659 | 0.034621227 | 0.016647479 | 0.003358708 | 0.077255411 | 0.087508867 |
| Sgta | S | 303 | -0.037160132 | -0.296661234 | -0.388449779 | -0.351289647 | -0.091788545 | 0.299419848 | 0.000129469 | 2.88695E-05 | 3.30341E-05 | 0.044906546 |
| Birc6 | S | 2245 | -0.172962315 | -0.280483791 | -0.416482003 | -0.243519689 | -0.135998213 | 0.00010676 | 0.00500778 | 0.010357961 | 0.04180202 | 0.117801159 |
| 2310022B05Rik | S | 176 | -0.451332521 | -0.319209846 | -0.425822907 | 0.025509614 | -0.106613061 | 0.003814636 | 0.049111221 | 0.020260446 | 0.832791305 | 0.464840316 |
| Mapk14 | Y | 182 | -0.375001695 | -0.352895649 | -0.582927283 | -0.207925587 | -0.230031634 | 0.015135048 | 0.077209228 | 0.009386933 | 0.185813641 | 0.227041554 |
| Prrc2a | S | 1667 | 0.003404099 | -0.421751319 | -0.394682472 | -0.398086571 | 0.027068847 | 0.920787762 | 6.0594E-05 | 0.000181055 | 0.002873251 | 0.565850467 |
| Kat6a | S | 447 | -0.037514172 | -0.529120368 | -0.521990169 | -0.484475997 | 0.007130199 | 0.746182684 | 2.25722E-05 | 0.027949297 | 0.029940945 | 0.95498582 |
| Synrg | S | 1067 | -0.0842716 | -0.305776657 | -0.34588286 | -0.261611259 | -0.040106202 | 0.061405453 | 0.002519038 | 8.30715E-06 | 0.002094805 | 0.396559444 |
| Gapvd1 | S | 892 | -0.308033512 | -0.169351283 | -0.520255789 | -0.212222277 | -0.350904506 | 0.013258619 | 0.020951095 | 0.001123931 | 0.054633895 | 0.007335158 |
| Sec31a | S | 801 | -0.128396469 | -0.256999022 | -0.328762917 | -0.200366447 | -0.071763895 | 0.001380523 | 0.000112925 | 0.002789712 | 0.02705126 | 0.18645911 |
| Atg2b | S | 1571 | -0.176825293 | -0.356809773 | -0.38001852 | -0.203193227 | -0.023208747 | 0.00075192 | 0.005902963 | 0.007974584 | 0.050228013 | 0.761747986 |
| Usp24 | S | 2558 | -0.118408545 | -0.46385868 | -0.459511324 | -0.341102779 | 0.004347355 | 0.131992079 | 0.003047488 | 0.001782561 | 0.008251536 | 0.952579187 |
| Dync1li1 | S | 398 | -0.062714403 | -0.323507381 | -0.356204341 | -0.293489938 | -0.03269696 | 0.112442509 | 7.33071E-06 | 0.001719061 | 0.00460159 | 0.4765277 |
| Eif4b | S | 219 | -0.102609754 | -0.262114308 | -0.411302263 | -0.30869251 | -0.149187955 | 0.024219407 | 0.000106969 | 0.001209736 | 0.000907243 | 0.013314017 |
| Mapre2 | S | 208 | -0.023603284 | -0.378934809 | -0.575049317 | -0.551446032 | -0.196114508 | 0.527540991 | 3.21676E-05 | 0.005681416 | 0.006690119 | 0.077403976 |
| Mthfd1 | S | 549 | -0.148450726 | -0.376156614 | -0.547291669 | -0.398840943 | -0.171135056 | 0.056542667 | 0.011424665 | 0.001623592 | 0.010998885 | 0.151322257 |
| Zcchc14 | S | 445 | -0.231426104 | -0.247800665 | -0.275344584 | -0.043918481 | -0.02754392 | 2.4909E-05 | 0.007941509 | 0.007407337 | 0.356132254 | 0.617170559 |
| Rpl29 | S | 138 | -0.181873262 | -0.489323828 | -0.377224149 | -0.195350887 | 0.112099679 | 0.074085394 | 0.005610164 | 0.001885576 | 0.059605855 | 0.186164129 |
| Bnip3 | S | 85 | -0.101212042 | -0.404451698 | -0.256902004 | -0.155689962 | 0.147549694 | 0.132549521 | 5.26383E-05 | 0.000688648 | 0.05488404 | 0.001530674 |
| Espl1 | S | 1394 | -0.200088844 | -0.335053613 | -0.527603178 | -0.327514334 | -0.192549565 | 0.006690026 | 0.001860745 | 0.029542556 | 0.075193011 | 0.195732131 |
| Mcph1 | S | 552 | -0.038386824 | -0.450309442 | -0.419332608 | -0.380945784 | 0.030976834 | 0.499234665 | 6.28792E-05 | 0.003044314 | 0.004022314 | 0.605601227 |
| Fer | Y | 402 | -0.548235647 | -0.369070766 | -0.51874039 | 0.029495257 | -0.149669624 | 0.007160406 | 0.167546781 | 0.029796741 | 0.851801853 | 0.538181308 |

| | | | | | | | | | | | | |
|---|---|---|---|---|---|---|---|---|---|---|---|---|
| Map1b | S | 1911 | -0.081072473 | -0.305176733 | -0.336837812 | -0.25576534 | -0.03166108 | 0.0195701 | 4.49655E-07 | 0.022627115 | 0.03465999 | 0.628551208 |
| C2cd5 | S | 323 | 0.008409153 | -0.333208186 | -0.453105585 | -0.461514739 | -0.119897399 | 0.84430251 | 3.91984E-05 | 0.000144009 | 0.000488623 | 0.060980767 |
| Dus2 | S | 445 | -0.102777852 | -0.366895382 | -0.447074612 | -0.34429676 | -0.08017923 | 0.173362698 | 0.001100525 | 0.000732714 | 0.007300259 | 0.265315079 |
| D10Wsu102e | S | 177 | 0.00098852 | -0.325319463 | -0.417595676 | -0.418584196 | -0.092276212 | 0.978826832 | 0.002572605 | 8.90027E-07 | 0.001035419 | 0.09992718 |
| Lmna | S | 406 | 0.051024486 | -0.360512103 | -0.376462012 | -0.427486498 | -0.015949909 | 0.45782471 | 2.87567E-06 | 0.000108712 | 0.006834135 | 0.613353094 |
| Baz1a | S | 1297 | -0.01301386 | -0.372738597 | -0.580651252 | -0.567637392 | -0.207912655 | 0.89125707 | 3.16752E-05 | 0.00393013 | 0.0060129 | 0.080663441 |
| Baz1a | S | 1299 | -0.01301386 | -0.372738597 | -0.580651252 | -0.567637392 | -0.207912655 | 0.89125707 | 3.16752E-05 | 0.00393013 | 0.0060129 | 0.080663441 |
| Hspb1 | S | 86 | 0.924112737 | -0.448210881 | -0.549879934 | -1.47399267 | -0.101669053 | 0.29762529 | 2.18545E-05 | 2.53166E-06 | 0.155954827 | 0.040004217 |
| Tmcc3 | S | 46 | -0.024242826 | -0.400833465 | -0.53303529 | -0.508792464 | -0.132201825 | 0.780850242 | 0.012051037 | 2.25115E-05 | 0.013336564 | 0.205146206 |
| Mia2 | S | 1240 | -0.363148877 | -0.275625638 | -0.515500768 | -0.152351891 | -0.23987513 | 0.010000267 | 0.094577891 | 0.003749042 | 0.210897646 | 0.142843942 |
| Tbc1d8b | S | 952 | -0.46103507 | -0.335261281 | -0.2365277 | 0.22450737 | 0.098733582 | 0.001998046 | 0.020841996 | 0.019693865 | 0.046050234 | 0.341719899 |
| Dvl3 | S | 48 | -0.474426881 | -0.28944008 | -0.556442255 | -0.082015374 | -0.267002175 | 0.006778965 | 0.10884921 | 0.022873012 | 0.607251041 | 0.189892385 |
| Ccm2 | S | 15 | -0.063046991 | -0.409441456 | -0.558214986 | -0.495167995 | -0.14877353 | 0.00913415 | 0.002961238 | 0.005720871 | 0.010929612 | 0.094569442 |
| Pnn | S | 96 | -0.108119381 | -0.463818933 | -0.515568831 | -0.407449451 | -0.051749898 | 0.114007982 | 0.00012799 | 0.045727472 | 0.07973496 | 0.730984384 |
| Usp15 | S | 225 | -0.133643558 | -0.400543881 | -0.431528338 | -0.297884779 | -0.030984457 | 0.148395812 | 0.027040903 | 4.55008E-05 | 0.035741786 | 0.756379978 |
| Larp4b | S | 500 | -0.069810996 | -0.328172978 | -0.555769878 | -0.485958882 | -0.2275969 | 0.116720754 | 0.000505558 | 0.00267435 | 0.00988761 | 0.043297537 |
| Taok1 | S | 9 | -0.49818809 | -0.410513418 | -0.572015085 | -0.073826995 | -0.161501666 | 0.017364964 | 0.046837607 | 0.068783654 | 0.735197924 | 0.502451562 |
| Slc1a5 | S | 518 | 0.08369931 | -0.402152893 | -0.488302311 | -0.572001621 | -0.086149418 | 0.141687608 | 0.000104278 | 2.853E-05 | 8.22465E-05 | 0.054746821 |
| Gpn1 | S | 314 | -0.1040861 | -0.326527877 | -0.327310916 | -0.223224816 | -0.000783039 | 0.05148412 | 6.05246E-06 | 0.009505962 | 0.024548016 | 0.988821904 |
| Chek1 | S | 280 | -0.137069335 | -0.385430131 | -0.476442858 | -0.339373522 | -0.091012726 | 0.278115096 | 0.002492283 | 0.000911797 | 0.037995115 | 0.178535389 |
| Tns3 | S | 946 | -0.175196421 | -0.311603694 | -0.521613668 | -0.346417247 | -0.210009974 | 0.005746033 | 0.054125946 | 0.001311843 | 0.015581345 | 0.129717268 |
| Ddx21 | S | 64 | 0.019930622 | -0.324084746 | -0.377501229 | -0.397431851 | -0.053416483 | 0.656541329 | 3.77934E-05 | 1.70258E-05 | 0.00076947 | 0.103681244 |
| Knop1 | S | 142 | -0.008061695 | -0.274900267 | -0.286659984 | -0.278598289 | -0.011759716 | 0.867026384 | 6.81455E-06 | 2.69171E-06 | 0.015944614 | 0.519708802 |
| Csde1 | T | 761 | -0.252136506 | -0.166913805 | -0.390447563 | -0.138311057 | -0.223533758 | 0.001893145 | 0.125764527 | 8.81435E-05 | 0.031217255 | 0.08349079 |
| Plec | S | 4389 | -0.135086863 | -0.233514324 | -0.57057636 | -0.435489496 | -0.337062035 | 0.059128305 | 0.01223866 | 0.000372828 | 0.001786762 | 0.004777009 |
| Rsrc2 | S | 45 | -0.023188894 | -0.261725499 | -0.325545158 | -0.302356264 | -0.063819659 | 0.385167524 | 0.000123752 | 1.96731E-06 | 0.000381946 | 0.066928847 |
| Stk39 | S | 397 | -0.240744122 | -0.280384844 | -0.284910865 | -0.044166742 | -0.00452602 | 0.0042672 | 0.03730661 | 7.51928E-05 | 0.353463107 | 0.95403595 |
| Arhgap5 | S | 1201 | -0.136544334 | -0.359803316 | -0.376063219 | -0.239518885 | -0.016259903 | 0.015929358 | 0.012236426 | 0.000323006 | 0.006119243 | 0.831295557 |
| Lrrc59 | S | 23 | 0.08180423 | -0.566905981 | -0.462341239 | -0.544145469 | 0.104564742 | 0.307487011 | 0.000159166 | 0.000706919 | 0.00094735 | 0.135653672 |
| Leo1 | S | 140 | -0.308088572 | -0.160839694 | -0.422470649 | -0.114382077 | -0.261630954 | 0.00648011 | 0.061791537 | 0.000354395 | 0.147022189 | 0.01836512 |
| Leo1 | S | 151 | -0.308088572 | -0.160839694 | -0.422470649 | -0.114382077 | -0.261630954 | 0.00648011 | 0.061791537 | 0.000354395 | 0.147022189 | 0.01836512 |
| Leo1 | S | 154 | -0.308088572 | -0.160839694 | -0.422470649 | -0.114382077 | -0.261630954 | 0.00648011 | 0.061791537 | 0.000354395 | 0.147022189 | 0.01836512 |
| Ddx24 | S | 92 | -0.015630287 | -0.400353924 | -0.281246987 | -0.2656167 | 0.119106936 | 0.768755999 | 1.35266E-05 | 0.000166684 | 0.008253955 | 0.015533521 |
| Bclaf3 | S | 600 | -0.09197377 | -0.394464083 | -0.564945186 | -0.472971416 | -0.170481103 | 0.327991612 | 0.001717388 | 0.002518834 | 0.008349358 | 0.131079422 |
| Plec | S | 4413 | 0.024633813 | -0.379054162 | -0.487807267 | -0.51244108 | -0.108753106 | 0.67372275 | 0.000316167 | 0.000124336 | 0.00116711 | 0.084535513 |
| Nab2 | S | 515 | -0.311383873 | -0.341614592 | -0.39714217 | -0.085758297 | -0.055527578 | 0.011936699 | 0.007850676 | 0.016006706 | 0.413072251 | 0.587049988 |
| Sh3pxd2a | S | 976 | -0.010843175 | -0.321212852 | -0.394468147 | -0.383624972 | -0.073255295 | 0.742405266 | 7.24737E-05 | 7.75204E-05 | 0.001085052 | 0.115823757 |
| Rtn4 | S | 892 | -0.226753056 | -0.366058217 | -0.323390154 | -0.096637098 | 0.042668063 | 0.010552085 | 0.000260092 | 0.050237654 | 0.408911083 | 0.692286365 |
| Nampt | S | 470 | -0.46148057 | -0.208886433 | -0.548605279 | -0.087124709 | -0.339718846 | 0.020968908 | 0.089036485 | 0.005451318 | 0.561198621 | 0.042289305 |
| Klhdc4 | S | 62 | -0.026829443 | -0.402809003 | -0.41220244 | -0.385372996 | -0.009393437 | 0.313428104 | 1.97664E-06 | 0.021578309 | 0.021807188 | 0.902028242 |
| Cad | T | 1824 | -0.013748399 | -0.31174448 | -0.321403139 | -0.307654741 | -0.009658659 | 0.706534974 | 0.000146748 | 3.7043E-06 | 0.00259699 | 0.739480144 |
| Sh3pxd2a | S | 514 | -0.172256124 | -0.318485688 | -0.494077377 | -0.321821253 | -0.175591689 | 0.001331558 | 0.017941674 | 0.008594112 | 0.037570393 | 0.126335795 |
| Sh3pxd2a | T | 513 | -0.172256124 | -0.318485688 | -0.494077377 | -0.321821253 | -0.175591689 | 0.001331558 | 0.017941674 | 0.008594112 | 0.037570393 | 0.126335795 |
| Nelfa | S | 225 | -0.189112317 | -0.273982121 | -0.295227501 | -0.106115184 | -0.02124538 | 0.005315049 | 0.000873277 | 0.002144362 | 0.100662379 | 0.691155679 |
| 2310033P09Rik | S | 242 | -0.441879975 | -0.3386896 | -0.213301991 | 0.228577983 | 0.125387609 | 0.006383719 | 0.001030282 | 0.094788968 | 0.102303302 | 0.256838789 |
| Tbc1d1 | T | 460 | -0.088566457 | -0.351059622 | -0.465865762 | -0.377299305 | -0.11480614 | 0.363688192 | 0.004463569 | 0.00011924 | 0.021013247 | 0.151503802 |
| Eif4b | S | 93 | -0.299921948 | -0.394177882 | -0.270787 | 0.029134949 | 0.123390882 | 0.003160162 | 0.000862288 | 0.128037593 | 0.830841808 | 0.400230652 |
| Pphln1 | S | 219 | -0.145747621 | -0.240209553 | -0.244512684 | -0.098765062 | -0.004303131 | 0.02402465 | 9.69099E-05 | 0.000100924 | 0.083407789 | 0.842992618 |
| Zfp771 | T | 198 | -0.290629569 | -0.312596236 | -0.984920176 | -0.694290607 | -0.67232394 | 0.032388804 | 0.017618755 | 0.074472069 | 0.140647333 | 0.149616401 |
| Gtse1 | S | 726 | -0.143157423 | -0.395910485 | -0.488703605 | -0.345546181 | -0.09279312 | 0.022373328 | 0.008256357 | 0.0051443 | 0.011077095 | 0.305663388 |
| Lmbrd2 | S | 610 | -0.311660381 | -0.33158905 | -0.461626546 | -0.149960165 | -0.130031497 | 0.005084132 | 0.011023306 | 0.041006359 | 0.341668522 | 0.409460009 |
| Riok2 | S | 481 | -0.109667462 | -0.28973705 | -0.342230712 | -0.23256325 | -0.052493662 | 0.029569074 | 0.006669796 | 3.03873E-05 | 0.00905504 | 0.35544559 |
| Ano1 | S | 106 | -0.22785676 | -0.505874382 | -0.199526914 | 0.028329846 | 0.306347468 | 0.003888384 | 0.000791438 | 0.094168911 | 0.744418919 | 0.034234036 |
| Sos1 | S | 1153 | -0.153112869 | -0.367494142 | -0.429565202 | -0.276452333 | -0.06207106 | 0.21672067 | 0.001058757 | 0.001908545 | 0.068055195 | 0.402304038 |
| Cep170 | S | 950 | -0.132187214 | -0.312586981 | -0.582768924 | -0.45058171 | -0.270181943 | 0.184779416 | 0.032190338 | 0.00024864 | 0.005159413 | 0.057415677 |
| Tagln2 | S | 163 | -0.411145202 | -0.364151884 | -0.583083129 | -0.171937927 | -0.218931245 | 0.026323965 | 0.087900454 | 0.016826309 | 0.324786448 | 0.287144859 |
| Lmna | S | 407 | 0.007210402 | -0.347700127 | -0.349233709 | -0.356444111 | -0.001533583 | 0.861148695 | 2.28459E-05 | 8.52236E-05 | 0.001409037 | 0.962260415 |
| Anxa1 | Y | 21 | -0.446307837 | -0.33700168 | -0.55162377 | -0.105315932 | -0.21462209 | 0.01756285 | 0.11356833 | 0.016843361 | 0.4500932 | 0.274154997 |
| Riok1 | S | 22 | -0.312292893 | -0.404967212 | -0.195263356 | 0.117029537 | 0.209703856 | 0.06459794 | 2.62193E-06 | 0.287593834 | 0.518474666 | 0.264104003 |
| Ric8a | T | 442 | -0.296011258 | -0.226014764 | -0.412600351 | -0.116589093 | -0.186585586 | 0.010817294 | 0.032560643 | 0.001235796 | 0.136710099 | 0.048092265 |
| Gpsm1 | T | 383 | 0.035699716 | -0.397978517 | -0.427221387 | -0.462921102 | -0.029242869 | 0.685988403 | 0.000502963 | 4.27288E-05 | 0.019098966 | 0.54766123 |
| Hps5 | S | 662 | -0.132143925 | -0.304334095 | -0.432938745 | -0.30079482 | -0.12860465 | 0.016203062 | 0.012023189 | 0.000582606 | 0.003651843 | 0.117482616 |
| Cdv3 | S | 63 | -0.177501715 | -0.343778661 | -0.551058429 | -0.373556714 | -0.207279767 | 0.101582255 | 0.02242801 | 0.001306251 | 0.016019033 | 0.107441633 |
| Eif4ebp1 | T | 45 | -0.256877742 | -0.343336645 | -0.549141672 | -0.29226393 | -0.205805028 | 0.027758527 | 0.008086027 | 0.022278318 | 0.112255464 | 0.215016537 |
| Ncl | S | 403 | -0.137804601 | -0.380528413 | -0.253299954 | -0.115495353 | 0.127228458 | 0.108604177 | 2.83739E-05 | 0.005818589 | 0.174742054 | 0.077095948 |
| Anln | S | 913 | -0.465189719 | -0.204470731 | -0.377042055 | 0.088147664 | -0.172571323 | 0.001795285 | 0.101821386 | 0.012008866 | 0.353237697 | 0.151399153 |
| Ftsj3 | S | 482 | -0.247473106 | -0.429371088 | -0.365063952 | -0.117590846 | 0.064307136 | 0.075834058 | 0.00089255 | 0.025979894 | 0.388890197 | 0.559988753 |
| Spag5 | S | 12 | -0.149742183 | -0.419670886 | -0.289050434 | -0.13930825 | 0.130620452 | 0.093928408 | 0.000507262 | 0.002672683 | 0.09906091 | 0.071466593 |
| Spag5 | S | 14 | -0.149742183 | -0.419670886 | -0.289050434 | -0.13930825 | 0.130620452 | 0.093928408 | 0.000507262 | 0.002672683 | 0.09906091 | 0.071466593 |
| Ptpn21 | S | 670 | -0.103958957 | -0.331433295 | -0.436498351 | -0.332539394 | -0.105065055 | 0.284080795 | 0.004517932 | 0.000116486 | 0.02860927 | 0.144975855 |
| Ilrun | S | 215 | -0.094478112 | -0.360798456 | -0.558179034 | -0.463700922 | -0.197380578 | 0.122403157 | 0.000469621 | 0.011456437 | 0.019621144 | 0.133677197 |
| 8030462N17Rik | S | 147 | 0.053260377 | -0.354578325 | -0.357745207 | -0.411005585 | -0.003166882 | 0.264497299 | 5.84031E-06 | 4.82553E-05 | 0.001475266 | 0.909898368 |
| Fam133b | S | 82 | -0.304103658 | -0.328951064 | -0.483404417 | -0.179300759 | -0.154453353 | 0.005569395 | 0.110859913 | 0.005898175 | 0.133695386 | 0.380106051 |
| Ythdf2 | S | 22 | -0.197836426 | -0.392416705 | -0.533295516 | -0.335459091 | -0.140878811 | 0.122867094 | 0.013339714 | 0.0035178 | 0.038302001 | 0.27477698 |
| Dync1li1 | S | 405 | -0.05008682 | -0.333218604 | -0.303585971 | -0.253499151 | 0.029632632 | 0.518131166 | 0.000381584 | 8.96026E-06 | 0.051724576 | 0.41678877 |
| Abl2 | S | 819 | -0.151814582 | -0.424439277 | -0.484920912 | -0.33310633 | -0.060481635 | 0.158620559 | 0.005227101 | 0.003490197 | 0.02637649 | 0.574741778 |
| Zc3h11a | S | 740 | -0.085780802 | -0.403693215 | -0.334437001 | -0.248656199 | 0.069256214 | 0.120375439 | 0.002755528 | 0.000165497 | 0.00366249 | 0.318203045 |
| Fxr1 | T | 398 | -0.131293907 | -0.347492403 | -0.475654799 | -0.344360891 | -0.128162396 | 0.113406642 | 0.030035119 | 0.000163069 | 0.004879337 | 0.271939399 |
| Fam207a | S | 38 | -0.000142359 | -0.243603303 | -0.272122103 | -0.271979744 | -0.0285188 | 0.996339646 | 4.05981E-06 | 1.05744E-06 | 0.005680352 | 0.10640407 |
| Tiam2 | S | 751 | 0.001431989 | -0.426330454 | -0.504499754 | -0.505931743 | -0.0781693 | 0.979076664 | 0.001065577 | 0.0004512 | 0.002654535 | 0.320896459 |
| Nufip2 | S | 626 | -0.042114076 | -0.418800425 | -0.322129687 | -0.280015612 | 0.096670738 | 0.415056479 | 0.000100847 | 0.000787581 | 0.004052043 | 0.106745935 |
| Afap1 | S | 279 | -0.099809265 | -0.362469218 | -0.407809167 | -0.307999902 | -0.045339949 | 0.184226927 | 0.001531581 | 0.000655622 | 0.002361111 | 0.388496865 |
| Prrc2c | T | 358 | -0.063234633 | -0.22830578 | -0.26679195 | -0.203557317 | -0.03848617 | 0.017953977 | 0.000338877 | 4.30057E-06 | 0.000150397 | 0.189908022 |
| Esf1 | S | 156 | -0.122823442 | -0.327852072 | -0.314508442 | -0.191685 | 0.013343629 | 0.02852498 | 0.000113059 | 0.005341557 | 0.039257003 | 0.819368295 |
| Eif4g1 | S | 1231 | -0.086563799 | -0.27948113 | -0.221017023 | -0.134453224 | 0.058464106 | 0.003621818 | 3.8354E-06 | 0.003975484 | 0.037061104 | 0.168145754 |
| Zmym4 | S | 161 | -0.397250362 | -0.221688437 | -0.571904447 | -0.174654085 | -0.35021601 | 0.014379705 | 0.290832762 | 0.002556649 | 0.078893049 | 0.128796908 |
| Usp36 | S | 512 | -0.040647099 | -0.531637651 | -0.297325132 | -0.256678033 | 0.234312518 | 0.430114807 | 9.53093E-05 | 0.004626712 | 0.010889286 | 0.014227794 |
| Rpl26 Gm15772 | T | 139 | -0.162506521 | -0.362860151 | -0.393398151 | -0.230891629 | -0.030538 | 0.080862141 | 0.000177203 | 0.017962641 | 0.071553989 | 0.728723494 |
| Cux1 | S | 1506 | -0.374260798 | -0.236085697 | -0.436752462 | -0.062491664 | -0.200666765 | 0.002706449 | 0.124588199 | 0.007025343 | 0.528350213 | 0.190682618 |
| Ranbp2 | S | 1599 | -0.253153968 | -0.512531987 | -0.376489094 | -0.123335126 | 0.136042893 | 0.214403536 | 0.001109881 | 0.026649326 | 0.506970922 | 0.246571805 |
| Foxk2 | S | 389 | 0.011368704 | -0.470331223 | -0.357705171 | -0.369073876 | 0.112626051 | 0.846308835 | 6.21636E-05 | 0.001024777 | 0.002869041 | 0.107214949 |
| Eml1 | S | 134 | -0.499983728 | -0.258605652 | -0.321750174 | 0.178233554 | -0.063144522 | 0.00018025 | 0.080077641 | 0.1544153 | 0.359893908 | 0.742613861 |
| Smc5 | S | 35 | -0.481820901 | -0.403506561 | -0.181920287 | 0.299900614 | 0.221586274 | 0.001280344 | 0.003278983 | 0.515363333 | 0.317204277 | 0.433291077 |
| Med14 | S | 1141 | -0.198809533 | -0.405839202 | -0.350104667 | -0.151295134 | 0.055734535 | 0.052927327 | 0.002944316 | 0.005537802 | 0.052998872 | 0.322264313 |

| | | | | | | | | | | | | |
|---|---|---|---|---|---|---|---|---|---|---|---|---|
| Med14 | S | 1147 | -0.198809533 | -0.405839202 | -0.350104667 | -0.151295134 | 0.055734535 | 0.052927327 | 0.002944316 | 0.005537802 | 0.052998872 | 0.322264313 |
| Zdhhc5 | Y | 637 | -0.320614027 | -0.587828117 | -1.544806567 | -1.22419254 | -0.95697845 | 0.249991638 | 0.264328396 | 0.057670894 | 0.119130437 | 0.189455894 |
| Adam19 | T | 771 | 0.025953596 | -0.3536583 | -0.39394858 | -0.419902176 | -0.040290279 | 0.223098288 | 0.002460965 | 5.34758E-07 | 2.18916E-06 | 0.364627126 |
| Cttn | S | 405 | -0.401696567 | -0.274402001 | -0.260572916 | 0.141123651 | 0.013829085 | 0.001428249 | 0.02657145 | 0.01788732 | 0.084313003 | 0.873078615 |
| Hsph1 | S | 810 | -0.120077719 | -0.205036446 | -0.262321807 | -0.142244088 | -0.057285361 | 0.005164091 | 0.000862762 | 3.34424E-05 | 0.005073237 | 0.116562632 |
| Fbl | S | 130 | 0.045129387 | -0.332241594 | -0.323637792 | -0.368767178 | 0.008603802 | 0.237719487 | 1.13277E-05 | 7.81118E-06 | 0.000639053 | 0.736364355 |
| Cdca2 | S | 306 | 0.001856517 | -0.316305429 | -0.427541366 | -0.429397883 | -0.111235937 | 0.957266415 | 0.000753724 | 2.20606E-05 | 0.000363464 | 0.05957927 |
| Strap | S | 340 | -0.040100648 | -0.309671384 | -0.337188768 | -0.29708812 | -0.027517384 | 0.366482496 | 0.000906935 | 1.07569E-05 | 0.003628049 | 0.484216415 |
| Nav3 | S | 543 | -0.104240655 | -0.346218887 | -0.424960036 | -0.320719381 | -0.078741149 | 0.167953964 | 0.002414732 | 0.000682009 | 0.010034396 | 0.158226202 |
| Stk3 | S | 412 | -0.356196926 | -0.260104036 | -0.371992836 | -0.01579591 | -0.111888799 | 0.000592227 | 0.021723776 | 0.065004003 | 0.905562497 | 0.453840167 |
| Rps15 | S | 55 | -0.134972617 | -0.327046021 | -0.294147219 | -0.159174602 | 0.032898802 | 0.032196841 | 0.000819151 | 0.000990746 | 0.001062212 | 0.129284308 |
| Larp7 | T | 251 | -0.195585591 | -0.24171275 | -0.311367606 | -0.115782015 | -0.069654856 | 0.008752669 | 0.01097782 | 0.000208303 | 0.0680975 | 0.266227928 |
| Irs1 | S | 526 | -0.46378505 | -0.386807891 | -0.424477699 | 0.039307351 | -0.037669808 | 0.021361834 | 0.041263298 | 0.036509866 | 0.802858187 | 0.809642904 |
| Fbxo38 | S | 624 | -0.019482427 | -0.34259441 | -0.409620566 | -0.390138139 | -0.067026157 | 0.716094972 | 0.00030517 | 0.000173085 | 0.000991497 | 0.188699685 |
| Map1a | S | 384 | -0.117486267 | -0.30929028 | -0.3005724 | -0.183086133 | 0.00871788 | 0.06022956 | 0.000380149 | 0.000596476 | 0.01688329 | 0.733118688 |
| Snap91 | T | 333 | -0.015350218 | -0.443985059 | -0.332627405 | -0.317277187 | 0.111357655 | 0.791835794 | 3.55727E-05 | 0.002084942 | 0.007247511 | 0.078879327 |
| Srp68 | S | 31 | -0.053918676 | -0.30712767 | -0.267090028 | -0.213171351 | 0.040037642 | 0.528532656 | 2.49586E-05 | 4.55264E-05 | 0.086544579 | 0.156509567 |
| Gtf2f1 | S | 381 | 0.050388591 | -0.499182141 | -0.47259045 | -0.522979041 | 0.02659169 | 0.341351333 | 7.68376E-05 | 0.002530827 | 0.00995451 | 0.730138822 |
| Ngrn | S | 239 | 0.073892378 | -0.498414848 | -0.397888824 | -0.471781202 | 0.100526024 | 0.212415209 | 0.000275733 | 7.8624E-05 | 0.00352128 | 0.135247999 |
| Tulp3 | S | 372 | -0.109715372 | -0.2993497 | -0.278206003 | -0.168490631 | 0.021143696 | 0.025728848 | 2.61383E-05 | 0.005274658 | 0.031232099 | 0.663731663 |
| Ube2t | S | 193 | -0.207997713 | -0.332481295 | -0.424279315 | -0.216281602 | -0.09179802 | 0.009224181 | 0.003542029 | 0.023948264 | 0.13821361 | 0.447672385 |
| E2f8 | S | 415 | 0.021696099 | -0.397377945 | -0.288088803 | -0.309784902 | 0.109289142 | 0.743058608 | 4.65027E-05 | 4.17591E-05 | 0.021563113 | 0.031747424 |
| Nup35 | T | 307 | -0.013690937 | -0.426563534 | -0.402465655 | -0.388774718 | 0.024097879 | 0.770355299 | 0.004199838 | 3.66577E-05 | 0.003038622 | 0.643157006 |
| Gpi1 | T | 109 | -0.291445465 | -0.343885646 | -0.414808393 | -0.123362928 | -0.070922747 | 0.01612862 | 0.004579798 | 0.040845169 | 0.356834314 | 0.569346096 |
| Suco | S | 1203 | -0.252356766 | -0.423081591 | -0.30200729 | -0.049650524 | 0.121074301 | 0.019657035 | 0.009159084 | 0.008233344 | 0.344152248 | 0.256517704 |
| Gja1 | S | 272 | 0.038362743 | -0.346713112 | -0.546165868 | -0.584528611 | -0.199452756 | 0.565354403 | 0.000920165 | 0.00014296 | 0.002616561 | 0.003220242 |
| Jpt1 | T | 54 | -0.081388507 | -0.287209684 | -0.474307769 | -0.392919262 | -0.187098085 | 0.305913997 | 0.009730031 | 6.21987E-05 | 0.000834085 | 0.043747603 |
| Nhsl1 | S | 466 | -0.148666804 | -0.481885859 | -0.247769338 | -0.099102533 | 0.234116521 | 0.107822303 | 0.000130838 | 0.025203348 | 0.286511816 | 0.03949261 |
| Anln | S | 182 | -0.297369748 | -0.19708562 | -0.206850953 | 0.090518795 | -0.009765333 | 3.44971E-05 | 0.081366755 | 0.001618909 | 0.052974949 | 0.901578597 |
| Ddx21 | T | 114 | 0.050747227 | -0.305669318 | -0.440492837 | -0.491240064 | -0.134823519 | 0.242979704 | 0.00111737 | 1.50519E-06 | 0.002947351 | 0.042220125 |
| Dbndd2 | S | 117 | -0.378296552 | -0.278325827 | -0.572366362 | -0.19406981 | -0.294040535 | 0.013619198 | 0.079041738 | 0.020909882 | 0.271967326 | 0.142872588 |
| Kif4 | S | 802 | -0.191814598 | -0.287909981 | -0.578611383 | -0.386796786 | -0.290701403 | 0.134823081 | 0.066645965 | 0.000491396 | 0.018686051 | 0.069416259 |
| Ddx24 | S | 91 | -0.058085511 | -0.346926404 | -0.359718829 | -0.301633318 | -0.012792424 | 0.450329708 | 0.000302585 | 0.000298273 | 0.026613464 | 0.734653669 |
| Ngdn | S | 214 | -0.102674715 | -0.327527677 | -0.452976878 | -0.350302164 | -0.125449201 | 0.181235484 | 0.001950506 | 0.00113869 | 0.007948593 | 0.138381144 |
| Disp2 | S | 1198 | -0.244820994 | -0.256062646 | -0.482300076 | -0.237479082 | -0.22623743 | 0.061055885 | 0.032814905 | 0.000922358 | 0.074274791 | 0.059502341 |
| Sub1 | S | 58 | -0.202921751 | -0.234332209 | -0.276323611 | -0.07340186 | -0.041991402 | 0.003952128 | 0.011330484 | 0.000250114 | 0.121329812 | 0.450140275 |
| Ncl | T | 121 | 0.074434705 | -0.310894216 | -0.420496949 | -0.494931655 | -0.109602734 | 0.009097619 | 0.000173284 | 1.9898E-07 | 6.35217E-06 | 0.021770423 |
| Cdv3 | S | 131 | -0.268934921 | -0.392475369 | -0.426601053 | -0.157666132 | -0.034125683 | 0.11291486 | 0.035365809 | 0.001286431 | 0.272863473 | 0.76162853 |
| Nfic | S | 337 | -0.090372251 | -0.197246548 | -0.266027425 | -0.175655174 | -0.068780877 | 0.016346434 | 1.97727E-05 | 0.000163245 | 0.002993541 | 0.058895 |
| Nav3 | S | 354 | -0.072701367 | -0.3723936 | -0.347950289 | -0.275248923 | 0.02444331 | 0.038141436 | 3.99918E-05 | 0.020480447 | 0.049869354 | 0.751348131 |
| Lmna | S | 423 | 0.031622562 | -0.390106113 | -0.324411283 | -0.356033846 | 0.065694829 | 0.471776129 | 3.1438E-05 | 9.3202E-05 | 0.000287969 | 0.085118946 |
| Usp24 | S | 1609 | -0.261179052 | -0.368612342 | -0.557446416 | -0.296267364 | -0.188834074 | 0.077178337 | 0.084183239 | 0.002659459 | 0.034330043 | 0.288465535 |
| Nup62 | S | 412 | -0.210238017 | -0.478111174 | -0.296000657 | -0.08576264 | 0.182110518 | 0.058021794 | 0.000184727 | 0.121873636 | 0.580573637 | 0.275874525 |
| Larp1 | S | 824 | -0.029253793 | -0.363157907 | -0.385705643 | -0.356451851 | -0.022547737 | 0.179799589 | 0.000288774 | 0.000899157 | 0.000483843 | 0.48917111 |
| Ell2 | S | 501 | -0.048056932 | -0.429598709 | -0.264479761 | -0.216422829 | 0.165118948 | 0.401900608 | 1.73426E-05 | 0.003957243 | 0.018842423 | 0.045151076 |
| Pex19 | S | 35 | -0.187348836 | -0.241057432 | -0.24058438 | -0.053235544 | 0.000473053 | 0.002725728 | 0.001166727 | 0.001355135 | 0.18986191 | 0.99083246 |
| Frmd4a | S | 712 | -0.014647223 | -0.364872684 | -0.352829547 | -0.338182324 | 0.012043137 | 0.722098961 | 1.51107E-06 | 0.007963673 | 0.004901531 | 0.814589087 |
| Map1b | S | 1815 | -0.129146916 | -0.292022353 | -0.31350754 | -0.184360624 | -0.021485186 | 0.041859528 | 0.000368278 | 0.001531425 | 0.022540269 | 0.662561816 |
| Psmf1 | S | 189 | 0.01509447 | -0.36902565 | -0.542686304 | -0.557780774 | -0.173660654 | 0.890236698 | 8.56942E-05 | 0.003150785 | 0.010055886 | 0.106419245 |
| Zfp36l2 | S | 57 | -0.108476707 | -0.443109543 | -0.52689284 | -0.418416134 | -0.083783298 | 0.416314508 | 0.00026233 | 0.029127831 | 0.056835519 | 0.552966229 |
| Sltm | S | 747 | -0.350282799 | -0.353808131 | -0.46081416 | -0.110531361 | -0.107006028 | 0.042028262 | 0.033821203 | 0.011370822 | 0.268285223 | 0.179159255 |
| Prrc2c | T | 617 | -0.23270924 | -0.483844193 | -0.242860151 | -0.010150912 | 0.240984042 | 0.013923617 | 0.000210356 | 0.270864431 | 0.956918177 | 0.274724425 |
| Gtf2f1 | T | 389 | 0.10182357 | -0.525911253 | -0.449637992 | -0.551461562 | 0.076273261 | 0.12206168 | 0.000215221 | 0.000233319 | 2.14165E-05 | 0.231097741 |
| Sdad1 | S | 585 | 0.026633632 | -0.326210716 | -0.383853655 | -0.410487288 | -0.05764294 | 0.733046089 | 0.000241233 | 2.26528E-05 | 0.023320827 | 0.174227208 |
| Fer | S | 434 | -0.087790206 | -0.232850889 | -0.361917583 | -0.274127376 | -0.129066694 | 0.022616024 | 0.006808834 | 3.57946E-05 | 0.001705644 | 0.054627218 |
| Ankle2 | S | 266 | -0.167889139 | -0.408747917 | -0.255740538 | -0.087851399 | 0.153007379 | 0.005312886 | 6.62165E-05 | 0.18399701 | 0.574017461 | 0.363995852 |
| Ankle2 | S | 275 | -0.167889139 | -0.408747917 | -0.255740538 | -0.087851399 | 0.153007379 | 0.005312886 | 6.62165E-05 | 0.18399701 | 0.574017461 | 0.363995852 |
| Eif3a | S | 895 | -0.097049206 | -0.33714455 | -0.469599034 | -0.372549828 | -0.132454484 | 0.263795554 | 0.00054287 | 0.003683517 | 0.013779364 | 0.173873244 |
| Cep250 | S | 1707 | 0.104359298 | -0.380130208 | -0.415185459 | -0.519544757 | -0.035055251 | 0.472803048 | 0.00103328 | 5.98311E-06 | 0.045077008 | 0.476521396 |
| Ranbp2 | T | 1104 | -0.1230184 | -0.426632772 | -0.475127978 | -0.352109578 | -0.048495206 | 0.106368641 | 0.000651402 | 0.032389092 | 0.076019365 | 0.71903073 |
| Ect2 | S | 367 | -0.266122456 | -0.300662459 | -0.552202367 | -0.286079911 | -0.251539907 | 0.027980774 | 0.045179372 | 0.008188016 | 0.083439647 | 0.124097096 |
| Ncbp1 | S | 22 | -0.018196964 | -0.309872678 | -0.453604887 | -0.435407923 | -0.143732209 | 0.789242142 | 2.02676E-05 | 0.002104897 | 0.00454239 | 0.068726607 |
| Gys1 | S | 700 | -0.335827303 | -0.292733158 | -0.455935945 | -0.120108642 | -0.163202788 | 0.004377114 | 0.064932603 | 0.021815552 | 0.361515832 | 0.308548023 |
| Arl14ep | S | 266 | -0.026111783 | -0.240211824 | -0.336046179 | -0.309934396 | -0.095834355 | 0.449010424 | 0.000220515 | 7.0121E-06 | 0.000559523 | 0.024700792 |
| Kif16b | S | 657 | -0.294540763 | -0.344269778 | -0.42106934 | -0.126528576 | -0.076799562 | 0.038168882 | 0.022458537 | 0.007302988 | 0.175010215 | 0.395920717 |
| Nampt | S | 472 | -0.349236936 | -0.325013226 | -0.499560698 | -0.150323762 | -0.174547472 | 0.026899325 | 0.110953983 | 0.006307619 | 0.273508253 | 0.3361935 |
| Zfp219 Zfp239 Zfp46 | T | 524;85;283;322;5 | -0.165250068 | -0.490588149 | -0.512410983 | -0.347160915 | -0.021822834 | 0.284534863 | 0.009161542 | 0.007350169 | 0.022632267 | 0.774561278 |
| Pnisr | S | 290 | 0.056654376 | -0.422633771 | -0.525102071 | -0.581756448 | -0.1024683 | 0.127922314 | 6.48963E-07 | 0.018615224 | 0.013190742 | 0.333116418 |
| Lemd3 | S | 338 | -0.040084633 | -0.299883697 | -0.435644879 | -0.395560246 | -0.135761182 | 0.470048248 | 0.001928822 | 9.00989E-05 | 0.001624987 | 0.055083496 |
| Gja1 | S | 306 | -0.042752981 | -0.330147942 | -0.450346911 | -0.407593929 | -0.120198969 | 0.376186072 | 0.000448271 | 0.001051479 | 0.010355754 | 0.133602031 |
| Map1b | S | 1403 | -0.180200414 | -0.314335173 | -0.201406557 | -0.021206144 | 0.112928616 | 0.007094156 | 0.000388688 | 0.004766241 | 0.413995 | 0.009606283 |
| Slc7a6os | T | 42 | -0.001220045 | -0.433200463 | -0.516990773 | -0.515770729 | -0.08379031 | 0.977808453 | 0.008185561 | 0.000190351 | 0.000584622 | 0.234774399 |
| Axl | Y | 696 | -0.503722886 | -0.347071288 | -0.462503575 | 0.041219311 | -0.115432287 | 0.011529503 | 0.11326967 | 0.050261613 | 0.81198963 | 0.580350233 |
| Bub1b | S | 535 | -0.075152368 | -0.200747876 | -0.33638733 | -0.261234962 | -0.135639454 | 0.340985144 | 8.74432E-05 | 2.47562E-05 | 0.038966666 | 0.009964658 |
| Phkg2 | S | 261 | -0.032743516 | -0.305366076 | -0.392097445 | -0.359353926 | -0.086731368 | 0.419858611 | 0.001455799 | 5.55887E-05 | 0.0016708 | 0.057882091 |
| Gtf2b | S | 76 | -0.258445744 | -0.275676341 | -0.393478733 | -0.135032988 | -0.117802392 | 0.009398163 | 0.096004862 | 0.001094146 | 0.03158631 | 0.378703731 |
| Alkbh1 | S | 378 | -0.139007353 | -0.353672376 | -0.373740322 | -0.234732969 | -0.020067947 | 0.030966 | 0.010085798 | 0.001373766 | 0.014606861 | 0.800671955 |
| Ccdc88a | S | 1653 | 0.092950871 | -0.346601403 | -0.364202516 | -0.457153387 | -0.017601113 | 0.021872339 | 1.58582E-05 | 2.30208E-06 | 0.00012444 | 0.511155083 |
| Pom121 | S | 450 | -0.179988286 | -0.421087569 | -0.321506704 | -0.141518418 | 0.099580865 | 0.025450099 | 0.033291807 | 0.00073267 | 0.057961376 | 0.42576953 |
| Cdc6 | S | 72 | -0.165413104 | -0.465208266 | -0.359643984 | -0.19423088 | 0.105564282 | 0.329058549 | 0.004008552 | 0.001896205 | 0.269580232 | 0.234167373 |
| Usp15 | S | 229 | -0.084975663 | -0.25466443 | -0.31797296 | -0.232997298 | -0.06330853 | 0.070309738 | 0.001093816 | 7.34716E-05 | 0.003769036 | 0.093555828 |
| Map1b | S | 1251 | -0.08831087 | -0.26373812 | -0.224727188 | -0.136416318 | 0.039010932 | 0.045562641 | 5.15795E-05 | 0.000172215 | 0.020105491 | 0.054110249 |
| Twistnb | S | 325 | -0.000504493 | -0.28353212 | -0.271274147 | -0.270769654 | 0.012257973 | 0.991539779 | 2.65578E-05 | 6.43434E-06 | 0.015312539 | 0.584061661 |
| Zbtb17 | S | 348 | -0.01106515 | -0.428269579 | -0.538443378 | -0.527378228 | -0.110173798 | 0.877944869 | 0.000390831 | 0.006080824 | 0.00340681 | 0.186852869 |
| Mia3 | S | 1733 | -0.237745732 | -0.324826699 | -0.439018158 | -0.201272426 | -0.114191459 | 0.030671967 | 0.029459831 | 0.004295946 | 0.08636504 | 0.34423074 |
| Grasp | S | 93 | -0.019446497 | -0.381982412 | -0.328905397 | -0.3094589 | 0.053077015 | 0.672705943 | 6.6536E-05 | 0.000872017 | 0.004420037 | 0.302437329 |
| Nudt9 | T | 67 | -0.294893072 | -0.379682983 | -0.35997281 | -0.065079738 | 0.019710173 | 0.036779333 | 0.016026059 | 0.010101255 | 0.518823217 | 0.835240142 |
| Strap | S | 335 | -0.050136214 | -0.305736703 | -0.320182867 | -0.270046653 | -0.014446164 | 0.141685402 | 0.001552382 | 3.78841E-05 | 0.000876611 | 0.696862695 |
| Rangap1 | S | 405 | -0.135587615 | -0.258742715 | -0.318281389 | -0.182693773 | -0.059538673 | 0.008432838 | 0.021711963 | 8.90061E-05 | 0.005682856 | 0.378917939 |
| Cmip | S | 349 | -0.170619402 | -0.390406611 | -0.300778312 | -0.130158911 | 0.089628299 | 0.048032047 | 0.001241831 | 0.00933783 | 0.123471089 | 0.229785575 |
| Trim47 | S | 591 | -0.091856323 | -0.24564764 | -0.344486155 | -0.252629833 | -0.098838516 | 0.13673785 | 0.000726992 | 0.000125599 | 0.00820911 | 0.05994932 |
| Arhgap22 | S | 424 | 0.00443729 | -0.228108953 | -0.309199454 | -0.313636743 | -0.0810905 | 0.789322225 | 0.00022364 | 2.65071E-07 | 1.01613E-05 | 0.028296428 |
| Tenm3 | S | 27 | 0.007146878 | -0.515471443 | -0.339928016 | -0.347074894 | 0.175543427 | 0.800831049 | 1.52287E-05 | 0.018848529 | 0.02241425 | 0.082920486 |
| Ankrd11 | S | 1799 | -0.220056927 | -0.299734804 | -0.511276075 | -0.291219148 | -0.211541271 | 0.008608006 | 0.050124445 | 0.009510269 | 0.071140788 | 0.162287287 |

| | | | | | | | | | | | | |
|---|---|---|---|---|---|---|---|---|---|---|---|---|
| Slain2 | S | 88 | -0.444184946 | -0.34564853 | -0.41504078 | 0.029144166 | -0.06939225 | 0.014000003 | 0.063294621 | 0.039391641 | 0.822149741 | 0.65219677 |
| Tbc1d1 | S | 564 | -0.087758818 | -0.344585248 | -0.400039106 | -0.312280288 | -0.055453858 | 0.183204573 | 0.001361459 | 0.00146085 | 0.00724923 | 0.377943004 |
| Prrc2b | S | 562 | 0.016363728 | -0.28968518 | -0.363567178 | -0.379930906 | -0.073881998 | 0.54474501 | 0.000566481 | 3.48523E-06 | 6.86581E-05 | 0.099890416 |
| Epn2 | T | 271 | -0.208531321 | -0.223796378 | -0.506740657 | -0.298209336 | -0.282944279 | 0.053844012 | 0.091052135 | 0.000385964 | 0.015518303 | 0.056272758 |
| Nes | S | 606 | -0.159303045 | -0.362557656 | -0.30543259 | -0.146129545 | 0.057125066 | 0.075116249 | 0.000148331 | 0.022311502 | 0.170810445 | 0.488988368 |
| Zfp24 | T | 330 | -0.146581801 | -0.288725567 | -0.357739428 | -0.211157627 | -0.069013862 | 0.049228402 | 0.002631978 | 0.001411041 | 0.018711621 | 0.239902083 |
| G3bp1 | T | 268 | -0.033898191 | -0.322105101 | -0.399932754 | -0.366034562 | -0.077827652 | 0.533639875 | 0.000724187 | 0.000223122 | 0.001693829 | 0.112831106 |
| Map7d1 | S | 519 | -0.057880933 | -0.369868958 | -0.311496528 | -0.253615594 | 0.05837243 | 0.587600239 | 0.000232196 | 0.000363166 | 0.092053805 | 0.201240725 |
| Myo5a | S | 1650 | -0.098199748 | -0.303093937 | -0.394646106 | -0.296446359 | -0.091552169 | 0.231574151 | 0.001959024 | 0.000423952 | 0.017468934 | 0.088415358 |
| Dab2 | T | 671 | -0.087854158 | -0.313049452 | -0.36447837 | -0.276624212 | -0.051428918 | 0.049333484 | 0.000196203 | 0.006482746 | 0.027078467 | 0.457574917 |
| Snap23 | T | 123 | -0.230052799 | -0.267495292 | -0.398956122 | -0.168903324 | -0.13146083 | 0.01882378 | 0.012590078 | 0.004569173 | 0.102049645 | 0.181194089 |
| C2cd2l | S | 468 | -0.010875516 | -0.300066463 | -0.254618837 | -0.243743321 | 0.045447626 | 0.714621046 | 1.10634E-05 | 4.80165E-05 | 0.000818341 | 0.107036825 |
| Peak1 | T | 1141 | -0.148176226 | -0.196052643 | -0.251700782 | -0.103524556 | -0.05564814 | 0.001112285 | 0.016681739 | 3.81971E-05 | 0.016942426 | 0.286642801 |
| Akt3 | T | 447 | -0.134333827 | -0.20773285 | -0.487566255 | -0.353232428 | -0.279833404 | 0.02958021 | 0.003445284 | 0.002209936 | 0.00751087 | 0.016090497 |
| Tpr | T | 2205 | -0.201712303 | -0.312586684 | -0.426496615 | -0.224784312 | -0.113909931 | 0.038672485 | 0.024187037 | 0.002275927 | 0.037875083 | 0.292262793 |
| Cspg4 | T | 2266 | -0.330406145 | -0.341144259 | -0.311392099 | 0.019014045 | 0.02975216 | 0.022699798 | 0.011887934 | 0.014448986 | 0.852726287 | 0.748163104 |
| Ubash3b | T | 12 | 0.001040586 | -0.481475631 | -0.484535197 | -0.485575783 | -0.003059567 | 0.980720508 | 0.000587092 | 0.004800822 | 0.002395596 | 0.963186552 |
| Cdk11b | S | 264 | -0.199975225 | -0.248767808 | -0.467173955 | -0.26719873 | -0.218406146 | 0.060927365 | 0.016335288 | 0.001600299 | 0.035322885 | 0.047557141 |
| Ahsa2 | S | 189 | -0.080042051 | -0.365222348 | -0.464532021 | -0.38448997 | -0.099309673 | 0.172987831 | 0.005152001 | 0.001590232 | 0.00384892 | 0.227874027 |
| Dnmt1 | S | 190 | -0.116638378 | -0.334297504 | -0.280752533 | -0.164114154 | 0.053544972 | 0.058018181 | 0.001947901 | 0.000449097 | 0.019076305 | 0.344928731 |
| Mybbp1a | T | 1232 | -0.143411254 | -0.430033217 | -0.355279345 | -0.211868092 | 0.074753872 | 0.347963661 | 0.000285993 | 0.014292462 | 0.211031445 | 0.420621652 |
| Alkbh5 | S | 375 | -0.22379925 | -0.31541064 | -0.514215934 | -0.290416683 | -0.198805294 | 0.018810345 | 0.061557681 | 0.006614166 | 0.035120761 | 0.174085345 |
| Slc12a2 | S | 74 | -0.309818709 | -0.255812585 | -0.424173059 | -0.11435435 | -0.168360474 | 0.022434152 | 0.058038682 | 0.003510251 | 0.210024708 | 0.126431031 |
| D10Wsu102e | S | 66 | -0.097218636 | -0.328260319 | -0.461815933 | -0.364597297 | -0.133555614 | 0.179033568 | 0.001034259 | 0.004659505 | 0.018270624 | 0.187459278 |
| Otud4 | S | 1000 | -0.022540744 | -0.238321825 | -0.316425228 | -0.293884483 | -0.078103402 | 0.374863488 | 0.001930179 | 7.14918E-07 | 0.000706083 | 0.052736657 |
| Dync1li2 | S | 446 | -0.047731171 | -0.185259297 | -0.303028948 | -0.255297777 | -0.117769651 | 0.17102677 | 6.96017E-05 | 1.06473E-05 | 0.001301266 | 0.007570426 |
| Sin3a | S | 431 | -0.128947032 | -0.319834952 | -0.36748404 | -0.238537008 | -0.047649088 | 0.175572953 | 0.00046003 | 0.003454348 | 0.048366609 | 0.482997489 |
| Hsp90ab1 | T | 446 | -0.019213154 | -0.306221921 | -0.328820458 | -0.309607304 | -0.022598537 | 0.719909122 | 5.64343E-06 | 0.001273304 | 0.006484984 | 0.557832603 |
| Clip2 | S | 295 | 0.002171723 | -0.377855701 | -0.363260054 | -0.365431778 | 0.014595647 | 0.954588422 | 2.59901E-05 | 0.002115302 | 0.004074221 | 0.779493216 |
| Ppp1r13l | S | 151 | 0.049596933 | -0.365611625 | -0.336362027 | -0.38595896 | 0.029249598 | 0.214545099 | 2.45084E-05 | 7.44657E-05 | 0.000352656 | 0.374295352 |
| Akt1s1 | S | 213 | -0.031435209 | -0.209497185 | -0.285656148 | -0.25422094 | -0.076158963 | 0.360054592 | 0.000132977 | 2.09056E-06 | 0.003786558 | 0.021614596 |
| H1f4 | T | 18 | 0.026293292 | -0.345243816 | -0.39099358 | -0.417286872 | -0.045749764 | 0.607063965 | 7.23938E-05 | 0.000306134 | 0.00108442 | 0.349313193 |
| Atg2b | S | 496 | -0.29966277 | -0.320689814 | -0.533900192 | -0.234237422 | -0.213210378 | 0.062568209 | 0.042745799 | 0.010423063 | 0.14430056 | 0.164103538 |
| Mpzl1 | S | 210 | -0.033749661 | -0.312201899 | -0.283215689 | -0.249466028 | 0.028986211 | 0.463504606 | 1.14946E-06 | 0.004043091 | 0.00719447 | 0.470536859 |
| Hif1a | S | 450 | 0.074624496 | -0.509762041 | -0.469518903 | -0.544143399 | 0.040243138 | 0.390002082 | 0.000934772 | 0.000653759 | 0.0034326 | 0.597311004 |
| Mtrex | S | 38 | -0.045585188 | -0.379399453 | -0.37602846 | -0.330443272 | 0.003370993 | 0.042475532 | 0.000529918 | 0.005357232 | 0.013398976 | 0.949497515 |
| Psma2 | S | 7 | 0.045937289 | -0.320280215 | -0.432778404 | -0.478715693 | -0.112498189 | 0.594221831 | 2.10243E-05 | 0.000714638 | 0.009522476 | 0.087452076 |
| Pard3 | Y | 1076 | -0.224166425 | -0.360401988 | -0.388097289 | -0.163930864 | -0.027695301 | 0.03835391 | 0.030829389 | 0.003089378 | 0.076925044 | 0.80670286 |
| Nav3 | S | 1248 | -0.04442664 | -0.340346897 | -0.335498217 | -0.291071576 | 0.00484868 | 0.327119991 | 3.27268E-05 | 0.003609987 | 0.00484515 | 0.91787572 |
| Tepsin | S | 525 | -0.375832819 | -0.51677747 | -0.877818967 | -0.501986147 | -0.361041496 | 0.133994751 | 0.071488115 | 0.096182304 | 0.241448059 | 0.373021145 |
| Cnot3 | S | 299 | -0.096787494 | -0.257419121 | -0.384907346 | -0.288119852 | -0.127488225 | 0.037658416 | 0.009898593 | 0.000191303 | 0.000898048 | 0.040307229 |
| Rpl4 | S | 295 | -0.053038871 | -0.245325695 | -0.316433685 | -0.263394814 | -0.071107991 | 0.040033724 | 0.013449549 | 2.30561E-06 | 0.000219751 | 0.163437145 |
| Zfp553 | S | 325 | -0.327261262 | -0.55636994 | -0.849355911 | -0.522094649 | -0.292985971 | 0.278899567 | 0.035061949 | 0.083018536 | 0.233834784 | 0.439961142 |
| Akt1 | S | 477 | -0.001933477 | -0.572113905 | -0.522154164 | -0.520220687 | 0.049959741 | 0.970759495 | 0.000793024 | 0.019235043 | 0.014777247 | 0.649172984 |
| Smarca5 | S | 136 | -0.320587908 | -0.283903318 | -0.542076955 | -0.221489047 | -0.258173637 | 0.058324801 | 0.141723567 | 0.003215837 | 0.116925173 | 0.157746663 |
| Anln | S | 653 | -0.20162264 | -0.274040745 | -0.485949224 | -0.284326585 | -0.211908479 | 0.044108713 | 0.030157029 | 0.002822959 | 0.046596063 | 0.086204703 |
| Cd3eap | S | 170 | 0.048695839 | -0.334904979 | -0.410376836 | -0.459072675 | -0.075471856 | 0.200541452 | 2.28277E-06 | 0.001268123 | 0.000717388 | 0.153775494 |
| Phactr4 | S | 457 | -0.09173555 | -0.38792089 | -0.575698258 | -0.483962709 | -0.187777368 | 0.380790069 | 0.001583164 | 0.013027672 | 0.013350597 | 0.12720141 |
| Inf2 | T | 377 | -0.140995037 | -0.389716186 | -0.463129384 | -0.322134347 | -0.073413198 | 0.043679541 | 0.000226085 | 0.122329286 | 0.223572 | 0.732442024 |
| Zswim8 | S | 1204 | -0.201267317 | -0.291136021 | -0.530394454 | -0.329127136 | -0.239258432 | 0.058698779 | 0.106759688 | 0.001131083 | 0.013446694 | 0.163126344 |
| Ssrp1 | S | 657 | -0.095306365 | -0.281680167 | -0.256896468 | -0.161590104 | 0.024783699 | 0.051022897 | 0.00094467 | 0.000116492 | 0.027964511 | 0.493538604 |
| Acaca | S | 79 | -0.348046291 | -0.273344724 | -0.380015773 | -0.031969482 | -0.106671049 | 0.000477132 | 0.104340881 | 0.058675413 | 0.801647829 | 0.525295844 |
| Dok1 | Y | 450 | -0.405680576 | -0.335991357 | -0.429932371 | -0.024251795 | -0.093941014 | 0.022822755 | 0.165637367 | 0.008266328 | 0.839126038 | 0.637887448 |
| Cc2d1a | S | 667 | -0.056919788 | -0.389283182 | -0.491454284 | -0.434534496 | -0.102171102 | 0.773544352 | 0.00129252 | 0.002428501 | 0.105917259 | 0.264475191 |
| Aspscr1 | S | 230 | -0.177204805 | -0.253194152 | -0.512814214 | -0.335609408 | -0.259620062 | 0.01145403 | 0.02850108 | 0.006338121 | 0.018805443 | 0.040865401 |
| Dnajc21 | S | 283 | -0.086070322 | -0.172004503 | -0.223501581 | -0.137431259 | -0.051497078 | 0.000986933 | 0.010095353 | 6.25068E-07 | 0.00043221 | 0.141381681 |
| Cobll1 | S | 647 | -0.071495854 | -0.214740919 | -0.294180342 | -0.222684488 | -0.079439423 | 0.143473763 | 6.3187E-05 | 0.000116566 | 0.006899279 | 0.049145265 |
| Tmpo | S | 384 | -0.017562292 | -0.230490958 | -0.512329234 | -0.494766942 | -0.281838276 | 0.765392774 | 0.033592282 | 5.39506E-06 | 0.007136172 | 0.035976672 |
| Tacc2 | S | 587 | -0.175323499 | -0.444334683 | -0.392706254 | -0.217382755 | 0.051628429 | 0.152725585 | 0.00058403 | 0.059977015 | 0.21864727 | 0.712039818 |
| Ftsj3 | S | 468 | -0.27466318 | -0.322343076 | -0.346207315 | -0.071544135 | -0.023864239 | 0.03333584 | 0.00262152 | 0.032339246 | 0.537812797 | 0.81234504 |
| 2310033P09Rik | S | 245 | -0.405332934 | -0.256664599 | -0.254714741 | 0.150618194 | 0.001949858 | 0.015104093 | 0.011221808 | 0.011977759 | 0.207469048 | 0.966947479 |
| Tcof1 | S | 311 | -0.146233611 | -0.281275015 | -0.496520551 | -0.35028694 | -0.215245537 | 0.086624936 | 0.082051732 | 0.000316226 | 0.001651848 | 0.154802607 |
| Eif4b | S | 504 | -0.034712963 | -0.161501485 | -0.227547851 | -0.192834888 | -0.066046366 | 0.038756432 | 3.69979E-05 | 5.01066E-07 | 7.50169E-05 | 0.009960523 |
| Hnrnpa0 | S | 282 | -0.186327812 | -0.23298711 | -0.45186414 | -0.265536328 | -0.21887703 | 0.033124321 | 0.063817328 | 0.000570226 | 0.009534118 | 0.079636868 |
| Nefm | S | 30 | 0.022708515 | -0.323311386 | -0.404036718 | -0.426745234 | -0.080725332 | 0.737877811 | 0.00042367 | 9.72828E-05 | 0.003619992 | 0.106969375 |
| Snapc1 | S | 268 | -0.041781928 | -0.277154159 | -0.293087037 | -0.251305108 | -0.015932878 | 0.472550303 | 0.000111957 | 7.53585E-05 | 0.016152885 | 0.600327628 |
| Gtse1 | S | 736 | -0.641770008 | -0.47958132 | -0.39312943 | 0.248640578 | 0.08645189 | 0.053697874 | 0.008048282 | 0.476368684 | 0.649298695 | 0.865992002 |
| Psma5 | S | 56 | -0.248995293 | -0.333438609 | -0.438908441 | -0.189913148 | -0.105469832 | 0.043340761 | 0.005556804 | 0.031713254 | 0.228778452 | 0.449441605 |
| Cdk17 | S | 122 | 0.002248252 | -0.304985346 | -0.34655361 | -0.348801862 | -0.041568264 | 0.94671067 | 0.002901879 | 2.84866E-06 | 0.002499507 | 0.354841027 |
| Med13 | S | 395 | -0.02821988 | -0.410527003 | -0.42061947 | -0.39239959 | -0.010092467 | 0.728064121 | 0.001836261 | 0.00085845 | 0.002794982 | 0.869941122 |
| Clasp2 | S | 1005 | -0.115802304 | -0.284098728 | -0.403391937 | -0.287589633 | -0.119293209 | 0.046442139 | 0.015149498 | 0.000566344 | 0.006676887 | 0.173635972 |
| Map1b | S | 1420 | -0.066810978 | -0.251309568 | -0.346779041 | -0.279968063 | -0.095469473 | 0.011995842 | 9.31829E-06 | 0.018962858 | 0.032072506 | 0.224019788 |
| Traf4 | S | 426 | -0.17657563 | -0.337101086 | -0.491872067 | -0.315296437 | -0.154770982 | 0.10265352 | 0.002130415 | 0.025058733 | 0.083123562 | 0.2926776 |
| Yap1 | S | 46 | -0.325709867 | -0.390994123 | -0.492812633 | -0.167102766 | -0.10181851 | 0.057864088 | 0.086689252 | 0.011853972 | 0.262429016 | 0.559593189 |
| Ddx39 | T | 171 | -0.076835748 | -0.278459669 | -0.422623369 | -0.345787621 | -0.144163701 | 0.491588485 | 0.002485616 | 0.00020653 | 0.047269042 | 0.022280833 |
| Nolc1 | T | 212 | 0.084707994 | -0.345263309 | -0.489024924 | -0.573732918 | -0.143761615 | 0.160527618 | 0.000202983 | 0.000123066 | 0.000409837 | 0.041651634 |
| Hmces | S | 302 | -0.228410622 | -0.276526603 | -0.235041784 | -0.006631162 | 0.041484819 | 0.057551255 | 9.59918E-05 | 0.013049305 | 0.938167086 | 0.445418794 |
| Rgs12 | S | 1233 | 0.03009591 | -0.408645814 | -0.41116583 | -0.44126174 | -0.002520016 | 0.741060609 | 0.000670906 | 0.000448603 | 0.01011381 | 0.942368296 |
| Cfl1 | S | 41 | -0.305939677 | -0.353009854 | -0.555523729 | -0.249584052 | -0.202513875 | 0.106623614 | 0.071833741 | 0.008843886 | 0.184494146 | 0.261815226 |
| Smg7 | S | 732 | -0.059391287 | -0.392379059 | -0.420948382 | -0.361557095 | -0.028569324 | 0.08690649 | 0.015840947 | 0.000677326 | 0.001429788 | 0.705900118 |
| Phldb1 | S | 503 | -0.006565088 | -0.436831554 | -0.501164238 | -0.49459915 | -0.064332684 | 0.9192491 | 0.004589862 | 0.001075518 | 0.003079909 | 0.497653887 |
| Bzw1 | S | 12 | -0.156888933 | -0.227698941 | -0.508638367 | -0.351749434 | -0.280939426 | 0.137878344 | 0.027707626 | 0.000584314 | 0.008973563 | 0.007485241 |
| Vps72 | T | 103 | 0.031189934 | -0.293396571 | -0.342692427 | -0.373882362 | -0.049295857 | 0.595584977 | 0.000366656 | 7.52288E-06 | 0.010175924 | 0.204071844 |
| Gja1 | S | 255 | 0.041519558 | -0.296004932 | -0.461915763 | -0.503435321 | -0.165910831 | 0.440830651 | 0.000723346 | 5.51466E-05 | 0.000226638 | 0.013896524 |
| Smg1 | S | 3585 | -0.093498377 | -0.278096478 | -0.340551521 | -0.247053144 | -0.062455043 | 0.200116684 | 1.35773E-05 | 0.010039195 | 0.0262801 | 0.339055276 |
| Ddx18 | S | 86 | 0.035890492 | -0.354804272 | -0.350040783 | -0.385931275 | 0.004763489 | 0.671670016 | 0.000796213 | 2.30533E-05 | 0.025707878 | 0.913732882 |
| Pdzrn3 | S | 840 | -0.125878296 | -0.270376066 | -0.436820963 | -0.310942667 | -0.166444897 | 0.091350989 | 0.02409786 | 0.000387826 | 0.000578185 | 0.107468254 |
| Myh9 | T | 1151 | -0.055582729 | -0.225655932 | -0.411885755 | -0.356303027 | -0.186229823 | 0.306859364 | 0.009240503 | 1.82665E-05 | 0.006278673 | 0.036890532 |
| Dnmt1 | S | 187 | -0.131598898 | -0.369165757 | -0.255985691 | -0.124386793 | 0.113180066 | 0.113056569 | 0.000830191 | 0.002296523 | 0.117012066 | 0.088916835 |
| Flnb | S | 983 | -0.152204155 | -0.311323227 | -0.337975418 | -0.185771263 | -0.026652191 | 0.063730816 | 0.003624985 | 0.002367228 | 0.005406962 | 0.539482265 |
| Prdx6 | T | 93 | -0.420036048 | -0.381270215 | -0.42319662 | -0.003160572 | -0.041926405 | 0.055182344 | 0.011142899 | 0.100361126 | 0.9886277 | 0.829099223 |
| Slc4a7 | S | 57 | -0.207979107 | -0.26990255 | -0.581734946 | -0.373755838 | -0.311832395 | 0.07095193 | 0.059575158 | 0.004230878 | 0.022285581 | 0.050579465 |
| Sdc3 | S | 428 | -0.251718796 | -0.263473752 | -0.393721578 | -0.142002781 | -0.130247825 | 0.004831608 | 0.122264639 | 0.003438998 | 0.126468712 | 0.373759207 |
| Rpl17 | S | 141 | -0.230537743 | -0.317701567 | -0.293788957 | -0.063251213 | 0.02391261 | 0.006022269 | 0.000237179 | 0.2045676 | 0.735078437 | 0.896092761 |

| | | | | | | | | | | | | |
|---|---|---|---|---|---|---|---|---|---|---|---|---|
| Zc3hav1 | S | 489 | -0.10103931 | -0.511685378 | -0.293628032 | -0.192588721 | 0.218057346 | 0.303362258 | 0.000462844 | 0.02080617 | 0.094771887 | 0.070656768 |
| Exoc1 | S | 487 | -0.341372641 | -0.206476729 | -0.451249788 | -0.109877146 | -0.244773059 | 0.030526593 | 0.057446712 | 0.005435603 | 0.378881694 | 0.06282992 |
| Snx17 | S | 409 | -0.179324932 | -0.222179659 | -0.43767627 | -0.258351337 | -0.215496611 | 0.03071135 | 0.079241182 | 0.000397176 | 0.00181656 | 0.097501525 |
| Pxn | S | 137 | -0.026204814 | -0.316954532 | -0.336123103 | -0.309918289 | -0.019168572 | 0.671603276 | 0.000189382 | 0.000329909 | 0.008409082 | 0.666622559 |
| Btf3 | S | 156 | -0.016586859 | -0.382906609 | -0.401279225 | -0.384692367 | -0.018372617 | 0.833358908 | 6.71889E-06 | 0.027425971 | 0.022585815 | 0.842585319 |
| Tpd52 | S | 170 | -0.107776684 | -0.291174253 | -0.521668254 | -0.41389157 | -0.230494001 | 0.252968316 | 0.025347421 | 0.000576943 | 0.00415654 | 0.056974786 |
| Map7d1 | S | 552 | -0.097572791 | -0.30212979 | -0.232197698 | -0.134624906 | 0.069932093 | 0.021465884 | 0.000224829 | 0.00233739 | 0.030221274 | 0.163966489 |
| Plcb3 | S | 537 | -0.086534105 | -0.428846822 | -0.257335284 | -0.170801179 | 0.171511538 | 0.13787412 | 0.00065415 | 0.003691395 | 0.028541821 | 0.043410997 |
| Eif3a | S | 1008 | -0.160500034 | -0.371695661 | -0.416587613 | -0.256087579 | -0.044891952 | 0.195270091 | 0.004259975 | 0.00645621 | 0.074503208 | 0.500545292 |
| Prrc2b | S | 226 | 0.014792154 | -0.346946148 | -0.331838782 | -0.346630936 | 0.015107366 | 0.717835591 | 0.000377132 | 5.67651E-05 | 0.002378265 | 0.690218388 |
| Ap3b1 | S | 761 | -0.027705835 | -0.358709264 | -0.382349873 | -0.354644038 | -0.023640609 | 0.629238379 | 0.001480909 | 0.000361159 | 0.002423462 | 0.699991303 |
| Gja1 | S | 282 | 0.026360944 | -0.322688639 | -0.482918364 | -0.509279309 | -0.160229726 | 0.672865219 | 0.001422869 | 0.000197444 | 0.002908448 | 0.000199241 |
| Map1b | S | 1621 | -0.178372543 | -0.38818364 | -0.308319672 | -0.129947129 | 0.079863968 | 0.06005583 | 0.000308009 | 0.071969918 | 0.341066127 | 0.516131107 |
| Mast3 | S | 370 | 0.020786913 | -0.520959025 | -0.409521968 | -0.43030888 | 0.111437057 | 0.834636582 | 0.029932254 | 2.4481E-05 | 0.030062211 | 0.387237089 |
| Pdlim5 | S | 111 | -0.083382507 | -0.320593349 | -0.289361445 | -0.205978938 | 0.031231904 | 0.031928008 | 0.003990879 | 0.000381726 | 0.00344122 | 0.531992215 |
| Rpl15 | T | 197 | 0.058455665 | -0.485322674 | -0.478283622 | -0.536739287 | 0.007039052 | 0.189040781 | 0.057315491 | 1.91349E-06 | 0.002378364 | 0.960640366 |
| Ppp1r14b | S | 143 | -0.208398611 | -0.338876421 | -0.569498274 | -0.361099663 | -0.230621853 | 0.042374683 | 0.020415784 | 0.030342078 | 0.103343792 | 0.228243074 |
| Eif4b | S | 542 | -0.09988065 | -0.298759135 | -0.439921483 | -0.340040832 | -0.141162348 | 0.258340141 | 0.01335878 | 0.000361478 | 0.010970748 | 0.133568607 |
| Dst | T | 2409 | -0.229028335 | -0.329314093 | -0.38152163 | -0.152493296 | -0.052207537 | 0.031987281 | 0.008269598 | 0.019567285 | 0.207305951 | 0.626470716 |
| Hnrnpab | S | 260 | -0.178203487 | -0.117804783 | -0.284462581 | -0.106259094 | -0.166657797 | 0.001005867 | 0.0468032 | 6.21243E-05 | 0.001923981 | 0.030008016 |
| Rsl1d1 | T | 360 | 0.090098121 | -0.336460309 | -0.488105246 | -0.578203367 | -0.151644937 | 0.345563367 | 0.002454834 | 2.93674E-05 | 0.013746788 | 0.071809801 |
| Tnks1bp1 | S | 568 | -0.157988143 | -0.243099834 | -0.191041955 | -0.033053812 | 0.052057879 | 0.003798725 | 0.000452374 | 0.00270761 | 0.325305946 | 0.195898276 |
| Acin1 | S | 583 | -0.105018339 | -0.252059924 | -0.208071113 | -0.103052774 | 0.043988811 | 0.00948269 | 0.000146412 | 0.001347368 | 0.043194968 | 0.248242599 |
| Ssrp1 | S | 621 | -0.067933346 | -0.24538094 | -0.306428211 | -0.238494866 | -0.061047271 | 0.080433761 | 6.5097E-05 | 0.001511226 | 0.005892858 | 0.201956403 |
| Heatr5b | S | 1737 | -0.097527676 | -0.31739598 | -0.36653742 | -0.269009743 | -0.04914144 | 0.031582258 | 0.002762072 | 0.004266465 | 0.013035139 | 0.455880852 |
| Nup98 | T | 546 | 0.08206466 | -0.276774208 | -0.422466886 | -0.504531546 | -0.145692678 | 0.3473528 | 2.71903E-05 | 0.000122613 | 0.00978281 | 0.012755495 |
| Arhgap31 | S | 829 | -0.186515452 | -0.29239643 | -0.4157458 | -0.229230347 | -0.123349369 | 0.129620588 | 0.010496154 | 0.002288649 | 0.088249963 | 0.192953776 |
| Rplp2 | S | 17 | -0.037393254 | -0.261034475 | -0.302580231 | -0.265186978 | -0.041545757 | 0.444763976 | 4.67214E-05 | 0.000284341 | 0.007467839 | 0.229330028 |
| Pard3 | S | 667 | -0.111507313 | -0.422890764 | -0.399638763 | -0.28813145 | 0.023252001 | 0.124600031 | 0.000799949 | 0.038564116 | 0.06360842 | 0.826458272 |
| Erf | S | 433 | -0.242769299 | -0.349824537 | -0.320789787 | -0.078020488 | 0.02903475 | 0.018134351 | 0.003558269 | 0.05414685 | 0.511643652 | 0.797688847 |
| Erf | S | 437 | -0.242769299 | -0.349824537 | -0.320789787 | -0.078020488 | 0.02903475 | 0.018134351 | 0.003558269 | 0.05414685 | 0.511643652 | 0.797688847 |
| Erf | S | 446 | -0.242769299 | -0.349824537 | -0.320789787 | -0.078020488 | 0.02903475 | 0.018134351 | 0.003558269 | 0.05414685 | 0.511643652 | 0.797688847 |
| Stim2 | S | 697 | -0.031386821 | -0.3951983 | -0.580620229 | -0.549233408 | -0.185421928 | 0.786425023 | 0.097477644 | 0.000100994 | 0.015358957 | 0.32526748 |
| Arhgef2 | S | 931 | -0.074190337 | -0.398245581 | -0.301426301 | -0.227235964 | 0.096819279 | 0.456582884 | 1.19636E-05 | 0.03832021 | 0.104117201 | 0.316396939 |
| Nhsl1 | S | 1235 | -0.052703848 | -0.23636779 | -0.296406986 | -0.243703138 | -0.060039196 | 0.068665431 | 3.74703E-05 | 0.001017964 | 0.0084217 | 0.170805944 |
| 2310057M21Rik | S | 435 | -0.093631497 | -0.383748468 | -0.360789519 | -0.267158022 | 0.022958948 | 0.1059504 | 0.000481582 | 0.020641609 | 0.049721875 | 0.798329852 |
| Spata5 | S | 18 | 0.035876592 | -0.382509934 | -0.365967204 | -0.401843797 | 0.01654273 | 0.3770145 | 0.00022945 | 0.000233338 | 0.000719745 | 0.729929773 |
| Dlgap4 | S | 716 | -0.145246863 | -0.210092441 | -0.366082505 | -0.220835643 | -0.155990064 | 0.025467098 | 0.017637081 | 0.000373751 | 0.001666701 | 0.065903436 |
| Prrc2a | S | 1666 | 0.007553471 | -0.361371726 | -0.359480072 | -0.367033543 | 0.001891654 | 0.758751487 | 1.66126E-06 | 0.017841054 | 0.022448547 | 0.978059637 |
| Cep170 | S | 1260 | -0.389155913 | -0.151235767 | -0.579996753 | -0.19084084 | -0.428760986 | 0.015483058 | 0.307881826 | 0.009127971 | 0.217059879 | 0.038587469 |
| Psme3 | S | 24 | -0.016703053 | -0.395162566 | -0.363673635 | -0.346970582 | 0.031488931 | 0.854317812 | 1.80034E-05 | 0.017038496 | 0.027680028 | 0.681255714 |
| Smarca4 | S | 1384 | -0.006337226 | -0.245355533 | -0.249312747 | -0.242975522 | -0.003957215 | 0.792528506 | 1.3475E-05 | 1.9548E-05 | 0.000226418 | 0.83666928 |
| Ank3 | T | 1803 | 0.040003541 | -0.272480923 | -0.437606452 | -0.477609992 | -0.165125529 | 0.434108515 | 0.000740741 | 3.44328E-05 | 0.000216928 | 0.010719487 |
| Phax | T | 80 | -0.136522462 | -0.233146272 | -0.348577484 | -0.212055022 | -0.115431212 | 0.322651781 | 0.00090951 | 0.000340157 | 0.172878614 | 0.062529799 |
| Dab2 | S | 323 | -0.154447062 | -0.271326595 | -0.365594411 | -0.211147349 | -0.094267816 | 0.099082089 | 0.000176526 | 0.018235308 | 0.078875243 | 0.296192248 |
| Cad | T | 1884 | 0.009469431 | -0.366826925 | -0.485829855 | -0.495299285 | -0.119002929 | 0.928544498 | 0.135125866 | 2.41505E-06 | 0.02788208 | 0.51840311 |
| Mybbp1a | T | 8 | -0.042485559 | -0.342345866 | -0.440273155 | -0.397787596 | -0.097927289 | 0.676139452 | 0.000462039 | 0.003281904 | 0.019163039 | 0.247211071 |
| Tasor2 | S | 1430 | -0.160284984 | -0.229435928 | -0.296859749 | -0.136574765 | -0.067423821 | 0.004451006 | 0.006172147 | 0.002751869 | 0.044017187 | 0.259652534 |
| Wdr43 | S | 588 | -0.026589444 | -0.242824703 | -0.26063499 | -0.234045546 | -0.017810287 | 0.470761836 | 2.87944E-05 | 5.46806E-05 | 0.010758965 | 0.253738297 |
| Slc5a3 | S | 594 | -0.232288957 | -0.266603363 | -0.364246799 | -0.131957842 | -0.097643436 | 0.063395939 | 0.016403741 | 0.002335825 | 0.225966947 | 0.265941285 |
| Cdv3 | S | 51 | -0.098704927 | -0.277372391 | -0.485235866 | -0.386530939 | -0.207863476 | 0.280692044 | 0.002848252 | 0.002549775 | 0.012558085 | 0.061899807 |
| Map1b | S | 2030 | -0.124775572 | -0.173102754 | -0.24059674 | -0.115821168 | -0.067493987 | 0.022011692 | 0.000870218 | 5.4788E-05 | 0.036075726 | 0.056594665 |
| Uba1 | S | 816 | -0.041094669 | -0.44921285 | -0.491186615 | -0.450091946 | -0.041973765 | 0.470862589 | 0.002362516 | 0.009968239 | 0.010250588 | 0.659462881 |
| Abl2 | S | 632 | -0.084434586 | -0.358262651 | -0.394416702 | -0.309982117 | -0.036154052 | 0.160514755 | 0.003638719 | 0.002932662 | 0.006459504 | 0.581494372 |
| Wfs1 | S | 236 | -0.373548629 | -0.232331722 | -0.458257522 | -0.084708893 | -0.225925799 | 0.003512423 | 0.26553046 | 0.022481769 | 0.513761111 | 0.296394483 |
| Pgd | S | 257 | -0.24689732 | -0.217759249 | -0.514916232 | -0.268018911 | -0.297156983 | 0.012200127 | 0.087382334 | 0.009172256 | 0.093601026 | 0.069835734 |
| Prune2 | S | 1750 | -0.0817407 | -0.329199367 | -0.294803523 | -0.213062823 | 0.034395844 | 0.148931659 | 9.00356E-06 | 0.039874477 | 0.073206617 | 0.673115885 |
| Zfp771 | T | 254 | -0.159220708 | -0.357991647 | -0.532784245 | -0.373563537 | -0.174792598 | 0.396591142 | 0.124929189 | 0.000304086 | 0.117958753 | 0.354164615 |
| Ap3d1 | S | 688 | -0.080024354 | -0.272571181 | -0.305820347 | -0.225795993 | -0.033249166 | 0.003556551 | 0.002354631 | 0.002217691 | 0.015258459 | 0.468835646 |
| Vps4b | S | 102 | -0.298556095 | -0.261169504 | -0.274358473 | 0.024197622 | -0.013188969 | 0.00471747 | 0.033958203 | 0.010626891 | 0.730580948 | 0.877348277 |
| Anln | S | 180 | -0.055491433 | -0.291142226 | -0.248978269 | -0.193486836 | 0.042163958 | 0.238450007 | 0.000290982 | 0.000124008 | 0.011091064 | 0.268886254 |
| 2810004N23Rik | S | 51 | -0.152761922 | -0.334998398 | -0.247779112 | -0.09501719 | 0.087219286 | 0.162139786 | 0.000552196 | 0.002834565 | 0.32590621 | 0.065555121 |
| Ftsj3 | T | 464 | -0.257797921 | -0.334664155 | -0.302559966 | -0.044762045 | 0.032104189 | 0.217977984 | 0.007122681 | 0.001471115 | 0.794331967 | 0.661894565 |
| Trim25 | S | 404 | -0.0663756 | -0.308385091 | -0.33568694 | -0.269311339 | -0.027301848 | 0.161736673 | 0.000203604 | 0.004378909 | 0.014058256 | 0.623175736 |
| Ptdss1 | S | 417 | -0.049141838 | -0.465369754 | -0.518002553 | -0.468860714 | -0.052632798 | 0.202266816 | 0.004659941 | 0.016853322 | 0.013725372 | 0.565596971 |
| Ezh2 | S | 380 | -0.05382829 | -0.51869483 | -0.286753478 | -0.232925188 | 0.231941352 | 0.629110443 | 0.00043101 | 0.009766206 | 0.075349317 | 0.015014222 |
| Eif4b | S | 352 | -0.122650986 | -0.29225342 | -0.399623669 | -0.276972683 | -0.107370249 | 0.034256418 | 0.030595934 | 0.001000212 | 0.010491733 | 0.277875602 |
| Hnrnpc | S | 166 | -0.084794223 | -0.277990372 | -0.567182548 | -0.482388324 | -0.289192176 | 0.401430093 | 0.040579422 | 0.00044864 | 0.010959031 | 0.05581064 |
| Errfi1 | S | 460 | -0.012791123 | -0.355531837 | -0.563978002 | -0.551186879 | -0.208446165 | 0.898689674 | 0.01555996 | 0.00054388 | 0.009679271 | 0.067428024 |
| Eif4b | S | 263 | -0.245822515 | -0.279935882 | -0.360957335 | -0.11513482 | -0.081021453 | 0.016703856 | 0.009565121 | 0.022875444 | 0.303580816 | 0.444441603 |
| Cdc6 | S | 75 | -0.168616086 | -0.417221435 | -0.35509201 | -0.186475924 | 0.062129426 | 0.239346391 | 0.000912869 | 0.003482912 | 0.203033241 | 0.402551241 |
| Eef1g | T | 46 | -0.149315733 | -0.186334653 | -0.338688584 | -0.189372851 | -0.152353931 | 0.034791774 | 0.00334693 | 0.000692691 | 0.023295132 | 0.036858442 |
| Osbpl6 | S | 9 | -0.032597772 | -0.219523546 | -0.314264379 | -0.281666607 | -0.094740833 | 0.283726433 | 0.000721252 | 1.97245E-05 | 0.000346322 | 0.036739355 |
| Peg10 | S | 30 | -0.120854889 | -0.311331507 | -0.277479787 | -0.156624897 | 0.033851721 | 0.108892336 | 0.000732311 | 0.002463031 | 0.051800111 | 0.487206642 |
| Pbk | S | 58 | -0.172720813 | -0.228260157 | -0.318302553 | -0.14558174 | -0.090042396 | 0.023577813 | 0.058792158 | 0.000116243 | 0.047976917 | 0.31720815 |
| Mycbp2 | S | 3550 | -0.209805668 | -0.196948527 | -0.44528267 | -0.235477003 | -0.248334144 | 0.082289035 | 0.071184386 | 0.000421902 | 0.082770942 | 0.057084867 |
| Bora | S | 137 | -0.134573708 | -0.297865763 | -0.400872892 | -0.266299184 | -0.103007129 | 0.10594778 | 0.001883589 | 0.008149938 | 0.027587525 | 0.217460203 |
| Mast3 | S | 750 | -0.062412611 | -0.309297984 | -0.402266299 | -0.339853688 | -0.092968315 | 0.285989115 | 0.001197223 | 0.002034473 | 0.005660515 | 0.195293518 |
| Adam19 | S | 820 | -0.008828846 | -0.539889544 | -0.525352644 | -0.516523798 | 0.0145369 | 0.860119412 | 0.010022388 | 0.004433374 | 0.001562366 | 0.868730112 |
| Pgm1 | S | 117 | -0.796836989 | -0.418204828 | -0.938029151 | -0.141192161 | -0.519824322 | 0.124984355 | 0.427642077 | 0.117715988 | 0.796866986 | 0.400566075 |
| Larp1 | S | 193 | -0.118113895 | -0.397004735 | -0.228273146 | -0.110159251 | 0.168731589 | 0.040913132 | 6.20823E-05 | 0.10071576 | 0.330679264 | 0.182501354 |
| Drosha | S | 374 | -0.257926824 | -0.335680648 | -0.204515728 | 0.053411096 | 0.13116492 | 0.005133911 | 0.001753784 | 0.108793137 | 0.581633933 | 0.248735496 |
| Cenpe | S | 2380 | -0.10529172 | -0.287930585 | -0.327076641 | -0.221784921 | -0.039146055 | 0.037877441 | 0.005575017 | 0.001159449 | 0.007305607 | 0.49074491 |
| Gja1 | S | 273 | 0.059546711 | -0.290088149 | -0.48863959 | -0.548186301 | -0.198551441 | 0.303084821 | 0.001247927 | 6.51605E-05 | 0.000281409 | 0.012277188 |
| Dync1li1 | S | 510 | -0.068248805 | -0.229691293 | -0.220901259 | -0.152652454 | 0.008790034 | 0.078591829 | 0.001147495 | 5.79949E-06 | 0.012578618 | 0.741588115 |
| Mybbp1a | S | 1280 | 0.060043616 | -0.28588856 | -0.401191227 | -0.461234843 | -0.115302667 | 0.039777471 | 5.31662E-05 | 2.59412E-05 | 0.000134463 | 0.015574603 |
| Parg | S | 135 | -0.153357086 | -0.172359304 | -0.200350776 | -0.04699369 | -0.027991472 | 0.001492942 | 0.008443647 | 6.75164E-05 | 0.120831586 | 0.451259234 |
| Luc7l | S | 331 | -0.241056734 | -0.221501878 | -0.518718242 | -0.277661508 | -0.297216364 | 0.055774821 | 0.059393177 | 0.004683526 | 0.055004168 | 0.047049297 |
| Snta1 | S | 195 | -0.124733961 | -0.198778757 | -0.159869394 | -0.035135432 | 0.038909363 | 0.009155027 | 0.000166775 | 0.000103796 | 0.25570567 | 0.130917795 |
| D1Ertd622e | S | 132 | -0.298377776 | -0.254903155 | -0.399215338 | -0.100837562 | -0.144312183 | 0.011336822 | 0.02467634 | 0.033745966 | 0.435802363 | 0.300985412 |
| Ttf1 | S | 437 | -0.040756194 | -0.335411151 | -0.347703152 | -0.306946958 | -0.012292002 | 0.406248345 | 0.000103605 | 0.005560358 | 0.014266515 | 0.843119299 |
| Szrd1 | S | 107 | -0.13766129 | -0.310671822 | -0.54093943 | -0.40327814 | -0.230267607 | 0.130754388 | 0.012911868 | 0.008968434 | 0.027093945 | 0.112952572 |
| Zyx | S | 336 | -0.107144552 | -0.32842343 | -0.415732649 | -0.308588097 | -0.087309219 | 0.058995297 | 0.007541252 | 0.005505612 | 0.010506138 | 0.277379656 |
| Fam133b | S | 86 | -0.454095231 | -0.160975968 | -0.49494201 | -0.040846779 | -0.333966042 | 0.002775124 | 0.38214044 | 0.040578971 | 0.799830139 | 0.15899843 |

| | | | | | | | | | | | | |
|---|---|---|---|---|---|---|---|---|---|---|---|---|
| Polr2m | S | 268 | -0.083889073 | -0.226410775 | -0.283504065 | -0.199614992 | -0.05709329 | 0.028322497 | 0.021333154 | 9.90687E-06 | 0.0021158 | 0.335983386 |
| Ufd1 | S | 247 | -0.032958403 | -0.255755221 | -0.254914985 | -0.221956582 | 0.000840236 | 0.08305911 | 1.48724E-06 | 0.005695686 | 0.016402296 | 0.981470502 |
| Rps9 | S | 163 | -0.225416908 | -0.319178069 | -0.502704741 | -0.277287833 | -0.183526672 | 0.046057742 | 0.053999933 | 0.012460776 | 0.085892172 | 0.254176239 |
| Irs1 | S | 307 | -0.115914484 | -0.322392179 | -0.340799886 | -0.224885402 | -0.018407707 | 0.468002043 | 0.002869858 | 0.000486857 | 0.218568503 | 0.76224067 |
| Prrc2c | S | 610 | -0.239548277 | -0.289491711 | -0.150144401 | 0.089403876 | 0.139347311 | 0.000651449 | 0.00030082 | 0.300690779 | 0.500393122 | 0.329141224 |
| Edc3 | S | 131 | -0.029334489 | -0.340505992 | -0.449124435 | -0.419789946 | -0.108618443 | 0.672283861 | 0.00172797 | 0.001590683 | 0.008909984 | 0.046112024 |
| Map1b | T | 1928 | -0.091797277 | -0.231551677 | -0.354053281 | -0.262256004 | -0.122501604 | 0.013358847 | 8.44121E-05 | 0.019925696 | 0.043754931 | 0.187415184 |
| Slc4a7 | S | 275 | -0.170896794 | -0.271590004 | -0.255709206 | -0.084812412 | 0.015880798 | 0.099602641 | 0.009662118 | 0.000105458 | 0.317356434 | 0.769935951 |
| Mbd3 | S | 56 | -0.427590756 | -0.215497895 | -0.502512936 | -0.07492218 | -0.287015041 | 0.033579807 | 0.319876617 | 0.006982316 | 0.61837658 | 0.206446597 |
| Hdac1 | S | 435 | -0.074722338 | -0.412115255 | -0.414487401 | -0.339765063 | -0.002372146 | 0.521815688 | 0.007443101 | 0.001544157 | 0.062673613 | 0.965816507 |
| Klhdc4 | S | 418 | -0.129330057 | -0.087498831 | -0.397904545 | -0.268574488 | -0.310405714 | 0.098090989 | 0.044772483 | 4.67125E-06 | 0.023488592 | 0.00062368 |
| Emd | S | 88 | -0.26594789 | -0.292844361 | -0.296438261 | -0.030490372 | -0.0035939 | 0.008612046 | 0.02109996 | 0.018287623 | 0.722978176 | 0.970405649 |
| Spag9 | T | 307 | -0.171413028 | -0.213365358 | -0.402995827 | -0.231582799 | -0.189630469 | 0.044599759 | 0.022008444 | 0.00123289 | 0.016515404 | 0.032145648 |
| Atad5 | S | 452 | -0.369652266 | -0.175651071 | -0.369112302 | 0.000539964 | -0.193461231 | 0.006070495 | 0.280551389 | 0.004025122 | 0.99342361 | 0.237087474 |
| Nsfl1c | S | 140 | -0.197112078 | -0.254598957 | -0.17749642 | 0.019615659 | 0.077102538 | 0.000586916 | 0.000201192 | 0.14104834 | 0.827790407 | 0.431945867 |
| Sipa1l3 | S | 1434 | 0.001118784 | -0.417299012 | -0.629568299 | -0.630687083 | -0.212269287 | 0.982074014 | 0.000111238 | 0.060790833 | 0.06527075 | 0.344734793 |
| Ccdc86 | S | 18 | 0.006391598 | -0.33334902 | -0.292200582 | -0.29859218 | 0.041148438 | 0.809437202 | 2.44959E-05 | 0.000939842 | 0.001072372 | 0.266945 |
| Tdrd3 | S | 627 | -0.190930781 | -0.239649093 | -0.345678565 | -0.154747784 | -0.106029473 | 0.03516166 | 0.000151691 | 0.049220982 | 0.210607228 | 0.343855969 |
| Knop1 | S | 57 | -0.043043613 | -0.32786743 | -0.22532859 | -0.182284977 | 0.10253884 | 0.448404819 | 7.00124E-05 | 0.000473625 | 0.030947009 | 0.021561266 |
| Rasa3 | S | 809 | -0.008848438 | -0.167051087 | -0.547018745 | -0.538170307 | -0.379967658 | 0.869489097 | 0.032085392 | 1.38324E-05 | 0.000169159 | 0.002747611 |
| Ddx21 | S | 118 | 0.00495078 | -0.241501427 | -0.328825135 | -0.333775914 | -0.087323708 | 0.920296064 | 0.001360357 | 3.84957E-06 | 0.008427475 | 0.059333204 |
| Ube2o | S | 406 | -0.268903742 | -0.181200975 | -0.346678183 | -0.077774441 | -0.165477208 | 0.173900277 | 0.022944867 | 0.000141479 | 0.620361847 | 0.037949022 |
| Rps10 | S | 146 | -0.013659608 | -0.303429077 | -0.41395544 | -0.400295832 | -0.110526363 | 0.781224321 | 6.20787E-05 | 0.004627221 | 0.003838222 | 0.155977083 |
| Srrm2 | S | 1515 | -0.076251987 | -0.323584316 | -0.270992196 | -0.19474021 | 0.05259212 | 0.076732946 | 2.32791E-05 | 0.032668575 | 0.071401636 | 0.488464143 |
| Kti12 | S | 157 | -0.113183139 | -0.280171453 | -0.372388069 | -0.259204931 | -0.092216616 | 0.153461931 | 0.000279653 | 0.011575086 | 0.037481478 | 0.284921637 |
| Mtmr14 | S | 528 | -0.108303899 | -0.418565031 | -0.409015146 | -0.300711247 | 0.009549885 | 0.180365862 | 0.006575974 | 0.009382493 | 0.019242984 | 0.90723531 |
| Rtkn | S | 106 | -0.040846273 | -0.264518764 | -0.449670487 | -0.408824214 | -0.185151724 | 0.495518312 | 0.000698957 | 0.001534042 | 0.00410609 | 0.054593602 |
| Utp23 | S | 220 | -0.02793952 | -0.332368235 | -0.444852425 | -0.416912905 | -0.112484189 | 0.654501294 | 0.000328143 | 0.00554606 | 0.004617692 | 0.149033455 |
| Map1b | S | 1255 | -0.113339759 | -0.216972955 | -0.195430071 | -0.082090313 | 0.021542884 | 0.02271709 | 0.000219822 | 0.000321491 | 0.066544012 | 0.293962584 |
| Arhgef5 | S | 851 | -0.204998938 | -0.202233941 | -0.379268085 | -0.174269147 | -0.177034144 | 0.037860861 | 0.020320295 | 0.001722471 | 0.07549797 | 0.050999743 |
| Tagln2 Tagln3 | S | 185;185 | -0.377811829 | -0.26754005 | -0.54751901 | -0.169707181 | -0.27997896 | 0.053836934 | 0.199427793 | 0.012289677 | 0.20526789 | 0.137275869 |
| Itprid2 | S | 758 | -0.042778032 | -0.221131533 | -0.256295977 | -0.213517944 | -0.035164444 | 0.534041907 | 0.001897233 | 5.84353E-07 | 0.064968816 | 0.193371295 |
| Zc3hav1 | S | 490 | 0.004253509 | -0.482896662 | -0.354107104 | -0.358360612 | 0.128789558 | 0.964824577 | 0.000810957 | 0.003488565 | 0.02183492 | 0.10842782 |
| Rsl1d1 | T | 399 | 0.050523487 | -0.318434031 | -0.429625843 | -0.48014933 | -0.111191812 | 0.33562628 | 0.00093783 | 6.19894E-05 | 0.000715229 | 0.075498073 |
| Lyar | S | 253 | -0.173683953 | -0.401030686 | -0.480458229 | -0.306774276 | -0.079427544 | 0.023428801 | 0.013250432 | 0.071044307 | 0.172422894 | 0.66929195 |
| Ccdc6 | S | 320 | -0.074617315 | -0.334297054 | -0.426545751 | -0.351928436 | -0.092248697 | 0.459316669 | 0.000880338 | 0.00602491 | 0.027488986 | 0.284858968 |
| Atxn2l | T | 507 | -0.12303043 | -0.379463104 | -0.289136602 | -0.166106172 | 0.090326502 | 0.077578509 | 0.000218099 | 0.060368247 | 0.161921936 | 0.377852769 |
| Map4 | S | 901 | -0.529365332 | -0.221737765 | -0.571620123 | -0.042254791 | -0.349882358 | 0.031464121 | 0.451875647 | 0.022053308 | 0.773187406 | 0.233090113 |
| Prune2 | S | 578 | -0.209200175 | -0.286051883 | -0.400785329 | -0.191585154 | -0.114733446 | 0.043367101 | 0.034353058 | 0.005071232 | 0.067730797 | 0.253400534 |
| BC055324 | S | 802 | -0.349018757 | -0.22296264 | -0.284326023 | 0.064692734 | -0.061363383 | 0.002146488 | 0.038766308 | 0.047258128 | 0.524752008 | 0.576841216 |
| Spag9 | S | 308 | -0.197221615 | -0.182593389 | -0.355271798 | -0.158050183 | -0.172678409 | 0.00615361 | 0.10813918 | 0.000692372 | 0.034431408 | 0.12481333 |
| Cobll1 | S | 1092 | -0.057888039 | -0.172732784 | -0.212483653 | -0.154595614 | -0.039750869 | 0.02627218 | 6.55677E-05 | 2.61992E-05 | 0.001097522 | 0.089840232 |
| Mettl6 | S | 15 | -0.344800471 | -0.288157268 | -0.670693536 | -0.325893065 | -0.382536268 | 0.015844008 | 0.135444823 | 0.083913746 | 0.286053019 | 0.234800887 |
| Atg2b | T | 1570 | -0.192499574 | -0.233366743 | -0.319894976 | -0.127395402 | -0.086528233 | 0.213167344 | 0.00093783 | 0.001789744 | 0.366505267 | 0.160686752 |
| Uvrag | S | 508 | -0.020167358 | -0.363717845 | -0.485540848 | -0.46537349 | -0.121823003 | 0.786470477 | 0.000104094 | 0.023914235 | 0.020533027 | 0.328610842 |
| Frmd4a | S | 318 | -0.042024767 | -0.304380254 | -0.334040716 | -0.29201595 | -0.029660463 | 0.398726125 | 0.000278451 | 0.001591248 | 0.007240103 | 0.567839905 |
| Tyms | S | 108 | -0.123476956 | -0.372287278 | -0.426155146 | -0.302678189 | -0.053867868 | 0.217494476 | 0.005026815 | 0.009774026 | 0.042926497 | 0.579436732 |
| Bin1 | S | 296 | -0.039610804 | -0.260664823 | -0.295203724 | -0.255592921 | -0.034538902 | 0.379856906 | 0.000178089 | 0.000306413 | 0.004518064 | 0.36032712 |
| Slc20a2 | S | 268 | -0.13483832 | -0.331724703 | -0.237155064 | -0.102316744 | 0.09456964 | 0.092347368 | 0.000114783 | 0.024200952 | 0.239813372 | 0.217889179 |
| Eif4ebp1 | Y | 33 | -0.258678525 | -0.286966714 | -0.36116668 | -0.102488155 | -0.074199966 | 0.01311509 | 0.016311403 | 0.036845364 | 0.399708092 | 0.542606993 |
| Ppm1g | S | 524 | -0.262105997 | -0.310228559 | -0.263449114 | -0.001343118 | 0.046779445 | 0.016439032 | 0.011871781 | 0.018322434 | 0.978766436 | 0.522858922 |
| Sh3pxd2a | S | 916 | -0.157960953 | -0.260854496 | -0.279796506 | -0.121835553 | -0.018942009 | 0.075045963 | 9.58346E-05 | 0.020427431 | 0.176792078 | 0.763764134 |
| Ppp1r14b | S | 142 | -0.171529929 | -0.296544644 | -0.489644218 | -0.31811429 | -0.193099575 | 0.058263843 | 0.008741826 | 0.023880916 | 0.086884142 | 0.213259303 |
| Ranbp2 | S | 778 | -0.080214732 | -0.259477648 | -0.257052551 | -0.176837819 | 0.002425097 | 0.068464612 | 0.000260608 | 0.001289074 | 0.012518617 | 0.945773862 |
| Gorasp1 | S | 277 | -0.003759754 | -0.467246709 | -0.370104839 | -0.366345085 | 0.09714187 | 0.954513528 | 0.023679291 | 5.88274E-05 | 0.009661331 | 0.385105831 |
| Pnn | S | 698 | 0.004907199 | -0.307286459 | -0.322165401 | -0.3270726 | -0.014878942 | 0.88583979 | 0.000766407 | 5.7397E-05 | 0.00034537 | 0.695116644 |
| Kirrel | S | 605 | -0.431906064 | -0.224112478 | -0.488638322 | -0.056732258 | -0.264525844 | 0.023771888 | 0.340756265 | 0.012161043 | 0.669402349 | 0.260680061 |
| Gnl3l | S | 571 | -0.085317234 | -0.267482006 | -0.337232337 | -0.251915103 | -0.069750331 | 0.038260311 | 0.00014839 | 0.01775262 | 0.054465139 | 0.388528418 |
| Sh3pxd2a | S | 1008 | -0.180979445 | -0.19654389 | -0.535683624 | -0.354704179 | -0.339139734 | 0.14107362 | 0.109043532 | 0.00069465 | 0.019298108 | 0.019485433 |
| Zfp36l2 | S | 75 | -0.06194502 | -0.338329437 | -0.481367947 | -0.419422927 | -0.14303851 | 0.600293512 | 0.009190864 | 0.001272176 | 0.022276795 | 0.166259281 |
| Eif3d | S | 161 | -0.000327199 | -0.218326095 | -0.31853723 | -0.318210031 | -0.100211135 | 0.995649029 | 4.82393E-07 | 0.001427234 | 0.016255883 | 0.029803659 |
| Ctcf | T | 289 | -0.154762884 | -0.192206108 | -0.403521846 | -0.248758961 | -0.211315737 | 0.016140912 | 0.009129304 | 0.004456216 | 0.019799884 | 0.031668665 |
| Arl14ep | S | 179 | -0.238052754 | -0.271399743 | -0.4959342 | -0.257881446 | -0.224534456 | 0.044875201 | 0.248331654 | 0.002068948 | 0.023039269 | 0.317227824 |
| Utp20 | S | 1735 | -0.080856268 | -0.267101145 | -0.372476085 | -0.291619817 | -0.10537494 | 0.178848319 | 0.001087074 | 0.002346717 | 0.006600345 | 0.09003156 |
| Sart1 | S | 84 | -0.210153542 | -0.346311046 | -0.366835886 | -0.156682343 | -0.02052484 | 0.020965731 | 0.075939983 | 0.005635987 | 0.109886824 | 0.883735531 |
| Itprid2 | S | 745 | -0.138892922 | -0.350391413 | -0.26615742 | -0.127264498 | 0.084233993 | 0.025843292 | 0.000275464 | 0.082686986 | 0.302815438 | 0.464361749 |
| Col4a3bp | S | 125 | -0.244797492 | -0.358465236 | -0.495121451 | -0.250323958 | -0.136656215 | 0.062446385 | 0.015992995 | 0.06036311 | 0.24931624 | 0.476726916 |
| Nes | S | 1419 | -0.162371415 | -0.244465015 | -0.216834088 | -0.054462673 | 0.027630927 | 0.00345299 | 0.00976776 | 0.001371754 | 0.185802491 | 0.583174526 |
| Heatr6 | S | 23 | -0.106528366 | -0.155792551 | -0.23179085 | -0.125262484 | -0.075998298 | 0.028460216 | 0.000151437 | 0.000144484 | 0.021260544 | 0.03696362 |
| Pebp1 | S | 52 | -0.348674718 | -0.22562423 | -0.430052145 | -0.081377427 | -0.204427915 | 0.009075155 | 0.116923601 | 0.02776759 | 0.524278859 | 0.212727123 |
| Ppp1r13l | S | 485 | -0.178878492 | -0.284100509 | -0.282296077 | -0.103417585 | 0.001804432 | 0.151183142 | 0.000183511 | 0.017073723 | 0.35877544 | 0.977312244 |
| Edc3 | T | 173 | 0.009424565 | -0.43197893 | -0.457848713 | -0.467273277 | -0.025869783 | 0.873115163 | 0.042053514 | 7.70507E-05 | 0.003196526 | 0.820112493 |
| Huwe1 | S | 3374 | -0.110207805 | -0.22560005 | -0.33275959 | -0.222551785 | -0.107159539 | 0.038608588 | 0.007166095 | 0.000656505 | 0.01433867 | 0.112097264 |
| Eif3a | S | 1028 | 0.053310741 | -0.276928924 | -0.378957085 | -0.432267826 | -0.102028161 | 0.611648408 | 0.000411793 | 6.74853E-05 | 0.029159436 | 0.055284144 |
| Npm1 | S | 258 | -0.083148426 | -0.318032624 | -0.356399782 | -0.273251356 | -0.038367157 | 0.111101877 | 0.001661891 | 0.005578257 | 0.008367606 | 0.512347284 |
| Prrc2b | S | 555 | -0.009910803 | -0.222062039 | -0.246084774 | -0.236173971 | -0.024022735 | 0.660145575 | 2.75119E-05 | 2.26729E-05 | 0.000702853 | 0.206111187 |
| Camk2d | S | 490 | -0.155317947 | -0.199603739 | -0.205799992 | -0.050482045 | -0.006196253 | 0.000188528 | 3.89391E-05 | 0.140644247 | 0.626708998 | 0.950861803 |
| Ehbp1l1 | S | 367 | -0.053499019 | -0.265825453 | -0.35896868 | -0.305469661 | -0.093143227 | 0.285446735 | 4.95401E-05 | 0.007209543 | 0.006662567 | 0.15880512 |
| Nup98 | T | 670 | -0.048445195 | -0.296985139 | -0.260942944 | -0.212497749 | 0.036042196 | 0.134798963 | 3.07514E-05 | 0.006651646 | 0.015964983 | 0.439976417 |
| Rai14 | S | 438 | -0.02490744 | -0.244114506 | -0.316325441 | -0.291418001 | -0.072210935 | 0.552715728 | 2.35371E-05 | 0.001012899 | 0.002820873 | 0.109392277 |
| Emd | S | 117 | -0.163525087 | -0.199503982 | -0.418825722 | -0.255300634 | -0.219321739 | 0.006640877 | 0.017475134 | 0.008850975 | 0.053239227 | 0.060296323 |
| Camsap2 | S | 818 | -0.258449495 | -0.241405846 | -0.290005901 | -0.031556406 | -0.048600055 | 0.003771069 | 0.037202669 | 0.013368894 | 0.672509074 | 0.608301495 |
| Picalm | S | 307 | -0.165471522 | -0.303603876 | -0.323092654 | -0.157621133 | -0.019488778 | 0.160894753 | 0.005983442 | 0.002068248 | 0.171319228 | 0.747082197 |
| Cdc42se1 | S | 75 | -0.062276616 | -0.47848466 | -0.42670737 | -0.364430755 | 0.05177729 | 0.154689804 | 0.014007413 | 0.007799912 | 0.002993471 | 0.518198854 |
| Nav1 | S | 510 | -0.317745848 | -0.226993061 | -0.471416449 | -0.153670601 | -0.244423388 | 0.011922701 | 0.117124459 | 0.02436054 | 0.301897847 | 0.165321276 |
| Slc25a3 | S | 27 | -0.104103661 | -0.362413883 | -0.451113001 | -0.34700934 | -0.088699118 | 0.132994797 | 0.002900604 | 0.024536786 | 0.047673602 | 0.46217526 |
| Iqsec1 | S | 104 | -0.352298593 | -0.147216588 | -0.518186289 | -0.165887697 | -0.370969701 | 0.025878087 | 0.408418853 | 0.00359503 | 0.172511565 | 0.080750509 |
| Pxn | S | 244 | -0.142079518 | -0.231173414 | -0.465192949 | -0.323113431 | -0.234019535 | 0.044345811 | 0.007203856 | 0.009567699 | 0.054041126 | 0.100327497 |
| Nop2 | S | 767 | -0.13005478 | -0.222433229 | -0.241467247 | -0.111412467 | -0.019034018 | 0.004826185 | 0.001440856 | 0.003693907 | 0.067113976 | 0.690681532 |
| Sfr1 | S | 192 | -0.205179508 | -0.29232392 | -0.522257698 | -0.317078191 | -0.229933778 | 0.054238707 | 0.103684121 | 0.006785203 | 0.039960482 | 0.181735793 |
| Sapcd2 | S | 283 | -0.208841226 | -0.226317074 | -0.450896858 | -0.242055632 | -0.224579785 | 0.266339572 | 0.048125508 | 0.000514601 | 0.216085709 | 0.052430891 |
| Gab1 | T | 639 | -0.074156572 | -0.212873445 | -0.413433206 | -0.339276634 | -0.200559761 | 0.333736572 | 0.001231303 | 0.000770445 | 0.014578207 | 0.007264011 |
| Eaf1 | T | 157 | 0.007179025 | -0.310967668 | -0.316247391 | -0.323426415 | -0.005279723 | 0.877715502 | 2.66893E-06 | 0.008243019 | 0.004978567 | 0.912556746 |
| Rpp30 | S | 251 | -0.099014109 | -0.252592363 | -0.194469919 | -0.09545581 | 0.058122444 | 0.079015551 | 4.30257E-05 | 0.001917663 | 0.096464278 | 0.13459791 |

| | | | | | | | | | | | | |
|---|---|---|---|---|---|---|---|---|---|---|---|---|
| Wnk1 | S | 1973 | -0.056504711 | -0.219341677 | -0.327101954 | -0.270597244 | -0.107760278 | 0.018742848 | 0.001110199 | 0.001046692 | 0.002333241 | 0.02832662 |
| Tex2 | S | 195 | -0.233306938 | -0.033723225 | -0.373902503 | -0.140595565 | -0.340179278 | 0.001414576 | 0.731121104 | 0.000101981 | 0.022212301 | 0.040518107 |
| Abl2 | S | 915 | -0.104652595 | -0.339831371 | -0.546889951 | -0.442237356 | -0.20705858 | 0.162563992 | 0.061911986 | 0.002986618 | 0.009225316 | 0.183693008 |
| Cdan1 | S | 282 | -0.102430486 | -0.421362912 | -0.55461186 | -0.452181374 | -0.133248948 | 0.221106359 | 0.048834544 | 0.006574272 | 0.008436948 | 0.36411407 |
| Dync1li2 | S | 391 | -0.028144487 | -0.279693664 | -0.354105879 | -0.325961392 | -0.074412214 | 0.601939296 | 0.000880873 | 0.000392856 | 0.003451147 | 0.04544042 |
| Rab12 | S | 15 | -0.196596749 | -0.203482722 | -0.2981441 | -0.101547351 | -0.094661378 | 0.00774521 | 0.012774999 | 0.004548443 | 0.155680592 | 0.19874124 |
| Crtc3 | S | 396 | -0.299800053 | -0.182175986 | -0.229811792 | 0.069988261 | -0.047635806 | 8.38316E-05 | 0.194295056 | 0.017625058 | 0.279091279 | 0.691249052 |
| Pnn | S | 68 | -0.189631813 | -0.26619674 | -0.404098741 | -0.214466929 | -0.137902001 | 0.091036526 | 0.055862408 | 0.00133071 | 0.085557931 | 0.23460506 |
| Jpt1 | S | 79 | -0.322831881 | -0.233955116 | -0.490861773 | -0.168029892 | -0.256906657 | 0.0145831 | 0.091960922 | 0.037317001 | 0.299430092 | 0.161495224 |
| Rftn2 | S | 429 | -0.335727802 | -0.225063899 | -0.503072876 | -0.167345074 | -0.278008978 | 0.038921945 | 0.148445947 | 0.012359066 | 0.279757718 | 0.118687543 |
| Mybbp1a | S | 1167 | -0.022123431 | -0.415668878 | -0.470822531 | -0.4486991 | -0.055153653 | 0.751971284 | 0.015484397 | 0.001119018 | 0.00607785 | 0.488750547 |
| Parp8 | S | 323 | -0.180047369 | -0.285274358 | -0.350179297 | -0.170131929 | -0.064904939 | 0.025666181 | 0.0578795 | 0.002091746 | 0.045297709 | 0.560257795 |
| Eif3c | S | 203 | 0.11174269 | -0.41275405 | -0.429710224 | -0.541452914 | -0.016956174 | 0.123062218 | 0.00145292 | 7.22449E-05 | 0.001159001 | 0.782543301 |
| Morf4l2 | S | 15 | -0.100496889 | -0.269407086 | -0.413841278 | -0.313344389 | -0.144434192 | 0.09393141 | 0.071394254 | 0.000253439 | 0.010582329 | 0.243565947 |
| Ahctf1 | S | 1931 | 0.004446406 | -0.233056298 | -0.353658991 | -0.358105396 | -0.120602693 | 0.884189376 | 6.15039E-07 | 0.00464437 | 0.001401129 | 0.06607386 |
| Rab1a | T | 195 | -0.080464831 | -0.342279329 | -0.245620497 | -0.165155666 | 0.096658832 | 0.12000922 | 0.000162975 | 0.012464676 | 0.060454658 | 0.174607859 |
| Dmwd | S | 487 | -0.158319301 | -0.271822495 | -0.39624905 | -0.237929749 | -0.124426555 | 0.056311528 | 0.013084843 | 0.006313833 | 0.044242637 | 0.213922991 |
| Rsrc2 | S | 46 | 0.02138526 | -0.240737817 | -0.328975146 | -0.350360406 | -0.08823733 | 0.794438914 | 0.001211503 | 7.24016E-06 | 0.034811007 | 0.034645318 |
| Pgam1 | S | 31 | -0.285137676 | -0.302427191 | -0.437617178 | -0.152479502 | -0.135189988 | 0.028148444 | 0.055400766 | 0.031745345 | 0.316998399 | 0.40415132 |
| Ctnnd1 | Y | 904 | -0.276792587 | -0.264721822 | -0.38899457 | -0.112201983 | -0.124272749 | 0.047831127 | 0.033530362 | 0.012336142 | 0.322661842 | 0.216421902 |
| Kdm4a | S | 570 | -0.238196911 | -0.295947071 | -0.333129761 | -0.094932851 | -0.037182691 | 0.024251515 | 0.006533949 | 0.046828651 | 0.447191381 | 0.750728179 |
| Unc5b | S | 479 | -0.271773699 | -0.225533238 | -0.441456059 | -0.16968236 | -0.215922821 | 0.12287371 | 0.155150665 | 0.00084313 | 0.276640168 | 0.174723644 |
| Tnks1bp1 | S | 1702 | -0.153770921 | -0.343076213 | -0.500598021 | -0.3468271 | -0.157521808 | 0.14006209 | 0.036774371 | 0.008038166 | 0.024777795 | 0.221091676 |
| Fscn1 | T | 403 | -0.280476753 | -0.225081044 | -0.36102622 | -0.080549467 | -0.135945177 | 0.016290819 | 0.034166679 | 0.016019366 | 0.422040726 | 0.205948993 |
| Txndc9 | S | 221 | -0.081043898 | -0.171056703 | -0.301916743 | -0.220872845 | -0.13086004 | 0.068179915 | 0.006858117 | 2.84419E-05 | 0.003206519 | 0.028772626 |
| Txndc9 | S | 223 | -0.081043898 | -0.171056703 | -0.301916743 | -0.220872845 | -0.13086004 | 0.068179915 | 0.006858117 | 2.84419E-05 | 0.003206519 | 0.028772626 |
| Prune2 | S | 2171 | -0.236583486 | -0.245659146 | -0.323854961 | -0.087271475 | -0.078195814 | 0.11143966 | 0.024640195 | 0.001026354 | 0.461284284 | 0.314183549 |
| Map3k4 | S | 207 | -0.24397505 | -0.204487805 | -0.421294579 | -0.177319529 | -0.216806775 | 0.026172846 | 0.054564728 | 0.00674815 | 0.119907325 | 0.076733941 |
| Nol7 | S | 214 | -0.025458333 | -0.337832587 | -0.281703686 | -0.256245352 | 0.056128901 | 0.749017446 | 0.000191975 | 0.001180156 | 0.037639217 | 0.272098055 |
| Atg2b | S | 379 | -0.045962661 | -0.398343541 | -0.549408988 | -0.503446328 | -0.151065447 | 0.350843263 | 0.004978929 | 0.017472394 | 0.028917361 | 0.265190595 |
| Topors | S | 780 | -0.036095446 | -0.395343295 | -0.30033761 | -0.264242164 | 0.095005684 | 0.541563753 | 3.62151E-05 | 0.034893654 | 0.049457403 | 0.326707818 |
| Gripap1 | S | 657 | -0.017746126 | -0.273581764 | -0.396261065 | -0.378514939 | -0.122679301 | 0.696249724 | 0.028668109 | 1.53088E-05 | 0.001430796 | 0.166631886 |
| Wasf2 | S | 284 | -0.260814332 | -0.252831884 | -0.57572323 | -0.314908898 | -0.322891346 | 0.072472399 | 0.2425303 | 0.005365149 | 0.060175386 | 0.161775067 |
| Map7d1 | S | 548 | -0.101586206 | -0.264092697 | -0.214933618 | -0.113347411 | 0.04915908 | 0.028189965 | 0.000461186 | 0.002530943 | 0.04227495 | 0.280628647 |
| Sqstm1 | S | 357 | 0.026329336 | -0.397418716 | -0.397053752 | -0.423383088 | 0.000364964 | 0.600566314 | 0.00797984 | 0.000123731 | 0.000790123 | 0.996050486 |
| Afap1 | S | 712 | 0.032696045 | -0.300304623 | -0.348258245 | -0.38095429 | -0.047953622 | 0.455329583 | 0.000434131 | 9.57564E-05 | 0.000820784 | 0.27779844 |
| Ncl | S | 67 | -0.053062134 | -0.245513453 | -0.309259829 | -0.256197695 | -0.063746377 | 0.193385951 | 2.76485E-05 | 0.005966112 | 0.009992572 | 0.261849116 |
| Dync1li1 | S | 516 | -0.049638977 | -0.250243627 | -0.259798482 | -0.210159505 | -0.009554855 | 0.284464227 | 0.00029699 | 0.000211338 | 0.014003394 | 0.692619306 |
| Tut1 | S | 133 | -0.053069293 | -0.297194581 | -0.444234048 | -0.391164755 | -0.147039466 | 0.520502231 | 0.062797096 | 6.87332E-05 | 0.018449919 | 0.213737601 |
| Snx29 | S | 815 | 0.052280504 | -0.37578841 | -0.402749661 | -0.455030145 | -0.026961231 | 0.503773162 | 4.77223E-05 | 0.007073626 | 0.005756879 | 0.686485496 |
| Spata5 | S | 291 | -0.058872081 | -0.196167111 | -0.3699134 | -0.311041319 | -0.173746289 | 0.16158135 | 0.000629607 | 0.000810295 | 0.003865795 | 0.02916488 |
| Sh3pxd2a | S | 1047 | 0.018872094 | -0.244506963 | -0.341784216 | -0.36065631 | -0.097277253 | 0.480112461 | 0.001228526 | 9.27214E-06 | 0.000140103 | 0.048038231 |
| Rab3ip | S | 215 | -0.098695785 | -0.272750002 | -0.409313629 | -0.310617844 | -0.136563627 | 0.048364315 | 0.000499933 | 0.036192238 | 0.081349803 | 0.2893675 |
| Ajuba | S | 146 | -0.094983874 | -0.21385264 | -0.356033836 | -0.261049962 | -0.142181196 | 0.016215381 | 0.000354036 | 0.015150668 | 0.044307445 | 0.134299593 |
| Dab2 | S | 325 | -0.044297594 | -0.183668627 | -0.292351754 | -0.24805416 | -0.108683127 | 0.084085779 | 0.002152322 | 2.58385E-05 | 0.000795271 | 0.023772126 |
| Hectd1 | S | 1390 | -0.055067977 | -0.245588229 | -0.497533023 | -0.442465046 | -0.251944794 | 0.651141756 | 0.036721129 | 0.000146387 | 0.031530337 | 0.041049509 |
| Akap10 | S | 187 | -0.044991244 | -0.206381069 | -0.230448286 | -0.185457043 | -0.024067217 | 0.376172528 | 0.000946177 | 1.92074E-06 | 0.037141406 | 0.317969797 |
| Cdca3 | S | 44 | -0.06674468 | -0.271740011 | -0.482345294 | -0.415600614 | -0.210605283 | 0.27978008 | 0.000432894 | 0.016488383 | 0.023871823 | 0.124349616 |
| Ifi211 | S | 158 | -0.374293027 | -0.183912676 | -0.572445574 | -0.198152547 | -0.388532899 | 0.0332383 | 0.326381164 | 0.013758837 | 0.239826778 | 0.084501711 |
| Rbsn | T | 637 | 0.003948682 | -0.449172883 | -0.480786807 | -0.484735489 | -0.031613924 | 0.976221169 | 0.000314533 | 0.037952276 | 0.040396136 | 0.820583067 |
| Lmna | S | 409 | 0.02953283 | -0.313540003 | -0.267586605 | -0.297119435 | 0.045953398 | 0.549261447 | 8.61122E-05 | 0.000103229 | 0.011907647 | 0.179417013 |
| Eif4b | S | 192 | -0.113924697 | -0.182311129 | -0.376276184 | -0.262351487 | -0.193965055 | 0.084188705 | 0.026990391 | 0.000176326 | 0.004517639 | 0.026352104 |
| Txlna | S | 515 | 0.007622724 | -0.194054426 | -0.263190053 | -0.270812777 | -0.069135627 | 0.693710744 | 1.68083E-05 | 1.04049E-05 | 0.000169148 | 0.018419159 |
| Gtf3c1 | S | 1477 | -0.038529647 | -0.398081575 | -0.287653603 | -0.249123957 | 0.110427972 | 0.632143935 | 1.09459E-05 | 0.068144304 | 0.086886567 | 0.323616013 |
| Trmt10c | S | 79 | 0.029561181 | -0.372016755 | -0.380290069 | -0.40985125 | -0.008273314 | 0.631799492 | 2.86815E-05 | 0.01274755 | 0.009048299 | 0.912352813 |
| Mtdh | S | 215 | -0.283992052 | -0.453575934 | -0.311418391 | -0.027426339 | 0.142157543 | 0.14192501 | 0.018136351 | 0.028876812 | 0.850751372 | 0.277643502 |
| Bub1b | S | 659 | -0.090376716 | -0.300732547 | -0.354909578 | -0.264532862 | -0.054177031 | 0.103275699 | 3.14421E-05 | 0.089885384 | 0.146080826 | 0.69363364 |
| Erbin | Y | 1097 | -0.16058458 | -0.227180215 | -0.389803378 | -0.229218799 | -0.162623164 | 0.049556793 | 0.044651373 | 0.00131319 | 0.021680504 | 0.118435157 |
| Efnb2 | Y | 307 | -0.262647266 | -0.259174761 | -0.348551058 | -0.085903792 | -0.089376296 | 0.076545053 | 0.0832213 | 0.001342432 | 0.451436042 | 0.44316238 |
| Cdca5 | S | 20 | -0.259187587 | -0.188638462 | -0.391649199 | -0.132461612 | -0.203010737 | 0.016276661 | 0.15886576 | 0.002745479 | 0.0036049 | 0.139492405 |
| Fosl1 | S | 267 | 0.033600011 | -0.364320125 | -0.456576803 | -0.490176814 | -0.092256678 | 0.440371573 | 0.001647969 | 0.000968741 | 0.000496459 | 0.07694045 |
| Prrc2c | S | 853 | -0.062843501 | -0.424880805 | -0.272275153 | -0.209431653 | 0.152605652 | 0.371621144 | 0.000130963 | 0.038191761 | 0.083044617 | 0.166654592 |
| Txlna | S | 19 | -0.02757771 | -0.286128474 | -0.4931842 | -0.46560649 | -0.207055726 | 0.707972946 | 4.65637E-05 | 0.024329879 | 0.017573185 | 0.148804383 |
| Stom | S | 8 | -0.484594377 | -0.058688227 | -0.772465845 | -0.287871469 | -0.713777619 | 0.002126123 | 0.775601711 | 0.115309106 | 0.443050206 | 0.127945472 |
| Sec16a | S | 2104 | -0.193442974 | -0.216727759 | -0.487113932 | -0.293670958 | -0.270386173 | 0.069624399 | 0.156241091 | 0.001411599 | 0.005396574 | 0.098424345 |
| Ddx21 | S | 13 | 0.070804949 | -0.302300865 | -0.363426594 | -0.434231543 | -0.061125729 | 0.054828808 | 1.51776E-05 | 0.000194714 | 0.000158706 | 0.098704605 |
| Tsc1 | S | 1074 | -0.015070576 | -0.332912961 | -0.454150754 | -0.439080178 | -0.121237793 | 0.866764754 | 0.001629803 | 0.002879872 | 0.015730773 | 0.077800615 |
| Orc6 | S | 206 | -0.065843828 | -0.308795308 | -0.298934003 | -0.233090175 | 0.009861305 | 0.02899921 | 0.001744019 | 0.005177461 | 0.004767117 | 0.795895619 |
| Slain2 | S | 303 | -0.274082961 | -0.260996198 | -0.510058028 | -0.235975067 | -0.24906183 | 0.043417094 | 0.100996279 | 0.017480677 | 0.154589491 | 0.166196065 |
| Ogfr | S | 403 | -0.114303043 | -0.226410366 | -0.338605677 | -0.224302634 | -0.11219531 | 0.330245002 | 0.000564255 | 0.001550289 | 0.113659311 | 0.094280735 |
| Ubap2l | S | 1012 | -0.367843366 | -0.248700146 | -0.460957767 | -0.093114401 | -0.212257621 | 0.0318428 | 0.209514303 | 0.015347949 | 0.345354186 | 0.229485358 |
| Map4 | T | 777 | -0.097937855 | -0.138685286 | -0.351084939 | -0.253147084 | -0.212399654 | 0.037774329 | 0.016488651 | 8.3211E-05 | 8.01676E-05 | 0.007272097 |
| Morc2a | S | 737 | -0.05339636 | -0.231299686 | -0.386666872 | -0.333270512 | -0.155367186 | 0.400941085 | 0.00184334 | 0.000522944 | 0.004336011 | 0.041872039 |
| Ddr2 | Y | 471 | -0.169267507 | -0.276227956 | -0.548967326 | -0.379699819 | -0.27273937 | 0.158194622 | 0.052468807 | 0.005998019 | 0.03264102 | 0.089073507 |
| Eif2ak2 | S | 163 | -0.163319596 | -0.43763446 | -0.326285947 | -0.162966351 | 0.111348513 | 0.172634579 | 0.006319914 | 0.022650515 | 0.008945024 | 0.080459312 |
| Flii | S | 436 | -0.074771676 | -0.185262122 | -0.444261106 | -0.369489431 | -0.258998984 | 0.098689971 | 0.006894183 | 0.000941657 | 0.009948194 | 0.01287029 |
| Tulp3 | S | 65 | -0.268277265 | -0.190346671 | -0.526808852 | -0.258531587 | -0.336462181 | 0.007356913 | 0.250632436 | 0.016344824 | 0.123134067 | 0.103515588 |
| D1Pas1 Ddx3x Ddx3y | S | 69;72;71 | -0.165425802 | -0.246497604 | -0.359720182 | -0.19429438 | -0.113222578 | 0.056606307 | 0.01866667 | 0.002840226 | 0.035193589 | 0.175048098 |
| Dlgap4 | T | 915 | -0.094533077 | -0.28176101 | -0.501711813 | -0.407178737 | -0.219950803 | 0.242313408 | 0.015591255 | 0.003548478 | 0.008733606 | 0.060383682 |
| Nab2 | S | 514 | -0.243586005 | -0.224164503 | -0.266080905 | -0.0224949 | -0.041916401 | 0.02146345 | 0.007100062 | 0.009718766 | 0.776594284 | 0.512692211 |
| Prrc2a | S | 931 | -0.15032693 | -0.284038503 | -0.559682661 | -0.409355731 | -0.275644158 | 0.243412488 | 0.034225925 | 0.006141252 | 0.028663065 | 0.078990333 |
| Csnk1e | S | 363 | -0.050901499 | -0.189946294 | -0.289670268 | -0.238768769 | -0.099723974 | 0.135253039 | 0.001093936 | 6.54561E-05 | 6.98247E-05 | 0.028488488 |
| Plxnb2 | S | 1248 | -0.391179326 | -0.178202182 | -0.476476781 | -0.085297455 | -0.298274599 | 0.027068549 | 0.264372915 | 0.012100783 | 0.275078268 | 0.033929266 |
| Cd3eap | S | 187 | -0.086889152 | -0.520707129 | -0.525481556 | -0.438592404 | -0.004774427 | 0.252205048 | 0.011825112 | 0.048896659 | 0.060013805 | 0.976480874 |
| Ftsj3 | S | 456 | -0.237156219 | -0.271719484 | -0.350919702 | -0.113763484 | -0.079200219 | 0.067402701 | 0.006204888 | 0.023042802 | 0.342736971 | 0.420885228 |
| Zyx | S | 300 | -0.084247806 | -0.24827184 | -0.244389697 | -0.160141891 | 0.003882143 | 0.024086144 | 7.69171E-05 | 0.012942141 | 0.042200569 | 0.937058394 |
| Mast4 | S | 2041 | -0.41990482 | -0.060014434 | -0.490565181 | -0.070660361 | -0.430550747 | 0.015540414 | 0.65606684 | 0.004092776 | 0.544196509 | 0.017534969 |
| Arfgap2 | S | 339 | -0.276402026 | -0.240914427 | -0.560509455 | -0.284107429 | -0.319595028 | 0.12856393 | 0.230860528 | 0.003612823 | 0.112091627 | 0.130823772 |
| Prrc2b | S | 387 | -0.048989505 | -0.219617458 | -0.204903145 | -0.15591364 | 0.014714314 | 0.095217456 | 4.9369E-05 | 0.000239088 | 0.003116708 | 0.507176462 |
| Ssrp1 | T | 642 | -0.103412826 | -0.266181179 | -0.281212116 | -0.17779929 | -0.015030937 | 0.05099844 | 0.011543276 | 0.000424372 | 0.004728103 | 0.806302771 |
| Nhsl1 | S | 1234 | -0.060570323 | -0.218271899 | -0.284074233 | -0.223503911 | -0.065802335 | 0.312859991 | 0.000655943 | 0.000136535 | 0.021096087 | 0.047394568 |
| Plekha2 | S | 314 | -0.209873643 | -0.353346558 | -0.550191448 | -0.340317804 | -0.19684489 | 0.092828822 | 0.090834186 | 0.017459398 | 0.080407813 | 0.325672859 |
| Dok1 | S | 290 | -0.084164452 | -0.231024513 | -0.320417348 | -0.236252897 | -0.089392835 | 0.023878659 | 0.00017489 | 0.016753866 | 0.038444197 | 0.239250021 |
| Gja1 | S | 257 | -0.036009503 | -0.297070874 | -0.540064106 | -0.504054604 | -0.242993233 | 0.649933506 | 0.006442804 | 0.003024253 | 0.009363071 | 0.064194706 |

| | | | | | | | | | | | | |
|---|---|---|---|---|---|---|---|---|---|---|---|---|
| Plekhg3 | S | 736 | -0.133532117 | -0.232318513 | -0.316411581 | -0.182879464 | -0.084093069 | 0.124032159 | 0.018295892 | 0.000233423 | 0.069764347 | 0.233941781 |
| Map1b | S | 1793 | -0.06672435 | -0.255342193 | -0.253101503 | -0.186377153 | 0.00224069 | 0.126859487 | 0.000577504 | 0.000618779 | 0.005837869 | 0.937545928 |
| Epha2 | T | 899 | -0.118565408 | -0.323696999 | -0.515306162 | -0.396740754 | -0.191609164 | 0.244432956 | 0.030685834 | 0.005164555 | 0.017439225 | 0.140698455 |
| Eif3k | S | 217 | 0.034238565 | -0.354781106 | -0.307384532 | -0.341623097 | 0.047396573 | 0.163025635 | 8.25451E-07 | 0.014833567 | 0.013005005 | 0.425809938 |
| Pikfyve | S | 1548 | -0.068674473 | -0.21735327 | -0.190573652 | -0.121899179 | 0.026779618 | 0.016407465 | 4.03228E-05 | 0.001685914 | 0.028425679 | 0.371202618 |
| Rprd2 | S | 845 | -0.213748062 | -0.422494267 | -0.408607144 | -0.194859082 | 0.013887123 | 0.021138016 | 0.030570665 | 0.08868645 | 0.308800451 | 0.94169563 |
| Crtc3 | S | 110 | -0.297478472 | -0.19433208 | -0.352140314 | -0.054661842 | -0.157808234 | 0.025253502 | 0.160831707 | 0.002096604 | 0.528774864 | 0.227107235 |
| Bora | S | 231 | 0.034394179 | -0.454017612 | -0.540230955 | -0.574625134 | -0.086213343 | 0.322870963 | 0.001601854 | 0.008678914 | 0.005278475 | 0.310027306 |
| Prpf4b | S | 180 | -0.075776749 | -0.286049591 | -0.32041762 | -0.244640871 | -0.034368028 | 0.196242684 | 0.00016734 | 0.012483255 | 0.019223452 | 0.566972565 |
| E2f8 | S | 20 | 0.010580534 | -0.329436281 | -0.252775234 | -0.263355768 | 0.076661046 | 0.810607717 | 5.48393E-05 | 0.000658186 | 0.004746719 | 0.086135269 |
| Tnk2 | S | 825 | -0.156264411 | -0.254084292 | -0.268101521 | -0.11183711 | -0.014017229 | 0.015768257 | 0.013007401 | 0.002790822 | 0.074197861 | 0.835660442 |
| Bub1 | S | 579 | -0.030546516 | -0.380256298 | -0.53829703 | -0.507750515 | -0.158040732 | 0.539564025 | 0.007191154 | 0.008404625 | 0.002768429 | 0.094481231 |
| Nav3 | S | 1539 | -0.1697962 | -0.352418785 | -0.520493231 | -0.350697031 | -0.168074445 | 0.168338414 | 0.019216633 | 0.025908636 | 0.065560936 | 0.2745039 |
| Strap | T | 345 | 0.055662582 | -0.340783389 | -0.398523274 | -0.454185855 | -0.057739884 | 0.262844182 | 3.73081E-06 | 0.01297931 | 0.005733317 | 0.439523261 |
| Ivns1abp | S | 338 | -0.04113212 | -0.355071223 | -0.364586975 | -0.323454854 | -0.009515751 | 0.495693487 | 0.000377784 | 0.012501799 | 0.015450594 | 0.899015501 |
| Epb41l2 | S | 580 | -0.346732618 | -0.164045232 | -0.579944405 | -0.233211787 | -0.415899174 | 0.011894014 | 0.429457346 | 0.021269345 | 0.210983728 | 0.115289384 |
| Bnip3 | S | 88 | -0.043803161 | -0.266898463 | -0.221616938 | -0.177813777 | 0.045281526 | 0.358237449 | 9.57749E-05 | 0.000376903 | 0.015948392 | 0.145507553 |
| Mast3 | S | 373 | -0.154219324 | -0.41552624 | -0.525390322 | -0.371170998 | -0.109864082 | 0.189648726 | 0.036335379 | 0.020983583 | 0.046687411 | 0.475890992 |
| Map1b | S | 1384 | -0.157029559 | -0.18810615 | -0.147119947 | 0.009909612 | 0.040986203 | 0.001100388 | 0.00034541 | 0.007707815 | 0.76944784 | 0.26691133 |
| Specc1 | S | 55 | -0.295164553 | -0.188938717 | -0.338457424 | -0.04329287 | -0.149518707 | 0.008141676 | 0.214538289 | 0.003836835 | 0.254777901 | 0.291719204 |
| Cttn | Y | 421 | -0.509197693 | -0.273137283 | -0.407101361 | 0.102096332 | -0.133964078 | 0.008585671 | 0.243351589 | 0.122131232 | 0.618768919 | 0.597934804 |
| Supt16 | S | 1026 | 0.094218667 | -0.398615052 | -0.48332701 | -0.577545677 | -0.084711958 | 0.101239861 | 0.002011528 | 0.000270575 | 0.000638614 | 0.265491342 |
| Cavin1 | S | 171 | -0.324138852 | -0.175760315 | -0.373111954 | -0.048973102 | -0.197351638 | 0.012409355 | 0.077778685 | 0.01644832 | 0.654655177 | 0.116329069 |
| Arhgap5 | S | 1194 | -0.111721505 | -0.164560119 | -0.293628776 | -0.181907271 | -0.129068656 | 0.225750236 | 0.004668122 | 2.86248E-05 | 0.100907087 | 0.017964334 |
| Psrc1 | S | 211 | -0.081281229 | -0.252736179 | -0.426245232 | -0.344964003 | -0.173509052 | 0.183541968 | 0.003675764 | 0.003913582 | 0.020700645 | 0.107459348 |
| Ago2 | S | 388 | -0.171627656 | -0.2561042 | -0.422556019 | -0.250928363 | -0.166451819 | 0.183122104 | 0.028024508 | 0.002309178 | 0.08777053 | 0.123084709 |
| Nf1 | S | 2498 | -0.093382914 | -0.228724428 | -0.317672709 | -0.224289795 | -0.08894828 | 0.117894259 | 0.041037966 | 3.85998E-05 | 0.014085232 | 0.25589498 |
| Samd4 | S | 168 | -0.167636634 | -0.277980624 | -0.466404949 | -0.298768315 | -0.188424325 | 0.258314574 | 0.010445044 | 0.008267243 | 0.093085075 | 0.143663624 |
| Pfas | T | 622 | 0.074364419 | -0.451412813 | -0.45675073 | -0.531115149 | -0.005337917 | 0.374188882 | 0.000685131 | 0.005029477 | 0.005158979 | 0.953831432 |
| Dync1li2 | S | 203 | -0.089942724 | -0.161513463 | -0.263168098 | -0.173225375 | -0.101654635 | 0.024528831 | 0.002706347 | 0.000117643 | 0.002954098 | 0.023501408 |
| Cmtm4 | S | 194 | -0.195655453 | -0.224186862 | -0.473553502 | -0.27789805 | -0.249366641 | 0.009321462 | 0.046790632 | 0.026277534 | 0.105027609 | 0.12169085 |
| Pde3b | S | 296 | -0.165126566 | -0.20896627 | -0.53024492 | -0.365118354 | -0.32127865 | 0.017840184 | 0.032639772 | 0.020562966 | 0.067850766 | 0.071298183 |
| Nolc1 | S | 563 | -0.134848673 | -0.248920827 | -0.198859977 | -0.064011303 | 0.050060851 | 0.081880977 | 0.000126624 | 0.005475974 | 0.33689624 | 0.262002936 |
| Trip12 | S | 1069 | -0.061484747 | -0.306818364 | -0.267272705 | -0.205787957 | 0.039545659 | 0.200878571 | 0.00360184 | 0.000326781 | 0.011555561 | 0.33764373 |
| Brf1 | S | 612 | -0.20024469 | -0.30278163 | -0.334940496 | -0.134695806 | -0.032158866 | 0.050403022 | 0.011092457 | 0.019256814 | 0.217458462 | 0.746027279 |
| Brf1 | S | 613 | -0.20024469 | -0.30278163 | -0.334940496 | -0.134695806 | -0.032158866 | 0.050403022 | 0.011092457 | 0.019256814 | 0.217458462 | 0.746027279 |
| Anln | S | 513 | -0.081517657 | -0.231875941 | -0.251804239 | -0.170286582 | -0.019928298 | 0.080852516 | 0.000969783 | 0.000582232 | 0.003113263 | 0.434387922 |
| Neurl4 | T | 841 | -0.201556096 | -0.26474183 | -0.350729584 | -0.149173489 | -0.085987755 | 0.06095771 | 0.021987632 | 0.006370843 | 0.040212516 | 0.108147503 |
| Jpt1 | S | 71 | -0.237088374 | -0.267241205 | -0.516330975 | -0.279242601 | -0.24908977 | 0.012140227 | 0.059327204 | 0.062579218 | 0.213068092 | 0.250408267 |
| Ciapin1 | S | 213 | -0.181538626 | -0.259307533 | -0.49281477 | -0.311276144 | -0.233507237 | 0.225015427 | 0.027898147 | 0.005496527 | 0.083545193 | 0.082049993 |
| Ncaph | S | 422 | -0.152970787 | -0.278166064 | -0.365819807 | -0.212849021 | -0.087653743 | 0.108686098 | 0.010587583 | 0.005646647 | 0.055758978 | 0.317727447 |
| Arfgef2 | T | 244 | -0.155512387 | -0.244094388 | -0.264952317 | -0.109439929 | -0.020857928 | 0.002285415 | 0.056656179 | 0.002186725 | 0.041331515 | 0.791714575 |
| Hdac1 | S | 434 | -0.04932358 | -0.411677064 | -0.467140132 | -0.417816552 | -0.055463068 | 0.783526868 | 0.016486486 | 0.002514715 | 0.098342012 | 0.612142454 |
| Gpr149 | S | 238 | -0.257347022 | -0.516958666 | -0.293564709 | -0.036217686 | 0.223393957 | 0.132435675 | 0.010251376 | 0.100311653 | 0.755546677 | 0.096073231 |
| Asap1 | S | 741 | -0.300882422 | -0.089118445 | -0.292023195 | 0.008859227 | -0.20290475 | 7.68304E-06 | 0.396695144 | 0.058341718 | 0.921740659 | 0.152696208 |
| Fyttd1 | S | 23 | -0.015752236 | -0.317313963 | -0.165301637 | -0.149549402 | 0.152012326 | 0.653793721 | 5.80109E-06 | 0.001397478 | 0.009960827 | 0.005609165 |
| Wdr44 | S | 563 | -0.077003827 | -0.234304592 | -0.279502364 | -0.202498537 | -0.045197772 | 0.133562764 | 5.54019E-05 | 0.007427368 | 0.020809809 | 0.390479883 |
| Bora | S | 279 | -0.197806933 | -0.23711606 | -0.222934021 | -0.025127088 | 0.014182039 | 0.157539504 | 0.000658546 | 0.002496654 | 0.816796421 | 0.737818696 |
| Strap | S | 338 | -0.058389557 | -0.316281706 | -0.309812839 | -0.251423283 | 0.006468867 | 0.164768116 | 0.003961292 | 0.001350513 | 0.002834474 | 0.877893467 |
| Gramd1a | T | 279 | 0.006658475 | -0.378280218 | -0.223296266 | -0.229954741 | 0.154983952 | 0.918605745 | 4.59493E-05 | 0.002493576 | 0.028729013 | 0.035445641 |
| Cep250 | S | 1223 | -0.100126752 | -0.227742248 | -0.378156658 | -0.278029906 | -0.15041441 | 0.121366878 | 0.022325923 | 0.000541916 | 0.004890484 | 0.084140103 |
| Sec16a | T | 2103 | -0.248689079 | -0.191955292 | -0.400989021 | -0.152299942 | -0.209033729 | 0.016662256 | 0.173085939 | 0.004047804 | 0.107763769 | 0.145989929 |
| Eif2b4 | S | 21 | -0.043265025 | -0.211458183 | -0.29401272 | -0.250747694 | -0.082554537 | 0.251561324 | 0.000129851 | 0.000744128 | 0.002896583 | 0.082286253 |
| Synrg | S | 517 | -0.4906714 | -0.134754759 | -0.362049558 | 0.128621842 | -0.227294799 | 0.007353014 | 0.336982324 | 0.02764053 | 0.140417448 | 0.041871542 |
| Cdv3 | Y | 213 | -0.428434179 | -0.261385453 | -0.333552213 | 0.094881966 | -0.072166761 | 0.016121521 | 0.105373949 | 0.057049629 | 0.368050274 | 0.543596648 |
| Rbsn | S | 635 | -0.022444881 | -0.448451379 | -0.314195339 | -0.291750458 | 0.13425604 | 0.644113712 | 0.000172827 | 0.041018848 | 0.061120031 | 0.239211339 |
| Phactr4 | S | 145 | 0.03553974 | -0.252239506 | -0.403134657 | -0.438674397 | -0.150895151 | 0.406806168 | 0.001112688 | 8.74009E-05 | 2.54869E-05 | 0.024988241 |
| Pnn | S | 700 | 0.039439286 | -0.28631619 | -0.357256761 | -0.396696047 | -0.070940571 | 0.212137642 | 1.16303E-05 | 0.001618136 | 0.001403626 | 0.16668854 |
| Zyx | S | 272 | -0.045789503 | -0.256287359 | -0.379059323 | -0.33326982 | -0.122771965 | 0.24142223 | 0.000206488 | 0.008824126 | 0.02271789 | 0.162992019 |
| Phldb2 | S | 359 | -0.158744025 | -0.358874815 | -0.225554234 | -0.066810209 | 0.133320581 | 0.143808466 | 0.000411167 | 0.044015452 | 0.526969253 | 0.16324615 |
| Mast3 | S | 1237 | -0.049607087 | -0.23724523 | -0.307133278 | -0.257526191 | -0.069888049 | 0.341417154 | 0.004925894 | 7.31262E-05 | 0.007008761 | 0.19350695 |
| Cad | T | 1833 | -0.049644098 | -0.278852317 | -0.273574565 | -0.223930467 | 0.005277752 | 0.134376687 | 0.000458725 | 0.002427191 | 0.003682728 | 0.887636353 |
| Ncbp1 | T | 21 | 0.010464746 | -0.239870751 | -0.362109704 | -0.37257445 | -0.122238953 | 0.832769667 | 0.000894324 | 0.00012 | 0.00273162 | 0.014128325 |
| Cd2ap | S | 510 | -0.026130302 | -0.1638911 | -0.283290974 | -0.257160673 | -0.119399875 | 0.46043053 | 8.27732E-05 | 4.81383E-05 | 0.002243715 | 0.010209542 |
| Adss Adssl1 | S | 452;451 | -0.16450516 | -0.177681087 | -0.33304489 | -0.16853973 | -0.155363803 | 0.031947985 | 0.060730566 | 0.000451507 | 0.033229387 | 0.091715011 |
| Zfyve19 | S | 219 | -0.070886101 | -0.446010344 | -0.496806239 | -0.425920138 | -0.050795895 | 0.317465254 | 0.004269147 | 0.052088767 | 0.069528167 | 0.75101078 |
| Esr1 | S | 170 | -0.469713468 | -0.412451208 | -0.309186203 | 0.160527265 | 0.103265004 | 0.054768181 | 0.042320301 | 0.165210492 | 0.432318234 | 0.526380662 |
| Esr1 | S | 171 | -0.469713468 | -0.412451208 | -0.309186203 | 0.160527265 | 0.103265004 | 0.054768181 | 0.042320301 | 0.165210492 | 0.432318234 | 0.526380662 |
| Esr1 | S | 172 | -0.469713468 | -0.412451208 | -0.309186203 | 0.160527265 | 0.103265004 | 0.054768181 | 0.042320301 | 0.165210492 | 0.432318234 | 0.526380662 |
| Map1b | T | 1629 | -0.059625985 | -0.284267108 | -0.344192605 | -0.28456622 | -0.059925497 | 0.127598943 | 0.0013108 | 0.00588668 | 0.005741129 | 0.278419558 |
| Raver1 Gm38431 | S | 576 | -0.013014431 | -0.234867 | -0.302969038 | -0.289954607 | -0.068102037 | 0.726732379 | 0.003105305 | 1.69983E-05 | 0.003532786 | 0.157185753 |
| Dennd4c | S | 1240 | -0.044361591 | -0.294421203 | -0.2953722 | -0.251010609 | -0.000950997 | 0.365849701 | 0.017971393 | 3.01165E-05 | 0.010379752 | 0.988147173 |
| Nup214 | S | 637 | 0.030338709 | -0.295516342 | -0.314708945 | -0.345047654 | -0.019192603 | 0.469244894 | 0.000286497 | 0.000172645 | 0.000226624 | 0.546352634 |
| Dennd4c | S | 1221 | 0.008813701 | -0.265346635 | -0.39365215 | -0.40246585 | -0.128305514 | 0.875476129 | 0.01520149 | 2.83573E-05 | 0.003472875 | 0.125241666 |
| Ubr5 | S | 321 | -0.390232374 | -0.156488902 | -0.522295337 | -0.132062962 | -0.365806435 | 0.044920343 | 0.442294815 | 0.007832322 | 0.3939683 | 0.117724231 |
| Ptpn2 | S | 298 | -0.05563103 | -0.180645393 | -0.277736952 | -0.222105922 | -0.097091559 | 0.072244144 | 0.000778756 | 0.000221219 | 0.00192257 | 0.032405637 |
| Ehmt2 | S | 289 | -0.254785021 | -0.275436326 | -0.436690884 | -0.181905863 | -0.161254558 | 0.062440353 | 0.138116779 | 0.007155537 | 0.042792697 | 0.302421192 |
| Rrp15 | S | 224 | -0.084048351 | -0.189124707 | -0.351152071 | -0.26710372 | -0.162027364 | 0.177105169 | 0.00038573 | 0.002572335 | 0.009937115 | 0.033249422 |
| Pnn | S | 666 | -0.088845181 | -0.165426047 | -0.151060592 | -0.062215411 | 0.014365455 | 0.100826113 | 2.53623E-05 | 2.23176E-05 | 0.19465438 | 0.242347835 |
| Ehd1 | S | 456 | -0.072005552 | -0.28712991 | -0.329616063 | -0.257610511 | -0.042486152 | 0.334186307 | 0.018008889 | 0.000175042 | 0.031969271 | 0.495310572 |
| Sos1 | S | 1120 | -0.142658702 | -0.285097683 | -0.489854068 | -0.347195365 | -0.204756385 | 0.190567738 | 0.060493739 | 0.003573558 | 0.02157172 | 0.129816256 |
| Rrp36 | S | 66 | -0.003022833 | -0.307028678 | -0.289776898 | -0.286754065 | 0.01725178 | 0.938221039 | 8.83056E-05 | 0.00218442 | 0.005516443 | 0.692670603 |
| Map1b | S | 1253 | -0.207122327 | -0.235705056 | -0.132375484 | 0.074746843 | 0.103329572 | 0.003969189 | 0.000629963 | 0.040740062 | 0.191753567 | 0.087228771 |
| Iqsec1 | S | 102 | -0.268735518 | -0.129586601 | -0.47393601 | -0.205200492 | -0.344349409 | 0.032212377 | 0.418849578 | 0.001661862 | 0.044120251 | 0.087256731 |
| Sfr1 | S | 140 | -0.215188319 | -0.248598332 | -0.472662436 | -0.257474117 | -0.224064104 | 0.064353781 | 0.038603403 | 0.018815327 | 0.121617424 | 0.160674064 |
| C530008M17Rik | S | 783 | -0.148245836 | -0.338478107 | -0.225165776 | -0.07691994 | 0.113312331 | 0.022026939 | 0.030381954 | 0.001702141 | 0.156354467 | 0.247933039 |
| Ybx1 | S | 207 | 0.061771822 | -0.363029251 | -0.427503419 | -0.489275241 | -0.064474168 | 0.210033936 | 0.000715679 | 0.000937745 | 0.000947783 | 0.304399351 |
| Gtse1 | S | 73 | -0.215926634 | -0.3270458 | -0.30387234 | -0.087945706 | 0.023173459 | 0.040622526 | 0.002781912 | 0.110379579 | 0.56197989 | 0.86695732 |
| Copg1 | T | 584 | -0.093819946 | -0.205341014 | -0.249477064 | -0.155657118 | -0.04413605 | 0.034522842 | 0.00019342 | 0.005003229 | 0.029844159 | 0.353153793 |
| Ldb3 | S | 179 | -0.186879685 | -0.25415354 | -0.15657395 | 0.030305735 | 0.09757959 | 0.007185487 | 0.001273013 | 0.021869912 | 0.499716204 | 0.073031154 |
| Map2k7 | S | 287 | -0.185753054 | -0.250179305 | -0.520937191 | -0.335184137 | -0.270757886 | 0.109412839 | 0.059867272 | 0.00964591 | 0.065290049 | 0.088845576 |
| Rif1 | S | 1439 | 0.035316727 | -0.271541709 | -0.20712144 | -0.242438166 | 0.06442027 | 0.261984932 | 6.0495E-06 | 4.10539E-05 | 0.001399601 | 0.020225029 |
| Tnks1bp1 | S | 876 | -0.081047686 | -0.18106289 | -0.435984028 | -0.354936342 | -0.254921137 | 0.223174238 | 0.037615852 | 0.000221944 | 0.000770206 | 0.011820217 |
| Afdn | S | 181 | -0.266307574 | -0.212034552 | -0.254726209 | 0.011581365 | -0.042691657 | 0.004790569 | 0.069883533 | 0.009475186 | 0.849038495 | 0.640509086 |

| | | | | | | | | | | | | |
|---|---|---|---|---|---|---|---|---|---|---|---|---|
| Mtor | S | 1261 | -0.35302033 | -0.232940491 | -0.30944388 | 0.043576449 | -0.076503389 | 0.023001325 | 0.133593869 | 0.010069302 | 0.660299834 | 0.533723445 |
| Map1b | S | 1260 | -0.052560244 | -0.217593957 | -0.200412383 | -0.147852139 | 0.017181574 | 0.401237974 | 8.73923E-05 | 6.8491E-05 | 0.087750051 | 0.42997052 |
| Tjp2 | S | 684 | -0.163177341 | -0.300069225 | -0.231236465 | -0.068059124 | 0.06883276 | 0.099792563 | 0.000386694 | 0.034652158 | 0.479021378 | 0.376536305 |
| Cavin2 | S | 274 | -0.250723603 | -0.20622873 | -0.42669424 | -0.175970637 | -0.22046551 | 0.050707675 | 0.102510539 | 0.005892307 | 0.058560518 | 0.040113957 |
| Zc3hav1 | S | 553 | -0.084330873 | -0.246030076 | -0.235023946 | -0.150693074 | 0.01100613 | 0.068717814 | 1.17799E-05 | 0.03602035 | 0.088995335 | 0.858264682 |
| Anln | S | 444 | 0.014471808 | -0.272921492 | -0.380221875 | -0.394693682 | -0.107300383 | 0.755588105 | 0.003396462 | 0.000125715 | 0.000264734 | 0.097865492 |
| Spag5 | S | 926 | -0.207002223 | -0.216300969 | -0.527517759 | -0.320515536 | -0.31121679 | 0.189117374 | 0.02987658 | 0.009306503 | 0.091145912 | 0.074035625 |
| Sqstm1 | S | 249 | 0.05968809 | -0.382431107 | -0.404084499 | -0.463772589 | -0.021653391 | 0.054680264 | 0.000198194 | 0.001302621 | 0.003843755 | 0.671882818 |
| Htt | S | 1179 | -0.051384259 | -0.205915801 | -0.290886493 | -0.239502234 | -0.084970692 | 0.141965599 | 0.003820524 | 9.63263E-05 | 0.001292561 | 0.091093136 |
| Sapcd2 | T | 218 | -0.231577906 | -0.15405188 | -0.546990373 | -0.315412466 | -0.392938493 | 0.07834167 | 0.355780294 | 0.001786765 | 0.018434362 | 0.058210053 |
| Nup214 | S | 660 | -0.049771722 | -0.266176449 | -0.364562477 | -0.314790755 | -0.098386027 | 0.245395493 | 0.000610913 | 0.006302365 | 0.008398351 | 0.165169222 |
| Plpp1 | S | 273 | -0.097619105 | -0.314537583 | -0.344746347 | -0.247127242 | -0.030208764 | 0.335593664 | 0.017086366 | 0.000809419 | 0.073026985 | 0.699370751 |
| Map1b | S | 1410 | -0.39750424 | -0.322034699 | -0.50105912 | -0.103554879 | -0.179024421 | 0.07649614 | 0.029869054 | 0.166464228 | 0.734330196 | 0.545733834 |
| Prkab1 | S | 25 | -0.415207033 | -0.180335925 | -0.307194955 | 0.108012078 | -0.12685903 | 0.003470883 | 0.053279257 | 0.153426986 | 0.546483343 | 0.477371526 |
| Ccdc91 | S | 417 | -0.167967521 | -0.324750854 | -0.395882525 | -0.227915003 | -0.07113167 | 0.117467312 | 0.02453932 | 0.012106417 | 0.088732341 | 0.551763842 |
| Utp14a | S | 567 | -0.153650163 | -0.24320097 | -0.293855766 | -0.140205603 | -0.050654796 | 0.029178197 | 0.003291495 | 0.015939817 | 0.115509523 | 0.491110228 |
| Gphn | S | 327 | -0.30069148 | -0.165749082 | -0.363180173 | -0.062488693 | -0.197431092 | 0.016837322 | 0.235678784 | 0.003874851 | 0.4171175 | 0.163094169 |
| Gja1 | S | 279 | 0.044768794 | -0.276818447 | -0.439950399 | -0.484719193 | -0.163131953 | 0.473182271 | 0.002504996 | 0.000247103 | 0.002816435 | 0.001145928 |
| Sqstm1 | S | 330 | -0.04918421 | -0.263528823 | -0.327426774 | -0.278242565 | -0.063897951 | 0.259161296 | 0.000610108 | 0.003474387 | 0.00818755 | 0.285165137 |
| Tgs1 | S | 152 | -0.076254988 | -0.162778274 | -0.401955401 | -0.325700413 | -0.239177127 | 0.452112398 | 0.015578997 | 8.25333E-05 | 0.04413273 | 0.006388741 |
| Specc1 | S | 9 | -0.021583416 | -0.327935548 | -0.356934927 | -0.335351512 | -0.02899938 | 0.783799518 | 0.002077538 | 0.001584265 | 0.015801461 | 0.503548559 |
| Bclaf1 | S | 752 | -0.073856833 | -0.430700793 | -0.467590417 | -0.393733584 | -0.036889624 | 0.171527117 | 0.004770186 | 0.063978744 | 0.078775049 | 0.802416339 |
| Pard3b | Y | 998 | -0.180624366 | -0.136115101 | -0.412434711 | -0.231810345 | -0.276319611 | 0.022451916 | 0.316228397 | 0.000432112 | 0.011659858 | 0.098729137 |
| Nop56 | S | 554 | 0.040690933 | -0.278521376 | -0.380696827 | -0.42138776 | -0.102175451 | 0.599931926 | 0.000589367 | 0.000441069 | 0.013435864 | 0.043407687 |
| Pacs1 | S | 779 | -0.155595688 | -0.31023752 | -0.249217546 | -0.093621858 | 0.061019975 | 0.17426368 | 0.002528174 | 0.006162201 | 0.346534109 | 0.23088262 |
| Rubcn | S | 389 | -0.112454677 | -0.219985064 | -0.209359589 | -0.096904911 | 0.010625475 | 0.05243992 | 0.000439134 | 0.002254629 | 0.092598085 | 0.776118322 |
| Camsap1 | S | 823 | -0.135209096 | -0.169298105 | -0.397146527 | -0.261937431 | -0.227848421 | 0.066386947 | 0.032464603 | 0.001129024 | 0.012858933 | 0.019839886 |
| Supv3l1 | S | 725 | 0.031448574 | -0.314425916 | -0.291691104 | -0.323139679 | 0.022734812 | 0.672743189 | 6.26272E-05 | 0.001523948 | 0.021057979 | 0.566325003 |
| Ankrd17 | S | 1605 | -0.051448273 | -0.283851678 | -0.31372078 | -0.262272507 | -0.029869102 | 0.366636411 | 9.17276E-05 | 0.014171503 | 0.023281259 | 0.651725529 |
| Ubap2 | S | 1020 | -0.180019977 | -0.345987313 | -0.466788666 | -0.286768689 | -0.120801353 | 0.315332548 | 0.030045654 | 0.011153544 | 0.119113374 | 0.260732179 |
| Vasp | S | 285 | -0.255283887 | -0.283155059 | -0.310287354 | -0.055003468 | -0.027132296 | 0.003133416 | 0.089081416 | 0.049724507 | 0.609128471 | 0.851060908 |
| Mphosph8 | S | 392 | -0.07326375 | -0.241661261 | -0.367218466 | -0.293954716 | -0.125557205 | 0.19489503 | 3.62756E-05 | 0.040107814 | 0.052101864 | 0.267603601 |
| Dbnl | S | 291 | -0.038372062 | -0.206843446 | -0.273112071 | -0.234740009 | -0.066268625 | 0.297231346 | 0.000747309 | 0.000129236 | 0.001737149 | 0.101880773 |
| Map1b | S | 614 | -0.184771914 | -0.211877826 | -0.213759109 | -0.028987195 | -0.001881283 | 0.001519864 | 0.02805004 | 0.006740319 | 0.505632702 | 0.97729069 |
| Sipa1l3 | S | 152 | -0.117233271 | -0.196345904 | -0.325290985 | -0.208057714 | -0.128945081 | 0.049715156 | 0.007329383 | 0.001993462 | 0.036151501 | 0.091552759 |
| Nufip2 | S | 113 | -0.026816563 | -0.282997008 | -0.300894926 | -0.274078363 | -0.017897918 | 0.53236345 | 0.000304719 | 0.002284866 | 0.004113566 | 0.667896677 |
| Ifitm3 | S | 38 | -0.262410617 | -0.361823151 | -0.507257916 | -0.244847299 | -0.145434765 | 0.089407529 | 0.127817133 | 0.029446399 | 0.186116013 | 0.508464963 |
| Zfp830 | S | 187 | -0.222068914 | -0.198490495 | -0.325650685 | -0.103581771 | -0.12716019 | 0.009090969 | 0.082985754 | 0.007072409 | 0.199420766 | 0.234283615 |
| Wdr43 | S | 658 | -0.022602253 | -0.404794901 | -0.247200072 | -0.224597819 | 0.157594829 | 0.694015495 | 0.000205329 | 0.011628735 | 0.023690152 | 0.070497086 |
| Parp8 | S | 345 | -0.131282684 | -0.224876762 | -0.436216214 | -0.30493353 | -0.211339452 | 0.135293502 | 0.005481695 | 0.011247657 | 0.039282055 | 0.105549913 |
| Nav1 | T | 543 | 0.068543014 | -0.324372472 | -0.419234396 | -0.48777741 | -0.094861924 | 0.537854321 | 0.000362388 | 0.00228581 | 0.016607871 | 0.196129099 |
| Myo1e | T | 1002 | -0.308137824 | -0.327812338 | -0.448333427 | -0.140195603 | -0.120521089 | 0.046286603 | 0.043889103 | 0.104777107 | 0.523660225 | 0.583180993 |
| Trim25 | S | 401 | -0.044673993 | -0.360556113 | -0.369588812 | -0.32491482 | -0.009032699 | 0.49141373 | 0.000760756 | 0.020086409 | 0.025706766 | 0.918845463 |
| Mthfr | S | 10 | -0.138609082 | -0.150820646 | -0.320765922 | -0.18215684 | -0.169945277 | 0.019398096 | 0.012987331 | 0.00156165 | 0.028698153 | 0.042660491 |
| D10Wsu102e | S | 143 | -0.130168516 | -0.336090295 | -0.410315716 | -0.2801472 | -0.074225421 | 0.228846872 | 0.007027321 | 0.021029562 | 0.075028287 | 0.527616647 |
| Prkd3 | S | 252 | -0.102446323 | -0.436424908 | -0.160311734 | -0.05786541 | 0.276113174 | 0.091500659 | 0.000558613 | 0.053707672 | 0.401505542 | 0.012584937 |
| Gpr39 | S | 271 | 0.015081396 | -0.34004842 | -0.255105761 | -0.270187157 | 0.084942659 | 0.795632654 | 0.000247768 | 0.000669351 | 0.012397317 | 0.084212995 |
| Edc3 | S | 161 | -0.051443583 | -0.268067262 | -0.462751066 | -0.411307483 | -0.194683804 | 0.449692713 | 0.008121038 | 0.002829574 | 0.005028547 | 0.047430042 |
| Dctn6 | T | 186 | -0.146404563 | -0.259543105 | -0.421018229 | -0.274613665 | -0.161475124 | 0.085196383 | 0.049879053 | 0.004650221 | 0.034216761 | 0.187623047 |
| Frmd4a | S | 696 | -0.119587899 | -0.204227603 | -0.387550895 | -0.267962997 | -0.183323292 | 0.055683238 | 0.103471608 | 0.000493548 | 0.016774213 | 0.135118589 |
| Zdhhc5 | S | 432 | -0.133605527 | -0.269498143 | -0.359367682 | -0.225762155 | -0.089869539 | 0.127166781 | 0.022068494 | 0.002988394 | 0.003629691 | 0.249273791 |
| Tnrc6a | S | 1254 | -0.28126832 | -0.214605507 | -0.501326782 | -0.220058462 | -0.286721274 | 0.081883489 | 0.178063603 | 0.009010952 | 0.136521673 | 0.081303592 |
| Pacsin2 | S | 387 | -0.152458159 | -0.270131569 | -0.405143724 | -0.252685565 | -0.135012156 | 0.081236455 | 0.020808837 | 0.011824122 | 0.025410611 | 0.129550544 |
| Map1b | S | 1627 | -0.060144491 | -0.286817689 | -0.378075789 | -0.317931298 | -0.0912581 | 0.424591688 | 0.002247635 | 0.004428149 | 0.014672039 | 0.262329451 |
| Rabl6 | S | 377 | -0.001218616 | -0.303769732 | -0.536928902 | -0.535710286 | -0.23315917 | 0.979258553 | 0.000426729 | 0.020960459 | 0.029762269 | 0.139583231 |
| Lin9 | S | 65 | -0.270030144 | -0.322137165 | -0.379897723 | -0.109867579 | -0.057760559 | 0.078538872 | 0.04126177 | 0.032945913 | 0.354246929 | 0.598218624 |
| Akap13 | S | 2527 | -0.327327621 | -0.219782664 | -0.521461917 | -0.194134296 | -0.301679253 | 0.026144014 | 0.297751161 | 0.027091134 | 0.267059376 | 0.206317693 |
| Sccpdh | S | 217 | 0.067829058 | -0.314274978 | -0.342892618 | -0.410721676 | -0.02861764 | 0.036711578 | 9.33812E-05 | 0.000139386 | 0.000137124 | 0.371248559 |
| Arhgef33 | S | 534 | -0.412267319 | -0.054589117 | -0.330963303 | 0.081304016 | -0.276374186 | 0.001032517 | 0.656416325 | 0.019221129 | 0.384451674 | 0.0854428 |
| Arhgef33 | S | 538 | -0.412267319 | -0.054589117 | -0.330963303 | 0.081304016 | -0.276374186 | 0.001032517 | 0.656416325 | 0.019221129 | 0.384451674 | 0.0854428 |
| Dact3 | S | 6 | -0.436163135 | -0.302330525 | -0.651906324 | -0.215743189 | -0.349575799 | 0.052566606 | 0.207562314 | 0.129475647 | 0.535836662 | 0.354022419 |
| Cers5 | S | 407 | -0.111584416 | -0.37435227 | -0.345026697 | -0.233442282 | 0.029325572 | 0.081799056 | 0.005765843 | 0.039219054 | 0.071334001 | 0.756003571 |
| Phldb2 | S | 510 | -0.195160221 | -0.239354624 | -0.337959703 | -0.142799482 | -0.098605079 | 0.032791387 | 0.025672563 | 0.012059808 | 0.160746188 | 0.32327205 |
| Parp8 | S | 387 | -0.112861916 | -0.280152804 | -0.238673746 | -0.12581183 | 0.041479058 | 0.021896506 | 0.000124698 | 0.10648261 | 0.295907291 | 0.694536079 |
| Ddx50 | S | 85 | -0.059714462 | -0.236603413 | -0.332385634 | -0.272671173 | -0.095782221 | 0.327090782 | 0.000131891 | 0.008272806 | 0.014941737 | 0.176351835 |
| Ect2 | S | 716 | -0.023029754 | -0.300839982 | -0.260829481 | -0.237799727 | 0.040010501 | 0.475995813 | 0.000183331 | 0.002889572 | 0.006083935 | 0.384653189 |
| Bdp1 | S | 1870 | -0.112737673 | -0.235692497 | -0.168346108 | -0.055608435 | 0.067346389 | 0.056962803 | 0.000350768 | 0.001707052 | 0.25637294 | 0.087204373 |
| Map1b | S | 1247 | -0.198081457 | -0.255744823 | -0.1268844 | 0.071197057 | 0.128860423 | 0.012881786 | 0.000528383 | 0.033882329 | 0.22969089 | 0.029140976 |
| Plekhg3 | S | 593 | -0.06029942 | -0.225071776 | -0.473381879 | -0.41308246 | -0.248310103 | 0.16861512 | 0.000532331 | 0.022173996 | 0.036006989 | 0.100722081 |
| Cep170 | T | 1405 | -0.062849706 | -0.362425341 | -0.361683338 | -0.298833632 | 0.000742002 | 0.557074928 | 0.002455428 | 0.009076773 | 0.038467577 | 0.992516102 |
| Specc1 | S | 360 | -0.028357327 | -0.340516379 | -0.204744805 | -0.176387478 | 0.135771574 | 0.522834921 | 0.00014161 | 0.002768655 | 0.004320249 | 0.011600183 |
| Tacc1 | S | 279 | -0.030042308 | -0.39199596 | -0.397743867 | -0.367701559 | -0.005747907 | 0.52606356 | 0.00496383 | 0.007803404 | 0.006947875 | 0.940092247 |
| Arhgef5 | S | 849 | -0.171366075 | -0.141915609 | -0.468400987 | -0.297034912 | -0.326485378 | 0.088997844 | 0.031002345 | 0.003598561 | 0.030268222 | 0.034549235 |
| Map4k4 | S | 708 | -0.019382974 | -0.252932639 | -0.210756491 | -0.191373517 | 0.042176148 | 0.359432794 | 0.001510333 | 1.18039E-05 | 0.000468985 | 0.213630944 |
| Nsd2 | S | 407 | -0.081862781 | -0.308231243 | -0.335689716 | -0.253826935 | -0.027458473 | 0.580461871 | 0.001278745 | 0.005428868 | 0.1598408 | 0.694978924 |
| Pxk | S | 452 | -0.233087299 | -0.422714145 | -0.522692889 | -0.28960559 | -0.099978744 | 0.180005695 | 0.069676228 | 0.045162517 | 0.179000633 | 0.641149125 |
| Tln1 | T | 144 | -0.35605261 | -0.198280984 | -0.518985661 | -0.162933051 | -0.320704677 | 0.040530656 | 0.434601138 | 0.013809848 | 0.247175459 | 0.230926926 |
| Mvb12a | S | 161 | -0.325301145 | -0.134680292 | -0.54197073 | -0.216669585 | -0.407290438 | 0.064950782 | 0.53113195 | 0.003985674 | 0.182559683 | 0.123769563 |
| Ric8b | S | 468 | -0.108564876 | -0.219629338 | -0.276134658 | -0.167569782 | -0.05650532 | 0.015674839 | 0.00041903 | 0.029431933 | 0.109197277 | 0.467365168 |
| Ric8b | T | 473 | -0.108564876 | -0.219629338 | -0.276134658 | -0.167569782 | -0.05650532 | 0.015674839 | 0.00041903 | 0.029431933 | 0.109197277 | 0.467365168 |
| Dnajb4 | S | 38 | -0.200055807 | -0.2885715 | -0.111836527 | 0.088219279 | 0.176734973 | 0.027844342 | 6.60731E-05 | 0.161797674 | 0.283206391 | 0.068050407 |
| Eif3a | S | 492 | -0.048435099 | -0.318985969 | -0.299822567 | -0.251387468 | 0.019163402 | 0.320136816 | 0.002171011 | 0.002730645 | 0.011248572 | 0.762459603 |
| Nmd3 | T | 470 | -0.08671651 | -0.202375568 | -0.311576073 | -0.224859563 | -0.109200505 | 0.066730746 | 0.010644764 | 0.000600855 | 0.010148055 | 0.098457883 |
| Ripor1 | S | 167 | -0.048733614 | -0.338775063 | -0.462423336 | -0.413689722 | -0.123648273 | 0.164083469 | 0.002822272 | 0.033022043 | 0.040997984 | 0.316950756 |
| Phldb2 | S | 465 | -0.090202902 | -0.233149337 | -0.208588112 | -0.11838521 | 0.024561226 | 0.033278316 | 0.00257932 | 0.000710624 | 0.020401742 | 0.555670799 |
| Stx18 | S | 180 | -0.109097598 | -0.302979502 | -0.276498783 | -0.167401185 | 0.026480718 | 0.05396019 | 0.00097985 | 0.038620877 | 0.127999166 | 0.759857243 |
| Tmem87b | T | 495 | -0.253857854 | -0.295764267 | -0.452948518 | -0.199090664 | -0.157184252 | 0.030343022 | 0.185876776 | 0.022071076 | 0.180363329 | 0.449434268 |
| Exoc8 | S | 28 | -0.305617428 | -0.310427761 | -0.508005605 | -0.202388178 | -0.197577845 | 0.119526796 | 0.134858513 | 0.025312337 | 0.141583259 | 0.200866169 |
| Zfp37 Zfp74 | T | 362;398 | -0.219688059 | -0.241673514 | -0.448319407 | -0.228631349 | -0.206645893 | 0.019319805 | 0.030109767 | 0.064235996 | 0.240811488 | 0.275255988 |
| Map1b | T | 2047 | 0.037302023 | -0.338644381 | -0.402835216 | -0.440137238 | -0.064190835 | 0.617436458 | 0.003493582 | 0.000700424 | 0.002944671 | 0.314214073 |
| Safb Safb2 | S | 306;300 | -0.2178066 | -0.266839706 | -0.135369351 | 0.082437249 | 0.131470355 | 0.004113778 | 0.005083545 | 0.033333668 | 0.037273274 | 0.071358017 |
| Rpp38 | S | 230 | 0.019160701 | -0.232321587 | -0.376313927 | -0.395474627 | -0.14399234 | 0.627494371 | 0.031842214 | 4.21211E-06 | 0.002644157 | 0.110508801 |
| Dbnl | S | 235 | -0.067359298 | -0.264982722 | -0.364574389 | -0.297215091 | -0.099591667 | 0.245396146 | 0.010809289 | 0.001313929 | 0.006565112 | 0.204833486 |
| Sipa1l1 | S | 1170 | -0.447905291 | -0.106642569 | -0.500172235 | -0.052266944 | -0.393529666 | 0.014875135 | 0.55326141 | 0.028515076 | 0.71237883 | 0.077073034 |

| | | | | | | | | | | | | |
|---|---|---|---|---|---|---|---|---|---|---|---|---|
| Plk1 | T | 210 | -0.016729853 | -0.25703408 | -0.404580188 | -0.387850334 | -0.147546108 | 0.309103583 | 0.002548537 | 0.002113625 | 0.005667568 | 0.026181971 |
| Pdzrn3 | S | 845 | -0.138813595 | -0.162492628 | -0.208762595 | -0.069949 | -0.046269967 | 0.001625561 | 0.075226701 | 0.000133435 | 0.019926044 | 0.482893603 |
| Lbh | S | 34 | -0.085605943 | -0.255869315 | -0.318672383 | -0.23306644 | -0.062803068 | 0.310563406 | 0.000777795 | 0.005515068 | 0.044998685 | 0.336672309 |
| Bora | S | 305 | -0.001669453 | -0.533293626 | -0.521285366 | -0.519615913 | 0.01200826 | 0.973034902 | 0.007734951 | 0.032107557 | 0.037574519 | 0.934534168 |
| Bora | S | 311 | -0.001669453 | -0.533293626 | -0.521285366 | -0.519615913 | 0.01200826 | 0.973034902 | 0.007734951 | 0.032107557 | 0.037574519 | 0.934534168 |
| Erbin | S | 912 | -0.387537886 | -0.253639042 | -0.40266059 | -0.015122704 | -0.149021549 | 0.014006 | 0.273805637 | 0.050798349 | 0.921629313 | 0.518876543 |
| Spag9 | S | 311 | -0.136357013 | -0.266073334 | -0.400301775 | -0.263944762 | -0.13422844 | 0.022545523 | 0.025760125 | 0.020349617 | 0.06251576 | 0.249479354 |
| Cdc5l | S | 303 | -0.139636269 | -0.426679882 | -0.476252382 | -0.336616112 | -0.049572499 | 0.219994249 | 0.042801697 | 0.023864248 | 0.056793599 | 0.74725739 |
| Nup35 | T | 265 | -0.006072951 | -0.279522159 | -0.43643073 | -0.43035778 | -0.156908571 | 0.698530757 | 0.001721787 | 0.003465536 | 0.007941261 | 0.035242201 |
| Dlgap4 | S | 683 | 0.010370451 | -0.393388006 | -0.291558866 | -0.301929317 | 0.101829139 | 0.940574287 | 0.001231768 | 0.001864827 | 0.122780628 | 0.115412151 |
| Ahctf1 | S | 1928 | -0.008924517 | -0.24897333 | -0.306470129 | -0.297545612 | -0.0574968 | 0.812834029 | 0.003655968 | 6.04305E-05 | 0.0016282 | 0.220282313 |
| Ampd2 | S | 12 | -0.277316939 | -0.380432602 | -0.276973159 | 0.00034378 | 0.103459443 | 0.039450925 | 0.011496514 | 0.171261801 | 0.998246439 | 0.52726435 |
| Ldha | T | 309 | -0.201179672 | -0.217378606 | -0.364597569 | -0.163417897 | -0.147218963 | 0.058247136 | 0.008088953 | 0.023484808 | 0.165536872 | 0.174218739 |
| Myef2 Gm9833 | S | 474 | -0.438766237 | -0.201464135 | -0.464924384 | -0.026158147 | -0.263460249 | 0.035175642 | 0.323426377 | 0.032905449 | 0.798064425 | 0.134379659 |
| Cdv3 | S | 70 | -0.22224389 | -0.355166084 | -0.727701628 | -0.505457738 | -0.372535544 | 0.116108807 | 0.168968966 | 0.052972754 | 0.111496525 | 0.235705803 |
| Psmg1 | T | 18 | -0.13786112 | -0.165324629 | -0.478098966 | -0.340237846 | -0.312774337 | 0.276168443 | 0.143646688 | 0.000324241 | 0.047083538 | 0.031483321 |
| Ubr4 | S | 362 | 0.010075958 | -0.244128943 | -0.39533605 | -0.405412008 | -0.151207108 | 0.851286437 | 0.000634034 | 0.00104819 | 0.00224871 | 0.056044708 |
| Syap1 | S | 283 | -0.066809892 | -0.279986413 | -0.443721949 | -0.376912058 | -0.163735536 | 0.342885822 | 0.000629599 | 0.030730339 | 0.043419724 | 0.234869778 |
| Med14 | S | 1138 | -0.17711281 | -0.36764311 | -0.175594412 | 0.001518397 | 0.192048698 | 0.00417983 | 0.008607154 | 0.09718103 | 0.984016705 | 0.100057663 |
| Naca | S | 114;2086 | -0.145242189 | -0.175948113 | -0.397381641 | -0.252139453 | -0.221433529 | 0.326833912 | 0.015884903 | 0.000755094 | 0.143706876 | 0.01706278 |
| Tardbp | S | 292 | -0.286570709 | -0.183484507 | -0.562922081 | -0.276351372 | -0.379437575 | 0.115874638 | 0.325874088 | 0.005634518 | 0.089562709 | 0.071667265 |
| Cdc6 | S | 129 | -0.090147054 | -0.204548097 | -0.321276021 | -0.231128968 | -0.116727925 | 0.194434208 | 0.00700889 | 0.000537812 | 0.01441143 | 0.031303713 |
| Arfgap3 | S | 241 | -0.403715301 | -0.130173398 | -0.556297145 | -0.152581844 | -0.426123747 | 0.046753612 | 0.503123615 | 0.014441914 | 0.360687485 | 0.064246702 |
| Osbpl6 | S | 319 | -0.051532782 | -0.231561688 | -0.537338172 | -0.48580539 | -0.305776484 | 0.622515423 | 0.137036293 | 0.000251317 | 0.008950335 | 0.083431108 |
| Brd1 | S | 416 | -0.104983117 | -0.307067316 | -0.210290328 | -0.105307211 | 0.096776988 | 0.090031264 | 0.001402699 | 0.006967608 | 0.014194177 | 0.096071178 |
| Brd1 | S | 422 | -0.104983117 | -0.307067316 | -0.210290328 | -0.105307211 | 0.096776988 | 0.090031264 | 0.001402699 | 0.006967608 | 0.014194177 | 0.096071178 |
| Brd1 | S | 424 | -0.104983117 | -0.307067316 | -0.210290328 | -0.105307211 | 0.096776988 | 0.090031264 | 0.001402699 | 0.006967608 | 0.014194177 | 0.096071178 |
| Brd1 | T | 419 | -0.104983117 | -0.307067316 | -0.210290328 | -0.105307211 | 0.096776988 | 0.090031264 | 0.001402699 | 0.006967608 | 0.014194177 | 0.096071178 |
| Brd1 | T | 423 | -0.104983117 | -0.307067316 | -0.210290328 | -0.105307211 | 0.096776988 | 0.090031264 | 0.001402699 | 0.006967608 | 0.014194177 | 0.096071178 |
| Vapa | T | 170 | -0.029521685 | -0.153092509 | -0.336841215 | -0.30731953 | -0.183748706 | 0.744020743 | 0.002189952 | 2.73415E-05 | 0.04838121 | 0.002472963 |
| Ddx24 | S | 80 | -0.028599811 | -0.284840168 | -0.187806124 | -0.159206313 | 0.097034044 | 0.327964272 | 0.000283705 | 0.000282804 | 0.002655684 | 0.035399687 |
| Prrc2b | S | 1007 | 0.049246725 | -0.433808986 | -0.333276444 | -0.382523169 | 0.100532542 | 0.39831356 | 0.016362223 | 4.8521E-05 | 0.005401282 | 0.275459296 |
| Braf | S | 464 | -0.178125997 | -0.414791241 | -0.473804592 | -0.295678595 | -0.059013351 | 0.17351101 | 0.200783176 | 0.005615704 | 0.047247823 | 0.824751404 |
| Caprin1 | S | 8 | -0.107277279 | -0.274992644 | -0.359376361 | -0.252099082 | -0.084383717 | 0.311184481 | 0.0088115 | 0.002955848 | 0.04612899 | 0.193457789 |
| Map1b | T | 2301 | -0.058958312 | -0.339604335 | -0.250017224 | -0.191058912 | 0.089587111 | 0.362058645 | 0.001136354 | 0.003847161 | 0.012178926 | 0.137646437 |
| Acap2 | S | 523 | -0.040311767 | -0.253195207 | -0.435123517 | -0.39481175 | -0.181928311 | 0.595293757 | 0.019531988 | 0.000593155 | 0.001747087 | 0.05513525 |
| Nsd2 | S | 8 | -0.036153683 | -0.21058614 | -0.168414523 | -0.132260841 | 0.042171617 | 0.381683863 | 1.63627E-05 | 0.000124063 | 0.049308929 | 0.032648708 |
| Anln | S | 68 | -0.153644693 | -0.201487945 | -0.278861475 | -0.125216782 | -0.07737353 | 0.023766084 | 0.005053873 | 0.008680531 | 0.115945649 | 0.266537933 |
| Abl2 | S | 785 | -0.125309576 | -0.234064274 | -0.470530527 | -0.34522095 | -0.236466253 | 0.303421315 | 0.066829946 | 0.001403273 | 0.027912488 | 0.059162493 |
| Jcad | S | 768 | -0.269594917 | -0.214273046 | -0.421977544 | -0.152382627 | -0.207704499 | 0.030571536 | 0.087865526 | 0.024845957 | 0.267502294 | 0.167875594 |
| Ptpre | S | 104 | -0.005653998 | -0.3176304 | -0.362346692 | -0.356692694 | -0.044716292 | 0.936412554 | 0.000116515 | 0.015923855 | 0.016490157 | 0.580657421 |
| Bcl11a Bcl11b Ikzf5 Z | T | 256;400;449;133 | -0.150265613 | -0.184392464 | -0.318662186 | -0.168396572 | -0.134269721 | 0.024323588 | 0.086146012 | 0.000726658 | 0.001715079 | 0.177094013 |
| Iqsec2 | S | 402 | -0.370865731 | -0.14496731 | -0.456143198 | -0.085277467 | -0.311175888 | 0.013004239 | 0.395568146 | 0.024395006 | 0.515268933 | 0.124839055 |
| Drosha | S | 356 | -0.045865262 | -0.228898383 | -0.336202611 | -0.290337349 | -0.107304227 | 0.417612423 | 0.000994262 | 0.00150619 | 0.007243081 | 0.078834816 |
| Adam19 | T | 781 | 0.025918601 | -0.290780188 | -0.269782836 | -0.295701437 | 0.020997352 | 0.561772237 | 9.98729E-05 | 0.000558897 | 0.004152978 | 0.522162731 |
| Acin1 | S | 605 | -0.120628803 | -0.273770112 | -0.287055116 | -0.166426313 | -0.013285004 | 0.079047161 | 0.030449928 | 0.001041843 | 0.03793402 | 0.867463286 |
| Washc2 | S | 798 | -0.039673622 | -0.280071139 | -0.281451178 | -0.241777556 | -0.001380039 | 0.463382647 | 2.86148E-05 | 0.020137317 | 0.018632621 | 0.97904081 |
| Ncoa3 | S | 562 | -0.022399106 | -0.354544437 | -0.401446369 | -0.379047263 | -0.046901932 | 0.604525898 | 0.000264898 | 0.044106615 | 0.041056892 | 0.666560599 |
| Blm | S | 274 | -0.066722961 | -0.318859391 | -0.316822164 | -0.250099204 | 0.002037227 | 0.375684795 | 0.003491971 | 0.003609143 | 0.004775049 | 0.962797947 |
| Map1b | S | 1271 | -0.170373719 | -0.281645983 | -0.402934548 | -0.232560828 | -0.121288565 | 0.341979561 | 0.020512765 | 0.005114779 | 0.222126913 | 0.235986044 |
| Nelfe | S | 255 | -0.222834593 | -0.170040317 | -0.372834009 | -0.149939416 | -0.202793692 | 0.027504458 | 0.152268688 | 0.003024772 | 0.045782384 | 0.089091737 |
| Rtn4 | S | 344 | -0.055829852 | -0.143985613 | -0.26608716 | -0.210257308 | -0.122101547 | 0.05429673 | 0.012805695 | 1.23793E-05 | 0.000814299 | 0.02257651 |
| Fam117b | T | 391 | -0.096766589 | -0.297561991 | -0.442042399 | -0.34527581 | -0.144480408 | 0.033724779 | 0.045048431 | 0.012293649 | 0.016944112 | 0.195192134 |
| Pttg1ip | Y | 171 | -0.224277558 | -0.200787907 | -0.427337589 | -0.203060031 | -0.226549682 | 0.07338343 | 0.169988463 | 0.003253505 | 0.062726277 | 0.11956017 |
| Carhsp1 | S | 53 | -0.203454766 | -0.231120728 | -0.561790225 | -0.358335459 | -0.330669497 | 0.117264545 | 0.138608772 | 0.011132653 | 0.058168901 | 0.087086613 |
| Sh3gl1 | S | 288 | -0.361897065 | -0.202019658 | -0.69497003 | -0.333072965 | -0.492950372 | 0.049849165 | 0.347492648 | 0.054753517 | 0.245363329 | 0.134555522 |
| Parp9 | S | 42 | -0.125503925 | -0.426656466 | -0.390580906 | -0.26507698 | 0.03607556 | 0.465517609 | 0.003128271 | 0.060822009 | 0.213919744 | 0.805416576 |
| Tuba1a Tuba1b Tuba | S | 48;48;48;48 | -0.255662448 | -0.251798191 | -0.459060087 | -0.203397639 | -0.207261897 | 0.031266384 | 0.061870492 | 0.057373656 | 0.273572797 | 0.274706987 |
| Cwc25 | S | 309 | -0.147797828 | -0.404246678 | -0.134844152 | 0.012953677 | 0.269402526 | 0.052857913 | 0.000743946 | 0.099681065 | 0.781608346 | 0.013829975 |
| Cdc5l | T | 355 | -0.147222022 | -0.184713312 | -0.219786635 | -0.072564613 | -0.035073323 | 0.002620587 | 0.003010537 | 0.01618307 | 0.239919376 | 0.541635002 |
| Nudt5 | S | 10 | -0.240233733 | -0.209960152 | -0.333850705 | -0.093616972 | -0.123890553 | 0.019468021 | 0.129297733 | 0.006663429 | 0.276026388 | 0.313743033 |
| Rif1 | S | 2296 | 0.140903033 | -0.32847357 | -0.340863498 | -0.481766531 | -0.012389928 | 0.473306286 | 0.000445128 | 0.000331117 | 0.090999806 | 0.746338378 |
| Ecm1 | S | 556 | -0.016885085 | -0.386706538 | -0.41838523 | -0.401500145 | -0.031678692 | 0.830845421 | 0.002777701 | 0.012382792 | 0.0124609 | 0.710811342 |
| Nvl | S | 194 | -0.133110737 | -0.235099961 | -0.359910459 | -0.226799722 | -0.124810498 | 0.320994416 | 0.00341761 | 0.005224872 | 0.142762256 | 0.130891654 |
| Dync1h1 | S | 4366 | -0.012350854 | -0.190506315 | -0.269049874 | -0.25669902 | -0.078543559 | 0.717241608 | 0.00511605 | 2.88399E-06 | 0.008630216 | 0.089198868 |
| Sipa1l3 | T | 1433 | -0.072184603 | -0.393711074 | -0.501563882 | -0.429379279 | -0.107852807 | 0.573299807 | 0.005750971 | 0.035711795 | 0.044615099 | 0.41082049 |
| Atp2b1 | S | 1139 | -0.288804221 | -0.226547361 | -0.574058063 | -0.285253842 | -0.347510702 | 0.06391678 | 0.236453274 | 0.025088863 | 0.167768113 | 0.131129453 |
| Eif4ebp2 | T | 46 | -0.18421177 | -0.215957528 | -0.231650857 | -0.047439086 | -0.015693329 | 0.030764589 | 0.00787251 | 0.005737494 | 0.38509966 | 0.580684075 |
| Caprin1 | S | 10 | -0.300413543 | -0.467972871 | -0.21149321 | 0.088920333 | 0.256479661 | 0.017504767 | 0.057850294 | 0.127928459 | 0.393373169 | 0.216011138 |
| Mid1ip1 | S | 109 | 0.076158239 | -0.258563905 | -0.488822382 | -0.564980621 | -0.230258477 | 0.28453379 | 0.002042435 | 0.000385462 | 0.000951306 | 0.030806344 |
| Ufd1 | S | 229 | -0.013195637 | -0.279956099 | -0.496193705 | -0.482998068 | -0.216237606 | 0.847027098 | 0.003908647 | 0.004928671 | 0.008156036 | 0.075489912 |
| Sacs | S | 1781 | -0.041796307 | -0.272926595 | -0.160030015 | -0.118233708 | 0.11289658 | 0.208078012 | 0.000300049 | 0.00019026 | 0.018968246 | 0.025858827 |
| Fxr2 | S | 455 | -0.052066165 | -0.215658778 | -0.246702301 | -0.194636136 | -0.031043523 | 0.358214861 | 0.001447551 | 0.000115678 | 0.038645857 | 0.365131705 |
| Hsf1 | S | 326 | -0.090123613 | -0.274564521 | -0.275538472 | -0.18541486 | -0.000973952 | 0.144190063 | 0.005161691 | 0.002301904 | 0.010767394 | 0.986191187 |
| Eif5 | S | 387 | -0.057431333 | -0.219090107 | -0.168598103 | -0.11116677 | 0.050492004 | 0.023383681 | 2.13462E-05 | 0.007072846 | 0.029697696 | 0.172295405 |
| Eif5 | S | 388 | -0.057431333 | -0.219090107 | -0.168598103 | -0.11116677 | 0.050492004 | 0.023383681 | 2.13462E-05 | 0.007072846 | 0.029697696 | 0.172295405 |
| Ddx21 | S | 89 | -0.070427053 | -0.250388985 | -0.452626368 | -0.382199315 | -0.202237383 | 0.327844878 | 0.020309771 | 0.00254598 | 0.008519413 | 0.070261339 |
| Dlgap4 | S | 946 | -0.174548248 | -0.193511066 | -0.291260183 | -0.116711935 | -0.097749117 | 0.007484671 | 0.07937086 | 0.003474235 | 0.094012112 | 0.299916411 |
| Slc7a1 | S | 457 | -0.077264157 | -0.352648864 | -0.202756531 | -0.125492374 | 0.149892333 | 0.246125725 | 0.001860042 | 0.002838881 | 0.107133521 | 0.076192354 |
| Setd2 | S | 345 | -0.370583208 | -0.213208204 | -0.472843418 | -0.10226021 | -0.259635214 | 0.050675296 | 0.298544167 | 0.021324214 | 0.264260197 | 0.154433377 |
| Gtf3c5 | Y | 331 | -0.17707041 | -0.27361242 | -0.24973406 | -0.07266365 | 0.02387836 | 0.038920444 | 0.006575006 | 0.021953647 | 0.308235536 | 0.711498812 |
| Map4k4 | S | 894 | -0.134481965 | -0.134231614 | -0.550537888 | -0.416055923 | -0.416306274 | 0.151192584 | 0.438843362 | 0.000416853 | 0.002062592 | 0.076906674 |
| Dcaf1 | S | 254 | 0.057324198 | -0.305758177 | -0.351169844 | -0.408494043 | -0.045411667 | 0.272084729 | 3.77939E-05 | 0.003358029 | 0.002408544 | 0.420558882 |
| Rpl22l1 | S | 118 | 0.043719801 | -0.313131476 | -0.29937836 | -0.343098161 | 0.013753115 | 0.242718121 | 8.94351E-06 | 0.005591226 | 0.003026124 | 0.756870569 |
| Pik3r4 | S | 905 | -0.172807965 | -0.256307856 | -0.24533965 | -0.072531684 | 0.010968206 | 0.070298739 | 0.000395483 | 0.068785264 | 0.494515884 | 0.902462433 |
| Map3k2 | S | 153 | -0.189602951 | -0.214140012 | -0.456799984 | -0.267197033 | -0.242659972 | 0.060281095 | 0.198118293 | 0.004063391 | 0.045193503 | 0.15924866 |
| Dock1 | T | 1844 | 0.029807067 | -0.3211913 | -0.477413752 | -0.507220819 | -0.156222452 | 0.498773336 | 0.002862592 | 0.0027961 | 0.000835068 | 0.044924168 |
| Cdc20 | S | 84 | -0.219266203 | -0.142386202 | -0.547133245 | -0.327867042 | -0.404747043 | 0.103131971 | 0.401506243 | 0.002102073 | 0.026444061 | 0.059138409 |
| Larp1 | S | 801 | -0.042800567 | -0.236830507 | -0.217434314 | -0.174633748 | 0.019396192 | 0.159527634 | 5.22808E-05 | 0.00411791 | 0.018267265 | 0.609809321 |
| Tsc2 | S | 981 | -0.084597034 | -0.409009665 | -0.379616588 | -0.295019555 | 0.029393077 | 0.180821344 | 0.029448499 | 0.010370058 | 0.031544836 | 0.813176409 |
| Esf1 | S | 306 | 0.0198476 | -0.202553603 | -0.205456431 | -0.22530403 | -0.002902828 | 0.513789758 | 3.40375E-05 | 6.33668E-06 | 0.004886592 | 0.864896845 |
| Esf1 | S | 307 | 0.0198476 | -0.202553603 | -0.205456431 | -0.22530403 | -0.002902828 | 0.513789758 | 3.40375E-05 | 6.33668E-06 | 0.004886592 | 0.864896845 |
| Esf1 | T | 305 | 0.0198476 | -0.202553603 | -0.205456431 | -0.22530403 | -0.002902828 | 0.513789758 | 3.40375E-05 | 6.33668E-06 | 0.004886592 | 0.864896845 |
| Smg9 | T | 118 | -0.07820451 | -0.430059884 | -0.291464047 | -0.213259538 | 0.138595836 | 0.103470506 | 0.021797522 | 0.007606481 | 0.012204095 | 0.17946286 |

| | | | | | | | | | | | | |
|---|---|---|---|---|---|---|---|---|---|---|---|---|
| Pom121 | T | 421 | 0.086406676 | -0.420921493 | -0.410919263 | -0.497325938 | 0.01000223 | 0.203154776 | 0.002464143 | 0.001073238 | 0.001459886 | 0.892003908 |
| Map4 | S | 667 | -0.047846668 | -0.146178939 | -0.259752173 | -0.211905505 | -0.113573234 | 0.204382003 | 0.000576174 | 7.10355E-05 | 0.003576083 | 0.023302259 |
| Trmt44 | S | 76 | -0.224967518 | -0.291791665 | -0.140707494 | 0.084260023 | 0.151084171 | 0.007412914 | 0.002699677 | 0.135837486 | 0.309523942 | 0.11627435 |
| C2cd2l | S | 464 | -0.029034369 | -0.282886333 | -0.242782999 | -0.21374863 | 0.040103335 | 0.469080495 | 0.000226037 | 0.002544681 | 0.009565357 | 0.353980704 |
| Brip1 | S | 209 | -0.304913359 | -0.371247132 | -0.361425822 | -0.056512463 | 0.00982131 | 0.083908819 | 0.046451178 | 0.079731034 | 0.64917624 | 0.936651904 |
| Exoc4 | T | 33 | -0.296520366 | -0.2407974 | -0.286924525 | 0.009595841 | -0.046127125 | 0.02584092 | 0.054577333 | 0.027876998 | 0.920356575 | 0.63438076 |
| Mical3 | S | 1993 | -0.323481419 | -0.287704017 | -0.239113016 | 0.084368403 | 0.048591001 | 0.067982157 | 0.022773488 | 0.030432862 | 0.534455877 | 0.613246531 |
| Pnn | S | 703 | 0.076759106 | -0.318866673 | -0.420396111 | -0.497155217 | -0.101529438 | 0.099636998 | 3.15923E-05 | 0.00499927 | 0.003565411 | 0.191460818 |
| Rpl37 Rpl37rt | S | 7 | -0.006015963 | -0.116276928 | -0.580421813 | -0.574405849 | -0.464144884 | 0.954824227 | 0.441047016 | 1.21955E-05 | 0.016950929 | 0.056178846 |
| Sqstm1 | S | 367 | -0.020431656 | -0.279612876 | -0.291567329 | -0.271135673 | -0.011954453 | 0.638575535 | 0.002751394 | 0.000409422 | 0.002030792 | 0.813576685 |
| Pea15a | S | 116 | -0.414655544 | -0.225910606 | -0.577163893 | -0.162508349 | -0.351253287 | 0.047177456 | 0.325394374 | 0.064639432 | 0.505596852 | 0.22933714 |
| Peak1 | S | 897 | -0.069524787 | -0.221859562 | -0.262842293 | -0.193317506 | -0.040982731 | 0.433336981 | 0.014520305 | 2.30035E-05 | 0.108429377 | 0.426757722 |
| Dnajb4 | S | 148 | -0.029141868 | -0.192720403 | -0.329455972 | -0.300314105 | -0.136735569 | 0.670953312 | 0.000926648 | 0.000358572 | 0.015439056 | 0.033041308 |
| Phldb2 | S | 410 | -0.19245339 | -0.258006521 | -0.442850204 | -0.250396814 | -0.184843683 | 0.059768674 | 0.050103443 | 0.026340239 | 0.091901969 | 0.188964501 |
| Phrf1 | S | 731 | 0.006351368 | -0.345568802 | -0.351764251 | -0.358115619 | -0.006195449 | 0.932300503 | 0.01431765 | 0.000294705 | 0.018989095 | 0.935759228 |
| Tbc1d15 | S | 662 | -0.092293036 | -0.34618534 | -0.319772194 | -0.227479158 | 0.026413145 | 0.343086936 | 0.005755711 | 0.008770128 | 0.037137541 | 0.591156242 |
| Fer | Y | 715 | -0.357717772 | -0.214749334 | -0.473256333 | -0.115538561 | -0.258506999 | 0.054402972 | 0.360768974 | 0.016605994 | 0.454115869 | 0.280305322 |
| Dync1li1 | T | 513 | -0.064019246 | -0.225492658 | -0.210241907 | -0.146222661 | 0.015250751 | 0.011703557 | 0.000127164 | 0.0170997 | 0.034348072 | 0.699993808 |
| Knop1 | S | 104 | 0.046171868 | -0.319456382 | -0.239753857 | -0.285925725 | 0.079702524 | 0.466506886 | 9.59054E-05 | 0.000511513 | 0.015467796 | 0.078154402 |
| Rps6kc1 | S | 599 | -0.263526922 | -0.183645983 | -0.285572029 | -0.022045107 | -0.101926046 | 0.022920868 | 0.097271001 | 0.005106997 | 0.779701149 | 0.282823019 |
| Pip5k1c | S | 555 | 0.00999596 | -0.520085784 | -0.421050616 | -0.431046576 | 0.099035168 | 0.881420867 | 0.014788791 | 0.007564761 | 0.007612504 | 0.423263838 |
| Wrnip1 | S | 65 | -0.000209671 | -0.17856066 | -0.181289826 | -0.181080155 | -0.002729165 | 0.993131366 | 2.18693E-06 | 4.18328E-05 | 0.003794169 | 0.846487621 |
| Trio | T | 2633 | -0.403009129 | -0.152879056 | -0.326629433 | 0.076379696 | -0.173750377 | 0.032585486 | 0.199853329 | 0.011543376 | 0.551683364 | 0.101696581 |
| Hells | S | 486 | 0.022781545 | -0.18616611 | -0.306313383 | -0.329094928 | -0.120147273 | 0.578400883 | 0.000293914 | 4.0924E-05 | 0.001403797 | 0.010682737 |
| Zfr | S | 1054 | -0.011086041 | -0.225221909 | -0.187616002 | -0.17652996 | 0.037605907 | 0.60209876 | 3.69092E-05 | 0.000147157 | 3.8826E-05 | 0.009299241 |
| Tacc2 | S | 933 | -0.243443009 | -0.235975743 | -0.214096617 | 0.029346392 | 0.021879126 | 0.072310293 | 0.014278101 | 0.004443655 | 0.759086475 | 0.727823242 |
| Pbdc1 | S | 184 | -0.200022157 | -0.184005641 | -0.345872999 | -0.145850842 | -0.161867358 | 0.011130911 | 0.072156978 | 0.0123799 | 0.147139609 | 0.150501549 |
| Esf1 | S | 686 | -0.07443106 | -0.321601027 | -0.206348451 | -0.131917391 | 0.115252576 | 0.19140128 | 0.00051579 | 0.011004094 | 0.057970642 | 0.092830881 |
| Esf1 | S | 688 | -0.07443106 | -0.321601027 | -0.206348451 | -0.131917391 | 0.115252576 | 0.19140128 | 0.00051579 | 0.011004094 | 0.057970642 | 0.092830881 |
| Esf1 | S | 689 | -0.07443106 | -0.321601027 | -0.206348451 | -0.131917391 | 0.115252576 | 0.19140128 | 0.00051579 | 0.011004094 | 0.057970642 | 0.092830881 |
| Pakap | Y | 439 | -0.198331256 | -0.271533293 | -0.420069374 | -0.221738118 | -0.148536081 | 0.079205092 | 0.033523488 | 0.031399176 | 0.167072085 | 0.304707734 |
| Usp24 | T | 2556 | -0.249182562 | -0.292138438 | -0.509717064 | -0.260534502 | -0.217578627 | 0.061234408 | 0.005272421 | 0.283003138 | 0.53891425 | 0.600218109 |
| Prune2 | S | 2540 | -0.17855955 | -0.286906101 | -0.225606133 | -0.047046583 | 0.061299968 | 0.033744632 | 0.004429952 | 0.04085497 | 0.517119302 | 0.422598916 |
| Arhgef17 | S | 389 | -0.201083809 | -0.308268993 | -0.278800645 | -0.077716836 | 0.029468348 | 0.07038043 | 0.064235313 | 0.00467622 | 0.381474782 | 0.801449093 |
| Btrc | S | 127 | -0.011817371 | -0.272087982 | -0.375867428 | -0.364050057 | -0.103779446 | 0.859402743 | 0.030465886 | 0.000104521 | 0.011438264 | 0.256945403 |
| Marchf7 | S | 302 | -0.194470812 | -0.180611066 | -0.325493796 | -0.131022983 | -0.14488273 | 0.032352809 | 0.057824574 | 0.004064287 | 0.004632583 | 0.06959557 |
| Iqsec1 | S | 398 | -0.254256806 | -0.098222616 | -0.456184353 | -0.201927547 | -0.357961737 | 0.04295414 | 0.473214867 | 0.001681248 | 0.094997878 | 0.055561158 |
| Sfswap | S | 821 | -0.121630848 | -0.137824924 | -0.191109331 | -0.069478483 | -0.053284407 | 0.001518273 | 0.000697642 | 0.006496147 | 0.142579379 | 0.214332179 |
| Zdhhc20 | S | 343 | 0.039012904 | -0.257110215 | -0.480240763 | -0.519253667 | -0.223130549 | 0.566077217 | 0.00770796 | 0.000539732 | 0.000676982 | 0.003726402 |
| Sqstm1 | S | 207 | -0.15422084 | -0.281010203 | -0.441003825 | -0.286782984 | -0.159993622 | 0.489236321 | 0.025479951 | 0.005259784 | 0.255037194 | 0.088497544 |
| Eml1 | S | 146 | -0.325541663 | -0.19585239 | -0.749581397 | -0.424039734 | -0.553729008 | 0.089214491 | 0.253435223 | 0.057592587 | 0.20552437 | 0.140119799 |
| Surf6 | T | 179 | -0.042572638 | -0.276716437 | -0.4076088 | -0.365036162 | -0.130892364 | 0.669506084 | 0.032629026 | 0.000486697 | 0.031402827 | 0.180149202 |
| H2bc1 H2bc11 H2bc1 | S | 113;113;113;114; | 0.108332011 | -0.328188585 | -0.242772339 | -0.351104351 | 0.085416246 | 0.544807709 | 1.94186E-05 | 0.001853811 | 0.138290252 | 0.097396302 |
| Afap1 | S | 711 | -0.013192754 | -0.275123539 | -0.518189137 | -0.504996382 | -0.243065597 | 0.789419601 | 0.000161456 | 0.046638046 | 0.045864123 | 0.185919297 |
| Slc20a2 | T | 318 | -0.202511691 | -0.177680048 | -0.529807724 | -0.327296033 | -0.352127676 | 0.062621241 | 0.216505199 | 0.008204316 | 0.057358977 | 0.058667192 |
| Phldb2 | S | 930 | -0.083543479 | -0.362958063 | -0.47308381 | -0.389540331 | -0.110125747 | 0.542261695 | 0.013670411 | 0.016736677 | 0.054800503 | 0.317562098 |
| Ctps | S | 571 | -0.137345154 | -0.193311083 | -0.271538241 | -0.134193088 | -0.078227158 | 0.043153548 | 0.001842498 | 0.011169569 | 0.10289633 | 0.251092323 |
| Nqo1 | S | 82 | -0.249011274 | -0.045581139 | -0.435441863 | -0.186430589 | -0.389860724 | 0.012231247 | 0.611332208 | 0.001690913 | 0.056785456 | 0.008008569 |
| Pdlim1 | S | 213 | -0.314357125 | -0.25369936 | -0.261769929 | 0.052587197 | -0.008070569 | 0.005518016 | 0.125258318 | 0.059647211 | 0.6216037 | 0.955655857 |
| Map1b | T | 1945 | -0.062169128 | -0.133128437 | -0.305441275 | -0.243272147 | -0.172312838 | 0.089973315 | 0.004917381 | 0.000157663 | 0.005374994 | 0.009589113 |
| Cit | S | 1966 | -0.028688063 | -0.344356622 | -0.347195479 | -0.318507416 | -0.002838857 | 0.769780027 | 0.013310396 | 0.000939488 | 0.034852515 | 0.973044418 |
| Lrrc8c | T | 198 | -0.168216238 | -0.305181566 | -0.329413719 | -0.161197481 | -0.024232153 | 0.114003192 | 0.041372757 | 0.007427887 | 0.104168915 | 0.820443475 |
| Phldb2 | S | 809 | -0.003403536 | -0.349933316 | -0.294612397 | -0.291208861 | 0.055320919 | 0.950427943 | 0.002441913 | 0.001335415 | 0.002215709 | 0.392363878 |
| Pdcd11 | S | 36 | 0.017703097 | -0.200815527 | -0.386689961 | -0.404393058 | -0.185874434 | 0.709285308 | 0.000797906 | 0.000394468 | 0.001299623 | 0.0106638 |
| Pom121 | S | 367 | 0.000100603 | -0.360568432 | -0.313735253 | -0.313835856 | 0.046833179 | 0.998902395 | 0.001436448 | 0.003765064 | 0.018955912 | 0.423320779 |
| Hmgn3 | S | 93 | -0.210599099 | -0.192295171 | -0.197644704 | 0.012954395 | -0.005349533 | 0.002913836 | 0.033424206 | 0.011288877 | 0.779582494 | 0.936061398 |
| Ybx1 | S | 165 | -0.328103262 | -0.116298943 | -0.506412191 | -0.178308929 | -0.390113248 | 0.006659485 | 0.278688007 | 0.060809891 | 0.369950374 | 0.114531188 |
| Hnrnpa2b1 | S | 236 | -0.175896307 | -0.304776031 | -0.551220234 | -0.375323927 | -0.246444203 | 0.350974231 | 0.143718599 | 0.00677731 | 0.079216641 | 0.187029346 |
| Ckb | S | 199 | -0.095351511 | -0.240058956 | -0.553273779 | -0.457922268 | -0.313214823 | 0.683139334 | 0.044628187 | 0.002818052 | 0.12521803 | 0.046539937 |
| Abl1 | S | 634 | -0.484768133 | -0.224526149 | -0.306253036 | 0.178515097 | -0.081726886 | 0.008204409 | 0.193154559 | 0.182041366 | 0.367124857 | 0.685138718 |
| Dock7 | S | 440 | -0.177637069 | -0.113341738 | -0.466320624 | -0.288683555 | -0.352978886 | 0.059613097 | 0.416209394 | 0.000745685 | 0.006556875 | 0.060189574 |
| Prrc2b | S | 762 | 0.008725523 | -0.424857518 | -0.408595385 | -0.417320908 | 0.016262132 | 0.862626811 | 0.003132527 | 0.013610533 | 0.005901707 | 0.831677939 |
| Rptor | S | 722 | -0.34879952 | -0.160668941 | -0.325218751 | 0.023580769 | -0.16454981 | 0.002289433 | 0.351685756 | 0.049692364 | 0.834036888 | 0.373144393 |
| Pla2g4a | S | 728 | -0.150856859 | -0.17013524 | -0.450669833 | -0.299812974 | -0.280534592 | 0.040592956 | 0.075094576 | 0.007062967 | 0.039513323 | 0.042301033 |
| Riok2 | T | 480 | -0.085160772 | -0.193719588 | -0.223158921 | -0.137998148 | -0.029439333 | 0.008568689 | 0.038465276 | 0.000119512 | 0.00366558 | 0.591595523 |
| Ppig | S | 671 | -0.327345436 | -0.480675199 | -0.384909899 | -0.057564464 | 0.0957653 | 0.123806132 | 0.183984032 | 0.038138131 | 0.758679303 | 0.748870995 |
| Ccdc6 | S | 410 | -0.137934797 | -0.148084587 | -0.311181959 | -0.173247162 | -0.163097372 | 0.011368808 | 0.026034413 | 0.003459284 | 0.05506159 | 0.047590269 |
| Arid1a | S | 1927 | -0.166099788 | -0.239186321 | -0.27586193 | -0.109762142 | -0.03667561 | 0.029557156 | 0.000599905 | 0.129715752 | 0.448549657 | 0.782787399 |
| Spred1 | S | 318 | -0.217679201 | -0.16468872 | -0.568136717 | -0.350457516 | -0.403447997 | 0.166408882 | 0.274524464 | 0.004312839 | 0.063476666 | 0.043364277 |
| Dda1 | S | 33 | 0.012453257 | -0.243382971 | -0.570620869 | -0.583074127 | -0.327237898 | 0.750187667 | 0.054837376 | 0.000571206 | 0.004235498 | 0.024198637 |
| Cd2ap | S | 514 | -0.024943106 | -0.163759276 | -0.290483354 | -0.265540247 | -0.126724078 | 0.461989674 | 4.53423E-05 | 0.000916209 | 0.001802979 | 0.029901899 |
| Dnm1l | S | 535 | -0.378528474 | -0.245530773 | -0.272026982 | 0.106501492 | -0.026496209 | 0.011131651 | 0.148844102 | 0.068132466 | 0.41035813 | 0.864393685 |
| Ccdc6 | S | 321 | -0.023880483 | -0.278780698 | -0.581386331 | -0.557505848 | -0.302605633 | 0.727471793 | 0.039418981 | 0.003641135 | 0.006589844 | 0.047635371 |
| Mapk6 | S | 189 | -0.091178015 | -0.307278218 | -0.294464495 | -0.20328648 | 0.012813723 | 0.091659275 | 0.011806392 | 0.007029619 | 0.02553501 | 0.861447369 |
| Nfic | S | 277 | 0.0409317 | -0.254670129 | -0.411094369 | -0.452026069 | -0.156424241 | 0.676706561 | 0.011996763 | 0.000144976 | 0.019161109 | 0.06353162 |
| Jcad | S | 1245 | -0.301172235 | -0.16942899 | -0.410669125 | -0.10949689 | -0.241240135 | 0.026858002 | 0.178385752 | 0.021416922 | 0.383428656 | 0.109560167 |
| Arnt | S | 77 | 0.008769576 | -0.216459337 | -0.292636226 | -0.301405803 | -0.076176889 | 0.892533582 | 0.000638171 | 0.00014367 | 0.015997632 | 0.055912748 |
| Kirrel | S | 606 | -0.453479428 | -0.143501043 | -0.48669971 | -0.033220282 | -0.343198667 | 0.061230048 | 0.450745918 | 0.020487573 | 0.840257153 | 0.062035734 |
| Ehd4 | S | 15 | -0.243704259 | -0.118314327 | -0.444789343 | -0.201085084 | -0.326475016 | 0.01342569 | 0.27808988 | 0.010401387 | 0.121003625 | 0.042713807 |
| Dnajb6 | S | 15 | -0.086125944 | -0.207547068 | -0.324474377 | -0.238348433 | -0.116927309 | 0.080943887 | 0.013846659 | 0.001810671 | 0.014114713 | 0.118234232 |
| Sec16a | S | 311 | -0.027143649 | -0.142815725 | -0.25583381 | -0.228690161 | -0.113018086 | 0.27113482 | 0.00083738 | 2.55487E-05 | 0.001356267 | 0.005271582 |
| Lrig1 | S | 1067 | 0.017832341 | -0.237417091 | -0.206920204 | -0.224752546 | 0.030496887 | 0.292948175 | 0.000142735 | 1.10688E-05 | 0.00013846 | 0.16066951 |
| Oprd1 | S | 249 | -0.077774924 | -0.227944327 | -0.28449454 | -0.206719617 | -0.056550214 | 0.193863676 | 0.006765345 | 0.000885499 | 0.006694537 | 0.271952409 |
| Svil | S | 1045 | -0.25183998 | -0.240768749 | -0.450650823 | -0.198810843 | -0.209882074 | 0.207245126 | 0.102998613 | 0.008355221 | 0.278018122 | 0.094624141 |
| Twistnb | T | 226 | 0.061381839 | -0.332922358 | -0.340284909 | -0.401666748 | -0.007362551 | 0.248586557 | 0.001127172 | 0.000419256 | 0.004108503 | 0.883240089 |
| Numa1 | S | 1951 | -0.216790608 | -0.228258068 | -0.368756636 | -0.151966028 | -0.140498567 | 0.023778782 | 0.032003312 | 0.05054622 | 0.283805673 | 0.321125731 |
| Mast4 | S | 1465 | -0.314367013 | -0.192910715 | -0.497556525 | -0.183189513 | -0.30464581 | 0.044462898 | 0.302124635 | 0.023583712 | 0.248248593 | 0.145236073 |
| Larp1 | S | 525 | -0.088486755 | -0.168285044 | -0.159602327 | -0.071115573 | 0.008682717 | 0.038954107 | 1.58963E-06 | 0.01822747 | 0.104039992 | 0.770391974 |
| Rps6kb1 | S | 394 | -0.178197019 | -0.314446652 | -0.479151226 | -0.300954207 | -0.164704574 | 0.062292228 | 0.04394849 | 0.066882637 | 0.178236316 | 0.41293174 |
| Eif4g1 | S | 1597 | -0.049700183 | -0.1705306 | -0.251904499 | -0.202204316 | -0.081373899 | 0.135029427 | 0.012535155 | 1.96189E-05 | 0.002270932 | 0.115427026 |
| Acaca | S | 29 | -0.110368993 | -0.225041611 | -0.217648917 | -0.107279923 | 0.007392694 | 0.063453126 | 0.009773409 | 0.000687613 | 0.074580615 | 0.882529834 |
| Eef1b2 | S | 106 | -0.104048917 | -0.313887481 | -0.250656722 | -0.146607805 | 0.063230759 | 0.260812995 | 0.002204119 | 0.010265914 | 0.148913888 | 0.373928737 |
| Rnf187 | S | 200 | -0.095998985 | -0.145899033 | -0.446480645 | -0.350481659 | -0.300581611 | 0.064506247 | 0.044989512 | 0.002335139 | 0.010036431 | 0.012205848 |
| Dennd4c | S | 1293 | 0.012319466 | -0.182289635 | -0.317890579 | -0.330210045 | -0.135600944 | 0.799048898 | 0.001805165 | 3.38406E-05 | 0.005048425 | 0.010279732 |

| | | | | | | | | | | | | |
|---|---|---|---|---|---|---|---|---|---|---|---|---|
| Nisch | S | 548 | 0.024452775 | -0.256118581 | -0.392654447 | -0.417107222 | -0.136535866 | 0.507345198 | 0.001180754 | 0.00103184 | 0.006385608 | 0.06207387 |
| Prrc2c | S | 613 | -0.212127746 | -0.306007438 | -0.126945863 | 0.085181884 | 0.179061575 | 0.002141629 | 0.003980772 | 0.310652456 | 0.467604022 | 0.19055093 |
| Pja1 | S | 175 | -0.016816797 | -0.257194438 | -0.432060212 | -0.415243415 | -0.174865774 | 0.896107857 | 0.010902342 | 0.00118643 | 0.039657236 | 0.064772026 |
| Map7d1 | S | 544 | -0.094385384 | -0.244198807 | -0.203873473 | -0.109488089 | 0.040325334 | 0.04223437 | 0.00055928 | 0.01465303 | 0.092102886 | 0.445566706 |
| Bicd2 | S | 602 | -0.159176067 | -0.194775468 | -0.398314622 | -0.239138555 | -0.203539154 | 0.136122866 | 0.028201772 | 0.005934623 | 0.065739484 | 0.069636987 |
| Fblim1 | S | 58 | -0.024756439 | -0.24598565 | -0.271363735 | -0.246607296 | -0.025378085 | 0.540567906 | 0.001248623 | 0.000591026 | 0.008963976 | 0.542320789 |
| Cdan1 | S | 270 | -0.065928993 | -0.257418362 | -0.394779855 | -0.328850862 | -0.137361493 | 0.021190831 | 0.055659673 | 0.004531896 | 0.007345781 | 0.168704839 |
| Anln | S | 209 | -0.36741877 | -0.734654903 | -0.849979029 | -0.482560259 | -0.115324127 | 0.342473286 | 0.363885732 | 0.122565984 | 0.347697404 | 0.883055287 |
| Cul4b | S | 250 | 0.081550046 | -0.287786148 | -0.530070094 | -0.61162014 | -0.242283946 | 0.807897832 | 0.046959119 | 0.000459135 | 0.166365581 | 0.080105383 |
| Tgs1 | S | 431 | -0.09323142 | -0.232262245 | -0.281246575 | -0.188015155 | -0.04898433 | 0.068036365 | 0.029171225 | 0.000793269 | 0.009600071 | 0.432650714 |
| Snx17 | S | 421 | 0.017538218 | -0.233094117 | -0.281247508 | -0.298785726 | -0.048153391 | 0.642745916 | 0.000233018 | 0.00028524 | 0.001008225 | 0.197083972 |
| Vcp | Y | 805 | -0.240364779 | -0.212838307 | -0.338528133 | -0.098163355 | -0.125689827 | 0.006369993 | 0.063181633 | 0.068541581 | 0.467701438 | 0.394604266 |
| Synrg | S | 576 | -0.132986701 | -0.263298251 | -0.321239799 | -0.188253097 | -0.057941548 | 0.002952436 | 0.031695599 | 0.048092212 | 0.144356431 | 0.601248424 |
| Pum2 | S | 587 | -0.091208568 | -0.171609811 | -0.391228301 | -0.300019733 | -0.219618489 | 0.161000506 | 0.068672151 | 0.000456746 | 0.005800072 | 0.036607232 |
| Pex19 | T | 41 | -0.223961601 | -0.192019077 | -0.184402858 | 0.039558743 | 0.007616219 | 0.078402422 | 7.40548E-05 | 0.063174366 | 0.684029268 | 0.903273793 |
| Gys1 | S | 701 | -0.195722167 | -0.185234212 | -0.506647243 | -0.310925076 | -0.321413031 | 0.040290483 | 0.144828162 | 0.017833245 | 0.074412761 | 0.074321563 |
| Adam19 | S | 804 | -0.243360445 | -0.232016275 | -0.485388109 | -0.242027665 | -0.253371834 | 0.045289675 | 0.113529398 | 0.041945483 | 0.198816313 | 0.197923691 |
| Ctnnd1 | T | 906 | -0.132634934 | -0.224453508 | -0.448647891 | -0.316012957 | -0.224194382 | 0.154631707 | 0.170264726 | 0.001714077 | 0.011196502 | 0.17101926 |
| Smg9 | S | 450 | -0.179502424 | -0.486310657 | -0.571771507 | -0.392269084 | -0.085460851 | 0.279124964 | 0.127003366 | 0.043685843 | 0.121937817 | 0.761128264 |
| Smg9 | S | 451 | -0.179502424 | -0.486310657 | -0.571771507 | -0.392269084 | -0.085460851 | 0.279124964 | 0.127003366 | 0.043685843 | 0.121937817 | 0.761128264 |
| Fbf1 | S | 172 | -0.419571218 | -0.107031442 | -0.54047123 | -0.120900012 | -0.433439788 | 0.012098574 | 0.547302392 | 0.068771754 | 0.578990332 | 0.126907979 |
| Ranbp3 | S | 277 | -0.166871526 | -0.254424997 | -0.324365607 | -0.157494081 | -0.06994061 | 0.012235964 | 0.020399571 | 0.056547892 | 0.212615481 | 0.544163501 |
| Ubr5 | S | 1543 | -0.05076237 | -0.195640379 | -0.192894788 | -0.142132418 | 0.002745591 | 0.029515304 | 0.008517848 | 2.60515E-05 | 0.001487115 | 0.934846364 |
| Syt17 | S | 129 | -0.590841753 | -0.097904818 | -0.830228788 | -0.239387035 | -0.73232397 | 0.071619777 | 0.802657944 | 0.088025755 | 0.543487754 | 0.174443892 |
| Bclaf1 | S | 756 | -0.013393663 | -0.261857963 | -0.329173721 | -0.315780058 | -0.067315758 | 0.841183738 | 0.000189903 | 0.007505876 | 0.013126172 | 0.318833499 |
| Mcph1 | S | 358 | 0.027271336 | -0.312367602 | -0.237102829 | -0.264374165 | 0.075264773 | 0.553087831 | 3.87001E-05 | 0.003471171 | 0.004985091 | 0.144225776 |
| Map1b | S | 1781 | -0.053389351 | -0.208046531 | -0.176435948 | -0.123046597 | 0.031610583 | 0.315794215 | 9.57047E-05 | 0.000390152 | 0.073278336 | 0.182040634 |
| Larp4b | S | 416 | -0.26778353 | -0.298540382 | -0.598401871 | -0.330618341 | -0.299861489 | 0.17305472 | 0.106889703 | 0.056844049 | 0.219391298 | 0.247293484 |
| Trim3 | S | 454 | -0.166145549 | -0.202429565 | -0.449795089 | -0.283649539 | -0.247365524 | 0.108540492 | 0.12199443 | 0.004719335 | 0.037832551 | 0.072789567 |
| Trim3 | S | 455 | -0.166145549 | -0.202429565 | -0.449795089 | -0.283649539 | -0.247365524 | 0.108540492 | 0.12199443 | 0.004719335 | 0.037832551 | 0.072789567 |
| Rtkn2 | S | 564 | -0.276690064 | -0.071328825 | -0.449250023 | -0.172559959 | -0.377921198 | 0.010368683 | 0.540876025 | 0.007729498 | 0.149021296 | 0.029854422 |
| D1Ertd622e | S | 133 | -0.20120829 | -0.249025847 | -0.360050055 | -0.158841765 | -0.111024208 | 0.086265915 | 0.096766393 | 0.006573515 | 0.09009087 | 0.351913917 |
| Tpd52l2 | S | 168 | -0.324098199 | -0.215114026 | -0.460631371 | -0.136533172 | -0.245517345 | 0.052111838 | 0.244778887 | 0.028385866 | 0.319258394 | 0.20781921 |
| Tcea1 | S | 97 | -0.161034396 | -0.199853689 | -0.262666077 | -0.101631681 | -0.062812388 | 0.021768437 | 0.006252692 | 0.01969123 | 0.206819781 | 0.391818797 |
| Nhsl1 | S | 512 | -0.114300975 | -0.34590999 | -0.684955173 | -0.570654198 | -0.339045183 | 0.266882488 | 0.05959737 | 0.050395943 | 0.05557107 | 0.165631991 |
| Nhsl1 | Y | 517 | -0.114300975 | -0.34590999 | -0.684955173 | -0.570654198 | -0.339045183 | 0.266882488 | 0.05959737 | 0.050395943 | 0.05557107 | 0.165631991 |
| Atxn2 | S | 836 | -0.055901977 | -0.19782846 | -0.347995281 | -0.292093304 | -0.150166821 | 0.39214734 | 0.01808764 | 0.00028221 | 0.00649765 | 0.048623021 |
| Carhsp1 | S | 42 | -0.348967138 | -0.244546179 | -0.438190377 | -0.089223239 | -0.193644198 | 0.034773032 | 0.100803571 | 0.114582974 | 0.686230834 | 0.404629564 |
| Art2b | T | 192 | -0.071258438 | -0.299914824 | -0.348037923 | -0.276779485 | -0.0481231 | 0.316024865 | 0.002315599 | 0.01960157 | 0.034672693 | 0.578928953 |
| Myc | S | 293 | -0.035020143 | -0.380712839 | -0.306551054 | -0.271530912 | 0.074161785 | 0.502314221 | 0.033804291 | 0.000564911 | 0.00087775 | 0.504881973 |
| Sppl2a | S | 203 | -0.283447189 | -0.257131822 | -0.176237662 | 0.107209527 | 0.080894161 | 0.056332873 | 0.001261994 | 0.137354973 | 0.405552696 | 0.416738218 |
| Iqgap1 | T | 1434 | -0.062295227 | -0.183087938 | -0.219749359 | -0.157454132 | -0.036661421 | 0.216341378 | 0.000235524 | 0.00074339 | 0.026521041 | 0.254036999 |
| Rsl1d1 | T | 407 | 0.017954058 | -0.313505814 | -0.464276389 | -0.482230447 | -0.150770575 | 0.564592257 | 0.000948445 | 0.012387329 | 0.01852466 | 0.151817842 |
| Tbc1d1 | T | 739 | 0.15168056 | -0.269406379 | -0.32327839 | -0.47495895 | -0.053872011 | 0.267162904 | 9.57817E-06 | 0.00107715 | 0.032717726 | 0.215071528 |
| Ebna1bp2 | S | 270 | -0.087907139 | -0.168228846 | -0.369200891 | -0.281293752 | -0.200972044 | 0.38794879 | 0.020399819 | 0.000379483 | 0.051422726 | 0.005084879 |
| Tle3 | T | 312 | -0.127303948 | -0.237113948 | -0.303691257 | -0.176387309 | -0.066577308 | 0.079823365 | 0.007874067 | 0.011556589 | 0.081022949 | 0.388207537 |
| Map1b | S | 1789 | -0.099624809 | -0.229009702 | -0.206189339 | -0.106564529 | 0.022820363 | 0.071492567 | 0.001824905 | 0.003207373 | 0.032213433 | 0.450232216 |
| Ddah1 | S | 33 | -0.150269616 | -0.142215604 | -0.323273831 | -0.173004215 | -0.181058227 | 0.020579015 | 0.07668913 | 0.001753591 | 0.03319331 | 0.046359706 |
| Aebp2 | S | 366 | -0.119690721 | -0.238725979 | -0.216853808 | -0.097163087 | 0.021872172 | 0.010456721 | 0.004348483 | 0.020072022 | 0.152399623 | 0.722482197 |
| Cep170b | S | 944 | -0.102528497 | -0.323431126 | -0.405061492 | -0.302532995 | -0.081630366 | 0.332709032 | 0.005591319 | 0.034595448 | 0.059625839 | 0.442008345 |
| Cep170b | S | 947 | -0.102528497 | -0.323431126 | -0.405061492 | -0.302532995 | -0.081630366 | 0.332709032 | 0.005591319 | 0.034595448 | 0.059625839 | 0.442008345 |
| Usp24 | S | 1282 | -0.280999403 | -0.128533657 | -0.475448565 | -0.194449163 | -0.346914908 | 0.051067218 | 0.445533977 | 0.00594071 | 0.119938609 | 0.081411724 |
| Rhbdf1 | S | 210 | -0.358932148 | -0.155587191 | -0.573756791 | -0.214824643 | -0.418169601 | 0.03464477 | 0.436036764 | 0.043883254 | 0.320379194 | 0.125169675 |
| Rps6kb1 | S | 401 | -0.087397734 | -0.5870376 | -0.735299584 | -0.647901849 | -0.148261983 | 0.561922697 | 0.053527669 | 0.115183279 | 0.147816271 | 0.691578267 |
| Casp8 | S | 213 | -0.151508129 | -0.236350441 | -0.373446405 | -0.221938275 | -0.137095963 | 0.005939141 | 0.079517631 | 0.030614419 | 0.096477903 | 0.285160673 |
| Ranbp1 | S | 60 | -0.195708985 | -0.191566318 | -0.337717738 | -0.142008753 | -0.146151419 | 0.020403496 | 0.072849661 | 0.012779418 | 0.153934626 | 0.187994937 |
| Wdr43 | S | 85 | 0.027038129 | -0.247135798 | -0.397623876 | -0.424662005 | -0.150488078 | 0.705351434 | 0.002383289 | 0.000914765 | 0.004648729 | 0.052746875 |
| Hnrnpa0 | S | 84 | -0.235616292 | -0.320272463 | -0.057309482 | 0.178306809 | 0.262962981 | 0.035658588 | 0.000191571 | 0.335241416 | 0.077048734 | 0.010787568 |
| Abcc5 | S | 815 | -0.104723103 | -0.23376322 | -0.192260216 | -0.087537113 | 0.041503004 | 0.023544447 | 0.000252378 | 0.04434667 | 0.233009499 | 0.513921228 |
| Cdc42ep1 | S | 39 | -0.150558932 | -0.18302761 | -0.240921182 | -0.09036225 | -0.057893572 | 0.002211673 | 0.010260507 | 0.030534376 | 0.245530234 | 0.434678693 |
| Arhgap1 | S | 51 | -0.209986175 | -0.22398367 | -0.418922433 | -0.208936259 | -0.194938764 | 0.059997976 | 0.113817031 | 0.015411023 | 0.12552632 | 0.191156401 |
| Arhgef5 | S | 169 | -0.13967356 | -0.175981961 | -0.307856577 | -0.168183018 | -0.131874616 | 0.006094196 | 0.025309165 | 0.014696203 | 0.079360576 | 0.135981166 |
| Sub1 | S | 92 | -0.155586539 | -0.185935912 | -0.357832406 | -0.202245867 | -0.171896495 | 0.095836747 | 0.003423485 | 0.023881423 | 0.106040078 | 0.140588941 |
| Zfp182 Zfp2 | T | 235;314 | -0.188779391 | -0.310971756 | -0.311229979 | -0.122450588 | -0.000258223 | 0.024150369 | 0.041897573 | 0.053298805 | 0.319024719 | 0.998465846 |
| Hsp90ab1 | S | 255 | -0.27061332 | -0.285268979 | -0.317918737 | -0.047305417 | -0.032649758 | 0.119418852 | 0.051998315 | 0.020768559 | 0.727927043 | 0.761369858 |
| Zfp24 | S | 274 | -0.095499845 | -0.245850709 | -0.315150053 | -0.219650208 | -0.069299344 | 0.065551034 | 0.001415637 | 0.040427009 | 0.104567836 | 0.489310166 |
| Ccdc86 | S | 161 | -0.108002469 | -0.375027145 | -0.260712851 | -0.152710381 | 0.114314294 | 0.151377272 | 0.001275365 | 0.113761324 | 0.283877509 | 0.399705509 |
| Rell1 | S | 245 | -0.178954387 | -0.19649793 | -0.357708742 | -0.178754355 | -0.161210813 | 0.071057215 | 0.058906215 | 0.006076222 | 0.051683078 | 0.064400114 |
| Tjp2 | S | 811 | -0.022416732 | -0.360812839 | -0.271029342 | -0.24861261 | 0.089783497 | 0.657892825 | 2.82128E-05 | 0.12992312 | 0.150153634 | 0.512509111 |
| Mical3 | S | 1216 | -0.092295399 | -0.284873411 | -0.254183955 | -0.161888556 | 0.030689456 | 0.233818804 | 0.007544032 | 0.00243128 | 0.081850469 | 0.588658312 |
| Casp3 | S | 12 | -0.204888929 | -0.188744671 | -0.370047992 | -0.165159063 | -0.181303321 | 0.090209274 | 0.088682389 | 0.004833589 | 0.148989158 | 0.098563434 |
| Pgk1 | S | 203 | -0.293688452 | -0.299006955 | -0.439764491 | -0.146076039 | -0.140757536 | 0.056121319 | 0.117879862 | 0.076343665 | 0.453871457 | 0.505644476 |
| Slc39a14 | S | 308 | 0.048213677 | -0.256295089 | -0.325756067 | -0.373969744 | -0.069460978 | 0.241038147 | 0.000493678 | 0.000241015 | 0.000863566 | 0.019412114 |
| Eif4b | S | 442 | 0.029724877 | -0.373279426 | -0.400619197 | -0.430344074 | -0.02733977 | 0.773819048 | 0.00302493 | 0.007642492 | 0.014168522 | 0.771868385 |
| Eif4b | S | 445 | 0.029724877 | -0.373279426 | -0.400619197 | -0.430344074 | -0.02733977 | 0.773819048 | 0.00302493 | 0.007642492 | 0.014168522 | 0.771868385 |
| Phactr4 | S | 148 | 0.057675812 | -0.273502058 | -0.333908447 | -0.391584259 | -0.06040639 | 0.132554252 | 0.000193238 | 0.000497623 | 0.001793897 | 0.200830011 |
| Tenm4 | S | 29 | -0.075676963 | -0.325650575 | -0.234492349 | -0.158815386 | 0.091158226 | 0.241296555 | 0.001806094 | 0.011919616 | 0.050228933 | 0.141802374 |
| Lima1 | S | 225 | -0.197814532 | -0.170122305 | -0.214924695 | -0.017110164 | -0.04480239 | 0.011695782 | 0.002017013 | 0.045389093 | 0.821458631 | 0.532955222 |
| Nolc1 | S | 116 | -0.02548383 | -0.256581724 | -0.212621962 | -0.187138133 | 0.043959762 | 0.491991946 | 0.000418353 | 0.000789459 | 0.002003261 | 0.224169189 |
| Igf2r | S | 2471 | -0.243736932 | -0.218430463 | -0.307228244 | -0.063491312 | -0.088797782 | 0.01727126 | 0.087189986 | 0.02676332 | 0.518414748 | 0.453124714 |
| Mcrs1 | S | 85 | -0.076187315 | -0.144008131 | -0.201526657 | -0.125339343 | -0.057518526 | 0.251022492 | 0.000448923 | 5.91827E-05 | 0.119426177 | 0.004001982 |
| Tbcb | S | 110 | -0.180629369 | -0.209392111 | -0.378643196 | -0.198013827 | -0.169251085 | 0.042622976 | 0.04874477 | 0.018677276 | 0.11524872 | 0.172057763 |
| Camsap2 | S | 573 | -0.100162928 | -0.177190521 | -0.23123429 | -0.131071362 | -0.054043769 | 0.075496366 | 0.000258837 | 0.007806542 | 0.05971227 | 0.274557625 |
| Serbp1 | S | 63 | 0.10662135 | -0.54675667 | -0.350963134 | -0.457584484 | 0.195793536 | 0.189247072 | 0.000126612 | 0.064545781 | 0.041198647 | 0.218848836 |
| Mcph1 | S | 327 | -0.080467331 | -0.319761188 | -0.358778721 | -0.27831139 | -0.039017533 | 0.399867739 | 0.003148299 | 0.025876615 | 0.057620552 | 0.678377088 |
| Srsf9 | S | 190 | 0.056097339 | -0.335695301 | -0.222017456 | -0.278114795 | 0.113677845 | 0.602547907 | 1.5602E-05 | 0.009739134 | 0.073597829 | 0.09148849 |
| Slc20a2 | S | 316 | -0.196169371 | -0.206704897 | -0.519386045 | -0.323216674 | -0.312681148 | 0.04575521 | 0.228614382 | 0.018236798 | 0.087425997 | 0.12999918 |
| Eif2ak3 | S | 551 | 0.007031356 | -0.381538346 | -0.299584636 | -0.306615992 | 0.081953709 | 0.863139247 | 0.000901453 | 0.01066876 | 0.009703034 | 0.251721278 |
| Nhsl1 | S | 867 | -0.007148907 | -0.31559765 | -0.281768594 | -0.274619687 | 0.033829056 | 0.930490833 | 6.50932E-05 | 0.0281973 | 0.042208337 | 0.639124764 |
| Slc38a2 | Y | 41 | -0.191120942 | -0.184928727 | -0.338986507 | -0.147865565 | -0.154057779 | 0.01100528 | 0.030441701 | 0.041903416 | 0.236634313 | 0.225073176 |
| Dvl2 | S | 618 | -0.232467621 | -0.140003874 | -0.284531234 | -0.052063613 | -0.14452736 | 0.035148613 | 0.216265673 | 0.000820211 | 0.498978909 | 0.206939701 |
| Zfand3 | S | 129 | 0.006856027 | -0.19705277 | -0.309163587 | -0.316019614 | -0.112110817 | 0.879339074 | 4.29888E-05 | 0.001945712 | 0.003300489 | 0.062515898 |
| Ivns1abp | S | 277 | -0.087678646 | -0.228512097 | -0.286112629 | -0.198433983 | -0.057600532 | 0.272690396 | 0.00019015 | 0.024481885 | 0.077389778 | 0.44013173 |
| Baz1b | S | 204 | -0.119249395 | -0.234405513 | -0.211270019 | -0.092020624 | 0.023135494 | 0.090861263 | 0.007596188 | 0.001690695 | 0.166409288 | 0.631478264 |

| | | | | | | | | | | | | |
|---|---|---|---|---|---|---|---|---|---|---|---|---|
| Mia3 | S | 274 | -0.24353441 | -0.04120506 | -0.417328729 | -0.173794318 | -0.376123668 | 0.00999425 | 0.755857262 | 0.002198693 | 0.069311032 | 0.051280122 |
| Cdv3 | S | 90 | -0.223644344 | -0.252905703 | -0.075017288 | 0.148627056 | 0.177888415 | 0.002209495 | 0.001334931 | 0.300876672 | 0.086176028 | 0.049817039 |
| Ctnna1 | S | 48 | -0.344221085 | -0.176448345 | -0.386865144 | -0.042644059 | -0.210416799 | 0.036285381 | 0.252429837 | 0.02389106 | 0.302986678 | 0.086178758 |
| Nucks1 | S | 50 | -0.194864577 | -0.099183215 | -0.35694989 | -0.162085313 | -0.257766675 | 0.009718429 | 0.172947508 | 0.004266752 | 0.091201207 | 0.022731647 |
| Hectd1 | S | 1576 | -0.308078008 | -0.280897934 | -0.179286435 | 0.128791573 | 0.101611499 | 0.015360698 | 0.021338329 | 0.14425962 | 0.188691246 | 0.262105114 |
| Dennd4c | S | 1321 | -0.351706619 | -0.19902929 | -0.525762115 | -0.174055495 | -0.326732824 | 0.060019968 | 0.409525861 | 0.031824472 | 0.322547093 | 0.20751397 |
| Iqsec1 | S | 179 | -0.319269797 | -0.148975104 | -0.268748424 | 0.050521373 | -0.11977332 | 0.011298838 | 0.270090235 | 0.009059222 | 0.543169528 | 0.348977177 |
| Plcg1 | S | 1221 | -0.096970298 | -0.266954762 | -0.347749166 | -0.250778868 | -0.080794404 | 0.171416158 | 0.009544417 | 0.012579716 | 0.052530876 | 0.393871253 |
| Rpl24 | S | 64 | 0.047137811 | -0.252746214 | -0.398156057 | -0.445293868 | -0.145409843 | 0.221117047 | 0.000267677 | 0.00180601 | 0.0049169 | 0.063908284 |
| Lysmd2 | S | 24 | -0.175213638 | -0.207013697 | -0.469022653 | -0.293809015 | -0.262008957 | 0.08416654 | 0.031993584 | 0.034483151 | 0.10214627 | 0.129664498 |
| Zdhhc5 | S | 246 | -0.047253755 | -0.302994293 | -0.231746151 | -0.184492397 | 0.071248142 | 0.461336113 | 5.04403E-05 | 0.049131787 | 0.086292076 | 0.375708545 |
| Sh3bp4 | S | 19 | -0.347397177 | -0.1467567 | -0.457921898 | -0.110524721 | -0.311165198 | 0.0116595 | 0.204445575 | 0.098660913 | 0.597051343 | 0.209235993 |
| Zc3h14 | S | 343 | -0.04511895 | -0.231887021 | -0.397210997 | -0.352092047 | -0.165323976 | 0.424478245 | 0.053543989 | 0.00051327 | 0.002890098 | 0.126939058 |
| Nedd1 | S | 397 | -0.519367638 | -0.179956183 | -0.488997371 | 0.030370267 | -0.309041188 | 0.025516384 | 0.390779722 | 0.149866234 | 0.905618067 | 0.304612863 |
| Srf | S | 97 | -0.158824353 | -0.189664445 | -0.234332433 | -0.075508079 | -0.044667988 | 0.032565887 | 0.06033217 | 0.00085242 | 0.20128291 | 0.541242553 |
| Cavin1 | S | 120 | -0.015535406 | -0.286739251 | -0.301329452 | -0.285794046 | -0.014590201 | 0.652994408 | 0.012659457 | 0.000337269 | 0.002588397 | 0.818546123 |
| Heatr3 | S | 144 | -0.055141039 | -0.227521046 | -0.345126597 | -0.289985558 | -0.117605551 | 0.130106083 | 0.002462014 | 0.010940931 | 0.006031666 | 0.076984596 |
| Tnks1bp1 | S | 977 | -0.246554493 | -0.142953217 | -0.291881615 | -0.045327123 | -0.148928398 | 0.090429282 | 0.124009122 | 0.000850299 | 0.665064463 | 0.117891655 |
| Syne1 | S | 6238 | 0.002412958 | -0.170040946 | -0.365246805 | -0.367659763 | -0.19520586 | 0.951861266 | 0.00545077 | 0.000113404 | 4.46008E-05 | 0.000168224 |
| Slc4a10 | S | 238 | -0.059735041 | -0.388847704 | -0.198771692 | -0.139036651 | 0.190076012 | 0.369714532 | 0.001208706 | 0.018353346 | 0.014254475 | 0.046766624 |
| Mcph1 | S | 357 | 0.016171747 | -0.317127463 | -0.269530631 | -0.285702378 | 0.047596832 | 0.72924486 | 0.000205661 | 0.005619355 | 0.007286593 | 0.408614577 |
| Cebpz | S | 670 | -0.033369041 | -0.351619159 | -0.470777206 | -0.437408165 | -0.119158047 | 0.838051896 | 0.017207088 | 0.009713113 | 0.058430787 | 0.360519119 |
| Btf3 Btf3l4 | T | 80;33 | -0.241171504 | -0.199997363 | -0.468391275 | -0.227219771 | -0.268393912 | 0.088108657 | 0.378687873 | 0.006312439 | 0.031507911 | 0.249212155 |
| Slc7a6os | S | 305 | 0.024721104 | -0.192470738 | -0.26582819 | -0.290549294 | -0.073357452 | 0.384638609 | 0.000750209 | 1.4263E-05 | 0.000191004 | 0.050987339 |
| Bub1 | S | 447 | 0.025283532 | -0.259464759 | -0.374067206 | -0.399350739 | -0.114602447 | 0.360870877 | 0.017811058 | 7.61722E-05 | 0.000143231 | 0.095180716 |
| Dclk1 | S | 352 | -0.177645702 | -0.163829971 | -0.236324652 | -0.05867895 | -0.072494682 | 0.02105685 | 0.023142472 | 0.004466843 | 0.20932412 | 0.074553787 |
| Arfgap1 | T | 135 | -0.074295051 | -0.208755495 | -0.186063339 | -0.111768288 | 0.022692155 | 0.086946995 | 0.003273443 | 0.000312866 | 0.03032375 | 0.364978854 |
| Cep170 | S | 878 | -0.067607603 | -0.292226659 | -0.339466617 | -0.271859013 | -0.047239958 | 0.100645117 | 0.002945861 | 0.04018855 | 0.058829501 | 0.61527007 |
| Hdac6 | S | 21 | -0.095417084 | -0.145526595 | -0.248063542 | -0.152646458 | -0.102536947 | 0.072730349 | 0.006648585 | 0.000418769 | 0.022345654 | 0.041291403 |
| Usp47 | S | 947 | -0.251735731 | -0.175911993 | -0.348520409 | -0.096784678 | -0.172608417 | 0.004838085 | 0.220116205 | 0.037834793 | 0.404912628 | 0.277744671 |
| Clasp2 | S | 1049 | -0.119349261 | -0.284937665 | -0.315294605 | -0.195945344 | -0.03035694 | 0.017541752 | 0.016793472 | 0.050942197 | 0.121815751 | 0.764242537 |
| Spty2d1 | S | 484 | 0.088820756 | -0.245046985 | -0.383195905 | -0.472016661 | -0.13814892 | 0.342665447 | 0.006076763 | 5.83968E-05 | 0.015359086 | 0.036968527 |
| Map1a | S | 1648 | -0.04611769 | -0.210944403 | -0.294018181 | -0.247900491 | -0.083073778 | 0.212683185 | 0.000864112 | 0.004775475 | 0.003011455 | 0.077782135 |
| Gclm | S | 59 | -0.163697112 | -0.255780927 | -0.453609402 | -0.28991229 | -0.197828475 | 0.140534028 | 0.030823537 | 0.03563703 | 0.117487395 | 0.232450236 |
| Txlna | S | 523 | 0.002162914 | -0.23034935 | -0.349396625 | -0.35155954 | -0.119047275 | 0.952446911 | 3.98198E-05 | 0.0139189 | 0.00570741 | 0.121278444 |
| Nav1 | S | 380 | -0.330588998 | -0.227936973 | -0.575820293 | -0.245231295 | -0.34788332 | 0.048189216 | 0.295080977 | 0.078478426 | 0.343322454 | 0.243068213 |
| Septin9 | S | 30 | -0.300993766 | -0.164493939 | -0.356097383 | -0.055103617 | -0.191603445 | 0.029418311 | 0.24686917 | 0.014415709 | 0.607107045 | 0.185779606 |
| Ywhae | S | 210 | -0.156774192 | -0.298045116 | -0.136963943 | 0.019810249 | 0.161081173 | 0.111110318 | 0.000477987 | 0.049372798 | 0.806894171 | 0.03591451 |
| Arhgap31 | S | 332 | -0.364732398 | -0.206193416 | -0.389771711 | -0.025039313 | -0.183578295 | 0.009046728 | 0.209930212 | 0.15159808 | 0.906079588 | 0.449040905 |
| Sec16a | S | 2101 | -0.240759666 | -0.150573378 | -0.288712693 | -0.047953027 | -0.138139319 | 0.011747386 | 0.169846614 | 0.007881319 | 0.470912443 | 0.192076143 |
| Ppfibp1 | T | 963 | -0.012373956 | -0.232936964 | -0.250228942 | -0.237854986 | -0.017291978 | 0.567887399 | 0.000518791 | 0.000926039 | 0.003236554 | 0.616808297 |
| Rbl1 | S | 350 | -0.408907125 | -0.15948288 | -0.55950277 | -0.150595645 | -0.400019891 | 0.070526498 | 0.579055321 | 0.030713939 | 0.431832486 | 0.202115192 |
| Farp1 | S | 516 | -0.185634371 | -0.215059185 | -0.187244513 | -0.001610142 | 0.027814672 | 0.004542625 | 0.063256716 | 0.007335653 | 0.969230672 | 0.736850993 |
| Carhsp1 | T | 46 | -0.355994114 | -0.217851925 | -0.470736024 | -0.11474191 | -0.2528841 | 0.042238053 | 0.146232568 | 0.108796541 | 0.624659201 | 0.314234836 |
| Cap1 | S | 307 | -0.235363317 | -0.316362035 | -0.158249005 | 0.077114312 | 0.15811303 | 0.03554077 | 0.008007317 | 0.09663709 | 0.249673538 | 0.027400216 |
| Nop2 | T | 599 | -0.001144415 | -0.233274639 | -0.313538346 | -0.312393931 | -0.080263707 | 0.980443976 | 0.002133641 | 0.000586127 | 0.000103153 | 0.065351915 |
| Uvrag | S | 507 | 0.09866614 | -0.363794177 | -0.448153571 | -0.546819711 | -0.084359394 | 0.06884181 | 4.55606E-05 | 0.018593015 | 0.014128302 | 0.41452778 |
| Jmjd1c | S | 1800 | 0.085283449 | -0.372261508 | -0.344985026 | -0.430268474 | 0.027276482 | 0.269152609 | 0.000292316 | 0.00639683 | 0.005151192 | 0.679740358 |
| Tacc2 | S | 778 | -0.209203653 | -0.346345326 | -0.369624656 | -0.160421003 | -0.02327933 | 0.392942787 | 0.028807487 | 0.023780003 | 0.506162568 | 0.851783376 |
| Ranbp2 | S | 2348 | 0.028227137 | -0.288072528 | -0.332290098 | -0.360517235 | -0.04421757 | 0.506556079 | 0.000338726 | 0.00424468 | 0.002204444 | 0.391380983 |
| Ssr3 | S | 105 | -0.018871475 | -0.180775863 | -0.322888269 | -0.304016794 | -0.142112406 | 0.499659024 | 5.19403E-05 | 0.006504243 | 0.007430873 | 0.066761932 |
| Heatr3 | S | 15 | -0.11182476 | -0.194336372 | -0.295650234 | -0.183825474 | -0.101313862 | 0.100932812 | 0.000920551 | 0.023656822 | 0.074742095 | 0.228737322 |
| Pnn | S | 66 | -0.138497178 | -0.23134595 | -0.361473631 | -0.222976453 | -0.130127681 | 0.231287133 | 0.037216881 | 0.004387325 | 0.078595711 | 0.159813517 |
| Cep55 | S | 423 | -0.039985495 | -0.217764812 | -0.339866163 | -0.299880668 | -0.122097751 | 0.280787794 | 0.000240325 | 0.020256817 | 0.028469273 | 0.18149862 |
| Oxr1 | S | 15 | -0.435436919 | -0.163859151 | -0.403281 | 0.032155918 | -0.23942185 | 0.030841002 | 0.371605019 | 0.052686112 | 0.823613629 | 0.177350184 |
| Dpp9 | S | 472 | -0.237731078 | -0.342226229 | -0.196916678 | 0.0408144 | 0.14530955 | 0.039685044 | 0.037390207 | 0.043538028 | 0.656319818 | 0.236381343 |
| Rrp15 | S | 19 | 0.00866551 | -0.215788844 | -0.25369571 | -0.26236122 | -0.037906866 | 0.873626784 | 2.10622E-06 | 0.010376817 | 0.013324343 | 0.387635895 |
| Atxn2 | S | 727 | -0.003587813 | -0.19805023 | -0.377129044 | -0.37354123 | -0.179078814 | 0.923381482 | 0.000896471 | 0.002084759 | 0.004871521 | 0.035582698 |
| Mns1 | S | 118 | -0.396668401 | -0.147069913 | -0.546510767 | -0.149842366 | -0.399440854 | 0.044245212 | 0.49217498 | 0.048600992 | 0.469277794 | 0.139149857 |
| Ybx1 | S | 174 | -0.04496551 | -0.219875892 | -0.352215868 | -0.307250357 | -0.132339976 | 0.485900731 | 0.005303565 | 0.002225149 | 0.008197215 | 0.107678652 |
| Fxr1 | S | 401 | -0.073941104 | -0.247198428 | -0.375067238 | -0.301126133 | -0.127868809 | 0.384367955 | 0.026622038 | 0.002346793 | 0.018773047 | 0.182415857 |
| Mcph1 | S | 260 | 0.018526675 | -0.240512362 | -0.196923291 | -0.215449966 | 0.043589071 | 0.483229574 | 5.28779E-05 | 0.000197081 | 0.000134403 | 0.016828651 |
| Tjap1 | S | 473 | 0.014984442 | -0.223248573 | -0.231992616 | -0.246977058 | -0.008744043 | 0.752246658 | 0.010067343 | 1.35428E-06 | 0.021760983 | 0.808002405 |
| Ranbp2 | S | 2117 | -0.055206601 | -0.327024526 | -0.206443491 | -0.15123689 | 0.120581034 | 0.343078283 | 0.00020463 | 0.042104736 | 0.103091765 | 0.16342507 |
| Morf4l2 | S | 69 | -0.200062554 | -0.219043452 | -0.382990681 | -0.182928127 | -0.163947229 | 0.091308413 | 0.088412975 | 0.012433295 | 0.109026593 | 0.163676069 |
| Ptpdc1 | S | 392 | -0.516858028 | -0.136517145 | -0.474984085 | 0.041873943 | -0.33846694 | 0.027954866 | 0.578437006 | 0.084274425 | 0.839555166 | 0.222542316 |
| Rnf4 | S | 98 | -0.122451801 | -0.230282796 | -0.214230641 | -0.09177884 | 0.016052155 | 0.139770698 | 0.000393496 | 0.026381515 | 0.285589805 | 0.779321654 |
| Rnf4 | S | 99 | -0.122451801 | -0.230282796 | -0.214230641 | -0.09177884 | 0.016052155 | 0.139770698 | 0.000393496 | 0.026381515 | 0.285589805 | 0.779321654 |
| Sh2b1 | S | 219 | -0.032117377 | -0.300950385 | -0.321449234 | -0.289331857 | -0.020498849 | 0.24563791 | 0.000719878 | 0.037126771 | 0.041551876 | 0.796134296 |
| Gnl3 | S | 101 | 0.109713041 | -0.363559011 | -0.336149481 | -0.445862522 | 0.02740953 | 0.040861859 | 0.000133614 | 0.001150819 | 0.001959602 | 0.604695288 |
| Nop56 | S | 522 | -0.231101544 | -0.10018741 | -0.224080299 | 0.007021245 | -0.123892889 | 0.002533219 | 0.032485628 | 0.017054862 | 0.899075035 | 0.080533376 |
| Nmd3 | S | 468 | -0.067683524 | -0.164382002 | -0.226912869 | -0.159229345 | -0.062530867 | 0.035498855 | 0.003757422 | 0.000706463 | 0.009606341 | 0.142967833 |
| Mast3 | S | 1289 | -0.114807316 | -0.19662064 | -0.387669884 | -0.272862567 | -0.191049244 | 0.123339261 | 0.049347236 | 0.004206329 | 0.024393785 | 0.080585645 |
| Klf12 Klf3 Klf5 Klf7 Zf | T | 155;342;151;594; | -0.152750728 | -0.11913849 | -0.387841948 | -0.235091221 | -0.268703458 | 0.116104098 | 0.105914363 | 0.001105202 | 0.03703319 | 0.009738547 |
| Tbc1d5 | S | 578 | -0.027374385 | -0.188215867 | -0.273676291 | -0.246301906 | -0.085460424 | 0.541425094 | 0.002977409 | 0.000173539 | 0.002722321 | 0.060377078 |
| Yap1 | S | 148 | -0.194054422 | -0.150370809 | -0.324083326 | -0.130028905 | -0.173712517 | 0.057142526 | 0.074634056 | 0.003760129 | 0.171601005 | 0.066755378 |
| Nol12 | S | 135 | -0.019196568 | -0.499622981 | -0.508053411 | -0.488856843 | -0.008430431 | 0.660950953 | 0.055894723 | 0.018436326 | 0.017070534 | 0.959209441 |
| Cdk4 | S | 300 | -0.051506851 | -0.168165597 | -0.321201532 | -0.269694681 | -0.153035934 | 0.196182169 | 0.006056313 | 0.000930606 | 0.00841641 | 0.029570832 |
| Cct4 | S | 381 | -0.037765404 | -0.217634026 | -0.333910915 | -0.296145511 | -0.116276889 | 0.175635553 | 0.000563225 | 0.01540858 | 0.018609895 | 0.135814564 |
| Dlg1 | S | 102 | -0.094589303 | -0.294976825 | -0.176136149 | -0.081546846 | 0.118840676 | 0.024291688 | 0.001851173 | 0.039037379 | 0.211019431 | 0.111924812 |
| Arhgap39 | S | 123 | -0.317240528 | -0.265555297 | -0.157466667 | 0.159773861 | 0.10808863 | 0.004300756 | 0.020241483 | 0.402754551 | 0.397655704 | 0.554042946 |
| Gtse1 | S | 68 | -0.115151089 | -0.307430012 | -0.292726688 | -0.177575599 | 0.014703324 | 0.180727802 | 0.002320106 | 0.062544365 | 0.190837356 | 0.89037637 |
| Abcc5 | S | 43 | -0.053745281 | -0.237853613 | -0.31163186 | -0.257886579 | -0.073778247 | 0.170388495 | 0.005548362 | 0.004818433 | 0.008572003 | 0.234502416 |
| Anln | S | 66 | -0.010331798 | -0.13929151 | -0.23005256 | -0.219720762 | -0.09076105 | 0.707109442 | 0.000409309 | 1.33757E-05 | 0.004204059 | 0.009302781 |
| Pcm1 | S | 1280 | -0.178706953 | -0.183345909 | -0.227464367 | -0.048757414 | -0.044118458 | 0.065408614 | 0.022570485 | 0.002286922 | 0.497751512 | 0.410879777 |
| Hnrnpc | S | 229 | -0.030553226 | -0.177432209 | -0.140189904 | -0.109636678 | 0.037242305 | 0.343981876 | 1.03745E-05 | 0.000178872 | 0.028416678 | 0.068058739 |
| Pum3 | S | 99 | -0.117992897 | -0.209013953 | -0.275734035 | -0.157741138 | -0.066720081 | 0.24563358 | 0.005406259 | 0.003303275 | 0.154192166 | 0.270590053 |
| H1f3 | T | 18 | 0.097843988 | -0.310377774 | -0.397396345 | -0.495240333 | -0.087018571 | 0.112685946 | 0.001713103 | 0.00040274 | 0.000728369 | 0.186492813 |
| Ppp1r12c | S | 403 | -0.107536245 | -0.20234722 | -0.263884603 | -0.156348358 | -0.061537383 | 0.182838895 | 0.002278298 | 0.005618357 | 0.090174214 | 0.260211356 |
| Prrc2c | S | 1965 | -0.016314522 | -0.282274715 | -0.246794933 | -0.230480412 | 0.035479781 | 0.694183048 | 0.000942768 | 0.002248254 | 0.009187642 | 0.272391152 |
| Pgam1 | S | 118 | -0.279547264 | -0.164553722 | -0.35475785 | -0.075210587 | -0.190204128 | 0.015894558 | 0.090579241 | 0.059289605 | 0.582641598 | 0.216562632 |
| Psen1 | S | 32 | -0.1807621 | -0.254172238 | -0.260081668 | -0.079319568 | -0.005909431 | 0.017906468 | 0.007678889 | 0.12141883 | 0.544072731 | 0.962865243 |
| H1f5 | S | 18 | -0.004721177 | -0.181597401 | -0.317938814 | -0.313217637 | -0.136341412 | 0.945369305 | 0.001518827 | 0.000306646 | 0.016379415 | 0.022405125 |
| Ppp1r12c | S | 455 | -0.39620079 | -0.12970867 | -0.562937364 | -0.166736574 | -0.433228694 | 0.05025292 | 0.598898813 | 0.037675168 | 0.385191244 | 0.13369133 |

| | | | | | | | | | | | | |
|---|---|---|---|---|---|---|---|---|---|---|---|---|
| Bicd1 | S | 548 | -0.134326543 | -0.079797372 | -0.264789875 | -0.130463332 | -0.184992503 | 0.005620877 | 0.145756934 | 0.000259639 | 0.013544886 | 0.016044702 |
| Slc16a1 | S | 461 | -0.099790303 | -0.206757893 | -0.282186517 | -0.182396214 | -0.075428624 | 0.212359142 | 0.000108622 | 0.044342028 | 0.115535254 | 0.393149468 |
| Usp24 | S | 1296 | -0.146434208 | -0.187914587 | -0.490651166 | -0.344216959 | -0.30273658 | 0.178024452 | 0.218506987 | 0.003468907 | 0.019693135 | 0.079253297 |
| Flna | S | 20 | -0.261806723 | -0.100346598 | -0.54710227 | -0.285295546 | -0.446755672 | 0.106372405 | 0.571180144 | 0.005045721 | 0.05953732 | 0.059778046 |
| Vcpip1 | S | 996 | -0.107014403 | -0.209429953 | -0.225545147 | -0.118530744 | -0.016115194 | 0.029902057 | 0.00132258 | 0.024934939 | 0.153763948 | 0.795376517 |
| Stim2 | S | 650 | -0.078878468 | -0.308296726 | -0.227596679 | -0.148718211 | 0.080700047 | 0.230015888 | 0.001135342 | 0.025092134 | 0.092465746 | 0.271281461 |
| Sh2b1 | S | 384 | -0.171525518 | -0.228139372 | -0.218514812 | -0.046989295 | 0.00962456 | 0.070583739 | 0.010598147 | 0.009498447 | 0.540013143 | 0.876286656 |
| Nf1 | S | 2525 | -0.26656352 | -0.213677481 | -0.367145985 | -0.100582465 | -0.153468504 | 0.145844429 | 0.052544474 | 0.021102948 | 0.535939656 | 0.193875873 |
| Cdc42bpb | S | 652 | 0.002387049 | -0.323454711 | -0.349437729 | -0.351824778 | -0.025983018 | 0.978621748 | 0.002371442 | 0.007424346 | 0.022456615 | 0.723422343 |
| Eif4ebp2 | T | 70 | 0.000983045 | -0.258586387 | -0.404713771 | -0.405696816 | -0.146127384 | 0.991547736 | 0.012317186 | 0.001341283 | 0.006793248 | 0.028047393 |
| Map1a | S | 1185 | -0.220974831 | -0.188524867 | -0.229737505 | -0.008762673 | -0.041212637 | 0.005414831 | 0.16006844 | 0.009207185 | 0.870487548 | 0.709739999 |
| Hnrnpm | S | 539 | -0.175419472 | -0.19456829 | -0.2207185 | -0.045299028 | -0.02615021 | 0.024156526 | 0.023136663 | 0.00793605 | 0.432675142 | 0.669288403 |
| Prune2 | S | 876 | -0.089497533 | -0.406434301 | -0.301218508 | -0.211720974 | 0.105215794 | 0.026812854 | 0.042370083 | 0.037616197 | 0.080526172 | 0.419660037 |
| Pi4kb | S | 275 | -0.148283203 | -0.212495528 | -0.137779607 | 0.010503596 | 0.074715921 | 0.002964144 | 0.000395925 | 0.191372337 | 0.899928165 | 0.416992915 |
| Anln | S | 318 | 0.00451686 | -0.184458648 | -0.27642582 | -0.28094268 | -0.091967172 | 0.897185856 | 0.00140599 | 7.7077E-05 | 0.000492794 | 0.034817679 |
| Nav1 | S | 952 | -0.292045151 | -0.171470356 | -0.4343912 | -0.142346049 | -0.262920844 | 0.04004626 | 0.288819184 | 0.025045618 | 0.338335118 | 0.168337068 |
| Kif5b | S | 933 | -0.082830195 | -0.161698037 | -0.293336861 | -0.210506665 | -0.131638824 | 0.197728485 | 0.000851024 | 0.004431191 | 0.023745509 | 0.048459441 |
| Bicd2 | S | 819 | -0.091291468 | -0.241047804 | -0.245576135 | -0.154284668 | -0.004528331 | 0.092807227 | 0.004310207 | 0.008306856 | 0.048885183 | 0.940014207 |
| Bag3 | S | 390 | -0.050971083 | -0.219483096 | -0.340153683 | -0.289182599 | -0.120670587 | 0.210435598 | 0.003776147 | 0.006574556 | 0.008369575 | 0.095118267 |
| Kpna3 | S | 56 | -0.078913803 | -0.22322968 | -0.274653275 | -0.195739473 | -0.051423595 | 0.047185872 | 0.003885858 | 0.013081788 | 0.032622003 | 0.409677437 |
| Map1s | S | 687 | -0.098140299 | -0.16955803 | -0.203212718 | -0.10507242 | -0.033654688 | 0.028767167 | 0.003049971 | 0.002225407 | 0.038578104 | 0.389050004 |
| Relch | T | 183 | -0.222080659 | -0.165049091 | -0.351936394 | -0.129855735 | -0.186887303 | 0.014485925 | 0.165504493 | 0.019125587 | 0.231567664 | 0.166590011 |
| Pno1 | S | 32 | -0.177214872 | -0.262859316 | -0.291787231 | -0.114572358 | -0.028927915 | 0.036373439 | 0.015912582 | 0.068354696 | 0.335135553 | 0.794755295 |
| Prrc2a | T | 995 | 0.258597886 | -0.294231464 | -0.413573245 | -0.672171131 | -0.119341781 | 0.499516066 | 3.03709E-06 | 0.03369296 | 0.160321342 | 0.291267843 |
| Gapvd1 | S | 964 | -0.292414407 | -0.225598983 | -0.222594845 | 0.069819562 | 0.003004138 | 0.024789634 | 0.054857378 | 0.040592756 | 0.429382945 | 0.972817782 |
| Araf | S | 157 | -0.30436084 | -0.164112301 | -0.573496685 | -0.269135845 | -0.409384383 | 0.085019382 | 0.340666627 | 0.031727423 | 0.214179271 | 0.096618842 |
| Kirrel | Y | 604 | -0.307743778 | -0.203714764 | -0.445391051 | -0.137647273 | -0.241676286 | 0.0842561 | 0.248802191 | 0.02416832 | 0.40370811 | 0.199469316 |
| Ppp1r12a | S | 994 | 0.006116036 | -0.323141482 | -0.359656182 | -0.365772218 | -0.036514701 | 0.830255978 | 0.006303315 | 0.002972913 | 0.003944135 | 0.556361385 |
| Eps8 | S | 40 | -0.253557095 | -0.247120677 | -0.922884708 | -0.669327613 | -0.675764031 | 0.166065315 | 0.40737633 | 0.075241667 | 0.144838564 | 0.153744814 |
| Syk | Y | 317 | -0.349507052 | -0.229261207 | -0.440432782 | -0.09092573 | -0.211171575 | 0.078623562 | 0.229089318 | 0.045754548 | 0.409143114 | 0.134624579 |
| Rbm42 | S | 133 | -0.191767994 | -0.279074082 | -0.260427813 | -0.068659819 | 0.018646269 | 0.123734089 | 0.003871895 | 0.08006363 | 0.589171057 | 0.857046542 |
| Zswim8 | S | 1831 | -0.167178618 | -0.154466861 | -0.479350441 | -0.312171823 | -0.324883581 | 0.053217702 | 0.043128976 | 0.030870305 | 0.070796367 | 0.066915636 |
| Dbnl | T | 278 | -0.501619456 | -0.092542584 | -0.741806227 | -0.240186772 | -0.649263643 | 0.071320223 | 0.748646483 | 0.078730916 | 0.473640107 | 0.122835443 |
| Znrf2 | S | 141 | -0.094571603 | -0.406600391 | -0.544926309 | -0.450354706 | -0.138325918 | 0.367277371 | 0.036715164 | 0.064977666 | 0.074531503 | 0.477921222 |
| Znrf2 | S | 147 | -0.094571603 | -0.406600391 | -0.544926309 | -0.450354706 | -0.138325918 | 0.367277371 | 0.036715164 | 0.064977666 | 0.074531503 | 0.477921222 |
| Ankrd11 | S | 153 | -0.078233231 | -0.301337643 | -0.307325315 | -0.229092083 | -0.005987672 | 0.124947634 | 0.035375162 | 0.004833161 | 0.038748153 | 0.950903903 |
| Cd2ap | S | 404 | -0.060650427 | -0.214767261 | -0.482368608 | -0.421718181 | -0.267601347 | 0.524304931 | 0.002589608 | 0.022062302 | 0.026081563 | 0.091832001 |
| Larp4 | S | 592 | -0.005335399 | -0.308207192 | -0.295496921 | -0.290161522 | 0.012710271 | 0.964313546 | 0.00013541 | 0.030875763 | 0.087438743 | 0.874651791 |
| Dlg5 | S | 900 | -0.402313561 | -0.316237972 | -0.331125783 | 0.071187777 | -0.014887811 | 0.037972978 | 0.148246506 | 0.175019852 | 0.740833931 | 0.949660552 |
| Dennd4c | S | 1310 | -0.056295966 | -0.126796447 | -0.294802527 | -0.238506561 | -0.16800608 | 0.278868323 | 0.005548429 | 0.000162461 | 0.007995111 | 0.006944437 |
| Lmna | S | 301 | 0.094405608 | -0.307769557 | -0.429730125 | -0.524135734 | -0.121960568 | 0.10153438 | 5.98478E-05 | 0.009299254 | 0.002458413 | 0.146992073 |
| Map1b | S | 1151 | -0.064498652 | -0.135366556 | -0.221761823 | -0.157263171 | -0.086395267 | 0.41744591 | 0.013321624 | 3.0221E-06 | 0.129113334 | 0.060335156 |
| Zfp609 | S | 1057 | -0.132321696 | -0.530081031 | -0.071497337 | 0.060824358 | 0.458583693 | 0.116763146 | 0.00109747 | 0.427171652 | 0.337609163 | 0.005105949 |
| Synj1 | S | 830 | -0.152419384 | -0.158616903 | -0.270067904 | -0.11764852 | -0.111451001 | 0.048002456 | 0.012557225 | 0.00648709 | 0.133162628 | 0.114211319 |
| Pinx1 | T | 228 | 0.01861281 | -0.273048666 | -0.339490381 | -0.358103191 | -0.066441715 | 0.697096343 | 0.000192742 | 0.011299774 | 0.009903107 | 0.361951135 |
| Tuba1a Tuba1b | S | 439;439 | -0.156696767 | -0.198352295 | -0.232520347 | -0.07582358 | -0.034168052 | 0.040551052 | 0.026950997 | 0.004351518 | 0.173407809 | 0.560385086 |
| Trip12 | S | 1460 | -0.100733338 | -0.183078187 | -0.225024585 | -0.124291248 | -0.041946399 | 0.148500042 | 0.002172027 | 0.002018191 | 0.097873486 | 0.288978731 |
| Arhgef2 | T | 152 | -0.224912169 | -0.162000946 | -0.515394081 | -0.290481911 | -0.353393135 | 0.095820183 | 0.24651776 | 0.014554057 | 0.097086448 | 0.06623294 |
| Ets2 | Y | 250 | -0.210946124 | -0.244098682 | -0.549362011 | -0.338415887 | -0.305263329 | 0.181196093 | 0.010636226 | 0.147492075 | 0.300440731 | 0.3357148 |
| Szrd1 | S | 39 | -0.140639452 | -0.202212852 | -0.338452962 | -0.197813511 | -0.13624011 | 0.100471132 | 0.093928245 | 0.002615322 | 0.050400463 | 0.207516946 |
| Sipa1l1 | S | 1117 | -0.385648451 | -0.155941445 | -0.298960922 | 0.086687529 | -0.143019477 | 0.010506877 | 0.254781499 | 0.070303841 | 0.45528036 | 0.301633764 |
| Otud5 | S | 447 | -0.118355534 | -0.280834399 | -0.28892735 | -0.170571816 | -0.008092951 | 0.031299854 | 0.016208237 | 0.040746068 | 0.138330919 | 0.935443346 |
| Nes | S | 623 | -0.06311647 | -0.250524497 | -0.164815483 | -0.101699013 | 0.085709014 | 0.13406784 | 0.000294066 | 0.007797268 | 0.047452692 | 0.074329573 |
| Rif1 | T | 2193 | -0.056597633 | -0.213640441 | -0.278901756 | -0.222304123 | -0.065261314 | 0.612367621 | 0.002496032 | 0.00108821 | 0.123063894 | 0.162381399 |
| Wdr43 | S | 666 | -0.031639853 | -0.259426373 | -0.217157002 | -0.185517149 | 0.042269372 | 0.42882019 | 0.001379258 | 0.001230629 | 0.00591418 | 0.322195989 |
| Wdr43 | S | 674 | -0.031639853 | -0.259426373 | -0.217157002 | -0.185517149 | 0.042269372 | 0.42882019 | 0.001379258 | 0.001230629 | 0.00591418 | 0.322195989 |
| Myh11 Myh14 Myh9 | S | 548;541;571 | -0.083869576 | -0.12015953 | -0.2582276 | -0.174358024 | -0.13806807 | 0.026050929 | 0.024340608 | 0.000272979 | 0.004616726 | 0.020998245 |
| Synrg | S | 744 | -0.244692639 | -0.219759003 | -0.140631618 | 0.104061021 | 0.079127385 | 0.004874982 | 0.009846917 | 0.138699706 | 0.222962556 | 0.333948323 |
| Tln1 | S | 425 | -0.061453994 | -0.166174548 | -0.318735449 | -0.257281455 | -0.152560901 | 0.367045078 | 0.001638005 | 0.002241811 | 0.020019212 | 0.029218612 |
| Cdk17 | S | 165 | -0.097658145 | -0.221685595 | -0.317940791 | -0.220282646 | -0.096255196 | 0.047777432 | 0.002146428 | 0.054656658 | 0.129642878 | 0.399415891 |
| Sec16a | S | 1342 | -0.35866991 | -0.271439654 | -0.218898984 | 0.139770927 | 0.052540671 | 0.021780871 | 0.057836285 | 0.171847201 | 0.258092112 | 0.627046238 |
| Hdgfl2 | S | 454 | -0.109926023 | -0.179224158 | -0.345602707 | -0.235676684 | -0.166378549 | 0.046272727 | 0.102076205 | 0.002282058 | 0.018270645 | 0.127224627 |
| Eif2b5 | S | 713 | -0.067773309 | -0.197629463 | -0.307730584 | -0.239957275 | -0.110101121 | 0.151419301 | 0.000715425 | 0.017543581 | 0.035350099 | 0.190414879 |
| Otud3 | S | 306 | -0.169802187 | -0.028963503 | -0.384722323 | -0.214920136 | -0.35575882 | 0.006542292 | 0.597873202 | 0.001069587 | 0.021157329 | 0.003549989 |
| Pcdh19 | S | 1134 | -0.119241085 | -0.209199855 | -0.198384139 | -0.079143055 | 0.010815715 | 0.013927216 | 0.034958 | 0.001913067 | 0.07001167 | 0.866790763 |
| Usp39 | S | 19 | -0.014463764 | -0.327994309 | -0.306572445 | -0.292108681 | 0.021421864 | 0.751646313 | 0.001241676 | 0.016776616 | 0.03130068 | 0.787028092 |
| Rtkn | S | 220 | -0.099505547 | -0.247332344 | -0.229804229 | -0.130298682 | 0.017528115 | 0.189933335 | 0.007152122 | 0.003067457 | 0.113661811 | 0.761650594 |
| Eif3b | S | 141 | -0.086536643 | -0.169698293 | -0.171318788 | -0.084782145 | -0.001620495 | 0.032740962 | 0.002199779 | 0.000834296 | 0.026636681 | 0.955431834 |
| Map1b | S | 1497 | -0.127897517 | -0.164283927 | -0.143137827 | -0.01524031 | 0.0211461 | 0.007354088 | 0.002377094 | 0.004694951 | 0.497199658 | 0.466549679 |
| Camsap2 | S | 391 | -0.288166282 | -0.149626853 | -0.350423842 | -0.06225756 | -0.200796989 | 0.008422533 | 0.277201773 | 0.042129611 | 0.597163314 | 0.219129471 |
| Tnks1bp1 | S | 713 | -0.067652907 | -0.204516419 | -0.223939434 | -0.156286527 | -0.019423016 | 0.326409508 | 0.000583769 | 0.002696798 | 0.078506013 | 0.580899783 |
| Sh3pxd2a | S | 766 | -0.293525777 | -0.152349655 | -0.321501121 | -0.027975344 | -0.169151466 | 0.014516013 | 0.316148838 | 0.019463866 | 0.722491692 | 0.262890001 |
| Hmgcs1 | S | 495 | -0.240351812 | -0.182116331 | -0.290555705 | -0.050203893 | -0.108439374 | 0.156190718 | 0.002027692 | 0.051702806 | 0.725312063 | 0.27522917 |
| Arhgap23 | S | 1184 | -0.315459091 | -0.188169227 | -0.361422008 | -0.045962916 | -0.17325278 | 0.050681948 | 0.240010697 | 0.022869621 | 0.66087639 | 0.211059234 |
| Anln | S | 656 | -0.038488845 | -0.225849544 | -0.354169842 | -0.315680997 | -0.128320298 | 0.351179702 | 0.004891865 | 0.005448614 | 0.027394509 | 0.12746968 |
| Micall2 | S | 153 | 0.009379392 | -0.194078626 | -0.294956058 | -0.30433545 | -0.100877432 | 0.793595433 | 0.002356325 | 0.000139134 | 0.00204984 | 0.02560058 |
| Gigyf2 | S | 594 | -0.061661902 | -0.257207479 | -0.152217046 | -0.090555145 | 0.104990433 | 0.182186803 | 9.27274E-05 | 0.017394969 | 0.111314729 | 0.065324552 |
| Fbf1 | S | 248 | -0.951554404 | -0.005787888 | -0.822859804 | 0.128694599 | -0.817071917 | 0.172329284 | 0.987305309 | 0.108409989 | 0.833120842 | 0.139645692 |
| Scamp4 | T | 194 | -0.302510495 | -0.181321865 | -0.349415687 | -0.046905192 | -0.168093822 | 0.053680039 | 0.180046935 | 0.021885386 | 0.68871033 | 0.170398663 |
| Klc1 | S | 459 | -0.114012167 | -0.153753732 | -0.220366333 | -0.106354166 | -0.066612601 | 0.0158647 | 0.003200516 | 0.008497548 | 0.088631842 | 0.219452594 |
| Nisch | S | 1373 | -0.007353733 | -0.387758291 | -0.410446416 | -0.403092684 | -0.022688125 | 0.817470447 | 0.017559407 | 0.010954651 | 0.022813095 | 0.821533903 |
| Gphn | S | 328 | -0.210853824 | -0.072839744 | -0.403285322 | -0.192431497 | -0.330445577 | 0.023994818 | 0.537885887 | 0.002550555 | 0.050288766 | 0.041665226 |
| Epha2 | S | 898 | -0.071461903 | -0.252130296 | -0.380594379 | -0.309132476 | -0.128464083 | 0.300883351 | 0.018181566 | 0.008170031 | 0.017154075 | 0.182265143 |
| Baiap2 | S | 326 | 0.012523633 | -0.261610589 | -0.282553785 | -0.295077419 | -0.020943196 | 0.728718524 | 0.000217486 | 0.004816061 | 0.003902206 | 0.654812829 |
| Lgals1 | S | 30 | -0.442939948 | -0.104885665 | -0.346337612 | 0.096602337 | -0.241451946 | 0.004478988 | 0.501793642 | 0.135029423 | 0.605202137 | 0.279954079 |
| Bcr | T | 304 | -0.394736164 | -0.012287541 | -0.492741741 | -0.098005577 | -0.480454199 | 0.005445007 | 0.953601882 | 0.056489917 | 0.588148587 | 0.106622885 |
| Rgl3 | S | 707 | -0.129581542 | -0.298143713 | -0.339832961 | -0.210251419 | -0.041689248 | 0.297278541 | 0.009458731 | 0.033934762 | 0.150513692 | 0.638460547 |
| Ston1 | S | 203 | -0.204183833 | -0.271863724 | -0.471752438 | -0.267568604 | -0.199888713 | 0.164916241 | 0.057840814 | 0.057737337 | 0.20377196 | 0.306284457 |
| Ranbp2 | S | 2113 | -0.015868961 | -0.297420449 | -0.23672879 | -0.220859828 | 0.060691659 | 0.741252663 | 0.000127202 | 0.023311648 | 0.033769986 | 0.37614253 |
| Mapre2 | S | 222 | -0.459581752 | -0.129188525 | -0.38109636 | 0.078485392 | -0.251907835 | 0.037450643 | 0.526256621 | 0.035323681 | 0.62482722 | 0.226698611 |
| Mapk1 | T | 183 | -0.12474613 | -0.26235091 | -0.350981933 | -0.226235802 | -0.088631023 | 0.130025019 | 0.016049818 | 0.031892047 | 0.112113739 | 0.450902971 |
| Arhgef1 | S | 905 | -0.016273645 | -0.299712111 | -0.232532166 | -0.216258521 | 0.067179945 | 0.595494962 | 0.00067398 | 0.007010741 | 0.022874692 | 0.243721009 |
| Hnrnpm | S | 411 | -0.078174309 | -0.104690495 | -0.389731367 | -0.311557058 | -0.285040872 | 0.251864648 | 0.147209296 | 0.000128179 | 0.006273829 | 0.00924399 |
| Nsun2 | S | 23 | -0.039654187 | -0.176955091 | -0.204702844 | -0.165048656 | -0.027747753 | 0.402579849 | 0.002763376 | 4.34509E-05 | 0.038973176 | 0.369002508 |

| | | | | | | | | | | | | |
|---|---|---|---|---|---|---|---|---|---|---|---|---|
| Cobll1 | S | 897 | -0.020164226 | -0.171303828 | -0.290328081 | -0.270163855 | -0.119024253 | 0.640957655 | 0.002130387 | 0.000338022 | 0.002275286 | 0.044168122 |
| Sec63 | S | 593 | -0.164204625 | -0.190418716 | -0.26312036 | -0.098915735 | -0.072701644 | 0.021776784 | 0.012509078 | 0.035352088 | 0.288858296 | 0.410804733 |
| Sec63 | S | 597 | -0.164204625 | -0.190418716 | -0.26312036 | -0.098915735 | -0.072701644 | 0.021776784 | 0.012509078 | 0.035352088 | 0.288858296 | 0.410804733 |
| Ostf1 | S | 203 | -0.219217747 | -0.204775102 | -0.52234135 | -0.303123603 | -0.317566248 | 0.190956962 | 0.213213906 | 0.01519672 | 0.120656735 | 0.106676658 |
| Arhgap1 | S | 49 | -0.248808892 | -0.247025616 | -0.179991848 | 0.068817045 | 0.067033768 | 0.022721816 | 0.016422777 | 0.086135409 | 0.401506969 | 0.348256105 |
| Hypk | S | 38 | 0.03617817 | -0.348941139 | -0.367937215 | -0.404115384 | -0.018996076 | 0.513973352 | 0.004956835 | 0.003668842 | 0.002894023 | 0.703432421 |
| Tjp2 | S | 960 | -0.322270393 | -0.214386273 | -0.140503823 | 0.181766569 | 0.073882449 | 0.001218255 | 0.040250766 | 0.440227592 | 0.338590379 | 0.674874258 |
| Gpatch4 | S | 258 | 0.006455051 | -0.223273526 | -0.251103818 | -0.257558869 | -0.027830292 | 0.877898782 | 0.000729715 | 0.000413818 | 0.001002843 | 0.355546976 |
| Maea | S | 311 | 0.000805653 | -0.204957213 | -0.302615794 | -0.303421447 | -0.097658581 | 0.984292988 | 0.005935955 | 0.0001759 | 0.000514149 | 0.082415127 |
| Rps6ka3 | S | 737 | -0.293389266 | -0.287893576 | -0.612018828 | -0.318629562 | -0.324125252 | 0.293941138 | 0.190832566 | 0.051122243 | 0.295890227 | 0.222194988 |
| Specc1 | S | 115 | -0.038565966 | -0.358421055 | -0.381924107 | -0.343358141 | -0.023503052 | 0.668780099 | 0.006624829 | 0.027376131 | 0.049331599 | 0.842117189 |
| Arrb1 | S | 412 | -0.179760428 | -0.097772632 | -0.378966153 | -0.199205726 | -0.281193521 | 0.012420958 | 0.263196508 | 0.006189833 | 0.081739357 | 0.030884387 |
| Ptpn12 | S | 748 | -0.096527261 | -0.245478538 | -0.28707623 | -0.190548969 | -0.041597692 | 0.047722843 | 0.013944088 | 0.018394148 | 0.032153302 | 0.516572381 |
| Hectd2 | S | 9 | -0.046488801 | -0.294719877 | -0.384622868 | -0.338134067 | -0.089902991 | 0.6767101 | 0.008089435 | 0.011515104 | 0.044088686 | 0.373525397 |
| Map1b | S | 1013 | -0.232156301 | -0.362931925 | -0.234604135 | -0.002447834 | 0.12832779 | 0.091091872 | 0.041765332 | 0.05812191 | 0.983696237 | 0.362600567 |
| Ibtk | S | 1005 | -0.083366912 | -0.171258101 | -0.463148358 | -0.379781445 | -0.291890257 | 0.418247782 | 0.023030252 | 0.005116264 | 0.021854979 | 0.040514616 |
| Cc2d1b | S | 502 | -0.150121307 | -0.179888604 | -0.24800801 | -0.097886702 | -0.068119406 | 0.026352841 | 0.049343583 | 0.004348482 | 0.132604188 | 0.350713624 |
| Msn | S | 407 | -0.340001622 | -0.110317615 | -0.367825275 | -0.027823653 | -0.25750766 | 0.073320435 | 0.385421354 | 0.006153795 | 0.83668909 | 0.070525913 |
| Kdm5c | S | 317 | -0.19486815 | -0.177890552 | -0.333891158 | -0.139023008 | -0.156000606 | 0.119443194 | 0.095555432 | 0.004458967 | 0.236070202 | 0.144469997 |
| Plekha3 | S | 211 | 0.049376873 | -0.408707705 | -0.412500274 | -0.461877147 | -0.003792569 | 0.888809945 | 0.011887123 | 0.011042928 | 0.265184363 | 0.972459578 |
| Gpatch8 | S | 1082 | -0.106017582 | -0.183603703 | -0.326370295 | -0.220352713 | -0.142766592 | 0.142435805 | 0.037539667 | 0.002423129 | 0.01647672 | 0.055207106 |
| Nop14 | S | 173 | 0.077839485 | -0.388966822 | -0.489284353 | -0.567123838 | -0.100317531 | 0.262560238 | 0.011037936 | 0.004353157 | 0.002459773 | 0.328017561 |
| Birc6 | S | 493 | -0.126051624 | -0.246007792 | -0.173731704 | -0.04768008 | 0.072276088 | 0.270629776 | 0.000816044 | 0.009114703 | 0.638239461 | 0.144921706 |
| Akt2 | T | 451 | -0.116342614 | -0.061829429 | -0.491839328 | -0.375496715 | -0.4300099 | 0.088765358 | 0.430787425 | 0.000800885 | 0.004901835 | 0.005217051 |
| Eif5b | S | 175 | -0.14360076 | -0.343870709 | -0.280886429 | -0.137285669 | 0.06298428 | 0.130657692 | 0.021124708 | 0.043372124 | 0.24327708 | 0.595879295 |
| Rrp15 | S | 242 | 0.035790095 | -0.209445782 | -0.338121162 | -0.373911256 | -0.128675379 | 0.280210585 | 0.000695712 | 0.000467818 | 0.003652983 | 0.056335473 |
| Trp53bp2 | S | 480 | -0.383615633 | -0.114438365 | -0.488582828 | -0.104967195 | -0.374144463 | 0.06766501 | 0.563227862 | 0.023063202 | 0.371329102 | 0.065360744 |
| Map7d1 | S | 78 | -0.046964099 | -0.130788576 | -0.191790504 | -0.144826405 | -0.061001928 | 0.365131666 | 0.000219761 | 6.25366E-05 | 0.068286545 | 0.007269986 |
| Pdlim5 | S | 339 | 0.00133447 | -0.289857916 | -0.381451419 | -0.382785889 | -0.091593503 | 0.996121771 | 0.011439285 | 0.002917049 | 0.245832101 | 0.130108897 |
| Tmem131 | S | 1742 | -0.332762563 | -0.230526789 | -0.520534546 | -0.187771983 | -0.290007758 | 0.040895162 | 0.324809609 | 0.103198113 | 0.457638887 | 0.336439679 |
| Reps1 | S | 708 | -0.135798662 | -0.137297805 | -0.290226199 | -0.154427537 | -0.152928394 | 0.041840419 | 0.077969403 | 0.001472131 | 0.018356515 | 0.047301846 |
| Myo9b | S | 1371 | -0.064927735 | -0.156316919 | -0.478862073 | -0.413934338 | -0.322545154 | 0.343109886 | 0.03461174 | 0.003761079 | 0.013896844 | 0.038576184 |
| Map1b | S | 1204 | -0.054302674 | -0.174036117 | -0.229663989 | -0.175361315 | -0.055627872 | 0.326448737 | 0.01598679 | 3.82984E-05 | 0.040337525 | 0.23655349 |
| Dock7 | S | 452 | -0.262325508 | -0.203342088 | -0.333596201 | -0.071270694 | -0.130254113 | 0.050686385 | 0.175727687 | 0.019821687 | 0.531981556 | 0.357936204 |
| Mast4 | S | 2478 | -0.044945434 | -0.213516756 | -0.2731851 | -0.228239666 | -0.059668344 | 0.430593241 | 0.069368349 | 3.54093E-05 | 0.031792897 | 0.462188539 |
| Pgm2 | S | 173 | -0.288767047 | -0.221747334 | -0.510207096 | -0.221440049 | -0.288459762 | 0.135294496 | 0.306267957 | 0.028249178 | 0.278009668 | 0.228006188 |
| Kpnb1 | S | 213 | -0.169861044 | -0.119958861 | -0.257866355 | -0.088005311 | -0.137907494 | 0.082863689 | 0.014807175 | 0.002141765 | 0.289489326 | 0.052549805 |
| Itprid2 | S | 1051 | -0.164103137 | -0.167155216 | -0.264748209 | -0.100645072 | -0.097592993 | 0.009399651 | 0.057491718 | 0.015129993 | 0.195085988 | 0.263269075 |
| Ndel1 | S | 198 | 0.023531937 | -0.227379638 | -0.247124182 | -0.270656119 | -0.019744544 | 0.573002147 | 0.000335435 | 0.000425105 | 0.004049325 | 0.540844978 |
| Eif1ad | S | 155 | 0.037993683 | -0.268031851 | -0.407656109 | -0.445649792 | -0.139624258 | 0.782666573 | 0.00185849 | 0.007279488 | 0.048842846 | 0.093368976 |
| Eif1ad | S | 159 | 0.037993683 | -0.268031851 | -0.407656109 | -0.445649792 | -0.139624258 | 0.782666573 | 0.00185849 | 0.007279488 | 0.048842846 | 0.093368976 |
| Itpkc | S | 212 | -0.216576733 | -0.085155559 | -0.298501972 | -0.081925239 | -0.213346412 | 0.009524648 | 0.362651357 | 0.002977879 | 0.154349656 | 0.060108184 |
| Usp37 | S | 165 | 0.025967268 | -0.213413999 | -0.31796173 | -0.343928999 | -0.104547731 | 0.60828102 | 0.000371819 | 0.001601135 | 0.002815702 | 0.093340247 |
| Rrp12 | S | 1050 | -0.054175979 | -0.197655582 | -0.272914784 | -0.218738805 | -0.075259202 | 0.156199139 | 0.000459271 | 0.01564883 | 0.037452474 | 0.278211288 |
| Sh3pxd2b | S | 769 | -0.085823493 | -0.187750577 | -0.264063323 | -0.17823983 | -0.076312746 | 0.208521126 | 0.008134902 | 0.001704783 | 0.03896815 | 0.016465016 |
| Lactb2 | T | 279 | -0.372754175 | -0.112146556 | -0.458406302 | -0.085652127 | -0.346259746 | 0.010292275 | 0.554858214 | 0.090002427 | 0.669171379 | 0.197077645 |
| Psmd13 | S | 323 | -0.337192254 | -0.164417925 | -0.312236011 | 0.024956243 | -0.147818086 | 0.007996817 | 0.26061141 | 0.089944027 | 0.859104013 | 0.396590064 |
| Zc3h12c | S | 470 | -0.407931094 | -0.108691622 | -0.443957957 | -0.036026863 | -0.335266335 | 0.061177949 | 0.563791945 | 0.022732616 | 0.839133573 | 0.138352173 |
| Slc4a7 | S | 271 | -0.045028229 | -0.198932462 | -0.184731256 | -0.139703026 | 0.014201206 | 0.394057067 | 0.009686056 | 1.20317E-05 | 0.078255066 | 0.662017262 |
| Tent4b | S | 545 | -0.259273846 | -0.214240654 | -0.317684691 | -0.058410845 | -0.103444037 | 0.035084944 | 0.178712343 | 0.026776341 | 0.486980324 | 0.455103154 |
| Mre11a | S | 674 | -0.226527271 | -0.211128675 | -0.340090873 | -0.113563601 | -0.128962197 | 0.061715468 | 0.087172961 | 0.028655475 | 0.276748923 | 0.240326495 |
| Cep170 | S | 1407 | -0.123340569 | -0.321505126 | -0.313029482 | -0.189688912 | 0.008475644 | 0.253262939 | 0.008652051 | 0.051767369 | 0.173462041 | 0.937598514 |
| Tnks1bp1 | S | 1136 | -0.092932344 | -0.124024445 | -0.139449831 | -0.046517487 | -0.015425387 | 0.014069774 | 0.00989913 | 2.17786E-05 | 0.107115293 | 0.538878808 |
| Cactin | S | 96 | -0.145382315 | -0.14852013 | -0.379092022 | -0.233709707 | -0.230571892 | 0.079195758 | 0.222829593 | 0.002081636 | 0.029120701 | 0.099971164 |
| Ubr4 | T | 2721 | -0.219581509 | -0.127237039 | -0.234427891 | -0.014846382 | -0.107190852 | 0.005192461 | 0.08617147 | 0.017225223 | 0.787282037 | 0.152143077 |
| Pkp4 | S | 232 | -0.374605668 | -0.171409951 | -0.641750662 | -0.267144994 | -0.47034071 | 0.127221327 | 0.485232528 | 0.05235106 | 0.344080522 | 0.155023342 |
| Knop1 | S | 102 | 0.02300864 | -0.210042015 | -0.213693354 | -0.236701994 | -0.003651339 | 0.670130322 | 7.50426E-05 | 0.000415851 | 0.01827869 | 0.887702948 |
| Utp18 | T | 30 | 0.103297064 | -0.322175739 | -0.413189084 | -0.516486149 | -0.091013345 | 0.097162008 | 0.000764578 | 0.002449244 | 0.002895076 | 0.240647193 |
| Map1b | S | 1438 | -0.122747685 | -0.143013128 | -0.142566896 | -0.019819212 | 0.000446232 | 0.002115298 | 0.000923442 | 0.022236872 | 0.608779524 | 0.990511731 |
| Tsnax | T | 165 | -0.184110011 | -0.211983371 | -0.191842306 | -0.007732295 | 0.020141064 | 0.012678489 | 0.013136156 | 0.053252684 | 0.91390315 | 0.786566968 |
| Fgf1 | S | 153 | -0.117800353 | -0.383555807 | -0.53533913 | -0.417538777 | -0.151783323 | 0.51122327 | 0.071067242 | 0.039178018 | 0.081652473 | 0.429236033 |
| Fgf1 | S | 154 | -0.117800353 | -0.383555807 | -0.53533913 | -0.417538777 | -0.151783323 | 0.51122327 | 0.071067242 | 0.039178018 | 0.081652473 | 0.429236033 |
| Hif1a | S | 679 | 0.104739701 | -0.353658092 | -0.453326392 | -0.558066093 | -0.0996683 | 0.033143314 | 1.90205E-05 | 0.033345693 | 0.023294532 | 0.413278311 |
| Ppp6r3 | S | 693 | -0.235169049 | -0.205682624 | -0.314740001 | -0.079570952 | -0.109057377 | 0.105234754 | 0.108795695 | 0.010314302 | 0.468777109 | 0.284478863 |
| Pprc1 | S | 232 | -0.192486159 | -0.425004199 | -0.275257145 | -0.082770986 | 0.149747054 | 0.258100876 | 0.042598498 | 0.04222509 | 0.607689284 | 0.344556185 |
| Pprc1 | S | 239 | -0.192486159 | -0.425004199 | -0.275257145 | -0.082770986 | 0.149747054 | 0.258100876 | 0.042598498 | 0.04222509 | 0.607689284 | 0.344556185 |
| Naxe | S | 43 | -0.277077704 | -0.190787185 | -0.262820158 | 0.014257546 | -0.072032973 | 0.140856079 | 0.034525501 | 0.013573639 | 0.923373961 | 0.348798566 |
| Arhgef18 | S | 1311 | -0.468984376 | -0.11818318 | -0.723960412 | -0.254976036 | -0.605777232 | 0.116649152 | 0.665419039 | 0.072738422 | 0.450409618 | 0.124114093 |
| Lnpk | S | 411 | -0.240159632 | -0.152104716 | -0.248166277 | -0.008006645 | -0.096061561 | 0.014254404 | 0.23559361 | 0.006314024 | 0.890999653 | 0.404776183 |
| Rrp15 | S | 229 | -0.012744743 | -0.173328158 | -0.269847029 | -0.257102286 | -0.096518871 | 0.815546506 | 0.002560108 | 0.000166701 | 0.012140323 | 0.036775529 |
| Prune2 | T | 2502 | 0.097646438 | -0.282399376 | -0.383555763 | -0.481202201 | -0.101156387 | 0.401273165 | 0.001806914 | 0.001760962 | 0.020475951 | 0.139280554 |
| Ggta1 | S | 20 | -0.387478443 | -0.084444767 | -0.425987786 | -0.038509343 | -0.341543019 | 0.023302793 | 0.671667144 | 0.032858695 | 0.787678498 | 0.146227917 |
| Adgre5 | S | 814 | -0.0789442 | -0.079498603 | -0.293540805 | -0.214596605 | -0.214042202 | 0.016047259 | 0.066040082 | 0.000250093 | 0.00476064 | 0.00384491 |
| Wnk1 | T | 17 | -0.059562979 | -0.123579581 | -0.221181861 | -0.161618883 | -0.097602281 | 0.164198835 | 0.024214566 | 1.08912E-05 | 0.023414163 | 0.062731646 |
| Osbpl5 | S | 88 | 0.049672448 | -0.291742217 | -0.46621493 | -0.515887378 | -0.174472713 | 0.253457216 | 0.000905389 | 0.01265843 | 0.009988292 | 0.116710536 |
| Ccdc86 | S | 253 | 0.028477456 | -0.427553272 | -0.419156037 | -0.447633493 | 0.008397235 | 0.86737365 | 0.006743972 | 0.03287424 | 0.063588632 | 0.951791875 |
| Nup35 | S | 73 | 0.082530902 | -0.296204917 | -0.41786265 | -0.500393552 | -0.121657733 | 0.050163252 | 3.9327E-05 | 0.012149918 | 0.010409851 | 0.195390601 |
| Map6 | S | 198 | -0.094752823 | -0.203034036 | -0.275274942 | -0.180522119 | -0.072240906 | 0.037361285 | 0.038795172 | 0.004395172 | 0.031158746 | 0.349643379 |
| Med1 | S | 671 | -0.307603163 | -0.189734847 | -0.487604681 | -0.180001518 | -0.297869834 | 0.088654586 | 0.401218407 | 0.029261882 | 0.248714367 | 0.198577772 |
| Map4k4 | S | 795 | -0.038948076 | -0.144410081 | -0.214943929 | -0.175995853 | -0.070533848 | 0.183299377 | 0.001131447 | 0.000222187 | 0.002240599 | 0.046848156 |
| Tcof1 | S | 154 | 0.003703079 | -0.244597994 | -0.28271079 | -0.286413868 | -0.038112795 | 0.940071424 | 0.002324422 | 0.001093514 | 0.001350639 | 0.233298478 |
| Sipa1l1 | S | 1249 | -0.010560286 | -0.23468788 | -0.207741837 | -0.197181551 | 0.026946042 | 0.787108351 | 0.007062927 | 3.38302E-05 | 0.014518316 | 0.506377322 |
| Tnks1bp1 | S | 912 | -0.254906986 | -0.108554217 | -0.350970476 | -0.09606349 | -0.242416259 | 0.031146464 | 0.294131441 | 0.009012379 | 0.012048035 | 0.016283346 |
| Nme1 | S | 120 | -0.258773052 | -0.058700231 | -0.483265041 | -0.224491988 | -0.42456481 | 0.1053401 | 0.7222798 | 0.002736538 | 0.129708673 | 0.059324917 |
| Bbx | S | 477 | 0.04194936 | -0.416120284 | -0.314057091 | -0.356006451 | 0.102063194 | 0.649745236 | 0.001243644 | 0.03271265 | 0.027625332 | 0.298937387 |
| Tbc1d1 | S | 231 | -0.135323831 | -0.167545108 | -0.164722847 | -0.029399016 | 0.002822261 | 0.017118408 | 0.007629131 | 0.004476988 | 0.492168438 | 0.945854074 |
| Ccdc88a | S | 1654 | 0.118690645 | -0.483436353 | -0.3937726 | -0.512463245 | 0.089663754 | 0.001946713 | 0.002406652 | 0.000447205 | 0.001163985 | 0.181654697 |
| Map4 | S | 519 | -0.163169011 | -0.110768086 | -0.252823771 | -0.089654759 | -0.142055684 | 0.008419693 | 0.054509271 | 0.006029793 | 0.157299483 | 0.057068626 |
| Farp1 | S | 335 | -0.157728934 | -0.314851557 | -0.625649253 | -0.467920319 | -0.310797695 | 0.121010394 | 0.117360605 | 0.104487886 | 0.176623481 | 0.32485165 |
| Bace1 | S | 498 | -0.478292687 | -0.239673549 | -0.314302245 | 0.163990443 | -0.074628696 | 0.221086541 | 0.143876304 | 0.028811493 | 0.612233148 | 0.580051321 |
| Spen | S | 647 | -0.115345114 | -0.220137593 | -0.304735784 | -0.189390671 | -0.084598191 | 0.167503395 | 0.025871154 | 0.006639076 | 0.00199489 | 0.085954815 |
| Tcaf3 | S | 460 | -0.096526851 | -0.160729657 | -0.220031433 | -0.123504582 | -0.059301776 | 0.030111758 | 0.011602686 | 0.002231677 | 0.028904539 | 0.220200236 |
| Tcaf3 | Y | 458 | -0.096526851 | -0.160729657 | -0.220031433 | -0.123504582 | -0.059301776 | 0.030111758 | 0.011602686 | 0.002231677 | 0.028904539 | 0.220200236 |
| Abcf1 | S | 194 | -0.191212462 | -0.168394842 | -0.204409748 | -0.013197286 | -0.036014907 | 0.048350267 | 0.006857824 | 0.017477695 | 0.857993976 | 0.528620432 |

| | | | | | | | | | | | | |
|---|---|---|---|---|---|---|---|---|---|---|---|---|
| Abcf1 | S | 613 | 0.000972017 | -0.190141921 | -0.254624816 | -0.255596832 | -0.064482895 | 0.980297133 | 0.000354638 | 0.000687183 | 0.003449183 | 0.111058375 |
| Cdk17 | S | 137 | -0.189961755 | -0.199583785 | -0.457313682 | -0.267351927 | -0.257729896 | 0.139565321 | 0.14385108 | 0.01836151 | 0.100416143 | 0.115570279 |
| Akt1s1 | S | 88 | -0.107889734 | -0.247606552 | -0.19249782 | -0.084608086 | 0.055108733 | 0.437254294 | 0.001378159 | 0.005003409 | 0.523807631 | 0.120662898 |
| Akt1s1 | S | 92 | -0.107889734 | -0.247606552 | -0.19249782 | -0.084608086 | 0.055108733 | 0.437254294 | 0.001378159 | 0.005003409 | 0.523807631 | 0.120662898 |
| Dhx29 | S | 198 | 0.067556168 | -0.291560993 | -0.289147214 | -0.356703381 | 0.002413779 | 0.043446165 | 1.41339E-05 | 0.004872947 | 0.000948492 | 0.9425186 |
| Adgra2 | S | 1324 | -0.275518908 | -0.132667513 | -0.367849993 | -0.092331085 | -0.23518248 | 0.031472483 | 0.456028779 | 0.011183043 | 0.30611331 | 0.213177759 |
| Bbx | S | 479 | 0.043069399 | -0.432531697 | -0.352463664 | -0.395533063 | 0.080068033 | 0.66830143 | 0.002845246 | 0.0278649 | 0.025027797 | 0.426518858 |
| Fxr1 | S | 449 | -0.112905037 | -0.217172475 | -0.178936405 | -0.066031368 | 0.03823607 | 0.120120669 | 0.005762653 | 0.002482069 | 0.299192099 | 0.401697031 |
| Lsm11 | S | 281 | 0.062117985 | -0.22421857 | -0.40909212 | -0.471210105 | -0.184873551 | 0.438442125 | 0.00105377 | 0.002461848 | 0.004635547 | 0.062989739 |
| Ets1 | S | 254 | -0.115961688 | -0.398211015 | -0.456656677 | -0.340694989 | -0.058445663 | 0.788077133 | 0.055953199 | 0.019257509 | 0.450843076 | 0.681839077 |
| Cdc42bpa | S | 1641 | -0.225633408 | -0.129871069 | -0.394093071 | -0.168459664 | -0.264222002 | 0.072557086 | 0.39763699 | 0.004317472 | 0.125477135 | 0.140067356 |
| Lig3 | S | 892 | 0.026224619 | -0.206008335 | -0.250882934 | -0.277107553 | -0.044874599 | 0.443903217 | 0.00019259 | 0.000410652 | 0.002268777 | 0.1047198 |
| Tnks1bp1 | S | 539 | -0.054906279 | -0.198367446 | -0.183732563 | -0.128826284 | 0.014634883 | 0.598431407 | 3.71737E-05 | 0.004923508 | 0.278037513 | 0.628704126 |
| Mtr | T | 1252 | 0.160976781 | -0.323016546 | -0.423987618 | -0.584964399 | -0.100971072 | 0.712627404 | 0.000453352 | 0.027437124 | 0.260384403 | 0.332264377 |
| Ptk2 | S | 910 | -0.072863903 | -0.151542977 | -0.297139935 | -0.224276032 | -0.145596958 | 0.161673497 | 0.000301632 | 0.016508801 | 0.020682275 | 0.077476769 |
| Prr14 | S | 140 | -0.098395003 | -0.283533159 | -0.256096101 | -0.157701098 | 0.027437058 | 0.19952463 | 0.007312666 | 0.020480813 | 0.095430668 | 0.685040201 |
| Nhsl1 | S | 832 | -0.053940736 | -0.252419603 | -0.208744843 | -0.154804107 | 0.04367476 | 0.395531096 | 0.00158663 | 0.003876217 | 0.045587473 | 0.104263939 |
| Espl1 | S | 1509 | -0.226539925 | -0.182090449 | -0.242982956 | -0.016443031 | -0.060892507 | 0.050547411 | 0.069057261 | 0.010187068 | 0.845925112 | 0.463910325 |
| Atxn2l | S | 453 | 0.046845612 | -0.18078134 | -0.22004516 | -0.266890772 | -0.039263819 | 0.006766697 | 0.000604575 | 7.25959E-08 | 2.85114E-05 | 0.043478367 |
| Tmsb4x | T | 29 | -0.472117616 | -0.101799543 | -0.332506721 | 0.139610895 | -0.230707177 | 0.003060926 | 0.606478302 | 0.233061228 | 0.565261981 | 0.419499243 |
| Adss Adssl1 | S | 449;448 | -0.263351921 | -0.159088461 | -0.37597366 | -0.112621739 | -0.216885199 | 0.138349863 | 0.014195513 | 0.059483154 | 0.507084002 | 0.165925505 |
| Tacc2 | S | 759 | -0.020562774 | -0.30941471 | -0.219459417 | -0.198896644 | 0.089955293 | 0.720991817 | 0.000712721 | 0.010983142 | 0.019442544 | 0.141934818 |
| Hsp90aa1 | S | 263 | -0.242013293 | -0.242688446 | -0.178518387 | 0.063494907 | 0.064170059 | 0.091704366 | 0.014432494 | 0.027931847 | 0.564559088 | 0.388070606 |
| Lrba | S | 1000 | -0.39198002 | -0.112929008 | -0.371464099 | 0.02051592 | -0.258535091 | 0.032871631 | 0.51558127 | 0.029675927 | 0.843531948 | 0.124433687 |
| Spag5 | S | 165 | 0.024497346 | -0.137297999 | -0.330759661 | -0.355257007 | -0.193461663 | 0.782354691 | 0.012459954 | 2.41595E-05 | 0.036306959 | 0.006166307 |
| Cryab | S | 59 | -0.146107655 | -0.23705279 | -0.202647058 | -0.056539403 | 0.034405732 | 0.069651973 | 0.005219438 | 0.032663963 | 0.441522042 | 0.575524028 |
| Rnf219 | S | 566 | 0.006658557 | -0.212737083 | -0.232517047 | -0.239175604 | -0.019779964 | 0.869556176 | 5.23699E-06 | 0.014207825 | 0.007391753 | 0.588101065 |
| Ncbp1 | S | 7 | 0.043478751 | -0.252901376 | -0.194351301 | -0.237830052 | 0.058550074 | 0.234417271 | 1.5926E-06 | 0.006750301 | 0.003017116 | 0.150241442 |
| Nav3 | S | 1042 | -0.126553079 | -0.249833075 | -0.257416233 | -0.130863154 | -0.007583158 | 0.107299367 | 0.029943077 | 0.0087743 | 0.04767736 | 0.921264848 |
| Phldb1 | T | 582 | -0.015978755 | -0.217129691 | -0.312760091 | -0.296781336 | -0.0956304 | 0.772186314 | 0.014462133 | 0.000468567 | 0.004143206 | 0.161952608 |
| Igfbp3 | S | 202 | -0.054335308 | -0.391232177 | -0.208297601 | -0.153962293 | 0.182934576 | 0.502547833 | 0.002209242 | 0.036142376 | 0.00673161 | 0.024967674 |
| Ahctf1 | S | 1381 | -0.081052561 | -0.236596033 | -0.225338335 | -0.144285774 | 0.011257699 | 0.055541482 | 0.002662608 | 0.02676779 | 0.096006307 | 0.865325065 |
| Top2b | T | 1307 | -0.185814561 | -0.261491814 | -0.148824418 | 0.036990143 | 0.112667395 | 0.165777199 | 0.000702848 | 0.078657576 | 0.740892807 | 0.141767486 |
| Fancd2 | S | 10 | -0.006993303 | -0.264306053 | -0.258251707 | -0.251258404 | 0.006054346 | 0.892436833 | 0.000984231 | 0.004176134 | 0.008913514 | 0.898416685 |
| Espl1 | S | 1497 | 0.022760031 | -0.237036108 | -0.444922078 | -0.467682108 | -0.207885969 | 0.909637818 | 0.039161877 | 0.001111686 | 0.102357049 | 0.045732272 |
| Gpatch4 | S | 410 | -0.008858432 | -0.268301206 | -0.415793029 | -0.406934597 | -0.147491824 | 0.783595726 | 0.012042158 | 0.007107088 | 0.001412616 | 0.04254798 |
| Abtb2 | T | 364 | -0.133583485 | -0.227171081 | -0.242051251 | -0.108467766 | -0.014880171 | 0.107112593 | 0.035370952 | 0.004161208 | 0.143151879 | 0.837017636 |
| Dennd4c | T | 1086 | -0.138723775 | -0.257503414 | -0.199365546 | -0.060641771 | 0.058137867 | 0.04706209 | 0.015358577 | 0.020353816 | 0.344055747 | 0.39695361 |
| Prrc2b | S | 168 | 0.067232089 | -0.458014047 | -0.47980555 | -0.547037638 | -0.021791503 | 0.06298075 | 0.005319288 | 0.012373177 | 0.006909447 | 0.790038098 |
| Nedd1 | S | 282 | -0.16111654 | -0.150092491 | -0.217741129 | -0.056624589 | -0.067648639 | 0.002998884 | 0.035414709 | 0.025922482 | 0.34484156 | 0.314004612 |
| Specc1 | S | 210 | -0.091287232 | -0.162805216 | -0.220622979 | -0.129335746 | -0.057817763 | 0.177502618 | 0.027467356 | 0.000130222 | 0.094361584 | 0.247260849 |
| Slc20a2 | S | 367 | 0.071668667 | -0.300798667 | -0.297074496 | -0.368743163 | 0.003724171 | 0.339286978 | 0.000156904 | 0.006577295 | 0.007894233 | 0.947838899 |
| Svil | S | 1046 | -0.273529943 | -0.104717927 | -0.383308402 | -0.109778458 | -0.278590475 | 0.06729371 | 0.362043553 | 0.006946732 | 0.320245395 | 0.029541121 |
| Plekhg3 | S | 502 | -0.238106608 | -0.119650625 | -0.534381955 | -0.296275348 | -0.414731331 | 0.098989147 | 0.436661213 | 0.014174866 | 0.099461276 | 0.040563468 |
| Rabl6 | S | 502 | -0.207474005 | -0.273267714 | -0.120134363 | 0.087339642 | 0.153133351 | 0.022088343 | 0.00257646 | 0.26599921 | 0.392771962 | 0.174851266 |
| Eif4b | S | 309 | -0.10966264 | -0.231431449 | -0.420331338 | -0.310668697 | -0.188899888 | 0.154922851 | 0.052712789 | 0.019443961 | 0.024551019 | 0.107601469 |
| Fra10ac1 | S | 243 | -0.037557476 | -0.200830762 | -0.363945553 | -0.326388077 | -0.163114792 | 0.30008455 | 0.047944631 | 0.001138675 | 0.007037362 | 0.084527882 |
| Wrnip1 | S | 75 | -0.129762182 | -0.283472336 | -0.120887632 | 0.00887455 | 0.162584704 | 0.029980709 | 0.000368148 | 0.281747406 | 0.928293395 | 0.183256532 |
| Ctnnd1 | S | 269 | -0.244088466 | -0.112912322 | -0.284665432 | -0.040576967 | -0.17175311 | 0.021926136 | 0.219199062 | 0.007306788 | 0.507833684 | 0.051137053 |
| Tpi1 | S | 130 | -0.11667743 | -0.317193292 | -0.33830348 | -0.22162605 | -0.021110187 | 0.176392521 | 0.027601415 | 0.039492979 | 0.10763064 | 0.859689944 |
| Nav1 | S | 454 | -0.107858118 | -0.109459857 | -0.323019059 | -0.215160941 | -0.213559201 | 0.10649209 | 0.131154749 | 0.000366625 | 0.020540285 | 0.032258127 |
| Tmem87a | S | 475 | 0.057086813 | -0.283501168 | -0.221377095 | -0.278463908 | 0.062124074 | 0.451563317 | 0.000237714 | 0.001002343 | 0.0312986 | 0.063793598 |
| Sipa1 | S | 77 | 0.014219289 | -0.413663276 | -0.345829656 | -0.360048945 | 0.067833619 | 0.854645982 | 0.033310769 | 0.002890987 | 0.022574482 | 0.487172597 |
| Mettl22 | S | 120 | 0.054999703 | -0.254357171 | -0.347781318 | -0.402781022 | -0.093424147 | 0.601711553 | 0.000726164 | 0.004459737 | 0.027341879 | 0.161959111 |
| Sccpdh | S | 215 | 0.036215822 | -0.362516966 | -0.494841252 | -0.531057075 | -0.132324286 | 0.374073194 | 0.001121088 | 0.060159362 | 0.054046377 | 0.430501852 |
| Crkl | Y | 251 | -0.105827698 | -0.231987001 | -0.267648861 | -0.161821162 | -0.035661859 | 0.161591547 | 0.000548571 | 0.091080361 | 0.226842143 | 0.743701971 |
| Aagab | S | 311 | -0.151878156 | -0.210969659 | -0.327549222 | -0.175671067 | -0.116579564 | 0.143083195 | 0.052521753 | 0.010029586 | 0.118786632 | 0.248343771 |
| Trmt13 | T | 385 | -0.022714496 | -0.246813674 | -0.292872187 | -0.270157691 | -0.046058513 | 0.659023304 | 0.003913875 | 0.003330322 | 0.008357538 | 0.459840566 |
| Eif4g1 | S | 1196 | 0.002622628 | -0.237423061 | -0.181884595 | -0.184507223 | 0.055538466 | 0.978889074 | 3.92607E-05 | 0.003480061 | 0.163752786 | 0.093737084 |
| Cpeb2 | S | 24 | -0.080772507 | -0.226461802 | -0.191782732 | -0.111010225 | 0.034679071 | 0.026995765 | 0.000253551 | 0.110843596 | 0.275024654 | 0.692755548 |
| Clspn | S | 785 | 0.015594404 | -0.367155452 | -0.259654447 | -0.275248851 | 0.107501005 | 0.878157285 | 0.000452853 | 0.037908038 | 0.066488044 | 0.259142905 |
| Fndc3b | S | 396 | -0.045940658 | -0.31690913 | -0.205213481 | -0.159272822 | 0.111695649 | 0.540808461 | 0.001577947 | 0.011358259 | 0.058187139 | 0.0127252 |
| Fgd6 | S | 697 | -0.322940869 | -0.158986796 | -0.368902471 | -0.045961602 | -0.209915675 | 0.092289402 | 0.352139164 | 0.012461893 | 0.759613723 | 0.231334414 |
| Nisch | S | 543 | -0.14322139 | -0.150705207 | -0.282081099 | -0.138859709 | -0.131375892 | 0.06143288 | 0.017679236 | 0.009632724 | 0.113082203 | 0.108077957 |
| Iws1 | S | 666 | 0.015031711 | -0.180344229 | -0.333827542 | -0.348859253 | -0.153483312 | 0.758236021 | 0.006218948 | 0.00025124 | 0.002758162 | 0.018431168 |
| Tab2 | S | 450 | -0.111210376 | -0.264775725 | -0.307409445 | -0.196199069 | -0.042633719 | 0.307949726 | 0.02037704 | 0.010906994 | 0.066715576 | 0.345205788 |
| Ankrd17 | S | 1562 | -0.096854001 | -0.210617762 | -0.470830315 | -0.373976314 | -0.260212553 | 0.362894679 | 0.192917943 | 0.003013693 | 0.006298228 | 0.117323443 |
| Eif4g1 | S | 1149 | 0.054451191 | -0.220007429 | -0.324557834 | -0.379009025 | -0.104550405 | 0.330643103 | 0.001920388 | 0.000291544 | 0.00129651 | 0.063135778 |
| Cdc42ep1 | T | 34 | -0.205693451 | -0.164869863 | -0.387433185 | -0.181739734 | -0.222563322 | 0.108370607 | 0.307768229 | 0.004884899 | 0.114534917 | 0.183745216 |
| Nvl | T | 149 | 0.088141038 | -0.345670694 | -0.370885046 | -0.459026084 | -0.025214351 | 0.342215126 | 0.00431135 | 0.002565783 | 0.005787746 | 0.750990997 |
| Dlgap4 | T | 975 | -0.329005049 | -0.158303759 | -0.597658713 | -0.268653664 | -0.439354954 | 0.015530881 | 0.389956343 | 0.190620514 | 0.482528003 | 0.296371576 |
| Srbd1 | S | 168 | -0.016212312 | -0.176768368 | -0.174353208 | -0.158140896 | 0.00241516 | 0.381670647 | 1.77693E-05 | 0.002116908 | 0.006970906 | 0.91873035 |
| Thrap3 | S | 552 | -0.082350858 | -0.151289075 | -0.164719077 | -0.082368219 | -0.013430002 | 0.003795341 | 0.00067097 | 0.016506707 | 0.099594379 | 0.721703417 |
| Ehmt2 | S | 287 | -0.130724648 | -0.300847483 | -0.187246199 | -0.056521552 | 0.113601284 | 0.134260179 | 0.003262199 | 0.060256246 | 0.526885626 | 0.204007477 |
| Akap10 | S | 96 | -0.287835541 | -0.107229084 | -0.448523891 | -0.160688351 | -0.341294808 | 0.059057126 | 0.505962128 | 0.014564063 | 0.239003372 | 0.075629007 |
| Sipa1 | S | 65 | 0.005367357 | -0.21529591 | -0.207628648 | -0.212996005 | 0.007667263 | 0.950335923 | 4.7796E-05 | 0.002862558 | 0.09070784 | 0.808600341 |
| Ube2e2 | S | 19 | -0.312404937 | -0.179507645 | -0.272104641 | 0.040300296 | -0.092596995 | 0.01809227 | 0.061591902 | 0.146877377 | 0.794548397 | 0.5503374 |
| Clasrp | S | 525 | -0.170462976 | -0.147468361 | -0.258556434 | -0.088093457 | -0.111088073 | 0.091671003 | 0.250076273 | 0.000269358 | 0.294884591 | 0.356280753 |
| Unc5b | S | 471 | -0.094228012 | -0.174639387 | -0.153878189 | -0.059650177 | 0.020761197 | 0.091280272 | 0.001004565 | 0.00208512 | 0.221281544 | 0.41893768 |
| Foxk1 | S | 243 | 0.067839254 | -0.235656371 | -0.202581272 | -0.270420526 | 0.033075099 | 0.152179298 | 8.99867E-05 | 4.55596E-05 | 0.009435766 | 0.193527319 |
| Ddx5 | S | 498 | -0.098511768 | -0.215581506 | -0.290730723 | -0.192218955 | -0.075149217 | 0.054080875 | 0.017227189 | 0.018869102 | 0.040230239 | 0.3042322 |
| Fam177a Fam177a2 | T | 60 | -0.462759222 | -0.22533718 | -0.308277447 | 0.154481776 | -0.082940267 | 0.097590424 | 0.064219387 | 0.18446247 | 0.548835175 | 0.671020931 |
| Acss2 | S | 267 | -0.065287817 | -0.202273163 | -0.298211913 | -0.232924095 | -0.09593875 | 0.281085189 | 0.017821005 | 0.00210965 | 0.012902619 | 0.181047344 |
| Slc6a6 | T | 603 | -0.278165736 | -0.264786478 | -0.568721691 | -0.290555956 | -0.303935213 | 0.055584862 | 0.222446319 | 0.16283507 | 0.401144125 | 0.400870463 |
| Srrm2 | S | 1315 | -0.082872837 | -0.245766032 | -0.305364066 | -0.22249123 | -0.059598035 | 0.209852714 | 0.001244257 | 0.069955288 | 0.136012755 | 0.604571486 |
| Atg16l1 | S | 269 | -0.266715 | -0.132785927 | -0.332800657 | -0.066085657 | -0.20001473 | 0.005826893 | 0.134655232 | 0.11134668 | 0.670904767 | 0.268600353 |
| Tln1 | S | 1641 | -0.141048107 | -0.145083185 | -0.168147481 | -0.027099374 | -0.023064295 | 1.70539E-05 | 0.140109245 | 0.031191351 | 0.515596124 | 0.764787136 |
| Sh3pxd2b | Y | 661 | -0.224892371 | -0.180474686 | -0.447025638 | -0.222133267 | -0.266550952 | 0.131915472 | 0.298508169 | 0.012500068 | 0.166175359 | 0.169764415 |
| Dab2 | S | 249 | -0.515405867 | -0.177314693 | -0.309393432 | 0.206012435 | -0.132078739 | 0.05467402 | 0.064337658 | 0.289006799 | 0.485856586 | 0.606868682 |
| Map7d1 | S | 454 | 0.027526477 | -0.268107498 | -0.344878947 | -0.372405424 | -0.076771449 | 0.440544606 | 0.002204482 | 0.003181179 | 0.00100181 | 0.137142874 |
| Macf1 | S | 1376 | 0.065383613 | -0.340284361 | -0.304454911 | -0.369838523 | 0.03582945 | 0.598736442 | 0.000401686 | 0.020198669 | 0.046818699 | 0.640054036 |
| Myh10 | S | 1952 | -0.394241653 | -0.148397715 | -0.426064554 | -0.031822901 | -0.277666839 | 0.031803328 | 0.471830675 | 0.079701714 | 0.860304589 | 0.248565853 |
| Ahctf1 | S | 1541 | 0.072152176 | -0.434553469 | -0.436319114 | -0.50847129 | -0.001765645 | 0.213431926 | 0.007503386 | 0.010253169 | 0.009642544 | 0.986635074 |
| Cd3eap | S | 315 | -0.004655719 | -0.266139567 | -0.28965946 | -0.285003741 | -0.023519893 | 0.874293101 | 0.001463137 | 0.007252776 | 0.014814325 | 0.682251067 |

| | | | | | | | | | | | | |
|---|---|---|---|---|---|---|---|---|---|---|---|---|
| Nes | T | 597 | -0.030071801 | -0.301805417 | -0.236890843 | -0.206819042 | 0.064914574 | 0.642641474 | 0.002539491 | 0.007336002 | 0.008049581 | 0.164147537 |
| Tpr | S | 1259 | -0.015361822 | -0.23390339 | -0.29597713 | -0.280615308 | -0.06207374 | 0.736948649 | 0.000551499 | 0.012053456 | 0.01797634 | 0.36539067 |
| Abcc1 | S | 289 | -0.087576804 | -0.142463747 | -0.147611657 | -0.060034853 | -0.00514791 | 0.001833906 | 0.001249767 | 0.010819542 | 0.137133768 | 0.876916418 |
| Cdk13 | S | 260 | -0.068193469 | -0.301090835 | -0.332019374 | -0.263825905 | -0.03092854 | 0.405852508 | 0.025775324 | 0.009847115 | 0.03533415 | 0.728226449 |
| Akap1 | S | 103 | -0.063763807 | -0.198712706 | -0.154981733 | -0.091217926 | 0.043730973 | 0.08332376 | 0.000453866 | 0.005309476 | 0.044425638 | 0.210847619 |
| Specc1 | S | 133 | -0.024812278 | -0.245801162 | -0.204297466 | -0.179485188 | 0.041503696 | 0.555622702 | 0.000605366 | 0.003886761 | 0.011995476 | 0.307719931 |
| Mink1 | S | 894 | -0.024022497 | -0.204787474 | -0.186096015 | -0.162073518 | 0.018691458 | 0.34147696 | 0.001589959 | 0.00031337 | 0.000377239 | 0.539383686 |
| Frmd4a | S | 675 | -0.207988902 | -0.133456659 | -0.349809607 | -0.141820704 | -0.216352948 | 0.024658196 | 0.247294883 | 0.013685908 | 0.176873065 | 0.114330483 |
| Rell1 | S | 158 | -0.104090188 | -0.297497639 | -0.325044204 | -0.220954016 | -0.027546565 | 0.097982594 | 0.085071784 | 0.013210047 | 0.058668154 | 0.835458341 |
| Rnf213 | T | 46 | -0.143211994 | -0.259200764 | -0.241879388 | -0.098667394 | 0.017321376 | 0.114992366 | 0.013402927 | 0.03021745 | 0.207772126 | 0.790837491 |
| Ctnnb1 | Y | 716 | -0.213626538 | -0.248560452 | -0.410075477 | -0.19644894 | -0.161515025 | 0.112711289 | 0.120565691 | 0.045557603 | 0.24527405 | 0.356929447 |
| Rbbp6 | S | 1705 | -0.064031946 | -0.342232046 | -0.343561342 | -0.279529396 | -0.001329296 | 0.519260616 | 0.003911585 | 0.07203659 | 0.117110845 | 0.992137368 |
| Sgta | S | 307 | 0.077939741 | -0.254278333 | -0.379684232 | -0.457623974 | -0.1254059 | 0.218346238 | 0.000121153 | 0.007517527 | 0.00338698 | 0.142123672 |
| Plekho2 | S | 211 | -0.057368811 | -0.158068161 | -0.215453681 | -0.158084869 | -0.05738552 | 0.405749205 | 0.019444181 | 2.33252E-05 | 0.097410486 | 0.216118901 |
| Mcrip2 | T | 86 | -0.060048829 | -0.263992446 | -0.315692051 | -0.255643222 | -0.051699605 | 0.396301234 | 0.001055857 | 0.056813092 | 0.07187833 | 0.604199038 |
| Tacc2 | T | 934 | -0.174930396 | -0.161799774 | -0.183816112 | -0.008885715 | -0.022016337 | 0.069430256 | 0.008814045 | 0.00572556 | 0.899816639 | 0.598423519 |
| Fxr1 | T | 430 | 0.017927295 | -0.215455916 | -0.319842144 | -0.337769438 | -0.104386227 | 0.887779571 | 0.069891264 | 3.14743E-05 | 0.088239131 | 0.263531368 |
| Ctnnd1 | S | 252 | -0.3045576 | -0.063438571 | -0.344601439 | -0.040043839 | -0.281162867 | 0.017670854 | 0.631366984 | 0.010612736 | 0.48994705 | 0.068194639 |
| Rplp0 | S | 307 | -0.076580674 | -0.210435979 | -0.139969714 | -0.06338904 | 0.070466265 | 0.095852253 | 0.00049701 | 0.006443287 | 0.12750565 | 0.058711586 |
| Pym1 | S | 138 | -0.033909982 | -0.385833975 | -0.091785829 | -0.057875847 | 0.294048146 | 0.732672543 | 5.62294E-05 | 0.078396319 | 0.561133349 | 0.001263922 |
| Sipa1l1 | S | 1412 | -0.024411947 | -0.132475764 | -0.156180642 | -0.131768694 | -0.023704877 | 0.174825371 | 0.000193516 | 3.93683E-05 | 0.000512614 | 0.180408925 |
| Crip2 | S | 115 | -0.436955948 | 0.010924326 | -0.548821215 | -0.111865267 | -0.559745541 | 0.030203932 | 0.961993344 | 0.043222247 | 0.548407457 | 0.065929071 |
| Atp2b1 | S | 17 | -0.237874832 | -0.116218174 | -0.492980081 | -0.25510525 | -0.376761907 | 0.091763791 | 0.479620305 | 0.011683334 | 0.114127931 | 0.075379985 |
| Slc25a3 | S | 31 | -0.153860346 | -0.300218377 | -0.212396731 | -0.058536385 | 0.087821646 | 0.181089233 | 0.070207716 | 0.002713557 | 0.546998868 | 0.447169844 |
| Wdr44 | S | 50 | -0.062726348 | -0.201468663 | -0.247399466 | -0.184673117 | -0.045930802 | 0.08353526 | 0.004692389 | 0.008134383 | 0.038398914 | 0.431275411 |
| Ywhaz | S | 207 | -0.168501324 | -0.194568871 | -0.225171612 | -0.056670288 | -0.030602741 | 0.031289668 | 0.043521861 | 0.014177366 | 0.394841247 | 0.68848641 |
| Wiz | S | 1184 | 0.065055253 | -0.19565759 | -0.255637731 | -0.320692984 | -0.05998014 | 0.54322978 | 0.000861106 | 8.5256E-05 | 0.06398748 | 0.094266803 |
| Wiz | T | 1186 | 0.065055253 | -0.19565759 | -0.255637731 | -0.320692984 | -0.05998014 | 0.54322978 | 0.000861106 | 8.5256E-05 | 0.06398748 | 0.094266803 |
| Klf10 | S | 248 | -0.049279627 | -0.204745475 | -0.235071171 | -0.185791544 | -0.030325696 | 0.379070852 | 0.00314728 | 0.001836783 | 0.021542542 | 0.186284036 |
| Frmd4a | S | 600 | -0.314496938 | -0.182794024 | -0.472973055 | -0.158476117 | -0.290179031 | 0.055248579 | 0.284977162 | 0.081840511 | 0.455720588 | 0.229544251 |
| Tuba1c | S | 439 | -0.077666597 | -0.201137952 | -0.231426767 | -0.15376017 | -0.030288815 | 0.174103902 | 0.00277353 | 0.007566551 | 0.044170337 | 0.549396823 |
| Flrt2 | S | 598 | -0.075294143 | -0.317568574 | -0.490259025 | -0.414964882 | -0.172690451 | 0.730439607 | 0.041191422 | 0.023386478 | 0.128790363 | 0.288561209 |
| Mcm3ap | S | 1368 | -0.370110769 | -0.192176727 | -0.165051726 | 0.205059043 | 0.027125001 | 0.001788881 | 0.204642216 | 0.318409924 | 0.234941136 | 0.87557895 |
| Map1a | T | 2064 | -0.103908743 | -0.120100873 | -0.133362411 | -0.029453668 | -0.013261539 | 0.02766224 | 0.00068177 | 0.000859528 | 0.384264723 | 0.490673372 |
| Cep170 | T | 875 | -0.051178551 | -0.25199678 | -0.280579435 | -0.229400884 | -0.028582655 | 0.190892798 | 0.002033269 | 0.033668715 | 0.043845376 | 0.693573999 |
| Hectd1 | S | 1350 | 0.016793962 | -0.271776066 | -0.255301097 | -0.272095059 | 0.016474969 | 0.731100855 | 0.000104034 | 0.018472867 | 0.01637513 | 0.786610575 |
| Mapk1 | Y | 185 | -0.122374162 | -0.233398984 | -0.339785861 | -0.217411699 | -0.106386878 | 0.132651251 | 0.024908851 | 0.030340202 | 0.108988044 | 0.359250751 |
| Hjurp A730008H23Ril | S | 499 | -0.092467608 | -0.203467636 | -0.166762835 | -0.074295227 | 0.0367048 | 0.06663101 | 1.87435E-05 | 0.189747675 | 0.484889692 | 0.712119234 |
| Tpd52 | S | 223 | -0.136247098 | -0.107479585 | -0.391491279 | -0.255244181 | -0.284011694 | 0.043778711 | 0.09250975 | 0.009112637 | 0.047402016 | 0.037171381 |
| Nufip2 | S | 376 | -0.069179386 | -0.285655885 | -0.3944531 | -0.325273714 | -0.108797215 | 0.36044661 | 0.013748774 | 0.036724218 | 0.049318941 | 0.375830106 |
| Hnrnpm | S | 85 | 0.004955285 | -0.463692109 | -0.417063191 | -0.422018475 | 0.046628918 | 0.972330521 | 0.003862097 | 0.131580308 | 0.131782475 | 0.827573949 |
| Rnaseh1 | S | 76 | -0.247873917 | -0.135174365 | -0.237981024 | 0.009892893 | -0.102806659 | 0.004797886 | 0.025607761 | 0.161604673 | 0.940980036 | 0.465087096 |
| Stip1 | T | 198 | -0.088221183 | -0.154069233 | -0.149758357 | -0.061537174 | 0.004310876 | 0.080640938 | 0.003428769 | 0.000282524 | 0.167093584 | 0.867999808 |
| Usp42 | S | 936 | -0.032439955 | -0.176040725 | -0.106207945 | -0.073767989 | 0.06983278 | 0.188669506 | 8.05646E-05 | 9.57202E-05 | 0.04100349 | 0.018548625 |
| Ssrp1 | S | 647 | -0.098171305 | -0.230501795 | -0.24710056 | -0.148929255 | -0.016598765 | 0.093503733 | 0.007832403 | 0.022006382 | 0.104575675 | 0.824152688 |
| Ticrr | S | 1778 | -0.09965017 | -0.173869967 | -0.343103471 | -0.243453301 | -0.169233505 | 0.194284832 | 0.045362025 | 0.00412105 | 0.028019945 | 0.061888412 |
| Cdk5rap2 | S | 485 | -0.284871134 | -0.139840994 | -0.377606823 | -0.09273569 | -0.23776583 | 0.07667315 | 0.290251056 | 0.017223019 | 0.413653454 | 0.040123003 |
| Tnks1bp1 | S | 1375 | -0.045118783 | -0.141952545 | -0.190450023 | -0.14533124 | -0.048497478 | 0.228143948 | 3.03302E-05 | 0.003574852 | 0.014738807 | 0.146455367 |
| Tbc1d15 | S | 205 | -0.11762371 | -0.141037008 | -0.473623823 | -0.356000113 | -0.332586815 | 0.298303792 | 0.242701475 | 0.002493305 | 0.007362912 | 0.016007247 |
| Snw1 | S | 518 | -0.063609234 | -0.203409249 | -0.346749467 | -0.283140232 | -0.143340218 | 0.37790598 | 0.015113131 | 0.005788588 | 0.018889937 | 0.091056075 |
| Map2k1 | T | 286 | -0.128562478 | -0.113474165 | -0.198027545 | -0.069465067 | -0.084553379 | 0.006347748 | 0.003877521 | 0.0155542 | 0.176376534 | 0.118850921 |
| Gtf3c5 | S | 333 | -0.141894199 | -0.19420869 | -0.247100444 | -0.105206245 | -0.052891754 | 0.055348855 | 0.018681462 | 0.019286759 | 0.178684964 | 0.443742003 |
| Gtf3c5 | Y | 329 | -0.141894199 | -0.19420869 | -0.247100444 | -0.105206245 | -0.052891754 | 0.055348855 | 0.018681462 | 0.019286759 | 0.178684964 | 0.443742003 |
| Tjp2 | S | 441 | -0.206256802 | -0.358706053 | -0.383227243 | -0.17697044 | -0.02452119 | 0.354814432 | 0.07375698 | 0.048228388 | 0.442490696 | 0.889448563 |
| Map1a | T | 1162 | -0.109202981 | -0.13409165 | -0.112783643 | -0.003580662 | 0.021308008 | 0.004346502 | 0.009686188 | 0.000347182 | 0.861861667 | 0.463053128 |
| P2ry2 | S | 356 | -0.044249918 | -0.126264067 | -0.494075087 | -0.449825169 | -0.36781102 | 0.608024046 | 0.27372473 | 0.000442013 | 0.001720705 | 0.026337358 |
| Fbxo41 | S | 540 | -0.372550027 | -0.124235073 | -0.498764443 | -0.126214416 | -0.37452937 | 0.054986001 | 0.561855525 | 0.056131412 | 0.526839896 | 0.155665666 |
| Fbxo41 | S | 545 | -0.372550027 | -0.124235073 | -0.498764443 | -0.126214416 | -0.37452937 | 0.054986001 | 0.561855525 | 0.056131412 | 0.526839896 | 0.155665666 |
| Abca1 | S | 2249 | -0.26305287 | -0.196527037 | -0.420122751 | -0.157069881 | -0.223595713 | 0.217319754 | 0.068817143 | 0.04385388 | 0.462542195 | 0.194472495 |
| Prkd1 | S | 243 | -0.166012703 | -0.201206334 | -0.135795451 | 0.030217252 | 0.065410883 | 0.014913435 | 0.007906694 | 0.03233476 | 0.26343327 | 0.147859204 |
| Fnip1 | S | 763 | -0.038852896 | -0.26163317 | -0.340103091 | -0.301250194 | -0.078469921 | 0.499512525 | 0.001781883 | 0.038565694 | 0.044614638 | 0.44193625 |
| Atg2b | T | 398 | -0.002643296 | -0.276129869 | -0.321023103 | -0.318379806 | -0.044893233 | 0.983589382 | 0.000922239 | 0.022167899 | 0.08059538 | 0.595503687 |
| Pdlim2 | S | 114 | 0.109172152 | -0.54479708 | -0.410118979 | -0.519291131 | 0.134678101 | 0.289672114 | 0.021225273 | 0.00729368 | 0.004857308 | 0.401188407 |
| Arhgap22 | T | 423 | -0.080813071 | -0.213265972 | -0.35312246 | -0.272309389 | -0.139856488 | 0.172127492 | 0.016480089 | 0.018299068 | 0.035975254 | 0.170061243 |
| Camsap2 | S | 836 | -0.036741994 | -0.267959662 | -0.271420851 | -0.234678857 | -0.003461189 | 0.477209599 | 0.00437602 | 0.009146468 | 0.013882464 | 0.945939906 |
| Map1b | S | 1401 | -0.11753416 | -0.218230753 | -0.222005022 | -0.104470862 | -0.003774269 | 0.045147813 | 0.005145104 | 0.047207663 | 0.243389896 | 0.961420864 |
| Zc3h8 | T | 66 | 0.043008743 | -0.285688846 | -0.260823368 | -0.303832111 | 0.024865479 | 0.294636128 | 0.007438052 | 0.00010624 | 0.001149839 | 0.642980563 |
| Zcchc8 | S | 423 | 0.08097406 | -0.260136439 | -0.330845908 | -0.411819969 | -0.070709469 | 0.07219283 | 0.000184614 | 0.0015276 | 0.001918865 | 0.201307863 |
| Tns3 | S | 1288 | -0.089568756 | -0.147842618 | -0.297323002 | -0.207754246 | -0.149480384 | 0.099195626 | 0.032385129 | 0.002566834 | 0.03210402 | 0.04975031 |
| Nufip2 | S | 213 | -0.0119778 | -0.16581357 | -0.143079546 | -0.131101746 | 0.022734024 | 0.545517992 | 0.000231277 | 2.33102E-05 | 0.004446624 | 0.235112165 |
| Nufip2 | S | 215 | -0.023247961 | -0.16986355 | -0.138890357 | -0.115642396 | 0.030973193 | 0.354051492 | 5.26038E-05 | 0.000360596 | 0.016489714 | 0.076314855 |
| Eml1 | S | 139 | -0.16131253 | -0.10616758 | -0.207773202 | -0.046460672 | -0.101605621 | 0.007208743 | 0.1131091 | 0.002262927 | 0.293861052 | 0.129025601 |
| Nop58 | S | 351 | -0.046306725 | -0.129350896 | -0.140866368 | -0.094559643 | -0.011515472 | 0.044401685 | 0.000338626 | 0.000182592 | 0.000973144 | 0.427520498 |
| Trp53bp2 | S | 479 | -0.310860263 | -0.129271028 | -0.371319202 | -0.060458939 | -0.242048174 | 0.051598647 | 0.421474789 | 0.020244657 | 0.501290041 | 0.123816567 |
| Dvl2 | S | 620 | -0.19126641 | -0.192880742 | -0.373395563 | -0.182129153 | -0.180514821 | 0.175236686 | 0.221037312 | 0.006106068 | 0.168927431 | 0.228793856 |
| Ctcf | T | 518 | -0.16362519 | -0.160627094 | -0.357752282 | -0.194127093 | -0.197125188 | 0.168583586 | 0.030530407 | 0.016931064 | 0.123956893 | 0.052926034 |
| Srrm2 | S | 1288 | -0.04197845 | -0.165681469 | -0.18718623 | -0.145207781 | -0.021504761 | 0.146136263 | 2.41838E-05 | 0.015265717 | 0.023565487 | 0.568333048 |
| Map3k4 | S | 492 | -0.161175075 | -0.133142906 | -0.203510612 | -0.042335537 | -0.070367706 | 0.013272561 | 0.112018906 | 0.002111271 | 0.372403795 | 0.326590738 |
| Ube2o | S | 836 | -0.018291939 | -0.19024595 | -0.334096868 | -0.315804929 | -0.143850918 | 0.77928752 | 0.001628273 | 0.005402375 | 0.011426469 | 0.075083445 |
| Plekhg3 | S | 76 | 0.103807188 | -0.321945098 | -0.242372006 | -0.346179194 | 0.079573091 | 0.022915655 | 4.45183E-05 | 0.00073009 | 0.000770463 | 0.071860495 |
| Rai14 | S | 374 | -0.137484582 | -0.286157134 | -0.156265628 | -0.018781047 | 0.129891506 | 0.121286998 | 0.000959368 | 0.150330112 | 0.850712753 | 0.210770284 |
| Mynn | S | 263 | 0.048298473 | -0.320935542 | -0.40207852 | -0.450376993 | -0.081142978 | 0.609724477 | 0.02943494 | 0.001970435 | 0.009105774 | 0.44444706 |
| Larp1 | S | 826 | 0.007013868 | -0.206564415 | -0.190491025 | -0.197504893 | 0.01607339 | 0.780020911 | 0.000369795 | 0.000253679 | 0.002261172 | 0.560800165 |
| Birc6 | S | 593 | -0.079838191 | -0.158532653 | -0.191613405 | -0.111775214 | -0.033080753 | 0.067649786 | 0.00472178 | 0.001682853 | 0.003049441 | 0.224847384 |
| Cdv3 | S | 58 | -0.072715812 | -0.212751185 | -0.388679678 | -0.315963866 | -0.175928493 | 0.622255786 | 0.033851392 | 0.004253795 | 0.106288102 | 0.093449872 |
| Zw10 | S | 438 | -0.131025674 | -0.165957728 | -0.138530143 | -0.007504469 | 0.027427585 | 0.001322848 | 0.005212061 | 0.059747845 | 0.879170478 | 0.617268889 |
| Elac2 | S | 815 | -0.315380369 | -0.153951407 | -0.258666648 | 0.05671372 | -0.104715241 | 0.096965508 | 0.068135253 | 0.022660466 | 0.685559693 | 0.245204781 |
| Twistnb | S | 306 | -0.018390138 | -0.174724247 | -0.22170929 | -0.203319153 | -0.046985043 | 0.49911461 | 0.007466619 | 7.37032E-05 | 0.001035708 | 0.239753511 |
| Arhgef2 | S | 174 | -0.054515474 | -0.245810275 | -0.314675825 | -0.260160352 | -0.068865551 | 0.496213537 | 0.016965975 | 0.004893277 | 0.003396849 | 0.216859779 |
| Ssh1 | S | 963 | -0.075181085 | -0.243745572 | -0.225597207 | -0.150416122 | 0.018148366 | 0.221680932 | 0.014726772 | 0.003220335 | 0.054600559 | 0.759755751 |
| Crtc2 | S | 461 | -0.058124123 | -0.256385059 | -0.155742843 | -0.09761872 | 0.100642216 | 0.212874827 | 0.000340899 | 0.025176539 | 0.103290454 | 0.095722931 |
| Hectd1 | S | 357 | -0.031304476 | -0.21281879 | -0.294531652 | -0.263227176 | -0.081712862 | 0.733640914 | 0.021764878 | 0.000550062 | 0.05926965 | 0.236239202 |
| Plec | S | 1443 | -0.102523707 | -0.162797115 | -0.209558701 | -0.107034993 | -0.046761585 | 0.090342589 | 0.008579378 | 0.002769336 | 0.078105132 | 0.00898304 |
| Zc3h8 | S | 67 | 0.034487974 | -0.334680927 | -0.300982471 | -0.335470445 | 0.033698456 | 0.523694418 | 0.006709484 | 0.002504079 | 0.004162119 | 0.590254137 |

| | | | | | | | | | | | | |
|---|---|---|---|---|---|---|---|---|---|---|---|---|
| Sipa1l3 | S | 140 | -0.204377247 | -0.145150709 | -0.262740074 | -0.058362827 | -0.117589364 | 0.018230429 | 0.061372704 | 0.033270056 | 0.492443956 | 0.220364787 |
| Eif2s2 | S | 67 | -0.045790555 | -0.174463951 | -0.223685115 | -0.17789456 | -0.049221164 | 0.205532474 | 0.002348714 | 0.001784507 | 0.006290936 | 0.120691377 |
| Aagab | S | 312 | -0.157745563 | -0.192683953 | -0.31188305 | -0.154137487 | -0.119199098 | 0.127955999 | 0.062920127 | 0.010905761 | 0.153411391 | 0.2275309 |
| Zfp507 | S | 376 | -0.039946976 | -0.211128767 | -0.202421933 | -0.162474957 | 0.008706834 | 0.051342763 | 0.007368715 | 0.001998404 | 0.007162884 | 0.819493432 |
| Abi1 | S | 225 | -0.099334137 | -0.150699184 | -0.406026822 | -0.306692685 | -0.255327638 | 0.294852685 | 0.238491089 | 0.001031363 | 0.017363615 | 0.086080546 |
| Map1b | S | 2068 | -0.094216212 | -0.124487232 | -0.242479769 | -0.148263557 | -0.117992537 | 0.030867479 | 0.017520292 | 0.002593128 | 0.046820121 | 0.051785371 |
| Cask | S | 577 | -0.036240939 | -0.339745314 | -0.190223613 | -0.153982674 | 0.149521702 | 0.693230619 | 0.000479438 | 0.049700449 | 0.167185552 | 0.106094173 |
| Pdlim5 | S | 137 | 0.040689341 | -0.270912905 | -0.303277986 | -0.343967327 | -0.032365082 | 0.422114185 | 0.00061843 | 0.005583275 | 0.007859817 | 0.592257734 |
| Eno1 Eno1b | S | 263 | -0.192730966 | -0.197232704 | -0.269222617 | -0.076491651 | -0.071989914 | 0.031054807 | 0.04444436 | 0.056296167 | 0.468706598 | 0.509785158 |
| Mtmr4 | S | 610 | 0.039850816 | -0.23675172 | -0.239679663 | -0.27953048 | -0.002927943 | 0.261612092 | 0.000195534 | 0.000954906 | 0.002625491 | 0.931446725 |
| Inpp5b | S | 156 | -0.023696533 | -0.169113768 | -0.282401914 | -0.258705381 | -0.113288146 | 0.646616958 | 0.001908385 | 0.001560185 | 0.007361269 | 0.07348771 |
| Tnrc6a | S | 1255 | -0.231094736 | -0.131617141 | -0.479656343 | -0.248561607 | -0.348039202 | 0.215597596 | 0.390941905 | 0.007386847 | 0.16996665 | 0.043787581 |
| Ppp1r18 | S | 140 | -0.462273069 | -0.08565085 | -0.373490666 | 0.088782403 | -0.287839816 | 0.026695795 | 0.666523959 | 0.068468626 | 0.423569549 | 0.128678648 |
| Clasp1 | S | 1193 | -0.088455048 | -0.183241639 | -0.118784787 | -0.030329739 | 0.064456852 | 0.010742483 | 0.000168617 | 0.055548056 | 0.499031391 | 0.207185993 |
| Snx33 | S | 187 | -0.085636298 | -0.194723458 | -0.310406659 | -0.22477036 | -0.115683201 | 0.195382764 | 0.024789492 | 0.006122252 | 0.025762795 | 0.146200885 |
| Snx33 | S | 192 | -0.085636298 | -0.194723458 | -0.310406659 | -0.22477036 | -0.115683201 | 0.195382764 | 0.024789492 | 0.006122252 | 0.025762795 | 0.146200885 |
| Nav3 | S | 1040 | -0.184631631 | -0.147786612 | -0.35109328 | -0.166461649 | -0.203306668 | 0.037426065 | 0.096289284 | 0.02937782 | 0.189337448 | 0.131360966 |
| Atad2 | S | 158 | -0.092185125 | -0.149020695 | -0.346458413 | -0.254273288 | -0.197437718 | 0.101839631 | 0.153659684 | 0.001631703 | 0.013506003 | 0.088551066 |
| Cwc22 | S | 60 | -0.123839978 | -0.178158982 | -0.204311582 | -0.080471604 | -0.0261526 | 0.014726227 | 0.003260328 | 0.066340682 | 0.342113074 | 0.735458624 |
| Zwilch | S | 536 | -0.139861512 | -0.161412196 | -0.270071066 | -0.130209554 | -0.10865887 | 0.023283732 | 0.057155763 | 0.014201411 | 0.128193411 | 0.216457629 |
| Lima1 | S | 602 | -0.077822322 | -0.221966523 | -0.282653099 | -0.204830777 | -0.060686576 | 0.246941946 | 0.010990147 | 0.009580589 | 0.044447861 | 0.363239076 |
| Myh10 | S | 1935 | -0.193493048 | -0.220598651 | -0.415490386 | -0.221997338 | -0.194891735 | 0.335413464 | 0.163085986 | 0.009940525 | 0.26372925 | 0.172388345 |
| Nrd1 | S | 103 | -0.070360569 | -0.21416508 | -0.328846006 | -0.258485437 | -0.114680926 | 0.401626738 | 0.00506446 | 0.016833062 | 0.04730976 | 0.222593545 |
| Lbr | S | 105 | -0.20603903 | -0.11863965 | -0.079769334 | 0.126269695 | 0.038870315 | 0.000453887 | 0.012185802 | 0.047578394 | 0.007541905 | 0.229612755 |
| Nifk | T | 261 | -0.022845165 | -0.164739363 | -0.246406153 | -0.223560988 | -0.08166679 | 0.607215863 | 0.021588422 | 4.31825E-05 | 0.013851721 | 0.119641056 |
| Map3k1 | S | 915 | -0.082327015 | -0.193514197 | -0.177492863 | -0.095165848 | 0.016021334 | 0.046042038 | 0.011548706 | 0.002194786 | 0.038820778 | 0.73486527 |
| Tnks1bp1 | T | 238 | -0.110999989 | -0.125350144 | -0.349932347 | -0.238932358 | -0.224582203 | 0.018038973 | 0.012110493 | 0.039507351 | 0.099836619 | 0.109979576 |
| Eps8 | S | 660 | -0.174209275 | -0.246375148 | -0.140410792 | 0.033798482 | 0.105964356 | 0.023474416 | 0.020545277 | 0.034552064 | 0.527379025 | 0.190097412 |
| Abi2 | S | 301 | -0.287204578 | -0.029636744 | -0.358985492 | -0.071780914 | -0.329348748 | 0.026216073 | 0.816017751 | 0.005253124 | 0.423539298 | 0.051813796 |
| Map1b | S | 1775 | -0.076061068 | -0.184353392 | -0.175519047 | -0.09945798 | 0.008834345 | 0.087814536 | 0.000749902 | 0.010881777 | 0.081349932 | 0.827102517 |
| Plec | T | 4037 | -0.505580026 | -0.01991236 | -0.353025604 | 0.152554422 | -0.333113244 | 0.033878491 | 0.918946412 | 0.028152608 | 0.366127284 | 0.145707614 |
| Map1a | S | 1789 | -0.094463121 | -0.127140372 | -0.110852822 | -0.016389701 | 0.01628755 | 0.002876896 | 0.00031452 | 0.009827526 | 0.539654139 | 0.523940876 |
| Ripor1 | S | 875 | -0.111617177 | -0.253628322 | -0.226650566 | -0.115033388 | 0.026977756 | 0.081094933 | 0.004637403 | 0.070966368 | 0.214748017 | 0.726758516 |
| Wiz | S | 1045 | -0.0335849 | -0.137159889 | -0.164244776 | -0.130659876 | -0.027084887 | 0.340723302 | 5.01022E-05 | 0.000511608 | 0.022719831 | 0.199937066 |
| Slc2a1 | S | 490 | 0.00691918 | -0.266228691 | -0.293886427 | -0.300805606 | -0.027657736 | 0.826247916 | 0.006831321 | 0.00195175 | 0.003192653 | 0.619577257 |
| Bicd2 | S | 604 | -0.137283771 | -0.170033089 | -0.319979814 | -0.182696042 | -0.149946725 | 0.160879025 | 0.023249518 | 0.013992813 | 0.116389927 | 0.131716937 |
| Nav3 | S | 275 | -0.082341887 | -0.136603666 | -0.196210499 | -0.113868612 | -0.059606833 | 0.101388942 | 0.007177992 | 0.00054672 | 0.049844935 | 0.094317129 |
| Vamp1 Vamp2 Vamp | S | 48;63;61 | -0.000651358 | -0.305998828 | -0.237725898 | -0.237074541 | 0.068272929 | 0.978516931 | 0.004656975 | 0.002769378 | 0.009691187 | 0.239101139 |
| Ogfr | S | 359 | -0.072345198 | -0.154441255 | -0.331940461 | -0.259595263 | -0.177499205 | 0.099026683 | 0.025611361 | 0.005821423 | 0.034280106 | 0.05339716 |
| Katnbl1 | S | 56 | -0.012089007 | -0.189163541 | -0.249535271 | -0.237446264 | -0.060371729 | 0.895067784 | 0.002676018 | 0.000529931 | 0.082328106 | 0.086917441 |
| Mtrex | S | 83 | -0.021388735 | -0.263110946 | -0.287001208 | -0.265612473 | -0.023890263 | 0.716890787 | 0.002369339 | 0.012508515 | 0.021380047 | 0.7383872 |
| Smg7 | S | 737 | 0.025743732 | -0.281941206 | -0.38519855 | -0.410942282 | -0.103257344 | 0.545793841 | 0.005160644 | 0.007592237 | 0.011034844 | 0.234924717 |
| Necap2 | S | 252 | -0.010750547 | -0.157197012 | -0.328710503 | -0.317959956 | -0.171513491 | 0.777348327 | 0.014841996 | 0.000418921 | 0.002730791 | 0.019456163 |
| Aldoa | S | 46 | -0.178622393 | -0.194755156 | -0.154735913 | 0.02388648 | 0.040019243 | 0.013567281 | 0.086580942 | 0.005474671 | 0.58003191 | 0.62449335 |
| Lrig1 | S | 957 | 0.00859453 | -0.270102372 | -0.28011687 | -0.2887114 | -0.010014498 | 0.808403199 | 0.003194673 | 0.003423421 | 0.002123453 | 0.808971068 |
| Phldb1 | S | 467 | -0.415530041 | 0.11094373 | -0.530476655 | -0.114946614 | -0.641420385 | 0.019307331 | 0.65452078 | 0.024950547 | 0.4589649 | 0.058069277 |
| Cluh | S | 1300 | -0.065271811 | -0.158980447 | -0.168862451 | -0.103590639 | -0.009882003 | 0.014354172 | 0.10667596 | 2.4141E-05 | 0.005361168 | 0.882826965 |
| Tubb3 Tubb4a Tubb4 | S | 172;172;172;172 | -0.415262784 | -0.365107319 | -0.048113395 | 0.367149388 | 0.316993923 | 0.016192094 | 0.045733088 | 0.838820092 | 0.18764754 | 0.227726747 |
| Dnaaf2 | S | 681 | -0.070934471 | -0.285615924 | -0.302875227 | -0.231940756 | -0.017259303 | 0.227386597 | 0.003276007 | 0.08774316 | 0.134510858 | 0.884008447 |
| Tln1 | S | 1643 | -0.091499068 | -0.146594836 | -0.203914847 | -0.11241578 | -0.057320011 | 0.202337939 | 0.001195053 | 0.003124124 | 0.142686298 | 0.16361844 |
| Uaca | S | 1342 | -0.014473309 | -0.228671475 | -0.255812062 | -0.241338753 | -0.027140587 | 0.745067654 | 0.002217928 | 0.003197663 | 0.007446152 | 0.599352914 |
| Myh9 | S | 1435 | -0.063094212 | -0.085754738 | -0.341928142 | -0.27883393 | -0.256173404 | 0.502905841 | 0.247680338 | 2.44805E-05 | 0.068340888 | 0.041612201 |
| Anapc1 | S | 343 | -0.195843474 | -0.172712838 | -0.323406185 | -0.127562712 | -0.150693347 | 0.075533443 | 0.205238643 | 0.009653646 | 0.094386817 | 0.220944123 |
| Golga3 | S | 385 | -0.022263204 | -0.262523834 | -0.108637495 | -0.08637429 | 0.153886339 | 0.23308982 | 1.39239E-06 | 0.234295671 | 0.316329795 | 0.140204889 |
| Cdca3 | T | 75 | 0.009856708 | -0.259979299 | -0.277177808 | -0.287034516 | -0.01719851 | 0.622945063 | 0.00609195 | 0.001092141 | 0.000320398 | 0.531813367 |
| Nav3 | T | 360 | 0.020587496 | -0.41351635 | -0.399688571 | -0.420276066 | 0.01382778 | 0.760523024 | 0.016356796 | 0.022872987 | 0.032964128 | 0.91654727 |
| Pwp2 | T | 262 | 0.07129632 | -0.278827234 | -0.391874919 | -0.463171239 | -0.113047685 | 0.154405232 | 0.00062515 | 0.007335821 | 0.008379866 | 0.194594885 |
| Atg2b | S | 885 | 0.121568012 | -0.355149083 | -0.445241474 | -0.566809487 | -0.090092391 | 0.262837335 | 0.000322749 | 0.046264385 | 0.019313213 | 0.518927181 |
| Eif2b5 | S | 540 | -0.111054568 | -0.182828694 | -0.18795537 | -0.076900802 | -0.005126676 | 0.025411536 | 0.028210386 | 0.004398717 | 0.118403697 | 0.92830295 |
| Pcdh19 | S | 1136 | -0.174905388 | -0.202210808 | -0.311895599 | -0.136990211 | -0.109684791 | 0.106471421 | 0.098911243 | 0.014862749 | 0.189045624 | 0.319180956 |
| Dbnl | S | 277 | -0.329012926 | -0.087418666 | -0.445104432 | -0.116091506 | -0.357685765 | 0.076262948 | 0.595984826 | 0.019637933 | 0.434972705 | 0.05813034 |
| Wiz | S | 1029 | -0.050035308 | -0.24892013 | -0.172580684 | -0.122545376 | 0.076339446 | 0.247998449 | 0.000448042 | 0.023291758 | 0.074214088 | 0.193543582 |
| Dab2 | S | 727 | -0.235990273 | -0.185425846 | -0.287070067 | -0.051079794 | -0.101644221 | 0.04516667 | 0.21120581 | 0.019506319 | 0.384685195 | 0.407441935 |
| Zc2hc1a | S | 212 | -0.247375496 | -0.231794124 | -0.180638589 | 0.066736907 | 0.051155535 | 0.028588961 | 0.041453462 | 0.091399406 | 0.503804823 | 0.611878548 |
| Slc39a10 | T | 542 | -0.167055457 | -0.248267718 | -0.514769996 | -0.347714539 | -0.266502278 | 0.248801135 | 0.206047977 | 0.029921007 | 0.097877056 | 0.224969312 |
| Rab6a | S | 184 | -0.166072113 | -0.286891189 | -0.367071362 | -0.20099925 | -0.080180173 | 0.130947303 | 0.072384713 | 0.064422652 | 0.199793955 | 0.594071316 |
| Eef2 | S | 279 | -0.229667544 | -0.251300671 | -0.423580629 | -0.193913085 | -0.172279959 | 0.118074284 | 0.183371917 | 0.057867309 | 0.285717852 | 0.394434008 |
| Clasp1 | S | 1220 | -0.002662873 | -0.26214217 | -0.174898827 | -0.172235953 | 0.087243343 | 0.950831435 | 0.001789467 | 0.000504074 | 0.021950574 | 0.095693755 |
| Csrp1 | S | 81 | -0.416127543 | -0.096976554 | -0.422562253 | -0.00643471 | -0.3255857 | 0.046569924 | 0.616424652 | 0.055623968 | 0.952740315 | 0.071451715 |
| Abl2 | S | 671 | -0.112107771 | -0.157167869 | -0.53463446 | -0.422526689 | -0.377466591 | 0.379219973 | 0.161795221 | 0.00905295 | 0.031881205 | 0.038573224 |
| Sik3 | S | 1160 | -0.01728033 | -0.273821997 | -0.216275005 | -0.198994725 | 0.057546942 | 0.742021012 | 0.000313413 | 0.025188673 | 0.038203673 | 0.383709219 |
| Ctdspl2 | S | 13 | -0.073413751 | -0.111854692 | -0.260480387 | -0.187066636 | -0.148625695 | 0.363867274 | 0.005954082 | 0.000431824 | 0.080254619 | 0.027397823 |
| Ufl1 | S | 789 | 0.019206471 | -0.205069265 | -0.201458845 | -0.220665316 | 0.00361042 | 0.683508533 | 0.000366458 | 0.000372331 | 0.013687818 | 0.874711051 |
| Syne1 | S | 6239 | -0.084972231 | -0.100936339 | -0.276611507 | -0.191639276 | -0.175675168 | 0.056613517 | 0.164666539 | 0.000215141 | 0.006508157 | 0.05749828 |
| Arhgap12 | S | 608 | -0.37491933 | -0.077931892 | -0.510094634 | -0.135175304 | -0.432162743 | 0.068272957 | 0.724068274 | 0.038978765 | 0.406973885 | 0.083301525 |
| Xkr5 | T | 652 | -0.067416928 | -0.189611044 | -0.29538274 | -0.227965812 | -0.105771696 | 0.167907461 | 0.07103717 | 0.001399628 | 0.021433688 | 0.240108331 |
| Rab11fip1 | S | 383 | -0.220872818 | 0.044622168 | -0.466319216 | -0.245446398 | -0.510941384 | 0.019290495 | 0.676271238 | 0.002991798 | 0.025013255 | 0.013868728 |
| Smarcc1 | T | 397 | -0.140028601 | -0.394973284 | -0.185795556 | -0.045766955 | 0.209177728 | 0.240110565 | 0.002964481 | 0.28472913 | 0.78191321 | 0.241338161 |
| Tarbp2 | S | 151 | -0.043793171 | -0.174536548 | -0.212564751 | -0.16877158 | -0.038028203 | 0.267119365 | 6.81681E-05 | 0.020088238 | 0.031619815 | 0.448362268 |
| Ythdf2 | S | 39 | -0.032423197 | -0.365646212 | -0.178601353 | -0.146178156 | 0.18704486 | 0.609176675 | 0.001406958 | 0.036355823 | 0.046351285 | 0.032516907 |
| Dennd4c | S | 720 | -0.121998221 | -0.089136408 | -0.325871989 | -0.203873767 | -0.23673558 | 0.085949884 | 0.17661172 | 0.000966819 | 0.011002413 | 0.005294675 |
| Srf | S | 99 | -0.22011903 | -0.195213766 | -0.384876941 | -0.164757911 | -0.189663175 | 0.049708154 | 0.243976768 | 0.041607049 | 0.261117363 | 0.302093644 |
| Znhit6 | S | 115 | -0.053372899 | -0.308550807 | -0.514852064 | -0.461479165 | -0.206301258 | 0.628001852 | 0.029649771 | 0.049572579 | 0.048767184 | 0.250697627 |
| Sik3 | T | 606 | -0.104449859 | -0.152170335 | -0.291247844 | -0.186797985 | -0.139077509 | 0.181630927 | 0.087022202 | 0.001075529 | 0.050155369 | 0.113455451 |
| Fam92a | S | 310 | -0.166601466 | -0.092815028 | -0.337542612 | -0.170941147 | -0.244727584 | 0.072396768 | 0.427480813 | 0.001275377 | 0.062190417 | 0.094440929 |
| Tnks1bp1 | S | 1290 | -0.062076027 | -0.157098938 | -0.333774659 | -0.271698632 | -0.176675721 | 0.20247307 | 0.010546817 | 0.007315784 | 0.022570618 | 0.062925911 |
| Smtn | T | 440 | 0.068887405 | -0.205706584 | -0.30413475 | -0.373022154 | -0.098428166 | 0.35118568 | 0.002263479 | 0.000244295 | 0.011925995 | 0.018090234 |
| Nedd4 | S | 456 | -0.134468366 | -0.258839731 | -0.227417061 | -0.092948695 | 0.03142267 | 0.165035779 | 0.003820567 | 0.085237241 | 0.423870437 | 0.740465247 |
| Eif4b | S | 498 | -0.058365574 | -0.099170891 | -0.189792858 | -0.131427283 | -0.090621966 | 0.010054272 | 0.023170359 | 0.000117126 | 0.000948458 | 0.024158688 |
| Socs6 | S | 18 | -0.348963072 | -0.161319324 | -0.815138078 | -0.466175006 | -0.653818755 | 0.203830153 | 0.658556745 | 0.080132467 | 0.248671135 | 0.186846419 |
| Eps15 | S | 798 | -0.175463086 | -0.128904308 | -0.221805419 | -0.046342333 | -0.092901112 | 0.008611641 | 0.198210481 | 0.005798638 | 0.34479547 | 0.321342193 |
| Phldb2 | S | 156 | -0.137541571 | -0.184562727 | -0.563693499 | -0.426151928 | -0.379130772 | 0.345321794 | 0.230131937 | 0.015260581 | 0.050959227 | 0.071537771 |
| Nav1 | S | 675 | -0.132496185 | -0.250510481 | -0.478412402 | -0.345916217 | -0.22790192 | 0.463217974 | 0.166744947 | 0.012452082 | 0.104937906 | 0.205959893 |
| Eif4b | T | 506 | 0.009019913 | -0.216393209 | -0.342349817 | -0.35136973 | -0.125956608 | 0.900597012 | 0.016372539 | 0.001120355 | 0.004607287 | 0.093053553 |

| | | | | | | | | | | | | |
|---|---|---|---|---|---|---|---|---|---|---|---|---|
| Med1 | S | 1449 | -0.042724671 | -0.184489788 | -0.211722145 | -0.168997475 | -0.027232357 | 0.511333305 | 0.069309446 | 9.31399E-06 | 0.082981933 | 0.662495812 |
| Zdhhc5 | S | 429 | -0.063073798 | -0.285077231 | -0.288250965 | -0.225177167 | -0.003173734 | 0.318267646 | 0.076457018 | 0.002547794 | 0.005697382 | 0.977688171 |
| Id4 | S | 5 | -0.002897754 | -0.24813886 | -0.255698262 | -0.252800508 | -0.007559401 | 0.896153332 | 0.007778877 | 0.001084782 | 0.001349332 | 0.85968485 |
| L3mbtl3 | S | 625 | -0.206199889 | -0.105721812 | -0.315543502 | -0.109343613 | -0.209821689 | 0.027717043 | 0.243026181 | 0.011260509 | 0.230048281 | 0.066527354 |
| L3mbtl3 | S | 626 | -0.206199889 | -0.105721812 | -0.315543502 | -0.109343613 | -0.209821689 | 0.027717043 | 0.243026181 | 0.011260509 | 0.230048281 | 0.066527354 |
| Prkab1 | S | 24 | -0.346460382 | -0.1242635 | -0.464766448 | -0.118306066 | -0.340502948 | 0.040083058 | 0.369737807 | 0.102788902 | 0.581344827 | 0.200948591 |
| Ppp2r5c | S | 497 | -0.181875794 | -0.112188911 | -0.432928575 | -0.251052781 | -0.320739664 | 0.145964854 | 0.378699804 | 0.005052402 | 0.040266183 | 0.030202161 |
| Cmtm4 | S | 183 | -0.411173902 | -0.086801846 | -0.518091205 | -0.106917303 | -0.431289359 | 0.05340698 | 0.734874302 | 0.070082504 | 0.595243091 | 0.15454381 |
| Knop1 | S | 261 | 0.08123136 | -0.297255374 | -0.292762517 | -0.373993877 | 0.004492856 | 0.194661837 | 0.001236011 | 0.001390564 | 0.00080223 | 0.918644324 |
| Tacc1 | S | 82 | -0.023918915 | -0.300982547 | -0.482702578 | -0.458783664 | -0.181720031 | 0.861794569 | 0.123084439 | 0.005677421 | 0.010834994 | 0.271022272 |
| Map1b | S | 1778 | -0.078400142 | -0.209813476 | -0.170076008 | -0.091675865 | 0.039737469 | 0.113801568 | 0.001303002 | 0.015971842 | 0.10984682 | 0.386686444 |
| Exo1 | S | 508 | -0.09702686 | -0.230195636 | -0.387299714 | -0.290272854 | -0.157104078 | 0.39211568 | 0.001370756 | 0.124169753 | 0.199769875 | 0.420825024 |
| Tom1l2 | S | 160 | -0.112204665 | -0.157600049 | -0.220012995 | -0.10780833 | -0.062412946 | 0.099571107 | 0.015673448 | 0.003551622 | 0.121776661 | 0.249876647 |
| Nbeal2 | T | 1855 | -0.177835223 | -0.260242902 | -0.129555549 | 0.048279675 | 0.130687353 | 0.088990359 | 0.000853457 | 0.256556619 | 0.673340222 | 0.254268699 |
| Mmachc | S | 276 | -0.069091075 | -0.14134456 | -0.34977525 | -0.280684175 | -0.20843069 | 0.430237509 | 0.152332249 | 0.000330729 | 0.033269808 | 0.070021805 |
| Rtn4 | S | 768 | -0.092146038 | -0.103630312 | -0.2249457 | -0.132799662 | -0.121315388 | 0.205796192 | 0.012963045 | 0.000215065 | 0.113306151 | 0.004640261 |
| Evi2a | S | 178 | -0.066650381 | -0.412840164 | -0.22753217 | -0.160881788 | 0.185307994 | 0.423561869 | 0.002123505 | 0.208563369 | 0.332457553 | 0.279530137 |
| Setd1b | S | 1301 | -0.159748928 | -0.164497738 | -0.291787026 | -0.132038098 | -0.127289287 | 0.138268509 | 0.121642324 | 0.003655264 | 0.185760072 | 0.186128174 |
| Srrm2 | S | 2335 | 0.008464254 | -0.226816803 | -0.13936081 | -0.147825064 | 0.087455993 | 0.867153882 | 4.85552E-05 | 0.001270827 | 0.055987786 | 0.015770931 |
| Psrc1 | S | 70 | -0.065450618 | -0.228873878 | -0.342795277 | -0.277344659 | -0.113921399 | 0.536194398 | 0.113487626 | 0.001159615 | 0.063767164 | 0.345435666 |
| Eif4b | S | 459 | -0.058929809 | -0.1882206 | -0.212134608 | -0.153204799 | -0.023914008 | 0.395920608 | 0.01857019 | 0.000239158 | 0.095034544 | 0.620048046 |
| Ppp1r2 | S | 23 | -0.185240562 | -0.137521396 | -0.292545839 | -0.107305277 | -0.155024443 | 0.034786798 | 0.081149899 | 0.022713469 | 0.256910525 | 0.130281868 |
| Zc3h4 | S | 1182 | -0.090199127 | -0.167708031 | -0.283633105 | -0.193433978 | -0.115925074 | 0.328181546 | 0.011214864 | 0.004279852 | 0.091274361 | 0.093941656 |
| Rassf8 | S | 105 | -0.174329567 | -0.169367274 | -0.405369811 | -0.231040243 | -0.236002537 | 0.066152751 | 0.197077332 | 0.026244102 | 0.125416294 | 0.147663557 |
| Setd1a | S | 1176 | -0.042795291 | -0.346361626 | -0.48897202 | -0.446176729 | -0.142610393 | 0.615720501 | 0.018541387 | 0.081497057 | 0.08154591 | 0.457015503 |
| Orc1 | S | 332 | -0.013961941 | -0.329276519 | -0.258391502 | -0.244429561 | 0.070885016 | 0.748864606 | 0.001052256 | 0.057014295 | 0.078578573 | 0.470678031 |
| Prrc2c | S | 30 | -0.360043406 | -0.129396391 | -0.427113229 | -0.067069823 | -0.297716837 | 0.04800442 | 0.496856273 | 0.065350148 | 0.670333476 | 0.164846514 |
| Slc7a6os | S | 299 | 0.039830806 | -0.195242438 | -0.278768873 | -0.318599679 | -0.083526434 | 0.313833007 | 0.003649701 | 0.0001106 | 0.001783938 | 0.092745516 |
| Ctdp1 | S | 830 | -0.144756133 | -0.308729808 | -0.211327926 | -0.066571793 | 0.097401881 | 0.091232623 | 0.013719154 | 0.108047771 | 0.54527068 | 0.408824961 |
| Mef2c | S | 240 | -0.028237396 | -0.225992997 | -0.252653243 | -0.224415847 | -0.026660246 | 0.531078256 | 0.004061764 | 0.00423472 | 0.010401242 | 0.621413547 |
| Mga | S | 848 | -0.039468702 | -0.277100623 | -0.24347976 | -0.204011059 | 0.033620862 | 0.536335229 | 0.00206963 | 0.022970915 | 0.048186738 | 0.641958143 |
| Ccdc88a | S | 1840 | 0.074858229 | -0.238145803 | -0.292633439 | -0.367491668 | -0.054487636 | 0.280357681 | 0.000751746 | 0.000982554 | 0.007580723 | 0.212788022 |
| Ranbp3 | S | 28 | -0.090300539 | -0.10893907 | -0.456871193 | -0.366570654 | -0.347932123 | 0.394022542 | 0.403059739 | 0.000714483 | 0.018631473 | 0.053226158 |
| Atxn2 | S | 860 | -0.002322687 | -0.247519322 | -0.349651862 | -0.347329175 | -0.102132541 | 0.973279078 | 0.000807963 | 0.033830281 | 0.032056035 | 0.344560299 |
| Bod1l | S | 1242 | -0.311048471 | -0.127287363 | -0.381017162 | -0.069968692 | -0.253729799 | 0.039670854 | 0.467613919 | 0.03948941 | 0.544867488 | 0.172689697 |
| Afap1 | S | 259 | -0.001794747 | -0.200354177 | -0.17829093 | -0.176496183 | 0.022063247 | 0.950994936 | 0.000338697 | 0.000619246 | 0.000821766 | 0.268429404 |
| Gja1 | S | 373 | 0.003917722 | -0.174104276 | -0.288959409 | -0.292877131 | -0.114855133 | 0.953520296 | 0.055352319 | 3.32973E-05 | 0.028272684 | 0.131122275 |
| Numa1 | T | 1836 | -0.446428366 | -0.12891116 | -0.31611004 | 0.130318326 | -0.18719888 | 0.006286517 | 0.549335563 | 0.274418469 | 0.613037105 | 0.534774412 |
| Npm1 | S | 220 | 0.067022682 | -0.244055236 | -0.310602474 | -0.377625156 | -0.066547238 | 0.472631985 | 0.000317233 | 0.006891031 | 0.017867069 | 0.306491452 |
| Osbpl3 | S | 191 | -0.252017992 | -0.146953633 | -0.293549026 | -0.041531033 | -0.146595393 | 0.03774167 | 0.192296411 | 0.029165724 | 0.607898766 | 0.141489229 |
| Mtmr14 | S | 20 | -0.562166936 | -0.400319303 | -0.282145737 | 0.280021199 | 0.118173566 | 0.116580688 | 0.3729222 | 0.183427482 | 0.351506493 | 0.767352795 |
| Rrbp1 | S | 191 | -0.079871815 | -0.101035955 | -0.326269909 | -0.246398094 | -0.225233954 | 0.409073402 | 0.070765301 | 0.000277623 | 0.072365885 | 0.001570462 |
| E2f8 | S | 357 | 0.03217735 | -0.288796093 | -0.179230155 | -0.211407505 | 0.109565938 | 0.74851701 | 2.54913E-05 | 0.03744284 | 0.116268762 | 0.129682936 |
| Tenm3 Tenm4 | S | 142;145 | -0.14979845 | -0.145935766 | -0.209467876 | -0.059669426 | -0.06353209 | 0.02269164 | 0.050923999 | 0.007988474 | 0.251332873 | 0.304884575 |
| Ubap2 | S | 236 | -0.252970492 | -0.46668774 | -0.054216405 | 0.198754086 | 0.412471334 | 0.07036039 | 0.00906851 | 0.776938263 | 0.297469055 | 0.082917249 |
| Phldb2 | S | 346 | -0.339872648 | -0.161396036 | -0.070122431 | 0.269750218 | 0.091273605 | 0.00256796 | 0.021054607 | 0.551733019 | 0.084790505 | 0.438491149 |
| Htt | S | 620 | -0.122449349 | -0.128106631 | -0.149297815 | -0.026848466 | -0.021191184 | 0.01082266 | 0.009088466 | 0.004627536 | 0.018348995 | 0.034775979 |
| Gphn | S | 270 | 0.023433637 | -0.301509858 | -0.335347519 | -0.358781156 | -0.033837661 | 0.645604708 | 0.011484622 | 0.004136344 | 0.003900496 | 0.645035062 |
| Rock2 | S | 1374 | -0.230089966 | 0.057299475 | -0.504819359 | -0.274729394 | -0.562118834 | 0.140170398 | 0.787367886 | 0.001052382 | 0.095323606 | 0.076495835 |
| Trim25 | S | 424 | -0.018824362 | -0.171434916 | -0.247637242 | -0.22881288 | -0.076202326 | 0.665216643 | 0.004267693 | 0.000564055 | 0.001196896 | 0.032792834 |
| Magi3 | S | 662 | -0.099578995 | -0.082890306 | -0.163198748 | -0.063619753 | -0.080308442 | 0.005005046 | 0.007990367 | 0.00111368 | 0.061278714 | 0.04600294 |
| Slc12a4 | S | 56 | -0.094755167 | -0.226094131 | -0.271651631 | -0.176896464 | -0.0455575 | 0.131678818 | 0.005772178 | 0.053069108 | 0.116258693 | 0.606431965 |
| Mdm2 | S | 163 | 0.00729803 | -0.203873146 | -0.406600216 | -0.413898247 | -0.202727071 | 0.90497821 | 0.013375831 | 0.003963803 | 0.004012689 | 0.032601846 |
| Wdr4 | S | 397 | -0.038247691 | -0.136409125 | -0.197903172 | -0.159655481 | -0.061494046 | 0.051951506 | 0.000636517 | 0.003014115 | 0.006115066 | 0.078032784 |
| Sipa1l1 | S | 1372 | -0.266511112 | -0.148954706 | -0.192358599 | 0.074152513 | -0.043403893 | 0.005824224 | 0.223713564 | 0.047164565 | 0.284606457 | 0.689806323 |
| Rasa3 | S | 818 | 0.013790994 | -0.243157406 | -0.38238174 | -0.396172735 | -0.139224334 | 0.772033081 | 0.000830313 | 0.0304906 | 0.021617615 | 0.210346621 |
| Snapin | S | 133 | -0.046095556 | -0.191338463 | -0.280181635 | -0.234086079 | -0.088843172 | 0.239982797 | 0.004099428 | 0.011038721 | 0.009950084 | 0.134848772 |
| Bnip3 | S | 79 | -0.106119105 | -0.228669898 | -0.086352797 | 0.019766308 | 0.142317101 | 0.058902405 | 0.000326485 | 0.053608869 | 0.643017158 | 0.009253435 |
| Ap2m1 | S | 236 | -0.076450268 | -0.242052371 | -0.486547246 | -0.410096978 | -0.244494875 | 0.54911959 | 0.172083491 | 0.007726276 | 0.001495188 | 0.132311059 |
| Ctps | S | 575 | -0.13001526 | -0.189160033 | -0.336271012 | -0.206255752 | -0.147110979 | 0.089772122 | 0.037111764 | 0.036344561 | 0.103147694 | 0.202325915 |
| Flvcr1 | S | 559 | -0.238259921 | -0.306651573 | -0.099568036 | 0.138691885 | 0.207083538 | 0.028119558 | 0.010084604 | 0.417613282 | 0.232913244 | 0.120642421 |
| Ctnnd1 | S | 244 | -0.218274599 | -0.161466594 | -0.235108462 | -0.016833862 | -0.073641868 | 0.023633136 | 0.081090981 | 0.042278795 | 0.842853858 | 0.422513477 |
| Apbb1ip | S | 507 | -0.075133612 | -0.169125178 | -0.341271491 | -0.266137879 | -0.172146313 | 0.274692243 | 0.036889384 | 0.004998145 | 0.019999281 | 0.069653212 |
| Mtpn | T | 31 | -0.342858044 | -0.168294936 | -0.447442509 | -0.104584465 | -0.279147573 | 0.054679636 | 0.364508881 | 0.110489395 | 0.646366706 | 0.286540031 |
| Vps4a | S | 378 | -0.278011901 | -0.299871001 | -0.389699559 | -0.111687657 | -0.089828558 | 0.342558978 | 0.078196792 | 0.08469356 | 0.696719852 | 0.622870777 |
| Ciapin1 | S | 207 | -0.166658175 | -0.240381076 | -0.428822309 | -0.262164134 | -0.188441233 | 0.115469826 | 0.024604691 | 0.18038842 | 0.354715982 | 0.48169673 |
| Per1 | T | 134 | -0.35757142 | -0.081749892 | -0.376890226 | -0.019318806 | -0.295140334 | 0.038876289 | 0.650826398 | 0.032707484 | 0.894994661 | 0.171063322 |
| Mier3 | S | 52 | 0.002742745 | -0.271078506 | -0.16375073 | -0.166493475 | 0.107327777 | 0.977761899 | 0.000784944 | 0.001832689 | 0.186823933 | 0.034932755 |
| Mier3 | S | 53 | 0.002742745 | -0.271078506 | -0.16375073 | -0.166493475 | 0.107327777 | 0.977761899 | 0.000784944 | 0.001832689 | 0.186823933 | 0.034932755 |
| Sh2b3 | S | 302 | -0.077171216 | -0.18495111 | -0.186747932 | -0.109576716 | -0.001796822 | 0.097980436 | 0.000890245 | 0.023514976 | 0.108749346 | 0.971817192 |
| Hsp90aa1 Hsp90ab1 | T | 294;285 | -0.032270745 | -0.212479825 | -0.2377196 | -0.205448854 | -0.025239775 | 0.64482877 | 0.0001315 | 0.032372243 | 0.061556267 | 0.695242759 |
| Ptges3 | S | 148 | -0.177937953 | -0.234242192 | -0.163122939 | 0.014815014 | 0.071119253 | 0.011602332 | 0.0437835 | 0.073524339 | 0.833766064 | 0.449080437 |
| Ptges3 | S | 151 | -0.177937953 | -0.234242192 | -0.163122939 | 0.014815014 | 0.071119253 | 0.011602332 | 0.0437835 | 0.073524339 | 0.833766064 | 0.449080437 |
| Casp8ap2 | S | 952 | -0.227264988 | -0.341575344 | -0.000803227 | 0.226461761 | 0.340772117 | 0.064888589 | 0.000517207 | 0.992886484 | 0.094673783 | 0.030053986 |
| Pds5a | S | 1174 | -0.327046664 | -0.463298905 | 0.029014681 | 0.356061345 | 0.492313585 | 0.032620283 | 0.013409489 | 0.830908018 | 0.020454469 | 0.012935024 |
| Nr2c2ap | S | 22 | -0.307921498 | -0.100315128 | -0.465224603 | -0.157303105 | -0.364909475 | 0.090185369 | 0.613832988 | 0.02429649 | 0.327870768 | 0.111752255 |
| Rgs12 | S | 1130 | 0.116352231 | -0.324067652 | -0.411085586 | -0.527437817 | -0.087017933 | 0.075020673 | 0.000949124 | 0.006270865 | 0.011860472 | 0.326244525 |
| Prkaa1 | S | 508 | -0.299434143 | -0.111896314 | -0.132030178 | 0.167403965 | -0.020133865 | 0.001955842 | 0.086033507 | 0.143948857 | 0.083843122 | 0.776643994 |
| Sh3pxd2b | S | 620 | -0.266595445 | -0.098509389 | -0.480215062 | -0.213619617 | -0.381705674 | 0.066074116 | 0.594029817 | 0.026751824 | 0.199841887 | 0.094531821 |
| Zdhhc8 | S | 603 | -0.091109995 | -0.172978642 | -0.388049593 | -0.296939598 | -0.215070952 | 0.274247994 | 0.341978513 | 0.001152861 | 0.01093145 | 0.259630733 |
| Bloc1s4 | S | 65 | -0.208132641 | -0.347263375 | -0.149323657 | 0.058808984 | 0.197939718 | 0.074640424 | 0.010218982 | 0.347979859 | 0.666594987 | 0.220073592 |
| Supt6 | T | 619 | 0.0105558 | -0.182142154 | -0.224523915 | -0.235079715 | -0.042381761 | 0.712439708 | 0.001696534 | 0.000219362 | 0.003969717 | 0.222357641 |
| Clasp2 | S | 1045 | -0.013689335 | -0.240082621 | -0.224196964 | -0.210507628 | 0.015885657 | 0.675251423 | 0.001483397 | 0.006081916 | 0.009562121 | 0.734671887 |
| Srsf6 | S | 301 | 0.009365345 | -0.21464535 | -0.179915165 | -0.18928051 | 0.034730185 | 0.622067624 | 3.32924E-05 | 0.005359908 | 0.013294956 | 0.31235777 |
| Wnk1 | T | 1972 | -0.099130085 | -0.146610231 | -0.256808284 | -0.1576782 | -0.110198053 | 0.155076665 | 0.014749762 | 0.004688251 | 0.057570108 | 0.098548708 |
| Ptprj | S | 910 | 0.017417835 | -0.183284961 | -0.321701493 | -0.339119328 | -0.138416532 | 0.726555686 | 0.030865852 | 0.000183802 | 0.001104991 | 0.077665783 |
| Ctps | S | 574 | -0.136756265 | -0.178672755 | -0.33537366 | -0.198617394 | -0.156700905 | 0.085271804 | 0.041854388 | 0.036413218 | 0.117201623 | 0.18678336 |
| Mybbp1a | S | 1160 | 0.063339008 | -0.19208207 | -0.29819666 | -0.361535668 | -0.10611459 | 0.125586847 | 2.54708E-05 | 0.002821273 | 0.000815404 | 0.051460942 |
| Pkn2 | S | 582 | -0.052148714 | -0.11901378 | -0.146808031 | -0.094659317 | -0.027794251 | 0.130086311 | 0.000837334 | 0.000140434 | 0.039703934 | 0.14537316 |
| Eif5b | S | 137 | -0.099941811 | -0.098080847 | -0.110673201 | -0.010731389 | -0.012592354 | 0.059804569 | 2.81675E-06 | 0.00760459 | 0.749847279 | 0.478209678 |
| Map2k4 | S | 392 | -0.132895103 | -0.12163214 | -0.344320396 | -0.211425293 | -0.222688256 | 0.306138441 | 0.116789043 | 0.001770319 | 0.143828955 | 0.005253858 |
| Hspa12a | S | 23 | -0.153089731 | -0.25449029 | -0.200808202 | -0.047718471 | 0.053682088 | 0.011790309 | 0.075526401 | 0.061613014 | 0.539872228 | 0.633000284 |
| Bcl7c | S | 126 | -0.036505281 | -0.189852496 | -0.223929002 | -0.187423721 | -0.034076506 | 0.192560159 | 0.034375526 | 0.000338036 | 0.012053563 | 0.557864493 |
| Ddx54 | S | 74 | 0.008860366 | -0.241738078 | -0.263159414 | -0.272019779 | -0.021421335 | 0.927768575 | 0.001652176 | 0.005811817 | 0.056342281 | 0.697415703 |

| | | | | | | | | | | | | |
|---|---|---|---|---|---|---|---|---|---|---|---|---|
| Ddx21 | S | 107 | 0.002190878 | -0.186967405 | -0.242994719 | -0.245185597 | -0.056027314 | 0.969863407 | 0.001511932 | 0.001075621 | 0.014985534 | 0.184537612 |
| Blvra | S | 229 | -0.276935374 | -0.175630573 | -0.387748717 | -0.110813343 | -0.212118144 | 0.029869816 | 0.336843158 | 0.093268896 | 0.526960744 | 0.34182512 |
| Clip1 | S | 311 | -0.188279133 | -0.149068215 | -0.294763849 | -0.106484717 | -0.145695634 | 0.001043612 | 0.275883557 | 0.109836462 | 0.437385088 | 0.385892447 |
| Trim28 | S | 828 | -0.151449292 | -0.201913388 | -0.470161697 | -0.318712405 | -0.268248309 | 0.20111912 | 0.127156148 | 0.037406193 | 0.112185247 | 0.157664219 |
| Ftsj3 | S | 532 | -0.081236028 | -0.249405051 | -0.177374973 | -0.096138945 | 0.072030078 | 0.180689669 | 0.00075392 | 0.080490472 | 0.268847248 | 0.362411868 |
| Akt1s1 | S | 203 | -0.032824774 | -0.122530023 | -0.208055005 | -0.175230231 | -0.085524981 | 0.294276904 | 0.01037078 | 5.28476E-05 | 0.008957082 | 0.057316927 |
| Zdhhc5 | T | 454 | -0.078916725 | -0.128904323 | -0.299989944 | -0.22107322 | -0.171085622 | 0.292830027 | 0.121480569 | 0.000342196 | 0.043013034 | 0.07613979 |
| Acap2 | S | 545 | -0.148262624 | -0.145255001 | -0.286810767 | -0.138548144 | -0.141555767 | 0.180430791 | 0.065706553 | 0.004429947 | 0.189707691 | 0.036530485 |
| Tjp1 | S | 1277 | -0.250054354 | -0.11931378 | -0.200254966 | 0.049799388 | -0.080941186 | 0.001739525 | 0.181078464 | 0.108399861 | 0.60584051 | 0.467825921 |
| Ncaph | T | 43 | -0.023622491 | -0.14808551 | -0.185379538 | -0.161757047 | -0.037294028 | 0.472492714 | 0.046441197 | 1.69357E-06 | 0.019603772 | 0.408888107 |
| Fgd1 | S | 135 | -0.140568134 | -0.18595878 | -0.170654894 | -0.03008676 | 0.015303886 | 0.046697164 | 0.079943697 | 0.001876729 | 0.557775257 | 0.836095638 |
| Bin1 | S | 324 | -0.120835972 | -0.130589755 | -0.23567555 | -0.114839578 | -0.105085795 | 0.067820141 | 0.022295774 | 0.005819778 | 0.103722531 | 0.097320702 |
| Farp1 | S | 875 | 0.041347664 | -0.299188444 | -0.288067497 | -0.329415161 | 0.011120948 | 0.226081422 | 0.000738393 | 0.009741624 | 0.008564166 | 0.84740305 |
| Otud5 | S | 503 | -0.071070847 | -0.207579679 | -0.18700847 | -0.115937623 | 0.020571209 | 0.227091872 | 0.00012477 | 0.081642011 | 0.209927813 | 0.777218468 |
| Rprd2 | T | 742 | -0.519451263 | -0.414484209 | -0.108099186 | 0.411352077 | 0.306385024 | 0.054139092 | 0.112142624 | 0.713020232 | 0.153591416 | 0.243548793 |
| Sgsm3 | S | 406 | -0.384638864 | -0.017589338 | -0.354777235 | 0.029861629 | -0.337187897 | 0.010398738 | 0.907496468 | 0.057215061 | 0.814023763 | 0.088633252 |
| Ddx39 Ddx39b | S | 40;41 | -0.134904386 | -0.147464234 | -0.160510545 | -0.025606159 | -0.013046312 | 0.003234618 | 0.011498419 | 0.054892051 | 0.650374715 | 0.824416172 |
| Hnrnpa1 | S | 285 | -0.023494494 | -0.137011991 | -0.342748545 | -0.319254051 | -0.205736553 | 0.597705029 | 0.04058476 | 0.000567635 | 0.003533208 | 0.010014895 |
| Arhgap5 | S | 765 | -0.082521613 | -0.229035096 | -0.252110324 | -0.169588712 | -0.023075228 | 0.220167103 | 0.000379083 | 0.163814963 | 0.292270684 | 0.866220335 |
| Rpl18a | S | 71 | -0.150157128 | -0.193720002 | -0.334986513 | -0.184829385 | -0.141266511 | 0.175127717 | 0.028375161 | 0.042205777 | 0.196335647 | 0.265595588 |
| Sacs | S | 4290 | 0.008633057 | -0.239269783 | -0.355468956 | -0.364102014 | -0.116199173 | 0.842667287 | 0.021463998 | 0.002667204 | 0.006710858 | 0.176955348 |
| Baz2a | S | 1174 | -0.051156362 | -0.251197014 | -0.199634706 | -0.148478345 | 0.051562308 | 0.366373747 | 0.042136164 | 0.000384508 | 0.056197269 | 0.504027787 |
| Nup133 | S | 56 | -0.19191854 | -0.190053453 | -0.34210506 | -0.15018652 | -0.152051607 | 0.205370404 | 0.136712254 | 0.014426947 | 0.249629241 | 0.113342502 |
| Samd4b | S | 172 | -0.071858958 | -0.200598695 | -0.218797675 | -0.146938718 | -0.018198981 | 0.018368909 | 0.012498245 | 0.025373606 | 0.060718593 | 0.751848083 |
| Rabgap1 | S | 486 | -0.235103864 | -0.276369709 | -0.153109664 | 0.0819942 | 0.123260045 | 0.028486117 | 0.016264364 | 0.37334433 | 0.591364785 | 0.443202029 |
| Zcchc8 | S | 426 | 0.05951449 | -0.228134163 | -0.344738308 | -0.404252798 | -0.116604145 | 0.238760971 | 0.000332543 | 0.006896471 | 0.00476264 | 0.143110379 |
| Capn2 | T | 464 | 0.101273265 | -0.375768718 | -0.467309885 | -0.56858315 | -0.091541167 | 0.229407913 | 0.003204271 | 0.028964246 | 0.008034596 | 0.411867687 |
| Apc2 | S | 2257 | -0.024534611 | -0.147818386 | -0.188043763 | -0.163509152 | -0.040225376 | 0.571410833 | 0.000272226 | 0.000892047 | 0.024226514 | 0.176278505 |
| Apc2 | T | 2254 | -0.024534611 | -0.147818386 | -0.188043763 | -0.163509152 | -0.040225376 | 0.571410833 | 0.000272226 | 0.000892047 | 0.024226514 | 0.176278505 |
| Snx8 | S | 450 | -0.000466319 | -0.186471128 | -0.172378591 | -0.171912272 | 0.014092537 | 0.985512895 | 1.02051E-05 | 0.008995944 | 0.008352306 | 0.657527872 |
| Bag3 | S | 200 | -0.24467895 | -0.281746325 | -0.509572972 | -0.264894022 | -0.227826647 | 0.314192485 | 0.23979087 | 0.058994333 | 0.074108548 | 0.06516304 |
| Matr3 | S | 188 | -0.047140991 | -0.129902867 | -0.194806187 | -0.147665196 | -0.06490332 | 0.060340701 | 0.000326583 | 0.007055862 | 0.021853458 | 0.14056839 |
| Nadk | S | 64 | -0.159425375 | -0.186233849 | -0.363158217 | -0.203732843 | -0.176924369 | 0.230681535 | 0.165673302 | 0.009195198 | 0.117225251 | 0.150847416 |
| Rbbp5 | S | 497 | -0.023774354 | -0.185005982 | -0.184937137 | -0.161162783 | 6.88455E-05 | 0.354556597 | 0.005189333 | 0.000306668 | 0.002233275 | 0.99841644 |
| Fnip1 | S | 760 | 0.057844628 | -0.271378912 | -0.252636808 | -0.310481436 | 0.018742105 | 0.558902362 | 0.00177112 | 0.002300852 | 0.047749696 | 0.617224894 |
| Frmd4a | S | 601 | -0.262301094 | -0.152058516 | -0.357686438 | -0.095385344 | -0.205627922 | 0.054914723 | 0.233955532 | 0.053305473 | 0.498713632 | 0.195409105 |
| Reep4 | S | 152 | -0.136880424 | -0.15856233 | -0.334940607 | -0.198060183 | -0.176378277 | 0.052421837 | 0.161257121 | 0.014341246 | 0.065466992 | 0.151913939 |
| Atxn7l3b | S | 92 | -0.239181247 | -0.172072265 | -0.349895102 | -0.110713855 | -0.177822837 | 0.032118383 | 0.149746029 | 0.105563748 | 0.51578964 | 0.336181654 |
| Ercc6l | S | 980 | 0.129118909 | -0.304809107 | -0.390646915 | -0.519765824 | -0.085837808 | 0.136397216 | 0.008841927 | 0.000783406 | 0.002722584 | 0.263712104 |
| Atxn2l | S | 234 | 0.030068651 | -0.159654866 | -0.199730981 | -0.229799632 | -0.040076114 | 0.597694243 | 3.20373E-05 | 0.00072046 | 0.029328402 | 0.145072827 |
| Nsfl1c | S | 99 | -0.168048214 | -0.179504849 | -0.44070971 | -0.272661496 | -0.261204861 | 0.23121809 | 0.236702864 | 0.015344957 | 0.070622963 | 0.09787314 |
| Thoc2 | S | 1417 | -0.135939307 | -0.125844509 | -0.123895933 | 0.012043374 | 0.001948577 | 0.004372333 | 0.010246684 | 0.011713288 | 0.714424708 | 0.954746649 |
| Cdh6 | S | 786 | 0.045195676 | -0.24757827 | -0.246535379 | -0.291731056 | 0.001042891 | 0.159407273 | 7.9975E-06 | 0.026938787 | 0.014157054 | 0.984925263 |
| Rplp2 | S | 105 | -0.049179982 | -0.224317967 | -0.15332543 | -0.104145448 | 0.070992536 | 0.596801966 | 0.000644582 | 0.00416364 | 0.302493276 | 0.054917578 |
| Map1a | T | 1797 | -0.103212014 | -0.145356555 | -0.124673901 | -0.021461887 | 0.020682655 | 0.07735662 | 0.000347578 | 0.008716045 | 0.640078214 | 0.456605483 |
| Mphosph8 | S | 379 | -0.024280825 | -0.253656694 | -0.196004164 | -0.171723339 | 0.057652531 | 0.738851377 | 0.000607982 | 0.015687451 | 0.088370847 | 0.290537611 |
| Flad1 | S | 468 | -0.058923467 | -0.198557771 | -0.204738348 | -0.145814881 | -0.006180577 | 0.191089549 | 0.001623472 | 0.016441332 | 0.051841589 | 0.906252968 |
| Habp4 | S | 108 | -0.027182515 | -0.224381654 | -0.271721852 | -0.244539337 | -0.047340198 | 0.43836324 | 0.010810796 | 0.004978998 | 0.009704451 | 0.436687819 |
| Trip12 | S | 975 | -0.092575729 | -0.14316708 | -0.178065783 | -0.085490054 | -0.034898703 | 0.00475293 | 0.001404347 | 0.072033234 | 0.26594681 | 0.603806909 |
| Pak1ip1 | S | 361 | -0.051968448 | -0.136434321 | -0.16787365 | -0.115905202 | -0.031439329 | 0.213212472 | 0.000701109 | 0.000985264 | 0.03858665 | 0.24513227 |
| Tmem123 | S | 187 | 0.05365538 | -0.326743847 | -0.295182904 | -0.348838284 | 0.031560943 | 0.10437791 | 0.007135375 | 0.000950489 | 0.000544093 | 0.547429994 |
| Ppfibp1 | S | 961 | -0.023142246 | -0.20363587 | -0.152104806 | -0.12896256 | 0.051531063 | 0.585678873 | 0.000126181 | 0.005368767 | 0.04278498 | 0.145904292 |
| Ripk2 | S | 364 | -0.203787374 | -0.140579396 | -0.148769384 | 0.055017991 | -0.008189987 | 0.011318367 | 0.062238212 | 0.017738126 | 0.328152058 | 0.886038943 |
| Eno1 Eno1b | S | 427 | -0.269807364 | -0.188639289 | -0.314847037 | -0.045039674 | -0.126207748 | 0.037959488 | 0.157488167 | 0.115375665 | 0.779703781 | 0.470608558 |
| Phldb2 | T | 570 | -0.022869546 | -0.252708451 | -0.23233433 | -0.209464784 | 0.020374121 | 0.508159811 | 0.000338365 | 0.060157536 | 0.076626844 | 0.796401096 |
| Racgap1 | S | 155 | -0.266041034 | -0.110201062 | -0.303715279 | -0.037674246 | -0.193514217 | 0.12707477 | 0.230458737 | 0.0080414 | 0.775859533 | 0.019704303 |
| Rplp2 | S | 86 | -0.029571423 | -0.146336252 | -0.232326283 | -0.202754859 | -0.085990031 | 0.593787542 | 0.006634758 | 0.000301312 | 0.034465844 | 0.051364991 |
| Nfx1 | S | 49 | 0.018414622 | -0.225551566 | -0.190660645 | -0.209075267 | 0.034890921 | 0.580611351 | 0.000350612 | 0.001500326 | 0.005784594 | 0.276349108 |
| Frmd4a | S | 642 | -0.006273908 | -0.205276962 | -0.184288627 | -0.178014719 | 0.020988335 | 0.902719379 | 0.000374895 | 0.002402297 | 0.046988601 | 0.33756612 |
| Ctps | S | 573 | -0.138096505 | -0.176884673 | -0.349809924 | -0.211713419 | -0.172925251 | 0.110534357 | 0.048662994 | 0.038120108 | 0.12166305 | 0.178780944 |
| Thada | S | 1006 | -0.248801989 | -0.181888406 | -0.313854675 | -0.065052685 | -0.131966268 | 0.098749673 | 0.243863656 | 0.023379109 | 0.532408946 | 0.316232492 |
| Trip12 | S | 161 | -0.011070311 | -0.170928736 | -0.262841814 | -0.251771503 | -0.091913078 | 0.51034033 | 0.112357779 | 3.04206E-05 | 0.000833002 | 0.289871194 |
| Srrm2 | S | 2666 | -0.10607117 | -0.132966981 | -0.181614726 | -0.075543556 | -0.048647744 | 0.070798864 | 0.019652269 | 0.001324074 | 0.15016762 | 0.232468661 |
| Gigyf2 | S | 202 | -0.303936319 | -0.042736006 | -0.391446484 | -0.087510164 | -0.348710478 | 0.03742661 | 0.805159897 | 0.017995668 | 0.454735977 | 0.100235228 |
| Prune2 | S | 1675 | -0.047088181 | -0.229129478 | -0.254803395 | -0.207715215 | -0.025673917 | 0.420803343 | 0.001904143 | 0.02907909 | 0.039057121 | 0.677094017 |
| Golga4 | S | 41 | 0.004935769 | -0.160819878 | -0.266075853 | -0.271011622 | -0.105255975 | 0.886999149 | 0.001014705 | 0.001423963 | 0.002741128 | 0.071959359 |
| Plec | S | 4391 | -0.035139306 | -0.137749155 | -0.268385949 | -0.233246642 | -0.130636793 | 0.369016506 | 0.015354153 | 0.000653617 | 0.003757692 | 0.027094526 |
| Chordc1 | T | 47 | -0.139054064 | -0.254451771 | -0.339623904 | -0.20056984 | -0.085172133 | 0.173669639 | 0.028200785 | 0.09198425 | 0.22003232 | 0.547658796 |
| Trim28 | S | 489 | -0.182337453 | -0.186835703 | -0.092164962 | 0.090172491 | 0.094670741 | 0.001603802 | 0.033994398 | 0.092538897 | 0.091222524 | 0.189113759 |
| Atxn2 | S | 636 | -0.072001645 | -0.201481874 | -0.343144973 | -0.271143328 | -0.141663099 | 0.173692607 | 0.042344535 | 0.015815481 | 0.038902247 | 0.180057833 |
| Dync1li2 | S | 407 | 0.020839864 | -0.204781071 | -0.267658943 | -0.288028808 | -0.062877872 | 0.703361446 | 0.015439938 | 0.000210209 | 0.010023932 | 0.274516136 |
| Srrm2 | T | 919 | -0.342859523 | -0.364521634 | 0.162081318 | 0.504940842 | 0.526602953 | 0.00853697 | 0.005151418 | 0.533841457 | 0.133383847 | 0.125169271 |
| Nolc1 | S | 646 | 0.003140203 | -0.210906661 | -0.309557206 | -0.312697409 | -0.098650545 | 0.950195319 | 0.046375967 | 0.000414872 | 0.000547425 | 0.24250623 |
| Fscn1 | S | 38 | -0.307881254 | -0.163988376 | -0.246175822 | 0.061705432 | -0.082187446 | 0.007171012 | 0.319735672 | 0.154170453 | 0.651476974 | 0.650749215 |
| Camsap2 | S | 439 | -0.209690669 | -0.140546283 | -0.242951359 | -0.03326069 | -0.102405076 | 0.01206706 | 0.272243295 | 0.023703482 | 0.616413223 | 0.409805808 |
| Setd2 | S | 731 | -0.071117252 | -0.328923791 | -0.23250459 | -0.161387338 | 0.096419201 | 0.504682058 | 0.0011171 | 0.184291714 | 0.340395425 | 0.5150444 |
| Rangap1 | T | 434 | -0.124875543 | -0.238074052 | -0.423710364 | -0.298834821 | -0.185636312 | 0.089251186 | 0.08435732 | 0.083626262 | 0.152780644 | 0.32543654 |
| Zzef1 | S | 1488 | -0.036025104 | -0.273420463 | -0.376044269 | -0.340019165 | -0.102623806 | 0.706087313 | 0.022445672 | 0.020276338 | 0.036191873 | 0.381393239 |
| Tulp4 | S | 1347 | -0.175045988 | -0.187598614 | -0.191117064 | -0.016071076 | -0.00351845 | 0.03126063 | 0.024963605 | 0.061418461 | 0.812179702 | 0.959293881 |
| Dnm1l | S | 622 | -0.106861706 | -0.173382858 | -0.246160562 | -0.139298856 | -0.072777704 | 0.058766974 | 0.008261411 | 0.043596933 | 0.14029065 | 0.371375377 |
| Srgap2 | S | 1013 | -0.029158397 | -0.273413918 | -0.48473004 | -0.455571642 | -0.211316122 | 0.81457244 | 0.030220754 | 0.033325914 | 0.043376889 | 0.215729267 |
| Slc16a1 | S | 213 | -0.074837576 | -0.317829264 | -0.128365618 | -0.053528042 | 0.189463646 | 0.125527695 | 0.001871358 | 0.111501253 | 0.43389004 | 0.04457128 |
| Aspscr1 | S | 279 | -0.007984492 | -0.185106357 | -0.372359962 | -0.364375469 | -0.187253605 | 0.907220285 | 0.032876767 | 0.002112305 | 0.008668905 | 0.038431369 |
| Nop58 | S | 407 | -0.077725554 | -0.073719366 | -0.267604297 | -0.189878743 | -0.19388493 | 0.070012981 | 0.173264917 | 0.000172593 | 0.002709102 | 0.014687733 |
| Csde1 | S | 276 | 0.006301732 | -0.185989684 | -0.180807756 | -0.187109488 | 0.005181928 | 0.760630295 | 0.003546284 | 4.04568E-05 | 0.000165255 | 0.859985953 |
| Kif23 | T | 796 | -0.080577338 | -0.24214217 | -0.224319999 | -0.14374266 | 0.017822171 | 0.267177153 | 0.001699664 | 0.072971258 | 0.190601404 | 0.836815754 |
| Dmwd | S | 537 | -0.171936564 | -0.161556221 | -0.269759426 | -0.097822862 | -0.108203205 | 0.156483356 | 0.077826008 | 0.008999594 | 0.372078735 | 0.204669761 |
| Strip1 | S | 59 | -0.132315768 | -0.235959153 | -0.272175383 | -0.139859615 | -0.036216229 | 0.156691205 | 0.016150571 | 0.069358658 | 0.257391343 | 0.729362323 |
| Ubr4 | T | 360 | -0.000739004 | -0.187852611 | -0.389210774 | -0.38847177 | -0.201358163 | 0.988099751 | 0.015397323 | 0.00467484 | 0.010644655 | 0.050334084 |
| Atf6 | T | 153 | -0.001423875 | -0.300760304 | -0.214340381 | -0.212916507 | 0.086419922 | 0.982246646 | 0.002723603 | 0.009837079 | 0.0069509 | 0.150704864 |
| E2f8 | T | 44 | 0.07705653 | -0.311448615 | -0.297458894 | -0.374515424 | 0.013989721 | 0.238806342 | 0.004318066 | 0.001938431 | 0.004671522 | 0.777843912 |
| Cdk2 Cdk3 | T | 160;160 | -0.019449198 | -0.137776579 | -0.198962591 | -0.179513393 | -0.061186012 | 0.513502855 | 0.001889276 | 0.000235741 | 0.000746452 | 0.027283321 |
| Igf2r | S | 2476 | -0.154218711 | -0.1781833 | -0.204215641 | -0.04999693 | -0.026032342 | 0.093884169 | 0.031372464 | 0.01305791 | 0.523818317 | 0.691929217 |
| Osbpl6 | S | 35 | -0.039855646 | -0.144782696 | -0.409893268 | -0.370037621 | -0.265110572 | 0.686460221 | 0.369246174 | 0.000287799 | 0.025519276 | 0.160027139 |

| | | | | | | | | | | | | |
|---|---|---|---|---|---|---|---|---|---|---|---|---|
| 8030462N17Rik | S | 145 | -0.047558228 | -0.208209657 | -0.163855822 | -0.116297594 | 0.044353835 | 0.001116116 | 0.006047868 | 0.038890166 | 0.079993183 | 0.360789894 |
| Dclk2 | S | 361 | -0.228926888 | -0.232533659 | -0.307970288 | -0.079043399 | -0.075436628 | 0.020404826 | 0.177305363 | 0.160426747 | 0.649675878 | 0.713090702 |
| Naca | S | 166;2138 | -0.058136341 | -0.151773161 | -0.265808243 | -0.207671902 | -0.114035082 | 0.231824558 | 0.001139458 | 0.019523887 | 0.024356174 | 0.113777579 |
| Sub1 | S | 57 | -0.238226899 | -0.183942338 | -0.213943127 | 0.024283771 | -0.030000789 | 0.044710536 | 0.075196471 | 0.059259288 | 0.796640372 | 0.748072441 |
| Eml3 | S | 199 | -0.041206053 | -0.132827407 | -0.332409782 | -0.29120373 | -0.199582376 | 0.460645513 | 0.018070262 | 0.002344565 | 0.007574033 | 0.026378659 |
| Rprd2 | S | 1086 | -0.309671837 | -0.144918666 | -0.313024117 | -0.00335228 | -0.168105451 | 0.051423664 | 0.29160833 | 0.057511767 | 0.980146442 | 0.247842886 |
| Sympk | S | 494 | -0.188033005 | -0.163725686 | -0.3376716 | -0.149638595 | -0.173945914 | 0.111382268 | 0.283037109 | 0.01136162 | 0.111673866 | 0.232036373 |
| Wasl | Y | 253 | -0.456710023 | -0.153345076 | -0.246562554 | 0.210147469 | -0.093217478 | 0.014987991 | 0.492918388 | 0.219171264 | 0.268462009 | 0.68888985 |
| Fermt2 | S | 666 | 0.005814398 | -0.155916101 | -0.394601403 | -0.400415801 | -0.238685302 | 0.918234483 | 0.009191268 | 0.003577373 | 0.004850195 | 0.030687783 |
| Oxr1 | S | 346 | -0.092384172 | -0.06516825 | -0.378153545 | -0.285769374 | -0.312985296 | 0.487618045 | 0.538375975 | 0.000101233 | 0.111361742 | 0.058296285 |
| Map1b | S | 828 | -0.164565779 | -0.244251631 | -0.104704813 | 0.059860966 | 0.139546818 | 0.065266828 | 0.002891077 | 0.13602563 | 0.412373095 | 0.047872553 |
| Map1b | S | 829 | -0.164565779 | -0.244251631 | -0.104704813 | 0.059860966 | 0.139546818 | 0.065266828 | 0.002891077 | 0.13602563 | 0.412373095 | 0.047872553 |
| Ccdc6 | S | 411 | -0.105711041 | -0.120204386 | -0.216891628 | -0.111180587 | -0.096687242 | 0.015150161 | 0.012430453 | 0.018644145 | 0.124894931 | 0.156695274 |
| Map7d1 | S | 805 | -0.054262793 | -0.198767747 | -0.291362358 | -0.237099565 | -0.092594611 | 0.289880246 | 0.019201394 | 0.00873425 | 0.009914029 | 0.116824992 |
| Mcm2 | S | 381 | -0.085466298 | -0.138814829 | -0.140815665 | -0.055349366 | -0.002000835 | 0.037368853 | 0.009320391 | 0.001015172 | 0.13280631 | 0.94471936 |
| Huwe1 | T | 2746 | -0.161265176 | -0.175512988 | -0.217042866 | -0.05577769 | -0.041529878 | 0.017243187 | 0.053371835 | 0.054591008 | 0.489127166 | 0.635930888 |
| Nlrp4e | S | 277 | -0.152196876 | -0.27686926 | -0.183377945 | -0.031181069 | 0.093491315 | 0.125052554 | 0.03779877 | 0.028975334 | 0.704403958 | 0.349957787 |
| Nlrp4e | T | 279 | -0.152196876 | -0.27686926 | -0.183377945 | -0.031181069 | 0.093491315 | 0.125052554 | 0.03779877 | 0.028975334 | 0.704403958 | 0.349957787 |
| Nlrp4e | T | 282 | -0.152196876 | -0.27686926 | -0.183377945 | -0.031181069 | 0.093491315 | 0.125052554 | 0.03779877 | 0.028975334 | 0.704403958 | 0.349957787 |
| Rsbn1 | S | 91 | -0.121990333 | -0.306976924 | -0.110093468 | 0.011896865 | 0.196883457 | 0.079585962 | 0.002912422 | 0.192416122 | 0.877238428 | 0.058268962 |
| Prr12 | S | 1070 | -0.313130811 | -0.033661764 | -0.497937284 | -0.184806473 | -0.464275519 | 0.035547505 | 0.837348553 | 0.050126132 | 0.342745881 | 0.075357013 |
| Cwc22 | S | 895 | -0.160255615 | -0.135543826 | -0.237843649 | -0.077588034 | -0.102299824 | 0.256781287 | 0.040095127 | 0.002513859 | 0.533630444 | 0.066701833 |
| Wfs1 | S | 237 | -0.334237539 | -0.130804544 | -0.308796125 | 0.025441414 | -0.17799158 | 0.025541074 | 0.453503021 | 0.076336059 | 0.858185138 | 0.360010594 |
| Srrm2 | S | 1878 | -0.224016431 | -0.165032839 | -0.156460822 | 0.067555609 | 0.008572017 | 0.015935347 | 0.045144866 | 0.071850182 | 0.330690117 | 0.889370986 |
| Srrm2 | T | 1880 | -0.224016431 | -0.165032839 | -0.156460822 | 0.067555609 | 0.008572017 | 0.015935347 | 0.045144866 | 0.071850182 | 0.330690117 | 0.889370986 |
| Ube2o | S | 833 | -0.023438113 | -0.170083074 | -0.332803578 | -0.309365464 | -0.162720503 | 0.65777228 | 0.00320062 | 0.0105581 | 0.015106427 | 0.096182923 |
| Trmt1 | S | 121 | -0.071090249 | -0.177257169 | -0.121394724 | -0.050304475 | 0.055862445 | 0.165771296 | 0.000228664 | 0.011131011 | 0.306019165 | 0.12950231 |
| Zc3hav1 | S | 324 | -0.022840902 | -0.143206959 | -0.210520643 | -0.187679741 | -0.067313684 | 0.546692995 | 0.000619107 | 0.001524489 | 0.007873709 | 0.106699325 |
| Sfswap | S | 903 | -0.009577406 | -0.226682151 | -0.181361935 | -0.171784529 | 0.045320215 | 0.811298254 | 0.001272099 | 0.002563607 | 0.0178986 | 0.211820696 |
| Frmd4a | S | 626 | -0.005750963 | -0.252133543 | -0.213420605 | -0.207669642 | 0.038712938 | 0.933560281 | 0.002027698 | 0.006112454 | 0.027058125 | 0.362721854 |
| Foxp1 | S | 468 | -0.028936596 | -0.297641495 | -0.277995286 | -0.24905869 | 0.019646209 | 0.840983729 | 0.0157606 | 0.010073345 | 0.153494119 | 0.806464256 |
| Prune2 | S | 704 | 0.076313902 | -0.143147062 | -0.223099953 | -0.299413855 | -0.079952891 | 0.24022269 | 8.17078E-05 | 5.32263E-05 | 0.017086512 | 0.012061259 |
| Map1b | S | 1877 | -0.162529223 | -0.17368394 | -0.237962008 | -0.075432785 | -0.064278069 | 0.092729836 | 0.017148056 | 0.047936827 | 0.437348629 | 0.445196487 |
| Diaph3 | S | 1070 | -0.041617091 | -0.293892147 | -0.231412545 | -0.189795454 | 0.062479602 | 0.248855697 | 0.009318213 | 0.030669037 | 0.048106218 | 0.408682752 |
| Sdc3 | T | 430 | -0.10760755 | -0.259763942 | -0.181747835 | -0.074140285 | 0.078016107 | 0.009316484 | 0.024196771 | 0.122212194 | 0.414717433 | 0.427832347 |
| Fblim1 | S | 193 | -0.245517191 | -0.218351845 | -0.198295158 | 0.047222033 | 0.020056686 | 0.070161206 | 0.061708209 | 0.071755625 | 0.6649814 | 0.839632272 |
| Tnrc6b | S | 840 | -0.086063797 | -0.189573552 | -0.134565385 | -0.048501588 | 0.055008167 | 0.26765448 | 0.000506256 | 0.011666595 | 0.499747689 | 0.164705484 |
| Cavin4 | S | 171 | -0.152178317 | -0.270629475 | -0.20960036 | -0.057422044 | 0.061029114 | 0.138352883 | 0.024946656 | 0.062679038 | 0.23933364 | 0.219143471 |
| Cavin4 | S | 172 | -0.152178317 | -0.270629475 | -0.20960036 | -0.057422044 | 0.061029114 | 0.138352883 | 0.024946656 | 0.062679038 | 0.23933364 | 0.219143471 |
| Dact3 | S | 237 | -0.017285256 | -0.211281231 | -0.268119869 | -0.250834614 | -0.056838638 | 0.835414231 | 0.000242773 | 0.040255551 | 0.060771152 | 0.48392715 |
| Map1a | T | 382 | -0.069523576 | -0.254254983 | -0.351709286 | -0.282185711 | -0.097454303 | 0.414395964 | 0.039190602 | 0.022764161 | 0.035324898 | 0.28871165 |
| Znhit6 | S | 78 | -0.063208287 | -0.29780782 | -0.176901221 | -0.113692935 | 0.120906598 | 0.529650904 | 0.004923272 | 0.016882245 | 0.271379332 | 0.107864175 |
| Fgd6 | S | 670 | -0.247408426 | -0.088212529 | -0.146312888 | 0.101095539 | -0.058100359 | 0.002650163 | 0.150918917 | 0.039444887 | 0.068365876 | 0.235476283 |
| Srsf9 | Y | 207 | -0.025847844 | -0.421793555 | -0.18720784 | -0.161359996 | 0.234585714 | 0.847950414 | 0.000425999 | 0.367382289 | 0.466059324 | 0.282083407 |
| Mvb12a | S | 168 | -0.295271 | -0.108730174 | -0.369391406 | -0.074120406 | -0.260661232 | 0.042286228 | 0.418830281 | 0.05583081 | 0.625293087 | 0.159613056 |
| Rock2 | T | 1365 | -0.166915278 | -0.073192995 | -0.439386554 | -0.272471276 | -0.366193559 | 0.078924953 | 0.399715504 | 0.011867719 | 0.078184875 | 0.038871702 |
| Tbc1d13 | S | 184 | 0.004957703 | -0.173767484 | -0.280822856 | -0.28578056 | -0.107055372 | 0.920663449 | 0.006274113 | 0.001063348 | 0.000401912 | 0.069945012 |
| Cenpe | S | 2414 | 0.018352603 | -0.226751292 | -0.198365274 | -0.216717876 | 0.028386019 | 0.721586401 | 0.002500325 | 0.000573758 | 0.018316944 | 0.477148173 |
| Tjap1 | T | 407 | -0.103211983 | -0.107904266 | -0.155163291 | -0.051951308 | -0.047259024 | 0.006816463 | 0.050490248 | 0.00124309 | 0.014664239 | 0.27861582 |
| Slc1a4 | T | 506 | -0.202122619 | -0.340371196 | -0.220445832 | -0.018323213 | 0.119925364 | 0.071207067 | 0.126976054 | 0.086389852 | 0.862961932 | 0.498652561 |
| Gprc5b | S | 369 | -0.112657888 | -0.215387814 | -0.352409089 | -0.2397512 | -0.137021275 | 0.31606012 | 0.157189609 | 0.007583335 | 0.08648614 | 0.329150636 |
| Msn | S | 576 | -0.41901169 | -0.04343725 | -0.19358135 | 0.22543034 | -0.1501441 | 0.010896572 | 0.638084435 | 0.040675342 | 0.0863263 | 0.118818032 |
| Samd4b | S | 585 | 0.047244014 | -0.327030292 | -0.133608791 | -0.180852805 | 0.193421501 | 0.493679253 | 0.000156202 | 0.009192665 | 0.062290614 | 0.005766338 |
| Samd4b | S | 587 | 0.047244014 | -0.327030292 | -0.133608791 | -0.180852805 | 0.193421501 | 0.493679253 | 0.000156202 | 0.009192665 | 0.062290614 | 0.005766338 |
| Pnisr | S | 597 | -0.197399827 | -0.154179369 | -0.250157594 | -0.052757766 | -0.095978224 | 0.024100982 | 0.187029235 | 0.030147484 | 0.470170447 | 0.377690039 |
| N4bp1 | S | 560 | -0.015913408 | -0.23386984 | -0.272759821 | -0.256846413 | -0.038889981 | 0.830248847 | 0.000752578 | 0.036706187 | 0.045659461 | 0.616895576 |
| Eif4b | S | 207 | -0.186131499 | -0.113324578 | -0.324134286 | -0.138002787 | -0.210809708 | 0.07419301 | 0.248712494 | 0.011109148 | 0.103114661 | 0.035787466 |
| Cc2d1a | S | 459 | -0.048532768 | -0.15875773 | -0.20748975 | -0.158956982 | -0.04873202 | 0.186799405 | 0.006416023 | 0.002384747 | 0.014946402 | 0.276141187 |
| Lmna | T | 548 | -0.119343236 | -0.169852327 | -0.257754325 | -0.138411089 | -0.087901998 | 0.267856845 | 0.03544757 | 0.005897181 | 0.198831605 | 0.064060499 |
| Rfx1 | T | 952 | -0.264966558 | -0.070971296 | -0.391983215 | -0.127016657 | -0.321011918 | 0.050306671 | 0.716199972 | 0.017881102 | 0.225012341 | 0.153175485 |
| Prrc2b | S | 415 | -0.089169665 | -0.12456812 | -0.173333667 | -0.084164002 | -0.048765546 | 0.014304781 | 0.001882828 | 0.025162645 | 0.153844379 | 0.339887522 |
| Npm1 | T | 217 | 0.125354743 | -0.295589388 | -0.396205112 | -0.521559854 | -0.100615723 | 0.003856213 | 0.034140875 | 1.02328E-05 | 4.21546E-05 | 0.273683892 |
| Tnks1bp1 | S | 866 | -0.13029765 | -0.145963494 | -0.161206711 | -0.030909061 | -0.015243218 | 0.046747467 | 0.004815716 | 0.022341058 | 0.587615318 | 0.744937115 |
| Eef1g | T | 43 | -0.006638197 | -0.109071135 | -0.266305854 | -0.259667657 | -0.157234719 | 0.868729721 | 0.011494446 | 9.75552E-05 | 0.003419521 | 0.000760399 |
| Hmga1 Hmga1b | S | 36 | -0.111324476 | -0.163282701 | -0.129219084 | -0.017894608 | 0.034063617 | 0.002735903 | 0.006161169 | 0.078159824 | 0.726121924 | 0.540289232 |
| Mia3 | S | 421 | -0.136190377 | -0.119751787 | -0.357617449 | -0.221427073 | -0.237865662 | 0.235450506 | 0.195011614 | 0.003866266 | 0.075405102 | 0.020317321 |
| Ddx3x | S | 131 | -0.010731659 | -0.240123927 | -0.220304504 | -0.209572845 | 0.019819423 | 0.874975857 | 0.002976704 | 0.005225573 | 0.056769051 | 0.583535561 |
| Dync1li1 | T | 408 | 0.036877653 | -0.190729498 | -0.292004767 | -0.32888242 | -0.101275269 | 0.400957416 | 6.87302E-05 | 0.012920319 | 0.005185768 | 0.14159767 |
| Slc12a4 | S | 104 | -0.141641242 | -0.267492983 | -0.262730726 | -0.121089485 | 0.004762256 | 0.061816664 | 0.012621926 | 0.271427989 | 0.56652728 | 0.981375113 |
| Sema6d | T | 956 | -0.029627183 | -0.174984006 | -0.161111591 | -0.131484408 | 0.013872415 | 0.243994789 | 9.86968E-06 | 0.06208436 | 0.088398866 | 0.792241363 |
| Mybbp1a | S | 1325 | -0.012693179 | -0.153102983 | -0.3717038 | -0.359010621 | -0.218600817 | 0.839946112 | 0.194497176 | 0.000275299 | 0.004011812 | 0.103036813 |
| Samd4 | S | 183 | -0.197917068 | -0.154029888 | -0.218089547 | -0.020172479 | -0.064059659 | 0.03317467 | 0.175756881 | 0.015314913 | 0.689457069 | 0.484063193 |
| Usp37 | S | 650 | -0.053809653 | -0.105136022 | -0.180036695 | -0.126227042 | -0.074900674 | 0.123976963 | 0.023479554 | 5.41677E-05 | 0.027836534 | 0.085711637 |
| Tbc1d1 | S | 700 | -0.22673628 | -0.174021558 | -0.070018814 | 0.156717466 | 0.104002745 | 0.00160129 | 0.045168882 | 0.173378526 | 0.019686839 | 0.163685314 |
| Map4k4 | S | 862 | -0.047232753 | -0.242076768 | -0.32695364 | -0.279720888 | -0.084876872 | 0.379973149 | 0.064669569 | 0.007517064 | 0.010847293 | 0.383350209 |
| Lgalsl | S | 25 | -0.144063205 | -0.205879996 | -0.246986665 | -0.10292346 | -0.041106669 | 0.123107773 | 0.020984919 | 0.053398594 | 0.320122681 | 0.641100776 |
| Nav1 | S | 762 | -0.009069058 | -0.187467382 | -0.098974812 | -0.089905754 | 0.08849257 | 0.859928449 | 6.13968E-06 | 0.003188864 | 0.17321512 | 0.00872654 |
| Usp8 | S | 603 | -0.373914152 | -0.13482419 | -0.368613329 | 0.005300822 | -0.233789139 | 0.048421302 | 0.429892554 | 0.117243107 | 0.975790488 | 0.24720776 |
| Vmn2r1 | S | 523 | -0.173045174 | -0.236178764 | -0.276067387 | -0.103022213 | -0.039888623 | 0.207286894 | 0.063396198 | 0.035181175 | 0.445571826 | 0.724886669 |
| Vmn2r1 | T | 524 | -0.173045174 | -0.236178764 | -0.276067387 | -0.103022213 | -0.039888623 | 0.207286894 | 0.063396198 | 0.035181175 | 0.445571826 | 0.724886669 |
| Zrsr2 | S | 387 | 0.099997957 | -0.331585501 | -0.232862211 | -0.332860167 | 0.098723291 | 0.101346988 | 0.00228668 | 0.000406292 | 0.005419753 | 0.129077757 |
| Zrsr2 | S | 389 | 0.099997957 | -0.331585501 | -0.232862211 | -0.332860167 | 0.098723291 | 0.101346988 | 0.00228668 | 0.000406292 | 0.005419753 | 0.129077757 |
| Med1 | T | 1217 | -0.059657862 | -0.119029551 | -0.081079949 | -0.021422087 | 0.037949602 | 0.005562981 | 0.00062306 | 0.000405214 | 0.141900064 | 0.069398531 |
| Slc7a6os | S | 32 | -0.067553682 | -0.148174788 | -0.29113733 | -0.223583649 | -0.142962542 | 0.317914841 | 0.05512834 | 0.00182815 | 0.019605489 | 0.070556594 |
| Alkbh5 | S | 384 | 0.094390538 | -0.373699883 | -0.271447993 | -0.365838531 | 0.10225189 | 0.224699839 | 0.001797381 | 0.009352475 | 0.002800413 | 0.106124842 |
| Smtn | S | 574 | -0.040889266 | -0.252826392 | -0.297927154 | -0.257037889 | -0.045100762 | 0.306375765 | 0.015743388 | 0.025945783 | 0.042998056 | 0.605916454 |
| Xpo4 | S | 521 | -0.013741664 | -0.215078394 | -0.195186752 | -0.181445088 | 0.019891642 | 0.721852642 | 0.000171824 | 0.02363567 | 0.022313883 | 0.663808816 |
| Stard13 | S | 572 | -0.117971715 | -0.284250586 | -0.469458796 | -0.351487081 | -0.18520821 | 0.385185256 | 0.148568116 | 0.043035702 | 0.10022781 | 0.373941774 |
| Camsap2 | S | 437 | -0.316489184 | -0.129676309 | -0.248559812 | 0.067929372 | -0.118883503 | 0.020315001 | 0.363383291 | 0.06956675 | 0.53283922 | 0.398137331 |
| Aldoa | S | 39 | -0.183342771 | -0.166090031 | -0.242559994 | -0.059217223 | -0.076469963 | 0.327936665 | 0.00639032 | 0.031128847 | 0.728873183 | 0.302605791 |
| Ddx54 | S | 774 | 0.022711011 | -0.224805951 | -0.279517149 | -0.30222816 | -0.054711198 | 0.844010191 | 0.017400943 | 0.000973271 | 0.087637309 | 0.377533538 |
| Arhgef7 | S | 673 | -0.144543316 | -0.119838128 | -0.19976128 | -0.055217964 | -0.079923152 | 0.013745 | 0.19267119 | 0.003900461 | 0.069766599 | 0.336305689 |
| Prune2 | S | 574 | -0.259380572 | -0.143392623 | -0.303247988 | -0.043867416 | -0.159855366 | 0.186350976 | 0.002601892 | 0.198935414 | 0.844102135 | 0.426248964 |
| Rbbp6 | S | 246 | -0.159652741 | -0.21710454 | -0.292697003 | -0.133044263 | -0.075592463 | 0.043192556 | 0.05006263 | 0.132686136 | 0.407765919 | 0.633284285 |

| | | | | | | | | | | | | |
|---|---|---|---|---|---|---|---|---|---|---|---|---|
| Rbbp6 | S | 247 | -0.159652741 | -0.21710454 | -0.292697003 | -0.133044263 | -0.075592463 | 0.043192556 | 0.05006263 | 0.132686136 | 0.407765919 | 0.633284285 |
| Rbbp6 | S | 248 | -0.159652741 | -0.21710454 | -0.292697003 | -0.133044263 | -0.075592463 | 0.043192556 | 0.05006263 | 0.132686136 | 0.407765919 | 0.633284285 |
| Cdc37 | S | 378 | -0.150879164 | -0.063423868 | -0.328670546 | -0.177791382 | -0.265246678 | 0.018435419 | 0.374229877 | 0.008137927 | 0.088348571 | 0.025140362 |
| Eloa | S | 82 | -0.131775056 | -0.142582611 | -0.123334844 | 0.008440213 | 0.019247767 | 0.028084358 | 0.004723545 | 0.014541367 | 0.844001821 | 0.557629145 |
| Arhgef2 | S | 646 | -0.046188727 | -0.232921876 | -0.211685879 | -0.165497153 | 0.021235997 | 0.498344714 | 0.005109509 | 0.009389705 | 0.032375243 | 0.580431384 |
| Rhpn2 | S | 654 | -0.343624673 | -0.101713252 | -0.386236115 | -0.042611442 | -0.284522863 | 0.028770604 | 0.206377724 | 0.209577432 | 0.867257516 | 0.316424948 |
| Satb2 | S | 39 | 0.005331167 | -0.229229813 | -0.257379062 | -0.262710229 | -0.028149249 | 0.869665316 | 0.000984887 | 0.013920539 | 0.021637898 | 0.637671233 |
| Dennd2a | S | 275 | -0.087956689 | -0.180670299 | -0.117887253 | -0.029930565 | 0.062783046 | 0.198396424 | 0.00196252 | 0.002424677 | 0.603309068 | 0.096249877 |
| D1Pas1 Ddx3x Ddx3y | T | 410;410;411 | -0.008642561 | -0.248316819 | -0.135460592 | -0.126818031 | 0.112856226 | 0.804224496 | 4.21952E-05 | 0.041939975 | 0.055341448 | 0.086542335 |
| Pimreg | S | 43 | -0.116296006 | -0.071981258 | -0.360370287 | -0.24407428 | -0.288389029 | 0.121006186 | 0.482547866 | 0.001210833 | 0.007225603 | 0.041260093 |
| Arhgap32 | S | 871 | -0.100126242 | -0.296457851 | -0.099728123 | 0.000398119 | 0.196729728 | 0.177960914 | 0.000370725 | 0.326658311 | 0.996734535 | 0.113752935 |
| Cobll1 | S | 1028 | -0.061089126 | -0.172791104 | -0.142638719 | -0.081549594 | 0.030152385 | 0.110425188 | 0.00206476 | 0.004419371 | 0.02930062 | 0.34170066 |
| Ppp1r12a | S | 507 | 0.003278302 | -0.161762676 | -0.469394284 | -0.472672987 | -0.307632008 | 0.975220915 | 0.130448373 | 0.001900738 | 0.003879916 | 0.010278528 |
| Rabep1 | T | 480 | -0.211863729 | -0.28824366 | -0.183255081 | 0.028608648 | 0.104988579 | 0.276966883 | 0.009819816 | 0.136666058 | 0.871035114 | 0.301130363 |
| Ctnnb1 | T | 551 | 0.021626317 | -0.158651862 | -0.267492685 | -0.289119002 | -0.108840823 | 0.739532579 | 0.005553236 | 0.000373865 | 0.01689183 | 0.039329496 |
| Canx | S | 553 | -0.159305484 | -0.121940136 | -0.255243229 | -0.095937745 | -0.133303093 | 0.045157654 | 0.239415631 | 0.005159782 | 0.093591592 | 0.190970046 |
| Kpna3 | S | 60 | -0.104051374 | -0.143365097 | -0.173268258 | -0.069216885 | -0.029903161 | 0.096356589 | 0.008589553 | 0.004958276 | 0.221263843 | 0.448534283 |
| Pacsin2 | S | 386 | -0.040975622 | -0.15425753 | -0.230615803 | -0.189640181 | -0.076358273 | 0.182035315 | 0.051275465 | 0.000435791 | 0.002762268 | 0.206767516 |
| Thrap3 | S | 860 | -0.45855772 | 0.086914334 | -0.272127798 | 0.186429922 | -0.359042132 | 0.002997978 | 0.657772524 | 0.077187664 | 0.160715201 | 0.133278885 |
| Hsf1 | S | 314 | -0.075845599 | -0.1986097 | -0.198109799 | -0.1222642 | 0.000499901 | 0.193219202 | 0.011166068 | 0.008148017 | 0.023295648 | 0.991094462 |
| Myo1e | S | 1008 | -0.174404845 | -0.261148872 | -0.135164143 | 0.039240701 | 0.125984728 | 0.053806862 | 0.015393632 | 0.139307393 | 0.561005105 | 0.141241679 |
| Ranbp3 | S | 257 | 0.001386484 | -0.162602497 | -0.269866056 | -0.27125254 | -0.107263559 | 0.977052938 | 0.014404754 | 0.00041811 | 0.003377782 | 0.067129419 |
| Med14 | S | 983 | 0.022048506 | -0.385401 | -0.303860659 | -0.325909166 | 0.08154034 | 0.919793542 | 0.007634852 | 0.052781225 | 0.204217501 | 0.452685415 |
| Abi1 | S | 296 | -0.350075053 | -0.025918942 | -0.450559288 | -0.100484235 | -0.424640346 | 0.06996803 | 0.859031885 | 0.032013536 | 0.574587901 | 0.054809382 |
| Pikfyve | S | 307 | 0.032165149 | -0.156750658 | -0.496216368 | -0.528381517 | -0.33946571 | 0.694692792 | 0.069076291 | 0.002338066 | 0.003814137 | 0.016513741 |
| Golm1 | S | 319 | -0.18668035 | -0.170525388 | -0.13780228 | 0.04887807 | 0.032723108 | 0.10306213 | 0.005317348 | 0.036127958 | 0.588699886 | 0.452235798 |
| Ppfibp1 | S | 355 | -0.120380356 | -0.296914299 | -0.127104049 | -0.006723693 | 0.16981025 | 0.032752754 | 0.005922303 | 0.290160434 | 0.948925067 | 0.191168839 |
| Vcl | T | 604 | -0.217597613 | -0.185634742 | -0.198305186 | 0.019292427 | -0.012670443 | 0.036353332 | 0.141383224 | 0.033802557 | 0.814098023 | 0.90461079 |
| Chd8 | S | 1999 | -0.046100064 | -0.2014356 | -0.167508669 | -0.121408605 | 0.033926931 | 0.186262711 | 0.000495334 | 0.03350674 | 0.097968558 | 0.52020359 |
| Ppp4r2 | S | 226 | -0.259219367 | -0.263647901 | -0.08758415 | 0.171635216 | 0.17606375 | 0.078004746 | 0.004128664 | 0.478216858 | 0.262000612 | 0.205958793 |
| Dst | S | 2410 | -0.14804947 | -0.269950555 | -0.235829658 | -0.087780188 | 0.034120897 | 0.14408557 | 0.024753927 | 0.102048892 | 0.416288743 | 0.743357115 |
| Plekho2 | S | 395 | -0.074124596 | -0.110266186 | -0.397957442 | -0.323832846 | -0.287691256 | 0.496119474 | 0.219656457 | 0.001247229 | 0.036956798 | 0.010135888 |
| Mtor | T | 1262 | -0.297444349 | -0.194654185 | -0.179392544 | 0.118051805 | 0.015261641 | 0.088759138 | 0.106340458 | 0.040676761 | 0.395083519 | 0.872172173 |
| Stk3 | S | 391 | 0.0343841 | -0.17239366 | -0.355930793 | -0.390314893 | -0.183537133 | 0.645786619 | 0.031225503 | 0.000649939 | 0.006631578 | 0.018828134 |
| Ptk2 | S | 29 | -0.028185746 | -0.088129705 | -0.196729569 | -0.168543823 | -0.108599864 | 0.670559736 | 0.001949252 | 3.63385E-05 | 0.088906684 | 0.003084364 |
| Dtnb | S | 394 | -0.080183661 | -0.110816782 | -0.141488936 | -0.061305275 | -0.030672155 | 0.005635247 | 0.014501243 | 0.002037218 | 0.075634352 | 0.356698805 |
| Crtc2 | S | 457 | -0.049997005 | -0.215681202 | -0.112894563 | -0.062897558 | 0.10278664 | 0.150559633 | 0.000196912 | 0.040696897 | 0.171401275 | 0.066689328 |
| Itprid2 | S | 442 | -0.276898621 | -0.048504816 | -0.422465453 | -0.145566832 | -0.373960636 | 0.04494014 | 0.723154523 | 0.030927343 | 0.319680146 | 0.057189639 |
| Cdk16 | S | 153 | -0.295872776 | -0.127012752 | -0.316721522 | -0.020848746 | -0.18970877 | 0.035148341 | 0.387557852 | 0.068556748 | 0.873626007 | 0.253338639 |
| Atg4b | S | 383 | -0.273857946 | -0.203252152 | -0.441963727 | -0.168105781 | -0.238711575 | 0.235621233 | 0.265632191 | 0.060055655 | 0.475912944 | 0.265344989 |
| Ubr4 | T | 1760 | -0.22209984 | -0.042207851 | -0.117066682 | 0.105033158 | -0.074858831 | 0.000310058 | 0.212944952 | 0.02209275 | 0.033697254 | 0.088972535 |
| Map7d1 | S | 728 | 0.002832049 | -0.151420277 | -0.286608408 | -0.289440457 | -0.135188131 | 0.949875642 | 0.004141586 | 0.00152204 | 0.003655509 | 0.03283337 |
| Tle3 | S | 245 | -0.145788247 | -0.228255615 | -0.207471866 | -0.061683619 | 0.020783749 | 0.218146838 | 0.016203495 | 0.038677503 | 0.578240484 | 0.788588475 |
| Ehd4 | S | 459 | -0.018536268 | -0.1801878 | -0.317761855 | -0.299225587 | -0.137574054 | 0.837989038 | 0.059153375 | 0.000924284 | 0.048518857 | 0.101574391 |
| Tbc1d1 | S | 559 | -0.020582581 | -0.314459489 | -0.414289729 | -0.393707148 | -0.09983024 | 0.864040188 | 0.022551388 | 0.050116987 | 0.057727995 | 0.483712376 |
| Abi1 | T | 297 | -0.573810343 | -0.150306834 | -0.427374443 | 0.1464359 | -0.27706761 | 0.037108741 | 0.576115174 | 0.433522719 | 0.766832301 | 0.59306744 |
| Srek1ip1 | S | 96 | 0.031027532 | -0.151127833 | -0.140732242 | -0.171759774 | 0.010395591 | 0.293525402 | 6.18028E-06 | 0.00016161 | 0.006218734 | 0.475459447 |
| Zc3h14 | S | 85 | -0.041022715 | -0.174703102 | -0.163180853 | -0.122158139 | 0.011522248 | 0.385628447 | 0.001958646 | 0.001756515 | 0.052962242 | 0.659268546 |
| Zbtb11 Rpl24 | S | 451 | -0.063524986 | -0.171370722 | -0.355086866 | -0.29156188 | -0.183716144 | 0.563026077 | 0.06222048 | 0.004799693 | 0.06265361 | 0.070371165 |
| Zbtb11 Rpl24 | S | 458 | -0.063524986 | -0.171370722 | -0.355086866 | -0.29156188 | -0.183716144 | 0.563026077 | 0.06222048 | 0.004799693 | 0.06265361 | 0.070371165 |
| Zbtb11 Rpl24 | T | 453 | -0.063524986 | -0.171370722 | -0.355086866 | -0.29156188 | -0.183716144 | 0.563026077 | 0.06222048 | 0.004799693 | 0.06265361 | 0.070371165 |
| Melk | S | 352 | -0.108412385 | -0.120977058 | -0.2490474 | -0.140635016 | -0.128070342 | 0.093682813 | 0.081812112 | 0.003074774 | 0.055162278 | 0.081360741 |
| Smap2 | S | 219 | -0.024411305 | -0.161191139 | -0.275149141 | -0.250737836 | -0.113958002 | 0.698690344 | 0.007901586 | 0.002619658 | 0.014212148 | 0.09109873 |
| Tarbp2 | S | 141 | -0.176862658 | -0.330707207 | -0.203775795 | -0.026913137 | 0.126931412 | 0.186819124 | 0.011101329 | 0.308357568 | 0.882717898 | 0.481014966 |
| Rhbdf1 | S | 8 | -0.182961855 | -0.13934259 | -0.488846388 | -0.305884533 | -0.349503798 | 0.227760163 | 0.44779652 | 0.018240925 | 0.090996402 | 0.108349019 |
| Hectd1 | S | 1389 | -0.014914729 | -0.15147058 | -0.317187036 | -0.302272306 | -0.165716455 | 0.810608819 | 0.031805433 | 0.001050355 | 0.002090002 | 0.005694593 |
| Acin1 | S | 838 | -0.086373087 | -0.146014749 | -0.20244229 | -0.116069203 | -0.056427541 | 0.044235326 | 0.113423514 | 0.001157695 | 0.005408243 | 0.443362661 |
| Larp1 | T | 765 | -0.012220902 | -0.165899152 | -0.235529124 | -0.223308222 | -0.069629972 | 0.688944385 | 0.000108327 | 0.02092597 | 0.022093862 | 0.257449127 |
| Chmp1a | S | 182 | -0.120589328 | -0.250036262 | -0.336980289 | -0.216390961 | -0.086944027 | 0.251234214 | 0.039171982 | 0.071442783 | 0.181088682 | 0.533954945 |
| Chordc1 | S | 139 | -0.088268242 | -0.29628679 | -0.317740127 | -0.229471885 | -0.021453337 | 0.336690841 | 0.03525604 | 0.059998216 | 0.13442907 | 0.87398665 |
| Nisch | S | 545 | 0.077096425 | -0.205915628 | -0.329521151 | -0.406617575 | -0.123605523 | 0.425536889 | 0.000628484 | 0.007206744 | 0.016671812 | 0.099590914 |
| Usp36 | S | 716 | -0.033192524 | -0.171793361 | -0.154462729 | -0.121270205 | 0.017330632 | 0.125498156 | 0.000991362 | 0.005421422 | 0.019487297 | 0.590537681 |
| Srsf5 | S | 153 | -0.185747746 | -0.298263458 | -0.144338591 | 0.041409155 | 0.153924868 | 0.022150137 | 0.061887415 | 0.179816129 | 0.630719274 | 0.268782532 |
| Golph3 | S | 35 | -0.093639959 | -0.119278744 | -0.185771811 | -0.092131852 | -0.066493067 | 0.021707572 | 0.003128456 | 0.026425693 | 0.146114105 | 0.253782746 |
| Crip2 | S | 74 | -0.270095852 | -0.015619662 | -0.308394962 | -0.038299109 | -0.292775299 | 0.019679781 | 0.921056455 | 0.010663268 | 0.507669386 | 0.13336117 |
| Cog3 | S | 525 | -0.164095497 | -0.17014243 | -0.160797674 | 0.003297823 | 0.009344757 | 0.021417644 | 0.110606096 | 0.012107342 | 0.949714205 | 0.908755127 |
| Mmachc | S | 272 | -0.086681165 | -0.14021499 | -0.302392682 | -0.215711517 | -0.162177692 | 0.302024071 | 0.125862885 | 0.001719553 | 0.041786253 | 0.089045179 |
| Ppp1r35 | S | 60 | -0.112348327 | -0.340168071 | -0.39858306 | -0.286234733 | -0.058414989 | 0.310378389 | 0.124591547 | 0.065035737 | 0.152989809 | 0.778870224 |
| Tnks1bp1 | S | 911 | -0.239031537 | -0.088907377 | -0.252667114 | -0.013635577 | -0.163759737 | 0.007338974 | 0.360853938 | 0.052981575 | 0.878392914 | 0.184531676 |
| Knl1 | S | 1477 | -0.189525376 | -0.549056164 | -0.18318203 | 0.006343346 | 0.365874134 | 0.246263735 | 0.053182336 | 0.31347215 | 0.971564355 | 0.157858694 |
| Apbb2 | S | 334 | -0.159041637 | -0.24922897 | -0.185024312 | -0.025982675 | 0.064204658 | 0.057616166 | 0.014327024 | 0.202993761 | 0.827710412 | 0.595260365 |
| Thoc2 | S | 1393 | -0.090680647 | -0.134403195 | -0.306924006 | -0.216243359 | -0.172520811 | 0.214221629 | 0.024342448 | 0.012050419 | 0.048306282 | 0.091974825 |
| Exoc1 | S | 565 | -0.315742728 | -0.043262082 | -0.35613821 | -0.040395482 | -0.312876127 | 0.024750379 | 0.713664807 | 0.047071689 | 0.727108177 | 0.068342172 |
| Eif4g3 | S | 1151 | -0.332335726 | -0.035875146 | -0.222480323 | 0.109855403 | -0.186605176 | 0.004955986 | 0.786183025 | 0.047500458 | 0.200129286 | 0.202176808 |
| Kdm3b | S | 357 | -0.280640816 | -0.183131345 | -0.230630902 | 0.050009914 | -0.047499557 | 0.113416295 | 0.258155961 | 0.017829597 | 0.711213827 | 0.73013949 |
| Hsp90ab1 | S | 445 | 0.012971406 | -0.147784076 | -0.215096749 | -0.228068154 | -0.067312673 | 0.789141443 | 4.87748E-05 | 0.004173901 | 0.014153977 | 0.110237925 |
| Ndrg3 | S | 361 | -0.194826424 | -0.168341011 | -0.150866938 | 0.043959486 | 0.017474073 | 0.085695387 | 0.017378118 | 0.031383465 | 0.610024825 | 0.669074176 |
| Dock8 | S | 2097 | -0.290878572 | -0.197100273 | -0.127616055 | 0.163262516 | 0.069484217 | 0.015219135 | 0.060536943 | 0.297322683 | 0.161256265 | 0.463572916 |
| Ehbp1 | S | 1061 | -0.196162239 | -0.132426691 | -0.139760281 | 0.056401958 | -0.00733359 | 0.001411725 | 0.016708598 | 0.376932442 | 0.693705427 | 0.958724346 |
| Mycbp2 | S | 2905 | -0.018869742 | -0.16417546 | -0.185370155 | -0.166500412 | -0.021194695 | 0.564296791 | 0.000873747 | 0.002320133 | 0.007702687 | 0.498524899 |
| Ttc4 | S | 51 | -0.624157144 | -0.414968892 | -0.187622553 | 0.436534591 | 0.227346339 | 0.372240341 | 0.039965736 | 0.623118215 | 0.534477524 | 0.538412789 |
| Plch1 | S | 924 | -0.049307734 | -0.182603803 | -0.214036495 | -0.164728761 | -0.031432692 | 0.332573205 | 0.013880853 | 0.002781324 | 0.016037769 | 0.539803415 |
| Dlg1 | S | 158 | 0.013287984 | -0.274935579 | -0.200023409 | -0.213311393 | 0.07491217 | 0.819368455 | 0.002110942 | 0.007522887 | 0.005341799 | 0.054162355 |
| Zranb1 | S | 216 | 0.009405895 | -0.186858988 | -0.280810484 | -0.290216379 | -0.093951496 | 0.830904532 | 0.002048593 | 0.005681537 | 0.0082733 | 0.164655924 |
| Pxn | S | 130 | -0.04677464 | -0.192958473 | -0.156214899 | -0.109440259 | 0.036743574 | 0.189996531 | 0.000845644 | 0.018236764 | 0.059120986 | 0.40976405 |
| Plcl1 | S | 77 | -0.340236936 | 0.012409895 | -0.679666951 | -0.339430016 | -0.692076847 | 0.116180193 | 0.96578931 | 0.055255963 | 0.239867768 | 0.082444846 |
| Hectd1 | S | 1777 | 0.036453682 | -0.202511052 | -0.365433723 | -0.401887405 | -0.16292267 | 0.359414231 | 0.019426809 | 0.001549587 | 0.003131865 | 0.05394226 |
| Carhsp1 | S | 31 | -0.306969733 | -0.151117415 | -0.35119708 | -0.044227347 | -0.200079665 | 0.044026446 | 0.267919344 | 0.153520204 | 0.827310075 | 0.363390967 |
| Erf | S | 131 | -0.197427116 | -0.237711549 | -0.332813775 | -0.135386659 | -0.095102226 | 0.212170584 | 0.065365639 | 0.098669148 | 0.461719214 | 0.553712379 |
| Scyl2 | S | 658 | -0.217246633 | -0.159145873 | -0.299727345 | -0.082480712 | -0.140581471 | 0.079787979 | 0.192928833 | 0.039672921 | 0.439822339 | 0.234544473 |
| Nop14 | S | 96 | 0.024864458 | -0.174663994 | -0.241058136 | -0.265922594 | -0.066394142 | 0.458132084 | 0.009637215 | 0.000120047 | 0.001416725 | 0.140571622 |
| Tanc1 | S | 445 | -0.020260753 | -0.200222851 | -0.289595503 | -0.269334749 | -0.089372651 | 0.600570293 | 0.010629185 | 0.008190964 | 0.003642281 | 0.072307511 |
| Bicd2 | S | 817 | -0.063003678 | -0.233355606 | -0.141016782 | -0.078013104 | 0.092338824 | 0.242709467 | 0.001548235 | 0.028827957 | 0.206698426 | 0.1047555 |
| Sbno1 | S | 791 | -0.050369973 | -0.21459287 | -0.355958809 | -0.305588836 | -0.141365939 | 0.599368451 | 0.122314185 | 0.003999191 | 0.04288657 | 0.239771089 |

| | | | | | | | | | | | | |
|---|---|---|---|---|---|---|---|---|---|---|---|---|
| Tex2 | S | 221 | -0.077680541 | -0.116135362 | -0.24698361 | -0.169303069 | -0.130848248 | 0.187731979 | 0.031493951 | 0.002010671 | 0.023615003 | 0.033974278 |
| Car13 | S | 5 | -0.332986995 | -0.128912655 | -0.329006372 | 0.003980623 | -0.200093717 | 0.033337252 | 0.307521293 | 0.158569327 | 0.983678993 | 0.346000277 |
| Nup155 | S | 992 | -0.127486411 | -0.148959747 | -0.156376337 | -0.028889926 | -0.00741659 | 0.093003766 | 0.001234664 | 0.050739763 | 0.681305181 | 0.88826598 |
| Stk24 | S | 369 | -0.235412367 | -0.220492244 | -0.194395296 | 0.041017071 | 0.026096948 | 0.021694763 | 0.09461293 | 0.198266948 | 0.750821167 | 0.856103836 |
| Ranbp2 | S | 2036 | 0.012748764 | -0.30737379 | -0.327715183 | -0.340463947 | -0.020341393 | 0.824100607 | 0.012696315 | 0.021472358 | 0.011338982 | 0.797050822 |
| Pnisr | S | 762 | -0.139689921 | -0.167631877 | -0.12993355 | 0.009756371 | 0.037698327 | 0.047148117 | 0.009589497 | 0.024368229 | 0.857839091 | 0.436657877 |
| Carhsp1 | S | 33 | -0.306156608 | -0.153839077 | -0.349330737 | -0.04317413 | -0.19549166 | 0.047078385 | 0.254500893 | 0.156977991 | 0.83299409 | 0.374624604 |
| Xrcc5 | S | 528 | -0.145943689 | -0.172331042 | -0.09808494 | 0.047858749 | 0.074246103 | 0.012040735 | 0.009409834 | 0.056999571 | 0.266172158 | 0.145138839 |
| Map1b | S | 1307 | -0.062488537 | -0.100520734 | -0.118767455 | -0.056278918 | -0.018246722 | 0.012317997 | 0.001078591 | 0.001539493 | 0.057891574 | 0.354151938 |
| Thoc2 | S | 1577 | -0.028622963 | -0.174872118 | -0.274694673 | -0.24607171 | -0.099822555 | 0.59142453 | 0.008384212 | 0.005715859 | 0.014890137 | 0.154643885 |
| Nvl | S | 190 | -0.053815022 | -0.141093467 | -0.118452552 | -0.06463753 | 0.022640915 | 0.037376859 | 0.000641445 | 0.00522845 | 0.065212899 | 0.357629624 |
| Map7d1 | S | 368 | -0.177864434 | -0.142259152 | -0.269422428 | -0.091557995 | -0.127163277 | 0.066604138 | 0.169446409 | 0.019411217 | 0.317008626 | 0.227636078 |
| Slc12a2 | S | 937 | -0.021103384 | -0.400924561 | -0.146630866 | -0.125527482 | 0.254293696 | 0.848458321 | 0.022425298 | 0.002595899 | 0.318248695 | 0.06708417 |
| Rin1 | S | 246 | -0.12737784 | -0.145716099 | -0.206281412 | -0.078903573 | -0.060565313 | 0.031225928 | 0.008628095 | 0.075779597 | 0.379776152 | 0.479333333 |
| Ikzf5 | T | 105 | -0.47206261 | -0.130160003 | -0.241910598 | 0.230152012 | -0.111750595 | 0.086379752 | 0.107715702 | 0.213608417 | 0.341624191 | 0.508806266 |
| Clcn6 | S | 684 | 0.072742295 | -0.223621557 | -0.247840686 | -0.320582981 | -0.024219129 | 0.394072824 | 0.000144153 | 0.009191175 | 0.022301814 | 0.604663145 |
| Atxn2l | S | 109 | 0.038650647 | -0.190542149 | -0.229020519 | -0.267671166 | -0.03847837 | 0.330719127 | 0.000117302 | 0.00427993 | 0.003179532 | 0.342386204 |
| Dclk1 | S | 353 | -0.158540898 | -0.109962324 | -0.233150301 | -0.074609403 | -0.123187977 | 0.122331794 | 0.094054471 | 0.003913962 | 0.388841713 | 0.052080887 |
| Dclk1 | S | 355 | -0.158540898 | -0.109962324 | -0.233150301 | -0.074609403 | -0.123187977 | 0.122331794 | 0.094054471 | 0.003913962 | 0.388841713 | 0.052080887 |
| Ctps | S | 568 | -0.139485378 | -0.156126098 | -0.326646147 | -0.187160769 | -0.170520049 | 0.094213799 | 0.074582951 | 0.043362459 | 0.145576046 | 0.174839497 |
| Gapvd1 | S | 914 | -0.177145362 | 0.011446899 | -0.215905523 | -0.038760161 | -0.227352422 | 0.000202672 | 0.672048022 | 0.005370658 | 0.319321934 | 0.004395127 |
| Rbm26 | S | 621 | 0.026593394 | -0.17857035 | -0.321597783 | -0.348191177 | -0.143027433 | 0.486451243 | 0.004039544 | 0.002956006 | 0.001866466 | 0.035304506 |
| Ncl | S | 616 | -0.126409876 | -0.123998536 | -0.08510667 | 0.041303206 | 0.038891866 | 0.002447275 | 0.009185082 | 0.017324828 | 0.069912011 | 0.22321614 |
| Ddx39b | T | 172 | 0.047416925 | -0.158210996 | -0.325594278 | -0.373011203 | -0.167383282 | 0.681417486 | 0.002518174 | 0.00280014 | 0.045727191 | 0.047419702 |
| Rplp2 | S | 102 | -0.049740021 | -0.211667897 | -0.139663983 | -0.089923961 | 0.072003914 | 0.583198942 | 0.000915747 | 0.006338378 | 0.346519981 | 0.055582421 |
| Rrp7a | T | 101 | 0.08984214 | -0.202421894 | -0.248155048 | -0.337997188 | -0.045733154 | 0.280447504 | 0.001723449 | 0.000283838 | 0.022445029 | 0.228466352 |
| Arhgap31 | T | 337 | -0.095239395 | -0.069370244 | -0.271148317 | -0.175908922 | -0.201778073 | 0.107188432 | 0.183100742 | 0.000616202 | 0.022438921 | 0.010935779 |
| Csnk1e | S | 389 | -0.104036201 | -0.176530314 | -0.554648807 | -0.450612607 | -0.378118494 | 0.464892261 | 0.197651848 | 0.028814831 | 0.059041624 | 0.08704632 |
| Atrx | S | 1103 | -0.187697686 | -0.189644671 | -0.080490422 | 0.107207265 | 0.10915425 | 0.058482065 | 0.005266978 | 0.0492202 | 0.188626599 | 0.048506974 |
| Fbf1 | S | 185 | -0.260634078 | -0.05159577 | -0.484935131 | -0.224301053 | -0.433339361 | 0.051525324 | 0.606696025 | 0.056616533 | 0.258979489 | 0.075208036 |
| Srrm2 | S | 1276 | -0.091796928 | -0.086060061 | -0.264508596 | -0.172711668 | -0.178448535 | 0.447070267 | 0.127668918 | 0.00014912 | 0.213286889 | 0.021304211 |
| Ube4b | S | 105 | -0.217129312 | -0.062142728 | -0.478620159 | -0.261490847 | -0.416477431 | 0.196799406 | 0.544494178 | 0.013911537 | 0.158947449 | 0.032386825 |
| Mtmr2 | S | 9 | 0.055684753 | -0.226294339 | -0.227573102 | -0.283257855 | -0.001278763 | 0.51965253 | 0.001564755 | 0.001775149 | 0.039250717 | 0.968157203 |
| Hnrnpc | S | 231 | -0.03252846 | -0.135622911 | -0.180564221 | -0.148035761 | -0.044941309 | 0.059177799 | 0.000817177 | 0.008038265 | 0.011262588 | 0.173608342 |
| Gatad2a | T | 185 | 0.03528523 | -0.182377542 | -0.219579852 | -0.254865082 | -0.037202309 | 0.332811966 | 0.000469668 | 0.000986428 | 0.001693424 | 0.283828803 |
| Ccsap | S | 222 | -0.222462032 | -0.12676293 | -0.353208198 | -0.130746166 | -0.226445268 | 0.06675096 | 0.352749416 | 0.037201115 | 0.286849106 | 0.141396911 |
| Rbbp8 | S | 710 | 0.064725197 | -0.250418755 | -0.29740901 | -0.362134207 | -0.046990255 | 0.060747907 | 6.75112E-05 | 0.018070747 | 0.016168902 | 0.482046278 |
| Cdca8 | T | 213 | -0.089256561 | -0.132387894 | -0.196920904 | -0.107664343 | -0.064533009 | 0.027685549 | 0.00596206 | 0.030805403 | 0.151266435 | 0.313754591 |
| Sec16a | S | 1371 | -0.216655575 | -0.017844582 | -0.410559965 | -0.19390439 | -0.392715384 | 0.078160819 | 0.880124067 | 0.007059586 | 0.097083202 | 0.017632711 |
| Vim | S | 7 | 0.003204576 | -0.156060989 | -0.264899789 | -0.268104365 | -0.108838801 | 0.96134923 | 0.006006095 | 0.001208956 | 0.026185264 | 0.041327661 |
| Nbea | S | 1311 | -0.066370566 | -0.15247375 | -0.248786237 | -0.182415671 | -0.096312487 | 0.32817915 | 0.012756624 | 0.005982867 | 0.049131669 | 0.132106595 |
| Nup93 | S | 364 | 0.025326602 | -0.180383632 | -0.238760996 | -0.264087598 | -0.058377363 | 0.204936171 | 0.023574545 | 2.44587E-05 | 0.000466439 | 0.249305679 |
| Rybp | S | 201 | 0.00917934 | -0.354530209 | -0.271074583 | -0.280253924 | 0.083455626 | 0.936572805 | 0.010902898 | 0.030508476 | 0.031641928 | 0.205795028 |
| 1810013L24Rik | T | 214 | -0.154399895 | -0.458698638 | -0.249793764 | -0.095393869 | 0.208904874 | 0.144020439 | 0.079214566 | 0.2511994 | 0.616078713 | 0.402528807 |
| Gas2l3 | S | 340 | -0.158003974 | -0.175134709 | -0.187486828 | -0.029482854 | -0.012352119 | 0.02134301 | 0.029522045 | 0.107312209 | 0.743835719 | 0.894100763 |
| Spen | S | 1267 | -0.199417197 | -0.134907125 | -0.345838583 | -0.146421386 | -0.210931459 | 0.243956751 | 0.453581377 | 0.004950254 | 0.354940875 | 0.267197398 |
| Gigyf2 | S | 23 | -0.070315226 | -0.36263815 | -0.215495794 | -0.145180569 | 0.147142356 | 0.628858145 | 0.007066252 | 0.108946022 | 0.334543182 | 0.187705766 |
| Srpk1 | S | 450 | -0.008030543 | -0.147818152 | -0.16019057 | -0.152160027 | -0.012372418 | 0.746198965 | 1.65332E-05 | 0.007330258 | 0.007094991 | 0.640962736 |
| Otud1 | S | 123 | -0.212198725 | -0.231719764 | -0.363915467 | -0.151716742 | -0.132195702 | 0.180789901 | 0.042808318 | 0.209469396 | 0.558521547 | 0.589772861 |
| Map1b | S | 2250 | 0.143892174 | -0.270717466 | -0.255967325 | -0.399859499 | 0.014750141 | 0.572878909 | 0.005585835 | 0.001237139 | 0.201498431 | 0.784779954 |
| Apc2 | S | 2259 | -0.084915327 | -0.23502163 | -0.103560462 | -0.018645135 | 0.131461169 | 0.046036985 | 0.002903183 | 0.067880096 | 0.670951795 | 0.036476724 |
| Rps6kb1 | S | 447 | -0.164880014 | -0.15556005 | -0.30494314 | -0.140063126 | -0.14938309 | 0.032039319 | 0.150649511 | 0.064221244 | 0.273397548 | 0.281260558 |
| Ets1 | S | 282 | -0.151411494 | -0.199454728 | -0.287771851 | -0.136360357 | -0.088317123 | 0.197140823 | 0.072035318 | 0.033194979 | 0.283945323 | 0.428533827 |
| Slc12a4 | S | 37 | -0.062614197 | -0.191404594 | -0.100009362 | -0.037395165 | 0.091395232 | 0.399986764 | 0.000779821 | 0.001656868 | 0.591003634 | 0.028933293 |
| Sipa1l1 | Y | 1133 | -0.160930247 | -0.215442688 | -0.52202101 | -0.361090763 | -0.306578322 | 0.323622716 | 0.140800041 | 0.085250764 | 0.187643229 | 0.24003009 |
| Mast4 | S | 1781 | -0.073127549 | -0.345094639 | -0.206126837 | -0.132999288 | 0.138967802 | 0.205855132 | 0.038765735 | 0.041272441 | 0.104358775 | 0.201352283 |
| Tbc1d32 | T | 495 | 0.082943555 | -0.296529077 | -0.309823021 | -0.392766576 | -0.013293944 | 0.311551445 | 0.019765732 | 0.001221541 | 0.007144133 | 0.868911444 |
| Tjp2 | S | 213 | -0.312228122 | -0.094904763 | -0.151969622 | 0.1602585 | -0.057064859 | 0.001773612 | 0.260597212 | 0.271174436 | 0.250950194 | 0.650294159 |
| Ska3 | S | 148 | -0.195096785 | -0.241078686 | -0.104451077 | 0.090645708 | 0.136627608 | 0.013096068 | 0.041865908 | 0.193011916 | 0.251678472 | 0.188575672 |
| Cdc42bpb | S | 970 | -0.069074407 | -0.212910287 | -0.125664999 | -0.056590592 | 0.087245288 | 0.377261407 | 0.001945293 | 0.006424897 | 0.453108196 | 0.068408394 |
| Uhrf1 | S | 519 | -0.144259737 | -0.317664315 | -0.100535926 | 0.043723812 | 0.217128389 | 0.172650289 | 0.004565935 | 0.208801341 | 0.583878128 | 0.008383153 |
| Strap | S | 343 | -0.052471564 | -0.437920333 | -0.586121279 | -0.533649715 | -0.148200945 | 0.703626452 | 0.07143832 | 0.183193264 | 0.211373813 | 0.688661856 |
| Tnfaip2 | S | 26 | -0.004871665 | -0.222291561 | -0.342577052 | -0.337705187 | -0.120285492 | 0.883127685 | 0.013373016 | 0.014543873 | 0.019940294 | 0.174499191 |
| Rbbp6 | S | 1078 | -0.099662535 | -0.233793645 | -0.234278956 | -0.134616421 | -0.000485311 | 0.626357478 | 0.049744843 | 0.003393737 | 0.518914161 | 0.99529826 |
| Myo1b | S | 548 | -0.034357837 | -0.15651511 | -0.221254524 | -0.186896687 | -0.064739414 | 0.479094437 | 0.006187438 | 0.002379556 | 0.010654518 | 0.151328551 |
| Srrm2 | S | 1478 | -0.029941067 | -0.170168853 | -0.243416174 | -0.213475107 | -0.073247321 | 0.541949096 | 0.013717251 | 0.002279738 | 0.007643435 | 0.118281005 |
| Evl | S | 343 | -0.137283511 | -0.253550499 | -0.396855451 | -0.25957194 | -0.143304951 | 0.439213162 | 0.123083113 | 0.038363598 | 0.184214688 | 0.363425067 |
| Xkr5 | S | 651 | -0.154964962 | -0.153732088 | -0.162059113 | -0.007094152 | -0.008327025 | 0.037829759 | 0.014104314 | 0.05745597 | 0.918400527 | 0.895494746 |
| Lsg1 | S | 58 | -0.100573086 | -0.205940567 | -0.611059695 | -0.510486609 | -0.405119129 | 0.212757652 | 0.231514074 | 0.074901985 | 0.103861232 | 0.157410985 |
| Map1a | S | 1159 | -0.10329175 | -0.10013193 | -0.101725215 | 0.001566535 | -0.001593286 | 0.006252409 | 0.015403902 | 0.001125672 | 0.943860154 | 0.950106607 |
| Eomes | S | 617 | -0.118889364 | -0.295651699 | -0.41281642 | -0.293927056 | -0.117164722 | 0.568079676 | 0.137617337 | 0.035097303 | 0.231982793 | 0.490342348 |
| Nav3 | T | 357 | 0.02662339 | -0.351274677 | -0.339452354 | -0.366075744 | 0.011822323 | 0.650126498 | 0.00849748 | 0.048517644 | 0.051551489 | 0.923015568 |
| Rai14 | S | 914 | -0.135249037 | -0.083224281 | -0.16324325 | -0.027994213 | -0.080018969 | 0.006808001 | 0.100913994 | 0.004406008 | 0.360490803 | 0.107150245 |
| Ibtk | S | 994 | -0.048643368 | -0.325825569 | -0.312312344 | -0.263668976 | 0.013513225 | 0.735335503 | 0.00427299 | 0.171905851 | 0.252834933 | 0.940455763 |
| Cdc25b | S | 372 | -0.117929784 | -0.184263397 | -0.420896502 | -0.302966718 | -0.236633105 | 0.323953089 | 0.079274172 | 0.043454297 | 0.109515183 | 0.17542244 |
| AI837181 | S | 36 | -0.024206242 | -0.218745468 | -0.426365783 | -0.402159541 | -0.207620315 | 0.805955361 | 0.182990419 | 0.004603708 | 0.006319874 | 0.188603547 |
| Pabpn1 | S | 146 | 0.010641884 | -0.183505324 | -0.144936873 | -0.155578757 | 0.038568451 | 0.792476547 | 0.000280066 | 0.000753704 | 0.029762121 | 0.118351193 |
| Ppme1 | S | 243 | -0.083969446 | -0.139988038 | -0.394330687 | -0.310361241 | -0.254342649 | 0.394192653 | 0.188464631 | 0.005472659 | 0.012348318 | 0.025290514 |
| Fhdc1 | S | 1018 | -0.172883737 | -0.26320998 | -0.193230101 | -0.020346364 | 0.069979879 | 0.318848743 | 0.010078598 | 0.111771144 | 0.902140449 | 0.499609226 |
| Plec | T | 2893 | -0.037432388 | -0.14616471 | -0.260487734 | -0.223055346 | -0.114323024 | 0.284738629 | 0.213698672 | 0.000171084 | 0.000233893 | 0.302959751 |
| Ubap2l | T | 864 | -0.009835179 | -0.173954151 | -0.363996318 | -0.35416114 | -0.190042167 | 0.851585714 | 0.008886471 | 0.013796815 | 0.024107822 | 0.107607911 |
| Ankrd17 | S | 1692 | -0.0685021 | -0.115242019 | -0.215634984 | -0.147132884 | -0.100392965 | 0.085231159 | 0.031612777 | 0.00210361 | 0.016459905 | 0.069461018 |
| Golga3 | S | 388 | -0.013027796 | -0.266819173 | -0.09796244 | -0.084934644 | 0.168856733 | 0.734583627 | 1.38913E-05 | 0.189602199 | 0.245641661 | 0.072350437 |
| Rps6ka3 | S | 17 | -0.095537383 | -0.13090904 | -0.279133893 | -0.18359651 | -0.148224852 | 0.112937151 | 0.014820231 | 0.028729532 | 0.087927571 | 0.151603653 |
| Wiz | S | 1370 | 0.030059935 | -0.179724053 | -0.265577959 | -0.295637895 | -0.085853906 | 0.276845995 | 0.000769341 | 0.002650412 | 0.01096209 | 0.108416724 |
| Cetn2 | S | 20 | -0.170428159 | -0.142877287 | -0.040045272 | 0.130382888 | 0.102832015 | 0.000457055 | 0.00359536 | 0.564412063 | 0.142744656 | 0.208554163 |
| Mybbp1a | S | 1253 | 0.130188162 | -0.221748938 | -0.35556788 | -0.485756042 | -0.133818943 | 0.059113689 | 0.000772067 | 0.001279562 | 0.000956861 | 0.047535974 |
| Atg2b | S | 1015 | -0.110608602 | -0.097905077 | -0.205809439 | -0.095200837 | -0.107904362 | 0.066644639 | 0.111186335 | 0.001130564 | 0.103357562 | 0.092830568 |
| Atg2b | S | 1017 | -0.110608602 | -0.097905077 | -0.205809439 | -0.095200837 | -0.107904362 | 0.066644639 | 0.111186335 | 0.001130564 | 0.103357562 | 0.092830568 |
| Gkap1 | S | 97 | -0.163764302 | -0.107999976 | -0.192066053 | -0.028301751 | -0.084066077 | 0.011378384 | 0.427099964 | 0.003861124 | 0.536915781 | 0.522915546 |
| Gkap1 | S | 98 | -0.163764302 | -0.107999976 | -0.192066053 | -0.028301751 | -0.084066077 | 0.011378384 | 0.427099964 | 0.003861124 | 0.536915781 | 0.522915546 |
| Usp24 | S | 75 | -0.154339211 | -0.194503263 | -0.204077541 | -0.04973833 | -0.009574278 | 0.067285806 | 0.058214262 | 0.042443182 | 0.548054057 | 0.914396839 |
| Ankrd10 | S | 232 | -0.120852396 | -0.161047094 | -0.326588842 | -0.205736446 | -0.165541749 | 0.160553048 | 0.12783521 | 0.017885412 | 0.083208534 | 0.162818185 |
| Ldlrap1 | S | 198 | -0.102752098 | -0.102901919 | -0.203253125 | -0.100501027 | -0.100351206 | 0.021316838 | 0.037305508 | 0.008880463 | 0.094281694 | 0.096259073 |

| | | | | | | | | | | | | |
|---|---|---|---|---|---|---|---|---|---|---|---|---|
| Cdc25b | S | 351 | -0.098444679 | -0.098547084 | -0.442318557 | -0.343873878 | -0.343771473 | 0.310466428 | 0.376213796 | 0.004675691 | 0.021625048 | 0.018280052 |
| Gys1 | S | 412 | -0.164095468 | -0.198529922 | -0.139775552 | 0.024319916 | 0.05875437 | 0.013374343 | 0.146853231 | 0.021848298 | 0.553309874 | 0.590735391 |
| Cd44 | S | 754 | 0.015136969 | -0.186469651 | -0.240516426 | -0.255653395 | -0.054046775 | 0.757246023 | 0.003244119 | 0.002056812 | 0.007193704 | 0.256307082 |
| Taf1d | S | 39 | -0.041609109 | -0.180401121 | -0.253892611 | -0.212283502 | -0.073491489 | 0.495456126 | 0.017912167 | 0.004736496 | 0.00838641 | 0.142107934 |
| Slk | S | 775 | -0.026764308 | -0.166356589 | -0.281325512 | -0.254561204 | -0.114968923 | 0.704930065 | 0.052503 | 0.001030011 | 0.026617068 | 0.137036263 |
| Scamp3 | S | 87 | -0.078706453 | -0.22390732 | -0.336356338 | -0.257649885 | -0.112449018 | 0.513596645 | 0.064542462 | 0.016520098 | 0.037189749 | 0.137003988 |
| Pxn | S | 126 | -0.021600318 | -0.188136677 | -0.179485228 | -0.157884911 | 0.008651449 | 0.65161162 | 0.001005519 | 0.006536074 | 0.034117867 | 0.820643773 |
| Fez1 | S | 58 | -0.202639369 | -0.016418438 | -0.232086974 | -0.029447605 | -0.215668536 | 0.019541162 | 0.876563279 | 0.001240068 | 0.592304642 | 0.130070739 |
| Rrp1 | S | 489 | -0.041511921 | -0.178247348 | -0.426021438 | -0.384509517 | -0.24777409 | 0.720559491 | 0.01390723 | 0.034860612 | 0.051739922 | 0.137341352 |
| Slc7a1 | T | 613 | -0.012134642 | -0.245591823 | -0.370920881 | -0.358786239 | -0.125329059 | 0.742043088 | 0.029310305 | 0.021544079 | 0.02875117 | 0.242347147 |
| Ccdc9b | S | 230 | 0.048498544 | -0.293258153 | -0.184188044 | -0.232686588 | 0.109070109 | 0.526170336 | 0.000158144 | 0.044009949 | 0.044629496 | 0.16560227 |
| Map7d1 | T | 49 | -0.062258451 | -0.25613963 | -0.087057643 | -0.024799192 | 0.169081987 | 0.393642673 | 3.75518E-05 | 0.291518186 | 0.783189673 | 0.111082327 |
| Bmp2k | S | 715 | 0.038078399 | -0.379500977 | -0.237153334 | -0.275231733 | 0.142347643 | 0.599976696 | 0.007164413 | 0.021687341 | 0.015850626 | 0.168462113 |
| Hax1 | S | 69 | -0.03340993 | -0.270060432 | -0.353123229 | -0.319713299 | -0.083062797 | 0.50330575 | 0.033168009 | 0.038863009 | 0.056597698 | 0.496739175 |
| Csnk2b | S | 205 | 0.0301361 | -0.243723803 | -0.247466594 | -0.277602694 | -0.003742792 | 0.737557192 | 0.00042182 | 0.037740123 | 0.04825784 | 0.959052976 |
| Prc1 | S | 195 | -0.222466997 | -0.193843739 | -0.125372009 | 0.097094988 | 0.068471731 | 0.089013847 | 0.004643301 | 0.236605735 | 0.427766826 | 0.47435604 |
| Cfdp1 | S | 127 | -0.016192292 | -0.146365156 | -0.223914326 | -0.207722034 | -0.07754917 | 0.499937419 | 0.103162409 | 4.56634E-05 | 0.000555158 | 0.295980794 |
| Dock5 | S | 1832 | -0.172570727 | -0.117824611 | -0.137582871 | 0.034987856 | -0.019758261 | 0.001598546 | 0.049569914 | 0.131290731 | 0.605605821 | 0.778603101 |
| Ube2e1 | T | 28 | 0.018645276 | -0.1872478 | -0.535201341 | -0.553846617 | -0.347953541 | 0.825294674 | 0.036535867 | 0.019682757 | 0.022253955 | 0.074711242 |
| Mprip | S | 270 | -0.25183035 | -0.167760972 | -0.124839032 | 0.126991318 | 0.04292194 | 0.008034325 | 0.173909638 | 0.102530684 | 0.123544963 | 0.687458112 |
| Foxk1 | S | 229 | 0.077486509 | -0.215661656 | -0.126143429 | -0.203629938 | 0.089518227 | 0.128099651 | 7.33024E-06 | 0.000500299 | 0.016053554 | 0.007628305 |
| Psma5 | T | 55 | -0.258026183 | -0.200996043 | -0.252664062 | 0.00536212 | -0.051668019 | 0.325385958 | 0.223287277 | 0.009143479 | 0.981324071 | 0.713116788 |
| Bbx | S | 476 | 0.038281601 | -0.360434684 | -0.172444842 | -0.210726443 | 0.187989842 | 0.324149212 | 0.000111298 | 0.149647102 | 0.111069012 | 0.130418914 |
| Plk4 | S | 400 | -0.179011427 | -0.072154122 | -0.276091113 | -0.097079686 | -0.20393699 | 0.073098897 | 0.318341896 | 0.005259447 | 0.232263198 | 0.011925896 |
| Dst | T | 2521 | -0.137972236 | -0.183488774 | -0.230391067 | -0.092418831 | -0.046902292 | 0.104451998 | 0.022498738 | 0.070107336 | 0.335921343 | 0.576661786 |
| Rnf213 | T | 293 | -0.082859537 | -0.379781292 | -0.188390895 | -0.105531359 | 0.191390397 | 0.487728448 | 0.011716464 | 0.120443601 | 0.175501927 | 0.005208781 |
| Anln | S | 790 | -0.076337018 | -0.262620994 | -0.224177057 | -0.147840039 | 0.038443937 | 0.315844379 | 0.026042724 | 0.027105744 | 0.126361359 | 0.639310574 |
| Atrx | S | 440 | -0.1684058 | -0.238014804 | -0.21776675 | -0.04936095 | 0.020248053 | 0.135440384 | 0.058334132 | 0.065020938 | 0.33467556 | 0.750682625 |
| Ice1 | S | 1677 | -0.023007276 | -0.209222909 | -0.204774223 | -0.181766948 | 0.004448685 | 0.493145691 | 0.002472826 | 0.015240426 | 0.044824698 | 0.934938582 |
| Prune2 | S | 698 | -0.141456358 | -0.124839032 | -0.18751649 | -0.046060132 | -0.062677458 | 0.061511698 | 0.02208045 | 0.020204971 | 0.496544581 | 0.279102302 |
| Phldb2 | S | 490 | -0.35530535 | -0.572090775 | -1.655189437 | -1.299884088 | -1.083098663 | 0.175958728 | 0.034738634 | NA | NA | NA |
| Cdc42bpa | S | 750 | -0.126025288 | -0.121243679 | -0.107154177 | 0.01887111 | 0.014089501 | 0.02843338 | 0.010498957 | 0.004643326 | 0.610167384 | 0.608195173 |
| Tbc1d9b | S | 464 | -0.037103263 | -0.088340758 | -0.312181404 | -0.275078141 | -0.223840646 | 0.49435304 | 0.171253117 | 0.000234702 | 0.009663982 | 0.025434524 |
| Thrap3 | S | 572 | -0.286521439 | -0.049251224 | -0.342457263 | -0.055935824 | -0.293206039 | 0.052053232 | 0.796378478 | 0.021666184 | 0.63432074 | 0.186737595 |
| 1110059G10Rik | S | 147 | -0.280230647 | -0.037970944 | -0.416792495 | -0.136561849 | -0.378821551 | 0.113052306 | 0.832156871 | 0.01676014 | 0.365868521 | 0.084232563 |
| Riok1 | T | 57 | -0.102642766 | -0.324563944 | -0.107600369 | -0.004957604 | 0.216963575 | 0.242747871 | 0.007964373 | 0.068829201 | 0.948245045 | 0.037079829 |
| Nf1 | S | 2545 | -0.35950904 | -0.020989343 | -0.370386676 | -0.010877637 | -0.349397334 | 0.03287301 | 0.908931274 | 0.061399617 | 0.939608348 | 0.115239247 |
| Zzef1 | S | 1475 | -0.068873277 | -0.190380202 | -0.215899312 | -0.147026036 | -0.025519111 | 0.19175883 | 0.026215961 | 0.00979205 | 0.045590152 | 0.698100553 |
| Akt1 | S | 129 | -0.150810218 | -0.080435955 | -0.424156874 | -0.273346656 | -0.343720919 | 0.15363082 | 0.42359983 | 0.011693548 | 0.062003111 | 0.033497118 |
| Prag1 | S | 209 | -0.009925434 | -0.218846427 | -0.186606828 | -0.176681393 | 0.0322396 | 0.790943709 | 0.001629631 | 0.009280149 | 0.014405528 | 0.462430575 |
| Map3k2 | S | 37 | -0.17011863 | -0.116224604 | -0.255569765 | -0.085451135 | -0.139345161 | 0.033829642 | 0.288983026 | 0.015888829 | 0.234391621 | 0.214998505 |
| Arhgef7 | S | 228 | -0.068318514 | -0.12676268 | -0.210001263 | -0.141682749 | -0.083238583 | 0.144152921 | 0.025662809 | 0.002477422 | 0.000501617 | 0.043260118 |
| Helb | T | 965 | -0.07373034 | -0.189297301 | -0.160304101 | -0.086573761 | 0.0289932 | 0.271829434 | 0.007851971 | 0.006480284 | 0.192842025 | 0.482269249 |
| Akirin2 | S | 18 | 0.001819726 | -0.146462206 | -0.224729593 | -0.226549319 | -0.078267387 | 0.93329507 | 0.019943414 | 0.00012899 | 0.005322339 | 0.115963111 |
| Tmem201 | S | 436 | 0.062333867 | -0.164121708 | -0.27816097 | -0.340494838 | -0.114039262 | 0.278091436 | 0.006620501 | 0.000201389 | 0.004566945 | 0.026405239 |
| Kif20b | S | 806 | -0.113189152 | -0.127519494 | -0.231057244 | -0.117868091 | -0.10353775 | 0.158554741 | 0.071241629 | 0.004164698 | 0.151421754 | 0.133295261 |
| Aptx | S | 142 | -0.200199816 | -0.125100623 | -0.405703485 | -0.20550367 | -0.280602863 | 0.334160638 | 0.381446533 | 0.011490857 | 0.328497031 | 0.113224647 |
| Rabgef1 | S | 132 | -0.22558343 | -0.103348748 | -0.258109938 | -0.032526509 | -0.15476119 | 0.030481321 | 0.377946128 | 0.028782663 | 0.62945976 | 0.184097084 |
| Flvcr1 | S | 558 | -0.297786444 | -0.254692363 | -0.094235689 | 0.203550755 | 0.160456673 | 0.007088916 | 0.129502271 | 0.582184158 | 0.274475713 | 0.41571387 |
| Ring1 | S | 324 | 0.049109853 | -0.232751577 | -0.401043247 | -0.4501531 | -0.16829167 | 0.50313797 | 0.060343322 | 0.00277412 | 0.006032259 | 0.144649303 |
| H2-D1 H2-L | S | 356;356;166 | -0.303452325 | 0.018133253 | -0.360940325 | -0.057488001 | -0.379073578 | 0.024039827 | 0.922389918 | 0.028046325 | 0.60920929 | 0.108012355 |
| Chic1 | S | 80 | 0.000691661 | -0.19953738 | -0.302344876 | -0.303036537 | -0.102807496 | 0.989755219 | 0.00819595 | 0.009719483 | 0.01264392 | 0.194576894 |
| Prdm6 | S | 408 | -0.087766332 | -0.214425253 | -0.255421983 | -0.167655651 | -0.04099673 | 0.588029379 | 0.003468154 | 0.062296521 | 0.35715892 | 0.662384071 |
| Prdm6 | S | 409 | -0.087766332 | -0.214425253 | -0.255421983 | -0.167655651 | -0.04099673 | 0.588029379 | 0.003468154 | 0.062296521 | 0.35715892 | 0.662384071 |
| Prdm6 | S | 414 | -0.087766332 | -0.214425253 | -0.255421983 | -0.167655651 | -0.04099673 | 0.588029379 | 0.003468154 | 0.062296521 | 0.35715892 | 0.662384071 |
| Mettl18 | S | 67 | -0.000785672 | -0.256445311 | -0.242469456 | -0.241683784 | 0.013975855 | 0.989854815 | 0.009243923 | 0.009698303 | 0.017238912 | 0.841210093 |
| Ndel1 | S | 197 | -0.237601828 | -0.146841257 | -0.253476633 | -0.015874805 | -0.106635376 | 0.072080876 | 0.043734125 | 0.153118933 | 0.918246064 | 0.466483435 |
| Nifk | T | 257 | -0.009097975 | -0.153033096 | -0.227449042 | -0.218351067 | -0.074415946 | 0.838284161 | 0.040744928 | 0.000145537 | 0.015030211 | 0.183219642 |
| Srrm1 | S | 735 | -0.316014551 | -0.244156937 | -0.045884567 | 0.270129983 | 0.19827237 | 0.026410387 | 0.027495376 | 0.616468045 | 0.055384448 | 0.08504967 |
| Zfp36l2 | S | 416 | 0.061084026 | -0.179001001 | -0.371038775 | -0.432122801 | -0.192037774 | 0.42174677 | 0.022241623 | 0.001400217 | 0.002373251 | 0.031716991 |
| Gja1 | S | 364 | 0.055001595 | -0.158790088 | -0.197580773 | -0.252582368 | -0.038790685 | 0.20182617 | 7.28816E-05 | 0.000513464 | 0.003965584 | 0.166070153 |
| Dkc1 | S | 481 | -0.022624143 | -0.106464376 | -0.22460375 | -0.201979607 | -0.118139373 | 0.55462085 | 0.013154947 | 0.000260779 | 0.002786308 | 0.003748966 |
| Setd2 | S | 753 | -0.089297669 | -0.212935496 | -0.046629658 | 0.042668011 | 0.166305838 | 0.020524444 | 0.000529983 | 0.146245188 | 0.057344559 | 0.009242916 |
| Tanc1 | S | 336 | -0.240296475 | 0.000711299 | -0.391845979 | -0.151549504 | -0.392557278 | 0.02612818 | 0.995726691 | 0.022205838 | 0.222915369 | 0.052786047 |
| Ybx3 | S | 328 | -0.01952309 | -0.082194736 | -0.464037464 | -0.444514375 | -0.381842728 | 0.801280305 | 0.356182743 | 0.000944985 | 0.007322539 | 0.002843767 |
| Riok2 | S | 437 | -0.036857822 | -0.169184881 | -0.240327858 | -0.203470037 | -0.071142978 | 0.290657033 | 0.011961147 | 0.008093915 | 0.010381195 | 0.18878165 |
| Brd1 | S | 814 | -0.00331072 | -0.139857398 | -0.150749611 | -0.147438891 | -0.010892213 | 0.908272463 | 0.002698821 | 4.60035E-05 | 0.01534876 | 0.6287264 |
| Ptpn12 | S | 673 | -0.246070859 | -0.154953691 | -0.117702018 | 0.128368841 | 0.037251673 | 0.064157781 | 0.02934947 | 0.050494401 | 0.21600362 | 0.513039266 |
| Tnks1bp1 | S | 976 | -0.111498447 | -0.112313128 | -0.26689522 | -0.155396772 | -0.154582092 | 0.057562402 | 0.185487483 | 0.00674646 | 0.059902637 | 0.104012263 |
| Ranbp2 | T | 779 | -0.016011208 | -0.147106346 | -0.181226044 | -0.165214836 | -0.034119698 | 0.691206895 | 0.001721343 | 0.001121429 | 0.01121917 | 0.261900309 |
| Kif20b | S | 1644 | 0.036086695 | -0.29308238 | -0.251060452 | -0.287147147 | 0.042021928 | 0.430424726 | 0.002400546 | 0.018809604 | 0.028914267 | 0.572426503 |
| Sugp2 | S | 206 | -0.198133059 | -0.094387706 | -0.520986799 | -0.322853739 | -0.426599092 | 0.135329581 | 0.53967519 | 0.042415284 | 0.143839425 | 0.08277697 |
| Nav3 | S | 1192 | -0.025950655 | -0.168461755 | -0.14469451 | -0.118743855 | 0.023767246 | 0.188391369 | 0.001217642 | 0.004635179 | 0.013521534 | 0.433701981 |
| Bmpr2 | S | 586 | -0.246553764 | -0.006534919 | -0.442169416 | -0.195615652 | -0.435634497 | 0.180412542 | 0.968163033 | 0.006698945 | 0.242064818 | 0.058813951 |
| Cyld | S | 556 | -0.25303297 | -0.049692838 | -0.255337279 | -0.00230436 | -0.205644441 | 0.020675053 | 0.538067995 | 0.022221733 | 0.971994138 | 0.036846045 |
| Dnttip2 | S | 16 | -0.014691684 | -0.259454554 | -0.296293194 | -0.28160151 | -0.03683864 | 0.717544722 | 0.068323782 | 0.005275322 | 0.007453582 | 0.70704441 |
| Dnttip2 | S | 17 | -0.014691684 | -0.259454554 | -0.296293194 | -0.28160151 | -0.03683864 | 0.717544722 | 0.068323782 | 0.005275322 | 0.007453582 | 0.70704441 |
| Ttc14 | S | 617 | -0.119445059 | -0.116427491 | -0.237435417 | -0.117990358 | -0.121007926 | 0.032154078 | 0.341941947 | 0.003574913 | 0.075258497 | 0.325536811 |
| Ampd2 | S | 53 | -0.240941372 | -0.154920731 | -0.489492148 | -0.248550776 | -0.334571417 | 0.146482253 | 0.448184279 | 0.077877465 | 0.287942137 | 0.213381937 |
| Cep170 | S | 443 | -0.148672525 | -0.180654538 | -0.473447034 | -0.324774509 | -0.292792496 | 0.249224124 | 0.246582127 | 0.049445026 | 0.12057508 | 0.167331911 |
| Ldha | S | 213 | -0.16079927 | -0.206125582 | -0.062349132 | 0.098450138 | 0.143776451 | 0.003475945 | 0.045499917 | 0.078701254 | 0.031491778 | 0.116102334 |
| Ythdc2 | S | 449 | 0.038144738 | -0.399028197 | -0.370133934 | -0.408278672 | 0.028894264 | 0.79031158 | 0.035591968 | 0.040097956 | 0.059646806 | 0.846014907 |
| Rnf169 | S | 410 | -0.136909217 | -0.090700097 | -0.423720916 | -0.286811699 | -0.33302082 | 0.022143746 | 0.406557304 | 0.046390499 | 0.115593703 | 0.078331357 |
| Rpl26 Gm15772 | S | 12 | -0.013388097 | -0.248760674 | -0.261365686 | -0.24797759 | -0.012605012 | 0.819333643 | 0.034143195 | 0.004866424 | 0.014177633 | 0.875088775 |
| Tubb2a Tubb2b Tubb | S | 338;338;338;338; | -0.05922052 | -0.167138547 | -0.326386358 | -0.267165838 | -0.15924781 | 0.401595075 | 0.02367475 | 0.020781295 | 0.04045141 | 0.135057157 |
| Brd1 | S | 852 | -0.075698864 | -0.134529653 | -0.167395972 | -0.091697108 | -0.032866319 | 0.019078758 | 0.019684718 | 0.010507455 | 0.07530853 | 0.479443544 |
| Dnmt1 | S | 150 | -0.031880541 | -0.095073321 | -0.192071586 | -0.160191046 | -0.096998265 | 0.1601657 | 0.017856371 | 0.000179147 | 0.001503829 | 0.020062744 |
| Gnl1 | T | 48 | -0.131281521 | -0.148928974 | -0.294813335 | -0.163531815 | -0.145884361 | 0.192535311 | 0.094599842 | 0.017812115 | 0.15684692 | 0.157953704 |
| Tes | S | 173 | -0.149173824 | -0.16730547 | -0.083242015 | 0.065931809 | 0.084063456 | 0.01067191 | 0.008035669 | 0.113209723 | 0.119246565 | 0.081491656 |
| Tmem230 | T | 25 | -0.282029169 | -0.112829526 | -0.246464241 | 0.035564928 | -0.133634715 | 0.024432491 | 0.324835568 | 0.091682116 | 0.771234248 | 0.338806999 |
| 4930579G24Rik | S | 43 | 0.035804182 | -0.394870052 | -0.459127222 | -0.494931404 | -0.06425717 | 0.810037609 | 0.025584705 | 0.099401966 | 0.087776047 | 0.767770571 |
| Larp4 | S | 500 | -0.019768549 | -0.313628417 | -0.39187744 | -0.372108891 | -0.078249023 | 0.781569664 | 0.042986177 | 0.050276746 | 0.040823362 | 0.56102883 |
| Tacc2 | S | 583 | -0.138960469 | -0.203713942 | -0.088949385 | 0.050011083 | 0.114764557 | 0.027178357 | 0.003985706 | 0.203638164 | 0.453901733 | 0.131948688 |
| Dok1 | Y | 361 | -0.315190649 | -0.172050383 | -0.188624361 | 0.126566288 | -0.016573978 | 0.013260343 | 0.30832573 | 0.228797869 | 0.366917107 | 0.923704472 |

| | | | | | | | | | | | | |
|---|---|---|---|---|---|---|---|---|---|---|---|---|
| Arcn1 | S | 259 | -0.283820089 | -0.266350009 | -0.280443342 | 0.003376747 | -0.014093333 | 0.075693218 | 0.13167483 | 0.335136488 | 0.989860879 | 0.95876799 |
| Tmf1 | S | 359 | -0.30754976 | -0.36674786 | -0.0759609 | 0.231588861 | 0.29078696 | 0.29991617 | 0.010764082 | 0.449536764 | 0.41188623 | 0.046035431 |
| Ttc3 | S | 1903 | -0.04640508 | -0.236044422 | -0.242038648 | -0.195633568 | -0.005994226 | 0.452841855 | 0.015780826 | 0.021748185 | 0.046625801 | 0.9313252 |
| Plec | S | 4392 | 0.064528817 | -0.171534891 | -0.345930512 | -0.410459329 | -0.174395621 | 0.127661984 | 0.003381098 | 0.001255681 | 0.006079797 | 0.032992871 |
| Frmd8 | S | 440 | -0.259459433 | -0.031392499 | -0.352164879 | -0.092705446 | -0.32077238 | 0.006675287 | 0.819599069 | 0.088494777 | 0.544068925 | 0.131689337 |
| Tbc1d5 | S | 544 | -0.252241089 | -0.112973848 | -0.240673644 | 0.011567445 | -0.127699796 | 0.056467304 | 0.259944943 | 0.034868718 | 0.890642197 | 0.06295345 |
| Phldb1 | S | 698 | -0.051160096 | -0.163890184 | -0.262770342 | -0.211610247 | -0.098880159 | 0.285991348 | 0.045148314 | 0.005243238 | 0.015645517 | 0.181543276 |
| Api5 | S | 462 | -0.098984059 | -0.141038728 | -0.313623714 | -0.214639655 | -0.172584986 | 0.126565427 | 0.052133931 | 0.031442611 | 0.069132052 | 0.11395631 |
| Peak1 | S | 852 | 0.017452743 | -0.246418767 | -0.310950911 | -0.328403654 | -0.064532144 | 0.213906145 | 0.014159279 | 0.005213649 | 0.009583542 | 0.26111037 |
| Otud7b | S | 501 | -0.345345255 | -0.063723646 | -0.404666596 | -0.059321341 | -0.340942951 | 0.033607964 | 0.676078152 | 0.136241042 | 0.781115325 | 0.191196107 |
| Cdc25b | S | 370 | -0.181228385 | -0.135943255 | -0.357469041 | -0.176240655 | -0.221525786 | 0.200320643 | 0.095198384 | 0.050700627 | 0.28550537 | 0.152342345 |
| Ints12 | S | 377 | -0.090474004 | -0.104019895 | -0.119764618 | -0.029290614 | -0.015744724 | 0.057642261 | 0.092190285 | 2.27503E-05 | 0.376863839 | 0.705464728 |
| Setd2 | S | 475 | -0.097963677 | -0.197110014 | -0.255209619 | -0.157245942 | -0.058099605 | 0.281394013 | 0.008160001 | 0.075725362 | 0.222437286 | 0.576775337 |
| Lsm14a | S | 373 | -0.155656026 | -0.121687932 | -0.342625976 | -0.18696995 | -0.220938045 | 0.138164107 | 0.222703371 | 0.020390878 | 0.106939302 | 0.081037066 |
| Sec62 | S | 356 | 0.031493478 | -0.213739368 | -0.218026544 | -0.249520021 | -0.004287175 | 0.481784164 | 0.005206681 | 0.00098411 | 0.00629535 | 0.915430239 |
| Aimp1 | T | 84 | -0.218940498 | -0.299536643 | -0.118566296 | 0.100374202 | 0.180970347 | 0.091135457 | 0.025279083 | 0.329690031 | 0.347326825 | 0.109966119 |
| Atxn2l | S | 236 | 0.0260624 | -0.149106245 | -0.17163107 | -0.19769347 | -0.022524825 | 0.242017867 | 0.00057055 | 7.92439E-05 | 0.000654717 | 0.300346671 |
| Prune2 | S | 1926 | -0.088851641 | -0.125356968 | -0.116242019 | -0.027390378 | 0.009114949 | 0.109926545 | 0.00031922 | 0.01706869 | 0.560348773 | 0.753596917 |
| Cebpz | S | 972 | -0.18479041 | -0.144748586 | -0.613420082 | -0.428629672 | -0.468671496 | 0.378340204 | 0.328394951 | 0.061326404 | 0.158792474 | 0.1147867 |
| Fip1l1 | S | 36 | -0.168585494 | -0.187525802 | -0.406140912 | -0.237555417 | -0.21861511 | 0.339132304 | 0.237089771 | 0.0311967 | 0.229034868 | 0.220300006 |
| Nt5c | T | 102 | -0.24149882 | -0.097678807 | -0.196662667 | 0.044836153 | -0.09898386 | 0.002814671 | 0.40206821 | 0.121867757 | 0.647925969 | 0.463109467 |
| Ripk2 | S | 411 | -0.153903435 | -0.097519067 | -0.201311786 | -0.047408351 | -0.103792719 | 0.004729057 | 0.192135016 | 0.034356512 | 0.458127565 | 0.22924526 |
| Afap1 | S | 278 | 0.037965078 | -0.264978258 | -0.179378309 | -0.217343387 | 0.085599949 | 0.282705542 | 0.001673143 | 0.002146371 | 0.005619414 | 0.10879293 |
| Ppp3ca | S | 462 | 0.073940697 | -0.20265007 | -0.247285533 | -0.321226231 | -0.044635463 | 0.444756136 | 0.023137557 | 0.000107768 | 0.047330109 | 0.406470138 |
| Sh3pxd2a | S | 993 | -0.073238182 | -0.15190437 | -0.291768475 | -0.218530293 | -0.139864104 | 0.556798644 | 0.0248221 | 0.008842573 | 0.152421572 | 0.098040044 |
| Eml3 | S | 162 | 0.039742155 | -0.198028744 | -0.312453573 | -0.352195728 | -0.114424829 | 0.666026262 | 0.015660455 | 0.002922885 | 0.01873847 | 0.120828368 |
| Ppp1r7 | S | 27 | -0.258556573 | -0.149818614 | -0.207479008 | 0.051077564 | -0.057660395 | 0.027732327 | 0.184366078 | 0.119541865 | 0.662103492 | 0.646779472 |
| Pum2 | S | 85 | -0.029769841 | -0.413350115 | -0.37118928 | -0.341419439 | 0.042160835 | 0.749554949 | 0.075391731 | 0.066627234 | 0.092092427 | 0.831862559 |
| Serpinh1 | S | 137 | -0.199180819 | -0.171983133 | -0.372428369 | -0.173247549 | -0.200445236 | 0.0971436 | 0.455564185 | 0.043711404 | 0.241407456 | 0.41099619 |
| Arhgap23 | S | 372 | -0.189269372 | -0.164589836 | -0.25644396 | -0.067174588 | -0.091854124 | 0.074131568 | 0.133150594 | 0.059471988 | 0.462752687 | 0.365995902 |
| Nsd2 | T | 115 | 0.057232257 | -0.259005014 | -0.29603544 | -0.353267697 | -0.037030425 | 0.115705985 | 0.001531284 | 0.008796527 | 0.007915661 | 0.542643703 |
| Ptpn12 | T | 519 | -0.100442505 | -0.083420873 | -0.155106163 | -0.054663658 | -0.07168529 | 0.022603499 | 0.032300231 | 0.002158617 | 0.105628846 | 0.030947021 |
| Ccm2 | S | 417 | -0.088011589 | -0.176971265 | -0.171095028 | -0.083083439 | 0.005876237 | 0.497955278 | 0.000937767 | 0.03151611 | 0.527526988 | 0.904116946 |
| Mapk8ip3 | S | 755 | 0.056464704 | -0.125763115 | -0.370072919 | -0.426537623 | -0.244309805 | 0.456697626 | 0.042072432 | 0.000382922 | 0.014343404 | 0.005105823 |
| Rpap3 | S | 429 | -0.089573452 | -0.075606386 | -0.147769541 | -0.058196088 | -0.072163154 | 0.020540271 | 0.004096598 | 0.004557908 | 0.121347137 | 0.064509226 |
| Flrt2 | S | 611 | -0.164931252 | -0.310160283 | -0.599025641 | -0.434094388 | -0.288865358 | 0.437426423 | 0.362047775 | 0.091881721 | 0.176806951 | 0.425607849 |
| Hspa12a | S | 24 | -0.121636998 | -0.292200852 | -0.152801227 | -0.031164229 | 0.139399625 | 0.194895925 | 0.013472773 | 0.130525935 | 0.753121144 | 0.17105324 |
| Map4 | T | 433 | -0.032591984 | -0.235286154 | -0.288818 | -0.256226016 | -0.053531845 | 0.756097284 | 0.001786451 | 0.103678973 | 0.143462736 | 0.67221812 |
| Samd4b | S | 243 | -0.012335686 | -0.116378085 | -0.225106567 | -0.212770881 | -0.108728482 | 0.842504722 | 0.013405853 | 0.000238623 | 0.039353478 | 0.020654307 |
| Akt1s1 | S | 204 | -0.016846072 | -0.099940288 | -0.213684791 | -0.196838719 | -0.113744503 | 0.654591611 | 0.010815999 | 0.000158267 | 0.010796772 | 0.007376506 |
| Cdca2 | S | 712 | 0.055633009 | -0.184147717 | -0.263986732 | -0.319619741 | -0.079839014 | 0.147686113 | 0.001784195 | 0.000808482 | 0.006089163 | 0.100931644 |
| Ampd2 | S | 87 | -0.130998494 | -0.159662758 | -0.279627164 | -0.14862867 | -0.119964407 | 0.104875519 | 0.107355163 | 0.031563241 | 0.161611616 | 0.267797904 |
| Triobp | S | 1591 | -0.218318736 | -0.057894462 | -0.270854702 | -0.052535966 | -0.21296024 | 0.035081554 | 0.488341482 | 0.015258165 | 0.203552881 | 0.001173732 |
| Clasp2 | S | 505 | -0.220734949 | 0.032350944 | -0.476126152 | -0.255391203 | -0.508477096 | 0.149707092 | 0.850904726 | 0.006543574 | 0.080543229 | 0.035713743 |
| Mpp1 | S | 261 | -0.277134478 | -0.124897913 | -0.213068643 | 0.064065835 | -0.08817073 | 0.003111089 | 0.224796584 | 0.427876273 | 0.795202579 | 0.728572422 |
| Rbm3 | S | 129 | -0.294122079 | -0.087227044 | -0.211898613 | 0.082223466 | -0.12467157 | 0.019680165 | 0.400873077 | 0.063756336 | 0.273366695 | 0.127248299 |
| Nav3 | S | 1140 | -0.099591793 | -0.156545141 | -0.412117901 | -0.312526108 | -0.25557276 | 0.40397236 | 0.140290976 | 0.020478268 | 0.062503208 | 0.092860977 |
| Errfi1 | S | 336 | -0.07490432 | -0.335386036 | -0.204440841 | -0.12953652 | 0.130945195 | 0.66680107 | 0.007403441 | 0.116589725 | 0.498413765 | 0.269504549 |
| Ndel1 | S | 194 | -0.171601245 | -0.124620415 | -0.181068445 | -0.009467199 | -0.056448029 | 0.052863312 | 0.009275106 | 0.10005421 | 0.917789281 | 0.49816395 |
| Cdc5l | T | 227 | 0.007434662 | -0.269117209 | -0.416384491 | -0.423819153 | -0.147267281 | 0.935878164 | 0.13844689 | 0.008329092 | 0.013609225 | 0.360527969 |
| Hsp90ab1 | S | 307 | -0.029291794 | -0.250770398 | -0.353236165 | -0.323944371 | -0.102465766 | 0.865749874 | 0.042913675 | 0.021386988 | 0.142487356 | 0.392238328 |
| Snip1 | S | 144 | -0.073790869 | -0.25732842 | -0.141653687 | -0.067862818 | 0.115674733 | 0.003556769 | 0.01872559 | 0.22172609 | 0.492834057 | 0.298113369 |
| Map1a | S | 1772 | -0.088222058 | -0.106135418 | -0.085859635 | 0.002362423 | 0.020275783 | 0.038002389 | 0.000373109 | 0.00570913 | 0.935614456 | 0.309466361 |
| Map4k1 | T | 58 | -0.168971631 | -0.214801554 | -0.333389806 | -0.164418175 | -0.118588552 | 0.451901325 | 0.044261138 | 0.073191603 | 0.488890703 | 0.417071351 |
| Tnks1bp1 | S | 1453 | -0.154527506 | -0.429012454 | -0.202144495 | -0.047616989 | 0.226867958 | 0.336294011 | 0.016289491 | 0.492651417 | 0.861228417 | 0.431182566 |
| Dnmt1 | S | 138 | -0.010775406 | -0.100165843 | -0.257937627 | -0.247162221 | -0.157771784 | 0.665043236 | 0.006428213 | 0.000946383 | 0.00814262 | 0.028035221 |
| Cep55 | S | 426 | 0.031706732 | -0.163866045 | -0.259280983 | -0.290987714 | -0.095414938 | 0.48951892 | 0.00237944 | 0.001939829 | 0.003072998 | 0.086755126 |
| Mapk6 | S | 386 | -0.078560394 | -0.126388253 | -0.16758351 | -0.089023116 | -0.041195257 | 0.205864492 | 0.011890284 | 0.002111778 | 0.168741266 | 0.275501733 |
| Mapk6 | T | 389 | -0.078560394 | -0.126388253 | -0.16758351 | -0.089023116 | -0.041195257 | 0.205864492 | 0.011890284 | 0.002111778 | 0.168741266 | 0.275501733 |
| Ide | S | 57 | -0.097235893 | -0.185043301 | -0.161468667 | -0.064232773 | 0.023574634 | 0.202184961 | 0.010216829 | 0.018710464 | 0.327888801 | 0.558396953 |
| Prkca | S | 319 | 0.022126306 | -0.323869691 | -0.302202563 | -0.324328869 | 0.021667128 | 0.671314733 | 0.003869856 | 0.082132752 | 0.07149641 | 0.858431006 |
| Zswim8 | S | 1042 | -0.024374368 | -0.121211246 | -0.347585162 | -0.323210794 | -0.226373916 | 0.842814301 | 0.38587695 | 0.000218128 | 0.085216128 | 0.170671427 |
| Phldb2 | S | 380 | -0.162076244 | -0.164547979 | -0.323020754 | -0.16094451 | -0.158472775 | 0.196821152 | 0.176109406 | 0.029369681 | 0.204384301 | 0.197197095 |
| Dph2 | S | 489 | -0.205822874 | -0.178151383 | -0.189103235 | 0.016719638 | -0.010951853 | 0.024781276 | 0.066684115 | 0.204220614 | 0.890893776 | 0.93101417 |
| Pdlim2 | S | 116 | 0.185995512 | -0.625319813 | -0.50639375 | -0.692389261 | 0.118926063 | 0.474167333 | 0.073513109 | 0.055607258 | 0.060094302 | 0.665884357 |
| Stk4 | S | 418 | -0.027208083 | -0.112575719 | -0.322371334 | -0.295163251 | -0.209795615 | 0.668663434 | 0.135976509 | 0.000669599 | 0.009670266 | 0.034169582 |
| Cobll1 | T | 1027 | -0.072486062 | -0.165416163 | -0.14623229 | -0.073746228 | 0.019183873 | 0.043423514 | 0.004536389 | 0.037898684 | 0.189335997 | 0.705243395 |
| Stmn1 | S | 25 | -0.118281107 | -0.243448037 | -0.342242813 | -0.223961706 | -0.098794776 | 0.262952608 | 0.106409186 | 0.061547656 | 0.166988967 | 0.52938441 |
| Osbpl5 | S | 82 | 0.046562119 | -0.202419432 | -0.267860707 | -0.314422826 | -0.065441275 | 0.517900521 | 0.00646744 | 0.002260985 | 0.012173717 | 0.25781184 |
| Ppfibp1 | S | 500 | -0.329546496 | 0.018804301 | -0.336894224 | -0.007347728 | -0.355698525 | 0.020876785 | 0.910167194 | 0.044215769 | 0.950396734 | 0.091367122 |
| Map1a | S | 2066 | -0.090027752 | -0.095913789 | -0.134553148 | -0.044525396 | -0.038639359 | 0.031779845 | 0.001557776 | 0.012595987 | 0.261548556 | 0.261286632 |
| Add1 | S | 12 | -0.082188846 | -0.13980662 | -0.286398584 | -0.204209738 | -0.146591964 | 0.297907148 | 0.092801973 | 0.00555441 | 0.011430414 | 0.026581227 |
| Bin1 | S | 298 | -0.076069426 | -0.124613004 | -0.136593793 | -0.060524367 | -0.011980789 | 0.082343102 | 0.006890697 | 0.003120808 | 0.152127654 | 0.700555572 |
| Nans | S | 275 | -0.208884683 | -0.082990895 | -0.278624662 | -0.069739979 | -0.195633767 | 0.014744504 | 0.442324065 | 0.050448937 | 0.501377952 | 0.166845116 |
| Ptpre | Y | 695 | -0.133402068 | -0.190357934 | -0.314950452 | -0.181548384 | -0.124592518 | 0.304121233 | 0.152804276 | 0.019383566 | 0.119038221 | 0.216847651 |
| Sync | S | 32 | -0.154294509 | -0.153841178 | -0.249010966 | -0.094716457 | -0.095169788 | 0.144872412 | 0.068900539 | 0.032599316 | 0.376653707 | 0.309493719 |
| Akap10 | S | 52 | -0.000949355 | -0.209711909 | -0.19160055 | -0.190651195 | 0.01811136 | 0.982103011 | 0.013650389 | 0.000921842 | 0.0126576 | 0.708475048 |
| Snd1 | T | 103 | -0.09855238 | -0.152946472 | -0.09804661 | -0.008191371 | 0.054899863 | 0.126754342 | 0.001015685 | 0.012991588 | 0.859637482 | 0.073418562 |
| Frmd4a | T | 627 | 0.065380419 | -0.18203526 | -0.30103627 | -0.366416689 | -0.11900101 | 0.416478694 | 0.00746244 | 0.001421491 | 0.009710323 | 0.069414452 |
| Lmnb1 | S | 285 | -0.050204967 | -0.169010413 | -0.104915806 | -0.05471084 | 0.064094606 | 0.061770183 | 0.000190909 | 0.067183799 | 0.248995841 | 0.195585697 |
| Tnks1bp1 | S | 980 | -0.073253746 | -0.18634153 | -0.273570422 | -0.200316676 | -0.087228892 | 0.225814064 | 0.050813015 | 0.020158652 | 0.034262472 | 0.26092323 |
| Rpl22 | S | 68 | -0.135461015 | -0.172913936 | -0.28606194 | -0.150600925 | -0.113148004 | 0.083695159 | 0.153267101 | 0.040047949 | 0.186759204 | 0.372805925 |
| Nfkb1 | S | 944 | -0.163976641 | -0.121723332 | -0.377026256 | -0.213049615 | -0.255302924 | 0.16685961 | 0.352591208 | 0.020521553 | 0.10263555 | 0.086556318 |
| Anln | S | 293 | -0.007109109 | -0.139818768 | -0.2160423 | -0.208933191 | -0.076223532 | 0.885482876 | 0.001989126 | 0.002284451 | 0.015670582 | 0.095531305 |
| Dync1li1 | T | 515 | 0.047342486 | -0.210214545 | -0.244974993 | -0.292317479 | -0.034760448 | 0.290663387 | 0.00138954 | 0.003614282 | 0.004963751 | 0.446256992 |
| Tjp1 | S | 1051 | -0.333517431 | -0.05811066 | -0.234520556 | 0.098996875 | -0.176409896 | 0.008882335 | 0.582194091 | 0.143314221 | 0.458034013 | 0.236683259 |
| Wiz | S | 1039 | -0.05078802 | -0.125549854 | -0.215134326 | -0.164346306 | -0.089584472 | 0.184559087 | 0.002761352 | 0.014597924 | 0.036740632 | 0.152862987 |
| Wiz | S | 1043 | -0.05078802 | -0.125549854 | -0.215134326 | -0.164346306 | -0.089584472 | 0.184559087 | 0.002761352 | 0.014597924 | 0.036740632 | 0.152862987 |
| Daxx | S | 515 | -0.131845688 | -0.146205423 | -0.259827991 | -0.127982303 | -0.113622568 | 0.297455191 | 0.004581959 | 0.075850082 | 0.362014503 | 0.288887791 |
| Pum1 | S | 247 | -0.071078786 | -0.239347168 | -0.194229406 | -0.12315062 | 0.045117762 | 0.387572884 | 0.005393196 | 0.065380111 | 0.214910083 | 0.564407226 |
| Rab11fip5 | S | 382 | -0.040003428 | -0.116628937 | -0.344147619 | -0.304144191 | -0.227518682 | 0.503647035 | 0.305502731 | 0.000795547 | 0.006958585 | 0.102204555 |
| Rif1 | T | 2190 | -0.028105722 | -0.348471374 | -0.26635424 | -0.238248518 | 0.082117134 | 0.811216677 | 0.013873508 | 0.08752011 | 0.114094187 | 0.502543161 |
| Map4 | S | 828 | -0.106179286 | -0.152051411 | -0.323186162 | -0.217006876 | -0.171134751 | 0.186193334 | 0.176448053 | 0.014815437 | 0.062954363 | 0.159956079 |
| Arhgap31 | S | 1059 | 0.043579165 | -0.198537002 | -0.327556356 | -0.371135522 | -0.129019355 | 0.390228326 | 0.014608952 | 0.002768728 | 0.004987237 | 0.102825239 |

| | | | | | | | | | | | | |
|---|---|---|---|---|---|---|---|---|---|---|---|---|
| Tcof1 | S | 456 | 0.043770296 | -0.254240744 | -0.347323947 | -0.391094243 | -0.093083204 | 0.319826304 | 0.001875463 | 0.038854073 | 0.042953185 | 0.399587924 |
| Racgap1 | T | 577 | -0.024199131 | -0.142173911 | -0.234703365 | -0.210504234 | -0.092529454 | 0.533745459 | 0.003597073 | 0.006668999 | 0.009981564 | 0.094805892 |
| Tnks1bp1 | S | 247 | -0.051437072 | -0.131404884 | -0.215817976 | -0.164380904 | -0.084413092 | 0.33809797 | 0.050431247 | 0.000812609 | 0.027761222 | 0.148503569 |
| Ccdc6 | S | 318 | -0.199231401 | -0.073139312 | -0.240274443 | -0.041043042 | -0.167135131 | 0.103527819 | 0.353497445 | 0.003970339 | 0.6664492 | 0.078245556 |
| Frmd4a | S | 785 | -0.162080165 | -0.135020033 | -0.34648803 | -0.184407864 | -0.211467996 | 0.075788913 | 0.312931985 | 0.039593838 | 0.183259019 | 0.19142374 |
| Pcbp1 | S | 173 | -0.068909569 | -0.172094126 | -0.227958498 | -0.159048929 | -0.055864372 | 0.279764119 | 0.014147663 | 0.019629236 | 0.073615918 | 0.394785717 |
| Son | S | 1623 | -0.22690242 | -0.223924509 | -0.109088544 | 0.117813875 | 0.114835965 | 0.025404589 | 0.031870631 | 0.42085065 | 0.400088961 | 0.41324389 |
| Spire1 | S | 520 | -0.219453114 | -0.07941167 | -0.118176652 | 0.101276462 | -0.038764982 | 0.016064152 | 0.17008453 | 0.006508645 | 0.14014997 | 0.427765021 |
| Gas7 | S | 108 | 0.06263789 | -0.27264574 | -0.221583998 | -0.284221888 | 0.051061742 | 0.235694227 | 0.000964642 | 0.00885505 | 0.006338607 | 0.332517594 |
| Akt1 | T | 450 | -0.056907531 | -0.108688377 | -0.323384259 | -0.266476728 | -0.214695882 | 0.375598262 | 0.081911334 | 0.003877074 | 0.014851563 | 0.025362195 |
| Zc3hc1 | S | 406 | -0.009829859 | -0.143379847 | -0.160070685 | -0.150240826 | -0.016690839 | 0.627016939 | 0.000422679 | 0.00267416 | 0.013404252 | 0.555211828 |
| Sipa1l1 | S | 1116 | -0.323971602 | -0.113974823 | -0.226159035 | 0.097812568 | -0.112184211 | 0.024158374 | 0.357617762 | 0.149265475 | 0.401992224 | 0.362578437 |
| Wnk1 | T | 70 | -0.022831747 | -0.200647482 | -0.300545273 | -0.277713526 | -0.099897791 | 0.778222873 | 0.039913647 | 0.007247531 | 0.006100253 | 0.138109168 |
| Srrm2 | T | 1074 | -0.18368698 | -0.284402717 | -0.036030335 | 0.147656645 | 0.248372381 | 0.136221958 | 0.001290556 | 0.69095854 | 0.249344956 | 0.058331841 |
| Tacc1 | S | 401 | 0.006204797 | -0.156208337 | -0.121083619 | -0.127288416 | 0.035124718 | 0.838749261 | 0.000134519 | 0.00059107 | 0.019799441 | 0.118945558 |
| Nav3 | S | 1220 | -0.093440199 | -0.204141813 | -0.194370807 | -0.100930608 | 0.009771006 | 0.099013471 | 0.013150331 | 0.086205466 | 0.281442578 | 0.90940304 |
| Esyt1 | S | 314 | -0.209489142 | -0.133423619 | -0.254829439 | -0.045340297 | -0.12140582 | 0.274067588 | 0.011909362 | 0.078350543 | 0.80001898 | 0.289619012 |
| Hspa8 | S | 254 | -0.16235462 | -0.11209169 | -0.19222214 | -0.02986752 | -0.08013045 | 0.030425037 | 0.135385846 | 0.019775977 | 0.6289467 | 0.283070232 |
| Pank4 | S | 393 | -0.064658809 | -0.262210625 | -0.169137208 | -0.104478398 | 0.093073417 | 0.023657779 | 0.031281348 | 0.082581882 | 0.194867883 | 0.296294936 |
| Rfc1 | T | 162 | 0.059774462 | -0.194721846 | -0.293692935 | -0.353467397 | -0.09897109 | 0.243596049 | 0.006336253 | 0.001361712 | 0.003767487 | 0.099029496 |
| Pttg1 | S | 162 | -0.24878801 | -0.137320433 | -0.256664025 | -0.007876015 | -0.119343592 | 0.192199107 | 0.093370946 | 0.047953175 | 0.961964915 | 0.258738047 |
| Map1b | S | 339 | 0.01207555 | -0.185077099 | -0.26770299 | -0.27977854 | -0.082625891 | 0.786024971 | 0.000432432 | 0.034185002 | 0.02274851 | 0.295363968 |
| Dock7 | S | 439 | -0.110617803 | -0.05456477 | -0.555026497 | -0.444408694 | -0.500461727 | 0.4685267 | 0.760785346 | 0.006126055 | 0.007651342 | 0.034272356 |
| Lig3 | S | 918 | 0.01357498 | -0.152984256 | -0.190765886 | -0.204340866 | -0.03778163 | 0.63965214 | 0.000201948 | 0.003837225 | 0.004141625 | 0.280990871 |
| Cep170b | S | 965 | -0.027760072 | -0.158817713 | -0.187640436 | -0.159880364 | -0.028822723 | 0.757041194 | 0.00134938 | 0.005981896 | 0.15880866 | 0.451592748 |
| Clasp1 | T | 727 | -0.089977371 | -0.186810875 | -0.262523176 | -0.172545805 | -0.075712301 | 0.189597014 | 0.026643026 | 0.051464126 | 0.144529242 | 0.455161629 |
| Pnkp | S | 143 | -0.025721595 | -0.132668854 | -0.269321816 | -0.24360022 | -0.136652962 | 0.724820045 | 0.0612081 | 0.000750745 | 0.038015336 | 0.060350212 |
| Rtkn2 | S | 565 | -0.145508264 | -0.136701505 | -0.318484395 | -0.172976131 | -0.18178289 | 0.225344042 | 0.183927257 | 0.016333388 | 0.189350443 | 0.133311763 |
| Ap3d1 | S | 686 | -0.04889461 | -0.237755324 | -0.103350417 | -0.054455806 | 0.134404907 | 0.100991553 | 0.002828255 | 0.040675563 | 0.190883845 | 0.030494301 |
| Prr11 | T | 361 | -0.194917488 | -0.177869588 | -0.299604396 | -0.104686908 | -0.121734809 | 0.133870967 | 0.139653641 | 0.078818335 | 0.481112013 | 0.407034467 |
| Eif3c | S | 39 | -0.002691579 | -0.106527434 | -0.121040383 | -0.118348804 | -0.014512949 | 0.848661527 | 0.000540782 | 1.13334E-05 | 0.00351333 | 0.217921555 |
| 8030462N17Rik | S | 66 | -0.035326606 | -0.153510006 | -0.145175733 | -0.109849127 | 0.008334273 | 0.303965134 | 0.000164912 | 0.025838841 | 0.056556981 | 0.818196852 |
| Atxn1l | S | 282 | -0.106128723 | -0.132621676 | -0.284710991 | -0.178582268 | -0.152089314 | 0.142397871 | 0.100800424 | 0.016228178 | 0.066490814 | 0.105580305 |
| Rbsn | S | 634 | -0.01789094 | -0.474883648 | -0.243356929 | -0.225465989 | 0.23152672 | 0.852004034 | 0.015560375 | 0.196244455 | 0.226979312 | 0.233351403 |
| Akt1 | S | 126 | -0.129392913 | -0.079463925 | -0.472970556 | -0.343577643 | -0.393506632 | 0.251373934 | 0.429560278 | 0.015209353 | 0.047907277 | 0.030176346 |
| Eno1 Eno1b Eno2 Enc | Y | 44;44;44 | -0.246697404 | -0.181049533 | -0.337723033 | -0.091025629 | -0.1566735 | 0.211766615 | 0.213894274 | 0.072462225 | 0.641140667 | 0.357655704 |
| Exoc4 | S | 32 | -0.182311145 | -0.117800395 | -0.293984147 | -0.111673002 | -0.176183752 | 0.09461632 | 0.164668229 | 0.04035867 | 0.337049363 | 0.151008118 |
| Map1b | S | 2252 | 0.016445068 | -0.156864001 | -0.313191329 | -0.329636397 | -0.156327328 | 0.894692548 | 0.047765379 | 0.001351807 | 0.07967901 | 0.057511033 |
| Magi3 | S | 237 | 0.03548324 | -0.21771714 | -0.167812739 | -0.203295979 | 0.049904401 | 0.227694919 | 0.00112669 | 0.000772176 | 0.001305575 | 0.181184675 |
| Jcad | S | 1223 | -0.258502273 | -0.078003318 | -0.278268439 | -0.019766166 | -0.200265121 | 0.016460987 | 0.458483711 | 0.096813 | 0.869860626 | 0.197128481 |
| Hat1 | S | 358 | -0.118552392 | -0.075082733 | -0.154254383 | -0.035701991 | -0.07917165 | 0.000871779 | 0.014878328 | 0.092293236 | 0.581934899 | 0.280706759 |
| Pola2 | S | 125 | -0.255933991 | -0.116887482 | -0.27732084 | -0.021386849 | -0.160433358 | 0.074889652 | 0.496891164 | 0.030927376 | 0.842878913 | 0.348902007 |
| Asph | S | 38 | -0.15621788 | -0.131609085 | -0.313270234 | -0.157052354 | -0.181661149 | 0.02201664 | 0.372767745 | 0.056247201 | 0.230492216 | 0.288323819 |
| Ralgapa1 | S | 714 | -0.361736257 | 0.100975406 | -0.358121881 | 0.003614376 | -0.459097287 | 0.025415174 | 0.613003392 | 0.024705817 | 0.961178135 | 0.07916527 |
| Mcm3ap | S | 550 | 0.02748495 | -0.170654731 | -0.123330568 | -0.150815518 | 0.047324164 | 0.499712504 | 7.96581E-05 | 0.000661408 | 0.037303663 | 0.021261283 |
| Nudc | S | 139 | -0.209905795 | -0.186068628 | -0.186038 | 0.023867796 | 3.06281E-05 | 0.067946372 | 0.123781448 | 0.068385429 | 0.804700607 | 0.999767661 |
| Phldb1 | S | 585 | 0.001126865 | -0.129191289 | -0.228527946 | -0.229654811 | -0.099336657 | 0.972040388 | 0.007741677 | 0.000656721 | 0.002073331 | 0.038895098 |
| Ctnnd1 | S | 899 | -0.060433098 | -0.135944892 | -0.197304432 | -0.136871333 | -0.061359539 | 0.131725599 | 0.08338396 | 0.001233103 | 0.012386881 | 0.340020943 |
| Prune2 | T | 2312 | -0.262467628 | -0.352180427 | -0.125911249 | 0.136556379 | 0.226269178 | 0.124622535 | 0.045948596 | 0.491208444 | 0.360851434 | 0.165139679 |
| Bag3 | S | 201 | -0.083035544 | -0.128080437 | -0.438298559 | -0.355263016 | -0.310218123 | 0.054851447 | 0.217287093 | 0.049911235 | 0.086897976 | 0.092964452 |
| Caml | S | 53 | -0.077055542 | -0.164191039 | -0.109427472 | -0.032371931 | 0.054763566 | 0.041979426 | 0.005295131 | 0.019256389 | 0.355277416 | 0.160361579 |
| Dnajc2 | S | 183 | 0.00354646 | -0.238474237 | -0.192320705 | -0.195867165 | 0.046153532 | 0.969470625 | 0.000168847 | 0.110869464 | 0.147972982 | 0.605809194 |
| Plekha2 | S | 184 | -0.180999252 | -0.124601812 | -0.543714929 | -0.362715677 | -0.419113117 | 0.24120655 | 0.453083855 | 0.054659482 | 0.145273208 | 0.113374126 |
| Arid4b | S | 656 | -0.101479947 | -0.136191106 | -0.122661697 | -0.021181751 | 0.013529409 | 0.00707657 | 0.04000934 | 0.016964481 | 0.538912077 | 0.778326025 |
| Rif1 | S | 2191 | 0.042236555 | -0.255407595 | -0.221824322 | -0.264060877 | 0.033583273 | 0.245866185 | 0.008830466 | 0.001195366 | 0.004592372 | 0.556997986 |
| Eif4g3 | S | 421 | -0.183471959 | -0.116167745 | -0.28737812 | -0.103906161 | -0.171210375 | 0.042435016 | 0.072856822 | 0.131119892 | 0.500171034 | 0.300252436 |
| Eif4g3 | T | 417 | -0.183471959 | -0.116167745 | -0.28737812 | -0.103906161 | -0.171210375 | 0.042435016 | 0.072856822 | 0.131119892 | 0.500171034 | 0.300252436 |
| Yap1 | S | 149 | -0.147633655 | -0.109995792 | -0.280553415 | -0.132919759 | -0.170557623 | 0.086875422 | 0.206185661 | 0.018293963 | 0.16906074 | 0.108290605 |
| Wipi2 | S | 395 | -0.140149427 | -0.102593572 | -0.264984727 | -0.1248353 | -0.162391155 | 0.080795537 | 0.226732301 | 0.011042911 | 0.093112763 | 0.066520415 |
| Lsm11 | S | 15 | -0.063817946 | -0.179540161 | -0.24149011 | -0.177672164 | -0.06194995 | 0.170025522 | 0.021800103 | 0.032175378 | 0.043469376 | 0.348712869 |
| Myo9b | S | 1266 | -0.047687064 | -0.170054371 | -0.253404272 | -0.205717208 | -0.083349901 | 0.5203042 | 0.033702704 | 0.005913159 | 0.018986125 | 0.075661644 |
| Upf1 | S | 1105 | -0.053308552 | -0.175309166 | -0.201541094 | -0.148232542 | -0.026231928 | 0.424858092 | 0.101810189 | 0.000554855 | 0.089173984 | 0.736304538 |
| Sra1 | S | 60 | -0.049365132 | -0.110991552 | -0.274570675 | -0.225205543 | -0.163578723 | 0.186399495 | 0.046767187 | 0.00509195 | 0.03082777 | 0.03952633 |
| Sema5a | S | 957 | -0.374622812 | -0.042238838 | -0.352847917 | 0.021774895 | -0.310609079 | 0.03266962 | 0.85760474 | 0.121667329 | 0.900696643 | 0.257196661 |
| Exoc8 | S | 390 | -0.254354292 | -0.356111257 | -0.296366409 | -0.042012117 | 0.059744848 | 0.184813248 | 0.196176739 | 0.232599634 | 0.860418199 | 0.831298998 |
| Mast3 | S | 168 | 0.022083038 | -0.260255011 | -0.220582697 | -0.242665735 | 0.039672314 | 0.661257184 | 0.004827905 | 0.011677927 | 0.013100455 | 0.542442903 |
| Pphln1 | S | 92 | -0.114388941 | -0.20509008 | -0.21931096 | -0.10492202 | -0.01422088 | 0.391082464 | 0.045814611 | 0.016321984 | 0.430934828 | 0.84205438 |
| Syde1 | S | 159 | -0.072321324 | -0.099569596 | -0.402446709 | -0.330125384 | -0.302877113 | 0.415901363 | 0.374067493 | 0.003513684 | 0.012839354 | 0.024410113 |
| Eif3a | T | 1334 | -0.078314577 | -0.176429939 | -0.189257793 | -0.110943216 | -0.012827854 | 0.085261547 | 0.003805987 | 0.11317423 | 0.280182685 | 0.884785164 |
| Twf1 | S | 143 | -0.099055944 | -0.090919765 | -0.220483787 | -0.121427843 | -0.129564022 | 0.02578953 | 0.203974359 | 0.004557563 | 0.051035804 | 0.108535575 |
| Arhgef17 | S | 414 | -0.16695114 | -0.21388437 | -0.171838705 | -0.004887565 | 0.042045665 | 0.053219716 | 0.079520731 | 0.095103243 | 0.9524858 | 0.690002918 |
| Map2k2 | S | 295 | -0.142316781 | -0.059275565 | -0.20490098 | -0.062584199 | -0.145625415 | 0.014410442 | 0.313289469 | 0.005507491 | 0.183585031 | 0.041952932 |
| Srrm2 | S | 812 | 0.183351293 | -0.355293449 | -0.30409231 | -0.487443604 | 0.051201139 | 0.007068946 | 0.000193316 | 0.004544006 | 0.002842165 | 0.412684291 |
| Relch | S | 182 | -0.244153014 | -0.159976274 | -0.310344529 | -0.066191515 | -0.150368254 | 0.086992314 | 0.178630056 | 0.145237295 | 0.716343921 | 0.420115083 |
| Crtc3 | S | 329 | -0.166327789 | -0.175125456 | -0.145459117 | 0.020868673 | 0.02966634 | 0.061824838 | 0.012832881 | 0.160100411 | 0.82445877 | 0.725073458 |
| Dnajc17 | S | 112 | -0.105690688 | -0.090640712 | -0.159704757 | -0.054014069 | -0.069064045 | 0.016991387 | 0.034348765 | 0.009730799 | 0.207338606 | 0.130019477 |
| Ogfr | S | 149 | -0.072573482 | -0.226696272 | -0.168408094 | -0.095834612 | 0.058288178 | 0.184736729 | 0.0037749 | 0.123068406 | 0.310339207 | 0.501344311 |
| Ogfr | S | 150 | -0.072573482 | -0.226696272 | -0.168408094 | -0.095834612 | 0.058288178 | 0.184736729 | 0.0037749 | 0.123068406 | 0.310339207 | 0.501344311 |
| Mycbp2 | S | 2904 | -0.033203834 | -0.185095741 | -0.278029478 | -0.244825644 | -0.092933737 | 0.453051125 | 0.001742482 | 0.088021224 | 0.117100148 | 0.435446113 |
| Kars | S | 468 | -0.0293413 | -0.34537381 | -0.233427118 | -0.204085818 | 0.111946692 | 0.800477328 | 0.008380031 | 0.120907519 | 0.20064736 | 0.399119253 |
| Atg2b | S | 255 | -0.055229356 | -0.214679796 | -0.302541024 | -0.247311668 | -0.087861229 | 0.552388874 | 0.045902525 | 0.022156667 | 0.067555273 | 0.371710616 |
| Ankrd11 | S | 2354 | -0.262260652 | -0.200721496 | -0.014141391 | 0.248119261 | 0.186580105 | 0.012128004 | 0.01257428 | 0.807852255 | 0.031098705 | 0.006820365 |
| Upf2 | T | 529 | -0.523179252 | -0.280311162 | -0.340128935 | 0.183050316 | -0.059817773 | 0.310395438 | 0.375802135 | 0.181837071 | 0.689462901 | 0.826587103 |
| Ythdc2 | S | 1292 | -0.281148315 | -0.192610635 | 0.019187208 | 0.300335523 | 0.211797843 | 0.003286317 | 0.021360614 | 0.822961747 | 0.024326794 | 0.048820646 |
| Ehbp1l1 | T | 244 | 0.085564144 | -0.382786958 | -0.42774876 | -0.513312904 | -0.044961802 | 0.038577761 | 0.017364922 | 0.014190537 | 0.007394485 | 0.628520543 |
| Nr1d1 | S | 115 | -0.068521656 | -0.108395387 | -0.217584905 | -0.149063249 | -0.109189517 | 0.337028511 | 0.099645989 | 0.000455929 | 0.099490636 | 0.097179186 |
| Cenpu | S | 182 | -0.174570478 | -0.097232604 | -0.252313765 | -0.077743287 | -0.155081161 | 0.051440126 | 0.200432166 | 0.027604867 | 0.335531245 | 0.111574462 |
| Ctdspl2 | T | 60 | -0.113574478 | -0.199330365 | -0.155857954 | -0.042283477 | 0.043472411 | 0.285636759 | 0.000591344 | 0.239827835 | 0.750728404 | 0.69602104 |
| Bclaf3 | S | 592 | -0.279448662 | -0.252872373 | -0.133523815 | 0.145924847 | 0.119348559 | 0.063184581 | 0.053325134 | 0.451658478 | 0.393674518 | 0.450996784 |
| Rps26 | S | 54 | -0.073584862 | -0.192947976 | -0.428975915 | -0.355391052 | -0.236027939 | 0.519070909 | 0.081857302 | 0.045828626 | 0.07451376 | 0.168131856 |
| Hspa14 | S | 62 | -0.155567432 | -0.167165772 | -0.099800427 | 0.055767005 | 0.067365345 | 0.019028006 | 0.022791047 | 0.095961255 | 0.323399584 | 0.273333051 |
| Kirrel | Y | 654 | -0.194351529 | -0.189010863 | -0.328615407 | -0.134263878 | -0.139604545 | 0.100959802 | 0.340504645 | 0.066064218 | 0.364459961 | 0.497532987 |
| Ggnbp2 | S | 360 | -0.172200658 | -0.236432563 | -0.182844499 | -0.010643841 | 0.053588065 | 0.103359352 | 0.109182446 | 0.061160937 | 0.856195905 | 0.6223333 |
| Ppp1r7 | S | 24 | -0.268681461 | -0.135063598 | -0.177536482 | 0.09114498 | -0.042472884 | 0.017762227 | 0.236992008 | 0.148600275 | 0.423535116 | 0.732298061 |

| | | | | | | | | | | | | |
|---|---|---|---|---|---|---|---|---|---|---|---|---|
| Diaph3 | S | 1072 | 0.069642585 | -0.144688825 | -0.166944191 | -0.236586776 | -0.022255366 | 0.108415701 | 7.70068E-07 | 0.001760775 | 0.004066248 | 0.225505184 |
| Mybbp1a | S | 1244 | 0.012954798 | -0.183526455 | -0.24939777 | -0.262352568 | -0.065871315 | 0.663562003 | 0.005372384 | 0.004717342 | 0.004110564 | 0.195868231 |
| Ncoa7 | S | 92 | -0.279549097 | -0.109933858 | -0.436951536 | -0.157402438 | -0.327017678 | 0.173890246 | 0.568483884 | 0.058776418 | 0.426205094 | 0.140887496 |
| Gtf2i | S | 695 | -0.062631747 | -0.234907865 | -0.262783714 | -0.200151967 | -0.027875849 | 0.282984389 | 0.095882242 | 0.01664353 | 0.041203234 | 0.795593335 |
| Gon4l | S | 784 | -0.025098725 | -0.17397943 | -0.178909597 | -0.153810872 | -0.004930167 | 0.587010119 | 0.002577663 | 0.008392823 | 0.023441693 | 0.900749819 |
| Eif4g3 | S | 1150 | -0.285658442 | -0.18851471 | -0.149640675 | 0.136017767 | 0.038874035 | 0.015788046 | 0.136801016 | 0.426521214 | 0.448088571 | 0.826509598 |
| Osbpl3 | S | 342 | -0.059588676 | -0.179828178 | -0.296043641 | -0.236454965 | -0.116215463 | 0.40012431 | 0.016357466 | 0.043261712 | 0.084127246 | 0.291728725 |
| Ric1 | S | 1188 | -0.142521782 | -0.153069243 | -0.201872833 | -0.059351051 | -0.04880359 | 0.037064455 | 0.065236605 | 0.073969981 | 0.498850543 | 0.598039402 |
| Mms19 | S | 1028 | -0.238682796 | -0.179534771 | -0.2013539 | 0.037328895 | -0.021819129 | 0.320363478 | 0.036823926 | 0.054623781 | 0.8609164 | 0.767237625 |
| Cul4b | S | 200 | -0.052053894 | -0.185240713 | -0.132577008 | -0.080523114 | 0.052663706 | 0.045389194 | 0.00422926 | 0.04187614 | 0.109009137 | 0.245804569 |
| Tut7 | S | 705 | -0.012627126 | -0.139709515 | -0.268811415 | -0.256184289 | -0.1291019 | 0.781095813 | 0.025088492 | 0.002205654 | 0.005994326 | 0.056125567 |
| Klf13 | S | 269 | -0.182430139 | -0.14878722 | -0.432496085 | -0.250065946 | -0.283708864 | 0.284268342 | 0.342101 | 0.040626596 | 0.214082491 | 0.155785575 |
| Anln | S | 139 | -0.109538472 | -0.092565427 | -0.214397302 | -0.104858831 | -0.121831876 | 0.023029574 | 0.179779222 | 0.008261371 | 0.091493886 | 0.117762535 |
| Bicd2 | S | 331 | -0.069852891 | -0.194231978 | -0.214564573 | -0.144711682 | -0.020332595 | 0.446621871 | 0.030694723 | 0.009878698 | 0.171824664 | 0.765458727 |
| Ddx21 | S | 236 | 0.058436584 | -0.295973607 | -0.233809023 | -0.292245607 | 0.062164584 | 0.799895589 | 0.003946901 | 0.033902795 | 0.275461712 | 0.450560417 |
| Larp7 | S | 227 | -0.272736201 | -0.074736 | -0.303966712 | -0.031230511 | -0.229230711 | 0.036917226 | 0.562483518 | 0.070240307 | 0.778412193 | 0.139069518 |
| Map3k3 | S | 337 | -0.019473518 | -0.211938856 | -0.380472075 | -0.360998557 | -0.168533219 | 0.861616121 | 0.150172195 | 0.007431948 | 0.026591822 | 0.205723513 |
| Chchd3 | S | 40 | 0.088218408 | -0.202303556 | -0.234151735 | -0.322370143 | -0.031848179 | 0.304195299 | 0.000946368 | 0.002350423 | 0.022216882 | 0.459742452 |
| Ccdc25 | S | 204 | -0.08610783 | -0.157875689 | -0.298777428 | -0.212669598 | -0.140901739 | 0.317750548 | 0.04321922 | 0.027871007 | 0.080313298 | 0.15217772 |
| Smtn | T | 359 | -0.039176145 | -0.241912272 | -0.220026106 | -0.18084996 | 0.021886167 | 0.394227102 | 0.001971698 | 0.161031308 | 0.224908515 | 0.858044351 |
| Mast4 | S | 2549 | -0.272379265 | -0.046898784 | -0.234840539 | 0.037538726 | -0.187941755 | 0.007701223 | 0.741519835 | 0.074238869 | 0.708740201 | 0.262087217 |
| Cd3eap | S | 86 | -0.005633714 | -0.144692492 | -0.206526285 | -0.200892571 | -0.061833794 | 0.91069538 | 0.042892819 | 0.000142832 | 0.026544201 | 0.255958837 |
| Cep350 | S | 2421 | -0.068086414 | -0.180919466 | -0.130788757 | -0.062702343 | 0.050130709 | 0.139401309 | 0.021515944 | 0.003972551 | 0.170797242 | 0.328554624 |
| Cdv3 | S | 78 | 0.001759888 | -0.353961176 | -0.432507046 | -0.434266934 | -0.078545869 | 0.992352602 | 0.134628442 | 0.033663381 | 0.036467172 | 0.670487853 |
| Kansl3 | S | 540 | -0.242755212 | -0.042702718 | -0.229728484 | 0.013026728 | -0.187025766 | 0.015260709 | 0.642839168 | 0.02837405 | 0.86472901 | 0.091594052 |
| Bclaf1 | S | 412 | -0.03697165 | -0.187923134 | -0.247444075 | -0.210472424 | -0.05952094 | 0.684187564 | 0.161496205 | 0.000662366 | 0.096624076 | 0.580860739 |
| Snx9 | S | 175 | -0.107451127 | -0.075859124 | -0.171957602 | -0.064506475 | -0.096098478 | 0.048529486 | 0.132834406 | 0.001039729 | 0.156389702 | 0.081604731 |
| Hmgn1 | S | 87 | -0.084072602 | -0.11146974 | -0.092484554 | -0.008411952 | 0.018985186 | 0.085869659 | 0.000528504 | 0.007055167 | 0.815674599 | 0.383551013 |
| Hspa4 | S | 415 | -0.085357493 | -0.108209134 | -0.149515166 | -0.064157673 | -0.041306032 | 0.382229253 | 0.017928321 | 0.00029635 | 0.492483007 | 0.21718156 |
| Sart1 | S | 42 | -0.19594852 | -0.230666861 | -0.157094395 | 0.038854125 | 0.073572465 | 0.033885094 | 0.036682388 | 0.43724294 | 0.834788579 | 0.701767977 |
| Ehbp1 | T | 657 | -0.093892842 | -0.065363246 | -0.432980841 | -0.339087999 | -0.367617595 | 0.475315232 | 0.54725217 | 0.00302456 | 0.042810019 | 0.008318988 |
| Tasor2 | S | 668 | 0.03813904 | -0.229064518 | -0.152960843 | -0.191099884 | 0.076103675 | 0.470197824 | 0.000480328 | 0.005365737 | 0.017824999 | 0.05004162 |
| Clasp1 | S | 724 | -0.134552528 | -0.121554547 | -0.195686974 | -0.061134446 | -0.074132427 | 0.043318346 | 0.085122892 | 0.025054599 | 0.300904174 | 0.260345764 |
| Mink1 | S | 892 | -0.104171923 | -0.299973033 | -0.318189323 | -0.2140174 | -0.01821629 | 0.418664163 | 0.183270532 | 0.035411427 | 0.175740855 | 0.923749872 |
| Ncapd2 | S | 1320 | -0.228321586 | -0.073654758 | -0.253912164 | -0.025590578 | -0.180257406 | 0.03292353 | 0.60408416 | 0.024627628 | 0.768715725 | 0.251672442 |
| Tln1 | T | 2270 | -0.188097237 | -0.222861656 | -0.125232829 | 0.062864408 | 0.097628827 | 0.049559959 | 0.084603021 | 0.08813492 | 0.429365567 | 0.357318995 |
| Pdlim1 | S | 144 | -0.015502814 | -0.152603385 | -0.223954717 | -0.208451904 | -0.071351333 | 0.453942472 | 0.001719741 | 0.018034626 | 0.015155785 | 0.168991478 |
| Ptpn21 | S | 637 | -0.083871336 | -0.173766646 | -0.171953147 | -0.088081811 | 0.001813499 | 0.047833163 | 0.02230069 | 0.046635534 | 0.202254053 | 0.978224117 |
| Pde4a | S | 786 | -0.433055816 | -0.222949164 | -0.379192212 | 0.053863604 | -0.156243049 | 0.255904601 | 0.298186683 | 0.221400842 | 0.887000947 | 0.600089766 |
| Bag3 | S | 297 | -0.029695114 | -0.091421574 | -0.232379202 | -0.202684087 | -0.140957628 | 0.481935899 | 0.011601935 | 0.001105137 | 0.007737944 | 0.025164905 |
| Flcn | S | 62 | -0.115629953 | -0.162386467 | -0.15826248 | -0.042632527 | 0.004123987 | 0.028424742 | 0.04030045 | 0.054215085 | 0.472505626 | 0.953004168 |
| Snrpa1 | S | 178 | -0.043601781 | -0.116584941 | -0.348872891 | -0.30527111 | -0.23228795 | 0.375205447 | 0.325380191 | 0.001972287 | 0.005377513 | 0.107435946 |
| Clasp2 | S | 947 | -0.129898777 | -0.141026268 | -0.210314303 | -0.080415526 | -0.069288036 | 0.033525794 | 0.277644378 | 0.013307336 | 0.210520865 | 0.561288495 |
| Heatr5a | S | 1704 | -0.104115583 | -0.113832932 | -0.169381558 | -0.065265975 | -0.055548626 | 0.005090816 | 0.084487048 | 0.030336302 | 0.239280218 | 0.384233452 |
| Ranbp2 | S | 781 | 0.008879587 | -0.112154106 | -0.166858488 | -0.175738075 | -0.054704382 | 0.772905224 | 0.001054117 | 0.000150628 | 0.012254781 | 0.008659383 |
| Zc3h3 | S | 460 | 0.034599739 | -0.200999813 | -0.228597568 | -0.263197307 | -0.027597755 | 0.329833023 | 0.025752314 | 0.000244503 | 0.00024497 | 0.620716423 |
| Map2k4 | S | 391 | -0.032356501 | -0.147565964 | -0.298056091 | -0.265699591 | -0.150490127 | 0.755030989 | 0.076970629 | 0.002647408 | 0.064553489 | 0.045923074 |
| Rsl1d1 | S | 403 | 0.067078308 | -0.180388523 | -0.436029823 | -0.50310813 | -0.255641299 | 0.588505971 | 0.130844565 | 0.002378433 | 0.014714773 | 0.057117031 |
| Rbbp6 | S | 1715 | -0.106474677 | -0.107448889 | -0.328594944 | -0.222120267 | -0.221146056 | 0.269438904 | 0.392355326 | 0.00394313 | 0.061232716 | 0.133418103 |
| Dnajc21 | S | 370 | -0.053120524 | -0.112038067 | -0.226294075 | -0.173173551 | -0.114256008 | 0.03720998 | 0.018836093 | 0.018065221 | 0.037912269 | 0.083067943 |
| Irs1 | S | 522 | -0.233890885 | -0.125857594 | -0.186867649 | 0.047023236 | -0.061010055 | 0.065728724 | 0.15755004 | 0.046516375 | 0.605173841 | 0.2935485 |
| Lmna | S | 94 | -0.280733719 | -0.063749925 | -0.257223076 | 0.023510643 | -0.193473151 | 0.034298462 | 0.642516805 | 0.04448984 | 0.665853173 | 0.149400239 |
| Morc2a | S | 775 | -0.010326574 | -0.106332922 | -0.13511617 | -0.124789596 | -0.028783248 | 0.656537585 | 0.00069384 | 0.000151228 | 0.004702721 | 0.064951115 |
| Arhgef7 | S | 236 | -0.013961404 | -0.132001499 | -0.30964821 | -0.295686806 | -0.177646711 | 0.761217688 | 0.04499726 | 0.003191381 | 0.005913348 | 0.034565369 |
| Tcof1 | S | 83 | 0.030563829 | -0.446097555 | -0.219673388 | -0.250237216 | 0.226424167 | 0.834747633 | 0.019243436 | 0.062192566 | 0.151829981 | 0.136811032 |
| Tcof1 | S | 84 | 0.030563829 | -0.446097555 | -0.219673388 | -0.250237216 | 0.226424167 | 0.834747633 | 0.019243436 | 0.062192566 | 0.151829981 | 0.136811032 |
| Tcof1 | S | 87 | 0.030563829 | -0.446097555 | -0.219673388 | -0.250237216 | 0.226424167 | 0.834747633 | 0.019243436 | 0.062192566 | 0.151829981 | 0.136811032 |
| Chd4 | S | 1342 | -0.083050962 | -0.246767961 | -0.242546536 | -0.159495574 | 0.004221425 | 0.245854 | 0.021849563 | 0.11725392 | 0.247597823 | 0.972866993 |
| Psrc1 | S | 185 | -0.083535 | -0.206858104 | -0.112685256 | -0.029150256 | 0.094172848 | 0.361393178 | 0.000676085 | 0.097292957 | 0.745043384 | 0.146760465 |
| Supt6 | S | 91 | -0.077561785 | -0.294035201 | -0.334604037 | -0.257042251 | -0.040568836 | 0.193428973 | 0.04209467 | 0.22107333 | 0.314672894 | 0.860240697 |
| Ncaph | T | 46 | -0.031145838 | -0.171504644 | -0.124295769 | -0.093149931 | 0.047208875 | 0.572662304 | 0.023896913 | 5.8815E-05 | 0.179860092 | 0.286256032 |
| Usp14 | S | 222 | -0.208784398 | -0.189367485 | -0.309600736 | -0.100816338 | -0.120233251 | 0.170381186 | 0.098776447 | 0.153333707 | 0.606160018 | 0.511876135 |
| Akt1s1 | S | 212 | -0.050309625 | -0.09726078 | -0.218822706 | -0.168513081 | -0.121561926 | 0.525326217 | 0.066987778 | 0.00027605 | 0.109395396 | 0.040660858 |
| Pdpk1 | T | 248 | -0.184418305 | -0.130548153 | -0.36605977 | -0.181641465 | -0.235511617 | 0.068353484 | 0.18717971 | 0.132287832 | 0.359052095 | 0.260176389 |
| Copa | S | 173 | -0.021507722 | -0.172908449 | -0.220099884 | -0.198592161 | -0.047191435 | 0.631839301 | 0.001590214 | 0.030655179 | 0.02797038 | 0.405808359 |
| Tlk1 | S | 133 | -0.247633026 | -0.085624616 | -0.139781074 | 0.107851952 | -0.054156458 | 0.036054212 | 0.181901626 | 0.019668685 | 0.233437608 | 0.30387678 |
| Rnf219 | S | 715 | 0.106723042 | -0.253042024 | -0.30348707 | -0.410210112 | -0.050445046 | 0.167532588 | 0.000714635 | 0.017036234 | 0.005149548 | 0.44851397 |
| Eif5b | S | 222 | -0.093465803 | -0.207315015 | -0.221474342 | -0.128008539 | -0.014159327 | 0.28041986 | 0.00940697 | 0.109177264 | 0.303716106 | 0.890191502 |
| Smtn | S | 439 | 0.031416215 | -0.232731753 | -0.31798439 | -0.349400605 | -0.085252637 | 0.673345028 | 0.015971027 | 0.014728078 | 0.017994677 | 0.337773542 |
| Dis3l2 | S | 12 | -0.095320227 | -0.187095476 | -0.274823024 | -0.179502797 | -0.087727548 | 0.23408257 | 0.04157837 | 0.05807016 | 0.160919624 | 0.424808854 |
| Map7d1 | S | 503 | -0.091947363 | -0.098500521 | -0.18720431 | -0.095256947 | -0.088703789 | 0.092121696 | 0.068634974 | 0.002911926 | 0.070433858 | 0.075062022 |
| Clcc1 | S | 433 | -0.095109865 | -0.167770721 | -0.087636314 | 0.007473552 | 0.080134408 | 0.155326979 | 0.000680423 | 0.058293622 | 0.895483377 | 0.087417019 |
| Clasp1 | S | 728 | -0.138994799 | -0.101314908 | -0.192409341 | -0.053414542 | -0.091094433 | 0.039279218 | 0.141701586 | 0.013523237 | 0.214755384 | 0.120508488 |
| Nfkbib | S | 346 | -0.19818005 | -0.047371751 | -0.110670462 | 0.087509587 | -0.063298712 | 0.000178168 | 0.306104503 | 0.123848051 | 0.200552474 | 0.338822772 |
| Cobll1 | S | 359 | -0.208655635 | -0.002504362 | -0.37928048 | -0.170624844 | -0.376776118 | 0.004895955 | 0.973327785 | 0.086383461 | 0.314987172 | 0.082401698 |
| Rbm25 | S | 678 | 0.084654638 | -0.205012503 | -0.328992129 | -0.413646767 | -0.123979625 | 0.042279692 | 0.002532809 | 0.00190852 | 0.004769085 | 0.070516658 |
| Rab3ip | S | 218 | -0.005099187 | -0.159738286 | -0.277622006 | -0.272522819 | -0.11788372 | 0.864645959 | 0.016633928 | 0.006166271 | 0.021292854 | 0.098938062 |
| Ctnnb1 | S | 552 | 0.034037019 | -0.06845105 | -0.242583121 | -0.27662014 | -0.174132071 | 0.1462454 | 0.004274598 | 1.02466E-05 | 0.000140856 | 0.001001385 |
| Fam53c | S | 234 | -0.166573921 | -0.035855281 | -0.135490783 | 0.031083138 | -0.099635502 | 0.001426777 | 0.672860631 | 0.004244027 | 0.150566208 | 0.291645175 |
| Parp3 | T | 21 | -0.234126213 | -0.10343748 | -0.291382732 | -0.057256519 | -0.187945252 | 0.061782739 | 0.411040769 | 0.057550491 | 0.557063623 | 0.150105156 |
| Dip2b | S | 99 | -0.022337426 | -0.171973225 | -0.36954655 | -0.347209124 | -0.197573325 | 0.720214893 | 0.051614987 | 0.017454051 | 0.014902927 | 0.076117936 |
| Thrap3 | S | 238 | -0.045395822 | -0.149649187 | -0.143193331 | -0.097797509 | 0.006455856 | 0.035893567 | 0.007069086 | 0.015584824 | 0.038853945 | 0.850394089 |
| Tsc2 | S | 1805 | -0.122117714 | -0.338734883 | -0.531008175 | -0.408890462 | -0.192273292 | 0.604788891 | 0.074790281 | 0.267952371 | 0.378803718 | 0.646600497 |
| Osbpl6 | S | 44 | -0.115149292 | -0.04397467 | -0.303544361 | -0.188395069 | -0.259569692 | 0.219016969 | 0.610825325 | 0.000930841 | 0.087860773 | 0.049251095 |
| Parp8 | S | 389 | -0.092775692 | -0.170646233 | -0.198424075 | -0.105648383 | -0.027777842 | 0.038079789 | 0.08826212 | 0.031950855 | 0.144032604 | 0.737311769 |
| Nes | S | 919 | -0.011633912 | -0.28198846 | -0.116019987 | -0.104386075 | 0.165968473 | 0.785366159 | 0.000656717 | 0.08861544 | 0.114606111 | 0.049201254 |
| Kirrel | S | 677 | -0.127307578 | -0.12784553 | -0.22204025 | -0.094732672 | -0.09419472 | 0.07406742 | 0.148403381 | 0.017027245 | 0.164535303 | 0.257363714 |
| 2410004B18Rik | S | 156 | -0.047602177 | -0.076181832 | -0.190618903 | -0.143016726 | -0.114437071 | 0.306685789 | 0.066710537 | 9.56641E-05 | 0.043322468 | 0.030295835 |
| Stmn1 | S | 38 | -0.192871944 | -0.112828218 | -0.404773831 | -0.211901886 | -0.291945612 | 0.141498 | 0.477015007 | 0.046006414 | 0.20915284 | 0.158303002 |
| Cdk16 Cdk17 | S | 119;146 | -0.263384099 | -0.070707107 | -0.357362162 | -0.093978063 | -0.286655055 | 0.098664856 | 0.65258006 | 0.04197147 | 0.457763262 | 0.089882346 |
| Luzp1 | S | 612 | -0.352916099 | -0.122666496 | -0.145101756 | 0.207814342 | -0.02243526 | 0.015965735 | 0.369146806 | 0.231614888 | 0.064846205 | 0.837617835 |
| Ralgapa1 | S | 710 | -0.318248316 | 0.047274929 | -0.356777428 | -0.038529112 | -0.404052357 | 0.039984642 | 0.808612421 | 0.032412474 | 0.682296102 | 0.089888312 |
| Ccm2 | T | 288 | -0.064660596 | -0.182400715 | -0.369392155 | -0.304731559 | -0.18699144 | 0.490180444 | 0.010775123 | 0.114932844 | 0.159381176 | 0.327346462 |
| Tasor2 | T | 697 | 0.134833626 | -0.395285766 | -0.422839173 | -0.557672799 | -0.027553407 | 0.017024635 | 0.002798546 | 0.02724336 | 0.01158355 | 0.771631112 |

| | | | | | | | | | | | | |
|---|---|---|---|---|---|---|---|---|---|---|---|---|
| Bmp2k | T | 714 | 0.003175 | -0.358707448 | -0.161297899 | -0.164472898 | 0.197409549 | 0.970822567 | 0.003230692 | 0.113329821 | 0.086060281 | 0.050438639 |
| Wiz | S | 1050 | -0.03445482 | -0.113169845 | -0.11781213 | -0.08335731 | -0.004642285 | 0.391307981 | 4.65781E-05 | 0.006200774 | 0.100598169 | 0.814803997 |
| Hjurp A730008H23Ril | S | 580 | 0.073544216 | -0.145956566 | -0.244371891 | -0.317916107 | -0.098415325 | 0.095511389 | 0.00151706 | 0.000166601 | 0.000928167 | 0.02518981 |
| Nasp | S | 188 | -0.070685843 | -0.212734839 | -0.201675413 | -0.13098957 | 0.011059426 | 0.417344233 | 0.00144319 | 0.180528677 | 0.356534872 | 0.924712851 |
| Mpp5 | S | 84 | -0.027876584 | -0.113627699 | -0.302504784 | -0.2746282 | -0.188877085 | 0.590536372 | 0.205554989 | 0.000787206 | 0.003343746 | 0.074504364 |
| Leo1 | S | 247 | -0.052876296 | -0.220763286 | -0.100859031 | -0.047982735 | 0.119904255 | 0.292134683 | 0.000429349 | 0.132028853 | 0.435305281 | 0.099274207 |
| Asap1 | S | 10 | -0.130932151 | -0.197765554 | -0.433359025 | -0.302426874 | -0.235593471 | 0.202975228 | 0.25246736 | 0.084317725 | 0.155675685 | 0.283943157 |
| Kif4 | S | 1218 | -0.202844081 | -0.132254823 | -0.286880225 | -0.084036144 | -0.154625401 | 0.16593655 | 0.320433303 | 0.026228834 | 0.472729343 | 0.200284407 |
| Myo9b | S | 1106 | -0.115753513 | -0.146949547 | -0.114117609 | 0.001635904 | 0.032831938 | 0.021520238 | 0.010075049 | 0.089675664 | 0.972960312 | 0.533705681 |
| Bod1l | S | 240 | -0.282098554 | -0.047685711 | -0.174144183 | 0.107954371 | -0.126458472 | 0.006663806 | 0.732761698 | 0.059084571 | 0.211401559 | 0.402611021 |
| Set | S | 30 | -0.10959237 | -0.133380732 | -0.106459238 | 0.003133132 | 0.026921494 | 0.058702645 | 0.045937056 | 0.00198228 | 0.936794788 | 0.544610642 |
| Ybx3 | S | 313 | -0.142146022 | -0.057953564 | -0.277311313 | -0.135165291 | -0.219357749 | 0.089529462 | 0.426167648 | 0.005385025 | 0.115456116 | 0.029330548 |
| 1700066M21Rik | S | 77 | -0.189566565 | -0.184912737 | -0.244791095 | -0.055224531 | -0.059878359 | 0.201164268 | 0.140952478 | 0.049688389 | 0.688597994 | 0.61722963 |
| Ccnf | S | 653 | -0.046223878 | -0.133333367 | -0.35306456 | -0.306840682 | -0.219731193 | 0.632433952 | 0.198526229 | 0.003877017 | 0.021515539 | 0.057647972 |
| Ccnf | S | 654 | -0.046223878 | -0.133333367 | -0.35306456 | -0.306840682 | -0.219731193 | 0.632433952 | 0.198526229 | 0.003877017 | 0.021515539 | 0.057647972 |
| Dnajb2 | S | 311 | 0.041286749 | -0.168193342 | -0.3253038 | -0.366590549 | -0.157110458 | 0.306177757 | 0.020813014 | 0.001966326 | 0.001233097 | 0.031697505 |
| Wdr77 | T | 5 | -0.053779181 | -0.172899681 | -0.161674904 | -0.107895724 | 0.011224777 | 0.25752184 | 0.04415573 | 0.001941832 | 0.057615704 | 0.840631025 |
| Dpysl3 | S | 472 | -0.153408848 | -0.038400254 | -0.250109878 | -0.096701031 | -0.211709624 | 0.003213839 | 0.495359902 | 0.037701186 | 0.247282327 | 0.052719996 |
| Pnisr | S | 726 | -0.191462718 | -0.066661158 | -0.396805861 | -0.205343143 | -0.330144704 | 0.076909743 | 0.721354955 | 0.033905196 | 0.175181102 | 0.161496219 |
| Eif5b | S | 108 | -0.269984652 | -0.092581193 | -0.262347959 | 0.007636693 | -0.169766766 | 0.017994387 | 0.498053119 | 0.145286108 | 0.957817507 | 0.346940633 |
| Pinx1 | S | 233 | 0.036978766 | -0.167253431 | -0.220343277 | -0.257322042 | -0.053089846 | 0.224310849 | 0.000244337 | 0.005961114 | 0.005592879 | 0.242677997 |
| Rab11fip5 | S | 640 | 0.033147076 | -0.231787934 | -0.201008203 | -0.234155278 | 0.030779731 | 0.473124396 | 0.007221014 | 0.002573274 | 0.011066061 | 0.426343717 |
| Irs2 | S | 556 | 0.04357592 | -0.183233669 | -0.29718742 | -0.34076334 | -0.113953751 | 0.659447581 | 0.023647185 | 0.003051724 | 0.034125146 | 0.130162549 |
| Dlgap5 | S | 328 | -0.042945331 | -0.094047843 | -0.18536328 | -0.142417949 | -0.091315437 | 0.300353851 | 0.051811905 | 0.000218723 | 0.027479046 | 0.069447252 |
| Hjurp A730008H23Ril | S | 503 | -0.114089171 | -0.257419235 | -0.086582483 | 0.027506687 | 0.170836752 | 0.069355374 | 0.003655034 | 0.483028161 | 0.817288676 | 0.225656376 |
| Rock2 | S | 1362 | -0.171294431 | -0.021739648 | -0.543740706 | -0.372446275 | -0.522001058 | 0.153715979 | 0.869163439 | 0.029698186 | 0.092443121 | 0.038865305 |
| Errfi1 | S | 251 | -0.059468062 | -0.198724352 | -0.158387425 | -0.098919363 | 0.040336927 | 0.343001419 | 0.005275483 | 0.034304869 | 0.184124052 | 0.488290922 |
| Polr1a | S | 1436 | 0.00249782 | -0.261496891 | -0.200348841 | -0.202846661 | 0.06114805 | 0.953703471 | 0.019052157 | 0.007034437 | 0.013896048 | 0.415218475 |
| Helb | S | 1026 | -0.099349553 | -0.116138879 | -0.279704562 | -0.18035501 | -0.163565684 | 0.098744481 | 0.04130817 | 0.050063279 | 0.140806077 | 0.172242685 |
| Dgcr8 | S | 109 | -0.027920472 | -0.163454002 | -0.23903133 | -0.211110858 | -0.075577328 | 0.682108239 | 0.002291402 | 0.033006756 | 0.052313325 | 0.265096479 |
| Sbno1 | S | 1059 | 0.052065982 | -0.301127227 | -0.178924407 | -0.230990389 | 0.12220282 | 0.314843025 | 0.003504369 | 0.006932034 | 0.010376164 | 0.062237882 |
| Cwc22 | S | 45 | -0.166275366 | -0.128246721 | -0.134111568 | 0.032163798 | -0.005864847 | 0.150078362 | 0.003142586 | 0.081372548 | 0.737739312 | 0.914320028 |
| Clasp2 | S | 461 | -0.182745136 | -0.105458335 | -0.284774218 | -0.102029082 | -0.179315883 | 0.100882558 | 0.319375263 | 0.028493437 | 0.353279343 | 0.14775365 |
| Tnks1bp1 | T | 533 | -0.086130391 | -0.164557164 | -0.086147989 | -1.75983E-05 | 0.078409175 | 0.307893746 | 9.53136E-05 | 0.097480251 | 0.999819642 | 0.126029479 |
| Plekha3 | S | 289 | -0.177496897 | -0.182015444 | -0.147455368 | 0.030041529 | 0.034560077 | 0.227933344 | 0.007665314 | 0.131701768 | 0.822526835 | 0.657561635 |
| Tns3 | T | 410 | -0.032362276 | -0.152178944 | -0.263940914 | -0.231578639 | -0.11176197 | 0.350272284 | 0.079053596 | 0.004507028 | 0.004783754 | 0.138755183 |
| Sh3kbp1 | T | 398 | -0.043862813 | -0.146201717 | -0.292747859 | -0.248885047 | -0.146546142 | 0.62860264 | 0.059565938 | 0.006491744 | 0.047139409 | 0.087829111 |
| Ncl | T | 76 | 0.189777778 | -0.308609691 | -0.371927277 | -0.561705055 | -0.063317586 | 0.039822218 | 0.011208919 | 0.000991769 | 0.001040507 | 0.434566635 |
| Frmd4a | S | 320 | -0.078692438 | -0.180675558 | -0.313011499 | -0.234319061 | -0.132335941 | 0.2478942 | 0.071205656 | 0.047808487 | 0.102253806 | 0.293277523 |
| Rrp1b | S | 336 | -0.003148912 | -0.191367456 | -0.421761185 | -0.418612273 | -0.23039373 | 0.980680739 | 0.066046604 | 0.022005573 | 0.037410855 | 0.127176584 |
| Tank | S | 377 | -0.147177637 | -0.077699569 | -0.276607804 | -0.129430166 | -0.198908235 | 0.09856291 | 0.347862712 | 0.010244332 | 0.034127817 | 0.013376275 |
| Limd1 | S | 372 | -0.101054863 | -0.258565122 | -0.300771325 | -0.199716461 | -0.042206202 | 0.337382938 | 0.026471029 | 0.200464595 | 0.354433684 | 0.827734243 |
| Limd1 | S | 376 | -0.101054863 | -0.258565122 | -0.300771325 | -0.199716461 | -0.042206202 | 0.337382938 | 0.026471029 | 0.200464595 | 0.354433684 | 0.827734243 |
| Eps15l1 | S | 229 | 0.020663805 | -0.192939264 | -0.205050549 | -0.225714354 | -0.012111285 | 0.386336359 | 0.00396334 | 0.00266738 | 0.00103876 | 0.682824788 |
| Baz1a | S | 1415 | -0.15117782 | -0.12739809 | -0.154242655 | -0.003064835 | -0.026844565 | 0.154567399 | 0.024008118 | 0.020491392 | 0.971644878 | 0.590540506 |
| Disp2 | S | 1121 | 0.017555021 | -0.311813733 | -0.079475417 | -0.097030437 | 0.232338316 | 0.660034506 | 0.000226963 | 0.067617753 | 0.014385722 | 0.006246677 |
| Nsd2 | S | 194 | -0.030475972 | -0.182884444 | -0.115469852 | -0.08499388 | 0.067414593 | 0.603947322 | 0.001139749 | 0.007006445 | 0.203493685 | 0.072997904 |
| Ralbp1 | S | 34 | -0.035507294 | -0.068552744 | -0.234996776 | -0.199489482 | -0.166444032 | 0.469215376 | 0.047615551 | 0.000328202 | 0.018209521 | 0.004457198 |
| Cdc25c | S | 180 | 0.045640558 | -0.21310615 | -0.366137662 | -0.41177822 | -0.153031512 | 0.472953791 | 0.004922445 | 0.033648227 | 0.028834436 | 0.222782429 |
| Pagr1a | S | 53 | -0.167196932 | -0.167907921 | -0.053211163 | 0.113985769 | 0.114696757 | 0.010707768 | 0.01012057 | 0.325447842 | 0.057675896 | 0.055740414 |
| Map1a | S | 1768 | -0.082600736 | -0.100684673 | -0.084524508 | -0.001923773 | 0.016160165 | 0.039740796 | 0.000908701 | 0.008231778 | 0.946520925 | 0.443262894 |
| Cep76 | S | 82 | -0.023559649 | -0.132055437 | -0.311872843 | -0.288313195 | -0.179817406 | 0.449172402 | 0.010979 | 0.023197989 | 0.044533029 | 0.090743329 |
| Usp9x | S | 2547 | -0.019061748 | -0.175160034 | -0.196115192 | -0.177053443 | -0.020955157 | 0.510173352 | 0.004184887 | 0.016259815 | 0.020613693 | 0.658168682 |
| Iqschfp Schip1 | S | 63;138 | -0.044528153 | -0.148791528 | -0.402899018 | -0.358370864 | -0.25410749 | 0.798981797 | 0.22887973 | 0.006598628 | 0.103929114 | 0.038280974 |
| Dlgap5 | S | 70 | -0.143682349 | -0.009096919 | -0.22901937 | -0.085337021 | -0.219922451 | 0.000205215 | 0.728055912 | 0.048556796 | 0.27458712 | 0.051021447 |
| 5031439G07Rik | S | 362 | -0.164366622 | -0.079844326 | -0.301293743 | -0.136927121 | -0.221449417 | 0.114894557 | 0.493034158 | 0.01186575 | 0.193630662 | 0.115674626 |
| Plcb3 | S | 1105 | -0.187529271 | -0.005039673 | -0.457419834 | -0.269890562 | -0.45238016 | 0.093909171 | 0.972765601 | 0.023295471 | 0.123797454 | 0.04061102 |
| Reps1 | S | 561 | -0.02296381 | -0.135679097 | -0.143914821 | -0.120951011 | -0.008235725 | 0.473565432 | 0.011102644 | 0.000318366 | 0.019005319 | 0.788912227 |
| Mast3 | S | 732 | -0.272043044 | -0.072738976 | -0.414844997 | -0.142801953 | -0.342106021 | 0.118906821 | 0.550378823 | 0.081684889 | 0.484992645 | 0.130805001 |
| Prune2 | S | 1757 | 0.062676628 | -0.288651495 | -0.244489069 | -0.307165697 | 0.044162426 | 0.681526158 | 0.035968898 | 0.003914259 | 0.134709795 | 0.623147944 |
| Epb41l2 | S | 698 | -0.036301436 | -0.114521264 | -0.117545153 | -0.081243717 | -0.003023889 | 0.251176394 | 0.012217346 | 7.87221E-05 | 0.058366256 | 0.900182745 |
| Cul4b | S | 82 | -0.177087891 | -0.096733672 | -0.296040037 | -0.118952146 | -0.199306364 | 0.063743992 | 0.377120256 | 0.038416484 | 0.283581139 | 0.143887189 |
| Mitf | S | 491 | -0.144198772 | -0.618121602 | -0.230417151 | -0.086218379 | 0.387704451 | 0.593080216 | 0.155752502 | 0.168219299 | 0.751217556 | 0.310688077 |
| Mitf | S | 504 | -0.144198772 | -0.618121602 | -0.230417151 | -0.086218379 | 0.387704451 | 0.593080216 | 0.155752502 | 0.168219299 | 0.751217556 | 0.310688077 |
| Rabep1 | T | 408 | -0.172611671 | -0.099218738 | -0.328693291 | -0.15608162 | -0.229474553 | 0.264054468 | 0.50855682 | 0.008725089 | 0.287085564 | 0.167120097 |
| Lima1 | S | 372 | 0.002958518 | -0.188523233 | -0.270786841 | -0.273745359 | -0.082263609 | 0.964494921 | 0.016838311 | 0.009903185 | 0.013797588 | 0.235654577 |
| Phrf1 | S | 841 | -0.374861427 | -0.417877065 | -0.052110114 | 0.322751312 | 0.365766951 | 0.378502243 | 0.044475208 | 0.328547381 | 0.436203117 | 0.064405672 |
| 2310022B05Rik | S | 172 | -0.504862889 | -0.031436423 | -0.174312359 | 0.33055053 | -0.142875935 | 0.016464808 | 0.877890114 | 0.265455593 | 0.089099184 | 0.506129976 |
| 5031439G07Rik | T | 293 | -0.055290496 | -0.117562756 | -0.246931188 | -0.191640692 | -0.129368432 | 0.239713661 | 0.138268044 | 0.002394566 | 0.012445745 | 0.118462602 |
| Cd44 | S | 733 | -0.026095377 | -0.056710589 | -0.574478796 | -0.548383418 | -0.517768207 | 0.848219461 | 0.672227612 | 0.004323972 | 0.004361795 | 0.005719111 |
| Ppp3cc | S | 462 | -0.241638685 | -0.039337223 | -0.372911697 | -0.131273012 | -0.333574474 | 0.096897691 | 0.809604845 | 0.030322469 | 0.352523176 | 0.102586918 |
| Map4 | S | 254 | -0.018322503 | -0.079832781 | -0.198155255 | -0.179832753 | -0.118322474 | 0.559256109 | 0.008420457 | 0.00032093 | 0.004184183 | 0.008086009 |
| Arfgap1 | S | 360 | -0.087004952 | -0.090060343 | -0.33575149 | -0.248746538 | -0.245691147 | 0.211336147 | 0.175512146 | 0.011972183 | 0.042473815 | 0.047273798 |
| Cwc25 | S | 307 | -0.101387314 | -0.272853481 | -0.052878225 | 0.048509089 | 0.219975257 | 0.097867873 | 0.002641216 | 0.387125445 | 0.226081062 | 0.014718786 |
| Aimp1 | S | 83 | -0.337603542 | -0.292383136 | -0.093644867 | 0.243958674 | 0.198738269 | 0.08717175 | 0.072250843 | 0.657999014 | 0.270858412 | 0.32290419 |
| Lap3 | S | 139 | 0.091876491 | -0.373471377 | -0.491608707 | -0.583485198 | -0.11813733 | 0.782051599 | 0.13466005 | 0.038279731 | 0.162935326 | 0.6092864 |
| Birc6 | S | 584 | -0.067104635 | -0.22903663 | -0.145774663 | -0.078670028 | 0.083261967 | 0.153616102 | 0.016603842 | 0.051254337 | 0.214519077 | 0.240635371 |
| Tcof1 | T | 395 | 0.108443001 | -0.199885784 | -0.256019217 | -0.364462218 | -0.056133433 | 0.146287938 | 0.005495084 | 0.000387712 | 0.01105351 | 0.220099243 |
| Map4k4 | T | 822 | -0.041080029 | -0.142374954 | -0.216003042 | -0.174923013 | -0.073628088 | 0.352558277 | 0.017859811 | 0.008298874 | 0.023257453 | 0.205543904 |
| Racgap1 | S | 158 | -0.133983412 | -0.084873894 | -0.380654021 | -0.246670608 | -0.295780127 | 0.142518309 | 0.343151224 | 0.027609739 | 0.103116325 | 0.068518286 |
| Pimreg | S | 58 | -0.133111377 | -0.08237956 | -0.483828198 | -0.350716821 | -0.401448638 | 0.251383288 | 0.577146742 | 0.024535059 | 0.071725835 | 0.063305776 |
| Nucks1 | T | 179 | -0.056029854 | -0.097565514 | -0.32327862 | -0.267248766 | -0.225713106 | 0.05469438 | 0.195189489 | 0.013709598 | 0.018392077 | 0.031936872 |
| Fhod1 | S | 502 | -0.009121641 | -0.108888873 | -0.134174488 | -0.125052847 | -0.025285616 | 0.802433239 | 0.000757145 | 0.00026683 | 0.037877584 | 0.187902766 |
| Huwe1 | T | 2747 | -0.046994236 | -0.372878691 | -0.431916206 | -0.384921971 | -0.059037516 | 0.764168785 | 0.038716676 | 0.257788778 | 0.300721297 | 0.855745671 |
| Rsbn1 | S | 92 | -0.049943731 | -0.173957318 | -0.091482311 | -0.04153858 | 0.082475007 | 0.120009141 | 0.001768241 | 0.018765961 | 0.129969065 | 0.045697976 |
| Qars | T | 118 | -0.113662084 | -0.254401604 | -0.197046545 | -0.083384461 | 0.057355058 | 0.462544383 | 0.063192702 | 0.026853039 | 0.582009138 | 0.579994451 |
| Cdr2 | S | 310 | -0.085590023 | -0.116993301 | -0.197015365 | -0.111425342 | -0.080022064 | 0.204198519 | 0.07825265 | 0.003229279 | 0.125465984 | 0.190431173 |
| Ube2e3 | S | 8 | -0.124929609 | -0.083019507 | -0.207541667 | -0.082612058 | -0.12452216 | 0.016690772 | 0.100874059 | 0.037480965 | 0.258183561 | 0.128736016 |
| Ripk2 | S | 538 | -0.164693565 | -0.177985562 | -0.232832085 | -0.06813852 | -0.054846523 | 0.08580479 | 0.154779725 | 0.081565408 | 0.506013523 | 0.65333443 |
| Ssh1 | S | 962 | -0.073386226 | -0.098632433 | -0.217598189 | -0.144211963 | -0.118965756 | 0.221122115 | 0.026916901 | 0.006577214 | 0.060872109 | 0.056254525 |
| Rab3gap1 | S | 579 | -0.235621862 | -0.195942954 | -0.002476117 | 0.233145745 | 0.193466836 | 0.007373003 | 0.017084941 | 0.981735392 | 0.099852739 | 0.147669953 |
| Rab3gap1 | S | 581 | -0.235621862 | -0.195942954 | -0.002476117 | 0.233145745 | 0.193466836 | 0.007373003 | 0.017084941 | 0.981735392 | 0.099852739 | 0.147669953 |
| Gps1 | S | 454 | -0.077577393 | -0.152661553 | -0.20745668 | -0.129879287 | -0.054795127 | 0.245808845 | 0.041693947 | 0.013062089 | 0.016722138 | 0.135631982 |
| Sh3pxd2a | S | 1029 | -0.179861674 | -0.071095223 | -0.273948542 | -0.094086868 | -0.202853319 | 0.079489352 | 0.550636405 | 0.012654335 | 0.202627837 | 0.122452876 |

| | | | | | | | | | | | | |
|---|---|---|---|---|---|---|---|---|---|---|---|---|
| Rbm28 | S | 97 | 0.030982048 | -0.266426542 | -0.295322364 | -0.326304412 | -0.028895822 | 0.764365899 | 0.001694133 | 0.139935405 | 0.114591544 | 0.847206568 |
| Nup35 | S | 258 | -0.02397054 | -0.318988802 | -0.197572831 | -0.173602291 | 0.121415971 | 0.835543639 | 0.009071967 | 0.093511419 | 0.232233627 | 0.265167971 |
| Peak1 | S | 1203 | -0.025239581 | -0.124611947 | -0.35383794 | -0.328598359 | -0.229225993 | 0.70913156 | 0.258990727 | 0.002208665 | 0.006208937 | 0.078987089 |
| Etl4 | S | 1868 | 0.037359922 | -0.162951473 | -0.370530484 | -0.407890406 | -0.207579011 | 0.373340997 | 0.010607945 | 0.010078163 | 0.005636657 | 0.038798594 |
| Map1a | T | 1156 | -0.162732757 | -0.147171408 | -0.149273469 | 0.013459287 | -0.002102062 | 0.026735602 | 0.11396569 | 0.06385628 | 0.834506114 | 0.979404821 |
| Arhgap1 | S | 223 | -0.172930504 | -0.13300768 | -0.099019808 | 0.073910697 | 0.033987873 | 0.001239089 | 0.111349716 | 0.277374558 | 0.388073069 | 0.71467379 |
| Cttnbp2nl | S | 522 | 0.000558367 | -0.160016532 | -0.178333866 | -0.178892233 | -0.018317334 | 0.992496436 | 0.001556912 | 0.005881955 | 0.043561193 | 0.625079643 |
| Rexo1 | S | 487 | -0.120962002 | -0.119231257 | -0.342419146 | -0.221457144 | -0.223187888 | 0.400941848 | 0.316401334 | 0.009060145 | 0.145991005 | 0.06500612 |
| Mtmr10 | S | 603 | -0.054566952 | -0.048041381 | -0.349233138 | -0.294666186 | -0.301191758 | 0.505684944 | 0.681169853 | 0.000439655 | 0.021960839 | 0.075141341 |
| Septin7 | S | 333 | -0.05860155 | -0.132384838 | -0.156317159 | -0.097715609 | -0.023932321 | 0.125302408 | 0.002358173 | 0.033779843 | 0.118243458 | 0.617273907 |
| Nisch | T | 1371 | -0.016347013 | -0.141659405 | -0.405486771 | -0.389139758 | -0.263827366 | 0.848855241 | 0.080914653 | 0.013352992 | 0.017699516 | 0.045914883 |
| Arhgap39 | S | 292 | 0.014859481 | -0.195053048 | -0.176991012 | -0.191850494 | 0.018062036 | 0.723106829 | 0.001077536 | 0.011822294 | 0.014229553 | 0.665230665 |
| Pla2g4a | S | 512 | -0.225904558 | -0.159561832 | -0.201154012 | 0.024750546 | -0.04159218 | 0.066009048 | 0.029957586 | 0.383363357 | 0.907718771 | 0.841960948 |
| Map1b | S | 2251 | 0.120563624 | -0.205651543 | -0.258671701 | -0.379235325 | -0.053020158 | 0.495867824 | 0.007519602 | 0.00220585 | 0.116626559 | 0.072812162 |
| Srsf9 | S | 217 | -0.016941276 | -0.293820872 | -0.118591033 | -0.101649757 | 0.175229839 | 0.779636245 | 0.001257298 | 0.117949266 | 0.158576978 | 0.059455259 |
| Tnks1bp1 | S | 500 | -0.131354738 | -0.10676644 | -0.133285668 | -0.001930929 | -0.026519228 | 0.060151055 | 0.158706676 | 0.001790186 | 0.967688152 | 0.661038198 |
| Baz2a | S | 1767 | 0.010907136 | -0.196769935 | -0.133649294 | -0.14455643 | 0.063120641 | 0.723204777 | 0.003512341 | 0.000829233 | 0.019282582 | 0.140971995 |
| Kif14 | S | 1290 | -0.187980423 | -0.211343774 | -0.127335661 | 0.060644762 | 0.084008113 | 0.023842179 | 0.099784677 | 0.22972179 | 0.513647226 | 0.486662552 |
| Dlgap4 | S | 973 | -0.125993655 | -0.094156455 | -0.15007257 | -0.024078915 | -0.055916115 | 0.00336291 | 0.022472055 | 0.160923165 | 0.771452978 | 0.520945836 |
| Strn4 | S | 167 | -0.080833398 | -0.10032329 | -0.239152148 | -0.15831875 | -0.138828859 | 0.289559255 | 0.094024336 | 0.003435261 | 0.090435901 | 0.046570173 |
| Aebp2 | S | 364 | -0.028649818 | -0.273504219 | -0.150039796 | -0.121389978 | 0.123464423 | 0.725791873 | 0.002676837 | 0.091577572 | 0.242572296 | 0.141436141 |
| Utp14a | S | 343 | 0.041659918 | -0.188687845 | -0.138267112 | -0.17992703 | 0.050420733 | 0.475545118 | 0.000411495 | 0.001915336 | 0.051580042 | 0.070110497 |
| Haus8 | S | 111 | -0.180039609 | -0.108264953 | -0.232578453 | -0.052538843 | -0.124313499 | 0.059458739 | 0.290115776 | 0.033654298 | 0.519140674 | 0.231302585 |
| Map1b | T | 527 | -0.190023946 | -0.206982592 | -0.146889227 | 0.043134719 | 0.060093365 | 0.024646599 | 0.127662753 | 0.214686252 | 0.653034141 | 0.643887112 |
| Rabep1 | S | 407 | -0.133820101 | -0.106589053 | -0.212828864 | -0.079008763 | -0.106239811 | 0.058161473 | 0.200829918 | 0.016326191 | 0.180353152 | 0.181300435 |
| Nedd4 | S | 385 | -0.161659441 | -0.334532596 | -0.560220772 | -0.398561332 | -0.225688177 | 0.427764259 | 0.104824545 | 0.389065902 | 0.520944143 | 0.705050638 |
| Pdcd11 | S | 1490 | -0.04783813 | -0.207719406 | -0.129851309 | -0.082013179 | 0.077868097 | 0.481983235 | 0.002941483 | 0.028661969 | 0.233610169 | 0.098976805 |
| Prkce | S | 368 | -0.237805971 | -0.073255116 | -0.319704955 | -0.081898984 | -0.246449839 | 0.064546586 | 0.542505831 | 0.060491894 | 0.501840497 | 0.109740164 |
| Pcdh19 | S | 1045 | 0.038728458 | -0.293834234 | -0.274125495 | -0.312853954 | 0.019708738 | 0.353592212 | 0.001441849 | 0.10877529 | 0.096962752 | 0.873592519 |
| Gapvd1 | S | 1017 | 0.02841867 | -0.214117604 | -0.164120361 | -0.192539031 | 0.049997242 | 0.718098568 | 0.001694893 | 0.00769136 | 0.082028756 | 0.277158996 |
| Bola1 | S | 81 | -0.2051245 | -0.157697551 | -0.177913824 | 0.027210676 | -0.020216273 | 0.02275573 | 0.223386505 | 0.13354718 | 0.780836188 | 0.87523191 |
| Map1b | S | 1797 | -0.143322935 | -0.155007074 | -0.161814098 | -0.018491163 | -0.006807024 | 0.089426825 | 0.051236705 | 0.048982219 | 0.733451375 | 0.862260162 |
| E2f8 | S | 102 | 0.035974071 | -0.224310398 | -0.207789221 | -0.243763292 | 0.016521177 | 0.433436012 | 0.000609934 | 0.031059718 | 0.01748584 | 0.776636676 |
| Sox4 | S | 213 | -0.123564322 | -0.171778587 | -0.136881161 | -0.013316839 | 0.034897426 | 0.008770996 | 0.049887518 | 0.183675325 | 0.871601252 | 0.707834891 |
| E2f8 | S | 416 | 0.089418898 | -0.276061716 | -0.228478212 | -0.31789711 | 0.047583504 | 0.028841114 | 9.47293E-05 | 0.02308924 | 0.008648077 | 0.410860935 |
| Slc1a4 | S | 507 | -0.041649661 | -0.309077713 | -0.14393128 | -0.102281619 | 0.165146433 | 0.448975142 | 0.014261962 | 0.060682867 | 0.147993732 | 0.098767503 |
| Golga3 | S | 391 | -0.032746676 | -0.254612931 | -0.065055816 | -0.032308842 | 0.189557413 | 0.262286641 | 7.40975E-05 | 0.5467712 | 0.757559688 | 0.166915979 |
| Rbm19 | S | 944 | 0.064192105 | -0.289564766 | -0.165961804 | -0.230153909 | 0.123602962 | 0.514231977 | 0.000390331 | 0.071317622 | 0.080722451 | 0.148212081 |
| Csnk1d | S | 382 | -0.056313311 | -0.049319863 | -0.283453301 | -0.22713999 | -0.234133439 | 0.504573385 | 0.391360607 | 0.000205734 | 0.06786429 | 0.015740431 |
| Gpbp1 | S | 260 | -0.022016388 | -0.127492607 | -0.228791229 | -0.206774841 | -0.101298622 | 0.643556131 | 0.044644074 | 0.001285453 | 0.003202355 | 0.074746632 |
| Lima1 | S | 230 | -0.207094101 | -0.158225516 | -0.124194192 | 0.082899909 | 0.034031323 | 0.009045343 | 0.106134822 | 0.314959362 | 0.462892229 | 0.774760067 |
| Smarcad1 | S | 81 | -0.07449464 | -0.107775398 | -0.233972189 | -0.159477549 | -0.126196791 | 0.13845609 | 0.206142275 | 0.003025264 | 0.02584508 | 0.151654682 |
| Frmd6 | S | 371 | -0.109857009 | -0.032818333 | -0.349461402 | -0.239604393 | -0.31664307 | 0.198871476 | 0.760373458 | 0.002540258 | 0.010355154 | 0.04406645 |
| Clic4 | S | 167 | -0.197740976 | -0.105796186 | -0.131666949 | 0.066074027 | -0.025870763 | 0.00477864 | 0.176259575 | 0.144624607 | 0.387897226 | 0.759232759 |
| Anln | S | 659 | 0.045470257 | -0.26189495 | -0.250001983 | -0.29547224 | 0.011892967 | 0.157606748 | 0.006724564 | 0.007478625 | 0.004015132 | 0.822169712 |
| Tpr | S | 711 | -0.080312457 | -0.364860824 | -0.415349588 | -0.335037131 | -0.050488764 | 0.524645174 | 0.211036195 | 0.099500309 | 0.166030746 | 0.856391127 |
| Helb | S | 1015 | -0.129092653 | -0.123336351 | -0.206607217 | -0.077514564 | -0.083270866 | 0.101647249 | 0.044545256 | 0.046108803 | 0.371956382 | 0.299846793 |
| Rab11fip5 | S | 416 | -0.124063428 | -0.218834681 | -0.156553444 | -0.032490017 | 0.062281236 | 0.233217361 | 0.003882203 | 0.302205998 | 0.82484867 | 0.643897036 |
| Ppm1b | S | 386 | -0.078382817 | -0.158556832 | -0.259394411 | -0.181011594 | -0.100837579 | 0.311982534 | 0.019127622 | 0.055684168 | 0.138467343 | 0.327622085 |
| Nek9 | S | 800 | -0.128834944 | -0.091421697 | -0.0636055 | 0.065229444 | 0.027816197 | 0.000119244 | 0.063734805 | 0.105461983 | 0.108128531 | 0.505450629 |
| Rrp8 | S | 103 | -0.019428913 | -0.141084251 | -0.183552482 | -0.164123569 | -0.042468231 | 0.80695831 | 0.083129052 | 9.85296E-05 | 0.134105045 | 0.475504187 |
| Dlg5 | S | 1222 | 0.151949366 | -0.462254562 | -0.296022884 | -0.44797225 | 0.166231678 | 0.190186564 | 0.029020856 | 0.00900472 | 0.013087057 | 0.256995305 |
| Kmt2d | S | 2208 | -0.162668407 | -0.300427701 | -0.053941329 | 0.108727078 | 0.246486372 | 0.147065447 | 0.005877764 | 0.633034613 | 0.383861898 | 0.081157871 |
| Ctcf | S | 402 | -0.1307294 | -0.124153656 | -0.193270653 | -0.062541253 | -0.069116997 | 0.235682269 | 0.00642714 | 0.062891625 | 0.564915286 | 0.3649923 |
| Igf2r | T | 2352 | -0.059750909 | -0.177365194 | -0.156293654 | -0.096542745 | 0.02107154 | 0.319461913 | 0.01206339 | 0.014863124 | 0.159714245 | 0.621863056 |
| Ncl | T | 92 | 0.170127758 | -0.256011144 | -0.343548248 | -0.513676006 | -0.087537104 | 0.051017564 | 0.009938894 | 0.000683888 | 0.001958509 | 0.198345244 |
| Cdc42bpa | S | 1638 | -0.145482957 | -0.059101403 | -0.261959073 | -0.116476116 | -0.20285767 | 0.09904182 | 0.619745955 | 0.003776464 | 0.135849263 | 0.157004453 |
| Arhgef40 | S | 1080 | -0.110489813 | -0.053059065 | -0.219332212 | -0.108842399 | -0.166273147 | 0.036259065 | 0.242265062 | 0.005065582 | 0.071956873 | 0.022229339 |
| Satb1 | S | 47 | -0.044193583 | -0.194740375 | -0.344733272 | -0.30053969 | -0.149992897 | 0.546208526 | 0.138405685 | 0.018731084 | 0.032596546 | 0.259372367 |
| Abcf1 | S | 22 | -0.031932201 | -0.139215796 | -0.11884239 | -0.086910189 | 0.020373406 | 0.321292968 | 0.000309986 | 0.015569955 | 0.059759236 | 0.50874128 |
| Rilpl1 | T | 348 | -0.092122345 | -0.149705775 | -0.098160194 | -0.006037849 | 0.051545581 | 0.27120755 | 0.000438317 | 0.056266343 | 0.935868056 | 0.226117514 |
| Nfx1 | S | 50 | 0.018889604 | -0.209130812 | -0.217827585 | -0.236717189 | -0.008696774 | 0.730058524 | 0.01055773 | 0.006020858 | 0.002199332 | 0.847772597 |
| Dock7 | S | 1382 | -0.008191177 | -0.162817758 | -0.252573144 | -0.244381967 | -0.089755386 | 0.905296793 | 0.083398431 | 0.001390518 | 0.022682196 | 0.260387508 |
| Nsd2 | T | 114 | 0.061926176 | -0.184145534 | -0.181734228 | -0.243660404 | 0.002411305 | 0.193835508 | 1.76066E-06 | 0.062653127 | 0.020571279 | 0.966230388 |
| Ehbp1l1 | S | 370 | -0.035061669 | -0.138535518 | -0.155218776 | -0.120157107 | -0.016683258 | 0.331399167 | 0.004874252 | 0.005597382 | 0.028421133 | 0.604034796 |
| Blm | S | 247 | -0.037907794 | -0.169781211 | -0.178027811 | -0.140120017 | -0.008246599 | 0.429818787 | 0.00211158 | 0.042315687 | 0.072260792 | 0.873686334 |
| Zfp800 | S | 593 | -0.06887101 | -0.311587603 | -0.061845659 | 0.007025351 | 0.249741945 | 0.360020405 | 0.00091504 | 0.492854036 | 0.938164693 | 0.047775037 |
| Erbin | S | 1013 | -0.311985243 | -0.004536069 | -0.200452297 | 0.111532946 | -0.195916228 | 0.004383543 | 0.969168553 | 0.132089439 | 0.323989801 | 0.187651264 |
| Srrm2 | S | 1194 | -0.193711523 | -0.246484963 | -0.125913999 | 0.067797524 | 0.120570964 | 0.04716248 | 0.062323579 | 0.371346206 | 0.620629044 | 0.42305166 |
| Sqstm1 | S | 363 | -0.067466511 | -0.180972514 | -0.17264964 | -0.105183129 | 0.008322874 | 0.190254392 | 0.036266017 | 0.016833053 | 0.071574679 | 0.894129375 |
| Srpk1 | T | 448 | 0.005455712 | -0.177706045 | -0.162449296 | -0.167905008 | 0.015256749 | 0.855042803 | 0.000225554 | 0.029603249 | 0.024264261 | 0.728379921 |
| Ect2 | T | 359 | -0.044802058 | -0.057100694 | -0.194155905 | -0.149353847 | -0.137055211 | 0.117234453 | 0.059170087 | 0.000378783 | 0.004551039 | 0.005632566 |
| Rplp0 | S | 304 | -0.076840285 | -0.166464506 | -0.090078928 | -0.013238642 | 0.076385578 | 0.096627983 | 0.002361526 | 0.048984514 | 0.708393785 | 0.054333561 |
| Vamp2 Vamp3 | S | 67;80 | -0.076313398 | -0.098143828 | -0.140144581 | -0.063831183 | -0.042000753 | 0.050100954 | 0.019080222 | 0.005976748 | 0.071631428 | 0.172610511 |
| Eif3a | T | 982 | -0.074286565 | -0.277901152 | -0.148040611 | -0.073754046 | 0.129860541 | 0.245975511 | 0.003223921 | 0.411122294 | 0.655010194 | 0.456485563 |
| Pdlim5 | S | 332 | -0.068562802 | -0.116453936 | -0.076688098 | -0.008125295 | 0.039765839 | 0.070499148 | 6.2821E-06 | 0.220380888 | 0.879168411 | 0.46178967 |
| Map1b | Y | 1405 | -0.09349695 | -0.174594667 | -0.235100844 | -0.141603895 | -0.060506178 | 0.30142512 | 0.046930946 | 0.036560312 | 0.17853532 | 0.481882145 |
| Cap1 | T | 306 | -0.070888837 | -0.132753999 | -0.282300818 | -0.211411981 | -0.149546818 | 0.178921896 | 0.152021063 | 0.01394386 | 0.031626756 | 0.138162174 |
| Gemin5 | S | 1414 | 0.047222229 | -0.190336789 | -0.235505658 | -0.282727887 | -0.045168869 | 0.36149584 | 0.004141695 | 0.003842321 | 0.003705933 | 0.326222005 |
| Hspa8 | S | 541 | -0.077132693 | -0.199829007 | -0.147447644 | -0.070314952 | 0.052381363 | 0.020330928 | 0.082495987 | 0.038807434 | 0.171059559 | 0.520006836 |
| Eif4a1 | S | 394 | 0.011963956 | -0.356121241 | -0.582857894 | -0.59482185 | -0.226736653 | 0.937706109 | 0.067613593 | 0.184449272 | 0.181356677 | 0.541754582 |
| Eno1 Eno1b | T | 272 | -0.203011429 | -0.142831473 | -0.293764514 | -0.090753085 | -0.150933041 | 0.203838084 | 0.390656391 | 0.031179788 | 0.541026401 | 0.383958989 |
| Add3 | S | 423 | 0.032418047 | -0.184555181 | -0.271979255 | -0.304397302 | -0.087424074 | 0.578632905 | 0.087429699 | 0.00069374 | 0.004278271 | 0.324233758 |
| Syvn1 | S | 608 | -0.088911049 | -0.092159612 | -0.144413058 | -0.055502009 | -0.052253446 | 0.07660593 | 0.015076908 | 0.007268201 | 0.233096481 | 0.159946333 |
| Serbp1 | S | 247 | -0.033123673 | -0.365595739 | -0.134443179 | -0.101319505 | 0.231152561 | 0.549976 | 0.005297754 | 0.273096775 | 0.375499918 | 0.107840549 |
| Pwp1 | S | 493 | -0.010305909 | -0.083939231 | -0.123786088 | -0.113480179 | -0.039846857 | 0.656246767 | 0.000786655 | 5.49793E-05 | 0.022975239 | 0.039967299 |
| Nfe2l2 | S | 425 | 0.062349698 | -0.305120898 | -0.269746503 | -0.332096201 | 0.035374395 | 0.496904221 | 0.040218359 | 0.006760886 | 0.033621202 | 0.662709797 |
| Eif2ak4 | S | 678 | -0.131481878 | -0.330444321 | -0.20624246 | -0.074760582 | 0.124201861 | 0.541595731 | 0.017631029 | 0.292975151 | 0.759193966 | 0.500089097 |
| Cast | T | 224 | 0.037475506 | -0.268569416 | -0.219127831 | -0.256603337 | 0.049441585 | 0.665726741 | 0.008067912 | 0.019823181 | 0.020792763 | 0.29329407 |
| Ranbp2 | S | 1605 | -0.047256671 | -0.132059895 | -0.168676449 | -0.121419777 | -0.036616554 | 0.28129938 | 0.014616843 | 0.004813491 | 0.005004425 | 0.127052336 |
| Yap1 | S | 274 | -0.14318212 | -0.205646599 | -0.103395679 | 0.039786441 | 0.10225092 | 0.008356008 | 0.076934352 | 0.237302007 | 0.591499216 | 0.337876785 |
| Mtmr6 | S | 557 | 0.045895687 | -0.201615798 | -0.187404619 | -0.233300306 | 0.014211179 | 0.433247652 | 0.000254235 | 0.022408231 | 0.019082442 | 0.772441664 |
| Rrp1 | T | 319 | 0.086439683 | -0.169951116 | -0.216738822 | -0.303178505 | -0.046787706 | 0.493419817 | 0.01554371 | 0.000236275 | 0.094734764 | 0.315288905 |
| Nop56 | T | 546 | 0.063603616 | -0.197279506 | -0.185596782 | -0.249200398 | 0.011682724 | 0.147448902 | 0.000863224 | 0.001333406 | 0.001534657 | 0.628723521 |

| | | | | | | | | | | | | |
|---|---|---|---|---|---|---|---|---|---|---|---|---|
| 2700062C07Rik | S | 136 | 0.056408001 | -0.130073656 | -0.233859323 | -0.290267324 | -0.103785668 | 0.324497927 | 0.001803132 | 0.000866922 | 0.007908031 | 0.044029033 |
| Nudt9 | S | 183 | -0.125199186 | -0.094165452 | -0.362962948 | -0.237763761 | -0.268797496 | 0.285720733 | 0.372713235 | 0.014687913 | 0.102076011 | 0.0643749 |
| Steap3 | S | 17 | 0.033860091 | -0.046020274 | -0.216300132 | -0.250160223 | -0.170279858 | 0.402039338 | 0.031026763 | 1.26479E-06 | 0.012488081 | 0.000537192 |
| Srgap2 | S | 206 | -0.021392542 | -0.096867028 | -0.225163495 | -0.203770953 | -0.128296467 | 0.636506597 | 0.034312873 | 0.000791458 | 0.014849975 | 0.009203626 |
| Prune2 | S | 878 | -0.074585201 | -0.153100113 | -0.150343043 | -0.075757843 | 0.00275707 | 0.227094881 | 0.01033986 | 0.01962499 | 0.232651971 | 0.944437743 |
| Erbin | S | 1150 | -0.209742088 | -0.100400793 | -0.251401526 | -0.041659438 | -0.151000733 | 0.035372985 | 0.422347516 | 0.075514085 | 0.696836211 | 0.295340404 |
| Raly | T | 175 | -0.067356688 | -0.140036194 | -0.253336008 | -0.18597932 | -0.113299814 | 0.593444058 | 0.049742976 | 0.008348627 | 0.196179049 | 0.107475179 |
| Satb2 | S | 594 | 0.011801213 | -0.196185955 | -0.14090286 | -0.152704073 | 0.055283095 | 0.768161269 | 0.000786626 | 0.009693308 | 0.013938149 | 0.160158546 |
| Zc3h18 | S | 742 | -0.009948428 | -0.242361618 | -0.158421808 | -0.148473379 | 0.08393981 | 0.886826891 | 0.001372162 | 0.076644082 | 0.134674099 | 0.260868934 |
| Bnip3 | T | 66 | 0.056089598 | -0.236412909 | -0.25144881 | -0.307538408 | -0.015035901 | 0.401859609 | 0.001923459 | 0.02864886 | 0.018510811 | 0.838575721 |
| Rasa3 | S | 833 | 0.065195258 | -0.112023829 | -0.285492952 | -0.35068821 | -0.173469123 | 0.158996437 | 0.033431811 | 6.15722E-05 | 0.007124356 | 0.026345384 |
| Tjp1 | S | 1576 | -0.305246933 | -0.03885338 | -0.273974844 | 0.031272088 | -0.235121464 | 0.016029544 | 0.661804373 | 0.180993306 | 0.854066322 | 0.233798062 |
| Cep170 | S | 985 | -0.119490458 | -0.026533925 | -0.543529062 | -0.424038604 | -0.516995137 | 0.521290212 | 0.890451424 | 0.009028625 | 0.044153418 | 0.035910483 |
| Psmc4 | S | 28 | -0.048772537 | -0.102604346 | -0.316980197 | -0.26820766 | -0.214375851 | 0.412230391 | 0.0693907 | 0.010437798 | 0.023759202 | 0.049892876 |
| Atg2a | S | 1243 | -0.160059503 | -0.007597892 | -0.328582581 | -0.168523078 | -0.320984689 | 0.314709734 | 0.926470725 | 0.001290745 | 0.295528015 | 0.019366105 |
| Orc1 | S | 254 | -0.002087809 | -0.269431769 | -0.121908954 | -0.119821145 | 0.147522815 | 0.975248847 | 0.001235415 | 0.062195118 | 0.153210958 | 0.044770978 |
| Luc7l | S | 363 | -0.101492635 | -0.110697398 | -0.103209277 | -0.001716642 | 0.007488121 | 0.03165708 | 0.004790784 | 0.042726325 | 0.967014927 | 0.836586863 |
| Nes | S | 644 | -0.169289128 | -0.037544043 | -0.190194637 | -0.02090551 | -0.152650595 | 0.005514997 | 0.162713503 | 0.059824815 | 0.759887016 | 0.110684443 |
| Fbxo28 | S | 235 | 0.049752109 | -0.223768359 | -0.231220939 | -0.280973047 | -0.007452579 | 0.389272313 | 0.082716662 | 0.000205872 | 0.012672956 | 0.930369333 |
| Rlim | S | 78 | -0.07267127 | -0.20329888 | -0.106494486 | -0.033823216 | 0.096804395 | 0.413401854 | 0.00178972 | 0.062575298 | 0.69595421 | 0.089990902 |
| Ago2 | S | 825 | -0.362666373 | -0.196664604 | -0.116154252 | 0.24651212 | 0.080510352 | 0.044852774 | 0.398369888 | 0.183383402 | 0.114117351 | 0.710488246 |
| Taf7 | S | 213 | -0.149629561 | -0.198173871 | -0.19840688 | -0.04877732 | -0.00023301 | 0.242696185 | 0.150431257 | 0.024713024 | 0.651969083 | 0.998274962 |
| Naca | T | 161;2133 | -0.068274683 | -0.106306826 | -0.199018686 | -0.130744003 | -0.09271186 | 0.062448094 | 0.009385199 | 0.042485717 | 0.085074834 | 0.168149422 |
| Tln1 | S | 2040 | -0.1541861 | -0.077942619 | -0.072253862 | 0.081932238 | 0.005688757 | 0.057939782 | 0.004173798 | 0.005482394 | 0.201274511 | 0.649534244 |
| Hspa9 | S | 162 | -0.029156863 | -0.358790496 | -0.024230266 | 0.004926597 | 0.33456023 | 0.567743722 | 0.000253666 | 0.780599658 | 0.948962871 | 0.026225842 |
| Alms1 Alms1-ps2 | T | 652 | 0.028562437 | -0.309276004 | -0.132465052 | -0.161027489 | 0.176810952 | 0.63896873 | 0.000470646 | 0.14857728 | 0.10901702 | 0.084466807 |
| Fam122a | S | 59 | -0.025940385 | -0.144902612 | -0.368304892 | -0.342364507 | -0.223402279 | 0.66554262 | 0.199590397 | 0.007537139 | 0.012511504 | 0.093084538 |
| Anapc1 | S | 547 | -0.248351012 | -0.237111804 | -0.222971201 | 0.025379811 | 0.014140603 | 0.033710058 | 0.420902298 | 0.249104858 | 0.879251272 | 0.961885679 |
| Eif2ak2 | S | 32 | -0.064744752 | -0.052681539 | -0.296510814 | -0.231766062 | -0.243829275 | 0.349279183 | 0.20204973 | 0.001708841 | 0.028418515 | 0.004915554 |
| Cct6a | S | 528 | -0.057234301 | -0.247062729 | -0.177808311 | -0.120574009 | 0.069254418 | 0.556417791 | 0.009385199 | 0.081090001 | 0.298257147 | 0.362802576 |
| H1f1 | S | 180 | 0.050643379 | -0.08507699 | -0.359162543 | -0.409805922 | -0.274085553 | 0.606943166 | 0.151050103 | 0.000374446 | 0.028500033 | 0.000740162 |
| Pdzd8 | S | 515 | 0.011668476 | -0.176966974 | -0.124679477 | -0.136347952 | 0.052287498 | 0.75873053 | 0.000283808 | 0.007607795 | 0.027614964 | 0.123344137 |
| Sema4b | S | 804 | 0.039612518 | -0.25753523 | -0.238631534 | -0.278244053 | 0.018903696 | 0.570351094 | 0.008341421 | 0.020559748 | 0.020570786 | 0.798580496 |
| Trip10 | S | 482 | -0.119074935 | -0.103148912 | -0.498936129 | -0.379861194 | -0.395787217 | 0.468370929 | 0.500639867 | 0.025057463 | 0.057339982 | 0.050052882 |
| Apbb2 | S | 160 | -0.033431022 | -0.221608476 | -0.255609797 | -0.222178775 | -0.034001321 | 0.441483002 | 0.016188847 | 0.080500382 | 0.097443805 | 0.73469975 |
| Per1 | S | 979 | -0.171496254 | -0.116605981 | -0.215775658 | -0.044279403 | -0.099169677 | 0.058454836 | 0.265292317 | 0.040352816 | 0.463926082 | 0.291511131 |
| Rrad | S | 39 | -0.302426251 | -0.153935825 | -0.012954338 | 0.289471913 | 0.140981487 | 0.002518837 | 0.148341915 | 0.909461582 | 0.076705322 | 0.307653907 |
| Myo9b | S | 1988 | -0.075901977 | -0.141152508 | -0.146864687 | -0.070962709 | -0.005712179 | 0.318867116 | 0.001501367 | 0.047165643 | 0.37561575 | 0.903874929 |
| Acaca | S | 25 | -0.259568436 | -0.088776006 | -0.171473697 | 0.088094739 | -0.082697691 | 0.023691873 | 0.395832577 | 0.085280233 | 0.294708832 | 0.37726166 |
| Ogfod1 | S | 371 | -0.172517202 | -0.056320392 | -0.401785933 | -0.229268731 | -0.345465541 | 0.349031408 | 0.808909588 | 0.008354283 | 0.24549758 | 0.223124027 |
| Trim3 | S | 427 | -0.023740404 | -0.14104935 | -0.189433405 | -0.165693001 | -0.048384055 | 0.675482217 | 0.01402538 | 0.00298487 | 0.037591809 | 0.143931104 |
| Usp47 | S | 911 | -0.072929833 | -0.120861466 | -0.178353351 | -0.105423518 | -0.057491885 | 0.287661123 | 0.041499883 | 0.003703979 | 0.164472037 | 0.232732692 |
| Eif4b | S | 406 | 0.080692657 | -0.20878034 | -0.447915365 | -0.528608021 | -0.239135025 | 0.112843469 | 0.001917219 | 0.040789605 | 0.034194403 | 0.155348904 |
| Tnks1bp1 | S | 496 | -0.144281896 | -0.117688572 | -0.114228484 | 0.030053412 | 0.003460089 | 0.051817554 | 0.14440149 | 0.004311968 | 0.558616921 | 0.955413147 |
| Tns3 | S | 1436 | -0.084132507 | -0.088607541 | -0.43059102 | -0.346458513 | -0.34198348 | 0.464255686 | 0.560234661 | 0.008179567 | 0.040587819 | 0.09044679 |
| Trip12 | S | 165 | 0.011003314 | -0.185716133 | -0.250881402 | -0.261884716 | -0.065165269 | 0.718341002 | 0.118695762 | 0.000698865 | 0.001485097 | 0.476397567 |
| Specc1 | S | 314 | -0.137800048 | -0.121038563 | -0.147461478 | -0.00966143 | -0.026422915 | 0.021077744 | 0.16284803 | 0.02948432 | 0.800521227 | 0.717647316 |
| Tex2 | S | 294 | -0.186564349 | -0.050812722 | -0.327596927 | -0.141032578 | -0.276784205 | 0.124793357 | 0.687041233 | 0.016737333 | 0.118619862 | 0.048279496 |
| Prpf40a | S | 34 | 0.01862954 | -0.30980883 | -0.358792538 | -0.377422078 | -0.048983708 | 0.667777599 | 0.022897772 | 0.092838353 | 0.094578487 | 0.754504516 |
| Bop1 | S | 112 | -0.040545021 | -0.179442387 | -0.127700426 | -0.087155405 | 0.051741962 | 0.47852707 | 0.003458631 | 0.014223127 | 0.14884523 | 0.123150987 |
| Bop1 | S | 113 | -0.040545021 | -0.179442387 | -0.127700426 | -0.087155405 | 0.051741962 | 0.47852707 | 0.003458631 | 0.014223127 | 0.14884523 | 0.123150987 |
| Smg9 | S | 53 | -0.216607355 | -0.10500409 | -0.185725445 | 0.03088191 | -0.080721355 | 0.087648725 | 0.302640284 | 0.022990252 | 0.748998964 | 0.396413169 |
| R3hdm2 | S | 37 | -0.351819655 | -0.132795334 | -0.047110549 | 0.304709106 | 0.085684785 | 0.004685122 | 0.280351557 | 0.688377989 | 0.05537371 | 0.530183059 |
| Clasp1 | S | 794 | -0.244239049 | -0.133269006 | -0.340142647 | -0.095903598 | -0.206873641 | 0.229030488 | 0.469993276 | 0.053044341 | 0.57184384 | 0.235043697 |
| Rgs12 | S | 879 | -0.099241408 | -0.102960202 | -0.440875239 | -0.341633831 | -0.337915037 | 0.379967388 | 0.422497707 | 0.020552394 | 0.066566296 | 0.052539944 |
| Fhod1 | T | 499 | 0.013400296 | -0.146876323 | -0.176096755 | -0.189497051 | -0.029220432 | 0.780217821 | 0.009593897 | 0.000564813 | 0.029023855 | 0.388533686 |
| Uimc1 | S | 689 | -0.105029446 | -0.163726069 | -0.104730064 | 0.000299382 | 0.058996005 | 0.177435254 | 0.008289211 | 0.030817251 | 0.996283491 | 0.188776786 |
| Marchf7 | S | 545 | -0.006218828 | -0.1162117 | -0.16029351 | -0.154074681 | -0.04408181 | 0.923625079 | 0.013431408 | 0.00015747 | 0.104884728 | 0.191152669 |
| Camsap2 | S | 1285 | -0.245661272 | -0.089340933 | -0.315498537 | -0.069837265 | -0.226157603 | 0.099176353 | 0.528855344 | 0.067368774 | 0.583093762 | 0.140135488 |
| Map1b | S | 1422 | -0.030980658 | -0.091428183 | -0.151145318 | -0.120164661 | -0.059717136 | 0.146995919 | 0.000269532 | 0.013964386 | 0.024513481 | 0.136818849 |
| Hsp90aa1 | S | 231 | -0.185746069 | -0.242969444 | -0.147691137 | 0.038054932 | 0.095278307 | 0.264133048 | 0.086287815 | 0.060073925 | 0.792998305 | 0.399891036 |
| N4bp1 | S | 286 | -0.163630469 | -0.045435008 | -0.271702073 | -0.108071604 | -0.226267065 | 0.077082983 | 0.714497721 | 0.008244787 | 0.185407385 | 0.13368213 |
| Usp14 | S | 393 | -0.128818053 | -0.114794079 | -0.080817444 | 0.048000609 | 0.033976635 | 0.012258603 | 0.004836015 | 0.171536205 | 0.381797597 | 0.497985579 |
| Ahctf1 | S | 1772 | -0.035564895 | -0.0908053 | -0.255800627 | -0.220235732 | -0.164995327 | 0.602857559 | 0.178111568 | 0.000568051 | 0.035159526 | 0.046781151 |
| Tcof1 | S | 794 | -0.083239658 | -0.172517993 | -0.107723597 | -0.024483939 | 0.064794396 | 0.30064001 | 0.003250428 | 0.034575652 | 0.727466795 | 0.078279219 |
| Htatsf1 | S | 388 | -0.268027624 | 0.010114498 | -0.239010376 | 0.029017248 | -0.249124874 | 0.006716888 | 0.931844323 | 0.078021439 | 0.764013162 | 0.115093033 |
| Nol8 | S | 300 | -0.052120941 | -0.149405409 | -0.068335617 | -0.016214677 | 0.081069792 | 0.215612443 | 0.00030735 | 0.016532824 | 0.63968957 | 0.002074736 |
| Champ1 | S | 298 | -0.35582793 | -0.195079114 | 0.121035744 | 0.476863674 | 0.316114859 | 0.00352651 | 0.043431149 | 0.502703592 | 0.070999317 | 0.155682626 |
| Atxn2l | T | 505 | 0.088398326 | -0.23511379 | -0.214398086 | -0.302796412 | 0.020715704 | 0.41667114 | 0.008282558 | 0.002661794 | 0.062565623 | 0.665345776 |
| Tfdp1 | S | 23 | -0.081071677 | -0.163041401 | -0.308539588 | -0.22746791 | -0.145498187 | 0.384094383 | 0.047482205 | 0.061681146 | 0.111021839 | 0.218650493 |
| Actr8 | S | 132 | -0.031813828 | -0.126094877 | -0.130006848 | -0.09819302 | -0.003911971 | 0.155295122 | 0.001123948 | 0.014685285 | 0.06104922 | 0.895187877 |
| Sh3gl1 Sh3gl2 Sh3gl3 | S | 2;2;2 | -0.151357399 | -0.064847888 | -0.469785533 | -0.318428133 | -0.404937645 | 0.291685577 | 0.691388566 | 0.02409817 | 0.092435998 | 0.068676771 |
| Arid5b | S | 264 | -0.125233746 | -0.081154649 | -0.187112291 | -0.061878545 | -0.105957642 | 0.012514721 | 0.442939487 | 0.011833212 | 0.227738195 | 0.338122392 |
| Mff | S | 131 | -0.003458144 | -0.108949279 | -0.259739146 | -0.256281003 | -0.150789867 | 0.953751424 | 0.014592849 | 0.003324227 | 0.017300298 | 0.025484117 |
| Abl2 | S | 1000 | 0.005093435 | -0.329157722 | -0.191591652 | -0.196685086 | 0.137566071 | 0.958112012 | 0.065599879 | 0.006884513 | 0.118930647 | 0.309316794 |
| Pgm3 | S | 64 | -0.369528572 | -0.066167207 | -0.389867252 | -0.02033868 | -0.323700045 | 0.142096998 | 0.798596643 | 0.104856838 | 0.921711573 | 0.229284497 |
| Map1b | T | 1962 | -0.036863915 | -0.068235966 | -0.176618083 | -0.139754168 | -0.108382117 | 0.422847278 | 0.026348134 | 0.000184465 | 0.044483328 | 0.002957465 |
| Eml1 | S | 165 | -0.079980448 | -0.195784832 | -0.42220086 | -0.342220413 | -0.226416028 | 0.652891709 | 0.280636338 | 0.028583324 | 0.139414658 | 0.243676416 |
| Fadd | S | 191 | -0.163982167 | -0.141415495 | -0.167750701 | -0.003768533 | -0.026335206 | 0.060321241 | 0.042151316 | 0.153601313 | 0.970133737 | 0.78424375 |
| Relch | S | 422 | -0.137727526 | -0.191687099 | -0.086152053 | 0.051575473 | 0.105535046 | 0.052727947 | 0.00954784 | 0.288535433 | 0.523677739 | 0.221437791 |
| Serbp1 | S | 25 | -0.127225586 | -0.16035393 | -0.08273483 | 0.044490755 | 0.077619099 | 0.023808773 | 0.008300584 | 0.244454471 | 0.47017359 | 0.250984558 |
| Farp1 | S | 433 | 0.058406211 | -0.176534719 | -0.143585176 | -0.201991387 | 0.032949543 | 0.145405089 | 0.014790534 | 1.7266E-06 | 0.013766117 | 0.339585144 |
| Mark2 | S | 40 | -0.064734823 | -0.103503392 | -0.244052366 | -0.179317542 | -0.140548974 | 0.388781059 | 0.07133736 | 0.005018676 | 0.061003486 | 0.047950048 |
| Yap1 | T | 48 | -0.087467471 | -0.172330421 | -0.191804582 | -0.104337111 | -0.019474161 | 0.413649327 | 0.030021268 | 0.022900334 | 0.353329844 | 0.766816438 |
| Pi4kb | T | 280 | -0.198292175 | -0.109448381 | -0.103447072 | 0.094845103 | 0.006001309 | 0.001491152 | 0.323017756 | 0.18394559 | 0.213908417 | 0.955330338 |
| Mphosph8 | S | 278 | -0.303793175 | -0.042633307 | -0.220813442 | 0.082979732 | -0.178180135 | 0.003561813 | 0.651238693 | 0.380154489 | 0.714981996 | 0.466334235 |
| Spp1 | S | 27 | -0.276369187 | -0.047315022 | -0.272842177 | 0.00352701 | -0.225527155 | 0.045501674 | 0.40276672 | 0.111056777 | 0.979524167 | 0.153977793 |
| Ddx21 | T | 387 | 0.03561961 | -0.222613459 | -0.294850709 | -0.330470318 | -0.07223725 | 0.603860566 | 0.024410603 | 0.01310404 | 0.014335523 | 0.406705917 |
| Rrp1 | S | 434 | -0.007826947 | -0.118548863 | -0.236149054 | -0.228322107 | -0.117600191 | 0.861569954 | 0.033724659 | 0.00152875 | 0.002212028 | 0.015992962 |
| Slc20a2 | S | 423 | 0.080953743 | -0.186548644 | -0.484637242 | -0.565590985 | -0.298088598 | 0.359913246 | 0.118339342 | 0.006517329 | 0.006152611 | 0.053761891 |
| Ripk2 | S | 168 | -0.265161737 | -0.090583749 | -0.149131628 | 0.116030109 | -0.058547879 | 0.009294829 | 0.293143965 | 0.245018205 | 0.320050814 | 0.596587461 |
| Kars | T | 458 | -0.179437827 | -0.235309709 | -0.187874481 | -0.008436653 | 0.047435228 | 0.109635024 | 0.182464051 | 0.111242448 | 0.937222136 | 0.759783176 |
| Dop1a | S | 1232 | 0.021920304 | -0.213971005 | -0.2965482 | -0.318468504 | -0.082577195 | 0.68157325 | 0.001992388 | 0.091064465 | 0.08034083 | 0.515225255 |
| Zcchc9 | S | 53 | 0.014343158 | -0.217477545 | -0.222058514 | -0.236401672 | -0.004580969 | 0.784184198 | 0.029274485 | 0.004886673 | 0.003701817 | 0.944557737 |

| | | | | | | | | | | | | |
|---|---|---|---|---|---|---|---|---|---|---|---|---|
| Ybx1 | S | 163 | 0.05515745 | -0.205894372 | -0.293968867 | -0.349126317 | -0.088074495 | 0.343204402 | 0.007846523 | 0.011365651 | 0.016435348 | 0.258682411 |
| Nap1l1 | T | 62 | -0.033990699 | -0.170659428 | -0.223637736 | -0.189647037 | -0.052978308 | 0.313045176 | 0.016316823 | 0.035130171 | 0.035318893 | 0.395427259 |
| Grb14 | S | 49 | -0.022478566 | -0.163399817 | -0.227029739 | -0.204551173 | -0.063629922 | 0.673663633 | 0.004555178 | 0.035889223 | 0.051634894 | 0.39067716 |
| Triobp | T | 1615 | -0.042280335 | -0.270903436 | -0.298344893 | -0.256064558 | -0.027441458 | 0.783293457 | 0.022696561 | 0.125155542 | 0.217019174 | 0.85556268 |
| Nufip2 | S | 649 | 0.045778769 | -0.222651144 | -0.38638036 | -0.432159129 | -0.163729216 | 0.576786076 | 0.071632147 | 0.013095726 | 0.023767513 | 0.191886939 |
| Otub1 | S | 16 | -0.163626143 | -0.068310386 | -0.226525155 | -0.062899012 | -0.158214769 | 0.041438194 | 0.261229051 | 0.033702142 | 0.437168453 | 0.100286025 |
| Rif1 | S | 1029 | -0.020284208 | -0.171349918 | -0.169034007 | -0.148749799 | 0.002315911 | 0.787251362 | 0.017579369 | 0.002738017 | 0.133792466 | 0.956449738 |
| Atxn2l | S | 333 | -0.104137797 | -0.11366129 | -0.236180531 | -0.132042734 | -0.122519241 | 0.191297138 | 0.122957419 | 0.014850184 | 0.0993822 | 0.092870325 |
| Plekhf1 | S | 197 | -0.216326908 | -0.146833317 | -0.219836366 | -0.003509458 | -0.073003049 | 0.033868142 | 0.195474896 | 0.246516863 | 0.982229153 | 0.667263988 |
| Nelfe | S | 49 | -0.288277557 | -0.028278376 | -0.126666042 | 0.161611516 | -0.098387666 | 0.003811761 | 0.778294875 | 0.092935951 | 0.006085923 | 0.31229103 |
| Tbc1d25 | S | 560 | -0.104552328 | -0.107328786 | -0.311357383 | -0.206805055 | -0.204028597 | 0.240985678 | 0.23759124 | 0.017633337 | 0.081645613 | 0.086609704 |
| Rai14 | S | 11 | -0.189140281 | -0.082735031 | -0.140776955 | 0.048363327 | -0.058041923 | 0.013166139 | 0.092078044 | 0.116284891 | 0.515207437 | 0.432421443 |
| Pdha1 | S | 232 | -0.138772936 | -0.143660929 | -0.079886581 | 0.058886355 | 0.063774347 | 0.001730799 | 0.032111295 | 0.437024903 | 0.54889301 | 0.533778735 |
| Gm13889 | S | 210 | -0.193734409 | -0.042772221 | -0.376696108 | -0.182961699 | -0.333923887 | 0.138611866 | 0.756170384 | 0.026313457 | 0.205624997 | 0.07161663 |
| Evl | S | 347 | -0.087410804 | -0.174703016 | -0.324142179 | -0.236731375 | -0.149439163 | 0.595552403 | 0.146429917 | 0.023030452 | 0.194946998 | 0.193728954 |
| Bzw2 | S | 412 | -0.053259851 | -0.136691586 | -0.171399454 | -0.118139603 | -0.034707869 | 0.188235538 | 0.005542198 | 0.035529377 | 0.087682471 | 0.515633354 |
| Bzw2 | S | 414 | -0.053259851 | -0.136691586 | -0.171399454 | -0.118139603 | -0.034707869 | 0.188235538 | 0.005542198 | 0.035529377 | 0.087682471 | 0.515633354 |
| Rgl1 | T | 631 | -0.349195293 | -0.123919574 | -0.183346111 | 0.165849182 | -0.059426537 | 0.011934753 | 0.49645348 | 0.495436702 | 0.528010042 | 0.830796167 |
| Iscu | S | 15 | -0.359166648 | -0.151555996 | -0.137564011 | 0.221602636 | 0.013991985 | 0.08882938 | 0.096810103 | 0.371561879 | 0.273190911 | 0.915075228 |
| Iscu | S | 21 | -0.359166648 | -0.151555996 | -0.137564011 | 0.221602636 | 0.013991985 | 0.08882938 | 0.096810103 | 0.371561879 | 0.273190911 | 0.915075228 |
| Prrc2a | S | 808 | -0.016095371 | -0.252921224 | -0.217466171 | -0.2013708 | 0.035455053 | 0.721979937 | 0.015494541 | 0.053748432 | 0.050423204 | 0.643156884 |
| Zfp93 | S | 257 | -0.075179197 | -0.210999838 | -0.160945121 | -0.085765924 | 0.050054717 | 0.35503823 | 0.027551779 | 0.03138343 | 0.322946759 | 0.469926478 |
| Pip4k2b | S | 315 | -0.391784653 | -0.004594574 | -0.204819321 | 0.186965332 | -0.200224747 | 0.009396866 | 0.96062606 | 0.253866141 | 0.296511788 | 0.269473784 |
| Gtpbp1 | S | 580 | -0.161419536 | -0.063979232 | -0.137523383 | 0.023896153 | -0.073544151 | 0.003949329 | 0.134525433 | 0.078146596 | 0.682399906 | 0.263122321 |
| Ranbp2 | S | 788 | 0.073625805 | -0.220281714 | -0.215581298 | -0.289207103 | 0.004700416 | 0.138413589 | 0.001122784 | 0.005792791 | 0.00387602 | 0.913105768 |
| Eme1 | T | 13 | -0.036575623 | -0.136851652 | -0.200770247 | -0.164194624 | -0.063918595 | 0.333133445 | 0.122504683 | 0.001244896 | 0.000363596 | 0.385142948 |
| Taf1c | S | 576 | -0.027676884 | -0.139747441 | -0.212866377 | -0.185189493 | -0.073118937 | 0.720275752 | 0.011406761 | 0.008362163 | 0.091862076 | 0.128247313 |
| Trpm7 | S | 1502 | 0.001717352 | -0.103945818 | -0.263609538 | -0.265326891 | -0.159663721 | 0.978490915 | 0.042569538 | 0.001241156 | 0.012991663 | 0.019459548 |
| Nfatc2 | S | 219 | -0.018059701 | -0.181970467 | -0.23174753 | -0.213687829 | -0.049777063 | 0.819059143 | 0.027315173 | 0.010708558 | 0.025459357 | 0.225996201 |
| Nfatc2 | S | 223 | -0.018059701 | -0.181970467 | -0.23174753 | -0.213687829 | -0.049777063 | 0.819059143 | 0.027315173 | 0.010708558 | 0.025459357 | 0.225996201 |
| Urgcp | S | 829 | 0.006997424 | -0.165121438 | -0.203553427 | -0.210550851 | -0.03843199 | 0.907712825 | 0.007995624 | 0.005583607 | 0.018912272 | 0.349628433 |
| Ppp1r12a | T | 508 | -0.082628492 | -0.114557603 | -0.181073866 | -0.098445374 | -0.066516263 | 0.084119355 | 0.082324965 | 0.010656285 | 0.082496652 | 0.273707645 |
| Ska3 | T | 120 | -0.081621833 | -0.097179656 | -0.24597676 | -0.164354927 | -0.148797105 | 0.075232423 | 0.182088101 | 0.013669128 | 0.036082643 | 0.086131199 |
| Srrm2 | T | 2421 | 0.023077485 | -0.146751391 | -0.137681956 | -0.160759441 | 0.009069436 | 0.515615585 | 2.19232E-05 | 0.014750115 | 0.014134339 | 0.700686262 |
| Recql5 | S | 728 | 0.070872818 | -0.203520253 | -0.178198118 | -0.249070936 | 0.025322135 | 0.207133657 | 0.000754064 | 0.003004501 | 0.009665166 | 0.48656459 |
| Sash1 | S | 500 | -0.040889827 | -0.111296032 | -0.141159305 | -0.100269478 | -0.029863272 | 0.456426301 | 0.002661694 | 0.003163668 | 0.138496601 | 0.282390538 |
| 5031439G07Rik | S | 279 | -0.078978037 | -0.141579469 | -0.189298861 | -0.110320824 | -0.047719392 | 0.179832395 | 0.010038859 | 0.066597585 | 0.214673878 | 0.519309248 |
| Psat1 | S | 344 | -0.124773565 | -0.129080316 | -0.313384187 | -0.188610622 | -0.184303871 | 0.09408862 | 0.195045939 | 0.08046431 | 0.220582908 | 0.235671833 |
| Rnf219 | S | 549 | -0.034687378 | -0.300126034 | -0.153079991 | -0.118392613 | 0.147046043 | 0.749481567 | 0.005393342 | 0.150531353 | 0.327310657 | 0.152398619 |
| Gab1 | S | 266 | -0.066944073 | -0.124443313 | -0.350451921 | -0.283507848 | -0.226008608 | 0.243819873 | 0.175690106 | 0.028339932 | 0.050674003 | 0.091006017 |
| Psmb6 | T | 55 | 0.074880717 | -0.024830684 | -0.31213838 | -0.387019097 | -0.287307696 | 0.529209533 | 0.545146755 | 3.28228E-06 | 0.056897187 | 0.005462661 |
| Nufip2 | S | 634 | -0.011382613 | -0.18878541 | -0.407983412 | -0.396600799 | -0.219198002 | 0.887839779 | 0.05535467 | 0.047815856 | 0.056835201 | 0.18630541 |
| Specc1l | S | 871 | -0.01907656 | -0.225443139 | -0.115921064 | -0.096844504 | 0.109522074 | 0.645649175 | 0.005003186 | 0.013662813 | 0.037252696 | 0.081458801 |
| Ctnnd1 | Y | 257 | -0.121029303 | -0.17039209 | -0.395256199 | -0.274226896 | -0.224864109 | 0.461067521 | 0.226882408 | 0.052183474 | 0.18768798 | 0.207014535 |
| Ulk1 | S | 637 | -0.180259519 | -0.079018231 | -0.101942811 | 0.078316708 | -0.02292458 | 0.000119585 | 0.357281656 | 0.347889022 | 0.450045246 | 0.841555977 |
| Mcmbp | S | 298 | -0.087976263 | -0.098948439 | -0.1138629 | -0.025886637 | -0.014914461 | 0.224909729 | 0.007027529 | 0.002490689 | 0.672608072 | 0.530294642 |
| Dyrk1a | S | 47 | -0.420952933 | -0.10291876 | -0.122630929 | 0.298322005 | -0.019712169 | 0.019750761 | 0.466383728 | 0.426629363 | 0.124091212 | 0.911722935 |
| Mrtfa | S | 370 | -0.20643272 | -0.100472362 | -0.444351193 | -0.237918473 | -0.343878831 | 0.180936892 | 0.513636352 | 0.089403294 | 0.282517464 | 0.158477311 |
| Mapkapk5 | T | 182 | -0.002083393 | -0.13240517 | -0.265279384 | -0.263195991 | -0.132874214 | 0.973050454 | 0.093051967 | 0.001399859 | 0.021069587 | 0.107057997 |
| Ank3 | S | 914 | -0.194122308 | -0.002790606 | -0.30507848 | -0.110956172 | -0.302287874 | 0.049060285 | 0.979924533 | 0.016938787 | 0.20039513 | 0.037006367 |
| Ank3 | S | 917 | -0.194122308 | -0.002790606 | -0.30507848 | -0.110956172 | -0.302287874 | 0.049060285 | 0.979924533 | 0.016938787 | 0.20039513 | 0.037006367 |
| Eif4b | S | 497 | -0.065702449 | -0.102115523 | -0.17156234 | -0.105859891 | -0.069446817 | 0.034311228 | 0.050675594 | 0.012751278 | 0.037301763 | 0.147043303 |
| Srrm2 | S | 1246 | 0.010290877 | -0.204406563 | -0.375994582 | -0.386285459 | -0.171588019 | 0.923392407 | 0.054740822 | 0.03055045 | 0.035615717 | 0.180914835 |
| Xrcc6 | S | 518 | -0.032702528 | -0.160584939 | -0.239069111 | -0.206366583 | -0.078484171 | 0.490457848 | 0.003542624 | 0.077140251 | 0.100272767 | 0.42047388 |
| Wiz | S | 1015 | -0.008784461 | -0.214886221 | -0.096175181 | -0.08739072 | 0.11871104 | 0.897593632 | 7.57259E-05 | 0.148336898 | 0.304161006 | 0.103950638 |
| Lrig1 | S | 1066 | 0.030453704 | -0.246701594 | -0.222023618 | -0.252477322 | 0.024677976 | 0.278458791 | 0.007090276 | 0.013293535 | 0.007755577 | 0.637975161 |
| Osbpl6 | S | 23 | -0.138919756 | -0.093904311 | -0.554107175 | -0.415187419 | -0.460202864 | 0.35046907 | 0.679984508 | 0.044232377 | 0.098733206 | 0.130713125 |
| Tapt1 | T | 472 | 0.050418977 | -0.319231806 | -0.299316829 | -0.349735806 | 0.019914977 | 0.329502848 | 0.086196958 | 0.005985284 | 0.002737795 | 0.866245448 |
| Kirrel | S | 648 | 0.024544479 | -0.170541358 | -0.297563052 | -0.322107531 | -0.127021694 | 0.904323354 | 0.043035885 | 0.007726913 | 0.200145862 | 0.110171975 |
| Adam10 | T | 720 | -0.153318386 | -0.107361719 | -0.41441801 | -0.261099624 | -0.307056291 | 0.075890468 | 0.513775686 | 0.094335964 | 0.218164566 | 0.200395582 |
| Acap2 | T | 548 | 0.037387923 | -0.154683064 | -0.276348803 | -0.313736726 | -0.121665739 | 0.538907686 | 0.035210069 | 0.001868038 | 0.00889426 | 0.076105078 |
| Abcf1 | S | 225 | -0.186523293 | -0.08062831 | -0.163484418 | 0.023038876 | -0.082856107 | 0.023872683 | 0.109625268 | 0.089429578 | 0.753418193 | 0.28558637 |
| H2-D1 H2-L | S | 350;350;160 | -0.246022958 | -0.013344631 | -0.225823352 | 0.020199606 | -0.212478721 | 0.013296147 | 0.927310174 | 0.05007634 | 0.794188001 | 0.209795163 |
| Clasp2 | S | 376 | -0.176764883 | -0.055696572 | -0.397195286 | -0.220430403 | -0.341498713 | 0.256577388 | 0.749281639 | 0.017485719 | 0.183970561 | 0.114867991 |
| Baz1b | S | 374 | -0.023460765 | -0.127025291 | -0.119097278 | -0.095636514 | 0.007928013 | 0.572537661 | 2.22013E-05 | 0.046832705 | 0.107130559 | 0.821068778 |
| Tmed8 | S | 21 | -0.19325681 | -0.220615671 | -0.192396765 | 0.000860045 | 0.028218906 | 0.073689646 | 0.04943678 | 0.510263009 | 0.997452469 | 0.916797744 |
| Arap2 | S | 986 | -0.071875237 | -0.140656459 | -0.209240244 | -0.137365007 | -0.068583784 | 0.170331257 | 0.049641562 | 0.025366379 | 0.088321562 | 0.344739628 |
| Arap2 | T | 985 | -0.071875237 | -0.140656459 | -0.209240244 | -0.137365007 | -0.068583784 | 0.170331257 | 0.049641562 | 0.025366379 | 0.088321562 | 0.344739628 |
| Arap2 | T | 993 | -0.071875237 | -0.140656459 | -0.209240244 | -0.137365007 | -0.068583784 | 0.170331257 | 0.049641562 | 0.025366379 | 0.088321562 | 0.344739628 |
| Arap2 | T | 995 | -0.071875237 | -0.140656459 | -0.209240244 | -0.137365007 | -0.068583784 | 0.170331257 | 0.049641562 | 0.025366379 | 0.088321562 | 0.344739628 |
| Pa2g4 | S | 360 | 0.037099014 | -0.120909471 | -0.255681371 | -0.292780384 | -0.134771899 | 0.474949522 | 0.015025661 | 0.00092204 | 0.003584685 | 0.023068308 |
| Zbtb41 Zfp12 Zfp281 | S | 690;342;337 | -0.064247552 | -0.207566277 | -0.205548309 | -0.141300757 | 0.002017967 | 0.450674478 | 0.202091349 | 0.003423236 | 0.152732935 | 0.987681903 |
| Mcph1 | S | 288 | 0.021585435 | -0.264469623 | -0.15278969 | -0.174375125 | 0.111679933 | 0.699970487 | 0.011364166 | 0.006141125 | 0.036957437 | 0.122971392 |
| Tsc2 | S | 1339 | -0.220820217 | -0.20065283 | -0.28892006 | -0.068099843 | -0.08826723 | 0.338514413 | 0.203918113 | 0.093034066 | 0.72465977 | 0.363693654 |
| Tsc2 | S | 1340 | -0.220820217 | -0.20065283 | -0.28892006 | -0.068099843 | -0.08826723 | 0.338514413 | 0.203918113 | 0.093034066 | 0.72465977 | 0.363693654 |
| Tsc2 | S | 1342 | -0.220820217 | -0.20065283 | -0.28892006 | -0.068099843 | -0.08826723 | 0.338514413 | 0.203918113 | 0.093034066 | 0.72465977 | 0.363693654 |
| Casp8ap2 | S | 1428 | -0.043704829 | -0.175117477 | -0.408335069 | -0.36463024 | -0.233217592 | 0.876661798 | 0.520697415 | 0.005094864 | 0.271739532 | 0.410202128 |
| Pola1 | T | 180 | -0.058712962 | -0.135859667 | -0.388986974 | -0.330274012 | -0.253127306 | 0.565892681 | 0.430752003 | 0.008549748 | 0.013156644 | 0.170429045 |
| Satb2 | S | 593 | -0.025858701 | -0.184294225 | -0.123323531 | -0.09746483 | 0.060970693 | 0.326680789 | 0.001460205 | 0.04660994 | 0.10408818 | 0.233086951 |
| Pdlim5 | S | 359 | -0.057452434 | -0.168447246 | -0.055013153 | 0.002439281 | 0.113434093 | 0.231370795 | 5.55904E-05 | 0.180352464 | 0.959213146 | 0.042521139 |
| Ahdc1 | S | 845 | -0.042449904 | -0.139193614 | -0.17486263 | -0.132412726 | -0.035669016 | 0.368450725 | 0.012868328 | 0.010047569 | 0.030376515 | 0.372745374 |
| Farp1 | S | 336 | -0.120898328 | -0.157743032 | -0.367820502 | -0.246922174 | -0.21007747 | 0.081614842 | 0.233195359 | 0.139332689 | 0.260199599 | 0.335594023 |
| Cep170b | S | 1029 | -0.214895288 | -0.074079107 | -0.196508295 | 0.018386992 | -0.122429189 | 0.035339938 | 0.320572571 | 0.060262715 | 0.83828387 | 0.201847857 |
| Cep170b | S | 1032 | -0.214895288 | -0.074079107 | -0.196508295 | 0.018386992 | -0.122429189 | 0.035339938 | 0.320572571 | 0.060262715 | 0.83828387 | 0.201847857 |
| Inpp5f | S | 714 | -0.231757558 | -0.153986641 | -0.361614875 | -0.129857316 | -0.207628234 | 0.165748965 | 0.316200225 | 0.159907064 | 0.5706032 | 0.381032836 |
| Arid1a | T | 301 | -0.026862399 | -0.231311971 | -0.2874021 | -0.260539701 | -0.056090129 | 0.82245613 | 0.195995225 | 0.007000697 | 0.1020061 | 0.698445474 |
| Dnaja1 | S | 112 | -0.083059118 | -0.208210934 | -0.122819679 | -0.039760561 | 0.085391255 | 0.004347944 | 0.049689943 | 0.241377017 | 0.65167227 | 0.414671097 |
| Trp53bp1 | S | 507 | -0.070231362 | -0.145521528 | -0.29863164 | -0.228400278 | -0.153110112 | 0.177023201 | 0.101297948 | 0.047564783 | 0.081520804 | 0.191848441 |
| Irf2bp2 | S | 226 | -0.040947176 | -0.13994309 | -0.074662754 | -0.033715578 | 0.065280336 | 0.217141563 | 0.001660099 | 0.00193124 | 0.288555143 | 0.057143764 |
| Ssh1 | S | 607 | -0.087227288 | -0.114429469 | -0.379858256 | -0.292630967 | -0.265428787 | 0.476472072 | 0.385226661 | 0.012111129 | 0.047482965 | 0.076820268 |
| Ahctf1 | S | 1980 | -0.086066592 | -0.192320395 | -0.173067102 | -0.08700051 | 0.019253292 | 0.201058898 | 0.017052041 | 0.106239524 | 0.349836596 | 0.816180765 |
| Rbm5 | S | 69 | -0.027513239 | -0.217658565 | -0.194463857 | -0.166950618 | 0.023194707 | 0.630346261 | 0.004516445 | 0.091621793 | 0.129647572 | 0.788864883 |
| Rbm5 | S | 72 | -0.027513239 | -0.217658565 | -0.194463857 | -0.166950618 | 0.023194707 | 0.630346261 | 0.004516445 | 0.091621793 | 0.129647572 | 0.788864883 |
| Ubap2l | S | 497 | 0.017997508 | -0.119589825 | -0.212123601 | -0.230121109 | -0.092533776 | 0.514087571 | 0.002423171 | 0.002201038 | 0.004417247 | 0.073638571 |

| | | | | | | | | | | | | |
|---|---|---|---|---|---|---|---|---|---|---|---|---|
| Hspa8 | S | 511 | -0.037265851 | -0.160758431 | -0.169342093 | -0.132076242 | -0.008583662 | 0.605713841 | 0.000766704 | 0.07216967 | 0.178096233 | 0.890279438 |
| Lima1 | S | 485 | -0.294959105 | -0.37465693 | -0.026490088 | 0.268469016 | 0.348166841 | 0.176934862 | 0.035247895 | 0.931406268 | 0.417511156 | 0.300399217 |
| Bicd2 | S | 570 | -0.017866988 | -0.127950258 | -0.250974704 | -0.233107716 | -0.123024446 | 0.743620135 | 0.064134358 | 0.002869565 | 0.006338413 | 0.068448978 |
| Ppfibp1 | S | 957 | -0.012892343 | -0.166032782 | -0.146029316 | -0.133136973 | 0.020003466 | 0.643726971 | 0.001528792 | 0.01823705 | 0.022221824 | 0.589470556 |
| Psmg1 | T | 32 | -0.11212057 | -0.283472347 | -0.243924753 | -0.131804183 | 0.039547595 | 0.285866486 | 0.119765589 | 0.114417291 | 0.342522395 | 0.811694491 |
| Ybx1 | S | 172 | 0.052748022 | -0.17753465 | -0.279271455 | -0.332019477 | -0.101736805 | 0.315834886 | 0.012073012 | 0.0041475 | 0.01154749 | 0.136276385 |
| Pard3b | S | 708 | -0.16110545 | -0.179904093 | -0.135725965 | 0.025379485 | 0.044178128 | 0.045717662 | 0.083396832 | 0.155334607 | 0.708926035 | 0.624572191 |
| Pard3b | S | 710 | -0.16110545 | -0.179904093 | -0.135725965 | 0.025379485 | 0.044178128 | 0.045717662 | 0.083396832 | 0.155334607 | 0.708926035 | 0.624572191 |
| Map4 | T | 776 | -0.113592269 | -0.052981165 | -0.189820598 | -0.076228329 | -0.136839433 | 0.152531199 | 0.150084556 | 0.001774197 | 0.290462246 | 0.014084263 |
| Srsf7 | S | 252 | 0.019009258 | -0.236893622 | -0.119911887 | -0.138921145 | 0.116981735 | 0.568206849 | 0.000213274 | 0.070622302 | 0.063031612 | 0.083910531 |
| Wdr59 | S | 603 | -0.013820101 | -0.141325104 | -0.188052641 | -0.17423254 | -0.046727538 | 0.591923032 | 0.001872186 | 0.021885316 | 0.039952153 | 0.369323413 |
| Camk2g | S | 384 | -0.211110436 | -0.023109106 | -0.367688743 | -0.156578307 | -0.344579637 | 0.124049123 | 0.871915068 | 0.026579056 | 0.205988603 | 0.048775517 |
| Polh | S | 670 | -0.080703147 | -0.195370366 | -0.16534121 | -0.084638063 | 0.030029156 | 0.175699678 | 0.031151637 | 0.062654208 | 0.267834816 | 0.701312772 |
| Scrib | S | 1490 | -0.251128528 | 0.000138809 | -0.410118705 | -0.158990177 | -0.410257514 | 0.097356592 | 0.999318576 | 0.049736915 | 0.354634288 | 0.085671387 |
| Pom121 | S | 440 | 0.062580964 | -0.22591346 | -0.179277755 | -0.24185872 | 0.046635705 | 0.198037718 | 0.00044722 | 0.014411323 | 0.00781706 | 0.340493833 |
| Ahctf1 | S | 1507 | -0.094542738 | -0.165389909 | -0.123397713 | -0.028854975 | 0.041992196 | 0.064321253 | 0.010105708 | 0.140459932 | 0.677280538 | 0.556086324 |
| Baz1a | S | 1412 | -0.019811643 | -0.119664213 | -0.139169907 | -0.119358264 | -0.019505693 | 0.674602037 | 0.006275084 | 0.000697685 | 0.0774731 | 0.44929591 |
| Pcm1 | S | 645 | -0.004949388 | -0.091496708 | -0.208335399 | -0.203386012 | -0.116838691 | 0.950540439 | 0.024541015 | 0.000409871 | 0.088164219 | 0.003098383 |
| Mast4 | S | 1776 | 0.010636851 | -0.139000709 | -0.135466936 | -0.146103787 | 0.003533773 | 0.819132804 | 0.004536401 | 0.000282562 | 0.054374164 | 0.891712639 |
| Tenm3 | S | 277 | -0.203993368 | -0.029686543 | -0.507940639 | -0.303947272 | -0.478254096 | 0.083842604 | 0.871020194 | 0.093578781 | 0.219306155 | 0.113741881 |
| Spdl1 | S | 518 | -0.042980281 | -0.137991879 | -0.115103298 | -0.072123017 | 0.022888581 | 0.310983418 | 0.003192276 | 0.007008958 | 0.134818144 | 0.462340665 |
| Mxra7 | S | 118 | -0.010597196 | -0.153293743 | -0.211286988 | -0.200689792 | -0.057993245 | 0.74092697 | 0.000924243 | 0.056638379 | 0.068661376 | 0.439251024 |
| Aup1 | S | 290 | -0.033328371 | -0.152361403 | -0.181175151 | -0.14784678 | -0.028813748 | 0.343299519 | 0.004299278 | 0.037983947 | 0.071937539 | 0.622525942 |
| Riok2 | S | 332 | -0.053473099 | -0.354062706 | -0.241851489 | -0.18837839 | 0.112211217 | 0.422577875 | 0.074502149 | 0.134869345 | 0.203190441 | 0.497872677 |
| Riok2 | S | 337 | -0.053473099 | -0.354062706 | -0.241851489 | -0.18837839 | 0.112211217 | 0.422577875 | 0.074502149 | 0.134869345 | 0.203190441 | 0.497872677 |
| Prkci | S | 576 | -0.101513333 | -0.19643878 | -0.175779312 | -0.074265979 | 0.020659468 | 0.092672944 | 0.112593346 | 0.051113178 | 0.313819326 | 0.839755881 |
| Phldb2 | S | 81 | 0.202751698 | -0.310623404 | -0.321866664 | -0.524618361 | -0.01124326 | 0.129240588 | 0.013048465 | 0.003435655 | 0.022069184 | 0.874357267 |
| Srsf6 | S | 297 | 0.022480924 | -0.186355172 | -0.140339346 | -0.16282027 | 0.046015826 | 0.343386809 | 9.05273E-05 | 0.028806727 | 0.024791345 | 0.293748487 |
| Srsf6 | S | 299 | 0.022480924 | -0.186355172 | -0.140339346 | -0.16282027 | 0.046015826 | 0.343386809 | 9.05273E-05 | 0.028806727 | 0.024791345 | 0.293748487 |
| Rps26 | T | 96 | 0.009041357 | -0.219960705 | -0.206066094 | -0.215107451 | 0.013894611 | 0.739640762 | 0.007734575 | 0.022298292 | 0.034703142 | 0.819904268 |
| Lig3 | S | 919 | 0.014413812 | -0.159735979 | -0.097882086 | -0.112295898 | 0.061853893 | 0.655847714 | 0.000121931 | 0.003432612 | 0.027558486 | 0.04368547 |
| Irf3 | S | 135 | -0.146789601 | -0.13662472 | -0.221912781 | -0.075123181 | -0.085288061 | 0.077979366 | 0.114028622 | 0.103050551 | 0.499381063 | 0.455607845 |
| Zc3hav1 | S | 342 | 0.423742311 | -0.422265185 | -1.70116488 | -2.124907191 | -1.278899695 | 0.013020127 | 0.012948793 | 0.128640628 | 0.090787947 | 0.204315219 |
| Ate1 | S | 166 | -0.192173674 | -0.024342666 | -0.257013041 | -0.064839368 | -0.232670375 | 0.011903409 | 0.770895212 | 0.054996649 | 0.499007795 | 0.082854063 |
| Actb Actg1 | S | 233;233 | -0.014889532 | -0.14915665 | -0.132928382 | -0.11803885 | 0.016228268 | 0.651725583 | 0.000287827 | 0.030300265 | 0.047307848 | 0.674719681 |
| Dync1li2 | S | 443 | -0.10576974 | -0.096138226 | -0.301944441 | -0.196174701 | -0.205806215 | 0.044491498 | 0.26971879 | 0.055055913 | 0.14204694 | 0.132812976 |
| Polk | S | 798 | -0.189615338 | -0.185108156 | -0.071892963 | 0.117722375 | 0.113215193 | 0.00948707 | 0.053621026 | 0.609882368 | 0.416170444 | 0.445420934 |
| Slc12a7 | S | 38 | -0.11339257 | -0.29109375 | -0.168616368 | -0.055223798 | 0.122477383 | 0.411690386 | 0.026110541 | 0.206158504 | 0.703888397 | 0.341040689 |
| Sik3 | S | 1049 | -0.016323067 | -0.11810846 | -0.283603171 | -0.267280104 | -0.16549471 | 0.864581716 | 0.058433944 | 0.004598852 | 0.05116724 | 0.04502297 |
| Tut7 | S | 57 | -0.051706393 | -0.183598567 | -0.233239277 | -0.181532884 | -0.049640711 | 0.363412754 | 0.106556114 | 0.014587372 | 0.042857201 | 0.597607394 |
| Frmd4a | S | 783 | -0.044226149 | -0.127508984 | -0.181927872 | -0.137701724 | -0.054418888 | 0.364131933 | 0.12817646 | 0.000793321 | 0.0384858 | 0.430267756 |
| Bin1 | S | 304 | -0.054985599 | -0.108387916 | -0.120821853 | -0.065836254 | -0.012433937 | 0.047109584 | 0.028274024 | 0.003293874 | 0.047755052 | 0.707178212 |
| Map1a | S | 908 | -0.16054268 | -0.120444094 | -0.085572887 | 0.074969793 | 0.034871208 | 0.005665942 | 0.033081401 | 0.325771197 | 0.381003034 | 0.668634699 |
| Sh3bp5 | S | 379 | -0.091125342 | -0.129021553 | -0.218005794 | -0.126880451 | -0.08898424 | 0.188489203 | 0.090163126 | 0.021566882 | 0.135362268 | 0.262710847 |
| Kdm2b | S | 768 | 0.015597351 | -0.129563687 | -0.255517541 | -0.271114892 | -0.125953854 | 0.716141716 | 0.002107931 | 0.014476453 | 0.01119482 | 0.103121487 |
| Fam92a | S | 312 | -0.036021707 | -0.099371028 | -0.171385335 | -0.135363628 | -0.072014307 | 0.294373271 | 0.029049309 | 0.001455174 | 0.001362012 | 0.052349726 |
| Rbbp6 | T | 1205 | -0.013471007 | -0.253118025 | -0.164043997 | -0.15057299 | 0.089074028 | 0.879808049 | 0.003202029 | 0.10082032 | 0.181697865 | 0.297559805 |
| Arid1a | S | 605 | -0.232600897 | -0.113257265 | -0.141458799 | 0.091142098 | -0.028201534 | 0.041460418 | 0.17362864 | 0.113490045 | 0.323055555 | 0.690366471 |
| Ice1 | S | 1820 | -0.039875644 | -0.165918917 | -0.110261278 | -0.070385633 | 0.055657639 | 0.312639433 | 0.000891106 | 0.054921752 | 0.177579964 | 0.23836411 |
| Huwe1 | S | 1084 | -0.185756903 | -0.200915303 | -0.009615262 | 0.176141641 | 0.191300041 | 0.003613948 | 0.035888543 | 0.95663282 | 0.37111342 | 0.341120227 |
| Kansl3 | S | 536 | -0.210670353 | -0.029625408 | -0.206015194 | 0.00465516 | -0.176389786 | 0.019454052 | 0.718137 | 0.02995904 | 0.947166309 | 0.080839165 |
| Nol8 | S | 306 | -0.048141668 | -0.144571208 | -0.0662756 | -0.018133932 | 0.078295607 | 0.257365708 | 0.000297588 | 0.016684982 | 0.613082908 | 0.006316544 |
| Eif4b | S | 597 | -0.161989117 | -0.088410117 | -0.134791727 | 0.02719739 | -0.04638161 | 0.014905973 | 0.183041386 | 0.04484637 | 0.610567423 | 0.464556932 |
| Dst | S | 2574 | -0.133517573 | -0.448270194 | 0.010604547 | 0.14412212 | 0.458874741 | 0.286873482 | 0.007237458 | 0.890470437 | 0.253274528 | 0.010697665 |
| Csde1 | S | 514 | -0.126639403 | -0.167861412 | -0.060383557 | 0.066255845 | 0.107477855 | 0.149750318 | 0.000887952 | 0.267256315 | 0.408100527 | 0.093957184 |
| Pakap | S | 486 | -0.03123562 | -0.116686559 | -0.192971084 | -0.161735464 | -0.076284525 | 0.45626678 | 0.038366872 | 0.002244597 | 0.01098702 | 0.133198974 |
| Zfp507 | S | 186 | -0.028015807 | -0.188299787 | -0.194365919 | -0.166350112 | -0.006066132 | 0.7877169 | 0.098707052 | 0.001646094 | 0.195525431 | 0.940573896 |
| Ahctf1 | S | 2132 | -0.088574695 | -0.173433198 | -0.188984773 | -0.100410079 | -0.015551576 | 0.023954591 | 0.085139271 | 0.123209534 | 0.319763008 | 0.880952022 |
| Zfp609 | S | 577 | -0.069622941 | -0.116656764 | -0.130010863 | -0.060387922 | -0.013354099 | 0.015460824 | 0.031181677 | 0.027781916 | 0.151254106 | 0.747233117 |
| Myo18a | S | 1994 | -0.091699385 | -0.14543009 | -0.124736726 | -0.033037342 | 0.020693364 | 0.04802677 | 0.02043732 | 0.068965924 | 0.534949152 | 0.704317967 |
| Myh9 | S | 1243 | -0.153061759 | -0.174962319 | -0.117257822 | 0.035803937 | 0.057704497 | 0.005846076 | 0.237076424 | 0.154809011 | 0.60038234 | 0.663713398 |
| Orc1 | S | 436 | -0.052563635 | -0.194539424 | -0.10723922 | -0.054675584 | 0.087300205 | 0.33960191 | 0.001247638 | 0.119052395 | 0.401013696 | 0.17907937 |
| Map4 | S | 345 | -0.059817851 | -0.086937889 | -0.139648323 | -0.079830472 | -0.052710434 | 0.086386191 | 0.023635847 | 0.002906424 | 0.016375331 | 0.071000434 |
| Ssrp1 | S | 667 | -0.048411632 | -0.125402724 | -0.065896352 | -0.01748472 | 0.059506372 | 0.020260078 | 0.000153427 | 0.121831264 | 0.595029544 | 0.159804571 |
| Ptpdc1 | S | 543 | -0.117418403 | -0.112218179 | -0.220544944 | -0.103126541 | -0.108326766 | 0.18327652 | 0.022454402 | 0.074454083 | 0.342645131 | 0.276154036 |
| Tdp1 | S | 118 | -0.287003766 | -0.164738359 | -0.180567917 | 0.106435849 | -0.015829559 | 0.088616178 | 0.17874172 | 0.265689923 | 0.506705795 | 0.902148291 |
| Ticrr | S | 1064 | -0.006612823 | -0.168123064 | -0.164536627 | -0.157923804 | 0.003586437 | 0.90601919 | 0.001032632 | 0.031268054 | 0.064387791 | 0.927288066 |
| Marchf7 | S | 301 | -0.058535003 | -0.275004026 | -0.291272564 | -0.232737561 | -0.016268537 | 0.648130145 | 0.191844245 | 0.027671977 | 0.076213209 | 0.922704449 |
| Dapk1 | S | 696 | -0.233828729 | -0.209421459 | -0.000983279 | 0.23284545 | 0.20843818 | 0.015257377 | 0.028572174 | 0.990107383 | 0.013706424 | 0.02371627 |
| Zfp593 | S | 124 | -0.06708846 | -0.239057981 | -0.353238021 | -0.286149561 | -0.11418004 | 0.536131368 | 0.095897274 | 0.100068185 | 0.159843036 | 0.523521417 |
| Kalrn Trio | S | 1779;1745 | 0.036021489 | -0.103029552 | -0.317888335 | -0.353909824 | -0.214858783 | 0.481157014 | 0.03219021 | 0.001726377 | 0.002676052 | 0.026266897 |
| Ranbp2 | T | 1526 | 0.052138484 | -0.162450046 | -0.218191456 | -0.270329941 | -0.05574141 | 0.522321567 | 8.02748E-05 | 0.052858541 | 0.038903264 | 0.433650545 |
| Lrrk2 | S | 910 | 0.002364501 | -0.113559218 | -0.120329992 | -0.122694494 | -0.006770774 | 0.931122999 | 0.000590667 | 0.000570312 | 0.012922195 | 0.710204233 |
| Wiz | S | 1550 | -0.006730607 | -0.120433193 | -0.16862977 | -0.161899163 | -0.048196577 | 0.751014748 | 0.024566514 | 0.000440629 | 0.003797617 | 0.228912377 |
| Plekha1 | S | 308 | -0.22056587 | 0.01241899 | -0.253663043 | -0.033097173 | -0.266082032 | 0.024125756 | 0.854149291 | 0.020793891 | 0.6871413 | 0.025114871 |
| Ppfibp1 | S | 959 | 0.02862345 | -0.159634937 | -0.198448678 | -0.227072128 | -0.03881374 | 0.090241928 | 0.000151652 | 0.012777786 | 0.013182128 | 0.334642548 |
| Pcm1 | S | 1332 | -0.046905661 | -0.145453745 | -0.114055582 | -0.067149921 | 0.031398163 | 0.226004211 | 0.004630751 | 0.012952427 | 0.053077566 | 0.189665465 |
| H1f3 H1f4 H1f5 | S | 89;89;90 | -0.054530623 | -0.135895406 | -0.299700138 | -0.245169515 | -0.163804733 | 0.630921697 | 0.272885643 | 0.004487401 | 0.086326144 | 0.191789186 |
| Clic4 | S | 157 | -0.191463452 | -0.10289924 | -0.181688644 | 0.009774808 | -0.078789404 | 0.053556172 | 0.36521626 | 0.035111447 | 0.895714338 | 0.461920012 |
| Gphn | S | 188 | -0.10163782 | -0.071045563 | -0.126359931 | -0.02472211 | -0.055314368 | 0.055721594 | 0.091540606 | 0.00172443 | 0.515064468 | 0.15841044 |
| Gphn | S | 194 | -0.10163782 | -0.071045563 | -0.126359931 | -0.02472211 | -0.055314368 | 0.055721594 | 0.091540606 | 0.00172443 | 0.515064468 | 0.15841044 |
| Map1b | S | 1373 | -0.080625735 | -0.091201853 | -0.095802712 | -0.015176977 | -0.004600859 | 0.043069197 | 0.000997112 | 0.041087645 | 0.675716346 | 0.877963909 |
| Prrc2a | S | 1302 | -0.173473907 | -0.128236014 | -0.290321708 | -0.116847801 | -0.162085694 | 0.34963527 | 0.396261988 | 0.018155837 | 0.507067516 | 0.296503189 |
| Uba1 | S | 803 | -0.148110133 | -0.090496167 | -0.379653266 | -0.231543132 | -0.289157099 | 0.326542489 | 0.285907444 | 0.039990198 | 0.200504335 | 0.106306882 |
| Zfand2b | S | 173 | -0.123241848 | -0.101898372 | -0.127421666 | -0.004179818 | -0.025523294 | 0.037184883 | 0.085437134 | 0.018838909 | 0.927152891 | 0.607789392 |
| Fbxo28 | S | 344 | -0.000701179 | -0.128626798 | -0.143144458 | -0.142443278 | -0.01451766 | 0.980194792 | 0.001951298 | 0.001872079 | 0.004162719 | 0.56624286 |
| Frmd4a | S | 590 | 0.174078417 | -0.198581646 | -0.358423646 | -0.532502063 | -0.159842 | 0.006759944 | 0.007243792 | 0.000130234 | 4.34696E-05 | 0.02306376 |
| Herc2 | S | 2455 | -0.125577133 | -0.239903231 | -0.123221195 | 0.002355937 | 0.116682035 | 0.16013389 | 0.051156626 | 0.108925813 | 0.977294028 | 0.228252602 |
| Cebpb | S | 184 | -0.010013117 | -0.137336051 | -0.186290195 | -0.176277078 | -0.048954143 | 0.793216732 | 0.013709436 | 0.00338749 | 0.008883195 | 0.277356993 |
| Inppl1 | S | 241 | -0.041995661 | -0.23551385 | -0.130557816 | -0.088562155 | 0.104956034 | 0.496363685 | 0.005522786 | 0.085318847 | 0.153578289 | 0.116650596 |
| Cep170b | T | 968 | 0.0377517 | -0.117767816 | -0.385622401 | -0.423374101 | -0.267854585 | 0.350120922 | 0.016012305 | 0.009100912 | 0.010950024 | 0.033237552 |
| Rps6ka3 | S | 19 | -0.1171182 | -0.277476315 | -0.145413634 | -0.028295433 | 0.132062682 | 0.252733317 | 0.017699008 | 0.35989981 | 0.843488545 | 0.386370659 |
| Micall2 | S | 172 | 0.060247434 | -0.191181381 | -0.329764047 | -0.39001148 | -0.138582666 | 0.472238006 | 0.042760547 | 0.006906233 | 0.004521382 | 0.104944543 |
| Rif1 | S | 1705 | -0.056728977 | -0.05715483 | -0.434707866 | -0.377978889 | -0.377553036 | 0.571439132 | 0.63892637 | 0.005420633 | 0.014468941 | 0.017009185 |

| | | | | | | | | | | | | |
|---|---|---|---|---|---|---|---|---|---|---|---|---|
| Ubr5 | S | 1302 | -0.063273629 | -0.085195845 | -0.144044833 | -0.080771204 | -0.058848988 | 0.027164854 | 0.005232196 | 0.034272437 | 0.126554049 | 0.21977585 |
| Vps4a | S | 97 | -0.112068284 | -0.113044817 | -0.152974649 | -0.040906365 | -0.039929832 | 0.11672849 | 0.163560528 | 0.004976962 | 0.490438556 | 0.559212366 |
| Sh3pxd2b | S | 291 | -0.033014294 | -0.15919706 | -0.210940661 | -0.177926367 | -0.051743601 | 0.341996262 | 0.019134672 | 0.029742381 | 0.031590881 | 0.379043267 |
| Mfap1a Mfap1b | S | 80;80 | -0.079931093 | -0.315200706 | -0.237627722 | -0.157696628 | 0.077572984 | 0.569172042 | 0.057982259 | 0.14037229 | 0.155561788 | 0.463146955 |
| Sipa1l3 | T | 395 | -0.129176743 | -0.13342795 | -0.164339992 | -0.03516325 | -0.030912042 | 0.204779842 | 0.059448205 | 0.025528377 | 0.696013638 | 0.618615941 |
| Klf16 | T | 51 | -0.104832495 | -0.168310735 | -0.338073163 | -0.233240668 | -0.169762428 | 0.633790284 | 0.028460927 | 0.118007434 | 0.373172188 | 0.342252114 |
| Map4k4 | S | 629 | -0.08784737 | -0.229986168 | -0.118014163 | -0.030166793 | 0.111972005 | 0.075078963 | 0.028672258 | 0.165041238 | 0.666180101 | 0.235457944 |
| Ralgapa1 | S | 796 | -0.049266529 | -0.090042019 | -0.207467632 | -0.158201104 | -0.117425613 | 0.236529058 | 0.179293578 | 0.001195339 | 0.01692013 | 0.122989381 |
| Nufip1 | S | 329 | 0.146909385 | -0.289042664 | -0.3460758 | -0.492985185 | -0.057033135 | 0.025557005 | 0.000963872 | 0.020681693 | 0.01301176 | 0.53221025 |
| Nufip1 | S | 331 | 0.146909385 | -0.289042664 | -0.3460758 | -0.492985185 | -0.057033135 | 0.025557005 | 0.000963872 | 0.020681693 | 0.01301176 | 0.53221025 |
| Nufip1 | S | 333 | 0.146909385 | -0.289042664 | -0.3460758 | -0.492985185 | -0.057033135 | 0.025557005 | 0.000963872 | 0.020681693 | 0.01301176 | 0.53221025 |
| Ubr4 | S | 364 | 0.035277681 | -0.167087758 | -0.281454278 | -0.316731959 | -0.11436652 | 0.539249132 | 0.079630856 | 0.00171716 | 0.000393065 | 0.176142411 |
| Dab2ip | S | 1168 | 0.020313262 | -0.276785073 | -0.315265598 | -0.33557886 | -0.038480526 | 0.819709684 | 0.023988247 | 0.081901721 | 0.071246022 | 0.780580467 |
| Rif1 | S | 1407 | -0.022583118 | -0.169237278 | -0.255011122 | -0.232428004 | -0.085773844 | 0.833000222 | 0.146144662 | 0.003385849 | 0.100607638 | 0.392274461 |
| Acap2 | S | 521 | -0.021358971 | -0.165761221 | -0.248031157 | -0.226672186 | -0.082269936 | 0.706929494 | 0.020287052 | 0.026482711 | 0.038299608 | 0.30842621 |
| Crip2 | T | 117 | -0.230582404 | 0.02322346 | -0.512768317 | -0.282185913 | -0.535991777 | 0.130652526 | 0.90523659 | 0.059773185 | 0.217602516 | 0.06734277 |
| Armc5 | S | 469 | -0.27577708 | -0.193401073 | -0.15401749 | 0.12175959 | 0.039383583 | 0.135186136 | 0.11923125 | 0.261403042 | 0.483075171 | 0.772998702 |
| Efr3a | S | 368 | -0.073717081 | -0.039989346 | -0.26753353 | -0.193816448 | -0.227544184 | 0.217297157 | 0.292118585 | 0.002159908 | 0.022452142 | 0.023163416 |
| Palm | S | 124 | -0.067485521 | -0.12936744 | -0.18037658 | -0.112891059 | -0.05100914 | 0.301528297 | 0.019209981 | 0.020174224 | 0.133560127 | 0.337310227 |
| Mbtps2 | S | 89 | -0.144162132 | -0.112241777 | -0.182888308 | -0.038726176 | -0.070646531 | 0.083086767 | 0.1788358 | 0.030206576 | 0.573335276 | 0.373621925 |
| Mbtps2 | S | 90 | -0.144162132 | -0.112241777 | -0.182888308 | -0.038726176 | -0.070646531 | 0.083086767 | 0.1788358 | 0.030206576 | 0.573335276 | 0.373621925 |
| Cfdp1 | S | 112 | 0.043892034 | -0.392925342 | -0.641207678 | -0.685099712 | -0.248282336 | 0.710663547 | 0.139969173 | 0.161157041 | 0.147504359 | 0.528572741 |
| Tra2b | T | 69 | -0.016173882 | -0.211162707 | -0.448738682 | -0.4325648 | -0.237575975 | 0.867833979 | 0.022378154 | 0.168071243 | 0.178954687 | 0.390788317 |
| Tra2b | Y | 68 | -0.016173882 | -0.211162707 | -0.448738682 | -0.4325648 | -0.237575975 | 0.867833979 | 0.022378154 | 0.168071243 | 0.178954687 | 0.390788317 |
| Xkr5 | S | 459 | -0.000139952 | -0.291103125 | -0.246695318 | -0.246555366 | 0.044407807 | 0.998549754 | 0.041214767 | 0.045373757 | 0.05530936 | 0.676469918 |
| Lypd6b | T | 108 | 0.105266769 | -0.276003051 | -0.200966025 | -0.306232794 | 0.075037026 | 0.31692745 | 0.00327936 | 0.013697577 | 0.045124546 | 0.059381138 |
| Lypd6b | T | 111 | 0.105266769 | -0.276003051 | -0.200966025 | -0.306232794 | 0.075037026 | 0.31692745 | 0.00327936 | 0.013697577 | 0.045124546 | 0.059381138 |
| Cd2ap | S | 458 | -0.147641921 | 0.025556494 | -0.346579728 | -0.198937807 | -0.372136222 | 0.064515173 | 0.810339054 | 0.009226685 | 0.070313027 | 0.024470936 |
| Hnrnpa1 | S | 312 | 0.016636087 | -0.184318169 | -0.525979123 | -0.54261521 | -0.341660954 | 0.900306246 | 0.164770416 | 0.040041865 | 0.033867496 | 0.10250542 |
| Sqstm1 | S | 24 | 0.044412398 | -0.142991384 | -0.302041151 | -0.346453549 | -0.159049767 | 0.49803582 | 0.053402792 | 0.00210905 | 0.005109169 | 0.05306459 |
| Specc1 | T | 357 | -0.090642178 | -0.278015731 | -0.229857431 | -0.139215253 | 0.0481583 | 0.270442636 | 0.138732897 | 0.086452469 | 0.245066101 | 0.76492449 |
| Ino80 | S | 238 | -0.07457291 | -0.103947282 | -0.111549956 | -0.036977047 | -0.007602675 | 0.235053174 | 0.001602989 | 0.014196231 | 0.519489759 | 0.774751255 |
| Sqstm1 | S | 344 | 0.04683078 | -0.238652408 | -0.242823361 | -0.289654141 | -0.004170953 | 0.162892271 | 0.00490036 | 0.016231096 | 0.015806081 | 0.94651487 |
| Cask | S | 575 | -0.092933995 | -0.273249207 | -0.134013762 | -0.041079767 | 0.139235445 | 0.356590875 | 0.036942915 | 0.08478856 | 0.681357687 | 0.166935632 |
| P2ry2 | T | 327 | -0.00948606 | -0.203209469 | -0.121930224 | -0.112444164 | 0.081279245 | 0.868902467 | 0.002130141 | 0.022703079 | 0.10030018 | 0.029166492 |
| Zswim8 | S | 1094 | -0.001215404 | -0.180366003 | -0.164871901 | -0.163656497 | 0.015494102 | 0.97124692 | 0.023827516 | 0.002144901 | 0.003452418 | 0.747855677 |
| Rab7 | S | 111 | -0.137832691 | -0.233788952 | -0.164734303 | -0.026901613 | 0.069054649 | 0.325569877 | 0.11855152 | 0.03796367 | 0.832586488 | 0.549850241 |
| Zdhhc5 | S | 634 | -0.014003771 | -0.155701645 | -0.402341238 | -0.388337467 | -0.246639593 | 0.83352633 | 0.07867208 | 0.036800977 | 0.036489467 | 0.105954167 |
| Arhgef17 | S | 516 | -0.033418881 | -0.146850954 | -0.155482158 | -0.122063276 | -0.008631204 | 0.221229541 | 0.082379333 | 0.001567659 | 0.009400988 | 0.884753854 |
| Mki67 | S | 250 | -0.121015185 | 0.03175873 | -0.216959986 | -0.095944801 | -0.248718716 | 0.009443814 | 0.474269055 | 0.000892998 | 0.039909816 | 0.003691948 |
| Efnb1 Efnb2 | S | 337;328 | 0.001949475 | -0.214990499 | -0.235422909 | -0.237372384 | -0.020432409 | 0.978702901 | 0.026212587 | 0.021116725 | 0.025093928 | 0.787950094 |
| Peak1 | S | 371 | 0.087429816 | -0.206513625 | -0.280470234 | -0.36790005 | -0.073956609 | 0.293501983 | 0.024999615 | 0.002752846 | 0.008122602 | 0.292269589 |
| Zc3h3 | S | 211 | 0.057320579 | -0.206471486 | -0.191331838 | -0.248652417 | 0.015139649 | 0.807732826 | 0.00794784 | 0.009661842 | 0.346705414 | 0.516007298 |
| Dennd4c | S | 1274 | -0.254991468 | -0.009532713 | -0.356287937 | -0.101296469 | -0.346755224 | 0.073035637 | 0.953895654 | 0.063046296 | 0.495834837 | 0.105022634 |
| Lin54 | S | 314 | -0.063164281 | -0.156577155 | -0.079307072 | -0.016142791 | 0.077270083 | 0.158545724 | 0.004115347 | 0.018670583 | 0.657447674 | 0.063309768 |
| Pcbp2 | S | 362 | -0.025401708 | -0.076382765 | -0.247806772 | -0.222405064 | -0.171424006 | 0.59774704 | 0.090910399 | 0.001239333 | 0.006499624 | 0.011906937 |
| Gnl3 | S | 493 | 0.051943093 | -0.185530278 | -0.174295666 | -0.226238759 | 0.011234612 | 0.418265807 | 0.000265821 | 0.02398395 | 0.02615992 | 0.797725032 |
| Zfp593 | S | 133 | -0.032284183 | -0.099282319 | -0.335727212 | -0.303443029 | -0.236444893 | 0.44317786 | 0.135207476 | 0.011741829 | 0.030607973 | 0.03766326 |
| Atxn3 | S | 219 | -0.012189844 | -0.131002856 | -0.241808916 | -0.229619072 | -0.11080606 | 0.788629094 | 0.047872154 | 0.004459113 | 0.014987297 | 0.096131319 |
| Prr11 | S | 315 | -0.047216603 | -0.311476506 | -0.078414495 | -0.031197892 | 0.233062011 | 0.535877374 | 0.001927507 | 0.41833928 | 0.764031304 | 0.07222172 |
| Afap1l2 | T | 345 | -0.228099782 | 0.01560808 | -0.373664229 | -0.145564447 | -0.38927231 | 0.106170934 | 0.912174977 | 0.028967939 | 0.175297958 | 0.019504455 |
| Brd8 | S | 460 | -0.064144579 | -0.241032223 | -0.031965943 | 0.032178636 | 0.20906628 | 0.136645887 | 0.000708028 | 0.503670064 | 0.405934599 | 0.005915502 |
| Bnip2 | T | 116 | -0.085120728 | -0.080995062 | -0.303018181 | -0.217897452 | -0.222023118 | 0.242546558 | 0.292226475 | 0.011394083 | 0.045673029 | 0.040557158 |
| Riok2 | S | 433 | -0.036673641 | -0.130284022 | -0.177820112 | -0.141146471 | -0.04753609 | 0.378228626 | 0.023203192 | 0.007097657 | 0.029024642 | 0.303116171 |
| Eps8 | S | 658 | -0.112751664 | -0.161679503 | -0.072632026 | 0.040119637 | 0.089047476 | 0.027293758 | 0.013947034 | 0.171064098 | 0.401627661 | 0.133100478 |
| Srrm2 | S | 344 | -0.081131131 | -0.256320757 | -0.078333477 | 0.002797654 | 0.177987279 | 0.409583421 | 0.025278284 | 0.012462091 | 0.974709498 | 0.074703165 |
| Ogfr | S | 361 | -0.033937352 | -0.138704751 | -0.311585834 | -0.277648481 | -0.172881083 | 0.508228632 | 0.125556023 | 0.015740741 | 0.033805786 | 0.115744221 |
| Hdac7 | S | 204 | -0.240619105 | -0.128785574 | -0.096776268 | 0.143842837 | 0.032009306 | 0.008955346 | 0.221478334 | 0.348503698 | 0.201910523 | 0.781886739 |
| Pdpk1 | S | 244 | -0.088161079 | -0.126955322 | -0.328275872 | -0.240114793 | -0.20132055 | 0.331809986 | 0.223099096 | 0.029242519 | 0.080529029 | 0.134228088 |
| 2310057M21Rik | S | 423 | -0.03634145 | -0.162636304 | -0.18855214 | -0.15221069 | -0.025915837 | 0.631119999 | 0.005571029 | 0.040695558 | 0.126784053 | 0.670278117 |
| Nudc | T | 108 | -0.097374489 | -0.096618416 | -0.233329006 | -0.135954517 | -0.13671059 | 0.281712264 | 0.043079713 | 0.026101068 | 0.192563979 | 0.119350066 |
| Usp47 | S | 934 | -0.040577267 | -0.122964786 | -0.115206147 | -0.07462888 | 0.007758639 | 0.260424867 | 0.000225871 | 0.056792009 | 0.160183065 | 0.841953401 |
| Tjp2 | S | 988 | -0.233309951 | -0.103810423 | -0.041306435 | 0.192003516 | 0.062503988 | 0.001761479 | 0.117290038 | 0.641231194 | 0.111530634 | 0.494316009 |
| Fxr1 | S | 641 | 0.013777399 | -0.260654024 | -0.171923143 | -0.185700542 | 0.088730881 | 0.723890799 | 0.034954845 | 0.004998184 | 0.008689184 | 0.282406461 |
| Pak2 | T | 143 | -0.23006436 | -0.031896981 | -0.217765039 | 0.012299321 | -0.185868058 | 0.024222232 | 0.677207609 | 0.058658639 | 0.895134296 | 0.105327963 |
| H1f5 | T | 135 | 0.043288444 | -0.088028395 | -0.264063024 | -0.307351468 | -0.176034629 | 0.625881345 | 0.055419905 | 0.000311181 | 0.045924496 | 0.001048565 |
| Nek9 | S | 983 | -0.074460226 | -0.191744826 | -0.194136129 | -0.119675903 | -0.002391303 | 0.313100022 | 0.046893482 | 0.050141386 | 0.191291392 | 0.977610092 |
| Adrm1 | T | 39 | -0.181082309 | -0.073111437 | -0.060126723 | 0.120955586 | 0.012984714 | 0.001852938 | 0.056897598 | 0.230435581 | 0.047387816 | 0.747291405 |
| Npm1 | S | 252 | -0.020732228 | -0.172426443 | -0.266873684 | -0.246141456 | -0.094447241 | 0.427969086 | 0.046575872 | 0.031325853 | 0.027541968 | 0.226988574 |
| Cldn12 | S | 231 | 0.037436108 | -0.215474041 | -0.350443643 | -0.387879751 | -0.134969602 | 0.743864474 | 0.127502399 | 0.010757499 | 0.030153076 | 0.309451462 |
| Map4 | S | 973 | -0.002578572 | -0.205453519 | -0.402005496 | -0.399426924 | -0.196551977 | 0.993192832 | 0.190589495 | 0.022813035 | 0.248965456 | 0.134288227 |
| Pum1 | S | 124 | -0.031947309 | -0.131533134 | -0.162397516 | -0.130450207 | -0.030864382 | 0.416103219 | 0.000337007 | 0.088963709 | 0.129073881 | 0.637615811 |
| Mast4 | S | 2055 | -0.242018044 | -0.035830645 | -0.244771821 | -0.002753777 | -0.208941176 | 0.045447915 | 0.754629486 | 0.050838759 | 0.96749216 | 0.064619112 |
| Dennd1b | S | 577 | -0.246742923 | -0.055644155 | -0.256203767 | -0.009460844 | -0.200559613 | 0.037713792 | 0.591554299 | 0.112857379 | 0.936982832 | 0.175829903 |
| Specc1 | T | 345 | -0.055305805 | -0.146528053 | -0.123591955 | -0.06828615 | 0.022936099 | 0.225279817 | 0.000659987 | 0.138383245 | 0.354891007 | 0.719165694 |
| Gripap1 | S | 620 | -0.142658939 | -0.238000945 | -0.493464954 | -0.350806015 | -0.255464009 | 0.655278556 | 0.59611363 | 0.048825374 | 0.330588346 | 0.578595951 |
| Mcm4 | S | 29 | -0.061703019 | -0.210001451 | -0.161196261 | -0.099493241 | 0.048805191 | 0.370262365 | 0.078587529 | 0.01273746 | 0.161676587 | 0.57392229 |
| Vapa | S | 214 | -0.069863168 | -0.103727189 | -0.121252977 | -0.051389809 | -0.017525788 | 0.022245502 | 0.042094609 | 0.012632948 | 0.153690687 | 0.661827419 |
| Ak5 | S | 113 | -0.377886999 | -0.138782379 | -0.164910349 | 0.21297665 | -0.02612797 | 0.013699191 | 0.666006919 | 0.567073731 | 0.458263938 | 0.94494179 |
| Map3k3 | S | 147 | -0.215311717 | -0.08306546 | -0.179858225 | 0.035453492 | -0.096792765 | 0.025078731 | 0.500604902 | 0.07316157 | 0.565243392 | 0.414052519 |
| Srrm2 | S | 204 | 0.008290339 | -0.25478622 | -0.217273602 | -0.225563941 | 0.037512617 | 0.907376039 | 0.052399799 | 0.01233475 | 0.027381804 | 0.693275613 |
| Utp14a | S | 341 | 0.042964281 | -0.18190078 | -0.133028204 | -0.175992486 | 0.048872576 | 0.439144087 | 0.00089931 | 0.003215607 | 0.051680201 | 0.125502727 |
| Ranbp2 | S | 2115 | 0.011475211 | -0.166110264 | -0.133145269 | -0.14462048 | 0.032964995 | 0.816443746 | 0.00093541 | 0.009657189 | 0.04126049 | 0.308964374 |
| Zfp106 | S | 431 | -0.043213746 | -0.094282493 | -0.136992689 | -0.093778943 | -0.042710196 | 0.090213412 | 0.003034309 | 0.016191833 | 0.054966192 | 0.255110459 |
| Tmem134 | S | 70 | -0.030412794 | -0.210567082 | -0.309789626 | -0.279376832 | -0.099222544 | 0.866854074 | 0.351705745 | 0.005136697 | 0.212379629 | 0.63312211 |
| Tcof1 | S | 918 | -0.221123427 | 0.003191663 | -0.226368362 | -0.005244936 | -0.229560026 | 0.030287657 | 0.979006422 | 0.019913779 | 0.923465537 | 0.122662355 |
| Pkp4 | S | 289 | -0.025063831 | -0.03214421 | -0.40568919 | -0.380625359 | -0.373544979 | 0.708570072 | 0.83343589 | 0.001287668 | 0.004338137 | 0.086838631 |
| Gatad2a | T | 184 | 0.060890431 | -0.233136728 | -0.214228231 | -0.275118662 | 0.018908497 | 0.317028592 | 0.021383285 | 0.002606419 | 0.013429266 | 0.758140262 |
| Mga | S | 531 | -0.040719048 | -0.085034154 | -0.161692732 | -0.120973684 | -0.076658578 | 0.383597051 | 0.002647558 | 0.006432951 | 0.059475005 | 0.055710738 |
| Usp37 | S | 652 | -0.001182401 | -0.106617737 | -0.200099986 | -0.198917584 | -0.093482248 | 0.97362916 | 0.011364654 | 0.0020166 | 0.005075099 | 0.049798816 |
| Ddi2 | S | 194 | -0.102433808 | -0.264630886 | -0.256319437 | -0.153885629 | 0.008311449 | 0.358179984 | 0.071442835 | 0.191857577 | 0.369789114 | 0.960291878 |
| Tnks1bp1 | S | 602 | -0.016292342 | -0.102050788 | -0.221307172 | -0.20501483 | -0.119256384 | 0.535024751 | 0.023293812 | 0.004665192 | 0.016318114 | 0.045118434 |
| Myo1e | S | 1004 | -0.120964663 | -0.269183892 | -0.144847006 | -0.023882343 | 0.124336886 | 0.527449675 | 0.01751572 | 0.192587884 | 0.895087267 | 0.19027167 |
| Lage3 | S | 50 | -0.075031607 | -0.093239587 | -0.404852037 | -0.32982043 | -0.31161245 | 0.392992717 | 0.375043382 | 0.022248667 | 0.069822437 | 0.058566466 |

| | | | | | | | | | | | | |
|---|---|---|---|---|---|---|---|---|---|---|---|---|
| Trim47 | S | 393 | -0.142490452 | -0.023298388 | -0.253712885 | -0.111222434 | -0.230414497 | 0.021295444 | 0.805394874 | 0.019939608 | 0.181555889 | 0.076807904 |
| Tjap1 | S | 506 | 0.152535617 | -0.220464195 | -0.24185825 | -0.394393867 | -0.021394055 | 0.005775007 | 0.000303172 | 0.000485634 | 0.000320913 | 0.503056115 |
| Tln1 | Y | 1777 | -0.942103133 | -0.325101577 | 0.034348505 | 0.976451638 | 0.359450082 | 0.301053646 | 0.128217381 | 0.85542134 | 0.287873163 | 0.133012648 |
| Phactr4 | S | 444 | -0.096161951 | -0.136935447 | -0.274808066 | -0.178646115 | -0.137872619 | 0.290881335 | 0.202420619 | 0.027416426 | 0.124382701 | 0.240712746 |
| Gja1 | S | 365 | 0.020558557 | -0.139764521 | -0.24895855 | -0.269517107 | -0.109194029 | 0.758206042 | 0.001326485 | 0.033540605 | 0.026415046 | 0.169963549 |
| Rbm12b2 | S | 665 | 0.023871881 | -0.143704675 | -0.108912804 | -0.132784684 | 0.034791871 | 0.740629613 | 0.000226409 | 0.003155246 | 0.157483547 | 0.154280617 |
| Myo9b | S | 1264 | -0.063547699 | -0.117938561 | -0.216946509 | -0.15339881 | -0.099007948 | 0.343867166 | 0.09017596 | 0.00890214 | 0.027041221 | 0.070983059 |
| Lmna | S | 637 | -0.251001398 | 0.004635643 | -0.229723497 | 0.021277901 | -0.23435914 | 0.028578848 | 0.972018506 | 0.036588647 | 0.710534119 | 0.123617659 |
| Iars | S | 521 | -0.275429339 | -0.015355728 | -0.212589977 | 0.062839362 | -0.19723425 | 0.046818295 | 0.851930811 | 0.037307861 | 0.55567059 | 0.059500055 |
| Pacs1 | S | 517 | -0.341579902 | -0.01152419 | -0.34564483 | -0.004064928 | -0.33412064 | 0.071286823 | 0.957356173 | 0.13527553 | 0.982171941 | 0.189655775 |
| Map1b | S | 1073 | -0.151604183 | -0.172651207 | -0.16845688 | -0.016852697 | 0.004194327 | 0.212056997 | 0.080792784 | 0.07577624 | 0.870502873 | 0.95661715 |
| Msl2 | S | 345 | -0.155329974 | -0.11858447 | 0.008839308 | 0.164169282 | 0.127423779 | 0.001026668 | 0.004634731 | 0.908665257 | 0.122640699 | 0.186913366 |
| Prune2 | S | 2329 | -0.294243354 | -0.190618879 | -0.292304369 | 0.001938985 | -0.101685491 | 0.157074948 | 0.521442727 | 0.176259674 | 0.992885922 | 0.741870022 |
| Snta1 | S | 194 | -0.218089184 | -0.078204693 | -0.028744574 | 0.18934461 | 0.049460119 | 0.002926184 | 0.037604043 | 0.375727602 | 0.01493597 | 0.045236905 |
| Tmem131 | S | 1744 | -0.113291109 | -0.12193588 | -0.678909877 | -0.565618768 | -0.556973997 | 0.674859178 | 0.625200622 | 0.067782584 | 0.141805575 | 0.133003481 |
| Ppp1r8 | S | 199 | -0.016350056 | -0.150767736 | -0.233306707 | -0.216956651 | -0.082538971 | 0.774251773 | 0.044236294 | 0.008804013 | 0.016158904 | 0.221292398 |
| Plekhh3 | S | 593 | -0.129616966 | -0.231072154 | -0.158567994 | -0.028951028 | 0.072504161 | 0.462797305 | 0.032473003 | 0.107305203 | 0.85278454 | 0.15682795 |
| Nolc1 | S | 626 | -0.016441811 | -0.218865168 | -0.179960015 | -0.163518204 | 0.038905153 | 0.78735412 | 0.035360627 | 0.013360388 | 0.065751696 | 0.560925777 |
| Hnrnpa1 | S | 316 | -0.282555067 | -0.529455898 | 0.021474726 | 0.304029794 | 0.550930624 | 0.257447942 | 0.059535705 | 0.92056763 | 0.093275835 | 0.03511592 |
| Osbpl3 | S | 405 | -0.359631432 | 0.057665443 | -0.272393309 | 0.087238123 | -0.330058751 | 0.021214245 | 0.69740464 | 0.116056804 | 0.521939602 | 0.080855196 |
| Bclaf1 | S | 759 | 0.013175962 | -0.150617789 | -0.222941364 | -0.236117327 | -0.072323576 | 0.756126514 | 0.001960715 | 0.02857671 | 0.024767994 | 0.291746092 |
| Map1b | T | 1784 | -0.038683179 | -0.171371556 | -0.117497037 | -0.078813857 | 0.05387452 | 0.376427135 | 0.004644595 | 0.02910508 | 0.056210394 | 0.12671025 |
| Phlpp2 | S | 1178 | -0.096516042 | -0.090872197 | -0.211720362 | -0.11520432 | -0.120848165 | 0.018693963 | 0.085996555 | 0.095367537 | 0.249646752 | 0.233211123 |
| Otud4 | S | 126 | -0.057743374 | -0.170733456 | -0.257228852 | -0.199485479 | -0.086495396 | 0.193793394 | 0.034016061 | 0.126237406 | 0.195115069 | 0.511777063 |
| Akap9 | S | 3694 | -0.071186875 | -0.227134149 | -0.033772834 | 0.037414041 | 0.193361314 | 0.159321317 | 0.000590259 | 0.579749415 | 0.535609812 | 0.03618438 |
| Dcbld2 | S | 582 | -0.226668214 | -0.113172053 | -0.288447163 | -0.061778949 | -0.17527511 | 0.163712148 | 0.408720969 | 0.087662623 | 0.692824116 | 0.261765771 |
| Impact | S | 137 | -0.0924528 | -0.101876683 | -0.289572691 | -0.197119891 | -0.187696008 | 0.362050957 | 0.241708904 | 0.014281104 | 0.091709432 | 0.06721099 |
| Hectd1 | S | 1493 | -0.12359945 | -0.204054396 | -0.104283396 | 0.019316054 | 0.099771 | 0.045649672 | 0.053213553 | 0.20956776 | 0.793008913 | 0.301495706 |
| Chaf1b | T | 401 | -0.096384633 | -0.137942577 | -0.121779527 | -0.025394894 | 0.01616305 | 0.056612999 | 0.015197062 | 0.119565161 | 0.671069051 | 0.778451616 |
| Glyr1 | S | 132 | -0.030867377 | -0.09268241 | -0.250116345 | -0.219248968 | -0.157433936 | 0.354689018 | 0.117242335 | 0.004202107 | 0.008218758 | 0.037515038 |
| Rpl12 | S | 126 | -0.004687115 | -0.270475357 | -0.055439626 | -0.05075251 | 0.215035731 | 0.879583144 | 0.000740985 | 0.080940899 | 0.146449475 | 0.003757862 |
| Gab1 | S | 264 | -0.123850967 | -0.08876894 | -0.365666976 | -0.24181601 | -0.276898036 | 0.119662056 | 0.328260675 | 0.077666041 | 0.187268022 | 0.143735961 |
| Rbm34 | S | 19 | 0.013988791 | -0.18324702 | -0.281507878 | -0.295496669 | -0.098260858 | 0.813816054 | 0.046350934 | 0.013923467 | 0.012290093 | 0.207648617 |
| Prrc2a | S | 166 | -0.243600587 | -0.085022389 | -0.176150016 | 0.067450572 | -0.091127627 | 0.028542719 | 0.518625464 | 0.108082684 | 0.324136796 | 0.451497431 |
| Bcar1 | Y | 391 | -0.187855973 | -0.080199933 | -0.228962994 | -0.041107022 | -0.148763062 | 0.104677362 | 0.567869671 | 0.023158715 | 0.67778825 | 0.326876645 |
| Akirin2 | S | 21 | 0.024585228 | -0.148678895 | -0.201975299 | -0.226560527 | -0.053296403 | 0.225565309 | 0.0001716 | 0.028346255 | 0.03244095 | 0.341023058 |
| Sos1 | S | 1261 | -0.044996306 | -0.095125764 | -0.165383333 | -0.120387028 | -0.07025757 | 0.521575344 | 0.023713029 | 0.00175296 | 0.159365551 | 0.084374836 |
| Cavin1 | T | 304 | -0.143521541 | -0.088424452 | -0.096805459 | 0.046716082 | -0.008381007 | 0.024220107 | 0.004504111 | 0.243643453 | 0.541293431 | 0.901946012 |
| Rbbp7 | S | 354 | -0.118254014 | -0.139598805 | -0.138982055 | -0.020728041 | 0.00061675 | 0.128265022 | 0.04778761 | 0.049272848 | 0.71883122 | 0.988503343 |
| Fra10ac1 | S | 283 | -0.042876661 | -0.117958364 | -0.222886513 | -0.180009852 | -0.104928149 | 0.299201278 | 0.085549145 | 0.00898099 | 0.017670808 | 0.125739408 |
| Zdhhc5 | S | 455 | -0.05980056 | -0.03287832 | -0.220232042 | -0.160431482 | -0.187353722 | 0.108608921 | 0.553312849 | 0.000542999 | 0.007091859 | 0.027899983 |
| Traf7 | S | 12 | 0.056901208 | -0.078099563 | -0.311021219 | -0.367922427 | -0.232921656 | 0.227884655 | 0.100218073 | 0.000153679 | 0.000314112 | 0.000792035 |
| Traf7 | S | 15 | 0.056901208 | -0.078099563 | -0.311021219 | -0.367922427 | -0.232921656 | 0.227884655 | 0.100218073 | 0.000153679 | 0.000314112 | 0.000792035 |
| Hdac5 | S | 651 | -0.413169066 | -0.160454352 | 0.032477739 | 0.445646805 | 0.192932091 | 0.011043212 | 0.189708169 | 0.751151755 | 0.012617668 | 0.110121711 |
| Bnip2 | S | 114 | -0.119403805 | -0.086948503 | -0.28819413 | -0.168790324 | -0.201245627 | 0.172301605 | 0.341881193 | 0.026954386 | 0.118002182 | 0.076136769 |
| Stag2 | S | 1161 | -0.238533272 | -0.105134897 | -0.214236323 | 0.024296949 | -0.109101425 | 0.162795276 | 0.089201996 | 0.164235178 | 0.882318582 | 0.406850677 |
| Rbbp6 | S | 1695 | -0.161849699 | -0.217115323 | -0.143827734 | 0.018021965 | 0.07328759 | 0.232183807 | 0.033309434 | 0.262668583 | 0.900388777 | 0.547233015 |
| Rps6ka1 Rps6ka2 Rp: | T | 577;562;570 | -0.150885867 | -0.068254075 | -0.16653178 | -0.015645913 | -0.098277705 | 0.207696066 | 0.134903902 | 0.005788475 | 0.871284092 | 0.004611914 |
| Nav3 | S | 1485 | -0.012572927 | -0.172841082 | -0.19960869 | -0.187035762 | -0.026767607 | 0.848810027 | 0.001999995 | 0.085476865 | 0.103010515 | 0.739792133 |
| Abtb2 | S | 365 | -0.158275382 | -0.165852309 | -0.145939042 | 0.012336341 | 0.019913268 | 0.141784015 | 0.161497741 | 0.043608691 | 0.879273977 | 0.828563177 |
| Als2 | S | 486 | -0.135913491 | -0.231308539 | -0.020357318 | 0.115556173 | 0.210951221 | 0.204350718 | 0.001213243 | 0.786599431 | 0.296915341 | 0.055826056 |
| Rsbn1 | S | 80 | 0.026999518 | -0.238834786 | -0.194242797 | -0.221242315 | 0.044591989 | 0.109195484 | 0.021189129 | 0.002894939 | 0.00531622 | 0.412335023 |
| Dlgap5 | S | 66 | -0.214665066 | 0.000257896 | -0.219261804 | -0.004596738 | -0.2195197 | 0.011383041 | 0.994646883 | 0.053494898 | 0.954711684 | 0.061676348 |
| Ubl7 | T | 243 | -0.168867569 | -0.099543029 | -0.159546764 | 0.009320806 | -0.060003735 | 0.019887289 | 0.0617249 | 0.297238591 | 0.944340522 | 0.655271255 |
| Nhsl1 | S | 1542 | -0.248150165 | -0.040527047 | -0.374314211 | -0.126164046 | -0.333787165 | 0.060713347 | 0.76602906 | 0.150671806 | 0.557036505 | 0.190398908 |
| Agps | S | 57 | -0.052977197 | -0.136571044 | -0.137876805 | -0.084899607 | -0.00130576 | 0.184386408 | 0.002441593 | 0.092068875 | 0.230558044 | 0.982297101 |
| Nucks1 | S | 181 | -0.046735286 | -0.06583218 | -0.244493492 | -0.197758205 | -0.178661312 | 0.107907444 | 0.105372544 | 0.009187701 | 0.019680887 | 0.020675471 |
| Samd4b | S | 238 | -0.144153244 | -0.075932029 | -0.270111925 | -0.125958681 | -0.194179896 | 0.152513502 | 0.434305201 | 0.023662466 | 0.234396215 | 0.105600484 |
| Prrg4 | S | 222 | -0.023180042 | -0.018819392 | -0.467037347 | -0.443857305 | -0.448217955 | 0.802528982 | 0.850443286 | 0.002913586 | 0.007673742 | 0.005564292 |
| Tnks1bp1 | S | 1131 | -0.098138523 | -0.149580065 | -0.050351216 | 0.047787307 | 0.099228849 | 0.029687366 | 0.003849952 | 0.215262718 | 0.148194267 | 0.018379588 |
| Gas2 | S | 283 | 0.036436255 | -0.135452198 | -0.19275498 | -0.229191235 | -0.057302782 | 0.411043557 | 0.020801907 | 0.00037572 | 0.013934415 | 0.197889769 |
| Map3k2 | S | 347 | 0.034686588 | -0.090568866 | -0.19906417 | -0.233750758 | -0.108495304 | 0.428442668 | 0.007423397 | 0.000263313 | 0.008323215 | 0.010254019 |
| Gtf3c2 | S | 230 | -0.161366371 | -0.17999058 | -0.186376114 | -0.025009743 | -0.006385534 | 0.247562732 | 0.107095213 | 0.092116284 | 0.832307644 | 0.939567744 |
| Map4 | S | 475 | -0.032928505 | -0.096177295 | -0.143599915 | -0.11067141 | -0.04742262 | 0.208005454 | 0.011796397 | 0.003816369 | 0.013421438 | 0.142692496 |
| Rnf213 | S | 252 | -0.106889628 | -0.260814179 | -0.110953778 | -0.00406415 | 0.149861201 | 0.251890183 | 0.023391027 | 0.23440939 | 0.881908327 | 0.003020726 |
| Plxna2 Plxna4 | S | 1633;1632 | -0.095875559 | -0.082022829 | -0.22776622 | -0.131890662 | -0.145743392 | 0.204839453 | 0.329956353 | 0.005802828 | 0.097153878 | 0.117742985 |
| Mafg | S | 124 | -0.043168438 | -0.197700136 | -0.166634005 | -0.123465567 | 0.031066132 | 0.430667187 | 0.043544041 | 0.022220343 | 0.035080689 | 0.656407879 |
| Ssh2 | S | 1216 | -0.092982758 | -0.098118615 | -0.186347179 | -0.093364421 | -0.088228564 | 0.094942792 | 0.084850731 | 0.028460941 | 0.169262177 | 0.188297222 |
| Ubtf | T | 388 | -0.101373286 | -0.096515605 | -0.214351222 | -0.112977936 | -0.117835617 | 0.212098459 | 0.097285521 | 0.022179483 | 0.207367875 | 0.125116117 |
| Esf1 | S | 651 | 0.016253793 | -0.226632157 | -0.143572966 | -0.159826759 | 0.083059191 | 0.715455814 | 0.0022702 | 0.042661608 | 0.045587559 | 0.154488007 |
| H2-Q8 | T | 199 | -0.133548988 | -0.353155622 | -0.222368171 | -0.088819182 | 0.130787451 | 0.320240684 | 0.153034037 | 0.232429969 | 0.61896057 | 0.573690068 |
| H2-Q8 | T | 203 | -0.133548988 | -0.353155622 | -0.222368171 | -0.088819182 | 0.130787451 | 0.320240684 | 0.153034037 | 0.232429969 | 0.61896057 | 0.573690068 |
| Hsf1 | T | 365 | -0.056758445 | -0.118908306 | -0.139005052 | -0.082246607 | -0.020096746 | 0.175439058 | 0.080605853 | 0.002522601 | 0.078273865 | 0.691424767 |
| Cep85 | S | 50 | -0.01210353 | -0.255510176 | -0.253079931 | -0.2409764 | 0.002430245 | 0.812397268 | 0.059182982 | 0.049279067 | 0.051072055 | 0.981913932 |
| Crhr1 Crhr2 | T | 298;302 | -0.084412044 | -0.159900048 | -0.093301324 | -0.00888928 | 0.066598724 | 0.371282123 | 0.004215234 | 0.03989167 | 0.914554214 | 0.019680849 |
| Crhr1 Crhr2 | Y | 305;309 | -0.084412044 | -0.159900048 | -0.093301324 | -0.00888928 | 0.066598724 | 0.371282123 | 0.004215234 | 0.03989167 | 0.914554214 | 0.019680849 |
| Acsl4 | S | 57 | 0.018855861 | -0.182204172 | -0.172484844 | -0.191340705 | 0.009719328 | 0.667738986 | 0.007892606 | 0.007347939 | 0.009910368 | 0.836369183 |
| Map4k4 | S | 799 | -0.03996585 | -0.231687169 | -0.115028546 | -0.075062697 | 0.116658622 | 0.466886471 | 0.002860392 | 0.195892796 | 0.341192896 | 0.183087644 |
| Usp47 | S | 921 | -0.16488504 | -0.124020947 | -0.227489231 | -0.06260419 | -0.103468284 | 0.208359783 | 0.304944978 | 0.033317783 | 0.603488987 | 0.400757583 |
| Plcb3 | S | 1107 | -0.235022995 | 0.074671925 | -0.434025098 | -0.199002103 | -0.508697023 | 0.109221341 | 0.682166022 | 0.032624438 | 0.20797282 | 0.042604966 |
| Gps1 | T | 459 | -0.042190697 | -0.098820355 | -0.200882608 | -0.15869191 | -0.102062253 | 0.194837401 | 0.014068522 | 0.025463453 | 0.06769924 | 0.15500511 |
| Srrm2 | S | 944 | -0.018896097 | -0.118213369 | -0.101345438 | -0.082449341 | 0.016867931 | 0.326401411 | 0.00596986 | 0.000847971 | 0.005342905 | 0.492070225 |
| Gpatch4 | T | 138 | 0.010714114 | -0.429265887 | -0.52680198 | -0.537516094 | -0.097536093 | 0.972133284 | 0.117680137 | 0.27232929 | 0.288125912 | 0.821085895 |
| Actn1 | S | 91 | -0.190042326 | -0.096961306 | -0.219208126 | -0.0291658 | -0.12224682 | 0.146993637 | 0.264039319 | 0.051726797 | 0.784514911 | 0.143171307 |
| Hivep2 | S | 1431 | -0.210626415 | -0.032491844 | -0.318743413 | -0.108116998 | -0.286251569 | 0.074977777 | 0.77306368 | 0.063499565 | 0.429086166 | 0.093419206 |
| Anp32a | S | 17 | 0.10255739 | -0.196623091 | -0.443134926 | -0.545692316 | -0.246511835 | 0.426933945 | 0.091993008 | 0.013865952 | 0.010629772 | 0.09253774 |
| Topbp1 | S | 1185 | -0.188685668 | -0.199710769 | -0.080803175 | 0.107882493 | 0.118907594 | 0.218548643 | 0.008551818 | 0.446708082 | 0.479557994 | 0.295603589 |
| Rbm45 | S | 464 | -0.044829549 | -0.131244192 | -0.214539863 | -0.169710314 | -0.083295671 | 0.455753124 | 0.018969582 | 0.032728265 | 0.073946483 | 0.265273633 |
| Mbd3 | S | 85 | -0.003422211 | -0.1541377 | -0.160974208 | -0.157551997 | -0.006836509 | 0.928188302 | 0.004436984 | 0.013109383 | 0.021439233 | 0.852185091 |
| Rock2 | S | 1366 | -0.196174265 | 0.038053655 | -0.805341217 | -0.609166952 | -0.843394872 | 0.3833671 | 0.895295517 | 0.089089511 | 0.15638938 | 0.088424273 |
| Fip1l1 | S | 508 | -0.322009847 | -0.058670326 | -0.005885937 | 0.31612391 | 0.052784389 | 0.001756897 | 0.176960771 | 0.95623669 | 0.055943588 | 0.620653315 |
| Lrrc59 | S | 6 | -0.075214534 | -0.110750512 | -0.138934686 | -0.063720152 | -0.028184174 | 0.484833247 | 0.026634258 | 0.003048686 | 0.54785947 | 0.424621334 |
| Mapre2 | S | 199 | 0.031014435 | -0.156593416 | -0.601813501 | -0.632827937 | -0.445220086 | 0.763574166 | 0.031057476 | 0.125435047 | 0.105280301 | 0.201428768 |
| Zfp609 | T | 745 | -0.026641216 | -0.12098707 | -0.149248696 | -0.12260748 | -0.028261625 | 0.342032443 | 0.003102489 | 0.021315764 | 0.051632197 | 0.500532446 |

| | | | | | | | | | | | | | |
|---|---|---|---|---|---|---|---|---|---|---|---|---|---|
| Bcr | S | | 303 | -0.174767522 | 0.001507028 | -0.297871199 | -0.123103677 | -0.299378227 | 0.0396939 | 0.989672698 | 0.031443185 | 0.242866651 | 0.06561849 |
| Srsf9 | S | | 205 | -0.05443365 | -0.229825488 | -0.095176913 | -0.040743262 | 0.134648576 | 0.468242074 | 0.002400965 | 0.217746395 | 0.639942682 | 0.117777286 |
| Tnip1 | S | | 73 | -0.082745315 | -0.288689778 | -0.346305772 | -0.263560457 | -0.057615993 | 0.549511338 | 0.218057669 | 0.109913574 | 0.202886607 | 0.804378856 |
| Ttf2 | S | | 436 | -0.088945583 | -0.331846029 | -0.221279222 | -0.132333638 | 0.110566807 | 0.413586816 | 0.134229073 | 0.13673981 | 0.321442599 | 0.56702569 |
| Birc6 | T | | 3954 | 0.005546837 | -0.124977284 | -0.365525211 | -0.371072047 | -0.240547927 | 0.957595908 | 0.288767029 | 0.005412914 | 0.030337811 | 0.093314707 |
| Sgpp1 | S | | 65 | 0.064410454 | -0.11974273 | -0.35461152 | -0.419021973 | -0.23486879 | 0.079254163 | 0.008083655 | 0.00384808 | 0.008079848 | 0.025488082 |
| Trpm7 | T | | 1503 | -0.182006452 | -0.015570211 | -0.468126176 | -0.286119724 | -0.452555965 | 0.297182906 | 0.926305988 | 0.034630343 | 0.072466252 | 0.021044958 |
| Top2a | S | | 1388 | -0.087824174 | -0.184000365 | -0.129350568 | -0.041526394 | 0.054649797 | 0.125274445 | 0.010760123 | 0.257886537 | 0.677185124 | 0.591497161 |
| Atp10a | S | | 519 | -0.222235279 | -0.106949619 | -0.042195685 | 0.180039595 | 0.064753934 | 0.004968818 | 0.056031932 | 0.71908272 | 0.197719117 | 0.576312127 |
| Atp10a | S | | 522 | -0.222235279 | -0.106949619 | -0.042195685 | 0.180039595 | 0.064753934 | 0.004968818 | 0.056031932 | 0.71908272 | 0.197719117 | 0.576312127 |
| Fbxo5 | S | | 85 | -0.20992791 | -0.185995723 | -0.199999559 | 0.009928351 | -0.014003836 | 0.264636847 | 0.211666013 | 0.090624257 | 0.953326449 | 0.918647446 |
| Fbxo5 | S | | 89 | -0.20992791 | -0.185995723 | -0.199999559 | 0.009928351 | -0.014003836 | 0.264636847 | 0.211666013 | 0.090624257 | 0.953326449 | 0.918647446 |
| Lmna | S | | 653 | -0.094413041 | -0.057782598 | -0.152843739 | -0.058430698 | -0.095061141 | 0.038989249 | 0.174774451 | 0.005808689 | 0.179052034 | 0.067640032 |
| Lats1 | S | | 612 | -0.071191702 | -0.085437473 | -0.191981827 | -0.120790125 | -0.106544354 | 0.16767161 | 0.119484402 | 0.007091381 | 0.060281213 | 0.088966921 |
| Plekhm2 | S | | 423 | -0.016964633 | -0.135576407 | -0.203449238 | -0.186484605 | -0.06787283 | 0.82379776 | 0.032415782 | 0.006145566 | 0.086993564 | 0.207495228 |
| Slc1a5 | S | | 506 | 0.026909618 | -0.14947428 | -0.184283072 | -0.211192691 | -0.034808792 | 0.128371915 | 0.000605734 | 0.008358164 | 0.016129228 | 0.34986495 |
| Stx7 | T | | 79 | -0.048476309 | -0.187500995 | -0.0427886 | 0.00568771 | 0.144712396 | 0.186304777 | 0.000227092 | 0.33413306 | 0.89082491 | 0.031873465 |
| Map4 | S | | 381 | -0.059042844 | -0.114728339 | -0.190987299 | -0.131944455 | -0.07625896 | 0.109009924 | 0.015068086 | 0.076687309 | 0.172576892 | 0.357596881 |
| Ddx19a Ddx25 | S | 85;92 | | -0.067689988 | -0.164403243 | -0.217765483 | -0.150075495 | -0.05336224 | 0.334742211 | 0.03443288 | 0.081407467 | 0.181055977 | 0.574848451 |
| Thoc2 | T | | 1385 | -0.059200454 | -0.106009688 | -0.217484266 | -0.158283812 | -0.111474578 | 0.181039728 | 0.018862564 | 0.057330922 | 0.106921342 | 0.204170606 |
| Anxa1 | S | | 37 | -0.194980194 | -0.08124888 | -0.204374557 | -0.009394363 | -0.123125676 | 0.025314092 | 0.427586991 | 0.129909241 | 0.927668767 | 0.342595681 |
| Rai14 | S | | 476 | -0.01274452 | -0.103038437 | -0.267856654 | -0.255112134 | -0.164818217 | 0.625119235 | 0.006615694 | 0.032348793 | 0.038218771 | 0.09177215 |
| Mtmr4 | S | | 629 | 0.086115541 | -0.247602908 | -0.268061795 | -0.354177335 | -0.020458887 | 0.052122511 | 0.002385842 | 0.018382848 | 0.012145531 | 0.764242828 |
| Prrc2c | S | | 389 | -0.008978705 | -0.193278741 | -0.104456254 | -0.095477549 | 0.088822486 | 0.784217091 | 0.002050234 | 0.027453876 | 0.040934842 | 0.05478961 |
| Nufip2 | S | | 217 | -0.082490056 | -0.121849548 | -0.169031264 | -0.086541208 | -0.047181716 | 0.253087358 | 0.00796544 | 0.082657547 | 0.335349431 | 0.510842155 |
| Gorasp1 | S | | 406 | -0.0680127 | -0.417930092 | -0.162016905 | -0.094004205 | 0.255913187 | 0.643398506 | 0.03859158 | 0.349223043 | 0.611993149 | 0.211131717 |
| Map3k2 | S | | 516 | -0.014384124 | -0.136678018 | -0.366025498 | -0.351641375 | -0.229347481 | 0.90111338 | 0.043294723 | 0.046933736 | 0.064177583 | 0.148207096 |
| Sorbs2 | S | | 1097 | -0.310347034 | -0.048949769 | -0.187053449 | 0.123293584 | -0.13810368 | 0.025561129 | 0.702183876 | 0.194319219 | 0.313943654 | 0.307237006 |
| Papola | S | | 545 | -0.208342421 | -0.097534832 | -0.170740805 | 0.037601616 | -0.073205973 | 0.140293329 | 0.150656076 | 0.065401044 | 0.74976928 | 0.362504668 |
| Fhdc1 | S | | 703 | -0.058292699 | -0.140842103 | -0.155666885 | -0.097374186 | -0.014824782 | 0.289611556 | 0.069151768 | 0.007771264 | 0.094650093 | 0.782677343 |
| Med24 | S | | 870 | -0.082908355 | -0.151509295 | -0.116182364 | -0.033274009 | 0.035326931 | 0.365985718 | 0.00979478 | 0.034468857 | 0.693096986 | 0.403639726 |
| Stk39 | T | | 366 | -0.03286028 | -0.183582979 | -0.145471348 | -0.112611068 | 0.038111631 | 0.480637567 | 0.007710016 | 0.054447674 | 0.107185802 | 0.514777253 |
| Hivep2 | S | | 1435 | -0.231419459 | 0.006640294 | -0.325385098 | -0.09396564 | -0.332025392 | 0.061406337 | 0.956147932 | 0.057325802 | 0.481134081 | 0.065227918 |
| Macf1 | S | | 4458 | -0.038228141 | -0.131945357 | -0.121642459 | -0.083414317 | 0.010302898 | 0.22521948 | 0.008903642 | 0.013686401 | 0.046094727 | 0.693211605 |
| Kif15 | S | | 676 | 0.057745275 | -0.214693197 | -0.465565847 | -0.523311122 | -0.25087265 | 0.58316307 | 0.247589598 | 0.016893955 | 0.011631268 | 0.200267621 |
| Kif15 | S | | 677 | 0.057745275 | -0.214693197 | -0.465565847 | -0.523311122 | -0.25087265 | 0.58316307 | 0.247589598 | 0.016893955 | 0.011631268 | 0.200267621 |
| Prpf4b | S | | 33 | -0.002448265 | -0.089643952 | -0.096333425 | -0.09388516 | -0.006689473 | 0.907219722 | 0.000366257 | 0.000261002 | 0.022489547 | 0.482426326 |
| Sh3bp5 | S | | 378 | -0.077467425 | -0.120632663 | -0.214324029 | -0.136856604 | -0.093691366 | 0.389219018 | 0.15684671 | 0.008894233 | 0.181876097 | 0.257405599 |
| Clasrp | S | | 294 | -0.011171129 | -0.201448427 | -0.173897128 | -0.162725999 | 0.027551298 | 0.684680887 | 0.033455328 | 0.014717371 | 0.012401158 | 0.621504752 |
| Gorab | S | | 347 | -0.033750842 | -0.329436275 | -0.108155816 | -0.074404974 | 0.22128046 | 0.795034459 | 0.009093122 | 0.202202019 | 0.526357344 | 0.051416322 |
| Fam53c | S | | 162 | 0.014305433 | -0.136459268 | -0.318975947 | -0.33328138 | -0.182516679 | 0.86259433 | 0.110615457 | 0.007733508 | 0.00813203 | 0.050798713 |
| Ripk2 | S | | 412 | -0.113152967 | -0.132384294 | -0.161584612 | -0.048431645 | -0.029200318 | 0.065136109 | 0.046058125 | 0.150586585 | 0.603348192 | 0.751505388 |
| Phldb1 | S | | 378 | -0.034488489 | -0.142469609 | -0.329323175 | -0.294834686 | -0.186853565 | 0.746242588 | 0.042820035 | 0.05517625 | 0.086159221 | 0.196669471 |
| Aimp1 | S | | 107 | -0.074795143 | -0.085758561 | -0.175120535 | -0.100325392 | -0.089361974 | 0.014987885 | 0.018324016 | 0.12775025 | 0.300533664 | 0.343037833 |
| Comt | S | | 261 | -0.1890216 | -0.028112469 | -0.41248734 | -0.22346574 | -0.384374871 | 0.129084927 | 0.844304236 | 0.067352137 | 0.238305915 | 0.094598795 |
| N4bp1 | S | | 299 | -0.197196913 | -0.054877073 | -0.266604299 | -0.069407385 | -0.211727226 | 0.130421259 | 0.645936655 | 0.032371963 | 0.530679297 | 0.113198568 |
| Vim | S | | 8 | -0.099744246 | -0.171267105 | -0.108150938 | -0.008406692 | 0.063116167 | 0.357511462 | 0.026491111 | 0.023892678 | 0.930362913 | 0.268740738 |
| Rps20 | T | | 9 | 0.052864443 | -0.206453916 | -0.250423438 | -0.30328788 | -0.043969522 | 0.623629601 | 0.010126071 | 0.032144204 | 0.055414844 | 0.585782916 |
| Rbbp8 | S | | 326 | 0.012585444 | -0.151186912 | -0.204503821 | -0.217089265 | -0.053316909 | 0.700432552 | 0.003033801 | 0.023928683 | 0.02532874 | 0.364099469 |
| Dvl2 | S | | 252 | -0.055658336 | -0.081791501 | -0.424707171 | -0.369048834 | -0.34291567 | 0.637031672 | 0.523940197 | 0.013328336 | 0.021855831 | 0.022352497 |
| Rfc1 | T | | 160 | 0.088195772 | -0.165548939 | -0.266929448 | -0.35512522 | -0.101380509 | 0.103792091 | 0.012364744 | 0.001239875 | 0.001373442 | 0.04311626 |
| Mapkap1 | S | | 510 | -0.198068708 | -0.005087919 | -0.335792087 | -0.137723379 | -0.330704168 | 0.098920729 | 0.976932773 | 0.036696206 | 0.277292605 | 0.130773442 |
| Fasn | S | | 2201 | -0.296603968 | 0.052302004 | -0.306295898 | -0.00969193 | -0.358597902 | 0.075440747 | 0.836474129 | 0.048617424 | 0.928543158 | 0.216526444 |
| Dock7 | S | | 1428 | -0.126260379 | -0.143799901 | -0.18624636 | -0.059985981 | -0.042446459 | 0.14759197 | 0.150611113 | 0.058145431 | 0.250150816 | 0.543314759 |
| Rbbp6 | S | | 1707 | -0.096045217 | -0.46774082 | -0.145380609 | -0.049335392 | 0.322360211 | 0.378142109 | 0.09020307 | 0.380592787 | 0.72769543 | 0.205733759 |
| Triobp | S | | 1633 | -0.060394471 | -0.213179505 | -0.16063808 | -0.100243608 | 0.052541425 | 0.5070039 | 0.013753421 | 0.120809259 | 0.354319879 | 0.549317096 |
| Apc | S | | 2290 | -0.068514257 | -0.146620309 | -0.187539548 | -0.119025291 | -0.040919239 | 0.210020033 | 0.09019429 | 0.027756266 | 0.107977195 | 0.596631741 |
| Erbin | S | | 910 | -0.25124732 | 0.027532059 | -0.266513938 | -0.015266618 | -0.294045997 | 0.009402427 | 0.822901659 | 0.13487636 | 0.911121456 | 0.129049943 |
| Kif20a | S | | 14 | -0.036779611 | -0.091583104 | -0.906318157 | -0.869538546 | -0.814735053 | 0.866371646 | 0.692743816 | 0.088144517 | 0.084941979 | 0.099120529 |
| Parva | S | | 28 | -0.224035184 | 0.004027096 | -0.435559779 | -0.211524595 | -0.439586875 | 0.141041485 | 0.984539619 | 0.067050696 | 0.291078082 | 0.126248656 |
| Afap1 | S | | 688 | 0.082681573 | -0.154453078 | -0.205018047 | -0.28769962 | -0.050564969 | 0.014867034 | 0.000779814 | 0.000348408 | 0.001216414 | 0.106889716 |
| Tnks1bp1 | S | | 763 | -0.038748493 | -0.262438519 | -0.107540445 | -0.068791952 | 0.154898074 | 0.751674788 | 0.00292735 | 0.232024618 | 0.591603804 | 0.107766078 |
| Racgap1 | T | | 576 | -0.01219892 | -0.313321569 | -0.254028342 | -0.241829422 | 0.059293227 | 0.95087927 | 0.008120956 | 0.394628049 | 0.458314531 | 0.827521431 |
| Uckl1 | S | | 56 | -0.008545286 | -0.124569525 | -0.16587768 | -0.157332394 | -0.041308155 | 0.750160174 | 0.024838802 | 0.002114435 | 0.005418664 | 0.285244664 |
| Dop1a | S | | 1166 | -0.056366069 | -0.228033813 | -0.141077512 | -0.084711442 | 0.086956301 | 0.362983178 | 0.018042871 | 0.123912371 | 0.29287843 | 0.321539724 |
| Dop1a | S | | 1169 | -0.056366069 | -0.228033813 | -0.141077512 | -0.084711442 | 0.086956301 | 0.362983178 | 0.018042871 | 0.123912371 | 0.29287843 | 0.321539724 |
| Scamp3 | S | | 78 | 0.026555219 | -0.160315446 | -0.095843856 | -0.122399076 | 0.06447159 | 0.619680155 | 0.000359481 | 0.004259372 | 0.097992682 | 0.037289329 |
| Kif18b | S | | 727 | -0.123821017 | -0.16370457 | -0.208039697 | -0.08421868 | -0.044335126 | 0.260659912 | 0.388537385 | 0.014751093 | 0.398516735 | 0.796002769 |
| Eef2k | S | | 473 | -0.160707697 | -0.078221368 | -0.056373702 | 0.104333996 | 0.021847666 | 8.37049E-05 | 0.256709451 | 0.551163419 | 0.316224885 | 0.828734771 |
| Ppp1r18 | S | | 224 | 0.02149456 | -0.177187021 | -0.134908503 | -0.156403063 | 0.042278518 | 0.460186761 | 0.000685449 | 0.019270893 | 0.015622877 | 0.302070943 |
| Tmem245 | S | | 323 | -0.045103184 | -0.085002655 | -0.176380412 | -0.131277228 | -0.091377757 | 0.371688898 | 0.05103137 | 0.002675043 | 0.049230064 | 0.001133873 |
| Nemf | S | | 738 | -0.080706268 | -0.130680075 | -0.145888852 | -0.065182584 | -0.015208777 | 0.095973473 | 0.066590597 | 0.032762622 | 0.238635229 | 0.803561099 |
| Nt5c | T | | 101 | -0.177900017 | -0.182370125 | -0.206072175 | -0.028172158 | -0.023702051 | 0.231163007 | 0.228506025 | 0.088796336 | 0.839882184 | 0.866666054 |
| C2cd3 | S | | 453 | -0.01575395 | -0.134389742 | -0.075456954 | -0.059703003 | 0.058932788 | 0.712922941 | 0.000507304 | 0.003523328 | 0.234412169 | 0.028586943 |
| Supt16 | S | | 1039 | 0.037066759 | -0.15336154 | -0.242735303 | -0.279802062 | -0.089373763 | 0.414795858 | 0.002330669 | 0.023960656 | 0.01444845 | 0.21579067 |
| Phldb1 | S | | 770 | 0.114786696 | -0.175225188 | -0.405532702 | -0.520319397 | -0.230307514 | 0.135754086 | 0.00924517 | 0.014714055 | 0.002157531 | 0.035167221 |
| Tmf1 | S | | 357 | -0.064247068 | -0.269087155 | -0.214264323 | -0.150017255 | 0.054822832 | 0.5380508 | 0.030653724 | 0.223060915 | 0.344974353 | 0.706522888 |
| Npm1 | S | | 125 | -0.128045052 | -0.142894728 | -0.048585988 | 0.079459064 | 0.094308739 | 0.019886302 | 0.012996986 | 0.311855926 | 0.064780418 | 0.035064423 |
| Atp1a2 | S | | 734 | -0.091106846 | -0.189217345 | -0.154588318 | -0.063481472 | 0.034629027 | 0.460597048 | 0.120832166 | 0.010967495 | 0.591388193 | 0.71073619 |
| Atp1a2 | S | | 736 | -0.091106846 | -0.189217345 | -0.154588318 | -0.063481472 | 0.034629027 | 0.460597048 | 0.120832166 | 0.010967495 | 0.591388193 | 0.71073619 |
| Mapk1 | S | | 358 | -0.1132933 | -0.144055721 | -0.27030913 | -0.15701583 | -0.12625341 | 0.167353753 | 0.18243988 | 0.100396161 | 0.269390742 | 0.381999976 |
| Pus7l | S | | 103 | -0.010934528 | -0.14667949 | -0.246527379 | -0.235592851 | -0.099847888 | 0.807048273 | 0.08696344 | 0.008341228 | 0.010679568 | 0.198310058 |
| Bag2 | S | | 20 | -0.147204138 | -0.047588179 | -0.115672096 | 0.031532042 | -0.068083918 | 0.00474092 | 0.453161755 | 0.025719484 | 0.400220612 | 0.300894811 |
| Sharpin | S | | 128 | -0.175322767 | -0.163649569 | -0.172368475 | 0.002954292 | -0.008718906 | 0.123241342 | 0.149749524 | 0.153063775 | 0.961124019 | 0.892941174 |
| Ets1 | S | | 285 | -0.151359684 | -0.148549708 | -0.259981433 | -0.108621749 | -0.111431725 | 0.205884932 | 0.247411122 | 0.092657385 | 0.426632085 | 0.436151148 |
| Snap23 | S | | 160 | -0.132590329 | -0.188153951 | -0.515718195 | -0.383127866 | -0.327564244 | 0.347544998 | 0.387510434 | 0.184672672 | 0.289652266 | 0.366016439 |
| Phf2 | S | | 701 | -0.290573587 | 0.020637367 | -0.241962647 | 0.04861094 | -0.262600013 | 0.035217531 | 0.896979494 | 0.075180391 | 0.491984666 | 0.13013333 |
| Anln | S | | 548 | 0.008647214 | -0.156474207 | -0.260171248 | -0.268818462 | -0.103697041 | 0.903203703 | 0.008729699 | 0.045120037 | 0.043768669 | 0.278781452 |
| Bicd2 | S | | 611 | 0.002333037 | -0.147803211 | -0.169822929 | -0.172155966 | -0.022019718 | 0.901377059 | 0.018788852 | 0.003559208 | 0.00175247 | 0.504040271 |
| Mrtfb | S | | 913 | -0.042853505 | -0.13204621 | -0.242307602 | -0.199454097 | -0.110261392 | 0.300090048 | 0.14958199 | 0.015961645 | 0.034461092 | 0.234139785 |
| Arhgef7 | T | | 596 | -0.079002485 | -0.182086736 | -0.182174042 | -0.103171557 | -8.73057E-05 | 0.319246059 | 0.030669809 | 0.111752754 | 0.33452675 | 0.999241619 |
| Nav3 | S | | 1189 | -0.041657504 | -0.140754915 | -0.134275685 | -0.09261818 | 0.00647923 | 0.348869887 | 0.0027466 | 0.026656457 | 0.110341009 | 0.883170152 |
| Map1a | S | | 319 | -0.159984389 | -0.038010993 | -0.154182092 | 0.005802298 | -0.116171099 | 0.015908379 | 0.632677789 | 0.01884221 | 0.815712129 | 0.177424101 |
| Tns3 | S | | 844 | -0.025172045 | -0.194863574 | -0.246021919 | -0.220849874 | -0.051158345 | 0.811919071 | 0.020673991 | 0.080944965 | 0.143955672 | 0.632123992 |

| | | | | | | | | | | | | |
|---|---|---|---|---|---|---|---|---|---|---|---|---|
| Cavin4 | S | 20 | -0.073062052 | -0.287989997 | -0.103822754 | -0.030760702 | 0.184167243 | 0.404459074 | 0.014064512 | 0.281310473 | 0.549672689 | 0.03715572 |
| Cep170 | S | 1384 | -0.2699069 | -0.169914175 | -0.270995742 | -0.001088842 | -0.101081567 | 0.19231479 | 0.531817886 | 0.137282645 | 0.995404309 | 0.704706219 |
| Col4a3bp | S | 380 | -0.220460268 | -0.066097197 | -0.195453128 | 0.02500714 | -0.12935593 | 0.01738091 | 0.548087448 | 0.17540884 | 0.832391178 | 0.374786433 |
| Zc3hav1 | S | 551 | 0.054369323 | -0.169127822 | -0.282758299 | -0.337127622 | -0.113630478 | 0.715194662 | 0.027293077 | 0.015866646 | 0.093647295 | 0.18849756 |
| Osbpl3 | S | 733 | -0.080776015 | -0.097872463 | -0.134291356 | -0.053515341 | -0.036418893 | 0.126943651 | 0.038073816 | 0.014529594 | 0.295815163 | 0.393309503 |
| Mnat1 | T | 51 | -0.01727037 | -0.044329893 | -0.243472373 | -0.226202003 | -0.19914248 | 0.824632141 | 0.222124552 | 0.000279043 | 0.062049877 | 0.002551284 |
| Lsm11 | S | 18 | -0.04679503 | -0.166513297 | -0.194785533 | -0.147990503 | -0.028272236 | 0.257845632 | 0.027403878 | 0.079784889 | 0.117694834 | 0.707869374 |
| Lsm11 | S | 21 | -0.04679503 | -0.166513297 | -0.194785533 | -0.147990503 | -0.028272236 | 0.257845632 | 0.027403878 | 0.079784889 | 0.117694834 | 0.707869374 |
| Top2b | S | 1563 | -0.081651964 | -0.150591334 | -0.171899499 | -0.090247535 | -0.021308165 | 0.20311955 | 0.026262725 | 0.102355264 | 0.32824013 | 0.796484504 |
| Ints3 | S | 1034 | 0.038054478 | -0.241901928 | -0.210882278 | -0.248936756 | 0.03101965 | 0.609072909 | 0.021481612 | 0.021424448 | 0.003248435 | 0.604961409 |
| Cry1 | S | 588 | -0.01564122 | -0.108096855 | -0.274550152 | -0.258908932 | -0.166453297 | 0.738809058 | 0.011394517 | 0.034273434 | 0.036915654 | 0.115119248 |
| Nova1 | S | 154 | -0.144078243 | -0.168697619 | -0.062049364 | 0.082028879 | 0.106648255 | 0.221124523 | 0.005421527 | 0.187884281 | 0.434000092 | 0.014099589 |
| Nop53 | S | 305 | 0.026212228 | -0.201225724 | -0.220700282 | -0.24691251 | -0.019474558 | 0.006835626 | 0.020546296 | 0.001413834 | 0.003117741 | 0.61833691 |
| Tom1 | S | 462 | 0.071585071 | -0.110498612 | -0.264691652 | -0.336276723 | -0.15419304 | 0.155798455 | 0.02339235 | 0.000373045 | 0.003514585 | 0.002546837 |
| Tnks1bp1 | T | 131 | -0.011979131 | -0.202078868 | -0.16659188 | -0.154612748 | 0.035486989 | 0.872023369 | 0.004246977 | 0.089795065 | 0.148748389 | 0.62406272 |
| Syap1 | T | 262 | -0.179141485 | -0.180456701 | -0.127539354 | 0.051602131 | 0.052917347 | 0.070742152 | 0.133828305 | 0.229535565 | 0.621096032 | 0.652609361 |
| 1810030O07Rik | T | 315 | -0.248700916 | -0.04277803 | -0.325647262 | -0.076946346 | -0.282869232 | 0.06123833 | 0.76101639 | 0.152414945 | 0.683727283 | 0.212960085 |
| Sipa1 | T | 82 | 0.027207066 | -0.372715613 | -0.274106203 | -0.301313269 | 0.098609409 | 0.672887028 | 0.017698822 | 0.266992729 | 0.239517436 | 0.656911472 |
| Exoc1 | S | 484 | -0.088013103 | -0.117213917 | -0.466977762 | -0.378964658 | -0.349763845 | 0.358483014 | 0.526941876 | 0.059139209 | 0.088778516 | 0.161073848 |
| Tmx1 | T | 277 | -0.042625092 | -0.170765143 | -0.158631807 | -0.116006715 | 0.012133336 | 0.339340533 | 0.046645858 | 0.020443775 | 0.06338176 | 0.825872172 |
| Wdr75 | T | 692 | 0.083468719 | -0.24927035 | -0.32984143 | -0.413310149 | -0.08057108 | 0.438672984 | 0.072328651 | 0.015060761 | 0.020870636 | 0.467651397 |
| Zfyve16 | S | 906 | 0.086322805 | -0.162054325 | -0.22561408 | -0.311936885 | -0.063559755 | 0.071754094 | 0.00437787 | 0.000861785 | 0.001503807 | 0.053624858 |
| Pdlim5 | S | 325 | -0.187557944 | 0.059395023 | -0.511241316 | -0.323683372 | -0.570636339 | 0.360978387 | 0.872300432 | 0.024776456 | 0.147108143 | 0.200058346 |
| Wnk1 | S | 2135 | -0.263990521 | -0.074663489 | -0.081780463 | 0.182210057 | -0.007116974 | 0.001861954 | 0.592859314 | 0.506041645 | 0.197931664 | 0.964513604 |
| Rplp2 | S | 79 | 0.014962655 | -0.142046775 | -0.107639685 | -0.12260234 | 0.03440709 | 0.347068955 | 0.000836574 | 0.001336864 | 0.007621634 | 0.165622533 |
| Mtbp | S | 697 | -0.309184925 | -0.122429235 | -0.460879839 | -0.151694914 | -0.338450604 | 0.327351995 | 0.46523026 | 0.226540595 | 0.697397838 | 0.345354013 |
| Scaf4 | S | 1167 | -0.120259879 | -0.11620669 | -0.061968682 | 0.058291197 | 0.054238008 | 0.140615847 | 0.002240024 | 0.082794009 | 0.396090161 | 0.106695052 |
| Ppp1r18 | T | 236 | -0.148919208 | -0.245728937 | -0.057216835 | 0.091702373 | 0.188512102 | 0.149550096 | 0.01293178 | 0.657685276 | 0.520088474 | 0.215862434 |
| Raly | S | 301 | 0.004477442 | -0.13418427 | -0.0882151 | -0.092692542 | 0.045969169 | 0.876775205 | 0.000339847 | 0.00332961 | 0.031039478 | 0.024335547 |
| Cavin2 | S | 264 | -0.373581742 | 0.044453587 | -0.389561986 | -0.015980245 | -0.434015574 | 0.104818959 | 0.857924857 | 0.134195368 | 0.934711856 | 0.123316533 |
| Btf3 | S | 199 | -0.016626067 | -0.150314687 | -0.212530985 | -0.195904919 | -0.062216299 | 0.554606137 | 0.032722897 | 0.021460983 | 0.033001942 | 0.338552284 |
| Ibtk | S | 1116 | 0.043250437 | -0.190087183 | -0.16389535 | -0.207145787 | 0.026191833 | 0.780351619 | 0.000283338 | 0.131496326 | 0.265540125 | 0.745575029 |
| Usp10 | S | 541 | -0.053963562 | -0.104115348 | -0.065093853 | -0.011130292 | 0.039021494 | 0.099613703 | 0.002797808 | 0.005318737 | 0.646549044 | 0.116119506 |
| Pcdh19 | S | 912 | 0.234123202 | -0.373624324 | -0.335813991 | -0.569937192 | 0.037810333 | 0.15049892 | 0.014756107 | 0.022360838 | 0.015301611 | 0.603798515 |
| Pcdh19 | S | 917 | 0.234123202 | -0.373624324 | -0.335813991 | -0.569937192 | 0.037810333 | 0.15049892 | 0.014756107 | 0.022360838 | 0.015301611 | 0.603798515 |
| Pcdh19 | T | 915 | 0.234123202 | -0.373624324 | -0.335813991 | -0.569937192 | 0.037810333 | 0.15049892 | 0.014756107 | 0.022360838 | 0.015301611 | 0.603798515 |
| Apbb1ip | S | 537 | 0.033210693 | -0.180510434 | -0.257487578 | -0.29069827 | -0.076977143 | 0.461494191 | 0.021779888 | 0.014133148 | 0.006461978 | 0.23716778 |
| Sik3 | S | 610 | -0.068326138 | -0.073986972 | -0.287559557 | -0.219233419 | -0.213572586 | 0.2731163 | 0.325601636 | 0.011591613 | 0.029834337 | 0.04733732 |
| Enc1 | S | 406 | 0.101828181 | -0.261706321 | -0.298647509 | -0.400475691 | -0.036941188 | 0.129579836 | 0.001837775 | 0.070414085 | 0.034402683 | 0.740373241 |
| Foxm1 | S | 521 | -0.052679358 | -0.10790046 | -0.213021214 | -0.160341856 | -0.105120755 | 0.151909883 | 0.02960812 | 0.054962597 | 0.114375549 | 0.225719957 |
| Tbc1d15 | Y | 215 | -0.257614202 | -0.159404679 | -0.446554636 | -0.188940434 | -0.287149957 | 0.34371276 | 0.414589368 | 0.223646572 | 0.602205596 | 0.404033235 |
| Usp6nl | S | 665 | -0.160158489 | -0.041872185 | -0.163294903 | -0.003136414 | -0.121422719 | 0.014332176 | 0.472432755 | 0.043642235 | 0.952182135 | 0.099835472 |
| Bag3 | S | 386 | -0.091152656 | -0.152038759 | -0.082477756 | 0.0086749 | 0.069561003 | 0.10696789 | 0.010701059 | 0.101000481 | 0.83262955 | 0.095074733 |
| Mink1 | S | 603 | -0.102873012 | -0.065222305 | -0.210688438 | -0.107815427 | -0.145466134 | 0.134798475 | 0.302022356 | 0.010195335 | 0.118857386 | 0.050221226 |
| Zkscan17 | S | 182 | -0.382989814 | -0.166020844 | -0.000310551 | 0.382679263 | 0.165710292 | 0.047489346 | 0.087903957 | 0.99861573 | 0.119809193 | 0.404382551 |
| Ctnnd1 | T | 916 | -0.10021915 | -0.048260108 | -0.212997289 | -0.112778139 | -0.164737181 | 0.107169159 | 0.521521426 | 0.004823765 | 0.068741498 | 0.092518407 |
| Rpl6 | S | 150 | -0.002334175 | -0.07919895 | -0.142557946 | -0.14022377 | -0.063358995 | 0.959424619 | 0.005531875 | 0.000351252 | 0.054925569 | 0.0197823 |
| Cr1l | T | 470 | -0.162364679 | -0.123176689 | -0.340387207 | -0.178022528 | -0.217210518 | 0.412690714 | 0.326212264 | 0.064590637 | 0.40559185 | 0.199320685 |
| Hdac9 | T | 448 | -0.206557833 | -0.225979612 | 0.031941594 | 0.238499426 | 0.257921206 | 0.034347477 | 0.018263333 | 0.858236631 | 0.248153623 | 0.223314213 |
| Arhgap18 | S | 68 | -0.289840887 | -0.111020885 | -0.164692123 | 0.125148764 | -0.053671239 | 0.040036604 | 0.41719201 | 0.311820524 | 0.429866209 | 0.742167764 |
| Myo18a | S | 1998 | -0.099091293 | -0.145235656 | -0.164672156 | -0.065580863 | -0.0194365 | 0.321247533 | 0.041164666 | 0.051919615 | 0.513716434 | 0.774234494 |
| Pkp4 | S | 235 | -0.198084471 | -0.041475226 | -0.307975502 | -0.109891032 | -0.266500276 | 0.110135975 | 0.772018339 | 0.053608738 | 0.407963981 | 0.139697075 |
| Larp4b | T | 629 | 0.093609508 | -0.220489879 | -0.176935348 | -0.270544856 | 0.043554531 | 0.081649787 | 0.00085618 | 0.006094415 | 0.002807104 | 0.261912093 |
| Pcgf6 | S | 118 | -0.07200598 | -0.118701737 | -0.152033923 | -0.080027943 | -0.033332186 | 0.191842701 | 0.085348063 | 0.010978951 | 0.170427919 | 0.536878305 |
| Rell1 | S | 110 | -0.029394296 | -0.135885051 | -0.25786789 | -0.228473593 | -0.121982839 | 0.798261349 | 0.161976355 | 0.006837722 | 0.12686428 | 0.209454905 |
| Cobll1 | T | 1091 | 0.029588074 | -0.113076652 | -0.143933674 | -0.173521748 | -0.030857022 | 0.403386598 | 0.000579309 | 0.001696805 | 0.00811888 | 0.224134844 |
| Spag9 | S | 733 | -0.062204571 | -0.065798468 | -0.2430062 | -0.180801629 | -0.177207732 | 0.457685206 | 0.377547614 | 0.001811847 | 0.092348207 | 0.066797755 |
| Gga1 | S | 267 | -0.054235016 | -0.177032422 | -0.304006345 | -0.249771329 | -0.126973923 | 0.494404307 | 0.133172206 | 0.062036774 | 0.110152267 | 0.362253146 |
| Bcr | S | 305 | -0.212346137 | -0.038648835 | -0.40825397 | -0.195907833 | -0.369605135 | 0.097449782 | 0.785667954 | 0.130127199 | 0.38817752 | 0.162503686 |
| Hadh | S | 13 | -0.167435097 | -0.192607718 | -0.142139013 | 0.025296084 | 0.050468705 | 0.129100215 | 0.129197722 | 0.174116194 | 0.522334104 | 0.526965306 |
| Ivns1abp | S | 336 | -0.051016461 | -0.13319609 | -0.252483885 | -0.201467424 | -0.119287796 | 0.474815225 | 0.019881317 | 0.082451993 | 0.133092122 | 0.300799012 |
| Cdc20 | T | 70 | 0.098400522 | -0.079614395 | -0.34758521 | -0.445985732 | -0.267970815 | 0.020865481 | 0.045226805 | 6.46134E-05 | 0.00075334 | 0.001812537 |
| Clasrp | S | 529 | -0.065736445 | -0.114541843 | -0.169888516 | -0.10415207 | -0.055346673 | 0.233243412 | 0.133424778 | 0.006830228 | 0.055453128 | 0.378220803 |
| Zkscan17 | S | 330 | -0.271116046 | -0.310791387 | -0.046793506 | 0.22432254 | 0.26399788 | 0.177335794 | 0.061335208 | 0.830320522 | 0.380788161 | 0.284248793 |
| Zkscan17 | S | 332 | -0.271116046 | -0.310791387 | -0.046793506 | 0.22432254 | 0.26399788 | 0.177335794 | 0.061335208 | 0.830320522 | 0.380788161 | 0.284248793 |
| Dnmt1 | S | 285 | 0.023994056 | -0.106279647 | -0.186523399 | -0.210517455 | -0.080243752 | 0.449433034 | 0.010810899 | 0.000792118 | 0.000356472 | 0.001166206 |
| Prmt3 | S | 22 | -0.076947406 | -0.119211949 | -0.237683178 | -0.160735772 | -0.118471228 | 0.228635285 | 0.058230931 | 0.072641037 | 0.164109215 | 0.265293688 |
| Peak1 | T | 1154 | -0.345945366 | 0.036802903 | -0.214936291 | 0.131009076 | -0.251739194 | 0.012226549 | 0.774657508 | 0.219136383 | 0.388283861 | 0.164925374 |
| Srrm2 | S | 522 | -0.263418665 | -0.176015882 | 0.089656533 | 0.353075198 | 0.265672415 | 0.009980955 | 0.019165793 | 0.561271648 | 0.09205956 | 0.168270017 |
| Srrm2 | S | 524 | -0.263418665 | -0.176015882 | 0.089656533 | 0.353075198 | 0.265672415 | 0.009980955 | 0.019165793 | 0.561271648 | 0.09205956 | 0.168270017 |
| Sdccag8 | S | 92 | -0.057957182 | -0.086808101 | -0.172156808 | -0.114199627 | -0.085348707 | 0.165724999 | 0.014160789 | 0.029900575 | 0.08984974 | 0.160140321 |
| Aptx | S | 143 | -0.01615407 | -0.254211268 | -0.182147362 | -0.165993293 | 0.072063905 | 0.800444387 | 0.065027641 | 0.026192687 | 0.008818582 | 0.481432507 |
| Map2k1 Map2k2 | S | 218;222 | -0.087124643 | -0.067207404 | -0.377555268 | -0.290430625 | -0.310347864 | 0.134932365 | 0.649338427 | 0.036187215 | 0.06973411 | 0.120265461 |
| Kif1b | S | 1645 | -0.444993512 | 0.000541078 | -0.372577971 | 0.072415541 | -0.373119048 | 0.080065606 | 0.998539974 | 0.32401683 | 0.819702318 | 0.337313084 |
| Srrm1 | S | 674 | -0.000857407 | -0.129627617 | -0.103696634 | -0.102839227 | 0.025930983 | 0.969126341 | 0.000355482 | 0.010524925 | 0.016297614 | 0.33089867 |
| Polr2m | S | 362 | -0.142180914 | -0.057423011 | -0.206848234 | -0.06466732 | -0.149425223 | 0.043203627 | 0.425783901 | 0.039626793 | 0.371318443 | 0.106191081 |
| Zfp703 | S | 257 | 0.034709474 | -0.125020755 | -0.261860614 | -0.296570088 | -0.136839859 | 0.463657645 | 0.01321871 | 0.007341165 | 0.005398648 | 0.054552936 |
| Akap12 | S | 273 | 0.025835931 | -0.091714324 | -0.369789491 | -0.395625422 | -0.278075167 | 0.743679577 | 0.146987551 | 0.006039372 | 0.008638031 | 0.039023827 |
| Eme1 | S | 9 | -0.036325488 | -0.10896125 | -0.160999239 | -0.124673751 | -0.05203799 | 0.285041677 | 0.101713935 | 0.002165082 | 0.003088609 | 0.343482669 |
| Rnpc3 | S | 21 | -0.106562168 | -0.12173006 | -0.088790165 | 0.017772004 | 0.032939895 | 0.023817492 | 0.009228388 | 0.315719489 | 0.822793077 | 0.680013244 |
| Rusc2 | S | 504 | -0.082198906 | -0.218548294 | -0.051885298 | 0.030313609 | 0.166662996 | 0.002202535 | 0.058107602 | 0.433360927 | 0.626977362 | 0.104542336 |
| Api5 | S | 469 | -0.05786674 | -0.110410425 | -0.223369781 | -0.165503041 | -0.112959355 | 0.398011104 | 0.072062784 | 0.019977591 | 0.075012282 | 0.102414374 |
| 1110008P14Rik | S | 82 | -0.158414069 | -0.068871241 | -0.216135189 | -0.05772112 | -0.147263948 | 0.044584221 | 0.512394619 | 0.056540596 | 0.485889442 | 0.222920318 |
| Taf1d | S | 206 | 0.025026562 | -0.25898443 | -0.217014135 | -0.242040698 | 0.041970295 | 0.60885832 | 0.002234569 | 0.237390797 | 0.206454001 | 0.784756711 |
| Adgre5 | S | 806 | -0.283393188 | 0.049787378 | -0.263795712 | 0.019597476 | -0.31358309 | 0.011845636 | 0.688026538 | 0.141009091 | 0.890732767 | 0.112886765 |
| Cdc42bpa | S | 1622 | -0.072346512 | -0.093585008 | -0.40152197 | -0.329175459 | -0.307936963 | 0.39484272 | 0.553083648 | 0.026424864 | 0.047627126 | 0.130600438 |
| Gpat3 | S | 432 | -0.030121584 | -0.1010461 | -0.161988098 | -0.131866514 | -0.060941998 | 0.370230825 | 0.002414747 | 0.031264804 | 0.04657476 | 0.228312746 |
| Phldb2 | S | 976 | -0.040595228 | -0.130252386 | -0.190445209 | -0.14984998 | -0.060192823 | 0.328234667 | 0.052728578 | 0.018760212 | 0.026300922 | 0.247321029 |
| 5031439G07Rik | S | 268 | -0.082949885 | -0.108387512 | -0.146479925 | -0.06353004 | -0.038092413 | 0.116995672 | 0.028543791 | 0.052402787 | 0.315972565 | 0.502700695 |
| Nmd3 | S | 490 | 0.004305727 | -0.207166676 | -0.463913979 | -0.468219707 | -0.256747303 | 0.953466616 | 0.132517481 | 0.086276262 | 0.08909884 | 0.2505846 |
| Fnip1 | S | 174 | -0.211302621 | 0.042807929 | -0.32247341 | -0.111170789 | -0.36528134 | 0.083948555 | 0.658003718 | 0.022054717 | 0.338980536 | 0.025651054 |
| Tnks1bp1 | S | 429 | -0.293026501 | 0.022012042 | -0.23484184 | 0.058184661 | -0.256853882 | 0.025210515 | 0.861650654 | 0.111776393 | 0.622096596 | 0.108480149 |
| Gtf3c2 | S | 231 | -0.172894385 | -0.203478261 | -0.23556177 | -0.062667385 | -0.032083509 | 0.140350847 | 0.179323708 | 0.311707644 | 0.764637755 | 0.883851334 |
| Kat5 | S | 90 | -0.067272532 | -0.119288497 | -0.199568476 | -0.132295944 | -0.080279979 | 0.528278604 | 0.074868397 | 0.012604634 | 0.253380657 | 0.190373785 |

| | | | | | | | | | | | | |
|---|---|---|---|---|---|---|---|---|---|---|---|---|
| Prr11 | S | 249 | -0.026047077 | -0.133676104 | -0.430939334 | -0.404892257 | -0.29726323 | 0.809491609 | 0.360310772 | 0.026082469 | 0.056944515 | 0.094833481 |
| Prmt3 | S | 24 | -0.076318344 | -0.11398656 | -0.243430154 | -0.16711181 | -0.129443594 | 0.224646386 | 0.086367265 | 0.058578372 | 0.137287975 | 0.216826155 |
| Larp1 | S | 81 | -0.083363408 | -0.050781012 | -0.073422754 | 0.009940653 | -0.022641742 | 0.002596343 | 0.055601593 | 0.006435575 | 0.464369424 | 0.279071144 |
| Baz1b | S | 189 | -0.003103297 | -0.185636557 | -0.164154164 | -0.161050866 | 0.021482393 | 0.920245407 | 0.029338848 | 0.009875652 | 0.019863652 | 0.713938418 |
| Cct3 | S | 252 | -0.043464373 | -0.164557477 | -0.146244826 | -0.102780453 | 0.018312651 | 0.253517981 | 0.117674352 | 0.006513966 | 0.000570918 | 0.813353797 |
| Klf13 | S | 229 | -0.308106714 | -0.062542952 | -0.224967997 | 0.083138718 | -0.162425045 | 0.091789569 | 0.374935657 | 0.212969674 | 0.65178239 | 0.332703486 |
| Ranbp9 | S | 392 | -0.092442467 | -0.049739094 | -0.158265821 | -0.065823354 | -0.108526728 | 0.053803595 | 0.372851613 | 0.003207471 | 0.092371955 | 0.094693323 |
| Rai14 | S | 449 | 0.012632679 | -0.031574498 | -0.328635397 | -0.341268076 | -0.297060899 | 0.790637513 | 0.639863715 | 0.000307426 | 0.000129561 | 0.015309282 |
| Smg6 | S | 331 | -0.285221663 | 0.0167696 | -0.255206749 | 0.030014914 | -0.271976349 | 0.038238784 | 0.893963099 | 0.096216887 | 0.811609927 | 0.099559467 |
| Txnrd3 | T | 15 | -0.110517351 | -0.132439258 | -0.269565131 | -0.15904778 | -0.137125873 | 0.265565895 | 0.191946777 | 0.071225784 | 0.228447237 | 0.286417252 |
| Marcks | S | 27 | -0.121001584 | -0.041851799 | -0.144629322 | -0.023627738 | -0.102777523 | 0.081983359 | 0.084698836 | 0.009288429 | 0.620870211 | 0.015674211 |
| Eif3b | S | 727 | -0.018811143 | -0.078824374 | -0.229344187 | -0.210533044 | -0.150519814 | 0.702504328 | 0.346002028 | 0.000482448 | 0.014090664 | 0.131590277 |
| Nmt1 | S | 69 | -0.112751604 | -0.053928034 | -0.207953659 | -0.095202056 | -0.154025625 | 0.031245653 | 0.23598136 | 0.049761436 | 0.255405319 | 0.119022957 |
| Bach1 | S | 419 | -0.180141294 | -0.195156781 | -0.03918849 | 0.140952804 | 0.15596829 | 0.031389583 | 0.045348265 | 0.661712903 | 0.155565164 | 0.143766686 |
| Nedd4l | S | 475 | -0.08289807 | -0.092307991 | -0.163424953 | -0.080526883 | -0.071116962 | 0.005839024 | 0.193467077 | 0.074711468 | 0.263116353 | 0.379750762 |
| Cdca3 | T | 69 | 0.082890998 | -0.172768274 | -0.2184074 | -0.301298398 | -0.045639125 | 0.15334314 | 0.021237145 | 0.00061038 | 0.005324307 | 0.380133654 |
| Notch2 | S | 2071 | -0.01990077 | -0.170982396 | -0.194206522 | -0.174305752 | -0.023224126 | 0.750467388 | 0.00832997 | 0.073456395 | 0.102504247 | 0.763643425 |
| Rnf20 | S | 525 | -0.137499304 | -0.240341581 | -0.140442505 | -0.002943201 | 0.099899076 | 0.356547084 | 0.042817896 | 0.243185751 | 0.984146951 | 0.397535614 |
| L3mbtl2 | S | 67 | -0.033386534 | -0.111355847 | -0.298251796 | -0.264865262 | -0.186895949 | 0.636470829 | 0.046643422 | 0.038602701 | 0.052784896 | 0.121204973 |
| Dmtn | S | 333 | -0.277145279 | -0.0564313 | -0.255737686 | 0.021407592 | -0.199306386 | 0.059806481 | 0.695350575 | 0.177365617 | 0.898489384 | 0.308629979 |
| Gga2 | S | 414 | -0.012922936 | -0.223546927 | -0.423088518 | -0.410165581 | -0.199541591 | 0.908300273 | 0.130580547 | 0.102654859 | 0.10053447 | 0.346688511 |
| Cobll1 | S | 896 | -0.031994403 | -0.084965377 | -0.234165902 | -0.202171499 | -0.149200525 | 0.608967607 | 0.161692687 | 0.002893451 | 0.017764506 | 0.016216055 |
| Plekhg3 | S | 737 | -0.029041826 | -0.095728531 | -0.284931491 | -0.255889665 | -0.18920296 | 0.608444951 | 0.428219791 | 0.003072999 | 0.009560847 | 0.174383975 |
| Ctnna1 Ctnna2 | S | 652;651 | -0.191833523 | -0.016780462 | -0.23901409 | -0.047180567 | -0.222233627 | 0.041847479 | 0.881513891 | 0.044537455 | 0.583426562 | 0.11006145 |
| Asap1 | S | 1045 | -0.133470718 | -0.027272064 | -0.151394282 | -0.017923564 | -0.124122217 | 0.002499766 | 0.474115972 | 0.060921219 | 0.741489334 | 0.093782398 |
| Zfp507 | S | 534 | -0.038175937 | -0.159087679 | -0.236737197 | -0.19856126 | -0.077649518 | 0.71927384 | 0.023955388 | 0.064706993 | 0.16242802 | 0.423283612 |
| Mlf2 | S | 237 | 0.01816883 | -0.178722357 | -0.12634342 | -0.144512249 | 0.052378937 | 0.700240545 | 0.00464774 | 0.00742223 | 0.041636682 | 0.19447468 |
| Smarcad1 | S | 79 | -0.078917932 | -0.121676311 | -0.270363717 | -0.191445785 | -0.148687406 | 0.142634993 | 0.206371718 | 0.066659503 | 0.14792486 | 0.248652027 |
| Suco | S | 1077 | -0.056822376 | -0.103782572 | -0.167938817 | -0.111116441 | -0.064156245 | 0.168114812 | 0.112956883 | 0.008785992 | 0.04953575 | 0.281933594 |
| Trip10 | S | 484 | -0.133257051 | -0.020326587 | -0.476946796 | -0.343689745 | -0.456620209 | 0.383423436 | 0.886077877 | 0.032442002 | 0.095101333 | 0.044089067 |
| Senp6 | S | 373 | -0.065911621 | -0.227191226 | -0.202886681 | -0.13697506 | 0.024304545 | 0.641954972 | 0.061923172 | 0.077989872 | 0.351635523 | 0.806416072 |
| Srgap3 | S | 858 | -0.215875522 | 0.024413537 | -0.228350834 | -0.012475312 | -0.252764372 | 0.124704157 | 0.706487301 | 0.004672136 | 0.904619119 | 0.021524532 |
| Larp4b | S | 246 | -0.206047345 | -0.201461128 | -0.115515854 | 0.090531491 | 0.085945275 | 0.157740866 | 0.120716002 | 0.224055118 | 0.48738385 | 0.459877846 |
| Lrrc47 | S | 530 | -0.03337849 | -0.166628436 | -0.197913125 | -0.164534635 | -0.031284689 | 0.837126593 | 0.014303946 | 0.05121369 | 0.363183919 | 0.651493215 |
| Sae1 | S | 189 | -0.0648424 | -0.112616066 | -0.115685455 | -0.050843056 | -0.00306939 | 0.104546651 | 0.038471648 | 0.015204333 | 0.178855051 | 0.928412876 |
| Nab1 | S | 405 | -0.117950834 | -0.095616985 | -0.145927912 | -0.027977078 | -0.050310928 | 0.098599495 | 0.122377326 | 0.030778151 | 0.562980714 | 0.191500204 |
| Specc1l | S | 882 | -0.070225443 | -0.115092779 | -0.306075237 | -0.235849793 | -0.190982457 | 0.273867133 | 0.172050783 | 0.058819922 | 0.11431045 | 0.168793425 |
| Rbm10 | T | 91 | -0.192638336 | 0.020252685 | -0.351540075 | -0.158901739 | -0.37179276 | 0.209447742 | 0.744763798 | 0.015399394 | 0.295977828 | 0.016215602 |
| Cttn | S | 113 | -0.344092937 | -0.039358694 | -0.264177093 | 0.079915844 | -0.224818399 | 0.081601323 | 0.821780837 | 0.186552641 | 0.481464381 | 0.112454049 |
| Med14 | S | 1149 | -0.135794717 | -0.110431254 | -0.02193974 | 0.113854977 | 0.088491514 | 0.003486138 | 0.010618734 | 0.649162094 | 0.069427013 | 0.117585623 |
| Usp42 | S | 960 | 0.033216762 | -0.186010815 | -0.208744152 | -0.241960915 | -0.022733338 | 0.433305652 | 0.004217578 | 0.034205635 | 0.036714696 | 0.731006427 |
| Usp42 | S | 962 | 0.033216762 | -0.186010815 | -0.208744152 | -0.241960915 | -0.022733338 | 0.433305652 | 0.004217578 | 0.034205635 | 0.036714696 | 0.731006427 |
| Myh9 | T | 1939 | -0.246980786 | -0.07361663 | -0.135842382 | 0.111138405 | -0.062225752 | 0.013891802 | 0.567209727 | 0.226493077 | 0.310662857 | 0.668486527 |
| Cdca2 | S | 165 | -0.016632662 | -0.254006834 | -0.140487754 | -0.123855092 | 0.11351908 | 0.773129151 | 0.003341949 | 0.301714873 | 0.345090639 | 0.379071795 |
| Ankrd17 | T | 728 | -0.035952311 | -0.06590795 | -0.22765315 | -0.191700839 | -0.1617452 | 0.482364005 | 0.436479144 | 0.000721063 | 0.023356715 | 0.128724389 |
| Pcm1 | S | 644 | -0.012753683 | -0.09211843 | -0.199545876 | -0.186792193 | -0.107427446 | 0.825990221 | 0.027660695 | 0.003808822 | 0.032396991 | 0.059929128 |
| Ppil2 | S | 520 | -0.13759543 | -0.440875614 | -0.227892386 | -0.090296956 | 0.212983228 | 0.571352213 | 0.121008989 | 0.434179839 | 0.785535353 | 0.510837867 |
| Cend1 | T | 15 | -0.019735764 | -0.160377918 | -0.209390605 | -0.189654841 | -0.049012687 | 0.769252256 | 0.055426126 | 0.016750097 | 0.003459356 | 0.308941227 |
| Ass1 | S | 219 | -0.181384454 | 0.043980012 | -0.410735398 | -0.229350944 | -0.45471541 | 0.481670247 | 0.638300455 | 0.005780148 | 0.391918017 | 0.009870466 |
| Apc | S | 1772 | -0.051275367 | -0.137102596 | -0.073170815 | -0.021895448 | 0.063931781 | 0.019726308 | 0.004679023 | 0.150827151 | 0.590141636 | 0.198858014 |
| Dpysl3 | S | 475 | -0.199667357 | -0.031360919 | -0.089142184 | 0.110525173 | -0.057781266 | 0.002786239 | 0.484070296 | 0.110773756 | 0.074991647 | 0.292002572 |
| Dock7 | S | 1420 | -0.123606448 | -0.072330988 | -0.215897095 | -0.092290648 | -0.143566107 | 0.071323227 | 0.325955561 | 0.043941574 | 0.265529274 | 0.134114721 |
| Kirrel | S | 772 | -0.321362073 | 0.012785917 | -0.169470584 | 0.151891489 | -0.182256501 | 0.03266521 | 0.912293506 | 0.074809562 | 0.199903724 | 0.146357629 |
| Kirrel | S | 775 | -0.321362073 | 0.012785917 | -0.169470584 | 0.151891489 | -0.182256501 | 0.03266521 | 0.912293506 | 0.074809562 | 0.199903724 | 0.146357629 |
| Kirrel | T | 771 | -0.321362073 | 0.012785917 | -0.169470584 | 0.151891489 | -0.182256501 | 0.03266521 | 0.912293506 | 0.074809562 | 0.199903724 | 0.146357629 |
| Nab1 | S | 401 | -0.191981572 | -0.002300181 | -0.170909247 | 0.021072325 | -0.168609066 | 0.009055418 | 0.976674488 | 0.048499691 | 0.739680992 | 0.09823372 |
| Gon4l | S | 414 | -0.060088552 | -0.264226533 | -0.293087597 | -0.232999044 | -0.028861063 | 0.737167134 | 0.135017483 | 0.104218978 | 0.29190733 | 0.865125008 |
| Ddx51 | S | 77 | -0.024449396 | -0.242442052 | -0.157782851 | -0.133333456 | 0.084659201 | 0.720499637 | 0.010630824 | 0.16010272 | 0.201772189 | 0.365168282 |
| Sltm | T | 74 | 0.038306635 | -0.339176144 | -0.23574082 | -0.274047455 | 0.103435324 | 0.653285598 | 0.058339378 | 0.05668555 | 0.038833479 | 0.467961398 |
| Ice2 | S | 587 | -0.07843635 | -0.110215871 | -0.194725595 | -0.116289246 | -0.084509725 | 0.024478033 | 0.264746616 | 0.053887685 | 0.167133842 | 0.411589814 |
| Ice2 | T | 589 | -0.07843635 | -0.110215871 | -0.194725595 | -0.116289246 | -0.084509725 | 0.024478033 | 0.264746616 | 0.053887685 | 0.167133842 | 0.411589814 |
| Arhgap23 | S | 655 | -0.238011884 | -0.301596543 | -0.092447855 | 0.145564029 | 0.209148688 | 0.266698303 | 0.064327912 | 0.633245409 | 0.557327655 | 0.343576162 |
| Sox2 | S | 39 | -0.093182362 | -0.243391302 | -0.076986586 | 0.016195776 | 0.166404716 | 0.228972223 | 0.009480021 | 0.471920658 | 0.863537525 | 0.164736294 |
| Gpalpp1 | S | 121 | -0.097054157 | -0.145935889 | -0.082742432 | 0.014311725 | 0.063193458 | 0.023435753 | 0.05385398 | 0.119701807 | 0.74973321 | 0.310904915 |
| Phrf1 | S | 1043 | -0.249485825 | -0.237718592 | -0.117587163 | 0.131898662 | 0.120131429 | 0.119317537 | 0.181680997 | 0.441167784 | 0.155707936 | 0.347831351 |
| Phrf1 | S | 1045 | -0.249485825 | -0.237718592 | -0.117587163 | 0.131898662 | 0.120131429 | 0.119317537 | 0.181680997 | 0.441167784 | 0.155707936 | 0.347831351 |
| Dzip1l | S | 722 | -0.20776287 | -0.0811862 | -0.007105637 | 0.200657233 | 0.074080563 | 0.000662961 | 0.11019286 | 0.956829375 | 0.219342525 | 0.586841328 |
| Naa38 | S | 25 | -0.037371608 | -0.106902459 | -0.12237889 | -0.085007282 | -0.015476431 | 0.281174559 | 0.006290633 | 0.014546067 | 0.060906791 | 0.60419039 |
| Pi4kb | T | 263 | 0.02847184 | -0.1607195 | -0.236462239 | -0.26493408 | -0.075742739 | 0.548337964 | 0.001503699 | 0.075406993 | 0.045439439 | 0.40195519 |
| Hsp90ab1 | S | 226 | -0.061740779 | -0.240605699 | -0.135905397 | -0.074164618 | 0.104700302 | 0.726726992 | 0.05965742 | 0.024834799 | 0.677946172 | 0.272488633 |
| Adar | S | 774 | -0.003896189 | -0.313633193 | -0.139194908 | -0.135298719 | 0.174438285 | 0.967620537 | 0.012238102 | 0.183157835 | 0.161258009 | 0.092578711 |
| Inpp5f | S | 936 | -0.01520624 | -0.107741252 | -0.096211481 | -0.081005241 | 0.011529771 | 0.639788227 | 0.000131383 | 0.021316059 | 0.077445259 | 0.639600073 |
| Msantd4 | S | 106 | 0.015350305 | -0.114706665 | -0.176194296 | -0.191544601 | -0.06148763 | 0.715422616 | 0.003942625 | 0.006185281 | 0.013201481 | 0.137451433 |
| Pkp1 | S | 156 | -0.120754245 | -0.081425476 | -0.139072486 | -0.018318241 | -0.057647011 | 0.092651976 | 0.032695879 | 0.070117849 | 0.791218021 | 0.338971388 |
| Samd4 | S | 254 | -0.161045421 | -0.116717861 | -0.128835534 | 0.032209887 | -0.012117673 | 0.117653944 | 0.064623388 | 0.123891343 | 0.721207996 | 0.839055705 |
| Cobll1 | S | 551 | -0.088382956 | -0.100693502 | -0.068640006 | 0.01974295 | 0.032053496 | 0.015702764 | 0.004739444 | 0.202461185 | 0.663123704 | 0.487896723 |
| Prrc2a | S | 2114 | 0.012060066 | -0.246640897 | -0.153655802 | -0.165715869 | 0.092985094 | 0.696889932 | 0.049745393 | 0.006741585 | 0.019259116 | 0.302330854 |
| Srrm2 | S | 1284 | -0.07639129 | -0.068856696 | -0.182212324 | -0.105821034 | -0.113355628 | 0.196392885 | 0.263497401 | 0.003502807 | 0.08460454 | 0.090722487 |
| Otud7b | S | 464 | -0.143330148 | -0.032780658 | -0.085985961 | 0.057344187 | -0.053205303 | 0.001484474 | 0.547392326 | 0.024888159 | 0.056479208 | 0.339296159 |
| Plekhh3 | S | 30 | -0.190661457 | -0.197319331 | -0.129404694 | 0.061256763 | 0.067914637 | 0.18344104 | 0.09595668 | 0.255210366 | 0.562658452 | 0.249334474 |
| Ulk1 | S | 622 | 0.039375471 | -0.227597143 | -0.230415289 | -0.26979076 | -0.002818146 | 0.589120085 | 0.092547592 | 0.006511028 | 0.021924473 | 0.977364581 |
| Golga1 | S | 30 | -0.140205663 | -0.07777961 | -0.07709412 | 0.063111543 | 0.000685489 | 0.009693277 | 0.264216012 | 0.022565619 | 0.099126218 | 0.99083993 |
| Nedd4l | S | 478 | -0.045787926 | -0.111066916 | -0.257788458 | -0.212000532 | -0.146721542 | 0.302902185 | 0.034785414 | 0.071866927 | 0.108803347 | 0.205738082 |
| Mtss2 | S | 419 | -0.35476371 | 0.054032783 | -0.349106536 | 0.005657174 | -0.403139319 | 0.131611593 | 0.814859398 | 0.086041042 | 0.973789921 | 0.108683122 |
| Nav3 | S | 1841 | 0.054115991 | -0.146048111 | -0.30398541 | -0.358101401 | -0.157937299 | 0.376813585 | 0.04200283 | 0.00760973 | 0.01252757 | 0.080841615 |
| Flnb | S | 1505 | -0.028825969 | -0.158598274 | -0.128700467 | -0.099874499 | 0.029897807 | 0.556174397 | 0.009293096 | 0.028457154 | 0.045839648 | 0.392347597 |
| Rai1 | S | 1312 | 0.017998832 | -0.133258986 | -0.210114237 | -0.228113069 | -0.076855251 | 0.608172258 | 0.007258272 | 0.013325761 | 0.022563564 | 0.196781641 |
| Phrf1 | S | 1157 | -0.149866725 | -0.150113344 | -0.122591318 | 0.027275407 | 0.027522027 | 0.067784461 | 0.172373579 | 0.110906249 | 0.414300269 | 0.736361852 |
| Jcad | S | 980 | -0.223681147 | -0.088082717 | -0.123846168 | 0.099834979 | -0.03576345 | 0.023629632 | 0.394365982 | 0.172732 | 0.12009291 | 0.671668184 |
| Ddr2 | S | 461 | 0.051127806 | -0.162108826 | -0.217471795 | -0.268599601 | -0.055362969 | 0.460719414 | 0.031530448 | 0.003196408 | 0.017304604 | 0.349217226 |
| Dpysl2 | S | 402 | -0.158824717 | -0.069089625 | -0.282587661 | -0.123762944 | -0.213498035 | 0.100734223 | 0.352571762 | 0.106976097 | 0.396121189 | 0.193291031 |
| Casc3 | S | 65 | -0.111535607 | -0.140411548 | -0.121619538 | -0.010083931 | 0.01879201 | 0.065081412 | 0.120147016 | 0.069564661 | 0.859401632 | 0.801631251 |
| Vim | S | 459 | -0.014157259 | -0.076286734 | -0.17338809 | -0.159230831 | -0.097101356 | 0.741033341 | 0.032736949 | 0.001046318 | 0.033466371 | 0.000620596 |
| Phlpp2 | S | 1179 | -0.080011345 | -0.093674039 | -0.155545045 | -0.0755337 | -0.061871005 | 0.039906052 | 0.045177992 | 0.084425179 | 0.284179829 | 0.369060357 |

| | | | | | | | | | | | | |
|---|---|---|---|---|---|---|---|---|---|---|---|---|
| Eya3 | S | 375 | 0.030475985 | -0.056856932 | -0.322968171 | -0.353444156 | -0.266111239 | 0.635016873 | 0.254929104 | 0.000899761 | 0.003764971 | 0.008302762 |
| Tulp3 | S | 166 | -0.004358259 | -0.195423662 | -0.208275051 | -0.203916792 | -0.012851389 | 0.970216435 | 0.15115551 | 0.005178341 | 0.169962508 | 0.903857451 |
| Vdac1 | S | 70 | -0.067309238 | -0.161068374 | -0.100270848 | -0.032961611 | 0.060797525 | 0.246656668 | 0.004454948 | 0.165578922 | 0.62732134 | 0.343396928 |
| Adrm1 | S | 213 | -0.022971436 | -0.102799753 | -0.210229162 | -0.187257727 | -0.107429409 | 0.725618828 | 0.143392003 | 0.002260106 | 0.043438152 | 0.128926126 |
| Usp45 | S | 432 | -0.281723611 | -0.152703779 | -0.185590336 | 0.096133275 | -0.032886557 | 0.082915252 | 0.337743942 | 0.387344531 | 0.653138547 | 0.880963879 |
| Zfyve16 | S | 888 | 0.006416607 | -0.307974338 | -0.232468274 | -0.23888488 | 0.075506065 | 0.95087403 | 0.045181264 | 0.117504066 | 0.135561052 | 0.589704805 |
| Bod1l | S | 1470 | 0.008222374 | -0.122589325 | -0.246351723 | -0.254574097 | -0.123762398 | 0.893361115 | 0.191374319 | 0.00187431 | 0.010329754 | 0.187773653 |
| Stat1 | S | 727 | -0.095340552 | -0.130541647 | -0.138698937 | -0.043358384 | -0.00815729 | 0.148473022 | 0.063144521 | 0.054061911 | 0.302466782 | 0.836805382 |
| Psmd11 | S | 14 | -0.21755825 | -0.019389787 | -0.215149428 | 0.002408821 | -0.195759642 | 0.034856414 | 0.83536436 | 0.073603532 | 0.979264223 | 0.104591212 |
| Ajuba | S | 129 | -0.04946929 | -0.179345339 | -0.135160973 | -0.085691683 | 0.044184365 | 0.421157536 | 0.003368742 | 0.231504793 | 0.424416367 | 0.646323484 |
| Ctnnd1 | T | 869 | -0.015934611 | -0.135043327 | -0.181173265 | -0.165238653 | -0.046129937 | 0.814485663 | 0.003573665 | 0.051720475 | 0.098101156 | 0.466190917 |
| Birc6 | S | 4047 | 0.082681968 | -0.203773494 | -0.206875474 | -0.289557443 | -0.00310198 | 0.341448091 | 0.000146968 | 0.157128296 | 0.07519923 | 0.977214186 |
| Rock1 | S | 1341 | -0.131394731 | -0.022597907 | -0.430054363 | -0.298659632 | -0.407456456 | 0.177606603 | 0.887102791 | 0.050578452 | 0.096843011 | 0.0864902 |
| Ppp1r12b | S | 941 | 0.006799454 | -0.280466516 | -0.196574526 | -0.20337398 | 0.08389199 | 0.876816685 | 0.0544886 | 0.04262447 | 0.053518965 | 0.447060539 |
| Tjap1 | S | 513 | 0.047235683 | -0.206874373 | -0.169338447 | -0.216574131 | 0.037535926 | 0.439843101 | 0.0079683 | 0.012933562 | 0.026188169 | 0.454967294 |
| Anapc1 | S | 355 | -0.255389009 | -0.170159067 | -0.082166092 | 0.173222917 | 0.087992975 | 0.054003335 | 0.165903597 | 0.484834468 | 0.058446953 | 0.255268067 |
| Ibtk | S | 1113 | -0.006970056 | -0.170183894 | -0.11454244 | -0.107572384 | 0.055641454 | 0.956628806 | 0.000660682 | 0.089441488 | 0.44489762 | 0.317137041 |
| Dnmt1 | T | 149 | 0.067762814 | -0.131289445 | -0.382089266 | -0.44985208 | -0.250799821 | 0.128895685 | 0.010118417 | 0.016134825 | 0.005085508 | 0.03092854 |
| R3hdm2 | S | 263 | -0.26364051 | 0.069915312 | -0.27689258 | -0.013252069 | -0.346807891 | 0.023696779 | 0.593053926 | 0.07056978 | 0.906024645 | 0.057496969 |
| Ivns1abp | S | 246 | 0.010438438 | -0.160312112 | -0.262388912 | -0.27282735 | -0.1020768 | 0.889918589 | 0.045670288 | 0.023900075 | 0.032479787 | 0.223309333 |
| Prrc2a | S | 1217 | 0.035769388 | -0.193298718 | -0.116046505 | -0.151815893 | 0.077252212 | 0.327293003 | 0.000976955 | 0.014627518 | 0.003992125 | 0.036494704 |
| Zyx | S | 144 | -0.140769755 | -0.156622478 | -0.241826537 | -0.101056782 | -0.085204059 | 0.252638299 | 0.195645927 | 0.123047297 | 0.496080965 | 0.554006916 |
| Cacnb2 | S | 485 | -0.05022765 | -0.099413422 | -0.228234662 | -0.178007012 | -0.12882124 | 0.302245241 | 0.085062419 | 0.025063739 | 0.051336996 | 0.114562764 |
| Cenpj | S | 603 | -0.052624324 | -0.165593688 | -0.206089358 | -0.153465034 | -0.04049567 | 0.383880506 | 0.024416236 | 0.129726508 | 0.219348652 | 0.705890497 |
| Adgrl2 | S | 1362 | -0.119991207 | -0.140653842 | -0.259987302 | -0.139996096 | -0.119333461 | 0.489904866 | 0.109377176 | 0.081918489 | 0.457773318 | 0.337794761 |
| Nmt1 | S | 47 | -0.026125607 | -0.048502766 | -0.175188453 | -0.149062846 | -0.126685687 | 0.297894824 | 0.036214716 | 0.001324048 | 0.004503817 | 0.008074174 |
| Map4 | S | 534 | 0.12043475 | -0.229746555 | -0.353319116 | -0.473753867 | -0.123572561 | 0.506836967 | 0.050539397 | 0.029735915 | 0.054955977 | 0.293009393 |
| Map4 | T | 536 | 0.12043475 | -0.229746555 | -0.353319116 | -0.473753867 | -0.123572561 | 0.506836967 | 0.050539397 | 0.029735915 | 0.054955977 | 0.293009393 |
| Utp14a | S | 570 | -0.129157738 | -0.191621581 | -0.26434537 | -0.135187632 | -0.072723789 | 0.410971362 | 0.158745414 | 0.139442979 | 0.468881793 | 0.653812149 |
| Gpbp1 | S | 381 | 0.08560182 | -0.279628819 | -0.268609638 | -0.354211458 | 0.011019181 | 0.216493481 | 0.004252961 | 0.103577327 | 0.045744835 | 0.920575989 |
| Zc3h3 | S | 380 | -0.003430111 | -0.179551973 | -0.198456389 | -0.195026278 | -0.018904415 | 0.935924229 | 0.04356442 | 0.018686761 | 0.03034351 | 0.792138124 |
| Cbx5 | S | 92 | 0.055287378 | -0.178896666 | -0.197435204 | -0.252722582 | -0.018538538 | 0.291255212 | 0.02367454 | 0.002263505 | 0.006495473 | 0.728522582 |
| Rassf8 | S | 385 | -0.206756906 | -0.12408588 | -0.146442397 | 0.060314509 | -0.022356517 | 0.044353123 | 0.407699623 | 0.160666627 | 0.371781537 | 0.866446774 |
| Specc1 | S | 359 | 0.020759894 | -0.334928346 | -0.191746106 | -0.212506 | 0.14318224 | 0.706391358 | 0.078101918 | 0.031558111 | 0.029196911 | 0.320992638 |
| Map4 | S | 517 | -0.043221231 | -0.047183322 | -0.190673699 | -0.147452468 | -0.143490377 | 0.398545321 | 0.311631195 | 0.000412775 | 0.043811678 | 0.03035557 |
| Nadk | S | 44 | -0.090851761 | -0.084153717 | -0.255287344 | -0.164435583 | -0.171133627 | 0.195750134 | 0.404371802 | 0.021300955 | 0.089567739 | 0.151672352 |
| Zfp36l1 | S | 54 | -0.246368332 | -0.052926612 | -0.230386793 | 0.015981539 | -0.177460181 | 0.061924466 | 0.690792168 | 0.131115757 | 0.901913733 | 0.263116634 |
| Jcad | S | 655 | -0.238338236 | -0.094222727 | -0.131851824 | 0.106486412 | -0.037629097 | 0.016734431 | 0.235045703 | 0.516577919 | 0.585049555 | 0.840538939 |
| Nono | T | 430 | -0.05186767 | -0.066068432 | -0.182188063 | -0.130320393 | -0.116119631 | 0.04566548 | 0.108140446 | 0.016027036 | 0.039374996 | 0.051474424 |
| Prrc2c | S | 189 | 0.016800492 | -0.142805443 | -0.156439619 | -0.17324011 | -0.013634175 | 0.638519717 | 0.002029749 | 0.017556397 | 0.016843042 | 0.724487205 |
| Iqsec1 | S | 88 | -0.194248189 | -0.036642339 | -0.169150032 | 0.025098157 | -0.132507693 | 0.010772188 | 0.55956171 | 0.152008069 | 0.794182141 | 0.238076134 |
| Cep170 | T | 1068 | -0.070915262 | -0.034083034 | -0.160516128 | -0.089600867 | -0.126433094 | 0.060890135 | 0.310005666 | 0.001898465 | 0.026343542 | 0.007966716 |
| Rpl28 | S | 115 | 0.025775783 | -0.143250234 | -0.370122714 | -0.395898497 | -0.22687248 | 0.782993478 | 0.257599986 | 0.011421311 | 0.017747797 | 0.12398292 |
| Rnf169 | S | 309 | -0.131667761 | -0.078405263 | -0.319634522 | -0.187966761 | -0.241229258 | 0.44696217 | 0.689186533 | 0.015812227 | 0.287419245 | 0.263599162 |
| Rnf169 | T | 306 | -0.131667761 | -0.078405263 | -0.319634522 | -0.187966761 | -0.241229258 | 0.44696217 | 0.689186533 | 0.015812227 | 0.287419245 | 0.263599162 |
| Cep170b | S | 1341 | -0.067336171 | -0.063313706 | -0.102620997 | -0.035284825 | -0.03930729 | 0.02043336 | 0.195181604 | 0.001562682 | 0.117243614 | 0.371073507 |
| Tom1l2 | S | 194 | -0.074226101 | -0.104263611 | -0.282559634 | -0.208333533 | -0.178296023 | 0.490199508 | 0.085961926 | 0.052727022 | 0.152762852 | 0.158244556 |
| Atg16l1 | S | 287 | -0.063958921 | -0.003714655 | -0.255555343 | -0.191596421 | -0.251840688 | 0.092217116 | 0.963557184 | 0.002343404 | 0.010785127 | 0.05086515 |
| Clpx | S | 618 | 0.028490247 | -0.1183657 | -0.251687295 | -0.280177541 | -0.133321595 | 0.598558318 | 0.057026971 | 0.003783953 | 0.008705496 | 0.050428586 |
| H2-D1 H2-L | S | 353;353;163 | -0.251218922 | 0.062677455 | -0.231920438 | 0.019298484 | -0.294597893 | 0.020680872 | 0.660035772 | 0.05494985 | 0.80046189 | 0.094088265 |
| Ess2 | S | 295 | -0.104625766 | -0.095141634 | -0.093119162 | 0.011506604 | 0.002022472 | 0.020109458 | 0.09650875 | 0.046241298 | 0.673212351 | 0.963593049 |
| Nfx1 | S | 147 | -0.068860792 | -0.073779367 | -0.110983632 | -0.04212284 | -0.037204265 | 0.083573581 | 0.017661014 | 0.014404704 | 0.270814384 | 0.247487669 |
| Prune2 | S | 1749 | 0.067284515 | -0.242548883 | -0.271152709 | -0.338437225 | -0.028603827 | 0.560682731 | 0.05509412 | 0.024333781 | 0.033808823 | 0.776967729 |
| Zeb1 | S | 663 | -0.10529335 | -0.143591031 | -0.16221669 | -0.05692334 | -0.018625659 | 0.344780647 | 0.369871415 | 0.004717629 | 0.57576995 | 0.895676171 |
| Spdl1 | S | 558 | 0.043555977 | -0.137432861 | -0.181906758 | -0.225462735 | -0.044473897 | 0.066323142 | 0.005435354 | 0.000769997 | 0.002583141 | 0.2045708 |
| Mdc1 | S | 476 | -0.071440288 | -0.470623031 | -0.201822588 | -0.1303823 | 0.268800443 | 0.854928482 | 0.077536567 | 0.379983241 | 0.747882273 | 0.309271525 |
| Prr12 | S | 1054 | -0.094569509 | -0.151691887 | -0.09733213 | -0.002762621 | 0.054359757 | 0.304406145 | 0.00294132 | 0.247301654 | 0.977641384 | 0.475051419 |
| Cast | S | 81 | -0.163120637 | -0.046971693 | -0.156025604 | 0.007095032 | -0.109053911 | 0.013531215 | 0.476506961 | 0.085171938 | 0.913899645 | 0.201303865 |
| Actb Actg1 | S | 234;234 | -0.005061661 | -0.219515585 | -0.07339783 | -0.068336169 | 0.146117755 | 0.909742019 | 0.000665057 | 0.179911107 | 0.221208172 | 0.053229475 |
| Ldlrap1 | S | 14 | 0.136545697 | -0.213442162 | -0.301454901 | -0.438000598 | -0.088012739 | 0.230056266 | 0.002362149 | 0.048337576 | 0.019785951 | 0.402639454 |
| E2f8 | S | 429 | 0.023622883 | -0.167675206 | -0.212208462 | -0.235831344 | -0.044533256 | 0.401947107 | 0.035422489 | 0.005982743 | 0.013992442 | 0.461033521 |
| Melk | S | 452 | -0.00920155 | -0.152894265 | -0.29128067 | -0.282079119 | -0.138386404 | 0.897900447 | 0.063827403 | 0.041266906 | 0.043626607 | 0.207818694 |
| Tom1 | S | 160 | -0.105052153 | -0.070554465 | -0.132313104 | -0.027260951 | -0.061758639 | 0.031946343 | 0.197111648 | 0.023797397 | 0.526609198 | 0.264911249 |
| Ahdc1 | S | 1180 | -0.248435665 | -0.07544846 | -0.282547677 | -0.034112013 | -0.207099217 | 0.161015177 | 0.628504543 | 0.114706526 | 0.813061729 | 0.17398577 |
| Rif1 | S | 1703 | -0.126578269 | -0.094228131 | -0.215014836 | -0.088436567 | -0.120786705 | 0.177020162 | 0.380682156 | 0.02894277 | 0.176705707 | 0.21763019 |
| Cttn | S | 298 | -0.157534449 | -0.099793183 | -0.285247516 | -0.127713067 | -0.185454333 | 0.226896444 | 0.504554615 | 0.059358521 | 0.198618616 | 0.180894596 |
| Tjp2 | S | 1133 | -0.140860243 | -0.128567908 | -0.055146438 | 0.085713805 | 0.07342147 | 0.039997354 | 0.034009097 | 0.176167184 | 0.145533581 | 0.139436714 |
| Mink1 | S | 903 | -0.080038981 | -0.152517511 | -0.26441355 | -0.184374569 | -0.111896039 | 0.146437819 | 0.158261938 | 0.141436402 | 0.252822257 | 0.471600964 |
| Pdxdc1 | S | 386 | -0.058838578 | -0.283485703 | -0.326768585 | -0.267930007 | -0.043282883 | 0.607726384 | 0.097285431 | 0.266286722 | 0.340326733 | 0.868622735 |
| Hsp90ab1 | S | 261 | -0.209324072 | -0.139430507 | -0.120553119 | 0.088770952 | 0.018877388 | 0.131954773 | 0.075599375 | 0.284108035 | 0.511123138 | 0.852046869 |
| Mid1ip1 | S | 71 | 0.014738869 | -0.152845054 | -0.274109232 | -0.288848101 | -0.121264178 | 0.859843532 | 0.02942553 | 0.036397421 | 0.040819542 | 0.214301522 |
| Mid1ip1 | T | 67 | 0.014738869 | -0.152845054 | -0.274109232 | -0.288848101 | -0.121264178 | 0.859843532 | 0.02942553 | 0.036397421 | 0.040819542 | 0.214301522 |
| Ppil4 | S | 178 | -0.074643533 | -0.131869007 | -0.077471889 | -0.002828356 | 0.054397118 | 0.005617917 | 0.021486927 | 0.298161265 | 0.964326955 | 0.450386166 |
| Vim | S | 18 | -0.02012623 | -0.098870833 | -0.117220269 | -0.097094039 | -0.018349436 | 0.242971435 | 0.001920769 | 0.018356765 | 0.056881124 | 0.551940082 |
| Txlna | S | 522 | 0.038833048 | -0.116393984 | -0.262210676 | -0.301043723 | -0.145816691 | 0.561299466 | 0.107417183 | 0.001944008 | 0.006790859 | 0.052388353 |
| Nfix | S | 265 | -0.095763841 | -0.082113511 | -0.181879767 | -0.086115925 | -0.099766256 | 0.160241103 | 0.212000359 | 0.016587176 | 0.163595637 | 0.112715619 |
| Fam207a | S | 76 | 0.029026793 | -0.22022378 | -0.253700949 | -0.282727742 | -0.033477168 | 0.758388203 | 0.045097934 | 0.040527127 | 0.020140511 | 0.621029699 |
| Fam207a | S | 79 | 0.029026793 | -0.22022378 | -0.253700949 | -0.282727742 | -0.033477168 | 0.758388203 | 0.045097934 | 0.040527127 | 0.020140511 | 0.621029699 |
| Bcar1 | S | 432 | -0.060320714 | -0.122441515 | -0.125165568 | -0.064844854 | -0.002724053 | 0.452229403 | 0.012760631 | 0.022643429 | 0.425246796 | 0.944609692 |
| Tbc1d5 | S | 608 | -0.005582109 | -0.21550375 | -0.30174502 | -0.296162911 | -0.086241269 | 0.967998192 | 0.294784158 | 0.0167553 | 0.114996916 | 0.644071322 |
| Ank3 | S | 1451 | -0.114864749 | -0.023108716 | -0.392984888 | -0.278120139 | -0.369876173 | 0.360296191 | 0.83640185 | 0.021204971 | 0.086757319 | 0.034970238 |
| Phldb1 | S | 697 | 0.001834773 | -0.13710964 | -0.2488126 | -0.250647373 | -0.11170296 | 0.978133599 | 0.096594325 | 0.009441902 | 0.003679146 | 0.09159486 |
| Phldb1 | T | 699 | 0.001834773 | -0.13710964 | -0.2488126 | -0.250647373 | -0.11170296 | 0.978133599 | 0.096594325 | 0.009441902 | 0.003679146 | 0.09159486 |
| Rubcn | S | 412 | -0.014213586 | -0.161605989 | -0.156855914 | -0.142642328 | 0.004750075 | 0.401926011 | 0.011255163 | 0.056499188 | 0.065196821 | 0.926764081 |
| Larp1 | S | 201 | -0.125757419 | -0.042646258 | -0.121538377 | 0.004219041 | -0.07889212 | 0.036895778 | 0.417763264 | 0.005231945 | 0.918325865 | 0.173418114 |
| Mast4 | S | 1273 | -0.185763799 | -0.282693767 | 0.040610654 | 0.226374454 | 0.323304421 | 0.031956071 | 0.048249918 | 0.821103406 | 0.277483574 | 0.159983326 |
| 1810037I17Rik | S | 19 | -0.246285921 | -0.225700776 | 0.096292583 | 0.342578504 | 0.321993359 | 0.016818792 | 0.021023869 | 0.390450276 | 0.028980328 | 0.043914943 |
| Osbpl6 | T | 47 | -0.029182377 | -0.092975224 | -0.311886156 | -0.282703779 | -0.218910932 | 0.786303655 | 0.334789849 | 0.00765163 | 0.075736221 | 0.077750584 |
| Chd9 | S | 610 | 0.006402606 | -0.138480371 | -0.131128241 | -0.137530847 | 0.00735213 | 0.930790481 | 0.000366451 | 0.058759636 | 0.156235831 | 0.867603899 |
| Chd9 | S | 615 | 0.006402606 | -0.138480371 | -0.131128241 | -0.137530847 | 0.00735213 | 0.930790481 | 0.000366451 | 0.058759636 | 0.156235831 | 0.867603899 |
| Ehbp1 | S | 764 | -0.085173293 | -0.132072439 | -0.209280989 | -0.124107695 | -0.077208549 | 0.176856676 | 0.063547572 | 0.134952638 | 0.310768177 | 0.503116509 |
| Nucks1 | S | 54 | -0.109046154 | -0.075308263 | -0.109607899 | -0.000561745 | -0.034299636 | 0.106322316 | 0.109620811 | 0.007496207 | 0.990927646 | 0.358928783 |
| Mib1 | S | 805 | -0.034215603 | -0.148121328 | -0.090954663 | -0.05673906 | 0.057166665 | 0.455697455 | 0.000677942 | 0.129907643 | 0.343295727 | 0.2832484 |
| Cobll1 | S | 794 | -0.145399965 | -0.072429288 | -0.220548499 | -0.075148534 | -0.148119211 | 0.075361338 | 0.378640723 | 0.075011119 | 0.419634087 | 0.176406716 |

| | | | | | | | | | | | | |
|---|---|---|---|---|---|---|---|---|---|---|---|---|
| Rbm15 | S | 768 | -0.03302486 | -0.126538698 | -0.161970063 | -0.128945203 | -0.035431365 | 0.140756705 | 0.002911264 | 0.188522698 | 0.263731673 | 0.716700416 |
| Zfp503 | S | 107 | 0.005812067 | -0.199159314 | -0.139901302 | -0.145713369 | 0.059258011 | 0.882747628 | 0.000763673 | 0.21300464 | 0.202707219 | 0.537298403 |
| Ercc6l | S | 1021 | -0.212393779 | -0.130400512 | -0.079698068 | 0.132695711 | 0.050702443 | 0.074658767 | 0.054852899 | 0.384421787 | 0.252575159 | 0.577755961 |
| Nop53 | S | 329 | 0.015380654 | -0.150364437 | -0.204634548 | -0.220015203 | -0.054270112 | 0.240335605 | 0.017791952 | 0.007364101 | 0.005124024 | 0.178372291 |
| Fhod1 Fhod3 | T | 1174;1555 | -0.281479335 | 0.104560225 | -0.461650491 | -0.180171156 | -0.566210716 | 0.090889278 | 0.691552769 | 0.110937621 | 0.450325387 | 0.117212305 |
| Tjp1 | S | 212 | -0.149346496 | -0.008847701 | -0.205952167 | -0.056605671 | -0.197104466 | 0.030839193 | 0.894002107 | 0.023234403 | 0.366275204 | 0.035280121 |
| Nav1 | S | 1380 | 0.155611873 | -0.220097825 | -0.319973551 | -0.475585425 | -0.099875727 | 0.179159548 | 0.075490245 | 0.002270747 | 0.012672734 | 0.330692603 |
| Zdhhc5 | S | 636 | -0.069380149 | -0.079822918 | -0.281161902 | -0.211781753 | -0.201338984 | 0.442063718 | 0.378691427 | 0.011918771 | 0.079916226 | 0.086843277 |
| Mef2c | S | 396 | -0.120108705 | -0.172038712 | -0.064176746 | 0.055931959 | 0.107861966 | 0.076675824 | 0.022933715 | 0.318683105 | 0.149955802 | 0.033134047 |
| Pcbp2 | S | 361 | -0.0495216 | -0.07075116 | -0.213885663 | -0.164364062 | -0.143134503 | 0.25977341 | 0.048395074 | 0.020658829 | 0.043343757 | 0.066549503 |
| Micall2 | S | 125 | -0.092251889 | -0.052563693 | -0.2070081 | -0.114756211 | -0.154444406 | 0.113066449 | 0.306053543 | 0.014804654 | 0.091676925 | 0.058861525 |
| Osbpl3 | S | 340 | -0.010530544 | -0.169343479 | -0.227610322 | -0.217079778 | -0.058266843 | 0.829213998 | 0.036675534 | 0.04541877 | 0.035201626 | 0.425016331 |
| Zfp553 | S | 229 | 0.054483178 | -0.238140766 | -0.251179384 | -0.305662563 | -0.013038618 | 0.169213884 | 0.002981313 | 0.096837106 | 0.070130858 | 0.902188233 |
| Smc6 | S | 11 | 0.00093497 | -0.096468366 | -0.138797358 | -0.139732328 | -0.042328992 | 0.978752333 | 0.009278547 | 0.001245626 | 0.016610289 | 0.14825462 |
| Hectd1 | S | 1514 | -0.177280951 | -0.07687937 | -0.211551236 | -0.034270284 | -0.134671866 | 0.029085777 | 0.274365466 | 0.273556455 | 0.834735374 | 0.447063479 |
| Fam129b | S | 650 | -0.022283013 | -0.094972823 | -0.22250976 | -0.200226747 | -0.127536937 | 0.725467085 | 0.094918018 | 0.005648537 | 0.023821191 | 0.046427551 |
| Flnb | T | 2554 | -0.009974992 | -0.110593147 | -0.105220951 | -0.095245959 | 0.005372196 | 0.696911574 | 0.00332847 | 0.00326104 | 0.005041406 | 0.780364666 |
| Mark2 | S | 400 | -0.07166112 | -0.06752457 | -0.269072255 | -0.197411135 | -0.201547684 | 0.144354292 | 0.477542502 | 0.018796512 | 0.031865867 | 0.104682845 |
| Jam2 | S | 286 | -0.132649749 | -0.091681282 | -0.172699479 | -0.040049731 | -0.081018197 | 0.108472122 | 0.253605144 | 0.044897537 | 0.4561804 | 0.189469107 |
| Dlc1 | S | 432 | -0.223184162 | -0.119335444 | -0.246697754 | -0.023513592 | -0.12736231 | 0.162325458 | 0.298851534 | 0.218780808 | 0.900849773 | 0.477339023 |
| Serbp1 | S | 252 | -0.085698853 | 0.024779655 | -0.390503739 | -0.304804885 | -0.415283394 | 0.166932315 | 0.917803983 | 0.014778548 | 0.02768397 | 0.179692039 |
| Cdca3 | S | 67 | 0.038544651 | -0.177375152 | -0.205361869 | -0.24390652 | -0.027986717 | 0.036000302 | 0.025223125 | 0.00116299 | 0.002318491 | 0.531593376 |
| Washc2 | S | 723 | -0.06620493 | -0.167517575 | -0.024883167 | 0.041321762 | 0.142634408 | 0.495877093 | 4.19001E-05 | 0.600254096 | 0.67498364 | 0.059055246 |
| Tln1 | S | 858 | -0.223398392 | -0.20318083 | -0.011686731 | 0.211711661 | 0.191494099 | 0.015699785 | 0.141613721 | 0.906379596 | 0.10260017 | 0.194044199 |
| Map3k1 | S | 142 | -0.024062564 | -0.16080976 | -0.194155875 | -0.170093311 | -0.033346116 | 0.80031443 | 0.176033244 | 0.00445198 | 0.150678029 | 0.727705044 |
| Eps8 | S | 684 | -0.012490372 | -0.126982169 | -0.266193179 | -0.253702807 | -0.13921101 | 0.857703012 | 0.134213855 | 0.012723154 | 0.019695414 | 0.120215366 |
| Eif3d | S | 529 | -0.016218958 | -0.120834063 | -0.12967462 | -0.113455662 | -0.008840557 | 0.597106657 | 0.005999681 | 0.013244161 | 0.027709217 | 0.760700078 |
| Atp2a2 | S | 661 | -0.004872491 | -0.228469119 | -0.120835167 | -0.115962677 | 0.107633952 | 0.904545849 | 0.00197857 | 0.245952018 | 0.260952779 | 0.286733266 |
| Smarca1 Smarca5 | S | 754;762 | -0.109481682 | -0.095252683 | -0.127745388 | -0.018263706 | -0.032492706 | 0.078854478 | 0.334366344 | 0.008971438 | 0.704714941 | 0.713951053 |
| Rc3h2 | S | 801 | -0.159733035 | -0.12943495 | -0.311971357 | -0.152238323 | -0.182536408 | 0.338264279 | 0.116206288 | 0.224909682 | 0.536164778 | 0.425708916 |
| Tsr1 | S | 49 | -0.10142752 | -0.040197367 | -0.462204802 | -0.360777282 | -0.422007435 | 0.413972886 | 0.73599349 | 0.040825878 | 0.097190559 | 0.07518565 |
| Sorbs3 | S | 459 | -0.176203767 | -0.005869194 | -0.130540523 | 0.045663245 | -0.124671328 | 0.007584145 | 0.947958348 | 0.028832253 | 0.349397686 | 0.233425955 |
| Rbbp6 | S | 1024 | -0.066981191 | -0.202009327 | -0.277975812 | -0.210994621 | -0.075966485 | 0.539892697 | 0.053279421 | 0.217398738 | 0.326550947 | 0.690799592 |
| Ubap1 | S | 146 | 0.161357992 | -0.18760924 | -0.414111274 | -0.575469267 | -0.226502034 | 0.08761683 | 0.057559297 | 0.003816692 | 0.009791261 | 0.046238768 |
| Spag7 | S | 114 | -0.094002574 | -0.093208752 | -0.228635919 | -0.134633344 | -0.135427166 | 0.070783131 | 0.232545297 | 0.0822266 | 0.220438101 | 0.23793224 |
| Ivns1abp | S | 326 | 0.116743021 | -0.285205456 | -0.256823365 | -0.373566385 | 0.028382091 | 0.041714535 | 0.006855603 | 0.014829889 | 0.004979619 | 0.689789583 |
| Ivns1abp | T | 330 | 0.116743021 | -0.285205456 | -0.256823365 | -0.373566385 | 0.028382091 | 0.041714535 | 0.006855603 | 0.014829889 | 0.004979619 | 0.689789583 |
| Snx7 | S | 379 | -0.119986728 | -0.08178414 | -0.229921308 | -0.10993458 | -0.148137167 | 0.093024452 | 0.194997709 | 0.102590716 | 0.348485817 | 0.23376438 |
| Map4k4 | S | 646 | -0.043301906 | -0.03043648 | -0.225981091 | -0.182679185 | -0.195544611 | 0.370588765 | 0.601899466 | 0.000628192 | 0.016494912 | 0.037542466 |
| Map6 | S | 386 | -0.090705911 | -0.008392245 | -0.356479893 | -0.265773982 | -0.348087647 | 0.242564991 | 0.790258983 | 0.015444599 | 0.032455394 | 0.022192193 |
| Nhsl1 | S | 328 | 0.048870206 | -0.139249945 | -0.237548241 | -0.286418447 | -0.098298296 | 0.571712666 | 0.000749074 | 0.074145873 | 0.049344309 | 0.310603733 |
| Trip12 | S | 1409 | 0.006804829 | -0.071421514 | -0.148785754 | -0.155590583 | -0.077364241 | 0.898271936 | 0.021659568 | 0.0001999 | 0.066584508 | 0.021346754 |
| Fmnl2 | S | 1015 | -0.036338581 | -0.144540021 | -0.162056382 | -0.1257178 | -0.017516361 | 0.53406188 | 0.033093062 | 0.026129638 | 0.107672424 | 0.744041605 |
| Matr3 | S | 164 | -0.162197937 | -0.170899713 | -0.046904028 | 0.115293909 | 0.123995685 | 0.044130848 | 0.036662915 | 0.586351548 | 0.160171814 | 0.145938379 |
| Atxn2 | S | 702 | -0.209403444 | 0.01838831 | -0.219966516 | -0.010563073 | -0.238354826 | 0.052445236 | 0.719687346 | 0.02254561 | 0.903017032 | 0.032972619 |
| Esf1 | S | 657 | 0.015714608 | -0.236441555 | -0.117951716 | -0.133666324 | 0.118489839 | 0.741579011 | 0.002572366 | 0.118033452 | 0.094131297 | 0.117739099 |
| Cep170 | S | 877 | 0.05146643 | -0.167982278 | -0.231857564 | -0.283323995 | -0.063875287 | 0.318290716 | 0.097820809 | 0.001345082 | 0.011284815 | 0.424270711 |
| Incenp | S | 218 | -0.0292013 | -0.083939902 | -0.109147807 | -0.079946508 | -0.025207905 | 0.307098018 | 0.000221544 | 0.03981237 | 0.082780473 | 0.446391025 |
| Cep170b | S | 478 | 0.004279178 | -0.165468339 | -0.210521722 | -0.2148009 | -0.045053384 | 0.774157219 | 0.119755268 | 0.004821341 | 0.012286101 | 0.570879689 |
| Rabep1 | S | 410 | -0.190241193 | -0.198371938 | -0.119930361 | 0.070310832 | 0.078441577 | 0.130029073 | 0.115750631 | 0.376825786 | 0.452058527 | 0.404290382 |
| Cacnb2 | S | 508 | -0.2224021 | 0.01958223 | -0.290922227 | -0.068520127 | -0.310504457 | 0.203410455 | 0.913972511 | 0.020149637 | 0.65377195 | 0.165427099 |
| Htt | S | 398 | -0.068910241 | -0.036301642 | -0.218689107 | -0.149778866 | -0.182387465 | 0.113596678 | 0.521469878 | 0.004958315 | 0.027785765 | 0.031826561 |
| Nav3 | S | 1116 | -0.069008009 | -0.117259998 | -0.256167772 | -0.187159763 | -0.138907775 | 0.280008305 | 0.240215965 | 0.038690928 | 0.101165813 | 0.229645969 |
| 2610507B11Rik | S | 253 | -0.200226784 | -0.11852998 | -0.157846405 | 0.042380379 | -0.039316425 | 0.032627733 | 0.544208581 | 0.178656277 | 0.652289048 | 0.838116144 |
| Tbc1d13 | S | 183 | 0.008260025 | -0.15141449 | -0.211162107 | -0.219422132 | -0.059747617 | 0.873777497 | 0.064108627 | 0.008274087 | 0.014180164 | 0.392594488 |
| Cct4 | S | 18 | -0.03992138 | -0.060854777 | -0.325611817 | -0.285690437 | -0.26475704 | 0.623638974 | 0.550735241 | 0.006042319 | 0.00499549 | 0.051025805 |
| Pkn2 | S | 127 | -0.046701313 | -0.139501124 | -0.135100893 | -0.08839958 | 0.004400231 | 0.209021687 | 0.003391167 | 0.214254533 | 0.369008352 | 0.96015389 |
| Ctps | S | 578 | -0.090885295 | -0.094573261 | -0.171950753 | -0.081065458 | -0.077377492 | 0.187196103 | 0.038626386 | 0.072317588 | 0.337708249 | 0.307119142 |
| Usp36 | S | 918 | 0.07198853 | -0.258293468 | -0.356784433 | -0.428772963 | -0.098490965 | 0.162886649 | 0.016396348 | 0.08090852 | 0.065147461 | 0.508525316 |
| Macf1 | S | 2077 | -0.118181882 | -0.108682072 | -0.097981228 | 0.020200655 | 0.010700844 | 0.193420472 | 0.014395146 | 0.080913278 | 0.796967226 | 0.789828098 |
| Macf1 | S | 2081 | -0.118181882 | -0.108682072 | -0.097981228 | 0.020200655 | 0.010700844 | 0.193420472 | 0.014395146 | 0.080913278 | 0.796967226 | 0.789828098 |
| Srsf7 | S | 254 | 0.005183601 | -0.234088233 | -0.08868642 | -0.09387002 | 0.145401814 | 0.89046289 | 0.001050668 | 0.207905835 | 0.18684152 | 0.076863069 |
| Akap12 | S | 1462 | -0.135909355 | 0.023381598 | -0.349992894 | -0.214083539 | -0.373374492 | 0.224362918 | 0.879744504 | 0.014311612 | 0.027998523 | 0.061657632 |
| Baz1a | T | 1413 | -0.016607321 | -0.196081664 | -0.146810159 | -0.130202837 | 0.049271505 | 0.893773875 | 0.039220848 | 0.016733448 | 0.353450689 | 0.457236631 |
| Tsc1 | S | 595 | -0.135847159 | 0.054576193 | -0.321823197 | -0.185976038 | -0.37639939 | 0.501055033 | 0.704894599 | 0.001500619 | 0.374410713 | 0.0787532 |
| Gga2 | S | 418 | 0.150148707 | -0.516445772 | -0.169414723 | -0.31956343 | 0.347031049 | 0.328882846 | 0.009778016 | 0.260664435 | 0.011776272 | 0.00727747 |
| Spag5 | S | 229 | 0.034871464 | -0.198442547 | -0.195478532 | -0.230349996 | 0.002964016 | 0.111163613 | 0.012483971 | 0.009115092 | 0.011923466 | 0.951108328 |
| Pafah1b2 | S | 64 | -0.000992408 | -0.12932625 | -0.235830984 | -0.234838576 | -0.106504734 | 0.986525974 | 0.150766927 | 0.00509565 | 0.005653171 | 0.202914598 |
| Eif3b | S | 79 | -0.006095526 | -0.094254434 | -0.109940217 | -0.103844692 | -0.015685784 | 0.816269071 | 0.002324445 | 0.002126183 | 0.01183121 | 0.388252097 |
| Hjurp A730008H23Ril | S | 388 | 0.114660125 | -0.195186994 | -0.274489013 | -0.389149138 | -0.079302019 | 0.000947561 | 0.001998456 | 0.000752858 | 0.001424653 | 0.084389089 |
| Kars | S | 393 | 0.111680987 | -0.083282537 | -0.342033176 | -0.453714163 | -0.258750639 | 0.060560668 | 0.181380882 | 8.06372E-05 | 0.005261307 | 0.024402119 |
| Morc2a | S | 777 | -0.002919462 | -0.099528065 | -0.11824255 | -0.115323089 | -0.018714485 | 0.914230067 | 0.004093651 | 0.002007725 | 0.006748915 | 0.384867206 |
| Sec16a | S | 1995 | -0.046452107 | -0.136531265 | -0.176447993 | -0.129995886 | -0.039916728 | 0.316210934 | 0.012198784 | 0.125011218 | 0.217211382 | 0.652936819 |
| Hspbp1 | S | 55 | -0.052018107 | -0.135061833 | -0.247759784 | -0.195741678 | -0.112697951 | 0.62639244 | 0.488576527 | 0.005017847 | 0.134473829 | 0.554735956 |
| Ctdspl2 | S | 134 | -0.019159482 | -0.123831484 | -0.238889036 | -0.219729554 | -0.115057552 | 0.636008304 | 0.040894958 | 0.036637086 | 0.062740031 | 0.196705061 |
| Map2k1 Map2k2 | S | 222;226 | -0.144682824 | -0.09846277 | -0.186549984 | -0.04186716 | -0.088087214 | 0.060988424 | 0.322706887 | 0.110622716 | 0.657722912 | 0.445542023 |
| Nefm | S | 44 | 0.021500438 | -0.195337995 | -0.148990036 | -0.170490474 | 0.046347959 | 0.636009818 | 0.003348095 | 0.056607524 | 0.047357922 | 0.419214843 |
| Syn1 | S | 67 | -0.024391791 | -0.057861764 | -0.181686488 | -0.157294697 | -0.123824724 | 0.844693144 | 0.071857703 | 0.000643626 | 0.282772774 | 0.000298914 |
| Dab2ip | S | 702 | -0.041015851 | -0.163443664 | -0.102604637 | -0.061588785 | 0.060839028 | 0.412394214 | 0.005204429 | 0.094811571 | 0.261582931 | 0.229776878 |
| Ube3a | S | 212 | -0.164922385 | -0.06008411 | -0.157686952 | 0.007235433 | -0.097602842 | 0.07187077 | 0.464817655 | 0.032354918 | 0.921823102 | 0.274545112 |
| Trip12 | S | 1030 | 0.098065824 | -0.171870659 | -0.230860346 | -0.32892617 | -0.058989687 | 0.016217446 | 0.001986508 | 0.001756023 | 0.002201913 | 0.187669105 |
| Prrc2b | S | 166 | 0.014872652 | -0.292214174 | -0.287349373 | -0.302222025 | 0.004864801 | 0.949000752 | 0.043125519 | 0.202399604 | 0.294543978 | 0.979526951 |
| Ric8a | T | 440 | -0.053617854 | -0.098734492 | -0.227140523 | -0.173522668 | -0.128406031 | 0.493286521 | 0.190889821 | 0.011371293 | 0.059332255 | 0.093707852 |
| Dennd5a | S | 1096 | -0.172599072 | -0.038191653 | -0.166765048 | 0.005834023 | -0.128573396 | 0.006249575 | 0.593118427 | 0.185347049 | 0.953538863 | 0.290569869 |
| N4bp1 | S | 287 | -0.080060399 | -0.085726744 | -0.309041645 | -0.228981246 | -0.223314901 | 0.374261178 | 0.291399927 | 0.038911715 | 0.097788178 | 0.097345855 |
| Fra10ac1 | S | 285 | -0.039541437 | -0.110310064 | -0.195931732 | -0.156390295 | -0.085621667 | 0.340416393 | 0.105122097 | 0.015685467 | 0.029754836 | 0.196920355 |
| Tprn | S | 274 | -0.060709543 | -0.239754291 | -0.097397324 | -0.03668778 | 0.142356967 | 0.577652055 | 0.006376427 | 0.3415018 | 0.769274847 | 0.198877164 |
| C530008M17Rik | S | 992 | -0.060740565 | -0.221829087 | -0.07627316 | -0.015532595 | 0.145555927 | 0.452456603 | 0.012289603 | 0.123288108 | 0.83552105 | 0.057306915 |
| Cdca2 | S | 390 | 0.007581487 | -0.171895657 | -0.224962454 | -0.23254394 | -0.053066797 | 0.900539335 | 0.080975653 | 0.014312039 | 0.026441457 | 0.480233106 |
| Eif3d | S | 528 | -0.013322506 | -0.122071139 | -0.129287474 | -0.115964968 | -0.007216335 | 0.654864495 | 0.005770791 | 0.014809045 | 0.025163718 | 0.804508896 |
| Mindy2 | S | 99 | -0.05143033 | -0.148807234 | -0.226578723 | -0.175148393 | -0.077771489 | 0.612542284 | 0.038032598 | 0.083532435 | 0.1932748 | 0.440263333 |
| Satb1 | T | 48 | 0.007018078 | -0.155753969 | -0.342531611 | -0.349549689 | -0.186777642 | 0.908424006 | 0.210470803 | 0.022668357 | 0.025785874 | 0.183775104 |
| Apbb1ip | S | 623 | -0.031378215 | -0.21284635 | -0.138094661 | -0.106716447 | 0.074751688 | 0.734271964 | 0.007356449 | 0.182536107 | 0.359571835 | 0.421086714 |
| Traf3ip1 | S | 177 | -0.202278401 | -0.078324725 | -0.206250121 | -0.00397172 | -0.127925395 | 0.03974495 | 0.373510463 | 0.269685896 | 0.979585706 | 0.450464986 |

| | | | | | | | | | | | | |
|---|---|---|---|---|---|---|---|---|---|---|---|---|
| Ccdc43 | T | 137 | -0.11816983 | -0.108428343 | -0.053692703 | 0.064477127 | 0.05473564 | 0.015904302 | 0.006345268 | 0.603835544 | 0.53987219 | 0.594593858 |
| Kif3b | S | 681 | -0.117718053 | -0.151248262 | -0.147581263 | -0.02986321 | 0.003666999 | 0.133809422 | 0.075433252 | 0.192643523 | 0.731296863 | 0.966936308 |
| Med1 | S | 988 | 0.044526259 | -0.090991437 | -0.47873381 | -0.523260069 | -0.387742373 | 0.597459417 | 0.276304805 | 0.01533988 | 0.016821653 | 0.039048305 |
| Arnt | S | 45 | -0.024584825 | -0.058019353 | -0.199919773 | -0.175334948 | -0.14190042 | 0.476030631 | 0.312349867 | 0.00066398 | 0.003440865 | 0.058196216 |
| Rab3gap1 | S | 536 | -0.015899845 | -0.098869676 | -0.147017608 | -0.131117762 | -0.048147932 | 0.70019258 | 0.027559535 | 0.002681553 | 0.022624513 | 0.148196003 |
| Map4k4 | S | 849 | -0.015917582 | -0.095797253 | -0.141394204 | -0.125476622 | -0.045596951 | 0.826199473 | 0.008058169 | 0.004772185 | 0.167573449 | 0.166183715 |
| Eif4a3 Eif4a3l2 | T | 163 | -0.106818048 | -0.051571455 | -0.166068293 | -0.059250245 | -0.114496838 | 0.082595625 | 0.299061661 | 0.014655421 | 0.312011707 | 0.07685441 |
| Rell1 | S | 248 | -0.063262403 | -0.086729679 | -0.246926178 | -0.183663774 | -0.160196498 | 0.4182308 | 0.261530388 | 0.013963158 | 0.016827204 | 0.024722318 |
| Lrrc47 | S | 519 | -0.102677343 | -0.071146323 | -0.175298145 | -0.072620803 | -0.104151822 | 0.057307932 | 0.183477517 | 0.055003846 | 0.314982981 | 0.186172408 |
| Cdt1 | S | 30 | -0.099040987 | -0.278344963 | -0.177304379 | -0.078263392 | 0.101040584 | 0.317735018 | 0.137081035 | 0.2157188 | 0.519642128 | 0.55683615 |
| Cdt1 | T | 28 | -0.099040987 | -0.278344963 | -0.177304379 | -0.078263392 | 0.101040584 | 0.317735018 | 0.137081035 | 0.2157188 | 0.519642128 | 0.55683615 |
| Rrp8 | S | 177 | -0.035390401 | -0.149346723 | -0.194465147 | -0.159074746 | -0.045118423 | 0.725329181 | 0.081880937 | 0.01800684 | 0.186648392 | 0.544407978 |
| Ccdc86 | S | 225 | 0.001505546 | -0.226076676 | -0.151811605 | -0.15331715 | 0.074265072 | 0.930441088 | 0.027283362 | 0.036237987 | 0.051674725 | 0.277003926 |
| Larp1 | S | 68 | -0.169273333 | -0.190123607 | -0.028125928 | 0.141147405 | 0.161997679 | 0.003480197 | 0.186887817 | 0.887322211 | 0.501599859 | 0.474047719 |
| Ncl | S | 303 | -0.04797608 | -0.178968923 | -0.110156969 | -0.06218089 | 0.068811953 | 0.550285156 | 0.011257021 | 0.073589801 | 0.445301593 | 0.227094167 |
| Incenp | T | 269 | -0.090668234 | -0.076171741 | -0.217864209 | -0.127195975 | -0.141692467 | 0.131462478 | 0.363869035 | 0.025327174 | 0.119515398 | 0.146625696 |
| Zdhhc5 | S | 621 | 0.038819591 | -0.246091528 | -0.114621842 | -0.153441433 | 0.131469686 | 0.141744454 | 0.00027258 | 0.164403577 | 0.109043623 | 0.129623536 |
| Ddx55 | S | 544 | 0.008237502 | -0.171749733 | -0.081045999 | -0.089283502 | 0.090703733 | 0.855157297 | 0.002796242 | 0.007277181 | 0.129854584 | 0.046928348 |
| Eif3b | S | 90 | -0.022243753 | -0.142400414 | -0.081679472 | -0.059435719 | 0.060720942 | 0.415937802 | 0.004157224 | 0.016968612 | 0.092942593 | 0.082591124 |
| Fyb | S | 203 | -0.329566303 | -0.174999852 | 0.057849265 | 0.387415568 | 0.232849118 | 0.00563208 | 0.239184593 | 0.476875756 | 0.004470074 | 0.14678371 |
| Pacs2 | S | 319 | -0.06860059 | -0.075870373 | -0.172950749 | -0.104350159 | -0.097080376 | 0.425696731 | 0.090938291 | 0.006645332 | 0.258344964 | 0.060903171 |
| Map6 | T | 634 | -0.105205889 | -0.104698456 | -0.185098583 | -0.079892694 | -0.080400127 | 0.05845719 | 0.34042438 | 0.062111001 | 0.301990886 | 0.478580344 |
| Ctnnbl1 | S | 389 | -0.044107105 | -0.002823579 | -0.218922982 | -0.174815877 | -0.216099403 | 0.25258111 | 0.963365249 | 0.000264206 | 0.00543984 | 0.043306356 |
| Spag9 | S | 730 | -0.082173306 | -0.056116151 | -0.201877642 | -0.119704335 | -0.145761491 | 0.299111054 | 0.437440793 | 0.003493338 | 0.16790606 | 0.103014071 |
| Fra10ac1 | S | 250 | -0.152325648 | -0.064607564 | -0.148235865 | 0.004089783 | -0.0836283 | 0.014467856 | 0.373441546 | 0.136552822 | 0.958864846 | 0.386151472 |
| Picalm | S | 303 | -0.065766792 | -0.232484504 | -0.052773703 | 0.012993089 | 0.179710802 | 0.421073172 | 0.006191315 | 0.244081378 | 0.86635271 | 0.021354041 |
| Cep63 | S | 304 | -0.0702532 | -0.18034849 | -0.4017673 | -0.331514099 | -0.22141881 | 0.692991205 | 0.419816054 | 0.066930983 | 0.099561425 | 0.317813986 |
| Srsf10 | T | 92 | -0.002052275 | -0.236284552 | -0.038495229 | -0.036442953 | 0.197789323 | 0.974270842 | 0.0001562 | 0.548767278 | 0.642532101 | 0.04828329 |
| Phlpp1 | S | 1488 | -0.832562445 | -0.263030433 | NA | NA | NA | 0.172507512 | 0.564239469 | NA | NA | NA |
| Phlpp1 | S | 1493 | -0.832562445 | -0.263030433 | NA | NA | NA | 0.172507512 | 0.564239469 | NA | NA | NA |
| Papola | S | 654 | -0.132311586 | -0.105610503 | -0.108970791 | 0.023340795 | -0.003360288 | 0.033342684 | 0.168723871 | 0.107622821 | 0.693109136 | 0.962894747 |
| Gpr39 | S | 397 | -0.068971858 | -0.032828876 | -0.707636881 | -0.638665023 | -0.674808005 | 0.409650025 | 0.864835474 | 0.098835043 | 0.116058743 | 0.096057622 |
| Usp15 | S | 961 | -0.075586511 | -0.077694281 | -0.089350557 | -0.013764046 | -0.011656276 | 0.057348467 | 0.003882033 | 0.080024921 | 0.742768417 | 0.744714026 |
| Mcl1 | T | 144 | 0.003176693 | -0.128775017 | -0.084182299 | -0.087358992 | 0.044592718 | 0.90740695 | 0.000649754 | 0.00750144 | 0.020024817 | 0.045113749 |
| Erf | T | 529 | -0.12503716 | -0.229663909 | -0.229563507 | -0.104526347 | 0.000100402 | 0.416571261 | 0.141379477 | 0.21424515 | 0.470133931 | 0.999387328 |
| Ube4b | S | 238 | -0.129686536 | -0.086580401 | -0.120925573 | 0.008760964 | -0.034345171 | 0.067516615 | 0.066046129 | 0.107616156 | 0.889829143 | 0.555987448 |
| Tns3 | S | 769 | -0.086672294 | -0.074890537 | -0.29664954 | -0.209977246 | -0.221759003 | 0.370986228 | 0.556378704 | 0.017918464 | 0.04862592 | 0.118923896 |
| Mga | S | 287 | -0.041477709 | -0.206413284 | -0.181824028 | -0.140346319 | 0.024589256 | 0.633001414 | 0.032439779 | 0.124892908 | 0.249942284 | 0.805994477 |
| Vcpip1 | T | 760 | -0.008565164 | -0.112170909 | -0.168706144 | -0.16014098 | -0.056535234 | 0.717443723 | 0.069933663 | 0.00287109 | 0.011472742 | 0.278597423 |
| Tle3 | S | 240 | -0.047125714 | -0.15935673 | -0.084755625 | -0.037629911 | 0.074601104 | 0.426662351 | 0.017666843 | 0.016302739 | 0.519241625 | 0.109480766 |
| Dffa | S | 110 | -0.14925125 | -0.117082586 | -0.265988945 | -0.116737695 | -0.148906359 | 0.249728548 | 0.184140387 | 0.164626433 | 0.503577663 | 0.371995337 |
| Myo9b | S | 1113 | -0.061986501 | -0.092819421 | -0.169245766 | -0.107259265 | -0.076426345 | 0.267878771 | 0.104441263 | 0.014119054 | 0.072774679 | 0.146620894 |
| Chd1 | S | 1094 | 0.005864634 | -0.065099426 | -0.132109494 | -0.137974128 | -0.067010068 | 0.893048357 | 0.034594672 | 4.49136E-05 | 0.061820469 | 0.0350446 |
| Chd1 | S | 1096 | 0.005864634 | -0.065099426 | -0.132109494 | -0.137974128 | -0.067010068 | 0.893048357 | 0.034594672 | 4.49136E-05 | 0.061820469 | 0.0350446 |
| Chd1 | S | 1098 | 0.005864634 | -0.065099426 | -0.132109494 | -0.137974128 | -0.067010068 | 0.893048357 | 0.034594672 | 4.49136E-05 | 0.061820469 | 0.0350446 |
| Ncl | S | 28 | -0.146496295 | -0.285930512 | 0.005932284 | 0.152428579 | 0.291862796 | 0.12206273 | 0.020797208 | 0.96183731 | 0.243305842 | 0.069692489 |
| Ncl | S | 34 | -0.146496295 | -0.285930512 | 0.005932284 | 0.152428579 | 0.291862796 | 0.12206273 | 0.020797208 | 0.96183731 | 0.243305842 | 0.069692489 |
| Ncl | S | 40 | -0.146496295 | -0.285930512 | 0.005932284 | 0.152428579 | 0.291862796 | 0.12206273 | 0.020797208 | 0.96183731 | 0.243305842 | 0.069692489 |
| Ncl | S | 41 | -0.146496295 | -0.285930512 | 0.005932284 | 0.152428579 | 0.291862796 | 0.12206273 | 0.020797208 | 0.96183731 | 0.243305842 | 0.069692489 |
| Gm14569 | S | 303 | -0.539857646 | -0.333262927 | 0.031281953 | 0.571139599 | 0.36454488 | 0.141671421 | 0.330487503 | 0.943859646 | 0.189094071 | 0.339665241 |
| Gm14569 | S | 308 | -0.539857646 | -0.333262927 | 0.031281953 | 0.571139599 | 0.36454488 | 0.141671421 | 0.330487503 | 0.943859646 | 0.189094071 | 0.339665241 |
| Gm14569 | T | 304 | -0.539857646 | -0.333262927 | 0.031281953 | 0.571139599 | 0.36454488 | 0.141671421 | 0.330487503 | 0.943859646 | 0.189094071 | 0.339665241 |
| Kif5b | S | 945 | -0.049665803 | -0.178083834 | -0.21912217 | -0.169456368 | -0.041038336 | 0.638370227 | 0.128720807 | 0.040434474 | 0.120089935 | 0.653828799 |
| Bud13 | T | 262 | -0.552825422 | -0.239700758 | 0.109691161 | 0.662516582 | 0.349391918 | 0.059877145 | 0.261376458 | 0.635875215 | 0.05906615 | 0.220294777 |
| Rif1 | S | 2121 | -0.012651031 | -0.141034223 | -0.12390025 | -0.11124922 | 0.017133973 | 0.817325093 | 0.003835826 | 0.031170089 | 0.114468045 | 0.674203527 |
| --- | S | 28 | -0.03056328 | -0.106674615 | -0.090874382 | -0.060311102 | 0.015800233 | 0.440985771 | 0.007721409 | 0.003739398 | 0.181539648 | 0.491494466 |
| Mical3 | S | 685 | 0.009720701 | -0.155629936 | -0.182107402 | -0.191828103 | -0.026477466 | 0.855163169 | 0.006906159 | 0.04569641 | 0.045193812 | 0.676588754 |
| H2-D1 H2-L | S | 352;352;162 | -0.171771072 | -0.045986529 | -0.140509888 | 0.031261184 | -0.094523358 | 0.02650223 | 0.561102889 | 0.055304699 | 0.333832165 | 0.190829033 |
| Lmna | S | 403 | 0.090544223 | -0.192153905 | -0.227997638 | -0.318541862 | -0.035843733 | 0.338103972 | 0.00503417 | 0.029707206 | 0.027526946 | 0.586018174 |
| Zdhhc5 | S | 274 | -0.12341952 | -0.152042104 | -0.135387712 | -0.011968192 | 0.016654392 | 0.057391584 | 0.145807934 | 0.216166664 | 0.89407657 | 0.880203906 |
| Polr2c | S | 124 | 0.012617446 | -0.090001754 | -0.091413865 | -0.104031311 | -0.001412112 | 0.737241953 | 0.000143921 | 0.001802899 | 0.070464191 | 0.911710906 |
| Usp39 | S | 24 | -0.083326861 | -0.133558005 | -0.118585312 | -0.035258451 | 0.014972693 | 0.487418105 | 0.011445995 | 0.06304003 | 0.761582368 | 0.759742555 |
| Map1a | S | 1190 | -0.127413791 | -0.054196682 | -0.069234088 | 0.058179703 | -0.015037406 | 0.011375644 | 0.087583791 | 0.040443035 | 0.134189485 | 0.600511064 |
| Herc2 | S | 1589 | -0.064069448 | -0.136703745 | -0.150532749 | -0.0864633 | -0.013829004 | 0.613579818 | 0.008776536 | 0.085841217 | 0.520999842 | 0.829008854 |
| Srrm2 | S | 1380 | -0.091382248 | -0.123092364 | -0.042265845 | 0.049116402 | 0.080826519 | 0.046756289 | 0.001474639 | 0.527455388 | 0.478001438 | 0.274626465 |
| Szrd1 | S | 37 | -0.069408059 | -0.13608587 | -0.212150456 | -0.142742397 | -0.076064586 | 0.146524883 | 0.168218596 | 0.080196116 | 0.177951904 | 0.485450623 |
| Cux1 | S | 427 | -0.017365507 | -0.134872902 | -0.123594472 | -0.106228965 | 0.01127843 | 0.643991586 | 0.026226897 | 0.005570661 | 0.028123554 | 0.78008927 |
| Bclaf1 | T | 339 | -0.035919072 | -0.184651946 | -0.069104267 | -0.033185195 | 0.115547679 | 0.420483312 | 0.003142612 | 0.122251535 | 0.26881224 | 0.003283594 |
| Fxr1 | S | 452 | -0.080070112 | -0.103878698 | -0.082495117 | -0.002425006 | 0.021383581 | 0.296354091 | 0.035241119 | 0.003258391 | 0.970497643 | 0.516120775 |
| Champ1 | S | 441 | -0.034402972 | -0.127079034 | -0.045079572 | -0.0106766 | 0.081999462 | 0.066145726 | 0.000208735 | 0.263200473 | 0.757818135 | 0.095363841 |
| Dync1li2 | S | 383 | 0.030169661 | -0.127960479 | -0.238423718 | -0.26859338 | -0.110463239 | 0.573633963 | 0.02182921 | 0.012887643 | 0.012851208 | 0.139392065 |
| Cdan1 | S | 280 | -0.0294275 | -0.093473812 | -0.23494785 | -0.20552035 | -0.141474039 | 0.676572403 | 0.183913416 | 0.006760835 | 0.057618164 | 0.078140827 |
| Adcy9 | S | 691 | -0.216756778 | 0.086534514 | -0.296217998 | -0.079461219 | -0.382752512 | 0.034576509 | 0.309294364 | 0.023855343 | 0.353766152 | 0.023334659 |
| H1f5 | S | 186 | -0.010752328 | -0.137026564 | -0.240248047 | -0.229495719 | -0.103221483 | 0.888250537 | 0.060023922 | 0.027203425 | 0.039165603 | 0.212614873 |
| Rai14 | S | 516 | -0.171571 | -0.011694821 | -0.192862444 | -0.021291444 | -0.181167623 | 0.017430324 | 0.872534883 | 0.071633653 | 0.791670468 | 0.105119603 |
| Cacnb2 | S | 510 | -0.214451205 | 0.03074017 | -0.2945466 | -0.080095395 | -0.32528677 | 0.162566234 | 0.853550208 | 0.026877046 | 0.564209212 | 0.126967594 |
| Npm1 | S | 70 | -0.053649846 | -0.297914658 | -0.053602124 | 4.77223E-05 | 0.244312534 | 0.505620289 | 0.006236578 | 0.601280722 | 0.999554513 | 0.06862694 |
| Srbd1 | S | 165 | -0.050353065 | -0.187803315 | -0.215010144 | -0.164657079 | -0.02720683 | 0.441663847 | 0.124719002 | 0.069048696 | 0.133154403 | 0.810131667 |
| Klc1 | S | 520 | -0.173910195 | -0.101748483 | -0.094579199 | 0.079330996 | 0.007169284 | 0.041173434 | 0.164806374 | 0.164925607 | 0.290793406 | 0.921387746 |
| Parp3 | S | 107 | -0.10790759 | -0.085759652 | -0.137042364 | -0.029134774 | -0.051282712 | 0.120299843 | 0.192244363 | 0.020207858 | 0.612267783 | 0.403851852 |
| Parp3 | T | 113 | -0.10790759 | -0.085759652 | -0.137042364 | -0.029134774 | -0.051282712 | 0.120299843 | 0.192244363 | 0.020207858 | 0.612267783 | 0.403851852 |
| Usp28 | S | 131 | -0.151948392 | -0.237193958 | -0.036356995 | 0.115591397 | 0.200836963 | 0.140620548 | 0.025182178 | 0.740004249 | 0.267881854 | 0.093872958 |
| Map4k5 | S | 433 | -0.157952137 | 0.064261194 | -0.311106767 | -0.153154631 | -0.375367962 | 0.093117317 | 0.610525402 | 0.014255421 | 0.112127572 | 0.035307385 |
| Pcyt1a | S | 323 | -0.07417527 | -0.070455819 | -0.210434818 | -0.136259548 | -0.139978999 | 0.236817628 | 0.310149087 | 0.011422969 | 0.001170422 | 0.047772147 |
| Bysl | T | 163 | -0.312984938 | -0.110150813 | -0.568560108 | -0.25557517 | -0.458409295 | 0.39852285 | 0.427923377 | 0.379960285 | 0.69115672 | 0.464257367 |
| Cpeb3 | S | 443 | -0.04985689 | -0.226291904 | -0.309077149 | -0.259220259 | -0.082785244 | 0.796158413 | 0.521914281 | 0.020884363 | 0.231930508 | 0.804021832 |
| Sec16a | S | 1368 | -0.102689516 | -0.039055006 | -0.165107498 | -0.062417982 | -0.126052492 | 0.078256775 | 0.502849514 | 0.006771057 | 0.177735515 | 0.078432737 |
| Sipa1 | S | 812 | -0.070743356 | -0.066576775 | -0.136874331 | -0.066130975 | -0.070297555 | 0.217541067 | 0.088216767 | 0.004918195 | 0.228825522 | 0.022851557 |
| Fam120a | T | 653 | -0.073772599 | -0.160132874 | -0.098368906 | -0.024596307 | 0.061763969 | 0.102255054 | 0.03746985 | 0.129903456 | 0.660256944 | 0.357947256 |
| Ubr4 | S | 178 | -0.034010942 | -0.114462903 | -0.184380361 | -0.150369419 | -0.069917459 | 0.220425276 | 0.027383517 | 0.062544248 | 0.109082166 | 0.344218911 |
| Dbnl | T | 279 | -0.274414872 | 0.0589256 | -0.318947161 | -0.044532289 | -0.377872761 | 0.041410938 | 0.678429375 | 0.134599652 | 0.787027449 | 0.098005992 |
| Stim2 | S | 661 | -0.248920104 | -0.016721449 | -0.189314906 | 0.059605198 | -0.172593457 | 0.00538918 | 0.915120854 | 0.421682575 | 0.783639359 | 0.505142185 |
| Pip5k1a | S | 56 | 0.033377197 | -0.144922171 | -0.343050026 | -0.376427223 | -0.198127854 | 0.646522799 | 0.173788579 | 0.015020679 | 0.012143973 | 0.113007186 |
| Dmxl1 | S | 915 | -0.168863307 | -0.141266344 | -0.096186855 | 0.072676453 | 0.045079489 | 0.173256086 | 0.033962185 | 0.284575121 | 0.535125844 | 0.568274721 |

| | | | | | | | | | | | | |
|---|---|---|---|---|---|---|---|---|---|---|---|---|
| Snta1 | S | 187 | 0.022793073 | -0.181957427 | -0.169953476 | -0.192746549 | 0.012003951 | 0.562255456 | 0.001953908 | 0.095698096 | 0.086050377 | 0.870194527 |
| Akap13 | T | 941 | -0.186172925 | -0.176791564 | -0.089733718 | 0.096439207 | 0.087057846 | 0.014591121 | 0.340747556 | 0.477370349 | 0.453396364 | 0.649008093 |
| Ccdc6 | S | 316 | -0.147042131 | -0.032889241 | -0.245498736 | -0.098456605 | -0.212609495 | 0.111096041 | 0.631019196 | 0.039875648 | 0.330672595 | 0.067321081 |
| Cfl1 | S | 24 | -0.138479068 | -0.092228913 | -0.363112304 | -0.224633236 | -0.270883391 | 0.418617997 | 0.673155094 | 0.05017037 | 0.140746934 | 0.224178617 |
| Ptpdc1 | S | 550 | -0.084470902 | -0.185061477 | -0.149660741 | -0.06518984 | 0.035400735 | 0.515094495 | 0.022176984 | 0.18272251 | 0.649066637 | 0.713502165 |
| Pdxdc1 | S | 652 | -0.215477821 | -0.065926562 | -0.315382859 | -0.099905038 | -0.249456297 | 0.192168243 | 0.651625057 | 0.120871136 | 0.598547125 | 0.231373479 |
| Ncl | S | 321 | -0.054389775 | -0.228557908 | -0.151782488 | -0.097392713 | 0.076775419 | 0.627028208 | 0.025077213 | 0.18865601 | 0.46217869 | 0.477263693 |
| Larp1 | T | 762 | -0.000250288 | -0.139270923 | -0.128066865 | -0.127816577 | 0.011204058 | 0.994658306 | 0.009210497 | 0.009632524 | 0.004081986 | 0.674922804 |
| Kdm5c | S | 1353 | -0.019854969 | -0.332751285 | -0.160386079 | -0.140531109 | 0.172365206 | 0.689575132 | 0.051158842 | 0.21142128 | 0.257584022 | 0.244998938 |
| Naa10 | S | 233 | -0.065214581 | -0.108201771 | -0.227909914 | -0.162695333 | -0.119708143 | 0.3531882 | 0.057881945 | 0.077402034 | 0.165849175 | 0.263135717 |
| Ppp1r14b | S | 32 | -0.041639573 | -0.076014933 | -0.218670457 | -0.177030884 | -0.142655524 | 0.502306117 | 0.133949185 | 0.008651464 | 0.043965712 | 0.048246911 |
| Sh2b1 | S | 376 | -0.049737426 | -0.151899434 | -0.244773653 | -0.195036228 | -0.092874219 | 0.528470503 | 0.050638919 | 0.11480562 | 0.180302147 | 0.457163328 |
| Sh3pxd2a | T | 995 | -0.095385084 | -0.077180462 | -0.359867437 | -0.264482353 | -0.282686975 | 0.494649504 | 0.376700201 | 0.047965463 | 0.153354627 | 0.088568197 |
| Anln | S | 531 | -0.085729627 | -0.221037416 | -0.048560836 | 0.037168791 | 0.17247658 | 0.066704225 | 0.029396527 | 0.388343536 | 0.521717192 | 0.051983526 |
| Amotl1 | S | 334 | 0.040097851 | -0.101906666 | -0.228829435 | -0.268927286 | -0.12692277 | 0.40964193 | 0.122830888 | 0.000582376 | 0.003654878 | 0.078998041 |
| Psmd9 | S | 128 | -0.136785387 | -0.038396067 | -0.161867156 | -0.025081769 | -0.12347109 | 0.050521831 | 0.652676809 | 0.017098711 | 0.54437144 | 0.195386417 |
| Lin54 | S | 310 | -0.033341313 | -0.145231054 | -0.072933401 | -0.039592088 | 0.072297652 | 0.225562615 | 0.013129272 | 0.013233921 | 0.122891639 | 0.107808689 |
| Usp15 | S | 965 | -0.09246899 | -0.075395795 | -0.07735279 | 0.0151162 | -0.001956996 | 0.097671241 | 0.001508645 | 0.109615969 | 0.754029507 | 0.955234703 |
| Tln1 | S | 1631 | -0.078116934 | -0.111439644 | -0.091110548 | -0.012993614 | 0.020329095 | 0.027815874 | 0.01658553 | 0.219456705 | 0.832999088 | 0.750773708 |
| Jpt1 | S | 77 | -0.21444369 | 0.037122422 | -0.456078448 | -0.241634758 | -0.49320087 | 0.161484713 | 0.812590361 | 0.09384346 | 0.296952913 | 0.083405062 |
| Toe1 | S | 288 | -0.001299999 | -0.136214226 | -0.195891736 | -0.194591737 | -0.05967751 | 0.982263726 | 0.029422955 | 0.019474625 | 0.032445409 | 0.337213895 |
| Orc1 | S | 197 | 0.029247103 | -0.202518431 | -0.219650765 | -0.248897868 | -0.017132334 | 0.54108877 | 0.047663316 | 0.020358265 | 0.016546635 | 0.829880251 |
| Rsrc1 | S | 59 | -0.10358954 | -0.100100929 | -0.03150328 | 0.072086259 | 0.068597649 | 0.023755652 | 0.001811432 | 0.452752614 | 0.155090745 | 0.161610424 |
| Ubtf | S | 484 | -0.01043384 | -0.123498289 | -0.143043693 | -0.132609852 | -0.019545403 | 0.623851582 | 0.001811727 | 0.066837061 | 0.08772439 | 0.709755029 |
| Smn1 | T | 84 | 0.012984758 | -0.089059305 | -0.240682677 | -0.253667434 | -0.151623372 | 0.75484595 | 0.134662035 | 0.002213194 | 0.011825607 | 0.031292087 |
| Cbl | S | 722 | -0.04973094 | -0.110439048 | -0.324039972 | -0.274309032 | -0.213600924 | 0.780663573 | 0.37481301 | 0.018805508 | 0.178189594 | 0.083954667 |
| Thrap3 | S | 187 | -0.112231742 | -0.054079012 | -0.192972139 | -0.080740397 | -0.138893127 | 0.159946529 | 0.437894045 | 0.013690806 | 0.24546375 | 0.061662489 |
| Wdr20rt | S | 432 | -0.057426828 | -0.124353273 | -0.216863247 | -0.159436419 | -0.092509974 | 0.203775959 | 0.080880953 | 0.095747547 | 0.161409418 | 0.366156193 |
| Chaf1a | S | 749 | -0.006255242 | -0.101307411 | -0.185736555 | -0.179481313 | -0.084429144 | 0.855813316 | 0.107939755 | 0.002147539 | 0.008260032 | 0.153651934 |
| Srrm1 | S | 500 | 0.10446895 | -0.317134873 | -0.250005009 | -0.354473959 | 0.067129864 | 0.520246233 | 0.053507154 | 0.047807321 | 0.081851379 | 0.551487724 |
| Pdxdc1 | T | 687 | 0.019323419 | -0.048348524 | -0.219055774 | -0.238379193 | -0.17070725 | 0.653955652 | 0.290422957 | 0.000102248 | 0.011042895 | 0.022679046 |
| Pkm | S | 100 | -0.141929859 | -0.074908856 | -0.173182294 | -0.031252434 | -0.098273438 | 0.107229246 | 0.341265639 | 0.047757205 | 0.678816514 | 0.221821142 |
| Tmem209 | S | 265 | 0.047558396 | -0.281796388 | -0.192770565 | -0.24032896 | 0.089025824 | 0.517897537 | 0.028324989 | 0.055322705 | 0.033665142 | 0.383821207 |
| Col4a3bp | S | 373 | -0.292419936 | -0.039379898 | -0.060603543 | 0.231816393 | -0.021223645 | 0.006791819 | 0.73846324 | 0.342683027 | 0.025455562 | 0.845743769 |
| Pcm1 | S | 988 | -0.069605792 | -0.289693181 | -0.11992688 | -0.050321088 | 0.169766301 | 0.521628729 | 0.034216064 | 0.318438697 | 0.645189726 | 0.182043304 |
| Cbl | S | 450 | -0.066664561 | -0.162107681 | -0.186513532 | -0.119848971 | -0.024405851 | 0.34275135 | 0.069178809 | 0.106433495 | 0.246075036 | 0.800670511 |
| Tjap1 | S | 527 | -0.127408081 | -0.022259423 | -0.1891854 | -0.061777319 | -0.166925977 | 0.071562241 | 0.643931442 | 0.014825986 | 0.34730416 | 0.03263633 |
| Prrc2a | S | 342 | -0.067791978 | -0.098640168 | -0.128924676 | -0.061132698 | -0.030284507 | 0.023678249 | 0.157784109 | 0.041522265 | 0.22024774 | 0.636263197 |
| Prrc2a | S | 350 | -0.067791978 | -0.098640168 | -0.128924676 | -0.061132698 | -0.030284507 | 0.023678249 | 0.157784109 | 0.041522265 | 0.22024774 | 0.636263197 |
| Bud13 | S | 122 | -0.015377168 | -0.126244182 | -0.117312246 | -0.101935078 | 0.008931936 | 0.719661867 | 0.012405803 | 0.007765698 | 0.075047925 | 0.796703779 |
| Crtc3 | S | 370 | -0.099231121 | -0.161532418 | -0.044831364 | 0.054399757 | 0.116701054 | 0.316537158 | 0.002733463 | 0.231573842 | 0.546534719 | 0.017188025 |
| Sh2b3 | S | 102 | -0.082659622 | -0.070234364 | -0.167130594 | -0.084470973 | -0.09689623 | 0.158203473 | 0.206599053 | 0.013750301 | 0.109661135 | 0.067235235 |
| Slc9a1 | S | 776 | 0.038911474 | -0.176782667 | -0.175346961 | -0.214258435 | 0.001435706 | 0.298105432 | 0.019320294 | 0.005748022 | 0.004810714 | 0.975781061 |
| Ilf3 | S | 73 | -0.07837508 | -0.058198019 | -0.21936678 | -0.1409917 | -0.161168761 | 0.315701349 | 0.602141074 | 0.004154949 | 0.111097136 | 0.211912053 |
| Eif3b | S | 143 | 0.02309117 | -0.123540436 | -0.131068554 | -0.154159724 | -0.007528118 | 0.440367718 | 0.005446827 | 0.001951293 | 0.001956777 | 0.746615164 |
| Map1a | S | 1580 | -0.096464352 | -0.060763269 | -0.162744526 | -0.066280174 | -0.101981256 | 0.013962971 | 0.221171396 | 0.091202756 | 0.360485797 | 0.2115815 |
| Nrbp1 | S | 422 | 0.044132467 | -0.126375654 | -0.343101404 | -0.387233871 | -0.216725751 | 0.638330638 | 0.022654042 | 0.048822272 | 0.033949991 | 0.144086613 |
| Ddx51 | S | 432 | -0.042942745 | -0.162608115 | -0.172370949 | -0.129428204 | -0.009762834 | 0.439761247 | 0.222264141 | 0.010474056 | 0.02313561 | 0.928349186 |
| Ip6k1 | S | 127 | -0.293596271 | -0.093146701 | -0.060790567 | 0.232805704 | 0.032356135 | 0.021697323 | 0.453561628 | 0.397513677 | 0.055826879 | 0.770881788 |
| Snx5 | S | 22 | -0.007294125 | -0.128595422 | -0.268831921 | -0.261537796 | -0.140236499 | 0.945122374 | 0.105901282 | 0.022335135 | 0.076182346 | 0.15195905 |
| Srgap2 | S | 994 | 0.014555249 | -0.187713662 | -0.125351065 | -0.139906314 | 0.062362597 | 0.553346195 | 0.025778017 | 0.004013734 | 0.005269176 | 0.254265139 |
| Eif5b | S | 584 | -0.068583038 | -0.13955701 | -0.178000299 | -0.109417261 | -0.038443289 | 0.582668257 | 0.046164881 | 0.054500671 | 0.409291646 | 0.585970663 |
| Eif5b | S | 585 | -0.068583038 | -0.13955701 | -0.178000299 | -0.109417261 | -0.038443289 | 0.582668257 | 0.046164881 | 0.054500671 | 0.409291646 | 0.585970663 |
| Eif5b | S | 587 | -0.068583038 | -0.13955701 | -0.178000299 | -0.109417261 | -0.038443289 | 0.582668257 | 0.046164881 | 0.054500671 | 0.409291646 | 0.585970663 |
| Eif5b | S | 591 | -0.068583038 | -0.13955701 | -0.178000299 | -0.109417261 | -0.038443289 | 0.582668257 | 0.046164881 | 0.054500671 | 0.409291646 | 0.585970663 |
| Pnkp | S | 142 | -0.07539046 | -0.12977386 | -0.200720337 | -0.125329877 | -0.070946477 | 0.237387499 | 0.059702918 | 0.135848129 | 0.291517105 | 0.51772448 |
| Tor1aip1 | Y | 243 | -0.019298672 | -0.197047132 | -0.316585714 | -0.297287042 | -0.119538582 | 0.862462007 | 0.1098452 | 0.100689166 | 0.121557699 | 0.424442084 |
| Bet1 | S | 50 | -0.225770584 | -0.029337508 | -0.065798198 | 0.159972386 | -0.03646069 | 0.000670375 | 0.822212444 | 0.528991214 | 0.196683918 | 0.810677189 |
| Utp14a | S | 37 | 0.089743674 | -0.230544432 | -0.740368642 | -0.830112316 | -0.50982421 | 0.844820078 | 0.562687293 | 0.093911287 | 0.071948876 | 0.036412957 |
| Pds5a | T | 1191 | -0.185048071 | -0.25002335 | -0.082445788 | 0.102602283 | 0.167577562 | 0.34675557 | 0.044613289 | 0.49746462 | 0.603928531 | 0.240189291 |
| Slc12a7 | T | 973 | -0.050709816 | -0.145433235 | -0.08987627 | -0.039166454 | 0.055556964 | 0.244646155 | 0.015447208 | 0.050600886 | 0.165937348 | 0.183216791 |
| Golm1 | Y | 332 | -0.180644646 | -0.12350739 | -0.097384524 | 0.083260122 | 0.026122867 | 0.091623097 | 0.13231592 | 0.182650975 | 0.374891438 | 0.732082168 |
| Nol8 | S | 870 | -0.088383523 | -0.27667931 | -0.096315065 | -0.007931542 | 0.180364245 | 0.469070308 | 0.015668448 | 0.571571422 | 0.962454858 | 0.306584623 |
| Nol8 | S | 872 | -0.088383523 | -0.27667931 | -0.096315065 | -0.007931542 | 0.180364245 | 0.469070308 | 0.015668448 | 0.571571422 | 0.962454858 | 0.306584623 |
| Cux1 | S | 909 | -0.046115736 | -0.0937915 | -0.212753094 | -0.166637358 | -0.118961594 | 0.287689876 | 0.03448653 | 0.065622821 | 0.109473951 | 0.202466408 |
| Larp4b | S | 628 | 0.014034236 | -0.182479104 | -0.160124655 | -0.174158891 | 0.022354449 | 0.805001002 | 0.018686832 | 0.025648706 | 0.025458039 | 0.695330511 |
| Fbxo4 | S | 11 | -0.174370769 | -0.031741878 | -0.174915973 | -0.000545204 | -0.143174095 | 0.075348753 | 0.667780064 | 0.031517503 | 0.994561339 | 0.12072315 |
| Dvl2 | S | 616 | -0.171101064 | -0.028205541 | -0.220457258 | -0.049356193 | -0.192251716 | 0.013390394 | 0.845708353 | 0.161205282 | 0.691964086 | 0.304102361 |
| Kmt5b | S | 427 | -0.046540444 | -0.113205633 | -0.092183605 | -0.045643161 | 0.021022028 | 0.094701836 | 0.014067958 | 0.044753594 | 0.218754896 | 0.580127705 |
| Prepl | S | 137 | -0.080856579 | -0.066795743 | -0.216697901 | -0.135841322 | -0.149902158 | 0.297098616 | 0.315573426 | 0.013056596 | 0.066496737 | 0.002307716 |
| Kank2 | T | 151 | 0.040442216 | -0.186301457 | -0.373801497 | -0.414243713 | -0.18750004 | 0.735787345 | 0.145791576 | 0.046222292 | 0.039998637 | 0.20160033 |
| Ddx10 | T | 587 | 0.03527988 | -0.148621332 | -0.128132493 | -0.163412373 | 0.020488839 | 0.516452516 | 0.004421649 | 0.005660291 | 0.041957634 | 0.515077568 |
| Arid1a | S | 608 | -0.162668565 | -0.136237067 | -0.178520637 | -0.015852072 | -0.04228357 | 0.241561851 | 0.103397594 | 0.216969609 | 0.915700035 | 0.721492097 |
| Rin1 | S | 198 | -0.083282407 | -0.12830646 | -0.15322614 | -0.069943733 | -0.02491968 | 0.174683527 | 0.070752677 | 0.099799966 | 0.39039472 | 0.751469377 |
| Pum2 | S | 87 | -0.049352347 | -0.192201353 | -0.23495475 | -0.185602403 | -0.042753396 | 0.533437371 | 0.150318499 | 0.075788359 | 0.134909943 | 0.740671153 |
| Ralbp1 | T | 27 | -0.006813448 | -0.092670062 | -0.266555813 | -0.259742365 | -0.173885751 | 0.803599681 | 0.261069851 | 0.006053884 | 0.017552753 | 0.084265582 |
| Itsn1 | S | 315 | 0.101085333 | -0.163052907 | -0.22189377 | -0.322979104 | -0.058840864 | 0.10636322 | 0.010626973 | 0.002032299 | 0.004828689 | 0.115333085 |
| Map6 | S | 905 | -0.0818793 | -0.086373557 | -0.18803547 | -0.10615617 | -0.101661912 | 0.162721522 | 0.069891398 | 0.078148915 | 0.245665746 | 0.258481042 |
| Tnks1bp1 | S | 1545 | -0.058866144 | -0.100977115 | -0.128237305 | -0.069371161 | -0.027260189 | 0.174855323 | 0.025421948 | 0.046195766 | 0.198325792 | 0.550344828 |
| Phlpp2 | S | 1207 | -0.036921043 | -0.134013274 | -0.10466012 | -0.067739076 | 0.029353154 | 0.234478197 | 0.014802791 | 0.042830796 | 0.133289427 | 0.508015067 |
| Tacc2 | S | 552 | -0.081546313 | -0.127289364 | -0.228269839 | -0.146723526 | -0.100980476 | 0.426127712 | 0.052376029 | 0.131487185 | 0.310643602 | 0.410772031 |
| Spp1 | S | 26 | 0.015444377 | -0.148819846 | -0.144230178 | -0.159674555 | 0.004589668 | 0.80051284 | 0.000211636 | 0.185600027 | 0.160668581 | 0.956201814 |
| Sipa1l1 | T | 1530 | -0.146046989 | -0.266895559 | -0.119410179 | 0.02663681 | 0.14748538 | 0.4509417 | 0.125701233 | 0.15566591 | 0.877801266 | 0.325582486 |
| Ercc5 | S | 704 | -0.12664264 | -0.033663318 | -0.115807943 | 0.010834697 | -0.082144625 | 0.01014709 | 0.572646094 | 0.024173055 | 0.725310687 | 0.201409105 |
| Srrm2 | S | 1414 | -0.061411553 | -0.170115873 | -0.115553525 | -0.054141972 | 0.054562348 | 0.086968164 | 0.055652626 | 0.151255659 | 0.423969983 | 0.505661644 |
| Mtdh | S | 179 | 0.006835756 | -0.218962233 | -0.213162602 | -0.219998359 | 0.005799631 | 0.938309256 | 0.010750849 | 0.206766641 | 0.20289141 | 0.96658653 |
| Thrap3 | S | 182 | -0.05520601 | -0.140809275 | -0.124572281 | -0.069366271 | 0.016236994 | 0.301631815 | 0.073121906 | 0.021825551 | 0.157061625 | 0.777135647 |
| Prrc2a | T | 609 | -0.015650178 | -0.137238096 | -0.06163676 | -0.045986582 | 0.075601337 | 0.457011285 | 0.00068945 | 0.040769221 | 0.08629852 | 0.030963147 |
| Erf | T | 193 | -0.156293584 | -0.064547992 | -0.142304967 | 0.013988616 | -0.077756976 | 0.157608008 | 0.285539255 | 0.02316794 | 0.870048861 | 0.132605152 |
| Rrp1 | S | 439 | 0.009055221 | -0.078799444 | -0.263343637 | -0.272398858 | -0.184544193 | 0.88960466 | 0.305499652 | 0.001990116 | 0.0177221 | 0.063754994 |
| Nes | S | 565 | 0.031190029 | -0.265042894 | -0.166857102 | -0.198047131 | 0.098185792 | 0.7712214 | 0.021601227 | 0.089599904 | 0.091728607 | 0.223200774 |
| Nes | T | 556 | 0.031190029 | -0.265042894 | -0.166857102 | -0.198047131 | 0.098185792 | 0.7712214 | 0.021601227 | 0.089599904 | 0.091728607 | 0.223200774 |
| Hdac4 Hdac5 Hdac9 | S | 245;220;250 | -0.196824652 | -0.083778267 | -0.081983435 | 0.114841217 | 0.001794831 | 0.053459362 | 0.311240849 | 0.059818888 | 0.178451307 | 0.980009095 |
| Fanci | S | 555 | -0.096070748 | -0.151670618 | -0.350743225 | -0.254672477 | -0.199072607 | 0.440133206 | 0.263858572 | 0.145882952 | 0.256434387 | 0.3588411 |

| | | | | | | | | | | | | |
|---|---|---|---|---|---|---|---|---|---|---|---|---|
| Arhgap17 | S | 698 | -0.142276401 | -0.106859878 | -0.08560273 | 0.056673671 | 0.021257148 | 0.066961774 | 0.069335466 | 0.1577545 | 0.400417584 | 0.69123025 |
| Hnrnpu | S | 4 | 0.687265518 | -0.283790538 | -0.286733171 | -0.973998689 | -0.002942633 | 0.36470577 | 0.001194203 | 0.030078275 | 0.240136401 | 0.967855568 |
| Mphosph8 | T | 163 | 0.017013512 | -0.159124025 | -0.133439431 | -0.150452943 | 0.025684593 | 0.352688242 | 0.000247831 | 0.116799205 | 0.100949925 | 0.678734505 |
| Kif23 | S | 160 | -0.068420862 | -0.100588437 | -0.136129413 | -0.067708551 | -0.035540976 | 0.037660202 | 0.028340315 | 0.185631842 | 0.440640187 | 0.674171479 |
| 4933427D14Rik | S | 603 | -0.088251778 | -0.16813279 | -0.113795078 | -0.0255433 | 0.054337711 | 0.310955991 | 0.043704988 | 0.106909471 | 0.766730844 | 0.450309717 |
| Akap12 | S | 767 | -0.091544951 | 0.007747517 | -0.305718914 | -0.214173963 | -0.313466431 | 0.421589554 | 0.912839884 | 0.004121978 | 0.112286488 | 0.000486787 |
| Lca5 | S | 90 | -0.219236065 | -0.041182831 | -0.266360412 | -0.047124347 | -0.225177581 | 0.153232949 | 0.737108533 | 0.092180993 | 0.751650113 | 0.153832821 |
| Fxr1 | S | 438 | -0.006633684 | -0.137036498 | -0.123497074 | -0.116863389 | 0.013539424 | 0.932352771 | 0.073943214 | 0.00089243 | 0.224105628 | 0.794100645 |
| Trex1 | S | 78 | -0.070564408 | -0.037920765 | -0.130564136 | -0.059999728 | -0.092643371 | 0.040025716 | 0.35517558 | 0.002827029 | 0.009711421 | 0.065157367 |
| Srrm2 | S | 946 | -0.020651904 | -0.099976959 | -0.08176923 | -0.061117326 | 0.018207729 | 0.320244841 | 0.006859493 | 0.002947092 | 0.01472598 | 0.418857636 |
| Dnaja4 | S | 79 | -0.066830681 | -0.13867164 | -0.496843969 | -0.430013288 | -0.358172329 | 0.814498343 | 0.709069566 | 0.051551437 | 0.2136503 | 0.382336224 |
| Kirrel | Y | 753 | -0.139443424 | -0.175403143 | -0.219726839 | -0.080283415 | -0.044323696 | 0.331411465 | 0.267547842 | 0.125200024 | 0.584015453 | 0.772817795 |
| Anxa2 | S | 89 | -0.207752395 | -0.268134582 | -0.14222756 | 0.065524835 | 0.125907022 | 0.580375948 | 0.028978818 | 0.671592026 | 0.885712797 | 0.707627307 |
| Anxa2 | S | 92 | -0.207752395 | -0.268134582 | -0.14222756 | 0.065524835 | 0.125907022 | 0.580375948 | 0.028978818 | 0.671592026 | 0.885712797 | 0.707627307 |
| Helz | S | 1785 | -0.139768414 | -0.094875087 | -0.01841392 | 0.121354494 | 0.076461167 | 0.012283129 | 0.010690345 | 0.534232788 | 0.030227981 | 0.01453579 |
| Champ1 | S | 276 | 0.035889392 | -0.178791659 | -0.101524549 | -0.137413942 | 0.07726711 | 0.326414457 | 0.00401346 | 0.003717755 | 0.018843572 | 0.080740761 |
| Kdm3a | S | 325 | -0.030543694 | -0.125609033 | -0.117304367 | -0.086760673 | 0.008304666 | 0.472178135 | 0.146336717 | 0.000948351 | 0.106928771 | 0.897959482 |
| Tut7 | S | 1468 | 0.042167612 | -0.144755489 | -0.298564237 | -0.340731849 | -0.153808748 | 0.569935448 | 0.059441952 | 0.023038206 | 0.014469501 | 0.082747777 |
| Rps3a1 | S | 263 | 0.010677814 | -0.103120812 | -0.121776626 | -0.13245444 | -0.018655814 | 0.831527958 | 4.46134E-05 | 0.066799655 | 0.080597902 | 0.649967788 |
| Ddx20 | S | 522 | -0.20080675 | -0.010263108 | -0.21916993 | -0.01836318 | -0.208906822 | 0.040607426 | 0.9317792 | 0.095885907 | 0.853950518 | 0.161734763 |
| Dok1 | Y | 408 | -0.165160191 | -0.084524854 | -0.127479305 | 0.037680886 | -0.042954451 | 0.041195617 | 0.222262469 | 0.185677122 | 0.617932162 | 0.56140727 |
| Jmjd6 | S | 396 | 0.056679142 | -0.161160384 | -0.377930799 | -0.43460994 | -0.216770414 | 0.513503821 | 0.14052814 | 0.025557591 | 0.022411364 | 0.132205249 |
| Atp9b | S | 525 | -0.042252495 | -0.062361673 | -0.518515287 | -0.476262793 | -0.456153614 | 0.850464481 | 0.717282237 | 0.034997906 | 0.111954046 | 0.073278206 |
| Cpne1 | S | 54 | -0.116045197 | -0.048968121 | -0.272490249 | -0.156445052 | -0.223522129 | 0.595230711 | 0.68337836 | 0.006977695 | 0.478696353 | 0.135952287 |
| Arhgap39 | S | 298 | 0.144287341 | -0.230487433 | -0.228997974 | -0.373285315 | 0.001489459 | 0.031008208 | 0.003752724 | 0.004266184 | 0.000686163 | 0.971573333 |
| Amfr | S | 542 | -0.010852471 | -0.233568755 | -0.230900908 | -0.220048438 | 0.002667847 | 0.906218439 | 0.119576111 | 0.059433713 | 0.055925814 | 0.982789488 |
| Txnrd3 | S | 50 | 0.075995663 | -0.079367463 | -0.283602904 | -0.359598567 | -0.204235441 | 0.287531362 | 0.119970799 | 0.000657475 | 0.012540054 | 0.000447577 |
| Pnkp | S | 110 | -0.196002595 | -0.064350124 | -0.162628238 | 0.033374357 | -0.098278114 | 0.031936865 | 0.543382679 | 0.185495874 | 0.737810151 | 0.42891311 |
| Stxbp1 | S | 594 | -0.018997161 | -0.143975959 | -0.208437094 | -0.189439933 | -0.064461135 | 0.903768795 | 0.130147795 | 0.012466319 | 0.2874312 | 0.407015234 |
| Snx17 | S | 336 | -0.021637936 | -0.095616517 | -0.127809711 | -0.106171775 | -0.032193194 | 0.483552989 | 0.023019571 | 0.005505676 | 0.012656471 | 0.302557081 |
| Setd1a | S | 1338 | -0.25163312 | -0.163200481 | -0.10631068 | 0.14532244 | 0.056889801 | 0.152676328 | 0.201387111 | 0.335017374 | 0.339761634 | 0.557156312 |
| Sec62 | S | 117 | -0.058872758 | -0.214209861 | -0.079144503 | -0.020271745 | 0.135065358 | 0.456455278 | 0.013085295 | 0.192091779 | 0.803863286 | 0.081495279 |
| Septin9 | S | 59 | -0.112540756 | -0.047520472 | -0.273617 | -0.161076244 | -0.226096528 | 0.137339359 | 0.63322032 | 0.045379066 | 0.165793606 | 0.103372931 |
| Clip1 | S | 146 | -0.164447773 | -0.159767644 | -0.093980023 | 0.07046775 | 0.065787622 | 0.132303168 | 0.107508474 | 0.239907717 | 0.394353548 | 0.338432317 |
| Zc3h4 | S | 159 | -0.107073201 | -0.146118819 | -0.03034519 | 0.07672801 | 0.115773628 | 0.009174981 | 0.048832539 | 0.399429087 | 0.04518787 | 0.094857157 |
| Plec | T | 119 | -0.209543186 | -0.126488635 | 0.000109921 | 0.209653107 | 0.126598556 | 0.036858209 | 0.018924794 | 0.999157665 | 0.126919971 | 0.284492748 |
| Cep170 | S | 1188 | 0.052816483 | -0.092588716 | -0.21045141 | -0.263267893 | -0.117862695 | 0.321025422 | 0.078474157 | 0.000328846 | 0.01028608 | 0.04536663 |
| Eef1d | S | 133 | -0.021653189 | -0.05555782 | -0.229233136 | -0.207579948 | -0.173675316 | 0.330015558 | 0.070484729 | 0.012746153 | 0.012805729 | 0.018334059 |
| H2bc11 H2bc13 H2bc | S | 39;39;39;39;39;39 | 0.185790196 | -0.254487629 | -0.280337279 | -0.466127475 | -0.02584965 | 0.306364099 | 0.082789156 | 0.005377081 | 0.064593443 | 0.812816088 |
| Als2 | S | 1329 | -0.098782185 | -0.071578289 | -0.11418194 | -0.015399755 | -0.042603651 | 0.054500771 | 0.217675401 | 0.01937991 | 0.700387952 | 0.432134753 |
| Dab2ip | S | 992 | -0.019731174 | -0.088568303 | -0.227105103 | -0.207373929 | -0.138536801 | 0.681526913 | 0.049282581 | 0.021468856 | 0.033533457 | 0.100090908 |
| Dab2ip | S | 995 | -0.019731174 | -0.088568303 | -0.227105103 | -0.207373929 | -0.138536801 | 0.681526913 | 0.049282581 | 0.021468856 | 0.033533457 | 0.100090908 |
| Mapre2 | S | 9 | -0.031940329 | -0.091952639 | -0.167159413 | -0.135219084 | -0.075206774 | 0.683355135 | 0.099519038 | 0.003749119 | 0.161314424 | 0.13711327 |
| Dlg1 | S | 575 | -0.001757913 | -0.099614522 | -0.107361622 | -0.105603709 | -0.0077471 | 0.959201915 | 0.004419589 | 0.002832441 | 0.054250271 | 0.628890488 |
| Senp6 | S | 344 | -0.057785535 | -0.183630467 | -0.364865356 | -0.307079822 | -0.18123489 | 0.681599976 | 0.440830123 | 0.064713648 | 0.098504288 | 0.45241692 |
| Senp6 | S | 355 | -0.057785535 | -0.183630467 | -0.364865356 | -0.307079822 | -0.18123489 | 0.681599976 | 0.440830123 | 0.064713648 | 0.098504288 | 0.45241692 |
| Senp6 | S | 356 | -0.057785535 | -0.183630467 | -0.364865356 | -0.307079822 | -0.18123489 | 0.681599976 | 0.440830123 | 0.064713648 | 0.098504288 | 0.45241692 |
| Msn | S | 468 | -0.094352043 | -0.076949844 | -0.166125603 | -0.07177356 | -0.089175759 | 0.314169699 | 0.228250269 | 0.010687593 | 0.426584378 | 0.187591652 |
| Kif1b | S | 1487 | -0.131694139 | -0.065514279 | -0.19242019 | -0.060726051 | -0.126905911 | 0.154947971 | 0.490455169 | 0.029258888 | 0.453410564 | 0.205411303 |
| Unc5b | S | 492 | -0.150926156 | -0.123153822 | -0.163861443 | -0.012935287 | -0.040707621 | 0.159724406 | 0.23845251 | 0.120814209 | 0.898738408 | 0.693168162 |
| Tob2 | S | 157 | 0.048981863 | -0.607448909 | -0.257991234 | -0.306973098 | 0.349457675 | 0.759968167 | 0.210377996 | 0.168640316 | 0.143923797 | 0.41827621 |
| Rabl6 | T | 526 | -0.144933412 | -0.119686673 | -0.106085634 | 0.038847777 | 0.013601039 | 0.233961304 | 0.081036807 | 0.072694844 | 0.712090252 | 0.819095794 |
| Tcf12 | S | 28 | -0.033443964 | -0.271026262 | -0.146901113 | -0.113457149 | 0.124125149 | 0.831322347 | 0.009479986 | 0.471500401 | 0.625035798 | 0.537832722 |
| Trip12 | T | 1410 | 0.018348446 | -0.096203959 | -0.144754599 | -0.163103045 | -0.04855064 | 0.740096131 | 0.063249729 | 0.000259766 | 0.067799976 | 0.25185656 |
| Mtdh | S | 297 | -0.154858155 | -0.037654021 | -0.284051435 | -0.12919328 | -0.246397414 | 0.190810163 | 0.768161849 | 0.048273922 | 0.222588901 | 0.082054783 |
| Casc4 | S | 365 | -0.030530304 | -0.065781358 | -0.190395842 | -0.159865538 | -0.124614484 | 0.765359406 | 0.20415565 | 0.001382591 | 0.203373995 | 0.050054456 |
| Gapdh | T | 81 | -0.141886764 | -0.078720615 | -0.168038738 | -0.026151974 | -0.089318122 | 0.147206982 | 0.404975255 | 0.037046927 | 0.748422945 | 0.369907557 |
| Rab11fip5 | S | 474 | -0.012459734 | -0.03171247 | -0.28159225 | -0.269132516 | -0.24987978 | 0.773796491 | 0.720401564 | 0.001234573 | 0.004185363 | 0.056957289 |
| Prune2 | S | 583 | -0.007640743 | -0.124026504 | -0.18516042 | -0.177519678 | -0.061133916 | 0.74806034 | 0.054109491 | 0.014923039 | 0.016967027 | 0.272810226 |
| Stk10 | S | 513 | -0.057729698 | -0.12570361 | -0.167215303 | -0.109485605 | -0.041511693 | 0.350154357 | 0.086306127 | 0.041352547 | 0.16225565 | 0.535786088 |
| Gab2 | S | 402 | -0.108121192 | -0.028944528 | -0.197929693 | -0.089808502 | -0.168985165 | 0.043565461 | 0.688231753 | 0.027247871 | 0.197197019 | 0.084587677 |
| Svil | S | 508 | -0.104045246 | -0.16266154 | -0.027988374 | 0.076056872 | 0.134673166 | 0.042917759 | 0.018455006 | 0.437689213 | 0.088102522 | 0.055783359 |
| Cep95 | S | 449 | 0.031560553 | -0.245178245 | -0.269804401 | -0.301364954 | -0.024626156 | 0.696757735 | 0.206900724 | 0.021958554 | 0.028273638 | 0.875611635 |
| Foxk1 | S | 199 | -0.086353059 | -0.163332901 | -0.043898591 | 0.042454467 | 0.11943431 | 0.21999226 | 0.003439139 | 0.381809548 | 0.541739432 | 0.070504133 |
| Ube3a | S | 8 | -0.083620869 | -0.026167647 | -0.208084227 | -0.124463358 | -0.18191658 | 0.144897297 | 0.670118996 | 0.006397872 | 0.012571009 | 0.032410992 |
| Dhx29 | S | 528 | -0.120329731 | -0.201725876 | -0.276484869 | -0.156155139 | -0.074758994 | 0.772417971 | 0.066321528 | 0.257189008 | 0.725508709 | 0.724826017 |
| Camsap1 | S | 1353 | -0.16810838 | -0.009359442 | -0.315035649 | -0.146927269 | -0.305676207 | 0.350321609 | 0.939754229 | 0.023873293 | 0.367547392 | 0.02932729 |
| Champ1 | S | 375 | 0.072305245 | -0.192489824 | -0.106820071 | -0.179125316 | 0.085669753 | 0.260868426 | 9.37451E-05 | 0.092904312 | 0.044140384 | 0.142564433 |
| Peak1 | S | 390 | -0.0017168 | -0.104590754 | -0.118040553 | -0.116323753 | -0.013449798 | 0.927111453 | 0.007144302 | 0.004589434 | 0.005539443 | 0.505776699 |
| Lpin3 | S | 218 | -0.037985716 | -0.081685316 | -0.135418275 | -0.097432559 | -0.05373296 | 0.351974221 | 0.104114773 | 0.002587395 | 0.051137134 | 0.229513209 |
| Zfp280c | S | 76 | 0.048909922 | -0.145944126 | -0.283017335 | -0.331927257 | -0.137073209 | 0.526972968 | 0.216915348 | 0.004698848 | 0.005113284 | 0.230234168 |
| Esyt2 | S | 667 | -0.100054101 | -0.112859988 | -0.048242784 | 0.051811317 | 0.064617204 | 0.062975147 | 0.002838629 | 0.484405293 | 0.476572443 | 0.368377378 |
| Nbeal2 | S | 756 | -0.177992493 | -0.155884417 | -0.12698096 | 0.051011532 | 0.028903457 | 0.205709819 | 0.189236074 | 0.155597608 | 0.686696425 | 0.791636213 |
| Tango6 | S | 787 | -0.029675914 | -0.150636685 | -0.148991678 | -0.119315764 | 0.001645008 | 0.705254951 | 0.015323988 | 0.069237069 | 0.215793267 | 0.979076328 |
| Kif4 | T | 800 | 0.010269011 | -0.045946403 | -0.251282866 | -0.261551877 | -0.205336463 | 0.898644399 | 0.547461498 | 0.000445163 | 0.049876814 | 0.065615095 |
| Sema4b | S | 780 | 0.092769359 | -0.202393352 | -0.309036096 | -0.401805455 | -0.106642744 | 0.229289606 | 0.067378095 | 0.013192788 | 0.006016692 | 0.255857054 |
| Rybp | S | 99 | -0.091042373 | -0.186433521 | -0.035154685 | 0.055887687 | 0.151278836 | 0.125846735 | 0.005518289 | 0.723694033 | 0.591743265 | 0.207097986 |
| Srrm1 | S | 450 | 0.001067452 | -0.16530505 | -0.073664895 | -0.074732347 | 0.091640155 | 0.977443624 | 0.000690445 | 0.078621772 | 0.103249672 | 0.041930992 |
| 4930579G24Rik | S | 18 | -0.236716209 | -0.130275658 | -0.124290053 | 0.112426156 | 0.005985605 | 0.174988458 | 0.209908033 | 0.223996195 | 0.464899452 | 0.948277818 |
| Tor1aip1 | S | 242 | -0.117789147 | -0.212961691 | -0.193025836 | -0.075236689 | 0.019935855 | 0.308995616 | 0.146725288 | 0.255542686 | 0.575643526 | 0.892906961 |
| Caskin2 | S | 940 | 0.025456368 | -0.179837809 | -0.171263709 | -0.196720076 | 0.0085741 | 0.817692856 | 0.034745897 | 0.020060533 | 0.153433132 | 0.865088337 |
| Cdk12 | S | 32 | -0.12569379 | -0.268773745 | -0.055135751 | 0.070558039 | 0.213637993 | 0.385844126 | 0.018924937 | 0.684033109 | 0.664389161 | 0.168985182 |
| Rtn4 | S | 223 | 0.014140898 | -0.184639282 | -0.098646737 | -0.112787635 | 0.085992545 | 0.762425429 | 0.002870777 | 0.048979457 | 0.032192281 | 0.032894116 |
| Ssrp1 | S | 535 | 0.083659451 | -0.279639248 | -0.229486992 | -0.313056443 | 0.050152256 | 0.641257327 | 0.067419141 | 0.043974822 | 0.162510579 | 0.675973147 |
| Il6st | S | 780 | -0.092652094 | -0.103514297 | -0.372706865 | -0.280054771 | -0.269192568 | 0.711731159 | 0.319359248 | 0.068030611 | 0.333425697 | 0.123372392 |
| Araf | S | 162 | 0.01877077 | -0.154472351 | -0.130906454 | -0.149677223 | 0.023565897 | 0.735623117 | 0.005631822 | 0.022819992 | 0.053783395 | 0.581336228 |
| Hnrnpu | S | 58 | -0.17006067 | -0.084504537 | -0.16836122 | 0.00169945 | -0.083856684 | 0.079494627 | 0.38680943 | 0.133306109 | 0.985005477 | 0.415602432 |
| Gprasp1 | S | 1306 | -0.01917218 | -0.076665432 | -0.252316771 | -0.23314459 | -0.175651339 | 0.886109602 | 0.206774125 | 0.006580741 | 0.162012848 | 0.030988225 |
| Trim28 | S | 471 | -0.103057972 | -0.07375736 | -0.171449488 | -0.068391516 | -0.097692128 | 0.133148523 | 0.33585036 | 0.028238746 | 0.0415989 | 0.140501322 |
| Mcph1 | S | 290 | -0.044297119 | -0.069712563 | -0.094590414 | -0.050293294 | -0.024877851 | 0.08753864 | 0.001666578 | 0.054206701 | 0.203381202 | 0.458843787 |
| Nck1 | S | 96 | -0.096185956 | -0.115899278 | -0.144478087 | -0.048292131 | -0.028578809 | 0.060996409 | 0.288820862 | 0.068086129 | 0.428985145 | 0.780069052 |
| Rps6ka3 | S | 369 | -0.109430683 | -0.034310026 | -0.247122026 | -0.137691342 | -0.212811999 | 0.155504806 | 0.666516048 | 0.025630661 | 0.120600534 | 0.044414717 |
| Rps6ka3 | S | 375 | -0.109430683 | -0.034310026 | -0.247122026 | -0.137691342 | -0.212811999 | 0.155504806 | 0.666516048 | 0.025630661 | 0.120600534 | 0.044414717 |
| Lysmd2 | S | 31 | -0.022256664 | -0.114248662 | -0.472211972 | -0.449955308 | -0.35796331 | 0.836638482 | 0.252717901 | 0.092112519 | 0.095592702 | 0.152090553 |

| | | | | | | | | | | | | |
|---|---|---|---|---|---|---|---|---|---|---|---|---|
| Rps6kc1 | S | 493 | 0.045171429 | -0.155925405 | -0.235522999 | -0.280694428 | -0.079597594 | 0.086488562 | 0.002649405 | 0.034585142 | 0.029940531 | 0.272483154 |
| Gpsm1 | S | 490 | -0.161925684 | -0.053549427 | -0.159940232 | 0.001985452 | -0.106390805 | 0.024871221 | 0.521152815 | 0.142050619 | 0.981230619 | 0.319497068 |
| Ddhd2 | S | 447 | -0.010805012 | -0.034138246 | -0.343050617 | -0.332245605 | -0.308912372 | 0.899138206 | 0.722038884 | 0.003896829 | 0.003387579 | 0.016413197 |
| Zc3h13 | S | 64 | -0.012696515 | -0.095838061 | -0.12297489 | -0.110278374 | -0.027136828 | 0.664701457 | 0.015333993 | 0.004567892 | 0.000833895 | 0.18466815 |
| Baiap2 | S | 367 | -0.081311895 | -0.037766356 | -0.193122537 | -0.111810642 | -0.155356181 | 0.287883268 | 0.617583168 | 0.002806464 | 0.173485442 | 0.108021086 |
| Cttn | S | 11 | -0.235440448 | -0.008877688 | -0.268144306 | -0.032703859 | -0.259266618 | 0.086207508 | 0.943341823 | 0.132699124 | 0.827737675 | 0.151163991 |
| Prag1 | T | 208 | -0.014278673 | -0.205664492 | -0.127821013 | -0.11354234 | 0.077843479 | 0.841218707 | 0.003415412 | 0.309524635 | 0.372929062 | 0.503418016 |
| Pml | S | 609 | -0.125352255 | 0.006335343 | -0.154176744 | -0.02882449 | -0.160512087 | 0.003020835 | 0.907532387 | 0.046252829 | 0.575951685 | 0.060593806 |
| Pam | S | 925 | -0.380014092 | 0.132217261 | -0.229893951 | 0.150120141 | -0.362111211 | 0.018140247 | 0.60947264 | 0.284867521 | 0.464606657 | 0.253012466 |
| Kmt2b | S | 1913 | -0.105591831 | -0.187358568 | -0.089959338 | 0.015632493 | 0.09739923 | 0.24636645 | 0.079829234 | 0.113180497 | 0.84671045 | 0.275926444 |
| Arhgef2 | S | 122 | -0.307351071 | -0.082810848 | -0.284242167 | 0.023108904 | -0.201431319 | 0.200959474 | 0.675718911 | 0.24526548 | 0.922805865 | 0.382301802 |
| Slbp | S | 182 | -0.039441715 | -0.086913679 | -0.177985246 | -0.138543531 | -0.091071567 | 0.497877877 | 0.353394844 | 0.002094402 | 0.080907242 | 0.336171905 |
| Ticrr | S | 919 | 0.10339331 | -0.176275148 | -0.223984788 | -0.327378098 | -0.04770964 | 0.06704655 | 0.009098185 | 0.003256929 | 0.00167079 | 0.212190517 |
| Hps1 | S | 282 | 0.004619806 | -0.084815664 | -0.361227367 | -0.365847173 | -0.276411704 | 0.963481814 | 0.291327132 | 0.018191492 | 0.025440181 | 0.062469888 |
| Lmna | S | 414 | -0.045529275 | -0.072974227 | -0.126113928 | -0.080584652 | -0.0531397 | 0.08424918 | 0.038141098 | 0.023138003 | 0.071348662 | 0.185465334 |
| Tfpt | T | 203 | -0.164285869 | -0.223488798 | -0.009247156 | 0.155038713 | 0.214241642 | 0.032267065 | 0.090707093 | 0.910486753 | 0.099935375 | 0.108023231 |
| Srrm2 | S | 202 | 0.021595832 | -0.187731009 | -0.164838431 | -0.186434263 | 0.022892577 | 0.699163002 | 0.096258389 | 0.004880623 | 0.024763071 | 0.781207959 |
| Samd4b | S | 550 | -0.001278056 | -0.100019844 | -0.188618364 | -0.187340308 | -0.08859852 | 0.979042958 | 0.003399353 | 0.050085385 | 0.047910931 | 0.223660957 |
| Prune2 | T | 695 | -0.028643609 | -0.075146933 | -0.150725772 | -0.122082163 | -0.075578839 | 0.376167982 | 0.05646213 | 0.006021782 | 0.02103293 | 0.088686778 |
| Cep170 | S | 1522 | -0.0687989 | -0.05269031 | -0.158103769 | -0.08930487 | -0.10541346 | 0.459381625 | 0.069490385 | 0.005782181 | 0.35745738 | 0.050511875 |
| Wdr62 | T | 1364 | 0.013599331 | -0.086243266 | -0.172893452 | -0.186492783 | -0.086650186 | 0.769242539 | 0.040825602 | 0.001858913 | 0.015738112 | 0.007798436 |
| Strip1 | S | 335 | -0.026260374 | -0.089239425 | -0.059311196 | -0.033050822 | 0.029928229 | 0.220974055 | 0.000235025 | 0.035364063 | 0.193361525 | 0.16584281 |
| Dkc1 | S | 422 | -0.060630187 | -0.168732147 | -0.156007499 | -0.095377312 | 0.012724647 | 0.500696577 | 0.172344578 | 0.025305187 | 0.286612634 | 0.897035609 |
| Ubr4 | S | 625 | -0.062192955 | -0.055831712 | -0.317950832 | -0.255757878 | -0.26211912 | 0.328545464 | 0.407422851 | 0.037969932 | 0.077410173 | 0.07029255 |
| Gng2 | T | 52 | -0.09916526 | -0.173778639 | -0.140877156 | -0.041711896 | 0.032901483 | 0.399211988 | 0.032724785 | 0.262375682 | 0.757804134 | 0.760145833 |
| Bdp1 | S | 491 | -0.011646777 | -0.197276206 | -0.137202447 | -0.12555567 | 0.060073759 | 0.800931051 | 0.074194777 | 0.018676246 | 0.051578373 | 0.464303349 |
| Pcdh18 | S | 1031 | -0.076926571 | -0.036488973 | -0.284533051 | -0.20760648 | -0.248044078 | 0.679607834 | 0.558879248 | 0.007019608 | 0.317505395 | 0.018076786 |
| Stard9 | S | 1164 | -0.18280491 | -0.110032253 | -0.130744503 | 0.052060407 | -0.02071225 | 0.015806209 | 0.481348013 | 0.388614366 | 0.703131422 | 0.909786349 |
| Usp42 | S | 1189 | -0.079930049 | -0.10836395 | -0.107274696 | -0.027344647 | 0.001089254 | 0.406303941 | 0.016777732 | 0.047389079 | 0.76121806 | 0.974312613 |
| Nab2 | S | 162 | -0.003604609 | -0.064729565 | -0.156001169 | -0.15239656 | -0.091271604 | 0.939621196 | 0.071134991 | 0.0005918 | 0.048816111 | 0.031911342 |
| Ctnnd1 | T | 98 | -0.149034613 | 0.033066416 | -0.264254501 | -0.115219888 | -0.297320917 | 0.268470943 | 0.816825366 | 0.007067231 | 0.364348818 | 0.115482698 |
| Mast4 | S | 1444 | -0.142477519 | -0.045400426 | -0.117495287 | 0.024982232 | -0.072094861 | 0.020322578 | 0.454138364 | 0.063395312 | 0.513591902 | 0.219725324 |
| Ptpn12 | S | 588 | -0.079539188 | -0.115812767 | -0.122842588 | -0.043303401 | -0.007029822 | 0.250708989 | 0.047640302 | 0.0658635 | 0.536871569 | 0.89752309 |
| Champ1 | T | 371 | 0.071199208 | -0.192255352 | -0.108711724 | -0.179910932 | 0.083543627 | 0.281024793 | 0.000131201 | 0.106761402 | 0.050337824 | 0.171860509 |
| Msantd4 | T | 100 | 0.059241886 | -0.191068846 | -0.214505402 | -0.273747288 | -0.023436556 | 0.486581059 | 0.004269918 | 0.109514086 | 0.067375812 | 0.812832473 |
| Ppfibp2 | S | 362 | 0.006284879 | -0.207472656 | -0.233542002 | -0.239826881 | -0.026069346 | 0.94783897 | 0.026112425 | 0.162273858 | 0.163956477 | 0.846712174 |
| Hdac5 | S | 650 | -0.278357963 | -0.075473684 | 0.007845792 | 0.286203755 | 0.083319475 | 0.00311456 | 0.396532396 | 0.898979064 | 0.00651636 | 0.346571136 |
| Nbea | S | 1519 | -0.099032585 | -0.071933328 | -0.127677253 | -0.028644668 | -0.055743925 | 0.129951737 | 0.133356546 | 0.028954945 | 0.59508757 | 0.181487322 |
| Tra2a | T | 200 | -0.217683341 | -0.129040164 | 0.080952091 | 0.298635433 | 0.209992256 | 0.005528125 | 0.049799132 | 0.659986879 | 0.191476487 | 0.307067631 |
| Tent4b | S | 586 | -0.225063775 | -0.071232487 | -0.219111252 | 0.005952523 | -0.147878765 | 0.128405395 | 0.627800996 | 0.153838528 | 0.965660161 | 0.352421867 |
| Sap30bp | S | 113 | -0.07267627 | -0.180342297 | -0.057795499 | 0.014880772 | 0.122546798 | 0.162226625 | 0.00509797 | 0.633301315 | 0.898817602 | 0.348774314 |
| Tbc1d1 | T | 590 | -0.079610273 | -0.037859884 | -0.20629753 | -0.126687258 | -0.168437646 | 0.277816553 | 0.654931724 | 0.004498347 | 0.110202166 | 0.110123701 |
| Zeb1 Zeb2 | S | 243;305 | 0.018932451 | -0.120844658 | -0.182995233 | -0.201927684 | -0.062150574 | 0.716818722 | 0.049834882 | 0.005313698 | 0.008093822 | 0.199688508 |
| Erbin | S | 1171 | -0.138256517 | -0.081712656 | -0.211002211 | -0.072745694 | -0.129289555 | 0.19809754 | 0.490340757 | 0.053063831 | 0.440162522 | 0.275779065 |
| Nbr1 | S | 487 | -0.058975436 | -0.007633954 | -0.359539804 | -0.300564368 | -0.35190585 | 0.637772378 | 0.945964456 | 0.007993177 | 0.049173163 | 0.017781034 |
| Rgl1 | S | 629 | -0.403211939 | 0.105309018 | -0.23302397 | 0.170187969 | -0.338332987 | 0.020272749 | 0.397187396 | 0.261378083 | 0.395201426 | 0.146423711 |
| Thrap3 | S | 184 | -0.137612297 | -0.111365343 | -0.119887446 | 0.017724851 | -0.008522102 | 0.08430739 | 0.157815016 | 0.165806079 | 0.811706537 | 0.910678695 |
| Mphosph9 | S | 482 | -0.24950425 | -0.058339675 | -0.198490807 | 0.051013443 | -0.140151131 | 0.099432292 | 0.443326392 | 0.25044056 | 0.764481387 | 0.384742903 |
| Trp53 | S | 312 | -0.045487956 | -0.08876318 | -0.140035429 | -0.094547473 | -0.051272249 | 0.431769781 | 0.056809556 | 0.010313638 | 0.125634021 | 0.06606844 |
| Ppfibp1 | S | 967 | 0.017316451 | -0.134759698 | -0.111165612 | -0.128482063 | 0.023594087 | 0.699291591 | 0.000536455 | 0.052944415 | 0.05892679 | 0.555976263 |
| Ccdc6 | S | 352 | -0.139360165 | -0.251372283 | -0.08123421 | 0.058125955 | 0.170138073 | 0.495036664 | 0.017095486 | 0.683924985 | 0.817781914 | 0.421898059 |
| Phf10 | S | 326 | 0.046141083 | -0.108419182 | -0.225471165 | -0.271612248 | -0.117051982 | 0.307649846 | 0.039612222 | 0.003092911 | 0.003833799 | 0.045298142 |
| Hnrnpc | S | 232 | 0.045559128 | -0.11511147 | -0.123477911 | -0.169037039 | -0.008366441 | 0.019594479 | 0.00080025 | 0.000229671 | 0.003813726 | 0.641735757 |
| Vim | Y | 38 | -0.066641588 | -0.145350658 | -0.167691085 | -0.101049497 | -0.022340426 | 0.501060197 | 0.132746479 | 0.037018784 | 0.288054242 | 0.755435553 |
| Huwe1 | S | 1298 | -0.323118189 | -0.106720761 | 0.176481076 | 0.499599264 | 0.283201837 | 0.001171434 | 0.156234458 | 0.364416139 | 0.077014696 | 0.195558622 |
| Prune2 | S | 696 | -0.01960675 | -0.058131985 | -0.19060122 | -0.170994469 | -0.132469235 | 0.577667191 | 0.269832615 | 0.001387542 | 0.005472829 | 0.04414273 |
| Asxl2 | S | 143 | -0.022435902 | -0.117975684 | -0.139905837 | -0.117469935 | -0.021930153 | 0.319286205 | 0.079530768 | 0.011190845 | 0.036802834 | 0.676341477 |
| Asxl2 | S | 150 | -0.022435902 | -0.117975684 | -0.139905837 | -0.117469935 | -0.021930153 | 0.319286205 | 0.079530768 | 0.011190845 | 0.036802834 | 0.676341477 |
| Kif18b | S | 814 | -0.084166219 | -0.035516507 | -0.129409079 | -0.04524286 | -0.093892572 | 0.031355016 | 0.534452773 | 0.006271169 | 0.107302347 | 0.151905451 |
| Ccdc6 | S | 345 | -0.098549393 | -0.086703626 | -0.174777293 | -0.076227901 | -0.088073667 | 0.323775587 | 0.064724663 | 0.071903147 | 0.472018187 | 0.27313496 |
| Eps8 | S | 25 | -0.15736791 | -0.088691912 | -0.165863914 | -0.008496003 | -0.077172002 | 0.062627908 | 0.598415367 | 0.095108065 | 0.921289342 | 0.654401817 |
| Rabgef1 | T | 389 | -0.132209745 | -0.181856343 | -0.136375878 | -0.004166132 | 0.045480465 | 0.130222934 | 0.187782037 | 0.262371876 | 0.969786188 | 0.743903593 |
| Ctnnd1 | S | 920 | -0.021955021 | -0.132643133 | -0.174440099 | -0.152485078 | -0.041796966 | 0.528223439 | 0.094573217 | 0.021159392 | 0.024378851 | 0.505277109 |
| Wdr3 | S | 249 | 0.021522273 | -0.102335029 | -0.121663072 | -0.143185345 | -0.019328043 | 0.598230711 | 0.005813188 | 0.001793405 | 0.035056558 | 0.324851856 |
| Wdr3 | S | 252 | 0.021522273 | -0.102335029 | -0.121663072 | -0.143185345 | -0.019328043 | 0.598230711 | 0.005813188 | 0.001793405 | 0.035056558 | 0.324851856 |
| Ccdc86 | S | 252 | 0.009514106 | -0.207456293 | -0.10719132 | -0.116705426 | 0.100264973 | 0.800988682 | 0.071117682 | 0.002435009 | 0.042638672 | 0.254856795 |
| Brca1 | S | 502 | -0.088481555 | -0.067249953 | -0.102167572 | -0.013686017 | -0.034917619 | 0.111801555 | 0.078817703 | 0.017505591 | 0.741952859 | 0.13606789 |
| Cherp | S | 743 | 0.011365466 | -0.155954612 | -0.233970226 | -0.245335692 | -0.078015613 | 0.815439209 | 0.054693603 | 0.038960984 | 0.035149674 | 0.370059354 |
| Nap1l1 | S | 10 | -0.103059655 | -0.158530732 | -0.180649319 | -0.077589665 | -0.022118588 | 0.352754081 | 0.126338246 | 0.140374655 | 0.544437322 | 0.845627154 |
| Cdk14 | S | 24 | -0.178582669 | 0.014401921 | -0.282932184 | -0.104349515 | -0.297334105 | 0.130846606 | 0.920105728 | 0.047728475 | 0.363611467 | 0.096585019 |
| Cdk4 | T | 172 | -0.036838301 | -0.116698631 | -0.229976666 | -0.193138365 | -0.113278035 | 0.549140666 | 0.077298043 | 0.061789968 | 0.0959946 | 0.255465795 |
| Aggf1 | S | 308 | -0.027182763 | -0.093253127 | -0.096563686 | -0.069380923 | -0.003310559 | 0.327690088 | 0.00363621 | 0.024881556 | 0.07838701 | 0.900875824 |
| Cbx5 | S | 95 | 0.054645448 | -0.218682253 | -0.234324126 | -0.288969574 | -0.015641873 | 0.347622311 | 0.032891558 | 0.039419162 | 0.019007342 | 0.850694554 |
| Macf1 | S | 6923 | -0.149727132 | -0.200202448 | -0.113337192 | 0.03638994 | 0.086865256 | 0.285005641 | 0.128387053 | 0.211727698 | 0.75083234 | 0.404301541 |
| Pdcl3 | S | 235 | -0.064381438 | -0.109795868 | -0.164218736 | -0.099837298 | -0.054422868 | 0.188280581 | 0.058557783 | 0.105400686 | 0.260259335 | 0.504722359 |
| Numa1 | S | 1874 | -0.141060243 | 0.043531206 | -0.357985347 | -0.216925103 | -0.401516553 | 0.257932401 | 0.722024469 | 0.021044474 | 0.134503317 | 0.040304867 |
| Pdap1 | S | 48 | -0.178879002 | -0.073400414 | -0.110952696 | 0.067926306 | -0.037552282 | 0.11525967 | 0.052018504 | 0.226351692 | 0.524399422 | 0.627800711 |
| Chd9 | S | 2078 | 0.008426131 | -0.149988516 | -0.151385699 | -0.159811829 | -0.001397183 | 0.722623165 | 0.008830775 | 0.038011898 | 0.027125647 | 0.975268584 |
| Sptan1 | Y | 1176 | -0.084002863 | -0.344638081 | -0.111862843 | -0.02785998 | 0.232775238 | 0.566308691 | 0.037095177 | 0.675699931 | 0.910127867 | 0.387279878 |
| Zc3h11a | S | 367 | -0.074599604 | -0.191371408 | -0.113813671 | -0.039214067 | 0.077557738 | 0.389989561 | 0.109711843 | 0.053134052 | 0.639215892 | 0.42112019 |
| Myo9b | S | 1218 | -0.030734998 | -0.06278143 | -0.156345647 | -0.125610648 | -0.093564217 | 0.294182186 | 0.200496273 | 0.002110338 | 0.011116268 | 0.094766972 |
| Nab2 | S | 147 | -0.0334755 | -0.028274651 | -0.19424274 | -0.16074924 | -0.165950089 | 0.137320121 | 0.104245041 | 0.006976162 | 0.00837392 | 0.010940355 |
| Raly | S | 177 | -0.020656998 | -0.065246872 | -0.144433588 | -0.123776589 | -0.079186716 | 0.739778041 | 0.050667722 | 0.001434667 | 0.127389698 | 0.020289851 |
| Atad2 | S | 892 | -0.025192591 | -0.193182314 | -0.225012142 | -0.199819551 | -0.031829828 | 0.779367602 | 0.070778081 | 0.114821248 | 0.141380418 | 0.773880209 |
| Nsd3 | S | 587 | 0.070502267 | -0.223553063 | -0.204612627 | -0.275114893 | 0.018940436 | 0.499929323 | 0.044614879 | 0.022792562 | 0.065149238 | 0.805855935 |
| Prpf38b | S | 478 | 0.085409069 | -0.160062707 | -0.172190054 | -0.257599123 | -0.012127347 | 0.370581609 | 0.00939673 | 0.007423001 | 0.057968521 | 0.741185589 |
| Fgd6 | S | 629 | -0.102928889 | -0.040342298 | -0.262072759 | -0.159143869 | -0.221730461 | 0.084758235 | 0.506924074 | 0.075864625 | 0.188640535 | 0.099985489 |
| Atrx | S | 2471 | -0.047817117 | -0.067836248 | -0.14270328 | -0.094886163 | -0.074867032 | 0.05596436 | 0.035496025 | 0.054505778 | 0.130975136 | 0.194506444 |
| Cdk11b | S | 270 | -0.02929154 | -0.132899504 | -0.157445594 | -0.128154055 | -0.024546091 | 0.330836737 | 0.045807172 | 0.057150692 | 0.101325708 | 0.70625111 |
| Cep170 | S | 1200 | 0.022282779 | -0.171795919 | -0.196763986 | -0.219046765 | -0.024968067 | 0.756489915 | 0.008517508 | 0.10569269 | 0.088943846 | 0.779889266 |
| Vim | T | 327 | -0.223111563 | -0.109377758 | -0.010515147 | 0.212596416 | 0.098862612 | 0.027041762 | 0.064889138 | 0.818570044 | 0.047404585 | 0.058379476 |
| Aak1 Gm38495 | S | 174 | -0.172742821 | -0.02748793 | -0.151746233 | 0.020996589 | -0.124258302 | 0.0588175 | 0.789996407 | 0.036008795 | 0.735494569 | 0.258829829 |
| Uevld | S | 365 | -0.105786881 | -0.094570982 | -0.138565623 | -0.032778743 | -0.043994641 | 0.09496035 | 0.303990144 | 0.044951144 | 0.268113473 | 0.569606819 |
| Map1b | S | 1153 | -0.108668653 | -0.117576892 | -0.108290943 | 0.00037771 | 0.009285949 | 0.193479101 | 0.089048971 | 0.071664183 | 0.995212635 | 0.827114511 |

| | | | | | | | | | | | | |
|---|---|---|---|---|---|---|---|---|---|---|---|---|
| Zc3h12c | S | 231 | -0.202976006 | -0.075598775 | -0.074624103 | 0.128351903 | 0.000974672 | 0.01479247 | 0.306730748 | 0.387300843 | 0.177569541 | 0.991025478 |
| Uhrf1bp1l | S | 1108 | -0.019204572 | -0.213938316 | -0.25955301 | -0.240348438 | -0.045614694 | 0.883781409 | 0.045586768 | 0.218937138 | 0.259114261 | 0.790686521 |
| Uhrf1bp1l | S | 1111 | -0.019204572 | -0.213938316 | -0.25955301 | -0.240348438 | -0.045614694 | 0.883781409 | 0.045586768 | 0.218937138 | 0.259114261 | 0.790686521 |
| Pcdh18 | S | 937 | -0.112017847 | -0.151316036 | -0.117832417 | -0.00581457 | 0.033483618 | 0.153485894 | 0.165246073 | 0.110847102 | 0.936236868 | 0.724181606 |
| Cdv3 | S | 95 | -0.0260918 | -0.015894047 | -0.212273152 | -0.186181352 | -0.196379105 | 0.644660841 | 0.814868592 | 0.000255468 | 0.04019087 | 0.06336128 |
| Aftph | S | 679 | -0.04265253 | -0.070258894 | -0.240434483 | -0.197781953 | -0.170175589 | 0.595881396 | 0.076214262 | 0.030817971 | 0.084096404 | 0.059705253 |
| Usp24 | S | 1940 | 0.103277163 | -0.052524369 | -0.265836288 | -0.369113451 | -0.213311918 | 0.618799442 | 0.401088419 | 0.000230838 | 0.169098666 | 0.027649275 |
| Add2 | S | 618 | -0.058444242 | -0.186945877 | -0.093626093 | -0.035181851 | 0.093319784 | 0.483210987 | 0.036781056 | 0.071896204 | 0.639100954 | 0.196939701 |
| Uhrf1bp1l | S | 1020 | -0.125946777 | 0.024549919 | -0.302072041 | -0.176125263 | -0.32662196 | 0.297472733 | 0.847725185 | 0.011552641 | 0.133734489 | 0.058538827 |
| Nedd4l | T | 331 | -0.111979871 | -0.086966737 | -0.044486 | 0.067493871 | 0.042480737 | 0.010571875 | 0.059404494 | 0.175600074 | 0.061596592 | 0.260544034 |
| Pus1 | T | 422 | -0.020775836 | -0.069107326 | -0.07387862 | -0.053102784 | -0.004771294 | 0.19866797 | 0.000111851 | 0.029106206 | 0.06293401 | 0.802929919 |
| Cpeb2 | S | 246 | -0.160548227 | -0.265363689 | -0.427260929 | -0.266712702 | -0.16189724 | 0.678384413 | 0.455158001 | 0.240458149 | 0.564194041 | 0.698536589 |
| Pdcd5 | Y | 126 | -0.073939643 | -0.017925476 | -0.177986888 | -0.104047245 | -0.160061412 | 0.092919913 | 0.696425785 | 0.00418929 | 0.04176632 | 0.01793042 |
| Cdc42ep4 | T | 161 | -0.034865586 | -0.17053601 | -0.248930066 | -0.21406448 | -0.078394056 | 0.785586856 | 0.171518999 | 0.056117946 | 0.084149876 | 0.327775544 |
| Nav1 | T | 940 | 0.008085226 | -0.087144977 | -0.234844666 | -0.242929892 | -0.147699689 | 0.894725534 | 0.13050054 | 0.006055028 | 0.008370853 | 0.042566241 |
| Myo1c | S | 864 | -0.022024417 | -0.108024475 | -0.085762313 | -0.063737896 | 0.022262161 | 0.665132574 | 0.00140068 | 0.029726647 | 0.271817603 | 0.432716257 |
| Etv3 | S | 29 | -0.085937514 | -0.242248363 | -0.018460655 | 0.067476859 | 0.223787708 | 0.214076877 | 0.009980504 | 0.79538109 | 0.187862388 | 0.011588625 |
| Eef1b2 | S | 174 | -0.126531147 | -0.047014114 | -0.253262896 | -0.126731749 | -0.206248782 | 0.262240126 | 0.624145704 | 0.034112851 | 0.291462924 | 0.100581048 |
| Lrch3 | S | 419 | -0.001961754 | -0.070343659 | -0.229726997 | -0.227765243 | -0.159383338 | 0.962141235 | 0.03532305 | 0.013868285 | 0.015126632 | 0.062323127 |
| Cavin1 | S | 173 | -0.065549985 | -0.079665046 | -0.189564691 | -0.124014706 | -0.109899645 | 0.519148895 | 0.182845988 | 0.012727588 | 0.245266579 | 0.015561186 |
| Pdlim2 | S | 210 | -0.268585972 | -0.056100962 | -0.210497435 | 0.058088537 | -0.154396474 | 0.098750226 | 0.771901702 | 0.207227547 | 0.623738712 | 0.400175177 |
| Mark3 | S | 469 | 0.074207119 | -0.119312636 | -0.146036792 | -0.220243911 | -0.026724157 | 0.106941433 | 0.005309867 | 0.000247167 | 0.010378046 | 0.310039201 |
| Raly | S | 254 | -0.056953939 | -0.318453839 | -0.100848267 | -0.043894328 | 0.217605571 | 0.260884156 | 0.061057739 | 0.501814093 | 0.754324773 | 0.237228898 |
| Psmf1 | S | 153 | -0.086047698 | -0.074178939 | -0.190019467 | -0.103971769 | -0.115840528 | 0.25447299 | 0.240162067 | 0.030740361 | 0.220834711 | 0.144059074 |
| Fam234b | S | 30 | -0.04774132 | -0.153973325 | -0.114522051 | -0.066780731 | 0.039451274 | 0.301392237 | 0.013580587 | 0.200659211 | 0.39683078 | 0.604028953 |
| Fam234b | S | 33 | -0.04774132 | -0.153973325 | -0.114522051 | -0.066780731 | 0.039451274 | 0.301392237 | 0.013580587 | 0.200659211 | 0.39683078 | 0.604028953 |
| Rps10 | S | 130 | 0.046890913 | -0.126466553 | -0.318421487 | -0.3653124 | -0.191954935 | 0.248984901 | 0.01945069 | 0.035558177 | 0.020069468 | 0.090570026 |
| Eif4enif1 | S | 212 | -0.049535712 | -0.245056379 | -0.190484425 | -0.140948712 | 0.054571955 | 0.736870547 | 0.084851603 | 0.170375747 | 0.426300539 | 0.681454035 |
| Tsc1 | S | 592 | -0.147202104 | -0.01346941 | -0.254041858 | -0.106839754 | -0.240572448 | 0.295511965 | 0.875275863 | 0.017327019 | 0.401850954 | 0.012240035 |
| Slain2 | S | 251 | -0.082981472 | -0.077812355 | -0.259414077 | -0.176432605 | -0.181601722 | 0.223252594 | 0.305720283 | 0.07287957 | 0.175109485 | 0.163994226 |
| Irf3 | S | 378 | -0.050159857 | -0.074535 | -0.23100631 | -0.180846453 | -0.15647131 | 0.296857072 | 0.157880707 | 0.03993184 | 0.081294471 | 0.108933669 |
| Nek7 | T | 302 | -0.084634265 | -0.110913396 | -0.130247856 | -0.045613591 | -0.01933446 | 0.282703098 | 0.074715035 | 0.054164879 | 0.487363531 | 0.550162881 |
| Zfp280c | S | 522 | -0.073853422 | -0.09549267 | -0.115472914 | -0.041619492 | -0.019980244 | 0.053879008 | 0.020856338 | 0.238275461 | 0.621722364 | 0.808306122 |
| S100a6 | S | 46 | -0.198640306 | -0.06024625 | -0.311967324 | -0.113327019 | -0.251721074 | 0.181761133 | 0.691194555 | 0.166400443 | 0.576431054 | 0.270733661 |
| Arhgef5 | S | 593 | -0.051521386 | -0.063384518 | -0.167501777 | -0.115980391 | -0.104117259 | 0.343322357 | 0.16736758 | 0.007355882 | 0.073394591 | 0.053576062 |
| Brwd1 | S | 1830 | -0.025017341 | -0.082947413 | -0.127891317 | -0.102873976 | -0.044943903 | 0.189482689 | 0.007379543 | 0.042211602 | 0.078354848 | 0.306313198 |
| Svil | S | 632 | -0.000160353 | -0.156999488 | -0.177164384 | -0.177004031 | -0.020164896 | 0.998072629 | 0.012658254 | 0.092650754 | 0.102503342 | 0.792581432 |
| Hivep2 | S | 2085 | -0.14637978 | 0.019952568 | -0.185039753 | -0.038659973 | -0.204992322 | 0.059886303 | 0.781858744 | 0.011835556 | 0.465571917 | 0.035921719 |
| Mxra7 | S | 79 | -0.090198744 | -0.100325925 | -0.054408933 | 0.035789812 | 0.045916992 | 0.05173585 | 0.012450344 | 0.197170301 | 0.386401521 | 0.218507897 |
| Tbx3 | S | 432 | -0.003774098 | -0.178697303 | -0.081998503 | -0.078224405 | 0.0966988 | 0.924452388 | 0.0017456 | 0.149500791 | 0.152436751 | 0.102029334 |
| Dnttip2 | S | 250 | -0.102909319 | -0.085646462 | -0.238036845 | -0.135127526 | -0.152390383 | 0.325881994 | 0.239807563 | 0.073716118 | 0.282887321 | 0.19338763 |
| Dhcr7 | S | 14 | -0.059973865 | -0.107663606 | -0.092474221 | -0.032500356 | 0.015189385 | 0.444372769 | 0.011554446 | 0.0322967 | 0.65802421 | 0.55538958 |
| Ngdn | S | 143 | 0.062376226 | -0.241041084 | -0.130383744 | -0.192759971 | 0.11065734 | 0.509177714 | 0.001445221 | 0.247561003 | 0.161578174 | 0.309028557 |
| Ngdn | S | 149 | 0.062376226 | -0.241041084 | -0.130383744 | -0.192759971 | 0.11065734 | 0.509177714 | 0.001445221 | 0.247561003 | 0.161578174 | 0.309028557 |
| Smarcc2 | S | 367 | 0.021106543 | -0.183770405 | -0.114321646 | -0.135428189 | 0.069448758 | 0.742480998 | 0.00103392 | 0.195517419 | 0.170084365 | 0.379732391 |
| Git2 | T | 536 | -0.270940482 | 0.056514945 | -0.333445132 | -0.06250465 | -0.389960077 | 0.103524116 | 0.6450308 | 0.114461733 | 0.735089389 | 0.084814913 |
| Ppp1r12a | S | 432 | 0.025958772 | -0.250822833 | -0.16105032 | -0.187009093 | 0.089772513 | 0.825235876 | 0.069476669 | 0.036776623 | 0.198454166 | 0.382046569 |
| Plec | S | 4381 | -0.217478541 | -0.026455305 | -0.342912352 | -0.12543381 | -0.316457047 | 0.257142047 | 0.874728822 | 0.108016937 | 0.549637849 | 0.180909406 |
| Poc5 | S | 496 | -0.015052105 | -0.166071238 | -0.321272983 | -0.306220878 | -0.155201745 | 0.926977506 | 0.077894716 | 0.142180735 | 0.200481025 | 0.39984412 |
| Lrch3 | S | 625 | 0.019095338 | -0.095955481 | -0.132744335 | -0.151839673 | -0.036788853 | 0.61521391 | 0.010046081 | 0.002002947 | 0.015549338 | 0.076876935 |
| Tbc1d1 | S | 497 | -0.057966322 | -0.138545968 | -0.22179752 | -0.163831198 | -0.083251552 | 0.380303431 | 0.052160946 | 0.197477963 | 0.306241862 | 0.563113698 |
| Map3k4 | S | 424 | -0.071159229 | -0.130336001 | -0.133117621 | -0.061958392 | -0.00278162 | 0.277868941 | 0.073714149 | 0.072574819 | 0.387102552 | 0.966880394 |
| Mtbp | S | 701 | -0.01384567 | -0.158940562 | -0.510952004 | -0.497106334 | -0.352011442 | 0.946744705 | 0.082862665 | 0.269545653 | 0.284926985 | 0.409489657 |
| Kpna2 | S | 55 | -0.090969749 | 0.021627996 | -0.315510551 | -0.224540802 | -0.337138547 | 0.416298629 | 0.765292298 | 0.00609581 | 0.099070832 | 0.008697638 |
| Map1a | S | 2637 | -0.128115384 | -0.096097981 | -0.105275649 | 0.022839735 | -0.009177668 | 0.043239403 | 0.101697158 | 0.271103108 | 0.778440431 | 0.908665512 |
| Lmna | S | 546 | 0.060651088 | -0.161289192 | -0.09896492 | -0.159616009 | 0.062324272 | 0.12929574 | 7.28635E-06 | 0.135852654 | 0.042364967 | 0.277460646 |
| Arhgap31 | S | 1144 | -0.226500432 | -0.121380905 | -0.180548002 | 0.04595243 | -0.059167097 | 0.21601792 | 0.327577696 | 0.229420353 | 0.799690631 | 0.698805355 |
| Nfx1 | S | 151 | 0.000412188 | -0.09841544 | -0.186881561 | -0.187293748 | -0.08846612 | 0.994336405 | 0.10379218 | 0.004651233 | 0.03167785 | 0.14220703 |
| Cdc27 | T | 367 | -0.092728284 | -0.095012054 | -0.574426597 | -0.481698313 | -0.479414543 | 0.565915126 | 0.463570886 | 0.192895375 | 0.248202407 | 0.249226869 |
| Brd1 | S | 808 | 0.026724134 | -0.088016346 | -0.094886058 | -0.121610192 | -0.006869711 | 0.543466786 | 0.00119625 | 0.00070094 | 0.072694842 | 0.55283976 |
| Palld | S | 1392 | -0.040651125 | -0.10456931 | -0.231539149 | -0.190888024 | -0.12696984 | 0.751222731 | 0.15390814 | 0.025676703 | 0.221431342 | 0.140784603 |
| Dicer1 | S | 1016 | -0.057393774 | -0.08768553 | -0.130840119 | -0.073446345 | -0.043154589 | 0.086589624 | 0.029040576 | 0.1087398 | 0.285945566 | 0.499704089 |
| Ahctf1 | S | 2188 | 0.093389809 | -0.174253942 | -0.166819193 | -0.260209002 | 0.00743475 | 0.242229583 | 0.007172656 | 0.007229241 | 0.041844621 | 0.802030832 |
| Srrm2 | S | 1287 | 0.039926815 | -0.132719856 | -0.150911707 | -0.190838522 | -0.018191851 | 0.414721786 | 0.000433589 | 0.064002113 | 0.032714304 | 0.728292376 |
| Lrrfip1 | T | 566 | -0.178582259 | -0.006707313 | -0.101265835 | 0.077316425 | -0.094558522 | 0.003383183 | 0.887231827 | 0.156524076 | 0.243489805 | 0.181632323 |
| Bora | S | 215 | 0.03060402 | -0.170795749 | -0.109978286 | -0.140582306 | 0.060817463 | 0.594801371 | 0.007808682 | 0.014071711 | 0.082200285 | 0.14603836 |
| Cdca3 | S | 197 | -0.040632193 | -0.091848727 | -0.15031679 | -0.109684597 | -0.058468063 | 0.359886382 | 0.022428961 | 0.048439899 | 0.115948472 | 0.311722544 |
| Cdca3 | S | 207 | -0.040632193 | -0.091848727 | -0.15031679 | -0.109684597 | -0.058468063 | 0.359886382 | 0.022428961 | 0.048439899 | 0.115948472 | 0.311722544 |
| Spg20 | S | 474 | 0.02915275 | -0.143695407 | -0.18264899 | -0.211801741 | -0.038953583 | 0.531983169 | 0.007152779 | 0.041725715 | 0.025379929 | 0.511685368 |
| Fhdc1 | S | 495 | -0.141352783 | -0.048019782 | -0.283151749 | -0.141798966 | -0.235131967 | 0.178132449 | 0.649772246 | 0.087602319 | 0.292996977 | 0.130238057 |
| Stim2 | S | 678 | -0.069887292 | -0.06920374 | -0.109659197 | -0.039771905 | -0.040455456 | 0.129550674 | 0.271027138 | 0.003419499 | 0.329533946 | 0.482070199 |
| Stim2 | S | 680 | -0.069887292 | -0.06920374 | -0.109659197 | -0.039771905 | -0.040455456 | 0.129550674 | 0.271027138 | 0.003419499 | 0.329533946 | 0.482070199 |
| Zfp516 | S | 116 | -0.048497828 | -0.150819735 | -0.023040698 | 0.02545713 | 0.127779037 | 0.23354795 | 0.000355092 | 0.607235365 | 0.604911455 | 0.060402399 |
| Wbp11 | S | 600 | -0.163799069 | 0.07060995 | -0.158369996 | 0.005429073 | -0.228979946 | 0.001541225 | 0.556482488 | 0.07480141 | 0.930240038 | 0.133779053 |
| Ppm1a | S | 377 | -0.203470667 | 0.008132003 | -0.241419399 | -0.037948732 | -0.249551402 | 0.156825053 | 0.939938899 | 0.041883085 | 0.760843912 | 0.06906198 |
| Lats1 | S | 181 | -0.070775663 | -0.093919896 | -0.194605954 | -0.123830291 | -0.100686058 | 0.205266008 | 0.252581575 | 0.046738743 | 0.147859474 | 0.272691439 |
| Stk10 | S | 20 | -0.146977264 | -0.140881648 | -0.100230116 | 0.046747148 | 0.040651532 | 0.132018475 | 0.045833668 | 0.505445941 | 0.754175014 | 0.771815983 |
| Vgll4 | S | 259 | -0.108531928 | -0.073016958 | -0.053770246 | 0.054761682 | 0.019246712 | 0.00433023 | 0.153169171 | 0.136332498 | 0.105309868 | 0.661633914 |
| Ckap4 | S | 17 | -0.123595472 | -0.018871895 | -0.087768966 | 0.035826505 | -0.068897071 | 0.041947837 | 0.691364686 | 0.00149257 | 0.383178497 | 0.221767942 |
| Srrm1 | S | 688 | -0.121404941 | -0.153697174 | -0.087736727 | 0.033668214 | 0.065960447 | 0.360166169 | 0.008767343 | 0.463318525 | 0.823139313 | 0.5695986 |
| Ajuba | S | 239 | -0.120399541 | -0.117982538 | -0.108954699 | 0.011444842 | 0.009027839 | 0.123477163 | 0.134896604 | 0.12380402 | 0.875170206 | 0.902645447 |
| Cap1 | S | 34 | -0.098041418 | -0.061165127 | -0.169313237 | -0.071271819 | -0.10814811 | 0.113704966 | 0.466446841 | 0.025693338 | 0.262226982 | 0.241180358 |
| Map1b | S | 343 | -0.073291973 | -0.091857416 | -0.032094033 | 0.04119794 | 0.059763383 | 0.012050026 | 0.002024936 | 0.539733949 | 0.438962246 | 0.29379028 |
| Rabl6 | S | 482 | -0.053730006 | -0.060278535 | -0.122459032 | -0.068729027 | -0.062180497 | 0.137338577 | 0.078007221 | 0.010673828 | 0.080324268 | 0.090595335 |
| Rabl6 | S | 483 | -0.053730006 | -0.060278535 | -0.122459032 | -0.068729027 | -0.062180497 | 0.137338577 | 0.078007221 | 0.010673828 | 0.080324268 | 0.090595335 |
| Dnajb4 | S | 122 | -0.155325616 | -0.204020495 | -0.134392683 | 0.020932933 | 0.069627812 | 0.221662978 | 0.15522657 | 0.373162439 | 0.872599746 | 0.625306031 |
| Samd4 | S | 272 | -0.14909824 | -0.082891482 | -0.108341229 | 0.040757011 | -0.025449747 | 0.100584356 | 0.117526283 | 0.14470618 | 0.612782535 | 0.651644179 |
| Tmem230 | S | 24 | -0.179268071 | -0.056449943 | -0.154042396 | 0.025225675 | -0.097592454 | 0.054118296 | 0.509420615 | 0.144792956 | 0.774754913 | 0.319750388 |
| Pakap | S | 120 | 0.031681231 | -0.118067562 | -0.168585394 | -0.200266625 | -0.050517832 | 0.126589846 | 0.012283968 | 0.002696983 | 0.003094957 | 0.170091553 |
| Utp14a | S | 29 | 0.080480484 | -0.264728621 | -0.485824473 | -0.566304958 | -0.221095853 | 0.774685597 | 0.352967228 | 0.089983368 | 0.04312717 | 0.23306681 |
| Utp14a | S | 31 | 0.080480484 | -0.264728621 | -0.485824473 | -0.566304958 | -0.221095853 | 0.774685597 | 0.352967228 | 0.089983368 | 0.04312717 | 0.23306681 |
| Ifih1 | S | 1022 | -0.168840159 | -0.148464589 | -0.103946279 | 0.064893881 | 0.04451831 | 0.079734482 | 0.228021318 | 0.327784502 | 0.528516922 | 0.717531778 |
| Nol6 | S | 65 | -0.021934047 | -0.228529895 | -0.153598485 | -0.131664437 | 0.07493141 | 0.673503249 | 0.056904717 | 0.129899538 | 0.17765969 | 0.482633731 |
| Armcx2 | T | 140 | 0.001177935 | -0.077314216 | -0.111517564 | -0.112695499 | -0.034203348 | 0.980026055 | 0.004731443 | 0.002275968 | 0.090950859 | 0.147695264 |

| | | | | | | | | | | | | |
|---|---|---|---|---|---|---|---|---|---|---|---|---|
| Gli3 | S | 93 | -0.126006487 | -0.115811771 | -0.0880671 | 0.037939387 | 0.02774467 | 0.215269643 | 0.096202759 | 0.057633118 | 0.660730047 | 0.620981635 |
| Msh6 | S | 77 | -0.053019994 | 0.001106561 | -0.200643205 | -0.147623212 | -0.201749766 | 0.335332596 | 0.98418757 | 0.000563306 | 0.05186415 | 0.034218675 |
| Uba1 | S | 835 | -0.121786328 | -0.077828264 | -0.175513967 | -0.053727638 | -0.097685702 | 0.05810898 | 0.349009626 | 0.141323763 | 0.582176993 | 0.389301346 |
| Usp47 | Y | 1020 | -0.121929744 | -0.161182968 | -0.103299209 | 0.018630535 | 0.057883759 | 0.162678301 | 0.137470295 | 0.174538731 | 0.715507918 | 0.478014301 |
| Pakap | S | 118 | 0.08356904 | -0.140740226 | -0.252035402 | -0.335604442 | -0.111295176 | 0.085207333 | 0.132959346 | 0.000518981 | 0.00036314 | 0.207460643 |
| Lima1 | S | 283 | -0.185531423 | -0.082656143 | -0.070226896 | 0.115304528 | 0.012429247 | 0.007688567 | 0.28482879 | 0.608389679 | 0.426635963 | 0.929611005 |
| Denr | S | 73 | -0.188212328 | -0.089974905 | -0.03879089 | 0.149421438 | 0.051184015 | 0.006339907 | 0.276419026 | 0.589943364 | 0.086074733 | 0.550539493 |
| Gnl1 | S | 51 | -0.091266969 | -0.086620766 | -0.209220973 | -0.117954004 | -0.122600207 | 0.3117915 | 0.275927493 | 0.047340274 | 0.260521987 | 0.213222495 |
| Sfswap | S | 601 | -0.148351396 | -0.181022424 | -0.007564277 | 0.140787119 | 0.173458147 | 0.197043218 | 0.007284868 | 0.92703162 | 0.246932587 | 0.106688282 |
| Ccnl1 | T | 331 | -0.100831212 | -0.126221778 | -0.158791895 | -0.057960683 | -0.032570116 | 0.449774126 | 0.064186617 | 0.115097553 | 0.672520591 | 0.696755979 |
| Syde1 | T | 698 | -0.091144715 | -0.015377338 | -0.286536217 | -0.195391502 | -0.27115888 | 0.555974159 | 0.916260732 | 0.007125137 | 0.24706581 | 0.142364038 |
| Sh3glb2 | S | 205 | -0.039513465 | -0.254531307 | -0.233510448 | -0.193996982 | 0.02102086 | 0.801037226 | 0.088851628 | 0.23761809 | 0.306611826 | 0.893645443 |
| Ctnnd1 | S | 320 | -0.099215556 | -0.118552268 | -0.169003586 | -0.06978803 | -0.050451318 | 0.260490404 | 0.210951465 | 0.074939371 | 0.258038553 | 0.454566083 |
| Tnk2 | Y | 284 | -0.1607138 | -0.153238924 | -0.104374161 | 0.056339639 | 0.048864762 | 0.076630953 | 0.128712876 | 0.54032354 | 0.726107235 | 0.766035318 |
| Cdca2 | S | 798 | 0.128696046 | -0.15299503 | -0.32579371 | -0.454489757 | -0.17279868 | 0.113410845 | 0.071953507 | 0.003683152 | 0.001523973 | 0.040438624 |
| Fermt2 | S | 181 | -0.074323527 | -0.30134722 | -0.178759727 | -0.1044362 | 0.122587494 | 0.469658822 | 0.118983723 | 0.375518026 | 0.567495161 | 0.568420852 |
| Plec | T | 4639 | -0.234912432 | 0.068033302 | -0.228903994 | 0.006008438 | -0.296937296 | 0.048454223 | 0.500456239 | 0.049538134 | 0.926697316 | 0.00651447 |
| Nasp | S | 489 | -0.083057094 | -0.027538725 | -0.322958459 | -0.239901365 | -0.295419734 | 0.598016248 | 0.871040519 | 0.012766044 | 0.171441885 | 0.147791727 |
| Tgfb1i1 | S | 186 | -0.059739734 | -0.077135954 | -0.153467806 | -0.093728072 | -0.076331852 | 0.09268637 | 0.103328486 | 0.058728023 | 0.173455039 | 0.252489394 |
| Zfp608 | S | 1269 | 0.008088483 | -0.242210565 | -0.039204729 | -0.047293211 | 0.203005836 | 0.858231645 | 0.000531714 | 0.612508772 | 0.546345107 | 0.068055845 |
| Myc | S | 282 | 0.054248612 | -0.180092772 | -0.139992126 | -0.194240738 | 0.040100646 | 0.225459201 | 0.001042385 | 0.060530209 | 0.028097479 | 0.467193106 |
| Zfp608 | S | 963 | -0.258393825 | -0.180917543 | 0.063114076 | 0.321507902 | 0.244031619 | 0.019565618 | 0.053463092 | 0.283280779 | 0.010669899 | 0.025699561 |
| Pard3 | S | 1112 | -0.07576009 | -0.172348591 | -0.186711052 | -0.110950962 | -0.014362461 | 0.312368195 | 0.07806169 | 0.270496538 | 0.473418507 | 0.923709241 |
| Flnb | S | 518 | -0.270408705 | -0.362035953 | 0.109666514 | 0.380075218 | 0.471702467 | 0.045957355 | 0.138544317 | 0.428418528 | 0.025647477 | 0.082037756 |
| Ddx20 | S | 679 | -0.091200371 | -0.089406168 | -0.058902164 | 0.032298207 | 0.030504004 | 0.072748585 | 0.003966674 | 0.316980808 | 0.585770539 | 0.571252818 |
| Ttll5 | S | 1301 | -0.241037335 | -0.116119567 | -0.112714649 | 0.128322686 | 0.003404917 | 0.133201754 | 0.285448582 | 0.290135752 | 0.383359973 | 0.975789607 |
| Ttll5 | S | 1308 | -0.241037335 | -0.116119567 | -0.112714649 | 0.128322686 | 0.003404917 | 0.133201754 | 0.285448582 | 0.290135752 | 0.383359973 | 0.975789607 |
| Ttll5 | T | 1306 | -0.241037335 | -0.116119567 | -0.112714649 | 0.128322686 | 0.003404917 | 0.133201754 | 0.285448582 | 0.290135752 | 0.383359973 | 0.975789607 |
| Mcrip1 | S | 18 | 0.020750154 | -0.073342817 | -0.236918771 | -0.257668925 | -0.163575954 | 0.694572965 | 0.167119871 | 0.002909642 | 0.002274547 | 0.022283109 |
| Flna | S | 1084 | -0.070038275 | -0.06463872 | -0.088174949 | -0.018136674 | -0.023536229 | 0.034438438 | 0.078266058 | 0.028689909 | 0.462539333 | 0.446343212 |
| Setd1b | S | 977 | -0.307131187 | 0.02213625 | -0.065524765 | 0.241606422 | -0.087661015 | 0.004782178 | 0.777404551 | 0.406017838 | 0.03839335 | 0.388406678 |
| Zc3hc1 | S | 407 | -0.094261516 | -0.121669944 | -0.111037548 | -0.016776031 | 0.010632397 | 0.037526705 | 0.188842287 | 0.172670061 | 0.802537488 | 0.90691691 |
| Ubash3b | S | 366 | -0.085701564 | -0.093596499 | -0.136005318 | -0.050303754 | -0.04240882 | 0.15016862 | 0.087595483 | 0.093507263 | 0.441137002 | 0.481472264 |
| Ftsj3 | S | 576 | 0.009196158 | -0.115103659 | -0.136275099 | -0.145471257 | -0.02117144 | 0.637813256 | 0.012429138 | 0.008965098 | 0.00476427 | 0.473102951 |
| Apc | S | 2350 | 0.072409216 | -0.170110936 | -0.175688773 | -0.248097989 | -0.005577837 | 0.407398874 | 0.005526364 | 0.036589735 | 0.045796045 | 0.923573276 |
| Uhrf1bp1l | S | 888 | -0.138788138 | -0.103256813 | -0.100333588 | 0.03845455 | 0.002923226 | 0.088907933 | 0.171261317 | 0.143593767 | 0.588055014 | 0.966025672 |
| Ehbp1l1 | S | 1444 | 0.054936769 | -0.109417564 | -0.280472098 | -0.335408867 | -0.171054534 | 0.346559909 | 0.067028981 | 0.009373024 | 0.006463194 | 0.049677367 |
| Las1l | S | 20 | -0.093647486 | -0.287479232 | 0.012147973 | 0.105795459 | 0.299627206 | 0.208431433 | 0.015383798 | 0.917873543 | 0.36208409 | 0.059114648 |
| Ppp1r2 | S | 88 | -0.212919768 | 4.78608E-05 | -0.211304143 | 0.001615625 | -0.211352003 | 0.054060876 | 0.999524886 | 0.128694656 | 0.989255093 | 0.136562922 |
| Vcl | S | 272 | -0.17009506 | -0.026255204 | -0.175677152 | -0.005582092 | -0.149421948 | 0.059697735 | 0.739864287 | 0.080778998 | 0.937130625 | 0.094926137 |
| Wee1 | S | 642 | -0.279042989 | -0.323973509 | -0.123573143 | 0.155469846 | 0.200400365 | 0.389739867 | 0.222262493 | 0.638674412 | 0.637886293 | 0.474875124 |
| Shroom2 | S | 1165 | -0.141633414 | -0.069147174 | -0.211939289 | -0.070305875 | -0.142792116 | 0.173800497 | 0.346880963 | 0.114594822 | 0.55601092 | 0.235351144 |
| Cfl1 | T | 25 | -0.189070707 | -0.00287933 | -0.233257061 | -0.044186354 | -0.230377731 | 0.054189665 | 0.979914841 | 0.125516135 | 0.702217527 | 0.145762714 |
| Tanc2 | S | 1480 | 0.026072869 | -0.132572194 | -0.40144543 | -0.427518299 | -0.268873235 | 0.443786744 | 0.105373361 | 0.065805259 | 0.060481439 | 0.127905801 |
| Hspa4 | S | 781 | -0.130591058 | -0.025366952 | -0.127162464 | 0.003428593 | -0.101795513 | 0.054015008 | 0.2812172 | 0.036514474 | 0.947919443 | 0.069616107 |
| Fam160b1 | T | 551 | -0.067614713 | -0.033002708 | -0.134926795 | -0.067312082 | -0.101924087 | 0.062814497 | 0.505340705 | 0.004040464 | 0.031290303 | 0.089959903 |
| Clasp2 | S | 374 | -0.095801208 | -0.046501712 | -0.318021757 | -0.222220549 | -0.271520045 | 0.474709363 | 0.667908249 | 0.033279585 | 0.154300516 | 0.065171173 |
| Atxn2l | S | 243 | 0.00324563 | -0.213356952 | -0.193778083 | -0.197023713 | 0.019578869 | 0.98574136 | 0.025057917 | 0.193275471 | 0.356026101 | 0.873999801 |
| Atxn2l | S | 246 | 0.00324563 | -0.213356952 | -0.193778083 | -0.197023713 | 0.019578869 | 0.98574136 | 0.025057917 | 0.193275471 | 0.356026101 | 0.873999801 |
| Tasor2 | S | 698 | -0.068844593 | -0.077559956 | -0.177554499 | -0.108709906 | -0.099994543 | 0.16753905 | 0.136832938 | 0.063162 | 0.189529545 | 0.216101067 |
| Cavin2 | S | 241 | -0.258071128 | 0.144444492 | -0.236175221 | 0.021895908 | -0.380619713 | 0.019183159 | 0.436750766 | 0.076417957 | 0.821898583 | 0.100926347 |
| Gorasp1 | S | 365 | -0.043891788 | -0.120147072 | -0.138011759 | -0.094119971 | -0.017864687 | 0.643347096 | 0.027186932 | 0.048514839 | 0.358672149 | 0.724494285 |
| Pum1 | S | 209 | -0.054635804 | -0.143154112 | -0.243631375 | -0.188995571 | -0.100477263 | 0.52251689 | 0.201572526 | 0.084298757 | 0.154501959 | 0.434199204 |
| Sntb2 | S | 373 | 0.019485374 | -0.143676159 | -0.16858303 | -0.188068404 | -0.024906871 | 0.638147582 | 0.034820761 | 0.014895186 | 0.01474116 | 0.592648757 |
| Serbp1 | S | 393 | -0.030561321 | -0.07914942 | -0.178692339 | -0.148131018 | -0.099542919 | 0.25481613 | 0.109331849 | 0.023217004 | 0.037066262 | 0.108714815 |
| Rpl22l1 | S | 120 | 0.053271776 | -0.152720486 | -0.143236029 | -0.196507805 | 0.009484457 | 0.264615035 | 0.000448621 | 0.066701134 | 0.029161266 | 0.855831107 |
| Trim56 | T | 413 | -0.003409518 | -0.147752872 | -0.131876309 | -0.128466791 | 0.015876564 | 0.933507118 | 0.024110389 | 0.028810653 | 0.044081623 | 0.729719419 |
| Atad3a | S | 41 | -0.001493472 | -0.092241875 | -0.172357398 | -0.170863925 | -0.080115523 | 0.973694196 | 0.060959982 | 0.006862947 | 0.023364488 | 0.120947806 |
| Gorasp2 | S | 418 | 0.008967167 | -0.122581281 | -0.149481318 | -0.158448486 | -0.026900037 | 0.830496452 | 0.031716296 | 0.009833923 | 0.013946292 | 0.548728844 |
| Uhrf1bp1l | S | 774 | -0.100954496 | 0.043458924 | -0.354639484 | -0.253684988 | -0.398098408 | 0.251848795 | 0.696434645 | 0.018076574 | 0.063512607 | 0.021656605 |
| Srrm2 | S | 220 | 0.034320909 | -0.180361075 | -0.164547063 | -0.198867972 | 0.015814013 | 0.679171206 | 0.11639237 | 0.005043631 | 0.087501751 | 0.852914874 |
| Numa1 | S | 1826 | -0.178387538 | -0.072477981 | -0.159033027 | 0.019354511 | -0.086555046 | 0.087125246 | 0.191835835 | 0.305186319 | 0.893093235 | 0.543099358 |
| Clasp2 | S | 465 | -0.133666743 | -0.03598495 | -0.237067187 | -0.103400444 | -0.201082237 | 0.200791447 | 0.737642231 | 0.040399473 | 0.331434757 | 0.125995849 |
| H1f3 | S | 189 | -0.017051721 | -0.169971877 | -0.251057033 | -0.234005312 | -0.081085157 | 0.858203373 | 0.100070355 | 0.0981769 | 0.115594392 | 0.499241422 |
| Kif23 | S | 860 | -0.084089561 | -0.108855407 | -0.10390005 | -0.019810488 | 0.004955358 | 0.187942638 | 0.042057933 | 0.111551253 | 0.759404963 | 0.927789866 |
| Frmd4a | S | 679 | -0.078291357 | -0.083254381 | -0.106689185 | -0.028397828 | -0.023434804 | 0.080381727 | 0.109250377 | 0.049698518 | 0.529084441 | 0.634957677 |
| Adgrl3 | S | 1240 | -0.112280244 | -0.135328323 | -0.216280563 | -0.104000319 | -0.08095224 | 0.239133376 | 0.25564227 | 0.180184555 | 0.476417933 | 0.597021225 |
| Lrrfip1 | S | 284 | -0.102033015 | -0.051866991 | -0.18036171 | -0.078328695 | -0.128494719 | 0.145581687 | 0.611388727 | 0.020742999 | 0.113990951 | 0.236840072 |
| Bicd2 | S | 308 | 0.035654631 | -0.179145066 | -0.147684 | -0.183338631 | 0.031461066 | 0.709827262 | 0.07148615 | 0.006515726 | 0.145075149 | 0.65369984 |
| Lima1 | S | 615 | -0.191660883 | -0.042974146 | -0.182350656 | 0.009310226 | -0.13937651 | 0.074816456 | 0.71925968 | 0.125557979 | 0.930439926 | 0.31741093 |
| Msrb3 | S | 244 | -0.146285683 | -0.034810129 | -0.191612462 | -0.045326778 | -0.156802333 | 0.031865032 | 0.626956409 | 0.151580372 | 0.665635516 | 0.219770851 |
| Nol7 | S | 129 | -0.096328708 | -0.187709868 | -0.063000045 | 0.033328663 | 0.124709824 | 0.168996266 | 0.035264386 | 0.423906856 | 0.68244626 | 0.185194025 |
| Zfp319 | S | 337 | -0.0841762 | -0.094543233 | -0.080486163 | 0.003690038 | 0.01405707 | 0.289238307 | 0.041941067 | 0.018318961 | 0.956085851 | 0.64649422 |
| Nfix | S | 268 | -0.032947535 | -0.063151482 | -0.145389257 | -0.112441723 | -0.082237775 | 0.557812913 | 0.142457171 | 0.002056795 | 0.111242616 | 0.06739817 |
| Rabgap1l | S | 292 | -0.100476098 | -0.155322716 | -0.115385144 | -0.014909046 | 0.039937572 | 0.218205925 | 0.131091434 | 0.130212015 | 0.744102498 | 0.586811803 |
| Nr3c1 | S | 275 | 0.020807319 | -0.123629294 | -0.208755075 | -0.229562394 | -0.085125781 | 0.499747597 | 0.048355182 | 0.013751942 | 0.020510552 | 0.191387698 |
| Ddx11 | S | 203 | 0.071302883 | -0.124706428 | -0.232869075 | -0.304171958 | -0.108162647 | 0.414276607 | 0.083784057 | 0.004064205 | 0.025567435 | 0.119009307 |
| Srek1 | S | 422 | -0.009600109 | -0.195731173 | -0.072281176 | -0.062681067 | 0.123449997 | 0.910904277 | 0.04420277 | 0.004400482 | 0.491631279 | 0.12286078 |
| Rc3h2 | S | 806 | -0.137030656 | -0.050077558 | -0.154734208 | -0.017703552 | -0.10465665 | 0.015276018 | 0.330296129 | 0.27864764 | 0.884878188 | 0.432001422 |
| Bcl6 | S | 396 | 0.104852786 | -0.326070278 | -0.291622133 | -0.39647492 | 0.034448145 | 0.342173024 | 0.099170695 | 0.06052171 | 0.031384245 | 0.829553135 |
| Kdm2a | S | 692 | -0.145221868 | -0.120885092 | -0.047155494 | 0.098066374 | 0.073729598 | 0.042145258 | 0.065788915 | 0.482980196 | 0.163734644 | 0.254506024 |
| Mier1 | S | 376 | 0.031165105 | -0.137369446 | -0.101212707 | -0.132377811 | 0.03615674 | 0.353922742 | 0.005780702 | 0.002634759 | 0.025315524 | 0.24558846 |
| Pacs2 | S | 408 | -0.130096186 | 0.006044679 | -0.220766297 | -0.090670111 | -0.226810976 | 0.106407169 | 0.951928483 | 0.022826955 | 0.195151623 | 0.070586577 |
| Sh3pxd2a | S | 764 | -0.10062804 | -0.01383266 | -0.34979526 | -0.24916722 | -0.3359626 | 0.411665728 | 0.926615694 | 0.030520955 | 0.062530111 | 0.060257472 |
| Dmd | S | 3616 | -0.186654485 | 0.016280019 | -0.243233319 | -0.056578834 | -0.259513338 | 0.158363069 | 0.914067009 | 0.039307105 | 0.579100184 | 0.13919943 |
| Zfyve16 | S | 310 | -0.08619973 | -0.192238155 | -0.223159734 | -0.136960004 | -0.030921578 | 0.198053984 | 0.306135641 | 0.216071823 | 0.399317388 | 0.879327303 |
| Tmem115 | T | 318 | -0.038406647 | -0.227478058 | -0.112819561 | -0.074412914 | 0.114658498 | 0.636661018 | 0.019357341 | 0.324811575 | 0.491904476 | 0.306848699 |
| Nf1 | S | 2523 | -0.094565068 | -0.065037045 | -0.227071038 | -0.13250597 | -0.162033993 | 0.25111932 | 0.464678598 | 0.041602053 | 0.161300888 | 0.12436769 |
| Rap2a Rap2b Rap2c | S | 66;66;66 | -0.057568752 | -0.239262169 | -0.13249164 | -0.074922888 | 0.106770529 | 0.720304213 | 0.04042159 | 0.263280546 | 0.674274046 | 0.380513623 |
| Cast | S | 219 | 0.001880331 | -0.109581587 | -0.134687431 | -0.136567763 | -0.025105845 | 0.95162401 | 0.06580994 | 0.002530294 | 0.011167153 | 0.570520009 |
| Asph | S | 27 | -0.13982364 | -0.058273349 | -0.160023601 | -0.020199961 | -0.101750252 | 0.151504886 | 0.158087533 | 0.111704256 | 0.836973058 | 0.251459074 |
| Otud5 | S | 64 | -0.030487611 | -0.110791066 | -0.138519782 | -0.108032171 | -0.027728716 | 0.582642966 | 0.066761547 | 0.015853226 | 0.117871636 | 0.586347371 |
| Pak2 | S | 141 | -0.163189379 | -0.052325765 | -0.217469083 | -0.054279704 | -0.165143319 | 0.079441186 | 0.614860276 | 0.160237398 | 0.679040068 | 0.287492133 |
| Srsf3 | S | 5 | -0.042801958 | -0.070306299 | -0.135960023 | -0.093158065 | -0.065653724 | 0.222428744 | 0.316060784 | 0.002765073 | 0.034686007 | 0.342575366 |

| | | | | | | | | | | | | | |
|---|---|---|---|---|---|---|---|---|---|---|---|---|---|
| Map2k4 | T | | 389 | -0.018974294 | -0.056612455 | -0.245403616 | -0.226429323 | -0.188791162 | 0.85768924 | 0.345267926 | 0.005314341 | 0.109543763 | 0.00032214 |
| Fermt2 | S | | 159 | -0.09309535 | -0.071276218 | -0.168444761 | -0.075349411 | -0.097168544 | 0.131955249 | 0.265157617 | 0.058753916 | 0.291120697 | 0.207559762 |
| Atxn2 | S | | 665 | -0.048065124 | -0.154100734 | -0.242024523 | -0.193959399 | -0.087923789 | 0.741734125 | 0.18007565 | 0.071697074 | 0.191491624 | 0.310376901 |
| Lats1 | S | | 216 | -0.088099311 | -0.058702886 | -0.064803845 | 0.023295466 | -0.006100958 | 0.003655392 | 0.038632707 | 0.268546841 | 0.646447846 | 0.903467174 |
| Taok1 | S | | 421 | -0.069291762 | -0.1800859 | -0.014718147 | 0.054573615 | 0.165367753 | 0.125216493 | 0.004653719 | 0.723152687 | 0.035657512 | 0.004862759 |
| Ice1 | S | | 676 | 0.041834574 | -0.145344556 | -0.094873194 | -0.136707768 | 0.050471362 | 0.36727903 | 0.001547316 | 0.009239714 | 0.048034099 | 0.017461751 |
| Npm3 | S | | 151 | -0.078958155 | -0.106824784 | -0.076197909 | 0.002760246 | 0.030626875 | 0.369665596 | 0.008036964 | 0.085213847 | 0.972646278 | 0.380190627 |
| Npm3 | S | | 155 | -0.078958155 | -0.106824784 | -0.076197909 | 0.002760246 | 0.030626875 | 0.369665596 | 0.008036964 | 0.085213847 | 0.972646278 | 0.380190627 |
| Rsl1d1 | S | | 10 | 0.031108832 | -0.324270915 | -0.247434904 | -0.278543736 | 0.076836011 | 0.805819304 | 0.157149341 | 0.110584997 | 0.115065676 | 0.701285523 |
| Tmx1 | S | | 260 | -0.023017251 | -0.174735577 | -0.135955074 | -0.112937823 | 0.038780503 | 0.770239846 | 0.049661409 | 0.054522834 | 0.239082281 | 0.562987904 |
| Cdc23 | T | | 596 | 0.038438098 | -0.142992195 | -0.244051445 | -0.282489543 | -0.10105925 | 0.805830857 | 0.107976008 | 0.02335757 | 0.154801645 | 0.272848968 |
| Eif3c | S | | 9 | 0.017634928 | -0.282416316 | -0.206716346 | -0.224351273 | 0.07569997 | 0.845273913 | 0.073738473 | 0.149694443 | 0.141538679 | 0.595420185 |
| Eif3c | S | | 13 | 0.017634928 | -0.282416316 | -0.206716346 | -0.224351273 | 0.07569997 | 0.845273913 | 0.073738473 | 0.149694443 | 0.141538679 | 0.595420185 |
| Eif3c | S | | 15 | 0.017634928 | -0.282416316 | -0.206716346 | -0.224351273 | 0.07569997 | 0.845273913 | 0.073738473 | 0.149694443 | 0.141538679 | 0.595420185 |
| Eif3c | S | | 16 | 0.017634928 | -0.282416316 | -0.206716346 | -0.224351273 | 0.07569997 | 0.845273913 | 0.073738473 | 0.149694443 | 0.141538679 | 0.595420185 |
| Cep170 | S | | 1529 | -0.040638224 | -0.153426108 | -0.116490345 | -0.07585212 | 0.036935763 | 0.417165261 | 0.025199482 | 0.127154348 | 0.295167783 | 0.569743186 |
| Skap2 | S | | 272 | -0.235837507 | -0.055033171 | -0.114673596 | 0.121163912 | -0.059640424 | 0.036801353 | 0.525467091 | 0.325071483 | 0.307137416 | 0.579431468 |
| Tom1l2 | T | | 164 | -0.083791265 | -0.12588586 | -0.14810751 | -0.064316245 | -0.02222165 | 0.195532917 | 0.249112906 | 0.062173693 | 0.282970915 | 0.816705403 |
| Mtbp | S | | 633 | 0.032598434 | -0.169780202 | -0.129839168 | -0.162437602 | 0.039941034 | 0.663361467 | 0.001875203 | 0.132590646 | 0.123055501 | 0.561732337 |
| Fnbp1 | S | | 296 | -0.166387598 | -0.016730696 | -0.128680451 | 0.037707147 | -0.111949754 | 0.013879795 | 0.828771578 | 0.113730698 | 0.555191715 | 0.221440138 |
| Tjp2 | S | | 186 | 0.065855451 | -0.223850456 | -0.193397701 | -0.259253151 | 0.030452756 | 0.329909843 | 0.042374496 | 0.022359827 | 0.00022466 | 0.671246407 |
| Npm3 | S | | 147 | -0.080046631 | -0.106470106 | -0.076779819 | 0.003266812 | 0.029690287 | 0.366532157 | 0.008941322 | 0.084273461 | 0.967812274 | 0.398344839 |
| Ncapd2 | S | | 757 | -0.01706303 | -0.147376805 | -0.160141342 | -0.143078312 | -0.012764537 | 0.836375422 | 0.017639499 | 0.104639992 | 0.195018696 | 0.866816396 |
| Jcad | S | | 1248 | -0.171297373 | -0.007612847 | -0.108895972 | 0.062401401 | -0.101283124 | 0.00507999 | 0.872181413 | 0.16343019 | 0.356462761 | 0.185217983 |
| Hdgf | S | | 128 | -0.306119404 | -0.066967132 | 0.025732097 | 0.331851501 | 0.09269923 | 0.005784703 | 0.431696665 | 0.845958009 | 0.072921118 | 0.496777539 |
| Zdhhc8 | S | | 679 | -0.081924436 | -0.166759245 | -0.106610185 | -0.02468575 | 0.06014906 | 0.157397644 | 0.043885046 | 0.393396529 | 0.832934744 | 0.616628781 |
| Sema4c | S | | 743 | -0.286648357 | -0.093533308 | -0.105433301 | 0.181215057 | -0.011899992 | 0.069186385 | 0.553683553 | 0.38720309 | 0.242049181 | 0.9445509 |
| Huwe1 | S | | 2743 | -0.034127144 | -0.127867155 | -0.15636235 | -0.122235206 | -0.028495195 | 0.516491252 | 0.017521052 | 0.137447418 | 0.216856502 | 0.726654372 |
| Bcas3 | S | | 570 | -0.075409574 | -0.077457816 | -0.170075787 | -0.094666213 | -0.092617971 | 0.127689747 | 0.176884314 | 0.073879023 | 0.243640361 | 0.252919799 |
| Osbpl3 | S | | 338 | -0.024228765 | -0.142336549 | -0.199942683 | -0.175713918 | -0.057606134 | 0.742491767 | 0.215502318 | 0.022109823 | 0.020295449 | 0.545080702 |
| Col4a3bp | S | | 379 | -0.157635979 | -0.068036728 | -0.179845791 | -0.022209811 | -0.111809063 | 0.095341775 | 0.455493896 | 0.149081865 | 0.837502244 | 0.35053291 |
| Cdc42bpb | T | | 423 | -0.03983098 | -0.093878162 | -0.093176321 | -0.053345341 | 0.00070184 | 0.34612921 | 0.013373448 | 0.026287247 | 0.222847505 | 0.980040554 |
| Srsf10 | S | | 219 | -0.101094392 | -0.071110217 | -0.151879236 | -0.050784844 | -0.080769019 | 0.132561258 | 0.539781588 | 0.02167478 | 0.377570989 | 0.484716993 |
| Srsf10 | S | | 221 | -0.101094392 | -0.071110217 | -0.151879236 | -0.050784844 | -0.080769019 | 0.132561258 | 0.539781588 | 0.02167478 | 0.377570989 | 0.484716993 |
| Dync1li1 | T | | 512 | -0.101742652 | -0.147674225 | -0.046797865 | 0.054944787 | 0.100876359 | 0.255435185 | 0.010019494 | 0.332426485 | 0.513146329 | 0.093744959 |
| Surf6 | T | | 223 | 0.083033135 | -0.200798958 | -0.221696663 | -0.304729798 | -0.020897706 | 0.159633072 | 0.029841493 | 0.013789657 | 0.014822744 | 0.76562862 |
| Tmod3 | S | | 71 | 0.125093623 | -0.166639134 | -0.246376576 | -0.371470198 | -0.079737441 | 0.015033948 | 0.0264549 | 0.000488662 | 0.000161891 | 0.178872874 |
| Atg2a | S | | 1338 | -0.310757886 | 0.002438703 | -0.077650824 | 0.233107062 | -0.080089527 | 0.020330544 | 0.978537924 | 0.244269221 | 0.052315875 | 0.408693168 |
| Atg2a | T | | 1335 | -0.310757886 | 0.002438703 | -0.077650824 | 0.233107062 | -0.080089527 | 0.020330544 | 0.978537924 | 0.244269221 | 0.052315875 | 0.408693168 |
| Kdm2b | S | | 991 | -0.00953429 | -0.140554596 | -0.132872422 | -0.123338131 | 0.007682174 | 0.833179057 | 0.009414231 | 0.085056486 | 0.107401277 | 0.894514015 |
| Gon4l | S | | 347 | -0.054458788 | -0.163092093 | -0.153358177 | -0.098899389 | 0.009733916 | 0.408734901 | 0.062091699 | 0.160558363 | 0.333030765 | 0.920392847 |
| Mta1 | S | | 639 | -0.082739028 | -0.073711025 | -0.207821033 | -0.125082004 | -0.134110008 | 0.256627887 | 0.415647925 | 0.035286344 | 0.119935561 | 0.165340378 |
| Snx2 | T | | 104 | 0.01069044 | -0.091571678 | -0.153575385 | -0.164265825 | -0.062003707 | 0.756209563 | 0.027943816 | 0.00478993 | 0.005973585 | 0.094258126 |
| Anapc1 | S | | 688 | -0.012741083 | -0.111058582 | -0.076355657 | -0.063614573 | 0.034702925 | 0.423480061 | 0.026105025 | 0.002268774 | 0.008914955 | 0.291733675 |
| Pla2g4a | S | | 437 | -0.210109961 | 0.035850407 | -0.190098162 | 0.020011799 | -0.225948569 | 0.068506903 | 0.561954903 | 0.030221739 | 0.821471604 | 0.027461485 |
| Evl | T | | 341 | -0.037538097 | -0.095855532 | -0.251428907 | -0.21389081 | -0.155573375 | 0.811974782 | 0.30911008 | 0.019000698 | 0.232330788 | 0.098708243 |
| Zfp36l2 | S | | 59 | -0.087260265 | -0.078605155 | -0.212254197 | -0.124993932 | -0.133649042 | 0.198328382 | 0.139734271 | 0.13435519 | 0.31414538 | 0.28283723 |
| Clasp2 | S | | 1004 | 0.036965243 | -0.186361307 | -0.320917124 | -0.357882366 | -0.134555817 | 0.654519654 | 0.175311651 | 0.051041235 | 0.030469679 | 0.338676861 |
| Map3k2 | S | | 135 | -0.022160284 | -0.155607851 | -0.117477198 | -0.095316913 | 0.038130653 | 0.760371456 | 0.033651422 | 0.039210726 | 0.24569819 | 0.499843443 |
| Add3 | S | | 402 | -0.043952008 | -0.251585739 | -0.348661879 | -0.304709871 | -0.09707614 | 0.82438608 | 0.354326624 | 0.146943855 | 0.139653205 | 0.69137665 |
| Ppp1r8 | T | | 202 | -0.066401025 | -0.10966601 | -0.186849998 | -0.120448973 | -0.077183988 | 0.297868236 | 0.096445216 | 0.119033092 | 0.258187834 | 0.428561848 |
| Ap3b1 | S | | 995 | -0.022156065 | -0.221640732 | -0.347988773 | -0.325832708 | -0.126348041 | 0.899754855 | 0.328868689 | 0.110624528 | 0.184518029 | 0.596062395 |
| Nosip | T | | 168 | -0.115127339 | -0.095101361 | -0.250532829 | -0.13540549 | -0.155431468 | 0.373791528 | 0.282227696 | 0.114770501 | 0.390201584 | 0.266235619 |
| Dtnb | S | | 528 | -0.125450415 | -0.188898704 | 0.011019808 | 0.136470224 | 0.199918512 | 0.035929924 | 0.032688078 | 0.921916286 | 0.282916504 | 0.15676462 |
| Prune2 | S | | 1876 | -0.049377863 | -0.066910122 | -0.057953271 | -0.008575407 | 0.008956851 | 0.109276367 | 0.003413468 | 0.02116648 | 0.739782375 | 0.613926377 |
| Cep55 | S | | 434 | -0.053139202 | -0.065100946 | -0.207273173 | -0.154133971 | -0.142172228 | 0.2687306 | 0.10932069 | 0.054814514 | 0.100459847 | 0.120065869 |
| Glcci1 | S | | 214 | -0.170711896 | -0.102136599 | -0.037835739 | 0.132876158 | 0.06430086 | 0.054431422 | 0.038541827 | 0.605305421 | 0.165588622 | 0.398526043 |
| Mycbp2 | S | | 2983 | -0.208547604 | -0.07149279 | -0.122626218 | 0.085921386 | -0.051133428 | 0.031183437 | 0.493492356 | 0.381016239 | 0.530202538 | 0.726864064 |
| Dagla | S | | 943 | -0.271005087 | -0.060069205 | -0.288438407 | -0.01743332 | -0.228369202 | 0.18623091 | 0.782742825 | 0.260675018 | 0.9266995 | 0.322881641 |
| Npm1 | S | | 106 | -0.131935719 | -0.111354599 | -0.221337047 | -0.089401328 | -0.109982448 | 0.149509027 | 0.351650294 | 0.222492067 | 0.577507335 | 0.528305451 |
| Efr3a | S | | 219 | -0.047127085 | -0.109430673 | -0.150515691 | -0.103388606 | -0.041085018 | 0.594318866 | 0.118682682 | 0.017736956 | 0.271672026 | 0.468042798 |
| Zzef1 | S | | 240 | -0.106914602 | -0.055360879 | -0.159226712 | -0.05231211 | -0.103865832 | 0.159045238 | 0.237874358 | 0.0489278 | 0.492593538 | 0.144340316 |
| Myo9b | T | | 1317 | -0.040738557 | -0.081722046 | -0.230216329 | -0.189477772 | -0.148494283 | 0.549508037 | 0.322888226 | 0.01858646 | 0.048886619 | 0.125756573 |
| Zc3h13 | S | | 1275 | -0.07902351 | -0.108156966 | -0.103674104 | -0.024650594 | 0.004482862 | 0.34716105 | 0.009912791 | 0.179305726 | 0.783189702 | 0.941434725 |
| Bcl10 | S | | 138 | -0.10278183 | -0.029312612 | -0.210754844 | -0.107973014 | -0.181442231 | 0.192533123 | 0.821303839 | 0.01657402 | 0.113877059 | 0.219714264 |
| Bcl10 | S | | 141 | -0.10278183 | -0.029312612 | -0.210754844 | -0.107973014 | -0.181442231 | 0.192533123 | 0.821303839 | 0.01657402 | 0.113877059 | 0.219714264 |
| Msh2 | S | | 921 | -0.093011723 | -0.104787034 | -0.275228171 | -0.182216448 | -0.170441137 | 0.419505494 | 0.316785857 | 0.106311098 | 0.226120186 | 0.244731741 |
| Gab1 | S | | 420 | -0.02510508 | -0.142011087 | -0.332962163 | -0.307857083 | -0.190951076 | 0.836016398 | 0.180097056 | 0.110973453 | 0.13807326 | 0.283349012 |
| Ranbp3 | S | | 282 | -0.094540878 | -0.158647743 | -0.204543137 | -0.110002259 | -0.045895394 | 0.289360791 | 0.139018703 | 0.275865855 | 0.527125768 | 0.788278769 |
| Myc Mycn | S | 62;62 | | 0.052802985 | -0.190913624 | -0.09210203 | -0.144905015 | 0.098811594 | 0.112788259 | 0.002916932 | 0.004990549 | 0.014427121 | 0.056922613 |
| Helb | S | | 946 | -0.085384625 | -0.045495342 | -0.092458976 | -0.007074351 | -0.046963634 | 0.127318657 | 0.096990628 | 0.007951288 | 0.866192012 | 0.003553129 |
| Scaf1 | S | | 710 | -0.104379009 | -0.14110444 | -0.168327444 | -0.063948434 | -0.027223004 | 0.283647686 | 0.214696776 | 0.130948362 | 0.518017659 | 0.800571104 |
| Etv6 | S | | 265 | 0.00065971 | -0.19541355 | -0.055073423 | -0.055733132 | 0.140340128 | 0.992743266 | 0.002120774 | 0.157822228 | 0.473456938 | 0.016957785 |
| Flrt2 | S | | 651 | 0.00148044 | -0.152853283 | -0.269824681 | -0.271305121 | -0.116971398 | 0.983233047 | 0.131051573 | 0.064289381 | 0.080618883 | 0.340397141 |
| Usp6nl | S | | 389 | -0.045394048 | -0.065997706 | -0.148237859 | -0.102843811 | -0.082240153 | 0.412920992 | 0.348526469 | 0.002310285 | 0.121435622 | 0.259810198 |
| Ppfibp1 | S | | 589 | -0.300243776 | 0.099270309 | -0.310147258 | -0.009903481 | -0.409417567 | 0.106716282 | 0.604452413 | 0.116473933 | 0.954788659 | 0.092186517 |
| Kank2 | S | | 157 | -0.062344789 | -0.032676765 | -0.16397119 | -0.101626401 | -0.131294425 | 0.094691877 | 0.600818456 | 0.007170654 | 0.056359699 | 0.095918835 |
| Pla2g4a | T | | 268 | -0.158916079 | 0.047241419 | -0.226688823 | -0.067772744 | -0.273930242 | 0.011878906 | 0.40074085 | 0.068066743 | 0.458939678 | 0.043933815 |
| Leo1 | S | | 262 | -0.004733851 | -0.1880222 | -0.127723056 | -0.122989205 | 0.060299145 | 0.903627696 | 0.006774559 | 0.250095726 | 0.264352555 | 0.544755713 |
| Dtnb | S | | 524 | -0.148617972 | -0.196051904 | 0.027027826 | 0.175645798 | 0.22307973 | 0.026282954 | 0.056406343 | 0.809658015 | 0.193406134 | 0.127695147 |
| U2surp | S | | 238 | 0.048667699 | -0.168516975 | -0.195680006 | -0.244347704 | -0.027163031 | 0.301796212 | 0.005561807 | 0.072426158 | 0.029028053 | 0.684940479 |
| Ring1 | S | | 38 | -0.244166401 | -0.018665673 | -0.129199823 | 0.114966579 | -0.11053415 | 0.101194002 | 0.806891173 | 0.062121529 | 0.345131053 | 0.201906644 |
| Ints10 | S | | 231 | -0.067163705 | -0.144110165 | -0.086380211 | -0.019216506 | 0.057729955 | 0.262140023 | 0.051630911 | 0.089484089 | 0.731154334 | 0.340656465 |
| Ccnl2 | S | | 335 | -0.042726401 | -0.131513607 | -0.041406851 | 0.00131955 | 0.090106755 | 0.32048082 | 0.000830776 | 0.298822228 | 0.976887743 | 0.070104921 |
| Bcl9l | S | | 1475 | -0.199047269 | 0.056489774 | -0.176292178 | 0.022755091 | -0.232781952 | 0.012626845 | 0.518476721 | 0.078719041 | 0.771693454 | 0.062054497 |
| Wdr91 | S | | 257 | 0.009977573 | -0.159751381 | -0.097817751 | -0.107795324 | 0.06193363 | 0.641644874 | 0.003532119 | 0.059014797 | 0.045303482 | 0.16336255 |
| 5031439G07Rik | S | | 351 | -0.144411795 | -0.069529862 | -0.142802004 | 0.001609791 | -0.073272142 | 0.119755031 | 0.529886283 | 0.054086106 | 0.980295953 | 0.476541401 |
| Maco1 | S | | 305 | 0.067963407 | -0.18546944 | -0.237419268 | -0.305382675 | -0.051949828 | 0.482629731 | 0.061325972 | 0.030143595 | 0.038055048 | 0.556046037 |
| Ubr5 | S | | 2120 | -0.005162846 | -0.201642207 | -0.080191433 | -0.075028588 | 0.121450773 | 0.950534746 | 0.018917631 | 0.045297954 | 0.398752669 | 0.092139945 |
| Iqsec2 | S | | 1133 | -0.127244272 | -0.062553595 | -0.040041337 | 0.087202935 | 0.022512258 | 0.000439593 | 0.3251489 | 0.563649105 | 0.268478559 | 0.780014004 |
| Tbc1d1 | T | | 1242 | -0.045353033 | -0.116620353 | -0.12868073 | -0.083327696 | -0.012060376 | 0.463098198 | 0.013953865 | 0.126697141 | 0.315512758 | 0.847922547 |
| Syn1 | S | | 568 | -0.200691035 | 0.084063634 | -0.496921011 | -0.296229976 | -0.580984645 | 0.263699325 | 0.670623972 | 0.098043377 | 0.26357052 | 0.075097847 |
| Tjp2 | S | | 395 | -0.150686213 | -0.173869534 | 0.076286928 | 0.226973142 | 0.250156462 | 0.004060563 | 0.017720031 | 0.219256713 | 0.021661755 | 0.01219674 |
| Tspyl2 | S | | 652 | -0.025970263 | -0.05874555 | -0.148200152 | -0.122229889 | -0.089454602 | 0.685659267 | 0.089572222 | 0.002828081 | 0.140858112 | 0.003894547 |

| | | | | | | | | | | | | |
|---|---|---|---|---|---|---|---|---|---|---|---|---|
| Tspyl2 | S | 655 | -0.025970263 | -0.05874555 | -0.148200152 | -0.122229889 | -0.089454602 | 0.685659267 | 0.089572222 | 0.002828081 | 0.140858112 | 0.003894547 |
| Pdcl3 | S | 237 | -0.058037678 | -0.101738271 | -0.156664411 | -0.098626734 | -0.05492614 | 0.243135103 | 0.063086897 | 0.107588558 | 0.251836085 | 0.484064636 |
| Gsk3a Gsk3b | Y | 279;216 | -0.137765183 | -0.055913778 | -0.107723436 | 0.030041747 | -0.051809657 | 0.023936179 | 0.359751892 | 0.150682896 | 0.594000448 | 0.436647079 |
| Arhgap21 | S | 1699 | -0.056811704 | -0.097016677 | -0.175079773 | -0.118268068 | -0.078063096 | 0.448021382 | 0.168721447 | 0.031465261 | 0.079544911 | 0.094624241 |
| Taf1d | T | 213 | 0.070719449 | -0.206268027 | -0.189210457 | -0.259929906 | 0.01705757 | 0.208422286 | 0.007572512 | 0.056978332 | 0.045887795 | 0.799166381 |
| Tle3 | S | 263 | -0.000341909 | -0.094420242 | -0.139187082 | -0.138845173 | -0.04476684 | 0.988919834 | 0.02787033 | 0.007693164 | 0.012684437 | 0.251964277 |
| Sh2b3 | S | 20 | -0.035948235 | -0.126726218 | -0.215548247 | -0.179600012 | -0.08882203 | 0.422019333 | 0.150821234 | 0.079007295 | 0.117464962 | 0.398651255 |
| Dync1li1 | S | 207 | -0.00121282 | -0.08246903 | -0.175560887 | -0.174348067 | -0.093091858 | 0.970074604 | 0.085240203 | 0.005970948 | 0.008580173 | 0.071360447 |
| Larp1 | S | 828 | -0.033061805 | -0.068272153 | -0.139052792 | -0.105990987 | -0.070780638 | 0.231191592 | 0.430235058 | 0.001634825 | 0.010593434 | 0.415330542 |
| Nedd4 | T | 838 | -0.203883015 | 0.013641355 | -0.272490608 | -0.068607594 | -0.286131964 | 0.199945253 | 0.80880713 | 0.056335123 | 0.640654081 | 0.045391494 |
| Hdgfl2 | S | 417 | -0.030470307 | -0.116896 | -0.126429624 | -0.095959316 | -0.009533623 | 0.492942228 | 0.049090713 | 0.030741666 | 0.041074804 | 0.806344084 |
| Rai14 | S | 424 | 0.020156694 | -0.04308547 | -0.222291282 | -0.242447976 | -0.179205812 | 0.477641946 | 0.137358947 | 0.001066033 | 0.002128587 | 0.006464994 |
| Kansl1 | S | 268 | -0.066492863 | -0.245330143 | -0.005723832 | 0.060769031 | 0.23960631 | 0.2417381 | 0.008776866 | 0.941067427 | 0.402921476 | 0.031760904 |
| Prkd1 | S | 427 | -0.102254048 | -0.0642688 | -0.144507545 | -0.042253496 | -0.080238745 | 0.111205855 | 0.278838744 | 0.05706252 | 0.521155676 | 0.263210181 |
| Incenp | S | 259 | -0.015845267 | -0.043818672 | -0.130456641 | -0.114611374 | -0.086637969 | 0.558487546 | 0.06561912 | 0.0008354 | 0.022069984 | 0.003030572 |
| Bok | S | 7 | -0.193419775 | 0.065799308 | -0.255839454 | -0.06241968 | -0.321638762 | 0.110104543 | 0.596730166 | 0.033152238 | 0.399685908 | 0.047002744 |
| Bok | S | 8 | -0.193419775 | 0.065799308 | -0.255839454 | -0.06241968 | -0.321638762 | 0.110104543 | 0.596730166 | 0.033152238 | 0.399685908 | 0.047002744 |
| Mapk8ip3 | T | 276 | -0.003904997 | -0.147006595 | -0.384119666 | -0.380214669 | -0.237113071 | 0.96624086 | 0.063349192 | 0.227698573 | 0.230140424 | 0.405365488 |
| Exoc6b | S | 264 | -0.082443869 | -0.421670541 | -0.266448697 | -0.184004828 | 0.155221843 | 0.757278261 | 0.198495142 | 0.507718069 | 0.638176542 | 0.703874081 |
| Cdv3 | S | 155 | -0.139190843 | -0.121966776 | -0.114735668 | 0.024455175 | 0.007231108 | 0.277122228 | 0.023257944 | 0.452587147 | 0.88299483 | 0.95877934 |
| Phrf1 | S | 838 | -0.029151075 | -0.295229708 | -0.11183239 | -0.082681315 | 0.183397318 | 0.882270097 | 0.027450132 | 0.421704929 | 0.673931327 | 0.165125757 |
| Phrf1 | S | 840 | -0.029151075 | -0.295229708 | -0.11183239 | -0.082681315 | 0.183397318 | 0.882270097 | 0.027450132 | 0.421704929 | 0.673931327 | 0.165125757 |
| Cavin2 | S | 247 | -0.253948695 | 0.041507526 | -0.376661251 | -0.122712556 | -0.418168777 | 0.24992188 | 0.842916585 | 0.120802202 | 0.487467522 | 0.060634856 |
| Itprid2 | S | 268 | -0.164902943 | -0.050122147 | -0.23811578 | -0.073212837 | -0.187993633 | 0.119510042 | 0.63338428 | 0.162373434 | 0.617282852 | 0.263768488 |
| Pip4p2 | S | 32 | -0.096967839 | -0.028660458 | -0.374891331 | -0.277923492 | -0.346230873 | 0.073835176 | 0.831433048 | 0.162361849 | 0.256505678 | 0.187724484 |
| U2surp | S | 236 | 0.103232747 | -0.139542991 | -0.297291304 | -0.400524052 | -0.157748313 | 0.195452734 | 0.054898844 | 0.009672986 | 0.006283168 | 0.082874934 |
| Prune2 | S | 1072 | -0.102319587 | -0.076307738 | -0.110519176 | -0.00819959 | -0.034211439 | 0.219759249 | 0.023995107 | 0.132703385 | 0.920032478 | 0.558209136 |
| Lats1 | S | 204 | -0.057055024 | -0.065914073 | -0.11512651 | -0.058071486 | -0.049212437 | 0.163889793 | 0.200751265 | 0.007265998 | 0.118851009 | 0.288323099 |
| Hivep2 | S | 1434 | -0.040148159 | -0.152758009 | -0.343803121 | -0.303654962 | -0.191045112 | 0.728556749 | 0.296219023 | 0.119343435 | 0.157071407 | 0.338397125 |
| Fhod1 | S | 1170;1551 | -0.256019686 | 0.162561934 | -0.443951729 | -0.187932043 | -0.606513662 | 0.108589809 | 0.531127208 | 0.122550021 | 0.432769378 | 0.094319435 |
| Pelo | S | 55 | -0.062020401 | -0.020609756 | -0.240379736 | -0.178359335 | -0.21976998 | 0.542170535 | 0.890410928 | 0.003897093 | 0.144552119 | 0.223275205 |
| Melk | S | 490 | -0.04742526 | -0.151462705 | -0.081468393 | -0.034043133 | 0.069994312 | 0.470995003 | 0.011116477 | 0.164548015 | 0.588408524 | 0.170294104 |
| Mtmr10 | T | 764 | 0.003587006 | -0.133923699 | -0.179990474 | -0.18357748 | -0.046066775 | 0.917844887 | 0.055078904 | 0.030777419 | 0.02587378 | 0.461211918 |
| Tuba1b Tuba4a | S | 340;340 | -0.175144383 | 0.018805831 | -0.16326356 | 0.011880823 | -0.182069391 | 0.032759669 | 0.829616341 | 0.05794218 | 0.804128926 | 0.068586413 |
| Spag9 | S | 109 | -0.058592245 | -0.004459992 | -0.19498878 | -0.136396535 | -0.190528788 | 0.271022715 | 0.922551047 | 0.002001685 | 0.034324351 | 0.006349896 |
| Pacs1 | S | 493 | 0.046983403 | -0.129915829 | -0.179096621 | -0.226080024 | -0.049180793 | 0.235476316 | 0.007169811 | 0.016745098 | 0.00700193 | 0.29166518 |
| Paxbp1 | S | 16 | 0.089310127 | -0.367496727 | -0.127040741 | -0.216350868 | 0.240455986 | 0.499126391 | 0.01072951 | 0.372599836 | 0.195460094 | 0.131875819 |
| Psd3 | S | 316 | -0.080376146 | -0.063909136 | -0.173636131 | -0.093259985 | -0.109726995 | 0.08920097 | 0.15963311 | 0.115060415 | 0.319800552 | 0.261404377 |
| Plekhm2 | S | 244 | 0.106297004 | -0.163032215 | -0.259046423 | -0.365343427 | -0.096014209 | 0.090952435 | 0.022901903 | 0.008182976 | 0.014491491 | 0.155864426 |
| Trappc12 | S | 309 | -0.09312128 | -0.04322784 | -0.085523058 | 0.007598223 | -0.042295218 | 0.010682074 | 0.40079769 | 0.030468434 | 0.779592542 | 0.421478432 |
| Sacs | S | 2513 | -0.145969953 | 0.035639579 | -0.263775063 | -0.11780511 | -0.299414642 | 0.23159235 | 0.803436386 | 0.019137129 | 0.32605756 | 0.109752209 |
| Prrc2c | S | 841 | -0.035980478 | -0.091302125 | -0.207925742 | -0.171945265 | -0.116623618 | 0.601355602 | 0.06569029 | 0.061735969 | 0.106532507 | 0.18137963 |
| Bcor | S | 1443 | -0.151882296 | -0.187308475 | 0.025677005 | 0.177559301 | 0.212985481 | 0.042222459 | 0.043612328 | 0.850363681 | 0.251502797 | 0.188944663 |
| Arhgap10 | S | 710 | -0.004675796 | -0.171501578 | -0.121790742 | -0.117114946 | 0.049710976 | 0.957974685 | 0.019235709 | 0.077970503 | 0.269141256 | 0.42594892 |
| Zc3h13 | S | 1069 | 0.037304284 | -0.116818146 | -0.12402107 | -0.161325354 | -0.007202924 | 0.196947116 | 0.016732342 | 0.000694879 | 0.00346792 | 0.809323187 |
| Phrf1 | T | 1376 | 0.036884565 | -0.182010546 | -0.192879091 | -0.229763656 | -0.010868545 | 0.734987426 | 0.14984184 | 0.013221126 | 0.117611528 | 0.913461622 |
| Pdcl3 | S | 65 | -0.017919911 | -0.08934482 | -0.153930649 | -0.136010737 | -0.064585829 | 0.543620816 | 0.019722205 | 0.042297404 | 0.064067579 | 0.255842046 |
| Mink1 | S | 729 | -0.08277304 | -0.064185557 | -0.201546729 | -0.118773688 | -0.137361172 | 0.195356842 | 0.313859397 | 0.059796477 | 0.192819388 | 0.149355489 |
| Map4 | S | 785 | 0.000510996 | -0.041279727 | -0.145302327 | -0.145813322 | -0.1040226 | 0.981501637 | 0.111741259 | 0.000257836 | 0.001042439 | 0.005401001 |
| Ppp6r1 | S | 529 | -0.02108212 | -0.067674542 | -0.118878168 | -0.097796048 | -0.051203626 | 0.859831425 | 0.041757383 | 0.001643642 | 0.449461866 | 0.104916727 |
| Ppp6r1 | S | 530 | -0.02108212 | -0.067674542 | -0.118878168 | -0.097796048 | -0.051203626 | 0.859831425 | 0.041757383 | 0.001643642 | 0.449461866 | 0.104916727 |
| Ppp6r1 | S | 531 | -0.02108212 | -0.067674542 | -0.118878168 | -0.097796048 | -0.051203626 | 0.859831425 | 0.041757383 | 0.001643642 | 0.449461866 | 0.104916727 |
| Thrap3 | S | 253 | 0.0293846 | -0.074588558 | -0.125749801 | -0.155134401 | -0.051161243 | 0.277548741 | 0.008407544 | 0.000413587 | 0.003547906 | 0.027487159 |
| Uhrf1 | S | 161 | -0.130419543 | -0.041074966 | -0.130476774 | -5.72308E-05 | -0.089401808 | 0.013902509 | 0.428948758 | 0.198492895 | 0.999436201 | 0.341815638 |
| Cdk16 | S | 95 | -0.189826686 | -0.071376997 | -0.089502276 | 0.10032441 | -0.018125279 | 0.024638604 | 0.359326571 | 0.408510876 | 0.368421898 | 0.867632843 |
| Bclaf1 | S | 448 | -0.06099243 | -0.108298053 | -0.133245549 | -0.072253119 | -0.024947496 | 0.209026432 | 0.18577505 | 0.03988612 | 0.215183577 | 0.7300806 |
| Fip1l1 | S | 483 | -0.050785696 | -0.071593157 | -0.140130662 | -0.089344966 | -0.068537505 | 0.302011989 | 0.04092815 | 0.042129017 | 0.164133974 | 0.210568472 |
| Phldb2 | S | 82 | 0.213646444 | -0.278973344 | -0.231387614 | -0.445034058 | 0.04758573 | 0.011052014 | 0.002647376 | 0.006169933 | 0.000727309 | 0.149528133 |
| Rpl13 | S | 106 | 0.032584507 | -0.116076391 | -0.183307971 | -0.215892478 | -0.067231581 | 0.354965072 | 0.00681518 | 0.027167468 | 0.023051579 | 0.259778897 |
| Hectd1 | T | 1805 | -0.159681284 | -0.188629062 | -0.293639441 | -0.133958157 | -0.105010379 | 0.401226877 | 0.367324543 | 0.326821418 | 0.653063128 | 0.728952719 |
| Pola2 | S | 126 | -0.171844703 | -0.080257518 | -0.140984694 | 0.030860009 | -0.060727176 | 0.088867865 | 0.518384216 | 0.152860856 | 0.635446979 | 0.577670332 |
| Map1a | T | 633 | -0.035340432 | -0.048259664 | -0.153870375 | -0.118529944 | -0.105610711 | 0.11355603 | 0.212234583 | 0.010381629 | 0.042249444 | 0.051149129 |
| Map1b | T | 945 | -0.084503101 | -0.116187587 | -0.088431209 | -0.003928108 | 0.027756379 | 0.213915876 | 0.043243193 | 0.131319511 | 0.943243855 | 0.45859021 |
| Trp53bp1 | S | 1615 | -0.063385525 | -0.095827679 | -0.229215522 | -0.165829997 | -0.133387843 | 0.495273213 | 0.101903053 | 0.111003863 | 0.231148473 | 0.278968021 |
| Eif3b | S | 111 | 0.01119382 | -0.078758188 | -0.168156749 | -0.179350569 | -0.089398561 | 0.754706823 | 0.076575324 | 0.002792522 | 0.002774425 | 0.035225384 |
| Ppp1r12c | S | 411 | -0.135029884 | -0.066784356 | -0.139379753 | -0.004349868 | -0.072595397 | 0.088761241 | 0.394150963 | 0.099829449 | 0.916629074 | 0.228762149 |
| Specc1 | S | 914 | -0.112678251 | -0.104882524 | -0.044272007 | 0.068406244 | 0.060610517 | 0.014633825 | 0.115087953 | 0.331810239 | 0.12466978 | 0.313659561 |
| Nedd4l | S | 477 | -0.021084494 | -0.010775138 | -0.15230518 | -0.131220686 | -0.141530043 | 0.323700466 | 0.712739447 | 0.000121036 | 0.000619648 | 0.013687787 |
| Csrp1 | S | 192 | -0.161002446 | -0.001055389 | -0.324752719 | -0.163750273 | -0.32369733 | 0.164578693 | 0.992722979 | 0.111230574 | 0.350951547 | 0.11833693 |
| Rps3 | T | 221 | -0.039865664 | -0.122123757 | -0.073713471 | -0.033847807 | 0.048410287 | 0.21118265 | 0.006803455 | 0.178157415 | 0.454558997 | 0.31806775 |
| Zc3h12c | S | 492 | -0.121153456 | -0.063123931 | -0.13304838 | -0.011894924 | -0.069924449 | 0.035673179 | 0.215928359 | 0.233911324 | 0.895914621 | 0.476991379 |
| Mcl1 | S | 81 | -0.039970145 | -0.306758139 | -0.160202205 | -0.12023206 | 0.146555934 | 0.649041892 | 0.096150655 | 0.356257818 | 0.485150591 | 0.450427121 |
| Epb41l2 | S | 38 | -0.132884871 | -0.0506473 | -0.079214946 | 0.053669926 | -0.028567646 | 0.010404489 | 0.172902353 | 0.301297494 | 0.450593789 | 0.664850222 |
| Eif5b | S | 9 | -0.059687902 | -0.19369324 | -0.157766018 | -0.098078117 | 0.035927221 | 0.40478178 | 0.053787513 | 0.362153349 | 0.546580975 | 0.822841464 |
| Eif5b | S | 12 | -0.059687902 | -0.19369324 | -0.157766018 | -0.098078117 | 0.035927221 | 0.40478178 | 0.053787513 | 0.362153349 | 0.546580975 | 0.822841464 |
| Map1a | S | 2603 | -0.144108414 | -0.038946895 | -0.055295777 | 0.088812636 | -0.016348882 | 0.005110935 | 0.417887867 | 0.141831609 | 0.026204734 | 0.65552258 |
| Ehbp1 | S | 446 | -0.094170619 | -0.125435138 | -0.115404779 | -0.021234159 | 0.010030359 | 0.210752494 | 0.118639919 | 0.130673557 | 0.670474791 | 0.849804432 |
| Mertk | T | 945 | 0.066961856 | -0.178566997 | -0.25516049 | -0.322122346 | -0.076593493 | 0.281504651 | 0.049093309 | 0.030860394 | 0.01798722 | 0.406169192 |
| Zswim8 | S | 1090 | 0.003203378 | -0.1664338 | -0.168964964 | -0.172168341 | -0.002531164 | 0.953659272 | 0.097754855 | 0.028707893 | 0.029908629 | 0.973574556 |
| Txnl1 | S | 113 | -0.19733187 | -0.059091856 | -0.128638008 | 0.068693862 | -0.069546152 | 0.021827561 | 0.51450844 | 0.459264401 | 0.668224579 | 0.677388302 |
| Akt1 | S | 124 | -0.151737955 | -0.014812899 | -0.234584281 | -0.082846326 | -0.219771382 | 0.10587241 | 0.867457439 | 0.088962423 | 0.47568789 | 0.118144253 |
| Clasp2 | S | 53 | -0.024904508 | -0.037117958 | -0.116456261 | -0.091551754 | -0.079338303 | 0.233300819 | 0.304430315 | 0.000254359 | 0.006635783 | 0.08444224 |
| Chd9 | S | 549 | -0.006101062 | -0.235990059 | -0.052415412 | -0.04631435 | 0.183574647 | 0.941460867 | 0.006375314 | 0.246798525 | 0.60634802 | 0.014376516 |
| Gorasp2 | S | 432 | -0.102736096 | -0.095348049 | -0.092527535 | 0.010208561 | 0.002820514 | 0.131880673 | 0.068854421 | 0.148071786 | 0.877715829 | 0.960535433 |
| Pcbp1 | S | 171 | -0.174906133 | -0.061872863 | -0.168307828 | 0.006598305 | -0.106434965 | 0.08516548 | 0.319124951 | 0.286143294 | 0.963627319 | 0.4653406 |
| Psmb6 | S | 57 | 0.064318878 | -0.123676224 | -0.227774084 | -0.292092962 | -0.10409786 | 0.331965414 | 0.050137222 | 0.008743902 | 0.008559791 | 0.121033702 |
| Gapdh | T | 97 | -0.120944648 | -0.094860353 | -0.108498133 | 0.012446515 | -0.01363778 | 0.03228783 | 0.290052468 | 0.239640387 | 0.871709812 | 0.890262943 |
| Vcl | Y | 822 | -0.156104063 | -0.098156989 | -0.124516014 | 0.03158805 | -0.026359024 | 0.188514682 | 0.106577859 | 0.274272936 | 0.80190655 | 0.792210927 |
| Ppig | S | 644 | -0.009552696 | -0.192575059 | -0.075263859 | -0.065711163 | 0.1173112 | 0.769640468 | 0.003861848 | 0.312914649 | 0.363968176 | 0.160322363 |
| Map1a | S | 1062 | -0.08491776 | -0.063646157 | -0.155631757 | -0.070713997 | -0.091985599 | 0.135149695 | 0.189019515 | 0.070907972 | 0.324730398 | 0.213980103 |
| Dock1 | S | 1858 | -0.266680929 | 0.012318188 | -0.15038451 | 0.116296419 | -0.162702697 | 0.032272743 | 0.916876022 | 0.251986959 | 0.320323201 | 0.225923172 |
| Arhgap5 | S | 1217 | -0.121097604 | -0.059901682 | -0.076738716 | 0.044358888 | -0.016837033 | 0.007680831 | 0.354721597 | 0.180627963 | 0.369558332 | 0.799777293 |
| Arhgap39 | S | 299 | -0.013181016 | -0.168453319 | -0.076669182 | -0.063488166 | 0.091784137 | 0.74113502 | 0.011992611 | 0.0686766 | 0.090435506 | 0.090378754 |
| Ccdc97 | S | 210 | -0.063000513 | -0.319491508 | 0.008734173 | 0.071734686 | 0.328225681 | 0.124539441 | 0.029505798 | 0.797783722 | 0.045713236 | 0.039828554 |

| | | | | | | | | | | | | |
|---|---|---|---|---|---|---|---|---|---|---|---|---|
| Mus81 | S | 95 | -0.184306163 | -0.209139967 | -0.054965187 | 0.129340976 | 0.15417478 | 0.190238888 | 0.162625432 | 0.453193771 | 0.3169814 | 0.26654629 |
| Ppp1r18 | S | 174 | -0.04931898 | -0.082272068 | -0.082168009 | -0.032849029 | 0.000104059 | 0.029946011 | 0.016983578 | 0.167916221 | 0.503466038 | 0.998284212 |
| Kif4 | S | 1224 | -0.027211326 | -0.115513667 | -0.095260396 | -0.068049069 | 0.020253271 | 0.494653769 | 0.02146079 | 0.031711453 | 0.163967234 | 0.596988936 |
| Dst | S | 2717 | -0.162794856 | -0.20999955 | -0.1377927 | 0.025002156 | 0.07220685 | 0.410463896 | 0.149636823 | 0.380413681 | 0.903639377 | 0.658751098 |
| Dst | S | 2721 | -0.162794856 | -0.20999955 | -0.1377927 | 0.025002156 | 0.07220685 | 0.410463896 | 0.149636823 | 0.380413681 | 0.903639377 | 0.658751098 |
| Zfp106 | S | 1218 | 0.005071402 | -0.167172811 | -0.105200441 | -0.110271843 | 0.06197237 | 0.907177698 | 0.027296236 | 0.024843082 | 0.077388343 | 0.223078222 |
| Ajuba | S | 272 | -0.08029769 | -0.064058076 | -0.189795909 | -0.109498219 | -0.125737833 | 0.214305227 | 0.132772731 | 0.09675512 | 0.2592387 | 0.194484203 |
| Tbc1d1 | S | 521 | -0.137949774 | -0.012917769 | -0.166356175 | -0.028406401 | -0.153438406 | 0.14560864 | 0.843036953 | 0.018558502 | 0.722658601 | 0.0792136 |
| Lrrc45 | S | 661 | -0.070879005 | -0.204212753 | -0.151103512 | -0.080224507 | 0.053109241 | 0.423429576 | 0.037971665 | 0.523314392 | 0.724418709 | 0.813041832 |
| Sipa1l3 | S | 1538 | -0.072745621 | -0.081577859 | -0.038354756 | 0.034390865 | 0.043223103 | 0.010604348 | 0.010385366 | 0.384907303 | 0.422333844 | 0.33433288 |
| Tlk1 | S | 134 | -0.222295996 | -0.040493774 | -0.024976405 | 0.197319591 | 0.015517368 | 0.002669218 | 0.626185389 | 0.716787237 | 0.043299636 | 0.856264092 |
| Osbpl11 | S | 194 | -0.127750979 | 0.026048577 | -0.299393594 | -0.171642615 | -0.325442171 | 0.225931744 | 0.80598044 | 0.031946224 | 0.156511657 | 0.034901381 |
| Spty2d1 | S | 596 | 0.040640879 | -0.149856119 | -0.134822038 | -0.175462917 | 0.015034081 | 0.649467948 | 0.00206214 | 0.109727467 | 0.13656977 | 0.811376001 |
| Dync1i2 | S | 81 | -0.039871515 | -0.058676666 | -0.106716027 | -0.066844512 | -0.048039361 | 0.219009953 | 0.125085173 | 0.003319157 | 0.082593363 | 0.189963746 |
| Rrp15 | S | 265 | -0.040823849 | -0.154802052 | -0.079393222 | -0.038569373 | 0.07540883 | 0.405936841 | 0.016587602 | 0.152282701 | 0.229213123 | 0.078607246 |
| Adam17 | S | 822 | -0.052360893 | -0.026439508 | -0.173536804 | -0.121175911 | -0.147097296 | 0.170772132 | 0.656441623 | 0.004658771 | 0.028038769 | 0.070653478 |
| Prcp | S | 238 | -0.07773408 | -0.093009605 | -0.135627884 | -0.057893803 | -0.042618279 | 0.112757339 | 0.119837435 | 0.137466035 | 0.444935589 | 0.582751922 |
| Sun1 | S | 139 | 0.045630207 | -0.095938081 | -0.159338014 | -0.204968221 | -0.063399933 | 0.302452715 | 0.00063656 | 0.02467343 | 0.01182653 | 0.190143996 |
| Map1a | T | 1794 | -0.069393905 | -0.084317874 | -0.121961802 | -0.052567897 | -0.037643927 | 0.215597744 | 0.052680138 | 0.083607943 | 0.412050157 | 0.489129027 |
| Nelfe | S | 115 | -0.217112459 | 0.017231477 | -0.178293954 | 0.038818505 | -0.195525431 | 0.050515799 | 0.897580938 | 0.121983058 | 0.698701052 | 0.209900608 |
| Cacnb3 | S | 152 | -0.075591688 | -0.113740142 | -0.055679405 | 0.019912284 | 0.058060737 | 0.036362715 | 0.052391828 | 0.210289677 | 0.615004813 | 0.27446621 |
| Ifih1 | S | 645 | -0.251109435 | -0.070791464 | -0.055876078 | 0.195233357 | 0.014915386 | 0.033054875 | 0.40934422 | 0.498461468 | 0.069482161 | 0.823292581 |
| Ifih1 | S | 648 | -0.251109435 | -0.070791464 | -0.055876078 | 0.195233357 | 0.014915386 | 0.033054875 | 0.40934422 | 0.498461468 | 0.069482161 | 0.823292581 |
| Ighmbp2 | S | 676 | -0.039411188 | -0.169158548 | -0.13349827 | -0.094087082 | 0.035660278 | 0.558651073 | 0.087942149 | 0.079685974 | 0.232522871 | 0.670053014 |
| Dkc1 | T | 485 | 0.054308409 | -0.139575931 | -0.121254656 | -0.175563066 | 0.018321275 | 0.24958175 | 0.016653371 | 0.001404918 | 0.026626873 | 0.603452052 |
| Brd7 | S | 289 | 0.002897404 | -0.175748551 | -0.10058107 | -0.103478474 | 0.075167481 | 0.967744593 | 0.011680784 | 0.084019946 | 0.169719869 | 0.037867949 |
| Vps13b | S | 3025 | 0.043145797 | -0.184837538 | -0.183563506 | -0.226709303 | 0.001274032 | 0.389948881 | 0.119698931 | 0.00742173 | 0.005701781 | 0.988808826 |
| Ppp1r11 | S | 82 | -0.130324886 | -0.039417643 | -0.196684147 | -0.066359261 | -0.157266503 | 0.077640252 | 0.649578199 | 0.105471581 | 0.504788792 | 0.19958782 |
| Atxn2l | S | 339 | -0.121713459 | -0.136918096 | -0.178636613 | -0.056923154 | -0.041718517 | 0.346489916 | 0.249053471 | 0.155080255 | 0.502304481 | 0.469836627 |
| Upf1 | S | 1102 | -0.003486597 | -0.121165878 | -0.07347542 | -0.069988822 | 0.047690458 | 0.886578988 | 0.001566805 | 0.051556495 | 0.070315057 | 0.158409944 |
| Phldb2 | S | 383 | -0.065362145 | -0.057459293 | -0.151450736 | -0.086088591 | -0.093991443 | 0.199688693 | 0.319231231 | 0.016075935 | 0.081011883 | 0.110585773 |
| Ppp1ccb Ppp1cc | T | 311 | -0.140101443 | -0.261699821 | 0.060125933 | 0.200227376 | 0.321825754 | 0.054418387 | 0.032594103 | 0.467424662 | 0.074587245 | 0.023056121 |
| Mast4 | S | 1521 | -0.094193939 | -0.125712409 | -0.052985669 | 0.041208271 | 0.07272674 | 0.079325022 | 0.05043911 | 0.253441041 | 0.428711462 | 0.210283103 |
| Piezo1 | S | 1864 | -0.109432044 | 0.04135538 | -0.389633805 | -0.28020176 | -0.430989185 | 0.356607726 | 0.753928385 | 0.035020851 | 0.092391543 | 0.03301469 |
| Ubn2 | S | 13 | 0.006536334 | -0.230433966 | -0.124994156 | -0.13153049 | 0.10543981 | 0.961868197 | 0.01334175 | 0.29779145 | 0.425050389 | 0.372652296 |
| Ppp1r11 | S | 79 | -0.130294576 | -0.038553356 | -0.196346212 | -0.066051636 | -0.157792856 | 0.076809961 | 0.655544874 | 0.105486697 | 0.505649026 | 0.197415496 |
| Svil | S | 220 | -0.161353205 | -0.040358223 | -0.145927678 | 0.015425527 | -0.105569455 | 0.053804148 | 0.557420668 | 0.145911088 | 0.845125634 | 0.229231348 |
| Pak4 | S | 148 | -0.127475087 | -0.10313735 | -0.113726938 | 0.013748149 | -0.010589588 | 0.085719724 | 0.180178398 | 0.262270926 | 0.884195989 | 0.914076981 |
| Flna | S | 2180 | -0.122868443 | -0.105379887 | -0.024193926 | 0.098674517 | 0.081185961 | 0.02717355 | 0.034651282 | 0.610864083 | 0.066199506 | 0.090552774 |
| Sh3pxd2a | S | 420 | -0.022045592 | -0.07904817 | -0.18713241 | -0.165086817 | -0.10808424 | 0.684582302 | 0.179982364 | 0.011938534 | 0.005670818 | 0.020531091 |
| Eef1g | S | 387 | -0.030466456 | -0.123469655 | -0.17155698 | -0.141090524 | -0.048087325 | 0.304845137 | 0.153633172 | 0.05860034 | 0.103794467 | 0.569406531 |
| Pcm1 | S | 93 | -0.153142689 | -0.098973788 | -0.029393697 | 0.123748991 | 0.069580091 | 0.01630143 | 0.130489934 | 0.583792639 | 0.035618076 | 0.209507836 |
| Fxr1 | S | 435 | -0.004897987 | -0.107148448 | -0.105884871 | -0.100986884 | 0.001263577 | 0.93027109 | 0.003666721 | 0.044232231 | 0.163194685 | 0.971467276 |
| Ercc6l | S | 957 | 0.075025267 | -0.183017315 | -0.158919781 | -0.233945048 | 0.024097535 | 0.138605789 | 0.001849263 | 0.054676784 | 0.02021698 | 0.682638803 |
| Prune2 | S | 2592 | 0.059329429 | -0.097001998 | -0.245203432 | -0.30453286 | -0.148201434 | 0.205725117 | 0.098356478 | 0.0028314 | 0.005015395 | 0.034116945 |
| Clint1 | S | 242 | -0.169810559 | -0.11240108 | -0.002694223 | 0.167116336 | 0.109706857 | 0.083246554 | 0.011293033 | 0.97767323 | 0.1824787 | 0.313380789 |
| Zzef1 | S | 1538 | -0.166210953 | 0.030148602 | -0.188822814 | -0.022611861 | -0.218971416 | 0.015221107 | 0.755604334 | 0.126306773 | 0.812799297 | 0.123406623 |
| Tbc1d5 | S | 811 | -0.128369899 | -0.269293544 | -0.158082377 | -0.029712478 | 0.111211167 | 0.487994668 | 0.113558339 | 0.569683436 | 0.919710715 | 0.694468263 |
| C2cd2l | S | 703 | 0.040958316 | -0.115191314 | -0.199692842 | -0.240651158 | -0.084501528 | 0.598978932 | 0.005061529 | 0.068592641 | 0.054828704 | 0.308190779 |
| Rbm15b | S | 559 | 0.036395887 | -0.136204819 | -0.151511276 | -0.187907163 | -0.015306457 | 0.283731907 | 0.000840053 | 0.093844795 | 0.055716497 | 0.80667957 |
| Clip1 | T | 144 | -0.189211749 | -0.162059111 | -0.049241707 | 0.139970042 | 0.112817404 | 0.056289664 | 0.311802324 | 0.465209921 | 0.143140005 | 0.461931952 |
| Myh9 | S | 1326 | -0.059518353 | 0.006939711 | -0.165111643 | -0.10559329 | -0.172051354 | 0.223332794 | 0.882212872 | 0.000684631 | 0.071340482 | 0.028242585 |
| Rnf213 | S | 274 | -0.086913455 | -0.207357023 | -0.044877644 | 0.042035811 | 0.162479379 | 0.284410074 | 0.02632778 | 0.539221163 | 0.34431509 | 0.002456839 |
| Pla2g4a | S | 515 | -0.242011528 | 0.044042105 | -0.166650213 | 0.075361314 | -0.210692319 | 0.017633849 | 0.738054658 | 0.233236674 | 0.540602129 | 0.215568527 |
| Bclaf1 | S | 758 | 0.044595445 | -0.109145921 | -0.159094343 | -0.203689788 | -0.049948421 | 0.546308687 | 0.02308053 | 0.007036532 | 0.056009941 | 0.225500295 |
| Bag3 | S | 274 | -0.049721391 | -0.051357953 | -0.104745857 | -0.055024466 | -0.053387905 | 0.240876978 | 0.080845291 | 0.005393137 | 0.187404091 | 0.015637738 |
| Inpp5f | S | 829 | -0.083887814 | -0.166840442 | -0.079457483 | 0.004430331 | 0.087382959 | 0.256225624 | 0.029006645 | 0.433239847 | 0.961016353 | 0.358389905 |
| Crtc1 | S | 421 | -0.099120035 | -0.066504913 | -0.186318925 | -0.08719889 | -0.119814012 | 0.044576939 | 0.331641561 | 0.211663655 | 0.495431969 | 0.382556563 |
| Sin3b | S | 670 | -0.129873384 | -0.2115685 | -0.134583921 | -0.004710537 | 0.076984578 | 0.494148876 | 0.072749715 | 0.474459675 | 0.983220005 | 0.668337972 |
| Mybl2 | S | 424 | -0.018908484 | -0.107402217 | -0.134440568 | -0.115532084 | -0.027038351 | 0.851651191 | 0.026375003 | 0.030830698 | 0.313631058 | 0.555009263 |
| Sorbs1 | S | 345 | -0.017063783 | -0.135238199 | -0.233931185 | -0.216867402 | -0.098692986 | 0.869816397 | 0.043443207 | 0.152193376 | 0.195359326 | 0.464339927 |
| Larp1 | S | 608 | 0.027481638 | -0.086126483 | -0.125706023 | -0.153187661 | -0.03957954 | 0.451117078 | 0.01145575 | 0.00181047 | 0.014811964 | 0.0433672 |
| Uaca | S | 280 | -0.122524898 | -0.065285679 | -0.224623211 | -0.102098313 | -0.159337532 | 0.203718199 | 0.56923026 | 0.093431072 | 0.354190693 | 0.233420555 |
| Rangap1 | S | 441 | 0.097181733 | -0.132644373 | -0.417254908 | -0.51443664 | -0.284610535 | 0.132738584 | 0.16539187 | 0.015625487 | 0.007019247 | 0.043620875 |
| Dyrk2 Dyrk4 | Y | 380;379 | 0.038697416 | -0.089075094 | -0.19231507 | -0.231012486 | -0.103239975 | 0.561516111 | 0.004792081 | 0.029711229 | 0.032083905 | 0.131801142 |
| Ccdc6 | S | 233 | -0.121562425 | -0.03927075 | -0.09508549 | 0.026476934 | -0.055814741 | 0.024492662 | 0.332218953 | 0.086112733 | 0.551019027 | 0.215507449 |
| Ccdc6 | S | 237 | -0.121562425 | -0.03927075 | -0.09508549 | 0.026476934 | -0.055814741 | 0.024492662 | 0.332218953 | 0.086112733 | 0.551019027 | 0.215507449 |
| Fosl1 | T | 269 | -0.251981383 | -0.661243752 | 0.151080089 | 0.403061472 | 0.81232384 | 0.095246973 | 0.232946811 | 0.366543761 | 0.041679776 | 0.172078025 |
| Chaf1b | S | 456 | 0.029459195 | -0.118264962 | -0.316170478 | -0.345629673 | -0.197905516 | 0.832551974 | 0.075219799 | 0.085358549 | 0.098798878 | 0.19840152 |
| Usp24 | S | 62 | -0.073285907 | -0.089380944 | -0.230041202 | -0.156755294 | -0.140660258 | 0.230517222 | 0.182843003 | 0.164728974 | 0.286650682 | 0.329819177 |
| Grb14 | S | 370 | -0.024113274 | -0.244409447 | -0.109235617 | -0.085122343 | 0.13517383 | 0.823003014 | 0.013110653 | 0.528814821 | 0.634149298 | 0.440471261 |
| Phf20 | S | 159 | 0.005163315 | -0.065280115 | -0.128403676 | -0.133566992 | -0.063123562 | 0.825010587 | 0.025759268 | 0.001714878 | 0.004198585 | 0.048149842 |
| Map1a | S | 1060 | -0.112154505 | -0.012156633 | -0.229772897 | -0.117618391 | -0.217616264 | 0.072714802 | 0.832162957 | 0.07548442 | 0.267403455 | 0.089108138 |
| Nfrkb | S | 351 | -0.27403518 | -0.169930006 | -0.045546951 | 0.228488228 | 0.124383055 | 0.18484216 | 0.159702867 | 0.723518117 | 0.251643783 | 0.28383212 |
| Rabgap1 | S | 987 | -0.109293195 | -0.037050776 | -0.075521086 | 0.033772109 | -0.03847031 | 0.011092731 | 0.471685544 | 0.04091629 | 0.152419885 | 0.417483807 |
| Ehbp1l1 | S | 1383 | -0.072529126 | -0.068611879 | -0.22071845 | -0.148189323 | -0.152106571 | 0.314078176 | 0.448334145 | 0.042539564 | 0.119607202 | 0.132580299 |
| Cobl | S | 294 | -0.095431867 | 0.021094556 | -0.14793166 | -0.052499793 | -0.169026216 | 0.01019855 | 0.669011929 | 0.014837356 | 0.21465877 | 0.031418392 |
| Camsap1 | S | 858 | 0.042833028 | -0.110888755 | -0.13648803 | -0.179321059 | -0.025599275 | 0.802494127 | 0.028277792 | 0.006670762 | 0.354255769 | 0.481963379 |
| Arhgef12 | S | 637 | 0.034340484 | -0.140197424 | -0.078475763 | -0.112816247 | 0.061721661 | 0.02682991 | 4.01021E-06 | 0.14361791 | 0.074422216 | 0.214099823 |
| Zc3h13 | S | 1305 | -0.114117603 | -0.045615491 | -0.104831959 | 0.009285643 | -0.059216469 | 0.035743365 | 0.236511915 | 0.106524677 | 0.866567762 | 0.300697228 |
| Ankrd17 | S | 2038 | 0.021884059 | -0.090266069 | -0.120940205 | -0.142824264 | -0.030674136 | 0.502446231 | 0.01157546 | 0.002706528 | 0.007398732 | 0.122349714 |
| Ankrd17 | S | 2040 | 0.021884059 | -0.090266069 | -0.120940205 | -0.142824264 | -0.030674136 | 0.502446231 | 0.01157546 | 0.002706528 | 0.007398732 | 0.122349714 |
| Cep97 | S | 797 | 0.031772308 | -0.109163618 | -0.375254039 | -0.407026347 | -0.266090421 | 0.760773426 | 0.257636434 | 0.048601545 | 0.043356117 | 0.118720206 |
| R3hcc1l | S | 165 | -0.269585666 | -0.239707336 | -0.190360772 | 0.079224895 | 0.049346565 | 0.433464221 | 0.481957831 | 0.323373518 | 0.80948933 | 0.88055347 |
| Jmjd6 | S | 401 | 0.086032277 | -0.160731557 | -0.374715635 | -0.460747912 | -0.213984078 | 0.29077982 | 0.088637616 | 0.050370407 | 0.045945595 | 0.183127761 |
| Phactr4 | S | 335 | -0.075721018 | -0.135679409 | -0.240458899 | -0.164737881 | -0.10477949 | 0.45898673 | 0.311309957 | 0.118920335 | 0.266861089 | 0.499942752 |
| Phactr4 | S | 337 | -0.075721018 | -0.135679409 | -0.240458899 | -0.164737881 | -0.10477949 | 0.45898673 | 0.311309957 | 0.118920335 | 0.266861089 | 0.499942752 |
| 4930579G24Rik | S | 11 | -0.189059903 | -0.090610455 | -0.118844403 | 0.0702155 | -0.028233948 | 0.218090886 | 0.144582223 | 0.258431884 | 0.62945688 | 0.762495954 |
| Ddx46 | S | 130 | -0.14060827 | -0.275558093 | -0.026385204 | 0.114223066 | 0.249172889 | 0.214891426 | 0.087452219 | 0.82274562 | 0.185779599 | 0.099934456 |
| Baiap2 | T | 361 | -0.053743317 | -0.055509283 | -0.278111394 | -0.224368077 | -0.222602111 | 0.631390969 | 0.679004711 | 0.018661916 | 0.068504462 | 0.139415029 |
| Gpsm1 | S | 543 | -0.125889431 | -0.076575387 | -0.091009866 | 0.034879565 | -0.01443448 | 0.201757071 | 0.015592402 | 0.276575842 | 0.726030803 | 0.838429667 |
| Ranbp2 | S | 796 | -0.044951372 | -0.185702573 | -0.07156349 | -0.026612119 | 0.114139083 | 0.385916146 | 0.00757326 | 0.584942331 | 0.837811938 | 0.411606929 |
| Sec61b | S | 13 | 0.019125172 | -0.12551219 | -0.23476746 | -0.253892631 | -0.109255269 | 0.792665887 | 0.252859344 | 0.013391454 | 0.016136155 | 0.303929292 |
| Dnttip2 | T | 495 | 0.063771842 | -0.241703298 | -0.183563034 | -0.247334877 | 0.058140264 | 0.288182174 | 0.004595714 | 0.261847883 | 0.166633761 | 0.682919615 |

| | | | | | | | | | | | | |
|---|---|---|---|---|---|---|---|---|---|---|---|---|
| Dst | S | 2241 | 0.013481889 | -0.145075956 | -0.169263707 | -0.182745595 | -0.02418775 | 0.851645585 | 0.037935274 | 0.049893652 | 0.077679299 | 0.723538867 |
| Cdc20 | T | 106 | 0.150247249 | -0.095707637 | -0.368478116 | -0.518725365 | -0.272770479 | 0.01388848 | 0.285853474 | 0.000137523 | 0.000460717 | 0.037388649 |
| Tprn | S | 272 | 0.119912442 | -0.161353017 | -0.252165168 | -0.372077609 | -0.09081215 | 0.411357932 | 0.068140512 | 0.015849258 | 0.063673908 | 0.131321944 |
| Baz1a | S | 1416 | 0.032449017 | -0.176607232 | -0.134257267 | -0.166706284 | 0.042349965 | 0.540841628 | 0.031054479 | 0.022895462 | 0.039362741 | 0.447421638 |
| Prrc2b | S | 418 | 0.019731782 | -0.114124589 | -0.131631097 | -0.151362879 | -0.017506508 | 0.561110922 | 0.001525881 | 0.064904029 | 0.045961802 | 0.717109115 |
| Fcho2 | S | 532 | -0.092285435 | -0.163032261 | -0.03527742 | 0.057008015 | 0.127754841 | 0.095063118 | 0.029936942 | 0.614455126 | 0.385229049 | 0.121369154 |
| Srrm2 | S | 2224 | 0.032791219 | -0.069283789 | -0.094841599 | -0.127632818 | -0.02555781 | 0.232579347 | 0.001504392 | 0.000192442 | 0.019420588 | 0.014882188 |
| Flna | S | 2158 | -0.144490688 | -0.127898974 | -0.145537854 | -0.001047166 | -0.01763888 | 0.391527689 | 0.168179345 | 0.170424209 | 0.994903074 | 0.864034054 |
| Tcof1 | T | 803 | 0.061280791 | -0.216593048 | -0.120425774 | -0.181706565 | 0.096167274 | 0.658737481 | 0.005005424 | 0.197930068 | 0.257036519 | 0.267293882 |
| Lima1 | S | 617 | -0.197912292 | -0.005763433 | -0.161854332 | 0.03605796 | -0.156090899 | 0.061971248 | 0.964020543 | 0.111037188 | 0.627165203 | 0.240852263 |
| Leo1 | S | 255 | -0.028647795 | -0.170032234 | -0.039397488 | -0.010749693 | 0.130634746 | 0.640123628 | 0.001274458 | 0.459127372 | 0.871033011 | 0.067214997 |
| Cebpz | S | 949 | -0.119954993 | -0.055897218 | -0.162819824 | -0.042864831 | -0.106922607 | 0.198128142 | 0.501667669 | 0.043056511 | 0.611063248 | 0.232885946 |
| Epb41l2 | S | 582 | 0.033185798 | -0.114121372 | -0.186883049 | -0.220068847 | -0.072761678 | 0.586447586 | 0.079122019 | 0.008220049 | 0.020843728 | 0.144098632 |
| Hspb6 | S | 157 | -0.019100374 | -0.181082435 | -0.181846636 | -0.162746261 | -0.000764201 | 0.887242214 | 0.186809632 | 0.044281715 | 0.262239439 | 0.994233097 |
| Med1 | S | 1455 | 0.097494698 | -0.317554683 | -0.209113477 | -0.306608175 | 0.108441206 | 0.262160328 | 0.117308356 | 0.023004375 | 0.015424505 | 0.497007649 |
| Med1 | S | 1465 | 0.097494698 | -0.317554683 | -0.209113477 | -0.306608175 | 0.108441206 | 0.262160328 | 0.117308356 | 0.023004375 | 0.015424505 | 0.497007649 |
| Inf2 | S | 624 | 0.114468858 | -0.389057389 | -0.176138059 | -0.290606916 | 0.212919331 | 0.397933224 | 0.015431799 | 0.39847108 | 0.187404808 | 0.295228091 |
| Oxr1 | S | 90 | -0.210702872 | 0.001550034 | -0.107572184 | 0.103130687 | -0.109122218 | 0.013632944 | 0.986052513 | 0.218035627 | 0.180915368 | 0.252708533 |
| Tbc1d1 | S | 608 | -0.016112741 | -0.177543773 | -0.167392461 | -0.15127972 | 0.010151312 | 0.77589838 | 0.182037222 | 0.038860548 | 0.049761307 | 0.927058885 |
| Map4 | T | 792 | 0.027340227 | -0.111972944 | -0.276374374 | -0.303714601 | -0.164401431 | 0.722410247 | 0.137888186 | 0.032361499 | 0.029167401 | 0.126650546 |
| Naa38 | S | 29 | -0.038394192 | -0.067638156 | -0.122715048 | -0.084320856 | -0.055076892 | 0.157589263 | 0.014131031 | 0.087086732 | 0.182569558 | 0.331275725 |
| Dab2 | T | 329 | 0.142720821 | -0.267942613 | -0.250360209 | -0.39308103 | 0.017582404 | 0.111773263 | 0.032085195 | 0.027349226 | 0.006501253 | 0.850193701 |
| Tra2a | S | 237 | 0.094641653 | -0.141925817 | -0.277763718 | -0.372405371 | -0.135837901 | 0.025515261 | 0.004419334 | 0.017292831 | 0.014235885 | 0.116658564 |
| Son | S | 961 | 0.18608866 | -0.295075376 | -0.32258757 | -0.50867623 | -0.027512194 | 0.022535294 | 0.036320632 | 0.011488068 | 0.002658141 | 0.780812831 |
| Fhdc1 | S | 645 | -0.168735181 | -0.076623925 | -0.112145332 | 0.056589849 | -0.035521407 | 0.08656177 | 0.270588257 | 0.257847901 | 0.54085946 | 0.634680482 |
| Rtn4 | T | 834 | 0.01527791 | -0.216519371 | -0.072924473 | -0.088202383 | 0.143594897 | 0.683571668 | 0.002424078 | 0.33316567 | 0.253342057 | 0.108076288 |
| Ctu1 | S | 416 | 0.062716123 | -0.112174378 | -0.260669146 | -0.32338527 | -0.148494768 | 0.236529557 | 0.350222443 | 0.00143522 | 0.000890233 | 0.241377497 |
| Itsn1 | S | 313 | 0.093770213 | -0.195834171 | -0.132944008 | -0.226714221 | 0.062890163 | 0.327023792 | 0.003313241 | 0.058208536 | 0.066069854 | 0.257915438 |
| Rps6ka4 | S | 347 | -0.008009894 | -0.080006516 | -0.153471633 | -0.14546174 | -0.073465118 | 0.81839071 | 0.007464181 | 0.048121243 | 0.052982353 | 0.219356605 |
| Srsf7 | S | 256 | 0.012873899 | -0.202566669 | -0.084589172 | -0.097463071 | 0.117977497 | 0.786530568 | 0.003500873 | 0.268191186 | 0.197775024 | 0.144390807 |
| Abl1 Abl2 | Y | 393;439 | -0.067302901 | -0.081635756 | -0.222809025 | -0.155506124 | -0.141173268 | 0.344538194 | 0.508878052 | 0.04047005 | 0.104158889 | 0.28570047 |
| Plxna1 | S | 1633 | -0.138748181 | 0.001123058 | -0.142582617 | -0.003834436 | -0.143705675 | 0.027767126 | 0.990694351 | 0.053569056 | 0.947032693 | 0.212475897 |
| Cic | T | 1489 | 0.016444454 | -0.183541973 | -0.19577472 | -0.212219174 | -0.012232747 | 0.682861004 | 0.169122445 | 0.02369354 | 0.037419239 | 0.912165907 |
| Bend3 | S | 90 | 0.054467743 | -0.302166916 | -0.194731571 | -0.249199314 | 0.107435345 | 0.833837098 | 0.053416682 | 0.268394265 | 0.381801438 | 0.49783061 |
| Bend3 | S | 91 | 0.054467743 | -0.302166916 | -0.194731571 | -0.249199314 | 0.107435345 | 0.833837098 | 0.053416682 | 0.268394265 | 0.381801438 | 0.49783061 |
| Kmt2d | S | 628 | -0.162748363 | -0.366500912 | -0.029161485 | 0.133586878 | 0.337339427 | 0.541820635 | 0.07770256 | 0.869193847 | 0.645257851 | 0.159874917 |
| Supt5 | S | 664 | -0.3160399 | -0.049498192 | 0.003289279 | 0.319329179 | 0.052787471 | 0.00968618 | 0.646918739 | 0.966838973 | 0.020297422 | 0.659509503 |
| Cttnbp2nl | S | 540 | 0.051852092 | -0.14461063 | -0.23968423 | -0.291536322 | -0.0950736 | 0.408418485 | 0.042313227 | 0.038030376 | 0.033270608 | 0.27833177 |
| Safb | S | 783 | -0.265961459 | -0.082306874 | -0.023079481 | 0.242881978 | 0.059227393 | 0.016496537 | 0.525705326 | 0.803300087 | 0.036338016 | 0.636708152 |
| Dnajc2 | S | 63 | 0.004112056 | -0.312188142 | -0.025521869 | -0.029633925 | 0.286666273 | 0.976892741 | 0.004822243 | 0.848190279 | 0.85767901 | 0.108929022 |
| Klc2 | S | 443 | 0.107023194 | -0.133954995 | -0.249851214 | -0.356874408 | -0.115896219 | 0.100664042 | 0.038626969 | 0.00379535 | 0.002852743 | 0.087141809 |
| Nufip2 | T | 88 | 0.131321689 | -0.132643489 | -0.348388626 | -0.479710315 | -0.215745137 | 0.297094446 | 0.267422252 | 0.006454182 | 0.016468638 | 0.113970405 |
| Cdca2 | S | 296 | -0.037090412 | -0.092831462 | -0.085995942 | -0.04890553 | 0.006835521 | 0.136891978 | 0.055272533 | 0.028023249 | 0.139206889 | 0.855979588 |
| H1f2 H1f3 H1f4 | S | 51;51;52 | 0.271125538 | -0.186517537 | -0.283774291 | -0.554899829 | -0.097256754 | 0.234865613 | 0.01521038 | 0.015481689 | 0.063767311 | 0.237494979 |
| Syde1 | T | 699 | -0.048706292 | -0.108558509 | -0.411970533 | -0.363264241 | -0.303412024 | 0.721821206 | 0.492995125 | 0.116438995 | 0.153375797 | 0.223630459 |
| Ccdc86 | S | 194 | -0.028615474 | -0.146473164 | -0.250641796 | -0.222026322 | -0.104168633 | 0.785817386 | 0.051374662 | 0.235827051 | 0.288516592 | 0.568252161 |
| Dock7 | S | 1422 | -0.124247489 | -0.071020119 | -0.203299857 | -0.079052369 | -0.132279738 | 0.184126591 | 0.474392946 | 0.121012989 | 0.443599786 | 0.257748202 |
| Prrc2a | T | 799 | 0.133400151 | -0.167152577 | -0.197821702 | -0.331221854 | -0.030669125 | 0.027606073 | 0.00581415 | 0.002011135 | 0.002950795 | 0.321654019 |
| Zeb1 | S | 664 | -0.093684268 | -0.1171331 | -0.144760536 | -0.051076269 | -0.027627436 | 0.28583317 | 0.35410523 | 0.052786607 | 0.478391518 | 0.801628092 |
| Ints8 | T | 18 | -0.059832543 | -0.343319662 | -0.226033312 | -0.166200769 | 0.117286349 | 0.795726646 | 0.178004077 | 0.3977715 | 0.570662276 | 0.68169115 |
| Gpatch2 | S | 283 | -0.032788704 | -0.080480472 | -0.206363736 | -0.173575033 | -0.125883264 | 0.651495474 | 0.325146464 | 0.016109493 | 0.082666541 | 0.177229774 |
| Ppp1r11 | T | 80 | -0.128199627 | -0.037693155 | -0.193970097 | -0.065770469 | -0.156276941 | 0.08405216 | 0.66495743 | 0.109956438 | 0.510619304 | 0.203422249 |
| Tnks1bp1 | S | 1063 | 0.068778688 | -0.137748164 | -0.174413572 | -0.24319226 | -0.036665408 | 0.258983054 | 0.011037929 | 0.014594843 | 0.01146146 | 0.412978429 |
| Ptpn12 | S | 632 | 0.020513377 | -0.11194248 | -0.258167152 | -0.278680525 | -0.146224672 | 0.851976059 | 0.346206116 | 0.013502001 | 0.079336066 | 0.251694472 |
| Ppp1cb | T | 316 | 0.001704219 | -0.06734479 | -0.23926641 | -0.240970629 | -0.17192162 | 0.981300241 | 0.248431542 | 0.010904937 | 0.024789968 | 0.042540191 |
| Rps9 | S | 160 | 0.077253546 | -0.178295333 | -0.4027155 | -0.479969045 | -0.224420167 | 0.246004043 | 0.107079701 | 0.068902649 | 0.045269559 | 0.20290088 |
| Coil | S | 234 | -0.013632739 | -0.134120906 | -0.094245137 | -0.080612398 | 0.039875769 | 0.602338308 | 0.005620498 | 0.149605806 | 0.200928023 | 0.477281925 |
| Cobll1 | S | 1007 | -0.081360591 | -0.143343031 | -0.142035411 | -0.06067482 | 0.00130762 | 0.436743101 | 0.17077516 | 0.096367513 | 0.577840205 | 0.988749627 |
| Dact3 | S | 456 | -0.158769784 | -0.008168102 | -0.239342517 | -0.080572733 | -0.231174415 | 0.179852976 | 0.942760367 | 0.071266591 | 0.506865465 | 0.117219052 |
| Col4a3bp | S | 377 | -0.11475505 | -0.072879332 | -0.086140231 | 0.028614819 | -0.013260899 | 0.135898707 | 0.28284234 | 0.027290467 | 0.63911254 | 0.825853357 |
| Noc2l | S | 674 | 0.038831791 | -0.12369876 | -0.115062982 | -0.153894774 | 0.008635777 | 0.412051703 | 0.007915336 | 0.007471412 | 0.034707689 | 0.77788567 |
| Ankrd28 | S | 1011 | -0.342231236 | 0.0753004 | -0.160744186 | 0.18148705 | -0.236044585 | 0.032284518 | 0.684161173 | 0.327831612 | 0.230917558 | 0.244371749 |
| Epb41l2 | S | 665 | -0.279640392 | -0.099746232 | -0.09074455 | 0.188895842 | 0.009001682 | 0.074611698 | 0.386853449 | 0.704234941 | 0.44259604 | 0.967421646 |
| Epb41l2 | S | 666 | -0.279640392 | -0.099746232 | -0.09074455 | 0.188895842 | 0.009001682 | 0.074611698 | 0.386853449 | 0.704234941 | 0.44259604 | 0.967421646 |
| Nckipsd | T | 174 | -0.07992547 | -0.093351672 | -0.144332364 | -0.064406894 | -0.050980692 | 0.340651327 | 0.040250317 | 0.180663403 | 0.546955744 | 0.568001962 |
| Ddx21 | T | 247 | -0.0624728 | -0.052180447 | -0.115146669 | -0.052673869 | -0.062966222 | 0.085222891 | 0.101576654 | 0.043853841 | 0.251863919 | 0.185467028 |
| Camsap1 | S | 1069 | 0.048790662 | -0.116192502 | -0.205992277 | -0.254782939 | -0.089799775 | 0.491267268 | 0.018205193 | 0.032359783 | 0.027832631 | 0.166870361 |
| Nhsl1 | S | 1459 | -0.022811005 | -0.096079308 | -0.162239437 | -0.139428432 | -0.066160128 | 0.680592062 | 0.106577141 | 0.023823066 | 0.079099742 | 0.247311602 |
| Soga1 | S | 1396 | -0.013362724 | -0.078716281 | -0.272738295 | -0.259375571 | -0.194022014 | 0.920065695 | 0.442557397 | 0.017630679 | 0.106993328 | 0.050799534 |
| Pik3c2a | S | 261 | -0.089591314 | -0.097011119 | -0.147882736 | -0.058291422 | -0.050871617 | 0.385129523 | 0.136983517 | 0.085134232 | 0.582103178 | 0.498188429 |
| Ncaph | S | 25 | 0.091727962 | -0.096404457 | -0.200421945 | -0.292149907 | -0.104017487 | 0.058331027 | 0.247467756 | 1.8472E-05 | 0.004435958 | 0.224757116 |
| Pagr1a | S | 233 | -0.221740836 | -0.143210445 | -0.065968519 | 0.155772316 | 0.077241926 | 0.189825206 | 0.106389507 | 0.676915535 | 0.418341192 | 0.608365807 |
| Arglu1 | S | 92 | 0.007924136 | -0.050306582 | -0.255048621 | -0.262972758 | -0.20474204 | 0.947014843 | 0.496610911 | 0.004808904 | 0.106360204 | 0.026253637 |
| Rbm27 | S | 129 | -0.023117418 | -0.040402392 | -0.128281473 | -0.105164055 | -0.087879081 | 0.754840181 | 0.050984857 | 0.00165888 | 0.236478542 | 0.021861122 |
| Ilrun | S | 265 | -0.145272845 | -0.066450056 | -0.300116301 | -0.154843457 | -0.233666246 | 0.31010799 | 0.467326233 | 0.199141514 | 0.475735775 | 0.288663278 |
| Ppil4 | S | 393 | -0.085101085 | -0.116527385 | -0.045147783 | 0.039953302 | 0.071379601 | 0.055785171 | 0.0269455 | 0.455129354 | 0.459139317 | 0.243762978 |
| Ubac1 | S | 98 | -0.097431882 | -0.012239267 | -0.178533366 | -0.081101484 | -0.166294098 | 0.158290668 | 0.840263308 | 0.015461537 | 0.144693621 | 0.023440207 |
| Gars | S | 643 | -0.092144886 | -0.170210119 | -0.135382633 | -0.043237747 | 0.034827486 | 0.390157563 | 0.190132472 | 0.150335973 | 0.696125897 | 0.769128125 |
| Ccdc14 | S | 781 | -0.018811447 | -0.179535136 | -0.071549678 | -0.052738231 | 0.107985458 | 0.683830446 | 0.00584357 | 0.317988409 | 0.435562405 | 0.16315704 |
| Ythdc1 | S | 118 | 0.075407817 | -0.245701356 | -0.2593238 | -0.334731617 | -0.013622444 | 0.462981927 | 0.043136534 | 0.156232941 | 0.095664666 | 0.929147156 |
| Tmem237 | S | 119 | -0.0276385 | -0.133749368 | -0.203672347 | -0.176033848 | -0.06992298 | 0.697936319 | 0.244903977 | 0.043797675 | 0.092724515 | 0.530315298 |
| Dock7 | S | 894 | 0.098960493 | -0.106716861 | -0.229646506 | -0.328606999 | -0.122929645 | 0.209635925 | 0.027465089 | 0.004885082 | 0.011016932 | 0.060974855 |
| Asph | S | 29 | -0.085790364 | -0.947820151 | -0.172838092 | -0.087047728 | 0.774982058 | 0.808820741 | 0.331655309 | 0.839549075 | 0.917843452 | 0.477106861 |
| Wdhd1 | S | 388 | 0.021457048 | -0.13652396 | -0.109938559 | -0.131395607 | 0.026585402 | 0.390781036 | 0.068149062 | 0.001061557 | 0.003230319 | 0.601612588 |
| Arhgap23 | S | 594 | -0.173565724 | 0.008298133 | -0.164926446 | 0.008639279 | -0.173224579 | 0.061725558 | 0.926276208 | 0.070851933 | 0.837061298 | 0.052484717 |
| Arpp21 R3hdm2 | S | 283;278 | -0.116402656 | 0.00282624 | -0.191371067 | -0.074968411 | -0.194197307 | 0.056735063 | 0.973473065 | 0.048350065 | 0.31095499 | 0.08791348 |
| Ppp1r11 | S | 78 | -0.128547795 | -0.035255355 | -0.195222632 | -0.066674837 | -0.159967277 | 0.0880403 | 0.686832947 | 0.109881405 | 0.508768656 | 0.19704712 |
| Zfp592 | S | 326 | -0.265911854 | -0.081818674 | 0.097093509 | 0.363005363 | 0.178912183 | 0.005168007 | 0.191916148 | 0.606751174 | 0.13976609 | 0.372664723 |
| Itprid2 | S | 736 | -0.222768405 | -0.005278344 | -0.154099947 | 0.068668458 | -0.148821604 | 0.051174295 | 0.97013444 | 0.192296866 | 0.507905086 | 0.33210449 |
| Asb3 | S | 35 | -0.230153208 | 0.060079621 | -0.205159589 | 0.024993619 | -0.26523921 | 0.072041696 | 0.67790007 | 0.088893986 | 0.807026479 | 0.119756562 |
| Abcf1 | T | 21 | 0.110762812 | -0.190285408 | -0.744022958 | -0.854785771 | -0.553737551 | 0.452667815 | 0.17861467 | 0.224893637 | 0.18650806 | 0.331962287 |
| Sart1 | S | 392 | -0.077369447 | -0.008600555 | -0.217197001 | -0.139827554 | -0.208596446 | 0.10124811 | 0.898784281 | 0.028661724 | 0.110885554 | 0.048484001 |
| Maz | S | 302 | 0.007482921 | -0.126654101 | -0.1634848 | -0.170967721 | -0.036830699 | 0.927218057 | 0.092176408 | 0.019201497 | 0.109724174 | 0.570962275 |
| Clasp1 | S | 572 | -0.146966374 | -0.0322357 | -0.02627824 | 0.120688134 | 0.00595746 | 0.001369326 | 0.261890403 | 0.488237295 | 0.028370245 | 0.837078966 |

| | | | | | | | | | | | | |
|---|---|---|---|---|---|---|---|---|---|---|---|---|
| Ubap2l | S | 628 | -0.015334518 | -0.083230031 | -0.123942018 | -0.1086075 | -0.040711986 | 0.440222842 | 0.046927053 | 0.01667989 | 0.051892715 | 0.314760398 |
| Ankrd17 | S | 1696 | 0.062303083 | -0.113616282 | -0.187965147 | -0.25026823 | -0.074348865 | 0.133935576 | 0.003098304 | 0.017404937 | 0.007717311 | 0.180717053 |
| Fbxo5 | S | 157 | 0.003695555 | -0.119094463 | -0.207979748 | -0.211675304 | -0.088885286 | 0.944036796 | 0.036894257 | 0.084697052 | 0.086109535 | 0.354425176 |
| Fkbp3 | S | 91 | -0.257062057 | -0.045566928 | -0.083131912 | 0.173930145 | -0.037564983 | 0.05447893 | 0.388360386 | 0.453834746 | 0.205819189 | 0.728493564 |
| Pus7 | S | 40 | 0.018675782 | -0.144238849 | -0.144290132 | -0.162965913 | -5.12829E-05 | 0.682450083 | 0.005719007 | 0.122792237 | 0.091878903 | 0.999413278 |
| Kif23 | S | 717 | -0.03865917 | -0.036427778 | -0.237683494 | -0.199024324 | -0.201255716 | 0.747502429 | 0.733248676 | 0.00530378 | 0.169264291 | 0.127919386 |
| Naf1 | S | 308 | -0.008442231 | -0.138383377 | -0.120893582 | -0.112451351 | 0.017489795 | 0.903879845 | 0.017941457 | 0.079572026 | 0.168535772 | 0.706634987 |
| Skap2 | S | 247 | -0.241593926 | -0.119982136 | 0.077751709 | 0.319345634 | 0.197733845 | 0.016730424 | 0.117833232 | 0.680700945 | 0.181344551 | 0.348050888 |
| Ncoa3 | S | 847 | -0.142613826 | 0.012961753 | -0.108728676 | 0.03388515 | -0.121690429 | 0.002513846 | 0.742820302 | 0.138337059 | 0.568894377 | 0.113628797 |
| Tle3 | S | 267 | 0.003558354 | -0.061914435 | -0.128524977 | -0.132083331 | -0.066610542 | 0.896598906 | 0.068494837 | 0.001066507 | 0.004650656 | 0.054907281 |
| Ikbkg | S | 380 | -0.046471709 | -0.081431104 | -0.143780636 | -0.097308927 | -0.062349532 | 0.36967094 | 0.146108427 | 0.028662292 | 0.035628803 | 0.091733512 |
| Kctd3 | S | 712 | 0.025151329 | -0.195651651 | -0.169754154 | -0.194905483 | 0.025897497 | 0.519490635 | 0.06929737 | 0.043283757 | 0.025864497 | 0.743308379 |
| Rnps1 | S | 141 | -0.010934267 | -0.170372818 | -0.055549749 | -0.044615482 | 0.11482307 | 0.83070357 | 0.001606284 | 0.378185756 | 0.50147141 | 0.122879323 |
| Peak1 | S | 825 | -0.30616768 | -0.00589859 | -0.229664181 | 0.076503499 | -0.223765591 | 0.107249364 | 0.969095423 | 0.361219637 | 0.758547474 | 0.401425475 |
| Tpr | S | 2111 | 0.023792096 | -0.025827801 | -0.230243139 | -0.254035235 | -0.204415338 | 0.780223854 | 0.441215022 | 0.001064687 | 0.056108534 | 0.013190046 |
| Bod1l | S | 2777 | -0.068323538 | -0.079825208 | -0.152124924 | -0.083801386 | -0.072299715 | 0.445479504 | 0.250420324 | 0.023354159 | 0.330962834 | 0.222720485 |
| Dennd4c | S | 1043 | -0.139087108 | 0.054394288 | -0.256725564 | -0.117638456 | -0.311119852 | 0.128614943 | 0.526414165 | 0.024609704 | 0.234094501 | 0.022960831 |
| Arhgap12 | S | 101 | -0.21648189 | 0.015554437 | -0.126810211 | 0.089671679 | -0.142364648 | 0.026845963 | 0.875489049 | 0.159856354 | 0.135995132 | 0.145580384 |
| Mettl2 | S | 140 | 0.064162991 | -0.071532348 | -0.244146097 | -0.308309088 | -0.172613749 | 0.521232092 | 0.14752266 | 0.0042568 | 0.043412782 | 0.024551006 |
| Mark2 | S | 592 | 0.085411501 | 0.317330805 | -0.577513792 | -0.662925293 | -0.894844597 | 0.803540061 | 0.415067622 | 0.001089698 | 0.149180295 | 0.098383839 |
| Prune2 | S | 710 | -0.049468218 | -0.061767249 | -0.100489179 | -0.051020962 | -0.038721931 | 0.115456798 | 0.268075068 | 0.006844945 | 0.046719975 | 0.440876337 |
| Ptpn12 | S | 434 | -0.108987686 | 0.075038046 | -0.253970061 | -0.144982375 | -0.329008107 | 0.329800548 | 0.658850937 | 0.005557286 | 0.207517526 | 0.13901862 |
| Sae1 | S | 16 | -0.073273866 | -0.0512897 | -0.164023271 | -0.090749405 | -0.112733571 | 0.367642757 | 0.499657744 | 0.011086551 | 0.262773225 | 0.171111784 |
| Zc3h13 | S | 408 | -0.026096307 | -0.04307551 | -0.130320879 | -0.104224572 | -0.087245369 | 0.375058123 | 0.425358582 | 0.000769032 | 0.02027479 | 0.169835171 |
| Canx | S | 563 | -0.160540284 | -0.021764331 | 0.039437521 | 0.199977805 | 0.061201852 | 1.65399E-05 | 0.486042642 | 0.614477654 | 0.093770965 | 0.460601226 |
| Mdc1 | T | 1298 | -0.055289286 | -0.083871319 | -0.146324156 | -0.09103487 | -0.062452837 | 0.224271143 | 0.053650645 | 0.143096146 | 0.296847013 | 0.441949241 |
| Smg6 | S | 478 | -0.150847352 | -0.10396666 | 0.043713403 | 0.194560754 | 0.147680063 | 0.021802401 | 0.001978524 | 0.244457785 | 0.010377785 | 0.029816041 |
| Rbbp6 | S | 1027 | -0.073738129 | -0.126627125 | -0.058945752 | 0.014792377 | 0.067681373 | 0.351496152 | 0.005240908 | 0.430908749 | 0.871277899 | 0.37813794 |
| Smc2 | T | 723 | -0.167966813 | -0.060506654 | -0.147327736 | 0.020639077 | -0.086821082 | 0.324172487 | 0.493202088 | 0.046039821 | 0.890751219 | 0.3299292 |
| Aimp2 | T | 82 | -0.00694024 | -0.164791294 | -0.132907008 | -0.125966768 | 0.031884286 | 0.910245699 | 0.051318848 | 0.072829933 | 0.054803169 | 0.611298158 |
| Tasor2 | S | 2012 | 0.013237052 | -0.151446583 | -0.065964725 | -0.079201777 | 0.085481858 | 0.840828499 | 0.002597456 | 0.074878293 | 0.275283005 | 0.009689127 |
| Mapk3 | T | 203 | -0.10908054 | -0.126757475 | -0.163129624 | -0.054049084 | -0.036372149 | 0.228930135 | 0.252631374 | 0.220824987 | 0.623180445 | 0.765050889 |
| Supt16 | S | 508 | -0.070534522 | -0.093889535 | -0.088729638 | -0.018195117 | 0.005159897 | 0.085515231 | 0.25446044 | 0.037393546 | 0.500872669 | 0.940537704 |
| Txndc5 | T | 160 | -0.080262015 | -0.132566535 | -0.334889445 | -0.25462743 | -0.20232291 | 0.624076651 | 0.254971601 | 0.236626328 | 0.362373259 | 0.432570918 |
| Ncor2 | S | 1963 | -0.018526032 | -0.058402528 | -0.179851657 | -0.161325625 | -0.121449129 | 0.731497818 | 0.338534634 | 0.004432073 | 0.024297206 | 0.076481883 |
| Mki67 | S | 159 | -0.261005649 | 0.094771143 | -0.292518929 | -0.03151328 | -0.387290072 | 0.104496674 | 0.519138068 | 0.114536857 | 0.784462483 | 0.038028525 |
| Cobll1 | S | 993 | 0.011110008 | -0.077859029 | -0.140105004 | -0.151215012 | -0.062245975 | 0.792570231 | 0.014508119 | 0.011465794 | 0.026015416 | 0.131362761 |
| Arhgef17 | S | 568 | -0.030180361 | -0.17491219 | -0.123297916 | -0.093117555 | 0.051614274 | 0.838207423 | 0.025226084 | 0.210915614 | 0.550622699 | 0.513630599 |
| Adar | S | 231 | -0.120043239 | -0.085501285 | -0.263718349 | -0.143675109 | -0.178217064 | 0.169666589 | 0.596506798 | 0.209911139 | 0.439415704 | 0.418173369 |
| Mta1 | S | 386 | 0.002526243 | -0.070091442 | -0.243820594 | -0.246346837 | -0.173729152 | 0.972384398 | 0.086815945 | 0.03406328 | 0.043808385 | 0.094990252 |
| Tln1 | Y | 436 | -0.221390582 | 0.062610958 | -0.151901637 | 0.069488946 | -0.214512595 | 0.03630211 | 0.491430752 | 0.053358733 | 0.385523447 | 0.060216516 |
| Map4 | T | 343 | -0.149408769 | -0.058648652 | -0.194732192 | -0.045323423 | -0.13608354 | 0.221211679 | 0.552229197 | 0.11299931 | 0.709206315 | 0.252353731 |
| Foxk2 | S | 364 | 0.060359422 | -0.197512637 | -0.243816108 | -0.30417553 | -0.046303471 | 0.363439558 | 0.06104039 | 0.064840466 | 0.037818714 | 0.666333547 |
| Ptpn23 | S | 1179 | -0.05490986 | -0.100486093 | -0.117597557 | -0.062687698 | -0.017111464 | 0.53646521 | 0.062606165 | 0.046774378 | 0.492740299 | 0.731716135 |
| Spry4 | S | 126 | 0.091614408 | -0.233585023 | -0.126269969 | -0.217884377 | 0.107315054 | 0.012172895 | 0.001446281 | 0.017998835 | 0.006872945 | 0.05190215 |
| Palld | S | 901 | 0.016585562 | -0.066163459 | -0.294095482 | -0.310681044 | -0.227932023 | 0.847019477 | 0.412726516 | 0.013775958 | 0.024679466 | 0.049786785 |
| Kif21b | S | 1217 | -0.128319743 | -0.044602283 | -0.111036858 | 0.017282885 | -0.066434576 | 0.105594799 | 0.390872732 | 0.053944257 | 0.758609595 | 0.05911908 |
| Trmt10a | S | 22 | -0.075297153 | -0.053555515 | -0.089491806 | -0.014194653 | -0.035936291 | 0.076019248 | 0.105736302 | 0.04662992 | 0.709624102 | 0.333496341 |
| Srebf1 | S | 446 | 0.074292789 | -0.141237235 | -0.1984035 | -0.27269629 | -0.057166265 | 0.271024642 | 0.013458534 | 0.027955407 | 0.015382359 | 0.363993596 |
| Peak1 | T | 778 | -0.046326468 | -0.14020177 | -0.120456495 | -0.074130027 | 0.019745275 | 0.4772965 | 0.099942275 | 0.075926786 | 0.329467366 | 0.781281942 |
| Phldb1 | S | 522 | 0.026433988 | -0.037057347 | -0.175402991 | -0.201836979 | -0.138345644 | 0.519096547 | 0.067255402 | 0.000728794 | 0.012350627 | 0.004657149 |
| Pacc1 | S | 14 | 0.023016005 | 0.037202931 | -0.393107378 | -0.416123383 | -0.430310309 | 0.879761935 | 0.789736747 | 0.004480748 | 0.056875259 | 0.037925862 |
| Cbl | S | 723 | -0.064056496 | -0.10792872 | -0.326627006 | -0.26257051 | -0.218698286 | 0.514047127 | 0.352820198 | 0.164485803 | 0.238737243 | 0.307778309 |
| Wdr4 | S | 392 | 0.003601274 | -0.087171018 | -0.083166862 | -0.086718135 | 0.004054156 | 0.864162824 | 0.006802352 | 0.00432286 | 0.003973987 | 0.808400631 |
| Ppp1r13l | T | 479 | -0.098758933 | -0.234306684 | -0.375911605 | -0.277152671 | -0.14160492 | 0.783064747 | 0.282648154 | 0.371990121 | 0.571767698 | 0.725348637 |
| Shc1 | Y | 423 | -0.225702077 | -0.100092382 | 0.121393453 | 0.347095531 | 0.221485835 | 0.001523035 | 0.047043483 | 0.183008275 | 0.020367079 | 0.065264847 |
| Brf1 | T | 365 | -0.03095984 | -0.068491348 | -0.078901985 | -0.047942146 | -0.010410638 | 0.398367595 | 0.015911594 | 0.009289066 | 0.212899043 | 0.108816289 |
| Helb | S | 945 | -0.143097046 | 0.0158771 | -0.114653307 | 0.028443739 | -0.130530407 | 0.002196723 | 0.787477274 | 0.193209735 | 0.692789652 | 0.17293383 |
| Cluh | S | 658 | -0.096419532 | -0.052942479 | -0.178449523 | -0.082029992 | -0.125507045 | 0.145062527 | 0.1195211 | 0.194027108 | 0.487002948 | 0.309796818 |
| Fam129b | S | 653 | 0.001771912 | -0.057504435 | -0.214235943 | -0.216007855 | -0.156731508 | 0.97903763 | 0.327593008 | 0.005254457 | 0.021893662 | 0.015770746 |
| Pfkfb2 | S | 496 | -0.153872346 | -0.005371118 | -0.132864587 | 0.021007758 | -0.12749347 | 0.027902446 | 0.950056701 | 0.10356422 | 0.762162299 | 0.233108704 |
| Dlg5 | S | 1021 | -0.074386803 | -0.081902402 | -0.214025961 | -0.139639159 | -0.132123559 | 0.273123038 | 0.357978157 | 0.09689189 | 0.224541364 | 0.27678332 |
| Spire1 | S | 365 | -0.101689803 | -0.033021619 | -0.140181103 | -0.0384913 | -0.107159483 | 0.059924569 | 0.434945564 | 0.072358194 | 0.533377917 | 0.139708931 |
| Rab40c | S | 268 | -0.010267618 | -0.122771181 | -0.126486593 | -0.116218975 | -0.003715412 | 0.771076371 | 0.14652312 | 0.009186145 | 0.002319991 | 0.954342182 |
| H1f4 | S | 187 | 0.034874125 | -0.094844026 | -0.1586445 | -0.193518625 | -0.063800473 | 0.654659988 | 0.037914636 | 0.008308992 | 0.07347083 | 0.126797867 |
| Dennd1b | S | 395 | -0.176348734 | -0.045601279 | -0.132323542 | 0.044025192 | -0.086722263 | 0.022177105 | 0.602886258 | 0.426675787 | 0.773245152 | 0.596625795 |
| Crtc2 | S | 623 | -0.013803989 | -0.095136663 | -0.20811805 | -0.194314061 | -0.112981387 | 0.584258789 | 0.132430141 | 0.052767282 | 0.076154982 | 0.194902965 |
| Eif4b | T | 610 | 0.002721852 | -0.082428721 | -0.083191733 | -0.085913585 | -0.000763012 | 0.935798413 | 0.003993867 | 0.006670593 | 0.068660071 | 0.968704346 |
| Tln1 | S | 1328 | -0.029458074 | -0.109999714 | -0.077170804 | -0.04771273 | 0.03282891 | 0.279955742 | 0.024833023 | 0.053335101 | 0.155034863 | 0.389165824 |
| Ifih1 | S | 302 | -0.153891384 | -0.092606035 | -0.103724993 | 0.050166391 | -0.011118958 | 0.089341301 | 0.258446778 | 0.322447768 | 0.628939782 | 0.913115489 |
| Ccdc88a | S | 1417 | 0.061130589 | -0.154845449 | -0.348962091 | -0.410092681 | -0.194116642 | 0.145753225 | 0.120594501 | 0.036039287 | 0.029379802 | 0.136831211 |
| Ak5 | S | 105 | -0.205357202 | -0.170311797 | -0.111027745 | 0.094329457 | 0.059284052 | 0.185662897 | 0.314445761 | 0.508900017 | 0.514341079 | 0.708670421 |
| Ak5 | S | 108 | -0.205357202 | -0.170311797 | -0.111027745 | 0.094329457 | 0.059284052 | 0.185662897 | 0.314445761 | 0.508900017 | 0.514341079 | 0.708670421 |
| Ppan | S | 240 | -0.023300007 | -0.145910935 | -0.270490133 | -0.247190126 | -0.124579198 | 0.945582381 | 0.253648332 | 0.082388126 | 0.502302286 | 0.401574489 |
| Ube4b | S | 124 | -0.094675842 | -0.060072755 | -0.140849928 | -0.046174086 | -0.080777173 | 0.104953585 | 0.186811897 | 0.140783017 | 0.564507679 | 0.33196679 |
| Ttk | S | 277 | 0.086274491 | -0.154996612 | -0.2210487 | -0.307323191 | -0.066052088 | 0.23919129 | 0.144373308 | 0.003854371 | 0.011661988 | 0.463026425 |
| Cmtr1 | S | 27 | -0.145704389 | 0.004570362 | -0.253649439 | -0.10794505 | -0.258219801 | 0.08751821 | 0.967664354 | 0.128575272 | 0.43852491 | 0.145531051 |
| Ddx20 | S | 496 | 0.087455236 | -0.175367471 | -0.271807184 | -0.35926242 | -0.096439713 | 0.139117611 | 0.075736681 | 0.017672512 | 0.022475636 | 0.315013207 |
| Prrc2a | S | 1012 | 0.038235822 | -0.100460054 | -0.118710084 | -0.156945906 | -0.01825003 | 0.696380036 | 0.001144279 | 0.049465332 | 0.195495971 | 0.640382471 |
| Ngef | S | 606 | -0.198394087 | -0.004691011 | -0.195942006 | 0.00245208 | -0.191250996 | 0.073965108 | 0.964917473 | 0.188236637 | 0.98428799 | 0.200861442 |
| Clasp1 | S | 600 | -0.093833794 | -0.071514652 | -0.165081199 | -0.071247405 | -0.093566547 | 0.181091002 | 0.415237545 | 0.076140399 | 0.29457249 | 0.285308064 |
| Tbc1d1 | S | 501 | -0.032018416 | -0.173659392 | -0.170963945 | -0.138945529 | 0.002695447 | 0.700983325 | 0.055924279 | 0.247226627 | 0.318788241 | 0.982418021 |
| Kifap3 | S | 102 | -0.109339035 | 0.020502529 | -0.255511478 | -0.146172444 | -0.276014007 | 0.216920214 | 0.836157084 | 0.028289351 | 0.102793801 | 0.022155613 |
| Fndc3b | S | 260 | 0.038991283 | -0.182320858 | -0.134049935 | -0.173041219 | 0.048270923 | 0.408951447 | 0.031692247 | 0.026944308 | 0.008966286 | 0.434636374 |
| Mnat1 | S | 279 | -0.174132977 | -0.03090517 | -0.054410901 | 0.119722076 | -0.023505731 | 0.021571899 | 0.356721024 | 0.177420455 | 0.064174544 | 0.54439395 |
| Zdhhc5 | S | 331 | -0.159438007 | -0.106440846 | -0.110570601 | 0.048867405 | -0.004129755 | 0.113815957 | 0.257652104 | 0.365016189 | 0.592874843 | 0.962277131 |
| Trmt1l | S | 607 | -0.037310178 | -0.061014663 | -0.141350114 | -0.104039936 | -0.080335451 | 0.362295068 | 0.061489733 | 0.032730455 | 0.07025415 | 0.080552166 |
| Setd2 | S | 132 | -0.06907003 | -0.093633477 | -0.000725417 | 0.068344613 | 0.09290806 | 0.00296537 | 0.009100735 | 0.975139315 | 0.043230366 | 0.018674063 |
| Sipa1l3 | S | 1528 | 0.004697993 | -0.169035288 | -0.219430138 | -0.224128131 | -0.05039485 | 0.910869306 | 0.085868128 | 0.123337983 | 0.132397768 | 0.674842612 |
| Nr3c1 | S | 212 | -0.065206761 | -0.052912543 | -0.123885884 | -0.058679123 | -0.070973341 | 0.211747995 | 0.123552269 | 0.033241369 | 0.286030751 | 0.149278488 |
| Rif1 | S | 1565 | -0.020813517 | -0.066754908 | -0.082333168 | -0.061519651 | -0.01557826 | 0.389343417 | 0.043823428 | 0.002468543 | 0.05679791 | 0.519047143 |
| Ccnd3 | S | 264 | -0.002273585 | -0.08194766 | -0.127625719 | -0.125352135 | -0.045678059 | 0.950086039 | 0.006215438 | 0.038752374 | 0.0477568 | 0.305153974 |
| Clint1 | S | 240 | -0.187849756 | -0.105530454 | -0.014079224 | 0.173770532 | 0.09145123 | 0.08370519 | 0.04372234 | 0.87962428 | 0.166597386 | 0.362972516 |
| Sh3pxd2a | S | 546 | -0.047731861 | -0.05257028 | -0.16238565 | -0.114653789 | -0.10981537 | 0.400039483 | 0.312898544 | 0.012533352 | 0.04567583 | 0.014081759 |

| | | | | | | | | | | | | |
|---|---|---|---|---|---|---|---|---|---|---|---|---|
| Ripk2 | S | 356 | -0.044231608 | -0.072065692 | -0.163771016 | -0.119539409 | -0.091705324 | 0.400062965 | 0.274237084 | 0.021551083 | 0.049165596 | 0.123029611 |
| Gfpt1 | T | 260 | -0.128483371 | -0.07853223 | -0.13053617 | -0.002052799 | -0.052003941 | 0.115819653 | 0.332786737 | 0.172260095 | 0.981691189 | 0.587385811 |
| Tep1 | S | 2529 | -0.094098529 | -0.067950939 | -0.292371713 | -0.198273184 | -0.224420774 | 0.643747665 | 0.437276496 | 0.080306678 | 0.386854006 | 0.147201566 |
| Bub1b | S | 1050 | -0.074728198 | -0.046772555 | -0.132689321 | -0.057961123 | -0.085916766 | 0.083013525 | 0.155820037 | 0.084386394 | 0.341661128 | 0.186757859 |
| Dmap1 | S | 455 | -0.000267651 | -0.138351616 | -0.027549104 | -0.027281453 | 0.110802511 | 0.990904993 | 6.28319E-05 | 0.493324993 | 0.521136728 | 0.067344085 |
| Relch | S | 243 | -0.122677457 | -0.062039839 | -0.100396424 | 0.022281033 | -0.038356584 | 0.082518546 | 0.304297473 | 0.107628802 | 0.574047255 | 0.222123269 |
| Pnn | S | 442 | -0.139680426 | -0.151387457 | -0.009158146 | 0.130522281 | 0.142229312 | 0.244110991 | 0.011223676 | 0.908109372 | 0.29745989 | 0.142343645 |
| Tjp2 | S | 386 | -0.024051292 | -0.294298093 | -0.334569735 | -0.310518443 | -0.040271642 | 0.960332007 | 0.290485554 | 0.268318289 | 0.556894147 | 0.899113306 |
| Prrc2b | S | 1070 | 0.131411882 | -0.186775531 | -0.270950567 | -0.402362449 | -0.084175035 | 0.286652852 | 0.110733487 | 0.015596781 | 0.02300336 | 0.391100901 |
| D1Pas1 Ddx3x Ddx3y | S | 68;71;70 | 0.001459396 | -0.022167399 | -0.277972282 | -0.279431678 | -0.255804883 | 0.984947963 | 0.810920121 | 0.004171443 | 0.01409064 | 0.044797338 |
| Kctd3 | T | 663 | -0.052884981 | -0.069171562 | -0.272264529 | -0.219379548 | -0.203092967 | 0.594090909 | 0.465565477 | 0.050557073 | 0.1044064 | 0.117099549 |
| D1Ertd622e | S | 25 | 0.018349379 | -0.100036949 | -0.099376635 | -0.117726014 | 0.000660313 | 0.414739955 | 0.005591454 | 0.006816349 | 0.006569808 | 0.979068933 |
| Mink1 | S | 537 | 0.041481195 | -0.130699591 | -0.087397648 | -0.128878842 | 0.043301943 | 0.304769989 | 0.002104194 | 0.012046653 | 0.038712085 | 0.086651144 |
| Ctnnd1 | S | 346 | -0.231781546 | -0.079074953 | -0.08187713 | 0.149904417 | -0.002802177 | 0.066100171 | 0.416244914 | 0.494434466 | 0.248631671 | 0.979295957 |
| Ostf1 | S | 214 | -0.186172383 | 0.006258248 | -0.215793843 | -0.02962146 | -0.222052091 | 0.076171253 | 0.956157681 | 0.167548434 | 0.824736117 | 0.187354819 |
| Ifih1 | S | 291 | -0.138751223 | -0.096447998 | -0.054123599 | 0.084627624 | 0.042324399 | 0.048850564 | 0.12358344 | 0.477319735 | 0.276039578 | 0.549232293 |
| Ctnna1 Ctnna2 | T | 658;657 | -0.019820286 | -0.132031293 | -0.076702966 | -0.05688268 | 0.055328327 | 0.583524982 | 0.02171374 | 0.04855658 | 0.068994757 | 0.198123004 |
| Dnmt1 | S | 182 | -0.036529903 | -0.058334162 | -0.213718315 | -0.177188412 | -0.155384153 | 0.461562159 | 0.074561145 | 0.077972 | 0.108542828 | 0.150021743 |
| Rps6kb1 | T | 444 | -0.099310912 | -0.036030347 | -0.259469184 | -0.160158272 | -0.223438837 | 0.241756125 | 0.665233861 | 0.085487229 | 0.227670501 | 0.125500945 |
| Jmjd6 | S | 400 | 0.095223321 | -0.16530497 | -0.34164087 | -0.436864191 | -0.1763359 | 0.177160736 | 0.040658618 | 0.062376703 | 0.055367004 | 0.242407598 |
| Camk1d | S | 358 | -0.105068205 | -0.073842393 | -0.154593844 | -0.049525639 | -0.08075145 | 0.080636588 | 0.301683855 | 0.221847151 | 0.649565829 | 0.490923156 |
| Rnf213 | S | 271 | -0.086827257 | -0.171842699 | -0.071695536 | 0.015131721 | 0.100147163 | 0.288205207 | 0.059093442 | 0.366271165 | 0.643123153 | 0.02248636 |
| Pla2g4a | S | 726 | -0.023814657 | 0.027288166 | -0.420829432 | -0.397014775 | -0.448117598 | 0.803286476 | 0.765656343 | 0.017260362 | 0.028359966 | 0.021758097 |
| Kmt2a | S | 2542 | -0.236862094 | -0.112186188 | 0.005315973 | 0.242178067 | 0.117502161 | 0.032986209 | 0.222615346 | 0.953851005 | 0.028432373 | 0.149342721 |
| Lpin3 | S | 198 | -0.027576019 | -0.08056771 | -0.160587048 | -0.133011029 | -0.080019338 | 0.591550541 | 0.12035285 | 0.02734759 | 0.058988186 | 0.172541512 |
| Cavin4 | T | 22 | -0.071923369 | -0.129062599 | -0.042802874 | 0.029120495 | 0.086259725 | 0.258497803 | 0.003781427 | 0.664302656 | 0.780233147 | 0.412284762 |
| Larp1 | S | 188 | -0.06215717 | -0.085545149 | -0.172363886 | -0.110206716 | -0.086818737 | 0.161776303 | 0.306481996 | 0.094226208 | 0.220229467 | 0.386401228 |
| Atp10a | S | 511 | -0.266901913 | -0.185539834 | 0.12917386 | 0.396075773 | 0.314713694 | 0.037402873 | 0.153182162 | 0.654757849 | 0.239292695 | 0.318171864 |
| Atp10a | T | 509 | -0.266901913 | -0.185539834 | 0.12917386 | 0.396075773 | 0.314713694 | 0.037402873 | 0.153182162 | 0.654757849 | 0.239292695 | 0.318171864 |
| Prrc2c | S | 192 | 0.103520632 | -0.148130802 | -0.158266305 | -0.261786937 | -0.010135503 | 0.544268937 | 0.009951551 | 0.038889941 | 0.203873258 | 0.808917588 |
| Eif5 | T | 153 | 0.097345516 | -0.152997276 | -0.172404317 | -0.269749833 | -0.019407041 | 0.32357905 | 0.005592007 | 0.044482813 | 0.047071518 | 0.73920766 |
| Ppp1r9b | S | 100 | -0.105230018 | -0.036947141 | -0.147930622 | -0.042700604 | -0.11098348 | 0.123889445 | 0.506255121 | 0.0505984 | 0.449280675 | 0.082189162 |
| Ppfia2 Ppfia3 | T | 307;342 | -0.158735275 | -0.121505723 | -0.062272254 | 0.096463021 | 0.059233469 | 0.283404365 | 0.103693041 | 0.194560686 | 0.476311231 | 0.359428358 |
| Lrp4 | S | 1887 | -0.11583003 | 0.086193373 | -0.221838845 | -0.106008816 | -0.308032218 | 0.083340295 | 0.462074367 | 0.009817707 | 0.076184559 | 0.061256614 |
| Ndel1 | S | 231 | 0.003822348 | -0.080925395 | -0.122029808 | -0.125852155 | -0.041104413 | 0.919406086 | 0.03579302 | 0.006011846 | 0.030899518 | 0.103524668 |
| Slc20a2 | S | 324 | -0.010772663 | -0.117707759 | -0.261920802 | -0.251148139 | -0.144213043 | 0.871146035 | 0.120041741 | 0.114862447 | 0.127756332 | 0.308093488 |
| N4bp1 | T | 289 | -0.019682506 | -0.05250974 | -0.246982032 | -0.227299527 | -0.194472292 | 0.782315598 | 0.526867626 | 0.012748729 | 0.009519137 | 0.028532315 |
| Dlgap4 | S | 666 | 0.092242436 | -0.170191761 | -0.23490295 | -0.327145385 | -0.064711189 | 0.114678705 | 0.133389547 | 0.003507202 | 0.001076476 | 0.48160997 |
| Sgip1 | S | 169 | -0.01516536 | -0.035724238 | -0.234083908 | -0.218918549 | -0.19835967 | 0.79276328 | 0.294402238 | 0.011510095 | 0.027527556 | 0.01065062 |
| Kif20a | S | 7 | -0.007823887 | -0.092793638 | -0.171734435 | -0.163910547 | -0.078940796 | 0.941390276 | 0.213915489 | 0.009883925 | 0.200901816 | 0.265591344 |
| Larp1 | T | 759 | -0.042969832 | -0.109776715 | -0.126028053 | -0.083058221 | -0.016251339 | 0.329091458 | 0.088889395 | 0.085776991 | 0.207226428 | 0.802405414 |
| Rrp7a | S | 99 | 0.088139398 | -0.21574157 | -0.163001109 | -0.251140507 | 0.052740461 | 0.277674805 | 0.027067252 | 0.033536038 | 0.031958502 | 0.437863856 |
| Zzef1 | T | 1519 | -0.092220105 | -0.168270351 | -0.105355738 | -0.013135633 | 0.062914613 | 0.282344239 | 0.182023965 | 0.197796369 | 0.88555692 | 0.581944849 |
| Ube2m | S | 50 | -0.039798236 | -0.092356194 | -0.210807329 | -0.171009093 | -0.118451135 | 0.69014165 | 0.163697839 | 0.064864461 | 0.17365689 | 0.22010297 |
| Msantd4 | S | 101 | 0.046300836 | -0.121778891 | -0.112092439 | -0.158393275 | 0.009686452 | 0.344633423 | 0.003123061 | 0.017574673 | 0.026607859 | 0.733664785 |
| Leo1 | S | 631 | -0.051947933 | -0.075317006 | -0.036769077 | 0.015178856 | 0.038547928 | 0.133622413 | 0.003432774 | 0.070887894 | 0.584528331 | 0.029117587 |
| Tuba1a Tuba1b Tuba | S | 277;277;277;277; | -0.106697718 | -0.028327016 | -0.27532741 | -0.168629692 | -0.247000393 | 0.578670435 | 0.800549155 | 0.033291902 | 0.37219062 | 0.021823581 |
| Iws1 | S | 209 | -0.02920715 | -0.145124691 | -0.04501445 | -0.015807299 | 0.100110241 | 0.261987891 | 0.006226023 | 0.301081835 | 0.676746761 | 0.069488761 |
| Eya3 | T | 206 | -0.037673596 | -0.045531921 | -0.235389317 | -0.19771572 | -0.189857396 | 0.352837044 | 0.615425576 | 0.023137262 | 0.048664923 | 0.112857913 |
| Pcdh19 | S | 830 | -0.071968853 | -0.06979649 | -0.194138981 | -0.122170128 | -0.124342491 | 0.312439757 | 0.376786823 | 0.060556404 | 0.166371537 | 0.178377552 |
| Dmap1 | S | 458 | 0.055486418 | -0.169076409 | -0.044229412 | -0.09971583 | 0.124846997 | 0.485964875 | 5.95489E-05 | 0.385268134 | 0.274545177 | 0.078030114 |
| Tmem237 | T | 118 | -0.046620574 | -0.057691945 | -0.169420582 | -0.122800008 | -0.111728637 | 0.33111501 | 0.314425309 | 0.022413739 | 0.073851881 | 0.118808924 |
| Zbtb7a | S | 331 | -0.075899866 | -0.110216063 | -0.058907378 | 0.016992488 | 0.051308685 | 0.098592779 | 0.039180168 | 0.288466162 | 0.752133014 | 0.373227609 |
| Ticrr | S | 1113 | -0.079148429 | -0.085110946 | -0.062749476 | 0.016398953 | 0.02236147 | 0.182889373 | 0.031335988 | 0.103277353 | 0.751718461 | 0.530839193 |
| Sgms1 | S | 60 | 0.070594746 | -0.158437038 | -0.272254961 | -0.342849707 | -0.113817923 | 0.257892911 | 0.051296504 | 0.047650778 | 0.0416546 | 0.291527154 |
| Fermt2 | Y | 179 | -0.310599187 | 0.075205977 | -0.183738239 | 0.126860948 | -0.258944216 | 0.068176347 | 0.636121369 | 0.183730016 | 0.367254326 | 0.139921873 |
| Dennd4b | S | 736 | 0.023019294 | -0.178988096 | -0.074375864 | -0.097395159 | 0.104612232 | 0.456143286 | 0.012896504 | 0.02018723 | 0.03247943 | 0.052891278 |
| Erc1 | S | 75 | -0.241093412 | -0.148376423 | -0.105186552 | 0.13590686 | 0.043189871 | 0.329248241 | 0.243973512 | 0.397512912 | 0.561215491 | 0.738365305 |
| Peak1 | T | 896 | 0.03581171 | -0.146668945 | -0.136971148 | -0.172782858 | 0.009697797 | 0.707798622 | 0.015205605 | 0.06376645 | 0.160098684 | 0.840831425 |
| Gpsm1 | S | 539 | -0.136209512 | 0.122860772 | -0.288099665 | -0.151890154 | -0.410960437 | 0.319819918 | 0.429949116 | 0.006041239 | 0.262955607 | 0.067745912 |
| Luzp1 | S | 982 | 0.042075957 | -0.105697193 | -0.143141529 | -0.185217486 | -0.037444337 | 0.34121415 | 0.009612014 | 0.011786181 | 0.010732825 | 0.312729627 |
| Slain2 | S | 248 | -0.060613509 | -0.061188358 | -0.236184498 | -0.175570989 | -0.17499614 | 0.271160972 | 0.458911588 | 0.073280072 | 0.149008595 | 0.160269448 |
| Tagln | S | 181 | -0.149751991 | -0.092736219 | -0.286900395 | -0.137148405 | -0.194164176 | 0.402777853 | 0.595136932 | 0.1841639 | 0.327252408 | 0.208240415 |
| Sirt6 | S | 317 | -0.143457489 | -0.053737932 | -0.062789105 | 0.080668384 | -0.009051173 | 0.032492956 | 0.168684061 | 0.300724533 | 0.235010161 | 0.864035973 |
| Map1a | S | 1177 | -0.135986007 | -0.090502861 | -0.211022888 | -0.075036881 | -0.120520027 | 0.36960217 | 0.668497507 | 0.082223979 | 0.623120583 | 0.583749044 |
| Nes | S | 645 | -0.076843507 | -0.084982762 | -0.16723588 | -0.090392373 | -0.082253119 | 0.352011197 | 0.200659361 | 0.091731271 | 0.316249823 | 0.298989479 |
| Gpsm1 | S | 410 | -0.123647501 | -0.042823378 | -0.174507203 | -0.050859702 | -0.131683825 | 0.09497198 | 0.49191839 | 0.155222826 | 0.621265137 | 0.251083404 |
| Zswim8 | S | 564 | -0.057435526 | -0.044525439 | -0.190690987 | -0.13325546 | -0.146165547 | 0.292048279 | 0.653722568 | 0.017148943 | 0.065341432 | 0.190802094 |
| Pkp4 | S | 460 | -0.152231684 | 0.133655155 | -0.388024235 | -0.235792551 | -0.521679389 | 0.517365748 | 0.489616591 | 0.015042273 | 0.323142724 | 0.05883587 |
| Chd1 | S | 1678 | -0.002241216 | -0.115400072 | -0.109323618 | -0.107082402 | 0.006076454 | 0.948252681 | 0.009977754 | 0.066861952 | 0.077375747 | 0.892393138 |
| Map1a | S | 895 | -0.112315748 | -0.06537932 | -0.095600883 | 0.016714865 | -0.030221563 | 0.055502491 | 0.130545566 | 0.279756726 | 0.836191213 | 0.702176957 |
| Sec16a | S | 1362 | -0.094022419 | -0.029079226 | -0.109348995 | -0.015326577 | -0.08026977 | 0.053919995 | 0.530282905 | 0.028562469 | 0.555218172 | 0.067033585 |
| Sec16a | S | 1365 | -0.094022419 | -0.029079226 | -0.109348995 | -0.015326577 | -0.08026977 | 0.053919995 | 0.530282905 | 0.028562469 | 0.555218172 | 0.067033585 |
| Dclk1 | S | 305 | -0.062488827 | 0.012159706 | -0.201370435 | -0.138881608 | -0.213530141 | 0.335097005 | 0.864516013 | 0.003634236 | 0.063028154 | 0.042591282 |
| Tspyl2 | S | 648 | -0.052883861 | -0.056881625 | -0.115485422 | -0.062601561 | -0.058603797 | 0.439341513 | 0.134489576 | 0.009136334 | 0.358035508 | 0.062798432 |
| Spty2d1 | S | 278 | -0.002044068 | -0.127226918 | -0.062503188 | -0.060459121 | 0.064723729 | 0.922489373 | 0.000391604 | 0.31155367 | 0.323760883 | 0.299010734 |
| Flna | S | 966 | -0.045955472 | -0.103247793 | -0.097288719 | -0.051333248 | 0.005959074 | 0.217700429 | 0.045430572 | 0.122407279 | 0.355349685 | 0.91269087 |
| Calm1 Calm2 Calm3 | Y | 100;100;100 | -0.051397915 | -0.077426559 | -0.100052813 | -0.048654898 | -0.022626254 | 0.142621552 | 0.224467441 | 0.020185449 | 0.203905595 | 0.685747368 |
| Prrc2b | S | 417 | -0.009115429 | -0.189712543 | -0.156161337 | -0.147045908 | 0.033551206 | 0.833328242 | 0.116847682 | 0.098545535 | 0.104071313 | 0.736416517 |
| Prrc2b | S | 421 | -0.009115429 | -0.189712543 | -0.156161337 | -0.147045908 | 0.033551206 | 0.833328242 | 0.116847682 | 0.098545535 | 0.104071313 | 0.736416517 |
| Plekhg3 | S | 1291 | -0.201610048 | -0.192037908 | -0.022820736 | 0.178789312 | 0.169217171 | 0.073265599 | 0.293968886 | 0.809317393 | 0.125667306 | 0.347635615 |
| Plekhg3 | S | 759 | -0.035410423 | -0.069704778 | -0.143357669 | -0.107947246 | -0.073652891 | 0.529542162 | 0.024039544 | 0.068287738 | 0.16506325 | 0.22843583 |
| Jcad | S | 888 | -0.013208927 | -0.132020619 | -0.04276221 | -0.029553284 | 0.089258408 | 0.553846624 | 0.000334711 | 0.534853995 | 0.65705631 | 0.25379628 |
| Nvl | S | 507 | 0.027058526 | -0.079920964 | -0.294711311 | -0.321769837 | -0.214790347 | 0.767449489 | 0.455786713 | 0.014870699 | 0.012735003 | 0.07569968 |
| Ppp3ca | S | 373 | -0.337294878 | -0.337671218 | 0.162509588 | 0.499804466 | 0.500180806 | 0.48307685 | 0.016455843 | 0.385793257 | 0.331879385 | 0.052959128 |
| Mybbp1a | T | 1162 | 0.008869172 | -0.077317625 | -0.131460963 | -0.140330134 | -0.054143338 | 0.834272848 | 0.057700893 | 0.004102548 | 0.035812399 | 0.10206736 |
| Nadk | T | 62 | -0.016161822 | -0.01110917 | -0.358915105 | -0.342753283 | -0.347805934 | 0.902377642 | 0.929753779 | 0.01690916 | 0.028565565 | 0.019877857 |
| Errfi1 | S | 214 | -0.025076466 | -0.157293401 | -0.095004262 | -0.069927796 | 0.062289139 | 0.528260517 | 0.00588314 | 0.507159776 | 0.618399001 | 0.653808598 |
| Tns3 | S | 1100 | -0.1842316 | 0.004510616 | -0.187516926 | -0.003285326 | -0.192027542 | 0.047094814 | 0.955240274 | 0.207151761 | 0.978407339 | 0.196595146 |
| 4930579G24Rik | S | 10 | -0.093484492 | -0.29754268 | -0.053198777 | 0.040285716 | 0.244343903 | 0.422849536 | 0.081284662 | 0.723774358 | 0.811838403 | 0.211356799 |
| Rgl1 | S | 632 | -0.362977386 | 0.160206456 | -0.226459127 | 0.136518259 | -0.386665583 | 0.014628319 | 0.216081276 | 0.283020911 | 0.491791075 | 0.117167691 |
| Scrib | S | 821 | 0.050572441 | -0.137034588 | -0.109699647 | -0.160272088 | 0.027334941 | 0.237032737 | 0.005995901 | 0.010114415 | 0.024301172 | 0.324746406 |
| Arhgap11a | S | 650 | -0.172289315 | -0.016386334 | -0.240965361 | -0.068676046 | -0.224579027 | 0.06296065 | 0.842522553 | 0.306389482 | 0.740141972 | 0.333076842 |
| Plk4 | S | 536 | -0.18229428 | -0.041130903 | -0.181032713 | 0.001261567 | -0.13990181 | 0.161224749 | 0.694122149 | 0.158398432 | 0.992383006 | 0.311524707 |

| | | | | | | | | | | | | |
|---|---|---|---|---|---|---|---|---|---|---|---|---|
| Dek | S | 55 | -0.015144261 | -0.174666324 | -0.132656972 | -0.117512711 | 0.042009352 | 0.853738478 | 0.037276358 | 0.190117701 | 0.25524635 | 0.627827148 |
| Camk1d | S | 359 | -0.128224464 | -0.033931126 | -0.351686157 | -0.223461693 | -0.31775503 | 0.313307997 | 0.608974939 | 0.193047667 | 0.36376861 | 0.226670403 |
| Drosha | S | 354 | -0.011345346 | -0.073777599 | -0.083833607 | -0.072488261 | -0.010056009 | 0.653105942 | 0.027461425 | 0.003406262 | 0.042026292 | 0.671761255 |
| Sipa1l1 | T | 1190 | -0.43073385 | 0.033537754 | 0.110899587 | 0.541633437 | 0.077361833 | 0.000240095 | 0.610716255 | 0.113209028 | 0.000784878 | 0.299461941 |
| Ralgapa1 | S | 720 | -0.128758284 | -0.008564067 | -0.177222458 | -0.048464174 | -0.168658391 | 0.066417523 | 0.930082112 | 0.086441923 | 0.524067765 | 0.159043567 |
| Tnks1bp1 | S | 699 | -0.074714915 | -0.118241016 | -0.228774602 | -0.154059687 | -0.110533586 | 0.381963157 | 0.248725047 | 0.206749391 | 0.35885897 | 0.503605639 |
| Rab11fip5 | S | 280 | -0.069210363 | -0.140515837 | -0.097445271 | -0.028234908 | 0.043070566 | 0.465707807 | 0.036025239 | 0.264398597 | 0.783178644 | 0.582311838 |
| Shb | S | 304 | -0.089790095 | -0.09960519 | -0.091225063 | -0.001434968 | 0.008380126 | 0.105868117 | 0.071314053 | 0.346632995 | 0.986259969 | 0.918876073 |
| Pkn1 | S | 382 | -0.096529706 | -0.023919904 | -0.140802992 | -0.044273286 | -0.116883088 | 0.077194793 | 0.796269999 | 0.030343754 | 0.372146217 | 0.263153645 |
| Pkn1 | S | 383 | -0.096529706 | -0.023919904 | -0.140802992 | -0.044273286 | -0.116883088 | 0.077194793 | 0.796269999 | 0.030343754 | 0.372146217 | 0.263153645 |
| Kdm2a | S | 422 | -0.016217556 | -0.149155332 | -0.164706627 | -0.148489071 | -0.015551295 | 0.697852255 | 0.069817313 | 0.143050289 | 0.171652391 | 0.870465143 |
| Amotl1 | S | 936 | 0.028260611 | -0.101711653 | -0.169092671 | -0.197353282 | -0.067381018 | 0.547007711 | 0.050267074 | 0.013736511 | 0.009916271 | 0.165618355 |
| Hpca Hpcal1 Ncald | S | 175;175;175 | -0.111555451 | -0.045480039 | -0.205047272 | -0.093491821 | -0.159567233 | 0.330540066 | 0.529368256 | 0.063934757 | 0.442422336 | 0.137176166 |
| Washc2 | S | 832 | -0.047711616 | -0.125602443 | -0.17621559 | -0.128503974 | -0.050613146 | 0.682727886 | 0.115272315 | 0.115671867 | 0.33382653 | 0.57934353 |
| Mapk6 | S | 703 | 0.050273346 | -0.125080662 | -0.162028438 | -0.212301784 | -0.036947776 | 0.245644791 | 0.041986602 | 0.006164335 | 0.000229133 | 0.374795735 |
| Usp10 | S | 208 | 0.027985947 | -0.126718466 | -0.100114196 | -0.128100142 | 0.02660427 | 0.424658905 | 0.017619176 | 0.006518995 | 0.016690929 | 0.443514067 |
| Phldb1 | S | 335 | 0.245679796 | -0.259514105 | -0.284234117 | -0.529913913 | -0.024720012 | 0.020308436 | 0.00688637 | 0.012894613 | 0.002471755 | 0.737900891 |
| Ptk2 | Y | 577 | -0.129446414 | -0.032921638 | -0.125106137 | 0.004340277 | -0.092184499 | 0.008815283 | 0.731517589 | 0.303641995 | 0.966954566 | 0.492830247 |
| Suv39h2 | S | 165 | 0.045646485 | -0.195626255 | -0.115726727 | -0.161373212 | 0.079899528 | 0.121325237 | 0.001591136 | 0.163846793 | 0.093894593 | 0.290541056 |
| Gapvd1 | S | 982 | -0.386385715 | 0.029417413 | -0.120587455 | 0.26579826 | -0.150004868 | 0.046655851 | 0.751011092 | 0.538295542 | 0.245385921 | 0.440952896 |
| Son | S | 94 | 0.171586146 | -0.208115687 | -0.241115482 | -0.412701628 | -0.032999795 | 0.573830026 | 0.07596237 | 0.051424661 | 0.239792447 | 0.48299484 |
| Cit | S | 1458 | 0.055732025 | -0.308031261 | -0.229488088 | -0.285220112 | 0.078543173 | 0.109074295 | 0.037553047 | 0.191949347 | 0.136691743 | 0.608429349 |
| Dmtn | S | 289 | -0.05188525 | -0.111051861 | -0.09860703 | -0.04672178 | 0.012444831 | 0.231940082 | 0.039752952 | 0.204634231 | 0.483167811 | 0.849005162 |
| Amer1 | S | 744 | -0.258226749 | -0.038900112 | -0.344687477 | -0.086460728 | -0.305787365 | 0.205949016 | 0.870692277 | 0.406480363 | 0.82491431 | 0.475445771 |
| Ibtk | S | 1111 | 0.009046581 | -0.116626562 | -0.15905011 | -0.168096691 | -0.042423547 | 0.911069684 | 0.096350581 | 0.022719231 | 0.117460585 | 0.503776467 |
| Cdca3 | S | 29 | 0.016949041 | -0.109493412 | -0.196061079 | -0.21301012 | -0.086567668 | 0.551541514 | 0.057862772 | 0.034918014 | 0.043005771 | 0.226423751 |
| Rreb1 | S | 970 | 0.009407242 | -0.202629341 | -0.044447656 | -0.053854898 | 0.158181685 | 0.848924681 | 0.002086612 | 0.443570705 | 0.383289205 | 0.04317164 |
| Gtpbp1 | S | 47 | -0.248795128 | 0.00270062 | -0.1354836 | 0.113311528 | -0.13818422 | 0.076025123 | 0.95168956 | 0.182690322 | 0.348767285 | 0.178023527 |
| Irs2 | S | 675 | -0.035387657 | -0.128916252 | -0.057054569 | -0.021666911 | 0.071861684 | 0.430153774 | 0.004149499 | 0.319127654 | 0.706856741 | 0.226330896 |
| Top1 | S | 396 | 0.046701415 | -0.135572075 | -0.111594551 | -0.158295966 | 0.023977524 | 0.409875365 | 0.008578104 | 0.01816954 | 0.04662201 | 0.52298402 |
| Sh3d19 | S | 564 | -0.113367795 | -0.046175922 | -0.139278525 | -0.02591073 | -0.093102603 | 0.099648554 | 0.484840242 | 0.094547248 | 0.620004253 | 0.163014875 |
| Setd3 | S | 12 | -0.136853093 | 0.006465346 | -0.280755607 | -0.143902514 | -0.287220953 | 0.171064767 | 0.958624961 | 0.107466686 | 0.326381883 | 0.117042699 |
| Klf16 | S | 103 | -0.023566491 | -0.169406488 | -0.202666547 | -0.179100056 | -0.033260058 | 0.771128288 | 0.331508987 | 0.059818354 | 0.119765287 | 0.837067317 |
| Rprd2 | S | 1117 | -0.043035232 | -0.194675524 | -0.116539261 | -0.073504029 | 0.078136264 | 0.470537674 | 0.138608327 | 0.155463085 | 0.369179616 | 0.484481758 |
| Nes | S | 883 | -0.106378291 | -0.027601229 | -0.098347872 | 0.008030419 | -0.070746643 | 0.075038518 | 0.748452066 | 0.013491444 | 0.856534109 | 0.437287129 |
| Frmd8 | S | 20 | -0.126297432 | 0.033949949 | -0.281750851 | -0.155453419 | -0.3157008 | 0.309812459 | 0.776470148 | 0.034822351 | 0.251758897 | 0.057965751 |
| Tbc1d9b | S | 997 | 0.145238147 | -0.051480919 | -0.354090247 | -0.499328394 | -0.302609327 | 0.327659941 | 0.102067221 | 0.007134841 | 0.032818429 | 0.018723737 |
| Amotl1 | T | 882 | -0.060532549 | -0.146377307 | -0.126111341 | -0.065578791 | 0.020265966 | 0.323207309 | 0.05200041 | 0.393576518 | 0.625394308 | 0.87827254 |
| Mdc1 | S | 1104 | -0.089783752 | 0.011571954 | -0.215081794 | -0.125298042 | -0.226653748 | 0.241921844 | 0.894672061 | 0.016271631 | 0.152630652 | 0.073811791 |
| Ednra | S | 245 | -0.158513789 | -0.199590618 | 0.020417651 | 0.178931441 | 0.220008269 | 0.116961417 | 0.067848978 | 0.866659251 | 0.153821583 | 0.101370958 |
| Ednra | T | 238 | -0.158513789 | -0.199590618 | 0.020417651 | 0.178931441 | 0.220008269 | 0.116961417 | 0.067848978 | 0.866659251 | 0.153821583 | 0.101370958 |
| Ednra | T | 244 | -0.158513789 | -0.199590618 | 0.020417651 | 0.178931441 | 0.220008269 | 0.116961417 | 0.067848978 | 0.866659251 | 0.153821583 | 0.101370958 |
| Ednra | Y | 251 | -0.158513789 | -0.199590618 | 0.020417651 | 0.178931441 | 0.220008269 | 0.116961417 | 0.067848978 | 0.866659251 | 0.153821583 | 0.101370958 |
| Epb41l2 | T | 584 | 0.069626875 | -0.166390443 | -0.268924113 | -0.338550988 | -0.10253367 | 0.503805535 | 0.10761892 | 0.059848668 | 0.043689958 | 0.390507717 |
| Map3k1 | S | 71 | -0.041931483 | -0.14323216 | -0.23987511 | -0.197943626 | -0.09664295 | 0.861260886 | 0.484815074 | 0.046089679 | 0.437554447 | 0.623506936 |
| Saal1 | S | 376 | -0.059348622 | -0.058235935 | -0.126064298 | -0.066715675 | -0.067828363 | 0.13289052 | 0.240536042 | 0.044644781 | 0.203587892 | 0.245125429 |
| Fam91a1 | S | 671 | -0.014062022 | -0.179349397 | -0.143883156 | -0.129821134 | 0.035466241 | 0.849879236 | 0.07281528 | 0.145868832 | 0.225833458 | 0.703034273 |
| Palb2 | S | 334 | -0.061514252 | -0.207667462 | -0.079635384 | -0.018121132 | 0.128032078 | 0.282580804 | 0.165269087 | 0.180386753 | 0.721590266 | 0.334975115 |
| Pde4d | S | 136 | -0.118337578 | -0.039499864 | -0.297928452 | -0.179590874 | -0.258428588 | 0.400563745 | 0.742032163 | 0.103587529 | 0.298417364 | 0.145167181 |
| Fgd6 | S | 494 | -0.109839318 | -0.048415082 | -0.032686437 | 0.077152882 | 0.015728645 | 0.002570755 | 0.199519825 | 0.409854583 | 0.10710636 | 0.701413451 |
| 2310039H08Rik | S | 15 | -0.073050479 | -0.095672383 | -0.23169952 | -0.158649041 | -0.136027157 | 0.567764573 | 0.197726831 | 0.150751746 | 0.34855051 | 0.338656209 |
| Dph2 | S | 188 | 0.009423865 | -0.072753769 | -0.149535087 | -0.158958952 | -0.076781318 | 0.912443053 | 0.070304284 | 0.008056311 | 0.146665473 | 0.077257444 |
| Ogfr | S | 616 | 0.10071732 | -0.064335429 | -0.190166944 | -0.290884265 | -0.125831515 | 0.594451643 | 0.268066367 | 0.000534322 | 0.20912422 | 0.07930197 |
| Arfgap1 | S | 359 | -0.048327409 | -0.06578186 | -0.249798123 | -0.201470714 | -0.184016263 | 0.408169035 | 0.309108113 | 0.091807176 | 0.14882234 | 0.172142403 |
| Coprs | S | 15 | -0.045671204 | -0.056553823 | -0.1630766 | -0.117405395 | -0.106522777 | 0.437204077 | 0.299640778 | 0.019220278 | 0.112521604 | 0.115069314 |
| Htt | S | 411 | -0.075738211 | -0.01101172 | -0.173814318 | -0.098076107 | -0.162802598 | 0.234187941 | 0.826667374 | 0.011577527 | 0.162928753 | 0.038245946 |
| Ermp1 | S | 34 | -0.035519601 | -0.113658118 | -0.089360803 | -0.053841202 | 0.024297315 | 0.431472441 | 0.156978642 | 0.012445384 | 0.244700242 | 0.701502542 |
| Ska3 | S | 110 | -0.066041686 | -0.015845134 | -0.160669926 | -0.09462824 | -0.144824791 | 0.273602411 | 0.807630317 | 0.005998784 | 0.13226288 | 0.091614849 |
| Eif5b | S | 139 | -0.087411553 | -0.065168651 | -0.081188405 | 0.006223148 | -0.016019754 | 0.096672356 | 0.104093417 | 0.113054704 | 0.893071803 | 0.705063605 |
| Eml4 | S | 138 | 0.21950698 | -0.22596889 | -0.175096877 | -0.394603858 | 0.050872013 | 0.446604942 | 0.007196572 | 0.111483668 | 0.226866428 | 0.565840708 |
| Plekhg3 | S | 762 | -0.051129467 | -0.097087036 | -0.009754777 | 0.041374689 | 0.087332258 | 0.129433686 | 0.00061556 | 0.483026075 | 0.200422283 | 0.006123433 |
| Cdk17 | S | 180 | -0.23229396 | 0.011526135 | -0.184714313 | 0.047579647 | -0.196240447 | 0.112839987 | 0.928499915 | 0.170093539 | 0.663814575 | 0.112600926 |
| Leo1 | S | 608 | 0.003565412 | -0.110328533 | -0.091862674 | -0.095428087 | 0.018465859 | 0.908309435 | 0.008182002 | 0.039780182 | 0.038966823 | 0.576928319 |
| Leo1 | S | 609 | 0.003565412 | -0.110328533 | -0.091862674 | -0.095428087 | 0.018465859 | 0.908309435 | 0.008182002 | 0.039780182 | 0.038966823 | 0.576928319 |
| Leo1 | S | 611 | 0.003565412 | -0.110328533 | -0.091862674 | -0.095428087 | 0.018465859 | 0.908309435 | 0.008182002 | 0.039780182 | 0.038966823 | 0.576928319 |
| Trim56 | S | 454 | -0.149289885 | -0.078279729 | 0.061359897 | 0.210649782 | 0.139639626 | 0.002536784 | 0.035741083 | 0.513544053 | 0.10330083 | 0.20694367 |
| Srrm1 | S | 260 | -0.113425404 | -0.153980972 | 0.034207432 | 0.147632837 | 0.188188404 | 0.13762693 | 0.003882974 | 0.537197485 | 0.090614666 | 0.02799614 |
| Axl | Y | 860 | -0.157027783 | 0.01115236 | -0.166380754 | -0.009352972 | -0.177533114 | 0.016162367 | 0.915613334 | 0.251164471 | 0.938467717 | 0.26651783 |
| Birc6 | S | 4850 | 0.173502041 | -0.374586069 | -0.311604621 | -0.485106662 | 0.062981448 | 0.095886021 | 0.122394796 | 0.039557507 | 0.011318265 | 0.739698369 |
| Sh3pxd2a | S | 656 | 0.026356977 | -0.145504104 | -0.10096689 | -0.127323867 | 0.044537214 | 0.445086427 | 0.005260424 | 0.065451997 | 0.051297235 | 0.2901838 |
| Map4k4 | T | 909 | -0.041100858 | -0.076102248 | -0.137216506 | -0.096115648 | -0.061114258 | 0.384934579 | 0.253654469 | 0.020328811 | 0.105628499 | 0.356795974 |
| Hook3 | S | 238 | -0.076592043 | -0.146030147 | -0.151761797 | -0.075169753 | -0.00573165 | 0.387706969 | 0.138225254 | 0.270746883 | 0.578161781 | 0.964663838 |
| Pom121 | S | 441 | -0.053011099 | -0.143830286 | -0.05238084 | 0.000630259 | 0.091449447 | 0.384763008 | 0.035174271 | 0.120459796 | 0.990367939 | 0.131834411 |
| Timm17a | S | 40 | -0.008563758 | -0.156460927 | -0.247892995 | -0.239329237 | -0.091432068 | 0.932443125 | 0.117319352 | 0.184999469 | 0.204812693 | 0.571589962 |
| Gopc | S | 442 | 0.061183984 | -0.121995794 | -0.156536175 | -0.217720159 | -0.034540381 | 0.488578212 | 0.026074002 | 0.018092071 | 0.071243796 | 0.474214102 |
| Akap12 | S | 270 | -0.00168147 | -0.021101484 | -0.22142908 | -0.21974761 | -0.200327596 | 0.978071512 | 0.698112589 | 0.002376293 | 0.022732464 | 0.0169194 |
| Mybbp1a | T | 1256 | 0.208033958 | -0.195196027 | -0.287667719 | -0.495701677 | -0.092471692 | 0.004344033 | 0.019393202 | 0.000770643 | 6.18338E-05 | 0.147715624 |
| Dock8 | S | 452 | -0.214118206 | -0.112525823 | -0.028634704 | 0.185483502 | 0.083891119 | 0.065464123 | 0.247321154 | 0.759369038 | 0.094258037 | 0.333498912 |
| Tor1aip1 | T | 235 | -0.087732377 | -0.000206853 | -0.092612332 | -0.004879956 | -0.09240548 | 0.015698906 | 0.996473029 | 0.011241047 | 0.841954259 | 0.119077276 |
| Srpk1 | S | 311 | -0.104034125 | -0.093238022 | -0.117124623 | -0.013090497 | -0.023886601 | 0.128717473 | 0.096595624 | 0.416182334 | 0.923140492 | 0.858157409 |
| Yju2 | S | 213 | -0.025413666 | -0.049129939 | -0.067908038 | -0.042494372 | -0.018778099 | 0.06901295 | 0.009156755 | 0.022298317 | 0.093272506 | 0.359930475 |
| Ddx46 | S | 129 | -0.106050042 | -0.158258909 | -0.087267549 | 0.018782493 | 0.07099136 | 0.291705018 | 0.11575478 | 0.348298268 | 0.856416944 | 0.483111845 |
| Ppan | S | 238 | -0.024708226 | -0.153460654 | -0.190877067 | -0.166168842 | -0.037416413 | 0.894699751 | 0.090741398 | 0.161956868 | 0.435929451 | 0.751180605 |
| Elavl1 | S | 202 | 0.032460198 | -0.100897579 | -0.166071095 | -0.198531293 | -0.065173517 | 0.44670505 | 0.056207249 | 0.010395927 | 0.013640767 | 0.138037803 |
| Sacs | S | 4266 | -0.061624439 | -0.06708614 | -0.087506945 | -0.025882507 | -0.020420805 | 0.203312958 | 0.050787124 | 0.066091541 | 0.585764059 | 0.579503615 |
| Pxn | S | 136 | -0.050177196 | -0.10617245 | -0.256416654 | -0.206239458 | -0.150244204 | 0.438789854 | 0.269005031 | 0.173378829 | 0.248977656 | 0.376274952 |
| Scmh1 | S | 574 | -0.045964447 | -0.071755441 | -0.090982715 | -0.045018269 | -0.019227275 | 0.095937888 | 0.086576796 | 0.064669458 | 0.265447999 | 0.650833249 |
| Tnrc6b | S | 1809 | -0.026890219 | -0.138067711 | -0.33154062 | -0.304650401 | -0.19347291 | 0.693494115 | 0.115153281 | 0.337471606 | 0.369827194 | 0.544389618 |
| Rabggta | S | 10 | -0.150549522 | -0.181434352 | 0.027202384 | 0.177751905 | 0.208636735 | 0.091724219 | 0.059800092 | 0.837556292 | 0.219723795 | 0.168056009 |
| Rabggta | T | 9 | -0.150549522 | -0.181434352 | 0.027202384 | 0.177751905 | 0.208636735 | 0.091724219 | 0.059800092 | 0.837556292 | 0.219723795 | 0.168056009 |
| Cep170 | S | 1069 | -0.042781703 | -0.016118728 | -0.145635866 | -0.102854162 | -0.129517138 | 0.182269106 | 0.750791588 | 0.003542219 | 0.033239354 | 0.059156641 |
| Tor1aip1 | S | 244 | -0.098704824 | -0.006898554 | -0.127867567 | -0.029162744 | -0.120969013 | 0.063364675 | 0.86323563 | 0.023987423 | 0.535411295 | 0.038337136 |
| Map4 | T | 260 | 0.002594422 | -0.158738399 | -0.195091981 | -0.197686403 | -0.036353582 | 0.9813983 | 0.039621549 | 0.230201568 | 0.257499067 | 0.791604778 |
| Dennd4b | S | 951 | 0.176616472 | -0.162613045 | -0.189614063 | -0.366230535 | -0.027001019 | 0.17344499 | 0.009093521 | 0.010867506 | 0.03830191 | 0.585865243 |

| | | | | | | | | | | | | |
|---|---|---|---|---|---|---|---|---|---|---|---|---|
| Plekhg3 | S | 1290 | -0.167874041 | -0.083166412 | -0.13631187 | 0.031562171 | -0.053145458 | 0.167462853 | 0.144672695 | 0.528301231 | 0.883578582 | 0.795610769 |
| Zc3h11a | S | 743 | 0.054510382 | -0.16900959 | -0.124163955 | -0.178674336 | 0.044845635 | 0.062619054 | 0.001305082 | 0.071228127 | 0.028005245 | 0.37313641 |
| Tbc1d15 | S | 203 | -0.275606715 | 0.00104217 | -0.338239206 | -0.06263249 | -0.339281376 | 0.305488198 | 0.991673569 | 0.260416068 | 0.844640075 | 0.259695922 |
| Cavin1 | T | 271 | -0.085803634 | -0.105807302 | -0.079908662 | 0.005894972 | 0.02589864 | 0.168536505 | 0.05353473 | 0.334797755 | 0.938251694 | 0.715675952 |
| Mavs | S | 220 | 0.000396485 | -0.080569534 | -0.164391832 | -0.164788318 | -0.083822298 | 0.994381269 | 0.011678511 | 0.079111208 | 0.086074667 | 0.264652323 |
| Sec62 | S | 115 | -0.102210917 | -0.1790713 | -0.076285266 | 0.025925651 | 0.102786034 | 0.354855698 | 0.079330916 | 0.459773449 | 0.838167769 | 0.379273435 |
| Srsf2 | S | 206 | 0.067154105 | -0.184624701 | -0.14482446 | -0.211978565 | 0.039800241 | 0.440247839 | 0.006386592 | 0.164872963 | 0.095778956 | 0.644069129 |
| Wdr44 | S | 172 | -0.073379104 | 0.018253898 | -0.28144625 | -0.208067146 | -0.299700148 | 0.052696343 | 0.739049753 | 0.07608059 | 0.144296365 | 0.060597562 |
| Map1a | S | 1762 | -0.054872803 | -0.101155404 | -0.067827493 | -0.01295469 | 0.033327911 | 0.162646217 | 0.050848364 | 0.122648104 | 0.680089586 | 0.416632565 |
| Gprc5a | S | 303 | 0.080887332 | -0.116906426 | -0.210670634 | -0.291557966 | -0.093764209 | 0.17519981 | 0.047250182 | 0.007252253 | 0.00587574 | 0.130904869 |
| Srrm2 | S | 1864 | -0.157219992 | -0.277166367 | -0.015626361 | 0.141593631 | 0.261540006 | 0.381818153 | 0.091055456 | 0.917540814 | 0.448020509 | 0.144109671 |
| Ncor2 | S | 2213 | -0.020297378 | -0.068735261 | -0.125468283 | -0.105170905 | -0.056733022 | 0.640558498 | 0.165618306 | 0.006580884 | 0.050256063 | 0.20524731 |
| Cask | S | 576 | -0.025113557 | -0.14489955 | -0.03020181 | -0.005088252 | 0.114697741 | 0.615431608 | 0.000982629 | 0.579310166 | 0.934985917 | 0.107185941 |
| Hcfc1 | S | 2029 | 0.004592463 | -0.046634759 | -0.138697323 | -0.143289786 | -0.092062563 | 0.80881811 | 0.036207941 | 0.003875035 | 0.004493756 | 0.01916609 |
| Ncapd2 | T | 1319 | 0.030041484 | -0.081859494 | -0.181088762 | -0.211130246 | -0.099229268 | 0.629348772 | 0.026776237 | 0.025912361 | 0.034519643 | 0.113987291 |
| Osbpl6 | S | 24 | -0.062046169 | -0.066703858 | -0.296478949 | -0.23443278 | -0.229775091 | 0.689488569 | 0.741216928 | 0.0489508 | 0.217204745 | 0.320962599 |
| Nono | T | 452 | 0.081932731 | -0.13623243 | -0.167777061 | -0.249709792 | -0.03154463 | 0.072866928 | 0.012188749 | 0.005979337 | 0.002059362 | 0.404905185 |
| Cep170 | T | 1249 | 0.146007949 | -0.218469275 | -0.33184451 | -0.477852459 | -0.113375235 | 0.098955649 | 0.022166353 | 0.083815323 | 0.037967115 | 0.441155394 |
| Clic4 | S | 236 | -0.205074149 | -0.088144583 | -0.202350932 | 0.002723217 | -0.114206349 | 0.291107446 | 0.480593584 | 0.319130178 | 0.989855585 | 0.53191056 |
| Gtpbp1 | S | 44 | -0.243100487 | 0.004197723 | -0.127474337 | 0.11562615 | -0.13167206 | 0.07024883 | 0.925382377 | 0.2007 | 0.327102949 | 0.192111921 |
| Nap1l1 | S | 69 | -0.005951963 | -0.067182369 | -0.181024468 | -0.175072505 | -0.113842099 | 0.93505611 | 0.488820724 | 0.004053477 | 0.079999574 | 0.277029528 |
| Dab2ip | S | 895 | -0.158590658 | 0.049754399 | -0.248379893 | -0.089789235 | -0.298134292 | 0.069769827 | 0.634990644 | 0.111451328 | 0.468845792 | 0.080142463 |
| Nav3 | S | 941 | 0.007028712 | -0.261519226 | -0.078274828 | -0.085303539 | 0.183244399 | 0.938697041 | 0.020219237 | 0.453749421 | 0.484587442 | 0.149512928 |
| Rplp1 | S | 104 | 0.007001987 | -0.105754366 | -0.091259025 | -0.098261012 | 0.014495341 | 0.792097981 | 0.011990417 | 0.021126029 | 0.021552576 | 0.645174597 |
| Ace2 | S | 623 | -0.011331774 | -0.045230336 | -0.212047424 | -0.20071565 | -0.166817088 | 0.761741229 | 0.400050087 | 0.011110074 | 0.017312639 | 0.040177005 |
| Ace2 | S | 626 | -0.011331774 | -0.045230336 | -0.212047424 | -0.20071565 | -0.166817088 | 0.761741229 | 0.400050087 | 0.011110074 | 0.017312639 | 0.040177005 |
| Ace2 | Y | 633 | -0.011331774 | -0.045230336 | -0.212047424 | -0.20071565 | -0.166817088 | 0.761741229 | 0.400050087 | 0.011110074 | 0.017312639 | 0.040177005 |
| Katnb1 | S | 362 | -0.146931442 | -0.093540145 | -0.097136809 | 0.049794632 | -0.003596664 | 0.113594447 | 0.176985646 | 0.477280176 | 0.709023361 | 0.976898509 |
| Cwc22 | S | 831 | -0.037091178 | -0.105012429 | -0.143212654 | -0.106121475 | -0.038200225 | 0.587565128 | 0.183076191 | 0.042265733 | 0.204848353 | 0.589703848 |
| Ssb | S | 219 | -0.02580299 | -0.035122035 | -0.161502568 | -0.135699579 | -0.126380534 | 0.612371108 | 0.303569225 | 0.0054093 | 0.045463349 | 0.029101779 |
| Nrbp1 | T | 433 | -0.088087796 | -0.056421979 | -0.068338337 | 0.019749459 | -0.011916358 | 0.014374453 | 0.302357446 | 0.144896475 | 0.584842538 | 0.825229456 |
| Cdc25b | S | 186 | -0.020962949 | -0.05437043 | -0.18714787 | -0.166184921 | -0.13277744 | 0.644050428 | 0.233433204 | 0.01981731 | 0.037395605 | 0.067347669 |
| Eif3b | S | 153 | -0.058133018 | -0.065376468 | -0.077965489 | -0.019832472 | -0.012589021 | 0.117861205 | 0.090815061 | 0.049580042 | 0.497477627 | 0.669957442 |
| Ap3d1 | T | 870 | -0.267382175 | -0.121983061 | -0.02953664 | 0.237845535 | 0.092446421 | 0.152214022 | 0.198072461 | 0.812748113 | 0.202432977 | 0.399957022 |
| Fosl2 | S | 120 | -0.031614526 | -0.125697891 | -0.078348952 | -0.046734426 | 0.04734894 | 0.517665818 | 0.021063326 | 0.141914336 | 0.28654562 | 0.222672277 |
| Gnl1 | T | 50 | -0.03003207 | -0.103458878 | -0.175467273 | -0.145435203 | -0.072008394 | 0.744012575 | 0.119313344 | 0.078435804 | 0.199374384 | 0.377779871 |
| Sipa1l1 | S | 187 | -0.174447424 | -0.074247326 | -0.026974215 | 0.147473209 | 0.047273111 | 0.018178813 | 0.267491195 | 0.777581494 | 0.174171339 | 0.605118505 |
| Noc2l | S | 673 | 0.040190077 | -0.119647215 | -0.108809789 | -0.148999866 | 0.010837426 | 0.416887413 | 0.011036289 | 0.013503299 | 0.04227169 | 0.739553533 |
| Mettl16 | S | 425 | 0.005983187 | -0.031729897 | -0.165738095 | -0.171721282 | -0.134008197 | 0.829836971 | 0.321734118 | 0.000944089 | 0.005301198 | 0.013907895 |
| Zfp800 | S | 337 | 0.067244986 | -0.213087036 | -0.06781034 | -0.135055326 | 0.145276695 | 0.12465062 | 0.000660883 | 0.154628962 | 0.021395118 | 0.016552106 |
| Tdp1 | S | 132 | -0.029633926 | -0.064936645 | -0.110203832 | -0.080569906 | -0.045267187 | 0.605512837 | 0.055201772 | 0.016020843 | 0.21602076 | 0.183286801 |
| Sh3d19 | S | 368 | -0.172620236 | 0.015631347 | -0.134698698 | 0.037921538 | -0.150330045 | 0.048423673 | 0.89255767 | 0.106586747 | 0.464718429 | 0.22404364 |
| Fmnl2 | T | 467 | -0.075776696 | -0.054693918 | -0.103997351 | -0.028220655 | -0.049303433 | 0.146328034 | 0.336607731 | 0.030571408 | 0.549974235 | 0.385902029 |
| Fam193a | S | 302 | -0.137164772 | -0.163560123 | -0.123551379 | 0.013613393 | 0.040008744 | 0.449636257 | 0.251413663 | 0.238472652 | 0.934221882 | 0.742034152 |
| Usp14 | S | 143 | -0.11021281 | -0.111796116 | -0.046528783 | 0.063684027 | 0.065267333 | 0.079106339 | 0.11308156 | 0.407120703 | 0.266457976 | 0.310179417 |
| Marcksl1 | S | 104 | 0.773913576 | -0.276807398 | -0.560869965 | -1.334783541 | -0.284062568 | 0.431441039 | 0.212862455 | 0.033738116 | 0.231503305 | 0.244677879 |
| Pum2 | S | 102 | -0.008301726 | -0.140443292 | -0.118747755 | -0.110446028 | 0.021695537 | 0.821070902 | 0.060410394 | 0.072540107 | 0.092096349 | 0.729788334 |
| Rab3gap1 | S | 590 | -0.153536747 | -0.177471023 | 0.021887089 | 0.175423836 | 0.199358112 | 0.10158293 | 0.068897462 | 0.869942768 | 0.221639936 | 0.180871423 |
| Map1a | S | 667 | -0.096049616 | -0.091978702 | -0.0790653 | 0.016984316 | 0.012913403 | 0.219578937 | 0.058185192 | 0.24571706 | 0.824031159 | 0.809522087 |
| Nol8 | S | 787 | -0.02695017 | -0.112282144 | -0.076649799 | -0.049699629 | 0.035632345 | 0.678748973 | 0.004474253 | 0.212312406 | 0.512219503 | 0.499641224 |
| Rps6kc1 | S | 426 | -0.098294216 | -0.000981809 | -0.099304454 | -0.001010238 | -0.098322645 | 0.019060531 | 0.98299007 | 0.027269324 | 0.966584861 | 0.075046659 |
| Spn | S | 347 | -0.300760456 | 0.000519312 | 0.006325315 | 0.307085771 | 0.005806003 | 0.005831276 | 0.995457499 | 0.952806831 | 0.046159921 | 0.959637569 |
| Zfp592 | S | 329 | -0.249438819 | -0.071255728 | 0.098358516 | 0.347797336 | 0.169614245 | 0.005949582 | 0.229516165 | 0.584372536 | 0.137190553 | 0.373037304 |
| Cdk7 | S | 164 | -0.024003508 | -0.056756395 | -0.16719776 | -0.143194252 | -0.110441365 | 0.48579544 | 0.113395034 | 0.037434477 | 0.055460334 | 0.101925385 |
| Gorab | S | 348 | -0.233156047 | -0.285320198 | -0.096320448 | 0.136835599 | 0.18899975 | 0.469705587 | 0.441013206 | 0.342913678 | 0.656748525 | 0.593508683 |
| Peak1 | S | 716 | -0.050285931 | -0.106620464 | -0.135100439 | -0.084814508 | -0.028479975 | 0.495050308 | 0.078217609 | 0.141272346 | 0.357107789 | 0.705821497 |
| Csde1 | S | 114 | 0.125415609 | -0.106575983 | -0.389613126 | -0.515028735 | -0.283037143 | 0.21017025 | 0.588914166 | 0.005355758 | 0.004188547 | 0.215073984 |
| P2ry2 | S | 329 | -0.025478696 | -0.134863279 | -0.086210357 | -0.060731661 | 0.048652921 | 0.620251422 | 0.02892972 | 0.128452503 | 0.162474135 | 0.24206943 |
| Fam13b | S | 696 | -0.01018006 | -0.147628923 | -0.095376149 | -0.085196088 | 0.052252774 | 0.875260323 | 0.005508448 | 0.367640623 | 0.449706111 | 0.593755804 |
| Spen | T | 1634 | -0.139865272 | -0.130797222 | -0.015789548 | 0.124075724 | 0.115007674 | 0.173618027 | 0.028575297 | 0.879911801 | 0.341277192 | 0.329637992 |
| Ets1 | T | 38 | 0.077211081 | -0.146135347 | -0.214323745 | -0.291534826 | -0.068188398 | 0.014351085 | 0.024856215 | 0.005320586 | 0.000684119 | 0.114946551 |
| Usp10 | S | 223 | 0.042181447 | -0.153686065 | -0.122156489 | -0.164337936 | 0.031529576 | 0.367425046 | 0.020309411 | 0.030428011 | 0.015051092 | 0.521726348 |
| Cdca2 | S | 238 | 0.114554225 | -0.148962613 | -0.134656726 | -0.249210951 | 0.014305886 | 0.751516083 | 0.048271135 | 0.019024157 | 0.511882041 | 0.795950405 |
| Prune2 | S | 1071 | -0.087606444 | -0.116997213 | -0.113561458 | -0.025955014 | 0.003435755 | 0.692300343 | 0.047624032 | 0.174783074 | 0.905669118 | 0.956595319 |
| Stard13 | S | 416 | -0.020783586 | 0.020259505 | -0.297762939 | -0.276979353 | -0.318022443 | 0.862495318 | 0.807277882 | 0.006129727 | 0.077229717 | 0.005579895 |
| Nupl2 | S | 104 | 0.003728808 | -0.148547369 | -0.141198089 | -0.144926898 | 0.00734928 | 0.914415201 | 0.130374071 | 0.034556243 | 0.039938215 | 0.926710914 |
| Nfix | S | 280 | 0.047373071 | -0.124931546 | -0.143065319 | -0.19043839 | -0.018133773 | 0.464416157 | 0.01789411 | 0.031770617 | 0.047201721 | 0.669978266 |
| Gmpr2 | S | 26 | -0.141398769 | 0.017792946 | -0.264341638 | -0.12294287 | -0.282134584 | 0.271193214 | 0.730292574 | 0.058631339 | 0.38113422 | 0.066421219 |
| Psmb7 | S | 277 | -0.262124657 | -0.050678874 | -0.11771505 | 0.144409607 | -0.067036176 | 0.073256554 | 0.688146015 | 0.575000107 | 0.508029044 | 0.753265429 |
| Nes | S | 886 | -0.078007085 | -0.04932782 | -0.097554445 | -0.01954736 | -0.048226626 | 0.14582031 | 0.307862244 | 0.027635181 | 0.632214868 | 0.257323927 |
| Map1b | S | 1649 | -0.070113061 | -0.121347108 | -0.032726497 | 0.037386564 | 0.088620612 | 0.127880262 | 0.023740454 | 0.429340267 | 0.060676435 | 0.000135073 |
| Ttc4 | S | 237 | -0.063436654 | -0.331693919 | -0.040946907 | 0.022489747 | 0.290747011 | 0.753295325 | 0.053774748 | 0.781490892 | 0.919974745 | 0.139484978 |
| Fosl1 | S | 261 | -0.189945266 | -0.07077024 | -0.221109494 | -0.031164229 | -0.150339254 | 0.27860603 | 0.657667327 | 0.249831419 | 0.843513245 | 0.345342526 |
| Cops3 | S | 423 | -0.057072006 | -0.032632726 | -0.097171795 | -0.040099788 | -0.064539069 | 0.27324508 | 0.088902315 | 0.010514563 | 0.419748709 | 0.057387672 |
| Ccnl2 | S | 391 | 0.003911132 | -0.098557437 | -0.176920624 | -0.180831756 | -0.078363187 | 0.930998783 | 0.156833385 | 0.025166049 | 0.031673128 | 0.275142147 |
| Slc20a2 | S | 259 | 0.018162181 | -0.152620471 | -0.071139552 | -0.089301732 | 0.081480919 | 0.809250464 | 0.004078091 | 0.15661674 | 0.312898666 | 0.118427765 |
| Upf2 | S | 8 | -0.008085682 | -0.060568174 | -0.238820857 | -0.230735175 | -0.178252683 | 0.897537583 | 0.356892593 | 0.025112149 | 0.035080888 | 0.071130636 |
| Fzr1 | S | 137 | 0.096152043 | -0.160417514 | -0.322543763 | -0.418695806 | -0.162126248 | 0.751734152 | 0.294331237 | 0.064637274 | 0.228051515 | 0.002134027 |
| Tbc1d1 | S | 256 | -0.07088277 | -0.071962058 | -0.153773209 | -0.082890439 | -0.081811151 | 0.374818816 | 0.156196928 | 0.105743284 | 0.383532613 | 0.300275363 |
| Ppp1r12a | T | 443 | -0.129639705 | 0.001428277 | -0.247338065 | -0.11769836 | -0.248766342 | 0.320667372 | 0.989800146 | 0.058924349 | 0.372488308 | 0.079909237 |
| Pank4 | T | 392 | -0.080427186 | -0.113784101 | -0.348225947 | -0.267798761 | -0.234441845 | 0.597845385 | 0.435999957 | 0.238770114 | 0.338613709 | 0.389252102 |
| Cd44 | S | 742 | 0.064183084 | -0.161708403 | -0.119451941 | -0.183635026 | 0.042256462 | 0.365212113 | 0.001051855 | 0.231619428 | 0.116178676 | 0.619208234 |
| Elob | S | 108 | -0.156965341 | -0.015738955 | -0.090361311 | 0.06660403 | -0.074622356 | 0.033216444 | 0.786809723 | 0.137423392 | 0.2052291 | 0.129815318 |
| Elob | S | 111 | -0.156965341 | -0.015738955 | -0.090361311 | 0.06660403 | -0.074622356 | 0.033216444 | 0.786809723 | 0.137423392 | 0.2052291 | 0.129815318 |
| Polk | S | 499 | 0.021523173 | -0.087408774 | -0.140331648 | -0.161854821 | -0.052922874 | 0.681194351 | 0.082889619 | 0.006531287 | 0.032821821 | 0.207885219 |
| Tnrc6a | S | 1524 | -0.049812729 | -0.195201657 | -0.448319538 | -0.398506809 | -0.253117881 | 0.894113766 | 0.6866511 | 0.184598308 | 0.332945267 | 0.605508324 |
| Ehbp1l1 | T | 448 | 0.022921595 | -0.123851032 | -0.061412527 | -0.084334122 | 0.062438504 | 0.522971313 | 0.001484785 | 0.043044996 | 0.061550079 | 0.02381632 |
| Map4 | T | 344 | -0.058129834 | -0.041806398 | -0.16821685 | -0.110087017 | -0.126410452 | 0.363812718 | 0.406173733 | 0.02709018 | 0.154413231 | 0.079521376 |
| Senp5 | S | 175 | 0.069138517 | -0.1219592 | -0.141245322 | -0.210383839 | -0.019286122 | 0.212846629 | 0.035564407 | 0.003048181 | 0.019357968 | 0.630174278 |
| Huwe1 | S | 3557 | -0.04628996 | -0.038536587 | -0.213692789 | -0.167402829 | -0.175156202 | 0.457237504 | 0.533865028 | 0.029101668 | 0.076527907 | 0.068771318 |
| Gfpt1 | S | 259 | -0.13111078 | 0.004591046 | -0.291824064 | -0.160713284 | -0.29641511 | 0.267639766 | 0.968055142 | 0.104592667 | 0.311632067 | 0.107386357 |
| Tacc1 | S | 376 | 0.009441636 | -0.200232155 | -0.209723351 | -0.219164987 | -0.009491196 | 0.926327314 | 0.082965113 | 0.254124463 | 0.245055985 | 0.953874196 |
| Atp11c | S | 1123 | -0.058261342 | -0.034284997 | -0.241134818 | -0.182873476 | -0.206849822 | 0.781449695 | 0.649744673 | 0.019684353 | 0.418988055 | 0.045200492 |
| Atp11c | S | 1126 | -0.058261342 | -0.034284997 | -0.241134818 | -0.182873476 | -0.206849822 | 0.781449695 | 0.649744673 | 0.019684353 | 0.418988055 | 0.045200492 |

| | | | | | | | | | | | | |
|---|---|---|---|---|---|---|---|---|---|---|---|---|
| Lig3 | S | 211 | 0.074981613 | -0.144953795 | -0.13751834 | -0.212499953 | 0.007435455 | 0.005521736 | 6.21424E-05 | 0.045000741 | 0.019477754 | 0.856845094 |
| Cobll1 | T | 382 | -0.074099961 | -0.030077185 | -0.159094781 | -0.08499482 | -0.129017597 | 0.219129349 | 0.550524033 | 0.030835645 | 0.216221895 | 0.080108277 |
| Yeats2 | S | 640 | 0.003479399 | -0.1655253 | -0.113083915 | -0.116563314 | 0.052441385 | 0.902286438 | 0.028121266 | 0.149775067 | 0.143904132 | 0.46592129 |
| Fbxo42 | S | 365 | 0.042974632 | -0.11402131 | -0.19702262 | -0.239997253 | -0.08300131 | 0.363270234 | 0.046542879 | 0.030258933 | 0.029939377 | 0.234430852 |
| Ube2e3 | S | 20 | -0.144946534 | -0.065493168 | -0.133088836 | 0.011857698 | -0.067595668 | 0.049139784 | 0.519769405 | 0.411183628 | 0.935814762 | 0.679050845 |
| Apbb1ip | S | 624 | 0.064243492 | -0.188873019 | -0.089820677 | -0.154064169 | 0.099052342 | 0.438260286 | 0.005185075 | 0.110570054 | 0.115276859 | 0.043787896 |
| Raf1 | S | 43 | -0.128965538 | 0.035821612 | -0.334891152 | -0.205925615 | -0.370712764 | 0.345380048 | 0.819115808 | 0.077604854 | 0.18467037 | 0.061572725 |
| Cd248 | S | 754 | -0.187283313 | 0.054626414 | -0.161183016 | 0.026100297 | -0.21580943 | 0.026129982 | 0.529637831 | 0.132756942 | 0.76435694 | 0.081047734 |
| Fzr1 | T | 121 | 0.061678788 | -0.047999283 | -0.317550187 | -0.379228974 | -0.269550903 | 0.121319273 | 0.442962058 | 0.002639882 | 0.011709827 | 0.012594407 |
| Dlg5 | S | 1797 | -0.081078256 | -0.122931152 | 0.016857114 | 0.09793537 | 0.139788266 | 0.049791716 | 0.005615191 | 0.674527632 | 0.070225159 | 0.025417123 |
| Gpsm1 | S | 491 | -0.051371417 | -0.046877764 | -0.187273513 | -0.135902096 | -0.140395749 | 0.476696874 | 0.566870051 | 0.020867554 | 0.056439429 | 0.10327174 |
| Ehmt2 | S | 172 | -0.045827471 | -0.18582907 | -0.048907038 | -0.003079568 | 0.136922032 | 0.496794172 | 0.018952691 | 0.546391961 | 0.966535471 | 0.121322993 |
| Lpin3 | S | 220 | -0.132169527 | -0.1261266 | -0.0060269 | 0.126142627 | 0.1200997 | 0.222931237 | 0.012498766 | 0.934277288 | 0.273225764 | 0.177648226 |
| Vcpip1 | S | 755 | -0.007777486 | -0.075046102 | -0.126980014 | -0.119202528 | -0.051933912 | 0.82430438 | 0.126457945 | 0.008719055 | 0.021413209 | 0.260832337 |
| Iws1 | S | 198 | -0.041090404 | -0.14301018 | -0.037296662 | 0.003793742 | 0.105713518 | 0.282465596 | 0.008049783 | 0.546615972 | 0.945169273 | 0.140405853 |
| Rybp | T | 215 | -0.118758674 | -0.245951174 | 0.036435543 | 0.155194217 | 0.282386717 | 0.318777719 | 0.02545514 | 0.716038375 | 0.229196279 | 0.035747009 |
| Tnik | S | 518 | -0.045776577 | -0.246648053 | -0.090813413 | -0.045036836 | 0.155834639 | 0.655815153 | 0.074087495 | 0.453248171 | 0.632636932 | 0.212561227 |
| Zc2hc1a | S | 222 | -0.00726374 | -0.080372195 | -0.111283429 | -0.104019688 | -0.030911234 | 0.891290352 | 0.002434224 | 0.122034261 | 0.18124942 | 0.568283553 |
| Nes | S | 1010 | -0.074573268 | -0.163239992 | -0.0336724 | 0.040900868 | 0.129567592 | 0.321067894 | 0.018567979 | 0.696174478 | 0.641466548 | 0.177253284 |
| Sh2b1 | S | 126 | 0.080009352 | -0.280126416 | -0.089680986 | -0.169690337 | 0.190445431 | 0.662764423 | 0.01044048 | 0.474819181 | 0.411211347 | 0.179898228 |
| Dock7 | S | 898 | 0.185907264 | -0.123149522 | -0.221936964 | -0.407844227 | -0.098787441 | 0.071317918 | 0.015233209 | 0.001698418 | 0.011725915 | 0.047411676 |
| Otud4 | S | 1016 | -0.051755776 | -0.041832158 | -0.046742572 | 0.005013204 | -0.004910414 | 0.005485608 | 0.338286044 | 0.008031483 | 0.663693002 | 0.89839166 |
| Pabpc1 | S | 315 | 0.033511659 | -0.233529051 | -0.138198626 | -0.171710284 | 0.095330426 | 0.565987056 | 0.126458039 | 0.047082779 | 0.043135606 | 0.439697324 |
| Rnps1 | S | 39 | 0.005954679 | -0.143903076 | -0.132927106 | -0.138881785 | 0.01097597 | 0.917561302 | 0.045589562 | 0.093010549 | 0.111833575 | 0.870933712 |
| Trim32 | S | 337 | 0.121801148 | -0.106497133 | -0.245395005 | -0.367196153 | -0.138897872 | 0.006153434 | 0.007215375 | 0.001420847 | 0.000960296 | 0.038464333 |
| Ankrd11 | S | 1832 | -0.119080904 | -0.152321078 | -0.048717504 | 0.0703634 | 0.103603574 | 0.145898192 | 0.262949572 | 0.209711536 | 0.328833124 | 0.409286542 |
| Zfp687 | S | 1191 | -0.033972454 | -0.073109225 | -0.185904362 | -0.151931908 | -0.112795137 | 0.767887751 | 0.263047709 | 0.032463577 | 0.230621138 | 0.09326274 |
| Sowahc | S | 82 | -0.151827393 | -0.000307387 | -0.142555896 | 0.009271497 | -0.142248509 | 0.074362913 | 0.997004937 | 0.097569856 | 0.840449329 | 0.064717607 |
| Map1b | S | 541 | -0.084118597 | -0.109241265 | -0.065567747 | 0.01855085 | 0.043673518 | 0.232047318 | 0.089460828 | 0.16973016 | 0.754310193 | 0.398718679 |
| Zc3hc1 | S | 410 | -0.036899055 | -0.116561035 | -0.09922882 | -0.062329765 | 0.017332216 | 0.309668325 | 0.172341162 | 0.050613958 | 0.188255172 | 0.805692238 |
| Rbmx2 | S | 266 | -0.026454663 | -0.123478114 | -0.066217742 | -0.039763079 | 0.057260372 | 0.677366274 | 0.05790318 | 0.019636018 | 0.541494082 | 0.250453164 |
| Larp1 | S | 302 | -0.029282323 | -0.10077757 | -0.074608832 | -0.045326508 | 0.026168738 | 0.330403444 | 0.025632353 | 0.097347714 | 0.242166554 | 0.51355343 |
| Slco3a1 | S | 15 | -0.145452791 | 0.034913503 | -0.205057424 | -0.059604634 | -0.239970927 | 0.109232103 | 0.82379244 | 0.08020315 | 0.550240247 | 0.214909672 |
| Ahctf1 | S | 2198 | 0.02897477 | -0.178594701 | -0.163635622 | -0.192610391 | 0.01495908 | 0.400444125 | 0.037735255 | 0.10890201 | 0.088567727 | 0.861283492 |
| Larp1 | T | 701 | -0.044009317 | -0.084751274 | -0.19744659 | -0.153437274 | -0.112695316 | 0.602670006 | 0.354070754 | 0.054383259 | 0.105149661 | 0.221276676 |
| Abcc1 | S | 1142 | 0.067268337 | -0.160174466 | -0.178417007 | -0.245685344 | -0.018242541 | 0.552740922 | 0.06538956 | 0.04537137 | 0.106111669 | 0.787185764 |
| Zc3hc1 | S | 353 | 0.111220511 | -0.144568639 | -0.234639394 | -0.345859905 | -0.090070755 | 0.21460888 | 0.03128116 | 0.029405265 | 0.014684331 | 0.235133665 |
| Bmp2k | S | 888 | -0.063635459 | -0.251101387 | -0.024036814 | 0.039598645 | 0.227064574 | 0.449942317 | 0.040062971 | 0.769430377 | 0.273865658 | 0.056690739 |
| Pum1 | S | 214 | -0.336538032 | -0.062415755 | -0.036343644 | 0.300194388 | 0.026072111 | 0.166363497 | 0.148310012 | 0.833028867 | 0.243724763 | 0.878580637 |
| Fhl3 | S | 232 | -0.121397895 | -0.043183395 | -0.145037637 | -0.023639743 | -0.101854242 | 0.06349034 | 0.638374583 | 0.197641481 | 0.800542525 | 0.392186874 |
| Plekhg3 | S | 606 | -0.114401103 | -0.147339615 | -0.002509546 | 0.111891557 | 0.144830069 | 0.20009554 | 0.019129265 | 0.962785592 | 0.223360398 | 0.057918908 |
| Fam98a | S | 285 | -0.016592863 | -0.194946791 | -0.145564243 | -0.128971379 | 0.049382549 | 0.861800644 | 0.073612981 | 0.265372062 | 0.263989163 | 0.635235659 |
| Fnip2 | S | 158 | 0.047845711 | -0.110318804 | -0.232444079 | -0.28028979 | -0.122125275 | 0.489535576 | 0.198186548 | 0.018309305 | 0.011058929 | 0.150809992 |
| Trps1 | S | 216 | -0.015273071 | -0.171091273 | -0.010478758 | 0.004794313 | 0.160612515 | 0.665998069 | 0.001251055 | 0.760032721 | 0.891815864 | 0.006520779 |
| Map3k2 | S | 239 | -0.108794729 | -0.052787247 | -0.145933211 | -0.037138482 | -0.093145963 | 0.233770588 | 0.522429727 | 0.072432766 | 0.664516454 | 0.294933275 |
| Larp7 | T | 335 | -0.088932899 | -0.066794807 | -0.031057925 | 0.057874974 | 0.035736882 | 0.033404002 | 0.028444844 | 0.423140611 | 0.200746295 | 0.360453785 |
| Mxra7 | S | 96 | -0.058381798 | -0.103323834 | -0.055991577 | 0.002390221 | 0.047332258 | 0.170986601 | 0.027191891 | 0.273692507 | 0.955141921 | 0.300721888 |
| Hook3 | S | 707 | 0.035857806 | -0.125537726 | -0.327566596 | -0.363424401 | -0.20202887 | 0.848558003 | 0.400616161 | 0.068014816 | 0.073539857 | 0.082554542 |
| 4833439L19Rik | S | 32 | -0.236257005 | -0.021198497 | -0.121074085 | 0.11518292 | -0.099875588 | 0.04099169 | 0.906276888 | 0.509042483 | 0.505654633 | 0.647393547 |
| Slc16a1 | T | 462 | -0.063653761 | -0.226595369 | -0.087689423 | -0.024035663 | 0.138905945 | 0.339157984 | 0.154752935 | 0.389994244 | 0.79682697 | 0.356584698 |
| Git1 | S | 507 | -0.23034046 | -0.084585757 | -0.121972869 | 0.108367591 | -0.037387112 | 0.140771586 | 0.532108856 | 0.485492939 | 0.537807883 | 0.821838584 |
| Eif3a | S | 1099 | 0.050175735 | -0.098276726 | -0.218504566 | -0.268680301 | -0.12022784 | 0.564030505 | 0.123526592 | 0.02161246 | 0.039896437 | 0.134610447 |
| Rabep2 | S | 180 | -0.158859807 | 0.001563191 | -0.100087826 | 0.058771981 | -0.101651017 | 0.022092206 | 0.986084879 | 0.164294897 | 0.344832488 | 0.303147302 |
| Rab13 | S | 178 | -0.0598312 | 0.005759623 | -0.312919249 | -0.253088048 | -0.318678872 | 0.661376825 | 0.970572821 | 0.028624364 | 0.084586134 | 0.087178662 |
| Ip6k1 | S | 125 | -0.264802801 | -0.088889586 | -0.016956207 | 0.247846594 | 0.07193338 | 0.04668691 | 0.557559105 | 0.749174359 | 0.069975044 | 0.623872293 |
| Rbm28 | S | 231 | 0.012661212 | -0.242540293 | -0.042088462 | -0.054749673 | 0.200451832 | 0.781163156 | 0.004810137 | 0.724580328 | 0.637372342 | 0.170869358 |
| Notch2 | S | 2082 | -0.021201963 | -0.228140402 | -0.117262056 | -0.096060094 | 0.110878346 | 0.818395438 | 0.074176194 | 0.325839336 | 0.355672338 | 0.354303469 |
| Espl1 | S | 239 | -0.165629256 | -0.012225933 | -0.140847955 | 0.024781302 | -0.128622022 | 0.26018201 | 0.793825867 | 0.040222153 | 0.844650998 | 0.070834559 |
| Map1a | S | 981 | -0.006777276 | -0.066543355 | -0.105139503 | -0.098362227 | -0.038596149 | 0.779957076 | 0.076487985 | 0.005258025 | 0.009225513 | 0.235392873 |
| Lrfn4 | S | 627 | 0.020284222 | -0.058101256 | -0.204168376 | -0.224452598 | -0.14606712 | 0.726422748 | 0.311127598 | 0.006286288 | 0.009465544 | 0.030494374 |
| Cdc42ep3 | S | 108 | -0.22227981 | 0.083639406 | -0.130598713 | 0.091681098 | -0.214238119 | 0.025928337 | 0.41764676 | 0.086469913 | 0.19185827 | 0.091119476 |
| Srp54a Srp54b | T | 21 | -0.181173842 | -0.285170814 | -0.049454198 | 0.131719644 | 0.235716616 | 0.48594754 | 0.155749451 | 0.795831865 | 0.54805459 | 0.078591446 |
| Vcpip1 | S | 1196 | -0.320480793 | 0.003163993 | -0.055525506 | 0.264955287 | -0.058689499 | 0.055980209 | 0.978737123 | 0.344961753 | 0.075637627 | 0.639869599 |
| Ftsj3 | S | 333 | 0.080267315 | -0.210038338 | -0.146587026 | -0.226854341 | 0.063451312 | 0.208634456 | 0.019986105 | 0.059014065 | 0.023633737 | 0.37022163 |
| Itprid2 | S | 664 | -0.149011189 | -0.043294151 | -0.084888306 | 0.064122883 | -0.041594154 | 0.083986157 | 0.332260123 | 0.200885237 | 0.413943686 | 0.501300377 |
| Tcea1 | S | 100 | -0.128622086 | 0.026299338 | -0.080686149 | 0.047935938 | -0.106985487 | 0.00081443 | 0.557556009 | 0.196262542 | 0.38937216 | 0.134185775 |
| Palld | S | 1129 | -0.056857329 | -0.012458128 | -0.139074239 | -0.082216911 | -0.126616112 | 0.164298635 | 0.767619057 | 0.008299649 | 0.077785182 | 0.033295894 |
| Frem1 | S | 3 | -0.40869059 | -0.261639098 | 0.084930567 | 0.493621157 | 0.346569665 | 0.236364667 | 0.265387252 | 0.731636045 | 0.187520675 | 0.218400694 |
| Nek4 | S | 629 | -0.200039282 | -0.035934305 | -0.176153445 | 0.023885838 | -0.14021914 | 0.116375669 | 0.730052731 | 0.347330367 | 0.892622817 | 0.444800835 |
| Slc16a1 | S | 210 | -0.052336443 | -0.228878171 | -0.061513393 | -0.00917695 | 0.167364779 | 0.603343299 | 0.054783163 | 0.468899886 | 0.931852849 | 0.148883228 |
| Stk17b | S | 362 | -0.001597876 | -0.121209151 | -0.227109673 | -0.225511797 | -0.105900521 | 0.983736366 | 0.156532912 | 0.105726912 | 0.109241423 | 0.368511831 |
| Fam110a | S | 7 | 0.042020657 | -0.072010754 | -0.263492156 | -0.305512813 | -0.191481402 | 0.592127647 | 0.34711183 | 0.013500429 | 0.009151917 | 0.037363739 |
| Ube4b | S | 88 | 0.047636599 | -0.127409778 | -0.12148529 | -0.169121889 | 0.005924489 | 0.314326703 | 0.016376097 | 0.017016543 | 0.01617431 | 0.806360484 |
| Nkap | S | 7 | -0.059382855 | -0.070821443 | -0.109289374 | -0.049906519 | -0.038467931 | 0.489033254 | 0.206685286 | 0.02049874 | 0.556929318 | 0.463321023 |
| Csnk1g3 | S | 346 | -0.104032964 | -0.05502047 | -0.084457749 | 0.019575214 | -0.029437279 | 0.011477493 | 0.499111427 | 0.310359967 | 0.790972369 | 0.764219452 |
| Eif4b | S | 16 | -0.104326231 | -0.146704374 | -0.178766454 | -0.074440223 | -0.032062079 | 0.457371747 | 0.149314439 | 0.453711445 | 0.760879613 | 0.887405575 |
| C2cd2l | S | 466 | 0.094024508 | -0.305268987 | -0.198170577 | -0.292195085 | 0.10709841 | 0.208218767 | 0.046154561 | 0.158617669 | 0.080722239 | 0.454505084 |
| Phf10 | S | 21 | 0.059111202 | -0.200603898 | -0.161091051 | -0.220202253 | 0.039512847 | 0.486537787 | 0.04118979 | 0.111629665 | 0.047930386 | 0.634082219 |
| Rhbdf1 | S | 11 | -0.192847516 | 0.055063418 | -0.340772851 | -0.147925335 | -0.39583627 | 0.342725239 | 0.710129555 | 0.108164082 | 0.506758016 | 0.086999961 |
| Rtn4 | S | 690 | 0.012708039 | -0.123623426 | -0.108092896 | -0.120800935 | 0.01553053 | 0.555501497 | 0.010140724 | 0.08182174 | 0.068514905 | 0.740706832 |
| Mcm3 | S | 522 | -0.143528605 | -0.074675332 | -0.190315001 | -0.046786395 | -0.115639669 | 0.174956882 | 0.42923314 | 0.369339005 | 0.813982636 | 0.571560878 |
| Mcm3 | S | 523 | -0.143528605 | -0.074675332 | -0.190315001 | -0.046786395 | -0.115639669 | 0.174956882 | 0.42923314 | 0.369339005 | 0.813982636 | 0.571560878 |
| Osbpl3 | S | 272 | -0.051116986 | -0.05799922 | -0.074861421 | -0.023744436 | -0.016862202 | 0.075164748 | 0.034856113 | 0.142664264 | 0.575121369 | 0.680799283 |
| Snip1 | S | 145 | -0.126301181 | -0.132706518 | -0.108613428 | 0.017687754 | 0.024093091 | 0.24828387 | 0.20456445 | 0.41669062 | 0.900614647 | 0.861944727 |
| Larp1 | T | 503 | 0.012329501 | -0.122627489 | -0.077462619 | -0.08979212 | 0.04516487 | 0.654607365 | 0.011701627 | 0.030422056 | 0.019106569 | 0.210595721 |
| Hnrnpc | S | 246 | -0.013931863 | -0.202525573 | -0.035187443 | -0.021255581 | 0.167338129 | 0.805210631 | 0.007127472 | 0.614861498 | 0.775746786 | 0.073300575 |
| Akap1 | S | 378 | -0.015760328 | -0.068160421 | -0.099746428 | -0.083986101 | -0.031586008 | 0.608719933 | 0.061550627 | 0.012443957 | 0.028037806 | 0.291949779 |
| Plekho2 | S | 279 | -0.066202214 | -0.026257178 | -0.099026907 | -0.032824693 | -0.072769729 | 0.046416756 | 0.332727704 | 0.039841494 | 0.359790133 | 0.105820085 |
| Itsn1 | S | 971 | -0.173495814 | -0.06288768 | -0.124179764 | 0.049316049 | -0.061292085 | 0.14684851 | 0.680510992 | 0.192445661 | 0.601753137 | 0.670394329 |
| Src | S | 74 | 0.064321873 | -0.086605326 | -0.181467884 | -0.245789757 | -0.094862559 | 0.519736664 | 0.016398898 | 0.031174101 | 0.079117022 | 0.118465272 |
| Anapc1 | S | 60 | -0.111690403 | -0.057434166 | -0.022759274 | 0.088931129 | 0.034674892 | 0.002956598 | 0.285505622 | 0.589090252 | 0.103018154 | 0.533969356 |
| 4933427D14Rik | S | 554 | -0.034408725 | -0.087916825 | -0.07937743 | -0.044968705 | 0.008539395 | 0.322117405 | 0.01469232 | 0.165318261 | 0.385267037 | 0.852493168 |
| 4933427D14Rik | S | 558 | -0.034408725 | -0.087916825 | -0.07937743 | -0.044968705 | 0.008539395 | 0.322117405 | 0.01469232 | 0.165318261 | 0.385267037 | 0.852493168 |
| Gapvd1 | S | 902 | 0.004753164 | -0.024950468 | -0.288272511 | -0.293025675 | -0.263322044 | 0.952051759 | 0.692354114 | 0.014741445 | 0.024167122 | 0.027110968 |

| | | | | | | | | | | | | |
|---|---|---|---|---|---|---|---|---|---|---|---|---|
| Rpl6 | S | 26 | 0.093193538 | -0.258520618 | -0.260789254 | -0.353982792 | -0.002268636 | 0.01766057 | 0.014271361 | 0.116854424 | 0.065433836 | 0.984614159 |
| Tbc1d12 | S | 691 | -0.019765559 | -0.120717499 | -0.089034799 | -0.06926924 | 0.0316827 | 0.807399998 | 0.019630023 | 0.126500987 | 0.445886917 | 0.5377788 |
| Cdc27 | S | 365 | 0.029998249 | -0.14348124 | -0.125312564 | -0.155310813 | 0.018168676 | 0.640747886 | 0.032919188 | 0.057455023 | 0.031474655 | 0.595452056 |
| Eme1 | S | 12 | -0.045175629 | -0.109408098 | -0.145561915 | -0.100386286 | -0.036153817 | 0.314982203 | 0.185986199 | 0.150412836 | 0.275490605 | 0.705750464 |
| Rad21 | S | 153 | -0.161965621 | -0.046482924 | -0.053929826 | 0.108035795 | -0.007446902 | 0.02934493 | 0.372587109 | 0.434132535 | 0.132213944 | 0.882672471 |
| Tom1l2 | S | 489 | -0.11431693 | -0.124643547 | -0.103585831 | 0.010731099 | 0.021057716 | 0.214693059 | 0.306973471 | 0.249408635 | 0.882883791 | 0.841882679 |
| Spats2 | S | 520 | 0.159445871 | -0.16703701 | -0.329243175 | -0.488689046 | -0.162206165 | 0.233981776 | 0.041067903 | 0.07109039 | 0.028726786 | 0.273323398 |
| Foxk1 | S | 225 | 0.054741678 | -0.142030474 | -0.044121968 | -0.098863647 | 0.097908506 | 0.367606735 | 0.000267036 | 0.078704102 | 0.164612811 | 0.002742879 |
| Btf3 | T | 180 | -0.127269281 | -0.167819202 | -0.114137181 | 0.0131321 | 0.053682022 | 0.367314533 | 0.248695294 | 0.359707498 | 0.907635185 | 0.644192486 |
| Crip2 | S | 104 | -0.195560838 | 0.096427765 | -0.242358752 | -0.046797914 | -0.338786517 | 0.1340815 | 0.361755423 | 0.044727818 | 0.687072618 | 0.025064035 |
| Mcm4 | S | 31 | -0.010535361 | -0.088610284 | -0.145072473 | -0.134537112 | -0.056462188 | 0.840780148 | 0.019639473 | 0.125746931 | 0.152790747 | 0.455626869 |
| Rrp1b | S | 705 | 0.109875292 | -0.13509187 | -0.179071683 | -0.288946975 | -0.043979813 | 0.01505898 | 0.003344804 | 0.00456097 | 0.001646775 | 0.25430374 |
| Arl6ip6 | S | 36 | 0.141796124 | -0.082895073 | -0.260496745 | -0.402292869 | -0.177601673 | 0.143826218 | 0.351755706 | 0.000479046 | 0.023690463 | 0.117254778 |
| Rtn4 | T | 488 | -0.026261765 | -0.071361447 | -0.06018302 | -0.033921255 | 0.011178427 | 0.460860644 | 0.014001947 | 0.019675377 | 0.332914783 | 0.517220265 |
| Bysl | S | 166 | -0.048176966 | -0.048812408 | -0.130960171 | -0.082783205 | -0.082147763 | 0.302120138 | 0.066538664 | 0.078766807 | 0.218329507 | 0.2002031 |
| Wdr44 | S | 344 | -0.12270857 | -0.121863302 | -0.103934307 | 0.018774262 | 0.017928995 | 0.227963062 | 0.215887895 | 0.368148576 | 0.879020175 | 0.882609755 |
| Uri1 | S | 439 | -0.137090582 | -0.036284757 | -0.128106264 | 0.008984318 | -0.091821507 | 0.092398073 | 0.72692544 | 0.140618447 | 0.843477403 | 0.348242038 |
| Anln | S | 334 | 0.058448252 | -0.156054631 | -0.107383236 | -0.165831488 | 0.048671395 | 0.053193696 | 0.000564869 | 0.08583697 | 0.043741777 | 0.336858062 |
| Vim | S | 9 | -0.069227717 | -0.121112865 | -0.049835736 | 0.019391981 | 0.071277128 | 0.284452598 | 0.06399736 | 0.14965726 | 0.733646608 | 0.202271931 |
| Sf3a1 | S | 449 | -0.051443511 | -0.107655294 | -0.12374224 | -0.072298729 | -0.016086946 | 0.403735972 | 0.216021098 | 0.081623448 | 0.274491023 | 0.838110922 |
| Dus1l | T | 462 | -0.185883965 | -0.14793386 | -0.14075956 | 0.045124405 | 0.0071743 | 0.352763988 | 0.291209142 | 0.501552286 | 0.854791393 | 0.973564799 |
| Eps15 | S | 561 | 0.058878343 | -0.118539186 | -0.18510342 | -0.243981763 | -0.066564234 | 0.391930921 | 0.0332924 | 0.037860187 | 0.026336244 | 0.32405767 |
| Pom121 | S | 244 | 0.107671907 | -0.177747156 | -0.153957734 | -0.261629641 | 0.023789421 | 0.103944589 | 0.037877852 | 0.00463504 | 0.013261275 | 0.678901428 |
| Ric8a | S | 435 | -0.00554494 | -0.072925111 | -0.200046663 | -0.194501723 | -0.127121552 | 0.94466044 | 0.309803197 | 0.023056619 | 0.061684086 | 0.115945417 |
| Nfx1 | S | 51 | 0.072283611 | -0.20153101 | -0.101036171 | -0.173319783 | 0.100494839 | 0.035021512 | 0.001131764 | 0.083106343 | 0.032927139 | 0.091305633 |
| Mcoln1 | T | 34 | -0.09096285 | -0.070401984 | -0.217617808 | -0.126654958 | -0.147215824 | 0.434067316 | 0.469856367 | 0.125897275 | 0.347713065 | 0.247938249 |
| Pagr1a | S | 236 | -0.065243884 | -0.10773037 | 0.000411179 | 0.065655063 | 0.108141549 | 0.048768664 | 0.006529277 | 0.979656777 | 0.066744822 | 0.01952986 |
| Wnk1 | T | 71 | -0.00191893 | -0.086923577 | -0.114193058 | -0.112274128 | -0.027269481 | 0.947130397 | 0.143133116 | 0.006743024 | 0.003830028 | 0.568113751 |
| Twf1 | T | 349 | -0.038375527 | -0.080570787 | -0.121305202 | -0.082929675 | -0.040734416 | 0.561572665 | 0.279275345 | 0.016206861 | 0.238776715 | 0.539345528 |
| Ndc1 | S | 343 | -0.082209637 | -0.628181499 | -0.106778962 | -0.024569325 | 0.521402537 | 0.886893563 | 0.254175702 | 0.811225211 | 0.970926009 | 0.400478953 |
| Pbk | S | 18 | -0.097819885 | 0.03071236 | -0.284504114 | -0.186684228 | -0.315216474 | 0.338336647 | 0.761158941 | 0.044291184 | 0.1445016 | 0.040377156 |
| Gps1 | S | 448 | -0.008981708 | -0.075375824 | -0.156946179 | -0.147964471 | -0.081570354 | 0.853483582 | 0.157876928 | 0.024089864 | 0.028001414 | 0.122852764 |
| Zrsr2 | S | 353 | -0.010746881 | -0.12230417 | -0.082029724 | -0.071282843 | 0.040274446 | 0.773997953 | 0.03049178 | 0.068133882 | 0.077238199 | 0.351350549 |
| Dync1li2 | S | 389 | -0.034111796 | -0.126141612 | -0.213935549 | -0.179823753 | -0.087793937 | 0.565698311 | 0.140719418 | 0.258474886 | 0.321666369 | 0.602081348 |
| Jade1 | S | 90 | -0.174550793 | -0.039500358 | -0.057650992 | 0.116899801 | -0.018150634 | 0.020008995 | 0.582311479 | 0.492377916 | 0.180833149 | 0.816634026 |
| Ybx3 | S | 33 | -0.015491518 | -0.103821566 | -0.068196492 | -0.052704974 | 0.035625073 | 0.501729989 | 0.003153182 | 0.27286165 | 0.369395253 | 0.521596301 |
| Rab11fip5 | S | 539 | 0.016875414 | -0.05999605 | -0.280847561 | -0.297722974 | -0.220851511 | 0.923213496 | 0.335007034 | 0.039117457 | 0.181380691 | 0.069309485 |
| Wdr3 | S | 333 | 0.073816461 | -0.12383552 | -0.246982945 | -0.320799406 | -0.123147425 | 0.196280812 | 0.035603638 | 0.044344557 | 0.030391669 | 0.215126664 |
| Map7d1 | T | 101 | 0.007826561 | -0.512680665 | -0.081233938 | -0.089060499 | 0.431446727 | 0.986225457 | 0.114094714 | 0.81719693 | 0.845149501 | 0.210722473 |
| Cdk5rap2 | S | 1435 | -0.206017117 | -0.136328799 | -0.070150191 | 0.135866927 | 0.066178608 | 0.175388455 | 0.357051422 | 0.568706035 | 0.399739326 | 0.684736299 |
| Ubn2 | S | 935 | 0.082949758 | -0.211401407 | -0.304312135 | -0.387261893 | -0.092910728 | 0.445322954 | 0.051464698 | 0.262196239 | 0.183474982 | 0.690598937 |
| Ubn2 | S | 936 | 0.082949758 | -0.211401407 | -0.304312135 | -0.387261893 | -0.092910728 | 0.445322954 | 0.051464698 | 0.262196239 | 0.183474982 | 0.690598937 |
| Mapk6 | S | 704 | 0.076924758 | -0.126232824 | -0.156693123 | -0.233617881 | -0.030460299 | 0.109965174 | 0.181080594 | 0.000343994 | 0.010080479 | 0.68561514 |
| Pum1 | S | 710 | 0.026905864 | -0.01670042 | -0.215796204 | -0.242702068 | -0.199095784 | 0.459421591 | 0.668645236 | 0.000882626 | 0.002693106 | 0.002353074 |
| Pwp2 | T | 891 | 0.02165232 | -0.14988666 | -0.058312554 | -0.079964875 | 0.091574105 | 0.672671846 | 0.004575043 | 0.104339715 | 0.183212591 | 0.04457465 |
| Pxn | S | 230 | 0.013282462 | 0.031331145 | -0.434472569 | -0.447755031 | -0.465803714 | 0.864979298 | 0.804328246 | 0.021099695 | 0.020490253 | 0.029035591 |
| Phactr4 | T | 146 | -0.052398349 | -0.093885065 | -0.197554052 | -0.145155703 | -0.103668987 | 0.531292734 | 0.225399995 | 0.149981053 | 0.260268765 | 0.384196712 |
| Ncoa6 | S | 1337 | -0.081427921 | -0.095323475 | -0.122501933 | -0.041074012 | -0.027178458 | 0.335085572 | 0.215738249 | 0.143758963 | 0.494965151 | 0.52814634 |
| Hjurp A730008H23Ril | S | 613 | 0.22947651 | -0.094005501 | -0.290412104 | -0.519888614 | -0.196406603 | 0.214250227 | 0.076571527 | 0.003840963 | 0.046023685 | 0.026993326 |
| Tax1bp1 | S | 619 | 0.051908316 | -0.126763316 | -0.149820002 | -0.201728318 | -0.023056686 | 0.144783534 | 0.003654931 | 0.064823227 | 0.038010083 | 0.677901749 |
| Hmga1 Hmga1b | T | 39 | -0.081420006 | -0.108374553 | -0.021641885 | 0.059778121 | 0.086732669 | 0.039737462 | 0.053121973 | 0.688480668 | 0.321909696 | 0.19901641 |
| Lrrc42 | S | 399 | -0.020934397 | -0.052406564 | -0.172265073 | -0.151330676 | -0.119858508 | 0.43627056 | 0.281143495 | 0.030148907 | 0.050442687 | 0.090359696 |
| Zfp638 | S | 1400 | -0.033714501 | -0.098407822 | -0.055759383 | -0.022044882 | 0.042648439 | 0.60800249 | 0.001818451 | 0.331218669 | 0.770544488 | 0.436027071 |
| Sipa1l2 | S | 1214 | -0.188946395 | -0.119810965 | 0.035930332 | 0.224876727 | 0.155741297 | 0.006456724 | 0.398422601 | 0.56476349 | 0.018596859 | 0.297780725 |
| Hnrnpf | T | 215 | -0.02930292 | -0.084349511 | -0.163060171 | -0.13375725 | -0.07871066 | 0.602260533 | 0.168679802 | 0.071938147 | 0.117213132 | 0.27305123 |
| Mak16 | S | 197 | 0.011170088 | -0.204605762 | -0.035436878 | -0.046606965 | 0.169168884 | 0.819975068 | 0.003569423 | 0.587427113 | 0.428092223 | 0.058350733 |
| Mak16 | S | 199 | 0.011170088 | -0.204605762 | -0.035436878 | -0.046606965 | 0.169168884 | 0.819975068 | 0.003569423 | 0.587427113 | 0.428092223 | 0.058350733 |
| Mrtfb | S | 535 | -0.006013927 | -0.015740805 | -0.153267318 | -0.147253391 | -0.137526513 | 0.912174091 | 0.750720101 | 0.000546544 | 0.071943227 | 0.062536063 |
| Mrtfb | S | 537 | -0.006013927 | -0.015740805 | -0.153267318 | -0.147253391 | -0.137526513 | 0.912174091 | 0.750720101 | 0.000546544 | 0.071943227 | 0.062536063 |
| Son | S | 256 | -0.053606019 | -0.097054938 | -0.101603723 | -0.047997704 | -0.004548785 | 0.40679998 | 0.106594115 | 0.107020778 | 0.500642449 | 0.937348387 |
| Rexo1 | S | 505 | -0.067158595 | -0.12102279 | -0.143643997 | -0.076485402 | -0.022621208 | 0.361386587 | 0.125418027 | 0.329585068 | 0.558970954 | 0.856851977 |
| Chd9 | S | 653 | 0.072377265 | -0.103136231 | -0.145543408 | -0.217920673 | -0.042407177 | 0.200757244 | 0.00163441 | 0.038309145 | 0.016108441 | 0.369197595 |
| Pakap | S | 482 | -0.084002039 | -0.026507893 | -0.174383264 | -0.090381225 | -0.147875371 | 0.349730045 | 0.729911435 | 0.032578779 | 0.326478146 | 0.115298032 |
| Plekhf2 | S | 239 | -0.056492484 | -0.109919374 | -0.127623342 | -0.071130858 | -0.017703968 | 0.637455315 | 0.093179003 | 0.152298552 | 0.577425326 | 0.815404519 |
| Ptk2 | S | 574 | -0.100727282 | -0.011065463 | -0.213984651 | -0.113257369 | -0.202919188 | 0.087366989 | 0.927143263 | 0.151547106 | 0.375151212 | 0.229074966 |
| Rbm27 | S | 187 | -0.044133812 | -0.161295791 | -0.02664697 | 0.017486842 | 0.134648821 | 0.428661219 | 0.007884139 | 0.775045977 | 0.848583864 | 0.214557888 |
| L3mbtl2 | S | 687 | -0.038129244 | -0.080602267 | -0.272302139 | -0.234172895 | -0.191699872 | 0.677377361 | 0.255808774 | 0.160573516 | 0.201603951 | 0.262659056 |
| Supt20 | S | 381 | -0.158838136 | -0.156019711 | -0.055207223 | 0.103630912 | 0.100812488 | 0.327362777 | 0.168309456 | 0.449704291 | 0.491886192 | 0.318637789 |
| Rrp9 | S | 470 | -0.010663745 | -0.071116175 | -0.086619178 | -0.075955433 | -0.015503004 | 0.593191419 | 0.038045725 | 0.015365508 | 0.030998854 | 0.588792385 |
| Nmt2 | S | 70 | -0.062883401 | -0.026199276 | -0.089145527 | -0.026262126 | -0.062946251 | 0.053402495 | 0.395792011 | 0.0251448 | 0.293646766 | 0.060321144 |
| Nktr | S | 390 | -0.060003547 | -0.175475326 | -0.035891373 | 0.024112174 | 0.139583953 | 0.373531617 | 0.048967354 | 0.377224867 | 0.692722193 | 0.095512037 |
| Arhgap18 | S | 65 | -0.252886689 | -0.028363954 | -0.084739861 | 0.168146828 | -0.056375907 | 0.055234161 | 0.79911177 | 0.561319664 | 0.294079311 | 0.707216668 |
| Erc1 | S | 37 | -0.082693002 | -0.026620104 | -0.167692967 | -0.084999965 | -0.141072863 | 0.241752576 | 0.787070455 | 0.040158417 | 0.018345614 | 0.179263607 |
| Nav1 | S | 455 | -0.122743823 | -0.038083525 | -0.243239246 | -0.120495423 | -0.205155721 | 0.270083722 | 0.720659544 | 0.182206934 | 0.422372306 | 0.227900339 |
| Uckl1 | T | 53 | 0.002603024 | -0.016930544 | -0.155934675 | -0.158537699 | -0.139004132 | 0.959506737 | 0.665376983 | 0.000538877 | 0.059120564 | 0.03409913 |
| Prpf4b | S | 37 | 0.018047189 | -0.10186594 | -0.197725363 | -0.215772552 | -0.095859423 | 0.6245008 | 0.064813321 | 0.072610739 | 0.048945001 | 0.248594519 |
| Eif3a | S | 1156 | -0.016039047 | -0.269364428 | -0.055391257 | -0.03935221 | 0.21397317 | 0.740794898 | 0.057684088 | 0.452559503 | 0.532262906 | 0.104216237 |
| Sympk | S | 1276 | -0.089760645 | -0.105985411 | -0.020801142 | 0.068959503 | 0.085184269 | 0.080765731 | 0.035123461 | 0.709502235 | 0.300581436 | 0.21204559 |
| Caml | S | 55 | -0.100546888 | -0.048078741 | -0.066339092 | 0.034207796 | -0.018260351 | 0.038102338 | 0.366957835 | 0.137747336 | 0.335965677 | 0.693149565 |
| Ttyh3 | S | 437 | -0.12993798 | -0.021068092 | -0.139356364 | -0.009418384 | -0.118288272 | 0.268050842 | 0.792072469 | 0.039447272 | 0.92491439 | 0.173359963 |
| Rhbdf2 | S | 60 | -0.093415682 | 0.013272816 | -0.165680282 | -0.0722646 | -0.178953098 | 0.013183049 | 0.71281961 | 0.126701892 | 0.408021946 | 0.112015537 |
| Eml3 | S | 890 | -0.238113953 | 0.106538129 | -0.211910741 | 0.026203212 | -0.31844887 | 0.068627826 | 0.308116744 | 0.093166391 | 0.826428422 | 0.044456562 |
| Eml3 | S | 894 | -0.238113953 | 0.106538129 | -0.211910741 | 0.026203212 | -0.31844887 | 0.068627826 | 0.308116744 | 0.093166391 | 0.826428422 | 0.044456562 |
| Micall1 | T | 311 | -0.003433243 | -0.072040247 | -0.094294456 | -0.090861214 | -0.022254209 | 0.95739631 | 0.209957311 | 0.001023249 | 0.246547894 | 0.639956232 |
| Parp8 | S | 344 | -0.071897084 | -0.056447681 | -0.075145575 | -0.003248491 | -0.018697895 | 0.203374945 | 0.218307596 | 0.026831627 | 0.942488322 | 0.612911676 |
| Tgfb1i1 | S | 194 | -0.145930103 | -0.011223272 | -0.070114439 | 0.075815664 | -0.058891166 | 0.010478771 | 0.84053654 | 0.294863269 | 0.263872486 | 0.416860289 |
| Zdhhc5 | S | 409 | 0.149621473 | -0.317773288 | -0.118317968 | -0.267939441 | 0.19945532 | 0.08536389 | 0.005862474 | 0.14194985 | 0.004941717 | 0.038145188 |
| Vim | S | 22 | -0.070963858 | -0.057416246 | -0.095045247 | -0.02408139 | -0.037629001 | 0.328218694 | 0.189279717 | 0.03634452 | 0.696599799 | 0.203280172 |
| Top2b | T | 1305 | -0.034489021 | -0.085163161 | -0.068316397 | -0.033827376 | 0.016846764 | 0.606936165 | 0.060606386 | 0.015967029 | 0.611284936 | 0.614336742 |
| Ube4b | S | 1136 | -0.089211654 | -0.022560612 | -0.07793347 | 0.011278184 | -0.055372858 | 0.122643752 | 0.569696803 | 0.008966418 | 0.794782011 | 0.207150411 |
| Strn4 | S | 276 | -0.138417089 | -0.043479939 | -0.05357733 | 0.084839759 | -0.010097391 | 0.13979015 | 0.300546339 | 0.049851123 | 0.296537054 | 0.778812995 |
| Rgl2 | T | 13 | -0.158971864 | -0.055225162 | -0.203790669 | -0.044818805 | -0.148565508 | 0.202907357 | 0.647702415 | 0.3044448 | 0.811739756 | 0.459994358 |
| Ska3 | S | 154 | 0.004318466 | -0.109140587 | -0.127157242 | -0.131475708 | -0.018016656 | 0.8753169 | 0.023164429 | 0.1071665 | 0.111040871 | 0.769196063 |
| Hnrnpa3 | S | 371 | -0.032801442 | -0.005716713 | -0.136238404 | -0.103436962 | -0.13052169 | 0.586477865 | 0.827184044 | 0.000813479 | 0.164359604 | 0.004069026 |

| | | | | | | | | | | | | | |
|---|---|---|---|---|---|---|---|---|---|---|---|---|---|
| Uhrf1 | S | | 91 | -0.081272579 | -0.098696545 | -0.058079985 | 0.023192594 | 0.04061656 | 0.173188793 | 0.069115609 | 0.316554566 | 0.68005016 | 0.429875067 |
| Champ1 | S | | 405 | 0.118384127 | -0.176953688 | -0.13839877 | -0.256782897 | 0.038554918 | 0.013512639 | 0.002400136 | 0.006950338 | 0.002254234 | 0.229627581 |
| Ncbp1 | S | | 680 | -0.223692356 | -0.079231766 | -0.114597236 | 0.10909512 | -0.03536547 | 0.093071886 | 0.693282426 | 0.58122673 | 0.595123288 | 0.8864925 |
| Ncbp1 | S | | 681 | -0.223692356 | -0.079231766 | -0.114597236 | 0.10909512 | -0.03536547 | 0.093071886 | 0.693282426 | 0.58122673 | 0.595123288 | 0.8864925 |
| Dlg1 | S | | 571 | -0.050030459 | -0.067452753 | -0.068060874 | -0.018030415 | -0.000608121 | 0.156800378 | 0.11227242 | 0.044884437 | 0.549394616 | 0.985785372 |
| Osbpl5 | S | | 749 | -0.126064858 | -0.049979948 | -0.081023776 | 0.045041082 | -0.031043828 | 0.050732255 | 0.51688702 | 0.214989016 | 0.448496985 | 0.68675791 |
| Osbpl5 | T | | 742 | -0.126064858 | -0.049979948 | -0.081023776 | 0.045041082 | -0.031043828 | 0.050732255 | 0.51688702 | 0.214989016 | 0.448496985 | 0.68675791 |
| Shcbp1 | S | | 630 | -0.154216578 | -0.024022659 | -0.164828372 | -0.010611795 | -0.140805714 | 0.061508002 | 0.7631628 | 0.359829178 | 0.948145346 | 0.42940325 |
| Srek1 | T | | 441 | -0.072448995 | -0.170028226 | -0.002744911 | 0.069704084 | 0.167283315 | 0.386682739 | 0.010949576 | 0.975736589 | 0.524429831 | 0.14673716 |
| Cmtr1 | S | | 28 | -0.057503239 | -0.050786307 | -0.127712387 | -0.070209147 | -0.076926079 | 0.1474022 | 0.241309728 | 0.097734348 | 0.285177642 | 0.254325526 |
| Cmtr1 | S | | 30 | -0.057503239 | -0.050786307 | -0.127712387 | -0.070209147 | -0.076926079 | 0.1474022 | 0.241309728 | 0.097734348 | 0.285177642 | 0.254325526 |
| Hk1 | T | | 608 | -0.151190381 | 0.001325401 | -0.113804334 | 0.037386047 | -0.115129735 | 0.204436654 | 0.972356115 | 0.021378451 | 0.704173682 | 0.031646851 |
| Cdc23 | S | | 593 | 0.104420786 | -0.157029428 | -0.247351921 | -0.351772708 | -0.090322493 | 0.549644183 | 0.143484126 | 0.056492061 | 0.119192933 | 0.433152771 |
| Bora | S | | 336 | -0.030875522 | -0.053385982 | -0.148423982 | -0.11754846 | -0.095038 | 0.340335679 | 0.09804361 | 0.081396525 | 0.141570449 | 0.204410762 |
| Fhod1 | S | | 370 | -0.163322932 | -0.058544993 | -0.000136131 | 0.163186801 | 0.058408862 | 0.010885314 | 0.250220221 | 0.998501309 | 0.10362097 | 0.475273321 |
| Sh3pxd2a | S | | 794 | -0.117175444 | -0.004715005 | -0.203571164 | -0.08639572 | -0.198856159 | 0.280091494 | 0.961211417 | 0.06576827 | 0.373554544 | 0.066433733 |
| Aspscr1 | S | | 535 | 0.008137231 | -0.096412278 | -0.215672737 | -0.223809968 | -0.119260459 | 0.859219075 | 0.382474162 | 0.028776123 | 0.033533814 | 0.314039676 |
| Jcad | S | | 1234 | -0.115168412 | -0.007293029 | -0.155940834 | -0.040772422 | -0.148647805 | 0.03501584 | 0.922552692 | 0.203838356 | 0.684508625 | 0.236365391 |
| Tpmt | S | | 34 | -0.20897254 | -0.083653145 | -0.117911792 | 0.091060748 | -0.034258647 | 0.133863004 | 0.648492167 | 0.456156736 | 0.438145633 | 0.841144909 |
| Ercc6l | S | | 1001 | -0.061503596 | -0.076901406 | -0.138427801 | -0.076924205 | -0.061526395 | 0.274573735 | 0.282459998 | 0.114499591 | 0.332181895 | 0.468711612 |
| Fus | S | | 270 | 0.039257174 | -0.323313142 | -0.1445061 | -0.183763274 | 0.178807042 | 0.774173259 | 0.10057258 | 0.368626958 | 0.258383868 | 0.338301234 |
| Map3k4 | S | | 1241 | -0.073910373 | -0.113667687 | -0.074851911 | -0.000941538 | 0.038815775 | 0.133844511 | 0.226661483 | 0.204572668 | 0.980344451 | 0.638462129 |
| Cdk11b | S | | 115 | -0.108316632 | -0.07673882 | -0.062846983 | 0.045469649 | 0.013891837 | 0.080081085 | 0.183825574 | 0.348527162 | 0.39696266 | 0.779934666 |
| Casc3 | T | | 13 | 0.073150957 | -0.233788785 | -0.147697399 | -0.220848357 | 0.086091386 | 0.371190003 | 0.069763446 | 0.073411538 | 0.023520552 | 0.380396642 |
| Ankrd11 | S | | 1837 | -0.233598698 | -0.165828012 | -0.013073521 | 0.220525176 | 0.152754491 | 0.158833298 | 0.308679489 | 0.865301727 | 0.177024607 | 0.348136729 |
| Ankrd11 | S | | 1844 | -0.233598698 | -0.165828012 | -0.013073521 | 0.220525176 | 0.152754491 | 0.158833298 | 0.308679489 | 0.865301727 | 0.177024607 | 0.348136729 |
| H1f2 H1f3 | T | 146;147 | | 0.139004451 | -0.086057357 | -0.24513791 | -0.384142361 | -0.159080553 | 0.057845222 | 0.142315669 | 0.000527219 | 0.005624929 | 0.029629257 |
| Map1a | S | | 1747 | -0.02883034 | -0.09389226 | -0.034762312 | -0.005931973 | 0.059129948 | 0.38984854 | 0.005904829 | 0.105106474 | 0.831520804 | 0.053526206 |
| Map1a | S | | 2281 | 0.029272189 | -0.090287511 | -0.131785686 | -0.161057874 | -0.041498175 | 0.596514401 | 0.091571544 | 0.008053765 | 0.058250553 | 0.320738639 |
| Rtn4 | S | | 838 | 0.01611466 | -0.138046213 | -0.057549946 | -0.073664606 | 0.080496267 | 0.519647414 | 0.004286405 | 0.088504717 | 0.040212178 | 0.046914363 |
| Golga3 | S | | 60 | -0.06037069 | -0.161807225 | 0.000320966 | 0.060691656 | 0.162128191 | 0.17478854 | 0.016468043 | 0.99329894 | 0.220258606 | 0.021556314 |
| Prrc2a | T | | 807 | 0.105053537 | -0.165959273 | -0.109079995 | -0.214133532 | 0.056879278 | 0.165332721 | 0.002252733 | 0.052409217 | 0.032645717 | 0.234149538 |
| Mxra7 | S | | 165 | -0.059098941 | -0.059379252 | -0.085692289 | -0.026593347 | -0.026313037 | 0.12437701 | 0.149420652 | 0.094057772 | 0.483904038 | 0.513214195 |
| Ubap2l | S | | 629 | 0.068102693 | -0.096528093 | -0.196689928 | -0.264792622 | -0.100161835 | 0.020183246 | 0.000674028 | 0.03473423 | 0.0172391 | 0.143392069 |
| Nol8 | S | | 783 | -0.013473221 | -0.100885892 | -0.072330338 | -0.058857117 | 0.028555554 | 0.80562064 | 0.004236349 | 0.202120989 | 0.385600063 | 0.549271702 |
| Map1a | T | | 1712 | 0.018449444 | -0.080642729 | -0.112104496 | -0.13055394 | -0.031461767 | 0.618015204 | 0.003520521 | 0.063641879 | 0.046062049 | 0.458408918 |
| Casp8ap2 | S | | 923 | -0.068240503 | -0.08949151 | -0.115761764 | -0.047521261 | -0.026270254 | 0.075578194 | 0.366247116 | 0.223641982 | 0.563459476 | 0.810984224 |
| Specc1 | T | | 148 | 0.061363086 | -0.159834495 | -0.304520795 | -0.365883882 | -0.1446863 | 0.464617109 | 0.083547622 | 0.180116661 | 0.139747756 | 0.455637646 |
| Srek1 | S | | 420 | -0.003394654 | -0.160462233 | -0.057462065 | -0.054067411 | 0.103000169 | 0.963497215 | 0.076176467 | 0.017788763 | 0.490363112 | 0.180851521 |
| Myo9b | S | | 1647 | -0.042907049 | -0.001297105 | -0.258780503 | -0.215873455 | -0.257483398 | 0.629245535 | 0.988768681 | 0.022596677 | 0.03288466 | 0.022933586 |
| Ercc6l2 | S | | 887 | -0.326362908 | 0.02423069 | -0.1016307 | 0.224732208 | -0.12586139 | 0.096739852 | 0.742882816 | 0.306714206 | 0.193015577 | 0.256815772 |
| Ercc6l2 | S | | 890 | -0.326362908 | 0.02423069 | -0.1016307 | 0.224732208 | -0.12586139 | 0.096739852 | 0.742882816 | 0.306714206 | 0.193015577 | 0.256815772 |
| Ercc6l2 | T | | 885 | -0.326362908 | 0.02423069 | -0.1016307 | 0.224732208 | -0.12586139 | 0.096739852 | 0.742882816 | 0.306714206 | 0.193015577 | 0.256815772 |
| Tnrc6a | S | | 1520 | -0.025479152 | -0.146654017 | -0.130417072 | -0.104937921 | 0.016236944 | 0.64441725 | 0.156840862 | 0.138642386 | 0.196244664 | 0.865358194 |
| Bclaf1 | S | | 275 | -0.143488684 | -0.158421641 | 0.057805496 | 0.201294179 | 0.216227137 | 0.107202045 | 0.02795979 | 0.555277139 | 0.11311118 | 0.092277941 |
| Denn2b | S | | 462 | -0.005355819 | -0.191294103 | -0.079457151 | -0.074101331 | 0.111836952 | 0.964682322 | 0.008177294 | 0.722771427 | 0.755152613 | 0.620375252 |
| Apc | S | | 2449 | -0.085844821 | -0.099359413 | 0.040957085 | 0.126801906 | 0.140316499 | 0.05167987 | 0.002136893 | 0.566520735 | 0.15301016 | 0.138367794 |
| Ep400 | T | | 314 | -0.051844985 | -0.013108905 | -0.295705197 | -0.243860212 | -0.282596292 | 0.581689137 | 0.835114754 | 0.058195205 | 0.105404994 | 0.076099034 |
| Phf11d | S | | 169 | -0.154815885 | -0.138061798 | -0.083863223 | 0.070952662 | 0.054198575 | 0.233288176 | 0.280735783 | 0.504204282 | 0.124326352 | 0.202109259 |
| Grk6 | S | | 566 | 0.043783654 | -0.119124051 | -0.150659553 | -0.194443207 | -0.031535501 | 0.652548547 | 0.008126357 | 0.175176311 | 0.155841923 | 0.720969826 |
| Grk6 | S | | 568 | 0.043783654 | -0.119124051 | -0.150659553 | -0.194443207 | -0.031535501 | 0.652548547 | 0.008126357 | 0.175176311 | 0.155841923 | 0.720969826 |
| Spen | S | | 1283 | 0.347497391 | -0.172745817 | -0.149340024 | -0.496837414 | 0.023405793 | 0.255909444 | 0.005029339 | 0.006695809 | 0.153346489 | 0.503641648 |
| Sh3pxd2a | S | | 266 | -0.063569492 | -0.069194333 | -0.190637517 | -0.127068025 | -0.121443184 | 0.497491851 | 0.505622701 | 0.073970988 | 0.069148614 | 0.154955531 |
| Champ1 | S | | 394 | 0.062274748 | -0.073480725 | -0.073129442 | -0.13540419 | 0.000351283 | 0.10065832 | 0.001677017 | 0.017214504 | 0.973736904 |
| Edc4 | S | | 680 | 0.015669192 | -0.120711554 | -0.06623228 | -0.081901472 | 0.054479274 | 0.702097592 | 0.008483731 | 0.048627117 | 0.102202889 | 0.1088968 |
| Atxn2l | S | | 337 | -0.028032261 | -0.046821104 | -0.128378848 | -0.100346587 | -0.081557744 | 0.430932453 | 0.228301241 | 0.018599671 | 0.046649563 | 0.082200711 |
| Rai14 | T | | 330 | -0.014522715 | -0.149801804 | -0.433550072 | -0.419027357 | -0.283748268 | 0.940372271 | 0.468951567 | 0.25956105 | 0.282382114 | 0.443692356 |
| Dclre1c | T | | 380 | -0.09015247 | 0.011017082 | -0.270707066 | -0.180554596 | -0.281724148 | 0.357730227 | 0.878099398 | 0.063281249 | 0.176518813 | 0.052039713 |
| Pgm3 | T | | 62 | -0.267833375 | 0.046422195 | -0.209617715 | 0.058215661 | -0.256039909 | 0.162858587 | 0.846698958 | 0.273159258 | 0.740559962 | 0.321180217 |
| Fntb | S | | 432 | -0.111028329 | -0.013697957 | -0.154890883 | -0.043862555 | -0.141192926 | 0.463698038 | 0.899693112 | 0.016008616 | 0.754741843 | 0.254040661 |
| Matr3 | S | | 596 | -0.046164963 | -0.102368343 | -0.001044843 | 0.045120119 | 0.1013235 | 0.178423309 | 0.000960449 | 0.983154016 | 0.413909799 | 0.133276296 |
| Smg1 | S | | 3567 | -0.028319359 | -0.141919135 | -0.085907833 | -0.057588474 | 0.056011303 | 0.705132513 | 0.056105067 | 0.169062841 | 0.425030037 | 0.345512605 |
| Esco1 | S | | 202 | 0.028235265 | -0.169373672 | -0.190191157 | -0.218426422 | -0.020817484 | 0.48035212 | 0.062915865 | 0.161694357 | 0.125054317 | 0.853010901 |
| Rftn1 | S | | 536 | 0.032797008 | -0.178851498 | -0.165355152 | -0.198152159 | 0.013496347 | 0.731312214 | 0.068778946 | 0.173396368 | 0.121313323 | 0.89171428 |
| Psmc4 | S | | 21 | -0.356637456 | -0.031226422 | -0.319079536 | 0.03755792 | -0.287853113 | 0.399748072 | 0.820407649 | 0.488225625 | 0.943736721 | 0.521681726 |
| Cldn12 | S | | 228 | 0.071185799 | -0.099728187 | -0.300573863 | -0.371759662 | -0.200845676 | 0.277303278 | 0.26920775 | 0.022341432 | 0.011141504 | 0.093478449 |
| Hmga1 Hmga1b | S | | 44 | -0.038465309 | -0.101086582 | -0.138940086 | -0.100474777 | -0.037853503 | 0.521198789 | 0.319495536 | 0.057428496 | 0.091695745 | 0.675160509 |
| Prrc2b | S | | 621 | 0.051798268 | -0.140710285 | -0.116193681 | -0.167991949 | 0.024516603 | 0.159574014 | 0.004688696 | 0.062651874 | 0.026842864 | 0.60728346 |
| Tasor | S | | 653 | -0.073703244 | -0.102324897 | -0.098281858 | -0.024578614 | 0.004043039 | 0.143130225 | 0.253911496 | 0.240113666 | 0.738067507 | 0.965831465 |
| Dbr1 | S | | 505 | -0.100707557 | 0.026422419 | -0.116996754 | -0.016289197 | -0.143419173 | 0.041359294 | 0.665808055 | 0.022900456 | 0.630918646 | 0.070543133 |
| Tom1l1 | S | | 472 | -0.180207134 | -0.149024913 | -0.046230823 | 0.133976311 | 0.10279409 | 0.167195356 | 0.288649968 | 0.697332557 | 0.146130363 | 0.341536606 |
| Dlc1 | S | | 420 | -0.297845458 | 0.056079004 | -0.209445283 | 0.088400176 | -0.265524286 | 0.12957302 | 0.73424266 | 0.335121036 | 0.680018561 | 0.244474084 |
| Tnip1 | S | | 279 | -0.07105044 | -0.053788996 | -0.04827606 | 0.02277438 | 0.005512936 | 0.184410038 | 0.106879652 | 0.020535533 | 0.604219398 | 0.829767572 |
| Ints3 | S | | 993 | 0.066926723 | -0.167306024 | -0.078155605 | -0.145082328 | 0.089150419 | 0.155279005 | 0.012098214 | 0.007299817 | 0.029176578 | 0.055977908 |
| Nes | S | | 894 | 0.031245306 | -0.166268518 | -0.076513702 | -0.107759009 | 0.089754816 | 0.60619584 | 0.011038538 | 0.143426815 | 0.091996563 | 0.007750855 |
| Fhdc1 | S | | 544 | -0.060310395 | -0.151919591 | -0.122844615 | -0.06253422 | 0.029074976 | 0.617308417 | 0.09563836 | 0.351721592 | 0.68209913 | 0.818282463 |
| Ank2 | S | | 31 | -0.058505168 | 0.042037272 | -0.193778244 | -0.135273076 | -0.235815516 | 0.300960395 | 0.425021666 | 0.003369661 | 0.037393617 | 0.004707686 |
| Ptpn12 | S | | 734 | -0.026759501 | -0.035703592 | -0.17199618 | -0.145236679 | -0.136292587 | 0.800716022 | 0.676706819 | 0.006521951 | 0.245781498 | 0.186121454 |
| Lima1 | S | | 735 | -0.038768759 | -0.0671345 | -0.096800145 | -0.058031386 | -0.029665645 | 0.316682146 | 0.087913547 | 0.067856701 | 0.205083582 | 0.461239207 |
| Fnbp4 | S | | 507 | -0.21977433 | -0.011588884 | -0.032266246 | 0.187508085 | -0.020677362 | 0.01137596 | 0.834971638 | 0.78760371 | 0.200734199 | 0.86520149 |
| Pphln1 | S | | 147 | -0.009696609 | -0.136817365 | -0.050700004 | -0.041003395 | 0.086117361 | 0.849750671 | 0.005346156 | 0.322464707 | 0.47416083 | 0.118504453 |
| Nosip | S | | 166 | -0.379892273 | -0.094758707 | -0.112925717 | 0.266966556 | -0.018167009 | 0.287212797 | 0.658538658 | 0.60583311 | 0.420069448 | 0.903017936 |
| Nfatc1 | S | | 263 | 0.068961065 | -0.229243805 | -0.231047148 | -0.300008212 | -0.001803342 | 0.17214971 | 0.147965271 | 0.057823183 | 0.041568548 | 0.9894091 |
| Sgk3 | S | | 126 | -0.293541449 | 0.085270041 | -0.067928684 | 0.225612764 | -0.153198725 | 0.006236279 | 0.294569919 | 0.468972927 | 0.061489894 | 0.106854662 |
| Sgk3 | S | | 129 | -0.293541449 | 0.085270041 | -0.067928684 | 0.225612764 | -0.153198725 | 0.006236279 | 0.294569919 | 0.468972927 | 0.061489894 | 0.106854662 |
| Anln | S | | 73 | 0.133581986 | -0.116134217 | -0.259065838 | -0.392647824 | -0.142931621 | 0.035372352 | 0.090610848 | 0.002086518 | 0.004385159 | 0.037476633 |
| Safb | S | | 623 | -0.138397407 | -0.097336658 | -0.041968287 | 0.09642912 | 0.055368371 | 0.046220626 | 0.40235983 | 0.484123146 | 0.02938517 | 0.597136726 |
| Rab11fip5 | S | | 278 | 0.056738565 | -0.129256871 | -0.208782982 | -0.265521547 | -0.079526112 | 0.338759503 | 0.026569194 | 0.104031786 | 0.054344102 | 0.419486674 |
| Prrc2b | S | | 1159 | 0.035872004 | -0.121028725 | -0.231514829 | -0.267386833 | -0.110486104 | 0.688615827 | 0.061253725 | 0.153935706 | 0.118726712 | 0.417760007 |
| Slain2 | S | | 134 | -0.081943662 | -0.184634389 | -0.073422279 | 0.008521383 | 0.11121211 | 0.391566308 | 0.158985715 | 0.387072557 | 0.905790564 | 0.326315291 |
| Cebpz | S | | 973 | 0.079508932 | -0.408331435 | -0.090283941 | -0.169792872 | 0.318047494 | 0.481466731 | 0.045202216 | 0.528111447 | 0.245600077 | 0.111501451 |
| Mtdh | S | | 423 | -0.208588246 | -0.123081257 | 0.000152153 | 0.208740398 | 0.123233409 | 0.164807504 | 0.121238783 | 0.998927598 | 0.207904878 | 0.330089928 |
| Sptan1 | S | | 1031 | 0.006770544 | 0.026271701 | -0.246742345 | -0.253512888 | -0.273014046 | 0.883857609 | 0.599411856 | 0.001491088 | 0.000469485 | 0.000663562 |
| Bcl9l | S | | 996 | -0.08886721 | -0.092460249 | -0.098178886 | -0.009311675 | -0.005718637 | 0.083184973 | 0.378224619 | 0.282206842 | 0.909170195 | 0.959816154 |

| | | | | | | | | | | | | |
|---|---|---|---|---|---|---|---|---|---|---|---|---|
| Mtdh | S | 491 | 0.07988042 | -0.122481249 | -0.263074355 | -0.342954775 | -0.140593107 | 0.146372726 | 0.06913169 | 0.03436654 | 0.013037624 | 0.133905536 |
| Fndc3b | S | 795 | -0.08722818 | -0.086513375 | -0.158309055 | -0.071080875 | -0.071795679 | 0.119287533 | 0.481539351 | 0.295485169 | 0.596630862 | 0.656927932 |
| Myo10 | S | 2056 | -0.234798409 | -0.194919208 | 0.022596735 | 0.257395144 | 0.217515943 | 0.16258103 | 0.201565248 | 0.695087686 | 0.146975524 | 0.174730951 |
| Ybx3 | S | 358 | -0.065078756 | -0.040018384 | -0.140891102 | -0.075812346 | -0.100872718 | 0.219807049 | 0.149977203 | 0.134628176 | 0.36074437 | 0.237638072 |
| Usp42 | S | 961 | 0.056390826 | -0.165865436 | -0.189273851 | -0.245664677 | -0.023408416 | 0.184844879 | 0.029680601 | 0.096712715 | 0.066946133 | 0.793510731 |
| Nkrf | S | 429 | 0.050212447 | -0.022457759 | -0.247564582 | -0.297777029 | -0.225106823 | 0.59951113 | 0.763931623 | 0.002921734 | 0.041725075 | 0.034145591 |
| Amot | S | 838 | -0.11242194 | -0.019076144 | -0.267439936 | -0.155017996 | -0.248363792 | 0.417640848 | 0.785567647 | 0.130357868 | 0.382805522 | 0.151995419 |
| Strn4 | S | 36 | -0.029033692 | -0.109732009 | -0.134078045 | -0.105044353 | -0.024346036 | 0.651943396 | 0.045734551 | 0.258585998 | 0.362614649 | 0.808794023 |
| Gtf2e2 | S | 31 | 0.030057394 | -0.131221763 | -0.112774079 | -0.142831474 | 0.018447684 | 0.503118604 | 0.036413606 | 0.039588288 | 0.000805438 | 0.601632634 |
| Bora | S | 344 | -0.07277155 | -0.029955464 | -0.170045966 | -0.097274416 | -0.140090503 | 0.096606013 | 0.448797074 | 0.167135584 | 0.365031387 | 0.23347788 |
| Mfhas1 | T | 1031 | 0.042364066 | -0.181972754 | -0.117647802 | -0.160011868 | 0.064324952 | 0.645243568 | 0.040102499 | 0.129905002 | 0.115133306 | 0.317870733 |
| Gigyf2 | S | 161 | -0.110316073 | 0.016499828 | -0.115582913 | -0.00526684 | -0.132082741 | 0.051376873 | 0.788008191 | 0.040536067 | 0.822126695 | 0.075922329 |
| Trip12 | S | 1350 | -0.001669371 | -0.082172835 | -0.115948718 | -0.114279346 | -0.033775883 | 0.947832643 | 0.021473521 | 0.076876751 | 0.10006526 | 0.4963477 |
| Trip12 | S | 1355 | -0.001669371 | -0.082172835 | -0.115948718 | -0.114279346 | -0.033775883 | 0.947832643 | 0.021473521 | 0.076876751 | 0.10006526 | 0.4963477 |
| Rin1 | S | 280 | -0.140258958 | 0.01321019 | -0.165291879 | -0.025032921 | -0.178502069 | 0.162118385 | 0.893172442 | 0.077587356 | 0.793508155 | 0.155712909 |
| Sik3 | S | 1114 | -0.102178249 | -0.130687836 | -0.063667742 | 0.038510507 | 0.067020095 | 0.165739216 | 0.167295499 | 0.520273077 | 0.638596142 | 0.495148047 |
| 1810013L24Rik | S | 266 | -0.055959192 | -0.024650627 | -0.197388282 | -0.14142909 | -0.172737654 | 0.088808837 | 0.624519497 | 0.125520061 | 0.216932705 | 0.15253079 |
| Zfyve21 | S | 17 | -0.003596122 | -0.076655382 | -0.209501261 | -0.205905139 | -0.132845879 | 0.955429618 | 0.101552107 | 0.107563883 | 0.110632224 | 0.23736996 |
| Setd2 | S | 242 | 0.022832962 | -0.098610439 | -0.105198649 | -0.128031611 | -0.00658821 | 0.497085739 | 0.009772745 | 0.047713473 | 0.032460601 | 0.863414178 |
| Srrm1 | S | 810 | 0.003126474 | -0.061756149 | -0.044365422 | -0.047491896 | 0.017390728 | 0.894848111 | 0.00024939 | 0.019587299 | 0.12609346 | 0.214004436 |
| Comt | S | 260 | -0.136370253 | 0.018868698 | -0.237021036 | -0.100650783 | -0.255889734 | 0.171317526 | 0.894018127 | 0.150471484 | 0.479495843 | 0.182014187 |
| Pkn2 | T | 121 | -0.031462945 | -0.096009509 | -0.068486294 | -0.03702335 | 0.027523215 | 0.524375727 | 0.032936692 | 0.097389031 | 0.441022711 | 0.43224515 |
| Clasrp | S | 448 | -0.220361143 | -0.020574554 | 0.031518092 | 0.251879234 | 0.052092646 | 0.003617165 | 0.705795539 | 0.800748911 | 0.130042326 | 0.673950626 |
| Clasrp | S | 450 | -0.220361143 | -0.020574554 | 0.031518092 | 0.251879234 | 0.052092646 | 0.003617165 | 0.705795539 | 0.800748911 | 0.130042326 | 0.673950626 |
| 0610010K14Rik | S | 96 | -0.048565118 | -0.072648744 | -0.008434508 | 0.04013061 | 0.064214236 | 0.021736743 | 0.003943144 | 0.755364393 | 0.182548712 | 0.072437487 |
| Nars | S | 493 | -0.087087769 | -0.064414617 | -0.186484562 | -0.099396793 | -0.122069945 | 0.284728066 | 0.404059108 | 0.207735062 | 0.437848431 | 0.356058279 |
| Sh3pxd2b | S | 550 | -0.044669252 | -0.105448409 | -0.08006637 | -0.035397118 | 0.025382039 | 0.48226627 | 0.077455985 | 0.118128794 | 0.51864632 | 0.538998251 |
| Slx4 | T | 325 | -0.016607891 | -0.13551039 | -0.106312974 | -0.089705083 | 0.029197416 | 0.811695797 | 0.0983263 | 0.100938958 | 0.176859931 | 0.630625913 |
| Stard13 | S | 334 | 0.021001205 | -0.108096999 | -0.092105593 | -0.113106798 | 0.015991406 | 0.62725308 | 0.024763518 | 0.026219027 | 0.041004919 | 0.588794564 |
| Myo18a | S | 2049 | 0.043879614 | -0.070070263 | -0.14263315 | -0.186512765 | -0.072562887 | 0.568217904 | 0.110771729 | 0.004544312 | 0.086548261 | 0.065700639 |
| Coprs | S | 16 | -0.077616852 | -0.103712148 | -0.098602351 | -0.020985499 | 0.005109797 | 0.410922766 | 0.094623432 | 0.278601817 | 0.838839459 | 0.948393632 |
| Fhod1 | S | 390 | -0.039844393 | -0.197757485 | -0.082845172 | -0.043000779 | 0.114912312 | 0.518852874 | 0.141056152 | 0.272934691 | 0.568249289 | 0.342216299 |
| Fam110a | S | 190 | -0.091529474 | -0.041464037 | -0.092874113 | -0.001344639 | -0.051410076 | 0.046398613 | 0.54060975 | 0.148162511 | 0.977090738 | 0.48051122 |
| Pclo | S | 2057 | -0.117493918 | -0.162337611 | -0.003001001 | 0.114492916 | 0.15933661 | 0.197449482 | 0.065742924 | 0.976929209 | 0.25938189 | 0.147660528 |
| Pclo | S | 2061 | -0.117493918 | -0.162337611 | -0.003001001 | 0.114492916 | 0.15933661 | 0.197449482 | 0.065742924 | 0.976929209 | 0.25938189 | 0.147660528 |
| Pclo | T | 2059 | -0.117493918 | -0.162337611 | -0.003001001 | 0.114492916 | 0.15933661 | 0.197449482 | 0.065742924 | 0.976929209 | 0.25938189 | 0.147660528 |
| Rai14 | S | 385 | 0.087943763 | -0.188398377 | -0.176068677 | -0.26401244 | 0.0123297 | 0.259810178 | 0.103090002 | 0.026498295 | 0.013086387 | 0.883761236 |
| Rai14 | S | 388 | 0.087943763 | -0.188398377 | -0.176068677 | -0.26401244 | 0.0123297 | 0.259810178 | 0.103090002 | 0.026498295 | 0.013086387 | 0.883761236 |
| Fancm | S | 1368 | 0.030776214 | -0.046145883 | -0.222722053 | -0.253498267 | -0.17657617 | 0.630969413 | 0.312037295 | 0.011556072 | 0.01588141 | 0.043252844 |
| Cobll1 | S | 1266 | -0.087567603 | 0.019621045 | -0.25404881 | -0.166481207 | -0.273669855 | 0.260297438 | 0.830428292 | 0.068030716 | 0.177933045 | 0.068087743 |
| Rgs12 | S | 454 | -0.138049576 | 0.152467488 | -0.413587135 | -0.275537559 | -0.566054624 | 0.342873914 | 0.374656746 | 0.050417653 | 0.12463963 | 0.020200046 |
| Iws1 | S | 211 | -0.044540831 | -0.141654283 | -0.012521354 | 0.032019478 | 0.129132929 | 0.211516511 | 0.011127185 | 0.834896436 | 0.574352484 | 0.094430499 |
| Chd9 | S | 611 | 0.006132689 | -0.097401895 | -0.098022194 | -0.104154884 | -0.000620299 | 0.94422662 | 0.006544246 | 0.154935945 | 0.322675202 | 0.990668793 |
| Amotl1 | S | 884 | -0.008033528 | -0.175695264 | -0.082549517 | -0.074515989 | 0.093145747 | 0.795063633 | 0.020756551 | 0.495946219 | 0.532353146 | 0.455021907 |
| Fam193a | T | 393 | 0.043809514 | -0.147006046 | -0.262549192 | -0.306358705 | -0.115543146 | 0.516644682 | 0.093231217 | 0.16477056 | 0.136996924 | 0.471190137 |
| Tjp1 | S | 353 | -0.019647756 | -0.123580776 | -0.146576774 | -0.126929017 | -0.022995997 | 0.749082704 | 0.092605648 | 0.193936332 | 0.249732591 | 0.815417937 |
| Ik | S | 402 | -0.050289419 | -0.151003814 | -0.020543064 | 0.029746355 | 0.13046075 | 0.327121045 | 0.013615664 | 0.814125371 | 0.723769858 | 0.197851344 |
| Ralbp1 | S | 638 | -0.169242283 | 0.008554467 | -0.059925992 | 0.109316291 | -0.068480458 | 0.005887682 | 0.840395991 | 0.467426828 | 0.227386535 | 0.406404902 |
| Chd1 | S | 1684 | 0.018236653 | -0.098749984 | -0.117152095 | -0.135388747 | -0.01840211 | 0.797060426 | 0.060746198 | 0.028600557 | 0.119496521 | 0.556347416 |
| Zc3h14 | S | 146 | -0.001446132 | -0.113841606 | -0.206115503 | -0.204669371 | -0.092273897 | 0.991976692 | 0.352972712 | 0.06118202 | 0.215444831 | 0.440068788 |
| Ptpn1 | S | 335 | -0.116071597 | 0.008318219 | -0.131755435 | -0.015683838 | -0.140073653 | 0.093011478 | 0.899602075 | 0.058811784 | 0.799702551 | 0.094264017 |
| Polr3c | S | 205 | -0.058554334 | -0.091613739 | -0.069427264 | -0.01087293 | 0.022186475 | 0.311316788 | 0.094122171 | 0.12568291 | 0.84727549 | 0.651328535 |
| Bod1l | S | 1071 | -0.011362216 | -0.119125667 | -0.174995316 | -0.1636331 | -0.055869649 | 0.895564524 | 0.253819347 | 0.07938508 | 0.009557205 | 0.421279928 |
| Zc3hc1 | S | 409 | -0.017585929 | -0.049720584 | -0.064772471 | -0.047186542 | -0.015051887 | 0.34247925 | 0.021199893 | 0.013036959 | 0.041402218 | 0.37538776 |
| Phldb1 | S | 1007 | 0.156335884 | -0.180858665 | -0.215035313 | -0.371371197 | -0.034176648 | 0.037234912 | 0.021713189 | 0.010474749 | 0.001987202 | 0.259453285 |
| Phldb1 | S | 1009 | 0.156335884 | -0.180858665 | -0.215035313 | -0.371371197 | -0.034176648 | 0.037234912 | 0.021713189 | 0.010474749 | 0.001987202 | 0.259453285 |
| Rbm10 | S | 797 | 0.060356364 | -0.111601482 | -0.0941992 | -0.154555563 | 0.017402283 | 0.498621377 | 0.000742153 | 0.163297843 | 0.161631853 | 0.740243193 |
| Nucks1 | S | 113 | -0.160915688 | -0.093792989 | -0.043805584 | 0.117110104 | 0.049987405 | 0.114512598 | 0.406708685 | 0.318294026 | 0.215461467 | 0.629412974 |
| Wiz | S | 1179 | 0.048079815 | -0.108452373 | -0.099773056 | -0.147852871 | 0.008679317 | 0.581192495 | 0.011877369 | 0.037251272 | 0.171206502 | 0.807746229 |
| Pno1 | S | 31 | -0.039592403 | -0.126148584 | -0.226133531 | -0.186541128 | -0.099984947 | 0.729846551 | 0.581599475 | 0.0940492 | 0.033508503 | 0.636796553 |
| Slc26a2 | S | 12 | -0.045601693 | -0.244724383 | 0.0313349 | 0.076936593 | 0.276059283 | 0.750459378 | 0.009014202 | 0.797831773 | 0.637946188 | 0.09724383 |
| Marcks | S | 171 | -0.245932068 | -0.084670998 | 0.054324923 | 0.300256991 | 0.138995921 | 0.016097341 | 0.396182128 | 0.535810486 | 0.029161557 | 0.253484211 |
| Espl1 | S | 1504 | 0.085173519 | -0.051507487 | -0.168968778 | -0.254142297 | -0.117461291 | 0.365683464 | 0.066759976 | 0.002048412 | 0.066971489 | 0.008825676 |
| Sin3a | S | 10 | 0.013637101 | -0.101747252 | -0.086827838 | -0.100464939 | 0.014919414 | 0.752398941 | 0.016119032 | 0.045945049 | 0.066355468 | 0.642765414 |
| Larp1 | S | 299 | -0.018024293 | -0.11846273 | -0.101645021 | -0.083620728 | 0.016817709 | 0.587849903 | 0.082948211 | 0.12271904 | 0.182211506 | 0.792940638 |
| Phf2 | S | 1057 | 0.007476076 | -0.073965784 | -0.153433617 | -0.160909693 | -0.079467833 | 0.907726225 | 0.119374481 | 0.030166011 | 0.073611287 | 0.178824344 |
| Mki67 | S | 337 | 0.040403189 | 0.016250133 | -0.254842543 | -0.295245732 | -0.271092676 | 0.590494834 | 0.803007017 | 0.001216571 | 0.02632284 | 0.018208955 |
| Ptpn12 | S | 670 | -0.071235844 | -0.059854626 | -0.148287335 | -0.077051491 | -0.08843271 | 0.338860902 | 0.471915514 | 0.079986165 | 0.270610837 | 0.277019908 |
| Phlpp1 | S | 1497 | -0.225666649 | 0.041196178 | -0.077827583 | 0.147839065 | -0.119023761 | 0.023735846 | 0.831432623 | 0.393187672 | 0.073044996 | 0.544138211 |
| Fnbp4 | S | 125 | -0.00941175 | -0.120843488 | -0.131729761 | -0.122318011 | -0.010886273 | 0.412271727 | 0.223543626 | 0.074975543 | 0.083824997 | 0.899805878 |
| Trio | S | 1655 | -0.00536854 | -0.192841385 | -0.199858557 | -0.194490017 | -0.007017172 | 0.982198977 | 0.231135504 | 0.20929481 | 0.43983167 | 0.961304926 |
| Emd | S | 29 | -0.055994344 | -0.072586305 | -0.076444583 | -0.020450239 | -0.003858278 | 0.590826573 | 0.032938466 | 0.088287168 | 0.840276373 | 0.90582614 |
| Rai1 | S | 856 | -0.196180048 | -0.124281898 | -0.141447716 | 0.054732332 | -0.017165819 | 0.40526566 | 0.483711983 | 0.36303327 | 0.811243702 | 0.924955512 |
| Tspyl1 | T | 27 | 0.04814162 | -0.192148637 | -0.113254953 | -0.161396573 | 0.078893684 | 0.446263825 | 0.017955044 | 0.213839882 | 0.097665173 | 0.304061729 |
| Scrib | T | 812 | 0.034001453 | -0.159464114 | -0.194894587 | -0.22889604 | -0.035430473 | 0.525843075 | 0.063632525 | 0.173741117 | 0.144585178 | 0.76925067 |
| Larp1 | S | 827 | -0.014329642 | -0.059417135 | -0.074870065 | -0.060540423 | -0.01545293 | 0.536709711 | 0.028531835 | 0.019167679 | 0.036035068 | 0.444598392 |
| Fam193a | S | 688 | 0.249608021 | -0.252229964 | -0.327250127 | -0.576858149 | -0.075020163 | 0.144385754 | 0.088765556 | 0.043867933 | 0.010234178 | 0.356512287 |
| Srrm2 | T | 1828 | -0.368829243 | -0.102886012 | 0.046957211 | 0.415786454 | 0.149843223 | 0.16322913 | 0.206824901 | 0.72470292 | 0.130255788 | 0.300630479 |
| Braf | S | 135 | 0.029525321 | -0.011905697 | -0.278326888 | -0.307852209 | -0.26642119 | 0.706472888 | 0.890466874 | 0.007900838 | 0.003001032 | 0.020166298 |
| Sec23ip | S | 735 | -0.028217401 | -0.040483776 | -0.130291185 | -0.102073784 | -0.089807409 | 0.630338807 | 0.209072688 | 0.016700231 | 0.148239895 | 0.06616703 |
| Dpysl3 | S | 518 | -0.201565401 | 0.120825752 | -0.286267509 | -0.084702107 | -0.407093261 | 0.15750065 | 0.569528279 | 0.143272241 | 0.585817933 | 0.120092775 |
| Topors | S | 82 | 0.151413823 | -0.206560818 | -0.162266147 | -0.31367997 | 0.044294671 | 0.023750553 | 0.004551561 | 0.02088766 | 0.003605315 | 0.418131498 |
| Bcl9l | S | 1007 | -0.104097076 | -0.107722483 | -0.118719897 | -0.014622822 | -0.010997414 | 0.092387266 | 0.480554722 | 0.430802334 | 0.91723483 | 0.952557359 |
| Golga3 | S | 389 | 0.042448032 | -0.198361483 | 0.006912243 | -0.035535789 | 0.205273726 | 0.187680463 | 5.54135E-05 | 0.85475928 | 0.400850896 | 0.010927275 |
| Rps6kc1 | S | 778 | 0.010789165 | -0.12044713 | -0.161491028 | -0.172280193 | -0.041043898 | 0.863359681 | 0.276038744 | 0.033454754 | 0.05549982 | 0.686072652 |
| Mycbp2 | S | 1621 | -0.183961178 | -0.008233702 | -0.114312165 | 0.069649013 | -0.106078463 | 0.074222682 | 0.912133401 | 0.291440366 | 0.470518822 | 0.263353631 |
| Phip | S | 659 | -0.13325491 | -0.084353674 | -0.131353395 | 0.001901516 | -0.046999721 | 0.409767941 | 0.56343583 | 0.118720476 | 0.98944668 | 0.72790371 |
| Jmjd6 | S | 402 | 0.101380415 | -0.169962479 | -0.313201185 | -0.4145816 | -0.143238706 | 0.150987075 | 0.075343069 | 0.090368702 | 0.059200997 | 0.343209538 |
| Ppp1r10 | S | 313 | 0.036164699 | -0.139879811 | -0.049658519 | -0.085823218 | 0.090221291 | 0.510496817 | 0.00202851 | 0.132746117 | 0.175278676 | 0.022475316 |
| Rcc2 | T | 519 | -0.103788868 | -0.048723477 | -0.177168723 | -0.073379855 | -0.128445246 | 0.170558331 | 0.616521845 | 0.230582941 | 0.577892857 | 0.39244593 |
| Snx2 | S | 97 | 0.07116889 | -0.102916179 | -0.137535557 | -0.208704447 | -0.034619378 | 0.084852114 | 0.058574597 | 0.001063533 | 0.005970332 | 0.399675013 |
| Gtf2f2 | S | 248 | -0.08396108 | -0.006270208 | -0.134092421 | -0.050131341 | -0.127822213 | 0.04637351 | 0.930814655 | 0.09454542 | 0.426549471 | 0.185408745 |
| Sar1a | T | 139 | -0.103027825 | 0.013763553 | -0.115010377 | -0.011982552 | -0.128773931 | 0.056506278 | 0.746357854 | 0.039137742 | 0.802483781 | 0.045925374 |
| Etv3 | S | 139 | -0.064412859 | -0.064507035 | -0.123986079 | -0.05957322 | -0.059479044 | 0.139073414 | 0.280480391 | 0.198696278 | 0.47643775 | 0.50360145 |

| | | | | | | | | | | | | |
|---|---|---|---|---|---|---|---|---|---|---|---|---|
| Map7d1 | S | 95 | 0.009434119 | -0.419524375 | -0.114964726 | -0.124398845 | 0.304559648 | 0.981411601 | 0.142324459 | 0.707972401 | 0.758579658 | 0.287461365 |
| Polr2a | S | 1815 | -0.036767039 | -0.123430289 | -0.080387173 | -0.043620134 | 0.043043117 | 0.61928521 | 0.043052933 | 0.248597336 | 0.555561762 | 0.423305207 |
| Atxn2l | S | 389 | 0.172553809 | -0.311469831 | -0.251414519 | -0.423968328 | 0.060055312 | 0.104514647 | 0.110474203 | 0.043985928 | 0.007389831 | 0.693873318 |
| Mast4 | S | 2206 | 0.123285042 | -0.181931162 | -0.270658713 | -0.393943755 | -0.088727551 | 0.062676202 | 0.013659637 | 0.099386336 | 0.05561483 | 0.481292215 |
| Snrk | S | 495 | -0.375682693 | -0.236695951 | 0.152331771 | 0.528014465 | 0.389027722 | 0.225666386 | 0.019744332 | 0.078721619 | 0.139580658 | 0.006553743 |
| Gas2 | S | 287 | -0.040845669 | -0.021351464 | -0.20896483 | -0.168119161 | -0.187613366 | 0.501794328 | 0.77342484 | 0.032199288 | 0.06874836 | 0.056136984 |
| Sh3bp4 | S | 295 | -0.151469831 | -0.059703576 | -0.094036387 | 0.057433444 | -0.034332811 | 0.129914722 | 0.413362726 | 0.376152315 | 0.609096509 | 0.736502861 |
| Dbn1 | T | 429 | 0.004447431 | -0.125837962 | -0.109526394 | -0.113973825 | 0.016311567 | 0.972978127 | 0.025329448 | 0.200365047 | 0.436025121 | 0.816454693 |
| Gnl3 | S | 485 | -0.103734724 | -0.164495129 | -0.035836966 | 0.067897758 | 0.128658163 | 0.398538388 | 0.07057362 | 0.677965321 | 0.580253008 | 0.185435204 |
| Ptbp3 | S | 29 | 0.037428643 | -0.104337775 | -0.110608343 | -0.148036986 | -0.006270568 | 0.593426859 | 0.003202407 | 0.157131295 | 0.132021772 | 0.916861008 |
| Atxn2l | S | 56 | -0.061154659 | -0.137088741 | -0.099260446 | -0.038105786 | 0.037828296 | 0.372402056 | 0.264730773 | 0.178003166 | 0.631340775 | 0.737863277 |
| Dab2ip | S | 910 | -0.247078775 | 0.107783746 | -0.207626153 | 0.039452622 | -0.315409899 | 0.062622208 | 0.460252418 | 0.235252133 | 0.7717715 | 0.102750365 |
| Vim | S | 29 | -0.000787611 | -0.013441532 | -0.222516896 | -0.221729285 | -0.209075364 | 0.991337023 | 0.742543273 | 0.009285518 | 0.041435467 | 0.020902152 |
| Prune2 | S | 1074 | 0.070941796 | -0.144911219 | -0.245486197 | -0.316427993 | -0.100574978 | 0.295504776 | 0.196751562 | 0.038419628 | 0.02278644 | 0.383749325 |
| Cflar | S | 185 | -0.149309077 | -0.03172174 | -0.081164351 | 0.068144726 | -0.049442611 | 0.100997808 | 0.653583697 | 0.162000909 | 0.366562378 | 0.449904013 |
| Cflar | S | 193 | -0.149309077 | -0.03172174 | -0.081164351 | 0.068144726 | -0.049442611 | 0.100997808 | 0.653583697 | 0.162000909 | 0.366562378 | 0.449904013 |
| Cflar | T | 189 | -0.149309077 | -0.03172174 | -0.081164351 | 0.068144726 | -0.049442611 | 0.100997808 | 0.653583697 | 0.162000909 | 0.366562378 | 0.449904013 |
| Kdm1a | S | 70 | -0.002997185 | -0.115916744 | -0.169048753 | -0.166051569 | -0.053132009 | 0.98063779 | 0.170901543 | 0.09948201 | 0.233813551 | 0.516731429 |
| Erf | T | 148 | -0.049449623 | -0.01926458 | -0.629476442 | -0.580026819 | -0.610211862 | 0.806545579 | 0.802835695 | 0.263264068 | 0.293016031 | 0.275163209 |
| Cep55 | T | 422 | -0.009555229 | -0.202551648 | -0.154660706 | -0.145105477 | 0.047890942 | 0.851764391 | 0.147434559 | 0.315080751 | 0.340910524 | 0.763109933 |
| Dicer1 | S | 1015 | -0.054527565 | -0.06060823 | -0.086791315 | -0.03226375 | -0.026183085 | 0.161075011 | 0.085618804 | 0.184615686 | 0.576617019 | 0.637565288 |
| Jam3 | S | 307 | 0.060994516 | -0.0231726 | -0.421664798 | -0.482659314 | -0.398492199 | 0.667721861 | 0.832815224 | 0.03781846 | 0.034482072 | 0.053905365 |
| Zc3h18 | S | 58 | 0.08886444 | -0.158501414 | -0.05059366 | -0.139458099 | 0.107907754 | 0.260030886 | 0.000246188 | 0.209304702 | 0.122022517 | 0.042955504 |
| Incenp | S | 867 | -0.055730439 | -0.09199256 | -0.070204623 | -0.014474184 | 0.021787938 | 0.196027787 | 0.09496793 | 0.240988369 | 0.781678136 | 0.704862782 |
| C77080 | S | 1003 | -0.092676834 | -0.011804006 | -0.184224004 | -0.09154717 | -0.172419998 | 0.442148345 | 0.896721809 | 0.039657062 | 0.461110063 | 0.147651272 |
| Nrd1 | S | 85 | -0.113281866 | 0.021313674 | -0.149096734 | -0.035814868 | -0.170410408 | 0.119067316 | 0.788795301 | 0.053160599 | 0.553679048 | 0.076453759 |
| Dst | S | 1970 | -0.068031814 | -0.137697143 | 0.0165399 | 0.084571714 | 0.154237043 | 0.133336067 | 0.010501479 | 0.672275232 | 0.023446753 | 0.000686664 |
| Clcc1 | S | 497 | -0.15003071 | -0.0650567 | -0.000117833 | 0.149912877 | 0.064938867 | 0.020986574 | 0.209221811 | 0.998547247 | 0.053813166 | 0.270032539 |
| Ankrd11 | S | 1676 | -0.062821762 | -0.056902967 | -0.128032031 | -0.065210269 | -0.071129063 | 0.368362067 | 0.439927749 | 0.052567525 | 0.319631347 | 0.317238852 |
| Rai14 | S | 341 | 0.064449746 | -0.129702776 | -0.11015473 | -0.174604476 | 0.019548046 | 0.169839165 | 0.048805169 | 0.003341944 | 0.022045186 | 0.659967564 |
| Farsa | S | 301 | -0.02913785 | 0.005255162 | -0.183263724 | -0.154125874 | -0.188518886 | 0.791708663 | 0.971606132 | 0.003844902 | 0.234816669 | 0.276620902 |
| Zeb2 | S | 784 | 0.097779311 | -0.177349003 | -0.124158427 | -0.221937738 | 0.053190576 | 0.039100415 | 0.002475925 | 0.049540355 | 0.010086417 | 0.302109334 |
| Nf1 | S | 659 | -0.181476323 | 0.050112547 | -0.10105055 | 0.080425773 | -0.151163097 | 0.012159126 | 0.643859421 | 0.333608935 | 0.412631908 | 0.264426721 |
| Ubr4 | S | 619 | -0.007096019 | -0.050604752 | -0.123893138 | -0.116797119 | -0.073288386 | 0.859403408 | 0.047542594 | 0.031900342 | 0.059918781 | 0.107468293 |
| Sorbs2 | S | 381 | -0.017958362 | -0.1214204 | -0.066877485 | -0.048919122 | 0.054542915 | 0.79410991 | 0.025690807 | 0.164271571 | 0.510207347 | 0.277726247 |
| Kif13a | S | 636 | 0.04286135 | -0.130139208 | -0.130960805 | -0.173822155 | -0.000821597 | 0.660022786 | 0.076218018 | 0.047051126 | 0.146204522 | 0.989155007 |
| Sh3kbp1 | S | 227 | -0.119788974 | -0.014460203 | -0.031312846 | 0.088476127 | -0.016852643 | 0.002330913 | 0.791323742 | 0.407186725 | 0.066874764 | 0.765141557 |
| Acta1 Acta2 Actb Act | S | 60;60;62;62;61;62 | 0.065082248 | -0.187613148 | -0.142649242 | -0.20773149 | 0.044963906 | 0.63218453 | 0.072057873 | 0.121698679 | 0.185254946 | 0.60265511 |
| Sh3rf1 | S | 745 | -0.073052716 | -0.139206577 | -0.012896201 | 0.060156515 | 0.126310375 | 0.109797297 | 0.059442543 | 0.828666422 | 0.348259763 | 0.123012671 |
| Cfap36 | S | 147 | 0.028270618 | -0.048023157 | -0.298886111 | -0.327156729 | -0.250862953 | 0.670508601 | 0.461311705 | 0.037393256 | 0.037495681 | 0.071556233 |
| Nmt2 | S | 68 | -0.054730891 | -0.024022615 | -0.071034558 | -0.016303667 | -0.047011943 | 0.164632158 | 0.599683323 | 0.002503378 | 0.613062208 | 0.341466879 |
| Nrcam | S | 1247 | 0.049114249 | -0.065718922 | -0.166382715 | -0.215496964 | -0.100663793 | 0.315451494 | 0.122994818 | 0.004714892 | 0.004910075 | 0.024799238 |
| Pakap | S | 740 | -0.042711065 | 0.029743236 | -0.223454493 | -0.180743429 | -0.253197729 | 0.50124959 | 0.68297953 | 0.010233544 | 0.0163414 | 0.012007546 |
| Ifih1 | S | 289 | -0.126842364 | -0.077980168 | -0.037435539 | 0.089406824 | 0.040544628 | 0.056673627 | 0.220607888 | 0.635559762 | 0.262563048 | 0.585157477 |
| Zdhhc18 | S | 53 | -0.085415153 | -0.113662649 | -0.006566159 | 0.078848994 | 0.10709649 | 0.097561546 | 0.039158464 | 0.915063363 | 0.194759297 | 0.114994112 |
| Arhgap11a | S | 708 | -0.005628205 | -0.16925056 | -0.090200492 | -0.084572288 | 0.079050067 | 0.857942384 | 0.078866959 | 0.182369858 | 0.205400927 | 0.336390811 |
| Phka1 | S | 759 | 0.029410895 | -0.119188328 | -0.294097282 | -0.323508177 | -0.174908954 | 0.779578115 | 0.290257566 | 0.127574164 | 0.106954366 | 0.315119499 |
| Mapk3 | Y | 205 | -0.162790142 | -0.041728883 | -0.139816328 | 0.022973814 | -0.098087445 | 0.151896373 | 0.719301088 | 0.309278249 | 0.863241939 | 0.505457741 |
| Kansl1 | S | 249 | -0.057627305 | -0.126521844 | -0.008652495 | 0.04897481 | 0.117869349 | 0.373307023 | 0.006086236 | 0.871994612 | 0.48897216 | 0.104516269 |
| Pola2 | S | 147 | -0.010627526 | -0.054532477 | -0.152489789 | -0.141862263 | -0.097957312 | 0.837025413 | 0.165401234 | 0.033799939 | 0.068800585 | 0.106602575 |
| Ythdc2 | S | 1105 | -0.067221084 | -0.060125809 | -0.101861973 | -0.034640889 | -0.041736164 | 0.146550922 | 0.214808434 | 0.189682975 | 0.613328069 | 0.553890472 |
| Vcl | S | 566 | -0.294920462 | 0.011226721 | -0.090566098 | 0.204354364 | -0.101792818 | 0.087497941 | 0.783552992 | 0.422667253 | 0.20248518 | 0.376648369 |
| Tnip1 | S | 416 | -0.085581682 | -0.038771441 | -0.075464866 | 0.010116816 | -0.036693424 | 0.104913891 | 0.198386664 | 0.140638936 | 0.841981652 | 0.417403657 |
| Anapc5 | S | 180 | -0.066634802 | -0.155573411 | -0.163830357 | -0.097195555 | -0.008256946 | 0.533106253 | 0.288915923 | 0.324206375 | 0.509072331 | 0.959700069 |
| Ccdc14 | S | 774 | 0.018505614 | -0.106414521 | -0.143945339 | -0.162450953 | -0.037530818 | 0.36780832 | 0.114341624 | 0.024848005 | 0.034690002 | 0.52557791 |
| Nes | S | 1188 | -0.011590811 | -0.104202927 | -0.064886023 | -0.053295212 | 0.039316904 | 0.557083831 | 0.014190108 | 0.195802386 | 0.266387321 | 0.403459074 |
| Dcaf10 | S | 67 | -0.047746282 | -0.051209065 | -0.056686167 | -0.008939885 | -0.005477102 | 0.076871921 | 0.169413319 | 0.043238587 | 0.422863796 | 0.843553323 |
| Rbmx Rbmxl1 | S | 313;316 | -0.065698895 | -0.010526408 | -0.187665889 | -0.121966994 | -0.177139481 | 0.619027306 | 0.917563826 | 0.0184058 | 0.382766228 | 0.154617129 |
| Cdan1 | T | 70 | -0.042840675 | -0.031256541 | -0.192589691 | -0.149749016 | -0.16133315 | 0.452011683 | 0.300962884 | 0.081385552 | 0.138835052 | 0.127575637 |
| Mtmr14 | S | 18 | -0.233127706 | -0.113949167 | -0.144839738 | 0.088287968 | -0.030890571 | 0.413174566 | 0.443222129 | 0.504645429 | 0.776157364 | 0.893638715 |
| Pom121 | S | 159 | 0.136050484 | -0.119393327 | -0.143757566 | -0.27980805 | -0.024364239 | 0.087001414 | 0.00884942 | 0.003653624 | 0.02350537 | 0.233284626 |
| Uhrf1bp1l | S | 890 | 0.026689814 | -0.122636723 | -0.187550879 | -0.214240693 | -0.064914156 | 0.830350276 | 0.048785488 | 0.217845156 | 0.230425205 | 0.616064431 |
| Rtn4 | T | 30 | -0.070289143 | -0.075262289 | -0.045968938 | 0.024320205 | 0.029293351 | 0.235620729 | 0.049355911 | 0.18644635 | 0.631543635 | 0.280085981 |
| Mcmbp | S | 154 | -0.031402298 | -0.061045211 | -0.088649989 | -0.057247691 | -0.027604778 | 0.531691412 | 0.060280889 | 0.048781157 | 0.307106783 | 0.438880137 |
| Zfp335 | S | 976 | -0.001954008 | -0.110411006 | -0.075306066 | -0.073352058 | 0.03510494 | 0.969963133 | 0.03024817 | 0.073927645 | 0.176839209 | 0.277204404 |
| Usp15 | S | 242 | -0.045935111 | -0.11989674 | -0.119051485 | -0.073116374 | 0.000845255 | 0.387459643 | 0.198988125 | 0.222227884 | 0.400697189 | 0.993246192 |
| Spout1 | T | 370 | -0.050883352 | -0.143553029 | -0.169793993 | -0.118910642 | -0.026240965 | 0.583584777 | 0.271778247 | 0.266567176 | 0.422831733 | 0.866812072 |
| Cfdp1 | S | 246 | -0.081243026 | -0.007203599 | -0.127839675 | -0.046596648 | -0.120636076 | 0.067033089 | 0.912965 | 0.073866141 | 0.416746304 | 0.166420479 |
| Ncapd2 | S | 1323 | 0.010726379 | -0.072255855 | -0.098924827 | -0.109651206 | -0.026668972 | 0.795315186 | 0.00862298 | 0.055535783 | 0.075249775 | 0.458699636 |
| Sos1 | S | 1196 | -0.062578489 | -0.03099857 | -0.120781241 | -0.058202751 | -0.089782671 | 0.051663419 | 0.614834327 | 0.108642733 | 0.349895409 | 0.264111973 |
| Pakap | T | 19 | 0.036656559 | -0.150090681 | -0.157056421 | -0.19371298 | -0.00696574 | 0.756975661 | 0.134865978 | 0.083919146 | 0.149088593 | 0.926491267 |
| Dek | S | 214 | -0.016429885 | 0.022698885 | -0.209060967 | -0.192631082 | -0.231759852 | 0.627380678 | 0.664710814 | 0.00379055 | 0.019153426 | 0.010337112 |
| Pus1 | S | 416 | 0.007809052 | -0.083720928 | -0.037633261 | -0.045442313 | 0.046087667 | 0.628657078 | 0.000901679 | 0.029259815 | 0.020440698 | 0.016056157 |
| Clk3 | S | 281 | -0.08262989 | -0.114030906 | -0.115527748 | -0.032897858 | -0.001496842 | 0.410737264 | 0.121143065 | 0.438651285 | 0.830315022 | 0.991510732 |
| Clk3 | S | 283 | -0.08262989 | -0.114030906 | -0.115527748 | -0.032897858 | -0.001496842 | 0.410737264 | 0.121143065 | 0.438651285 | 0.830315022 | 0.991510732 |
| Kank2 | S | 153 | -0.018990255 | -0.044549437 | -0.252818171 | -0.233827916 | -0.208268734 | 0.818514252 | 0.412567882 | 0.074738166 | 0.092278393 | 0.099449528 |
| Slc4a7 | S | 960 | 0.073005761 | -0.156948318 | -0.106630821 | -0.179636582 | 0.050317497 | 0.039230161 | 0.00178714 | 0.05316381 | 0.018172726 | 0.266363438 |
| Csnk2b | S | 209 | 0.116296201 | -0.144822536 | -0.299239728 | -0.415535928 | -0.154417192 | 0.132130422 | 0.018166692 | 0.122337925 | 0.058719804 | 0.328779489 |
| Zfp710 | S | 544 | 0.044446466 | -0.107872943 | -0.182328201 | -0.226774666 | -0.074455257 | 0.292985364 | 0.034053767 | 0.068669589 | 0.048640963 | 0.341358346 |
| Eif4enif1 | S | 76 | 0.022371044 | -0.102599794 | -0.111235018 | -0.133606062 | -0.008635223 | 0.493609113 | 0.013018196 | 0.082235416 | 0.054264819 | 0.856910199 |
| Prpf40a | T | 886 | -0.068957631 | -0.136090457 | -0.011410054 | 0.057547577 | 0.124680403 | 0.236157419 | 0.021985797 | 0.911342049 | 0.576771868 | 0.276692194 |
| Trappc12 | T | 313 | -0.085592723 | -0.103004943 | -0.018178367 | 0.067414355 | 0.084826576 | 0.060444409 | 0.081636378 | 0.764052246 | 0.323471501 | 0.251947378 |
| Specc1 | S | 343 | -0.05072003 | -0.088363853 | -0.005140922 | 0.045579107 | 0.08322293 | 0.224034702 | 0.001230427 | 0.90192109 | 0.387795846 | 0.139875864 |
| Ndc1 | S | 473 | 0.05038974 | -0.037751454 | -0.15765663 | -0.208046369 | -0.119905175 | 0.403904836 | 0.205745593 | 0.001086495 | 0.034299889 | 0.005988345 |
| Heatr3 | T | 31 | -0.090429253 | -0.073089433 | -0.10158646 | -0.011157207 | -0.028497026 | 0.313990913 | 0.116564319 | 0.305872604 | 0.917409385 | 0.748872308 |
| Src | T | 73 | -0.019160565 | -0.121430289 | -0.103204204 | -0.084043639 | 0.018226084 | 0.729684729 | 0.05159534 | 0.23077044 | 0.327097002 | 0.813522773 |
| Lima1 | S | 371 | -0.035357509 | -0.164321908 | -0.06914414 | -0.033786631 | 0.095177768 | 0.554744796 | 0.103941879 | 0.244118925 | 0.607985264 | 0.288927747 |
| Farp1 | S | 373 | -0.05419453 | -0.10966063 | -0.046243006 | 0.007951523 | 0.063417624 | 0.154528406 | 0.093117828 | 0.293998333 | 0.808585655 | 0.273706323 |
| Jcad | S | 193 | -0.180227809 | -0.066984152 | -0.098721696 | 0.081506114 | -0.031737543 | 0.111283809 | 0.694649345 | 0.449627505 | 0.470378911 | 0.852528674 |
| Ptpra | S | 204 | -0.029258905 | -0.169800466 | -0.033478711 | -0.004219806 | 0.136321754 | 0.388706091 | 0.060036189 | 0.272308827 | 0.85022333 | 0.115342824 |
| Mettl1 | T | 262 | 0.20245308 | -0.271755436 | -0.181182587 | -0.383635667 | 0.090572849 | 0.160518485 | 0.0029684 | 0.384600336 | 0.130477345 | 0.63981098 |
| Gemin5 | S | 1410 | 0.068424615 | -0.108773454 | -0.194999834 | -0.263424448 | -0.086226379 | 0.373168134 | 0.124417172 | 0.02494723 | 0.010452352 | 0.134909081 |
| Map3k20 | S | 593 | -0.114083774 | -0.074743078 | -0.097565205 | 0.016518569 | -0.022822128 | 0.072971177 | 0.518517753 | 0.405538958 | 0.876090066 | 0.868416065 |
| Afdn | S | 1140 | 7.02729E-05 | -0.124440265 | -0.069201849 | -0.069272122 | 0.055238416 | 0.998258377 | 0.031573336 | 0.089233941 | 0.120831725 | 0.240645243 |

| | | | | | | | | | | | | |
|---|---|---|---|---|---|---|---|---|---|---|---|---|
| Ampd2 | S | 37 | -0.104617808 | 0.018342218 | -0.336899897 | -0.232282089 | -0.355242115 | 0.437638148 | 0.893191485 | 0.138918205 | 0.262861383 | 0.127219549 |
| Hnrnpc | S | 249 | -0.047146505 | -0.116307909 | -0.065402572 | -0.018256067 | 0.050905337 | 0.512206723 | 0.042851038 | 0.299123584 | 0.79889418 | 0.359901854 |
| Kif18a | S | 695 | -0.007505473 | -0.042589213 | -0.095943375 | -0.088437903 | -0.053354162 | 0.920159754 | 0.178227967 | 0.002145645 | 0.305613038 | 0.10699309 |
| Dab2 | S | 328 | 0.07894067 | -0.172825814 | -0.28430119 | -0.36324186 | -0.111475376 | 0.599893986 | 0.157961621 | 0.178644895 | 0.128585078 | 0.548626619 |
| Sox13 | S | 377 | -0.202121413 | -0.086736823 | -0.059734109 | 0.142387304 | 0.027002714 | 0.135539431 | 0.463550169 | 0.618109324 | 0.07792839 | 0.511420929 |
| Trmo | S | 220 | -0.237700361 | -0.106043753 | -0.117641208 | 0.120059154 | -0.011597454 | 0.364113302 | 0.458510185 | 0.499864116 | 0.651352452 | 0.949260057 |
| Trmo | T | 219 | -0.237700361 | -0.106043753 | -0.117641208 | 0.120059154 | -0.011597454 | 0.364113302 | 0.458510185 | 0.499864116 | 0.651352452 | 0.949260057 |
| Gga3 | S | 159 | 0.075349464 | -0.178598627 | -0.122413686 | -0.19776315 | 0.056184941 | 0.532716869 | 0.071765538 | 0.055421375 | 0.172864725 | 0.447655222 |
| Dock1 | T | 1767 | -0.077460938 | -0.034788534 | -0.095678935 | -0.018217997 | -0.060890401 | 0.050425628 | 0.517723315 | 0.14447407 | 0.715812809 | 0.357810881 |
| Myo1c | S | 408 | -0.056109456 | -0.039246503 | -0.081645842 | -0.025536386 | -0.04239934 | 0.156781565 | 0.359801005 | 0.028159783 | 0.492041496 | 0.344769413 |
| Cwc22 | S | 809 | -0.168687219 | -0.126214985 | -0.037870972 | 0.130816247 | 0.088344013 | 0.122331101 | 0.368728215 | 0.718120682 | 0.223710656 | 0.514817885 |
| Arhgap12 | Y | 241 | -0.158612253 | -0.041276508 | -0.090313528 | 0.068298725 | -0.04903702 | 0.049635559 | 0.758252678 | 0.461507028 | 0.578737401 | 0.763305273 |
| Dido1 | S | 887 | -0.14458835 | -0.196199582 | -0.029013132 | 0.115575218 | 0.16718645 | 0.257017013 | 0.228433766 | 0.803651408 | 0.245582731 | 0.255537106 |
| Flnb | S | 2481 | -0.098633595 | -0.016070975 | -0.104366443 | -0.005732847 | -0.088295468 | 0.111681443 | 0.741633162 | 0.070518239 | 0.891692457 | 0.038171391 |
| Anapc4 | S | 777 | -0.052090607 | -0.08312675 | -0.017541806 | 0.034548801 | 0.065584944 | 0.019590247 | 0.093018478 | 0.221928999 | 0.080112372 | 0.147668198 |
| Zeb1 | S | 299 | 0.066106026 | -0.07233265 | -0.214618012 | -0.280724038 | -0.142285362 | 0.505860451 | 0.279533132 | 0.012707965 | 0.046461714 | 0.067673393 |
| Miip | S | 307 | -0.061485405 | -0.071591341 | -0.104873886 | -0.043388481 | -0.033282545 | 0.430705934 | 0.184411717 | 0.107731999 | 0.565253813 | 0.508990935 |
| Zfyve19 | S | 280 | -0.130500423 | -0.053665388 | -0.125936796 | 0.004563627 | -0.072271408 | 0.151149308 | 0.626489705 | 0.27451987 | 0.965621445 | 0.572173499 |
| Cux1 | S | 1265 | 0.012666012 | -0.119151937 | -0.052273773 | -0.064939785 | 0.066878164 | 0.599884714 | 0.003350136 | 0.150866525 | 0.094319189 | 0.077928674 |
| Tacc2 | S | 771 | -0.053616666 | -0.074521935 | -0.042872132 | 0.010744535 | 0.031649803 | 0.335573906 | 0.010204637 | 0.287297652 | 0.846716279 | 0.402947003 |
| Otud4 | S | 1017 | -0.042584725 | -0.03859539 | -0.032141529 | 0.010443196 | 0.006453861 | 0.006264415 | 0.146437668 | 0.041152103 | 0.331748121 | 0.764657878 |
| Fam193a | S | 672 | 0.200391104 | -0.219282112 | -0.268492596 | -0.4688837 | -0.049210483 | 0.038647446 | 0.038239571 | 0.020955859 | 0.003791472 | 0.545107431 |
| Fam193a | S | 683 | 0.200391104 | -0.219282112 | -0.268492596 | -0.4688837 | -0.049210483 | 0.038647446 | 0.038239571 | 0.020955859 | 0.003791472 | 0.545107431 |
| Rab12 | S | 20 | -0.020027869 | -0.048450121 | -0.117642199 | -0.09761433 | -0.069192078 | 0.589909365 | 0.223270241 | 0.018600891 | 0.016981092 | 0.038528669 |
| Fkbp15 | S | 342 | -0.081582382 | -0.045085123 | -0.208703973 | -0.12712159 | -0.16361885 | 0.74104255 | 0.670449843 | 0.060061223 | 0.60353844 | 0.092635547 |
| Gpr176 | S | 364 | -0.048078114 | -0.385683474 | 0.126626334 | 0.174704448 | 0.512309808 | 0.538075574 | 0.00919613 | 0.186995423 | 0.057100578 | 0.005656871 |
| Usp5 | T | 623 | -0.102271416 | -0.01302349 | -0.196233712 | -0.093962295 | -0.183210222 | 0.169065591 | 0.836389223 | 0.178286024 | 0.45218743 | 0.208110192 |
| Prkab1 | S | 182 | -0.040579769 | -0.008952506 | -0.177058616 | -0.136478847 | -0.168106111 | 0.459259769 | 0.77464818 | 0.020302337 | 0.073969501 | 0.042319242 |
| Ubr4 | S | 2719 | -0.15578097 | -0.086031399 | -0.066389542 | 0.089391427 | 0.019641857 | 0.156764184 | 0.437030219 | 0.38603543 | 0.399377833 | 0.86243231 |
| Fam129b | S | 628 | -0.079520046 | 0.028954493 | -0.154961919 | -0.075441873 | -0.183916412 | 0.102129187 | 0.473744054 | 0.020443238 | 0.155139163 | 0.019531145 |
| Arhgap17 | S | 671 | -0.149921515 | -0.083460172 | -0.075739725 | 0.07418179 | 0.007720447 | 0.164288747 | 0.391881528 | 0.425157851 | 0.250610964 | 0.862539992 |
| Map4k4 Mink1 | T | 186;187 | -0.21930843 | -0.122611696 | -0.063723931 | 0.155584499 | 0.058887765 | 0.352631657 | 0.245697132 | 0.664219977 | 0.506944902 | 0.648747735 |
| Araf | S | 214 | -0.175709852 | -0.038154645 | 0.009467254 | 0.185177106 | 0.047621899 | 0.006227183 | 0.552145525 | 0.834593876 | 0.013195095 | 0.475402514 |
| Supt6 | T | 1530 | -0.024158081 | -0.11638718 | -0.195900795 | -0.171742714 | -0.079513615 | 0.621953689 | 0.223929248 | 0.239689428 | 0.287246323 | 0.603525393 |
| Tsc1 | S | 1094 | -0.20543769 | -0.097815875 | -0.117288373 | 0.088149316 | -0.019472499 | 0.261369351 | 0.535618571 | 0.510163839 | 0.631856611 | 0.907039957 |
| Dag1 | T | 788 | -0.16730105 | 0.060143803 | -0.263481026 | -0.096179976 | -0.32362483 | 0.084587981 | 0.73746464 | 0.323281525 | 0.688535831 | 0.277823852 |
| Jade3 | S | 578 | -0.019612974 | -0.209341386 | -0.011033075 | 0.008579899 | 0.198308311 | 0.798346627 | 0.014365735 | 0.838990921 | 0.884965509 | 0.033484844 |
| Srsf10 | S | 209 | -0.101245816 | -0.178040474 | -0.078305207 | 0.022940609 | 0.099735267 | 0.410212834 | 0.149554418 | 0.688875511 | 0.897988567 | 0.583801145 |
| Arhgap5 | S | 1088 | 0.067076452 | -0.341461885 | -0.140182739 | -0.20725919 | 0.201279147 | 0.427500072 | 0.128717052 | 0.229745243 | 0.107174972 | 0.310452576 |
| Clspn | S | 990 | 0.057171797 | -0.146356274 | -0.169081223 | -0.22625302 | -0.022724949 | 0.424377809 | 0.092465468 | 0.063901531 | 0.034958378 | 0.778701709 |
| Clspn | S | 996 | 0.057171797 | -0.146356274 | -0.169081223 | -0.22625302 | -0.022724949 | 0.424377809 | 0.092465468 | 0.063901531 | 0.034958378 | 0.778701709 |
| Kifap3 | S | 60 | -0.03880426 | -0.032427117 | -0.117971085 | -0.079166824 | -0.085543968 | 0.670142801 | 0.462131232 | 0.006777391 | 0.411730206 | 0.09714061 |
| Lrba | S | 2489 | 0.206752348 | -0.402097471 | -0.437417811 | -0.644170159 | -0.03532034 | 0.320012562 | 0.249637096 | 0.241209869 | 0.124956904 | 0.928816174 |
| Arhgap23 | S | 550 | 0.012043389 | -0.241086059 | -0.110574677 | -0.122618065 | 0.130511382 | 0.928390483 | 0.171394271 | 0.191853185 | 0.378364162 | 0.398101876 |
| Spire1 | S | 577 | -0.107824155 | 0.028003125 | -0.206262238 | -0.098438083 | -0.234265363 | 0.19101756 | 0.769468265 | 0.08525954 | 0.309031591 | 0.071954704 |
| Rrp1 | S | 438 | 0.016400241 | -0.183403025 | -0.139665389 | -0.156065629 | 0.043737636 | 0.853625823 | 0.163807432 | 0.139741246 | 0.166226607 | 0.711087496 |
| Hecw2 | T | 50 | 0.084683809 | -0.182611037 | -0.099345407 | -0.184029216 | 0.08326563 | 0.57824101 | 0.092451964 | 0.025146019 | 0.285327119 | 0.331739346 |
| Tax1bp1 | S | 693 | 0.075724316 | -0.097274064 | -0.145280959 | -0.221005275 | -0.048006896 | 0.340711873 | 0.005059826 | 0.072725479 | 0.049281825 | 0.418641511 |
| Rtn4 | S | 836 | 0.010871453 | -0.115369613 | -0.05083484 | -0.061706293 | 0.064534772 | 0.715984712 | 0.004100657 | 0.146653021 | 0.072169531 | 0.059705591 |
| Gas2l3 | S | 377 | 0.016862279 | -0.032104963 | -0.216537013 | -0.233399292 | -0.18443205 | 0.743363544 | 0.901389848 | 0.00551373 | 0.006245077 | 0.501207038 |
| Mga | S | 604 | 0.057808619 | -0.126024207 | -0.169138278 | -0.226946897 | -0.043114071 | 0.366792254 | 0.103788483 | 0.034486798 | 0.025631811 | 0.532329671 |
| Chd1 | S | 214 | 0.004289684 | -0.123309899 | -0.041435826 | -0.04572551 | 0.081874073 | 0.889721866 | 0.009663571 | 0.092607699 | 0.209208023 | 0.03987217 |
| Chd1 | S | 215 | 0.004289684 | -0.123309899 | -0.041435826 | -0.04572551 | 0.081874073 | 0.889721866 | 0.009663571 | 0.092607699 | 0.209208023 | 0.03987217 |
| Bclaf1 | T | 305 | -0.01737246 | -0.050114418 | -0.09837812 | -0.08100566 | -0.048263702 | 0.451906076 | 0.190097429 | 0.015152147 | 0.038722903 | 0.222091775 |
| Slain2 | S | 252 | -0.038245546 | -0.044564361 | -0.21295492 | -0.174709374 | -0.168390559 | 0.462491488 | 0.697858281 | 0.071585404 | 0.122628742 | 0.233633266 |
| Atxn1l | S | 62 | -0.004061205 | -0.217061451 | 0.04036519 | 0.044426395 | 0.257426642 | 0.948759755 | 0.002538718 | 0.756345366 | 0.735086528 | 0.13800209 |
| Lrrtm2 | S | 458 | -0.056627812 | -0.034542354 | -0.227426213 | -0.170798401 | -0.192883858 | 0.572262722 | 0.738778914 | 0.075213209 | 0.198861633 | 0.168240157 |
| Cfap36 | S | 85 | 0.029101713 | -0.165884741 | -0.14846064 | -0.177562353 | 0.017424101 | 0.749637176 | 0.113680582 | 0.142955916 | 0.057526827 | 0.821203661 |
| Abcf1 | S | 90 | -0.086670962 | -0.008507967 | -0.122237746 | -0.035566783 | -0.113729779 | 0.059014711 | 0.627427984 | 0.142556406 | 0.591328617 | 0.160531863 |
| Trp53bp1 | S | 1675 | 0.000221566 | -0.089649764 | -0.138945295 | -0.139166861 | -0.049295531 | 0.996374248 | 0.207410143 | 0.038949326 | 0.060599408 | 0.479410199 |
| Map1b | S | 888 | -0.062790809 | -0.032885297 | -0.105939145 | -0.043148336 | -0.073053848 | 0.25256937 | 0.590482989 | 0.02642917 | 0.421668991 | 0.283944609 |
| Ist1 | S | 238 | 0.106131782 | -0.201513946 | -0.157839553 | -0.263971336 | 0.043674392 | 0.363964382 | 0.244167975 | 0.011391226 | 0.089079348 | 0.762422108 |
| Mtdh | S | 308 | 0.091726697 | -0.108728803 | -0.190364727 | -0.282091424 | -0.081635924 | 0.124414399 | 0.12736061 | 0.004716945 | 0.003365991 | 0.197025309 |
| Micall2 | S | 143 | 0.006776526 | -0.073061808 | -0.080602849 | -0.087379375 | -0.007541041 | 0.652565423 | 0.002614045 | 0.09022791 | 0.091980211 | 0.82432439 |
| Ube3a | S | 213 | -0.003819952 | -0.140587952 | -0.028679745 | -0.024859793 | 0.111908207 | 0.899647143 | 0.003168066 | 0.577305305 | 0.643720433 | 0.102332969 |
| Tln1 | S | 2162 | -0.087948404 | -0.171889649 | -0.004529717 | 0.083418687 | 0.167359932 | 0.160298473 | 0.098680239 | 0.966074095 | 0.468666536 | 0.216338082 |
| Ank3 | S | 1813 | -0.000187993 | -0.103598107 | -0.139149213 | -0.138961219 | -0.035551106 | 0.997046763 | 0.060677778 | 0.160249943 | 0.163752635 | 0.649904472 |
| Nes | S | 1837 | -0.022367225 | -0.16115309 | -0.072566585 | -0.05019936 | 0.088586505 | 0.759034494 | 0.055486705 | 0.331707924 | 0.232800337 | 0.047963137 |
| Ralgapa1 | S | 863 | -0.136731371 | 0.020539777 | -0.077850207 | 0.058881164 | -0.098389984 | 0.005357261 | 0.696386358 | 0.352911047 | 0.456358264 | 0.272445256 |
| Zfyve16 | S | 884 | 0.004870476 | -0.089527181 | -0.103110894 | -0.10798137 | -0.013583713 | 0.906305816 | 0.012580234 | 0.153886333 | 0.147559271 | 0.808300213 |
| Nop56 | S | 543 | 0.05065053 | -0.115822851 | -0.098811853 | -0.149462383 | 0.017010997 | 0.182239543 | 0.00794816 | 0.030088338 | 0.00871545 | 0.58051067 |
| Rabgap1 | T | 991 | -0.155197641 | 0.002739503 | -0.043678443 | 0.111519198 | -0.046417946 | 0.006113699 | 0.964959733 | 0.543317928 | 0.171760251 | 0.559532002 |
| Trp53bp1 | S | 654 | -0.137433369 | -0.073618077 | -0.082175721 | 0.055257648 | -0.008557644 | 0.271243082 | 0.181752332 | 0.426038709 | 0.672016653 | 0.924600378 |
| Zc3h11a | S | 132 | -0.053317205 | -0.12571508 | -0.111371681 | -0.058054476 | 0.014343399 | 0.502934752 | 0.152267306 | 0.303090449 | 0.508601158 | 0.868121143 |
| Plekhg3 | S | 862 | -0.025056409 | -0.026579987 | -0.229420807 | -0.204364399 | -0.202840821 | 0.543374544 | 0.807971776 | 0.04507434 | 0.073482588 | 0.154353349 |
| Rtkn2 | S | 196 | -0.034594797 | -0.005534433 | -0.211660978 | -0.17706618 | -0.206126545 | 0.48685889 | 0.939012026 | 0.029029424 | 0.068578166 | 0.050772712 |
| Mepce | T | 188 | -0.157425778 | 0.030612411 | -0.042512067 | 0.114913711 | -0.073124478 | 0.004750043 | 0.681039348 | 0.280145184 | 0.017892144 | 0.348709608 |
| Pikfyve | S | 1189 | -0.190719259 | 0.035832926 | -0.052544936 | 0.138174323 | -0.088377862 | 0.005547943 | 0.535914968 | 0.481359202 | 0.10953918 | 0.251569046 |
| Bloc1s3 | T | 59 | -0.065141497 | -0.070584498 | 0.001303086 | 0.066444583 | 0.071887584 | 0.088991723 | 0.002422424 | 0.961205984 | 0.106525564 | 0.069296997 |
| Sipa1l3 | S | 1541 | -0.002767117 | -0.100971089 | -0.080887924 | -0.078120807 | 0.020083165 | 0.944741384 | 0.055804203 | 0.051914062 | 0.055445011 | 0.585120145 |
| Nedd4 | S | 309 | -0.098987468 | -0.040911071 | -0.14589604 | -0.046908573 | -0.10498497 | 0.247907245 | 0.679698146 | 0.131691263 | 0.35713212 | 0.215557188 |
| Bin1 | S | 332 | -0.054098443 | -0.062589354 | -0.108699507 | -0.054601064 | -0.046110153 | 0.530425835 | 0.270749708 | 0.050462679 | 0.518139268 | 0.375876456 |
| Gphn | S | 200 | -0.030317922 | -0.155527305 | -0.04483003 | -0.014512109 | 0.110697275 | 0.512517266 | 0.050795417 | 0.347822461 | 0.618078879 | 0.12321258 |
| Srrm2 | S | 1473 | 0.039518046 | -0.086629132 | -0.114539551 | -0.154057597 | -0.027910419 | 0.390302041 | 0.024458137 | 0.020722537 | 0.020928732 | 0.380044509 |
| Adnp | S | 607 | -0.044923784 | -0.075112609 | -0.089344016 | -0.044420232 | -0.014231407 | 0.268577887 | 0.092444315 | 0.208051486 | 0.471250778 | 0.809667463 |
| Ehbp1l1 | S | 1135 | -0.080008606 | -0.041074118 | -0.266522452 | -0.186513846 | -0.225448334 | 0.288281227 | 0.686571949 | 0.258763667 | 0.394083488 | 0.324872314 |
| Rtn4 | S | 839 | 0.012920328 | -0.113545045 | -0.045088128 | -0.058008456 | 0.068456917 | 0.692321526 | 0.003066709 | 0.139013823 | 0.116490108 | 0.038835075 |
| Pacs2 | S | 424 | -0.094069095 | 0.044728361 | -0.171856105 | -0.077787011 | -0.216584466 | 0.167173177 | 0.570388246 | 0.026930838 | 0.131340172 | 0.032126481 |
| Chd4 | T | 1682 | -0.1105123 | -0.135408112 | -0.066189391 | 0.044322909 | 0.069218722 | 0.432330022 | 0.106068436 | 0.598920635 | 0.77727887 | 0.568361545 |
| Nbea | S | 2565 | -0.040823952 | -0.158033471 | -0.190072285 | -0.149248333 | -0.032038815 | 0.768331494 | 0.157634114 | 0.438750166 | 0.536409237 | 0.883456252 |
| Sphk2 | S | 364 | 0.013103092 | -0.102142418 | -0.15703548 | -0.170138571 | -0.054893062 | 0.763702253 | 0.03457617 | 0.187430602 | 0.16660149 | 0.579420463 |
| Sphk2 | S | 379 | 0.013103092 | -0.102142418 | -0.15703548 | -0.170138571 | -0.054893062 | 0.763702253 | 0.03457617 | 0.187430602 | 0.16660149 | 0.579420463 |
| Otud7b | S | 467 | -0.118348575 | -0.015096101 | -0.051916434 | 0.066432141 | -0.036820333 | 0.007534087 | 0.830299997 | 0.374367049 | 0.263841686 | 0.6421481 |
| Mast4 | S | 2353 | 0.129851727 | -0.095730788 | -0.222927809 | -0.352779536 | -0.12719702 | 0.224812051 | 0.034043678 | 0.024228203 | 0.024145351 | 0.085168583 |

| | | | | | | | | | | | | |
|---|---|---|---|---|---|---|---|---|---|---|---|---|
| Sox2 | S | 253 | 0.109403679 | -0.177563684 | -0.218566839 | -0.327970518 | -0.041003155 | 0.180756624 | 0.072487309 | 0.058380687 | 0.026309628 | 0.636921529 |
| Slc20a2 | S | 253 | 0.130095598 | -0.173082281 | -0.052570913 | -0.182666511 | 0.120511368 | 0.336859147 | 0.000286466 | 0.378744011 | 0.214586381 | 0.110911005 |
| Frmd8 | S | 21 | -0.185396274 | 0.042418204 | -0.154677095 | 0.03071918 | -0.197095298 | 0.026137032 | 0.617750493 | 0.435728925 | 0.863777846 | 0.338125707 |
| Stom | S | 7 | -0.111940864 | 0.074286862 | -0.231723044 | -0.11978218 | -0.306009906 | 0.425106199 | 0.417244089 | 0.015503445 | 0.404292485 | 0.030483828 |
| Ice1 | T | 674 | -0.00578732 | -0.142364051 | -0.160957428 | -0.155170109 | -0.018593377 | 0.851806449 | 0.142091294 | 0.239750247 | 0.252003362 | 0.882417104 |
| Ift43 | S | 77 | -0.36659636 | 0.114799153 | -0.00522912 | 0.36136724 | -0.120028274 | 0.011888755 | 0.583743036 | 0.978077437 | 0.128878076 | 0.616969119 |
| Arhgef7 | S | 795 | -0.120914279 | -0.133505368 | -0.011620417 | 0.109293863 | 0.121884951 | 0.148470015 | 0.125921755 | 0.92721604 | 0.437848823 | 0.394099731 |
| Map1b | S | 825 | -0.02510463 | -0.128947806 | -0.020845084 | 0.004259546 | 0.108102722 | 0.674702674 | 0.003589454 | 0.713327023 | 0.954653263 | 0.144741501 |
| Kirrel | S | 674 | -0.106256409 | -0.006523489 | -0.121212849 | -0.014956439 | -0.11468936 | 0.060826865 | 0.952005898 | 0.159314204 | 0.837745613 | 0.368198165 |
| Srsf6 | S | 295 | 0.016652149 | -0.152977005 | -0.083539849 | -0.100191998 | 0.069437156 | 0.669521169 | 0.024868939 | 0.20373595 | 0.142682123 | 0.295314231 |
| Rsrc1 | S | 61 | -0.050920487 | -0.044419014 | -0.053958758 | -0.003038271 | -0.009539745 | 0.144818632 | 0.066094923 | 0.075884182 | 0.92085288 | 0.660832164 |
| Chaf1a | S | 696 | -0.095987273 | -0.119406938 | -0.108187973 | -0.0122007 | 0.011218964 | 0.226395552 | 0.232672165 | 0.590345723 | 0.950675507 | 0.955588118 |
| Sipa1l1 | S | 1195 | -0.199848634 | 0.03919432 | -0.044265975 | 0.155582659 | -0.083460296 | 0.013697406 | 0.510024805 | 0.192892252 | 0.030922883 | 0.216361544 |
| Thoc2 | S | 1448 | -0.040026902 | -0.06141947 | -0.079673124 | -0.039646222 | -0.018253654 | 0.278597348 | 0.111449091 | 0.088207289 | 0.351266527 | 0.647722818 |
| Tmbim1 | S | 81 | -0.003462549 | -0.081539897 | -0.07623625 | -0.072773701 | 0.005303647 | 0.937880717 | 0.061450354 | 0.021551062 | 0.164056728 | 0.867064292 |
| Map4k4 | S | 706 | 0.007460116 | -0.086562697 | -0.087137805 | -0.094597921 | -0.000575109 | 0.901450834 | 0.041943769 | 0.036832503 | 0.183560526 | 0.985494566 |
| Usp37 | S | 212 | 0.091597151 | -0.092066897 | -0.149619168 | -0.241216319 | -0.057552271 | 0.030882239 | 0.009004448 | 0.002788241 | 0.001504192 | 0.104510652 |
| Srek1ip1 | S | 94 | 0.079218159 | -0.107154226 | -0.165264607 | -0.244482765 | -0.058110381 | 0.106742574 | 0.034933551 | 0.012171095 | 0.005140842 | 0.236032501 |
| Nufip2 | T | 569 | 0.031586925 | -0.109167577 | -0.09118618 | -0.122773105 | 0.017981397 | 0.391478839 | 0.015586319 | 0.046050416 | 0.012226414 | 0.519517007 |
| Nav1 | S | 1010 | -0.05918112 | -0.123943263 | -0.020009263 | 0.039171857 | 0.103933999 | 0.249654281 | 0.021742116 | 0.855824307 | 0.719476678 | 0.378578857 |
| Otud7b | S | 449 | 0.039210836 | -0.152950497 | -0.148558378 | -0.187769213 | 0.004392119 | 0.492265788 | 0.013494184 | 0.365985822 | 0.277281423 | 0.976082297 |
| Med8 | S | 82 | 0.025197423 | -0.053386306 | -0.148387102 | -0.173584525 | -0.095000796 | 0.432541415 | 0.260888073 | 0.003835561 | 0.005799089 | 0.093782042 |
| Bloc1s3 | S | 61 | -0.070811315 | -0.073699564 | 0.002235754 | 0.073047068 | 0.075935318 | 0.06861176 | 0.00654488 | 0.920760126 | 0.068962826 | 0.029098164 |
| Tnrc6c | S | 714 | 0.022674319 | -0.073992522 | -0.118217001 | -0.140891321 | -0.044224479 | 0.748374577 | 0.09772393 | 0.016496101 | 0.118901058 | 0.169178963 |
| Cep41 | S | 357 | -0.082295166 | -0.127493712 | -0.13718076 | -0.054885595 | -0.009687048 | 0.669573202 | 0.210472262 | 0.307112181 | 0.790202027 | 0.940636779 |
| Cavin1 | S | 169 | 0.045446673 | -0.05976891 | -0.181370193 | -0.226816866 | -0.121601283 | 0.433876593 | 0.284831408 | 0.006858915 | 0.009826259 | 0.04755408 |
| Ranbp2 | T | 1602 | -0.112535522 | -0.166777798 | -0.001211496 | 0.111324026 | 0.165566302 | 0.109442899 | 0.190201344 | 0.985292857 | 0.038713793 | 0.192452356 |
| Apex1 | S | 18 | -0.117457603 | 0.019874988 | -0.074509987 | 0.042947616 | -0.094384975 | 0.01193262 | 0.722026938 | 0.132046766 | 0.2783017 | 0.147165072 |
| Uri1 | S | 440 | -0.210026418 | -0.024191043 | -0.202739992 | 0.007286426 | -0.178548949 | 0.336930612 | 0.831967741 | 0.314906906 | 0.975758139 | 0.374084608 |
| Mki67 | T | 1363 | -0.018274673 | -0.03467956 | -0.082631106 | -0.064356433 | -0.047951547 | 0.688879687 | 0.220075772 | 0.001993918 | 0.23307438 | 0.128958315 |
| Bag6 | S | 986 | -0.140733277 | -0.048055746 | -0.042302367 | 0.09843091 | 0.005753379 | 0.049791627 | 0.347606105 | 0.574093726 | 0.265816955 | 0.942282142 |
| Cttnbp2nl | S | 562 | 0.002557819 | -0.127282919 | -0.04049765 | -0.04305547 | 0.086785269 | 0.938351092 | 0.005492044 | 0.280037308 | 0.283979715 | 0.060937864 |
| Nedd4l | T | 347 | -0.083658454 | -0.046971178 | -0.09109848 | -0.007440026 | -0.044127302 | 0.263672401 | 0.342878556 | 0.08871709 | 0.908548836 | 0.321080472 |
| Map7d1 | S | 315 | 0.035725311 | -0.014408899 | -0.389794796 | -0.425520107 | -0.375385897 | 0.805685387 | 0.940404171 | 0.048604487 | 0.046087878 | 0.12070092 |
| Racgap1 | T | 561 | 0.037663133 | -0.052865072 | -0.164845428 | -0.202508562 | -0.111980357 | 0.549807118 | 0.156633158 | 0.010742016 | 0.03215584 | 0.053460004 |
| Nolc1 | S | 696 | -0.002519227 | -0.077792082 | -0.196897425 | -0.194378198 | -0.119105343 | 0.959860194 | 0.546658758 | 0.038595931 | 0.043306079 | 0.392129422 |
| Srrm2 | S | 1495 | -0.005624973 | -0.123995129 | -0.113210153 | -0.10758518 | 0.010784976 | 0.937116593 | 0.071256733 | 0.187131833 | 0.262299705 | 0.889814909 |
| Klf16 | S | 113 | -0.021206587 | -0.132130856 | -0.200095843 | -0.178889256 | -0.067964987 | 0.794429645 | 0.431956076 | 0.143363128 | 0.191340444 | 0.699450555 |
| Lysmd3 | S | 53 | -0.139003358 | -0.051930358 | -0.002503236 | 0.136500122 | 0.049427122 | 0.007634513 | 0.500782595 | 0.931900713 | 0.014232221 | 0.513151121 |
| Rnps1 | S | 52 | -0.061729843 | -0.069888888 | -0.036323653 | 0.02540619 | 0.033565235 | 0.051879633 | 0.049812622 | 0.5829588 | 0.683201381 | 0.602013654 |
| Anln | S | 259 | 0.014254767 | -0.054061438 | -0.197518903 | -0.21177367 | -0.143457464 | 0.811858135 | 0.364173945 | 0.026678514 | 0.024656404 | 0.070195597 |
| Ptpn12 | S | 331 | -0.057167775 | -0.048305341 | -0.112715716 | -0.055547941 | -0.064410375 | 0.196423747 | 0.256921487 | 0.148208066 | 0.404147915 | 0.343162399 |
| Zfand2b | T | 185 | 0.003280896 | -0.207665105 | -0.116183908 | -0.119464804 | 0.091481197 | 0.974357357 | 0.075468023 | 0.457094655 | 0.458397427 | 0.558587297 |
| Larp7 | T | 260 | -0.070419631 | -0.050944125 | -0.052437018 | 0.017982613 | -0.001492893 | 0.019734431 | 0.46443058 | 0.177260186 | 0.589170006 | 0.982369391 |
| Magi3 | S | 1445 | -0.133244236 | -0.050326324 | -0.098026189 | 0.035218046 | -0.047699866 | 0.158073348 | 0.574315633 | 0.263174115 | 0.667446459 | 0.574218549 |
| Phf23 | S | 315 | -0.088100069 | -0.002196262 | -0.111753781 | -0.023653711 | -0.109557518 | 0.019956661 | 0.939757545 | 0.197980418 | 0.736152354 | 0.212149199 |
| Fmr1 | S | 602 | -0.1291147 | 0.031596678 | -0.09991929 | 0.02919541 | -0.131515967 | 0.020330489 | 0.50619704 | 0.118560289 | 0.53029064 | 0.053636636 |
| Nav1 | S | 582 | -0.050559223 | -0.083910219 | -0.172585094 | -0.122025871 | -0.088674875 | 0.700681105 | 0.174861093 | 0.22487244 | 0.457697307 | 0.478137266 |
| Map1b | S | 992 | -0.019284045 | -0.096031644 | -0.086432268 | -0.067148223 | 0.009599376 | 0.731652786 | 0.125265003 | 0.05576039 | 0.276503092 | 0.853293566 |
| Prrc2a | T | 782 | 0.019248681 | -0.16864927 | -0.049551906 | -0.068800586 | 0.119097364 | 0.53421881 | 0.010163234 | 0.272333432 | 0.142004707 | 0.053994956 |
| Map3k4 | S | 1240 | 0.035398283 | -0.124994014 | -0.1327044 | -0.168102683 | -0.007710385 | 0.726091378 | 0.08892667 | 0.079547133 | 0.178520846 | 0.912042511 |
| Lrrfip1 | S | 658 | -0.044139243 | -0.192363135 | -0.188600173 | -0.14446093 | 0.003762962 | 0.861617819 | 0.424304353 | 0.218161366 | 0.591904729 | 0.987244826 |
| Bag6 | T | 985 | -0.122830273 | -0.027343457 | -0.054929973 | 0.0679003 | -0.027586516 | 0.030609654 | 0.555651631 | 0.34614628 | 0.27951021 | 0.645747539 |
| Osbpl3 | S | 33 | -0.162915941 | -0.005503817 | -0.074878402 | 0.088037539 | -0.069374585 | 0.041887374 | 0.945743549 | 0.336476906 | 0.151085044 | 0.343144322 |
| Sp100 | T | 313 | -0.12123716 | -0.084721009 | -0.115821333 | 0.005415827 | -0.031100324 | 0.231324223 | 0.537346238 | 0.304058601 | 0.951494352 | 0.810721032 |
| Atad2 | S | 780 | -0.073876094 | -0.121532163 | -0.09309506 | -0.019218967 | 0.028437103 | 0.349368734 | 0.211852656 | 0.359039299 | 0.819235283 | 0.768486575 |
| Dennd5a | S | 193 | -0.239121106 | 0.032540026 | -0.072134822 | 0.166986284 | -0.104674848 | 0.034085028 | 0.776881272 | 0.534328609 | 0.133280468 | 0.362954826 |
| Dck | T | 73 | -0.145164613 | -0.145268252 | -0.018256892 | 0.126907721 | 0.12701136 | 0.112548458 | 0.336871654 | 0.820113793 | 0.132327176 | 0.384387726 |
| Pcyt1a | S | 322 | -0.093932484 | 0.038179495 | -0.095834829 | -0.001902345 | -0.134014324 | 0.024881527 | 0.228905582 | 0.009394274 | 0.950098896 | 0.008699424 |
| Stim2 | S | 719 | -0.102014431 | -0.071705171 | -0.088165584 | 0.013848846 | -0.016460414 | 0.075157222 | 0.539125893 | 0.369542323 | 0.879227845 | 0.901257627 |
| Ubap2l | S | 436 | 0.080764542 | -0.119475907 | -0.220211952 | -0.300976494 | -0.100736045 | 0.166221153 | 0.240354946 | 0.011920725 | 0.011672091 | 0.303371346 |
| Fnip1 | S | 170 | -0.158012926 | -0.084656859 | -0.040745291 | 0.117267635 | 0.043911568 | 0.273983498 | 0.082131079 | 0.755620735 | 0.488782407 | 0.730126385 |
| Rrp1 | S | 490 | 0.001619106 | 0.000913296 | -0.223080474 | -0.22469958 | -0.22399377 | 0.980112463 | 0.987434634 | 0.008762824 | 0.005825261 | 0.009803583 |
| Apc | S | 2773 | -0.127507362 | -0.421587798 | 0.185302577 | 0.31280994 | 0.606890376 | 0.437723412 | 0.063358648 | 0.448435074 | 0.207741774 | 0.053132455 |
| Eif5b | S | 132 | -0.000666881 | -0.0764258 | -0.123489791 | -0.12282291 | -0.047063991 | 0.988013282 | 0.243906151 | 0.022059469 | 0.012373266 | 0.405042824 |
| Layn | Y | 380 | -0.158787183 | -0.063955251 | -0.210004965 | -0.051217782 | -0.146049715 | 0.412843102 | 0.517416184 | 0.402264555 | 0.848436705 | 0.546540878 |
| Layn | Y | 381 | -0.158787183 | -0.063955251 | -0.210004965 | -0.051217782 | -0.146049715 | 0.412843102 | 0.517416184 | 0.402264555 | 0.848436705 | 0.546540878 |
| Anxa5 | S | 35 | -0.131691443 | 0.009229427 | -0.20292891 | -0.071237467 | -0.212158337 | 0.106282273 | 0.919988335 | 0.289171816 | 0.669793126 | 0.273222166 |
| Ep400 | S | 321 | -0.02993099 | -0.077623346 | -0.073123908 | -0.043192918 | 0.004499438 | 0.425158775 | 0.018251684 | 0.293776304 | 0.518784192 | 0.940910138 |
| Ripor1 | S | 347 | -0.008952317 | -0.034081719 | -0.088078686 | -0.079126369 | -0.053996967 | 0.776552422 | 0.369745191 | 0.000921025 | 0.078873415 | 0.199231732 |
| Setd2 | S | 1862 | -0.215534785 | -0.082349468 | -0.002447975 | 0.213086809 | 0.079901492 | 0.130487413 | 0.234212978 | 0.974856168 | 0.138885783 | 0.322111577 |
| Ddx20 | S | 569 | -0.026011494 | -0.064690967 | -0.101177272 | -0.075165778 | -0.036486305 | 0.368532889 | 0.085691414 | 0.128194643 | 0.215317875 | 0.507210606 |
| Anxa2 | T | 19 | -0.106235355 | -0.102609708 | 0.009130178 | 0.115365534 | 0.111739887 | 0.072738672 | 0.063895474 | 0.814198227 | 0.066421451 | 0.059809127 |
| Nup188 | S | 1719 | -0.03615481 | -0.077108601 | -0.041250726 | -0.005095916 | 0.035857875 | 0.130008709 | 0.018230511 | 0.425350414 | 0.915997033 | 0.4860294 |
| Dpm1 | S | 13 | 0.03311733 | -0.106906099 | -0.112530542 | -0.145647871 | -0.005624443 | 0.753810651 | 0.032272894 | 0.102892785 | 0.23735373 | 0.909790793 |
| Thrap3 | S | 695 | -0.045856541 | -0.106605843 | -0.031769999 | 0.014086541 | 0.074835844 | 0.208986243 | 0.044134723 | 0.402354189 | 0.718298914 | 0.140689842 |
| Ift172 | S | 342 | -0.29601471 | -0.446779775 | 0.239001838 | 0.535016548 | 0.685781613 | 0.415715428 | 0.22746066 | 0.670682278 | 0.324967927 | 0.23941162 |
| Ift172 | S | 343 | -0.29601471 | -0.446779775 | 0.239001838 | 0.535016548 | 0.685781613 | 0.415715428 | 0.22746066 | 0.670682278 | 0.324967927 | 0.23941162 |
| Ift172 | T | 345 | -0.29601471 | -0.446779775 | 0.239001838 | 0.535016548 | 0.685781613 | 0.415715428 | 0.22746066 | 0.670682278 | 0.324967927 | 0.23941162 |
| L3mbtl2 | S | 13 | -0.369297465 | -0.060660403 | 0.108725864 | 0.478023329 | 0.169386266 | 0.022278124 | 0.770266804 | 0.444271646 | 0.018809018 | 0.439194598 |
| Ehbp1 | S | 659 | 0.178077111 | -0.138075575 | -0.284031581 | -0.462108692 | -0.145956006 | 0.333895296 | 0.305387398 | 0.020549282 | 0.063558193 | 0.249574001 |
| Clip1 | S | 194 | 0.007705193 | -0.04784531 | -0.213366259 | -0.221071451 | -0.165520948 | 0.931111415 | 0.578554245 | 0.02770972 | 0.038233274 | 0.066789083 |
| Pcdh19 | S | 983 | -0.06478423 | -0.085136146 | -0.050457751 | 0.01432648 | 0.034678396 | 0.11223409 | 0.119494409 | 0.412627348 | 0.812643814 | 0.597894796 |
| Phf10 | S | 269 | -0.076698603 | -0.03636145 | -0.083192804 | -0.006494201 | -0.046831354 | 0.066231967 | 0.500862909 | 0.149040334 | 0.882609745 | 0.431317099 |
| Ssr1 | S | 281 | 0.060225124 | -0.124924914 | -0.128779742 | -0.189004866 | -0.003854828 | 0.368859211 | 0.025443869 | 0.073194922 | 0.039291103 | 0.936337016 |
| Ssr1 | S | 284 | 0.060225124 | -0.124924914 | -0.128779742 | -0.189004866 | -0.003854828 | 0.368859211 | 0.025443869 | 0.073194922 | 0.039291103 | 0.936337016 |
| Polr2a | S | 1892 | 0.044972447 | -0.104422849 | -0.127813872 | -0.17278632 | -0.023391023 | 0.404603511 | 0.019004344 | 0.078398305 | 0.043722596 | 0.654422571 |
| Slc39a6 | S | 488 | 0.132886797 | -0.106390661 | -0.258630056 | -0.391516853 | -0.152239394 | 0.094650961 | 0.061674094 | 0.016925862 | 0.006221511 | 0.091805377 |
| Rtn4 | S | 244 | -0.046226708 | -0.100993306 | -0.087040141 | -0.040813432 | 0.013953166 | 0.061342414 | 0.265357923 | 0.389802928 | 0.659903989 | 0.899913035 |
| Akap1 | S | 104 | 0.119362052 | -0.130500676 | -0.197246464 | -0.316608516 | -0.066745788 | 0.097609147 | 0.063051782 | 0.011569933 | 0.002840083 | 0.136315998 |
| Esrra | S | 19 | 0.027201914 | -0.103395842 | -0.142231553 | -0.169433467 | -0.038835711 | 0.270524178 | 0.011785473 | 0.147068694 | 0.117457418 | 0.614301447 |
| Esrra | S | 22 | 0.027201914 | -0.103395842 | -0.142231553 | -0.169433467 | -0.038835711 | 0.270524178 | 0.011785473 | 0.147068694 | 0.117457418 | 0.614301447 |
| Ak2 | T | 195 | -0.064526123 | -0.082210442 | -0.092586113 | -0.02805999 | -0.010375671 | 0.381187182 | 0.480388254 | 0.056428625 | 0.658288294 | 0.922024198 |
| Cep85 | S | 49 | -0.062264728 | -0.128680762 | -0.094037497 | -0.031772768 | 0.034643265 | 0.296922495 | 0.274748154 | 0.318148969 | 0.719098664 | 0.773262381 |

| | | | | | | | | | | | | |
|---|---|---|---|---|---|---|---|---|---|---|---|---|
| Fam110a | S | 188 | -0.11006879 | -0.00172317 | -0.124562907 | -0.014494117 | -0.122839737 | 0.208328006 | 0.975532733 | 0.061888396 | 0.854641754 | 0.100953424 |
| Rptor | S | 877 | -0.020324097 | -0.07142345 | -0.118813125 | -0.098489028 | -0.047389674 | 0.566617535 | 0.220261602 | 0.062299872 | 0.102944468 | 0.441665555 |
| Hspa9 | S | 164 | -0.014357133 | -0.132538166 | -0.135196929 | -0.120839796 | -0.002658764 | 0.857914492 | 0.165863075 | 0.188149485 | 0.160746034 | 0.973358875 |
| Elac2 | S | 200 | -0.010684274 | 0.002482157 | -0.152859397 | -0.142175122 | -0.155341554 | 0.708618276 | 0.937758675 | 0.002349895 | 0.007090168 | 0.00483519 |
| Maml1 | S | 45 | 0.015460723 | -0.126046702 | -0.078588795 | -0.094049518 | 0.047457908 | 0.644941257 | 0.015408863 | 0.180058524 | 0.130190364 | 0.388753051 |
| Pard6g | S | 4 | -0.110694755 | 0.05815853 | -0.190012567 | -0.079317812 | -0.248171098 | 0.140523954 | 0.601152462 | 0.067515705 | 0.322193794 | 0.080583404 |
| Pard6g | S | 8 | -0.110694755 | 0.05815853 | -0.190012567 | -0.079317812 | -0.248171098 | 0.140523954 | 0.601152462 | 0.067515705 | 0.322193794 | 0.080583404 |
| Avl9 | S | 324 | -0.100122719 | -0.174848827 | 0.094764952 | 0.194887671 | 0.269613779 | 0.306947974 | 0.010548028 | 0.581904603 | 0.312196954 | 0.197056096 |
| Acta1 Acta2 Actb Act | S | 52;52;54;54;53;54 | -0.235325094 | -0.045036247 | -0.004498016 | 0.230827079 | 0.040538231 | 0.053538104 | 0.552686443 | 0.94926195 | 0.068448203 | 0.399724502 |
| Cavin1 | S | 239 | -0.325735716 | 0.138555894 | -0.187491132 | 0.138244584 | -0.326047026 | 0.087937209 | 0.459833617 | 0.346116802 | 0.423486701 | 0.122922528 |
| Farp2 | S | 1058 | 0.03061446 | -0.185792461 | -0.00573616 | -0.03635062 | 0.180056301 | 0.658263523 | 0.001502228 | 0.921665118 | 0.633986416 | 0.042510263 |
| Dennd4c | S | 1045 | -0.018183312 | 0.000432712 | -0.28017389 | -0.261990578 | -0.280606602 | 0.795423968 | 0.996182967 | 0.04148866 | 0.05779079 | 0.053566 |
| Kif20a | S | 21 | 0.015534788 | -0.070140669 | -0.185851932 | -0.201386719 | -0.115711263 | 0.868338131 | 0.510822792 | 0.023076513 | 0.094925606 | 0.279689877 |
| Morc2a | S | 741 | 0.019744349 | -0.103847906 | -0.058073391 | -0.07781774 | 0.045774516 | 0.389291869 | 0.005844271 | 0.053326039 | 0.03305729 | 0.128216406 |
| Cnst | S | 279 | 0.031575994 | -0.084745747 | -0.183718723 | -0.215294717 | -0.098972976 | 0.627420743 | 0.250126048 | 0.034825895 | 0.038053831 | 0.23635308 |
| Lrfn4 | S | 585 | 0.027880985 | -0.114488337 | -0.231068677 | -0.258949662 | -0.11658034 | 0.384086721 | 0.256958069 | 0.074299024 | 0.072424103 | 0.329232223 |
| Mcm3ap | S | 1917 | -0.071176231 | -0.077990128 | -0.091809887 | -0.020633655 | -0.013819759 | 0.296789913 | 0.220462915 | 0.227676099 | 0.749798114 | 0.816040832 |
| Mbd3 | S | 86 | 0.003402904 | -0.246545713 | -0.040490309 | -0.043893213 | 0.206055404 | 0.932450188 | 0.065607856 | 0.365744266 | 0.394718254 | 0.085214201 |
| Clint1 | S | 235 | -0.091661776 | -0.08526 | 0.000235846 | 0.091897622 | 0.085495846 | 0.257571555 | 0.006747441 | 0.997025412 | 0.316237905 | 0.253758071 |
| Smarcc1 | S | 800 | -0.142373516 | -0.25423921 | 0.142249028 | 0.284622545 | 0.396488238 | 0.262897608 | 0.026048771 | 0.334432023 | 0.100075944 | 0.051105164 |
| Iqsec1 | S | 159 | -0.095301762 | -0.031321848 | -0.24455391 | -0.149252147 | -0.213232061 | 0.490481331 | 0.699014246 | 0.189790628 | 0.429437255 | 0.237605737 |
| Frk | S | 503 | -0.262086465 | -0.12032034 | 0.027535881 | 0.289622346 | 0.147856222 | 0.214492581 | 0.213372912 | 0.865658393 | 0.224260216 | 0.391295034 |
| Hdac4 Hdac5 Hdac9 2 | S | 265;89;240;270 | -0.143952769 | -0.056543831 | 0.032303867 | 0.176256637 | 0.088847699 | 0.002849107 | 0.445693111 | 0.594667513 | 0.061774432 | 0.304803024 |
| Kmt2d | S | 2597 | -0.041488652 | -0.112921946 | -0.031190994 | 0.010297658 | 0.081730952 | 0.364278209 | 0.027366164 | 0.438771965 | 0.705437806 | 0.023305009 |
| Tor1aip1 | S | 94 | -0.120506041 | 0.026355679 | -0.129610088 | -0.009104047 | -0.155965766 | 0.046812644 | 0.750493551 | 0.167076555 | 0.906219769 | 0.164598913 |
| Son | S | 1870 | 0.067849022 | -0.103547605 | -0.246234733 | -0.314083755 | -0.142687128 | 0.809213237 | 0.506092622 | 0.050989843 | 0.322804273 | 0.381010184 |
| Phax | S | 355 | -0.024132413 | -0.156895563 | -0.06040415 | -0.036271737 | 0.096491413 | 0.852677348 | 0.044201495 | 0.384905292 | 0.766628409 | 0.052233624 |
| Atad3a | S | 320 | -0.039461577 | -0.153088995 | -3.12243E-05 | 0.039430353 | 0.15305777 | 0.620501311 | 0.007631847 | 0.999399544 | 0.600510827 | 0.007603389 |
| Evl | S | 150 | -0.106417087 | 0.008571006 | -0.202028892 | -0.095611805 | -0.210599899 | 0.482485572 | 0.941072789 | 0.063964006 | 0.485027796 | 0.086486676 |
| Fbxw7 | S | 375 | -0.117872025 | -0.073611036 | -0.000899784 | 0.116972241 | 0.072711252 | 0.062836432 | 0.085548198 | 0.98204231 | 0.074435994 | 0.170406494 |
| Chd4 | S | 1301 | -0.214056105 | -0.09028412 | 0.047336415 | 0.26139252 | 0.137620535 | 0.03501633 | 0.279648121 | 0.48253345 | 0.022742977 | 0.105668498 |
| Sirt2 | S | 23 | -0.129345978 | -0.038273357 | -0.037191581 | 0.092154397 | 0.001081775 | 0.050422894 | 0.311874528 | 0.469902285 | 0.169396119 | 0.983193591 |
| Mtmr2 | S | 6 | -0.016713577 | -0.080598205 | -0.098147969 | -0.081434393 | -0.017549764 | 0.823257548 | 0.14035136 | 0.048219856 | 0.298534905 | 0.622322362 |
| Rad51ap1 | S | 276 | -0.016254773 | -0.07617753 | -0.084282672 | -0.068027899 | -0.008105142 | 0.394723043 | 0.053377763 | 0.142887032 | 0.208722305 | 0.865482945 |
| Armcx2 | T | 354 | -0.205847986 | 0.024734349 | -0.076705347 | 0.129142639 | -0.101439697 | 0.036800531 | 0.820375814 | 0.456288161 | 0.177782867 | 0.358675454 |
| Rem1 | S | 18 | -0.21879193 | 0.135249893 | -0.253540635 | -0.034748705 | -0.388790528 | 0.16089718 | 0.327518714 | 0.104887585 | 0.809046019 | 0.037489184 |
| Lipe | S | 559 | -0.334912565 | 0.058122366 | 0.027086051 | 0.361998616 | -0.031036315 | 0.009948346 | 0.585686893 | 0.84921066 | 0.073330842 | 0.841597986 |
| Bmp2k | S | 908 | -0.104805247 | -0.038759784 | -0.054052534 | 0.050752712 | -0.01529275 | 0.039752831 | 0.6559877 | 0.210094854 | 0.104703817 | 0.848680784 |
| Pard3 | S | 695 | -0.016950272 | -0.051593567 | -0.134746214 | -0.117795942 | -0.083152647 | 0.419878609 | 0.288501158 | 0.055408611 | 0.073746737 | 0.18640669 |
| Lrrc8a | S | 217 | 0.173593744 | -0.124608191 | -0.350781233 | -0.524374977 | -0.226173042 | 0.155666666 | 0.111148609 | 0.04482898 | 0.01589791 | 0.137910774 |
| Clspn | S | 998 | 0.072944689 | -0.159056669 | -0.164104306 | -0.237048995 | -0.005047636 | 0.277771849 | 0.052831983 | 0.095977415 | 0.039979164 | 0.950703361 |
| Maged1 | S | 166 | 0.035633446 | -0.108504963 | -0.176443302 | -0.212076748 | -0.06793834 | 0.752909447 | 0.092954181 | 0.137967595 | 0.166329948 | 0.487910231 |
| Csnk1d | T | 355 | -0.064455849 | -0.058351057 | -0.114218825 | -0.049762976 | -0.055867768 | 0.421580091 | 0.17200104 | 0.18193616 | 0.596735311 | 0.456959611 |
| Aarsd1 | S | 409 | -0.064302356 | -0.074751303 | -0.120543038 | -0.056240681 | -0.045791735 | 0.406020243 | 0.214812245 | 0.235046198 | 0.568267006 | 0.603027819 |
| Ring1 | S | 229 | -0.221111034 | -0.046093765 | -0.223483402 | -0.002372367 | -0.177389637 | 0.463297334 | 0.691326395 | 0.398557283 | 0.994493236 | 0.501368736 |
| Tars | S | 8 | -0.183800842 | 0.10732206 | -0.406241458 | -0.222440616 | -0.513563518 | 0.168439428 | 0.397061874 | 0.260753407 | 0.489924632 | 0.190693688 |
| Bclaf3 | S | 210 | -0.115754043 | -0.038618751 | -0.079852381 | 0.035901662 | -0.04123363 | 0.229273036 | 0.494904016 | 0.112423673 | 0.662300017 | 0.416066842 |
| Exosc9 | S | 306 | 0.023544667 | -0.133337659 | -0.122092881 | -0.145637549 | 0.011244778 | 0.178150776 | 0.050949827 | 0.071925924 | 0.061275138 | 0.842832876 |
| Ppp1ca Ppp1cb Ppp1c | S | 129;128;129 | -0.02918456 | -0.166449067 | -0.050856639 | -0.021672079 | 0.115592429 | 0.604220688 | 0.078638253 | 0.375401157 | 0.606792192 | 0.172948671 |
| Rab18 | S | 144 | 0.028221275 | -0.040254971 | -0.167039969 | -0.195261244 | -0.126784999 | 0.563262938 | 0.171029123 | 0.012493615 | 0.018781868 | 0.03646498 |
| Hjurp A730008H23Ril | S | 505 | -0.044128993 | -0.08232342 | -0.200829927 | -0.156700934 | -0.118506506 | 0.70528974 | 0.141520557 | 0.33362584 | 0.456639764 | 0.535656212 |
| Lmnb1 Lmnb2 | S | 394;385 | -0.059371432 | -0.051165286 | -0.032524301 | 0.026847131 | 0.018640984 | 0.233893408 | 0.015199301 | 0.174569448 | 0.546044775 | 0.342720983 |
| Rnps1 | S | 27 | -0.039349923 | -0.086953981 | -0.033259987 | 0.006089936 | 0.053693994 | 0.443917044 | 0.058514332 | 0.047046825 | 0.897827267 | 0.157090891 |
| 8030462N17Rik | S | 34 | -0.066452462 | -0.09854202 | -0.030675045 | 0.035774416 | 0.067866974 | 0.188556273 | 0.078530292 | 0.430800679 | 0.464819478 | 0.201173692 |
| Strn4 | S | 35 | 0.01129211 | -0.111125752 | -0.116637864 | -0.127929974 | -0.005512111 | 0.81146847 | 0.060055741 | 0.134377719 | 0.114958927 | 0.933488863 |
| Nes | S | 728 | -0.079629275 | -0.002935976 | -0.048458822 | 0.031170453 | -0.045522846 | 0.001356633 | 0.905673651 | 0.270201456 | 0.439050051 | 0.306684788 |
| Map1a | S | 1606 | -0.08539638 | 0.009921035 | -0.18142544 | -0.096029059 | -0.191346475 | 0.167366177 | 0.787771236 | 0.096227824 | 0.296033907 | 0.084115967 |
| Ptpn21 | S | 777 | -0.022268938 | -0.110054736 | -0.102770663 | -0.080501725 | 0.007284074 | 0.734935351 | 0.372215744 | 0.037530841 | 0.244778762 | 0.946849914 |
| Hjurp A730008H23Ril | S | 449 | 0.116097472 | -0.080182994 | -0.188236906 | -0.304334378 | -0.108053912 | 0.034826377 | 0.044666241 | 0.001300333 | 0.007481935 | 0.000190862 |
| Wdr46 | S | 41 | 0.09663319 | -0.098128927 | -0.143441801 | -0.240074992 | -0.045312875 | 0.258501066 | 0.130707454 | 0.003210279 | 0.050207206 | 0.409640935 |
| Cdan1 | S | 165 | -0.012534348 | -0.049032091 | -0.136388367 | -0.123854019 | -0.087356276 | 0.772376529 | 0.362523453 | 0.024107854 | 0.027111754 | 0.107846296 |
| Nab2 | S | 159 | -0.010482106 | -0.02412898 | -0.093941928 | -0.083459822 | -0.069812949 | 0.770708689 | 0.387364434 | 0.001333134 | 0.091932747 | 0.053732388 |
| Pak1 | S | 223 | 0.009072467 | -0.045788978 | -0.175268183 | -0.184340649 | -0.129479204 | 0.884898222 | 0.324218933 | 0.026357573 | 0.03769406 | 0.083658957 |
| Plcb3 | S | 1009 | -0.006625436 | -0.064919414 | -0.163007196 | -0.156381761 | -0.098087782 | 0.89312465 | 0.266660348 | 0.062858204 | 0.079761499 | 0.198889044 |
| Plekhg3 | S | 550 | -0.013321391 | -0.152588359 | -0.069495432 | -0.05617404 | 0.083092928 | 0.845903559 | 0.051188415 | 0.347253306 | 0.502534451 | 0.308662679 |
| Pdlim5 | T | 208 | -0.045843847 | -0.016437588 | -0.195120549 | -0.149276703 | -0.178682961 | 0.594503139 | 0.875849897 | 0.040725143 | 0.134044166 | 0.15272152 |
| Osbpl11 | S | 179 | -0.079333588 | 0.038118693 | -0.2088754 | -0.129541813 | -0.246994093 | 0.258395654 | 0.560176958 | 0.04247393 | 0.132769873 | 0.036046669 |
| Flna | S | 2128 | -0.084576808 | -0.101462246 | -0.074496619 | 0.010080189 | 0.026965627 | 0.298464427 | 0.209362473 | 0.367277661 | 0.890956041 | 0.708463038 |
| Tmem45a | S | 269 | -0.023482245 | 0.040870192 | -0.222238406 | -0.198756161 | -0.263108598 | 0.742090822 | 0.339121357 | 0.003625651 | 0.046438784 | 0.012253398 |
| Pkn1 | S | 346 | 0.025870806 | -0.101720654 | -0.228469556 | -0.254340361 | -0.126748902 | 0.711866102 | 0.094303528 | 0.212813767 | 0.180389989 | 0.432747481 |
| Plec | S | 2047 | -0.029132326 | -0.030093743 | -0.083505257 | -0.054372931 | -0.053411514 | 0.479766685 | 0.405434947 | 0.003964437 | 0.229005749 | 0.179381858 |
| Mast1 Mast2 Mast4 | S | 1331;1430;1254 | 0.061881785 | -0.109129639 | -0.174078533 | -0.235960319 | -0.064948894 | 0.48739918 | 0.080814309 | 0.071537381 | 0.060730055 | 0.404711657 |
| Szt2 | S | 3230 | 0.000400967 | -0.063188155 | -0.207683098 | -0.208084065 | -0.144494944 | 0.995362207 | 0.397457431 | 0.066396514 | 0.074549649 | 0.172397736 |
| Phldb1 | S | 220 | 0.025378262 | -0.064571254 | -0.087157686 | -0.112535948 | -0.022586432 | 0.342190289 | 0.020174275 | 0.007451283 | 0.009574538 | 0.329007893 |
| Nfib | T | 327 | -0.060569825 | -0.068594473 | -0.011593796 | 0.048976028 | 0.057000676 | 0.008569983 | 0.131092515 | 0.574113133 | 0.047104347 | 0.186942217 |
| U2surp | S | 951 | -0.167447833 | -0.152926706 | 0.127771223 | 0.295219055 | 0.280697929 | 0.043168046 | 0.062380318 | 0.31072098 | 0.069409941 | 0.076810258 |
| Git2 | S | 359 | 0.199083846 | -0.059248954 | -0.217807713 | -0.41689156 | -0.15855876 | 0.384356949 | 0.215682731 | 0.002990899 | 0.143322446 | 0.014318154 |
| Ptpn21 | S | 679 | -0.092897145 | -0.032190636 | -0.05908498 | 0.033812165 | -0.026894344 | 0.052021018 | 0.420081408 | 0.220668918 | 0.408899031 | 0.481679412 |
| Eps15 | T | 779 | -0.034261153 | -0.100245343 | -0.078079908 | -0.043818755 | 0.022165435 | 0.506886977 | 0.068822363 | 0.270849736 | 0.461623809 | 0.682954901 |
| Kank2 | S | 503 | -0.025347827 | 0.017699307 | -0.325806329 | -0.300458502 | -0.343505636 | 0.848093838 | 0.849850577 | 0.052992857 | 0.110775295 | 0.046532519 |
| Dst | S | 2862 | 0.023644959 | -0.227494588 | -0.029639928 | -0.053284888 | 0.19785466 | 0.741956807 | 0.016759067 | 0.67192568 | 0.303383266 | 0.020754356 |
| Ubr4 | S | 620 | 0.005173731 | -0.0676079 | -0.112434342 | -0.117608072 | -0.044826442 | 0.909480333 | 0.068365944 | 0.047632288 | 0.079674186 | 0.309419611 |
| Usp47 | S | 1014 | -0.09302122 | -0.028834548 | -0.06568944 | 0.02733178 | -0.036854891 | 0.079184909 | 0.582487842 | 0.113381302 | 0.508463275 | 0.456650355 |
| Frmd6 | S | 375 | -0.125784408 | 0.124880844 | -0.296561932 | -0.170777523 | -0.421442775 | 0.198937988 | 0.433399348 | 0.082162435 | 0.249023063 | 0.062074027 |
| Anks1 | S | 677 | -0.091634882 | -0.008194533 | -0.122074802 | -0.03043992 | -0.113880269 | 0.0292237 | 0.915132461 | 0.284060213 | 0.757999358 | 0.35172188 |
| Tbc1d1 | S | 255 | -0.150959517 | -0.007648482 | -0.108953424 | 0.042006094 | -0.101304942 | 0.115677385 | 0.948341429 | 0.235246828 | 0.56862599 | 0.384069069 |
| Serpinh1 | S | 138 | -0.066192896 | -0.141767166 | -0.131272027 | -0.065079132 | 0.010495139 | 0.749055135 | 0.301585858 | 0.23315803 | 0.760544758 | 0.939122918 |
| Wnk1 | S | 2046 | -0.08276186 | -0.063215435 | -0.226323265 | -0.143561405 | -0.16310783 | 0.716456034 | 0.785365223 | 0.116311141 | 0.506949479 | 0.468870315 |
| Wnk1 | S | 2048 | -0.08276186 | -0.063215435 | -0.226323265 | -0.143561405 | -0.16310783 | 0.716456034 | 0.785365223 | 0.116311141 | 0.506949479 | 0.468870315 |
| Smarcad1 | T | 99 | -0.024885028 | -0.05536261 | -0.075244701 | -0.050359673 | -0.019882091 | 0.420732194 | 0.125441906 | 0.035478205 | 0.174308577 | 0.556565045 |
| Map4 | T | 511 | -0.113548012 | -0.07867274 | -0.141013522 | -0.02746551 | -0.062340782 | 0.386209455 | 0.556132459 | 0.248330223 | 0.696941451 | 0.462163646 |
| Rabgef1 | S | 386 | -0.039807188 | -0.096172069 | -0.165000314 | -0.125193126 | -0.068828245 | 0.647096929 | 0.228718884 | 0.249882989 | 0.362849747 | 0.584892223 |
| Flna | S | 2152 | -0.14056428 | -0.000533831 | -0.097158033 | 0.043406247 | -0.096624202 | 0.063553766 | 0.98800266 | 0.260417202 | 0.608211041 | 0.263207016 |
| Ubr4 | S | 181 | -0.033561825 | 0.001257126 | -0.144343116 | -0.11078129 | -0.145600241 | 0.436144716 | 0.946488314 | 0.008670321 | 0.061897464 | 0.019451077 |

| | | | | | | | | | | | | |
|---|---|---|---|---|---|---|---|---|---|---|---|---|
| Helb | S | 942 | -0.166280815 | 0.068987772 | -0.070481489 | 0.095799326 | -0.139469261 | 0.009091849 | 0.177196958 | 0.058415384 | 0.077446885 | 0.043827802 |
| Lima1 | S | 360 | 0.006449375 | -0.071759072 | -0.077567113 | -0.084016489 | -0.005808041 | 0.849062656 | 0.057671444 | 0.016127585 | 0.061282698 | 0.813293474 |
| Zc3h13 | S | 317 | -0.027609527 | -0.09214027 | -0.092994948 | -0.065385421 | -0.000854679 | 0.670058767 | 0.238008886 | 0.05652852 | 0.333739752 | 0.989829046 |
| Rabep2 | S | 176 | -0.193929188 | 0.048770344 | -0.0466472 | 0.147281988 | -0.095417544 | 0.009231278 | 0.625671149 | 0.486649813 | 0.055458226 | 0.364257286 |
| Tmem200a | S | 323 | -0.08608295 | -0.085896769 | -0.059262773 | 0.026820177 | 0.026633996 | 0.329154032 | 0.168576381 | 0.254451693 | 0.731412644 | 0.59843585 |
| Arhgap21 | S | 874 | -0.122854497 | -0.00997745 | -0.082068582 | 0.040785915 | -0.072091132 | 0.063531467 | 0.849990891 | 0.202114509 | 0.507124544 | 0.249474702 |
| Rab8a | S | 181 | -0.003524671 | -0.144382393 | -0.093551318 | -0.090026647 | 0.050831076 | 0.918003069 | 0.067861353 | 0.283500158 | 0.298585293 | 0.562980785 |
| Diaph1 | S | 22 | -0.050408651 | -0.037128945 | -0.078950356 | -0.028541705 | -0.04182141 | 0.113930881 | 0.243513972 | 0.102874412 | 0.486430141 | 0.336989091 |
| Eif4g1 | T | 212 | 0.098022934 | -0.106688884 | -0.174763933 | -0.272786867 | -0.068075049 | 0.057453377 | 0.146398968 | 0.001617228 | 0.002570458 | 0.301742017 |
| Eno1 Eno1b | S | 419 | -0.170916724 | 0.000892658 | -0.082985681 | 0.087931044 | -0.083878339 | 0.036633728 | 0.992801155 | 0.519104026 | 0.492414718 | 0.553257651 |
| Myo9b | S | 1992 | -0.113935109 | -0.10197772 | -0.280349774 | -0.166414664 | -0.178372054 | 0.413677762 | 0.67371551 | 0.463215682 | 0.653456329 | 0.659745293 |
| Cpne3 | S | 14 | -0.13888784 | 0.004066178 | -0.110244768 | 0.028643072 | -0.114310945 | 0.121592619 | 0.958647657 | 0.153895795 | 0.681279963 | 0.142884049 |
| Dnaja1 | S | 188 | -0.089367175 | -0.023927739 | -0.02279226 | 0.066574916 | 0.00113548 | 0.00276747 | 0.546967419 | 0.443174971 | 0.087173127 | 0.977788822 |
| Ripor1 | S | 838 | 0.005947126 | -0.074296786 | -0.238142163 | -0.244089289 | -0.163845377 | 0.944482464 | 0.41095367 | 0.098684754 | 0.10527468 | 0.196697784 |
| Ripor1 | S | 844 | 0.005947126 | -0.074296786 | -0.238142163 | -0.244089289 | -0.163845377 | 0.944482464 | 0.41095367 | 0.098684754 | 0.10527468 | 0.196697784 |
| Tnks1bp1 | S | 300 | 0.011458672 | -0.068655571 | -0.115756761 | -0.127215433 | -0.04710119 | 0.750725591 | 0.098603427 | 0.029148976 | 0.029577202 | 0.196159112 |
| Nmt2 | S | 38 | -0.097612445 | -0.059876597 | -0.054951631 | 0.042660815 | 0.004924966 | 0.057771598 | 0.564220291 | 0.269289313 | 0.358529594 | 0.960596938 |
| Snd1 | T | 240 | -0.008003796 | -0.080733707 | -0.152505761 | -0.144501965 | -0.071772054 | 0.94169604 | 0.292961677 | 0.065740966 | 0.27032326 | 0.393670967 |
| Mdm2 | S | 161 | 0.007640364 | -0.019271326 | -0.120042925 | -0.127683289 | -0.100771599 | 0.803470675 | 0.494666834 | 0.001082805 | 0.020777635 | 0.020987135 |
| Ttc28 | S | 24 | -0.185876877 | -0.081266426 | -0.035249639 | 0.150627238 | 0.046016787 | 0.08919516 | 0.546625895 | 0.798279089 | 0.26623149 | 0.752936048 |
| Phactr4 | T | 134 | 0.065574781 | -0.052899798 | -0.244551419 | -0.3101262 | -0.191651621 | 0.338654745 | 0.533952632 | 0.009660754 | 0.003656837 | 0.048563899 |
| Trp53bp1 | T | 434 | -0.012138864 | -0.084966453 | -0.063602551 | -0.051463688 | 0.021363901 | 0.873485039 | 0.026013855 | 0.099731695 | 0.515695325 | 0.479475146 |
| Prkab1 | S | 108 | -0.116423102 | -0.005900572 | -0.119956591 | -0.003533489 | -0.114056018 | 0.111406843 | 0.948200526 | 0.176112012 | 0.958635922 | 0.256307276 |
| Atrx | S | 84 | 0.007234861 | -0.130769008 | -0.094896099 | -0.10213096 | 0.035872909 | 0.825289252 | 0.089545657 | 0.112792886 | 0.105763028 | 0.592431612 |
| Bud13 | S | 297 | -0.310722891 | -0.100329122 | 0.006889485 | 0.317612376 | 0.107218607 | 0.249654705 | 0.281363973 | 0.843671587 | 0.244224779 | 0.258943423 |
| Srrm2 | S | 2360 | -0.078404412 | -0.033894924 | -0.08854265 | -0.010138238 | -0.054647725 | 0.108904269 | 0.414147345 | 0.181077633 | 0.853966831 | 0.345372869 |
| Anln | S | 887 | -0.030867991 | -0.053902781 | -0.10404019 | -0.073172199 | -0.050137409 | 0.155537297 | 0.395412254 | 0.078082383 | 0.151699853 | 0.459585167 |
| Leo1 | S | 171 | -0.015321241 | -0.047287714 | -0.132895207 | -0.117573966 | -0.085607493 | 0.804986019 | 0.074421907 | 0.086885301 | 0.169763156 | 0.184474285 |
| Leo1 | S | 179 | -0.015321241 | -0.047287714 | -0.132895207 | -0.117573966 | -0.085607493 | 0.804986019 | 0.074421907 | 0.086885301 | 0.169763156 | 0.184474285 |
| Camkk2 | S | 132 | -0.23144541 | -0.0424547 | -0.137613376 | 0.093832034 | -0.095158676 | 0.232388444 | 0.783018869 | 0.531839855 | 0.697492946 | 0.680040089 |
| Camkk2 | S | 136 | -0.23144541 | -0.0424547 | -0.137613376 | 0.093832034 | -0.095158676 | 0.232388444 | 0.783018869 | 0.531839855 | 0.697492946 | 0.680040089 |
| Mdfic | S | 129 | 0.015567945 | -0.095084674 | -0.158827343 | -0.174395289 | -0.063742669 | 0.76302719 | 0.390302196 | 0.030791801 | 0.028232728 | 0.537802381 |
| Ogfr | S | 614 | 0.032820524 | -0.127908443 | -0.138742924 | -0.171563449 | -0.010834482 | 0.847982789 | 0.122243898 | 0.129642748 | 0.375497344 | 0.898920332 |
| Ticrr | S | 934 | -0.031183755 | -0.05040701 | -0.156726248 | -0.125542493 | -0.106319238 | 0.434176565 | 0.146844939 | 0.196967279 | 0.272697063 | 0.334717752 |
| Tbc1d20 | S | 388 | 0.127372796 | -0.179805368 | -0.17305316 | -0.300425956 | 0.006752208 | 0.039918105 | 0.084902672 | 0.005347199 | 0.003079419 | 0.928357866 |
| Aldoa | T | 37 | -0.064974915 | -0.196922257 | -0.050955128 | 0.014019787 | 0.145967129 | 0.676698087 | 0.085569406 | 0.748447936 | 0.943600646 | 0.4050532 |
| Ndc1 | S | 470 | 0.015574911 | -0.040502523 | -0.195885002 | -0.211459913 | -0.155382479 | 0.853848408 | 0.284042016 | 0.038140872 | 0.081097723 | 0.073377726 |
| Plcb3 | S | 481 | -0.210806283 | 0.065098481 | -0.134933506 | 0.075872777 | -0.200031987 | 0.076160463 | 0.701749806 | 0.322151867 | 0.556733094 | 0.305369703 |
| Map1a | S | 991 | -0.071934884 | -0.042843058 | -0.075779558 | -0.003844673 | -0.032936499 | 0.112219862 | 0.278879231 | 0.207471216 | 0.944607685 | 0.558647951 |
| Nhsl1 | S | 720 | -0.019020416 | -0.096548779 | -0.157468236 | -0.13844782 | -0.060919457 | 0.831510842 | 0.108687461 | 0.270798904 | 0.34822749 | 0.629803982 |
| Slx4 | S | 922 | -0.003498497 | -0.169423908 | -0.096423488 | -0.092924991 | 0.07300042 | 0.958346413 | 0.135363859 | 0.222089822 | 0.245387966 | 0.47370976 |
| Lmnb1 | S | 279 | 0.048580165 | -0.103841473 | -0.0892285 | -0.137808665 | 0.014612973 | 0.270527822 | 0.011482835 | 0.035447483 | 0.022843741 | 0.61646623 |
| Tra2a | S | 18 | 0.023781382 | -0.093050301 | -0.11275507 | -0.136536453 | -0.019704769 | 0.62484469 | 0.078463657 | 0.043733889 | 0.014427319 | 0.105817692 |
| Prpf38b | S | 523 | 0.123104587 | -0.295278352 | -0.075604427 | -0.198709015 | 0.219673925 | 0.12366077 | 0.070500363 | 0.009070779 | 0.048158308 | 0.118347893 |
| Svil | S | 510 | 0.024031216 | -0.259515978 | -0.077036994 | -0.101068211 | 0.182478983 | 0.846683399 | 0.089768041 | 0.457538524 | 0.497035094 | 0.213278848 |
| Epb41 | S | 703 | -0.139966656 | -0.01156686 | -0.20872751 | -0.068760854 | -0.19716065 | 0.24412507 | 0.944888841 | 0.297220056 | 0.681456422 | 0.353435192 |
| Cep41 | S | 358 | 0.032891597 | -0.058307141 | -0.285171991 | -0.318063588 | -0.22686485 | 0.887164441 | 0.726600459 | 0.058751044 | 0.220253858 | 0.164260225 |
| Prr14 | S | 29 | 0.045178631 | -0.053748816 | -0.224829404 | -0.270008036 | -0.171080588 | 0.697168849 | 0.300636601 | 0.041496848 | 0.087171748 | 0.102785992 |
| Ncor2 | S | 1309 | -0.005729201 | -0.060281015 | -0.073166597 | -0.067437396 | -0.012885582 | 0.786451923 | 0.013425692 | 0.08426812 | 0.104754089 | 0.683962648 |
| Setd2 | S | 1326 | -0.00888914 | -0.124635296 | -0.043119375 | -0.034230235 | 0.081515921 | 0.87780688 | 0.011667564 | 0.423264256 | 0.585654257 | 0.148718119 |
| Smarcd2 | S | 15 | -0.051694947 | -0.233606261 | -0.22116064 | -0.169465693 | 0.012445621 | 0.848722511 | 0.355119314 | 0.507480259 | 0.648946266 | 0.971399655 |
| Snx24 | S | 116 | -0.011981814 | -0.017151851 | -0.159598079 | -0.147616265 | -0.142446228 | 0.76056591 | 0.706653847 | 0.012441124 | 0.022979964 | 0.0329356 |
| Arfgef1 | S | 664 | -0.065405261 | -0.133811267 | -0.14358311 | -0.07817785 | -0.009771843 | 0.480881885 | 0.414062316 | 0.31245026 | 0.533138121 | 0.954202842 |
| Nbn | S | 398 | -0.037078883 | -0.0913867 | -0.07913684 | -0.042057957 | 0.01224986 | 0.508347678 | 0.149234007 | 0.138532434 | 0.483273186 | 0.833105192 |
| Osbpl6 | S | 290 | -0.014771557 | -0.090492849 | -0.142869111 | -0.128097554 | -0.052376261 | 0.776553114 | 0.239716819 | 0.120308326 | 0.151992464 | 0.545273147 |
| Map3k2 | S | 349 | -0.00434056 | -0.133225804 | -0.03344735 | -0.02910679 | 0.099778455 | 0.889058292 | 0.014662827 | 0.309876459 | 0.421731403 | 0.047987548 |
| Nup98 | S | 1027 | -0.057854637 | -0.201040223 | 0.019383748 | 0.077238385 | 0.220423972 | 0.295137584 | 0.04897714 | 0.754165452 | 0.149624365 | 0.04377587 |
| Top1 | S | 167 | -0.102910422 | -0.034488385 | -0.097356442 | 0.00555398 | -0.062868057 | 0.197274492 | 0.21635902 | 0.324182967 | 0.955825775 | 0.493230376 |
| Scrib | S | 1209 | -0.007554996 | -0.077425899 | -0.15940247 | -0.151847474 | -0.081976571 | 0.911359164 | 0.3301299 | 0.070489016 | 0.115979502 | 0.362925076 |
| Scrib | S | 1212 | -0.007554996 | -0.077425899 | -0.15940247 | -0.151847474 | -0.081976571 | 0.911359164 | 0.3301299 | 0.070489016 | 0.115979502 | 0.362925076 |
| Med12 | T | 642 | -0.025839023 | -0.041806414 | -0.1437984 | -0.117959376 | -0.101991986 | 0.697271614 | 0.702715927 | 0.019743962 | 0.142595394 | 0.387965229 |
| Sh3rf1 | S | 741 | -0.047557746 | -0.140571782 | -0.020597444 | 0.026960302 | 0.119974338 | 0.417435493 | 0.03878179 | 0.680227073 | 0.632039581 | 0.074530069 |
| Snx11 | S | 168 | -0.054727589 | -0.001209334 | -0.198144986 | -0.143417398 | -0.196935653 | 0.670616775 | 0.978868908 | 0.03510899 | 0.319169349 | 0.050751184 |
| Relch | S | 193 | -0.057905437 | -0.094376955 | -0.120020313 | -0.062114876 | -0.025643358 | 0.586204953 | 0.195468673 | 0.266633477 | 0.604444353 | 0.780138652 |
| Dpy19l1 | S | 22 | -0.078001005 | -0.017762692 | -0.041079088 | 0.036921916 | -0.023316396 | 0.011319244 | 0.646124546 | 0.134424923 | 0.191693644 | 0.566836745 |
| Arhgef40 | S | 226 | -0.035366074 | -0.083725855 | -0.064225541 | -0.028859467 | 0.019500314 | 0.652273585 | 0.071511302 | 0.114082706 | 0.716036666 | 0.639012182 |
| Ccnd1 | T | 286 | -0.022866738 | -0.014336688 | -0.099262128 | -0.07639539 | -0.084925441 | 0.295814739 | 0.629182016 | 0.005590115 | 0.02366103 | 0.044818073 |
| Cit | S | 439 | -0.021702185 | -0.067123421 | -0.111444329 | -0.089742143 | -0.044320907 | 0.750210755 | 0.089060761 | 0.122245219 | 0.290407128 | 0.458803314 |
| Knop1 | T | 314 | 0.162402188 | -0.107817363 | -0.177544417 | -0.339946606 | -0.069727054 | 0.461189601 | 0.014099626 | 0.097935274 | 0.189592153 | 0.405330453 |
| Kif20b | S | 950 | -0.124331054 | -0.030256869 | -0.052562427 | 0.071768627 | -0.022305558 | 0.038873656 | 0.555810176 | 0.478999975 | 0.367402907 | 0.773016442 |
| Eif3b | S | 75 | -0.000791281 | -0.053832698 | -0.052114468 | -0.051323187 | 0.00171823 | 0.9722496 | 0.012866921 | 0.012968285 | 0.073346588 | 0.88351525 |
| Prrx1 | S | 74 | -0.133766017 | 0.047543239 | -0.101635293 | 0.032130724 | -0.149178532 | 0.022968784 | 0.413393415 | 0.126324675 | 0.568985063 | 0.063412007 |
| Prrx1 | T | 71 | -0.133766017 | 0.047543239 | -0.101635293 | 0.032130724 | -0.149178532 | 0.022968784 | 0.413393415 | 0.126324675 | 0.568985063 | 0.063412007 |
| Hnrnpm | S | 636 | -0.082626572 | 0.0265717 | -0.160884067 | -0.078257495 | -0.187455767 | 0.114664917 | 0.801246136 | 0.093253613 | 0.32629331 | 0.16276186 |
| Fam83h | S | 970 | -0.065237583 | 0.016674717 | -0.125346012 | -0.06010843 | -0.142020729 | 0.221960792 | 0.780818823 | 0.01699629 | 0.199739634 | 0.067877873 |
| Cit | S | 581 | -0.052717908 | -0.089308347 | -0.083874517 | -0.031156609 | 0.00543383 | 0.429864838 | 0.197674956 | 0.191850154 | 0.462854664 | 0.892299728 |
| Cemip2 | S | 24 | 0.055867306 | -0.182667478 | -0.053758658 | -0.109625964 | 0.12890882 | 0.154379805 | 0.001900939 | 0.418133722 | 0.153931024 | 0.113876707 |
| Rtn4 | S | 489 | -0.00868539 | -0.041395694 | -0.081268722 | -0.072583332 | -0.039873027 | 0.824590608 | 0.097685732 | 0.010005306 | 0.129480045 | 0.068416222 |
| Pml | T | 504 | -0.165528502 | -0.031967259 | 0.035793138 | 0.20132164 | 0.067760397 | 0.003582241 | 0.579312507 | 0.602170437 | 0.064454169 | 0.382145222 |
| Taok3 | S | 324 | -0.168179946 | 0.095370798 | -0.187867837 | -0.019687891 | -0.283238635 | 0.160767103 | 0.346250379 | 0.060425022 | 0.811000663 | 0.02501986 |
| Ppp1r2 | S | 90 | -0.166457269 | 0.09191106 | -0.21965997 | -0.053202701 | -0.31157103 | 0.059366617 | 0.289800084 | 0.179696239 | 0.69476197 | 0.093292917 |
| Smg1 | T | 3570 | -0.039369311 | -0.104348363 | -0.065581843 | -0.026212532 | 0.03876652 | 0.520577364 | 0.096229018 | 0.236366682 | 0.649765487 | 0.470194744 |
| Kars | S | 592 | -0.049884692 | -0.016906612 | -0.059390169 | -0.009505477 | -0.042483557 | 0.042129366 | 0.498291303 | 0.030103153 | 0.57124335 | 0.10102364 |
| Tpm1 | S | 206 | -0.142059339 | -0.090753344 | 0.006389572 | 0.148448911 | 0.097142916 | 0.029194382 | 0.432859844 | 0.924253558 | 0.102054049 | 0.430893967 |
| Ect2 | S | 879 | -0.10225798 | -0.423267922 | NA | NA | NA | 0.707024034 | 0.243490351 | NA | NA | NA |
| Incenp | T | 452 | 0.036433933 | -0.098406336 | -0.102122002 | -0.138555935 | -0.003715666 | 0.337842216 | 0.011881943 | 0.086734452 | 0.042480092 | 0.933539199 |
| Mlh1 | S | 481 | -0.058546745 | -0.035042541 | -0.118152609 | -0.059605864 | -0.083110068 | 0.168117838 | 0.568341967 | 0.124211381 | 0.367380231 | 0.297390652 |
| Specc1 | S | 847 | 0.136756324 | -0.234276372 | -0.154848554 | -0.291604878 | 0.079427818 | 0.094549617 | 0.011270157 | 0.19647713 | 0.052252184 | 0.43197409 |
| Sh2b3 | S | 17 | -0.030878646 | -0.073850408 | -0.083248072 | -0.052369426 | -0.009397664 | 0.324441184 | 0.247135286 | 0.086517797 | 0.2058758 | 0.871816164 |
| Soga1 | S | 924 | -0.122636642 | 0.022487664 | -0.098878943 | 0.023757699 | -0.121366607 | 0.02338558 | 0.661422433 | 0.23840422 | 0.744181117 | 0.174987638 |
| Prune2 | S | 1424 | -0.07246527 | -0.023035668 | -0.143824775 | -0.071359506 | -0.120789107 | 0.460124788 | 0.828577875 | 0.050497425 | 0.472838968 | 0.3108845 |
| Numa1 | S | 1835 | -0.169717499 | 0.048596156 | -0.139597771 | 0.030119728 | -0.188193928 | 0.050595717 | 0.667591811 | 0.342735151 | 0.82436989 | 0.275821445 |
| Tex2 | S | 791 | -0.176877251 | -0.074463587 | 0.04755843 | 0.224435681 | 0.122022017 | 0.028234228 | 0.243754764 | 0.581695709 | 0.062034863 | 0.221539842 |
| Gemin5 | T | 1416 | 0.077156619 | -0.145620464 | -0.196842794 | -0.273999414 | -0.05122233 | 0.413184471 | 0.19898256 | 0.06464129 | 0.019197616 | 0.58860935 |

| | | | | | | | | | | | | |
|---|---|---|---|---|---|---|---|---|---|---|---|---|
| Cxxc1 | S | 114 | -0.089075997 | -0.133355531 | -0.052759275 | 0.036316721 | 0.080596255 | 0.479947468 | 0.121089815 | 0.572298419 | 0.79303611 | 0.449782957 |
| Rap1gap | S | 499 | -0.144698052 | 0.161314727 | -0.301548414 | -0.156850362 | -0.462863141 | 0.153482679 | 0.344803324 | 0.09035632 | 0.299725383 | 0.054037137 |
| Irf2bp2 | S | 406 | 0.058509935 | -0.172796863 | -0.146975876 | -0.20548581 | 0.025820988 | 0.191725707 | 0.084455808 | 0.076370323 | 0.036769171 | 0.761936367 |
| Exoc2 | S | 435 | -0.150906514 | -0.082716187 | 0.031382279 | 0.182288793 | 0.114098466 | 0.060047827 | 0.106137182 | 0.608739847 | 0.049641902 | 0.10842664 |
| Sin3b | S | 667 | -0.121077536 | -0.11120149 | -0.160800457 | -0.039722922 | -0.049598968 | 0.609244465 | 0.424796869 | 0.36315125 | 0.876045091 | 0.783524291 |
| Sin3b | T | 666 | -0.121077536 | -0.11120149 | -0.160800457 | -0.039722922 | -0.049598968 | 0.609244465 | 0.424796869 | 0.36315125 | 0.876045091 | 0.783524291 |
| Ing5 | T | 152 | -0.002752989 | -0.124123734 | -0.065813162 | -0.063060173 | 0.058310571 | 0.947340323 | 0.014308641 | 0.483273322 | 0.490400419 | 0.516696136 |
| Cavin4 | S | 325 | -0.214060881 | -0.118227404 | 0.015779007 | 0.229839887 | 0.134006411 | 0.136245667 | 0.369658887 | 0.906444654 | 0.028292113 | 0.108689776 |
| Eif3c | S | 11 | 0.079296895 | -0.105289304 | -0.232802886 | -0.31209978 | -0.127513582 | 0.393598522 | 0.325132107 | 0.034670236 | 0.022441699 | 0.25815419 |
| Zfyve16 | T | 908 | 0.089334296 | -0.12163509 | -0.234496324 | -0.323830621 | -0.112861234 | 0.277649416 | 0.092139604 | 0.087152451 | 0.030178035 | 0.275169852 |
| Bag3 | S | 382 | -0.022239365 | -0.088032447 | -0.09856103 | -0.076321665 | -0.010528583 | 0.805594838 | 0.0714669 | 0.188852736 | 0.462994571 | 0.870599908 |
| Fxr1 | S | 643 | 0.043060139 | -0.117595094 | -0.072662563 | -0.115722701 | 0.044932531 | 0.098122531 | 0.002028656 | 0.089466675 | 0.047763484 | 0.226954662 |
| Fam117b | S | 405 | 0.051253404 | -0.040960557 | -0.230673173 | -0.281926576 | -0.189712615 | 0.84574841 | 0.775119822 | 0.020539213 | 0.339624956 | 0.235490967 |
| Cep350 | S | 447 | 0.034074806 | -0.102387364 | -0.126568714 | -0.16064352 | -0.02418135 | 0.755285515 | 0.087890163 | 0.087720892 | 0.202923149 | 0.620003725 |
| Apbb2 | T | 34 | -0.052288665 | -0.113420509 | -0.00335809 | 0.048930575 | 0.11006242 | 0.430005271 | 0.008779716 | 0.950514014 | 0.506523332 | 0.110293934 |
| Polh | S | 659 | -0.020236992 | -0.053796902 | -0.110751592 | -0.0905146 | -0.05695469 | 0.64359307 | 0.124795223 | 0.081555952 | 0.1576647 | 0.285495386 |
| Tox4 | S | 178 | -0.009928981 | -0.078239114 | -0.028263871 | -0.01833489 | 0.049975243 | 0.580961948 | 0.004406931 | 0.153901012 | 0.001911338 | 0.011006879 |
| Cast | S | 165 | -0.078827563 | 0.026541767 | -0.072903784 | 0.005923779 | -0.099445551 | 0.025213856 | 0.713959313 | 0.028062538 | 0.721791883 | 0.241217444 |
| Smchd1 | T | 1936 | 0.049018281 | -0.107590102 | -0.064911276 | -0.113929556 | 0.042678826 | 0.11049793 | 0.000503427 | 0.109312323 | 0.024908978 | 0.2225537 |
| Scrib | S | 1284 | 0.054052978 | -0.045511715 | -0.133172384 | -0.187225362 | -0.087660669 | 0.346030713 | 0.2157322 | 0.001548895 | 0.037795303 | 0.054215228 |
| Atrx | S | 1978 | 0.046609002 | -0.305046953 | -0.141603663 | -0.188212665 | 0.16344329 | 0.420136498 | 0.137705336 | 0.368773348 | 0.260710288 | 0.417222481 |
| Cep170 | S | 1048 | -0.18419926 | 0.026986097 | -0.010314874 | 0.173884386 | -0.037300971 | 0.004196816 | 0.786504678 | 0.922689473 | 0.186244273 | 0.775953744 |
| Smarca5 | T | 112 | 0.033130839 | -0.082974724 | -0.065368918 | -0.098499757 | 0.017605806 | 0.313232889 | 0.051580212 | 0.001488998 | 0.048676703 | 0.537246304 |
| Ddx1 | T | 346 | -0.010605396 | -0.083291714 | -0.099040058 | -0.088434663 | -0.015748344 | 0.89920639 | 0.021999966 | 0.280613602 | 0.425196856 | 0.842064049 |
| Srrm2 | S | 1822 | -0.145194927 | -0.089174741 | 0.02271126 | 0.167906187 | 0.111886001 | 0.193212312 | 0.042348906 | 0.817868857 | 0.217048149 | 0.314279798 |
| Nemf | S | 737 | -0.038871252 | -0.088318329 | -0.10270991 | -0.063838657 | -0.014391581 | 0.295449248 | 0.226920168 | 0.260681111 | 0.445898802 | 0.874260874 |
| Fam53c | S | 232 | -0.096488736 | -0.015723395 | -0.078036676 | 0.018452061 | -0.062313281 | 0.081459047 | 0.831989903 | 0.122429528 | 0.699296489 | 0.426435808 |
| Ssh2 | S | 36 | -0.31168538 | 0.012244729 | -0.086074112 | 0.225611267 | -0.098318842 | 0.166985987 | 0.911538141 | 0.520644396 | 0.298747924 | 0.51495172 |
| Ssh2 | S | 40 | -0.31168538 | 0.012244729 | -0.086074112 | 0.225611267 | -0.098318842 | 0.166985987 | 0.911538141 | 0.520644396 | 0.298747924 | 0.51495172 |
| Dennd4c | S | 1058 | -0.125419491 | 0.027755598 | -0.106078919 | 0.019340572 | -0.133834517 | 0.037251255 | 0.535945373 | 0.162174086 | 0.759727875 | 0.103961398 |
| Srrm2 | S | 1506 | 0.056977836 | -0.182321834 | -0.038496112 | -0.095473949 | 0.143825722 | 0.230212187 | 0.002025694 | 0.447517378 | 0.105005454 | 0.052506115 |
| Cdca7l | S | 136 | 0.076902846 | -0.055618011 | -0.196221761 | -0.273124607 | -0.14060375 | 0.091832384 | 0.234215725 | 0.002235055 | 0.001443537 | 0.007907376 |
| Magi3 | S | 954 | -0.043848858 | -0.091012886 | -0.126827059 | -0.082978201 | -0.035814174 | 0.67959909 | 0.307542612 | 0.148744965 | 0.488499801 | 0.709225568 |
| Paf1 | S | 456 | -0.056324053 | -0.112856217 | -0.115480754 | -0.059156701 | -0.002624537 | 0.596627262 | 0.257060817 | 0.27301625 | 0.609956983 | 0.97987051 |
| Paf1 | S | 460 | -0.056324053 | -0.112856217 | -0.115480754 | -0.059156701 | -0.002624537 | 0.596627262 | 0.257060817 | 0.27301625 | 0.609956983 | 0.97987051 |
| Paf1 | S | 466 | -0.056324053 | -0.112856217 | -0.115480754 | -0.059156701 | -0.002624537 | 0.596627262 | 0.257060817 | 0.27301625 | 0.609956983 | 0.97987051 |
| Cep131 | S | 76 | -0.10896852 | -0.034154641 | -0.120265427 | -0.011296907 | -0.086110786 | 0.428133541 | 0.598122973 | 0.112463401 | 0.930918447 | 0.26344084 |
| Nes | S | 235 | -0.092481874 | -0.127751612 | 0.042931498 | 0.135413371 | 0.170683109 | 0.195324725 | 0.021137602 | 0.627216924 | 0.217057153 | 0.139731878 |
| Hid1 | S | 593 | -0.104036759 | -0.00335482 | -0.124619271 | -0.020582512 | -0.12126445 | 0.18520812 | 0.977167352 | 0.11357196 | 0.759517814 | 0.334636159 |
| Otud4 | S | 204 | -0.165528967 | -0.173613279 | 0.010065047 | 0.175594013 | 0.183678326 | 0.288072314 | 0.218232346 | 0.96026155 | 0.452016049 | 0.42130716 |
| Nav1 | S | 998 | 0.006518509 | -0.014175978 | -0.207458278 | -0.213976787 | -0.1932823 | 0.776784736 | 0.578479134 | 0.020139879 | 0.026799565 | 0.029234464 |
| Fam160b1 | S | 552 | -0.032674485 | -0.006463532 | -0.115772407 | -0.083097922 | -0.109308875 | 0.337793746 | 0.891037483 | 0.009779303 | 0.015134834 | 0.068090208 |
| Trim47 | T | 100 | -0.054881324 | -0.065284147 | -0.055240854 | -0.000359531 | 0.010043293 | 0.326772663 | 0.137101165 | 0.124310652 | 0.993639031 | 0.724531571 |
| Slmap | S | 148 | -0.032544414 | -0.089112864 | -0.103020908 | -0.070476494 | -0.013908044 | 0.66086627 | 0.079514669 | 0.29208534 | 0.501422026 | 0.873133267 |
| Ehbp1 | S | 1038 | -0.09187314 | -0.03236838 | -0.093135475 | -0.001262335 | -0.060767095 | 0.10501127 | 0.584743422 | 0.24813741 | 0.98402875 | 0.405661293 |
| Osbpl3 | S | 184 | 0.000743674 | 0.04313476 | -0.247431083 | -0.248174757 | -0.290565844 | 0.992952682 | 0.534628804 | 0.006669701 | 0.050778313 | 0.014695166 |
| Osbpl3 | S | 185 | 0.000743674 | 0.04313476 | -0.247431083 | -0.248174757 | -0.290565844 | 0.992952682 | 0.534628804 | 0.006669701 | 0.050778313 | 0.014695166 |
| Osbpl3 | S | 187 | 0.000743674 | 0.04313476 | -0.247431083 | -0.248174757 | -0.290565844 | 0.992952682 | 0.534628804 | 0.006669701 | 0.050778313 | 0.014695166 |
| Rps18 | T | 60 | 0.031184041 | -0.053613378 | -0.143107127 | -0.174291169 | -0.08949375 | 0.266560523 | 0.021630705 | 0.033783902 | 0.019128865 | 0.112075429 |
| Stim2 | S | 631 | -0.147194394 | -0.01414768 | -0.044536724 | 0.102657671 | -0.030389044 | 0.054127416 | 0.779538513 | 0.299282188 | 0.133640581 | 0.580287846 |
| Adam17 | T | 764 | -0.144686975 | -0.097866819 | -0.011630162 | 0.133056813 | 0.086236657 | 0.097473668 | 0.331168835 | 0.890160445 | 0.186737247 | 0.427173253 |
| Trp53 | S | 389 | -0.14936488 | 0.083265446 | -0.127706792 | 0.021658088 | -0.210972238 | 0.042099722 | 0.240260622 | 0.058289625 | 0.542522225 | 0.029329987 |
| Papolg | S | 524 | -0.042107841 | -0.10061234 | 0.003258644 | 0.045366485 | 0.103870984 | 0.083256418 | 0.018290801 | 0.954013239 | 0.442665271 | 0.148821537 |
| Adam17 | S | 794 | -0.053349931 | -0.064546357 | -0.111786971 | -0.05843704 | -0.047240614 | 0.398364479 | 0.305725708 | 0.166859816 | 0.471934323 | 0.550389814 |
| Arhgap31 | S | 1242 | 0.206059538 | -0.115360217 | -0.348499737 | -0.554559276 | -0.23313952 | 0.018020399 | 0.113835942 | 0.005172318 | 0.002804026 | 0.040588579 |
| Trim28 | S | 473 | -0.051464132 | -0.043903914 | -0.081038479 | -0.029574348 | -0.037134565 | 0.151008199 | 0.395171687 | 0.100604626 | 0.413151802 | 0.468520894 |
| Atxn2l | S | 562 | 0.034170389 | -0.077617936 | -0.100142266 | -0.134312655 | -0.022524329 | 0.395296497 | 0.030113512 | 0.026898459 | 0.027996451 | 0.511285703 |
| Tjap1 | S | 305 | -0.069406581 | -0.076694264 | -0.039609149 | 0.029797432 | 0.037085115 | 0.07440874 | 0.214573039 | 0.455869984 | 0.5171827 | 0.551044817 |
| Ndel1 | S | 215 | -0.061308083 | -0.049737433 | -0.095880389 | -0.034572306 | -0.046142956 | 0.196092669 | 0.371317677 | 0.178595131 | 0.565124144 | 0.489743102 |
| 8030462N17Rik | S | 143 | 0.010448833 | -0.08167632 | -0.068024382 | -0.078473215 | 0.013651938 | 0.628890587 | 0.002506519 | 0.234631222 | 0.188333785 | 0.77569339 |
| Pnn | S | 100 | -0.200704548 | -0.041588257 | 0.01572006 | 0.216424608 | 0.057308318 | 0.033956914 | 0.503399492 | 0.857359013 | 0.073739782 | 0.52807589 |
| Palld | S | 1126 | 0.012036181 | 0.106582866 | -0.386990417 | -0.399026598 | -0.493573283 | 0.933809697 | 0.550795272 | 0.020771223 | 0.016827149 | 0.039064371 |
| Bysl | S | 169 | -0.05912628 | -0.025042085 | -0.176169589 | -0.117043309 | -0.151127504 | 0.260905457 | 0.571420801 | 0.187891412 | 0.33417742 | 0.238706885 |
| Ranbp2 | S | 2729 | -0.035581309 | -0.133824322 | -0.006036245 | 0.029545064 | 0.127788076 | 0.408295749 | 0.01690305 | 0.900931837 | 0.39167738 | 0.03570066 |
| Axl | S | 509 | -0.069351896 | -0.083351887 | -0.131580176 | -0.06222828 | -0.048228289 | 0.357243653 | 0.392818528 | 0.303222486 | 0.561551482 | 0.682384638 |
| Cep170 | T | 1230 | 0.000748084 | -0.055927069 | -0.182287954 | -0.183036038 | -0.126360886 | 0.988622358 | 0.269965505 | 0.082332139 | 0.090596329 | 0.18508676 |
| Ttc28 | S | 41 | -0.115740054 | 0.005305125 | -0.065402112 | 0.050337943 | -0.070707236 | 0.033174132 | 0.928936808 | 0.182603139 | 0.102188385 | 0.217827786 |
| Adamtsl1 | S | 787 | -0.110789715 | 0.023990223 | -0.06443393 | 0.046355784 | -0.088424154 | 0.024205023 | 0.603413583 | 0.074776584 | 0.194140167 | 0.103233861 |
| Trim28 | S | 501 | -0.163632063 | -0.221353198 | 0.099518839 | 0.263150902 | 0.320872038 | 0.140162277 | 0.117804903 | 0.370086758 | 0.062720415 | 0.056041473 |
| Sgce | S | 363 | -0.139840215 | 0.08123761 | -0.233515579 | -0.093675364 | -0.314753189 | 0.064503328 | 0.572985197 | 0.295144386 | 0.635447101 | 0.206366012 |
| Tacc1 | S | 253 | 0.202045438 | -0.288532068 | -0.238721226 | -0.440766665 | 0.049810842 | 0.002055339 | 0.067604339 | 0.003036928 | 0.001617695 | 0.646565733 |
| Pak1 | S | 144 | -0.209723947 | 0.154015345 | -0.120752576 | 0.088971371 | -0.274767921 | 0.024603696 | 0.30359494 | 0.140545257 | 0.264227797 | 0.118802014 |
| Nolc1 | S | 701 | 0.070227287 | -0.086590426 | -0.125269608 | -0.195496894 | -0.038679182 | 0.066058175 | 0.008182313 | 0.013966141 | 0.00357456 | 0.236954866 |
| Dstn | S | 24 | -0.142475073 | 0.175466698 | -0.348310201 | -0.205835128 | -0.523776899 | 0.285309381 | 0.393694626 | 0.087407559 | 0.232842796 | 0.055757382 |
| Spp1 | S | 176 | 0.149522198 | -0.157892353 | -0.187299516 | -0.336821714 | -0.029407163 | 0.015940799 | 0.003607235 | 0.037630542 | 0.006755857 | 0.622902614 |
| Nsmce2 | S | 16 | -0.081508404 | -0.208673773 | -0.100055445 | -0.018547041 | 0.108618329 | 0.497064809 | 0.330319831 | 0.616921369 | 0.918542766 | 0.648485854 |
| Naca | S | 203;2175 | -0.12735304 | -0.07601804 | -0.133371789 | -0.006018749 | -0.057353749 | 0.309615155 | 0.668440177 | 0.34294606 | 0.94703908 | 0.721144317 |
| Eif4g2 | T | 505 | -0.116815616 | -0.057811043 | 0.02695945 | 0.143775066 | 0.084770493 | 0.002764929 | 0.231057688 | 0.293837781 | 0.000101901 | 0.125591353 |
| Map1a | S | 119 | -0.012455446 | -0.076102532 | -0.180131603 | -0.167676157 | -0.104029071 | 0.76541518 | 0.282888131 | 0.163587932 | 0.183821634 | 0.372929367 |
| Kat7 | T | 87 | -0.038430382 | -0.054193725 | -0.084859231 | -0.046428849 | -0.030665505 | 0.545729189 | 0.120934973 | 0.094189529 | 0.496923512 | 0.47759722 |
| Prr11 | S | 362 | -0.155323761 | -0.085229445 | -0.076198458 | 0.079125302 | 0.009030987 | 0.230737868 | 0.507107323 | 0.528064525 | 0.560738483 | 0.947906405 |
| Slc35a5 | S | 407 | 0.21387264 | -0.185211966 | -0.539729608 | -0.753602248 | -0.354517641 | 0.119641795 | 0.195867586 | 0.123392822 | 0.075905154 | 0.256430867 |
| Mical3 | S | 977 | -0.029562243 | -0.015237144 | -0.199666544 | -0.170104302 | -0.184429401 | 0.729602219 | 0.826603072 | 0.044829798 | 0.11354695 | 0.064068917 |
| Cc2d1a | S | 312 | -0.120658862 | 0.041976888 | -0.179345968 | -0.058687105 | -0.221322856 | 0.294644147 | 0.365344428 | 0.040199077 | 0.593228458 | 0.024807519 |
| Slc19a2 | S | 492 | -0.051558444 | -0.035580231 | -0.120107962 | -0.068549518 | -0.084527731 | 0.673609866 | 0.66271628 | 0.022909324 | 0.572447875 | 0.320352816 |
| Slc4a8 | S | 1076 | -0.07333186 | -0.000188322 | -0.21213909 | -0.13880723 | -0.211950769 | 0.472921973 | 0.997917642 | 0.097950541 | 0.288443307 | 0.104101033 |
| Nbeal2 | S | 2727 | -0.089216898 | -0.073234801 | -0.057505198 | 0.031711699 | 0.015729602 | 0.369093209 | 0.134975675 | 0.320565628 | 0.743639326 | 0.786790518 |
| Sntb2 | S | 202 | -0.050020707 | -0.036782072 | -0.118107886 | -0.068087179 | -0.081325813 | 0.345953185 | 0.730701345 | 0.047913087 | 0.240144416 | 0.470013436 |
| Sntb2 | S | 211 | -0.050020707 | -0.036782072 | -0.118107886 | -0.068087179 | -0.081325813 | 0.345953185 | 0.730701345 | 0.047913087 | 0.240144416 | 0.470013436 |
| Ywhaz | T | 232 | -0.019390875 | -0.033449462 | -0.082624321 | -0.063233446 | -0.049174859 | 0.389321993 | 0.363170715 | 0.009964225 | 0.031971107 | 0.208780782 |
| Tor1aip1 | S | 241 | -0.087394938 | 0.007114333 | -0.075237297 | 0.012157641 | -0.08235163 | 0.015230367 | 0.840990434 | 0.167455556 | 0.776203586 | 0.14526592 |
| Zc3hc1 | T | 371 | 0.105449289 | -0.247778834 | -0.292845718 | -0.398295006 | -0.045066884 | 0.143184151 | 0.140341713 | 0.195551879 | 0.111664813 | 0.822316351 |
| Clspn | S | 417 | 0.031590781 | -0.078499062 | -0.076953906 | -0.108544687 | 0.001545156 | 0.170174983 | 0.057093202 | 0.001766441 | 0.008764799 | 0.955361935 |
| Kif20a | S | 823 | 0.03864986 | -0.039674031 | -0.188945931 | -0.22759579 | -0.1492719 | 0.436976889 | 0.370908771 | 0.011766554 | 0.011393512 | 0.035323353 |

| | | | | | | | | | | | | |
|---|---|---|---|---|---|---|---|---|---|---|---|---|
| Cdk2 Cdk3 | Y | 15;15 | -0.018904304 | -0.009614033 | -0.19815491 | -0.179250606 | -0.188540877 | 0.762796919 | 0.903415582 | 0.030223657 | 0.060042318 | 0.086852402 |
| Uhrf1bp1l | S | 414 | -0.034729284 | 0.011721999 | -0.167940411 | -0.133211127 | -0.17966241 | 0.083619136 | 0.758835666 | 0.054706778 | 0.092023291 | 0.036759717 |
| Uhrf1bp1l | S | 418 | -0.034729284 | 0.011721999 | -0.167940411 | -0.133211127 | -0.17966241 | 0.083619136 | 0.758835666 | 0.054706778 | 0.092023291 | 0.036759717 |
| Pphln1 | S | 222 | 0.000316046 | -0.177813668 | -0.08779436 | -0.088110406 | 0.090019308 | 0.996599933 | 0.146160216 | 0.24482313 | 0.327311345 | 0.415322781 |
| Rnmt | S | 100 | -0.128166037 | 0.019077681 | -0.08124293 | 0.046923107 | -0.100320611 | 0.039962122 | 0.816021185 | 0.211905797 | 0.360304148 | 0.255560625 |
| Reps1 | S | 482 | -0.071789039 | -0.087416824 | -0.032245221 | 0.039543819 | 0.055171603 | 0.128648262 | 0.10662632 | 0.691537696 | 0.630545567 | 0.518185235 |
| Srrm1 | T | 409 | -0.056073596 | -0.042390126 | -0.134797997 | -0.0787244 | -0.092407871 | 0.310668263 | 0.748882734 | 0.091216652 | 0.225016009 | 0.500574123 |
| Uvrag | S | 497 | 0.01876765 | -0.07995483 | -0.139256281 | -0.158023931 | -0.059301451 | 0.809471853 | 0.114632664 | 0.088404472 | 0.131933726 | 0.387215583 |
| Polk | S | 607 | -0.083567558 | -0.028402933 | -0.241510515 | -0.157942957 | -0.213107582 | 0.575662181 | 0.650316634 | 0.215317926 | 0.441165905 | 0.259669104 |
| Pdpk1 | S | 25 | -0.024261033 | -0.042226068 | -0.126928847 | -0.102667814 | -0.084702779 | 0.632247273 | 0.302550022 | 0.05682065 | 0.108455026 | 0.155807322 |
| Ubr4 | S | 2716 | -0.084234023 | -0.024288277 | -0.060057609 | 0.024176414 | -0.035769332 | 0.177987849 | 0.511488062 | 0.049759902 | 0.639387221 | 0.321695554 |
| Srrm2 | S | 279 | 0.002398493 | -0.027754102 | -0.206410089 | -0.208808582 | -0.178655987 | 0.970709702 | 0.81597276 | 0.026342601 | 0.028961865 | 0.199688282 |
| Srrm2 | T | 955 | 0.014191755 | -0.228401676 | -0.001700652 | -0.015892407 | 0.226701024 | 0.873914496 | 0.016095136 | 0.984815445 | 0.858028125 | 0.040042579 |
| Tjap1 | T | 529 | -0.057482328 | -0.083167206 | -0.074538229 | -0.017055901 | 0.008628977 | 0.386005073 | 0.367062363 | 0.105650512 | 0.779741644 | 0.917488856 |
| Ctnnd1 | T | 59 | -0.040179735 | -0.007263975 | -0.145115275 | -0.104935541 | -0.137851301 | 0.68655691 | 0.936151846 | 0.014052864 | 0.337860276 | 0.209007956 |
| Usp45 | S | 452 | -0.150357367 | 0.038057553 | -0.131701798 | 0.018655569 | -0.169759351 | 0.115675481 | 0.79133116 | 0.205675411 | 0.837263356 | 0.289673637 |
| Rab12 | S | 24 | 0.012373609 | -0.019596268 | -0.155068156 | -0.167441765 | -0.135471889 | 0.801506254 | 0.646651382 | 0.00601263 | 0.029597151 | 0.017136331 |
| Stim2 | S | 640 | -0.058619956 | -0.154902658 | 0.000210641 | 0.058830597 | 0.155113299 | 0.494191322 | 0.032684427 | 0.997907669 | 0.509652567 | 0.095441477 |
| Cep170b | S | 840 | -0.086658478 | -0.081783783 | -4.29594E-05 | 0.086615519 | 0.081740824 | 0.103764216 | 0.055354572 | 0.998257361 | 0.096304996 | 0.050883773 |
| Prune2 | S | 1678 | -0.068071224 | -0.050871261 | -0.079676992 | -0.011605767 | -0.02880573 | 0.252832656 | 0.218943453 | 0.222897132 | 0.864231324 | 0.633387532 |
| Topbp1 | S | 839 | -0.163966294 | 0.048027195 | -0.242365992 | -0.078399698 | -0.290393186 | 0.282451537 | 0.734786566 | 0.23127195 | 0.693261702 | 0.196959781 |
| Steap1 | T | 29 | -0.05534672 | -0.14363565 | -0.049074223 | 0.006272496 | 0.094561426 | 0.486470136 | 0.095878044 | 0.636232905 | 0.946566358 | 0.348756028 |
| Srrm1 | S | 676 | -0.011963851 | -0.037001926 | -0.157470659 | -0.145506807 | -0.120468732 | 0.815644969 | 0.578443375 | 0.031790866 | 0.050051543 | 0.133906324 |
| Chmp2b | S | 199 | -0.04109472 | -0.047060062 | -0.187189709 | -0.14609499 | -0.140129647 | 0.714724926 | 0.629056342 | 0.095685126 | 0.266869772 | 0.235050361 |
| Fnip1 | S | 272 | -0.133782065 | -0.094352166 | -0.089731469 | 0.044050595 | 0.004620697 | 0.194515198 | 0.691611396 | 0.436029977 | 0.723372459 | 0.984881258 |
| Supt20 | S | 520 | -0.271595438 | 0.046078639 | -0.137163739 | 0.1344317 | -0.183242378 | 0.121176399 | 0.59849298 | 0.465638765 | 0.51856112 | 0.351414531 |
| Prpf4b | S | 366 | -0.081173536 | -0.059847519 | -0.026027608 | 0.055145928 | 0.033819911 | 0.054912684 | 0.153897144 | 0.645074496 | 0.379980806 | 0.58336595 |
| Pja1 | S | 317 | -0.04933425 | -0.068433387 | -0.033030254 | 0.016303996 | 0.035403133 | 0.16460781 | 0.043187638 | 0.442870461 | 0.692295157 | 0.389200921 |
| Jcad | S | 774 | -0.093210265 | -0.014600406 | -0.126205488 | -0.032995223 | -0.111605082 | 0.35247565 | 0.792947247 | 0.086408776 | 0.733099252 | 0.134802156 |
| Elac2 | S | 195 | 0.028705903 | -0.022632462 | -0.20598331 | -0.234689213 | -0.183350848 | 0.433829574 | 0.548615532 | 0.01098022 | 0.019544996 | 0.029455956 |
| Topors | S | 538 | 0.058386906 | -0.181049314 | -0.097099715 | -0.155486621 | 0.0839496 | 0.24646924 | 0.011648794 | 0.362595905 | 0.192180721 | 0.412813673 |
| Fam53c | S | 273 | -0.096790388 | 0.011164526 | -0.126912662 | -0.030122274 | -0.138077188 | 0.105485434 | 0.847881343 | 0.137228698 | 0.648692596 | 0.109870112 |
| Ctdspl2 | S | 132 | -0.040466496 | -0.047275589 | -0.141065689 | -0.100599193 | -0.0937901 | 0.774579856 | 0.47786737 | 0.056261578 | 0.488332606 | 0.146837883 |
| Sox13 | S | 378 | -0.19998256 | -0.062524913 | -0.017808219 | 0.182174342 | 0.044716694 | 0.098687216 | 0.555536948 | 0.85475886 | 0.067760755 | 0.50228365 |
| Dclk1 | S | 330 | -0.028693573 | -0.018737756 | -0.116216455 | -0.087522882 | -0.097478698 | 0.239614009 | 0.511930888 | 0.040644383 | 0.087600136 | 0.067235561 |
| Emd | S | 177 | -0.056174012 | -0.029374035 | -0.190054611 | -0.133880599 | -0.160680576 | 0.488571753 | 0.696127805 | 0.131783881 | 0.249142121 | 0.181059484 |
| Rbm28 | S | 239 | -0.00436198 | -0.195338013 | 0.01809695 | 0.02245893 | 0.213434964 | 0.92014856 | 0.009849625 | 0.894490776 | 0.866719069 | 0.203827781 |
| Cndp2 | S | 354 | -0.075453925 | -0.08797899 | -0.094020352 | -0.018566427 | -0.006041362 | 0.401196513 | 0.164950238 | 0.477985996 | 0.887581067 | 0.959351119 |
| Cndp2 | S | 359 | -0.075453925 | -0.08797899 | -0.094020352 | -0.018566427 | -0.006041362 | 0.401196513 | 0.164950238 | 0.477985996 | 0.887581067 | 0.959351119 |
| Cndp2 | Y | 360 | -0.075453925 | -0.08797899 | -0.094020352 | -0.018566427 | -0.006041362 | 0.401196513 | 0.164950238 | 0.477985996 | 0.887581067 | 0.959351119 |
| Kif14 | S | 1462 | -0.188668383 | -0.04128975 | -0.15998484 | 0.028683543 | -0.118695089 | 0.464602909 | 0.690478046 | 0.333603476 | 0.910820731 | 0.429343411 |
| Kif14 | T | 1466 | -0.188668383 | -0.04128975 | -0.15998484 | 0.028683543 | -0.118695089 | 0.464602909 | 0.690478046 | 0.333603476 | 0.910820731 | 0.429343411 |
| Nfkb1 | S | 897 | 0.04977313 | -0.12423856 | -0.127930975 | -0.177704105 | -0.003692415 | 0.539698498 | 0.209048389 | 0.034242407 | 0.085702019 | 0.963520479 |
| Vps50 | S | 559 | -0.128177265 | -0.028651198 | -0.062524551 | 0.065652714 | -0.033873353 | 0.162047987 | 0.491635396 | 0.245026518 | 0.427177523 | 0.528858273 |
| Ankrd17 | S | 2043 | 0.013954598 | -0.079342742 | -0.071092178 | -0.085046776 | 0.008250563 | 0.843845912 | 0.078971692 | 0.021720581 | 0.295329888 | 0.810480313 |
| Srrm2 | S | 1378 | -0.030438449 | -0.087067267 | -0.003205521 | 0.027232929 | 0.083861747 | 0.373180734 | 0.00200452 | 0.947009529 | 0.607908546 | 0.175092099 |
| 4930550C14Rik | S | 272 | -0.161005739 | -0.141851411 | -0.031460001 | 0.129545738 | 0.110391411 | 0.239652264 | 0.457989775 | 0.684333273 | 0.320292755 | 0.550395369 |
| 4930550C14Rik | S | 275 | -0.161005739 | -0.141851411 | -0.031460001 | 0.129545738 | 0.110391411 | 0.239652264 | 0.457989775 | 0.684333273 | 0.320292755 | 0.550395369 |
| Pard3b | Y | 953 | -0.107343343 | -0.049104263 | -0.160512378 | -0.053169036 | -0.111408115 | 0.377879679 | 0.674438437 | 0.269576583 | 0.727595934 | 0.480962466 |
| Hdgfl2 | S | 640 | -0.050304091 | -0.036341833 | -0.081897682 | -0.031593591 | -0.045555849 | 0.135000531 | 0.529024434 | 0.083030809 | 0.435201409 | 0.463163754 |
| Fbxo42 | S | 488 | -0.033534114 | 0.059896022 | -0.297471885 | -0.263937771 | -0.357367907 | 0.782747705 | 0.669942843 | 0.038310272 | 0.019147731 | 0.031279848 |
| Tmem245 | S | 327 | -0.006281355 | -0.034841005 | -0.073477239 | -0.067195884 | -0.038636235 | 0.785968701 | 0.103174327 | 0.007380865 | 0.050690206 | 0.104089085 |
| Kmt2a | S | 181 | 0.094970509 | -0.139333632 | -0.246694324 | -0.341664833 | -0.107360692 | 0.495792724 | 0.285782867 | 0.095607883 | 0.073964217 | 0.458303825 |
| Slc12a2 | T | 259 | -0.066114614 | -0.136692743 | 0.03819448 | 0.104309094 | 0.174887222 | 0.324251661 | 0.007729174 | 0.477802788 | 0.174114444 | 0.027364234 |
| Lpar6 | S | 317 | -0.122696821 | -0.102317336 | 0.018790075 | 0.141486896 | 0.121107411 | 0.050554577 | 0.250441606 | 0.817545166 | 0.135615613 | 0.238493789 |
| Map3k2 | S | 344 | 0.056814073 | -0.089454313 | -0.170799157 | -0.22761323 | -0.081344844 | 0.283064957 | 0.044872719 | 0.077825611 | 0.032443529 | 0.276095484 |
| Tjp1 | T | 354 | 0.057565951 | -0.110175189 | -0.130525391 | -0.188091342 | -0.020350202 | 0.26401227 | 0.070785714 | 0.037027022 | 0.005342002 | 0.597106858 |
| Zc3h13 | S | 1273 | -0.047218369 | -0.07332031 | -0.045743971 | 0.001474399 | 0.02757634 | 0.48919847 | 0.028614933 | 0.330864683 | 0.982956176 | 0.500627339 |
| Cep170b | S | 1186 | 0.083296455 | -0.092339379 | -0.252690182 | -0.335986636 | -0.160350803 | 0.056225809 | 0.023842285 | 0.069248917 | 0.037691457 | 0.164353465 |
| Scarf2 | S | 638 | 0.039727053 | -0.150206165 | -0.106159318 | -0.14588637 | 0.044046848 | 0.585021184 | 0.065963662 | 0.181141828 | 0.025409397 | 0.308239147 |
| Tsr1 | S | 795 | -0.126454517 | -0.003212952 | -0.057898741 | 0.068555776 | -0.054685789 | 0.051748046 | 0.917919419 | 0.233094796 | 0.244047955 | 0.229377105 |
| Tsr1 | S | 798 | -0.126454517 | -0.003212952 | -0.057898741 | 0.068555776 | -0.054685789 | 0.051748046 | 0.917919419 | 0.233094796 | 0.244047955 | 0.229377105 |
| Prr14 | S | 142 | 0.081621926 | -0.101101857 | -0.359531247 | -0.441153173 | -0.25842939 | 0.397767741 | 0.251383572 | 0.154420226 | 0.106900331 | 0.261710437 |
| Vim | T | 361 | -0.056579513 | -0.031103938 | -0.144316335 | -0.087736822 | -0.113212397 | 0.417852955 | 0.682318979 | 0.091650881 | 0.243119599 | 0.183790767 |
| Samd4 | S | 658 | -0.093418484 | -0.066820141 | -0.05697707 | 0.036441414 | 0.009843071 | 0.168944815 | 0.288452156 | 0.41915949 | 0.518841454 | 0.848603158 |
| Tcof1 | S | 1124 | -0.159654448 | 0.024555467 | -0.114909556 | 0.044744892 | -0.139465023 | 0.042322023 | 0.676107701 | 0.444549025 | 0.745726653 | 0.358766387 |
| Arl6ip4 | S | 140 | -0.124730907 | -0.080833698 | -0.041977558 | 0.082753349 | 0.038856141 | 0.165049026 | 0.392291941 | 0.508550625 | 0.281473721 | 0.632395329 |
| Gapdh | S | 208 | 0.003860939 | -0.086666263 | -0.159300943 | -0.163161882 | -0.07263468 | 0.945730799 | 0.359462599 | 0.08025899 | 0.079801575 | 0.464674988 |
| Irak1 | S | 642 | 0.004512397 | -0.009804687 | -0.161457487 | -0.165969884 | -0.1516528 | 0.971414419 | 0.833752366 | 0.007781651 | 0.261464688 | 0.007521691 |
| Camk2d | S | 280 | -0.136529615 | 0.058977977 | -0.156238069 | -0.019708454 | -0.215216046 | 0.060045082 | 0.332699074 | 0.142928084 | 0.816929579 | 0.079317784 |
| Camk2b Camk2g | S | 280;280 | -0.136529615 | 0.058977977 | -0.156238069 | -0.019708454 | -0.215216046 | 0.060045082 | 0.332699074 | 0.142928084 | 0.816929579 | 0.079317784 |
| Trps1 | S | 157 | -0.077518299 | -0.121594651 | 0.007940493 | 0.085458792 | 0.129535144 | 0.160185656 | 0.072027804 | 0.914916037 | 0.258975333 | 0.132714276 |
| Nmt2 | S | 61 | -0.193334691 | -0.005273944 | -0.077485965 | 0.115848726 | -0.072212021 | 0.12732805 | 0.960594865 | 0.390545449 | 0.339364182 | 0.545809213 |
| Kif7 | S | 662 | 0.037877905 | -0.097539927 | -0.123401859 | -0.161279764 | -0.025861932 | 0.118794011 | 0.033309434 | 0.042743199 | 0.017952748 | 0.536947183 |
| Fhdc1 | S | 681 | 0.007906196 | -0.038585174 | -0.161506805 | -0.169413001 | -0.122921632 | 0.870933265 | 0.161194453 | 0.055549057 | 0.053075283 | 0.117275993 |
| Phldb2 | S | 389 | 0.092533109 | -0.088605393 | -0.231648506 | -0.324181614 | -0.143043113 | 0.163598577 | 0.120084231 | 0.032407551 | 0.014260603 | 0.122293176 |
| Rtn4 | S | 291 | -0.063176548 | -0.097823464 | 0.009904015 | 0.073080563 | 0.107727478 | 0.134419142 | 0.020019163 | 0.699488042 | 0.104740559 | 0.020180555 |
| Brwd1 | S | 1857 | -0.026983608 | -0.064697028 | -0.041975613 | -0.014992005 | 0.022721415 | 0.072146756 | 0.067667897 | 0.280444637 | 0.658881904 | 0.562123944 |
| Vcl | S | 913 | -0.113976437 | -0.022380187 | -0.104619572 | 0.009356865 | -0.082239385 | 0.421036931 | 0.698976384 | 0.082535898 | 0.940678533 | 0.084278338 |
| Vcl | S | 914 | -0.113976437 | -0.022380187 | -0.104619572 | 0.009356865 | -0.082239385 | 0.421036931 | 0.698976384 | 0.082535898 | 0.940678533 | 0.084278338 |
| Leng8 | S | 352 | -0.03554339 | -0.028187831 | -0.044106819 | -0.008563428 | -0.015918987 | 0.051618061 | 0.057169458 | 0.120477261 | 0.709010164 | 0.48091818 |
| Dnaja2 | S | 187 | -0.027079161 | -0.086966238 | -0.231530263 | -0.204451102 | -0.144564025 | 0.667571863 | 0.327749432 | 0.342729243 | 0.390068801 | 0.527703 |
| Ncor2 | S | 921 | 0.030659604 | -0.055530686 | -0.105674122 | -0.136333726 | -0.050143437 | 0.590269018 | 0.434886572 | 0.00157845 | 0.095266855 | 0.470948377 |
| Map1b | S | 1293 | 0.006567765 | -0.073391385 | -0.07685465 | -0.083422415 | -0.003463265 | 0.786379325 | 0.036458476 | 0.06350765 | 0.06037613 | 0.915588357 |
| Bcl2l13 | S | 35 | -0.112179721 | -0.106763386 | -0.081441258 | 0.030738463 | 0.025322128 | 0.513957462 | 0.220772838 | 0.497141624 | 0.856989129 | 0.792884985 |
| Wdr55 | S | 380 | 0.027521697 | -0.253605457 | 0.004295673 | -0.023226025 | 0.257901129 | 0.82602847 | 0.020120975 | 0.978773103 | 0.894174697 | 0.181444875 |
| Wdr55 | S | 387 | 0.027521697 | -0.253605457 | 0.004295673 | -0.023226025 | 0.257901129 | 0.82602847 | 0.020120975 | 0.978773103 | 0.894174697 | 0.181444875 |
| Map7d1 | S | 127 | 0.092910286 | -0.037485818 | -0.182591668 | -0.275501954 | -0.14510585 | 0.421688488 | 0.12810212 | 0.009835405 | 0.084209908 | 0.032746155 |
| Lmna | S | 398 | 0.143591115 | -0.11806234 | -0.228139541 | -0.371730656 | -0.110077201 | 0.010487259 | 0.027635274 | 0.006570336 | 0.003282301 | 0.085809504 |
| Tns3 | S | 937 | -0.063673601 | 0.004438822 | -0.174274214 | -0.110600613 | -0.178713036 | 0.457641868 | 0.961930024 | 0.058838905 | 0.154917118 | 0.08005303 |
| Arhgap17 | S | 669 | -0.096134358 | -0.092321333 | -0.089235245 | 0.006899113 | 0.003086088 | 0.390858664 | 0.334603883 | 0.379415539 | 0.930424247 | 0.949563185 |
| Epha2 | S | 572 | -0.116175904 | 0.001544316 | -0.067018433 | 0.049157471 | -0.068562749 | 0.060374928 | 0.986306119 | 0.168042781 | 0.310052615 | 0.45817143 |
| Map1a | S | 118 | 0.016828119 | -0.07234555 | -0.124014731 | -0.14084285 | -0.051669181 | 0.822978448 | 0.316251734 | 0.023886016 | 0.132852019 | 0.446735896 |
| Nrip1 | S | 519 | -0.00356905 | -0.058592718 | -0.130310253 | -0.126741203 | -0.071717535 | 0.952356261 | 0.33767842 | 0.040579936 | 0.055856364 | 0.19120858 |

| | | | | | | | | | | | | |
|---|---|---|---|---|---|---|---|---|---|---|---|---|
| Akt1s1 | S | 117 | -0.141262191 | -0.100893001 | -0.111125514 | 0.030136678 | -0.010232513 | 0.566784436 | 0.458474213 | 0.337034512 | 0.902264038 | 0.944893268 |
| Peak1 | S | 1050 | 0.075995933 | -0.088722431 | -0.158305888 | -0.234301821 | -0.069583458 | 0.183331711 | 0.177093293 | 0.00764119 | 0.011361648 | 0.235895066 |
| Poldip2 | T | 292 | 0.058000927 | -0.210722407 | -0.089031118 | -0.147032045 | 0.121691289 | 0.178723718 | 0.012293521 | 0.456769318 | 0.264543505 | 0.337845745 |
| Nrbp1 | T | 431 | -0.02002264 | -0.155034767 | -0.089804922 | -0.069782282 | 0.065229845 | 0.796599298 | 0.189025347 | 0.286632756 | 0.312185772 | 0.520621375 |
| Tnks1bp1 | S | 1466 | -0.041822379 | -0.013758993 | -0.128757519 | -0.08693514 | -0.114998526 | 0.378306546 | 0.79638802 | 0.036453813 | 0.144175218 | 0.102784203 |
| Tns3 | S | 934 | -0.09146744 | -0.007981589 | -0.133395098 | -0.041927658 | -0.125413508 | 0.203164853 | 0.894356375 | 0.156642461 | 0.601371371 | 0.165211269 |
| Ppp4r3b | S | 811 | 0.043058813 | -0.072397849 | -0.135334174 | -0.178392988 | -0.062936326 | 0.19783887 | 0.120826955 | 0.010919782 | 0.008322788 | 0.177606469 |
| Epb41l1 | S | 564 | -0.039514595 | 0.066665787 | -0.183203958 | -0.143689363 | -0.249869745 | 0.186937023 | 0.142500685 | 0.003757339 | 0.018335296 | 0.003468357 |
| Polr2m | S | 363 | -0.095178481 | -0.03518578 | -0.060609974 | 0.034568507 | -0.025424193 | 0.092980888 | 0.474240754 | 0.298425352 | 0.378300679 | 0.484140091 |
| Klc2 | S | 575 | -0.076004974 | -0.099926035 | -0.109213526 | -0.033208552 | -0.00928749 | 0.554690144 | 0.439897346 | 0.214845608 | 0.799435169 | 0.942039506 |
| Topors | S | 568 | 0.000427568 | -0.073953532 | -0.074939723 | -0.075367292 | -0.000986191 | 0.993302543 | 0.126505186 | 0.027766601 | 0.222700332 | 0.979946542 |
| Erf | S | 161 | -0.036811205 | -0.025079454 | -0.120965158 | -0.084153953 | -0.095885704 | 0.41048482 | 0.386183383 | 0.067304068 | 0.180073198 | 0.118898607 |
| Plekhg3 | S | 1129 | 0.056483276 | -0.111831562 | -0.09779427 | -0.154277546 | 0.014037292 | 0.082218733 | 0.010951097 | 0.035394581 | 0.011026614 | 0.697723363 |
| Zmpste24 | S | 298 | -0.070803067 | -0.079132458 | -0.124874093 | -0.054071026 | -0.045741635 | 0.270745394 | 0.638366537 | 0.254271371 | 0.593941032 | 0.79679969 |
| Map3k4 | T | 1243 | -0.035816833 | -0.099480359 | -0.058265898 | -0.022449065 | 0.04121446 | 0.580251406 | 0.037884684 | 0.519458727 | 0.819867204 | 0.645133693 |
| Ncapd2 | S | 466 | -0.014997482 | -0.087510175 | -0.133962967 | -0.118965484 | -0.046452792 | 0.868176635 | 0.233242796 | 0.151739588 | 0.300154623 | 0.601516475 |
| Srsf7 | S | 210 | -0.037656258 | -0.093095564 | -0.032596193 | 0.005060065 | 0.060499371 | 0.359118107 | 0.031755217 | 0.53778366 | 0.916183353 | 0.250557938 |
| Tasor2 | S | 2014 | 0.015596657 | -0.140778732 | -0.076440761 | -0.092037418 | 0.064337971 | 0.847180887 | 0.040196059 | 0.349787168 | 0.364661134 | 0.432598925 |
| Wnk1 | S | 10 | -0.017009385 | -0.073076194 | -0.181656044 | -0.16464666 | -0.10857985 | 0.877273834 | 0.561034247 | 0.096149284 | 0.199322182 | 0.411542311 |
| Arhgap31 | S | 814 | 0.011154015 | -0.045524783 | -0.159631035 | -0.170785051 | -0.114106252 | 0.729862306 | 0.464976017 | 0.022737743 | 0.01810667 | 0.141801631 |
| Bag6 | S | 1103 | -0.114242757 | -0.002794454 | -0.117055998 | -0.002813241 | -0.114261543 | 0.157861528 | 0.974904841 | 0.196030116 | 0.965122841 | 0.205451637 |
| Rab3gap1 | T | 535 | 0.062358826 | -0.12632623 | -0.170395735 | -0.232754561 | -0.044069505 | 0.151155692 | 0.097665178 | 0.053982751 | 0.03105862 | 0.566553122 |
| Kif4 | S | 788 | -0.02431729 | 0.013093845 | -0.086325768 | -0.062008478 | -0.099419613 | 0.196913272 | 0.610978213 | 0.000788618 | 0.020707877 | 0.026722818 |
| Map4k5 | S | 304 | -0.053170229 | -0.035954631 | -0.17188764 | -0.118717411 | -0.135933009 | 0.424089634 | 0.668956785 | 0.151412465 | 0.282159656 | 0.258966793 |
| Mpdz | S | 496 | 0.046570914 | -0.123920656 | -0.089711713 | -0.136282627 | 0.034208943 | 0.270041322 | 0.019879438 | 0.100993356 | 0.033574541 | 0.36426726 |
| Adrm1 | S | 405 | 0.038612915 | -0.123051308 | -0.051288338 | -0.089901253 | 0.07176297 | 0.377408603 | 0.019550295 | 0.043347122 | 0.102005458 | 0.090766167 |
| Pold3 | S | 419 | -0.034917838 | -0.086244969 | -0.14243227 | -0.107514432 | -0.0561873 | 0.700159877 | 0.493207411 | 0.125112162 | 0.24917714 | 0.640128695 |
| Osbp | S | 236 | -0.02741121 | -0.090134526 | -0.115893626 | -0.088482416 | -0.0257591 | 0.787626349 | 0.460261168 | 0.068899948 | 0.38486876 | 0.812464775 |
| Lats1 | T | 261 | -0.011198615 | -0.074906223 | -0.165838014 | -0.1546394 | -0.090931792 | 0.880311244 | 0.438774508 | 0.101843516 | 0.110619663 | 0.369855053 |
| Lats1 | T | 262 | -0.011198615 | -0.074906223 | -0.165838014 | -0.1546394 | -0.090931792 | 0.880311244 | 0.438774508 | 0.101843516 | 0.110619663 | 0.369855053 |
| Tln1 | S | 1201 | -0.051051356 | -0.076234001 | -0.034267376 | 0.01678398 | 0.041966625 | 0.233679549 | 0.098918989 | 0.279003173 | 0.622662049 | 0.271136549 |
| Mast4 | S | 1393 | -0.096691393 | -0.079463321 | -0.018266938 | 0.078424456 | 0.061196383 | 0.0438978 | 0.382853268 | 0.731838739 | 0.149147738 | 0.492156991 |
| Psen2 | S | 52 | -0.135745717 | -0.083480769 | 0.071037641 | 0.206783358 | 0.15451841 | 0.062443667 | 0.065157809 | 0.60242464 | 0.204049055 | 0.307363859 |
| Cep170b | S | 1101 | -0.0388583 | 0.003383267 | -0.28567495 | -0.24681665 | -0.289058217 | 0.610736467 | 0.96766446 | 0.123292104 | 0.165492082 | 0.122351908 |
| Nucb2 | S | 332 | -0.19795957 | -0.033965952 | 0.099369692 | 0.297329262 | 0.133335643 | 5.00923E-05 | 0.461460547 | 0.012302555 | 0.000735737 | 0.044760294 |
| Gja1 | S | 368 | 0.051698352 | -0.081429562 | -0.194543073 | -0.246241425 | -0.113113511 | 0.604599974 | 0.077961675 | 0.151146487 | 0.113672902 | 0.331490613 |
| Tfeb | S | 454 | -0.465655592 | 0.004040732 | 0.233102745 | 0.698758337 | 0.229062013 | 0.023190317 | 0.96766446 | 0.475889647 | 0.099664561 | 0.513967011 |
| Eme1 | S | 15 | -0.027686634 | -0.078067748 | -0.052065812 | -0.024379178 | 0.026001936 | 0.396095464 | 0.032499417 | 0.384804623 | 0.651935335 | 0.627390289 |
| Myo5a | S | 600 | -0.079701413 | -0.051077747 | -0.009088987 | 0.070612426 | 0.04198876 | 0.029797383 | 0.114529947 | 0.850969036 | 0.190955609 | 0.379545684 |
| Rabl6 | S | 480 | 0.017481573 | -0.105359021 | -0.092090534 | -0.109572107 | 0.013268487 | 0.427222942 | 0.053010501 | 0.0948676 | 0.067699106 | 0.785750201 |
| Bcl9l | S | 118 | -0.179242466 | 0.048128684 | -0.05332424 | 0.125918226 | -0.101452924 | 0.016080953 | 0.5894537 | 0.475335136 | 0.108466445 | 0.291956794 |
| Cfl1 | S | 3 | -0.137267108 | -0.031804632 | -0.091869833 | 0.045397276 | -0.060065201 | 0.112154929 | 0.673553138 | 0.534686542 | 0.741748516 | 0.663028127 |
| Mcm4 | S | 23 | -0.172208134 | 0.110992356 | -0.189953657 | -0.017745523 | -0.300946013 | 0.132383536 | 0.394634639 | 0.123342759 | 0.879828513 | 0.085012502 |
| Chd7 | S | 2548 | 0.002189886 | -0.082076434 | -0.110435011 | -0.112624897 | -0.028358577 | 0.97795208 | 0.112419102 | 0.12171281 | 0.216260618 | 0.586387933 |
| Srrm2 | S | 1216 | -0.076025637 | -0.118569603 | 0.021867657 | 0.097893294 | 0.14043726 | 0.173613273 | 0.037022366 | 0.682767805 | 0.077797376 | 0.023775511 |
| Nav1 | S | 202 | 0.025845241 | -0.105869209 | -0.107829039 | -0.133674279 | -0.00195983 | 0.612451295 | 0.132989713 | 0.061068044 | 0.044905094 | 0.973846871 |
| Usp20 | S | 374 | 0.002142698 | -0.08279546 | -0.08145769 | -0.083600388 | 0.001337771 | 0.976624953 | 0.122573054 | 0.053611522 | 0.311184011 | 0.97644872 |
| Prrx1 | S | 72 | -0.128177546 | 0.04236295 | -0.09037491 | 0.037802635 | -0.13273786 | 0.019525081 | 0.462309075 | 0.222698561 | 0.565367977 | 0.121676592 |
| Mphosph9 | S | 815 | 0.047760104 | -0.045065396 | -0.327741458 | -0.375501563 | -0.282676063 | 0.699680867 | 0.469287598 | 0.114019604 | 0.084605788 | 0.137750721 |
| Mphosph9 | S | 816 | 0.047760104 | -0.045065396 | -0.327741458 | -0.375501563 | -0.282676063 | 0.699680867 | 0.469287598 | 0.114019604 | 0.084605788 | 0.137750721 |
| Prrc2a | S | 1113 | 0.009247934 | -0.103399974 | -0.080445657 | -0.089693591 | 0.022954317 | 0.698663523 | 0.05698528 | 0.122643766 | 0.101291287 | 0.64475651 |
| Tnks1bp1 | S | 1370 | -0.036196692 | -0.035509469 | -0.304963168 | -0.268766477 | -0.2694537 | 0.792837925 | 0.720924778 | 0.20285544 | 0.257276647 | 0.247529049 |
| Tnks1bp1 | S | 1371 | -0.036196692 | -0.035509469 | -0.304963168 | -0.268766477 | -0.2694537 | 0.792837925 | 0.720924778 | 0.20285544 | 0.257276647 | 0.247529049 |
| Serbp1 | S | 329 | -0.243250744 | -0.017964754 | 0.03621962 | 0.279470363 | 0.054184374 | 0.02437229 | 0.818842209 | 0.700593871 | 0.035579532 | 0.555720913 |
| Mast4 | T | 1388 | -0.088211938 | -0.052987806 | -0.069931373 | 0.018280565 | -0.016943567 | 0.06035182 | 0.636801066 | 0.425108856 | 0.81301937 | 0.890500139 |
| Ptpn21 | S | 799 | -0.042340048 | -0.030382551 | -0.108028585 | -0.065688537 | -0.077646033 | 0.406531835 | 0.395498014 | 0.071641005 | 0.269739609 | 0.153092082 |
| Erf | S | 154 | -0.034165527 | -0.023532573 | -0.123740603 | -0.089575076 | -0.10020803 | 0.442667952 | 0.328170233 | 0.073849726 | 0.170937884 | 0.124816203 |
| Tacc3 | T | 59 | 0.043336056 | -0.002752419 | -0.200597833 | -0.243933888 | -0.197845413 | 0.451964295 | 0.933110055 | 0.003380925 | 0.016941331 | 0.005558052 |
| Mcrip1 | S | 21 | 0.013535994 | -0.031708557 | -0.104431212 | -0.117967206 | -0.072722655 | 0.653251058 | 0.218220691 | 0.004403424 | 0.014102745 | 0.001514619 |
| Ralgapb | S | 502 | -0.120734911 | -0.141636963 | -0.009035623 | 0.111699288 | 0.13260134 | 0.318453831 | 0.180082993 | 0.892884573 | 0.333081773 | 0.187906531 |
| Gtpbp1 | S | 25 | -0.073779606 | -0.029933524 | -0.073513932 | 0.000265673 | -0.043580408 | 0.077936366 | 0.632689002 | 0.197212975 | 0.995687962 | 0.533811041 |
| Cdr2l | S | 178 | -0.048361374 | -0.033448441 | -0.091568443 | -0.043207069 | -0.058120002 | 0.130304617 | 0.299124867 | 0.204715323 | 0.485679416 | 0.372313132 |
| Dab2 | S | 759 | 0.010825091 | -0.142573403 | -0.08899176 | -0.099816851 | 0.053581643 | 0.831584022 | 0.131710521 | 0.161286793 | 0.143682493 | 0.520310097 |
| Gpbp1 | S | 398 | -0.063946681 | -0.094688844 | -0.109407284 | -0.045460603 | -0.01471844 | 0.363421152 | 0.206557733 | 0.55300111 | 0.797442997 | 0.933458528 |
| Tpd52l2 | S | 209 | -0.071944042 | -0.030235414 | -0.04411505 | 0.027828991 | -0.013879637 | 0.174148318 | 0.25918945 | 0.074236158 | 0.526771949 | 0.470400318 |
| Tnk2 | S | 622 | -0.00157569 | -0.0465755 | -0.088945137 | -0.087369447 | -0.042369637 | 0.976769128 | 0.176658694 | 0.016384335 | 0.179952069 | 0.190844521 |
| Tnk2 | T | 618 | -0.00157569 | -0.0465755 | -0.088945137 | -0.087369447 | -0.042369637 | 0.976769128 | 0.176658694 | 0.016384335 | 0.179952069 | 0.190844521 |
| Aldh3a1 | S | 446 | 0.022140015 | 0.028545634 | -0.244592166 | -0.266732181 | -0.2731378 | 0.709743392 | 0.62362251 | 0.007269155 | 0.007657583 | 0.00772728 |
| Zmym4 | S | 1065 | -0.096442388 | -0.118613718 | -0.075427764 | 0.021014624 | 0.043185954 | 0.530203785 | 0.195106622 | 0.567985573 | 0.904186063 | 0.746242583 |
| Zmym4 | S | 1069 | -0.096442388 | -0.118613718 | -0.075427764 | 0.021014624 | 0.043185954 | 0.530203785 | 0.195106622 | 0.567985573 | 0.904186063 | 0.746242583 |
| Zmym4 | T | 1071 | -0.096442388 | -0.118613718 | -0.075427764 | 0.021014624 | 0.043185954 | 0.530203785 | 0.195106622 | 0.567985573 | 0.904186063 | 0.746242583 |
| Tbc1d10b | S | 650 | 0.083942066 | 0.015043868 | -0.327754889 | -0.411696955 | -0.342798757 | 0.492191927 | 0.876349815 | 0.014014365 | 0.032070861 | 0.03078578 |
| Polk | S | 605 | -0.07281416 | -0.083996164 | -0.153505314 | -0.080691154 | -0.06950915 | 0.587023403 | 0.524914047 | 0.245566832 | 0.502260676 | 0.550668606 |
| Map4k4 | S | 556 | -0.032968956 | -0.06918507 | -0.100126649 | -0.067157693 | -0.030941579 | 0.458723808 | 0.253967232 | 0.16405835 | 0.298103755 | 0.647721094 |
| Spindoc | S | 313 | -0.039900296 | -0.119566826 | -0.048958074 | -0.009057778 | 0.070608751 | 0.583912251 | 0.072598267 | 0.489384152 | 0.911481764 | 0.346340191 |
| Klf16 | S | 225 | -0.011474074 | -0.129740831 | -0.030257926 | -0.018783852 | 0.099482904 | 0.920764254 | 0.010671492 | 0.741341689 | 0.887967229 | 0.319863494 |
| Stx12 | S | 218 | -0.020587628 | -0.157700289 | -0.139553875 | -0.118966247 | 0.018146414 | 0.91392929 | 0.174373379 | 0.468046477 | 0.629854512 | 0.924198102 |
| Eif2s2 | S | 105 | -0.073015547 | -0.02740404 | -0.064320071 | 0.008695476 | -0.036916031 | 0.121372677 | 0.537339264 | 0.119339893 | 0.773768194 | 0.293824056 |
| Mki67 | T | 2538 | -0.10534288 | -0.067214184 | -0.075176811 | 0.030166069 | -0.007962627 | 0.236233572 | 0.423844327 | 0.399860144 | 0.701042221 | 0.916110758 |
| Hdac1 | S | 409 | -0.031363746 | -0.139234899 | 0.037216575 | 0.068580321 | 0.176451474 | 0.563233463 | 0.002299597 | 0.495825102 | 0.312251575 | 0.039848349 |
| Vps50 | S | 561 | -0.141476815 | -0.031679892 | -0.0592717 | 0.082205115 | -0.027591808 | 0.199020916 | 0.485793428 | 0.311057131 | 0.416743257 | 0.641546985 |
| Samd4 | S | 573 | -0.031081468 | -0.019412717 | -0.117003337 | -0.085921869 | -0.09759062 | 0.593705566 | 0.612569757 | 0.022252926 | 0.192985841 | 0.050496614 |
| Nsd3 | S | 547 | -0.055290868 | -0.092952923 | -0.045063097 | 0.010227771 | 0.047889826 | 0.269107065 | 0.171097386 | 0.35082244 | 0.837312953 | 0.450143549 |
| Rgs17 | S | 55 | -0.057660891 | -0.2555078 | 0.129564243 | 0.187225134 | 0.385072043 | 0.423132589 | 0.011002893 | 0.282425077 | 0.152582272 | 0.030352355 |
| Lbr | S | 103 | -0.10842594 | -0.058061976 | 0.011846879 | 0.120272819 | 0.069908856 | 0.036564162 | 0.077814632 | 0.482033952 | 0.03520367 | 0.056502463 |
| Zc3h3 | S | 118 | 0.019321374 | -0.068719588 | -0.263489265 | -0.282810639 | -0.194769677 | 0.692339558 | 0.271857449 | 0.176520482 | 0.161290522 | 0.275647277 |
| Nol7 | T | 120 | -0.00297128 | -0.085253337 | -0.075785278 | -0.072813998 | 0.009468059 | 0.96916372 | 0.035162875 | 0.205217984 | 0.410505304 | 0.85288857 |
| Apex1 | S | 53 | -0.042492771 | -0.045362199 | -0.150357415 | -0.107864644 | -0.104995216 | 0.479737779 | 0.448182948 | 0.162855113 | 0.289838119 | 0.29979959 |
| Cdc42ep4 | S | 154 | -0.086228244 | 0.016735819 | -0.119160369 | -0.032932125 | -0.135896188 | 0.131678643 | 0.803635407 | 0.093285006 | 0.602929738 | 0.134891938 |
| Ptpn21 | S | 822 | -0.082809452 | -0.039287493 | 0.003864218 | 0.08667367 | 0.043151711 | 0.008999216 | 0.122635709 | 0.875269393 | 0.018518857 | 0.126234316 |
| Dcp1a | S | 200 | 0.026900584 | -0.083622251 | -0.141060246 | -0.16796083 | -0.057437995 | 0.565939557 | 0.093888642 | 0.101801593 | 0.07158112 | 0.421594298 |
| Gas7 | S | 107 | 0.153868581 | -0.269950432 | -0.141597109 | -0.29546569 | 0.128353323 | 0.053593379 | 0.02561306 | 0.104913924 | 0.01082733 | 0.196766577 |
| Ubap2l | S | 116 | -0.023140934 | -0.051837312 | -0.07421566 | -0.051074725 | -0.022378347 | 0.450726619 | 0.137962238 | 0.080534263 | 0.185247417 | 0.532680327 |
| Clp1 | T | 96 | -0.103281515 | -0.089631912 | -0.056196463 | 0.047085051 | 0.033435448 | 0.299004778 | 0.300715484 | 0.465807584 | 0.570139669 | 0.618982222 |

| | | | | | | | | | | | | | |
|---|---|---|---|---|---|---|---|---|---|---|---|---|---|
| Clp1 | T | | 98 | -0.103281515 | -0.089631912 | -0.056196463 | 0.047085051 | 0.033435448 | 0.299004778 | 0.300715484 | 0.465807584 | 0.570139669 | 0.618982222 |
| Phldb1 | S | | 405 | 0.227162164 | -0.180984115 | -0.279937312 | -0.507099476 | -0.098953197 | 0.000499094 | 0.077311449 | 0.000255829 | 0.000107831 | 0.232335047 |
| Clk3 | S | | 226 | -0.102819553 | -0.055748798 | -0.059292253 | 0.0435273 | -0.003543455 | 0.161414078 | 0.351982184 | 0.462848871 | 0.61863393 | 0.965538506 |
| Cobll1 | S | | 541 | -0.121654944 | 0.051376427 | -0.102141638 | 0.019513306 | -0.153518065 | 0.025833517 | 0.18882372 | 0.07553796 | 0.691968549 | 0.028057403 |
| Map3k11 | S | | 549 | -0.086136705 | -0.00815906 | -0.149095145 | -0.062958441 | -0.140936085 | 0.20313475 | 0.920531696 | 0.198146016 | 0.512570095 | 0.223667479 |
| Champ1 | S | | 421 | 0.009136447 | -0.130487944 | 0.0117122 | 0.002575752 | 0.142200144 | 0.715021911 | 0.000384228 | 0.658504634 | 0.920349057 | 0.004993253 |
| Champ1 | S | | 425 | 0.009136447 | -0.130487944 | 0.0117122 | 0.002575752 | 0.142200144 | 0.715021911 | 0.000384228 | 0.658504634 | 0.920349057 | 0.004993253 |
| Dennd1b | S | | 571 | -0.246267861 | 0.167840718 | -0.356764516 | -0.110496655 | -0.524605234 | 0.238675295 | 0.361826234 | 0.248110342 | 0.694677417 | 0.125943036 |
| Bmp2k | S | | 701 | 0.06840991 | -0.044351693 | -0.214528463 | -0.282938373 | -0.170176771 | 0.366295577 | 0.609879595 | 0.009457044 | 0.01440169 | 0.107102564 |
| Map7d1 | T | | 74 | -0.047077098 | -0.104697059 | -0.143245459 | -0.09616836 | -0.0385484 | 0.718714893 | 0.406500943 | 0.236854554 | 0.5258611 | 0.781962717 |
| Esyt2 | S | | 359 | 0.006832401 | -0.12434574 | -0.049791663 | -0.056624064 | 0.074554076 | 0.87139869 | 0.037402283 | 0.213589604 | 0.106196026 | 0.136043238 |
| Spag9 | T | | 217 | 0.030842574 | -0.053088611 | -0.099284607 | -0.130127181 | -0.046195996 | 0.375603052 | 0.020524402 | 0.02500897 | 0.021064639 | 0.173870582 |
| Kifc5b Kifc1 | S | | 492 | 0.019958536 | -0.084134148 | -0.079600904 | -0.09955944 | 0.004533244 | 0.650211099 | 0.014266302 | 0.163131706 | 0.126173617 | 0.917802509 |
| Med1 | S | | 1042 | -0.102013038 | -0.155479212 | -0.008523414 | 0.093489624 | 0.146955798 | 0.539464128 | 0.089140434 | 0.950182109 | 0.629442518 | 0.338832924 |
| Erc1 Erc2 | S | 191;187 | | -0.03617771 | -0.051877102 | -0.091635455 | -0.055457745 | -0.039758353 | 0.509817178 | 0.381127028 | 0.061900879 | 0.343754958 | 0.501016835 |
| Hnrnpm | S | | 466 | -0.130100858 | 0.025778426 | -0.179308689 | -0.049207831 | -0.205087115 | 0.177604048 | 0.73450021 | 0.248521984 | 0.71370287 | 0.203336402 |
| Mastl | S | | 587 | -0.123868465 | 0.032100208 | -0.132984234 | -0.009115769 | -0.165084442 | 0.073388036 | 0.616043618 | 0.232660072 | 0.924952689 | 0.164410414 |
| Eprs | S | | 713 | -0.077541355 | 0.013576313 | -0.100434322 | -0.022892967 | -0.114010635 | 0.174013883 | 0.683450596 | 0.031965748 | 0.636651213 | 0.021296643 |
| Phtf1 | S | | 254 | -0.128818815 | -0.007961877 | -0.023789494 | 0.105029322 | -0.015827617 | 0.023449745 | 0.678975897 | 0.476654891 | 0.052356459 | 0.587588135 |
| Pcm1 | S | | 110 | -0.118064956 | -0.096829643 | -0.011833088 | 0.106231868 | 0.084996555 | 0.343091015 | 0.084481117 | 0.87981232 | 0.422117929 | 0.34532883 |
| Tbc1d12 | S | | 205 | -0.096017812 | -0.019848009 | -0.116219769 | -0.020201956 | -0.09637176 | 0.169014191 | 0.799298555 | 0.246417936 | 0.792204549 | 0.306992707 |
| Psma2 | T | | 222 | 0.012912177 | -0.0930194 | -0.147577593 | -0.16048977 | -0.054558193 | 0.848978431 | 0.143521177 | 0.179251144 | 0.166965316 | 0.570992918 |
| Rell1 | S | | 162 | 0.045818855 | -0.266876818 | -0.16163422 | -0.207453076 | 0.105242598 | 0.628502599 | 0.215585215 | 0.393319164 | 0.285970526 | 0.643779013 |
| Baz1a | T | | 732 | 0.040327708 | -0.169235151 | -0.080407116 | -0.120734824 | 0.088828035 | 0.453095304 | 0.054914826 | 0.199120759 | 0.105397926 | 0.256245469 |
| Sec16a | S | | 1323 | 0.003784772 | -0.053749439 | -0.14460589 | -0.148390662 | -0.090856451 | 0.961044609 | 0.405832039 | 0.043097926 | 0.088249689 | 0.09006987 |
| Aebp2 | S | | 21 | 0.028939209 | -0.130825141 | -0.070059035 | -0.098998244 | 0.060766106 | 0.440979489 | 0.010220668 | 0.377563573 | 0.239475347 | 0.419289163 |
| Aebp2 | S | | 24 | 0.028939209 | -0.130825141 | -0.070059035 | -0.098998244 | 0.060766106 | 0.440979489 | 0.010220668 | 0.377563573 | 0.239475347 | 0.419289163 |
| Cc2d1b | S | | 484 | -0.062598901 | -0.021335367 | -0.109083056 | -0.046484155 | -0.087747689 | 0.257197783 | 0.642593814 | 0.107848275 | 0.451948758 | 0.171017804 |
| Acss2 | S | | 30 | -0.143859458 | -0.015911588 | -0.087436977 | 0.056422481 | -0.071525389 | 0.098514023 | 0.863172185 | 0.473496448 | 0.63800927 | 0.581278507 |
| Fam117b | S | | 409 | -0.023309057 | -0.053022143 | -0.090782511 | -0.067473454 | -0.037760368 | 0.744724337 | 0.163513015 | 0.072095016 | 0.374313144 | 0.317229443 |
| Pfas | T | | 619 | 0.002349339 | -0.062550414 | -0.136635571 | -0.13898491 | -0.074085157 | 0.971341804 | 0.385513865 | 0.047620358 | 0.058480983 | 0.266675087 |
| Map1b | T | | 1277 | 0.04736476 | -0.08609208 | -0.203956181 | -0.251320941 | -0.117864101 | 0.537385906 | 0.134270675 | 0.137425822 | 0.090937511 | 0.322510086 |
| Brd4 | S | | 1237 | -0.062126573 | -0.084963081 | -0.09843821 | -0.036311637 | -0.013475129 | 0.490854912 | 0.3578594 | 0.240216091 | 0.619148126 | 0.853324041 |
| Brd4 | S | | 1240 | -0.062126573 | -0.084963081 | -0.09843821 | -0.036311637 | -0.013475129 | 0.490854912 | 0.3578594 | 0.240216091 | 0.619148126 | 0.853324041 |
| Trim32 | S | | 341 | 0.117376334 | -0.08293818 | -0.269913857 | -0.387290191 | -0.186975677 | 0.033543573 | 0.077013415 | 0.016759285 | 0.010041341 | 0.072515255 |
| Nap1l1 | S | | 237 | -0.137209839 | 0.008815526 | -0.156351986 | -0.019142146 | -0.165167512 | 0.295510141 | 0.970865581 | 0.218918216 | 0.889086724 | 0.523560716 |
| Elac2 | S | | 191 | 0.001579153 | -0.026601115 | -0.163474609 | -0.165053762 | -0.136873494 | 0.969697197 | 0.351531294 | 0.041911652 | 0.042068962 | 0.074719811 |
| Kctd10 | S | | 23 | -0.101953838 | 0.003437689 | -0.138372583 | -0.036418744 | -0.141810272 | 0.201036324 | 0.961570285 | 0.180198394 | 0.658998521 | 0.149218076 |
| Eif3h | S | | 183 | -0.111687506 | -0.11768267 | 0.036441135 | 0.148128641 | 0.154123805 | 0.141962372 | 0.075868686 | 0.592511252 | 0.096949635 | 0.068578146 |
| Thoc5 | T | | 19 | 0.004643432 | -0.04009128 | -0.098098497 | -0.102741929 | -0.058007217 | 0.936240268 | 0.249560917 | 0.011598336 | 0.159258819 | 0.078074085 |
| Golga1 | S | | 41 | -0.085687742 | -0.094197619 | -0.019975825 | 0.065711918 | 0.074221795 | 0.198092393 | 0.222166547 | 0.426747287 | 0.293922618 | 0.309974862 |
| Med12 | S | | 699 | -0.104952979 | -0.132952637 | 0.051644792 | 0.156597771 | 0.184597428 | 0.116378728 | 0.075252961 | 0.524212246 | 0.111647179 | 0.078537575 |
| Anapc1 | S | | 522 | -0.067790538 | -0.01601949 | -0.324371122 | -0.256580584 | -0.308351632 | 0.398459342 | 0.902618435 | 0.343908893 | 0.435038087 | 0.368498338 |
| Add1 | S | | 366 | -0.098242964 | 0.008485479 | -0.058842646 | 0.039400318 | -0.067328125 | 0.011348111 | 0.771545905 | 0.310843163 | 0.454096341 | 0.252098636 |
| Rtkn | S | | 448 | -0.272591677 | 0.199820036 | -0.824984016 | -0.552392339 | -1.024804052 | 0.649018505 | 0.417586837 | 0.332579785 | 0.500239758 | 0.262598577 |
| Cyb5r3 | T | | 264 | -0.022327821 | -0.227903926 | -0.037011237 | -0.014683416 | 0.190892689 | 0.851070625 | 0.078753419 | 0.916473812 | 0.967417609 | 0.604966461 |
| Srrm1 | S | | 655 | -0.070135628 | -0.09909655 | 0.041320489 | 0.111456117 | 0.14041704 | 0.069013699 | 0.021692649 | 0.489927453 | 0.123875962 | 0.076675452 |
| Lap3 | S | | 238 | -0.051417527 | -0.077884067 | -0.051517949 | -0.000100422 | 0.026366118 | 0.329217367 | 0.06610396 | 0.540001266 | 0.999033915 | 0.737977097 |
| Ppfibp1 | S | | 40 | 0.102394672 | -0.10513453 | -0.127596167 | -0.229990839 | -0.022461637 | 0.004675505 | 0.007595851 | 0.00170027 | 0.000240726 | 0.359040623 |
| Sh2b3 | S | | 13 | -0.050037638 | -0.053434937 | -0.067795432 | -0.017757794 | -0.014360496 | 0.191564631 | 0.394670938 | 0.138907739 | 0.582568787 | 0.802913101 |
| Lsm12 | S | | 194 | 0.117234296 | 0.023093873 | -0.26906027 | -0.386294566 | -0.292154143 | 0.312585177 | 0.602277152 | 0.001102877 | 0.035872517 | 0.002613525 |
| Lsm12 | S | | 195 | 0.117234296 | 0.023093873 | -0.26906027 | -0.386294566 | -0.292154143 | 0.312585177 | 0.602277152 | 0.001102877 | 0.035872517 | 0.002613525 |
| Pdlim5 | S | | 322 | -0.056482874 | -0.050899327 | -0.07305346 | -0.016570586 | -0.022154133 | 0.297465235 | 0.097985016 | 0.367328648 | 0.838036291 | 0.76475378 |
| Myh9 | S | | 1943 | -0.134177644 | 0.033440591 | 0.006889154 | 0.141066798 | -0.026551437 | 0.00018174 | 0.482480301 | 0.763217373 | 0.007990354 | 0.571124969 |
| Prrc2b | S | | 793 | 0.026852017 | -0.191398691 | -0.006926945 | -0.033778962 | 0.184471745 | 0.860396128 | 0.013193323 | 0.886105447 | 0.82001639 | 0.027521736 |
| Gpam | S | | 668 | -0.167689685 | 0.1672823 | -0.406253988 | -0.238564303 | -0.573536288 | 0.516720312 | 0.264954181 | 0.079544238 | 0.407324044 | 0.034326686 |
| Filip1l | S | | 1045 | -0.21022279 | -0.005043782 | -0.031653859 | 0.178568931 | -0.026610077 | 0.07409977 | 0.94840443 | 0.637906432 | 0.116377229 | 0.754928384 |
| Zfp638 | S | | 606 | -0.11022491 | -0.153456943 | 0.095984636 | 0.206209546 | 0.249441579 | 0.240680891 | 0.007766143 | 0.214539466 | 0.078269296 | 0.035699699 |
| Stat3 | S | | 727 | 0.029217135 | -0.027728646 | -0.233184085 | -0.26240122 | -0.205455439 | 0.787823074 | 0.65435324 | 0.045070083 | 0.076372706 | 0.077427716 |
| Myo18a | S | | 164 | 0.020558036 | -0.045515831 | -0.153355095 | -0.173913131 | -0.107839264 | 0.820837547 | 0.592766116 | 0.018122568 | 0.128981536 | 0.243343444 |
| Rfc1 | S | | 244 | 0.00329196 | -0.067902183 | -0.0646818 | -0.06797376 | 0.003220382 | 0.925224731 | 0.02881104 | 0.084324657 | 0.148659704 | 0.915890614 |
| Camk2d | S | | 330 | -0.112325779 | -0.052228005 | -0.065334077 | 0.046991703 | -0.013106072 | 0.11840236 | 0.565585184 | 0.502133353 | 0.617165378 | 0.903459296 |
| Pwp1 | S | | 49 | -0.011390737 | -0.049125766 | -0.046056837 | -0.0346661 | 0.003068929 | 0.502933216 | 0.006970459 | 0.157602787 | 0.256967831 | 0.904004378 |
| Yju2 | S | | 211 | -0.013368756 | -0.03087819 | -0.05327843 | -0.039909674 | -0.02240024 | 0.161208427 | 0.06895758 | 0.031011126 | 0.056915589 | 0.221971989 |
| Zfp106 | S | | 1296 | 0.089404633 | -0.115389493 | -0.157628215 | -0.247032848 | -0.042238722 | 0.143352182 | 0.277107709 | 0.005576642 | 0.010266132 | 0.65094827 |
| Gtse1 | S | | 139 | 0.111625331 | -0.113599232 | -0.285937876 | -0.397563207 | -0.172338643 | 0.00954397 | 0.231856514 | 0.007341374 | 0.004976268 | 0.112722541 |
| Gigyf2 | S | | 19 | -0.169950921 | -0.010321705 | -0.147009457 | 0.022941463 | -0.136687752 | 0.182194778 | 0.934643 | 0.526770534 | 0.918890263 | 0.563596689 |
| Syde1 | S | | 647 | -0.065438851 | -0.080633517 | -0.086334585 | -0.020895734 | -0.005701068 | 0.4126749 | 0.442027441 | 0.198797736 | 0.769694645 | 0.952518698 |
| Ranbp1 | T | | 13 | -0.17168277 | 0.045717125 | -0.236628383 | -0.064945614 | -0.282345509 | 0.207273113 | 0.611151159 | 0.346677643 | 0.783611011 | 0.279847707 |
| Ifi211 | S | | 190 | -0.334371998 | -0.034387707 | 0.080411528 | 0.414783526 | 0.114799235 | 0.054422078 | 0.685017351 | 0.51705529 | 0.037541761 | 0.313252671 |
| Osbpl3 | S | | 271 | -0.066754419 | -0.001780959 | -0.142791558 | -0.076037139 | -0.141010599 | 0.169737737 | 0.977839563 | 0.150587551 | 0.37723394 | 0.17024444 |
| Stk11 | S | | 31 | -0.127313264 | -0.074351916 | -0.030072056 | 0.097241208 | 0.04427986 | 0.122260266 | 0.430227141 | 0.705468454 | 0.292138614 | 0.661899084 |
| Ppp1r2 | S | | 123 | -0.152969023 | 0.082898563 | -0.189167156 | -0.036198133 | -0.272065719 | 0.092001649 | 0.369852159 | 0.198833646 | 0.771907338 | 0.100180516 |
| Ddx54 | S | | 38 | 0.131595462 | -0.110794273 | -0.18462822 | -0.316223682 | -0.073833947 | 0.187490667 | 0.243671403 | 0.008054251 | 0.032508039 | 0.391467556 |
| Ddx54 | S | | 40 | 0.131595462 | -0.110794273 | -0.18462822 | -0.316223682 | -0.073833947 | 0.187490667 | 0.243671403 | 0.008054251 | 0.032508039 | 0.391467556 |
| Srsf5 | S | | 86 | 0.064604091 | -0.07209485 | -0.230885083 | -0.295489174 | -0.158790234 | 0.430027388 | 0.309705376 | 0.057367235 | 0.034868157 | 0.145787278 |
| Paxip1 | S | | 230 | -0.019771867 | -0.105356379 | -0.045739342 | -0.025967474 | 0.059617037 | 0.67717962 | 0.100714751 | 0.160586009 | 0.606537952 | 0.272830834 |
| Camsap1 | S | | 1133 | 0.016446671 | -0.038716674 | -0.138016977 | -0.154463648 | -0.099300303 | 0.745827716 | 0.397120029 | 0.017276955 | 0.026239209 | 0.057967497 |
| Tnks1bp1 | S | | 246 | -0.027366307 | -0.045377856 | -0.195528143 | -0.168161836 | -0.150150287 | 0.716787972 | 0.59479419 | 0.141155601 | 0.187517406 | 0.234647018 |
| Serbp1 | S | | 219 | 0.116803056 | -0.144777616 | -0.152874678 | -0.269677735 | -0.008097062 | 0.376867048 | 0.077603096 | 0.097456963 | 0.099488253 | 0.918304759 |
| Tox4 | T | | 176 | -0.010689506 | -0.057790515 | -0.045051692 | -0.034362186 | 0.012738823 | 0.708599669 | 0.016945122 | 0.101335156 | 0.273381093 | 0.538172357 |
| Ahctf1 | S | | 1218 | 0.052706349 | -0.153187538 | -0.112983388 | -0.165689736 | 0.04020415 | 0.361963115 | 0.03314612 | 0.286259153 | 0.154377077 | 0.670544956 |
| Maf1 | S | | 60 | -0.061376522 | -0.062734714 | -0.055863393 | 0.005513129 | 0.006871321 | 0.401266502 | 0.148915015 | 0.235698965 | 0.933192406 | 0.832177534 |
| Arhgef40 | S | | 415 | -0.004923396 | -0.069313462 | -0.051526763 | -0.046603367 | 0.017786698 | 0.850948954 | 0.043820307 | 0.067868842 | 0.175159968 | 0.502766446 |
| Rif1 | S | | 1683 | 0.017497184 | -0.125901081 | -0.037792721 | -0.055289905 | 0.088108359 | 0.725191853 | 0.010313234 | 0.415989261 | 0.294748729 | 0.07611422 |
| Nup35 | S | | 137 | -0.234198904 | 0.004415162 | -0.035225235 | 0.198973669 | -0.039640397 | 0.103707563 | 0.979164752 | 0.542701221 | 0.148484602 | 0.814189892 |
| Zc3h13 | S | | 1344 | -0.056515462 | -0.136555862 | -0.017852023 | 0.038663439 | 0.118703839 | 0.170571866 | 0.167583751 | 0.841928462 | 0.670560807 | 0.309188022 |
| Srrm2 | S | | 1399 | 0.06660004 | -0.083939048 | -0.102306781 | -0.168906821 | -0.018367734 | 0.137103684 | 0.004315052 | 0.050536921 | 0.015361582 | 0.607693748 |
| Foxm1 | T | | 659 | -0.060277159 | -0.033356213 | -0.320361565 | -0.260084406 | -0.287005352 | 0.719077576 | 0.922718863 | 0.244160052 | 0.320730963 | 0.461771682 |
| Cmc2 | S | | 71 | 0.00507938 | -0.09927422 | -0.097061482 | -0.102140861 | 0.002212738 | 0.916651059 | 0.064932216 | 0.253704272 | 0.227085383 | 0.975034866 |
| Irs1 | S | | 369 | -0.173867126 | 0.061256785 | -0.147071971 | 0.026795154 | -0.208328756 | 0.150569498 | 0.574939567 | 0.216074706 | 0.811146784 | 0.114074739 |
| Nfib | S | | 312 | -0.108978142 | -0.075475498 | 0.056332562 | 0.165310703 | 0.131808059 | 0.011175565 | 0.064281261 | 0.222973771 | 0.016667086 | 0.031893873 |
| Lig1 | S | | 65 | -0.179209516 | 0.046692855 | -0.016870411 | 0.162339104 | -0.063563266 | 0.003688612 | 0.380258899 | 0.718888337 | 0.009325386 | 0.210941914 |
| Ptpn14 | S | | 594 | -0.030784005 | -0.038895609 | -0.044525811 | -0.013741806 | -0.005630201 | 0.053461502 | 0.084308604 | 0.235864783 | 0.665651748 | 0.864935629 |

| | | | | | | | | | | | | |
|---|---|---|---|---|---|---|---|---|---|---|---|---|
| Sirt2 | S | 25 | -0.124896753 | -0.031639531 | -0.019560703 | 0.105336051 | 0.012078829 | 0.054442595 | 0.393283312 | 0.669857982 | 0.113836959 | 0.803301421 |
| Slc4a2 | S | 112 | 0.035201663 | -0.07015858 | -0.139339451 | -0.174541115 | -0.069180871 | 0.617078596 | 0.29189081 | 0.029831873 | 0.080512162 | 0.313139219 |
| Kat5 | T | 77 | 0.001242829 | -0.143272779 | -0.075886015 | -0.077128844 | 0.067386764 | 0.989316569 | 0.063689169 | 0.474453051 | 0.543467048 | 0.536517259 |
| Rangap1 | T | 411 | 0.186172455 | -0.131450065 | -0.231846003 | -0.418018457 | -0.100395938 | 0.090042152 | 0.106907951 | 0.01463132 | 0.012572199 | 0.063726942 |
| Tob2 | S | 255 | 0.024539189 | -0.104663748 | -0.079069197 | -0.103608386 | 0.025594551 | 0.748224371 | 0.039177486 | 0.186690355 | 0.265611399 | 0.638273291 |
| Hdac4 | S | 349 | -0.115967525 | -0.077815728 | -0.09587884 | 0.020088685 | -0.018063112 | 0.462845506 | 0.38716209 | 0.41252833 | 0.903890818 | 0.879500525 |
| Phf2 | S | 848 | -0.162857332 | 0.047528215 | -0.062760025 | 0.100097307 | -0.11028824 | 0.037531559 | 0.570514263 | 0.252859553 | 0.120918589 | 0.212599455 |
| Cbl | S | 480 | -0.027101381 | -0.032182225 | -0.11111838 | -0.084016999 | -0.078936155 | 0.620359989 | 0.43809075 | 0.046116002 | 0.179440735 | 0.118149209 |
| Anln | S | 556 | 0.020516421 | -0.081407091 | -0.065724804 | -0.086241225 | 0.015682287 | 0.645963284 | 0.02017245 | 0.096932875 | 0.112885354 | 0.595466679 |
| Rps17 | S | 113 | -0.070840476 | -0.134269598 | 0.00914462 | 0.079985096 | 0.143414218 | 0.332285538 | 0.060573326 | 0.887093827 | 0.204951339 | 0.026067912 |
| Srsf7 | S | 260 | 0.013579751 | -0.079519181 | -0.047003798 | -0.060583549 | 0.032515383 | 0.815325814 | 0.00788644 | 0.163415456 | 0.350988981 | 0.288848501 |
| Stambpl1 | S | 242 | -0.01217973 | -0.023785172 | -0.080472648 | -0.068292918 | -0.056687476 | 0.759765214 | 0.356979014 | 0.005676425 | 0.158126996 | 0.050670775 |
| Mki67 | S | 162 | 0.036418009 | -0.057982269 | -0.090860652 | -0.127278661 | -0.032878383 | 0.560133401 | 0.073977919 | 0.0153466 | 0.122736325 | 0.027319196 |
| Trio | S | 2481 | -0.05837377 | -0.009457902 | -0.093766003 | -0.035392233 | -0.084308101 | 0.179155185 | 0.786464905 | 0.073480967 | 0.447481226 | 0.101204857 |
| Ehd2 | S | 468 | -0.071781306 | -0.06604451 | -0.023608416 | 0.04817289 | 0.042436094 | 0.268887187 | 0.098566872 | 0.334138469 | 0.424958605 | 0.243287748 |
| Zc3hc1 | S | 358 | -0.02784498 | -0.084109015 | -0.181654206 | -0.153809225 | -0.097545191 | 0.763410798 | 0.341205183 | 0.293942113 | 0.377367041 | 0.55159894 |
| Sap30bp | T | 35 | 0.075413489 | -0.158160693 | -0.432294208 | -0.507707697 | -0.274133516 | 0.780080929 | 0.430178183 | 0.399038235 | 0.349094653 | 0.577080956 |
| Stx16 | S | 74 | -0.113517039 | -0.041173106 | -0.020039025 | 0.093478014 | 0.021134081 | 0.017020493 | 0.675647219 | 0.876393952 | 0.490493583 | 0.886517942 |
| Ppp1r18 | S | 208 | -0.068301821 | -0.026078793 | -0.073644831 | -0.00534301 | -0.047566038 | 0.184804997 | 0.58098543 | 0.114125951 | 0.86660137 | 0.179258084 |
| Smpd4 | S | 749 | -0.005821867 | -0.087834272 | -0.091189491 | -0.085367624 | -0.003355218 | 0.897875265 | 0.12190474 | 0.168086256 | 0.216056308 | 0.95715037 |
| Lysmd1 | S | 99 | -0.172853639 | 0.043187109 | -0.089764108 | 0.083089532 | -0.132951217 | 0.041744608 | 0.618195623 | 0.467346487 | 0.462329039 | 0.293753288 |
| Srrm2 | S | 785 | -0.140917289 | -0.096241126 | 0.12758847 | 0.268505759 | 0.223829596 | 0.042497006 | 0.006720928 | 0.137693388 | 0.020524305 | 0.046421107 |
| Rabgap1 | S | 42 | -0.068592973 | 0.04336147 | -0.148022202 | -0.079429229 | -0.191383671 | 0.227029168 | 0.616133701 | 0.042403199 | 0.137811743 | 0.07726458 |
| Capg | S | 341 | -0.205122553 | 0.028674946 | -0.072389031 | 0.132733522 | -0.101063977 | 0.069682578 | 0.810064695 | 0.586659166 | 0.360889175 | 0.518040196 |
| Filip1l | S | 1047 | -0.278253663 | 0.123348348 | -0.019508822 | 0.258744841 | -0.142857169 | 0.006525952 | 0.325778261 | 0.895118581 | 0.158015632 | 0.403154929 |
| Prrc2c | T | 1217 | 0.06222383 | -0.12128805 | -0.070613931 | -0.132837761 | 0.050674119 | 0.084698251 | 0.006209425 | 0.047592467 | 0.005447297 | 0.004924294 |
| Arhgap31 | S | 1355 | -0.048900772 | -0.086989293 | -0.0340111 | 0.014889672 | 0.052978193 | 0.41071255 | 0.035167269 | 0.70774403 | 0.876567035 | 0.563751875 |
| Acap2 | S | 378 | -0.15417641 | 0.021195523 | -0.079052286 | 0.075124124 | -0.100247809 | 0.083457963 | 0.877388224 | 0.343899116 | 0.390025782 | 0.493702866 |
| Map11 | S | 495 | 0.017768867 | -0.188224644 | 0.00804947 | -0.009719397 | 0.196274114 | 0.723586861 | 0.007867648 | 0.938745111 | 0.926499941 | 0.147625627 |
| Phf10 | S | 26 | 0.117419686 | -0.207540828 | -0.08474254 | -0.202162227 | 0.122798288 | 0.58086422 | 0.026218338 | 0.507851523 | 0.385208213 | 0.337143921 |
| Tbc1d1 | S | 621 | -0.107698771 | 0.012542493 | -0.157097657 | -0.049398886 | -0.16964015 | 0.399449922 | 0.83868034 | 0.110161783 | 0.69976773 | 0.09693374 |
| Myo9a | S | 2458 | -0.107971751 | -0.027286642 | -0.138282624 | -0.030310873 | -0.110995981 | 0.234360122 | 0.802394667 | 0.347344416 | 0.813987741 | 0.453593088 |
| Mast3 | S | 107 | -0.057948148 | 0.003972527 | -0.150833455 | -0.092885307 | -0.154805982 | 0.344698775 | 0.939780916 | 0.07636759 | 0.247483376 | 0.080812146 |
| Twist2 | S | 55 | -0.000591186 | -0.152290537 | -0.045377661 | -0.044786475 | 0.106912876 | 0.988807606 | 0.042139956 | 0.574616842 | 0.573879914 | 0.251849467 |
| Pogz | S | 421 | -0.034443908 | -0.165869272 | -0.029407252 | 0.005036657 | 0.13646202 | 0.684343384 | 0.070549496 | 0.812697138 | 0.97024106 | 0.33507933 |
| Dlg5 | S | 1254 | -0.099917433 | -0.114923902 | -0.044856794 | 0.055060639 | 0.070067108 | 0.28173305 | 0.370732596 | 0.556844288 | 0.484661543 | 0.55111957 |
| Itga3 | S | 1044 | -0.087553909 | -0.003180495 | -0.127764363 | -0.040210455 | -0.124583868 | 0.245123712 | 0.974644261 | 0.14700772 | 0.435259914 | 0.211146308 |
| Nifk | S | 253 | -0.047620455 | -0.083451191 | -0.07431667 | -0.026696215 | 0.009134521 | 0.447848789 | 0.427441896 | 0.127385538 | 0.638481886 | 0.924073386 |
| Tep1 | S | 2532 | -0.051154706 | -0.049587253 | -0.202536243 | -0.151381537 | -0.152948989 | 0.784212339 | 0.551262555 | 0.201333487 | 0.484964494 | 0.311376737 |
| Hspa4 | S | 816 | -0.00922426 | -0.094475575 | -0.080115542 | -0.070891282 | 0.014360033 | 0.84995753 | 0.266683353 | 0.070828627 | 0.142142922 | 0.838760369 |
| Setd2 | T | 734 | -0.077268251 | -0.145790868 | -0.07189776 | 0.005370492 | 0.073893108 | 0.476868719 | 0.253790173 | 0.681084871 | 0.973699276 | 0.669985569 |
| Ccdc93 | S | 305 | -0.071653828 | 0.017673341 | -0.087780575 | -0.016126747 | -0.105453916 | 0.173424045 | 0.423570076 | 0.012296543 | 0.713264958 | 0.019515612 |
| Eapp | S | 55 | -0.040696049 | -0.244671095 | -0.037906179 | 0.00278987 | 0.206764915 | 0.568793861 | 0.272440467 | 0.611445095 | 0.974227382 | 0.336023694 |
| Eapp | S | 56 | -0.040696049 | -0.244671095 | -0.037906179 | 0.00278987 | 0.206764915 | 0.568793861 | 0.272440467 | 0.611445095 | 0.974227382 | 0.336023694 |
| Smn1 | T | 59 | -0.031536456 | -0.039133674 | -0.075494337 | -0.043957881 | -0.036360662 | 0.339826437 | 0.320537019 | 0.063336448 | 0.186510669 | 0.334814302 |
| Ccdc9 | S | 60 | -0.164086227 | 0.043895677 | -0.011817355 | 0.152268872 | -0.055713031 | 0.003375939 | 0.523930837 | 0.781634762 | 0.013627977 | 0.413315716 |
| Jcad | S | 1243 | -0.167352655 | 0.003946025 | -0.044614477 | 0.122738178 | -0.048560502 | 0.03698511 | 0.962224257 | 0.719034271 | 0.335839642 | 0.691841495 |
| Kmt2b | S | 941 | -0.09258689 | -0.063580911 | -0.032860344 | 0.059726546 | 0.030720567 | 0.270008416 | 0.141388666 | 0.503579048 | 0.461842113 | 0.531620247 |
| Strn4 | S | 206 | -0.121070278 | 0.065823805 | -0.117915894 | 0.003154384 | -0.183739699 | 0.089383197 | 0.279421274 | 0.047340808 | 0.953645282 | 0.020013088 |
| Mepce | S | 229 | 0.032689069 | -0.044108138 | -0.147180857 | -0.179869926 | -0.103072719 | 0.320590652 | 0.272357674 | 0.012392108 | 0.024626506 | 0.050448927 |
| Eipr1 | S | 320 | -0.225169668 | 0.05434895 | -0.069252311 | 0.155917357 | -0.123601261 | 0.052199796 | 0.615009602 | 0.568754329 | 0.239691817 | 0.356595827 |
| Srrm2 | T | 2362 | -0.069580894 | -0.029865885 | -0.071701509 | -0.002120615 | -0.041835624 | 0.125414078 | 0.45268395 | 0.24826004 | 0.967147168 | 0.434000739 |
| Sipa1l1 | S | 1127 | -0.09771292 | -0.054511645 | -0.047551218 | 0.050161701 | 0.006960426 | 0.120937547 | 0.507195674 | 0.421016878 | 0.383531409 | 0.92891217 |
| Lrrfip1 | S | 348 | -0.109828191 | -0.00044339 | -0.003434591 | 0.1063936 | -0.002991201 | 0.001801043 | 0.9925341 | 0.940442156 | 0.105858089 | 0.961041582 |
| Bag2 | S | 25 | -0.188302049 | 0.040657879 | -0.019852378 | 0.168449671 | -0.060510257 | 0.012443267 | 0.573750491 | 0.661622316 | 0.026235443 | 0.418869134 |
| Tmcc3 | T | 194 | 0.030254057 | -0.237316927 | -0.002865238 | -0.033119295 | 0.234451689 | 0.845734961 | 0.029552961 | 0.986999136 | 0.868785445 | 0.239188388 |
| Mdc1 | S | 929 | -0.014451773 | -0.030365648 | -0.083263501 | -0.068811729 | -0.052897853 | 0.721191034 | 0.276518352 | 0.017062798 | 0.140832317 | 0.007706063 |
| Nufip2 | S | 570 | 0.058483214 | -0.113363808 | -0.110743627 | -0.169226841 | 0.002620182 | 0.254344463 | 0.058641931 | 0.011767536 | 0.952137499 | 0.952137499 |
| Prkca Prkcb Prkcg | T | 501;504;518 | -0.050086675 | -0.162699737 | 0.011016106 | 0.06110278 | 0.173715843 | 0.298179285 | 0.07860133 | 0.856612317 | 0.342126653 | 0.079693556 |
| Brd2 | S | 297 | -0.046511198 | -0.061975644 | 0.005734013 | 0.052245211 | 0.067709657 | 0.012608827 | 0.040125631 | 0.653708784 | 0.010392427 | 0.051763252 |
| Tnrc6b | S | 600 | -0.2527513 | 0.068043845 | -0.040853145 | 0.211898156 | -0.10889699 | 0.037551657 | 0.65666897 | 0.757977614 | 0.172326098 | 0.535390881 |
| Cobl | S | 655 | -0.067920218 | -0.060155354 | -0.087621783 | -0.019701565 | -0.02746643 | 0.34201693 | 0.477732319 | 0.215411535 | 0.749392051 | 0.721985554 |
| Bmp2k | S | 685 | -0.008770784 | -0.098198923 | -0.09359158 | -0.084820796 | 0.004607344 | 0.934369092 | 0.129246939 | 0.223462311 | 0.452559466 | 0.941028795 |
| Helz | S | 1317 | -0.093943073 | -0.033588503 | -0.038660265 | 0.055282808 | -0.005071762 | 0.130160185 | 0.127763996 | 0.578112188 | 0.481962414 | 0.938724685 |
| Cyren | S | 122 | -0.017249677 | -0.058987277 | -0.136845936 | -0.119596259 | -0.07785866 | 0.68675421 | 0.301295527 | 0.16144952 | 0.20358683 | 0.383945856 |
| Prrc2a | S | 1104 | -0.01858317 | -0.135531707 | 0.043465281 | 0.062048451 | 0.178996989 | 0.573299262 | 0.000217341 | 0.088344126 | 0.11635013 | 0.001376534 |
| Ccdc88a | S | 1738 | 0.009587894 | -0.017131411 | -0.291206007 | -0.300793901 | -0.274074596 | 0.937887958 | 0.859780517 | 0.099376028 | 0.102520652 | 0.119838721 |
| Zcchc8 | S | 660 | -0.021923853 | -0.032363642 | -0.127620887 | -0.105697034 | -0.095257245 | 0.687221252 | 0.573954471 | 0.048623836 | 0.016602974 | 0.043799221 |
| Golim4 | Y | 604 | -0.054386397 | -0.20226116 | 0.014167402 | 0.068553799 | 0.216428562 | 0.374744877 | 0.113530685 | 0.844285985 | 0.356469723 | 0.103257632 |
| Tab1 | S | 16 | -0.305293054 | 0.001025719 | 0.022061637 | 0.327354691 | 0.021035918 | 0.071132348 | 0.995008163 | 0.890335157 | 0.037284156 | 0.8719477 |
| Dlgap5 | S | 679 | -0.083177828 | 0.044108246 | -0.162668987 | -0.079491159 | -0.206777234 | 0.282681937 | 0.322989235 | 0.033153021 | 0.326505352 | 0.021494337 |
| Tbc1d4 | T | 273 | -0.023579742 | -0.088501453 | -0.14637814 | -0.122798398 | -0.057876687 | 0.886994678 | 0.598377876 | 0.101359045 | 0.486700006 | 0.728014488 |
| Kmt2d | T | 2197 | 0.038198529 | -0.13365768 | -0.063800046 | -0.101998575 | 0.069857634 | 0.360125746 | 0.015308099 | 0.255039607 | 0.08714064 | 0.184147352 |
| Ctnna1 | S | 453 | -0.105128159 | -0.208020034 | 0.05066098 | 0.15578914 | 0.258681014 | 0.600346983 | 0.052806362 | 0.56954814 | 0.437723362 | 0.007789001 |
| Irs1 | S | 422 | -0.101890484 | -0.272277572 | 0.007114252 | 0.109004736 | 0.279391824 | 0.743782407 | 0.178717859 | 0.973607825 | 0.759454671 | 0.302687263 |
| Mllt3 | S | 484 | -0.011764928 | -0.157919045 | -0.039400476 | -0.027635548 | 0.118518569 | 0.89158422 | 0.057333913 | 0.626095892 | 0.7124373 | 0.098218572 |
| Mff | S | 115 | -0.092021663 | -0.180679702 | 0.038205083 | 0.130226746 | 0.218884785 | 0.374460103 | 0.0593384 | 0.466787631 | 0.232734093 | 0.054436154 |
| Camsap1 | S | 734 | -0.110462562 | -0.029243835 | -0.183953469 | -0.073490907 | -0.154709634 | 0.420690894 | 0.866370096 | 0.300471969 | 0.672768404 | 0.460800811 |
| Vcl | Y | 692 | -0.152391542 | 0.026623856 | -0.068933212 | 0.08345833 | -0.095557068 | 0.03673735 | 0.743281458 | 0.485292888 | 0.378364221 | 0.366005326 |
| Uqcrc1 | S | 159 | 0.111849235 | -0.241381991 | -0.115599426 | -0.227448662 | 0.125782565 | 0.606546006 | 0.132875792 | 0.282714253 | 0.317974641 | 0.332282196 |
| Arhgef10 | S | 355 | -0.123620597 | 0.172673444 | -0.257492366 | -0.133871768 | -0.43016581 | 0.246630853 | 0.34444137 | 0.057964009 | 0.17340173 | 0.06396356 |
| Ranbp2 | S | 837 | -0.107397915 | -0.151533366 | 0.071833932 | 0.179231847 | 0.223367298 | 0.142554445 | 0.056198275 | 0.349849102 | 0.050594755 | 0.027129455 |
| Basp1 | S | 6 | -0.149976491 | 0.046650474 | -0.081242468 | 0.068734024 | -0.127892941 | 0.050847519 | 0.635453351 | 0.283512924 | 0.333384073 | 0.240842325 |
| Washc2 | S | 783 | 0.060674712 | -0.10317227 | -0.13352764 | -0.194202351 | -0.030355369 | 0.135695309 | 0.013814361 | 0.125604177 | 0.053223655 | 0.648178862 |
| Cdca8 | S | 175 | 0.036968395 | -0.033575068 | -0.12230909 | -0.159277485 | -0.088734022 | 0.288704323 | 0.268727666 | 0.002841849 | 0.00596141 | 0.016833747 |
| Rplp1 | S | 101 | 0.013387149 | -0.082359478 | -0.063287951 | -0.076675099 | 0.019071527 | 0.63588846 | 0.037338389 | 0.080168593 | 0.048853005 | 0.561915365 |
| Gpn1 | S | 338 | -0.012947405 | -0.009479388 | -0.120145077 | -0.107197672 | -0.110665689 | 0.725722746 | 0.710297908 | 0.013343374 | 0.038683324 | 0.03933481 |
| Zfp503 | S | 232 | -0.094015722 | -0.006127338 | -0.120625087 | -0.026609365 | -0.114497749 | 0.116250033 | 0.927896047 | 0.320902535 | 0.804882873 | 0.3495637 |
| Zfp503 | S | 234 | -0.094015722 | -0.006127338 | -0.120625087 | -0.026609365 | -0.114497749 | 0.116250033 | 0.927896047 | 0.320902535 | 0.804882873 | 0.3495637 |
| Gm4070 Gvin1 | S | 851 | -0.284336001 | 0.045169806 | 0.027305335 | 0.311641335 | -0.017864471 | 0.012549783 | 0.517307692 | 0.683101387 | 0.01493982 | 0.72857728 |
| Lima1 | S | 539 | -0.064924146 | -0.104914048 | -0.203174708 | -0.138250562 | -0.09826066 | 0.657282783 | 0.494599303 | 0.447740662 | 0.602938681 | 0.709976577 |
| Tor1aip1 | S | 328 | 0.023389477 | -0.250795521 | 0.001420405 | -0.021969072 | 0.252215926 | 0.800018364 | 0.035392334 | 0.992888806 | 0.874126279 | 0.167123523 |
| Mtcl1 | S | 1501 | -0.113552186 | 0.057559156 | -0.129619451 | -0.016067266 | -0.187178607 | 0.236587409 | 0.452664515 | 0.038802246 | 0.833443531 | 0.06976694 |
| Aak1 Gm38495 | S | 115 | -0.066761063 | 0.008551688 | -0.139036748 | -0.072275685 | -0.147588436 | 0.262011855 | 0.928589856 | 0.101154395 | 0.344583899 | 0.209470465 |

| | | | | | | | | | | | | |
|---|---|---|---|---|---|---|---|---|---|---|---|---|
| Nop58 | S | 521 | 0.039961544 | -0.074028429 | -0.065491556 | -0.1054531 | 0.008536872 | 0.179550509 | 0.011514499 | 0.014799292 | 0.015623362 | 0.62359949 |
| Arhgap31 | S | 343 | -0.040490221 | 0.004465769 | -0.21112896 | -0.170638739 | -0.215594729 | 0.599369373 | 0.963812998 | 0.093956516 | 0.148000725 | 0.096938033 |
| Snta1 | S | 183 | 0.02235506 | -0.075270404 | -0.115780785 | -0.138135846 | -0.040510382 | 0.462796941 | 0.047548974 | 0.125645335 | 0.105856294 | 0.504807386 |
| Incenp | S | 284 | 0.056141718 | -0.035181511 | -0.180404145 | -0.236545863 | -0.145222634 | 0.118786606 | 0.613922381 | 0.002160693 | 0.0035763 | 0.106441692 |
| Ctcf | T | 317 | -0.172955619 | -0.010281968 | -0.168533846 | 0.004421773 | -0.158251878 | 0.547484787 | 0.858069666 | 0.262320557 | 0.98772213 | 0.286029974 |
| Plekho2 | S | 292 | -0.088817202 | -0.074367451 | -0.02074565 | 0.068071552 | 0.053621801 | 0.183837845 | 0.150051437 | 0.770697259 | 0.425227587 | 0.494897664 |
| C2cd2l | S | 613 | 0.002534295 | -0.149142853 | -0.085300028 | -0.087834323 | 0.063842825 | 0.977268678 | 0.082998855 | 0.522668983 | 0.553048879 | 0.638564003 |
| Gorasp1 | T | 363 | 0.015839812 | -0.174491768 | -0.111270252 | -0.127110063 | 0.063221517 | 0.907807195 | 0.222412381 | 0.304756545 | 0.406450389 | 0.645517425 |
| Inka2 | S | 281 | 0.142371965 | -0.115937321 | -0.081353474 | -0.223725439 | 0.034583847 | 0.082311015 | 0.000892453 | 0.0314561 | 0.027873804 | 0.237793832 |
| Lima1 | S | 607 | 0.001425731 | -0.100641061 | -0.090578964 | -0.092004695 | 0.010062097 | 0.982779414 | 0.138308823 | 0.179193575 | 0.065508969 | 0.663610124 |
| Fosl2 | S | 320 | -0.100982436 | -0.05724827 | -0.005781462 | 0.095200974 | 0.051466807 | 0.024001058 | 0.508523176 | 0.895698915 | 0.058649569 | 0.546202625 |
| Smcr8 | S | 491 | -0.020176103 | -0.040704434 | -0.157505985 | -0.137329881 | -0.11680155 | 0.906425219 | 0.548772835 | 0.077520452 | 0.445514692 | 0.105700751 |
| Arfgef1 | S | 661 | 0.106499566 | -0.224188616 | -0.245542386 | -0.352041952 | -0.021353769 | 0.66950046 | 0.263800474 | 0.368862755 | 0.274932915 | 0.933757654 |
| Dact3 | S | 327 | -0.119965915 | -0.039622367 | -0.131298528 | -0.011332613 | -0.091676161 | 0.412864602 | 0.685412399 | 0.321444624 | 0.943402123 | 0.495958784 |
| Dact3 | S | 330 | -0.119965915 | -0.039622367 | -0.131298528 | -0.011332613 | -0.091676161 | 0.412864602 | 0.685412399 | 0.321444624 | 0.943402123 | 0.495958784 |
| Mphosph10 | S | 244 | -0.022636424 | -0.200575126 | -0.025046191 | -0.002409767 | 0.175528935 | 0.903941473 | 0.067339785 | 0.904615736 | 0.992126444 | 0.415590295 |
| Mphosph10 | S | 247 | -0.022636424 | -0.200575126 | -0.025046191 | -0.002409767 | 0.175528935 | 0.903941473 | 0.067339785 | 0.904615736 | 0.992126444 | 0.415590295 |
| Slc4a2 | S | 172 | -0.04937939 | -0.184224563 | -0.010584939 | 0.03879445 | 0.173639624 | 0.683749367 | 0.091987987 | 0.89968107 | 0.69593641 | 0.068671682 |
| Cwc22 | S | 558 | 0.076195368 | -0.017093343 | -0.205198445 | -0.281393812 | -0.188105101 | 0.098738814 | 0.607745112 | 0.001402078 | 0.001628607 | 0.005742276 |
| Cwc22 | S | 562 | 0.076195368 | -0.017093343 | -0.205198445 | -0.281393812 | -0.188105101 | 0.098738814 | 0.607745112 | 0.001402078 | 0.001628607 | 0.005742276 |
| Cwc22 | T | 556 | 0.076195368 | -0.017093343 | -0.205198445 | -0.281393812 | -0.188105101 | 0.098738814 | 0.607745112 | 0.001402078 | 0.001628607 | 0.005742276 |
| Cwc22 | Y | 555 | 0.076195368 | -0.017093343 | -0.205198445 | -0.281393812 | -0.188105101 | 0.098738814 | 0.607745112 | 0.001402078 | 0.001628607 | 0.005742276 |
| Hnrnpa3 | S | 367 | 0.070457522 | -0.019017986 | -0.138805735 | -0.209263258 | -0.11978775 | 0.608618922 | 0.755125044 | 0.001564097 | 0.212833064 | 0.134825659 |
| Fam117b | S | 407 | 0.057230544 | -0.086386 | -0.172135315 | -0.229365859 | -0.085749315 | 0.720444304 | 0.4934595 | 0.048064734 | 0.217346236 | 0.470844447 |
| Sipa1l1 | S | 1234 | -0.122206053 | -0.104237374 | 0.080304421 | 0.202510474 | 0.184541795 | 0.086041343 | 0.023939199 | 0.23979204 | 0.038146425 | 0.045838511 |
| Rtf1 | S | 702 | 0.084090696 | -0.03630238 | -0.198650875 | -0.282741571 | -0.162348495 | 0.038735981 | 0.573158719 | 0.000605669 | 0.000402405 | 0.071928535 |
| Cavin4 | S | 361 | -0.108972078 | -0.148683255 | -0.016215913 | 0.092756165 | 0.132467341 | 0.37844029 | 0.236093026 | 0.889363622 | 0.280021849 | 0.1401871 |
| Cavin4 | S | 362 | -0.108972078 | -0.148683255 | -0.016215913 | 0.092756165 | 0.132467341 | 0.37844029 | 0.236093026 | 0.889363622 | 0.280021849 | 0.1401871 |
| Evi2a | S | 197 | 0.167222639 | -0.139330204 | -0.162366351 | -0.32958899 | -0.023036147 | 0.037402185 | 0.042121574 | 0.00616513 | 0.009209 | 0.627571519 |
| Esf1 | S | 75 | 0.030836788 | -0.199889724 | -0.09424651 | -0.125083298 | 0.105643215 | 0.573078955 | 0.111242049 | 0.383713628 | 0.279077367 | 0.411818332 |
| Esf1 | S | 77 | 0.030836788 | -0.199889724 | -0.09424651 | -0.125083298 | 0.105643215 | 0.573078955 | 0.111242049 | 0.383713628 | 0.279077367 | 0.411818332 |
| Esf1 | S | 79 | 0.030836788 | -0.199889724 | -0.09424651 | -0.125083298 | 0.105643215 | 0.573078955 | 0.111242049 | 0.383713628 | 0.279077367 | 0.411818332 |
| Esf1 | S | 82 | 0.030836788 | -0.199889724 | -0.09424651 | -0.125083298 | 0.105643215 | 0.573078955 | 0.111242049 | 0.383713628 | 0.279077367 | 0.411818332 |
| Dhx30 | S | 249 | -0.093446222 | -0.022587537 | -0.132902901 | -0.039456678 | -0.110315363 | 0.197006056 | 0.740234248 | 0.37774124 | 0.765498509 | 0.438929273 |
| Srrm2 | S | 1077 | -0.059425451 | -0.128909761 | 0.015409747 | 0.074835198 | 0.144319508 | 0.298901068 | 0.056074998 | 0.873580309 | 0.43045135 | 0.189685842 |
| Mcm4 | S | 141 | -0.088339958 | -0.105727961 | 0.010169284 | 0.098509241 | 0.115897244 | 0.103150547 | 0.176656137 | 0.818487471 | 0.093422029 | 0.153134419 |
| Add1 | S | 586 | -0.115150096 | 0.112312643 | -0.2266566 | -0.111506504 | -0.338969243 | 0.140259226 | 0.18158933 | 0.062388267 | 0.253662435 | 0.023059097 |
| Tut7 | S | 1471 | 0.023944638 | -0.080954648 | -0.228316376 | -0.252261014 | -0.147361727 | 0.764378434 | 0.407027046 | 0.165095516 | 0.137779316 | 0.341045415 |
| Stub1 | S | 274 | -0.06186436 | -0.011805472 | -0.145218595 | -0.083354235 | -0.133413123 | 0.33537849 | 0.773541065 | 0.16029577 | 0.385497424 | 0.183367113 |
| Ptges3 | S | 113 | -0.121087312 | 0.070076065 | -0.12315009 | -0.002062778 | -0.193226155 | 0.111192311 | 0.44392471 | 0.08315583 | 0.976518876 | 0.100397247 |
| Dlgap5 | S | 680 | -0.097361815 | 0.039328482 | -0.145746065 | -0.04838425 | -0.185074547 | 0.184868529 | 0.358004936 | 0.068193069 | 0.520211506 | 0.056914568 |
| Baz1a | T | 1366 | 0.223567263 | -0.172850417 | -0.248472912 | -0.472040176 | -0.075622496 | 0.150861896 | 0.09975344 | 0.055027108 | 0.025557035 | 0.495150891 |
| Tcf20 | S | 1550 | -0.029391392 | -0.038881607 | -0.093615914 | -0.064224522 | -0.054734307 | 0.661912193 | 0.592152092 | 0.028580795 | 0.369627945 | 0.46183808 |
| Pum1 | S | 228 | -0.14987841 | -0.121377046 | -0.060926932 | 0.088951478 | 0.060450114 | 0.448286652 | 0.68050984 | 0.310938702 | 0.634899862 | 0.833995763 |
| Wdr70 | S | 641 | 0.008733361 | -0.012861835 | -0.176431701 | -0.185165062 | -0.163569866 | 0.808386678 | 0.800879702 | 0.022269554 | 0.021078592 | 0.044411087 |
| Gab1 | S | 251 | -0.113865764 | -0.053548554 | -0.071450389 | 0.042415376 | -0.017901835 | 0.211963792 | 0.507013912 | 0.51236372 | 0.681204583 | 0.854687073 |
| Nedd4 | T | 432 | 0.026689481 | -0.065678364 | -0.133521768 | -0.160211249 | -0.067843404 | 0.642690706 | 0.293916738 | 0.04038209 | 0.029459773 | 0.25397778 |
| Msrb3 | S | 242 | -0.138802316 | -0.037548801 | -0.197787431 | -0.058985115 | -0.16023863 | 0.497162276 | 0.754989064 | 0.418440624 | 0.824321415 | 0.498474877 |
| Nlk | S | 522 | -0.026732092 | -0.094590615 | -0.040227432 | -0.01349534 | 0.054363182 | 0.764054906 | 0.05386091 | 0.304828018 | 0.87872104 | 0.226926307 |
| Map1a | S | 117 | 0.018746957 | -0.124340972 | -0.140342952 | -0.159089909 | -0.016001979 | 0.710359457 | 0.162453365 | 0.261987865 | 0.218146304 | 0.89177319 |
| Cul4b | S | 149 | -0.071753715 | -0.04171212 | -0.092307761 | -0.020554045 | -0.050595641 | 0.416786878 | 0.219595137 | 0.314735021 | 0.845379893 | 0.548556018 |
| Hlcs | S | 73 | -0.004556165 | -0.019007298 | -0.162116062 | -0.157559897 | -0.143108763 | 0.965870325 | 0.721641316 | 0.034995798 | 0.198378515 | 0.055896051 |
| Slc20a2 | S | 256 | 0.033847604 | -0.128185002 | -0.010030017 | -0.043877621 | 0.118154986 | 0.322061104 | 0.000618388 | 0.770760086 | 0.320588354 | 0.037905416 |
| Iws1 | S | 474 | 0.032613723 | -0.121533035 | -0.284168416 | -0.316782139 | -0.16263538 | 0.8021599 | 0.407658397 | 0.302391265 | 0.263791394 | 0.534297381 |
| Nfib | S | 276 | -0.08758693 | -0.172394699 | 0.031024599 | 0.118611529 | 0.203419298 | 0.101972711 | 0.267911676 | 0.745362045 | 0.275799435 | 0.227460166 |
| Ddx21 | S | 245 | -0.041632097 | -0.003425106 | -0.129096673 | -0.087464577 | -0.125671567 | 0.565252857 | 0.957000864 | 0.033732172 | 0.227516011 | 0.092555467 |
| Ptk2 | Y | 576 | -0.12445828 | 0.011022635 | -0.049805311 | 0.074652968 | -0.060827946 | 0.019091001 | 0.891657394 | 0.592928024 | 0.447032265 | 0.5970716 |
| Ehmt1 | S | 381 | -0.161761968 | -0.044057965 | 0.011967838 | 0.173729807 | 0.056025804 | 0.050464906 | 0.48005927 | 0.829679789 | 0.046207584 | 0.346082324 |
| Gemin2 | T | 37 | 0.019529978 | -0.237506187 | -0.060286533 | -0.079816512 | 0.177219654 | 0.805471873 | 0.119883796 | 0.577328738 | 0.452131321 | 0.241550946 |
| Rev1 | S | 1082 | 0.029453171 | -0.108111195 | -0.114681691 | -0.144134862 | -0.006570496 | 0.66743773 | 0.206611901 | 0.092110819 | 0.059019699 | 0.924951139 |
| Rsl1d1 | S | 12 | -0.06843534 | -0.131529483 | -0.091036941 | -0.022601601 | 0.040492542 | 0.392981337 | 0.481310529 | 0.465792512 | 0.844124774 | 0.832444935 |
| Lrrfip1 | S | 605 | -0.089429497 | 0.00463899 | -0.079645459 | 0.009784038 | -0.084284449 | 0.110025204 | 0.94557144 | 0.136429404 | 0.623747062 | 0.213820905 |
| Kif1c | S | 1031 | -0.095861898 | 0.028398791 | -0.102555822 | -0.006693924 | -0.130954613 | 0.071594774 | 0.612713642 | 0.149579276 | 0.905145392 | 0.093143521 |
| Hcfc1 | S | 598 | -0.035905706 | -0.046122298 | -0.061363034 | -0.025457329 | -0.015240736 | 0.305035218 | 0.258861001 | 0.107338381 | 0.234171296 | 0.600362392 |
| Inpp5e | S | 236 | -0.231095956 | 0.180911302 | -0.154608339 | 0.076487616 | -0.335519641 | 0.152962991 | 0.357594226 | 0.098468657 | 0.569237965 | 0.146976418 |
| Cep350 | S | 563 | -0.057419716 | -0.054942727 | -0.049243985 | 0.008175731 | 0.005698742 | 0.142776764 | 0.347972348 | 0.278608335 | 0.786254551 | 0.913320081 |
| Camsap2 | S | 1240 | -0.202175774 | 0.054447832 | -0.11021232 | 0.091963454 | -0.164660152 | 0.119079604 | 0.639672844 | 0.381972518 | 0.439441652 | 0.194467896 |
| Klhl40 | S | 425 | -0.13865711 | 0.019825508 | -0.0890015 | 0.049655611 | -0.108827008 | 0.074564853 | 0.745633935 | 0.360534445 | 0.604405954 | 0.284518849 |
| Klhl40 | S | 428 | -0.13865711 | 0.019825508 | -0.0890015 | 0.049655611 | -0.108827008 | 0.074564853 | 0.745633935 | 0.360534445 | 0.604405954 | 0.284518849 |
| Klhl40 | S | 431 | -0.13865711 | 0.019825508 | -0.0890015 | 0.049655611 | -0.108827008 | 0.074564853 | 0.745633935 | 0.360534445 | 0.604405954 | 0.284518849 |
| Klhl40 | Y | 435 | -0.13865711 | 0.019825508 | -0.0890015 | 0.049655611 | -0.108827008 | 0.074564853 | 0.745633935 | 0.360534445 | 0.604405954 | 0.284518849 |
| Usp24 | S | 2585 | -0.191455427 | 0.103776919 | -0.326106576 | -0.134651149 | -0.429883495 | 0.278506149 | 0.313187777 | 0.245446645 | 0.611447005 | 0.157265433 |
| Mrtfa | S | 606 | 0.082285059 | -0.085848006 | -0.109825248 | -0.192110307 | -0.023977241 | 0.31616647 | 0.126391878 | 0.010055092 | 0.079051777 | 0.607625848 |
| Polr3e | S | 162 | 0.027952838 | -0.096107305 | -0.060173211 | -0.088126049 | 0.035934094 | 0.44491116 | 0.024236351 | 0.09454655 | 0.077910631 | 0.314502596 |
| Brf1 | S | 432 | -0.04249276 | 0.017596624 | -0.103318339 | -0.060825579 | -0.120914963 | 0.662183562 | 0.552444644 | 0.00224476 | 0.542340643 | 0.016508262 |
| Ddhd1 | S | 370 | 0.127461941 | -0.077128216 | -0.278220181 | -0.405682122 | -0.201091965 | 0.598442544 | 0.594152888 | 0.06210217 | 0.172881774 | 0.247991288 |
| Ddhd1 | S | 374 | 0.127461941 | -0.077128216 | -0.278220181 | -0.405682122 | -0.201091965 | 0.598442544 | 0.594152888 | 0.06210217 | 0.172881774 | 0.247991288 |
| Ints3 | S | 1033 | 0.140712615 | -0.138167336 | -0.19489688 | -0.335609494 | -0.056729544 | 0.057729172 | 0.083156581 | 0.020513834 | 0.004097411 | 0.306100716 |
| Tox4 | S | 182 | -0.007783761 | -0.061738476 | -0.031413764 | -0.023630003 | 0.030324713 | 0.68180865 | 0.014163609 | 0.124829048 | 0.056492187 | 0.05025046 |
| Usp47 | S | 1018 | -0.080071017 | -0.026270909 | -0.052578606 | 0.027492411 | -0.026307697 | 0.110130606 | 0.615750573 | 0.20266714 | 0.492075714 | 0.584402726 |
| Rabl6 | S | 436 | -0.083239845 | -0.001956346 | -0.023290037 | 0.059949808 | -0.021333691 | 0.002646413 | 0.933025465 | 0.60506925 | 0.241318682 | 0.634304217 |
| Rabl6 | S | 438 | -0.083239845 | -0.001956346 | -0.023290037 | 0.059949808 | -0.021333691 | 0.002646413 | 0.933025465 | 0.60506925 | 0.241318682 | 0.634304217 |
| Vapa | S | 164 | -0.037923255 | -0.018347417 | -0.104237243 | -0.066313988 | -0.085889825 | 0.419788684 | 0.751704215 | 0.04510206 | 0.166357965 | 0.180291915 |
| Map1s | S | 684 | 0.016286646 | -0.067659677 | -0.088881355 | -0.105168 | -0.021221678 | 0.678508187 | 0.066501966 | 0.08073601 | 0.056141237 | 0.563852272 |
| Thrap3 | S | 379 | -0.124046196 | 0.014561306 | -0.043788341 | 0.080257855 | -0.058349647 | 0.033315052 | 0.853798963 | 0.331406665 | 0.073487169 | 0.444017643 |
| Sipa1 | S | 879 | -0.093216137 | -0.100063249 | -0.018174301 | 0.075041835 | 0.081888947 | 0.376074819 | 0.137146687 | 0.741602356 | 0.435348596 | 0.102516381 |
| Znrf2 | S | 18 | -0.044211397 | -0.02772434 | -0.079758858 | -0.035547462 | -0.052034519 | 0.303457916 | 0.603711383 | 0.060520542 | 0.406146499 | 0.35728631 |
| Vapa | S | 153 | 0.045215614 | -0.061996802 | -0.112596467 | -0.157812082 | -0.050599665 | 0.358411359 | 0.138115017 | 0.015320751 | 0.031977392 | 0.236480831 |
| Leo1 | T | 189 | -0.032916645 | -0.039928966 | -0.072431286 | -0.039514641 | -0.032502591 | 0.107747803 | 0.455732645 | 0.147088647 | 0.359199329 | 0.583117289 |
| Camkk2 | S | 91 | -0.063842093 | -0.011574758 | -0.098585344 | -0.034743251 | -0.087010585 | 0.226945785 | 0.776746507 | 0.11942937 | 0.522091871 | 0.143326987 |
| Trank1 | S | 2386 | -0.024525644 | -0.049852967 | -0.083501235 | -0.058975591 | -0.033648268 | 0.692753658 | 0.244039956 | 0.079986138 | 0.316729153 | 0.098347757 |
| Frmd4a | T | 878 | 0.077219954 | -0.085005827 | -0.170939366 | -0.24815932 | -0.085933539 | 0.152471807 | 0.124199954 | 0.028736752 | 0.01238643 | 0.163919151 |
| Flna | S | 860 | -0.076390089 | 0.023665486 | -0.203072439 | -0.126682351 | -0.226737926 | 0.36592359 | 0.860694365 | 0.169096431 | 0.34346528 | 0.201532862 |
| Dclk1 | S | 310 | 0.005470741 | -0.012429143 | -0.2065601 | -0.212030841 | -0.194130958 | 0.902738538 | 0.843366779 | 0.046762622 | 0.063960623 | 0.064305003 |

| | | | | | | | | | | | | |
|---|---|---|---|---|---|---|---|---|---|---|---|---|
| Ccny | S | 21 | -0.116194403 | 0.016144718 | -0.114315651 | 0.001878753 | -0.130460369 | 0.137742666 | 0.371010202 | 0.151952345 | 0.980657436 | 0.129278333 |
| Prpf3 | S | 133 | 0.160815824 | -0.1646598 | -0.203081102 | -0.363896926 | -0.038421302 | 0.005801453 | 0.11990588 | 0.001740191 | 0.000515535 | 0.640373137 |
| Zdhhc5 | S | 584 | -0.104868769 | 0.041680843 | -0.160556531 | -0.055687761 | -0.202237374 | 0.232315678 | 0.757329098 | 0.170392045 | 0.505406257 | 0.18212987 |
| Brd2 | S | 631 | 0.009210368 | -0.084922358 | -0.036877423 | -0.046087791 | 0.048044935 | 0.754209513 | 0.011534299 | 0.18033165 | 0.123266337 | 0.064885082 |
| Pola2 | T | 127 | -0.033769867 | -0.017234796 | -0.101827785 | -0.068057917 | -0.084592989 | 0.268368794 | 0.685284997 | 0.067165829 | 0.169238533 | 0.144738559 |
| Map1a | S | 2211 | -0.111873022 | 0.018461739 | -0.333420815 | -0.221547793 | -0.351882554 | 0.546411522 | 0.919458371 | 0.34593527 | 0.526103552 | 0.338938507 |
| Add1 Add2 Add3 | T | 691;710;722 | -0.144405092 | 0.142857442 | -0.247261069 | -0.102855977 | -0.390118511 | 0.268122682 | 0.296882646 | 0.077108055 | 0.455241297 | 0.040530378 |
| Snrk | S | 351 | -0.032979537 | -0.04696923 | -0.058504499 | -0.025524961 | -0.011535269 | 0.368155813 | 0.290705788 | 0.071600242 | 0.371839184 | 0.740167337 |
| Sufu | S | 301 | -0.151100046 | 0.196543018 | -0.285493597 | -0.134393551 | -0.482036615 | 0.125845174 | 0.079284404 | 0.039333556 | 0.222217854 | 0.008917764 |
| Kmt2b | S | 694 | -0.104263061 | 0.246211621 | -0.356334151 | -0.25207109 | -0.602545773 | 0.754786318 | 0.375656308 | 0.027138516 | 0.462474652 | 0.097164005 |
| Pdap1 | S | 57 | -0.065799382 | -0.024867843 | -0.047824197 | 0.017975184 | -0.022956355 | 0.06127531 | 0.423507896 | 0.302773391 | 0.639636295 | 0.555496887 |
| Prrc2a | T | 1106 | -0.012169685 | -0.088308649 | -0.01641216 | -0.004242476 | 0.071896489 | 0.6073521 | 0.007678343 | 0.677934144 | 0.909191989 | 0.133922672 |
| Anxa7 | S | 311 | -0.094848535 | -0.050010739 | -0.044387393 | 0.050461142 | 0.005623346 | 0.136487846 | 0.443293661 | 0.493058823 | 0.379331538 | 0.925062431 |
| Mmp24 | S | 144 | -0.198896192 | 0.075586187 | 0.062894307 | 0.261790499 | -0.012691879 | 0.000475575 | 0.55469565 | 0.498573289 | 0.070879516 | 0.92749522 |
| Mmp24 | T | 142 | -0.198896192 | 0.075586187 | 0.062894307 | 0.261790499 | -0.012691879 | 0.000475575 | 0.55469565 | 0.498573289 | 0.070879516 | 0.92749522 |
| Mmp24 | T | 149 | -0.198896192 | 0.075586187 | 0.062894307 | 0.261790499 | -0.012691879 | 0.000475575 | 0.55469565 | 0.498573289 | 0.070879516 | 0.92749522 |
| Mmp24 | Y | 148 | -0.198896192 | 0.075586187 | 0.062894307 | 0.261790499 | -0.012691879 | 0.000475575 | 0.55469565 | 0.498573289 | 0.070879516 | 0.92749522 |
| Son | S | 1975 | 0.003873831 | -0.071568173 | -0.064835243 | -0.068709074 | 0.00673293 | 0.947359087 | 0.015545207 | 0.294278955 | 0.371036382 | 0.899944476 |
| Son | S | 1977 | 0.003873831 | -0.071568173 | -0.064835243 | -0.068709074 | 0.00673293 | 0.947359087 | 0.015545207 | 0.294278955 | 0.371036382 | 0.899944476 |
| Kmt2d | S | 1815 | -0.138085937 | -0.143623429 | 0.061801317 | 0.199887253 | 0.205424745 | 0.274813053 | 0.077205712 | 0.470607948 | 0.147380565 | 0.041998309 |
| Tjap1 | S | 214 | 0.074558704 | -0.169287569 | -0.131594325 | -0.206153029 | 0.037693245 | 0.107856107 | 0.110874146 | 0.059243651 | 0.018205632 | 0.658174229 |
| Cers5 | S | 350 | -0.069420296 | -0.133243215 | -0.016088086 | 0.05333221 | 0.117155129 | 0.431638793 | 0.133162789 | 0.848491694 | 0.417712984 | 0.086329737 |
| Cers5 | S | 354 | -0.069420296 | -0.133243215 | -0.016088086 | 0.05333221 | 0.117155129 | 0.431638793 | 0.133162789 | 0.848491694 | 0.417712984 | 0.086329737 |
| Cers5 | S | 355 | -0.069420296 | -0.133243215 | -0.016088086 | 0.05333221 | 0.117155129 | 0.431638793 | 0.133162789 | 0.848491694 | 0.417712984 | 0.086329737 |
| Cers5 | S | 356 | -0.069420296 | -0.133243215 | -0.016088086 | 0.05333221 | 0.117155129 | 0.431638793 | 0.133162789 | 0.848491694 | 0.417712984 | 0.086329737 |
| Pdlim7 | S | 217 | 0.050820268 | -0.120668395 | -0.068485325 | -0.119305593 | 0.05218307 | 0.376547151 | 0.028443868 | 0.123306769 | 0.076530939 | 0.19085289 |
| Tnks1bp1 | S | 1373 | -0.076289554 | 0.014108342 | -0.090987378 | -0.014697824 | -0.10509572 | 0.05004868 | 0.62858168 | 0.152114654 | 0.779995877 | 0.119444441 |
| Phldb2 | S | 203 | 0.053164972 | -0.058697643 | -0.137033832 | -0.190198804 | -0.078336189 | 0.199480021 | 0.117522495 | 0.016669508 | 0.007650984 | 0.095251431 |
| Ccdc6 | S | 344 | -0.165700646 | 0.0239085 | -0.176310862 | -0.010610215 | -0.200219361 | 0.247382062 | 0.76297916 | 0.386384227 | 0.957706421 | 0.330348377 |
| Myg1 | S | 283 | -0.104190874 | 0.025348672 | -0.136167158 | -0.031976284 | -0.16151583 | 0.287615454 | 0.403189847 | 0.070186815 | 0.731425007 | 0.052614781 |
| Aff4 | S | 207 | -0.070671495 | -0.109740561 | -0.010509031 | 0.060162464 | 0.099231529 | 0.347762896 | 0.117204356 | 0.775046581 | 0.421653962 | 0.147810021 |
| Rrbp1 | S | 1162 | -0.153288666 | 0.021026915 | -0.118466472 | 0.034822194 | -0.139493387 | 0.161927115 | 0.831778117 | 0.368698976 | 0.776039061 | 0.2907341 |
| Ptpn12 | S | 603 | -0.092976836 | -0.00448424 | -0.067787047 | 0.025189789 | -0.063302808 | 0.247560907 | 0.885859826 | 0.068688206 | 0.718138969 | 0.125445373 |
| Ptpn12 | S | 606 | -0.092976836 | -0.00448424 | -0.067787047 | 0.025189789 | -0.063302808 | 0.247560907 | 0.885859826 | 0.068688206 | 0.718138969 | 0.125445373 |
| Ptpn12 | S | 608 | -0.092976836 | -0.00448424 | -0.067787047 | 0.025189789 | -0.063302808 | 0.247560907 | 0.885859826 | 0.068688206 | 0.718138969 | 0.125445373 |
| Ankra2 | Y | 114 | -0.106100851 | -0.01881164 | -0.093440847 | 0.012660004 | -0.074629207 | 0.251046462 | 0.643760884 | 0.300039056 | 0.90061519 | 0.38604392 |
| Son | S | 2147 | -0.025288227 | -0.157767813 | 0.039616464 | 0.064904691 | 0.197384277 | 0.737194228 | 0.007099584 | 0.513249192 | 0.429560709 | 0.040527987 |
| Cep170 | T | 1047 | -0.216599497 | 0.09265584 | -0.096796527 | 0.11980297 | -0.189452367 | 0.090136094 | 0.600952462 | 0.387637262 | 0.36527546 | 0.346824186 |
| Zc3h14 | S | 579 | 0.095700691 | -0.085084456 | -0.287256751 | -0.382957442 | -0.202172295 | 0.243251929 | 0.409923387 | 0.059663965 | 0.032460763 | 0.16113767 |
| Sash1 | S | 400 | -0.229254755 | 0.224594896 | -0.220034039 | 0.009220716 | -0.444628935 | 0.040686351 | 0.150981924 | 0.192378354 | 0.948138327 | 0.047964706 |
| Git2 | S | 535 | -0.108157531 | 0.06313267 | -0.130823465 | -0.022665934 | -0.193956136 | 0.051432169 | 0.323511874 | 0.13962088 | 0.735595675 | 0.058919722 |
| Rpl30 | S | 10 | -0.018835603 | -0.001865222 | -0.16201758 | -0.143181977 | -0.160152358 | 0.767604565 | 0.968830361 | 0.037677133 | 0.07423731 | 0.051866894 |
| Clasp1 | S | 598 | -0.092827866 | -0.010804299 | -0.071539601 | 0.021288265 | -0.060735302 | 0.10283302 | 0.841414307 | 0.27416922 | 0.710035705 | 0.33665677 |
| Cdk14 | S | 120 | -0.116105619 | -0.030774946 | -0.14468014 | -0.028574522 | -0.113905194 | 0.398804684 | 0.758480664 | 0.353234842 | 0.848599796 | 0.403235031 |
| Trip12 | S | 236 | 0.009541947 | -0.099910244 | -0.058227761 | -0.067769708 | 0.041682482 | 0.816457823 | 0.031068298 | 0.310283932 | 0.227889312 | 0.411061278 |
| Nasp | S | 712 | -0.119869609 | -0.042646457 | -0.015115577 | 0.104754032 | 0.02753088 | 0.121647212 | 0.264911451 | 0.755403223 | 0.177656339 | 0.591962996 |
| Plec | S | 2586 | -0.115858863 | 0.046247828 | -0.140452908 | -0.024594044 | -0.186700736 | 0.27935084 | 0.445090532 | 0.067999692 | 0.789084698 | 0.022123438 |
| Arhgef40 | S | 397 | -0.027538533 | -0.06514532 | -0.055347041 | -0.027808508 | 0.009798279 | 0.516588434 | 0.232905891 | 0.093443195 | 0.455315486 | 0.825441194 |
| Sema4b | S | 777 | 0.176921445 | -0.255722353 | -0.148715404 | -0.325636848 | 0.107006949 | 0.003703265 | 0.007111521 | 0.046516578 | 0.006722816 | 0.155104535 |
| Tut7 | S | 830 | -0.050133653 | -0.11396704 | 0.013763399 | 0.063897052 | 0.127730439 | 0.148019807 | 0.071997338 | 0.819779372 | 0.337053852 | 0.119366018 |
| Hnrnpc | S | 171 | -0.087874223 | -0.170218692 | -0.016958609 | 0.070915614 | 0.153260083 | 0.57415723 | 0.213222838 | 0.743536101 | 0.6464559 | 0.249926718 |
| Rai14 | S | 480 | 0.039708851 | -0.033872002 | -0.201305312 | -0.241014163 | -0.16743331 | 0.532094191 | 0.449878892 | 0.04222186 | 0.029952745 | 0.064958494 |
| Tmcc3 | S | 216 | 0.139932367 | -0.168114506 | -0.165846811 | -0.305779177 | 0.002267695 | 0.026763372 | 0.0130313 | 0.071289003 | 0.022410262 | 0.972303815 |
| Nefm | S | 610 | 0.018487117 | -0.229915847 | 0.03516738 | 0.016680263 | 0.265083227 | 0.751916894 | 0.009625158 | 0.498620087 | 0.65780874 | 0.016484971 |
| Golga3 | S | 1479 | -0.035813575 | -0.014206853 | -0.147262076 | -0.111448501 | -0.133055223 | 0.6333912 | 0.799026032 | 0.073613552 | 0.21205921 | 0.100003622 |
| 2700049A03Rik | S | 1220 | 0.00259688 | -0.114704515 | -0.171830481 | -0.174427361 | -0.057125967 | 0.971598076 | 0.266663874 | 0.353480951 | 0.34703141 | 0.745088816 |
| Myo18a | S | 2034 | -0.097071968 | -0.021913163 | -0.079601572 | 0.017470396 | -0.057688409 | 0.373479737 | 0.500535711 | 0.177298584 | 0.865393611 | 0.290425624 |
| Ripk2 | S | 362 | -0.083386639 | -0.025421217 | -0.050913097 | 0.032473542 | -0.02549188 | 0.073632199 | 0.562365377 | 0.400555036 | 0.536985445 | 0.632298897 |
| Srbd1 | S | 174 | 0.016427353 | -0.141737065 | -0.055752054 | -0.072179407 | 0.085985012 | 0.781375136 | 0.045966624 | 0.467307656 | 0.418944934 | 0.315649239 |
| Mrpl30 | S | 96 | -0.082605734 | -0.058027157 | -0.015151105 | 0.067454629 | 0.042876052 | 0.069237548 | 0.267730148 | 0.809114798 | 0.301136611 | 0.51483689 |
| Trim2 | S | 428 | -0.100201888 | 0.058016797 | -0.155386731 | -0.055184843 | -0.213403528 | 0.142743789 | 0.415181958 | 0.109535944 | 0.44358407 | 0.04592989 |
| Ybx3 | S | 359 | -0.043701403 | -0.032053713 | -0.070602212 | -0.026900808 | -0.038548498 | 0.194400375 | 0.132227991 | 0.33152676 | 0.689532681 | 0.563319141 |
| Rab3ip | T | 217 | -0.089295755 | -0.045501222 | -0.056307959 | 0.032987796 | -0.010806737 | 0.259955888 | 0.586830109 | 0.204899281 | 0.64661597 | 0.895121196 |
| Pard3 | S | 692 | 0.018487259 | -0.020007624 | -0.223883106 | -0.242370365 | -0.203875482 | 0.870282089 | 0.536408307 | 0.086429887 | 0.127102832 | 0.119754508 |
| Copa | S | 704 | -0.105566521 | 0.024593097 | -0.13965805 | -0.034091529 | -0.164251146 | 0.264708009 | 0.806337501 | 0.16759818 | 0.567762951 | 0.081029231 |
| Copa | Y | 707 | -0.105566521 | 0.024593097 | -0.13965805 | -0.034091529 | -0.164251146 | 0.264708009 | 0.806337501 | 0.16759818 | 0.567762951 | 0.081029231 |
| Sgta | S | 309 | 0.084241622 | -0.009788338 | -0.200655574 | -0.284897195 | -0.190867236 | 0.445825389 | 0.889656652 | 0.005148659 | 0.076108056 | 0.053037536 |
| Ctu2 | T | 508 | 0.022559151 | -0.12597973 | -0.059905944 | -0.082465094 | 0.066073786 | 0.796869587 | 0.115629908 | 0.127410067 | 0.376819837 | 0.33061342 |
| Kat7 | S | 275 | -0.0755384 | 0.01430555 | -0.044004115 | 0.031534285 | -0.058309665 | 0.031405995 | 0.823285613 | 0.069664308 | 0.249476397 | 0.40076372 |
| Fmn1 | S | 532 | -0.005143501 | -0.168023468 | -0.086503331 | -0.08135983 | 0.081520138 | 0.958745997 | 0.181592087 | 0.484555964 | 0.389179125 | 0.459570377 |
| Naa15 | S | 854 | -0.072555825 | -0.026686784 | -0.030389682 | 0.042166143 | -0.003702897 | 0.116248141 | 0.13264657 | 0.374389026 | 0.346344627 | 0.900416965 |
| Gtpbp4 | S | 468 | 0.095921411 | -0.098627996 | -0.127279186 | -0.223200597 | -0.028651189 | 0.036512735 | 0.155632811 | 0.001040807 | 0.003240814 | 0.602524145 |
| Gtpbp4 | S | 470 | 0.095921411 | -0.098627996 | -0.127279186 | -0.223200597 | -0.028651189 | 0.036512735 | 0.155632811 | 0.001040807 | 0.003240814 | 0.602524145 |
| Gtpbp4 | S | 472 | 0.095921411 | -0.098627996 | -0.127279186 | -0.223200597 | -0.028651189 | 0.036512735 | 0.155632811 | 0.001040807 | 0.003240814 | 0.602524145 |
| Vim | S | 26 | -0.035474584 | -0.01764211 | -0.082014707 | -0.046540122 | -0.064372597 | 0.390580698 | 0.669943156 | 0.031399921 | 0.214000124 | 0.134444639 |
| Brd1 | S | 128 | 0.01395216 | -0.039434855 | -0.159686269 | -0.173638429 | -0.120251413 | 0.851896697 | 0.538689725 | 0.050805908 | 0.090512636 | 0.150501389 |
| Myof | S | 340 | -0.036389007 | -0.10704717 | -0.019322585 | 0.017066422 | 0.087724586 | 0.432412407 | 0.070628448 | 0.649069635 | 0.70546906 | 0.128526529 |
| Srpk2 | S | 10 | -0.191599331 | 0.077811276 | 0.015990579 | 0.207589909 | -0.061820698 | 0.000700062 | 0.246058654 | 0.850953943 | 0.099830983 | 0.530137161 |
| Uhrf1bp1l | S | 886 | -0.120657189 | -0.038577563 | -0.024050794 | 0.096606395 | 0.014526769 | 0.052009504 | 0.716005906 | 0.727621 | 0.165482544 | 0.890889291 |
| Itpr3 | S | 1832 | -0.10857758 | 0.136293572 | -0.320409379 | -0.211831799 | -0.456702952 | 0.028427846 | 0.017324555 | 0.071693456 | 0.160866724 | 0.036001468 |
| Ice1 | S | 1616 | -0.040591793 | -0.073227454 | -0.125225443 | -0.084633649 | -0.051997988 | 0.800048837 | 0.335817074 | 0.246062229 | 0.630491596 | 0.622576177 |
| Tgif1 | S | 20 | 0.023333148 | -0.135782412 | -0.148283514 | -0.171616661 | -0.012501102 | 0.919970403 | 0.203814143 | 0.386786722 | 0.505557923 | 0.931865242 |
| Add2 | S | 528 | -0.035947817 | -0.13025286 | 0.017397751 | 0.053345568 | 0.147650611 | 0.316830338 | 0.029308045 | 0.655378057 | 0.213926163 | 0.026997826 |
| Tfip11 | S | 60 | -0.154639514 | -0.067262837 | 0.099836496 | 0.25447601 | 0.167099333 | 0.008018081 | 0.108731788 | 0.126128865 | 0.011013533 | 0.051378375 |
| Ttc7b | S | 678 | -0.109353605 | -0.175684332 | -0.026985321 | 0.082368284 | 0.148699011 | 0.397476569 | 0.427627882 | 0.732989946 | 0.475939041 | 0.486076407 |
| Zc3h13 | S | 681 | -0.003100618 | 0.070613605 | -0.334001643 | -0.330901025 | -0.404615249 | 0.966407546 | 0.3369075 | 0.034131024 | 0.052134083 | 0.039178328 |
| Spg20 | S | 126 | -0.164334173 | 0.056967696 | -0.077630323 | 0.08670385 | -0.13459802 | 0.031114272 | 0.416877659 | 0.410693572 | 0.331281198 | 0.176619757 |
| Cttnbp2nl | S | 488 | -0.088981569 | -0.109656771 | 0.061459964 | 0.150441533 | 0.171116735 | 0.058320261 | 0.031908474 | 0.183319329 | 0.019542995 | 0.013412684 |
| Anln | S | 238 | -0.108452545 | -0.161419927 | 0.025002885 | 0.133455429 | 0.186422812 | 0.479026868 | 0.079577313 | 0.523764837 | 0.39827686 | 0.059460909 |
| Macf1 | S | 3889 | -0.026367054 | -0.02972319 | -0.068356905 | -0.041989852 | -0.038633716 | 0.246252026 | 0.418930399 | 0.057871531 | 0.177803185 | 0.333786591 |
| Ktn1 | S | 75 | -0.019734091 | -0.086261045 | -0.023717156 | -0.003983065 | 0.062543888 | 0.608843904 | 0.03397878 | 0.361832917 | 0.914603483 | 0.094334444 |
| Pla2g4a | S | 435 | -0.12441151 | 0.069354982 | -0.122365122 | 0.002046388 | -0.191720103 | 0.201604357 | 0.29770377 | 0.035997107 | 0.979907705 | 0.032366689 |
| Cdca2 | S | 626 | 0.010462572 | -0.147912431 | -0.069592577 | -0.080055149 | 0.078319854 | 0.807286069 | 0.134605669 | 0.281926305 | 0.209753851 | 0.38634489 |
| Phf2 | S | 893 | 0.009378453 | -0.052650956 | -0.076071503 | -0.085449956 | -0.023420547 | 0.758011344 | 0.044861728 | 0.077762809 | 0.07479408 | 0.490249889 |

| | | | | | | | | | | | | |
|---|---|---|---|---|---|---|---|---|---|---|---|---|
| Dido1 | S | 886 | -0.158939821 | -0.116987604 | 0.00609973 | 0.165039551 | 0.123087334 | 0.244953101 | 0.377126922 | 0.955578718 | 0.279744999 | 0.410251712 |
| Hprt | S | 11 | -0.394895526 | 0.104129965 | -0.034320347 | 0.360575179 | -0.138450312 | 0.059816598 | 0.353843587 | 0.875561756 | 0.187314184 | 0.53477719 |
| Raly | S | 252 | -0.027583579 | -0.072614846 | -0.05904994 | -0.03146636 | 0.013564906 | 0.578961651 | 0.22445357 | 0.139906295 | 0.485322499 | 0.78442345 |
| Plekhg2 | S | 1277 | -0.117572614 | -0.041370369 | 0.087615462 | 0.205188076 | 0.128985831 | 0.004916507 | 0.170617958 | 0.439293911 | 0.150819605 | 0.291423976 |
| Map4 | S | 760 | 0.053290637 | -0.030566561 | -0.151708818 | -0.204999455 | -0.121142256 | 0.129910332 | 0.318110582 | 0.003144227 | 0.001827982 | 0.021340766 |
| Trim24 | S | 110 | -0.27686043 | -0.007448792 | -0.002436212 | 0.274424218 | 0.005012581 | 0.116944633 | 0.968660061 | 0.985960582 | 0.073675464 | 0.973572999 |
| Sin3b | S | 698 | -0.028695533 | -0.16825973 | 0.015528834 | 0.044224367 | 0.183788564 | 0.659949572 | 0.037653763 | 0.792368482 | 0.326670099 | 0.024338347 |
| Lmnb1 | S | 280 | 0.03208899 | -0.046429732 | -0.107571039 | -0.13966003 | -0.061141307 | 0.314263991 | 0.54551797 | 0.002022945 | 0.01034896 | 0.43673316 |
| Ttc4 | S | 235 | 0.109311248 | -0.023612098 | -0.402937279 | -0.512248527 | -0.379325181 | 0.502319709 | 0.662601488 | 0.098050643 | 0.059589576 | 0.112196364 |
| Abca2 | T | 2411 | -0.03211924 | -0.067480037 | -0.062842529 | -0.030723289 | 0.004637507 | 0.311579848 | 0.228799982 | 0.281556762 | 0.552573097 | 0.940094769 |
| Morc2a | S | 854 | 0.032752702 | -0.090735128 | -0.065259098 | -0.0980118 | 0.02547603 | 0.422568581 | 0.013128064 | 0.184580672 | 0.101639016 | 0.55598815 |
| Resf1 | S | 1102 | -0.296443637 | 0.100404173 | -0.152987089 | 0.143456548 | -0.253391262 | 0.178694182 | 0.52751846 | 0.49119836 | 0.566241151 | 0.29604741 |
| Phf12 | S | 131 | -0.053250271 | -0.068348318 | -0.022173065 | 0.031077206 | 0.046175253 | 0.307435661 | 0.104735187 | 0.374974555 | 0.519937375 | 0.227178236 |
| Mrtfa | S | 41 | -0.035609446 | -0.092773141 | -0.0001166 | 0.035492846 | 0.092656541 | 0.537271847 | 0.011855373 | 0.997041201 | 0.527369185 | 0.017243236 |
| Prpf4b | S | 233 | -0.034112546 | -0.176766007 | 1.55029E-07 | 0.034112701 | 0.176766162 | 0.642264433 | 0.080894904 | 0.999997464 | 0.629999223 | 0.087563246 |
| Pard3 | S | 728 | -0.072511827 | 0.03219225 | -0.127341471 | -0.054829643 | -0.159533721 | 0.504975577 | 0.449928623 | 0.015854097 | 0.600228222 | 0.006216384 |
| Ccdc9b | S | 241 | -0.075363295 | -0.110637495 | 0.042700504 | 0.1180638 | 0.153337999 | 0.225007127 | 0.021527534 | 0.396239664 | 0.118546167 | 0.038707641 |
| Setd2 | S | 1819 | 0.023404168 | -0.007472786 | -0.129492167 | -0.152896335 | -0.12201938 | 0.767602323 | 0.793912909 | 0.005197048 | 0.143958245 | 0.013115385 |
| Nkap | S | 5 | -0.073619157 | -0.044284029 | -0.064800457 | 0.0088187 | -0.020516428 | 0.291351348 | 0.504107004 | 0.223748474 | 0.8926043 | 0.754840633 |
| Ppfibp1 | T | 562 | -0.186800283 | 0.151598022 | -0.223347912 | -0.036547629 | -0.374945934 | 0.123661598 | 0.250882437 | 0.179116175 | 0.807654136 | 0.067836362 |
| Arhgap31 | S | 941 | -0.136602742 | 0.053457161 | -0.10143573 | 0.035167012 | -0.154892891 | 0.09188916 | 0.557909917 | 0.220505099 | 0.609111075 | 0.130347717 |
| Herc2 | S | 1949 | -0.078198964 | -0.010710134 | -0.094642831 | -0.016443868 | -0.083932697 | 0.235171074 | 0.896024475 | 0.163130348 | 0.673118117 | 0.274405646 |
| Cpox | S | 99 | -0.039207326 | -0.080909441 | -0.071392932 | -0.032185606 | 0.00951651 | 0.594387907 | 0.146332726 | 0.424274205 | 0.726315877 | 0.903185718 |
| Crtc2 | S | 70 | -0.092767554 | -0.106426705 | -0.067519814 | 0.02524774 | 0.038906891 | 0.488087147 | 0.332718843 | 0.606901607 | 0.858652954 | 0.750017375 |
| Crtc2 | S | 86 | -0.092767554 | -0.106426705 | -0.067519814 | 0.02524774 | 0.038906891 | 0.488087147 | 0.332718843 | 0.606901607 | 0.858652954 | 0.750017375 |
| Crtc2 | S | 90 | -0.092767554 | -0.106426705 | -0.067519814 | 0.02524774 | 0.038906891 | 0.488087147 | 0.332718843 | 0.606901607 | 0.858652954 | 0.750017375 |
| Tnks1bp1 | S | 1442 | 0.006569951 | -0.048481193 | -0.171786651 | -0.178356603 | -0.123305459 | 0.92141689 | 0.438017938 | 0.117165108 | 0.108707071 | 0.215011187 |
| Cspg4 | S | 1300 | -0.040267645 | -0.035549643 | -0.16396155 | -0.123693904 | -0.128411906 | 0.761969805 | 0.646775398 | 0.15366201 | 0.421246588 | 0.261819266 |
| Nes | S | 907 | -0.034226804 | -0.191840436 | 0.00538748 | 0.039614285 | 0.197227917 | 0.592687014 | 0.101660829 | 0.968536594 | 0.759354394 | 0.22654249 |
| Actl6a | S | 233 | -0.080527868 | -0.021648601 | -0.044376916 | 0.036150952 | -0.022728315 | 0.203081085 | 0.711698741 | 0.075282811 | 0.504550272 | 0.690509467 |
| Jakmip3 | S | 710 | -0.107551192 | -0.048517155 | -0.031408004 | 0.076143188 | 0.017109151 | 0.139670293 | 0.41389442 | 0.658487433 | 0.244610989 | 0.73075026 |
| Ddx20 | S | 673 | -0.07586274 | -0.133532051 | 0.042209771 | 0.118072511 | 0.175741822 | 0.501450496 | 0.038008208 | 0.756503487 | 0.472421573 | 0.267290448 |
| Ddx20 | S | 678 | -0.07586274 | -0.133532051 | 0.042209771 | 0.118072511 | 0.175741822 | 0.501450496 | 0.038008208 | 0.756503487 | 0.472421573 | 0.267290448 |
| Tpr | S | 453 | 0.008454162 | -0.055283122 | -0.074401247 | -0.08285541 | -0.019118125 | 0.841109438 | 0.061903999 | 0.086143008 | 0.121340004 | 0.551718984 |
| Rbmx2 | S | 144 | -0.0475873 | -0.075372879 | 0.027422846 | 0.075010147 | 0.102795725 | 0.144994163 | 0.009463237 | 0.617926904 | 0.239226703 | 0.145005036 |
| Rbmx2 | S | 149 | -0.0475873 | -0.075372879 | 0.027422846 | 0.075010147 | 0.102795725 | 0.144994163 | 0.009463237 | 0.617926904 | 0.239226703 | 0.145005036 |
| Rbmx2 | T | 140 | -0.0475873 | -0.075372879 | 0.027422846 | 0.075010147 | 0.102795725 | 0.144994163 | 0.009463237 | 0.617926904 | 0.239226703 | 0.145005036 |
| Nck1 | S | 166 | 0.018982367 | 0.002838369 | -0.288687157 | -0.307669524 | -0.291525526 | 0.846866644 | 0.968351013 | 0.083582326 | 0.078187 | 0.090310579 |
| Adat1 | S | 191 | 0.024322919 | -0.017919063 | -0.125375399 | -0.149698317 | -0.107456336 | 0.41751252 | 0.574852109 | 0.004281247 | 0.002020549 | 0.007956687 |
| Gle1 | S | 368 | -0.01619041 | -0.097724696 | -0.039915787 | -0.023725377 | 0.057808908 | 0.683282713 | 0.066724986 | 0.41310097 | 0.617597785 | 0.287905528 |
| Htatsf1 | S | 441 | -0.22598567 | -0.039141132 | 0.059283804 | 0.285269474 | 0.098424936 | 0.04405086 | 0.432892772 | 0.377470103 | 0.023948096 | 0.19103319 |
| Dvl1 | S | 89 | -0.166072347 | -0.070033407 | -0.012531146 | 0.153541201 | 0.05750226 | 0.157019566 | 0.577262486 | 0.930740105 | 0.343798087 | 0.724783513 |
| Dvl1 | S | 93 | -0.166072347 | -0.070033407 | -0.012531146 | 0.153541201 | 0.05750226 | 0.157019566 | 0.577262486 | 0.930740105 | 0.343798087 | 0.724783513 |
| Tgs1 | S | 849 | -0.113092807 | -0.029997523 | -0.039474937 | 0.07361787 | -0.009477414 | 0.112756568 | 0.575218942 | 0.545162101 | 0.353246731 | 0.898865569 |
| Phf2 | S | 677 | -0.211152457 | -0.034112739 | 0.07142391 | 0.282576367 | 0.105536649 | 0.038191993 | 0.506354256 | 0.562398828 | 0.087835 | 0.406210111 |
| Tjp1 | S | 617 | -0.039686843 | -0.036873778 | -0.063840639 | -0.024153796 | -0.026966861 | 0.454522479 | 0.340955638 | 0.077761302 | 0.639689166 | 0.481932843 |
| Rin1 | S | 238 | -0.115065646 | 0.037485701 | -0.091644375 | 0.023421272 | -0.129130076 | 0.07173139 | 0.465738908 | 0.176758782 | 0.645281782 | 0.071968276 |
| Vapa | S | 166 | 0.008603788 | -0.077622318 | -0.103877172 | -0.11248096 | -0.026254854 | 0.811588596 | 0.416641292 | 0.048163859 | 0.039416817 | 0.769226765 |
| Kpna4 | S | 72 | -0.107947987 | 0.030279332 | -0.175588896 | -0.067640908 | -0.205868228 | 0.397864249 | 0.636343205 | 0.146145068 | 0.620893342 | 0.112700589 |
| Tmem115 | T | 329 | 0.065172049 | -0.093698068 | -0.301499874 | -0.366671924 | -0.207801806 | 0.60463392 | 0.314798601 | 0.266979456 | 0.200555882 | 0.408736537 |
| Atxn2l | S | 407 | 0.08753456 | -0.060234513 | -0.311670725 | -0.399205284 | -0.251436212 | 0.240275983 | 0.364409322 | 0.080790213 | 0.056763359 | 0.14573538 |
| 9530068E07Rik | S | 165 | -0.138335702 | -0.040596083 | 0.060509962 | 0.198845664 | 0.101106045 | 0.009858116 | 0.304234292 | 0.383609471 | 0.048976937 | 0.183618201 |
| Flcn | S | 302 | -0.040335088 | 0.021146447 | -0.173834454 | -0.133499366 | -0.194980901 | 0.74964218 | 0.691272229 | 0.036388795 | 0.336267064 | 0.030892652 |
| Ubr4 | S | 1761 | -0.034904148 | -0.002103142 | -0.119929721 | -0.085025573 | -0.117826579 | 0.453109229 | 0.951999771 | 0.048868283 | 0.162037064 | 0.057841365 |
| Mcm6 | S | 802 | -0.065577323 | 0.093377355 | -0.263854562 | -0.19827724 | -0.357231918 | 0.672888214 | 0.371371399 | 0.040198482 | 0.272914309 | 0.029065551 |
| Cep55 | T | 25 | -0.126491033 | 0.038234976 | -0.113555156 | 0.012935877 | -0.151790131 | 0.177184139 | 0.739984144 | 0.216016784 | 0.859073833 | 0.21033135 |
| Stim2 | S | 28 | -0.059420887 | 0.034495156 | -0.246001113 | -0.186580226 | -0.280496269 | 0.469639324 | 0.613243814 | 0.139950775 | 0.229408006 | 0.127008127 |
| Iqsec1 | S | 362 | 0.067315143 | -0.011433811 | -0.187766695 | -0.255081839 | -0.176332884 | 0.363864748 | 0.809854148 | 0.005462259 | 0.026789382 | 0.018427604 |
| Hectd1 | S | 1492 | -0.098130187 | -0.055596635 | -0.038634152 | 0.059496035 | 0.016962483 | 0.204397553 | 0.384355291 | 0.55227312 | 0.345565503 | 0.706672672 |
| Safb | S | 554 | -0.23936227 | 0.07610189 | -0.089283078 | 0.150079192 | -0.165384968 | 0.110327282 | 0.698374026 | 0.594249523 | 0.407461317 | 0.468903863 |
| Pi4ka | S | 268 | 0.02408356 | -0.04418766 | -0.083432844 | -0.107516403 | -0.039245184 | 0.419904951 | 0.210553348 | 0.00558065 | 0.019823407 | 0.250869393 |
| Etv3 | S | 427 | 0.071750968 | -0.250032785 | -0.107083639 | -0.178834607 | 0.142949145 | 0.429304096 | 0.123535962 | 0.43494126 | 0.211242976 | 0.383425969 |
| Dcaf5 | S | 531 | 0.10596106 | -0.260580252 | -0.198289097 | -0.304250156 | 0.062291156 | 0.44783613 | 0.302567923 | 0.2716039 | 0.110001967 | 0.794209667 |
| Dcaf5 | S | 533 | 0.10596106 | -0.260580252 | -0.198289097 | -0.304250156 | 0.062291156 | 0.44783613 | 0.302567923 | 0.2716039 | 0.110001967 | 0.794209667 |
| Stard13 | S | 133 | -0.097029178 | 0.04553389 | -0.128398577 | -0.031369399 | -0.173932467 | 0.185530351 | 0.497854159 | 0.09595158 | 0.471289806 | 0.01012718 |
| Brd2 | S | 300 | -0.034430487 | -0.05577031 | 0.005776789 | 0.040207276 | 0.061547099 | 0.023762108 | 0.021017178 | 0.664196903 | 0.01827089 | 0.022783127 |
| Mast2 | S | 796 | -0.118681587 | 0.051597609 | -0.091490511 | 0.027191076 | -0.14308812 | 0.059665717 | 0.593917212 | 0.243113174 | 0.696296714 | 0.212932914 |
| Arhgap17 | S | 574 | -0.063252604 | -0.125675339 | -0.093340792 | -0.030088187 | 0.032334547 | 0.412821379 | 0.449201783 | 0.594537112 | 0.859666319 | 0.87725772 |
| Nap1l4 | T | 51 | -0.049227954 | -0.010594326 | -0.067245713 | -0.018017759 | -0.056651387 | 0.094148354 | 0.686742487 | 0.131598895 | 0.633429668 | 0.195710908 |
| Akirin2 | S | 39 | -0.201531642 | -0.039967142 | 0.020821265 | 0.222352907 | 0.060788407 | 0.086236458 | 0.616205613 | 0.79682179 | 0.063706252 | 0.264171188 |
| Avil | S | 622 | -0.0472736 | -0.01540677 | -0.089829362 | -0.042555762 | -0.074422592 | 0.140215055 | 0.795492656 | 0.155568264 | 0.434293363 | 0.320696003 |
| Itprid2 | S | 326 | -0.072909948 | 0.030347249 | -0.171234361 | -0.098324414 | -0.201581611 | 0.263761358 | 0.599562948 | 0.135055818 | 0.338373193 | 0.096360315 |
| Sh2b1 | S | 127 | -0.033441584 | -0.124771633 | 0.01743121 | 0.050872794 | 0.142202843 | 0.621157949 | 0.019953072 | 0.854132259 | 0.637066885 | 0.217559446 |
| Atxn2 | S | 753 | 0.054521736 | -0.127915534 | -0.05636437 | -0.110886105 | 0.071551164 | 0.172742758 | 0.003016982 | 0.425018246 | 0.17562182 | 0.328861117 |
| Camsap1 | T | 559 | -0.027176384 | -0.023152367 | -0.139718849 | -0.112542465 | -0.116566482 | 0.743225024 | 0.768592047 | 0.074185278 | 0.235288554 | 0.20335915 |
| Snrpc | S | 17 | -0.00682062 | -0.063426049 | -0.109946036 | -0.103125416 | -0.046519987 | 0.921464893 | 0.335122499 | 0.119029988 | 0.13667456 | 0.391200602 |
| Pdlim7 | T | 215 | 0.064059339 | -0.117172735 | -0.077417479 | -0.141476818 | 0.039755256 | 0.220698803 | 0.024710827 | 0.098829649 | 0.030642371 | 0.31727652 |
| Pkn2 | S | 110 | -0.011855784 | -0.011848065 | -0.10249074 | -0.090634956 | -0.090642675 | 0.71208995 | 0.7443153 | 0.016621149 | 0.046307634 | 0.065057038 |
| Clk3 | S | 215 | 0.10593818 | -0.093485469 | -0.09631826 | -0.20225644 | -0.002832791 | 0.482774411 | 0.01915478 | 0.133999082 | 0.237779704 | 0.955329347 |
| Arhgap23 | T | 549 | -0.065115587 | -0.010135286 | -0.143936171 | -0.078820584 | -0.133800886 | 0.629796524 | 0.873827445 | 0.113356022 | 0.566045026 | 0.119380869 |
| Vim | S | 412 | 0.049936074 | -0.056629926 | -0.170326038 | -0.220262112 | -0.113696112 | 0.235090591 | 0.272531129 | 0.031405078 | 0.029060129 | 0.107732141 |
| Micall1 | T | 314 | -0.002594746 | -0.017585941 | -0.119852666 | -0.11725792 | -0.102266725 | 0.957332297 | 0.741111591 | 0.019499609 | 0.069771444 | 0.12088144 |
| Notch2 | S | 1723 | -0.018363939 | 0.02556708 | -0.408091806 | -0.389727867 | -0.433658887 | 0.901120652 | 0.874273389 | 0.22108274 | 0.238288882 | 0.202538014 |
| Notch2 | S | 1724 | -0.018363939 | 0.02556708 | -0.408091806 | -0.389727867 | -0.433658887 | 0.901120652 | 0.874273389 | 0.22108274 | 0.238288882 | 0.202538014 |
| Clk3 | S | 224 | 0.029126725 | -0.116773364 | -0.080926357 | -0.110053082 | 0.035847007 | 0.434705883 | 0.124099052 | 0.099612726 | 0.047598401 | 0.577425352 |
| Pbrm1 | S | 39 | -0.04205843 | -0.038700408 | -0.08439588 | -0.042337451 | -0.045695473 | 0.317634402 | 0.220683457 | 0.324029259 | 0.598623313 | 0.560216371 |
| Zbtb11 Rpl24 | S | 493 | -0.035255609 | -0.043870552 | -0.243532841 | -0.208277232 | -0.199662289 | 0.808994341 | 0.871915315 | 0.221669182 | 0.331954142 | 0.519504866 |
| Specc1l | S | 982 | -0.040664964 | -0.095025084 | -0.006903602 | 0.033761361 | 0.088121482 | 0.326289991 | 0.050908485 | 0.919939592 | 0.606162214 | 0.242236836 |
| Thrap3 | S | 502 | -0.000303498 | -0.118722072 | -0.069223552 | -0.068920054 | 0.04949852 | 0.996012031 | 0.083414342 | 0.480961476 | 0.492647525 | 0.611559535 |
| Nol8 | S | 364 | -0.005192554 | -0.001931455 | -0.244185296 | -0.238992742 | -0.24225384 | 0.977334694 | 0.989546356 | 0.097889043 | 0.196292688 | 0.072912055 |
| Nol8 | S | 365 | -0.005192554 | -0.001931455 | -0.244185296 | -0.238992742 | -0.24225384 | 0.977334694 | 0.989546356 | 0.097889043 | 0.196292688 | 0.072912055 |
| Aen | S | 64 | 0.040688852 | -0.158549147 | -0.077025565 | -0.117714416 | 0.081523583 | 0.627067715 | 0.07184125 | 0.41667232 | 0.26940724 | 0.400660974 |
| Wiz | S | 1576 | 0.071913105 | -0.156348744 | -0.081820502 | -0.153733607 | 0.074528242 | 0.421389473 | 0.051952219 | 0.249264792 | 0.102425975 | 0.132550248 |
| Patl1 | S | 177 | -0.011745809 | -0.04101375 | -0.100235538 | -0.088489729 | -0.059221789 | 0.658014523 | 0.302828966 | 0.103941113 | 0.14454116 | 0.292003814 |

| | | | | | | | | | | | | |
|---|---|---|---|---|---|---|---|---|---|---|---|---|
| Hectd1 | S | 1522 | -0.204429249 | 0.098240403 | -0.132735821 | 0.071693428 | -0.230976224 | 0.077332226 | 0.384139471 | 0.39531696 | 0.643097778 | 0.201635246 |
| Cep170b | T | 541 | -0.018543881 | -0.050010898 | -0.062940455 | -0.044396574 | -0.012929557 | 0.755625358 | 0.080276778 | 0.159275546 | 0.506146451 | 0.734361899 |
| Dis3l2 | S | 864 | -0.101478433 | -0.014367207 | -0.036951978 | 0.064526455 | -0.022584771 | 0.041921174 | 0.81536921 | 0.563458058 | 0.308892342 | 0.750024903 |
| Golga2 | T | 916 | -0.102638235 | -0.222950473 | -0.052884276 | 0.049753958 | 0.170066197 | 0.348087292 | 0.59443072 | 0.836520318 | 0.842980535 | 0.70569763 |
| Cic | S | 1208 | -0.132123774 | -0.118562629 | 0.139262833 | 0.271386607 | 0.257825463 | 0.018029619 | 0.156972776 | 0.264845442 | 0.086549191 | 0.088016022 |
| Srrm2 | S | 775 | 0.002781451 | -0.037065652 | -0.032350282 | -0.035131733 | 0.00471537 | 0.923428014 | 0.032179596 | 0.003820183 | 0.296365156 | 0.671769111 |
| Gsn | S | 307 | -0.07277171 | -0.087442048 | -0.039684501 | 0.033087209 | 0.047757547 | 0.223304532 | 0.255327552 | 0.815135395 | 0.845566396 | 0.783759674 |
| Prcp | S | 47 | -0.266404909 | -0.064987106 | 0.139043876 | 0.405448784 | 0.204030982 | 0.015926838 | 0.526186664 | 0.179536917 | 0.010826317 | 0.115472065 |
| Prcp | Y | 46 | -0.266404909 | -0.064987106 | 0.139043876 | 0.405448784 | 0.204030982 | 0.015926838 | 0.526186664 | 0.179536917 | 0.010826317 | 0.115472065 |
| Prcp | Y | 50 | -0.266404909 | -0.064987106 | 0.139043876 | 0.405448784 | 0.204030982 | 0.015926838 | 0.526186664 | 0.179536917 | 0.010826317 | 0.115472065 |
| Flna | S | 1734 | -0.025875646 | -0.029140722 | -0.075215131 | -0.049339485 | -0.046074409 | 0.721754333 | 0.213720349 | 0.058684146 | 0.520147929 | 0.183876449 |
| Srrm2 | S | 945 | -0.018262278 | -0.04626637 | -0.041573257 | -0.023310979 | 0.004693114 | 0.33894849 | 0.081945527 | 0.142490994 | 0.311304067 | 0.846607394 |
| Afap1 | T | 338 | 0.058380395 | -0.078055101 | -0.222459725 | -0.28084012 | -0.144404624 | 0.745387178 | 0.353559119 | 0.186872125 | 0.214476258 | 0.341844667 |
| Golim4 | S | 540 | -0.082606243 | -0.162179319 | 0.078884212 | 0.161490455 | 0.241063531 | 0.228249312 | 0.075326336 | 0.503661386 | 0.220314318 | 0.106320565 |
| Grasp | S | 384 | -0.030711053 | -0.021847208 | -0.157306629 | -0.126595576 | -0.135459421 | 0.640467714 | 0.734904794 | 0.13152975 | 0.213901045 | 0.188685901 |
| Ptges3 | S | 118 | -0.107910254 | -0.004594815 | -0.109545218 | -0.001634964 | -0.104950403 | 0.504896935 | 0.946953634 | 0.126135886 | 0.991640104 | 0.239497771 |
| Eno1 Eno1b | S | 176 | -0.051227915 | -0.192314932 | -0.054860839 | -0.003632924 | 0.137454093 | 0.692227197 | 0.297983619 | 0.686518488 | 0.965248653 | 0.389314227 |
| Lyst | S | 2118 | -0.260752416 | -0.07976196 | 0.048934137 | 0.309686553 | 0.128696097 | 0.217533268 | 0.34158335 | 0.592374953 | 0.163725108 | 0.224781877 |
| Bend3 | S | 486 | -0.025897267 | -0.096248492 | -0.037632228 | -0.011734961 | 0.058616265 | 0.881996842 | 0.020787294 | 0.785262123 | 0.954941026 | 0.675184485 |
| Tmem200a | S | 487 | -0.003765143 | -0.102954811 | -0.051081761 | -0.047316618 | 0.051873051 | 0.958214582 | 0.091541402 | 0.28336756 | 0.512513571 | 0.322449738 |
| Ddx27 | S | 23 | 0.018896902 | -0.058344328 | -0.0514249 | -0.070321802 | 0.006919428 | 0.339216481 | 0.019260765 | 0.029680017 | 0.001004354 | 0.261456988 |
| Ddx27 | S | 25 | 0.018896902 | -0.058344328 | -0.0514249 | -0.070321802 | 0.006919428 | 0.339216481 | 0.019260765 | 0.029680017 | 0.001004354 | 0.261456988 |
| Lmna | S | 22 | 0.002793995 | -0.069740711 | -0.102699286 | -0.105493281 | -0.032958575 | 0.961879127 | 0.214828509 | 0.064011848 | 0.127726296 | 0.696233653 |
| Jmjd1c | S | 135 | -0.178318832 | -0.015253348 | 0.009966242 | 0.188285074 | 0.02521959 | 0.048157406 | 0.770412325 | 0.874403575 | 0.059729368 | 0.728953444 |
| Atp10a | S | 470 | 0.02619247 | -0.118053882 | -0.053332416 | -0.079524886 | 0.064721466 | 0.575725574 | 0.052311817 | 0.20291445 | 0.08234621 | 0.185811731 |
| Tbc1d17 | S | 20 | -0.048365817 | -0.015296725 | -0.124312889 | -0.075947072 | -0.109016164 | 0.491372079 | 0.789684703 | 0.117484369 | 0.358219351 | 0.17317581 |
| Exoc2 | S | 431 | -0.152137308 | -0.036412998 | 0.042366196 | 0.194503504 | 0.078779194 | 0.022558944 | 0.414315351 | 0.47640092 | 0.022417959 | 0.182572281 |
| Sipa1l2 | S | 1030 | 0.04806987 | -0.080141699 | -0.127489946 | -0.175559816 | -0.047348247 | 0.674616821 | 0.409702941 | 0.043877099 | 0.196285125 | 0.582006134 |
| 5031439G07Rik | S | 324 | -0.073380589 | 0.061838421 | -0.244638907 | -0.171258318 | -0.306477328 | 0.405519146 | 0.365310571 | 0.09858135 | 0.211477957 | 0.068480047 |
| Ncl | S | 605 | -0.072427246 | -0.079207253 | -0.021521418 | 0.050905828 | 0.057685835 | 0.233966231 | 0.214828509 | 0.703455794 | 0.303562537 | 0.280498833 |
| Wnk1 | T | 12 | -0.02081753 | -0.059854033 | -0.12631514 | -0.10549761 | -0.066461107 | 0.692430568 | 0.496424457 | 0.176480382 | 0.229014474 | 0.500828438 |
| Rai14 | S | 340 | 0.143218998 | -0.062548515 | -0.216893664 | -0.360112662 | -0.154345149 | 0.045640774 | 0.489027633 | 0.001128371 | 0.00555269 | 0.156037464 |
| Sf3b1 | T | 142 | -0.052926862 | -0.030245225 | -0.0364778 | 0.016449062 | -0.006232575 | 0.036853191 | 0.51738236 | 0.332033973 | 0.617448147 | 0.899633268 |
| Tjp2 | S | 111 | -0.182983585 | -0.030961458 | -0.066177071 | 0.116806514 | -0.035215612 | 0.281871603 | 0.68317149 | 0.6472577 | 0.536920426 | 0.803243379 |
| Gpbp1l1 | S | 289 | -0.047782085 | -0.145510429 | -0.00142966 | 0.046352425 | 0.144080769 | 0.408578447 | 0.12718243 | 0.983839723 | 0.55263174 | 0.165283669 |
| Pkn2 | S | 301 | -0.026512597 | -0.083291923 | -0.100294382 | -0.073781785 | -0.017002459 | 0.718690141 | 0.399806226 | 0.221096138 | 0.392678059 | 0.864790106 |
| Fam117b | S | 188 | -0.074676462 | -0.010765475 | -0.054059056 | 0.020617406 | -0.043293581 | 0.080145271 | 0.678354883 | 0.286164468 | 0.677563482 | 0.387358725 |
| Morc3 | S | 505 | -0.113548927 | -0.143448764 | -0.022554792 | 0.090994135 | 0.120893972 | 0.670109863 | 0.19535214 | 0.847382299 | 0.741373194 | 0.394973318 |
| Mcmbp | S | 156 | -0.062886976 | -0.063755917 | -0.030629897 | 0.032257079 | 0.03312602 | 0.285803475 | 0.154007141 | 0.565063478 | 0.590610271 | 0.499285353 |
| Mapk8ip3 | S | 366 | 0.042667857 | -0.01192647 | -0.245609007 | -0.288276864 | -0.233682537 | 0.627729361 | 0.879622175 | 0.049377707 | 0.034687382 | 0.061351811 |
| Sik2 | S | 587 | -0.143833252 | 0.062814479 | -0.240161679 | -0.096328427 | -0.302976158 | 0.332521138 | 0.566574155 | 0.286740722 | 0.660993192 | 0.206636456 |
| Ehbp1 | S | 438 | -0.240232784 | 0.121145088 | -0.049450856 | 0.190781928 | -0.170595944 | 0.034070225 | 0.287208019 | 0.298917335 | 0.068974672 | 0.173523934 |
| Tmcc2 | S | 435 | 0.13660244 | -0.161420231 | -0.156216311 | -0.292818751 | 0.00520392 | 0.026036813 | 0.013133127 | 0.081451664 | 0.024244498 | 0.93542385 |
| Pds5b | S | 1191 | -0.033191312 | -0.092250124 | -0.048055941 | -0.014864629 | 0.044194183 | 0.603804337 | 0.107848818 | 0.53659866 | 0.8414202 | 0.521717328 |
| Ppp1r2 | S | 122 | -0.138767806 | 0.09012002 | -0.180113863 | -0.041346056 | -0.270233882 | 0.110326115 | 0.311805401 | 0.214741524 | 0.739248754 | 0.100836465 |
| Trim56 | S | 376 | -0.047169498 | -0.010666018 | -0.057599405 | -0.010429907 | -0.046933387 | 0.213604257 | 0.825990668 | 0.034610187 | 0.756972582 | 0.379937696 |
| Hdgf | S | 132 | -0.205918119 | -0.066336972 | 0.043455061 | 0.24937318 | 0.109792033 | 0.107044488 | 0.441259757 | 0.569962899 | 0.073055686 | 0.234706294 |
| Ccdc88a | S | 1677 | 0.047472531 | -0.083783362 | -0.12564148 | -0.173114011 | -0.041858118 | 0.175811583 | 0.051445298 | 0.095872646 | 0.046334278 | 0.484416049 |
| Cip2a | T | 586 | -0.032216767 | -0.004188938 | -0.259471357 | -0.22725459 | -0.255282419 | 0.490397511 | 0.933153531 | 0.267336635 | 0.314648684 | 0.272169504 |
| Twistnb | S | 318 | -0.03649774 | -0.053160938 | -0.032899471 | 0.003598268 | 0.020261466 | 0.675027333 | 0.154509149 | 0.045748523 | 0.966145018 | 0.510497221 |
| Numa1 | S | 1751 | -0.150001051 | 0.016023229 | -0.004744915 | 0.145256136 | -0.020768144 | 0.013180173 | 0.708363631 | 0.905844236 | 0.027459079 | 0.67245112 |
| Helq | S | 28 | 0.031270541 | -0.029205415 | -0.112746445 | -0.144016985 | -0.08354103 | 0.292569222 | 0.507874276 | 0.003044818 | 0.004611415 | 0.120449989 |
| Prrc2a | S | 1687 | 0.057486782 | -0.09944428 | -0.065223386 | -0.122710168 | 0.034220894 | 0.247180621 | 0.018412209 | 0.079850263 | 0.049555877 | 0.241218618 |
| Ripor1 | S | 841 | 0.001954961 | -0.055142355 | -0.157923393 | -0.159878354 | -0.102781038 | 0.969370926 | 0.298272339 | 0.202302466 | 0.203518146 | 0.359607833 |
| Ripor1 | S | 842 | 0.001954961 | -0.055142355 | -0.157923393 | -0.159878354 | -0.102781038 | 0.969370926 | 0.298272339 | 0.202302466 | 0.203518146 | 0.359607833 |
| Rock2 | S | 1052 | -0.067211393 | 0.038285061 | -0.123854856 | -0.056643463 | -0.162139917 | 0.241023009 | 0.462713215 | 0.047077717 | 0.268707438 | 0.016587048 |
| Rap1gap | S | 498 | -0.098868601 | 0.147930339 | -0.277864017 | -0.178995416 | -0.425794356 | 0.367061026 | 0.388276889 | 0.093280342 | 0.227868165 | 0.063719183 |
| Zc3h13 | S | 365 | 0.044137775 | 0.032599158 | -0.216085708 | -0.260223484 | -0.248684866 | 0.690934767 | 0.672317393 | 0.010332401 | 0.087994886 | 0.031260399 |
| Jmjd1c | S | 138 | -0.202247053 | -0.013949139 | 0.037329818 | 0.23957687 | 0.051278957 | 0.030802996 | 0.808586666 | 0.587512008 | 0.032388995 | 0.522943617 |
| Cast | S | 444 | 0.00654585 | -0.048031395 | -0.10185578 | -0.108401631 | -0.053824385 | 0.937354865 | 0.345208565 | 0.05297471 | 0.251129343 | 0.26732148 |
| Lgalsl | S | 22 | 0.027300062 | -0.034879418 | -0.162592148 | -0.18989221 | -0.12771273 | 0.706464787 | 0.72065733 | 0.033711835 | 0.049352704 | 0.242955877 |
| Add3 | S | 677 | 0.044340547 | -0.021701614 | -0.179981023 | -0.22432157 | -0.158279409 | 0.281444367 | 0.438828713 | 0.017186107 | 0.010707283 | 0.051578935 |
| Sav1 | S | 27 | -0.0518717 | -0.030439911 | -0.062631747 | -0.010760047 | -0.032191837 | 0.208113781 | 0.625145143 | 0.146769553 | 0.795077486 | 0.615133459 |
| Mast4 | S | 1375 | 0.038000512 | -0.027564119 | -0.221326858 | -0.259327369 | -0.193762738 | 0.699255606 | 0.727454302 | 0.066818291 | 0.073140032 | 0.110568728 |
| Cadm1 | T | 436 | 0.088866888 | -0.311378011 | -0.233439918 | -0.322306806 | 0.077938092 | 0.26596697 | 0.196951377 | 0.534565906 | 0.410810219 | 0.839914692 |
| Glipr2 | S | 55 | -0.000794992 | -0.085363815 | -0.049821617 | -0.049026626 | 0.035542198 | 0.972422749 | 0.052121035 | 0.303704849 | 0.311233356 | 0.472844972 |
| Limd1 | S | 397 | 0.008500815 | -0.047238816 | -0.127002918 | -0.135503733 | -0.079764102 | 0.846898761 | 0.197421734 | 0.128384884 | 0.113775527 | 0.275425095 |
| Lsm8 | S | 49 | 0.027742356 | -0.063589145 | -0.13110154 | -0.158843896 | -0.067512395 | 0.729227794 | 0.501948108 | 0.046708994 | 0.104051834 | 0.439979316 |
| Actr10 | T | 415 | -0.024824011 | 0.08356015 | -0.336936882 | -0.312112871 | -0.420497032 | 0.837083974 | 0.59419675 | 0.094561418 | 0.115697975 | 0.068034011 |
| Znfx1 | S | 83 | 0.017206446 | -0.030933547 | -0.165201213 | -0.18240766 | -0.134267666 | 0.751206454 | 0.632702097 | 0.051300117 | 0.044586725 | 0.117536083 |
| Mcm3ap | S | 295 | 0.020722381 | -0.103958657 | -0.05345066 | -0.074173041 | 0.050507997 | 0.782672712 | 0.036598185 | 0.359068465 | 0.377660635 | 0.36107557 |
| Trmt1 | S | 35 | 0.012437632 | -0.070949601 | -0.067237771 | -0.079675404 | 0.003711829 | 0.768001122 | 0.055917365 | 0.155213861 | 0.144568556 | 0.924274643 |
| Hdgf | S | 133 | -0.20378161 | -0.068989566 | 0.046375964 | 0.250157574 | 0.11536553 | 0.107562111 | 0.439609901 | 0.562303807 | 0.071232942 | 0.230697191 |
| Ubr4 | S | 1759 | -0.04124423 | -0.007440003 | -0.068704996 | -0.027460766 | -0.061264992 | 0.14186804 | 0.807735545 | 0.072252087 | 0.33177656 | 0.097446256 |
| Knop1 | S | 51 | 0.023824921 | -0.091869404 | -0.075646656 | -0.099471577 | 0.016222748 | 0.707690271 | 0.055778124 | 0.249417707 | 0.213820833 | 0.773728647 |
| Dtd1 | S | 196 | 0.027811093 | -0.067381137 | -0.123005512 | -0.150816605 | -0.055624374 | 0.74166975 | 0.217552485 | 0.112209485 | 0.143902048 | 0.379677202 |
| Rps2 | S | 77 | -0.000681202 | -0.160099213 | -0.064651283 | -0.063970081 | 0.095447931 | 0.995191254 | 0.13119526 | 0.619899579 | 0.654853932 | 0.477955162 |
| Usp16 | S | 414 | -0.047181464 | -0.027358164 | -0.057773865 | -0.010592401 | -0.030415701 | 0.186326412 | 0.437596039 | 0.178313961 | 0.704197806 | 0.338176635 |
| Mapre3 | S | 162 | 0.022500375 | 0.008422073 | -0.188560318 | -0.211060694 | -0.196982391 | 0.762890049 | 0.910077761 | 0.020954763 | 0.022494115 | 0.028115887 |
| Mcph1 | S | 162 | 0.10437118 | -0.21173012 | -0.150698084 | -0.255069264 | 0.061032037 | 0.0822145 | 0.101629193 | 0.123047903 | 0.040848613 | 0.589866839 |
| Kif21a | S | 1241 | 0.042352691 | -0.068897877 | -0.174429903 | -0.216782594 | -0.105532026 | 0.042342715 | 0.041001244 | 0.09294312 | 0.068643703 | 0.216039908 |
| Dtna | T | 501 | -0.010955749 | 0.02019874 | -0.232473269 | -0.221517519 | -0.252672008 | 0.852900452 | 0.732655282 | 0.071012357 | 0.087040386 | 0.065477761 |
| Clasp1 | S | 559 | -0.162180352 | 0.086142165 | -0.109193291 | 0.052987062 | -0.195335456 | 0.044795008 | 0.401097035 | 0.397521013 | 0.644349571 | 0.186708724 |
| Jcad | S | 405 | -0.001655169 | -0.049700672 | -0.135778332 | -0.134123163 | -0.08607766 | 0.981836645 | 0.660626769 | 0.072545301 | 0.152484536 | 0.476323349 |
| Gsk3a Gsk3b | S | 278;215 | -0.092008516 | -0.039491876 | -0.020213705 | 0.071794811 | 0.019278171 | 0.069902277 | 0.462719246 | 0.748082649 | 0.244970854 | 0.741702489 |
| Tmem209 | S | 149 | -0.039127413 | -0.111480523 | -0.012207682 | 0.026919732 | 0.099272841 | 0.629521121 | 0.061635563 | 0.790301086 | 0.723983846 | 0.073540209 |
| Zfp326 | S | 270 | -0.078539504 | -0.095145015 | 0.015142056 | 0.09368156 | 0.110287072 | 0.287194358 | 0.08478799 | 0.834728946 | 0.288451697 | 0.178840096 |
| Acin1 | S | 595 | -0.014586248 | -0.053748327 | -0.071209894 | -0.056623646 | -0.017461567 | 0.794939747 | 0.215282775 | 0.105809038 | 0.288217812 | 0.449982379 |
| Pml | S | 503 | -0.169032571 | 0.028764972 | 0.038754676 | 0.207787247 | 0.009989704 | 0.00205898 | 0.733867392 | 0.421019575 | 0.024068439 | 0.906626161 |
| Rrbp1 | S | 135 | 0.027968456 | 0.062264683 | -0.250479594 | -0.27844805 | -0.312744278 | 0.864030847 | 0.400064648 | 0.012501787 | 0.174135809 | 0.000496516 |
| Zc3h12c | S | 690 | -0.031831001 | 0.064784534 | -0.308548405 | -0.276717405 | -0.37333294 | 0.567366841 | 0.49003983 | 0.109894738 | 0.148057029 | 0.071188033 |
| Fam122a | S | 17 | -0.177263597 | -0.005873071 | -0.074288933 | 0.102974664 | -0.068415862 | 0.14529201 | 0.942650321 | 0.764568594 | 0.68154276 | 0.776584292 |
| Ankrd13a | S | 584 | 0.041039259 | -0.066480848 | -0.356309583 | -0.397348843 | -0.289828735 | 0.74110447 | 0.715812759 | 0.246508769 | 0.211500541 | 0.343138529 |
| Stard13 | S | 314 | -0.043488025 | -0.028661712 | -0.100608414 | -0.05712039 | -0.071946702 | 0.444986676 | 0.618237278 | 0.147465439 | 0.229484468 | 0.157088675 |

| | | | | | | | | | | | | |
|---|---|---|---|---|---|---|---|---|---|---|---|---|
| Ube4b | S | 84 | 0.088729135 | -0.084741363 | -0.206537288 | -0.295266422 | -0.121795925 | 0.248933988 | 0.276853411 | 0.058734128 | 0.024016291 | 0.205732562 |
| Tbc1d17 | S | 633 | -0.428545752 | 0.27469796 | -0.132363724 | 0.296182028 | -0.407061684 | 0.022766278 | 0.054123513 | 0.364323027 | 0.114195581 | 0.0466739 |
| Tbc1d17 | S | 634 | -0.428545752 | 0.27469796 | -0.132363724 | 0.296182028 | -0.407061684 | 0.022766278 | 0.054123513 | 0.364323027 | 0.114195581 | 0.0466739 |
| Ppfibp1 | S | 560 | -0.150998428 | 0.088526673 | -0.135152511 | 0.015845917 | -0.223679184 | 0.121423463 | 0.451105464 | 0.231254457 | 0.883986036 | 0.140145258 |
| Apbb1ip | S | 532 | 0.154272274 | -0.05988145 | -0.41758607 | -0.571858345 | -0.357704621 | 0.032621297 | 0.37246188 | 0.027161667 | 0.011044754 | 0.033563754 |
| Ccz1 | S | 264 | -0.057147417 | -0.018907632 | -0.059893132 | -0.002745715 | -0.0409855 | 0.274030724 | 0.547880223 | 0.108388328 | 0.951443135 | 0.165183073 |
| Rtn4 | S | 686 | -0.153016855 | 0.019542144 | -0.041253885 | 0.11176297 | -0.060796029 | 0.067532959 | 0.62731557 | 0.392046528 | 0.146272856 | 0.251119319 |
| Lig3 | S | 906 | 0.044824632 | -0.072881583 | -0.085824276 | -0.130648908 | -0.012942693 | 0.181684603 | 0.077914447 | 0.017748531 | 0.010701405 | 0.701623604 |
| Zc3h13 | S | 355 | -0.091106731 | -0.060935538 | -0.016994311 | 0.074112419 | 0.043941226 | 0.168293597 | 0.257069764 | 0.851653281 | 0.470429341 | 0.65189811 |
| Zc3h13 | T | 353 | -0.091106731 | -0.060935538 | -0.016994311 | 0.074112419 | 0.043941226 | 0.168293597 | 0.257069764 | 0.851653281 | 0.470429341 | 0.65189811 |
| Rad21 | T | 186 | -0.285691984 | 0.062276031 | 0.006522902 | 0.292214886 | -0.055753128 | 0.041126799 | 0.548897734 | 0.946881357 | 0.041341742 | 0.532020925 |
| Cpsf2 | S | 419 | -0.005286228 | -0.12210943 | -0.005974777 | -0.000688549 | 0.116134653 | 0.895397339 | 0.01985039 | 0.893151449 | 0.979666249 | 0.019199677 |
| Cpsf2 | S | 420 | -0.005286228 | -0.12210943 | -0.005974777 | -0.000688549 | 0.116134653 | 0.895397339 | 0.01985039 | 0.893151449 | 0.979666249 | 0.019199677 |
| Cpsf2 | S | 423 | -0.005286228 | -0.12210943 | -0.005974777 | -0.000688549 | 0.116134653 | 0.895397339 | 0.01985039 | 0.893151449 | 0.979666249 | 0.019199677 |
| Klf6 | S | 151 | -0.302707569 | -0.020472416 | -0.067971271 | 0.234736298 | -0.047498854 | 0.401618209 | 0.821799725 | 0.709008351 | 0.52081398 | 0.7951346 |
| Acin1 | S | 561 | 0.091493895 | -0.046172408 | -0.185530644 | -0.277024539 | -0.139358236 | 0.210189045 | 0.330606957 | 0.013601773 | 0.014516041 | 0.04693379 |
| Map1a | T | 1806 | 0.077129236 | -0.094035939 | -0.118872997 | -0.196002233 | -0.024837058 | 0.326920909 | 0.070056339 | 0.090307959 | 0.06253729 | 0.67111215 |
| Eif3b | S | 88 | 0.021065802 | -0.123751926 | -0.060506555 | -0.081572357 | 0.063245371 | 0.780976826 | 0.12503212 | 0.213304004 | 0.309273598 | 0.361159308 |
| Map1a | S | 527 | 0.010285656 | -0.032662708 | -0.082461124 | -0.092746781 | -0.049798416 | 0.811936872 | 0.483397637 | 0.007754344 | 0.108507624 | 0.31179634 |
| Cobl | S | 826 | -0.069975045 | -0.035184966 | -0.03192131 | 0.038053735 | 0.003263656 | 0.060968765 | 0.483054849 | 0.549468999 | 0.496069522 | 0.959024302 |
| Nap1l4 | S | 121 | -0.090733589 | 0.024357131 | -0.103618515 | -0.012884926 | -0.127975646 | 0.177641299 | 0.739125283 | 0.167470392 | 0.856032818 | 0.167060168 |
| Cep170 | S | 1231 | -0.016377046 | -0.030647399 | -0.096600967 | -0.080223921 | -0.065953568 | 0.795395631 | 0.583795444 | 0.043340397 | 0.228183152 | 0.222975281 |
| Nap1l1 | S | 242 | -0.023194741 | -0.03115634 | -0.1806202 | -0.157425459 | -0.14946386 | 0.847802372 | 0.871939871 | 0.125662871 | 0.123239461 | 0.431904075 |
| Inf2 | S | 351 | 0.067620508 | -0.150339015 | -0.091585593 | -0.159206101 | 0.058753422 | 0.097515001 | 0.01309962 | 0.307531118 | 0.136528303 | 0.486479812 |
| Nes | T | 627 | 0.048506575 | -0.187312434 | -0.057456139 | -0.105962713 | 0.129856295 | 0.607158035 | 0.071381786 | 0.528819808 | 0.148681584 | 0.079029592 |
| Tbc1d1 | S | 1159 | -0.061577317 | -0.024784064 | -0.029844014 | 0.031733303 | -0.00505995 | 0.01935684 | 0.518297119 | 0.625308677 | 0.597051474 | 0.936407605 |
| Rb1cc1 | S | 257 | -0.123510455 | 0.016269889 | -0.022335501 | 0.101174954 | -0.038605389 | 0.016494905 | 0.780520076 | 0.695638529 | 0.155743963 | 0.602625214 |
| Ubtf | S | 389 | -0.028488885 | -0.004078419 | -0.103139387 | -0.074650501 | -0.099060967 | 0.724276462 | 0.896238966 | 0.02363885 | 0.387706855 | 0.028643878 |
| Gatad2a | S | 335 | -0.05514211 | -0.015761816 | -0.043387284 | 0.011754827 | -0.027625467 | 0.070889971 | 0.673956 | 0.16983042 | 0.558246868 | 0.428019878 |
| Inka2 | S | 98 | -0.026609238 | -0.092302392 | -0.147977212 | -0.121367974 | -0.05567482 | 0.780022462 | 0.241256706 | 0.556230797 | 0.623411438 | 0.813798514 |
| Inka2 | S | 101 | -0.026609238 | -0.092302392 | -0.147977212 | -0.121367974 | -0.05567482 | 0.780022462 | 0.241256706 | 0.556230797 | 0.623411438 | 0.813798514 |
| Cdc42bpa | S | 1706 | -0.079880062 | -0.066940415 | -0.049431434 | 0.030448628 | 0.01750898 | 0.382036996 | 0.277197076 | 0.566188071 | 0.752860401 | 0.814213008 |
| Srrm2 | S | 1376 | -0.058885247 | -0.095202728 | 0.008500001 | 0.067385248 | 0.103702729 | 0.424021077 | 0.046451348 | 0.913890628 | 0.487151744 | 0.255657177 |
| Cwc22 | S | 776 | -0.061462854 | -0.098523108 | 0.013397494 | 0.074860348 | 0.111920602 | 0.306064662 | 0.056954479 | 0.738432624 | 0.25497364 | 0.064228442 |
| Tjp2 | S | 1008 | -0.087306741 | -0.03596688 | -0.081199177 | 0.006107564 | -0.045232297 | 0.231495747 | 0.744366275 | 0.389438382 | 0.950008898 | 0.727679531 |
| Ppp1r12c | T | 560 | -0.063084233 | 0.254360936 | -0.305315889 | -0.242231656 | -0.559676825 | 0.698697186 | 0.408812771 | 0.024581868 | 0.177567127 | 0.140853555 |
| Eif3c | S | 530 | 0.008252243 | -0.069457887 | -0.112959586 | -0.121211829 | -0.043501699 | 0.826737012 | 0.401187714 | 0.087336489 | 0.079376514 | 0.60476883 |
| Trim3 | S | 437 | -0.009989499 | -0.00874723 | -0.102179733 | -0.092190233 | -0.093432503 | 0.763424342 | 0.777311027 | 0.018644018 | 0.014121625 | 0.023129754 |
| Ralbp1 | S | 30 | -0.003582636 | -0.015810761 | -0.149220578 | -0.145637941 | -0.133409817 | 0.976760675 | 0.82689902 | 0.048387424 | 0.293006115 | 0.116904868 |
| Fgd1 | S | 714 | -0.041614518 | -0.036973769 | -0.04382053 | -0.002206013 | -0.006846761 | 0.219643029 | 0.168282205 | 0.304475043 | 0.955813446 | 0.848040632 |
| Slc38a10 | S | 441 | -0.034346741 | -0.207523449 | -0.032987174 | 0.001359567 | 0.174536276 | 0.910992306 | 0.162747035 | 0.877417858 | 0.996898096 | 0.466122904 |
| Baz1b | S | 1315 | -0.035941122 | -0.040713127 | -0.085984833 | -0.050043711 | -0.045271706 | 0.558729169 | 0.497655084 | 0.124982924 | 0.46637115 | 0.496208357 |
| Nup214 | S | 1139 | -0.121539128 | 0.024024091 | -0.05279143 | 0.068747697 | -0.076815521 | 0.03110204 | 0.652916903 | 0.512954715 | 0.386959598 | 0.352712211 |
| Pkm | S | 406 | -0.117479913 | -0.000478989 | -0.056332941 | 0.061146973 | -0.055853952 | 0.088754737 | 0.996762201 | 0.48933743 | 0.472350672 | 0.667808337 |
| Klc1 | T | 461 | -0.077265439 | -0.049863877 | -0.053572321 | 0.023693118 | -0.003708445 | 0.338057863 | 0.285547996 | 0.489542286 | 0.791449848 | 0.95733716 |
| Exoc7 | S | 242 | -0.0067669 | -0.014003397 | -0.141218946 | -0.134452046 | -0.127215549 | 0.916190508 | 0.744335564 | 0.052073034 | 0.106931838 | 0.076005449 |
| Zdhhc8 | S | 659 | -0.045332718 | -0.017118347 | -0.259921016 | -0.214588299 | -0.242802669 | 0.647303663 | 0.900864086 | 0.30986447 | 0.382140598 | 0.345370953 |
| Chmp5 | S | 30 | -0.09481069 | 0.033594335 | -0.125811563 | -0.031000872 | -0.159405898 | 0.321151348 | 0.724238062 | 0.126076303 | 0.731313847 | 0.153491384 |
| Cast | T | 23 | -0.203548886 | 0.119250082 | -0.21449654 | -0.010947654 | -0.333746622 | 0.258800885 | 0.520391426 | 0.327951579 | 0.94776122 | 0.123374513 |
| Ehbp1l1 | S | 170 | 0.050871139 | -0.135271095 | -0.12193983 | -0.172810969 | 0.013331265 | 0.50180631 | 0.234114589 | 0.152931537 | 0.043738782 | 0.89054309 |
| Samd4 | T | 256 | -0.127229553 | 0.006299582 | -0.071544213 | 0.05568534 | -0.077843795 | 0.099542882 | 0.907618091 | 0.517258685 | 0.621952938 | 0.487974123 |
| Usp6nl | S | 669 | -0.017160426 | 0.02979158 | -0.159234819 | -0.142074394 | -0.1890264 | 0.749246193 | 0.537679023 | 0.018008401 | 0.037744061 | 0.011343491 |
| Daxx | S | 669 | -0.007547597 | -0.004807399 | -0.10801287 | -0.100465273 | -0.103205472 | 0.937131312 | 0.946392338 | 0.01145769 | 0.345012029 | 0.220533722 |
| Cdr2l | S | 396 | -0.106816614 | 0.029432535 | -0.082079007 | 0.024737606 | -0.111511542 | 0.110377757 | 0.679217409 | 0.216156947 | 0.485702647 | 0.097493505 |
| Smg7 | S | 519 | 0.042666834 | -0.050645583 | -0.233817347 | -0.276484182 | -0.183171765 | 0.696835792 | 0.648798573 | 0.11677133 | 0.0641749 | 0.145881994 |
| Rpl7l1 | S | 230 | 0.170688147 | 0.037768166 | -0.27232412 | -0.443012267 | -0.310092287 | 0.129353123 | 0.419931568 | 0.000246567 | 0.017506065 | 0.002528078 |
| Pum2 | S | 82 | 0.03564891 | -0.094618671 | -0.043570366 | -0.079219276 | 0.051048305 | 0.447655935 | 0.010042893 | 0.261865659 | 0.160688331 | 0.199158142 |
| Fam83d | S | 458 | 0.059244844 | -0.038636768 | -0.116942463 | -0.176187307 | -0.078305695 | 0.517829182 | 0.283277646 | 0.013638031 | 0.13243127 | 0.056916196 |
| Sh3bp2 | S | 225 | -0.217102549 | 0.16237223 | -0.167507704 | 0.049594845 | -0.329879934 | 0.089703386 | 0.377373079 | 0.373631531 | 0.767427247 | 0.164251761 |
| Erc1 | S | 415 | 0.022869806 | -0.093702389 | -0.048687617 | -0.071557423 | 0.045014772 | 0.482638351 | 0.015515447 | 0.319388496 | 0.161723738 | 0.320562447 |
| Eme1 | S | 7 | -0.035323264 | -0.074509809 | -0.044783567 | -0.009460304 | 0.029726241 | 0.482879279 | 0.089224716 | 0.618369175 | 0.914404901 | 0.727931872 |
| Arhgap10 | S | 591 | 0.05062469 | -0.069177953 | -0.095520254 | -0.146144944 | -0.026342301 | 0.446716395 | 0.125557889 | 0.040121827 | 0.093300328 | 0.301368339 |
| Apeh | S | 185 | -0.119439223 | -0.014412041 | -0.064984935 | 0.054454288 | -0.050572894 | 0.305598895 | 0.799426604 | 0.25498266 | 0.606093013 | 0.351040899 |
| Asxl2 | S | 653 | -0.251294361 | 0.035054603 | -0.389597927 | -0.138303566 | -0.42465253 | 0.656464543 | 0.944208833 | 0.593826462 | 0.86669245 | 0.600971514 |
| Gpbp1l1 | S | 76 | -0.082566198 | 0.01439866 | -0.1117001 | -0.029133901 | -0.12609876 | 0.218775346 | 0.851293863 | 0.198745349 | 0.693170449 | 0.185189693 |
| Afap1l2 | S | 344 | -0.178922781 | 0.19413572 | -0.244808675 | -0.065885895 | -0.438944396 | 0.159486209 | 0.241500547 | 0.149116957 | 0.586725338 | 0.038947916 |
| Tbc1d1 | S | 1158 | -0.111853009 | -0.080065235 | 0.011647342 | 0.123500352 | 0.091712577 | 0.278427098 | 0.141993181 | 0.848391936 | 0.240196299 | 0.125106568 |
| Map1b | T | 1499 | -0.030279164 | -0.104470342 | -0.072252228 | -0.041973065 | 0.032218114 | 0.656474928 | 0.285989955 | 0.406844742 | 0.506516463 | 0.712773225 |
| Arhgap35 | Y | 1087 | -0.056191615 | -0.015483169 | -0.062698632 | -0.006507016 | -0.047215462 | 0.215786524 | 0.684060272 | 0.128285888 | 0.878555579 | 0.269272498 |
| Supt6 | T | 1532 | 0.0459354 | -0.049698433 | -0.183966448 | -0.229901848 | -0.134268014 | 0.457521525 | 0.410551741 | 0.065463225 | 0.037973598 | 0.143816189 |
| Trip12 | T | 83 | -0.131949321 | -0.06073541 | -0.006734267 | 0.125215055 | 0.054001144 | 0.170236352 | 0.425117383 | 0.95583219 | 0.355524122 | 0.657776081 |
| Nav3 | S | 1280 | -0.047492778 | -0.161474819 | 0.012580945 | 0.060073723 | 0.174055763 | 0.334633337 | 0.180769096 | 0.951629674 | 0.774624323 | 0.452868031 |
| Ctr9 | S | 970 | -0.062887113 | -0.10948303 | 0.009785856 | 0.072672969 | 0.119268886 | 0.432979223 | 0.079667069 | 0.923360472 | 0.508191266 | 0.278584364 |
| Zc3hav1 | S | 564 | -0.045646589 | -0.128414663 | 0.002547428 | 0.048194016 | 0.130962091 | 0.391385155 | 0.113209054 | 0.973804905 | 0.567474505 | 0.195779543 |
| Lats1 | S | 463 | -0.052254154 | 0.008487314 | -0.076310399 | -0.024056245 | -0.084797714 | 0.195945661 | 0.825914047 | 0.052030053 | 0.54441062 | 0.099164622 |
| Psmd2 | S | 363 | -0.049457377 | 0.034056319 | -0.137014323 | -0.087556946 | -0.171070642 | 0.471519478 | 0.492226657 | 0.031315413 | 0.189459692 | 0.008085684 |
| Nup160 | S | 1123 | -0.005842748 | -0.062640834 | -0.025257649 | -0.019414901 | 0.037383185 | 0.768825485 | 0.012072676 | 0.358523918 | 0.425731916 | 0.181707875 |
| Rpgr | S | 396 | -0.152597156 | 0.045106398 | -0.086704189 | 0.065892967 | -0.131810587 | 0.078241092 | 0.458781417 | 0.379301302 | 0.519440089 | 0.220604846 |
| Arhgef17 | S | 538 | -0.104604215 | -0.038834555 | -0.024531251 | 0.080072964 | 0.014303304 | 0.103867248 | 0.603456575 | 0.673690979 | 0.193688772 | 0.841519209 |
| Tdp1 | S | 61 | 0.123680218 | -0.132478839 | -0.206123812 | -0.32980403 | -0.073644973 | 0.00677331 | 0.004921775 | 0.111828491 | 0.049157764 | 0.458719001 |
| Larp1 | Y | 839 | 0.028130808 | 0.027312769 | -0.279764422 | -0.30789523 | -0.307077191 | 0.885696948 | 0.741081345 | 0.065263501 | 0.209488256 | 0.043911097 |
| Irf2bp2 | S | 71 | 0.05446958 | -0.058436771 | -0.090405767 | -0.144875347 | -0.031968996 | 0.039158295 | 0.063458771 | 0.002214642 | 0.001802823 | 0.23770815 |
| Rsl1d1 | S | 8 | 0.125748372 | -0.060204249 | -0.171866319 | -0.297614692 | -0.111662071 | 0.207786126 | 0.290644352 | 0.00933076 | 0.036717545 | 0.075932472 |
| Zc3h13 | S | 682 | 0.058222454 | -0.105231797 | -0.089134616 | -0.14735707 | 0.016097181 | 0.165825187 | 0.036538303 | 0.095175795 | 0.026714431 | 0.72748075 |
| Gab2 | S | 143 | -0.065472135 | -0.083386705 | -0.012740252 | 0.052731884 | 0.070646453 | 0.124108795 | 0.324320185 | 0.831389407 | 0.414368855 | 0.435886139 |
| Rbm45 | S | 199 | -0.059565869 | -0.065063639 | 0.036292732 | 0.095858601 | 0.101356371 | 0.21378576 | 0.004916903 | 0.465071169 | 0.143299314 | 0.1228536 |
| Dcp2 | S | 284 | -0.0595165 | -0.014281895 | -0.103337244 | -0.043820745 | -0.089055349 | 0.332585755 | 0.749404544 | 0.205637363 | 0.565650506 | 0.250771384 |
| Lyn | T | 502 | -0.316208894 | -0.251434351 | 0.192722586 | 0.50893148 | 0.444156937 | 0.435925742 | 0.329221415 | 0.594083993 | 0.291549154 | 0.256522704 |
| Zc3h11a | S | 741 | 0.050176791 | -0.103413462 | -0.083126608 | -0.133303399 | 0.020286854 | 0.206918172 | 0.015667469 | 0.229943196 | 0.09766145 | 0.735138065 |
| Mki67 | S | 810 | -0.070964974 | -0.015528311 | -0.101842983 | -0.030878008 | -0.086314672 | 0.321446212 | 0.837887278 | 0.229992359 | 0.664349385 | 0.296961153 |
| Cblb | T | 527 | 0.031639639 | -0.08844818 | -0.123972282 | -0.155611921 | -0.035524102 | 0.677913669 | 0.139445202 | 0.248585827 | 0.194249342 | 0.709819383 |
| Rbmxl1 | S | 172 | -0.16028946 | -0.068148858 | 0.019430661 | 0.179720121 | 0.087579519 | 0.196686083 | 0.303549919 | 0.755780155 | 0.164000212 | 0.237366863 |
| Arl2 | S | 45 | -0.068404689 | -0.028195899 | -0.113890752 | -0.045486063 | -0.085694854 | 0.528817239 | 0.731747625 | 0.214718629 | 0.682284842 | 0.359665475 |
| Hcfc1 | S | 666 | -0.009792329 | -0.098477753 | -0.052270988 | -0.042478659 | 0.046206766 | 0.895035025 | 0.120036749 | 0.359558332 | 0.567183772 | 0.415494487 |

| | | | | | | | | | | | | |
|---|---|---|---|---|---|---|---|---|---|---|---|---|
| Bcl7b | S | 122 | -0.004414611 | -0.074423062 | -0.16141733 | -0.157002719 | -0.086994268 | 0.918623816 | 0.470955081 | 0.262343331 | 0.272596185 | 0.554478366 |
| Mga | S | 2023 | 0.007971683 | -0.044076121 | -0.126048255 | -0.134019938 | -0.081972134 | 0.901395333 | 0.497015352 | 0.073434237 | 0.024139335 | 0.0843979 |
| Anln | S | 512 | 0.033289878 | -0.071092714 | -0.110967517 | -0.144257395 | -0.039874802 | 0.244212814 | 0.072683473 | 0.100092811 | 0.06313789 | 0.469423761 |
| Clcc1 | S | 429 | -0.117661667 | -0.074326668 | 0.046870491 | 0.164532158 | 0.121197159 | 0.127477062 | 0.065082635 | 0.347696784 | 0.062819596 | 0.062731079 |
| Tpd52l2 | S | 180 | -0.042724066 | 0.204538886 | -0.316189096 | -0.273465031 | -0.520727983 | 0.742671671 | 0.262052798 | 0.017519934 | 0.099686109 | 0.040675068 |
| Zcchc8 | S | 334 | -0.09265358 | 0.019224686 | -0.16859006 | -0.075936479 | -0.178714746 | 0.311910611 | 0.702362292 | 0.263904833 | 0.597155916 | 0.227726672 |
| Dlc1 | T | 404 | 0.027197884 | -0.068460426 | -0.168647282 | -0.195845166 | -0.100186855 | 0.668805473 | 0.301065334 | 0.180000645 | 0.147508421 | 0.361783717 |
| Hdlbp | S | 31 | -0.017227181 | -0.005248029 | -0.078697914 | -0.061470734 | -0.073449886 | 0.407728108 | 0.820335765 | 0.017761072 | 0.04834034 | 0.034466255 |
| Plekha8 | S | 189 | -0.349573034 | 0.099596977 | -0.252821351 | 0.096751682 | -0.352418328 | 0.43542662 | 0.820346662 | 0.665337998 | 0.881775103 | 0.599946265 |
| Rps6 Rps6-ps4 | S | 148 | 0.084690596 | -0.117668156 | -0.136850203 | -0.221540799 | -0.019182048 | 0.093099502 | 0.228288509 | 0.016303757 | 0.005516808 | 0.813860682 |
| Cdca2 | S | 886 | 0.072080633 | -0.109007721 | -0.142299947 | -0.21438058 | -0.033292226 | 0.454206049 | 0.171351517 | 0.150996221 | 0.086267889 | 0.696040315 |
| Cdca2 | S | 888 | 0.072080633 | -0.109007721 | -0.142299947 | -0.21438058 | -0.033292226 | 0.454206049 | 0.171351517 | 0.150996221 | 0.086267889 | 0.696040315 |
| Ccdc6 | S | 45 | -0.07959778 | 0.026479217 | -0.127184658 | -0.047586878 | -0.153663875 | 0.237834651 | 0.790910409 | 0.190298715 | 0.597496241 | 0.22956159 |
| Usp24 | S | 1923 | -0.062715026 | -0.024832651 | -0.125703834 | -0.062988808 | -0.100871183 | 0.50522963 | 0.717523903 | 0.24435908 | 0.573829064 | 0.325605367 |
| Tnik | S | 678 | 0.01381036 | -0.07446645 | -0.064004972 | -0.077815332 | 0.010461479 | 0.854666513 | 0.095005243 | 0.153456224 | 0.331659238 | 0.735806193 |
| Afap1 | S | 680 | 0.12955297 | -0.074960151 | -0.113306821 | -0.242859791 | -0.038346671 | 0.353911503 | 0.220512385 | 0.007119936 | 0.153443328 | 0.468558701 |
| Rnf169 | S | 279 | -0.235278715 | 0.128198346 | -0.076528284 | 0.158750431 | -0.20472663 | 0.047599459 | 0.308621452 | 0.415034689 | 0.109413996 | 0.120779577 |
| Lpar6 | S | 314 | -0.128872143 | -0.092773722 | 0.072439144 | 0.201311287 | 0.165212866 | 0.09311438 | 0.190189511 | 0.487645856 | 0.103001679 | 0.15237784 |
| Dst | S | 1381 | -0.056543035 | -0.006981293 | -0.072664942 | -0.016121907 | -0.065683649 | 0.268293861 | 0.864915663 | 0.096071116 | 0.650044036 | 0.05361148 |
| Arhgap31 | S | 384 | -0.072531968 | -0.10726968 | -0.028894294 | 0.043637674 | 0.078375386 | 0.332409426 | 0.351137912 | 0.710109208 | 0.378137019 | 0.445663338 |
| Ank3 | S | 1883 | 0.044181268 | -0.009296174 | -0.162885103 | -0.207066371 | -0.153588929 | 0.260924791 | 0.764962037 | 0.006151559 | 0.004970581 | 0.01506514 |
| Akap13 | S | 1532 | -0.159789442 | 0.062711399 | 0.001216391 | 0.161005833 | -0.061495008 | 0.004241016 | 0.484966119 | 0.98415359 | 0.060084067 | 0.517742646 |
| Anapc1 | S | 518 | -0.05183011 | 0.014036101 | -0.148035513 | -0.096205403 | -0.162071614 | 0.602859666 | 0.872799907 | 0.088892002 | 0.323443889 | 0.094033089 |
| Sp2 | S | 78 | 0.045878233 | -0.101535165 | -0.07367746 | -0.119555693 | 0.027857705 | 0.577981089 | 0.047659791 | 0.250373093 | 0.215624588 | 0.618890198 |
| Dvl3 | S | 192 | 0.098351584 | 0.003778517 | -0.295336993 | -0.393688576 | -0.29911551 | 0.435844434 | 0.966166996 | 0.030783355 | 0.033363377 | 0.037675623 |
| Src | S | 21 | -0.184496698 | -0.063497525 | 0.096733127 | 0.281229825 | 0.160230653 | 0.059271418 | 0.507097957 | 0.62502953 | 0.22391207 | 0.441794959 |
| Psd3 | S | 1000 | -0.108597157 | 0.083987867 | -0.148448357 | -0.039851201 | -0.232436225 | 0.075847384 | 0.466365246 | 0.272339121 | 0.735687949 | 0.162314152 |
| Ptpn21 | S | 377 | -0.045608769 | 0.009347734 | -0.097828554 | -0.052219785 | -0.107176289 | 0.258258524 | 0.809574879 | 0.068983654 | 0.24386098 | 0.053587749 |
| Mzt2 | S | 127 | -0.188713861 | 0.004346697 | 0.018755299 | 0.20746916 | 0.014408603 | 0.036538335 | 0.956771721 | 0.820460045 | 0.023313551 | 0.822377582 |
| Rad54l2 | S | 1154 | -0.088872383 | -0.112546805 | 0.010568114 | 0.099440496 | 0.123114918 | 0.304254131 | 0.214561143 | 0.937951322 | 0.514207633 | 0.42752667 |
| Ercc5 | S | 705 | -0.075457439 | 0.003697522 | -0.052747436 | 0.022710002 | -0.056444958 | 0.083791807 | 0.937222888 | 0.186111957 | 0.565190756 | 0.326591263 |
| Hdgfl2 | S | 659 | 0.072106562 | -0.076634123 | -0.150414356 | -0.222520918 | -0.073780233 | 0.084398006 | 0.066465527 | 0.045877449 | 0.022481664 | 0.224600443 |
| Ralgapa1 | S | 859 | -0.108387889 | 0.040911833 | -0.081834106 | 0.026553783 | -0.122745939 | 0.090358255 | 0.549997592 | 0.204649448 | 0.670999275 | 0.145481278 |
| Micall2 | S | 249 | -0.033262118 | -0.083336677 | -0.093214527 | -0.059952409 | -0.00987785 | 0.557141486 | 0.342515057 | 0.441714569 | 0.597229076 | 0.935518409 |
| Serbp1 | S | 327 | -0.152072161 | 0.019996382 | -0.026080551 | 0.12599161 | -0.046076933 | 0.04428348 | 0.794008006 | 0.714632401 | 0.070116009 | 0.50720545 |
| Xrn1 | S | 1382 | -0.000626005 | -0.080951859 | -0.035548358 | -0.034922353 | 0.045403501 | 0.98254229 | 0.051676196 | 0.2556429 | 0.360142503 | 0.218115254 |
| Ppp1r10 | S | 591 | -0.03245061 | -0.095112836 | -0.172595641 | -0.140145031 | -0.077482805 | 0.820551247 | 0.490506049 | 0.44745756 | 0.5219604 | 0.711299097 |
| Med1 | S | 1483 | 0.027906369 | -0.122868066 | -0.038751252 | -0.066657621 | 0.084116814 | 0.510073103 | 0.022795567 | 0.482872088 | 0.224033437 | 0.152256044 |
| Apobr | S | 484 | -0.14006616 | 0.076419396 | -0.031836412 | 0.108229748 | -0.108255808 | 0.00341378 | 0.224245122 | 0.454495138 | 0.05501658 | 0.128594061 |
| Tmem115 | S | 340 | 0.135208307 | -0.139445117 | -0.3576333 | -0.492841607 | -0.218188183 | 0.376293783 | 0.458275379 | 0.210940608 | 0.122740939 | 0.410467982 |
| Tomm20 | S | 138 | -0.050125044 | -0.052541315 | -0.067985193 | -0.017860148 | -0.015443877 | 0.407373916 | 0.4972525 | 0.246550589 | 0.794616701 | 0.851149816 |
| R3hdm2 | S | 347 | -0.12401516 | 0.055258526 | -0.045639376 | 0.078375785 | -0.100897902 | 0.01494022 | 0.483488607 | 0.451079031 | 0.209552245 | 0.256693084 |
| Ahctf1 | S | 1725 | -0.054966237 | -0.100639669 | -0.09570741 | -0.040741173 | 0.004932259 | 0.59365415 | 0.40996973 | 0.538599163 | 0.80371652 | 0.97687748 |
| Pard3 | S | 221 | 0.022211913 | 0.099372027 | -0.25237665 | -0.274588563 | -0.351748676 | 0.689784504 | 0.187747324 | 0.003006825 | 0.00239953 | 0.013863451 |
| Clasrp | S | 641 | 0.046167744 | -0.177421108 | -0.016968781 | -0.063136525 | 0.160452327 | 0.730035074 | 0.038246451 | 0.741506286 | 0.641630617 | 0.059689471 |
| Clasrp | S | 643 | 0.046167744 | -0.177421108 | -0.016968781 | -0.063136525 | 0.160452327 | 0.730035074 | 0.038246451 | 0.741506286 | 0.641630617 | 0.059689471 |
| Pgrmc2 | S | 202 | -0.016742886 | -0.078020637 | -0.117429618 | -0.100686732 | -0.039408981 | 0.751632039 | 0.427890981 | 0.282728695 | 0.352328003 | 0.744947795 |
| Arhgap21 | S | 1108 | -0.052626678 | -0.074822198 | 0.020377302 | 0.07300398 | 0.095199499 | 0.145898567 | 0.021123749 | 0.359312633 | 0.071455455 | 0.014127192 |
| Ppfibp1 | S | 595 | -0.071197389 | 0.072904949 | -0.181932007 | -0.110734618 | -0.254836956 | 0.424298567 | 0.41786886 | 0.059642613 | 0.28068994 | 0.047350714 |
| Smg6 | S | 483 | -0.098373933 | -0.047212458 | 0.02660289 | 0.124976823 | 0.073815348 | 0.053826162 | 0.167076248 | 0.604123981 | 0.070162584 | 0.195493293 |
| Sipa1l3 | S | 1553 | -0.139866184 | -0.078228285 | -0.094957404 | 0.04490878 | -0.016729119 | 0.544542665 | 0.58090404 | 0.61945849 | 0.860957853 | 0.932034505 |
| Rpp25l | S | 147 | -0.053999044 | 0.000959888 | -0.131266484 | -0.07726744 | -0.132226371 | 0.454886283 | 0.989754515 | 0.140391864 | 0.31668152 | 0.136118164 |
| Camkk2 | S | 89 | -0.114081661 | -0.003204387 | -0.090703131 | 0.023378531 | -0.087498743 | 0.324914265 | 0.947674645 | 0.281089256 | 0.837548846 | 0.279437571 |
| Ppfibp1 | S | 37 | 0.014884088 | -0.064829433 | -0.069349747 | -0.084233835 | -0.004520314 | 0.545152009 | 0.040597602 | 0.171579863 | 0.122191396 | 0.914958837 |
| Aatf | T | 281 | -0.044095424 | -0.08093828 | -0.030413538 | 0.013681886 | 0.050524743 | 0.610909149 | 0.10660997 | 0.539531968 | 0.863369726 | 0.152770247 |
| Scrib | T | 810 | 0.096790896 | -0.135751 | -0.112998991 | -0.209789888 | 0.022752009 | 0.158249201 | 0.072077253 | 0.079528818 | 0.017240562 | 0.615311641 |
| Pcbp2 | S | 185 | -0.068884452 | -0.009956537 | -0.072878285 | -0.003993832 | -0.062921748 | 0.176783016 | 0.800912219 | 0.250910448 | 0.948767539 | 0.332207597 |
| Supt20 | S | 505 | -0.066287943 | -0.03357498 | -0.100501231 | -0.034213287 | -0.06692625 | 0.405149621 | 0.602733777 | 0.328526515 | 0.752032321 | 0.52522076 |
| Erf | S | 491 | -0.077203263 | -0.028048134 | -0.065463218 | 0.011740046 | -0.037415084 | 0.220400664 | 0.62016339 | 0.379752944 | 0.831667886 | 0.499091393 |
| Dmrta2 | S | 204 | -0.025198738 | 0.01229045 | -0.121468599 | -0.096269861 | -0.133759049 | 0.593189864 | 0.896899647 | 0.037695015 | 0.001858337 | 0.221193494 |
| Wnk1 | S | 2052 | -0.33588074 | -0.020155249 | 0.051534419 | 0.387415159 | 0.071689668 | 0.188382068 | 0.874566464 | 0.848658344 | 0.244561607 | 0.77777046 |
| Ptk2 | S | 390 | -0.126332652 | -0.02146798 | -0.046141259 | 0.080191393 | -0.024673279 | 0.310616919 | 0.680969511 | 0.306981866 | 0.487352108 | 0.601260954 |
| Prpf4b | S | 368 | -0.06261499 | -0.041739563 | -0.008802352 | 0.053812638 | 0.032937211 | 0.065515474 | 0.200544218 | 0.861290537 | 0.351463234 | 0.554074988 |
| Slc7a1 | S | 455 | 0.025594971 | -0.026486765 | -0.138157519 | -0.16375249 | -0.111670754 | 0.632563378 | 0.545655957 | 0.031899157 | 0.032013943 | 0.063967656 |
| Nap1l1 | S | 246 | -0.064182943 | -0.071918902 | -0.108674341 | -0.044491398 | -0.03675544 | 0.533105257 | 0.314223188 | 0.643747804 | 0.851956499 | 0.873481532 |
| Smcr8 | S | 488 | -0.02472984 | 0.021416568 | -0.225633727 | -0.200903887 | -0.247050295 | 0.88437142 | 0.786325129 | 0.098167843 | 0.302110032 | 0.086180216 |
| Gorab | S | 82 | -0.040906702 | -0.085268625 | -0.004050084 | 0.036856618 | 0.081218541 | 0.556034327 | 0.036397497 | 0.9059724 | 0.592430056 | 0.059529711 |
| Tmf1 | T | 360 | -0.030700805 | -0.14881355 | -0.029962142 | 0.000738663 | 0.118851408 | 0.776567453 | 0.158436022 | 0.741129111 | 0.994410429 | 0.247202668 |
| Eif3e | S | 399 | 0.032225673 | -0.08376248 | -0.102859911 | -0.135085583 | -0.01909743 | 0.392422741 | 0.10020538 | 0.151084856 | 0.087018639 | 0.760036166 |
| Arid4b | S | 778 | -0.073758371 | -0.057062914 | 0.013292914 | 0.087051284 | 0.070355827 | 0.184618279 | 0.060894516 | 0.848746574 | 0.29276253 | 0.353811764 |
| Slc30a1 | S | 502 | 0.056248316 | -0.085960493 | -0.123765985 | -0.180014301 | -0.037805492 | 0.091013427 | 0.033990419 | 0.100363286 | 0.045309816 | 0.520251438 |
| Cdk1 | Y | 15 | 0.008706329 | 0.013266468 | -0.179356966 | -0.188063295 | -0.192623434 | 0.875383363 | 0.863212743 | 0.032614955 | 0.039770669 | 0.076819016 |
| Zdhhc5 | S | 299 | -0.169921148 | 0.09135797 | -0.149972208 | 0.01994894 | -0.241330179 | 0.166093042 | 0.358007949 | 0.245165621 | 0.86743423 | 0.083599829 |
| H1f5 | S | 51 | 0.031559558 | -0.031653387 | -0.15016271 | -0.181722268 | -0.118509323 | 0.62049129 | 0.568343001 | 0.040442932 | 0.021217442 | 0.052427425 |
| Ell | S | 300 | -0.033002618 | -0.019495316 | -0.076186014 | -0.043183396 | -0.056690698 | 0.41399418 | 0.572980101 | 0.088190608 | 0.212086895 | 0.111202923 |
| Nfat5 | S | 560 | 0.037254175 | -0.052747739 | -0.061602195 | -0.09885637 | -0.008854456 | 0.392860519 | 0.02332193 | 0.039534746 | 0.081447185 | 0.681974743 |
| Cars | S | 102 | -0.03060347 | -0.029151832 | -0.066452857 | -0.035849387 | -0.037301026 | 0.375335957 | 0.165013388 | 0.254273021 | 0.517306284 | 0.473716428 |
| Pi4k2a | S | 51 | 0.000159146 | -0.040235478 | -0.093519094 | -0.09367824 | -0.053283616 | 0.996200269 | 0.251982 | 0.094979378 | 0.098581051 | 0.260476467 |
| Eif4g3 | S | 314 | -0.201796479 | -0.070694747 | 0.068833548 | 0.270630027 | 0.139528295 | 0.104647978 | 0.451597903 | 0.531123152 | 0.066900404 | 0.222085895 |
| Marcks | S | 141 | 0.106830853 | -0.04497952 | -0.142234895 | -0.249065748 | -0.097255375 | 0.369444002 | 0.293697996 | 0.011818468 | 0.106376913 | 0.079195879 |
| Dok1 | S | 48 | -0.201000372 | 0.04859602 | -0.044986992 | 0.156013381 | -0.093583012 | 0.057241447 | 0.515882347 | 0.710576139 | 0.264955054 | 0.46122463 |
| Dock4 | S | 1640 | -0.009508162 | -0.007159294 | -0.062815418 | -0.053307256 | -0.055656125 | 0.829375718 | 0.849886236 | 0.001930711 | 0.298837404 | 0.229169211 |
| Zc3hc1 | S | 68 | 0.030410244 | -0.154494827 | -0.0510874 | -0.081497644 | 0.103407427 | 0.463397039 | 0.103310239 | 0.256841151 | 0.08166528 | 0.222140234 |
| Irs2 | S | 161 | -0.151956772 | 0.083888401 | -0.220383308 | -0.068426536 | -0.304271709 | 0.515935415 | 0.627321746 | 0.215075684 | 0.702535128 | 0.027546595 |
| Prkra | S | 18 | -0.26425601 | 0.097503057 | 0.054561145 | 0.318817155 | -0.042941912 | 0.002868467 | 0.280458389 | 0.512970269 | 0.017440708 | 0.639227459 |
| Arhgef12 | S | 1492 | -0.078020063 | -0.037161547 | -0.121063452 | -0.043043388 | -0.083901905 | 0.643092832 | 0.806695573 | 0.224980009 | 0.787617488 | 0.572034632 |
| Mycbp2 | S | 2884 | -0.086456438 | -0.059612512 | 0.017645404 | 0.104101842 | 0.077275916 | 0.116395416 | 0.157923609 | 0.792032343 | 0.191653646 | 0.288105735 |
| Kmt2d | S | 1627 | -0.129105617 | -0.039046473 | 0.032410523 | 0.16151614 | 0.071456996 | 0.042125969 | 0.366153361 | 0.551947286 | 0.03152168 | 0.175017911 |
| Ncdn | S | 448 | -0.1063913 | -0.138261164 | 0.004934924 | 0.111326224 | 0.143196088 | 0.423700193 | 0.305362026 | 0.984714999 | 0.655905508 | 0.572483241 |
| Rab3ip | S | 240 | -0.113396979 | 0.034373006 | -0.08960853 | 0.023788449 | -0.123981536 | 0.064834115 | 0.482104734 | 0.367143205 | 0.799831389 | 0.249576939 |
| Zfhx4 | S | 478 | -0.249662668 | 0.018885068 | -0.04509108 | 0.204571588 | -0.063976148 | 0.187122931 | 0.841724439 | 0.7706418 | 0.324162066 | 0.679829218 |
| AU040320 | S | 977 | 0.077814827 | -0.124637302 | -0.068531377 | -0.146346204 | 0.056105925 | 0.264848991 | 0.008666871 | 0.402028717 | 0.155086871 | 0.482834242 |
| Kmt2e | S | 522 | 0.10222076 | -0.056567962 | -0.081886173 | -0.184106933 | -0.02531821 | 0.401454278 | 0.061863639 | 0.011256423 | 0.19459253 | 0.351747847 |
| Satb2 | S | 20 | -0.011854061 | -0.104085105 | -0.050740739 | -0.038886678 | 0.053344366 | 0.82758693 | 0.142274926 | 0.456913022 | 0.471259133 | 0.410710798 |

| | | | | | | | | | | | | |
|---|---|---|---|---|---|---|---|---|---|---|---|---|
| Wnk1 | S | 2265 | -0.131907097 | 0.132958193 | -0.28619211 | -0.154285013 | -0.419150304 | 0.426888272 | 0.438385688 | 0.186653884 | 0.386758257 | 0.068406178 |
| Gigyf2 | S | 26 | -0.004535827 | 0.000625289 | -0.17123707 | -0.166701242 | -0.171862359 | 0.926373087 | 0.99018938 | 0.06671686 | 0.065990067 | 0.062814336 |
| Srsf9 | S | 212 | 0.021000905 | -0.138719035 | -0.007212048 | -0.028212953 | 0.131506987 | 0.668013578 | 0.015627723 | 0.887203189 | 0.511455409 | 0.037100574 |
| Cobll1 | S | 546 | -0.051416592 | -0.041162178 | -0.003567808 | 0.047848784 | 0.03759437 | 0.13622503 | 0.044749453 | 0.908453524 | 0.243335875 | 0.292573952 |
| Axin1 | S | 217 | -0.03187588 | -0.017914127 | -0.100460865 | -0.068584985 | -0.082546738 | 0.540021369 | 0.739696511 | 0.098618571 | 0.133754502 | 0.103084234 |
| Rbm12 | S | 413 | -0.146006219 | 0.019371988 | -0.014164375 | 0.131841844 | -0.033536363 | 0.027739041 | 0.727013531 | 0.831981014 | 0.074692002 | 0.552283261 |
| Nup93 | S | 767 | -0.018129716 | -0.077794093 | 0.003477728 | 0.021607444 | 0.081271821 | 0.382400817 | 0.011672719 | 0.841129409 | 0.21673367 | 0.018525873 |
| Akap12 | T | 583 | -0.102238673 | 0.097743518 | -0.201987094 | -0.099748421 | -0.299730612 | 0.310243115 | 0.257853599 | 0.080271315 | 0.337955796 | 0.026490516 |
| Arhgap11a | S | 640 | -0.027153181 | -0.010867355 | -0.171499037 | -0.144345856 | -0.160631682 | 0.796633959 | 0.88040821 | 0.14201723 | 0.281305862 | 0.171824949 |
| Mia3 | S | 1458 | -0.038011134 | -0.134894471 | -0.016290808 | 0.021720326 | 0.118603663 | 0.559670744 | 0.178603412 | 0.779463388 | 0.722136594 | 0.223635186 |
| 5730455P16Rik | S | 55 | -0.164970555 | 0.129707932 | -0.074005524 | 0.090965031 | -0.203713456 | 0.011317082 | 0.193849387 | 0.30714311 | 0.204285295 | 0.082455803 |
| Dnttip2 | S | 173 | -0.002404713 | -0.066964256 | -0.067116276 | -0.064711563 | -0.00015202 | 0.895292329 | 0.10249788 | 0.294155504 | 0.308207044 | 0.997946739 |
| Zc3h13 | S | 316 | -0.02332304 | -0.082668629 | -0.075758655 | -0.052435615 | 0.006909974 | 0.693090194 | 0.382501433 | 0.260747235 | 0.48158066 | 0.942065695 |
| Nup188 | S | 1727 | -0.075558766 | -0.094848907 | -0.061380471 | 0.014178295 | 0.033468436 | 0.507581465 | 0.382288469 | 0.645158863 | 0.913051007 | 0.789583683 |
| Pds5b | S | 1404 | -0.1941415 | 0.087916096 | -0.092480948 | 0.101660552 | -0.180397044 | 0.024089016 | 0.218572059 | 0.630089938 | 0.589002099 | 0.374091267 |
| Pds5b | S | 1414 | -0.1941415 | 0.087916096 | -0.092480948 | 0.101660552 | -0.180397044 | 0.024089016 | 0.218572059 | 0.630089938 | 0.589002099 | 0.374091267 |
| Pds5b | S | 1418 | -0.1941415 | 0.087916096 | -0.092480948 | 0.101660552 | -0.180397044 | 0.024089016 | 0.218572059 | 0.630089938 | 0.589002099 | 0.374091267 |
| H2ax | S | 140 | 0.033738283 | -0.103620535 | -0.027055336 | -0.060793619 | 0.076565199 | 0.429214402 | 0.007274983 | 0.454026455 | 0.190237641 | 0.062269997 |
| Cavin4 | S | 19 | -0.028451217 | -0.102776393 | 0.010621636 | 0.039072852 | 0.113398029 | 0.538780759 | 0.035344624 | 0.930726848 | 0.756104301 | 0.400855648 |
| Sf3b1 | S | 194 | 0.181641769 | -0.074670944 | -0.169831358 | -0.351473127 | -0.095160414 | 0.233965513 | 0.081855386 | 0.030324617 | 0.069847449 | 0.146886853 |
| Eif4enif1 | S | 77 | -0.018350066 | -0.041634426 | -0.027415539 | -0.009065473 | 0.014218888 | 0.297017533 | 0.072545005 | 0.202355229 | 0.607705146 | 0.493001616 |
| Prrc2a | T | 1081 | -0.002136621 | -0.127449532 | -0.02177042 | -0.019633799 | 0.105679112 | 0.950560726 | 0.056126801 | 0.785295067 | 0.814417122 | 0.269358563 |
| Edc4 | T | 731 | -0.105479166 | -0.080818856 | -0.083573865 | 0.021905301 | -0.002755009 | 0.611914357 | 0.440201845 | 0.608933824 | 0.921342191 | 0.984873619 |
| Itfg2 | S | 219 | -0.169684323 | 0.03441666 | -0.036305461 | 0.133378862 | -0.070722121 | 0.046818938 | 0.552106062 | 0.713112601 | 0.216895609 | 0.44617015 |
| Eif5b | S | 510 | -0.025705211 | 0.027586606 | -0.129029112 | -0.103323901 | -0.156615718 | 0.891898342 | 0.883806746 | 0.024353882 | 0.599861376 | 0.445775242 |
| Rpl22l1 | S | 112 | 0.009339973 | -0.083206066 | -0.097062792 | -0.106402764 | -0.013856726 | 0.895860681 | 0.3469849 | 0.158392472 | 0.206490921 | 0.870798939 |
| Aftph | S | 395 | 0.056727185 | -0.112728223 | -0.090075246 | -0.146802431 | 0.022652977 | 0.208812102 | 0.125081132 | 0.061218602 | 0.018433795 | 0.707010238 |
| Gpat3 | S | 77 | 0.043547756 | -0.057152451 | -0.123887332 | -0.167435088 | -0.066734881 | 0.447745183 | 0.310355356 | 0.046948157 | 0.011714191 | 0.119040485 |
| Map1b | S | 1321 | -0.096601711 | 0.01881202 | -0.076217015 | 0.020384696 | -0.095029035 | 0.098454855 | 0.697486338 | 0.317354747 | 0.773054291 | 0.226773592 |
| Ripk2 | S | 392 | -0.164254232 | 0.023413611 | -0.014632011 | 0.149622221 | -0.038045622 | 0.037660716 | 0.703308127 | 0.878752738 | 0.177861022 | 0.680234346 |
| Als2 | T | 1031 | -0.129583732 | -0.038644968 | -0.011365349 | 0.118218383 | 0.027279619 | 0.107677464 | 0.795737908 | 0.765912458 | 0.139019063 | 0.85267775 |
| Als2 | T | 1036 | -0.129583732 | -0.038644968 | -0.011365349 | 0.118218383 | 0.027279619 | 0.107677464 | 0.795737908 | 0.765912458 | 0.139019063 | 0.85267775 |
| Als2 | Y | 1030 | -0.129583732 | -0.038644968 | -0.011365349 | 0.118218383 | 0.027279619 | 0.107677464 | 0.795737908 | 0.765912458 | 0.139019063 | 0.85267775 |
| Als2 | Y | 1033 | -0.129583732 | -0.038644968 | -0.011365349 | 0.118218383 | 0.027279619 | 0.107677464 | 0.795737908 | 0.765912458 | 0.139019063 | 0.85267775 |
| Zdhhc8 | Y | 538 | -0.112817161 | -0.022218875 | -0.001026782 | 0.111790379 | 0.021192093 | 0.035861088 | 0.801426362 | 0.988194097 | 0.180648269 | 0.83320785 |
| Zdhhc5 | Y | 533 | -0.112817161 | -0.022218875 | -0.001026782 | 0.111790379 | 0.021192093 | 0.035861088 | 0.801426362 | 0.988194097 | 0.180648269 | 0.83320785 |
| Nop56 | S | 536 | 0.046185229 | -0.090137908 | -0.05155852 | -0.097743749 | 0.038579388 | 0.202782536 | 0.011261798 | 0.134075064 | 0.0340542 | 0.201897436 |
| Lyst | S | 2122 | -0.157766838 | -0.008308537 | 0.00927934 | 0.167046178 | 0.017587878 | 0.051469732 | 0.880300132 | 0.896998944 | 0.098748583 | 0.830128398 |
| Aatf | S | 283 | -0.042432307 | -0.123144364 | -0.017813198 | 0.024619109 | 0.105331166 | 0.727133043 | 0.124720629 | 0.805787148 | 0.819456136 | 0.040311208 |
| Uhrf1bp1l | S | 988 | -0.18931283 | -0.044051898 | 0.093759196 | 0.283072026 | 0.137811094 | 0.009178977 | 0.601034153 | 0.169512453 | 0.009499077 | 0.17822073 |
| Traf6 | S | 291 | 0.009287156 | -0.053963966 | -0.054725704 | -0.064012861 | -0.000761738 | 0.920165418 | 0.10545479 | 0.089458957 | 0.509935415 | 0.975356565 |
| Srrm2 | S | 964 | -0.030207925 | -0.103657356 | 0.020485656 | 0.050693581 | 0.124143012 | 0.473196058 | 0.019494319 | 0.581479355 | 0.220721859 | 0.012477883 |
| Lnpk | S | 194 | 0.091449551 | -0.096549234 | -0.099809298 | -0.191258849 | -0.003260064 | 0.268996149 | 0.196477943 | 0.017968238 | 0.078168307 | 0.957456708 |
| Myo9b | S | 1396 | -0.089940313 | -0.144960401 | 0.078227111 | 0.168167424 | 0.223187511 | 0.207473323 | 0.149454112 | 0.544253237 | 0.249864575 | 0.162445208 |
| Kdm4a | S | 450 | -0.150948105 | -0.030805572 | -0.036427545 | 0.11452056 | -0.005621973 | 0.177810487 | 0.775539478 | 0.824992771 | 0.476259349 | 0.9708582 |
| Fntb | T | 436 | -0.506151793 | 0.080202137 | 0.040028283 | 0.546180076 | -0.040173853 | 0.174035442 | 0.690919682 | 0.831836538 | 0.154447176 | 0.859948022 |
| Ptger4 | S | 420 | 0.044856285 | -0.01486386 | -0.367163702 | -0.412019988 | -0.352299842 | 0.661109591 | 0.922635858 | 0.187644299 | 0.162290777 | 0.209077534 |
| Srrm2 | S | 1434 | -0.03624507 | -0.074075995 | 0.054756862 | 0.091001932 | 0.128832857 | 0.10255342 | 0.000910574 | 0.144727424 | 0.047470585 | 0.028656573 |
| Mrto4 | S | 225 | 0.02184649 | -0.099619629 | -0.066310199 | -0.088156688 | 0.033309431 | 0.539524602 | 0.105858677 | 0.203056049 | 0.134596794 | 0.555694552 |
| Cby1 | S | 26 | -0.023010288 | 0.008063152 | -0.172358946 | -0.149348658 | -0.180422098 | 0.534213561 | 0.812409168 | 0.121596462 | 0.169186083 | 0.128568812 |
| Etfa | T | 93 | -0.048432445 | -0.084024964 | 0.005841151 | 0.054273596 | 0.089866115 | 0.389699934 | 0.053387246 | 0.873876444 | 0.330993291 | 0.045562048 |
| Srrm2 | S | 1518 | 0.119243713 | -0.096446872 | -0.079049161 | -0.198292874 | 0.017397712 | 0.271558286 | 0.019398808 | 0.069853199 | 0.121781365 | 0.602409318 |
| Arhgef2 | S | 959 | -0.021387577 | -0.093299119 | 0.000901264 | 0.022288841 | 0.094200383 | 0.438828064 | 0.038052183 | 0.980253396 | 0.471676245 | 0.054735226 |
| Srrm2 | S | 222 | 0.095085001 | -0.220823396 | -0.114392948 | -0.209477949 | 0.106430448 | 0.46296662 | 0.243046328 | 0.260311947 | 0.119554063 | 0.515229534 |
| Tsc2 | S | 1421 | 0.017926891 | 0.010916101 | -0.154259431 | -0.172186322 | -0.165175532 | 0.705040392 | 0.748652397 | 0.013372124 | 0.020843501 | 0.018206138 |
| Map1a | S | 526 | 0.006171576 | -0.029440556 | -0.078552422 | -0.084723999 | -0.049111867 | 0.882768119 | 0.517232482 | 0.016381639 | 0.11906828 | 0.312932531 |
| Arhgef40 | S | 215 | -0.079580176 | 0.049235403 | -0.135385618 | -0.055805441 | -0.18462102 | 0.310305908 | 0.709773684 | 0.152527215 | 0.552193825 | 0.243338375 |
| Pkd2 | S | 164 | 0.105775108 | -0.151946116 | -0.134237509 | -0.240012617 | 0.017708606 | 0.105034255 | 0.122345226 | 0.066793703 | 0.0102148 | 0.815406427 |
| Etl4 | S | 1647 | 0.04232989 | -0.033734069 | -0.149786017 | -0.192115907 | -0.116051948 | 0.462313512 | 0.463026775 | 0.038530304 | 0.032181662 | 0.0915752 |
| Cep170 | S | 925 | 0.002490564 | -0.062707262 | -0.041349995 | -0.043840558 | 0.021357268 | 0.919694051 | 0.060246498 | 0.164381391 | 0.103391537 | 0.43167055 |
| Samd4b | S | 600 | 0.044387373 | -0.049939104 | -0.13797314 | -0.182360513 | -0.088034036 | 0.391758777 | 0.453070384 | 0.034021729 | 0.026766381 | 0.248211844 |
| Camsap1 | S | 661 | -0.187745541 | -0.199744004 | -1.178641229 | -0.990895688 | -0.978897225 | 0.485229449 | 0.619380329 | NA | NA | NA |
| Uba1 | S | 46 | -0.062196026 | 0.004971375 | -0.075514623 | -0.013318597 | -0.080485998 | 0.190779383 | 0.900555382 | 0.148615761 | 0.72583432 | 0.09148991 |
| Dnmt1 | S | 125 | 0.056343817 | -0.040808573 | -0.138861827 | -0.195205643 | -0.098053253 | 0.131535284 | 0.321903946 | 0.009906028 | 0.00499053 | 0.070295972 |
| Flna | T | 2599 | -0.008343685 | -0.050054433 | -0.081707515 | -0.07336383 | -0.031653083 | 0.720628081 | 0.21967662 | 0.224365175 | 0.262680768 | 0.608670739 |
| Vcpip1 | S | 767 | 0.011184609 | -0.022426807 | -0.071314945 | -0.082499554 | -0.048888138 | 0.724969418 | 0.066667977 | 0.034508767 | 0.07379805 | 0.081201223 |
| Snap23 | S | 23 | 0.029952589 | -0.082268677 | -0.059661235 | -0.089613824 | 0.022607442 | 0.405496474 | 0.063033385 | 0.109242675 | 0.011506594 | 0.415314315 |
| Mast4 | S | 255 | -0.130379623 | -0.094909921 | 0.069772911 | 0.200152534 | 0.164682832 | 0.104417171 | 0.305705243 | 0.555260537 | 0.144013527 | 0.212378455 |
| Psma1 | S | 247 | 0.058338535 | -0.054967536 | -0.111910496 | -0.170249031 | -0.05694296 | 0.139156535 | 0.25650855 | 0.007379506 | 0.005389943 | 0.226917898 |
| Pogz | S | 292 | -0.077352823 | -0.204460169 | -0.014346306 | 0.063006517 | 0.190113863 | 0.626236593 | 0.369202009 | 0.923812717 | 0.517294538 | 0.352051324 |
| Zzef1 | S | 2407 | -0.05256009 | 0.024752917 | -0.097418408 | -0.044858318 | -0.122171325 | 0.26740486 | 0.600570268 | 0.059469465 | 0.381551277 | 0.072217433 |
| Uri1 | S | 415 | -0.343237575 | 0.247376919 | -0.13995964 | 0.203277935 | -0.387336559 | 0.054503516 | 0.202143701 | 0.516054768 | 0.30090639 | 0.108469184 |
| Pi4kb | S | 277 | -0.037747073 | -0.01210097 | -0.152014651 | -0.114267578 | -0.139913681 | 0.671954476 | 0.900137522 | 0.17295153 | 0.32697746 | 0.26519561 |
| Mphosph9 | S | 746 | 0.002868728 | 0.027272902 | -0.148853105 | -0.151721833 | -0.176126007 | 0.966919454 | 0.585248291 | 0.012827143 | 0.099507825 | 0.021936683 |
| Dlgap4 | S | 665 | 0.034570383 | -0.033775907 | -0.168290675 | -0.202861058 | -0.134514768 | 0.481180688 | 0.467367059 | 0.067671203 | 0.050979722 | 0.125903892 |
| Macf1 | S | 7231 | -0.090155606 | -0.040262753 | 0.001510701 | 0.091666307 | 0.041773454 | 0.093379168 | 0.299876813 | 0.957502964 | 0.085819771 | 0.335131629 |
| Psmb8 | S | 242 | -0.058932922 | -0.044145876 | -0.12098558 | -0.062052658 | -0.076839704 | 0.683455139 | 0.74885412 | 0.246389799 | 0.691114164 | 0.614238423 |
| Psmb8 | Y | 241 | -0.058932922 | -0.044145876 | -0.12098558 | -0.062052658 | -0.076839704 | 0.683455139 | 0.74885412 | 0.246389799 | 0.691114164 | 0.614238423 |
| Psmb8 | Y | 249 | -0.058932922 | -0.044145876 | -0.12098558 | -0.062052658 | -0.076839704 | 0.683455139 | 0.74885412 | 0.246389799 | 0.691114164 | 0.614238423 |
| Acap2 | T | 382 | -0.097033172 | 0.088107494 | -0.116233793 | -0.019200621 | -0.204341287 | 0.128011617 | 0.316025003 | 0.076895196 | 0.756268281 | 0.071674569 |
| Eif3j1 Eif3j2 | T | 112;114 | -0.042482031 | -0.014117912 | -0.101242413 | -0.058760382 | -0.087124501 | 0.271092613 | 0.862212191 | 0.218117444 | 0.419873112 | 0.383024053 |
| Jun Jund | S | 73;100 | 0.105911863 | -0.090926402 | -0.116508539 | -0.222420402 | -0.025582136 | 0.036558375 | 0.034044193 | 0.013011455 | 0.004197384 | 0.198284764 |
| Zc3h18 | S | 528 | 0.021316579 | -0.063513437 | -0.057152983 | -0.078469562 | 0.006360454 | 0.70816508 | 0.040891568 | 0.188685006 | 0.247646458 | 0.863848936 |
| Itpr2 | T | 339 | 0.01960218 | -0.021180836 | -0.140842641 | -0.160444821 | -0.119661805 | 0.640769653 | 0.54218394 | 0.04839078 | 0.038633773 | 0.082888118 |
| Mrto4 | S | 229 | 0.02903689 | -0.084134224 | -0.080239189 | -0.109276079 | 0.003895035 | 0.472758286 | 0.149181949 | 0.110063871 | 0.073114149 | 0.941721975 |
| Mrto4 | S | 233 | 0.02903689 | -0.084134224 | -0.080239189 | -0.109276079 | 0.003895035 | 0.472758286 | 0.149181949 | 0.110063871 | 0.073114149 | 0.941721975 |
| Sipa1l1 | S | 114 | -0.039451427 | -0.074988135 | -0.025845848 | 0.013605579 | 0.049142287 | 0.471230861 | 0.101198812 | 0.748113555 | 0.871932234 | 0.549485418 |
| Asap1 | S | 858 | 0.02402569 | -0.032088077 | -0.092798093 | -0.116823782 | -0.060710016 | 0.518768961 | 0.135653201 | 0.040432317 | 0.041630434 | 0.092064125 |
| Mxra7 | S | 94 | -0.077093208 | -0.070401669 | 0.037662901 | 0.114756109 | 0.108064571 | 0.106454144 | 0.097607433 | 0.561258619 | 0.129465757 | 0.15068505 |
| Oga | T | 709 | -0.028597672 | -0.04467061 | -0.056930374 | -0.028332702 | -0.012259765 | 0.446472883 | 0.258769455 | 0.265597023 | 0.469011737 | 0.74482867 |
| Afdn | S | 655 | -0.030592762 | -0.09284075 | -0.014042984 | 0.016549778 | 0.078797766 | 0.595959491 | 0.081010154 | 0.751485971 | 0.797124075 | 0.182835407 |
| Camkk2 | S | 495 | -0.126338928 | 0.11962906 | -0.2349937 | -0.108654772 | -0.35462276 | 0.288710923 | 0.289530125 | 0.155698798 | 0.430629261 | 0.06181404 |
| Ccdc88a | S | 1588 | 0.121675157 | -0.060590499 | -0.160593264 | -0.282268421 | -0.100002765 | 0.393029354 | 0.468154867 | 0.01959186 | 0.117605189 | 0.253214349 |
| Nhlrc2 | Y | 403 | -0.097461849 | 0.051042906 | -0.099429272 | -0.001967423 | -0.150472178 | 0.131923228 | 0.408046367 | 0.140724601 | 0.975026663 | 0.068426587 |

| | | | | | | | | | | | | |
|---|---|---|---|---|---|---|---|---|---|---|---|---|
| Rnps1 | S | 53 | -0.016213692 | -0.028552388 | -0.101512042 | -0.08529835 | -0.072959654 | 0.7346103 | 0.561797213 | 0.111340487 | 0.115421513 | 0.147057534 |
| Irs1 | S | 632 | -0.006239096 | -0.038423373 | -0.129657462 | -0.123418366 | -0.091234088 | 0.922664749 | 0.591659211 | 0.138447055 | 0.192377597 | 0.329181898 |
| Hivep1 | S | 1776 | 0.003374886 | 0.021349253 | -0.1586911 | -0.162065986 | -0.180040353 | 0.961920414 | 0.79850354 | 0.028518671 | 0.0516902 | 0.08598115 |
| Sf3b1 | T | 434 | -0.038927734 | -0.064712435 | 0.014267775 | 0.053195508 | 0.078980209 | 0.146683132 | 0.03376214 | 0.700805648 | 0.181219389 | 0.086593774 |
| Casp8ap2 | S | 1763 | -0.071813302 | -0.036823261 | -0.010169354 | 0.061643948 | 0.026653907 | 0.134084669 | 0.324209753 | 0.482284562 | 0.187729458 | 0.444322827 |
| Sh3bp2 | S | 277 | -0.06366903 | -0.049412498 | -0.044370848 | 0.019298182 | 0.00504165 | 0.336180047 | 0.262984694 | 0.605920715 | 0.837280996 | 0.952827486 |
| Ttc28 | S | 2370 | 0.023241195 | -0.090553591 | -0.027286649 | -0.050527845 | 0.063266942 | 0.740567804 | 0.00941444 | 0.6780268 | 0.56688346 | 0.36997196 |
| Kansl2 | S | 147 | -0.094475559 | -0.063137236 | 0.014073575 | 0.108549134 | 0.07721081 | 0.106938437 | 0.350932607 | 0.852592819 | 0.178036852 | 0.332395265 |
| Kansl2 | S | 149 | -0.094475559 | -0.063137236 | 0.014073575 | 0.108549134 | 0.07721081 | 0.106938437 | 0.350932607 | 0.852592819 | 0.178036852 | 0.332395265 |
| Cdk14 | S | 95 | -0.121450035 | 0.01045528 | -0.075172144 | 0.046277891 | -0.085627424 | 0.201737725 | 0.914715229 | 0.413700451 | 0.47002775 | 0.290224864 |
| Baz1b | S | 330 | 0.077940643 | -0.007734836 | -0.146670272 | -0.224610915 | -0.138935437 | 0.507097646 | 0.823298192 | 0.00653783 | 0.137386195 | 0.012237561 |
| Ehbp1l1 | S | 239 | -0.139456871 | -0.156189221 | 0.099464829 | 0.2389217 | 0.25565405 | 0.34656037 | 0.094285317 | 0.345106878 | 0.168243468 | 0.065111617 |
| Ehbp1l1 | S | 240 | -0.139456871 | -0.156189221 | 0.099464829 | 0.2389217 | 0.25565405 | 0.34656037 | 0.094285317 | 0.345106878 | 0.168243468 | 0.065111617 |
| Map7d1 | S | 313 | 0.09851696 | 0.114065455 | -0.370647237 | -0.469164197 | -0.484712691 | 0.557367654 | 0.591224819 | 0.027991699 | 0.042641856 | 0.083335995 |
| Wdr41 | S | 459 | 0.057452578 | -0.007126149 | -0.172700785 | -0.230153363 | -0.165574636 | 0.612254397 | 0.872960289 | 0.024007391 | 0.115393827 | 0.046087867 |
| Rsrc1 | S | 12 | -0.078508937 | -0.037289154 | -0.032659204 | 0.045849733 | 0.004629949 | 0.17187808 | 0.412315849 | 0.682278429 | 0.58657462 | 0.953270225 |
| Eif2ak2 | T | 31 | -0.044244965 | -0.096442586 | -0.034308231 | 0.009936734 | 0.062134355 | 0.682629077 | 0.177927263 | 0.624229058 | 0.915888737 | 0.13302578 |
| BC005624 | S | 17 | -0.050289023 | 0.023003377 | -0.089087609 | -0.038798586 | -0.112090986 | 0.072565073 | 0.46163214 | 0.119294505 | 0.411087385 | 0.072664831 |
| Eif4g2 | S | 17 | -0.06177368 | -0.031744596 | -0.061424969 | 0.000348711 | -0.029680373 | 0.215778674 | 0.572887259 | 0.441954995 | 0.995853278 | 0.686940178 |
| Rapgef2 | S | 501 | -0.181477576 | 0.124419619 | -0.136194384 | 0.045283192 | -0.260614003 | 0.122913893 | 0.278508249 | 0.231373458 | 0.295120883 | 0.007541165 |
| Rbmxl1 | S | 186 | -0.055324083 | -0.136271129 | 0.068664042 | 0.123988125 | 0.204935171 | 0.240126768 | 0.021234293 | 0.182277023 | 0.064894782 | 0.012501494 |
| Cdc42bpb | S | 363 | -0.024896281 | -0.073257799 | -0.083522959 | -0.058626678 | -0.01026516 | 0.740061716 | 0.407822353 | 0.29381406 | 0.519051979 | 0.914262485 |
| Arhgef5 | S | 782 | -0.131160163 | 0.047520635 | -0.011676281 | 0.119483883 | -0.059196915 | 0.008519892 | 0.484224538 | 0.751187 | 0.018097224 | 0.387659095 |
| Septin7 | S | 76 | -0.198908624 | -0.012256841 | -0.005196618 | 0.193712006 | 0.007060222 | 0.142801424 | 0.921655216 | 0.985418314 | 0.522703514 | 0.980612615 |
| Ncoa6 | S | 1748 | -0.059346185 | -0.038271605 | -0.026405631 | 0.032940554 | 0.011865973 | 0.650071146 | 0.026140279 | 0.535047263 | 0.801475476 | 0.766290094 |
| Oxsr1 | S | 339 | -0.112784659 | 0.024360384 | -0.067231298 | 0.045553361 | -0.091591682 | 0.089844159 | 0.321533222 | 0.247288461 | 0.481341529 | 0.15399092 |
| Pom121 | S | 116 | -0.028723075 | -0.119350392 | -0.099944948 | -0.071221872 | 0.019405445 | 0.781311975 | 0.478497697 | 0.454621013 | 0.614031763 | 0.914704533 |
| Vps13a | T | 831 | -0.262038425 | -0.101345859 | 0.04782347 | 0.309861895 | 0.14916933 | 0.329365147 | 0.682582526 | 0.889152717 | 0.353229993 | 0.62614177 |
| Irs2 | S | 728 | -0.082885766 | -0.072674933 | 0.046904069 | 0.129789835 | 0.119579002 | 0.156540973 | 0.021028486 | 0.234053309 | 0.061448939 | 0.030723868 |
| Dennd4c | S | 1637 | 0.005483715 | -0.122811553 | -0.018079827 | -0.023563542 | 0.104731726 | 0.969561336 | 0.043278414 | 0.857683267 | 0.88630385 | 0.356839464 |
| Osbp | S | 238 | -0.02431398 | -0.108037578 | -0.010763894 | 0.013550086 | 0.097273684 | 0.748846859 | 0.070134474 | 0.85685231 | 0.843788343 | 0.090377027 |
| Nfic | S | 323 | 0.007076643 | -0.060125139 | -0.073744073 | -0.080820716 | -0.013618934 | 0.876831729 | 0.184955431 | 0.153876399 | 0.065762872 | 0.635095207 |
| Prune2 | S | 915 | -0.099290085 | 0.006515988 | -0.136388904 | -0.037098819 | -0.142904892 | 0.556637611 | 0.958935554 | 0.252147741 | 0.819534245 | 0.286980737 |
| Erf | S | 185 | -0.091307246 | 0.011615237 | -0.045229216 | 0.046078029 | -0.056844453 | 0.07825537 | 0.801761107 | 0.322021893 | 0.072657804 | 0.068530216 |
| Lipe | S | 557 | -0.146577038 | 0.040668629 | -0.08692059 | 0.059656447 | -0.12758922 | 0.104921421 | 0.354852913 | 0.383998678 | 0.573072901 | 0.236486827 |
| Taf3 | S | 428 | -0.066751848 | -0.016320888 | -0.051571212 | 0.015180637 | -0.035250324 | 0.164514738 | 0.76307811 | 0.314062512 | 0.601011307 | 0.434831166 |
| Etv4 | S | 148 | 0.071536434 | -0.130748131 | -0.156752847 | -0.22828928 | -0.026004715 | 0.250820919 | 0.219317317 | 0.169973415 | 0.082492707 | 0.822537629 |
| Map3k3 | S | 340 | -0.07719268 | 0.002977264 | -0.060365144 | 0.016827537 | -0.063342407 | 0.074870271 | 0.946320034 | 0.439395711 | 0.805986836 | 0.409101606 |
| Atad2 | S | 893 | -0.040569159 | -0.103787019 | -0.058941901 | -0.018372742 | 0.044845118 | 0.712339274 | 0.358014476 | 0.463000992 | 0.865109733 | 0.674153517 |
| Atad2 | S | 894 | -0.040569159 | -0.103787019 | -0.058941901 | -0.018372742 | 0.044845118 | 0.712339274 | 0.358014476 | 0.463000992 | 0.865109733 | 0.674153517 |
| Pde12 | S | 101 | 0.169067014 | -0.099571561 | -0.156474414 | -0.325541428 | -0.056902853 | 0.4379203 | 0.144399267 | 0.107044043 | 0.194775981 | 0.433007914 |
| Pde12 | S | 102 | 0.169067014 | -0.099571561 | -0.156474414 | -0.325541428 | -0.056902853 | 0.4379203 | 0.144399267 | 0.107044043 | 0.194775981 | 0.433007914 |
| Pds5a | S | 1300 | 0.040400659 | -0.031082541 | -0.127762033 | -0.168162692 | -0.096679492 | 0.352199877 | 0.439561563 | 0.021104524 | 0.006981568 | 0.039927349 |
| Reps1 | S | 390 | -0.144784277 | 0.02303437 | -0.024092193 | 0.120692084 | -0.047126563 | 0.04549295 | 0.542871672 | 0.626172696 | 0.087354783 | 0.278225463 |
| Nf1 | S | 2821 | -0.07940256 | 0.100666413 | -0.217647767 | -0.138245207 | -0.31831418 | 0.170825946 | 0.151006401 | 0.093604838 | 0.226215344 | 0.04057031 |
| Pheta1 | S | 144 | -0.025834647 | -0.014629869 | -0.15988654 | -0.134051892 | -0.14525667 | 0.766287629 | 0.788711665 | 0.191072912 | 0.287678494 | 0.225249064 |
| Atg2b | S | 1780 | -0.018196764 | 0.019510379 | -0.252306076 | -0.234109312 | -0.271816455 | 0.833954506 | 0.882590744 | 0.189429468 | 0.216013164 | 0.193360479 |
| Gtf2f1 | S | 217 | -0.043488973 | 0.019448408 | -0.052203461 | -0.008714488 | -0.071651868 | 0.1366675 | 0.661840748 | 0.020361474 | 0.705834238 | 0.185652782 |
| Cep131 | S | 144 | 0.036746452 | -0.073871322 | -0.075819184 | -0.112565637 | -0.001947862 | 0.656129624 | 0.094931075 | 0.176960081 | 0.228618467 | 0.961559998 |
| Vim | S | 5 | -0.034252771 | -0.082153465 | -0.066510108 | -0.032257337 | 0.015643357 | 0.817668232 | 0.260053598 | 0.42285046 | 0.835463172 | 0.856120759 |
| Thrap3 | S | 505 | 0.007061404 | -0.083478862 | -0.116056427 | -0.123117832 | -0.032577566 | 0.906114036 | 0.215721972 | 0.405988621 | 0.371462912 | 0.794247454 |
| Nav1 | S | 394 | -0.052480429 | -0.054188727 | -0.00556439 | 0.046916038 | 0.048624337 | 0.220790956 | 0.112416131 | 0.859551129 | 0.312240968 | 0.233268448 |
| Anapc1 | S | 51 | -0.014484985 | -0.067298377 | -0.02598845 | -0.011503465 | 0.041309928 | 0.446012587 | 0.065369634 | 0.585305299 | 0.794345064 | 0.417731687 |
| Fubp1 | S | 411 | -0.022299997 | -0.056358565 | -0.005727882 | 0.016572115 | 0.050630683 | 0.580555546 | 0.012611713 | 0.677792406 | 0.677738032 | 0.027326699 |
| Snx2 | S | 94 | -0.062178425 | 0.010404218 | -0.1419771 | -0.079798675 | -0.152381318 | 0.597770826 | 0.927200131 | 0.168819055 | 0.549362528 | 0.283440693 |
| Lsm8 | S | 48 | 0.059392481 | -0.010057041 | -0.131074268 | -0.190466749 | -0.121017227 | 0.344845566 | 0.783702053 | 0.003766738 | 0.043016143 | 0.027743386 |
| Ftsj3 | S | 465 | -0.0988296 | -0.195803672 | -0.039259313 | 0.059570287 | 0.156544359 | 0.796373884 | 0.419599529 | 0.811878122 | 0.871433833 | 0.47362446 |
| Arhgef17 | S | 519 | -0.035256071 | -0.040270487 | -0.00588693 | 0.029369141 | 0.034383557 | 0.202906154 | 0.026649421 | 0.825669828 | 0.362292877 | 0.246267153 |
| Ahctf1 | S | 1138 | -0.054656489 | -0.088576126 | -0.007694754 | 0.046961735 | 0.080881372 | 0.254430025 | 0.251167212 | 0.895274185 | 0.494303047 | 0.345441295 |
| Nup133 | S | 492 | -0.037946651 | -0.054454075 | -0.019919859 | 0.018026792 | 0.034534216 | 0.085281829 | 0.233503403 | 0.806815987 | 0.82362651 | 0.689507004 |
| Sgcd | S | 7 | -0.05056875 | 0.009134944 | -0.168488853 | -0.117920103 | -0.177623797 | 0.500229861 | 0.943422084 | 0.24361788 | 0.388801342 | 0.303085008 |
| Mki67 | T | 2054 | -0.113342321 | -0.053244064 | -0.01532908 | 0.098013241 | 0.037914984 | 0.201508361 | 0.500244422 | 0.930944579 | 0.578712826 | 0.822201665 |
| Bag3 | T | 420 | -0.011364125 | 0.021916809 | -0.134431603 | -0.123067478 | -0.156348412 | 0.835244309 | 0.704557903 | 0.027935356 | 0.044064775 | 0.029979023 |
| Brd4 | S | 602 | -0.206101207 | -0.076256538 | 0.12463983 | 0.330741037 | 0.200896368 | 0.028075636 | 0.208530781 | 0.085960287 | 0.007319098 | 0.015640355 |
| Adrm1 | S | 211 | -0.091235476 | -0.05732577 | 0.02453978 | 0.115775257 | 0.08186555 | 0.063027064 | 0.32980399 | 0.64411528 | 0.098012853 | 0.254452512 |
| Anln | S | 96 | 0.139240322 | -0.106945875 | -0.172227767 | -0.311468089 | -0.065281892 | 0.021774956 | 0.10033887 | 0.010672009 | 0.00570473 | 0.223600799 |
| Rcor2 | S | 63 | 0.004649486 | -0.110092235 | -0.109607971 | -0.114257457 | 0.000484263 | 0.912180308 | 0.292063332 | 0.434210573 | 0.413556403 | 0.997345326 |
| Zc3h13 | S | 242 | -0.038372669 | -0.057234106 | 0.015198276 | 0.053570946 | 0.072432382 | 0.194119759 | 0.016306662 | 0.61066422 | 0.146087788 | 0.075028176 |
| Vrk1 | T | 378 | 0.052777409 | 0.026115338 | -0.1766389 | -0.229416309 | -0.202754238 | 0.577202607 | 0.551357537 | 0.006422379 | 0.088366822 | 0.002616068 |
| Nsrp1 | S | 323 | -0.157571759 | 0.083157279 | -0.11125966 | 0.046312099 | -0.194416939 | 0.217127928 | 0.546946906 | 0.268531264 | 0.68825197 | 0.205147891 |
| Rb1 | T | 364 | 0.009765207 | 0.003919029 | -0.107566019 | -0.117331226 | -0.111485048 | 0.800311581 | 0.885533045 | 0.008354415 | 0.039197347 | 0.013069778 |
| Tbc1d10a | S | 445 | 0.083147896 | -0.064461249 | -0.139522465 | -0.222670361 | -0.075061216 | 0.241715092 | 0.285428654 | 0.027492116 | 0.023052392 | 0.187988226 |
| Bop1 | S | 84 | -0.020449861 | 0.011535482 | -0.137487923 | -0.117038062 | -0.149023405 | 0.809971713 | 0.859716139 | 0.058570866 | 0.255686522 | 0.105029738 |
| Fhod1 | S | 527 | -0.096204031 | -0.047309343 | 0.022203957 | 0.118407988 | 0.0695133 | 0.042891986 | 0.343718688 | 0.480767946 | 0.029496256 | 0.206399089 |
| Map1b | S | 933 | -0.066204496 | -0.04503353 | -0.051465118 | 0.014739377 | -0.006431588 | 0.228671448 | 0.617505598 | 0.507298932 | 0.849712983 | 0.950144678 |
| Map1b | S | 934 | -0.066204496 | -0.04503353 | -0.051465118 | 0.014739377 | -0.006431588 | 0.228671448 | 0.617505598 | 0.507298932 | 0.849712983 | 0.950144678 |
| Luc7l3 | S | 425 | -0.073265491 | 0.081257204 | -0.039098512 | 0.034166978 | -0.120355717 | 0.003427632 | 0.249274637 | 0.054038819 | 0.039514544 | 0.139854212 |
| Wdr62 | S | 33 | -0.039112358 | 0.022268919 | -0.129172314 | -0.090059956 | -0.151441233 | 0.405943077 | 0.328510331 | 0.052842041 | 0.158807203 | 0.045754364 |
| Lrfn4 | S | 575 | 0.068504412 | -0.221860749 | -0.058269201 | -0.126773613 | 0.163591548 | 0.533971529 | 0.100523614 | 0.714658926 | 0.458451124 | 0.358914336 |
| Srsf7 | S | 212 | -0.015048811 | -0.064820494 | -0.025415881 | -0.01036707 | 0.039404612 | 0.523373244 | 0.058418327 | 0.579116484 | 0.801539125 | 0.40103816 |
| Cdr2l | T | 154 | 0.042422881 | 0.065963328 | -0.204678133 | -0.247101014 | -0.270641461 | 0.332132037 | 0.421213596 | 0.002690659 | 0.004539605 | 0.038552319 |
| Nsrp1 | S | 432 | 0.012040328 | -0.076549631 | -0.095377161 | -0.107417488 | -0.018827529 | 0.817234552 | 0.296012607 | 0.204252156 | 0.184976601 | 0.815889426 |
| Tjp2 | S | 393 | -0.197450244 | -0.109290284 | 0.087701085 | 0.285151329 | 0.196991369 | 0.030122229 | 0.499295891 | 0.087760261 | 0.020911149 | 0.272910483 |
| Xpo6 | S | 208 | -0.075013331 | -0.032554021 | -0.099554942 | -0.024541611 | -0.067000921 | 0.456317416 | 0.736482988 | 0.399061308 | 0.845384804 | 0.599374936 |
| Xpo6 | T | 204 | -0.075013331 | -0.032554021 | -0.099554942 | -0.024541611 | -0.067000921 | 0.456317416 | 0.736482988 | 0.399061308 | 0.845384804 | 0.599374936 |
| Kif11 | T | 925 | -0.061339416 | 0.068050604 | -0.121019386 | -0.059679971 | -0.18906999 | 0.008191056 | 0.094098302 | 0.123851719 | 0.350746576 | 0.042621515 |
| Rnf149 | T | 327 | 0.038046735 | -0.105291508 | -0.093813501 | -0.131860236 | 0.011478007 | 0.714213036 | 0.185285625 | 0.299264386 | 0.243976356 | 0.875724175 |
| Ank3 | S | 1569 | 0.106836543 | -0.163737497 | -0.175122177 | -0.28195872 | -0.01138468 | 0.646257352 | 0.366729839 | 0.306607096 | 0.218373461 | 0.913869405 |
| Phf12 | S | 768 | -0.101318273 | -0.037727736 | 0.055464178 | 0.156782451 | 0.093191915 | 0.024471503 | 0.324104679 | 0.575295923 | 0.191459153 | 0.369052775 |
| Thada | S | 861 | -0.001998704 | -0.110991959 | -0.054456156 | -0.052457453 | 0.056535803 | 0.934820431 | 0.148499557 | 0.579574074 | 0.590218316 | 0.596235476 |
| Smc1a | S | 970 | 0.037202713 | -0.065131526 | -0.11833792 | -0.155540632 | -0.053206394 | 0.671882525 | 0.275715277 | 0.148254493 | 0.142171003 | 0.427518554 |
| Incenp | S | 433 | 0.01316056 | -0.016582695 | -0.177163099 | -0.19032366 | -0.160580405 | 0.777857978 | 0.808989 | 0.098206108 | 0.085769689 | 0.140089736 |
| Rtn4 | S | 832 | -0.022479521 | -0.199017869 | 0.002240732 | 0.024720253 | 0.201258601 | 0.667658765 | 0.216050242 | 0.989948553 | 0.889165847 | 0.359403647 |
| Braf | S | 766 | -0.023247771 | -0.017098778 | -0.0935477 | -0.070299929 | -0.076448922 | 0.694212399 | 0.745196154 | 0.085840539 | 0.235195778 | 0.148606049 |

| | | | | | | | | | | | | |
|---|---|---|---|---|---|---|---|---|---|---|---|---|
| Fndc3a | S | 202 | -0.099443257 | -0.08632616 | 0.033217892 | 0.132661149 | 0.119544052 | 0.180645399 | 0.159666521 | 0.425833058 | 0.106850488 | 0.075189474 |
| Sgce | S | 362 | -0.176516773 | 0.14456476 | -0.153010093 | 0.02350668 | -0.297574853 | 0.1651098 | 0.498494262 | 0.360037438 | 0.87982091 | 0.234086121 |
| Emd | S | 57 | -0.052514811 | -0.035863735 | -0.037766941 | 0.01474787 | -0.001903206 | 0.248390226 | 0.433107348 | 0.353361342 | 0.694918608 | 0.961096196 |
| Hectd1 | S | 251 | 0.011671359 | -0.125078962 | -0.081254815 | -0.092926174 | 0.043824147 | 0.843173985 | 0.322790364 | 0.305809019 | 0.25989745 | 0.718696055 |
| Esrra | S | 26 | 0.043154214 | -0.097030943 | -0.144024764 | -0.187178978 | -0.046993822 | 0.107893923 | 0.103770496 | 0.198487944 | 0.131315901 | 0.623388489 |
| Stag2 | T | 1160 | -0.193419788 | 0.043216489 | -0.085449757 | 0.10797003 | -0.128666247 | 0.223285649 | 0.455379675 | 0.380845067 | 0.476800054 | 0.219059104 |
| Cog3 | S | 655 | -0.201317906 | -0.01475146 | -0.017569485 | 0.183748421 | -0.002818026 | 0.209474772 | 0.91389755 | 0.898026334 | 0.219649719 | 0.981045802 |
| Pi4k2a | S | 468 | -0.333539364 | 0.323104981 | -0.303479498 | 0.030059866 | -0.626584479 | 0.072750079 | 0.073918344 | 0.278268211 | 0.896662762 | 0.079119528 |
| Acin1 | S | 606 | -0.011944156 | -0.091939872 | -0.069209829 | -0.057265673 | 0.022730044 | 0.854441529 | 0.274518416 | 0.399559171 | 0.524299233 | 0.810111866 |
| Ddb1 | S | 766 | -0.085574537 | -0.057414678 | 0.004077513 | 0.08965205 | 0.061492191 | 0.259906715 | 0.182118335 | 0.961598165 | 0.381641445 | 0.488753842 |
| Myo18a | S | 865 | -0.08110494 | 0.181120913 | -0.270478726 | -0.189373786 | -0.451599639 | 0.672679013 | 0.345298524 | 0.050394347 | 0.363390462 | 0.070126048 |
| Cherp | S | 811 | 0.006900874 | -0.096712001 | -0.025776532 | -0.032677405 | 0.070935469 | 0.852367225 | 0.0345422 | 0.668381123 | 0.546260124 | 0.251980122 |
| Cherp | S | 813 | 0.006900874 | -0.096712001 | -0.025776532 | -0.032677405 | 0.070935469 | 0.852367225 | 0.0345422 | 0.668381123 | 0.546260124 | 0.251980122 |
| Cherp | S | 815 | 0.006900874 | -0.096712001 | -0.025776532 | -0.032677405 | 0.070935469 | 0.852367225 | 0.0345422 | 0.668381123 | 0.546260124 | 0.251980122 |
| Virma | S | 1577 | -0.002726838 | -0.043255417 | -0.065095638 | -0.0623688 | -0.021840221 | 0.956692543 | 0.267676799 | 0.097381362 | 0.231709686 | 0.457966963 |
| Dst | S | 2121 | -0.094904656 | -0.121191867 | 0.123775696 | 0.218680352 | 0.244967564 | 0.122673022 | 0.039870677 | 0.220895855 | 0.073608537 | 0.058597873 |
| Ccdc61 | S | 471 | -0.110114513 | -0.004561466 | -0.078525 | 0.031589513 | -0.073963534 | 0.156983079 | 0.959207876 | 0.682180036 | 0.869185864 | 0.712080562 |
| Bcar3 | S | 420 | -0.135083007 | -0.043614609 | -0.069408135 | 0.065674872 | -0.025793526 | 0.427737447 | 0.791859633 | 0.56785424 | 0.712000959 | 0.886405681 |
| Rai14 | S | 483 | 0.0604274 | -0.025440197 | -0.188587691 | -0.249015091 | -0.163147494 | 0.549615543 | 0.771933072 | 0.051970926 | 0.075692555 | 0.156148072 |
| Amot | T | 836 | -0.00335044 | -0.026587955 | -0.184122076 | -0.180771637 | -0.157534122 | 0.968938856 | 0.798101273 | 0.193711789 | 0.204044186 | 0.280244449 |
| Irf2 | S | 256 | -0.075874057 | 0.02511207 | -0.104952824 | -0.029078767 | -0.130064894 | 0.312296252 | 0.827201093 | 0.204131236 | 0.647858546 | 0.287915278 |
| Sp1 | S | 58 | -0.120329085 | 0.000316169 | -0.044876789 | 0.075452296 | -0.045192958 | 0.138950308 | 0.997309669 | 0.606923807 | 0.382079189 | 0.651758549 |
| Mndal | T | 232 | 0.037339833 | -0.08337324 | -0.059293866 | -0.096633699 | 0.024079374 | 0.440131657 | 0.042751663 | 0.222715794 | 0.102905381 | 0.543267042 |
| Sbf2 | T | 1103 | -0.02327801 | 0.038527294 | -0.152484016 | -0.129206006 | -0.19101131 | 0.704816503 | 0.521917852 | 0.038778816 | 0.009050279 | 0.001165664 |
| Bag3 | T | 291 | 0.031860289 | -0.039980979 | -0.122556066 | -0.154416355 | -0.082575086 | 0.395787173 | 0.233369791 | 0.073271012 | 0.047407729 | 0.182595831 |
| Sh3pxd2a | S | 659 | -0.175956817 | -0.14398281 | 0.099913267 | 0.275870084 | 0.243896076 | 0.258559759 | 0.102991112 | 0.164499518 | 0.128628601 | 0.020589615 |
| Sh3pxd2a | S | 661 | -0.175956817 | -0.14398281 | 0.099913267 | 0.275870084 | 0.243896076 | 0.258559759 | 0.102991112 | 0.164499518 | 0.128628601 | 0.020589615 |
| Map2k5 | S | 149 | -0.145597842 | 0.11562752 | -0.149318968 | -0.003721125 | -0.264946488 | 0.179408417 | 0.32839181 | 0.195231603 | 0.942360177 | 0.025584351 |
| Ilf3 | S | 382 | -0.061964057 | -0.045437993 | 0.007989849 | 0.069953906 | 0.053427843 | 0.089125818 | 0.137714532 | 0.708799253 | 0.071702685 | 0.113411361 |
| Dbnl | S | 24 | -0.006703503 | -0.033705424 | -0.17689865 | -0.170195147 | -0.143193226 | 0.932512359 | 0.675474206 | 0.246077527 | 0.256270242 | 0.321931052 |
| Chd8 | S | 1993 | -0.10974497 | -0.050426612 | 0.027584992 | 0.137329962 | 0.078011604 | 0.188832626 | 0.201738162 | 0.805513952 | 0.302502123 | 0.505719157 |
| Zc3hav1 | S | 538 | -0.013704274 | -0.092533226 | 0.00708261 | 0.020786884 | 0.099615836 | 0.630615575 | 0.020124276 | 0.738920131 | 0.402156084 | 0.028622911 |
| Phf14 | S | 828 | -0.048050121 | -0.042273027 | -0.012636904 | 0.035413217 | 0.029636123 | 0.199416336 | 0.15800345 | 0.627846109 | 0.260188967 | 0.099922207 |
| Polr3c | T | 210 | 0.003674401 | -0.084067079 | -0.037566749 | -0.04124115 | 0.04650033 | 0.887917826 | 0.129973649 | 0.213241015 | 0.102873683 | 0.334910436 |
| Tiam2 | S | 292 | -0.11366025 | -0.032535685 | 0.012634697 | 0.126294947 | 0.045170383 | 0.074453206 | 0.522125482 | 0.789605855 | 0.071295396 | 0.415539461 |
| Runx2 | S | 426 | -0.026001308 | 0.019968385 | -0.176017028 | -0.150015719 | -0.195985412 | 0.720574277 | 0.769830148 | 0.117518917 | 0.176717967 | 0.098428164 |
| Tbc1d10a | S | 18 | 0.008235564 | -0.028592129 | -0.091544138 | -0.099779702 | -0.062952009 | 0.886469067 | 0.644470275 | 0.035479215 | 0.14200597 | 0.316674364 |
| Nfib | S | 332 | -0.055173039 | -0.034436432 | 0.002599182 | 0.057772221 | 0.037035614 | 0.123508397 | 0.054868851 | 0.822221647 | 0.127399605 | 0.052090527 |
| Ankib1 | S | 738 | -0.034098601 | -0.1171226 | 0.009050724 | 0.043149325 | 0.126173325 | 0.574786653 | 0.09094859 | 0.879947248 | 0.054628718 | 0.001836952 |
| Ddx20 | S | 568 | -0.023160324 | -0.087549635 | -0.055942176 | -0.032781853 | 0.031607459 | 0.706831713 | 0.271342945 | 0.470377836 | 0.698513271 | 0.728123903 |
| Kank2 | T | 170 | -0.06593597 | -0.097660742 | 0.000639141 | 0.06657511 | 0.098299882 | 0.412490905 | 0.205011779 | 0.992932702 | 0.244524814 | 0.052577535 |
| Rgl1 | S | 627 | -0.038504749 | -0.10705126 | -0.056507942 | -0.018003194 | 0.050543317 | 0.65988176 | 0.386247682 | 0.538157332 | 0.839293854 | 0.672253187 |
| Mapk6 | S | 363 | -0.044061226 | -0.045361399 | -0.031770307 | 0.012290919 | 0.013591092 | 0.557358909 | 0.184491524 | 0.300802288 | 0.866219478 | 0.687329794 |
| Map1b | S | 2205 | 0.030577116 | -0.037842799 | -0.086130919 | -0.116708034 | -0.048288119 | 0.671273825 | 0.200026492 | 0.057377539 | 0.179849065 | 0.197060204 |
| Cic | S | 2247 | -0.292347948 | 0.132439144 | 0.064613717 | 0.356961665 | -0.067825427 | 0.011947656 | 0.345373284 | 0.769097525 | 0.191000981 | 0.772887884 |
| Zfpm2 | S | 904 | -0.03868209 | -0.00671027 | -0.06535993 | -0.026677841 | -0.058649661 | 0.217252334 | 0.815592417 | 0.150706259 | 0.463832881 | 0.170660188 |
| Reps1 | S | 555 | -0.053849149 | -0.076109861 | -0.085709283 | -0.031860134 | -0.009599422 | 0.814422452 | 0.439552596 | 0.410685484 | 0.889948409 | 0.925648074 |
| Scnm1 | S | 182 | -0.055596674 | -0.017072634 | -0.04323282 | 0.012363855 | -0.026160186 | 0.153146491 | 0.772255311 | 0.258314375 | 0.740500745 | 0.664879804 |
| Rps3 | S | 224 | 0.048661927 | -0.041705161 | -0.154487261 | -0.203149187 | -0.1127821 | 0.513481571 | 0.6164388 | 0.055485558 | 0.047430336 | 0.232562717 |
| Rif1 | S | 405 | 0.150197905 | -0.179223636 | -0.170531859 | -0.320729764 | 0.008691777 | 0.002559649 | 0.010928887 | 0.067926914 | 0.0173946 | 0.899005235 |
| Svil | S | 1181 | -0.001729412 | -0.067262505 | -0.078264888 | -0.076535476 | -0.011002382 | 0.967269019 | 0.172151913 | 0.372068472 | 0.380435353 | 0.89230446 |
| Zeb2 | T | 782 | 0.111809692 | -0.158761335 | -0.063098051 | -0.174907743 | 0.095663284 | 0.096146632 | 0.014021476 | 0.201198455 | 0.034355332 | 0.109374445 |
| Pkn2 | T | 124 | 0.054115266 | -0.054654423 | -0.09237129 | -0.146486556 | -0.037716868 | 0.424529974 | 0.303750087 | 0.023226463 | 0.096652392 | 0.460523968 |
| Cdk14 | S | 122 | -0.032590512 | -0.126774981 | -0.09035974 | -0.057769228 | 0.036415241 | 0.700893023 | 0.479948683 | 0.594936796 | 0.724251277 | 0.863221165 |
| E2f3 | S | 155 | 0.02404013 | -0.061463732 | -0.116375695 | -0.140415825 | -0.054911963 | 0.747030262 | 0.540804278 | 0.099287215 | 0.080623199 | 0.530778046 |
| Ccdc61 | S | 374 | -0.08834709 | -0.011407464 | -0.051603627 | 0.036743462 | -0.040196163 | 0.134855146 | 0.908738052 | 0.551242939 | 0.652256344 | 0.722637364 |
| Yap1 | S | 404 | -0.036629923 | -0.182162824 | -0.082188103 | -0.04555818 | 0.099974721 | 0.824594434 | 0.45188562 | 0.719375304 | 0.849581445 | 0.725388947 |
| Tjp2 | S | 390 | -0.162704082 | -0.096197359 | 0.103547594 | 0.266251676 | 0.199744953 | 0.023518315 | 0.41217896 | 0.148552815 | 0.0133128 | 0.153671934 |
| Ctr9 | S | 1041 | 0.035668903 | -0.087905842 | -0.050320868 | -0.08598977 | 0.037584975 | 0.274808132 | 0.012654469 | 0.371467736 | 0.17610848 | 0.486425106 |
| Bcl9l | S | 116 | -0.122887765 | 0.096933657 | -0.102716902 | 0.020170864 | -0.199650559 | 0.061455845 | 0.31842237 | 0.253840628 | 0.798452985 | 0.106998646 |
| Washc2 | S | 5 | -0.046964946 | -0.12080576 | 0.02592502 | 0.072889966 | 0.14673078 | 0.490459462 | 0.11748937 | 0.854234269 | 0.590588152 | 0.328039758 |
| Washc2 | T | 4 | -0.046964946 | -0.12080576 | 0.02592502 | 0.072889966 | 0.14673078 | 0.490459462 | 0.11748937 | 0.854234269 | 0.590588152 | 0.328039758 |
| Gpatch8 | S | 809 | 0.351229339 | -0.139919234 | -0.161930642 | -0.513159981 | -0.022011408 | 0.408672262 | 0.068801813 | 0.128892523 | 0.266535692 | 0.804297775 |
| Gpatch8 | S | 810 | 0.351229339 | -0.139919234 | -0.161930642 | -0.513159981 | -0.022011408 | 0.408672262 | 0.068801813 | 0.128892523 | 0.266535692 | 0.804297775 |
| Gpatch8 | S | 815 | 0.351229339 | -0.139919234 | -0.161930642 | -0.513159981 | -0.022011408 | 0.408672262 | 0.068801813 | 0.128892523 | 0.266535692 | 0.804297775 |
| S1pr2 | T | 341 | 0.005777975 | -0.023966808 | -0.252498941 | -0.258276917 | -0.228532134 | 0.970983815 | 0.819056569 | 0.274821927 | 0.297338856 | 0.313468751 |
| Psmb4 | S | 127 | -0.15563812 | -0.102408523 | 0.033367986 | 0.189006105 | 0.135776508 | 0.260395041 | 0.505839138 | 0.762149924 | 0.215479806 | 0.410646053 |
| Psip1 | T | 141 | -0.012801252 | -0.035104571 | -0.053426636 | -0.040625384 | -0.018322065 | 0.673100183 | 0.112004816 | 0.235880388 | 0.370164923 | 0.635731528 |
| Capg | S | 327 | 0.008538369 | -0.014214196 | -0.121460227 | -0.129998597 | -0.107246031 | 0.794030059 | 0.683150049 | 0.053361946 | 0.061331519 | 0.083551183 |
| Tacc2 | S | 419 | 0.048124534 | -0.034877711 | -0.075803543 | -0.123928077 | -0.040925832 | 0.618620757 | 0.272875665 | 0.014867051 | 0.266571209 | 0.193892818 |
| Lamtor1 | S | 27 | -0.01925263 | -0.069664431 | -0.040606757 | -0.021354128 | 0.029057674 | 0.619190085 | 0.238712116 | 0.325189908 | 0.585177819 | 0.592889532 |
| Hivep3 | S | 553 | -0.002707806 | -0.12903204 | -0.059148447 | -0.056440641 | 0.069883593 | 0.973286309 | 0.212408049 | 0.614335954 | 0.622683769 | 0.573415687 |
| Akap10 | S | 280 | 0.107849666 | 0.04461942 | -0.289214856 | -0.397064522 | -0.333834276 | 0.246580087 | 0.557268902 | 0.008421597 | 0.010961125 | 8.45636E-05 |
| Cd3eap | S | 242 | 0.095311164 | -0.083308158 | -0.227967879 | -0.323279043 | -0.144659721 | 0.403753056 | 0.205763833 | 0.257204468 | 0.148265022 | 0.431546423 |
| Mapkbp1 | S | 1089 | -0.050320656 | 0.091254027 | -0.215652973 | -0.165332317 | -0.306907001 | 0.631826027 | 0.541319461 | 0.096722455 | 0.234035874 | 0.117457329 |
| Usp34 | S | 1495 | 0.06234328 | -0.149895223 | -0.066290077 | -0.128633357 | 0.083605145 | 0.741411365 | 0.143933298 | 0.473754426 | 0.490591121 | 0.177769235 |
| Usp34 | S | 1498 | 0.06234328 | -0.149895223 | -0.066290077 | -0.128633357 | 0.083605145 | 0.741411365 | 0.143933298 | 0.473754426 | 0.490591121 | 0.177769235 |
| Usp34 | S | 1506 | 0.06234328 | -0.149895223 | -0.066290077 | -0.128633357 | 0.083605145 | 0.741411365 | 0.143933298 | 0.473754426 | 0.490591121 | 0.177769235 |
| Usp34 | S | 1509 | 0.06234328 | -0.149895223 | -0.066290077 | -0.128633357 | 0.083605145 | 0.741411365 | 0.143933298 | 0.473754426 | 0.490591121 | 0.177769235 |
| Tob2 | S | 155 | -0.119404427 | 0.036665482 | -0.120683134 | -0.001278707 | -0.157348616 | 0.25819615 | 0.77707588 | 0.430175358 | 0.993173065 | 0.375571045 |
| Cep76 | S | 83 | 0.016149246 | -0.044580361 | -0.165572452 | -0.181721698 | -0.120992091 | 0.844530699 | 0.345233301 | 0.280093612 | 0.256377026 | 0.400193275 |
| C2cd5 | S | 643 | -0.078990245 | -0.050823568 | -0.06909595 | 0.009894295 | -0.018272383 | 0.384533205 | 0.455124893 | 0.723609729 | 0.959615213 | 0.923395037 |
| Ap3d1 | S | 760 | 0.03608363 | -0.045108528 | -0.051897799 | -0.08798143 | -0.006789271 | 0.618705158 | 0.008433991 | 0.194929938 | 0.290641329 | 0.83293957 |
| Apeh | S | 25 | -0.176868119 | -0.021641308 | 0.082063368 | 0.258931486 | 0.103704676 | 0.011662066 | 0.908450028 | 0.312066639 | 0.02807329 | 0.599161782 |
| Rai1 | S | 1520 | -0.023655615 | -0.099077356 | 0.025847733 | 0.049503348 | 0.12492509 | 0.573633889 | 0.018305242 | 0.569020521 | 0.318701347 | 0.041376897 |
| Rab10 | T | 73 | 0.016815249 | -0.13597177 | -0.046021037 | -0.062836286 | 0.089950732 | 0.841177342 | 0.16683486 | 0.479228722 | 0.471754223 | 0.329064997 |
| Retreg3 | S | 26 | -0.042245226 | -0.07740667 | 0.042492837 | 0.084738062 | 0.119899506 | 0.3187974 | 0.013363431 | 0.489343735 | 0.232083973 | 0.128326795 |
| Dnmt1 | T | 308 | 0.019770852 | 0.004546859 | -0.112086772 | -0.131857624 | -0.116633631 | 0.767232203 | 0.918346401 | 0.010394974 | 0.1346881 | 0.065907852 |
| Gps1 | S | 220 | -0.04932096 | -0.018575066 | -0.062652936 | -0.013331975 | -0.044077869 | 0.21627019 | 0.732930159 | 0.306029215 | 0.814122292 | 0.52332708 |
| Srrm1 | S | 843 | 0.019284458 | -0.07960711 | 0.008690679 | -0.010593779 | 0.088297789 | 0.313058955 | 0.000255929 | 0.841998469 | 0.812163967 | 0.140412295 |
| Arhgef10 | S | 21 | 0.014870511 | 0.048280175 | -0.310840946 | -0.325711457 | -0.359121121 | 0.940614848 | 0.585591384 | 0.119092767 | 0.207000017 | 0.09185962 |
| Arhgef10 | S | 34 | 0.014870511 | 0.048280175 | -0.310840946 | -0.325711457 | -0.359121121 | 0.940614848 | 0.585591384 | 0.119092767 | 0.207000017 | 0.09185962 |
| Wnk1 | S | 2022 | -0.064137223 | 0.025184542 | -0.11914443 | -0.055007208 | -0.144328973 | 0.597519634 | 0.59692682 | 0.075719597 | 0.651891072 | 0.049922927 |
| Ercc6 | S | 482 | 0.036290854 | -0.190631164 | 0.034440953 | -0.001849901 | 0.225072117 | 0.469002797 | 0.011355944 | 0.583660879 | 0.972173301 | 0.016548174 |

| | | | | | | | | | | | | |
|---|---|---|---|---|---|---|---|---|---|---|---|---|
| Ercc6 | S | 485 | 0.036290854 | -0.190631164 | 0.034440953 | -0.001849901 | 0.225072117 | 0.469002797 | 0.011355944 | 0.583660879 | 0.972173301 | 0.016548174 |
| Pard3 | T | 579 | 0.031770148 | -0.134559858 | -0.048148079 | -0.079918227 | 0.086411779 | 0.579711811 | 0.108483512 | 0.411971446 | 0.295181995 | 0.280054217 |
| Tuba1a Tuba1b Tuba | S | 54;54;54;54 | -0.062190204 | 0.152590294 | -0.162170179 | -0.099979975 | -0.314760473 | 0.249511686 | 0.314920165 | 0.031812225 | 0.12669602 | 0.098712057 |
| Xaf1 | S | 128 | -0.037176316 | -0.112201488 | 0.016273552 | 0.053449868 | 0.12847504 | 0.549595199 | 0.08306126 | 0.82547459 | 0.424726289 | 0.106587475 |
| Srrm2 | S | 1647 | 0.018823874 | -0.097664367 | -0.005683606 | -0.024507479 | 0.091980762 | 0.444780867 | 0.004628718 | 0.928184341 | 0.693600542 | 0.224292165 |
| Osbp | S | 349 | -0.180821537 | -0.038545933 | 0.091647839 | 0.272469376 | 0.130193772 | 0.107140973 | 0.239061226 | 0.507182233 | 0.124544757 | 0.369916842 |
| 4933427D14Rik | T | 553 | 0.009572191 | -0.123672633 | 0.008126462 | -0.001445729 | 0.131799095 | 0.899083057 | 0.022836763 | 0.888265884 | 0.984926826 | 0.075543048 |
| Hdac7 | S | 344 | -0.066912667 | -0.034132373 | -0.037523739 | 0.029388928 | -0.003391366 | 0.221301451 | 0.550853584 | 0.505612709 | 0.648407714 | 0.960677021 |
| Tcf12 | S | 583 | 0.104667014 | -0.07220112 | -0.274293193 | -0.378960208 | -0.202092073 | 0.050467424 | 0.152595036 | 0.091006664 | 0.049050181 | 0.16183473 |
| E2f8 | S | 412 | -0.04671138 | -0.03486579 | -0.038198983 | 0.008512396 | -0.003333193 | 0.328226049 | 0.370908857 | 0.330478409 | 0.848275134 | 0.927960384 |
| Nop56 | T | 545 | 0.100304414 | -0.1178427 | -0.071105938 | -0.171410352 | 0.046736761 | 0.044382729 | 0.030975976 | 0.01690497 | 0.022081759 | 0.25356909 |
| Git2 | S | 507 | -0.277724338 | 0.116428732 | -0.516400038 | -0.238675701 | -0.632828771 | 0.710523334 | 0.365825505 | 0.437599175 | 0.790334389 | 0.358445409 |
| Plec | S | 4372 | -0.06657106 | -0.237253655 | 0.016863081 | 0.083434141 | 0.254116736 | 0.740564296 | 0.34098957 | 0.956280586 | 0.781206151 | 0.447939847 |
| Dtna | S | 567 | -0.519808054 | -0.041608632 | 0.615240585 | 1.135048639 | 0.656849216 | 0.01087976 | 0.853613191 | 0.199886783 | 0.075333404 | 0.181180997 |
| Dtl | S | 694 | -0.047604809 | -0.018981096 | -0.113439373 | -0.065834564 | -0.094458277 | 0.57404802 | 0.839955247 | 0.246257294 | 0.513327929 | 0.397731438 |
| Phb2 | S | 267 | 0.047589787 | -0.113588125 | -0.028591161 | -0.076180949 | 0.084996963 | 0.471163513 | 0.014042771 | 0.645770303 | 0.334804048 | 0.214322571 |
| Ctr9 | S | 1039 | 0.035875859 | -0.088226621 | -0.050231593 | -0.086107452 | 0.037995028 | 0.283035825 | 0.013692437 | 0.39467568 | 0.193132984 | 0.502969027 |
| Nudcd3 | S | 357 | 0.242495374 | -0.218541562 | -0.356283483 | -0.598778857 | -0.13774192 | 0.105618051 | 0.176253128 | 0.214207848 | 0.08240508 | 0.590255805 |
| Ahctf1 | S | 1129 | -0.11570185 | 0.017240142 | -0.07579136 | 0.039910489 | -0.093031503 | 0.170275335 | 0.824488139 | 0.517153168 | 0.704274618 | 0.402319616 |
| Fbxo30 | T | 147 | -0.035359503 | 0.049704711 | -0.187781791 | -0.152422288 | -0.237486502 | 0.464849662 | 0.34626184 | 0.08314634 | 0.141080771 | 0.055033332 |
| Mtmr2 | S | 630 | 0.067667842 | -0.087063503 | -0.063811874 | -0.131479716 | 0.023251629 | 0.386601631 | 0.060225076 | 0.10765019 | 0.15508693 | 0.405146935 |
| Fam129b | S | 647 | 0.016593713 | 0.003814524 | -0.158847762 | -0.175441476 | -0.162662286 | 0.737513267 | 0.927321284 | 0.043376251 | 0.037873738 | 0.062417249 |
| Psmd1 | T | 270 | -0.023598075 | -0.082945989 | -0.00572986 | 0.017868215 | 0.077216129 | 0.618489823 | 0.053848434 | 0.937707552 | 0.80740594 | 0.323604412 |
| Apobr | S | 520 | -0.133864976 | 0.095376685 | -0.089928589 | 0.043936387 | -0.185305275 | 0.031765156 | 0.139382037 | 0.214764123 | 0.477595502 | 0.040323781 |
| Glmn | S | 430 | 0.048585536 | -0.086333826 | -0.083938024 | -0.13252356 | 0.002395802 | 0.280734066 | 0.048516676 | 0.223771685 | 0.099079981 | 0.968297692 |
| Phf23 | S | 313 | -0.050253036 | 0.012415862 | -0.110512128 | -0.060259092 | -0.122927991 | 0.19506556 | 0.683717837 | 0.236832488 | 0.470519147 | 0.2099772 |
| U2surp | S | 485 | -0.015880306 | -0.086535471 | -0.023358239 | -0.007477933 | 0.063177233 | 0.696854253 | 0.109787818 | 0.631069692 | 0.84351281 | 0.216879179 |
| Kdm7a | S | 634 | -0.023648314 | -0.114891059 | 0.004399195 | 0.028047508 | 0.119290254 | 0.738202981 | 0.07270161 | 0.936982893 | 0.684853628 | 0.074728841 |
| Sipa1l1 | S | 1345 | -0.11456315 | -0.056914871 | 0.006774258 | 0.121337407 | 0.063689129 | 0.324789152 | 0.244148151 | 0.905462497 | 0.300304969 | 0.20208411 |
| Shroom4 | S | 1050 | -0.035551961 | -0.107811582 | -0.104545255 | -0.068993294 | 0.003266327 | 0.743268995 | 0.365035617 | 0.724721819 | 0.817995041 | 0.991277143 |
| Irs2 | T | 771 | -0.059343361 | -0.05292424 | -0.054770446 | 0.004572915 | -0.001846206 | 0.38922947 | 0.538742783 | 0.488505693 | 0.94720357 | 0.982751186 |
| Usp42 | S | 858 | 0.038410612 | -0.082935034 | -0.104413619 | -0.142824231 | -0.021478585 | 0.389712592 | 0.183074175 | 0.169565093 | 0.091634677 | 0.765150151 |
| Map1a | S | 2048 | -0.042370355 | 0.014648049 | -0.118053114 | -0.075682759 | -0.132701163 | 0.357517387 | 0.870281971 | 0.177715399 | 0.348567981 | 0.251919109 |
| Map1a | T | 504 | -0.013783 | -0.007390682 | -0.07162698 | -0.05784398 | -0.064236298 | 0.66155771 | 0.536006231 | 0.044696909 | 0.157705902 | 0.064588147 |
| Dock9 | S | 936 | -0.156129512 | 0.002358723 | 0.011204683 | 0.167334196 | 0.008845961 | 0.057562787 | 0.95801883 | 0.850550062 | 0.067282528 | 0.882145247 |
| Ect2 | T | 444 | 0.013192118 | -0.045586596 | -0.072110887 | -0.085303006 | -0.026524291 | 0.743811896 | 0.329722201 | 0.058633407 | 0.108990038 | 0.567803165 |
| Rbm12b1 | S | 667 | -0.053744866 | -0.029158158 | -0.009010184 | 0.044734682 | 0.020147975 | 0.075309832 | 0.304761757 | 0.708562454 | 0.077742908 | 0.356091605 |
| Tcof1 | T | 453 | 0.247458308 | -0.205043545 | -0.222242464 | -0.469700772 | -0.017198919 | 0.039751159 | 0.034424496 | 0.127758749 | 0.019353243 | 0.878561611 |
| Ep400 | S | 164 | -0.128292513 | -0.107570915 | 0.045588517 | 0.173881029 | 0.153159432 | 0.359935357 | 0.249541511 | 0.67405145 | 0.238223863 | 0.153476078 |
| Epn1 | S | 419 | -0.080774199 | -0.039661475 | -0.039295134 | 0.041479065 | 0.000366341 | 0.270468601 | 0.543164684 | 0.63489615 | 0.655257499 | 0.996745039 |
| Arhgef10 | S | 107 | -0.2324442 | 0.051586312 | -0.122853306 | 0.109590894 | -0.174439618 | 0.423720357 | 0.883501762 | 0.608104148 | 0.735686302 | 0.659244739 |
| Eif4b | S | 489 | -0.092726177 | 0.03675079 | -0.086423176 | 0.006303001 | -0.123173966 | 0.302619035 | 0.614108543 | 0.136601497 | 0.930552593 | 0.121270602 |
| Gga2 | S | 286 | -0.071645385 | -0.187114477 | -0.024696324 | 0.046949061 | 0.162418153 | 0.675730576 | 0.416408344 | 0.93481828 | 0.88002647 | 0.632031317 |
| Dnajc2 | S | 60 | 0.008910095 | -0.242635372 | 0.092867922 | 0.083957827 | 0.335503293 | 0.939550504 | 0.062597757 | 0.696545459 | 0.723454353 | 0.226470289 |
| Osbpl6 | S | 195 | 0.013472392 | -0.003867552 | -0.16072307 | -0.174195462 | -0.156855518 | 0.842628145 | 0.950849639 | 0.074153977 | 0.082945502 | 0.097014838 |
| Pkn1 | S | 536 | -0.141984219 | -0.049513816 | -0.029710947 | 0.112273272 | 0.01980287 | 0.356712956 | 0.660254353 | 0.777811538 | 0.479912642 | 0.875547659 |
| Pkn1 | S | 540 | -0.141984219 | -0.049513816 | -0.029710947 | 0.112273272 | 0.01980287 | 0.356712956 | 0.660254353 | 0.777811538 | 0.479912642 | 0.875547659 |
| Slc38a10 | S | 792 | -0.061137148 | -0.011850731 | -0.033614276 | 0.027522872 | -0.021763545 | 0.103887818 | 0.744853478 | 0.377525469 | 0.306094068 | 0.459417545 |
| Arid1a | S | 365 | -0.115584456 | -0.038514105 | 0.004958408 | 0.120542864 | 0.043472513 | 0.168276505 | 0.404944732 | 0.845121621 | 0.164519551 | 0.33703531 |
| Zc3hc1 | S | 369 | 0.054037736 | -0.060806146 | -0.084745285 | -0.138783021 | -0.023939139 | 0.110197432 | 0.128953801 | 0.013570813 | 0.008280341 | 0.495288061 |
| Rsrp1 | S | 17 | -0.039326884 | -0.118723475 | 0.00281282 | 0.042139704 | 0.121536295 | 0.488696175 | 0.178092893 | 0.96884185 | 0.576759956 | 0.217564919 |
| Anks1 | S | 679 | -0.125501606 | 0.020311439 | 0.001661996 | 0.127163602 | -0.018649443 | 0.026063879 | 0.842231751 | 0.97279326 | 0.049964767 | 0.856300737 |
| Specc1 | T | 149 | 0.103723224 | -0.069198624 | -0.245418271 | -0.349141496 | -0.176219647 | 0.277938465 | 0.253388394 | 0.163499142 | 0.080287079 | 0.276596423 |
| Vim | S | 47 | 0.012863138 | -0.027204225 | -0.076526467 | -0.089389605 | -0.049322242 | 0.831709124 | 0.394303911 | 0.038542593 | 0.218891516 | 0.185635174 |
| Map4k4 | S | 648 | -0.136993317 | 0.065851033 | -0.132805087 | 0.00418823 | -0.19865612 | 0.237257891 | 0.591229564 | 0.394040958 | 0.97484537 | 0.215440157 |
| Map4k4 | T | 647 | -0.136993317 | 0.065851033 | -0.132805087 | 0.00418823 | -0.19865612 | 0.237257891 | 0.591229564 | 0.394040958 | 0.97484537 | 0.215440157 |
| Zc3h13 | S | 1513 | -0.200052989 | 0.124743701 | -0.184622243 | 0.015430746 | -0.309365945 | 0.351357434 | 0.273211147 | 0.189282828 | 0.939089389 | 0.056333003 |
| Zc3h13 | S | 1516 | -0.200052989 | 0.124743701 | -0.184622243 | 0.015430746 | -0.309365945 | 0.351357434 | 0.273211147 | 0.189282828 | 0.939089389 | 0.056333003 |
| Zc3h13 | S | 1526 | -0.200052989 | 0.124743701 | -0.184622243 | 0.015430746 | -0.309365945 | 0.351357434 | 0.273211147 | 0.189282828 | 0.939089389 | 0.056333003 |
| Actb Actg1 | T | 203;203 | 0.047764389 | -0.037634052 | -0.123129647 | -0.170894036 | -0.085495596 | 0.645866786 | 0.508788221 | 0.056677025 | 0.175612917 | 0.197393643 |
| Cdan1 | S | 162 | -0.006408591 | -0.002598262 | -0.126620127 | -0.120211535 | -0.124021864 | 0.890902766 | 0.961008403 | 0.075691828 | 0.085678359 | 0.074205674 |
| Tmpo | T | 153;153 | -0.143760425 | 0.022377119 | -0.149258872 | -0.005498447 | -0.171635991 | 0.530669426 | 0.916503095 | 0.437635491 | 0.97344663 | 0.28983205 |
| Map3k3 | S | 250 | -0.096092148 | 0.064883971 | -0.129214291 | -0.033122143 | -0.194098262 | 0.318725927 | 0.530729955 | 0.167996765 | 0.704671774 | 0.103385638 |
| Rbbp8 | S | 233 | 0.021635991 | -0.112412102 | -0.070666769 | -0.09230276 | 0.041745332 | 0.830651066 | 0.169918707 | 0.487668763 | 0.484765974 | 0.700301517 |
| Frmd6 | S | 542 | -0.125725724 | 0.037129174 | -0.020323661 | 0.105402063 | -0.057452835 | 0.02157086 | 0.48968393 | 0.750710737 | 0.17034156 | 0.434961669 |
| Dido1 | S | 1336 | -0.056404943 | -0.014849154 | -0.000612379 | 0.055792564 | 0.014236775 | 0.011165326 | 0.510814059 | 0.968898673 | 0.003846237 | 0.446426134 |
| Gapvd1 | S | 569 | -0.100064904 | -0.033802292 | -0.002801457 | 0.097263446 | 0.031000835 | 0.100310396 | 0.669488569 | 0.955401031 | 0.160677541 | 0.717054075 |
| Dhx29 | S | 27 | 0.120985082 | -0.105839485 | -0.2284622 | -0.349447283 | -0.122622716 | 0.050870042 | 0.073879696 | 0.130590303 | 0.066492078 | 0.341028473 |
| Fam160a2 | S | 510 | -0.030178514 | -0.016378358 | -0.113161497 | -0.082982983 | -0.096783139 | 0.625985066 | 0.874514935 | 0.17971852 | 0.249772288 | 0.381158325 |
| Ppp4r1 | S | 482 | -0.111515541 | -0.013632246 | -0.022870405 | 0.088645136 | -0.009238159 | 0.153520284 | 0.674364212 | 0.777744998 | 0.37058119 | 0.911156508 |
| Map1a | S | 2212 | -0.057400298 | 0.075689616 | -0.177638316 | -0.120238018 | -0.253327932 | 0.584278055 | 0.367838664 | 0.060608075 | 0.290623762 | 0.031201735 |
| Eri1 | S | 30 | -0.066627733 | -0.029691288 | -0.011690938 | 0.054936795 | 0.018000349 | 0.072952319 | 0.490428474 | 0.855083161 | 0.423342465 | 0.792269982 |
| Cttnbp2nl | S | 555 | -0.127580561 | -0.089576859 | 0.032629522 | 0.160210083 | 0.122206382 | 0.243686835 | 0.467918607 | 0.824426363 | 0.350687199 | 0.487720382 |
| Smg6 | S | 1190 | -0.083599057 | -0.014513502 | -0.02762298 | 0.055976077 | -0.013109478 | 0.11852245 | 0.719542366 | 0.623434454 | 0.371721387 | 0.812792046 |
| Smg6 | S | 1195 | -0.083599057 | -0.014513502 | -0.02762298 | 0.055976077 | -0.013109478 | 0.11852245 | 0.719542366 | 0.623434454 | 0.371721387 | 0.812792046 |
| Cwc25 | S | 258 | 0.066609487 | -0.084567089 | -0.140921261 | -0.207530748 | -0.056354172 | 0.306142188 | 0.280131693 | 0.109727095 | 0.049807715 | 0.523365487 |
| Cwc25 | S | 260 | 0.066609487 | -0.084567089 | -0.140921261 | -0.207530748 | -0.056354172 | 0.306142188 | 0.280131693 | 0.109727095 | 0.049807715 | 0.523365487 |
| Naa38 | S | 24 | 0.018224146 | -0.077980263 | -0.043286505 | -0.06151065 | 0.034693758 | 0.712055565 | 0.071768506 | 0.292709797 | 0.189028307 | 0.161398836 |
| Rnf214 | S | 48 | -0.085214096 | -0.032897703 | -0.064699515 | 0.020514581 | -0.031801813 | 0.432566356 | 0.673674086 | 0.46079121 | 0.84855502 | 0.703885114 |
| Samd4 | S | 579 | 0.07335271 | -0.190201813 | -0.005868561 | -0.079221271 | 0.184333251 | 0.467035398 | 0.028203488 | 0.948555668 | 0.473630556 | 0.093187185 |
| Phldb2 | S | 333 | 0.093849972 | -0.092458175 | -0.090428898 | -0.18427887 | 0.002029277 | 0.351109028 | 0.0843981 | 0.117075219 | 0.122920893 | 0.96226446 |
| Sorbs1 | T | 286 | -0.080719168 | 0.01351403 | -0.09445943 | -0.013740263 | -0.10797346 | 0.335068111 | 0.824190809 | 0.256214284 | 0.869633112 | 0.177073837 |
| Soga1 | S | 926 | -0.062676455 | 0.038084612 | -0.099189647 | -0.036513192 | -0.137274259 | 0.162748307 | 0.705673599 | 0.228766812 | 0.59901222 | 0.255985345 |
| Cactin | S | 94 | -0.043925945 | -0.01190988 | -0.09938687 | -0.055460924 | -0.087476989 | 0.468056023 | 0.879981216 | 0.228777006 | 0.475869566 | 0.358000387 |
| Ctnna1 Ctnna2 | S | 655;654 | -0.021196295 | -0.021302737 | -0.051106818 | -0.029910522 | -0.02980408 | 0.445118663 | 0.421542492 | 0.107384953 | 0.274515015 | 0.255700012 |
| Ncoa6 | S | 2047 | -0.045876377 | -0.04317633 | -0.071606893 | -0.025730516 | -0.028430563 | 0.498672414 | 0.435119123 | 0.461117007 | 0.789577109 | 0.753468741 |
| Naa38 | S | 23 | 0.025133789 | -0.083971024 | -0.040595131 | -0.06572892 | 0.043375893 | 0.562283905 | 0.050052551 | 0.295791113 | 0.103574137 | 0.076730999 |
| Cdk14 | S | 123 | -0.094117936 | -0.012995109 | -0.01444027 | 0.079677666 | -0.001445161 | 0.072185711 | 0.796791434 | 0.83359068 | 0.252006146 | 0.981152095 |
| Mindy4 | S | 143 | -0.21592288 | 0.165701081 | -0.133342856 | 0.082580024 | -0.299043937 | 0.037663254 | 0.163156867 | 0.488613302 | 0.64769089 | 0.179274879 |
| Sfr1 | S | 120 | 0.086684714 | -0.051228492 | -0.170202362 | -0.256887076 | -0.11897387 | 0.090159387 | 0.297358297 | 0.021268494 | 0.008783455 | 0.068800616 |
| Msn | S | 74 | -0.154826412 | 0.008165143 | -0.030269713 | 0.1245567 | -0.038434856 | 0.166841189 | 0.918603964 | 0.711125474 | 0.278026865 | 0.68254792 |
| Srrm2 | S | 1418 | 0.031793402 | -0.091358276 | -0.059270272 | -0.091063673 | 0.032088004 | 0.393367554 | 0.029806333 | 0.483822651 | 0.317686482 | 0.694478768 |
| Kif1b | S | 1658 | -0.08458139 | 0.124277573 | -0.175318431 | -0.090737041 | -0.299596004 | 0.28251357 | 0.184885155 | 0.046769096 | 0.287092784 | 0.023252661 |
| Irf9 | S | 393 | -0.160770664 | 0.021260869 | -0.0037746 | 0.156996064 | -0.025035469 | 0.054950476 | 0.672177448 | 0.971711267 | 0.211828882 | 0.804509973 |

| | | | | | | | | | | | | |
|---|---|---|---|---|---|---|---|---|---|---|---|---|
| Ubr5 | T | 2264 | 0.047108459 | 0.039677934 | -0.178634174 | -0.225742632 | -0.218312108 | 0.52476234 | 0.480480049 | 0.006665036 | 0.042021004 | 0.009300716 |
| Palld | S | 1004 | -0.01156004 | -0.047738099 | -0.054059059 | -0.042499019 | -0.00632096 | 0.764228061 | 0.386533179 | 0.129694965 | 0.189922362 | 0.889587621 |
| Epb41l2 | S | 116 | -0.029375253 | -0.086682757 | 0.00739798 | 0.036773233 | 0.094080737 | 0.43277504 | 0.078433668 | 0.915206555 | 0.59107326 | 0.236343016 |
| Pakap | S | 768 | 0.082510291 | -0.063888462 | -0.104021274 | -0.186531565 | -0.040132811 | 0.063708353 | 0.108525745 | 0.008892188 | 0.006366816 | 0.279492565 |
| Atxn2 | S | 653 | 0.116513246 | -0.115374105 | -0.156841478 | -0.273354724 | -0.041467373 | 0.042062138 | 0.036291495 | 0.10875855 | 0.031217355 | 0.592026201 |
| Uhrf1 | S | 93 | -0.01139519 | -0.067342635 | -0.068567213 | -0.057172023 | -0.001224578 | 0.883128967 | 0.207076411 | 0.439457854 | 0.562507483 | 0.987749396 |
| Fxr2 | S | 534 | -0.271261268 | 0.179573611 | -0.051422773 | 0.219838496 | -0.230996383 | 0.023383164 | 0.156455599 | 0.579533195 | 0.052979078 | 0.088027758 |
| Specc1l | S | 113 | -0.173101466 | -0.008574366 | 0.009333412 | 0.182434878 | 0.017907778 | 0.121814115 | 0.945615911 | 0.93014939 | 0.072300817 | 0.870265343 |
| Chd4 | T | 1538 | -0.120431471 | 0.018139558 | -0.025125651 | 0.09530582 | -0.043265209 | 0.076912454 | 0.729540532 | 0.567601986 | 0.14799072 | 0.453089026 |
| Paf1 | S | 439 | 0.072579518 | -0.226629725 | -0.078574925 | -0.151154443 | 0.1480548 | 0.648593624 | 0.238594453 | 0.582596915 | 0.439549991 | 0.461118579 |
| Paf1 | S | 440 | 0.072579518 | -0.226629725 | -0.078574925 | -0.151154443 | 0.1480548 | 0.648593624 | 0.238594453 | 0.582596915 | 0.439549991 | 0.461118579 |
| Zc3h14 | S | 309 | 0.063533189 | -0.110978641 | -0.085798997 | -0.149332187 | 0.025179643 | 0.224534276 | 0.04236005 | 0.316705998 | 0.132816485 | 0.749162273 |
| Usp6nl | S | 433 | -0.143189074 | 0.068760614 | -0.056391014 | 0.08679806 | -0.125151628 | 0.136164822 | 0.435240079 | 0.197310939 | 0.305201732 | 0.204964776 |
| Map3k2 | S | 331 | 0.016959122 | 0.009762165 | -0.111565306 | -0.128524427 | -0.12132747 | 0.707143467 | 0.742765082 | 0.008679884 | 0.042334013 | 0.005436752 |
| Mtss2 | S | 542 | -0.103899991 | 0.112411539 | -0.171928361 | -0.06802837 | -0.2843399 | 0.30011316 | 0.244818438 | 0.091503535 | 0.406876396 | 0.013900856 |
| Arhgap17 | T | 740 | -0.041841312 | -0.022852163 | -0.048188321 | -0.00634701 | -0.025336159 | 0.331276036 | 0.544860932 | 0.234724832 | 0.883524464 | 0.538191317 |
| Gnl3 | S | 95 | -0.039500074 | -0.082169618 | 0.022305634 | 0.061805708 | 0.104475252 | 0.393737342 | 0.030783717 | 0.466618634 | 0.188862113 | 0.001234846 |
| Smarca4 | S | 1389 | 0.011670729 | -0.103064718 | -0.077715374 | -0.089386103 | 0.025349343 | 0.934903421 | 0.450978774 | 0.230741288 | 0.557858039 | 0.847199923 |
| Crk | S | 83 | -0.02660785 | -0.02704942 | -0.040443099 | -0.013835249 | -0.013393678 | 0.663824768 | 0.294658067 | 0.093181938 | 0.811625764 | 0.43407189 |
| Map1b | S | 1334 | -0.142091292 | 0.052920454 | -0.071205479 | 0.070885813 | -0.124125933 | 0.04464641 | 0.142734668 | 0.409433899 | 0.430288613 | 0.203435307 |
| Srrm2 | S | 224 | 0.141097302 | -0.256221388 | -0.091580889 | -0.23267819 | 0.1646405 | 0.13371855 | 0.151464996 | 0.129022549 | 0.052968855 | 0.30129007 |
| Larp1 | S | 498 | 0.096303176 | -0.031815142 | -0.150079865 | -0.246383041 | -0.118264723 | 0.238963051 | 0.484077188 | 0.011985622 | 0.036752969 | 0.017310771 |
| Ap1g1 | S | 764 | 0.036152577 | -0.104837736 | -0.035319667 | -0.071472244 | 0.069518069 | 0.197341107 | 0.008111811 | 0.632366645 | 0.364925659 | 0.376696845 |
| Mxra7 | T | 102 | 0.095702231 | -0.076545333 | -0.102916296 | -0.198618527 | -0.026370963 | 0.085242866 | 0.031273141 | 0.04456697 | 0.012551324 | 0.478558649 |
| Nes | S | 632 | 0.056295644 | -0.129453182 | -0.05534451 | -0.111640154 | 0.074108672 | 0.563460592 | 0.087741876 | 0.437772338 | 0.250712441 | 0.182581289 |
| Safb2 | T | 593 | 0.021254372 | -0.060584668 | -0.061818803 | -0.083073175 | -0.001234136 | 0.703147529 | 0.12955041 | 0.172552674 | 0.183230083 | 0.970877253 |
| Rps29 | S | 9 | -0.056919801 | -0.130044409 | 0.081973417 | 0.138893218 | 0.212017826 | 0.239687842 | 0.027524561 | 0.181276984 | 0.047982287 | 0.013918649 |
| Trip12 | S | 310 | 0.016497273 | -0.08311932 | -0.067284982 | -0.083782255 | 0.015834338 | 0.835125148 | 0.133131701 | 0.427509709 | 0.432774054 | 0.849841139 |
| Dnttip2 | S | 178 | -0.011006755 | -0.039901859 | -0.083198699 | -0.072191945 | -0.04329684 | 0.72897973 | 0.397705637 | 0.244022362 | 0.300156904 | 0.535601176 |
| Cobll1 | S | 366 | -0.073634548 | 0.023768747 | -0.019276313 | 0.054358235 | -0.04304506 | 0.007977245 | 0.442085139 | 0.382182135 | 0.017185203 | 0.179381246 |
| Arhgap11a | S | 758 | -0.010341769 | 0.003121801 | -0.079796672 | -0.069454903 | -0.082918473 | 0.76680076 | 0.954101106 | 0.021656271 | 0.078148166 | 0.195563495 |
| Ptpn1 | S | 50 | 0.173494278 | -0.138747042 | -0.143586923 | -0.317081201 | -0.004839881 | 0.012001545 | 0.011942931 | 0.037919824 | 0.003753051 | 0.921658581 |
| Nde1 | S | 330 | 0.113058232 | -0.05825741 | -0.18627492 | -0.299333152 | -0.12801751 | 0.008360411 | 0.100297864 | 0.00691372 | 0.003793025 | 0.038583674 |
| Ssh2 | T | 1422 | 0.007103854 | -0.031475789 | -0.117286236 | -0.12439009 | -0.085810447 | 0.888237065 | 0.569689439 | 0.130671391 | 0.127460138 | 0.26473298 |
| Senp3 | S | 67 | 0.008914269 | 0.001221863 | -0.183816539 | -0.192730808 | -0.185038403 | 0.862006782 | 0.974558871 | 0.111884672 | 0.102026206 | 0.119842603 |
| Fam76a | S | 306 | 0.035446369 | -0.051890156 | -0.135267522 | -0.170713891 | -0.083377366 | 0.603909855 | 0.376935233 | 0.156626059 | 0.115948927 | 0.351207171 |
| Thoc2 | S | 1578 | 0.080892422 | -0.083526907 | -0.217896491 | -0.298788913 | -0.134369584 | 0.639092085 | 0.637436646 | 0.204254558 | 0.174286618 | 0.506355814 |
| Clip1 | Y | 1315 | -0.036857698 | -0.06850728 | -0.083001586 | -0.046143887 | -0.014494305 | 0.572978726 | 0.622154141 | 0.419991843 | 0.593298839 | 0.919066042 |
| Macroh2a1 | S | 138 | 0.122391899 | -0.125571551 | -0.224668399 | -0.347060298 | -0.099096848 | 0.424549277 | 0.403035889 | 0.226374982 | 0.059703453 | 0.453413165 |
| Zfp592 | S | 1089 | -0.051971644 | -0.067954402 | -0.033065656 | 0.018905988 | 0.034888746 | 0.462416688 | 0.384806055 | 0.568080098 | 0.694000767 | 0.558595697 |
| Abca1 | S | 2234 | -0.060056953 | -0.080172174 | 0.021144023 | 0.081200975 | 0.101316197 | 0.282109905 | 0.146726455 | 0.75872923 | 0.245157093 | 0.16560115 |
| Ppig | S | 672 | 0.038197259 | -0.168192115 | -0.0619851 | -0.100182359 | 0.106207015 | 0.805790338 | 0.142322204 | 0.824446817 | 0.728251415 | 0.697934102 |
| Ahctf1 | T | 1979 | -0.031537128 | -0.10626279 | -0.015050057 | 0.016487071 | 0.091212733 | 0.658610648 | 0.177299225 | 0.859777898 | 0.83793014 | 0.304712401 |
| Ptpn21 | S | 590 | -0.028151062 | -0.064960724 | -0.012071353 | 0.016079709 | 0.052889371 | 0.441853094 | 0.105597923 | 0.761820695 | 0.59059315 | 0.136447453 |
| Gpbp1 | S | 276 | -0.033794679 | -0.06838057 | -0.002148439 | 0.03164624 | 0.066232131 | 0.46370725 | 0.074744779 | 0.96515431 | 0.557926214 | 0.211974478 |
| Top2a | T | 1245 | -0.028543571 | 0.006792593 | -0.078245748 | -0.049702178 | -0.085038341 | 0.269734368 | 0.777539676 | 0.08530968 | 0.218150248 | 0.075829842 |
| Lmnb1 Lmnb2 | S | 392;383 | -0.051396574 | -0.022811169 | -0.014309213 | 0.037087362 | 0.008501956 | 0.225563929 | 0.178070232 | 0.394404667 | 0.34821869 | 0.616238689 |
| Cit | S | 1999 | 0.041563834 | -0.026238603 | -0.082908587 | -0.124472422 | -0.056669984 | 0.578492435 | 0.105318572 | 0.050803242 | 0.176066632 | 0.138892928 |
| Map1b | S | 2083 | -0.13569878 | -0.04969419 | 0.034031352 | 0.169730132 | 0.083725542 | 0.116448332 | 0.411967152 | 0.466314465 | 0.067783445 | 0.242461748 |
| Wdr37 | S | 30 | -0.10017882 | 0.03076473 | -0.062539306 | 0.037639514 | -0.093304036 | 0.137396436 | 0.646339686 | 0.343272042 | 0.307242516 | 0.090091533 |
| Vim | S | 42 | -0.003641736 | -0.051300094 | -0.072913559 | -0.069271823 | -0.021613464 | 0.960298356 | 0.314154825 | 0.221305516 | 0.33724541 | 0.569090547 |
| Cdca2 | S | 133 | 0.002043559 | -0.032134133 | -0.084299713 | -0.086343272 | -0.05216558 | 0.958620176 | 0.473653263 | 0.098963994 | 0.081013333 | 0.265085656 |
| Lamtor1 | T | 28 | 0.005702762 | -0.074067909 | -0.035264273 | -0.040967035 | 0.038803636 | 0.8050745 | 0.070351347 | 0.404780543 | 0.317748956 | 0.383317146 |
| Cpne1 | T | 525 | -0.058902819 | -0.006684279 | -0.077162609 | -0.018259791 | -0.07047833 | 0.247906739 | 0.868554787 | 0.392610328 | 0.830111919 | 0.425684525 |
| Fam122a | S | 267 | -0.089172265 | -0.005664853 | -0.000959388 | 0.088212878 | 0.004705466 | 0.031700118 | 0.879892476 | 0.981829396 | 0.066837977 | 0.900373537 |
| Exoc7 | S | 243 | -0.009585518 | -0.035618511 | -0.093068189 | -0.08348267 | -0.057449678 | 0.852026256 | 0.50147208 | 0.195266231 | 0.240332409 | 0.396301412 |
| Casc3 | S | 10 | 0.085206567 | -0.168353405 | -0.119573271 | -0.204779838 | 0.048780134 | 0.29309494 | 0.202696583 | 0.279107634 | 0.08965216 | 0.694311013 |
| Rnaseh2b | S | 33 | -0.045905313 | -0.005494435 | -0.092131245 | -0.046225933 | -0.08663681 | 0.392482036 | 0.888541179 | 0.258784662 | 0.558924754 | 0.283743063 |
| Cttnbp2nl | S | 548 | -0.153114487 | -0.091109893 | 0.121657589 | 0.274772076 | 0.212767482 | 0.001499541 | 0.132984133 | 0.002985425 | 0.000408771 | 0.024138015 |
| Arhgap17 | T | 730 | -0.039496472 | -0.025090975 | -0.04791126 | -0.008414788 | -0.022820285 | 0.36756197 | 0.516190898 | 0.249769578 | 0.855342886 | 0.602493311 |
| Ddx24 | S | 857 | 0.048650438 | -0.074147902 | -0.065693171 | -0.11434361 | 0.008454731 | 0.582144869 | 0.147038566 | 0.123705566 | 0.253647782 | 0.851583504 |
| Irf2bp2 | S | 250 | -0.016083481 | -0.050730147 | -0.037949821 | -0.021866341 | 0.012780325 | 0.595779613 | 0.219437229 | 0.290286305 | 0.568295473 | 0.762461447 |
| Pak1 | S | 115 | -0.072456819 | 0.110086911 | -0.173763582 | -0.101306763 | -0.283850492 | 0.384262841 | 0.302477501 | 0.067359862 | 0.181135959 | 0.035556722 |
| Scrib | S | 506 | -0.013427489 | -0.046364587 | -0.059187778 | -0.045760289 | -0.012823192 | 0.783508466 | 0.332673496 | 0.214583401 | 0.411019446 | 0.798248851 |
| Atrx | S | 912 | -0.080824616 | -0.066013275 | 0.047024281 | 0.127848897 | 0.113037555 | 0.135631793 | 0.21151567 | 0.670083735 | 0.300570446 | 0.348760433 |
| Jph2 | S | 162 | -0.140093153 | -0.124765735 | 0.10776534 | 0.247858493 | 0.232531075 | 0.200730178 | 0.183729232 | 0.32215415 | 0.046477708 | 0.049747549 |
| Dab2 | S | 326 | 0.151262787 | -0.105899118 | -0.236181576 | -0.387444364 | -0.130282458 | 0.065897973 | 0.358810395 | 0.033455236 | 0.0089174 | 0.294815451 |
| Anapc4 | S | 757 | -0.083888961 | -0.05696906 | 0.013414177 | 0.097303138 | 0.070383237 | 0.158048407 | 0.396749198 | 0.890625605 | 0.355875787 | 0.502044866 |
| Anapc4 | S | 758 | -0.083888961 | -0.05696906 | 0.013414177 | 0.097303138 | 0.070383237 | 0.158048407 | 0.396749198 | 0.890625605 | 0.355875787 | 0.502044866 |
| Phip | S | 136 | -0.06567263 | -0.016704282 | -0.022913427 | 0.042759203 | -0.006209144 | 0.041153657 | 0.842155923 | 0.796946456 | 0.632664695 | 0.955571278 |
| Prkd1 | S | 234 | -0.05764387 | -0.009502634 | -0.10058862 | -0.04294475 | -0.091085986 | 0.404222501 | 0.926191175 | 0.343791886 | 0.676258936 | 0.480694168 |
| Pkn2 | S | 305 | 0.008788442 | -0.023081708 | -0.143947424 | -0.152735866 | -0.120865716 | 0.881677397 | 0.622336678 | 0.162277276 | 0.149028356 | 0.222044966 |
| Ppp1r9b | S | 192 | 0.149305476 | -0.114954264 | -0.249174173 | -0.398479649 | -0.134219909 | 0.567692284 | 0.603761797 | 0.20149884 | 0.199362507 | 0.579404644 |
| Ppp1r9b | S | 205 | 0.149305476 | -0.114954264 | -0.249174173 | -0.398479649 | -0.134219909 | 0.567692284 | 0.603761797 | 0.20149884 | 0.199362507 | 0.579404644 |
| Alkbh5 | S | 65 | -0.010199026 | -0.106679232 | 0.019459726 | 0.029658752 | 0.126138958 | 0.756048603 | 0.024700465 | 0.598914389 | 0.224047813 | 0.010945163 |
| Alkbh5 | S | 70 | -0.010199026 | -0.106679232 | 0.019459726 | 0.029658752 | 0.126138958 | 0.756048603 | 0.024700465 | 0.598914389 | 0.224047813 | 0.010945163 |
| Syne1 | S | 5890 | -0.069958682 | -0.046276351 | -0.022712742 | 0.04724594 | 0.023563609 | 0.291508083 | 0.328882393 | 0.847780742 | 0.699798832 | 0.839021905 |
| Miga2 | S | 113 | 0.071489232 | -0.04073468 | -0.253412107 | -0.324901339 | -0.212677426 | 0.10842296 | 0.600109875 | 0.063418442 | 0.053027558 | 0.102910738 |
| Fez2 | S | 200 | -0.069167588 | 0.079129968 | -0.095297551 | -0.026129963 | -0.174427519 | 0.073688697 | 0.115052772 | 0.033046898 | 0.482133193 | 0.015031551 |
| Arhgap23 | S | 515 | -0.059905526 | -0.087073817 | 0.071392395 | 0.131297921 | 0.158466213 | 0.057889736 | 0.032147736 | 0.133313853 | 0.039366899 | 0.01716241 |
| H1f2 H1f3 H1f4 H1f5 | S | 102;102;104;102; | -0.038866034 | 0.06250349 | -0.178190472 | -0.139324438 | -0.240693961 | 0.760190292 | 0.659882093 | 0.101287159 | 0.258752939 | 0.133991816 |
| Stx16 | T | 71 | -0.079695539 | -0.012898451 | -0.10765175 | -0.02795621 | -0.094753298 | 0.327589009 | 0.886650157 | 0.579211261 | 0.883189214 | 0.63338271 |
| Leo1 | S | 198 | -0.050571691 | -0.05999931 | 0.059215997 | 0.109787689 | 0.119215308 | 0.178679388 | 0.017247174 | 0.479066874 | 0.243961295 | 0.22041427 |
| Sh3bp4 | S | 636 | -0.146942654 | -0.023609796 | -0.063070047 | 0.083872607 | -0.039460251 | 0.428772767 | 0.826991781 | 0.667647602 | 0.644983096 | 0.731439794 |
| Specc1 | T | 809 | -0.002213789 | -0.088503239 | -0.010247492 | -0.008033703 | 0.078255747 | 0.958472489 | 0.047865669 | 0.777305171 | 0.769305519 | 0.010446634 |
| Prkab2 | S | 183 | -0.009314816 | -0.02850148 | -0.090034164 | -0.080719348 | -0.061532683 | 0.792384189 | 0.474237752 | 0.189928536 | 0.232616635 | 0.346735793 |
| Slf2 | S | 719 | -0.091120159 | -0.028085435 | -0.093534669 | -0.00241451 | -0.065449235 | 0.628074656 | 0.708790333 | 0.448726758 | 0.990252135 | 0.582399086 |
| Tacc2 | S | 510 | -0.043750097 | -0.058296384 | 0.04307469 | 0.086824787 | 0.101371074 | 0.066976456 | 0.018570147 | 0.238649187 | 0.061817578 | 0.053508853 |
| Tacc2 | S | 514 | -0.043750097 | -0.058296384 | 0.04307469 | 0.086824787 | 0.101371074 | 0.066976456 | 0.018570147 | 0.238649187 | 0.061817578 | 0.053508853 |
| Cep170 | S | 1528 | -0.02192633 | 0.013382136 | -0.119630826 | -0.097704496 | -0.133012962 | 0.879444339 | 0.772354054 | 0.058260387 | 0.526809866 | 0.064606369 |
| Numb | S | 440 | -0.019207281 | -0.024849914 | -0.0798133 | -0.060606019 | -0.054963385 | 0.749276043 | 0.524201711 | 0.169795361 | 0.377621411 | 0.29886628 |
| Med12 | S | 701 | -0.090863781 | -0.115701703 | 0.092365356 | 0.183229137 | 0.208067059 | 0.24314739 | 0.088585989 | 0.351327286 | 0.11053482 | 0.085748129 |
| Clasrp | S | 527 | 0.038990086 | -0.081615324 | -0.081220597 | -0.120210683 | 0.000394727 | 0.542192381 | 0.19121234 | 0.195302703 | 0.037620446 | 0.988233584 |
| Tab3 | S | 126 | 0.048101495 | -0.074656434 | -0.23177349 | -0.279874985 | -0.157117056 | 0.561688341 | 0.398519644 | 0.368380114 | 0.296259735 | 0.520868015 |

| | | | | | | | | | | | | |
|---|---|---|---|---|---|---|---|---|---|---|---|---|
| Lmna | T | 394 | 0.091042943 | -0.031903868 | -0.127798226 | -0.21884117 | -0.095894358 | 0.069388203 | 0.331544449 | 0.002290018 | 0.006775104 | 0.033013167 |
| Zc3h11a | S | 677 | 0.031811171 | -0.149168843 | -0.019973085 | -0.051784255 | 0.129195758 | 0.70305466 | 0.083150241 | 0.765968199 | 0.561144261 | 0.146771655 |
| Epha2 | S | 893 | 0.00363852 | 0.01826533 | -0.233748317 | -0.237386838 | -0.252013647 | 0.88950886 | 0.728633317 | 0.14425602 | 0.136788574 | 0.113588136 |
| Mkrn2 | S | 139 | -0.007592804 | -0.040584492 | -0.045817538 | -0.038224734 | -0.005233046 | 0.734164448 | 0.248448722 | 0.161088664 | 0.256229748 | 0.884500149 |
| Mybl1 | S | 5 | -0.131041144 | -0.005099849 | 0.026513995 | 0.157555139 | 0.031613844 | 0.027126321 | 0.95005405 | 0.535843825 | 0.017238175 | 0.692673109 |
| Mybl1 | S | 7 | -0.131041144 | -0.005099849 | 0.026513995 | 0.157555139 | 0.031613844 | 0.027126321 | 0.95005405 | 0.535843825 | 0.017238175 | 0.692673109 |
| Clasp2 | S | 884 | 0.207640077 | -0.031779798 | -0.285114407 | -0.492754484 | -0.253334609 | 0.143585903 | 0.633194641 | 0.015585285 | 0.01697914 | 0.048537425 |
| Ppm1a | S | 375 | -0.149085162 | 0.049126481 | -0.047867701 | 0.101217462 | -0.096994182 | 0.084867866 | 0.617966282 | 0.731978098 | 0.453627696 | 0.495426448 |
| Ctr9 | S | 941 | -0.014355335 | -0.067515972 | -0.011078041 | 0.003277294 | 0.056437931 | 0.758073537 | 0.046904589 | 0.744247836 | 0.948001055 | 0.166860807 |
| Ctr9 | S | 943 | -0.014355335 | -0.067515972 | -0.011078041 | 0.003277294 | 0.056437931 | 0.758073537 | 0.046904589 | 0.744247836 | 0.948001055 | 0.166860807 |
| Incenp | S | 796 | -0.05087646 | -0.020054869 | -0.005360666 | 0.045515794 | 0.014694203 | 0.02644333 | 0.474133233 | 0.909863954 | 0.376085256 | 0.766650245 |
| Incenp | S | 799 | -0.05087646 | -0.020054869 | -0.005360666 | 0.045515794 | 0.014694203 | 0.02644333 | 0.474133233 | 0.909863954 | 0.376085256 | 0.766650245 |
| Incenp | T | 800 | -0.05087646 | -0.020054869 | -0.005360666 | 0.045515794 | 0.014694203 | 0.02644333 | 0.474133233 | 0.909863954 | 0.376085256 | 0.766650245 |
| Ncoa5 | S | 96 | -0.069952299 | -0.04773307 | 0.014905555 | 0.084857854 | 0.062638625 | 0.153109941 | 0.24490237 | 0.863711802 | 0.38338256 | 0.501321243 |
| Dtd1 | S | 197 | 0.045253968 | 0.05697813 | -0.236016875 | -0.281270844 | -0.292995006 | 0.895737929 | 0.665310128 | 0.067500825 | 0.450613651 | 0.083948973 |
| Camk1 | S | 324 | 0.015393688 | -0.062518481 | -0.164301972 | -0.179695659 | -0.10178349 | 0.924095301 | 0.533310211 | 0.357267555 | 0.390667822 | 0.541344717 |
| Dmap1 | T | 446 | -0.105297999 | -0.067628702 | 0.007489243 | 0.112787242 | 0.075117945 | 0.408160372 | 0.275786455 | 0.898857235 | 0.367053981 | 0.053760667 |
| Npm1 | S | 88 | 0.036193934 | -0.116365778 | -0.114005901 | -0.150199836 | 0.002359876 | 0.739343238 | 0.467182324 | 0.279909171 | 0.276265172 | 0.988411485 |
| Ambra1 | S | 1206 | -0.033015408 | -0.04229614 | -0.056080721 | -0.023065313 | -0.013784581 | 0.485704863 | 0.376161443 | 0.435428412 | 0.726717426 | 0.833136703 |
| Ccdc12 | S | 165 | -0.021907972 | -0.020093088 | -0.041692716 | -0.019784744 | -0.021599628 | 0.630942542 | 0.125213802 | 0.176529454 | 0.683501273 | 0.415868665 |
| Epb41 | T | 704 | -0.106753516 | 0.078102647 | -0.182545253 | -0.075791737 | -0.2606479 | 0.412324979 | 0.673217868 | 0.343349101 | 0.653631307 | 0.252028065 |
| Lemd2 | S | 145 | -0.150339265 | -0.053575431 | 0.091325588 | 0.241664852 | 0.144901018 | 0.135686692 | 0.131339341 | 0.36262125 | 0.078600381 | 0.199934669 |
| Sec62 | T | 158 | -0.038995825 | -0.093339696 | 0.034792308 | 0.073788134 | 0.128132004 | 0.452655994 | 0.054315084 | 0.641390171 | 0.376830476 | 0.160566634 |
| Cep170b | S | 509 | 0.133328107 | -0.054246483 | -0.201752205 | -0.335080312 | -0.147505722 | 0.295601073 | 0.577605007 | 0.040798259 | 0.046271421 | 0.121191143 |
| Zswim8 | S | 53 | -0.090153689 | 0.028124015 | -0.047707635 | 0.042446054 | -0.07583165 | 0.150128835 | 0.649795969 | 0.218425928 | 0.445070593 | 0.272731033 |
| Stau2 | S | 489 | 0.001817519 | 0.019095225 | -0.120787284 | -0.122604803 | -0.139882509 | 0.967296753 | 0.770638728 | 0.030092496 | 0.064318822 | 0.11449947 |
| Pdlim2 | S | 209 | -0.135612249 | -0.042109832 | -0.042286617 | 0.093325632 | -0.000176785 | 0.469437987 | 0.678519997 | 0.696888272 | 0.613559922 | 0.998680231 |
| Sh3kbp1 | S | 454 | -0.033265044 | 0.003835552 | -0.09182008 | -0.058555035 | -0.095655632 | 0.572063273 | 0.945267232 | 0.110424784 | 0.342782455 | 0.143703726 |
| Iws1 | S | 196 | -0.038843833 | -0.095656814 | 0.049404297 | 0.08824813 | 0.145061111 | 0.330722457 | 0.042858326 | 0.500378572 | 0.256319873 | 0.112550058 |
| Ptbp2 | S | 26 | -0.090479438 | -0.236224517 | 0.112793467 | 0.203272906 | 0.349017984 | 0.657996396 | 0.220822933 | 0.649082399 | 0.360680016 | 0.166390329 |
| Trps1 | S | 1081 | -0.018270078 | -0.092758951 | 0.015077869 | 0.033347947 | 0.10783682 | 0.662894946 | 0.033892722 | 0.680786909 | 0.308520993 | 0.005504362 |
| Trps1 | S | 1085 | -0.018270078 | -0.092758951 | 0.015077869 | 0.033347947 | 0.10783682 | 0.662894946 | 0.033892722 | 0.680786909 | 0.308520993 | 0.005504362 |
| Dnajc2 | S | 47 | -0.082987283 | -0.064745453 | -0.006295388 | 0.076691894 | 0.058450064 | 0.488642292 | 0.236716707 | 0.928630409 | 0.543789281 | 0.452026958 |
| Ywhaq | S | 232 | -0.02846127 | -0.004257567 | -0.067937027 | -0.039475757 | -0.06367946 | 0.379048631 | 0.90053652 | 0.114944412 | 0.315669646 | 0.155092699 |
| Cdk13 | T | 495 | 0.000939911 | -0.116303728 | -0.017685724 | -0.018625635 | 0.098618005 | 0.980859076 | 0.109073169 | 0.776745003 | 0.752639902 | 0.218674869 |
| Dtl | S | 655 | 0.018037672 | -0.007122199 | -0.112198246 | -0.130235918 | -0.105076047 | 0.628550857 | 0.808307325 | 0.031609626 | 0.026261917 | 0.052220248 |
| Wbp1l | S | 177 | -0.214265784 | 0.028756109 | 0.040836868 | 0.255102652 | 0.01208076 | 0.03981858 | 0.605905952 | 0.613787853 | 0.036543287 | 0.8585873 |
| Erbin | T | 914 | -0.309093714 | 0.131436756 | -0.130405215 | 0.1786885 | -0.261841971 | 0.339078204 | 0.407324719 | 0.422481261 | 0.565108844 | 0.194162391 |
| Tmem131 | S | 1596 | -0.012255445 | -0.073501997 | -0.066115056 | -0.053859611 | 0.007386941 | 0.880524885 | 0.190148116 | 0.610720305 | 0.696605946 | 0.953015225 |
| Stim2 | S | 626 | -0.108733871 | 0.091829196 | -0.121486899 | -0.012753029 | -0.213316096 | 0.043120349 | 0.125761712 | 0.232498466 | 0.878977744 | 0.081607449 |
| Arfgap2 | S | 145 | -0.026918526 | -0.00849242 | -0.135207066 | -0.10828854 | -0.126714646 | 0.749561055 | 0.78897082 | 0.244865286 | 0.381732984 | 0.271843955 |
| Rpl13 | S | 139 | -0.084954838 | -0.018798732 | -0.040000851 | 0.044953987 | -0.021202119 | 0.168680653 | 0.814037358 | 0.707448814 | 0.670821491 | 0.854038 |
| Denn2b | S | 446 | 0.011845737 | -0.114126509 | -0.032978439 | -0.044824176 | 0.081148069 | 0.868761811 | 0.079464456 | 0.855852118 | 0.804447134 | 0.654052874 |
| Cep170b | T | 1124 | -0.018951842 | 0.029943551 | -0.124126114 | -0.105174272 | -0.154069665 | 0.633412579 | 0.496455392 | 0.045122856 | 0.086347994 | 0.033724527 |
| Nfix | S | 361 | 0.010400339 | -0.051949886 | -0.098831431 | -0.10923177 | -0.046881545 | 0.846444058 | 0.291255798 | 0.275761505 | 0.24482606 | 0.569556298 |
| Wasf2 | S | 473 | -0.006595016 | -0.043723122 | -0.087014678 | -0.080419662 | -0.043291556 | 0.947590703 | 0.541008567 | 0.18168424 | 0.433901498 | 0.51045383 |
| Ddx55 | S | 594 | 0.055269227 | -0.078837449 | -0.080222654 | -0.135491881 | -0.001385205 | 0.024496855 | 0.118939418 | 0.009545365 | 0.002722473 | 0.971730947 |
| Kctd15 | S | 38 | 0.069235823 | -0.168470355 | -0.078086187 | -0.147322011 | 0.090384168 | 0.487501385 | 0.217550853 | 0.357092345 | 0.117009279 | 0.43667026 |
| Phf8-ps Phf8 | S | 820 | -0.060887426 | -0.028381525 | 0.001180874 | 0.0620683 | 0.029562399 | 0.09576101 | 0.258869242 | 0.96588315 | 0.126276955 | 0.367298413 |
| AU022252 | T | 47 | -0.086051727 | -0.021402561 | -0.051788017 | 0.034263711 | -0.030385456 | 0.241171904 | 0.814416371 | 0.646002795 | 0.735907611 | 0.790527942 |
| Psmc1 | T | 434 | 0.017591285 | -0.067858957 | -0.064800616 | -0.082391901 | 0.00305834 | 0.704917675 | 0.196279535 | 0.221452539 | 0.187074161 | 0.9559309 |
| Aktip | T | 29 | 0.019923437 | -0.063088874 | -0.07708551 | -0.097008947 | -0.013996635 | 0.859184768 | 0.359791999 | 0.173457833 | 0.429296581 | 0.839817411 |
| Rps6ka1 Rps6ka3 Rps | T | 231;225;256 | -0.155033102 | -0.044269786 | -0.005162054 | 0.149871048 | 0.039107732 | 0.278291942 | 0.793414957 | 0.929789755 | 0.28937607 | 0.812904952 |
| Dus3l | S | 264 | -0.165264566 | -0.026833026 | -0.076085652 | 0.089178914 | -0.049252626 | 0.497816572 | 0.870967653 | 0.698953262 | 0.749462331 | 0.832785635 |
| Smg5 | S | 415 | 0.127159466 | -0.127679002 | -0.239220209 | -0.366379675 | -0.111541207 | 0.353185619 | 0.484190863 | 0.195417563 | 0.094816425 | 0.600749767 |
| Smg5 | T | 418 | 0.127159466 | -0.127679002 | -0.239220209 | -0.366379675 | -0.111541207 | 0.353185619 | 0.484190863 | 0.195417563 | 0.094816425 | 0.600749767 |
| Ahctf1 | S | 1930 | 0.055068677 | -0.074419023 | -0.124583878 | -0.179652555 | -0.050164855 | 0.566948254 | 0.550233272 | 0.103805925 | 0.119440914 | 0.679209991 |
| Nucks1 | S | 30 | -0.080758916 | 0.057030902 | -0.165632473 | -0.084873556 | -0.222663375 | 0.461640804 | 0.513279272 | 0.207461027 | 0.496427397 | 0.109773624 |
| Nolc1 | T | 504 | 0.139897375 | -0.102232202 | -0.162024311 | -0.301921687 | -0.05979211 | 0.170502448 | 0.216279281 | 0.064463863 | 0.035816521 | 0.1770367 |
| Prkd3 | S | 414 | 0.03919763 | -0.090500376 | -0.067876868 | -0.107074498 | 0.022623508 | 0.467195054 | 0.056824686 | 0.46549574 | 0.280635819 | 0.786034248 |
| Kmt2d | S | 1107 | -0.094523983 | -0.124702293 | 0.054498036 | 0.149022019 | 0.179200329 | 0.424516979 | 0.157503382 | 0.479480648 | 0.217450871 | 0.036495503 |
| Gpsm2 | S | 578 | -0.089309717 | -0.09483297 | 0.05823114 | 0.147540857 | 0.153064111 | 0.187734231 | 0.119979238 | 0.284797862 | 0.045686281 | 0.013174714 |
| Tmpo | S | 288 | -0.06514104 | -0.042444403 | 0.012885907 | 0.078026947 | 0.055330309 | 0.051349305 | 0.419552233 | 0.702361429 | 0.041832776 | 0.300936351 |
| Prrc2a | S | 1083 | 0.065370787 | -0.049044295 | -0.097060933 | -0.162431719 | -0.048016638 | 0.013263178 | 0.098473506 | 0.002670334 | 0.002242577 | 0.110999332 |
| Tlr7 | S | 301 | -0.088917321 | -0.022015122 | -0.034920723 | 0.053996598 | -0.012905601 | 0.184675577 | 0.813322364 | 0.713562516 | 0.584201076 | 0.911287911 |
| Hspd1 | S | 70 | 0.021927402 | -0.032620609 | -0.066576539 | -0.088503941 | -0.033955931 | 0.350680699 | 0.213639329 | 0.026879782 | 0.018112456 | 0.23187752 |
| Atad5 | S | 451 | -0.126084499 | 0.167489373 | -0.333622707 | -0.207538208 | -0.50111208 | 0.370735526 | 0.30236732 | 0.273834729 | 0.467063524 | 0.143238829 |
| Bcl9l | S | 702 | -0.06949116 | 0.042154153 | -0.139692072 | -0.070200912 | -0.181846224 | 0.250783958 | 0.457357344 | 0.258246444 | 0.522945628 | 0.173253203 |
| Nkap Nkapl | S | 147;141 | -0.045931905 | -0.099308595 | 0.067346199 | 0.113278103 | 0.166654794 | 0.34327172 | 0.050005979 | 0.539530865 | 0.328613274 | 0.19993704 |
| Osbpl6 | S | 45 | -0.00451469 | 0.070644114 | -0.228906896 | -0.224392206 | -0.299551009 | 0.951562098 | 0.488476263 | 0.07352384 | 0.084532211 | 0.045851476 |
| Piezo1 | S | 1389 | 0.04510295 | -0.09734323 | -0.102161234 | -0.147264184 | -0.004818004 | 0.40258222 | 0.228235277 | 0.232528335 | 0.116677986 | 0.955781365 |
| Rnaseh2b | S | 34 | -0.12289539 | 0.024123769 | -0.063864622 | 0.059030768 | -0.087988391 | 0.134279433 | 0.519817581 | 0.565278649 | 0.616614363 | 0.442301375 |
| Nfat5 | S | 618 | -0.05176247 | -0.057018591 | -0.034041624 | 0.017720846 | 0.022976967 | 0.444570476 | 0.467157456 | 0.523096807 | 0.771776512 | 0.750457139 |
| Osbpl8 | T | 39 | 0.028373055 | -0.072956175 | -0.093031672 | -0.121404726 | -0.020075496 | 0.701109806 | 0.467227452 | 0.141698496 | 0.186256611 | 0.839842667 |
| Pik3cb | S | 318 | -0.034133958 | -0.007534411 | -0.162109841 | -0.127975883 | -0.15457543 | 0.722675359 | 0.883562782 | 0.327384992 | 0.447949504 | 0.345893988 |
| Tgfb1i1 | S | 203 | 0.011865313 | -0.10964613 | -0.110498836 | -0.122364149 | -0.000852706 | 0.917043739 | 0.346467974 | 0.562050342 | 0.532254214 | 0.996373 |
| Sorbs1 | S | 470 | -0.080259569 | -0.067911307 | 0.0009818 | 0.081241368 | 0.068893107 | 0.424758814 | 0.260515577 | 0.989360075 | 0.452094032 | 0.379324318 |
| Nes | S | 813 | -0.060246973 | -0.042712069 | 0.011421741 | 0.071668714 | 0.054133809 | 0.09479123 | 0.273805002 | 0.793729034 | 0.168320605 | 0.290382362 |
| Atp5a1 | S | 76 | -0.020036181 | -0.070442841 | -0.053789468 | -0.033753286 | 0.016653373 | 0.701448859 | 0.445519455 | 0.349970846 | 0.416403868 | 0.841113294 |
| Igf2r | S | 2401 | -0.0433685 | -0.057986054 | -0.038116554 | 0.005251946 | 0.0198695 | 0.604967359 | 0.333680501 | 0.508653493 | 0.937252771 | 0.354518674 |
| Rlim | S | 212 | 0.133731183 | -0.100700337 | -0.067154467 | -0.20088565 | 0.03354587 | 0.200254162 | 0.013761245 | 0.134519927 | 0.09198538 | 0.405901623 |
| Rlim | S | 214 | 0.133731183 | -0.100700337 | -0.067154467 | -0.20088565 | 0.03354587 | 0.200254162 | 0.013761245 | 0.134519927 | 0.09198538 | 0.405901623 |
| Tbcel | S | 41 | 0.035436957 | -0.033819646 | -0.208495977 | -0.243932934 | -0.17467633 | 0.507685336 | 0.634109756 | 0.187531631 | 0.147959846 | 0.249785953 |
| Apc | S | 1370 | 0.020638916 | -0.034996398 | -0.12586959 | -0.146508506 | -0.090873192 | 0.781770805 | 0.648571607 | 0.12871362 | 0.008125928 | 0.061025234 |
| Prrc2a | S | 1108 | 0.006157686 | -0.077956861 | -0.024378329 | -0.030536016 | 0.053578532 | 0.843985288 | 0.069051042 | 0.471055463 | 0.410619258 | 0.207037665 |
| Mki67 | T | 492 | 0.190491146 | -0.097426072 | -0.224415264 | -0.41490641 | -0.126989192 | 0.07751271 | 0.18319352 | 0.039717771 | 0.007961386 | 0.167551699 |
| Atrx | S | 590 | -0.052336793 | -0.073154856 | 0.04118761 | 0.093524403 | 0.114342466 | 0.142181837 | 0.033559443 | 0.196732385 | 0.01930657 | 0.011439989 |
| Snx5 | S | 20 | -0.032082666 | 0.02299904 | -0.163837318 | -0.131754652 | -0.186836358 | 0.584114291 | 0.70932117 | 0.19640771 | 0.272516703 | 0.160629493 |
| Ptrh2 | S | 59 | -0.027861235 | -0.013748575 | -0.06333118 | -0.035469945 | -0.049582605 | 0.689081331 | 0.805647944 | 0.077602867 | 0.601127406 | 0.38153263 |
| Flna | S | 2327 | 0.013507975 | -0.044117941 | -0.042914692 | -0.056422667 | 0.001203249 | 0.590460947 | 0.09251396 | 0.093892892 | 0.083872019 | 0.959848167 |
| Sap30bp | Y | 17 | 0.203056621 | -0.199540832 | -0.298721826 | -0.501778447 | -0.099180994 | 0.402232235 | 0.331915943 | 0.40566176 | 0.221989138 | 0.7758575 |
| Tpx2 | S | 486 | 0.068903057 | -0.030052896 | -0.108347932 | -0.177250989 | -0.078295036 | 0.20652721 | 0.455939633 | 0.006904353 | 0.02868676 | 0.094023157 |
| Zfhx4 | S | 507 | -0.090881796 | -0.050115054 | 0.018930132 | 0.109811928 | 0.069045186 | 0.138965656 | 0.376044126 | 0.671351219 | 0.114861732 | 0.295448764 |
| Atxn2 | S | 678 | 0.014592647 | -0.192531523 | 0.008066502 | -0.006526145 | 0.200598025 | 0.949223031 | 0.151551267 | 0.962786598 | 0.978651205 | 0.261948139 |

| | | | | | | | | | | | | |
|---|---|---|---|---|---|---|---|---|---|---|---|---|
| Crtc2 | S | 434 | 0.010152452 | -0.095468895 | -0.032172873 | -0.042325325 | 0.063296023 | 0.880635513 | 0.091070542 | 0.662616669 | 0.622330459 | 0.413504516 |
| Gnl1 | S | 324 | -0.04685267 | 0.008042974 | -0.070648992 | -0.023796322 | -0.078691966 | 0.282204467 | 0.824151279 | 0.170761282 | 0.620948615 | 0.13584529 |
| Gsn | T | 384;385 | -0.080684337 | -0.018268873 | -0.047141999 | 0.033542338 | -0.028873126 | 0.266475928 | 0.696427353 | 0.623071033 | 0.733130978 | 0.743825886 |
| Eif4b | T | 500 | 0.018033398 | -0.013985725 | -0.080354092 | -0.09838749 | -0.066368367 | 0.579766351 | 0.560493462 | 0.018458608 | 0.043347181 | 0.058128553 |
| Cep170 | S | 872 | -0.032295876 | -0.037698893 | -0.03856286 | -0.006266984 | -0.000863967 | 0.259499401 | 0.358814474 | 0.549884207 | 0.918664195 | 0.989502993 |
| Prkab1 | S | 181 | -0.023970853 | -0.075106485 | -0.048218968 | -0.024248115 | 0.026887517 | 0.600073094 | 0.369991441 | 0.536828565 | 0.736917218 | 0.773414205 |
| Trps1 | S | 1066 | 0.004210961 | -0.063497148 | -0.039665982 | -0.043876943 | 0.023831166 | 0.881169779 | 0.127664809 | 0.300175407 | 0.303502582 | 0.578651288 |
| Utp18 | S | 202 | 0.052696203 | -0.087782044 | -0.055119308 | -0.107815512 | 0.032662736 | 0.042174379 | 0.02219938 | 0.067565762 | 0.009577477 | 0.278849611 |
| Phldb1 | S | 983 | 0.074771234 | -0.066877547 | -0.091160984 | -0.165932219 | -0.024283437 | 0.188006491 | 0.128141381 | 0.041852954 | 0.029284271 | 0.472759458 |
| Uhrf1bp1l | S | 1106 | -0.033371062 | -0.042992409 | -0.118789622 | -0.08541856 | -0.075797214 | 0.798348678 | 0.609031064 | 0.417122338 | 0.592093842 | 0.563720084 |
| Mis18bp1 | T | 983 | 0.04559517 | -0.15567131 | -0.10162995 | -0.147225119 | 0.054041361 | 0.704091208 | 0.431689539 | 0.367392116 | 0.31280852 | 0.781410337 |
| Alkbh5 | S | 253 | -0.032446579 | 0.020383692 | -0.199278462 | -0.166831883 | -0.219662154 | 0.807033263 | 0.770625549 | 0.227488017 | 0.359517554 | 0.197382194 |
| Top2b | S | 1537 | -0.063138473 | -0.068740834 | 0.066919829 | 0.130058302 | 0.135660664 | 0.241012507 | 0.017784854 | 0.330201205 | 0.122843258 | 0.120124071 |
| Rps3a1 | S | 237 | -0.054579052 | -0.030767243 | 0.020991748 | 0.0755708 | 0.051758991 | 0.042891152 | 0.245837661 | 0.798316258 | 0.396818556 | 0.542195614 |
| Dcps | S | 100 | -0.048465353 | -0.008180278 | -0.072244802 | -0.023779449 | -0.064064525 | 0.465994687 | 0.889093135 | 0.221192811 | 0.732156263 | 0.349542036 |
| Map1s | S | 658 | -0.030903922 | -0.037106706 | -0.020527143 | 0.010376779 | 0.016579563 | 0.405117397 | 0.134661821 | 0.533638253 | 0.7882634 | 0.571561261 |
| Mast3 | S | 731 | -0.074145968 | -0.207538638 | 0.044880808 | 0.119026776 | 0.252419446 | 0.539732763 | 0.333319951 | 0.653966404 | 0.230764758 | 0.253308776 |
| Vat1 | T | 122 | -0.169538736 | 0.085612045 | -0.053075002 | 0.116463734 | -0.138687047 | 0.054970225 | 0.442537341 | 0.776372579 | 0.54916548 | 0.49997447 |
| Pcnx3 | S | 521 | -0.220512214 | 0.022946133 | -0.014035458 | 0.206476756 | -0.036981591 | 0.217112465 | 0.816968055 | 0.890059319 | 0.255128154 | 0.763299846 |
| Srrm1 | S | 461 | -0.032634472 | -0.06489798 | 0.032186643 | 0.064821115 | 0.097084623 | 0.405904445 | 0.005063173 | 0.271403805 | 0.169992001 | 0.029077313 |
| Prune2 | T | 2642 | -0.017866256 | -0.022406342 | -0.133028743 | -0.115162487 | -0.110622401 | 0.851911829 | 0.71109282 | 0.280874083 | 0.380593404 | 0.34481782 |
| Mga | S | 56 | -0.049045409 | 0.058129815 | -0.200018536 | -0.150973127 | -0.258148351 | 0.637595218 | 0.515634407 | 0.165789882 | 0.248329323 | 0.106244547 |
| Arhgap17 | S | 575 | -0.028100173 | -0.010038304 | -0.134681823 | -0.10658165 | -0.124643519 | 0.748057139 | 0.840115036 | 0.272493073 | 0.412605649 | 0.302811757 |
| Pcm1 | S | 774 | 0.038581345 | 0.028399643 | -0.246697416 | -0.28527876 | -0.275097058 | 0.776990877 | 0.710252562 | 0.103722932 | 0.132386139 | 0.077694593 |
| Pcm1 | S | 777 | 0.038581345 | 0.028399643 | -0.246697416 | -0.28527876 | -0.275097058 | 0.776990877 | 0.710252562 | 0.103722932 | 0.132386139 | 0.077694593 |
| Dennd2a | S | 497 | -0.072474126 | -0.051569523 | -0.001153301 | 0.071320825 | 0.050416222 | 0.196286754 | 0.451502586 | 0.985585464 | 0.351025027 | 0.547159742 |
| Mia3 | S | 1917 | -0.068426162 | -0.100702587 | 0.063072727 | 0.131498889 | 0.163775315 | 0.333603199 | 0.041288635 | 0.259643852 | 0.113108828 | 0.026651796 |
| Nfatc1 | S | 239 | 0.040616744 | -0.109809996 | -0.128053886 | -0.16867063 | -0.01824389 | 0.766570905 | 0.319719727 | 0.497998612 | 0.423342788 | 0.921407102 |
| Nfatc1 | S | 241 | 0.040616744 | -0.109809996 | -0.128053886 | -0.16867063 | -0.01824389 | 0.766570905 | 0.319719727 | 0.497998612 | 0.423342788 | 0.921407102 |
| Ccdc9b | S | 237 | -0.041939494 | -0.022868432 | -0.166834955 | -0.12489546 | -0.143966522 | 0.813090055 | 0.822140787 | 0.40448751 | 0.579687921 | 0.448792214 |
| Mcrip2 | S | 82 | 0.048550237 | -0.100473521 | -0.055599052 | -0.104149289 | 0.044874469 | 0.328712964 | 0.073490844 | 0.27310394 | 0.057368367 | 0.33295211 |
| Alkbh5 | S | 385 | -0.018849067 | -0.052320972 | -0.022285219 | -0.003436152 | 0.030035753 | 0.663934838 | 0.119575346 | 0.48830026 | 0.924471468 | 0.137731385 |
| Nfu1 | S | 247 | -0.258536948 | -0.033980992 | -0.002575511 | 0.255961437 | 0.031405482 | 0.479430869 | 0.728384674 | 0.978365326 | 0.479221268 | 0.441176792 |
| Nexn | S | 436 | -0.089662083 | -0.026243954 | -0.010188815 | 0.079473268 | 0.016055139 | 0.236230775 | 0.457096437 | 0.825862959 | 0.284633591 | 0.647568496 |
| Runx1 | S | 212 | -0.057375944 | -0.033596943 | -0.030615471 | 0.026760474 | 0.002981472 | 0.255574899 | 0.508931917 | 0.642654054 | 0.603931427 | 0.954152229 |
| Xrn1 | S | 1664 | -0.098946539 | -0.049064792 | -0.116892631 | -0.017946093 | -0.06782784 | 0.745427727 | 0.769899711 | 0.554312901 | 0.957005279 | 0.768534147 |
| Parn | S | 605 | -0.21969559 | 0.179352464 | -0.137666504 | 0.082029086 | -0.317018969 | 0.13913889 | 0.292399631 | 0.39495486 | 0.614506316 | 0.133792079 |
| Micall2 | T | 831 | -0.029355346 | -0.077306788 | 0.006683467 | 0.036038813 | 0.083990255 | 0.448165659 | 0.090307147 | 0.799277137 | 0.350759546 | 0.079452243 |
| Bub1b | S | 567 | 0.004703688 | -0.012161098 | -0.163695236 | -0.168398925 | -0.151534139 | 0.971071638 | 0.847019952 | 0.200816306 | 0.30375683 | 0.229694486 |
| Sap30bp | T | 26 | -0.399476983 | 0.096934289 | -0.152173797 | 0.247303186 | -0.249108086 | 0.251267562 | 0.101391352 | 0.611810041 | 0.527235858 | 0.430229838 |
| Cwc22 | S | 105 | -0.15308459 | -0.055616242 | 0.067676255 | 0.220760845 | 0.123292498 | 0.08311122 | 0.179274843 | 0.116923674 | 0.044051699 | 0.009110172 |
| Atf7ip | S | 951 | -0.044971248 | -0.041244784 | -0.051658859 | -0.006687611 | -0.010414076 | 0.460543529 | 0.478860854 | 0.522395699 | 0.92470713 | 0.880062741 |
| Emd | S | 124 | -0.045627569 | -0.002008081 | -0.071648396 | -0.026020826 | -0.069640314 | 0.349767965 | 0.965473203 | 0.247436308 | 0.617895238 | 0.226217102 |
| Taf3 | S | 278 | -0.118370484 | -0.020190757 | -0.065688394 | 0.05268209 | -0.045497637 | 0.343210778 | 0.873030563 | 0.756130701 | 0.781147462 | 0.812837992 |
| Afap1 | S | 283 | 0.034623345 | -0.045942445 | -0.083034858 | -0.117658203 | -0.037092412 | 0.286551894 | 0.196315225 | 0.063792973 | 0.029161046 | 0.272866223 |
| Zcrb1 | S | 210 | -0.050348954 | -0.037163071 | -0.000524295 | 0.049824659 | 0.036638776 | 0.133600913 | 0.26485662 | 0.987424176 | 0.078106336 | 0.170068891 |
| Zcrb1 | S | 216 | -0.050348954 | -0.037163071 | -0.000524295 | 0.049824659 | 0.036638776 | 0.133600913 | 0.26485662 | 0.987424176 | 0.078106336 | 0.170068891 |
| Spag9 | S | 183 | 0.072452967 | 0.044987153 | -0.197444786 | -0.269897753 | -0.242431939 | 0.33140179 | 0.395094222 | 0.004622937 | 0.024408471 | 0.002084741 |
| Tor1aip1 | S | 201 | 0.109979876 | -0.008557403 | -0.149964785 | -0.259944662 | -0.141407382 | 0.144873301 | 0.722784052 | 0.002395667 | 0.022235468 | 0.006034591 |
| Syne1 | T | 8305 | -0.074664398 | -0.026136902 | -0.033828907 | 0.040835491 | -0.007692005 | 0.351421851 | 0.736631053 | 0.408808398 | 0.57953901 | 0.917529968 |
| Cux1 | S | 1332 | -0.15850971 | 0.040347133 | -0.102302734 | 0.056206977 | -0.142649867 | 0.388519006 | 0.337740058 | 0.272075467 | 0.750426818 | 0.162256377 |
| Runx2 | S | 107 | 0.080971289 | -0.080263125 | -0.073385776 | -0.154357065 | 0.006877349 | 0.047292146 | 0.005674337 | 0.11894503 | 0.017895314 | 0.849815724 |
| Med26 | S | 361 | 0.0794195 | -0.16196896 | -0.135868242 | -0.215287742 | 0.026100718 | 0.009600271 | 0.084826804 | 0.139086494 | 0.066602314 | 0.764558059 |
| Trip12 | S | 1049 | -0.050434431 | -0.058066506 | 0.045489815 | 0.095924247 | 0.103556321 | 0.074327207 | 0.037256301 | 0.234651107 | 0.048162232 | 0.045401277 |
| Srrm2 | T | 1782 | -0.235581766 | -0.065026949 | 0.134700038 | 0.370281805 | 0.199726987 | 0.014330915 | 0.483491967 | 0.033907826 | 0.004614719 | 0.094683656 |
| Dclre1a | S | 204 | 0.10065893 | -0.133635885 | -0.204005667 | -0.304664597 | -0.070369781 | 0.194355637 | 0.421651454 | 0.163849852 | 0.088192126 | 0.688205833 |
| Dbf4 | S | 543 | -0.001825863 | -0.074245398 | -0.035459766 | -0.033633903 | 0.038785632 | 0.958104831 | 0.204253102 | 0.36598522 | 0.426589612 | 0.486044726 |
| Dbf4 | T | 539 | -0.001825863 | -0.074245398 | -0.035459766 | -0.033633903 | 0.038785632 | 0.958104831 | 0.204253102 | 0.36598522 | 0.426589612 | 0.486044726 |
| Mgrn1 | S | 380 | -0.070300286 | -0.120086325 | -0.049507788 | 0.020792498 | 0.070578537 | 0.604120492 | 0.660476845 | 0.712672883 | 0.850715874 | 0.787240912 |
| Lmna | S | 575 | -0.178054459 | -0.228027518 | 0.213311912 | 0.391366371 | 0.44133943 | 0.271316556 | 0.033244672 | 0.042016516 | 0.070856784 | 0.000226444 |
| S1pr2 | S | 342 | 0.0375838 | 0.020416453 | -0.183556278 | -0.221140079 | -0.203972731 | 0.756916218 | 0.876915925 | 0.072099348 | 0.105571633 | 0.160235418 |
| Rnf34 | S | 258 | -0.185334536 | -0.159892005 | 0.093755195 | 0.279089731 | 0.2536472 | 0.306257072 | 0.346971049 | 0.292903638 | 0.16615758 | 0.176776979 |
| Ktn1 | T | 905 | 0.006771983 | -0.091184778 | -0.057748523 | -0.064520506 | 0.033436255 | 0.97263497 | 0.213362781 | 0.577802921 | 0.751790229 | 0.709045699 |
| Pkm | S | 346 | -0.12160609 | 0.000503212 | -0.057472474 | 0.064133616 | -0.057975686 | 0.464359466 | 0.994547136 | 0.38003227 | 0.691706804 | 0.512581343 |
| Tubgcp4 | S | 437 | -0.243918303 | 0.093680008 | -0.01258005 | 0.231338253 | -0.106260058 | 0.08528534 | 0.463185302 | 0.916416312 | 0.124425708 | 0.432588506 |
| Tubgcp4 | S | 448 | -0.243918303 | 0.093680008 | -0.01258005 | 0.231338253 | -0.106260058 | 0.08528534 | 0.463185302 | 0.916416312 | 0.124425708 | 0.432588506 |
| Tubgcp4 | S | 456 | -0.243918303 | 0.093680008 | -0.01258005 | 0.231338253 | -0.106260058 | 0.08528534 | 0.463185302 | 0.916416312 | 0.124425708 | 0.432588506 |
| Uba2 | S | 548 | -0.140979318 | 0.02682018 | 0.035649865 | 0.176629184 | 0.008829685 | 0.00418773 | 0.380999728 | 0.410019878 | 0.010277625 | 0.792759035 |
| Mprip | S | 364 | -0.030981912 | 0.005543803 | -0.110734376 | -0.079751564 | -0.116277279 | 0.594836452 | 0.924066675 | 0.183233258 | 0.343748548 | 0.196975557 |
| Setd2 | S | 1926 | -0.068426789 | -0.043026159 | -0.02994212 | 0.038484669 | 0.013084039 | 0.333840338 | 0.546233326 | 0.697213306 | 0.674915906 | 0.889048803 |
| Mapk8ip3 | S | 585 | 0.082066083 | -0.049794227 | -0.164228986 | -0.246295069 | -0.114434759 | 0.273024565 | 0.603270232 | 0.047738029 | 0.022873054 | 0.277867605 |
| Slc35e1 | S | 363 | -0.000328524 | -0.147661835 | -0.011712839 | -0.011384315 | 0.135948996 | 0.99717702 | 0.195402856 | 0.828812635 | 0.892780386 | 0.224087641 |
| Ccnb2 | S | 397 | -0.010166332 | -0.053447211 | -0.056087759 | -0.045921427 | -0.002640548 | 0.866616489 | 0.216586599 | 0.442597804 | 0.570874366 | 0.968980473 |
| Tbc1d10b | S | 664 | -0.10138237 | -0.005092472 | 0.025076956 | 0.126459326 | 0.030169429 | 0.028265202 | 0.899522006 | 0.741489232 | 0.169785666 | 0.680224461 |
| Ube2e3 | T | 15 | -0.079814318 | 0.07057736 | -0.16024301 | -0.080428691 | -0.23082037 | 0.464545756 | 0.336400272 | 0.134458299 | 0.467378093 | 0.066107765 |
| Gtf2f1 | T | 331 | -0.028392028 | -0.036153988 | -0.038865669 | -0.010473641 | -0.002711681 | 0.472274821 | 0.247069116 | 0.490926224 | 0.858911881 | 0.960472309 |
| Ckap2 | T | 561 | -0.080281038 | -0.17640891 | 0.113995575 | 0.194276613 | 0.290404485 | 0.473013964 | 0.080207102 | 0.290576725 | 0.15479595 | 0.039620424 |
| Brf2 | S | 386 | 0.016405965 | 0.000180631 | -0.111038043 | -0.127444008 | -0.111218674 | 0.84227353 | 0.997412069 | 0.04327017 | 0.187625572 | 0.083950106 |
| Rgs12 | S | 934 | -0.024448665 | -0.009518259 | -0.165498052 | -0.141049387 | -0.155979793 | 0.765603685 | 0.909221548 | 0.334600135 | 0.393390958 | 0.35395788 |
| Frs2 | S | 327 | 0.083558632 | -0.021203024 | -0.118124698 | -0.20168333 | -0.096921675 | 0.625129962 | 0.653039497 | 0.031334427 | 0.295384645 | 0.034376974 |
| Thrap3 | S | 531 | 0.047043343 | -0.117361169 | -0.081172485 | -0.128215828 | 0.036188684 | 0.216601733 | 0.078109248 | 0.442301035 | 0.266687605 | 0.727273812 |
| Ybx1 | S | 311 | -0.092037981 | -0.033790666 | -0.032230843 | 0.059807138 | 0.001559823 | 0.503722782 | 0.686414471 | 0.401044435 | 0.652017336 | 0.984649281 |
| Cdr2l | S | 179 | -0.013388054 | -0.019317388 | -0.084056554 | -0.0706685 | -0.064739166 | 0.624472831 | 0.530993538 | 0.241974018 | 0.306169562 | 0.34148294 |
| Pnkp | S | 114 | -0.052376365 | 0.064993277 | -0.110862994 | -0.058486629 | -0.175856271 | 0.253094944 | 0.516106438 | 0.131327928 | 0.360026638 | 0.150526991 |
| Ezh2 | T | 487 | 0.011850395 | -0.071280191 | -0.012422687 | -0.024273082 | 0.058857505 | 0.712826473 | 0.020105045 | 0.668453741 | 0.444193111 | 0.060845345 |
| Prrc2c | S | 2646 | -0.015359401 | -0.023314157 | -0.037395802 | -0.022036402 | -0.014081645 | 0.433016671 | 0.305204998 | 0.174832527 | 0.369724935 | 0.579197446 |
| Hcfc1 | S | 2030 | 0.032487614 | 0.061537133 | -0.174691504 | -0.207179119 | -0.236228637 | 0.71612748 | 0.4051108 | 0.01248521 | 0.084468541 | 0.027775046 |
| Vps35 | S | 7 | 0.063993975 | -0.055351075 | -0.183189273 | -0.247183249 | -0.127838198 | 0.042108202 | 0.197185995 | 0.076921428 | 0.045712508 | 0.151438493 |
| Emsy | S | 1085 | -0.071927663 | -0.045163114 | -0.066199752 | 0.005727912 | -0.021036637 | 0.520512236 | 0.722051695 | 0.557579215 | 0.965712688 | 0.886017164 |
| Lima1 | S | 488 | -0.040992775 | -0.080694595 | 0.013780044 | 0.054772818 | 0.094474639 | 0.340272132 | 0.148748429 | 0.71388913 | 0.09476006 | 0.100920298 |
| Etv6 | S | 319 | -0.010597661 | -0.12084694 | 0.02757767 | 0.038175331 | 0.148424611 | 0.529063053 | 0.07917152 | 0.686976914 | 0.579966196 | 0.11266393 |
| Srrm2 | S | 1774 | -0.061454736 | -0.033973926 | -0.035907416 | 0.02554732 | -0.00193349 | 0.338081724 | 0.525571247 | 0.634898808 | 0.703442609 | 0.973751542 |
| Ptk2 | S | 705 | -0.047289182 | 0.061134384 | -0.177497297 | -0.130208115 | -0.238631681 | 0.27096486 | 0.15001639 | 0.110699826 | 0.204420396 | 0.076720577 |
| Ptk2 | S | 708 | -0.047289182 | 0.061134384 | -0.177497297 | -0.130208115 | -0.238631681 | 0.27096486 | 0.15001639 | 0.110699826 | 0.204420396 | 0.076720577 |

| | | | | | | | | | | | | |
|---|---|---|---|---|---|---|---|---|---|---|---|---|
| Sacs | T | 4265 | -0.02046026 | -0.029748656 | -0.147214827 | -0.126754567 | -0.117466171 | 0.71753102 | 0.698216254 | 0.441503987 | 0.499390952 | 0.535055891 |
| Fip1l1 | T | 481 | 0.009434723 | -0.011452809 | -0.08600144 | -0.095436163 | -0.074548631 | 0.704231394 | 0.628000154 | 0.044618744 | 0.043733778 | 0.084304349 |
| Kif13a | S | 1274 | -0.071391697 | -0.130164114 | 0.066798042 | 0.138189739 | 0.196962156 | 0.075783266 | 0.194559364 | 0.146001428 | 0.026467063 | 0.09387529 |
| Kif13a | T | 1272 | -0.071391697 | -0.130164114 | 0.066798042 | 0.138189739 | 0.196962156 | 0.075783266 | 0.194559364 | 0.146001428 | 0.026467063 | 0.09387529 |
| Specc1 | S | 311 | -0.021304543 | -0.173428633 | 0.073239073 | 0.094543616 | 0.246667706 | 0.771564306 | 0.062817162 | 0.469985755 | 0.291765579 | 0.045144989 |
| Tbc1d1 | S | 607 | -0.00957708 | -0.015762772 | -0.039850085 | -0.030273006 | -0.024087313 | 0.748137527 | 0.613745989 | 0.021374023 | 0.344514608 | 0.448065357 |
| Vps13b | T | 3024 | 0.138562039 | -0.080539268 | -0.200984578 | -0.339546617 | -0.12044531 | 0.084770368 | 0.424021042 | 0.026641223 | 0.004982179 | 0.244525284 |
| Sync | S | 314 | -0.26080227 | 0.157049966 | -0.012317205 | 0.248485065 | -0.169367171 | 0.022800006 | 0.213103969 | 0.923240138 | 0.097979254 | 0.237203942 |
| Helb | T | 992 | -0.060896995 | -0.044801831 | 0.002424216 | 0.063321211 | 0.047226046 | 0.232440596 | 0.260530844 | 0.942276432 | 0.20356385 | 0.191053187 |
| Smpd4 | S | 283 | -0.041844954 | -0.023922459 | -0.074430207 | -0.032585253 | -0.050507747 | 0.564614429 | 0.768296948 | 0.304622302 | 0.649884136 | 0.539374587 |
| Larp1 | S | 198 | -0.069575729 | -0.052136846 | 0.001362468 | 0.070938197 | 0.053499314 | 0.191551214 | 0.473359658 | 0.978708068 | 0.267468453 | 0.497768686 |
| Ktn1 | S | 90 | 0.054406896 | -0.094893968 | -0.043579301 | -0.097986197 | 0.051314667 | 0.686042391 | 0.087488734 | 0.423921299 | 0.482885527 | 0.312196197 |
| Med15 | S | 657 | -0.074079045 | -0.020837052 | -0.009992597 | 0.064086448 | 0.010844455 | 0.157346091 | 0.5270702 | 0.822857012 | 0.260982029 | 0.800298881 |
| Agap3 | S | 573 | 0.031998066 | -0.044526069 | -0.084272015 | -0.116270081 | -0.039745946 | 0.793938567 | 0.383936924 | 0.137078177 | 0.383265496 | 0.441534769 |
| Sltm | S | 549 | -0.059765344 | -0.073067369 | -6.20252E-05 | 0.059703319 | 0.073005344 | 0.271058914 | 0.42453761 | 0.999460278 | 0.506783212 | 0.507675645 |
| Sqstm1 | S | 345 | 0.056765801 | -0.137961743 | -0.14875388 | -0.205519681 | -0.010792137 | 0.532561495 | 0.549977529 | 0.287207331 | 0.159553066 | 0.962639462 |
| Reps1 | S | 274 | -0.071268681 | 0.030073295 | -0.058376068 | 0.012892613 | -0.088449363 | 0.171661345 | 0.613069429 | 0.223786143 | 0.754850556 | 0.164092604 |
| Fam110b | S | 257 | -0.12128976 | 0.07831045 | -0.090522021 | 0.030767738 | -0.168832471 | 0.117777875 | 0.450868221 | 0.449504736 | 0.79427458 | 0.253712497 |
| Heatr5a | S | 1647 | -0.071789377 | 0.036481268 | -0.052245087 | 0.019544291 | -0.088726355 | 0.209105272 | 0.481609934 | 0.087768443 | 0.690158047 | 0.151425807 |
| Fhod1 | S | 514 | -0.21360126 | 0.139293661 | -0.174549908 | 0.039051352 | -0.313843569 | 0.247187871 | 0.381956358 | 0.499593957 | 0.871791973 | 0.245128134 |
| Mus81 | S | 94 | -0.127059572 | 0.036175097 | -0.026317429 | 0.100742143 | -0.062492525 | 0.079734734 | 0.683611277 | 0.790339378 | 0.317397021 | 0.564784555 |
| Arhgap17 | T | 734 | -0.035715535 | -0.02519984 | -0.044695088 | -0.008979553 | -0.019495248 | 0.43961312 | 0.545626584 | 0.298880745 | 0.856256347 | 0.679301781 |
| Abi1 | S | 267 | -0.012894212 | -0.056853792 | -0.059829221 | -0.046935009 | -0.002975429 | 0.770658137 | 0.244730136 | 0.561067962 | 0.642676836 | 0.976050763 |
| Ahctf1 | T | 1769 | 0.094681142 | -0.111669527 | -0.088424076 | -0.183105218 | 0.023245451 | 0.065508148 | 0.058675396 | 0.080134218 | 0.001174494 | 0.534390397 |
| Tln1 | S | 979 | -0.080256115 | -0.020707607 | -0.021294755 | 0.05896136 | -0.000587147 | 0.298707223 | 0.696755342 | 0.45916712 | 0.419389875 | 0.990890336 |
| Zfp687 | S | 1184 | -0.075675248 | -0.093415671 | 0.016219369 | 0.091894617 | 0.10963504 | 0.443444628 | 0.319148258 | 0.867278598 | 0.268923287 | 0.163998796 |
| Ash1l | S | 897 | -0.000294861 | -0.094946547 | -0.019716326 | -0.019421465 | 0.075230221 | 0.994891363 | 0.152385307 | 0.58553144 | 0.695307026 | 0.236198631 |
| Pelp1 | S | 483 | -0.028273184 | -0.112439825 | 0.025431008 | 0.053704192 | 0.137870833 | 0.729088185 | 0.072101624 | 0.611862679 | 0.481773439 | 0.023025931 |
| Sh3kbp1 | S | 225 | 0.041620147 | -0.038196124 | -0.091570152 | -0.133190299 | -0.053374028 | 0.772565323 | 0.410143017 | 0.114455858 | 0.398997422 | 0.341962375 |
| Map1s | S | 786 | -0.000880407 | -0.058383292 | -0.051339153 | -0.050458746 | 0.007044139 | 0.988464377 | 0.277527953 | 0.297491052 | 0.328898546 | 0.812569854 |
| Ppp1r9b | T | 193 | -0.047324534 | -0.032179324 | -0.048131005 | -0.00080647 | -0.01595168 | 0.565551784 | 0.339170141 | 0.528753343 | 0.993479069 | 0.829133553 |
| Stim2 | S | 718 | -0.056438525 | -0.016525598 | -0.036506723 | 0.019931802 | -0.019981125 | 0.118638638 | 0.839112801 | 0.653534367 | 0.805732727 | 0.84947594 |
| Serinc3 | S | 367 | -0.158089584 | -0.010412525 | 0.02265509 | 0.180744674 | 0.033067615 | 0.128310223 | 0.947328775 | 0.812166881 | 0.013494334 | 0.814045038 |
| Serinc3 | T | 365 | -0.158089584 | -0.010412525 | 0.02265509 | 0.180744674 | 0.033067615 | 0.128310223 | 0.947328775 | 0.812166881 | 0.013494334 | 0.814045038 |
| Rps6kc1 | S | 281 | 0.04292769 | -0.030330354 | -0.124053303 | -0.166980992 | -0.093722948 | 0.319090962 | 0.390871982 | 0.068180304 | 0.033227857 | 0.13196353 |
| Map2k1 | T | 386 | -0.046827453 | -0.002546676 | -0.10266162 | -0.055834167 | -0.100114944 | 0.609405485 | 0.971139097 | 0.27703234 | 0.586653491 | 0.2835061 |
| Ppig | S | 656 | -0.033899354 | -0.070117732 | 0.011220346 | 0.0451197 | 0.081338078 | 0.434262408 | 0.107160407 | 0.8644462 | 0.538014315 | 0.289362072 |
| Osbpl3 | S | 288 | 0.236043791 | -0.175772383 | -0.01050922 | -0.246553011 | 0.165263163 | 0.529079574 | 0.012949757 | 0.878152494 | 0.512958279 | 0.050366756 |
| Osbpl3 | S | 291 | 0.236043791 | -0.175772383 | -0.01050922 | -0.246553011 | 0.165263163 | 0.529079574 | 0.012949757 | 0.878152494 | 0.512958279 | 0.050366756 |
| Ccs | S | 267 | -0.065714717 | 0.027327962 | -0.109866221 | -0.044151504 | -0.137194183 | 0.25835453 | 0.795216662 | 0.430752575 | 0.72746852 | 0.379900035 |
| Hdac7 | S | 398 | -0.132424282 | 0.005663279 | 0.022662032 | 0.155086314 | 0.016998753 | 0.062445345 | 0.918842471 | 0.849075885 | 0.253021602 | 0.881688951 |
| Kpna4 | S | 60 | 0.024436516 | -0.032529364 | -0.085582134 | -0.11001865 | -0.05305277 | 0.695799483 | 0.618248235 | 0.058931947 | 0.140679936 | 0.393673261 |
| Anln | T | 903 | 0.180931599 | -0.115631591 | -0.283482566 | -0.464414165 | -0.167850976 | 0.069825219 | 0.237031671 | 0.099313441 | 0.042494399 | 0.260387236 |
| Nol8 | S | 716 | 0.020283526 | -0.024136748 | -0.077898618 | -0.098182143 | -0.053761869 | 0.722291936 | 0.445626023 | 0.062668934 | 0.149943511 | 0.122344374 |
| Sfswap | S | 812 | -0.137386289 | 0.029694796 | -0.090711619 | 0.046674669 | -0.120406416 | 0.232237 | 0.773327959 | 0.748091921 | 0.870758989 | 0.680781268 |
| Hivep1 | S | 1735 | -0.311124465 | 0.162549104 | 0.039272845 | 0.35039731 | -0.123276259 | 0.013917014 | 0.14508976 | 0.881423045 | 0.25637148 | 0.639583595 |
| Tpx2 | S | 125 | -0.06150711 | 0.02829659 | -0.063054523 | -0.001547413 | -0.091351113 | 0.316332328 | 0.340955276 | 0.052537982 | 0.976940015 | 0.013924097 |
| Sh3pxd2a | S | 641 | -0.051400333 | 0.011780298 | -0.077049409 | -0.025649076 | -0.088829707 | 0.341276158 | 0.888698327 | 0.254765476 | 0.581503008 | 0.309566584 |
| Trim25 | T | 84 | 0.065561926 | -0.103187097 | -0.132728296 | -0.198290222 | -0.029541198 | 0.61654691 | 0.317604758 | 0.384275523 | 0.278079674 | 0.843693134 |
| Phactr4 | S | 282 | -0.228127272 | -0.036719656 | -0.040226004 | 0.187901268 | -0.003506348 | 0.578512488 | 0.833183681 | 0.829849172 | 0.646771064 | 0.985277113 |
| Rfc1 | S | 189 | 0.090506886 | -0.233566129 | 0.123671013 | 0.033164127 | 0.357237142 | 0.553784986 | 0.00240063 | 0.217226021 | 0.832184055 | 0.025513208 |
| Dhx33 | T | 43 | 0.014408638 | -0.036680077 | -0.104658075 | -0.119066713 | -0.067977998 | 0.563189125 | 0.635688358 | 0.10804816 | 0.097466836 | 0.43475367 |
| Ppfibp1 | S | 558 | -0.086582982 | 0.026980829 | -0.068955248 | 0.017627734 | -0.095936077 | 0.093011806 | 0.619761066 | 0.6235687 | 0.894719567 | 0.502666506 |
| Nufip2 | T | 221 | 0.140793334 | -0.137268353 | -0.115450866 | -0.2562442 | 0.021817488 | 0.013899713 | 0.043379357 | 0.028820666 | 0.000665624 | 0.628634703 |
| Pald1 | S | 347 | 0.02831937 | -0.177926096 | 0.007445799 | -0.020873571 | 0.185371895 | 0.692216655 | 0.097241019 | 0.938548591 | 0.828594683 | 0.143056 |
| Afap1 | S | 683 | 0.267941934 | -0.071986878 | -0.107677535 | -0.375619468 | -0.035690657 | 0.330141326 | 0.10417417 | 0.017193663 | 0.215865977 | 0.248572453 |
| Pafah1b1 | S | 109 | -0.089913439 | 0.034889891 | -0.064038049 | 0.02587539 | -0.09892794 | 0.238618473 | 0.680009441 | 0.30167384 | 0.662974531 | 0.244816674 |
| Vcl | S | 383 | -0.175200587 | 0.113658 | -0.08504769 | 0.090152897 | -0.19870569 | 0.088262763 | 0.228342758 | 0.404362802 | 0.332122894 | 0.069962221 |
| Srrm2 | S | 1395 | 0.071300779 | -0.070963021 | -0.070150994 | -0.141451773 | 0.000812027 | 0.111622626 | 0.027287336 | 0.091671115 | 0.020530798 | 0.977713905 |
| Plekhg3 | S | 1151 | 0.087361603 | -0.156561095 | -0.016892907 | -0.10425451 | 0.139668188 | 0.064316959 | 0.003991225 | 0.689786398 | 0.077661306 | 0.031656601 |
| BC055324 | S | 247 | -0.171355365 | 0.053839471 | -0.061295863 | 0.110059502 | -0.115135334 | 0.194743454 | 0.584352408 | 0.659608083 | 0.477050668 | 0.42081547 |
| Maco1 | S | 332 | 0.092703628 | -0.033105127 | -0.213887866 | -0.306591494 | -0.180782739 | 0.156187378 | 0.596719898 | 0.04912587 | 0.021707257 | 0.084080426 |
| Add1 Add2 Add3 | S | 693;712;724 | -0.128746858 | 0.131506659 | -0.127937369 | 0.00080949 | -0.259444028 | 0.157244256 | 0.330776758 | 0.266417557 | 0.993571269 | 0.113472398 |
| Cdk12 | S | 1079 | 0.032347405 | -0.095955862 | 0.001565876 | -0.030781529 | 0.097521738 | 0.458749096 | 0.007637684 | 0.966537242 | 0.505803412 | 0.060316308 |
| Limk1 | S | 298 | 0.012095697 | -0.021440146 | -0.114213066 | -0.126308763 | -0.09277262 | 0.7573896 | 0.561729279 | 0.151970827 | 0.127408569 | 0.219848993 |
| Limk1 | S | 310 | 0.012095697 | -0.021440146 | -0.114213066 | -0.126308763 | -0.09277262 | 0.7573896 | 0.561729279 | 0.151970827 | 0.127408569 | 0.219848993 |
| Phactr4 | S | 264 | 0.042098602 | 0.002516827 | -0.170633342 | -0.212731943 | -0.173150169 | 0.359872286 | 0.962734905 | 0.042479895 | 0.03154271 | 0.048864198 |
| AI597479 | S | 193 | -0.081166777 | -0.015038701 | -0.075847085 | 0.005319692 | -0.060808384 | 0.507424566 | 0.881624498 | 0.473170408 | 0.969045711 | 0.631532037 |
| Tmpo | T | 449 | -0.054309211 | -0.002365965 | -0.107816446 | -0.053507235 | -0.105450481 | 0.583377198 | 0.978761683 | 0.346560386 | 0.618337666 | 0.322147508 |
| Arcn1 | S | 493 | 0.005845153 | -0.127830824 | -0.014866937 | -0.02071209 | 0.112963887 | 0.959552736 | 0.158708007 | 0.86495376 | 0.831296634 | 0.077351644 |
| Srsf7 | S | 208 | -0.012179324 | -0.100401539 | 0.01249796 | 0.024677284 | 0.112899499 | 0.809212553 | 0.073884701 | 0.838459884 | 0.630988748 | 0.088971678 |
| Cwc25 | S | 254 | 0.078988357 | -0.105054325 | -0.138741291 | -0.217729648 | -0.033686966 | 0.15679727 | 0.147892445 | 0.232875963 | 0.112920905 | 0.7442613 |
| Cwc25 | S | 259 | 0.078988357 | -0.105054325 | -0.138741291 | -0.217729648 | -0.033686966 | 0.15679727 | 0.147892445 | 0.232875963 | 0.112920905 | 0.7442613 |
| Elac2 | T | 194 | 0.023890669 | 0.022531046 | -0.118944859 | -0.142835529 | -0.141475905 | 0.569234279 | 0.512059069 | 0.008409119 | 0.022227329 | 0.008411813 |
| Mfap1a Mfap1b | S | 94;94 | -0.019515739 | -0.044447854 | -0.031251759 | -0.01173602 | 0.013196095 | 0.572892587 | 0.292706021 | 0.366684264 | 0.757896205 | 0.754864579 |
| Slain2 | T | 86 | -0.011391717 | 0.016615113 | -0.121601179 | -0.110209462 | -0.138216292 | 0.831129338 | 0.765987637 | 0.094485975 | 0.127746085 | 0.079162411 |
| Pias1 | S | 503 | -0.136844776 | 0.020898091 | -0.003417909 | 0.133426867 | -0.024316001 | 0.078069086 | 0.691606062 | 0.959695702 | 0.144132288 | 0.742258987 |
| Ubxn1 | S | 200 | 0.037842239 | -0.018818149 | -0.187601359 | -0.225443599 | -0.16878321 | 0.075147497 | 0.520067568 | 0.064703934 | 0.048180797 | 0.072801814 |
| Knl1 | S | 32 | 0.04189383 | -0.096225418 | -0.007251667 | -0.049145497 | 0.088973752 | 0.368085555 | 0.007791921 | 0.8413858 | 0.319013785 | 0.068572609 |
| Slc39a6 | S | 476 | -0.043165879 | -0.079868786 | -0.015202619 | 0.02796326 | 0.064666167 | 0.720818635 | 0.230984339 | 0.830750567 | 0.80770361 | 0.249329933 |
| Xkr5 | S | 597 | -0.013607209 | -0.060060803 | -0.077772736 | -0.064165527 | -0.017711934 | 0.87178904 | 0.413975417 | 0.474893317 | 0.559893575 | 0.85861009 |
| Map4k4 | S | 836 | -0.1329835 | -0.009388224 | 0.033747918 | 0.166731418 | 0.043136141 | 0.074224176 | 0.908302785 | 0.724797935 | 0.127955344 | 0.647133268 |
| Akap13 | S | 1869 | -0.154127016 | 0.128180091 | -0.090158865 | 0.063968151 | -0.218338956 | 0.108652064 | 0.299603959 | 0.3168311 | 0.483382942 | 0.12356794 |
| Tor4a | T | 84 | 0.107330938 | -0.063127663 | -0.155886796 | -0.263217734 | -0.092759133 | 0.294255161 | 0.564872789 | 0.048805005 | 0.049920713 | 0.409303294 |
| Ube4b | S | 1149 | 0.005242787 | 0.071955199 | -0.167621063 | -0.17286385 | -0.239576262 | 0.934624651 | 0.636697747 | 0.048589661 | 0.051683133 | 0.190511069 |
| Prpf40a | S | 884 | -0.035742033 | -0.089870416 | 0.03629985 | 0.072041883 | 0.126170267 | 0.486799397 | 0.082678459 | 0.667519377 | 0.427002158 | 0.204217233 |
| Apaf1 | S | 790 | -0.088879281 | 0.042708241 | -0.094550186 | -0.005670905 | -0.137258427 | 0.217673353 | 0.608741379 | 0.410822088 | 0.953863496 | 0.252279944 |
| Ncor2 | S | 955 | 0.009487707 | -0.076052453 | 0.008642427 | -0.00084528 | 0.08469488 | 0.64322883 | 0.00705638 | 0.854920805 | 0.985013341 | 0.15869868 |
| Topors | S | 734 | 0.06561282 | -0.142356063 | -0.026989258 | -0.092602079 | 0.115366804 | 0.222144905 | 0.03394109 | 0.523514015 | 0.078350881 | 0.071637064 |
| Topors | S | 735 | 0.06561282 | -0.142356063 | -0.026989258 | -0.092602079 | 0.115366804 | 0.222144905 | 0.03394109 | 0.523514015 | 0.078350881 | 0.071637064 |
| Srrm2 | T | 789 | -0.116301487 | -0.033987659 | 0.040474107 | 0.156775594 | 0.074461767 | 0.140562863 | 0.426459136 | 0.633362253 | 0.155277084 | 0.410437379 |
| Mkrn2 | S | 365 | -0.195952709 | 0.221223788 | -0.135727081 | 0.060225628 | -0.356950869 | 0.028115248 | 0.101196893 | 0.33154284 | 0.638183744 | 0.060695837 |
| Csnk1d | S | 356 | 0.021552368 | -0.000952607 | -0.09597751 | -0.117529878 | -0.095024903 | 0.642631717 | 0.989886154 | 0.021852592 | 0.053176537 | 0.265838171 |
| Cdk11b | S | 47 | 0.028166777 | -0.017464545 | -0.082593081 | -0.110759858 | -0.065128536 | 0.532302392 | 0.565436219 | 0.027571666 | 0.068875785 | 0.088886458 |

| | | | | | | | | | | | | |
|---|---|---|---|---|---|---|---|---|---|---|---|---|
| Crtc3 | S | 62 | 0.001937545 | -0.013062297 | -0.112467299 | -0.114404843 | -0.099405002 | 0.98998595 | 0.749549412 | 0.159873479 | 0.495348261 | 0.20040708 |
| Ckap4 | S | 26 | -0.111934481 | -0.044316621 | 0.026986056 | 0.138920537 | 0.071302677 | 0.329643533 | 0.220837497 | 0.68939287 | 0.264005992 | 0.328030716 |
| Srrm2 | T | 963 | 0.02752339 | -0.031280225 | -0.05021118 | -0.077734569 | -0.018930955 | 0.35804313 | 0.128722005 | 0.027088138 | 0.067071113 | 0.131551014 |
| Mapk8ip3 | S | 586 | -0.002540093 | 0.021636735 | -0.124772863 | -0.122232769 | -0.146409598 | 0.964506614 | 0.673038807 | 0.059313209 | 0.038791344 | 0.0253361 |
| Jam3 | S | 284 | 0.119687871 | -0.127148698 | -0.095946639 | -0.215634509 | 0.031202059 | 0.082984507 | 0.020929544 | 0.268559178 | 0.055073576 | 0.681136243 |
| Ubap2l | T | 515 | -0.006562631 | 0.069169982 | -0.180133978 | -0.173571348 | -0.249303961 | 0.928467939 | 0.355227556 | 0.041902701 | 0.061769225 | 0.020009103 |
| Bsdc1 | S | 214 | -0.140077377 | -0.004414168 | -0.017242204 | 0.122835173 | -0.012828036 | 0.217308346 | 0.978826674 | 0.91833227 | 0.472331619 | 0.94909446 |
| BC005624 | S | 15 | -0.038351483 | 0.036718875 | -0.096208638 | -0.057857155 | -0.132927513 | 0.102094676 | 0.224174468 | 0.124454404 | 0.280867034 | 0.05888239 |
| Nedd4l | Y | 514 | -0.05361499 | -0.004511652 | -0.079869299 | -0.026254309 | -0.075357647 | 0.237937414 | 0.944530557 | 0.559811178 | 0.843811879 | 0.598117611 |
| Tmem41b | T | 18 | 0.042921679 | -0.132085047 | -0.065408867 | -0.108330545 | 0.06667618 | 0.76194261 | 0.216574529 | 0.652981567 | 0.552725721 | 0.663886893 |
| Chd4 | Y | 103 | -0.022518965 | -0.08200749 | -0.026884034 | -0.004365068 | 0.055123457 | 0.694570149 | 0.333968643 | 0.596251018 | 0.932106544 | 0.485750693 |
| Kif2a | S | 75 | -0.129455066 | -0.065058254 | 0.043034958 | 0.172490023 | 0.108093212 | 0.255255361 | 0.34591653 | 0.491115407 | 0.157425853 | 0.08556674 |
| Rab3gap1 | S | 5 | -0.051160944 | 0.009831718 | -0.109052367 | -0.057891423 | -0.118884085 | 0.710103949 | 0.76477379 | 0.187570095 | 0.69002303 | 0.167734194 |
| Ubr5 | T | 1220 | 0.016866622 | 0.042329658 | -0.147876621 | -0.164743242 | -0.190206279 | 0.86353627 | 0.19896458 | 0.010748154 | 0.180786405 | 0.012812775 |
| Fam117b | S | 51 | 0.372707534 | -0.189727424 | -0.186576648 | -0.559284182 | 0.003150776 | 0.493185402 | 0.243329958 | 0.257794175 | 0.333515184 | 0.977209215 |
| Gpatch1 | S | 200 | 0.029413839 | -0.070948536 | -0.111792946 | -0.141206786 | -0.040844411 | 0.550346137 | 0.50237211 | 0.208018884 | 0.137820081 | 0.719230485 |
| Gpatch1 | S | 203 | 0.029413839 | -0.070948536 | -0.111792946 | -0.141206786 | -0.040844411 | 0.550346137 | 0.50237211 | 0.208018884 | 0.137820081 | 0.719230485 |
| Gpatch1 | Y | 210 | 0.029413839 | -0.070948536 | -0.111792946 | -0.141206786 | -0.040844411 | 0.550346137 | 0.50237211 | 0.208018884 | 0.137820081 | 0.719230485 |
| Trp53bp1 | S | 571 | -0.030987573 | -0.026464202 | -0.062419093 | -0.03143152 | -0.035954891 | 0.66013961 | 0.558083143 | 0.312002522 | 0.657493876 | 0.465233478 |
| Ahctf1 | S | 1792 | 0.10125909 | -0.065178108 | -0.069736211 | -0.170995301 | -0.004558102 | 0.077427039 | 0.002587919 | 0.109445921 | 0.016199433 | 0.885089562 |
| Dok1 | S | 269 | -0.170067278 | 0.132773133 | -0.117446544 | 0.052620734 | -0.250219677 | 0.126645896 | 0.248920283 | 0.365824547 | 0.662726118 | 0.102071835 |
| Nes | S | 586 | 0.058207078 | -0.062755271 | -0.045814355 | -0.104021432 | 0.016940917 | 0.352370699 | 0.006701111 | 0.447483751 | 0.204902178 | 0.764043129 |
| Nav1 | S | 1860 | -0.093444276 | -0.094964267 | 0.045195454 | 0.13863973 | 0.140159721 | 0.403547309 | 0.282437117 | 0.680846957 | 0.279288622 | 0.217916146 |
| Osbpl5 | S | 67 | 0.047782106 | -0.08359999 | -0.046985896 | -0.094768003 | 0.036614093 | 0.11232434 | 0.017142003 | 0.276367421 | 0.071915622 | 0.348739477 |
| Ppp1ca | T | 320 | 0.084682372 | -0.004220657 | -0.224372144 | -0.309054516 | -0.220151487 | 0.237779722 | 0.94374659 | 0.043576859 | 0.019860876 | 0.051424453 |
| Otud4 | S | 197 | -0.044749505 | -0.143590232 | 0.026957439 | 0.071706944 | 0.170547671 | 0.748320298 | 0.252670644 | 0.898861492 | 0.756876917 | 0.462122678 |
| Otud4 | S | 199 | -0.044749505 | -0.143590232 | 0.026957439 | 0.071706944 | 0.170547671 | 0.748320298 | 0.252670644 | 0.898861492 | 0.756876917 | 0.462122678 |
| Otud4 | S | 202 | -0.044749505 | -0.143590232 | 0.026957439 | 0.071706944 | 0.170547671 | 0.748320298 | 0.252670644 | 0.898861492 | 0.756876917 | 0.462122678 |
| Pcyt1a | T | 342 | -0.01057221 | 0.013374871 | -0.106845489 | -0.096273279 | -0.12022036 | 0.609832791 | 0.589640994 | 0.083051237 | 0.11444758 | 0.067070272 |
| Nup214 | S | 930 | 0.015975234 | -0.053970078 | -0.033180751 | -0.049155986 | 0.020789327 | 0.5665169 | 0.078406969 | 0.207396901 | 0.062547514 | 0.240815019 |
| Mki67 | S | 588 | 0.007547458 | 0.043226712 | -0.170533708 | -0.178081165 | -0.21376042 | 0.844581499 | 0.477446206 | 0.04857493 | 0.043907257 | 0.034170214 |
| Pakap | T | 738 | -0.015868919 | 0.014705436 | -0.121507297 | -0.105638378 | -0.136212733 | 0.808095991 | 0.861334972 | 0.136295709 | 0.158181061 | 0.157374203 |
| Ppp1r2 | S | 131 | -0.184399056 | 0.133899024 | -0.102559388 | 0.081839668 | -0.236458411 | 0.062927633 | 0.16484545 | 0.464443044 | 0.562150574 | 0.157725304 |
| Pip4k2b | S | 326 | 0.020652044 | -0.05532022 | -0.051132072 | -0.071784115 | 0.004188149 | 0.698773193 | 0.265799536 | 0.137479775 | 0.22004307 | 0.918611564 |
| Knop1 | S | 312 | 0.021954002 | -0.084353641 | -0.012292563 | -0.034246566 | 0.072061078 | 0.588444569 | 0.033343912 | 0.629683711 | 0.388490323 | 0.057793303 |
| Gpsm2 | S | 535 | -0.226023486 | 0.016497719 | -0.035066062 | 0.190957424 | -0.051563781 | 0.386025845 | 0.947591254 | 0.87293344 | 0.490085895 | 0.849317776 |
| Gpsm2 | S | 540 | -0.226023486 | 0.016497719 | -0.035066062 | 0.190957424 | -0.051563781 | 0.386025845 | 0.947591254 | 0.87293344 | 0.490085895 | 0.849317776 |
| Xaf1 | S | 127 | -0.060051828 | -0.072176993 | -0.020680085 | 0.039371742 | 0.051496907 | 0.509793306 | 0.413005845 | 0.889599172 | 0.773308062 | 0.704718134 |
| Otud3 | S | 46 | -0.097051131 | -0.042577189 | -0.035877383 | 0.061173747 | 0.006699806 | 0.407210013 | 0.721588837 | 0.801903492 | 0.670957065 | 0.963329928 |
| Shc4 | S | 132 | 0.015278138 | -0.072776 | -0.054160733 | -0.069438871 | 0.018615267 | 0.88343047 | 0.324790702 | 0.288468694 | 0.510135316 | 0.766237113 |
| Ppp1r12a | S | 299 | 0.065452028 | -0.101720146 | -0.116696249 | -0.182148276 | -0.014976103 | 0.174201573 | 0.156909944 | 0.243762072 | 0.121327277 | 0.871073576 |
| Erc1 | S | 55 | 0.011282734 | -0.034229491 | -0.111914924 | -0.123197658 | -0.077685433 | 0.854258691 | 0.367235119 | 0.317033392 | 0.289564029 | 0.456055218 |
| Brwd1 | S | 1902 | 0.063523876 | 0.013328689 | -0.187433432 | -0.250957308 | -0.20076212 | 0.374628438 | 0.769097836 | 0.030065091 | 0.024482323 | 0.031275712 |
| Srrm2 | S | 1820 | -0.081609169 | -0.021119813 | -0.051369705 | 0.030239464 | -0.030249891 | 0.412594944 | 0.624202491 | 0.709278594 | 0.841879951 | 0.822280183 |
| Wdr44 | S | 346 | -0.052195858 | 0.014833763 | -0.104252595 | -0.052056736 | -0.119086357 | 0.537482259 | 0.85002085 | 0.268729746 | 0.602440833 | 0.253393045 |
| Itsn1 | S | 895;904 | -0.204072795 | 0.11249832 | 0.015466114 | 0.21953891 | -0.097032205 | 0.004108979 | 0.09985808 | 0.857425369 | 0.071449754 | 0.300095427 |
| Hsp90ab1 | S | 718 | 0.065834502 | -0.014087292 | -0.205380273 | -0.271214775 | -0.191292981 | 0.571424832 | 0.864296128 | 0.130652326 | 0.105978336 | 0.164776123 |
| Rps7 | T | 121 | 0.125866425 | -0.061056412 | -0.213616754 | -0.339483178 | -0.152560341 | 0.190531378 | 0.558898161 | 0.063150206 | 0.023618643 | 0.232983486 |
| Ice1 | T | 1607 | -0.001516289 | -0.057931066 | -0.071481109 | -0.069964821 | -0.013550043 | 0.993281934 | 0.468195684 | 0.316421402 | 0.692378451 | 0.798863895 |
| Iws1 | S | 321 | -0.120118678 | -0.036039406 | 0.04909735 | 0.169216027 | 0.085136756 | 0.079474279 | 0.431339778 | 0.353482256 | 0.042671316 | 0.184939856 |
| Pgk1 | S | 415 | 0.139369415 | -0.189747432 | -0.107277314 | -0.246646729 | 0.082470119 | 0.587266849 | 0.26695215 | 0.503477852 | 0.334830699 | 0.498872606 |
| Gramd1b | S | 550 | -0.076438745 | -0.008303647 | 0.023067943 | 0.099506688 | 0.031371159 | 0.005211378 | 0.714339214 | 0.217530313 | 0.003382419 | 0.186125263 |
| Psip1 | S | 441 | -0.09966319 | 0.037403599 | -0.070988652 | 0.028674537 | -0.108392251 | 0.185685012 | 0.698562219 | 0.553638123 | 0.792901231 | 0.405142186 |
| Steap1 | S | 28 | -0.039174201 | -0.04537521 | -0.003767448 | 0.035406753 | 0.041607762 | 0.272382165 | 0.228363878 | 0.949531607 | 0.538944883 | 0.480945673 |
| Relch | T | 143 | -0.047231952 | 0.008469084 | -0.046377586 | 0.000854366 | -0.05484667 | 0.410225962 | 0.877834198 | 0.100996236 | 0.986508909 | 0.350252313 |
| Psmd1 | T | 273 | -0.03078023 | 0.009432875 | -0.051496801 | -0.020716571 | -0.060929677 | 0.054137339 | 0.727577568 | 0.28807707 | 0.627043928 | 0.237839186 |
| Cttn | T | 323 | -0.059609355 | 0.031226381 | -0.174167971 | -0.114558616 | -0.205394352 | 0.650882722 | 0.792613949 | 0.358405385 | 0.538207569 | 0.28807988 |
| Zc3h13 | S | 318 | -0.027840199 | -0.047542978 | -0.056364391 | -0.028524192 | -0.008821413 | 0.675735357 | 0.584924853 | 0.380634959 | 0.722984141 | 0.925752001 |
| Araf | S | 267 | -0.008430745 | 0.017283331 | -0.160595804 | -0.152165059 | -0.177879135 | 0.949149757 | 0.852120933 | 0.175050574 | 0.273743852 | 0.103011947 |
| Zfp516 | S | 387 | -0.039681378 | -0.010073202 | -0.090218334 | -0.050536956 | -0.080145132 | 0.683066356 | 0.824976771 | 0.29833778 | 0.655156486 | 0.353090516 |
| Lmod2 | S | 407 | -0.021792525 | 0.003004998 | -0.104401808 | -0.082609283 | -0.107406807 | 0.824579283 | 0.97015492 | 0.154148012 | 0.449711593 | 0.268477624 |
| Rsrc2 | S | 16 | 0.068370217 | -0.134638567 | -0.059562379 | -0.127932597 | 0.075076187 | 0.489404585 | 0.157410041 | 0.447063338 | 0.184088018 | 0.318379375 |
| Nsrp1 | S | 255 | -0.080270254 | -0.054535935 | 0.040285962 | 0.120556216 | 0.094821897 | 0.118788989 | 0.234411255 | 0.376592261 | 0.019398408 | 0.020805096 |
| Atp13a1 | S | 896 | -0.033116526 | -0.073586988 | -0.012860272 | 0.020256254 | 0.060726716 | 0.432520175 | 0.341072791 | 0.834341928 | 0.702032168 | 0.450392362 |
| Kmt2b | T | 2079 | -0.013642342 | -0.086218946 | -0.015664212 | -0.00202187 | 0.070554734 | 0.629961611 | 0.205398512 | 0.866844831 | 0.982175487 | 0.500435298 |
| Kdm2b | S | 880 | -0.004656702 | -0.042751585 | -0.033336208 | -0.028679506 | 0.009415377 | 0.897232028 | 0.224569654 | 0.231324443 | 0.424266472 | 0.755067968 |
| Numa1 | S | 1844 | -0.054155376 | -0.007219401 | -0.040751165 | 0.013404211 | -0.033531763 | 0.238007861 | 0.585957115 | 0.553984611 | 0.85196322 | 0.618411766 |
| Wnk1 | S | 1803 | 0.056610502 | -0.197607167 | -0.060062643 | -0.116673144 | 0.137544524 | 0.752758737 | 0.370401745 | 0.583860234 | 0.550615084 | 0.527062106 |
| Add3 | S | 64 | -0.037579055 | -0.047234846 | -0.109032147 | -0.071453092 | -0.061797301 | 0.744165205 | 0.648807685 | 0.570460229 | 0.706471129 | 0.737137687 |
| Rmdn3 | S | 212 | -0.063227126 | -0.039234113 | -0.008803124 | 0.054424001 | 0.030430989 | 0.425566382 | 0.234695493 | 0.933854667 | 0.652764961 | 0.772965271 |
| Champ1 | S | 726 | 0.037899363 | -0.004930155 | -0.134482209 | -0.172381572 | -0.129552054 | 0.787333445 | 0.968859338 | 0.083422406 | 0.276767888 | 0.34489624 |
| Fndc3a | Y | 536 | -0.07404632 | -0.015873655 | -0.033870724 | 0.040175596 | -0.017997068 | 0.365402531 | 0.731438579 | 0.510069056 | 0.623229906 | 0.749952255 |
| Wdr33 | S | 1204 | -0.029146238 | -0.071105071 | 0.023772969 | 0.052919207 | 0.094878039 | 0.699487896 | 0.036578675 | 0.694575609 | 0.549073786 | 0.189399973 |
| Npdc1 | S | 236 | 0.046930692 | -0.204913833 | 0.01957087 | -0.027359822 | 0.224484703 | 0.605023361 | 0.095642796 | 0.895715261 | 0.831227105 | 0.176722373 |
| Fndc3b | S | 257 | 0.078674501 | -0.093691842 | -0.085908193 | -0.164582694 | 0.007783649 | 0.074751823 | 0.036943967 | 0.172176724 | 0.04084249 | 0.880773154 |
| Tjap1 | S | 212 | -0.02571648 | -0.07929733 | -0.015601427 | 0.010115053 | 0.063695903 | 0.659975144 | 0.252472292 | 0.803707497 | 0.86697625 | 0.366099919 |
| Pcm1 | S | 380 | 0.027355436 | -0.080506651 | 0.003194104 | -0.024161332 | 0.083700755 | 0.660254435 | 0.009726572 | 0.953675202 | 0.749704163 | 0.212210613 |
| Pcm1 | S | 384 | 0.027355436 | -0.080506651 | 0.003194104 | -0.024161332 | 0.083700755 | 0.660254435 | 0.009726572 | 0.953675202 | 0.749704163 | 0.212210613 |
| Srrm2 | S | 962 | 0.001636314 | -0.105175679 | 0.023687018 | 0.022050705 | 0.128862697 | 0.983580003 | 0.036117837 | 0.687860382 | 0.790609856 | 0.076261183 |
| Ranbp2 | T | 2578 | -0.004091718 | -0.065844587 | -0.009456344 | -0.005364626 | 0.056388243 | 0.875518835 | 0.06245474 | 0.83733508 | 0.909336062 | 0.289220386 |
| Gja1 | S | 341 | 0.073286346 | -0.038457278 | -0.206764881 | -0.280051227 | -0.168307603 | 0.214272138 | 0.50472293 | 0.117810214 | 0.058264004 | 0.169430284 |
| Msl1 | S | 207 | -0.027140176 | -0.047778957 | 0.010754937 | 0.037895112 | 0.058533894 | 0.477506235 | 0.034784702 | 0.783357353 | 0.441192824 | 0.206560042 |
| Socs5 | S | 39 | 0.081654441 | -0.11412007 | -0.063110127 | -0.144764569 | 0.051009943 | 0.366343784 | 0.140387477 | 0.231269912 | 0.159889456 | 0.459655896 |
| Fam120c | S | 436 | 0.183065432 | -0.107610595 | -0.190049992 | -0.373115423 | -0.082439397 | 0.019198746 | 0.137300224 | 0.017126944 | 0.000500604 | 0.125074785 |
| Gtf2i | S | 207 | -0.004648238 | -0.106742225 | -0.041229847 | -0.036581609 | 0.065512377 | 0.955922034 | 0.349719775 | 0.61602189 | 0.666266512 | 0.550854926 |
| Gtf2i | S | 210 | -0.004648238 | -0.106742225 | -0.041229847 | -0.036581609 | 0.065512377 | 0.955922034 | 0.349719775 | 0.61602189 | 0.666266512 | 0.550854926 |
| Usp47 | T | 1016 | -0.053443141 | -0.017406978 | -0.022470657 | 0.030972484 | -0.005063679 | 0.287366327 | 0.42740068 | 0.579491006 | 0.566685628 | 0.895471859 |
| Epb41l1 | S | 86 | -0.03721123 | 0.010258997 | -0.154470819 | -0.117259589 | -0.164729816 | 0.735808022 | 0.946165583 | 0.341497923 | 0.389838105 | 0.331511488 |
| Ppp6r1 | S | 739 | -0.032832434 | -0.024158983 | -0.049978166 | -0.017145732 | -0.025819183 | 0.592751549 | 0.332125523 | 0.463693376 | 0.830393001 | 0.690257017 |
| Ankib1 | S | 886 | 0.010298558 | -0.076388726 | -0.055515502 | -0.06581406 | 0.020873223 | 0.939182016 | 0.238484479 | 0.543421644 | 0.659413427 | 0.818085564 |
| Fndc3a | S | 203 | -0.06753753 | -0.105281801 | 0.051085232 | 0.118622762 | 0.156367033 | 0.22458927 | 0.130494965 | 0.189175607 | 0.068923101 | 0.060786436 |
| Tln2 | S | 428 | 0.043965249 | -0.001741539 | -0.149687033 | -0.193652282 | -0.147945494 | 0.442453371 | 0.96878474 | 0.04950887 | 0.034701093 | 0.058120417 |
| Zc3h13 | S | 324 | -0.022038739 | -0.073804323 | -0.033255442 | -0.011216703 | 0.040548881 | 0.684855531 | 0.412891628 | 0.546653036 | 0.865308873 | 0.658598455 |
| Cc2d1a | T | 91 | -0.011229992 | 0.028135918 | -0.091264717 | -0.080034725 | -0.119400635 | 0.900567555 | 0.628531623 | 0.028991838 | 0.406720629 | 0.104535779 |

| | | | | | | | | | | | | |
|---|---|---|---|---|---|---|---|---|---|---|---|---|
| Usp36 | S | 554 | 0.01448214 | 0.004592968 | -0.155679776 | -0.170161916 | -0.160272744 | 0.850162972 | 0.951708263 | 0.141715712 | 0.145311032 | 0.162961262 |
| Prpf40a | T | 754 | -0.080375777 | -0.17681807 | 0.048346142 | 0.128721919 | 0.225164211 | 0.616837825 | 0.431932089 | 0.831282633 | 0.553074713 | 0.394443797 |
| Sun2 | S | 147 | -0.086226364 | -0.00220001 | -0.128202045 | -0.041975681 | -0.126002035 | 0.476532529 | 0.987108525 | 0.664069436 | 0.887649868 | 0.682129185 |
| Vcl | S | 755 | -0.075454663 | 0.041224486 | -0.033414814 | 0.042039849 | -0.0746393 | 0.019027538 | 0.391086949 | 0.572804744 | 0.472996796 | 0.284954683 |
| Srrm2 | S | 1922 | -0.072819468 | -0.039791078 | -0.010797153 | 0.062022315 | 0.028993925 | 0.361938695 | 0.484960955 | 0.831231099 | 0.427297573 | 0.600756742 |
| Ssh1 | S | 781 | 0.078810784 | -0.122935507 | -0.00494402 | -0.083754805 | 0.117991486 | 0.398325118 | 0.011826206 | 0.943787551 | 0.416652992 | 0.165627597 |
| Thrap3 | S | 935 | -0.106179366 | -0.061301677 | 0.05911915 | 0.165298516 | 0.120420826 | 0.169477926 | 0.135984964 | 0.221569882 | 0.068427905 | 0.037645868 |
| Tjp2 | T | 915 | -0.009285056 | -0.026748681 | -0.109525834 | -0.100240778 | -0.082777153 | 0.938435112 | 0.748537809 | 0.280797718 | 0.454630315 | 0.415224923 |
| Ypel5 | S | 118 | -0.211561325 | 0.055794365 | -0.049236355 | 0.16232497 | -0.10503072 | 0.229003296 | 0.343263546 | 0.552004726 | 0.334339996 | 0.237894641 |
| Kmt2a | S | 1053 | -0.084750156 | -0.008526434 | -0.076371952 | 0.008378204 | -0.067845518 | 0.564972281 | 0.940821719 | 0.464504285 | 0.943373009 | 0.387700574 |
| Bag3 | S | 380 | 0.192993787 | -0.096542121 | -0.176725889 | -0.369719676 | -0.080183768 | 0.219336642 | 0.229867433 | 0.092233345 | 0.059840048 | 0.403254317 |
| Sec61a1 | S | 408 | 0.000556682 | -0.078826362 | -0.009704958 | -0.010261641 | 0.069121403 | 0.991206753 | 0.083691889 | 0.814360425 | 0.844146389 | 0.153375208 |
| Dph1 | S | 418 | 0.030332116 | -0.075609732 | -0.047526333 | -0.077858449 | 0.028083399 | 0.553427392 | 0.117515971 | 0.374662438 | 0.207019358 | 0.578907461 |
| Mtor | S | 2448 | 0.127911918 | -0.07542262 | -0.205700606 | -0.333612524 | -0.130277986 | 0.090486101 | 0.195477032 | 0.100363304 | 0.027547309 | 0.224613067 |
| Polr2a | S | 1845 | 0.040944083 | -0.093433815 | -0.148080839 | -0.189024921 | -0.054647023 | 0.603470975 | 0.555156247 | 0.35179269 | 0.260707053 | 0.77458747 |
| Setd1b | S | 1283 | -0.151102006 | -0.040301949 | 0.066929076 | 0.218031082 | 0.107231024 | 0.115242151 | 0.670512228 | 0.503304779 | 0.049588467 | 0.263300529 |
| Cbx5 | S | 110 | 0.029185949 | -0.041161032 | -0.074270159 | -0.103456107 | -0.033109127 | 0.450716183 | 0.274467207 | 0.108998207 | 0.049871004 | 0.40382669 |
| Soga1 | T | 922 | -0.123493556 | -0.077725322 | 0.062068619 | 0.185562175 | 0.139793941 | 0.139023183 | 0.649650092 | 0.54930055 | 0.14481233 | 0.458905218 |
| Filip1l | S | 1008 | -0.071369918 | -0.092204172 | -0.007966345 | 0.063403574 | 0.084237827 | 0.636045516 | 0.414080045 | 0.955912406 | 0.66749705 | 0.464265396 |
| Filip1l | S | 1010 | -0.071369918 | -0.092204172 | -0.007966345 | 0.063403574 | 0.084237827 | 0.636045516 | 0.414080045 | 0.955912406 | 0.66749705 | 0.464265396 |
| Filip1l | S | 1011 | -0.071369918 | -0.092204172 | -0.007966345 | 0.063403574 | 0.084237827 | 0.636045516 | 0.414080045 | 0.955912406 | 0.66749705 | 0.464265396 |
| Mark2 | S | 616 | 0.056273288 | -0.069098254 | -0.068064339 | -0.124337627 | 0.001033915 | 0.401215125 | 0.091562603 | 0.258139941 | 0.130995494 | 0.982937188 |
| Sirt2 | S | 368 | 0.024407738 | -0.037598585 | -0.086596363 | -0.111004101 | -0.048997777 | 0.767446632 | 0.553304613 | 0.138353708 | 0.226617054 | 0.422772656 |
| Pou2f1 | S | 365 | -0.178496506 | 0.076116766 | -0.043278995 | 0.13521751 | -0.119395761 | 0.123893656 | 0.40399425 | 0.657766811 | 0.160607685 | 0.109164988 |
| Usp42 | S | 71 | 0.015224616 | -0.064173897 | -0.073307619 | -0.088532235 | -0.009133722 | 0.701659567 | 0.156929245 | 0.544224393 | 0.471637106 | 0.936814403 |
| Nefm | S | 669 | 0.009201534 | -0.069187812 | -0.008335326 | -0.017536859 | 0.060852486 | 0.861785142 | 0.037236706 | 0.808048851 | 0.747397637 | 0.11581905 |
| Axin1 | S | 485 | -0.060317444 | 0.03241521 | -0.091355111 | -0.031037668 | -0.123770321 | 0.194151982 | 0.738739249 | 0.460565141 | 0.785076139 | 0.387833151 |
| Stat6 | S | 11 | -0.078582446 | 0.095540131 | -0.141709906 | -0.063127461 | -0.237250037 | 0.360095672 | 0.327646918 | 0.143334733 | 0.210165242 | 0.018207438 |
| Kpna4 | T | 71 | -0.083982223 | 0.089194099 | -0.237600657 | -0.153618433 | -0.326794756 | 0.828700132 | 0.787441193 | 0.346698796 | 0.715039007 | 0.397577713 |
| Akap12 | T | 1191 | -0.053754024 | -0.042636006 | 0.040706977 | 0.094461001 | 0.083342983 | 0.107095424 | 0.048587175 | 0.290102222 | 0.062075326 | 0.08504578 |
| Stat3 | T | 714 | 0.012506233 | -0.064379011 | -0.073072008 | -0.08557824 | -0.008692997 | 0.829191454 | 0.316728127 | 0.403955093 | 0.329726761 | 0.91614568 |
| Pard3 | S | 824 | 0.027496652 | 0.076554793 | -0.231826975 | -0.259323627 | -0.308381767 | 0.722041673 | 0.429103624 | 0.051213058 | 0.042862722 | 0.026090524 |
| Vcp | S | 770 | -0.072521075 | 0.019770638 | -0.061222278 | 0.011298796 | -0.080992916 | 0.358642465 | 0.777883742 | 0.281399142 | 0.874673451 | 0.271549615 |
| Nfkb1 | S | 755 | 0.026666913 | -0.066018566 | -0.103992279 | -0.130659193 | -0.037973713 | 0.797233532 | 0.5836458 | 0.268664317 | 0.135830297 | 0.679502818 |
| Fmnl3 | S | 174 | -0.040807947 | -0.020704686 | -0.022180625 | 0.018627322 | -0.001475939 | 0.248555149 | 0.617674225 | 0.401871347 | 0.496528806 | 0.966960048 |
| Usp24 | S | 1922 | -0.053046297 | -0.041038488 | -0.061266407 | -0.00822011 | -0.02022792 | 0.698265359 | 0.762092445 | 0.401660435 | 0.949612595 | 0.876197754 |
| Mthfsd | S | 285 | -0.013550085 | -0.001240197 | -0.095739838 | -0.082189752 | -0.094499641 | 0.936476647 | 0.990426224 | 0.125214075 | 0.634602185 | 0.383752274 |
| Top2b | T | 1418 | -0.130804906 | -0.039967987 | 0.033969111 | 0.164774017 | 0.073937098 | 0.246078089 | 0.569666272 | 0.7495271 | 0.243601957 | 0.520020257 |
| Marcks | S | 46 | -0.056648494 | -0.032760671 | -0.029404771 | 0.027243723 | 0.0033559 | 0.536793845 | 0.308581995 | 0.735839206 | 0.812372414 | 0.969511721 |
| Chd8 | S | 1997 | 0.032713505 | -0.128586845 | 0.0106759 | -0.022037605 | 0.139262745 | 0.447683979 | 0.030553409 | 0.925204641 | 0.844553745 | 0.287039409 |
| Dennd4c | S | 971 | -0.032413132 | -0.020610823 | -0.021737734 | 0.010675398 | -0.001126912 | 0.355277141 | 0.333675931 | 0.380539091 | 0.737934363 | 0.949909947 |
| Map4 | S | 617 | -0.018181708 | -0.021904291 | -0.041775321 | -0.023593613 | -0.01987103 | 0.454749079 | 0.446079191 | 0.295306696 | 0.481872719 | 0.571193108 |
| Sacs | T | 4263 | -0.010006578 | -0.043409269 | -0.079116323 | -0.069109746 | -0.035707055 | 0.747957618 | 0.53691797 | 0.425146481 | 0.472281022 | 0.731695616 |
| Napg | T | 287 | -0.110900223 | -0.027635318 | -0.030136092 | 0.08076413 | -0.002500774 | 0.371442147 | 0.908392793 | 0.739523812 | 0.549802001 | 0.991942706 |
| Actb Actg1 | S | 199;199 | 0.326498505 | -0.105525539 | -0.125143018 | -0.451641523 | -0.019617479 | 0.40987256 | 0.115745887 | 0.096875026 | 0.287780675 | 0.757096503 |
| Gtpbp1 | S | 8 | -0.048291787 | 0.000826567 | -0.043495854 | 0.004795933 | -0.044322421 | 0.234544313 | 0.985711347 | 0.34062597 | 0.918464447 | 0.429571464 |
| Pds5b | S | 1176 | -0.061235063 | -0.024755547 | -0.001545926 | 0.059689137 | 0.023209621 | 0.176892591 | 0.437501846 | 0.951264862 | 0.194366447 | 0.508390343 |
| Pip4k2b | T | 322 | 0.019378552 | -0.060413521 | -0.044732771 | -0.064111323 | 0.01568075 | 0.731355847 | 0.22457677 | 0.233950716 | 0.275850607 | 0.697720484 |
| D830031N03Rik | S | 468 | -0.062283389 | -0.041607392 | 0.02291003 | 0.085193419 | 0.064517422 | 0.121964787 | 0.368152789 | 0.708011185 | 0.22401382 | 0.349085515 |
| Ddx10 | S | 576 | -0.019037561 | -0.07664991 | -0.025640562 | -0.006603001 | 0.051009348 | 0.83894413 | 0.222753911 | 0.780405126 | 0.956120214 | 0.610964089 |
| Bcl7b | S | 114 | 0.14757078 | -0.080852098 | -0.150346085 | -0.297916865 | -0.069493987 | 0.040692716 | 0.337211306 | 0.006706953 | 0.006492619 | 0.394946629 |
| Arid4b | S | 839 | -0.046561121 | -0.056153293 | -0.007508263 | 0.039052858 | 0.04864503 | 0.449383071 | 0.3218329 | 0.881813145 | 0.57395912 | 0.457625043 |
| Pde4d | S | 129 | -0.198618956 | 0.086471016 | 0.055210561 | 0.253829517 | -0.031260455 | 0.007344568 | 0.359429698 | 0.434560843 | 0.016746125 | 0.727781073 |
| Pard3b | S | 337 | -0.224665618 | -0.038991292 | 0.106060599 | 0.330726217 | 0.145051891 | 0.116611368 | 0.767609488 | 0.379607257 | 0.021812391 | 0.161289079 |
| Prr11 | S | 363 | 0.018667954 | -0.067773468 | -0.066556618 | -0.085224572 | 0.00121685 | 0.847548167 | 0.163951609 | 0.610409286 | 0.562799004 | 0.992071543 |
| Arhgap39 | S | 295 | -0.196282709 | -0.006604546 | 0.009115396 | 0.205398104 | 0.015719942 | 0.300738652 | 0.934094401 | 0.899212074 | 0.283581086 | 0.704626554 |
| Dab2ip | S | 1108 | 0.068090713 | -0.027192813 | -0.170102854 | -0.238193567 | -0.142910041 | 0.4784445 | 0.748622769 | 0.111459021 | 0.034453952 | 0.105075572 |
| Nes | S | 321 | 0.020759064 | -0.026721255 | -0.041272772 | -0.062031836 | -0.014551517 | 0.613798158 | 0.260241208 | 0.026804912 | 0.207248086 | 0.464154236 |
| Sltm | S | 999 | -0.12118655 | -0.0646106 | 0.068606993 | 0.189793544 | 0.133217594 | 0.169301168 | 0.147811917 | 0.184601231 | 0.067530847 | 0.040147654 |
| Anapc1 | S | 1347 | -0.028578226 | 0.013625402 | -0.047331951 | -0.018753725 | -0.060957353 | 0.374183443 | 0.83365297 | 0.067634804 | 0.50776923 | 0.385942509 |
| Huwe1 | S | 3922 | -0.014506199 | -0.013135777 | -0.027275871 | -0.012769671 | -0.014140094 | 0.337529982 | 0.192912673 | 0.224147583 | 0.553269424 | 0.478599666 |
| Cdc42bpa | S | 1532 | -0.005492574 | -0.007363981 | -0.062148735 | -0.056656161 | -0.054784754 | 0.895671027 | 0.761486109 | 0.07172121 | 0.231584896 | 0.088736162 |
| Anks1 | S | 461 | 0.023830974 | -0.080934777 | -0.03022503 | -0.054056004 | 0.050709747 | 0.321259395 | 0.044719751 | 0.540205884 | 0.297489119 | 0.33544115 |
| Smc1a | S | 360 | 0.017897504 | -0.099655923 | 0.007456414 | -0.01044109 | 0.107112337 | 0.590933118 | 0.024836275 | 0.834932708 | 0.792109154 | 0.043881519 |
| Tbc1d10a | S | 39 | -0.029653379 | -0.068374964 | -0.009771267 | 0.019882113 | 0.058603697 | 0.535748378 | 0.252429875 | 0.910845112 | 0.824048915 | 0.538657673 |
| Anln | S | 787 | -0.000631783 | -0.128620648 | -0.030685426 | -0.030053643 | 0.097935221 | 0.995817568 | 0.300749372 | 0.819232685 | 0.812188589 | 0.451131791 |
| Ino80 | S | 242 | 0.021800969 | -0.084217593 | -0.049640336 | -0.071441304 | 0.034577258 | 0.676521302 | 0.166835363 | 0.534575797 | 0.361992048 | 0.648546045 |
| Setd2 | S | 890 | 0.07390219 | -0.06449577 | -0.090126923 | -0.164029113 | -0.025631152 | 0.416956214 | 0.251735945 | 0.130781069 | 0.125224249 | 0.234927089 |
| Supt6 | S | 73 | 0.025360015 | -0.144049733 | 0.024114597 | -0.001245418 | 0.16816433 | 0.716178576 | 0.048779233 | 0.685525002 | 0.976635256 | 0.000206987 |
| Supt6 | S | 75 | 0.025360015 | -0.144049733 | 0.024114597 | -0.001245418 | 0.16816433 | 0.716178576 | 0.048779233 | 0.685525002 | 0.976635256 | 0.000206987 |
| Supt6 | S | 78 | 0.025360015 | -0.144049733 | 0.024114597 | -0.001245418 | 0.16816433 | 0.716178576 | 0.048779233 | 0.685525002 | 0.976635256 | 0.000206987 |
| Vim | S | 39 | 0.120224644 | -0.064444838 | -0.170399608 | -0.290624252 | -0.10595477 | 0.560808223 | 0.723733987 | 0.128371315 | 0.212437167 | 0.543101588 |
| Ap3b1 | S | 276 | -0.052215685 | 0.065527747 | -0.054673062 | -0.002457377 | -0.120200809 | 0.199158315 | 0.390099449 | 0.033979468 | 0.941656186 | 0.179441777 |
| Anks1 | S | 692 | -0.06275642 | 0.022009801 | -0.062491928 | 0.000264493 | -0.084501728 | 0.258144706 | 0.668531214 | 0.299765879 | 0.996744533 | 0.247500641 |
| Elp2 | S | 483 | -0.13874674 | 0.042302298 | -0.082893917 | 0.055852823 | -0.125196215 | 0.351559679 | 0.753396774 | 0.61629509 | 0.704041716 | 0.3901449 |
| 4932438A13Rik | S | 2755 | 0.003371668 | -0.037051837 | -0.06470053 | -0.068072198 | -0.027648693 | 0.964952662 | 0.53073756 | 0.186815584 | 0.4298497 | 0.665613869 |
| Rlim | S | 227 | -0.045291152 | -0.023362753 | -0.022302315 | 0.022988837 | 0.001060437 | 0.338694252 | 0.545253103 | 0.46328826 | 0.622878352 | 0.979245625 |
| Uhrf1 | S | 656 | -0.048608294 | 0.009591418 | -0.058668791 | -0.010060497 | -0.068260208 | 0.464263661 | 0.789323602 | 0.16365848 | 0.86291873 | 0.042457059 |
| Itch | S | 62 | 0.074244906 | -0.04636996 | -0.15569241 | -0.229937316 | -0.10932245 | 0.368551582 | 0.413675335 | 0.152324826 | 0.073737189 | 0.271659779 |
| Srrm2 | S | 814 | 0.129052557 | -0.133449556 | -0.092545941 | -0.221598498 | 0.040903615 | 0.015830725 | 0.024696939 | 0.067362394 | 0.002892828 | 0.356912499 |
| Specc1 | S | 424 | 0.084149579 | -0.042547536 | -0.216872972 | -0.301022552 | -0.174325437 | 0.46640074 | 0.442890244 | 0.26305711 | 0.161889811 | 0.344887106 |
| Nes | T | 158 | -0.036937008 | 0.005629833 | -0.041256239 | -0.00431923 | -0.046886072 | 0.176622058 | 0.756768719 | 0.201902222 | 0.89013435 | 0.16349015 |
| Aptx | S | 123 | 0.091521972 | -0.097335301 | -0.085007405 | -0.176529377 | 0.012327896 | 0.148214726 | 0.313166354 | 0.027986756 | 0.042857715 | 0.882465494 |
| Aptx | S | 150 | 0.091521972 | -0.097335301 | -0.085007405 | -0.176529377 | 0.012327896 | 0.148214726 | 0.313166354 | 0.027986756 | 0.042857715 | 0.882465494 |
| Anln | S | 557 | 0.134436752 | -0.069652525 | -0.149124383 | -0.283561135 | -0.079471857 | 0.045434862 | 0.182312743 | 0.018931104 | 0.004141169 | 0.102943962 |
| Zfp462 | S | 1346 | -0.203820822 | -0.000535621 | 0.026499356 | 0.230320178 | 0.027034978 | 0.218436379 | 0.994143971 | 0.737306102 | 0.180490453 | 0.69782197 |
| Rpl37 Rpl37rt | S | 97 | 0.081310882 | -0.016642793 | -0.157070036 | -0.238380918 | -0.140427242 | 0.065804482 | 0.684900693 | 0.009057181 | 0.005644366 | 0.015466345 |
| Prrc2a | S | 1087 | 0.032673706 | -0.000281089 | -0.058229313 | -0.090903019 | -0.057948225 | 0.348817846 | 0.995746626 | 0.000662206 | 0.071876486 | 0.337101276 |
| Ugdh | T | 474 | -0.016416896 | 0.000200158 | -0.069981972 | -0.053565077 | -0.07018213 | 0.667005999 | 0.997632634 | 0.116919459 | 0.071288527 | 0.314328811 |
| Mycbp2 | S | 3577 | -0.041905343 | 0.007783785 | -0.112174807 | -0.070269464 | -0.119958592 | 0.674047344 | 0.875689798 | 0.292235953 | 0.560681404 | 0.261952586 |
| Foxm1 | S | 635 | -0.093036478 | -0.040643889 | 0.056040993 | 0.149077471 | 0.096684881 | 0.078893025 | 0.378371073 | 0.431237122 | 0.086253359 | 0.191473314 |
| Dlg1 | S | 122 | 0.065651902 | -0.066294873 | -0.112367138 | -0.17801904 | -0.046072265 | 0.06629839 | 0.174149772 | 0.068164683 | 0.023145517 | 0.406971264 |
| Map1b | S | 1325 | -0.121795588 | 0.062994451 | -0.073462988 | 0.0483326 | -0.136457438 | 0.073593567 | 0.12321856 | 0.378961528 | 0.56495924 | 0.160975491 |
| Rgs19 | S | 80 | -0.013925875 | -0.06985996 | 0.019195074 | 0.03312095 | 0.089055034 | 0.391578588 | 0.067602591 | 0.712074929 | 0.529568566 | 0.169750671 |

| | | | | | | | | | | | | |
|---|---|---|---|---|---|---|---|---|---|---|---|---|
| Triobp | S | 1598 | 0.122752483 | -0.123298518 | -0.120879964 | -0.243632447 | 0.002418554 | 0.037489205 | 0.091728686 | 0.075237898 | 0.011319653 | 0.970755164 |
| Rbbp7 | S | 95 | -0.041784842 | -0.042655142 | -0.028696364 | 0.013088478 | 0.013958778 | 0.513592698 | 0.466633581 | 0.606883955 | 0.851688893 | 0.831603275 |
| Nes | S | 1444 | 0.006865381 | -0.0406506 | -0.084138254 | -0.091003636 | -0.043487654 | 0.920813073 | 0.477833889 | 0.302181788 | 0.331060252 | 0.5897405 |
| Epb41l2 | S | 555 | 0.01535199 | 0.004421367 | -0.075876554 | -0.091228544 | -0.080297921 | 0.746308847 | 0.928643757 | 0.01824101 | 0.128942846 | 0.176847691 |
| Nes | S | 688 | -0.054847899 | -0.077956844 | 0.035796131 | 0.09064403 | 0.113752976 | 0.341673269 | 0.179907079 | 0.567968288 | 0.090807236 | 0.071474519 |
| Csnk1g3 | S | 345 | 0.000391943 | -0.029483444 | -0.101367756 | -0.101759699 | -0.071884311 | 0.996990501 | 0.785877789 | 0.235708776 | 0.275972214 | 0.442235661 |
| Scrib | S | 1331 | -0.027966147 | 0.011045521 | -0.062904166 | -0.03493802 | -0.073949687 | 0.305405703 | 0.64994462 | 0.138297387 | 0.347085013 | 0.102353918 |
| Dab2ip | S | 957 | 0.034882748 | -0.024499515 | -0.142792633 | -0.177675381 | -0.118293118 | 0.805346679 | 0.749936908 | 0.216643881 | 0.291644846 | 0.29944053 |
| Srrm2 | S | 1068 | -0.031546367 | -0.096994908 | 0.064102438 | 0.095648805 | 0.161097346 | 0.53891561 | 0.042727801 | 0.4416658 | 0.276406993 | 0.127391026 |
| Sdcbp | Y | 57 | -0.125707755 | 0.073930039 | -0.127656783 | -0.001949028 | -0.201586822 | 0.378565345 | 0.473955458 | 0.384749339 | 0.990890613 | 0.234785091 |
| Trip12 | S | 312 | 0.01796402 | -0.047356446 | -0.019813636 | -0.037777657 | 0.027542809 | 0.516214639 | 0.028775581 | 0.263191231 | 0.202999569 | 0.089826497 |
| Ahdc1 | S | 842 | -0.083436586 | 0.008983467 | -0.020332575 | 0.063104011 | -0.029316041 | 0.125844103 | 0.843465765 | 0.69085482 | 0.293883785 | 0.596298451 |
| Synj1 | S | 1049 | 0.038449127 | -0.07798482 | -0.107702472 | -0.146151599 | -0.029717652 | 0.424436359 | 0.483174491 | 0.215120497 | 0.124440679 | 0.801526505 |
| Synj1 | S | 1053 | 0.038449127 | -0.07798482 | -0.107702472 | -0.146151599 | -0.029717652 | 0.424436359 | 0.483174491 | 0.215120497 | 0.124440679 | 0.801526505 |
| Ubr4 | T | 4459 | -0.078548837 | -0.028048624 | -0.079059052 | -0.000510215 | -0.051010428 | 0.48348049 | 0.842321483 | 0.738566082 | 0.998191721 | 0.8307469 |
| Ubr4 | T | 4460 | -0.078548837 | -0.028048624 | -0.079059052 | -0.000510215 | -0.051010428 | 0.48348049 | 0.842321483 | 0.738566082 | 0.998191721 | 0.8307469 |
| Zfp638 | S | 128 | 0.027684741 | -0.046265531 | -0.103227289 | -0.130912031 | -0.056961758 | 0.664799469 | 0.453134513 | 0.253159631 | 0.143950671 | 0.440729168 |
| Filip1l | S | 1017 | -0.094357905 | -0.027638249 | -0.025610827 | 0.068747078 | 0.002027422 | 0.355775753 | 0.753045654 | 0.866720444 | 0.631850504 | 0.987905776 |
| Rnps1 | S | 139 | 0.006353161 | -0.119109192 | 0.051511037 | 0.045157875 | 0.170620229 | 0.912452157 | 0.017548282 | 0.430131394 | 0.539685455 | 0.056779159 |
| Lmf2 | S | 683 | 0.067712387 | -0.079422378 | -0.109374634 | -0.17708702 | -0.029952255 | 0.127868472 | 0.18248212 | 0.136319245 | 0.05296684 | 0.652831416 |
| Tgfbrap1 | S | 430 | -0.12331241 | -0.039831854 | 0.009293081 | 0.132605491 | 0.049124935 | 0.292505079 | 0.748183267 | 0.871960659 | 0.264103381 | 0.690250058 |
| Flnb | T | 519 | -0.062726161 | -0.093330365 | -0.015808748 | 0.046917413 | 0.077521617 | 0.573107655 | 0.565323181 | 0.887568019 | 0.728269294 | 0.658479183 |
| Spast | S | 99 | 0.020099678 | -0.082835051 | -0.029824331 | -0.049924009 | 0.053010721 | 0.836276424 | 0.158454413 | 0.568892043 | 0.59683736 | 0.235587027 |
| Prkca Prkcb Prkcg | T | 497;500;514 | -0.030591841 | -0.005744897 | -0.08390412 | -0.053312279 | -0.078159223 | 0.468921255 | 0.853124259 | 0.398663297 | 0.578936367 | 0.426336928 |
| Larp7 | S | 334 | -0.078914652 | -0.018586615 | 0.031057457 | 0.10997211 | 0.049644073 | 0.03200416 | 0.377689094 | 0.292003071 | 0.018146632 | 0.11503685 |
| Wdr44 | S | 405 | -0.093989381 | 0.033524516 | -0.026934841 | 0.06705454 | -0.060459357 | 0.084641012 | 0.567391689 | 0.60677013 | 0.26964453 | 0.376948475 |
| Zmynd11 | S | 420 | 0.144249228 | -0.191349754 | -0.061540796 | -0.205790023 | 0.129808959 | 0.333140178 | 0.090668786 | 0.590828746 | 0.239986216 | 0.336517182 |
| Pelp1 | S | 482 | 0.014898707 | -0.179153334 | 0.047676819 | 0.032778112 | 0.226830153 | 0.787784093 | 0.047015003 | 0.358796307 | 0.386369956 | 0.037733177 |
| Srrm2 | T | 973 | 0.016258707 | -0.049313299 | -0.020283802 | -0.036542509 | 0.029029497 | 0.659673654 | 0.06347528 | 0.230633688 | 0.347412685 | 0.194584397 |
| Slc23a2 | T | 19 | 0.14943926 | -0.102954926 | -0.153980075 | -0.303419335 | -0.05102515 | 0.047898133 | 0.241522514 | 0.025874416 | 0.004924517 | 0.528306546 |
| Fosl1 | S | 101 | 0.14382988 | -0.137166946 | -0.209159498 | -0.352989378 | -0.071992552 | 0.002932345 | 0.038034992 | 0.093513248 | 0.04184785 | 0.45413834 |
| Thrap3 | S | 53 | -0.121918711 | -0.054890863 | 0.040483099 | 0.162401811 | 0.095373962 | 0.2445052 | 0.518754841 | 0.588870145 | 0.121365339 | 0.161595731 |
| 8030462N17Rik | S | 69 | 0.051435822 | -0.104697907 | -0.061643195 | -0.113079016 | 0.043054713 | 0.524582404 | 0.154283001 | 0.531799681 | 0.288564631 | 0.639029611 |
| Map1b | T | 905 | -0.077919502 | 0.001534394 | -0.012211841 | 0.065707661 | -0.013746235 | 0.123415558 | 0.980567842 | 0.750536299 | 0.214261069 | 0.836499096 |
| Fnbp4 | S | 487 | -0.013920307 | -0.078347132 | -0.015686816 | -0.001766509 | 0.062660316 | 0.922085397 | 0.173013011 | 0.836165603 | 0.990657416 | 0.450191306 |
| Pcm1 | S | 1936 | 0.148803967 | -0.204353976 | 0.031668997 | -0.117134969 | 0.236022973 | 0.260101338 | 0.005462566 | 0.59811434 | 0.353815315 | 0.010056554 |
| Kdm4a | S | 616 | -0.085971311 | -0.081009803 | 0.029654457 | 0.115625768 | 0.11066426 | 0.339689442 | 0.363734799 | 0.543944318 | 0.217007934 | 0.229909593 |
| Nup188 | S | 1718 | -0.006109598 | -0.048715824 | -0.016563187 | -0.01045359 | 0.032152637 | 0.646306127 | 0.090021493 | 0.791880988 | 0.865598962 | 0.622133087 |
| Anapc1 | S | 680 | 0.173768239 | -0.16027483 | -0.123719029 | -0.297487268 | 0.0365558 | 0.28302154 | 0.05708744 | 0.610601001 | 0.291274227 | 0.875465659 |
| Retreg2 | S | 339 | -0.00632213 | -0.020823243 | -0.063901427 | -0.057579298 | -0.043078184 | 0.93857913 | 0.69869085 | 0.147742176 | 0.481535081 | 0.358915928 |
| Slc38a10 | S | 991 | 0.029099677 | -0.029402362 | -0.132171696 | -0.161271373 | -0.102769334 | 0.394498866 | 0.331929041 | 0.230794752 | 0.17229356 | 0.322783414 |
| Nin | S | 1540 | -0.198354149 | -0.116998812 | 0.056512595 | 0.254866744 | 0.173511407 | 0.446608467 | 0.649565223 | 0.603767881 | 0.341841695 | 0.504049418 |
| Ophn1 | T | 736 | -0.109896423 | 0.064756855 | -0.050850471 | 0.059045951 | -0.115607326 | 0.018328677 | 0.083485324 | 0.478604568 | 0.423900711 | 0.174285451 |
| Prkaa1 | S | 531 | -0.06492353 | -0.062187252 | -0.008302021 | 0.056621509 | 0.053885231 | 0.404742322 | 0.608482558 | 0.846143692 | 0.428628468 | 0.644449826 |
| Plec | S | 728 | -0.098607783 | 0.017457299 | -0.025225104 | 0.07338268 | -0.042682402 | 0.143960095 | 0.733171807 | 0.718177795 | 0.353700586 | 0.546435174 |
| Utp18 | S | 206 | 0.062756939 | -0.084305157 | -0.057742806 | -0.120499745 | 0.026562352 | 0.022825423 | 0.027945174 | 0.069331866 | 0.008508178 | 0.389518056 |
| Lrrk1 | S | 1854 | -0.145892751 | 0.008302188 | -0.077970193 | 0.067922558 | -0.086272382 | 0.532200611 | 0.94515562 | 0.682014993 | 0.794710882 | 0.655188912 |
| Mxra8 | S | 423 | -0.041702563 | -0.084607463 | 0.036910215 | 0.078612778 | 0.121517678 | 0.327326119 | 0.147508592 | 0.482012506 | 0.194138658 | 0.095600773 |
| Rab8a | S | 185 | -0.004004061 | -0.018007289 | -0.044350746 | -0.040346685 | -0.026343457 | 0.902210376 | 0.533666959 | 0.089340581 | 0.27634226 | 0.392759062 |
| Ranbp2 | S | 2576 | 0.004059453 | -0.05834092 | -0.008372156 | -0.01243161 | 0.049968764 | 0.875545671 | 0.035904036 | 0.85219712 | 0.789010477 | 0.319099384 |
| Erf | S | 327 | -0.078219817 | 0.027447215 | -0.055517434 | 0.022702384 | -0.082964649 | 0.088740272 | 0.325078018 | 0.508514213 | 0.78075267 | 0.347697154 |
| Micall2 | S | 122 | -0.02733811 | 0.05533587 | -0.130218904 | -0.102880794 | -0.185554774 | 0.599967172 | 0.376133499 | 0.084091689 | 0.125764509 | 0.031652183 |
| Ankrd11 | S | 276 | -0.013420675 | -0.039950986 | -0.02578913 | -0.012368455 | 0.014161856 | 0.581208466 | 0.202100546 | 0.599439503 | 0.794117125 | 0.773825782 |
| Mon2 | S | 534 | -0.092654309 | 0.056624968 | -0.049393536 | 0.043260773 | -0.106018503 | 0.095206054 | 0.271000653 | 0.259781294 | 0.388022373 | 0.082965421 |
| Osbpl8 | S | 120 | -0.133671985 | 0.002613138 | 0.048245516 | 0.181917501 | 0.045632378 | 0.043661242 | 0.964088502 | 0.493534005 | 0.037334929 | 0.452397302 |
| Pex10 | S | 258 | 0.061958219 | 0.061509289 | -0.169481094 | -0.231439313 | -0.230990383 | 0.472593875 | 0.515088531 | 0.011483561 | 0.061742372 | 0.080212793 |
| Telo2 | S | 486 | -0.049098908 | -0.007415534 | -0.04248086 | 0.006618048 | -0.035065326 | 0.341612426 | 0.632342084 | 0.528721056 | 0.928069673 | 0.594374653 |
| Telo2 | S | 488 | -0.049098908 | -0.007415534 | -0.04248086 | 0.006618048 | -0.035065326 | 0.341612426 | 0.632342084 | 0.528721056 | 0.928069673 | 0.594374653 |
| Telo2 | S | 492 | -0.049098908 | -0.007415534 | -0.04248086 | 0.006618048 | -0.035065326 | 0.341612426 | 0.632342084 | 0.528721056 | 0.928069673 | 0.594374653 |
| Exo1 | S | 691 | 0.032602826 | -0.045063622 | -0.169946705 | -0.202549531 | -0.124883083 | 0.761896011 | 0.535183955 | 0.430052361 | 0.369495324 | 0.54643138 |
| Erbin | S | 1099 | -0.008013317 | -0.069842698 | -0.077848379 | -0.069835062 | -0.008005681 | 0.97965211 | 0.507342288 | 0.532146506 | 0.823215403 | 0.929316915 |
| Parp3 | S | 167 | 0.009236052 | -0.010080275 | -0.05831791 | -0.067553962 | -0.048237635 | 0.82770108 | 0.672620512 | 0.042022174 | 0.173464695 | 0.020039741 |
| Spen | S | 747 | -0.025271285 | -0.092662014 | 0.018662479 | 0.043933764 | 0.111324494 | 0.748145942 | 0.145994891 | 0.842372362 | 0.697330542 | 0.309754896 |
| Top2b | S | 1428 | -0.116389036 | -0.005437191 | -0.058692092 | 0.057696944 | -0.053254901 | 0.423554645 | 0.951643971 | 0.787210745 | 0.810716922 | 0.813744503 |
| Efr3a | S | 363 | 0.056787573 | -0.021084463 | -0.20739808 | -0.264185653 | -0.186313617 | 0.22006311 | 0.647071412 | 0.131661382 | 0.092142927 | 0.166461336 |
| Gorasp1 | T | 439 | -0.000334657 | -0.045550712 | -0.032531371 | -0.032196714 | 0.013019341 | 0.993200362 | 0.271838565 | 0.27165072 | 0.469903916 | 0.74708818 |
| Anp32b | T | 265 | -0.104297799 | 0.127285719 | -0.059698097 | 0.044599702 | -0.186983816 | 0.016407931 | 0.169415058 | 0.290620508 | 0.417231144 | 0.075316742 |
| Rad51c | S | 11 | -0.128007224 | 0.045983535 | -0.076144948 | 0.051862276 | -0.122128483 | 0.523980901 | 0.797834032 | 0.397013002 | 0.778267528 | 0.491752555 |
| Lpin3 | S | 106 | 0.16993774 | -0.31467317 | 0.124196868 | -0.045740873 | 0.438870037 | 0.275019927 | 0.015379472 | 0.660733443 | 0.873233213 | 0.201080447 |
| Magi3 | S | 1351 | -0.133472531 | -0.01937209 | 0.047014463 | 0.180486995 | 0.066386553 | 0.127141618 | 0.908126243 | 0.747523508 | 0.254230843 | 0.729731897 |
| Nmnat1 | S | 116 | 0.046922646 | -0.004877102 | -0.125510101 | -0.172432747 | -0.120632999 | 0.560741421 | 0.909357256 | 0.060390996 | 0.092642661 | 0.069840455 |
| Cttnbp2nl | S | 443 | -0.110974564 | -0.050361151 | 0.043986089 | 0.154960653 | 0.09434724 | 0.329057996 | 0.192405245 | 0.443831745 | 0.210312012 | 0.144924271 |
| Zfpm2 | S | 846 | -0.145929642 | 0.006943292 | 0.042912441 | 0.188842083 | 0.035969149 | 0.023802971 | 0.880433009 | 0.210323349 | 0.03144626 | 0.38157753 |
| Marcks | S | 140 | -0.037926042 | -0.05417874 | -0.004390091 | 0.033535951 | 0.049788649 | 0.423777349 | 0.311309459 | 0.930591637 | 0.571217683 | 0.431593822 |
| Ncor2 | S | 750 | -0.059758976 | -0.033301723 | 0.018186348 | 0.077945323 | 0.051488071 | 0.11303815 | 0.564727842 | 0.874862096 | 0.52210277 | 0.675578999 |
| Mki67 | S | 1366 | -0.004943361 | -0.009129153 | -0.048898179 | -0.043954818 | -0.039769026 | 0.93314888 | 0.800234451 | 0.047622808 | 0.480151023 | 0.306352842 |
| Eif4g2 | T | 507 | -0.087258266 | -0.021575417 | 0.032039693 | 0.119297959 | 0.05361511 | 0.046763714 | 0.602917501 | 0.396896085 | 0.021372512 | 0.217828738 |
| Plekhf2 | S | 247 | 0.039793517 | -0.069589051 | -0.154998914 | -0.194792431 | -0.085409863 | 0.429020676 | 0.505749902 | 0.329591107 | 0.245862397 | 0.599794935 |
| Plekhf2 | S | 248 | 0.039793517 | -0.069589051 | -0.154998914 | -0.194792431 | -0.085409863 | 0.429020676 | 0.505749902 | 0.329591107 | 0.245862397 | 0.599794935 |
| Flnb | S | 2465 | 0.04872961 | -0.05232364 | -0.119698163 | -0.168427773 | -0.067374523 | 0.324950787 | 0.376485612 | 0.17488651 | 0.095670605 | 0.397702293 |
| Fhod1 | S | 495 | -0.091431814 | -0.012314721 | -0.013747936 | 0.077683878 | -0.001433215 | 0.247046458 | 0.854553974 | 0.849612948 | 0.377319705 | 0.985595045 |
| Mtmr1 | S | 656 | -0.016542445 | -0.03257675 | -0.039117398 | -0.022574953 | -0.006540647 | 0.849620739 | 0.450579782 | 0.243020732 | 0.788013706 | 0.834652138 |
| Topors | S | 585 | 0.094178727 | -0.087468359 | -0.058146259 | -0.152324985 | 0.0293221 | 0.167043729 | 0.012710432 | 0.368835169 | 0.089176395 | 0.617320755 |
| Proc | S | 446 | -0.191828724 | 0.08461683 | 0.055980935 | 0.247809658 | -0.028635895 | 0.007910317 | 0.456383952 | 0.305023715 | 0.00123872 | 0.779786122 |
| Fam120a | S | 382 | 0.040747401 | -0.066201718 | -0.133149642 | -0.173897042 | -0.066947924 | 0.635943296 | 0.500179839 | 0.3582599 | 0.231169284 | 0.596134859 |
| Plec | S | 4573 | 0.000401237 | -0.063141932 | -0.037633613 | -0.03803485 | 0.025508319 | 0.994625223 | 0.206637703 | 0.621631502 | 0.616150931 | 0.705127435 |
| Inpp5f | S | 1103 | 0.008670158 | -0.018135149 | -0.077258943 | -0.085929101 | -0.059123794 | 0.821877328 | 0.760888325 | 0.103675002 | 0.039007206 | 0.308352555 |
| Spidr | S | 840 | -0.002930453 | -0.0880472 | -0.008360397 | -0.005429944 | 0.079686802 | 0.934183396 | 0.1662859 | 0.85617113 | 0.875614824 | 0.205156876 |
| Lpin2 | S | 174 | -0.030166276 | 0.026775239 | -0.121416384 | -0.091250108 | -0.148191623 | 0.583511759 | 0.430745413 | 0.16195116 | 0.28194032 | 0.123611231 |
| Lpin2 | S | 243 | -0.029045882 | -0.037215205 | -0.046388168 | -0.017342286 | -0.009172963 | 0.59046841 | 0.622960588 | 0.458148243 | 0.778131812 | 0.907905597 |
| Fhl3 | T | 229 | -0.095912777 | -0.002170305 | 0.015726055 | 0.111638832 | 0.01789636 | 0.088885068 | 0.983395481 | 0.89774658 | 0.389271659 | 0.899777205 |
| Mier1 | S | 52 | -0.125643545 | 0.019393828 | 0.03030571 | 0.155949255 | 0.010911881 | 0.033734687 | 0.703778545 | 0.603500814 | 0.031410597 | 0.824288421 |
| Mier1 | S | 53 | -0.125643545 | 0.019393828 | 0.03030571 | 0.155949255 | 0.010911881 | 0.033734687 | 0.703778545 | 0.603500814 | 0.031410597 | 0.824288421 |
| Zeb2 | S | 360 | 0.014008311 | -0.017462922 | -0.061969698 | -0.075978009 | -0.044506776 | 0.806718367 | 0.254649299 | 0.144357906 | 0.269211013 | 0.254668174 |

| | | | | | | | | | | | | |
|---|---|---|---|---|---|---|---|---|---|---|---|---|
| Zfp830 | S | 342 | -0.036046765 | -0.081946847 | 0.051970068 | 0.088016833 | 0.133916916 | 0.36300465 | 0.065356198 | 0.400527644 | 0.179664633 | 0.081025033 |
| Washc2 | S | 768 | 0.007694735 | -0.132091982 | -0.006543745 | -0.01423848 | 0.125548237 | 0.918964569 | 0.213572214 | 0.94488071 | 0.826013506 | 0.214075965 |
| Bclaf1 | S | 523 | -0.100731406 | -0.066775362 | 0.084820916 | 0.185552321 | 0.151596278 | 0.000128464 | 0.324058816 | 0.000777301 | 0.00015339 | 0.094903916 |
| Ubap2l | S | 624 | 0.016109136 | -0.100406827 | -0.0168068 | -0.032915936 | 0.083600027 | 0.778766876 | 0.124700376 | 0.856534079 | 0.681629645 | 0.350089863 |
| Ubap2l | S | 627 | 0.016109136 | -0.100406827 | -0.0168068 | -0.032915936 | 0.083600027 | 0.778766876 | 0.124700376 | 0.856534079 | 0.681629645 | 0.350089863 |
| Ssrp1 | S | 668 | -0.025092277 | -0.067872544 | 0.044203221 | 0.069295498 | 0.112075765 | 0.207353595 | 0.013202096 | 0.137600859 | 0.056443202 | 0.011173528 |
| Golga5 | S | 150 | 0.003678726 | -0.040713482 | -0.047005616 | -0.050684343 | -0.006292135 | 0.952534647 | 0.257123162 | 0.348099313 | 0.493212638 | 0.900367516 |
| Nup133 | S | 491 | -0.040015355 | -0.055900482 | 0.005965451 | 0.045980806 | 0.061865933 | 0.296663736 | 0.353253629 | 0.951076172 | 0.645085375 | 0.562312363 |
| Zfp652 | S | 203 | -0.114470383 | 0.009117731 | 0.044024916 | 0.158495299 | 0.034907185 | 0.038119394 | 0.889714714 | 0.662858069 | 0.189105432 | 0.735048417 |
| Gpatch2 | S | 381 | 0.058919154 | -0.059567859 | -0.094075695 | -0.152994849 | -0.034507835 | 0.461559291 | 0.289586509 | 0.205487526 | 0.101238128 | 0.527273839 |
| Scrib | T | 735 | -0.13840091 | -0.009928184 | 0.038557453 | 0.176958364 | 0.048485638 | 0.131740156 | 0.943590872 | 0.73215681 | 0.190561962 | 0.763252261 |
| Mknk1 | S | 39 | -0.043389366 | -0.027769858 | -0.062545751 | -0.019156385 | -0.034775893 | 0.653312246 | 0.706819295 | 0.494268849 | 0.860990827 | 0.710296106 |
| Ncor2 | T | 2350 | -0.021241696 | -0.024982541 | -0.089357671 | -0.068115975 | -0.06437513 | 0.645783038 | 0.684651475 | 0.499663578 | 0.603760281 | 0.631786721 |
| Hdgfl2 | S | 620 | -0.090140624 | -0.015661557 | 0.025281778 | 0.115422402 | 0.040943335 | 0.072881013 | 0.69385242 | 0.571384851 | 0.072134334 | 0.444608503 |
| Rest | S | 848 | 0.010438386 | 0.01037737 | -0.121928783 | -0.132367169 | -0.132306154 | 0.895512112 | 0.811315858 | 0.106243185 | 0.183977801 | 0.092919363 |
| Nsd2 | S | 121 | 0.061905529 | -0.065647723 | -0.09069345 | -0.152598979 | -0.025045726 | 0.003669347 | 0.008004862 | 0.089280008 | 0.037040367 | 0.525075134 |
| Trp53bp1 | S | 1110 | -0.100789386 | -0.089336343 | 0.110547616 | 0.211337002 | 0.199883959 | 0.14818376 | 0.18869338 | 0.265411018 | 0.075207345 | 0.085523259 |
| Mcm4 | S | 130 | 0.037421854 | -0.058056989 | -0.012463683 | -0.049885538 | 0.045593306 | 0.471475814 | 0.005082141 | 0.780513367 | 0.429181672 | 0.353332071 |
| Senp1 | S | 412 | -0.114227235 | -0.059813102 | 0.041692909 | 0.155920144 | 0.101506011 | 0.281557794 | 0.431931983 | 0.52666371 | 0.185187167 | 0.269631392 |
| Marcks | S | 138 | -0.044633561 | -0.020571848 | 0.027977121 | 0.072610682 | 0.048548969 | 0.027621383 | 0.117203244 | 0.359069401 | 0.071989614 | 0.16949578 |
| Myo18a | S | 1966 | -0.050799662 | 0.008090366 | -0.012805703 | 0.037993959 | -0.020896069 | 0.046076808 | 0.835062372 | 0.581036838 | 0.054668901 | 0.575719995 |
| Zfpm2 | S | 434 | -0.144842043 | -0.015076733 | 0.039023918 | 0.183865961 | 0.054100651 | 0.039877683 | 0.688256211 | 0.073603874 | 0.034192986 | 0.208792947 |
| Fam117b | S | 176 | -0.091099957 | 0.060262636 | -0.068175725 | 0.022924232 | -0.128438361 | 0.254674651 | 0.30952369 | 0.650070427 | 0.185150436 | |
| Pkp4 | T | 507 | -0.028802464 | 0.008641267 | -0.090434385 | -0.061631921 | -0.099075652 | 0.65270357 | 0.918777473 | 0.248688301 | 0.415417028 | 0.309658518 |
| Cttnbp2nl | S | 444 | -0.031673795 | -0.020432386 | -0.036415907 | -0.004742112 | -0.015983522 | 0.262661767 | 0.606697426 | 0.621820775 | 0.944898241 | 0.827204591 |
| Gtf3c1 | S | 845 | -0.051689035 | -0.061963682 | 0.029512083 | 0.081201118 | 0.091475765 | 0.203995162 | 0.232642622 | 0.47040796 | 0.047697788 | 0.093965158 |
| Tbl2 | S | 415 | -0.090548336 | -0.015406556 | -0.009453747 | 0.081094589 | 0.00595281 | 0.292010783 | 0.741665324 | 0.870798145 | 0.338952625 | 0.897604172 |
| Clasp1 | S | 594 | -0.012534781 | 0.069584868 | -0.173140952 | -0.160606172 | -0.24272582 | 0.889828555 | 0.574112056 | 0.12854124 | 0.087940139 | 0.079935031 |
| Cep170 | S | 920 | 0.034560212 | -0.107884648 | -0.040641732 | -0.075201944 | 0.067242916 | 0.546365111 | 0.120162083 | 0.767481363 | 0.581896821 | 0.623845576 |
| Eif4g3 | S | 281 | 0.053679837 | -0.055895726 | -0.153436403 | -0.20711624 | -0.097540678 | 0.607293017 | 0.640163717 | 0.278387986 | 0.183582749 | 0.51555249 |
| Dennd4c | S | 740 | -0.034461334 | -0.095644908 | 0.061197985 | 0.095659319 | 0.156842893 | 0.359409431 | 0.060928444 | 0.3094628 | 0.16103366 | 0.053309204 |
| Zfp462 | S | 681 | 0.026822558 | -0.120512331 | 0.043695481 | 0.016872923 | 0.164207812 | 0.577654295 | 0.005212377 | 0.27527626 | 0.72105986 | 0.009450685 |
| Snip1 | S | 48 | -0.10329259 | 0.076535418 | -0.044467025 | 0.058825565 | -0.121002442 | 0.025079197 | 0.272558917 | 0.661382999 | 0.565478949 | 0.303367222 |
| Tsc22d4 | S | 62 | 0.079425819 | 0.06903394 | -0.218540323 | -0.297966142 | -0.287574263 | 0.381918016 | 0.267430271 | 0.008947499 | 0.033486045 | 0.00030365 |
| Rab34 | S | 241 | 0.017095541 | -0.043706198 | -0.075776256 | -0.092871798 | -0.032070059 | 0.661695315 | 0.585811519 | 0.167587959 | 0.113830578 | 0.696875948 |
| Tln2 | S | 474 | -0.009070068 | 0.011713096 | -0.197985782 | -0.188915714 | -0.209698878 | 0.899790154 | 0.8588372 | 0.363074266 | 0.384473024 | 0.342086766 |
| Eif3c | S | 907 | 0.002827111 | -0.072251773 | -0.013387867 | -0.016214977 | 0.058863906 | 0.963427677 | 0.144116677 | 0.673882169 | 0.780283414 | 0.193544203 |
| Dab2ip | S | 763 | 0.020684778 | -0.024103447 | -0.161488616 | -0.182173394 | -0.137385168 | 0.910204058 | 0.833398115 | 0.346049737 | 0.408813537 | 0.4183214 |
| Tjp2 | S | 410 | -0.062157071 | -0.029600977 | 0.036634619 | 0.09879169 | 0.066235596 | 0.105691011 | 0.321242206 | 0.669408983 | 0.300573361 | 0.453620408 |
| Fhod1 | S | 524 | -0.106453511 | -0.033347283 | 0.039038152 | 0.145491663 | 0.072385436 | 0.089177206 | 0.508485971 | 0.2840544 | 0.045107319 | 0.192633188 |
| Map2k3 Map2k6 | S | 218;207 | -0.085471605 | 0.01733171 | -0.047204747 | 0.038266857 | -0.064536457 | 0.212207658 | 0.7756226 | 0.667559902 | 0.736005822 | 0.575916585 |
| Cd44 | T | 756 | -0.01576341 | -0.033873163 | -0.034459535 | -0.018696126 | -0.000586373 | 0.683174042 | 0.32074997 | 0.467090774 | 0.69957801 | 0.989423762 |
| Dennd4b | S | 735 | 0.034813887 | -0.156851009 | 0.049406739 | 0.014592851 | 0.206257747 | 0.515897758 | 0.029511209 | 0.611505606 | 0.866129007 | 0.095620024 |
| Srrm1 | S | 832 | 0.020101999 | -0.054570473 | -0.002032653 | -0.022134652 | 0.05253782 | 0.438709723 | 0.003791269 | 0.927304244 | 0.481921335 | 0.101478933 |
| Fzr1 | S | 138 | 0.034925495 | 0.015336057 | -0.143004528 | -0.177930023 | -0.158340585 | 0.83094496 | 0.86974263 | 0.109954029 | 0.332046677 | 0.164263384 |
| Fzr1 | S | 151 | 0.034925495 | 0.015336057 | -0.143004528 | -0.177930023 | -0.158340585 | 0.83094496 | 0.86974263 | 0.109954029 | 0.332046677 | 0.164263384 |
| Specc1 | S | 863 | 0.029384843 | -0.111871145 | -0.013938555 | -0.043323398 | 0.097932591 | 0.440411556 | 0.080864205 | 0.77277322 | 0.405901707 | 0.148532034 |
| Phf11d | S | 152 | -0.137488286 | 0.018807803 | 0.046355907 | 0.183844193 | 0.027548104 | 0.016042819 | 0.746429041 | 0.273975251 | 0.006084947 | 0.582486721 |
| Phldb2 | S | 202 | 0.012509518 | 0.003892113 | -0.112961511 | -0.125471029 | -0.116853625 | 0.822048621 | 0.936689421 | 0.109582264 | 0.068733764 | 0.091954234 |
| Prkd1 | S | 231 | -0.091942532 | -0.01878133 | -0.059601824 | 0.032340708 | -0.040820494 | 0.469232407 | 0.936891378 | 0.736115341 | 0.852683275 | 0.875457055 |
| Itprid2 | S | 90 | -0.05171807 | -0.014133501 | -0.008033274 | 0.043684796 | 0.006100227 | 0.189083673 | 0.558892058 | 0.738761622 | 0.242857391 | 0.759518789 |
| Rrp8 | S | 64 | 0.022581312 | -0.056444688 | -0.036337271 | -0.058918582 | 0.020107417 | 0.247942803 | 0.075003295 | 0.237857562 | 0.099840699 | 0.532050643 |
| Slc11a2 | S | 564 | -0.055580477 | -0.029502989 | -0.052313844 | 0.003266633 | -0.022810855 | 0.597967129 | 0.68922888 | 0.614940013 | 0.976871724 | 0.797434275 |
| Hif1a | S | 244 | -0.053035999 | -0.037600038 | -0.005768485 | 0.047267514 | 0.031831552 | 0.399808456 | 0.418502251 | 0.84086969 | 0.442950283 | 0.477737482 |
| Hif1a | S | 247 | -0.053035999 | -0.037600038 | -0.005768485 | 0.047267514 | 0.031831552 | 0.399808456 | 0.418502251 | 0.84086969 | 0.442950283 | 0.477737482 |
| Hif1a | T | 241 | -0.053035999 | -0.037600038 | -0.005768485 | 0.047267514 | 0.031831552 | 0.399808456 | 0.418502251 | 0.84086969 | 0.442950283 | 0.477737482 |
| Hif1a | Y | 254 | -0.053035999 | -0.037600038 | -0.005768485 | 0.047267514 | 0.031831552 | 0.399808456 | 0.418502251 | 0.84086969 | 0.442950283 | 0.477737482 |
| Kif1b | S | 1613 | 0.039001533 | -0.064549926 | -0.136588114 | -0.175589647 | -0.072038189 | 0.751466255 | 0.33644369 | 0.579880124 | 0.505133248 | 0.764406727 |
| Hecw2 | S | 48 | -0.027041405 | -0.054856727 | -0.019693788 | 0.007347617 | 0.035162939 | 0.622653394 | 0.340854251 | 0.744836992 | 0.896298186 | 0.549367398 |
| Epha2 | S | 571 | -0.008492491 | -0.031952233 | -0.033833565 | -0.025341074 | -0.001881331 | 0.842757906 | 0.269455593 | 0.348340866 | 0.610261777 | 0.959264977 |
| Rpl18a | T | 24 | -0.061478822 | 0.026163885 | -0.137331731 | -0.075852908 | -0.163495616 | 0.818577609 | 0.849794427 | 0.337938981 | 0.759853253 | 0.056878978 |
| Ireb2 Kmt2a | S | 277;158 | 0.039664707 | -0.015185407 | -0.081012432 | -0.120677139 | -0.065827026 | 0.59900274 | 0.574716488 | 0.056470653 | 0.188783488 | 0.111534902 |
| Lats1 | S | 278 | -0.05241278 | -0.006625398 | -0.04721446 | 0.00519832 | -0.040589062 | 0.283209911 | 0.897758432 | 0.630112058 | 0.956690745 | 0.682800396 |
| Mef2a | S | 98 | 0.004660997 | -0.052557279 | -0.018462794 | -0.023123791 | 0.034094485 | 0.918223233 | 0.19614541 | 0.255325823 | 0.625652124 | 0.352786484 |
| Slk | S | 189 | -0.115889875 | 0.033882649 | 0.022184666 | 0.138074541 | -0.011697983 | 0.014264014 | 0.411286585 | 0.65338213 | 0.046000303 | 0.788046485 |
| Pak1 | T | 212 | 0.004493766 | 0.029878605 | -0.131083628 | -0.135577393 | -0.160962233 | 0.970138326 | 0.721462149 | 0.106951075 | 0.315982316 | 0.123587643 |
| Git2 | S | 583 | 0.106755226 | -0.006287149 | -0.367733919 | -0.474489146 | -0.36144677 | 0.422919178 | 0.947613129 | 0.24846519 | 0.164294941 | 0.253155156 |
| Scaf1 | S | 716 | -0.074148649 | -0.037822355 | 0.032891825 | 0.107040474 | 0.07071418 | 0.157178471 | 0.574302375 | 0.803935997 | 0.447744415 | 0.612414395 |
| Scrn1 | S | 360 | -0.060622058 | 0.08744844 | -0.194690301 | -0.134068243 | -0.282138741 | 0.524957987 | 0.419344707 | 0.264941729 | 0.40530856 | 0.141215482 |
| Srrm1 | S | 838 | 0.018549357 | -0.054535558 | 0.000177456 | -0.018371901 | 0.054713013 | 0.50049564 | 0.004377555 | 0.993751141 | 0.575372536 | 0.097147165 |
| Map1a | S | 1720 | 0.067459037 | -0.009567446 | -0.139606639 | -0.207065676 | -0.130039194 | 0.062632355 | 0.775006303 | 0.008814773 | 0.003540536 | 0.020456435 |
| Tut7 | S | 172 | -0.011116552 | -0.059153202 | -0.010982403 | 0.000134149 | 0.048170798 | 0.865899429 | 0.162012272 | 0.709946014 | 0.998445612 | 0.270644801 |
| Papola | T | 544 | 0.003595238 | -0.032422091 | -0.070438655 | -0.074033893 | -0.038016565 | 0.950548928 | 0.476222403 | 0.309141196 | 0.322300515 | 0.544509434 |
| Dido1 | S | 1840 | 0.02919864 | -0.092926671 | -0.017886561 | -0.047085201 | 0.07504011 | 0.590816748 | 0.120730253 | 0.5099233 | 0.407971617 | 0.182741884 |
| Aftph | S | 151 | -0.035889924 | -0.017949593 | -0.016217488 | 0.019672436 | 0.001732105 | 0.27384167 | 0.46086156 | 0.571833439 | 0.554285776 | 0.948417515 |
| Rnpc3 | S | 108 | 0.044688232 | 0.021532415 | -0.190851342 | -0.235539574 | -0.212383757 | 0.720170283 | 0.815954669 | 0.137306409 | 0.158659417 | 0.135993083 |
| Ankrd27 | S | 634 | 0.034255548 | -0.064666801 | -0.096794363 | -0.131049912 | -0.032127562 | 0.659032657 | 0.586657033 | 0.25832866 | 0.043415122 | 0.754920529 |
| Ar | S | 630 | -0.047431233 | 0.033720954 | -0.07478768 | -0.027356447 | -0.108508635 | 0.415201068 | 0.669602263 | 0.212914071 | 0.631730079 | 0.220926854 |
| Inpp5k | S | 298 | -0.19179111 | -0.058953054 | 0.131599487 | 0.323390597 | 0.190552541 | 0.213243218 | 0.70099274 | 0.594194836 | 0.234871776 | 0.448003462 |
| Rbm10 | S | 723 | -0.018429854 | -0.054551524 | 0.024039327 | 0.042469181 | 0.078590851 | 0.416609556 | 0.016049469 | 0.281947418 | 0.148038074 | 0.020436832 |
| Itprid2 | S | 271 | -0.119064702 | 0.121000306 | -0.112428647 | 0.006636055 | -0.233428954 | 0.124197263 | 0.134371763 | 0.205802052 | 0.938589442 | 0.048745437 |
| Itprid2 | S | 274 | -0.119064702 | 0.121000306 | -0.112428647 | 0.006636055 | -0.233428954 | 0.124197263 | 0.134371763 | 0.205802052 | 0.938589442 | 0.048745437 |
| Washc2 | T | 428 | 0.013570731 | -0.060951203 | -0.066329994 | -0.079900725 | -0.005378791 | 0.81174662 | 0.297537717 | 0.520649847 | 0.45749629 | 0.957584647 |
| Rnf6 | S | 559 | -0.024044876 | 0.024560635 | -0.150708952 | -0.126664076 | -0.175269587 | 0.921302026 | 0.935891424 | 0.305947336 | 0.625192162 | 0.58763346 |
| Pip4p2 | T | 22 | -0.009398868 | -0.025066845 | -0.062214545 | -0.052815678 | -0.037147701 | 0.910291831 | 0.565908408 | 0.290429196 | 0.570594143 | 0.537572389 |
| Srrm2 | S | 1872 | -0.014324419 | -0.184666275 | 0.112205292 | 0.126529711 | 0.296871566 | 0.878239986 | 0.079556262 | 0.44763988 | 0.409781553 | 0.106591584 |
| Fnip1 | S | 261 | -0.058808773 | -0.076149092 | 0.006221511 | 0.065030284 | 0.082370603 | 0.42721006 | 0.475577267 | 0.820673322 | 0.387487244 | 0.445141779 |
| Cul4b | S | 110 | 0.111270234 | 0.005496505 | -0.203437022 | -0.314707256 | -0.208933527 | 0.065059233 | 0.823306116 | 0.006706593 | 0.003029078 | 0.019168718 |
| Slk | S | 543 | -0.105558245 | 0.033069213 | -0.037611743 | 0.067946502 | -0.070680956 | 0.304009853 | 0.588711185 | 0.335762067 | 0.470137361 | 0.254101351 |
| Cdr2l | S | 407 | 0.087453147 | -0.033831466 | -0.138314246 | -0.225767393 | -0.104482781 | 0.403728195 | 0.583123478 | 0.094453601 | 0.089131494 | 0.189835467 |
| Ogfr | S | 563 | -0.011610329 | -0.003065708 | -0.097134243 | -0.085523914 | -0.094068536 | 0.844865097 | 0.933672031 | 0.252622025 | 0.344217592 | 0.2666097 |
| Epn2 | S | 180 | 0.148631267 | -0.053719924 | -0.177189995 | -0.325821262 | -0.123470071 | 0.069242556 | 0.070532103 | 0.075572761 | 0.012424905 | 0.154106125 |
| Tjp2 | S | 411 | -0.054855607 | -0.020707185 | 0.012475004 | 0.067330611 | 0.033182188 | 0.120321089 | 0.49417605 | 0.888731085 | 0.469429687 | 0.704577798 |
| Ptcd3 | S | 680 | 0.028780164 | -0.081531868 | -0.025511276 | -0.05429144 | 0.056020592 | 0.539539227 | 0.09402691 | 0.583403253 | 0.351510538 | 0.306042649 |

| | | | | | | | | | | | | |
|---|---|---|---|---|---|---|---|---|---|---|---|---|
| Camsap1 | S | 553 | 0.079036685 | -0.017211668 | -0.096985028 | -0.176021713 | -0.079773359 | 0.401254009 | 0.597760658 | 0.025183406 | 0.132717892 | 0.02906529 |
| Nap1l4 | S | 125 | -0.111598761 | 0.077690475 | -0.071372038 | 0.040226723 | -0.149062512 | 0.148803526 | 0.287898245 | 0.35468724 | 0.63689731 | 0.138390452 |
| Arhgap24 | S | 390 | -0.064789149 | -0.020298917 | 0.025699676 | 0.090488825 | 0.045998593 | 0.048759289 | 0.462252016 | 0.438267419 | 0.026661315 | 0.139392148 |
| Kdm5c | T | 286 | -0.128928006 | -0.022123106 | 0.072948621 | 0.201876628 | 0.095071727 | 0.015844009 | 0.346021839 | 0.053316517 | 0.004868401 | 0.03526458 |
| Parp10 | T | 959 | -0.145721631 | 0.012813541 | -0.011634078 | 0.134087553 | -0.024447619 | 0.289617209 | 0.896856045 | 0.910248268 | 0.286211724 | 0.740433232 |
| Gtf2i | T | 558 | 0.163162261 | -0.085542498 | -0.205079483 | -0.368241744 | -0.119536986 | 0.027277397 | 0.116020342 | 0.060710887 | 0.01408941 | 0.20232362 |
| Map1s | S | 670 | -0.029306066 | -0.02966903 | -0.040784774 | -0.011478708 | -0.011115744 | 0.544090103 | 0.603126221 | 0.514746206 | 0.861529578 | 0.876310235 |
| Nbr1 | S | 812 | 0.198736277 | -0.322452075 | -0.127821113 | -0.326557391 | 0.194630962 | 0.440712196 | 0.246158046 | 0.835496415 | 0.613960427 | 0.759570765 |
| Cast | T | 467 | -0.004799798 | 0.028180662 | -0.13336442 | -0.128564622 | -0.161545083 | 0.942320325 | 0.438447098 | 0.09884619 | 0.157034319 | 0.079155934 |
| Map1a | S | 1208 | 0.019080697 | 0.108708566 | -0.216074873 | -0.23515557 | -0.324783438 | 0.899887931 | 0.24915892 | 0.034366981 | 0.188012077 | 0.01113853 |
| Afap1 | S | 684 | 0.370849789 | -0.06707136 | -0.120791365 | -0.491641155 | -0.053720005 | 0.408760828 | 0.223067398 | 0.022116061 | 0.302808789 | 0.263355984 |
| Afap1 | T | 687 | 0.370849789 | -0.06707136 | -0.120791365 | -0.491641155 | -0.053720005 | 0.408760828 | 0.223067398 | 0.022116061 | 0.302808789 | 0.263355984 |
| Tjp2 | T | 907 | 0.164562338 | 0.11584115 | -0.696868805 | -0.861431143 | -0.812709955 | 0.351018987 | 0.6331267 | 0.196579106 | 0.147024894 | 0.151827679 |
| Abitram | T | 192 | -0.005607818 | -0.035540829 | -0.022146026 | -0.016538208 | 0.013394803 | 0.845688089 | 0.194242741 | 0.376171266 | 0.534772459 | 0.553863573 |
| Diaph2 | S | 69 | -0.088814313 | 0.063622468 | -0.042543424 | 0.04627089 | -0.106165892 | 0.050656052 | 0.244455647 | 0.377757839 | 0.249224578 | 0.081774504 |
| Snrnp70 | S | 408 | 0.021996426 | -0.046268119 | -0.03798244 | -0.059978866 | 0.008285678 | 0.315266837 | 0.025181635 | 0.48761373 | 0.307389289 | 0.869386104 |
| Cdca7l | S | 183 | -0.004553871 | -0.040742059 | -0.022591074 | -0.018037203 | 0.018150985 | 0.923647798 | 0.155805596 | 0.504785261 | 0.701169433 | 0.484341876 |
| Gpatch8 | S | 503 | -0.012648524 | 0.016401436 | -0.092126724 | -0.0794782 | -0.10852816 | 0.873913582 | 0.826150151 | 0.130582755 | 0.293396433 | 0.158144286 |
| Kdm4a | S | 523 | -0.12424657 | 0.028640944 | 0.034971812 | 0.159218382 | 0.006330868 | 0.060067109 | 0.77104563 | 0.806651141 | 0.313691937 | 0.966919384 |
| Fam83d | S | 360 | 0.114388518 | -0.092123201 | -0.090751146 | -0.205139664 | 0.001372055 | 0.110363317 | 0.051545636 | 0.175246254 | 0.034182741 | 0.980522088 |
| Fam83d | S | 363 | 0.114388518 | -0.092123201 | -0.090751146 | -0.205139664 | 0.001372055 | 0.110363317 | 0.051545636 | 0.175246254 | 0.034182741 | 0.980522088 |
| Fam83d | S | 366 | 0.114388518 | -0.092123201 | -0.090751146 | -0.205139664 | 0.001372055 | 0.110363317 | 0.051545636 | 0.175246254 | 0.034182741 | 0.980522088 |
| Fam83d | T | 364 | 0.114388518 | -0.092123201 | -0.090751146 | -0.205139664 | 0.001372055 | 0.110363317 | 0.051545636 | 0.175246254 | 0.034182741 | 0.980522088 |
| Trafd1 | S | 450 | -0.04432635 | -0.018037432 | -0.004404454 | 0.039921896 | 0.013632979 | 0.139607388 | 0.513003327 | 0.96275638 | 0.674573486 | 0.883312548 |
| Ppp6r3 | S | 824 | 0.057827337 | -0.037080843 | -0.079708683 | -0.13753602 | -0.04262784 | 0.045790032 | 0.085199476 | 0.026797106 | 0.006108902 | 0.146711036 |
| H2ax | Y | 143 | 0.031442105 | -0.097656696 | 0.013137057 | -0.018305049 | 0.110793753 | 0.484056072 | 0.017151275 | 0.745386678 | 0.687365301 | 0.036448769 |
| Dpysl3 | T | 524 | -0.097708834 | 0.183060026 | -0.237673624 | -0.13996479 | -0.42073365 | 0.423879692 | 0.338031924 | 0.251511863 | 0.450362312 | 0.105084463 |
| Nucb1 | S | 85 | -0.259452232 | 0.071378955 | 0.10756343 | 0.367015662 | 0.036184475 | 0.000602177 | 0.145476193 | 0.035448439 | 0.000754783 | 0.233602127 |
| Bcar3 | S | 271 | -0.0663968 | 0.063941592 | -0.123899515 | -0.057502715 | -0.187841106 | 0.361744455 | 0.313483658 | 0.226052558 | 0.563869534 | 0.107978652 |
| Spred1 | S | 239 | -0.094167483 | 0.075467328 | -0.085554959 | 0.008612524 | -0.161022287 | 0.119682858 | 0.234436281 | 0.34918916 | 0.917961481 | 0.129536352 |
| Pif1 | S | 641 | 0.03939172 | -0.107874877 | -0.03546672 | -0.07485844 | 0.072408157 | 0.656810295 | 0.166566193 | 0.7341787 | 0.457388463 | 0.427418454 |
| Cdk14 | S | 112 | -0.077651881 | 0.08551566 | -0.10453743 | -0.026885549 | -0.190053089 | 0.294576046 | 0.387809037 | 0.229600384 | 0.689778156 | 0.095607666 |
| Alad | S | 214 | -0.016597613 | 0.008691609 | -0.125378622 | -0.108781009 | -0.134070231 | 0.891160975 | 0.881664614 | 0.2713045 | 0.458553653 | 0.247019652 |
| Kdm3b | S | 515 | -0.125370509 | -0.081397505 | 0.086237381 | 0.21160789 | 0.167634885 | 0.305838608 | 0.463817594 | 0.568951948 | 0.254628643 | 0.347094243 |
| Nudt4 | T | 149 | -0.014662875 | 0.096891219 | -0.379249881 | -0.364587005 | -0.476141099 | 0.931686561 | 0.465600595 | 0.322965886 | 0.34465618 | 0.24041954 |
| Golga2 | S | 789 | 0.10977791 | -0.021833245 | -0.229908338 | -0.339686248 | -0.208075093 | 0.014959692 | 0.592172577 | 0.014469736 | 0.016806757 | 0.02633513 |
| Sf3b3 | T | 1200 | -0.031652161 | -0.006498101 | -0.027867361 | 0.0037848 | -0.02136926 | 0.141571238 | 0.866200381 | 0.519564293 | 0.928632262 | 0.684822135 |
| Bahcc1 | S | 1658 | -0.053546656 | -0.061875274 | -0.030166642 | 0.023380014 | 0.031708633 | 0.570102165 | 0.630055981 | 0.843157909 | 0.866358386 | 0.84282456 |
| Bahcc1 | S | 1662 | -0.053546656 | -0.061875274 | -0.030166642 | 0.023380014 | 0.031708633 | 0.570102165 | 0.630055981 | 0.843157909 | 0.866358386 | 0.84282456 |
| Fermt2 | S | 351 | 0.133203565 | -0.269721801 | -0.04574006 | -0.178943625 | 0.223981741 | 0.054940665 | 0.048961292 | 0.67745107 | 0.176672238 | 0.133830091 |
| Ppfibp1 | T | 39 | 0.086001912 | -0.100152159 | -0.076946491 | -0.162948402 | 0.023205668 | 0.22949327 | 0.043077591 | 0.56971229 | 0.288211831 | 0.859210863 |
| Camsap2 | S | 585 | -0.046672791 | -0.066157252 | -0.005871166 | 0.040801625 | 0.060286086 | 0.559339989 | 0.439178927 | 0.960930123 | 0.756429205 | 0.653364487 |
| Sgip1 | S | 287 | 0.01774855 | -0.049808575 | -0.144479544 | -0.162228094 | -0.094670969 | 0.935744143 | 0.75360706 | 0.473263543 | 0.532437519 | 0.651284253 |
| Emsy | S | 174 | 0.000201171 | -0.132627411 | 0.086673576 | 0.086472404 | 0.219300987 | 0.995306537 | 0.003292344 | 0.090186151 | 0.09492813 | 0.008562425 |
| Dhx32 | S | 24 | -0.077824261 | -0.174219001 | 0.054369165 | 0.132193426 | 0.228588166 | 0.790496296 | 0.505090854 | 0.92040736 | 0.813274175 | 0.686115641 |
| Dhx32 | S | 25 | -0.077824261 | -0.174219001 | 0.054369165 | 0.132193426 | 0.228588166 | 0.790496296 | 0.505090854 | 0.92040736 | 0.813274175 | 0.686115641 |
| Prpf4b | Y | 141 | -0.061460858 | -0.014496299 | -0.026488746 | 0.034972112 | -0.011992447 | 0.31071329 | 0.652546477 | 0.848611088 | 0.807053825 | 0.930609127 |
| Mtmr14 | S | 622 | 0.011017744 | -0.090419987 | -0.082203894 | -0.093221639 | 0.008216092 | 0.947054463 | 0.525639567 | 0.627068739 | 0.644809158 | 0.963589053 |
| Cd80 | S | 290 | 0.090175747 | -0.005778211 | -0.167148115 | -0.257323862 | -0.161369904 | 0.063155657 | 0.952437166 | 0.008323401 | 0.002488386 | 0.176358739 |
| Zfp800 | S | 455 | -0.046561348 | -0.07706439 | 0.035083851 | 0.081645199 | 0.112148242 | 0.23352703 | 0.109425504 | 0.143649902 | 0.083905943 | 0.059374546 |
| Itsn2 | S | 843 | 0.000490456 | -0.039687426 | -0.023869249 | -0.024359705 | 0.015818177 | 0.979594642 | 0.226564232 | 0.286776568 | 0.178586375 | 0.580211609 |
| Tab3 | S | 389 | 0.026965831 | -0.050529243 | -0.079599243 | -0.106565074 | -0.02907 | 0.69062357 | 0.44887297 | 0.297195287 | 0.111815974 | 0.586349222 |
| Ank3 | S | 897 | 0.122105324 | -0.017646406 | -0.34398633 | -0.466091654 | -0.326339924 | 0.36271128 | 0.8889165 | 0.223198208 | 0.135177445 | 0.242308656 |
| Dock4 | S | 1709 | -0.035242026 | -0.064545874 | 0.00626904 | 0.041511066 | 0.070814914 | 0.544363115 | 0.290462442 | 0.954742878 | 0.698084484 | 0.523021854 |
| Vim | S | 51 | -0.008423266 | -0.008973453 | -0.064508247 | -0.056084981 | -0.055534794 | 0.865051089 | 0.841369851 | 0.171031126 | 0.264312648 | 0.218014968 |
| Dgkq | S | 17 | 0.003430595 | -0.038800469 | -0.0380705 | -0.041501096 | 0.000729969 | 0.943138078 | 0.397514598 | 0.229359202 | 0.396183186 | 0.985588986 |
| Dennd4c | S | 1057 | -0.096781787 | 0.073123157 | -0.059364855 | 0.037416932 | -0.132488012 | 0.065513687 | 0.164780452 | 0.316721962 | 0.524638843 | 0.078345857 |
| Cog5 | S | 408 | 0.219881086 | -0.028021069 | -0.302561622 | -0.522442708 | -0.274540553 | 0.134827496 | 0.887150495 | 0.032676333 | 0.013264457 | 0.229051281 |
| Nmt2 | T | 64 | -0.114414122 | 0.021221533 | -0.024119572 | 0.09029455 | -0.045341105 | 0.259329046 | 0.845308764 | 0.775717784 | 0.426374892 | 0.717442443 |
| Rbm15 | S | 782 | -0.098677983 | 0.01855148 | -0.026248234 | 0.072429749 | -0.044799714 | 0.224277484 | 0.813646747 | 0.739861674 | 0.179886019 | 0.403866572 |
| Arhgef12 | S | 190 | 0.050360278 | -0.056108623 | -0.125787498 | -0.176147776 | -0.069678875 | 0.281587867 | 0.332908623 | 0.254951143 | 0.148457774 | 0.50090091 |
| Pdzrn3 | S | 775 | -0.160295817 | -0.008787646 | 0.048202604 | 0.208498421 | 0.05699025 | 0.157748735 | 0.887709567 | 0.563965848 | 0.104237569 | 0.478774409 |
| Top2b | S | 1448 | -0.083540007 | -0.050882948 | 0.057763434 | 0.141303441 | 0.108646381 | 0.136434801 | 0.410159029 | 0.46391226 | 0.122734536 | 0.185714843 |
| Srrm2 | S | 821 | 0.005476879 | -0.068420203 | -0.042393899 | -0.047870778 | 0.026026304 | 0.974694839 | 0.255419321 | 0.726515905 | 0.803837826 | 0.821201908 |
| Ranbp3 | Y | 47 | -0.081521888 | -0.099351294 | 0.119888911 | 0.201410799 | 0.219240205 | 0.110038344 | 0.065456148 | 0.092242223 | 0.026109251 | 0.019707192 |
| Fhdc1 | S | 540 | -0.084942055 | -0.074535473 | 0.013919647 | 0.098861702 | 0.08845512 | 0.396597018 | 0.71186874 | 0.93785571 | 0.578278609 | 0.70989503 |
| Brms1l | S | 197 | -0.117820452 | -0.079246573 | 0.106585068 | 0.22440552 | 0.185831641 | 0.166274226 | 0.040452986 | 0.055875805 | 0.040629229 | 0.017805736 |
| Asxl2 | S | 477 | -0.014754753 | -0.07895143 | -0.019669564 | -0.004914811 | 0.059281866 | 0.885437992 | 0.314407097 | 0.809108049 | 0.966437585 | 0.53936582 |
| Dpysl3 | S | 541 | -0.05983255 | 0.05393579 | -0.110565064 | -0.050732513 | -0.164500853 | 0.545892074 | 0.5959696 | 0.261044813 | 0.641185733 | 0.187143595 |
| Alkbh3 | S | 28 | 0.133865804 | 0.004035937 | -0.250398601 | -0.384264405 | -0.254434538 | 0.094244822 | 0.959830701 | 0.020989615 | 0.007120744 | 0.035282993 |
| Macroh2a1 | T | 129 | 0.076058257 | -0.058563927 | -0.062321093 | -0.13837935 | -0.003757166 | 0.217803505 | 0.174982255 | 0.051922531 | 0.075953312 | 0.91393808 |
| Zcchc8 | S | 697 | 0.184949897 | -0.086250791 | -0.220754256 | -0.405704153 | -0.134503464 | 0.093211532 | 0.42398708 | 0.055580359 | 0.000196443 | 0.094389954 |
| Pan2 | S | 1188 | -0.048673128 | 0.002405873 | -0.032095358 | 0.01657777 | -0.034501231 | 0.307340209 | 0.944843004 | 0.364516725 | 0.733174166 | 0.436752998 |
| Mfap1a Mfap1b | S | 118;118 | -0.030110654 | -0.087271112 | 0.049487308 | 0.079597961 | 0.13675842 | 0.431662024 | 0.051028886 | 0.253021809 | 0.040869164 | 0.008917854 |
| Myo18a | S | 1065 | -0.038401017 | 0.028070671 | -0.086819815 | -0.048418797 | -0.114890486 | 0.754582741 | 0.536968448 | 0.112118777 | 0.687777307 | 0.025728712 |
| Piezo1 | S | 1646 | 0.022063497 | -0.028847672 | -0.052877065 | -0.074940563 | -0.024029393 | 0.634569478 | 0.408302271 | 0.102416652 | 0.174352628 | 0.497106822 |
| Trim28 | T | 599 | -0.106633818 | 0.026603843 | -0.078904903 | 0.027728916 | -0.105508746 | 0.347420248 | 0.730545351 | 0.713656163 | 0.901426981 | 0.63518872 |
| Pcnp | S | 119 | -0.087028399 | 0.054981669 | -0.059682085 | 0.027346314 | -0.114663754 | 0.125469651 | 0.36905885 | 0.458935131 | 0.735950114 | 0.228996807 |
| Srrm1 | S | 731 | -0.067781265 | -0.06086933 | 0.062293293 | 0.130074558 | 0.123162623 | 0.159085031 | 0.108859245 | 0.205410781 | 0.050742722 | 0.047798247 |
| Epb41l1 | T | 741 | -0.033635639 | 0.016911079 | -0.072876726 | -0.039241087 | -0.089787805 | 0.542632155 | 0.842136233 | 0.252341716 | 0.57023123 | 0.365864521 |
| Nucks1 | S | 214 | -0.09393512 | -0.064041869 | 0.064259486 | 0.158194606 | 0.128301356 | 0.243660911 | 0.362175389 | 0.491406864 | 0.160016636 | 0.220384845 |
| Prpf3 | S | 619 | -0.086116389 | -0.038535362 | 0.045548232 | 0.131664622 | 0.084083594 | 0.177525663 | 0.38572762 | 0.521762523 | 0.142864125 | 0.277324576 |
| Inpp5e | S | 103 | 0.051883881 | -0.042142857 | -0.070667258 | -0.122551139 | -0.028524401 | 0.200184661 | 0.254178858 | 0.054278439 | 0.030299035 | 0.265318798 |
| Dst | T | 2418 | -0.059408951 | -0.055394569 | 0.051600812 | 0.111009763 | 0.106995381 | 0.1781265 | 0.188041617 | 0.392498086 | 0.106121198 | 0.116515515 |
| Heatr3 | S | 28 | 0.043129376 | -0.062161313 | -0.126890584 | -0.170019961 | -0.064729271 | 0.705128146 | 0.500122354 | 0.44643102 | 0.346622921 | 0.684588472 |
| Npm1 | S | 216 | 0.222627357 | -0.17018631 | -0.113001336 | -0.335628694 | 0.057184974 | 0.039885841 | 0.024028806 | 0.160100758 | 0.013846038 | 0.359359675 |
| Map3k1 | S | 270 | -0.01576561 | -0.039092554 | -0.04067651 | -0.0249109 | -0.001583956 | 0.766444062 | 0.3395322 | 0.641973498 | 0.77589735 | 0.984453427 |
| Map1b | S | 967 | -0.049868752 | -0.085118235 | 0.084738704 | 0.134607456 | 0.169856939 | 0.403861001 | 0.080822323 | 0.420749644 | 0.241266679 | 0.170071982 |
| Epb41l2 | T | 187 | -0.118240379 | 0.134437609 | -0.047832095 | 0.070408284 | -0.182269704 | 0.004702673 | 0.044090328 | 0.253746847 | 0.124611824 | 0.020202762 |
| Exoc4 | S | 226 | -0.084493914 | -0.047711151 | 0.065305485 | 0.149799399 | 0.113016636 | 0.040622933 | 0.534045931 | 0.278204328 | 0.057369321 | 0.220584436 |
| Gtf2f1 | S | 218 | -0.048455482 | 0.033608894 | -0.029472847 | 0.018982635 | -0.063081741 | 0.10124576 | 0.434488484 | 0.146752557 | 0.433030683 | 0.196583294 |
| Gtf2f1 | S | 221 | -0.048455482 | 0.033608894 | -0.029472847 | 0.018982635 | -0.063081741 | 0.10124576 | 0.434488484 | 0.146752557 | 0.433030683 | 0.196583294 |
| Gtf2f1 | S | 224 | -0.048455482 | 0.033608894 | -0.029472847 | 0.018982635 | -0.063081741 | 0.10124576 | 0.434488484 | 0.146752557 | 0.433030683 | 0.196583294 |
| Pacsin2 | S | 305 | -0.029768248 | -0.041645253 | -0.040830371 | -0.011062123 | 0.000814883 | 0.689518134 | 0.634155709 | 0.53174117 | 0.901116015 | 0.993384276 |

| | | | | | | | | | | | | | |
|---|---|---|---|---|---|---|---|---|---|---|---|---|---|
| Jmjd1c | S | | 1815 | 0.068406025 | -0.119580958 | -0.061391779 | -0.129797804 | 0.05818918 | 0.346016469 | 0.317868001 | 0.225535996 | 0.111657011 | 0.590158856 |
| Cry2 | S | | 557 | -0.02883886 | 0.006961555 | -0.043740479 | -0.014901619 | -0.050702034 | 0.523440506 | 0.889320844 | 0.137249274 | 0.71626084 | 0.337168643 |
| Cdk5rap2 | S | | 530 | 0.006237781 | -0.028387257 | -0.072079025 | -0.078316806 | -0.043691768 | 0.911853928 | 0.652189306 | 0.248899423 | 0.115278939 | 0.402424891 |
| Trmt10a | T | | 21 | 0.002810006 | 0.008036278 | -0.129460685 | -0.132270691 | -0.137496963 | 0.94345931 | 0.877674251 | 0.212860771 | 0.200982135 | 0.19543304 |
| Col3a1 | S | | 735 | -0.025737692 | -0.053941815 | 0.008078569 | 0.033816261 | 0.062020383 | 0.685834714 | 0.115449704 | 0.840680748 | 0.600084063 | 0.135314564 |
| Gpatch2l | S | | 31 | 0.020185296 | 0.014341596 | -0.210949922 | -0.231135218 | -0.225291518 | 0.90773373 | 0.909224256 | 0.330299415 | 0.33940526 | 0.304810266 |
| Ahctf1 | T | | 1224 | -0.037703692 | -0.099036264 | 0.06670125 | 0.104404942 | 0.165737514 | 0.46210652 | 0.041509838 | 0.195547854 | 0.071049503 | 0.023877573 |
| Prpsap2 | S | | 227 | -0.045835586 | 0.009846333 | -0.045760622 | 7.4964E-05 | -0.055606955 | 0.183565356 | 0.707019592 | 0.489076308 | 0.999044239 | 0.405027835 |
| Topors | S | | 718 | 0.111201954 | -0.038381385 | -0.215855397 | -0.327057351 | -0.177474012 | 0.15303802 | 0.596086682 | 0.092938888 | 0.034149988 | 0.149215822 |
| Map2k4 | S | | 255 | -0.068927959 | 0.036197726 | -0.071111893 | -0.002183934 | -0.107309619 | 0.228764575 | 0.552489607 | 0.482040056 | 0.982549534 | 0.337919408 |
| Rfc1 | T | | 350 | 0.014870662 | 0.023027383 | -0.093661998 | -0.108532661 | -0.116689382 | 0.758795344 | 0.623906137 | 0.031772782 | 0.092362787 | 0.069214335 |
| Map1b | S | | 2291 | 0.037753483 | 0.004467189 | -0.117144014 | -0.154897498 | -0.121611204 | 0.751475763 | 0.95650085 | 0.101222585 | 0.270673772 | 0.23382457 |
| Scrib | S | | 1457 | -0.076380804 | 0.00856497 | -0.008034821 | 0.068345983 | -0.016599791 | 0.128651968 | 0.76444615 | 0.714989967 | 0.166788113 | 0.510012984 |
| Rusc2 | S | | 534 | -0.009304904 | 0.009289659 | -0.076254023 | -0.06694912 | -0.085543682 | 0.900345678 | 0.789069653 | 0.104489542 | 0.403232769 | 0.072924337 |
| Eif3b | S | | 436 | 0.15576499 | -0.139082306 | -0.103300663 | -0.259065654 | 0.035781643 | 0.103736384 | 0.060955301 | 0.266475154 | 0.043802421 | 0.668072077 |
| Map4k5 | S | | 335 | -0.073047129 | 0.032931337 | -0.037205345 | 0.035841785 | -0.070136682 | 0.109047025 | 0.563909571 | 0.616255899 | 0.641579293 | 0.425013456 |
| Ubr4 | S | | 2715 | -0.017091077 | -0.025452614 | -0.054522198 | -0.037431121 | -0.029069584 | 0.800803607 | 0.671223641 | 0.357980789 | 0.423337134 | 0.289269854 |
| Arhgap21 | S | | 471 | -0.01717117 | -0.015391216 | -0.050773188 | -0.033602018 | -0.035381972 | 0.730460176 | 0.740824268 | 0.271768096 | 0.546642752 | 0.505919619 |
| Nde1 | S | | 326 | 0.008976347 | -0.005700306 | -0.077763262 | -0.086739608 | -0.072062955 | 0.90302859 | 0.931205283 | 0.116794289 | 0.265239307 | 0.277327439 |
| Asap1 | S | | 1146 | 0.049468223 | -0.15419103 | 0.01222406 | -0.037244164 | 0.16641509 | 0.749181624 | 0.143804725 | 0.934425984 | 0.826210358 | 0.266663248 |
| Supt6 | S | | 125 | -0.003409883 | -0.098887271 | 0.010801639 | 0.014211521 | 0.10968891 | 0.926235048 | 0.167393475 | 0.869022155 | 0.813591359 | 0.197511549 |
| Tmem87b | S | | 534 | 0.091779119 | 0.048901529 | -0.453565464 | -0.545344583 | -0.502466993 | 0.609263172 | 0.503546735 | 0.249398119 | 0.187836558 | 0.219396797 |
| Tax1bp1 | S | | 138 | 0.176755895 | 0.075748745 | -0.404409253 | -0.581165148 | -0.480157998 | 0.619522734 | 0.708525494 | 0.167771965 | 0.182067377 | 0.125880622 |
| Tax1bp1 | S | | 124 | 0.151498165 | 0.099191123 | -0.399186582 | -0.550684747 | -0.498377705 | 0.311094065 | 0.456916509 | 0.049228125 | 0.019945772 | 0.032258901 |
| Tcp11l1 | S | | 44 | -0.041711025 | 0.064622794 | -0.169790802 | -0.128079777 | -0.234413596 | 0.778067024 | 0.504059358 | 0.222832884 | 0.44876565 | 0.120264581 |
| Gpatch8 | S | | 1013 | -0.111640175 | -0.020882982 | 0.046206199 | 0.157846374 | 0.067089181 | 0.084138605 | 0.65891569 | 0.340500376 | 0.040921597 | 0.235496531 |
| Arhgap10 | S | | 706 | -0.064605093 | -0.022559955 | -0.017345774 | 0.047259319 | 0.005214181 | 0.359850981 | 0.774869871 | 0.796903022 | 0.338329773 | 0.932610146 |
| Srrm2 | S | | 887 | -0.043652649 | -0.061359145 | 0.033710456 | 0.077363106 | 0.095069601 | 0.318953838 | 0.126150415 | 0.364571682 | 0.123651423 | 0.05103533 |
| Pqbp1 | S | | 245 | 0.112723622 | -0.062313586 | -0.091723198 | -0.20444682 | -0.029409613 | 0.065428928 | 0.122520022 | 0.023070609 | 0.022644184 | 0.30264173 |
| Peak1 | S | | 877 | -0.032831111 | -0.024570309 | -0.010180864 | 0.022650248 | 0.014389445 | 0.336624232 | 0.360039133 | 0.800572483 | 0.605690076 | 0.717333873 |
| Otud5 | S | | 165 | -0.049015186 | -0.049062126 | -0.011469865 | 0.037545322 | 0.037592261 | 0.515089635 | 0.500805181 | 0.923770659 | 0.740109304 | 0.73716051 |
| Tbx3 | S | | 369 | 0.078632916 | -0.142735175 | -0.012148468 | -0.090781384 | 0.130586707 | 0.11630061 | 0.017952446 | 0.827062555 | 0.108343852 | 0.06139533 |
| Kif22 | S | | 407 | -0.035882909 | 0.091184164 | -0.086700251 | -0.050817342 | -0.177884415 | 0.600919827 | 0.292854302 | 0.017197394 | 0.459370135 | 0.108553679 |
| Kif20b | S | | 1711 | 0.021822169 | -0.19411732 | 0.077029833 | 0.055207665 | 0.271147154 | 0.853022355 | 0.059013702 | 0.339920793 | 0.581454979 | 0.027242843 |
| Nab2 | S | | 157 | 0.030118868 | -0.006743634 | -0.184322766 | -0.214441635 | -0.177579132 | 0.431420515 | 0.924391289 | 0.21793156 | 0.175778967 | 0.234811328 |
| Sec62 | S | | 307 | -0.091407165 | -0.028565184 | 0.061095339 | 0.152502504 | 0.089660523 | 0.092883189 | 0.605018835 | 0.546436242 | 0.192466451 | 0.385547089 |
| Srrm2 | S | | 281 | 0.106210935 | -0.044765521 | -0.12303892 | -0.229249855 | -0.078273399 | 0.547938812 | 0.621919909 | 0.130449525 | 0.246158743 | 0.326525374 |
| Eif4g1 | T | | 218 | 0.046008791 | -0.014160213 | -0.124967819 | -0.170976611 | -0.110807606 | 0.393040459 | 0.821783342 | 0.090426941 | 0.039381878 | 0.11667799 |
| Atf7 | T | | 190 | -0.112921822 | -0.01460427 | 0.014417756 | 0.127339578 | 0.029022026 | 0.255434909 | 0.916871738 | 0.916887884 | 0.344720511 | 0.85297338 |
| Traf7 | S | | 32 | 0.061175078 | -0.086636253 | -0.060413794 | -0.121588872 | 0.026222459 | 0.452426483 | 0.091267687 | 0.66377482 | 0.435877172 | 0.849243479 |
| Anapc1 | S | | 686 | 0.031325259 | -0.04210257 | -0.111445627 | -0.142770886 | -0.069343058 | 0.71585322 | 0.519434923 | 0.379657873 | 0.304682859 | 0.56878463 |
| Actl7a | S | | 37 | -0.085044489 | -0.025048585 | 0.011874256 | 0.096918746 | 0.036922841 | 0.235476299 | 0.727954169 | 0.826077445 | 0.144271469 | 0.545369702 |
| Actl7a | T | | 35 | -0.085044489 | -0.025048585 | 0.011874256 | 0.096918746 | 0.036922841 | 0.235476299 | 0.727954169 | 0.826077445 | 0.144271469 | 0.545369702 |
| Ercc4 | S | | 363 | -0.052537152 | -0.008389226 | 0.029405455 | 0.081942608 | 0.037794681 | 0.005675592 | 0.692547186 | 0.313595827 | 0.055267897 | 0.238885734 |
| Mga | S | | 414 | 0.0895291 | -0.103557863 | -0.042141625 | -0.131670725 | 0.061416238 | 0.353310274 | 0.143606829 | 0.231427132 | 0.21390254 | 0.328315129 |
| Mga | T | | 406 | 0.0895291 | -0.103557863 | -0.042141625 | -0.131670725 | 0.061416238 | 0.353310274 | 0.143606829 | 0.231427132 | 0.21390254 | 0.328315129 |
| Mga | Y | | 413 | 0.0895291 | -0.103557863 | -0.042141625 | -0.131670725 | 0.061416238 | 0.353310274 | 0.143606829 | 0.231427132 | 0.21390254 | 0.328315129 |
| Urb1 | S | | 17 | 0.268900658 | -0.128327897 | -0.212238462 | -0.481139121 | -0.083910565 | 0.045644474 | 0.100305879 | 0.06634898 | 0.005052929 | 0.331168332 |
| Irf2bpl | S | | 644 | -0.044518935 | -0.028551207 | -0.039092495 | 0.00542644 | -0.010541288 | 0.408249772 | 0.718642883 | 0.797636709 | 0.970561361 | 0.945831082 |
| Rrp8 | S | | 62 | 0.028405228 | -0.052606903 | -0.04357786 | -0.071983088 | 0.009029044 | 0.171283203 | 0.113527774 | 0.173762845 | 0.061319209 | 0.783175137 |
| Chfr | S | | 214 | -0.079244113 | -0.06053016 | 0.044069015 | 0.123313128 | 0.104599175 | 0.239526568 | 0.389281894 | 0.458847284 | 0.14017071 | 0.219026678 |
| Zc3hc1 | S | | 58 | -0.103347382 | 0.048882695 | -0.031870517 | 0.071476865 | -0.080753212 | 0.183769859 | 0.658828321 | 0.662609235 | 0.14160071 | 0.431369006 |
| Dnttip2 | S | | 476 | -0.026277761 | -0.06548851 | 0.017546553 | 0.043824314 | 0.083035063 | 0.717790435 | 0.109085032 | 0.664548219 | 0.549570665 | 0.062525702 |
| Cdk12 | S | | 340 | -0.020351535 | -0.038309215 | -0.011766523 | 0.008585011 | 0.026542692 | 0.352828786 | 0.321703165 | 0.902609601 | 0.928629971 | 0.789034333 |
| Cdk12 | S | | 342 | -0.020351535 | -0.038309215 | -0.011766523 | 0.008585011 | 0.026542692 | 0.352828786 | 0.321703165 | 0.902609601 | 0.928629971 | 0.789034333 |
| Cdk12 | S | | 344 | -0.020351535 | -0.038309215 | -0.011766523 | 0.008585011 | 0.026542692 | 0.352828786 | 0.321703165 | 0.902609601 | 0.928629971 | 0.789034333 |
| Cdc5l | S | | 427 | 0.099233397 | -0.124203297 | -0.112399922 | -0.21163332 | 0.011803374 | 0.212093551 | 0.232302629 | 0.32583055 | 0.119911548 | 0.922671768 |
| Ncor2 | S | | 747 | -0.021906582 | -0.023918448 | -0.041313999 | -0.019407417 | -0.017395551 | 0.62363197 | 0.388434685 | 0.637779704 | 0.826109624 | 0.834802125 |
| Ncor2 | T | | 745 | -0.021906582 | -0.023918448 | -0.041313999 | -0.019407417 | -0.017395551 | 0.62363197 | 0.388434685 | 0.637779704 | 0.826109624 | 0.834802125 |
| Cep55 | S | | 428 | 0.223746925 | 0.036667809 | -0.333221892 | -0.556968818 | -0.369889701 | 0.104094379 | 0.604144305 | 0.010236926 | 0.008633753 | 0.019562817 |
| Xrn2 | S | | 448 | 0.01174857 | -0.023259118 | -0.081588567 | -0.093337137 | -0.058329449 | 0.872920776 | 0.660016629 | 0.263678834 | 0.264371548 | 0.356947801 |
| Mex3d | S | | 495 | -0.073673757 | 0.040665968 | -0.040143141 | 0.033530616 | -0.080809109 | 0.077277354 | 0.269455542 | 0.427461434 | 0.430146103 | 0.136808059 |
| Ralgapa1 | S | | 772 | -0.005398128 | -0.005936469 | -0.065970434 | -0.060572307 | -0.060033966 | 0.885577297 | 0.874923213 | 0.183814303 | 0.213136423 | 0.217413596 |
| Fam122a | S | | 170 | -0.253045962 | -0.046588114 | 0.170575453 | 0.423621415 | 0.217163567 | 0.5515912 | NA | NA | NA | NA |
| Fam122a | S | | 174 | -0.253045962 | -0.046588114 | 0.170575453 | 0.423621415 | 0.217163567 | 0.5515912 | NA | NA | NA | NA |
| Rbmx Rbmxl1 | S | 329;332 | | 0.047677672 | -0.045783556 | -0.099883033 | -0.147560705 | -0.054099477 | 0.466385564 | 0.464404092 | 0.216059849 | 0.072728927 | 0.391307075 |
| Tcf3 | S | | 140 | -0.103765779 | -0.016360369 | 0.017609169 | 0.121374948 | 0.033969538 | 0.287583596 | 0.665034808 | 0.850431067 | 0.326681907 | 0.721224415 |
| Hivep3 | S | | 492 | -0.030142002 | -0.018942886 | -0.043887031 | -0.013745029 | -0.024944144 | 0.53020872 | 0.715929296 | 0.519438777 | 0.794564417 | 0.661636433 |
| Mprip | T | | 623 | 0.028935795 | -0.05606495 | -0.080445041 | -0.109380836 | -0.024380091 | 0.595935048 | 0.398426528 | 0.372372959 | 0.227656961 | 0.764516683 |
| Atrx | S | | 1224 | -0.095551014 | -0.031090646 | 0.047681306 | 0.14323232 | 0.078771952 | 0.097343145 | 0.405958903 | 0.260588584 | 0.040138288 | 0.131974834 |
| Map1a | S | | 2209 | -0.03642813 | 0.093908384 | -0.144422039 | -0.107993908 | -0.238330423 | 0.635336789 | 0.13297332 | 0.036062216 | 0.164221366 | 0.000697655 |
| Ticrr | S | | 861 | 0.077419606 | 0.063493532 | -0.22427772 | -0.301697326 | -0.287771252 | 0.336513642 | 0.428463945 | 0.025280676 | 0.017269086 | 0.020829033 |
| Lmnb1 | T | | 26 | -0.080485531 | 0.014393679 | -0.035440239 | 0.045045292 | -0.049833918 | 0.333058007 | 0.824673851 | 0.575281764 | 0.595662665 | 0.507063597 |
| Zfp292 | T | | 1850 | 0.0910382 | -0.200756674 | -0.047609199 | -0.138647399 | 0.153147475 | 0.155637662 | 0.117496154 | 0.662339158 | 0.242241796 | 0.268378781 |
| Myh10 | S | | 1956 | -0.116704071 | 0.001287678 | 0.031631815 | 0.148335885 | 0.030344137 | 0.12002714 | 0.984879076 | 0.686083144 | 0.125764033 | 0.715298574 |
| Hdac7 | S | | 178 | -0.139249406 | 0.036049998 | 0.053340891 | 0.192590297 | 0.017290893 | 0.007412465 | 0.41231864 | 0.270270762 | 0.014401286 | 0.740776097 |
| Hdgfl2 | S | | 661 | 0.040608491 | -0.01726326 | -0.107418254 | -0.148026744 | -0.090154994 | 0.219580453 | 0.610389031 | 0.066726759 | 0.046130049 | 0.108124853 |
| Trim24 | S | | 688 | -0.238870684 | 0.063654685 | 0.165690276 | 0.40456096 | 0.102035591 | 0.011611303 | 0.578235955 | 0.276370699 | 0.065502167 | 0.509170718 |
| Vcl | T | | 603 | -0.014411168 | -0.019667987 | -0.024253248 | -0.009842079 | -0.004585261 | 0.702681161 | 0.245508863 | 0.416736909 | 0.81044264 | 0.857744021 |
| Eif4g3 | S | | 436 | 0.058628239 | -0.072203695 | -0.072351928 | -0.130980167 | -0.000148233 | 0.40203239 | 0.163014765 | 0.415554072 | 0.19776505 | 0.998491193 |
| Prpf38a | S | | 209 | -0.002664861 | -0.112541105 | 0.065732058 | 0.06839692 | 0.178273163 | 0.960981668 | 0.040553157 | 0.451714483 | 0.42577893 | 0.108716153 |
| Mier1 | S | | 483 | 0.01649434 | -0.100901736 | -0.003665592 | -0.020159932 | 0.097236144 | 0.743588973 | 0.144823807 | 0.970352208 | 0.839531297 | 0.380150639 |
| Kif20a | S | | 239 | -0.012531785 | -0.052714137 | -0.053515812 | -0.040984026 | -0.000801675 | 0.847036893 | 0.600967356 | 0.57307882 | 0.65287239 | 0.994372623 |
| Nsd2 | S | | 406 | 0.142404513 | 0.017730954 | -0.256169776 | -0.39857429 | -0.273900731 | 0.080037641 | 0.571678972 | 0.010433356 | 0.004039844 | 0.01776641 |
| Brca1 | S | | 1479 | -0.059161151 | -0.06890121 | 0.070028397 | 0.129189548 | 0.138929608 | 0.391071375 | 0.070054284 | 0.32144968 | 0.158002944 | 0.109488061 |
| Add1 | S | | 364 | -0.146609495 | 0.04954635 | -0.037580057 | 0.109029438 | -0.087126407 | 0.250662876 | 0.431551726 | 0.613930293 | 0.362959919 | 0.171999419 |
| Nap1l4 | S | | 294 | 0.079978868 | -0.033626608 | -0.150805385 | -0.230784252 | -0.117178777 | 0.523315398 | 0.61145407 | 0.216192831 | 0.151953335 | 0.302909262 |
| Ints9 | T | | 563 | -0.051670567 | -0.005623295 | -0.046663832 | 0.005006735 | -0.041040537 | 0.358336077 | 0.967879742 | 0.666048079 | 0.962904705 | 0.800480722 |
| Rbbp6 | S | | 1329 | -0.137654555 | -0.027261698 | 0.115255668 | 0.252910223 | 0.142517366 | 0.019458463 | 0.592723136 | 0.14204406 | 0.026347393 | 0.088768887 |
| Stk10 | T | | 950 | 0.043769125 | -0.042157991 | -0.043428807 | -0.087197932 | -0.001270816 | 0.057435239 | 0.046216072 | 0.039298785 | 0.013538536 | 0.874355251 |
| Casp8ap2 | T | | 1531 | 0.005887237 | -0.075673098 | -0.019734474 | -0.025621711 | 0.055938624 | 0.898178964 | 0.212965166 | 0.810918792 | 0.764001858 | 0.538444519 |
| Pnpla2 | S | | 406 | -0.013982014 | -0.018537449 | -0.074721231 | -0.060739217 | -0.056183782 | 0.750559567 | 0.765284003 | 0.440778435 | 0.501499963 | 0.556932764 |
| Fcho2 | S | | 387 | 0.030804222 | -0.055372599 | -0.094340838 | -0.12514506 | -0.038968239 | 0.824062175 | 0.588190204 | 0.417796602 | 0.449272613 | 0.765164782 |
| Ncor1 | S | | 1206 | -0.090381888 | -0.045594695 | 0.073970076 | 0.164351964 | 0.119564772 | 0.11600125 | 0.35028211 | 0.349631356 | 0.083905933 | 0.161715739 |

| | | | | | | | | | | | | |
|---|---|---|---|---|---|---|---|---|---|---|---|---|
| Prrc2b | S | 806 | 0.031626867 | -0.047701318 | -0.074466796 | -0.106093663 | -0.026765478 | 0.562620802 | 0.402988239 | 0.281888663 | 0.132221275 | 0.656694836 |
| Prpsap1 | S | 215 | -0.054742081 | 0.026494754 | -0.045391097 | 0.009350983 | -0.071885851 | 0.225697118 | 0.507717238 | 0.31726854 | 0.846268411 | 0.176703168 |
| Slc11a2 | S | 567 | -0.125671008 | 0.019579544 | -0.028790644 | 0.096880364 | -0.048370188 | 0.35317707 | 0.839295108 | 0.815558413 | 0.490035391 | 0.658713578 |
| Rpl29 | S | 31 | 0.183333543 | -0.111876543 | -0.192291136 | -0.375624679 | -0.080414593 | 0.024100669 | 0.123444313 | 0.066671276 | 0.011355077 | 0.350637267 |
| Kdm5c | S | 287 | -0.096767634 | -0.028242862 | 0.046037712 | 0.142805346 | 0.074280574 | 0.039662329 | 0.482547173 | 0.101602777 | 0.016235507 | 0.121574888 |
| Wac | S | 64 | -0.102966249 | 0.019671773 | 0.017266994 | 0.120233243 | -0.002404779 | 0.055171463 | 0.656715182 | 0.706505193 | 0.062596814 | 0.962167135 |
| Rbm12b1 Rbm12b2 | S | 502;502 | -0.044271136 | 0.040874312 | -0.08609982 | -0.041828684 | -0.126974133 | 0.474771157 | 0.69885469 | 0.311084427 | 0.620577822 | 0.312609802 |
| Ahi1 | S | 26 | -0.063869319 | 0.052393606 | -0.051969717 | 0.011899602 | -0.104363323 | 0.057643808 | 0.128281408 | 0.224441915 | 0.761903362 | 0.056479389 |
| Hdac5 | T | 486 | -0.18226851 | 0.0763478 | 0.015770825 | 0.198039335 | -0.060576975 | 0.041123426 | 0.208730456 | 0.905888311 | 0.212761087 | 0.647333756 |
| Lrrfip1 | T | 91 | -0.018274075 | 0.040528577 | -0.11567595 | -0.097401875 | -0.156204527 | 0.86552209 | 0.548919357 | 0.13444307 | 0.385664479 | 0.052565465 |
| Tasor2 | S | 2015 | 0.018933555 | -0.040672041 | -0.073012698 | -0.091946253 | -0.032340657 | 0.779014638 | 0.616241413 | 0.279845145 | 0.295202938 | 0.723230475 |
| Nup35 | T | 272 | -0.019974654 | -0.042344739 | 0.00713464 | 0.027109294 | 0.049479379 | 0.494250888 | 0.134817198 | 0.854091558 | 0.480254154 | 0.228983533 |
| Strn4 | S | 53 | -0.189635459 | 0.160341661 | -0.31213353 | -0.122498071 | -0.472475191 | 0.694172563 | 0.312561817 | 0.386639486 | 0.819094792 | 0.229153197 |
| Srsf7 | S | 184 | 0.136529945 | -0.170472571 | -0.089353803 | -0.225883748 | 0.081118768 | 0.548776961 | 0.40248268 | 0.447090021 | 0.336368775 | 0.657362474 |
| Srsf7 | S | 186 | 0.136529945 | -0.170472571 | -0.089353803 | -0.225883748 | 0.081118768 | 0.548776961 | 0.40248268 | 0.447090021 | 0.336368775 | 0.657362474 |
| Srsf7 | S | 188 | 0.136529945 | -0.170472571 | -0.089353803 | -0.225883748 | 0.081118768 | 0.548776961 | 0.40248268 | 0.447090021 | 0.336368775 | 0.657362474 |
| Slc45a4 | T | 765 | -0.048085852 | -0.024738775 | -0.02298345 | 0.025102402 | 0.001755325 | 0.337389036 | 0.76749342 | 0.827266708 | 0.799840382 | 0.987846834 |
| Pard3 | S | 174 | -0.167234207 | 0.226182224 | -0.194370257 | -0.02713605 | -0.420552481 | 0.294133408 | 0.194038625 | 0.234838984 | 0.828404626 | 0.031683792 |
| Rbm12b1 | S | 555 | -0.053458005 | 0.007644171 | -0.030956256 | 0.022501749 | -0.038600427 | 0.231632837 | 0.872688438 | 0.595947818 | 0.721691742 | 0.575330502 |
| Pask | T | 1002 | -0.279266028 | 0.076473329 | 0.00227401 | 0.281540038 | -0.074199319 | 0.293458931 | 0.573761707 | 0.987445391 | 0.305335846 | 0.66728301 |
| Spn | S | 394 | 0.088610757 | 0.002978554 | -0.162369337 | -0.250980093 | -0.16534789 | 0.059699035 | 0.943354109 | 0.007922825 | 0.002671914 | 0.012253286 |
| Slc41a2 | S | 137 | -0.003326963 | -0.085993986 | -0.003814494 | -0.000487531 | 0.082179491 | 0.945368923 | 0.249091351 | 0.951172584 | 0.993342097 | 0.304715016 |
| Slc41a2 | S | 138 | -0.003326963 | -0.085993986 | -0.003814494 | -0.000487531 | 0.082179491 | 0.945368923 | 0.249091351 | 0.951172584 | 0.993342097 | 0.304715016 |
| Acta1 Acta2 Actc1 Ac | S | 235;235;234;235 | -0.180307037 | 0.0300051 | 0.077001819 | 0.257308856 | 0.046996719 | 0.067746873 | 0.71631443 | 0.522791235 | 0.097336137 | 0.703556524 |
| Nfic | S | 284 | -0.052238434 | 0.019837414 | -0.050124281 | 0.002114153 | -0.069961694 | 0.391508271 | 0.768087029 | 0.369380997 | 0.974690315 | 0.376648293 |
| Acap2 | S | 379 | -0.1297399 | 0.151159176 | -0.040401876 | 0.089338024 | -0.191561052 | 0.026914946 | 0.192533948 | 0.422531481 | 0.04221873 | 0.133932942 |
| Rfc1 | T | 504 | 0.017457798 | -0.088976156 | -0.003228358 | -0.020686156 | 0.085747798 | 0.725152479 | 0.121760952 | 0.942904626 | 0.689037697 | 0.152087585 |
| Gpatch2l | S | 248 | 0.032317246 | -0.075531712 | -0.056888024 | -0.08920527 | 0.018643688 | 0.637867012 | 0.252626268 | 0.60217704 | 0.40889934 | 0.84905033 |
| Mia2 | S | 1188 | -0.059281462 | -0.016207714 | 0.024508727 | 0.083790189 | 0.040716441 | 0.065604575 | 0.585994234 | 0.536661342 | 0.091159484 | 0.334436487 |
| Psmc6 | S | 131 | 0.024445498 | 0.007338756 | -0.177317604 | -0.201763102 | -0.184656361 | 0.68285901 | 0.960343541 | 0.265257068 | 0.22076365 | 0.347175155 |
| Srrm2 | S | 1876 | -0.015988878 | -0.081972564 | 0.004558836 | 0.020547714 | 0.0865314 | 0.830609245 | 0.260856309 | 0.95103322 | 0.754279572 | 0.196642469 |
| Foxp1 | T | 641 | -0.028811731 | -0.118183537 | 0.024165741 | 0.052977473 | 0.142349278 | 0.813153295 | 0.269081601 | 0.659716771 | 0.664109705 | 0.202499132 |
| Specc1l | S | 10 | -0.008520718 | -0.038987491 | -0.059122961 | -0.050602243 | -0.020135471 | 0.833918262 | 0.639116863 | 0.514127249 | 0.576892649 | 0.852413456 |
| Tnks1bp1 | S | 811 | 0.123282552 | -0.01762121 | -0.158344371 | -0.281626923 | -0.140723161 | 0.301833928 | 0.806251773 | 0.048586047 | 0.066473511 | 0.077983908 |
| Ank | T | 328 | -0.130226862 | 0.002126807 | -0.001606953 | 0.128619909 | -0.003733761 | 0.353484748 | 0.985658545 | 0.975393085 | 0.355213593 | 0.973788235 |
| Bud13 | S | 371 | -0.055517441 | -0.016608698 | 0.035134734 | 0.090652175 | 0.051743432 | 0.01933804 | 0.533765943 | 0.314767343 | 0.056321498 | 0.194500483 |
| Bud13 | S | 376 | -0.055517441 | -0.016608698 | 0.035134734 | 0.090652175 | 0.051743432 | 0.01933804 | 0.533765943 | 0.314767343 | 0.056321498 | 0.194500483 |
| Trappc10 | S | 708 | -0.051438545 | 0.020403953 | -0.04582954 | 0.005609005 | -0.066233493 | 0.277415553 | 0.671355489 | 0.411505813 | 0.918696915 | 0.28591793 |
| Smg9 | S | 32 | -0.049260067 | -0.010705327 | -0.002841928 | 0.046418139 | 0.007863398 | 0.155854418 | 0.762134513 | 0.95384469 | 0.399297333 | 0.884441923 |
| Mki67 | S | 1772 | -0.044481791 | 0.036381756 | -0.086963138 | -0.042481347 | -0.123344894 | 0.408407292 | 0.223267804 | 0.155836222 | 0.505454731 | 0.086379724 |
| Tut7 | S | 929 | -0.10839987 | 0.123310653 | -0.184198006 | -0.075798136 | -0.307508659 | 0.617562648 | 0.592673958 | 0.431637727 | 0.627200213 | 0.129990847 |
| Nmnat1 | S | 136 | -0.007224476 | -0.095217052 | 0.038511901 | 0.045736378 | 0.133728954 | 0.867485161 | 0.074121321 | 0.505750836 | 0.460405957 | 0.080417767 |
| Mphosph8 | S | 164 | 0.028836309 | 0.032818555 | -0.089216129 | -0.118052438 | -0.122034683 | 0.317535012 | 0.303265977 | 0.001906901 | 0.0174162 | 0.023563346 |
| Apbb1ip | S | 531 | 0.157624488 | -0.091712351 | -0.194168705 | -0.351793192 | -0.102456354 | 0.068570141 | 0.495678773 | 0.049556878 | 0.009244928 | 0.43010567 |
| Arhgap35 | S | 1174 | 0.018755009 | 0.047581131 | -0.16640967 | -0.185164679 | -0.2139908 | 0.846522498 | 0.473353071 | 0.10523541 | 0.122491908 | 0.062783848 |
| Arhgap35 | S | 1176 | 0.018755009 | 0.047581131 | -0.16640967 | -0.185164679 | -0.2139908 | 0.846522498 | 0.473353071 | 0.10523541 | 0.122491908 | 0.062783848 |
| Arhgap21 | S | 248 | -0.029581384 | -0.031802109 | -0.035227934 | -0.005646549 | -0.003425825 | 0.755192751 | 0.532722377 | 0.597983868 | 0.955729536 | 0.959144446 |
| Shb | S | 301 | 0.00411846 | -0.085443772 | -0.075431554 | -0.079550014 | 0.010012218 | 0.96849771 | 0.54029422 | 0.744269521 | 0.723874364 | 0.965925307 |
| Arid1a | S | 302 | -0.051625071 | -0.010851361 | -0.023889794 | 0.027735276 | -0.013038434 | 0.374569567 | 0.888963449 | 0.6240296 | 0.469737663 | 0.844750503 |
| Tex2 | S | 265 | 0.050609928 | 0.005561519 | -0.133543925 | -0.184153854 | -0.139105444 | 0.212958706 | 0.928087965 | 0.035934068 | 0.016029082 | 0.099256743 |
| Nek3 | T | 477 | -0.158650989 | 0.145393064 | -0.029398317 | 0.129252673 | -0.174791381 | 0.047015542 | 0.33510356 | 0.852681334 | 0.441217521 | 0.379675172 |
| Zbed3 | S | 111 | -0.028590955 | -0.020531057 | -0.051635385 | -0.02304443 | -0.031104328 | 0.517234212 | 0.801830888 | 0.603757136 | 0.80927294 | 0.783893479 |
| Sipa1l3 | S | 1248 | 0.104967122 | -0.14614023 | -0.072399024 | -0.177366145 | 0.073741207 | 0.292264015 | 0.175490349 | 0.509700627 | 0.093171675 | 0.426207645 |
| Dek | S | 247 | -0.156642229 | 0.079477537 | -0.051120127 | 0.105522102 | -0.130597664 | 0.350657174 | 0.586009201 | 0.581189889 | 0.508582395 | 0.385934922 |
| Dek | S | 248 | -0.156642229 | 0.079477537 | -0.051120127 | 0.105522102 | -0.130597664 | 0.350657174 | 0.586009201 | 0.581189889 | 0.508582395 | 0.385934922 |
| Dek | S | 255 | -0.156642229 | 0.079477537 | -0.051120127 | 0.105522102 | -0.130597664 | 0.350657174 | 0.586009201 | 0.581189889 | 0.508582395 | 0.385934922 |
| Vcl | S | 456 | -0.095104721 | 0.026750069 | -0.032281009 | 0.062823712 | -0.059031078 | 0.329965026 | 0.812424578 | 0.662933849 | 0.509410246 | 0.608396205 |
| Mllt10 | S | 370 | 0.136208419 | -0.101835011 | -0.230618065 | -0.366826484 | -0.128783054 | 0.067873491 | 0.158881045 | 0.286049733 | 0.14148359 | 0.512071916 |
| Casp8ap2 | S | 798 | -0.017928561 | -0.072836131 | 0.043149487 | 0.061078047 | 0.115985618 | 0.720145269 | 0.045855751 | 0.445867218 | 0.345470436 | 0.105901374 |
| Sltm | S | 959 | -0.012810298 | -0.071225638 | 0.028599768 | 0.041410066 | 0.099825406 | 0.769469158 | 0.119465769 | 0.784813018 | 0.691058387 | 0.376641342 |
| Tbc1d5 | S | 565 | -0.083628133 | 0.054161546 | -0.03192084 | 0.051707293 | -0.086082386 | 0.121925232 | 0.522936272 | 0.583112296 | 0.361201 | 0.341181922 |
| Tbc1d5 | S | 568 | -0.083628133 | 0.054161546 | -0.03192084 | 0.051707293 | -0.086082386 | 0.121925232 | 0.522936272 | 0.583112296 | 0.361201 | 0.341181922 |
| Ftsj3 | S | 335 | 0.035717855 | -0.101650191 | 0.011962996 | -0.023754859 | 0.113613187 | 0.520782475 | 0.038686767 | 0.746000625 | 0.684449531 | 0.047085972 |
| Ftsj3 | S | 336 | 0.035717855 | -0.101650191 | 0.011962996 | -0.023754859 | 0.113613187 | 0.520782475 | 0.038686767 | 0.746000625 | 0.684449531 | 0.047085972 |
| Abraxas1 | S | 48 | 0.057955222 | 0.017145272 | -0.13297654 | -0.190931763 | -0.150121812 | 0.503382671 | 0.84038488 | 0.057247721 | 0.093557256 | 0.163112777 |
| Thrap3 | S | 243 | 0.020632055 | -0.044289695 | -0.020247756 | -0.040879811 | 0.024041939 | 0.260552048 | 0.054065535 | 0.240951914 | 0.019059136 | 0.167374589 |
| Syn1 | S | 62 | 0.013584447 | -0.039184671 | -0.077968061 | -0.091552508 | -0.038783391 | 0.808738976 | 0.468234788 | 0.466608995 | 0.404820512 | 0.705167174 |
| Brip1 | S | 1103 | -0.041665666 | 0.094385551 | -0.147704402 | -0.106038737 | -0.242089954 | 0.468589361 | 0.084802968 | 0.049266418 | 0.151370728 | 0.013435248 |
| Zfp800 | S | 460 | -0.028782797 | -0.068474037 | 0.03155297 | 0.060335767 | 0.100027007 | 0.594129962 | 0.119982682 | 0.528478999 | 0.268016665 | 0.080206798 |
| Hnrnpf | S | 77 | -0.062827046 | -0.021136109 | -0.000936639 | 0.061890407 | 0.02019947 | 0.259379184 | 0.813100893 | 0.983603612 | 0.298465235 | 0.82461464 |
| Raly | S | 229 | -0.114264263 | -0.092172655 | 0.004635849 | 0.118900112 | 0.096808503 | 0.61395854 | 0.823121442 | 0.982865112 | 0.194957715 | 0.801617209 |
| Lbr | S | 140 | -0.068029219 | 0.051380212 | -0.106167331 | -0.038138112 | -0.157547543 | 0.475893796 | 0.305652515 | 0.251099948 | 0.727552634 | 0.129539438 |
| Taf1 | S | 1802 | -0.104932543 | 0.025400463 | 0.006205631 | 0.111138175 | -0.019194832 | 0.098578672 | 0.613059543 | 0.933153508 | 0.172786435 | 0.766947654 |
| Arhgef2 | S | 885 | -0.099228314 | 0.064189147 | -0.060157125 | 0.039071189 | -0.124346271 | 0.184290289 | 0.428979946 | 0.569374117 | 0.669097729 | 0.250639053 |
| Aatk | S | 1263 | -0.055325616 | 0.065745333 | -0.109343817 | -0.054016862 | -0.17508915 | 0.618495526 | 0.502041086 | 0.214571363 | 0.564237619 | 0.073415521 |
| Luc7l3 | S | 393 | -0.135272067 | -0.01158887 | 0.069790227 | 0.205062294 | 0.081379097 | 0.03772857 | 0.782650551 | 0.157745801 | 0.013438614 | 0.114774377 |
| Luc7l3 | S | 395 | -0.135272067 | -0.01158887 | 0.069790227 | 0.205062294 | 0.081379097 | 0.03772857 | 0.782650551 | 0.157745801 | 0.013438614 | 0.114774377 |
| Lima1 | S | 367 | -0.212123532 | 0.154378241 | 0.019538432 | 0.231661963 | -0.13483981 | 0.047015195 | 0.359816848 | 0.888011187 | 0.148946227 | 0.460564086 |
| Cnp | S | 22 | -0.226488736 | 0.354518541 | -0.114629281 | 0.111859455 | -0.469147822 | 0.042210618 | 0.105442942 | 0.237041641 | 0.224517363 | 0.06298247 |
| Rai14 | S | 627 | 0.145836918 | -0.021890217 | -0.21328048 | -0.359117398 | -0.191390263 | 0.096291334 | 0.752409944 | 0.029988683 | 0.009080363 | 0.066310931 |
| Cep89 | S | 104 | -0.148506085 | 0.047413174 | 0.036710047 | 0.185216132 | -0.010703128 | 0.015882299 | 0.356395987 | 0.322866229 | 0.011906851 | 0.809740408 |
| Sars | S | 506 | 0.074742943 | -0.046150974 | -0.148121807 | -0.22286475 | -0.101970833 | 0.620779725 | 0.80687662 | 0.257071553 | 0.228023806 | 0.619047573 |
| Inpp5f | S | 882 | 0.120612261 | -0.001214886 | -0.316072953 | -0.436685214 | -0.314858067 | 0.498653655 | 0.993964823 | 0.267814094 | 0.167371615 | 0.280292375 |
| Ect2 | S | 865 | -0.02513765 | -0.047653096 | -0.016830689 | 0.008306961 | 0.030822407 | 0.442255565 | 0.609874978 | 0.798407275 | 0.900910425 | 0.76728665 |
| Usp37 | S | 928 | 0.046476159 | -0.032009972 | -0.054870004 | -0.101346163 | -0.022860032 | 0.553774893 | 0.384880702 | 0.109855319 | 0.250061497 | 0.404131498 |
| H1f2 H1f5 | S | 173;170 | 0.019674856 | 0.04705671 | -0.129007574 | -0.14868243 | -0.176064283 | 0.855275143 | 0.339293503 | 0.042769492 | 0.23531969 | 0.014562187 |
| Dpysl3 | S | 522 | -0.071828351 | 0.122227351 | -0.126174177 | -0.054345826 | -0.248401529 | 0.241977429 | 0.057699996 | 0.044175922 | 0.355959867 | 0.00728768 |
| Cdc5l | T | 435 | -0.040863342 | -0.039546209 | -0.017753733 | 0.023109609 | 0.021792476 | 0.529499174 | 0.624106355 | 0.830269445 | 0.720785669 | 0.783411228 |
| Cnnm4 | S | 684 | -0.206849521 | 0.0626173 | -0.017000007 | 0.189849514 | -0.079617306 | 0.346459266 | 0.657081665 | 0.868075386 | 0.36852251 | 0.50390268 |
| Wdr44 | T | 221 | 0.064418709 | -0.036318421 | -0.078823786 | -0.143242496 | -0.042505366 | 0.469761631 | 0.633376104 | 0.079705262 | 0.173466545 | 0.55421779 |
| Ktn1 | S | 87 | 0.08110289 | -0.074466584 | -0.117554468 | -0.198657358 | -0.043087884 | 0.540526994 | 0.53960505 | 0.324921632 | 0.207176366 | 0.743862355 |
| Pds5b | S | 1405 | -0.042661561 | -0.236103387 | 0.092006593 | 0.134668154 | 0.32810998 | 0.603833515 | 0.108065028 | 0.05465381 | 0.169851436 | 0.067340888 |
| Hnrnpc | S | 261 | 0.023055118 | -0.133711573 | 0.055487395 | 0.032432276 | 0.189198968 | 0.623335544 | 0.02862726 | 0.383092626 | 0.503115541 | 0.024942997 |
| Amotl1 | S | 820 | 0.055387479 | -0.063547863 | -0.117270984 | -0.172658463 | -0.05372312 | 0.285541278 | 0.257096394 | 0.408904509 | 0.26035087 | 0.680554283 |

| | | | | | | | | | | | | |
|---|---|---|---|---|---|---|---|---|---|---|---|---|
| Nup35 | T | 48 | 0.102383647 | -0.212350727 | 0.035590861 | -0.066792786 | 0.247941588 | 0.146824448 | 0.016357804 | 0.588530918 | 0.319401544 | 0.015408481 |
| Zcchc7 | S | 222 | 0.116786779 | -0.274092084 | 0.071431162 | -0.045355617 | 0.345523245 | 0.4911276 | 0.077706785 | 0.587771571 | 0.739906165 | 0.019153671 |
| Arhgef40 | S | 1472 | -0.019183827 | -0.06121765 | 0.004985301 | 0.024169128 | 0.066202951 | 0.644057433 | 0.246699909 | 0.826306967 | 0.573881074 | 0.220966679 |
| Cmtr1 | S | 50 | -0.032451432 | -0.073246097 | 0.041751578 | 0.07420301 | 0.114997675 | 0.288282973 | 0.061780745 | 0.117480393 | 0.051401995 | 0.01991294 |
| Cmtr1 | S | 52 | -0.032451432 | -0.073246097 | 0.041751578 | 0.07420301 | 0.114997675 | 0.288282973 | 0.061780745 | 0.117480393 | 0.051401995 | 0.01991294 |
| Rock2 | S | 439 | -0.181534888 | 0.066427055 | 0.014450742 | 0.19598563 | -0.051976313 | 0.138366337 | 0.54143374 | 0.874759176 | 0.087501224 | 0.474018762 |
| Mff | T | 112 | -0.043120635 | -0.12412886 | 0.069315945 | 0.112436581 | 0.193444805 | 0.708814035 | 0.186793594 | 0.482194158 | 0.352932445 | 0.079000423 |
| Tmpo | S | 221 | -0.051709699 | -0.002071253 | -0.04588787 | 0.005821829 | -0.043816617 | 0.427477631 | 0.973648869 | 0.667979472 | 0.958469399 | 0.700024036 |
| Map3k20 | S | 638 | 0.00731598 | 0.004792573 | -0.088191602 | -0.095507582 | -0.092984175 | 0.849533249 | 0.931047323 | 0.154596629 | 0.146001795 | 0.215933153 |
| Spag9 | S | 185 | 0.076698884 | -0.03251125 | -0.211683434 | -0.288382319 | -0.179172184 | 0.393357724 | 0.699238646 | 0.290109766 | 0.184236976 | 0.356419994 |
| Arhgap19 | S | 31 | 0.132715869 | -0.133237856 | -0.15682332 | -0.289539189 | -0.023585463 | 0.056188931 | 0.0960899 | 0.339843811 | 0.139939557 | 0.874091323 |
| Kif20a | S | 876 | 0.033399928 | -0.017659577 | -0.103316603 | -0.136716531 | -0.085657026 | 0.390789597 | 0.52300007 | 0.179238752 | 0.106957418 | 0.247100162 |
| Kif1b | S | 1655 | -0.11973586 | 0.148514427 | -0.13321473 | -0.01347887 | -0.281729158 | 0.302023375 | 0.195164642 | 0.197481122 | 0.911281466 | 0.060407665 |
| Garem1 | T | 659 | -0.11176718 | -0.015872128 | 0.008772073 | 0.120539254 | 0.024644201 | 0.386808283 | 0.806207103 | 0.877620038 | 0.352790199 | 0.673570004 |
| Setd2 | S | 246 | -0.001425975 | -0.060704176 | -0.016973903 | -0.015547929 | 0.043730272 | 0.976288195 | 0.306352941 | 0.708112016 | 0.721951019 | 0.425892377 |
| Arhgap10 | S | 715 | 0.137112716 | -0.102685221 | -0.123586043 | -0.260698759 | -0.020900821 | 0.012816317 | 0.288250143 | 0.00872158 | 0.002103287 | 0.804221147 |
| Erbin | S | 854 | 0.004915884 | -0.023060725 | -0.054945274 | -0.059861158 | -0.031884548 | 0.903264092 | 0.583394928 | 0.29016486 | 0.25406721 | 0.521167292 |
| Cast | S | 324 | -0.160427472 | -0.003973932 | -0.010999608 | 0.149427864 | -0.007025676 | 0.533873146 | 0.984276971 | 0.947337408 | 0.586895556 | 0.975595675 |
| Pdxdc1 | S | 387 | 0.152323559 | 0.006077588 | -0.437975563 | -0.590299122 | -0.444053151 | 0.169398905 | 0.962460009 | 0.167294192 | 0.09909293 | 0.161908706 |
| Clspn | S | 932 | 0.014029401 | 0.107511685 | -0.188948109 | -0.202977509 | -0.296459794 | 0.846533476 | 0.401577241 | 0.068913664 | 0.066478393 | 0.075901463 |
| Mga | S | 2837 | -0.231768257 | -0.054806605 | 0.157076658 | 0.388844914 | 0.211883263 | 0.297495048 | 0.70109032 | 0.520134524 | 0.210914291 | 0.419577778 |
| Mga | S | 2849 | -0.231768257 | -0.054806605 | 0.157076658 | 0.388844914 | 0.211883263 | 0.297495048 | 0.70109032 | 0.520134524 | 0.210914291 | 0.419577778 |
| Arhgef2 | S | 955 | 0.00022365 | -0.076930039 | 0.034944818 | 0.034721169 | 0.111874857 | 0.99519789 | 0.039419751 | 0.531590019 | 0.539533799 | 0.110231181 |
| Sfpq | S | 33 | -0.00647088 | -0.042472619 | -0.076538548 | -0.070067668 | -0.034065929 | 0.957902291 | 0.785610129 | 0.497687639 | 0.635140555 | 0.843409201 |
| Wdr81 | S | 1570 | -0.003680669 | -0.064626793 | -0.009726864 | -0.006046195 | 0.054899929 | 0.943300059 | 0.263081042 | 0.84572147 | 0.925795027 | 0.411007414 |
| Klc1 | S | 523 | -0.131336984 | -0.01319243 | 0.043916914 | 0.175253898 | 0.057109344 | 0.216279503 | 0.854678006 | 0.618815775 | 0.170934478 | 0.580740231 |
| Pard3 | S | 201 | 0.036087013 | -0.152562326 | -0.008674216 | -0.044761229 | 0.14388811 | 0.572218242 | 0.241484244 | 0.931777235 | 0.68378358 | 0.324692414 |
| Cttnbp2nl | S | 559 | 0.097496527 | -0.107692063 | -0.027051005 | -0.124547532 | 0.080641058 | 0.091407007 | 0.03446214 | 0.144535774 | 0.053621176 | 0.065225526 |
| Ubr2 | S | 1009 | -0.00053004 | 0.00702922 | -0.093288428 | -0.092758388 | -0.100317648 | 0.983827169 | 0.898241327 | 0.226032171 | 0.230721021 | 0.236909112 |
| Abhd15 | S | 442 | 0.010146531 | -0.138619358 | 0.038668226 | 0.028521695 | 0.177287585 | 0.905648165 | 0.178533157 | 0.734827582 | 0.798004983 | 0.184809797 |
| Pgrmc2 | T | 110 | -0.019265147 | -0.053469543 | -0.029179657 | -0.00991451 | 0.024289885 | 0.861864361 | 0.560924694 | 0.618152584 | 0.929793762 | 0.792627643 |
| Lrrfip1 | S | 649 | -0.061697602 | -0.058529594 | 0.087743295 | 0.149440897 | 0.146272888 | 0.126291128 | 0.421437475 | 0.533131217 | 0.330374135 | 0.347700494 |
| Eps15 | S | 562 | 0.025804579 | -0.09599654 | -0.019097783 | -0.044902363 | 0.076898757 | 0.772244916 | 0.255259412 | 0.792839083 | 0.671316294 | 0.427190991 |
| Tmpo | S | 115;115 | -0.037638461 | 0.024682015 | -0.060565118 | -0.022926657 | -0.085247132 | 0.426058686 | 0.573882793 | 0.263805043 | 0.693895574 | 0.183781824 |
| Cnbp | T | 173 | -0.054534042 | -0.001446264 | -0.019186381 | 0.03534766 | -0.017740118 | 0.251042175 | 0.954475146 | 0.810766748 | 0.678793822 | 0.823824637 |
| Prpf31 | S | 439 | -0.004833104 | 0.01551789 | -0.073702626 | -0.068869522 | -0.089220516 | 0.954350366 | 0.839283749 | 0.136286128 | 0.420217771 | 0.271057887 |
| Prpf31 | T | 440 | -0.004833104 | 0.01551789 | -0.073702626 | -0.068869522 | -0.089220516 | 0.954350366 | 0.839283749 | 0.136286128 | 0.420217771 | 0.271057887 |
| Sorbs2 | S | 27 | -0.083329559 | 0.153739537 | -0.201948484 | -0.118618925 | -0.355688022 | 0.5820653 | 0.371806091 | 0.257164845 | 0.425064096 | 0.075801168 |
| Ankrd17 | S | 2041 | 0.038440587 | -0.035903947 | -0.039218276 | -0.077658864 | -0.003314329 | 0.564383478 | 0.220410598 | 0.175782242 | 0.297722775 | 0.914252671 |
| Snip1 | S | 50 | -0.089194573 | 0.080793153 | -0.04509095 | 0.044103623 | -0.125884103 | 0.040814854 | 0.251829899 | 0.626550171 | 0.63295771 | 0.256882152 |
| Srrm1 | S | 434 | 0.005019705 | -0.065900476 | -0.026106038 | -0.031125743 | 0.039794438 | 0.927015984 | 0.387135564 | 0.602737401 | 0.450228466 | 0.547221767 |
| Washc2 | S | 1169 | 0.010565047 | -0.006861614 | -0.082284756 | -0.092849803 | -0.075423142 | 0.697395003 | 0.836563391 | 0.186502179 | 0.152760507 | 0.223261935 |
| Washc2 | S | 1172 | 0.010565047 | -0.006861614 | -0.082284756 | -0.092849803 | -0.075423142 | 0.697395003 | 0.836563391 | 0.186502179 | 0.152760507 | 0.223261935 |
| Ythdc2 | S | 1104 | -0.092604012 | 0.044749361 | -0.083643936 | 0.008960075 | -0.128393298 | 0.456895161 | 0.251606151 | 0.28519345 | 0.943245803 | 0.145575126 |
| Dst | S | 237 | 0.004070258 | -0.013355879 | -0.054655413 | -0.058725672 | -0.041299535 | 0.911379731 | 0.802912291 | 0.172385894 | 0.014622122 | 0.392892177 |
| Rnf2 | S | 168 | -0.001504473 | 0.013536642 | -0.337184014 | -0.335679541 | -0.350720656 | 0.995149675 | 0.955503149 | 0.671217826 | 0.67520726 | 0.661856981 |
| C2cd5 | S | 637 | -0.039202697 | 0.049590497 | -0.103291178 | -0.06408848 | -0.152881674 | 0.711372587 | 0.654485407 | 0.258349287 | 0.509118928 | 0.185365322 |
| Polh | S | 668 | 0.032138021 | -0.088795121 | 0.009247978 | -0.022890043 | 0.098043099 | 0.718459739 | 0.061810938 | 0.759833607 | 0.789683288 | 0.055588959 |
| Zc3hav1 | S | 805 | 0.025395768 | -0.089161686 | -0.049028172 | -0.07442394 | 0.040133514 | 0.868046241 | 0.567795516 | 0.520970076 | 0.650989593 | 0.800725216 |
| Sh3pxd2a | S | 566 | -0.007571486 | -0.001334768 | -0.049719787 | -0.042148301 | -0.048385019 | 0.787360302 | 0.946325895 | 0.181559054 | 0.287492428 | 0.19527224 |
| Ap2m1 | S | 234 | -0.024516363 | -0.030893006 | -0.020687174 | 0.003829189 | 0.010205832 | 0.549748596 | 0.571683722 | 0.679048784 | 0.926374826 | 0.850345284 |
| Hps4 | S | 482 | -0.040422691 | -0.0005648 | -0.033410854 | 0.007011837 | -0.032846054 | 0.416656882 | 0.994745755 | 0.500301945 | 0.882552741 | 0.703814807 |
| Smg8 | S | 742 | -0.008036407 | -0.06074433 | 0.008959537 | 0.016995944 | 0.069703867 | 0.900907566 | 0.134318833 | 0.862474397 | 0.806596587 | 0.206181925 |
| Ddx47 | S | 9 | 0.037024584 | -0.06777613 | -0.023379346 | -0.06040393 | 0.044396784 | 0.351761739 | 0.104261545 | 0.396870107 | 0.176996485 | 0.254669979 |
| Arhgap17 | T | 736 | -0.038529111 | 0.005244126 | -0.029610909 | 0.008918202 | -0.034855035 | 0.324032479 | 0.886162249 | 0.42813791 | 0.826447849 | 0.410065381 |
| Lyst | S | 1503 | -0.010999472 | 0.057177814 | -0.1321894 | -0.121189928 | -0.189367214 | 0.864579914 | 0.552431014 | 0.167640987 | 0.217371522 | 0.136901143 |
| Map1a | S | 999 | -0.03679126 | -0.014498192 | -0.03058044 | 0.00621082 | -0.016082248 | 0.446909659 | 0.794038501 | 0.665165483 | 0.926873082 | 0.82534761 |
| Map1a | S | 1008 | -0.03679126 | -0.014498192 | -0.03058044 | 0.00621082 | -0.016082248 | 0.446909659 | 0.794038501 | 0.665165483 | 0.926873082 | 0.82534761 |
| Champ1 | S | 108 | 0.09148973 | -0.064589455 | -0.086807942 | -0.178297672 | -0.022218487 | 0.135406481 | 0.324651676 | 0.056676967 | 0.032195476 | 0.686424429 |
| Rin1 | S | 446 | -0.034541312 | -0.003281967 | -0.059335473 | -0.024794161 | -0.056053506 | 0.664859325 | 0.964679976 | 0.470910533 | 0.73616394 | 0.428491348 |
| Baz1b | S | 326 | -0.014788649 | -0.008796564 | -0.046315225 | -0.031526577 | -0.037518661 | 0.697678063 | 0.79961855 | 0.339862052 | 0.534140061 | 0.448113349 |
| Baz1b | S | 327 | -0.014788649 | -0.008796564 | -0.046315225 | -0.031526577 | -0.037518661 | 0.697678063 | 0.79961855 | 0.339862052 | 0.534140061 | 0.448113349 |
| Dclre1c | S | 527 | -0.015122566 | -0.025592683 | -0.035768799 | -0.020646232 | -0.010176116 | 0.782747115 | 0.396017274 | 0.642990793 | 0.806839426 | 0.890919456 |
| Srrm2 | S | 200 | 0.022675761 | -0.081046741 | -0.037129899 | -0.05980566 | 0.043916841 | 0.551785335 | 0.438373979 | 0.347867243 | 0.01918723 | 0.649663395 |
| Egln1 | S | 131 | 0.03730621 | -0.02235885 | -0.114474794 | -0.151781003 | -0.092115944 | 0.233701814 | 0.408942225 | 0.19626719 | 0.126568268 | 0.274515289 |
| Brwd1 | S | 1851 | 0.036157522 | -0.033427616 | -0.069699563 | -0.105857085 | -0.036271947 | 0.44521588 | 0.297238379 | 0.252155192 | 0.141625857 | 0.515145497 |
| Brwd1 | T | 1853 | 0.036157522 | -0.033427616 | -0.069699563 | -0.105857085 | -0.036271947 | 0.44521588 | 0.297238379 | 0.252155192 | 0.141625857 | 0.515145497 |
| Srrm2 | S | 1810 | -0.03889865 | -0.093607908 | 0.104685631 | 0.142984282 | 0.198293539 | 0.423301635 | 0.077759339 | 0.352394415 | 0.234498206 | 0.14341436 |
| Nefm | S | 715 | -0.076533528 | -0.089609804 | 0.088319799 | 0.164853327 | 0.177929603 | 0.298693382 | 0.195784354 | 0.255863645 | 0.03742008 | 0.032708435 |
| Map1a | S | 1316 | -0.066455078 | 0.024649097 | -0.026066693 | 0.040388385 | -0.05071579 | 0.154415284 | 0.544711953 | 0.631904725 | 0.512592596 | 0.428082431 |
| Gnl1 | S | 32 | -0.061226704 | -0.019431215 | 0.072173142 | 0.133399846 | 0.091604357 | 0.036278373 | 0.454077853 | 0.464863217 | 0.236193851 | 0.370373997 |
| Gnl1 | S | 33 | -0.061226704 | -0.019431215 | 0.072173142 | 0.133399846 | 0.091604357 | 0.036278373 | 0.454077853 | 0.464863217 | 0.236193851 | 0.370373997 |
| Gnl1 | S | 34 | -0.061226704 | -0.019431215 | 0.072173142 | 0.133399846 | 0.091604357 | 0.036278373 | 0.454077853 | 0.464863217 | 0.236193851 | 0.370373997 |
| Sltm | T | 997 | -0.062014358 | -0.026618314 | 0.00340339 | 0.065417748 | 0.030021703 | 0.413629406 | 0.556187218 | 0.891573503 | 0.389251945 | 0.487535905 |
| Fbxo46 | S | 240 | -0.00615915 | -0.061250328 | -0.034215222 | -0.028056072 | 0.027035105 | 0.92221992 | 0.558306515 | 0.62059942 | 0.682690659 | 0.79536397 |
| Cdk1 | T | 161 | 0.04086574 | -0.035460813 | -0.053204604 | -0.094070345 | -0.017743791 | 0.18619432 | 0.200599697 | 0.084957696 | 0.007572056 | 0.232586474 |
| Eps8 | S | 624 | -0.022241507 | -0.149099552 | 0.095880038 | 0.118121545 | 0.244979589 | 0.674604483 | 0.038983848 | 0.120561782 | 0.039749473 | 0.007367296 |
| Svil | T | 1183 | -0.071491878 | -0.005297974 | 0.042981636 | 0.114473513 | 0.04827961 | 0.023098338 | 0.864436226 | 0.364956774 | 0.070630884 | 0.340505012 |
| Cep57 | S | 53 | 0.172012452 | -0.112903641 | -0.129322566 | -0.301335018 | -0.016418925 | 4.93807E-05 | 0.000522446 | 0.017965954 | 0.003147097 | 0.603932146 |
| Rab11fip1 | S | 243 | -0.04346364 | -0.036146973 | 0.026826042 | 0.070289682 | 0.062973015 | 0.242756514 | 0.274609403 | 0.555317708 | 0.202583696 | 0.233956833 |
| Per1 | S | 978 | 0.001175826 | -0.045765636 | -0.088131616 | -0.089307443 | -0.042365981 | 0.992855334 | 0.686683842 | 0.607226277 | 0.576333499 | 0.768885021 |
| Dyrk1a | S | 49 | -0.058139274 | -0.00157608 | -0.055176151 | 0.002963124 | -0.05360007 | 0.685489429 | 0.990254282 | 0.53538897 | 0.983870695 | 0.695636767 |
| Map3k4 | S | 65 | -0.12646686 | 0.058159552 | -0.057422369 | 0.069044491 | -0.115581922 | 0.286991289 | 0.581783336 | 0.788611561 | 0.758989555 | 0.611553607 |
| Slc9a8 | S | 566 | -0.008896132 | -0.073723336 | -0.006044116 | 0.002852016 | 0.06767922 | 0.914082135 | 0.332814096 | 0.913234014 | 0.970840415 | 0.348165043 |
| Slc9a8 | S | 568 | -0.008896132 | -0.073723336 | -0.006044116 | 0.002852016 | 0.06767922 | 0.914082135 | 0.332814096 | 0.913234014 | 0.970840415 | 0.348165043 |
| Pi4kb | S | 428 | -0.061392721 | 0.090818318 | -0.083765405 | -0.022372684 | -0.174583723 | 0.197100263 | 0.259152755 | 0.193758007 | 0.675779644 | 0.077146194 |
| Rbl2 | T | 398 | -0.086278317 | 0.083882322 | -0.119611007 | -0.03333269 | -0.203493329 | 0.454143592 | 0.478190158 | 0.417061595 | 0.828748106 | 0.2387584 |
| Nit2 | T | 125 | -0.074249664 | 0.017883305 | -0.006292042 | 0.067957622 | -0.024175346 | 0.140932841 | 0.736036281 | 0.882572692 | 0.181219114 | 0.648200688 |
| Phtf1 | S | 181 | 0.010050562 | -0.075101034 | -0.004474961 | -0.014525523 | 0.070626073 | 0.837379735 | 0.173778153 | 0.956892062 | 0.867983091 | 0.442153631 |
| Slc16a1 | S | 482 | 0.040769589 | -0.061270345 | -0.087014545 | -0.127784134 | -0.0257442 | 0.655142776 | 0.505953522 | 0.456219106 | 0.34484836 | 0.83844052 |
| Rpsa | Y | 139 | -0.028622647 | 0.026446112 | -0.112963986 | -0.084341339 | -0.139410098 | 0.825840724 | 0.894470468 | 0.418616374 | 0.57283313 | 0.518185912 |
| Mapre3 | T | 161 | 0.021839195 | 0.028289577 | -0.123346715 | -0.145185911 | -0.151636292 | 0.628527384 | 0.637385684 | 0.0886021 | 0.06909436 | 0.06115703 |
| Wdr43 | S | 432 | -0.054790901 | -0.101067042 | 0.078993923 | 0.133784824 | 0.180060965 | 0.26465579 | 0.061955198 | 0.062983082 | 0.036964195 | 0.012934958 |
| Numa1 | S | 1812 | -0.000488992 | 0.014677164 | -0.071561946 | -0.071072954 | -0.08623911 | 0.989276234 | 0.778036706 | 0.117955809 | 0.067183206 | 0.125905735 |

| | | | | | | | | | | | | | |
|---|---|---|---|---|---|---|---|---|---|---|---|---|---|
| Ppp1r9b | S | | 99 | 0.023124469 | 0.051497991 | -0.175453825 | -0.198578294 | -0.226951817 | 0.878373368 | 0.574286835 | 0.171233124 | 0.279313896 | 0.116575078 |
| Larp1 | S | | 298 | 0.026900266 | -0.058417902 | -0.054828464 | -0.081728731 | 0.003589437 | 0.602126238 | 0.394829118 | 0.427438298 | 0.199637302 | 0.956681282 |
| Ap3d1 | S | | 755 | 0.052863392 | -0.042544803 | -0.054477061 | -0.107340453 | -0.011932258 | 0.249681091 | 0.115524106 | 0.194028004 | 0.072003758 | 0.735949672 |
| Ranbp3 | S | | 31 | -0.043371164 | 0.022609067 | -0.075884966 | -0.032513802 | -0.098494033 | 0.595627237 | 0.677675573 | 0.370117298 | 0.743562508 | 0.279722408 |
| Med1 | S | | 1209 | 0.08477406 | -0.133106138 | -0.02739895 | -0.11217301 | 0.105707188 | 0.125533497 | 0.045862183 | 0.645665572 | 0.148470145 | 0.167515536 |
| Mprip | S | | 502 | -0.110037888 | 0.071925475 | -0.046399413 | 0.063638475 | -0.118324888 | 0.167178543 | 0.295275863 | 0.486362307 | 0.24499681 | 0.012445123 |
| Specc1 | S | | 371 | 0.14647235 | -0.12516297 | -0.075200063 | -0.221672413 | 0.049962907 | 0.226840401 | 0.083726586 | 0.466975779 | 0.137268221 | 0.62756431 |
| Tnks1bp1 | S | | 1024 | 0.114229585 | -0.100813216 | -0.094964228 | -0.209193813 | 0.005848988 | 0.384987687 | 0.19074108 | 0.51293505 | 0.252986869 | 0.96687726 |
| Flna | S | | 1533 | 0.094335484 | -0.145175758 | -0.038654681 | -0.132990165 | 0.106521077 | 0.296351184 | 0.127488454 | 0.680223617 | 0.080069511 | 0.13017524 |
| Srgap2 | S | | 1061 | 0.004038075 | -0.063102462 | -0.008468444 | -0.01250652 | 0.054634018 | 0.905696514 | 0.192283765 | 0.914837499 | 0.867662606 | 0.50804074 |
| Srsf1 | S | | 199 | -0.053173776 | 0.003794513 | 0.001607617 | 0.054781394 | -0.002186896 | 0.097510784 | 0.946228597 | 0.964210775 | 0.135636811 | 0.968966589 |
| Tpx2 | S | | 645 | -0.076364095 | -0.015461762 | -0.029351618 | 0.047012477 | -0.013889856 | 0.545914713 | 0.892191074 | 0.835876747 | 0.67418912 | 0.888680645 |
| Cdca2 | S | | 572 | 0.071836521 | -0.047568676 | -0.135329637 | -0.207166158 | -0.087760961 | 0.362928432 | 0.393510466 | 0.327660135 | 0.177770312 | 0.493513683 |
| Cr1l | S | | 477 | -0.000636039 | 0.005533223 | -0.059892477 | -0.059256438 | -0.0654257 | 0.994126906 | 0.925283671 | 0.132203603 | 0.51962072 | 0.327355721 |
| Rgs12 | S | | 171 | -0.089724924 | 0.073618356 | -0.072056436 | 0.017668488 | -0.145674792 | 0.19841964 | 0.295602536 | 0.426179257 | 0.832724626 | 0.150143635 |
| Raly | S | | 249 | -0.009671407 | -0.12180746 | -0.037283035 | -0.027611628 | 0.084524425 | 0.969752124 | 0.578332825 | 0.926207547 | 0.947540421 | 0.834400586 |
| Anxa2 | S | | 127 | -0.026178575 | 0.118586954 | -0.257573048 | -0.231394474 | -0.376160003 | 0.871513771 | 0.354044262 | 0.225102395 | 0.285183265 | 0.111020553 |
| Fam122a | S | | 73 | -0.033934034 | 0.169471352 | -0.252251047 | -0.218317013 | -0.421722399 | 0.550600368 | 0.249659541 | 0.167398955 | 0.211248467 | 0.061992708 |
| Kdm2a | S | | 731 | 0.012619711 | -0.00230916 | -0.057204049 | -0.06982376 | -0.054894889 | 0.715735796 | 0.924858122 | 0.079810406 | 0.096789144 | 0.076015015 |
| Mia3 | S | | 1693 | -0.022124417 | 0.017137591 | -0.065672752 | -0.043548335 | -0.082810342 | 0.656391985 | 0.779454231 | 0.259692734 | 0.371146968 | 0.205150719 |
| Trip4 | S | | 142 | 0.114909599 | -0.110340446 | -0.145940495 | -0.260850094 | -0.035600049 | 0.416562393 | 0.510588236 | 0.384630755 | 0.188571915 | 0.854308541 |
| Rspry1 | S | | 50 | -0.101617693 | 0.005786667 | 0.03855671 | 0.140174402 | 0.032770043 | 0.079982896 | 0.866469448 | 0.560530319 | 0.10064018 | 0.603762993 |
| Son | S | | 1807 | 0.086576475 | -0.149514452 | 0.016990169 | -0.069586307 | 0.166504621 | 0.063335979 | 0.004760021 | 0.702697372 | 0.124932659 | 0.011403566 |
| Gmps | S | | 332 | -0.051400469 | 0.023412284 | -0.045516365 | 0.005884104 | -0.068928649 | 0.259556047 | 0.634799769 | 0.538943534 | 0.930556661 | 0.364731992 |
| Elk3 | S | | 115 | 0.122518686 | -0.09244596 | -0.056565422 | -0.179084108 | 0.035880538 | 0.618924318 | 0.256517029 | 0.441247322 | 0.47635713 | 0.445847375 |
| Incenp | S | | 72 | 0.150751358 | -0.00977578 | -0.147752766 | -0.298504124 | -0.137976986 | 0.505854225 | 0.711519883 | 0.094693758 | 0.247755074 | 0.122651508 |
| Larp1 | S | | 1023 | -0.037273942 | 0.021418394 | -0.078496299 | -0.041222357 | -0.099914694 | 0.608185791 | 0.574545464 | 0.318548751 | 0.655698079 | 0.234722819 |
| Etl4 | S | | 1734 | 0.05396882 | 0.024933174 | -0.213073295 | -0.267042115 | -0.238006468 | 0.748274682 | 0.799721172 | 0.272472171 | 0.258531238 | 0.239437095 |
| Slc12a4 | S | | 967 | 0.045588944 | -0.066629313 | -0.038331622 | -0.083920566 | 0.02829769 | 0.080114732 | 0.034072436 | 0.385261776 | 0.122070212 | 0.494319887 |
| Arhgap31 | S | | 667 | -0.039319239 | -0.04366039 | -0.034088575 | 0.005230664 | 0.009571816 | 0.73912239 | 0.733056985 | 0.741810021 | 0.970969306 | 0.949490555 |
| Arhgap31 | T | | 666 | -0.039319239 | -0.04366039 | -0.034088575 | 0.005230664 | 0.009571816 | 0.73912239 | 0.733056985 | 0.741810021 | 0.970969306 | 0.949490555 |
| Sumo2 Sumo3 | T | 38;37 | | -0.124429917 | 0.147576845 | -0.055376112 | 0.069053805 | -0.202952957 | 0.052835573 | 0.26876745 | 0.624259359 | 0.544491163 | 0.208430458 |
| Rin1 | S | | 341 | -0.088143253 | 0.024696207 | -0.005148463 | 0.08299479 | -0.02984467 | 0.156682327 | 0.691709381 | 0.945123806 | 0.284943378 | 0.688221931 |
| Otud4 | S | | 442 | 0.150294477 | -0.081521122 | -0.239909744 | -0.390204221 | -0.158388622 | 0.019702384 | 0.120899721 | 0.146678837 | 0.06369181 | 0.28419971 |
| Phf20 | S | | 362 | 0.108025574 | -0.089957019 | -0.081285845 | -0.18931142 | 0.008671173 | 0.56348005 | 0.341596142 | 0.427145658 | 0.332228149 | 0.907792355 |
| Sik3 | S | | 856 | 0.006646147 | 0.001033971 | -0.093548854 | -0.100195001 | -0.094582825 | 0.867423471 | 0.981707399 | 0.250292856 | 0.223585061 | 0.244133715 |
| Rhbdf1 | S | | 76 | -0.100871985 | 0.087366974 | -0.002767349 | 0.098104636 | -0.090134323 | 0.00143477 | 0.053094212 | 0.88329008 | 0.00343157 | 0.061025431 |
| Map7d1 | T | | 99 | -0.113025341 | 0.031977625 | -0.031696982 | 0.081328358 | -0.063674607 | 0.361826248 | 0.73152272 | 0.79430903 | 0.554856749 | 0.590230335 |
| Abraxas2 | S | | 280 | 0.053603348 | -0.12183758 | -0.020181337 | -0.073784685 | 0.101656243 | 0.388684578 | 0.178765893 | 0.663751528 | 0.292906304 | 0.24664929 |
| Ash1l | S | | 958 | 0.004106867 | -0.051742271 | -0.039191774 | -0.043298641 | 0.012550497 | 0.941038226 | 0.410453054 | 0.671710514 | 0.656646583 | 0.897773681 |
| Samd4b | S | | 252 | -0.028340421 | 0.009583462 | -0.052910031 | -0.024569609 | -0.062493492 | 0.431459351 | 0.80021394 | 0.417607264 | 0.69763112 | 0.362767263 |
| Rrp36 | T | | 64 | -0.055125099 | 0.016966627 | -0.047723102 | 0.007401997 | -0.064689729 | 0.364127848 | 0.730322494 | 0.585330566 | 0.931218138 | 0.453650929 |
| Ikbkb | S | | 672 | -0.103855982 | 0.008472547 | 0.042083471 | 0.145939452 | 0.033610924 | 0.123334892 | 0.895623656 | 0.738314298 | 0.304570224 | 0.794170806 |
| Tbrg1 | S | | 400 | -0.003475824 | -0.07260281 | 0.009085788 | 0.012561612 | 0.081688598 | 0.947504875 | 0.186345045 | 0.835837969 | 0.840363551 | 0.197899994 |
| Tnks1bp1 | S | | 1611 | -0.03981403 | -0.005734615 | 0.018423003 | 0.058237033 | 0.024157618 | 0.034756714 | 0.822540703 | 0.726890006 | 0.323109022 | 0.658363169 |
| Dock1 | S | | 1704 | 0.010963092 | -0.006477314 | -0.040282104 | -0.051245196 | -0.03380479 | 0.941621379 | 0.835622883 | 0.086431605 | 0.735127161 | 0.275581746 |
| Bmp2k | S | | 1010 | -0.015670311 | -0.07753601 | 0.039763858 | 0.055434169 | 0.117299868 | 0.759128233 | 0.046067676 | 0.258390801 | 0.280280838 | 0.002711638 |
| Fam110a | S | | 253 | 0.006123409 | -0.042991651 | -0.031573977 | -0.037697386 | 0.011417674 | 0.910423403 | 0.276707186 | 0.676654758 | 0.65816959 | 0.881082177 |
| Slc30a6 | S | | 381 | 0.033963128 | -0.134782235 | 0.017212574 | -0.016750554 | 0.151994808 | 0.619643536 | 0.150669885 | 0.835135575 | 0.857247383 | 0.178691503 |
| Phldb2 | S | | 486 | 0.027234967 | -0.05458214 | 0.00359418 | -0.023640787 | 0.05817632 | 0.57791725 | 0.02102718 | 0.936952742 | 0.696769064 | 0.272329657 |
| Tcp1 | S | | 544 | -0.04537193 | -0.016574854 | 0.000275689 | 0.045647619 | 0.016850543 | 0.371603866 | 0.476560212 | 0.994681389 | 0.442120434 | 0.703688617 |
| Nipbl | S | | 1071 | 0.062059196 | -0.13638606 | 0.020631769 | -0.041427427 | 0.157017829 | 0.268786917 | 0.02350205 | 0.567337405 | 0.42086141 | 0.020269066 |
| Tjp2 | S | | 902 | 0.116738807 | -0.043692679 | -0.228230661 | -0.344969468 | -0.184537982 | 0.370275761 | 0.838940435 | 0.235621635 | 0.100246799 | 0.435549846 |
| Ruvbl2 | S | | 463 | 0.050658497 | -0.059790592 | -0.090230515 | -0.140889012 | -0.030439922 | 0.796486389 | 0.65690607 | 0.445411235 | 0.503462863 | 0.833142373 |
| 2210016L21Rik | S | | 139 | 0.200114478 | -0.058947924 | -0.328786237 | -0.528900715 | -0.269838313 | 0.033070882 | 0.486664945 | 0.067767366 | 0.019066873 | 0.100281843 |
| Farp1 | T | | 527 | 0.030717719 | -0.008370486 | -0.087190028 | -0.117907747 | -0.078819542 | 0.410229626 | 0.861085035 | 0.097318023 | 0.052704856 | 0.138369524 |
| Tmem209 | S | | 131 | -0.029688345 | 0.032449685 | -0.111714818 | -0.082026473 | -0.144164503 | 0.832504663 | 0.81383676 | 0.380381345 | 0.614128636 | 0.386085574 |
| Dffa | S | | 107 | 0.156676866 | -0.03990705 | -0.163847387 | -0.320524253 | -0.123940336 | 0.432700162 | 0.832743258 | 0.11377864 | 0.171019089 | 0.534572893 |
| Spice1 | S | | 765 | -0.019278025 | -0.014190085 | -0.08406589 | -0.064787866 | -0.069875805 | 0.903992887 | 0.91031272 | 0.506642623 | 0.682594522 | 0.574668257 |
| Lrch3 | S | | 395 | 0.000771742 | -0.086765686 | -0.026367933 | -0.027139675 | 0.060397753 | 0.995422113 | 0.499704615 | 0.803702172 | 0.861219982 | 0.680659326 |
| Eif5b | S | | 165 | -0.014788556 | -0.008904807 | -0.040591214 | -0.025802658 | -0.031686407 | 0.751284875 | 0.790030851 | 0.342995007 | 0.640475299 | 0.496959722 |
| Ska3 | S | | 34 | 0.059744745 | 0.0108969 | -0.16274876 | -0.222493505 | -0.17364566 | 0.514675457 | 0.860068147 | 0.146892197 | 0.100117774 | 0.130032791 |
| Cspp1 | S | | 915 | 0.12522121 | -0.103591015 | -0.080077824 | -0.205299034 | 0.023513192 | 0.529460371 | 0.404917741 | 0.337400775 | 0.332789197 | 0.836587135 |
| Ccm2 | S | | 296 | -0.225542958 | 0.143213466 | -0.069018803 | 0.156524155 | -0.21223227 | 0.217009203 | 0.30866001 | 0.748920861 | 0.50735817 | 0.354545112 |
| Kctd12 | S | | 187 | 0.046605612 | -0.10229215 | -0.083286261 | -0.129891873 | 0.019005888 | 0.649346211 | 0.528374037 | 0.606016719 | 0.465907007 | 0.924856336 |
| Zfyve16 | S | | 927 | 0.153068428 | -0.134501045 | -0.104163658 | -0.257232086 | 0.030337387 | 0.18340525 | 0.111966536 | 0.461433721 | 0.145506157 | 0.825582997 |
| Hjurp A730008H23Ril | S | | 356 | 0.093834245 | -0.108064504 | -0.083028947 | -0.176863193 | 0.025035556 | 0.454353977 | 0.244464395 | 0.621034039 | 0.359867338 | 0.879845426 |
| Zswim8 | T | | 1154 | -0.212951786 | -0.008520306 | 0.070333298 | 0.283285084 | 0.078853604 | 0.435606767 | 0.951546831 | 0.802729742 | 0.432492404 | 0.786099114 |
| Btbd7 | S | | 722 | -0.062273265 | -0.04465283 | 0.024502669 | 0.086775933 | 0.069155498 | 0.463676542 | 0.493103432 | 0.771864099 | 0.415470299 | 0.462221731 |
| Ccm2 | T | | 294 | 0.004530332 | 0.025978574 | -0.202640073 | -0.207170405 | -0.228618647 | 0.975203979 | 0.837247535 | 0.462622822 | 0.440426682 | 0.396722697 |
| Fra10ac1 | S | | 248 | -0.072667723 | 0.026098555 | -0.031562667 | 0.041105056 | -0.057661222 | 0.405452905 | 0.759423158 | 0.513466524 | 0.628532078 | 0.521089402 |
| Mycbp2 | S | | 2992 | 0.02538734 | -0.075184363 | -0.019219452 | -0.044606792 | 0.055964911 | 0.694733938 | 0.207343451 | 0.782708248 | 0.52158989 | 0.386112964 |
| Etl4 | S | | 1843 | -0.094286192 | 0.081284824 | -0.116998273 | -0.022712081 | -0.198283097 | 0.426843557 | 0.210156645 | 0.274275785 | 0.85618326 | 0.111225476 |
| Atxn3 | S | | 272 | 0.021136473 | -0.036706188 | -0.05711551 | -0.078251983 | -0.020409322 | 0.641790371 | 0.560268967 | 0.286351821 | 0.108554683 | 0.726055043 |
| Pias1 | S | | 510 | -0.046677945 | 0.01190429 | -0.065088122 | -0.018410177 | -0.076992412 | 0.481156987 | 0.798709877 | 0.597866474 | 0.878863168 | 0.525626953 |
| Aldh1l2 | T | | 213 | -0.16043443 | -0.024119429 | 0.112675026 | 0.273109455 | 0.136794455 | 0.088917868 | 0.572793699 | 0.203028957 | 0.033294114 | 0.14916033 |
| Sh3kbp1 | S | | 156 | -0.12653307 | 0.040755514 | 0.031782033 | 0.158315103 | -0.008973481 | 0.117289558 | 0.701787798 | 0.794192173 | 0.25461355 | 0.949011457 |
| Srrm1 | S | | 714 | -0.032274034 | -0.028101725 | 0.022986587 | 0.055260621 | 0.051088313 | 0.102474936 | 0.066809986 | 0.096075631 | 0.034309225 | 0.006136766 |
| Kif23 | T | | 906 | 0.03836403 | -0.00237342 | -0.125693899 | -0.164057929 | -0.123320479 | 0.214955229 | 0.942954648 | 0.093649333 | 0.051800588 | 0.095149713 |
| Rbm3 | S | | 143 | -0.054659448 | 0.08394247 | -0.136135112 | -0.081475664 | -0.220077582 | 0.369764356 | 0.137511167 | 0.172153192 | 0.384090643 | 0.062267965 |
| Atrx | S | | 111 | 0.022897254 | -0.088658199 | -0.016062178 | -0.038959432 | 0.072596021 | 0.507826397 | 0.201219039 | 0.787614257 | 0.53804836 | 0.358034969 |
| Cbx8 | S | | 238 | 0.001937938 | 0.018863356 | -0.128667758 | -0.130605696 | -0.147531114 | 0.97724511 | 0.686110021 | 0.27884598 | 0.280169574 | 0.233014266 |
| Inpp5b | S | | 287 | 0.027226773 | -0.049695935 | -0.010486656 | -0.037713429 | 0.039209279 | 0.385817185 | 0.042755931 | 0.547837906 | 0.246332286 | 0.077974789 |
| Vcl | S | | 721 | -0.113280283 | 0.010370569 | 0.018155336 | 0.131435618 | 0.007784767 | 0.234453498 | 0.939754491 | 0.787181681 | 0.187040491 | 0.954691759 |
| Nbn | S | | 433 | 0.044597121 | -0.072179682 | -0.025741319 | -0.07033844 | 0.046438363 | 0.293127132 | 0.093178204 | 0.497852634 | 0.125006496 | 0.24270213 |
| Amacr | S | | 323 | -0.052428899 | -0.149311645 | 0.137574373 | 0.190003271 | 0.286886017 | 0.488773255 | 0.148891753 | 0.289112648 | 0.183939568 | 0.080605822 |
| Ehbp1l1 | S | | 1331 | 0.031243014 | -0.11871115 | -0.011256399 | -0.042499413 | 0.10745475 | 0.731970803 | 0.279641763 | 0.95253628 | 0.823242141 | 0.589864948 |
| Morc4 | S | | 541 | 0.117746184 | -0.16227567 | 0.01691557 | -0.100830614 | 0.17919124 | 0.183726244 | 0.014999211 | 0.775052601 | 0.263389589 | 0.038203845 |
| Atrx | S | | 766 | -0.01854553 | -0.044442032 | 0.014195675 | 0.032741205 | 0.058637708 | 0.71218759 | 0.118087951 | 0.676703516 | 0.541802009 | 0.131264685 |
| Stat5a | S | | 127 | -0.090195135 | 0.068961949 | -0.066842392 | 0.023352743 | -0.135804341 | 0.452359483 | 0.532588299 | 0.381513408 | 0.83156562 | 0.242383714 |
| Uba3 | S | | 377 | -0.099715311 | -0.032312725 | 0.063226655 | 0.162941965 | 0.095539379 | 0.083322557 | 0.6802521 | 0.248775367 | 0.018718744 | 0.256740348 |
| Pdlim2 | T | | 142 | -0.140758377 | 0.073924513 | 0.043295106 | 0.184053483 | -0.030629408 | 0.006513353 | 0.238130116 | 0.383393711 | 0.019121245 | 0.628027432 |
| Dync1li2 | S | | 194 | -0.025245281 | 0.054569883 | -0.076683739 | -0.051438458 | -0.131253622 | 0.336669579 | 0.049166714 | 0.01881781 | 0.064592718 | 0.002853189 |
| Cops3 | S | | 410 | -0.021693724 | 0.011589611 | -0.054994425 | -0.033300701 | -0.066584036 | 0.451217392 | 0.861809659 | 0.408507228 | 0.590326969 | 0.431690801 |

| | | | | | | | | | | | | |
|---|---|---|---|---|---|---|---|---|---|---|---|---|
| Irs1 | S | 1096 | -0.065957605 | -0.008471837 | -0.006817894 | 0.059139711 | 0.001653944 | 0.345403903 | 0.903366772 | 0.956332649 | 0.651497565 | 0.98982329 |
| Rbm10 | S | 89 | 0.058358441 | -0.008847026 | -0.049410751 | -0.107769192 | -0.040563726 | 0.839382826 | 0.845047414 | 0.058902271 | 0.711838805 | 0.385137842 |
| Ctnna1 | S | 455 | -0.142435117 | -0.132474626 | 0.250417362 | 0.392852479 | 0.382891988 | 0.032106573 | 0.151793876 | 0.042877141 | 0.012790296 | 0.013777388 |
| Prkar2a | S | 392 | -0.050078719 | -0.086826869 | 0.018925692 | 0.069004412 | 0.105752562 | 0.676890466 | 0.563248467 | 0.868705042 | 0.599998172 | 0.506210815 |
| Huwe1 | S | 649 | -0.110877986 | 0.149290081 | -0.227492303 | -0.116614317 | -0.376782384 | 0.323222392 | 0.080973471 | 0.239796744 | 0.514757507 | 0.11301349 |
| Napa | S | 195 | -0.065768018 | -0.021884094 | -0.00690022 | 0.058867799 | 0.014983875 | 0.492800403 | 0.766978617 | 0.952792596 | 0.613039357 | 0.881933495 |
| Pcm1 | S | 1432 | -0.021248268 | -0.121157872 | 0.107517235 | 0.128765503 | 0.228675108 | 0.59755366 | 0.020135331 | 0.114774902 | 0.082381855 | 0.019310407 |
| Rc3h1 | S | 770 | 0.076133186 | -0.135415212 | -0.046618824 | -0.12275201 | 0.088796387 | 0.138423055 | 0.125413362 | 0.547974711 | 0.186846523 | 0.357518406 |
| Rc3h1 | S | 777 | 0.076133186 | -0.135415212 | -0.046618824 | -0.12275201 | 0.088796387 | 0.138423055 | 0.125413362 | 0.547974711 | 0.186846523 | 0.357518406 |
| Rmdn2 | S | 121 | 0.102857685 | 0.04936544 | -0.150752436 | -0.253610121 | -0.200117876 | 0.185236701 | 0.487441006 | 0.00392875 | 0.033940195 | 0.059041036 |
| Vcpip1 | S | 993 | -0.052354546 | 0.023267634 | -0.019084589 | 0.033269957 | -0.042352224 | 0.134916071 | 0.359421322 | 0.360856661 | 0.276908482 | 0.092427992 |
| Srf | S | 251 | 0.062552974 | -0.042500193 | -0.122183991 | -0.184736965 | -0.079683798 | 0.501968297 | 0.408637024 | 0.429395143 | 0.284964993 | 0.592071511 |
| Smcr8 | S | 512 | 0.040926059 | -0.057932501 | -0.116753781 | -0.15767984 | -0.05882128 | 0.566380343 | 0.582640187 | 0.500516954 | 0.377749708 | 0.73792504 |
| Nectin3 | S | 469 | -0.012483563 | -0.111413156 | 0.049339874 | 0.061823437 | 0.16075303 | 0.84008668 | 0.161613742 | 0.484550115 | 0.191591554 | 0.037887263 |
| Rrp1b | S | 405 | -0.048853139 | 0.020371224 | -0.047296946 | 0.001556193 | -0.067668171 | 0.502544451 | 0.699124193 | 0.393549463 | 0.981832956 | 0.216500523 |
| Mfap1a Mfap1b | S | 116;116 | -0.028370659 | -0.076121509 | 0.055405269 | 0.083775928 | 0.131526779 | 0.403945249 | 0.052912084 | 0.147763776 | 0.016070092 | 0.004403043 |
| Atg2b | S | 898 | -0.06403894 | -0.058981333 | 0.038542085 | 0.102581025 | 0.097523418 | 0.230571881 | 0.508581764 | 0.359491252 | 0.068517457 | 0.297749273 |
| Zfp609 | S | 252 | 0.102706488 | 0.050081626 | -0.333783365 | -0.436489853 | -0.383864991 | 0.58986727 | 0.539775864 | 0.21507393 | 0.153694068 | 0.178477345 |
| Slc35c2 | S | 335 | 0.004627735 | -0.029432357 | -0.029116974 | -0.033744708 | 0.000315383 | 0.901467251 | 0.363072513 | 0.429865595 | 0.312707757 | 0.989095722 |
| Myo9b | S | 1949 | -0.11763043 | 0.010519166 | 0.036454686 | 0.154085116 | 0.02593552 | 0.086703234 | 0.655667027 | 0.485579144 | 0.058908474 | 0.605866724 |
| Npm1 | S | 241 | 0.060089095 | -0.034827261 | -0.105205995 | -0.165295091 | -0.070378734 | 0.011583721 | 0.142588894 | 0.048529218 | 0.015579736 | 0.109074315 |
| Nfrkb | S | 1288 | -0.093570368 | 0.089180117 | -0.057085676 | 0.036484692 | -0.146265793 | 0.193496486 | 0.347261698 | 0.421290984 | 0.537053702 | 0.147496034 |
| Sec23ip | S | 737 | -0.040479375 | 0.094058672 | -0.132561682 | -0.092082307 | -0.226620355 | 0.538408934 | 0.582044152 | 0.097324698 | 0.112804198 | 0.022002211 |
| Nes | S | 731 | 0.03370105 | -0.012582644 | -0.052130104 | -0.085831154 | -0.03954746 | 0.141387314 | 0.425005081 | 0.017427638 | 0.009664897 | 0.03776618 |
| Med13 | S | 889 | -0.121220569 | -0.039921343 | 0.129927936 | 0.251148505 | 0.16984928 | 0.04236439 | 0.157125092 | 0.071939564 | 0.011658126 | 0.053225287 |
| D1Ertd622e | S | 24 | 0.031507873 | -0.061635504 | -0.02968853 | -0.061196402 | 0.031946974 | 0.517006537 | 0.20965903 | 0.517086706 | 0.251373941 | 0.505637897 |
| Nol9 | T | 115 | 0.065084983 | 0.013888336 | -0.192681228 | -0.257766211 | -0.206569564 | 0.633537274 | 0.929406459 | 0.258937514 | 0.098832323 | 0.174848067 |
| Asap1 | S | 1059 | -0.12504936 | 0.095248744 | -0.026219423 | 0.098829937 | -0.121468167 | 0.031251857 | 0.122360792 | 0.710289237 | 0.191522627 | 0.136916592 |
| Arhgap31 | S | 1365 | 0.029672495 | -0.516897367 | 0.279319446 | 0.249646951 | 0.796216812 | 0.837035285 | 0.274271221 | 0.500940798 | 0.534546791 | 0.1757524 |
| Plekho2 | S | 245 | 0.057738994 | -0.003498667 | -0.114174347 | -0.171913341 | -0.11067568 | 0.582174473 | 0.973117095 | 0.141412047 | 0.171847776 | 0.344446235 |
| Six4 | S | 44 | 0.060346879 | -0.136371698 | -0.005011408 | -0.065358286 | 0.13136029 | 0.310047729 | 0.09919545 | 0.980817747 | 0.759420011 | 0.553944108 |
| Map4k4 | S | 561 | 0.038521067 | -0.027834168 | -0.059314829 | -0.097835896 | -0.031480661 | 0.237125758 | 0.16214103 | 0.186212654 | 0.07070151 | 0.424117601 |
| Ninl | S | 896 | 0.023533353 | -0.073975357 | 0.02407555 | 0.000542197 | 0.098050907 | 0.458588379 | 0.021045146 | 0.694180952 | 0.992751631 | 0.187608881 |
| Cttnbp2nl | S | 440 | -0.143792929 | -0.010353858 | 0.033028242 | 0.176821171 | 0.0433821 | 0.375248722 | 0.872984152 | 0.703628287 | 0.305352504 | 0.63408464 |
| Rnaseh2a | S | 258 | -0.099852687 | -0.018894728 | 0.036895716 | 0.136748403 | 0.055790444 | 0.207943833 | 0.801024888 | 0.515334188 | 0.116827089 | 0.473821131 |
| Srrm1 | S | 790 | 0.044216665 | -0.055606365 | -0.029137336 | -0.073354 | 0.026469029 | 0.153186757 | 0.032032812 | 0.438921648 | 0.107346269 | 0.451886539 |
| Ankrd11 | T | 278 | -0.046490999 | -0.121016621 | 0.106237386 | 0.152728385 | 0.227254007 | 0.646761411 | 0.208246289 | 0.493118711 | 0.351286212 | 0.193720137 |
| Mki67 | S | 1246 | 0.010711792 | 0.025187245 | -0.130521752 | -0.141233544 | -0.155708997 | 0.805027704 | 0.506357345 | 0.167595661 | 0.144106071 | 0.119633293 |
| Cep131 | S | 78 | -0.28689022 | 0.145860171 | 0.151928293 | 0.438818513 | 0.006068122 | 0.005116635 | 0.108138336 | 0.432242362 | 0.096228579 | 0.97274195 |
| Ddx18 | T | 101 | 0.041288689 | -0.040061335 | -0.073217256 | -0.114505945 | -0.033155921 | 0.252489219 | 0.421508874 | 0.178626458 | 0.073293118 | 0.571354448 |
| Rc3h1 | S | 460 | -0.050149108 | 0.010379133 | -0.020246566 | 0.029902542 | -0.030625699 | 0.257235304 | 0.834506229 | 0.703439357 | 0.47385509 | 0.527562902 |
| Septin9 | S | 41 | -0.013836246 | -0.09537576 | 0.003275699 | 0.017111945 | 0.098651459 | 0.71226993 | 0.525409676 | 0.93903421 | 0.52143649 | 0.509889473 |
| Ppfibp1 | S | 535 | 0.055171379 | -0.037660485 | -0.104840461 | -0.16001184 | -0.067179977 | 0.345602745 | 0.383869161 | 0.30300175 | 0.162828792 | 0.485219141 |
| Gmpr2 | S | 28 | 0.026874285 | -0.030363086 | -0.074637183 | -0.101511468 | -0.044274097 | 0.561809487 | 0.762241442 | 0.221783533 | 0.094951071 | 0.660234381 |
| Mastl | S | 683 | 0.054958727 | -0.067949102 | -0.046481565 | -0.101440292 | 0.021467537 | 0.446553808 | 0.269497812 | 0.376136123 | 0.207524349 | 0.717779676 |
| Tnrc18 | S | 1789 | -0.054583035 | 0.059195542 | -0.080979296 | -0.026396261 | -0.140174838 | 0.497006837 | 0.428681549 | 0.261732437 | 0.686419858 | 0.060336015 |
| Afdn | S | 1143 | 0.006771943 | -0.070670215 | 0.003117287 | -0.003654556 | 0.073787602 | 0.767199579 | 0.150692373 | 0.849152143 | 0.865962724 | 0.143807034 |
| Map1s | S | 660 | -0.043328232 | -0.011381013 | -0.005453277 | 0.037874954 | 0.005927735 | 0.32901755 | 0.731063903 | 0.918292692 | 0.498693288 | 0.902311386 |
| Mvp | S | 837 | -0.033767118 | 0.012962573 | -0.080604934 | -0.046837816 | -0.093567507 | 0.832882178 | 0.915431046 | 0.443957368 | 0.790330364 | 0.527661066 |
| Plekhm1 | S | 404 | -0.006971912 | 0.015980225 | -0.091491555 | -0.084519643 | -0.10747178 | 0.901638957 | 0.784373657 | 0.292604725 | 0.315791231 | 0.226316651 |
| Zfhx4 | S | 2700 | 0.008750738 | -0.008023187 | -0.086466647 | -0.095217384 | -0.07844346 | 0.957154315 | 0.966022963 | 0.346413372 | 0.582544323 | 0.68902234 |
| Rps8 | S | 160 | 0.052810133 | -0.017034782 | -0.106837845 | -0.159647978 | -0.089803063 | 0.320336688 | 0.756230011 | 0.11993477 | 0.050830038 | 0.208283851 |
| Champ1 | S | 416 | 0.007318964 | -0.084902083 | 0.052162552 | 0.044843588 | 0.137064635 | 0.742914123 | 0.002666511 | 0.07990977 | 0.11941896 | 0.008494108 |
| Tom1 | S | 313 | -0.140247999 | -0.198077415 | 0.10897433 | 0.249222329 | 0.307051745 | NA | 0.63551649 | NA | NA | NA |
| Tom1 | S | 314 | -0.140247999 | -0.198077415 | 0.10897433 | 0.249222329 | 0.307051745 | NA | 0.63551649 | NA | NA | NA |
| Gpr149 | S | 517 | 0.055894453 | -0.251967382 | 0.1663956 | 0.110501147 | 0.418362982 | 0.469601225 | 0.000812868 | 0.015275946 | 0.192080962 | 0.0024534 |
| Braf | S | 483 | -0.085916759 | 0.027635881 | -0.014561159 | 0.0713556 | -0.04219704 | 0.24218455 | 0.681554268 | 0.8260293 | 0.384709476 | 0.595761648 |
| Kdm5c | S | 301 | -0.019449664 | -0.012543345 | -0.028296677 | -0.008847013 | -0.015753332 | 0.616471597 | 0.668921465 | 0.546474536 | 0.854235413 | 0.713173408 |
| Huwe1 | S | 3363 | -0.070487254 | -0.026480149 | 0.016417522 | 0.086904776 | 0.042897671 | 0.342498418 | 0.725983299 | 0.752998762 | 0.21993201 | 0.529153808 |
| Farp1 | S | 340 | 0.020712181 | -0.037955306 | -0.054695799 | -0.07540798 | -0.016740493 | 0.495820212 | 0.454642702 | 0.330368633 | 0.218039363 | 0.791196451 |
| Nbeal2 | S | 758 | -0.119446257 | 0.038400693 | -0.081032496 | 0.038413761 | -0.119433189 | 0.456083143 | 0.638506482 | 0.828882185 | 0.920852379 | 0.751958644 |
| Ercc4 | S | 522 | -0.131537708 | 0.030107713 | 0.050080642 | 0.18161835 | 0.019972929 | 0.042739228 | 0.587742975 | 0.37938548 | 0.03466756 | 0.766362762 |
| Npat | T | 1264 | 0.07610776 | -0.140112718 | -0.000805469 | -0.076913229 | 0.139307249 | 0.245269704 | 0.063754724 | 0.990070901 | 0.079689259 | 0.022962472 |
| Slain2 | S | 302 | -0.189828256 | -0.184663474 | 0.227903808 | 0.417732064 | 0.412567281 | 0.240871689 | 0.323194553 | 0.131532468 | 0.026434352 | 0.060731086 |
| Raly | S | 135 | 0.022907605 | -0.05792127 | -0.01715941 | -0.040067015 | 0.040761859 | 0.586564226 | 0.15402966 | 0.703711834 | 0.479106484 | 0.437550104 |
| Stip1 | S | 481 | -0.029155737 | 0.059015238 | -0.070322345 | -0.041166608 | -0.129337583 | 0.47176761 | 0.166569128 | 0.041272631 | 0.270039459 | 0.030677519 |
| Pi4ka | T | 253 | -0.115742194 | 0.045916806 | -0.022552337 | 0.093189857 | -0.068469144 | 0.241244101 | 0.484613256 | 0.790600112 | 0.340934034 | 0.349443535 |
| Srrm2 | S | 773 | -0.004411672 | -0.026973212 | 0.001489678 | 0.00590135 | 0.02846289 | 0.882827992 | 0.051531035 | 0.930962454 | 0.849171156 | 0.145569581 |
| Ralgapb | T | 379 | 0.016252083 | -0.012727444 | -0.068453444 | -0.084705526 | -0.055725999 | 0.732635524 | 0.806366612 | 0.236030816 | 0.204600411 | 0.398141043 |
| Bag3 | S | 404 | 0.115015825 | -0.009614001 | -0.195442203 | -0.310458028 | -0.185828202 | 0.080047353 | 0.850983517 | 0.039502529 | 0.012250007 | 0.058572841 |
| Rad54l2 | S | 1171 | -0.055983757 | -0.027955703 | 0.037835021 | 0.093818779 | 0.065790724 | 0.191990729 | 0.670454826 | 0.707077338 | 0.387705596 | 0.554364899 |
| Abl1 | S | 658 | 0.031790729 | -0.100488899 | 0.014694798 | -0.017095931 | 0.115183696 | 0.495086491 | 0.065496314 | 0.655305725 | 0.696167602 | 0.051081449 |
| Arhgap12 | S | 199 | -0.031590031 | -0.013101063 | -0.024120563 | 0.007469468 | -0.0110195 | 0.484832956 | 0.844982587 | 0.67852195 | 0.887023207 | 0.877353677 |
| Mki67 | S | 2545 | 0.020188362 | 0.077229376 | -0.142545009 | -0.162733371 | -0.219774385 | 0.801684284 | 0.148394708 | 0.020287047 | 0.117996951 | 0.008832236 |
| Adcy9 | S | 613 | -0.180794833 | 0.026604033 | 0.080001188 | 0.260796021 | 0.053397155 | 0.176737518 | 0.836186735 | 0.616461494 | 0.112678807 | 0.671752904 |
| Mki67 | S | 2016 | 0.006900077 | 0.011227515 | -0.046400678 | -0.053300755 | -0.057628193 | 0.926586108 | 0.669172769 | 0.041661578 | 0.501803932 | 0.079091796 |
| Soga1 | S | 927 | -0.068703325 | 0.088666226 | -0.149285788 | -0.080582462 | -0.237952014 | 0.497270681 | 0.356390248 | 0.385669862 | 0.625206904 | 0.206261659 |
| Filip1l | S | 1033 | -0.058009892 | -0.007809442 | 0.009583241 | 0.067593133 | 0.017392683 | 0.20576325 | 0.813010683 | 0.773722029 | 0.123793949 | 0.05768989 |
| 1700030K09Rik | S | 357 | -0.081119992 | 0.010002438 | -0.008117154 | 0.073002837 | -0.018119592 | 0.352980599 | 0.94612821 | 0.919862197 | 0.14953017 | 0.892041513 |
| Mfhas1 | S | 1034 | 0.011996949 | 0.045163776 | -0.091084681 | -0.10308163 | -0.136248457 | 0.879946089 | 0.549441592 | 0.06678565 | 0.253391057 | 0.136092877 |
| Rad23a | S | 92 | -0.039954297 | 0.018677665 | -0.058675734 | -0.018721437 | -0.077353399 | 0.431758385 | 0.829073297 | 0.616246199 | 0.873532064 | 0.569777551 |
| Elf2 | S | 107 | 0.004151029 | -0.052177548 | -0.011679533 | -0.015830562 | 0.040498016 | 0.945721788 | 0.319512866 | 0.710282555 | 0.778379822 | 0.385910326 |
| Rreb1 | S | 1583 | -0.176910992 | -0.16070158 | 0.1762937 | 0.353204691 | 0.33699528 | 0.568150625 | 0.223417934 | 0.222033183 | 0.303079107 | 0.063619982 |
| Cdc42ep3 | S | 100 | -0.00375469 | 0.028976964 | -0.135755381 | -0.132000691 | -0.164732346 | 0.976423437 | 0.665235755 | 0.324461976 | 0.435225753 | 0.257599817 |
| Epha2 | S | 902 | 0.1787518 | -0.149023898 | -0.124145121 | -0.302896921 | 0.024878777 | 0.03629669 | 0.181180903 | 0.082316823 | 0.008284938 | 0.792100936 |
| Huwe1 | S | 2594 | -0.1020671 | 0.001058957 | -0.033419417 | 0.068647683 | -0.034478374 | 0.698158484 | 0.992282478 | 0.730621545 | 0.790085861 | 0.734729652 |
| Epb41l1 | S | 540 | -0.032770207 | 0.018701495 | -0.04130256 | -0.008532353 | -0.060004055 | 0.494424317 | 0.816950531 | 0.37754031 | 0.835567419 | 0.465864259 |
| Pdap1 | S | 63 | -0.026825187 | 0.019347051 | -0.032950145 | -0.006124957 | -0.052297195 | 0.31339693 | 0.599780612 | 0.249092659 | 0.836527155 | 0.23146662 |
| Myo9a | S | 1259 | -0.056590683 | -0.013323958 | 0.010690035 | 0.067280717 | 0.024013993 | 0.240748926 | 0.825031746 | 0.837360708 | 0.205488989 | 0.695313351 |
| Glcci1 | S | 136 | -0.106270207 | 0.006952601 | -0.010191646 | 0.096078562 | -0.017144247 | 0.486735212 | 0.907391914 | 0.853609615 | 0.527125464 | 0.788583164 |
| Arhgap24 | S | 374 | -0.11113623 | -0.00745768 | 0.041137296 | 0.152273527 | 0.048594976 | 0.170478282 | 0.842760062 | 0.422107593 | 0.093844757 | 0.312261268 |
| Vcp | S | 702 | -0.075237747 | 0.029788422 | 0.039324348 | 0.114562094 | 0.009535926 | 0.00380007 | 0.216811512 | 0.592911946 | 0.204218448 | 0.893134966 |
| Srrm2 | S | 1852 | 0.100950252 | -0.197207316 | 0.029773683 | -0.071176569 | 0.226980999 | 0.30358877 | 0.076551234 | 0.829915356 | 0.576690942 | 0.146932567 |
| Kcmf1 | S | 282 | -0.27158233 | 0.083949417 | 0.034582065 | 0.306164395 | -0.049367352 | 0.145637052 | 0.251502682 | 0.780593223 | 0.131577159 | 0.698560543 |

| | | | | | | | | | | | | |
|---|---|---|---|---|---|---|---|---|---|---|---|---|
| Mcph1 | T | 394 | -0.147064325 | 0.034098071 | -0.01418258 | 0.132881745 | -0.04828065 | 0.46508034 | 0.807333112 | 0.894035692 | 0.512179084 | 0.742768804 |
| Zmym2 | S | 112 | 0.181880652 | -0.11401034 | -0.155890014 | -0.337770666 | -0.041879675 | 0.118113266 | 0.228904737 | 0.197491556 | 0.038653784 | 0.717619274 |
| Bcor | S | 423 | -0.029307905 | -0.017654423 | -0.010519077 | 0.018788827 | 0.007135346 | 0.581017619 | 0.532388602 | 0.728380887 | 0.716998094 | 0.79394709 |
| Cep170 | T | 906 | -0.014297449 | -0.060848698 | 0.003829653 | 0.018127102 | 0.06467835 | 0.760257104 | 0.378731896 | 0.932313024 | 0.684115349 | 0.347506076 |
| Pde4d | S | 287 | 0.022857967 | 0.02425472 | -0.158683597 | -0.181541565 | -0.182938317 | 0.778162507 | 0.710934505 | 0.270793077 | 0.233916359 | 0.222593726 |
| Rrp9 | S | 50 | 0.035638915 | -0.077736976 | -0.015738829 | -0.051377744 | 0.061998147 | 0.07278389 | 0.059796977 | 0.366288242 | 0.031766524 | 0.103451419 |
| Rrp9 | S | 51 | 0.035638915 | -0.077736976 | -0.015738829 | -0.051377744 | 0.061998147 | 0.07278389 | 0.059796977 | 0.366288242 | 0.031766524 | 0.103451419 |
| Rrp9 | S | 53 | 0.035638915 | -0.077736976 | -0.015738829 | -0.051377744 | 0.061998147 | 0.07278389 | 0.059796977 | 0.366288242 | 0.031766524 | 0.103451419 |
| Nudt4 | S | 147 | -0.01052284 | -0.038123677 | -0.055045379 | -0.044522538 | -0.016921701 | 0.925931083 | 0.670424564 | 0.714130307 | 0.778583458 | 0.906713946 |
| Peak1 | T | 507 | 0.011926199 | -0.071511907 | -0.031489863 | -0.043416063 | 0.040022043 | 0.867207636 | 0.475214702 | 0.734308503 | 0.63509369 | 0.716511905 |
| Jcad | S | 1165 | -0.040473495 | 0.008664084 | -0.081496967 | -0.041023471 | -0.09016105 | 0.805520784 | 0.904542867 | 0.540525898 | 0.824787888 | 0.487248666 |
| Tbc1d15 | S | 201 | -0.170677993 | 0.138069872 | -0.07060785 | 0.100070143 | -0.208677722 | 0.244204806 | 0.323227302 | 0.592468216 | 0.547889692 | 0.243601792 |
| Ppp1r3f | S | 545 | 6.68532E-05 | -0.135790391 | 0.076315094 | 0.076248241 | 0.212105485 | 0.999232861 | 0.050068071 | 0.201523159 | 0.360386751 | 0.021106926 |
| Trmt61a | S | 264 | -0.034194343 | 0.071842329 | -0.184632892 | -0.150438549 | -0.256475221 | 0.738463962 | 0.425082272 | 0.361038528 | 0.46066837 | 0.240427937 |
| Anks1 | S | 668 | 0.000812706 | -0.015678143 | -0.080817708 | -0.081630414 | -0.065139565 | 0.983151753 | 0.805225871 | 0.510774976 | 0.505729955 | 0.60529946 |
| Srrm2 | T | 1409 | 0.026446531 | -0.056116205 | -0.038688255 | -0.065134786 | 0.01742795 | 0.654253585 | 0.266903428 | 0.687712198 | 0.528260441 | 0.855319974 |
| Oga | S | 364 | -0.114182589 | -0.000185483 | 0.067300849 | 0.181483438 | 0.067486333 | 0.042580782 | 0.995955646 | 0.229063332 | 0.019506713 | 0.198875506 |
| Bin1 | T | 328 | -0.064302353 | -0.019283422 | 0.012429913 | 0.076732266 | 0.031713336 | 0.538521283 | 0.49110784 | 0.927378115 | 0.634131199 | 0.817302088 |
| Med19 | S | 226 | -0.127346983 | 0.016205966 | 0.036277367 | 0.16362435 | 0.020071401 | 0.233612638 | 0.87567819 | 0.766504233 | 0.273355652 | 0.88738662 |
| Arhgap23 | S | 1221 | -0.005043358 | -0.03417537 | -0.008400173 | -0.003356815 | 0.025775197 | 0.862023228 | 0.335965003 | 0.60307181 | 0.900149858 | 0.438717358 |
| R3hdm2 | S | 143 | 0.005946514 | 0.014111495 | -0.107220982 | -0.113167496 | -0.121332477 | 0.709191274 | 0.422456073 | 0.162958462 | 0.155124712 | 0.136785693 |
| Supt16 | S | 1015 | 0.090110994 | 0.124233822 | -0.341139914 | -0.431250908 | -0.465373736 | 0.454383638 | 0.064986405 | 0.022973082 | 0.025592202 | 0.013394823 |
| Tbc1d10b | S | 673 | -0.080711956 | 0.165668358 | -0.16770598 | -0.086994024 | -0.333374338 | 0.308832441 | 0.077510773 | 0.095163611 | 0.243725186 | 0.009276847 |
| Nup133 | S | 488 | -0.005479849 | -0.064663891 | -0.002504511 | 0.002975338 | 0.06215938 | 0.805911314 | 0.401411307 | 0.978218125 | 0.973681881 | 0.571137496 |
| Traf3ip1 | S | 409 | -0.018624604 | -0.010334435 | -0.026402386 | -0.007777782 | -0.016067951 | 0.651021719 | 0.785520657 | 0.450113207 | 0.842360997 | 0.659671649 |
| Ep300 | S | 500 | 0.077218655 | 0.046628748 | -0.124467563 | -0.201686218 | -0.171096311 | 0.514114088 | 0.36963179 | 0.014453601 | 0.170696981 | 0.033024879 |
| Kif18b | S | 679 | 0.090067627 | -0.024488474 | -0.089016599 | -0.179084225 | -0.064528125 | 0.062522526 | 0.318426903 | 0.014107531 | 0.008243137 | 0.06104223 |
| Clasp2 | S | 333 | -0.131240047 | 0.095509693 | -0.0180724 | 0.113167647 | -0.113582093 | 0.071338002 | 0.242341772 | 0.803639587 | 0.095392934 | 0.152402309 |
| Iws1 | S | 200 | -0.010099616 | -0.113934583 | 0.073955544 | 0.08405516 | 0.187890127 | 0.831023176 | 0.122592043 | 0.432939976 | 0.381560024 | 0.11231689 |
| Kmt2e | S | 1364 | 0.04618417 | -0.045882088 | -0.075722301 | -0.121906471 | -0.029840212 | 0.38350048 | 0.493896438 | 0.273792311 | 0.098459073 | 0.669874899 |
| Vps13a | S | 845 | 0.088320815 | -0.070780362 | -0.120375887 | -0.208696703 | -0.049595525 | 0.493789902 | 0.649704746 | 0.346339425 | 0.107162383 | 0.724751736 |
| Atg4c | T | 452 | -0.066463317 | 0.002450666 | -0.040406219 | 0.026057098 | -0.042856885 | 0.65049051 | 0.976735738 | 0.697808343 | 0.855831808 | 0.614717672 |
| Rif1 | S | 1028 | -0.020872193 | -0.016634323 | -0.013523619 | 0.007348575 | 0.003110704 | 0.474331263 | 0.543041351 | 0.790738023 | 0.873464936 | 0.945131748 |
| Nfic | S | 286 | -0.131200565 | -0.019126656 | 0.098479709 | 0.229680274 | 0.117606365 | 0.084141481 | 0.86363767 | 0.306757089 | 0.05375026 | 0.349170054 |
| Trip10 | S | 298 | -0.062651588 | 0.022778066 | -0.056358347 | 0.006293241 | -0.079136413 | 0.591505519 | 0.76182142 | 0.568053369 | 0.962730286 | 0.482206358 |
| Nub1 | S | 488 | -0.112814869 | -0.096862469 | 0.077809693 | 0.190624562 | 0.174672163 | 0.651588092 | 0.371051506 | 0.451891258 | 0.459222878 | 0.112970498 |
| Dclk1 | S | 337 | -0.043135009 | 0.05138664 | -0.059413907 | -0.016278897 | -0.110800546 | 0.133592115 | 0.136700348 | 0.207073389 | 0.687971928 | 0.057453658 |
| Apc | S | 2713 | -0.065690285 | 0.007782453 | -0.006545947 | 0.059144338 | -0.0143284 | 0.287467926 | 0.856291458 | 0.876879351 | 0.372602622 | 0.794831062 |
| Zfp367 | S | 55 | -0.085038097 | -0.105507446 | 0.142513785 | 0.227551881 | 0.248021231 | 0.490259307 | 0.524881112 | 0.698189467 | 0.539466738 | 0.515096279 |
| Brca2 | S | 487 | -0.004739929 | -0.057939575 | -0.013332401 | -0.008592472 | 0.044607175 | 0.884746323 | 0.486347066 | 0.815585106 | 0.868406635 | 0.61135337 |
| Prpf4b | S | 518 | -0.137028757 | 0.019013134 | 0.008897598 | 0.145926355 | -0.010115536 | 0.380761084 | 0.797663913 | 0.964287565 | 0.529904796 | 0.959356891 |
| Prpf4b | S | 519 | -0.137028757 | 0.019013134 | 0.008897598 | 0.145926355 | -0.010115536 | 0.380761084 | 0.797663913 | 0.964287565 | 0.529904796 | 0.959356891 |
| Prpf4b | S | 520 | -0.137028757 | 0.019013134 | 0.008897598 | 0.145926355 | -0.010115536 | 0.380761084 | 0.797663913 | 0.964287565 | 0.529904796 | 0.959356891 |
| Exosc2 | S | 124 | 0.008326182 | 0.062988919 | -0.120073877 | -0.128400058 | -0.183062795 | 0.851479897 | 0.238327836 | 0.042454332 | 0.043618547 | 0.019361065 |
| Adcy6 | S | 573 | -0.038842715 | 0.001270054 | -0.021471689 | 0.017371027 | -0.022741743 | 0.357777205 | 0.977247093 | 0.717989591 | 0.778691102 | 0.724201249 |
| Gtf2i | S | 830 | -0.017054127 | -0.056455675 | 0.012923068 | 0.029977195 | 0.069378743 | 0.71370337 | 0.350844086 | 0.878360175 | 0.695381415 | 0.422283004 |
| Msl1 | S | 364 | -0.011748736 | -0.039388996 | 0.02555552 | 0.037304256 | 0.064944516 | 0.811902308 | 0.005357031 | 0.136986989 | 0.48061167 | 0.014478397 |
| Junb | S | 256 | 0.029621114 | -0.005515306 | -0.080020899 | -0.109642013 | -0.074505593 | 0.394061262 | 0.897723577 | 0.111029441 | 0.059411997 | 0.199374409 |
| Junb | T | 252 | 0.029621114 | -0.005515306 | -0.080020899 | -0.109642013 | -0.074505593 | 0.394061262 | 0.897723577 | 0.111029441 | 0.059411997 | 0.199374409 |
| Pja2 | S | 306 | 0.039159998 | 0.084860617 | -0.228635164 | -0.267795161 | -0.313495781 | 0.462574398 | 0.214136533 | 0.055585866 | 0.055982046 | 0.027580201 |
| Nemp1 | S | 366 | -0.016781144 | -0.022131923 | -0.019468675 | -0.002687531 | 0.002663247 | 0.818062234 | 0.615329677 | 0.503248394 | 0.969020287 | 0.941983303 |
| Jade3 | S | 774 | -0.030628205 | -0.048537287 | 0.011232319 | 0.041860524 | 0.059769606 | 0.751456033 | 0.332678059 | 0.818153578 | 0.682572255 | 0.339153209 |
| Jade3 | S | 776 | -0.030628205 | -0.048537287 | 0.011232319 | 0.041860524 | 0.059769606 | 0.751456033 | 0.332678059 | 0.818153578 | 0.682572255 | 0.339153209 |
| Dact3 | T | 267 | -0.093122472 | 0.099610577 | -0.065765867 | 0.027356605 | -0.165376444 | 0.296437033 | 0.266975665 | 0.247220006 | 0.707836254 | 0.107698704 |
| Emd | S | 197 | -0.098694113 | -0.031726405 | 0.038894473 | 0.137588586 | 0.070620878 | 0.262243548 | 0.081000434 | 0.030214751 | 0.162014025 | 0.00947412 |
| Zeb2 | S | 1203 | -0.024931417 | -0.033810992 | 0.030773321 | 0.055704738 | 0.064584313 | 0.172893936 | 0.039206184 | 0.105110916 | 0.021490421 | 0.015987394 |
| Ptbp2 | S | 27 | 0.03626009 | -0.0458157 | -0.093956068 | -0.130216158 | -0.048140368 | 0.553417463 | 0.385216238 | 0.61361077 | 0.494667938 | 0.7878783 |
| Lcor | S | 42 | 0.069845712 | -0.095294592 | 0.017406171 | -0.052439541 | 0.112700763 | 0.403139723 | 0.025118006 | 0.791177907 | 0.568207886 | 0.159421798 |
| Atp11a | S | 459 | -0.107639141 | 0.057932531 | 0.007278657 | 0.114917798 | -0.050653874 | 0.055122253 | 0.337469044 | 0.875394922 | 0.004046989 | 0.291226899 |
| Emsy | S | 371 | -0.012218658 | 0.010198694 | -0.046322694 | -0.034104036 | -0.056521387 | 0.840459056 | 0.826871974 | 0.197250124 | 0.559535773 | 0.227426129 |
| 5730455P16Rik | S | 8 | -0.15377683 | 0.11939127 | -0.057536574 | 0.096240256 | -0.176927844 | 0.338394086 | 0.384962191 | 0.480237573 | 0.513775736 | 0.217876785 |
| Dclk1 | S | 298 | 0.00391256 | 0.091878674 | -0.169917644 | -0.173830204 | -0.261796317 | 0.963896953 | 0.399969821 | 0.139844778 | 0.096420946 | 0.042136077 |
| Ticrr | S | 1531 | -0.094825247 | -0.024235718 | 0.032286098 | 0.127111345 | 0.056521816 | 0.277942815 | 0.681909787 | 0.41287703 | 0.179843115 | 0.338894414 |
| Usp7 | S | 54 | -0.16432696 | 0.12716939 | -0.057852427 | 0.106474533 | -0.185021817 | 0.224063966 | 0.21963473 | 0.43677633 | 0.403226167 | 0.116529581 |
| Mast4 | S | 2053 | -0.034016873 | 0.038932135 | -0.099616515 | -0.065599642 | -0.13854865 | 0.759295296 | 0.672127151 | 0.397307317 | 0.549116619 | 0.190602931 |
| Map3k7 | S | 412 | -0.009267783 | 0.07462032 | -0.134186565 | -0.124918782 | -0.208806885 | 0.86261182 | 0.187046659 | 0.07328043 | 0.081791064 | 0.021201434 |
| Habp4 | S | 7 | 0.033826456 | -0.037099206 | -0.065051786 | -0.098878242 | -0.027952581 | 0.486707983 | 0.49690633 | 0.324946695 | 0.160347125 | 0.663368337 |
| Atrx | S | 770 | -0.048802116 | -0.01979758 | 0.008278734 | 0.057080851 | 0.028076314 | 0.399949538 | 0.579819428 | 0.823582218 | 0.344940016 | 0.477902239 |
| Cmtr1 | S | 54 | -0.026456137 | -0.072339649 | 0.043006295 | 0.069462432 | 0.115345944 | 0.399966306 | 0.064715149 | 0.084747699 | 0.070663995 | 0.022118877 |
| Hsf1 | S | 307 | -0.024798768 | 0.023386839 | -0.04187132 | -0.017072551 | -0.065258159 | 0.348166498 | 0.438774256 | 0.219796936 | 0.512812838 | 0.075268081 |
| Amotl2 | S | 716 | -0.133537423 | 0.128453686 | -0.140299021 | -0.006761598 | -0.268752707 | 0.56367094 | 0.457531713 | 0.493586987 | 0.975488774 | 0.162117787 |
| Zfp609 | S | 575 | -0.001692756 | -0.057151803 | 0.016565198 | 0.018257954 | 0.073717001 | 0.958505523 | 0.100038446 | 0.568408977 | 0.598321707 | 0.067084627 |
| Gnl3 | S | 505 | 0.066716934 | -0.087491826 | -0.025886744 | -0.092603678 | 0.061605082 | 0.181965843 | 0.06090451 | 0.590430202 | 0.077188532 | 0.149518031 |
| Enkd1 | S | 91 | 0.037527065 | -0.084278275 | -0.026933953 | -0.064461018 | 0.057344322 | 0.481742608 | 0.205849858 | 0.811982008 | 0.567014839 | 0.616924549 |
| Dock1 | S | 1842 | -0.05460081 | -0.042867531 | 0.02427726 | 0.07887807 | 0.067144791 | 0.608904927 | 0.333924942 | 0.622052088 | 0.464926051 | 0.112774707 |
| Apc | S | 1036 | -0.003759085 | -0.034485162 | -0.020745161 | -0.016986076 | 0.013740001 | 0.946807808 | 0.403465615 | 0.720299959 | 0.801132724 | 0.807142955 |
| Speg | S | 2476 | 0.017096775 | 0.099994894 | -0.195112263 | -0.212209038 | -0.295107157 | 0.845324041 | 0.292999094 | 0.095060652 | 0.057631577 | 0.021932653 |
| Prune2 | S | 1682 | -0.055996143 | -0.014934279 | 0.01570148 | 0.071697622 | 0.030635758 | 0.211072504 | 0.743514451 | 0.608937391 | 0.148192127 | 0.535403364 |
| Slc39a6 | S | 593 | 0.113299371 | -0.18575578 | 0.032799241 | -0.080500131 | 0.21855502 | 0.076393434 | 0.010851674 | 0.721317505 | 0.410922072 | 0.083308331 |
| Tnks1bp1 | S | 1612 | -0.039687436 | -0.003665694 | 0.019981696 | 0.059669132 | 0.02364739 | 0.057337279 | 0.874063069 | 0.691538566 | 0.29325244 | 0.645226086 |
| Epb41l2 | T | 745 | -0.07680148 | 0.02827009 | -0.038863274 | 0.037938206 | -0.067133363 | 0.497415008 | 0.620735002 | 0.529928858 | 0.710782429 | 0.103280088 |
| Afap1 | S | 666 | 0.055048132 | -0.036413681 | -0.041443847 | -0.096491979 | -0.005030166 | 0.087623195 | 0.024673498 | 0.168844903 | 0.028441583 | 0.833734389 |
| Enkd1 | S | 51 | 0.040084767 | -0.047801471 | -0.040610076 | -0.080694842 | 0.007191395 | 0.123329282 | 0.236552018 | 0.109708564 | 0.024438027 | 0.841869686 |
| Dkc1 | S | 451 | 0.046525918 | -0.021961695 | -0.042912225 | -0.089438143 | -0.02095053 | 0.499704058 | 0.394680556 | 0.108287085 | 0.247574378 | 0.303551837 |
| Ccdc61 | S | 371 | -0.062743149 | -0.022292292 | 0.028729677 | 0.091472826 | 0.051021968 | 0.303670326 | 0.726428552 | 0.77395417 | 0.373540762 | 0.605841269 |
| Mcm2 | T | 59 | -0.191984275 | 0.030020776 | -0.000914661 | 0.191069614 | -0.030935437 | 0.409334489 | 0.497622372 | 0.993191443 | 0.426301099 | 0.779833569 |
| 2700049A03Rik | S | 333 | 0.046996012 | -0.077895656 | -0.004851599 | -0.051847611 | 0.073044057 | 0.843290724 | 0.217166297 | 0.946613325 | 0.8285042 | 0.32614843 |
| Trim56 | S | 450 | -0.186269856 | 0.035279941 | 0.160387156 | 0.346657012 | 0.125107216 | 0.006462215 | 0.463901128 | 0.170344635 | 0.046771446 | 0.257498501 |
| Svil | S | 234 | -0.012972701 | -0.029612531 | -0.012892439 | 8.02622E-05 | 0.016720093 | 0.754550401 | 0.473493902 | 0.739508905 | 0.998464256 | 0.68472649 |
| Myof | S | 950 | 0.097462935 | -0.074126305 | -0.204572218 | -0.302035153 | -0.130445912 | 0.4601926 | 0.714320206 | 0.478414553 | 0.319643619 | 0.668468267 |
| Setx | S | 1667 | -0.037190613 | -0.002714705 | -0.044426819 | -0.007236205 | -0.041712114 | 0.694001309 | 0.960612043 | 0.627805465 | 0.950270839 | 0.65988185 |
| Mef2a | S | 108 | -0.038072672 | -0.022215261 | 0.025720269 | 0.063792941 | 0.04793553 | 0.101244536 | 0.287422895 | 0.24228426 | 0.024455332 | 0.051156853 |
| Nucks1 | S | 75 | -0.155598673 | 0.05430289 | 0.022537336 | 0.17813601 | -0.031765553 | 0.078122368 | 0.474532397 | 0.330199363 | 0.056257033 | 0.663580066 |
| Arrdc1 | S | 292 | -0.105917629 | 0.070508976 | -0.084240848 | 0.021676781 | -0.154749824 | 0.468518086 | 0.59748518 | 0.687146567 | 0.919027939 | 0.477533775 |

| | | | | | | | | | | | | |
|---|---|---|---|---|---|---|---|---|---|---|---|---|
| Bag3 | S | 270 | 0.006427457 | 0.012871261 | -0.070575961 | -0.077003418 | -0.083447222 | 0.8719344 | 0.608851568 | 0.129023742 | 0.142887242 | 0.104331292 |
| Mta1 | S | 522 | -0.070873414 | 0.019109769 | -0.004398626 | 0.066474787 | -0.023508395 | 0.180890977 | 0.60839843 | 0.913961455 | 0.218307277 | 0.551116737 |
| Cuedc1 | S | 8 | -0.165470591 | 0.114207876 | 0.048199767 | 0.213670359 | -0.066008108 | 0.008946152 | 0.275286246 | 0.336728155 | 0.003894628 | 0.487573329 |
| Calcoco1 | S | 563 | 0.066237386 | -0.060499111 | -0.126574357 | -0.192811743 | -0.066075246 | 0.090317611 | 0.272313811 | 0.302924826 | 0.168499115 | 0.562381968 |
| 5031439G07Rik | T | 267 | -0.032000467 | -0.042230251 | 0.026434721 | 0.058435188 | 0.068664972 | 0.612159673 | 0.152027699 | 0.490557594 | 0.412282301 | 0.150932524 |
| Jmjd1c | S | 1789 | -0.054756028 | 0.005406702 | -0.008181941 | 0.046574087 | -0.013588643 | 0.310576101 | 0.864107983 | 0.782645752 | 0.380239657 | 0.678659832 |
| Gprc5b | S | 361 | 0.017766273 | -0.046826036 | -0.03976557 | -0.057531843 | 0.007060467 | 0.807801993 | 0.523559601 | 0.537097136 | 0.510459107 | 0.932255493 |
| Pip4p2 | S | 47 | -0.023848562 | 0.012553345 | -0.091846704 | -0.067998142 | -0.104400049 | 0.791486707 | 0.805464106 | 0.514353788 | 0.65581949 | 0.471372199 |
| Phldb2 | S | 199 | 0.034685197 | 0.049227691 | -0.126638214 | -0.16132341 | -0.175865904 | 0.523005448 | 0.295700891 | 0.030663187 | 0.023575032 | 0.008378593 |
| Hsf1 | S | 303 | -0.025880339 | 0.023767449 | -0.041098741 | -0.015218402 | -0.06486619 | 0.328069625 | 0.418181321 | 0.236619742 | 0.57714098 | 0.081899202 |
| Eif4g1 | S | 230 | 0.020456734 | -0.034046207 | -0.038325889 | -0.058782623 | -0.004279682 | 0.754647143 | 0.341771938 | 0.540044696 | 0.471521784 | 0.94327591 |
| Prdx3 | S | 238 | 0.026974463 | -0.071220964 | -0.028830694 | -0.055805157 | 0.04239027 | 0.335045439 | 0.338203086 | 0.401808413 | 0.176337956 | 0.552133332 |
| Mink1 | S | 699 | -0.073336357 | 0.016963729 | 0.020531859 | 0.093868216 | 0.00356813 | 0.042865885 | 0.581140821 | 0.516244445 | 0.007600305 | 0.856750369 |
| Kdm2a | S | 739 | 0.164744426 | -0.078499247 | -0.10872106 | -0.273465486 | -0.030221813 | 0.239937699 | 0.197093524 | 0.2365469 | 0.097372605 | 0.705599814 |
| Srrm2 | S | 531 | 0.060509255 | -0.0907172 | -0.033891021 | -0.094400275 | 0.05682618 | 0.518607266 | 0.361354932 | 0.454800333 | 0.340883204 | 0.54580925 |
| Srrm2 | S | 540 | 0.060509255 | -0.0907172 | -0.033891021 | -0.094400275 | 0.05682618 | 0.518607266 | 0.361354932 | 0.454800333 | 0.340883204 | 0.54580925 |
| Ttc28 | S | 1366 | 0.039744668 | 0.070206 | -0.17362333 | -0.213367999 | -0.24382933 | 0.654274546 | 0.167853374 | 0.038671229 | 0.081485975 | 0.016411619 |
| Tmem184b | S | 402 | 0.043961669 | -0.093647227 | -0.003754993 | -0.047716662 | 0.089892234 | 0.165404372 | 0.047247848 | 0.948032877 | 0.445089334 | 0.205310552 |
| Tcof1 | S | 1128 | -0.047834455 | -0.001565291 | -0.01192442 | 0.035910035 | -0.010359129 | 0.514702626 | 0.964672814 | 0.570841474 | 0.615710947 | 0.768232099 |
| Uprt | S | 23 | 0.072672124 | -0.045420321 | -0.146693038 | -0.219365162 | -0.101272717 | 0.333440368 | 0.507509582 | 0.384872833 | 0.232843068 | 0.530039494 |
| Sipa1l1 | S | 1421 | -0.029200263 | -0.274075792 | 0.220125062 | 0.249325325 | 0.494200854 | 0.688002651 | 0.127078975 | 0.206318924 | 0.178005511 | 0.044061329 |
| Sipa1l1 | S | 1422 | -0.029200263 | -0.274075792 | 0.220125062 | 0.249325325 | 0.494200854 | 0.688002651 | 0.127078975 | 0.206318924 | 0.178005511 | 0.044061329 |
| Vcpip1 | S | 756 | -0.03350591 | -0.035655288 | 0.014257013 | 0.047762923 | 0.049912301 | 0.777396321 | 0.23947362 | 0.832353532 | 0.711809882 | 0.485506515 |
| Nfib | S | 292 | -0.031394755 | 0.016281819 | -0.031283874 | 0.000110881 | -0.047565693 | 0.30182774 | 0.763642423 | 0.541319355 | 0.99818164 | 0.474946916 |
| Srrm2 | S | 1982 | 0.007949177 | -0.004094841 | -0.042517275 | -0.050466451 | -0.038424234 | 0.901117086 | 0.888064277 | 0.173852033 | 0.469888537 | 0.298244375 |
| Arhgap11a | S | 847 | -0.039574807 | 0.053069829 | -0.093550514 | -0.053975707 | -0.146620343 | 0.538437018 | 0.306084394 | 0.250105854 | 0.477767908 | 0.110259748 |
| Gon4l | S | 1414 | 0.05208714 | -0.102154188 | -0.021803443 | -0.073890583 | 0.080350745 | 0.325055474 | 0.161742682 | 0.764728416 | 0.317935348 | 0.331027616 |
| Brix1 | S | 261 | -0.015720753 | -0.040134752 | -0.000482533 | 0.015238221 | 0.039652219 | 0.834071983 | 0.331390198 | 0.991925259 | 0.834458455 | 0.312095281 |
| Prkd3 | S | 30 | -0.026937218 | 0.079661662 | -0.230235851 | -0.203298632 | -0.309897513 | 0.812162077 | 0.126880683 | 0.213135732 | 0.282133052 | 0.137513882 |
| Chd4 | S | 312 | -0.142477147 | -0.070161902 | 0.175095869 | 0.317573016 | 0.24525777 | 0.033653425 | 0.214801888 | 0.037613906 | 0.010022307 | 0.027792135 |
| Ralgapa1 | S | 999 | -0.041825705 | 0.022649754 | -0.033633489 | 0.008192216 | -0.056283243 | 0.361296377 | 0.719887367 | 0.532227819 | 0.883925529 | 0.444168058 |
| Smg1 | S | 3553 | -0.013060839 | -0.025454881 | -0.005996846 | 0.007063992 | 0.019458035 | 0.800326853 | 0.30948926 | 0.777987064 | 0.885257676 | 0.27827703 |
| Scaf8 | S | 273 | -0.041341202 | -0.049944736 | 0.040403993 | 0.081745195 | 0.090348729 | 0.231808762 | 0.35784098 | 0.346293127 | 0.107453731 | 0.160885047 |
| Slc20a1 | S | 420 | 0.078472553 | -0.070533218 | -0.068437837 | -0.14691039 | 0.00209538 | 0.454637911 | 0.364612668 | 0.409326043 | 0.188610512 | 0.975545084 |
| Rrbp1 | S | 154 | 0.021126904 | -0.040981664 | -0.086604 | -0.107730903 | -0.045622336 | 0.912408598 | 0.789689022 | 0.599265045 | 0.624628478 | 0.806431736 |
| Prpf40a | S | 904 | 0.022214905 | -0.158605164 | 0.087273655 | 0.06505875 | 0.245878819 | 0.881260135 | 0.12186425 | 0.459905668 | 0.654628919 | 0.067255288 |
| Inpp5e | S | 89 | -0.076169768 | 0.000971364 | 0.063805396 | 0.139975164 | 0.062834032 | 0.036014764 | 0.982752672 | 0.421987991 | 0.148489498 | 0.441476576 |
| Fam120a | T | 386 | 0.040479695 | -0.015861116 | -0.094139544 | -0.134619239 | -0.078278428 | 0.336287673 | 0.743759496 | 0.174385832 | 0.086962077 | 0.269738824 |
| Pes1 | T | 384 | 0.037474643 | -0.046292616 | -0.057277272 | -0.094751915 | -0.010984657 | 0.507794717 | 0.682631703 | 0.231129957 | 0.14904103 | 0.921287978 |
| Tbc1d10a | S | 40 | -0.027569492 | -0.005903802 | -0.006508018 | 0.021061474 | -0.000604217 | 0.201368699 | 0.824545876 | 0.944832373 | 0.823248517 | 0.994903984 |
| Tbc1d10a | S | 43 | -0.027569492 | -0.005903802 | -0.006508018 | 0.021061474 | -0.000604217 | 0.201368699 | 0.824545876 | 0.944832373 | 0.823248517 | 0.994903984 |
| Spice1 | S | 766 | 0.015351456 | 0.065952689 | -0.191144194 | -0.20649565 | -0.257096883 | 0.777809194 | 0.588919184 | 0.254008257 | 0.227620422 | 0.182725018 |
| Phf20 | S | 364 | -0.137918724 | 0.095770845 | -0.101538199 | 0.036380525 | -0.197309044 | 0.616148683 | 0.572677713 | 0.579323674 | 0.891054979 | 0.263743189 |
| Peak1 | T | 675 | 0.172324831 | -0.057179495 | -0.173479023 | -0.345803854 | -0.116299528 | 0.056376451 | 0.538391662 | 0.030567323 | 0.006354791 | 0.215707803 |
| Reep3 | S | 150 | 0.000798306 | 0.012321337 | -0.07620275 | -0.077001056 | -0.088524087 | 0.984450409 | 0.832276859 | 0.276885461 | 0.286556507 | 0.280100201 |
| Sacs | S | 4259 | 0.014067885 | -0.035413631 | -0.054036438 | -0.068104323 | -0.018622807 | 0.658026706 | 0.59450331 | 0.467340834 | 0.362656352 | 0.824104297 |
| Tkt | Y | 275 | 0.071637572 | 0.027978392 | -0.210531807 | -0.282169379 | -0.2385102 | 0.563961044 | 0.790494121 | 0.244075683 | 0.13772361 | 0.189536555 |
| Dlg5 | S | 841 | -0.031257602 | -0.013354095 | -0.003712166 | 0.027545436 | 0.009641929 | 0.404059297 | 0.651404658 | 0.92467334 | 0.478963043 | 0.760956757 |
| Ncbp3 | S | 563 | -0.014848415 | -0.075342954 | 0.041765887 | 0.056614302 | 0.117108841 | 0.71766589 | 0.225227641 | 0.607216716 | 0.514171283 | 0.236094105 |
| Msl2 | S | 343 | -0.092612555 | -0.042220491 | 0.112564506 | 0.205177061 | 0.154784998 | 0.185950505 | 0.60293789 | 0.498354107 | 0.262011798 | 0.372167173 |
| Cacnb2 | S | 530 | -0.16591259 | 0.004250668 | 0.04256043 | 0.20847302 | 0.038309762 | 0.348479355 | 0.95472916 | 0.614627532 | 0.259755045 | 0.596979382 |
| Msi2 | S | 279 | -0.052984133 | -0.058436191 | 0.069295728 | 0.122279861 | 0.127731919 | 0.403634481 | 0.354844373 | 0.544584333 | 0.333507762 | 0.315074913 |
| Gigyf1 | S | 157 | -0.166424387 | 0.072397718 | 0.034718448 | 0.201142835 | -0.03767927 | 0.13872852 | 0.513429633 | 0.774918588 | 0.150359517 | 0.762746039 |
| Shb | S | 311 | 0.031607844 | -0.146537898 | 0.059682293 | 0.028074449 | 0.206220191 | 0.810269528 | 0.144562502 | 0.541767781 | 0.822565347 | 0.056558851 |
| Hnrnph1 Hnrnph2 | S | 104;104 | 0.020688225 | -0.027880596 | -0.036380573 | -0.057068797 | -0.008499977 | 0.530203711 | 0.392146492 | 0.289854471 | 0.0320341 | 0.622770342 |
| Dkc1 | S | 455 | 0.051889952 | -0.032078103 | -0.035992002 | -0.087881955 | -0.003913899 | 0.487443373 | 0.247784318 | 0.199996221 | 0.280173784 | 0.822261153 |
| Stard13 | S | 171 | 0.177822478 | 0.015638697 | -0.246675137 | -0.424497615 | -0.262313834 | 0.116319588 | 0.869353511 | 0.031299108 | 0.00521836 | 0.006357719 |
| Gigyf2 | T | 383 | -0.045014244 | -0.032951997 | 0.03765939 | 0.082673633 | 0.070611387 | 0.132517296 | 0.066170082 | 0.044893654 | 0.04368741 | 0.004764372 |
| Rabgef1 | S | 390 | 0.06498718 | 0.039151131 | -0.196843055 | -0.261830235 | -0.235994186 | 0.519112128 | 0.35087465 | 0.103916709 | 0.077214814 | 0.063509907 |
| Cluap1 | S | 409 | -0.148851034 | 0.069036943 | 0.036754838 | 0.185605872 | -0.032282104 | 0.029753513 | 0.159480776 | 0.725154322 | 0.16161268 | 0.760004713 |
| Palld | S | 1009 | -0.003839662 | 0.038486061 | -0.085255567 | -0.081395905 | -0.123721629 | 0.907031593 | 0.270665955 | 0.081009934 | 0.09511005 | 0.03220763 |
| Add3 | T | 654 | -0.108201343 | 0.122299638 | -0.05150012 | 0.056701223 | -0.173799758 | 0.273461164 | 0.403176922 | 0.402085482 | 0.497213419 | 0.263452415 |
| Map1a | T | 2605 | -0.07691749 | -0.052609986 | 0.046982475 | 0.123899965 | 0.099592461 | 0.366032421 | 0.536941024 | 0.422797043 | 0.186117945 | 0.280392768 |
| Rnps1 | S | 35 | -0.065927474 | -0.074525733 | 0.096334164 | 0.162261638 | 0.170859897 | 0.396239576 | 0.436229612 | 0.555067574 | 0.343174495 | 0.330539755 |
| Anapc1 | S | 50 | -0.059566673 | 0.013658425 | 0.005600373 | 0.065167047 | -0.008058051 | 0.150009739 | 0.771641974 | 0.93293604 | 0.374572592 | 0.911270932 |
| Ctr9 | T | 925 | 0.008502953 | -0.058411837 | 0.015478062 | 0.006975109 | 0.073889898 | 0.821735224 | 0.066948939 | 0.522798748 | 0.863768467 | 0.051297464 |
| Sik3 | S | 534 | -0.073883742 | 0.047030113 | -0.044187552 | 0.02969619 | -0.091217666 | 0.279766766 | 0.358588967 | 0.525222754 | 0.688633149 | 0.210098236 |
| Atad5 | S | 351 | 0.000369356 | -0.035234488 | -0.043046511 | -0.043415867 | -0.007812023 | 0.994515913 | 0.794621123 | 0.523914057 | 0.48148186 | 0.954045109 |
| Nup214 | S | 1142 | -0.007237965 | -0.018981284 | -0.02523372 | -0.017995755 | -0.006252437 | 0.790998954 | 0.59570375 | 0.594126355 | 0.681162403 | 0.895356428 |
| Ptdss2 | S | 16 | 0.058987339 | -0.055381122 | -0.050993644 | -0.109980983 | 0.004387478 | 0.168500497 | 0.140641671 | 0.295085612 | 0.0608557 | 0.913555779 |
| Dcbld2 | S | 719 | -0.061917938 | 0.05571556 | -0.083995417 | -0.022077479 | -0.139710977 | 0.615207929 | 0.483774313 | 0.386503543 | 0.868461983 | 0.211611197 |
| Dcbld2 | S | 721 | -0.061917938 | 0.05571556 | -0.083995417 | -0.022077479 | -0.139710977 | 0.615207929 | 0.483774313 | 0.386503543 | 0.868461983 | 0.211611197 |
| Skp1a | T | 131 | -0.091451233 | 0.04165389 | 0.003834597 | 0.09528583 | -0.037819293 | 0.139703588 | 0.565204499 | 0.940289108 | 0.089028913 | 0.558744624 |
| Pdap1 | S | 60 | -0.026977092 | 0.023821637 | -0.031465926 | -0.004488834 | -0.055287563 | 0.329777397 | 0.532566468 | 0.266455715 | 0.882156747 | 0.217564459 |
| Lmf2 | S | 695 | 0.082085729 | -0.00464987 | -0.132972322 | -0.215058052 | -0.128322452 | 0.292954044 | 0.941766021 | 0.098568113 | 0.040517074 | 0.11514535 |
| Rif1 | S | 1540 | 0.037322179 | -0.036853117 | -0.025979951 | -0.06330213 | 0.010873165 | 0.543006214 | 0.117824429 | 0.550598787 | 0.363317716 | 0.78473222 |
| Pja1 | S | 319 | 0.003611553 | -0.053036651 | -0.0096359 | -0.013247453 | 0.043400752 | 0.950448673 | 0.364842476 | 0.870475566 | 0.672602373 | 0.176396571 |
| Stxbp5 | S | 760 | -0.137918291 | 0.196656989 | -0.139840854 | -0.001922563 | -0.336497843 | 0.282862039 | 0.26436401 | 0.420382218 | 0.989568146 | 0.118069998 |
| Otud5 | S | 177 | -0.056902862 | 0.001888966 | 0.007643515 | 0.064546378 | 0.00575455 | 0.229108215 | 0.970694738 | 0.900257586 | 0.298978521 | 0.924764773 |
| Rpl13 | S | 140 | -0.049295195 | 0.042904646 | -0.026812147 | 0.022483048 | -0.069716794 | 0.083158024 | 0.314971717 | 0.552091672 | 0.626336914 | 0.23334123 |
| Atad5 | S | 740 | 0.020778672 | -0.012520725 | -0.044341339 | -0.065120011 | -0.031820614 | 0.653553745 | 0.721742155 | 0.162748044 | 0.210294454 | 0.367721316 |
| Nme1 Nme2 | T | 94;94 | -0.06522754 | 0.085248627 | -0.103685549 | -0.038458009 | -0.188934176 | 0.360233729 | 0.317267506 | 0.399747848 | 0.732787304 | 0.173967449 |
| Cdc23 | T | 562 | -0.022208904 | 0.002820966 | -0.043262907 | -0.021054003 | -0.046083873 | 0.613071088 | 0.965418207 | 0.556415753 | 0.769314366 | 0.590350277 |
| Lrrc42 | S | 398 | 0.030039523 | 0.058542585 | -0.200557862 | -0.230597385 | -0.259100447 | 0.386129771 | 0.405046518 | 0.119954128 | 0.099653414 | 0.065728434 |
| Clasp2 | S | 1021 | 0.183730046 | -0.058253249 | -0.133187359 | -0.316917405 | -0.07493411 | 0.008872243 | 0.052517676 | 0.011620688 | 0.001296431 | 0.074420181 |
| Tmem200a | S | 397 | 0.061685609 | -0.078824646 | -0.008890075 | -0.070575684 | 0.069934571 | 0.207095041 | 0.033469656 | 0.784401343 | 0.148967232 | 0.036435293 |
| Atp7a | S | 356 | 0.012668366 | -0.096506997 | 0.014799092 | 0.002130726 | 0.11130609 | 0.739942969 | 0.222835752 | 0.739839588 | 0.947708915 | 0.174823914 |
| Sbno1 | S | 687 | 0.052939976 | 0.035184732 | -0.139107374 | -0.19204735 | -0.174292106 | 0.781796644 | 0.678576795 | 0.187286939 | 0.35712813 | 0.103342233 |
| Sde2 | S | 269 | 0.00921121 | -0.002856553 | -0.051968224 | -0.061179434 | -0.049111671 | 0.671519883 | 0.904352243 | 0.187033336 | 0.141947142 | 0.224094912 |
| Son | S | 1021 | 0.047793538 | -0.084802573 | 0.007418707 | -0.040374831 | 0.09222128 | 0.504102955 | 0.073219076 | 0.817328944 | 0.568971762 | 0.068645233 |
| Rab11fip1 | S | 358 | -0.038483822 | 0.051153377 | -0.1240802 | -0.085596378 | -0.175233576 | 0.553505551 | 0.246422853 | 0.335868204 | 0.494304956 | 0.214665646 |
| Tbc1d9b | T | 397 | -0.098965876 | 0.100581484 | -0.054532448 | 0.044433428 | -0.155113932 | 0.052823818 | 0.06411441 | 0.356544129 | 0.36385986 | 0.033987329 |
| Dhx9 | S | 136 | 0.036073436 | 0.049223078 | -0.108526999 | -0.144600435 | -0.157750076 | 0.806382979 | 0.802158404 | 0.163716595 | 0.366687258 | 0.451364715 |
| Cdc42bpb | S | 972 | -0.071290391 | 0.077423717 | -0.023244151 | 0.04804624 | -0.100667868 | 0.032131532 | 0.155183059 | 0.458143951 | 0.090422208 | 0.093137721 |

| | | | | | | | | | | | | |
|---|---|---|---|---|---|---|---|---|---|---|---|---|
| Cdk14 | S | 78 | -0.134713151 | 0.182730841 | -0.117535693 | 0.017177458 | -0.300266534 | 0.261347265 | 0.224946204 | 0.359711461 | 0.899607666 | 0.105744197 |
| Tnks1bp1 | S | 697 | -0.063143946 | 0.008456082 | -0.041716813 | 0.021427133 | -0.050172895 | 0.743607094 | 0.9421752 | 0.646925028 | 0.911490925 | 0.681004812 |
| Lima1 | S | 594 | 0.042982147 | 0.103906381 | -0.252513192 | -0.29549534 | -0.356419573 | 0.735336791 | 0.459811885 | 0.190152351 | 0.128326355 | 0.081678893 |
| Srsf7 | S | 262 | 0.024997663 | -0.031268268 | -0.026204835 | -0.051202498 | 0.005063433 | 0.338546809 | 0.198981861 | 0.259168445 | 0.113184946 | 0.837517982 |
| Sap130 | S | 13 | 0.113556114 | -0.081065047 | -0.14212401 | -0.255680124 | -0.061058962 | 0.123619421 | 0.221765691 | 0.351589221 | 0.150930517 | 0.652831091 |
| Eif5b | S | 1164 | -0.052479827 | -0.006567898 | 0.012625902 | 0.065105729 | 0.019193799 | 0.229943455 | 0.893197451 | 0.769927919 | 0.239630752 | 0.740921125 |
| Rtn4 | S | 240 | 0.031664092 | -0.014542266 | -0.07582432 | -0.107488413 | -0.061282054 | 0.798706258 | 0.835835457 | 0.353646734 | 0.440296851 | 0.498869765 |
| Polr2a | S | 1878 | 0.067180705 | -0.020650772 | -0.078180015 | -0.14536072 | -0.057529243 | 0.465016581 | 0.474941774 | 0.208312148 | 0.183439743 | 0.320167026 |
| Cnot2 | S | 165 | 0.053471137 | -0.02820595 | -0.08840517 | -0.141876308 | -0.06019922 | 0.449574795 | 0.615461162 | 0.276713048 | 0.107754531 | 0.373046725 |
| Trim3 | S | 7 | -0.116845771 | -0.018876524 | 0.052682733 | 0.169528504 | 0.071559258 | 0.36426566 | 0.910232671 | 0.693249724 | 0.103166949 | 0.623320695 |
| Cdc42ep4 | S | 165 | 0.112955569 | -0.047493305 | -0.173306121 | -0.28626169 | -0.125812816 | 0.37279383 | 0.638525417 | 0.336957189 | 0.172548904 | 0.482174386 |
| Als2 | T | 504 | -0.043551406 | -0.035672895 | 0.009658082 | 0.053209488 | 0.045330977 | 0.522178699 | 0.709626162 | 0.895879091 | 0.536813731 | 0.67305762 |
| Tfpt | S | 255 | -0.02619653 | -0.091906013 | 0.076662776 | 0.102859306 | 0.168568789 | 0.458612421 | 0.034502412 | 0.068837021 | 0.023905175 | 0.00513246 |
| Micall1 | S | 651 | 0.01211605 | -0.005977385 | -0.034262395 | -0.046378445 | -0.028285011 | 0.681773104 | 0.830375069 | 0.099892458 | 0.17371201 | 0.332861303 |
| Ctnnd1 | S | 230 | 0.022966281 | 0.000464581 | -0.090313781 | -0.113280062 | -0.090778362 | 0.319947225 | 0.990012763 | 0.155663892 | 0.106319804 | 0.164972538 |
| Aarsd1 | S | 88 | -0.026293064 | 0.069545577 | -0.099399395 | -0.073106331 | -0.168944972 | 0.782052441 | 0.142228826 | 0.05713068 | 0.456271265 | 0.007488427 |
| Ap2a1 | S | 655 | 0.0155327 | -0.019048456 | -0.022076187 | -0.037608886 | -0.003027731 | 0.763950499 | 0.351080486 | 0.275522607 | 0.483425058 | 0.851372615 |
| Zfp800 | S | 645 | -0.009733905 | -0.05513188 | 0.031190458 | 0.040924363 | 0.086322338 | 0.893072839 | 0.113417386 | 0.559631683 | 0.627406876 | 0.181779356 |
| Cit | S | 698 | -0.066237993 | 0.019538084 | 0.018146837 | 0.08438483 | -0.001391247 | 0.071714424 | 0.737764073 | 0.602870743 | 0.077127258 | 0.9818313 |
| Rif1 | S | 2144 | 0.023367449 | -0.014527917 | -0.078404702 | -0.101772151 | -0.063876785 | 0.673779407 | 0.782697701 | 0.365310527 | 0.263947413 | 0.449089596 |
| Prkd1 | T | 406 | -0.070290185 | -0.029219786 | 0.037617145 | 0.107907331 | 0.066836932 | 0.304691384 | 0.550033159 | 0.434978676 | 0.162982945 | 0.249940333 |
| Prkd3 | T | 394 | -0.070290185 | -0.029219786 | 0.037617145 | 0.107907331 | 0.066836932 | 0.304691384 | 0.550033159 | 0.434978676 | 0.162982945 | 0.249940333 |
| Rab34 | T | 76 | -0.038866349 | 0.009211996 | -0.029568051 | 0.009298298 | -0.038780047 | 0.561099561 | 0.734363473 | 0.47852953 | 0.889170365 | 0.318769123 |
| Nsd3 | S | 561 | -0.079287123 | -0.03484901 | 0.07999213 | 0.159279253 | 0.11484114 | 0.08446865 | 0.432201807 | 0.181045782 | 0.039737842 | 0.102065209 |
| Camkk2 | S | 85 | -0.105577506 | 0.052715181 | -0.011437198 | 0.094140309 | -0.064152379 | 0.10880278 | 0.225493787 | 0.870649274 | 0.270719527 | 0.402009071 |
| Suv39h1 | S | 391 | 0.048918439 | -0.024999877 | -0.049522332 | -0.098440771 | -0.024522455 | 0.144693489 | 0.321516611 | 0.052110972 | 0.027088673 | 0.315478937 |
| Ppp4r3a | S | 685 | 0.066169064 | -0.115437278 | -0.000105135 | -0.066274199 | 0.115332143 | 0.587031397 | 0.200848316 | 0.998994922 | 0.584942892 | 0.21295658 |
| Bmp2k | S | 1012 | -0.005279671 | -0.051882277 | 0.006969559 | 0.012249229 | 0.058851836 | 0.924451415 | 0.297755501 | 0.892560192 | 0.786965571 | 0.147196322 |
| Bmp2k | S | 1013 | -0.005279671 | -0.051882277 | 0.006969559 | 0.012249229 | 0.058851836 | 0.924451415 | 0.297755501 | 0.892560192 | 0.786965571 | 0.147196322 |
| Herc2 | S | 1939 | -0.172502333 | 0.027727692 | 0.014793275 | 0.187295608 | -0.012934417 | 0.388239279 | 0.592188515 | 0.890577934 | 0.371304349 | 0.896751929 |
| Srrm2 | S | 203 | 0.067078401 | -0.05492453 | -0.077737204 | -0.144815605 | -0.022812674 | 0.079733789 | 0.521226345 | 0.062365053 | 0.00597394 | 0.775235757 |
| Raly | S | 236 | 0.077866605 | -0.095537765 | -0.090865239 | -0.168731845 | 0.004672526 | 0.631883999 | 0.619971454 | 0.595480033 | 0.369833577 | 0.981626397 |
| Cep170 | S | 1155 | -0.043944874 | 0.061247276 | -0.103716682 | -0.059771808 | -0.164963958 | 0.611770604 | 0.332705568 | 0.29834418 | 0.584509722 | 0.142456743 |
| Nav3 | S | 1212 | -0.084980602 | 0.044417181 | -0.021144301 | 0.063836301 | -0.065561481 | 0.277721371 | 0.613405966 | 0.849147667 | 0.53661406 | 0.552452858 |
| Pa2g4 | S | 383 | 0.056274632 | -0.021438357 | -0.047366439 | -0.103641071 | -0.025928082 | 0.714928863 | 0.767428429 | 0.14313809 | 0.517397976 | 0.715574949 |
| Kdm2b | S | 886 | 0.0242683 | -0.024601555 | -0.056757895 | -0.081026196 | -0.03215634 | 0.760066344 | 0.573556565 | 0.473659631 | 0.403180359 | 0.6494096 |
| Kdm2b | S | 887 | 0.0242683 | -0.024601555 | -0.056757895 | -0.081026196 | -0.03215634 | 0.760066344 | 0.573556565 | 0.473659631 | 0.403180359 | 0.6494096 |
| Kdm2b | T | 877 | 0.0242683 | -0.024601555 | -0.056757895 | -0.081026196 | -0.03215634 | 0.760066344 | 0.573556565 | 0.473659631 | 0.403180359 | 0.6494096 |
| Mical2 | S | 738 | 0.130559734 | -0.035077412 | -0.156810567 | -0.2873703 | -0.121733154 | 0.216279677 | 0.747934738 | 0.110892772 | 0.018829004 | 0.205862408 |
| Nectin1 | S | 509 | -0.059202162 | 0.210369779 | -0.276130739 | -0.216928577 | -0.486500518 | 0.654147555 | 0.030568176 | 0.041023991 | 0.175746257 | 0.009536725 |
| Cdk9 | S | 347 | -0.022201659 | 0.006706004 | -0.024354476 | -0.002152818 | -0.03106048 | 0.367038635 | 0.938349476 | 0.596283663 | 0.962491003 | 0.739833018 |
| Wiz | S | 1112 | -0.001678084 | -0.114061819 | 0.027305139 | 0.028983223 | 0.141366957 | 0.993081824 | 0.455729472 | 0.867846004 | 0.900024395 | 0.484769414 |
| Rif1 | S | 2125 | 0.096032053 | -0.080011152 | -0.100107407 | -0.196139459 | -0.020096254 | 0.003554526 | 0.198472817 | 0.022388369 | 0.004092569 | 0.710762508 |
| Helq | T | 30 | 0.1309428 | -0.052084653 | -0.188129555 | -0.319072356 | -0.136044902 | 0.250662064 | 0.696012815 | 0.226361693 | 0.088540352 | 0.437465989 |
| Gclc | S | 5 | -0.075975561 | 0.096228518 | -0.084775904 | -0.008800343 | -0.181004421 | 0.329926126 | 0.14964647 | 0.18689035 | 0.901298988 | 0.028735238 |
| Psme3 | T | 23 | 0.027087121 | -0.014041337 | -0.105141698 | -0.132228819 | -0.091100361 | 0.898994542 | 0.964295073 | 0.497787179 | 0.563473204 | 0.777191525 |
| Atf7ip | S | 539 | -0.00979237 | -0.093054366 | 0.046779039 | 0.056571409 | 0.139833406 | 0.843864939 | 0.276704752 | 0.629385458 | 0.584001522 | 0.253589079 |
| Polr3c | S | 204 | 0.021599106 | -0.045102886 | -0.009256678 | -0.030855784 | 0.035846208 | 0.281347284 | 0.059230185 | 0.781903134 | 0.409445764 | 0.345791047 |
| Spen | S | 762 | -0.052110858 | -0.057112594 | 0.084042414 | 0.136153272 | 0.141155008 | 0.069032048 | 0.050139833 | 0.028163555 | 0.008021479 | 0.007373822 |
| Rbmx2 | S | 274 | 0.084526537 | 0.086157129 | -0.400615127 | -0.485141664 | -0.486772256 | 0.556720391 | 0.563436711 | 0.31132279 | 0.242450874 | 0.240765788 |
| Rai14 | S | 448 | 0.008871531 | 0.119873736 | -0.155348114 | -0.164219645 | -0.27522185 | 0.909782073 | 0.195577471 | 0.038936638 | 0.070675394 | 0.035663709 |
| Srsf5 | S | 245 | 0.070809017 | -0.072809234 | -0.052375932 | -0.123184949 | 0.020433302 | 0.058552507 | 0.086760061 | 0.261128404 | 0.039950027 | 0.634483659 |
| Arhgef6 | S | 224 | -0.044715923 | 0.119295298 | -0.150573005 | -0.105857082 | -0.269868302 | 0.706092171 | 0.310298936 | 0.206351071 | 0.21151913 | 0.011768492 |
| Pikfyve | S | 1753 | -0.043823796 | -0.007769536 | -0.010167886 | 0.033655911 | -0.00239835 | 0.637825433 | 0.779066988 | 0.753151615 | 0.710726056 | 0.903016516 |
| Pak4 | S | 146 | -0.109264153 | 0.050514202 | -0.067188152 | 0.042076002 | -0.117702354 | 0.489809143 | 0.257473916 | 0.48531327 | 0.798529716 | 0.272114178 |
| Mphosph8 | S | 85 | 0.048890718 | -0.059196914 | -0.008824374 | -0.057715093 | 0.05037254 | 0.349131793 | 0.055878799 | 0.743564492 | 0.28535852 | 0.115044207 |
| Sh3pxd2a | S | 592 | -0.022114044 | 0.01151326 | -0.038270165 | -0.016156121 | -0.049783426 | 0.638797113 | 0.775719272 | 0.381417798 | 0.641576314 | 0.093601738 |
| Rnps1 | S | 155 | -0.022910541 | -0.033592423 | 0.040080692 | 0.062991234 | 0.073673116 | 0.404257187 | 0.077381684 | 0.368362077 | 0.204367292 | 0.164593116 |
| Rnps1 | S | 157 | -0.022910541 | -0.033592423 | 0.040080692 | 0.062991234 | 0.073673116 | 0.404257187 | 0.077381684 | 0.368362077 | 0.204367292 | 0.164593116 |
| Trp53bp1 | S | 1653 | 0.007337614 | -0.052479256 | -0.005490144 | -0.012827759 | 0.046989111 | 0.879757404 | 0.297324801 | 0.961190187 | 0.904814009 | 0.668537151 |
| Eps15l1 | Y | 562 | -0.311703177 | 0.102757193 | 0.11310813 | 0.424811307 | 0.010350938 | 0.058075003 | 0.456219516 | 0.20646116 | 0.019978932 | 0.940479013 |
| Rapgef2 | S | 1094 | -0.063789893 | 0.026270479 | -0.026080585 | 0.037709308 | -0.052351064 | 0.218942671 | 0.404226656 | 0.789888003 | 0.707663582 | 0.59465333 |
| 1110038F14Rik | S | 31 | -0.070359928 | -0.004371676 | 0.041485398 | 0.111845326 | 0.045857074 | 0.034822043 | 0.923523359 | 0.173953718 | 0.015863752 | 0.370371859 |
| Asxl1 | S | 500 | -0.0849155 | -0.064578554 | 0.109046647 | 0.193962146 | 0.173625201 | 0.160270367 | 0.164751548 | 0.100693299 | 0.021957698 | 0.047014279 |
| Itprid2 | S | 351 | -0.07808894 | -0.04306211 | 0.09061295 | 0.16870189 | 0.13367506 | 0.145081556 | 0.554481977 | 0.338522134 | 0.129265118 | 0.222781975 |
| Med1 | S | 1403 | 0.063263762 | 0.016454552 | -0.114804742 | -0.178068504 | -0.131259295 | 0.046976894 | 0.540420461 | 0.005390733 | 0.002125248 | 0.006270413 |
| Pak1 | T | 230 | 0.00115948 | 0.024644443 | -0.075917852 | -0.077077332 | -0.100562295 | 0.980266584 | 0.662503501 | 0.219986097 | 0.158184809 | 0.114181055 |
| Lmtk2 | S | 545 | 0.180509652 | -0.230625059 | -0.184728567 | -0.365238219 | 0.045896491 | 0.14123129 | 0.444036459 | 0.429376468 | 0.175615448 | 0.88556461 |
| Lmtk2 | S | 548 | 0.180509652 | -0.230625059 | -0.184728567 | -0.365238219 | 0.045896491 | 0.14123129 | 0.444036459 | 0.429376468 | 0.175615448 | 0.88556461 |
| Srsf7 | S | 250 | 0.094885249 | -0.159883084 | 0.005608276 | -0.089276973 | 0.16549136 | 0.320319392 | 0.124446192 | 0.95913635 | 0.291496032 | 0.103111484 |
| Ppp1r35 | S | 46 | 0.043193122 | -0.041975103 | -0.075281211 | -0.118474333 | -0.033306107 | 0.675105266 | 0.586616134 | 0.546282026 | 0.433856975 | 0.800831878 |
| Dtl | S | 674 | -0.030520799 | -0.01899999 | 0.000442937 | 0.030963736 | 0.019442927 | 0.444899865 | 0.730795119 | 0.991899958 | 0.504493335 | 0.741555547 |
| Map3k11 | S | 796 | 0.010859359 | 0.041339849 | -0.091946195 | -0.102805553 | -0.133286043 | 0.768408596 | 0.509072883 | 0.107330912 | 0.085575661 | 0.098451398 |
| Spen | S | 771 | -0.078117079 | -0.028917767 | 0.070010216 | 0.148127295 | 0.098927983 | 0.065236982 | 0.439151102 | 0.122187299 | 0.010025468 | 0.034306611 |
| Hirip3 | T | 167 | -0.065864303 | 0.026221465 | 0.01307604 | 0.078940343 | -0.013145426 | 0.098472566 | 0.696867476 | 0.751653254 | 0.095094813 | 0.845050092 |
| Thrap3 | S | 530 | 0.041481006 | -0.053559171 | -0.063305538 | -0.104786543 | -0.009746367 | 0.076039745 | 0.236913632 | 0.287635261 | 0.132614662 | 0.869122223 |
| Rnf216 | S | 707 | -0.132161448 | 0.002486748 | 0.076779615 | 0.208941063 | 0.074292867 | 0.069952803 | 0.962619274 | 0.19963983 | 0.020929652 | 0.193471945 |
| Ptk2b | S | 746 | -0.010869906 | -0.034397128 | -0.061543309 | -0.050673403 | -0.027146181 | 0.947729593 | 0.787613023 | 0.804506196 | 0.853914316 | 0.916392767 |
| Ptk2b | S | 758 | -0.010869906 | -0.034397128 | -0.061543309 | -0.050673403 | -0.027146181 | 0.947729593 | 0.787613023 | 0.804506196 | 0.853914316 | 0.916392767 |
| Ncor2 | S | 1816 | 0.140726392 | -0.038314619 | -0.122216009 | -0.2629424 | -0.08390139 | 0.216847015 | 0.497689182 | 0.099923949 | 0.062229289 | 0.179001989 |
| Ncor2 | S | 1819 | 0.140726392 | -0.038314619 | -0.122216009 | -0.2629424 | -0.08390139 | 0.216847015 | 0.497689182 | 0.099923949 | 0.062229289 | 0.179001989 |
| Nav1 | S | 197 | 0.036339541 | -0.040647911 | -0.030861965 | -0.067201506 | 0.009785947 | 0.431636353 | 0.283564779 | 0.355959806 | 0.178551418 | 0.768732849 |
| Dnmt1 | S | 122 | 0.033962259 | 0.006425818 | -0.074084595 | -0.108046854 | -0.080510414 | 0.249731894 | 0.886607041 | 0.0476638 | 0.022560963 | 0.159751278 |
| Acin1 | S | 729 | -0.085493646 | -0.040006064 | 0.051595931 | 0.137089577 | 0.091601995 | 0.24248345 | 0.284100946 | 0.1027618 | 0.107282141 | 0.061480105 |
| Thrap3 | S | 924 | -0.101483693 | -0.037753216 | 0.091315868 | 0.192799561 | 0.129069083 | 0.098709542 | 0.218595769 | 0.068469108 | 0.019556345 | 0.034059342 |
| Pak4 | S | 293 | 0.011008096 | 0.004345035 | -0.032563266 | -0.043571362 | -0.036908301 | 0.687868604 | 0.755043348 | 0.037043153 | 0.183270415 | 0.020043281 |
| Ranbp2 | S | 1154 | -0.003740773 | -0.067915507 | 0.010620543 | 0.014361317 | 0.07853605 | 0.969849941 | 0.401595283 | 0.912435446 | 0.90234331 | 0.458741297 |
| Emc4 | S | 36 | 0.019077954 | -0.025117659 | -0.053724445 | -0.072802399 | -0.028606786 | 0.633529076 | 0.487426874 | 0.520798928 | 0.396806397 | 0.716596278 |
| Atp10a | S | 961 | 0.212746166 | 0.20946083 | -0.261618487 | -0.474364653 | -0.471079317 | 0.222375613 | 0.358243375 | 0.00237202 | 0.053971829 | 0.113499529 |
| Pxn | T | 132 | -0.093164727 | -0.078906782 | 0.092107521 | 0.185272248 | 0.171014303 | 0.291323751 | 0.288843269 | 0.142286368 | 0.081291867 | 0.059515051 |
| Rpl26 Gm15772 | S | 23 | 0.043029375 | -0.019424753 | -0.091238722 | -0.134268097 | -0.071813969 | 0.444219193 | 0.697603058 | 0.300065321 | 0.162420502 | 0.389171079 |
| Arhgef40 | T | 1072 | -0.049345817 | 0.053267961 | -0.064138548 | -0.014792731 | -0.117406509 | 0.447397005 | 0.220266894 | 0.218537079 | 0.82947372 | 0.074054253 |
| Setd1a | S | 1483 | -0.085526503 | -0.008796007 | 0.046731066 | 0.132257568 | 0.055527072 | 0.198938819 | 0.921634138 | 0.535732169 | 0.133571076 | 0.574838826 |
| Zeb1 | S | 624 | 0.001170692 | -0.099475249 | 0.036841591 | 0.035670899 | 0.13631684 | 0.986666832 | 0.279891626 | 0.657269316 | 0.712484377 | 0.232193956 |

| | | | | | | | | | | | | |
|---|---|---|---|---|---|---|---|---|---|---|---|---|
| Aimp1 | S | 99 | -0.014199371 | -0.007880727 | -0.024103428 | -0.009904057 | -0.0162227 | 0.571551923 | 0.871424476 | 0.632917088 | 0.829604556 | 0.78673624 |
| Papola | S | 648 | -0.092811586 | 0.045338712 | 0.017872345 | 0.110683931 | -0.027466366 | 0.040998754 | 0.437257762 | 0.423172121 | 0.038385655 | 0.614689463 |
| Dock7 | S | 182 | -0.111738137 | 0.053597201 | -0.006446936 | 0.105291201 | -0.060044137 | 0.215021559 | 0.44369441 | 0.93332446 | 0.299495154 | 0.49615881 |
| Cpox | S | 101 | -0.023311908 | -0.030039606 | -0.00282383 | 0.020488078 | 0.027215776 | 0.713187322 | 0.591752882 | 0.947571439 | 0.759979003 | 0.651714503 |
| Nfkb2 | S | 715 | 0.070427146 | -0.058416604 | -0.033404355 | -0.103831501 | 0.025012249 | 0.388579751 | 0.085725357 | 0.719214722 | 0.370317834 | 0.786977565 |
| Arhgef10 | S | 108 | 0.088143519 | -0.00997415 | -0.14673504 | -0.234878559 | -0.136760889 | 0.665226885 | 0.982176494 | 0.332833315 | 0.267378191 | 0.759828824 |
| Itprid2 | S | 330 | -0.030347442 | 0.041402118 | -0.078828033 | -0.048480591 | -0.120230151 | 0.580097972 | 0.360940018 | 0.314709902 | 0.550802894 | 0.171524302 |
| Msl1 | T | 404 | -0.007762883 | -0.062152055 | 0.013495254 | 0.021258137 | 0.075647309 | 0.92847671 | 0.385782561 | 0.8752649 | 0.811553943 | 0.34256155 |
| Tra2b | S | 22 | -0.006788719 | -0.084296705 | 0.014427775 | 0.021216495 | 0.09872448 | 0.972617201 | 0.447618435 | 0.845887763 | 0.913765309 | 0.36992777 |
| Arhgap35 | S | 985 | -0.059051356 | 0.009720361 | -0.011443592 | 0.047607765 | -0.021163953 | 0.431459662 | 0.871302646 | 0.878879255 | 0.494576743 | 0.697623591 |
| Cenpu | S | 111 | -0.007407851 | 0.00993653 | -0.064345645 | -0.056937794 | -0.074282175 | 0.836144046 | 0.766703602 | 0.40996462 | 0.463645241 | 0.353963032 |
| Kmt2a | S | 2560 | -0.24822804 | 0.061781839 | 0.033069718 | 0.281297757 | -0.028712121 | 0.432366664 | 0.721179371 | 0.784683652 | 0.383020485 | 0.867999206 |
| Caskin2 | S | 799 | -0.09354243 | 0.052253486 | -0.024547908 | 0.068994522 | -0.076801394 | 0.16985742 | 0.299120369 | 0.830785265 | 0.557999947 | 0.505669528 |
| Rplp1 | S | 38 | 0.075168509 | -0.08522418 | -0.089429462 | -0.164597971 | -0.004205282 | 0.409787162 | 0.523329535 | 0.504277692 | 0.24038688 | 0.977778721 |
| Tti2 | S | 21 | 0.04785282 | 0.001359069 | -0.075648385 | -0.123501205 | -0.077007454 | 0.387733042 | 0.965875469 | 0.074707041 | 0.075699103 | 0.061239068 |
| Lrrfip2 | S | 111 | 0.032747136 | -0.014116627 | -0.059490634 | -0.09223777 | -0.045374006 | 0.359510008 | 0.661011455 | 0.16095785 | 0.053142343 | 0.182474881 |
| H3c13 H3c14 H3c15 H | S | 29;29 | 0.122408584 | 0.043051793 | -0.134922746 | -0.25733133 | -0.17797454 | 0.176965512 | 0.656149069 | 0.007007135 | 0.044789403 | 0.151500028 |
| Zfp462 | T | 1000 | -0.035206789 | -0.146478499 | 0.124005267 | 0.159212055 | 0.270483766 | 0.620879747 | 0.080612927 | 0.118815847 | 0.101795217 | 0.020778305 |
| Arhgap1 | S | 47 | -0.20968979 | 0.023452334 | 0.106266571 | 0.315956361 | 0.082814237 | 0.123155281 | 0.629820517 | 0.346094803 | 0.062122629 | 0.435430609 |
| Smarcc1 | S | 309 | -0.01291205 | -0.064757312 | 0.042935277 | 0.055847328 | 0.107692589 | 0.818155741 | 0.176358399 | 0.535683046 | 0.459860206 | 0.1722603 |
| Fgf13 | S | 32 | -0.039720604 | -0.047161215 | 0.068350523 | 0.108071127 | 0.115511738 | 0.304667711 | 0.50236013 | 0.638102881 | 0.47053122 | 0.456343369 |
| Pclaf | S | 71 | -0.085723189 | -0.004514915 | 0.049588386 | 0.135311575 | 0.054103301 | 0.124778682 | 0.941434564 | 0.387015294 | 0.028484029 | 0.346756342 |
| Zdhhc5 | T | 659 | 0.074906679 | -0.052780177 | -0.0730092 | -0.147915879 | -0.020229023 | 0.514139277 | 0.525160424 | 0.394697476 | 0.186962247 | 0.601731087 |
| Syde1 | S | 66 | 0.020544791 | -0.016344149 | -0.052014829 | -0.072559621 | -0.03567068 | 0.667964822 | 0.726146214 | 0.32938029 | 0.106512479 | 0.344153162 |
| Dmtn | S | 26 | -0.020033469 | -0.004320792 | -0.059487324 | -0.039453854 | -0.055166532 | 0.88304513 | 0.93171352 | 0.670787117 | 0.82234113 | 0.68602666 |
| Cdc42bpb | S | 1692 | -0.111651651 | -0.064724634 | 0.130729243 | 0.242380894 | 0.195453877 | 0.473883347 | 0.655311668 | 0.660344873 | 0.429515856 | 0.511100582 |
| Tbx3 | S | 456 | -0.057773021 | -0.006373469 | 0.01894406 | 0.07671708 | 0.025317529 | 0.193846613 | 0.867408968 | 0.481879662 | 0.118796662 | 0.545809698 |
| Zeb2 | S | 1124 | -0.022665384 | -0.013053257 | -0.010347562 | 0.012317822 | 0.002705695 | 0.444455919 | 0.769656957 | 0.938266733 | 0.925535279 | 0.983945396 |
| Wdr76 | S | 195 | -0.021332943 | 0.006063904 | -0.026162497 | -0.004829553 | -0.032226401 | 0.618878864 | 0.896402277 | 0.453603483 | 0.891773151 | 0.452805263 |
| Basp1 | T | 36 | 0.065038718 | -0.099953088 | -0.011550383 | -0.076589101 | 0.088402705 | 0.347196602 | 0.160618811 | 0.777757843 | 0.298205052 | 0.211852398 |
| Map3k1 | S | 97 | 0.04312406 | -0.236112261 | 0.022596012 | -0.020528048 | 0.258708273 | 0.879331665 | 0.594388807 | 0.864689117 | 0.943948333 | 0.565471293 |
| Ercc6 | S | 428 | 0.06296502 | -0.114250247 | -0.035015495 | -0.097980516 | 0.079234751 | 0.084059446 | 0.151772624 | 0.529986987 | 0.153150328 | 0.318266908 |
| Ercc6 | S | 429 | 0.06296502 | -0.114250247 | -0.035015495 | -0.097980516 | 0.079234751 | 0.084059446 | 0.151772624 | 0.529986987 | 0.153150328 | 0.318266908 |
| Lbr | S | 101 | -0.051200047 | -0.019154368 | 0.034549501 | 0.085749548 | 0.053703869 | 0.192998286 | 0.43699317 | 0.322231052 | 0.086381239 | 0.185839918 |
| Ints3 | S | 992 | 0.100953206 | -0.129148002 | -0.058807378 | -0.159760585 | 0.070340624 | 0.193003188 | 0.240349933 | 0.520159891 | 0.161649436 | 0.552379128 |
| Mesd | S | 211 | -0.176741824 | 0.025845849 | 0.141396616 | 0.31813844 | 0.115550767 | 0.031364481 | 0.724718038 | 0.204159725 | 0.037516633 | 0.266952987 |
| Aatf | S | 287 | -0.038783452 | -0.037095688 | 0.018842174 | 0.057625625 | 0.055937861 | 0.665740033 | 0.436400621 | 0.686583696 | 0.499822567 | 0.014123818 |
| Aatf | S | 288 | -0.038783452 | -0.037095688 | 0.018842174 | 0.057625625 | 0.055937861 | 0.665740033 | 0.436400621 | 0.686583696 | 0.499822567 | 0.014123818 |
| Jmy | S | 94 | -0.044957748 | 0.04134635 | -0.066252651 | -0.021294903 | -0.107599001 | 0.586512082 | 0.579912388 | 0.508978966 | 0.807721426 | 0.249242389 |
| Srrm2 | S | 1397 | 0.096496895 | -0.043335735 | -0.100544479 | -0.197041374 | -0.057208744 | 0.02109977 | 0.058006045 | 0.105807248 | 0.019060097 | 0.272963429 |
| Tenm3 | S | 215 | 0.100957936 | -0.192263229 | 0.054187525 | -0.046770411 | 0.246450753 | 0.471501153 | 0.128811268 | 0.710625953 | 0.76962143 | 0.147100287 |
| Zranb2 | S | 290 | 0.094781257 | 0.070209107 | -0.298183452 | -0.392964709 | -0.368392559 | 0.311013544 | 0.457223016 | 0.128303993 | 0.075113034 | 0.085436771 |
| Zranb2 | S | 293 | 0.094781257 | 0.070209107 | -0.298183452 | -0.392964709 | -0.368392559 | 0.311013544 | 0.457223016 | 0.128303993 | 0.075113034 | 0.085436771 |
| Afdn | S | 1211 | 0.008397195 | -0.007275199 | -0.030628144 | -0.039025339 | -0.023352945 | 0.809202668 | 0.653238046 | 0.266274306 | 0.33779197 | 0.368976244 |
| Pkd2 | S | 810 | -0.023779368 | 0.049997977 | -0.078713078 | -0.054933709 | -0.128711055 | 0.607096813 | 0.470944242 | 0.31997404 | 0.45605173 | 0.174156905 |
| Ehmt2 | S | 193 | 0.018456678 | -0.047614728 | -0.005403981 | -0.023860659 | 0.042210747 | 0.614810659 | 0.190071038 | 0.867200517 | 0.326307223 | 0.061120048 |
| N4bp1 | T | 241 | -0.062348732 | 0.056998153 | -0.028859683 | 0.033489049 | -0.085857836 | 0.20718025 | 0.433482077 | 0.612573038 | 0.579457987 | 0.303091859 |
| Pcyt1a | S | 346 | 0.059138136 | 0.060128274 | -0.136646997 | -0.195785133 | -0.196775271 | 0.051409661 | 0.113258456 | 0.00072053 | 0.001207008 | 0.004785696 |
| Zc2hc1a | T | 243 | 0.029606443 | 0.021107516 | -0.06778525 | -0.097391694 | -0.088892766 | 0.718406005 | 0.78699402 | 0.114213001 | 0.282829823 | 0.297836878 |
| Phkb | S | 693 | -0.06519833 | 0.079298313 | -0.043443059 | 0.021755271 | -0.122741371 | 0.13056364 | 0.184595743 | 0.3484524 | 0.521094987 | 0.066496709 |
| Prr12 | T | 1053 | -0.005096802 | -0.022765416 | -0.025311972 | -0.020215171 | -0.002546556 | 0.868073668 | 0.741780267 | 0.662089095 | 0.724078987 | 0.974645971 |
| Foxc2 | S | 231 | 0.05822919 | -0.014095694 | -0.104546255 | -0.162775445 | -0.090450561 | 0.265320187 | 0.806846975 | 0.164564247 | 0.065574754 | 0.195169507 |
| Foxc2 | S | 239 | 0.05822919 | -0.014095694 | -0.104546255 | -0.162775445 | -0.090450561 | 0.265320187 | 0.806846975 | 0.164564247 | 0.065574754 | 0.195169507 |
| Rad18 | S | 103 | -0.041755945 | -0.007716553 | -0.000618977 | 0.041136968 | 0.007097576 | 0.530144159 | 0.773015612 | 0.974872914 | 0.530544614 | 0.749133477 |
| Ell | S | 299 | -0.032397174 | 0.000300794 | -0.007928484 | 0.02446869 | -0.008229278 | 0.365025568 | 0.993288889 | 0.884740418 | 0.647204949 | 0.876764924 |
| Champ1 | S | 605 | 0.110451738 | -0.104639404 | -0.012978852 | -0.12343059 | 0.091660552 | 0.035479214 | 0.004589334 | 0.707582015 | 0.034200939 | 0.04907585 |
| Rock1 | S | 456 | 0.00383237 | 0.022997129 | -0.123334146 | -0.127166516 | -0.146331274 | 0.964626513 | 0.762657381 | 0.480460305 | 0.459402791 | 0.399755478 |
| Fam172a | S | 400 | 0.045236728 | -0.072107587 | 0.010519931 | -0.034716796 | 0.082627519 | 0.635328191 | 0.089255922 | 0.674595165 | 0.710258227 | 0.069608133 |
| Phf8-ps Phf8 | S | 843 | 0.054416551 | -0.00703094 | -0.086168855 | -0.140585406 | -0.079137915 | 0.178014083 | 0.831223047 | 0.068909099 | 0.023368741 | 0.10147714 |
| Gtf3c1 | S | 1063 | -0.02216421 | -0.041351724 | 0.026530857 | 0.048695068 | 0.067882581 | 0.517323263 | 0.409803199 | 0.666939277 | 0.435305025 | 0.336510791 |
| Mdc1 | T | 875 | -0.089357894 | 0.022207162 | 0.00058692 | 0.089944814 | -0.021620242 | 0.337323942 | 0.633373038 | 0.993915554 | 0.389544906 | 0.767823819 |
| Srsf5 | S | 247 | 0.056491827 | -0.065900752 | 0.000283651 | -0.056208176 | 0.066184403 | 0.307067621 | 0.035838211 | 0.991732252 | 0.303676674 | 0.027410282 |
| Tssc4 | S | 57 | 0.015705572 | 0.020565676 | -0.103291233 | -0.118996805 | -0.123856909 | 0.800710325 | 0.72512488 | 0.307842607 | 0.276321444 | 0.254873482 |
| Tssc4 | S | 64 | 0.015705572 | 0.020565676 | -0.103291233 | -0.118996805 | -0.123856909 | 0.800710325 | 0.72512488 | 0.307842607 | 0.276321444 | 0.254873482 |
| Lpin2 | S | 186 | -0.033808233 | 0.043063152 | -0.053194499 | -0.019386266 | -0.096257651 | 0.217264167 | 0.144970677 | 0.254893099 | 0.652454557 | 0.087188221 |
| Lpin2 | S | 187 | -0.033808233 | 0.043063152 | -0.053194499 | -0.019386266 | -0.096257651 | 0.217264167 | 0.144970677 | 0.254893099 | 0.652454557 | 0.087188221 |
| Cenpu | S | 106 | -0.007018287 | 0.004883283 | -0.045793809 | -0.038775522 | -0.050677091 | 0.864499116 | 0.893679057 | 0.418250289 | 0.51141262 | 0.384428159 |
| Atp5a1 | S | 451 | -0.022877883 | -0.117602685 | 0.01897095 | 0.041848833 | 0.136573634 | 0.819402188 | 0.691499943 | 0.786708206 | 0.666061239 | 0.646236962 |
| Pcm1 | S | 1855 | -0.14321023 | -0.010269802 | 0.095087346 | 0.238297576 | 0.105357147 | 0.096442219 | 0.864896312 | 0.170963886 | 0.022954405 | 0.087107162 |
| Cd2bp2 | S | 49 | -0.081307611 | 0.003416095 | 0.014566772 | 0.095874383 | 0.011150677 | 0.3748889 | 0.920527388 | 0.789165238 | 0.329152457 | 0.836834109 |
| Htatsf1 | S | 697 | -0.101166998 | 0.044759673 | 0.006230238 | 0.107397235 | -0.038529435 | 0.26890599 | 0.648370116 | 0.946565364 | 0.105269411 | 0.587323767 |
| Iws1 | S | 261 | -0.010797166 | -0.0450504 | 0.027927563 | 0.038724729 | 0.072977963 | 0.511014753 | 0.105106338 | 0.248648247 | 0.123949983 | 0.036968691 |
| Mastl | S | 389 | -0.092092959 | 0.124982332 | -0.091273841 | 0.000819118 | -0.216256173 | 0.473124389 | 0.355325585 | 0.364504021 | 0.994107789 | 0.132759398 |
| Git1 | S | 371 | 0.025084369 | 0.028217154 | -0.092698953 | -0.117783322 | -0.120916107 | 0.533868045 | 0.480876411 | 0.087387286 | 0.039619501 | 0.038375494 |
| Atp5a1 | S | 22 | 0.080820032 | 0.134781919 | -0.305215591 | -0.386035624 | -0.439997511 | 0.531130804 | 0.379047105 | 0.126970903 | 0.07507074 | 0.055589022 |
| Ctdp1 | S | 530 | -0.092246118 | 0.015863723 | -0.015571826 | 0.076674292 | -0.031435549 | 0.625576387 | 0.708254873 | 0.71779782 | 0.678937984 | 0.167722504 |
| Frmd8 | S | 24 | -0.026363524 | 0.07404059 | -0.120774533 | -0.09441101 | -0.194815123 | 0.71015334 | 0.399559944 | 0.319394985 | 0.406067673 | 0.150992955 |
| Usp24 | S | 2077 | 0.149559733 | -0.001086636 | -0.149489505 | -0.299049237 | -0.148402869 | 0.066170288 | 0.987799804 | 0.009162663 | 0.024693435 | 0.115160867 |
| Rbm15 | S | 256 | -0.002531039 | -0.053593401 | 0.024177409 | 0.026708447 | 0.077770809 | 0.971759305 | 0.25932551 | 0.797973235 | 0.790366952 | 0.416335763 |
| Rbm15 | S | 258 | -0.002531039 | -0.053593401 | 0.024177409 | 0.026708447 | 0.077770809 | 0.971759305 | 0.25932551 | 0.797973235 | 0.790366952 | 0.416335763 |
| Mki67 | S | 2193 | 0.081403243 | -0.020750548 | -0.12543675 | -0.206839993 | -0.104686203 | 0.02369432 | 0.562840188 | 0.036052478 | 0.017851581 | 0.068919938 |
| Mki67 | T | 2190 | 0.081403243 | -0.020750548 | -0.12543675 | -0.206839993 | -0.104686203 | 0.02369432 | 0.562840188 | 0.036052478 | 0.017851581 | 0.068919938 |
| Axin1 | S | 495 | 0.00320471 | 0.053260148 | -0.096605676 | -0.099810385 | -0.149865823 | 0.957118243 | 0.406884299 | 0.143160804 | 0.185185117 | 0.078917892 |
| Sdhaf4 | S | 56 | 0.082565196 | -0.012573525 | -0.277913441 | -0.360478637 | -0.265339916 | 0.376706561 | 0.902469794 | 0.482677861 | 0.381674107 | 0.501412699 |
| Slc4a7 | S | 238 | 0.011913765 | -0.044737638 | -0.023287552 | -0.035201317 | 0.021450086 | 0.60249224 | 0.566986076 | 0.538024395 | 0.337974337 | 0.782814282 |
| Pkn1 | S | 920 | -0.061680037 | 0.029800137 | -0.010312337 | 0.0513677 | -0.040112473 | 0.240764737 | 0.470438351 | 0.746497503 | 0.320769813 | 0.359238207 |
| Srsf2 | S | 208 | 0.057229774 | -0.058324686 | -0.016730863 | -0.073960637 | 0.041593823 | 0.495465785 | 0.129985671 | 0.826615646 | 0.474143386 | 0.594011006 |
| Son | S | 1621 | -0.10919934 | -0.038496349 | 0.059703244 | 0.168902583 | 0.098199592 | 0.448196585 | 0.840020086 | 0.590961274 | 0.271440272 | 0.615053502 |
| Luc7l2 | S | 370 | -0.049160045 | 0.14842787 | -0.168914245 | -0.1197542 | -0.317342115 | 0.388932827 | 0.048161212 | 0.078335873 | 0.173226789 | 0.012950524 |
| Flnb | S | 2478 | -0.012784342 | 0.038561034 | -0.069961425 | -0.057177084 | -0.10852246 | 0.76466947 | 0.283753844 | 0.154238923 | 0.25086106 | 0.052321624 |
| Aak1 Gm38495 | S | 292 | -0.053077694 | 0.096511644 | -0.065665229 | -0.012587535 | -0.162176873 | 0.073698093 | 0.090972645 | 0.234219584 | 0.787314574 | 0.038952628 |
| Tut1 | S | 744 | 0.067045428 | -0.120092439 | -0.042066767 | -0.109112195 | 0.078025672 | 0.307636725 | 0.358831004 | 0.66885573 | 0.323278743 | 0.581707863 |
| Phactr4 | S | 557 | -0.012717968 | -0.097632123 | 0.165299087 | 0.178017055 | 0.26293121 | 0.669252556 | 0.001128441 | 0.058059773 | 0.044187013 | 0.027876619 |
| Topors | S | 99 | 0.06388541 | -0.04218869 | -0.065393785 | -0.129279195 | -0.023205094 | 0.108720195 | 0.150449548 | 0.268945477 | 0.076794452 | 0.662050786 |

| | | | | | | | | | | | | |
|---|---|---|---|---|---|---|---|---|---|---|---|---|
| Hirip3 | S | 163 | -0.104196972 | 0.062875451 | 0.025041284 | 0.129238257 | -0.037834166 | 0.045561735 | 0.427577838 | 0.472071596 | 0.028273055 | 0.606842422 |
| Rrp8 | S | 58 | -0.022306086 | -0.004779687 | -0.006071765 | 0.01623432 | -0.001292079 | 0.422469512 | 0.918309437 | 0.767169381 | 0.529153351 | 0.977419686 |
| Cd2bp2 | S | 196 | -0.037620149 | 0.035258578 | -0.020297213 | 0.017322936 | -0.055555792 | 0.033122989 | 0.020343744 | 0.110489417 | 0.192472981 | 0.002551317 |
| Hmox1 | S | 174 | 0.146109579 | 0.059854677 | -0.256644278 | -0.402753858 | -0.316498955 | 0.180529762 | 0.372159675 | 0.031458867 | 0.015643254 | 0.023158849 |
| Tcof1 | S | 593 | -0.018147578 | -0.0150231 | -0.00017704 | 0.017970538 | 0.014846061 | 0.624495613 | 0.482270869 | 0.993637684 | 0.601068453 | 0.293731159 |
| Zfp462 | S | 2158 | -0.170434872 | 0.082557373 | 0.038107655 | 0.208542527 | -0.044449718 | 0.120569684 | 0.336413386 | 0.759015936 | 0.140241822 | 0.681388844 |
| Prkar2a | S | 96 | -0.000462239 | 0.023252393 | -0.035379298 | -0.03491706 | -0.058631691 | 0.980371316 | 0.633044106 | 0.081568709 | 0.136516505 | 0.287476996 |
| Psip1 | S | 520 | -0.169164427 | 0.007268918 | 0.111288461 | 0.280452888 | 0.104019543 | 0.184819535 | 0.897829109 | 0.424141684 | 0.119565235 | 0.442552754 |
| Syne1 | S | 8281 | -0.006249538 | -0.032462117 | -0.016234573 | -0.009985035 | 0.016227544 | 0.957567946 | 0.526513145 | 0.920316501 | 0.956373051 | 0.918283257 |
| Syne1 | S | 8284 | -0.006249538 | -0.032462117 | -0.016234573 | -0.009985035 | 0.016227544 | 0.957567946 | 0.526513145 | 0.920316501 | 0.956373051 | 0.918283257 |
| Anxa6 | S | 173 | 0.208018812 | 0.103924417 | -0.601786515 | -0.809805327 | -0.705710932 | 0.312517555 | 0.667565507 | 0.265011117 | 0.170471806 | 0.210873438 |
| Chp1 | S | 6 | -0.050503259 | -0.016246694 | 0.025820729 | 0.076323988 | 0.042067423 | 0.315778479 | 0.554959065 | 0.457003737 | 0.165926504 | 0.194902229 |
| Polr2a | S | 1917 | 0.07055426 | -0.045624046 | -0.056397704 | -0.126951964 | -0.010773658 | 0.321475196 | 0.600011714 | 0.15010389 | 0.129030798 | 0.900433046 |
| Mdc1 | S | 1017 | -0.024393595 | -0.080369041 | 0.061403675 | 0.08579727 | 0.141772715 | 0.683683498 | 0.369051239 | 0.580870024 | 0.446866053 | 0.275704573 |
| Cc2d1a | S | 435 | -0.058795896 | 0.0281578 | 0.000154558 | 0.058950454 | -0.028003242 | 0.136586106 | 0.475318442 | 0.996857609 | 0.079428519 | 0.36867476 |
| Tenm2 Tenm3 | S | 43;42 | -0.012185648 | -0.022684236 | -0.005636154 | 0.006549494 | 0.017048082 | 0.760614659 | 0.58124596 | 0.872010758 | 0.783131723 | 0.519671863 |
| Lig1 | S | 49 | -0.119613721 | 0.228029985 | -0.113866934 | 0.005746786 | -0.341896919 | 0.341143524 | 0.137458843 | 0.101482507 | 0.958106214 | 0.068426534 |
| Kif18b | S | 499 | 0.080837787 | -0.022155107 | -0.094186269 | -0.175024056 | -0.072031162 | 0.505164944 | 0.777972232 | 0.259188181 | 0.173153827 | 0.095628868 |
| Camk2d | S | 494 | 0.009514552 | 0.117876089 | -0.126567333 | -0.136081885 | -0.244443423 | 0.86212925 | 0.208873077 | 0.030991249 | 0.08026783 | 0.047153415 |
| Nol7 | S | 122 | -0.060926671 | -0.038075233 | 0.054211534 | 0.115138205 | 0.092286767 | 0.319892243 | 0.466521514 | 0.351401926 | 0.080217114 | 0.092660827 |
| Dclk1 | S | 332 | -0.01535891 | 0.045346175 | -0.074951591 | -0.059592681 | -0.120297766 | 0.655060755 | 0.172022561 | 0.121543906 | 0.206820391 | 0.040388668 |
| Cdk13 | S | 224 | 0.014564439 | 0.011122641 | -0.074757843 | -0.089322281 | -0.085880484 | 0.791178138 | 0.875525633 | 0.319651625 | 0.190481267 | 0.260467463 |
| Ticrr | S | 1439 | 0.029721544 | -0.053188219 | -0.02772799 | -0.057449533 | 0.02546023 | 0.310607798 | 0.491770538 | 0.283041354 | 0.078334293 | 0.728432807 |
| Ercc6l | S | 998 | -0.036272031 | 0.037436052 | -0.063269987 | -0.026997956 | -0.100706039 | 0.587038212 | 0.48004004 | 0.46037003 | 0.754325975 | 0.257886553 |
| Akt1 | S | 122 | -0.033766916 | 0.074492204 | -0.104800577 | -0.071033661 | -0.17929278 | 0.789700864 | 0.443574565 | 0.307492875 | 0.593375758 | 0.132257555 |
| Rprd2 | S | 504 | 0.115128856 | -0.049507666 | -0.12754726 | -0.242676116 | -0.078039594 | 0.052811057 | 0.209598827 | 0.142495835 | 0.032780551 | 0.310883927 |
| Srrm2 | S | 1554 | -0.009468986 | 0.007331462 | -0.036181006 | -0.02671202 | -0.043512468 | 0.712185333 | 0.824541671 | 0.426778678 | 0.562620837 | 0.395637905 |
| Dst | S | 2560 | -0.03361022 | -0.058736647 | 0.065483093 | 0.099093312 | 0.12421974 | 0.447746925 | 0.149825571 | 0.244130743 | 0.11730017 | 0.065405505 |
| Dst | S | 2563 | -0.03361022 | -0.058736647 | 0.065483093 | 0.099093312 | 0.12421974 | 0.447746925 | 0.149825571 | 0.244130743 | 0.11730017 | 0.065405505 |
| Ppp6r3 | T | 820 | 0.016142224 | 0.00832673 | -0.086479695 | -0.10262192 | -0.094806425 | 0.790118358 | 0.893270013 | 0.387277103 | 0.305609134 | 0.33936787 |
| Pwp2 | S | 902 | 0.044467466 | -0.074114528 | 0.013793713 | -0.030673753 | 0.087908241 | 0.367530492 | 0.041624791 | 0.624269805 | 0.504183729 | 0.023859776 |
| Dgcr8 | S | 238 | 0.14892663 | -0.060560394 | -0.112202959 | -0.26112959 | -0.051642565 | 0.425159388 | 0.687356432 | 0.240263596 | 0.215151579 | 0.742027385 |
| Sik3 | T | 411 | -0.044223437 | 0.165438357 | -0.194227475 | -0.150004038 | -0.359665833 | 0.567044414 | 0.06930873 | 0.080574545 | 0.125888726 | 0.013841906 |
| N4bp3 | S | 172 | -0.027307523 | 0.005389163 | -0.028290359 | -0.000982836 | -0.033679522 | 0.756150769 | 0.943856592 | 0.584046663 | 0.991549994 | 0.690404027 |
| Eef2k | S | 469 | -0.11236252 | 0.081558588 | 0.021259323 | 0.133621843 | -0.060299266 | 0.003306724 | 0.044383838 | 0.599755485 | 0.03631982 | 0.214082739 |
| Nhsl1 | S | 713 | -0.009470051 | -0.010076456 | -0.043637746 | -0.034167695 | -0.03356129 | 0.918585364 | 0.801184725 | 0.730002075 | 0.810507628 | 0.785766302 |
| Nhsl1 | S | 717 | -0.009470051 | -0.010076456 | -0.043637746 | -0.034167695 | -0.03356129 | 0.918585364 | 0.801184725 | 0.730002075 | 0.810507628 | 0.785766302 |
| Gsn | T | 651 | -0.121387672 | 0.250314854 | -0.129663826 | -0.008276154 | -0.37997868 | 0.182814924 | 0.111287354 | 0.169220021 | 0.929508249 | 0.04155816 |
| Oxr1 | S | 488 | -0.024188338 | 0.023026145 | -0.043348062 | -0.019159724 | -0.066374207 | 0.697472585 | 0.761655338 | 0.480808784 | 0.782447966 | 0.434031873 |
| Lasp1 | S | 134 | 0.187762888 | -0.061799261 | -0.222251968 | -0.410014857 | -0.160452707 | 0.041605666 | 0.413171881 | 0.090195932 | 0.028766379 | 0.182508633 |
| Sash1 | S | 241 | -0.064023708 | 0.014359142 | 0.015030321 | 0.079054029 | 0.000671179 | 0.21683232 | 0.773485268 | 0.786665314 | 0.116352687 | 0.987064541 |
| Chd4 | S | 302 | -0.143608982 | -0.051718108 | 0.149751106 | 0.293360088 | 0.201469214 | 0.064054435 | 0.450663528 | 0.06522125 | 0.005620966 | 0.008527025 |
| Ticrr | S | 1187 | -0.056129754 | -0.038687987 | 0.129026411 | 0.185156165 | 0.167714398 | 0.172929065 | 0.181233214 | 0.327916055 | 0.202204235 | 0.237262132 |
| Eif4g2 | S | 16 | -0.05967171 | -0.019521608 | 0.016229408 | 0.075901118 | 0.035751016 | 0.574385557 | 0.846904904 | 0.871103885 | 0.151829944 | 0.210636002 |
| Chaf1b | T | 426 | 0.116832261 | -0.083859811 | -0.065051036 | -0.181883297 | 0.018808775 | 0.204869004 | 0.262584009 | 0.281890186 | 0.082669612 | 0.751709971 |
| Ddx21 | T | 90 | 0.202279252 | -0.061039553 | -0.214292684 | -0.416571936 | -0.153253131 | 0.042257629 | 0.318748491 | 0.084840635 | 0.014733873 | 0.184794617 |
| Pafah1b2 | T | 220 | -0.00642704 | 0.029191451 | -0.112494229 | -0.106067188 | -0.14168568 | 0.961084207 | 0.75792946 | 0.536151608 | 0.571913864 | 0.425057269 |
| Bsg | T | 358 | -0.086797147 | -0.146935873 | 0.182938054 | 0.2697352 | 0.329873927 | 0.568955487 | 0.209997729 | 0.241620391 | 0.141754481 | 0.064960091 |
| Ncor2 | S | 2161 | -0.077050323 | -0.061585974 | 0.089070668 | 0.166120991 | 0.150656641 | 0.068696338 | 0.255314729 | 0.031885636 | 0.012911747 | 0.054647657 |
| Tbx3 | S | 705 | -0.005770608 | -0.081535904 | 0.052142135 | 0.057912743 | 0.133678039 | 0.91554538 | 0.181500265 | 0.445218026 | 0.28314508 | 0.059415398 |
| Dst | S | 1080 | 0.067514548 | -0.077207637 | -0.011310688 | -0.078825236 | 0.065896949 | 0.269293958 | 0.08057051 | 0.823937556 | 0.273301542 | 0.274443434 |
| Mtmr2 | S | 51 | -0.061976875 | -0.018693763 | 0.049921535 | 0.111898411 | 0.068615299 | 0.33687829 | 0.755183693 | 0.666806099 | 0.380843037 | 0.57287822 |
| Pakap | S | 494 | 0.024061565 | -0.019265725 | -0.039980785 | -0.06404235 | -0.02071506 | 0.80373182 | 0.687594031 | 0.456470147 | 0.535118187 | 0.712802491 |
| Amotl1 | S | 253 | -0.037418981 | 0.063164465 | -0.07885563 | -0.041436649 | -0.142020095 | 0.520552661 | 0.217775261 | 0.224382821 | 0.553174197 | 0.073554239 |
| Arap1 | S | 232 | -0.12877772 | 0.082941605 | 0.009907091 | 0.138684811 | -0.073034514 | 0.062823937 | 0.202662189 | 0.821084934 | 0.056164125 | 0.223039797 |
| Usp1 | S | 474 | -0.083572026 | 0.076299201 | -0.03544084 | 0.048131186 | -0.111740041 | 0.238123696 | 0.25612752 | 0.465803653 | 0.484377094 | 0.142346678 |
| Wac | S | 470 | -0.014764147 | -0.06333362 | 0.022652969 | 0.037417117 | 0.085986589 | 0.869087188 | 0.515099614 | 0.84284008 | 0.731193348 | 0.462711866 |
| Clk1 | S | 137 | 0.095190958 | -0.057500236 | -0.079311119 | -0.174502078 | -0.021810883 | 0.66889582 | 0.666146342 | 0.510284676 | 0.434619544 | 0.838961966 |
| Clk1 | S | 139 | 0.095190958 | -0.057500236 | -0.079311119 | -0.174502078 | -0.021810883 | 0.66889582 | 0.666146342 | 0.510284676 | 0.434619544 | 0.838961966 |
| Lamb1 | S | 1666 | -0.030199061 | 0.022263336 | -0.025577892 | 0.004621169 | -0.047841228 | 0.325997905 | 0.678095636 | 0.692553102 | 0.94200722 | 0.532120612 |
| Tanc2 | T | 293 | 0.018277398 | 0.156564477 | -0.290385998 | -0.308663395 | -0.446950475 | 0.829714166 | 0.046764912 | 0.048100726 | 0.043317036 | 0.013522803 |
| Rfc1 | S | 309 | -0.020352565 | 0.014103411 | -0.045846787 | -0.025494222 | -0.059950197 | 0.820244541 | 0.894122737 | 0.556168214 | 0.774058155 | 0.57778232 |
| Smg1 | T | 3547 | 0.009549776 | -0.034282252 | -0.027634661 | -0.037184436 | 0.006647592 | 0.896690826 | 0.53806598 | 0.820948402 | 0.763407987 | 0.953862088 |
| Cep170 | S | 1521 | 0.062274759 | -0.027820215 | -0.058847302 | -0.121122061 | -0.031027087 | 0.618662925 | 0.741964723 | 0.293286927 | 0.353907952 | 0.671914265 |
| Slx4 | S | 426 | 0.105019739 | -0.05936318 | -0.087084184 | -0.192103923 | -0.027721003 | 0.259766083 | 0.304216321 | 0.362794732 | 0.127392848 | 0.76005416 |
| Tnrc6a | S | 1804 | -0.01015145 | -0.07812509 | 0.115507175 | 0.125658624 | 0.193632265 | 0.86358813 | 0.162521606 | 0.635030445 | 0.610590085 | 0.450966321 |
| Irs1 | S | 887 | -0.043016473 | 0.030666064 | -0.010767309 | 0.032249164 | -0.041433374 | 0.231378473 | 0.683589554 | 0.839730587 | 0.514339901 | 0.607118689 |
| Brca1 | T | 1126 | -0.026946292 | 0.013525173 | -0.063465924 | -0.036519631 | -0.076991096 | 0.794036761 | 0.902853979 | 0.719269453 | 0.847868244 | 0.694126244 |
| Toe1 | T | 411 | -0.044250053 | 0.01556157 | -0.056898144 | -0.012648091 | -0.072459714 | 0.722394606 | 0.733351728 | 0.723308926 | 0.945020655 | 0.649065738 |
| Gpatch2 | S | 114 | -0.089835157 | 0.041536713 | 0.016802194 | 0.106637352 | -0.024734519 | 0.104196022 | 0.403570309 | 0.743839951 | 0.095115383 | 0.637175741 |
| Gpatch2 | S | 116 | -0.089835157 | 0.041536713 | 0.016802194 | 0.106637352 | -0.024734519 | 0.104196022 | 0.403570309 | 0.743839951 | 0.095115383 | 0.637175741 |
| Spcs2 | S | 29 | 0.067675937 | -0.07475638 | -0.00913409 | -0.076810027 | 0.06562229 | 0.698446951 | 0.270761638 | 0.91323688 | 0.668221292 | 0.436762325 |
| Vim | S | 56 | -0.003884062 | -0.017931544 | -0.013028469 | -0.009144406 | 0.004903075 | 0.906505388 | 0.617430476 | 0.667242506 | 0.757122553 | 0.878878613 |
| Phkg2 | T | 402 | -0.066979389 | 0.164676251 | -0.18331163 | -0.116332241 | -0.347987881 | 0.420491384 | 0.113983732 | 0.201153295 | 0.383701055 | 0.053280334 |
| Fam129b | T | 655 | 0.028250067 | -0.000616561 | -0.060803433 | -0.0890535 | -0.060186873 | 0.637993876 | 0.992717771 | 0.237813731 | 0.232095983 | 0.441526189 |
| Mpdz | T | 911 | -0.029528015 | -0.021309558 | 0.006310272 | 0.035838287 | 0.02761983 | 0.714065791 | 0.55355168 | 0.88888359 | 0.662044509 | 0.499435947 |
| Srrm2 | T | 487 | 0.005788502 | -0.042081351 | -0.017150369 | -0.022938872 | 0.024930981 | 0.959643306 | 0.578643247 | 0.873775691 | 0.870969307 | 0.829513667 |
| Zc3h13 | S | 953 | -0.052644609 | -0.003067256 | 0.021508447 | 0.074153056 | 0.024575702 | 0.246042982 | 0.88072948 | 0.586547167 | 0.17445156 | 0.51184967 |
| Nasp | S | 617 | -0.106713385 | 0.084289963 | -0.013551955 | 0.09316143 | -0.097841918 | 0.074769746 | 0.174239706 | 0.878314946 | 0.304524671 | 0.287978553 |
| Aqp1 | S | 247 | 0.049116747 | -0.039349517 | -0.043785172 | -0.092901919 | -0.004435655 | 0.058423018 | 0.115447056 | 0.198243208 | 0.036250923 | 0.877196188 |
| Peak1 | T | 501 | 0.104299944 | -0.094336797 | -0.085272831 | -0.189572775 | 0.009063966 | 0.366786668 | 0.530290859 | 0.444915685 | 0.10371145 | 0.94683965 |
| Tomm70a | S | 94 | -0.074450917 | 0.014170941 | 0.052855347 | 0.127306264 | 0.038684405 | 0.057591653 | 0.754591942 | 0.420739345 | 0.117861232 | 0.577040102 |
| Rab11fip1 | S | 357 | -0.097962289 | 0.037710762 | -0.018417611 | 0.079544678 | -0.056128373 | 0.391195742 | 0.454866121 | 0.859277636 | 0.562772298 | 0.60393319 |
| Nckap5l | S | 573 | 0.124590426 | -0.101987118 | -0.17519928 | -0.299789705 | -0.073212161 | 0.35896699 | 0.603373429 | 0.570005252 | 0.354283212 | 0.816075878 |
| Trio | S | 2458 | -0.068726287 | 0.062809341 | -0.029685042 | 0.039041245 | -0.092494383 | 0.129019955 | 0.170462832 | 0.512387166 | 0.429440753 | 0.111443763 |
| Nhsl1 | S | 1079 | -0.069743974 | 0.046145794 | -0.048398157 | 0.021345817 | -0.094543951 | 0.473566317 | 0.677187013 | 0.851923795 | 0.932388429 | 0.715890686 |
| Mcm6 | S | 804 | -0.214889241 | 0.12493466 | 0.099371461 | 0.314260702 | -0.025563199 | 0.019544012 | 0.131357738 | 0.576814414 | 0.159873006 | 0.877702817 |
| Mphosph8 | S | 267 | 0.03750168 | -0.047218416 | -0.04138624 | -0.078887921 | 0.005832176 | 0.441119892 | 0.524932067 | 0.446003895 | 0.191884539 | 0.937985377 |
| Naa15 | S | 855 | -0.013855057 | -0.014861391 | -0.006935528 | 0.006919529 | 0.007925863 | 0.738818571 | 0.69113243 | 0.771882242 | 0.870008765 | 0.835921403 |
| Eif4g1 | T | 214 | 0.035238173 | -0.015166618 | -0.047167067 | -0.08240524 | -0.032000448 | 0.149913957 | 0.734331298 | 0.053816725 | 0.006812131 | 0.472329568 |
| Ctps2 | S | 568 | -0.129568783 | 0.043088094 | 0.087299357 | 0.21686814 | 0.044211263 | 0.022280743 | 0.512132561 | 0.243508075 | 0.037055998 | 0.587527262 |
| Ctps2 | S | 574 | -0.129568783 | 0.043088094 | 0.087299357 | 0.21686814 | 0.044211263 | 0.022280743 | 0.512132561 | 0.243508075 | 0.037055998 | 0.587527262 |
| Dcaf10 | S | 100 | 0.098299805 | -0.091311456 | -0.141966612 | -0.240266417 | -0.050655156 | 0.128346897 | 0.516089675 | 0.353390465 | 0.17027271 | 0.774407166 |
| Cic | T | 1643 | 0.055001661 | -0.03449022 | -0.027100263 | -0.082101924 | 0.007389957 | 0.432163118 | 0.143663097 | 0.503839161 | 0.28864713 | 0.847587071 |

| | | | | | | | | | | | | |
|---|---|---|---|---|---|---|---|---|---|---|---|---|
| Tgif1 | S | 11 | -0.035056217 | -0.012864838 | -0.011367143 | 0.023689074 | 0.001497695 | 0.812284322 | 0.796375902 | 0.870168312 | 0.872267176 | 0.97766476 |
| Tgif1 | S | 13 | -0.035056217 | -0.012864838 | -0.011367143 | 0.023689074 | 0.001497695 | 0.812284322 | 0.796375902 | 0.870168312 | 0.872267176 | 0.97766476 |
| Tgif1 | S | 15 | -0.035056217 | -0.012864838 | -0.011367143 | 0.023689074 | 0.001497695 | 0.812284322 | 0.796375902 | 0.870168312 | 0.872267176 | 0.97766476 |
| Rpsa | S | 43 | 0.047525018 | -0.031943676 | -0.099075392 | -0.14660041 | -0.067131716 | 0.405710847 | 0.617454771 | 0.500159023 | 0.339403626 | 0.633062901 |
| Pcyt1a | S | 321 | 0.079623742 | 0.102807354 | -0.226925203 | -0.306548946 | -0.329732557 | 0.434703125 | 0.309446754 | 0.066973364 | 0.02007184 | 0.015960814 |
| Limk2 | S | 297 | 0.012314767 | -0.024634019 | -0.025891295 | -0.038206062 | -0.001257276 | 0.880028355 | 0.363883015 | 0.869011966 | 0.820437452 | 0.993538974 |
| Wnk1 | S | 2007 | -0.06042275 | 0.041575137 | -0.014738569 | 0.045684181 | -0.056313706 | 0.161247459 | 0.303088791 | 0.724657382 | 0.379037099 | 0.298938478 |
| Arhgap39 | S | 362 | 0.042547107 | -0.035610277 | -0.034926452 | -0.077473559 | 0.000683826 | 0.305863325 | 0.302646291 | 0.331627228 | 0.059519198 | 0.967341279 |
| Zbtb33 | S | 188 | -0.042130411 | -0.063876831 | 0.107259145 | 0.149389557 | 0.171135976 | 0.612206682 | 0.37350154 | 0.645979921 | 0.540299869 | 0.485609189 |
| Cav1 | S | 9 | -0.066476926 | 0.020041833 | 0.035185168 | 0.101662094 | 0.015143335 | 0.066668322 | 0.58618876 | 0.559149967 | 0.156101997 | 0.79000167 |
| Mki67 | S | 286 | -0.000642767 | 0.048672597 | -0.075833372 | -0.075190606 | -0.12450597 | 0.993412395 | 0.244671001 | 0.080428899 | 0.357186231 | 0.005779475 |
| Pard3b | T | 810 | -0.050821166 | -0.007950334 | 0.051355845 | 0.102177011 | 0.059306179 | 0.211725337 | 0.93979114 | 0.733945263 | 0.51452254 | 0.728550011 |
| Ppp1r12a | T | 596 | 0.080204977 | -0.018264076 | -0.107494235 | -0.187699212 | -0.089230159 | 0.272008697 | 0.795511421 | 0.171454651 | 0.012450531 | 0.054019907 |
| Itprid2 | S | 667 | -0.02263778 | 0.00292573 | -0.008547418 | 0.014090362 | -0.011473148 | 0.412572827 | 0.872623673 | 0.700487405 | 0.60282636 | 0.550256447 |
| Fam91a1 | S | 355 | -0.107121112 | 0.020234591 | 0.014862736 | 0.121983848 | -0.005371856 | 0.417851592 | 0.759557078 | 0.819572497 | 0.354754104 | 0.913306955 |
| Wipf2 | S | 267 | 0.073111665 | 0.005651906 | -0.073010281 | -0.146121946 | -0.078662187 | 0.460706668 | 0.868536412 | 0.068274482 | 0.203419926 | 0.077298841 |
| Qrich1 | S | 658 | 0.010081147 | -0.025973844 | -0.021217189 | -0.031298336 | 0.004756655 | 0.877274699 | 0.517232997 | 0.729015074 | 0.702165331 | 0.941633049 |
| Hectd1 | S | 1572 | -0.092255621 | -0.125833215 | 0.172095851 | 0.264351471 | 0.297929065 | 0.427909765 | 0.197064543 | 0.166407708 | 0.06259736 | 0.040254034 |
| Aftph | S | 680 | 0.067035828 | 0.072286792 | -0.190876848 | -0.257912676 | -0.26316364 | 0.131108075 | 0.417047029 | 0.035984862 | 0.014956926 | 0.043655285 |
| Afap1 | T | 342 | 0.050004658 | 0.004902348 | -0.068173863 | -0.118178521 | -0.073076212 | 0.744124574 | 0.921468852 | 0.227991737 | 0.472103134 | 0.273749379 |
| Pdlim1 | S | 90 | -0.021813237 | 0.04737062 | -0.051477752 | -0.029664515 | -0.098848372 | 0.637819259 | 0.593096027 | 0.345289976 | 0.448013346 | 0.290954196 |
| Stk39 | S | 382 | 0.052175517 | -0.034899529 | -0.036690716 | -0.088866233 | -0.001791187 | 0.649198812 | 0.55194466 | 0.406865675 | 0.455952839 | 0.973992907 |
| Stk39 | S | 383 | 0.052175517 | -0.034899529 | -0.036690716 | -0.088866233 | -0.001791187 | 0.649198812 | 0.55194466 | 0.406865675 | 0.455952839 | 0.973992907 |
| 0610010F05Rik | S | 600 | 0.035650204 | 0.023403284 | -0.090612985 | -0.12626319 | -0.114016269 | 0.544929274 | 0.552675146 | 0.117642798 | 0.101624482 | 0.072876988 |
| Epb41l2 | S | 153 | 0.001922342 | 0.049928945 | -0.110030607 | -0.111952949 | -0.159959552 | 0.935339983 | 0.194700852 | 0.131752222 | 0.124542898 | 0.05391292 |
| Thoc5 | S | 28 | 0.004550674 | -0.045149941 | -0.004617674 | -0.009168348 | 0.040532267 | 0.943598942 | 0.50046636 | 0.957084422 | 0.911558301 | 0.633518609 |
| Gpam | S | 694 | 0.019474485 | -0.03814534 | 0.01048053 | -0.008993955 | 0.048625869 | 0.655060553 | 0.076280469 | 0.801284027 | 0.864474565 | 0.290321078 |
| Atf7ip | S | 591 | -0.07939577 | 0.079902501 | -0.028387812 | 0.051007958 | -0.108290313 | 0.061929231 | 0.134244175 | 0.678817439 | 0.483536515 | 0.199864367 |
| Tex2 | T | 268 | 0.034921175 | 0.022097987 | -0.124461149 | -0.159382325 | -0.146559136 | 0.664920306 | 0.889254285 | 0.385431161 | 0.300747923 | 0.456984952 |
| Zdhhc5 | S | 554 | 0.018123106 | -0.193525411 | 0.109343409 | 0.091220303 | 0.30286882 | 0.670548077 | 0.136168767 | 0.301294072 | 0.376825584 | 0.056921186 |
| Arhgap39 | S | 683 | -0.007875649 | 0.035116572 | -0.041603018 | -0.033727369 | -0.07671959 | 0.886486831 | 0.481767502 | 0.10981848 | 0.560149382 | 0.188484398 |
| Dennd1b | S | 516 | -0.160958592 | 0.11213171 | 0.036505353 | 0.197463945 | -0.075626358 | 0.018041775 | 0.132094226 | 0.523253634 | 0.015322334 | 0.233139122 |
| Hipk1 | S | 871 | -0.04192028 | -0.022986166 | -0.008478433 | 0.033441847 | 0.014507734 | 0.777509165 | 0.900346514 | 0.946030943 | 0.849438579 | 0.943702633 |
| Hnrnpc | S | 247 | 0.130780733 | -0.078882539 | -0.040818115 | -0.171598848 | 0.038064424 | 0.451229786 | 0.237940125 | 0.536561497 | 0.340003163 | 0.436646426 |
| Cbx8 | S | 164 | 0.013486215 | -0.024817558 | -0.031575407 | -0.045061621 | -0.006757849 | 0.707236024 | 0.57708064 | 0.611688842 | 0.491577886 | 0.918960862 |
| Srrm2 | T | 1448 | 0.034154045 | -0.071462664 | 0.012343498 | -0.021810547 | 0.083806162 | 0.402000202 | 0.073006745 | 0.567841936 | 0.557142304 | 0.060538521 |
| Pdcd5 | S | 119 | 0.025359588 | 0.056603299 | -0.132687797 | -0.158047386 | -0.189291096 | 0.521576001 | 0.432919234 | 0.131793641 | 0.096094031 | 0.078941829 |
| Jmjd1c | S | 471 | -0.067923447 | -0.016323939 | 0.04014479 | 0.108068237 | 0.056468729 | 0.257114603 | 0.615505249 | 0.297924002 | 0.117796851 | 0.190451424 |
| Mki67 | T | 1243 | 0.031740105 | -0.003905571 | -0.094142998 | -0.125883103 | -0.090237427 | 0.117174594 | 0.876481892 | 0.167560168 | 0.106844942 | 0.179327417 |
| Pcyt1a | S | 331 | -0.105576509 | 0.094473946 | -0.029628384 | 0.075948126 | -0.124102329 | 0.027923251 | 0.034528896 | 0.609971098 | 0.219567743 | 0.089309059 |
| Ubn2 | S | 678 | 0.045835545 | -0.109659756 | 0.038106383 | -0.007729162 | 0.147766138 | 0.360674679 | 0.04707774 | 0.382825088 | 0.87278034 | 0.025531593 |
| Psen1 | S | 353 | -0.129333918 | -0.005076197 | 0.073170659 | 0.202504576 | 0.078246856 | 0.265066665 | 0.942476335 | 0.395907424 | 0.140085828 | 0.434496882 |
| Ubl7 | S | 242 | 0.023495346 | -0.039911009 | -0.050420713 | -0.073916059 | -0.010509704 | 0.551696488 | 0.549457397 | 0.696854114 | 0.572949258 | 0.936818895 |
| Tprn | S | 196 | -0.02905337 | -0.024069194 | 0.039698571 | 0.068751942 | 0.063767766 | 0.300305171 | 0.527921917 | 0.610985977 | 0.400732325 | 0.438572602 |
| Rbm4b Rbm4 Gm21S | S | 86;86 | 0.049676672 | 0.028637255 | -0.166255182 | -0.215931854 | -0.194892437 | 0.594237897 | 0.648293224 | 0.307570125 | 0.21603541 | 0.249880466 |
| Bcl9l | S | 1018 | -0.039439208 | 0.000459841 | -0.025948166 | 0.013491042 | -0.026408007 | 0.666828868 | 0.997349942 | 0.924359839 | 0.961053103 | 0.927346594 |
| Sox13 | S | 326 | -0.07427511 | -0.001603746 | 0.043083825 | 0.117358935 | 0.044687571 | 0.357588979 | 0.985475969 | 0.75221909 | 0.383683567 | 0.727937077 |
| Kdm4a | S | 357 | -0.094823188 | 0.008702092 | 0.035353664 | 0.130176853 | 0.026651572 | 0.404519457 | 0.927166724 | 0.766443362 | 0.293169504 | 0.792678391 |
| Palld | S | 779 | -0.023902809 | 0.038927594 | -0.028598639 | -0.004695829 | -0.067526233 | 0.219494373 | 0.230452189 | 0.188372251 | 0.806567243 | 0.08041512 |
| Huwe1 | S | 2918 | -0.0652424 | -0.00573769 | 0.02370601 | 0.08894841 | 0.0294437 | 0.49316395 | 0.936961692 | 0.836193892 | 0.455466417 | 0.772059378 |
| Spag9 | T | 292 | 0.182754934 | 0.212611964 | -0.284769001 | -0.467523935 | -0.497380965 | 0.265764361 | 0.004944424 | 2.96558E-05 | 0.058500433 | 0.00130955 |
| Map4 | S | 598 | 0.028131876 | 0.008117934 | -0.058507506 | -0.086639382 | -0.06662544 | 0.558588247 | 0.82936074 | 0.146208006 | 0.098683911 | 0.018532033 |
| Clic1 | S | 156 | -0.050871356 | 0.053823915 | -0.06656804 | -0.015696685 | -0.120391956 | 0.154990966 | 0.047999437 | 0.349433556 | 0.810142234 | 0.157458722 |
| Ctr9 | S | 1037 | 0.051783161 | -0.064400714 | -0.003061658 | -0.054844819 | 0.061339056 | 0.101834843 | 0.020781821 | 0.951317182 | 0.324771936 | 0.278663578 |
| Arhgap12 | S | 163 | -0.043570989 | -0.020973188 | 0.024995055 | 0.068566044 | 0.045968243 | 0.469118596 | 0.732867151 | 0.699582872 | 0.405267014 | 0.579118564 |
| Mast2 Mast4 | S | 747;862 | 0.006422569 | -0.038405813 | -0.006276163 | -0.012698732 | 0.03212965 | 0.927724114 | 0.487642833 | 0.915898731 | 0.869687463 | 0.617757758 |
| Eef2k | S | 444 | -0.250856766 | 0.14563935 | 0.123433043 | 0.374289809 | -0.022206306 | 0.012846289 | 0.241115767 | 0.235513191 | 0.018634076 | 0.85894114 |
| Ankzf1 | S | 385 | -0.071728451 | 0.025624427 | -0.015462915 | 0.056265536 | -0.041087342 | 0.498699893 | 0.702621928 | 0.907795158 | 0.693698511 | 0.740745557 |
| Golga5 | S | 153 | 0.016834759 | -0.021921167 | -0.026796205 | -0.043630964 | -0.004875038 | 0.796715524 | 0.575340481 | 0.591321917 | 0.52595522 | 0.909025333 |
| Atp2b1 Atp2b2 | Y | 1129;1107 | -0.15354808 | 0.109848429 | -0.025584538 | 0.127963543 | -0.135432967 | 0.135440443 | 0.104569812 | 0.618909118 | 0.194986147 | 0.047241317 |
| Champ1 | S | 603 | 0.079953259 | -0.08527407 | -0.027692717 | -0.107645976 | 0.057581352 | 0.007799626 | 0.004923729 | 0.612946378 | 0.132038017 | 0.327760629 |
| Prune2 | S | 2638 | -0.058023397 | 0.024263892 | 0.015188378 | 0.073211775 | -0.009075513 | 0.118273888 | 0.594259202 | 0.704319227 | 0.147293819 | 0.861690036 |
| Ccdc102a | S | 26 | 0.093956506 | 0.066427903 | -0.165098788 | -0.259055294 | -0.231526691 | 0.230400147 | 0.772377641 | 0.066622092 | 0.02717669 | 0.364780975 |
| Ccdc102a | S | 28 | 0.093956506 | 0.066427903 | -0.165098788 | -0.259055294 | -0.231526691 | 0.230400147 | 0.772377641 | 0.066622092 | 0.02717669 | 0.364780975 |
| Eif3b | T | 31 | -0.169003095 | 0.081839981 | 0.015958405 | 0.1849615 | -0.065881575 | 0.222835691 | 0.337226652 | 0.90718042 | 0.296868071 | 0.65439056 |
| Zcchc7 | S | 221 | 0.157462857 | -0.269916504 | 0.081330273 | -0.076132583 | 0.351246777 | 0.32574487 | 0.09811465 | 0.593806019 | 0.442564466 | 0.016550324 |
| Sema6d | S | 734 | 0.154822283 | -0.088340993 | -0.066205476 | -0.221027758 | 0.022135517 | 0.235493602 | 0.224043922 | 0.319954318 | 0.131540433 | 0.651165728 |
| Agfg2 | S | 451 | -0.112463895 | 0.123702219 | -0.083285836 | 0.029178059 | -0.206988055 | 0.556084413 | 0.237378868 | 0.270789701 | 0.87602567 | 0.097998681 |
| Abcf1 | S | 138 | 0.028915934 | 0.014291958 | -0.053700843 | -0.082616778 | -0.067992801 | 0.493965821 | 0.572594499 | 0.053851524 | 0.122082616 | 0.059220811 |
| Srrm2 | S | 2671 | 0.049038004 | -0.035313663 | -0.086568537 | -0.135606541 | -0.051254874 | 0.153248817 | 0.405784596 | 0.381497535 | 0.218522967 | 0.585108635 |
| Pkd1l3 | S | 1033 | 0.010986439 | -0.003284195 | -0.064134637 | -0.075121076 | -0.060850442 | 0.816956729 | 0.955168185 | 0.540205937 | 0.465191238 | 0.557436174 |
| Add2 | S | 614 | -0.01853912 | -0.094953556 | 0.074050397 | 0.092589518 | 0.169003954 | 0.717622803 | 0.065042158 | 0.088238901 | 0.102508697 | 0.011100767 |
| Msh6 | S | 285 | 0.096908505 | -0.144725372 | -0.02361736 | -0.120525865 | 0.121108012 | 0.076635455 | 0.087773728 | 0.817497367 | 0.29584956 | 0.31002987 |
| Msh6 | S | 287 | 0.096908505 | -0.144725372 | -0.02361736 | -0.120525865 | 0.121108012 | 0.076635455 | 0.087773728 | 0.817497367 | 0.29584956 | 0.31002987 |
| Reps1 | S | 536 | -0.137692622 | 0.059813435 | -0.005617447 | 0.132075175 | -0.065430882 | 0.292565516 | 0.305049977 | 0.941272953 | 0.318514186 | 0.367395641 |
| Dixdc1 | S | 287 | -0.158631512 | -0.006642678 | 0.117535273 | 0.276166785 | 0.124177951 | 0.224115984 | 0.915123119 | 0.356217744 | 0.10969321 | 0.330976613 |
| Riok3 | S | 125 | -0.056399701 | 0.03798843 | -0.023935145 | 0.032464556 | -0.061923575 | 0.235854292 | 0.261161906 | 0.640223043 | 0.5546607 | 0.24821035 |
| Riok3 | S | 127 | -0.056399701 | 0.03798843 | -0.023935145 | 0.032464556 | -0.061923575 | 0.235854292 | 0.261161906 | 0.640223043 | 0.5546607 | 0.24821035 |
| Riok3 | S | 128 | -0.056399701 | 0.03798843 | -0.023935145 | 0.032464556 | -0.061923575 | 0.235854292 | 0.261161906 | 0.640223043 | 0.5546607 | 0.24821035 |
| Inpp5f | S | 908 | -0.105999552 | 0.076173012 | -0.063700867 | 0.042298685 | -0.13987388 | 0.455170997 | 0.346125867 | 0.675752849 | 0.804140601 | 0.362709031 |
| Flna | S | 1055 | 0.092394122 | -0.092759646 | -0.083503402 | -0.175897524 | 0.009256244 | 0.563552821 | 0.557594679 | 0.736628304 | 0.446579428 | 0.965418873 |
| Gtf2e2 | S | 29 | 0.163087327 | -0.09632656 | -0.149588538 | -0.312675865 | -0.053261979 | 0.023085782 | 0.246086161 | 0.106559779 | 0.027932478 | 0.534951013 |
| Noa1 | S | 421 | -0.049927678 | -0.033484325 | 0.034993916 | 0.084921594 | 0.068478241 | 0.567330049 | 0.450507692 | 0.438600939 | 0.339494553 | 0.023641749 |
| Apc | S | 849 | -0.038687851 | -0.113431217 | 0.088286336 | 0.126974187 | 0.201717553 | 0.708906468 | 0.240540954 | 0.238703671 | 0.302565734 | 0.088319166 |
| Mavs | S | 186 | 0.043030559 | 0.023489858 | -0.113458495 | -0.156489054 | -0.136948354 | 0.551511902 | 0.751841816 | 0.238033972 | 0.161041722 | 0.214962082 |
| Bod1l | S | 2867 | -0.085710834 | 0.010329399 | 0.035572693 | 0.121283526 | 0.025243294 | 0.331212141 | 0.86367545 | 0.722517932 | 0.280055845 | 0.775132058 |
| Lemd3 | S | 136 | 0.07509665 | -0.066792689 | -0.105050522 | -0.180147172 | -0.038257834 | 0.050662808 | 0.22677317 | 0.378193174 | 0.192883966 | 0.732076077 |
| Asap1 | T | 799 | 0.044426549 | -0.017519813 | -0.048070733 | -0.092497283 | -0.03055092 | 0.306455613 | 0.600422936 | 0.162167933 | 0.07053565 | 0.326497516 |
| Golga2 | S | 306 | 0.064291521 | -0.082030851 | -0.027241543 | -0.091533064 | 0.054789308 | 0.171333136 | 0.120255961 | 0.787661125 | 0.386731624 | 0.58385129 |
| Hcfc1 | S | 1506 | 0.016136038 | 0.025097639 | -0.079660439 | -0.095796477 | -0.104758078 | 0.893568199 | 0.836833597 | 0.400012616 | 0.477523652 | 0.44355197 |
| Etl4 | S | 1796 | 0.009339883 | 0.03339887 | -0.101980232 | -0.111320115 | -0.135379102 | 0.752244348 | 0.178715874 | 0.135002547 | 0.108467677 | 0.075563999 |
| Mia3 | T | 1929 | 0.174400654 | -0.039326001 | -0.094533037 | -0.268933691 | -0.055207037 | 0.108913964 | 0.357385963 | 0.028263641 | 0.056637823 | 0.142070708 |
| Slk | S | 347 | -0.038776297 | 0.026564769 | -0.032086626 | 0.006689671 | -0.058651395 | 0.570080222 | 0.475337915 | 0.483115091 | 0.91694335 | 0.153188956 |
| Micall1 | S | 647 | 0.013612242 | 0.012943962 | -0.04481658 | -0.058428822 | -0.057760542 | 0.563395396 | 0.598425437 | 0.088397707 | 0.042132393 | 0.051178928 |

| | | | | | | | | | | | | |
|---|---|---|---|---|---|---|---|---|---|---|---|---|
| Wdr11 | S | 401 | 0.003932016 | 0.012272412 | -0.050071631 | -0.054003647 | -0.062344043 | 0.974056015 | 0.832282739 | 0.390373392 | 0.667599896 | 0.359788848 |
| Usp10 | S | 570 | 0.056835001 | -0.060986018 | -0.001243393 | -0.058078394 | 0.059742625 | 0.129969443 | 0.018329722 | 0.971861225 | 0.205563759 | 0.166091712 |
| Fam102b | S | 267 | -0.296438863 | 0.263849853 | 0.050236913 | 0.346675776 | -0.21361294 | 0.077402286 | 0.266828498 | 0.744025269 | 0.062809639 | 0.353922375 |
| Reep3 | S | 224 | -0.069516672 | 0.07162174 | -0.044520625 | 0.024996047 | -0.116142365 | 0.138194247 | 0.008206393 | 0.043168784 | 0.508967408 | 0.001844 |
| Reep3 | S | 227 | -0.069516672 | 0.07162174 | -0.044520625 | 0.024996047 | -0.116142365 | 0.138194247 | 0.008206393 | 0.043168784 | 0.508967408 | 0.001844 |
| Epha2 | S | 374 | -0.014492966 | -0.009823216 | -0.00362387 | 0.010869096 | 0.006199347 | 0.639605877 | 0.816551474 | 0.744429082 | 0.71647597 | 0.881633913 |
| Lrsam1 | S | 11 | -0.022696784 | -0.009492257 | -0.009930712 | 0.012766072 | -0.000438454 | 0.749028821 | 0.858009082 | 0.86218165 | 0.828783236 | 0.989536359 |
| Chmp3 | S | 200 | -0.025231246 | 0.02515176 | -0.038747989 | -0.013516744 | -0.063899749 | 0.527412535 | 0.431800578 | 0.450637886 | 0.787096618 | 0.229928434 |
| Champ1 | S | 532 | 0.045267068 | -0.071278456 | 0.025776969 | -0.019490099 | 0.097055425 | 0.354658426 | 0.038043593 | 0.631214178 | 0.74438972 | 0.139895767 |
| Bcor | S | 1131 | 0.022469383 | 0.021561124 | -0.070461934 | -0.092931317 | -0.092023058 | 0.658498693 | 0.726482623 | 0.199613475 | 0.179519439 | 0.235207598 |
| Ncam1 | T | 788 | -0.01292446 | -0.038984054 | 0.022385959 | 0.035310418 | 0.061370013 | 0.875327247 | 0.262613002 | 0.566480845 | 0.676222876 | 0.150204974 |
| Ak1 | S | 192 | -0.167410709 | -0.021459092 | 0.135172915 | 0.302583624 | 0.156632007 | 0.180915429 | 0.773936921 | 0.219248 | 0.048192676 | 0.148311217 |
| Ppfibp1 | S | 923 | 0.073504382 | -0.093757987 | -0.012240018 | -0.0857444 | 0.081517968 | 0.028013177 | 0.016815328 | 0.824551699 | 0.19598659 | 0.20316471 |
| Ctdspl2 | S | 165 | -0.071821707 | 0.088714993 | -0.017011808 | 0.054809899 | -0.105726801 | 0.089391766 | 0.258973699 | 0.65729574 | 0.043331938 | 0.197554911 |
| Zc3hc1 | S | 337 | 0.045781113 | -0.03518699 | -0.03499873 | -0.080779843 | 0.00018826 | 0.413705568 | 0.560968973 | 0.273586715 | 0.204772731 | 0.997511698 |
| Ppp1r12a | S | 445 | -0.14135782 | 0.212757934 | -0.116718193 | 0.024639627 | -0.329476126 | 0.294531979 | 0.211646347 | 0.430394785 | 0.86287745 | 0.10146053 |
| Srgap2 | S | 796 | -0.030098071 | 0.051246349 | -0.088069413 | -0.057971342 | -0.139315762 | 0.736470217 | 0.547482328 | 0.537710706 | 0.684326519 | 0.354103407 |
| Pgls | S | 46 | -0.293596651 | 0.12167238 | 0.078609489 | 0.37220614 | -0.04306289 | 0.182982192 | 0.400034593 | 0.542451828 | 0.121497677 | 0.783518267 |
| Spen | S | 766 | -0.059334274 | -0.055445945 | 0.104744476 | 0.16407875 | 0.160190421 | 0.144936038 | 0.06660815 | 0.043464443 | 0.014559604 | 0.022976909 |
| Asap1 | S | 540 | 0.093381421 | -0.044999927 | -0.106652887 | -0.200034308 | -0.06165296 | 0.046467339 | 0.293850417 | 0.145349477 | 0.043022442 | 0.332657227 |
| Dab2 | T | 221 | -0.106597546 | 0.03334444 | 0.060195878 | 0.166793423 | 0.026851438 | 0.033982281 | 0.458760889 | 0.256253256 | 0.026688471 | 0.518284825 |
| Akirin2 | S | 55 | 0.048036133 | -0.007497278 | -0.05787149 | -0.105907623 | -0.050374213 | 0.431481502 | 0.878002958 | 0.165705458 | 0.135629819 | 0.323852298 |
| Plekho2 | S | 293 | -0.010865908 | 0.01124364 | -0.03749386 | -0.026627952 | -0.0487375 | 0.901180849 | 0.890445027 | 0.525911652 | 0.756737949 | 0.550733548 |
| Ppp1r37 | S | 56 | -0.012179683 | 0.101557958 | -0.162800636 | -0.150620953 | -0.264358594 | 0.882900626 | 0.189528235 | 0.184404377 | 0.234200824 | 0.069383275 |
| Cdc5l | T | 430 | -0.077218408 | -0.002758558 | 0.048227482 | 0.12544589 | 0.05098604 | 0.367784883 | 0.982813318 | 0.694231584 | 0.285236744 | 0.712483982 |
| Specc1l | S | 869 | 0.038511348 | -0.080400624 | 0.032236791 | -0.006274557 | 0.112637415 | 0.418692599 | 0.028796467 | 0.292527437 | 0.873200237 | 0.000931768 |
| Apc | S | 2129 | 0.025651108 | -0.044330637 | 0.000627784 | -0.025023324 | 0.044958421 | 0.80186087 | 0.307317081 | 0.994355832 | 0.833678356 | 0.598212018 |
| Afdn | S | 557 | 0.083574896 | -0.115004996 | 0.006514959 | -0.077059937 | 0.121519955 | 0.27140043 | 0.10713609 | 0.906825468 | 0.293538234 | 0.093773985 |
| Topors | S | 1016 | 0.061538167 | 0.053374133 | -0.173118485 | -0.234656652 | -0.226492618 | 0.205862841 | 0.175407412 | 0.033246001 | 0.0169936 | 0.037223066 |
| Zeb2 | S | 1122 | -0.068989992 | 0.015222188 | 0.003971753 | 0.072961745 | -0.011250434 | 0.244256384 | 0.24047042 | 0.954184907 | 0.387691326 | 0.870727907 |
| Phc3 | S | 312 | 0.298835555 | -0.153980117 | -0.049152221 | -0.347987776 | 0.104827896 | 0.211863979 | 0.071284244 | 0.466176707 | 0.166044378 | 0.210203614 |
| Brd1 | S | 803 | 0.040046351 | -0.04479993 | -0.016456128 | -0.056502479 | 0.028343802 | 0.158018396 | 0.101618094 | 0.530717959 | 0.056888744 | 0.222247346 |
| Arhgef12 | S | 41 | -0.144561001 | 0.079405847 | -0.006736898 | 0.137824103 | -0.086142745 | 0.331613047 | 0.407773592 | 0.958044772 | 0.390436536 | 0.477460139 |
| Nrf1 | S | 136 | -0.011221228 | -0.002504629 | -0.012208606 | -0.000987378 | -0.009703978 | 0.678208319 | 0.922324852 | 0.675043644 | 0.961422538 | 0.615026863 |
| Incenp | S | 147 | 0.033585106 | -0.062222735 | 0.011373963 | -0.022211144 | 0.073596698 | 0.55439536 | 0.173514036 | 0.860867717 | 0.733718464 | 0.260965991 |
| Srrm2 | S | 2618 | -0.033704655 | -0.033543701 | 0.026597188 | 0.060301843 | 0.060140888 | 0.768149965 | 0.548354566 | 0.745923268 | 0.621127602 | 0.433352429 |
| Srrm2 | S | 1630 | -0.01078314 | 0.002432489 | -0.02246333 | -0.01168019 | -0.024895818 | 0.780412564 | 0.956254419 | 0.604488959 | 0.812234379 | 0.645135058 |
| Srpk1 | S | 51 | 0.062577054 | -0.028218271 | -0.067782917 | -0.130359972 | -0.039564647 | 0.234843051 | 0.651271997 | 0.184480121 | 0.036208083 | 0.510107862 |
| Srrm1 | S | 463 | -0.021684239 | -0.025372051 | 0.025070409 | 0.046754648 | 0.050442461 | 0.678006545 | 0.423505862 | 0.709834344 | 0.55747997 | 0.486029192 |
| Mast4 | S | 1368 | -0.072400559 | 0.153498492 | -0.126210551 | -0.053809992 | -0.279709043 | 0.578491293 | 0.213341741 | 0.224840896 | 0.632729355 | 0.041644328 |
| Nfrkb | S | 180 | -0.112047654 | -0.035693843 | 0.056231739 | 0.168279393 | 0.091925582 | 0.596048339 | 0.698739409 | 0.462468797 | 0.434557362 | 0.227736209 |
| Rad51ap1 | S | 309 | 0.152522118 | -0.050645133 | -0.134436475 | -0.286958593 | -0.083791342 | 0.022271514 | 0.285387149 | 0.041045569 | 0.003419827 | 0.116042351 |
| Ccnl2 | S | 366 | -0.004943259 | -0.014619596 | -0.001565678 | 0.003377581 | 0.013053917 | 0.851134542 | 0.460151849 | 0.930383255 | 0.87295516 | 0.236653509 |
| Tns2 | S | 648 | -0.128314472 | 0.157994467 | -0.084650543 | 0.043663929 | -0.24264501 | 0.153580506 | 0.098233415 | 0.449814665 | 0.710686499 | 0.098419619 |
| Racgap1 | S | 250 | 0.042908476 | 0.05766427 | -0.185638481 | -0.228546958 | -0.243302751 | 0.472639673 | 0.318186216 | 0.162343667 | 0.114746369 | 0.104543652 |
| Flot1 | S | 14 | 0.032011339 | -0.161252211 | 0.104740093 | 0.072728754 | 0.265992304 | 0.72354326 | 0.217926292 | 0.55263386 | 0.674343679 | 0.20388798 |
| Numa1 | S | 200 | 0.019843247 | -0.071696321 | -0.0189724 | -0.038815647 | 0.05272392 | 0.886904223 | 0.759674389 | 0.863226855 | 0.803761956 | 0.827004993 |
| Numa1 | S | 201 | 0.019843247 | -0.071696321 | -0.0189724 | -0.038815647 | 0.05272392 | 0.886904223 | 0.759674389 | 0.863226855 | 0.803761956 | 0.827004993 |
| Atg2a | S | 1799 | 0.040184918 | -0.050018189 | -0.033871961 | -0.074056878 | 0.016146228 | 0.817105261 | 0.812522689 | 0.598617866 | 0.673092563 | 0.938302997 |
| Mterf4 | S | 328 | -0.088529363 | 0.055180739 | -0.004362839 | 0.084166524 | -0.059543578 | 0.291431774 | 0.486325291 | 0.96692448 | 0.368606165 | 0.501009184 |
| Mterf4 | S | 330 | -0.088529363 | 0.055180739 | -0.004362839 | 0.084166524 | -0.059543578 | 0.291431774 | 0.486325291 | 0.96692448 | 0.368606165 | 0.501009184 |
| Mterf4 | S | 333 | -0.088529363 | 0.055180739 | -0.004362839 | 0.084166524 | -0.059543578 | 0.291431774 | 0.486325291 | 0.96692448 | 0.368606165 | 0.501009184 |
| Limd1 | S | 659 | 0.002341583 | -0.008909979 | -0.023961528 | -0.026303111 | -0.015051549 | 0.973095386 | 0.878929213 | 0.564267217 | 0.7249815 | 0.814716496 |
| Parg | S | 256 | -0.021140374 | 0.082705049 | -0.073891925 | -0.052751552 | -0.156596974 | 0.570159398 | 0.120616708 | 0.062965729 | 0.14772932 | 0.025185671 |
| Parg | S | 259 | -0.021140374 | 0.082705049 | -0.073891925 | -0.052751552 | -0.156596974 | 0.570159398 | 0.120616708 | 0.062965729 | 0.14772932 | 0.025185671 |
| Phip | S | 1598 | 0.006618671 | -0.009929356 | -0.01954517 | -0.026163841 | -0.009615814 | 0.903655491 | 0.65815002 | 0.57655924 | 0.658008085 | 0.76953207 |
| Gpalpp1 | S | 104 | -0.06860583 | -0.007297575 | 0.066890209 | 0.135496039 | 0.074187784 | 0.027783614 | 0.867076509 | 0.107865267 | 0.018282229 | 0.183978116 |
| Zc3hc1 | S | 394 | 0.047000329 | -0.049418664 | -0.029206641 | -0.07620697 | 0.020212024 | 0.004408341 | 0.018496754 | 0.245545014 | 0.040356633 | 0.405267525 |
| Ccnh | T | 315 | 0.00412594 | 0.021883986 | -0.075965259 | -0.080091199 | -0.097849245 | 0.927445502 | 0.439152893 | 0.322964083 | 0.3231349 | 0.231420938 |
| Smtn | T | 309 | 0.126893956 | -0.143240827 | -0.041222031 | -0.168115987 | 0.102018796 | 0.162685382 | 0.249112066 | 0.478194987 | 0.089898096 | 0.373122673 |
| Tjp2 | S | 895 | -0.064094021 | -0.072326708 | 0.096509311 | 0.160603331 | 0.168836019 | 0.633899569 | 0.64721909 | 0.638511682 | 0.449274898 | 0.451264621 |
| Srrm2 | S | 559 | -0.191739784 | 0.048661878 | 0.009632453 | 0.201372237 | -0.039029425 | 0.615335971 | 0.648075355 | 0.960197381 | 0.61379992 | 0.833795465 |
| Srrm2 | S | 561 | -0.191739784 | 0.048661878 | 0.009632453 | 0.201372237 | -0.039029425 | 0.615335971 | 0.648075355 | 0.960197381 | 0.61379992 | 0.833795465 |
| Srrm2 | S | 563 | -0.191739784 | 0.048661878 | 0.009632453 | 0.201372237 | -0.039029425 | 0.615335971 | 0.648075355 | 0.960197381 | 0.61379992 | 0.833795465 |
| Eif2ak3 | T | 978 | -0.16453255 | -0.046759326 | 0.127578269 | 0.292110819 | 0.174337595 | 0.383831375 | 0.88363136 | 0.419127133 | 0.202506464 | 0.609782105 |
| Ptbp3 | S | 27 | 0.227502607 | -0.083913759 | -0.006159067 | -0.233661674 | 0.077754692 | 0.415710294 | 0.028009621 | 0.780922567 | 0.405460623 | 0.041388751 |
| Atn1 | S | 118 | -0.020412639 | -0.042494334 | 0.023014039 | 0.043426677 | 0.065508373 | 0.826904231 | 0.528262162 | 0.718018106 | 0.581599802 | 0.092252662 |
| Trps1 | S | 178 | -0.033979349 | -0.012150904 | 0.027788232 | 0.061767581 | 0.039939137 | 0.348360327 | 0.643051683 | 0.578917541 | 0.278033231 | 0.434052196 |
| Bard1 | T | 291 | -0.040517894 | 0.008140328 | -0.010395358 | 0.030122536 | -0.018535685 | 0.622500458 | 0.868231728 | 0.892026588 | 0.764177997 | 0.819180085 |
| Cep170 | S | 404 | -0.162844019 | 0.096407394 | -0.01073702 | 0.152106999 | -0.107144415 | 0.47154969 | 0.557825618 | 0.937188764 | 0.515745354 | 0.555077292 |
| Srrm2 | T | 823 | 0.041922667 | -0.056466071 | -0.001986578 | -0.043909245 | 0.054479493 | 0.265164821 | 0.095302098 | 0.953577248 | 0.273550731 | 0.150019908 |
| Mdc1 | S | 1286 | 0.093525196 | -0.057063125 | -0.1039342 | -0.197459395 | -0.046871075 | 0.214465903 | 0.406509343 | 0.443424024 | 0.19242515 | 0.704934598 |
| Prpf38b | S | 291 | -0.054162448 | -0.008035972 | 0.039699895 | 0.093862344 | 0.047735868 | 0.425139574 | 0.892692097 | 0.746473599 | 0.470360995 | 0.697706463 |
| Tnks1bp1 | S | 807 | 0.085037802 | 0.005514436 | -0.149133795 | -0.234171597 | -0.154648232 | 0.582393896 | 0.938051288 | 0.397653395 | 0.283633679 | 0.386154806 |
| Pcdh19 | S | 938 | -0.012829153 | -0.049323855 | 0.004172721 | 0.017001874 | 0.053496575 | 0.894742834 | 0.77149131 | 0.958767555 | 0.878177014 | 0.761964298 |
| Pcdh19 | T | 940 | -0.012829153 | -0.049323855 | 0.004172721 | 0.017001874 | 0.053496575 | 0.894742834 | 0.77149131 | 0.958767555 | 0.878177014 | 0.761964298 |
| Arid4b | S | 1192 | -0.008289786 | -0.025687961 | 0.011016497 | 0.019306284 | 0.036704459 | 0.769155548 | 0.471726958 | 0.810474199 | 0.669219529 | 0.463574036 |
| Fosl2 | S | 200 | -0.004226228 | -0.076113423 | 0.051594554 | 0.055820783 | 0.127707977 | 0.896234226 | 0.048652286 | 0.076782582 | 0.14335621 | 0.012211443 |
| Vim | S | 261 | -0.015666826 | -0.030008466 | -0.009752896 | 0.00591393 | 0.02025557 | 0.920333504 | 0.849246672 | 0.905645862 | 0.968647732 | 0.893739559 |
| Zc3h6 | S | 182 | 0.000956695 | -0.014110575 | -0.015855223 | -0.016811917 | -0.001744647 | 0.988837434 | 0.698123262 | 0.73949276 | 0.803976691 | 0.960409911 |
| Zc3h6 | S | 184 | 0.000956695 | -0.014110575 | -0.015855223 | -0.016811917 | -0.001744647 | 0.988837434 | 0.698123262 | 0.73949276 | 0.803976691 | 0.960409911 |
| Melk | S | 497 | 0.072581187 | 0.045699273 | -0.100299921 | -0.172881109 | -0.145999194 | 0.12685178 | 0.162535833 | 0.001394687 | 0.028755413 | 0.017552888 |
| Ski | S | 498 | 0.066294267 | 0.014429222 | -0.107769259 | -0.174063526 | -0.12219848 | 0.20003976 | 0.864955636 | 0.090539633 | 0.030591301 | 0.229074297 |
| Iws1 | S | 263 | -0.017006365 | -0.035439134 | 0.036487884 | 0.05349425 | 0.071927019 | 0.385846505 | 0.181991587 | 0.215505075 | 0.097244263 | 0.048813677 |
| Tnrc18 | S | 2403 | -0.107266512 | 0.01154186 | 0.080592693 | 0.187859204 | 0.069050833 | 0.012520966 | 0.449955346 | 0.078239472 | 0.006853417 | 0.126699934 |
| Mdc1 | T | 943 | 0.026576851 | -0.003390289 | -0.04385378 | -0.070430631 | -0.040463491 | 0.486255466 | 0.919790343 | 0.219815694 | 0.049583285 | 0.049182781 |
| Tlk1 | T | 38 | -0.023101918 | -0.016238555 | 0.007156133 | 0.03025805 | 0.023394688 | 0.63862818 | 0.676254926 | 0.749945003 | 0.53064081 | 0.526218624 |
| Irs2 | S | 762 | -0.00233702 | -0.020022106 | -0.01272598 | -0.01038896 | 0.007296126 | 0.932053178 | 0.750114646 | 0.838316001 | 0.863724682 | 0.92622708 |
| Afdn | S | 1275 | 0.01573216 | 0.058833415 | -0.08273764 | -0.0984698 | -0.141571056 | 0.704235688 | 0.400439726 | 0.093946799 | 0.017510362 | 0.093162188 |
| Phldb1 | S | 325 | 0.076361369 | -0.032464572 | -0.081308513 | -0.157669882 | -0.048843941 | 0.205795302 | 0.494382634 | 0.257733319 | 0.060162187 | 0.413909545 |
| Traf5 | S | 265 | -0.034790104 | 0.079331083 | -0.103615677 | -0.068825573 | -0.182946759 | 0.718028322 | 0.562378383 | 0.568027764 | 0.715116856 | 0.393245957 |
| Traf5 | S | 270 | -0.034790104 | 0.079331083 | -0.103615677 | -0.068825573 | -0.182946759 | 0.718028322 | 0.562378383 | 0.568027764 | 0.715116856 | 0.393245957 |
| Usp36 | S | 547 | -0.034678636 | 0.070852563 | -0.034106599 | 0.000572037 | -0.104959162 | 0.737016547 | 0.628311457 | 0.260163469 | 0.995491289 | 0.488537912 |
| Tbc1d2b | S | 951 | -0.004781844 | 0.051859275 | -0.080400782 | -0.075618938 | -0.132260058 | 0.946243823 | 0.484490418 | 0.295015311 | 0.051542486 | 0.010621861 |

| | | | | | | | | | | | | | |
|---|---|---|---|---|---|---|---|---|---|---|---|---|---|
| Spen | S | | 773 | 0.029371326 | -0.166401318 | 0.067534328 | 0.038163002 | 0.233935646 | 0.65594901 | 0.21986013 | 0.379717545 | 0.527976036 | 0.119564686 |
| Cad | S | | 1406 | -0.040572615 | 0.192000733 | -0.225163361 | -0.184590746 | -0.417164094 | 0.662422478 | 0.103166804 | 0.139466774 | 0.190853396 | 0.029465085 |
| Kif20a | S | | 881 | 0.056147798 | -0.031825326 | -0.073260059 | -0.129407857 | -0.041434734 | 0.223776297 | 0.416136642 | 0.404668931 | 0.190507147 | 0.612630796 |
| Cenpa | S | | 51 | 0.054481429 | 0.127429047 | -0.220622196 | -0.275103625 | -0.348051243 | 0.640460649 | 0.405052682 | 0.165907574 | 0.107541146 | 0.08537382 |
| Cenpa | T | | 52 | 0.054481429 | 0.127429047 | -0.220622196 | -0.275103625 | -0.348051243 | 0.640460649 | 0.405052682 | 0.165907574 | 0.107541146 | 0.08537382 |
| Adgra2 | S | | 1271 | -0.145368178 | 0.113162332 | 0.013960338 | 0.159328516 | -0.099201994 | 0.085578701 | 0.222285274 | 0.829517575 | 0.047316187 | 0.228632706 |
| Arhgap21 | S | | 850 | -0.108991618 | 0.100734059 | -0.075229841 | 0.033761777 | -0.1759639 | 0.293683358 | 0.060376055 | 0.301225248 | 0.742313105 | 0.063944884 |
| Armcx2 | T | | 93 | -0.023471194 | -0.07549292 | 0.086124705 | 0.109595899 | 0.161617625 | 0.839756932 | 0.304201448 | 0.541120775 | 0.485887637 | 0.278981499 |
| Farp2 | S | | 36 | 0.015975116 | 0.030997581 | -0.084078705 | -0.10005382 | -0.115076285 | 0.774129707 | 0.513171463 | 0.250295603 | 0.202905088 | 0.142265167 |
| Zmym2 | S | | 837 | 0.036583551 | 0.005513727 | -0.056427713 | -0.093011265 | -0.06194144 | 0.382494241 | 0.920088612 | 0.125534157 | 0.050692275 | 0.283026573 |
| Farp1 | T | | 34 | 0.075800863 | -0.075850778 | -0.028808512 | -0.104609376 | 0.047042266 | 0.198771801 | 0.211363316 | 0.507451928 | 0.086635293 | 0.332299193 |
| Pogz | S | | 422 | -0.075104482 | -0.035615421 | 0.13187499 | 0.206979471 | 0.16749041 | 0.065511093 | 0.587959621 | 0.205385748 | 0.101116906 | 0.143483998 |
| Pcm1 | S | | 69 | 0.01550285 | -0.024901231 | -0.016648432 | -0.032151282 | 0.008252799 | 0.661132415 | 0.42616895 | 0.763871699 | 0.583278825 | 0.880941461 |
| Eps15 | S | | 324 | 0.061409355 | 0.015776487 | -0.120156643 | -0.181565998 | -0.13593313 | 0.191780682 | 0.838085588 | 0.120923432 | 0.047843201 | 0.17817816 |
| Myo5a | S | | 1450 | 0.016972808 | 0.027132155 | -0.081591908 | -0.098564716 | -0.108724063 | 0.722672292 | 0.517396146 | 0.241427201 | 0.164731083 | 0.142494106 |
| Med26 | S | | 232 | -0.007891595 | -0.144413852 | 0.085358822 | 0.093250417 | 0.229772674 | 0.942926975 | 0.296059916 | 0.328440848 | 0.375257063 | 0.131533874 |
| Zc3h18 | S | | 838 | -0.077637984 | 0.023441539 | 0.041068497 | 0.118706481 | 0.017626958 | 0.094814037 | 0.572011841 | 0.450699317 | 0.06607759 | 0.707174365 |
| Katnb1 | T | | 395 | 0.098428063 | 0.066407727 | -0.160438991 | -0.258867054 | -0.226846718 | 0.260576585 | 0.225005561 | 0.016697087 | 0.040400982 | 0.001046117 |
| Ctps2 | S | | 571 | -0.130622824 | 0.047965996 | 0.086897549 | 0.217520373 | 0.038931553 | 0.034104064 | 0.474656254 | 0.257072277 | 0.039109639 | 0.639379497 |
| Lemd3 | S | | 137 | 0.067741549 | -0.048013594 | -0.093654177 | -0.161395727 | -0.045640583 | 0.060690068 | 0.253107454 | 0.357496737 | 0.174184984 | 0.628657585 |
| Aldoa | T | | 9 | 0.039394816 | -0.005388942 | -0.047688827 | -0.087083644 | -0.042299885 | 0.578569729 | 0.944932585 | 0.240148959 | 0.269294232 | 0.600835088 |
| Eif3a | S | | 1159 | 0.036663067 | -0.040087247 | -0.026831135 | -0.063494203 | 0.013256111 | 0.571446882 | 0.488664551 | 0.665248608 | 0.399105958 | 0.841185608 |
| Arhgap21 | S | | 435 | -0.126191696 | 0.138786929 | -0.10948434 | 0.016707355 | -0.248271269 | 0.230456062 | 0.09905331 | 0.52200798 | 0.919400578 | 0.205980148 |
| Srrm1 | S | | 412 | 0.002934228 | -0.015466541 | -0.014419697 | -0.017353926 | 0.001046844 | 0.943909971 | 0.770504348 | 0.643483162 | 0.631565625 | 0.982676498 |
| Tec | S | | 307 | 0.010471084 | 0.042884744 | -0.072151018 | -0.082622102 | -0.115035762 | 0.905396382 | 0.569659463 | 0.246931373 | 0.404536501 | 0.204621655 |
| Uhrf1 | S | | 95 | 0.038969242 | -0.059089535 | 0.003920887 | -0.035048354 | 0.063010423 | 0.460369652 | 0.182018121 | 0.955548492 | 0.627037047 | 0.378312176 |
| Camkk2 | S | | 572 | 0.038690879 | -0.006247707 | -0.083458825 | -0.122149704 | -0.077211118 | 0.482814763 | 0.797341719 | 0.444492468 | 0.304342426 | 0.473835138 |
| Srrm1 | S | | 822 | 0.006529814 | -0.032294355 | 0.011978531 | 0.005448717 | 0.044272886 | 0.756945964 | 0.096135959 | 0.414011671 | 0.75079089 | 0.04334694 |
| Chaf1a | S | | 310 | 0.043314002 | -0.002592536 | -0.073201351 | -0.116515353 | -0.070608814 | 0.501431856 | 0.963967006 | 0.310288588 | 0.077483842 | 0.209127431 |
| Edc4 | S | | 849 | -0.031305629 | 0.037189747 | -0.026884007 | 0.004421622 | -0.064073755 | 0.420586582 | 0.470467147 | 0.522432833 | 0.845884884 | 0.187234485 |
| Npat | S | | 209 | 0.004978851 | 0.003829901 | -0.035426153 | -0.040405003 | -0.039256053 | 0.942261386 | 0.916756472 | 0.465520785 | 0.592688038 | 0.433129926 |
| Zhx1 | S | | 648 | -0.079279644 | -7.53046E-05 | 0.049929539 | 0.129209183 | 0.050004844 | 0.274702831 | 0.998957361 | 0.480505949 | 0.125485935 | 0.438365965 |
| Atad5 | S | | 1592 | 0.027607004 | 0.026730587 | -0.068936859 | -0.096543862 | -0.095667445 | 0.589907179 | 0.43752168 | 0.100171709 | 0.141428815 | 0.064160151 |
| Ccdc61 | S | | 472 | -0.009928185 | 0.003436577 | -0.024126464 | -0.014198279 | -0.027563041 | 0.833936458 | 0.94825136 | 0.66156826 | 0.827923499 | 0.692272534 |
| Prkab2 | S | | 38 | -0.24263602 | 0.10651778 | 0.175727088 | 0.418363108 | 0.069209308 | 0.036126095 | 0.43513328 | 0.2824406 | 0.053548883 | 0.67152842 |
| Sipa1l1 | S | | 1487 | -0.004838831 | 0.039627327 | -0.061615414 | -0.056776584 | -0.101242742 | 0.960202054 | 0.578873017 | 0.329749885 | 0.517454099 | 0.107161024 |
| Hif1a | S | | 499 | 0.034631579 | -0.118982044 | 0.04667859 | 0.012047011 | 0.165660635 | 0.720244682 | 0.292809197 | 0.626743993 | 0.902372063 | 0.178505439 |
| Gak | S | | 827 | -0.074167738 | 0.058824564 | 0.016375666 | 0.090543405 | -0.042448898 | 0.001667453 | 0.019629775 | 0.676868247 | 0.102339156 | 0.325481277 |
| Chd9 | S | | 2750 | 0.124878999 | -0.109723132 | -0.102126407 | -0.227005406 | 0.007596725 | 0.042184004 | 0.21262558 | 0.312791541 | 0.08049155 | 0.94107707 |
| Med1 | T | | 998 | 0.008569384 | -0.027222287 | 0.000198628 | -0.008370756 | 0.027420916 | 0.822063089 | 0.346737948 | 0.994688546 | 0.832623116 | 0.389545353 |
| Ncoa6 | S | | 1812 | -0.035474688 | -0.016969411 | 0.028292247 | 0.063766936 | 0.045261658 | 0.407569165 | 0.648007419 | 0.514224515 | 0.246355564 | 0.366864863 |
| Map1a | S | | 2082 | -0.074969263 | 0.048667462 | -0.009063516 | 0.065905747 | -0.057730978 | 0.138280187 | 0.168334259 | 0.864124907 | 0.307552552 | 0.3333996 |
| Cdca2 | S | | 210 | 0.027081265 | -0.064822232 | 0.027404406 | 0.000323142 | 0.092226638 | 0.300397311 | 0.038449497 | 0.428172074 | 0.990168597 | 0.038082942 |
| Foxm1 | T | | 180 | 0.047171996 | 0.039239463 | -0.122583347 | -0.169755344 | -0.16182281 | 0.823111277 | 0.733984166 | 0.406327191 | 0.439129409 | 0.218634678 |
| Pcnt | T | | 1958 | -0.13471241 | -0.079975715 | 0.101869445 | 0.236581855 | 0.18184516 | 0.373469811 | 0.647536472 | 0.170644326 | 0.174298812 | 0.332590814 |
| Tns2 | S | | 120 | -0.11532791 | 0.128040545 | -0.063027076 | 0.052300834 | -0.191067621 | 0.259770897 | 0.226773072 | 0.639608854 | 0.694096973 | 0.208615607 |
| Tdp1 | S | | 119 | -0.107425163 | 0.072778654 | -0.023665066 | 0.083760097 | -0.09644372 | 0.170381739 | 0.068908062 | 0.482538653 | 0.258401531 | 0.007762264 |
| Septin5 | S | | 225 | -0.044467103 | 0.013448437 | 0.007685437 | 0.05215254 | -0.005763 | 0.348926808 | 0.738434207 | 0.912919207 | 0.44804372 | 0.9253149 |
| Pcyt1a | S | | 345 | 0.008850167 | 0.053713607 | -0.085340507 | -0.094190674 | -0.139054114 | 0.838767289 | 0.533746289 | 0.240618523 | 0.19664941 | 0.183337202 |
| Racgap1 | T | | 564 | 0.137004059 | 0.019781688 | -0.141672497 | -0.278676556 | -0.161454185 | 0.098190303 | 0.512343326 | 0.010417392 | 0.017959469 | 0.020281409 |
| Sap30 | S | | 131 | 0.025725281 | -0.060722819 | 0.013188186 | -0.012537096 | 0.073911004 | 0.479710982 | 0.155673318 | 0.663781361 | 0.63609385 | 0.087997913 |
| Ccnl2 | S | | 328 | -0.052309251 | -0.051426732 | 0.081812315 | 0.134121566 | 0.133239047 | 0.404762244 | 0.100724374 | 0.09770413 | 0.095557748 | 0.030574009 |
| Cic | S | | 1644 | 0.015364712 | -0.029528647 | -0.009072326 | -0.024437038 | 0.020456321 | 0.79169108 | 0.477197449 | 0.870777708 | 0.738849749 | 0.741509588 |
| Hcfc1 | S | | 411 | 0.053840589 | -0.05782364 | -0.004896557 | -0.058737146 | 0.052927083 | 0.64104745 | 0.319601849 | 0.91386 | 0.60878186 | 0.335294963 |
| Arhgef5 | S | | 1109 | -0.080470358 | 0.088581527 | -0.041377948 | 0.03909241 | -0.129959475 | 0.237529844 | 0.243273609 | 0.63614039 | 0.684769979 | 0.23565837 |
| Dtl | S | | 680 | -0.004932156 | 0.01592374 | -0.049639547 | -0.044707391 | -0.065563287 | 0.966380395 | 0.788857998 | 0.552846279 | 0.714067303 | 0.387124476 |
| Ppp1r18 | S | | 125 | -0.007619651 | 0.041938324 | -0.099601357 | -0.091981706 | -0.141539681 | 0.915162993 | 0.378178839 | 0.40978722 | 0.463951219 | 0.272905098 |
| Khdrbs1 | T | | 33 | -0.04059905 | -0.097945669 | 0.094088004 | 0.134687054 | 0.192033674 | 0.684221984 | 0.390801366 | 0.370722439 | 0.26301791 | 0.160302498 |
| Tcof1 | S | | 1127 | -0.071822127 | 0.042041674 | -0.015660035 | 0.056162092 | -0.057701709 | 0.370365122 | 0.375752821 | 0.808755992 | 0.491074927 | 0.33076522 |
| Ppfibp1 | T | | 561 | -0.123412152 | 0.194570882 | -0.076997076 | 0.046415076 | -0.271567958 | 0.223031593 | 0.227691285 | 0.509541895 | 0.711912425 | 0.144156848 |
| Cggbp1 | S | | 94 | 0.020471196 | -0.012699664 | -0.04637649 | -0.066847686 | -0.033676826 | 0.478285836 | 0.835306987 | 0.435327583 | 0.281114901 | 0.648041488 |
| Leo1 | S | | 278 | 0.039907908 | 0.009084947 | -0.056939172 | -0.09684708 | -0.066024119 | 0.161434494 | 0.769853361 | 0.033459299 | 0.012298061 | 0.081113824 |
| Gapvd1 | S | | 466 | 0.132714565 | -0.093049591 | -0.094180686 | -0.226895251 | -0.001131094 | 0.007841737 | 0.060916273 | 0.163331316 | 0.022320099 | 0.984656156 |
| Mark3 | S | | 400 | 0.092814717 | -0.073691568 | -0.044971205 | -0.137785922 | 0.028720363 | 0.418533627 | 0.408210535 | 0.612498425 | 0.277668634 | 0.765245464 |
| Tasor2 | S | | 1213 | -0.009176676 | -0.004122222 | -0.030888502 | -0.021711827 | -0.02676628 | 0.874998305 | 0.946068669 | 0.850875747 | 0.897198553 | 0.874033994 |
| Calu | S | | 261 | 0.014894687 | 0.059323132 | -0.173923205 | -0.188817892 | -0.233246336 | 0.896427235 | 0.336981093 | 0.371367416 | 0.359994191 | 0.261555267 |
| Svil | S | | 248 | -0.006849589 | -0.020384299 | 0.002430604 | 0.009280193 | 0.022814903 | 0.875761633 | 0.609300706 | 0.952151384 | 0.846262538 | 0.610282697 |
| Fcho2 | S | | 391 | -0.055738141 | 0.010804336 | 0.007594009 | 0.06333215 | -0.003210327 | 0.552600258 | 0.84705904 | 0.949113482 | 0.650642277 | 0.979173089 |
| Srgap2 | S | | 990 | 0.039956623 | -0.066684363 | 0.036624829 | -0.003331794 | 0.103309191 | 0.538541403 | 0.089151909 | 0.653204125 | 0.970696752 | 0.262232115 |
| Sorbs1 | S | | 1209 | 0.028624521 | -0.064506451 | -0.000874811 | -0.029499331 | 0.06363164 | 0.818566753 | 0.563041004 | 0.992768463 | 0.795564738 | 0.522465774 |
| Chordc1 | S | | 200 | -0.056600978 | 0.055981372 | -0.030321504 | 0.026279474 | -0.086302876 | 0.386913826 | 0.410420342 | 0.672240938 | 0.735770375 | 0.310896805 |
| Ripk2 | S | | 178 | -0.053556894 | 0.065186727 | -0.055427973 | -0.001871079 | -0.120614701 | 0.389728441 | 0.112874987 | 0.266775097 | 0.975876115 | 0.049273254 |
| Fhdc1 | S | | 948 | -0.062480611 | -0.029992753 | 0.049954099 | 0.11243471 | 0.079946852 | 0.458945125 | 0.425212195 | 0.251452613 | 0.22721807 | 0.065142311 |
| Ncoa3 | S | | 720 | -0.014786731 | 0.014012762 | -0.018304124 | -0.003517393 | -0.032316887 | 0.691704209 | 0.629112341 | 0.426474157 | 0.92007348 | 0.266718276 |
| Kri1 | S | | 177 | 0.015106133 | -0.048838315 | 0.01744278 | 0.002336647 | 0.066281096 | 0.71953876 | 0.150824056 | 0.464924182 | 0.953079585 | 0.073403149 |
| Kri1 | S | | 178 | 0.015106133 | -0.048838315 | 0.01744278 | 0.002336647 | 0.066281096 | 0.71953876 | 0.150824056 | 0.464924182 | 0.953079585 | 0.073403149 |
| Dnttip2 | S | | 133 | -0.033944713 | 0.062648424 | -0.057176877 | -0.023232164 | -0.119825301 | 0.569979326 | 0.354153913 | 0.381890789 | 0.711788917 | 0.131926327 |
| Dnttip2 | S | | 137 | -0.033944713 | 0.062648424 | -0.057176877 | -0.023232164 | -0.119825301 | 0.569979326 | 0.354153913 | 0.381890789 | 0.711788917 | 0.131926327 |
| Dnttip2 | S | | 140 | -0.033944713 | 0.062648424 | -0.057176877 | -0.023232164 | -0.119825301 | 0.569979326 | 0.354153913 | 0.381890789 | 0.711788917 | 0.131926327 |
| Pwp2 | S | | 895 | 0.045577658 | -0.066525046 | 0.018999236 | -0.026578421 | 0.085524282 | 0.37338335 | 0.054497042 | 0.525896211 | 0.578882031 | 0.027671158 |
| Pwp2 | S | | 898 | 0.045577658 | -0.066525046 | 0.018999236 | -0.026578421 | 0.085524282 | 0.37338335 | 0.054497042 | 0.525896211 | 0.578882031 | 0.027671158 |
| Tbc1d1 | S | | 782 | 0.019472789 | -0.081581443 | 0.013613426 | -0.005859363 | 0.095194869 | 0.544680432 | 0.028314718 | 0.532634354 | 0.846404512 | 0.235405046 |
| Wdtc1 | S | | 511 | 0.034395996 | 0.033952897 | -0.085564217 | -0.119960213 | -0.119517114 | 0.496515249 | 0.530831903 | 0.12872414 | 0.08937184 | 0.10256103 |
| Macf1 | S | | 1367 | 0.034319708 | -0.011313327 | -0.040131575 | -0.074451283 | -0.028818248 | 0.428334998 | 0.780058497 | 0.255129929 | 0.155009799 | 0.525496563 |
| Ociad1 | S | | 193 | 0.27130165 | -0.227377776 | -0.057388465 | -0.328690115 | 0.169989311 | 0.33588475 | 0.254471377 | 0.83197558 | 0.354747849 | 0.56925673 |
| Baiap3 Mepce | S | 191;214 | | -0.227746497 | 0.253227994 | 0.042616355 | 0.270362852 | -0.210611639 | 0.006631564 | 0.082574408 | 0.558009243 | 0.014316548 | 0.127458701 |
| Wrnip1 | S | | 133 | 0.054866862 | -0.0251375 | -0.073714935 | -0.128581796 | -0.048577434 | 0.630328744 | 0.772179496 | 0.597647653 | 0.365181913 | 0.680351206 |
| Elac2 | S | | 797 | -0.057648834 | 0.01734781 | 0.013198713 | 0.070847547 | -0.004149097 | 0.268782003 | 0.493323788 | 0.83438184 | 0.344653372 | 0.945444494 |
| Map4 Mapt | S | 585;983 | | -0.049751779 | 0.047301881 | -0.023608825 | 0.026142954 | -0.070910706 | 0.448828452 | 0.485926168 | 0.651733143 | 0.692443598 | 0.330367313 |
| Sipa1l1 | S | | 1544 | -0.049221973 | 0.015823896 | 0.04585062 | 0.095072593 | 0.030026725 | 0.025164746 | 0.446365795 | 0.42719951 | 0.171241619 | 0.584823284 |
| Nol9 | S | | 128 | 0.010503812 | -0.043312608 | 0.036224056 | 0.025720244 | 0.079536665 | 0.884336324 | 0.105488194 | 0.489340122 | 0.752213256 | 0.190726801 |
| Itsn2 | S | | 838 | 0.091529507 | -0.063107361 | -0.05663709 | -0.148166596 | 0.006470271 | 0.008688252 | 0.132295598 | 0.046372373 | 0.002333793 | 0.847515381 |
| Champ1 | S | | 434 | -0.041756554 | -0.092613202 | 0.108891326 | 0.150647881 | 0.201504529 | 0.302826958 | 0.042680286 | 0.02097884 | 0.007636712 | 0.002586604 |
| Tnks1bp1 | T | | 1275 | -0.00618994 | -0.036749645 | 0.029163229 | 0.035353169 | 0.065912874 | 0.789273489 | 0.153716648 | 0.307148087 | 0.164909938 | 0.039576771 |

| | | | | | | | | | | | | |
|---|---|---|---|---|---|---|---|---|---|---|---|---|
| Sipa1l1 | S | 310 | -0.058040908 | 0.00842044 | 0.019058566 | 0.077099475 | 0.010638126 | 0.370198009 | 0.851949711 | 0.662310833 | 0.214720321 | 0.532052915 |
| Sipa1l1 | S | 311 | -0.058040908 | 0.00842044 | 0.019058566 | 0.077099475 | 0.010638126 | 0.370198009 | 0.851949711 | 0.662310833 | 0.214720321 | 0.532052915 |
| Srrm1 | S | 657 | -0.010391085 | -0.02053446 | 0.024974059 | 0.035365144 | 0.045508518 | 0.483507056 | 0.207040484 | 0.395510305 | 0.252452355 | 0.170351823 |
| Rpl37 Rpl37rt | S | 96 | 0.022090116 | 0.041618133 | -0.098716914 | -0.12080703 | -0.140335048 | 0.672949257 | 0.374656272 | 0.182073691 | 0.143694689 | 0.094718481 |
| Champ1 | S | 265 | -0.068079129 | 0.02357254 | 0.023177842 | 0.091256972 | -0.000394698 | 0.113377075 | 0.46205175 | 0.450469816 | 0.061229418 | 0.973828891 |
| Gatad2a | S | 545 | 0.139044083 | -0.113726811 | -0.068688269 | -0.207732352 | 0.045038542 | 0.064381577 | 0.173688857 | 0.306008995 | 0.001464136 | 0.437573848 |
| Qk | S | 69 | 0.000937254 | 0.017413227 | -0.037287722 | -0.038224976 | -0.054700949 | 0.993864977 | 0.65302438 | 0.333640235 | 0.754452383 | 0.147976677 |
| Snx24 | S | 113 | 0.138242396 | -0.003827302 | -0.182232092 | -0.320474488 | -0.17840479 | 0.017197086 | 0.936377782 | 0.019449139 | 0.00324729 | 0.030104499 |
| Tln1 | S | 729 | -0.012367014 | -0.013365014 | 0.007600367 | 0.019967381 | 0.020965381 | 0.668483211 | 0.726644375 | 0.936067953 | 0.830705143 | 0.826405674 |
| Chchd3 | S | 112 | 0.127911162 | -0.046002393 | -0.111944238 | -0.2398554 | -0.065941845 | 0.033731042 | 0.250773277 | 0.089948355 | 0.010964453 | 0.240723715 |
| Hnrnpl | S | 182 | 0.013459158 | -0.048438784 | 0.03898822 | 0.025529062 | 0.087427004 | 0.600548086 | 0.075277378 | 0.526375212 | 0.661365491 | 0.218836206 |
| Depdc1b | S | 448 | -0.193370463 | -0.098871064 | 0.287938756 | 0.481309219 | 0.38680982 | 0.300509257 | 0.545873263 | 0.269473163 | 0.110090563 | 0.166229024 |
| Nedd4 | T | 287 | -0.009909404 | 0.038307922 | -0.060397791 | -0.050488387 | -0.098705712 | 0.872866256 | 0.468599611 | 0.367504277 | 0.480082378 | 0.167168354 |
| 0610010K14Rik | T | 160 | -0.065928974 | 0.037738228 | -0.010701493 | 0.055227482 | -0.048439721 | 0.227769157 | 0.225966376 | 0.886126893 | 0.508989641 | 0.531577487 |
| Numa1 | S | 1771 | -0.038687162 | 0.081173614 | -0.088422151 | -0.04973499 | -0.169595766 | 0.376364704 | 0.107890318 | 0.268553 | 0.48333385 | 0.081276935 |
| Prdm16 | S | 66 | 0.040215063 | -0.02567873 | -0.066031732 | -0.106246795 | -0.040353002 | 0.68933303 | 0.748203288 | 0.724281421 | 0.591556486 | 0.8289315 |
| Srrm2 | S | 1778 | -0.041319171 | -0.015370062 | 0.03617966 | 0.077498831 | 0.051549722 | 0.394830247 | 0.721313726 | 0.506482724 | 0.218249579 | 0.366183671 |
| Zranb2 | S | 153 | -0.063144495 | 0.026422238 | 0.028964184 | 0.092108679 | 0.002541946 | 0.082228331 | 0.679000362 | 0.30142742 | 0.038250908 | 0.968551664 |
| Rhbdf1 | S | 391 | -0.087394537 | 0.061515892 | 0.013223904 | 0.100618441 | -0.048291988 | 0.131022144 | 0.338966395 | 0.802711729 | 0.161999507 | 0.507340998 |
| Kmt2b | S | 1170 | -0.00169241 | -0.016305254 | -0.003505068 | -0.001812658 | 0.012800186 | 0.97648528 | 0.60633879 | 0.953873004 | 0.981238985 | 0.840936707 |
| Ptpn14 | S | 646 | -0.048307588 | 0.05550139 | -0.033067072 | 0.015240516 | -0.088568462 | 0.300834072 | 0.190222033 | 0.408793535 | 0.726029383 | 0.069358089 |
| Tmbim1 | S | 79 | -0.039880929 | -0.039347389 | 0.096715264 | 0.136596193 | 0.136062653 | 0.088754981 | 0.135943551 | 0.090659325 | 0.05001019 | 0.04432391 |
| Terf2 | S | 367 | 0.051213702 | -0.056641914 | -0.011233227 | -0.062446929 | 0.045408687 | 0.367027957 | 0.235651074 | 0.806083699 | 0.198721899 | 0.109163011 |
| Zfp646 | S | 1409 | -0.338597965 | 0.046390589 | 0.192776562 | 0.531374526 | 0.146385973 | 0.041471759 | 0.478330678 | 0.056455185 | 0.010942314 | 0.123322525 |
| Prkd2 | S | 711 | -0.017200081 | 0.084637146 | -0.086660155 | -0.069460074 | -0.171297301 | 0.833326336 | 0.415244075 | 0.294516518 | 0.362999199 | 0.130922893 |
| Incenp | S | 454 | 0.01058061 | -0.011031377 | -0.024731959 | -0.035312569 | -0.013700581 | 0.93644477 | 0.886000334 | 0.677457369 | 0.797571492 | 0.87219375 |
| Parp12 | S | 273 | 0.056442331 | -0.008276035 | -0.06888978 | -0.125332111 | -0.060613745 | 0.731338394 | 0.899652892 | 0.52263845 | 0.493912074 | 0.571093195 |
| Sik3 | S | 493 | -0.057054854 | 0.149430139 | -0.134821576 | -0.077766722 | -0.284251715 | 0.535604353 | 0.151057864 | 0.22195965 | 0.428237454 | 0.037485909 |
| Pja1 | S | 32 | 0.023833305 | -0.035994202 | -0.004009958 | -0.027843263 | 0.031984244 | 0.484527399 | 0.253477848 | 0.93202573 | 0.540955832 | 0.472262206 |
| Ankrd52 | S | 1075 | 0.056250743 | 0.008980451 | -0.083914947 | -0.140165691 | -0.092895399 | 0.317174196 | 0.817989645 | 0.142730852 | 0.061769745 | 0.113437357 |
| Miga2 | T | 206 | 0.087923972 | 0.025306996 | -0.082584178 | -0.170508149 | -0.107891173 | 0.399536963 | 0.475743491 | 0.034047253 | 0.171838668 | 0.011522555 |
| Spry1 | S | 50 | -0.085487823 | 0.024516866 | 0.053231738 | 0.138719561 | 0.028714872 | 0.217463074 | 0.707595266 | 0.611082344 | 0.224074993 | 0.769475101 |
| Cdan1 | S | 148 | 0.004155694 | -0.035374476 | -0.004673141 | -0.008828836 | 0.030701335 | 0.957315277 | 0.739345152 | 0.941394212 | 0.919751204 | 0.784827094 |
| Akap12 | T | 632 | -0.007424968 | 0.018432585 | -0.03760992 | -0.030184952 | -0.056042505 | 0.826226017 | 0.700195602 | 0.550707779 | 0.601453844 | 0.402786709 |
| Ccny | S | 83 | -0.016880455 | 0.079829109 | -0.101996639 | -0.085116184 | -0.181825748 | 0.853192082 | 0.205550292 | 0.193297945 | 0.41994743 | 0.061365604 |
| Synj1 | S | 1084 | 0.06297819 | -0.035784425 | -0.052176197 | -0.115154388 | -0.016391772 | 0.388257329 | 0.573555514 | 0.434010269 | 0.17826641 | 0.812494135 |
| Map3k20 | S | 275 | 0.075619995 | -0.027836913 | -0.083217202 | -0.158837197 | -0.05538029 | 0.410728947 | 0.773947152 | 0.39064376 | 0.146540398 | 0.589614847 |
| Tex2 | S | 269 | 0.06265535 | -0.016824585 | -0.06667681 | -0.12933216 | -0.049852225 | 0.143458092 | 0.695713 | 0.120965095 | 0.001736387 | 0.134125856 |
| Ash1l | S | 1787 | -0.031527542 | 0.012147755 | -0.002505786 | 0.029021756 | -0.014653541 | 0.513694684 | 0.819292311 | 0.968984028 | 0.662178865 | 0.832435384 |
| Mast1 Mast2 Mast3 | S | 177;239;161 | -0.028284809 | -0.029978055 | 0.052720231 | 0.08100504 | 0.082698286 | 0.483432874 | 0.359973113 | 0.418787178 | 0.244369976 | 0.235596079 |
| Mast1 Mast2 Mast3 | S | 179;241;163 | -0.028284809 | -0.029978055 | 0.052720231 | 0.08100504 | 0.082698286 | 0.483432874 | 0.359973113 | 0.418787178 | 0.244369976 | 0.235596079 |
| Gkap1 | S | 27 | 0.013401199 | 0.014743657 | -0.061590379 | -0.074991578 | -0.076334036 | 0.832900981 | 0.73706182 | 0.453682055 | 0.418410962 | 0.36900057 |
| Fam174c | S | 93 | 0.025956839 | -0.00034134 | -0.0529839 | -0.078940739 | -0.05264256 | 0.2891238 | 0.993324246 | 0.222962334 | 0.117783166 | 0.293376149 |
| Dpy19l1 | S | 23 | -0.055486886 | 0.029909779 | 0.032735594 | 0.08822248 | 0.002825816 | 0.026424659 | 0.3558962 | 0.416593334 | 0.092353408 | 0.945486585 |
| H2ax | S | 137 | 0.040232999 | -0.057504704 | 0.017210886 | -0.023022113 | 0.074715589 | 0.423503204 | 0.065092863 | 0.515073997 | 0.603784846 | 0.000659213 |
| D830031N03Rik | S | 276 | 0.027540806 | 0.006251197 | -0.05038357 | -0.077924377 | -0.056634768 | 0.182711087 | 0.723413943 | 0.071161972 | 0.022514038 | 0.056651347 |
| Lrch4 | S | 513 | 0.06132783 | 0.082056205 | -0.17947252 | -0.240800349 | -0.261528725 | 0.429162388 | 0.203199751 | 0.068907195 | 0.033744213 | 0.036883658 |
| Prpf4b | S | 431 | -0.008436071 | -0.01415497 | 0.008127886 | 0.016563957 | 0.022282856 | 0.806530856 | 0.372127493 | 0.666578172 | 0.641110925 | 0.246333451 |
| Phf20 | S | 485 | -0.006230482 | -0.068324896 | 0.040107529 | 0.046338011 | 0.108432424 | 0.94002223 | 0.386575715 | 0.480755837 | 0.604404058 | 0.223458631 |
| Map4 | S | 99 | -0.024788871 | -0.025003202 | 0.023086702 | 0.047875573 | 0.048089905 | 0.796921071 | 0.733033784 | 0.855159747 | 0.726799139 | 0.701872887 |
| Trp53bp1 | S | 652 | -0.087408179 | 0.016570018 | 0.012314471 | 0.099722651 | -0.004255546 | 0.500075266 | 0.492293447 | 0.906316734 | 0.51729654 | 0.96711542 |
| Araf Raf1 | S | 580;621 | -0.006691179 | 0.036129029 | -0.05625726 | -0.049566081 | -0.092386289 | 0.882098822 | 0.455665635 | 0.351933526 | 0.415093247 | 0.175205853 |
| Arhgef17 | S | 456 | -0.055900252 | -0.114688507 | 0.11906367 | 0.174963922 | 0.233752178 | 0.714838815 | 0.081787399 | 0.065124668 | 0.305498432 | 0.005580471 |
| Nucks1 | S | 73 | -0.085500028 | 0.021092173 | 0.018392572 | 0.1038926 | -0.002699601 | 0.351397984 | 0.773328213 | 0.440199734 | 0.279595253 | 0.969705 |
| Srebf1 | S | 448 | 0.168035063 | -0.164922393 | -0.007675685 | -0.175710748 | 0.157246708 | 0.123635204 | 0.070007024 | 0.870475066 | 0.12666443 | 0.093408154 |
| Kif7 | S | 1337 | -0.042032331 | 0.013476849 | 0.000120495 | 0.042152826 | -0.013356354 | 0.578730416 | 0.835138226 | 0.998427009 | 0.469695948 | 0.727170967 |
| Rfc1 | S | 535 | 0.079895486 | -0.006742233 | -0.072340614 | -0.1522361 | -0.065598381 | 0.204652163 | 0.877554219 | 0.079323212 | 0.065188646 | 0.120663031 |
| Fyb | S | 222 | 0.025980399 | -0.04013809 | -0.015918564 | -0.041898963 | 0.024219526 | 0.710881309 | 0.578116504 | 0.868039031 | 0.62577637 | 0.777418038 |
| U2surp | S | 969 | 0.012176893 | -0.070146931 | 0.028945823 | 0.016768931 | 0.099092755 | 0.818995474 | 0.266979952 | 0.441989691 | 0.737874011 | 0.14743932 |
| Cast | T | 479 | -0.076724337 | 0.092904838 | -0.030126282 | 0.046598055 | -0.12303112 | 0.05788281 | 0.051274788 | 0.433900824 | 0.2294271 | 0.028989326 |
| Atf7 | S | 82 | -0.107139547 | -0.003537655 | 0.109281804 | 0.216421351 | 0.112819459 | 0.074703502 | 0.907042631 | 0.163139639 | 0.034636206 | 0.166472754 |
| Camsap1 | S | 560 | 0.056020724 | 0.011616705 | -0.076549611 | -0.132570335 | -0.088166316 | 0.218003295 | 0.638447092 | 0.059382388 | 0.033795843 | 0.045926436 |
| Srrm2 | S | 1377 | -0.080254027 | -0.05857613 | 0.145006734 | 0.225260761 | 0.203582864 | 0.396378742 | 0.246408304 | 0.234314832 | 0.119386428 | 0.145442623 |
| Tra2b | T | 201 | -0.031710566 | -0.018134572 | 0.042457025 | 0.07416759 | 0.060591597 | 0.466314677 | 0.508269232 | 0.518433649 | 0.317411017 | 0.37770072 |
| Ptbp2 | S | 308 | 0.117846897 | -0.081731767 | -0.036480763 | -0.15432766 | 0.045251004 | 0.186995455 | 0.13395319 | 0.59440835 | 0.12916215 | 0.497407958 |
| Sgo1 | S | 497 | 0.017862323 | 0.10246658 | -0.085681724 | -0.103544047 | -0.188148304 | 0.883426157 | 0.428204816 | 0.104005179 | 0.423006344 | 0.203007683 |
| Bod1l | S | 2968 | 0.012909839 | -0.010852566 | -0.026018083 | -0.038927922 | -0.015165518 | 0.736583868 | 0.813657678 | 0.608126964 | 0.454226125 | 0.784348313 |
| Srrm2 | S | 1780 | -0.045929954 | -0.011149012 | 0.040487678 | 0.086417631 | 0.051636689 | 0.381519524 | 0.86391749 | 0.562357859 | 0.235476825 | 0.501725999 |
| Srrm1 | S | 400 | 0.055300363 | -0.041337024 | -0.034536626 | -0.08983699 | 0.006800398 | 0.223460747 | 0.446835156 | 0.302089175 | 0.08573586 | 0.894406379 |
| Incenp | T | 215 | 0.053977086 | -0.108829371 | 0.038596415 | -0.015380671 | 0.147425785 | 0.452768076 | 0.183803572 | 0.66811067 | 0.874744155 | 0.190677348 |
| Myo18a | S | 1063 | -0.011506332 | 0.133800874 | -0.158781409 | -0.147275077 | -0.292582283 | 0.89055174 | 0.221825094 | 0.19447488 | 0.236009766 | 0.062850032 |
| Slc2a1 | T | 478 | 0.016600652 | 0.014369052 | -0.049902115 | -0.066502767 | -0.064271167 | 0.674083893 | 0.633032643 | 0.259496328 | 0.190262425 | 0.161333859 |
| Eif4enif1 | S | 586 | -0.039096245 | 0.017186194 | 0.003009533 | 0.042105778 | -0.01417666 | 0.256409145 | 0.42468905 | 0.950500669 | 0.428475795 | 0.75937833 |
| Arhgap10 | T | 703 | -0.038348217 | -0.006036762 | 0.014482305 | 0.052830522 | 0.020519068 | 0.655483874 | 0.940935976 | 0.843875744 | 0.384359057 | 0.678760981 |
| Entr1 | S | 241 | 0.058987123 | -0.036210344 | -0.103388971 | -0.162376094 | -0.067178628 | 0.249054333 | 0.655881974 | 0.523074462 | 0.344658642 | 0.678615136 |
| Entr1 | S | 245 | 0.058987123 | -0.036210344 | -0.103388971 | -0.162376094 | -0.067178628 | 0.249054333 | 0.655881974 | 0.523074462 | 0.344658642 | 0.678615136 |
| Tacc1 | S | 480 | -0.000529447 | -0.021354607 | -0.002777677 | -0.00224823 | 0.01857693 | 0.99208086 | 0.711471796 | 0.976586793 | 0.979118476 | 0.833502085 |
| Ofd1 | S | 670 | -0.059674161 | 0.070467744 | -0.041048264 | 0.018625897 | -0.111516009 | 0.480619419 | 0.509324143 | 0.767355021 | 0.893784552 | 0.475292979 |
| Ofd1 | T | 663 | -0.059674161 | 0.070467744 | -0.041048264 | 0.018625897 | -0.111516009 | 0.480619419 | 0.509324143 | 0.767355021 | 0.893784552 | 0.475292979 |
| Tnrc6c | T | 925 | -0.138084908 | 0.052310774 | 0.055489569 | 0.193574478 | 0.003178795 | 0.218318081 | 0.596991028 | 0.490396724 | 0.120059827 | 0.974305632 |
| Usp8 | S | 680 | -0.013543049 | 0.07188951 | -0.08171504 | -0.068171991 | -0.153604551 | 0.782010364 | 0.336187727 | 0.291370548 | 0.373342314 | 0.125991237 |
| Kmt2d | S | 1907 | -0.039111944 | 0.006473764 | 0.004328273 | 0.043440217 | -0.002145492 | 0.668086539 | 0.960734083 | 0.953750355 | 0.545683132 | 0.985709792 |
| Ankrd11 | S | 1808 | 0.037020356 | -0.218109138 | 0.18814126 | 0.151120904 | 0.406250398 | 0.719211751 | 0.053825339 | 0.154299511 | 0.219568792 | 0.019967155 |
| Wdr44 | S | 473 | -0.020893969 | 0.014467354 | -0.016053276 | 0.004840694 | -0.030520629 | 0.60346 | 0.525991493 | 0.613509466 | 0.904272592 | 0.291929658 |
| Wdr44 | S | 474 | -0.020893969 | 0.014467354 | -0.016053276 | 0.004840694 | -0.030520629 | 0.60346 | 0.525991493 | 0.613509466 | 0.904272592 | 0.291929658 |
| Zfp516 | S | 16 | 0.016086634 | -0.057739619 | 0.043989076 | 0.027902442 | 0.101728695 | 0.789155337 | 0.077103106 | 0.305878232 | 0.655097465 | 0.061484071 |
| Exoc1 | S | 481 | 0.044698229 | 0.022592251 | -0.107154569 | -0.151852798 | -0.12974682 | 0.359077561 | 0.677051309 | 0.221115037 | 0.119563368 | 0.16685343 |
| Kif4 | S | 1230 | -0.036320847 | 0.018248798 | -0.011794966 | 0.02452588 | -0.030043765 | 0.73943835 | 0.901043141 | 0.911657243 | 0.829122073 | 0.841056556 |
| Dck | S | 11 | -0.087787943 | 0.039304762 | 0.036570838 | 0.124358781 | -0.002733923 | 0.073627636 | 0.306078786 | 0.458860342 | 0.058169746 | 0.952914982 |
| Ywhab | S | 216 | -0.15385976 | 0.016126215 | 0.136086505 | 0.289946265 | 0.119960291 | 0.083298008 | 0.891169767 | 0.182283511 | 0.025113079 | 0.323066959 |
| Ywhab | T | 217 | -0.15385976 | 0.016126215 | 0.136086505 | 0.289946265 | 0.119960291 | 0.083298008 | 0.891169767 | 0.182283511 | 0.025113079 | 0.323066959 |
| Ywhab | T | 228 | -0.15385976 | 0.016126215 | 0.136086505 | 0.289946265 | 0.119960291 | 0.083298008 | 0.891169767 | 0.182283511 | 0.025113079 | 0.323066959 |
| Ctnnd1 | S | 47 | 0.028908903 | 0.022084986 | -0.064659061 | -0.093567964 | -0.086744047 | 0.365242462 | 0.357180861 | 0.069383377 | 0.037322224 | 0.034050468 |
| Trappc10 | S | 710 | -0.034336178 | 0.11902256 | -0.141878359 | -0.107542181 | -0.260900919 | 0.661164591 | 0.080548253 | 0.146333544 | 0.287212622 | 0.036437465 |

| | | | | | | | | | | | | |
|---|---|---|---|---|---|---|---|---|---|---|---|---|
| Garnl3 | S | 893 | 0.033123226 | 0.043863001 | -0.125258494 | -0.158381719 | -0.169121495 | 0.73618856 | 0.566323198 | 0.371420925 | 0.276628938 | 0.240227281 |
| Map1a | T | 1003 | -0.020642572 | 0.009189985 | -0.014450104 | 0.006192468 | -0.02364009 | 0.696678508 | 0.864991637 | 0.88105179 | 0.949247296 | 0.810132417 |
| Mbd1 | S | 440 | 0.063626174 | -0.0799532 | 0.009687486 | -0.053938688 | 0.089640686 | 0.396380845 | 0.166206868 | 0.822408104 | 0.443577441 | 0.110889845 |
| Slc12a2 | S | 127 | 0.002637058 | -0.172150721 | 0.136013144 | 0.133376086 | 0.308163865 | 0.977145325 | 0.086601722 | 0.109260113 | 0.181430043 | 0.016879451 |
| C2cd2l | S | 662 | 0.058867242 | -0.041105068 | -0.040089134 | -0.098956376 | 0.001015934 | 0.336614335 | 0.428283101 | 0.534252237 | 0.160859647 | 0.984928652 |
| Ube4b | S | 674 | -0.082994757 | 0.093493119 | -0.07111121 | 0.011883547 | -0.164604329 | 0.293736701 | 0.150684143 | 0.590228348 | 0.928619161 | 0.265715088 |
| Ubap2 | S | 634 | 0.049870158 | 0.041930461 | -0.064554466 | -0.114424624 | -0.106484926 | 0.038351202 | 0.417741756 | 0.000677883 | 0.006421952 | 0.12050545 |
| Cdca5 | S | 164 | -0.090114132 | 0.09637665 | -0.022356624 | 0.067757508 | -0.118733274 | 0.4063951 | 0.399236726 | 0.614719358 | 0.505618566 | 0.312358804 |
| Hspe1 | S | 57 | 0.005273226 | -0.032782443 | 0.019838505 | 0.014565279 | 0.052620948 | 0.883928942 | 0.328313857 | 0.758366574 | 0.830340656 | 0.45227581 |
| Flnb | S | 789 | 0.008702149 | -0.03393245 | 0.003051073 | -0.005651076 | 0.036983524 | 0.845275876 | 0.54759876 | 0.942314077 | 0.854092488 | 0.44942248 |
| Wdr7 | S | 1455 | 0.06083341 | -0.037888593 | -0.05252169 | -0.113355099 | -0.014633097 | 0.123927131 | 0.227426898 | 0.429585872 | 0.147176467 | 0.807009632 |
| Myo1e | S | 1001 | 0.126943619 | 0.024207136 | -0.177066469 | -0.304010088 | -0.201273605 | 0.34168793 | 0.782203008 | 0.195865381 | 0.084135231 | 0.148351226 |
| Cic | T | 2283 | -0.112140306 | -0.012758991 | 0.089650076 | 0.201790382 | 0.102409067 | 0.180503394 | 0.873946377 | 0.185391642 | 0.037259478 | 0.185382289 |
| Numa1 | T | 1770 | -0.031543614 | 0.03688359 | -0.022585574 | 0.00895804 | -0.059469165 | 0.318321813 | 0.407741596 | 0.741660402 | 0.890539427 | 0.423334275 |
| Ssh2 | S | 1217 | 0.053951123 | -0.03938792 | -0.026359088 | -0.080310211 | 0.013028832 | 0.465548004 | 0.545599843 | 0.481729539 | 0.312862817 | 0.841742248 |
| Map7d1 | S | 118 | -0.023520762 | 0.016880935 | -0.012229415 | 0.011291348 | -0.029110349 | 0.497429106 | 0.527271139 | 0.701029384 | 0.728544622 | 0.284132701 |
| Tsen2 | T | 147 | 0.01512019 | 0.069676449 | -0.105530678 | -0.120650867 | -0.175207127 | 0.789298432 | 0.511164773 | 0.2810794 | 0.21647375 | 0.176764653 |
| Rrp12 | T | 88 | -0.03427457 | 0.004931983 | 0.01725957 | 0.051534139 | 0.012327586 | 0.379756767 | 0.899208601 | 0.756581235 | 0.357927957 | 0.813910015 |
| Cep170b | S | 534 | -0.009674564 | 0.014142999 | -0.03776523 | -0.028090665 | -0.051908229 | 0.81424606 | 0.433600882 | 0.53363905 | 0.675643462 | 0.413946477 |
| Kmt2e | S | 1002 | -0.019243213 | 0.00614429 | -0.005594603 | 0.01364861 | -0.011738894 | 0.591146373 | 0.878961516 | 0.948681214 | 0.871604115 | 0.891299678 |
| Ppan | S | 362 | 0.208876771 | 0.016692326 | -0.339442339 | -0.548319109 | -0.356134665 | 0.125329487 | 0.795835595 | 0.144871834 | 0.046965173 | 0.133191536 |
| Slk | S | 348 | -0.017671681 | 0.044353614 | -0.053482911 | -0.03581123 | -0.097836525 | 0.829632747 | 0.332706802 | 0.302449747 | 0.645090036 | 0.055713187 |
| Retreg3 | S | 435 | -0.026590517 | 0.016115233 | 0.010056536 | 0.036647052 | -0.006058698 | 0.187728714 | 0.728742515 | 0.779181659 | 0.345309283 | 0.908361381 |
| Rif1 | S | 376 | 0.096960454 | -0.120837829 | -0.001632542 | -0.098592996 | 0.119205287 | 0.538885649 | 0.405299514 | 0.991356815 | 0.605612247 | 0.512456938 |
| Hmg20a | S | 104 | -0.050407347 | 0.007187663 | 0.028600364 | 0.079007711 | 0.021412701 | 0.228895029 | 0.84687332 | 0.460560362 | 0.016432189 | 0.155150258 |
| Srebf1 | S | 96 | 0.073337961 | -0.075836616 | -0.038224663 | -0.111562624 | 0.037611953 | 0.023085854 | 0.151630852 | 0.232155303 | 0.016670473 | 0.414635305 |
| Huwe1 | S | 648 | -0.045039835 | 0.043206984 | -0.018855814 | 0.026184021 | -0.062062799 | 0.602148677 | 0.642502043 | 0.740932032 | 0.745585069 | 0.497535445 |
| Lsm14a | S | 182 | -0.023580079 | -0.007590341 | 0.008721922 | 0.032302001 | 0.016312263 | 0.711349995 | 0.897164314 | 0.899550632 | 0.67562157 | 0.823715187 |
| Psen1 | S | 327 | 0.100670675 | -0.043556878 | -0.049767837 | -0.150438512 | -0.006210959 | 0.120381408 | 0.34376953 | 0.107132689 | 0.05899528 | 0.875578367 |
| Peak1 | T | 568 | -0.103773337 | 0.11049129 | -0.035592576 | 0.068180761 | -0.146083866 | 0.183871884 | 0.161162861 | 0.681016783 | 0.353419587 | 0.096796148 |
| Zc3h12c | S | 233 | -0.001419163 | -0.11580084 | 0.103973708 | 0.105392871 | 0.219774549 | 0.985889903 | 0.340258073 | 0.493339329 | 0.502532578 | 0.232400875 |
| Plec | S | 114 | 0.036440224 | 0.000759357 | -0.065576114 | -0.102016338 | -0.06633547 | 0.274287911 | 0.978389421 | 0.23546378 | 0.104242889 | 0.228251446 |
| Specc1 | S | 810 | -0.03239829 | -0.028305658 | 0.037162463 | 0.069560753 | 0.065468121 | 0.556154715 | 0.588863846 | 0.442880648 | 0.128973693 | 0.102776199 |
| Sort1 | S | 819 | 0.056383799 | -0.031175177 | -0.046130576 | -0.102514375 | -0.0149554 | 0.259234569 | 0.309826091 | 0.533118028 | 0.225918646 | 0.824644385 |
| Atxn3 | S | 273 | -0.029414664 | 0.049493164 | -0.051699722 | -0.022285058 | -0.101192886 | 0.519567675 | 0.217063678 | 0.387928812 | 0.719101676 | 0.146474396 |
| Med1 | S | 1481 | -0.001364595 | -0.034433426 | 0.027328624 | 0.028693219 | 0.06176205 | 0.985241345 | 0.320017139 | 0.602002209 | 0.713464767 | 0.243944404 |
| Srrm1 | S | 672 | 0.022815137 | -0.021121264 | -0.01979909 | -0.042614227 | 0.001322173 | 0.187847074 | 0.197291649 | 0.636359793 | 0.342194393 | 0.97298481 |
| Ncor2 | S | 1509 | -0.137279727 | 0.106557969 | 0.031998215 | 0.169277942 | -0.074559754 | 0.063680909 | 0.226384077 | 0.574612383 | 0.025279359 | 0.328222037 |
| Hdac4 | S | 630 | -0.054374467 | 0.050822475 | -0.015617982 | 0.038756485 | -0.066440457 | 0.419524543 | 0.454796621 | 0.752820956 | 0.580221305 | 0.369211327 |
| Abi1 | T | 215 | -0.013282969 | 0.058340095 | -0.082339063 | -0.069056094 | -0.140679158 | 0.732721716 | 0.222618564 | 0.298637281 | 0.354458833 | 0.12057083 |
| Suz12 | S | 543 | 0.065712248 | -0.033184131 | -0.047396874 | -0.113109122 | -0.014212743 | 0.399236364 | 0.458718624 | 0.551072573 | 0.278861103 | 0.861277537 |
| Birc6 | S | 3960 | -0.095462059 | -0.148564182 | 0.233868309 | 0.329330369 | 0.382432492 | 0.294796771 | 0.183464875 | 0.084495437 | 0.046035982 | 0.020649807 |
| Tpx2 | S | 737 | 0.067444671 | 0.010130959 | -0.078898119 | -0.14634279 | -0.089029078 | 0.054574899 | 0.630061603 | 0.011454637 | 0.005816742 | 0.014154463 |
| Smc3 | S | 1083 | -0.0621248 | 0.053596115 | 0.00015396 | 0.062278761 | -0.053442155 | 0.231331333 | 0.408694502 | 0.998624819 | 0.501245759 | 0.58118018 |
| Kri1 | S | 182 | 0.015375423 | -0.048514194 | 0.023452802 | 0.008077379 | 0.071966995 | 0.732592537 | 0.151048018 | 0.371641748 | 0.851909967 | 0.057428508 |
| Chd8 | S | 2204 | -0.141925471 | 0.001919999 | 0.093848757 | 0.235774228 | 0.091928758 | 0.304066564 | 0.982280914 | 0.32664879 | 0.145433401 | 0.424690061 |
| Snapc2 | S | 356 | 0.079359184 | -0.153396875 | 0.028295215 | -0.051063968 | 0.18169209 | 0.553535364 | 0.301954343 | 0.546095941 | 0.694336335 | 0.242101409 |
| Son | S | 1866 | 0.175673362 | -0.018093178 | -0.101422224 | -0.277095586 | -0.083329046 | 0.372201435 | 0.907885632 | 0.129289201 | 0.208648249 | 0.601415876 |
| Taf6 | S | 673 | -0.05716994 | 0.025147592 | -0.004214219 | 0.052955721 | -0.029361811 | 0.246676587 | 0.129375024 | 0.86746313 | 0.283184813 | 0.267558997 |
| Setd1a | T | 553 | -0.032999812 | 0.106150518 | -0.071510785 | -0.038510972 | -0.177661303 | 0.488025667 | 0.298794675 | 0.321730689 | 0.597411718 | 0.140241298 |
| Tjp2 | S | 209 | -0.018423772 | -0.002205085 | -0.005205406 | 0.013218366 | -0.00300032 | 0.855655278 | 0.958600848 | 0.94424661 | 0.909677909 | 0.969497816 |
| Noc2l | S | 679 | 0.051911327 | -0.006532299 | -0.041727684 | -0.093639011 | -0.035195385 | 0.352415048 | 0.882681004 | 0.144271978 | 0.149253907 | 0.418619887 |
| Srrt | S | 4 | 0.023337931 | -0.026439965 | -0.008908537 | -0.032246468 | 0.017531428 | 0.723423306 | 0.541608564 | 0.838009664 | 0.638041647 | 0.708097618 |
| Clcc1 | S | 498 | 0.002359268 | -0.033350467 | 0.010075239 | 0.007715972 | 0.043425706 | 0.973130388 | 0.635149715 | 0.883276159 | 0.84712309 | 0.303191478 |
| Safb | S | 636 | -0.129741401 | -0.0293017 | 0.156070293 | 0.285811694 | 0.185371993 | 0.047140357 | 0.641862463 | 0.059241664 | 0.011429499 | 0.055063866 |
| Rbmxl1 | T | 178 | 0.081448782 | -0.123297208 | 0.040600717 | -0.040848065 | 0.163897925 | 0.136249261 | 0.037527314 | 0.555971528 | 0.489593369 | 0.072072341 |
| Tom1 | S | 472 | 0.037955925 | -0.014046913 | -0.042457652 | -0.080413577 | -0.028410739 | 0.207802409 | 0.809685829 | 0.18313811 | 0.012563928 | 0.61145766 |
| Nol9 | T | 496 | 0.138043661 | -0.09220234 | -0.119092499 | -0.257136161 | -0.02689016 | 0.254661879 | 0.620829966 | 0.461204934 | 0.174678771 | 0.898745906 |
| Rbm26 | S | 127 | 0.038958332 | -0.02747026 | -0.021574996 | -0.060533328 | 0.005895264 | 0.399433092 | 0.275973622 | 0.746458155 | 0.423461078 | 0.927290595 |
| Chrac1 | S | 122 | -0.034052862 | 0.043347718 | -0.03034662 | 0.003706242 | -0.073694338 | 0.144886224 | 0.13344002 | 0.637417651 | 0.951321032 | 0.301288345 |
| Anxa5 | S | 133 | -0.034223617 | -0.000892277 | 0.014028574 | 0.048252191 | 0.014920851 | 0.628954769 | 0.989967065 | 0.827145338 | 0.564661827 | 0.857872457 |
| Zfp746 | S | 360 | -0.169262962 | 0.144572931 | -0.061537007 | 0.107725954 | -0.206109938 | 0.225174596 | 0.080372768 | 0.596134064 | 0.469402473 | 0.137672416 |
| Pkp4 | S | 775 | -0.136388804 | -0.031191618 | 0.134531225 | 0.270920029 | 0.165722843 | 0.142317564 | 0.575099989 | 0.089039374 | 0.02602102 | 0.064959332 |
| Trim56 | S | 448 | -0.127246903 | 0.003488946 | 0.208396586 | 0.335643489 | 0.20490764 | 0.018743703 | 0.938426176 | 0.158107254 | 0.075255449 | 0.16962532 |
| Fadd | T | 204 | -0.023890295 | 0.10729981 | -0.132890629 | -0.108994334 | -0.240190439 | 0.838294308 | 0.268526044 | 0.323682681 | 0.409302443 | 0.103748766 |
| Dbr1 | S | 489 | -0.011651545 | 0.047453406 | -0.062984935 | -0.05133339 | -0.110438341 | 0.858818327 | 0.299399703 | 0.304876201 | 0.448185737 | 0.09992544 |
| Zfp106 | T | 1383 | -0.047395885 | 0.033204416 | 0.011074401 | 0.058470286 | -0.022130015 | 0.113131206 | 0.255937292 | 0.910191014 | 0.563539344 | 0.819975948 |
| Tjp2 | S | 1044 | 0.015502434 | 0.024855763 | -0.066151372 | -0.081653807 | -0.091007136 | 0.860200391 | 0.746618954 | 0.513788856 | 0.403601358 | 0.321424461 |
| Dennd1a | S | 536 | 0.058468773 | 0.015841585 | -0.072142163 | -0.130610936 | -0.087983748 | 0.250897117 | 0.694162333 | 0.071332497 | 0.04630094 | 0.04237113 |
| Snrpd2 | T | 12 | -0.02541613 | -0.025413061 | 0.042372825 | 0.067788955 | 0.067785887 | 0.646803486 | 0.677248346 | 0.673956111 | 0.506816299 | 0.514654002 |
| Fhdc1 | S | 655 | 0.100959779 | -0.027630491 | -0.08149089 | -0.182450668 | -0.053860398 | 0.081043947 | 0.228022268 | 0.17490832 | 0.029399964 | 0.321118715 |
| Ar | S | 75 | -0.069580769 | 0.118044053 | -0.090730685 | -0.021149916 | -0.208774738 | 0.469750424 | 0.203856022 | 0.423492898 | 0.868096351 | 0.141737201 |
| Scrib | T | 1531 | -0.090521504 | -0.112090703 | 0.17294371 | 0.263465213 | 0.285034413 | 0.511809075 | 0.369764715 | 0.227642034 | 0.159752869 | 0.114074432 |
| 4932438A13Rik | S | 4367 | -0.099831869 | 0.191037778 | -0.131374099 | -0.031542229 | -0.322411877 | 0.378761782 | 0.143329396 | 0.3487319 | 0.801416474 | 0.061985859 |
| Zzef1 | S | 1518 | 0.093750004 | 0.068263183 | -0.136934943 | -0.230684947 | -0.205198126 | 0.299173531 | 0.464562739 | 0.050595064 | 0.049125244 | 0.081389176 |
| Septin9 | T | 42 | -0.003075644 | -0.022632111 | 0.013283167 | 0.016358811 | 0.035915278 | 0.936192537 | 0.547855783 | 0.782055302 | 0.662815556 | 0.3634811 |
| Matr3 | S | 22 | 0.012917044 | -0.047193026 | -0.003706016 | -0.016623061 | 0.043487009 | 0.977886257 | 0.858219552 | 0.992458325 | 0.973025873 | 0.893585048 |
| Tle1 | S | 284 | -0.133886709 | -0.00395151 | 0.097058077 | 0.230944786 | 0.101009587 | 0.285022251 | 0.971988428 | 0.288167748 | 0.0932576 | 0.332389127 |
| Isg15 | T | 104 | -0.175948162 | -0.021363282 | 0.157335371 | 0.333283533 | 0.178698653 | 0.385716659 | 0.826105196 | 0.361076879 | 0.176235481 | 0.297571116 |
| Anp32a | T | 15 | -0.059764409 | 0.027998944 | 0.00761765 | 0.067382058 | -0.020381294 | 0.528537195 | 0.713229737 | 0.919095835 | 0.453410307 | 0.761069688 |
| Akap13 | S | 1905 | -0.262939341 | 0.190897711 | 0.051256617 | 0.314195957 | -0.139641095 | 0.043950708 | 0.087961178 | 0.632624604 | 0.025452447 | 0.152691702 |
| Iffo2 | S | 343 | 0.04683247 | -0.070415842 | 0.020046511 | -0.02678596 | 0.090462353 | 0.775605628 | 0.463150962 | 0.874770338 | 0.887820381 | 0.533089752 |
| Sik3 | S | 533 | -0.059871916 | 0.068797948 | -0.050774168 | 0.009097748 | -0.119572117 | 0.386787647 | 0.191855755 | 0.566915036 | 0.917193392 | 0.216598667 |
| Sp3 | S | 72 | 0.043944285 | -0.08516573 | 0.031352801 | -0.012591484 | 0.116518531 | 0.358364149 | 0.130304917 | 0.567788756 | 0.818643828 | 0.097900739 |
| Wiz | S | 1016 | 0.067635303 | -0.119467178 | 0.035221867 | -0.032413435 | 0.154689045 | 0.492329476 | 0.244722504 | 0.663353734 | 0.667223942 | 0.109705592 |
| Samd4 | S | 644 | -0.035340121 | 0.029228631 | -0.026104368 | 0.009235753 | -0.055333 | 0.693335472 | 0.479041072 | 0.661752625 | 0.920286544 | 0.337125617 |
| Ep400 | S | 315 | -0.011024981 | 0.017501914 | -0.020584875 | -0.009559894 | -0.038086789 | 0.849550968 | 0.90089734 | 0.783314924 | 0.902135025 | 0.79672024 |
| Phldb1 | S | 382 | 0.222618628 | -0.117762176 | -0.205449094 | -0.428067722 | -0.087686918 | 0.001251661 | 0.247088915 | 0.022179764 | 0.003391864 | 0.38232269 |
| Srrm2 | S | 870 | -0.025024054 | 0.008262085 | -0.003741479 | 0.021282575 | -0.012003564 | 0.704935113 | 0.816102788 | 0.923777603 | 0.753219485 | 0.76640172 |
| Rbbp6 | T | 1194 | -0.140838293 | -0.04104859 | 0.211832326 | 0.352670619 | 0.252880915 | 0.117115398 | 0.53050022 | 0.143406764 | 0.047626025 | 0.114943472 |
| Edc4 | S | 745 | -0.0174921 | -0.000623039 | -0.004517006 | 0.012975095 | -0.003893967 | 0.837087633 | 0.993021773 | 0.971559538 | 0.92320334 | 0.975777326 |
| Top2b | T | 1562 | -0.066424986 | 0.038747019 | 0.032195373 | 0.098620359 | -0.006551646 | 0.225468534 | 0.661227657 | 0.645048476 | 0.184729858 | 0.943489528 |
| Spp1 | T | 175 | 0.050691932 | 0.024492282 | -0.09335915 | -0.144051083 | -0.117851433 | 0.475001257 | 0.718088512 | 0.262782241 | 0.105054842 | 0.155701487 |
| Tsc1 | S | 562 | 0.030486664 | -0.030421906 | -0.02131516 | -0.051801824 | 0.009106746 | 0.497062096 | 0.441856971 | 0.857697972 | 0.675522311 | 0.939568774 |

| | | | | | | | | | | | | | |
|---|---|---|---|---|---|---|---|---|---|---|---|---|---|
| Baz1a | S | | 963 | 0.198011384 | -0.124387978 | -0.133179809 | -0.331191192 | -0.008791831 | 0.130615522 | 0.33335607 | 0.433028688 | 0.116674393 | 0.959894227 |
| Deaf1 | T | | 313 | 0.007839719 | -0.085363172 | 0.059424209 | 0.05158449 | 0.14478738 | 0.878016104 | 0.31766216 | 0.468151364 | 0.542480716 | 0.189648985 |
| Spp1 | S | | 183 | 0.022457066 | 0.122453038 | -0.156366792 | -0.178823858 | -0.27881983 | 0.799531723 | 0.21237328 | 0.121100777 | 0.018512401 | 0.005969375 |
| Rbmx Rbmxl1 | S | 212;215 | | -0.033011685 | -0.036091702 | 0.050773029 | 0.083784714 | 0.086864731 | 0.452855052 | 0.376912778 | 0.209996012 | 0.085085066 | 0.056603309 |
| Lmnb1 | S | | 24 | -0.042780438 | 0.035768343 | -0.005731802 | 0.037048636 | -0.041500146 | 0.162793247 | 0.186991592 | 0.817010924 | 0.143999642 | 0.050389164 |
| Rassf8 | S | | 387 | -0.000754466 | 0.019222928 | -0.036547603 | -0.035793138 | -0.055770531 | 0.988340234 | 0.81887351 | 0.644177971 | 0.680519699 | 0.602428478 |
| Akap1 | S | | 101 | 0.123701652 | -0.029287649 | -0.118824656 | -0.242526308 | -0.089537007 | 0.018619796 | 0.445663935 | 0.041202119 | 0.004388113 | 0.093970541 |
| Kars | T | | 589 | -0.070997465 | -0.004069636 | 0.033969886 | 0.104967351 | 0.038039522 | 0.57165366 | 0.929960472 | 0.518779493 | 0.421893347 | 0.438791696 |
| Acin1 | S | | 427 | -0.073263713 | -0.024081128 | 0.14947613 | 0.222739843 | 0.173557258 | 0.057874351 | 0.461700102 | 0.156266232 | 0.083322226 | 0.13154016 |
| Melk | S | | 521 | 0.0009356 | 0.069295725 | -0.097895651 | -0.098831251 | -0.167191376 | 0.990857048 | 0.369098561 | 0.323673558 | 0.358166115 | 0.142072244 |
| Kif1c | S | | 680 | -0.066172264 | 0.043362482 | 0.026055762 | 0.092228026 | -0.01730672 | 0.096762184 | 0.504595023 | 0.400111554 | 0.040930063 | 0.765482768 |
| Vps13a | S | | 835 | -0.054226768 | -0.019537675 | 0.045790043 | 0.100016812 | 0.065327719 | 0.554845338 | 0.804292525 | 0.501959474 | 0.301153931 | 0.419626146 |
| Cobll1 | S | | 884 | 0.107759867 | -0.102302705 | -0.033263845 | -0.141023711 | 0.06903886 | 0.103329253 | 0.12786452 | 0.730705959 | 0.215486708 | 0.501471202 |
| Srrm1 | T | | 414 | 0.110106053 | -0.047841171 | -0.093689754 | -0.203795807 | -0.045848583 | 0.096758993 | 0.600470412 | 0.156302993 | 0.004183265 | 0.574492445 |
| Ndc1 | T | | 439 | 0.098565409 | -0.147650277 | 0.107283945 | 0.008718536 | 0.254934223 | 0.569384692 | 0.103598056 | 0.489354105 | 0.964492102 | 0.159136513 |
| Ndc1 | T | | 448 | 0.098565409 | -0.147650277 | 0.107283945 | 0.008718536 | 0.254934223 | 0.569384692 | 0.103598056 | 0.489354105 | 0.964492102 | 0.159136513 |
| Relch | S | | 56 | -0.012372611 | -0.015496771 | 0.005060592 | 0.017433203 | 0.020557363 | 0.902506636 | 0.875051606 | 0.95501385 | 0.80741881 | 0.761166438 |
| Usp20 | T | | 378 | 0.058711957 | -0.066131941 | 0.00291425 | -0.055797707 | 0.069046191 | 0.344663877 | 0.196560147 | 0.950195625 | 0.325484698 | 0.123649112 |
| Otud4 | S | | 544 | 0.063391156 | 0.096115868 | -0.261638025 | -0.325029181 | -0.357753892 | 0.679357269 | 0.523598095 | 0.4276044 | 0.334840792 | 0.29785594 |
| Snx2 | T | | 101 | 0.122403745 | -0.046692685 | -0.125870082 | -0.248273827 | -0.079177398 | 0.026392772 | 0.624188055 | 0.066488142 | 0.011441756 | 0.428288155 |
| Ppp6r1 | S | | 633 | 0.033103823 | -0.046750687 | 0.008900616 | -0.024203207 | 0.055651303 | 0.643816321 | 0.314832116 | 0.782173517 | 0.731908482 | 0.247933812 |
| Abca2 | S | | 2418 | 0.096283664 | -0.09812946 | -0.020741548 | -0.117025212 | 0.077387912 | 0.343237587 | 0.313525155 | 0.893291981 | 0.431067194 | 0.582510307 |
| Dync1li2 | S | | 205 | 0.063493839 | -0.022409682 | -0.066586086 | -0.130079925 | -0.044176404 | 0.256059897 | 0.698164086 | 0.336724424 | 0.086291817 | 0.487297659 |
| Itsn1 | S | | 203 | -0.029343651 | 0.032542165 | -0.019040756 | 0.010302894 | -0.051582921 | 0.621181596 | 0.525797981 | 0.646346379 | 0.833191387 | 0.237583269 |
| Ankrd11 | S | | 1114 | -0.00418192 | -0.076806433 | 0.083136953 | 0.087318873 | 0.159943385 | 0.956227473 | 0.19900643 | 0.335580942 | 0.399322651 | 0.123005914 |
| Ankrd11 | T | | 1111 | -0.00418192 | -0.076806433 | 0.083136953 | 0.087318873 | 0.159943385 | 0.956227473 | 0.19900643 | 0.335580942 | 0.399322651 | 0.123005914 |
| Aff4 | S | | 1040 | -0.04090136 | 0.015790608 | 0.010976666 | 0.051878026 | -0.004813942 | 0.448982614 | 0.704435462 | 0.849902777 | 0.427563699 | 0.930186238 |
| Spen | S | | 758 | -0.044259049 | -0.044353654 | 0.088724825 | 0.132983875 | 0.133078479 | 0.081907705 | 0.080625052 | 0.016635754 | 0.011419879 | 0.012392747 |
| Calu | Y | | 47 | -0.106964546 | 0.041237135 | 0.069467636 | 0.176432182 | 0.028230501 | 0.129322174 | 0.390217826 | 0.540721794 | 0.186786324 | 0.791924029 |
| Palld | S | | 782 | 0.022166128 | 0.001114181 | -0.037379688 | -0.059545816 | -0.038493869 | 0.400160281 | 0.954707211 | 0.294567217 | 0.138374194 | 0.270768703 |
| Phactr4 | S | | 436 | -0.012820675 | 0.006785195 | -0.009053607 | 0.003767068 | -0.015838802 | 0.773353591 | 0.848589595 | 0.838490677 | 0.93184121 | 0.662110555 |
| Phactr4 | S | | 118 | 0.034142128 | -0.00877918 | -0.032453632 | -0.06659576 | -0.023674452 | 0.321508217 | 0.75950617 | 0.251845617 | 0.091668712 | 0.403954358 |
| Arhgap23 | S | | 619 | -0.03497923 | 0.023129527 | 0.008617925 | 0.043597155 | -0.014511601 | 0.32328779 | 0.678988545 | 0.826776676 | 0.289550195 | 0.79962309 |
| Als2 | S | | 1458 | -0.12877383 | 0.103969419 | 0.011580526 | 0.140354357 | -0.092388893 | 0.080008881 | 0.116230911 | 0.906909463 | 0.208644514 | 0.36305135 |
| Phf2 | S | | 1060 | 0.03322404 | 0.005831316 | -0.042511003 | -0.075735043 | -0.048342318 | 0.491872398 | 0.796506894 | 0.220920117 | 0.180756534 | 0.163886467 |
| Ston2 | S | | 267 | -0.151124783 | 0.039785114 | 0.095464708 | 0.246589492 | 0.055679595 | 0.145365023 | 0.667844631 | 0.276560541 | 0.058204201 | 0.615723282 |
| Nat10 | S | | 934 | 0.197578352 | -0.06763943 | -0.083013431 | -0.280591783 | -0.015374001 | 0.194493825 | 0.08715038 | 0.544090581 | 0.144214538 | 0.905012016 |
| Prpf4b | T | | 439 | -0.029316418 | -0.012434374 | 0.016601367 | 0.045917786 | 0.029035741 | 0.567481656 | 0.722965585 | 0.359982921 | 0.392253114 | 0.422164837 |
| G3bp2 | S | | 149 | -0.032200863 | -0.028821114 | 0.052843822 | 0.085044685 | 0.081664936 | 0.393097896 | 0.393470961 | 0.237873426 | 0.067431067 | 0.084193307 |
| Cdc42ep1 | S | | 368 | -0.077728202 | 0.076749183 | -0.001745212 | 0.075982989 | -0.078494395 | 0.227510644 | 0.350903666 | 0.979061948 | 0.252250831 | 0.3433998 |
| Acap2 | S | | 384 | -0.053116044 | 0.140287243 | -0.098537791 | -0.045421747 | -0.238825034 | 0.443890048 | 0.092296008 | 0.151122768 | 0.351061365 | 0.017283292 |
| Fzr1 | S | | 120 | 0.182037115 | 0.049982332 | -0.396398399 | -0.578435515 | -0.446380731 | 0.111956958 | 0.596239944 | 0.159478792 | 0.085910225 | 0.143648996 |
| Stard13 | T | | 313 | 0.005590179 | 0.035058577 | -0.063004803 | -0.068594982 | -0.09806338 | 0.964318236 | 0.670565162 | 0.552306611 | 0.602561234 | 0.322456827 |
| Srrm2 | S | | 908 | -0.140057426 | -0.204274013 | 0.217081125 | 0.357138551 | 0.421355139 | 0.676229043 | 0.775462575 | 0.476245498 | 0.399896625 | 0.581392141 |
| Nop2 | T | | 169 | 0.091020949 | -0.037237172 | -0.079063323 | -0.170084272 | -0.041826151 | 0.00726698 | 0.317542495 | 0.031332777 | 0.003217433 | 0.269197396 |
| Cast | S | | 163 | -0.050809118 | 0.078782643 | -0.063465588 | -0.01265647 | -0.142248231 | 0.442894298 | 0.26182564 | 0.595677366 | 0.909290321 | 0.26950357 |
| Bod1l | S | | 2554 | -0.039415068 | 0.014688378 | 0.017769619 | 0.057184687 | 0.003081241 | 0.268559368 | 0.695080522 | 0.603391525 | 0.005616313 | 0.877377962 |
| Ppp1r12a Ppp1r12b | S | 644;693 | | 0.009564173 | 0.202884112 | -0.144012118 | -0.153576291 | -0.34689623 | 0.943589747 | 0.331514094 | 0.116331223 | 0.275203248 | 0.155215693 |
| Mdh1 | S | | 242 | -0.134306146 | 0.196116652 | -0.071831315 | 0.062474831 | -0.267947967 | 0.278082245 | 0.242971143 | 0.662937993 | 0.675626049 | 0.179463346 |
| Prkra | T | | 20 | -0.213971852 | 0.058339156 | 0.174251237 | 0.388223088 | 0.115912081 | 0.007488435 | 0.594420907 | 0.027952839 | 0.002297111 | 0.324239548 |
| Clk3 | S | | 197 | -0.011206116 | -0.055293341 | 0.051650329 | 0.062856445 | 0.10694367 | 0.786623575 | 0.587390998 | 0.595025798 | 0.517867189 | 0.407464111 |
| Clk3 | S | | 199 | -0.011206116 | -0.055293341 | 0.051650329 | 0.062856445 | 0.10694367 | 0.786623575 | 0.587390998 | 0.595025798 | 0.517867189 | 0.407464111 |
| Slc35c2 | S | | 336 | 0.027117113 | -0.024032817 | -0.016747588 | -0.043864701 | 0.007285228 | 0.514837237 | 0.564658423 | 0.705147005 | 0.299448084 | 0.853378221 |
| Srrm2 | T | | 1812 | -0.033353586 | -0.044189095 | 0.087846264 | 0.121199849 | 0.132035359 | 0.427869624 | 0.413450931 | 0.343362242 | 0.221704996 | 0.192636727 |
| Ilf3 | S | | 57 | -0.184296486 | 0.089581382 | 0.086974348 | 0.271270835 | -0.002607034 | 0.086271959 | 0.416980268 | 0.27976422 | 0.035966695 | 0.980543612 |
| Ilf3 | S | | 60 | -0.184296486 | 0.089581382 | 0.086974348 | 0.271270835 | -0.002607034 | 0.086271959 | 0.416980268 | 0.27976422 | 0.035966695 | 0.980543612 |
| Fkbp15 | S | | 386 | -0.038285495 | 0.001314239 | 0.01756397 | 0.055849465 | 0.016249731 | 0.701167632 | 0.990470406 | 0.847824107 | 0.59721776 | 0.886801541 |
| Evl | S | | 244 | -0.003640923 | -0.018773026 | 0.007583309 | 0.011224232 | 0.026356335 | 0.965180289 | 0.753204847 | 0.917604684 | 0.905473251 | 0.731463966 |
| Armcx2 | S | | 104 | 0.001354459 | -0.044606371 | 0.028683064 | 0.027328605 | 0.073289435 | 0.977273762 | 0.550849443 | 0.711070137 | 0.747203577 | 0.466388324 |
| Cadm4 | S | | 361 | 0.09885749 | -0.027649511 | -0.120645272 | -0.219502763 | -0.092995761 | 0.126423426 | 0.617350106 | 0.273996291 | 0.096204989 | 0.367810631 |
| Ankib1 | S | | 907 | 0.152883788 | -0.015882304 | -0.189356065 | -0.342239853 | -0.173473761 | 0.285185342 | 0.941102726 | 0.318864333 | 0.117659522 | 0.483346182 |
| Cobll1 | S | | 553 | -0.011954395 | -0.033064273 | 0.019184315 | 0.03113871 | 0.052248588 | 0.815790733 | 0.548754179 | 0.403842328 | 0.539095394 | 0.359709616 |
| Tpr | S | | 2141 | 0.033969195 | -0.047403736 | -0.000436723 | -0.034405918 | 0.046967012 | 0.336008608 | 0.274191838 | 0.982175139 | 0.337660101 | 0.280358524 |
| Itga4 | Y | | 1031 | -0.027557854 | 0.01065993 | 0.000407996 | 0.02796585 | -0.010251935 | 0.614981044 | 0.74957182 | 0.986469883 | 0.60309434 | 0.740436061 |
| Chd4 | S | | 301 | -0.131733345 | 0.018560493 | 0.092080319 | 0.223813663 | 0.073519826 | 0.115188633 | 0.761174193 | 0.169221163 | 0.034402015 | 0.304736331 |
| Nfkb1 | S | | 940 | 0.000410079 | 0.128807553 | -0.142330644 | -0.142740723 | -0.271138197 | 0.996168764 | 0.198192468 | 0.154287824 | 0.169717319 | 0.040111297 |
| Ung | S | | 43 | -0.061100644 | 0.005157325 | 0.038108005 | 0.099208649 | 0.032950679 | 0.452034909 | 0.912104949 | 0.594134815 | 0.305478471 | 0.638478008 |
| Srrm2 | S | | 1866 | -0.002031228 | -0.060292428 | 0.08621455 | 0.088245779 | 0.146506978 | 0.970373279 | 0.17167834 | 0.412778825 | 0.404936134 | 0.214762064 |
| Foxk1 | S | | 239 | 0.136802332 | -0.152776872 | -0.042865734 | -0.179668066 | 0.109911138 | 0.14301645 | 0.273506405 | 0.681180294 | 0.168595475 | 0.458213888 |
| Tgfbrap1 | S | | 852 | 0.045876172 | -0.07684177 | 0.042509569 | -0.003366603 | 0.119351339 | 0.653365644 | 0.279118228 | 0.72908253 | 0.979362166 | 0.341027807 |
| Cenpe | S | | 815 | 0.061846662 | 0.012166698 | -0.051571798 | -0.11341846 | -0.063738497 | 0.678801231 | 0.891564426 | 0.298753091 | 0.466478831 | 0.48596119 |
| Lbr | S | | 368 | 0.115431708 | -0.071379395 | -0.072652811 | -0.188084518 | -0.001273415 | 0.416805118 | 0.742455327 | 0.486700289 | 0.243261556 | 0.995386044 |
| Crtc1 | S | | 64 | -0.044581505 | 0.021191257 | 4.50389E-05 | 0.044626543 | -0.021146218 | 0.641825872 | 0.714492572 | 0.999542199 | 0.674120992 | 0.785266 |
| Map3k1 | S | | 261 | -0.035687775 | -0.001859882 | 0.036373709 | 0.072061485 | 0.038233592 | 0.282487821 | 0.948254618 | 0.467061672 | 0.190063722 | 0.425769796 |
| Gnl3 | S | | 375 | -0.027042861 | 0.019620785 | -0.014642364 | 0.012400496 | -0.03426315 | 0.648464009 | 0.418812035 | 0.65745087 | 0.839627616 | 0.337905854 |
| Rtn4 | T | | 430 | 0.188472266 | -0.081338849 | -0.157280672 | -0.345752938 | -0.075941823 | 0.083910473 | 0.411623131 | 0.244710334 | 0.043608859 | 0.477822804 |
| Baiap2 | T | | 341 | 0.031879023 | 0.002182038 | -0.037633035 | -0.069512058 | -0.039815074 | 0.416632192 | 0.980717516 | 0.248796332 | 0.085152112 | 0.657788688 |
| Fip1l1 | S | | 541 | -0.141257076 | -0.005939267 | 0.197431849 | 0.338688925 | 0.203371116 | 0.005947371 | 0.888785356 | 0.032606209 | 0.020520911 | 0.035731928 |
| Etv6 | T | | 18 | 0.003757722 | 0.02409278 | -0.034354119 | -0.038111841 | -0.058446899 | 0.915579675 | 0.472546893 | 0.285741887 | 0.329987847 | 0.140042987 |
| Ranbp3 | S | | 138 | 0.01101372 | 0.012549173 | -0.04309224 | -0.05410596 | -0.055641413 | 0.802352582 | 0.788993422 | 0.558987119 | 0.466856699 | 0.461991088 |
| Zfp219 | S | | 16 | -0.05543071 | 0.061850945 | -0.013661766 | 0.041768944 | -0.075512711 | 0.372412864 | 0.484004664 | 0.92622704 | 0.785250117 | 0.64380852 |
| Zranb2 | S | | 305 | -0.122504414 | -0.014312714 | 0.086421966 | 0.208926381 | 0.10073468 | 0.218822864 | 0.602283965 | 0.084350347 | 0.080384151 | 0.063231186 |
| Zranb2 | S | | 307 | -0.122504414 | -0.014312714 | 0.086421966 | 0.208926381 | 0.10073468 | 0.218822864 | 0.602283965 | 0.084350347 | 0.080384151 | 0.063231186 |
| Lmnb1 | S | | 407 | -0.004242404 | -0.031527159 | 0.021696527 | 0.02593893 | 0.053223686 | 0.967554865 | 0.547341299 | 0.65641996 | 0.809829877 | 0.386119384 |
| Tmem184b | S | | 388 | 0.061369878 | 0.174820546 | -0.316164384 | -0.377534262 | -0.49098493 | 0.548916718 | 0.115543645 | 0.096599513 | 0.070889289 | 0.038765708 |
| Bnip3 | S | | 60 | 0.028091064 | -0.087184408 | 0.106780245 | 0.078689181 | 0.193964654 | 0.815436844 | 0.084713285 | 0.349819802 | 0.593650099 | 0.148586049 |
| Yeats2 | S | | 118 | 0.011453217 | -0.006444921 | -0.016423905 | -0.027877122 | -0.009978984 | 0.680868187 | 0.872716639 | 0.612191709 | 0.408084387 | 0.816105062 |
| Srsf6 | S | | 303 | 0.004091837 | -0.010347396 | -0.00472068 | -0.008812517 | 0.005626716 | 0.813337701 | 0.728852157 | 0.876897161 | 0.749181126 | 0.872989636 |
| Pak4 | S | | 192 | -0.133306001 | 0.082048917 | 0.011382319 | 0.14468832 | -0.070666597 | 0.28908315 | 0.296018548 | 0.879062171 | 0.236435521 | 0.090504052 |
| Prpf4b | S | | 437 | 0.011998967 | -0.006797216 | -0.008734606 | -0.020733573 | -0.00193739 | 0.668788664 | 0.641753755 | 0.598983006 | 0.470454892 | 0.889178588 |
| Ulk1 | S | | 638 | -0.02666098 | 0.10700394 | -0.207931918 | -0.181270938 | -0.314935858 | 0.870156767 | 0.417488088 | 0.654205174 | 0.696234956 | 0.509087654 |
| Rai1 | S | | 1085 | -0.083063784 | -0.022880494 | 0.066843661 | 0.149907444 | 0.089724155 | 0.451150697 | 0.6082311 | 0.233734486 | 0.228522695 | 0.162987432 |
| Lrrk2 | S | | 974 | -0.246384319 | 0.074012193 | 0.119702523 | 0.366086842 | 0.04569033 | 0.096912811 | 0.223809758 | 0.311840942 | 0.046168577 | 0.663972906 |
| Smarca5 | S | | 115 | 0.106296468 | -0.018184703 | -0.095641144 | -0.201937612 | -0.077456441 | 0.243404636 | 0.811461583 | 0.22543793 | 0.042604744 | 0.071328603 |

| | | | | | | | | | | | | |
|---|---|---|---|---|---|---|---|---|---|---|---|---|
| Mga | S | 55 | 0.059070033 | 0.101445497 | -0.171140848 | -0.230210881 | -0.272586345 | 0.635283185 | 0.304122087 | 0.157538207 | 0.089430496 | 0.041789959 |
| Zfp646 | S | 601 | -0.058767682 | 0.042395613 | 0.003415163 | 0.062182845 | -0.038980449 | 0.138566215 | 0.17025947 | 0.926959743 | 0.145226112 | 0.258684312 |
| Foxm1 | S | 714 | 0.034588346 | -0.046727411 | 0.005351087 | -0.029237259 | 0.052078498 | 0.665100897 | 0.47052511 | 0.938233915 | 0.711120189 | 0.427267286 |
| Prrc2a | S | 455 | 0.206649951 | 0.067892527 | -0.283845625 | -0.490495577 | -0.351738153 | 0.074945353 | 0.501285545 | 0.029813902 | 0.002562956 | 0.008866799 |
| Fgd6 | S | 557 | -0.020110002 | -0.008460344 | 0.018127692 | 0.038237693 | 0.026588036 | 0.561226167 | 0.834569146 | 0.595809122 | 0.25101091 | 0.492953744 |
| Zfp638 | S | 869 | -0.041874078 | -0.029768608 | 0.061156364 | 0.103030441 | 0.090924972 | 0.224576935 | 0.53762348 | 0.157651076 | 0.043508744 | 0.123322253 |
| Hsp90aa1 | T | 726 | 0.325262567 | 0.112825377 | -0.812229405 | -1.137491972 | -0.925054782 | 0.323742981 | 0.550433218 | 0.356739651 | 0.232822788 | 0.308142459 |
| Sbf2 | S | 1710 | 0.180209014 | -0.034281388 | -0.102660588 | -0.282869602 | -0.068379199 | 0.146241143 | 0.58206731 | 0.085745895 | 0.071195215 | 0.188795503 |
| Slc4a2 | S | 144 | 0.019639409 | -0.040136433 | 0.005212111 | -0.014427299 | 0.045348544 | 0.781741958 | 0.619840426 | 0.930245224 | 0.829367765 | 0.563486121 |
| Mief1 | S | 94 | -0.012447497 | 0.011856775 | -0.006760379 | 0.005687118 | -0.018617153 | 0.4215496 | 0.494839593 | 0.816185199 | 0.834414523 | 0.523632855 |
| Mastl | T | 508 | -0.073948164 | 0.073278383 | -0.039363019 | 0.034585145 | -0.112641403 | 0.380846349 | 0.290519952 | 0.838855117 | 0.86202143 | 0.578338468 |
| Bud13 | S | 272 | -0.018892271 | 0.008512501 | 0.006154254 | 0.025046525 | -0.002358248 | 0.550060971 | 0.922789693 | 0.856343794 | 0.332141799 | 0.97812095 |
| Cdk11b | Y | 751 | -0.0701461 | -0.043886958 | 0.114037248 | 0.184183347 | 0.157924206 | 0.200208726 | 0.397919977 | 0.123872928 | 0.040360903 | 0.059076267 |
| Nkrf | S | 428 | 0.253621153 | -0.070458025 | -0.107642237 | -0.36126339 | -0.037184211 | 0.33898233 | 0.492802404 | 0.321713521 | 0.21109356 | 0.612552536 |
| Zbtb5 | S | 127 | 0.024917076 | 0.01103154 | -0.052949504 | -0.077866579 | -0.063981044 | 0.526331717 | 0.853784546 | 0.410781014 | 0.247473471 | 0.398743398 |
| Art4 | T | 82 | 0.074003314 | -0.096170365 | -0.006309603 | -0.080312917 | 0.089860762 | 0.166359009 | 0.211574083 | 0.901601686 | 0.051637388 | 0.212445572 |
| Tanc1 | S | 1611 | -0.029237365 | 0.078855193 | -0.09938081 | -0.070143445 | -0.178236004 | 0.810182996 | 0.207585408 | 0.354368346 | 0.616037568 | 0.144098207 |
| Myh10 | S | 1975 | -0.082351196 | -0.006718188 | 0.084798429 | 0.167149625 | 0.091516617 | 0.246110932 | 0.909818918 | 0.304349323 | 0.073739071 | 0.229643074 |
| Ctnna1 | S | 44 | -0.080329862 | 0.052021939 | -0.022790209 | 0.057539653 | -0.074812148 | 0.508268138 | 0.326433811 | 0.87477482 | 0.732027001 | 0.604720237 |
| Klf6 | S | 150 | -0.18265199 | 0.076140816 | 0.016369832 | 0.199021821 | -0.059770984 | 0.537746422 | 0.425433294 | 0.92863221 | 0.536755378 | 0.747105641 |
| Tjp2 | S | 378 | -0.105609631 | 0.003813208 | 0.147424464 | 0.253034095 | 0.143611256 | 0.002183161 | 0.918821303 | 0.024310327 | 0.01511244 | 0.031998265 |
| Tjp2 | S | 380 | -0.105609631 | 0.003813208 | 0.147424464 | 0.253034095 | 0.143611256 | 0.002183161 | 0.918821303 | 0.024310327 | 0.01511244 | 0.031998265 |
| Zmym2 | S | 622 | -0.023876264 | 0.11402147 | -0.097754746 | -0.073877482 | -0.211775216 | 0.713515349 | 0.328307244 | 0.368019598 | 0.446106065 | 0.139325697 |
| Adgrl3 | S | 1509 | -0.113065079 | 0.058980367 | 0.024457706 | 0.137522786 | -0.034522661 | 0.316641846 | 0.396476501 | 0.763139543 | 0.27659638 | 0.703974386 |
| Srrm2 | S | 1493 | 0.152871122 | -0.016223938 | -0.251237406 | -0.404108529 | -0.235013468 | 0.065785405 | 0.822252823 | 0.201668195 | 0.098225036 | 0.229593914 |
| Rilpl1 | S | 349 | 0.064255497 | 0.06557963 | -0.126179129 | -0.190434626 | -0.191758759 | 0.36251903 | 0.372722738 | 0.086927121 | 0.014534107 | 0.025459424 |
| Map1b | S | 1141 | -0.099711031 | 0.137140427 | -0.006928489 | 0.092782542 | -0.144068916 | 0.316823077 | 0.476679868 | 0.909435621 | 0.336687653 | 0.455566456 |
| Vcl | S | 637 | 0.177463328 | -0.077544772 | -0.049740968 | -0.227204296 | 0.027803804 | 0.502816362 | 0.466433637 | 0.630275098 | 0.412572553 | 0.824747823 |
| Il1rl1 | T | 16 | 0.018896809 | -0.04627286 | 0.027811216 | 0.008914407 | 0.074084076 | 0.876327066 | 0.381344733 | 0.616333623 | 0.939418364 | 0.110211533 |
| Il1rl1 | T | 22 | 0.018896809 | -0.04627286 | 0.027811216 | 0.008914407 | 0.074084076 | 0.876327066 | 0.381344733 | 0.616333623 | 0.939418364 | 0.110211533 |
| Il1rl1 | Y | 20 | 0.018896809 | -0.04627286 | 0.027811216 | 0.008914407 | 0.074084076 | 0.876327066 | 0.381344733 | 0.616333623 | 0.939418364 | 0.110211533 |
| Srrm2 | S | 1791 | -0.016073008 | -0.003082204 | 0.002862664 | 0.018935673 | 0.005944868 | 0.890658641 | 0.975763245 | 0.984005037 | 0.9052267 | 0.968348975 |
| Zranb2 | S | 188 | -0.057956786 | -0.029845441 | 0.10167046 | 0.159627246 | 0.131515901 | 0.245384025 | 0.491674756 | 0.229923798 | 0.101595606 | 0.150069345 |
| Srrt | S | 26 | -0.046195156 | 0.04995026 | -0.002438912 | 0.043756244 | -0.052389173 | 0.277535565 | 0.417429552 | 0.966314787 | 0.454530432 | 0.460983457 |
| Akap8 | S | 596 | -0.022255251 | -0.016189645 | 0.021103477 | 0.043358728 | 0.037293122 | 0.777969812 | 0.856835498 | 0.728480835 | 0.474893859 | 0.617175931 |
| Specc1 | T | 214 | 0.069734202 | -0.009774249 | -0.071984987 | -0.141719189 | -0.062210737 | 0.64184607 | 0.918649815 | 0.606199365 | 0.444109255 | 0.670776513 |
| Rps2 | S | 281 | 0.046979991 | 0.07597717 | -0.141189135 | -0.188169127 | -0.217166305 | 0.604239341 | 0.163905742 | 0.102808191 | 0.107921818 | 0.04109393 |
| Ncaph | S | 191 | -0.058543996 | 0.018233137 | 0.041026543 | 0.099570539 | 0.022793406 | 0.065858525 | 0.614905976 | 0.185752954 | 0.005291813 | 0.458371447 |
| Sipa1l1 | S | 286 | 0.00232818 | 0.102192621 | -0.118608516 | -0.120936696 | -0.220801137 | 0.957820226 | 0.188993555 | 0.161957084 | 0.160630619 | 0.046733673 |
| Patl1 | S | 184 | 0.148694431 | -0.085978798 | -0.081815428 | -0.230509859 | 0.00416337 | 0.081404299 | 0.25124589 | 0.314156901 | 0.015255369 | 0.938062395 |
| Ppp1r12a Ppp1r12b | T | 645;694 | 0.006962178 | 0.245459453 | -0.15589989 | -0.162862068 | -0.401359343 | 0.964147903 | 0.314089281 | 0.106803415 | 0.312554245 | 0.158640885 |
| Smcr8 | S | 497 | -0.005408724 | 0.078315779 | -0.088221478 | -0.082812754 | -0.166537256 | 0.945291338 | 0.328154489 | 0.316028015 | 0.420604246 | 0.13506982 |
| Zeb1 | S | 1018 | -0.017865203 | 0.010374111 | 0.010022361 | 0.027887564 | -0.000351749 | 0.549890614 | 0.902009372 | 0.91468064 | 0.764277565 | 0.997564135 |
| Srrm1 | S | 448 | -0.007578123 | -0.029182425 | 0.029102177 | 0.0366803 | 0.058284602 | 0.808448277 | 0.30712622 | 0.30744349 | 0.30225325 | 0.100368404 |
| Depdc1b | S | 160 | 0.023505611 | 0.120243818 | -0.152423755 | -0.175929366 | -0.272667573 | 0.80880557 | 0.260340731 | 0.177538447 | 0.151710165 | 0.055410606 |
| Arhgap35 | S | 1236 | -0.065135965 | 0.046105654 | 0.013107181 | 0.078243146 | -0.032998473 | 0.481475636 | 0.641524739 | 0.911620233 | 0.525069928 | 0.791472343 |
| Mcm3 | S | 681 | -0.026400049 | -0.025289918 | 0.033841224 | 0.060241273 | 0.059131142 | 0.6020067 | 0.772718737 | 0.485739261 | 0.066844447 | 0.477098744 |
| Klf4 | S | 251 | 0.048219498 | -0.036624249 | -0.01976918 | -0.067988678 | 0.016855068 | 0.502364038 | 0.497266806 | 0.821362983 | 0.482715089 | 0.840788304 |
| Tle3 | S | 286 | -0.001156585 | 0.021287559 | -0.022686612 | -0.021530027 | -0.043974171 | 0.976093234 | 0.372789525 | 0.241061439 | 0.590765699 | 0.1171406 |
| Morc3 | S | 563 | 0.124345897 | -0.089561061 | -0.028476227 | -0.152822124 | 0.061084834 | 0.206564678 | 0.184206542 | 0.735668956 | 0.171016712 | 0.452517357 |
| Sik3 | S | 514 | -0.072297121 | 0.082665291 | -0.055308989 | 0.016988131 | -0.13797428 | 0.293133237 | 0.127464447 | 0.650431477 | 0.887052787 | 0.301242471 |
| Ccnb3 | S | 509 | -0.040638797 | -0.020012737 | 0.029798526 | 0.070437322 | 0.049811262 | 0.614796848 | 0.676282111 | 0.299115797 | 0.403939551 | 0.311879962 |
| Araf | S | 186 | 0.107788816 | -0.007160222 | -0.111199043 | -0.21898786 | -0.104038821 | 0.445438848 | 0.944872978 | 0.411038303 | 0.152366866 | 0.339478381 |
| Ino80b | S | 370 | -0.016816335 | -0.051026046 | 0.102619111 | 0.119435445 | 0.153645156 | 0.719396025 | 0.063253338 | 0.193558337 | 0.156725195 | 0.107801956 |
| Tecta | S | 1907 | -0.043292055 | 0.101373148 | -0.037200683 | 0.006091373 | -0.138573831 | 0.449566174 | 0.419896923 | 0.547779876 | 0.905902429 | 0.292647895 |
| Tecta | S | 1910 | -0.043292055 | 0.101373148 | -0.037200683 | 0.006091373 | -0.138573831 | 0.449566174 | 0.419896923 | 0.547779876 | 0.905902429 | 0.292647895 |
| Zcchc7 | S | 117 | 0.011112427 | 0.017174608 | -0.040714602 | -0.051827029 | -0.05788921 | 0.857798538 | 0.752523626 | 0.574866844 | 0.430206779 | 0.350028371 |
| Psmd12 | S | 434 | -0.126052747 | 0.092978205 | 0.04798272 | 0.174035466 | -0.044995485 | 0.512675207 | 0.761478946 | 0.836524257 | 0.386880236 | 0.88144705 |
| Trim28 | S | 624 | -0.080861485 | -0.024657623 | 0.084766125 | 0.16562761 | 0.109423748 | 0.025055521 | 0.873947163 | 0.023044055 | 0.002512797 | 0.506791659 |
| Dcaf10 | S | 96 | 0.071813491 | -0.018040109 | -0.076068768 | -0.147882259 | -0.058028659 | 0.267400411 | 0.783586666 | 0.357917722 | 0.095426432 | 0.428660152 |
| Ifnar2 | S | 403 | -0.127120613 | 0.03597766 | 0.09086499 | 0.217985603 | 0.05488713 | 0.02664289 | 0.279308911 | 0.135042691 | 0.014165069 | 0.315378046 |
| Zkscan1 | S | 13 | -0.069866633 | 0.037790408 | 0.035185362 | 0.105051995 | -0.002605046 | 0.184840827 | 0.468800527 | 0.58999421 | 0.169297056 | 0.968876511 |
| Aspscr1 | S | 193 | -0.017959193 | 0.019280259 | -0.026322667 | -0.008363474 | -0.045602926 | 0.675878712 | 0.432682651 | 0.778824925 | 0.930359975 | 0.629158882 |
| Ranbp10 | S | 489 | -0.006774895 | -0.137732889 | 0.125162598 | 0.131937494 | 0.262895487 | 0.97013033 | 0.388624331 | 0.402979948 | 0.47326849 | 0.134812967 |
| Hdac4 | S | 629 | -0.077646002 | 0.069907226 | 0.008388358 | 0.08603436 | -0.061518868 | 0.220248133 | 0.333499666 | 0.911366733 | 0.332337914 | 0.500577804 |
| Atxn2 | S | 741 | -0.002004268 | -0.038397426 | 0.057052201 | 0.059056468 | 0.095449627 | 0.920797441 | 0.514102087 | 0.725707038 | 0.716158791 | 0.576104836 |
| Sipa1l1 | S | 1507 | -0.045018465 | -0.062633366 | 0.088966938 | 0.133985403 | 0.151600304 | 0.555286037 | 0.402077104 | 0.225117583 | 0.053293776 | 0.02608324 |
| Zc3h13 | S | 704 | 0.001803094 | -0.012456249 | 0.003165092 | 0.001361999 | 0.015621341 | 0.968312848 | 0.766557002 | 0.950772626 | 0.974506363 | 0.693568838 |
| Rplp1 | S | 83 | 0.011751911 | -0.034436819 | 0.022907411 | 0.0111555 | 0.05734423 | 0.83094826 | 0.436481507 | 0.727989173 | 0.866459642 | 0.369668685 |
| Jak1 | S | 568 | -0.096760292 | 0.128968314 | -0.014797415 | 0.081962877 | -0.143765729 | 0.172915496 | 0.278619981 | 0.937584529 | 0.663932102 | 0.48783855 |
| Mical1 | S | 948 | -0.065047437 | 0.159989984 | -0.120628767 | -0.05558133 | -0.280618752 | 0.471652833 | 0.141095329 | 0.310512402 | 0.573990097 | 0.05009749 |
| Pou2f1 | S | 451 | -0.007815113 | -0.057299958 | 0.06666966 | 0.074484774 | 0.123969618 | 0.759292187 | 0.105501551 | 0.12978522 | 0.105602178 | 0.02939877 |
| Pds5b | T | 1121 | 0.039914723 | 0.023773698 | -0.09265675 | -0.132571473 | -0.116430448 | 0.57202606 | 0.368056552 | 0.288488621 | 0.206527481 | 0.209263842 |
| Npm1 | S | 240 | -0.038635281 | -0.024397105 | 0.056205866 | 0.094841147 | 0.080602971 | 0.507932733 | 0.652126458 | 0.395977208 | 0.244937608 | 0.30168278 |
| Prdm10 | S | 418 | -0.121413004 | -0.030897046 | 0.142138874 | 0.263551877 | 0.17303592 | 0.069980969 | 0.462832101 | 0.036196812 | 0.008789828 | 0.025660941 |
| Phf2 | S | 622 | -0.035370739 | -0.006698402 | 0.040150967 | 0.075521706 | 0.046849368 | 0.249337724 | 0.787158102 | 0.280150753 | 0.103301606 | 0.247288193 |
| Srrm1 | S | 733 | -0.038294372 | -0.038388102 | 0.067824508 | 0.10611888 | 0.106212609 | 0.556822226 | 0.294663684 | 0.19942186 | 0.18061911 | 0.081153855 |
| Pcbp1 | S | 189 | -0.009384199 | 0.024457932 | -0.02397712 | -0.014592921 | -0.048435052 | 0.940354856 | 0.779368807 | 0.715209652 | 0.898610691 | 0.497527632 |
| Tnfaip3 | S | 571 | -0.003547752 | 0.083377677 | -0.088755218 | -0.085207467 | -0.172132895 | 0.931284225 | 0.013380005 | 0.013266703 | 0.095955044 | 0.001974358 |
| Lrrk2 | S | 955 | -0.202365208 | 0.092889441 | 0.152012021 | 0.354377229 | 0.059122581 | 0.001857637 | 0.186450932 | 0.040051446 | 0.006765363 | 0.400346351 |
| Lrpprc | S | 1021 | 0.025113786 | -0.060131103 | 0.041821311 | 0.016707525 | 0.101952414 | 0.755212114 | 0.421371442 | 0.74692997 | 0.891303928 | 0.426693939 |
| Fam193a | S | 1136 | 0.000693448 | -0.004599939 | -0.010561512 | -0.01125496 | -0.005961573 | 0.99375048 | 0.964739332 | 0.953676354 | 0.953458039 | 0.976021292 |
| Spp1 | S | 180 | -0.0501668 | 0.098929158 | -0.050619918 | -0.000453118 | -0.149549077 | 0.412206564 | 0.110594731 | 0.192047258 | 0.993071704 | 0.048718825 |
| Srrm2 | S | 1628 | -0.003332194 | 0.016402952 | -0.018723053 | -0.015390859 | -0.035126005 | 0.907858282 | 0.673602265 | 0.614351536 | 0.689162272 | 0.452793698 |
| Map2k5 | S | 148 | -0.067256473 | 0.180013899 | -0.124203086 | -0.056946613 | -0.304216985 | 0.560749626 | 0.214309978 | 0.348152486 | 0.489604423 | 0.043414926 |
| Nfrkb | S | 298 | 0.013217767 | 0.06050133 | -0.129608656 | -0.142826422 | -0.190109985 | 0.788984921 | 0.313398734 | 0.381169852 | 0.341886065 | 0.236981049 |
| Haus1 | S | 247 | -0.009204639 | -0.052576061 | 0.04131207 | 0.05051671 | 0.093888131 | 0.911374217 | 0.675582008 | 0.619375939 | 0.152961227 | 0.408769209 |
| Cdca8 | S | 228 | 0.065484257 | -0.005043275 | -0.085666403 | -0.151150659 | -0.080623128 | 0.020837961 | 0.82733302 | 0.051247109 | 0.013561775 | 0.067589214 |
| Trio | S | 2462 | 0.058437154 | 0.003490717 | -0.046371889 | -0.104809043 | -0.049862607 | 0.223068459 | 0.918510449 | 0.076873417 | 0.076281335 | 0.198912389 |
| Mrtfb | S | 209 | -0.123846797 | 0.053298001 | 0.04443144 | 0.168278237 | -0.008866561 | 0.467233004 | 0.606276151 | 0.752684371 | 0.363422744 | 0.942415239 |
| Akap12 | T | 635 | -0.01082104 | 0.03497056 | -0.02704244 | -0.016221401 | -0.062013001 | 0.626397068 | 0.571152282 | 0.690743849 | 0.80388424 | 0.462475493 |
| Pbrm1 | S | 10 | -0.022786072 | 0.250067938 | -0.199687214 | -0.176901142 | -0.449755151 | 0.882381859 | 0.159172751 | 0.116370725 | 0.291796668 | 0.042547772 |
| Setdb1 | S | 1042 | -0.053466633 | -0.039196001 | 0.098121481 | 0.151588114 | 0.137317482 | 0.614470334 | 0.402178509 | 0.353257533 | 0.271810659 | 0.22636092 |
| Prkci | S | 590 | -0.057259115 | 0.096608037 | -0.035422667 | 0.021836448 | -0.132030704 | 0.392395855 | 0.283851058 | 0.539019072 | 0.683896918 | 0.160614028 |

| | | | | | | | | | | | | |
|---|---|---|---|---|---|---|---|---|---|---|---|---|
| Srrm2 | T | 2241 | -0.038416812 | -0.002760801 | 0.057287439 | 0.095704252 | 0.06004824 | 0.373663111 | 0.979662716 | 0.597668469 | 0.394247789 | 0.662420983 |
| Specc1 | S | 355 | -0.035150862 | -0.054861603 | 0.083095662 | 0.118246524 | 0.137957266 | 0.415834541 | 0.207697164 | 0.098332341 | 0.019063777 | 0.013046254 |
| Eps8 | S | 661 | -0.016041228 | -0.069576023 | 0.070697616 | 0.086738844 | 0.140273639 | 0.703359301 | 0.190342259 | 0.122579325 | 0.018887208 | 0.026553489 |
| Slk | S | 647 | -0.025903323 | -0.001236793 | 0.020402944 | 0.046306267 | 0.021639737 | 0.415483755 | 0.972044673 | 0.452568598 | 0.224665605 | 0.585617989 |
| Edc4 | S | 727 | 0.000990926 | -0.006214507 | -0.001921018 | -0.002911944 | 0.004293489 | 0.965273751 | 0.892268347 | 0.960198392 | 0.93452069 | 0.933693766 |
| Spp1 | S | 228 | 0.0807832 | -0.069189446 | -0.025454065 | -0.106237265 | 0.043735381 | 0.129228643 | 0.181006884 | 0.716384735 | 0.162820254 | 0.468323037 |
| Map4k4 | S | 621 | 0.011275979 | 0.073981947 | -0.089838601 | -0.10111458 | -0.163820548 | 0.753063344 | 0.118788781 | 0.074606444 | 0.052936608 | 0.015722175 |
| Fkbp15 | S | 310 | -0.036352053 | -0.023142803 | 0.043002565 | 0.079354618 | 0.066145368 | 0.239489467 | 0.663826501 | 0.129467477 | 0.045654739 | 0.27198072 |
| Pcyt1a | S | 329 | -0.088323207 | 0.101922483 | -0.042998242 | 0.045324965 | -0.144920726 | 0.150598313 | 0.034115216 | 0.370359831 | 0.410522235 | 0.018930465 |
| Ankrd44 | S | 983 | -0.112353724 | 0.052976279 | 0.02984698 | 0.142200704 | -0.023129299 | 0.316120471 | 0.627683418 | 0.362952681 | 0.234699419 | 0.825149287 |
| Mcm3ap | S | 502 | 0.073224873 | -0.04224755 | -0.030936632 | -0.104161504 | 0.011310918 | 0.432788778 | 0.550704091 | 0.639730228 | 0.229823363 | 0.742556982 |
| Slain2 | S | 87 | -0.000747757 | 0.062002392 | -0.093265293 | -0.092517535 | -0.155267685 | 0.988434767 | 0.257817622 | 0.345006249 | 0.333027669 | 0.159389572 |
| Cobll1 | T | 550 | -0.085324006 | 0.003340649 | 0.081079901 | 0.166403906 | 0.077739252 | 0.082193733 | 0.941220219 | 0.114008093 | 0.017284474 | 0.155931681 |
| Wac | T | 531 | -0.140765052 | -0.047612444 | 0.179898654 | 0.320663706 | 0.227511098 | 0.039061849 | 0.62595005 | 0.029254812 | 0.006917857 | 0.072951846 |
| Brca1 | S | 1178 | 0.050548845 | 0.006674726 | -0.038575982 | -0.089124828 | -0.045250708 | 0.658241165 | 0.959909859 | 0.399867235 | 0.444912044 | 0.729943781 |
| Ubqln2 | S | 25 | -0.007815698 | 0.015157866 | -0.013889562 | -0.006073865 | -0.029047428 | 0.791069747 | 0.685810063 | 0.780239966 | 0.89331089 | 0.569060613 |
| Pnisr | S | 764 | 0.016779866 | 0.044987288 | -0.062379945 | -0.079159811 | -0.107367233 | 0.83291618 | 0.548752466 | 0.380272792 | 0.384093748 | 0.230521096 |
| Mki67 | S | 1342 | 0.049640889 | 0.010078547 | -0.063575439 | -0.113216329 | -0.073653987 | 0.529901683 | 0.889961198 | 0.411374016 | 0.048110269 | 0.068389591 |
| Tle5 | S | 196 | -0.171514797 | 0.111441216 | 0.012616856 | 0.184131653 | -0.09882436 | 0.1194222 | 0.120014481 | 0.512228487 | 0.112905211 | 0.161644689 |
| Ehd2 Ehd4 | S | 373;376 | -0.123618452 | 0.053656433 | 0.03434911 | 0.157967562 | -0.019307323 | 0.183740463 | 0.263770467 | 0.454574871 | 0.126925808 | 0.38241401 |
| Etl4 | S | 1905 | -0.022920871 | 0.078803065 | -0.049215845 | -0.026294974 | -0.128018909 | 0.515174099 | 0.461627276 | 0.638383338 | 0.792167697 | 0.354318851 |
| Mast3 | S | 69 | -0.082184835 | 0.011805165 | 0.052802513 | 0.134987348 | 0.040997348 | 0.275524794 | 0.856027355 | 0.284059727 | 0.109164079 | 0.490029743 |
| Eif4g3 | T | 420 | -0.096366722 | 0.018843132 | 0.097941142 | 0.194307863 | 0.079098009 | 0.519467812 | 0.918364805 | 0.701568904 | 0.484331091 | 0.781539248 |
| Dbn1 | S | 387 | 0.056489333 | -0.036651744 | -0.002353848 | -0.05884318 | 0.034297897 | 0.430228848 | 0.228455746 | 0.956473355 | 0.424043147 | 0.388480239 |
| Zdhhc5 | S | 694 | 0.065728506 | -0.010119362 | -0.03877979 | -0.104508296 | -0.028660428 | 0.087059854 | 0.546517361 | 0.045046465 | 0.045525922 | 0.024658126 |
| 4930579G24Rik | S | 19 | -0.02104328 | 0.073519591 | -0.080352411 | -0.059309131 | -0.153872002 | 0.931403257 | 0.493162149 | 0.569825167 | 0.818481799 | 0.316192969 |
| Arhgap31 | S | 1058 | 0.043488262 | -0.009484358 | -0.043919998 | -0.08740826 | -0.03443564 | 0.476738213 | 0.901181532 | 0.513296573 | 0.095350967 | 0.590001354 |
| Eif5b | S | 191 | 0.003777885 | 0.03234674 | -0.058375856 | -0.062153741 | -0.090722596 | 0.852516044 | 0.150948903 | 0.2257584 | 0.206679378 | 0.106254239 |
| Rap1gap | S | 515 | -0.055839471 | 0.037387606 | 0.038874991 | 0.094714462 | 0.001487385 | 0.197328615 | 0.527423892 | 0.675523964 | 0.348692213 | 0.987805256 |
| Tmem184b | S | 358 | -0.007182151 | -0.015473631 | 0.015310827 | 0.022492979 | 0.030784459 | 0.973884753 | 0.870636335 | 0.913320421 | 0.927546351 | 0.847394425 |
| Cdca7l | S | 185 | -0.025801539 | -0.074160626 | 0.116064715 | 0.141866254 | 0.190225341 | 0.60496882 | 0.049032561 | 0.064356573 | 0.054380528 | 0.028257409 |
| Gatad2b | S | 487 | 0.041256516 | -0.027288325 | -0.03851072 | -0.079767236 | -0.011222395 | 0.022758395 | 0.204139378 | 0.321753684 | 0.103489302 | 0.757999907 |
| Tomm20 | S | 135 | -0.064458995 | 0.102576704 | -0.04097378 | 0.023485216 | -0.143550484 | 0.444150355 | 0.392787751 | 0.771845074 | 0.875028813 | 0.404984758 |
| Zfp800 | S | 453 | 0.02268667 | 0.075398546 | -0.078133042 | -0.100819712 | -0.153531588 | 0.784880853 | 0.477889771 | 0.244959634 | 0.203040871 | 0.182701796 |
| Ssrp1 | S | 672 | -0.009186102 | -0.037382529 | 0.054593466 | 0.063779568 | 0.091975995 | 0.644106032 | 0.135111032 | 0.161082775 | 0.12263758 | 0.052752826 |
| Cuedc1 | S | 12 | -0.108769832 | 0.096871401 | -0.003262232 | 0.1055076 | -0.100133632 | 0.391158785 | 0.375448987 | 0.973265939 | 0.418854711 | 0.38507791 |
| Sltm | S | 919 | -0.019525767 | -0.036560378 | 0.04460046 | 0.064126227 | 0.081160838 | 0.845847915 | 0.653775845 | 0.558715561 | 0.453426343 | 0.168938152 |
| Etv6 | S | 22 | 0.019639936 | 0.024124794 | -0.040665326 | -0.060305261 | -0.064790119 | 0.616172202 | 0.476088436 | 0.21028774 | 0.169613237 | 0.096851373 |
| Syt6 | S | 217 | 0.075822548 | 0.047249385 | -0.165937333 | -0.241759881 | -0.213186718 | 0.550387208 | 0.586710259 | 0.396742987 | 0.266725793 | 0.298093719 |
| Hdac6 | S | 43 | -0.025297099 | 0.022755492 | -0.005260085 | 0.020037014 | -0.028015577 | 0.630087449 | 0.599111144 | 0.927217527 | 0.744067523 | 0.614047403 |
| Plekhg3 | S | 947 | 0.064107555 | -0.116586945 | 0.049775897 | -0.014331658 | 0.166362842 | 0.250494896 | 0.054291194 | 0.243744009 | 0.735966032 | 0.023481661 |
| Sf3b1 | S | 129 | -0.034837819 | -0.00606668 | 0.03736183 | 0.072199649 | 0.04342851 | 0.247788633 | 0.857355303 | 0.238679474 | 0.020527346 | 0.131831998 |
| Galnt7 | S | 103 | 0.015970278 | -0.016748489 | -0.000111431 | -0.016081709 | 0.016637058 | 0.675441657 | 0.562310981 | 0.997314103 | 0.658753721 | 0.536379629 |
| Dmap1 | S | 457 | 0.060320089 | -0.087317127 | 0.032808345 | -0.027511743 | 0.120125473 | 0.356391943 | 0.152198459 | 0.566750008 | 0.710579298 | 0.127334121 |
| Phf14 | S | 270 | -0.232724218 | 0.10247345 | 0.152658639 | 0.385382857 | 0.050185189 | 0.055423679 | 0.327063235 | 0.245370789 | 0.027313985 | 0.573894941 |
| Cavin2 | S | 390 | 0.002602622 | 0.064680684 | -0.06423326 | -0.066835882 | -0.128913944 | 0.967071148 | 0.470288594 | 0.38040015 | 0.342858182 | 0.198362316 |
| Larp7 | S | 253 | -0.00691699 | -0.004996252 | 0.00770897 | 0.01462596 | 0.012705221 | 0.803989042 | 0.759278127 | 0.652253051 | 0.633758422 | 0.550305666 |
| Trim2 | S | 440 | -0.047080735 | 0.064609269 | -0.013978791 | 0.033101944 | -0.07858806 | 0.098264634 | 0.102123375 | 0.694719018 | 0.369894011 | 0.102827249 |
| Rgs12 | T | 669 | -0.240313551 | 0.14317376 | 0.045643188 | 0.285956739 | -0.097530572 | 0.10953217 | 0.072583019 | 0.808936044 | 0.213693255 | 0.602263693 |
| Rai1 | S | 863 | -0.018571645 | 0.048317095 | -0.025285223 | -0.006713579 | -0.073602318 | 0.710356709 | 0.610417121 | 0.713701348 | 0.916894506 | 0.475544203 |
| Mcm6 | S | 689 | -0.055677112 | 0.016483184 | 0.022147233 | 0.077824345 | 0.00566405 | 0.289714187 | 0.516632868 | 0.405495074 | 0.172968327 | 0.76160797 |
| Ybx1 | S | 312 | 0.049553161 | -0.017871707 | -0.039481235 | -0.089034396 | -0.021609528 | 0.060852705 | 0.413997207 | 0.172131661 | 0.01790557 | 0.355296562 |
| Scaf4 | T | 149 | 0.005702066 | 0.085733206 | -0.140641453 | -0.146343519 | -0.226374659 | 0.986111449 | 0.547452924 | 0.635120481 | 0.718656362 | 0.470196678 |
| Htatsf1 | S | 613 | -0.082492849 | 0.011451285 | 0.077583484 | 0.160076333 | 0.066132199 | 0.274191473 | 0.776685398 | 0.428373924 | 0.17748649 | 0.493550146 |
| Riok2 | S | 538 | 0.231007049 | -0.09026127 | -0.128727162 | -0.359734211 | -0.038465892 | 0.26739439 | 0.385971434 | 0.555026802 | 0.204547056 | 0.857311172 |
| Ppp6r3 | S | 588 | -0.043985582 | 0.043534732 | -0.012613456 | 0.031372127 | -0.056148188 | 0.405974477 | 0.272693338 | 0.786468411 | 0.605279444 | 0.30631334 |
| Arhgef17 | S | 2005 | -0.031340302 | -0.067792106 | 0.104536314 | 0.135876616 | 0.172328419 | 0.739991921 | 0.312607996 | 0.286153266 | 0.247186918 | 0.114446877 |
| Pds5b | S | 1140 | 0.037809686 | -0.013388044 | -0.053314885 | -0.09112457 | -0.03992684 | 0.528558934 | 0.930402804 | 0.69151173 | 0.519507874 | 0.831827149 |
| Sacs | T | 4262 | 0.058341858 | -0.016138085 | -0.037103944 | -0.095445802 | -0.020965859 | 0.637306488 | 0.876441299 | 0.626831853 | 0.458256482 | 0.84182974 |
| Sh3kbp1 | S | 274 | -0.135613541 | 0.075754232 | 0.067846181 | 0.203459723 | -0.007908051 | 0.03250898 | 0.257705417 | 0.170682427 | 0.011403042 | 0.894577853 |
| Slc9a3r1 | S | 294 | -0.143279363 | 0.140535315 | 0.008102636 | 0.151381998 | -0.132432679 | 0.098070185 | 0.125894518 | 0.939869675 | 0.242823904 | 0.301875689 |
| Wnk1 | S | 2024 | -0.025181122 | 0.065916276 | -0.060405745 | -0.035224623 | -0.126322021 | 0.745695346 | 0.14631762 | 0.261766523 | 0.667173574 | 0.058574291 |
| Gpatch8 | S | 699 | -0.060121692 | 0.047713123 | 0.028571496 | 0.088693188 | -0.019141627 | 0.37486208 | 0.627694784 | 0.802511884 | 0.428642479 | 0.875792135 |
| Dhx9 | T | 792 | -0.041896489 | -0.05563402 | 0.112468592 | 0.154365081 | 0.168102612 | 0.59661414 | 0.331384995 | 0.275563255 | 0.202935994 | 0.145431462 |
| Ccdc186 | S | 759 | -0.032883829 | 0.037394666 | -0.00585547 | 0.027028359 | -0.043250136 | 0.383008308 | 0.376382206 | 0.878226264 | 0.349673807 | 0.238732368 |
| Sppl2a | T | 508 | 0.007954777 | 0.05131163 | -0.046845349 | -0.054800126 | -0.09815698 | 0.841107287 | 0.40491998 | 0.219490858 | 0.280393468 | 0.173954538 |
| Lbr | S | 67 | -0.085597828 | 0.032526753 | 0.075832919 | 0.161430747 | 0.043306166 | 0.000266208 | 0.320892863 | 0.005032491 | 0.000735803 | 0.218354848 |
| Srrm2 | S | 1193 | -0.03124972 | -0.029445356 | 0.089726566 | 0.120976286 | 0.119171922 | 0.765125449 | 0.764029639 | 0.732423195 | 0.643179819 | 0.646087436 |
| Apc | S | 1359 | 0.010569972 | -0.083147282 | 0.066206064 | 0.055636092 | 0.149353346 | 0.804070023 | 0.02350926 | 0.02393261 | 0.239099632 | 0.012049412 |
| Apc | S | 1361 | 0.010569972 | -0.083147282 | 0.066206064 | 0.055636092 | 0.149353346 | 0.804070023 | 0.02350926 | 0.02393261 | 0.239099632 | 0.012049412 |
| Ubr5 | S | 1695 | 0.275480609 | -0.076156604 | -0.181252278 | -0.456732887 | -0.105095674 | 0.235239757 | 0.627590692 | 0.317015261 | 0.099720478 | 0.573969995 |
| Mki67 | T | 1247 | 0.047718397 | -0.043408687 | -0.005477785 | -0.053196182 | 0.037930902 | 0.565842119 | 0.544385128 | 0.957027046 | 0.640849544 | 0.72285745 |
| Mef2d | S | 201 | 0.007689883 | 0.006961568 | -0.021323716 | -0.029013599 | -0.028285284 | 0.947858563 | 0.965918888 | 0.89706368 | 0.876155753 | 0.894497663 |
| Tjp1 | S | 311 | 0.013269657 | -0.037909657 | 0.018292035 | 0.005022378 | 0.056201693 | 0.818385178 | 0.646551867 | 0.793745058 | 0.921224317 | 0.481566453 |
| Srrm2 | S | 2646 | -0.004148987 | -0.0108714 | 0.012923667 | 0.017072654 | 0.023795067 | 0.906152108 | 0.641074791 | 0.625899031 | 0.595421214 | 0.210696261 |
| Tcf12 | S | 564 | -0.038570552 | 0.024790704 | 0.026519779 | 0.065090332 | 0.001729075 | 0.335390753 | 0.761982042 | 0.626084692 | 0.27587283 | 0.984208011 |
| Arhgef12 | S | 1389 | -0.049303697 | 0.017360878 | 0.025940553 | 0.07524425 | 0.008579674 | 0.479411041 | 0.71909206 | 0.66244168 | 0.26932024 | 0.826863027 |
| Fam117b | S | 268 | -0.029696747 | 0.06694853 | -0.072471797 | -0.04277505 | -0.139420327 | 0.74453629 | 0.124133252 | 0.30977343 | 0.672421903 | 0.102219389 |
| Ash1l | S | 569 | 0.106994127 | -0.048011967 | -0.062538932 | -0.169533058 | -0.014526965 | 0.739248823 | 0.790674602 | 0.891266849 | 0.737486287 | 0.974522198 |
| Sipa1l1 | S | 1547 | -0.043006133 | 0.01220729 | 0.050142039 | 0.093148172 | 0.037934749 | 0.126346138 | 0.504088475 | 0.46542049 | 0.23029858 | 0.566008638 |
| Nucks1 | S | 204 | -0.067332921 | 0.023366125 | 0.069205438 | 0.136538359 | 0.045839313 | 0.049013589 | 0.523184678 | 0.234658968 | 0.070223357 | 0.403435529 |
| Sh3pxd2b | S | 551 | 0.060719396 | 0.116761388 | -0.276539659 | -0.337259056 | -0.393301047 | 0.547185544 | 0.110360401 | 0.162029755 | 0.111803611 | 0.09639825 |
| Tmx4 | S | 304 | 0.060050607 | -0.061023394 | -0.011134953 | -0.071185561 | 0.04988844 | 0.048331925 | 0.095477717 | 0.734075197 | 0.063172711 | 0.170226559 |
| Ralbp1 | S | 116 | -0.055941277 | 0.141939169 | -0.058351332 | -0.002410055 | -0.2002905 | 0.454390623 | 0.179054308 | 0.22617831 | 0.970537735 | 0.101606646 |
| Ybx3 | T | 355 | 0.019873122 | -0.045279281 | 0.027696752 | 0.00782363 | 0.072976033 | 0.603354401 | 0.256853561 | 0.502882711 | 0.872823657 | 0.176704606 |
| Hmga1 Hmga1b | S | 99 | -0.001514035 | -0.03845559 | 0.062457937 | 0.063971973 | 0.100913528 | 0.9616615 | 0.230172173 | 0.411154469 | 0.392216382 | 0.229511383 |
| Scrib | S | 943 | -0.016801459 | 0.016354293 | -0.00558228 | 0.011219178 | -0.021936574 | 0.744154084 | 0.652061893 | 0.912105235 | 0.834635066 | 0.602030608 |
| Nup133 | S | 44 | 0.030022375 | -0.011148529 | -0.020425032 | -0.050447407 | -0.009276502 | 0.765100724 | 0.91014241 | 0.805219138 | 0.590437756 | 0.918085524 |
| Nup133 | S | 49 | 0.030022375 | -0.011148529 | -0.020425032 | -0.050447407 | -0.009276502 | 0.765100724 | 0.91014241 | 0.805219138 | 0.590437756 | 0.918085524 |
| Ubr5 | S | 630 | 0.082020277 | 0.062110076 | -0.158738411 | -0.240758688 | -0.220848486 | 0.302095477 | 0.398515073 | 0.137143696 | 0.060182832 | 0.072641056 |
| H2bc1 H2bc11 H2bc1 | S | 56;56;56;57;56;5 | 0.764458769 | -0.157940855 | -0.06416632 | -0.828625089 | 0.093774535 | 0.310846007 | 0.03183389 | 0.484972552 | 0.28183894 | 0.286278717 |
| Bclaf1 | S | 299 | 0.016084938 | -0.092541216 | 0.0502074 | 0.034122461 | 0.142748616 | 0.889546343 | 0.187217663 | 0.096709996 | 0.769575435 | 0.087234205 |
| Bclaf1 | T | 303 | 0.016084938 | -0.092541216 | 0.0502074 | 0.034122461 | 0.142748616 | 0.889546343 | 0.187217663 | 0.096709996 | 0.769575435 | 0.087234205 |
| Bclaf1 | S | 221 | -0.016391298 | 0.002039841 | 0.014021573 | 0.030412871 | 0.011981732 | 0.728758249 | 0.943182739 | 0.816013805 | 0.65566132 | 0.838871 |

| | | | | | | | | | | | | | |
|---|---|---|---|---|---|---|---|---|---|---|---|---|---|
| 2310022A10Rik | S | | 115 | -0.017444383 | -0.045812854 | 0.06283246 | 0.080276843 | 0.108645314 | 0.63060214 | 0.522666375 | 0.356962211 | 0.268308847 | 0.240279747 |
| Slbp | S | | 7 | -0.08583951 | 0.058528194 | 0.062968366 | 0.148807876 | 0.004440172 | 0.029726614 | 0.277415036 | 0.319903576 | 0.073007148 | 0.946908791 |
| Mlh1 | S | | 469 | -0.037789129 | 0.008973706 | 0.040234614 | 0.078023743 | 0.031260908 | 0.130265905 | 0.696376187 | 0.321557234 | 0.106152003 | 0.404493177 |
| Srsf4 | S | | 78 | 0.007289439 | 0.03890308 | -0.037730215 | -0.045019654 | -0.076633295 | 0.805330538 | 0.452336721 | 0.270150204 | 0.232944495 | 0.197828986 |
| Gapvd1 | S | | 566 | -0.044639423 | 0.022285582 | 0.042968348 | 0.087607771 | 0.020682766 | 0.119114915 | 0.630148928 | 0.394408783 | 0.14228271 | 0.724065037 |
| H1f3 | S | | 174 | 0.115103922 | 0.062161893 | -0.147385592 | -0.262489515 | -0.209547485 | 0.375689037 | 0.29084113 | 0.073977351 | 0.100997548 | 0.0390686 |
| Camkk2 | S | | 99 | 0.07012224 | 0.074150768 | -0.135745178 | -0.205867418 | -0.209895946 | 0.405996669 | 0.370888468 | 0.126973372 | 0.009762155 | 0.001995994 |
| Cyfip1 | S | | 1233 | 0.013570973 | 0.028408604 | -0.055542228 | -0.069113201 | -0.083950832 | 0.829314259 | 0.818435716 | 0.729339356 | 0.65757441 | 0.642517065 |
| Recql4 | S | | 256 | 0.052376493 | -0.004366236 | -0.037179208 | -0.0895557 | -0.032812972 | 0.595755811 | 0.946508914 | 0.521235186 | 0.365446039 | 0.54366923 |
| Akap13 | S | | 583 | 0.080480549 | -0.012624516 | -0.075416199 | -0.155896748 | -0.062791683 | 0.141582568 | 0.791675014 | 0.199408401 | 0.027594413 | 0.221707582 |
| Ddx20 | S | | 472 | 0.059863955 | -0.019972051 | -0.051214687 | -0.111078642 | -0.031242636 | 0.017026694 | 0.40602672 | 0.074016798 | 0.008233645 | 0.249438527 |
| Cep152 | S | | 1575 | 0.027457532 | -0.074589269 | 0.030920191 | 0.00346266 | 0.10550946 | 0.59838568 | 0.447404679 | 0.624497861 | 0.946462654 | 0.304305542 |
| Slk | S | | 542 | -0.061031755 | -0.052171093 | 0.123367503 | 0.184399258 | 0.175538596 | 0.077195173 | 0.347317684 | 0.04213415 | 0.01933376 | 0.032502395 |
| Daxx | S | | 687 | -0.016478008 | 0.017641276 | -0.013003325 | 0.003474683 | -0.030644601 | 0.87658356 | 0.684168546 | 0.882205612 | 0.977330634 | 0.715538942 |
| Ddx41 | S | | 23 | -0.001156998 | -0.029872042 | 0.042952753 | 0.044109752 | 0.072824795 | 0.968262674 | 0.367747554 | 0.494786753 | 0.467846014 | 0.275570493 |
| Pphln1 | S | | 154 | 0.050744642 | -0.081544927 | 0.040972777 | -0.009771865 | 0.122517704 | 0.486791165 | 0.250399756 | 0.62053461 | 0.90583128 | 0.180778876 |
| Crkl | S | | 107 | -0.012163227 | -0.018388032 | 0.036136078 | 0.048299305 | 0.05452411 | 0.783250255 | 0.566760929 | 0.529159831 | 0.432647813 | 0.345518895 |
| Tbc1d10b | S | | 22 | 0.085242217 | -0.021426424 | -0.065136923 | -0.15037914 | -0.043710499 | 0.210604619 | 0.664163403 | 0.311578741 | 0.078739733 | 0.468135172 |
| Slx4 | S | | 988 | -0.057111947 | -0.061571306 | 0.15999583 | 0.217107778 | 0.221567136 | 0.337531168 | 0.111030261 | 0.095993275 | 0.047682138 | 0.055876357 |
| Safb | S | | 611 | -0.136350527 | 0.023575017 | 0.152166427 | 0.288516953 | 0.12859141 | 0.19869771 | 0.794129841 | 0.365547856 | 0.136872315 | 0.425219104 |
| Adgra2 | S | | 973 | -0.014495168 | 0.057574054 | -0.039242664 | -0.024747496 | -0.096816719 | 0.745113046 | 0.548731553 | 0.672084469 | 0.786510344 | 0.427581089 |
| Lrrc8a | T | | 215 | 0.176469002 | -0.003201699 | -0.202945528 | -0.37941453 | -0.199743829 | 0.11884238 | 0.967514731 | 0.162952466 | 0.0382115 | 0.16941904 |
| Tnip1 | S | | 448 | 0.023906903 | 0.006149336 | -0.02473478 | -0.048641684 | -0.030884116 | 0.810988975 | 0.91410721 | 0.679597231 | 0.634680506 | 0.608073714 |
| Itprid2 | S | | 109 | 0.005469763 | 0.022756352 | -0.032691849 | -0.038161612 | -0.055448201 | 0.916471918 | 0.593121443 | 0.577094929 | 0.494260957 | 0.291976975 |
| Dyrk1a | S | | 758 | 0.12124543 | 0.038245575 | -0.102139787 | -0.223385217 | -0.140385362 | 0.393734592 | 0.463902506 | 0.077796151 | 0.177494457 | 0.014950321 |
| Flywch1 | S | | 294 | 0.051328749 | -0.032579966 | -0.04772537 | -0.099054119 | -0.015145404 | 0.170077626 | 0.523615207 | 0.587961282 | 0.307369419 | 0.866964843 |
| Tanc1 | S | | 1436 | -0.006851116 | -0.009776685 | 0.020144022 | 0.026995137 | 0.029920707 | 0.889632238 | 0.679022872 | 0.68484862 | 0.658112742 | 0.531816535 |
| Srek1ip1 | S | | 97 | 0.02727411 | -0.019535233 | -0.001796887 | -0.029070997 | 0.017738346 | 0.150445313 | 0.094703952 | 0.883047673 | 0.139949719 | 0.16728871 |
| Osbpl3 | S | | 286 | -0.001803392 | -0.020130672 | 0.021355514 | 0.023158905 | 0.041486186 | 0.981596597 | 0.722038537 | 0.715261806 | 0.746314896 | 0.376881014 |
| Cobl | S | | 676 | -0.060743326 | 0.009467132 | 0.059796505 | 0.120539831 | 0.050329374 | 0.123189766 | 0.802482356 | 0.202748868 | 0.033410647 | 0.243360571 |
| Clasrp | S | | 541 | 0.101343174 | -0.072686028 | -0.021582086 | -0.122925259 | 0.051103943 | 0.055030402 | 0.071297786 | 0.596217754 | 0.040025156 | 0.20891033 |
| Rad18 | S | | 158 | 0.073007089 | -0.030578084 | -0.02194449 | -0.094951579 | 0.008633594 | 0.242682185 | 0.229895035 | 0.569509734 | 0.167833292 | 0.821862288 |
| Srrm1 | S | | 743 | -0.023522777 | -0.002125199 | 0.02332346 | 0.046846236 | 0.025448659 | 0.638871244 | 0.904619187 | 0.563157432 | 0.442305353 | 0.547004569 |
| Clk2 | S | | 17 | -0.05356408 | -0.031814783 | 0.079443286 | 0.133007366 | 0.111258069 | 0.563099478 | 0.663363833 | 0.351161218 | 0.240965921 | 0.254035762 |
| Cbx1 Cbx3 | S | 133;129 | | 0.002201327 | -0.013810213 | 0.012075169 | 0.009873842 | 0.025885382 | 0.97847552 | 0.887909306 | 0.912679772 | 0.927799719 | 0.829797411 |
| Kmt2d | T | | 1822 | -0.069285717 | -0.044809542 | 0.07790143 | 0.147187147 | 0.122710972 | 0.272596761 | 0.401560374 | 0.039242777 | 0.070089007 | 0.076214257 |
| Rtf1 | S | | 58 | -0.354175458 | -0.569313546 | -0.090682583 | 0.263492875 | 0.478630963 | NA | NA | NA | NA | NA |
| Rtf1 | S | | 63 | -0.354175458 | -0.569313546 | -0.090682583 | 0.263492875 | 0.478630963 | NA | NA | NA | NA | NA |
| Rtf1 | S | | 65 | -0.354175458 | -0.569313546 | -0.090682583 | 0.263492875 | 0.478630963 | NA | NA | NA | NA | NA |
| Mdm2 | S | | 284 | -0.23111659 | NA | NA | NA | NA | NA | NA | NA | NA | NA |
| Mdm2 | S | | 288 | -0.23111659 | NA | NA | NA | NA | NA | NA | NA | NA | NA |
| Emc2 | S | | 297 | NA | 0.681797573 | NA | NA | NA | NA | NA | NA | NA | NA |
| Rtf1 | T | | 60 | -0.354175458 | -0.569313546 | -0.090682583 | 0.263492875 | 0.478630963 | NA | NA | NA | NA | NA |
| Mdm2 | T | | 286 | -0.23111659 | NA | NA | NA | NA | NA | NA | NA | NA | NA |
| Ahctf1 | T | | 1190 | NA | NA | NA | NA | NA | NA | NA | NA | NA | NA |
| Abi1 | Y | | 306 | -1.169362711 | -1.357445575 | NA | NA | NA | NA | NA | NA | NA | NA |
| Ccnl1 | S | | 344 | 0.035092029 | 0.506130702 | 0.564885903 | 0.529793875 | 0.058755201 | 0.651578555 | 8.30996E-08 | 1.25307E-07 | 0.014574707 | 0.004834253 |
| Tmpo | T | 159;159 | | 0.067838892 | 0.439601625 | 0.520038896 | 0.452200004 | 0.080437272 | 0.060933576 | 3.87164E-06 | 5.13866E-08 | 0.000266253 | 0.025370145 |
| Synpo | S | | 833 | 0.155638316 | 0.567140912 | 0.535065112 | 0.379426796 | -0.0320758 | 0.05167558 | 4.58869E-06 | 4.53313E-05 | 0.003677758 | 0.479090722 |
| Synpo | S | | 765 | 0.155935254 | 0.396250996 | 0.4710049 | 0.315069647 | 0.074753904 | 0.044091473 | 2.99376E-06 | 1.15313E-06 | 0.014112522 | 0.009440613 |
| Srrm2 | S | | 1467 | 0.074867087 | 0.494342789 | 0.527979341 | 0.453112254 | 0.033636552 | 0.227363675 | 9.13505E-06 | 1.17712E-06 | 0.004656643 | 0.254848734 |
| Synpo | S | | 512 | 0.307287876 | 0.560318019 | 0.494193147 | 0.186905271 | -0.066124872 | 0.000287733 | 0.004692631 | 0.000139479 | 0.02555382 | 0.476782556 |
| Cdc42ep1 | S | | 121 | 0.268527472 | 0.512258795 | 0.450019459 | 0.181491987 | -0.062239336 | 0.002824325 | 6.64444E-05 | 0.000248914 | 0.012768298 | 0.173650254 |
| Arvcf | S | | 337 | 0.087544247 | 0.468135327 | 0.526621697 | 0.439077451 | 0.05848637 | 0.00342312 | 1.96926E-06 | 0.000799346 | 0.004930402 | 0.233548184 |
| Plec | S | | 4625 | -0.004710953 | 0.51915139 | 0.560814921 | 0.565525874 | 0.041663531 | 0.867590125 | 9.60169E-06 | 3.77986E-07 | 1.12049E-05 | 0.225716079 |
| Specc1l | S | | 390 | 0.141015397 | 0.423226141 | 0.569291451 | 0.428276054 | 0.14606531 | 0.087116347 | 5.80869E-05 | 3.51415E-06 | 0.011484751 | 0.02699777 |
| Smc1a | S | | 956 | 0.187688948 | 0.530718997 | 0.494858026 | 0.307169077 | -0.035860972 | 0.044890828 | 2.82066E-08 | 0.018796279 | 0.034900095 | 0.683800538 |
| Tmpo | S | 166;166 | | 0.0450457 | 0.449361847 | 0.552655617 | 0.507609917 | 0.10329377 | 0.226432989 | 0.001807001 | 2.0467E-08 | 0.000821313 | 0.072012168 |
| Pdlim2 | S | | 129 | 0.012169761 | 0.352357745 | 0.499338959 | 0.487169198 | 0.146981214 | 0.681749686 | 2.2624E-06 | 8.23473E-08 | 0.000345791 | 0.002055154 |
| Sipa1l1 | S | | 1712 | 0.054369154 | 0.311094213 | 0.406806127 | 0.352436973 | 0.095711913 | 0.047354261 | 9.65627E-08 | 1.97757E-06 | 9.38795E-05 | 0.010293621 |
| Becn1 | S | | 29 | 0.194903614 | 0.548196656 | 0.505148593 | 0.310244979 | -0.043048063 | 0.000784379 | 0.00013287 | 0.007819613 | 0.030933511 | 0.597302742 |
| Kat14 | S | | 284 | 0.157044081 | 0.527970158 | 0.429902732 | 0.272858652 | -0.098067426 | 0.02211728 | 2.06336E-05 | 0.000731501 | 0.010097323 | 0.155881908 |
| Ttc7 | S | | 690 | 0.352810644 | 0.487972893 | 0.526488244 | 0.173677599 | 0.038515351 | 0.003234983 | 0.010585486 | 0.000608321 | 0.032597314 | 0.715137763 |
| Mettl16 | T | | 459 | 0.013558138 | 0.393252193 | 0.391147375 | 0.377589237 | -0.002104818 | 0.438086904 | 1.91986E-06 | 2.06294E-07 | 1.71081E-06 | 0.904956013 |
| Uhrf2 | S | | 668 | 0.124262349 | 0.344563048 | 0.32479128 | 0.200528932 | -0.019771768 | 0.001595219 | 2.20693E-06 | 4.75882E-05 | 0.001894441 | 0.473797707 |
| Elf1 | S | | 187 | 0.030201258 | 0.356593398 | 0.571685969 | 0.541484711 | 0.215092571 | 0.513602252 | 3.04595E-05 | 1.96914E-06 | 0.000256673 | 0.005442218 |
| Rb1cc1 | S | | 623 | 0.062269142 | 0.492368624 | 0.48064766 | 0.418378517 | -0.011720965 | 0.406930877 | 9.72559E-06 | 6.40951E-05 | 0.00737048 | 0.744990555 |
| Cavin2 | S | | 293 | 0.099120256 | 0.364502865 | 0.388724799 | 0.289604543 | 0.024221934 | 0.014467012 | 5.939E-06 | 2.7577E-05 | 0.000827174 | 0.418547924 |
| Zfp219 | S | | 118 | 0.049413934 | 0.49301287 | 0.524363591 | 0.474949656 | 0.03135072 | 0.721583919 | 3.65869E-05 | 1.89295E-05 | 0.04905529 | 0.512581288 |
| Cars | S | | 390 | 0.065235576 | 0.503937203 | 0.48707441 | 0.421838834 | -0.016862793 | 0.374947355 | 0.00017876 | 7.49754E-06 | 0.008319425 | 0.723536959 |
| Cavin2 | S | | 287 | 0.23632497 | 0.330863693 | 0.311559679 | 0.075234709 | -0.019304014 | 0.00043943 | 8.42751E-05 | 0.000216801 | 0.091705598 | 0.557352868 |
| Specc1l | S | | 386 | 0.083256466 | 0.344977636 | 0.372637862 | 0.289381396 | 0.027660227 | 0.0125273 | 0.000125065 | 2.88159E-07 | 0.000362733 | 0.33941991 |
| Sipa1l1 | T | | 239 | 0.29485416 | 0.370117629 | 0.562101499 | 0.267247339 | 0.19198387 | 0.007277432 | 0.002985548 | 0.000519743 | 0.02773943 | 0.058452507 |
| Tns3 | Y | | 354 | 0.125998301 | 0.510254422 | 0.562218477 | 0.436220176 | 0.051964055 | 0.213642021 | 0.000198256 | 0.000191681 | 0.017541267 | 0.180703922 |
| Irs2 | S | | 723 | -0.023899413 | 0.393934117 | 0.508781095 | 0.532680508 | 0.114846979 | 0.503089821 | 2.66991E-06 | 2.12992E-06 | 8.02541E-05 | 0.013394943 |
| Sipa1l1 | S | | 1708 | 0.092436356 | 0.439515682 | 0.509678752 | 0.417242397 | 0.07016307 | 0.259991041 | 7.015E-06 | 0.000340907 | 0.009656958 | 0.111766851 |
| Marf1 | S | | 1083 | 0.064319605 | 0.346602393 | 0.412489132 | 0.348169527 | 0.065886739 | 0.376559829 | 7.22255E-06 | 1.93371E-06 | 0.019188593 | 0.029795102 |
| Dtna | S | | 455 | 0.084126249 | 0.407050847 | 0.357450027 | 0.273323778 | -0.04960082 | 0.084308781 | 3.3473E-06 | 5.76776E-05 | 0.002271628 | 0.173471549 |
| Cavin2 | S | | 288 | 0.104245669 | 0.366086656 | 0.387624432 | 0.283378763 | 0.021537776 | 0.017446843 | 6.91501E-06 | 0.000171037 | 0.002224442 | 0.552947547 |
| Acin1 | S | | 210 | 0.007684844 | 0.457187937 | 0.443894584 | 0.436209739 | -0.013293353 | 0.7916006 | 8.31109E-07 | 0.000158722 | 0.001275957 | 0.715897097 |
| Cep131 | T | | 483 | 0.078829454 | 0.501944087 | 0.473799835 | 0.394970381 | -0.028144252 | 0.599766027 | 2.21328E-05 | 0.000178368 | 0.075369504 | 0.593844943 |
| Gmnn | S | | 19 | 0.316882825 | 0.40871121 | 0.437095109 | 0.120212284 | 0.028383899 | 0.001320388 | 0.050278967 | 0.000156403 | 0.032605969 | 0.829181346 |
| Ptpn22 | S | | 518 | 0.205052937 | 0.351898745 | 0.479861752 | 0.274808815 | 0.127963007 | 0.04804638 | 1.24883E-05 | 0.002012451 | 0.015903049 | 0.061319627 |
| Sun2 | S | | 120 | 0.036580101 | 0.300877497 | 0.311224552 | 0.274644451 | 0.010347055 | 0.129910706 | 3.90364E-09 | 2.73555E-05 | 0.000353771 | 0.410352115 |
| Spry2 | S | | 111 | 0.288615038 | 0.438643753 | 0.360495201 | 0.071880163 | -0.078148552 | 0.050626566 | 0.000754217 | 0.00014681 | 0.479687058 | 0.231188702 |
| Zfp318 | S | | 89 | 0.137663419 | 0.150426202 | 0.411382682 | 0.273719263 | 0.26095648 | 0.002120166 | 0.000269172 | 4.39171E-07 | 0.000475529 | 0.000122641 |
| Acin1 | S | | 206 | -0.000663278 | 0.403283874 | 0.408747409 | 0.409410687 | 0.005463535 | 0.978616587 | 9.73983E-07 | 1.79255E-05 | 0.000708432 | 0.843744594 |
| Acin1 | S | | 207 | -0.000663278 | 0.403283874 | 0.408747409 | 0.409410687 | 0.005463535 | 0.978616587 | 9.73983E-07 | 1.79255E-05 | 0.000708432 | 0.843744594 |
| Foxk1 | S | | 427 | -0.003528 | 0.386655556 | 0.412344709 | 0.415872709 | 0.025689153 | 0.872850858 | 2.63579E-05 | 3.99193E-07 | 8.66875E-06 | 0.343156057 |
| Nusap1 | S | | 261 | 0.181389482 | 0.512845765 | 0.471897391 | 0.290507909 | -0.040948374 | 0.019353625 | 6.27633E-05 | 0.017104186 | 0.070902142 | 0.706985288 |
| Zc3h4 | S | | 1270 | 0.05105895 | 0.409559998 | 0.477371323 | 0.426312374 | 0.067811325 | 0.24813624 | 6.88224E-06 | 0.000286014 | 0.001091232 | 0.201011084 |
| Chd4 | S | | 1524 | 0.022800443 | 0.412026427 | 0.54315013 | 0.520349687 | 0.131123703 | 0.630047075 | 4.74021E-06 | 0.000347964 | 0.001442812 | 0.010477683 |
| Lmna | S | | 633 | 0.064588839 | 0.292292164 | 0.38389913 | 0.319310291 | 0.091606966 | 0.101700179 | 0.000205626 | 1.43197E-07 | 0.004327398 | 0.034463332 |
| Ahdc1 | S | | 1498 | 0.032108141 | 0.375296117 | 0.43833323 | 0.406225089 | 0.063037112 | 0.418807522 | 2.32428E-05 | 1.15378E-05 | 0.001345546 | 0.092165745 |

| | | | | | | | | | | | | |
|---|---|---|---|---|---|---|---|---|---|---|---|---|
| Irs2 | S | | 1165 | 0.357741554 | 0.519997499 | 0.5842046 | 0.226463046 | 0.064207101 | 0.295540113 | 0.000225558 | 0.010092999 | 0.471489064 | 0.456792316 |
| Synpo | S | | 672 | 0.11848318 | 0.362447123 | 0.409394571 | 0.290911391 | 0.046947448 | 0.124311937 | 3.83236E-05 | 7.8191E-05 | 0.012841835 | 0.258055536 |
| Ptprk | S | | 868 | 0.201885521 | 0.444313491 | 0.359956469 | 0.158070948 | -0.084357021 | 0.010386255 | 0.000221854 | 0.002456953 | 0.068965114 | 0.247107916 |
| Sun2 | T | | 117 | 0.045826836 | 0.290161716 | 0.311083281 | 0.265256444 | 0.020921565 | 0.089461028 | 2.36029E-09 | 0.000215524 | 0.000285358 | 0.234958528 |
| Pi4ka | S | | 1439 | 0.355333385 | 0.491911727 | 0.528563826 | 0.173230441 | 0.036652099 | 0.008419365 | 0.035691319 | 0.002873304 | 0.151436763 | 0.813629976 |
| Cic | S | | 766 | 0.115227398 | 0.291749894 | 0.409860189 | 0.294632791 | 0.118110295 | 0.002887798 | 4.24348E-06 | 0.00183806 | 0.005720379 | 0.063171248 |
| Sorbs2 | S | | 325 | 0.06017188 | 0.256180439 | 0.450543468 | 0.390371588 | 0.194363029 | 0.107518959 | 0.000453281 | 5.61259E-07 | 0.000354576 | 0.004207103 |
| Pard3b | S | | 730 | -0.045594853 | 0.355614318 | 0.487078985 | 0.532673839 | 0.131464667 | 0.404395693 | 4.08429E-06 | 6.68522E-06 | 0.001291151 | 0.013490437 |
| Irs2 | S | | 388 | 0.03156798 | 0.442467472 | 0.514957286 | 0.483389306 | 0.072489814 | 0.456539171 | 4.22665E-06 | 0.001740055 | 0.003420103 | 0.286076953 |
| Zc3h13 | S | | 215 | 0.504570154 | 0.047503087 | 0.510900997 | 0.006330843 | 0.46339791 | 0.000382137 | 0.468509707 | 0.000151396 | 0.919400242 | 0.002526607 |
| Zc3h13 | S | | 218 | 0.504570154 | 0.047503087 | 0.510900997 | 0.006330843 | 0.46339791 | 0.000382137 | 0.468509707 | 0.000151396 | 0.919400242 | 0.002526607 |
| Bbx | S | | 811 | 0.006609662 | 0.345649883 | 0.459435933 | 0.452826271 | 0.11378605 | 0.798487435 | 5.31777E-06 | 2.31043E-05 | 0.000982125 | 0.031543495 |
| Plec | S | | 4629 | 0.016285392 | 0.44231782 | 0.364674789 | 0.348389398 | -0.077643031 | 0.607023233 | 0.000146183 | 1.16257E-06 | 0.000670926 | 0.077722816 |
| Ctnnd1 | S | | 349 | 0.124998795 | 0.291810136 | 0.344830313 | 0.219831519 | 0.053020177 | 0.011586149 | 0.000114477 | 2.99912E-05 | 0.001263916 | 0.046102857 |
| Ncor1 | S | | 2150 | 0.12714942 | 0.317372074 | 0.5169079 | 0.38975848 | 0.199535825 | 0.094968662 | 0.00116696 | 5.60266E-05 | 0.002641817 | 0.00830316 |
| Ctnnd1 | S | | 352 | 0.098760241 | 0.304202237 | 0.361173071 | 0.26241283 | 0.056970834 | 0.040440467 | 6.03827E-05 | 1.9732E-05 | 0.001345883 | 0.073030597 |
| Wac | T | | 457 | 0.200307564 | 0.419199683 | 0.538871156 | 0.338563592 | 0.119671473 | 0.092890491 | 0.017336994 | 8.63762E-05 | 0.027544662 | 0.293574558 |
| Epb41l3 | S | | 96 | 0.07739737 | 0.283464272 | 0.492335106 | 0.414937737 | 0.208870834 | 0.04713168 | 9.02223E-06 | 0.000492213 | 0.001171855 | 0.016101129 |
| Nectin2 | T | | 401 | 0.135370042 | 0.345406312 | 0.409338945 | 0.273968902 | 0.063932632 | 0.023937826 | 0.000779264 | 9.65778E-05 | 0.000180049 | 0.2270959 |
| Cdkn1b | S | | 10 | 0.084645414 | 0.365718757 | 0.424369508 | 0.339724094 | 0.058650751 | 0.106790298 | 0.000151516 | 8.96749E-05 | 0.006813669 | 0.196680582 |
| Tmpo | T | 74;74 | | 0.052493001 | 0.260518465 | 0.405477823 | 0.352984822 | 0.144959358 | 0.324431949 | 1.07078E-05 | 3.89983E-06 | 0.012447574 | 0.006409131 |
| Fundc1 | S | | 13 | 0.09701383 | 0.281816408 | 0.352418389 | 0.255404559 | 0.07060198 | 0.157688386 | 2.60298E-05 | 5.53925E-06 | 0.026503887 | 0.008725554 |
| Peak1 | Y | | 632 | 0.20509748 | 0.383447846 | 0.404757827 | 0.199660347 | 0.021309981 | 0.016897003 | 0.002530438 | 0.000394087 | 0.015439332 | 0.752913249 |
| Afdn | T | | 1232 | 0.185010873 | 0.459570226 | 0.544705711 | 0.359694838 | 0.085135485 | 0.089847602 | 0.000258391 | 0.010283378 | 0.015169455 | 0.308319923 |
| Ubl4a Gm44504 | S | | 90 | 0.364289396 | 0.08785548 | 0.539086776 | 0.174797379 | 0.451231295 | 0.000599513 | 0.458310498 | 6.20408E-05 | 0.009223574 | 0.031937959 |
| Slain2 | S | | 354 | 0.246394106 | 0.569557319 | 0.5276996 | 0.281305494 | -0.041857719 | 0.045220917 | 0.003609532 | 0.013255624 | 0.088840556 | 0.760398608 |
| Cdc42ep1 | S | | 77 | 0.106834316 | 0.411169763 | 0.442464843 | 0.335630526 | 0.03129508 | 0.072789563 | 0.001791288 | 9.13358E-05 | 0.001498085 | 0.607239455 |
| Eif4enif1 | S | | 137 | 0.277113314 | 0.397962413 | 0.343202296 | 0.066088982 | -0.054760117 | 0.000500058 | 0.005350387 | 0.00887469 | 0.373575909 | 0.531496596 |
| Edc4 | T | | 922 | 0.329310791 | 0.548220367 | 0.236688589 | -0.092622202 | -0.311531777 | 0.00924194 | 7.65116E-05 | 0.175109339 | 0.541570544 | 0.112252344 |
| Plec | S | | 4627 | -0.023255533 | 0.461481437 | 0.396283391 | 0.419538924 | -0.065198046 | 0.56030555 | 1.32438E-05 | 4.13279E-05 | 0.000422216 | 0.058500584 |
| Hp1bp3 | S | | 158 | 0.147638052 | 0.327496345 | 0.371881891 | 0.224243838 | 0.044385545 | 0.212272879 | 2.01496E-05 | 0.00018614 | 0.102439289 | 0.279051656 |
| Sptbn1 | T | | 2327 | 0.140156131 | 0.270113583 | 0.235600771 | 0.09544464 | -0.034512812 | 0.001977875 | 1.21314E-05 | 0.000138202 | 0.022206456 | 0.203540476 |
| Synpo | S | | 567 | 0.217291907 | 0.312091512 | 0.292443484 | 0.075151577 | -0.019648029 | 0.00335815 | 0.000450497 | 0.00067962 | 0.010379066 | 0.347533715 |
| Thrap3 | S | | 743 | 0.147285377 | 0.445723815 | 0.527090491 | 0.379805113 | 0.081366676 | 0.020951875 | 0.00918131 | 0.000907489 | 0.002954957 | 0.351331571 |
| Mcoln1 | S | | 10 | 0.127129915 | 0.421059577 | 0.428202133 | 0.301072218 | 0.007142556 | 0.228539254 | 0.000416643 | 0.00026786 | 0.038776586 | 0.839901698 |
| Unk | S | | 240 | 0.018077517 | 0.291857964 | 0.333670549 | 0.315593032 | 0.041812585 | 0.446505752 | 3.77589E-05 | 2.17212E-07 | 0.000375384 | 0.106488078 |
| Synpo | S | | 854 | 0.16122081 | 0.298989378 | 0.267390074 | 0.106169264 | -0.031599303 | 0.003373373 | 0.000119672 | 0.000283319 | 0.0227585 | 0.305229792 |
| Tmpo | T | | 278 | 0.119777754 | 0.269283038 | 0.384880376 | 0.265102622 | 0.115597338 | 0.163277412 | 3.82651E-05 | 4.85134E-05 | 0.030365646 | 0.027024267 |
| Palm | T | | 145 | 0.117045753 | 0.308747847 | 0.38961967 | 0.272573917 | 0.080871822 | 0.005408557 | 1.91287E-05 | 0.004612934 | 0.013520038 | 0.196118047 |
| Zfp740 | S | | 19 | -0.010241583 | 0.432381342 | 0.515770807 | 0.52601239 | 0.083389465 | 0.788401448 | 7.56109E-07 | 0.005413044 | 0.002623477 | 0.252169192 |
| Cobll1 | S | | 12 | 0.373556311 | 0.288984388 | 0.458370298 | 0.084813987 | 0.16938591 | 0.002277699 | 0.081681227 | 0.001815159 | 0.279894721 | 0.242184867 |
| H2bc11 H2bc13 H2bc | S | 7;7;7;7 | | 0.799566965 | 0.571075919 | 0.372231961 | -0.427335003 | -0.198843957 | 0.210154759 | 0.000292379 | 0.190474656 | 0.444846245 | 0.412742353 |
| D830031N03Rik | S | | 180 | 0.092931817 | 0.262096886 | 0.351887527 | 0.25895571 | 0.089790641 | 0.044829729 | 8.98914E-05 | 2.32234E-05 | 0.003967821 | 0.009172691 |
| Foxk1 | S | | 431 | -0.014598733 | 0.282040725 | 0.285559886 | 0.300158618 | 0.00351916 | 0.445067806 | 6.12156E-07 | 1.22321E-07 | 0.000252155 | 0.784823055 |
| Spata6 | S | | 354 | 0.243662351 | 0.401849945 | 0.516377119 | 0.272714768 | 0.114527174 | 0.065043617 | 0.056315424 | 0.000142283 | 0.052116681 | 0.421955953 |
| Synpo | S | | 488 | 0.45923392 | 0.51972245 | 0.279296173 | -0.179937747 | -0.240426277 | 0.000311446 | 0.009212302 | 0.365724921 | 0.533949493 | 0.429170786 |
| Senp7 | S | | 12 | 0.113942742 | 0.370502274 | 0.423861246 | 0.309918504 | 0.053358972 | 0.001954129 | 0.001058605 | 0.002816194 | 0.01690049 | 0.364998781 |
| Irs2 | S | | 1163 | 0.334377082 | 0.261697185 | 0.411890944 | 0.077513863 | 0.150193759 | 0.399548914 | 0.00103052 | 2.04657E-05 | 0.828718711 | 0.019187008 |
| Tor1aip1 | S | | 60 | 0.038611392 | 0.413621414 | 0.518843051 | 0.480231659 | 0.105221637 | 0.630468662 | 0.002386969 | 4.13645E-05 | 0.008746524 | 0.135040264 |
| Trpm4 | S | | 527 | 0.126578424 | 0.340612409 | 0.497287175 | 0.370708751 | 0.156674767 | 0.466003275 | 0.000311542 | 0.00025582 | 0.108046011 | 0.046634656 |
| Pold1 | S | | 29 | 0.203117506 | 0.413221058 | 0.427695933 | 0.224578428 | 0.014474875 | 0.285664515 | 0.000709713 | 0.000745274 | 0.252529099 | 0.742047525 |
| Brca1 | S | | 1206 | 0.217181114 | 0.312907871 | 0.372850352 | 0.155669238 | 0.059942481 | 0.004486167 | 0.000603831 | 0.006357185 | 0.110389802 | 0.432587608 |
| Brca1 | S | | 1207 | 0.217181114 | 0.312907871 | 0.372850352 | 0.155669238 | 0.059942481 | 0.004486167 | 0.000603831 | 0.006357185 | 0.110389802 | 0.432587608 |
| Ltbp3 | S | | 1164 | -0.004174255 | 0.510650744 | 0.39870372 | 0.402877975 | -0.111947023 | 0.950382573 | 0.000208676 | 0.000104058 | 0.005993668 | 0.091495706 |
| Mtfr2 | S | | 317 | 0.093387128 | 0.316603341 | 0.333776221 | 0.240389093 | 0.01717288 | 0.157274275 | 9.87918E-06 | 0.000282567 | 0.015542797 | 0.613114723 |
| Mdm1 | S | | 315 | 0.103214309 | 0.473845657 | 0.431702686 | 0.328488377 | -0.042142972 | 0.150073507 | 3.08096E-05 | 0.019632517 | 0.041782626 | 0.677114529 |
| Atrx | S | | 586 | 0.112433193 | 0.454324041 | 0.482399823 | 0.369966631 | 0.028075782 | 0.000646696 | 0.003182071 | 0.014418956 | 0.029818401 | 0.743479334 |
| Nop56 | T | | 467 | 0.116588692 | 0.301361503 | 0.355133255 | 0.238544563 | 0.053771752 | 0.125053658 | 1.5098E-06 | 0.002651126 | 0.019519099 | 0.256914276 |
| Macf1 | T | | 5286 | 0.136948692 | 0.394669793 | 0.514504626 | 0.377555933 | 0.119834832 | 0.154792063 | 0.000124788 | 0.008883944 | 0.022606673 | 0.268652084 |
| Pml | S | | 515 | 0.05184053 | 0.298920331 | 0.367930844 | 0.316090315 | 0.069010514 | 0.374529819 | 5.22273E-05 | 1.83608E-05 | 0.007958372 | 0.066200771 |
| Hnrnpul2 | S | | 191 | 0.082484983 | 0.269181878 | 0.401342291 | 0.318857308 | 0.132160413 | 0.008200362 | 4.25572E-06 | 0.004413002 | 0.006972607 | 0.061195675 |
| Rps6ka5 | S | | 814 | 0.225921173 | 0.375863144 | 0.354158168 | 0.128236995 | -0.021704976 | 0.008593799 | 0.006781439 | 0.00143909 | 0.096867994 | 0.782863644 |
| Ezh2 | S | | 366 | 0.25856054 | 0.508514274 | 0.493191526 | 0.234630985 | -0.015322748 | 0.006325416 | 0.000919364 | 0.199830237 | 0.469852308 | 0.959728211 |
| Nipbl | S | | 280 | 0.14201911 | 0.275556623 | 0.304516102 | 0.162496991 | 0.028959479 | 0.032020447 | 9.21739E-05 | 0.000188514 | 0.023381734 | 0.402008815 |
| Klf3 | S | | 249 | 0.148402223 | 0.200235799 | 0.402082273 | 0.25368005 | 0.201846474 | 0.39444729 | 0.00018602 | 4.33899E-06 | 0.206669583 | 0.000556149 |
| Uqcrc1 | S | | 212 | 0.315078694 | 0.534550255 | 0.163683832 | -0.151394862 | -0.370866422 | 0.007032895 | 0.000501707 | 0.078756136 | 0.006241606 | 0.010185552 |
| Thrap3 | S | | 740 | 0.134016404 | 0.40103056 | 0.578377638 | 0.444361234 | 0.177347078 | 0.144561862 | 0.058753674 | 8.18089E-05 | 0.007494055 | 0.234067065 |
| Rcc2 | S | | 48 | 0.085516912 | 0.401416348 | 0.393361289 | 0.307844377 | -0.008055059 | 0.131443774 | 0.000163234 | 0.00135974 | 0.020862835 | 0.892959412 |
| Rcc2 | S | | 49 | 0.085516912 | 0.401416348 | 0.393361289 | 0.307844377 | -0.008055059 | 0.131443774 | 0.000163234 | 0.00135974 | 0.020862835 | 0.892959412 |
| Cic | S | | 1338 | 0.031724093 | 0.296302888 | 0.405280879 | 0.373556786 | 0.108977991 | 0.221688675 | 0.001886719 | 1.77817E-06 | 7.79293E-05 | 0.043741658 |
| Pxn | S | | 258 | 0.058506984 | 0.21409593 | 0.253949249 | 0.195442265 | 0.039853319 | 0.109906067 | 4.46268E-06 | 5.61244E-07 | 0.009114556 | 0.035617973 |
| Arel1 | S | | 337 | 0.175812559 | 0.335925997 | 0.544842823 | 0.369030264 | 0.208916826 | 0.008013173 | 0.002817914 | 0.011144509 | 0.033895274 | 0.109024638 |
| Synpo | S | | 227 | 0.223608221 | 0.538740941 | 0.429984356 | 0.206376135 | -0.108756585 | 0.016324708 | 0.001034479 | 0.094687065 | 0.315439504 | 0.569084791 |
| Tent4b | S | | 58 | 0.143304974 | 0.400960703 | 0.496083241 | 0.352778267 | 0.095122538 | 0.221836802 | 0.020527829 | 7.6724E-05 | 0.036616646 | 0.385361333 |
| Cpsf6 | S | | 525 | 0.51802511 | 0.409319121 | 0.207715725 | -0.310309385 | -0.201603395 | 0.002157681 | 0.001119308 | 0.410348502 | 0.256367614 | 0.418904855 |
| Tulp4 | S | | 577 | 0.113874928 | 0.193098639 | 0.315675899 | 0.201800971 | 0.12257259 | 0.000544857 | 0.002256966 | 1.73846E-05 | 0.001101745 | 0.014970479 |
| Pml | S | | 514 | 0.050905578 | 0.305192516 | 0.378769827 | 0.327864249 | 0.073577311 | 0.445874101 | 0.000110018 | 2.7614E-05 | 0.013424811 | 0.050106397 |
| Pik3r4 | S | | 865 | 0.006368085 | 0.410679664 | 0.427086226 | 0.420718141 | 0.016406562 | 0.956953438 | 5.52485E-07 | 0.009623988 | 0.035746269 | 0.799406937 |
| Smarca2 | S | | 1499 | -0.0278733 | 0.525136531 | 0.425780225 | 0.453653526 | -0.099356306 | 0.589086705 | 8.63376E-05 | 0.000708177 | 0.001789139 | 0.164625662 |
| Smarca2 | S | | 1503 | -0.0278733 | 0.525136531 | 0.425780225 | 0.453653526 | -0.099356306 | 0.589086705 | 8.63376E-05 | 0.000708177 | 0.001789139 | 0.164625662 |
| Smarca2 | S | | 1515 | -0.0278733 | 0.525136531 | 0.425780225 | 0.453653526 | -0.099356306 | 0.589086705 | 8.63376E-05 | 0.000708177 | 0.001789139 | 0.164625662 |
| Syne1 | S | | 8308 | 0.091220217 | 0.303082658 | 0.356237536 | 0.265017319 | 0.053154878 | 0.109160095 | 0.000243042 | 0.000103618 | 0.003430443 | 0.162420313 |
| Arl6ip6 | T | | 19 | 0.150282316 | 0.346764757 | 0.295371984 | 0.145089667 | -0.051392774 | 0.178169626 | 1.58425E-06 | 0.009039403 | 0.187040199 | 0.343107062 |
| Gtf2f1 | S | | 436 | 0.280944391 | 0.2502652 | 0.480684542 | 0.199740151 | 0.230419343 | 0.002099915 | 0.049292618 | 0.003589342 | 0.078820331 | 0.08253578 |
| Mia2 | S | | 1131 | 0.465812723 | 0.262253342 | 0.467904257 | 0.002091534 | 0.205650915 | 0.004358086 | 0.248091696 | 0.003421548 | 0.966100736 | 0.333736693 |
| Mia2 | S | | 1143 | 0.465812723 | 0.262253342 | 0.467904257 | 0.002091534 | 0.205650915 | 0.004358086 | 0.248091696 | 0.003421548 | 0.966100736 | 0.333736693 |
| Zfp318 | S | | 91 | 0.188483758 | 0.18793453 | 0.326188829 | 0.137705071 | 0.138254299 | 0.00170163 | 0.006386823 | 8.03221E-05 | 0.003144104 | 0.033338232 |
| Epb41l3 | S | | 599 | 0.116288945 | 0.275978042 | 0.335085979 | 0.218797034 | 0.059107937 | 0.019862833 | 0.000111828 | 0.000671694 | 0.008285828 | 0.223210804 |
| Peak1 | S | | 1363 | 0.40672528 | 0.245509671 | 0.352349318 | -0.054375961 | 0.106839647 | 0.05772669 | 0.005979355 | 0.000597563 | 0.695811853 | 0.109878119 |
| Lemd2 | T | | 158 | 0.01515501 | 0.356135137 | 0.521335037 | 0.506180028 | 0.165199901 | 0.802811974 | 0.000114708 | 0.000672866 | 0.001532875 | 0.052795702 |
| Gas2l1 | S | | 489 | 0.304212227 | 0.361910804 | 0.217217922 | -0.086994305 | -0.144692882 | 0.000879453 | 9.74721E-05 | 0.297893636 | 0.637590035 | 0.453609849 |
| Sun2 | S | | 39 | 0.10517973 | 0.353157312 | 0.305343583 | 0.200163852 | -0.04781373 | 0.111832925 | 3.21796E-05 | 0.001560777 | 0.022647825 | 0.296362345 |
| Atp2b1 | T | | 1165 | 0.151004636 | 0.251990339 | 0.322430654 | 0.171426018 | 0.070440315 | 0.01333505 | 0.001011192 | 0.000180044 | 0.009797514 | 0.021891052 |
| Ank3 | S | | 1795 | 0.169636139 | 0.23442485 | 0.254953058 | 0.085316919 | 0.020528208 | 0.004313377 | 0.001639806 | 4.22992E-05 | 0.035019043 | 0.473404714 |

| | | | | | | | | | | | | | |
|---|---|---|---|---|---|---|---|---|---|---|---|---|---|
| Epb41l3 | S | | 486 | 0.032803598 | 0.199907974 | 0.309424842 | 0.276621244 | 0.109516868 | 0.255664704 | 1.08209E-05 | 1.08663E-06 | 0.003056358 | 0.001534665 |
| Srsf4 | S | | 443 | 0.2712505 | 0.325516742 | 0.423711406 | 0.152460905 | 0.098194664 | 0.330129545 | 4.22595E-05 | 0.00827025 | 0.553685351 | 0.250313015 |
| Stk10 | S | | 416 | 0.229951922 | 0.284808133 | 0.239446888 | 0.009494967 | -0.045361245 | 0.012280851 | 0.000908344 | 0.000449727 | 0.848189101 | 0.283212771 |
| Thrap3 | S | | 679 | 0.371900235 | 0.410739932 | 0.150033453 | -0.221866782 | -0.260706479 | 0.000605449 | 0.000647136 | 0.383538504 | 0.237378013 | 0.185409907 |
| Synpo | T | | 547 | 0.093882094 | 0.310194666 | 0.470019077 | 0.376136982 | 0.15982441 | 0.059375908 | 0.00113684 | 0.00105423 | 0.014445237 | 0.065595887 |
| Frs2 | S | | 365 | 0.02639597 | 0.430994701 | 0.463732061 | 0.437336092 | 0.032737361 | 0.60655634 | 0.004515897 | 6.83362E-05 | 0.001536643 | 0.667777518 |
| Szt2 | S | | 1650 | 0.12011223 | 0.424716296 | 0.292485156 | 0.172372926 | -0.132231141 | 0.182492977 | 8.82634E-05 | 0.002932424 | 0.088903544 | 0.07296815 |
| Hspa5 | T | | 519 | 0.281010764 | 0.309876957 | 0.505516201 | 0.224505437 | 0.195639244 | 0.04678237 | 0.042352895 | 0.001319484 | 0.088684054 | 0.141377421 |
| Acin1 | S | | 491 | 0.077010912 | 0.285099222 | 0.301613736 | 0.224602824 | 0.016514514 | 0.062649855 | 0.000160773 | 6.56303E-05 | 0.00251771 | 0.534221724 |
| Thrap3 | S | | 737 | 0.131189106 | 0.30470856 | 0.387544221 | 0.256355115 | 0.082835661 | 0.000887427 | 0.002177316 | 0.005708272 | 0.01044068 | 0.115134828 |
| Stk24 Stk25 | T | 178;174 | | 0.141607522 | 0.422598827 | 0.466176912 | 0.32456939 | 0.043578085 | 0.023519187 | 0.001643634 | 0.018243937 | 0.051629876 | 0.689354953 |
| Flnb | T | | 1609 | 0.095545156 | 0.316057053 | 0.424920489 | 0.329375333 | 0.108863435 | 0.422395721 | 0.000241316 | 0.000384154 | 0.055370309 | 0.099152945 |
| Cobll1 | T | | 16 | 0.503978916 | 0.3678333 | 0.388710322 | -0.115268594 | 0.020877022 | 0.008264832 | 0.059704689 | 0.030568336 | 0.383714259 | 0.889421829 |
| Gas2l1 | S | | 352 | 0.035934377 | 0.388612594 | 0.502284443 | 0.466350067 | 0.113671849 | 0.544860973 | 0.000104084 | 0.003224264 | 0.002929029 | 0.15530696 |
| Sipa1l1 | S | | 174 | -0.061003655 | 0.370506309 | 0.482378214 | 0.543381869 | 0.111871905 | 0.143540915 | 3.97226E-05 | 7.8646E-05 | 0.001442095 | 0.060908929 |
| Ltbp3 | S | | 1163 | 0.057088041 | 0.493519969 | 0.308161852 | 0.25107381 | -0.185358117 | 0.190004933 | 0.001317808 | 0.00022866 | 0.003241725 | 0.023897173 |
| Tmpo | T | | 207 | 0.250297145 | 0.210172781 | 0.288061421 | 0.037764277 | 0.077888641 | 0.114129936 | 0.00076009 | 2.72954E-05 | 0.735606869 | 0.055616596 |
| Cx3cl1 | S | | 262 | 0.12467457 | 0.311373735 | 0.396741211 | 0.272066641 | 0.085367476 | 0.109101695 | 0.001525094 | 0.000416215 | 0.015049241 | 0.013049718 |
| Wdr60 | S | | 28 | 0.010913847 | 0.25195108 | 0.29239194 | 0.281478093 | 0.04044086 | 0.847758547 | 2.46159E-05 | 1.01412E-06 | 0.025802663 | 0.094924731 |
| Stim1 | S | | 575 | 0.241973438 | 0.25771377 | 0.343836966 | 0.101863528 | 0.086123196 | 0.01941114 | 0.014061773 | 0.000327006 | 0.191202911 | 0.243321234 |
| Edc4 | S | | 919 | 0.197015162 | 0.397897269 | 0.257861893 | 0.060846731 | -0.140035377 | 0.015677772 | 0.000161184 | 0.038292296 | 0.496423185 | 0.175943486 |
| Tmpo | S | | 183 | 0.066731001 | 0.22413856 | 0.279480572 | 0.212749571 | 0.055342012 | 0.172212449 | 2.97875E-05 | 8.23844E-06 | 0.014215954 | 0.034184076 |
| Klf3 | S | | 100 | 0.096525194 | 0.283139418 | 0.424754445 | 0.328229251 | 0.141615027 | 0.222346429 | 7.61618E-06 | 0.006151385 | 0.012333087 | 0.093208914 |
| Ccar2 | S | | 677 | 0.036326633 | 0.153573078 | 0.230673319 | 0.194346686 | 0.077100241 | 0.207397682 | 4.70327E-07 | 8.29622E-08 | 0.007047594 | 0.000924552 |
| Top2b | S | | 1453 | 0.005690586 | 0.223468678 | 0.353117021 | 0.347426435 | 0.129648343 | 0.902745088 | 5.39628E-05 | 1.67518E-06 | 0.006579859 | 0.003487691 |
| Cntln | T | | 1334 | 0.023586192 | 0.300672715 | 0.428887285 | 0.405301093 | 0.12821457 | 0.853544847 | 0.000382361 | 4.69045E-05 | 0.057701558 | 0.019158319 |
| Lmna | T | | 224 | 0.11131368 | 0.305902258 | 0.456301545 | 0.344987865 | 0.150399287 | 0.197884783 | 0.003774999 | 0.000271697 | 0.018253491 | 0.064918095 |
| Edc4 | S | | 923 | 0.167755986 | 0.37291078 | 0.255881259 | 0.088125274 | -0.117029521 | 0.016625865 | 9.99345E-05 | 0.023591146 | 0.280013943 | 0.179319486 |
| Synpo | S | | 134 | 0.098555702 | 0.328950829 | 0.362820146 | 0.264264445 | 0.033869317 | 0.036586555 | 5.94844E-05 | 0.009502601 | 0.03009514 | 0.624474026 |
| Sash1 | S | | 714 | 0.11232594 | 0.519035074 | 0.372916919 | 0.260590979 | -0.146118154 | 0.016310716 | 0.013702705 | 0.0027298 | 0.022065392 | 0.212321612 |
| Pold1 | S | | 31 | 0.381989846 | 0.435672351 | 0.208900127 | -0.173089718 | -0.226772223 | 0.028097585 | 0.000439325 | 0.136591074 | 0.257067177 | 0.121673674 |
| Prkaca Prkacb | T | 196;196 | | 0.102404791 | 0.224080684 | 0.267438945 | 0.165034154 | 0.043358261 | 0.006614199 | 0.000148596 | 0.000185349 | 0.01053208 | 0.188274043 |
| Sin3a | S | | 833 | 0.061461197 | 0.211697643 | 0.350705832 | 0.289244635 | 0.139008189 | 0.172129319 | 0.000146734 | 2.47045E-05 | 0.002197576 | 0.007438026 |
| Rapgef3 | T | | 380 | 0.281425348 | 0.335122292 | 0.283069624 | 0.001644277 | -0.052052668 | 0.038068023 | 0.000123448 | 0.047462742 | 0.988272364 | 0.570374837 |
| Dab2ip | S | | 978 | 0.043748253 | 0.299077899 | 0.309050752 | 0.265302499 | 0.009972853 | 0.410503979 | 6.82161E-05 | 6.04628E-05 | 0.00814486 | 0.695259531 |
| Zfp318 | S | | 732 | 0.083349772 | 0.199277602 | 0.444375264 | 0.361025492 | 0.245097662 | 0.173157643 | 0.000971519 | 6.7937E-05 | 0.004670692 | 0.001091842 |
| Abcf1 | S | | 103 | -0.036244881 | 0.48680784 | 0.415310915 | 0.451555795 | -0.071496926 | 0.523209356 | 3.52406E-06 | 0.016988673 | 0.004297719 | 0.342037244 |
| Ccar2 | S | | 674 | 0.056150053 | 0.156360109 | 0.256093263 | 0.19994321 | 0.099733153 | 0.025193798 | 2.61507E-05 | 1.01053E-06 | 0.000511266 | 0.000918528 |
| Oxsr1 | S | | 325 | 0.222926206 | 0.310030431 | 0.278394712 | 0.055468506 | -0.031635719 | 0.037074013 | 0.001416082 | 0.001897068 | 0.467347417 | 0.384028607 |
| Il16 | S | | 583 | 0.012394802 | 0.539249562 | 0.496111534 | 0.483716733 | -0.043138028 | 0.872874352 | 0.004031232 | 0.001164182 | 0.003533951 | 0.683310725 |
| Spp1 | S | | 158 | 0.251581617 | 0.424247352 | 0.275642613 | 0.024060997 | -0.148604739 | 0.006982434 | 0.004673916 | 0.033115251 | 0.738224316 | 0.110234367 |
| Thrap3 | S | | 405 | 0.171430745 | 0.386350058 | 0.312405286 | 0.140974541 | -0.073944772 | 0.014955621 | 0.000235564 | 0.056490772 | 0.269339702 | 0.514383287 |
| Fxr2 | S | | 477 | 0.099114008 | 0.303218304 | 0.387819885 | 0.288705877 | 0.084601581 | 0.284188938 | 0.000217602 | 0.001015882 | 0.03049698 | 0.178391946 |
| Acin1 | S | | 1175 | 0.247581953 | 0.352266745 | 0.369939015 | 0.122357062 | 0.017672269 | 0.002075051 | 0.026756183 | 0.018263671 | 0.202846332 | 0.86702868 |
| Acin1 | S | | 1177 | 0.247581953 | 0.352266745 | 0.369939015 | 0.122357062 | 0.017672269 | 0.002075051 | 0.026756183 | 0.018263671 | 0.202846332 | 0.86702868 |
| Patl1 | S | | 179 | 0.100484361 | 0.218855348 | 0.319860374 | 0.219376013 | 0.101005026 | 0.010687926 | 0.01053457 | 1.14035E-05 | 0.000508405 | 0.1065768 |
| Ints1 | S | | 1329 | 0.066252676 | 0.276146125 | 0.304660236 | 0.23840756 | 0.028514111 | 0.100209571 | 0.00080405 | 2.92822E-05 | 0.002641083 | 0.385276277 |
| Srek1 | S | | 240 | 0.526576504 | 0.5428402 | 0.114737624 | -0.41183888 | -0.428102576 | 0.004576501 | 0.004336416 | 0.771018169 | 0.343470874 | 0.329399664 |
| Tmpo | S | | 291 | 0.064672555 | 0.320799114 | 0.218283671 | 0.153611117 | -0.102515443 | 0.030306793 | 0.000221407 | 6.10881E-05 | 0.000268369 | 0.043696222 |
| Arhgef12 | T | | 736 | 0.264607649 | 0.330021489 | 0.317696531 | 0.053088882 | -0.012324959 | 0.0050555 | 0.008260388 | 0.017630071 | 0.546545243 | 0.897647662 |
| Ssbp3 | S | | 387 | 0.062355933 | 0.332993642 | 0.34479523 | 0.282439297 | 0.011801588 | 0.351806728 | 0.000530883 | 0.000169851 | 0.011200691 | 0.792922508 |
| Fxyd1 | S | | 83 | 0.373229048 | 0.330042146 | 0.186568261 | -0.186660786 | -0.143473885 | 0.002235046 | 0.003831057 | 0.072019465 | 0.064205592 | 0.117397926 |
| Kansl3 | S | | 515 | 0.247855237 | 0.342677126 | 0.435227401 | 0.187372164 | 0.092550276 | 0.021749451 | 0.001474808 | 0.063850085 | 0.296874882 | 0.56305987 |
| Dcaf8 | S | | 118 | 0.151924222 | 0.502955243 | 0.428205709 | 0.276281487 | -0.074749534 | 0.661236171 | 0.004580324 | 0.445138998 | 0.353366492 | |
| Rrbp1 | S | | 788 | 0.181770428 | 0.35555817 | 0.227084355 | 0.045313927 | -0.128473816 | 0.048288386 | 0.000181431 | 0.007071178 | 0.531514318 | 0.057830215 |
| H3c13 H3c14 H3c15 H | T | 81;81;81;81 | | 0.133118402 | 0.250594443 | 0.311793047 | 0.178674645 | 0.061198604 | 0.06543323 | 0.009570716 | 1.33816E-05 | 0.04491431 | 0.277254235 |
| Syn1 | S | | 551 | 0.433062942 | 0.434881223 | 0.10439425 | -0.328668691 | -0.330486973 | 0.002673231 | 0.001310408 | 0.546251207 | 0.123459957 | 0.130060495 |
| Syn1 | S | | 553 | 0.433062942 | 0.434881223 | 0.10439425 | -0.328668691 | -0.330486973 | 0.002673231 | 0.001310408 | 0.546251207 | 0.123459957 | 0.130060495 |
| Rere | S | | 675 | -0.015917499 | 0.455408051 | 0.479510114 | 0.495427613 | 0.024102062 | 0.814037997 | 0.000152133 | 0.004703899 | 0.003153784 | 0.67737239 |
| Tnrc18 | T | | 2077 | 0.204286021 | 0.327136957 | 0.343532427 | 0.139246407 | 0.01639547 | 2.64765E-05 | 0.017711516 | 0.08066879 | 0.32196522 | 0.899804101 |
| C77080 | S | | 340 | 0.293532131 | 0.396315673 | 0.366344656 | 0.072812525 | -0.029971017 | 0.022517956 | 0.040195243 | 0.006356198 | 0.366446047 | 0.817971043 |
| Ccm2 | S | | 393 | 0.060536522 | 0.237906547 | 0.311625238 | 0.251088717 | 0.073718691 | 0.098426147 | 0.000207763 | 5.62201E-05 | 0.001674021 | 0.049067928 |
| Vim | S | | 87 | 0.345622203 | 0.277268152 | 0.490461789 | 0.144839586 | 0.213193638 | 0.021677714 | 0.063453668 | 0.009479866 | 0.279871953 | 0.155927544 |
| Stag2 | S | | 1061 | 0.145102496 | 0.25405211 | 0.203941977 | 0.058839481 | -0.050110133 | 0.024685511 | 2.02748E-05 | 0.001648932 | 0.224463117 | 0.163024743 |
| Stag2 | S | | 1064 | 0.145102496 | 0.25405211 | 0.203941977 | 0.058839481 | -0.050110133 | 0.024685511 | 2.02748E-05 | 0.001648932 | 0.224463117 | 0.163024743 |
| Pcnt | S | | 1437 | 0.059125437 | 0.392272448 | 0.226276056 | 0.16715062 | -0.165996392 | 0.288282047 | 9.35937E-06 | 0.00353646 | 0.0324041 | 0.025448407 |
| Wwc2 | T | | 999 | 0.189059983 | 0.301145444 | 0.321050411 | 0.131990428 | 0.019904967 | 0.268364623 | 0.032794251 | 3.14951E-06 | 0.402214427 | 0.788393023 |
| Hdgf | S | | 200 | 0.266150559 | 0.343778682 | 0.279269161 | 0.013118602 | -0.064509521 | 0.01416816 | 0.002698807 | 0.021143392 | 0.878896976 | 0.448499478 |
| Tmpo | S | | 179 | 0.075639362 | 0.212011656 | 0.326937626 | 0.251298264 | 0.11492597 | 0.143347146 | 0.000282142 | 3.65989E-05 | 0.005513078 | 0.013747473 |
| Bbx | T | | 466 | 0.142772631 | 0.299223751 | 0.431593804 | 0.288821174 | 0.132370053 | 0.145500551 | 0.009818054 | 0.000451355 | 0.022111854 | 0.127937324 |
| Ccdc82 | S | | 167 | 0.161920985 | 0.408423809 | 0.436861608 | 0.274940622 | 0.028437799 | 0.146078707 | 0.005880831 | 0.005148209 | 0.024993441 | 0.724226866 |
| Ccdc82 | S | | 170 | 0.161920985 | 0.408423809 | 0.436861608 | 0.274940622 | 0.028437799 | 0.146078707 | 0.005880831 | 0.005148209 | 0.024993441 | 0.724226866 |
| Ccdc82 | S | | 173 | 0.161920985 | 0.408423809 | 0.436861608 | 0.274940622 | 0.028437799 | 0.146078707 | 0.005880831 | 0.005148209 | 0.024993441 | 0.724226866 |
| Sun2 | S | | 123 | 0.018729193 | 0.292084579 | 0.315633379 | 0.296904186 | 0.0235488 | 0.393077025 | 2.26534E-07 | 0.005755409 | 0.002365844 | 0.508627331 |
| L3mbtl3 | S | | 605 | 0.108980699 | 0.213226806 | 0.301516155 | 0.192535455 | 0.088289349 | 0.048015018 | 5.29842E-05 | 0.00062488 | 0.010735502 | 0.070425333 |
| Numa1 | S | | 1739 | 0.015686799 | 0.252550614 | 0.306062999 | 0.2903762 | 0.053512385 | 0.712197494 | 3.32435E-05 | 1.83012E-05 | 0.003814379 | 0.071706053 |
| Wdr47 | S | | 312 | 0.023976254 | 0.453871073 | 0.422084028 | 0.398107774 | -0.031787045 | 0.740476421 | 0.002811613 | 0.000494282 | 0.00102553 | 0.685178092 |
| Arhgap12 | T | | 229 | 0.068706513 | 0.200463182 | 0.297045992 | 0.228339479 | 0.09658281 | 0.019441277 | 0.033616798 | 1.47899E-07 | 0.001549457 | 0.147842202 |
| Ccar2 | S | | 680 | 0.079303996 | 0.172859709 | 0.284811018 | 0.205507022 | 0.111951309 | 0.00889664 | 0.002320609 | 5.02607E-06 | 0.00013714 | 0.024488754 |
| Spp1 | S | | 162 | 0.177171261 | 0.334715194 | 0.132406103 | -0.044765159 | -0.202309091 | 0.01679268 | 4.17124E-06 | 0.059062471 | 0.411513346 | 0.022813487 |
| Sipa1l3 | T | | 1700 | 0.151035831 | 0.284177226 | 0.504440204 | 0.353404373 | 0.220262979 | 0.117247067 | 0.021607783 | 0.000864832 | 0.000766322 | 0.04172483 |
| Palm | T | | 141 | 0.136854727 | 0.381420161 | 0.472854389 | 0.336009662 | 0.091434227 | 0.05450097 | 0.000972903 | 0.039087467 | 0.068760649 | 0.479207529 |
| Pip5k1c | S | | 554 | 0.037901955 | 0.439662301 | 0.281587107 | 0.243685153 | -0.158075194 | 0.221259266 | 5.6124E-05 | 0.007026759 | 0.016799563 | 0.040122822 |
| Hnrnpul1 | S | | 195 | 0.045798363 | 0.211602876 | 0.363288288 | 0.317489925 | 0.151685412 | 0.180385554 | 0.000205059 | 6.32615E-05 | 0.000893248 | 0.010856265 |
| Cttnbp2nl | S | | 567 | 0.027599848 | 0.261195251 | 0.311380469 | 0.283780621 | 0.050185218 | 0.174022243 | 4.24555E-07 | 0.003853793 | 0.004364943 | 0.227934717 |
| Cdc23 | S | | 588 | 0.012760526 | 0.26402047 | 0.370333536 | 0.357573009 | 0.106313066 | 0.432670239 | 2.26703E-07 | 0.005967164 | 0.006066839 | 0.092004321 |
| Svil | T | | 811 | 0.10742244 | 0.339827841 | 0.239212292 | 0.131789852 | -0.100615549 | 0.009968621 | 0.009383408 | 6.56213E-05 | 0.005348187 | 0.18033775 |
| Rbm15 | S | | 20 | 0.042982819 | 0.172654045 | 0.222361355 | 0.179378536 | 0.04970731 | 0.254105722 | 8.80581E-08 | 1.70456E-05 | 0.015560678 | 0.017410689 |
| Smarcc2 | S | | 347 | 0.029331578 | 0.189813545 | 0.291151448 | 0.261819871 | 0.101337903 | 0.288576381 | 1.96441E-05 | 8.06425E-06 | 0.000788376 | 0.014897048 |
| Sorbs2 | S | | 358 | 0.267960144 | 0.392775992 | 0.200406763 | -0.067553382 | -0.19236923 | 0.00445133 | 0.000601223 | 0.212987511 | 0.616679071 | 0.229113651 |
| Zeb1 | S | | 292 | 0.512255827 | 0.203272901 | 0.464109578 | -0.048146249 | 0.260836677 | 0.188460287 | 0.12536788 | 0.000458635 | 0.873708224 | 0.075388562 |
| Camk2d | T | | 336 | 0.046769362 | 0.452787602 | 0.386935967 | 0.340166605 | -0.065851635 | 0.559758436 | 0.023601443 | 3.37854E-05 | 0.02431169 | 0.540993092 |
| Atn1 | S | | 42 | 0.213804133 | 0.233920445 | 0.408143602 | 0.194339469 | 0.174223157 | 0.002680119 | 0.00158546 | 0.049427536 | 0.210744187 | 0.248596789 |
| Lmna | S | | 573 | 0.028259581 | 0.237059157 | 0.373229116 | 0.344969535 | 0.136169959 | 0.457346666 | 6.88132E-06 | 0.000785563 | 0.001055319 | 0.021131877 |

| | | | | | | | | | | | | |
|---|---|---|---|---|---|---|---|---|---|---|---|---|
| Pacsin3 | S | 347 | 0.23631078 | 0.551378957 | 0.302332759 | 0.066021979 | -0.249046198 | 0.048782024 | 0.001583175 | 0.17845351 | 0.720794304 | 0.246195966 |
| Top2b | S | 1509 | 0.014931467 | 0.199910655 | 0.306616903 | 0.291685436 | 0.106706248 | 0.756526146 | 8.58045E-05 | 1.69458E-06 | 0.015310238 | 0.007364376 |
| 1110051M20Rik | S | 304 | 0.253956753 | 0.252863917 | 0.206403476 | -0.047553276 | -0.04646044 | 0.216948022 | 8.26419E-07 | 0.01404042 | 0.773890781 | 0.286427912 |
| Adgra2 | S | 1292 | 0.117372523 | 0.304890782 | 0.249694026 | 0.132321502 | -0.055196756 | 0.016924496 | 0.000159625 | 0.006830047 | 0.067581141 | 0.330684549 |
| Son | S | 1854 | 0.419202783 | 0.258527041 | 0.517163216 | 0.097960433 | 0.258636176 | 0.088707589 | 0.023766204 | 0.017597624 | 0.626465309 | 0.121247609 |
| Zrsr1 | S | 389 | -0.079633848 | 0.501484914 | 0.39783559 | 0.477469438 | -0.103649324 | 0.348568714 | 6.56406E-06 | 0.016448814 | 0.008023883 | 0.284276934 |
| Hdgf | S | 206 | 0.15242893 | 0.267651142 | 0.233345819 | 0.08091689 | -0.034305323 | 0.003307698 | 0.000581974 | 0.006398416 | 0.162993671 | 0.514403019 |
| Mre11a | S | 2 | 0.131386054 | 0.253493282 | 0.289151244 | 0.157765189 | 0.035657961 | 0.013084669 | 0.000968188 | 0.00193781 | 0.044641921 | 0.493077742 |
| Champ1 | S | 272 | 0.007906082 | 0.195458609 | 0.293872901 | 0.285966819 | 0.098414292 | 0.797688127 | 4.82367E-05 | 1.45524E-06 | 0.005019682 | 0.008554469 |
| Rbl2 | S | 659 | 0.041233524 | 0.358326924 | 0.456064427 | 0.414830904 | 0.097737503 | 0.388223703 | 0.000181611 | 0.008828147 | 0.002657279 | 0.161427522 |
| Kat6b | S | 866 | 0.213151449 | 0.314167679 | 0.426758125 | 0.213606676 | 0.112590446 | 0.008825537 | 0.005425486 | 0.048715942 | 0.183493921 | 0.424490416 |
| Srrm2 | S | 1335 | 0.039587809 | 0.335303473 | 0.336015139 | 0.29642733 | 0.000711666 | 0.086332211 | 0.000320196 | 0.001681183 | 0.004116437 | 0.985765641 |
| Stim1 | S | 257 | 0.128811327 | 0.268930727 | 0.311540618 | 0.182729291 | 0.042609891 | 0.329835359 | 0.000341202 | 0.000349813 | 0.206100984 | 0.28302104 |
| Lzts3 | T | 675 | 0.174433823 | 0.261522434 | 0.291719171 | 0.117285348 | 0.030196737 | 0.007329354 | 0.009135527 | 0.001498686 | 0.073950154 | 0.642711035 |
| Rbbp6 | S | 874 | 0.290000739 | 0.313931442 | 0.41185849 | 0.121857751 | 0.097927048 | 0.027598489 | 0.046111791 | 0.008178986 | 0.17329366 | 0.402473422 |
| Cdc45 | S | 480 | 0.224224108 | 0.326982936 | 0.262796057 | 0.03857195 | -0.064186879 | 0.011491015 | 0.004002906 | 0.013494281 | 0.59942538 | 0.413051722 |
| Erbin | S | 444 | 0.130534341 | 0.383249732 | 0.284649372 | 0.154115031 | -0.098600361 | 0.091626348 | 0.000289071 | 0.016468742 | 0.115153027 | 0.240405789 |
| Ahdc1 | S | 1463 | 0.059686264 | 0.276720563 | 0.342846377 | 0.283160113 | 0.066125814 | 0.295945598 | 0.003476043 | 3.20183E-05 | 0.009150627 | 0.211305102 |
| Sipa1l1 | S | 321 | 0.09793418 | 0.230855721 | 0.412276836 | 0.314342656 | 0.181421115 | 0.393674241 | 0.002274061 | 0.000144841 | 0.069029934 | 0.004194617 |
| Dnttip1 | S | 160 | 0.232179814 | 0.319566499 | 0.439083612 | 0.206903799 | 0.119517113 | 0.069344006 | 0.034615064 | 0.003739587 | 0.049759099 | 0.262981307 |
| Vps9d1 | S | 116 | 0.129450519 | 0.366117084 | 0.376899114 | 0.247448595 | 0.01078203 | 0.046174851 | 0.002040752 | 0.015688949 | 0.053686961 | 0.905643102 |
| Lig1 | T | 77 | 0.05058517 | 0.317034032 | 0.3340712 | 0.28348603 | 0.017037168 | 0.342188426 | 0.000633778 | 0.000371238 | 0.000514094 | 0.520075717 |
| Pcm1 | S | 1777 | 0.050749932 | 0.217074864 | 0.348456861 | 0.297706929 | 0.131381997 | 0.339519484 | 0.000474573 | 4.91203E-05 | 0.013646139 | 0.01511155 |
| Hmga1 Hmga1b | S | 103 | 0.031717897 | 0.224967829 | 0.302646047 | 0.27092815 | 0.077678219 | 0.299999008 | 1.12116E-05 | 0.000256328 | 0.000919571 | 0.062988832 |
| Gatad2b | S | 335 | 0.07452793 | 0.200740695 | 0.342570828 | 0.268042898 | 0.141830133 | 0.154148175 | 0.024008642 | 1.65015E-06 | 0.009397133 | 0.062565322 |
| Phf2 | S | 876 | 0.083844193 | 0.2160183 | 0.278516922 | 0.194672729 | 0.062498622 | 0.086351283 | 5.08278E-05 | 0.000513806 | 0.008727075 | 0.11857218 |
| Srrm2 | S | 1465 | 0.061849444 | 0.348312684 | 0.42533731 | 0.363487867 | 0.077024626 | 0.275681706 | 0.000101793 | 0.01952083 | 0.032567062 | 0.448681288 |
| Akap8l | S | 284 | 0.062894013 | 0.270728362 | 0.32423827 | 0.261344257 | 0.053509908 | 0.100803219 | 0.000125998 | 0.001897815 | 0.001784458 | 0.165270612 |
| Dcaf8 | S | 100 | -0.091805609 | 0.326515752 | 0.349958079 | 0.441763688 | 0.023442327 | 0.009402097 | 9.59315E-06 | 4.45184E-06 | 4.21175E-05 | 0.358937919 |
| Arhgap12 | S | 238 | 0.055961143 | 0.194561121 | 0.278120843 | 0.2221597 | 0.083559722 | 0.072106103 | 0.021200745 | 1.05427E-07 | 0.004326576 | 0.130955997 |
| Tbc1d10b | S | 643 | 0.081331356 | 0.189102428 | 0.28682282 | 0.205491464 | 0.097720392 | 0.063033815 | 0.000552463 | 4.42538E-05 | 0.003932145 | 0.022814381 |
| Mex3c | S | 530 | 0.101088475 | 0.383940031 | 0.292154704 | 0.191066229 | -0.091785327 | 0.291676263 | 4.53314E-05 | 0.023314829 | 0.117456402 | 0.288362218 |
| Synpo | S | 685 | 0.179081059 | 0.277640251 | 0.208327723 | 0.029246664 | -0.069312528 | 0.012230751 | 0.000162773 | 0.018254964 | 0.622950651 | 0.245551256 |
| Irs1 | S | 343 | 0.006082088 | 0.369631285 | 0.428889637 | 0.422807549 | 0.059258352 | 0.787976858 | 0.000845247 | 0.001205591 | 0.000833857 | 0.204509108 |
| Map2 | S | 1799 | 0.3406517 | 0.484403937 | 0.176544863 | -0.164106837 | -0.307859074 | 0.056171549 | 0.000689997 | 0.267697389 | 0.334666082 | 0.107137544 |
| Map2 | S | 1800 | 0.3406517 | 0.484403937 | 0.176544863 | -0.164106837 | -0.307859074 | 0.056171549 | 0.000689997 | 0.267697389 | 0.334666082 | 0.107137544 |
| Map2 | S | 1801 | 0.3406517 | 0.484403937 | 0.176544863 | -0.164106837 | -0.307859074 | 0.056171549 | 0.000689997 | 0.267697389 | 0.334666082 | 0.107137544 |
| Zc3h13 | S | 379 | 0.264432874 | 0.407989476 | 0.477844502 | 0.213411628 | 0.069855026 | 0.032532398 | 0.017244131 | 0.075656775 | 0.278353948 | 0.695972445 |
| Prkaa1 | S | 524 | 0.070251573 | 0.308206785 | 0.290948655 | 0.220697082 | -0.01725813 | 0.212053659 | 1.1539E-06 | 0.040835878 | 0.053372093 | 0.812676736 |
| Top2b | S | 1568 | 0.041579108 | 0.17224584 | 0.325556689 | 0.283977581 | 0.153310849 | 0.30834209 | 0.00072682 | 4.1848E-06 | 0.000721988 | 0.007705478 |
| Cstf2 | S | 120 | 0.181875761 | 0.251133571 | 0.371800841 | 0.18992508 | 0.12066727 | 0.331425454 | 0.003033491 | 0.000479334 | 0.315377714 | 0.012507863 |
| Bcl2l13 | S | 347 | 0.172837941 | 0.328768285 | 0.241146983 | 0.068309042 | -0.087621303 | 0.009720702 | 0.000385212 | 0.056629662 | 0.452217012 | 0.359843416 |
| Fam76b | S | 193 | 0.050567211 | 0.242204694 | 0.329438515 | 0.278871304 | 0.087233821 | 0.35923789 | 0.000128788 | 0.00030237 | 0.007870736 | 0.063947649 |
| Cxxc1 | S | 100 | 0.147943543 | 0.275445218 | 0.355877418 | 0.207933874 | 0.080432199 | 0.109293777 | 0.010867604 | 0.000505893 | 0.051256467 | 0.280505721 |
| Lmna | S | 277 | -0.01204599 | 0.285266095 | 0.407578583 | 0.419624572 | 0.122312488 | 0.849665021 | 0.000848974 | 8.53185E-05 | 0.002757414 | 0.017262462 |
| Mecp2 | S | 360 | 0.179518658 | 0.259062921 | 0.48368086 | 0.304162203 | 0.22461794 | 0.089888293 | 0.005276772 | 0.009218935 | 0.043229464 | 0.084940624 |
| Hmga1 Hmga1b | S | 102 | 0.023902371 | 0.215170305 | 0.298138101 | 0.27423573 | 0.082967796 | 0.406699738 | 1.53705E-05 | 0.000125449 | 0.000668246 | 0.045102614 |
| Cdkn2aip | S | 169 | 0.121675006 | 0.241293867 | 0.232371935 | 0.110696929 | -0.008921932 | 0.03855155 | 0.000219082 | 0.000954033 | 0.062218288 | 0.802814461 |
| Prkce | T | 710 | 0.120897784 | 0.288748088 | 0.471906104 | 0.35100832 | 0.183158016 | 0.388647517 | 0.006739555 | 0.001171283 | 0.05802405 | 0.064325364 |
| Bclaf1 | S | 263 | 0.01431937 | 0.403401528 | 0.296297277 | 0.281977907 | -0.107104251 | 0.786654131 | 0.000175334 | 0.001395802 | 0.004663145 | 0.097755767 |
| Aplf | S | 344 | 0.216799465 | 0.279310397 | 0.330324467 | 0.113525002 | 0.05101407 | 0.028069449 | 0.004127574 | 0.012459739 | 0.241152171 | 0.541676591 |
| Aplf | S | 347 | 0.216799465 | 0.279310397 | 0.330324467 | 0.113525002 | 0.05101407 | 0.028069449 | 0.004127574 | 0.012459739 | 0.241152171 | 0.541676591 |
| Ctnnb1 | S | 191 | 0.081803013 | 0.155882182 | 0.283082619 | 0.201279606 | 0.127200437 | 0.055608586 | 0.000541233 | 1.93153E-05 | 0.00414955 | 0.004125119 |
| Top2b | S | 1511 | 0.008405565 | 0.19870177 | 0.309533484 | 0.301127919 | 0.110831714 | 0.858191312 | 0.000129909 | 4.68137E-06 | 0.013425192 | 0.008862679 |
| Gatad2b | S | 123 | 0.067331531 | 0.150491474 | 0.301112577 | 0.233781046 | 0.150621102 | 0.019252651 | 0.002658067 | 1.06287E-05 | 0.003806752 | 0.005680414 |
| Tor1aip1 | T | 85 | -0.002760716 | 0.388780566 | 0.31462871 | 0.317389425 | -0.074151857 | 0.975024111 | 0.01158906 | 2.73592E-06 | 0.053600371 | 0.296214857 |
| Bckdk | S | 31 | 0.046221949 | 0.236725737 | 0.313054913 | 0.266832964 | 0.076329176 | 0.108393208 | 0.005556104 | 1.2014E-05 | 0.000318555 | 0.134525001 |
| Cic | S | 1343 | 0.033238311 | 0.241004158 | 0.346221909 | 0.312983598 | 0.105217751 | 0.407718933 | 2.72889E-05 | 0.001007492 | 0.001815138 | 0.051825306 |
| Thrap3 | S | 248 | 0.170907134 | 0.208345108 | 0.201228849 | 0.030321715 | -0.007116258 | 0.000136028 | 0.000260001 | 0.055532846 | 0.633803792 | 0.909388218 |
| Sun2 | S | 122 | 0.248408512 | 0.407468661 | 0.354730855 | 0.106322343 | -0.052737806 | 0.055871067 | 0.004335058 | 0.071023599 | 0.489193184 | 0.710736849 |
| Gigyf1 | S | 235 | 0.276657418 | 0.261475153 | 0.225428301 | -0.051229117 | -0.036046852 | 0.009932975 | 0.008694088 | 0.006810739 | 0.462119688 | 0.571680826 |
| Ncor1 | S | 1772 | 0.229699384 | 0.23889848 | 0.43208085 | 0.202381467 | 0.19318237 | 0.017008395 | 0.041262962 | 0.007025165 | 0.067455524 | 0.097657637 |
| Magi3 | S | 1286 | 0.219869213 | 0.230768736 | 0.463592601 | 0.243723387 | 0.232823865 | 0.008774795 | 0.041562324 | 0.012424291 | 0.078533151 | 0.092527961 |
| Prkaca Prkacb | T | 198;198 | 0.148927079 | 0.231552134 | 0.245696472 | 0.096769392 | 0.014144338 | 0.059380226 | 0.00032158 | 0.001170652 | 0.165572693 | 0.713411557 |
| Bcl2l13 | S | 343 | 0.029677334 | 0.369019549 | 0.389359618 | 0.359682284 | 0.020340069 | 0.304282076 | 0.001506785 | 0.001874197 | 0.003659433 | 0.706579894 |
| Zc3h13 | S | 1329 | 0.191176117 | 0.398230476 | 0.430434108 | 0.239257991 | 0.032203632 | 0.118917849 | 0.008413785 | 0.022589201 | 0.121051815 | 0.792500639 |
| Irs1 | S | 1214 | 0.026286679 | 0.519581062 | 0.351181236 | 0.324894556 | -0.168399826 | 0.742027267 | 0.000592193 | 0.012752576 | 0.027970378 | 0.128497813 |
| Anapc2 | S | 514 | 0.284539141 | 0.511959575 | 0.378543181 | 0.09400404 | -0.133416394 | 0.0353841 | 0.052587916 | 0.043803171 | 0.511215362 | 0.499049635 |
| Carmil1 | T | 920 | 0.132120401 | 0.433276982 | 0.236628729 | 0.104508327 | -0.196648254 | 0.049346861 | 0.000278258 | 0.079293618 | 0.321737881 | 0.114091438 |
| Hmga2 | S | 100 | 0.057660317 | 0.167992053 | 0.214216653 | 0.156556336 | 0.0462246 | 0.018371871 | 5.387E-05 | 2.00636E-05 | 0.000889462 | 0.042642586 |
| Atp2b1 | S | 1155 | 0.103406683 | 0.223693593 | 0.401507846 | 0.298101163 | 0.177814253 | 0.15688612 | 0.003887835 | 0.000599774 | 0.008145523 | 0.042456576 |
| Thrap3 | S | 403 | 0.171400635 | 0.406709769 | 0.350098833 | 0.178698199 | -0.056610935 | 0.04700241 | 0.001564531 | 0.08429974 | 0.283887732 | 0.703707163 |
| Map2 | S | 1791 | 0.183096141 | 0.289251745 | 0.215685676 | 0.032589535 | -0.073566069 | 0.300692424 | 3.44994E-05 | 0.004388962 | 0.830100453 | 0.07462093 |
| Arhgap31 | T | 776 | 0.361816918 | 0.28296963 | 0.334763056 | -0.027053861 | 0.051793426 | 0.002002409 | 0.104759168 | 0.058883717 | 0.821050676 | 0.751770107 |
| Zfp219 | S | 688 | 0.00897725 | 0.273210347 | 0.305803487 | 0.296826236 | 0.03259314 | 0.805381592 | 2.23793E-05 | 0.000541249 | 0.001585942 | 0.38293225 |
| Pitpnm1 | S | 593 | 0.093228351 | 0.254701091 | 0.287275828 | 0.194047477 | 0.032574736 | 0.087645403 | 0.008509943 | 4.81515E-05 | 0.022675319 | 0.521295888 |
| Stx4a | S | 117 | 0.067821444 | 0.197580758 | 0.35296698 | 0.285145536 | 0.155386222 | 0.32575537 | 0.000314336 | 0.000211671 | 0.016540342 | 0.028001196 |
| Bbx | S | 242 | 0.062724158 | 0.237508195 | 0.301828615 | 0.239104457 | 0.06432042 | 0.346003858 | 0.000649436 | 8.0003E-05 | 0.030418363 | 0.04489616 |
| Rbl1 | S | 959 | 0.332632404 | 0.198894919 | 0.351735435 | 0.019103031 | 0.152840516 | 0.005704702 | 0.21022015 | 0.004129701 | 0.594222574 | 0.297717485 |
| Dcp1a | S | 543 | 0.059036222 | 0.185845434 | 0.264350137 | 0.205313915 | 0.078504703 | 0.342489456 | 0.000101834 | 1.85691E-05 | 0.041078048 | 0.016677914 |
| Zfp318 | T | 93 | 0.117941141 | 0.133903562 | 0.507961244 | 0.390020102 | 0.374057682 | 0.044910857 | 0.027362862 | 0.000535418 | 0.002506322 | 0.002925479 |
| Irs1 | S | 1213 | -0.010292962 | 0.480588382 | 0.432764116 | 0.443057079 | -0.047824265 | 0.886110676 | 0.000303817 | 0.018387151 | 0.027623117 | 0.654951025 |
| Pdlim2 | S | 124 | 0.008838022 | 0.181019836 | 0.36547999 | 0.356641968 | 0.184460154 | 0.811842674 | 0.000678025 | 9.19508E-06 | 0.000750318 | 0.00039981 |
| Pura | S | 255 | 0.098109305 | 0.409184468 | 0.253542991 | 0.155433685 | -0.155641477 | 0.052608466 | 0.001067932 | 0.014738921 | 0.076331193 | 0.084004226 |
| R3hdm2 | S | 142 | 0.162891997 | 0.214417593 | 0.274048855 | 0.111156859 | 0.059631262 | 0.008075715 | 0.00066738 | 0.011858384 | 0.145155208 | 0.358696934 |
| Tent4b | S | 50 | 0.038362089 | 0.36003356 | 0.411186656 | 0.372824566 | 0.051153095 | 0.567002105 | 0.021346572 | 0.000132781 | 0.003324069 | 0.588195938 |
| Septin7 | T | 425 | 0.447522699 | 0.260958323 | 0.262333386 | -0.185189314 | 0.001375063 | 0.003855798 | 0.128855548 | 0.036902117 | 0.013348698 | 0.991248355 |
| Map9 | S | 414 | 0.26316387 | 0.240474284 | 0.323235102 | 0.060071232 | 0.082760817 | 0.027367708 | 0.007822348 | 0.013073459 | 0.546949943 | 0.329609336 |
| Tns2 | Y | 483 | -0.103585634 | 0.36944463 | 0.454183816 | 0.55776945 | 0.084739186 | 0.164530939 | 0.000657984 | 0.000128551 | 0.001137578 | 0.187029554 |
| Ptpra | S | 176 | 0.047761002 | 0.453084859 | 0.326007313 | 0.278246311 | -0.127077547 | 0.561936917 | 0.000621311 | 0.009555325 | 0.021543261 | 0.148450305 |
| Msh6 | S | 256 | 0.014123494 | 0.214749251 | 0.222605561 | 0.208482067 | 0.007856311 | 0.573084901 | 3.71532E-05 | 4.36248E-06 | 0.002123629 | 0.659754761 |
| Ahdc1 | S | 1392 | 0.010912017 | 0.311921252 | 0.300260563 | 0.289348545 | -0.01166069 | 0.52963362 | 4.43779E-07 | 0.034833896 | 0.042200627 | 0.870245768 |
| Arl6ip6 | S | 64 | 0.120933755 | 0.411752451 | 0.295333756 | 0.174400001 | -0.116418695 | 0.283974316 | 0.000190338 | 0.047562005 | 0.211968014 | 0.296795567 |
| Setd2 | T | 2074 | 0.730255621 | 0.305749729 | 0.343590004 | -0.386665618 | 0.037840274 | 0.092539464 | 0.224244338 | 0.013309589 | 0.2716891 | 0.856395928 |

| | | | | | | | | | | | | | |
|---|---|---|---|---|---|---|---|---|---|---|---|---|---|
| Klf3 | S | | 91 | 0.026093515 | 0.219455486 | 0.36625105 | 0.340157536 | 0.146795565 | 0.566674187 | 1.35333E-05 | 0.001826939 | 0.002415779 | 0.041424989 |
| Cdk13 | S | | 526 | 0.059996702 | 0.248428847 | 0.307487947 | 0.247491245 | 0.0590591 | 0.019874491 | 1.70611E-05 | 0.023941679 | 0.048618664 | 0.4094669 |
| Samhd1 | S | | 55 | 0.06750751 | 0.289428793 | 0.221706699 | 0.154199189 | -0.067722094 | 0.200356584 | 1.16591E-05 | 0.006368782 | 0.03583027 | 0.159054718 |
| Sash1 | S | | 83 | -0.091167374 | 0.503965268 | 0.455827985 | 0.546995359 | -0.048137283 | 0.233629456 | 0.00031694 | 0.005199178 | 0.002389494 | 0.534568075 |
| Tbx2 | S | | 704 | 0.085802102 | 0.284679968 | 0.300804209 | 0.215002107 | 0.016124241 | 0.269736528 | 0.001560611 | 0.000523933 | 0.044991796 | 0.731179887 |
| Tbx2 | S | | 709 | 0.085802102 | 0.284679968 | 0.300804209 | 0.215002107 | 0.016124241 | 0.269736528 | 0.001560611 | 0.000523933 | 0.044991796 | 0.731179887 |
| Ccpg1 | S | | 189 | 0.105479373 | 0.370485145 | 0.443925097 | 0.338445724 | 0.073439953 | 0.040545382 | 0.004335131 | 0.041069123 | 0.087182538 | 0.585880017 |
| Rbmx Rbmxl1 | S | 73;73 | | 0.119733221 | 0.283456095 | 0.287127143 | 0.167393922 | 0.003671048 | 0.050342057 | 0.008836429 | 0.000765941 | 0.017048521 | 0.953960369 |
| Syne2 | S | | 4096 | -0.047539301 | 0.259348671 | 0.300307162 | 0.347846463 | 0.040958491 | 0.277472367 | 3.3696E-05 | 1.99661E-05 | 0.004753217 | 0.049877622 |
| Top2a | S | | 1472 | 0.261021855 | 0.649323491 | 0.175608488 | -0.085413367 | -0.473715004 | 0.074826644 | 0.050502747 | 0.0045446 | 0.418973465 | 0.092582522 |
| Usp34 | S | | 3443 | 0.163529282 | 0.126026635 | 0.237451788 | 0.073922506 | 0.111425153 | 0.001541663 | 0.00160577 | 0.001266869 | 0.089138557 | 0.028423702 |
| Terf2ip | S | | 151 | 0.060730322 | 0.267331404 | 0.195528948 | 0.134798627 | -0.071802456 | 0.277481375 | 6.23023E-05 | 0.000178433 | 0.06675046 | 0.045262397 |
| Psip1 | S | | 118 | 0.025558029 | 0.2092477 | 0.362299677 | 0.336741648 | 0.153051977 | 0.685187506 | 4.34531E-05 | 0.000641885 | 0.009829251 | 0.026164379 |
| Ahdc1 | S | | 630 | 0.231375015 | 0.445084332 | 0.211252306 | -0.02012271 | -0.233832026 | 0.027057434 | 0.001424821 | 0.22969703 | 0.89038874 | 0.20184973 |
| Frmd6 | S | | 525 | 0.091658882 | 0.334027455 | 0.124617811 | 0.032958929 | -0.209409644 | 0.031039503 | 3.70364E-05 | 0.008904385 | 0.083497678 | 0.000445413 |
| Frmd6 | S | | 522 | 0.114221124 | 0.371392445 | 0.122429219 | 0.008208095 | -0.248963226 | 0.029098785 | 7.32316E-05 | 0.02843467 | 0.7653047 | 0.001588869 |
| Prom1 | S | | 721 | 0.287007072 | 0.411874478 | 0.145778246 | -0.141228826 | -0.266096232 | 0.079862823 | 0.000273005 | 0.198227445 | 0.316591348 | 0.061320487 |
| Prom1 | S | | 724 | 0.287007072 | 0.411874478 | 0.145778246 | -0.141228826 | -0.266096232 | 0.079862823 | 0.000273005 | 0.198227445 | 0.316591348 | 0.061320487 |
| Prom1 | S | | 734 | 0.287007072 | 0.411874478 | 0.145778246 | -0.141228826 | -0.266096232 | 0.079862823 | 0.000273005 | 0.198227445 | 0.316591348 | 0.061320487 |
| Prom1 | T | | 730 | 0.287007072 | 0.411874478 | 0.145778246 | -0.141228826 | -0.266096232 | 0.079862823 | 0.000273005 | 0.198227445 | 0.316591348 | 0.061320487 |
| Slc35a5 | S | | 432 | 0.005804062 | 0.219208191 | 0.311432026 | 0.305627964 | 0.092223835 | 0.730560145 | 0.00026485 | 3.5191E-05 | 0.001343033 | 0.020419766 |
| Sgms2 | S | | 364 | 0.125648277 | 0.228813964 | 0.286506831 | 0.160858554 | 0.057692867 | 0.058502431 | 0.001804234 | 0.001441913 | 0.037820374 | 0.272743272 |
| Trim33 | T | | 1102 | 0.190956247 | 0.390108151 | 0.300064937 | 0.10910869 | -0.090043214 | 0.134495095 | 0.001521765 | 0.047126631 | 0.398352691 | 0.394571143 |
| Ptpra | S | | 180 | -0.008217598 | 0.373826553 | 0.279493621 | 0.287711219 | -0.094332932 | 0.79178673 | 0.001171881 | 8.41662E-05 | 0.000538222 | 0.109108391 |
| Pxn | S | | 261 | 0.09670945 | 0.270640278 | 0.296591839 | 0.199882388 | 0.02595156 | 0.050473736 | 5.89781E-05 | 0.034706771 | 0.081115988 | 0.738162292 |
| Lmna | T | | 24 | 0.21883317 | 0.223942718 | 0.325612441 | 0.106779271 | 0.101669723 | 0.000162444 | 0.045445674 | 0.065129785 | 0.368551796 | 0.414083748 |
| Ncoa2 | S | | 736 | 0.01571779 | 0.208254922 | 0.303717624 | 0.287999834 | 0.095462702 | 0.462343765 | 1.11329E-05 | 0.000579713 | 0.001815259 | 0.042580017 |
| Fam122b | S | | 159 | 0.387942294 | 0.529389752 | -0.02537042 | -0.413312714 | -0.554760172 | 0.007253858 | 0.001993965 | 0.930083487 | 0.234460144 | 0.146163593 |
| Ahdc1 | S | | 99 | 0.474218948 | 0.456503212 | 0.200692511 | -0.273526436 | -0.255810701 | 0.090034752 | 0.003019808 | 0.353794595 | 0.310556157 | 0.262247163 |
| Pard3b | S | | 746 | -0.034574462 | 0.237602363 | 0.28608566 | 0.320660122 | 0.048483297 | 0.234857739 | 8.80447E-05 | 3.96911E-06 | 0.000304588 | 0.093676792 |
| Cdk13 | T | | 871 | 0.170008956 | 0.191556884 | 0.436178226 | 0.266169271 | 0.244621342 | 0.233117885 | 0.014767931 | 0.000853862 | 0.104676334 | 0.033792935 |
| Gpbp1l1 | T | | 353 | -0.017242685 | 0.153484089 | 0.383742379 | 0.400985064 | 0.23025829 | 0.530469045 | 0.000905465 | 2.80731E-06 | 1.69798E-05 | 0.000257176 |
| Tbc1d4 | S | | 794 | 0.114581192 | 0.389716568 | 0.131478736 | 0.016897543 | -0.258237832 | 0.11719575 | 1.77241E-06 | 0.335246438 | 0.892881871 | 0.129492687 |
| Meis1 | S | | 196 | 0.037197516 | 0.236541879 | 0.434515973 | 0.397318457 | 0.197974094 | 0.574716824 | 0.013888645 | 0.000105459 | 0.013466011 | 0.053539856 |
| Tab2 | S | | 527 | 0.190615444 | 0.521289717 | 0.118020129 | -0.072595315 | -0.403269588 | 0.056266772 | 0.000576016 | 0.210428545 | 0.264551219 | 0.003029006 |
| Vps45 | S | | 556 | 0.064778014 | 0.361278961 | 0.320939945 | 0.256161931 | -0.040339016 | 0.505600032 | 0.000866763 | 0.004098103 | 0.050064044 | 0.553001284 |
| Baz1b | S | | 1468 | 0.046551617 | 0.159283312 | 0.260236268 | 0.213684651 | 0.100952956 | 0.165221079 | 0.000123658 | 3.13705E-05 | 0.00199718 | 0.011978933 |
| Atrx | S | | 1088 | -0.060863375 | 0.18669928 | 0.246776387 | 0.307639762 | 0.060077107 | 0.140429974 | 7.41154E-06 | 1.10911E-07 | 0.00530196 | 0.00652791 |
| Zdhhc20 | S | | 320 | 0.052510902 | 0.247586348 | 0.265034211 | 0.21252331 | 0.017447863 | 0.022131125 | 1.13482E-05 | 0.023271912 | 0.041750671 | 0.755056155 |
| Rictor | S | | 1576 | 0.040618563 | 0.352914274 | 0.4340011 | 0.393382537 | 0.081086825 | 0.717942226 | 0.004829001 | 0.002091776 | 0.017129334 | 0.303033617 |
| Pi4ka | S | | 259 | 0.025511105 | 0.367035632 | 0.312487667 | 0.286976562 | -0.054547965 | 0.660741835 | 0.00141964 | 0.00100724 | 0.000792136 | 0.355062313 |
| Ncor2 | S | | 1023 | 0.166285961 | 0.259992344 | 0.281675188 | 0.115389227 | 0.021682844 | 0.048313346 | 0.01137293 | 0.001835051 | 0.134020327 | 0.735513967 |
| Sltm | S | | 374 | 0.081216596 | 0.389194056 | 0.324474037 | 0.243257441 | -0.064720018 | 0.316469272 | 0.000493173 | 0.025905626 | 0.060838597 | 0.480525941 |
| Hp1bp3 | S | | 157 | -0.037553312 | 0.303871646 | 0.328209398 | 0.365762711 | 0.024337752 | 0.585084722 | 4.49052E-05 | 0.000940765 | 0.007797754 | 0.57094435 |
| Hmga2 | S | | 104 | 0.046366044 | 0.168665806 | 0.205740866 | 0.159374822 | 0.03707506 | 0.057346786 | 8.86005E-05 | 2.05285E-05 | 0.000459936 | 0.053484525 |
| Arhgef17 | S | | 728 | -0.006838339 | 0.160162042 | 0.382678786 | 0.389517125 | 0.222516744 | 0.797126132 | 0.001103831 | 9.48193E-06 | 0.001235516 | 0.001173232 |
| Map3k5 | S | | 993 | 0.1017262 | 0.36260473 | 0.476785861 | 0.375059661 | 0.114181131 | 0.618289019 | 0.016100528 | 0.003006657 | 0.135899631 | 0.294975661 |
| Pbrm1 | T | | 377 | 0.029421675 | 0.446370855 | 0.451243354 | 0.421821679 | 0.004872498 | 0.720949776 | 0.016124522 | 0.002365388 | 0.01019046 | 0.967467654 |
| Dzip1l | S | | 651 | -0.081910778 | 0.327301743 | 0.402246914 | 0.484157692 | 0.07494517 | 0.273198094 | 0.001102147 | 8.68776E-05 | 0.003329055 | 0.197148484 |
| Ccdc61 | S | | 446 | 0.126473452 | 0.239343843 | 0.284022043 | 0.157548592 | 0.044678201 | 0.005516732 | 0.016183269 | 0.002562784 | 0.023558863 | 0.469750662 |
| Hdgfl2 | S | | 391 | 0.082509785 | 0.195253673 | 0.220432341 | 0.137922556 | 0.025178669 | 0.106310641 | 7.71223E-06 | 0.002232869 | 0.029778229 | 0.404673816 |
| Hdgfl2 | S | | 392 | 0.082509785 | 0.195253673 | 0.220432341 | 0.137922556 | 0.025178669 | 0.106310641 | 7.71223E-06 | 0.002232869 | 0.029778229 | 0.404673816 |
| Hdgfl2 | S | | 393 | 0.082509785 | 0.195253673 | 0.220432341 | 0.137922556 | 0.025178669 | 0.106310641 | 7.71223E-06 | 0.002232869 | 0.029778229 | 0.404673816 |
| Hdgfl2 | S | | 395 | 0.082509785 | 0.195253673 | 0.220432341 | 0.137922556 | 0.025178669 | 0.106310641 | 7.71223E-06 | 0.002232869 | 0.029778229 | 0.404673816 |
| Hdgfl2 | T | | 389 | 0.082509785 | 0.195253673 | 0.220432341 | 0.137922556 | 0.025178669 | 0.106310641 | 7.71223E-06 | 0.002232869 | 0.029778229 | 0.404673816 |
| Gatad2b | S | | 130 | 0.076613733 | 0.132769063 | 0.269007253 | 0.192393521 | 0.136238191 | 0.008441246 | 0.00072482 | 0.000183538 | 0.006246995 | 0.008811538 |
| Rpl36a Rpl36al | S | | 32 | 0.417983747 | 0.27273936 | 0.1897511 | -0.228232647 | -0.082988261 | 0.004674936 | 0.037533358 | 0.099394967 | 0.011585133 | 0.295671529 |
| Speg | S | | 439 | 0.171827398 | 0.317639925 | 0.227974848 | 0.05614745 | -0.089665076 | 0.013073078 | 0.000990841 | 0.079712612 | 0.512122834 | 0.325334979 |
| Rprd1b | S | | 166 | 0.072870462 | 0.189299599 | 0.271361376 | 0.198490914 | 0.082061777 | 0.100431385 | 0.000870229 | 0.000160032 | 0.016760262 | 3.19932E-06 |
| Tle3 | S | | 217 | 0.161077156 | 0.146219129 | 0.160937857 | -0.000139299 | 0.014718728 | 0.002139565 | 0.000740884 | 0.000734011 | 0.995444228 | 0.347009555 |
| Tomm70a | T | | 88 | 0.137403963 | 0.436928927 | 0.366377248 | 0.228973284 | -0.070551679 | 0.234244172 | 0.008875248 | 0.017918364 | 0.06079931 | 0.523427837 |
| Nfia | S | | 310 | 0.124894218 | 0.178871359 | 0.286942187 | 0.162047969 | 0.108070828 | 0.016891151 | 0.000193824 | 0.010308411 | 0.041864767 | 0.126806336 |
| Atrx | S | | 588 | 0.22344478 | 0.269857892 | 0.33945955 | 0.11601477 | 0.069601658 | 0.029235678 | 0.008744013 | 0.033223623 | 0.316847668 | 0.516542007 |
| Baz1b | S | | 1338 | 0.121313616 | 0.380910989 | 0.234764371 | 0.113450755 | -0.146146618 | 0.215261879 | 0.00104443 | 0.009738898 | 0.206220529 | 0.019761033 |
| Taok1 | T | | 669 | 0.20538825 | 0.257776239 | 0.227963615 | 0.022575365 | -0.029812624 | 0.017616212 | 0.005500912 | 0.010353217 | 0.565346591 | 0.381015811 |
| Phc3 | S | | 840 | 0.037006916 | 0.305883885 | 0.444026751 | 0.407019836 | 0.138142867 | 0.536560044 | 0.032983439 | 0.000256817 | 0.006105251 | 0.140328891 |
| Trp53bp1 | S | | 1632 | 0.129667451 | 0.188842362 | 0.225448927 | 0.095781477 | 0.036606565 | 0.042686283 | 0.003947108 | 0.000126559 | 0.099265461 | 0.338772542 |
| Akap12 | S | | 637 | 0.149191406 | 0.509099983 | 0.320333447 | 0.171142041 | -0.188766537 | 0.049859007 | 0.008512711 | 0.101224653 | 0.292761028 | 0.278497475 |
| Cep68 | S | | 419 | 0.020975308 | 0.18686339 | 0.464136492 | 0.443161184 | 0.277273102 | 0.730331579 | 0.010751589 | 8.92432E-05 | 0.001334274 | 0.000553462 |
| Jun | S | | 246 | 0.105446587 | 0.203706154 | 0.25732168 | 0.151875092 | 0.053615526 | 0.051014712 | 0.00330513 | 0.000319448 | 0.020326161 | 0.200510313 |
| Dpysl3 | T | | 509 | -0.107177719 | 0.578597177 | 0.449676474 | 0.556854193 | -0.128920703 | 0.032997914 | 0.000184216 | 0.009307811 | 0.007323261 | 0.185962401 |
| Tmpo | S | | 547 | 0.145951735 | 0.384629115 | 0.32176522 | 0.175813485 | -0.062863895 | 0.286569926 | 0.003591979 | 0.01364451 | 0.197135692 | 0.418649713 |
| Gtf2ird1 | S | | 448 | 0.065696432 | 0.22708542 | 0.270625293 | 0.204928861 | 0.043539872 | 0.259885748 | 0.002037454 | 8.87367E-05 | 0.037280895 | 0.27433676 |
| Spp1 | S | | 250 | 0.010893028 | 0.345300654 | 0.177433567 | 0.166540539 | -0.167867088 | 0.762426456 | 0.002100116 | 1.13908E-06 | 0.028188556 | 0.016974806 |
| Pitpnm1 | S | | 342 | 0.072886703 | 0.27947602 | 0.25987787 | 0.186991167 | -0.01959815 | 0.103796637 | 0.000286164 | 0.005940774 | 0.045034448 | 0.704684454 |
| Pitpnm1 | S | | 345 | 0.072886703 | 0.27947602 | 0.25987787 | 0.186991167 | -0.01959815 | 0.103796637 | 0.000286164 | 0.005940774 | 0.045034448 | 0.704684454 |
| Pitpnm1 | S | | 346 | 0.072886703 | 0.27947602 | 0.25987787 | 0.186991167 | -0.01959815 | 0.103796637 | 0.000286164 | 0.005940774 | 0.045034448 | 0.704684454 |
| Gpr149 | S | | 572 | 0.312530153 | -0.029842654 | 0.479324667 | 0.166794514 | 0.50916732 | 0.004682235 | 0.724414233 | 0.000661499 | 0.051781372 | 0.008200163 |
| Phc3 | S | | 269 | -0.039477268 | 0.299826221 | 0.354779636 | 0.394256904 | 0.054953415 | 0.237212854 | 0.001229478 | 4.33349E-05 | 0.000229346 | 0.193712236 |
| Psip1 | S | | 176 | 0.036838297 | 0.235892193 | 0.214160581 | 0.177322284 | -0.021731612 | 0.244627007 | 1.67134E-06 | 0.005142672 | 0.006308177 | 0.478912383 |
| Gapdh | T | | 182 | 0.176329585 | 0.322306047 | 0.302822369 | 0.126492784 | -0.019483678 | 0.067482316 | 0.015704569 | 0.007685287 | 0.087590792 | 0.815545454 |
| Dtna | S | | 568 | -0.052418043 | 0.313701677 | 0.440736269 | 0.493154312 | 0.127034592 | 0.509468621 | 0.002239014 | 0.000373231 | 0.00225234 | 0.09135393 |
| Srrm2 | S | | 1453 | 0.099069677 | 0.238219168 | 0.212745413 | 0.113675736 | -0.025473756 | 0.006412136 | 2.32737E-05 | 0.066869782 | 0.205169144 | 0.726316559 |
| Prr12 | S | | 1562 | 0.011420082 | 0.178360906 | 0.261525949 | 0.250105867 | 0.083165043 | 0.710067991 | 9.86749E-05 | 1.33052E-05 | 0.001651456 | 0.01334693 |
| Prr12 | T | | 1555 | 0.011420082 | 0.178360906 | 0.261525949 | 0.250105867 | 0.083165043 | 0.710067991 | 9.86749E-05 | 1.33052E-05 | 0.001651456 | 0.01334693 |
| Chd4 | S | | 1530 | 0.135587806 | 0.197385903 | 0.37988967 | 0.244301864 | 0.182503767 | 0.330678108 | 0.002066994 | 0.001650673 | 0.136202914 | 0.030873763 |
| Caskin2 | S | | 403 | 0.012604934 | 0.266027404 | 0.288153058 | 0.275548124 | 0.022125654 | 0.589705927 | 0.00013033 | 0.000850445 | 0.006081595 | 0.560054702 |
| Magi3 | S | | 1285 | 0.232887304 | 0.196506924 | 0.280040279 | 0.047152974 | 0.083533355 | 0.004563533 | 0.022710876 | 0.017556025 | 0.518694912 | 0.331246773 |
| Irs2 | S | | 796 | 0.056773676 | 0.336229372 | 0.317602403 | 0.260828727 | -0.018626968 | 0.211085191 | 0.012917609 | 0.000627873 | 0.003763987 | 0.786605526 |
| Pitpnm1 | S | | 358 | -0.061457084 | 0.401819691 | 0.292180915 | 0.353637998 | -0.109638776 | 0.5867092 | 0.000135631 | 0.002441172 | 0.043685679 | 0.095420612 |
| Zfp609 | S | | 803 | 0.128981852 | 0.218300057 | 0.304872939 | 0.175891087 | 0.086572882 | 0.005436265 | 0.000456522 | 0.063483142 | 0.17960378 | 0.431312583 |
| Klf3 | S | | 71 | -0.037903536 | 0.214437986 | 0.400684669 | 0.438588206 | 0.186246683 | 0.227451283 | 0.000365933 | 6.58965E-05 | 0.002066574 | 0.006883998 |
| Cobll1 | T | | 328 | 0.078936635 | 0.227558439 | 0.254499097 | 0.175562462 | 0.026940658 | 0.163619219 | 0.000964365 | 0.000494067 | 0.018463087 | 0.44872322 |
| Acap2 | T | | 244 | 0.265840279 | 0.340620781 | 0.103348687 | -0.162491592 | -0.237272094 | 0.004085593 | 0.00398614 | 0.146784626 | 0.017109915 | 0.02486968 |

| | | | | | | | | | | | | |
|---|---|---|---|---|---|---|---|---|---|---|---|---|
| Retreg3 | S | 320 | 0.051775045 | 0.154054213 | 0.262719133 | 0.210944088 | 0.10866492 | 0.157709302 | 0.002590663 | 7.67357E-06 | 0.004709582 | 0.014112537 |
| Map3k7 | S | 432 | 0.154531088 | 0.270225965 | 0.128546093 | -0.025984996 | -0.141679873 | 0.001627079 | 0.005867291 | 0.003622897 | 0.087507128 | 0.066427291 |
| Nfic | S | 339 | 0.064514359 | 0.208888275 | 0.204125933 | 0.139611574 | -0.004762342 | 0.061792642 | 0.000243661 | 0.000330631 | 0.003988966 | 0.845962042 |
| Thrap3 | S | 734 | 0.084032163 | 0.199983727 | 0.287691206 | 0.203659043 | 0.087707479 | 0.0557047 | 0.000889023 | 0.001922913 | 0.002583098 | 0.023784193 |
| Cavin1 | S | 368 | 0.107702374 | 0.186206007 | 0.300969655 | 0.19326728 | 0.114763647 | 0.082427781 | 0.006485513 | 0.00041547 | 0.017888368 | 0.045159065 |
| Arhgap31 | T | 769 | 0.389546516 | 0.287368488 | 0.397395456 | 0.00784894 | 0.110026968 | 0.028059373 | 0.152180212 | 0.046999104 | 0.960634649 | 0.567774828 |
| Srsf4 | S | 401 | 0.043980374 | 0.206280266 | 0.32764731 | 0.283666936 | 0.121367044 | 0.466414065 | 0.006259451 | 4.53547E-05 | 0.013620131 | 0.052284632 |
| Ahdc1 | S | 1467 | 0.002351495 | 0.264804941 | 0.287918277 | 0.285566782 | 0.023113335 | 0.964752173 | 0.001236975 | 5.86366E-05 | 0.012084121 | 0.54368038 |
| Eif3g | T | 38 | 0.314037298 | 0.09336477 | 0.355055042 | 0.041017744 | 0.261690272 | 0.047875773 | 0.252230702 | 0.00036218 | 0.696024744 | 0.023408049 |
| Ttc7b | S | 630 | 0.100399275 | 0.195754615 | 0.254885307 | 0.154486033 | 0.059130692 | 0.167798323 | 0.000602938 | 0.000545616 | 0.070029439 | 0.139851303 |
| Mast4 | S | 351 | 0.026996011 | 0.167837375 | 0.204566149 | 0.177570138 | 0.036728774 | 0.140259439 | 7.79188E-05 | 1.24997E-05 | 1.8005E-05 | 0.009660051 |
| Rai14 | S | 318 | 0.109256689 | 0.245257066 | 0.290928543 | 0.181671853 | 0.045671477 | 0.139737757 | 0.000884856 | 0.005380885 | 0.052849753 | 0.4321895 |
| Nktr | S | 1148 | 0.09249283 | 0.265808291 | 0.211938761 | 0.119445931 | -0.05386953 | 0.157742654 | 4.6444E-05 | 0.012232839 | 0.114818774 | 0.303284219 |
| Zbtb8a | S | 186 | 0.03382507 | 0.25999211 | 0.371103818 | 0.337278748 | 0.111111708 | 0.400810721 | 0.022040434 | 0.000118299 | 0.000962318 | 0.154806 |
| Marcks | T | 143 | 0.119431817 | 0.266757302 | 0.22830074 | 0.108868923 | -0.038456562 | 0.019212795 | 0.001320974 | 0.01443426 | 0.134224591 | 0.538243637 |
| Spp1 | T | 170 | 0.132138601 | 0.244159845 | 0.077280314 | -0.054858287 | -0.166879531 | 0.001567605 | 5.74476E-06 | 0.190158731 | 0.314569282 | 0.048850065 |
| Gpatch8 | S | 1039 | 0.23680839 | 0.403026425 | 0.157600411 | -0.079207979 | -0.245426014 | 0.075825158 | 0.015828039 | 0.004554216 | 0.416719721 | 0.061030676 |
| Arhgap23 | S | 421 | 0.222125899 | 0.165363438 | 0.246612884 | 0.024486985 | 0.081249446 | 0.001810848 | 0.071510386 | 0.004647091 | 0.599178958 | 0.30536605 |
| Gab1 | S | 455 | 0.103831265 | 0.287947601 | 0.246165961 | 0.142334696 | -0.04178164 | 0.023740637 | 0.009543164 | 0.002490784 | 0.028187975 | 0.511120967 |
| Spata6 | S | 424 | 0.03907332 | 0.318594274 | 0.274968492 | 0.235895172 | -0.043625783 | 0.465341684 | 0.004011432 | 0.000291757 | 0.01098912 | 0.443247562 |
| Pbrm1 | S | 371 | -0.017723609 | 0.305655511 | 0.507836395 | 0.525560004 | 0.202180884 | 0.829867457 | 0.036547214 | 0.000276471 | 0.002866926 | 0.124522068 |
| Pbrm1 | S | 375 | -0.017723609 | 0.305655511 | 0.507836395 | 0.525560004 | 0.202180884 | 0.829867457 | 0.036547214 | 0.000276471 | 0.002866926 | 0.124522068 |
| Nf1 | T | 2516 | 0.185678758 | 0.23123937 | 0.27541385 | 0.089735092 | 0.04417448 | 0.027221343 | 0.010657576 | 0.008725427 | 0.189663171 | 0.444325908 |
| Safb | S | 579 | 0.175444656 | 0.214833794 | 0.296436269 | 0.120991613 | 0.081602475 | 0.12106478 | 0.027848162 | 0.000506721 | 0.233019803 | 0.255928628 |
| Stk10 | T | 185 | 0.089885321 | 0.231915204 | 0.248957726 | 0.159072405 | 0.017042521 | 0.084982589 | 0.005105277 | 0.00034681 | 0.017135358 | 0.701002452 |
| Amot | S | 693 | 0.072007155 | 0.167214627 | 0.220613695 | 0.14860654 | 0.053399068 | 0.021070971 | 0.007773695 | 1.73585E-05 | 0.003140891 | 0.182995288 |
| Dpysl2 | S | 517 | -0.038670252 | 0.519223967 | 0.30809441 | 0.346764662 | -0.211129556 | 0.303314054 | 0.003560892 | 0.000777374 | 0.001077109 | 0.0365503 |
| Dlg5 | S | 1700 | 0.101704203 | 0.437676775 | 0.518495758 | 0.416791555 | 0.080818982 | 0.570131958 | 0.014407278 | 0.02573102 | 0.070255935 | 0.587495807 |
| Dlg5 | S | 1702 | 0.101704203 | 0.437676775 | 0.518495758 | 0.416791555 | 0.080818982 | 0.570131958 | 0.014407278 | 0.02573102 | 0.070255935 | 0.587495807 |
| Tcf20 | S | 671 | 0.10987634 | 0.283850762 | 0.284130958 | 0.174254618 | 0.000280196 | 0.050684536 | 0.007883091 | 0.004769725 | 0.051243108 | 0.996880171 |
| Zfp687 | S | 243 | 0.090104344 | 0.299083251 | 0.359646693 | 0.269542349 | 0.060563441 | 0.1476907 | 0.002913642 | 0.014080279 | 0.048986711 | 0.51133382 |
| Rnf213 | S | 298 | -0.063029926 | 0.299168732 | 0.463003669 | 0.526033595 | 0.163834937 | 0.383785181 | 0.003604596 | 0.00034409 | 0.000241038 | 0.019441844 |
| Zbtb44 | S | 161 | 0.041627737 | 0.229736964 | 0.309000438 | 0.267372701 | 0.079263474 | 0.339799877 | 0.005503407 | 0.000125317 | 0.001727145 | 0.158477107 |
| Srrm2 | S | 1455 | 0.080797123 | 0.234422122 | 0.210862426 | 0.130065303 | -0.023559696 | 0.013359065 | 1.70496E-05 | 0.057314249 | 0.148429 | 0.728595075 |
| Srsf10 | S | 197 | 0.334702424 | 0.16094906 | 0.306918763 | -0.02778366 | 0.145969703 | 0.001798685 | 0.054629956 | 0.105017038 | 0.842384428 | 0.347780527 |
| Tmpo | S | 189 | 0.164660072 | 0.197211221 | 0.218777414 | 0.054117342 | 0.021566193 | 0.033408297 | 0.006424689 | 0.001080186 | 0.334022606 | 0.585714758 |
| Top2b | S | 1600 | 0.056454497 | 0.157065612 | 0.195260229 | 0.138805731 | 0.038194617 | 0.138798738 | 0.004727595 | 2.78255E-07 | 0.028544181 | 0.160031248 |
| Coq8a | S | 300 | 0.002369331 | 0.451104644 | 0.264059601 | 0.26169027 | -0.187045043 | 0.963721373 | 2.62382E-05 | 0.101961007 | 0.104784001 | 0.192329991 |
| Mast4 | S | 1382 | 0.058195053 | 0.270930625 | 0.224887486 | 0.166692433 | -0.046043139 | 0.418655361 | 5.12549E-05 | 0.004345586 | 0.076957725 | 0.303671979 |
| Atp2b1 | S | 1182 | 0.161027263 | 0.151377951 | 0.295856898 | 0.134829636 | 0.144478948 | 0.086817638 | 0.00165469 | 0.002062426 | 0.116066354 | 0.020738657 |
| Epha2 | S | 911 | 0.222835429 | 0.350015548 | 0.443752424 | 0.220916995 | 0.093736876 | 0.233439814 | 0.0118951 | 0.050920298 | 0.301986345 | 0.566872147 |
| Bclaf1 | S | 122 | 0.076093425 | 0.211884512 | 0.327762332 | 0.251668907 | 0.11587782 | 0.154053569 | 0.006307498 | 0.000676941 | 0.005665685 | 0.078949115 |
| Bclaf1 | S | 125 | 0.076093425 | 0.211884512 | 0.327762332 | 0.251668907 | 0.11587782 | 0.154053569 | 0.006307498 | 0.000676941 | 0.005665685 | 0.078949115 |
| Med13l | S | 542 | 0.250176055 | 0.226252939 | 0.259572987 | 0.009396932 | 0.033320048 | 0.03623505 | 0.041817804 | 0.003903554 | 0.904174591 | 0.658385065 |
| Med13l | S | 545 | 0.250176055 | 0.226252939 | 0.259572987 | 0.009396932 | 0.033320048 | 0.03623505 | 0.041817804 | 0.003903554 | 0.904174591 | 0.658385065 |
| Med13l | S | 548 | 0.250176055 | 0.226252939 | 0.259572987 | 0.009396932 | 0.033320048 | 0.03623505 | 0.041817804 | 0.003903554 | 0.904174591 | 0.658385065 |
| Med13l | S | 555 | 0.250176055 | 0.226252939 | 0.259572987 | 0.009396932 | 0.033320048 | 0.03623505 | 0.041817804 | 0.003903554 | 0.904174591 | 0.658385065 |
| Fnbp4 | S | 19 | 0.07632334 | 0.114335509 | 0.203762261 | 0.127438922 | 0.089426752 | 0.007772093 | 0.001040804 | 2.77408E-05 | 0.004798251 | 0.004837383 |
| Micall2 | S | 766 | 0.215086389 | 0.165814492 | 0.116847485 | -0.098238904 | -0.048967007 | 0.000422607 | 0.002424879 | 0.022950526 | 0.042337996 | 0.208445847 |
| Dpysl2 | T | 512 | 0.065537573 | 0.353589832 | 0.213088947 | 0.147551374 | -0.140500885 | 0.223504577 | 0.000707589 | 0.006420071 | 0.040172506 | 0.017293268 |
| Prpf4b | S | 249 | 0.346852056 | 0.418212535 | 0.140252048 | -0.206600008 | -0.277960487 | 0.021390363 | 0.005903803 | 0.456206719 | 0.281598616 | 0.182724231 |
| Ubl4a Gm44504 | T | 93 | 0.338029854 | 0.089483066 | 0.481249346 | 0.143219491 | 0.39176628 | 0.033058799 | 0.67053618 | 0.002678681 | 0.268796163 | 0.143570556 |
| Magi3 | S | 1287 | 0.248057896 | 0.191206445 | 0.242595469 | -0.005462427 | 0.051389024 | 0.003324869 | 0.029094215 | 0.026988316 | 0.93541804 | 0.533795307 |
| Tmpo | S | 176 | 0.045931635 | 0.190603795 | 0.328015098 | 0.282083463 | 0.137411303 | 0.009560718 | 0.005653205 | 0.011954269 | 0.0371888 | |
| 3110082I17Rik | S | 48 | 0.083808387 | 0.182296638 | 0.172707791 | 0.088899404 | -0.009588846 | 0.008481404 | 0.000141108 | 0.001853901 | 0.041835833 | 0.732703905 |
| Prc1 | T | 599 | 0.220743361 | 0.24239976 | 0.304437413 | 0.083694053 | 0.062037653 | 0.035662908 | 0.034031707 | 0.010311041 | 0.074874489 | 0.342106369 |
| Dapk3 | S | 306 | 0.18362726 | 0.256751037 | 0.408642169 | 0.225014909 | 0.151891132 | 0.105788116 | 0.063660191 | 0.005443498 | 0.075164206 | 0.226476849 |
| Ttc7 | T | 696 | 0.055187645 | 0.252775557 | 0.384858595 | 0.329670951 | 0.132083038 | 0.551935767 | 0.000222661 | 0.012124132 | 0.030002433 | 0.169130571 |
| Sptbn1 | T | 2186 | 0.125563393 | 0.265597738 | 0.242704925 | 0.117141532 | -0.022892813 | 0.028644507 | 0.000780403 | 0.036711602 | 0.203606233 | 0.760057162 |
| Prickle2 | S | 754 | -0.000328168 | 0.395545029 | 0.410187217 | 0.410515385 | 0.014642188 | 0.996960838 | 0.001199775 | 0.015092085 | 0.016642113 | 0.879622382 |
| Tut7 | S | 193 | 0.155229445 | 0.241146829 | 0.249538893 | 0.094309448 | 0.008392064 | 0.000579294 | 0.006937828 | 0.092504738 | 0.387579755 | 0.934765053 |
| Ttc7 | S | 697 | 0.222224845 | 0.317893176 | 0.446210675 | 0.22398583 | 0.128317499 | 0.107702819 | 0.156926866 | 0.007910127 | 0.027046951 | 0.484086747 |
| Bnip3l | S | 64 | 0.107901407 | 0.254495087 | 0.323728632 | 0.215827225 | 0.069233545 | 0.141252785 | 0.051273904 | 0.000332657 | 0.024587849 | 0.455897856 |
| Sipa1l2 | S | 1406 | 0.123446126 | 0.182737533 | 0.149292968 | 0.025846843 | -0.033444564 | 0.002465665 | 0.00202977 | 0.001056036 | 0.177631646 | 0.297529017 |
| Sgo2a | T | 974 | 0.310540459 | 0.258672647 | 0.224622183 | -0.085918277 | -0.034050464 | 0.008528046 | 0.056120942 | 0.038167358 | 0.351527838 | 0.751263541 |
| Fas | S | 204 | 0.345656884 | 0.435918263 | 0.423284295 | 0.077627411 | -0.012633968 | 0.216211007 | 0.051442846 | 0.066705419 | 0.726764656 | 0.915402734 |
| Tjp1 | S | 315 | -0.052691061 | 0.235036545 | 0.259397227 | 0.312088288 | 0.024360683 | 0.262456509 | 1.75566E-05 | 5.59151E-05 | 0.004039572 | 0.332880713 |
| U2surp | S | 67 | 0.392998455 | 0.21295789 | 0.212502557 | -0.180495898 | -0.000455333 | 0.003158059 | 0.129596459 | 0.062175476 | 0.093384631 | 0.996919388 |
| Smarcd3 | S | 175 | -0.012184386 | 0.201690017 | 0.245515525 | 0.257699911 | 0.043825508 | 0.600332783 | 6.47745E-05 | 2.3961E-05 | 2.23668E-05 | 0.047103894 |
| Xkr5 | S | 559 | 0.188867217 | 0.157443602 | 0.206485754 | 0.017618537 | 0.049042152 | 0.000877478 | 0.001328763 | 0.062255949 | 0.809270225 | 0.513919191 |
| Inf2 | T | 1203 | 0.126548396 | 0.245461861 | 0.235855894 | 0.109307498 | -0.009605967 | 0.10322402 | 0.002449124 | 0.002785335 | 0.128029152 | 0.70258359 |
| Jun | T | 242 | 0.025441899 | 0.404579359 | 0.56023437 | 0.534792471 | 0.15565501 | 0.747974017 | 0.022824371 | 0.01011499 | 0.00768064 | 0.245711471 |
| Rapgef2 | S | 1079 | 0.201620362 | 0.162756392 | 0.283188739 | 0.081568377 | 0.120432347 | 0.000961622 | 0.060200199 | 0.018284356 | 0.269737977 | 0.179211084 |
| Sipa1 | S | 907 | 0.077185841 | 0.264111264 | 0.312923943 | 0.235738102 | 0.048812679 | 0.222675475 | 0.008771167 | 0.001047615 | 0.009015785 | 0.461378684 |
| Atrx | S | 717 | 0.006646808 | 0.128640246 | 0.261201288 | 0.25455448 | 0.132561042 | 0.871272902 | 0.00030082 | 1.51316E-06 | 0.013103841 | 0.001127586 |
| Wdr90 | S | 16 | 0.154956364 | 0.213775832 | 0.22764929 | 0.072692925 | 0.013873457 | 0.10452856 | 0.013360849 | 0.000266553 | 0.346077531 | 0.775885492 |
| Ei24 | S | 56 | 0.091912587 | 0.19817008 | 0.227159208 | 0.135246621 | 0.028989128 | 0.066351382 | 0.00219411 | 0.000483641 | 0.017764014 | 0.403517267 |
| Atrx | S | 1089 | -0.007550165 | 0.163528 | 0.248842157 | 0.256392322 | 0.085314157 | 0.882592769 | 0.000101025 | 6.76858E-06 | 0.024312268 | 0.001140935 |
| Pnisr | S | 611 | 0.112732476 | 0.230125261 | 0.299982077 | 0.187249601 | 0.069856817 | 0.074792906 | 0.010955483 | 0.00215993 | 0.02235673 | 0.30275926 |
| Arid1a | S | 1601 | -0.126713184 | 0.366634609 | 0.278532954 | 0.405246139 | -0.088101654 | 0.345733363 | 0.000434771 | 0.00010383 | 0.056962998 | 0.10892619 |
| Ralgapb | S | 359 | 0.03629799 | 0.199444608 | 0.279453056 | 0.243155065 | 0.080008448 | 0.643014011 | 0.000911277 | 9.8046E-05 | 0.055477063 | 0.034102367 |
| Erc1 | S | 15 | 0.307034369 | 0.441849404 | 0.080360745 | -0.226673623 | -0.361488659 | 0.01539176 | 0.002204947 | 0.789239596 | 0.472108788 | 0.29316611 |
| Parn | S | 573 | 0.15480904 | 0.257977843 | 0.452896861 | 0.298087821 | 0.194919019 | 0.020950294 | 0.040737186 | 0.035099259 | 0.078478825 | 0.181276832 |
| Chd8 | S | 2519 | 0.280184195 | 0.293426651 | 0.292552586 | 0.012368391 | -0.000874065 | 0.035640737 | 0.069444831 | 0.021203797 | 0.901709729 | 0.994376737 |
| Stag1 | S | 1062 | 0.125251727 | 0.262252408 | 0.315324568 | 0.19007284 | 0.053072159 | 0.14022578 | 0.007625204 | 0.059395898 | 0.44853011 | |
| Git1 | Y | 607 | 0.182294794 | 0.478966747 | 0.193424599 | 0.011129805 | -0.285542148 | 0.056337368 | 0.006476932 | 0.128598979 | 0.917795888 | 0.064756916 |
| Uimc1 | S | 171 | 0.087047768 | 0.295702936 | 0.434647195 | 0.347599427 | 0.138944259 | 0.319435855 | 0.011955396 | 0.008025546 | 0.038191983 | 0.20879934 |
| Thoc2 | T | 1289 | 0.018227914 | 0.327015673 | 0.214244922 | 0.196017009 | -0.112770751 | 0.495833692 | 1.59624E-05 | 0.024691432 | 0.040725472 | 0.130364414 |
| Slc12a6 | S | 1032 | 0.120154542 | 0.129245389 | 0.180969323 | 0.06081478 | 0.051723934 | 0.037965247 | 0.000661767 | 8.28424E-05 | 0.162050389 | 0.048826803 |
| Retreg2 | S | 133 | 0.060351253 | 0.239503704 | 0.319516634 | 0.25916538 | 0.08001293 | 0.226210664 | 0.001472818 | 0.003260571 | 0.01825176 | 0.219107327 |
| Sorbs2 | T | 1171 | 0.119972513 | 0.416870442 | 0.182427953 | 0.06245544 | -0.234442489 | 0.160229737 | 0.001069322 | 0.048000657 | 0.301593299 | 0.013345794 |
| Crebbp | S | 1764 | 0.116820037 | 0.279247077 | 0.45572297 | 0.338902932 | 0.176475893 | 0.073473066 | 0.03437347 | 0.015499862 | 0.028273705 | 0.156091344 |
| Zhx2 | S | 824 | 0.09800678 | 0.185424168 | 0.193900111 | 0.095893331 | 0.008475943 | 0.113922999 | 0.000362212 | 0.000438078 | 0.124415054 | 0.703670699 |
| Zhx2 | S | 826 | 0.09800678 | 0.185424168 | 0.193900111 | 0.095893331 | 0.008475943 | 0.113922999 | 0.000362212 | 0.000438078 | 0.124415054 | 0.703670699 |

| | | | | | | | | | | | | |
|---|---|---|---|---|---|---|---|---|---|---|---|---|
| Farp1 | S | 23 | 0.108819838 | 0.174072762 | 0.283395969 | 0.174576131 | 0.109323207 | 0.077657689 | 0.031804019 | 0.000125278 | 0.042099698 | 0.12803719 |
| Esco2 | S | 200 | 0.116572729 | 0.200293705 | 0.283152953 | 0.166580224 | 0.082859248 | 0.036633179 | 0.002686491 | 0.006854511 | 0.046525125 | 0.211148147 |
| Lzts2 | S | 572 | -0.006549808 | 0.145508625 | 0.363090799 | 0.369640607 | 0.217582174 | 0.914192031 | 0.004318448 | 1.15885E-05 | 0.011487755 | 0.001402302 |
| Mia3 | S | 1766 | 0.209711098 | 0.337032033 | 0.177129292 | -0.032581806 | -0.159902742 | 0.024004795 | 0.008700109 | 0.042436058 | 0.47074932 | 0.101579872 |
| Marf1 | S | 715 | 0.131923335 | 0.221203342 | 0.340143078 | 0.208219743 | 0.118939735 | 0.057720506 | 0.030703869 | 0.003009961 | 0.029657179 | 0.185536981 |
| Sipa1l1 | S | 97 | 0.013132378 | 0.274898303 | 0.25767915 | 0.244546772 | -0.017219153 | 0.673646292 | 1.8295E-05 | 0.009818983 | 0.013671733 | 0.724401187 |
| Rbm14 | T | 206 | 0.077976432 | 0.24524163 | 0.318460306 | 0.240483874 | 0.073218676 | 0.225714935 | 0.00625146 | 0.001501138 | 0.00419375 | 0.145494842 |
| Atxn1 | S | 382 | -0.03582595 | 0.197776392 | 0.339841595 | 0.375667545 | 0.142065204 | 0.468616177 | 0.000354237 | 6.8224E-05 | 0.001924931 | 0.018563923 |
| Parn | S | 569 | 0.146725875 | 0.23992508 | 0.468567775 | 0.3218419 | 0.228642695 | 0.03306026 | 0.028175738 | 0.033685233 | 0.068460673 | 0.133509073 |
| Fyco1 | T | 372 | 0.013330858 | 0.234215142 | 0.372060764 | 0.358729907 | 0.137845622 | 0.691187733 | 7.36728E-05 | 0.00864148 | 0.006083164 | 0.086215229 |
| Chd2 | S | 1085 | 0.02991325 | 0.203565906 | 0.245120417 | 0.215207167 | 0.04155451 | 0.486102936 | 0.000446657 | 0.000114248 | 0.007840407 | 0.144273132 |
| Chd2 | S | 1087 | 0.02991325 | 0.203565906 | 0.245120417 | 0.215207167 | 0.04155451 | 0.486102936 | 0.000446657 | 0.000114248 | 0.007840407 | 0.144273132 |
| Cobl | S | 332 | 0.30699636 | 0.33760039 | 0.166829928 | -0.140166432 | -0.170770462 | 0.007973224 | 0.005465876 | 0.486566584 | 0.543709842 | 0.469675188 |
| Sipa1l1 | S | 234 | 0.106492963 | 0.347793137 | 0.579443584 | 0.47295062 | 0.231650446 | 0.393942564 | 0.094617023 | 0.008115424 | 0.033150371 | 0.232107628 |
| Pard3b | S | 635 | 0.083323804 | 0.297023154 | 0.234471407 | 0.151147603 | -0.062551747 | 0.181382758 | 0.000481392 | 0.013379569 | 0.069168301 | 0.316494321 |
| Slu7 | S | 514 | -0.000548419 | 0.274327788 | 0.243929525 | 0.244477944 | -0.030398263 | 0.984872944 | 1.01725E-05 | 0.006112002 | 0.008793642 | 0.486801507 |
| Cavin1 | S | 391 | 0.104860652 | 0.19580727 | 0.302038128 | 0.197177476 | 0.106230858 | 0.071364635 | 0.019934702 | 0.000679733 | 0.012868456 | 0.098021369 |
| Lmna | S | 282 | 0.115531564 | 0.193317266 | 0.215826725 | 0.100295161 | 0.02250946 | 0.04125638 | 0.00197899 | 0.001573652 | 0.068370124 | 0.291209014 |
| Irf2bpl | S | 526 | 0.057003307 | 0.281790503 | 0.268133709 | 0.211130403 | -0.013656794 | 0.172845441 | 0.000405059 | 0.012756283 | 0.020361439 | 0.80038042 |
| Mki67 | T | 981 | 0.359254258 | 0.218747779 | 0.152029276 | -0.207224982 | -0.066718503 | 0.002638478 | 0.048360692 | 0.089332426 | 0.027807489 | 0.421474594 |
| Smc1a | S | 957 | 0.034551313 | 0.240064291 | 0.327045897 | 0.292494584 | 0.086981607 | 0.53459707 | 0.005885125 | 0.000322732 | 0.001961611 | 0.144748479 |
| Crocc | S | 501 | 0.047363093 | 0.211720067 | 0.407797266 | 0.360434173 | 0.1960772 | 0.241101923 | 0.021478588 | 0.000724112 | 0.009573753 | 0.037328447 |
| Marf1 | S | 1676 | 0.033463547 | 0.292749949 | 0.287676056 | 0.25421251 | -0.005073893 | 0.675513803 | 5.14938E-05 | 0.01966664 | 0.044469693 | 0.938054775 |
| Decr2 | S | 223 | 0.063905829 | 0.235945517 | 0.256214009 | 0.19230818 | 0.020268492 | 0.138627306 | 0.000655498 | 0.00348546 | 0.015922508 | 0.672860581 |
| Cavin1 | S | 389 | 0.106497388 | 0.197220276 | 0.303359892 | 0.196862504 | 0.106139616 | 0.067042091 | 0.020015494 | 0.00090946 | 0.012662021 | 0.099439702 |
| Sipa1l1 | S | 211 | 0.161956461 | 0.144131029 | 0.377495719 | 0.215539258 | 0.233364691 | 0.055983024 | 0.07737147 | 0.001328204 | 0.00525358 | 0.002387575 |
| Ncor1 | S | 2198 | -0.00308913 | 0.222078019 | 0.359228082 | 0.362317212 | 0.137150063 | 0.947570752 | 0.002846106 | 0.000212788 | 0.001441425 | 0.023682875 |
| Adgra2 | S | 1116 | 0.090656618 | 0.118144336 | 0.17782666 | 0.087170042 | 0.059682324 | 0.001969144 | 0.006772156 | 3.62232E-05 | 0.00236149 | 0.078049762 |
| Irf2bpl | S | 638 | 0.078752608 | 0.199049321 | 0.222344758 | 0.14359215 | 0.023295437 | 0.284074991 | 6.85755E-06 | 0.009774122 | 0.101972879 | 0.538276026 |
| Cep131 | S | 222 | 0.14336885 | 0.208117636 | 0.287323054 | 0.143954204 | 0.079205418 | 0.31170942 | 0.002805131 | 0.001717532 | 0.310897082 | 0.164131159 |
| Rnf168 | S | 469 | 0.197794803 | 0.219743762 | 0.240851153 | 0.04305635 | 0.02110739 | 0.004230992 | 0.016705746 | 0.040502773 | 0.584123089 | 0.808245965 |
| Sorbs2 | T | 357 | 0.166029799 | 0.325824137 | 0.211532312 | 0.045502513 | -0.114291825 | 0.027522184 | 0.000928846 | 0.197845798 | 0.73606899 | 0.426177119 |
| Afap1l2 | S | 142 | 0.010791608 | 0.359801606 | 0.213499841 | 0.202708233 | -0.146301765 | 0.773532891 | 4.77403E-05 | 0.019028568 | 0.03380462 | 0.066161711 |
| Afap1l2 | S | 143 | 0.010791608 | 0.359801606 | 0.213499841 | 0.202708233 | -0.146301765 | 0.773532891 | 4.77403E-05 | 0.019028568 | 0.03380462 | 0.066161711 |
| Afap1l2 | S | 147 | 0.010791608 | 0.359801606 | 0.213499841 | 0.202708233 | -0.146301765 | 0.773532891 | 4.77403E-05 | 0.019028568 | 0.03380462 | 0.066161711 |
| Add3 | S | 679 | -0.042241383 | 0.259313593 | 0.406114298 | 0.44835568 | 0.146800705 | 0.696686359 | 2.28214E-05 | 0.013484283 | 0.020675351 | 0.133695286 |
| Map2 | S | 1654 | 0.320809905 | 0.273857517 | 0.02377873 | -0.297031175 | -0.250078787 | 0.257147396 | 0.07971534 | 0.096402556 | 0.283631689 | 1.09431E-06 |
| Psmd4 | T | 250 | 0.134749226 | 0.192153418 | 0.184171928 | 0.049422702 | -0.00798149 | 0.07971534 | 0.001244018 | 0.000817402 | 0.409706222 | 0.785676686 |
| Atf7 | S | 97 | -0.029339527 | 0.235723915 | 0.288944169 | 0.318283695 | 0.053220253 | 0.809810562 | 0.000494453 | 0.000141247 | 0.089335238 | 0.065440091 |
| Atf7 | T | 101 | -0.029339527 | 0.235723915 | 0.288944169 | 0.318283695 | 0.053220253 | 0.809810562 | 0.000494453 | 0.000141247 | 0.089335238 | 0.065440091 |
| Lmna | S | 12 | 0.108770711 | 0.199840146 | 0.296872831 | 0.188102121 | 0.097032685 | 0.151734186 | 0.003222796 | 0.002511015 | 0.044352695 | 0.127749584 |
| Zswim8 | S | 48 | 0.006928983 | 0.41211414 | 0.296995639 | 0.290066656 | -0.115118501 | 0.92351953 | 0.002701182 | 0.00434855 | 0.002948907 | 0.175469325 |
| Ccnk | S | 325 | -0.007015016 | 0.2219307 | 0.269285883 | 0.2763009 | 0.047355183 | 0.867428992 | 0.000337344 | 0.000126963 | 0.002369401 | 0.179147238 |
| Pds5b | S | 1177 | 0.115991967 | 0.142878951 | 0.310100294 | 0.194108327 | 0.167221342 | 0.14829474 | 0.027650868 | 0.000126813 | 0.051358129 | 0.021524352 |
| Rbms1 | S | 40 | 0.085567943 | 0.24348643 | 0.384990893 | 0.29942295 | 0.141504463 | 0.157343824 | 0.051934802 | 0.001217236 | 0.008280618 | 0.187221275 |
| Filip1l | T | 996 | 0.084926636 | 0.248390069 | 0.389919669 | 0.304993033 | 0.1415296 | 0.145467862 | 0.008809103 | 0.008580927 | 0.035691189 | 0.164422779 |
| Usp6nl | S | 413 | -0.009315916 | 0.272643985 | 0.303981573 | 0.313297489 | 0.031337588 | 0.943305565 | 0.008667132 | 4.92883E-05 | 0.107088559 | 0.573952941 |
| Cpsf7 | S | 423 | 0.066237585 | 0.436309218 | 0.393943435 | 0.32770585 | -0.042365783 | 0.527179937 | 0.007065065 | 0.030094897 | 0.066202224 | 0.743164019 |
| Palm | S | 152 | 0.104691276 | 0.255953789 | 0.248898701 | 0.144207425 | -0.007505088 | 0.208785225 | 0.001886192 | 0.003776438 | 0.114882026 | 0.893431754 |
| Atrx | T | 1092 | -0.069326 | 0.305533849 | 0.297545421 | 0.366871421 | -0.007988428 | 0.608732592 | 9.14371E-05 | 0.002263952 | 0.071876207 | 0.854425412 |
| Ttc14 | S | 661 | 0.209718033 | 0.151871256 | 0.227666648 | 0.017948615 | 0.075795392 | 0.002610875 | 0.009051179 | 0.03252619 | 0.797015399 | 0.319528258 |
| Lmna | S | 463 | 0.083150245 | 0.217889064 | 0.414732384 | 0.331582139 | 0.19684332 | 0.387765382 | 0.001353395 | 0.011278958 | 0.031034684 | 0.097735788 |
| Spag9 | S | 732 | -0.075441167 | 0.229312713 | 0.325178098 | 0.400619265 | 0.095865385 | 0.059510353 | 0.016924313 | 5.0993E-08 | 0.001388386 | 0.094783912 |
| Sash1 | S | 805 | -0.117500115 | 0.400340174 | 0.266996561 | 0.384496676 | -0.133343612 | 0.031292957 | 1.41284E-05 | 0.000956904 | 0.000789213 | 0.027498805 |
| Prr12 | S | 1128 | 0.079876474 | 0.204919031 | 0.279129731 | 0.199253257 | 0.0742107 | 0.050429859 | 0.00061939 | 0.011219308 | 0.040579581 | 0.262543631 |
| Atrx | S | 719 | 0.00942346 | 0.132411551 | 0.271603535 | 0.262180075 | 0.139191983 | 0.828074029 | 0.000118971 | 2.36665E-05 | 0.012923904 | 0.001163566 |
| Prrx1 | S | 21 | 0.16069251 | 0.208850588 | 0.243647878 | 0.082955368 | 0.03479729 | 0.08162282 | 0.009171403 | 0.002412961 | 0.2757945 | 0.441480846 |
| Son | S | 1934 | 0.505275262 | 0.183726951 | 0.342712806 | -0.162562456 | 0.158985854 | 0.019828217 | 0.285047957 | 0.072730662 | 0.370558833 | 0.40296292 |
| Pml | T | 527 | 0.034662437 | 0.232267847 | 0.323081269 | 0.288418832 | 0.090813423 | 0.487605086 | 0.014934961 | 0.000148243 | 0.004804707 | 0.191268739 |
| Gba2 | S | 893 | 0.101034898 | 0.175460907 | 0.129939088 | 0.02890419 | -0.045521819 | 0.040196281 | 8.84719E-05 | 0.000592077 | 0.407639976 | 0.014533986 |
| Adgra2 | S | 1117 | 0.181662753 | 0.186337191 | 0.283010825 | 0.101348072 | 0.096673635 | 0.190429553 | 0.003509297 | 0.003760278 | 0.407134221 | 0.121200122 |
| Lin37 | S | 138 | 0.076516191 | 0.198691426 | 0.194935263 | 0.118419071 | -0.003756163 | 0.27462826 | 0.000210177 | 0.000516635 | 0.136633963 | 0.87968681 |
| H2bc3 | S | 5 | 0.209926575 | 0.249950307 | 0.298604372 | 0.088677797 | 0.048654066 | 0.114615966 | 0.019084135 | 0.01130454 | 0.415959026 | 0.450114263 |
| Srrm2 | S | 1064 | 0.264924937 | 0.307781617 | -0.003204621 | -0.268129558 | -0.310986237 | 0.002087546 | 0.000388676 | 0.982743274 | 0.162851322 | 0.132436674 |
| Ccar1 | S | 326 | 0.148498731 | 0.248437231 | 0.360188145 | 0.211689414 | 0.111750914 | 0.342380658 | 0.020194109 | 0.00310925 | 0.213983487 | 0.193244068 |
| Ccar1 | S | 330 | 0.148498731 | 0.248437231 | 0.360188145 | 0.211689414 | 0.111750914 | 0.342380658 | 0.020194109 | 0.00310925 | 0.213983487 | 0.193244068 |
| Dlc1 | S | 89 | 0.083176953 | 0.368028878 | 0.304394383 | 0.22121743 | -0.063634495 | 0.162885621 | 0.007143017 | 0.020684197 | 0.071074311 | 0.527612138 |
| Wnk1 | S | 2367 | 0.097067626 | 0.433087265 | 0.148799746 | 0.051732121 | -0.284287519 | 0.174469564 | 0.000555265 | 0.082097171 | 0.349401013 | 0.0126588 |
| Sorbs3 | S | 436 | -0.094040798 | 0.530085127 | 0.322745541 | 0.416786339 | -0.207339586 | 0.266447743 | 9.07677E-05 | 0.059426097 | 0.034495817 | 0.16551897 |
| Plpp7 | S | 62 | -0.01011195 | 0.432641181 | 0.214588734 | 0.224700684 | -0.218052447 | 0.844518204 | 4.54222E-05 | 0.056929941 | 0.06145813 | 0.068337912 |
| Bclaf1 | S | 712 | 0.052143985 | 0.249939885 | 0.381315788 | 0.329171802 | 0.131375902 | 0.418353938 | 0.002808015 | 0.005981213 | 0.009312938 | 0.098420024 |
| Dst | S | 739 | 0.155100713 | 0.439889102 | 0.129751516 | -0.025349196 | -0.310137586 | 0.088660627 | 0.001181139 | 0.160774926 | 0.697136437 | 0.017594421 |
| Pml | S | 404 | 0.054005063 | 0.174325288 | 0.252771121 | 0.198766058 | 0.078445833 | 0.118255086 | 0.003199343 | 0.000170841 | 0.002479552 | 0.073317905 |
| Srrm2 | S | 1498 | 0.01720009 | 0.203884851 | 0.241284453 | 0.224084363 | 0.037399602 | 0.66545084 | 6.94088E-05 | 0.000586626 | 0.005312688 | 0.250677221 |
| Map4 | S | 914 | 0.099852595 | 0.195821982 | 0.137771449 | 0.037918854 | -0.058050533 | 0.010309401 | 0.000253702 | 0.004060566 | 0.246961338 | 0.103567269 |
| Prkd3 | S | 41 | 0.055338548 | 0.348081535 | 0.304624775 | 0.249286227 | -0.04345676 | 0.538029424 | 0.002776656 | 0.008637017 | 0.035500141 | 0.571863982 |
| Nf1 | S | 881 | 0.144855501 | 0.204732476 | 0.215637134 | 0.070781633 | 0.010904658 | 0.111540763 | 0.00262887 | 0.00210228 | 0.353613123 | 0.73754173 |
| Cenpt | S | 296 | 0.248389281 | 0.495183132 | 0.104178755 | -0.144210526 | -0.391004377 | 0.231960072 | 0.002476367 | 0.098057564 | 0.436503032 | 0.012537059 |
| Zzz3 | S | 388 | 0.291939098 | 0.371672859 | 0.180834187 | -0.11110491 | -0.190838672 | 0.022384242 | 0.007468294 | 0.384860614 | 0.559503717 | 0.349903131 |
| Brd3 | S | 262 | 0.001669467 | 0.142725273 | 0.233475089 | 0.231805623 | 0.090749817 | 0.956248206 | 0.000150647 | 4.99363E-06 | 0.004506055 | 0.004847361 |
| Pds5b | S | 1162 | -0.02081513 | 0.406163469 | 0.359744692 | 0.380559822 | -0.046418776 | 0.719150518 | 0.002620385 | 0.006612737 | 0.005050346 | 0.525052751 |
| Mki67 | S | 212 | 0.228029248 | 0.300417106 | 0.267518578 | 0.03948933 | -0.032898528 | 0.100054384 | 0.012088176 | 0.037188027 | 0.746588811 | 0.742564306 |
| Aff4 | S | 387 | 0.400640403 | 0.168701635 | 0.268898901 | -0.131741502 | 0.100197266 | 0.061287395 | 0.022035632 | 0.033752987 | 0.386719761 | 0.276404446 |
| Jcad | S | 954 | 0.112354301 | 0.199671322 | 0.238505654 | 0.126151353 | 0.038834332 | 0.003881222 | 0.014390656 | 0.006153084 | 0.03172966 | 0.429902934 |
| Sidt2 | S | 401 | 0.054249812 | 0.261108649 | 0.249966664 | 0.195716851 | -0.011141985 | 0.374475702 | 0.000381703 | 0.005162143 | 0.032261708 | 0.814160165 |
| Sidt2 | S | 403 | 0.054249812 | 0.261108649 | 0.249966664 | 0.195716851 | -0.011141985 | 0.374475702 | 0.000381703 | 0.005162143 | 0.032261708 | 0.814160165 |
| Zc3h13 | T | 363 | 0.544159514 | 0.356058069 | -0.035521658 | -0.579681172 | -0.391579727 | 0.004166702 | 0.02643704 | 0.923238308 | 0.203198517 | 0.338654577 |
| Srsf4 | S | 399 | 0.029180572 | 0.20251639 | 0.302096104 | 0.272915532 | 0.099579715 | 0.549886009 | 0.014201747 | 3.93511E-05 | 0.011304688 | 0.122651015 |
| Cir1 | S | 202 | -0.001175261 | 0.28077237 | 0.195539816 | 0.196715077 | -0.085232553 | 0.958164006 | 0.000153825 | 0.000365727 | 0.002111859 | 0.039630383 |
| Rnf139 | T | 667 | 0.087459323 | 0.179349403 | 0.303556621 | 0.216097298 | 0.124207218 | 0.07806562 | 0.016496983 | 0.000762556 | 0.007719537 | 0.069840736 |
| Gigyf1 | S | 228 | 0.258258369 | 0.270833938 | 0.148076839 | -0.11018153 | -0.122757099 | 0.495086391 | 0.000902022 | 0.001138939 | 0.757792394 | 0.037228692 |
| Dyrk1a | S | 529 | -0.116423474 | 0.476017949 | 0.296655552 | 0.413079027 | -0.179362397 | 0.023710328 | 9.99309E-06 | 0.01415612 | 0.011845554 | 0.07882984 |
| Pdzd8 | S | 532 | 0.337876014 | 0.204926942 | 0.211685961 | -0.126190053 | 0.006759019 | 0.001262657 | 0.036029399 | 0.340492699 | 0.53794801 | 0.972708331 |
| Psip1 | S | 129 | -0.039365805 | 0.244746795 | 0.451047706 | 0.490413512 | 0.206300911 | 0.611686227 | 0.000851166 | 0.002822518 | 0.004067474 | 0.046220199 |

| | | | | | | | | | | | | |
|---|---|---|---|---|---|---|---|---|---|---|---|---|
| Sidt2 | S | 404 | 0.069162683 | 0.260425766 | 0.247332838 | 0.178170155 | -0.013092928 | 0.334758159 | 0.000995466 | 0.003388137 | 0.060303198 | 0.786916194 |
| Rprd2 | S | 738 | 0.099415024 | 0.21883666 | 0.206353174 | 0.10693815 | -0.012483486 | 0.084152114 | 0.032148848 | 4.35564E-05 | 0.073886115 | 0.814746408 |
| Ahdc1 | S | 1396 | 0.04967887 | 0.342182934 | 0.339493071 | 0.289814201 | -0.002689863 | 0.345911851 | 0.001439009 | 0.032870929 | 0.030625419 | 0.973436554 |
| Oxsr1 | S | 324 | 0.141204588 | 0.316195644 | 0.25695537 | 0.115750781 | -0.059240274 | 0.154386454 | 0.017204754 | 0.00555012 | 0.197753679 | 0.487504815 |
| Zc3h4 | S | 1115 | 0.106206166 | 0.145470706 | 0.224041714 | 0.117835548 | 0.078571008 | 0.072693292 | 4.85228E-05 | 0.005944772 | 0.059354421 | 0.100391131 |
| Cnnm3 | S | 706 | 0.098430641 | 0.217490228 | 0.299949005 | 0.201518364 | 0.082458777 | 0.137850971 | 0.003166604 | 0.005966732 | 0.035575434 | 0.235904709 |
| Zfx | S | 269 | 0.055716018 | 0.192313293 | 0.250066478 | 0.19435046 | 0.057753185 | 0.238566597 | 0.001817194 | 0.000332535 | 0.011930896 | 0.156084777 |
| Pcm1 | S | 814 | 0.066931333 | 0.22022368 | 0.332913198 | 0.265981865 | 0.112689519 | 0.063119034 | 0.030472014 | 0.001256889 | 0.01492687 | 0.160553474 |
| Rbm5 | S | 78 | -0.015709771 | 0.206814983 | 0.303813765 | 0.319523537 | 0.096998783 | 0.727506044 | 0.000169208 | 0.000466336 | 0.002273658 | 0.062454924 |
| Ncor1 | S | 1292 | 0.102003964 | 0.167416419 | 0.289298026 | 0.187294062 | 0.121881606 | 0.185537766 | 0.018494037 | 0.00021234 | 0.054415491 | 0.058812929 |
| Baz2a | S | 1042 | 0.129340527 | 0.17820989 | 0.159200792 | 0.029860265 | -0.019009098 | 0.000913597 | 0.062470395 | 0.000287076 | 0.056863485 | 0.755836657 |
| Map4k3 | S | 329 | 0.190621389 | 0.32861388 | 0.276373941 | 0.085752552 | -0.052239939 | 0.098739101 | 0.029742274 | 0.018196481 | 0.279987061 | 0.592655566 |
| Map4k3 | T | 327 | 0.190621389 | 0.32861388 | 0.276373941 | 0.085752552 | -0.052239939 | 0.098739101 | 0.029742274 | 0.018196481 | 0.279987061 | 0.592655566 |
| Uqcrfs1 | S | 138 | 0.11840925 | 0.333170388 | 0.246784957 | 0.128375708 | -0.086385431 | 0.277587894 | 0.001702767 | 0.027183057 | 0.26502292 | 0.306684354 |
| Uqcrfs1 | S | 139 | 0.11840925 | 0.333170388 | 0.246784957 | 0.128375708 | -0.086385431 | 0.277587894 | 0.001702767 | 0.027183057 | 0.26502292 | 0.306684354 |
| Uqcrfs1 | S | 141 | 0.11840925 | 0.333170388 | 0.246784957 | 0.128375708 | -0.086385431 | 0.277587894 | 0.001702767 | 0.027183057 | 0.26502292 | 0.306684354 |
| Atl1 | S | 10 | -0.006286692 | 0.332512089 | 0.321190634 | 0.327477325 | -0.011321456 | 0.842119058 | 7.45515E-05 | 0.03341918 | 0.035342046 | 0.890102061 |
| Ldb1 | S | 302 | 0.010798884 | 0.216772416 | 0.254718981 | 0.243920097 | 0.037946565 | 0.899294484 | 0.017165603 | 2.3834E-06 | 0.079642922 | 0.424050256 |
| Nsrp1 | S | 292 | 0.119469983 | 0.219920196 | 0.207086481 | 0.087616497 | -0.012833715 | 0.011478524 | 0.000533058 | 0.056898924 | 0.289989674 | 0.855583744 |
| Ptpn21 | S | 492 | 0.281218746 | 0.303692883 | 0.085032838 | -0.196185909 | -0.218660045 | 0.027542964 | 0.000423036 | 0.489431465 | 0.184640014 | 0.155590824 |
| Myof | S | 174 | 0.092500548 | 0.224671079 | 0.406254757 | 0.313754209 | 0.181583679 | 0.161929022 | 0.002534507 | 0.025995848 | 0.053013685 | 0.168163682 |
| Trp53bp1 | S | 1104 | 0.051794814 | 0.301425189 | 0.286105759 | 0.234310945 | -0.01531943 | 0.573810692 | 0.001422819 | 0.006030211 | 0.067671885 | 0.805853539 |
| Arl13b | S | 328 | 0.026629416 | 0.17585163 | 0.266970878 | 0.240341461 | 0.091119248 | 0.273898473 | 2.10229E-05 | 0.003716457 | 0.004944901 | 0.077934039 |
| Rai14 | S | 335 | 0.207639233 | 0.258305721 | 0.289373263 | 0.081734031 | 0.031067542 | 0.052453222 | 0.04592621 | 0.01434047 | 0.173319538 | 0.708994974 |
| Nexn | T | 502 | 0.123121247 | 0.136512361 | 0.277293263 | 0.154172016 | 0.140780902 | 0.046171134 | 0.014432782 | 0.000798322 | 0.026964994 | 0.022860749 |
| Nelfe | S | 181 | 0.017985749 | 0.146907624 | 0.201289613 | 0.183303864 | 0.054381989 | 0.388666547 | 9.16386E-05 | 1.80661E-05 | 0.001477234 | 0.016873162 |
| Svil | S | 857 | 0.066589572 | 0.517887449 | 0.179499546 | 0.112909974 | -0.338387903 | 0.482298435 | 0.000956042 | 0.085868067 | 0.081230339 | 0.006059423 |
| Fundc1 | S | 17 | 0.353640963 | 0.492275815 | 0.303416552 | -0.050224411 | -0.188859263 | 0.120441364 | 0.055430809 | 0.178956007 | 0.623730607 | 0.226711097 |
| Cast | S | 120 | 0.162311307 | 0.316968388 | 0.156156301 | -0.006155006 | -0.160812088 | 0.042863105 | 0.00248654 | 0.048633627 | 0.759357064 | 0.027024532 |
| Ccdc57 | T | 572 | 0.252887233 | 0.456739923 | 0.37870214 | 0.125814907 | -0.078037783 | 0.280033476 | 0.037135908 | 0.076545875 | 0.554152436 | 0.645239309 |
| Smarca2 | S | 1382 | 0.040284515 | 0.325310722 | 0.247599556 | 0.20731504 | -0.077711166 | 0.340849984 | 8.49594E-05 | 0.064227452 | 0.102247261 | 0.416954035 |
| Pam | S | 952 | 0.020293859 | 0.347850128 | 0.347416196 | 0.327122337 | -0.000433932 | 0.633262261 | 0.001050767 | 0.024094364 | 0.018216073 | 0.995413108 |
| Pam | S | 962 | 0.020293859 | 0.347850128 | 0.347416196 | 0.327122337 | -0.000433932 | 0.633262261 | 0.001050767 | 0.024094364 | 0.018216073 | 0.995413108 |
| Pam | S | 964 | 0.020293859 | 0.347850128 | 0.347416196 | 0.327122337 | -0.000433932 | 0.633262261 | 0.001050767 | 0.024094364 | 0.018216073 | 0.995413108 |
| Pola2 | T | 156 | 0.1572963 | 0.280572474 | 0.197479274 | 0.040182974 | -0.0830932 | 0.07515714 | 0.003176608 | 0.022127193 | 0.577345341 | 0.212855648 |
| Wrnip1 | T | 116 | 0.219321703 | 0.145120049 | 0.342303003 | 0.122981299 | 0.197182954 | 0.051539476 | 0.193399834 | 0.001670826 | 0.194464364 | 0.102321773 |
| Pard3 | T | 943 | 0.333640672 | 0.270002953 | 0.227120252 | -0.10652042 | -0.042882701 | 0.155058952 | 0.027767293 | 0.012119535 | 0.564167885 | 0.611259154 |
| Vim | S | 214 | 0.110170587 | 0.212235347 | 0.231212038 | 0.121041451 | 0.018976691 | 0.022849204 | 0.000934208 | 0.028015162 | 0.146731854 | 0.763332818 |
| Aplf | S | 131 | 0.146231865 | 0.216137322 | 0.337682051 | 0.191450186 | 0.121544728 | 0.033759209 | 0.022291976 | 0.018851808 | 0.115395827 | 0.247766733 |
| Ahdc1 | S | 1088 | 0.170580933 | 0.212873606 | 0.20319218 | 0.032611247 | -0.009681426 | 0.15978657 | 0.003618403 | 0.002210228 | 0.733414621 | 0.826707192 |
| Iws1 | S | 375 | 0.024843288 | 0.193736277 | 0.290216651 | 0.265373363 | 0.096480374 | 0.473917213 | 0.001159236 | 0.000489807 | 0.002245993 | 0.058486175 |
| Pard3b | S | 338 | -0.000258002 | 0.239186689 | 0.276322759 | 0.27658076 | 0.03713607 | 0.990693457 | 7.87681E-05 | 0.003225845 | 0.013557742 | 0.399632465 |
| Srrm2 | S | 1477 | -0.064021841 | 0.348464514 | 0.279105653 | 0.343127493 | -0.069358862 | 0.393886151 | 0.003864674 | 8.9279E-05 | 0.027208627 | 0.251401071 |
| Caap1 | S | 306 | 0.060798115 | 0.158055068 | 0.330751195 | 0.269953081 | 0.172696127 | 0.433177883 | 0.017188044 | 0.000115105 | 0.03084912 | 0.017355209 |
| Stx4a | S | 15 | 0.057266498 | 0.165052206 | 0.186154159 | 0.128887661 | 0.021101953 | 0.203970621 | 0.000363336 | 9.57363E-05 | 0.039863565 | 0.237069628 |
| Tmpo | S | 544 | 0.344447807 | 0.438819692 | 0.158271948 | -0.186175859 | -0.280547743 | 0.023902661 | 0.052614573 | 0.224148955 | 0.194042704 | 0.135353028 |
| Sec16a | S | 1245 | 0.135448765 | 0.252651454 | 0.142600843 | 0.007152078 | -0.110050611 | 0.114187377 | 0.000157599 | 0.024691619 | 0.918615213 | 0.063234042 |
| Mcm10 | S | 61 | 0.576968642 | 0.313972592 | 0.621973604 | 0.045004962 | 0.308001012 | 0.027031686 | 0.34152794 | 0.423549977 | 0.944158872 | 0.656984767 |
| Tmpo | S | 277 | 0.070850602 | 0.310468762 | 0.509007838 | 0.438157236 | 0.198539076 | 0.546045508 | 0.062141514 | 0.00584615 | 0.014962351 | 0.17065334 |
| Mprip | S | 992 | -0.046395268 | 0.198352155 | 0.319974673 | 0.366369941 | 0.121622517 | 0.172788023 | 0.001315127 | 1.39553E-05 | 0.00028778 | 0.014974265 |
| Sorbs2 | T | 1001 | 0.031686363 | 0.197839963 | 0.2853006 | 0.253614237 | 0.087460637 | 0.295864701 | 0.001434288 | 0.000865854 | 0.001932505 | 0.06523859 |
| Lrp1 | S | 4524 | 0.123780356 | 0.196810778 | 0.22294829 | 0.099167935 | 0.026137513 | 0.026676008 | 0.001547405 | 0.016908899 | 0.159781033 | 0.64358771 |
| Son | S | 1935 | 0.48395279 | 0.182918129 | 0.338488641 | -0.145464149 | 0.155570512 | 0.026028738 | 0.289509065 | 0.070719591 | 0.418905608 | 0.411306041 |
| Marcksl1 | T | 85 | 0.091328165 | 0.160943069 | 0.545391212 | 0.454063048 | 0.384448143 | 0.232417884 | 0.09565088 | 0.003228055 | 0.016133265 | 0.018327314 |
| Washc2 | S | 614 | 0.083483333 | 0.225866256 | 0.167703128 | 0.084219795 | -0.058163128 | 0.072072566 | 0.000788101 | 0.00230142 | 0.048598618 | 0.068109674 |
| Map2 | S | 1783 | 0.048686648 | 0.237047181 | 0.265537475 | 0.216850827 | 0.028490295 | 0.239835538 | 0.000263222 | 0.011898555 | 0.029381526 | 0.62459184 |
| Dvl1 | S | 676 | 0.026157863 | 0.138176379 | 0.399006596 | 0.372848732 | 0.260830217 | 0.615543644 | 0.074531701 | 3.13475E-05 | 0.001652888 | 0.018862917 |
| Dcaf8 | S | 135 | -0.049228989 | 0.229087227 | 0.357274422 | 0.406503411 | 0.128187195 | 0.108530555 | 0.01221161 | 6.08821E-06 | 0.000823687 | 0.07644092 |
| Cavin3 | S | 165 | -0.025333532 | 0.189717645 | 0.370280344 | 0.395613876 | 0.180562699 | 0.449370709 | 0.00032052 | 0.000633187 | 0.000897725 | 0.028773226 |
| Ralgapa2 | Y | 1015 | 0.294439426 | 0.405246685 | 0.186720298 | -0.107719127 | -0.218526387 | 0.098053494 | 0.003573998 | 0.418916106 | 0.647001871 | 0.355058639 |
| Smarcc2 | S | 302 | 0.013370965 | 0.17537437 | 0.225926541 | 0.212555577 | 0.050552171 | 0.709885716 | 0.004855387 | 2.56685E-06 | 0.013873031 | 0.116713156 |
| Zcchc24 | S | 65 | 0.094406228 | 0.182194667 | 0.209470398 | 0.115064169 | 0.02727573 | 0.040558989 | 0.001829238 | 0.002582259 | 0.02098574 | 0.358277074 |
| Pitpnm1 | S | 300 | 0.082854855 | 0.303939096 | 0.167894754 | 0.085039898 | -0.136044343 | 0.20738015 | 0.000109122 | 0.049132016 | 0.26548928 | 0.081332477 |
| Vrk2 | S | 344 | 0.135246261 | 0.268390355 | 0.201606329 | 0.066360068 | -0.066784026 | 0.083178823 | 0.000840033 | 0.049576658 | 0.435094785 | 0.376903988 |
| Gas2l1 | S | 403 | 0.138761915 | 0.194349236 | 0.267033508 | 0.128271593 | 0.072684272 | 0.012870798 | 0.011627115 | 0.019138661 | 0.152209107 | 0.355924214 |
| Ppp6r2 | S | 746 | 0.01733588 | 0.303001484 | 0.238990057 | 0.221654176 | -0.064011427 | 0.775674436 | 0.00061401 | 0.003223324 | 0.01586185 | 0.197948545 |
| Tmpo | T | 320 | 0.186934908 | 0.216904168 | 0.217359037 | 0.030424129 | 0.000454869 | 0.10763748 | 0.00025228 | 0.078026895 | 0.775156078 | 0.995498397 |
| Phrf1 | S | 919 | -0.00534437 | 0.175429873 | 0.353281444 | 0.358625814 | 0.177851571 | 0.872002797 | 0.038175256 | 1.20308E-05 | 0.0003673 | 0.055236243 |
| Sap130 | S | 927 | 0.031304755 | 0.403889845 | 0.36746302 | 0.336158265 | -0.036426825 | 0.727996674 | 0.030308423 | 0.00350787 | 0.014412394 | 0.767115027 |
| Gigyf1 | S | 24 | 0.061820139 | 0.242307929 | 0.300231659 | 0.23841152 | 0.05792373 | 0.235814671 | 0.000141266 | 0.045270974 | 0.065581196 | 0.521015676 |
| Cavin3 | S | 166 | -0.023817726 | 0.201118221 | 0.384816926 | 0.408634652 | 0.183698705 | 0.505985427 | 0.000153094 | 0.002348217 | 0.001748681 | 0.040372364 |
| Phf2 | S | 536 | 0.087359835 | 0.186596435 | 0.232206211 | 0.144846376 | 0.045609776 | 0.261031757 | 0.005058576 | 0.000237902 | 0.116351049 | 0.268556522 |
| Rbm15b | S | 10 | 0.437462434 | 0.364895765 | 0.110238263 | -0.327224171 | -0.254657503 | 0.017712465 | 0.031402016 | 0.504752084 | 0.065784787 | 0.112879943 |
| Amotl1 | S | 918 | 0.125300217 | 0.192010191 | 0.273792121 | 0.148491904 | 0.08178193 | 0.002966208 | 0.035459274 | 0.015087061 | 0.060463882 | 0.256230801 |
| Zbtb22 | S | 602 | -0.0252855 | 0.384424334 | 0.415181631 | 0.440467131 | 0.030757298 | 0.828568002 | 0.004862067 | 0.011303228 | 0.016030152 | 0.738707257 |
| Itprid2 | T | 85 | -0.10450201 | 0.363589207 | 0.296977456 | 0.401479466 | -0.066611751 | 0.05017812 | 0.000136618 | 0.000589625 | 0.00056553 | 0.076344321 |
| Pak2 | T | 134 | 0.254029329 | 0.231629012 | 0.165587416 | -0.088441913 | -0.066041596 | 0.005400645 | 0.047145986 | 0.039471671 | 0.223033972 | 0.461435764 |
| Fam131b | S | 313 | 0.094951815 | 0.221046737 | 0.227488738 | 0.132536923 | 0.006442001 | 0.152645174 | 0.001252695 | 0.005592672 | 0.081252603 | 0.890892505 |
| Ccdc92 | S | 261 | 0.318267253 | 0.031406275 | 0.561557927 | 0.243290675 | 0.530151653 | 0.051089929 | 0.93259902 | 0.00567849 | 0.003512639 | 0.233107317 |
| Sin3a | S | 1113 | 0.041258618 | 0.19919781 | 0.324535529 | 0.283276911 | 0.125337719 | 0.461047439 | 7.70307E-05 | 0.014591268 | 0.014372593 | 0.13205986 |
| Pkp4 | S | 220 | 0.016157038 | 0.248360496 | 0.296978029 | 0.280820991 | 0.048617533 | 0.736409904 | 0.019653907 | 8.84297E-05 | 0.006603221 | 0.428284694 |
| Pde2a | S | 911 | 0.551905401 | 0.301621912 | 0.219320502 | -0.332584899 | -0.082301409 | 0.014266188 | 0.114484833 | 0.512862484 | 0.327683421 | 0.784038398 |
| Rapgef3 | S | 531 | 0.018306479 | 0.247206593 | 0.225397442 | 0.207090964 | -0.02180915 | 0.623609559 | 0.001188034 | 0.000356505 | 0.001132527 | 0.558360447 |
| Sik3 | T | 1208 | -0.105422831 | 0.543139256 | 0.428979112 | 0.534401943 | -0.114160144 | 0.37757285 | 0.01234832 | 0.005424537 | 0.009387862 | 0.40992784 |
| Palm | S | 122 | -0.002053234 | 0.154407266 | 0.325953286 | 0.328006521 | 0.17154602 | 0.962921489 | 0.006714767 | 2.74797E-05 | 0.002594164 | 0.008459256 |
| Flna | T | 1750 | 0.071790879 | 0.154534827 | 0.222645713 | 0.150854834 | 0.068110886 | 0.015490758 | 0.040630539 | 6.12774E-05 | 0.002028417 | 0.224500395 |
| Mrps26 | S | 183 | 0.196168475 | 0.178568924 | 0.2445632 | 0.048394725 | 0.065994276 | 0.013029645 | 0.001104099 | 0.139219753 | 0.700298339 | 0.597782095 |
| Mrps26 | S | 186 | 0.196168475 | 0.178568924 | 0.2445632 | 0.048394725 | 0.065994276 | 0.013029645 | 0.001104099 | 0.139219753 | 0.700298339 | 0.597782095 |
| Mrps26 | Y | 187 | 0.196168475 | 0.178568924 | 0.2445632 | 0.048394725 | 0.065994276 | 0.013029645 | 0.001104099 | 0.139219753 | 0.700298339 | 0.597782095 |
| Ncoa1 | S | 702 | 0.020492082 | 0.226901083 | 0.378312386 | 0.357820304 | 0.151411303 | 0.638412072 | 0.001635648 | 0.005071819 | 0.019658918 | 0.099393391 |
| Wdr60 | S | 26 | -0.032911758 | 0.230077972 | 0.257803429 | 0.290715188 | 0.027725457 | 0.602901367 | 0.002205959 | 2.34826E-05 | 0.03037374 | 0.415108822 |
| Secisbp2l | S | 272 | 0.084962363 | 0.356299721 | 0.398807625 | 0.313845262 | 0.042507904 | 0.251118589 | 0.028039757 | 0.020769622 | 0.039458874 | 0.72957326 |
| Amer1 | S | 872 | 0.253903186 | 0.452543174 | 0.093189776 | -0.16071341 | -0.359353398 | 0.013318759 | 0.008286317 | 0.671548514 | 0.483226848 | 0.184457115 |
| Cdk12 | T | 889 | 0.177843129 | 0.206364174 | 0.3155427 | 0.13769957 | 0.109178525 | 0.181961113 | 0.015980883 | 0.007810161 | 0.281235757 | 0.178901538 |

| | | | | | | | | | | | | |
|---|---|---|---|---|---|---|---|---|---|---|---|---|
| Trio | S | 1900 | 0.046128106 | 0.267474474 | 0.217694119 | 0.171566014 | -0.049780355 | 0.36069643 | 0.00079495 | 0.003467842 | 0.011948626 | 0.228033498 |
| Fam83h | S | 871 | 0.002370948 | 0.247208008 | 0.203278381 | 0.200907433 | -0.043929626 | 0.948063826 | 0.002418927 | 2.17506E-05 | 0.0166266 | 0.248869959 |
| Numa1 | S | 2059 | 0.038715982 | 0.202823589 | 0.219769526 | 0.181053544 | 0.016945937 | 0.457822599 | 0.001366139 | 0.000176719 | 0.046056815 | 0.558645715 |
| Zfp608 | T | 480 | 0.073361091 | 0.161128942 | 0.277757932 | 0.20439684 | 0.116628989 | 0.279075071 | 0.008484304 | 0.000231907 | 0.034990052 | 0.029984104 |
| Ptk2b | S | 375 | -0.075750821 | 0.568641964 | 0.29427908 | 0.3700299 | -0.274362884 | 0.342776831 | 0.00168802 | 0.018344126 | 0.012090523 | 0.044690571 |
| Psip1 | S | 345 | 0.021897637 | 0.211583393 | 0.256648975 | 0.234751338 | 0.045065582 | 0.468043996 | 0.035787498 | 7.93E-06 | 0.002136773 | 0.458587731 |
| Peak1 | S | 1206 | 0.304737964 | 0.145894508 | 0.259414255 | -0.045323709 | 0.113519747 | 0.011378188 | 0.136828358 | 0.02153924 | 0.509276051 | 0.12397148 |
| Vim | S | 205 | 0.075443952 | 0.125414816 | 0.198573159 | 0.123129207 | 0.073158343 | 0.073982496 | 0.001352646 | 0.000102608 | 0.021785054 | 0.022993427 |
| Elk3 | T | 284 | 0.079089352 | 0.186737925 | 0.22073826 | 0.141648908 | 0.034000335 | 0.118130354 | 7.52337E-05 | 0.015351964 | 0.053725424 | 0.486370606 |
| Trp53bp1 | S | 267 | -0.009279588 | 0.185872863 | 0.272185408 | 0.281464996 | 0.086312545 | 0.82381417 | 0.000373752 | 0.000184476 | 0.002565877 | 0.053034946 |
| Osbpl8 | S | 807 | 0.084273044 | 0.164533946 | 0.20981952 | 0.125546476 | 0.045285574 | 0.135777983 | 1.97012E-05 | 0.013370208 | 0.049994817 | 0.223757576 |
| Osbpl8 | S | 808 | 0.084273044 | 0.164533946 | 0.20981952 | 0.125546476 | 0.045285574 | 0.135777983 | 1.97012E-05 | 0.013370208 | 0.049994817 | 0.223757576 |
| Osbpl8 | S | 810 | 0.084273044 | 0.164533946 | 0.20981952 | 0.125546476 | 0.045285574 | 0.135777983 | 1.97012E-05 | 0.013370208 | 0.049994817 | 0.223757576 |
| Mdc1 | S | 438 | 0.134892227 | 0.199426741 | 0.207590627 | 0.0726984 | 0.008163886 | 0.025471637 | 0.055799557 | 0.000662011 | 0.143687023 | 0.903794753 |
| Myof | T | 172 | 0.160245841 | 0.232392164 | 0.416425058 | 0.256179217 | 0.184032894 | 0.133409337 | 0.013049576 | 0.048817112 | 0.150505584 | 0.255891302 |
| Sipa1l1 | S | 208 | 0.107731437 | 0.125112926 | 0.328235639 | 0.220504202 | 0.203122713 | 0.077690318 | 0.052226593 | 0.000598188 | 0.005836125 | 0.00253437 |
| Rarg | S | 451 | 0.05401362 | 0.26090686 | 0.290397944 | 0.236384324 | 0.029491084 | 0.25831102 | 0.004361699 | 0.005374792 | 0.006892518 | 0.429114038 |
| Zfp318 | S | 41 | 0.01096279 | 0.206672343 | 0.354065538 | 0.343102747 | 0.147393195 | 0.871202267 | 0.001734437 | 0.001628026 | 0.011364256 | 0.031861968 |
| Irs1 | S | 1074 | -0.028642703 | 0.279655067 | 0.250437926 | 0.279080629 | -0.029217141 | 0.549701728 | 0.000494163 | 0.000595901 | 0.001937686 | 0.410140415 |
| Bclaf1 | S | 383 | 0.360457024 | 0.331710959 | 0.074504948 | -0.285952077 | -0.257206011 | 0.011963242 | 0.006537005 | 0.779540183 | 0.336492714 | 0.376762316 |
| Zmym3 | T | 807 | 0.187420682 | 0.141816297 | 0.253236447 | 0.065815765 | 0.11142015 | 0.040747912 | 0.075613682 | 0.001125524 | 0.343481574 | 0.134705408 |
| Vim | Y | 319 | 0.30038683 | 0.397507104 | 0.023589997 | -0.276796834 | -0.373917107 | 0.012583484 | 0.003390308 | 0.931857377 | 0.362155588 | 0.252656315 |
| Pbrm1 | S | 498 | 0.118706287 | 0.201486063 | 0.301902308 | 0.183196021 | 0.100416245 | 0.119622037 | 0.028989216 | 0.002533613 | 0.019389356 | 0.153690261 |
| Atrx | S | 1090 | -0.064290374 | 0.141384131 | 0.239324551 | 0.303614925 | 0.097940419 | 0.151007189 | 2.32917E-05 | 1.13723E-06 | 0.005891189 | 0.002377813 |
| Top2b | S | 1513 | -0.005638013 | 0.167148375 | 0.269448482 | 0.275086495 | 0.102300107 | 0.874835408 | 0.000629646 | 6.92657E-05 | 0.00225823 | 0.015934304 |
| Lmna | S | 17 | 0.065601562 | 0.153389145 | 0.198608647 | 0.133007085 | 0.045219502 | 0.042538969 | 0.003600225 | 0.00019323 | 0.003212441 | 0.176264339 |
| Cavin1 | S | 48 | 0.270392926 | 0.445397882 | 0.276064352 | 0.005671427 | -0.16933353 | 0.433573566 | 0.007457547 | 0.14168005 | 0.986154158 | 0.294760652 |
| Slc39a6 | S | 502 | 0.182357658 | 0.206651836 | 0.236392823 | 0.054035164 | 0.029740986 | 0.024053085 | 0.048719274 | 0.007749466 | 0.206037998 | 0.678155602 |
| Ncor2 | S | 1021 | 0.125787451 | 0.240248583 | 0.182640263 | 0.056852813 | -0.05760832 | 0.037873968 | 0.009075337 | 0.005103602 | 0.266657877 | 0.335829257 |
| Slc9a1 | S | 790 | -0.012664787 | 0.156497821 | 0.229936488 | 0.242601276 | 0.073438667 | 0.679105178 | 0.000246219 | 1.16991E-05 | 0.002680127 | 0.014431054 |
| Prkd2 | S | 197 | 0.103100479 | 0.197021109 | 0.20161372 | 0.098513241 | 0.004592611 | 0.242985145 | 0.011111056 | 9.99681E-05 | 0.257893546 | 0.895527153 |
| Ckap4 | S | 68 | 0.144548165 | 0.291802738 | 0.164237484 | 0.019689318 | -0.127565254 | 0.053538572 | 0.004797432 | 0.023393968 | 0.687876146 | 0.089017653 |
| Hsf1 | S | 319 | 0.500246857 | 0.722469026 | 0.349771228 | -0.150475628 | -0.372697798 | 0.18142484 | 0.086641215 | 0.589151969 | 0.792072556 | 0.540273442 |
| Hsf1 | S | 320 | 0.500246857 | 0.722469026 | 0.349771228 | -0.150475628 | -0.372697798 | 0.18142484 | 0.086641215 | 0.589151969 | 0.792072556 | 0.540273442 |
| Plekhg2 | S | 465 | 0.143440143 | 0.190507654 | 0.347457929 | 0.204017786 | 0.156950274 | 0.236448722 | 0.015651509 | 0.005796836 | 0.129912554 | 0.068055148 |
| Chd8 | S | 2040 | 0.127701871 | 0.134039507 | 0.317245121 | 0.18954325 | 0.183205614 | 0.00311242 | 0.006751597 | 0.041648337 | 0.119879901 | 0.123819238 |
| Chd8 | T | 2039 | 0.127701871 | 0.134039507 | 0.317245121 | 0.18954325 | 0.183205614 | 0.00311242 | 0.006751597 | 0.041648337 | 0.119879901 | 0.123819238 |
| Chd8 | Y | 2043 | 0.127701871 | 0.134039507 | 0.317245121 | 0.18954325 | 0.183205614 | 0.00311242 | 0.006751597 | 0.041648337 | 0.119879901 | 0.123819238 |
| Ptk2 | S | 722 | 0.01089784 | 0.348830069 | 0.303718016 | 0.292820176 | -0.045112052 | 0.796578082 | 0.019118303 | 0.000957031 | 0.002888557 | 0.586992074 |
| Caskin2 | S | 694 | 0.205719982 | 0.223685667 | 0.248745153 | 0.043025172 | 0.025059486 | 0.037377072 | 0.058718659 | 0.011320326 | 0.545136826 | 0.773995249 |
| Dnajc5 | S | 15 | 0.265374536 | 0.172216752 | 0.246412189 | -0.018962347 | 0.074195438 | 0.002001091 | 0.275942574 | 0.032511382 | 0.794408219 | 0.615434443 |
| Dcaf6 | S | 866 | -0.063921072 | 0.242770394 | 0.306288498 | 0.370209571 | 0.063518104 | 0.046748355 | 0.000155346 | 0.000130979 | 0.003462334 | 0.113239538 |
| Brd3 | S | 564 | -0.005933092 | 0.189635421 | 0.281627669 | 0.287560761 | 0.091992249 | 0.87010282 | 0.001312975 | 0.000143984 | 0.001260081 | 0.045003548 |
| Cavin1 | S | 42 | 0.032641886 | 0.166416105 | 0.341258327 | 0.308616441 | 0.174842222 | 0.440169111 | 0.005512663 | 0.000682002 | 0.002117326 | 0.012420899 |
| Nes | S | 963 | 0.006130233 | 0.190471356 | 0.252299936 | 0.246169704 | 0.061828581 | 0.854411804 | 0.000277883 | 0.000466262 | 0.00180485 | 0.11314655 |
| Pid1 | S | 214 | 0.034686238 | 0.207946119 | 0.219218108 | 0.18453187 | 0.011271989 | 0.446784685 | 0.000358136 | 0.001133104 | 0.021619717 | 0.655757166 |
| Atrx | S | 855 | 0.068076059 | 0.172589684 | 0.21464881 | 0.146572751 | 0.042059126 | 0.141289288 | 0.000246801 | 0.003258889 | 0.025311689 | 0.282602887 |
| Timeless | S | 1165 | 0.018201748 | 0.204824119 | 0.155787462 | 0.137585715 | -0.049036657 | 0.314236952 | 2.07514E-06 | 0.006330481 | 0.004073831 | 0.091175688 |
| Pcm1 | S | 813 | 0.101181728 | 0.210951134 | 0.273146306 | 0.171964578 | 0.062195172 | 0.042526709 | 0.037513651 | 0.002660102 | 0.024011002 | 0.410478934 |
| Map2 | S | 1787 | 0.117304297 | 0.287784714 | 0.240266112 | 0.122961815 | -0.047518603 | 0.515463204 | 0.000923035 | 0.019540913 | 0.502525265 | 0.487748853 |
| Map2 | S | 1788 | 0.117304297 | 0.287784714 | 0.240266112 | 0.122961815 | -0.047518603 | 0.515463204 | 0.000923035 | 0.019540913 | 0.502525265 | 0.487748853 |
| Map2 | T | 1781 | 0.117304297 | 0.287784714 | 0.240266112 | 0.122961815 | -0.047518603 | 0.515463204 | 0.000923035 | 0.019540913 | 0.502525265 | 0.487748853 |
| Nsrp1 | S | 291 | 0.122701676 | 0.223588334 | 0.189242598 | 0.066540921 | -0.034345737 | 0.014660483 | 0.000680009 | 0.092191481 | 0.441559469 | 0.674882832 |
| Tmpo | S | 167;167 | 0.00578386 | 0.224385222 | 0.232411531 | 0.226627671 | 0.008026309 | 0.829870138 | 8.39562E-05 | 0.00243217 | 0.006923339 | 0.811632892 |
| Retreg2 | S | 276 | 0.033470199 | 0.131933614 | 0.198045204 | 0.164575005 | 0.066111591 | 0.366053606 | 0.000302501 | 2.56194E-05 | 0.019959269 | 0.00464087 |
| Retreg2 | T | 274 | 0.033470199 | 0.131933614 | 0.198045204 | 0.164575005 | 0.066111591 | 0.366053606 | 0.000302501 | 2.56194E-05 | 0.019959269 | 0.00464087 |
| Runx1t1 | S | 244 | 0.114965952 | 0.231127599 | 0.383373033 | 0.268407081 | 0.152245434 | 0.024830786 | 0.013966256 | 0.066899201 | 0.137153603 | 0.312973125 |
| Incenp | S | 93 | 0.320251093 | 0.22460451 | 0.28208654 | -0.038164553 | 0.05748203 | 0.021005416 | 0.101045428 | 0.085024121 | 0.733392299 | 0.650124371 |
| Rpain | S | 110 | 0.219020981 | 0.399499497 | 0.367476787 | 0.148455806 | -0.032022709 | 0.409254319 | 0.013135943 | 0.107345294 | 0.591636728 | 0.856484008 |
| Pcnx3 | S | 96 | 0.175910354 | 0.195133021 | 0.201926386 | 0.026016032 | 0.006793365 | 0.003482048 | 0.003847586 | 0.146865408 | 0.806430855 | 0.94903259 |
| Iws1 | S | 372 | 0.067468248 | 0.130855623 | 0.156496538 | 0.089028291 | 0.025640916 | 0.055707941 | 0.000314796 | 0.000112037 | 0.030547212 | 0.203734666 |
| Trp53bp1 | S | 1423 | 0.007694739 | 0.347326566 | 0.201330463 | 0.193635725 | -0.145996102 | 0.864587358 | 0.000234008 | 0.014550609 | 0.020978402 | 0.042370036 |
| Zc3h18 | S | 113 | 0.07221619 | 0.231681467 | 0.258483107 | 0.186266917 | 0.02680164 | 0.402143955 | 0.002892205 | 0.002626727 | 0.094153133 | 0.573742155 |
| Bclaf1 | T | 839 | 0.343129913 | 0.255426027 | 0.176390453 | -0.16673946 | -0.079035574 | 0.018025418 | 0.186277469 | 0.02022151 | 0.110456881 | 0.61577195 |
| Mib2 | S | 251 | 0.106718951 | 0.354499677 | 0.210629545 | 0.103910594 | -0.143870131 | 0.099894533 | 0.001939722 | 0.1155654 | 0.354322784 | 0.238310939 |
| Srrm2 | T | 1894 | 0.179401574 | 0.206212017 | 0.191651677 | 0.012250103 | -0.01456034 | 0.021252821 | 0.015187295 | 0.014964866 | 0.716750746 | 0.731134784 |
| Foxj3 | S | 223 | 0.088580216 | 0.136035665 | 0.196110983 | 0.107530767 | 0.060075318 | 0.023521888 | 0.004316656 | 0.000518009 | 0.019502974 | 0.107100517 |
| Errfi1 | T | 126 | 0.447904077 | 0.171959266 | 0.287987536 | -0.159916541 | 0.11602827 | 0.001547666 | 0.318399018 | 0.407169625 | 0.621116263 | 0.728848419 |
| Vim | S | 201 | 0.082884574 | 0.152902844 | 0.257264545 | 0.174379971 | 0.1043617 | 0.377776751 | 0.017062905 | 6.09286E-05 | 0.13662242 | 0.071358048 |
| Kif18b | T | 669 | 0.257354779 | 0.265206848 | 0.214283279 | -0.0430715 | -0.050923569 | 0.018786248 | 0.037379104 | 0.096664665 | 0.634410317 | 0.641246338 |
| Tlk1 | S | 22 | 0.02322897 | 0.312195987 | 0.294870738 | 0.271641768 | -0.017325249 | 0.471527474 | 0.001808297 | 0.014733068 | 0.029155291 | 0.802588305 |
| Msantd2 | S | 27 | 0.049597411 | 0.160616361 | 0.258675718 | 0.209078307 | 0.098059357 | 0.056832257 | 0.114856012 | 1.7492E-05 | 0.003083293 | 0.258364709 |
| Arid4b | S | 790 | 0.054819394 | 0.128774832 | 0.159678333 | 0.104858939 | 0.030903501 | 0.178036389 | 0.001583768 | 2.85351E-06 | 0.055630275 | 0.097089888 |
| Arid4b | T | 793 | 0.054819394 | 0.128774832 | 0.159678333 | 0.104858939 | 0.030903501 | 0.178036389 | 0.001583768 | 2.85351E-06 | 0.055630275 | 0.097089888 |
| Hexim1 | T | 233 | 0.054057585 | 0.216203333 | 0.176156026 | 0.122098441 | -0.040047307 | 0.070226498 | 0.010011164 | 0.00010639 | 0.005058058 | 0.375469418 |
| Cir1 | S | 425 | 0.011195825 | 0.224019529 | 0.266181204 | 0.254985379 | 0.042161675 | 0.77459504 | 0.004143082 | 0.00026089 | 0.003422036 | 0.331441195 |
| Zzz3 | S | 613 | -0.040323254 | 0.162476815 | 0.351400902 | 0.391724156 | 0.188924086 | 0.631373303 | 0.003488211 | 8.05398E-05 | 0.023631839 | 0.003784243 |
| Atxn2l | S | 683 | 0.018826132 | 0.229965566 | 0.212329394 | 0.193503262 | -0.017636173 | 0.539455152 | 0.00537304 | 5.85602E-05 | 0.003961211 | 0.632836903 |
| Invs | S | 807 | 0.240571252 | 0.214999739 | 0.29747221 | 0.056900958 | 0.082472471 | 0.031601064 | 0.056497207 | 0.050100838 | 0.624010561 | 0.49499247 |
| Ibtk | S | 1046 | -0.111304784 | 0.247008809 | 0.333936156 | 0.445240941 | 0.086927348 | 0.147966561 | 0.001670099 | 3.73655E-05 | 0.006458438 | 0.090552718 |
| Sash1 | S | 694 | -0.025543256 | 0.385670466 | 0.297655609 | 0.323198866 | -0.088014856 | 0.66230361 | 0.009049853 | 0.001437564 | 0.000981601 | 0.330101957 |
| Add3 | S | 681 | 0.004499542 | 0.183960326 | 0.129923265 | 0.125423723 | -0.054037061 | 0.747864339 | 0.00016509 | 0.001170388 | 0.018226639 | |
| Srrm1 | T | 882 | 0.05233383 | 0.248088942 | 0.153269725 | 0.100935894 | -0.094819217 | 0.324763002 | 1.23782E-05 | 0.034833301 | 0.138970748 | 0.110795719 |
| Pcm1 | S | 116 | 0.093765224 | 0.092274768 | 0.164727789 | 0.070962565 | 0.07245302 | 0.133876736 | 0.000482139 | 1.31929E-05 | 0.216580369 | 0.002476905 |
| Htatsf1 | S | 455 | 0.010242292 | 0.238045817 | 0.235995543 | 0.225753252 | -0.002050274 | 0.874901514 | 0.000425006 | 0.001519785 | 0.028400341 | 0.952901825 |
| Sntb2 | S | 369 | 0.019318722 | 0.297092605 | 0.352686589 | 0.333367867 | 0.055593984 | 0.783697623 | 0.008322689 | 0.004797236 | 0.01125736 | 0.495562143 |
| Trp53bp1 | S | 1459 | -0.020152579 | 0.198845921 | 0.278203851 | 0.29835643 | 0.07935793 | 0.802658166 | 0.000854616 | 0.000266569 | 0.034131064 | 0.066499889 |
| Rprd2 | S | 749 | 0.033998242 | 0.108098446 | 0.301247432 | 0.26724919 | 0.193148986 | 0.397431006 | 0.007909479 | 4.81253E-05 | 0.007204256 | 0.00123827 |
| Lig1 | S | 911 | 0.024171834 | 0.204215963 | 0.197091645 | 0.172919812 | -0.007124318 | 0.213172718 | 6.56969E-06 | 0.02030556 | 0.022016712 | 0.8519051 |
| Fv4 Gm42102 Gm46( | S | 639;677 | 0.712610829 | 0.04595063 | 0.211697212 | -0.500913617 | 0.165746582 | 0.334609222 | 0.115671751 | 6.24706E-05 | 0.469112381 | 0.00127189 |
| Map2 | T | 1609 | 0.157063173 | 0.231595023 | 0.172709499 | 0.015646326 | -0.058885524 | 0.310512488 | 0.019852272 | 0.000120849 | 0.905978071 | 0.312109527 |
| Farp1 | T | 24 | 0.128651753 | 0.201058162 | 0.2103208 | 0.081669047 | 0.009262638 | 0.054009124 | 0.007248359 | 0.007026763 | 0.120571216 | 0.819920376 |
| Grk5 | S | 484 | 0.133652049 | 0.245164713 | 0.303579612 | 0.169927563 | 0.058414899 | 0.166494751 | 0.082361195 | 0.00233883 | 0.10211998 | 0.587023456 |
| Hmmr | S | 20 | 0.149137464 | 0.169794026 | 0.205736132 | 0.056598668 | 0.035942106 | 0.030774659 | 0.008079656 | 0.00762337 | 0.318811246 | 0.440154286 |

| | | | | | | | | | | | | |
|---|---|---|---|---|---|---|---|---|---|---|---|---|
| Prr12 | S | 909 | 0.13032229 | 0.269611264 | 0.171248395 | 0.040926105 | -0.098362869 | 0.03975836 | 0.000446989 | 0.187933413 | 0.704680107 | 0.389841347 |
| Jak2 | S | 523 | 0.247080493 | 0.256996412 | 0.140792772 | -0.106287721 | -0.116203641 | 0.288840473 | 0.000334278 | 0.057237911 | 0.606835938 | 0.102577711 |
| Luzp1 | Y | 264 | 0.192569203 | 0.52329667 | -0.003286653 | -0.195855855 | -0.526583323 | 0.068370593 | 0.000936811 | 0.973587134 | 0.038852767 | 0.002870004 |
| Fnbp4 | S | 472 | 0.041921229 | 0.143198458 | 0.216073453 | 0.174152224 | 0.072874995 | 0.320235519 | 0.001564965 | 6.89433E-05 | 0.02113736 | 0.033046334 |
| Mki67 | S | 1734 | 0.17683903 | 0.314031722 | 0.282640574 | 0.105801544 | -0.031391148 | 0.230270285 | 0.111383001 | 0.003069111 | 0.430067748 | 0.827546035 |
| Gas2l1 | S | 405 | 0.130226412 | 0.36136608 | 0.159850541 | 0.029624129 | -0.20151554 | 0.167196855 | 0.000285194 | 0.336359213 | 0.84897766 | 0.251322002 |
| Kmt2a | S | 1056 | 0.103652845 | 0.323611213 | 0.462522216 | 0.358869371 | 0.138911003 | 0.4539588 | 0.036810554 | 0.026289513 | 0.070682022 | 0.31905386 |
| Zfp687 | S | 66 | 0.104877635 | 0.137077055 | 0.253699134 | 0.1488215 | 0.116622079 | 0.141535724 | 0.055814746 | 9.22133E-05 | 0.071527555 | 0.082773589 |
| Tmem104 | S | 90 | 0.043593447 | 0.178415813 | 0.322408516 | 0.278815069 | 0.143992703 | 0.501192964 | 0.009016306 | 0.000725464 | 0.009290597 | 0.034857233 |
| Tmem104 | S | 91 | 0.043593447 | 0.178415813 | 0.322408516 | 0.278815069 | 0.143992703 | 0.501192964 | 0.009016306 | 0.000725464 | 0.009290597 | 0.034857233 |
| Tmem104 | S | 94 | 0.043593447 | 0.178415813 | 0.322408516 | 0.278815069 | 0.143992703 | 0.501192964 | 0.009016306 | 0.000725464 | 0.009290597 | 0.034857233 |
| Tmem104 | T | 92 | 0.043593447 | 0.178415813 | 0.322408516 | 0.278815069 | 0.143992703 | 0.501192964 | 0.009016306 | 0.000725464 | 0.009290597 | 0.034857233 |
| Lmna | T | 505 | 0.146550713 | 0.184059343 | 0.201067129 | 0.054516417 | 0.017007787 | 0.018592013 | 0.007215481 | 0.017880048 | 0.33472597 | 0.748326358 |
| Srrm1 | T | 666 | 0.257558815 | 0.279292682 | 0.176700482 | -0.080858334 | -0.1025922 | 0.044020728 | 0.003514938 | 0.297519211 | 0.617098461 | 0.503964269 |
| Rfx5 | S | 538 | 0.029349491 | 0.292484521 | 0.150241348 | 0.120891857 | -0.142243173 | 0.303937585 | 6.92743E-05 | 0.022481176 | 0.065171192 | 0.053268111 |
| Bcl9 | S | 62 | 0.164099056 | 0.241178022 | 0.404077083 | 0.239978027 | 0.162899062 | 0.192258307 | 0.112172805 | 0.009671644 | 0.104583908 | 0.258426971 |
| Srrm2 | S | 1058 | 0.239241807 | 0.227750194 | 0.073736064 | -0.165505743 | -0.15401413 | 0.001890449 | 0.002579351 | 0.501442024 | 0.192121875 | 0.212322269 |
| Trp53bp1 | S | 268 | -0.028328028 | 0.127280799 | 0.20134463 | 0.229672658 | 0.074063831 | 0.339186653 | 9.18869E-05 | 5.65797E-07 | 0.005984003 | 0.003580078 |
| Son | S | 1856 | 0.357618248 | 0.068473467 | 0.355068792 | -0.002549456 | 0.286595325 | 0.014437729 | 0.153744356 | 0.051963081 | 0.984024078 | 0.095572209 |
| Thrap3 | S | 532 | 0.064146834 | 0.128245997 | 0.235714707 | 0.171567874 | 0.10746871 | 0.029444521 | 0.004358287 | 0.000745661 | 0.006829936 | 0.026531573 |
| Dhx57 | S | 301 | -0.003763912 | 0.257778029 | 0.27201938 | 0.275783292 | 0.014241351 | 0.953521303 | 0.001428101 | 0.001595183 | 0.008570515 | 0.737690385 |
| Atf2 | S | 94 | 0.038881228 | 0.16036154 | 0.202499966 | 0.163618738 | 0.042138426 | 0.427586626 | 0.002980166 | 2.63039E-05 | 0.041367358 | 0.173516269 |
| Dlg1 Dlg2 | S | 301;175 | 0.099565093 | 0.135485141 | 0.313016824 | 0.213451731 | 0.177531683 | 0.343264654 | 0.019399719 | 0.000463322 | 0.101299431 | 0.019231291 |
| Spcs2 | S | 31 | 0.151081593 | 0.246528003 | 0.466515759 | 0.315434166 | 0.219987755 | 0.076444713 | 0.256630441 | 0.014648579 | 0.056027363 | 0.311920502 |
| Map2 | T | 1650 | 0.273738668 | 0.281775164 | 0.059880068 | -0.213858601 | -0.221895096 | 0.304947931 | 0.000142475 | 0.031060225 | 0.397398309 | 0.000348649 |
| Srrm2 | S | 1062 | 0.249390601 | 0.275429542 | 0.091269711 | -0.15812089 | -0.184159831 | 0.007696358 | 0.003609923 | 0.485993819 | 0.256867024 | 0.207214759 |
| Srrm2 | S | 1458 | 0.061479501 | 0.204709048 | 0.200110198 | 0.138630696 | -0.004598851 | 0.100747476 | 3.5443E-05 | 0.04876205 | 0.094567415 | 0.937229446 |
| Sp4 | S | 768 | 0.152826457 | 0.350265127 | 0.382005908 | 0.229179451 | 0.031740781 | 0.300542169 | 0.197559772 | 0.005260417 | 0.152341221 | 0.885875222 |
| Sp4 | T | 772 | 0.152826457 | 0.350265127 | 0.382005908 | 0.229179451 | 0.031740781 | 0.300542169 | 0.197559772 | 0.005260417 | 0.152341221 | 0.885875222 |
| Nipbl | S | 2652 | -0.022185901 | 0.218447719 | 0.3319748 | 0.3541607 | 0.11352708 | 0.555804914 | 0.00073433 | 0.001676686 | 0.007363849 | 0.087595493 |
| Ckap4 | S | 367 | 0.010267405 | 0.212146078 | 0.310772773 | 0.300505368 | 0.098626695 | 0.725884507 | 0.000189048 | 0.01040151 | 0.006422335 | 0.123090077 |
| Tns2 | S | 455 | -0.008354667 | 0.230505649 | 0.269594243 | 0.27794891 | 0.039088594 | 0.855427955 | 0.001168126 | 0.000832661 | 0.002322128 | 0.374786356 |
| Tns2 | S | 466 | -0.008354667 | 0.230505649 | 0.269594243 | 0.27794891 | 0.039088594 | 0.855427955 | 0.001168126 | 0.000832661 | 0.002322128 | 0.374786356 |
| Clip2 | S | 205 | 0.067499822 | 0.201347547 | 0.402034748 | 0.334534926 | 0.200687201 | 0.509403095 | 0.011780595 | 0.00605035 | 0.030565496 | 0.081419235 |
| Cep131 | T | 473 | -0.052532812 | 0.253146996 | 0.338784158 | 0.391316971 | 0.085637162 | 0.110586506 | 0.000909636 | 0.000533398 | 0.00268439 | 0.112846807 |
| Fip1l1 | S | 280 | 0.136140531 | 0.170845101 | 0.168908512 | 0.032767981 | -0.001936589 | 0.009960122 | 0.001706643 | 0.035465097 | 0.560678693 | 0.968952042 |
| Larp4b | S | 721 | 0.304762901 | 0.132081919 | 0.133318164 | -0.171444737 | 0.001236245 | 0.366877511 | 0.000353592 | 0.000490906 | 0.582171405 | 0.949365048 |
| Map7d1 | S | 125 | 0.156264554 | 0.246924347 | 0.182355279 | 0.026090725 | -0.064569068 | 0.077163885 | 0.000653701 | 0.132950124 | 0.802010919 | 0.506459995 |
| Zfp37 | S | 590 | -0.033996289 | 0.293596931 | 0.357551514 | 0.391547803 | 0.063954583 | 0.46095818 | 6.0078E-05 | 0.043814029 | 0.033632634 | 0.542507597 |
| Xrn1 | S | 1328 | 0.056324156 | 0.170801671 | 0.179917936 | 0.123593779 | 0.009116264 | 0.316506206 | 0.000232696 | 0.000782744 | 0.08116318 | 0.721749592 |
| Xrn2 | S | 471 | 0.127409823 | 0.199283561 | 0.171443245 | 0.044033423 | -0.027840316 | 0.025187256 | 0.003475279 | 0.016862808 | 0.332012051 | 0.53394619 |
| Ralgapa1 | S | 1003 | 0.086553599 | 0.135516041 | 0.119235976 | 0.032682377 | -0.016280065 | 0.093240953 | 5.64493E-05 | 0.000328814 | 0.413518168 | 0.284222595 |
| Ppig | S | 694 | 0.046925414 | 0.203782673 | 0.149304363 | 0.102378949 | -0.05447831 | 0.244162398 | 0.000155025 | 0.001455708 | 0.06137616 | 0.061464728 |
| Atxn1 | S | 751 | -0.03409645 | 0.15820821 | 0.227252049 | 0.261348499 | 0.069043839 | 0.18050785 | 7.4807E-05 | 1.34946E-05 | 0.000237513 | 0.013012675 |
| Ap2b1 | S | 497 | 0.294977819 | 0.277330579 | 0.203525805 | -0.091452014 | -0.073804775 | 0.01262049 | 0.059268642 | 0.215108036 | 0.513988421 | 0.633490472 |
| Lmnb2 | S | 403 | 0.200962915 | 0.165743953 | 0.131616778 | -0.069346137 | -0.034127175 | 0.015489222 | 0.001501722 | 0.050907791 | 0.28852409 | 0.478892892 |
| Hexim1 | S | 234 | 0.054996595 | 0.209072536 | 0.176297744 | 0.121301149 | -0.032774792 | 0.116346958 | 0.011845338 | 0.000103732 | 0.024531831 | 0.452046051 |
| Stim2 | S | 523 | 0.229486684 | 0.136288256 | 0.235696479 | 0.006209795 | 0.099408223 | 0.006814812 | 0.306991796 | 0.005506141 | 0.82116498 | 0.422711393 |
| Tcf20 | S | 447 | 0.451113726 | 0.345193169 | 0.381918048 | -0.069195678 | 0.036724879 | 0.055051275 | 0.184455973 | 0.310137744 | 0.836107938 | 0.91673804 |
| Tcf20 | S | 455 | 0.451113726 | 0.345193169 | 0.381918048 | -0.069195678 | 0.036724879 | 0.055051275 | 0.184455973 | 0.310137744 | 0.836107938 | 0.91673804 |
| Mdc1 | T | 623 | 0.001702419 | 0.222474402 | 0.220627562 | 0.218925143 | -0.001846839 | 0.979617517 | 0.000381803 | 0.000874067 | 0.040310855 | 0.95657064 |
| Mier1 | S | 487 | 0.002867988 | 0.280949427 | 0.305873568 | 0.30300558 | 0.024924141 | 0.957179913 | 0.000348592 | 0.024139118 | 0.025510493 | 0.742998388 |
| Cdc42ep2 | S | 141 | 0.075919435 | 0.221355987 | 0.179267805 | 0.10334837 | -0.042088182 | 0.046971543 | 0.000293384 | 0.04342107 | 0.160322397 | 0.490414491 |
| Fnbp4 | S | 440 | 0.02869918 | 0.121865405 | 0.219598133 | 0.190898953 | 0.097732727 | 0.161281708 | 0.000241893 | 0.002083057 | 0.015093766 |
| Nexn | S | 16 | -0.012833369 | 0.181763671 | 0.284404083 | 0.297237452 | 0.102640412 | 0.746580492 | 0.001726913 | 0.000193872 | 0.003506269 | 0.00029179 |
| Camk2d | S | 333 | -0.043962988 | 0.302836806 | 0.219035714 | 0.262998702 | -0.083801092 | 0.563580293 | 0.000312358 | 0.001971283 | 0.03667293 | 0.037556687 |
| Pdlim7 | T | 251 | 0.121725878 | 0.204752672 | 0.265070246 | 0.143344368 | 0.060317573 | 0.134713797 | 0.008218644 | 0.011596736 | 0.114125039 | 0.363206993 |
| Spp1 | S | 271 | 0.028715249 | 0.301661241 | 0.084245022 | 0.055529773 | -0.217416219 | 0.621903051 | 9.10602E-07 | 0.140354605 | 0.420849971 | 0.022416162 |
| Sipa1l1 | S | 1528 | 0.030481846 | 0.138165111 | 0.285086162 | 0.254604316 | 0.146921051 | 0.476820588 | 0.007119806 | 0.000135301 | 0.001755918 | 0.001953437 |
| Bag3 | S | 138 | 0.322244448 | 0.349053159 | 0.00612786 | -0.316116588 | -0.342925299 | 0.009486522 | 0.00624378 | 0.979093772 | 0.250238162 | 0.22291259 |
| Sptbn1 | S | 2160 | 0.008748479 | 0.212247587 | 0.184544411 | 0.175795933 | -0.027703176 | 0.796313702 | 3.66749E-05 | 0.0025408 | 0.007380421 | 0.381935731 |
| Larp4b | S | 595 | 0.364744118 | 0.516472126 | 0.128771437 | -0.235972681 | -0.387700689 | 0.217148533 | 0.015019694 | 0.390392189 | 0.392040007 | 0.062566257 |
| Arhgap23 | S | 1388 | 0.124104733 | 0.148734007 | 0.128092959 | 0.003988226 | -0.020641048 | 0.012551168 | 0.000180996 | 0.02237638 | 0.918433345 | 0.550449351 |
| Sorbs2 | S | 40 | 0.022623889 | 0.104827569 | 0.22834388 | 0.205719991 | 0.123516311 | 0.161564998 | 0.004905815 | 1.08747E-05 | 0.000382504 | 0.00459636 |
| Srrm2 | S | 375 | 0.139536714 | 0.14083821 | 0.181113513 | 0.0415768 | 0.040275303 | 0.001500566 | 0.003488635 | 0.057462499 | 0.520248355 | 0.537550195 |
| Cep131 | S | 481 | -0.068541452 | 0.260395953 | 0.351049986 | 0.419591438 | 0.090654033 | 0.156753804 | 0.003134362 | 0.000279007 | 0.000634235 | 0.140537101 |
| Srrm2 | S | 1334 | 0.009180443 | 0.169737724 | 0.372098218 | 0.362917775 | 0.202360493 | 0.81672309 | 0.000130326 | 0.011659804 | 0.006239251 | 0.057161935 |
| Lsm14a | S | 183 | -0.000281994 | 0.153309856 | 0.189215283 | 0.189497277 | 0.035905427 | 0.993575404 | 0.00047021 | 9.52272E-06 | 0.016585514 | 0.12380842 |
| Sun2 | S | 49 | 0.131959087 | 0.241227073 | 0.24791504 | 0.115955953 | 0.006687967 | 0.127892799 | 0.022704126 | 0.008685295 | 0.18806264 | 0.929484538 |
| Itgb3bp | S | 70 | 0.078878048 | 0.13880329 | 0.266686755 | 0.187808706 | 0.127883465 | 0.183242889 | 0.108208223 | 4.42965E-05 | 0.032568979 | 0.137320882 |
| Adnp | S | 961 | 0.047088681 | 0.151396379 | 0.274399393 | 0.227310712 | 0.123003014 | 0.260834954 | 0.006864527 | 0.000543637 | 0.001655403 | 0.009035962 |
| Tjp1 | S | 313 | -0.041580184 | 0.198227842 | 0.202289853 | 0.243870037 | 0.004062011 | 0.512125278 | 0.000100669 | 9.18303E-05 | 0.033560496 | 0.849392564 |
| Mia2 | S | 1228 | 0.175424097 | 0.179832282 | 0.376591754 | 0.201167657 | 0.196759471 | 0.117624882 | 0.136434987 | 0.008413632 | 0.069896586 | 0.096047283 |
| Arhgef6 | S | 487 | -0.032606073 | 0.418281833 | 0.168783128 | 0.201389201 | -0.249498705 | 0.556040005 | 0.00027083 | 0.015862618 | 0.012966148 | 0.008036049 |
| Ppp1r18 | S | 146 | 0.038729676 | 0.131422447 | 0.146422215 | 0.107692539 | 0.014999768 | 0.020756324 | 0.00232451 | 1.62001E-05 | 0.000925051 | 0.466293897 |
| Fam171b | S | 808 | -0.021410726 | 0.256795917 | 0.344800275 | 0.366211001 | 0.088004358 | 0.456292633 | 0.022457222 | 0.000247157 | 0.000436398 | 0.158935553 |
| Mki67 | S | 503 | 0.137432583 | 0.237388717 | 0.184694995 | 0.047262412 | -0.052693722 | 0.069564536 | 0.01277049 | 0.009530001 | 0.380364548 | 0.380118443 |
| Thrap3 | S | 915 | 0.429858798 | 0.129457577 | 0.393373291 | -0.036485507 | 0.263915714 | 0.130101231 | 0.320381607 | 0.02150767 | 0.869181517 | 0.111713118 |
| Rps27l Rps27 Rps27r | S | 78;78 | 0.558806979 | 0.282682915 | 0.207205649 | -0.351601331 | -0.075477267 | 0.028862194 | 0.237244569 | 0.295428686 | 0.068671375 | 0.616422284 |
| Ahdc1 | T | 1394 | -0.027854835 | 0.281659571 | 0.261028054 | 0.28888289 | -0.020631517 | 0.654571983 | 1.31024E-05 | 0.053083934 | 0.032860319 | 0.79616695 |
| Baiap2 | S | 493 | 0.131923158 | 0.328414104 | 0.184690488 | 0.05276733 | -0.143723616 | 0.136178729 | 0.02509905 | 0.008996509 | 0.474008553 | 0.184181864 |
| Aplf | S | 238 | 0.184659324 | 0.320050428 | 0.197042539 | 0.012383215 | -0.123007889 | 0.021627136 | 0.013626547 | 0.240974134 | 0.928921445 | 0.436927942 |
| Lmnb2 | T | 494 | 0.074687514 | 0.179351263 | 0.178789521 | 0.104102007 | -0.000561743 | 0.02823113 | 0.000744473 | 0.011836142 | 0.082039721 | 0.989328121 |
| Zfp280c | S | 531 | -0.006983836 | 0.226929196 | 0.375324991 | 0.382308826 | 0.148395794 | 0.777825552 | 0.010409924 | 0.001471661 | 0.005570067 | 0.046795998 |
| Erf | T | 3 | 0.343361765 | 0.171485692 | 0.091514182 | -0.251847583 | -0.07997151 | 0.00087056 | 0.15013772 | 0.155773986 | 0.004770889 | 0.421100739 |
| Exoc1 | S | 501 | 0.214818154 | 0.169009046 | 0.220646951 | 0.005828797 | 0.051637905 | 0.012457563 | 0.112055361 | 0.01588681 | 0.905849869 | 0.54682322 |
| Pds5b | S | 1165 | 0.113015162 | 0.423807771 | 0.293237702 | 0.18022254 | -0.130570069 | 0.353655712 | 0.012784429 | 0.095176032 | 0.267154304 | 0.405862075 |
| Phrf1 | S | 98 | 0.114305768 | 0.132376792 | 0.16560385 | 0.051298083 | 0.033227059 | 0.018102008 | 0.005523396 | 0.001752207 | 0.15165696 | 0.201047019 |
| Mki67 | S | 498 | 0.176255362 | 0.229962379 | 0.042408524 | -0.133846838 | -0.187553855 | 0.00410733 | 0.000471482 | 0.318417693 | 0.029971161 | 0.009412288 |
| Akap12 | S | 498 | 0.238949964 | 0.233217343 | 0.116711306 | -0.122238658 | -0.116506036 | 0.012297825 | 0.013374219 | 0.114434798 | 0.069859261 | 0.074970441 |
| Gatad2b | T | 121 | 0.093318074 | 0.210042189 | 0.28067941 | 0.187361335 | 0.070637221 | 0.027076731 | 0.0144083 | 0.027117123 | 0.046701854 | 0.288109858 |
| Bclaf1 | S | 718 | 0.11671636 | 0.210376036 | 0.378133394 | 0.261417034 | 0.167757358 | 0.342257994 | 0.069677155 | 0.004552212 | 0.081199457 | 0.139864907 |
| Cbx5 | S | 132 | 0.016974995 | 0.142765338 | 0.359008444 | 0.342033449 | 0.216243106 | 0.736198872 | 0.037733462 | 0.000156197 | 0.000812822 | 0.010450691 |
| Atrx | S | 92 | 0.032242095 | 0.150620693 | 0.266273278 | 0.234031183 | 0.115652585 | 0.390654707 | 0.003704136 | 0.000429183 | 0.002734415 | 0.02802597 |

| | | | | | | | | | | | | |
|---|---|---|---|---|---|---|---|---|---|---|---|---|
| Mcph1 | S | 273 | 0.284644216 | 0.238491607 | 0.131856602 | -0.152787615 | -0.106635005 | 0.009888536 | 0.009037468 | 0.455040832 | 0.394432806 | 0.529733003 |
| Cep290 | S | 1783 | 0.046809454 | 0.416855733 | 0.384970145 | 0.338160692 | -0.031885588 | 0.778113153 | 0.027094599 | 0.024958748 | 0.124429392 | 0.816205733 |
| Atrx | S | 854 | 0.04968074 | 0.156455285 | 0.201271636 | 0.151590896 | 0.044816351 | 0.331085583 | 0.001158568 | 0.000359117 | 0.039699185 | 0.167429636 |
| Fam122a | S | 186 | 0.04704178 | 0.158434596 | 0.245872168 | 0.198830387 | 0.087437572 | 0.23348188 | 0.000662376 | 0.003247466 | 0.010030583 | 0.09980379 |
| Smc3 | S | 1067 | -0.011034615 | 0.143187527 | 0.241454667 | 0.252489281 | 0.09826714 | 0.798972023 | 0.000143689 | 0.000167279 | 0.008579636 | 0.016164713 |
| Srrm2 | T | 1636 | 0.278494506 | 0.113897757 | 0.261557703 | -0.016936803 | 0.147659946 | 0.00423898 | 0.055197918 | 0.140540552 | 0.901248075 | 0.330366484 |
| Atn1 | S | 77 | 0.057790199 | 0.143451231 | 0.164538306 | 0.106748107 | 0.021087075 | 0.064871498 | 0.000700936 | 0.00059065 | 0.013102938 | 0.410255843 |
| Sec16a | S | 1238 | 0.202438225 | 0.191303782 | 0.203974051 | 0.001535826 | 0.012670269 | 0.066357689 | 0.070982376 | 0.00366241 | 0.983487278 | 0.86230971 |
| Lsm14a | S | 192 | -0.002890416 | 0.15030799 | 0.183224683 | 0.1861151 | 0.032916694 | 0.933343188 | 0.000462808 | 8.61049E-06 | 0.016505153 | 0.138486046 |
| Tor1aip1 | S | 79 | 0.092173976 | 0.38329037 | 0.215639682 | 0.123465706 | -0.167650688 | 0.511085541 | 0.004214357 | 0.042956364 | 0.347922979 | 0.005422765 |
| Cep55 | S | 96 | 0.061975878 | 0.355508882 | 0.17304874 | 0.111072862 | -0.182460143 | 0.126068162 | 0.000686296 | 0.171280036 | 0.323657145 | 0.155512029 |
| Itpr3 | S | 934 | 0.091143109 | 0.364983107 | 0.127636711 | 0.036493602 | -0.237346397 | 0.148877226 | 0.001185034 | 0.106192546 | 0.489155416 | 0.016866834 |
| Ythdc1 | S | 309 | 0.075981991 | 0.10435959 | 0.224632567 | 0.148650576 | 0.120272977 | 0.092088074 | 0.033895333 | 4.62677E-05 | 0.02340812 | 0.035239151 |
| Ythdc1 | S | 316 | 0.075981991 | 0.10435959 | 0.224632567 | 0.148650576 | 0.120272977 | 0.092088074 | 0.033895333 | 4.62677E-05 | 0.02340812 | 0.035239151 |
| Ythdc1 | S | 318 | 0.075981991 | 0.10435959 | 0.224632567 | 0.148650576 | 0.120272977 | 0.092088074 | 0.033895333 | 4.62677E-05 | 0.02340812 | 0.035239151 |
| Ythdc1 | S | 319 | 0.075981991 | 0.10435959 | 0.224632567 | 0.148650576 | 0.120272977 | 0.092088074 | 0.033895333 | 4.62677E-05 | 0.02340812 | 0.035239151 |
| Ythdc1 | S | 321 | 0.075981991 | 0.10435959 | 0.224632567 | 0.148650576 | 0.120272977 | 0.092088074 | 0.033895333 | 4.62677E-05 | 0.02340812 | 0.035239151 |
| Sbno2 | S | 1256 | 0.146320497 | 0.218385211 | 0.180300435 | 0.033979938 | -0.038084776 | 0.042149445 | 0.004653468 | 0.043334643 | 0.610560837 | 0.531229854 |
| Tmpo | S | 291 | 0.21638752 | 0.21459972 | 0.160536303 | -0.055851217 | -0.054063417 | 0.003970006 | 0.048265906 | 0.095059497 | 0.453944129 | 0.569086652 |
| Dpysl2 | S | 518 | 0.054200984 | 0.306726917 | 0.097710548 | 0.043509563 | -0.209016369 | 0.386528206 | 2.37708E-05 | 0.094507334 | 0.515373792 | 0.015624299 |
| Nsrp1 | S | 288 | 0.148588002 | 0.114078865 | 0.229224751 | 0.080636748 | 0.115145886 | 0.00302872 | 0.010943071 | 0.037532321 | 0.298926447 | 0.184866047 |
| Smarcc2 | S | 264 | 0.01640851 | 0.309375626 | 0.334556615 | 0.318148105 | 0.025180989 | 0.78813415 | 0.000962828 | 0.055027304 | 0.070188957 | 0.826119947 |
| Rb1cc1 | S | 222 | 0.097019089 | 0.247168932 | 0.128263251 | 0.031244162 | -0.118905681 | 0.251860666 | 0.001434368 | 0.00257085 | 0.668146268 | 0.031785792 |
| Zbtb22 | S | 203 | 0.010176153 | 0.263894743 | 0.309354293 | 0.29917814 | 0.04545955 | 0.918682757 | 0.013928852 | 0.001469318 | 0.057855784 | 0.489440658 |
| Bicral | S | 1037 | 0.157763317 | 0.531340159 | 0.197579925 | 0.039816608 | -0.333760234 | 0.565972589 | 0.003237874 | 0.331789856 | 0.888525016 | 0.137807824 |
| Atrx | S | 665 | 0.042993457 | 0.194557706 | 0.219027819 | 0.176034362 | 0.024470113 | 0.207080367 | 0.00427504 | 0.001103145 | 0.005437687 | 0.484226638 |
| Trim24 | S | 1026 | 0.018990224 | 0.233885993 | 0.268849632 | 0.249859408 | 0.034963639 | 0.657093922 | 0.003643049 | 0.002388462 | 0.00726823 | 0.526594852 |
| Trim24 | S | 1029 | 0.018990224 | 0.233885993 | 0.268849632 | 0.249859408 | 0.034963639 | 0.657093922 | 0.003643049 | 0.002388462 | 0.00726823 | 0.526594852 |
| Ift140 | S | 1445 | 0.042350784 | 0.192850698 | 0.141391604 | 0.099040819 | -0.051459094 | 0.158262235 | 9.19437E-06 | 0.022097781 | 0.052595948 | 0.203993068 |
| Rbm33 | S | 1002 | 0.100675556 | 0.177651056 | 0.208182577 | 0.107507021 | 0.030531521 | 0.055457006 | 0.017101609 | 0.002440362 | 0.021386293 | 0.528828444 |
| Cnot2 | S | 170 | 0.173632683 | 0.288766275 | 0.332516804 | 0.158884121 | 0.043750529 | 0.30270708 | 0.011577357 | 0.080441923 | 0.405438743 | 0.755820038 |
| Dennd2a | S | 358 | 0.09697188 | 0.199138569 | 0.206275043 | 0.109303163 | 0.007136474 | 0.028494263 | 0.021425287 | 0.005145134 | 0.054330827 | 0.899096899 |
| Rb1cc1 | S | 237 | -0.032882406 | 0.296034028 | 0.227332581 | 0.260214987 | -0.068701447 | 0.58532644 | 0.000549369 | 0.003094532 | 0.008856855 | 0.185453152 |
| Zfp609 | S | 841 | 0.253890411 | 0.18586212 | 0.28856576 | 0.034675349 | 0.10270364 | 0.065535633 | 0.028944487 | 0.076773594 | 0.802518115 | 0.415568081 |
| Zfp609 | T | 839 | 0.253890411 | 0.18586212 | 0.28856576 | 0.034675349 | 0.10270364 | 0.065535633 | 0.028944487 | 0.076773594 | 0.802518115 | 0.415568081 |
| Pou3f2 | S | 431 | 0.124633442 | 0.224741708 | 0.320283295 | 0.195649853 | 0.095541587 | 0.254439582 | 0.044444903 | 0.007424049 | 0.081544584 | 0.25265367 |
| Rbbp6 | S | 1501 | 0.046359845 | 0.134767337 | 0.153590026 | 0.107230181 | 0.018822689 | 0.053015815 | 0.000249666 | 0.000537078 | 0.00723389 | 0.377891657 |
| Acin1 | S | 657 | -0.013060007 | 0.148760961 | 0.254066097 | 0.267126104 | 0.105305136 | 0.373970316 | 0.002025495 | 2.10413E-05 | 9.3761E-05 | 0.007645759 |
| Zeb1 | S | 293 | 0.55998857 | 0.268919726 | 0.293079904 | -0.266908666 | 0.024160178 | 0.188895889 | 0.041792927 | 0.280377485 | 0.499177029 | 0.918298181 |
| Irs1 | S | 1097 | -0.092260072 | 0.284327474 | 0.269521591 | 0.361781663 | -0.014805883 | 0.323224018 | 0.000604588 | 0.000934091 | 0.023205534 | 0.660632678 |
| Mdc1 | S | 442 | 0.07378393 | 0.161791118 | 0.157841328 | 0.084057398 | -0.00394979 | 0.009960543 | 0.019285215 | 0.000368088 | 0.013575194 | 0.919418075 |
| Bcar3 | S | 312 | 0.033184688 | 0.246089488 | 0.24858633 | 0.215401642 | 0.002496842 | 0.691362797 | 0.001082628 | 0.008284629 | 0.063385219 | 0.964168878 |
| Il16 | S | 680 | 0.068187006 | 0.23139006 | 0.346507197 | 0.278320192 | 0.115117138 | 0.532521568 | 0.138715013 | 0.00054069 | 0.078521015 | 0.367451666 |
| Il16 | S | 686 | 0.068187006 | 0.23139006 | 0.346507197 | 0.278320192 | 0.115117138 | 0.532521568 | 0.138715013 | 0.00054069 | 0.078521015 | 0.367451666 |
| Pds5b | T | 1253 | 0.343880647 | 0.355846014 | 0.011483864 | -0.332396783 | -0.34436215 | 0.021262003 | 0.007194823 | 0.962926408 | 0.253075284 | 0.241328593 |
| 4932438A13Rik | S | 3835 | 0.153426149 | 0.123088821 | 0.245008189 | 0.09158204 | 0.121919368 | 0.152973648 | 0.028305843 | 0.001049152 | 0.333067212 | 0.008297692 |
| Pds5b | S | 1367 | 0.173964602 | 0.159981422 | 0.14957175 | -0.024392852 | -0.010409671 | 0.051173014 | 0.009708349 | 0.003696049 | 0.689111021 | 0.782602478 |
| Bclaf1 | S | 529 | 0.239546982 | 0.320379278 | 0.074963559 | -0.164583423 | -0.245415719 | 0.07521986 | 0.000675808 | 0.655211887 | 0.382295672 | 0.220037021 |
| Lin54 | S | 635 | 0.022520176 | 0.161353109 | 0.266891939 | 0.244371763 | 0.10553883 | 0.438893355 | 0.021251203 | 9.82906E-05 | 0.001522656 | 0.080137472 |
| Baz1b | S | 312 | 0.03398504 | 0.31095231 | 0.168847736 | 0.134862695 | -0.142104574 | 0.550242139 | 0.000432256 | 0.023108199 | 0.053620663 | 0.040453046 |
| Srsf10 | S | 199 | 0.314386776 | 0.083037498 | 0.366348366 | 0.05196159 | 0.283310868 | 0.029141453 | 0.426119757 | 0.023676845 | 0.678897456 | 0.064760903 |
| Ralgapb | T | 363 | 0.167536768 | 0.14374842 | 0.254678676 | 0.087141908 | 0.110930257 | 0.217902897 | 0.001135446 | 0.019952387 | 0.470795568 | 0.117851938 |
| Smarca4 | S | 699 | 0.017729726 | 0.141207112 | 0.205562065 | 0.18783234 | 0.064354953 | 0.345881256 | 7.73646E-05 | 0.000952113 | 0.004556357 | 0.071137881 |
| Tram2 | S | 332 | 0.187790304 | 0.570764907 | 0.102640321 | -0.085149984 | -0.468124586 | 0.201335522 | 0.00625155 | 0.574105202 | 0.546018019 | 0.05630082 |
| Ifngr1 | T | 373 | 0.063287716 | 0.147558042 | 0.176040523 | 0.112752807 | 0.028482481 | 0.273547365 | 0.001809257 | 0.000166867 | 0.10682032 | 0.291647962 |
| Cherp | S | 746 | 0.186324388 | 0.268293115 | 0.236477559 | 0.050153172 | -0.031815555 | 0.030314866 | 0.009489566 | 0.262521146 | 0.777657722 | 0.859293951 |
| Cherp | S | 748 | 0.186324388 | 0.268293115 | 0.236477559 | 0.050153172 | -0.031815555 | 0.030314866 | 0.009489566 | 0.262521146 | 0.777657722 | 0.859293951 |
| Sipa1l1 | S | 162 | 0.248386808 | 0.209550949 | 0.175572919 | -0.072813889 | -0.03397803 | 0.018043035 | 0.039266125 | 0.080535804 | 0.258410708 | 0.600347784 |
| Senp7 | S | 434 | 0.056427131 | 0.145538364 | 0.40176009 | 0.345332959 | 0.256221726 | 0.067788571 | 0.023575064 | 0.010078015 | 0.009683749 | 0.014794138 |
| Senp7 | S | 435 | 0.056427131 | 0.145538364 | 0.40176009 | 0.345332959 | 0.256221726 | 0.067788571 | 0.023575064 | 0.010078015 | 0.009683749 | 0.014794138 |
| Mecp2 | S | 80 | 0.034789639 | 0.289573943 | 0.338403184 | 0.303613545 | 0.048829241 | 0.650497781 | 0.000531979 | 0.09821021 | 0.112044469 | 0.729102592 |
| Ctnnal1 | S | 537 | -0.077029122 | 0.31267711 | 0.246391039 | 0.323420161 | -0.066286071 | 0.103009778 | 0.002095123 | 9.0901E-05 | 0.002301822 | 0.111489672 |
| Hexim2 | S | 75 | 0.057761425 | 0.257797611 | 0.284073754 | 0.226312329 | 0.026276144 | 0.504511538 | 0.008502066 | 0.007632479 | 0.032739057 | 0.667952399 |
| Setd2 | S | 532 | 0.149642502 | 0.167628338 | 0.199204029 | 0.049561528 | 0.031575691 | 0.024984557 | 0.022026683 | 0.011704325 | 0.315756054 | 0.548243741 |
| Senp3 | S | 206 | 0.039016591 | 0.248266651 | 0.227508607 | 0.188492016 | -0.020758044 | 0.101605777 | 0.002881406 | 0.014661502 | 0.021724515 | 0.66338296 |
| Kdm1a | S | 167 | 0.09613133 | 0.122827734 | 0.109698519 | 0.013567189 | -0.013129215 | 0.002274497 | 0.002413686 | 0.001285732 | 0.38997299 | 0.567214912 |
| Phrf1 | S | 102 | 0.121734447 | 0.105882995 | 0.111568742 | -0.010165705 | 0.005685747 | 0.003984069 | 0.002041892 | 0.001569481 | 0.661849013 | 0.680630854 |
| Ttc7b | S | 629 | 0.08228678 | 0.147697834 | 0.245268675 | 0.162981895 | 0.097570841 | 0.157451149 | 0.013092271 | 0.001165078 | 0.027745315 | 0.071661499 |
| Mre11a | S | 648 | 0.080638299 | 0.138855397 | 0.169176841 | 0.088538542 | 0.030321444 | 0.014735217 | 0.000371576 | 0.014455453 | 0.086601774 | 0.443230939 |
| Itgb5 | S | 770 | 0.083712193 | 0.140188056 | 0.159493142 | 0.075780949 | 0.019305086 | 0.063378172 | 0.001461642 | 0.001140955 | 0.08390895 | 0.420836559 |
| Cavin1 | S | 367 | 0.049275297 | 0.163228112 | 0.225708181 | 0.176432885 | 0.06248007 | 0.31474961 | 3.85989E-05 | 0.018513101 | 0.028662289 | 0.231901718 |
| Kmt2d | S | 688 | -0.050764965 | 0.386315209 | 0.526227013 | 0.576991978 | 0.139911804 | 0.693843031 | 0.012536362 | 0.038397705 | 0.031125077 | 0.435895458 |
| Glyr1 | S | 122 | 0.360515088 | 0.260255051 | 0.266261609 | -0.094253479 | 0.006006559 | 0.062967124 | 0.194173926 | 0.078768364 | 0.504043079 | 0.969659099 |
| Srrm2 | S | 1340 | 0.025984014 | 0.128499961 | 0.187909215 | 0.161925201 | 0.059409254 | 0.178614931 | 5.67995E-05 | 0.000975934 | 0.002485795 | 0.057029677 |
| Nipbl | T | 735 | 0.046170182 | 0.230270243 | 0.266559427 | 0.220389245 | 0.036289184 | 0.26324682 | 0.0008 | 0.03566033 | 0.077064097 | 0.648720688 |
| Srrm2 | S | 1804 | 0.053217123 | 0.224911336 | 0.1243233 | 0.071106177 | -0.100588035 | 0.033771852 | 1.65165E-05 | 0.164817521 | 0.352881866 | 0.228431218 |
| Cdc42ep3 | S | 144 | -0.076140356 | 0.313924686 | 0.240172388 | 0.316312744 | -0.073752298 | 0.077900015 | 0.001904782 | 7.66345E-05 | 0.001293206 | 0.101674042 |
| Mta3 | S | 427 | 0.061047693 | 0.113014907 | 0.185997441 | 0.124949748 | 0.072982534 | 0.075563985 | 0.002998166 | 0.000190151 | 0.006874596 | 0.014329233 |
| Bclaf1 | S | 119 | 0.120518774 | 0.21969488 | 0.274264369 | 0.153745596 | 0.054569489 | 0.078326005 | 0.02866284 | 0.020651891 | 0.119365789 | 0.543155192 |
| Rfc1 | S | 107 | 0.090092332 | 0.127308325 | 0.233755902 | 0.14366357 | 0.106447577 | 0.077892226 | 0.030652976 | 0.000563522 | 0.016216373 | 0.053800034 |
| Fam117b | T | 53 | 0.37177715 | 0.182555463 | 0.25109306 | -0.12068409 | 0.068537597 | 0.021621583 | 0.154884106 | 0.149943825 | 0.411449638 | 0.603735381 |
| Mki67 | S | 287 | 0.197384913 | 0.201441301 | 0.069919696 | -0.127465217 | -0.131521605 | 0.000713665 | 0.012078866 | 0.215372062 | 0.061950949 | 0.073896106 |
| Slc12a2 | T | 210 | 0.184375379 | 0.160359246 | 0.26071991 | 0.076344531 | 0.100360664 | 0.070783345 | 0.164969124 | 0.003066702 | 0.337872556 | 0.326650848 |
| Kif16b | S | 575 | 0.110555375 | 0.084640836 | 0.378637256 | 0.268081881 | 0.29399642 | 0.398266025 | 0.187872356 | 0.000257654 | 0.10877779 | 0.002766406 |
| Nhej1 | S | 245 | -0.095479221 | 0.335456201 | 0.341873121 | 0.437352343 | 0.00641692 | 0.199840181 | 5.90864E-05 | 0.040088881 | 0.016281236 | 0.946469174 |
| Lmna | S | 611 | 0.438471877 | 0.128800407 | 0.235087732 | -0.203384145 | 0.106287325 | 0.011259172 | 0.470176457 | 0.09346913 | 0.107054079 | 0.509012776 |
| Cdk12 | S | 677 | 0.117445887 | 0.168271981 | 0.234572499 | 0.117126612 | 0.066300518 | 0.02359614 | 0.042151347 | 0.007854306 | 0.060472576 | 0.300913033 |
| Srrm2 | S | 1457 | 0.127678453 | 0.203270644 | 0.18757434 | 0.059895887 | -0.015696304 | 0.032011051 | 0.001195661 | 0.123235 | 0.516136851 | 0.855361025 |
| Pml | S | 528 | 0.001514078 | 0.175690638 | 0.245273654 | 0.243759576 | 0.069583016 | 0.967608324 | 0.007036033 | 8.38938E-05 | 0.003634807 | 0.135199602 |
| Magi1 | S | 1359 | -0.053557032 | 0.236103859 | 0.257590823 | 0.311147855 | 0.021486964 | 0.215735023 | 3.44953E-05 | 0.00425267 | 0.001823547 | 0.551380731 |
| Sec16a | S | 1239 | 0.199191279 | 0.264164434 | 0.179326133 | -0.019865146 | -0.084838302 | 0.033017442 | 0.036256955 | 0.06690454 | 0.80981221 | 0.389348269 |
| Itsn1 | S | 979 | 0.137759209 | 0.095996843 | 0.223359349 | 0.08560014 | 0.127362506 | 0.011313618 | 0.092596475 | 0.001750441 | 0.078894284 | 0.040950137 |
| Brca2 | S | 706 | 0.332526941 | 0.081873677 | 0.381494918 | 0.048967977 | 0.299621241 | 0.045569399 | 0.548353931 | 0.020315209 | 0.721805402 | 0.094721008 |
| Phf2 | S | 1059 | 0.118314947 | 0.235391768 | 0.169746702 | 0.051431755 | -0.065645066 | 0.305043881 | 0.004625312 | 0.004744605 | 0.620469997 | 0.223241741 |

| | | | | | | | | | | | | |
|---|---|---|---|---|---|---|---|---|---|---|---|---|
| Lig1 | S | 907 | 0.02437538 | 0.222236616 | 0.203887785 | 0.179512404 | -0.018348832 | 0.351699571 | 0.000132569 | 0.021587713 | 0.019319949 | 0.666292154 |
| Afdn | S | 246 | 0.33693103 | 0.177264744 | -0.009540204 | -0.346471234 | -0.186804948 | 0.000482842 | 0.014803517 | 0.962244845 | 0.183740961 | 0.397398612 |
| Mki67 | S | 1152 | 0.316534571 | 0.268674079 | 0.114566747 | -0.201967824 | -0.154107331 | 0.018405008 | 0.07351825 | 0.126582551 | 0.067378108 | 0.213431396 |
| Ralbp1 | S | 29 | -0.032269537 | 0.204104719 | 0.231586301 | 0.263855838 | 0.027481582 | 0.524813876 | 0.002093879 | 8.92396E-05 | 0.021828072 | 0.39909674 |
| Epn2 | S | 433 | -0.01995333 | 0.170825277 | 0.369443702 | 0.389397032 | 0.198618424 | 0.736358913 | 0.127810435 | 4.12645E-05 | 0.004406946 | 0.101871216 |
| Acin1 | S | 479 | -0.004266254 | 0.149313276 | 0.238481403 | 0.242747657 | 0.089168127 | 0.920480382 | 0.001876541 | 8.37373E-05 | 0.009488051 | 0.011874186 |
| Lemd2 | T | 156 | -0.041958762 | 0.255932499 | 0.487815592 | 0.529774354 | 0.231883093 | 0.66958547 | 0.014611255 | 0.007978987 | 0.008777023 | 0.088446009 |
| Rbl2 | S | 935 | 0.115828561 | 0.52263444 | 0.485726223 | 0.369897662 | -0.036908217 | 0.5340213 | 0.058211184 | 0.152061304 | 0.239680091 | 0.900962516 |
| Zmpste24 | S | 310 | 0.128872847 | 0.203203804 | 0.152992973 | 0.024120126 | -0.050210831 | 0.028619218 | 0.00398162 | 0.03794319 | 0.630193867 | 0.355491189 |
| Safb | S | 616 | 0.396740906 | 0.102329497 | 0.317738404 | -0.079002502 | 0.215408907 | 0.091220922 | 0.492832707 | 0.01084149 | 0.646666084 | 0.205351346 |
| Safb | S | 618 | 0.396740906 | 0.102329497 | 0.317738404 | -0.079002502 | 0.215408907 | 0.091220922 | 0.492832707 | 0.01084149 | 0.646666084 | 0.205351346 |
| Fnbp4 | S | 443 | 0.027092434 | 0.175631644 | 0.246599699 | 0.219507265 | 0.070968055 | 0.635017162 | 0.002103943 | 0.001160437 | 0.016579647 | 0.121766843 |
| Chd2 | S | 156 | 0.242501873 | 0.145936455 | 0.401195262 | 0.158693389 | 0.255258807 | 0.200709475 | 0.167577941 | 0.014203989 | 0.358512224 | 0.058807704 |
| Chd2 | S | 165 | 0.242501873 | 0.145936455 | 0.401195262 | 0.158693389 | 0.255258807 | 0.200709475 | 0.167577941 | 0.014203989 | 0.358512224 | 0.058807704 |
| Chd2 | S | 168 | 0.242501873 | 0.145936455 | 0.401195262 | 0.158693389 | 0.255258807 | 0.200709475 | 0.167577941 | 0.014203989 | 0.358512224 | 0.058807704 |
| Chd2 | T | 164 | 0.242501873 | 0.145936455 | 0.401195262 | 0.158693389 | 0.255258807 | 0.200709475 | 0.167577941 | 0.014203989 | 0.358512224 | 0.058807704 |
| Hivep3 | S | 928 | 0.108644539 | 0.083829769 | 0.179217581 | 0.070573042 | 0.095387813 | 0.007984377 | 0.012518201 | 0.00103812 | 0.064011456 | 0.033872385 |
| Pdlim2 | S | 204 | 0.020089142 | 0.161539692 | 0.182174357 | 0.162085215 | 0.020634665 | 0.401682155 | 6.39236E-05 | 0.002217286 | 0.005390909 | 0.450613501 |
| Map3k11 | S | 751 | 0.193345312 | 0.137015697 | 0.111039824 | -0.082305488 | -0.025975873 | 0.012326961 | 0.019150664 | 0.004125385 | 0.141033136 | 0.488254344 |
| Wdr90 | S | 20 | 0.102861057 | 0.140254095 | 0.184895579 | 0.082034522 | 0.044641484 | 0.134103837 | 0.007622055 | 0.000715802 | 0.203319723 | 0.233118248 |
| Tnrc18 | S | 1849 | 0.066828062 | 0.236576466 | 0.236797862 | 0.1699698 | 0.000221396 | 0.379560496 | 0.004371285 | 0.009497235 | 0.063589498 | 0.996849129 |
| Luc7l3 | S | 431 | 0.020343903 | 0.178552934 | 0.13827707 | 0.117933167 | -0.040275864 | 0.401615711 | 4.59311E-05 | 0.001166282 | 0.007526716 | 0.114702429 |
| H2bc1 H2bc11 H2bc1 | S | 92;92;92;93;92;92 | 0.103227857 | 0.159038471 | 0.150187022 | 0.046959165 | -0.008851449 | 0.010043895 | 0.002603962 | 0.018200054 | 0.286759487 | 0.833881061 |
| Taf3 | S | 297 | 0.098877383 | 0.092281271 | 0.135718733 | 0.03684135 | 0.043437462 | 0.02518576 | 8.52739E-05 | 0.002424375 | 0.237948876 | 0.096833618 |
| Rbmxl1 | T | 163 | 0.020880002 | 0.154595867 | 0.212985633 | 0.192105631 | 0.058389766 | 0.4637872 | 0.001111729 | 0.000431862 | 0.003892741 | 0.096123773 |
| Scaf1 | S | 302 | 0.104476395 | 0.16309773 | 0.227466592 | 0.122990197 | 0.064368862 | 0.107237158 | 0.022191675 | 0.002552845 | 0.059971647 | 0.190723659 |
| Naa15 | T | 858 | 0.390570925 | 0.306156725 | 0.157736541 | -0.232834384 | -0.148420184 | 0.014914366 | 0.173599344 | 0.323039996 | 0.19254627 | 0.491304567 |
| Ebag9 | T | 41 | 0.048643907 | 0.15574604 | 0.240095932 | 0.191452025 | 0.084349891 | 0.31590082 | 0.004181479 | 0.001141936 | 0.014883319 | 0.078059369 |
| Mki67 | T | 1344 | 0.13669425 | 0.184526299 | 0.246397553 | 0.109703303 | 0.061871254 | 0.238131083 | 0.043710598 | 0.002109649 | 0.318046295 | 0.356831687 |
| Tsc1 | S | 502 | 0.076749249 | 0.262288351 | 0.221802113 | 0.145052863 | -0.040486238 | 0.3678774 | 0.002869624 | 0.022228156 | 0.15416919 | 0.506034284 |
| Brd4 | S | 471 | 0.091718335 | 0.164542203 | 0.191231959 | 0.099513623 | 0.026689755 | 0.060884359 | 0.005364137 | 0.005322075 | 0.061275407 | 0.518765519 |
| Nrcam | S | 1178 | 0.109286896 | 0.237599691 | 0.182465405 | 0.07317851 | -0.055134286 | 0.238783172 | 0.01062713 | 0.004201559 | 0.38603122 | 0.345169692 |
| Kif27 | S | 988 | -0.007857838 | 0.179237124 | 0.169131024 | 0.176988862 | -0.0101061 | 0.840481106 | 0.001609441 | 9.73038E-06 | 0.030021017 | 0.653883883 |
| Svil | S | 227 | 0.024126626 | 0.173342749 | 0.247169496 | 0.22304287 | 0.073826747 | 0.629608623 | 0.003713979 | 0.000680388 | 0.006178235 | 0.074485799 |
| Dcp1a | S | 545 | 0.028641353 | 0.146463867 | 0.18788115 | 0.159239797 | 0.041417283 | 0.251474097 | 0.001569136 | 0.000211331 | 0.002081574 | 0.166136189 |
| Retreg3 | S | 258 | 0.024467103 | 0.123501727 | 0.193083665 | 0.168616562 | 0.069581938 | 0.204352876 | 0.000324252 | 0.000354128 | 0.003704424 | 0.035398135 |
| Retreg3 | S | 260 | 0.024467103 | 0.123501727 | 0.193083665 | 0.168616562 | 0.069581938 | 0.204352876 | 0.000324252 | 0.000354128 | 0.003704424 | 0.035398135 |
| Adgrl3 | S | 1449 | 0.161701458 | 0.226139686 | 0.337282575 | 0.175581117 | 0.111142889 | 0.34435746 | 0.014543284 | 0.041040048 | 0.339315169 | 0.355349246 |
| Cavin2 | S | 204 | -0.068488895 | 0.23432665 | 0.352532811 | 0.421021706 | 0.118206161 | 0.147793548 | 0.008419927 | 0.00019707 | 0.000504852 | 0.08802082 |
| Lig3 | S | 244 | 0.092446634 | 0.109207367 | 0.338452368 | 0.246005733 | 0.229245001 | 0.184108928 | 0.10215236 | 0.000957967 | 0.004107819 | 0.00686855 |
| Dpy19l1 | S | 46 | 0.187550389 | 0.146928548 | 0.163611158 | -0.023939231 | 0.01668261 | 0.102619204 | 0.00859505 | 0.004650834 | 0.772028976 | 0.534076722 |
| Larp1 | S | 830 | -0.057410497 | 0.289537639 | 0.194817799 | 0.252228296 | -0.09471984 | 0.110630592 | 6.55445E-05 | 0.00175239 | 0.002293293 | 0.050180917 |
| Tmem237 | S | 126 | 0.041694417 | 0.139029553 | 0.234699035 | 0.193004618 | 0.095669482 | 0.266553049 | 0.017818791 | 0.000140535 | 0.0061711 | 0.055909339 |
| Dxo | S | 390 | 0.094668827 | 0.34810883 | 0.217647526 | 0.1229787 | -0.130461304 | 0.084776478 | 0.021929906 | 0.057562295 | 0.189832516 | 0.240071334 |
| Trrap | T | 1272 | 0.227773692 | 0.053299214 | 0.351979262 | 0.12420557 | 0.298680048 | 0.027610091 | 0.669764259 | 0.004762827 | 0.084935107 | 0.0622626 |
| Srsf10 | S | 160 | 0.236214966 | 0.152704744 | 0.129909276 | -0.10630569 | -0.022795469 | 0.002899607 | 0.081706792 | 0.048096074 | 0.09345457 | 0.741094231 |
| Pdlim2 | S | 205 | 0.04708897 | 0.126424498 | 0.1642579 | 0.11716893 | 0.037833402 | 0.011184721 | 0.012245046 | 0.000128186 | 0.005999541 | 0.229658554 |
| Bud13 | S | 148 | 0.294229568 | 0.045884527 | 0.156941759 | -0.137287809 | 0.111057231 | 0.000183839 | 0.626352162 | 0.052178566 | 0.080857239 | 0.302313385 |
| Bud13 | T | 144 | 0.294229568 | 0.045884527 | 0.156941759 | -0.137287809 | 0.111057231 | 0.000183839 | 0.626352162 | 0.052178566 | 0.080857239 | 0.302313385 |
| Incenp | S | 94 | 0.238463454 | 0.218049627 | 0.221469402 | -0.016994051 | 0.003419775 | 0.030822684 | 0.073694014 | 0.077304797 | 0.828223043 | 0.972614526 |
| Epb41l3 | S | 451 | 0.107806916 | 0.189908456 | 0.267095884 | 0.159288968 | 0.077187427 | 0.123861306 | 0.022842851 | 0.011144836 | 0.06855153 | 0.29814981 |
| Cic | T | 470 | -0.003884763 | 0.147258426 | 0.256667812 | 0.260552576 | 0.109409386 | 0.936442809 | 0.004540069 | 9.46202E-05 | 0.015912152 | 0.026568386 |
| Dgcr8 | S | 373 | -0.021274148 | 0.200587906 | 0.263330595 | 0.284604743 | 0.062742689 | 0.478039532 | 7.91429E-05 | 0.00574919 | 0.003679734 | 0.195222939 |
| Zfp639 | T | 75 | 0.263288249 | 0.243718809 | 0.120298534 | -0.142989715 | -0.123420275 | 0.00869795 | 0.011590635 | 0.580936635 | 0.510030536 | 0.563949744 |
| Sun2 | S | 48 | 0.100426542 | 0.210396498 | 0.220731747 | 0.120305204 | 0.010335249 | 0.273400979 | 0.040634365 | 0.001021593 | 0.209074015 | 0.87776131 |
| Spp1 | S | 290 | 0.069568597 | 0.208653848 | 0.108324032 | 0.038755435 | -0.100329816 | 0.000392118 | 0.004719694 | 0.029480732 | 0.223636476 | 0.031762389 |
| Nexn | S | 495 | 0.107306498 | 0.111289116 | 0.287286104 | 0.179979606 | 0.175996988 | 0.071089381 | 0.08367368 | 0.001688247 | 0.018479832 | 0.019158676 |
| Ptpn14 | S | 486 | 0.133208905 | 0.260920667 | 0.145980787 | 0.012911882 | -0.11493988 | 0.096345619 | 0.01329275 | 0.013523014 | 0.838373539 | 0.128289496 |
| Srrm2 | S | 1787 | 0.143221301 | 0.2340533 | 0.170232701 | 0.027011399 | -0.0638206 | 0.094930142 | 0.001111013 | 0.149470483 | 0.793947687 | 0.505215082 |
| Pip4p2 | T | 89 | 0.06094105 | 0.165647829 | 0.199661892 | 0.138720842 | 0.034014062 | 0.095988242 | 0.001501067 | 0.007086368 | 0.0414224 | 0.437804569 |
| Lmna | S | 629 | 0.069733175 | 0.20519408 | 0.209127953 | 0.139394778 | 0.003933873 | 0.145408417 | 0.003457458 | 0.011688698 | 0.052943579 | 0.941517217 |
| Runx1t1 | T | 565 | 0.029393422 | 0.2753319 | 0.263459812 | 0.23406639 | -0.011872088 | 0.323493708 | 0.002840191 | 0.03095045 | 0.056272857 | 0.874237475 |
| Retreg2 | S | 278 | 0.035732308 | 0.127739186 | 0.224018285 | 0.188285976 | 0.096279099 | 0.337695503 | 0.001684336 | 0.000481835 | 0.005235266 | 0.037644766 |
| Bclaf1 | S | 102 | 0.273261128 | 0.129843392 | 0.228958997 | -0.04430213 | 0.099115605 | 0.022465135 | 0.255882244 | 0.0166141 | 0.603969478 | 0.375559998 |
| Bclaf1 | S | 104 | 0.273261128 | 0.129843392 | 0.228958997 | -0.04430213 | 0.099115605 | 0.022465135 | 0.255882244 | 0.0166141 | 0.603969478 | 0.375559998 |
| Ano8 | S | 637 | 0.190966248 | 0.108139486 | 0.184616993 | -0.006349255 | 0.076477508 | 0.002194987 | 0.008535617 | 0.134813204 | 0.945357424 | 0.438799731 |
| Mia3 | S | 1770 | 0.100143596 | 0.272074996 | 0.330166396 | 0.2300228 | 0.0580914 | 0.055449362 | 0.05267562 | 0.056985746 | 0.1301146 | 0.662331368 |
| Spidr | S | 80 | 0.024138961 | 0.116495017 | 0.145378349 | 0.121239387 | 0.028883331 | 0.279245528 | 0.000200505 | 2.86699E-05 | 0.005440289 | 0.087444189 |
| Cavin2 | S | 203 | -0.066201535 | 0.243562142 | 0.3578686 | 0.424070135 | 0.114306458 | 0.195068828 | 0.004876843 | 0.000822522 | 0.001177624 | 0.101910813 |
| Magi3 | S | 702 | 0.028793947 | 0.142699964 | 0.208987821 | 0.180193875 | 0.066287858 | 0.320939564 | 0.000140699 | 0.003121371 | 0.006000077 | 0.115163728 |
| Pcif1 | S | 116 | 0.007523941 | 0.184233072 | 0.227448386 | 0.219924445 | 0.043215314 | 0.90991851 | 0.001898748 | 0.000574931 | 0.042142713 | 0.139942614 |
| Gigyf1 | S | 413 | 0.108458377 | 0.201400765 | 0.13730102 | 0.028842643 | -0.064099745 | 0.014656128 | 0.00274323 | 0.052127675 | 0.574918386 | 0.275742412 |
| Nexn | S | 22 | 0.109334265 | 0.297558037 | 0.292206855 | 0.18287259 | -0.005351182 | 0.304771628 | 0.007105596 | 0.094858199 | 0.247942696 | 0.967461671 |
| Rbms2 | T | 38 | 0.238079373 | 0.212920965 | 0.198476406 | -0.039602968 | -0.014444559 | 0.015614668 | 0.148685421 | 0.050389446 | 0.520097513 | 0.903901274 |
| Epb41l3 | T | 518 | 0.13837146 | 0.140410287 | 0.191925895 | 0.053554435 | 0.051515608 | 0.017195327 | 0.020889369 | 0.012749265 | 0.329418933 | 0.357689755 |
| Ccny | S | 326 | -0.024173241 | 0.157144211 | 0.239022771 | 0.263196012 | 0.08187856 | 0.343904823 | 0.002042771 | 3.07707E-05 | 9.79537E-05 | 0.038276458 |
| Camk2a Camk2d | T | 287;286 | 0.034396585 | 0.316961187 | 0.165037102 | 0.130640517 | -0.151924085 | 0.329455791 | 0.000931938 | 0.043764219 | 0.0748345 | 0.051593573 |
| Aqp1 | S | 262 | 0.076800156 | 0.158070826 | 0.216102378 | 0.139302221 | 0.058031552 | 0.178439879 | 0.003508491 | 0.004007195 | 0.051615074 | 0.196875043 |
| Pcm1 | S | 1766 | 0.126422832 | 0.121047315 | 0.179819339 | 0.053396507 | 0.058772023 | 0.00641961 | 0.005465529 | 0.026512207 | 0.340182345 | 0.298990758 |
| Cenpc1 | S | 60 | 0.141836859 | 0.082062902 | 0.215217111 | 0.073380251 | 0.133154208 | 0.014176229 | 0.114147143 | 0.001215697 | 0.095781772 | 0.02280532 |
| Cdk7 | T | 170 | 0.212058602 | 0.174329966 | 0.24277024 | 0.030711638 | 0.068440274 | 0.041998452 | 0.045332316 | 0.052326017 | 0.760991177 | 0.482870064 |
| C77080 | S | 512 | -0.053922764 | 0.202426907 | 0.234628396 | 0.288551159 | 0.032201489 | 0.248055581 | 0.000435547 | 0.000179178 | 0.003906988 | 0.178432036 |
| Xpc | S | 93 | 0.030483188 | 0.088498424 | 0.203646823 | 0.173163635 | 0.115148399 | 0.464825292 | 0.001770149 | 1.6294E-05 | 0.024451906 | 0.004817829 |
| Srrm2 | S | 984 | 0.164648754 | 0.121663531 | 0.242200278 | 0.077551524 | 0.120536747 | 0.024455903 | 0.153371135 | 0.004656077 | 0.195535021 | 0.153634952 |
| Taf12 | S | 51 | 0.050586595 | 0.109967826 | 0.179207571 | 0.128620976 | 0.069239745 | 0.056931423 | 4.88669E-05 | 0.005321111 | 0.008423525 | 0.060930494 |
| Cttnbp2nl | S | 484 | 0.037670617 | 0.371471289 | 0.323314154 | 0.285643537 | -0.048157135 | 0.783503029 | 0.017040381 | 0.02962393 | 0.059075815 | 0.300885497 |
| Kank2 | S | 173 | 0.015187164 | 0.229587728 | 0.233618351 | 0.218431187 | 0.004030623 | 0.729481036 | 0.002233107 | 0.003871783 | 0.01182947 | 0.907466623 |
| Cnpy4 | S | 64 | 0.036529958 | 0.162488197 | 0.274369266 | 0.237839308 | 0.111881069 | 0.241242508 | 0.001382191 | 0.009679606 | 0.035192032 | 0.130225151 |
| Nradd | S | 226 | 0.178706117 | 0.244144965 | 0.287974696 | 0.109268579 | 0.043829731 | 0.153250743 | 0.011344684 | 0.130645644 | 0.506240094 | 0.762424103 |
| Lyn | Y | 508 | 0.014471196 | 0.219934455 | 0.263430701 | 0.248959505 | 0.043496246 | 0.660628157 | 0.000762571 | 0.014735168 | 0.012493878 | 0.430802436 |
| Tmx1 | S | 223 | 0.066887912 | 0.117959052 | 0.123153067 | 0.056265155 | 0.005194015 | 0.004881713 | 0.003556204 | 0.000445147 | 0.024172267 | 0.818085423 |
| Bag3 | S | 136 | 0.345998466 | 0.340045277 | 0.033192766 | -0.3128057 | -0.306852511 | 0.020482233 | 0.020765726 | 0.828933721 | 0.077206319 | 0.095689253 |
| Fam131b | S | 117 | 0.035817812 | 0.198651611 | 0.1651609 | 0.129343088 | -0.033490711 | 0.332241654 | 0.000368702 | 0.005051401 | 0.024527968 | 0.331612276 |
| Kmt2b | S | 1101 | 0.176034663 | 0.207024216 | 0.193344303 | 0.01730964 | -0.013679913 | 0.025125483 | 0.156751275 | 0.008389799 | 0.765288889 | 0.904028055 |

| | | | | | | | | | | | | | |
|---|---|---|---|---|---|---|---|---|---|---|---|---|---|
| Bicra | S | | 1427 | 0.015386739 | 0.123576289 | 0.243421992 | 0.228035253 | 0.119845703 | 0.592743661 | 0.001626608 | 0.000300009 | 0.001600744 | 0.020721171 |
| Cfap97 | S | | 230 | 0.099133619 | 0.167872832 | 0.129757924 | 0.030624305 | -0.038114908 | 0.078855237 | 0.001239369 | 0.005401097 | 0.486561298 | 0.190262192 |
| Ppig | S | | 354 | 0.098867054 | 0.155258225 | 0.204040907 | 0.105173852 | 0.048782682 | 0.094953144 | 0.009319139 | 0.004525106 | 0.090554314 | 0.278582207 |
| Pard3 Pard3b | S | 920;780 | | -0.037713939 | 0.177296934 | 0.25789788 | 0.295611819 | 0.080600946 | 0.44428754 | 0.003905508 | 7.37385E-05 | 0.006727247 | 0.075344772 |
| Fam111a | S | | 59 | 0.065235074 | 0.153960135 | 0.191208959 | 0.125973886 | 0.037248824 | 0.174751683 | 0.008488684 | 0.000616385 | 0.04571348 | 0.175559631 |
| Phc3 | S | | 860 | 0.052711016 | 0.210053539 | 0.221822619 | 0.169111603 | 0.01176908 | 0.519000221 | 0.010368134 | 0.001256649 | 0.106597691 | 0.817387371 |
| Cdkl5 | S | | 306 | 0.113233434 | 0.130636423 | 0.337930768 | 0.224697334 | 0.207294345 | 0.13879755 | 0.038983241 | 0.009194074 | 0.040223636 | 0.042242865 |
| Zc3hav1 | S | | 804 | 0.26606981 | 0.202244113 | 0.081960831 | -0.184108979 | -0.120283282 | 0.000970439 | 0.081779122 | 0.304011552 | 0.075424771 | 0.248742113 |
| Cobll1 | S | | 266 | 0.109676311 | 0.348774821 | 0.196701578 | 0.087025267 | -0.152073243 | 0.455300523 | 0.000915083 | 0.244220003 | 0.632902681 | 0.340243067 |
| Golga3 | S | | 381 | 0.070580921 | 0.082529391 | 0.387606757 | 0.317025836 | 0.305077366 | 0.665457486 | 0.163774214 | 0.0001838 | 0.141522688 | 0.001714829 |
| Plcd3 | S | | 492 | 0.090672232 | 0.146895411 | 0.210381826 | 0.119709594 | 0.063486416 | 0.11646637 | 0.008951125 | 0.002949693 | 0.065614523 | 0.162624352 |
| Pkn2 | T | | 815 | 0.083109746 | 0.189183437 | 0.241592158 | 0.158482411 | 0.05240872 | 0.29492696 | 0.019344527 | 0.002649832 | 0.080555781 | 0.338752382 |
| Cbfa2t2 | S | | 24 | 0.122632101 | 0.164885228 | 0.193694959 | 0.071062857 | 0.028809731 | 0.053223302 | 0.001837785 | 0.050354826 | 0.355355867 | 0.65658954 |
| Trp53bp1 | S | | 424 | 0.023877855 | 0.222746546 | 0.291862583 | 0.267984728 | 0.069116037 | 0.767599665 | 0.002665572 | 0.010798914 | 0.036005208 | 0.328911056 |
| Hnrnpul2 | S | | 186 | 0.079047923 | 0.26085 | 0.414351875 | 0.335303952 | 0.153501875 | 0.244528469 | 0.031739377 | 0.055424426 | 0.090974188 | 0.333773906 |
| Mysm1 | S | | 330 | 0.056765713 | 0.092810057 | 0.163764058 | 0.106998345 | 0.070954002 | 0.011680215 | 0.008232667 | 0.000136841 | 0.003583159 | 0.030045441 |
| AI597479 | S | | 189 | -0.044358986 | 0.23318553 | 0.33926594 | 0.383624926 | 0.10608041 | 0.043050617 | 0.007851554 | 0.000226522 | 0.002145305 | 0.065482071 |
| Arhgap23 | T | | 1391 | 0.11486243 | 0.293483935 | 0.129534255 | 0.014671824 | -0.163949681 | 0.063057316 | 0.004359501 | 0.096578864 | 0.821216521 | 0.068857818 |
| Epb41l3 | S | | 601 | 0.112693639 | 0.091848663 | 0.177098759 | 0.06440512 | 0.085250096 | 0.111371746 | 0.014439182 | 0.000139838 | 0.287200233 | 0.024381318 |
| Sipa1l3 | T | | 1694 | 0.101366173 | 0.141114045 | 0.359044107 | 0.257677934 | 0.217930062 | 0.110156315 | 0.09693885 | 0.007366586 | 0.048168534 | 0.054462102 |
| Sipa1l3 | S | | 94 | 0.049702596 | 0.165304237 | 0.25990706 | 0.210204463 | 0.094602823 | 0.396979042 | 0.000300435 | 0.020987268 | 0.032815498 | 0.171391832 |
| Fzr1 | S | | 36 | 0.010133899 | 0.79407902 | 1.113022721 | 1.102888822 | 0.318943701 | 0.96150501 | 0.232050508 | 0.231170673 | 0.236150287 | 0.718277566 |
| Fzr1 | S | | 39 | 0.010133899 | 0.79407902 | 1.113022721 | 1.102888822 | 0.318943701 | 0.96150501 | 0.232050508 | 0.231170673 | 0.236150287 | 0.718277566 |
| Fzr1 | S | | 40 | 0.010133899 | 0.79407902 | 1.113022721 | 1.102888822 | 0.318943701 | 0.96150501 | 0.232050508 | 0.231170673 | 0.236150287 | 0.718277566 |
| Pten | S | | 385 | 0.462027086 | 0.418363267 | 0.395415458 | -0.066611628 | -0.022947809 | 0.197083863 | 0.235929764 | 0.259938582 | 0.161120141 | 0.56796356 |
| Lysmd1 | S | | 192 | 0.174598476 | 0.296605743 | 0.075917045 | -0.098681431 | -0.220688698 | 0.003629806 | 0.01878331 | 0.314420828 | 0.215069006 | 0.052881914 |
| Myc Mycn | T | 58;58 | | 0.07772668 | 0.110114924 | 0.203408416 | 0.125681736 | 0.093293492 | 0.137483766 | 0.055674128 | 5.34341E-05 | 0.059463426 | 0.098597398 |
| Srrm1 | S | | 784 | 0.057935428 | 0.129971426 | 0.179288404 | 0.121352975 | 0.049316978 | 0.157900564 | 0.002382202 | 0.000571436 | 0.021825708 | 0.085568007 |
| Xpc | S | | 875 | 0.052856649 | 0.238978122 | 0.197302796 | 0.144446147 | -0.041675325 | 0.334557423 | 0.004637183 | 0.006546947 | 0.026399727 | 0.432653367 |
| Xpc | S | | 876 | 0.052856649 | 0.238978122 | 0.197302796 | 0.144446147 | -0.041675325 | 0.334557423 | 0.004637183 | 0.006546947 | 0.026399727 | 0.432653367 |
| Foxk2 | S | | 415 | 0.170226046 | 0.159610859 | 0.251453761 | 0.081227715 | 0.091842902 | 0.04057059 | 0.056139263 | 0.027126454 | 0.306029067 | 0.265555246 |
| Foxk2 | S | | 419 | 0.170226046 | 0.159610859 | 0.251453761 | 0.081227715 | 0.091842902 | 0.04057059 | 0.056139263 | 0.027126454 | 0.306029067 | 0.265555246 |
| Zmynd11 | S | | 447 | 0.092252461 | 0.127017217 | 0.179227729 | 0.086975268 | 0.052210512 | 0.193906845 | 0.000462873 | 0.00415633 | 0.215913679 | 0.167471972 |
| Nelfb | S | | 557 | 0.027965679 | 0.15504404 | 0.2830364 | 0.255070721 | 0.12799236 | 0.469900789 | 0.049761296 | 0.000229402 | 0.001777306 | 0.090194814 |
| Bod1l | S | | 480 | 0.006770273 | 0.111442776 | 0.237434709 | 0.230664436 | 0.125991933 | 0.817649225 | 0.005743676 | 3.02957E-05 | 0.00150141 | 0.006732906 |
| Bod1l | S | | 482 | 0.006770273 | 0.111442776 | 0.237434709 | 0.230664436 | 0.125991933 | 0.817649225 | 0.005743676 | 3.02957E-05 | 0.00150141 | 0.006732906 |
| Rnf168 | S | | 508 | 0.306206176 | 0.267586946 | 0.217557039 | -0.088649137 | -0.050029907 | 0.054295645 | 0.080978338 | 0.190028646 | 0.484152949 | 0.681449385 |
| Hexim1 | S | | 249 | 0.038223604 | 0.189826879 | 0.160995399 | 0.122771795 | -0.02883148 | 0.424648745 | 0.001023224 | 0.001200996 | 0.054163646 | 0.366555724 |
| Chd2 | S | | 207 | 0.098626766 | 0.137295412 | 0.262396673 | 0.163769906 | 0.12510126 | 0.385835148 | 0.002209231 | 0.006964613 | 0.201216692 | 0.047584046 |
| Chd2 | S | | 208 | 0.098626766 | 0.137295412 | 0.262396673 | 0.163769906 | 0.12510126 | 0.385835148 | 0.002209231 | 0.006964613 | 0.201216692 | 0.047584046 |
| Slc4a7 | T | | 951 | 0.203185765 | 0.092116468 | 0.13207812 | -0.071107644 | 0.039961652 | 0.004479014 | 0.135345148 | 0.002614495 | 0.126190929 | 0.441828108 |
| Mki67 | S | | 125 | 0.133821657 | 0.132165334 | 0.10997851 | -0.023843147 | -0.022186824 | 0.004951775 | 0.005831767 | 0.012235701 | 0.317563542 | 0.256864878 |
| Nfya | S | | 325 | -0.003269793 | 0.14660515 | 0.264406858 | 0.267676651 | 0.117801708 | 0.929487983 | 0.005720987 | 0.000182974 | 0.000277899 | 0.011070771 |
| Afap1 | S | | 662 | 0.001605683 | 0.216602997 | 0.218200935 | 0.216595252 | 0.001597939 | 0.97208438 | 0.010855889 | 0.000195214 | 0.014605962 | 0.97136306 |
| Tmpo | S | 67;67 | | 0.069704723 | 0.138206254 | 0.162156137 | 0.092451413 | 0.023949882 | 0.202320597 | 0.001809437 | 0.000642261 | 0.123532244 | 0.211129948 |
| Ppig | T | | 356 | 0.077722919 | 0.151542431 | 0.165964125 | 0.088241206 | 0.014421694 | 0.111304053 | 0.004370942 | 0.00149339 | 0.085119588 | 0.632713968 |
| Rbm33 | S | | 243 | 0.005024325 | 0.161371604 | 0.194566566 | 0.189542241 | 0.033194962 | 0.921257899 | 8.84506E-05 | 0.001742678 | 0.034186638 | 0.2335746 |
| Gatad2b | S | | 136 | 0.095142062 | 0.210215885 | 0.261394684 | 0.166252622 | 0.051178799 | 0.059913231 | 0.01970616 | 0.036104172 | 0.068221827 | 0.465323333 |
| Mki67 | S | | 2980 | 0.266957373 | 0.351091885 | 0.144853808 | -0.122103565 | -0.206238077 | 0.149288305 | 0.024050238 | 0.178901318 | 0.452840878 | 0.130712771 |
| Ice1 | S | | 1076 | 0.201895164 | 0.069667436 | 0.299235356 | 0.097340193 | 0.22956792 | 0.128242829 | 0.245696688 | 0.00146214 | 0.382690858 | 0.003625622 |
| Ice1 | S | | 1078 | 0.201895164 | 0.069667436 | 0.299235356 | 0.097340193 | 0.22956792 | 0.128242829 | 0.245696688 | 0.00146214 | 0.382690858 | 0.003625622 |
| Arhgef7 | S | | 830 | 0.168018101 | 0.413210025 | 0.116719948 | -0.051298153 | -0.296490077 | 0.142495902 | 0.005880651 | 0.405649981 | 0.65033578 | 0.08061961 |
| 4932438A13Rik | S | | 3894 | 0.133350055 | 0.219767328 | 0.092073414 | -0.04127664 | -0.127693914 | 0.013449188 | 0.000897056 | 0.257718996 | 0.575767606 | 0.153345627 |
| Exoc1 | T | | 471 | 0.271133504 | 0.104749293 | 0.253351182 | -0.017782322 | 0.148601888 | 0.018766231 | 0.333072995 | 0.02338625 | 0.355367299 | 0.130950209 |
| Ptpn13 | S | | 1620 | 0.063198677 | 0.229760574 | 0.215390013 | 0.152191336 | -0.014370561 | 0.638776449 | 0.003162629 | 0.006949983 | 0.310819975 | 0.786877016 |
| Phip | S | | 1783 | 0.063008494 | 0.124528766 | 0.186101785 | 0.123093291 | 0.061573019 | 0.007450384 | 0.002992888 | 0.008114016 | 0.035885794 | 0.144048099 |
| Smarca4 | S | | 1419 | 0.052798517 | 0.11560294 | 0.201340453 | 0.148541937 | 0.085737514 | 0.221275761 | 0.000866784 | 0.001153927 | 0.020240031 | 0.044643963 |
| Rb1cc1 | S | | 281 | 0.034026843 | 0.354785056 | 0.328847464 | 0.294820621 | -0.025937591 | 0.806235164 | 0.032857761 | 0.016128275 | 0.119577807 | 0.794086852 |
| Phf6 | S | | 155 | 0.006894298 | 0.114855166 | 0.165499929 | 0.15860563 | 0.050644763 | 0.809494752 | 0.000241804 | 2.10447E-05 | 0.010980598 | 0.02026858 |
| Kmt2a | S | | 349 | 0.177334099 | 0.189962923 | 0.145129786 | -0.032204313 | -0.044833138 | 0.03651777 | 0.023988781 | 0.022278744 | 0.600575571 | 0.456917835 |
| Marf1 | S | | 1680 | 0.030190079 | 0.281393093 | 0.28196204 | 0.251771962 | 0.000568947 | 0.664792403 | 0.011819443 | 0.011720888 | 0.029630789 | 0.993076081 |
| Nsrp1 | S | | 290 | 0.122314599 | 0.163475283 | 0.177769592 | 0.055454993 | 0.014294309 | 0.011394993 | 0.006913321 | 0.060268647 | 0.423928926 | 0.832421305 |
| AW554918 | S | | 474 | 0.017652656 | 0.129761114 | 0.287787174 | 0.270134518 | 0.15802606 | 0.79232873 | 0.097650615 | 2.88885E-05 | 0.034479913 | 0.077279228 |
| Slc9a1 | S | | 697 | -0.001208679 | 0.156547344 | 0.183166182 | 0.184374861 | 0.026618838 | 0.966239852 | 0.000509406 | 0.000161489 | 0.007309535 | 0.221478082 |
| Plekhg5 | S | | 938 | -0.008625275 | 0.236285919 | 0.254745205 | 0.26337048 | 0.018459286 | 0.795312182 | 0.00199704 | 0.004226194 | 0.013589488 | 0.725301734 |
| Cenpc1 | S | | 145 | 0.04543106 | 0.116191804 | 0.144748712 | 0.099317652 | 0.028556908 | 0.093088151 | 0.000771444 | 0.000212703 | 0.00716999 | 0.108851971 |
| Rbm12b1 Rbm12b2 | S | 256;256 | | 0.207003696 | 0.086139801 | 0.169755355 | -0.037248342 | 0.083615553 | 0.00029849 | 0.397629853 | 0.030950525 | 0.46164633 | 0.423162389 |
| Tle4 | S | | 208 | -0.012523502 | 0.218087256 | 0.324597374 | 0.337120875 | 0.106510118 | 0.888240171 | 0.000596031 | 0.022450562 | 0.028729066 | 0.242709742 |
| Cavin1 | T | | 40 | -0.006705249 | 0.150681384 | 0.238899609 | 0.245604858 | 0.088218225 | 0.675805093 | 4.6747E-05 | 0.003280969 | 0.010576419 | 0.071999949 |
| Bclaf1 | S | | 121 | 0.174216696 | 0.256466589 | 0.261596858 | 0.087380161 | 0.005130269 | 0.065903713 | 0.084995387 | 0.059760278 | 0.396008419 | 0.968561781 |
| Tnrc18 | S | | 1847 | 0.024322531 | 0.274720854 | 0.252623876 | 0.228301345 | -0.022096978 | 0.748152523 | 0.017253665 | 0.00339769 | 0.039824682 | 0.764436738 |
| Mki67 | S | | 1836 | 0.100276166 | 0.134564796 | 0.166834023 | 0.066557857 | 0.032269227 | 0.025083816 | 0.01886464 | 0.002538916 | 0.035186912 | 0.391567126 |
| Brca1 | S | | 710 | 0.091530448 | 0.106659779 | 0.240187076 | 0.148656627 | 0.133527296 | 0.014806441 | 0.134491323 | 0.001442829 | 0.016412277 | 0.085811885 |
| Ing4 | S | | 150 | -0.111934379 | 0.282639743 | 0.435220702 | 0.547155081 | 0.152580958 | 0.045200914 | 0.00169643 | 0.003172721 | 0.000209506 | 0.020026625 |
| Mboat1 | S | | 483 | 0.141791019 | 0.235013593 | 0.158216698 | 0.016425679 | -0.076796895 | 0.029875709 | 0.0025353 | 0.288129009 | 0.897910575 | 0.561351535 |
| Kmt2b | S | | 1098 | 0.175238653 | 0.2007964 | 0.181481837 | 0.006243185 | -0.019314562 | 0.019350175 | 0.155204128 | 0.012683012 | 0.910234059 | 0.86042846 |
| Erbin | S | | 599 | 0.058834007 | 0.106284678 | 0.237251678 | 0.178417671 | 0.130967 | 0.056960114 | 0.006306017 | 0.002607374 | 0.019628507 | 0.033348811 |
| Erbin | S | | 600 | 0.058834007 | 0.106284678 | 0.237251678 | 0.178417671 | 0.130967 | 0.056960114 | 0.006306017 | 0.002607374 | 0.019628507 | 0.033348811 |
| Mier1 | S | | 159 | -0.00188933 | 0.158982671 | 0.220773237 | 0.222662567 | 0.061790567 | 0.968070611 | 0.039159696 | 6.61537E-06 | 0.024957861 | 0.246306315 |
| Rpain | S | | 18 | -0.029889101 | 0.226779318 | 0.197301054 | 0.227190155 | -0.029478264 | 0.5052668 | 0.000163385 | 0.003097423 | 0.006162661 | 0.426358236 |
| Zfp768 | S | | 23 | 0.135159097 | 0.2153214 | 0.407253932 | 0.272094835 | 0.191932532 | 0.163373765 | 0.140089402 | 0.031698249 | 0.063529366 | 0.197726211 |
| Zfp768 | S | | 27 | 0.135159097 | 0.2153214 | 0.407253932 | 0.272094835 | 0.191932532 | 0.163373765 | 0.140089402 | 0.031698249 | 0.063529366 | 0.197726211 |
| Fbxw8 | S | | 84 | 0.071477032 | 0.246788649 | 0.115761055 | 0.044284023 | -0.131027594 | 0.169300037 | 0.000159164 | 0.111188633 | 0.493510249 | 0.089851495 |
| Kmt2a | T | | 3024 | 0.057480357 | 0.210893594 | 0.395065756 | 0.337585398 | 0.184172162 | 0.620494836 | 0.040541594 | 0.010538835 | 0.050726652 | 0.108031127 |
| Srsf10 | S | | 156 | 0.171436961 | 0.125329906 | 0.144524768 | -0.026912193 | 0.019194862 | 0.006731944 | 0.052069371 | 0.012745453 | 0.533243061 | 0.700473607 |
| Adgra2 | S | | 1104 | 0.020840773 | 0.148868732 | 0.200147684 | 0.179306912 | 0.051278953 | 0.579514923 | 0.006132143 | 0.000110774 | 0.01386508 | 0.164349689 |
| Iqgap1 | T | | 178 | 0.190544268 | 0.260435264 | 0.174551425 | -0.015992843 | -0.085883839 | 0.135711948 | 0.003067775 | 0.262956269 | 0.916152162 | 0.538361743 |
| Rnf168 | S | | 506 | 0.198244927 | 0.312865697 | 0.144754921 | -0.053490006 | -0.168110776 | 0.043782927 | 0.024904347 | 0.224955764 | 0.611426908 | 0.203818449 |
| Fnbp4 | S | | 439 | 0.023004839 | 0.260933797 | 0.173140691 | 0.150135852 | -0.087793106 | 0.751591208 | 3.33621E-05 | 0.119935302 | 0.182768045 | 0.335717617 |
| Pacs2 | S | | 387 | -0.138849254 | 0.516670181 | 0.440607307 | 0.579456561 | -0.076062874 | 0.000119509 | 0.000315744 | 0.011250358 | 0.007922641 | 0.314542649 |
| Mecp2 | S | | 229 | 0.174064609 | 0.165653544 | 0.182052204 | 0.007987594 | 0.01639866 | 0.03324519 | 0.032346031 | 0.029195875 | 0.876786753 | 0.706260634 |
| Dgcr8 | S | | 271 | 0.134118457 | 0.118672907 | 0.12822062 | -0.005897837 | 0.009547713 | 0.011465787 | 0.022908629 | 0.002314071 | 0.851984907 | 0.77540657 |
| Ebag9 | S | | 36 | 0.017672304 | 0.18118001 | 0.212953328 | 0.195281024 | 0.031773319 | 0.325404656 | 0.000782994 | 0.00588315 | 0.008691643 | 0.374150437 |
| Srrm2 | S | | 726 | 0.295581397 | 0.029316753 | 0.317858552 | 0.022277155 | 0.288541799 | 0.092287975 | 0.848142898 | 0.001821596 | 0.861153431 | 0.146346701 |

| | | | | | | | | | | | | |
|---|---|---|---|---|---|---|---|---|---|---|---|---|
| Fhdc1 | S | 568 | 0.336164507 | 0.119049488 | 0.224053294 | -0.112111213 | 0.105003806 | 0.030482172 | 0.070559691 | 0.127333146 | 0.411651844 | 0.388474487 |
| Ece1 | T | 311 | -0.145774184 | 0.210762104 | 0.397750151 | 0.543524334 | 0.186988047 | 0.05683323 | 1.39632E-05 | 0.00561067 | 0.000641841 | 0.033398851 |
| Sash1 | S | 692 | -0.067065121 | 0.351412669 | 0.256806001 | 0.323871122 | -0.094606668 | 0.194928512 | 0.009773788 | 0.0006061 | 0.001130288 | 0.253679735 |
| Dbndd2 | T | 137 | 0.063403755 | 0.318521066 | 0.247853153 | 0.184449397 | -0.070667913 | 0.655755452 | 0.001340794 | 0.132129016 | 0.313636549 | 0.586137881 |
| Agfg1 | S | 181 | 0.07468466 | 0.12361263 | 0.078005808 | 0.003321148 | -0.045606822 | 0.004282505 | 0.000249097 | 0.00306558 | 0.710023161 | 0.019468418 |
| Xrn2 | S | 475 | 0.165480916 | 0.171208179 | 0.092292646 | -0.07318827 | -0.078915533 | 0.003866144 | 0.0029664 | 0.235512575 | 0.319012908 | 0.290549219 |
| Atp2b1 | S | 1178 | 0.08803405 | 0.183697082 | 0.251247286 | 0.163213237 | 0.067550204 | 0.290724662 | 0.002192504 | 0.031528611 | 0.117137478 | 0.327715459 |
| Daxx | S | 626 | 0.145466875 | 0.279837288 | 0.06156616 | -0.083900716 | -0.218271129 | 0.009977078 | 0.005633479 | 0.234521284 | 0.092604402 | 0.020287386 |
| Pnp | S | 289 | 0.330442506 | 0.287926329 | 0.227156433 | -0.103286073 | -0.060769895 | 0.173447494 | 0.074284232 | 0.119209437 | 0.617930011 | 0.64798176 |
| Prr12 | S | 859 | 0.15319799 | 0.178838435 | 0.277379597 | 0.124181607 | 0.098541162 | 0.126928467 | 0.062297945 | 0.0182716 | 0.235551138 | 0.301016927 |
| Gcfc2 | S | 418 | 0.061739356 | 0.081519743 | 0.161937401 | 0.100198045 | 0.080417658 | 0.018512425 | 0.00860672 | 0.000134268 | 0.000537025 | 0.011686045 |
| Gcfc2 | S | 419 | 0.061739356 | 0.081519743 | 0.161937401 | 0.100198045 | 0.080417658 | 0.018512425 | 0.00860672 | 0.000134268 | 0.000537025 | 0.011686045 |
| Ank3 | S | 27 | 0.072794409 | 0.17861125 | 0.118298253 | 0.045503844 | -0.060312998 | 0.048156133 | 0.000796829 | 0.012338906 | 0.154076358 | 0.086471251 |
| Jund | S | 249 | 0.076488507 | 0.105184619 | 0.235525876 | 0.15903737 | 0.130341257 | 0.018380048 | 0.003781269 | 0.014240028 | 0.053694123 | 0.086799842 |
| Lin37 | S | 135 | 0.073256452 | 0.194692074 | 0.180731005 | 0.107474552 | -0.01396107 | 0.306616396 | 0.000776939 | 0.01961466 | 0.18648773 | 0.77316391 |
| Itgb3bp | S | 19 | 0.145508607 | 0.132733383 | 0.242273176 | 0.096764569 | 0.109539792 | 0.01278675 | 0.082611762 | 0.024920608 | 0.226598184 | 0.213431092 |
| Rai14 | S | 293 | 0.009300392 | 0.218290847 | 0.213317046 | 0.204016653 | -0.004973802 | 0.853147072 | 0.001861915 | 0.003240508 | 0.008223643 | 0.902421828 |
| Setd1a | S | 1160 | -0.00814934 | 0.08463881 | 0.243631079 | 0.251780419 | 0.158992269 | 0.794376793 | 0.001449173 | 2.24648E-05 | 0.003632631 | 0.000767222 |
| Bcl2l13 | T | 342 | -0.003792633 | 0.393079153 | 0.425911486 | 0.429704119 | 0.032832332 | 0.973550155 | 0.04792207 | 0.028342928 | 0.033026255 | 0.828533478 |
| Ddx46 | S | 105 | 0.282697155 | 0.239787403 | 0.234875543 | -0.047821612 | -0.00491186 | 0.071774028 | 0.058266342 | 0.188495766 | 0.772339819 | 0.97468793 |
| Taf1 | S | 1820 | 0.116280281 | 0.098914297 | 0.227209754 | 0.110929473 | 0.128295457 | 0.008604819 | 0.009495165 | 0.03020336 | 0.156665984 | 0.127800892 |
| Snrnp200 | S | 2135 | 0.102561153 | 0.212508728 | 0.165121857 | 0.062560704 | -0.047386871 | 0.29470991 | 0.000331865 | 0.074587849 | 0.532779351 | 0.479329946 |
| Snrnp200 | T | 2131 | 0.102561153 | 0.212508728 | 0.165121857 | 0.062560704 | -0.047386871 | 0.29470991 | 0.000331865 | 0.074587849 | 0.532779351 | 0.479329946 |
| Gpatch8 | S | 980 | 0.578655577 | 0.101883999 | 0.163415298 | -0.415240279 | 0.061531298 | 0.300243951 | 0.007768019 | 0.02554395 | 0.425071239 | 0.186693739 |
| Cyb5r3 | T | 171 | 0.091226638 | 0.353288604 | 0.185548958 | 0.09432232 | -0.167739646 | 0.237920575 | 0.018555208 | 0.043832581 | 0.279384157 | 0.114205016 |
| Iffo2 | S | 328 | 0.170020526 | 0.209789695 | 0.207739287 | 0.037718762 | -0.002050407 | 0.122071854 | 0.050667118 | 0.017036672 | 0.682975846 | 0.980028194 |
| Susd6 | S | 283 | 0.023164758 | 0.196277293 | 0.281392243 | 0.258227485 | 0.08511495 | 0.684750062 | 0.023851176 | 0.001416735 | 0.003688837 | 0.203719719 |
| Eea1 Rab11fip3 | S | 1265;765 | 0.091996354 | 0.219769593 | 0.18529141 | 0.093295056 | -0.034478183 | 0.096703451 | 0.002302564 | 0.077388292 | 0.283421745 | 0.65793699 |
| Scap | S | 934 | 0.089481343 | 0.063162808 | 0.15618313 | 0.066701787 | 0.093020322 | 0.021583425 | 0.009056711 | 0.000142152 | 0.080956185 | 8.54069E-06 |
| Med26 | S | 435 | 0.157984202 | 0.114715699 | 0.230822363 | 0.072838162 | 0.116106665 | 0.112089266 | 0.040774775 | 0.004676268 | 0.392193812 | 0.062552509 |
| Cyb5b | S | 33 | 0.15071868 | 0.029927756 | 0.217513714 | 0.066795034 | 0.187585958 | 0.00508309 | 0.456849578 | 0.000667359 | 0.052240308 | 0.001948658 |
| Dpysl2 | T | 514 | -0.058986239 | 0.45401479 | 0.213453379 | 0.272439618 | -0.240561411 | 0.212088409 | 0.001768626 | 0.020211111 | 0.011420537 | 0.021675504 |
| Sipa1l1 | S | 1370 | 0.12035919 | 0.038401228 | 0.216035505 | 0.095676315 | 0.177634277 | 0.006536753 | 0.370138869 | 0.000268801 | 0.008365277 | 0.010887462 |
| Hivep3 | S | 1922 | 0.001021658 | 0.201643574 | 0.191550595 | 0.190528937 | -0.010092979 | 0.98809899 | 0.001802216 | 0.000751627 | 0.072469358 | 0.744493594 |
| Zbtb18 | S | 333 | 0.040872802 | 0.205119506 | 0.259468773 | 0.218595972 | 0.054349267 | 0.379360483 | 0.000990439 | 0.040089134 | 0.065336136 | 0.506545766 |
| Ncor2 | S | 1929 | 0.19481856 | 0.051918116 | 0.198032803 | 0.003214243 | 0.146114687 | 0.004067331 | 0.531250821 | 0.002593112 | 0.919615115 | 0.147926521 |
| Srsf4 | S | 407 | 0.084063805 | 0.428892726 | 0.223855711 | 0.139791906 | -0.205037014 | 0.502617145 | 0.004529627 | 0.271113082 | 0.459749837 | 0.296158543 |
| Rtp4 | S | 179 | 0.016061424 | 0.149015977 | 0.536756769 | 0.520695345 | 0.387740792 | 0.882911722 | 0.203110984 | 0.002930205 | 0.003126434 | 0.005076012 |
| Ppig | S | 751 | 0.096414892 | 0.182079163 | 0.201975723 | 0.105560831 | 0.019896561 | 0.107267295 | 0.025353136 | 0.005881064 | 0.104352797 | 0.722990987 |
| Luc7l2 | S | 281 | 0.136463064 | 0.271860943 | 0.243217501 | 0.106754436 | -0.028643443 | 0.233756626 | 0.029355326 | 0.043135023 | 0.167813807 | 0.487981791 |
| Luc7l2 | S | 283 | 0.136463064 | 0.271860943 | 0.243217501 | 0.106754436 | -0.028643443 | 0.233756626 | 0.029355326 | 0.043135023 | 0.167813807 | 0.487981791 |
| Luc7l2 | S | 285 | 0.136463064 | 0.271860943 | 0.243217501 | 0.106754436 | -0.028643443 | 0.233756626 | 0.029355326 | 0.043135023 | 0.167813807 | 0.487981791 |
| Ssbp2 | S | 360 | -0.014104348 | 0.133922143 | 0.546346251 | 0.560450598 | 0.412424107 | 0.921315262 | 0.269884346 | 0.001224762 | 0.017650072 | 0.014284358 |
| Acin1 | S | 454 | 0.202780831 | 0.234625902 | 0.091822215 | -0.110958616 | -0.142803688 | 0.023744766 | 0.00510062 | 0.33950062 | 0.279779742 | 0.178231599 |
| Cenpc1 | S | 517 | 0.014643542 | 0.190810466 | 0.26311116 | 0.248467618 | 0.072300693 | 0.764709561 | 0.000743161 | 0.013619107 | 0.017875552 | 0.275812184 |
| Larp1 | S | 604 | 0.057121155 | 0.174300568 | 0.113208633 | 0.056087477 | -0.061091936 | 0.11599965 | 0.00019709 | 0.009285909 | 0.139954336 | 0.084920773 |
| Rbm25 | S | 672 | 0.097384065 | 0.084161518 | 0.162313135 | 0.06492907 | 0.078151617 | 0.008941197 | 0.016039117 | 0.001459971 | 0.059047733 | 0.042456179 |
| Chaf1b | S | 417 | -0.021839968 | 0.166622229 | 0.162955155 | 0.184795123 | -0.003667074 | 0.443541177 | 0.000196947 | 8.30733E-05 | 0.006279238 | 0.832396224 |
| Cybc1 | S | 178 | 0.021011075 | 0.156668693 | 0.285128195 | 0.26411712 | 0.128459502 | 0.539778287 | 0.013547246 | 0.001507379 | 0.01082416 | 0.052178355 |
| Arid4a | S | 275 | 0.076685831 | 0.180149899 | 0.282939339 | 0.206253509 | 0.10278944 | 0.178977798 | 0.016184716 | 0.017639144 | 0.063607003 | 0.232217114 |
| Arid4a | S | 276 | 0.076685831 | 0.180149899 | 0.282939339 | 0.206253509 | 0.10278944 | 0.178977798 | 0.016184716 | 0.017639144 | 0.063607003 | 0.232217114 |
| Arid4a | S | 277 | 0.076685831 | 0.180149899 | 0.282939339 | 0.206253509 | 0.10278944 | 0.178977798 | 0.016184716 | 0.017639144 | 0.063607003 | 0.232217114 |
| Pgrmc1 | S | 181 | 0.191201307 | 0.131013486 | 0.147368946 | -0.043832361 | 0.01635546 | 0.260005129 | 0.000318171 | 0.020074982 | 0.759625308 | 0.651657198 |
| Bcr | S | 202 | 0.015498681 | 0.177680522 | 0.222329622 | 0.206830942 | 0.044649101 | 0.661673122 | 0.002843123 | 0.001664869 | 0.007938034 | 0.293409325 |
| Bcr | S | 203 | 0.015498681 | 0.177680522 | 0.222329622 | 0.206830942 | 0.044649101 | 0.661673122 | 0.002843123 | 0.001664869 | 0.007938034 | 0.293409325 |
| Cpsf7 | S | 429 | 0.076424057 | 0.286870203 | 0.487683405 | 0.411259348 | 0.200813202 | 0.550828376 | 0.068706636 | 0.05144094 | 0.071800055 | 0.282876912 |
| Zfp638 | S | 560 | 0.045019195 | 0.141696148 | 0.176456831 | 0.131437636 | 0.034760683 | 0.360604491 | 0.017712377 | 4.99121E-05 | 0.06374242 | 0.330740185 |
| Zfp638 | S | 562 | 0.045019195 | 0.141696148 | 0.176456831 | 0.131437636 | 0.034760683 | 0.360604491 | 0.017712377 | 4.99121E-05 | 0.06374242 | 0.330740185 |
| Tcf20 | T | 1699 | -0.003936836 | 0.154759413 | 0.191268743 | 0.195205579 | 0.03650933 | 0.898137082 | 0.0007255 | 0.000272061 | 0.003577001 | 0.041125722 |
| Cic | T | 1342 | -0.014940674 | 0.225980668 | 0.426580787 | 0.441521461 | 0.200600119 | 0.904663399 | 0.054859947 | 0.004516722 | 0.022514183 | 0.003461254 |
| Zfp219 | S | 702 | 0.03443153 | 0.137944927 | 0.234539141 | 0.200107611 | 0.096594214 | 0.054074148 | 0.001665157 | 0.010662157 | 0.024616948 | 0.084749699 |
| Cstf2 | S | 527 | 0.029667646 | 0.101615603 | 0.16701266 | 0.137345014 | 0.065397057 | 0.406272013 | 0.007780812 | 5.51188E-06 | 0.032404268 | 0.04431986 |
| Nsd2 | S | 437 | 0.349575287 | 0.113802175 | -0.023982389 | -0.373557677 | -0.137784565 | 0.000119458 | 0.063142619 | 0.896317663 | 0.142083469 | 0.481752838 |
| Sipa1l1 | S | 1533 | 0.123244499 | 0.108653907 | 0.321563309 | 0.19831881 | 0.212909402 | 0.179662697 | 0.135667868 | 0.003171188 | 0.067472672 | 0.029193364 |
| Brca1 | T | 1157 | 0.147964146 | 0.148251289 | 0.148889461 | 0.000925316 | 0.000638173 | 0.126102841 | 0.007728935 | 0.00529875 | 0.989752533 | 0.981294913 |
| Purb | S | 316 | 0.071741333 | 0.218983137 | 0.110230289 | 0.038488956 | -0.108752847 | 0.069645627 | 0.000113584 | 0.175412853 | 0.57247265 | 0.180379292 |
| Cttn | S | 407 | -0.029747867 | 0.157084964 | 0.21135909 | 0.241106957 | 0.054274126 | 0.065396712 | 0.000154257 | 9.88299E-05 | 0.000191702 | 0.033817428 |
| Numbl | S | 411 | -0.004447595 | 0.254297042 | 0.238020994 | 0.242468589 | -0.016276048 | 0.932337875 | 0.002382328 | 0.008652365 | 0.02038643 | 0.748592674 |
| Ctnna1 Ctnna2 | T | 654;653 | 0.103965362 | 0.188179375 | 0.190031046 | 0.086065685 | 0.001851671 | 0.075540679 | 0.009489643 | 0.026680349 | 0.189661043 | 0.974354593 |
| Prrc2c | T | 2634 | 0.154603432 | 0.086921183 | 0.217251324 | 0.062647892 | 0.130330141 | 0.010059522 | 0.407000251 | 0.00205056 | 0.111990265 | 0.246046856 |
| Micall2 | S | 768 | 0.291115108 | 0.171613121 | 0.047014968 | -0.24410014 | -0.124598153 | 0.002016701 | 0.021582542 | 0.714717536 | 0.142186071 | 0.356065624 |
| Prr14 | S | 124 | 0.079706838 | 0.193235285 | 0.223549257 | 0.143842419 | 0.030313972 | 0.254880832 | 0.008743476 | 0.012036306 | 0.096215341 | 0.610336943 |
| Map2 | S | 1796 | 0.403756206 | 0.427108235 | 0.114182718 | -0.289573488 | -0.312925517 | 0.18857637 | 0.039474918 | 0.490514401 | 0.319439634 | 0.145995693 |
| Pcnt | S | 2806 | 0.038332769 | 0.243763324 | 0.170607368 | 0.132274599 | -0.073155956 | 0.284822671 | 0.000546191 | 0.053485058 | 0.119077188 | 0.289588364 |
| Slbp | S | 23 | 0.022725051 | 0.171586729 | 0.137524414 | 0.114799363 | -0.034062315 | 0.392402319 | 5.81869E-05 | 0.005939786 | 0.016852847 | 0.259038483 |
| Htatsf1 | S | 621 | 0.056588435 | 0.070738637 | 0.200281037 | 0.143692602 | 0.1295424 | 0.040450711 | 0.220661622 | 9.08455E-06 | 0.006329849 | 0.085404899 |
| Cenpc1 | S | 404 | 0.155575054 | 0.131492784 | 0.294410173 | 0.13883512 | 0.16291739 | 0.026524303 | 0.247891135 | 0.016699637 | 0.146892954 | 0.192350286 |
| Insyn1 | S | 242 | 0.014667826 | 0.38084392 | 0.174975048 | 0.160307222 | -0.205868872 | 0.716733981 | 0.003836943 | 0.039685957 | 0.076299278 | 0.045484637 |
| Pola2 | T | 130 | 0.216028453 | 0.193553589 | 0.168348434 | -0.047680019 | -0.025205156 | 0.036875102 | 0.049743501 | 0.069455282 | 0.570784544 | 0.757192928 |
| Clip1 | S | 1314 | 0.165691572 | 0.240706579 | 0.078247362 | -0.08744421 | -0.162459218 | 0.008809172 | 0.005693851 | 0.37885042 | 0.319752611 | 0.120118768 |
| Mre11a | S | 686 | 0.008531269 | 0.131126815 | 0.137277953 | 0.128746684 | 0.006151138 | 0.632433066 | 8.80971E-05 | 0.000137294 | 0.003183549 | 0.533793702 |
| Casp8ap2 | S | 1844 | 0.206306114 | 0.135571015 | 0.096810657 | -0.109495457 | -0.038760358 | 0.001314228 | 0.012508902 | 0.320198209 | 0.271399586 | 0.656669288 |
| Bclaf1 | S | 198 | 0.024271972 | 0.200302774 | 0.1698847 | 0.145612728 | -0.030418074 | 0.565715015 | 0.00016348 | 0.016802769 | 0.036913937 | 0.480608576 |
| Mga | T | 1486 | 0.125374728 | 0.160811747 | 0.111354462 | -0.014020266 | -0.049457285 | 0.020162667 | 0.000245821 | 0.210538458 | 0.851798833 | 0.512463308 |
| Map7d1 | S | 401 | 0.266141342 | 0.154672437 | 0.185944706 | -0.080196636 | 0.031272269 | 0.004398521 | 0.185929205 | 0.186282189 | 0.502924771 | 0.816443155 |
| Srrm2 | S | 1650 | 0.088851236 | 0.158992086 | 0.133736822 | 0.044885586 | -0.025255264 | 0.012155194 | 0.000207107 | 0.174686364 | 0.57353664 | 0.741020114 |
| Epb41l3 | T | 64 | 0.047359869 | 0.127993024 | 0.243028557 | 0.195668688 | 0.115035534 | 0.303117325 | 0.004420807 | 0.002703255 | 0.012626609 | 0.047492283 |
| Ttc33 | S | 200 | -0.035778722 | 0.284716938 | 0.369858228 | 0.40563695 | 0.08514129 | 0.801585096 | 0.013283028 | 0.014835533 | 0.057885262 | 0.400314308 |
| Palm | S | 345 | 0.085197468 | 0.177420918 | 0.192117904 | 0.106920436 | 0.014696986 | 0.201369122 | 0.008837639 | 0.00658791 | 0.106408946 | 0.709084809 |
| Mki67 | T | 1150 | 0.302076895 | 0.222920819 | 0.102177407 | -0.199899488 | -0.120743412 | 0.02822423 | 0.052436041 | 0.179749051 | 0.098117986 | 0.208367403 |
| Pnisr | S | 211 | 0.015404824 | 0.133810892 | 0.178165988 | 0.162761165 | 0.044355096 | 0.25926976 | 8.09335E-05 | 0.003644746 | 0.011979862 | 0.153870578 |
| Fam117b | S | 10 | 0.452660962 | 0.194153966 | -0.081635033 | -0.534295996 | -0.275789 | 0.004857095 | 0.017200865 | 0.688562546 | 0.073745266 | 0.248428667 |
| Osbp | T | 375 | 0.251469487 | 0.183288664 | 0.064603878 | -0.186865609 | -0.118684786 | 0.016315011 | 0.007687971 | 0.238115295 | 0.05584418 | 0.017441284 |
| Tcof1 | S | 852 | 0.081561779 | 0.142110499 | 0.140872249 | 0.05931047 | -0.00123825 | 0.02919996 | 0.004583203 | 0.005652769 | 0.124153978 | 0.970651078 |

| | | | | | | | | | | | | |
|---|---|---|---|---|---|---|---|---|---|---|---|---|
| Srpr | S | 295 | 0.058632251 | 0.103791881 | 0.207606211 | 0.14897396 | 0.10381433 | 0.095179702 | 0.022419407 | 0.000443499 | 0.005962287 | 0.02118665 |
| Srpr | S | 296 | 0.058632251 | 0.103791881 | 0.207606211 | 0.14897396 | 0.10381433 | 0.095179702 | 0.022419407 | 0.000443499 | 0.005962287 | 0.02118665 |
| Srpr | S | 297 | 0.058632251 | 0.103791881 | 0.207606211 | 0.14897396 | 0.10381433 | 0.095179702 | 0.022419407 | 0.000443499 | 0.005962287 | 0.02118665 |
| Swap70 | S | 312 | -0.04685528 | 0.36329392 | 0.294158846 | 0.341014126 | -0.069135074 | 0.395973839 | 0.010866454 | 0.008026317 | 0.006913386 | 0.437782037 |
| Trpm7 | S | 1858 | 0.252314031 | 0.143640169 | 0.174559446 | -0.077754585 | 0.030919277 | 0.027509262 | 0.092187588 | 0.04842928 | 0.314602905 | 0.458552357 |
| Pola2 | T | 115 | 0.217890623 | 0.221635498 | 0.2628983 | 0.045007676 | 0.041262801 | 0.042307539 | 0.099146273 | 0.1409817 | 0.741029808 | 0.788196165 |
| Spp1 | S | 288 | 0.052713117 | 0.178699732 | 0.098540519 | 0.045827402 | -0.080159213 | 0.047445971 | 0.000144139 | 0.024359218 | 0.165375559 | 0.04365106 |
| Cdk13 | T | 1147 | 0.10903433 | 0.160506127 | 0.263321269 | 0.154286939 | 0.102815143 | 0.241872099 | 0.043300633 | 0.006154555 | 0.113993643 | 0.02371694 |
| Dxo | T | 392 | 0.05313679 | 0.333891674 | 0.28493974 | 0.231802951 | -0.048951933 | 0.413063361 | 0.008088985 | 0.122102898 | 0.166859766 | 0.721509599 |
| Srrt | S | 543 | 0.034236767 | 0.159613137 | 0.20524229 | 0.171005523 | 0.045629153 | 0.452723915 | 0.00412771 | 0.001476073 | 0.008412408 | 0.171689522 |
| Hnrnpul2 | T | 163 | -0.003603648 | 0.149246091 | 0.260666802 | 0.26427045 | 0.111420711 | 0.866206353 | 0.000166642 | 0.006793899 | 0.009823925 | 0.071362246 |
| Nup93 | S | 52 | 0.0143722 | 0.204457787 | 0.126868589 | 0.112496389 | -0.077589198 | 0.639049898 | 0.000207603 | 0.002942181 | 0.005007331 | 0.008103948 |
| Ncoa1 | S | 22 | -0.042315804 | 0.165859287 | 0.26632823 | 0.308644034 | 0.100468943 | 0.630536364 | 0.006026582 | 0.000280862 | 0.050833829 | 0.065369556 |
| Chtf18 | S | 91 | 0.117860021 | 0.108615332 | 0.132539353 | 0.014679332 | 0.023924022 | 0.004531567 | 0.049630016 | 0.002687216 | 0.541710932 | 0.554285498 |
| Srrm2 | S | 1649 | 0.040627348 | 0.136432324 | 0.163894314 | 0.123266966 | 0.02746199 | 0.216767002 | 2.58333E-05 | 0.019475375 | 0.03266572 | 0.448668031 |
| Spcs2 | S | 30 | 0.328506548 | 0.116631576 | 0.264037781 | -0.064468766 | 0.147406205 | 0.020458184 | 0.24501778 | 0.142533092 | 0.676366565 | 0.35509337 |
| Wbp1l | T | 245 | 0.124211223 | 0.078704468 | 0.272752092 | 0.148540869 | 0.194047624 | 0.035535988 | 0.296904701 | 0.002047988 | 0.036647719 | 0.051614549 |
| Supt6 | S | 1703 | 0.144813682 | 0.168973956 | 0.302088366 | 0.157274684 | 0.13311441 | 0.116757868 | 0.04296064 | 0.045752631 | 0.211633112 | 0.260944189 |
| Anapc4 | S | 779 | 0.070656554 | 0.104745116 | 0.199045252 | 0.128388698 | 0.094300135 | 0.107875286 | 0.000366025 | 0.011351827 | 0.037838318 | 0.088588704 |
| Clip1 | S | 199 | 0.173951508 | 0.238799778 | 0.087357389 | -0.086594118 | -0.151442389 | 0.086541978 | 0.001881044 | 0.20134135 | 0.327441822 | 0.051707415 |
| Pcdh18 | S | 890 | 0.133445549 | 0.133287175 | 0.323461013 | 0.190015464 | 0.190173837 | 0.07012593 | 0.219035033 | 0.011286086 | 0.08403805 | 0.122111837 |
| Pik3r1 | Y | 14 | 0.044380157 | 0.188247857 | 0.21025861 | 0.165878453 | 0.022010753 | 0.409419619 | 0.038459532 | 0.000487826 | 0.03398596 | 0.710589674 |
| Tcof1 | S | 853 | 0.070071239 | 0.167357878 | 0.171932451 | 0.101861212 | 0.004574573 | 0.109131948 | 0.001916222 | 0.018199853 | 0.095400909 | 0.915316801 |
| Dhx29 | S | 579 | 0.243046327 | 0.238761275 | 0.041460836 | -0.20158549 | -0.197300439 | 0.025708715 | 0.003219826 | 0.652302694 | 0.097079377 | 0.10903986 |
| Pdlim2 | T | 138 | -0.075979377 | 0.220657651 | 0.307728158 | 0.383707535 | 0.087070507 | 0.084482713 | 0.001747945 | 0.000691608 | 0.000502304 | 0.064162272 |
| Rbl1 | T | 385 | 0.133566664 | 0.213472049 | 0.084849667 | -0.048716997 | -0.128622382 | 0.045764119 | 0.000941792 | 0.174032016 | 0.416060821 | 0.075469208 |
| Ctnnd1 | T | 177 | 0.257637287 | 0.31360866 | 0.129586455 | -0.128050832 | -0.184022205 | 0.149758013 | 0.009757357 | 0.405292762 | 0.482628076 | 0.245222741 |
| Hdgf | S | 202 | 0.009330174 | 0.261638858 | 0.203917949 | 0.194587775 | -0.057720909 | 0.837601504 | 0.001109943 | 0.024119557 | 0.047041684 | 0.354018306 |
| Coil | T | 169 | 0.168700311 | 0.201143153 | 0.211035315 | 0.042335004 | 0.009892162 | 0.179535803 | 0.023451425 | 0.035649074 | 0.698709891 | 0.885805997 |
| Ythdc1 | S | 152 | 0.139285562 | 0.209682889 | 0.166735949 | 0.027450387 | -0.04294694 | 0.046401915 | 0.002208501 | 0.284442427 | 0.839994857 | 0.747532129 |
| Dgcr8 | S | 275 | 0.118794603 | 0.116889942 | 0.12317061 | 0.004376007 | 0.006280668 | 0.010217251 | 0.023639568 | 0.00307072 | 0.881522991 | 0.855332929 |
| Brca1 | S | 1152 | 0.054461191 | 0.109063889 | 0.229047034 | 0.174585844 | 0.119983145 | 0.179074671 | 0.032475909 | 0.000472636 | 0.008082041 | 0.031382138 |
| Brca1 | S | 1154 | 0.054461191 | 0.109063889 | 0.229047034 | 0.174585844 | 0.119983145 | 0.179074671 | 0.032475909 | 0.000472636 | 0.008082041 | 0.031382138 |
| Parn | S | 575 | 0.05850078 | 0.21555206 | 0.271526149 | 0.213025368 | 0.055974089 | 0.065393247 | 0.044393555 | 0.021205305 | 0.04632073 | 0.503912567 |
| Rab11fip5 | S | 188 | 0.29842904 | 0.110473206 | 0.098216759 | -0.20021228 | -0.012256447 | 0.000519105 | 0.304809577 | 0.235279726 | 0.056851407 | 0.907180651 |
| Dido1 | S | 802 | 0.105481067 | 0.158048958 | 0.171895899 | 0.066414832 | 0.013846942 | 0.063482953 | 0.012156757 | 0.012706442 | 0.202505053 | 0.76210011 |
| Psip1 | S | 272 | -0.048653587 | 0.199498203 | 0.270801696 | 0.319455283 | 0.071303494 | 0.282642358 | 0.003048104 | 0.000626778 | 0.0011016 | 0.144177584 |
| Psip1 | S | 274 | -0.048653587 | 0.199498203 | 0.270801696 | 0.319455283 | 0.071303494 | 0.282642358 | 0.003048104 | 0.000626778 | 0.0011016 | 0.144177584 |
| Abtb2 | S | 67 | 0.138573806 | 0.233608755 | 0.357251624 | 0.218677818 | 0.123642869 | 0.4993401 | 0.13567103 | 0.012908518 | 0.302228507 | 0.347062502 |
| Tor1aip1 | S | 252 | 0.439330858 | 0.345730433 | -0.104488557 | -0.543819416 | -0.450218991 | 0.016435374 | 0.042681688 | 0.810853242 | 0.280613099 | 0.349532898 |
| Sipa1l1 | S | 1564 | -0.014031278 | 0.127114193 | 0.200421024 | 0.214452302 | 0.073306831 | 0.561763365 | 0.000663204 | 9.17018E-05 | 0.0004572 | 0.023867748 |
| Arid1a | T | 1874 | 0.052963369 | 0.099909897 | 0.23787682 | 0.184913451 | 0.137966923 | 0.241831191 | 0.003990788 | 0.001876097 | 0.012712086 | 0.021498508 |
| Cobl | S | 372 | 0.217157855 | 0.188839747 | 0.219890698 | 0.002732843 | 0.031050951 | 0.012653797 | 0.057402222 | 0.278546947 | 0.987452992 | 0.861714661 |
| Srsf10 | S | 158 | 0.129964603 | 0.113664355 | 0.135719316 | 0.005754713 | 0.022054961 | 0.004040074 | 0.052145139 | 0.00771004 | 0.857227797 | 0.61328651 |
| Ercc6l2 | S | 1006 | 0.203493823 | 0.031587116 | 0.17626013 | -0.027233693 | 0.144673014 | 0.001792942 | 0.653882793 | 0.003931454 | 0.356561233 | 0.102716838 |
| Cep89 | S | 221 | 0.127636713 | 0.109792418 | 0.1513523 | 0.023715588 | 0.041559882 | 0.003263646 | 0.115551247 | 0.004937352 | 0.472062375 | 0.474742462 |
| Hdgfl2 | S | 366 | -0.009029518 | 0.125846706 | 0.170694789 | 0.179724307 | 0.044848082 | 0.666105396 | 0.000411144 | 4.4848E-05 | 0.000322705 | 0.025484832 |
| Hdgfl2 | S | 367 | -0.009029518 | 0.125846706 | 0.170694789 | 0.179724307 | 0.044848082 | 0.666105396 | 0.000411144 | 4.4848E-05 | 0.000322705 | 0.025484832 |
| Mboat2 | T | 471 | 0.298486104 | 0.332139752 | 0.024996294 | -0.273489809 | -0.307143457 | 0.072882231 | 0.006676405 | 0.853090921 | 0.128118384 | 0.083799196 |
| Kars | T | 585 | 0.04735579 | 0.173954145 | 0.142817721 | 0.095461931 | -0.031136424 | 0.286893694 | 0.001325986 | 0.002662769 | 0.082321678 | 0.31296837 |
| Phf14 | T | 280 | 0.11331897 | 0.199176189 | 0.211406626 | 0.098087657 | 0.012230437 | 0.201141245 | 0.001538893 | 0.116188806 | 0.408401791 | 0.90065904 |
| Xrn2 | S | 473 | 0.155358528 | 0.240752517 | 0.141670154 | -0.013688374 | -0.099082363 | 0.02470447 | 0.054780896 | 0.05459353 | 0.818512036 | 0.31360457 |
| Samhd1 | T | 52 | 0.026823042 | 0.154960078 | 0.155662109 | 0.128839067 | 0.000702031 | 0.229245647 | 0.000288971 | 0.004698748 | 0.032034449 | 0.979710255 |
| Scaf1 | T | 796 | 0.14667516 | 0.187386301 | 0.202793929 | 0.056118769 | 0.015407628 | 0.106532644 | 0.032723697 | 0.025567383 | 0.368225509 | 0.681686026 |
| Trim24 | S | 655 | 0.035535148 | 0.133103437 | 0.198417057 | 0.162881909 | 0.065313619 | 0.326655955 | 0.024158426 | 0.000131769 | 0.009556225 | 0.161363102 |
| Senp3 | S | 175 | 0.077821246 | 0.232789966 | 0.224225254 | 0.146404008 | -0.008564712 | 0.409427026 | 0.011475502 | 0.017816596 | 0.173396672 | 0.903596054 |
| Mki67 | S | 2424 | 0.234247777 | 0.486196974 | -0.02278009 | -0.257027867 | -0.508977064 | 0.049856224 | 0.013170841 | 0.871090706 | 0.153383181 | 0.030027763 |
| Rbl1 | S | 1004 | 0.115306507 | 0.1573563 | 0.185270574 | 0.069964067 | 0.027914273 | 0.052824444 | 0.03591394 | 0.009751735 | 0.189245996 | 0.625316447 |
| Mysm1 | S | 332 | 0.043784702 | 0.098570222 | 0.157772449 | 0.113987748 | 0.059202228 | 0.023842138 | 0.000740308 | 0.002144702 | 0.024724987 | 0.069318143 |
| Snrnp200 | S | 2133 | 0.097968741 | 0.202704306 | 0.156825947 | 0.058857206 | -0.045878359 | 0.305431665 | 0.00040146 | 0.07893541 | 0.548841798 | 0.483130306 |
| Ssr1 | T | 260 | 0.004081813 | 0.126218694 | 0.113301313 | 0.1092195 | -0.012917381 | 0.801102775 | 1.58763E-05 | 0.000147545 | 0.002831587 | 0.316407751 |
| Ano8 | S | 986 | 0.173153154 | 0.148285403 | 0.10421514 | -0.068938014 | -0.044070263 | 0.024722249 | 0.001233864 | 0.170583007 | 0.355306761 | 0.48737161 |
| Sorbs2 | S | 328 | 0.070043466 | 0.205036434 | 0.168333606 | 0.09829014 | -0.036702828 | 0.133537274 | 0.006825834 | 0.015195948 | 0.088131353 | 0.499578606 |
| Kmt2a | S | 748 | 0.111105282 | 0.216029076 | 0.206496413 | 0.09539113 | -0.009532663 | 0.385710564 | 0.004264786 | 0.040192206 | 0.462407174 | 0.890350553 |
| Srsf5 | S | 264 | 0.261002724 | 0.094726803 | 0.321829925 | 0.060827201 | 0.227103123 | 0.023950805 | 0.433013679 | 0.058185677 | 0.628550207 | 0.15932832 |
| Srsf5 | S | 267 | 0.261002724 | 0.094726803 | 0.321829925 | 0.060827201 | 0.227103123 | 0.023950805 | 0.433013679 | 0.058185677 | 0.628550207 | 0.15932832 |
| Fam122b | S | 25 | 0.082771075 | 0.240930732 | 0.220050422 | 0.137279346 | -0.02088031 | 0.353112083 | 0.021531837 | 0.013897943 | 0.122208652 | 0.733477967 |
| Fosl2 | S | 230 | 0.145084698 | 0.172235379 | 0.180996781 | 0.035912083 | 0.008761402 | 0.031297546 | 0.171966838 | 0.005813751 | 0.464843643 | 0.930337625 |
| Med14 | S | 992 | 0.015686457 | 0.112887605 | 0.203889424 | 0.188202967 | 0.091001819 | 0.724696905 | 0.013213463 | 3.11007E-05 | 0.025947899 | 0.035383276 |
| Med1 | T | 1051 | 0.042668899 | 0.098991623 | 0.214694076 | 0.172025177 | 0.115702453 | 0.128476653 | 0.055749262 | 0.000117244 | 0.0001927 | 0.056890763 |
| Nelfb | T | 559 | -0.005357977 | 0.174899868 | 0.248800012 | 0.254157989 | 0.073900144 | 0.921143539 | 0.043257373 | 0.000113032 | 0.019117757 | 0.261368357 |
| Ociad1 | S | 147 | 0.334185513 | 0.131037545 | 0.266065236 | -0.068120277 | 0.13502769 | 0.046640255 | 0.347951543 | 0.07093797 | 0.632376705 | 0.377303172 |
| Atn1 | S | 34 | 0.06821504 | 0.156643804 | 0.206057126 | 0.137842085 | 0.049413321 | 0.026187336 | 0.002036291 | 0.071209621 | 0.163486611 | 0.52823958 |
| Ncor2 | S | 2411 | 0.029074004 | 0.226467941 | 0.300488492 | 0.271414487 | 0.07402055 | 0.508576446 | 0.021484612 | 0.012420381 | 0.007753754 | 0.261219745 |
| Rai14 | S | 317 | 0.152080797 | 0.103451532 | 0.170955087 | 0.01887429 | 0.067503555 | 0.067416515 | 0.036898306 | 0.00301281 | 0.755076872 | 0.116336913 |
| Ltbp1 | S | 1607 | 0.085735308 | 0.252668469 | 0.13756794 | 0.051832632 | -0.11510053 | 0.149191879 | 0.00061606 | 0.229234767 | 0.608735061 | 0.29611513 |
| Rbms2 | S | 33 | 0.474792528 | 0.102879729 | 0.134446196 | -0.340346332 | 0.031566467 | 0.004771028 | 0.619689543 | 0.296253865 | 0.022151473 | 0.868328808 |
| Ifngr1 | S | 402 | 0.070008807 | 0.273638018 | 0.159896551 | 0.089887744 | -0.113741467 | 0.293621666 | 0.012760971 | 0.013254424 | 0.164586538 | 0.157511572 |
| Arhgef6 Arhgef7 | S | 648;682 | 0.035955078 | 0.126560255 | 0.364662845 | 0.328707766 | 0.238102589 | 0.629121842 | 0.268199532 | 0.000441251 | 0.009812986 | 0.091656627 |
| Cic | S | 2299 | -0.021746653 | 0.094326821 | 0.233656421 | 0.255403074 | 0.1393296 | 0.559619876 | 0.000424046 | 9.49238E-05 | 0.005267761 | 0.003328863 |
| Jmjd1c | S | 871 | 0.051567071 | 0.114229168 | 0.156235372 | 0.104668301 | 0.042006204 | 0.295848218 | 0.021020819 | 2.0217E-05 | 0.100636593 | 0.2172239 |
| Sipa1l1 | S | 1629 | 0.018397905 | 0.132881832 | 0.275990531 | 0.257592626 | 0.143108699 | 0.602880116 | 0.025919645 | 0.000665303 | 0.002876557 | 0.029983161 |
| Specc1l | S | 963 | 0.101983557 | 0.432867672 | 0.183815015 | 0.081831458 | -0.249052657 | 0.712832105 | 0.016862667 | 0.070183095 | 0.765672277 | 0.091948053 |
| Scrib | S | 1523 | 0.180951219 | 0.398895332 | 0.467964486 | 0.287013267 | 0.069069154 | 0.592889001 | 0.148896594 | 0.101534703 | 0.321261669 | 0.270942758 |
| Poldip3 | S | 127 | 0.203073881 | 0.20679408 | 0.232678514 | 0.029604633 | 0.025884434 | 0.00949442 | 0.109036146 | 0.254608841 | 0.862588898 | 0.888761948 |
| Bcl7c | S | 97 | -0.030691312 | 0.192966985 | 0.434398381 | 0.465089693 | 0.241431396 | 0.587312515 | 0.115941937 | 0.001325875 | 0.001760673 | 0.070616232 |
| Bcl7c | S | 100 | -0.030691312 | 0.192966985 | 0.434398381 | 0.465089693 | 0.241431396 | 0.587312515 | 0.115941937 | 0.001325875 | 0.001760673 | 0.070616232 |
| Mki67 | S | 213 | 0.11112659 | 0.203140626 | 0.178072746 | 0.066946156 | -0.02506788 | 0.234013457 | 0.007336259 | 0.024534769 | 0.451148541 | 0.669803946 |
| Kat7 | S | 113 | 0.049451182 | 0.213255013 | 0.171106802 | 0.12165562 | -0.042148211 | 0.271216916 | 0.00790042 | 0.005539798 | 0.003666328 | 0.381175796 |
| Smarca4 | S | 695 | 0.0183483 | 0.116236742 | 0.175239443 | 0.156891144 | 0.059002701 | 0.449257792 | 0.000536487 | 0.000486903 | 0.002881234 | 0.045707094 |
| Mecom | S | 1041 | 0.24495616 | 0.174904636 | 0.213440209 | -0.031515951 | 0.038535573 | 0.27217622 | 0.005390117 | 0.105844523 | 0.875268787 | 0.693362633 |
| Git1 | S | 394 | 0.036272124 | 0.146731438 | 0.083097951 | 0.046825828 | -0.063633487 | 0.026134295 | 8.87044E-05 | 0.00390337 | 0.045806618 | 0.019035601 |
| Mtcl1 | S | 74 | 0.333704035 | 0.080661415 | 0.178299272 | -0.155404763 | 0.097637857 | 0.004608306 | 0.46067247 | 0.114785578 | 0.134711962 | 0.377716901 |
| Top2a | S | 1465 | 0.255409563 | 0.577739335 | 0.042113233 | -0.21329633 | -0.535626102 | 0.097502027 | 0.073949795 | 0.338941215 | 0.132762705 | 0.083513349 |

| | | | | | | | | | | | | | |
|---|---|---|---|---|---|---|---|---|---|---|---|---|---|
| Map4 | S | | 256 | 0.169459325 | 0.167652133 | 0.251041141 | 0.081581816 | 0.083389008 | 0.082323403 | 0.016142285 | 0.120654119 | 0.542513102 | 0.510887866 |
| Baz1b | S | | 1464 | 0.08892921 | 0.105346249 | 0.129738195 | 0.040808985 | 0.024391946 | 0.203464491 | 9.71118E-05 | 0.003948493 | 0.49673795 | 0.295145416 |
| Ramac | S | | 36 | 0.00989081 | 0.118686528 | 0.147187401 | 0.137296591 | 0.028500874 | 0.726108654 | 0.000330152 | 8.12016E-05 | 0.014269316 | 0.050834516 |
| Ifi204 Ifi211 | T | 106;106 | | 0.061948278 | 0.320296556 | 0.196984037 | 0.135035759 | -0.123312519 | 0.517301523 | 0.00692762 | 0.059802231 | 0.164922313 | 0.164775349 |
| Gtf2f1 | S | | 433 | -0.037216816 | 0.239178767 | 0.23892591 | 0.276142726 | -0.000252856 | 0.399918526 | 0.012279812 | 0.000402542 | 0.003012122 | 0.996141443 |
| Prr12 | S | | 911 | -0.058787163 | 0.247713389 | 0.286074675 | 0.344861839 | 0.038361287 | 0.363374988 | 0.002022726 | 0.005880667 | 0.006477126 | 0.475199696 |
| Khdrbs1 | S | | 58 | 0.078742008 | 0.112710568 | 0.197956639 | 0.119214632 | 0.085246071 | 0.089562804 | 0.010151188 | 0.003722622 | 0.041096149 | 0.083686874 |
| Cdk13 | S | | 361 | 0.08547758 | 0.10330014 | 0.200860434 | 0.115382855 | 0.097560295 | 0.000702814 | 0.047178345 | 0.025676989 | 0.092290543 | 0.118582768 |
| Pum2 | S | | 136 | 0.014907999 | 0.136684072 | 0.13891929 | 0.124011291 | 0.002235218 | 0.420431313 | 9.8786E-05 | 0.001135209 | 0.004689262 | 0.911554273 |
| Arl6ip6 | S | | 65 | 0.004288297 | 0.31872396 | 0.471792128 | 0.467503831 | 0.153068168 | 0.970259484 | 0.144124488 | 0.014944044 | 0.022046548 | 0.437763668 |
| Fosl1 | S | | 16 | 0.135117265 | 0.12281675 | 0.297932382 | 0.162815116 | 0.175115632 | 0.128225736 | 0.151137272 | 0.008175657 | 0.015375925 | 0.001900757 |
| Adgra2 | S | | 1280 | 0.038193438 | 0.171159275 | 0.232460858 | 0.19426742 | 0.061301582 | 0.028286693 | 0.007233314 | 0.027010804 | 0.040551167 | 0.294297607 |
| Mdc1 | T | | 444 | 0.047642229 | 0.142284972 | 0.184492764 | 0.136850535 | 0.042207792 | 0.269459188 | 0.005273611 | 0.001607787 | 0.023606961 | 0.265014996 |
| Suds3 | S | | 53 | 0.076341886 | 0.093787355 | 0.164744499 | 0.088402612 | 0.070957144 | 0.028003352 | 0.027846229 | 0.000641943 | 0.016861519 | 0.069201123 |
| Sap130 | T | | 862 | 0.054084553 | 0.194554414 | 0.357144551 | 0.303059999 | 0.162590137 | 0.240270247 | 0.019815575 | 0.045690037 | 0.058650611 | 0.192424109 |
| Septin9 | S | | 325 | 0.041959179 | 0.21757485 | 0.126942636 | 0.084983457 | -0.090632215 | 0.803233943 | 1.86598E-05 | 0.080387464 | 0.628211071 | 0.153348298 |
| Spata13 | T | | 83 | 0.001799394 | 0.306868703 | 0.252983403 | 0.251184009 | -0.0538853 | 0.983657489 | 0.108330396 | 0.000607084 | 0.064401251 | 0.689122688 |
| Parp1 | S | | 781 | 0.086154485 | 0.252246806 | 0.182606973 | 0.096452488 | -0.069639833 | 0.38200806 | 0.000354372 | 0.259594466 | 0.537510883 | 0.617367755 |
| Lmna | S | | 458 | 0.046008717 | 0.228326429 | 0.2203207 | 0.174311983 | -0.00800573 | 0.501688688 | 0.006301227 | 0.017271179 | 0.042346443 | 0.881635228 |
| Esyt2 | S | | 661 | 0.144206181 | 0.19954891 | 0.218525389 | 0.074319208 | 0.018976479 | 0.092251893 | 0.051136094 | 0.038957562 | 0.40379192 | 0.830765829 |
| Tmpo | S | 66;66 | | 0.078105889 | 0.130551092 | 0.147762896 | 0.069657008 | 0.017211804 | 0.179619325 | 0.003423134 | 0.001694841 | 0.215488263 | 0.358692069 |
| Nes | S | | 977 | -0.058334048 | 0.190028438 | 0.292478158 | 0.350812206 | 0.10244972 | 0.266533736 | 0.003326754 | 0.000881672 | 0.001414429 | 0.077927654 |
| Anapc1 | S | | 377 | 0.053457632 | 0.135220049 | 0.162686363 | 0.109228731 | 0.027466313 | 0.173863812 | 0.013419529 | 0.000399556 | 0.032848324 | 0.429699041 |
| Pcnx3 | S | | 287 | 0.047182083 | 0.217612449 | 0.133978007 | 0.086795924 | -0.083634442 | 0.194661223 | 0.000599132 | 0.043515447 | 0.138954094 | 0.15485132 |
| Eif4g3 | S | | 470 | 0.097642363 | 0.241344987 | 0.152826799 | 0.055184436 | -0.088518188 | 0.14860797 | 0.012259062 | 0.029549296 | 0.387003656 | 0.219404308 |
| Sap130 | S | | 864 | 0.007838086 | 0.232093833 | 0.221019431 | 0.213181345 | -0.011074402 | 0.800654025 | 0.001842109 | 0.017592464 | 0.024506228 | 0.842425225 |
| Sap130 | T | | 865 | 0.007838086 | 0.232093833 | 0.221019431 | 0.213181345 | -0.011074402 | 0.800654025 | 0.001842109 | 0.017592464 | 0.024506228 | 0.842425225 |
| Glyr1 | S | | 113 | 0.148024195 | 0.109642212 | 0.206906315 | 0.05888212 | 0.097264103 | 0.187770113 | 0.060926444 | 0.001796468 | 0.535505913 | 0.076470715 |
| Sltm | S | | 344 | -0.006574467 | 0.163452532 | 0.199994693 | 0.20656916 | 0.036542161 | 0.86637216 | 0.000354956 | 0.002738713 | 0.00680941 | 0.311203967 |
| Susd6 | S | | 286 | 0.000633955 | 0.287058646 | 0.373030318 | 0.372396363 | 0.085971672 | 0.992959041 | 0.150843342 | 0.003406478 | 0.006914292 | 0.598322822 |
| Dgcr8 | T | | 279 | 0.220079826 | 0.079445965 | 0.187473719 | -0.032606107 | 0.108027753 | 0.023278649 | 0.163662745 | 0.012611409 | 0.634524784 | 0.100251615 |
| Sntb2 | S | | 88 | 0.122281258 | 0.234854022 | 0.132266789 | 0.009985531 | -0.102587233 | 0.012549281 | 0.016533173 | 0.175121313 | 0.896830785 | 0.267219443 |
| Cenpc1 | S | | 552 | 0.057151773 | 0.140154572 | 0.16107913 | 0.103927357 | 0.020924559 | 0.416132012 | 0.002920999 | 0.000881966 | 0.19415469 | 0.4779179 |
| Retreg2 | T | | 324 | 0.191233307 | 0.208840638 | 0.188360587 | -0.00287272 | -0.020480051 | 0.370841918 | 0.016236499 | 0.026028261 | 0.987866964 | 0.599968134 |
| Mki67 | S | | 2255 | 0.17293547 | 0.135424256 | 0.196259395 | 0.023323925 | 0.060835139 | 0.036783749 | 0.050984343 | 0.037861505 | 0.749209082 | 0.383847554 |
| Bclaf3 | S | | 410 | 0.128353487 | 0.201970147 | 0.198409893 | 0.070056406 | -0.003560254 | 0.290347806 | 0.004745546 | 0.057499208 | 0.557802334 | 0.960672992 |
| Rbm15 | S | | 725 | 0.155595356 | 0.164571729 | 0.124334632 | -0.031260724 | -0.040237097 | 0.011012654 | 0.008572488 | 0.171881313 | 0.667989469 | 0.585664295 |
| Lrrk2 | S | | 898 | -0.049873586 | 0.245697828 | 0.303078713 | 0.352952299 | 0.057380886 | 0.188375895 | 5.7484E-05 | 0.05759567 | 0.038870444 | 0.555053721 |
| Prkaa1 | T | | 526 | 0.006202846 | 0.260676322 | 0.199532285 | 0.193329439 | -0.061144037 | 0.916248351 | 0.002208351 | 0.016715195 | 0.021245524 | 0.288172678 |
| Sorbs3 | S | | 437 | -0.20464728 | 0.494526485 | 0.350140873 | 0.554788153 | -0.144385612 | 0.033106463 | 0.001136806 | 0.017128528 | 0.00626284 | 0.208863395 |
| Afdn | S | | 1083 | -0.030120296 | 0.159468211 | 0.2203318 | 0.250452096 | 0.060863589 | 0.465255583 | 0.000994219 | 0.000525566 | 0.002748793 | 0.096482014 |
| Klc4 | T | | 612 | -0.014960783 | 0.23210475 | 0.275375769 | 0.290336552 | 0.043271019 | 0.847947064 | 0.044900783 | 0.000913751 | 0.028944852 | 0.582766627 |
| Rtn4 | S | | 167 | 0.2423536 | 0.21976594 | 0.068869496 | -0.173484105 | -0.150896444 | 0.00345775 | 0.038154154 | 0.66887335 | 0.334493309 | 0.393569954 |
| Rps8 | T | | 130 | 0.007969695 | 0.273246579 | 0.244833978 | 0.236864283 | -0.028412601 | 0.911799433 | 0.050361598 | 0.001620168 | 0.036177531 | 0.753284489 |
| Mia2 | S | | 1124 | 0.187299175 | 0.094933473 | 0.228745644 | 0.041446469 | 0.133812172 | 0.071438391 | 0.164995957 | 0.006839049 | 0.580555722 | 0.023377 |
| Acin1 | S | | 216 | -0.003605068 | 0.116232536 | 0.231565291 | 0.235170359 | 0.115332755 | 0.912229854 | 0.003257218 | 0.000248525 | 0.001376488 | 0.022987066 |
| Smarcc2 | S | | 283 | 0.038320041 | 0.168538564 | 0.115401086 | 0.077081045 | -0.053137478 | 0.040683743 | 0.006145835 | 0.000798212 | 0.009814442 | 0.132754413 |
| Kmt2a | T | | 335 | 0.047910497 | 0.214673863 | 0.261757554 | 0.213847057 | 0.047083692 | 0.580260306 | 0.003880162 | 0.036712934 | 0.089270326 | 0.566210077 |
| Clasp1 | S | | 826 | 0.365792754 | 0.149114499 | 0.278566956 | -0.087225798 | 0.129452457 | 0.042324797 | 0.324976605 | 0.174436495 | 0.644416172 | 0.514733562 |
| Vps50 | S | | 492 | 0.016406422 | 0.200290983 | 0.165061011 | 0.148654589 | -0.035229972 | 0.775970927 | 0.0031018 | 0.001393031 | 0.068868783 | 0.366540806 |
| Fez1 | S | | 316 | -0.221256349 | 0.49598865 | 0.320322596 | 0.541578946 | -0.175666054 | 0.013357021 | 0.000367668 | 0.013188667 | 0.010584716 | 0.097650878 |
| Crebbp | T | | 1076 | -0.049024266 | 0.105587485 | 0.287550128 | 0.336574394 | 0.181962643 | 0.632089414 | 0.014813364 | 6.59657E-05 | 0.055512932 | 0.001325878 |
| Ptk2b | S | | 747 | -0.069470933 | 0.351649308 | 0.293922946 | 0.363393879 | -0.057726362 | 0.544769487 | 0.001073099 | 0.093083998 | 0.063191301 | 0.65666384 |
| Ptk2b | S | | 762 | -0.069470933 | 0.351649308 | 0.293922946 | 0.363393879 | -0.057726362 | 0.544769487 | 0.001073099 | 0.093083998 | 0.063191301 | 0.65666384 |
| Srrm2 | S | | 1703 | 0.103185102 | 0.200225348 | 0.200365226 | 0.097180124 | 0.000139879 | 0.340812478 | 0.023007668 | 0.008510065 | 0.357971689 | 0.998151472 |
| Srrm2 | S | | 1705 | 0.103185102 | 0.200225348 | 0.200365226 | 0.097180124 | 0.000139879 | 0.340812478 | 0.023007668 | 0.008510065 | 0.357971689 | 0.998151472 |
| Srrm2 | S | | 1706 | 0.103185102 | 0.200225348 | 0.200365226 | 0.097180124 | 0.000139879 | 0.340812478 | 0.023007668 | 0.008510065 | 0.357971689 | 0.998151472 |
| Brd7 | S | | 279 | 0.050993136 | 0.198773595 | 0.153206004 | 0.102212868 | -0.04556759 | 0.453638441 | 0.000666181 | 0.020383138 | 0.189934227 | 0.323995616 |
| U2surp | S | | 930 | 0.195054147 | 0.205047117 | 0.217613172 | 0.022559025 | 0.012566054 | 0.255773011 | 0.016283732 | 0.078574478 | 0.88681513 | 0.891576436 |
| U2surp | S | | 932 | 0.195054147 | 0.205047117 | 0.217613172 | 0.022559025 | 0.012566054 | 0.255773011 | 0.016283732 | 0.078574478 | 0.88681513 | 0.891576436 |
| U2surp | S | | 934 | 0.195054147 | 0.205047117 | 0.217613172 | 0.022559025 | 0.012566054 | 0.255773011 | 0.016283732 | 0.078574478 | 0.88681513 | 0.891576436 |
| Srrm2 | S | | 1888 | 0.148127121 | 0.174404789 | 0.152620164 | 0.004493043 | -0.021784625 | 0.043917693 | 0.028182343 | 0.03715944 | 0.877982453 | 0.578412591 |
| Srrm2 | T | | 1892 | 0.148127121 | 0.174404789 | 0.152620164 | 0.004493043 | -0.021784625 | 0.043917693 | 0.028182343 | 0.03715944 | 0.877982453 | 0.578412591 |
| Msh6 | S | | 252 | 0.025220605 | 0.139220616 | 0.184350215 | 0.15912961 | 0.045129599 | 0.446927008 | 0.00355666 | 0.000772364 | 0.002415581 | 0.111336986 |
| Wnk1 | S | | 165 | -0.000483656 | 0.219255684 | 0.159946821 | 0.160430476 | -0.059308863 | 0.99013214 | 0.000646792 | 0.005621474 | 0.010389454 | 0.158685976 |
| Rbmx Rbmxl1 | S | 88;88 | | 0.059922906 | 0.096841198 | 0.191131241 | 0.131208335 | 0.094290043 | 0.423803463 | 0.003650644 | 0.000469071 | 0.154781881 | 0.007436556 |
| Erc1 | S | | 21 | 0.206969432 | 0.104161643 | 0.092042821 | -0.114926612 | -0.012118822 | 0.000524706 | 0.142406589 | 0.088205668 | 0.051786734 | 0.844497654 |
| Ttf1 | S | | 137 | 0.303662646 | 0.236183787 | -0.050430618 | -0.354093265 | -0.286614406 | 0.000641853 | 0.056755023 | 0.661374662 | 0.0557202 | 0.076925205 |
| Sap130 | S | | 681 | 0.070101513 | 0.217474134 | 0.217650989 | 0.147549477 | 0.000176856 | 0.291381669 | 0.007772676 | 0.035994958 | 0.117671584 | 0.998092459 |
| Pml | T | | 513 | -0.069649366 | 0.284341023 | 0.254497193 | 0.324146559 | -0.02984383 | 0.462696648 | 0.0023422 | 0.009292657 | 0.028510378 | 0.647034787 |
| Rere | S | | 681 | 0.101168928 | 0.070132986 | 0.225168956 | 0.123485028 | 0.15503597 | 0.081083453 | 0.104334618 | 0.000803379 | 0.045667985 | 0.00475537 |
| Pcif1 | S | | 30 | -0.013938271 | 0.14996965 | 0.396381689 | 0.41031996 | 0.246412039 | 0.920899575 | 0.136184146 | 0.000985223 | 0.061623495 | 0.036734996 |
| Mki67 | S | | 767 | 0.211753987 | 0.206565431 | 0.124552148 | -0.087201839 | -0.082013283 | 0.056484948 | 0.020055304 | 0.144742174 | 0.368184627 | 0.329021911 |
| Nfib | S | | 333 | 0.046512646 | 0.114841593 | 0.245199013 | 0.198686367 | 0.13035742 | 0.199146164 | 0.023115257 | 0.001955361 | 0.022684135 | 0.034882702 |
| Ppp1r12c | S | | 647 | 0.130623586 | 0.194695856 | 0.129269699 | -0.001353887 | -0.065426157 | 0.071637459 | 0.002833909 | 0.131124054 | 0.985751655 | 0.367937143 |
| Mettl14 | S | | 399 | 0.055005438 | 0.109116431 | 0.162037736 | 0.107032298 | 0.052921305 | 0.017110307 | 0.002346396 | 0.008243527 | 0.040470265 | 0.165989312 |
| Pml | S | | 557 | 0.076280969 | 0.199642641 | 0.12757935 | 0.051298381 | -0.072063291 | 0.146460094 | 0.000742856 | 0.068151421 | 0.397159738 | 0.22497582 |
| Sf1 | S | | 80 | 0.167156552 | 0.168437778 | 0.117384947 | -0.049771605 | -0.051052831 | 0.014462086 | 0.013716364 | 0.146265243 | 0.44190965 | 0.423796698 |
| Mki67 | S | | 2333 | 0.183380884 | 0.316653053 | 0.074370831 | -0.109010053 | -0.242282223 | 0.19834151 | 0.002771571 | 0.415768029 | 0.41123362 | 0.035188815 |
| Ssh1 | S | | 980 | 0.057944968 | 0.267056052 | 0.260708496 | 0.202763527 | -0.006347556 | 0.582321244 | 0.019765737 | 0.028140641 | 0.134933586 | 0.943657606 |
| Sf1 | S | | 82 | 0.166983864 | 0.168321947 | 0.11737466 | -0.049609204 | -0.050947287 | 0.014540205 | 0.013773876 | 0.14608434 | 0.442809598 | 0.424208118 |
| Retreg3 | S | | 256 | 0.021666816 | 0.129714352 | 0.201891324 | 0.180224508 | 0.072176972 | 0.404457815 | 0.000715443 | 0.004065092 | 0.008688721 | 0.101175978 |
| Msh6 | S | | 62 | -0.00419957 | 0.20359513 | 0.134071939 | 0.138271509 | -0.069523192 | 0.933601173 | 0.001341657 | 0.000427216 | 0.073280584 | 0.07256132 |
| Serhl | S | | 210 | 0.170258202 | 0.204975278 | 0.264580397 | 0.094322195 | 0.059605118 | 0.235460378 | 0.04722126 | 0.055062074 | 0.51323599 | 0.584941043 |
| Dvl1 | S | | 679 | -0.01634524 | 0.096924461 | 0.329902461 | 0.346247701 | 0.232978 | 0.620333375 | 0.064834327 | 0.000108025 | 0.000449969 | 0.005195484 |
| Rbm15 | S | | 655 | -0.02017536 | 0.185179099 | 0.216183551 | 0.236358912 | 0.031004453 | 0.286909267 | 5.67432E-05 | 0.012475445 | 0.007700956 | 0.421807262 |
| Fnbp4 | S | | 446 | 0.066862514 | 0.247584668 | 0.204993052 | 0.138130538 | -0.042591616 | 0.351189035 | 0.003305459 | 0.087719898 | 0.197629906 | 0.616860539 |
| Pitpnm1 | T | | 287 | 0.142087105 | 0.268517766 | 0.143150351 | 0.001063246 | -0.125367415 | 0.483717837 | 0.001638785 | 0.144885782 | 0.995670943 | 0.188865978 |
| Fam120a | S | | 1021 | 0.083782732 | 0.175823851 | 0.19811092 | 0.114328187 | 0.022287068 | 0.172053506 | 0.014558876 | 0.014887863 | 0.083353428 | 0.669500151 |
| Dido1 | S | | 806 | 0.111893872 | 0.15955865 | 0.152470531 | 0.040576659 | -0.007088119 | 0.066463025 | 0.012044157 | 0.020785136 | 0.432753142 | 0.876888125 |
| Tnrc6c | T | | 776 | 0.000874468 | 0.082374105 | 0.237100882 | 0.236226414 | 0.154726777 | 0.983824888 | 0.008324183 | 5.43019E-05 | 0.01015959 | 0.002581634 |
| Srrm2 | T | | 252 | 0.122596004 | 0.112676231 | 0.165396988 | 0.042800984 | 0.052720757 | 0.047599963 | 0.002144987 | 0.046208864 | 0.490809555 | 0.355797926 |
| Rbl1 | T | | 369 | 0.024195878 | 0.187048419 | 0.117076898 | 0.09288102 | -0.069971521 | 0.345500788 | 0.000230665 | 0.008835366 | 0.027413039 | 0.056108012 |
| Myo1c | S | | 736 | 0.126925749 | 0.351870708 | 0.047177133 | -0.079748617 | -0.304693576 | 0.122249958 | 0.002191665 | 0.543687971 | 0.145825876 | 0.003001057 |

| | | | | | | | | | | | | |
|---|---|---|---|---|---|---|---|---|---|---|---|---|
| Bag3 | S | 179 | 0.231340733 | 0.293440823 | -0.112093996 | -0.34343473 | -0.405534819 | 0.003150814 | 0.001042278 | 0.205828871 | 0.018037887 | 0.017954798 |
| Mycbp2 | S | 3057 | 0.252288215 | 0.174613585 | 0.141951991 | -0.110336223 | -0.032661594 | 0.023222034 | 0.18667266 | 0.057459798 | 0.207674711 | 0.769725286 |
| Brd8 | S | 284 | 0.138267939 | 0.114980546 | 0.218484241 | 0.080216302 | 0.103503695 | 0.045240515 | 0.111310863 | 0.010750966 | 0.201877813 | 0.14844736 |
| Hirip3 | S | 153 | 0.012997681 | 0.102243272 | 0.1238823 | 0.110884619 | 0.021639028 | 0.698366313 | 0.000306499 | 1.49114E-05 | 0.057172338 | 0.13476032 |
| Itsn1 | S | 558;561 | -0.020070265 | 0.145877643 | 0.253563255 | 0.273633521 | 0.107685612 | 0.730058613 | 0.004159167 | 0.000873985 | 0.011981721 | 0.041314838 |
| Ssbp3 | S | 381 | 0.0018939 | 0.307776449 | 0.276534582 | 0.274640682 | -0.031241867 | 0.986475701 | 0.004424447 | 0.07142435 | 0.102000957 | 0.779120054 |
| Palm | T | 153 | 0.057515212 | 0.201794543 | 0.225381022 | 0.167865811 | 0.02358648 | 0.201307708 | 0.002397592 | 0.086751893 | 0.168820654 | 0.797803321 |
| Atm | S | 1987 | 0.120819185 | 0.205832046 | 0.107611953 | -0.013207232 | -0.098220093 | 0.394635478 | 0.000137765 | 0.140063864 | 0.921411224 | 0.173849464 |
| Prpf38a Srrm1 Thrap | S | 250;609;134 | 0.076989542 | 0.079812348 | 0.159053039 | 0.082063498 | 0.079240691 | 0.000674029 | 0.023842953 | 0.01044819 | 0.061297162 | 0.059481733 |
| Prpf38a Srrm1 Thrap | S | 252;611;136 | 0.076989542 | 0.079812348 | 0.159053039 | 0.082063498 | 0.079240691 | 0.000674029 | 0.023842953 | 0.01044819 | 0.061297162 | 0.059481733 |
| Pdlim2 | S | 199 | 0.033023474 | 0.118271724 | 0.14789571 | 0.114872236 | 0.029623986 | 0.029247916 | 0.004312182 | 0.001105697 | 0.004765043 | 0.225324215 |
| Cep89 | T | 112 | 0.069610376 | 0.21606431 | 0.180043431 | 0.110433055 | -0.036020879 | 0.212921652 | 0.060531632 | 0.00245097 | 0.092007574 | 0.640904523 |
| Eif2ak3 | T | 980 | 0.101016958 | 0.255652132 | 0.231306765 | 0.130289807 | -0.024345367 | 0.244546596 | 0.039568967 | 0.04078061 | 0.122464777 | 0.787961969 |
| Sipa1l1 | S | 1626 | 0.009337733 | 0.1221057 | 0.241552215 | 0.232214482 | 0.119446516 | 0.881202668 | 0.014712457 | 0.000324039 | 0.031308911 | 0.010736475 |
| Atxn2 | T | 710 | 0.054952311 | 0.109371732 | 0.184467638 | 0.129515327 | 0.075095906 | 0.113440817 | 0.023810665 | 0.000680054 | 0.002383076 | 0.07013839 |
| Prpf4b | S | 292 | 0.108037209 | 0.110053395 | 0.171099235 | 0.063062027 | 0.06104584 | 0.030247534 | 0.01242784 | 0.017046521 | 0.23897171 | 0.235059291 |
| Map3k6 | S | 32 | 0.192777366 | 0.298842602 | 0.262931999 | 0.070154633 | -0.035910603 | 0.234627448 | 0.06216948 | 0.152072151 | 0.691624229 | 0.826820205 |
| Suds3 | S | 45 | 0.069648956 | 0.092869825 | 0.17046133 | 0.100812375 | 0.077591505 | 0.031434807 | 0.027889345 | 0.001125541 | 0.018327077 | 0.062336376 |
| Suds3 | T | 49 | 0.069648956 | 0.092869825 | 0.17046133 | 0.100812375 | 0.077591505 | 0.031434807 | 0.027889345 | 0.001125541 | 0.018327077 | 0.062336376 |
| Tns3 | S | 332 | -0.036704513 | 0.185746471 | 0.274220237 | 0.31092475 | 0.088473766 | 0.218102696 | 0.000784881 | 0.004129399 | 0.002573666 | 0.092973368 |
| Map4 | T | 915 | 0.122706804 | 0.19206647 | 0.139807255 | 0.017100451 | -0.052259215 | 0.080782505 | 0.021545695 | 0.022637245 | 0.769267363 | 0.385778415 |
| Lpgat1 | S | 233 | 0.035939047 | 0.215291871 | 0.144538566 | 0.108599519 | -0.070753305 | 0.432561827 | 0.000251448 | 0.06321926 | 0.13404362 | 0.261994727 |
| Scaf1 | S | 1139 | 0.057560826 | 0.147714731 | 0.198764198 | 0.141203372 | 0.051049467 | 0.235347413 | 0.003242139 | 0.012614825 | 0.05006657 | 0.321694326 |
| Scaf1 | S | 1140 | 0.057560826 | 0.147714731 | 0.198764198 | 0.141203372 | 0.051049467 | 0.235347413 | 0.003242139 | 0.012614825 | 0.05006657 | 0.321694326 |
| Scaf1 | S | 1142 | 0.057560826 | 0.147714731 | 0.198764198 | 0.141203372 | 0.051049467 | 0.235347413 | 0.003242139 | 0.012614825 | 0.05006657 | 0.321694326 |
| Amer1 | S | 871 | 0.090015502 | 0.197443089 | 0.243075426 | 0.153059924 | 0.045632337 | 0.157988975 | 0.093308632 | 0.010831509 | 0.056315178 | 0.63100628 |
| Cdc42ep4 | S | 140 | -0.085895731 | 0.343044776 | 0.336214702 | 0.422110433 | -0.006830074 | 0.007263044 | 0.026775761 | 0.000236009 | 0.000178515 | 0.925710784 |
| Bclaf1 | S | 510 | 0.051436518 | 0.186304516 | 0.213113536 | 0.161677018 | 0.02680902 | 0.408194678 | 0.012997487 | 0.007200756 | 0.022805934 | 0.396605829 |
| Lsm14a | S | 216 | 0.00099712 | 0.127731674 | 0.15374755 | 0.152750429 | 0.026015875 | 0.968782556 | 0.000273349 | 0.000334191 | 0.004534174 | 0.229046376 |
| Ctnnd1 | S | 312 | 0.40677889 | 0.393867941 | -0.070728237 | -0.477507127 | -0.464596178 | 0.052509019 | 0.02981292 | 0.778491692 | 0.119688243 | 0.130443891 |
| Wdr81 | S | 829 | 0.133693203 | 0.105944173 | 0.198073645 | 0.064380442 | 0.092129472 | 0.040846493 | 0.010468608 | 0.043344334 | 0.344688308 | 0.187992324 |
| Ddx46 | S | 102 | 0.253842821 | 0.182026988 | 0.248711091 | -0.005131731 | 0.066684103 | 0.08782028 | 0.083384979 | 0.157045362 | 0.973933707 | 0.650375876 |
| Ddx46 | S | 104 | 0.254974192 | 0.176723291 | 0.247561779 | -0.007412413 | 0.070838489 | 0.086447174 | 0.084070491 | 0.150145547 | 0.961565812 | 0.62136609 |
| Mphosph6 | S | 111 | 0.191569606 | 0.120774087 | 0.092756893 | -0.098812713 | -0.028017194 | 0.001102871 | 0.016595398 | 0.402759348 | 0.37469371 | 0.781949807 |
| Ncor2 | T | 1350 | 0.117597912 | 0.131974881 | 0.235424201 | 0.117826289 | 0.10344932 | 0.019497459 | 0.02940108 | 0.079150572 | 0.274552977 | 0.324704417 |
| Srsf4 | S | 405 | 0.064076468 | 0.418345994 | 0.214726931 | 0.150650463 | -0.203619063 | 0.661167858 | 0.005599848 | 0.31241929 | 0.472881752 | 0.32289822 |
| Msh6 | S | 137 | 0.06773746 | 0.164868897 | 0.283641201 | 0.215903741 | 0.118772304 | 0.585094452 | 0.110455081 | 0.001767079 | 0.154704674 | 0.209729455 |
| Dock7 | S | 900 | 0.052119291 | 0.13882217 | 0.153178411 | 0.10105912 | 0.01435624 | 0.054610011 | 0.003425324 | 0.008270316 | 0.010465444 | 0.526765791 |
| Srrm2 | S | 1638 | 0.177975182 | 0.058286393 | 0.224400534 | 0.046425352 | 0.166114141 | 0.008366193 | 0.194525246 | 0.026575263 | 0.493433347 | 0.063492758 |
| Srrm2 | T | 1640 | 0.177975182 | 0.058286393 | 0.224400534 | 0.046425352 | 0.166114141 | 0.008366193 | 0.194525246 | 0.026575263 | 0.493433347 | 0.063492758 |
| Lins1 | S | 646 | 0.183725722 | 0.131887562 | 0.181260403 | -0.002465319 | 0.049372841 | 0.0176824 | 0.265791127 | 0.017846207 | 0.946897051 | 0.626090212 |
| Lins1 | S | 647 | 0.183725722 | 0.131887562 | 0.181260403 | -0.002465319 | 0.049372841 | 0.0176824 | 0.265791127 | 0.017846207 | 0.946897051 | 0.626090212 |
| Lins1 | S | 650 | 0.183725722 | 0.131887562 | 0.181260403 | -0.002465319 | 0.049372841 | 0.0176824 | 0.265791127 | 0.017846207 | 0.946897051 | 0.626090212 |
| Atn1 | S | 79 | 0.106001596 | 0.102200699 | 0.113996286 | 0.00799469 | 0.011795587 | 0.009524763 | 0.01575295 | 0.004934096 | 0.701377212 | 0.628415035 |
| Ubr4 | T | 2712 | 0.039209729 | 0.174086982 | 0.275925005 | 0.236715276 | 0.101838023 | 0.719262678 | 0.033822392 | 0.003486295 | 0.105427333 | 0.1711244 |
| Dlgap1 | T | 531 | 0.129691292 | 0.209451143 | 0.180517678 | 0.050826386 | -0.028933465 | 0.152811069 | 0.067800732 | 0.012845835 | 0.504282779 | 0.729535146 |
| Sun1 | S | 97 | -0.054624306 | 0.133418425 | 0.300026022 | 0.354650328 | 0.166607597 | 0.182330487 | 0.02818887 | 1.29237E-05 | 0.001038531 | 0.049201517 |
| Slc35a5 | T | 436 | -0.017788639 | 0.20385198 | 0.298194379 | 0.315983017 | 0.094342399 | 0.657793003 | 0.019999366 | 0.002809539 | 0.003391183 | 0.176969576 |
| Rai14 | S | 414 | 0.046432125 | 0.193773345 | 0.140382914 | 0.093950789 | -0.053390431 | 0.035341419 | 0.003396229 | 0.034813169 | 0.089892371 | 0.259867754 |
| Akap12 | S | 371 | 0.25055136 | 0.441162784 | 0.061632874 | -0.188918485 | -0.37952991 | 0.050166133 | 0.048352604 | 0.828569493 | 0.518273749 | 0.257799607 |
| Tmem115 | S | 320 | 0.043421094 | 0.166380191 | 0.194926291 | 0.151505197 | 0.0285461 | 0.015388434 | 0.017416822 | 0.018335012 | 0.045377839 | 0.55715798 |
| Baz1b | S | 361 | 0.076509652 | 0.131939386 | 0.219433549 | 0.142923898 | 0.087494163 | 0.425103928 | 0.019087405 | 0.002393365 | 0.192758521 | 0.010203731 |
| Ppig | S | 513 | 0.076398669 | 0.096168477 | 0.160196812 | 0.083798143 | 0.064028335 | 0.025165203 | 0.010681 | 0.004109399 | 0.074419361 | 0.10326573 |
| Dcaf8 | S | 123 | -0.034519334 | 0.197901459 | 0.290641131 | 0.325160474 | 0.092739672 | 0.10808741 | 0.010784418 | 0.000658651 | 0.000160735 | 0.048531595 |
| Cdk9 | T | 186 | 0.246670769 | 0.239898172 | 0.063399887 | -0.183270882 | -0.176498285 | 0.038483453 | 0.011338052 | 0.546974482 | 0.142673796 | 0.140646821 |
| Luzp1 | S | 424 | 0.427121157 | 0.133867673 | 0.13589332 | -0.291227837 | 0.002025647 | 0.009271265 | 0.487633833 | 0.300312988 | 0.016656561 | 0.990274116 |
| Scaf1 | S | 312 | 0.084395177 | 0.103449121 | 0.190774571 | 0.106379394 | 0.08732545 | 0.076143011 | 0.042051735 | 0.002015174 | 0.019895685 | 0.044119657 |
| Cobll1 | T | 319 | 0.13424162 | 0.296692103 | 0.199961866 | 0.065720246 | -0.096730237 | 0.325800326 | 0.141879523 | 0.008509227 | 0.589721574 | 0.541417286 |
| Vim | S | 73 | 0.230827183 | 0.248204979 | -0.064149096 | -0.29497628 | -0.312354075 | 0.004607632 | 0.003463383 | 0.619739091 | 0.100816786 | 0.088151467 |
| Bclaf1 | Y | 283 | 0.352107613 | 0.193063612 | 0.040617893 | -0.311489721 | -0.152445719 | 0.003316673 | 0.126539078 | 0.870638406 | 0.280720251 | 0.559132331 |
| Cavin3 | T | 14 | 0.010078378 | 0.211476438 | 0.478967663 | 0.468889285 | 0.267491225 | 0.927997633 | 0.152560678 | 0.011227722 | 0.023154429 | 0.09313909 |
| Xrcc1 | S | 445 | 0.149000963 | 0.113367792 | 0.16667654 | 0.017675577 | 0.053308747 | 0.009831957 | 0.06364181 | 0.039328368 | 0.755468148 | 0.410843031 |
| Ppp6r2 | S | 745 | -0.061870484 | 0.252493466 | 0.302488514 | 0.364358998 | 0.049995048 | 0.504628415 | 0.019827398 | 0.002809805 | 0.0170062 | 0.520938345 |
| Ulk1 | S | 450 | 0.021521376 | 0.216586958 | 0.196727892 | 0.175206516 | -0.019859066 | 0.845384262 | 0.003900032 | 0.008766589 | 0.185031774 | 0.612886444 |
| Itga5 | S | 154 | 0.022200935 | 0.246395856 | 0.19661954 | 0.174418605 | -0.049776316 | 0.789188845 | 0.000632556 | 0.081805162 | 0.145964404 | 0.542066596 |
| Itga5 | S | 155 | 0.022200935 | 0.246395856 | 0.19661954 | 0.174418605 | -0.049776316 | 0.789188845 | 0.000632556 | 0.081805162 | 0.145964404 | 0.542066596 |
| Itga5 | Y | 163 | 0.022200935 | 0.246395856 | 0.19661954 | 0.174418605 | -0.049776316 | 0.789188845 | 0.000632556 | 0.081805162 | 0.145964404 | 0.542066596 |
| Thoc2 | T | 1285 | 0.019381286 | 0.192946717 | 0.12630345 | 0.106923065 | -0.066642366 | 0.616912937 | 7.52336E-05 | 0.028332681 | 0.066491865 | 0.147664429 |
| Men1 | S | 544 | 0.047709793 | 0.119476366 | 0.215467035 | 0.167757242 | 0.095990669 | 0.044597104 | 0.000753185 | 0.048025357 | 0.072369152 | 0.195292004 |
| Numa1 | S | 398 | -0.119124513 | 0.188437833 | 0.434732733 | 0.553857246 | 0.2462949 | 0.01666396 | 0.002960847 | 0.000733433 | 0.003978012 | 0.013506821 |
| Trim28 | S | 752 | -0.011068656 | 0.10622896 | 0.237441706 | 0.248510362 | 0.131212746 | 0.735529394 | 0.064987244 | 1.20614E-05 | 0.003293322 | 0.047400014 |
| Zc3h18 | S | 92 | 0.016027503 | 0.086988917 | 0.154290597 | 0.138263094 | 0.06730168 | 0.411743547 | 0.001150745 | 5.24425E-05 | 0.001215806 | 0.009358745 |
| Cavin1 | S | 38 | -0.076294008 | 0.191250507 | 0.320186907 | 0.396480915 | 0.1289364 | 0.188297944 | 0.00394528 | 0.001445083 | 0.001798197 | 0.058124841 |
| Ehmt2 | T | 608 | 0.16634255 | 0.167393221 | 0.485955152 | 0.319612602 | 0.318561931 | 0.129528281 | 0.32157626 | 0.082980095 | 0.158139609 | 0.186527662 |
| Rsrc1 | S | 5 | 0.011138604 | 0.206176905 | 0.167957001 | 0.156818398 | -0.038219903 | 0.851779129 | 0.000543213 | 0.016412364 | 0.060622091 | 0.400742193 |
| Srsf4 | S | 403 | -0.004818329 | 0.198256434 | 0.241365638 | 0.246183967 | 0.043109204 | 0.933604431 | 0.044335032 | 0.000524322 | 0.015304277 | 0.52880358 |
| Pimreg | S | 128 | 0.067357956 | 0.081392655 | 0.114111328 | 0.046753372 | 0.032718673 | 0.008193001 | 0.005306433 | 0.000887262 | 0.057225553 | 0.131726778 |
| Sipa1l3 | T | 1698 | 0.090435257 | 0.085206728 | 0.359438843 | 0.269003586 | 0.274232115 | 0.111111459 | 0.292027357 | 0.006460338 | 0.040234574 | 0.027853957 |
| Nrcam | S | 1203 | 0.123753877 | 0.253047885 | 0.17933117 | 0.055577293 | -0.073716715 | 0.3098357 | 0.020796351 | 0.045518623 | 0.631988981 | 0.390883584 |
| Zc3h4 | S | 1276 | 0.10783746 | 0.160058663 | 0.180736321 | 0.07289886 | 0.020677658 | 0.071293304 | 0.018386404 | 0.03404578 | 0.248721065 | 0.697443442 |
| Scaf1 | S | 520 | -0.01887301 | 0.1649651 | 0.276860276 | 0.295733285 | 0.111895176 | 0.758644374 | 0.004941128 | 0.003505821 | 0.00871466 | 0.091014821 |
| Bod1l | S | 2989 | 0.022548899 | 0.097333703 | 0.166255775 | 0.143706875 | 0.068922071 | 0.232949709 | 0.000127544 | 0.002485586 | 0.004607316 | 0.053291736 |
| Rcor3 | S | 212 | 0.034143674 | 0.195367841 | 0.241261792 | 0.207118117 | 0.045893951 | 0.698522513 | 0.008235435 | 0.011799634 | 0.071391101 | 0.433642875 |
| Rbbp6 | S | 1053 | 0.034927093 | 0.110562089 | 0.188577113 | 0.153650019 | 0.078015024 | 0.29496963 | 0.003390646 | 0.00139954 | 0.007885697 | 0.061643593 |
| Lyst | S | 2099 | 0.230473738 | 0.282737405 | 0.051825843 | -0.178647895 | -0.230911562 | 0.053707502 | 0.006777768 | 0.815460959 | 0.453374065 | 0.348175519 |
| Cbx5 | S | 11 | 0.026721673 | 0.137184339 | 0.157203682 | 0.130482009 | 0.020019344 | 0.345781521 | 0.003118415 | 0.000860499 | 0.001503528 | 0.418450748 |
| Cdk12 | S | 333 | 0.061415956 | 0.107892397 | 0.126530432 | 0.065114476 | 0.018638035 | 0.006866269 | 0.007330102 | 0.004362962 | 0.045600356 | 0.496025814 |
| Jmjd1c | S | 191 | 0.144874234 | 0.236641035 | 0.213775344 | 0.06890111 | -0.022865691 | 0.279093194 | 0.005919119 | 0.208664554 | 0.680991411 | 0.868273824 |
| Jmjd1c | S | 194 | 0.144874234 | 0.236641035 | 0.213775344 | 0.06890111 | -0.022865691 | 0.279093194 | 0.005919119 | 0.208664554 | 0.680991411 | 0.868273824 |
| Jmjd1c | S | 197 | 0.144874234 | 0.236641035 | 0.213775344 | 0.06890111 | -0.022865691 | 0.279093194 | 0.005919119 | 0.208664554 | 0.680991411 | 0.868273824 |
| Sh3kbp1 | S | 108 | 0.024562488 | 0.190509514 | 0.191424482 | 0.166861994 | 0.000914967 | 0.687904937 | 0.004731542 | 0.005159454 | 0.055281825 | 0.935498298 |
| Epb41l1 | S | 466 | -0.094979462 | 0.290707082 | 0.481589748 | 0.57656921 | 0.190882666 | 0.254413217 | 0.005124808 | 0.042737247 | 0.018600192 | 0.222196908 |
| Thrap3 | S | 264 | 0.016969301 | 0.119454486 | 0.248399734 | 0.231430433 | 0.128945248 | 0.783161891 | 0.096023486 | 9.55271E-05 | 0.034539722 | 0.092657585 |
| Mdc1 | S | 168 | -0.006304085 | 0.204359738 | 0.162464574 | 0.16876866 | -0.041895163 | 0.879077825 | 0.001027068 | 0.003773446 | 0.005454628 | 0.203120188 |

| | | | | | | | | | | | | |
|---|---|---|---|---|---|---|---|---|---|---|---|---|
| Kmt2a | S | 2862 | 0.04295641 | 0.118293783 | 0.23198337 | 0.189026961 | 0.113689587 | 0.615694331 | 0.14041801 | 7.65427E-05 | 0.112054269 | 0.158329686 |
| Pknox1 | S | 41 | 0.096286828 | 0.191771407 | 0.255834784 | 0.159547955 | 0.064063377 | 0.435840881 | 0.222903589 | 0.001508595 | 0.239311239 | 0.630881876 |
| Cobll1 | S | 1143 | 0.456779818 | 0.397816909 | -0.276360208 | -0.733140026 | -0.674177117 | 0.021266661 | 0.034755771 | 0.599891241 | 0.235275106 | 0.263580668 |
| Bod1l | S | 2714 | 0.17890208 | 0.222715184 | 0.137112427 | -0.041789653 | -0.085602757 | 0.036483199 | 0.013570801 | 0.393325631 | 0.77328116 | 0.566326669 |
| Bod1l | S | 2715 | 0.17890208 | 0.222715184 | 0.137112427 | -0.041789653 | -0.085602757 | 0.036483199 | 0.013570801 | 0.393325631 | 0.77328116 | 0.566326669 |
| Bod1l | S | 2716 | 0.17890208 | 0.222715184 | 0.137112427 | -0.041789653 | -0.085602757 | 0.036483199 | 0.013570801 | 0.393325631 | 0.77328116 | 0.566326669 |
| Ncoa1 | S | 570 | 0.086701605 | 0.102132415 | 0.240164877 | 0.153463272 | 0.138032462 | 0.153090833 | 0.142525174 | 0.001403934 | 0.036437774 | 0.073400219 |
| Smarcc2 | S | 304 | 0.011867227 | 0.158287655 | 0.218777076 | 0.206909849 | 0.060489422 | 0.757898775 | 0.014581721 | 0.000995697 | 0.013275833 | 0.107414456 |
| Ralbp1 | S | 92 | -0.004064467 | 0.165985475 | 0.182683957 | 0.186748424 | 0.016698482 | 0.86580321 | 0.002787488 | 0.000810175 | 0.00390497 | 0.615874039 |
| Ralbp1 | S | 93 | -0.004064467 | 0.165985475 | 0.182683957 | 0.186748424 | 0.016698482 | 0.86580321 | 0.002787488 | 0.000810175 | 0.00390497 | 0.615874039 |
| Bicral | S | 930 | 0.107767207 | 0.179465044 | 0.218581168 | 0.110813961 | 0.039116124 | 0.114596463 | 0.008077849 | 0.112884144 | 0.341805963 | 0.702459429 |
| Synpo | T | 69 | 0.123501362 | 0.298215428 | 0.217333792 | 0.093832431 | -0.080881635 | 0.241053751 | 0.044975502 | 0.090893415 | 0.434219844 | 0.527847587 |
| Sipa1l1 | S | 1687 | -0.00916335 | 0.077994638 | 0.257425875 | 0.266589225 | 0.179431237 | 0.68773006 | 0.025498564 | 3.46924E-05 | 3.62401E-06 | 0.005285797 |
| Zfp608 | S | 1429 | 0.111369499 | 0.070600455 | 0.410764615 | 0.299395116 | 0.34016416 | 0.079321369 | 0.436649626 | 0.013582534 | 0.038815107 | 0.028711751 |
| Ank3 | S | 830 | 0.125831179 | 0.12696276 | 0.126832583 | 0.001001404 | -0.000130177 | 0.082821848 | 0.015266937 | 0.005452117 | 0.984955427 | 0.997095232 |
| Gemin8 | S | 181 | 0.06158164 | 0.492056784 | 0.03779287 | -0.02378877 | -0.454263914 | 0.503916061 | 0.001523335 | 0.731159069 | 0.750808241 | 0.010922542 |
| Dnm1 | S | 774 | 0.033230222 | 0.28193865 | 0.151540326 | 0.118310104 | -0.130398324 | 0.598967773 | 0.001832059 | 0.068726391 | 0.158999247 | 0.110875813 |
| Ddx46 | S | 106 | 0.246760946 | 0.162325236 | 0.244696585 | -0.002064361 | 0.082371349 | 0.072674866 | 0.096389894 | 0.151339603 | 0.988797237 | 0.563120162 |
| Nipbl | T | 2661 | 0.106449019 | 0.219503774 | 0.157761221 | 0.051312202 | -0.061742552 | 0.115226989 | 0.004279494 | 0.173274196 | 0.600664539 | 0.523508923 |
| Cenpc1 | S | 699 | -0.000608854 | 0.191326729 | 0.234008685 | 0.234617539 | 0.042681956 | 0.98856871 | 0.002333479 | 0.011206179 | 0.011832074 | 0.432126556 |
| Lmod1 | S | 135 | -0.043039797 | 0.17041415 | 0.422217243 | 0.465257041 | 0.251803093 | 0.509284371 | 0.024519178 | 0.005812498 | 0.010340255 | 0.053968082 |
| Tcf20 | S | 612 | -0.044737142 | 0.219537885 | 0.225548796 | 0.270285938 | 0.006010912 | 0.194637207 | 0.003157523 | 0.001051498 | 0.001257105 | 0.880179911 |
| Cbx5 | S | 12 | 0.02589986 | 0.13609868 | 0.15652394 | 0.13062408 | 0.020425259 | 0.36075422 | 0.003188569 | 0.000896635 | 0.00159837 | 0.40771253 |
| Lysmd1 | S | 193 | 0.067070257 | 0.388614455 | 0.19836413 | 0.131293873 | -0.190250325 | 0.239619934 | 0.031600366 | 0.139626338 | 0.274563317 | 0.203058439 |
| Vim | S | 278 | 0.140108336 | 0.106863544 | 0.202714155 | 0.062605819 | 0.095850611 | 0.128442484 | 0.032865419 | 0.011367532 | 0.442906998 | 0.11989611 |
| Rbl1 | S | 1036 | 0.133712353 | 0.100162224 | 0.21020066 | 0.076488307 | 0.110038436 | 0.033762036 | 0.317865997 | 0.003857123 | 0.081432717 | 0.272505457 |
| Nfib | S | 295 | 0.07886444 | 0.125480441 | 0.179289193 | 0.100424753 | 0.053808753 | 0.041560366 | 0.079904329 | 0.00278565 | 0.012387321 | 0.291692614 |
| Rbm14 | S | 571 | 0.029924524 | 0.123683391 | 0.227344821 | 0.197420297 | 0.103661429 | 0.456452277 | 0.020559934 | 0.001102106 | 0.00617026 | 0.047020737 |
| Pi4ka | S | 263 | 0.242611495 | 0.217724295 | -0.01031555 | -0.252927045 | -0.228039845 | 0.000958403 | 0.042359478 | 0.867794253 | 0.01695319 | 0.042926497 |
| Nop58 | S | 509 | 0.044526373 | 0.096383477 | 0.144226717 | 0.099700344 | 0.04784324 | 0.212556034 | 0.002922372 | 0.000320122 | 0.046259184 | 0.026632231 |
| Spp1 | S | 289 | 0.048709738 | 0.15936298 | 0.095098355 | 0.046388617 | -0.064264625 | 0.015367844 | 0.002876402 | 0.008662891 | 0.072223592 | 0.046639884 |
| Ilvbl | S | 369 | 0.087879649 | 0.269627684 | 0.13880272 | 0.050923071 | -0.130824965 | 0.217787448 | 0.004135293 | 0.158123092 | 0.530418353 | 0.173357445 |
| Sp110 | S | 214 | -0.026624023 | 0.153229522 | 0.332403678 | 0.359027701 | 0.179174156 | 0.575945426 | 0.014011213 | 0.002731385 | 0.008331704 | 0.042017413 |
| Sp110 | S | 226 | -0.026624023 | 0.153229522 | 0.332403678 | 0.359027701 | 0.179174156 | 0.575945426 | 0.014011213 | 0.002731385 | 0.008331704 | 0.042017413 |
| Mki67 | S | 2425 | 0.16363461 | 0.252422197 | 0.131840078 | -0.031794532 | -0.12058212 | 0.180048726 | 0.032610444 | 0.04992027 | 0.754449882 | 0.194231437 |
| Sipa1l1 | S | 255 | 0.012721939 | 0.190842948 | 0.260180736 | 0.247458797 | 0.069337788 | 0.751004806 | 0.002783934 | 0.027382202 | 0.047507734 | 0.376181866 |
| Rcc1 | S | 11 | -0.002546875 | 0.098104882 | 0.201087685 | 0.203634559 | 0.102982803 | 0.955020999 | 0.005898173 | 6.86803E-05 | 0.020932889 | 0.007363406 |
| Pml | S | 490 | 0.034585463 | 0.191966768 | 0.152350014 | 0.117764551 | -0.039616753 | 0.497615183 | 0.000400411 | 0.038130796 | 0.103255735 | 0.43563284 |
| Chd4 | S | 1528 | 0.027303646 | 0.12070055 | 0.166726223 | 0.139422577 | 0.046025672 | 0.414677627 | 0.001365221 | 0.001545497 | 0.019004496 | 0.061187856 |
| Ctnnd1 | T | 310 | 0.181568456 | 0.191994651 | 0.071092075 | -0.110476381 | -0.120902576 | 0.056543359 | 0.0053996 | 0.159513749 | 0.187565444 | 0.019070605 |
| Eif5b | S | 209 | 0.169550033 | 0.163323444 | 0.125002404 | -0.044547629 | -0.03832104 | 0.052981623 | 0.032512293 | 0.041614814 | 0.477896002 | 0.443786321 |
| Msl3 | S | 371 | 0.200395725 | 0.073681637 | 0.320975207 | 0.120579482 | 0.24729357 | 0.130614874 | 0.595574665 | 0.007685346 | 0.281196671 | 0.126554898 |
| Wdr26 | S | 101 | 0.059336301 | 0.117224991 | 0.131405355 | 0.072069054 | 0.014180364 | 0.011971212 | 0.010421666 | 0.004676443 | 0.056665903 | 0.654948854 |
| Usp6nl | S | 631 | 0.243406373 | 0.397899136 | 0.057917122 | -0.18548925 | -0.339982014 | 0.101571642 | 0.021486042 | 0.837945998 | 0.525726216 | 0.288235662 |
| Irf2bpl | S | 313 | 0.000194308 | 0.125767948 | 0.271523125 | 0.271328817 | 0.145755177 | 0.994143291 | 0.001613157 | 0.005800757 | 0.009001821 | 0.043214969 |
| Sp100 | S | 190 | 0.065693741 | 0.188149446 | 0.261019978 | 0.195326237 | 0.072870533 | 0.481912591 | 0.037984043 | 0.011542238 | 0.055439898 | 0.200696997 |
| Eif4enif1 | S | 949 | 0.007066545 | 0.1887933 | 0.160324601 | 0.153258056 | -0.028468699 | 0.841359354 | 0.02014194 | 0.000153494 | 0.023688592 | 0.530384492 |
| Acin1 | S | 710 | 0.003437388 | 0.175183467 | 0.286049075 | 0.282611687 | 0.110865608 | 0.947036132 | 0.002945509 | 0.020420009 | 0.022872854 | 0.197603446 |
| Trp53bp1 | S | 622 | 0.053080457 | 0.155000074 | 0.119011581 | 0.065931124 | -0.035988493 | 0.139966778 | 0.001956061 | 0.006764128 | 0.031448011 | 0.08696445 |
| Paxbp1 | S | 559 | 0.053279676 | 0.076692806 | 0.197898528 | 0.144618851 | 0.121205722 | 0.24899583 | 0.179370797 | 2.74282E-05 | 0.04681082 | 0.088754811 |
| Paxbp1 | S | 560 | 0.053279676 | 0.076692806 | 0.197898528 | 0.144618851 | 0.121205722 | 0.24899583 | 0.179370797 | 2.74282E-05 | 0.04681082 | 0.088754811 |
| Ccdc9b | T | 146 | 0.033915072 | 0.188694469 | 0.180848283 | 0.146933211 | -0.007846186 | 0.453447474 | 0.000740198 | 0.046037061 | 0.082336827 | 0.893430124 |
| Brca1 | S | 1150 | 0.040733532 | 0.097583443 | 0.267314812 | 0.22658128 | 0.169731369 | 0.256452107 | 0.054386184 | 0.001675108 | 0.006187935 | 0.017501176 |
| Mycbp2 | S | 2893 | 0.066446414 | 0.177729734 | 0.246578412 | 0.180131998 | 0.068848679 | 0.134410326 | 0.0284121 | 0.033662192 | 0.078089037 | 0.40855047 |
| Smarcc2 | T | 322 | 0.214213563 | 0.196487364 | 0.219990758 | 0.005777195 | 0.023503394 | 0.137469065 | 0.120695623 | 0.06518989 | 0.959156646 | 0.805677611 |
| Rai14 | T | 297 | 0.294452286 | 0.111609088 | 0.123366555 | -0.171085731 | 0.011757467 | 0.006187961 | 0.315842017 | 0.143911496 | 0.025635832 | 0.896178353 |
| Cbx5 | S | 13 | 0.024379754 | 0.13170348 | 0.151981327 | 0.127601574 | 0.020277847 | 0.368179373 | 0.002875919 | 0.000844017 | 0.002101733 | 0.393671533 |
| Xrcc6 | S | 6 | 0.08283165 | 0.187932226 | 0.185675204 | 0.102843555 | -0.002257022 | 0.525828609 | 0.003453629 | 0.029717528 | 0.446953781 | 0.966707436 |
| Xrcc6 | Y | 7 | 0.08283165 | 0.187932226 | 0.185675204 | 0.102843555 | -0.002257022 | 0.525828609 | 0.003453629 | 0.029717528 | 0.446953781 | 0.966707436 |
| Xrcc6 | Y | 8 | 0.08283165 | 0.187932226 | 0.185675204 | 0.102843555 | -0.002257022 | 0.525828609 | 0.003453629 | 0.029717528 | 0.446953781 | 0.966707436 |
| Arhgef17 | S | 822 | 0.049400327 | 0.057188301 | 0.201417408 | 0.152017081 | 0.144229107 | 0.255844036 | 0.050701093 | 6.08204E-05 | 0.02617377 | 0.001506049 |
| Nipbl | S | 2666 | 0.07556398 | 0.206846457 | 0.200255175 | 0.124691195 | -0.006591282 | 0.183079981 | 0.006315762 | 0.093605826 | 0.232854834 | 0.941181036 |
| Ncor1 | S | 2165 | -0.016893255 | 0.104114442 | 0.26745956 | 0.284352815 | 0.163345119 | 0.7027499 | 0.074718976 | 5.02822E-05 | 0.004646943 | 0.028045516 |
| Smarca4 | S | 613 | 0.002717496 | 0.161739743 | 0.228594113 | 0.225876617 | 0.066854371 | 0.969201328 | 0.011130702 | 0.001701382 | 0.03763564 | 0.183255625 |
| Klf16 | S | 233 | 0.082095364 | 0.202895149 | 0.360691594 | 0.27859623 | 0.157796445 | 0.551804735 | 0.093956353 | 0.024649672 | 0.109554532 | 0.231546797 |
| Slc38a10 | S | 887 | -0.034425406 | 0.150953389 | 0.294509975 | 0.328935381 | 0.143556586 | 0.298020731 | 0.00022239 | 0.011513682 | 0.005669969 | 0.066417688 |
| Eif4g1 | S | 1146 | 0.053783011 | 0.350444034 | 0.07915539 | 0.025372378 | -0.271288645 | 0.484999475 | 0.000611437 | 0.421397675 | 0.790427307 | 0.057544389 |
| Bclaf1 | S | 258 | 0.027894961 | 0.27116162 | 0.323244888 | 0.295349927 | 0.052083268 | 0.580847865 | 0.020470216 | 0.073656203 | 0.097073928 | 0.691934083 |
| Ulk2 | S | 781 | 0.047646767 | 0.229245109 | 0.238661482 | 0.191014716 | 0.009416373 | 0.466362019 | 0.01582384 | 0.033108468 | 0.075875663 | 0.906603895 |
| Ttc7 | S | 183 | 0.062317385 | 0.143938444 | 0.256711656 | 0.194394271 | 0.112773211 | 0.334980813 | 0.052030718 | 0.005606832 | 0.001258717 | 0.004412321 |
| Sfswap | S | 863 | 0.198069312 | 0.099595334 | 0.128689179 | -0.069380133 | 0.029093845 | 0.002495729 | 0.16437647 | 0.092794679 | 0.27954277 | 0.68612307 |
| Cdk13 | S | 359 | 0.079679223 | 0.108124312 | 0.210852906 | 0.131173683 | 0.102728594 | 0.061513019 | 0.04249095 | 0.007862767 | 0.028689324 | 0.067007808 |
| Fgd1 | S | 82 | 0.37022656 | 0.09404415 | 0.21204171 | -0.15818485 | 0.11799756 | 0.044850067 | 0.424466637 | 0.09299502 | 0.294333982 | 0.374747963 |
| Palm | T | 367 | 0.095702493 | 0.405387408 | 0.235324976 | 0.139622482 | -0.170062432 | 0.589451738 | 0.057556258 | 0.083460466 | 0.439076849 | 0.325884308 |
| Srsf4 | S | 315 | 0.011911414 | 0.351889253 | 0.162739785 | 0.150828371 | -0.189149468 | 0.905541751 | 0.001812711 | 0.166471769 | 0.229702426 | 0.130064935 |
| Srsf4 | S | 317 | 0.011911414 | 0.351889253 | 0.162739785 | 0.150828371 | -0.189149468 | 0.905541751 | 0.001812711 | 0.166471769 | 0.229702426 | 0.130064935 |
| Srsf4 | S | 321 | 0.011911414 | 0.351889253 | 0.162739785 | 0.150828371 | -0.189149468 | 0.905541751 | 0.001812711 | 0.166471769 | 0.229702426 | 0.130064935 |
| Top2b | S | 1376 | -0.027662969 | 0.146105255 | 0.18583454 | 0.21349751 | 0.039729285 | 0.268134856 | 0.006131089 | 2.97091E-05 | 0.000684663 | 0.221602949 |
| Nsd3 | S | 560 | 0.11971156 | 0.106153633 | 0.195969559 | 0.076257999 | 0.089815927 | 0.097725519 | 0.065318501 | 0.006223015 | 0.200905257 | 0.032173092 |
| Vim | S | 438 | 0.125968619 | 0.105094071 | 0.25504872 | 0.129080101 | 0.149954649 | 0.106060821 | 0.238883292 | 0.00595681 | 0.049083654 | 0.103347323 |
| Naa15 | S | 588 | 0.075902954 | 0.154471602 | 0.225258952 | 0.149355999 | 0.07078735 | 0.101257659 | 0.019629243 | 0.035892333 | 0.100033901 | 0.35491368 |
| Ptpra | Y | 825 | -0.077349976 | 0.236763277 | 0.30171308 | 0.379063056 | 0.064949803 | 0.300537767 | 0.031025404 | 0.000856113 | 0.00665919 | 0.404779063 |
| Tnrc18 | Y | 1844 | -0.018182999 | 0.312865242 | 0.26844989 | 0.286632889 | -0.044415351 | 0.842996352 | 0.045886171 | 0.00823461 | 0.031391227 | 0.687153246 |
| C77080 | S | 325 | 0.205341306 | 0.275895173 | 0.215110384 | 0.009769079 | -0.060784789 | 0.113317322 | 0.107805552 | 0.178796158 | 0.945662776 | 0.71391771 |
| Fndc3a | S | 213 | 0.040272176 | 0.086283661 | 0.270041026 | 0.229768851 | 0.183757365 | 0.779811321 | 0.067823774 | 0.000304111 | 0.201314787 | 0.003566474 |
| Hmga2 | T | 99 | 0.040129514 | 0.100017809 | 0.111431283 | 0.071301769 | 0.011413474 | 0.076252896 | 0.001266193 | 0.000482768 | 0.009711639 | 0.429141475 |
| Larp4b | T | 735 | -0.031216732 | 0.146779753 | 0.127481333 | 0.158698065 | -0.019298421 | 0.22058468 | 5.1623E-06 | 0.000966541 | 0.003008405 | 0.26746456 |
| Zbtb11 Rpl24 | T | 341 | 0.132393813 | 0.19606461 | 0.042498451 | -0.089895362 | -0.153566158 | 0.015717579 | 0.00152012 | 0.425325354 | 0.115058352 | 0.031857282 |
| Hmgn3 | S | 6 | 0.014970631 | 0.182084807 | 0.229186602 | 0.214215972 | 0.047101795 | 0.790485063 | 0.00241467 | 0.019037587 | 0.030559334 | 0.440374816 |
| Chaf1a | T | 297 | 0.207028813 | 0.200868976 | 0.176511312 | -0.030517501 | -0.024357664 | 0.056744514 | 0.107516737 | 0.113722343 | 0.622839712 | 0.788791444 |
| Snrnp27 | S | 61 | 0.043281128 | 0.10625324 | 0.179185956 | 0.135904828 | 0.072932715 | 0.175132421 | 0.002724096 | 0.004089692 | 0.016931222 | 0.097438285 |
| Clcc1 | S | 58 | 0.02045731 | 0.148524549 | 0.132911724 | 0.112454414 | -0.015612825 | 0.707220699 | 0.000469462 | 0.002032824 | 0.125270511 | 0.502889511 |
| Msh6 | S | 261 | 0.020303833 | 0.13048374 | 0.171506632 | 0.151202799 | 0.041022892 | 0.512481082 | 0.004151301 | 0.000905557 | 0.001861041 | 0.099737437 |
| Sdc1 | S | 288 | 0.004323964 | 0.141380532 | 0.503411982 | 0.499088019 | 0.36203145 | 0.960708569 | 0.146259444 | 0.009926025 | 0.02629654 | 0.050556177 |

| | | | | | | | | | | | | |
|---|---|---|---|---|---|---|---|---|---|---|---|---|
| Dxo | S | 394 | 0.06145005 | 0.320007209 | 0.200320065 | 0.138870014 | -0.119687144 | 0.216909775 | 0.027528298 | 0.100404881 | 0.199358478 | 0.309989931 |
| Hells | S | 815 | 0.305303459 | 0.18474095 | 0.068546234 | -0.236757226 | -0.116194716 | 0.003216752 | 0.142572727 | 0.771046152 | 0.360407471 | 0.634807253 |
| Vps50 | S | 494 | 0.013103946 | 0.231291709 | 0.173744684 | 0.160640738 | -0.057547026 | 0.762928976 | 0.033133295 | 0.000852557 | 0.022111556 | 0.399981036 |
| Arhgef17 | S | 324 | 0.11870945 | 0.223515949 | 0.151684391 | 0.032974941 | -0.071831558 | 0.044596088 | 0.012132414 | 0.256467822 | 0.773699528 | 0.554606847 |
| Arhgef17 | S | 330 | 0.11870945 | 0.223515949 | 0.151684391 | 0.032974941 | -0.071831558 | 0.044596088 | 0.012132414 | 0.256467822 | 0.773699528 | 0.554606847 |
| Lig1 | S | 906 | 0.018423339 | 0.193851795 | 0.11394004 | 0.095516701 | -0.079911755 | 0.566331922 | 9.42342E-05 | 0.037399741 | 0.06800153 | 0.090246683 |
| Sipa1l1 | S | 210 | 0.035031458 | 0.116047804 | 0.319022735 | 0.283991277 | 0.202974931 | 0.542117286 | 0.188459634 | 0.001163091 | 0.003465451 | 0.05230029 |
| Dbr1 | S | 491 | 0.083485495 | 0.174682447 | 0.124810595 | 0.0413251 | -0.049871851 | 0.094863858 | 0.005514421 | 0.031764666 | 0.284186379 | 0.194441318 |
| Prune2 | S | 2634 | 0.102589388 | 0.197609148 | 0.16923424 | 0.066644851 | -0.028374909 | 0.311011402 | 0.022563872 | 0.015317193 | 0.488540113 | 0.620550161 |
| Srrm2 | S | 1230 | 0.001953674 | 0.088761532 | 0.213276658 | 0.211322984 | 0.124515126 | 0.958649934 | 0.008357924 | 0.000114449 | 0.006172927 | 0.005144945 |
| Prpf6 | S | 279 | 0.20852983 | 0.141838808 | 0.268989728 | 0.060459898 | 0.127150921 | 0.156156824 | 0.153293345 | 0.044834192 | 0.654922607 | 0.258944555 |
| Rrbp1 | S | 786 | 0.023730286 | 0.221928732 | 0.130863301 | 0.107133015 | -0.091065431 | 0.806137817 | 0.001088571 | 0.013646903 | 0.313700743 | 0.029825864 |
| Gtf3c4 | S | 607 | 0.003275419 | 0.109667571 | 0.190540297 | 0.187264879 | 0.080872726 | 0.930277383 | 0.004755705 | 0.000210499 | 0.011075738 | 0.008594969 |
| Gtf2i | S | 674 | 0.086988718 | 0.070218393 | 0.119148503 | 0.032159785 | 0.04893011 | 0.002155102 | 0.010078244 | 0.008670549 | 0.263781175 | 0.132189647 |
| Herpud2 | S | 90 | -0.028354953 | 0.224488742 | 0.279914021 | 0.308268975 | 0.055425279 | 0.502447483 | 0.003055635 | 0.019526231 | 0.008259477 | 0.380956071 |
| Prkaca | S | 339 | 0.068339144 | 0.189807527 | 0.137562441 | 0.069223297 | -0.052245086 | 0.115631456 | 0.007814685 | 0.026769426 | 0.162457261 | 0.32940271 |
| Scaf1 | S | 510 | 0.003684346 | 0.123042985 | 0.120968388 | 0.117284042 | -0.002074597 | 0.878808107 | 0.000751125 | 6.06111E-05 | 0.017775874 | 0.895329861 |
| Septin10 | S | 413 | 0.109653394 | 0.266074293 | 0.146719863 | 0.037066468 | -0.119354431 | 0.286780102 | 0.012059914 | 0.091925506 | 0.615968931 | 0.004103123 |
| Cenpc1 | S | 59 | 0.144666205 | 0.077935061 | 0.202830539 | 0.058164334 | 0.124895478 | 0.011364267 | 0.253863037 | 0.016328469 | 0.304476969 | 0.125127887 |
| Carmil1 | S | 1335 | 0.066196661 | 0.276314182 | 0.037797868 | -0.028398793 | -0.238516313 | 0.213513579 | 0.00024306 | 0.316596474 | 0.521746501 | 0.001978951 |
| Gatad2b | S | 334 | -0.081193844 | 0.208191009 | 0.422058523 | 0.503252367 | 0.213867514 | 0.655278524 | 0.040506327 | 0.009053257 | 0.064187341 | 0.079581714 |
| Dab2ip | S | 747 | 0.065016342 | 0.126919102 | 0.1525275 | 0.087511158 | 0.025608398 | 0.085237758 | 0.007387533 | 0.008621558 | 0.077112961 | 0.457778071 |
| Mki67 | S | 2392 | 0.170346665 | 0.258333987 | 0.120469426 | -0.049877238 | -0.137864561 | 0.143479723 | 0.013619808 | 0.24404853 | 0.664025193 | 0.206310646 |
| Raf1 | S | 642 | 0.020148075 | 0.175403683 | 0.201969433 | 0.181821357 | 0.02656575 | 0.474145446 | 0.069982816 | 0.000526736 | 0.005898123 | 0.692464569 |
| Nf1 | S | 878 | 0.123565986 | 0.158864472 | 0.132326754 | 0.008760768 | -0.026537718 | 0.071395762 | 0.016194835 | 0.037341814 | 0.878155363 | 0.606593963 |
| Cep89 | T | 35 | 0.059051274 | 0.138556439 | 0.167845032 | 0.108793758 | 0.029288593 | 0.156691601 | 0.023544185 | 0.002988565 | 0.007716949 | 0.465692993 |
| Bclaf1 | S | 261 | 0.158895764 | 0.222960559 | 0.308498117 | 0.149602353 | 0.085537558 | 0.103311121 | 0.238209623 | 0.084656267 | 0.31023514 | 0.651088534 |
| Arfgef2 | S | 627 | 0.112422842 | 0.174046948 | 0.148170356 | 0.035747514 | -0.025876592 | 0.061000056 | 0.045392186 | 0.023868668 | 0.154207353 | 0.657172505 |
| Zc3h13 | S | 371 | 0.18641019 | 0.185744216 | 0.242941123 | 0.056530933 | 0.057196907 | 0.052616875 | 0.129396921 | 0.148303366 | 0.667970064 | 0.689072207 |
| Ccnl1 | S | 347 | 0.000746561 | 0.145580506 | 0.185563838 | 0.184817277 | 0.039983332 | 0.976748945 | 0.01070873 | 0.00032457 | 0.003442824 | 0.211019031 |
| Zfp687 | S | 242 | -0.047780983 | 0.153359758 | 0.300986786 | 0.348767769 | 0.147627028 | 0.573531153 | 0.03287447 | 0.00071165 | 0.024648099 | 0.032804583 |
| Srrm2 | S | 762 | 0.129962936 | 0.127133853 | 0.123238332 | -0.006724604 | -0.003895521 | 0.009077502 | 0.023628409 | 0.074878968 | 0.89413135 | 0.941572648 |
| Mki67 | T | 2213 | 0.255304094 | 0.20054245 | 0.196969878 | -0.058334216 | -0.003572572 | 0.086449042 | 0.099345533 | 0.197064436 | 0.669627983 | 0.975295058 |
| Depdc5 | S | 525 | 0.086772353 | 0.160499969 | 0.187383288 | 0.100610935 | 0.026883319 | 0.186907622 | 0.016282142 | 0.022883161 | 0.172726736 | 0.62692566 |
| Atf7 | S | 188 | 0.320382182 | 0.251474443 | 0.217556068 | -0.102826114 | -0.033918375 | 0.052517363 | 0.277274569 | 0.322641621 | 0.591390413 | 0.887591981 |
| Jph2 | S | 593 | 0.04377102 | 0.162584017 | 0.282413021 | 0.238642001 | 0.119829005 | 0.67107745 | 0.008721498 | 0.024179207 | 0.095439804 | 0.142811226 |
| Jph2 | S | 597 | 0.04377102 | 0.162584017 | 0.282413021 | 0.238642001 | 0.119829005 | 0.67107745 | 0.008721498 | 0.024179207 | 0.095439804 | 0.142811226 |
| Mapkbp1 | S | 19 | 0.242033636 | 0.147535122 | 0.090930482 | -0.151103154 | -0.056604639 | 0.004332282 | 0.093666794 | 0.380946425 | 0.180094087 | 0.582894036 |
| Cpsf6 | S | 494 | 0.226189722 | 0.171820958 | 0.169479151 | -0.056710571 | -0.002341807 | 0.036760094 | 0.312217783 | 0.045848829 | 0.497704129 | 0.98755597 |
| Cpsf6 | S | 500 | 0.226189722 | 0.171820958 | 0.169479151 | -0.056710571 | -0.002341807 | 0.036760094 | 0.312217783 | 0.045848829 | 0.497704129 | 0.98755597 |
| Top2b | S | 1363 | -0.024296603 | 0.142137322 | 0.197964923 | 0.222261527 | 0.055827601 | 0.318504288 | 0.022380377 | 1.92393E-05 | 0.000946308 | 0.177232038 |
| Nipbl | T | 757 | 0.108097135 | 0.1548232 | 0.270391862 | 0.162294727 | 0.115568662 | 0.21546647 | 0.120821549 | 0.016515871 | 0.078259298 | 0.206001628 |
| Tfe3 | S | 565 | 0.012835617 | 0.15016199 | 0.17921464 | 0.166379023 | 0.02905265 | 0.738147631 | 0.004755141 | 0.001614997 | 0.003814304 | 0.349335707 |
| Sipa1l3 | T | 1381 | 0.159836627 | 0.080396247 | 0.19279666 | 0.032960033 | 0.112400414 | 0.170390711 | 0.351581443 | 0.000502943 | 0.717927289 | 0.230368025 |
| Vamp4 | S | 30 | 0.060956501 | 0.142573287 | 0.185050919 | 0.124094418 | 0.042477632 | 0.230868819 | 0.020549282 | 0.005149227 | 0.017571207 | 0.279575049 |
| Iws1 | S | 295 | 0.201726639 | 0.09234016 | 0.218407935 | 0.016681296 | 0.126067775 | 0.032301771 | 0.184674949 | 0.052051703 | 0.851387086 | 0.194883557 |
| Hmbox1 | S | 148 | 0.107760318 | 0.129666234 | 0.230449034 | 0.122688717 | 0.100782801 | 0.168781678 | 0.04029371 | 0.02056663 | 0.171238998 | 0.187804712 |
| Peak1 | S | 584 | 0.399399795 | 0.074069596 | 0.004217511 | -0.395182284 | -0.069852085 | 0.000263598 | 0.578529318 | 0.958890494 | 0.013813358 | 0.624402735 |
| Gpatch8 | S | 1179 | 0.067987081 | 0.085788124 | 0.125325682 | 0.057338601 | 0.039537557 | 0.020216866 | 0.01729862 | 0.001129412 | 0.005089797 | 0.139929575 |
| Zrsr1 | S | 50 | 0.040221249 | 0.163628241 | 0.098860745 | 0.058639496 | -0.064767496 | 0.173998432 | 0.001015145 | 0.007050931 | 0.033185145 | 0.062113821 |
| Dennd1a | S | 592 | -0.034663166 | 0.210449831 | 0.169513808 | 0.204176973 | -0.040936024 | 0.482396029 | 0.001014591 | 0.003407495 | 0.009779472 | 0.199277216 |
| Marf1 | S | 1677 | 0.026916611 | 0.270036238 | 0.276248024 | 0.249331414 | 0.006211787 | 0.717180625 | 0.084886615 | 0.011040312 | 0.027739525 | 0.958300784 |
| Atad2 | S | 870 | -0.204434286 | 0.571599736 | 0.27632103 | 0.480755316 | -0.295278706 | 0.64641633 | 0.011796135 | 0.145403398 | 0.322390625 | 0.084239539 |
| Atad2 | T | 867 | -0.204434286 | 0.571599736 | 0.27632103 | 0.480755316 | -0.295278706 | 0.64641633 | 0.011796135 | 0.145403398 | 0.322390625 | 0.084239539 |
| Carmil1 | S | 972 | 0.081584399 | 0.291859883 | 0.137188949 | 0.05560455 | -0.154670934 | 0.308129952 | 0.009894797 | 0.107372625 | 0.1130078 | 0.066104925 |
| Mfsd11 | S | 204 | 0.091616574 | 0.281800509 | 0.149417091 | 0.057800517 | -0.132383418 | 0.224253485 | 0.014063264 | 0.125181866 | 0.454937009 | 0.179911241 |
| Srrm2 | S | 1229 | 0.000456454 | 0.08621832 | 0.209889221 | 0.209432767 | 0.123670901 | 0.990441153 | 0.008990994 | 0.00010923 | 0.007206981 | 0.004562962 |
| Cdca2 | S | 758 | 0.082603389 | 0.153441299 | 0.113187325 | 0.030583936 | -0.040253974 | 0.098395792 | 0.001366044 | 0.052547105 | 0.543942343 | 0.360367816 |
| Snrnp27 | S | 59 | 0.044755704 | 0.106642238 | 0.165507825 | 0.120752121 | 0.058865588 | 0.170914933 | 0.003422263 | 0.003785436 | 0.019983401 | 0.13699935 |
| Sbno2 | T | 1237 | -0.032209733 | 0.217493027 | 0.179310874 | 0.211520608 | -0.038182153 | 0.72634491 | 0.003745137 | 0.00310783 | 0.100580891 | 0.40824883 |
| Frmd6 | S | 349 | -0.025824266 | 0.288227066 | 0.296244144 | 0.32206841 | 0.008017077 | 0.869566659 | 0.029395485 | 0.021173461 | 0.102907215 | 0.905754589 |
| Lemd3 | S | 140 | 0.05065234 | 0.145414035 | 0.145697313 | 0.095044973 | 0.000283279 | 0.168336317 | 0.006180955 | 0.006562663 | 0.049918362 | 0.992889383 |
| Chaf1a | T | 50 | 0.018325976 | 0.216419194 | 0.169801832 | 0.151475856 | -0.046617362 | 0.740156502 | 0.000361962 | 0.095141084 | 0.126403101 | 0.540408168 |
| Msh6 | S | 63 | -0.000469201 | 0.23562843 | 0.201885386 | 0.202354587 | -0.033743044 | 0.98532347 | 0.008577348 | 0.010516366 | 0.007235925 | 0.479891722 |
| Numa1 | S | 1782 | 0.159137689 | 0.202446649 | 0.216460977 | 0.057323288 | 0.014014328 | 0.119975578 | 0.045876833 | 0.14745923 | 0.639272162 | 0.903308175 |
| Zgrf1 | S | 462 | 0.090321578 | 0.094666586 | 0.173777218 | 0.08345564 | 0.079110632 | 0.019662671 | 0.001582577 | 0.0966546 | 0.30283421 | 0.322749173 |
| Pikfyve | S | 475 | 0.152432661 | 0.239359629 | 0.106871789 | -0.045560872 | -0.13248784 | 0.057120318 | 0.018958156 | 0.261823053 | 0.620201446 | 0.202367741 |
| Brpf3 | S | 712 | 0.085767386 | 0.217105782 | 0.114789076 | 0.02902169 | -0.102316705 | 0.197519507 | 0.026995886 | 0.007031915 | 0.610928898 | 0.14082758 |
| Pdlim2 | S | 137 | -0.0364169 | 0.094698421 | 0.259019413 | 0.295436313 | 0.164320992 | 0.361014877 | 0.034851483 | 3.03156E-05 | 0.00261845 | 0.008185311 |
| Trp53bp1 | S | 1115 | 0.004676169 | 0.097173176 | 0.270747359 | 0.26607119 | 0.173574183 | 0.911051475 | 0.018431813 | 0.000987503 | 0.003133722 | 0.016417175 |
| Heatr6 | S | 673 | 0.090605623 | 0.330976015 | 0.242807038 | 0.152201415 | -0.088168978 | 0.472712282 | 0.006228296 | 0.426069682 | 0.604541276 | 0.752178012 |
| Heatr6 | S | 675 | 0.090605623 | 0.330976015 | 0.242807038 | 0.152201415 | -0.088168978 | 0.472712282 | 0.006228296 | 0.426069682 | 0.604541276 | 0.752178012 |
| Heatr6 | S | 679 | 0.090605623 | 0.330976015 | 0.242807038 | 0.152201415 | -0.088168978 | 0.472712282 | 0.006228296 | 0.426069682 | 0.604541276 | 0.752178012 |
| Septin9 | S | 85 | 0.156147684 | 0.110635464 | 0.085268295 | -0.070879389 | -0.025367169 | 0.003194172 | 0.133799841 | 0.016175331 | 0.063514458 | 0.656676559 |
| Zc3h13 | S | 1343 | -0.074066074 | 0.214945913 | 0.467977075 | 0.54204315 | 0.253031162 | 0.28111243 | 0.008274313 | 0.026536534 | 0.025243776 | 0.131588149 |
| Rnmt | S | 15 | 0.02333605 | 0.118081887 | 0.148546891 | 0.12521084 | 0.030465004 | 0.084333008 | 0.001440113 | 0.004690503 | 0.017998047 | 0.267724778 |
| Acly | S | 325 | 0.106583715 | 0.153837134 | 0.16158789 | 0.055004175 | 0.007750756 | 0.245538044 | 0.036860132 | 0.005837778 | 0.50243616 | 0.871597326 |
| Acly | T | 322 | 0.106583715 | 0.153837134 | 0.16158789 | 0.055004175 | 0.007750756 | 0.245538044 | 0.036860132 | 0.005837778 | 0.50243616 | 0.871597326 |
| Septin9 | T | 143 | 0.382415442 | 0.106689053 | 0.022578647 | -0.359836795 | -0.084110406 | 0.000764083 | 0.435715561 | 0.867869074 | 0.078922888 | 0.612000235 |
| Lrrfip1 | S | 343 | 0.008215245 | 0.288450906 | 0.212162705 | 0.203947459 | -0.076288201 | 0.909508683 | 0.006070973 | 0.061491447 | 0.080569023 | 0.413144368 |
| Atn1 | T | 615 | 0.029308024 | 0.112857072 | 0.215259122 | 0.185951098 | 0.10240205 | 0.50614766 | 0.013854654 | 0.001654622 | 0.0089251 | 0.042792203 |
| Nsd1 | S | 2129 | 0.134920492 | 0.111125143 | 0.193104754 | 0.058184262 | 0.081979611 | 0.00974943 | 0.024201297 | 0.178520263 | 0.613146381 | 0.489715771 |
| Arl6 | S | 8 | -0.061519689 | 0.185253066 | 0.208856845 | 0.270376534 | 0.023603779 | 0.199164256 | 0.003392578 | 0.000310552 | 0.01425187 | 0.472861076 |
| Cit | S | 2021 | 0.28527931 | 0.281004867 | -0.106172691 | -0.391452001 | -0.387177558 | 0.023603949 | 0.003248091 | 0.446462584 | 0.04759217 | 0.061102018 |
| Svil | S | 50 | 0.216698384 | 0.046376714 | 0.255880618 | 0.039182235 | 0.209503904 | 0.015482044 | 0.770450949 | 0.029625887 | 0.6391498 | 0.262866273 |
| Aplf | S | 330 | -0.028627897 | 0.139257664 | 0.200172323 | 0.228800219 | 0.060914658 | 0.346641016 | 0.003040696 | 0.000265115 | 0.001473912 | 0.074326352 |
| Rbl1 | S | 983 | 0.149093981 | 0.187022896 | 0.179017788 | 0.029923806 | -0.008005109 | 0.070822678 | 0.115305864 | 0.045532324 | 0.584445337 | 0.930553742 |
| Rbl1 | T | 992 | 0.149093981 | 0.187022896 | 0.179017788 | 0.029923806 | -0.008005109 | 0.070822678 | 0.115305864 | 0.045532324 | 0.584445337 | 0.930553742 |
| Fam171a2 | S | 785 | -0.114714356 | 0.281466872 | 0.332013685 | 0.446728041 | 0.050546813 | 0.262528416 | 0.042895628 | 0.001668341 | 0.010809039 | 0.607862375 |
| Ralbp1 | S | 99 | 0.231077083 | 0.207760879 | 0.277949129 | 0.046872046 | 0.07018825 | 0.126896148 | 0.153865664 | 0.178593033 | 0.796215861 | 0.698287331 |
| Pard3b | S | 346 | 0.065510854 | 0.108090114 | 0.245149657 | 0.179638802 | 0.137059543 | 0.054791896 | 0.037492191 | 0.020863572 | 0.054505957 | 0.087960293 |
| Sipa1 | S | 53 | 0.105680948 | 0.070118024 | 0.109679736 | 0.003998787 | 0.039561712 | 0.004775805 | 0.024604985 | 0.00638795 | 0.839957417 | 0.137009204 |
| AW554918 | S | 20 | 0.071153565 | 0.096866484 | 0.0654415 | -0.005712065 | -0.031424983 | 0.00471342 | 0.00057214 | 0.012030931 | 0.745960134 | 0.119488374 |
| Eftud2 | S | 361 | -0.020197966 | 0.131834804 | 0.294259042 | 0.314457008 | 0.162424239 | 0.55470921 | 0.006833768 | 0.003100936 | 0.008780276 | 0.036362836 |

| | | | | | | | | | | | | |
|---|---|---|---|---|---|---|---|---|---|---|---|---|
| Son | S | 2027 | 0.083801915 | 0.030467559 | 0.15781988 | 0.074017965 | 0.127352321 | 0.005088893 | 0.431261263 | 0.00014944 | 0.006846039 | 0.036300898 |
| Son | S | 2029 | 0.083801915 | 0.030467559 | 0.15781988 | 0.074017965 | 0.127352321 | 0.005088893 | 0.431261263 | 0.00014944 | 0.006846039 | 0.036300898 |
| Son | S | 2031 | 0.083801915 | 0.030467559 | 0.15781988 | 0.074017965 | 0.127352321 | 0.005088893 | 0.431261263 | 0.00014944 | 0.006846039 | 0.036300898 |
| Srbd1 | S | 139 | 0.017942023 | 0.26093676 | 0.384684769 | 0.366742746 | 0.123748009 | 0.77577782 | 0.044255079 | 0.06303046 | 0.091870763 | 0.438840204 |
| Yap1 | S | 112 | 0.240846284 | 0.068547533 | 0.228806844 | -0.01203944 | 0.160259311 | 0.015860442 | 0.430500081 | 0.079502097 | 0.899108489 | 0.17305501 |
| Cast | S | 11 | 0.182625958 | 0.160901389 | 0.094081228 | -0.08854473 | -0.066820161 | 0.020597989 | 0.072664372 | 0.064080937 | 0.160694946 | 0.345565329 |
| Atad5 | S | 353 | 0.099320278 | 0.180204148 | 0.13584546 | 0.036525182 | -0.044358688 | 0.125492843 | 0.011092029 | 0.046646377 | 0.552761024 | 0.41754283 |
| Pmf1 | S | 19 | -0.004754614 | 0.216972126 | 0.148815733 | 0.153570346 | -0.068156394 | 0.855401132 | 0.001960514 | 0.006838154 | 0.00717559 | 0.094838869 |
| Setd1a | S | 483 | 0.104207901 | 0.13580597 | 0.155557384 | 0.051349483 | 0.019751414 | 0.058850048 | 0.0237344 | 0.02944923 | 0.311766768 | 0.680667619 |
| Srsf5 | S | 262 | 0.241776408 | 0.103211818 | 0.310401462 | 0.068625054 | 0.207189644 | 0.028665941 | 0.393575234 | 0.156049289 | 0.690495485 | 0.303439702 |
| Safb | S | 639 | -0.03415164 | 0.193362227 | 0.222306441 | 0.25645808 | 0.028944214 | 0.702725916 | 0.044735159 | 0.000352153 | 0.070345787 | 0.628040813 |
| Sh3kbp1 | S | 159 | 0.234263326 | 0.255672711 | 0.170743378 | -0.063519948 | -0.084929333 | 0.2459994 | 0.024217474 | 0.311579771 | 0.755746206 | 0.56563498 |
| Farsb | T | 380 | 0.173478801 | 0.071761811 | 0.337733227 | 0.164254426 | 0.265971416 | 0.379051645 | 0.429605992 | 0.005020382 | 0.391665682 | 0.013832894 |
| Nusap1 | S | 152 | -0.081827013 | 0.254190562 | 0.243225052 | 0.325052065 | -0.01096551 | 0.137835508 | 0.000253533 | 0.025368554 | 0.012100337 | 0.864910118 |
| Sh3bp1 | S | 535 | 0.090875302 | 0.229158463 | 0.075668653 | -0.015206649 | -0.153489809 | 0.109774514 | 0.000497854 | 0.51365232 | 0.890948191 | 0.242783689 |
| Cast | S | 121 | 0.116733914 | 0.283690655 | 0.11709576 | 0.000361845 | -0.166594895 | 0.359940344 | 0.007627755 | 0.150089575 | 0.997235578 | 0.018083841 |
| Lpar1 | S | 344 | 0.060316722 | 0.344918761 | 0.139763524 | 0.079446802 | -0.205155237 | 0.690447169 | 0.006827109 | 0.132434058 | 0.600403886 | 0.062234249 |
| Lpar1 | S | 346 | 0.060316722 | 0.344918761 | 0.139763524 | 0.079446802 | -0.205155237 | 0.690447169 | 0.006827109 | 0.132434058 | 0.600403886 | 0.062234249 |
| Lpar1 | S | 347 | 0.060316722 | 0.344918761 | 0.139763524 | 0.079446802 | -0.205155237 | 0.690447169 | 0.006827109 | 0.132434058 | 0.600403886 | 0.062234249 |
| Samhd1 | T | 56 | 0.024762815 | 0.128211539 | 0.128614564 | 0.103851748 | 0.000403024 | 0.25335115 | 0.000657864 | 0.004205372 | 0.032999683 | 0.986181677 |
| Sorbs2 | S | 109 | 0.121821518 | 0.080466277 | 0.228009095 | 0.106187577 | 0.147542818 | 0.04330625 | 0.222814515 | 0.009945232 | 0.07845404 | 0.068399091 |
| Ess2 | T | 389 | 0.336752837 | 0.219700305 | -0.070858997 | -0.407611834 | -0.290559302 | 0.008481543 | 0.032498298 | 0.744292436 | 0.144610615 | 0.249270747 |
| Cavin3 | T | 24 | 0.165497416 | 0.240522256 | 0.281621176 | 0.11612376 | 0.04109892 | 0.444341987 | 0.076851381 | 0.082992224 | 0.600022834 | 0.768570332 |
| Mki67 | S | 517 | 0.260821867 | 0.219053404 | 0.098350414 | -0.162471453 | -0.120702989 | 0.097934625 | 0.027405158 | 0.346839085 | 0.265903564 | 0.255807839 |
| Srrm2 | T | 1654 | -0.048521262 | 0.236501467 | 0.19906529 | 0.247586551 | -0.037436177 | 0.11972112 | 0.000147978 | 0.023389774 | 0.025064482 | 0.48376544 |
| Tasor | S | 921 | 0.040146928 | 0.200994381 | 0.109585007 | 0.069438079 | -0.091409374 | 0.358741241 | 0.00053498 | 0.060658907 | 0.21589262 | 0.100331832 |
| Foxp1 | S | 643 | 0.012414721 | 0.092873744 | 0.269443368 | 0.257028647 | 0.176569624 | 0.871212748 | 0.332128293 | 5.4639E-05 | 0.058625223 | 0.137294742 |
| Pold3 | S | 306 | -0.016325202 | 0.096921644 | 0.169080893 | 0.185406095 | 0.072159249 | 0.419917556 | 0.001385377 | 4.06293E-05 | 0.000400555 | 0.002593059 |
| Rapgef2 | T | 1401 | 0.250271788 | 0.310254824 | 0.067857973 | -0.182413815 | -0.242396851 | 0.171811709 | 0.019605607 | 0.495886901 | 0.289906491 | 0.066658098 |
| Nr2c1 | S | 325 | -0.122775294 | 0.296861009 | 0.338799168 | 0.461574462 | 0.041938158 | 0.153471184 | 0.002441308 | 0.023262821 | 0.011266899 | 0.646223485 |
| Nr2c1 | S | 326 | -0.122775294 | 0.296861009 | 0.338799168 | 0.461574462 | 0.041938158 | 0.153471184 | 0.002441308 | 0.023262821 | 0.011266899 | 0.646223485 |
| Nr2c1 | S | 330 | -0.122775294 | 0.296861009 | 0.338799168 | 0.461574462 | 0.041938158 | 0.153471184 | 0.002441308 | 0.023262821 | 0.011266899 | 0.646223485 |
| Rap1gap | S | 541 | 0.426453577 | 0.041545606 | 0.134944948 | -0.291508629 | 0.093399342 | 0.01671723 | 0.631715606 | 0.129461284 | 0.060308087 | 0.279786885 |
| Rap1gap | S | 542 | 0.426453577 | 0.041545606 | 0.134944948 | -0.291508629 | 0.093399342 | 0.01671723 | 0.631715606 | 0.129461284 | 0.060308087 | 0.279786885 |
| Bclaf1 | S | 494 | 0.001626016 | 0.093199361 | 0.165197452 | 0.163571436 | 0.07199809 | 0.967299244 | 0.00264788 | 0.000101277 | 0.026236999 | 0.009643043 |
| Septin9 | S | 82 | 0.14589549 | 0.106263277 | 0.062025602 | -0.083869887 | -0.044237674 | 0.003505031 | 0.010259989 | 0.100179319 | 0.036030126 | 0.124868921 |
| Bud13 | T | 157 | 0.148184706 | 0.108903719 | 0.211970119 | 0.063785413 | 0.1030664 | 0.070953262 | 0.154208533 | 0.020936928 | 0.13662794 | 0.041867391 |
| Bud13 | T | 160 | 0.148184706 | 0.108903719 | 0.211970119 | 0.063785413 | 0.1030664 | 0.070953262 | 0.154208533 | 0.020936928 | 0.13662794 | 0.041867391 |
| Abcf1 | S | 107 | 0.048100521 | 0.203077256 | 0.095967687 | 0.047867165 | -0.107109569 | 0.413589494 | 8.89031E-05 | 0.188780065 | 0.523720673 | 0.158755533 |
| Mast4 | S | 353 | 0.00763882 | 0.11015286 | 0.159448973 | 0.151810154 | 0.049296114 | 0.608399608 | 0.000267118 | 0.002821704 | 0.009119096 | 0.123627865 |
| Micall2 | S | 655 | 0.138324477 | 0.113459741 | 0.115264409 | -0.023060068 | 0.001804668 | 0.005666113 | 0.048632347 | 0.073452082 | 0.635630789 | 0.973042321 |
| Hdac1 | S | 421 | 0.063051875 | 0.165431126 | 0.176109638 | 0.113057763 | 0.010678512 | 0.324600435 | 0.019265734 | 0.009538787 | 0.086760704 | 0.808321013 |
| Zc3h4 | S | 1109 | 0.098940086 | 0.058077662 | 0.177830711 | 0.078890625 | 0.119753049 | 0.00484183 | 0.144430877 | 0.008732939 | 0.061117561 | 0.028673334 |
| Brca1 | S | 185 | 0.065772824 | 0.14959728 | 0.115507544 | 0.049734721 | -0.034089736 | 0.241930888 | 0.036993836 | 0.000281616 | 0.346175846 | 0.442556937 |
| Slc12a6 | T | 31 | 0.011232405 | 0.115657488 | 0.291556253 | 0.280323848 | 0.175898764 | 0.754459076 | 0.000383079 | 0.03988741 | 0.032748635 | 0.108541476 |
| Strn | S | 204 | 0.110493634 | 0.242967564 | 0.189699359 | 0.079205726 | -0.053268205 | 0.51491538 | 0.031452458 | 0.03677448 | 0.628916809 | 0.527834213 |
| Cluap1 | S | 314 | 0.038097466 | 0.148833886 | 0.119024994 | 0.080927528 | -0.029808892 | 0.579869686 | 0.001291914 | 0.002671373 | 0.285219369 | 0.093146253 |
| Cluap1 | S | 324 | 0.038097466 | 0.148833886 | 0.119024994 | 0.080927528 | -0.029808892 | 0.579869686 | 0.001291914 | 0.002671373 | 0.285219369 | 0.093146253 |
| Pds5b | S | 1257 | 0.053837831 | 0.131987177 | 0.099054839 | 0.045217008 | -0.032932338 | 0.039336943 | 0.000582626 | 0.037648451 | 0.228344062 | 0.345843552 |
| Uba1 | S | 24 | -0.017745263 | 0.169420879 | 0.146430652 | 0.164175915 | -0.022990227 | 0.462026365 | 5.5827E-06 | 0.049189399 | 0.029626508 | 0.601193584 |
| Ankrd55 | S | 474 | 0.041560037 | 0.317646906 | 0.011466191 | -0.030093846 | -0.306180715 | 0.623135994 | 3.97904E-05 | 0.842666111 | 0.737059784 | 0.014771781 |
| Ibtk | S | 993 | 0.040413067 | 0.20372762 | 0.18771646 | 0.147303393 | -0.016011161 | 0.468132948 | 0.002347647 | 0.077853079 | 0.137639314 | 0.834884715 |
| Cnksr3 | S | 383 | 0.008323501 | 0.208039325 | 0.241514043 | 0.233190543 | 0.033474718 | 0.877046121 | 0.003016611 | 0.051612985 | 0.054437439 | 0.685571203 |
| Srrm2 | S | 2381 | 0.186371884 | 0.149510211 | 0.059652581 | -0.126719303 | -0.08985763 | 0.004598224 | 0.012730019 | 0.623304294 | 0.332477532 | 0.464629278 |
| Arhgap39 | S | 580 | 0.022912727 | 0.235257001 | 0.315409083 | 0.292496357 | 0.080152083 | 0.873771079 | 0.080817296 | 0.0148932 | 0.127590922 | 0.477042463 |
| Sp4 | S | 134 | 0.067871653 | 0.177928066 | 0.278591047 | 0.210719395 | 0.100662981 | 0.566528827 | 0.014006191 | 0.051996079 | 0.17005554 | 0.345389707 |
| Med13 Med13l | T | 1013;1048 | 0.040939209 | 0.103623961 | 0.346929639 | 0.305990429 | 0.243305678 | 0.777856095 | 0.051230757 | 0.006646013 | 0.11671948 | 0.036416782 |
| Med13 Med13l | T | 1016;1051 | 0.040939209 | 0.103623961 | 0.346929639 | 0.305990429 | 0.243305678 | 0.777856095 | 0.051230757 | 0.006646013 | 0.11671948 | 0.036416782 |
| Asph | S | 32 | 0.576330715 | -0.013408071 | 0.459654278 | -0.116676437 | 0.473062349 | 0.169088591 | 0.969525113 | 0.121950677 | 0.735068987 | 0.204747901 |
| Fcho2 | S | 495 | 0.039533046 | 0.125087711 | 0.133042326 | 0.09350928 | 0.007954615 | 0.229906008 | 0.003073047 | 0.00283416 | 0.036519194 | 0.76885295 |
| Zfp768 | S | 128 | 0.09279412 | 0.116266726 | 0.109649384 | 0.016855264 | -0.006617342 | 0.333885268 | 0.00157336 | 0.003389352 | 0.842191284 | 0.773865339 |
| Rgl1 | S | 508 | -0.031200304 | 0.283881944 | 0.191507432 | 0.222707736 | -0.092374512 | 0.590568547 | 0.000605807 | 0.108563606 | 0.081487683 | 0.351662785 |
| Zc3h4 | S | 816 | 0.005585138 | 0.096454734 | 0.193614733 | 0.188029595 | 0.09716 | 0.84053442 | 0.006559872 | 0.000326465 | 0.005771092 | 0.00288524 |
| Zc3h4 | S | 817 | 0.005585138 | 0.096454734 | 0.193614733 | 0.188029595 | 0.09716 | 0.84053442 | 0.006559872 | 0.000326465 | 0.005771092 | 0.00288524 |
| Sipa1l1 | S | 1690 | -0.015696145 | 0.054800031 | 0.261303306 | 0.276999451 | 0.206503275 | 0.497763137 | 0.082717086 | 1.41561E-05 | 2.55161E-05 | 0.001718035 |
| Glis2 | S | 51 | 0.133285227 | 0.031795409 | 0.301095125 | 0.167809898 | 0.269299716 | 0.110775919 | 0.789071446 | 0.002484934 | 0.041483969 | 0.093869007 |
| Hdac1 | S | 423 | 0.062942666 | 0.163931665 | 0.174422717 | 0.11148005 | 0.010491052 | 0.321062075 | 0.019355985 | 0.009621226 | 0.088443072 | 0.812041621 |
| Sash1 | S | 697 | 0.008650982 | 0.281248492 | 0.154300492 | 0.14564951 | -0.126948 | 0.863032492 | 0.009218078 | 0.018006617 | 0.031855263 | 0.111202712 |
| Usp7 | S | 19 | 0.047312297 | 0.122406951 | 0.156607803 | 0.109295506 | 0.034200852 | 0.132850706 | 0.005044817 | 0.009166262 | 0.028729016 | 0.333378177 |
| Wnk1 | S | 172 | 0.040306531 | 0.224343854 | 0.1339739 | 0.093667369 | -0.090369954 | 0.326294193 | 0.012016569 | 0.013052579 | 0.049930521 | 0.15954166 |
| Zfp768 | S | 135 | 0.067510531 | 0.147264493 | 0.120338949 | 0.052828418 | -0.026925545 | 0.075094035 | 0.002988442 | 0.03858465 | 0.244421341 | 0.49759942 |
| Zfp768 | S | 142 | 0.067510531 | 0.147264493 | 0.120338949 | 0.052828418 | -0.026925545 | 0.075094035 | 0.002988442 | 0.03858465 | 0.244421341 | 0.49759942 |
| Ints13 | S | 715 | 0.181004477 | 0.095371327 | 0.112146838 | -0.06885764 | 0.016775511 | 0.007848059 | 0.053980518 | 0.104005763 | 0.272039369 | 0.750800475 |
| Zcchc8 | S | 425 | 0.009313331 | 0.208172332 | 0.256447752 | 0.247134421 | 0.04827542 | 0.8787976 | 0.020272458 | 0.015645462 | 0.023527998 | 0.404020834 |
| Pax3 | S | 201 | 0.073281456 | 0.154155084 | 0.178903993 | 0.105622537 | 0.024748909 | 0.114129807 | 0.003437552 | 0.105988045 | 0.258909957 | 0.75808009 |
| Pax3 | S | 205 | 0.073281456 | 0.154155084 | 0.178903993 | 0.105622537 | 0.024748909 | 0.114129807 | 0.003437552 | 0.105988045 | 0.258909957 | 0.75808009 |
| Pax3 | S | 209 | 0.073281456 | 0.154155084 | 0.178903993 | 0.105622537 | 0.024748909 | 0.114129807 | 0.003437552 | 0.105988045 | 0.258909957 | 0.75808009 |
| Snrnp27 | S | 63 | 0.041978138 | 0.104923769 | 0.160712547 | 0.118734409 | 0.055788778 | 0.205315019 | 0.003721469 | 0.003989574 | 0.021074478 | 0.145006196 |
| Gpatch8 | S | 1002 | 0.076344347 | 0.206997513 | 0.177299834 | 0.100955486 | -0.02969768 | 0.272217253 | 0.09636879 | 0.005853635 | 0.154041476 | 0.744636902 |
| Gpatch8 | S | 1004 | 0.076344347 | 0.206997513 | 0.177299834 | 0.100955486 | -0.02969768 | 0.272217253 | 0.09636879 | 0.005853635 | 0.154041476 | 0.744636902 |
| Slf2 | S | 712 | 0.190878339 | 0.296396138 | 0.223141871 | 0.032263532 | -0.073254267 | 0.270917258 | 0.133710951 | 0.115889112 | 0.83231937 | 0.65580617 |
| Tmem131l | S | 1229 | 0.115173762 | 0.160237116 | 0.252921437 | 0.137747674 | 0.092684321 | 0.265894267 | 0.040536674 | 0.054751086 | 0.259619426 | 0.354680451 |
| Mdc1 | T | 782 | 0.013718905 | 0.107975691 | 0.080140156 | 0.066421251 | -0.027835534 | 0.6633266 | 3.58239E-05 | 0.000161059 | 0.123681772 | 0.031084934 |
| Brca1 | S | 183 | 0.076150138 | 0.198648582 | 0.065270211 | -0.010879927 | -0.133378371 | 0.15480592 | 0.000697755 | 0.104344007 | 0.812127451 | 0.013051923 |
| Brca1 | S | 186 | 0.076150138 | 0.198648582 | 0.065270211 | -0.010879927 | -0.133378371 | 0.15480592 | 0.000697755 | 0.104344007 | 0.812127451 | 0.013051923 |
| Zc3h14 | S | 515 | 0.01344016 | 0.135795416 | 0.247783289 | 0.234343129 | 0.111987872 | 0.803817643 | 0.021610273 | 0.003423834 | 0.018214793 | 0.050889567 |
| Xrn2 | S | 445 | 0.226954333 | 0.287915889 | 0.243196007 | 0.016241674 | -0.044719882 | 0.221979212 | 0.095576256 | 0.281928375 | 0.943286911 | 0.835953663 |
| Chd2 | S | 1365 | 0.05375006 | 0.099775668 | 0.128309994 | 0.074559934 | 0.028534327 | 0.056109173 | 0.026496914 | 0.000694358 | 0.026009859 | 0.367283294 |
| Pard3b | T | 695 | 0.014853175 | 0.191956184 | 0.189564863 | 0.174711689 | -0.002391321 | 0.683915217 | 0.003137026 | 0.025441665 | 0.039641783 | 0.96592243 |
| Afap1 | S | 343 | 0.110589733 | 0.068399798 | 0.148208283 | 0.037618551 | 0.079808485 | 0.000914381 | 0.077162824 | 0.051689705 | 0.455519822 | 0.18449129 |
| Rbm33 | S | 41 | 0.076259168 | 0.115806668 | 0.120153827 | 0.04389466 | 0.004347159 | 0.045293929 | 0.008829457 | 0.011177419 | 0.147377903 | 0.863948045 |
| Efr3a | S | 692 | 0.037905663 | 0.108414051 | 0.170450473 | 0.13254481 | 0.062036422 | 0.105389491 | 0.055856256 | 0.0007337 | 0.003910623 | 0.173686973 |
| Washc2 | S | 1067 | 0.074701442 | 0.139713671 | 0.064622289 | -0.010079153 | -0.075091382 | 0.0268148 | 0.00105384 | 0.036927439 | 0.580783824 | 0.004389334 |
| Bcar3 | S | 378 | -0.036882698 | 0.121237736 | 0.231178958 | 0.268061656 | 0.109941222 | 0.583280572 | 0.075580287 | 3.48424E-05 | 0.034318395 | 0.105094192 |

| | | | | | | | | | | | | |
|---|---|---|---|---|---|---|---|---|---|---|---|---|
| Zc3hc1 | T | 28 | 0.14744222 | 0.181658993 | 0.188031906 | 0.040589686 | 0.006372913 | 0.077485687 | 0.045688693 | 0.152228957 | 0.687198523 | 0.950194633 |
| Smarca4 | S | 610 | -0.030709654 | 0.097512831 | 0.207957637 | 0.23866729 | 0.110444806 | 0.429204339 | 0.004769116 | 8.61413E-05 | 0.004921019 | 0.011816811 |
| Brd1 | S | 1055 | -0.01032437 | 0.077478452 | 0.217057894 | 0.227382264 | 0.139579442 | 0.667378446 | 0.015947621 | 5.87046E-05 | 0.000470029 | 0.001467168 |
| Ptpn21 | S | 577 | 0.213558676 | 0.158759968 | 0.034690088 | -0.178868588 | -0.12406988 | 0.005353603 | 0.025448706 | 0.591485451 | 0.025254842 | 0.070175726 |
| Hcfc1 | S | 1848 | -0.023697694 | 0.088383895 | 0.201900301 | 0.225597996 | 0.113516407 | 0.603445867 | 0.005983857 | 6.83778E-05 | 0.015552141 | 0.004449355 |
| Wnk1 | S | 183 | 0.135089931 | 0.411848846 | 0.1233326 | -0.011757331 | -0.288516246 | 0.298124048 | 0.014271019 | 0.65769838 | 0.963280938 | 0.328015247 |
| Palm | S | 116 | -0.005042608 | 0.106138655 | 0.250979573 | 0.25602218 | 0.144840918 | 0.910630044 | 0.015578076 | 0.001054412 | 0.004968628 | 0.017145823 |
| Nop56 | S | 465 | 0.068883224 | 0.132179882 | 0.181875827 | 0.112992603 | 0.049695945 | 0.286284644 | 0.060977691 | 0.002254857 | 0.127828917 | 0.367025472 |
| Madd | S | 828 | 0.240986243 | 0.094002154 | 0.16699865 | -0.073987593 | 0.072996497 | 0.014491334 | 0.279344856 | 0.118480201 | 0.397560231 | 0.424448972 |
| Ercc6l | S | 994 | -0.03109144 | 0.382686254 | 0.224693971 | 0.255785411 | -0.157992283 | 0.747418358 | 0.015103762 | 0.065976109 | 0.060743532 | 0.215627712 |
| Srrm2 | S | 757 | 0.133215738 | 0.12079644 | 0.127686256 | -0.005529483 | 0.006889815 | 0.017592589 | 0.037688868 | 0.06560781 | 0.915446983 | 0.899070383 |
| Rprd2 | T | 751 | 0.039581917 | 0.143370052 | 0.220811693 | 0.181229776 | 0.077441641 | 0.62372049 | 0.011883625 | 0.009091955 | 0.090737762 | 0.192153642 |
| Pacsin3 | S | 383 | 0.082491071 | 0.161418804 | 0.226274458 | 0.143783387 | 0.064855654 | 0.131795491 | 0.039156955 | 0.051855657 | 0.149755484 | 0.467616782 |
| Jmjd1c | S | 460 | 0.024846278 | 0.152475008 | 0.0994149 | 0.074568622 | -0.053060109 | 0.269344809 | 5.15227E-05 | 0.051900806 | 0.104672594 | 0.198607005 |
| Cep170b | T | 529 | 0.237179803 | 0.11316934 | 0.141514465 | -0.095665338 | 0.028345125 | 0.011565737 | 0.449874472 | 0.067688506 | 0.156705263 | 0.836134917 |
| Atn1 | S | 101 | 0.05106873 | 0.114196398 | 0.136072599 | 0.085003869 | 0.021876201 | 0.141924041 | 0.005190328 | 0.004615755 | 0.036728877 | 0.430753238 |
| Flna | T | 2336 | 0.098753964 | 0.130124488 | 0.063996725 | -0.034757239 | -0.066127763 | 0.018775593 | 0.000369645 | 0.232319722 | 0.487244954 | 0.223980084 |
| Rbm14 | T | 572 | 0.084306767 | 0.096283244 | 0.208115726 | 0.123808959 | 0.111832483 | 0.193070554 | 0.2401537 | 0.000956043 | 0.089191955 | 0.190074333 |
| Cdk13 | S | 1224 | 0.079471176 | 0.117455634 | 0.215043594 | 0.135572418 | 0.09758796 | 0.236853739 | 0.036115713 | 0.011215732 | 0.096776313 | 0.133275789 |
| Dcaf8 | S | 124 | -0.071564586 | 0.162412694 | 0.219607249 | 0.291171834 | 0.057194555 | 0.162805995 | 0.011292617 | 6.81888E-05 | 0.00619097 | 0.174048157 |
| Ralgapb | S | 357 | 0.002060245 | 0.182512212 | 0.281434472 | 0.279374226 | 0.09892226 | 0.985487549 | 0.063959425 | 0.004609028 | 0.08531357 | 0.249864305 |
| Phactr1 Phactr4 | S | 582;467 | 0.136724837 | 0.2393322 | 0.113666678 | -0.023058159 | -0.125665522 | 0.272939708 | 0.022995218 | 0.056525965 | 0.825490623 | 0.148503653 |
| Macf1 | S | 280 | 0.155286758 | 0.105238223 | 0.099700629 | -0.055586129 | -0.005537594 | 0.034556301 | 0.007245129 | 0.081710773 | 0.328076428 | 0.895370294 |
| Sf3b1 | T | 296 | 0.071758095 | 0.098685606 | 0.17064818 | 0.098890084 | 0.071962574 | 0.094866638 | 0.040499992 | 0.004090382 | 0.02766435 | 0.069190817 |
| Trip12 | S | 77 | 0.119696106 | 0.157912758 | 0.126586175 | 0.006890069 | -0.031326583 | 0.037921755 | 0.03101016 | 0.072987489 | 0.904410396 | 0.617769465 |
| Srrm2 | S | 1927 | 0.177514974 | 0.166426162 | 0.119135439 | -0.058379535 | -0.047290723 | 0.021538908 | 0.023468254 | 0.409353291 | 0.660159493 | 0.71702845 |
| Srrt | S | 492 | 0.066962179 | 0.171489638 | 0.117503283 | 0.050541103 | -0.053986356 | 0.163293662 | 0.001765741 | 0.07578737 | 0.383765311 | 0.323284896 |
| Brip1 | S | 128 | 0.290265375 | 0.231053505 | -0.089324163 | -0.379589538 | -0.320377668 | 0.006586844 | 0.01731672 | 0.540046096 | 0.077490867 | 0.096683664 |
| Zfp768 | S | 17 | 0.107753652 | 0.2093902 | 0.247369513 | 0.139615861 | 0.037979313 | 0.253971743 | 0.109929664 | 0.044833209 | 0.203589088 | 0.722801656 |
| Rbm33 | S | 816 | -0.008946776 | 0.142313523 | 0.256752054 | 0.265698829 | 0.114438531 | 0.86787299 | 0.002758898 | 0.012767024 | 0.013516969 | 0.117874456 |
| Grk6 | S | 484 | 0.252695413 | 0.115050609 | 0.092586635 | -0.160108778 | -0.022463974 | 0.004284157 | 0.384343394 | 0.132820826 | 0.034854572 | 0.851250883 |
| Pitpnm1 | S | 621 | 0.045768836 | 0.131591088 | 0.164673504 | 0.118904668 | 0.033082416 | 0.210174669 | 0.000388253 | 0.074550548 | 0.137423755 | 0.593926048 |
| Mia3 | S | 1765 | 0.01017288 | 0.130707395 | 0.10416326 | 0.09399038 | -0.026544135 | 0.859472641 | 0.000303063 | 0.000723216 | 0.194704737 | 0.132441829 |
| Fam171b | S | 777 | 0.002439404 | 0.228193233 | 0.275924179 | 0.273484775 | 0.047730946 | 0.967117543 | 0.001390393 | 0.167840254 | 0.170172868 | 0.757430696 |
| Khsrp | T | 101 | 0.149115405 | 0.097691911 | 0.196609877 | 0.047494472 | 0.098917966 | 0.016860881 | 0.0965838 | 0.096909516 | 0.596747243 | 0.317391476 |
| Map3k7 | S | 389 | 0.027226167 | 0.217117885 | 0.071728039 | 0.044501872 | -0.145389846 | 0.376492029 | 0.000251689 | 0.07567304 | 0.151283098 | 0.006685765 |
| Nf1 | S | 2599 | -0.003864346 | 0.129296536 | 0.184329892 | 0.188194237 | 0.055033356 | 0.92775046 | 0.007503679 | 0.000712682 | 0.020789493 | 0.110279438 |
| Washc2 | S | 1066 | 0.105146356 | 0.353117571 | 0.154282988 | 0.049136632 | -0.198834583 | 0.220160515 | 0.047041362 | 0.202857279 | 0.664985044 | 0.193319856 |
| Ccar2 | S | 626 | 0.013323715 | 0.138782767 | 0.191354339 | 0.178030624 | 0.052571572 | 0.731757431 | 0.005654108 | 0.004266397 | 0.017407009 | 0.097185732 |
| Sipa1l1 | S | 1087 | 0.379713868 | 0.164733445 | 0.0386355 | -0.341078368 | -0.126097945 | 0.222878014 | 0.001780927 | 0.525795612 | 0.255631614 | 0.106126802 |
| Bclaf1 | S | 646 | 0.033996488 | 0.184281667 | 0.162157068 | 0.12816058 | -0.022124599 | 0.371539211 | 0.039025258 | 0.002773705 | 0.026655354 | 0.681684508 |
| Gripap1 | S | 653 | 0.045478545 | 0.180469771 | 0.291930256 | 0.246451712 | 0.111460485 | 0.451643013 | 0.084333686 | 0.019030979 | 0.05901738 | 0.286173621 |
| Gripap1 | S | 656 | 0.045478545 | 0.180469771 | 0.291930256 | 0.246451712 | 0.111460485 | 0.451643013 | 0.084333686 | 0.019030979 | 0.05901738 | 0.286173621 |
| Gsk3b | S | 389 | 0.300505214 | 0.208536531 | -0.073872514 | -0.374377728 | -0.282409045 | 0.008115748 | 0.012867324 | 0.51188633 | 0.03133267 | 0.068526543 |
| Prune2 | S | 2850 | 0.016274621 | 0.205847124 | 0.167690896 | 0.151416275 | -0.038156228 | 0.656508063 | 0.002962687 | 0.033499286 | 0.055159021 | 0.511111633 |
| Llgl1 | S | 982 | 0.147128655 | 0.135046428 | 0.114968477 | -0.032160178 | -0.020077952 | 0.002244704 | 0.084193311 | 0.238610236 | 0.695583194 | 0.827378864 |
| Llgl1 | S | 989 | 0.147128655 | 0.135046428 | 0.114968477 | -0.032160178 | -0.020077952 | 0.002244704 | 0.084193311 | 0.238610236 | 0.695583194 | 0.827378864 |
| Llgl1 | S | 992 | 0.147128655 | 0.135046428 | 0.114968477 | -0.032160178 | -0.020077952 | 0.002244704 | 0.084193311 | 0.238610236 | 0.695583194 | 0.827378864 |
| 4932438A13Rik | S | 3837 | 0.14534728 | 0.089441651 | 0.159864538 | 0.014517258 | 0.070422887 | 0.045968886 | 0.054252251 | 0.029306333 | 0.810029505 | 0.218769683 |
| Klf13 | S | 163 | -0.037635019 | 0.182789519 | 0.199751941 | 0.23738696 | 0.016962422 | 0.174099569 | 0.000143153 | 0.015107623 | 0.006064315 | 0.661795392 |
| Cep350 | S | 1259 | 0.01443018 | 0.148841057 | 0.164426853 | 0.149996673 | 0.015585796 | 0.620697823 | 0.047995779 | 0.000225226 | 0.004449444 | 0.747846794 |
| Cep350 | T | 1255 | 0.01443018 | 0.148841057 | 0.164426853 | 0.149996673 | 0.015585796 | 0.620697823 | 0.047995779 | 0.000225226 | 0.004449444 | 0.747846794 |
| Appl1 | S | 401 | 0.025329228 | 0.111354694 | 0.147295408 | 0.121966179 | 0.035940714 | 0.288549069 | 0.002720695 | 0.002448726 | 0.006745765 | 0.111234906 |
| Sash1 | S | 1014 | 0.026516247 | 0.163189911 | 0.20396788 | 0.17780541 | 0.040779968 | 0.593779254 | 0.015331881 | 0.008449171 | 0.020747713 | 0.453907569 |
| Add1 | T | 614 | 0.043238298 | 0.114677596 | 0.087296644 | 0.044058346 | -0.027380952 | 0.22184074 | 0.000277891 | 0.003382958 | 0.217112453 | 0.167060759 |
| Ints3 | S | 500 | -0.014468953 | 0.083082448 | 0.285311563 | 0.299780516 | 0.202229115 | 0.732689196 | 0.068310643 | 0.000293274 | 0.000732378 | 0.00816265 |
| Clip2 | S | 208 | 0.037312848 | 0.113176929 | 0.230554355 | 0.193241506 | 0.117377425 | 0.48514164 | 0.010015899 | 0.009164779 | 0.026901359 | 0.088379641 |
| Cdk12 | S | 322 | 0.08663457 | 0.114146883 | 0.168188729 | 0.08155416 | 0.054041846 | 0.210437837 | 0.023457068 | 0.007743736 | 0.241473002 | 0.246109717 |
| Cdk12 | S | 324 | 0.08663457 | 0.114146883 | 0.168188729 | 0.08155416 | 0.054041846 | 0.210437837 | 0.023457068 | 0.007743736 | 0.241473002 | 0.246109717 |
| Ifi205 Ifi211 Mndal | T | 442;356;335 | -0.0664321 | 0.225955914 | 0.253914289 | 0.320346389 | 0.027958375 | 0.238665026 | 0.000497094 | 0.034837757 | 0.014933282 | 0.694945223 |
| Mrtfa | S | 492 | 0.059352166 | 0.215585291 | 0.095891429 | 0.036539263 | -0.119693862 | 0.320117707 | 0.003020008 | 0.044223986 | 0.525331235 | 0.041893002 |
| Mrtfa | T | 488 | 0.059352166 | 0.215585291 | 0.095891429 | 0.036539263 | -0.119693862 | 0.320117707 | 0.003020008 | 0.044223986 | 0.525331235 | 0.041893002 |
| Ubn1 | S | 493 | -0.01474254 | 0.131888341 | 0.270662141 | 0.285404681 | 0.1387738 | 0.568626141 | 0.000528528 | 0.021620076 | 0.01901212 | 0.097313911 |
| Spag9 | T | 226 | 0.073571908 | 0.111587364 | 0.175829173 | 0.102257265 | 0.064241809 | 0.219534814 | 0.032429707 | 0.004615625 | 0.100939147 | 0.033131089 |
| Palld | S | 771 | 0.383156893 | -0.164666706 | 0.271852832 | -0.111304061 | 0.436519538 | 0.078283379 | 0.572236844 | 0.003497009 | 0.475059807 | 0.213903217 |
| Scarf2 | S | 640 | -0.029247374 | 0.263623567 | 0.206898477 | 0.236145851 | -0.05672509 | 0.567292032 | 0.020237019 | 0.005402752 | 0.007866767 | 0.445340611 |
| Rapgef2 | S | 1114 | 0.070778946 | 0.255722763 | 0.158554461 | 0.087775516 | -0.097168301 | 0.390647727 | 0.006724725 | 0.163739526 | 0.431061466 | 0.344224513 |
| Rapgef2 | S | 1119 | 0.070778946 | 0.255722763 | 0.158554461 | 0.087775516 | -0.097168301 | 0.390647727 | 0.006724725 | 0.163739526 | 0.431061466 | 0.344224513 |
| Erbin | S | 595 | 0.078475663 | 0.242843012 | 0.146231454 | 0.067755791 | -0.096611557 | 0.308606781 | 0.014029328 | 0.083877509 | 0.094750483 | 0.029937361 |
| Dpysl2 | S | 522 | 0.0076677 | 0.348640329 | 0.082617721 | 0.074950022 | -0.266022608 | 0.896587771 | 0.000590938 | 0.374791026 | 0.393448217 | 0.040553817 |
| Hivep1 | S | 774 | 0.174859043 | 0.166945045 | 0.067794754 | -0.107064289 | -0.099150291 | 0.041425691 | 0.004121386 | 0.490320259 | 0.322310791 | 0.333062706 |
| Hivep1 | S | 777 | 0.174859043 | 0.166945045 | 0.067794754 | -0.107064289 | -0.099150291 | 0.041425691 | 0.004121386 | 0.490320259 | 0.322310791 | 0.333062706 |
| Osbp | S | 191 | 0.028710801 | 0.160116041 | 0.122570448 | 0.093859647 | -0.037545593 | 0.471772101 | 0.001739025 | 0.00817916 | 0.063352431 | 0.275598947 |
| Spag9 | S | 203 | 0.065988435 | 0.202173554 | 0.186046722 | 0.120058287 | -0.016126832 | 0.34765809 | 0.024734872 | 0.033713316 | 0.164185521 | 0.808831796 |
| Zfp687 | S | 71 | 0.078180056 | 0.121307942 | 0.234206844 | 0.156026788 | 0.112898903 | 0.017169941 | 0.226800036 | 0.023942111 | 0.080507702 | 0.271586716 |
| Dcaf6 | S | 863 | -0.072133813 | 0.179513924 | 0.32429671 | 0.396430523 | 0.144782786 | 0.137060255 | 0.008891709 | 0.002799232 | 0.0020697 | 0.053590071 |
| Sipa1l1 | S | 176 | -0.030850355 | 0.143957734 | 0.201024925 | 0.231875281 | 0.057067191 | 0.24192597 | 0.006463911 | 0.000288595 | 0.000869406 | 0.138453515 |
| Med15 | T | 604 | -0.007832264 | 0.107339581 | 0.186541773 | 0.194374037 | 0.079202192 | 0.849631102 | 0.003319092 | 0.000613534 | 0.011475587 | 0.034729817 |
| Ccdc9b | T | 496 | 0.063485179 | 0.150090391 | 0.246832509 | 0.18334733 | 0.096742118 | 0.335749452 | 0.059444971 | 0.016549643 | 0.053505586 | 0.202195294 |
| Wee1 | T | 173 | 0.016512153 | 0.138556028 | 0.152349344 | 0.135837191 | 0.013793315 | 0.728396513 | 0.004230268 | 0.001884811 | 0.051614723 | 0.559513534 |
| Fnbp1 | S | 299 | 0.081978671 | 0.157980558 | 0.110857739 | 0.028879067 | -0.047122819 | 0.112868667 | 0.012796005 | 0.018124118 | 0.522987397 | 0.278302991 |
| Ncor2 | S | 149 | -0.019049031 | 0.10790479 | 0.158508437 | 0.177557468 | 0.050603647 | 0.447827816 | 0.000807902 | 0.000187448 | 0.002857216 | 0.040794572 |
| Mcm3 | T | 719 | -0.024414885 | 0.126261945 | 0.215640769 | 0.240055654 | 0.089378823 | 0.463501918 | 0.007679447 | 0.00055277 | 3.83988E-05 | 0.001262578 |
| Ctnna1 | S | 117 | 0.114338694 | 0.125975056 | 0.11710159 | 0.002762895 | -0.008873466 | 0.012483897 | 0.005506263 | 0.253996846 | 0.974506629 | 0.91776848 |
| Sp3 | S | 568 | -0.006195879 | 0.220093119 | 0.389649706 | 0.395845585 | 0.169556587 | 0.949330533 | 0.218396401 | 0.008843632 | 0.017517862 | 0.325452763 |
| Mki67 | S | 2268 | 0.240899196 | 0.388888483 | 0.134436446 | -0.10646275 | -0.254452037 | 0.505533204 | 0.03203282 | 0.389894932 | 0.753154746 | 0.077078587 |
| Pcm1 | S | 1366 | 0.252754364 | 0.17318515 | 0.135728153 | -0.117026211 | -0.037456997 | 0.042294824 | 0.076284515 | 0.392859286 | 0.452898408 | 0.789806198 |
| Spata6 | S | 217 | 0.115996108 | 0.118092867 | 0.039131011 | -0.076865097 | -0.078961856 | 0.00184175 | 0.001829008 | 0.360627899 | 0.124814298 | 0.118165043 |
| Axin1 | S | 77 | 0.168415265 | 0.152287607 | -0.010947206 | -0.179362471 | -0.163234813 | 0.002260204 | 0.003424705 | 0.936906173 | 0.273103196 | 0.307636809 |
| Washc2 | S | 157 | 0.02356187 | 0.158982025 | 0.154633738 | 0.131071867 | -0.004348287 | 0.439958577 | 0.033057796 | 0.000910268 | 0.011873198 | 0.921316585 |
| Washc2 | S | 159 | 0.02557167 | 0.160217182 | 0.154180108 | 0.128608438 | -0.006037075 | 0.402844282 | 0.033574971 | 0.001084673 | 0.012350858 | 0.892521301 |
| Suds3 | S | 234 | 0.024792193 | 0.092767074 | 0.200456355 | 0.175664163 | 0.107689281 | 0.550680418 | 0.023276056 | 0.00072949 | 0.00813036 | 0.015539501 |
| Eml4 | S | 146 | 0.005137066 | 0.170307907 | 0.180535416 | 0.17539835 | 0.010227509 | 0.800623433 | 0.003584003 | 0.011013465 | 0.022286804 | 0.805709674 |
| Slc9a9 | S | 498 | 0.195304547 | 0.417393113 | 0.064405917 | -0.13089863 | -0.352987196 | 0.362307581 | 0.013013948 | 0.788144895 | 0.638591113 | 0.21186122 |

| | | | | | | | | | | | | |
|---|---|---|---|---|---|---|---|---|---|---|---|---|
| Slc9a3r1 | S | 289 | 0.062641889 | 0.14779278 | 0.167781943 | 0.105140055 | 0.019989163 | 0.43558057 | 0.059716713 | 0.001557688 | 0.23773167 | 0.713963818 |
| Syne2 | S | 6348 | -0.035425979 | 0.213006728 | 0.172366393 | 0.207792372 | -0.040640335 | 0.492162833 | 0.001243339 | 0.011236555 | 0.01146525 | 0.340341863 |
| Cic | S | 20 | -0.032159011 | 0.198926137 | 0.237823546 | 0.269982557 | 0.038897409 | 0.198675401 | 0.041480384 | 0.000485419 | 0.001254899 | 0.525732165 |
| Carmil1 | S | 971 | -0.032327599 | 0.265659221 | 0.123582992 | 0.155910591 | -0.14207623 | 0.517093011 | 0.000729724 | 0.024910753 | 0.023796466 | 0.025067252 |
| Myo1c | S | 1041 | 0.037468785 | 0.278959559 | 0.345522643 | 0.308053858 | 0.066563084 | 0.88314861 | 0.08028076 | 0.053017905 | 0.303893719 | 0.623081409 |
| Megf10 | S | 1059 | 0.096836303 | 0.144752363 | 0.221066488 | 0.124230185 | 0.076314125 | 0.281187195 | 0.118212851 | 0.010128998 | 0.194793777 | 0.361886383 |
| Bud13 | S | 226 | 0.126840011 | 0.066029488 | 0.153200813 | 0.026360802 | 0.087171325 | 0.058442992 | 0.080018914 | 0.004746811 | 0.60517995 | 0.044259003 |
| Bud13 | T | 222 | 0.126840011 | 0.066029488 | 0.153200813 | 0.026360802 | 0.087171325 | 0.058442992 | 0.080018914 | 0.004746811 | 0.60517995 | 0.044259003 |
| Lrrk2 | S | 935 | -0.090288285 | 0.302447114 | 0.322831936 | 0.413120221 | 0.020384822 | 0.28193019 | 0.008617153 | 0.032272459 | 0.010540376 | 0.808755547 |
| Brd1 | S | 1052 | -0.033135736 | 0.104235976 | 0.241064511 | 0.274200247 | 0.136828535 | 0.350769787 | 0.016064086 | 0.000175676 | 0.000241576 | 0.002820622 |
| Gng12 | S | 10 | 0.089433473 | 0.115418428 | 0.227037152 | 0.137603679 | 0.111618724 | 0.351501983 | 0.211846003 | 0.002076979 | 0.189247227 | 0.221924305 |
| C77080 | S | 402 | -0.051893961 | 0.223134116 | 0.226054049 | 0.27794801 | 0.002919933 | 0.139282728 | 0.003942019 | 0.00384783 | 0.001415372 | 0.940355131 |
| Glg1 | S | 710 | -0.101238628 | 0.233327123 | 0.415557582 | 0.51679621 | 0.18223046 | 0.578149242 | 0.006448844 | 0.066431668 | 0.060658333 | 0.290163001 |
| Surf2 | S | 93 | -0.011793192 | 0.286689576 | 0.168567755 | 0.180360947 | -0.118121822 | 0.908031601 | 0.001157148 | 0.164309776 | 0.198311914 | 0.288474419 |
| Ctnnal1 | S | 373 | -0.132237469 | 0.214356854 | 0.346810579 | 0.479048049 | 0.132453725 | 0.228816447 | 0.020819735 | 0.002570491 | 0.009244915 | 0.065991758 |
| Mis18bp1 | S | 907 | 0.068911413 | 0.177771087 | 0.100026618 | 0.031115205 | -0.077744469 | 0.531427307 | 0.007075873 | 0.00308025 | 0.766998886 | 0.091200981 |
| Brd4 | S | 470 | 0.024856074 | 0.079700156 | 0.197334268 | 0.172478193 | 0.117634112 | 0.612678453 | 0.014898559 | 0.00054298 | 0.034326953 | 0.00952565 |
| Srrm2 | S | 759 | 0.116726971 | 0.105243139 | 0.096205298 | -0.020521673 | -0.009037841 | 0.003146154 | 0.028961902 | 0.09433196 | 0.63418536 | 0.844512204 |
| Epb41l3 | T | 725 | 0.097724017 | 0.111118825 | 0.375866306 | 0.278142289 | 0.264747481 | 0.20753501 | 0.130226232 | 0.05663332 | 0.103994271 | 0.116250308 |
| Stag1 | S | 1065 | 0.018422808 | 0.156316691 | 0.241393466 | 0.222970658 | 0.085076775 | 0.796914747 | 0.006033117 | 0.02417097 | 0.04796133 | 0.258780822 |
| Mcm7 | S | 500 | 0.168442592 | 0.146533577 | 0.171567462 | 0.00312487 | 0.025033886 | 0.099736814 | 0.073894525 | 0.075981789 | 0.973078349 | 0.755305131 |
| Caskin2 | S | 406 | -0.015865583 | 0.257024963 | 0.175393319 | 0.191258902 | -0.081631644 | 0.878284961 | 0.001745842 | 0.075763129 | 0.156707518 | 0.314573887 |
| Smarcc2 | S | 286 | 0.032031099 | 0.190639829 | 0.184459454 | 0.152428355 | -0.006180375 | 0.615166898 | 0.017324179 | 0.012357437 | 0.084496708 | 0.901187544 |
| Slc30a4 | S | 32 | 0.138055028 | 0.115819529 | 0.215879788 | 0.07782476 | 0.100060258 | 0.338372956 | 0.08741905 | 0.01118412 | 0.555947935 | 0.076697434 |
| Baiap3 Mepce | S | 192;215 | 0.03605972 | 0.120975209 | 0.146389004 | 0.110329284 | 0.025413795 | 0.305457048 | 0.035846914 | 0.000455324 | 0.028544075 | 0.514804684 |
| Edc4 | S | 884 | 0.095966429 | 0.105475744 | 0.110717315 | 0.014750886 | 0.005241571 | 0.014469396 | 0.019027481 | 0.031660459 | 0.68175632 | 0.892215459 |
| Foxk1 | S | 414 | -0.010011422 | 0.133863615 | 0.136731538 | 0.14674296 | 0.002867923 | 0.736230031 | 0.000503433 | 0.001374655 | 0.005895948 | 0.888493479 |
| Ccny | S | 99 | -0.015655277 | 0.204377803 | 0.160121801 | 0.175777078 | -0.044256002 | 0.674210717 | 0.001089675 | 0.018732664 | 0.028219356 | 0.336811044 |
| Knl1 | S | 397 | 0.247810058 | 0.045423148 | 0.126781429 | -0.121028629 | 0.081358282 | 0.003376017 | 0.568181773 | 0.09012752 | 0.083361094 | 0.308174016 |
| Ttf1 | S | 439 | 0.039829478 | 0.092518516 | 0.168960964 | 0.129131486 | 0.076442448 | 0.195718015 | 0.014781587 | 0.002108013 | 0.012402554 | 0.056845314 |
| Triobp | S | 2011 | 0.066012999 | 0.147745259 | 0.086617993 | 0.020604993 | -0.061127266 | 0.225481454 | 0.001031165 | 0.021226638 | 0.659658825 | 0.046138681 |
| Pml | S | 519 | -0.024255213 | 0.138692346 | 0.224452159 | 0.248707372 | 0.085759813 | 0.51014689 | 0.000337599 | 0.014622937 | 0.008066404 | 0.145108116 |
| Arhgap21 | S | 913 | 0.093034401 | 0.285087364 | 0.154999903 | 0.061965502 | -0.13008746 | 0.461748361 | 0.007678454 | 0.266703962 | 0.671136013 | 0.317676662 |
| Spp1 | S | 291 | 0.038702939 | 0.184362084 | 0.088144191 | 0.049441252 | -0.096217893 | 0.103998263 | 0.001009691 | 0.077549074 | 0.219426665 | 0.05624357 |
| Akap12 | S | 682 | 0.016433771 | 0.412033284 | 0.049237505 | 0.032803734 | -0.362795779 | 0.812835998 | 0.001343981 | 0.482902281 | 0.175816488 | 0.000319475 |
| Rbm39 | S | 129 | 0.113301177 | 0.089289378 | 0.319915726 | 0.206614548 | 0.230626348 | 0.172935911 | 0.158545361 | 0.032811989 | 0.111126198 | 0.104250813 |
| Ncoa5 | S | 378 | 0.069592132 | 0.097813952 | 0.192529313 | 0.122937181 | 0.094715361 | 0.017036437 | 0.013419166 | 0.013419166 | 0.043330419 | 0.143145115 |
| Dpy19l1 | T | 43 | 0.125354656 | 0.104039802 | 0.142447733 | 0.017093077 | 0.038407931 | 0.227281449 | 0.035522118 | 0.004590738 | 0.842898266 | 0.318958805 |
| Rbm39 | S | 136 | 0.050238378 | 0.071398732 | 0.108225148 | 0.05798677 | 0.036826416 | 0.032284809 | 0.006248553 | 0.000798219 | 0.015842261 | 0.062541713 |
| Lnpk | S | 373 | 0.044739914 | 0.18586307 | 0.188859646 | 0.144119732 | 0.002996576 | 0.542959895 | 0.01903842 | 0.017810012 | 0.061273948 | 0.914964918 |
| Prpf4b | S | 294 | 0.098604545 | 0.089840518 | 0.152774357 | 0.054169812 | 0.062933839 | 0.050153835 | 0.015633886 | 0.027457445 | 0.316774683 | 0.226697169 |
| Rere | S | 679 | 0.007319544 | 0.129328392 | 0.174454379 | 0.167134835 | 0.045125987 | 0.885007244 | 0.003812625 | 0.002697402 | 0.031468574 | 0.209322574 |
| Slc39a3 | S | 125 | 0.001351566 | 0.145879914 | 0.181685828 | 0.180334262 | 0.035805914 | 0.985710633 | 0.075924486 | 0.000128477 | 0.104191881 | 0.539023596 |
| Sfpq | T | 679 | 0.324446804 | 0.076986033 | 0.151779822 | -0.172666981 | 0.07479379 | 0.01388609 | 0.626730527 | 0.16210218 | 0.038889204 | 0.597455324 |
| Sptan1 | S | 1217 | -0.011948765 | 0.187437283 | 0.167086961 | 0.179035726 | -0.020350322 | 0.704759988 | 0.000895876 | 0.020133268 | 0.011311262 | 0.593301977 |
| Casc3 | S | 145 | 0.036345144 | 0.112326471 | 0.122209891 | 0.085864746 | 0.00988342 | 0.182723608 | 0.000168762 | 0.030479606 | 0.07305103 | 0.763097473 |
| Bclaf1 | S | 656 | 0.063227013 | 0.145845076 | 0.165324248 | 0.102097235 | 0.019479172 | 0.219956822 | 0.019140122 | 0.017043161 | 0.057791342 | 0.599399043 |
| Cdk12 | T | 1365 | 0.083180571 | 0.096367527 | 0.182997367 | 0.099816796 | 0.08662984 | 0.068365396 | 0.030256473 | 0.021861596 | 0.124923887 | 0.166309656 |
| Zfp827 | S | 681 | 0.135630829 | 0.071276378 | 0.170232126 | 0.034601298 | 0.098955748 | 0.053130845 | 0.265973504 | 0.004958993 | 0.51615591 | 0.143161722 |
| Ppp4r3a | S | 761 | 0.173109923 | 0.109499309 | 0.126151487 | -0.046958435 | 0.016652178 | 0.033785809 | 0.031078889 | 0.13108823 | 0.546658801 | 0.793954237 |
| Retreg3 | T | 440 | 0.107024255 | 0.135985442 | 0.10996367 | 0.002939416 | -0.026021772 | 0.011434756 | 0.055397993 | 0.053815303 | 0.938714879 | 0.652427645 |
| Ccp110 | S | 176 | 0.041008748 | 0.150898306 | 0.147400593 | 0.106391844 | -0.003497713 | 0.226582796 | 0.004436063 | 0.025714775 | 0.094773929 | 0.93749492 |
| Rab11fip5 | S | 307 | 0.154549572 | 0.167564078 | 0.064750124 | -0.089799448 | -0.102813954 | 0.012878188 | 0.019647054 | 0.371798329 | 0.251646782 | 0.207133012 |
| Cdk13 | S | 364 | -0.006537591 | 0.171895625 | 0.274498424 | 0.281036015 | 0.102602799 | 0.959380204 | 0.095244952 | 0.002947662 | 0.111282364 | 0.262429204 |
| Prpf40a | S | 908 | 0.403598009 | -0.076703648 | 0.18579609 | -0.217801919 | 0.262499738 | 0.008412624 | 0.657219098 | 0.086413592 | 0.067609929 | 0.184886043 |
| Gmip | S | 436 | 0.016848164 | 0.128345287 | 0.168409767 | 0.151561603 | 0.04006448 | 0.691243838 | 0.01230628 | 0.001378317 | 0.018212926 | 0.235783434 |
| Sgta | T | 82 | -0.082755258 | 0.274765115 | 0.156958604 | 0.239713862 | -0.11780651 | 0.260141212 | 0.000521496 | 0.022065812 | 0.022758585 | 0.064790603 |
| Cavin2 | S | 218 | -0.080232749 | 0.204272595 | 0.30429471 | 0.384527458 | 0.100022114 | 0.250634814 | 0.008394589 | 0.007158154 | 0.00535504 | 0.192753978 |
| Nsd1 | S | 2267 | 0.014110571 | 0.161247445 | 0.128625103 | 0.114514533 | -0.032622342 | 0.760275867 | 0.000101927 | 0.060374544 | 0.103279565 | 0.488695292 |
| Pde7a | S | 58 | 0.108374938 | 0.453036205 | 0.045170018 | -0.06320492 | -0.407866187 | 0.466897522 | 0.007820811 | 0.763484604 | 0.673926098 | 0.035107928 |
| Dido1 | S | 1256 | 0.065173716 | 0.032223197 | 0.215440759 | 0.150267043 | 0.183217562 | 0.362056815 | 0.216415861 | 0.000110963 | 0.109846372 | 0.001231259 |
| Rbl1 | S | 970 | 0.066272261 | 0.192964189 | 0.078558706 | 0.012286444 | -0.114405483 | 0.313572664 | 0.001189942 | 0.065430792 | 0.840520826 | 0.027177138 |
| Setd1a | S | 1159 | 0.0225438 | 0.074277936 | 0.24093841 | 0.21839461 | 0.166660474 | 0.463690236 | 0.075322773 | 0.000807361 | 0.002785449 | 0.009305491 |
| Dst | S | 7336 | 0.118313619 | 0.128424728 | 0.144629912 | 0.026316293 | 0.016205184 | 0.108921361 | 0.002994547 | 0.1732512 | 0.772600944 | 0.842943286 |
| Thrap3 | S | 320 | -0.04803844 | 0.052771654 | 0.308646801 | 0.356685242 | 0.255875147 | 0.140256974 | 0.149231486 | 1.21352E-05 | 0.000109191 | 0.001062756 |
| Git1 | S | 397 | 0.024482536 | 0.143843743 | 0.084749327 | 0.060266791 | -0.059094415 | 0.102304584 | 0.000715294 | 0.010467713 | 0.054319291 | 0.051099654 |
| Adnp | S | 959 | 0.006573074 | 0.09590769 | 0.174556151 | 0.167983078 | 0.078648461 | 0.796933586 | 0.011636026 | 0.000316462 | 0.000275373 | 0.020879571 |
| Lin52 | S | 28 | -0.070457585 | 0.160577604 | 0.282508894 | 0.352966479 | 0.121931291 | 0.178126553 | 0.001974003 | 0.004870376 | 0.003201741 | 0.087904675 |
| Hnrnpul2 | S | 166 | -0.008311803 | 0.137563579 | 0.263532416 | 0.271844219 | 0.125968837 | 0.777186217 | 0.003287912 | 0.017077044 | 0.011581878 | 0.084979232 |
| Sipa1l1 | S | 318 | -0.022155764 | 0.102381478 | 0.244725204 | 0.266880968 | 0.142343726 | 0.714338757 | 0.001401206 | 0.0048274 | 0.017502783 | 0.030458973 |
| Trap1 | S | 228 | 0.12389142 | 0.222149561 | 0.139925743 | 0.016034323 | -0.082223818 | 0.423192919 | 0.00359944 | 0.202421452 | 0.918494905 | 0.410195029 |
| Pacc1 | S | 7 | 0.50027721 | 0.241789659 | 0.139420947 | -0.360856264 | -0.102368713 | 0.09413707 | 0.240671724 | 0.747344938 | 0.438532752 | 0.807429627 |
| Nek1 | S | 997 | -0.000788524 | 0.191462584 | 0.18279126 | 0.183579784 | -0.008671324 | 0.98627878 | 0.009565994 | 0.008356189 | 0.011163304 | 0.862258825 |
| Atrx | S | 991 | -0.00192127 | 0.10892372 | 0.138289733 | 0.140211003 | 0.029366012 | 0.961063653 | 0.004203955 | 0.000132189 | 0.040647394 | 0.222241069 |
| Tnik | S | 651 | 0.457981251 | 0.485710274 | -0.050147372 | -0.508128623 | -0.535857646 | 0.26720339 | 0.055134889 | 0.853020793 | 0.235834179 | 0.084908406 |
| Pard3b | S | 350 | -0.039552706 | 0.227851324 | 0.246457079 | 0.286009786 | 0.018605755 | 0.476889115 | 0.003203655 | 0.036936863 | 0.034990762 | 0.810772242 |
| Cdt1 | S | 403 | 0.093806326 | 0.290070533 | 0.038876003 | -0.054930323 | -0.25119453 | 0.470928085 | 0.000882306 | 0.434163001 | 0.649601783 | 0.005096457 |
| Srrm2 | S | 869 | 0.022794393 | 0.071706114 | 0.139915284 | 0.117120891 | 0.06820917 | 0.514826247 | 0.004746456 | 0.000109324 | 0.044378739 | 0.003941614 |
| Eef1d | T | 147 | 0.044605036 | 0.142213031 | 0.168405683 | 0.123800647 | 0.026192652 | 0.326843226 | 0.034382643 | 0.004421924 | 0.035295277 | 0.598627473 |
| Rb1cc1 | S | 229 | 0.003524253 | 0.14007146 | 0.242918671 | 0.239394418 | 0.102847211 | 0.957428934 | 0.026574653 | 0.004520512 | 0.01983127 | 0.104159656 |
| Cenpc1 | S | 158 | 0.052460903 | 0.088790527 | 0.128030301 | 0.075569397 | 0.039239773 | 0.063518487 | 0.004264696 | 0.007395916 | 0.055173 | 0.215337801 |
| Strn | S | 245 | 0.029202189 | 0.103279893 | 0.113471613 | 0.084269424 | 0.01019172 | 0.231121107 | 0.008867725 | 0.000279707 | 0.021686564 | 0.634882076 |
| Fam117a | S | 213 | -0.113851691 | 0.284308151 | 0.393213028 | 0.507064718 | 0.108904877 | 0.043947026 | 0.025027508 | 0.005073666 | 0.012234324 | 0.28745705 |
| Rgs12 | S | 671 | 0.164309396 | 0.123545714 | 0.2222356 | 0.057926204 | 0.098689886 | 0.01978138 | 0.089348288 | 0.297901852 | 0.755230804 | 0.605948034 |
| Lrrc8a | S | 202 | 0.220993533 | 0.245379284 | 0.162424832 | -0.058568701 | -0.082954452 | 0.314904135 | 0.132727103 | 0.069422715 | 0.758949314 | 0.523684146 |
| Nup133 | S | 498 | 0.096678426 | 0.218416539 | 0.292562853 | 0.195884427 | 0.074146314 | 0.335552551 | 0.062033247 | 0.136489401 | 0.269717344 | 0.642483978 |
| Tmpo | S | 308 | 0.144661795 | 0.09559968 | 0.175096463 | 0.030434668 | 0.079496783 | 0.095521877 | 0.121448053 | 0.018564475 | 0.637609958 | 0.077766838 |
| Nfat5 | T | 135 | 0.196607789 | 0.082370822 | 0.171859381 | -0.024748408 | 0.089488559 | 0.025252144 | 0.377624869 | 0.043999559 | 0.498370434 | 0.273215309 |
| Zc3h18 | S | 106 | 0.018076853 | 0.063470195 | 0.167519059 | 0.149442205 | 0.104048864 | 0.378842654 | 0.040683974 | 5.39989E-05 | 0.000485256 | 0.011350578 |
| Arl6ip6 | S | 80 | 0.03078177 | 0.125508477 | 0.130095283 | 0.099313513 | 0.004586805 | 0.43414864 | 0.00362218 | 0.003136726 | 0.04222147 | 0.829810554 |
| Brd8 | S | 617 | 0.004372768 | 0.189906544 | 0.15468899 | 0.150316222 | -0.035217554 | 0.924402602 | 0.002378293 | 0.020497406 | 0.029623425 | 0.414975861 |
| Afap1 | S | 344 | 0.103281343 | 0.07254988 | 0.146203279 | 0.042921936 | 0.073653399 | 0.001067219 | 0.136311111 | 0.058561389 | 0.414623298 | 0.241205798 |
| Cntln | S | 508 | -0.066273197 | 0.210312515 | 0.169748427 | 0.236021624 | -0.040564088 | 0.307009566 | 0.000306499 | 0.015613554 | 0.021107772 | 0.34363394 |
| Cntln | S | 510 | -0.066273197 | 0.210312515 | 0.169748427 | 0.236021624 | -0.040564088 | 0.307009566 | 0.000306499 | 0.015613554 | 0.021107772 | 0.34363394 |

| | | | | | | | | | | | | |
|---|---|---|---|---|---|---|---|---|---|---|---|---|
| Ubn2 | T | 238 | 0.085831706 | 0.103990703 | 0.156874927 | 0.071043221 | 0.052884224 | 0.174339456 | 0.001444374 | 0.06658712 | 0.333189713 | 0.383937865 |
| Aak1 | T | 389 | 0.208538489 | 0.164455643 | -0.101530024 | -0.310068513 | -0.265985668 | 0.000259013 | 0.003164807 | 0.173674171 | 0.016959488 | 0.021045426 |
| Nucb1 | S | 368 | -0.04594225 | 0.129273474 | 0.383558224 | 0.429500474 | 0.25428475 | 0.431801134 | 0.051800207 | 0.004452749 | 0.008049913 | 0.03170952 |
| Zc3h13 | S | 1090 | -0.014061898 | 0.130325941 | 0.18916915 | 0.203231048 | 0.058843209 | 0.72559474 | 0.006957434 | 0.001253685 | 0.009138093 | 0.12844035 |
| Zc3h13 | S | 1093 | -0.014061898 | 0.130325941 | 0.18916915 | 0.203231048 | 0.058843209 | 0.72559474 | 0.006957434 | 0.001253685 | 0.009138093 | 0.12844035 |
| Phf14 | S | 273 | 0.094192716 | 0.286724452 | 0.303145379 | 0.208952663 | 0.016420927 | 0.599221239 | 0.054710663 | 0.173283751 | 0.380212497 | 0.929195937 |
| Setd2 | S | 614 | 0.035350375 | 0.119866394 | 0.172868235 | 0.13751786 | 0.05300184 | 0.057730297 | 0.002257225 | 0.064168619 | 0.106705299 | 0.400773881 |
| Cep89 | S | 44 | 0.052384217 | 0.114623321 | 0.152559009 | 0.100174793 | 0.037935689 | 0.147994501 | 0.038467319 | 0.00331314 | 0.003142901 | 0.33549574 |
| Akap13 | T | 1342 | 0.135997722 | 0.113200952 | 0.109909445 | -0.026088277 | -0.003291507 | 0.019485603 | 0.068166335 | 0.048454609 | 0.354609817 | 0.937900535 |
| Pimreg | S | 72 | 0.166826023 | 0.056175444 | 0.130673855 | -0.036152169 | 0.074498411 | 0.009499905 | 0.218767826 | 0.02722756 | 0.434453208 | 0.169824405 |
| Epb41l2 | T | 642 | 0.227458355 | 0.133121275 | 0.112473149 | -0.114985205 | -0.020648126 | 0.020812257 | 0.129275568 | 0.244022281 | 0.163696411 | 0.772135114 |
| 4932438A13Rik | S | 1312 | 0.02066679 | 0.090352722 | 0.209858519 | 0.189191729 | 0.119505797 | 0.603644342 | 0.184832386 | 0.000137733 | 0.013910589 | 0.121828444 |
| Dync1h1 | S | 1228 | 0.088421702 | 0.268980314 | 0.143948474 | 0.055526772 | -0.12503184 | 0.304360428 | 0.007581247 | 0.351109957 | 0.696450951 | 0.401520689 |
| Phtf1 | S | 299 | 0.076355774 | 0.157663053 | 0.174691115 | 0.098335342 | 0.017028062 | 0.205689398 | 0.051394207 | 0.020004826 | 0.144162157 | 0.76452572 |
| Wdr48 | S | 335 | -0.005792424 | 0.173103228 | 0.13683429 | 0.142626714 | -0.036268938 | 0.855955662 | 0.004948968 | 0.001812382 | 0.00484944 | 0.324929952 |
| Pcm1 | S | 1769 | 0.088141451 | 0.094630333 | 0.115380076 | 0.027238626 | 0.020749743 | 0.019248726 | 0.007318769 | 0.051258581 | 0.532655021 | 0.615075501 |
| Nav3 | S | 989 | 0.208811406 | 0.098918461 | 0.06039076 | -0.148420646 | -0.038527702 | 0.001792049 | 0.092370889 | 0.443567992 | 0.119944498 | 0.609461983 |
| Epb41l2 | S | 201 | 0.03333417 | 0.180752033 | 0.105019733 | 0.071685563 | -0.0757323 | 0.520043371 | 0.001713453 | 0.021735053 | 0.175516517 | 0.013639002 |
| Dock7 | S | 190 | 0.076884887 | 0.222390671 | 0.26891172 | 0.192026833 | 0.046521049 | 0.458414461 | 0.110316899 | 0.046107217 | 0.157220928 | 0.71249423 |
| Nfic | S | 333 | 0.072853869 | 0.11437334 | 0.133790177 | 0.060936309 | 0.019416838 | 0.09260661 | 0.026541959 | 0.008991437 | 0.056131467 | 0.534141383 |
| Ccnk | S | 329 | -0.003770984 | 0.115912749 | 0.148317713 | 0.152088698 | 0.032404964 | 0.902753796 | 0.00098748 | 0.001604108 | 0.007587847 | 0.226509397 |
| Trim28 | S | 600 | 0.002294818 | 0.170360116 | 0.174954757 | 0.172659939 | 0.004594641 | 0.935772401 | 0.060134642 | 0.000535912 | 0.001167028 | 0.938867781 |
| Acin1 | S | 208 | -0.009351122 | 0.166581939 | 0.318972786 | 0.328323908 | 0.152390847 | 0.883620315 | 0.030568665 | 0.019067606 | 0.015578053 | 0.11696695 |
| Nsd1 | S | 844 | 0.046002324 | 0.082666074 | 0.146732134 | 0.10072981 | 0.06406606 | 0.214029687 | 0.002428553 | 0.00516423 | 0.041175863 | 0.059411325 |
| Sltm | S | 371 | 0.026318644 | 0.144846819 | 0.184842912 | 0.158524268 | 0.039996093 | 0.654828623 | 0.016834481 | 0.005775579 | 0.032270839 | 0.241054976 |
| Thrap3 | S | 190 | 0.289834121 | -0.031249372 | 0.12904408 | -0.160790041 | 0.160293452 | 0.000968131 | 0.651861861 | 0.098124509 | 0.061202039 | 0.089477097 |
| Ppp4r3a | S | 764 | 0.030666155 | 0.120100146 | 0.12884385 | 0.098177695 | 0.008743704 | 0.249499267 | 0.014376909 | 0.00123647 | 0.002833037 | 0.764676554 |
| Ppig | S | 254 | 0.05556481 | 0.082835862 | 0.121483929 | 0.065919119 | 0.038648067 | 0.155226004 | 0.013005376 | 0.000841632 | 0.111740608 | 0.120548848 |
| Ppig | S | 255 | 0.05556481 | 0.082835862 | 0.121483929 | 0.065919119 | 0.038648067 | 0.155226004 | 0.013005376 | 0.000841632 | 0.111740608 | 0.120548848 |
| Ppig | S | 257 | 0.05556481 | 0.082835862 | 0.121483929 | 0.065919119 | 0.038648067 | 0.155226004 | 0.013005376 | 0.000841632 | 0.111740608 | 0.120548848 |
| Epb41l3 | T | 495 | 0.111366817 | 0.082290846 | 0.164640935 | 0.053274118 | 0.082350089 | 0.024543732 | 0.112426804 | 0.024359731 | 0.322236193 | 0.17965776 |
| Sbf1 | S | 1120 | 0.3360476 | 0.034412704 | 0.074671585 | -0.261376015 | 0.040258881 | 0.002223199 | 0.520527445 | 0.389135296 | 0.031770763 | 0.591487787 |
| Bclaf3 | S | 445 | 0.142464765 | 0.091302591 | 0.120232684 | -0.022232082 | 0.028930093 | 0.007182742 | 0.155169825 | 0.051752235 | 0.594506502 | 0.626742514 |
| Hnrnpul2 | S | 183 | -0.077942742 | 0.202090374 | 0.422173246 | 0.500115988 | 0.220082873 | 0.123018885 | 0.003354413 | 0.037246086 | 0.025363967 | 0.137448235 |
| Bcar3 | S | 333 | 0.02088367 | 0.147603781 | 0.168255415 | 0.147371745 | 0.020651634 | 0.277329959 | 0.008271059 | 0.014736183 | 0.041783669 | 0.633478954 |
| Ank | T | 8 | 0.000989885 | 0.071796142 | 0.202724557 | 0.201734672 | 0.130928415 | 0.988053524 | 0.074694941 | 5.33598E-05 | 0.065667179 | 0.01683442 |
| Mark2 | S | 483 | 0.169444646 | 0.410071177 | -0.049227011 | -0.218671657 | -0.459298188 | 0.371817743 | 0.001766679 | 0.534587291 | 0.26989169 | 0.002752247 |
| Atrx | S | 663 | -0.010185928 | 0.156702141 | 0.141428507 | 0.151614435 | -0.015273634 | 0.639375095 | 0.011920823 | 0.000260597 | 0.002027945 | 0.633663983 |
| Tcof1 | S | 169 | 0.063953027 | 0.097011416 | 0.190793978 | 0.126840951 | 0.093782562 | 0.294594504 | 0.033652604 | 0.00572115 | 0.080868945 | 0.07595113 |
| Cobll1 | S | 1145 | 0.449223701 | 0.412142038 | -0.343905644 | -0.793129344 | -0.756047681 | 0.037746602 | 0.046826765 | 0.496082515 | 0.186912364 | 0.201766562 |
| Pxn | S | 272 | -0.029281637 | 0.232590927 | 0.369694327 | 0.398975964 | 0.1371034 | 0.69376282 | 0.021620982 | 0.065357687 | 0.071257544 | 0.37085375 |
| Scaf1 | S | 912 | 0.044921822 | 0.140312144 | 0.20420025 | 0.159278428 | 0.063888106 | 0.264998814 | 0.00420184 | 0.076480204 | 0.111645906 | 0.421948079 |
| Ncor2 | S | 152 | -0.015008453 | 0.107484102 | 0.156905215 | 0.171913667 | 0.049421113 | 0.527134664 | 0.001453478 | 0.000416989 | 0.002475092 | 0.053970827 |
| Ubn2 | S | 236 | 0.071966051 | 0.097959667 | 0.106509079 | 0.034543028 | 0.008549413 | 0.043394231 | 0.000994941 | 0.060307382 | 0.427544236 | 0.820074412 |
| Rbm15 | S | 701 | 0.20243131 | 0.311918365 | 0.024235367 | -0.178195942 | -0.287682997 | 0.163934475 | 0.0110604 | 0.885592773 | 0.371582154 | 0.1725743 |
| Senp7 | S | 25 | -0.013399503 | 0.138956313 | 0.208312512 | 0.221712015 | 0.069356199 | 0.694783875 | 0.000415164 | 0.025520204 | 0.018470209 | 0.247515479 |
| Rif1 | S | 391 | 0.137022695 | 0.068308329 | 0.575657303 | 0.438634608 | 0.507348974 | 0.173989958 | 0.635813765 | 0.089605189 | 0.14897613 | 0.108642519 |
| Acin1 | S | 354 | 0.28938003 | 0.209218796 | 0.002308555 | -0.287071475 | -0.206910241 | 0.006262142 | 0.152501592 | 0.991122322 | 0.244405258 | 0.378129092 |
| Srrm2 | S | 2642 | 0.048222492 | 0.08714382 | 0.16633127 | 0.118108778 | 0.07918745 | 0.050607513 | 0.001918647 | 0.042221894 | 0.075665731 | 0.165318438 |
| Epb41l2 | S | 193 | 0.050289693 | 0.216460327 | 0.258797077 | 0.208507384 | 0.04233675 | 0.560246542 | 0.014393859 | 0.139240616 | 0.208629535 | 0.757189239 |
| Spen | S | 636 | 0.048412311 | 0.053453917 | 0.263044687 | 0.214632377 | 0.20959077 | 0.663979528 | 0.275063663 | 0.000380188 | 0.143948985 | 0.010339343 |
| Sav1 | S | 95 | -0.023345771 | 0.248384064 | 0.117699382 | 0.141045153 | -0.130684683 | 0.573101178 | 0.000822736 | 0.035162472 | 0.021947145 | 0.015780652 |
| Stt3b | S | 17 | 0.184771537 | 0.144013871 | 0.186479297 | 0.00170776 | 0.042465426 | 0.124409005 | 0.089341243 | 0.118447872 | 0.988289998 | 0.675540328 |
| Lemd3 | S | 263 | 0.06695041 | 0.129792553 | 0.171429653 | 0.104479243 | 0.0416371 | 0.156214689 | 0.00747103 | 0.061759193 | 0.184404185 | 0.525770905 |
| Spp1 | S | 24 | 0.154794234 | 0.12172662 | 0.235697249 | 0.080903015 | 0.113970629 | 0.180155026 | 0.212381254 | 0.034412267 | 0.344477942 | 0.074738369 |
| Epb41l1 | S | 544 | 0.082836724 | 0.095780229 | 0.082244738 | -0.000591985 | -0.013535491 | 0.023334287 | 0.00909759 | 0.010698726 | 0.977589213 | 0.492720532 |
| Nfic | S | 294 | 0.02871102 | 0.091194029 | 0.103524843 | 0.074813823 | 0.012330814 | 0.225440408 | 0.00349689 | 0.000309131 | 0.024749304 | 0.48382769 |
| Sorbs2 | S | 339 | -0.062145385 | 0.371472306 | 0.209122101 | 0.271267485 | -0.162350205 | 0.510656828 | 0.00755812 | 0.098426486 | 0.052092178 | 0.121265388 |
| Frmd6 | S | 609 | 0.000185549 | 0.280266007 | 0.09952129 | 0.099335741 | -0.180744717 | 0.99751893 | 0.000702086 | 0.182025102 | 0.200871123 | 0.046881083 |
| Mtch1 | S | 381 | -0.008769953 | 0.154633583 | 0.116205986 | 0.124975939 | -0.038427597 | 0.817101953 | 0.000713045 | 0.002975153 | 0.0302273 | 0.071084857 |
| Tra2b | T | 33 | 0.151113445 | 0.145225784 | 0.180435325 | 0.02932188 | 0.035209541 | 0.061116909 | 0.025205245 | 0.313623887 | 0.853513329 | 0.819826929 |
| Usp9x | S | 1600 | 0.059665334 | 0.169180618 | 0.082088267 | 0.022422933 | -0.087092351 | 0.258989363 | 0.00070298 | 0.088569561 | 0.662825062 | 0.082840021 |
| Pola1 | S | 215 | -0.017021534 | 0.191417051 | 0.149387775 | 0.166409309 | -0.042029276 | 0.473533524 | 0.001164911 | 0.011871846 | 0.007738518 | 0.258319075 |
| Mki67 | T | 1400 | 0.216111401 | 0.252414702 | 0.179746994 | -0.036364407 | -0.072667708 | 0.169592052 | 0.216989062 | 0.137204852 | 0.798132426 | 0.688689663 |
| Zranb2 | S | 294 | 0.026170304 | 0.121255757 | 0.26241443 | 0.236244126 | 0.141158674 | 0.651568123 | 0.040329009 | 0.009311593 | 0.020051762 | 0.080287428 |
| Nsd2 | S | 408 | 0.150583025 | 0.130923253 | 0.097581597 | -0.053001428 | -0.033341656 | 0.070454124 | 0.002339397 | 0.363781312 | 0.618560277 | 0.731568932 |
| Mki67 | S | 1393 | 0.189039854 | 0.168591439 | 0.096566163 | -0.092473691 | -0.072025276 | 0.106881169 | 0.013026334 | 0.323477105 | 0.422635584 | 0.447145204 |
| Ints12 | S | 388 | 0.253336657 | 0.071842652 | 0.139391647 | -0.11394501 | 0.067548995 | 0.002088887 | 0.383849727 | 0.475255629 | 0.55070013 | 0.72274381 |
| Tcf20 | S | 730 | 0.036986808 | 0.151551469 | 0.19302987 | 0.156043062 | 0.041478401 | 0.556838476 | 0.03221488 | 0.007973825 | 0.032646514 | 0.362296157 |
| Pkn1 | S | 777 | 0.202033701 | 0.402782205 | 0.052367003 | -0.149666698 | -0.350415201 | 0.193851338 | 0.041108389 | 0.743854041 | 0.372421382 | 0.092384253 |
| Aqp1 | T | 246 | 0.199506547 | 0.170688914 | -0.006944079 | -0.206450625 | -0.177632992 | 0.010246116 | 0.008928216 | 0.937703513 | 0.085980084 | 0.121187146 |
| Irf2bpl | S | 618 | 0.014717124 | 0.086689986 | 0.231187515 | 0.216470391 | 0.144497529 | 0.799671006 | 0.077565741 | 0.00064295 | 0.023060422 | 0.009825996 |
| Rock1 | S | 1328 | 0.200546936 | 0.193256708 | -0.00805923 | -0.208606166 | -0.201315938 | 0.00593128 | 0.027949247 | 0.968512247 | 0.360911025 | 0.375349555 |
| Ttc33 | S | 197 | -0.062477755 | 0.192817877 | 0.27872479 | 0.341202544 | 0.085906912 | 0.206710184 | 0.004994499 | 0.010064896 | 0.004975928 | 0.201951044 |
| Rcor3 | S | 227 | 0.060495751 | 0.157350836 | 0.216336262 | 0.15584051 | 0.058985426 | 0.488041231 | 0.017857343 | 0.043073864 | 0.156175665 | 0.436231686 |
| Caap1 | S | 15 | -0.013402961 | 0.396271455 | 0.157196166 | 0.170599127 | -0.239075289 | 0.924733857 | 0.005950531 | 0.298000507 | 0.297552541 | 0.129652876 |
| Ociad1 | S | 108 | 0.031563272 | 0.123177161 | 0.146230183 | 0.114666911 | 0.023053023 | 0.402000866 | 0.009761139 | 0.003756539 | 0.015766538 | 0.338975983 |
| Nipbl | S | 305 | 0.029483042 | 0.124563264 | 0.177976387 | 0.148493345 | 0.053413123 | 0.54698736 | 0.019598963 | 0.003843093 | 0.02777246 | 0.091455296 |
| Mier1 | S | 165 | 0.014330061 | 0.174073472 | 0.20385986 | 0.189529799 | 0.029786388 | 0.673788675 | 0.030964468 | 0.009437673 | 0.008185437 | 0.526698836 |
| Zdhhc8 | S | 523 | 0.163401345 | 0.10410302 | 0.127023874 | -0.036377471 | 0.022920854 | 0.045146669 | 0.086022033 | 0.052901843 | 0.576468134 | 0.692326348 |
| Irf2bp1 | S | 371 | 0.150692406 | 0.143062091 | 0.204631584 | 0.053939178 | 0.061569493 | 0.03347277 | 0.145294539 | 0.186961703 | 0.673800517 | 0.655244679 |
| Fbxw8 | T | 93 | 0.063550433 | 0.245582276 | 0.047679723 | -0.01587071 | -0.197902553 | 0.543057739 | 0.000196233 | 0.500743825 | 0.885962532 | 0.058465172 |
| Pimreg | S | 193 | 0.069546151 | 0.075377728 | 0.128178854 | 0.058632703 | 0.052801126 | 0.064852486 | 0.022663738 | 0.003340215 | 0.106854473 | 0.092101008 |
| Mki67 | S | 523 | 0.142925045 | 0.178897616 | 0.138671205 | -0.00425384 | -0.040226412 | 0.037161916 | 0.06817338 | 0.238849769 | 0.966734532 | 0.719667725 |
| Clasp1 | S | 1088 | 0.148033805 | 0.206196125 | 0.073711843 | -0.074321962 | -0.132484282 | 0.065786792 | 0.01215142 | 0.470262808 | 0.444486954 | 0.215079996 |
| Srrm2 | S | 2084 | 0.115770058 | 0.09168604 | 0.122779531 | 0.007009473 | 0.031093491 | 0.015390646 | 0.05168853 | 0.04956814 | 0.874870269 | 0.517382434 |
| Suds3 | S | 236 | -0.003771888 | 0.075062575 | 0.216771497 | 0.220543386 | 0.141708922 | 0.919784712 | 0.039596162 | 0.000226278 | 0.004007938 | 0.000866399 |
| Pacc1 | T | 8 | 0.14256243 | 0.192177356 | 0.181784675 | 0.039222245 | -0.010392681 | 0.200295037 | 0.143201342 | 0.051872733 | 0.670897464 | 0.920954201 |
| Magi3 | S | 1221 | 0.32042157 | 0.173074325 | 0.091075956 | -0.229345614 | -0.081998369 | 0.067813192 | 0.152146942 | 0.316525359 | 0.159052864 | 0.359552737 |
| Magi3 | S | 1223 | 0.32042157 | 0.173074325 | 0.091075956 | -0.229345614 | -0.081998369 | 0.067813192 | 0.152146942 | 0.316525359 | 0.159052864 | 0.359552737 |
| Ifi204 | S | 190 | -0.178452688 | 0.184541737 | 0.33824729 | 0.516699978 | 0.153705553 | 0.089569519 | 0.011305481 | 0.000687793 | 0.006060981 | 0.044096972 |
| Tcof1 | S | 1267 | 0.039113045 | 0.155164094 | 0.207771431 | 0.168658386 | 0.052607336 | 0.122414701 | 0.01159552 | 0.091721041 | 0.14278513 | 0.552694829 |
| Add1 | T | 610 | 0.005226088 | 0.136246021 | 0.13650417 | 0.131278082 | 0.000258149 | 0.802937691 | 0.00030753 | 0.01828065 | 0.021703502 | 0.993602365 |
| Trpm7 | S | 1255 | -0.115615912 | 0.145586754 | 0.285577044 | 0.401192956 | 0.139990291 | 0.244550014 | 0.002425923 | 0.002956002 | 0.015881288 | 0.058242579 |

| | | | | | | | | | | | | |
|---|---|---|---|---|---|---|---|---|---|---|---|---|
| Zfp37 | S | 69 | -0.025628147 | 0.225112249 | 0.151559895 | 0.177188041 | -0.073552355 | 0.659998847 | 0.010249386 | 0.004244531 | 0.050066424 | 0.21565004 |
| Fxr2 | S | 449 | 0.224341064 | 0.292094313 | 0.107685544 | -0.11665552 | -0.184408769 | 0.126138239 | 0.044733198 | 0.721006707 | 0.691070647 | 0.537563619 |
| Slc39a3 | S | 129 | -0.002449589 | 0.153392261 | 0.180356186 | 0.182805776 | 0.026963925 | 0.973509195 | 0.064071308 | 0.000445089 | 0.093129294 | 0.635027769 |
| Srsf4 | S | 441 | 0.436067104 | 0.048695794 | 0.242198047 | -0.193869056 | 0.193502254 | 0.183409434 | 0.471111491 | 0.087232701 | 0.481224934 | 0.140274622 |
| Uimc1 | S | 665 | 0.024696656 | 0.098828993 | 0.15610278 | 0.131406125 | 0.057273787 | 0.260155917 | 0.002047976 | 0.009297778 | 0.027242103 | 0.163479487 |
| Arhgap35 | S | 970 | 0.137546658 | 0.158139872 | 0.041968577 | -0.095578081 | -0.116171296 | 0.194301507 | 6.60157E-05 | 0.748056251 | 0.523590634 | 0.413567527 |
| Epb41l2 | S | 641 | 0.206492643 | 0.188678007 | 0.075232927 | -0.131259716 | -0.11344508 | 0.050208079 | 0.043112512 | 0.403305621 | 0.133408386 | 0.12683769 |
| Rrbp1 | S | 766 | 0.049427401 | 0.215991081 | 0.108191728 | 0.058764328 | -0.107799352 | 0.6097319 | 0.002270996 | 0.086836392 | 0.543055491 | 0.076513319 |
| Trp53bp1 | S | 1096 | -0.031248175 | 0.12088773 | 0.21638311 | 0.247631285 | 0.09549538 | 0.595515464 | 0.001019461 | 0.007293041 | 0.01906936 | 0.065398072 |
| Ercc5 | S | 572 | -0.026681885 | 0.132207971 | 0.154103376 | 0.180785261 | 0.021895405 | 0.4346306 | 0.000241548 | 0.004290124 | 0.006507822 | 0.440515624 |
| Prkacb | S | 339 | 0.013072382 | 0.142045266 | 0.224029654 | 0.210957271 | 0.081984388 | 0.709174587 | 0.010018397 | 0.019192396 | 0.047823904 | 0.228199152 |
| Trerf1 | S | 490 | -0.030364714 | 0.195502385 | 0.253275857 | 0.28364057 | 0.057773472 | 0.712944267 | 0.007500558 | 0.030595381 | 0.03622566 | 0.402171238 |
| Anln | S | 637 | 0.320941205 | 0.105199911 | -0.003627371 | -0.324568576 | -0.108827282 | 0.001771813 | 0.216067505 | 0.965491026 | 0.013793574 | 0.22525694 |
| Rps6ka4 | S | 343 | 0.06344615 | 0.186970626 | 0.127014167 | 0.063568018 | -0.059956459 | 0.283933916 | 0.008255769 | 0.067076696 | 0.315316297 | 0.313324246 |
| Kif13a | S | 1481 | 0.029088027 | 0.183563834 | 0.19914444 | 0.170056414 | 0.015580607 | 0.761030766 | 0.008091016 | 0.04516141 | 0.159256359 | 0.823756192 |
| Rbm15 | T | 567 | 0.039069312 | 0.074939653 | 0.117820309 | 0.078750997 | 0.042880656 | 0.038877723 | 0.001235863 | 0.008443355 | 0.036244079 | 0.153054797 |
| Phf2 | S | 726 | 0.024319413 | 0.140796889 | 0.20996343 | 0.185644017 | 0.069166541 | 0.781210103 | 0.001245876 | 0.060947786 | 0.130437817 | 0.376166039 |
| Phf2 | S | 729 | 0.024319413 | 0.140796889 | 0.20996343 | 0.185644017 | 0.069166541 | 0.781210103 | 0.001245876 | 0.060947786 | 0.130437817 | 0.376166039 |
| Phf2 | S | 730 | 0.024319413 | 0.140796889 | 0.20996343 | 0.185644017 | 0.069166541 | 0.781210103 | 0.001245876 | 0.060947786 | 0.130437817 | 0.376166039 |
| Mdc1 | S | 591 | 0.213227417 | 0.148116341 | -0.016658862 | -0.229886278 | -0.164775203 | 0.001765736 | 0.030192937 | 0.748712032 | 0.009259062 | 0.027343738 |
| Ccdc88a | S | 1439 | -0.139038439 | 0.451695815 | 0.225223242 | 0.364261681 | -0.226472573 | 0.154352037 | 0.002415118 | 0.134736772 | 0.048760967 | 0.132225783 |
| Safb | S | 577 | 0.132201857 | 0.079608521 | 0.249723674 | 0.117521817 | 0.170115153 | 0.300520574 | 0.347338062 | 0.006398336 | 0.350540476 | 0.100234227 |
| Tns2 | S | 820 | 0.125849313 | 0.120560134 | 0.183429246 | 0.057579933 | 0.062869111 | 0.061772491 | 0.263143757 | 0.025983996 | 0.379543754 | 0.534233401 |
| Sipa1l2 | S | 1488 | 0.054671051 | 0.108933655 | 0.10403347 | 0.049362418 | -0.004900185 | 0.090323529 | 0.006395729 | 0.008223241 | 0.025681502 | 0.677249197 |
| Cenpu | S | 164 | 0.050467524 | 0.15720588 | 0.177056846 | 0.126589322 | 0.019850966 | 0.262874567 | 0.009294908 | 0.0662085 | 0.107718862 | 0.73412684 |
| Bclaf3 | S | 78 | 0.09598885 | 0.044505296 | 0.162387123 | 0.066398273 | 0.117881828 | 0.052620644 | 0.106299197 | 0.003320534 | 0.153658365 | 0.027349247 |
| Bclaf3 | S | 80 | 0.09598885 | 0.044505296 | 0.162387123 | 0.066398273 | 0.117881828 | 0.052620644 | 0.106299197 | 0.003320534 | 0.153658365 | 0.027349247 |
| Plekhm3 | S | 132 | 0.04923751 | 0.198779007 | 0.150339549 | 0.101102039 | -0.048439458 | 0.733780221 | 0.013265688 | 0.021643297 | 0.493568669 | 0.322536704 |
| Zc3h4 | S | 1104 | 0.087801244 | 0.059591943 | 0.147197138 | 0.059395894 | 0.087605194 | 0.089757746 | 0.001190704 | 0.030956599 | 0.225514128 | 0.087448495 |
| Akap12 | T | 910 | -0.20495511 | 0.43881891 | 0.279922063 | 0.484877173 | -0.158896847 | 0.468091812 | 0.014473123 | 0.080542177 | 0.15117448 | 0.272512131 |
| S1pr2 | S | 332 | -0.032654961 | 0.214706882 | 0.243263771 | 0.275918732 | 0.028556889 | 0.277040403 | 0.003974949 | 0.030042334 | 0.021151375 | 0.657104784 |
| Cbx1 Cbx3 | S | 132;128 | 0.106075946 | 0.132270898 | 0.298519729 | 0.192443783 | 0.16624883 | 0.044633741 | 0.305542245 | 0.096284053 | 0.217676032 | 0.318416607 |
| Arhgap23 | S | 351 | 0.24712025 | 0.144953363 | 0.246158226 | -0.000962024 | 0.101204864 | 0.343361945 | 0.318868726 | 0.041481408 | 0.996727495 | 0.485465406 |
| Eif5b | S | 114 | 0.209803099 | 0.024070253 | 0.209503163 | -0.000299936 | 0.185432911 | 0.025810141 | 0.779618496 | 0.031082792 | 0.995007177 | 0.041227766 |
| Srrm2 | S | 743 | 0.1181272 | 0.047905727 | 0.139400501 | 0.021273301 | 0.091494774 | 0.00158901 | 0.1172571 | 0.068232935 | 0.684137191 | 0.165102827 |
| Emd | T | 30 | -0.026664912 | 0.124594447 | 0.198315841 | 0.224980754 | 0.073721395 | 0.393258252 | 0.001876439 | 0.002859505 | 0.002462506 | 0.068873546 |
| Zfp768 | T | 35 | 0.063701506 | 0.156342165 | 0.267344808 | 0.203643303 | 0.111002643 | 0.560225413 | 0.074286953 | 0.028797924 | 0.143846574 | 0.169210646 |
| Mdc1 | S | 595 | 0.060477197 | 0.117394953 | 0.092932357 | 0.03245516 | -0.024462596 | 0.035629885 | 0.001822528 | 0.072482856 | 0.407275435 | 0.51594474 |
| Zfp318 | S | 99 | -0.009513648 | 0.080104027 | 0.38788584 | 0.397399488 | 0.307781813 | 0.858124364 | 0.07712779 | 0.006942158 | 0.005443334 | 0.016911081 |
| Zfp318 | S | 101 | -0.009513648 | 0.080104027 | 0.38788584 | 0.397399488 | 0.307781813 | 0.858124364 | 0.07712779 | 0.006942158 | 0.005443334 | 0.016911081 |
| Safb Safb2 | S | 366;387 | 0.020158402 | 0.072835903 | 0.155962044 | 0.135803641 | 0.083126141 | 0.567139298 | 0.013324845 | 0.000258452 | 0.020459449 | 0.019526427 |
| Akap12 | S | 652 | 0.070358489 | 0.26630278 | 0.049052554 | -0.021305935 | -0.217250225 | 0.299687971 | 0.001685119 | 0.421539578 | 0.724456313 | 0.023621645 |
| Rfc1 | S | 365 | 0.062504173 | 0.108814236 | 0.098178271 | 0.035674097 | -0.010635966 | 0.160322999 | 0.000572017 | 0.040860947 | 0.424188097 | 0.724995065 |
| Vrk2 | T | 449 | 0.077709728 | 0.087963269 | 0.151748432 | 0.074038704 | 0.063785163 | 0.159033563 | 0.200505691 | 0.000740227 | 0.173507007 | 0.313236555 |
| Mast2 | S | 815 | -0.122051674 | 0.31370953 | 0.281350498 | 0.403402172 | -0.032359032 | 0.16099112 | 0.005950894 | 0.027856881 | 0.01537837 | 0.690423211 |
| Eif2ak4 | T | 666 | 0.050652213 | 0.148276208 | 0.129657322 | 0.079005108 | -0.018618887 | 0.209446689 | 0.028983186 | 0.00788335 | 0.082929207 | 0.690891196 |
| Cic | S | 1403 | 0.037963958 | 0.043577612 | 0.279891199 | 0.241927241 | 0.236313587 | 0.456692302 | 0.001097192 | 0.007338266 | 0.008532203 | |
| Zhx2 | T | 353 | 0.510356284 | 0.045187177 | 0.063292625 | -0.447063659 | 0.018105448 | 0.00712137 | 0.817760489 | 0.814668855 | 0.171604323 | 0.948283145 |
| Ctnnb1 | S | 675 | 0.029897303 | 0.181396796 | 0.130947633 | 0.10105033 | -0.050449163 | 0.422282571 | 0.002224114 | 0.073598048 | 0.145356856 | 0.383969195 |
| Cltc | T | 394 | -0.013147547 | 0.093348401 | 0.178982132 | 0.192129679 | 0.085633731 | 0.553957321 | 0.050946 | 3.35807E-05 | 0.000593864 | 0.072494408 |
| Rbbp6 | S | 1179 | 0.019844857 | 0.075457228 | 0.166693521 | 0.146848664 | 0.091236292 | 0.634653382 | 0.045338712 | 0.000135647 | 0.036916433 | 0.030943523 |
| Resf1 | S | 1480 | 0.086913708 | 0.160682067 | 0.170810703 | 0.083896996 | 0.010128637 | 0.176680242 | 0.055458109 | 0.045477661 | 0.186957799 | 0.87423574 |
| Zzz3 | S | 89 | 0.079806163 | 0.091027544 | 0.170663067 | 0.090856904 | 0.079635523 | 0.154824835 | 0.073090354 | 0.006834934 | 0.080917415 | 0.049017452 |
| Vim | S | 339 | 0.071475126 | 0.194532007 | 0.133524004 | 0.062048878 | -0.061008003 | 0.054501794 | 0.033941008 | 0.122997659 | 0.374190417 | 0.428830169 |
| Tcf20 | S | 1047 | 0.031375421 | 0.117153005 | 0.228596576 | 0.197221156 | 0.111443571 | 0.621408303 | 0.066571646 | 0.004546849 | 0.029125835 | 0.090744603 |
| Ctnnd1 | T | 308 | 0.190352418 | 0.202167557 | 0.013071102 | -0.177281316 | -0.189096455 | 0.009010483 | 0.044594554 | 0.785587543 | 0.023836402 | 0.052925708 |
| Micall1 | S | 628 | 0.195626933 | 0.17624958 | -0.005685054 | -0.201311987 | -0.181934634 | 0.017793541 | 0.00816515 | 0.97290734 | 0.299676794 | 0.337792414 |
| Fam117a | S | 316 | -0.084506657 | 0.526990671 | 0.233858339 | 0.318364996 | -0.293132333 | 0.703302453 | 0.007873869 | 0.557173989 | 0.456499126 | 0.471399212 |
| Syne2 | S | 3673 | -0.064926265 | 0.281093744 | 0.223063944 | 0.287990209 | -0.0580298 | 0.561699179 | 0.053870188 | 0.005240014 | 0.070716545 | 0.567628758 |
| C77080 | S | 543 | 0.028958818 | 0.171661454 | 0.167892729 | 0.138933912 | -0.003768725 | 0.435240578 | 0.001487706 | 0.132924098 | 0.169812938 | 0.961721311 |
| Safb Safb2 | S | 298;292 | 0.026798305 | 0.192836732 | 0.189540188 | 0.162741882 | -0.003296544 | 0.497916245 | 0.039972596 | 0.01851238 | 0.031307218 | 0.96044579 |
| Hnrnpf | S | 104 | 0.144896119 | 0.111322241 | 0.123057604 | -0.021838514 | 0.011735364 | 0.03282586 | 0.156765544 | 0.037359115 | 0.672228092 | 0.860060624 |
| Taf1 | T | 333 | 0.121654929 | 0.08913727 | 0.136444463 | 0.014789534 | 0.047307193 | 0.012388321 | 0.083554285 | 0.076975787 | 0.786408523 | 0.450863716 |
| Osbp | S | 387 | 0.122138023 | 0.203028126 | 0.091573717 | -0.030564305 | -0.111454409 | 0.172174246 | 0.011096203 | 0.219879906 | 0.696505252 | 0.113953232 |
| Zc3h13 | S | 1297 | 0.099463919 | 0.160315139 | 0.175946818 | 0.076482899 | 0.015631679 | 0.149351879 | 0.150967002 | 0.024864177 | 0.138707441 | 0.858570038 |
| Phf20l1 | S | 599 | 0.106759543 | 0.081456307 | 0.172897143 | 0.0661376 | 0.091440836 | 0.032895102 | 0.061924121 | 0.053456823 | 0.336118572 | 0.222168549 |
| Lrrk2 | S | 895 | -0.046763201 | 0.222813368 | 0.279567207 | 0.326330408 | 0.056753839 | 0.146139892 | 0.007036448 | 0.020849997 | 0.013843267 | 0.41047666 |
| Myo9a | S | 1243 | 0.108458445 | 0.201663558 | 0.102469041 | -0.005989404 | -0.099194517 | 0.166087392 | 0.019682676 | 0.127040108 | 0.917210741 | 0.124267317 |
| Safb | S | 411 | 0.262254753 | 0.149396807 | 0.040311365 | -0.221943388 | -0.109085442 | 0.008433055 | 0.103785664 | 0.820458793 | 0.283407936 | 0.56218383 |
| Setd2 | S | 1387 | 0.044083792 | 0.153290333 | 0.106246822 | 0.06216303 | -0.047043511 | 0.035211753 | 0.059235644 | 0.005242207 | 0.051595654 | 0.400528598 |
| Setd2 | S | 1389 | 0.044083792 | 0.153290333 | 0.106246822 | 0.06216303 | -0.047043511 | 0.035211753 | 0.059235644 | 0.005242207 | 0.051595654 | 0.400528598 |
| Setd2 | S | 1391 | 0.044083792 | 0.153290333 | 0.106246822 | 0.06216303 | -0.047043511 | 0.035211753 | 0.059235644 | 0.005242207 | 0.051595654 | 0.400528598 |
| Reps1 | S | 166 | 0.135875812 | 0.110211997 | 0.069731959 | -0.066143853 | -0.040480038 | 0.014964128 | 0.014256223 | 0.174585303 | 0.196543257 | 0.34355833 |
| Bud13 | S | 200 | 0.070745946 | 0.116998183 | 0.089013197 | 0.018267251 | -0.027984985 | 0.075690791 | 0.003335041 | 0.035914486 | 0.616382373 | 0.391977959 |
| Bud13 | T | 196 | 0.070745946 | 0.116998183 | 0.089013197 | 0.018267251 | -0.027984985 | 0.075690791 | 0.003335041 | 0.035914486 | 0.616382373 | 0.391977959 |
| Nipbl | S | 889 | 0.021948839 | 0.142084692 | 0.289796361 | 0.267847522 | 0.147711669 | 0.656015952 | 0.118663516 | 0.01113153 | 0.025717467 | 0.140536689 |
| Pcm1 | S | 1260 | 0.06229579 | 0.065767672 | 0.179152249 | 0.116856459 | 0.113384577 | 0.038630081 | 0.173662159 | 0.003853763 | 0.022541666 | 0.056956325 |
| Gpatch8 | S | 439 | -0.023227961 | 0.11691048 | 0.219053274 | 0.242281235 | 0.102142794 | 0.575767822 | 0.005905309 | 0.003069233 | 0.005546868 | 0.038242267 |
| H1f4 | S | 36 | 0.050631242 | 0.186565676 | 0.154362169 | 0.103730927 | -0.032203507 | 0.518303951 | 0.016128846 | 0.032643462 | 0.17563997 | 0.304425325 |
| Ppfibp1 | S | 435 | -0.029066267 | 0.150655669 | 0.172678652 | 0.201744918 | 0.022022983 | 0.205561699 | 0.010857352 | 0.000334482 | 0.000905309 | 0.415027611 |
| Als2 | S | 460 | 0.048688235 | 0.346557919 | 0.092139206 | 0.043450971 | -0.254418713 | 0.469781101 | 0.009568659 | 0.319747917 | 0.544903198 | 0.044151997 |
| Huwe1 | S | 2362 | -0.00419985 | 0.148333198 | 0.084283664 | 0.088483515 | -0.064049534 | 0.835381235 | 4.9589E-05 | 0.014055166 | 0.017140665 | 0.045548926 |
| Mast2 | S | 88 | -0.031119898 | 0.215195536 | 0.127134147 | 0.158254044 | -0.08806139 | 0.664920647 | 0.000832632 | 0.03193977 | 0.090662534 | 0.091276143 |
| Aak1 | T | 672 | 0.107462166 | 0.209758738 | 0.021177216 | -0.08628495 | -0.188581522 | 0.044245156 | 0.002662914 | 0.683747121 | 0.086000125 | 0.018719508 |
| Mcoln1 | S | 547 | 0.096662493 | 0.127033701 | 0.114883315 | 0.018220821 | -0.012150387 | 0.077294998 | 0.001898054 | 0.259521066 | 0.839140582 | 0.885637038 |
| Dlg1 | S | 570 | 0.052325164 | 0.092678655 | 0.336830288 | 0.284505123 | 0.244151632 | 0.106467761 | 0.190512276 | 0.038398022 | 0.05852051 | 0.06450608 |
| Atrx | S | 1891 | 0.038274268 | 0.110697113 | 0.203169854 | 0.164895585 | 0.092472741 | 0.244553282 | 0.008636721 | 0.035187592 | 0.076711702 | 0.208611328 |
| Atrx | S | 1898 | 0.038274268 | 0.110697113 | 0.203169854 | 0.164895585 | 0.092472741 | 0.244553282 | 0.008636721 | 0.035187592 | 0.076711702 | 0.208611328 |
| Irf2bpl | S | 195 | 0.020355375 | 0.112280035 | 0.24211084 | 0.221755465 | 0.129830805 | 0.591549462 | 0.014249023 | 0.017855006 | 0.024509831 | 0.093503394 |
| Zfp609 | S | 845 | 0.178727605 | 0.09919866 | 0.17144862 | -0.007278985 | 0.07224996 | 0.047344978 | 0.053405839 | 0.228802311 | 0.953422858 | 0.555658999 |
| Zfp609 | S | 848 | 0.178727605 | 0.09919866 | 0.17144862 | -0.007278985 | 0.07224996 | 0.047344978 | 0.053405839 | 0.228802311 | 0.953422858 | 0.555658999 |
| Hmga2 | S | 101 | 0.033744908 | 0.087809097 | 0.109581765 | 0.075836878 | 0.021772688 | 0.147627672 | 0.004672106 | 0.001606218 | 0.006893163 | 0.180206483 |
| Stat5a | S | 128 | 0.004494573 | 0.283348376 | 0.101461529 | 0.096966956 | -0.181886846 | 0.897426736 | 0.00091429 | 0.299029197 | 0.317065242 | 0.117843118 |
| Tnrc6b | S | 604 | -0.151068285 | 0.310872066 | 0.33850274 | 0.489571024 | 0.027630674 | 0.256607513 | 0.021261244 | 0.021629717 | 0.017811658 | 0.80528693 |

| | | | | | | | | | | | | |
|---|---|---|---|---|---|---|---|---|---|---|---|---|
| Senp3 | S | 226 | 0.089650185 | 0.076609489 | 0.103681394 | 0.01403121 | 0.027071905 | 0.021995104 | 0.019031997 | 0.018321647 | 0.660253735 | 0.353107556 |
| Sin3a | S | 941 | 0.041050203 | 0.144937748 | 0.232471868 | 0.191421665 | 0.08753412 | 0.332155276 | 0.039090709 | 0.035392165 | 0.049511721 | 0.261252576 |
| Sipa1l1 | S | 1149 | 0.105909311 | 0.09242398 | 0.225618266 | 0.119708955 | 0.133194287 | 0.040823426 | 0.052665191 | 0.132428766 | 0.342719665 | 0.304039695 |
| Plec | S | 4399 | 0.177627483 | 0.024680489 | 0.49726909 | 0.319641608 | 0.472588601 | 0.33434904 | 0.894552543 | 0.036063932 | 0.146032656 | 0.063444204 |
| Epn2 | S | 173 | 0.055456057 | 0.120180885 | 0.154066294 | 0.098610237 | 0.033885409 | 0.100403811 | 0.077068807 | 0.007637854 | 0.049668196 | 0.538888522 |
| Sfswap | S | 867 | 0.127174143 | 0.043141255 | 0.093238119 | -0.033936025 | 0.050096864 | 0.002837982 | 0.439774579 | 0.003497035 | 0.190536993 | 0.373304436 |
| Repin1 | S | 341 | 0.215718409 | 0.313264162 | 0.109367397 | -0.106351011 | -0.203896765 | 0.122733798 | 0.078124299 | 0.665062975 | 0.662260515 | 0.439918731 |
| Hnrnpu | S | 243 | 0.193280987 | 0.010038647 | 0.205257356 | 0.011976369 | 0.19521871 | 0.008368856 | 0.894492386 | 0.050774826 | 0.868839815 | 0.076894928 |
| Scaf1 | S | 304 | -0.004098628 | 0.200664815 | 0.292436065 | 0.296534693 | 0.09177125 | 0.967999298 | 0.027481716 | 0.049024195 | 0.065982509 | 0.397939099 |
| Il10 | S | 136 | -0.110489828 | 0.160871716 | 0.314670901 | 0.425160729 | 0.153799185 | 0.047317243 | 0.001396291 | 0.003696502 | 0.001661831 | 0.054593857 |
| 2310022A10Rik | S | 269 | 0.067236568 | 0.181490172 | 0.109829619 | 0.042593051 | -0.071660553 | 0.304579824 | 0.020195897 | 0.021197622 | 0.497605164 | 0.191439128 |
| Itprid2 | S | 1056 | 0.189289321 | 0.018370344 | 0.4175599 | 0.228270579 | 0.399189557 | 0.16894476 | 0.899916462 | 0.042067625 | 0.198649248 | 0.070134513 |
| Gse1 | S | 840 | 0.172613096 | 0.106423248 | 0.034348662 | -0.138264435 | -0.072074587 | 0.001324255 | 0.06451589 | 0.748548973 | 0.266730167 | 0.516243626 |
| Afdn | S | 1107 | 0.010909924 | 0.076655936 | 0.1394375 | 0.128527576 | 0.062781564 | 0.466716763 | 9.35494E-05 | 0.006410418 | 0.005483099 | 0.05636818 |
| Mki67 | S | 765 | 0.119564818 | 0.224795725 | 0.019314968 | -0.10024985 | -0.205480756 | 0.092860039 | 0.002412185 | 0.739713091 | 0.178274387 | 0.02068851 |
| Safb | S | 569 | 0.095721483 | 0.321220284 | 0.579966907 | 0.484245424 | 0.258746623 | 0.396925963 | 0.288508844 | 0.292869947 | 0.360033733 | 0.618589853 |
| Cav1 | Y | 14 | -0.082449064 | 0.073018433 | 0.314013947 | 0.396463011 | 0.240995515 | 0.509717439 | 0.153099924 | 0.000193479 | 0.0540991 | 0.001930654 |
| Fnbp1 | T | 294 | 0.072380486 | 0.150441519 | 0.103068737 | 0.030688251 | -0.047372782 | 0.215704178 | 0.012020519 | 0.022888737 | 0.563242364 | 0.252171403 |
| Mybl2 | T | 490 | 0.237374132 | 0.106433638 | 0.171536683 | -0.065837449 | 0.065103045 | 0.081385291 | 0.134371909 | 0.184160946 | 0.618323119 | 0.538686112 |
| Smarcc2 | T | 308 | 0.104090512 | 0.089726439 | 0.133243162 | 0.02915265 | 0.043516723 | 0.066097944 | 0.063881651 | 0.015989032 | 0.442848689 | 0.103117086 |
| Chd4 | S | 96 | 0.014689935 | 0.139952407 | 0.273629428 | 0.258939493 | 0.133677021 | 0.840676587 | 0.080754273 | 0.009669754 | 0.024235362 | 0.12758147 |
| Chd4 | S | 98 | 0.014689935 | 0.139952407 | 0.273629428 | 0.258939493 | 0.133677021 | 0.840676587 | 0.080754273 | 0.009669754 | 0.024235362 | 0.12758147 |
| Chd4 | S | 101 | 0.014689935 | 0.139952407 | 0.273629428 | 0.258939493 | 0.133677021 | 0.840676587 | 0.080754273 | 0.009669754 | 0.024235362 | 0.12758147 |
| Dab2 | S | 393 | -0.094872199 | 0.140276458 | 0.429769903 | 0.524642102 | 0.289493445 | 0.276099774 | 0.035015759 | 0.00836344 | 0.005017988 | 0.035150177 |
| Dab2 | S | 394 | -0.094872199 | 0.140276458 | 0.429769903 | 0.524642102 | 0.289493445 | 0.276099774 | 0.035015759 | 0.00836344 | 0.005017988 | 0.035150177 |
| Pds5b | S | 1383 | 0.035811746 | 0.114902541 | 0.1874788 | 0.151667054 | 0.072576259 | 0.564301399 | 0.033475702 | 0.004721057 | 0.071787775 | 0.158056783 |
| Sun2 | S | 12 | 0.101114139 | 0.187541599 | 0.123108703 | 0.021994564 | -0.064432897 | 0.537293905 | 0.006344417 | 0.089952344 | 0.888520608 | 0.269344381 |
| Srrm1 | S | 683 | 0.179193613 | -0.014380892 | 0.205162508 | 0.025968895 | 0.2195434 | 0.023756855 | 0.827242622 | 0.007442375 | 0.64407372 | 0.037125771 |
| Zfp692 | S | 261 | 0.280403829 | 0.126317686 | 0.252138796 | -0.028265034 | 0.12582111 | 0.124117167 | 0.585357776 | 0.111907799 | 0.828954477 | 0.55693012 |
| Lmna | S | 51 | 0.079249413 | 0.181266294 | 0.103839983 | 0.02459057 | -0.07742631 | 0.201935974 | 0.017632195 | 0.050829484 | 0.669465157 | 0.193635698 |
| Zfp414 | S | 283 | 0.170830186 | 0.227913675 | 0.145531397 | -0.025298789 | -0.082382278 | 0.0828398 | 0.196888598 | 0.171768588 | 0.720300514 | 0.587260368 |
| Fam117a | S | 29 | 0.010711996 | 0.268612993 | 0.255020968 | 0.244308973 | -0.013592024 | 0.910187598 | 0.081915584 | 0.037400241 | 0.019308109 | 0.903755429 |
| Cep170 | S | 226 | 0.038235468 | 0.111787468 | 0.209885667 | 0.171650198 | 0.098098199 | 0.51777819 | 0.044579063 | 0.007131682 | 0.032996493 | 0.095511097 |
| Usp28 | S | 1027 | 0.065090298 | 0.17398331 | 0.219091823 | 0.154001525 | 0.045108513 | 0.361858481 | 0.08233286 | 0.036846056 | 0.103286489 | 0.567370034 |
| Gm3414 | S | 294 | -0.002567051 | 0.208631087 | 0.066417264 | 0.068984315 | -0.142213823 | 0.957891951 | 0.000269881 | 0.037020304 | 0.215283464 | 4.52542E-05 |
| Irf2bpl | S | 601 | 0.073753863 | 0.214605575 | 0.162342125 | 0.088588262 | -0.05226345 | 0.29338178 | 0.059965452 | 0.054002345 | 0.269455929 | 0.567116078 |
| Brca1 | T | 1199 | 0.108276972 | 0.071664288 | 0.131137148 | 0.022860176 | 0.05947286 | 0.02037524 | 0.113135386 | 0.01885732 | 0.515310918 | 0.171090451 |
| Pnkp | T | 122 | 0.062722707 | 0.104715829 | 0.146020255 | 0.083297548 | 0.041304426 | 0.093332856 | 0.036187757 | 0.013828055 | 0.090386261 | 0.364797317 |
| Gpatch8 | S | 735 | 0.028480628 | 0.088764953 | 0.168019943 | 0.139539316 | 0.079254991 | 0.406911314 | 0.002871644 | 0.009472824 | 0.020193912 | 0.077619041 |
| Rnpc3 | S | 379 | 0.056551927 | 0.125423888 | 0.210744421 | 0.154192494 | 0.085320533 | 0.248821546 | 0.027700758 | 0.043120449 | 0.111635302 | 0.286487263 |
| Carmil1 | S | 1150 | -0.08671773 | 0.299677857 | 0.263572227 | 0.350289957 | -0.03610563 | 0.232473932 | 0.001675116 | 0.127665349 | 0.074848522 | 0.783987888 |
| Arhgap12 | S | 213 | 0.07693912 | 0.22155579 | 0.073069281 | -0.003869838 | -0.148486508 | 0.203577732 | 0.004108886 | 0.265481377 | 0.939257921 | 0.043496315 |
| Pard3 | Y | 1123 | 0.183963355 | 0.05774079 | 0.185891747 | 0.001928392 | 0.128150956 | 0.015399771 | 0.297854887 | 0.123254684 | 0.983763267 | 0.240749662 |
| Ppp1r18 | S | 510 | 0.011841102 | 0.148320776 | 0.16295848 | 0.151117378 | 0.014637704 | 0.764980018 | 0.021887586 | 0.003735824 | 0.013364382 | 0.745439925 |
| Srsf1 | S | 231 | 0.009857381 | 0.093961969 | 0.170610604 | 0.160753223 | 0.076648635 | 0.804886974 | 0.015782066 | 0.000919685 | 0.012157574 | 0.032089049 |
| Terf2 | S | 399 | 0.019586942 | 0.236230292 | 0.153297458 | 0.133710516 | -0.082932834 | 0.756949067 | 0.004862307 | 0.123005046 | 0.157894834 | 0.323201805 |
| Pimreg | S | 191 | 0.066911792 | 0.071670507 | 0.117785187 | 0.050873394 | 0.04611468 | 0.061277695 | 0.032417157 | 0.002716571 | 0.126616605 | 0.124232009 |
| Tra2a Tra2b | S | 266;261 | 0.048123007 | 0.098910515 | 0.141305584 | 0.093182576 | 0.042395068 | 0.213615464 | 0.002665139 | 0.024874222 | 0.09143914 | 0.316314632 |
| Pdha1 | S | 293 | 0.017302851 | 0.122373312 | 0.246371909 | 0.229069058 | 0.123998597 | 0.601734832 | 0.042060364 | 0.012002742 | 0.008467209 | 0.060626386 |
| Zfyve26 | S | 646 | 0.042548169 | 0.218003753 | 0.096768892 | 0.054220723 | -0.121234861 | 0.36274606 | 0.002309423 | 0.188227035 | 0.378719841 | 0.117483793 |
| Brca1 | S | 717 | 0.04921262 | 0.096083533 | 0.186885693 | 0.137673073 | 0.09080216 | 0.18575043 | 0.103842013 | 0.004463374 | 0.027139767 | 0.128875076 |
| Rere | T | 593 | 0.023862437 | 0.120097482 | 0.148400728 | 0.124538291 | 0.028303246 | 0.42906672 | 0.001446133 | 0.027122058 | 0.048644072 | 0.507333046 |
| Kif18b | S | 561 | 0.184637121 | 0.131072969 | 0.045175599 | -0.139461523 | -0.085897371 | 0.023194538 | 0.009747716 | 0.6211922 | 0.204251751 | 0.379760525 |
| Rapgef2 | S | 1022 | 0.031701791 | 0.143072935 | 0.169697034 | 0.137995243 | 0.026624099 | 0.542963358 | 0.074998022 | 0.00241127 | 0.060164629 | 0.646592197 |
| Smarca4 | T | 1390 | 0.030769112 | 0.113544868 | 0.230267654 | 0.199498543 | 0.116722787 | 0.536715728 | 0.159506367 | 0.002775234 | 0.004037593 | 0.140820841 |
| Tenm4 | S | 40 | 0.049368718 | 0.117951564 | 0.141896789 | 0.092528071 | 0.023945225 | 0.273156344 | 0.028024184 | 0.005673972 | 0.055663577 | 0.517118864 |
| Tenm4 | S | 42 | 0.049368718 | 0.117951564 | 0.141896789 | 0.092528071 | 0.023945225 | 0.273156344 | 0.028024184 | 0.005673972 | 0.055663577 | 0.517118864 |
| Tenm4 | T | 44 | 0.049368718 | 0.117951564 | 0.141896789 | 0.092528071 | 0.023945225 | 0.273156344 | 0.028024184 | 0.005673972 | 0.055663577 | 0.517118864 |
| Tenm4 | Y | 39 | 0.049368718 | 0.117951564 | 0.141896789 | 0.092528071 | 0.023945225 | 0.273156344 | 0.028024184 | 0.005673972 | 0.055663577 | 0.517118864 |
| Fam126a | S | 451 | -0.016303872 | 0.398361029 | 0.191249454 | 0.207553326 | -0.207111575 | 0.890034502 | 0.026383821 | 0.153202489 | 0.133547665 | 0.177401679 |
| Nf1 | T | 2567 | -0.03766788 | 0.182259988 | 0.167619876 | 0.205287757 | -0.014640111 | 0.370139425 | 0.003846477 | 0.00485277 | 0.007441293 | 0.714532226 |
| Pkp4 | S | 75 | 0.120368946 | 0.038428931 | 0.181261066 | 0.06089212 | 0.142832135 | 0.043654665 | 0.494153309 | 0.005150527 | 0.248058164 | 0.058989041 |
| Pik3r4 | S | 813 | 0.027317501 | 0.246281005 | 0.162317988 | 0.135000488 | -0.083963017 | 0.692550566 | 0.015658821 | 0.078157842 | 0.11130616 | 0.312149309 |
| Prpf6 | T | 275 | 0.0811112 | 0.094278993 | 0.18125698 | 0.100145779 | 0.086977986 | 0.234637275 | 0.068885255 | 0.010438278 | 0.162656142 | 0.109930642 |
| Mov10 | S | 970 | 0.014478159 | 0.097534328 | 0.199958086 | 0.185479927 | 0.102423758 | 0.741290504 | 0.040135086 | 0.00172089 | 0.005165394 | 0.012836025 |
| Gcfc2 | S | 625 | 0.097955727 | 0.167023717 | 0.068603825 | -0.029351903 | -0.098419893 | 0.130534938 | 0.004518454 | 0.156933999 | 0.57584396 | 0.034635369 |
| Ifi203 | S | 173 | 0.044998682 | 0.217045095 | 0.31622891 | 0.271230228 | 0.099183815 | 0.726143546 | 0.21857675 | 0.029074038 | 0.033963955 | 0.481755258 |
| Sp110 | S | 277 | 0.025091334 | 0.096668442 | 0.265965405 | 0.240874071 | 0.169296963 | 0.705077436 | 0.099418554 | 0.005076185 | 0.017926687 | 0.036714384 |
| Glyr1 | S | 166 | 0.222427073 | 0.133115398 | 0.068595978 | -0.153831095 | -0.064519419 | 0.011126432 | 0.065660751 | 0.724838298 | 0.450509768 | 0.734245052 |
| Jcad | S | 486 | 0.08093167 | 0.127356746 | 0.200139904 | 0.119208235 | 0.072783159 | 0.227704655 | 0.007486392 | 0.151655978 | 0.333807704 | 0.513263596 |
| Jcad | T | 484 | 0.08093167 | 0.127356746 | 0.200139904 | 0.119208235 | 0.072783159 | 0.227704655 | 0.007486392 | 0.151655978 | 0.333807704 | 0.513263596 |
| Srrm2 | S | 1869 | 0.117919039 | 0.16349541 | 0.117190659 | -0.00072838 | -0.046304751 | 0.049570385 | 0.014795181 | 0.406569505 | 0.995371917 | 0.718267892 |
| Ptprg | S | 1179 | -0.006392408 | 0.177675261 | 0.332626001 | 0.339018408 | 0.15495074 | 0.950120706 | 0.016503445 | 0.081365463 | 0.079717465 | 0.307470388 |
| Ccnl1 | S | 348 | 0.046473805 | 0.067971855 | 0.179215642 | 0.132741837 | 0.111243786 | 0.216409842 | 0.009980531 | 0.008678138 | 0.027724263 | 0.052459142 |
| Trp53bp1 | S | 382 | 0.033047221 | 0.084414464 | 0.230786227 | 0.197739006 | 0.146371763 | 0.38413265 | 0.012104942 | 0.018381864 | 0.027890323 | 0.078239918 |
| Kdm5a | S | 204 | 0.075033119 | 0.156789866 | 0.12187151 | 0.04683839 | -0.034918357 | 0.148243333 | 0.004799084 | 0.182696431 | 0.553280557 | 0.637511445 |
| Rad9a | S | 373 | 0.054176644 | 0.17145106 | 0.158608007 | 0.104431363 | -0.012843053 | 0.034215105 | 0.049621867 | 0.101323818 | 0.19973805 | 0.865530249 |
| Pdlim2 | S | 143 | -0.033635955 | 0.12179203 | 0.197342035 | 0.23097799 | 0.075550004 | 0.286458947 | 0.004479644 | 0.001364901 | 0.001752616 | 0.058561248 |
| Esyt1 | S | 807 | -0.012178514 | 0.2232804 | 0.133640304 | 0.145818818 | -0.089640097 | 0.859190863 | 0.000449929 | 0.180849444 | 0.178384227 | 0.322963258 |
| Prpf6 | T | 212 | 0.275089854 | 0.237489715 | 0.211823206 | -0.063266648 | -0.025666509 | 0.138838769 | 0.335758598 | 0.30712683 | 0.765996216 | 0.921337779 |
| Mef2d | S | 98 | 0.036761541 | 0.11810567 | 0.171360761 | 0.13459922 | 0.053255091 | 0.346769117 | 0.032971073 | 0.007657386 | 0.028478172 | 0.267148955 |
| Mdc1 | S | 1128 | 0.053739446 | 0.164993126 | 0.344071234 | 0.290331788 | 0.179078108 | 0.474649077 | 0.031226181 | 0.163415884 | 0.207808756 | 0.385160617 |
| Phf14 | S | 774 | 0.140726868 | 0.239531525 | 0.127258665 | -0.013468203 | -0.11227286 | 0.256797121 | 0.043065438 | 0.21864542 | 0.909574733 | 0.304141474 |
| Ddx39b | S | 184 | 0.135489354 | 0.111125908 | 0.144305421 | 0.008816067 | 0.033179513 | 0.162383883 | 0.095761215 | 0.02290171 | 0.911949017 | 0.553809915 |
| Srrm1 | T | 913 | 0.065090984 | 0.072161022 | 0.090891142 | 0.025800157 | 0.01873012 | 0.020660657 | 0.014765688 | 0.00523368 | 0.022385905 | 0.046621886 |
| Mki67 | T | 1175 | 0.086314593 | 0.188548388 | 0.156596972 | 0.070282379 | -0.031951416 | 0.39516839 | 0.030970185 | 0.066049556 | 0.437527454 | 0.551289445 |
| Mprip | S | 485 | -0.008238538 | 0.143100448 | 0.163841754 | 0.172080292 | 0.020741306 | 0.872598251 | 0.003514449 | 0.008979826 | 0.035559116 | 0.590119891 |
| Cacnb2 | S | 214 | 0.116353075 | 0.194314243 | 0.116792862 | 0.000439787 | -0.07752138 | 0.082798213 | 0.015422887 | 0.449409659 | 0.99753952 | 0.600941277 |
| Ncapd2 | S | 1367 | -0.022733958 | 0.120816538 | 0.180444181 | 0.203178139 | 0.059627643 | 0.612619677 | 0.005731069 | 0.00197552 | 0.008746067 | 0.1084394 |
| Slc12a7 | T | 980 | 0.142252576 | 0.099357179 | 0.254795169 | 0.112542593 | 0.15543799 | 0.208777206 | 0.238594335 | 0.043603887 | 0.324634511 | 0.1427311 |
| Rai1 | S | 1091 | 0.082395017 | 0.057852053 | 0.109293233 | 0.026898216 | 0.05144118 | 0.161174432 | 0.005723209 | 0.002544687 | 0.573906306 | 0.052058496 |
| Rb1cc1 | S | 261 | 0.035577303 | 0.16033365 | 0.114599775 | 0.079022472 | -0.045733875 | 0.503772352 | 0.017227105 | 0.005002926 | 0.185584397 | 0.311790258 |
| Cenpc1 | S | 737 | 0.090879399 | 0.12427995 | 0.131912098 | 0.041032699 | 0.007632148 | 0.032629353 | 0.010800691 | 0.241488043 | 0.668963397 | 0.935347535 |
| Setd1a | S | 481 | 0.070013876 | 0.305506623 | 0.235789652 | 0.165775776 | -0.069716971 | 0.625846292 | 0.08031028 | 0.139190753 | 0.313457483 | 0.659930124 |

| | | | | | | | | | | | | |
|---|---|---|---|---|---|---|---|---|---|---|---|---|
| Gak | T | 847 | 0.127825312 | 0.155976182 | 0.104170797 | -0.023654515 | -0.051805385 | 0.067765004 | 0.02873361 | 0.199960037 | 0.712980763 | 0.424876873 |
| Hnrnpul2 | S | 653 | 0.110844889 | 0.140553296 | 0.203842718 | 0.092997829 | 0.063289422 | 0.146352226 | 0.094537304 | 0.088805964 | 0.330214042 | 0.494280056 |
| Dst | S | 4678 | 0.174647915 | 0.14427047 | 0.181348465 | 0.00670055 | 0.037077995 | 0.049453953 | 0.203680302 | 0.205260552 | 0.953040329 | 0.779660576 |
| Gzf1 | T | 196 | 0.252991358 | 0.029080713 | 0.144918786 | -0.108072571 | 0.115838073 | 0.010815903 | 0.746642225 | 0.100655455 | 0.094205408 | 0.172504571 |
| Sh3kbp1 | Y | 109 | 0.068413579 | 0.23311493 | 0.147547307 | 0.079133729 | -0.085567623 | 0.268364248 | 0.079160199 | 0.047748116 | 0.180671078 | 0.414936027 |
| Rprd1a | S | 156 | -0.000611453 | 0.144043009 | 0.219659879 | 0.220271332 | 0.07561687 | 0.985852701 | 0.073470237 | 0.003055501 | 0.004642642 | 0.230728679 |
| Plec | S | 4637 | -0.074874721 | 0.37215421 | 0.094763557 | 0.169638279 | -0.277390653 | 0.317223612 | 0.000962965 | 0.151205599 | 0.051901516 | 0.017499755 |
| Mast4 | S | 359 | -0.027105352 | 0.200620416 | 0.176896725 | 0.204002078 | -0.02372369 | 0.766403449 | 0.002745299 | 0.047340408 | 0.096071835 | 0.705902093 |
| Atn1 | S | 649 | 0.058826733 | 0.071694303 | 0.144189299 | 0.085362566 | 0.072494997 | 0.112010836 | 0.029040078 | 0.004978985 | 0.052635734 | 0.072326039 |
| Atn1 | T | 638 | 0.058826733 | 0.071694303 | 0.144189299 | 0.085362566 | 0.072494997 | 0.112010836 | 0.029040078 | 0.004978985 | 0.052635734 | 0.072326039 |
| Gapvd1 | S | 742 | 0.009884428 | 0.119579246 | 0.304592436 | 0.294708008 | 0.18501319 | 0.948411432 | 0.318143992 | 0.00269651 | 0.14447783 | 0.162014729 |
| Gapvd1 | S | 746 | 0.009884428 | 0.119579246 | 0.304592436 | 0.294708008 | 0.18501319 | 0.948411432 | 0.318143992 | 0.00269651 | 0.14447783 | 0.162014729 |
| Ncor2 | S | 1928 | 0.18840518 | 0.056369409 | 0.221434218 | 0.033029038 | 0.165064809 | 0.051466098 | 0.562404381 | 0.052644857 | 0.679668514 | 0.128605595 |
| Ifi203 Mndal | S | 110;110 | 0.102710252 | 0.105071774 | 0.159389698 | 0.056679446 | 0.054317924 | 0.253666862 | 0.019347298 | 0.040216813 | 0.517695121 | 0.34134118 |
| Slc20a2 | T | 346 | 0.175432776 | 0.189413234 | 0.056637353 | -0.118795423 | -0.132775881 | 0.206504724 | 0.007117676 | 0.358673399 | 0.352404901 | 0.066881118 |
| Edc4 | T | 879 | -0.014084723 | 0.113341247 | 0.233125899 | 0.247210622 | 0.119784652 | 0.702962206 | 0.135491839 | 0.000511445 | 0.00110363 | 0.125894011 |
| Thrap3 | T | 870 | 0.388776997 | 0.033223317 | -0.036436858 | -0.425213855 | -0.069660174 | 0.00011624 | 0.79845459 | 0.237430374 | 0.003027058 | 0.599226003 |
| Hirip3 | S | 152 | 0.003004372 | 0.101333349 | 0.110619754 | 0.107615382 | 0.009286405 | 0.93022519 | 0.009199885 | 6.80793E-05 | 0.064704436 | 0.643081524 |
| Sptbn1 | S | 2164 | -0.00288408 | 0.121636579 | 0.111145361 | 0.114029441 | -0.010491218 | 0.911498709 | 0.001164194 | 0.001718555 | 0.005597038 | 0.520677951 |
| Cbx5 | S | 14 | 0.00505884 | 0.112583842 | 0.118452131 | 0.113393291 | 0.005868288 | 0.809602744 | 0.003330029 | 0.00111988 | 0.002961657 | 0.771999648 |
| Hnrnpd | S | 83 | -2.44762E-05 | 0.06060046 | 0.302380723 | 0.302405199 | 0.241780263 | 0.99975495 | 0.282939407 | 0.000935392 | 0.026606313 | 0.00942431 |
| Scarf1 | S | 612 | -0.011585632 | 0.330983868 | 0.07279669 | 0.084382322 | -0.258187178 | 0.855984144 | 0.000701409 | 0.511992967 | 0.440331845 | 0.089473652 |
| Sorbs2 | S | 382 | 0.018067771 | 0.124770608 | 0.297491964 | 0.279424194 | 0.172721356 | 0.7168629 | 0.04228369 | 0.030060082 | 0.059198221 | 0.13688272 |
| Fam131b | S | 114 | 0.024524751 | 0.171184685 | 0.082374215 | 0.057849464 | -0.08881047 | 0.473619866 | 0.000593966 | 0.073637189 | 0.203016023 | 0.066337071 |
| Acin1 | S | 1283 | 0.219640998 | 0.081476816 | 0.169682548 | -0.04995845 | 0.088205732 | 0.251342543 | 0.145559018 | 0.028195101 | 0.758354757 | 0.18690404 |
| Med13l | S | 918 | 0.038243135 | 0.132675682 | 0.309042623 | 0.270799489 | 0.176366941 | 0.581336212 | 0.124766351 | 0.027212623 | 0.045459832 | 0.132800701 |
| Bud13 | T | 131 | 0.07071277 | 0.1454241 | 0.153423107 | 0.082710337 | 0.007999006 | 0.24188561 | 0.015942745 | 0.074098559 | 0.281961473 | 0.901326978 |
| Pard6a | S | 278 | 0.005547473 | 0.140157522 | 0.22638728 | 0.220839807 | 0.086229759 | 0.952540623 | 0.015222704 | 0.019829876 | 0.091846178 | 0.205312785 |
| Fam92a | S | 231 | 0.030269175 | 0.228049356 | 0.155607788 | 0.125338613 | -0.072441569 | 0.419923657 | 0.031588868 | 0.053807828 | 0.077521312 | 0.313333426 |
| Pds5b | S | 1182 | 0.029850982 | 0.117055435 | 0.098552886 | 0.068701904 | -0.018502549 | 0.28553547 | 8.34448E-05 | 0.094741493 | 0.18528067 | 0.654302622 |
| Tti1 | S | 282 | 0.026775127 | 0.137465685 | 0.165913989 | 0.139138861 | 0.028448303 | 0.489894093 | 0.044770483 | 0.005322358 | 0.023333168 | 0.574861956 |
| Tti1 | T | 283 | 0.026775127 | 0.137465685 | 0.165913989 | 0.139138861 | 0.028448303 | 0.489894093 | 0.044770483 | 0.005322358 | 0.023333168 | 0.574861956 |
| Spp1 | S | 62 | 0.192386276 | 0.187183071 | 0.096365379 | -0.096020898 | -0.090817692 | 0.081032706 | 0.048478808 | 0.454869182 | 0.457596992 | 0.455660923 |
| Zfp318 | S | 1988 | -0.03003951 | 0.108809895 | 0.304177751 | 0.334217261 | 0.195367856 | 0.337934691 | 0.023150177 | 0.005256822 | 0.013946326 | 0.027249429 |
| Rbck1 | S | 270 | 0.065814788 | 0.488907581 | 0.091804191 | 0.025989404 | -0.39710339 | 0.501329365 | 0.03658147 | 0.540124172 | 0.841345077 | 0.081057361 |
| Srsf4 | S | 289 | 0.225585668 | 0.03641036 | 0.195181074 | -0.030404593 | 0.158770714 | 0.021221687 | 0.719144554 | 0.09300425 | 0.752878208 | 0.220194876 |
| Mef2a | S | 253 | 0.013757118 | 0.087248839 | 0.146343398 | 0.13258628 | 0.059094559 | 0.577581201 | 0.01266941 | 0.000836123 | 0.005033956 | 0.02442632 |
| Vim | S | 325 | -0.029111555 | 0.100241156 | 0.188737442 | 0.217848997 | 0.088496286 | 0.512688513 | 0.000794203 | 0.004208824 | 0.009980356 | 0.059006489 |
| Plce1 | S | 680 | 0.020635735 | 0.20647015 | 0.208441542 | 0.187805806 | 0.001971391 | 0.75005149 | 0.025046265 | 0.057263795 | 0.079865662 | 0.980105125 |
| Plce1 | S | 681 | 0.020635735 | 0.20647015 | 0.208441542 | 0.187805806 | 0.001971391 | 0.75005149 | 0.025046265 | 0.057263795 | 0.079865662 | 0.980105125 |
| Plce1 | Y | 684 | 0.020635735 | 0.20647015 | 0.208441542 | 0.187805806 | 0.001971391 | 0.75005149 | 0.025046265 | 0.057263795 | 0.079865662 | 0.980105125 |
| Usp34 | S | 2438 | 0.119030431 | 0.249272227 | 0.126504149 | 0.007473718 | -0.122768077 | 0.371070488 | 0.104639749 | 0.044472245 | 0.950058158 | 0.32975762 |
| Phf2 | S | 734 | 0.061016289 | 0.284186934 | 0.2606895 | 0.199673211 | -0.023497434 | 0.653335106 | 0.125822845 | 0.090058498 | 0.247730089 | 0.885254347 |
| Prr12 | S | 1924 | 0.007431336 | 0.212829872 | 0.29788688 | 0.290455545 | 0.085057009 | 0.940211964 | 0.181193969 | 0.01770845 | 0.017423432 | 0.528935023 |
| Srrm2 | S | 761 | 0.130813061 | 0.104675091 | 0.098590827 | -0.032222234 | -0.006084264 | 0.017152599 | 0.061787179 | 0.12734548 | 0.547582679 | 0.912523471 |
| Trim44 | T | 346 | 0.050688343 | 0.179439474 | 0.160187081 | 0.109498738 | -0.019252393 | 0.362964557 | 0.008166333 | 0.136908563 | 0.265660057 | 0.820371425 |
| Map1b | T | 1329 | 0.113218647 | 0.149068604 | 0.178666411 | 0.065447764 | 0.029597807 | 0.261718581 | 0.11037427 | 0.040530189 | 0.485668816 | 0.708466406 |
| Tjp1 | S | 131 | -0.007113711 | 0.081974944 | 0.158667345 | 0.165781056 | 0.076692402 | 0.716069029 | 0.015907572 | 0.000142739 | 0.001514032 | 0.043443866 |
| Trim33 | S | 895 | 0.056602374 | 0.155478305 | 0.253508281 | 0.196905907 | 0.098029976 | 0.65185325 | 0.022993028 | 0.08151081 | 0.222480627 | 0.386328221 |
| Mon1a | S | 56 | 0.035891292 | 0.084402403 | 0.091932291 | 0.056041 | 0.007529889 | 0.109956259 | 0.012670631 | 0.000564989 | 0.033504501 | 0.712878901 |
| Usp6nl | S | 704 | -0.007719114 | 0.146200145 | 0.186932688 | 0.194651802 | 0.040732543 | 0.793640465 | 0.002348061 | 0.027635618 | 0.013147646 | 0.3688161 |
| Polrmt | S | 1163 | -0.250801529 | 0.249493369 | 0.325749267 | 0.576550797 | 0.076255898 | 0.219037796 | 0.04685943 | 0.001429246 | 0.048789006 | 0.415317048 |
| Polrmt | S | 1165 | -0.250801529 | 0.249493369 | 0.325749267 | 0.576550797 | 0.076255898 | 0.219037796 | 0.04685943 | 0.001429246 | 0.048789006 | 0.415317048 |
| Kat5 | S | 199 | -0.01885203 | 0.112421999 | 0.178640504 | 0.197492533 | 0.066218505 | 0.765817359 | 0.057738719 | 0.000234145 | 0.056290513 | 0.187153595 |
| Ppp1r12a | S | 422 | 0.014151769 | 0.146314607 | 0.118918161 | 0.104766392 | -0.027396446 | 0.67256313 | 0.001135969 | 0.029831078 | 0.052096162 | 0.468198852 |
| Dnttip2 | S | 37 | 0.097978486 | 0.114309849 | 0.23494613 | 0.136967643 | 0.120636281 | 0.220651908 | 0.199331956 | 0.031096888 | 0.155402072 | 0.217806664 |
| Dnttip2 | S | 39 | 0.097978486 | 0.114309849 | 0.23494613 | 0.136967643 | 0.120636281 | 0.220651908 | 0.199331956 | 0.031096888 | 0.155402072 | 0.217806664 |
| Dnttip2 | S | 42 | 0.097978486 | 0.114309849 | 0.23494613 | 0.136967643 | 0.120636281 | 0.220651908 | 0.199331956 | 0.031096888 | 0.155402072 | 0.217806664 |
| Mafk | S | 25 | -0.020323568 | 0.122049867 | 0.191627009 | 0.211950577 | 0.069577142 | 0.705189642 | 0.049521701 | 0.000576095 | 0.023478067 | 0.185591841 |
| Usp6nl | S | 452 | 0.041488716 | 0.063467858 | 0.239967649 | 0.198478933 | 0.176499791 | 0.289371296 | 0.064379391 | 0.008269414 | 0.018508674 | 0.034183693 |
| Anapc1 | T | 902 | -0.015252645 | 0.133895838 | 0.160737162 | 0.175989808 | 0.026841325 | 0.391721192 | 0.000181347 | 0.027203125 | 0.037395076 | 0.527756444 |
| Bod1l | S | 2001 | 0.16330378 | 0.116621376 | 0.098615051 | -0.06468873 | -0.018006326 | 0.31911097 | 0.065563097 | 0.005217429 | 0.655645447 | 0.671342531 |
| Pcm1 | S | 119 | 0.107610322 | 0.084300722 | 0.123192404 | 0.015582082 | 0.038891683 | 0.137058651 | 0.078882326 | 0.006279601 | 0.781894045 | 0.341635848 |
| Tfip11 | S | 211 | 0.071610961 | 0.0994806 | 0.09908879 | 0.027477829 | -0.000391809 | 0.09793537 | 0.000999441 | 0.099179287 | 0.585410365 | 0.992684381 |
| Taf1 | S | 1823 | 0.101979805 | 0.058716923 | 0.188005763 | 0.086025958 | 0.12928884 | 0.014409408 | 0.167032373 | 0.057359245 | 0.260405509 | 0.13270491 |
| Tcf20 | S | 588 | -0.080222186 | 0.177906608 | 0.267940585 | 0.348162771 | 0.090033977 | 0.172751097 | 0.003747857 | 0.011594074 | 0.006754416 | 0.211180471 |
| Mki67 | S | 1704 | 0.156129609 | 0.118415814 | 0.057948307 | -0.098181302 | -0.060467507 | 0.004633921 | 0.069802123 | 0.381447972 | 0.176612436 | 0.388854921 |
| Exosc5 | S | 23 | 0.092162029 | 0.039386203 | 0.139818981 | 0.047656952 | 0.100432778 | 0.014248827 | 0.060334967 | 0.016910673 | 0.211002709 | 0.041832104 |
| Lmna | T | 19 | 0.137863933 | 0.129749508 | 0.119148334 | -0.018715599 | -0.010601173 | 0.076679522 | 0.059784585 | 0.108821892 | 0.749410317 | 0.81887036 |
| Srrm2 | S | 1338 | 0.027824877 | 0.070853518 | 0.151169588 | 0.123344711 | 0.08031607 | 0.274958933 | 0.003682518 | 0.005066814 | 0.012626889 | 0.063921495 |
| Kat5 | S | 202 | 0.110825933 | 0.135335356 | 0.18202219 | 0.071196257 | 0.046686834 | 0.221557836 | 0.145561082 | 0.03181995 | 0.424054592 | 0.577381245 |
| Ppig | S | 515 | 0.085019324 | 0.066405008 | 0.149120635 | 0.064101311 | 0.082715627 | 0.019821417 | 0.077653005 | 0.030050501 | 0.226032244 | 0.14251919 |
| Gas2l1 | S | 493 | -0.05253273 | 0.155085053 | 0.250432219 | 0.302964949 | 0.095347166 | 0.119620976 | 0.002476943 | 0.009382519 | 0.014920489 | 0.141286091 |
| Nexn | S | 23 | 0.020137077 | 0.074214525 | 0.184164072 | 0.164026995 | 0.109949547 | 0.557677224 | 0.033360372 | 0.00139 | 0.005739378 | 0.027136303 |
| Scrib | S | 1548 | 0.004690652 | 0.110129543 | 0.130219804 | 0.125529152 | 0.020090261 | 0.899602426 | 0.002366377 | 0.002877182 | 0.030934415 | 0.443471178 |
| Esyt2 | S | 662 | 0.098770548 | 0.121833136 | 0.194076097 | 0.095305549 | 0.072242961 | 0.187040861 | 0.184109162 | 0.02413682 | 0.238475609 | 0.413552278 |
| Septin5 | T | 336 | 0.136169417 | 0.194508186 | 0.077851935 | -0.058317483 | -0.116656252 | 0.13555071 | 0.010353267 | 0.45826304 | 0.603293621 | 0.296231248 |
| Als2 | S | 1454 | 0.124837503 | 0.407824621 | 0.077083429 | -0.047754073 | -0.330741192 | 0.725411263 | 0.017818177 | 0.506811115 | 0.893884546 | 0.054524614 |
| Foxj2 | S | 172 | -0.00541105 | 0.120173281 | 0.133379469 | 0.138790519 | 0.013206188 | 0.821019262 | 0.000799511 | 0.006897811 | 0.010724435 | 0.626222792 |
| Wasl | S | 426 | -0.126295851 | 0.305184887 | 0.226535426 | 0.352831277 | -0.078649461 | 0.094042972 | 0.004753092 | 0.014882779 | 0.002859448 | 0.264374718 |
| Phf14 | S | 523 | -0.006939878 | 0.066085342 | 0.156322377 | 0.163262255 | 0.090237034 | 0.881399906 | 0.002052399 | 0.000451604 | 0.042735425 | 0.012809478 |
| Raly | S | 274 | 0.128660995 | 0.098189275 | 0.092300761 | -0.036360234 | -0.005888514 | 0.008659353 | 0.043458177 | 0.226536044 | 0.580854269 | 0.92821562 |
| Whamm | S | 779 | 0.065552467 | 0.111231883 | 0.10687855 | 0.041326083 | -0.004353333 | 0.126137725 | 0.025692764 | 0.01021122 | 0.304432775 | 0.901556719 |
| Grk6 | T | 485 | 0.281307534 | 0.182582522 | 0.052206545 | -0.22910099 | -0.130375978 | 0.031584089 | 0.140332666 | 0.75236768 | 0.244165371 | 0.479916068 |
| Ppp1r12a | T | 872 | 0.034158339 | 0.125790817 | 0.17468732 | 0.140528982 | 0.048896503 | 0.616604401 | 0.007480541 | 0.027349904 | 0.114625008 | 0.369126026 |
| Zfpm2 | S | 472 | -0.190555144 | 0.318114736 | 0.250921705 | 0.441476849 | -0.067193031 | 0.018342387 | 0.001591277 | 0.005283111 | 0.000781356 | 0.185606313 |
| Mpp7 | S | 409 | 0.079873864 | 0.044957606 | 0.246226242 | 0.166352378 | 0.201268636 | 0.35660404 | 0.671536745 | 0.001230237 | 0.109000755 | 0.138033282 |
| Foxk1 | S | 402 | -0.048125717 | 0.291030286 | 0.236120814 | 0.284246531 | -0.054909472 | 0.441795373 | 0.089249823 | 0.008220482 | 0.005880001 | 0.662038954 |
| Prkca | T | 494 | 0.107904855 | 0.229039824 | 0.085770147 | -0.022134708 | -0.143269677 | 0.221177099 | 0.006501998 | 0.547775468 | 0.875003256 | 0.339640661 |
| Ints10 | S | 381 | 0.008192102 | 0.123569602 | 0.167548857 | 0.159356756 | 0.043979255 | 0.892369 | 0.014582036 | 0.004514266 | 0.070508815 | 0.287678785 |
| Lats2 | T | 533 | -0.0890243 | 0.169781705 | 0.207861248 | 0.296885547 | 0.038079543 | 0.019076416 | 8.71673E-05 | 0.005583934 | 0.001480354 | 0.302219657 |
| 1110004F10Rik | S | 15 | 0.055773164 | 0.176545862 | 0.066667589 | 0.010894425 | -0.109878273 | 0.652146751 | 0.000539929 | 0.094387063 | 0.927966541 | 0.023087237 |
| Kmt2a | T | 1841 | 0.095861031 | 0.105645704 | 0.124553645 | 0.028692614 | 0.018907942 | 0.194126628 | 0.177638239 | 0.00200675 | 0.638774262 | 0.762845371 |
| Srsf10 | S | 256 | -0.004996441 | 0.161666766 | 0.279124952 | 0.284121393 | 0.117458186 | 0.898969208 | 0.041794254 | 0.025955079 | 0.040404428 | 0.212477486 |

| | | | | | | | | | | | | | |
|---|---|---|---|---|---|---|---|---|---|---|---|---|---|
| Zfp148 | S | | 150 | 0.122315283 | 0.134207593 | 0.088048123 | -0.034267161 | -0.046159471 | 0.056076132 | 0.071419855 | 0.050012672 | 0.497789523 | 0.43182941 |
| Sart1 | S | | 602 | 0.019019054 | 0.175587907 | 0.134587681 | 0.115568627 | -0.041000226 | 0.762810487 | 0.001359504 | 0.102421557 | 0.186301424 | 0.521531211 |
| Pask | S | | 1347 | 0.034279953 | 0.188504488 | 0.159825441 | 0.125545488 | -0.028679047 | 0.659315434 | 0.020866049 | 0.03613965 | 0.106763077 | 0.454099851 |
| Rab31 | S | | 35 | 0.044359901 | 0.133829045 | 0.175836152 | 0.131476251 | 0.042007107 | 0.524249889 | 0.003271478 | 0.085667356 | 0.188773474 | 0.569104889 |
| Arhgef17 | S | | 611 | 0.143115516 | 0.013564586 | 0.104601893 | -0.038513623 | 0.091037307 | 0.001768677 | 0.785666612 | 0.01038687 | 0.141634472 | 0.130280713 |
| Zmym4 | S | | 1548 | 0.051102539 | 0.079584295 | 0.121786615 | 0.070684076 | 0.04220232 | 0.246786092 | 0.0110691 | 0.003327896 | 0.139584476 | 0.124977583 |
| Lsm14a | T | | 194 | 0.01547428 | 0.157193586 | 0.203869022 | 0.188394742 | 0.046675436 | 0.835264504 | 0.062636333 | 0.008414145 | 0.060146154 | 0.49624918 |
| Pcm1 | S | | 1254 | 0.054778476 | 0.067127117 | 0.162775011 | 0.107996535 | 0.095647894 | 0.065073971 | 0.192740807 | 0.00273244 | 0.019350992 | 0.099969647 |
| Pcm1 | S | | 1257 | 0.054778476 | 0.067127117 | 0.162775011 | 0.107996535 | 0.095647894 | 0.065073971 | 0.192740807 | 0.00273244 | 0.019350992 | 0.099969647 |
| Uso1 | S | | 940 | -0.000880526 | 0.2231451 | 0.279971544 | 0.280852071 | 0.056826444 | 0.984687491 | 0.049985404 | 0.061705675 | 0.068473557 | 0.622431265 |
| Bud13 | T | | 134 | 0.066254134 | 0.147201042 | 0.162723301 | 0.096469167 | 0.015522259 | 0.433351144 | 0.024435384 | 0.040443465 | 0.281655018 | 0.780527869 |
| Mrgbp | S | | 195 | 0.259835985 | 0.158352437 | 0.101562105 | -0.15827388 | -0.056790332 | 0.076766666 | 0.172077008 | 0.296039041 | 0.221284759 | 0.570784135 |
| Phip | S | | 1281 | 0.08517715 | 0.096826353 | 0.182051383 | 0.096874233 | 0.08522503 | 0.409977706 | 0.083664695 | 0.008601315 | 0.343746591 | 0.006382973 |
| Phip | S | | 1283 | 0.08517715 | 0.096826353 | 0.182051383 | 0.096874233 | 0.08522503 | 0.409977706 | 0.083664695 | 0.008601315 | 0.343746591 | 0.006382973 |
| Mnx1 | S | | 277 | 0.062436366 | 0.112067173 | 0.156595546 | 0.09415918 | 0.044528373 | 0.129538775 | 0.025255268 | 0.045020228 | 0.170174335 | 0.43566465 |
| Mnx1 | T | | 276 | 0.062436366 | 0.112067173 | 0.156595546 | 0.09415918 | 0.044528373 | 0.129538775 | 0.025255268 | 0.045020228 | 0.170174335 | 0.43566465 |
| Mnx1 | T | | 281 | 0.062436366 | 0.112067173 | 0.156595546 | 0.09415918 | 0.044528373 | 0.129538775 | 0.025255268 | 0.045020228 | 0.170174335 | 0.43566465 |
| Mnx1 | T | | 283 | 0.062436366 | 0.112067173 | 0.156595546 | 0.09415918 | 0.044528373 | 0.129538775 | 0.025255268 | 0.045020228 | 0.170174335 | 0.43566465 |
| Phf6 | S | | 138 | 0.086362216 | 0.084898538 | 0.102916909 | 0.016554693 | 0.018018371 | 0.013368779 | 0.063202453 | 0.029798534 | 0.622217231 | 0.65300971 |
| Mastl | S | | 545 | 0.037542754 | 0.253868979 | 0.078223988 | 0.040681233 | -0.175644992 | 0.396029782 | 0.004334966 | 0.239888699 | 0.504751142 | 0.046334015 |
| Axl | S | | 845 | -0.016898059 | 0.166524677 | 0.193820189 | 0.210718248 | 0.027295512 | 0.851183498 | 0.020376648 | 0.010205431 | 0.080967382 | 0.432014251 |
| Spp1 | S | | 61 | 0.191808622 | 0.185591386 | 0.091899617 | -0.099909005 | -0.09369177 | 0.080982737 | 0.051303839 | 0.482651939 | 0.448223903 | 0.452554678 |
| Osbp | S | | 384 | 0.071261518 | 0.150040363 | 0.077761711 | 0.006500192 | -0.072278652 | 0.04609105 | 0.011577862 | 0.116227776 | 0.853279546 | 0.164620482 |
| Pak4 | S | | 181 | -6.62313E-05 | 0.172228079 | 0.12365009 | 0.123716321 | -0.048577989 | 0.998632122 | 0.001224601 | 0.044384165 | 0.051893323 | 0.301308668 |
| Mef2d | S | | 180 | 0.127890189 | 0.135478128 | 0.125533461 | -0.002356728 | -0.009944667 | 0.055842147 | 0.120313886 | 0.088392048 | 0.968286932 | 0.895529947 |
| Sub1 | S | | 17 | 0.206266468 | 0.104401453 | 0.17785356 | -0.028412908 | 0.073452107 | 0.150176772 | 0.297524404 | 0.057256969 | 0.803440915 | 0.388312855 |
| Tmpo | S | 82;82 | | 0.04555503 | 0.181401324 | 0.089784981 | 0.044229951 | -0.091616344 | 0.246965329 | 0.006157339 | 0.076395951 | 0.326023464 | 0.099965806 |
| Exosc5 | S | | 19 | 0.16275975 | 0.113755088 | 0.085064387 | -0.077695363 | -0.028690701 | 0.0271291 | 0.023329369 | 0.414616009 | 0.458743557 | 0.76367251 |
| Phf20 | S | | 418 | 0.126270491 | 0.071821707 | 0.17433828 | 0.048067789 | 0.102516573 | 0.053964479 | 0.135573388 | 0.056657852 | 0.508776977 | 0.186663084 |
| Prkar2b | S | | 112 | 0.013842302 | 0.096190008 | 0.118867329 | 0.105025027 | 0.022677321 | 0.370285114 | 0.001366181 | 0.006035588 | 0.018516477 | 0.3661099 |
| Ubxn1 | S | | 188 | -0.00945963 | 0.21670767 | 0.150872941 | 0.160332572 | -0.065834729 | 0.563468949 | 0.005006967 | 0.040700077 | 0.048062007 | 0.23476803 |
| Trp53bp1 | S | | 533 | 0.004692099 | 0.081576519 | 0.197246611 | 0.192554512 | 0.115670092 | 0.869678443 | 0.035647969 | 0.001296276 | 0.014283715 | 0.020115929 |
| Ubxn1 | S | | 187 | 0.027625854 | 0.168680954 | 0.11851619 | 0.090890336 | -0.050164764 | 0.404128272 | 0.004685057 | 0.056983017 | 0.112874943 | 0.326884996 |
| Dtnbp1 | T | | 336 | -0.034380101 | 0.235914563 | 0.1802489 | 0.214629001 | -0.055665663 | 0.511016202 | 0.017680835 | 0.015281061 | 0.017303031 | 0.391171058 |
| Afap1l2 | S | | 645 | 0.029721249 | 0.151169341 | 0.21515858 | 0.185437332 | 0.06398924 | 0.599936874 | 0.196603599 | 0.00475776 | 0.033883817 | 0.523490051 |
| Cdk12 | S | | 305 | 0.092592943 | 0.058584169 | 0.063760079 | -0.028832864 | 0.00517591 | 0.012854427 | 0.020922589 | 0.005043315 | 0.236942418 | 0.773662358 |
| Pid1 | S | | 204 | 0.069218475 | 0.175250961 | 0.097792097 | 0.028573623 | -0.077458863 | 0.245431328 | 0.010358575 | 0.090337843 | 0.481134304 | 0.040087156 |
| Akap13 | T | | 1906 | 0.104255276 | 0.367308754 | 0.129636516 | 0.02538124 | -0.237672238 | 0.238983381 | 0.050172008 | 0.601486077 | 0.913270003 | 0.380660377 |
| Pds5b | S | | 1381 | 0.029355392 | 0.129344473 | 0.097568019 | 0.068212627 | -0.031776454 | 0.597431508 | 0.002352587 | 0.008423145 | 0.265100411 | 0.243323674 |
| Mki67 | S | | 2390 | 0.122969335 | 0.230876404 | 0.057117391 | -0.065851944 | -0.173759013 | 0.076298578 | 0.023588504 | 0.504518515 | 0.443714225 | 0.111372376 |
| Pds5b | S | | 1281 | -0.013920319 | 0.123979893 | 0.146043991 | 0.159964309 | 0.022064098 | 0.541753221 | 0.010641404 | 0.000679994 | 0.001634379 | 0.476971352 |
| Ddx20 | S | | 465 | 0.093734266 | 0.086796473 | 0.101519205 | 0.007784939 | 0.014722732 | 0.063133181 | 0.01186144 | 0.046286081 | 0.862266341 | 0.672919882 |
| Arhgap21 | S | | 917 | 0.101534766 | 0.162494391 | 0.077554206 | -0.02398056 | -0.084940185 | 0.03989497 | 0.050656476 | 0.103941786 | 0.58693214 | 0.213082682 |
| Incenp | S | | 245 | 0.101524906 | 0.129795809 | 0.208930456 | 0.10740555 | 0.079134647 | 0.011078125 | 0.16901431 | 0.260772825 | 0.510867324 | 0.634474202 |
| Ssh3 | S | | 639 | -0.049300387 | 0.150068167 | 0.343450302 | 0.392750689 | 0.193382135 | 0.598102913 | 0.135693266 | 0.007452933 | 0.001039976 | 0.008742965 |
| G3bp1 | S | | 149 | 0.082992389 | 0.173620687 | 0.089007922 | 0.006015534 | -0.084612764 | 0.313177196 | 0.003018138 | 0.213102348 | 0.943891761 | 0.229111368 |
| Sap130 | S | | 884 | -0.01240227 | 0.139153963 | 0.249079119 | 0.261481389 | 0.109925156 | 0.876797869 | 0.001244638 | 0.097971368 | 0.083989786 | 0.340415826 |
| Cpd | T | | 1365 | 0.037090423 | 0.127491604 | 0.156427153 | 0.11933673 | 0.028935548 | 0.490780887 | 0.019127507 | 0.014718909 | 0.058883337 | 0.434208029 |
| Gramd1a | S | | 265 | 0.20809189 | 0.102543225 | 0.074631677 | -0.133460213 | -0.027911548 | 0.008748672 | 0.199430533 | 0.336913434 | 0.135091096 | 0.744112794 |
| Gramd1a | S | | 269 | 0.20809189 | 0.102543225 | 0.074631677 | -0.133460213 | -0.027911548 | 0.008748672 | 0.199430533 | 0.336913434 | 0.135091096 | 0.744112794 |
| Baz1b | S | | 359 | 0.072354366 | 0.125614728 | 0.091679396 | 0.019319629 | -0.033940732 | 0.111682806 | 0.023369783 | 0.019956028 | 0.61349618 | 0.397555873 |
| Prpf38a | S | | 277 | 0.180891936 | 0.108430677 | 0.127585131 | -0.053306805 | 0.019154454 | 0.084410733 | 0.231037205 | 0.050897323 | 0.518693045 | 0.81498814 |
| Parn | S | | 556 | 0.066211369 | 0.112207597 | 0.05946241 | -0.006748959 | -0.052745187 | 0.031382046 | 0.000553024 | 0.249100278 | 0.880738965 | 0.293814228 |
| Scrib | S | | 750 | 0.026108492 | 0.092611761 | 0.180633411 | 0.154524919 | 0.088021649 | 0.345892893 | 0.032302671 | 0.006552168 | 0.026321351 | 0.084933132 |
| Atp2a2 | S | | 663 | 0.026174843 | 0.145349948 | 0.170762957 | 0.144588114 | 0.025413009 | 0.470843381 | 0.022818488 | 0.023322461 | 0.061765027 | 0.65238632 |
| Rragc | S | | 94 | 0.092038226 | 0.259224908 | 0.068037964 | -0.024000261 | -0.191186944 | 0.19953802 | 0.020985121 | 0.275689382 | 0.720844639 | 0.05988456 |
| Pnisr | S | | 601 | 0.116616072 | 0.195202215 | 0.01315425 | -0.103461822 | -0.182047966 | 0.050404362 | 0.003421251 | 0.891117199 | 0.321291796 | 0.143586558 |
| Hjurp A730008H23Ril | S | | 602 | 0.394731281 | 0.035090217 | 0.043133895 | -0.351597385 | 0.008043678 | 0.004628829 | 0.784103981 | 0.668030612 | 0.003955809 | 0.939511506 |
| Dnttip2 | T | | 43 | 0.030058046 | 0.180721073 | 0.288181379 | 0.258123333 | 0.107460306 | 0.866819637 | 0.049302643 | 0.069105929 | 0.240451036 | 0.386035883 |
| Bap1 | S | | 520 | -0.052254234 | 0.169158682 | 0.276126068 | 0.328380302 | 0.106967386 | 0.407295347 | 0.005001679 | 0.038095985 | 0.024132884 | 0.268755053 |
| Rhbdf2 | S | | 246 | 0.417944603 | -0.114382606 | 0.25545011 | -0.162494493 | 0.369832716 | 0.006664611 | 0.323654841 | 0.253688695 | 0.419770371 | 0.1396475 |
| Wnk1 | S | | 181 | 0.016009624 | 0.165116305 | 0.152405524 | 0.1363959 | -0.012710782 | 0.80409858 | 0.009540089 | 0.026413327 | 0.088234956 | 0.790001528 |
| Khsrp | S | | 182 | 0.054118502 | 0.093964978 | 0.102846787 | 0.048728286 | 0.008881809 | 0.257352116 | 0.007408632 | 0.005681341 | 0.286374706 | 0.61032497 |
| Brpf3 | S | | 17 | -0.012516803 | 0.111572337 | 0.2149151 | 0.227431904 | 0.103342763 | 0.693751485 | 0.001903644 | 0.021091563 | 0.016473631 | 0.1333165 |
| Brpf3 | S | | 19 | -0.012516803 | 0.111572337 | 0.2149151 | 0.227431904 | 0.103342763 | 0.693751485 | 0.001903644 | 0.021091563 | 0.016473631 | 0.1333165 |
| Cep112 | S | | 240 | 0.025855715 | 0.342143719 | 0.163463873 | 0.137608158 | -0.178679846 | 0.833935761 | 0.010683499 | 0.445046907 | 0.512938104 | 0.397524202 |
| Cep112 | S | | 241 | 0.025855715 | 0.342143719 | 0.163463873 | 0.137608158 | -0.178679846 | 0.833935761 | 0.010683499 | 0.445046907 | 0.512938104 | 0.397524202 |
| Msh6 | S | | 14 | 0.094872469 | 0.124367884 | 0.086701657 | -0.008170813 | -0.037666228 | 0.037851658 | 0.012564599 | 0.184185133 | 0.872942318 | 0.48325145 |
| Jmy | S | | 969 | 0.023515389 | 0.11542741 | 0.128046968 | 0.104531579 | 0.012619559 | 0.522531859 | 0.008157838 | 0.006049344 | 0.036582097 | 0.428554562 |
| Clasp2 | S | | 531 | -0.024347582 | 0.210339506 | 0.201106819 | 0.225454401 | -0.009232686 | 0.689388104 | 0.062126524 | 0.006462126 | 0.023139747 | 0.906189521 |
| Ap2a1 | T | | 653 | 0.086530592 | 0.09996547 | 0.137026539 | 0.050495947 | 0.037061069 | 0.105591031 | 0.012504664 | 0.090828916 | 0.45389179 | 0.542169061 |
| Heatr5a | T | | 1527 | -0.005914557 | 0.144652371 | 0.169067085 | 0.174981642 | 0.024414714 | 0.769681602 | 0.068331648 | 0.000861952 | 0.010883045 | 0.657995447 |
| Antxr1 | S | | 360 | 0.037951915 | 0.07693214 | 0.153062514 | 0.115110599 | 0.076130374 | 0.565311108 | 0.016234307 | 0.002292793 | 0.157620331 | 0.044972386 |
| Cadm1 | S | | 448 | 0.017017402 | 0.113896851 | 0.126547355 | 0.109529953 | 0.012650505 | 0.751240127 | 0.004706855 | 0.004848366 | 0.123546062 | 0.656187556 |
| H1f2 H1f3 | S | 36;37 | | 0.155467193 | 0.165537988 | 0.008064774 | -0.147402419 | -0.157473214 | 0.017659299 | 0.012812204 | 0.896918019 | 0.05160525 | 0.043269469 |
| Fam117b | S | | 399 | -0.178474477 | 0.173773565 | 0.375918158 | 0.554392635 | 0.202144593 | 0.006165911 | 0.006247725 | 0.000525034 | 0.003613104 | 0.022944119 |
| Erbin | S | | 849 | -0.062894837 | 0.229099393 | 0.187313867 | 0.250208704 | -0.041785527 | 0.180522903 | 0.009289251 | 0.00701582 | 0.008703984 | 0.466009914 |
| Cask | S | | 582 | 0.038021738 | 0.091609444 | 0.188945486 | 0.150923749 | 0.097336042 | 0.392731442 | 0.049160872 | 0.007941677 | 0.024704364 | 0.093514856 |
| Zfat | S | | 153 | 0.020045899 | 0.336892105 | 0.189198292 | 0.169152393 | -0.147693813 | 0.88404841 | 0.028203736 | 0.235703666 | 0.227608419 | 0.247983042 |
| Tmem184b | S | | 403 | 0.039579126 | 0.100450874 | 0.108631147 | 0.069052021 | 0.008180273 | 0.253765491 | 0.02077862 | 0.002324823 | 0.077544725 | 0.769264831 |
| Dock8 | S | | 905 | 0.054748031 | -0.050900929 | 0.381930622 | 0.327182591 | 0.432831551 | 0.562576916 | 0.430665358 | 0.000527237 | 0.027900962 | 0.000313849 |
| Atrx | S | | 626 | 0.019837413 | 0.105736067 | 0.123125648 | 0.103288235 | 0.01738958 | 0.373724383 | 0.034007038 | 0.000884764 | 0.007735491 | 0.593647433 |
| Usp6nl | S | | 405 | 0.059714412 | 0.199447754 | 0.214099958 | 0.154385546 | 0.014652204 | 0.435754736 | 0.144246938 | 0.042192617 | 0.127697746 | 0.892770115 |
| Mtmr3 | T | | 611 | 0.02808693 | 0.124382907 | 0.142410029 | 0.1143231 | 0.018027122 | 0.60202438 | 0.006134481 | 0.018879535 | 0.102295796 | 0.656589273 |
| Setd2 | T | | 616 | 0.027228135 | 0.144930865 | 0.195245038 | 0.168016903 | 0.050314173 | 0.54768852 | 0.131703034 | 0.00618984 | 0.017973791 | 0.52587405 |
| Stxbp5 | S | | 724 | 0.029629566 | 0.198909528 | 0.164107733 | 0.134478167 | -0.034801794 | 0.516862341 | 0.015649058 | 0.098304709 | 0.165637196 | 0.671217288 |
| Zmym4 | S | | 121 | 0.010201136 | 0.098024421 | 0.123399425 | 0.113198288 | 0.025375004 | 0.69296538 | 0.002854117 | 0.003000467 | 0.008811068 | 0.292597293 |
| Exosc5 | S | | 20 | 0.133093554 | 0.125996096 | 0.066447284 | -0.06664627 | -0.059548812 | 0.017361739 | 0.023998006 | 0.403558791 | 0.388716374 | 0.435800695 |
| Sltm | S | | 289 | 0.006224338 | 0.069190168 | 0.106258186 | 0.100033848 | 0.037068018 | 0.802898329 | 0.001638886 | 0.000164471 | 0.031327437 | 0.006491543 |
| Retreg3 | T | | 307 | 0.09614693 | 0.078315225 | 0.195960306 | 0.099813376 | 0.117645081 | 0.257350969 | 0.073943169 | 0.024660954 | 0.256256287 | 0.075116593 |
| Dlg5 | T | | 1011 | 0.237161896 | 0.146131198 | 0.007740396 | -0.2294215 | -0.138390802 | 0.01175466 | 0.070267264 | 0.971282363 | 0.339145027 | 0.531896377 |
| Syne1 | T | | 8363 | 0.016610389 | 0.106412978 | 0.104193716 | 0.087583327 | -0.002219262 | 0.665274619 | 0.001882809 | 0.003642806 | 0.059128148 | 0.919685067 |
| Rbmxl1 | S | | 246 | 0.004495028 | 0.093553874 | 0.200279197 | 0.195784168 | 0.106725322 | 0.933628495 | 0.07475403 | 0.00129547 | 0.039449386 | 0.069588234 |
| Cux1 | S | | 1316 | 0.229104263 | 0.147195799 | 0.169026012 | -0.060078251 | 0.021830213 | 0.211631819 | 0.079993173 | 0.291808167 | 0.750907911 | 0.878679935 |

| | | | | | | | | | | | | |
|---|---|---|---|---|---|---|---|---|---|---|---|---|
| Anks1 | S | 642 | 0.093074379 | 0.193261078 | 0.055606015 | -0.037468363 | -0.137655063 | 0.138731122 | 0.0074803 | 0.294647223 | 0.371624099 | 0.003773414 |
| Slc12a7 | S | 62 | 0.058056584 | 0.057070142 | 0.319883823 | 0.261827238 | 0.262813681 | 0.589628928 | 0.488260945 | 0.005752604 | 0.060079038 | 0.003380477 |
| Atad5 | S | 969 | 0.045517607 | 0.166813338 | 0.119141488 | 0.07362388 | -0.04767185 | 0.282731624 | 0.02054349 | 0.040994942 | 0.173567675 | 0.368028945 |
| Klf6 | T | 147 | -0.072488459 | 0.358973114 | 0.337738352 | 0.410226811 | -0.021234762 | 0.8104891 | 0.048952764 | 0.194056709 | 0.262679153 | 0.923784043 |
| Psen1 | S | 57 | 0.137697151 | 0.148473115 | 0.029455014 | -0.108242137 | -0.119018101 | 0.020012135 | 0.014819959 | 0.530747562 | 0.049854453 | 0.038422381 |
| Osbp | S | 188 | -0.011565518 | 0.145346727 | 0.136292631 | 0.147858149 | -0.009054096 | 0.785040628 | 0.004691786 | 0.006315206 | 0.017078279 | 0.774025841 |
| Zfp687 | S | 1107 | 0.101883561 | 0.181503722 | 0.153372697 | 0.051489136 | -0.028131025 | 0.403081834 | 0.051729437 | 0.079542714 | 0.629626104 | 0.628357364 |
| Exoc7 | S | 250 | 0.094589289 | 0.078595063 | 0.070444211 | -0.024145079 | -0.008150852 | 0.000721746 | 0.126782926 | 0.072715123 | 0.421595716 | 0.853176486 |
| Purb | S | 310 | -0.018976529 | 0.159001178 | 0.145727247 | 0.164703777 | -0.01327393 | 0.685634015 | 0.006125598 | 0.007516196 | 0.014436505 | 0.703122521 |
| Lemd2 | S | 149 | -0.082028633 | 0.100543132 | 0.291458051 | 0.373486684 | 0.190914919 | 0.247717507 | 0.020468816 | 0.001557738 | 0.005456252 | 0.025347905 |
| Yap1 | S | 352 | 0.189350528 | 0.133619775 | 0.168833875 | -0.020516653 | 0.0352141 | 0.132568317 | 0.057771181 | 0.346206498 | 0.906151751 | 0.8264495 |
| Psen2 | T | 49 | 0.002229401 | 0.206131237 | 0.148540605 | 0.146311204 | -0.057590632 | 0.980492623 | 0.006438593 | 0.067818324 | 0.179012155 | 0.376071659 |
| Prpf38a | S | 279 | 0.201596731 | 0.041204254 | 0.130707701 | -0.07088903 | 0.089503446 | 0.007612495 | 0.671500668 | 0.108469679 | 0.325335215 | 0.403293541 |
| Prpf38a | S | 281 | 0.201596731 | 0.041204254 | 0.130707701 | -0.07088903 | 0.089503446 | 0.007612495 | 0.671500668 | 0.108469679 | 0.325335215 | 0.403293541 |
| Srp14 | S | 44 | 0.056248476 | 0.283518481 | -0.010792885 | -0.067041361 | -0.294311366 | 0.258036929 | 0.00083784 | 0.890936741 | 0.379207623 | 0.025034288 |
| Svil | S | 666 | -0.027905235 | 0.129654735 | 0.145507576 | 0.173412811 | 0.015852841 | 0.24273602 | 0.035460366 | 5.16023E-05 | 0.00192311 | 0.667409562 |
| Rbbp6 | S | 1274 | 0.073054692 | 0.150890663 | 0.101919676 | 0.028864984 | -0.048970987 | 0.178322505 | 0.017655154 | 0.068929758 | 0.29760154 | 0.085728031 |
| Srsf4 | S | 439 | 0.279944764 | 0.049889912 | 0.272351301 | -0.007593463 | 0.222461388 | 0.119088543 | 0.71078421 | 0.123560296 | 0.965421459 | 0.226077085 |
| Tmem245 | S | 16 | 0.226106319 | 0.179813039 | 0.022676128 | -0.203430191 | -0.157136911 | 0.081526512 | 0.019897147 | 0.831417293 | 0.154942162 | 0.19477442 |
| Paxbp1 | S | 297 | 0.101091731 | 0.076965038 | 0.122043457 | 0.020951726 | 0.045078419 | 0.210305869 | 0.008117338 | 0.03113848 | 0.758097148 | 0.221712802 |
| Gatad2b | S | 339 | -0.022144657 | 0.091343972 | 0.170183537 | 0.192328194 | 0.078839565 | 0.319455027 | 0.049742696 | 5.52355E-05 | 0.000311451 | 0.08750037 |
| Clip1 | S | 203 | 0.208073309 | 0.251194443 | -0.005585398 | -0.213658706 | -0.256779841 | 0.089708746 | 0.024697025 | 0.953672018 | 0.067260913 | 0.02479991 |
| Nav3 | S | 1511 | 0.181126334 | 0.162274054 | 0.061195753 | -0.119930581 | -0.101078301 | 0.183698586 | 0.005131699 | 0.600522767 | 0.423795715 | 0.415901069 |
| Cdk13 | S | 350 | 0.070959409 | 0.112746397 | 0.173901022 | 0.102941614 | 0.061154625 | 0.353243022 | 0.024978445 | 0.043619746 | 0.242885937 | 0.340173058 |
| Rhbdf1 | S | 61 | 0.075352616 | 0.055252756 | 0.064726136 | -0.01062648 | 0.00947338 | 0.001268803 | 0.137721163 | 0.002951875 | 0.165656542 | 0.739864554 |
| Pdlim1 | S | 130 | 0.215249295 | 0.050496023 | 0.047695997 | -0.167553298 | -0.002800026 | 0.00060071 | 0.589521976 | 0.341369083 | 0.02957563 | 0.975612359 |
| Tra2b | S | 37 | 0.201179341 | 0.129790041 | 0.116717251 | -0.08446209 | -0.01307279 | 0.0300008 | 0.098111779 | 0.549937908 | 0.653240087 | 0.942465738 |
| Zfp367 | S | 300 | -0.052459933 | 0.145456674 | 0.195752934 | 0.248212867 | 0.05029626 | 0.211122036 | 0.004703091 | 0.003386342 | 0.003363325 | 0.194548648 |
| Pbrm1 | S | 1405 | 0.123709046 | 0.10357754 | 0.126288361 | 0.002579315 | 0.022710821 | 0.175575736 | 0.003937442 | 0.229876085 | 0.980122089 | 0.792558687 |
| Pacrgl | S | 47 | 0.287758402 | 0.157391198 | -0.12400817 | -0.411766572 | -0.281399367 | 0.003339922 | 0.039688559 | 0.390575358 | 0.054452327 | 0.119849898 |
| Ptpra | S | 202 | 0.085012918 | 0.081472435 | 0.127106148 | 0.04209323 | 0.045633713 | 0.022848692 | 0.0508931 | 0.063223309 | 0.415889931 | 0.395953017 |
| Map4 | T | 658 | 0.017179595 | 0.165003989 | 0.119545875 | 0.102366279 | -0.045458114 | 0.492347835 | 0.004125197 | 0.054681701 | 0.072924273 | 0.328428257 |
| Bclaf1 | S | 286 | 0.164035971 | 0.129090973 | 0.082729642 | -0.081306329 | -0.046361331 | 0.01131903 | 0.070318277 | 0.591143091 | 0.590968223 | 0.757305209 |
| Ints10 | S | 382 | 0.024515352 | 0.101450573 | 0.156634206 | 0.132118854 | 0.055183633 | 0.464643692 | 0.009846032 | 0.012355867 | 0.025646023 | 0.190531283 |
| Arhgef17 | S | 305 | 0.08970564 | 0.156195999 | 0.149842134 | 0.060136494 | -0.006353865 | 0.430466491 | 0.009706253 | 0.158496501 | 0.629920774 | 0.939470582 |
| Xrcc1 | T | 452 | 0.025286763 | 0.084500486 | 0.146797162 | 0.121510399 | 0.062296677 | 0.49543797 | 0.022442706 | 0.002096621 | 0.028349337 | 0.009214185 |
| Gpatch8 | S | 1185 | 0.051595679 | 0.067197372 | 0.125497902 | 0.073902223 | 0.058300529 | 0.097721198 | 0.051953624 | 0.002835499 | 0.005734912 | 0.03705798 |
| Epb41l2 | S | 541 | -0.040164976 | 0.141413607 | 0.193387636 | 0.233552612 | 0.051974029 | 0.664535961 | 0.148479568 | 0.000277146 | 0.092644857 | 0.509068681 |
| Tanc2 | S | 1583 | 0.096653587 | 0.129346039 | 0.009323382 | -0.087330205 | -0.120022657 | 0.019557261 | 0.000439721 | 0.890563209 | 0.27041974 | 0.172365363 |
| Fkbp15 | S | 1091 | 0.036358789 | 0.132241006 | 0.103546834 | 0.067188045 | -0.028694172 | 0.420717809 | 0.005914479 | 0.017417892 | 0.137158455 | 0.132454927 |
| Plekhg3 | S | 1136 | 0.317557927 | 0.091678811 | -0.078972059 | -0.396529986 | -0.17065087 | 0.001452146 | 0.187972669 | 0.653319497 | 0.108790594 | 0.363737072 |
| Plekhg3 | S | 1141 | 0.317557927 | 0.091678811 | -0.078972059 | -0.396529986 | -0.17065087 | 0.001452146 | 0.187972669 | 0.653319497 | 0.108790594 | 0.363737072 |
| Zbtb1 | T | 356 | 0.022041048 | 0.106446118 | 0.077104507 | 0.055063459 | -0.029341611 | 0.408427771 | 0.004900898 | 0.0006104 | 0.112753846 | 0.197602246 |
| Srrm2 | S | 741 | 0.112851785 | 0.035855081 | 0.129072139 | 0.016220354 | 0.093217058 | 0.001433327 | 0.242653089 | 0.0855261 | 0.757188295 | 0.162996894 |
| U2af2 | S | 79 | 0.033961172 | 0.04791548 | 0.107671068 | 0.073709896 | 0.059755588 | 0.343831055 | 0.009621212 | 0.000119981 | 0.109576084 | 0.001418324 |
| Atf7ip | S | 112 | 0.053554724 | 0.150643842 | 0.114455699 | 0.060900975 | -0.036188143 | 0.277775638 | 0.015821841 | 0.04509818 | 0.040713871 | 0.154884686 |
| Klf3 | S | 110 | 0.033449827 | 0.268835237 | 0.120569461 | 0.087119633 | -0.148265776 | 0.625477053 | 0.008565678 | 0.282466686 | 0.419634264 | 0.211247341 |
| Tbc1d14 | S | 296 | 0.006575974 | 0.359565955 | -0.013109944 | -0.019685918 | -0.372675899 | 0.947768845 | 0.000491416 | 0.946138541 | 0.922493142 | 0.152186254 |
| Zzz3 | S | 96 | -0.067579879 | 0.065482472 | 0.312409287 | 0.379989166 | 0.246926815 | 0.266613461 | 0.102535759 | 0.000416598 | 0.00470648 | 0.002154938 |
| Bclaf1 | S | 703 | 0.235190478 | 0.139517852 | 0.099605863 | -0.135584615 | -0.03991199 | 0.075331297 | 0.063430485 | 0.543769287 | 0.441887339 | 0.793919751 |
| Cic | S | 1080 | 0.046568256 | 0.137302893 | 0.188660197 | 0.142091941 | 0.051357304 | 0.546386602 | 0.044965319 | 0.027046654 | 0.107858387 | 0.377024275 |
| Bcl9l | S | 747 | -0.029512909 | 0.10960224 | 0.222659101 | 0.252172011 | 0.113056862 | 0.468951469 | 0.046224107 | 0.001330767 | 0.00023417 | 0.04366633 |
| Bclaf1 | S | 715 | 0.151026922 | 0.076913817 | 0.321684065 | 0.170657143 | 0.244770248 | 0.339589699 | 0.530099917 | 0.039512422 | 0.310076673 | 0.116762054 |
| Scaf1 | T | 751 | 0.13438786 | 0.138324026 | 0.12679423 | -0.00759363 | -0.011529796 | 0.066200772 | 0.115084457 | 0.146843171 | 0.906137116 | 0.882869461 |
| Arhgap31 | S | 765 | 0.083345999 | 0.10292546 | 0.100384847 | 0.017038847 | -0.002540613 | 0.027597964 | 0.062631363 | 0.042516839 | 0.611601129 | 0.955552435 |
| Mtmr3 | S | 633 | 0.023300554 | 0.09876103 | 0.151049446 | 0.127748892 | 0.052288416 | 0.405170408 | 0.002623655 | 0.028803686 | 0.049224592 | 0.273869999 |
| Sec23ip | S | 748 | 0.075458407 | 0.330640806 | -0.001453012 | -0.076911419 | -0.332093818 | 0.323524493 | 0.00386391 | 0.987272353 | 0.346231015 | 0.014164865 |
| Mast4 | S | 355 | 0.012975563 | 0.08875462 | 0.148055056 | 0.135079493 | 0.059300436 | 0.574979562 | 0.003129873 | 0.008276736 | 0.0168129 | 0.138170935 |
| Ccar2 | S | 625 | 0.025764521 | 0.070993642 | 0.117230027 | 0.091465506 | 0.046236385 | 0.327661677 | 0.015455935 | 0.000635671 | 0.024144862 | 0.071889082 |
| Brd8 | T | 264 | 0.041056106 | 0.085302853 | 0.158533568 | 0.117477462 | 0.073230716 | 0.286667885 | 0.086211906 | 0.00307183 | 0.006204878 | 0.100104425 |
| Abraxas1 | S | 355 | 0.094911219 | 0.148414518 | 0.209821993 | 0.114910774 | 0.061407475 | 0.117205192 | 0.354868462 | 0.047996948 | 0.19752645 | 0.686784977 |
| Mbd2 | S | 410 | 0.05002875 | 0.063714141 | 0.136052965 | 0.086024214 | 0.072338823 | 0.141927692 | 0.096327089 | 0.001457264 | 0.024114706 | 0.062103656 |
| Myo18a | T | 2041 | 0.042800878 | 0.09168634 | 0.116679861 | 0.073878983 | 0.02499352 | 0.254661147 | 0.012475807 | 0.006638249 | 0.070029357 | 0.299314156 |
| Dido1 | S | 1035 | 0.012886099 | 0.040265428 | 0.130674944 | 0.117788845 | 0.090409516 | 0.422832692 | 0.233603622 | 2.7575E-06 | 0.007934419 | 0.063439454 |
| Ncor2 | S | 743 | 0.060095834 | 0.159859861 | 0.132382981 | 0.072287147 | -0.02747688 | 0.344993982 | 0.002170924 | 0.303441078 | 0.552338496 | 0.805335204 |
| Kdm5a | S | 1111 | -0.050743728 | 0.124377178 | 0.330996194 | 0.381739922 | 0.206619016 | 0.143882507 | 0.007957139 | 0.015095919 | 0.012000162 | 0.04886981 |
| Setd1b | S | 210 | 0.13300923 | 0.213015897 | 0.273464714 | 0.140455484 | 0.060448817 | 0.442698621 | 0.225643748 | 0.129665415 | 0.433189904 | 0.720115287 |
| Nphp4 | S | 145 | -0.129928466 | 0.271160918 | 0.177224493 | 0.307152959 | -0.093936425 | 0.015587236 | 0.005292476 | 0.000372847 | 0.003930632 | 0.127288036 |
| Trip12 | S | 239 | 0.04157298 | 0.054994686 | 0.130704391 | 0.089131411 | 0.075709705 | 0.051900378 | 0.014342342 | 0.007076625 | 0.037092886 | 0.059427728 |
| Ank2 | T | 3785 | -0.031550331 | 0.312862673 | 0.0948029 | 0.126353232 | -0.218059773 | 0.701888268 | 0.002841099 | 0.20188784 | 0.108507319 | 0.003450537 |
| Nrcam | S | 1206 | 0.024551504 | 0.219128735 | 0.215431185 | 0.190879681 | -0.00369755 | 0.808605175 | 0.021456486 | 0.142495059 | 0.212824523 | 0.97491789 |
| Eps15 | S | 107 | 0.06853938 | 0.164396127 | 0.252103234 | 0.183563854 | 0.087707107 | 0.598042084 | 0.119289401 | 0.053854845 | 0.232576144 | 0.437738344 |
| Git1 | T | 401 | 0.030680008 | 0.189990282 | 0.101443374 | 0.070763365 | -0.088546908 | 0.077919736 | 0.016162807 | 0.111256811 | 0.214054058 | 0.162051786 |
| Atrx | S | 1290 | -0.006589241 | 0.26450202 | 0.432297214 | 0.438886455 | 0.167795194 | 0.910520233 | 0.084127579 | 0.186418691 | 0.179856885 | 0.536991279 |
| Zgrf1 | S | 663 | 0.055238539 | 0.083991459 | 0.091049833 | 0.035811294 | 0.007058374 | 0.037537045 | 0.006523 | 0.031755852 | 0.261867078 | 0.801405968 |
| Ptpn14 | S | 314 | 0.037508685 | 0.079821403 | 0.12860045 | 0.091091765 | 0.048779047 | 0.11488933 | 0.007352663 | 0.016809341 | 0.034803821 | 0.151995923 |
| Sh3bp4 | S | 278 | 0.04259924 | 0.167380649 | 0.087884605 | 0.045285364 | -0.079496044 | 0.301222961 | 0.016507963 | 0.02173701 | 0.275313087 | 0.128941693 |
| Kdm5c | S | 896 | 0.125035852 | 0.123225813 | 0.174174555 | 0.049138703 | 0.050948742 | 0.105186042 | 0.13982134 | 0.116892472 | 0.597212113 | 0.598810447 |
| Kars | S | 594 | 0.081609554 | 0.218742084 | 0.04575924 | -0.035850314 | -0.172982844 | 0.220989423 | 0.002997391 | 0.588606433 | 0.66730777 | 0.105948816 |
| Tarbp2 | S | 155 | 0.262258996 | 0.050374338 | 0.155894945 | -0.106364051 | 0.105520607 | 0.018530966 | 0.587227406 | 0.276803538 | 0.399504539 | 0.408586385 |
| Pml | S | 17 | -0.051676363 | 0.156152818 | 0.255049256 | 0.306725619 | 0.098896438 | 0.237831897 | 0.007884413 | 0.014877254 | 0.021943939 | 0.187250473 |
| Papolg | S | 556 | 0.060643756 | 0.100902058 | 0.100674887 | 0.040031131 | -0.000227171 | 0.173205689 | 0.038152478 | 0.004842723 | 0.313992297 | 0.994452738 |
| Cep89 | S | 40 | 0.032388968 | 0.045271777 | 0.252684207 | 0.220295239 | 0.20741243 | 0.613623129 | 0.518237428 | 0.000873681 | 0.030271048 | 0.045372129 |
| Rbm15b | S | 263 | 0.046125474 | 0.062627861 | 0.120685214 | 0.07455974 | 0.058057353 | 0.018236292 | 0.047628114 | 0.007780415 | 0.029495293 | 0.07670763 |
| Dnajc1 | S | 477 | 0.027169981 | 0.112573217 | 0.128056035 | 0.100886054 | 0.015482818 | 0.628882235 | 0.01121195 | 0.006118784 | 0.135495665 | 0.318061379 |
| Tbc1d10a | S | 407 | -0.006223285 | 0.197250208 | 0.142221321 | 0.148444607 | -0.055028887 | 0.888923604 | 0.026295052 | 0.009811464 | 0.036410168 | 0.325411774 |
| Brsk1 | S | 311 | 0.159679759 | 0.275303252 | 0.383526001 | 0.223846242 | 0.108222749 | 0.564116831 | 0.366686323 | 0.186277435 | 0.495636824 | 0.743549359 |
| Atn1 | S | 617 | -0.013188917 | 0.103595079 | 0.161223481 | 0.174412398 | 0.057628402 | 0.6906391 | 0.001857418 | 0.007086506 | 0.008582106 | 0.133124285 |
| Tra2b | S | 97 | -0.020630263 | 0.07949376 | 0.223468739 | 0.244099002 | 0.143974979 | 0.620956654 | 0.106816275 | 0.000475645 | 0.006946811 | 0.031056748 |
| Sfswap | T | 866 | 0.150377552 | 0.052127435 | 0.106347553 | -0.044029998 | 0.054220118 | 0.00883531 | 0.430375227 | 0.043704144 | 0.298500375 | 0.403649877 |
| Tulp4 | S | 1409 | -0.01640509 | 0.134157906 | 0.192676795 | 0.209081885 | 0.058518889 | 0.557112999 | 0.039029646 | 0.002679908 | 0.003299839 | 0.250723504 |
| Tulp4 | S | 1410 | -0.01640509 | 0.134157906 | 0.192676795 | 0.209081885 | 0.058518889 | 0.557112999 | 0.039029646 | 0.002679908 | 0.003299839 | 0.250723504 |
| Tulp4 | S | 1412 | -0.01640509 | 0.134157906 | 0.192676795 | 0.209081885 | 0.058518889 | 0.557112999 | 0.039029646 | 0.002679908 | 0.003299839 | 0.250723504 |
| Aak1 | S | 668 | 0.238318236 | 0.296549991 | -0.092914946 | -0.331233182 | -0.389464937 | 0.120843669 | 0.002117268 | 0.135072845 | 0.064043808 | 0.00196566 |

| | | | | | | | | | | | | |
|---|---|---|---|---|---|---|---|---|---|---|---|---|
| Aak1 | S | 674 | 0.238318236 | 0.296549991 | -0.092914946 | -0.331233182 | -0.389464937 | 0.120843669 | 0.002117268 | 0.135072845 | 0.064043808 | 0.00196566 |
| Ppp1r37 | S | 583 | 0.382479028 | 0.090619336 | -0.077897039 | -0.460376067 | -0.168516375 | 0.000990748 | 0.425095005 | 0.376685106 | 0.005911297 | 0.174178204 |
| Kif4 | S | 887 | 0.136942118 | 0.105227467 | 0.127350792 | -0.009591326 | 0.022123325 | 0.061209261 | 0.08682045 | 0.13654515 | 0.896380588 | 0.75884391 |
| Kif4 | S | 888 | 0.136942118 | 0.105227467 | 0.127350792 | -0.009591326 | 0.022123325 | 0.061209261 | 0.08682045 | 0.13654515 | 0.896380588 | 0.75884391 |
| Madd | S | 1237 | -0.11256875 | 0.12068452 | 0.354232665 | 0.466801414 | 0.233548144 | 0.038362284 | 0.048553079 | 0.000552677 | 0.001014722 | 0.00890236 |
| Rbfox2 | T | 65 | 0.065987345 | 0.054047776 | 0.154834345 | 0.088847 | 0.10078657 | 0.069111092 | 0.374814499 | 0.002080768 | 0.039312241 | 0.153295404 |
| Mphosph8 | S | 136 | 0.177519791 | 0.119718027 | 0.166416061 | -0.01110373 | 0.046698035 | 0.087786345 | 0.280978712 | 0.131636366 | 0.913139302 | 0.691689727 |
| Mphosph8 | S | 138 | 0.177519791 | 0.119718027 | 0.166416061 | -0.01110373 | 0.046698035 | 0.087786345 | 0.280978712 | 0.131636366 | 0.913139302 | 0.691689727 |
| Ercc5 | S | 355 | 0.143394232 | 0.061599288 | 0.17712913 | 0.033734898 | 0.115529842 | 0.046068211 | 0.307707762 | 0.066517354 | 0.63718387 | 0.169250956 |
| Ercc5 | S | 356 | 0.143394232 | 0.061599288 | 0.17712913 | 0.033734898 | 0.115529842 | 0.046068211 | 0.307707762 | 0.066517354 | 0.63718387 | 0.169250956 |
| Cul4b | T | 140 | 0.216775026 | 0.166581683 | 0.163440564 | -0.053334462 | -0.003141118 | 0.092098074 | 0.304210048 | 0.270671083 | 0.636763715 | 0.983263261 |
| Pdlim2 | S | 134 | -0.070729607 | 0.16530695 | 0.23459266 | 0.305322267 | 0.06928571 | 0.063532628 | 0.004739015 | 0.004445379 | 0.001463811 | 0.14017393 |
| Mief1 | S | 55 | -0.0541388 | 0.095073774 | 0.188423878 | 0.242562679 | 0.093350104 | 0.461922805 | 0.135022221 | 3.30129E-05 | 0.053426767 | 0.15066247 |
| Glyr1 | S | 114 | 0.127755087 | 0.089118887 | 0.187361993 | 0.059606906 | 0.098243106 | 0.181642595 | 0.219606873 | 0.035438257 | 0.509616879 | 0.222276516 |
| Git1 | S | 601 | 0.049128058 | 0.260125452 | 0.04478987 | -0.004338188 | -0.215335582 | 0.554044952 | 0.002016202 | 0.469104412 | 0.955694772 | 0.012257114 |
| Ttc28 | S | 2073 | -0.076654864 | 0.125482367 | 0.260475519 | 0.337130383 | 0.134993152 | 0.209869352 | 0.018131443 | 0.002223628 | 0.003246035 | 0.0599457 |
| Dact3 | S | 129 | 0.041692767 | 0.002733025 | 0.333900504 | 0.292207737 | 0.331167479 | 0.679731206 | 0.968805533 | 0.001317447 | 0.047302412 | 0.002460731 |
| Myl6 | S | 57 | 0.082763654 | 0.122177886 | 0.131153879 | 0.048390225 | 0.008975993 | 0.151848714 | 0.021475459 | 0.104575674 | 0.492677107 | 0.88580942 |
| Vim | T | 436 | 0.095919576 | 0.073876536 | 0.163331633 | 0.067412057 | 0.089455097 | 0.105880244 | 0.167058642 | 0.020654065 | 0.202328241 | 0.097941472 |
| Fbxo10 | S | 338 | -0.077054725 | 0.222227248 | 0.208721127 | 0.285775852 | -0.013506121 | 0.413005581 | 0.00725624 | 0.034511565 | 0.038674636 | 0.850354125 |
| Ccser2 | T | 429 | 0.044786756 | 0.116313069 | 0.208405709 | 0.163618953 | 0.09209264 | 0.183717538 | 0.077655969 | 0.045824821 | 0.063097827 | 0.218851837 |
| Nsfl1c | S | 176 | 0.108998107 | 0.260648485 | -0.050066003 | -0.15906411 | -0.310714488 | 0.06875689 | 0.001602632 | 0.403298615 | 0.024892935 | 0.002698401 |
| Septin9 | S | 228 | 0.107517579 | 0.156514627 | 0.057530712 | -0.049986867 | -0.098983915 | 0.088341718 | 0.031232477 | 0.087085515 | 0.332308231 | 0.119965142 |
| Rbmx2 | S | 225 | 0.114987208 | 0.098278984 | 0.189132222 | 0.074145014 | 0.090853238 | 0.077487701 | 0.241774092 | 0.071663676 | 0.382307537 | 0.355190072 |
| Anapc2 | S | 485 | -0.071370247 | 0.142605569 | 0.157177283 | 0.228547531 | 0.014571714 | 0.090149956 | 0.002237821 | 0.000433256 | 0.004146569 | 0.575955635 |
| Atxn1 | S | 87 | 0.057191797 | 0.179185598 | 0.157829137 | 0.100637339 | -0.021356462 | 0.373925005 | 0.054983408 | 0.061707251 | 0.158767797 | 0.778193251 |
| Rmdn3 | S | 46 | 0.053772965 | 0.166085697 | 0.034337757 | -0.019435208 | -0.13174794 | 0.093610036 | 0.001075863 | 0.317470104 | 0.412450645 | 0.013374497 |
| Samhd1 | S | 49 | 0.022533477 | 0.105144602 | 0.096192673 | 0.073659196 | -0.008951929 | 0.314103791 | 0.001498543 | 0.015658877 | 0.048161409 | 0.721006034 |
| Macroh2a1 | T | 178 | 0.116436507 | 0.136077064 | 0.134216706 | 0.017780199 | -0.001860357 | 0.230643633 | 0.138689837 | 0.029192668 | 0.827059365 | 0.979429145 |
| Atn1 | S | 107 | 0.036906121 | 0.092036275 | 0.126659563 | 0.089753441 | 0.034623287 | 0.292063296 | 0.016583576 | 0.007094379 | 0.031524386 | 0.223931251 |
| Ccny | S | 100 | 0.058697383 | 0.06184373 | 0.213623993 | 0.154926611 | 0.151780264 | 0.055200522 | 0.091402727 | 0.04241722 | 0.087659329 | 0.085801222 |
| Ppp1r10 | S | 542 | 0.085376364 | 0.182288794 | 0.214116282 | 0.128739918 | 0.031827488 | 0.325338995 | 0.034604105 | 0.275993762 | 0.478348606 | 0.850236568 |
| Clasp2 | S | 469 | 0.084979019 | 0.2785896 | 0.177374725 | 0.092395706 | -0.101214875 | 0.347765778 | 0.06927702 | 0.310207041 | 0.563520319 | 0.565203424 |
| Pias1 | S | 466 | -0.038342573 | 0.14508653 | 0.230627224 | 0.268969797 | 0.085540694 | 0.289161463 | 0.015670808 | 0.007827804 | 0.003295368 | 0.114069136 |
| Pias1 | S | 467 | -0.038342573 | 0.14508653 | 0.230627224 | 0.268969797 | 0.085540694 | 0.289161463 | 0.015670808 | 0.007827804 | 0.003295368 | 0.114069136 |
| Pias1 | S | 468 | -0.038342573 | 0.14508653 | 0.230627224 | 0.268969797 | 0.085540694 | 0.289161463 | 0.015670808 | 0.007827804 | 0.003295368 | 0.114069136 |
| Specc1l | S | 836 | 0.094283041 | 0.145381656 | 0.0898869 | -0.00439614 | -0.055494755 | 0.053819634 | 0.011199767 | 0.416286024 | 0.964466815 | 0.591404085 |
| Nexn | S | 296 | -0.087957139 | 0.101520729 | 0.221881312 | 0.309838451 | 0.120360583 | 0.163508065 | 0.057694026 | 5.59879E-05 | 0.0125208 | 0.046319072 |
| Ubr4 | S | 1633 | -0.076350571 | 0.187321397 | 0.178234454 | 0.254585025 | -0.009086943 | 0.004727221 | 0.01028594 | 5.20905E-05 | 0.000216103 | 0.802170681 |
| Mdc1 | S | 939 | -0.03522056 | 0.301438515 | 0.050529682 | 0.085750241 | -0.250908834 | 0.776486724 | 0.000621533 | 0.401310687 | 0.502474518 | 0.006218129 |
| Dus3l | S | 263 | 0.168136423 | 0.169257604 | 0.164116603 | -0.00401982 | -0.005141002 | 0.132586122 | 0.119634816 | 0.324386524 | 0.9797542 | 0.973889736 |
| Dus3l | T | 260 | 0.168136423 | 0.169257604 | 0.164116603 | -0.00401982 | -0.005141002 | 0.132586122 | 0.119634816 | 0.324386524 | 0.9797542 | 0.973889736 |
| Parg | S | 318 | 0.085530974 | 0.162737749 | 0.185029068 | 0.099498094 | 0.022291319 | 0.33950389 | 0.102158062 | 0.07140974 | 0.335750616 | 0.81464072 |
| Ubn1 | S | 173 | 0.043043523 | 0.165073156 | 0.128572412 | 0.08552889 | -0.036500744 | 0.469934871 | 0.007118376 | 0.114611744 | 0.274727996 | 0.577185006 |
| Ubn1 | S | 175 | 0.043043523 | 0.165073156 | 0.128572412 | 0.08552889 | -0.036500744 | 0.469934871 | 0.007118376 | 0.114611744 | 0.274727996 | 0.577185006 |
| Pxn | Y | 118 | -0.117095224 | 0.197650144 | 0.211039005 | 0.328134229 | 0.013388861 | 0.017716096 | 0.013873788 | 0.000121736 | 0.001888858 | 0.762464828 |
| C77080 | S | 502 | 0.082840894 | 0.135979222 | 0.104436325 | 0.021595431 | -0.031542898 | 0.12424097 | 0.026983432 | 0.095951637 | 0.657484539 | 0.532773383 |
| Rbms2 | S | 41 | 0.269522665 | 0.133540863 | 0.005795952 | -0.263726713 | -0.127744911 | 0.087788483 | 0.01070948 | 0.912273077 | 0.087570033 | 0.063125272 |
| Wac | S | 511 | 0.031330384 | 0.151200027 | 0.088777654 | 0.05744727 | -0.062422373 | 0.398577473 | 0.025132136 | 0.004365463 | 0.163034202 | 0.193966185 |
| Macf1 | S | 7258 | 0.05239511 | 0.076348961 | 0.300025445 | 0.247630335 | 0.223676484 | 0.418961792 | 0.256712205 | 0.02139571 | 0.048825313 | 0.061640863 |
| Rock1 | S | 1108 | 0.071285957 | 0.126444385 | 0.127645356 | 0.056359399 | 0.001200971 | 0.233203403 | 0.062696256 | 0.022572488 | 0.263865011 | 0.979222458 |
| Daxx | S | 736 | 0.051976512 | 0.165600667 | 0.147762676 | 0.095786164 | -0.017837991 | 0.050942552 | 0.16901479 | 0.046452894 | 0.120403023 | 0.853953973 |
| Daxx | S | 738 | 0.051976512 | 0.165600667 | 0.147762676 | 0.095786164 | -0.017837991 | 0.050942552 | 0.16901479 | 0.046452894 | 0.120403023 | 0.853953973 |
| Ap1m1 | T | 152 | 0.162679512 | 0.127814864 | 0.044505689 | -0.118173823 | -0.083309175 | 0.011132707 | 0.077746332 | 0.472710115 | 0.094859398 | 0.246812649 |
| Atad1 | S | 322 | 0.06297168 | 0.122102579 | 0.118360612 | 0.055388932 | -0.003741967 | 0.460945513 | 0.017147037 | 0.019408936 | 0.493521312 | 0.837985202 |
| Llgl1 | S | 964 | 0.002047038 | 0.14967352 | 0.173293417 | 0.17124638 | 0.023619898 | 0.978760982 | 0.0653278 | 0.004751259 | 0.099406886 | 0.698451904 |
| Mki67 | T | 2106 | 0.051381499 | 0.163052358 | 0.050878004 | -0.000503495 | -0.112174354 | 0.505198336 | 0.000338395 | 0.215635091 | 0.994685126 | 0.039977227 |
| Arvcf | S | 345 | -0.004749784 | 0.214643923 | 0.243018631 | 0.247768416 | 0.028374708 | 0.95270116 | 0.035624455 | 0.08354437 | 0.095917462 | 0.77637253 |
| Carmil1 | T | 1254 | 0.100701791 | 0.13030593 | 0.036099393 | -0.064602399 | -0.094206537 | 0.132548012 | 0.001006413 | 0.305362884 | 0.285364006 | 0.039657036 |
| Add3 | S | 683 | -0.071061267 | 0.357978444 | 0.185792881 | 0.256854148 | -0.172185563 | 0.545801847 | 0.012226396 | 0.187454746 | 0.129471467 | 0.23230442 |
| Nr2c1 | S | 338 | -0.086160953 | 0.154462224 | 0.226621004 | 0.312781957 | 0.07215878 | 0.247544267 | 0.012202148 | 0.003691629 | 0.010197978 | 0.149104512 |
| Hnrnpa2b1 | S | 344 | 0.07463143 | 0.071016704 | 0.133991365 | 0.059359935 | 0.062974661 | 0.3503022 | 0.050048077 | 0.003815145 | 0.438894499 | 0.065369482 |
| Nexn | T | 501 | 0.056691942 | 0.10746763 | 0.173471455 | 0.116779513 | 0.066003825 | 0.222576418 | 0.23597733 | 0.007691019 | 0.027162493 | 0.422015748 |
| Srsf4 | S | 291 | 0.173066893 | 0.054301822 | 0.142024847 | -0.031042046 | 0.087723025 | 0.006200042 | 0.542113298 | 0.19360166 | 0.731389677 | 0.447338836 |
| Stx5a | S | 178 | 0.062341371 | 0.074238907 | 0.081321777 | 0.018980406 | 0.007082869 | 0.047316028 | 0.030006034 | 0.004999043 | 0.391614843 | 0.743778571 |
| Stx5a | S | 181 | 0.062341371 | 0.074238907 | 0.081321777 | 0.018980406 | 0.007082869 | 0.047316028 | 0.030006034 | 0.004999043 | 0.391614843 | 0.743778571 |
| Rb1 | T | 246 | 0.075381038 | 0.2104195 | 0.083902201 | 0.008521163 | -0.126517299 | 0.23995561 | 0.008883428 | 0.398892234 | 0.920268785 | 0.21963106 |
| Sorbs3 | S | 669 | 0.22087243 | 0.208239916 | -0.000224232 | -0.221096662 | -0.208464148 | 0.22014606 | 0.006440176 | 0.998762238 | 0.282489534 | 0.229697091 |
| Fzr1 | S | 42 | 0.298427142 | 0.17080509 | 0.048732134 | -0.249695007 | -0.122072955 | 0.057272357 | 0.165664993 | 0.728945916 | 0.165163105 | 0.434654291 |
| Fzr1 | T | 30 | 0.298427142 | 0.17080509 | 0.048732134 | -0.249695007 | -0.122072955 | 0.057272357 | 0.165664993 | 0.728945916 | 0.165163105 | 0.434654291 |
| Fzr1 | T | 32 | 0.298427142 | 0.17080509 | 0.048732134 | -0.249695007 | -0.122072955 | 0.057272357 | 0.165664993 | 0.728945916 | 0.165163105 | 0.434654291 |
| Ttc4 | S | 244 | 0.23903223 | -0.021457651 | 0.431396196 | 0.192363966 | 0.452853848 | 0.093246968 | 0.843413469 | 0.147636477 | 0.443758436 | 0.147923969 |
| Srsf4 | T | 442 | 0.438418107 | 0.035263933 | 0.253009692 | -0.185408415 | 0.217745759 | 0.29603951 | 0.602359394 | 0.117032914 | 0.621026523 | 0.149258669 |
| Fam117b | S | 444 | 0.076823863 | 0.206027849 | 0.057513239 | -0.019310624 | -0.14851461 | 0.159235727 | 0.005446787 | 0.538007094 | 0.813000594 | 0.164990321 |
| Sash1 | S | 892 | -0.146344774 | 0.364976374 | 0.324297063 | 0.470641836 | -0.040679311 | 0.004351413 | 0.011588431 | 0.010375437 | 0.007820966 | 0.641793248 |
| Hirip3 | S | 205 | -0.001345427 | 0.127994708 | 0.107625223 | 0.10897065 | -0.020369485 | 0.920081072 | 0.000211848 | 0.048521532 | 0.044442596 | 0.541545641 |
| Hirip3 | S | 207 | -0.001345427 | 0.127994708 | 0.107625223 | 0.10897065 | -0.020369485 | 0.920081072 | 0.000211848 | 0.048521532 | 0.044442596 | 0.541545641 |
| Hirip3 | S | 208 | -0.001345427 | 0.127994708 | 0.107625223 | 0.10897065 | -0.020369485 | 0.920081072 | 0.000211848 | 0.048521532 | 0.044442596 | 0.541545641 |
| H1f3 H1f4 H1f5 | S | 113;113;114 | 0.270255564 | 0.12410547 | 0.055366302 | -0.214889261 | -0.068739167 | 0.190622713 | 0.032110349 | 0.248509241 | 0.26861342 | 0.085127979 |
| Dact3 | S | 370 | -0.03859241 | 0.157703374 | 0.242689338 | 0.281281748 | 0.084985964 | 0.196250821 | 0.004341637 | 0.029516926 | 0.030748215 | 0.25540674 |
| Dact3 | S | 374 | -0.03859241 | 0.157703374 | 0.242689338 | 0.281281748 | 0.084985964 | 0.196250821 | 0.004341637 | 0.029516926 | 0.030748215 | 0.25540674 |
| Cdk13 | T | 588 | 0.014204526 | 0.080336751 | 0.204402818 | 0.190198292 | 0.124066066 | 0.641442715 | 0.013146986 | 0.014458387 | 0.022007393 | 0.079227969 |
| Sh3bp5l | S | 361 | 0.076735906 | 0.185564206 | 0.118937262 | 0.042201356 | -0.066626944 | 0.265312618 | 0.02089886 | 0.20589563 | 0.639865994 | 0.441144408 |
| Srrm2 | S | 739 | 0.113718631 | 0.031090151 | 0.121825737 | 0.008107105 | 0.090735586 | 0.002060289 | 0.266431999 | 0.077658461 | 0.866455039 | 0.14884083 |
| Specc1l | S | 974 | 0.085596103 | 0.124556436 | 0.203777242 | 0.11818114 | 0.079220806 | 0.512351275 | 0.037291207 | 0.078590199 | 0.413876817 | 0.373985776 |
| Ino80 | S | 51 | 0.087934932 | 0.026599335 | 0.14784927 | 0.059914338 | 0.121249935 | 0.20728766 | 0.707369024 | 0.000174569 | 0.346513782 | 0.178451061 |
| Tgfbrap1 | S | 853 | 0.168420694 | 0.252947018 | -0.014603539 | -0.183024234 | -0.267550557 | 0.058062209 | 0.025431699 | 0.796896157 | 0.055691827 | 0.019555266 |
| Wdcp | S | 460 | 0.168906063 | 0.130182277 | 0.073205765 | -0.095700298 | -0.056976512 | 0.013587302 | 0.209150872 | 0.304647123 | 0.163918465 | 0.54805002 |
| Nsd2 | S | 421 | 0.366671808 | 0.343693151 | -0.176649837 | -0.543321644 | -0.520342988 | 0.039505819 | 0.013044295 | 0.056965445 | 0.009807284 | 0.00293025 |
| Epb41l1 | S | 461 | -0.110520476 | 0.194117105 | 0.350441823 | 0.460962299 | 0.156324718 | 0.333336656 | 0.004448145 | 0.088690746 | 0.037199143 | 0.307884734 |
| Ppfibp1 | S | 538 | 0.054970203 | 0.105930407 | 0.069048493 | 0.01407829 | -0.036881914 | 0.111349296 | 0.000403184 | 0.187969663 | 0.761275066 | 0.418945034 |
| Prrc2c | S | 1236 | 0.1517422 | 0.254212061 | 0.031619675 | -0.120122526 | -0.222592386 | 0.120375129 | 0.031740899 | 0.758875915 | 0.228364859 | 0.070581684 |
| Prrc2c | T | 1238 | 0.1517422 | 0.254212061 | 0.031619675 | -0.120122526 | -0.222592386 | 0.120375129 | 0.031740899 | 0.758875915 | 0.228364859 | 0.070581684 |
| Prrc2c | T | 1240 | 0.1517422 | 0.254212061 | 0.031619675 | -0.120122526 | -0.222592386 | 0.120375129 | 0.031740899 | 0.758875915 | 0.228364859 | 0.070581684 |

| | | | | | | | | | | | | |
|---|---|---|---|---|---|---|---|---|---|---|---|---|
| Cic | S | 1809 | 0.043300687 | 0.082971491 | 0.179949699 | 0.136649012 | 0.096978208 | 0.493295947 | 0.129484868 | 0.003496831 | 0.099390408 | 0.098266004 |
| Pitpnc1 | S | 274 | 0.085389113 | 0.393635987 | 0.091085993 | 0.00569688 | -0.302549994 | 0.658454401 | 0.030429101 | 0.547921446 | 0.969993082 | 0.013923971 |
| Gtf3c1 | S | 1890 | 0.017341284 | 0.095604451 | 0.170590825 | 0.153249542 | 0.074986374 | 0.661269266 | 0.034363224 | 0.005440725 | 0.012929834 | 0.094626059 |
| Tmpo | S | 660 | -0.046290219 | 0.172686338 | 0.219023241 | 0.26531346 | 0.046336903 | 0.298452399 | 0.008366121 | 0.020488601 | 0.007675914 | 0.398724212 |
| D830031N03Rik | S | 25 | 0.031052484 | 0.102642111 | 0.191501029 | 0.160448545 | 0.088858917 | 0.535347761 | 0.133774725 | 0.005349003 | 0.021625299 | 0.181502865 |
| Tpbg | S | 424 | 0.101157899 | 0.121427592 | 0.175655732 | 0.074497833 | 0.05422814 | 0.212789931 | 0.129862961 | 0.061140826 | 0.336116945 | 0.45351502 |
| Klf16 | S | 241 | 0.051216922 | 0.140186704 | 0.151891221 | 0.100674299 | 0.011704517 | 0.686919947 | 0.027668515 | 0.025559485 | 0.445958962 | 0.800842319 |
| Bhlhe41 | S | 122 | 0.094494224 | 0.118595675 | 0.117038504 | 0.02254428 | -0.001557171 | 0.064595175 | 0.120550691 | 0.054214063 | 0.65024658 | 0.981045876 |
| Foxp1 | S | 649 | -0.060384286 | 0.08312899 | 0.19048114 | 0.250865426 | 0.10735215 | 0.093267585 | 0.026298464 | 2.44907E-05 | 0.004445061 | 0.023127709 |
| Tnrc6b | S | 1807 | 0.411785596 | 0.352925709 | 0.270208476 | -0.14157712 | -0.082717233 | 0.505443233 | 0.329887742 | 0.491925743 | 0.799452889 | 0.751913658 |
| Trappc12 | S | 125 | -0.000897989 | 0.073478601 | 0.128505746 | 0.129403735 | 0.055027145 | 0.959540975 | 0.018295628 | 0.000167456 | 0.001071896 | 0.054649267 |
| Lpin3 | S | 94 | 0.027744326 | 0.139128175 | 0.082764292 | 0.055019966 | -0.056363883 | 0.647948714 | 0.005485683 | 0.007976948 | 0.401648857 | 0.059827996 |
| Arfgef2 | S | 624 | 0.034430242 | 0.106287756 | 0.144351362 | 0.10992112 | 0.038063606 | 0.553868334 | 0.059066742 | 0.003600748 | 0.119802699 | 0.387694482 |
| Hirip3 | S | 389 | 0.123357576 | 0.244104838 | 0.174634758 | 0.051277181 | -0.06947008 | 0.380149767 | 0.110268304 | 0.221369985 | 0.276135401 | 0.141069416 |
| Nol9 | S | 93 | 0.180683447 | 0.171102538 | -0.047563565 | -0.228247012 | -0.218666103 | 0.016662624 | 0.020240117 | 0.859000175 | 0.43074887 | 0.447256868 |
| Srrm2 | S | 907 | 0.003652378 | 0.260737024 | 0.159706782 | 0.156054405 | -0.101030242 | 0.987517757 | 0.020107764 | 0.125829454 | 0.516007983 | 0.160243732 |
| Srrm2 | S | 909 | 0.003652378 | 0.260737024 | 0.159706782 | 0.156054405 | -0.101030242 | 0.987517757 | 0.020107764 | 0.125829454 | 0.516007983 | 0.160243732 |
| Ttc7 | S | 648 | 0.018597619 | 0.107660768 | 0.233845536 | 0.215247917 | 0.126184768 | 0.686356299 | 0.162812313 | 0.00778413 | 0.014151449 | 0.129349929 |
| 4833420G17Rik | T | 544 | 0.05330367 | 0.111240928 | 0.266470766 | 0.213167096 | 0.155229838 | 0.572984234 | 0.177813405 | 0.027091388 | 0.09856826 | 0.13913949 |
| Brd4 | S | 1153 | 0.047011998 | 0.099057526 | 0.179308055 | 0.132296057 | 0.080250528 | 0.24233306 | 0.127665099 | 0.013174506 | 0.057683337 | 0.217982505 |
| Zfp608 | T | 635 | 0.186454191 | 0.018164977 | 0.134449313 | -0.052004877 | 0.116284336 | 0.03554248 | 0.56965636 | 0.025941547 | 0.419520817 | 0.049561619 |
| Zc3h18 | S | 530 | 0.07559714 | 0.130259099 | 0.145935243 | 0.070338102 | 0.015676143 | 0.235592349 | 0.072272148 | 0.044141283 | 0.093450027 | 0.705166354 |
| Ppp4r3a | S | 767 | 0.028571626 | 0.129455486 | 0.107959783 | 0.079388158 | -0.021495703 | 0.243285622 | 0.030392491 | 0.008682086 | 0.035079579 | 0.600701474 |
| Ank2 | S | 3786 | 0.001474272 | 0.348576056 | 0.058150188 | 0.056675916 | -0.290425868 | 0.98650553 | 0.004412959 | 0.473186319 | 0.274081238 | 0.008837673 |
| Septin7 | T | 227 | -0.012286559 | 0.133245119 | 0.142367653 | 0.154654212 | 0.009122533 | 0.598154917 | 0.01523925 | 0.002595039 | 0.005778362 | 0.800749205 |
| Zc3h18 | S | 114 | -0.000219959 | 0.087150596 | 0.213511333 | 0.213731292 | 0.126360736 | 0.994609066 | 0.044167339 | 0.004835509 | 0.018093494 | 0.04328436 |
| Git1 | S | 605 | 0.046677427 | 0.253725127 | 0.043157111 | -0.003520316 | -0.210568016 | 0.646954287 | 0.001369862 | 0.589652287 | 0.974095463 | 0.061965569 |
| Tbc1d10b | T | 136 | 0.002627937 | 0.115589314 | 0.289500552 | 0.286872614 | 0.173911237 | 0.944192813 | 0.090471734 | 0.022719017 | 0.042012775 | 0.091832473 |
| Ino80b | S | 149 | 0.033540571 | 0.076010752 | 0.152714174 | 0.119173603 | 0.076703422 | 0.221569797 | 0.018952199 | 0.013349628 | 0.027620019 | 0.08727362 |
| Add3 | S | 461 | 0.197301743 | 0.208756652 | 0.036147188 | -0.161154555 | -0.172609464 | 0.084022455 | 0.045723348 | 0.822115499 | 0.341540002 | 0.308786007 |
| Cntrob | S | 63 | 0.12631009 | 0.157899082 | 0.063907337 | -0.062402752 | -0.093991745 | 0.057478783 | 0.025033749 | 0.463936862 | 0.427296744 | 0.267598236 |
| Srrm1 | S | 387 | 0.073337659 | 0.057257503 | 0.079425728 | 0.006088069 | 0.022168225 | 0.000763021 | 0.032685701 | 0.112204757 | 0.863188724 | 0.567647312 |
| Mtss2 | S | 564 | 0.202977928 | 0.07738703 | 0.050408778 | -0.152569151 | -0.026978252 | 0.00442795 | 0.219360617 | 0.476300484 | 0.073482981 | 0.684236405 |
| Ccdc9b | S | 138 | 0.042749848 | 0.071603911 | 0.181411586 | 0.138661737 | 0.109807674 | 0.34028521 | 0.439676838 | 0.000908803 | 0.025133246 | 0.272294838 |
| Pdlim4 | S | 111 | 0.05140188 | 0.182659653 | 0.050785536 | -0.000616344 | -0.131874117 | 0.266119813 | 0.002537635 | 0.219068419 | 0.985229189 | 0.009877149 |
| Tra2b | S | 29 | 0.0125155 | 0.124456767 | 0.320405733 | 0.307890233 | 0.195948966 | 0.903028328 | 0.032761335 | 0.081995342 | 0.089248464 | 0.19463885 |
| Khsrp | S | 481 | 0.058252178 | 0.070028214 | 0.083080295 | 0.024828117 | 0.013052081 | 0.025358295 | 0.046988747 | 0.005500329 | 0.035481409 | 0.58618765 |
| Tmpo | S | 422 | 0.049140812 | 0.112626884 | 0.078145836 | 0.029005023 | -0.034481048 | 0.099697628 | 0.005075978 | 0.055966447 | 0.32785536 | 0.278220272 |
| Osbp | S | 380 | 0.046478306 | 0.12777324 | 0.075958945 | 0.02948064 | -0.051814295 | 0.139531781 | 0.010422064 | 0.029893929 | 0.088350926 | 0.153634078 |
| Rbm12b2 | S | 658 | 0.112491686 | 0.098289377 | 0.225782665 | 0.113290979 | 0.127493288 | 0.517382704 | 0.061698747 | 0.064747992 | 0.53015965 | 0.194307808 |
| Rbm12b1 | S | 660 | 0.112491686 | 0.098289377 | 0.225782665 | 0.113290979 | 0.127493288 | 0.517382704 | 0.061698747 | 0.064747992 | 0.53015965 | 0.194307808 |
| Tnrc18 | S | 1573 | 0.032567137 | 0.120353408 | 0.164867543 | 0.132300406 | 0.044514135 | 0.62221681 | 0.049041643 | 0.011907393 | 0.090422888 | 0.35027726 |
| Rnf169 | S | 234 | 0.030206843 | 0.098737039 | 0.193385068 | 0.163178226 | 0.09464803 | 0.410449577 | 0.037290027 | 0.024417948 | 0.029325308 | 0.116258145 |
| Ncoa7 | S | 353 | 0.142257054 | 0.219500579 | 0.002257155 | -0.140269899 | -0.217243424 | 0.162414138 | 0.005768568 | 0.978369811 | 0.217362214 | 0.061339893 |
| Spata13 | S | 82 | -0.018330114 | 0.319084772 | 0.163615808 | 0.181945921 | -0.155468965 | 0.842949654 | 0.02571682 | 0.152355119 | 0.171727514 | 0.201466424 |
| Bcl9 | S | 278 | 0.090344967 | 0.03033486 | 0.211804933 | 0.121459966 | 0.181470073 | 0.063293776 | 0.581008327 | 0.013922789 | 0.087330766 | 0.034929829 |
| Sf3b1 | T | 235 | 0.137354312 | 0.197373238 | 0.010712644 | -0.126821668 | -0.186660594 | 0.153032144 | 0.004062332 | 0.926811981 | 0.352153056 | 0.193467519 |
| Slbp | S | 20 | 0.088696267 | 0.135185599 | 0.094882932 | 0.006186665 | -0.040302668 | 0.033988189 | 0.02945226 | 0.293929886 | 0.937004318 | 0.631839019 |
| Sf3a3 | S | 365 | 0.076978135 | 0.08698774 | 0.222942632 | 0.145964497 | 0.135954892 | 0.118583411 | 0.213208236 | 0.054264989 | 0.137577305 | 0.173419535 |
| Sf3a3 | S | 367 | 0.076978135 | 0.08698774 | 0.222942632 | 0.145964497 | 0.135954892 | 0.118583411 | 0.213208236 | 0.054264989 | 0.137577305 | 0.173419535 |
| Sf3a3 | S | 369 | 0.076978135 | 0.08698774 | 0.222942632 | 0.145964497 | 0.135954892 | 0.118583411 | 0.213208236 | 0.054264989 | 0.137577305 | 0.173419535 |
| Tra2b | S | 26 | 0.130047916 | 0.015828085 | 0.260686743 | 0.130638827 | 0.244858659 | 0.585464129 | 0.895691384 | 0.002024815 | 0.580764611 | 0.13046508 |
| Ehd2 | S | 438 | -0.069044137 | 0.13221771 | 0.13799992 | 0.207044057 | 0.005782209 | 0.145377639 | 0.005663543 | 0.00012383 | 0.016638092 | 0.816721585 |
| Pkn2 | S | 621 | 0.030738382 | 0.183056718 | 0.211333765 | 0.180595383 | 0.028277047 | 0.833405741 | 0.033130695 | 0.091298163 | 0.282654483 | 0.753536326 |
| Tra2b | S | 95 | -0.024269449 | 0.076080449 | 0.257915045 | 0.282184494 | 0.181834596 | 0.747480236 | 0.251257729 | 0.000994806 | 0.031701568 | 0.038770771 |
| Peak1 | S | 785 | 0.079705543 | 0.128990442 | 0.085072387 | 0.005366845 | -0.043918055 | 0.203798199 | 0.002253005 | 0.273003315 | 0.944750381 | 0.529843911 |
| Srrm1 | S | 391 | 0.071509812 | 0.05681059 | 0.07898016 | 0.007470348 | 0.02216957 | 0.000748709 | 0.031727717 | 0.11346662 | 0.833001933 | 0.566806923 |
| Scaf1 | S | 491 | 0.092609896 | 0.018996048 | 0.101507042 | 0.008897147 | 0.082510994 | 0.006145425 | 0.399195228 | 0.003600464 | 0.679700454 | 0.021482307 |
| Scaf1 | S | 493 | 0.092609896 | 0.018996048 | 0.101507042 | 0.008897147 | 0.082510994 | 0.006145425 | 0.399195228 | 0.003600464 | 0.679700454 | 0.021482307 |
| Pitpnm1 | S | 895 | -0.036148172 | 0.078574138 | 0.219470577 | 0.255618749 | 0.140896439 | 0.435243154 | 0.130910331 | 0.000254226 | 0.008016839 | 0.034618963 |
| Haspin | T | 379 | 0.161716337 | 0.079334112 | 0.094655441 | -0.067060896 | 0.015321328 | 0.146865718 | 0.314044266 | 0.004586199 | 0.449825923 | 0.824368139 |
| Mia3 | S | 1769 | -0.009983201 | 0.286007284 | 0.618182599 | 0.6281658 | 0.332175315 | 0.887805965 | 0.308072335 | 0.180846337 | 0.177834205 | 0.432339066 |
| Phf14 | S | 775 | 0.036937146 | 0.166956545 | 0.107633586 | 0.07069644 | -0.059322959 | 0.608595812 | 0.002623035 | 0.163097199 | 0.414034331 | 0.375285613 |
| Brca1 | S | 706 | 0.07950994 | 0.09965742 | 0.167438524 | 0.087928583 | 0.067781103 | 0.279845611 | 0.131871583 | 0.020793216 | 0.210820841 | 0.229319881 |
| Tra2a Tra2b | S | 264;259 | 0.029564921 | 0.094338915 | 0.131034532 | 0.101469611 | 0.036695618 | 0.429909833 | 0.006450271 | 0.018143045 | 0.060773425 | 0.329132154 |
| Phldb1 | S | 223 | 0.103584102 | 0.080004979 | 0.076009592 | -0.027574511 | -0.003995387 | 0.015057277 | 0.03445326 | 0.129444301 | 0.493978289 | 0.912649028 |
| Erbin | S | 1077 | 0.090761829 | 0.090894937 | 0.100698922 | 0.009937093 | 0.009803985 | 0.242790062 | 0.005209434 | 0.066339686 | 0.887907776 | 0.799550221 |
| Cbx2 | S | 326 | 0.037184283 | 0.194106528 | 0.138576721 | 0.101392439 | -0.055529807 | 0.67873667 | 0.003816747 | 0.292596237 | 0.462340356 | 0.635995878 |
| Srsf10 | S | 253 | -0.011872516 | 0.1465988 | 0.257753027 | 0.269625543 | 0.111154228 | 0.735383388 | 0.029475129 | 0.039147092 | 0.04985638 | 0.231609741 |
| Srrm2 | S | 1646 | 0.066182994 | 0.059199395 | 0.114896048 | 0.048713054 | 0.055696653 | 0.08297725 | 0.028712389 | 0.012852709 | 0.207984253 | 0.113574608 |
| Zc3h13 | S | 369 | 0.121544079 | 0.115039141 | 0.152501727 | 0.030957648 | 0.037462587 | 0.052820467 | 0.141953515 | 0.194339371 | 0.74838893 | 0.716491892 |
| Mprip | S | 225 | 0.005186356 | 0.082036365 | 0.13524475 | 0.130058394 | 0.053208385 | 0.792778271 | 0.044445426 | 0.000385373 | 0.001889556 | 0.134321989 |
| Zfp638 | S | 1569 | 0.136628532 | 0.024506286 | 0.130977306 | -0.005651227 | 0.106471019 | 0.047132284 | 0.227958254 | 0.016431488 | 0.906107526 | 0.039356201 |
| Ano8 | S | 665 | 0.093241305 | 0.051101725 | 0.179934108 | 0.086692803 | 0.128832383 | 0.148408115 | 0.394277885 | 0.008281733 | 0.171907317 | 0.071205369 |
| Fyco1 | S | 196 | -0.01963394 | 0.143759293 | 0.216130057 | 0.235763997 | 0.072370765 | 0.678667032 | 0.12254624 | 0.003469244 | 0.000148539 | 0.352852013 |
| Cic | S | 1514 | 0.324693545 | 0.179054169 | 0.138058331 | -0.186635214 | -0.040995838 | 0.118632856 | 0.299688403 | 0.587265732 | 0.431935387 | 0.843461462 |
| Washc2 | S | 533 | 0.073012345 | 0.05043525 | 0.169071318 | 0.096058973 | 0.118636068 | 0.092815587 | 0.035409044 | 0.039114151 | 0.13688618 | 0.098499209 |
| Tjp1 | S | 125 | 0.000753254 | 0.101011378 | 0.161445832 | 0.160692578 | 0.060434453 | 0.982359386 | 0.046499844 | 0.001872337 | 0.013160224 | 0.151740891 |
| Ankzf1 | S | 555 | 0.009641856 | 0.126902902 | 0.426958436 | 0.41731658 | 0.300055534 | 0.863298885 | 0.088931818 | 0.107768717 | 0.115935905 | 0.19594271 |
| Safb Safb2 | S | 465;488 | 0.166164779 | 0.267135502 | -0.067069266 | -0.233234045 | -0.334204769 | 0.093313892 | 0.008376977 | 0.580154282 | 0.107116459 | 0.055508568 |
| Rere | S | 600 | 0.005948545 | 0.107496214 | 0.141455271 | 0.135506727 | 0.033959057 | 0.658770831 | 0.000334346 | 0.08599853 | 0.093529556 | 0.539771287 |
| Huwe1 | S | 2377 | 0.152737664 | 0.123712232 | 0.111128865 | -0.0415188 | -0.012493368 | 0.377375896 | 0.075274484 | 0.033373956 | 0.79083045 | 0.820909395 |
| Lmna | S | 613 | 0.091859524 | 0.078640704 | 0.211540791 | 0.119681268 | 0.132900087 | 0.539483446 | 0.271269715 | 0.008245955 | 0.435724387 | 0.097010609 |
| Lmna | S | 616 | 0.091859524 | 0.078640704 | 0.211540791 | 0.119681268 | 0.132900087 | 0.539483446 | 0.271269715 | 0.008245955 | 0.435724387 | 0.097010609 |
| Smarcad1 | S | 103 | 0.044532373 | 0.120583519 | 0.094184629 | 0.049652256 | -0.02639889 | 0.062285396 | 0.035254331 | 0.02848126 | 0.161807217 | 0.536654761 |
| Nipbl | T | 713 | -0.023395686 | 0.090224211 | 0.256153319 | 0.279549005 | 0.165929107 | 0.617191209 | 0.0734198 | 0.004119826 | 0.006925342 | 0.034422297 |
| Lmnb1 Lmnb2 | S | 396;387 | 0.080191073 | 0.141881639 | 0.124225909 | 0.044034836 | -0.01765573 | 0.239332463 | 0.011236893 | 0.22845294 | 0.649491782 | 0.839617996 |
| Zc3hc1 | S | 62 | 0.023404378 | 0.091881215 | 0.12713568 | 0.103731302 | 0.035254464 | 0.380292077 | 0.011672685 | 0.008898735 | 0.039927879 | 0.266928581 |
| Arfgef2 | S | 227 | 0.014241999 | 0.116911869 | 0.09790953 | 0.083667531 | -0.019002339 | 0.770458423 | 0.009855343 | 0.002400963 | 0.169223579 | 0.484626672 |
| Rreb1 | S | 1659 | 0.01541176 | -0.009216393 | 0.355073917 | 0.339662157 | 0.36429031 | 0.783160123 | 0.922867771 | 0.001368612 | 0.008269562 | 0.020342956 |
| Wdhd1 | T | 956 | 0.179445666 | -0.005488307 | 0.108165817 | -0.071279849 | 0.113654124 | 0.003364812 | 0.956651407 | 0.0509957 | 0.104991785 | 0.311431733 |
| Luc7l2 | S | 383 | 0.092629092 | 0.110019728 | 0.122189144 | 0.029560052 | 0.012169416 | 0.196442936 | 0.037565444 | 0.068070096 | 0.665113945 | 0.813611292 |
| Afap1 | S | 665 | 0.036244875 | 0.161342454 | 0.09967526 | 0.063430385 | -0.061667194 | 0.315682616 | 0.061648537 | 0.008868031 | 0.124204746 | 0.325906278 |
| Tra2b | S | 39 | 0.121825303 | 0.084022858 | 0.254107296 | 0.132281993 | 0.170084438 | 0.177661517 | 0.268480189 | 0.097260076 | 0.316938087 | 0.214746192 |

| | | | | | | | | | | | | |
|---|---|---|---|---|---|---|---|---|---|---|---|---|
| Pask | S | 1345 | 0.056229579 | 0.158202324 | 0.166891442 | 0.110661863 | 0.008689118 | 0.60444135 | 0.021487581 | 0.105364378 | 0.371341018 | 0.912151926 |
| Zdhhc8 | S | 335 | 0.018799019 | 0.116642629 | 0.081501065 | 0.062702046 | -0.035141564 | 0.569927654 | 0.001223479 | 0.01776567 | 0.107260351 | 0.162426249 |
| Prrc2c | T | 2625 | 0.119013593 | 0.054260877 | 0.16586586 | 0.046852267 | 0.111604982 | 0.076507146 | 0.394474473 | 0.025072925 | 0.118293436 | 0.034407205 |
| Gmip | S | 439 | -0.000766434 | 0.173524038 | 0.173266606 | 0.17403304 | -0.000257432 | 0.988641358 | 0.040460997 | 0.021398167 | 0.039398963 | 0.996830302 |
| Tle3 | S | 203 | 0.144709477 | 0.200477267 | -0.020788574 | -0.16549805 | -0.221265841 | 0.05958826 | 0.007739198 | 0.803910761 | 0.111802615 | 0.057349978 |
| Iqsec2 | S | 363 | 0.037861425 | 0.072948708 | 0.113744058 | 0.075882633 | 0.04079535 | 0.114724227 | 0.055338138 | 0.003047337 | 0.024175783 | 0.216073756 |
| Olfr867 | S | 313 | 0.121800308 | 0.075983407 | 0.080814506 | -0.040985802 | 0.004831099 | 0.009076086 | 0.356672389 | 0.034849026 | 0.246525186 | 0.948028871 |
| Epb41 | S | 86 | 0.023358824 | 0.286928681 | 0.244412497 | 0.221053673 | -0.042516184 | 0.756360991 | 0.101339561 | 0.162931624 | 0.192357531 | 0.802084702 |
| Ctr9 | S | 1083 | 0.27061316 | 0.117112586 | -0.011303839 | -0.281916999 | -0.128416426 | 0.009570633 | 0.164829652 | 0.948302557 | 0.186438898 | 0.473584969 |
| Trappc12 | T | 130 | 0.004392442 | 0.07526162 | 0.138898494 | 0.134506052 | 0.063636874 | 0.828564429 | 0.042032474 | 0.000419741 | 0.003098261 | 0.066473511 |
| Htt | S | 1841 | 0.003706416 | 0.415255169 | -0.067205519 | -0.070911935 | -0.482460688 | 0.953495489 | 0.000653144 | 0.544070159 | 0.49323138 | 0.028676577 |
| Ank3 | S | 1406 | 0.084506672 | 0.201046318 | 0.056631701 | -0.027874971 | -0.144414617 | 0.045342224 | 0.036027966 | 0.403024688 | 0.641206672 | 0.118721579 |
| Cdc42ep1 | S | 371 | 0.018649439 | 0.175572334 | 0.092301562 | 0.073652124 | -0.083270772 | 0.843553896 | 0.001801054 | 0.116267417 | 0.468528043 | 0.149688293 |
| Bclaf1 | S | 709 | 0.104801516 | -0.017510257 | 0.224835357 | 0.12003384 | 0.242345613 | 0.298814455 | 0.703620939 | 0.000704422 | 0.250007612 | 0.008346614 |
| Rbm15b | S | 261 | 0.041201662 | 0.061119931 | 0.114020834 | 0.072819172 | 0.052900903 | 0.003834274 | 0.096845761 | 0.013978267 | 0.05111869 | 0.145632017 |
| Eif4g3 | T | 168 | 0.102066674 | 0.11048666 | 0.062293095 | -0.039773578 | -0.048193565 | 0.060722842 | 0.017605554 | 0.137996708 | 0.399689442 | 0.250671014 |
| Ankib1 | S | 938 | 0.116343961 | 0.081805169 | 0.060343684 | -0.056000278 | -0.021461485 | 0.009281608 | 0.084853738 | 0.107387445 | 0.036719162 | 0.530962791 |
| Akap13 | S | 1567 | 0.063867732 | 0.12230176 | 0.162498447 | 0.098630715 | 0.040196687 | 0.444320172 | 0.066233258 | 0.031373693 | 0.283256382 | 0.523744209 |
| Hexim1 | S | 230 | 0.20359737 | 0.28306907 | 0.028927643 | -0.174669726 | -0.254141426 | 0.120458958 | 0.095574087 | 0.793172998 | 0.112275529 | 0.117299614 |
| Arfgef2 | S | 1520 | 0.050356327 | 0.143527779 | 0.115801284 | 0.065444957 | -0.027726495 | 0.181458826 | 0.071753609 | 0.028465522 | 0.123094887 | 0.648633735 |
| Frmd6 | S | 352 | -0.049196478 | 0.163258267 | 0.188628115 | 0.237824594 | 0.025369849 | 0.397353211 | 0.043060181 | 0.003009487 | 0.011722693 | 0.655345542 |
| Katnb1 | S | 360 | -0.113764685 | 0.223103374 | 0.344313643 | 0.458078327 | 0.121210268 | 0.314481925 | 0.004540724 | 0.144933116 | 0.070186594 | 0.500565998 |
| Hdac1 | S | 393 | 0.03859515 | 0.058116779 | 0.119851018 | 0.081255868 | 0.061734239 | 0.089469607 | 0.010917911 | 0.011413654 | 0.045414288 | 0.093250573 |
| Trappc12 | S | 128 | -0.00291393 | 0.073986272 | 0.131667718 | 0.134581648 | 0.057681447 | 0.873450136 | 0.034534072 | 0.000166035 | 0.001062273 | 0.076311159 |
| C2cd3 | S | 2039 | -0.06508658 | 0.143875865 | 0.249995065 | 0.315081645 | 0.1061192 | 0.631085228 | 0.269398547 | 0.001309061 | 0.100515475 | 0.386507142 |
| Dido1 | T | 148 | -0.027894388 | 0.108283109 | 0.130158139 | 0.158052527 | 0.02187503 | 0.30055229 | 0.003094955 | 0.001102127 | 0.000783506 | 0.23163021 |
| Foxk1 | S | 406 | 0.011516339 | 0.07619437 | 0.104413244 | 0.092896905 | 0.028218874 | 0.629240174 | 0.005984754 | 0.000965535 | 0.018326106 | 0.180383622 |
| Herpud2 | S | 88 | -0.016604752 | 0.183163713 | 0.201247642 | 0.217852394 | 0.018083929 | 0.708437261 | 0.017908489 | 0.054196489 | 0.04825645 | 0.814424088 |
| Kmt2d | S | 4789 | -0.01844086 | 0.09742578 | 0.251463395 | 0.269904255 | 0.154037615 | 0.734121358 | 0.118837109 | 0.004426953 | 0.007676446 | 0.02798651 |
| Sun2 | S | 55 | 0.219701788 | 0.125945698 | 0.042454454 | -0.177247334 | -0.083491244 | 0.027360708 | 0.148735478 | 0.512821301 | 0.056573019 | 0.242127793 |
| Magi3 | S | 700 | -0.01753648 | 0.133257168 | 0.176713865 | 0.194250344 | 0.043456697 | 0.788694853 | 0.002126062 | 0.059552093 | 0.062916374 | 0.502642445 |
| Trp53bp2 | S | 735 | 0.120715502 | 0.179477096 | 0.13162791 | 0.010912408 | -0.047849186 | 0.175325822 | 0.11852911 | 0.187030117 | 0.911848311 | 0.659350267 |
| Myo1c | S | 536 | 0.068612663 | 0.129197128 | 0.065573442 | -0.003039221 | -0.063623686 | 0.134412861 | 0.006341018 | 0.128451733 | 0.935921877 | 0.100021526 |
| Nucks1 | S | 19 | 0.001469563 | 0.190157225 | 0.10182446 | 0.100354897 | -0.088332765 | 0.959683644 | 0.005817712 | 0.05179747 | 0.058471108 | 0.093929808 |
| Ercc5 | S | 357 | 0.126084971 | 0.054167106 | 0.153404135 | 0.027319164 | 0.099237029 | 0.042396993 | 0.402869391 | 0.043727882 | 0.614018275 | 0.166793972 |
| Kat6a | S | 786 | 0.028872811 | 0.077774983 | 0.180297948 | 0.151425137 | 0.102522965 | 0.598837781 | 0.027261592 | 0.011753392 | 0.054288427 | 0.095072301 |
| Alad | S | 215 | 0.192159182 | 0.19165751 | 0.041206608 | -0.150952574 | -0.150450902 | 0.054140554 | 0.088944766 | 0.722152561 | 0.196317482 | 0.218838213 |
| Pcm1 | S | 859 | -0.091555276 | 0.225886649 | 0.200322539 | 0.291877815 | -0.02556411 | 0.125760936 | 0.003089635 | 0.02993452 | 0.013304656 | 0.666367938 |
| Peak1 | Y | 638 | 0.046084787 | 0.163742225 | 0.130137135 | 0.084052348 | -0.03360509 | 0.355324204 | 0.045446316 | 0.054438818 | 0.167053789 | 0.612868953 |
| Pi4ka | S | 260 | 0.183115513 | 0.165140136 | -0.025984865 | -0.209100378 | -0.191125002 | 0.008955716 | 0.043773491 | 0.665976959 | 0.019598565 | 0.039499558 |
| Hmga2 | S | 61 | 0.290767448 | 0.348827476 | -0.141987035 | -0.432754483 | -0.490814511 | 0.109970133 | 0.053372989 | 0.60421824 | 0.165618218 | 0.143759427 |
| Smarcc2 | S | 306 | 0.048398723 | 0.077701753 | 0.146097236 | 0.097698512 | 0.068395482 | 0.239588647 | 0.0205311 | 0.025611021 | 0.092037357 | 0.16321728 |
| Jdp2 | T | 148 | 0.303584922 | 0.083538713 | -0.033457474 | -0.337042396 | -0.116996187 | 0.002088216 | 0.41287592 | 0.757141206 | 0.042770187 | 0.360965531 |
| Slc25a4 | S | 149 | 0.054049394 | 0.047789465 | 0.190943722 | 0.136894329 | 0.143154257 | 0.196441778 | 0.271931397 | 0.005469803 | 0.028833134 | 0.028160315 |
| Septin9 | T | 49 | 0.334022526 | 0.063157379 | 0.078331458 | -0.255691068 | 0.015174079 | 0.012584064 | 0.728296594 | 0.62272197 | 0.15306373 | 0.939119502 |
| Trerf1 | S | 764 | 0.113934238 | 0.20543572 | 0.14862868 | 0.034694443 | -0.056807039 | 0.501204468 | 0.021862852 | 0.386028934 | 0.863357602 | 0.713007201 |
| Champ1 | S | 641 | 0.009449299 | 0.104978462 | 0.095526927 | 0.086077628 | -0.009451535 | 0.80063369 | 0.004481 | 0.003075058 | 0.09466863 | 0.620689604 |
| Chd8 | S | 2225 | 0.116991668 | 0.035787861 | 0.184732127 | 0.067740458 | 0.148944266 | 0.001766731 | 0.494708914 | 0.227844203 | 0.596473959 | 0.302388967 |
| Nipbl | T | 724 | 0.052910244 | 0.138203835 | 0.14232639 | 0.089416145 | 0.004122555 | 0.353943781 | 0.027220822 | 0.07880644 | 0.199608566 | 0.940447027 |
| Atrx | S | 1315 | 0.054165133 | 0.121593306 | 0.126368861 | 0.072203727 | 0.004775554 | 0.504305005 | 0.026547447 | 0.025181536 | 0.3651475 | 0.855238779 |
| Pde4a | S | 152 | 0.063815021 | 0.21873042 | 0.050321938 | -0.013493083 | -0.168408482 | 0.576761278 | 0.001323698 | 0.649759092 | 0.923594814 | 0.204615184 |
| Pde4a | Y | 149 | 0.063815021 | 0.21873042 | 0.050321938 | -0.013493083 | -0.168408482 | 0.576761278 | 0.001323698 | 0.649759092 | 0.923594814 | 0.204615184 |
| Kif1b | S | 1211 | 0.193143119 | -0.042461882 | 0.224441545 | 0.031298426 | 0.266903428 | 0.166052866 | 0.725820528 | 0.006321572 | 0.775080787 | 0.109458668 |
| Srrm2 | S | 453 | 0.039964291 | 0.063436219 | 0.115248044 | 0.075283753 | 0.051811824 | 0.084233419 | 0.102266045 | 0.002086361 | 0.021321856 | 0.161244733 |
| Osbpl11 | S | 186 | 0.056264335 | 0.028277285 | 0.209858429 | 0.153594094 | 0.181581144 | 0.271116498 | 0.460764929 | 0.002566924 | 0.024150079 | 0.011606433 |
| Rpl27a | S | 68 | 0.215406634 | 0.005448696 | 0.522436572 | 0.307029939 | 0.516987877 | 0.377757334 | 0.985009821 | 0.111436069 | 0.26367442 | 0.135155605 |
| Srrm2 | S | 2404 | 0.036770588 | 0.050985303 | 0.087995458 | 0.05122487 | 0.037010156 | 0.041738287 | 0.060726098 | 0.000498306 | 0.01174167 | 0.135434386 |
| Cavin1 | S | 204 | -0.036989384 | 0.112258794 | 0.213413031 | 0.250402415 | 0.101154237 | 0.477828831 | 0.010847435 | 0.010168475 | 0.010401282 | 0.088144159 |
| Cavin1 | S | 205 | -0.036989384 | 0.112258794 | 0.213413031 | 0.250402415 | 0.101154237 | 0.477828831 | 0.010847435 | 0.010168475 | 0.010401282 | 0.088144159 |
| Nab2 | T | 8 | 0.309779339 | 0.21624691 | -0.239527425 | -0.549306764 | -0.455774335 | 0.008280461 | 0.017676704 | 0.205106115 | 0.039776665 | 0.074873721 |
| Luc7l2 | S | 271 | 0.251517169 | 0.161444921 | 0.128125632 | -0.123391537 | -0.033319289 | 0.099368527 | 0.26441912 | 0.469550515 | 0.51764516 | 0.860876859 |
| Luc7l2 | S | 273 | 0.251517169 | 0.161444921 | 0.128125632 | -0.123391537 | -0.033319289 | 0.099368527 | 0.26441912 | 0.469550515 | 0.51764516 | 0.860876859 |
| Csf1 | S | 530 | 0.057491483 | 0.150486952 | 0.170575705 | 0.113084223 | 0.020088753 | 0.58160269 | 0.132991247 | 0.022034026 | 0.324843519 | 0.808056129 |
| Sipa1l2 | S | 1461 | 0.124611152 | 0.040177956 | 0.10204764 | -0.022563512 | 0.061869685 | 0.010482923 | 0.270120489 | 0.050934196 | 0.509892929 | 0.15136059 |
| Reps1 | T | 173 | 0.103478052 | 0.103797015 | 0.044328408 | -0.059149644 | -0.059468607 | 0.017690752 | 0.017161094 | 0.277380307 | 0.102659146 | 0.101100263 |
| Slx4 | S | 909 | 0.161782365 | 0.088710825 | 0.108149408 | -0.053632957 | 0.019438584 | 0.012702518 | 0.365075062 | 0.23700462 | 0.492669263 | 0.851978255 |
| Sptbn1 | S | 2102 | -0.014203966 | 0.066060328 | 0.137150215 | 0.151354181 | 0.071089887 | 0.46998447 | 0.003486414 | 0.000467305 | 0.001242393 | 0.025796125 |
| Rbm25 | S | 371 | 0.106042077 | 0.057471089 | 0.132454378 | 0.026412302 | 0.07498329 | 0.120714765 | 0.301688684 | 0.00825823 | 0.634313244 | 0.209055799 |
| Rc3h2 | T | 803 | 0.208125714 | 0.376109671 | -0.029419737 | -0.237545451 | -0.405529407 | 0.514438179 | 0.017717526 | 0.841675402 | 0.475690753 | 0.057634892 |
| Coil | S | 222 | 0.363688881 | 0.152476189 | -0.175999987 | -0.539688868 | -0.328476175 | 0.006936113 | 0.139072048 | 0.477134022 | 0.103718854 | 0.235966907 |
| Clta | S | 104 | 0.033808291 | 0.066043356 | 0.206164992 | 0.172356702 | 0.140121636 | 0.46970955 | 0.31902962 | 0.002907146 | 0.014349354 | 0.080601633 |
| Mlh1 | T | 419 | 0.135214963 | 0.235971454 | 0.003958865 | -0.131256098 | -0.232012589 | 0.312780999 | 0.004670681 | 0.975297086 | 0.421241595 | 0.1617141 |
| Phf6 | S | 154 | -0.004630869 | 0.104236194 | 0.168002767 | 0.172633636 | 0.063766573 | 0.917092155 | 0.010504599 | 0.011937922 | 0.020427937 | 0.189440706 |
| Chd8 | S | 2222 | 0.156308268 | 0.015300336 | 0.215136126 | 0.058827859 | 0.19983579 | 0.032726557 | 0.685167575 | 0.087255606 | 0.508395616 | 0.087348933 |
| Mis18bp1 | S | 215 | 0.117001316 | 0.164815713 | 0.112871397 | -0.004129919 | -0.051944316 | 0.241101283 | 0.034395162 | 0.282347048 | 0.970866377 | 0.599549219 |
| Zc3h18 | T | 105 | 0.033325431 | 0.082809727 | 0.176569335 | 0.143243904 | 0.093759608 | 0.297869357 | 0.141731498 | 0.007659127 | 0.012821854 | 0.107558076 |
| Epb41l3 | T | 493 | 0.129520088 | 0.041567091 | 0.243003989 | 0.113483901 | 0.201436898 | 0.069184152 | 0.484081644 | 0.087992334 | 0.330018686 | 0.14146171 |
| Rbl1 | T | 332 | 0.172797308 | 0.100727353 | 0.049713356 | -0.123083952 | -0.051013997 | 0.052211762 | 0.062115465 | 0.195956308 | 0.132427729 | 0.228230221 |
| Zfp687 | T | 246 | -0.065821692 | 0.111264116 | 0.281396525 | 0.347218218 | 0.170132409 | 0.319677961 | 0.154417525 | 0.001600736 | 0.005194313 | 0.062815899 |
| Cbx2 | S | 328 | 0.015507034 | 0.226481313 | 0.174960565 | 0.159453531 | -0.051520748 | 0.922689743 | 0.022526538 | 0.160739216 | 0.36647951 | 0.589023928 |
| Mprip | S | 976 | -0.018346676 | 0.141415337 | 0.203607093 | 0.221953768 | 0.062191756 | 0.749350226 | 0.013910854 | 0.036045825 | 0.033641316 | 0.371113191 |
| Cit | S | 1331 | 0.249062814 | -0.03019096 | 0.207618651 | -0.041444163 | 0.237809611 | 0.05059678 | 0.73852336 | 0.061555155 | 0.679727074 | 0.039313642 |
| Nck2 | S | 94 | 0.04445452 | 0.155614467 | 0.104658887 | 0.060204368 | -0.050955579 | 0.322177766 | 0.006931281 | 0.180970764 | 0.403565306 | 0.452822459 |
| Trpm7 | T | 1466 | 0.07674476 | 0.233761695 | 0.167251986 | 0.090507226 | -0.066509709 | 0.388647818 | 0.176153224 | 0.10004644 | 0.322475815 | 0.65209885 |
| Prkaa1 | S | 527 | -0.023209196 | 0.223144252 | 0.292178786 | 0.315387982 | 0.069034534 | 0.777428063 | 0.012905791 | 0.051814951 | 0.047931715 | 0.626621614 |
| Poll | S | 175 | -0.009042177 | 0.19643985 | 0.129192462 | 0.138234639 | -0.067247389 | 0.772925601 | 0.010891646 | 0.045102226 | 0.038466917 | 0.238131259 |
| Bcl9l | S | 21 | -0.023050707 | 0.110544354 | 0.171680057 | 0.194730764 | 0.061135703 | 0.392425328 | 0.02425324 | 0.002019571 | 0.00233554 | 0.136355945 |
| Kdm4b | S | 629 | 0.041455964 | 0.020547831 | 0.258702639 | 0.217246675 | 0.238154807 | 0.458120566 | 0.633677573 | 0.002387756 | 0.013748347 | 0.008320449 |
| Sbf2 | S | 1085 | 0.151604559 | 0.189482408 | 0.067777973 | -0.083826586 | -0.121704435 | 0.382579456 | 0.009522978 | 0.536302381 | 0.634531168 | 0.292171129 |
| Vps51 | S | 643 | 0.192380577 | 0.100705316 | 0.206040386 | 0.013659809 | 0.10533507 | 0.298988362 | 0.462427359 | 0.058330171 | 0.929728661 | 0.387341123 |
| Arid1a | S | 1756 | 0.003487787 | 0.099407695 | 0.187362879 | 0.183875092 | 0.087955184 | 0.916962741 | 0.009535695 | 0.025698007 | 0.030629741 | 0.162641387 |
| Pelp1 | T | 751 | 0.025454703 | 0.062885168 | 0.147955441 | 0.122500737 | 0.085070273 | 0.107550039 | 0.068357062 | 0.005563872 | 0.008987569 | 0.034681346 |
| Srsf2 | S | 212 | 0.026622732 | 0.124305104 | 0.180736262 | 0.15411353 | 0.056431158 | 0.734571704 | 0.031736466 | 0.034881872 | 0.135832344 | 0.372432067 |
| Srsf2 | S | 220 | 0.026622732 | 0.124305104 | 0.180736262 | 0.15411353 | 0.056431158 | 0.734571704 | 0.031736466 | 0.034881872 | 0.135832344 | 0.372432067 |

| | | | | | | | | | | | | |
|---|---|---|---|---|---|---|---|---|---|---|---|---|
| Mfsd6 | S | 646 | 0.363793441 | -0.18962926 | 0.338792078 | -0.025001363 | 0.528421338 | 0.043444804 | 0.304920211 | 0.077597993 | 0.816067971 | 0.019331781 |
| Sf3b1 | T | 303 | 0.088227174 | 0.079515828 | 0.10197827 | 0.013751096 | 0.022462442 | 0.056048112 | 0.078995043 | 0.041068369 | 0.597160454 | 0.416498892 |
| Stat4 | S | 188 | 0.120065588 | 0.184063448 | 0.122142555 | 0.002076966 | -0.061920893 | 0.464242736 | 0.060024367 | 0.128492934 | 0.989198486 | 0.449843594 |
| Stat4 | S | 205 | 0.120065588 | 0.184063448 | 0.122142555 | 0.002076966 | -0.061920893 | 0.464242736 | 0.060024367 | 0.128492934 | 0.989198486 | 0.449843594 |
| Stat4 | T | 197 | 0.120065588 | 0.184063448 | 0.122142555 | 0.002076966 | -0.061920893 | 0.464242736 | 0.060024367 | 0.128492934 | 0.989198486 | 0.449843594 |
| Bckdha | S | 344 | 0.048330953 | 0.060265704 | 0.292016974 | 0.243686021 | 0.23175127 | 0.394904359 | 0.118014008 | 0.044961314 | 0.058520024 | 0.076348148 |
| Cdk1 | T | 14 | 0.199108787 | 0.104807787 | 0.048096291 | -0.151012495 | -0.056711496 | 0.024373352 | 0.13091642 | 0.432543629 | 0.055789915 | 0.148753825 |
| Net1 | S | 508 | 0.107008805 | 0.081943987 | 0.215699022 | 0.108690218 | 0.133755035 | 0.207874073 | 0.454350615 | 0.032921092 | 0.216769157 | 0.252338574 |
| Rbm12b1 Rbm12b2 | S | 250;250 | 0.10372076 | 0.036613092 | 0.111902077 | 0.008181317 | 0.075288985 | 0.008891444 | 0.550101687 | 0.017295778 | 0.802324587 | 0.272935131 |
| Tns2 | S | 835 | 0.067621986 | 0.083022842 | 0.094183938 | 0.026561952 | 0.011161096 | 0.119375538 | 0.012678436 | 0.045169018 | 0.532987706 | 0.739895612 |
| Mtfr2 | S | 318 | -0.074429417 | 0.252376648 | 0.220878109 | 0.295307526 | -0.031498538 | 0.184366008 | 0.002064552 | 0.168738974 | 0.10941411 | 0.803630656 |
| Prrc2c | S | 191 | 0.097363405 | 0.189724921 | 0.061616832 | -0.035746573 | -0.128108089 | 0.23070338 | 0.011663063 | 0.443274873 | 0.662542375 | 0.14069974 |
| Cavin3 | T | 202 | 0.092711489 | 0.065959173 | 0.198010904 | 0.105299415 | 0.132051731 | 0.351455051 | 0.51950102 | 0.006311094 | 0.287557441 | 0.235502653 |
| 4932438A13Rik | S | 1355 | 0.064648355 | 0.064379609 | 0.139721205 | 0.07507285 | 0.075341596 | 0.116933981 | 0.082959076 | 0.018307877 | 0.127753457 | 0.116969345 |
| Cstf2t | S | 579 | 0.028851766 | 0.126850757 | 0.099995233 | 0.071143468 | -0.026855524 | 0.455679557 | 0.000711509 | 0.184727433 | 0.314209766 | 0.659057498 |
| Twist1 | S | 68 | -0.01416392 | 0.114715812 | 0.148239765 | 0.162403684 | 0.033523953 | 0.758866291 | 0.036816864 | 0.001820947 | 0.029645307 | 0.414858678 |
| Tardbp | S | 183 | 0.051791793 | 0.093108619 | 0.110002914 | 0.058211122 | 0.016894295 | 0.087424595 | 0.097362945 | 0.011874216 | 0.104931164 | 0.708519189 |
| Mcm9 | S | 709 | -0.009619081 | 0.125974475 | 0.253778804 | 0.263397884 | 0.127804329 | 0.815993132 | 0.121938548 | 0.013161314 | 0.009143936 | 0.131632423 |
| Prkci | T | 411 | 0.174531197 | 0.008633113 | 0.247601559 | 0.073070361 | 0.238968446 | 0.076641465 | 0.880472313 | 0.067850686 | 0.512830791 | 0.092091732 |
| Prx | S | 381 | 0.045349957 | 0.052126777 | 0.163751682 | 0.118401725 | 0.111624905 | 0.328899427 | 0.027919056 | 0.010689767 | 0.060822565 | 0.046366474 |
| Mta3 | S | 516 | 0.079377131 | 0.086707447 | 0.071473154 | -0.007903977 | -0.015234292 | 0.021016378 | 0.014077981 | 0.16854349 | 0.842835991 | 0.709127625 |
| Rftn2 | T | 408 | 0.093770453 | 0.203052077 | 0.05024305 | -0.043527403 | -0.152809026 | 0.126309747 | 0.011231122 | 0.73989026 | 0.767916243 | 0.353854264 |
| Xrcc1 | S | 446 | 0.027444829 | 0.077338819 | 0.1025292 | 0.07508437 | 0.02519038 | 0.43301134 | 0.010626672 | 0.002910038 | 0.078235316 | 0.170321293 |
| Sppl2a | S | 507 | 0.070038428 | 0.125804156 | 0.077091318 | 0.00705289 | -0.048712839 | 0.196165078 | 0.008152306 | 0.123119962 | 0.894954677 | 0.292225259 |
| Rif1 | S | 2181 | 0.102201987 | 0.044192967 | 0.122480835 | 0.020278847 | 0.078287868 | 0.038524276 | 0.267277163 | 0.018538885 | 0.494492579 | 0.043656612 |
| Nes | S | 1858 | 0.01736656 | 0.068116335 | 0.134754337 | 0.117387776 | 0.066638001 | 0.536655846 | 0.068780584 | 0.000733586 | 0.007675765 | 0.075071692 |
| Sufu | S | 346 | 0.044362704 | 0.21220858 | 0.044407851 | 4.51471E-05 | -0.167800728 | 0.400497023 | 0.002195613 | 0.498242342 | 0.999392769 | 0.05932789 |
| Cdk13 | T | 1245 | 0.047070518 | 0.054374356 | 0.172220945 | 0.125150426 | 0.117846589 | 0.175309305 | 0.019987746 | 0.032185827 | 0.062174722 | 0.084193263 |
| Huwe1 | S | 2394 | 0.128077765 | 0.092633321 | 0.208588846 | 0.080511081 | 0.115955526 | 0.067388139 | 0.486364223 | 0.115974093 | 0.451011022 | 0.441785907 |
| Arhgap31 | S | 1054 | 0.052936746 | 0.255580821 | 0.145939895 | 0.093003149 | -0.109640926 | 0.458900643 | 0.098743717 | 0.1317785 | 0.227451846 | 0.399229868 |
| Cbx2 | S | 245 | 0.203644002 | 0.045318317 | 0.200479181 | -0.003164821 | 0.155160864 | 0.097846484 | 0.497925286 | 0.121483955 | 0.978795164 | 0.199113949 |
| Zfp768 | S | 36 | 0.103813399 | 0.215058523 | 0.181201484 | 0.077388085 | -0.033857039 | 0.353455482 | 0.122015876 | 0.231968586 | 0.607585628 | 0.819685677 |
| Dennd1a | T | 707 | -0.06103394 | 0.151568009 | 0.170747553 | 0.231781493 | 0.019179544 | 0.169074327 | 0.001494313 | 0.014256515 | 0.007207483 | 0.649799747 |
| Pnisr | S | 600 | 0.253066125 | 0.157497818 | 0.026838285 | -0.226227839 | -0.130659533 | 0.031824182 | 0.188557722 | 0.83982915 | 0.129818406 | 0.342304919 |
| Mb21d2 | S | 370 | 0.02946161 | 0.177654755 | 0.093394085 | 0.063932475 | -0.08426067 | 0.573316161 | 0.004251675 | 0.178440283 | 0.361387945 | 0.218271845 |
| Setd2 | S | 2053 | 0.078022128 | 0.068200315 | 0.13080493 | 0.052782803 | 0.062604615 | 0.035784222 | 0.052895389 | 0.093716913 | 0.392735698 | 0.323630241 |
| Pimreg | S | 189 | 0.05826116 | 0.069214589 | 0.064026527 | 0.005765366 | -0.005188063 | 0.07127122 | 0.000490769 | 0.088747942 | 0.856341696 | 0.842284118 |
| Raly | T | 304 | 0.047815269 | 0.083141025 | 0.13457485 | 0.086759581 | 0.051433825 | 0.258792087 | 0.06457077 | 0.010794912 | 0.01615522 | 0.030176458 |
| Kbtbd12 | S | 311 | 0.067214657 | 0.039947419 | 0.150081847 | 0.082867191 | 0.110134428 | 0.098064571 | 0.266073505 | 0.005287458 | 0.076550903 | 0.031963317 |
| Rnf169 | S | 236 | -0.026166449 | 0.069968724 | 0.283438842 | 0.309605291 | 0.213470118 | 0.804120344 | 0.098508732 | 0.007107336 | 0.060831991 | 0.013981815 |
| Dok1 | S | 355 | -0.061242737 | 0.261656089 | 0.16870328 | 0.229946016 | -0.092952809 | 0.201536809 | 0.011545689 | 0.038155956 | 0.018167122 | 0.230490466 |
| Rhbdf1 | S | 51 | 0.047625443 | 0.085300688 | 0.052384516 | 0.004759073 | -0.032916172 | 0.044694688 | 0.003267333 | 0.031464124 | 0.731357124 | 0.05987006 |
| Uaca | S | 514 | 0.099447749 | 0.182843142 | 0.108248442 | 0.008800693 | -0.074594699 | 0.365033128 | 0.059077718 | 0.126494584 | 0.932655884 | 0.367863785 |
| Uaca | Y | 512 | 0.099447749 | 0.182843142 | 0.108248442 | 0.008800693 | -0.074594699 | 0.365033128 | 0.059077718 | 0.126494584 | 0.932655884 | 0.367863785 |
| Nhsl1 | S | 1373 | -0.037390375 | 0.240931724 | 0.090267852 | 0.127658227 | -0.150663872 | 0.297881972 | 0.006914977 | 0.01292579 | 0.019186393 | 0.028272442 |
| Nhsl1 | S | 1375 | -0.037390375 | 0.240931724 | 0.090267852 | 0.127658227 | -0.150663872 | 0.297881972 | 0.006914977 | 0.01292579 | 0.019186393 | 0.028272442 |
| Cic | S | 2275 | -0.011177058 | 0.059998639 | 0.175714813 | 0.18689187 | 0.115716174 | 0.842740398 | 0.053927999 | 0.000668407 | 0.041652791 | 0.014964511 |
| Rif1 | S | 1868 | 0.076573349 | 0.063479876 | 0.103664904 | 0.027091555 | 0.040185028 | 0.068079058 | 0.125427891 | 0.009944427 | 0.372089129 | 0.236594643 |
| Magi3 | S | 833 | 0.030661827 | 0.083143304 | 0.106477657 | 0.07581583 | 0.023334353 | 0.413962023 | 0.209734222 | 0.000114996 | 0.119490369 | 0.665817333 |
| Mast4 | S | 1389 | -0.004509405 | 0.17358829 | 0.141157172 | 0.145666576 | -0.032431118 | 0.890840122 | 0.029652435 | 0.018873862 | 0.02896669 | 0.577032428 |
| Dcaf8 | S | 22 | 0.096760577 | 0.420360588 | 0.074927028 | -0.021833549 | -0.34543356 | 0.58641946 | 0.066311953 | 0.549355931 | 0.885321823 | 0.11514044 |
| Dcaf8 | S | 23 | 0.096760577 | 0.420360588 | 0.074927028 | -0.021833549 | -0.34543356 | 0.58641946 | 0.066311953 | 0.549355931 | 0.885321823 | 0.11514044 |
| Rbmx Rbmxl1 | S | 58;58 | 0.095238821 | 0.173446941 | 0.099543737 | 0.004304916 | -0.073903204 | 0.190871354 | 0.031426588 | 0.3244143 | 0.958370738 | 0.401271787 |
| Dcp1a | S | 162 | 0.011635748 | 0.119953662 | 0.11257574 | 0.100939992 | -0.007377923 | 0.717012673 | 0.015231226 | 0.00815393 | 0.011998535 | 0.81588075 |
| Mta3 | S | 425 | 0.061088784 | 0.075048877 | 0.13135349 | 0.070264706 | 0.056304613 | 0.141923086 | 0.056500491 | 0.02485886 | 0.148001098 | 0.205202421 |
| Raly | T | 292 | 0.034774894 | 0.074787908 | 0.125300594 | 0.0905257 | 0.050512685 | 0.298370923 | 0.040835671 | 0.005195982 | 0.011651265 | 0.055656996 |
| Pard3b | S | 990 | 0.016447983 | 0.123228877 | 0.131530369 | 0.115082386 | 0.008301491 | 0.64323148 | 0.009098227 | 0.03654692 | 0.06594332 | 0.839249332 |
| Pard3 | S | 143 | -0.004756517 | 0.120225333 | 0.117883791 | 0.122640308 | -0.002341543 | 0.931627633 | 0.009776675 | 0.005860841 | 0.115331202 | 0.930653754 |
| Ercc4 | S | 521 | 0.091926718 | 0.115121121 | 0.057679817 | -0.034246901 | -0.057441303 | 0.029938046 | 0.045419274 | 0.133252462 | 0.363867746 | 0.237737738 |
| Hnrnpd | S | 80 | -0.07426714 | 0.097349154 | 0.284206582 | 0.358473722 | 0.186857427 | 0.369927952 | 0.056873714 | 0.004162915 | 0.014063669 | 0.021063856 |
| Ptpn4 | S | 403 | 0.22827497 | 0.032495245 | 0.036453323 | -0.191821647 | 0.003958078 | 0.001058916 | 0.596137469 | 0.657465167 | 0.088049421 | 0.963686953 |
| Macf1 | S | 7245 | 0.229470823 | 0.021974609 | 0.161026153 | -0.06844467 | 0.139051544 | 0.034883418 | 0.909821241 | 0.123704087 | 0.430645497 | 0.490674219 |
| Iws1 | S | 460 | 0.058985958 | 0.067983283 | 0.122723708 | 0.063737749 | 0.054740424 | 0.136785991 | 0.066255079 | 0.013023259 | 0.116388471 | 0.132360561 |
| Samd4b | S | 621 | 0.080796577 | 0.118776184 | 0.032182365 | -0.048614212 | -0.086593819 | 0.26078591 | 0.000505353 | 0.216133274 | 0.457876119 | 0.017099968 |
| Xpc | S | 397 | -0.01304785 | 0.105492056 | 0.183286137 | 0.196333988 | 0.077794082 | 0.907488009 | 0.077612745 | 0.003564963 | 0.17073772 | 0.12895493 |
| Scaf1 | S | 1141 | 0.034237527 | 0.112017594 | 0.16579561 | 0.131558083 | 0.053778016 | 0.484083466 | 0.013883243 | 0.078241982 | 0.134940387 | 0.448856993 |
| Fcho2 | T | 494 | 0.063072962 | 0.060636795 | 0.129091971 | 0.066019009 | 0.068455176 | 0.336474899 | 0.169447557 | 0.001694414 | 0.319703943 | 0.137653284 |
| Trp53bp1 | S | 1756 | 0.102489675 | 0.077490364 | 0.080010588 | -0.022479087 | 0.002520225 | 0.005428786 | 0.070405774 | 0.293845249 | 0.733403544 | 0.970520387 |
| Foxp1 | S | 646 | 0.013938905 | 0.063627355 | 0.238765629 | 0.224826724 | 0.175138275 | 0.824947149 | 0.39179299 | 0.002371613 | 0.031352069 | 0.075354243 |
| Foxp1 | S | 648 | 0.013938905 | 0.063627355 | 0.238765629 | 0.224826724 | 0.175138275 | 0.824947149 | 0.39179299 | 0.002371613 | 0.031352069 | 0.075354243 |
| Trim33 | S | 1120 | 0.046543759 | 0.054562134 | 0.182849071 | 0.136305312 | 0.128286937 | 0.296919712 | 0.226166172 | 0.005630637 | 0.03420009 | 0.039626731 |
| Tsc22d4 | T | 183 | 0.046554588 | 0.076434151 | 0.164712247 | 0.118157659 | 0.088278097 | 0.116165064 | 0.161818644 | 0.019387812 | 0.070021996 | 0.160272328 |
| Nf1 | S | 884 | 0.069832358 | 0.127982708 | 0.086572281 | 0.016739923 | -0.041410427 | 0.164925211 | 0.022662103 | 0.101969931 | 0.682486096 | 0.324460446 |
| Ttc14 | T | 660 | 0.16099668 | 0.05031791 | 0.191846457 | 0.030849777 | 0.141528547 | 0.063851481 | 0.599413526 | 0.096694049 | 0.733109164 | 0.252665041 |
| Ovol1 | S | 20 | 0.005726593 | 0.280660382 | 0.243035414 | 0.23730882 | -0.037624968 | 0.985423758 | 0.042850102 | 0.29358543 | 0.511138414 | 0.856862014 |
| Psip1 | T | 271 | -0.041625526 | 0.15519716 | 0.178178478 | 0.219804004 | 0.022981317 | 0.424214055 | 0.028434974 | 0.006969919 | 0.003192013 | 0.617370762 |
| Srrm2 | S | 1972 | 0.070463929 | 0.095579892 | 0.124120526 | 0.053656597 | 0.028540635 | 0.130874263 | 0.024583744 | 0.112362327 | 0.419261861 | 0.638063571 |
| Srrm2 | T | 1974 | 0.070463929 | 0.095579892 | 0.124120526 | 0.053656597 | 0.028540635 | 0.130874263 | 0.024583744 | 0.112362327 | 0.419261861 | 0.638063571 |
| Champ1 | S | 652 | 0.016949253 | 0.199590147 | 0.100254161 | 0.083304908 | -0.099335985 | 0.829421603 | 0.0105892 | 0.095405611 | 0.277274786 | 0.072764065 |
| Cic | S | 2284 | -0.003994298 | 0.071770828 | 0.168218151 | 0.172212449 | 0.096447323 | 0.950995479 | 0.055524166 | 0.001371534 | 0.071744168 | 0.014049624 |
| Mta1 | S | 449 | 0.089926105 | 0.152064544 | 0.041505521 | -0.048420584 | -0.110559023 | 0.102001301 | 0.006456078 | 0.517961853 | 0.450805334 | 0.14287074 |
| Nes | T | 630 | 0.248009838 | 0.078216606 | 0.072106724 | -0.175903113 | -0.006109881 | 0.019843464 | 0.406825609 | 0.442577882 | 0.05368007 | 0.935922423 |
| Ino80 | S | 48 | 0.092043772 | 0.014019616 | 0.14582029 | 0.053776519 | 0.131800675 | 0.227098412 | 0.859923322 | 0.00034629 | 0.429374275 | 0.192069618 |
| Dennd2a | T | 356 | 0.088198067 | 0.149324364 | 0.125690684 | 0.037492617 | -0.023633679 | 0.299750316 | 0.12694335 | 0.050840013 | 0.615467108 | 0.760236309 |
| Armc10 | S | 41 | -0.00313751 | 0.236572802 | 0.120934604 | 0.124072114 | -0.115638198 | 0.968742081 | 0.004480186 | 0.320724819 | 0.331162163 | 0.335684872 |
| Wdr13 | S | 79 | 0.16105052 | 0.103519546 | -0.007117483 | -0.168168003 | -0.11063703 | 0.002389349 | 0.061304032 | 0.864719648 | 0.012098388 | 0.056992055 |
| Srp68 | S | 26 | 0.128014129 | 0.080036496 | 0.081477535 | -0.046536594 | 0.001441039 | 0.027239539 | 0.273938112 | 0.052513882 | 0.303665837 | 0.982113393 |
| Arhgap31 | S | 1093 | 0.004218721 | 0.129841713 | 0.107124793 | 0.102906072 | -0.02271692 | 0.919588349 | 0.01153462 | 0.008988456 | 0.056677153 | 0.481819997 |
| Retreg3 | S | 313 | 0.010574515 | 0.09456327 | 0.225758064 | 0.215183549 | 0.131194794 | 0.786629159 | 0.057515654 | 0.023487056 | 0.017261695 | 0.068418348 |
| Kmt2d | S | 691 | 0.052151632 | 0.263164384 | 0.164892154 | 0.112740522 | -0.09827223 | 0.564335114 | 0.040994706 | 0.377791854 | 0.517567731 | 0.580769281 |
| Kmt2d | S | 709 | 0.052151632 | 0.263164384 | 0.164892154 | 0.112740522 | -0.09827223 | 0.564335114 | 0.040994706 | 0.377791854 | 0.517567731 | 0.580769281 |
| Trp53bp1 | S | 429 | 0.016000328 | 0.102000015 | 0.210622375 | 0.194622046 | 0.10862236 | 0.54957668 | 0.031400515 | 0.049382892 | 0.041424212 | 0.145719651 |
| Cdk11b | S | 234 | 0.104513805 | 0.121150507 | 0.01968808 | -0.084825725 | -0.101462427 | 0.019770436 | 0.005550663 | 0.861952879 | 0.479167948 | 0.40864207 |
| Xpc | S | 395 | -0.034408768 | 0.119406094 | 0.168403752 | 0.20281252 | 0.048997658 | 0.750125365 | 0.034479467 | 0.004560328 | 0.150234095 | 0.211137316 |

| | | | | | | | | | | | | | |
|---|---|---|---|---|---|---|---|---|---|---|---|---|---|
| Pkp4 | S | | 334 | 0.011863182 | 0.134616389 | 0.232257029 | 0.220393847 | 0.09764064 | 0.90681735 | 0.080125393 | 0.03836079 | 0.108433715 | 0.282733972 |
| Zfp638 | S | | 490 | 0.109608578 | 0.196206034 | 0.200358032 | 0.090749454 | 0.004151998 | 0.546440399 | 0.147418015 | 0.149594293 | 0.633668878 | 0.975850643 |
| Necap2 | S | | 181 | -0.011765298 | 0.358363242 | 0.046328563 | 0.058093861 | -0.312034679 | 0.908587425 | 0.005631163 | 0.582412772 | 0.613611013 | 0.022385078 |
| Pkn2 | S | | 619 | -0.056058098 | 0.131651905 | 0.213726015 | 0.269784113 | 0.08207411 | 0.055669425 | 0.055503771 | 0.000517833 | 0.001095722 | 0.161340985 |
| Ogfrl1 | S | | 65 | -0.073870638 | 0.221664185 | 0.228478753 | 0.302349391 | 0.006814568 | 0.305128486 | 0.028027682 | 0.030784901 | 0.017207718 | 0.935766933 |
| Ankrd17 | S | | 2401 | 0.201971729 | 0.01152176 | 0.033930565 | -0.168041163 | 0.022408805 | 0.000404396 | 0.868434119 | 0.349101269 | 0.004269973 | 0.745287844 |
| Rab11fip2 | S | | 277 | 0.025623452 | 0.157043243 | 0.138038472 | 0.112415019 | -0.019004772 | 0.600449578 | 0.013136192 | 0.115493925 | 0.174466992 | 0.772201364 |
| Cnot2 | S | | 169 | 0.112474412 | 0.168608868 | 0.131100554 | 0.018626142 | -0.037508313 | 0.196538247 | 0.078118567 | 0.292416505 | 0.845475388 | 0.705008004 |
| Pnisr | S | | 613 | 0.077670735 | 0.104630771 | 0.095008081 | 0.017337345 | -0.009622691 | 0.08459355 | 0.028004566 | 0.136116489 | 0.742758589 | 0.853055899 |
| Bclaf1 | T | | 256 | 0.025534729 | 0.152179753 | 0.078931346 | 0.053396617 | -0.073248407 | 0.680275874 | 0.004609799 | 0.053527451 | 0.381566171 | 2.59321E-05 |
| Tjp2 | S | | 968 | -0.029041417 | -0.009628611 | 0.338684797 | 0.367726214 | 0.348313408 | 0.535832071 | 0.76576323 | 0.000265484 | 0.001113966 | 0.006795317 |
| Msn | T | | 469 | -0.03755774 | 0.221254059 | 0.111379932 | 0.148937673 | -0.109874126 | 0.377608978 | 0.003060347 | 0.070209255 | 0.033980155 | 0.086866625 |
| Sntb2 | S | | 90 | -0.000916919 | 0.113781473 | 0.13221507 | 0.13313199 | 0.018433598 | 0.929974068 | 0.013627778 | 0.008568257 | 0.002308159 | 0.409897318 |
| Mast2 | S | | 817 | -0.101722875 | 0.287753054 | 0.231019975 | 0.33274285 | -0.056733078 | 0.076760137 | 0.010561709 | 0.043727628 | 0.036080576 | 0.542834825 |
| Trim33 | T | | 1117 | 0.046057653 | 0.055544556 | 0.195200592 | 0.149142939 | 0.139656035 | 0.349903736 | 0.209197023 | 0.008166997 | 0.038616171 | 0.036252563 |
| Rb1 | S | | 804 | -0.031469415 | 0.044333886 | 0.235183769 | 0.266653185 | 0.190849883 | 0.414937808 | 0.349934986 | 0.000160983 | 0.002405686 | 0.016944011 |
| Rprd2 | S | | 392 | -0.010512537 | 0.054000694 | 0.181955414 | 0.192467952 | 0.12795472 | 0.800142256 | 0.063612962 | 0.000742785 | 0.013941124 | 0.007567512 |
| Washc2 | S | | 613 | 0.116981724 | 0.074811092 | 0.102362219 | -0.014619505 | 0.027551127 | 0.123350251 | 0.102092294 | 0.040015887 | 0.798731888 | 0.237136307 |
| Flna | S | | 16 | 0.030964792 | 0.097068552 | 0.161364718 | 0.130399926 | 0.064296166 | 0.44453752 | 0.030592091 | 0.033093106 | 0.071378363 | 0.263042179 |
| Ezh2 | T | | 367 | 0.114888824 | 0.07281056 | 0.116872391 | 0.001983567 | 0.044061832 | 0.004272343 | 0.150625779 | 0.418982812 | 0.987865365 | 0.744732357 |
| Retreg2 | S | | 342 | 0.141186541 | 0.135966734 | 0.064798725 | -0.076387815 | -0.071168008 | 0.134949598 | 0.033474992 | 0.323652343 | 0.375043623 | 0.248297715 |
| Hipk2 | T | | 838 | 0.025594306 | 0.134946743 | 0.105194166 | 0.07959986 | -0.029752576 | 0.364791204 | 0.118701506 | 0.002868105 | 0.021478954 | 0.643617813 |
| Clip2 | S | | 203 | 0.014176767 | 0.055581949 | 0.150529998 | 0.136353231 | 0.094948049 | 0.728871269 | 0.08715719 | 0.000697005 | 0.046646728 | 0.025783155 |
| Dennd4c | T | | 966 | 0.071674595 | 0.11603658 | 0.067205072 | -0.004469523 | -0.048831508 | 0.12953048 | 0.014843575 | 0.093981901 | 0.901005297 | 0.150702206 |
| Atn1 | S | | 646 | 0.036686632 | 0.123029188 | 0.083340154 | 0.046653523 | -0.039689033 | 0.241374058 | 0.001225972 | 0.242576317 | 0.470453693 | 0.526165208 |
| Srrm2 | S | | 1037 | 0.056843461 | 0.128152009 | 0.147399519 | 0.090556058 | 0.01924751 | 0.185348031 | 0.204480839 | 0.029932862 | 0.123528023 | 0.825185839 |
| E2f1 | S | | 329 | -0.07223373 | 0.213418676 | 0.221210397 | 0.293444127 | 0.007791721 | 0.170398121 | 0.007361197 | 0.050490441 | 0.027183874 | 0.922214132 |
| Rbbp6 | S | | 773 | 0.064981357 | 0.118831956 | 0.287115313 | 0.222133956 | 0.168283357 | 0.139433366 | 0.369091162 | 0.122187552 | 0.194622487 | 0.340576167 |
| Cdc42ep2 | S | | 145 | 0.07370792 | 0.143348961 | 0.117369508 | 0.043661587 | -0.025979453 | 0.092902076 | 0.079483692 | 0.170403075 | 0.540546497 | 0.740894347 |
| Prrc2c | S | | 1215 | 0.047671612 | 0.033609157 | 0.09989014 | 0.052218528 | 0.066280983 | 0.113761732 | 0.2278686 | 0.00016834 | 0.100756697 | 0.063892306 |
| Havcr2 | S | | 229 | 0.700079128 | -0.439575785 | 0.460596506 | -0.239482623 | 0.900172291 | 0.094636415 | 0.476700597 | 0.283067374 | 0.39460988 | 0.179096384 |
| Peak1 | S | | 691 | 0.08605163 | 0.023518781 | 0.112458653 | 0.026407023 | 0.088939872 | 0.022994682 | 0.532076797 | 0.004177288 | 0.299656078 | 0.057438148 |
| Cenpv | S | | 105 | 0.06542439 | 0.072870723 | 0.15044305 | 0.08501866 | 0.077572327 | 0.498296012 | 0.181587594 | 0.0040493 | 0.389080928 | 0.146278551 |
| Cep350 | T | | 1253 | 0.009784797 | 0.111412932 | 0.174691695 | 0.164906898 | 0.063278763 | 0.774611711 | 0.017150878 | 0.002280224 | 0.006418928 | 0.463950804 |
| Mki67 | S | | 128 | 0.100771579 | 0.096651262 | 0.096102367 | -0.004669212 | -0.000548895 | 0.048652833 | 0.049238098 | 0.215624506 | 0.940802385 | 0.992815103 |
| Tns2 | S | | 1087 | -0.029063683 | 0.130990921 | 0.191931999 | 0.220995682 | 0.060941079 | 0.497327604 | 0.004118483 | 0.04566558 | 0.02853915 | 0.356798572 |
| Ppfibp1 | S | | 753 | 0.106657833 | 0.150213673 | 0.100631133 | -0.0060267 | -0.049582541 | 0.394213808 | 0.035125913 | 0.132961452 | 0.956342828 | 0.200553661 |
| Prpf4b | S | | 383 | 0.124109624 | 0.103045468 | 0.201891445 | 0.077781821 | 0.098845977 | 0.330877494 | 0.242934993 | 0.075907604 | 0.541474134 | 0.318867825 |
| Srsf2 | S | | 26 | 0.109262024 | 0.141243908 | 0.189587531 | 0.080325507 | 0.048343624 | 0.621675125 | 0.069550459 | 0.122124389 | 0.722733126 | 0.622558241 |
| Cdc42ep1 | T | | 378 | 0.128827993 | 0.092681468 | 0.153411518 | 0.024583525 | 0.06073005 | 0.486850231 | 0.338706787 | 0.008891385 | 0.887240023 | 0.507831605 |
| Numa1 | S | | 1710 | 0.095452094 | 0.199088702 | -0.033812151 | -0.129264245 | -0.232900853 | 0.204069763 | 0.001165927 | 0.840533286 | 0.477596774 | 0.250941796 |
| Ncam1 | S | | 774 | 0.139858861 | 0.200916094 | 0.037529066 | -0.102329795 | -0.163387028 | 0.120050765 | 0.039401477 | 0.673586738 | 0.135628642 | 0.054538342 |
| Sh3kbp1 | S | | 631 | 0.009961907 | 0.134261824 | 0.188264949 | 0.178303043 | 0.054003126 | 0.891782243 | 0.088427026 | 0.018365725 | 0.036434723 | 0.308808946 |
| Prr12 | S | | 441 | 0.033103468 | 0.122922211 | 0.196878957 | 0.16377549 | 0.073956746 | 0.700324758 | 0.015457893 | 0.119115934 | 0.202457243 | 0.456616397 |
| Nipbl | S | | 2503 | 0.013080489 | 0.186354096 | 0.105056424 | 0.091975935 | -0.081297671 | 0.838092443 | 0.002922105 | 0.252099309 | 0.34530572 | 0.351872234 |
| Nipbl | S | | 2505 | 0.013080489 | 0.186354096 | 0.105056424 | 0.091975935 | -0.081297671 | 0.838092443 | 0.002922105 | 0.252099309 | 0.34530572 | 0.351872234 |
| Nipbl | S | | 2507 | 0.013080489 | 0.186354096 | 0.105056424 | 0.091975935 | -0.081297671 | 0.838092443 | 0.002922105 | 0.252099309 | 0.34530572 | 0.351872234 |
| Tra2b | S | | 99 | -0.044208013 | 0.079341179 | 0.251051081 | 0.295259094 | 0.171709902 | 0.504459235 | 0.23239262 | 0.00113308 | 0.013485241 | 0.039771536 |
| Sun1 | S | | 99 | 0.057927408 | 0.082553438 | 0.217832359 | 0.159904951 | 0.135278922 | 0.459093787 | 0.229340062 | 0.02230747 | 0.108369637 | 0.117587363 |
| Ppp4r3a | S | | 126 | 0.069659278 | 0.088046859 | 0.181616649 | 0.111957371 | 0.09356979 | 0.234298822 | 0.091380514 | 0.070526138 | 0.201965063 | 0.253959115 |
| Tgfbr2 | S | | 578 | -0.073429086 | 0.292542787 | 0.232433627 | 0.305862712 | -0.06010916 | 0.122573784 | 0.017150878 | 0.0769691 | 0.036725853 | 0.537555392 |
| Nes | S | | 297 | 0.059397272 | 0.152477259 | 0.031760039 | -0.027637233 | -0.120717221 | 0.117710081 | 0.002548652 | 0.452137148 | 0.530220991 | 0.042173302 |
| Ptpn14 | S | | 811 | 0.091622697 | 0.130545644 | 0.05877809 | -0.032844608 | -0.071767554 | 0.097846366 | 0.025577979 | 0.18346421 | 0.516203902 | 0.171067285 |
| Oip5 | S | | 215 | 0.022710481 | 0.174464691 | 0.130413071 | 0.10770253 | -0.044051168 | 0.800676071 | 0.009742845 | 0.148314763 | 0.3239083 | 0.563179997 |
| Ilf3 | T | | 67 | 0.093912939 | 0.075321173 | 0.131833197 | 0.037920258 | 0.056512024 | 0.094935953 | 0.10964326 | 0.071119728 | 0.540071504 | 0.351346569 |
| Lrrk2 | S | | 973 | -0.179232393 | 0.193806506 | 0.353757072 | 0.532989466 | 0.159950567 | 0.107642403 | 0.053293003 | 0.005931426 | 0.004901732 | 0.113664064 |
| Cactin | S | | 80 | -0.001767617 | 0.12462274 | 0.102067618 | 0.103835234 | -0.022555122 | 0.961843486 | 0.002987659 | 0.021591574 | 0.044669937 | 0.476604463 |
| Ripor1 | S | | 748 | 0.069019177 | 0.310184548 | 0.081816896 | 0.012797718 | -0.228367652 | 0.524480617 | 0.026918369 | 0.618958294 | 0.939748658 | 0.231721034 |
| Rere | S | | 613 | 0.011628509 | 0.168081938 | 0.096776385 | 0.085147875 | -0.071305554 | 0.755695823 | 0.00410035 | 0.123698658 | 0.159927942 | 0.206598975 |
| Mki67 | S | | 2254 | 0.23706428 | 0.205156023 | 0.015141614 | -0.221922666 | -0.190014409 | 0.106329124 | 0.094918308 | 0.881198188 | 0.147094858 | 0.161489047 |
| Phc3 | S | | 261 | -0.029207835 | 0.056087335 | 0.190086721 | 0.219294556 | 0.133999386 | 0.637889316 | 0.153776282 | 0.00024626 | 0.041368965 | 0.018391098 |
| Osbpl6 | S | | 229 | 0.154978538 | 0.066628167 | 0.051352439 | -0.103626098 | -0.015275728 | 0.003786543 | 0.194630617 | 0.436447926 | 0.164597478 | 0.810218492 |
| Cyld | S | | 414 | 0.195519379 | 0.281216674 | 0.058628882 | -0.136890497 | -0.222587792 | 0.423404068 | 0.047194301 | 0.752599297 | 0.616927197 | 0.301447512 |
| Sugp2 | S | | 838 | 0.022457512 | 0.135621937 | 0.141687567 | 0.119230055 | 0.00606563 | 0.738140835 | 0.015008617 | 0.061618478 | 0.15201774 | 0.906067413 |
| Mki67 | S | | 2103 | 0.037511651 | 0.155824332 | 0.045575429 | 0.008063778 | -0.110248903 | 0.574086643 | 0.000429061 | 0.338803464 | 0.909325641 | 0.075309454 |
| Puf60 | S | | 117 | 0.275984997 | 0.092591641 | 0.098136483 | -0.177848514 | 0.005544842 | 0.015175716 | 0.632484278 | 0.70752997 | 0.504394859 | 0.984715185 |
| Tcof1 | T | | 171 | 0.202647342 | 0.057328186 | 0.146480485 | -0.056166857 | 0.089152299 | 0.41137841 | 0.078590785 | 0.0517582 | 0.804470706 | 0.175355776 |
| Spen | S | | 1397 | 0.058784689 | 0.052274767 | 0.088879192 | 0.030094503 | 0.036604425 | 0.067835388 | 0.124587345 | 0.00226121 | 0.25218181 | 0.233447735 |
| Ncor1 | S | | 779 | -0.000983483 | 0.116473848 | 0.233510015 | 0.234493498 | 0.117036167 | 0.980473061 | 0.075966692 | 0.026680684 | 0.022189767 | 0.14174828 |
| Cbx8 | S | | 284 | -0.024310606 | 0.057900422 | 0.221765973 | 0.24607658 | 0.163865551 | 0.612087137 | 0.358155243 | 0.000348688 | 0.00752604 | 0.057404766 |
| Sh3bp2 | S | | 414 | 0.078425831 | 0.137866142 | 0.083207902 | 0.004782071 | -0.054658241 | 0.0997104 | 0.049991145 | 0.15271471 | 0.927697387 | 0.372685623 |
| Scaf1 | S | | 577 | 0.047018842 | 0.071893841 | 0.146715791 | 0.099696949 | 0.074821949 | 0.343669767 | 0.091264941 | 0.009788802 | 0.087863541 | 0.10154714 |
| Scaf1 | S | | 579 | 0.047018842 | 0.071893841 | 0.146715791 | 0.099696949 | 0.074821949 | 0.343669767 | 0.091264941 | 0.009788802 | 0.087863541 | 0.10154714 |
| Cavin1 | S | | 381 | 0.123274666 | 0.119318012 | 0.006217052 | -0.117057614 | -0.11310096 | 0.028322814 | 0.00638623 | 0.915145571 | 0.130674851 | 0.14028564 |
| Htatsf1 | S | | 705 | -0.029938956 | 0.106013871 | 0.177018106 | 0.206957062 | 0.071004234 | 0.41990172 | 0.03580754 | 0.002513327 | 0.002528285 | 0.109064688 |
| Gzf1 | S | | 198 | 0.227279298 | 0.052647171 | 0.150739316 | -0.076539982 | 0.098092146 | 0.071862124 | 0.579078773 | 0.167826058 | 0.381488219 | 0.146781568 |
| Spp1 | Y | | 166 | 0.089477173 | 0.112797112 | 0.023667031 | -0.065810142 | -0.089130081 | 0.171978795 | 0.000275203 | 0.638000654 | 0.342219743 | 0.164475525 |
| Rpl18a | Y | | 10 | 0.027839694 | 0.11315523 | 0.216958464 | 0.18911877 | 0.103803234 | 0.519938658 | 0.038036664 | 0.092848306 | 0.107391405 | 0.293126336 |
| Ndufs4 | S | | 9 | 0.014699563 | 0.116478525 | 0.126639124 | 0.111939561 | 0.010160598 | 0.766015637 | 0.002067358 | 0.092332058 | 0.142822531 | 0.848961181 |
| Nup88 | S | | 170 | 0.053008408 | 0.114349138 | 0.306428832 | 0.253420424 | 0.192079694 | 0.63627236 | 0.431736505 | 0.039394915 | 0.104981155 | 0.247091814 |
| Mcm2 | S | | 139 | 0.018998487 | 0.089774584 | 0.08432928 | 0.065330793 | -0.005445304 | 0.306180734 | 0.001415066 | 0.030564616 | 0.054371538 | 0.816260955 |
| Matr3 | S | | 619 | -0.031262489 | 0.095102817 | 0.203109848 | 0.234372337 | 0.108007031 | 0.452362983 | 0.019794983 | 0.006846332 | 0.0065403 | 0.06445213 |
| Cbx3 | S | | 102 | 0.024610788 | 0.095243442 | 0.228838203 | 0.204227415 | 0.133594761 | 0.618755369 | 0.081573445 | 0.039133193 | 0.068298757 | 0.155187981 |
| Luc7l2 | S | | 354 | 0.095114247 | 0.08091164 | 0.282913017 | 0.18779877 | 0.202001377 | 0.552024719 | 0.643673696 | 0.023240646 | 0.295024739 | 0.307997343 |
| Ppig | S | | 252 | 0.009853307 | 0.05790491 | 0.181263979 | 0.171410672 | 0.123359069 | 0.879732505 | 0.061716257 | 0.003192998 | 0.071726173 | 0.021974344 |
| Nck2 | S | | 90 | 0.031700081 | 0.115153716 | 0.128273903 | 0.096573823 | 0.013120188 | 0.226994239 | 0.029807289 | 0.057800609 | 0.113844557 | 0.797877383 |
| Cdk11b | S | | 65 | 0.106650099 | 0.107239347 | 0.067388802 | -0.039261296 | -0.039850545 | 0.031640615 | 0.04989015 | 0.303522675 | 0.510168243 | 0.520966969 |
| Ercc5 | S | | 384 | 0.127578001 | 0.182712073 | 0.020493034 | -0.107084967 | -0.162219039 | 0.091920023 | 0.019535266 | 0.853990298 | 0.353542626 | 0.207379173 |
| Nop56 | S | | 466 | 0.005716434 | 0.099616555 | 0.082541115 | 0.076824681 | -0.01707544 | 0.941825621 | 0.00312365 | 0.003701407 | 0.377138032 | 0.381126716 |
| Fosl2 | S | | 19 | -0.237257777 | 0.698419748 | 1.103683596 | 1.340941373 | 0.405263848 | 0.193424956 | 0.336624164 | 0.266313516 | 0.205318962 | 0.682803024 |
| Dock7 | S | | 864 | 0.112571238 | 0.097889463 | 0.036827754 | -0.075743483 | -0.061061709 | 0.022395149 | 0.011964275 | 0.5703945 | 0.285326118 | 0.361616978 |
| Proser3 | S | | 588 | -0.119517908 | 0.280983967 | 0.319391593 | 0.438909501 | 0.038407626 | 0.07294157 | 0.061776854 | 0.012209507 | 0.008939769 | 0.764120926 |
| Mfap1a Mfap1b | S | 52;52 | | 0.042607247 | 0.042867895 | 0.125309215 | 0.082701968 | 0.08244132 | 0.145532222 | 0.148188527 | 0.001994175 | 0.006347938 | 0.008063882 |
| Mfap1a Mfap1b | S | 53;53 | | 0.042607247 | 0.042867895 | 0.125309215 | 0.082701968 | 0.08244132 | 0.145532222 | 0.148188527 | 0.001994175 | 0.006347938 | 0.008063882 |

| | | | | | | | | | | | | |
|---|---|---|---|---|---|---|---|---|---|---|---|---|
| Lrrc8c | S | 215 | -0.106872633 | 0.075687071 | 0.264454656 | 0.371327289 | 0.188767584 | 0.025204849 | 0.0323707 | 8.67438E-05 | 0.000415372 | 0.004156848 |
| Kmt2a | S | 2865 | 0.137081428 | 0.180485582 | 0.14205282 | 0.004971392 | -0.038432762 | 0.239743215 | 0.148881957 | 0.27862652 | 0.969929475 | 0.77448905 |
| Kmt2a | S | 2866 | 0.137081428 | 0.180485582 | 0.14205282 | 0.004971392 | -0.038432762 | 0.239743215 | 0.148881957 | 0.27862652 | 0.969929475 | 0.77448905 |
| Kmt2a | S | 2867 | 0.137081428 | 0.180485582 | 0.14205282 | 0.004971392 | -0.038432762 | 0.239743215 | 0.148881957 | 0.27862652 | 0.969929475 | 0.77448905 |
| Kmt2a | S | 2868 | 0.137081428 | 0.180485582 | 0.14205282 | 0.004971392 | -0.038432762 | 0.239743215 | 0.148881957 | 0.27862652 | 0.969929475 | 0.77448905 |
| Lig1 | S | 51 | 0.01658265 | 0.187098233 | 0.047988565 | 0.031405914 | -0.139109668 | 0.615718359 | 0.000949754 | 0.351652674 | 0.474444016 | 0.047622224 |
| Setd2 | S | 2055 | 0.081598174 | 0.058283596 | 0.136214685 | 0.05461651 | 0.077931088 | 0.044728873 | 0.246281311 | 0.038756842 | 0.283371581 | 0.199461665 |
| Akap9 | S | 2807 | 0.195341495 | 0.115929764 | 0.058884595 | -0.1364569 | -0.057045169 | 0.146430392 | 0.043811485 | 0.376424563 | 0.269196742 | 0.350983557 |
| Tcf20 | T | 1790 | -0.070843362 | 0.188738129 | 0.204626965 | 0.275470327 | 0.015888836 | 0.071610534 | 0.001192921 | 0.059633023 | 0.047337351 | 0.826451802 |
| Brd8 | S | 268 | 0.071690849 | 0.04969081 | 0.124990772 | 0.053299923 | 0.075299962 | 0.077696039 | 0.208900991 | 0.01144178 | 0.00252201 | 0.028468818 |
| Ufl1 | S | 753 | 0.115290954 | 0.054487757 | 0.139229904 | 0.023938951 | 0.084742148 | 0.605010788 | 0.434055836 | 0.00075731 | 0.910916725 | 0.263868769 |
| Nelfe | S | 357 | 0.166677541 | 0.08040565 | 0.063304335 | -0.103373205 | -0.017101314 | 0.015020246 | 0.19054458 | 0.366625809 | 0.18480815 | 0.810170199 |
| Tcof1 | S | 592 | 0.048383752 | 0.100807092 | 0.209775103 | 0.161391352 | 0.108968011 | 0.386116147 | 0.077156784 | 0.078554476 | 0.127050672 | 0.248065039 |
| Cpeb4 | S | 99 | -0.003564048 | 0.135906802 | 0.113026013 | 0.11659006 | -0.022880789 | 0.933734327 | 0.011380796 | 0.015136803 | 0.040503419 | 0.53631131 |
| Wapl | S | 77 | 0.063051941 | 0.051340029 | 0.167383912 | 0.104331971 | 0.116043883 | 0.127782408 | 0.30627551 | 0.013987833 | 0.072744921 | 0.075966551 |
| Ppfibp1 | S | 428 | 0.18442188 | 0.029995533 | 0.266181736 | 0.081759856 | 0.236186203 | 0.186378884 | 0.751730714 | 0.088938929 | 0.574119614 | 0.119218916 |
| Ppfibp1 | T | 427 | 0.18442188 | 0.029995533 | 0.266181736 | 0.081759856 | 0.236186203 | 0.186378884 | 0.751730714 | 0.088938929 | 0.574119614 | 0.119218916 |
| Frmd6 | T | 523 | 0.095442665 | 0.148365712 | 0.069950008 | -0.025492657 | -0.078415704 | 0.035314244 | 0.073546738 | 0.368797234 | 0.710173044 | 0.370236829 |
| Zbtb1 | S | 355 | 0.017484596 | 0.094059221 | 0.072753571 | 0.055268974 | -0.02130565 | 0.590278849 | 0.004355398 | 0.003606413 | 0.154595755 | 0.298940257 |
| Kmt2a | S | 1839 | 0.074191742 | 0.154396833 | 0.113044285 | 0.038852542 | -0.041352548 | 0.142837792 | 0.193621659 | 0.058062404 | 0.464490197 | 0.679092235 |
| Carmil1 | S | 1295 | 0.095855129 | 0.196054812 | 0.017273557 | -0.078581572 | -0.178781255 | 0.158061693 | 0.009001649 | 0.757918621 | 0.166003359 | 0.018690347 |
| Cdk11b | S | 578 | 0.069638707 | 0.148643667 | 0.095300913 | 0.025662206 | -0.053342754 | 0.304812017 | 0.033823204 | 0.115698712 | 0.705540752 | 0.32668412 |
| Ep400 | T | 320 | 0.098601842 | 0.072581241 | 0.170451491 | 0.071849649 | 0.097870249 | 0.150539952 | 0.222674034 | 0.062211755 | 0.360395609 | 0.220814842 |
| Hp1bp3 | T | 51 | -0.007237731 | 0.100665518 | 0.150713272 | 0.157951003 | 0.050047754 | 0.804708226 | 0.005526679 | 0.020306897 | 0.024159761 | 0.267795479 |
| Arvcf | S | 347 | 0.054367022 | 0.179981844 | 0.142200663 | 0.087833641 | -0.037781181 | 0.434373702 | 0.07733265 | 0.111619199 | 0.239971756 | 0.660439977 |
| Exoc2 | T | 440 | 0.107207799 | 0.08002995 | 0.102786167 | -0.004421632 | 0.022756217 | 0.263311049 | 0.00418757 | 0.195127065 | 0.962969139 | 0.719753928 |
| Rprd1b | S | 134 | 0.014648585 | 0.092225336 | 0.131484034 | 0.116835449 | 0.039258698 | 0.258784094 | 0.021069706 | 0.021122045 | 0.041924575 | 0.293392285 |
| Ncoa5 | S | 126 | -0.019761833 | 0.119905868 | 0.156822507 | 0.17658434 | 0.036916639 | 0.763197338 | 0.005234494 | 0.033913393 | 0.064658297 | 0.463902131 |
| Mki67 | S | 2017 | 0.122575722 | 0.133699088 | 0.040033326 | -0.082542396 | -0.093665762 | 0.150981933 | 0.017320821 | 0.265784632 | 0.284595855 | 0.061789812 |
| Srsf2 | T | 25 | 0.28226264 | 0.12327095 | 0.161044794 | -0.121217846 | 0.037773845 | 0.472133773 | 0.163182003 | 0.193228118 | 0.746207058 | 0.733779202 |
| Ppp2r5e | S | 33 | -0.033486693 | 0.233348642 | 0.131282118 | 0.164768811 | -0.102066524 | 0.371528466 | 0.058726284 | 0.006265434 | 0.006844204 | 0.276336018 |
| Tnrc18 | S | 2622 | -0.088031634 | 0.18645989 | 0.192209156 | 0.28024079 | 0.005749266 | 0.314282268 | 0.014484664 | 0.01815242 | 0.031094056 | 0.920919481 |
| Rnf20 | S | 138 | 0.053074234 | 0.053752094 | 0.18297777 | 0.129903535 | 0.129225675 | 0.156408543 | 0.269992125 | 0.016289553 | 0.066758287 | 0.063111527 |
| Arid4a | S | 1167 | -0.040339013 | 0.136677848 | 0.207668751 | 0.248007765 | 0.070990904 | 0.678121015 | 0.01150242 | 0.049294521 | 0.076000179 | 0.34410698 |
| Bcr | S | 1263 | -0.079504835 | 0.310238854 | 0.149202133 | 0.228706968 | -0.16103672 | 0.102798944 | 0.017136298 | 0.023366571 | 0.007523293 | 0.100020166 |
| Nfia | S | 281 | 0.119357513 | 0.227477132 | 0.120660567 | 0.001303054 | -0.106816565 | 0.348486142 | 0.076650327 | 0.41126065 | 0.993616125 | 0.489347889 |
| Abcb1b | S | 665 | 0.008059013 | 0.197570242 | 0.086729207 | 0.078670195 | -0.110841034 | 0.859289895 | 0.005110017 | 0.173250034 | 0.222859422 | 0.113726569 |
| Abcb1b | S | 669 | 0.008059013 | 0.197570242 | 0.086729207 | 0.078670195 | -0.110841034 | 0.859289895 | 0.005110017 | 0.173250034 | 0.222859422 | 0.113726569 |
| Ncoa5 | S | 381 | 0.022629295 | 0.120554619 | 0.165663936 | 0.143034641 | 0.045109317 | 0.61290727 | 0.017650983 | 0.083165634 | 0.110561474 | 0.52308119 |
| Mprip | S | 221 | -0.012141527 | 0.085359309 | 0.133219453 | 0.14536098 | 0.047860143 | 0.594283899 | 0.010267822 | 0.002318524 | 0.003607917 | 0.116130432 |
| Tjp1 | Y | 132 | 0.034942608 | 0.200787338 | 0.210568647 | 0.17562604 | 0.00978131 | 0.803601171 | 0.087254407 | 0.128694564 | 0.313214835 | 0.935385765 |
| Eloa | S | 123 | 0.096539353 | 0.11614367 | 0.128498539 | 0.031959186 | 0.012354869 | 0.073587083 | 0.218583014 | 0.120588789 | 0.636733408 | 0.893168741 |
| Aff4 | T | 669 | 0.112185512 | -0.001826447 | 0.102514601 | -0.009670911 | 0.104341048 | 0.002025314 | 0.947136819 | 0.023715049 | 0.749589472 | 0.029342211 |
| Oser1 | S | 82 | 0.101482241 | 0.084859999 | 0.079917555 | -0.021564685 | -0.004942444 | 0.245384627 | 0.015571805 | 0.061898641 | 0.775240021 | 0.866403577 |
| Oser1 | S | 84 | 0.101482241 | 0.084859999 | 0.079917555 | -0.021564685 | -0.004942444 | 0.245384627 | 0.015571805 | 0.061898641 | 0.775240021 | 0.866403577 |
| Snrnp27 | T | 60 | 0.042828859 | 0.112680495 | 0.117049264 | 0.074220404 | 0.004368769 | 0.335277224 | 0.071143153 | 0.018951557 | 0.121727057 | 0.928036863 |
| Epn2 | S | 192 | 0.067864912 | 0.107299369 | 0.098291919 | 0.030427007 | -0.009007451 | 0.198061939 | 0.01103471 | 0.171458217 | 0.645092316 | 0.874362102 |
| Epn2 | S | 195 | 0.067864912 | 0.107299369 | 0.098291919 | 0.030427007 | -0.009007451 | 0.198061939 | 0.01103471 | 0.171458217 | 0.645092316 | 0.874362102 |
| Rbmx2 | S | 223 | 0.121086865 | 0.090857298 | 0.176528074 | 0.055441209 | 0.085670776 | 0.132896097 | 0.27434584 | 0.125118601 | 0.582214143 | 0.425614509 |
| Psmc1 | T | 53 | 0.089727721 | 0.031744246 | 0.222845449 | 0.133117728 | 0.191101203 | 0.259932255 | 0.644580619 | 0.013786246 | 0.078757355 | 0.011286847 |
| Dnajc1 | S | 478 | -0.004902377 | 0.105462287 | 0.140377811 | 0.145280188 | 0.034915524 | 0.94601636 | 0.014451182 | 0.011630049 | 0.121562231 | 0.331338307 |
| Arfgef2 | S | 1521 | 0.079348982 | 0.110149484 | 0.062436237 | -0.016912745 | -0.047713247 | 0.102798476 | 0.029517023 | 0.082548611 | 0.680893601 | 0.240564472 |
| Casp8ap2 | S | 951 | 0.214760763 | 0.249409704 | -0.0366339 | -0.251394663 | -0.286043604 | 0.109016758 | 0.067890939 | 0.884854197 | 0.348489545 | 0.301069618 |
| Ptpn11 | S | 558 | 0.037194143 | 0.153336398 | 0.036828501 | -0.000365642 | -0.116507897 | 0.358728544 | 0.000692024 | 0.373450422 | 0.993813619 | 0.04758961 |
| Brca1 | S | 867 | 0.039139624 | 0.043778737 | 0.157665538 | 0.118525913 | 0.113886801 | 0.239553665 | 0.420191639 | 0.001542924 | 0.013670925 | 0.09550037 |
| Ap2a1 | S | 652 | 0.049465614 | 0.087850008 | 0.117482201 | 0.068016587 | 0.029632193 | 0.120440913 | 0.05242549 | 0.042763759 | 0.165581583 | 0.521018076 |
| Crebbp | S | 1077 | -0.061299534 | 0.199467483 | 0.252919366 | 0.3142189 | 0.053451883 | 0.264530376 | 0.01773247 | 0.065949496 | 0.025540861 | 0.53110862 |
| Reps1 | S | 272 | 0.01902761 | 0.107436315 | 0.040366054 | 0.021338444 | -0.067070261 | 0.189172565 | 0.000892103 | 0.021368882 | 0.13050676 | 0.01232144 |
| Borcs6 | S | 130 | 0.049642891 | 0.088884117 | 0.182377838 | 0.132734947 | 0.093493721 | 0.500156016 | 0.171392304 | 0.017597931 | 0.082985554 | 0.069512978 |
| Pgm2l1 | S | 175 | -0.046312652 | 0.094932445 | 0.413064219 | 0.459376871 | 0.318131774 | 0.756707454 | 0.101365812 | 0.057327816 | 0.057037744 | 0.113824484 |
| Srrm2 | S | 1984 | 0.095052878 | 0.070849083 | 0.105051936 | 0.009999058 | 0.034202853 | 0.040603596 | 0.095954968 | 0.111833794 | 0.850372075 | 0.529151262 |
| Acin1 | S | 380 | 0.154341854 | 0.013104609 | 0.170444181 | 0.016102328 | 0.157339573 | 0.057835692 | 0.879792946 | 0.0423742 | 0.688730443 | 0.08696465 |
| Zc3hc1 | S | 334 | 0.012936149 | 0.06291834 | 0.092435254 | 0.079499105 | 0.029516915 | 0.57438236 | 0.011117796 | 0.000675783 | 0.029478134 | 0.134295683 |
| Ppig | T | 746 | 0.012884362 | 0.094372436 | 0.113064093 | 0.100179731 | 0.018691657 | 0.737919246 | 0.00585705 | 0.016972476 | 0.061197247 | 0.545240196 |
| Phc3 | S | 614 | -0.012494281 | 0.052768683 | 0.257809387 | 0.270303669 | 0.205040704 | 0.787475472 | 0.252341903 | 0.003028906 | 0.006701261 | 0.02075081 |
| Scaf1 | S | 518 | 0.008078284 | 0.105387608 | 0.099671125 | 0.091592841 | -0.005716483 | 0.827609642 | 0.005822236 | 0.012324744 | 0.050660018 | 0.798035107 |
| Vrk3 | S | 59 | 0.019586117 | 0.096455782 | 0.143379055 | 0.123792938 | 0.046923273 | 0.636663172 | 0.033298359 | 0.016173682 | 0.048547373 | 0.288270538 |
| Arid1a | S | 1185 | 0.000569813 | 0.146320825 | 0.159265022 | 0.158695209 | 0.012944197 | 0.993593894 | 0.010522101 | 0.091818776 | 0.121745059 | 0.849914194 |
| Sipa1l1 | S | 1384 | -0.019511079 | 0.09078205 | 0.170149403 | 0.189660482 | 0.079367353 | 0.708420156 | 0.02129464 | 0.006494692 | 0.02894203 | 0.079375156 |
| Zc3hc1 | S | 343 | 0.037544695 | 0.074771416 | 0.086142624 | 0.048597929 | 0.011371207 | 0.182434374 | 0.06764845 | 0.001951447 | 0.119768549 | 0.689605696 |
| Dido1 | S | 153 | -0.018159974 | 0.09189189 | 0.129675498 | 0.147835472 | 0.037783608 | 0.491263251 | 0.00909567 | 0.002205061 | 0.00243323 | 0.133073383 |
| Nup88 | S | 235 | 0.246610721 | 0.103032057 | 0.186666701 | -0.05994402 | 0.083634644 | 0.204285807 | 0.599229914 | 0.166413114 | 0.726950531 | 0.665331279 |
| Cbx1 | S | 89 | 0.002577923 | 0.102927198 | 0.180571681 | 0.177993757 | 0.077644483 | 0.972873794 | 0.082236918 | 0.009479536 | 0.075627292 | 0.154221509 |
| Iws1 | S | 368 | 0.039651527 | 0.039986343 | 0.102899147 | 0.063247619 | 0.062912804 | 0.120509807 | 0.119648047 | 0.000872321 | 0.031075797 | 0.032459398 |
| Mki67 | S | 226 | 0.215303445 | 0.079457107 | 0.059390849 | -0.155912596 | -0.020066259 | 0.022650424 | 0.331495928 | 0.400341696 | 0.05450274 | 0.7539163 |
| Aaas | S | 495 | 0.027727994 | 0.107493632 | 0.17447129 | 0.146743296 | 0.066977658 | 0.617551302 | 0.072682162 | 0.029138175 | 0.048125158 | 0.237723103 |
| Frs2 | S | 177 | 0.104994485 | 0.1144843 | 0.224383133 | 0.119388648 | 0.109898833 | 0.465663887 | 0.123157951 | 0.144447736 | 0.475813962 | 0.39728119 |
| Snip1 | Y | 94 | 0.026317022 | 0.064901131 | 0.108771637 | 0.082454614 | 0.043870506 | 0.3575273 | 0.077889895 | 0.001113072 | 0.039860875 | 0.183338404 |
| Cbx3 | S | 99 | -0.003502896 | 0.060112644 | 0.13251081 | 0.136013706 | 0.072398166 | 0.888037135 | 0.020553067 | 0.000840158 | 0.007040708 | 0.02027202 |
| Ncbp3 | S | 410 | 0.090066433 | 0.026737787 | 0.062467099 | -0.027599334 | 0.035729312 | 0.001260953 | 0.249946325 | 0.0320939 | 0.236169016 | 0.189048062 |
| Herpud2 | S | 93 | -0.033340758 | 0.183842192 | 0.193043474 | 0.226384232 | 0.009201282 | 0.371429916 | 0.007119899 | 0.105042509 | 0.076857131 | 0.915376147 |
| Cast | S | 323 | 0.235119214 | 0.14328573 | 0.073061271 | -0.162057943 | -0.07022446 | 0.041258972 | 0.378191755 | 0.604038098 | 0.30150611 | 0.701048195 |
| Slc45a4 | S | 41 | 0.05115769 | 0.139078273 | 0.091937554 | 0.040779864 | -0.047140719 | 0.498051541 | 0.029649014 | 0.040685179 | 0.578284866 | 0.33236868 |
| Slc45a4 | S | 44 | 0.05115769 | 0.139078273 | 0.091937554 | 0.040779864 | -0.047140719 | 0.498051541 | 0.029649014 | 0.040685179 | 0.578284866 | 0.33236868 |
| Kank2 | S | 174 | -0.018199228 | 0.174312862 | 0.173389249 | 0.191588477 | -0.000923613 | 0.845177152 | 0.034806714 | 0.049916972 | 0.078190926 | 0.985173419 |
| Zc3h18 | S | 771 | 0.050424813 | 0.128634302 | 0.058861752 | 0.008436939 | -0.06977255 | 0.095716905 | 0.00316291 | 0.419508824 | 0.896522328 | 0.346966671 |
| Ric8a | S | 522 | 0.168233671 | 0.237341318 | 0.312674212 | 0.144440542 | 0.075332894 | 0.377910134 | 0.521150925 | 0.278310879 | 0.605428978 | 0.849761908 |
| Ric8a | S | 523 | 0.168233671 | 0.237341318 | 0.312674212 | 0.144440542 | 0.075332894 | 0.377910134 | 0.521150925 | 0.278310879 | 0.605428978 | 0.849761908 |
| Ric8a | S | 527 | 0.168233671 | 0.237341318 | 0.312674212 | 0.144440542 | 0.075332894 | 0.377910134 | 0.521150925 | 0.278310879 | 0.605428978 | 0.849761908 |
| Spen | S | 1395 | 0.036560476 | 0.073651199 | 0.070432365 | 0.033871889 | -0.003218833 | 0.162135495 | 0.026390291 | 0.002265095 | 0.180621793 | 0.877491189 |
| Dst | S | 7345 | 0.158795401 | 0.113596707 | 0.128167615 | -0.030627786 | 0.014570908 | 0.112006942 | 0.300546184 | 0.177151798 | 0.754079091 | 0.895572616 |
| 3110082I17Rik | S | 180 | 0.014604971 | 0.096992732 | 0.122284891 | 0.107679921 | 0.02529216 | 0.577961038 | 0.003718242 | 0.050619328 | 0.077885583 | 0.554054023 |
| Washc2 | S | 747 | 0.029447176 | 0.118106269 | 0.06441728 | 0.034970105 | -0.053688988 | 0.145327459 | 0.006039197 | 0.069558913 | 0.242954785 | 0.138307737 |
| Zfp687 | S | 228 | -0.055446311 | 0.122318256 | 0.237036763 | 0.292483074 | 0.114718507 | 0.450310739 | 0.100114823 | 0.005653333 | 0.018071887 | 0.082314079 |

| | | | | | | | | | | | | | |
|---|---|---|---|---|---|---|---|---|---|---|---|---|---|
| Eps15l1 | S | | 255 | -0.123537233 | 0.134007597 | 0.297415249 | 0.420952483 | 0.163407653 | 0.062537994 | 0.092120299 | 0.000870204 | 0.001747268 | 0.064290581 |
| C530008M17Rik | S | | 487 | 0.041533381 | 0.120589115 | 0.200609478 | 0.159076097 | 0.080020363 | 0.704835857 | 0.295350009 | 0.014211791 | 0.206483442 | 0.457925374 |
| Srrm2 | S | | 2075 | 0.128376392 | 0.103712411 | 0.046204196 | -0.082172195 | -0.057508214 | 0.051127018 | 0.019330783 | 0.535337377 | 0.317672736 | 0.43279001 |
| Casp12 | S | | 106 | -0.18310646 | 0.238161086 | 0.192080209 | 0.37518667 | -0.046080876 | 0.107781567 | 0.000882756 | 0.044945329 | 0.016829502 | 0.489299454 |
| Ppp2r5b | S | | 44 | 0.311970922 | 0.244456981 | -0.168023798 | -0.47999472 | -0.41248078 | 0.090408769 | 0.021521307 | 0.372497595 | 0.063986934 | 0.091946346 |
| Eif4enif1 | S | | 563 | 0.001055317 | 0.08707434 | 0.105910091 | 0.104854774 | 0.018835751 | 0.979626832 | 0.01128396 | 0.002466666 | 0.073730268 | 0.426189653 |
| Rab1a Rab1b | T | 43;40 | | 0.186023061 | 0.216989137 | -0.01041674 | -0.196439801 | -0.227405877 | 0.110707335 | 0.045380072 | 0.90569734 | 0.07077047 | 0.004771196 |
| Fbxo38 | S | | 599 | 0.018765821 | 0.118181822 | 0.184472941 | 0.16570712 | 0.066291119 | 0.656020378 | 0.031882668 | 0.070325167 | 0.068946542 | 0.343731785 |
| Fbxo38 | S | | 601 | 0.018765821 | 0.118181822 | 0.184472941 | 0.16570712 | 0.066291119 | 0.656020378 | 0.031882668 | 0.070325167 | 0.068946542 | 0.343731785 |
| Washc2 | S | | 684 | -0.051080281 | 0.074500331 | 0.161728984 | 0.212809265 | 0.087228653 | 0.326220734 | 0.03691751 | 0.000213069 | 0.024425334 | 0.025837117 |
| Washc2 | T | | 683 | -0.051080281 | 0.074500331 | 0.161728984 | 0.212809265 | 0.087228653 | 0.326220734 | 0.03691751 | 0.000213069 | 0.024425334 | 0.025837117 |
| Zfp318 | S | | 1032 | 0.10478704 | 0.059884866 | 0.250944332 | 0.146157292 | 0.191059467 | 0.393334563 | 0.648774125 | 0.027057118 | 0.250994251 | 0.196660503 |
| Zfp318 | S | | 1038 | 0.10478704 | 0.059884866 | 0.250944332 | 0.146157292 | 0.191059467 | 0.393334563 | 0.648774125 | 0.027057118 | 0.250994251 | 0.196660503 |
| Specc1l | S | | 949 | 0.023921971 | 0.158009298 | 0.14086626 | 0.116944289 | -0.017143038 | 0.60363953 | 0.013098673 | 0.185076827 | 0.248860606 | 0.844794795 |
| Kank2 | S | | 83 | -0.017912308 | 0.152160055 | 0.139608988 | 0.157521296 | -0.012551067 | 0.70979549 | 0.005761181 | 0.065149319 | 0.052910255 | 0.80408692 |
| Syde1 | S | | 683 | 0.107584465 | 0.114227136 | 0.079102476 | -0.028481989 | -0.03512466 | 0.176929672 | 0.027732339 | 0.206596756 | 0.697855634 | 0.494045378 |
| Syde1 | S | | 685 | 0.107584465 | 0.114227136 | 0.079102476 | -0.028481989 | -0.03512466 | 0.176929672 | 0.027732339 | 0.206596756 | 0.697855634 | 0.494045378 |
| Fam120a | T | | 483 | 0.007399226 | 0.206682433 | 0.114618864 | 0.107219638 | -0.092063569 | 0.949181969 | 0.004392605 | 0.352105439 | 0.471435566 | 0.431698269 |
| Atn1 | S | | 103 | -0.019375153 | 0.107690266 | 0.181704812 | 0.201079965 | 0.074014545 | 0.741155901 | 0.027658623 | 0.014284638 | 0.024415072 | 0.177055331 |
| Pnpla7 | S | | 379 | 0.067943665 | 0.164296345 | 0.072996651 | 0.005052987 | -0.091299694 | 0.212743495 | 0.016419768 | 0.348396935 | 0.933828192 | 0.225385213 |
| Tnrc6a | S | | 1876 | 0.079720667 | 0.326908286 | 0.039353005 | -0.040367662 | -0.287555281 | 0.624955554 | 0.020188283 | 0.775105111 | 0.827012427 | 0.106831138 |
| Tnrc6a | S | | 1878 | 0.079720667 | 0.326908286 | 0.039353005 | -0.040367662 | -0.287555281 | 0.624955554 | 0.020188283 | 0.775105111 | 0.827012427 | 0.106831138 |
| Fbxo38 | S | | 798 | 0.095730547 | 0.195067374 | 0.031692608 | -0.064037939 | -0.163374766 | 0.41390206 | 0.003407058 | 0.817915716 | 0.69246957 | 0.299251277 |
| Safb Safb2 | S | 296;290 | | 0.090529309 | 0.09028466 | 0.284684333 | 0.194155024 | 0.194399673 | 0.261144605 | 0.373986806 | 0.120236769 | 0.236211045 | 0.247517167 |
| Tns2 | S | | 991 | 0.045882387 | 0.077181796 | 0.28909588 | 0.243213493 | 0.211914085 | 0.602345153 | 0.232033377 | 0.048676576 | 0.087102184 | 0.116942202 |
| Junb | S | | 248 | -0.000357858 | 0.126067208 | 0.187740228 | 0.188098085 | 0.06167302 | 0.990964658 | 0.023941692 | 0.054758601 | 0.0739973 | 0.398354971 |
| Brwd1 | S | | 1782 | 0.004638703 | 0.215463416 | 0.142078254 | 0.137439551 | -0.073385161 | 0.92362347 | 0.012313921 | 0.272401243 | 0.285269399 | 0.538046594 |
| Brwd1 | S | | 1784 | 0.004638703 | 0.215463416 | 0.142078254 | 0.137439551 | -0.073385161 | 0.92362347 | 0.012313921 | 0.272401243 | 0.285269399 | 0.538046594 |
| Brwd1 | S | | 1789 | 0.004638703 | 0.215463416 | 0.142078254 | 0.137439551 | -0.073385161 | 0.92362347 | 0.012313921 | 0.272401243 | 0.285269399 | 0.538046594 |
| Terf1 | S | | 21 | 0.149701404 | 0.180934313 | 0.250567977 | 0.100866572 | 0.069633664 | 0.246861188 | 0.273278756 | 0.398401603 | 0.713594235 | 0.805886317 |
| Dst | S | | 7333 | 0.020168416 | 0.144149259 | 0.11627842 | 0.096110004 | -0.027870838 | 0.671689918 | 0.01942917 | 0.055973469 | 0.069236294 | 0.526220647 |
| Epc1 | S | | 119 | -0.001708884 | 0.2834147 | 0.261436091 | 0.263144975 | -0.021978608 | 0.986562961 | 0.111186268 | 0.21117379 | 0.209807714 | 0.912392556 |
| Ccp110 | S | | 426 | 0.20205513 | 0.146081649 | 0.104335408 | -0.097719721 | -0.041746241 | 0.081518174 | 0.286390921 | 0.465929443 | 0.508694191 | 0.795097846 |
| Rest | S | | 85 | -0.016760513 | 0.068230326 | 0.181287424 | 0.198047938 | 0.113057099 | 0.758531889 | 0.130112941 | 0.001039275 | 0.028008987 | 0.029829303 |
| Raly | T | | 268 | 0.13674367 | 0.080409099 | 0.058236822 | -0.078506847 | -0.022172277 | 0.007982557 | 0.151061402 | 0.442074482 | 0.311211179 | 0.765740152 |
| Ppig | S | | 743 | 0.01089722 | 0.092841915 | 0.111386216 | 0.100488997 | 0.018544301 | 0.776539672 | 0.005392936 | 0.017401576 | 0.061012057 | 0.541768761 |
| Taf3 | S | | 243 | 0.057027777 | 0.026018812 | 0.164505131 | 0.107477354 | 0.138486319 | 0.301783942 | 0.520194682 | 0.001608535 | 0.114111423 | 0.035727472 |
| Rai14 | S | | 413 | 0.081658173 | 0.174265145 | 0.048335695 | -0.033322478 | -0.12592945 | 0.051968056 | 0.078287006 | 0.239662392 | 0.302022489 | 0.159292724 |
| Scaf1 | T | | 923 | 0.03198281 | 0.060431045 | 0.19939953 | 0.16741672 | 0.138968485 | 0.214220771 | 0.111426714 | 0.030477592 | 0.039199331 | 0.056064863 |
| Llgl1 | S | | 986 | 0.044456556 | 0.120956206 | 0.167847795 | 0.123391239 | 0.046891589 | 0.21315925 | 0.121216976 | 0.079590844 | 0.14181718 | 0.562826975 |
| Tmx4 | S | | 302 | 0.027758458 | 0.100401884 | 0.100230211 | 0.072471754 | -0.000171673 | 0.497704451 | 0.044978999 | 0.005609725 | 0.129169896 | 0.996096541 |
| Zzz3 | S | | 321 | -0.111754217 | 0.162369856 | 0.114705718 | 0.226459935 | -0.047664138 | 0.32333027 | 0.001030571 | 0.011415144 | 0.112238971 | 0.15564097 |
| Trp53bp2 | S | | 121 | 0.132201532 | 0.049088837 | 0.120033125 | -0.012168407 | 0.070944288 | 0.044683946 | 0.370748975 | 0.076314214 | 0.770790616 | 0.138643019 |
| Osbp | S | | 383 | 0.046976052 | 0.111703626 | 0.073701925 | 0.026725873 | -0.0380017 | 0.068022775 | 0.063657081 | 0.030431205 | 0.266654964 | 0.410121069 |
| Mtcl1 | S | | 1407 | 0.069487706 | 0.041815187 | 0.104976208 | 0.035488502 | 0.063161021 | 0.042826814 | 0.184137755 | 0.011686765 | 0.233205703 | 0.069542459 |
| Mtcl1 | T | | 1403 | 0.069487706 | 0.041815187 | 0.104976208 | 0.035488502 | 0.063161021 | 0.042826814 | 0.184137755 | 0.011686765 | 0.233205703 | 0.069542459 |
| Arhgap31 | S | | 272 | -0.027272355 | 0.227870755 | 0.170387491 | 0.197659846 | -0.057483264 | 0.839590426 | 0.024806299 | 0.150083254 | 0.202101564 | 0.512731841 |
| Zfp687 | S | | 254 | -0.054676919 | 0.098468915 | 0.186523783 | 0.241200701 | 0.088054868 | 0.245758809 | 0.024549846 | 0.00171465 | 0.005289045 | 0.055384005 |
| Spindoc | S | | 249 | 0.009962867 | 0.083718767 | 0.213937024 | 0.203974157 | 0.130218257 | 0.854527093 | 0.008274002 | 0.075358404 | 0.074621335 | 0.190246722 |
| Spindoc | S | | 252 | 0.009962867 | 0.083718767 | 0.213937024 | 0.203974157 | 0.130218257 | 0.854527093 | 0.008274002 | 0.075358404 | 0.074621335 | 0.190246722 |
| Srf | S | | 220 | 0.036837737 | 0.040084139 | 0.067382884 | 0.030545148 | 0.027298745 | 0.274627356 | 0.005154365 | 0.00030128 | 0.345466838 | 0.032594467 |
| Mki67 | T | | 527 | 0.068863064 | 0.100970993 | 0.048383798 | -0.020479266 | -0.052587195 | 0.032736489 | 0.00464852 | 0.455397933 | 0.733998605 | 0.414518437 |
| Zhx3 | S | | 718 | 0.060302991 | 0.099783134 | 0.079662008 | 0.019359017 | -0.020121126 | 0.198341354 | 0.006562017 | 0.140470337 | 0.70589533 | 0.63707676 |
| Cic | S | | 2311 | -0.057825652 | 0.083152711 | 0.218541029 | 0.276366681 | 0.135388318 | 0.057575378 | 0.004632559 | 0.002269886 | 0.000607412 | 0.008726068 |
| Spidr | T | | 257 | 0.171508538 | 0.145153286 | 0.006547061 | -0.164961477 | -0.138606226 | 0.011260899 | 0.163838998 | 0.902723398 | 0.020985102 | 0.180748322 |
| Chaf1a | S | | 294 | 0.055614739 | 0.124896119 | 0.139702197 | 0.084087458 | 0.014806078 | 0.371149055 | 0.039485722 | 0.120579966 | 0.295828446 | 0.821512711 |
| Mdc1 | S | | 593 | 0.133083164 | 0.145003004 | 0.074901442 | -0.058181722 | -0.070101562 | 0.114290201 | 0.073799384 | 0.402940085 | 0.546628444 | 0.459270055 |
| Atn1 | S | | 630 | -0.000282968 | 0.119210764 | 0.105800029 | 0.106082997 | -0.013410735 | 0.987786857 | 0.000354744 | 0.158265141 | 0.164874628 | 0.814657547 |
| Hnrnpll | S | | 37 | -0.016531988 | 0.091431124 | 0.098357866 | 0.114889854 | 0.006926742 | 0.611742514 | 0.027837431 | 0.000149426 | 0.044123418 | 0.77867999 |
| Chaf1a | S | | 173 | 0.067119713 | 0.09524051 | 0.093452386 | 0.026332673 | -0.001788124 | 0.073312606 | 0.137194977 | 0.035591858 | 0.386607236 | 0.972490878 |
| Cavin3 | S | | 4 | 0.124809749 | 0.036244409 | 0.137430851 | 0.012621102 | 0.101186442 | 0.105025816 | 0.429754578 | 0.026428122 | 0.843440692 | 0.095431471 |
| Pard3 | S | | 869 | 0.042440238 | 0.201660421 | 0.058996328 | 0.01655609 | -0.142664093 | 0.389899332 | 0.007619755 | 0.442409483 | 0.798902735 | 0.107205995 |
| Atxn7l3 | S | | 163 | 0.085164144 | 0.061418226 | 0.138355986 | 0.053191842 | 0.07693776 | 0.136613999 | 0.090111511 | 0.065446977 | 0.41546976 | 0.22909212 |
| Prpf4b | S | | 61 | 0.041322029 | 0.087213148 | 0.163524509 | 0.12220248 | 0.076311361 | 0.475484541 | 0.049308801 | 0.040980054 | 0.121638358 | 0.227929942 |
| Lmnb1 | S | | 405 | 0.121663352 | 0.134396225 | -0.001408156 | -0.123071508 | -0.135804382 | 0.026308033 | 0.015374874 | 0.987849159 | 0.246856327 | 0.214384088 |
| Zfp768 | S | | 86 | 0.029537474 | 0.07064802 | 0.195449141 | 0.165911668 | 0.124801122 | 0.292351921 | 0.346341333 | 0.010284071 | 0.042067745 | 0.151922958 |
| Arid4a | S | | 867 | 0.015533876 | 0.070445834 | 0.102460052 | 0.086926176 | 0.032014218 | 0.516697823 | 0.00067492 | 0.033106404 | 0.048935837 | 0.311304183 |
| Rprd2 | S | | 927 | 0.002139354 | 0.054150425 | 0.228651585 | 0.226512231 | 0.17450116 | 0.953136414 | 0.172849028 | 0.005520237 | 0.012585709 | 0.024266791 |
| Ammecr1l | S | | 74 | -0.061607992 | 0.072251888 | 0.233983041 | 0.295591033 | 0.161731153 | 0.112245519 | 0.049940502 | 0.000650903 | 0.000953677 | 0.005590907 |
| Scaf1 | S | | 447 | 0.039268709 | 0.187152725 | 0.120930346 | 0.081661637 | -0.06622238 | 0.444904897 | 0.022359846 | 0.29124437 | 0.438807049 | 0.535701725 |
| Ccdc112 | S | | 260 | 0.152715015 | 0.102041222 | 0.10506267 | -0.047652345 | 0.003021448 | 0.06758749 | 0.219601414 | 0.265663328 | 0.597222032 | 0.973932531 |
| Osbp | S | | 377 | 0.029420889 | 0.127476736 | 0.075112708 | 0.045691819 | -0.052364028 | 0.237266562 | 0.030255007 | 0.022414396 | 0.079234532 | 0.23546443 |
| Hdac4 | T | | 463 | 0.088809573 | 0.056266327 | 0.129558637 | 0.040749064 | 0.07329231 | 0.352669667 | 0.046213111 | 0.026218648 | 0.648179526 | 0.107028157 |
| Fam117b | S | | 132 | 0.202092151 | 0.029012586 | 0.179353698 | -0.022738454 | 0.150341112 | 0.188930563 | 0.881906742 | 0.037861565 | 0.858429059 | 0.467702222 |
| Palld | S | | 1141 | 0.011589286 | 0.182781421 | 0.072831805 | 0.061242519 | -0.109949616 | 0.745886171 | 0.001795198 | 0.359171425 | 0.415033235 | 0.202147473 |
| Pcm1 | S | | 121 | 0.045550266 | 0.067554912 | 0.138589708 | 0.093039442 | 0.071034796 | 0.600084489 | 0.003920565 | 0.051122709 | 0.341257346 | 0.19716135 |
| Lig1 | T | | 186 | -0.057828034 | 0.143735149 | 0.11562571 | 0.173453743 | -0.02810944 | 0.43380375 | 0.000394118 | 0.063594042 | 0.078908966 | 0.527966109 |
| Ino80d | T | | 353 | 0.039102382 | 0.037010003 | 0.248402706 | 0.209300324 | 0.211392703 | 0.580310219 | 0.724766172 | 0.004824919 | 0.031320702 | 0.11582595 |
| Trp53bp1 | S | | 552 | -0.030896035 | 0.098366308 | 0.206102314 | 0.236998349 | 0.107736006 | 0.435459964 | 0.013315806 | 0.017343662 | 0.011930437 | 0.103221846 |
| Atp11c | S | | 1105 | 0.068612913 | 0.109817683 | 0.222865282 | 0.154252369 | 0.113047599 | 0.693320824 | 0.112951955 | 0.076254904 | 0.416696467 | 0.277458474 |
| Irf2bpl | S | | 69 | 0.116541345 | 0.029968308 | 0.077381836 | -0.03915951 | 0.047413528 | 0.113604316 | 0.261282098 | 0.001745958 | 0.480399563 | 0.115095036 |
| Gpatch8 | S | | 438 | 0.006270268 | 0.075120885 | 0.193488222 | 0.187217954 | 0.118367337 | 0.896462925 | 0.021430438 | 0.025735292 | 0.032108701 | 0.098991022 |
| Smad5 | S | | 465 | 0.221842298 | 0.126019817 | 0.208520031 | -0.013322268 | 0.082500214 | 0.338013613 | 0.290176576 | 0.275985487 | 0.956232847 | 0.650473168 |
| Erbin | S | | 829 | 0.104096682 | 0.038739822 | 0.113163283 | 0.009066601 | 0.074423461 | 0.009321557 | 0.30928773 | 0.1175741 | 0.869134238 | 0.25382477 |
| Nck2 | T | | 92 | -0.008198141 | 0.072202127 | 0.20969255 | 0.217890691 | 0.137490423 | 0.906984824 | 0.207249562 | 0.00333094 | 0.044016835 | 0.053834209 |
| Nck2 | Y | | 110 | -0.008198141 | 0.072202127 | 0.20969255 | 0.217890691 | 0.137490423 | 0.906984824 | 0.207249562 | 0.00333094 | 0.044016835 | 0.053834209 |
| Arfgef2 | S | | 1518 | 0.056123904 | 0.110701355 | 0.071860511 | 0.015736607 | -0.038840845 | 0.047585517 | 0.038986446 | 0.120847327 | 0.672307546 | 0.410636206 |
| Prkar1a | S | | 83 | 0.108225238 | 0.221186401 | -0.033508928 | -0.141734165 | -0.254695329 | 0.105408816 | 0.006906654 | 0.556468243 | 0.015888994 | 0.002250338 |
| Abi1 | S | | 183 | 0.027193903 | 0.15038634 | 0.063983787 | 0.036789884 | -0.086402553 | 0.452761875 | 0.00303941 | 0.198433961 | 0.378935955 | 0.09767561 |
| Srrm2 | S | | 1687 | 0.06029594 | 0.016276266 | 0.10642798 | 0.04613204 | 0.090151714 | 0.064720987 | 0.571622694 | 0.000494395 | 0.129238709 | 0.041029763 |
| Ncoa5 | S | | 29 | 0.026974944 | 0.104007437 | 0.113485287 | 0.086510343 | 0.009477849 | 0.553840555 | 0.002510569 | 0.124527765 | 0.225820598 | 0.862799858 |
| Scrib | S | | 431 | 0.115967862 | 0.230313796 | 0.017290758 | -0.098677104 | -0.213023038 | 0.227803866 | 0.025725689 | 0.758106117 | 0.28978985 | 0.039485268 |
| Jmy | S | | 108 | -0.039138451 | 0.129948175 | 0.149170163 | 0.188308613 | 0.019221987 | 0.186722783 | 0.001190952 | 0.030430086 | 0.022159946 | 0.654213268 |
| Cdk12 | S | | 300 | 0.044031991 | 0.057389886 | 0.058611731 | 0.01457974 | 0.001221845 | 0.105563715 | 0.001310408 | 0.023920598 | 0.545831942 | 0.940229527 |

| | | | | | | | | | | | | |
|---|---|---|---|---|---|---|---|---|---|---|---|---|
| Aff4 | S | 544 | 0.064143274 | 0.097724037 | 0.104020725 | 0.039877451 | 0.006296688 | 0.055601139 | 0.09917113 | 0.095906207 | 0.426656564 | 0.913562488 |
| Rps6ka3 | S | 715 | 0.061991666 | 0.168145054 | 0.087622761 | 0.025631095 | -0.080522293 | 0.295725658 | 0.036270017 | 0.196680751 | 0.635778408 | 0.240345248 |
| Srrm2 | T | 1986 | 0.090856568 | 0.068377691 | 0.093956623 | 0.003100055 | 0.025578931 | 0.030491676 | 0.082671614 | 0.149822305 | 0.953211352 | 0.636723085 |
| Srrm2 | S | 744 | 0.098446084 | 0.022368027 | 0.119585193 | 0.021139109 | 0.097217166 | 0.005854317 | 0.426623561 | 0.080515733 | 0.669068056 | 0.129080212 |
| Dcp1a | S | 164 | 0.030616376 | 0.17905264 | 0.136121347 | 0.105504971 | -0.042931294 | 0.53272489 | 0.054448049 | 0.11972911 | 0.20358009 | 0.609011323 |
| Rb1 | S | 243 | -0.013029534 | 0.199712953 | 0.092487726 | 0.10551726 | -0.107225227 | 0.767920924 | 0.008277149 | 0.08438075 | 0.051715951 | 0.083677578 |
| Ercc5 | S | 341 | 0.072168669 | 0.104019383 | 0.113063789 | 0.04089512 | 0.009044407 | 0.166007547 | 0.100373996 | 0.070225849 | 0.352535433 | 0.857851161 |
| Farp2 | S | 863 | 0.12165151 | 0.065812963 | 0.068205948 | -0.053445562 | 0.002392986 | 0.009862838 | 0.150631521 | 0.281100735 | 0.381015313 | 0.968000714 |
| Fxr2 | T | 611 | 0.099741354 | 0.112122854 | 0.158659604 | 0.05891825 | 0.04653675 | 0.09235463 | 0.16789575 | 0.260651305 | 0.632159002 | 0.716689379 |
| Itsn1 | S | 897;906 | 0.008860149 | 0.112751385 | 0.13755156 | 0.128691411 | 0.024800175 | 0.696633953 | 0.052876257 | 0.014314111 | 0.022372839 | 0.588719627 |
| Mcoln1 | S | 557 | 0.045799033 | 0.118883222 | 0.110408536 | 0.064609503 | -0.008474686 | 0.310580424 | 0.029091473 | 0.089508464 | 0.252091156 | 0.868943601 |
| Map1a | T | 1928 | 0.317809112 | 0.097491117 | 0.032046543 | -0.28576257 | -0.065444575 | 0.0381338 | 0.445766784 | 0.748854 | 0.056965131 | 0.502187979 |
| Chaf1a | S | 296 | 0.137102209 | 0.133813162 | 0.148339751 | 0.011237542 | 0.014526589 | 0.191536384 | 0.244831819 | 0.203725719 | 0.861798513 | 0.861052039 |
| Fip1l1 | S | 84 | 0.091563966 | 0.113553122 | 0.085347902 | -0.006216064 | -0.02820522 | 0.148216904 | 0.218611768 | 0.02466658 | 0.89771841 | 0.710794326 |
| Fip1l1 | S | 86 | 0.091563966 | 0.113553122 | 0.085347902 | -0.006216064 | -0.02820522 | 0.148216904 | 0.218611768 | 0.02466658 | 0.89771841 | 0.710794326 |
| Fip1l1 | S | 88 | 0.091563966 | 0.113553122 | 0.085347902 | -0.006216064 | -0.02820522 | 0.148216904 | 0.218611768 | 0.02466658 | 0.89771841 | 0.710794326 |
| Irf2bpl | S | 637 | 0.064221537 | 0.098167302 | 0.061201452 | -0.003020085 | -0.03696585 | 0.090089647 | 0.003244603 | 0.354339046 | 0.960561844 | 0.544612745 |
| Ddx59 | S | 149 | 0.041472189 | 0.145225278 | 0.013577349 | -0.027894841 | -0.131647929 | 0.497195222 | 7.26014E-05 | 0.841174398 | 0.735621606 | 0.151529111 |
| Ubr5 | T | 1723 | 0.001487351 | 0.126815831 | 0.153236678 | 0.151749327 | 0.026420847 | 0.973522156 | 0.040716759 | 0.025118447 | 0.03067163 | 0.617026825 |
| Haspin | T | 311 | 0.083081329 | 0.090518132 | 0.109059581 | 0.025978252 | 0.018541448 | 0.201784738 | 0.155362109 | 0.029749758 | 0.608892917 | 0.69387948 |
| Slu7 | S | 215 | 0.043327925 | 0.071543303 | 0.124735502 | 0.081407577 | 0.0531922 | 0.112433994 | 0.019346193 | 0.0847751 | 0.201535085 | 0.350752476 |
| Ubap2l | S | 496 | 0.068473118 | 0.068853921 | 0.045490587 | -0.022982531 | -0.023363334 | 0.012433783 | 0.010743077 | 0.1636473 | 0.41279885 | 0.401765981 |
| Zfp638 | S | 1846 | 0.055089581 | 0.101679319 | 0.080616383 | 0.025526802 | -0.021062936 | 0.388738328 | 0.020232447 | 0.028417751 | 0.657500666 | 0.446665069 |
| Add3 | S | 673 | 0.15150479 | 0.07638378 | 0.049484837 | -0.102019953 | -0.026898943 | 0.022872692 | 0.085459372 | 0.434625279 | 0.165068015 | 0.64379071 |
| Rnmt | S | 64 | -0.009046458 | 0.069347202 | 0.112777264 | 0.121823721 | 0.043430062 | 0.583964281 | 0.00305216 | 0.002272641 | 0.003832729 | 0.078807948 |
| Pds5b | S | 1255 | 0.03913325 | 0.035365008 | 0.061073975 | 0.021940725 | 0.025708967 | 0.153425674 | 0.006489578 | 0.000331374 | 0.355759056 | 0.022276419 |
| Rusc2 | S | 557 | -0.044627923 | 0.093176339 | 0.174029499 | 0.218657422 | 0.08085316 | 0.049053699 | 0.023673142 | 0.000390948 | 0.000482293 | 0.040805889 |
| Bclaf1 | S | 395 | -0.032788286 | 0.08504019 | 0.135755692 | 0.168543977 | 0.050715501 | 0.197461675 | 0.008433418 | 0.000781862 | 0.000410457 | 0.026294232 |
| Spen | S | 1276 | 0.04568052 | 0.068511224 | 0.158859269 | 0.113178749 | 0.090348044 | 0.581589776 | 0.19711548 | 0.006043876 | 0.222445255 | 0.087253541 |
| Dcaf1 | S | 203 | 0.15627799 | -0.041864433 | 0.249011038 | 0.092733048 | 0.290875471 | 0.154888269 | 0.662045551 | 0.020594172 | 0.27866923 | 0.014441025 |
| Cd248 | S | 764 | 0.114961271 | 0.099919508 | 0.106714248 | -0.008247023 | 0.00679474 | 0.094498585 | 0.257012028 | 0.093980913 | 0.871306581 | 0.927178476 |
| Bcl7c | T | 118 | 0.027115493 | 0.133145727 | 0.213275532 | 0.186160039 | 0.080129805 | 0.744230321 | 0.111465292 | 0.065925817 | 0.124266366 | 0.398631032 |
| Dtx2 | S | 217 | -0.098937101 | 0.068270707 | 0.195069798 | 0.294006899 | 0.126799092 | 0.138646589 | 0.181971846 | 1.10377E-05 | 0.020039691 | 0.067309972 |
| Myo1c | S | 1039 | 0.035474292 | 0.101350916 | 0.145114192 | 0.1096399 | 0.043763276 | 0.807538139 | 0.135382409 | 0.0065843 | 0.475096749 | 0.431994253 |
| Srrm2 | T | 1692 | 0.059287298 | 0.014182548 | 0.125965334 | 0.066678036 | 0.111782786 | 0.03219782 | 0.643359777 | 0.002798952 | 0.044033035 | 0.019596138 |
| Sptbn1 | S | 2168 | 0.005897182 | 0.096107827 | 0.084117917 | 0.078220735 | -0.01198991 | 0.802516149 | 0.003424908 | 0.011035164 | 0.014779744 | 0.515763382 |
| Rb1 | S | 781 | 0.037385254 | 0.111866542 | 0.100965361 | 0.063580107 | -0.010901181 | 0.295320314 | 0.013481051 | 0.099984431 | 0.235141982 | 0.807014778 |
| Gorasp2 | S | 448 | 0.162781717 | 0.02993606 | 0.156080398 | -0.006701319 | 0.126144338 | 0.391666825 | 0.479594608 | 0.009717868 | 0.968589677 | 0.016659889 |
| Ncam1 | S | 270 | 0.094150414 | 0.043509264 | 0.146320288 | 0.052169874 | 0.102811024 | 0.133450005 | 0.213676833 | 0.036979297 | 0.410132192 | 0.119139204 |
| Taf3 | S | 183 | 0.010544434 | 0.102608097 | 0.169962299 | 0.159417864 | 0.067354202 | 0.693727409 | 0.140515534 | 0.010721367 | 0.031829278 | 0.305260818 |
| Irf2bp2 | S | 169 | 0.054832812 | 0.060599295 | 0.105932109 | 0.051099297 | 0.045332814 | 0.062053357 | 0.03641539 | 0.05833157 | 0.252059249 | 0.295496862 |
| Prpf38b | S | 460 | 0.237963808 | -0.052274231 | 0.147819662 | -0.090144146 | 0.200093894 | 0.081142672 | 0.69259976 | 0.021687666 | 0.390981376 | 0.208641641 |
| Fam126a | S | 453 | -0.082627554 | 0.40641512 | 0.148399591 | 0.231027145 | -0.25801553 | 0.412741295 | 0.022042943 | 0.31885775 | 0.139957099 | 0.135644029 |
| Cdc42ep2 | S | 137 | 0.060146072 | 0.129259807 | 0.087889389 | 0.027743317 | -0.041370417 | 0.208881755 | 0.01600357 | 0.251143946 | 0.696136324 | 0.548850041 |
| Hmga2 | S | 59 | 0.252613041 | 0.333150041 | -0.163660825 | -0.416273867 | -0.496810867 | 0.151951312 | 0.046425869 | 0.545438602 | 0.175458997 | 0.14049774 |
| Kmt2a | S | 516 | -0.046324066 | 0.207269788 | 0.141767927 | 0.188091993 | -0.06550186 | 0.720955184 | 0.0030728 | 0.299371025 | 0.277509404 | 0.591974058 |
| Dlg1 | S | 598 | -0.040886234 | 0.178881782 | 0.1561789 | 0.197065134 | -0.022702882 | 0.58400969 | 0.042595669 | 0.019916832 | 0.047512519 | 0.707984488 |
| Wdr44 | T | 403 | 0.024326919 | 0.211042659 | 0.073537563 | 0.049210644 | -0.137505097 | 0.674234126 | 0.007123406 | 0.383951173 | 0.549620136 | 0.149900995 |
| Rras | S | 201 | 0.084879645 | 0.109783302 | 0.026594874 | -0.058284771 | -0.083188429 | 0.045497494 | 0.004544952 | 0.670735352 | 0.384991912 | 0.243896933 |
| Actn4 | S | 910 | 0.101379139 | 0.08213424 | 0.248007721 | 0.146628583 | 0.165873481 | 0.594302249 | 0.343125583 | 0.051647802 | 0.468495415 | 0.162507762 |
| Retreg3 | S | 436 | 0.049094333 | 0.061718838 | 0.088209509 | 0.039115176 | 0.026490671 | 0.017741639 | 0.232533666 | 0.009946309 | 0.129971408 | 0.565498822 |
| Srrm2 | S | 1097 | 0.066450739 | 0.048933813 | 0.093698779 | 0.02724804 | 0.044764966 | 0.059470297 | 0.04228536 | 0.036951512 | 0.450851809 | 0.214596236 |
| Tmx2 | S | 211 | 0.035900247 | 0.088064692 | 0.134817166 | 0.098916919 | 0.046752473 | 0.366462129 | 0.171888598 | 0.008889958 | 0.019352268 | 0.399402591 |
| Srrm2 | S | 1538 | 0.050122461 | 0.049127558 | 0.095085476 | 0.036963015 | 0.045957918 | 0.004678576 | 0.370356245 | 0.022042993 | 0.214250129 | 0.407642609 |
| Ppp1r18 | S | 134 | 0.102139643 | 0.210335213 | -0.00433004 | -0.106469683 | -0.214665253 | 0.173735821 | 0.011633789 | 0.977038148 | 0.493311835 | 0.227968263 |
| Hspa4 | S | 76 | 0.341298591 | 0.168482278 | -0.021039626 | -0.362338217 | -0.189521905 | 0.056776898 | 0.320292246 | 0.900934048 | 0.041595307 | 0.213430367 |
| Srrm2 | S | 2644 | 0.012641713 | 0.056124406 | 0.095761426 | 0.083119714 | 0.039637021 | 0.580805878 | 0.009900814 | 0.001511933 | 0.017018617 | 0.070782625 |
| Ppig | S | 742 | 0.010199358 | 0.092670825 | 0.112074375 | 0.101875017 | 0.01940355 | 0.801192276 | 0.007614951 | 0.02038498 | 0.067524639 | 0.545450891 |
| Mki67 | S | 2319 | 0.101822811 | 0.178684934 | 0.019763536 | -0.082059274 | -0.158921397 | 0.250535612 | 0.008434406 | 0.717661005 | 0.323447759 | 0.02380263 |
| Aff4 | S | 614 | 0.096465518 | 0.113918318 | 0.11561235 | 0.019146832 | 0.001694032 | 0.200602374 | 0.02713876 | 0.356414104 | 0.872201537 | 0.987647497 |
| Son | S | 1805 | 0.05737884 | 0.208110468 | 0.283553023 | 0.226174183 | 0.075442555 | 0.681840031 | 0.094336094 | 0.37040336 | 0.469475621 | 0.794106187 |
| Rapgef2 | S | 959 | -0.076808139 | 0.1559734 | 0.163296447 | 0.240104586 | 0.007323047 | 0.141331605 | 0.02676295 | 0.001822765 | 0.007329829 | 0.873242193 |
| Pole | T | 2020 | 0.053380524 | 0.101781838 | 0.123361235 | 0.069980711 | 0.021579398 | 0.261414843 | 0.15814265 | 0.024142663 | 0.0632056 | 0.701342764 |
| Peak1 | S | 504 | 0.097780913 | 0.060469093 | 0.092462907 | -0.005318005 | 0.031993815 | 0.095114188 | 0.289955154 | 0.014703292 | 0.905658917 | 0.544772106 |
| Trp53bp1 | S | 719 | 0.001900113 | 0.075719007 | 0.110587312 | 0.108687199 | 0.034868304 | 0.920372569 | 0.077260985 | 0.000440497 | 0.002533529 | 0.312563244 |
| Tuba1a Tuba1b Tuba | S | 287;287;287;287 | -0.011401876 | 0.100450409 | 0.13804019 | 0.149442066 | 0.037589781 | 0.691699506 | 0.054182612 | 0.002815596 | 0.000993956 | 0.339030129 |
| Svil | S | 241 | 0.022667028 | 0.097563785 | 0.086356449 | 0.063689421 | -0.011207336 | 0.451250449 | 0.025134698 | 0.008352124 | 0.0601139 | 0.698542069 |
| Tmem201 | S | 441 | 0.058839963 | 0.067903806 | 0.095701427 | 0.036861465 | 0.027797621 | 0.094333903 | 0.011590896 | 0.116485403 | 0.462314249 | 0.551121736 |
| Tmpo | S | 411 | 0.173999053 | 0.085069606 | 0.047426834 | -0.126572219 | -0.037642773 | 0.324296172 | 0.011887987 | 0.117417937 | 0.446321148 | 0.097599471 |
| Lmna | S | 107 | 0.006322578 | 0.123747719 | 0.216086057 | 0.209763479 | 0.092338338 | 0.910421051 | 0.037091625 | 0.095771715 | 0.107457465 | 0.37470745 |
| Trim28 | S | 594 | -0.025145005 | 0.09686594 | 0.130398132 | 0.155543137 | 0.033532192 | 0.143667063 | 0.030335325 | 0.000377957 | 0.000983495 | 0.291080744 |
| Ahdc1 | S | 1540 | 0.044850724 | 0.041457293 | 0.232631056 | 0.187780332 | 0.191173763 | 0.384951893 | 0.683111385 | 0.009546571 | 0.045457259 | 0.125533727 |
| Denn2b | S | 230 | 0.032580749 | 0.177387406 | 0.07305946 | 0.040478711 | -0.104327946 | 0.65009879 | 0.004799242 | 0.339713062 | 0.629964937 | 0.196494713 |
| Denn2b | S | 233 | 0.032580749 | 0.177387406 | 0.07305946 | 0.040478711 | -0.104327946 | 0.65009879 | 0.004799242 | 0.339713062 | 0.629964937 | 0.196494713 |
| Rxylt1 | S | 64 | 0.11684219 | 0.04573369 | 0.187123356 | 0.070281166 | 0.141389666 | 0.430291707 | 0.533791831 | 0.014627848 | 0.611107098 | 0.075055969 |
| Ckap4 | S | 9 | 0.067318254 | 0.183608184 | 0.066840602 | -0.000477652 | -0.116767583 | 0.579504387 | 0.031440866 | 0.110887502 | 0.996648521 | 0.123859696 |
| Notch2 | S | 1846 | 0.039545481 | 0.139687772 | 0.234466556 | 0.194921075 | 0.094778784 | 0.669153668 | 0.284044435 | 0.054252588 | 0.113306402 | 0.471045515 |
| Cobll1 | S | 396 | 0.161776929 | 0.167586901 | -0.039244629 | -0.201021557 | -0.206831529 | 0.050409524 | 0.015323633 | 0.516047046 | 0.027661499 | 0.011892994 |
| Mtmr2 | S | 58 | 0.071157887 | 0.109514687 | 0.054185371 | -0.016972516 | -0.055329316 | 0.084504887 | 0.006120434 | 0.464738557 | 0.809053944 | 0.447506943 |
| Rnf169 | S | 390 | 0.083345833 | 0.057346346 | 0.099047032 | 0.015701199 | 0.041700686 | 0.071044126 | 0.137378104 | 0.040822186 | 0.653310495 | 0.206228848 |
| Fam83h | S | 1041 | -0.215044542 | 0.185663038 | 0.292735762 | 0.507780304 | 0.107072724 | 0.025648573 | 0.03865625 | 0.00056414 | 0.008428746 | 0.164727899 |
| Kmt2a | S | 740 | 0.121594261 | 0.135467838 | 0.103922693 | -0.017671567 | -0.031545144 | 0.206520322 | 0.154413703 | 0.171308911 | 0.835825144 | 0.703840369 |
| Akap12 | S | 684 | 0.023517425 | 0.340895003 | -0.031821301 | -0.055338726 | -0.372716304 | 0.736699479 | 0.003183711 | 0.65338374 | 0.066485692 | 0.000438424 |
| Ptpn21 | S | 602 | 0.10536357 | 0.19050658 | 0.07442258 | -0.03094099 | -0.116084 | 0.456330849 | 0.015635835 | 0.612922851 | 0.860423277 | 0.442069893 |
| Ptpn21 | S | 616 | 0.10536357 | 0.19050658 | 0.07442258 | -0.03094099 | -0.116084 | 0.456330849 | 0.015635835 | 0.612922851 | 0.860423277 | 0.442069893 |
| Nsd3 | S | 671 | 0.099581901 | 0.147806722 | 0.095476823 | -0.004105078 | -0.052329899 | 0.330977858 | 0.038310601 | 0.293747967 | 0.970370045 | 0.54338178 |
| Scrib | S | 1361 | 0.001893455 | 0.137635146 | 0.023603109 | 0.021709654 | -0.114032037 | 0.936027336 | 1.21121E-05 | 0.606471931 | 0.650307213 | 0.092789864 |
| Lima1 | T | 629 | 0.131012436 | 0.084649586 | 0.181841236 | 0.0508288 | 0.09719165 | 0.120898287 | 0.211164183 | 0.282958114 | 0.734209482 | 0.522001625 |
| Cdk11b | T | 584 | 0.065708324 | 0.063385597 | 0.092310319 | 0.026601995 | 0.028924722 | 0.106059454 | 0.115035389 | 0.016883889 | 0.394172429 | 0.356072915 |
| Syne2 | S | 6761 | 0.002145015 | 0.065820677 | 0.127388294 | 0.125243279 | 0.061567617 | 0.978349135 | 0.032429076 | 0.002117349 | 0.206021493 | 0.000621187 |
| Clk3 | S | 157 | 0.091625351 | 0.047532906 | 0.020442935 | -0.071182416 | -0.027089971 | 0.000205005 | 0.106470974 | 0.290198301 | 0.024329259 | 0.311855226 |
| Tprn | S | 501 | 0.133767972 | 0.179981733 | 0.065193997 | -0.068573976 | -0.114787736 | 0.193942888 | 0.076097665 | 0.483291469 | 0.413751811 | 0.163529194 |
| Nktr | S | 379 | 0.026956231 | -0.006248981 | 0.133417187 | 0.106460956 | 0.139666168 | 0.396115995 | 0.803418208 | 8.87188E-06 | 0.048013108 | 0.019200886 |
| Iws1 | S | 384 | 0.031127007 | 0.063842183 | 0.175879136 | 0.144752129 | 0.112036953 | 0.203950049 | 0.224505145 | 0.018901108 | 0.055542209 | 0.097783755 |

| | | | | | | | | | | | | |
|---|---|---|---|---|---|---|---|---|---|---|---|---|
| Iws1 | S | 386 | 0.031127007 | 0.063842183 | 0.175879136 | 0.144752129 | 0.112036953 | 0.203950049 | 0.224505145 | 0.018901108 | 0.055542209 | 0.097783755 |
| Nlk | T | 298 | -0.017235101 | 0.093347079 | 0.136470927 | 0.153706028 | 0.043123848 | 0.64643006 | 0.02399869 | 0.00425021 | 0.017819427 | 0.221187207 |
| Fxr2 | S | 602 | 0.079447898 | 0.072946207 | 0.089821181 | 0.010373283 | 0.016874974 | 0.064678931 | 0.172688301 | 0.038402729 | 0.757750128 | 0.709815237 |
| Nes | S | 1021 | -0.016853055 | 0.041305059 | 0.219647893 | 0.236500948 | 0.178342834 | 0.730551775 | 0.307126163 | 0.001179701 | 0.004549613 | 0.006398665 |
| Tmpo | S | 377 | 0.115655147 | 0.111648475 | 0.075706092 | -0.039949055 | -0.035942383 | 0.145832846 | 0.027102672 | 0.415777404 | 0.685041951 | 0.684601711 |
| Lmbrd1 | S | 525 | -0.015441387 | 0.125828968 | 0.123088544 | 0.138529931 | -0.002740424 | 0.39739303 | 0.18775431 | 0.000198685 | 0.001032767 | 0.970287905 |
| Avl9 | S | 368 | 0.045437744 | 0.186290491 | 0.088220033 | 0.042782289 | -0.098070458 | 0.62739259 | 0.015409405 | 0.278367567 | 0.665517015 | 0.222052403 |
| Plxna1 Plxna2 Plxna3 | S | 1793;1793;1792;1 | 0.076950513 | 0.165793659 | 0.122036289 | 0.045085776 | -0.043757369 | 0.385264211 | 0.058625123 | 0.25846638 | 0.660468309 | 0.647562375 |
| Rb1 | T | 367 | -0.003167311 | 0.136594667 | 0.079444156 | 0.082611467 | -0.057150511 | 0.907721683 | 0.002401833 | 0.060074913 | 0.064218468 | 0.119240263 |
| Prkce | S | 729 | -0.094103513 | 0.150698057 | 0.228777473 | 0.322880985 | 0.078079415 | 0.512532402 | 0.021788902 | 0.041407479 | 0.089989057 | 0.346229237 |
| Bclaf1 | S | 284 | 0.03429276 | 0.055852265 | 0.087460204 | 0.053167444 | 0.031607939 | 0.298400326 | 0.014107787 | 0.005947761 | 0.145592023 | 0.158946904 |
| Prim2 | T | 470 | 0.267844152 | 0.041803113 | 0.06643858 | -0.201405573 | 0.024635466 | 0.019223958 | 0.756372557 | 0.470276981 | 0.024422353 | 0.835659653 |
| Farp1 | S | 881 | 0.07360499 | 0.15732492 | 0.188155792 | 0.114550802 | 0.030830872 | 0.338724524 | 0.112583482 | 0.277520166 | 0.468959102 | 0.841416593 |
| Cers2 | T | 354 | 0.033127809 | 0.105088093 | 0.27286366 | 0.239735851 | 0.167775566 | 0.680061296 | 0.457319305 | 0.034193254 | 0.061538815 | 0.286346381 |
| Msl3 | T | 409 | 0.072675626 | 0.153705722 | 0.081329094 | 0.008653468 | -0.072376629 | 0.285458647 | 0.047484332 | 0.18044227 | 0.88558846 | 0.265121017 |
| Zfhx3 | S | 2634 | 0.057157738 | 0.100886611 | 0.159840299 | 0.102682561 | 0.058953688 | 0.505137284 | 0.111476187 | 0.050598899 | 0.289404102 | 0.395758977 |
| Slc30a4 | S | 36 | 0.027405579 | 0.117832027 | 0.167374171 | 0.139968593 | 0.049542144 | 0.633205844 | 0.081056069 | 0.052428885 | 0.085689195 | 0.404046474 |
| Kank2 | S | 92 | 0.184653825 | -0.049749693 | 0.070544558 | -0.114109267 | 0.120294251 | 0.001076345 | 0.536141029 | 0.09108378 | 0.01326588 | 0.196297903 |
| Piezo1 | S | 1631 | 0.113933955 | 0.095572291 | 0.00432948 | -0.109604475 | -0.091242811 | 0.003069858 | 0.050758387 | 0.965207508 | 0.336245484 | 0.409212472 |
| Spp1 | S | 283 | -0.022968639 | 0.145105253 | 0.065013433 | 0.087982073 | -0.08009182 | 0.302095632 | 0.00073628 | 0.021392757 | 0.004130268 | 0.016634783 |
| Fnbp4 | S | 128 | 0.143068999 | 0.078559976 | 0.130664003 | -0.012404995 | 0.052104027 | 0.237258949 | 0.144356763 | 0.133063495 | 0.911772666 | 0.463078076 |
| Prkd3 | S | 364 | 0.020546032 | 0.114445267 | 0.078813655 | 0.058267623 | -0.035631612 | 0.606923379 | 0.009832235 | 0.029122312 | 0.157344399 | 0.193407634 |
| Kdm4b | S | 510 | -0.002947889 | 0.078265733 | 0.125198354 | 0.128146243 | 0.04693262 | 0.960429067 | 0.045794574 | 0.002202958 | 0.115243584 | 0.173169415 |
| Pard3 | Y | 377 | 0.217287042 | 0.09158317 | 0.003231297 | -0.214055745 | -0.088351873 | 0.02059065 | 0.135082633 | 0.977269203 | 0.146011491 | 0.469961209 |
| Ccny | S | 324 | 0.146565287 | 0.102823783 | 0.066320849 | -0.080244437 | -0.036502933 | 0.100410575 | 0.034028748 | 0.598353779 | 0.547286751 | 0.761994181 |
| Foxc1 | S | 237 | 0.104863687 | 0.119216467 | 0.085661535 | -0.019202152 | -0.033554932 | 0.297371194 | 0.040890429 | 0.184921765 | 0.839767595 | 0.580621316 |
| Foxc1 | S | 243 | 0.104863687 | 0.119216467 | 0.085661535 | -0.019202152 | -0.033554932 | 0.297371194 | 0.040890429 | 0.184921765 | 0.839767595 | 0.580621316 |
| Ppp6r2 | S | 670 | -0.011873129 | 0.172794221 | 0.0729406 | 0.084813729 | -0.099853621 | 0.710230391 | 0.002362397 | 0.117183974 | 0.076809043 | 0.0543091 |
| Pacs1 | T | 502 | 0.288615095 | 0.029258514 | -0.042723095 | -0.33133819 | -0.071981609 | 0.000557835 | 0.682570053 | 0.315001757 | 8.0988E-05 | 0.318372549 |
| Taf2 | S | 1090 | 0.021664166 | 0.094362255 | 0.103610706 | 0.081946541 | 0.009248451 | 0.326884426 | 0.005146074 | 0.095612477 | 0.169614 | 0.834226722 |
| Nhsl1 | T | 179 | 0.237353197 | 0.100218154 | 0.109583212 | -0.127769986 | 0.009365057 | 0.297244481 | 0.151419371 | 0.264127124 | 0.542776895 | 0.899572169 |
| Setd1b | S | 1678 | 0.147618379 | 0.097215365 | 0.041411718 | -0.106206662 | -0.055803647 | 0.014169808 | 0.192649952 | 0.537648591 | 0.175988609 | 0.495211899 |
| Zbed6 | S | 365 | 0.079995248 | 0.090842352 | 0.124513874 | 0.044518626 | 0.033671522 | 0.113017352 | 0.139202407 | 0.122132404 | 0.508613988 | 0.636966221 |
| Zbed6 | S | 367 | 0.079995248 | 0.090842352 | 0.124513874 | 0.044518626 | 0.033671522 | 0.113017352 | 0.139202407 | 0.122132404 | 0.508613988 | 0.636966221 |
| Ncor2 | S | 2215 | 0.026665901 | 0.062687317 | 0.096969245 | 0.070303343 | 0.034281927 | 0.430255455 | 0.020366857 | 0.005650187 | 0.091890266 | 0.172834259 |
| Lats2 | S | 534 | -0.050128789 | 0.122098269 | 0.184102733 | 0.234231522 | 0.062004464 | 0.16054303 | 0.001876357 | 0.036799344 | 0.016595059 | 0.288061982 |
| Zbtb7c | S | 537 | -0.061583146 | 0.169827462 | 0.216479881 | 0.278063027 | 0.046652419 | 0.160557572 | 0.176967886 | 0.002334417 | 0.00214854 | 0.650439202 |
| Bap1 | S | 327 | 0.187435763 | 0.079658549 | 0.017973738 | -0.169462025 | -0.06168481 | 0.01273412 | 0.187499396 | 0.686180686 | 0.025637889 | 0.300474544 |
| Nipbl | S | 318 | -0.005849882 | 0.091403835 | 0.142449696 | 0.148299578 | 0.05104586 | 0.863532821 | 0.060741615 | 0.004720606 | 0.001833222 | 0.180136168 |
| 2310022A10Rik | S | 141 | -0.009125882 | 0.156348063 | 0.111487198 | 0.120613081 | -0.044860865 | 0.762399905 | 0.012716763 | 0.052493994 | 0.058569063 | 0.38307493 |
| 2310022A10Rik | T | 145 | -0.009125882 | 0.156348063 | 0.111487198 | 0.120613081 | -0.044860865 | 0.762399905 | 0.012716763 | 0.052493994 | 0.058569063 | 0.38307493 |
| Atg2a | S | 1241 | 0.185595395 | -0.010384139 | 0.146311758 | -0.039283636 | 0.156695898 | 0.045410033 | 0.91448022 | 0.071067384 | 0.602704574 | 0.168156557 |
| Trp53bp1 | T | 796 | 0.019224922 | 0.145096612 | 0.157692501 | 0.138467579 | 0.012595889 | 0.668543774 | 0.015999154 | 0.224288882 | 0.269940371 | 0.9070538 |
| Gprc5b | S | 355 | 0.028529567 | 0.104638998 | 0.097551255 | 0.069021689 | -0.007087742 | 0.45440985 | 0.030827783 | 0.026642455 | 0.145253873 | 0.825081684 |
| Nsd2 | T | 422 | 0.086862413 | 0.065041615 | 0.063893864 | -0.022968549 | -0.001147751 | 0.207261523 | 0.012275342 | 0.046231056 | 0.69222645 | 0.957897801 |
| Rad18 | S | 485 | 0.069714709 | 0.006284759 | 0.137002068 | 0.067287359 | 0.130717309 | 0.055284952 | 0.832129923 | 0.004269317 | 0.067010719 | 0.00841932 |
| Tut4 | S | 245 | 0.009712357 | 0.17203353 | 0.156060836 | 0.146348478 | -0.015972694 | 0.919228884 | 0.004824229 | 0.433572411 | 0.474240516 | 0.929931528 |
| Tut4 | S | 248 | 0.009712357 | 0.17203353 | 0.156060836 | 0.146348478 | -0.015972694 | 0.919228884 | 0.004824229 | 0.433572411 | 0.474240516 | 0.929931528 |
| Tut4 | S | 250 | 0.009712357 | 0.17203353 | 0.156060836 | 0.146348478 | -0.015972694 | 0.919228884 | 0.004824229 | 0.433572411 | 0.474240516 | 0.929931528 |
| Pacc1 | Y | 10 | 0.088626144 | 0.199337391 | 0.338567494 | 0.24994135 | 0.139230103 | 0.642607059 | 0.397050522 | 0.21628917 | 0.376084238 | 0.631042159 |
| Zc3h18 | S | 71 | 0.051064954 | 0.000570089 | 0.194222915 | 0.143157961 | 0.193652826 | 0.475937378 | 0.9858072 | 0.00111569 | 0.113748405 | 0.003165178 |
| Frmd6 | S | 450 | -0.109560919 | 0.126540838 | 0.237223424 | 0.346784344 | 0.110682587 | 0.067845786 | 0.08310982 | 0.000588035 | 0.004128195 | 0.122611063 |
| Ccnl1 | S | 341 | 0.004506401 | 0.080861193 | 0.139158811 | 0.13465241 | 0.058297618 | 0.811287303 | 0.007338245 | 0.034270062 | 0.033748219 | 0.194174935 |
| Foxj2 | S | 164 | -0.025396923 | 0.084948987 | 0.113600007 | 0.13899693 | 0.02865102 | 0.426382009 | 0.003253052 | 0.003793818 | 0.011278463 | 0.241582167 |
| Rbbp6 | S | 1278 | 0.018124346 | 0.057787719 | 0.125266903 | 0.107142557 | 0.067479184 | 0.510330592 | 0.228038796 | 0.000821104 | 0.010129157 | 0.179121699 |
| Kdm4b | S | 630 | 0.034835914 | 0.047467789 | 0.217950246 | 0.183114332 | 0.170482458 | 0.591232929 | 0.190426538 | 0.018913794 | 0.057635362 | 0.041265073 |
| Rtn4 | T | 171 | 0.130092486 | 0.067902251 | 0.144538666 | 0.01444618 | 0.076636415 | 0.064224673 | 0.492376169 | 0.144508585 | 0.852846643 | 0.479552932 |
| Bclaf1 | T | 400 | -0.006773466 | 0.085629699 | 0.143362613 | 0.150136079 | 0.057732914 | 0.839406347 | 0.065192266 | 0.00368306 | 0.001302022 | 0.130670527 |
| Tnk2 | Y | 842 | 0.198464395 | 0.128303013 | 0.019247 | -0.179217395 | -0.109056013 | 0.122927844 | 0.034372745 | 0.911486028 | 0.370198388 | 0.543921136 |
| Mphosph10 | S | 168 | 0.157617073 | 0.004800916 | 0.13454444 | -0.023072633 | 0.129743524 | 0.180486956 | 0.930229471 | 0.00798541 | 0.805762914 | 0.074336776 |
| Wapl | S | 468 | 0.028335075 | 0.055609447 | 0.094318418 | 0.065983342 | 0.038708971 | 0.454252264 | 0.051679769 | 0.001368329 | 0.145041754 | 0.13287171 |
| Wapl | S | 470 | 0.028335075 | 0.055609447 | 0.094318418 | 0.065983342 | 0.038708971 | 0.454252264 | 0.051679769 | 0.001368329 | 0.145041754 | 0.13287171 |
| Dido1 | S | 149 | -0.061950322 | 0.162961268 | 0.130077929 | 0.192028252 | -0.032883339 | 0.219137613 | 0.001015642 | 0.068251761 | 0.027083349 | 0.527555992 |
| Lmna | S | 570 | 0.001757901 | 0.026182115 | 0.295940884 | 0.294182982 | 0.269758769 | 0.990765703 | 0.630785615 | 0.006056331 | 0.145216042 | 0.012079824 |
| Rbms1 Rbms3 | S | 112;106 | -0.050683094 | 0.203772977 | 0.180209585 | 0.230892679 | -0.023563392 | 0.346846175 | 0.07687727 | 0.017894871 | 0.010260968 | 0.78487287 |
| Camk1d | T | 180 | 0.019019516 | 0.18211145 | 0.049329596 | 0.03031008 | -0.132781854 | 0.735199612 | 0.001417108 | 0.544373895 | 0.723459226 | 0.172591403 |
| Cdk13 | S | 343 | 0.014917052 | 0.087962155 | 0.120477123 | 0.105560071 | 0.032514968 | 0.762515591 | 0.030606028 | 0.012713217 | 0.079338578 | 0.252725215 |
| Mark1 Mark2 Mark3 | T | 211;214;215;208 | 0.142956208 | 0.052138996 | 0.113906033 | -0.029050175 | 0.061767037 | 0.11348759 | 0.380151543 | 0.063595266 | 0.665649977 | 0.156228912 |
| Brca2 | S | 231 | 0.055224251 | 0.089443766 | 0.187232995 | 0.132008744 | 0.097789229 | 0.1887584 | 0.548649111 | 0.030780432 | 0.059597562 | 0.521538638 |
| Stub1 | S | 24 | -0.019324478 | 0.134803148 | 0.108738711 | 0.128063189 | -0.026064437 | 0.514535702 | 0.004937045 | 0.03298614 | 0.025879895 | 0.505278979 |
| Prr12 | S | 1299 | 0.078766695 | 0.079113403 | 0.172847155 | 0.09408046 | 0.093733752 | 0.333761675 | 0.200284535 | 0.06325458 | 0.317145624 | 0.250957304 |
| Ttf1 | S | 845 | 0.090845919 | 0.071388453 | 0.071341149 | -0.01950477 | -4.73036E-05 | 0.075319724 | 0.155699037 | 0.033512462 | 0.612345099 | 0.99905742 |
| Nucb2 | S | 85 | -0.066499562 | 0.137363834 | 0.246947933 | 0.313447496 | 0.109584099 | 0.224133449 | 0.113350854 | 0.006854765 | 0.004527888 | 0.188678683 |
| Reps1 | T | 538 | 0.056342725 | 0.150951517 | 0.069486778 | 0.013144053 | -0.08146474 | 0.556137519 | 0.018674648 | 0.121497155 | 0.885918815 | 0.130098587 |
| Lrp1 | T | 4525 | 0.136849258 | 0.100794656 | 0.087158365 | -0.049690893 | -0.013636291 | 0.140246463 | 0.121593408 | 0.225403798 | 0.56947174 | 0.847469444 |
| Sec16a | S | 1028 | 0.069049317 | 0.092733661 | 0.102694236 | 0.033644919 | 0.009960575 | 0.123578468 | 0.179635083 | 0.04684788 | 0.462893355 | 0.869786433 |
| Gys1 | S | 672 | -0.009004581 | 0.074326882 | 0.18074789 | 0.189752471 | 0.106421007 | 0.735619853 | 0.0636757 | 0.007556464 | 0.010560448 | 0.05139062 |
| Arhgef40 | S | 959 | 0.046325516 | 0.063433787 | 0.082802249 | 0.036476734 | 0.019368462 | 0.027463327 | 0.0487757 | 0.052908787 | 0.271143518 | 0.564611663 |
| Larp4b | S | 603 | 0.026451765 | 0.063714644 | 0.069411163 | 0.042959398 | 0.005696519 | 0.438708848 | 0.005543805 | 0.004214324 | 0.241556784 | 0.669121087 |
| Iws1 | S | 458 | 0.060095914 | 0.055696363 | 0.110191789 | 0.050095876 | 0.054495427 | 0.126328879 | 0.111418616 | 0.025484274 | 0.217230207 | 0.162123115 |
| Ksr1 | S | 392 | 0.000209024 | 0.117552994 | 0.140289381 | 0.140080357 | 0.022736388 | 0.996550424 | 0.012214734 | 0.066362381 | 0.075949998 | 0.67429521 |
| Nes | S | 1541 | 0.191011503 | 0.195756829 | 0.15979668 | -0.031214823 | -0.035960149 | 0.425716041 | 0.183546528 | 0.450137528 | 0.900150859 | 0.838515625 |
| Hdgfl2 | S | 629 | -0.046262048 | 0.112952112 | 0.211866204 | 0.258128253 | 0.098914093 | 0.277523926 | 0.125716779 | 0.003432374 | 0.005341377 | 0.167595258 |
| Mtcl1 | S | 768 | 0.006754055 | 0.09483014 | 0.094683062 | 0.087929007 | -0.000147078 | 0.740880079 | 0.005155487 | 0.027052879 | 0.054842948 | 0.995990823 |
| Twistnb | S | 299 | 0.107097776 | 0.132820933 | 0.079216549 | -0.027881228 | -0.053604384 | 0.304013119 | 0.054989715 | 0.197385209 | 0.763914125 | 0.344513976 |
| Twistnb | S | 301 | 0.107097776 | 0.132820933 | 0.079216549 | -0.027881228 | -0.053604384 | 0.304013119 | 0.054989715 | 0.197385209 | 0.763914125 | 0.344513976 |
| Twistnb | S | 302 | 0.107097776 | 0.132820933 | 0.079216549 | -0.027881228 | -0.053604384 | 0.304013119 | 0.054989715 | 0.197385209 | 0.763914125 | 0.344513976 |
| Mki67 | T | 3021 | 0.181719011 | 0.215229088 | 0.015702796 | -0.166016215 | -0.199526292 | 0.242131896 | 0.063036025 | 0.811753762 | 0.275440774 | 0.083139506 |
| Kmt2d | S | 3122 | 0.05745501 | 0.124792735 | 0.101589507 | 0.044134497 | -0.023203228 | 0.352418451 | 0.068269004 | 0.075338142 | 0.382501192 | 0.626791041 |
| Amot | S | 1083 | -0.07012728 | 0.380523725 | 0.037810064 | 0.107937344 | -0.342713661 | 0.286728328 | 0.002630787 | 0.650077224 | 0.178211227 | 0.010469418 |
| Tcof1 | S | 1242 | 0.007264846 | 0.100146103 | 0.08915367 | 0.081888824 | -0.010992433 | 0.824216699 | 0.002116039 | 0.052700714 | 0.093093863 | 0.734190769 |
| Bcl9 | S | 18 | 0.098940952 | 0.095585683 | 0.140904483 | 0.041963531 | 0.0453188 | 0.223091494 | 0.266894296 | 0.081564887 | 0.607353805 | 0.599810568 |
| Spen | T | 2373 | 0.045044011 | 0.159465703 | 0.209396346 | 0.164352334 | 0.049930643 | 0.696298942 | 0.094511941 | 0.185896042 | 0.32180247 | 0.715431522 |
| Ino80b | S | 114 | 0.035963033 | 0.02963787 | 0.234232036 | 0.198269003 | 0.204594166 | 0.595495358 | 0.587079519 | 0.007381493 | 0.030574443 | 0.017438186 |

| | | | | | | | | | | | | |
|---|---|---|---|---|---|---|---|---|---|---|---|---|
| Ino80b | S | 116 | 0.035963033 | 0.02963787 | 0.234232036 | 0.198269003 | 0.204594166 | 0.595495358 | 0.587079519 | 0.007381493 | 0.030574443 | 0.017438186 |
| Ncoa5 | S | 34 | 0.022143512 | 0.099845481 | 0.115418525 | 0.093275012 | 0.015573044 | 0.629525099 | 0.004238037 | 0.120648882 | 0.200496747 | 0.779216322 |
| Arhgap32 | S | 952 | -0.002921847 | 0.085930275 | 0.126111494 | 0.129033341 | 0.040181218 | 0.908503105 | 0.177608074 | 0.000623745 | 0.004151467 | 0.459122879 |
| Prc1 | S | 4 | 0.416188456 | 0.110398628 | -0.270670033 | -0.686858489 | -0.381068661 | 0.004285024 | 0.426934532 | 0.427121362 | 0.123334731 | 0.293645386 |
| Rbmx Rbmxl1 | S | 205;208 | 0.094397614 | 0.116873479 | 0.018927771 | -0.075469843 | -0.097945709 | 0.032716396 | 0.017914471 | 0.71988286 | 0.23231214 | 0.14796175 |
| Kmt2a | S | 2152 | 0.080820591 | 0.082738422 | 0.06577798 | -0.015042611 | -0.016960442 | 0.041275703 | 0.161016519 | 0.055471599 | 0.636478155 | 0.726330701 |
| Atxn2 | S | 697 | 0.087118284 | 0.06185705 | 0.154441982 | 0.067323698 | 0.092584933 | 0.270716998 | 0.265177732 | 0.038792802 | 0.395641216 | 0.171556653 |
| Srp14 | S | 45 | -0.002647625 | 0.33690927 | -0.011811669 | -0.009164043 | -0.348720939 | 0.974615481 | 0.003776944 | 0.924257474 | 0.934502406 | 0.054933095 |
| Srsf3 | S | 140 | -0.002006774 | 0.094441552 | 0.120898514 | 0.122905288 | 0.026456962 | 0.946122194 | 0.00064094 | 0.117615587 | 0.107455902 | 0.637848406 |
| Igf2r | S | 2336 | 0.083397404 | 0.273141489 | -0.028195529 | -0.111592933 | -0.301337018 | 0.346455597 | 0.010182222 | 0.770645856 | 0.219804892 | 0.021684591 |
| Clk2 | S | 50 | 0.062051571 | 0.052330703 | 0.125552449 | 0.063500878 | 0.073221746 | 0.163122256 | 0.467168592 | 0.006470662 | 0.164045053 | 0.337663464 |
| Msh6 | S | 254 | -0.006036488 | 0.078982068 | 0.12073869 | 0.126775179 | 0.041756622 | 0.867667941 | 0.024056446 | 0.003969223 | 0.017332056 | 0.086319119 |
| Rbmx Rbmxl1 | S | 349;352 | -0.004634667 | 0.068367185 | 0.136249054 | 0.140883721 | 0.067881869 | 0.834047524 | 0.145387099 | 0.000691275 | 0.002724647 | 0.149666261 |
| Srsf1 | S | 234 | 0.129038251 | 0.091319211 | 0.02774842 | -0.101289831 | -0.063570791 | 0.017133671 | 0.050682588 | 0.779655818 | 0.345229925 | 0.523543909 |
| Mtres1 | S | 106 | 0.046703899 | 0.044934416 | 0.143663381 | 0.096959483 | 0.098728965 | 0.206734971 | 0.230511772 | 0.011467366 | 0.0498791 | 0.047290377 |
| Man1b1 | S | 102 | 0.021007104 | 0.078322156 | 0.171748993 | 0.150741889 | 0.093426838 | 0.511655733 | 0.273516986 | 0.009783439 | 0.02515741 | 0.21799235 |
| Agrn | S | 571 | -0.000173186 | 0.362334573 | 0.13436968 | 0.134542866 | -0.227964893 | 0.99879352 | 0.28243869 | 0.042188286 | 0.304258989 | 0.457841124 |
| Nsd3 | S | 150 | 0.04212918 | 0.054540085 | 0.083059456 | 0.040930275 | 0.02851937 | 0.128867673 | 0.012873429 | 0.027453439 | 0.20407572 | 0.292798267 |
| Cic | S | 662 | -0.003544598 | 0.126068534 | 0.125319236 | 0.128863834 | -0.000749298 | 0.923569273 | 0.030774791 | 0.024520111 | 0.026710646 | 0.986386705 |
| Brpf1 | S | 858 | -0.001534477 | 0.212646296 | 0.137719066 | 0.139253543 | -0.07492723 | 0.981446118 | 0.020390482 | 0.272952764 | 0.25762733 | 0.501344707 |
| Dda1 | S | 95 | 0.082704921 | 0.082849564 | 0.022097552 | -0.060607369 | -0.060752013 | 0.044851915 | 0.002409793 | 0.560013119 | 0.193957189 | 0.183904668 |
| Rbbp6 | S | 781 | -0.015688635 | 0.15005794 | 0.173610506 | 0.189299141 | 0.023552567 | 0.803787096 | 0.006965327 | 0.208396574 | 0.181569191 | 0.833550365 |
| Rbm15 | S | 108 | 0.282643614 | 0.128365029 | 0.03453091 | -0.248112705 | -0.09383412 | 0.231198466 | 0.103077071 | 0.553860794 | 0.280115271 | 0.147292467 |
| Slx4 | S | 940 | -0.003458273 | 0.065028342 | 0.166131452 | 0.169589725 | 0.10110311 | 0.924190297 | 0.280413383 | 0.001227581 | 0.006231128 | 0.134876623 |
| Bod1l | S | 3000 | 0.07249497 | 0.122965894 | 0.093037261 | 0.020542291 | -0.029928634 | 0.078682704 | 0.129526458 | 0.18670443 | 0.732814055 | 0.699630826 |
| Med13l | T | 393 | 0.052190805 | 0.095974501 | 0.300574213 | 0.248383408 | 0.204599711 | 0.526242272 | 0.216344512 | 0.146833159 | 0.166191552 | 0.225367159 |
| Irf2bpl | S | 641 | 0.082203465 | 0.113700949 | 0.019070644 | -0.063132821 | -0.094630305 | 0.326555385 | 0.000746569 | 0.485020815 | 0.430918419 | 0.027769691 |
| H1f3 | S | 172 | 0.233022366 | 0.109851917 | 0.090194219 | -0.142828148 | -0.019657698 | 0.238721265 | 0.256766868 | 0.235300255 | 0.423512052 | 0.779355012 |
| Taf3 | S | 291 | 0.019060428 | 0.088035224 | 0.136960983 | 0.117900555 | 0.048925759 | 0.676107287 | 0.041106343 | 0.025859077 | 0.054491598 | 0.268675588 |
| Hnrnph1 Hnrnph2 | S | 63;63 | 0.018063557 | 0.054599512 | 0.166772662 | 0.148709106 | 0.112173151 | 0.698515876 | 0.127713011 | 0.007124129 | 0.027802475 | 0.04959709 |
| Ccnl2 | S | 348 | 0.017540887 | 0.033855084 | 0.129496475 | 0.111955588 | 0.095641391 | 0.576433636 | 0.123854308 | 0.000818273 | 0.020542935 | 0.017907438 |
| Slx4 | S | 926 | -0.002105905 | 0.071315114 | 0.168879069 | 0.170984975 | 0.097563955 | 0.957023611 | 0.266707302 | 0.001957848 | 0.009831811 | 0.16143428 |
| Specc1 | S | 935 | -0.080679088 | 0.198166572 | 0.112834916 | 0.193514004 | -0.085331656 | 0.204689333 | 0.002449692 | 0.045955062 | 0.023425203 | 0.086119748 |
| Champ1 | S | 617 | 0.029090872 | 0.060272889 | 0.125125016 | 0.096034143 | 0.064852127 | 0.352114458 | 0.088777361 | 0.009090447 | 0.040140879 | 0.06976777 |
| Esyt2 | S | 685 | -0.017910748 | 0.050169561 | 0.135571822 | 0.15348257 | 0.08540226 | 0.699405434 | 0.029423357 | 0.000807927 | 0.042969891 | 0.016251658 |
| Dennd5a | S | 1085 | -0.053389781 | 0.266779415 | 0.147814162 | 0.201203943 | -0.118965253 | 0.626804952 | 0.010828188 | 0.372216152 | 0.257523447 | 0.446427301 |
| Eif4ebp1 | S | 99 | -0.105203425 | 0.238028467 | 0.259749875 | 0.364953299 | 0.021721408 | 0.258932962 | 0.024382731 | 0.108301835 | 0.054080484 | 0.860631117 |
| Cdc42ep1 | S | 205 | -0.033578847 | 0.022662025 | 0.329385685 | 0.362964532 | 0.30672366 | 0.575137684 | 0.709311511 | 0.003322306 | 0.00618648 | 0.009368851 |
| Irf9 | S | 139 | 0.058482744 | 0.093973649 | 0.198300836 | 0.139818091 | 0.104327187 | 0.506608928 | 0.275261424 | 0.048383341 | 0.03091507 | 0.0264011 |
| Peak1 | S | 500 | 0.04531767 | 0.084407239 | 0.083877122 | 0.038559452 | -0.000530117 | 0.142876705 | 0.052956906 | 0.03629925 | 0.266111825 | 0.988310384 |
| Tnrc18 | S | 1921 | -0.052991209 | 0.22907077 | 0.2668333 | 0.319824509 | 0.03776253 | 0.271239428 | 0.073129393 | 0.084910973 | 0.073679806 | 0.771354051 |
| Rb1 | S | 773 | 0.028299574 | 0.119864533 | 0.090839676 | 0.062540101 | -0.029024858 | 0.530740441 | 0.014167672 | 0.084240659 | 0.198595558 | 0.454261632 |
| Sptbn1 | S | 2163 | 0.00342472 | 0.112995007 | 0.106152638 | 0.102727918 | -0.006842369 | 0.910593685 | 0.009722176 | 0.041795714 | 0.062333642 | 0.835909177 |
| Tra2b | T | 103 | 0.037878518 | 0.161690914 | 0.191668895 | 0.153790378 | 0.029977981 | 0.719089948 | 0.097502796 | 0.156046658 | 0.270185889 | 0.795513688 |
| Srrm2 | S | 1537 | 0.058428823 | 0.043005432 | 0.08728683 | 0.028858007 | 0.044281398 | 0.004984277 | 0.427662686 | 0.022866013 | 0.278845921 | 0.424875937 |
| Zfat | S | 648 | 0.087670533 | 0.081578628 | 0.064683754 | -0.022986779 | -0.016894875 | 0.067516563 | 0.020734549 | 0.317283672 | 0.707863141 | 0.762429515 |
| Met | T | 975 | 0.023255045 | 0.112954179 | 0.105603907 | 0.082348863 | -0.007350272 | 0.672249715 | 0.01689233 | 0.059507301 | 0.185975701 | 0.846775655 |
| Cenpc1 | S | 302 | 0.037705406 | 0.13892108 | 0.081789452 | 0.044084046 | -0.057131628 | 0.453702958 | 0.009349472 | 0.23809898 | 0.502793676 | 0.368818275 |
| Nucks1 | S | 220 | 0.071038918 | 0.094489244 | 0.141040639 | 0.07000172 | 0.046551394 | 0.296661204 | 0.110705438 | 0.099629772 | 0.392471191 | 0.514884583 |
| Zfat | S | 647 | 0.078891582 | 0.156405121 | 0.072085639 | -0.006805942 | -0.084319481 | 0.208292331 | 0.076248769 | 0.213196679 | 0.911081763 | 0.27851007 |
| Golim4 | Y | 633 | -0.00626677 | 0.05343645 | 0.172402741 | 0.178669511 | 0.118966292 | 0.916107025 | 0.1278148 | 0.002636594 | 0.045065205 | 0.023679573 |
| Numa1 | S | 270 | 0.04043853 | 0.114139146 | 0.105182314 | 0.064743784 | -0.008956832 | 0.235133432 | 0.043018799 | 0.113164751 | 0.267230438 | 0.872588503 |
| Gmpr2 | S | 279 | 0.023730231 | 0.164253606 | 0.140436234 | 0.116706003 | -0.023817372 | 0.758934183 | 0.072261446 | 0.089688276 | 0.106252008 | 0.71092705 |
| Sap30 | S | 138 | 0.000959259 | 0.029808314 | 0.212376157 | 0.211416897 | 0.182567843 | 0.981612481 | 0.667258598 | 0.001065172 | 0.001692785 | 0.063947637 |
| Cep170 | S | 1123 | -0.025689739 | 0.1698588 | 0.12068754 | 0.146377279 | -0.049171261 | 0.269096645 | 0.010247499 | 0.040868064 | 0.035618887 | 0.326063179 |
| Dgcr8 | S | 95 | 0.079806187 | 0.084453379 | 0.059716613 | -0.020089574 | -0.024736766 | 0.052000801 | 0.011731463 | 0.44111154 | 0.784021173 | 0.729313679 |
| Ppig | S | 413 | 0.016882795 | 0.074370046 | 0.091391476 | 0.074508682 | 0.017021431 | 0.46993352 | 0.004463883 | 0.027826581 | 0.054057364 | 0.521307082 |
| Lrrfip1 | S | 88 | 0.05300156 | 0.11459969 | 0.027554101 | -0.025447459 | -0.087045589 | 0.079277317 | 0.004772979 | 0.349130453 | 0.186990976 | 0.016384566 |
| Mtmr3 | S | 632 | -0.027698656 | 0.098604886 | 0.122444119 | 0.150142775 | 0.023839233 | 0.441250986 | 0.021886161 | 0.002517653 | 0.013834273 | 0.449992562 |
| Tgfbr2 | S | 576 | -0.061502332 | 0.266524655 | 0.239567864 | 0.301070196 | -0.026956791 | 0.105404011 | 0.046112716 | 0.076035602 | 0.043219382 | 0.795857536 |
| Ddx46 | S | 804 | 0.005088591 | 0.108065392 | 0.092091194 | 0.087002603 | -0.015974198 | 0.815794898 | 0.007616437 | 0.032828168 | 0.040433019 | 0.609986311 |
| Caskin2 | S | 743 | 0.117818107 | 0.107379782 | -0.026495718 | -0.144313825 | -0.1338755 | 0.009709176 | 0.009733881 | 0.553632188 | 0.034706152 | 0.043169526 |
| Sipa1l1 | S | 1611 | -0.012284802 | 0.156627918 | 0.093843941 | 0.106128743 | -0.062783977 | 0.812122596 | 0.009980489 | 0.05635937 | 0.103585025 | 0.19756157 |
| Arhgap39 | S | 282 | 0.157759475 | 0.076134001 | 0.005665682 | -0.152093793 | -0.070468319 | 0.001723564 | 0.308811761 | 0.949564908 | 0.181022831 | 0.50123069 |
| Pds5b | S | 1190 | 0.020280907 | 0.171654611 | 0.163498517 | 0.14321761 | -0.008156095 | 0.752063115 | 0.240612582 | 0.032812256 | 0.044139451 | 0.947201101 |
| Ilf3 | S | 864 | 0.034937267 | 0.118433953 | 0.098161116 | 0.063223848 | -0.020272837 | 0.45035192 | 0.003753938 | 0.31352904 | 0.49771583 | 0.81154896 |
| Pde4a | S | 147 | -0.098420409 | 0.159765213 | 0.233096811 | 0.33151722 | 0.073331598 | 0.336635461 | 0.083435854 | 0.012030324 | 0.029459051 | 0.270006131 |
| Epb41l1 | T | 475 | 0.054833333 | 0.077430217 | 0.04973811 | -0.005095223 | -0.027692108 | 0.063008433 | 0.015640578 | 0.074249735 | 0.733612919 | 0.088671952 |
| Nes | S | 1565 | 0.087096044 | 0.116533327 | 0.029388098 | -0.057707946 | -0.087145229 | 0.274137761 | 0.001839056 | 0.593517684 | 0.481452358 | 0.185307446 |
| Casp12 | S | 104 | -0.127500414 | 0.269203027 | 0.169521568 | 0.297021981 | -0.099681459 | 0.022611297 | 0.000512825 | 0.134430391 | 0.050558091 | 0.313603397 |
| Jmjd1c | S | 996 | 0.023863839 | 0.006358998 | 0.207671831 | 0.183807993 | 0.201312834 | 0.675510864 | 0.942065322 | 0.000978176 | 0.035679086 | 0.1046085 |
| Pdlim1 | S | 185 | 0.059764573 | 0.202526547 | -0.021178626 | -0.080943199 | -0.223705173 | 0.420454454 | 0.001036832 | 0.673216525 | 0.302780205 | 0.010035148 |
| Rbbp6 | S | 1535 | 0.002819117 | 0.050187975 | 0.189244122 | 0.186425005 | 0.139056147 | 0.912620005 | 0.125455918 | 0.006459759 | 0.010707506 | 0.021764442 |
| Pole | S | 2019 | 0.088996879 | 0.10812826 | 0.137368035 | 0.048371156 | 0.029239776 | 0.191074643 | 0.2245991 | 0.129802086 | 0.511415799 | 0.738174357 |
| Gcfc2 | S | 169 | -0.008576847 | 0.087721739 | 0.120614204 | 0.129191051 | 0.032892465 | 0.778244124 | 0.039377321 | 0.003364413 | 0.006543458 | 0.285627457 |
| Brd9 | S | 587 | 0.035408297 | 0.159994502 | 0.073717034 | 0.038308737 | -0.086277468 | 0.544905083 | 0.005445121 | 0.413592138 | 0.679668514 | 0.352373667 |
| Setd5 | S | 569 | 0.16379635 | 0.168843291 | 0.049468581 | -0.114327769 | -0.119374709 | 0.303939297 | 0.046305548 | 0.624634272 | 0.472021353 | 0.261056937 |
| Smarcad1 | S | 127 | 0.079997338 | 0.028953487 | 0.128820169 | 0.048822831 | 0.099866682 | 0.047082055 | 0.387176656 | 0.036075331 | 0.299340936 | 0.076281167 |
| Lats2 | S | 171 | 0.024216023 | 0.239414654 | 0.103227494 | 0.079011471 | -0.13618716 | 0.729797274 | 0.036326563 | 0.335161818 | 0.435935225 | 0.250557142 |
| Srrm2 | S | 1552 | 0.219902017 | 0.187355276 | -0.038514235 | -0.258416252 | -0.225869511 | 0.079691444 | 0.116608091 | 0.890494204 | 0.387533063 | 0.439465231 |
| Srrm2 | S | 1548 | 0.219902017 | 0.187355276 | -0.038514235 | -0.258416252 | -0.225869511 | 0.079691444 | 0.116608091 | 0.890494204 | 0.387533063 | 0.439465231 |
| Hdac2 | S | 394 | 0.035110062 | 0.039059475 | 0.077979606 | 0.042869544 | 0.038920131 | 0.018107041 | 0.057203281 | 0.009329332 | 0.074500737 | 0.102701552 |
| Cdc42ep4 | S | 285 | 0.031746638 | 0.141174295 | 0.228300684 | 0.196554046 | 0.087126389 | 0.739594516 | 0.097646328 | 0.164492747 | 0.21653013 | 0.517412686 |
| Nr2f6 | S | 35 | 0.0049345 | 0.091165896 | 0.129694869 | 0.124760369 | 0.038528973 | 0.9280389 | 0.158760653 | 0.002419471 | 0.097559566 | 0.476021536 |
| Pwwp3a | S | 105 | 0.123079739 | 0.10752776 | 0.041904319 | -0.08117542 | -0.065623442 | 0.05482603 | 0.06934992 | 0.466048907 | 0.239976471 | 0.320918158 |
| Vps13c | S | 871 | 0.097653339 | 0.09517595 | 0.072842958 | -0.024810381 | -0.022332992 | 0.471358095 | 0.029752358 | 0.05307144 | 0.843246311 | 0.424569235 |
| Lurap1l | S | 74 | 0.022194623 | 0.063068358 | 0.108903515 | 0.086708892 | 0.045835157 | 0.77295236 | 0.282054465 | 0.000237671 | 0.323904502 | 0.403715262 |
| Trp53bp2 | S | 736 | 0.076519763 | 0.155270642 | 0.13838769 | 0.061867927 | -0.016882952 | 0.528675693 | 0.140387825 | 0.123341515 | 0.619482142 | 0.861572902 |
| Mki67 | S | 2649 | 0.124590557 | 0.198868074 | 0.091549863 | -0.033040693 | -0.107318211 | 0.43039104 | 0.088145737 | 0.394999999 | 0.834185383 | 0.369399597 |
| Zcchc8 | S | 429 | 0.104020835 | 0.074657941 | 0.152312314 | 0.048291479 | 0.077654373 | 0.120843334 | 0.26182311 | 0.163188068 | 0.589549056 | 0.411158625 |
| Ralgapb | S | 1310 | 0.066628926 | 0.15477561 | 0.067836574 | 0.001207648 | -0.086939036 | 0.261883204 | 0.020050682 | 0.446413106 | 0.987898139 | 0.320428491 |
| Pip5k1c | T | 552 | 0.026731401 | 0.177940752 | 0.059619526 | 0.032888125 | -0.118321226 | 0.498915234 | 0.006160788 | 0.438076252 | 0.642055963 | 0.174152236 |
| Srrm2 | S | 2691 | 0.067049849 | 0.038778255 | 0.096588863 | 0.029539014 | 0.057810608 | 0.013255949 | 0.320496913 | 0.038296674 | 0.323700753 | 0.199293924 |
| Suv39h2 | S | 448 | 0.077036658 | 0.13107001 | 0.03743399 | -0.039602668 | -0.09363602 | 0.037827459 | 0.079567841 | 0.239929466 | 0.180614044 | 0.165731351 |

| | | | | | | | | | | | | |
|---|---|---|---|---|---|---|---|---|---|---|---|---|
| Suv39h2 | S | 451 | 0.077036658 | 0.13107001 | 0.03743399 | -0.039602668 | -0.09363602 | 0.037827459 | 0.079567841 | 0.239929466 | 0.180614044 | 0.165731351 |
| Suv39h2 | S | 455 | 0.077036658 | 0.13107001 | 0.03743399 | -0.039602668 | -0.09363602 | 0.037827459 | 0.079567841 | 0.239929466 | 0.180614044 | 0.165731351 |
| Baz1b | S | 706 | 0.072515538 | 0.087862076 | 0.13836072 | 0.065845181 | 0.050498643 | 0.166010998 | 0.055709074 | 0.239291767 | 0.53050766 | 0.618083568 |
| Vps13b | S | 998 | 0.059053134 | 0.138607828 | 0.183563871 | 0.124510737 | 0.044956043 | 0.189688788 | 0.247547109 | 0.185750961 | 0.318551087 | 0.744463642 |
| Vps13b | S | 1001 | 0.059053134 | 0.138607828 | 0.183563871 | 0.124510737 | 0.044956043 | 0.189688788 | 0.247547109 | 0.185750961 | 0.318551087 | 0.744463642 |
| Gas2l1 | S | 297 | 0.021182327 | 0.175317003 | 0.170993429 | 0.149811102 | -0.004323574 | 0.647533568 | 0.046335161 | 0.260990112 | 0.309676986 | 0.974154173 |
| Mecom | S | 1124 | -0.009590504 | 0.132269489 | 0.152393165 | 0.161983669 | 0.020123676 | 0.881948696 | 0.112193907 | 0.014107063 | 0.057637287 | 0.760928557 |
| Supt7l | S | 321 | 0.037634849 | 0.110417616 | 0.194450494 | 0.156815645 | 0.084032878 | 0.545715154 | 0.083229025 | 0.126624014 | 0.191328752 | 0.427313278 |
| Taf4b | S | 312 | 0.05947631 | 0.225576206 | 0.016442162 | -0.043034148 | -0.209134044 | 0.529531381 | 0.005866943 | 0.917810636 | 0.798233326 | 0.262023438 |
| Tmpo | T | 354 | 0.1665067 | 0.091623968 | 0.079645467 | -0.086861233 | -0.011978502 | 0.142105962 | 0.249029963 | 0.160529176 | 0.36934779 | 0.843612955 |
| Rap1gap | S | 484 | 0.026364285 | 0.238943944 | -0.027200826 | -0.053565111 | -0.26614477 | 0.8174119 | 0.000485649 | 0.616933539 | 0.648022652 | 0.010490378 |
| Slc33a1 | S | 42 | 0.107588105 | 0.130321967 | 0.064067246 | -0.043520859 | -0.066254722 | 0.04987626 | 0.140721767 | 0.42312816 | 0.563120006 | 0.477488704 |
| Srsf4 | S | 325 | 0.042440659 | 0.086812802 | 0.102130903 | 0.059690244 | 0.015318101 | 0.605540318 | 0.102485619 | 0.006477936 | 0.476771871 | 0.70840248 |
| Gpatch8 | S | 915 | 0.04597332 | 0.050004006 | 0.194177949 | 0.148204629 | 0.144173943 | 0.261366985 | 0.242500599 | 0.039082971 | 0.096555874 | 0.09786491 |
| Gpatch8 | S | 918 | 0.04597332 | 0.050004006 | 0.194177949 | 0.148204629 | 0.144173943 | 0.261366985 | 0.242500599 | 0.039082971 | 0.096555874 | 0.09786491 |
| Ofd1 | S | 791 | 0.144407168 | 0.111878592 | 0.019089995 | -0.125317173 | -0.092788597 | 0.11541781 | 0.017335134 | 0.739513422 | 0.169625994 | 0.154243635 |
| Lrrfip1 | T | 82 | -0.066629561 | 0.316611942 | 0.080520778 | 0.147150338 | -0.236091164 | 0.483844871 | 0.008035036 | 0.347705689 | 0.079215361 | 0.008761403 |
| Usp34 | S | 3395 | 0.040755699 | 0.076750873 | 0.09352798 | 0.052772281 | 0.016777107 | 0.05980295 | 0.0874 | 0.046480095 | 0.167911595 | 0.680461106 |
| Cep120 | S | 380 | 0.083192749 | 0.010457005 | 0.136346587 | 0.053153838 | 0.125889583 | 0.042828898 | 0.791242908 | 0.017013399 | 0.209281583 | 0.031507328 |
| Cdk13 | S | 341 | 0.008758413 | 0.071832108 | 0.116699652 | 0.107941239 | 0.044867544 | 0.857190314 | 0.033333819 | 0.005817478 | 0.088962892 | 0.114318889 |
| Srsf1 | S | 238 | 0.100299615 | 0.081917299 | 0.058427904 | -0.04187171 | -0.023489394 | 0.029426605 | 0.050606869 | 0.446824822 | 0.560267462 | 0.731646895 |
| Trp53bp1 | S | 851 | 0.053263878 | 0.228207661 | 0.141968899 | 0.088705021 | -0.086238763 | 0.553451898 | 0.093295958 | 0.337492095 | 0.549722837 | 0.577017091 |
| Scfd1 | S | 300 | 0.066998793 | 0.129552176 | 0.029101718 | -0.037897075 | -0.100450458 | 0.050282732 | 0.015020368 | 0.60653053 | 0.507245966 | 0.149370976 |
| Cavin3 | S | 62 | 0.074335697 | 0.219251283 | 0.047958438 | -0.026377259 | -0.171292844 | 0.435534998 | 0.015971001 | 0.795864775 | 0.891454985 | 0.399526258 |
| Srrm2 | S | 884 | 0.059236636 | 0.096341274 | 0.127074784 | 0.067838149 | 0.03073351 | 0.339837857 | 0.113903703 | 0.061537975 | 0.160278737 | 0.374312019 |
| Palld | S | 1146 | -0.003197158 | 0.163151181 | 0.084039114 | 0.087236272 | -0.079112067 | 0.920277283 | 0.002196127 | 0.290518804 | 0.275672973 | 0.310598743 |
| Ino80 | S | 47 | 0.076450853 | 0.082310951 | 0.187090857 | 0.110640004 | 0.104779905 | 0.359426029 | 0.272507981 | 0.073810577 | 0.281400973 | 0.274437719 |
| Fnip1 | S | 945 | 0.174341466 | 0.097641182 | 0.0264815 | -0.147859966 | -0.071159682 | 0.03392351 | 0.146996208 | 0.664312032 | 0.038106214 | 0.035336329 |
| Cic | S | 2195 | 0.102452536 | 0.131210028 | 0.233930698 | 0.131478162 | 0.10272067 | 0.469754619 | 0.449214847 | 0.121278052 | 0.430282416 | 0.581635779 |
| Trim24 | S | 812 | -0.05082842 | 0.088343222 | 0.247426228 | 0.298254648 | 0.159083006 | 0.502475557 | 0.284722236 | 0.004405665 | 0.026561995 | 0.099595729 |
| Spop | S | 6 | -0.013133252 | 0.228182455 | 0.073669589 | 0.086802841 | -0.154512867 | 0.795641973 | 0.004698373 | 0.419864038 | 0.366078355 | 0.153185537 |
| Srrm2 | S | 1533 | 0.069768431 | 0.043497125 | 0.086447521 | 0.01667909 | 0.042950396 | 0.006151751 | 0.387597942 | 0.05244617 | 0.593282228 | 0.419643601 |
| Irf2bpl | S | 316 | -0.000429512 | 0.064023459 | 0.199283783 | 0.199713294 | 0.135260324 | 0.985734616 | 0.023364457 | 0.035429972 | 0.044140933 | 0.097984746 |
| Ddi2 | S | 120 | 0.100191844 | 0.160021663 | 0.047575823 | -0.05261602 | -0.112445839 | 0.250182281 | 0.029126437 | 0.510046045 | 0.555843989 | 0.170143462 |
| Zfp768 | S | 191 | -0.055059781 | 0.137931488 | 0.174160336 | 0.229220117 | 0.036228848 | 0.443556496 | 0.074625896 | 0.008056976 | 0.03107823 | 0.530766854 |
| Rrbp1 | S | 155 | 0.428258021 | 0.107975398 | 0.007396681 | -0.420861341 | -0.100578717 | 0.078885735 | 0.577265119 | 0.964448408 | 0.09498343 | 0.615602203 |
| Emsy | S | 173 | 0.054143413 | -0.011389843 | 0.178767927 | 0.124624514 | 0.19015777 | 0.298883595 | 0.696975111 | 0.001009922 | 0.059385108 | 0.002895496 |
| Ncor1 | S | 2449 | -0.025617104 | 0.28276975 | 0.109854484 | 0.135471588 | -0.172915266 | 0.733842578 | 0.015281065 | 0.432631651 | 0.350756462 | 0.258358631 |
| Ncor1 | S | 2451 | -0.025617104 | 0.28276975 | 0.109854484 | 0.135471588 | -0.172915266 | 0.733842578 | 0.015281065 | 0.432631651 | 0.350756462 | 0.258358631 |
| Topbp1 | S | 890 | 0.088940048 | 0.179837415 | 0.067556676 | -0.021384373 | -0.112281739 | 0.53063585 | 0.031774541 | 0.322220287 | 0.870673824 | 0.094980023 |
| Pxn | S | 340 | 0.039376128 | 0.199603032 | 0.102435067 | 0.063058939 | -0.097167964 | 0.632848059 | 0.096723466 | 0.101350061 | 0.413235744 | 0.329911137 |
| Gcfc2 | S | 66 | 0.516759405 | -0.010663417 | 0.101962704 | -0.414796702 | 0.112626121 | 0.389712897 | 0.630280228 | 0.011680978 | 0.474113218 | 0.017517591 |
| Mllt10 | S | 688 | 0.074251749 | 0.122678849 | 0.017825051 | -0.056426699 | -0.104853798 | 0.062540165 | 0.010012821 | 0.59674134 | 0.006091283 | 0.010593508 |
| Mllt10 | S | 691 | 0.074251749 | 0.122678849 | 0.017825051 | -0.056426699 | -0.104853798 | 0.062540165 | 0.010012821 | 0.59674134 | 0.006091283 | 0.010593508 |
| Vps26b | S | 304 | 0.16553853 | 0.245258201 | 0.032187238 | -0.133351292 | -0.213070962 | 0.260106723 | 0.099196485 | 0.83528135 | 0.27998566 | 0.130637033 |
| Kif7 | S | 898 | -0.015357045 | 0.233128978 | 0.094919557 | 0.110276601 | -0.138209421 | 0.881200344 | 0.010961504 | 0.34150002 | 0.377194795 | 0.197699461 |
| Cdk13 | S | 338 | 0.040319883 | 0.157775241 | 0.127227635 | 0.086907751 | -0.030547606 | 0.674926979 | 0.049895755 | 0.15482495 | 0.371896747 | 0.639770563 |
| Sipa1l1 | S | 1624 | 0.021009969 | 0.11281339 | 0.175488747 | 0.154478778 | 0.062675357 | 0.879726857 | 0.096875076 | 0.045814368 | 0.312050751 | 0.300394263 |
| Sipa1l1 | S | 1628 | 0.021009969 | 0.11281339 | 0.175488747 | 0.154478778 | 0.062675357 | 0.879726857 | 0.096875076 | 0.045814368 | 0.312050751 | 0.300394263 |
| Klhl22 | T | 605 | 0.133879783 | 0.153448979 | -0.00758542 | -0.141465203 | -0.161034399 | 0.177234065 | 0.00956217 | 0.91530838 | 0.188861373 | 0.087253018 |
| Usp1 | S | 67 | 0.055496568 | 0.098373218 | 0.035869351 | -0.019627217 | -0.062503867 | 0.040006956 | 0.007657443 | 0.38484614 | 0.602218657 | 0.172446391 |
| Srrm2 | S | 454 | 0.023862398 | 0.056806967 | 0.094534835 | 0.070672437 | 0.037727868 | 0.282192836 | 0.027134728 | 0.008321293 | 0.033433736 | 0.15700536 |
| Senp3 | S | 163 | 0.105912515 | 0.14565289 | 0.043655652 | -0.062256864 | -0.101997238 | 0.083948537 | 0.060747377 | 0.604479451 | 0.424110914 | 0.254029953 |
| Atrx | S | 322 | -0.008501921 | 0.134802479 | 0.186035675 | 0.194537596 | 0.051233196 | 0.939088802 | 0.076280081 | 0.055587551 | 0.174281745 | 0.505478113 |
| Ep400 | S | 923 | -0.004499058 | 0.075547019 | 0.124655626 | 0.129154684 | 0.049108607 | 0.865209067 | 0.047145256 | 0.003345493 | 0.005552722 | 0.153468574 |
| Baz1b | S | 709 | 0.078910274 | 0.06740334 | 0.135450123 | 0.05653985 | 0.068046783 | 0.06579842 | 0.114011755 | 0.221788138 | 0.554188092 | 0.485171998 |
| Ccdc71 | S | 202 | 0.120014237 | 0.077904221 | 0.081401719 | -0.038612517 | 0.003497498 | 0.077085028 | 0.193741926 | 0.161439842 | 0.527006411 | 0.951953281 |
| Sltm | S | 550 | 0.024071966 | 0.025199471 | 0.117795383 | 0.093723417 | 0.092595912 | 0.342586847 | 0.205292392 | 0.000579837 | 0.025956293 | 0.002460398 |
| Srrm1 | S | 220 | 0.035107101 | 0.035696661 | 0.113918496 | 0.078811395 | 0.078221835 | 0.134349716 | 0.231949247 | 0.003463407 | 0.02337984 | 0.045695208 |
| Nup35 | S | 121 | 0.397355065 | 0.311013069 | -0.037380281 | -0.434735347 | -0.348393351 | 0.177593988 | 0.398775898 | 0.887921726 | 0.039959997 | 0.274073225 |
| Nup35 | T | 127 | 0.397355065 | 0.311013069 | -0.037380281 | -0.434735347 | -0.348393351 | 0.177593988 | 0.398775898 | 0.887921726 | 0.039959997 | 0.274073225 |
| Nup35 | T | 128 | 0.397355065 | 0.311013069 | -0.037380281 | -0.434735347 | -0.348393351 | 0.177593988 | 0.398775898 | 0.887921726 | 0.039959997 | 0.274073225 |
| Ifngr1 | S | 370 | -0.034087877 | 0.146021259 | 0.198526687 | 0.232614564 | 0.052505428 | 0.155768977 | 0.055358905 | 0.017156238 | 0.009468212 | 0.370532802 |
| Fbxw8 | S | 86 | 0.033379476 | 0.141551815 | 0.085635044 | 0.052255568 | -0.055916771 | 0.417177421 | 0.021785514 | 0.169722376 | 0.377916319 | 0.358973999 |
| Rbbp6 | S | 862 | 0.154301006 | 0.078408352 | 0.059410387 | -0.094890618 | -0.018997965 | 0.119252136 | 0.044023246 | 0.415737888 | 0.328399397 | 0.768146557 |
| Srsf3 | S | 138 | -0.006303057 | 0.093399013 | 0.117212168 | 0.123515225 | 0.023813155 | 0.836609735 | 0.000401636 | 0.146425994 | 0.125809349 | 0.693591834 |
| Srrm2 | T | 1574 | 0.021140173 | 0.116393825 | 0.067519774 | 0.046379601 | -0.04887405 | 0.57576417 | 0.002661437 | 0.152308829 | 0.342793552 | 0.264700707 |
| Mecp2 | S | 421 | 0.000912373 | 0.131791196 | 0.156491388 | 0.155579015 | 0.024700192 | 0.98466656 | 0.048264737 | 0.062879892 | 0.084863866 | 0.698402722 |
| Atrx | S | 993 | 0.001456264 | 0.086981194 | 0.091198643 | 0.089742379 | 0.004217449 | 0.97421316 | 0.022502301 | 0.00379046 | 0.129143669 | 0.861440687 |
| Tsc22d4 | S | 298 | 0.061935173 | 0.108036038 | 0.055002778 | -0.006932395 | -0.05303326 | 0.083990035 | 0.023072755 | 0.284328621 | 0.865469394 | 0.288784388 |
| Myo18a | S | 2039 | 0.000729255 | 0.067561608 | 0.09403651 | 0.093307256 | 0.026474902 | 0.980673789 | 0.00434826 | 0.007041048 | 0.045019077 | 0.236131385 |
| Cttn | S | 417 | -0.056406675 | 0.186915277 | 0.176053642 | 0.232460317 | -0.010861635 | 0.018052209 | 0.005656598 | 0.030293437 | 0.019752197 | 0.827009518 |
| Toe1 | S | 5 | -0.002387029 | 0.086810125 | 0.061817216 | 0.064204245 | -0.024992909 | 0.884089734 | 0.014834955 | 0.00032573 | 0.026668482 | 0.234327397 |
| Meox2 | S | 154 | -0.174771337 | 0.108002451 | 0.237752277 | 0.412523614 | 0.129749826 | 0.018126026 | 0.045576447 | 4.00354E-05 | 0.002994279 | 0.035292691 |
| Ehmt1 | S | 393 | 0.098850353 | 0.086032253 | 0.242492701 | 0.143642348 | 0.156460448 | 0.25927097 | 0.252967432 | 0.246887978 | 0.45318687 | 0.415050482 |
| Ehmt1 | S | 399 | 0.098850353 | 0.086032253 | 0.242492701 | 0.143642348 | 0.156460448 | 0.25927097 | 0.252967432 | 0.246887978 | 0.45318687 | 0.415050482 |
| Ehmt1 | T | 402 | 0.098850353 | 0.086032253 | 0.242492701 | 0.143642348 | 0.156460448 | 0.25927097 | 0.252967432 | 0.246887978 | 0.45318687 | 0.415050482 |
| Nucks1 | S | 225 | 0.053535653 | 0.092947392 | 0.116773537 | 0.063237884 | 0.023826144 | 0.266347515 | 0.051649322 | 0.117314618 | 0.348283216 | 0.691259812 |
| Nucks1 | S | 230 | 0.053535653 | 0.092947392 | 0.116773537 | 0.063237884 | 0.023826144 | 0.266347515 | 0.051649322 | 0.117314618 | 0.348283216 | 0.691259812 |
| Nucks1 | S | 231 | 0.053535653 | 0.092947392 | 0.116773537 | 0.063237884 | 0.023826144 | 0.266347515 | 0.051649322 | 0.117314618 | 0.348283216 | 0.691259812 |
| Akap8 | S | 336 | 0.061312811 | 0.007223451 | 0.07815399 | 0.01684118 | 0.070930539 | 0.006644486 | 0.658332777 | 0.002613115 | 0.2617208 | 0.00582159 |
| Cuedc1 | S | 9 | -0.121263253 | 0.27823173 | 0.150044894 | 0.271308146 | -0.128186836 | 0.237656402 | 0.032080599 | 0.029281676 | 0.053545092 | 0.206380703 |
| Dtl | S | 601 | 0.162972227 | 0.098581083 | 0.125219723 | -0.037752503 | 0.02663864 | 0.133307422 | 0.205225801 | 0.440395684 | 0.815290381 | 0.86437366 |
| Dkc1 | S | 508 | 0.097816543 | 0.070880422 | 0.052792319 | -0.045024224 | -0.018088103 | 0.125339209 | 0.017066343 | 0.180848667 | 0.421480336 | 0.576265462 |
| Matr3 | S | 604 | -0.072420994 | 0.099595308 | 0.232248473 | 0.304669467 | 0.132653165 | 0.194899875 | 0.036059046 | 0.008145988 | 0.005279703 | 0.070837664 |
| Sec62 | S | 341 | 0.099242995 | 0.029466467 | 0.220526111 | 0.121283116 | 0.191059644 | 0.361156589 | 0.656047205 | 0.035379027 | 0.305276515 | 0.067151209 |
| Akap12 | S | 1285 | -0.003412018 | 0.472680964 | -0.082831145 | -0.079419127 | -0.555512109 | 0.981073713 | 0.006665753 | 0.488916026 | 0.447473234 | 0.000557633 |
| Myo18a | S | 2016 | 0.100643124 | 0.254797102 | 0.062958148 | -0.037684976 | -0.191838954 | 0.305456474 | 0.248405288 | 0.148966827 | 0.669926353 | 0.351002192 |
| Zc3hc1 | S | 357 | 0.006705633 | 0.093067882 | 0.110632491 | 0.103926858 | 0.017564609 | 0.822600776 | 0.001463381 | 0.133805597 | 0.142378189 | 0.746282581 |
| Tra2a | S | 36 | 0.063027158 | 0.079898643 | 0.177749522 | 0.114722364 | 0.097850879 | 0.543828046 | 0.2110746 | 0.046385783 | 0.31005118 | 0.197999862 |
| Tra2a | S | 38 | 0.063027158 | 0.079898643 | 0.177749522 | 0.114722364 | 0.097850879 | 0.543828046 | 0.2110746 | 0.046385783 | 0.31005118 | 0.197999862 |
| Syne1 | S | 8726 | 0.164369304 | 0.110407984 | 0.104189623 | -0.060179681 | -0.006218361 | 0.138000967 | 0.221835669 | 0.400469528 | 0.591357971 | 0.948258494 |
| Syne1 | S | 8729 | 0.164369304 | 0.110407984 | 0.104189623 | -0.060179681 | -0.006218361 | 0.138000967 | 0.221835669 | 0.400469528 | 0.591357971 | 0.948258494 |
| Chaf1b | S | 442 | 0.0420195 | 0.085462884 | 0.098869172 | 0.056849672 | 0.013406288 | 0.412930544 | 0.052806731 | 0.028686193 | 0.281918732 | 0.708624909 |

| | | | | | | | | | | | | |
|---|---|---|---|---|---|---|---|---|---|---|---|---|
| Mtcl1 | S | 276 | 0.152152317 | 0.131840658 | 0.010078699 | -0.142073618 | -0.121761959 | 0.043264007 | 0.101398811 | 0.809798607 | 0.062888282 | 0.127777205 |
| Mtcl1 | S | 277 | 0.152152317 | 0.131840658 | 0.010078699 | -0.142073618 | -0.121761959 | 0.043264007 | 0.101398811 | 0.809798607 | 0.062888282 | 0.127777205 |
| Ing5 | T | 147 | 0.002586775 | 0.06072269 | 0.118685682 | 0.116098906 | 0.057962992 | 0.934976978 | 0.09119684 | 0.001237942 | 0.025666175 | 0.102760083 |
| Ddx46 | T | 305 | 0.048572333 | 0.047913929 | 0.351341876 | 0.302769543 | 0.303427947 | 0.729521432 | 0.553715398 | 0.060213395 | 0.127621358 | 0.076152829 |
| Il6st | S | 665 | -0.050795971 | 0.057788595 | 0.205200787 | 0.255996758 | 0.147412193 | 0.515648449 | 0.175136931 | 0.001558862 | 0.040317865 | 0.015771765 |
| Pkp4 | S | 313 | 0.036033372 | 0.046916681 | 0.125445684 | 0.089412313 | 0.078529003 | 0.300513424 | 0.185299638 | 0.006400849 | 0.028830035 | 0.041152905 |
| Ap1m1 | T | 154 | 0.180108821 | 0.066021265 | -0.036874105 | -0.216982927 | -0.10289537 | 0.000279083 | 0.464355401 | 0.525676961 | 0.035700487 | 0.308890072 |
| Fbf1 | S | 206 | 0.038033833 | 0.182546503 | 0.209588335 | 0.171554502 | 0.027041832 | 0.619502976 | 0.180351924 | 0.189385375 | 0.261507243 | 0.862140459 |
| Add1 | S | 465 | -0.055957112 | 0.151307696 | 0.16480844 | 0.220765552 | 0.013500744 | 0.095309151 | 0.026651874 | 0.006821271 | 0.001933665 | 0.738464672 |
| Spag17 | S | 1845 | 0.116663862 | 0.087907503 | 0.092853945 | -0.023809917 | 0.004946442 | 0.092993764 | 0.264449192 | 0.155482753 | 0.713040633 | 0.947795869 |
| Fmr1 | T | 462 | 0.195311856 | 0.223307682 | -0.061979247 | -0.257291103 | -0.285286928 | 0.15052107 | 0.048197879 | 0.704249199 | 0.195648144 | 0.153977407 |
| Lrrc8c | S | 212 | -0.116971826 | 0.073659858 | 0.250818274 | 0.3677901 | 0.177158415 | 0.015758291 | 0.043944667 | 0.000126794 | 0.000333474 | 0.002718665 |
| Huwe1 | S | 2365 | -0.012745931 | 0.132753126 | 0.119829596 | 0.132575527 | -0.012923531 | 0.622576377 | 0.036527987 | 0.017154867 | 0.02521729 | 0.78267452 |
| Nr2f6 | T | 29 | 0.170071365 | 0.137514861 | 0.102928535 | -0.067142831 | -0.034586327 | 0.086020141 | 0.443248294 | 0.41302353 | 0.531054626 | 0.844384689 |
| Nr2f6 | T | 40 | 0.170071365 | 0.137514861 | 0.102928535 | -0.067142831 | -0.034586327 | 0.086020141 | 0.443248294 | 0.41302353 | 0.531054626 | 0.844384689 |
| Scap | S | 821 | 0.009618702 | 0.151631194 | 0.070377775 | 0.060759073 | -0.081253418 | 0.699944501 | 0.030822441 | 0.029925608 | 0.060618092 | 0.135374473 |
| Wwc2 | S | 1017 | -0.089996937 | 0.176636371 | 0.253711092 | 0.343708029 | 0.077074721 | 0.000117893 | 0.023024434 | 0.000627905 | 0.000935462 | 0.122059634 |
| Arhgef6 | Y | 643 | -0.002413558 | 0.497246797 | -0.034236229 | -0.031822671 | -0.531483026 | 0.98867092 | 0.024359947 | 0.85630838 | 0.773453115 | 0.012286649 |
| Stt3b | S | 18 | 0.141898058 | 0.054923358 | 0.089552299 | -0.052345759 | 0.034628941 | 0.087846321 | 0.213302247 | 0.164566953 | 0.475843712 | 0.557307703 |
| Zfp148 | S | 784 | 0.029737741 | 0.127005901 | 0.099685317 | 0.069947576 | -0.027320584 | 0.717308129 | 0.011005096 | 0.164228419 | 0.449438422 | 0.630676865 |
| Ppp4r3a | S | 758 | 0.019008683 | 0.11477749 | 0.089275826 | 0.070267143 | -0.025501664 | 0.455554419 | 0.031103932 | 0.044878807 | 0.092157485 | 0.543714711 |
| Cdk13 | S | 396 | 0.022342661 | 0.067779722 | 0.154531762 | 0.132189101 | 0.08675204 | 0.720689361 | 0.012603422 | 0.065137013 | 0.135309635 | 0.199001223 |
| Cdk13 | S | 398 | 0.022342661 | 0.067779722 | 0.154531762 | 0.132189101 | 0.08675204 | 0.720689361 | 0.012603422 | 0.065137013 | 0.135309635 | 0.199001223 |
| Sipa1l1 | S | 193 | 0.022165709 | 0.062548942 | 0.124070067 | 0.101904358 | 0.061521125 | 0.471042638 | 0.025335276 | 0.026544497 | 0.053518052 | 0.158482582 |
| Ctr9 | S | 1079 | 0.239486409 | 0.078919195 | -0.020651289 | -0.260137698 | -0.099570484 | 0.013011167 | 0.297991262 | 0.889417645 | 0.158027807 | 0.503840772 |
| Maml1 | S | 127 | 0.330737047 | 0.014760075 | 0.104573399 | -0.226163648 | 0.089813324 | 0.062643464 | 0.897757448 | 0.466670331 | 0.155269091 | 0.392242182 |
| Rb1 | S | 848 | 0.045600394 | 0.136170281 | 0.09219335 | 0.046592956 | -0.043976931 | 0.42916843 | 0.042445885 | 0.138715489 | 0.17288272 | 0.161597257 |
| Trim24 | S | 1043 | 0.040158194 | 0.04745871 | 0.239385286 | 0.199227092 | 0.191926576 | 0.795828298 | 0.690709439 | 0.010000462 | 0.26279205 | 0.172276618 |
| Taf3 | S | 301 | 0.070736848 | 0.003834124 | 0.107931278 | 0.03719443 | 0.104097155 | 0.046204374 | 0.818555216 | 0.00335959 | 0.227653065 | 0.013234649 |
| Kmt2a | S | 3032 | 0.078961623 | 0.076963228 | 0.069965608 | -0.008996015 | -0.00699762 | 0.145372832 | 0.019694685 | 0.196385634 | 0.872324265 | 0.877696009 |
| Rbl1 | S | 650 | -0.043204561 | 0.119965782 | 0.111117392 | 0.154321953 | -0.00884839 | 0.392442824 | 0.058235739 | 0.000746878 | 0.059240911 | 0.826147487 |
| Pard3 | S | 144 | 0.110310814 | 0.077684974 | 0.110973249 | 0.000662434 | 0.033288275 | 0.21596897 | 0.084172922 | 0.192358215 | 0.994210114 | 0.644394129 |
| Ccdc88a | S | 237 | -0.03678657 | 0.131072135 | 0.172636237 | 0.209422807 | 0.041564101 | 0.616345289 | 0.004672274 | 0.150569376 | 0.107481112 | 0.654965615 |
| Zranb2 | S | 120 | 0.047805586 | 0.015720552 | 0.10989576 | 0.062090174 | 0.094175208 | 0.066676572 | 0.503607017 | 0.002445858 | 0.020939332 | 0.010901954 |
| Chaf1a | S | 51 | -0.028742747 | 0.106319971 | 0.164931769 | 0.193674516 | 0.058611798 | 0.489031734 | 0.029557735 | 0.019111221 | 0.015827132 | 0.259958542 |
| Kank2 | S | 246 | -0.024459237 | 0.202364289 | 0.139161782 | 0.163621019 | -0.063202507 | 0.728352602 | 0.062260843 | 0.066646297 | 0.022794291 | 0.43129386 |
| Hivep1 | S | 148 | 0.009169804 | 0.093626289 | 0.183546039 | 0.174376234 | 0.08991975 | 0.900176454 | 0.125461698 | 0.029869099 | 0.077607854 | 0.202403281 |
| Cenpc1 | S | 280 | 0.062509318 | 0.066489491 | 0.089051582 | 0.026542264 | 0.022562091 | 0.144352966 | 0.083539937 | 0.047696808 | 0.463133994 | 0.455217003 |
| Raly | S | 294 | 0.016420174 | 0.066665967 | 0.100999703 | 0.084579529 | 0.034333736 | 0.545017065 | 0.038138535 | 0.007053273 | 0.00319371 | 0.050913203 |
| Zdhhc5 | S | 593 | 0.045847798 | 0.149295061 | 0.154226795 | 0.108378997 | 0.004931734 | 0.566971617 | 0.086413396 | 0.188183748 | 0.327116519 | 0.960863828 |
| Epb41l1 | S | 541 | 0.097667854 | 0.108875115 | 0.059323096 | -0.038344758 | -0.049552019 | 0.068952468 | 0.158529092 | 0.189333256 | 0.262660425 | 0.437527556 |
| Map3k4 | S | 449 | 0.103156973 | 0.06922063 | 0.096778395 | -0.006378578 | 0.027557765 | 0.102722928 | 0.460908237 | 0.040604404 | 0.898703672 | 0.7592371 |
| Map3k4 | S | 454 | 0.103156973 | 0.06922063 | 0.096778395 | -0.006378578 | 0.027557765 | 0.102722928 | 0.460908237 | 0.040604404 | 0.898703672 | 0.7592371 |
| Erbin | S | 440 | -0.035965028 | 0.220919681 | 0.122398169 | 0.158363197 | -0.098521512 | 0.778779059 | 0.005090856 | 0.509109457 | 0.449475483 | 0.583402041 |
| Rcor1 | S | 121 | 0.176175336 | 0.177854217 | -0.079776195 | -0.255951531 | -0.257630413 | 0.109250185 | 0.022835696 | 0.690387556 | 0.268609848 | 0.266290043 |
| Scrib | S | 1292 | -0.01751391 | 0.080641964 | 0.083446277 | 0.100960187 | 0.002804314 | 0.23999812 | 0.000258763 | 0.010906814 | 0.008684801 | 0.875845011 |
| Sox4 | S | 301 | -0.297634907 | 0.017188923 | 0.947598803 | 1.245233709 | 0.93040988 | 0.632008927 | 0.921903606 | 0.066225057 | 0.159501233 | 0.083100914 |
| Fam219b | S | 125 | 0.055567844 | 0.137915165 | 0.112193511 | 0.056625667 | -0.025721653 | 0.374542275 | 0.114851572 | 0.116081784 | 0.437193154 | 0.743110853 |
| Fam219b | S | 127 | 0.055567844 | 0.137915165 | 0.112193511 | 0.056625667 | -0.025721653 | 0.374542275 | 0.114851572 | 0.116081784 | 0.437193154 | 0.743110853 |
| Cep131 | S | 47 | -0.084076555 | 0.353420879 | 0.197366113 | 0.281442667 | -0.156054766 | 0.468204568 | 0.047187833 | 0.303570904 | 0.187250439 | 0.428766121 |
| Smarcad1 | S | 124 | 0.084686191 | 0.027385285 | 0.126236235 | 0.041550043 | 0.09885095 | 0.059643889 | 0.437803505 | 0.042133576 | 0.397486087 | 0.087592208 |
| Zfp462 | S | 2161 | 0.067480417 | 0.07538622 | 0.097712758 | 0.030232341 | 0.022326538 | 0.119922633 | 0.142729249 | 0.070029237 | 0.518921274 | 0.659085976 |
| Bcl9l | S | 1473 | -0.134678125 | 0.282019025 | 0.253487674 | 0.388165799 | -0.028531352 | 0.133024997 | 0.007794592 | 0.217105056 | 0.102950456 | 0.867303852 |
| Alkbh5 | S | 372 | -0.114483064 | 0.140129299 | 0.193395322 | 0.307878386 | 0.053265383 | 0.164466523 | 0.001705999 | 0.042837776 | 0.015507758 | 0.405292459 |
| Tada3 | S | 298 | 0.037904989 | 0.038858602 | 0.143855968 | 0.105950979 | 0.104997365 | 0.405356436 | 0.363552513 | 0.004183725 | 0.071955562 | 0.057856608 |
| Prpf4b | S | 394 | -0.022960689 | 0.086710606 | 0.167051946 | 0.190012635 | 0.08034134 | 0.169617422 | 0.000981827 | 0.054420676 | 0.046748922 | 0.208745688 |
| Prpf4b | S | 396 | -0.022960689 | 0.086710606 | 0.167051946 | 0.190012635 | 0.08034134 | 0.169617422 | 0.000981827 | 0.054420676 | 0.046748922 | 0.208745688 |
| Kdm1a | S | 138 | 0.036398125 | 0.036270862 | 0.111745238 | 0.075347113 | 0.075474376 | 0.250711452 | 0.148206999 | 0.004173047 | 0.061847677 | 0.030546851 |
| Creb1 | S | 340 | -0.020283505 | 0.056816993 | 0.171291268 | 0.191574773 | 0.114474275 | 0.408674088 | 0.013385065 | 0.007989663 | 0.001957115 | 0.010189794 |
| Ubap2l | S | 491 | 0.051212812 | 0.064295895 | 0.133780739 | 0.082567927 | 0.069484844 | 0.18260696 | 0.204435696 | 0.040437081 | 0.134440048 | 0.233944209 |
| Kmt2d | S | 2318 | 0.052594946 | 0.036091752 | 0.154377518 | 0.101782572 | 0.118285767 | 0.056935183 | 0.50239834 | 0.033945285 | 0.115482376 | 0.108985288 |
| Taf2 | S | 1093 | 0.028777871 | 0.144537695 | 0.106330255 | 0.077552384 | -0.03820744 | 0.676252155 | 0.008793368 | 0.348078389 | 0.504419581 | 0.710798753 |
| Snx33 | S | 169 | -0.016149617 | 0.210156931 | 0.082344451 | 0.098494068 | -0.12781248 | 0.756766804 | 0.024787346 | 0.083028718 | 0.082327906 | 0.111442566 |
| Mdc1 | S | 918 | 0.086647765 | 0.130304062 | 0.04157589 | -0.045071875 | -0.088728173 | 0.257734388 | 0.010073214 | 0.582008824 | 0.608306126 | 0.273148089 |
| Atxn2l | S | 687 | 0.353949963 | -0.008487076 | -0.062477855 | -0.416427818 | -0.053990779 | 0.001328992 | 0.957595198 | 0.382121344 | 0.001930945 | 0.733939254 |
| Phc3 | T | 607 | -0.028485569 | 0.057058881 | 0.216780198 | 0.245265766 | 0.159721316 | 0.481435784 | 0.156390725 | 0.004360051 | 0.007452006 | 0.038281628 |
| Bcl9 | S | 104 | -0.014435723 | 0.119069808 | 0.269379139 | 0.283814862 | 0.15030933 | 0.676577647 | 0.270405076 | 0.03595446 | 0.030470282 | 0.213986508 |
| Fam118a | S | 311 | 0.029551303 | 0.115212356 | 0.020843818 | -0.008707485 | -0.094368539 | 0.192191374 | 0.000430657 | 0.618681822 | 0.830328856 | 0.108697829 |
| Trp53bp1 | S | 1427 | -0.016083787 | 0.164435167 | 0.062625322 | 0.078709109 | -0.101809845 | 0.768651684 | 0.001707754 | 0.222415768 | 0.259651902 | 0.090469811 |
| Zc3h3 | S | 317 | 0.20810535 | 0.058099519 | 0.057853178 | -0.150252173 | -0.000246341 | 0.019198983 | 0.724588162 | 0.42792478 | 0.019664141 | 0.998735916 |
| Ccdc9b | S | 225 | 0.03081465 | 0.053877259 | 0.159604558 | 0.128790007 | 0.105727298 | 0.305326762 | 0.076097198 | 0.045773318 | 0.058025914 | 0.093555776 |
| Htatsf1 | S | 724 | 0.014130334 | 0.066073668 | 0.15504651 | 0.140916176 | 0.088972842 | 0.519271038 | 0.117253574 | 0.017434178 | 0.046734053 | 0.102637637 |
| Igf2r | S | 2337 | 0.045659762 | 0.376248507 | -0.058115626 | -0.103775388 | -0.434364132 | 0.691199775 | 0.014907099 | 0.770780346 | 0.571063965 | 0.100766891 |
| Srrm2 | S | 1683 | 0.052060261 | 0.011976963 | 0.092340255 | 0.040279994 | 0.080363292 | 0.05582885 | 0.641685246 | 0.000908658 | 0.112142495 | 0.043753629 |
| Pde4d | S | 141 | 0.000835985 | 0.124167235 | 0.151140226 | 0.150304241 | 0.026972991 | 0.972850406 | 0.015781324 | 0.143737459 | 0.150243239 | 0.736386454 |
| Prrc2a | S | 1321 | 0.162711292 | 0.062601398 | 0.121542103 | -0.04116919 | 0.058940704 | 0.059627171 | 0.702949185 | 0.20651934 | 0.622524498 | 0.722403933 |
| Prrc2a | S | 1323 | 0.162711292 | 0.062601398 | 0.121542103 | -0.04116919 | 0.058940704 | 0.059627171 | 0.702949185 | 0.20651934 | 0.622524498 | 0.722403933 |
| Prrc2a | S | 1324 | 0.162711292 | 0.062601398 | 0.121542103 | -0.04116919 | 0.058940704 | 0.059627171 | 0.702949185 | 0.20651934 | 0.622524498 | 0.722403933 |
| Prrc2a | T | 1319 | 0.162711292 | 0.062601398 | 0.121542103 | -0.04116919 | 0.058940704 | 0.059627171 | 0.702949185 | 0.20651934 | 0.622524498 | 0.722403933 |
| Ywhag | S | 71 | -0.003796403 | 0.252389136 | 0.1543462 | 0.158142603 | -0.098042936 | 0.970480235 | 0.119213941 | 0.149572992 | 0.240953393 | 0.461811561 |
| Mia2 | S | 1271 | 0.008055992 | 0.200355781 | 0.318566099 | 0.310510108 | 0.118210318 | 0.76199494 | 0.216569258 | 0.251675032 | 0.260456434 | 0.641043426 |
| Dnajc5 | S | 10 | 0.057839938 | 0.067093871 | 0.03256212 | -0.025277817 | -0.03453175 | 0.01143874 | 0.007162358 | 0.373831219 | 0.467879541 | 0.345059583 |
| Srrm2 | T | 1365 | 0.109644934 | 0.081893681 | 0.034345158 | -0.075299776 | -0.047548523 | 0.015788633 | 0.066966884 | 0.669973771 | 0.385015051 | 0.566719957 |
| Rbm15 | S | 51 | -0.033529674 | 0.111401466 | 0.151224415 | 0.18475409 | 0.039822949 | 0.606282617 | 0.020696868 | 0.02832467 | 0.05864408 | 0.331353765 |
| Acap2 | S | 381 | 0.0133687 | 0.362045876 | -0.066445538 | -0.079814238 | -0.428491414 | 0.910778736 | 0.004889329 | 0.675496115 | 0.622623372 | 0.063510161 |
| Nfib | S | 265 | 0.004295843 | 0.05806624 | 0.073253779 | 0.068957936 | 0.015187539 | 0.828351862 | 0.008262467 | 0.001121538 | 0.026827626 | 0.271442666 |
| Ppfibp1 | S | 433 | -5.48856E-05 | 0.10630148 | 0.235349966 | 0.235404852 | 0.129048486 | 0.999392884 | 0.138214718 | 0.064630362 | 0.063845319 | 0.203651754 |
| Rest | S | 87 | -0.023663371 | 0.071790276 | 0.171591223 | 0.195254594 | 0.099800947 | 0.674647145 | 0.107849627 | 0.004761509 | 0.024784241 | 0.049348514 |
| Nhsl1 | S | 1218 | 0.153958823 | 0.11562926 | 0.108230988 | -0.045727835 | -0.007398272 | 0.168198386 | 0.275850909 | 0.321586875 | 0.415582429 | 0.878643478 |
| Cep152 | S | 47 | 0.155903117 | 0.107925063 | 0.110577496 | -0.045325621 | 0.002652433 | 0.143990412 | 0.369091778 | 0.266593262 | 0.670834997 | 0.982675217 |
| Cep152 | S | 48 | 0.155903117 | 0.107925063 | 0.110577496 | -0.045325621 | 0.002652433 | 0.143990412 | 0.369091778 | 0.266593262 | 0.670834997 | 0.982675217 |
| Cep152 | S | 54 | 0.155903117 | 0.107925063 | 0.110577496 | -0.045325621 | 0.002652433 | 0.143990412 | 0.369091778 | 0.266593262 | 0.670834997 | 0.982675217 |
| Cep152 | S | 57 | 0.155903117 | 0.107925063 | 0.110577496 | -0.045325621 | 0.002652433 | 0.143990412 | 0.369091778 | 0.266593262 | 0.670834997 | 0.982675217 |
| Sympk | S | 1275 | 0.079096271 | 0.123875636 | 0.115632858 | 0.036536588 | -0.008242778 | 0.37391226 | 0.056339191 | 0.279172872 | 0.736407842 | 0.927943696 |

| | | | | | | | | | | | | |
|---|---|---|---|---|---|---|---|---|---|---|---|---|
| Mef2a | S | 406 | 0.023859866 | 0.090119129 | 0.027570363 | 0.003710497 | -0.062548766 | 0.587561024 | 9.05452E-05 | 0.143847661 | 0.931125088 | 0.027962224 |
| Hira | T | 554 | 0.096645709 | 0.005820692 | 0.120067037 | 0.023421328 | 0.114246345 | 0.091395007 | 0.864917087 | 0.009313049 | 0.582194469 | 0.003258252 |
| Epb41l3 | S | 428 | 0.040211789 | 0.152817209 | 0.071661548 | 0.031449759 | -0.081155661 | 0.397831978 | 0.04293061 | 0.143671023 | 0.383281213 | 0.1897514 |
| Nr3c1 | T | 161 | 0.012708988 | 0.153113429 | 0.098904573 | 0.086195585 | -0.054208856 | 0.797478248 | 0.014982683 | 0.193475628 | 0.235834326 | 0.421850519 |
| Ankzf1 | S | 86 | -0.014728601 | 0.112295127 | 0.12945137 | 0.14417997 | 0.017156243 | 0.638952294 | 0.027164532 | 0.021876711 | 0.020026257 | 0.692651162 |
| Sipa1l1 | S | 1692 | 0.121270268 | 0.079890575 | 0.15679 | 0.035519732 | 0.076899426 | 0.068086558 | 0.492362318 | 0.275692424 | 0.778624953 | 0.618399384 |
| Palm | T | 263 | 0.300312618 | 0.206904894 | 0.010368792 | -0.289943826 | -0.196536103 | 0.165519291 | 0.301893507 | 0.939467196 | 0.194609379 | 0.353045449 |
| Wdcp | S | 463 | 0.041777861 | 0.124730439 | 0.193183734 | 0.151405873 | 0.068453296 | 0.657147219 | 0.524620221 | 0.025464164 | 0.142070931 | 0.713740068 |
| Abca1 | S | 1138 | 0.101794927 | 0.06212034 | 0.100251361 | -0.001543566 | 0.038131021 | 0.05422229 | 0.308462614 | 0.141363462 | 0.978736376 | 0.58444515 |
| Rap1gap | S | 624 | 0.269084356 | 0.347866258 | -0.204177458 | -0.473261814 | -0.552043716 | 0.014620842 | 0.047175628 | 0.031749345 | 0.004950209 | 0.029290029 |
| Parp12 | S | 268 | 0.021271557 | 0.18344019 | 0.114587374 | 0.093315817 | -0.068852816 | 0.771490405 | 0.043420307 | 0.207208697 | 0.312452603 | 0.440790083 |
| Reps1 | S | 162 | 0.073995928 | 0.080483007 | 0.059594979 | -0.014400949 | -0.020888028 | 0.18280835 | 0.047692033 | 0.061216763 | 0.759054682 | 0.526530656 |
| Myo18a | S | 140 | 0.004904785 | 0.081312016 | 0.213394247 | 0.208489462 | 0.132082231 | 0.918806166 | 0.059903586 | 0.067737058 | 0.073742112 | 0.187373321 |
| Prpf4b | S | 379 | 0.106580846 | 0.036471905 | 0.166763572 | 0.060182727 | 0.130291668 | 0.420146976 | 0.593443423 | 0.018697729 | 0.624511845 | 0.08078589 |
| Glce | S | 73 | -0.002759148 | 0.071978536 | 0.120532499 | 0.123291647 | 0.048553963 | 0.916089349 | 0.061431261 | 0.003584369 | 0.000929967 | 0.125974666 |
| Prkacb | T | 341 | 0.0438909 | 0.077062333 | 0.126474252 | 0.082583352 | 0.049411919 | 0.284915851 | 0.315355427 | 0.016108229 | 0.025996538 | 0.477435158 |
| Fam111a | T | 68 | 0.073558279 | 0.079077687 | 0.158155785 | 0.084597506 | 0.079078098 | 0.125310246 | 0.201152743 | 0.195071097 | 0.4288507 | 0.467277075 |
| Copg1 | T | 583 | 0.24833659 | 0.104369298 | -0.131688587 | -0.380025177 | -0.236057886 | 0.005935747 | 0.128147337 | 0.423969369 | 0.093047429 | 0.207750791 |
| Strn4 | S | 223 | 0.116308638 | 0.060976821 | 0.075696763 | -0.040611874 | 0.014719942 | 0.035175263 | 0.262811018 | 0.206053445 | 0.379685596 | 0.76602704 |
| Sec13 | S | 254 | 0.046772867 | 0.118537337 | 0.150773443 | 0.104000576 | 0.032236106 | 0.514627011 | 0.160949104 | 0.082571843 | 0.237679553 | 0.700412715 |
| Yeats2 | S | 371 | 0.111187157 | 0.145865517 | 0.030328835 | -0.080858322 | -0.115536682 | 0.092369026 | 0.061068939 | 0.714255619 | 0.368669895 | 0.235688494 |
| Srrm2 | S | 1508 | 0.073266556 | 0.053495018 | 0.109673925 | 0.036407369 | 0.056178907 | 0.101739841 | 0.202158282 | 0.062022716 | 0.375098134 | 0.21905607 |
| Rexo1 | S | 514 | 0.179663334 | -0.004712573 | 0.145015654 | -0.03464768 | 0.149728228 | 0.042795077 | 0.941761652 | 0.156949481 | 0.68147257 | 0.161716019 |
| Mis18bp1 | T | 741 | 0.188250773 | 0.13232172 | 0.028699867 | -0.159550906 | -0.103621853 | 0.112119871 | 0.115748384 | 0.810103456 | 0.254259484 | 0.387648283 |
| Erc1 | S | 17 | 0.015503597 | 0.059545317 | 0.138405549 | 0.122901953 | 0.078860233 | 0.509397969 | 0.309953034 | 0.003383273 | 0.012266871 | 0.20885136 |
| Eef1d | S | 162 | 0.133012558 | -0.003386372 | 0.124460107 | -0.008552451 | 0.127846479 | 0.042749209 | 0.951315779 | 0.046565311 | 0.767353197 | 0.029518093 |
| Srrm2 | S | 1688 | 0.052869575 | 0.01085265 | 0.091988038 | 0.039118463 | 0.081135387 | 0.055660109 | 0.682908958 | 0.00100262 | 0.122256188 | 0.047825395 |
| Eif2ak3 | S | 715 | 0.052808077 | 0.180275676 | -0.002075612 | -0.054883689 | -0.182351288 | 0.574500283 | 0.001434109 | 0.986051903 | 0.694318037 | 0.215803072 |
| Msn | S | 491 | 0.030915491 | 0.177452348 | 0.053867101 | 0.02295161 | -0.123585247 | 0.643720058 | 0.013173153 | 0.290242971 | 0.750107729 | 0.070362589 |
| Stim1 | S | 667 | 0.007598377 | 0.11866042 | 0.066283623 | 0.058685246 | -0.052376797 | 0.832781901 | 0.006537294 | 0.053297687 | 0.113253188 | 0.075026496 |
| Snd1 | S | 909 | 0.197982706 | 0.017757854 | 0.106386913 | -0.091595793 | 0.088629059 | 0.039693388 | 0.842095244 | 0.229125521 | 0.116276364 | 0.217168529 |
| Hmga2 | S | 14 | 0.076694299 | 0.144082706 | 0.123812051 | 0.047117752 | -0.020270655 | 0.172300787 | 0.209432274 | 0.260558878 | 0.636007403 | 0.867691264 |
| Map4 | T | 447 | 0.145944337 | 0.161895004 | -0.032905816 | -0.178850153 | -0.19480082 | 0.084725982 | 0.037684565 | 0.827621735 | 0.291499274 | 0.260238575 |
| Kif4 | S | 816 | 0.158857708 | 0.108450368 | 0.126178407 | -0.032679301 | 0.017728039 | 0.360643436 | 0.240310538 | 0.195551002 | 0.840115989 | 0.848814657 |
| Sos1 | S | 1147 | 0.049421552 | 0.22375223 | 0.104753242 | 0.05533169 | -0.118998989 | 0.805154416 | 0.054287636 | 0.335812767 | 0.788930882 | 0.315664701 |
| Zfp608 | S | 1427 | -0.010709187 | 0.078947283 | 0.267233215 | 0.277942402 | 0.188285932 | 0.910559878 | 0.352379646 | 0.024894513 | 0.048526792 | 0.096292645 |
| Rps6ka4 | S | 745 | 0.03633719 | 0.149859202 | 0.08459114 | 0.04825395 | -0.065268062 | 0.524124482 | 0.032056624 | 0.18561501 | 0.299311072 | 0.202602075 |
| Fam102b | S | 299 | 0.064786377 | 0.167622176 | 0.160178084 | 0.095391707 | -0.007444092 | 0.618193531 | 0.30794322 | 0.085393799 | 0.45149997 | 0.958517985 |
| Fam102b | S | 300 | 0.064786377 | 0.167622176 | 0.160178084 | 0.095391707 | -0.007444092 | 0.618193531 | 0.30794322 | 0.085393799 | 0.45149997 | 0.958517985 |
| Aebp2 | S | 136 | 0.136604577 | 0.160275392 | 0.011837994 | -0.124766583 | -0.148437398 | 0.169378055 | 0.041692093 | 0.858637269 | 0.175560331 | 0.010068952 |
| Ubxn7 | S | 328 | 0.019426448 | 0.075866737 | 0.148784979 | 0.129358531 | 0.072918242 | 0.835298932 | 0.125087503 | 0.014721328 | 0.230000927 | 0.109317266 |
| Mafk | T | 127 | -0.003634892 | 0.077888742 | 0.173299514 | 0.176934406 | 0.095410772 | 0.954580617 | 0.282916378 | 0.005616378 | 0.054195459 | 0.20951629 |
| Esyt2 | S | 679 | 0.048320781 | 0.038927735 | 0.114754405 | 0.066433624 | 0.07582667 | 0.098392276 | 0.499544319 | 0.007556728 | 0.068134886 | 0.241856009 |
| Akap13 | S | 1253 | 0.080716103 | 0.146142863 | 0.04745672 | -0.033259383 | -0.098686142 | 0.134741837 | 0.03025762 | 0.700556361 | 0.778167913 | 0.440898035 |
| Dpysl2 | T | 521 | 0.040193341 | 0.285657493 | -0.035265704 | -0.075459046 | -0.320923197 | 0.608204138 | 0.005728369 | 0.714110797 | 0.418376795 | 0.023615302 |
| Pdha1 | S | 295 | -0.017477955 | 0.071291137 | 0.294376373 | 0.311854328 | 0.223085236 | 0.875416767 | 0.233449049 | 0.040185184 | 0.062306227 | 0.095882024 |
| Pxn | S | 83 | 0.168814795 | 0.075510207 | 0.024134482 | -0.144680313 | -0.051375725 | 0.023248122 | 0.22181945 | 0.592465279 | 0.046455821 | 0.379028693 |
| Ep400 | T | 922 | 0.042601874 | 0.108682528 | 0.135658717 | 0.093056843 | 0.026976189 | 0.580172995 | 0.29404533 | 0.019840103 | 0.282297518 | 0.7702383 |
| Ssrp1 | S | 444 | 0.039425087 | 0.029882258 | 0.078104439 | 0.038679352 | 0.04822218 | 0.170872357 | 0.082975264 | 0.001597388 | 0.183156264 | 0.007419568 |
| Rere | S | 656 | 0.009312295 | 0.107959534 | 0.086798631 | 0.077486336 | -0.021160902 | 0.683372301 | 0.040655744 | 0.016211359 | 0.036578968 | 0.53762026 |
| Git2 | S | 508 | -0.108326824 | 0.132432159 | 0.261207276 | 0.3695341 | 0.128775117 | 0.191243427 | 0.121528446 | 0.00689594 | 0.007814904 | 0.147764564 |
| Vim | T | 458 | 0.124223453 | 0.092365827 | 0.078215349 | -0.046008104 | -0.014150478 | 0.251270633 | 0.056410916 | 0.280115182 | 0.657678387 | 0.83107169 |
| Ttc7b | S | 202 | -0.002311879 | 0.105017141 | 0.254137413 | 0.256449292 | 0.149120272 | 0.954260993 | 0.121721263 | 0.088584949 | 0.080269731 | 0.217123002 |
| Aven | S | 82 | -0.036010512 | 0.120245234 | 0.078096794 | 0.114107306 | -0.04214844 | 0.504351216 | 5.49511E-05 | 0.279889025 | 0.178136599 | 0.513362042 |
| Cap2 | S | 311 | 0.120284576 | 0.044580642 | 0.083742132 | -0.036542444 | 0.03916149 | 0.028484603 | 0.258417599 | 0.242296811 | 0.572024671 | 0.528118134 |
| Aak1 | S | 621 | 0.103228272 | 0.170783377 | 0.046362081 | -0.056866191 | -0.124421295 | 0.187027088 | 0.063130282 | 0.651255663 | 0.583825172 | 0.276701479 |
| Thrap3 | S | 315 | 0.106095795 | 0.162294393 | 0.142675907 | 0.036580112 | -0.019618486 | 0.397128653 | 0.382396372 | 0.133671153 | 0.771368856 | 0.91021453 |
| Mier1 | Y | 171 | 0.017290556 | 0.052546746 | 0.196294489 | 0.179003933 | 0.143747744 | 0.818446319 | 0.385255497 | 0.009053428 | 0.05907119 | 0.020230816 |
| Tulp3 | S | 184 | 0.108497226 | 0.102601639 | 0.117422564 | 0.008925338 | 0.014820925 | 0.414599271 | 0.365073846 | 0.040773391 | 0.941398625 | 0.884568449 |
| Caap1 | S | 183 | -0.02041953 | 0.091505315 | 0.157530875 | 0.177950405 | 0.06602556 | 0.413870656 | 0.017693983 | 0.023174845 | 0.02859305 | 0.194198027 |
| Kank2 | S | 120 | 0.195595366 | 0.132283591 | -0.060659977 | -0.256255343 | -0.192943568 | 0.030471664 | 0.08282821 | 0.666332772 | 0.14867082 | 0.237317057 |
| Ppp1r18 | S | 194 | 0.109693947 | 0.023384297 | 0.079171382 | -0.030522564 | 0.055787085 | 0.017329985 | 0.487837566 | 0.080440376 | 0.34131273 | 0.133395703 |
| Sart1 | S | 48 | 0.351464365 | 0.285465926 | -0.269235481 | -0.620699846 | -0.554701407 | 0.091618692 | 0.136584008 | 0.506118238 | 0.186971811 | 0.22364713 |
| Mpdz | S | 231 | 0.062571976 | 0.05863243 | 0.061514506 | -0.00105747 | 0.002882076 | 0.264307928 | 0.028272264 | 0.01826759 | 0.982281124 | 0.878353569 |
| Crebbp | S | 1762 | -0.011434855 | 0.184948958 | 0.170713474 | 0.182148329 | -0.014235484 | 0.837509247 | 0.162188842 | 0.053509918 | 0.054143249 | 0.897113953 |
| Prex2 | S | 1049 | -0.233335946 | 0.30423023 | 0.344719959 | 0.578055905 | 0.040489729 | 0.203457974 | 0.112235704 | 0.04499059 | 0.016096066 | 0.774423322 |
| Kmt2e | S | 1449 | 0.287665108 | -0.036339504 | 0.104359559 | -0.183305549 | 0.140699062 | 0.139103291 | 0.343353814 | 0.011443467 | 0.272788794 | 0.013509409 |
| Spen | S | 852 | -0.013673339 | 0.047682529 | 0.120096292 | 0.13376963 | 0.072413763 | 0.813944997 | 0.13236705 | 0.000329804 | 0.110784734 | 0.055620716 |
| Spen | S | 855 | -0.013673339 | 0.047682529 | 0.120096292 | 0.13376963 | 0.072413763 | 0.813944997 | 0.13236705 | 0.000329804 | 0.110784734 | 0.055620716 |
| Spen | S | 858 | -0.013673339 | 0.047682529 | 0.120096292 | 0.13376963 | 0.072413763 | 0.813944997 | 0.13236705 | 0.000329804 | 0.110784734 | 0.055620716 |
| Spen | S | 869 | -0.013673339 | 0.047682529 | 0.120096292 | 0.13376963 | 0.072413763 | 0.813944997 | 0.13236705 | 0.000329804 | 0.110784734 | 0.055620716 |
| Abca1 | S | 879 | 0.104361074 | 0.045279409 | 0.164426872 | 0.060065798 | 0.119147462 | 0.187812688 | 0.415001996 | 0.096116194 | 0.504307884 | 0.18816167 |
| Sash1 | S | 699 | -0.115890932 | 0.454801627 | 0.260759041 | 0.376649973 | -0.194042586 | 0.623047311 | 0.165594282 | 0.290952316 | 0.198593832 | 0.531491393 |
| Git1 | Y | 392 | 0.019206864 | 0.109362424 | 0.127811803 | 0.108604939 | 0.018449379 | 0.850143829 | 0.038640926 | 0.069093818 | 0.348793886 | 0.741188593 |
| Supt6 | T | 1718 | 0.039788773 | 0.067418218 | 0.135426338 | 0.095637565 | 0.06800812 | 0.317683428 | 0.159261808 | 0.035910674 | 0.103172773 | 0.225228296 |
| Ube2j1 | S | 266 | 0.067323912 | 0.134522949 | 0.124589511 | 0.057265599 | -0.009933438 | 0.249096518 | 0.063330755 | 0.438312873 | 0.70709385 | 0.947412982 |
| Zeb2 | T | 203 | 0.008564707 | 0.148607004 | 0.380562832 | 0.371998125 | 0.231955827 | 0.931012389 | 0.250070054 | 0.221794256 | 0.225502855 | 0.409094132 |
| Ccdc102a | S | 12 | 0.017832849 | 0.146847487 | 0.055295392 | 0.037462543 | -0.091552095 | 0.553842411 | 0.009970052 | 0.175584257 | 0.315521066 | 0.074719015 |
| Erbb2 | S | 1175 | 0.095268761 | 0.05701199 | 0.071073409 | -0.024195352 | 0.014061419 | 0.073746675 | 0.163770896 | 0.08774868 | 0.509176283 | 0.504150774 |
| Lemd3 | S | 296 | 0.032714053 | 0.086131588 | 0.167016322 | 0.134302269 | 0.080884734 | 0.652842448 | 0.470697216 | 0.011628777 | 0.147342054 | 0.498419 |
| Tra2a | S | 84 | 0.619150896 | 0.009212319 | 0.08551227 | -0.533638626 | 0.076299951 | 0.314508855 | 0.83095269 | 0.243247085 | 0.370036123 | 0.274803965 |
| Tra2a | T | 86 | 0.619150896 | 0.009212319 | 0.08551227 | -0.533638626 | 0.076299951 | 0.314508855 | 0.83095269 | 0.243247085 | 0.370036123 | 0.274803965 |
| Nrcam | S | 1254 | 0.174626443 | 0.103849695 | 0.008040149 | -0.166586294 | -0.095809546 | 0.047709456 | 0.120942909 | 0.841117659 | 0.046352156 | 0.154963199 |
| Eps8 | S | 573 | 0.030566435 | 0.049967954 | 0.174296705 | 0.143730269 | 0.124328751 | 0.670892801 | 0.326445797 | 0.010891112 | 0.09677062 | 0.033212391 |
| Spen | S | 749 | 0.03988668 | 0.05233645 | 0.121109844 | 0.081223163 | 0.068773394 | 0.425618768 | 0.147564231 | 0.011857718 | 0.134358123 | 0.057625278 |
| Cdyl | S | 116 | 0.016543349 | 0.0638874 | 0.212048656 | 0.195505307 | 0.148161256 | 0.885851288 | 0.16942597 | 0.032743156 | 0.174962942 | 0.101180971 |
| Kmt2a | S | 514 | -0.010280361 | 0.105776787 | 0.084988793 | 0.095269154 | -0.020787994 | 0.842604798 | 0.006488842 | 0.030355836 | 0.133747447 | 0.419874609 |
| Kif3a | S | 381 | 0.152301196 | -0.016612166 | 0.299314593 | 0.147013397 | 0.315926759 | 0.343284778 | 0.918203029 | 0.08159475 | 0.373498596 | 0.110997193 |
| Kif3a | S | 383 | 0.152301196 | -0.016612166 | 0.299314593 | 0.147013397 | 0.315926759 | 0.343284778 | 0.918203029 | 0.08159475 | 0.373498596 | 0.110997193 |
| Kif3a | S | 386 | 0.152301196 | -0.016612166 | 0.299314593 | 0.147013397 | 0.315926759 | 0.343284778 | 0.918203029 | 0.08159475 | 0.373498596 | 0.110997193 |
| Kif3a | S | 388 | 0.152301196 | -0.016612166 | 0.299314593 | 0.147013397 | 0.315926759 | 0.343284778 | 0.918203029 | 0.08159475 | 0.373498596 | 0.110997193 |
| Atad5 | T | 1200 | 0.127087544 | 0.186613696 | -0.073910482 | -0.200998026 | -0.260524179 | 0.170411681 | 0.007164871 | 0.573777488 | 0.197732702 | 0.130929877 |
| Purb | S | 113 | 0.015081703 | 0.105172546 | 0.133436841 | 0.118355138 | 0.028264295 | 0.645780018 | 0.028694613 | 0.113383697 | 0.142010545 | 0.666106723 |
| Ccdc102a | S | 536 | -0.048423586 | 0.118891701 | 0.165144515 | 0.213568102 | 0.046252814 | 0.434505539 | 0.03483665 | 0.020782405 | 0.027241582 | 0.390970861 |

| | | | | | | | | | | | | |
|---|---|---|---|---|---|---|---|---|---|---|---|---|
| Crhr2 | S | 404 | -0.007595052 | 0.075468601 | 0.220703905 | 0.228298957 | 0.145235304 | 0.915664642 | 0.147133133 | 0.030845519 | 0.041952157 | 0.111040015 |
| Vim | S | 54 | 0.088153839 | 0.090930095 | 0.101976787 | 0.013822948 | 0.011046692 | 0.250012048 | 0.11875124 | 0.149438557 | 0.859585715 | 0.864610373 |
| Srpk1 | S | 37 | 0.165556841 | 0.018215764 | 0.133319767 | -0.032237074 | 0.115104004 | 0.067713375 | 0.732371805 | 0.169095631 | 0.719050699 | 0.214061852 |
| Dennd1a | S | 893 | 0.531246425 | 0.212780187 | 0.453013335 | -0.07823309 | 0.240233148 | NA | 0.620243523 | 0.160350688 | NA | 0.592695693 |
| Vim | T | 417 | 0.158629337 | 0.010275512 | 0.135942632 | -0.022686706 | 0.125667119 | 0.036162301 | 0.914500568 | 0.192564909 | 0.798662425 | 0.296567806 |
| U2af2 | S | 53 | 0.119091566 | 0.033533344 | 0.144281812 | 0.025190246 | 0.110748468 | 0.157273237 | 0.775995861 | 0.045671775 | 0.681040909 | 0.362061521 |
| U2af2 | S | 55 | 0.119091566 | 0.033533344 | 0.144281812 | 0.025190246 | 0.110748468 | 0.157273237 | 0.775995861 | 0.045671775 | 0.681040909 | 0.362061521 |
| Scamp1 | T | 45 | 0.194000678 | 0.069624798 | 0.09409918 | -0.099901498 | 0.024474382 | 0.08534419 | 0.456173798 | 0.376249889 | 0.389753913 | 0.820659322 |
| Hivep2 | S | 565 | -0.019590671 | 0.092853945 | 0.082623353 | 0.102214024 | -0.010230592 | 0.217628117 | 0.013864245 | 0.000761506 | 0.001388534 | 0.640251713 |
| Fnbp1l | S | 295 | -0.040005757 | 0.097564334 | 0.156867377 | 0.196873134 | 0.059303044 | 0.203453548 | 0.051294363 | 0.003540891 | 0.004280224 | 0.18725107 |
| Ppig | S | 411 | 0.010463106 | 0.071101108 | 0.083980873 | 0.073517767 | 0.012879765 | 0.696432825 | 0.002799684 | 0.04068774 | 0.076090754 | 0.631078708 |
| Ehbp1 | S | 177 | 0.098466223 | 0.128249631 | -0.032759822 | -0.131226045 | -0.161009453 | 0.124049872 | 0.003406274 | 0.820063696 | 0.409330581 | 0.327666356 |
| Atad5 | S | 1194 | 0.121578268 | 0.09474763 | -0.019935906 | -0.141514174 | -0.114683536 | 0.014985395 | 0.021630826 | 0.649841034 | 0.031425734 | 0.05466556 |
| Furin | S | 772 | 0.0852069 | 0.02434693 | 0.066389769 | -0.018817131 | 0.042042839 | 0.011831714 | 0.360730658 | 0.046328458 | 0.474122086 | 0.153310213 |
| Furin | S | 774 | 0.0852069 | 0.02434693 | 0.066389769 | -0.018817131 | 0.042042839 | 0.011831714 | 0.360730658 | 0.046328458 | 0.474122086 | 0.153310213 |
| Arhgef6 | S | 639 | 0.044222834 | 0.329223338 | -0.05506938 | -0.099292214 | -0.384292718 | 0.610039747 | 0.008482156 | 0.564130991 | 0.135225304 | 0.002975072 |
| Zfp280c | S | 526 | -0.027919499 | 0.123044203 | 0.190234314 | 0.218153813 | 0.067190111 | 0.665725942 | 0.043922604 | 0.070764555 | 0.055958112 | 0.415533827 |
| Nipbl | S | 274 | 0.069261535 | 0.061530615 | 0.064038383 | -0.005223152 | 0.002507768 | 0.052591121 | 0.084542901 | 0.102821815 | 0.820649365 | 0.922879128 |
| Acin1 | S | 655 | -0.043906771 | 0.080412029 | 0.152980214 | 0.196886985 | 0.072568185 | 0.250887515 | 0.115784429 | 0.00071215 | 0.008436199 | 0.136618037 |
| Sorbs2 | S | 239 | -0.073146279 | 0.172062744 | 0.165984935 | 0.239131214 | -0.006077808 | 0.369046242 | 0.029885577 | 0.045662039 | 0.022787805 | 0.907554492 |
| Trim28 | S | 598 | -0.036928093 | 0.120940843 | 0.136311099 | 0.173239191 | 0.015370255 | 0.363548554 | 0.063567642 | 0.005755626 | 0.006119165 | 0.748834077 |
| Setd1a | S | 1110 | 0.0086119 | 0.038343403 | 0.136590644 | 0.127978745 | 0.098247241 | 0.836742685 | 0.203159844 | 0.001661861 | 0.046478092 | 0.004923125 |
| Hivep3 | S | 1933 | -0.085097815 | 0.103104725 | 0.344144234 | 0.42924205 | 0.241039509 | 0.213098604 | 0.151449452 | 0.022510974 | 0.023340111 | 0.077415258 |
| Zfp369 | S | 476 | 0.02509994 | 0.187807114 | 0.128379475 | 0.103279535 | -0.059427639 | 0.874426324 | 0.064720999 | 0.212909185 | 0.506114636 | 0.352200248 |
| Tjp2 | S | 107 | -0.004709797 | 0.063024258 | 0.210173294 | 0.214883091 | 0.147149036 | 0.92089913 | 0.026617124 | 0.063050076 | 0.045183602 | 0.119418451 |
| Dok1 | S | 415 | 0.179317595 | 0.025584611 | 0.121799595 | -0.057518 | 0.096214984 | 0.088528603 | 0.776058634 | 0.150466577 | 0.429085205 | 0.194407811 |
| Vasp | S | 235 | 0.021601375 | 0.44420315 | -0.079640619 | -0.101241994 | -0.523843769 | 0.863491692 | 0.015081634 | 0.599604857 | 0.379593745 | 0.011954539 |
| Sap30bp | S | 22 | 0.170260261 | 0.042781459 | 0.045544049 | -0.124716211 | 0.00276259 | 0.016190459 | 0.435728362 | 0.441643883 | 0.087657818 | 0.962933192 |
| Yeats2 | T | 1204 | -0.13409316 | 0.062045051 | 0.257596081 | 0.391689242 | 0.19555103 | 0.207982608 | 0.290153456 | 0.000491133 | 0.026749423 | 0.019633826 |
| Atad2 | T | 807 | 0.155197636 | 0.061234587 | 0.018742515 | -0.136455121 | -0.042492071 | 0.004278781 | 0.455682081 | 0.715543912 | 0.056157859 | 0.602372272 |
| Irs2 | S | 1190 | 0.036956938 | 0.076277441 | 0.123923359 | 0.086966421 | 0.047645918 | 0.20451781 | 0.127083191 | 0.06644053 | 0.162480156 | 0.411841543 |
| Tjp1 | S | 912 | -0.048854828 | 0.082913751 | 0.225929432 | 0.27478426 | 0.143015681 | 0.420799379 | 0.180848943 | 0.007776989 | 0.003005251 | 0.030403652 |
| Map1a | S | 899 | 0.106817801 | 0.074775932 | 0.238834676 | 0.132016875 | 0.164058743 | 0.359087327 | 0.246015587 | 0.264537057 | 0.51448046 | 0.407413654 |
| Adcy6 | S | 53 | 0.059536812 | 0.046242823 | 0.081999135 | 0.022462323 | 0.035756312 | 0.05885233 | 0.127521598 | 0.048087576 | 0.424657462 | 0.250431101 |
| Sptbn1 | S | 2159 | -0.004212404 | 0.10147567 | 0.088507819 | 0.092720223 | -0.012967851 | 0.874165077 | 0.010654807 | 0.027959153 | 0.037633822 | 0.628916235 |
| Nfib | S | 410 | 0.023709031 | 0.186707508 | 0.083866769 | 0.060157738 | -0.102840739 | 0.591665666 | 0.014995428 | 0.561108882 | 0.669895132 | 0.48768009 |
| Pou3f2 | S | 343 | -0.090615151 | 0.043692701 | 0.283641952 | 0.374257102 | 0.239949251 | 0.451330973 | 0.474535166 | 0.002466825 | 0.0456248 | 0.003407783 |
| Gtse1 | S | 398 | 0.086775762 | 0.102350494 | 0.260923002 | 0.174147239 | 0.158572507 | 0.276095555 | 0.340275131 | 0.313511335 | 0.471738246 | 0.515040831 |
| Ehbp1 | S | 335 | 0.04753868 | 0.123747149 | 0.086255867 | 0.038717186 | -0.037491282 | 0.345884505 | 0.076692717 | 0.119559001 | 0.379663535 | 0.502842346 |
| Ranbp3 | S | 32 | 0.033267667 | 0.065867714 | 0.052103639 | 0.018835973 | -0.013764074 | 0.110977194 | 0.001164616 | 0.224996935 | 0.614668963 | 0.700054619 |
| Setd2 | S | 322 | 0.066363904 | 0.100722461 | 0.08670631 | 0.020342406 | -0.014016152 | 0.079929322 | 0.081463975 | 0.346247282 | 0.807120037 | 0.870849603 |
| Setd2 | S | 324 | 0.066363904 | 0.100722461 | 0.08670631 | 0.020342406 | -0.014016152 | 0.079929322 | 0.081463975 | 0.346247282 | 0.807120037 | 0.870849603 |
| Cox7b | S | 32 | 0.003429597 | 0.185486397 | 0.190845536 | 0.187415939 | 0.005359139 | 0.959286433 | 0.124517142 | 0.169881302 | 0.178282866 | 0.967644088 |
| Thumpd2 | S | 172 | -0.101929236 | 0.136685182 | 0.222605232 | 0.324534468 | 0.08592005 | 0.360386514 | 0.170370234 | 0.007343037 | 0.045202342 | 0.348684306 |
| Stk38l | S | 282 | 0.030623034 | 0.196923888 | 0.017515815 | -0.013107219 | -0.179408073 | 0.484479994 | 0.006134301 | 0.789446928 | 0.823797397 | 0.047128134 |
| Peak1 | S | 1239 | 0.056691918 | 0.106847327 | 0.063170248 | 0.00647833 | -0.043677079 | 0.080767318 | 0.041214561 | 0.362116657 | 0.916402666 | 0.524049461 |
| Ncor1 | S | 2329 | -0.047060265 | 0.05889969 | 0.143861203 | 0.190921468 | 0.084961513 | 0.46004028 | 0.009633599 | 0.00443783 | 0.05015431 | 0.042072826 |
| Fmnl1 | S | 1021 | -0.165761795 | 0.118018751 | 0.260713606 | 0.426475401 | 0.142694855 | 0.010915219 | 0.012274283 | 0.000838487 | 0.000545986 | 0.017347492 |
| Spen | T | 2376 | 0.070121454 | 0.007137104 | 0.186862622 | 0.116741168 | 0.179725518 | 0.408702057 | 0.890327322 | 0.009334176 | 0.191388778 | 0.008028978 |
| Itprid2 | S | 1036 | 0.068066971 | 0.02601883 | 0.116670591 | 0.04860362 | 0.09065176 | 0.079974895 | 0.408219235 | 0.027844397 | 0.22592876 | 0.081113434 |
| Supt7l | S | 332 | 0.020526956 | 0.104515461 | 0.244853161 | 0.224326205 | 0.1403377 | 0.818869771 | 0.179577348 | 0.116330299 | 0.148400133 | 0.299375517 |
| Syne1 | S | 5892 | 0.007635915 | 0.14329908 | 0.177541334 | 0.169905419 | 0.034224254 | 0.878862016 | 0.038185336 | 0.247723349 | 0.262231401 | 0.793540654 |
| Cdk12 | S | 331 | 0.021431077 | 0.050342792 | 0.150310554 | 0.128879477 | 0.099967762 | 0.707293118 | 0.22915262 | 0.007751803 | 0.069896371 | 0.009489911 |
| G3bp2 | T | 227 | -0.00693427 | 0.078714717 | 0.114731103 | 0.121665373 | 0.036016387 | 0.762706112 | 0.003351595 | 0.049817613 | 0.035589748 | 0.358756658 |
| Lin37 | S | 182 | 0.090927799 | 0.122916359 | 0.03574798 | -0.055179819 | -0.08716838 | 0.112126956 | 0.036854643 | 0.624165173 | 0.454103464 | 0.262241189 |
| Dyrk1a Dyrk1b | Y | 111;63 | 0.06401339 | 0.155190778 | 0.028714443 | -0.035298947 | -0.126476335 | 0.261956556 | 0.016925735 | 0.584882018 | 0.553072828 | 0.067241165 |
| Dyrk1a Dyrk1b | Y | 112;64 | 0.06401339 | 0.155190778 | 0.028714443 | -0.035298947 | -0.126476335 | 0.261956556 | 0.016925735 | 0.584882018 | 0.553072828 | 0.067241165 |
| Champ1 | S | 309 | 0.068919052 | 0.026016263 | 0.196046765 | 0.127127713 | 0.170030502 | 0.455785306 | 0.771427201 | 0.01643037 | 0.186478136 | 0.115520012 |
| Hsp90b1 | S | 42 | -0.055084089 | 0.180946317 | 0.132001377 | 0.187085466 | -0.04894494 | 0.605787424 | 0.033475206 | 0.068845656 | 0.143311351 | 0.325235219 |
| Slain2 | S | 324 | 0.091678618 | 0.111282188 | 0.073172398 | -0.018506221 | -0.03810979 | 0.020340327 | 0.400189893 | 0.303742698 | 0.760431788 | 0.766733003 |
| Mga | S | 1250 | 0.043364038 | 0.203352478 | 0.167957163 | 0.124593125 | -0.035395315 | 0.784366425 | 0.271302444 | 0.122107766 | 0.455277968 | 0.830297246 |
| Mga | S | 1254 | 0.043364038 | 0.203352478 | 0.167957163 | 0.124593125 | -0.035395315 | 0.784366425 | 0.271302444 | 0.122107766 | 0.455277968 | 0.830297246 |
| Slc39a3 | S | 131 | 0.024116403 | 0.157736764 | 0.126839156 | 0.102722753 | -0.030897608 | 0.810278275 | 0.125337385 | 0.078534884 | 0.369948265 | 0.714113709 |
| Slc7a6os | T | 155 | 0.057335983 | 0.324808547 | -0.038881532 | -0.096217515 | -0.363690079 | 0.589586503 | 0.018701469 | 0.775950979 | 0.397783426 | 0.037444941 |
| Rictor | S | 1301 | 0.113997702 | 0.388957643 | -0.10387497 | -0.217872672 | -0.492832613 | 0.478846917 | 0.035994418 | 0.638172461 | 0.38184534 | 0.09421553 |
| Hps5 | S | 693 | 0.126266047 | 0.182593289 | -0.049489217 | -0.175755264 | -0.232082506 | 0.083455591 | 0.02487416 | 0.556020488 | 0.09656228 | 0.048241316 |
| Slc9a3r1 | S | 297 | -0.100792556 | 0.156996863 | 0.173519919 | 0.274312475 | 0.016523056 | 0.003470226 | 0.000290691 | 0.017939372 | 0.011346172 | 0.690419049 |
| Ncf4 | Y | 63 | 0.034014812 | 0.122732348 | 0.091647214 | 0.057632401 | -0.031085134 | 0.430904298 | 0.031988427 | 0.188532145 | 0.380518024 | 0.624140613 |
| Kmt2d | S | 1136 | 0.225018648 | 0.08841323 | 0.034742175 | -0.190276473 | -0.053671055 | 0.049143044 | 0.358098383 | 0.810422869 | 0.209152462 | 0.663443796 |
| Srrm2 | S | 231 | 0.032235486 | 0.065789715 | 0.150335979 | 0.118100493 | 0.084546264 | 0.663378433 | 0.047862512 | 0.062200954 | 0.207371133 | 0.193539092 |
| Gad1 | T | 53 | 0.318436474 | 0.140276185 | -0.166160855 | -0.48459733 | -0.30643704 | 0.02361671 | 0.278150185 | 0.566053941 | 0.172875589 | 0.321876139 |
| Ssx2ip | S | 313 | 0.230384193 | 0.064341434 | -0.007780467 | -0.23816466 | -0.072121901 | 0.024695093 | 0.242790455 | 0.958307505 | 0.196530649 | 0.634958825 |
| Pcm1 | S | 190 | 0.068106 | 0.208849197 | 0.120510365 | 0.052404364 | -0.088338832 | 0.332484071 | 0.247987616 | 0.276916994 | 0.582025648 | 0.600947113 |
| Slc20a2 | S | 436 | 0.172711587 | 0.087354884 | -0.070058497 | -0.242770084 | -0.157413381 | 0.003198185 | 0.203985766 | 0.686970184 | 0.243256819 | 0.405305409 |
| C1qbp | S | 197 | 0.066219296 | 0.055226244 | 0.096220714 | 0.030001418 | 0.04099447 | 0.247745399 | 0.019605941 | 0.138056648 | 0.640832674 | 0.436682368 |
| Snw1 | S | 232 | 0.043536902 | 0.031850425 | 0.082141312 | 0.03860441 | 0.050290887 | 0.049469344 | 0.209275507 | 0.009119871 | 0.1034161 | 0.08101472 |
| 3110082I17Rik | S | 177 | 0.11825872 | 0.082879918 | 0.068197604 | -0.050061115 | -0.014682314 | 0.047012183 | 0.149779842 | 0.530300583 | 0.621116463 | 0.882651564 |
| Parva | T | 16 | 0.045496823 | 0.212971645 | 0.16068641 | 0.115189587 | -0.052285235 | 0.58606155 | 0.167775512 | 0.310613821 | 0.461550785 | 0.758381539 |
| Pias2 | S | 498 | 0.209193108 | 0.070158666 | 0.067311092 | -0.141882016 | -0.002847574 | 0.336817478 | 0.059233775 | 0.303091036 | 0.489790195 | 0.958826483 |
| Sart1 | S | 604 | 0.060840376 | 0.06440355 | 0.096876251 | 0.036035874 | 0.032472701 | 0.340438042 | 0.034874053 | 0.08848508 | 0.584288697 | 0.460473876 |
| Fam219a | S | 87 | 0.023779975 | 0.018502554 | 0.140053278 | 0.116273303 | 0.121550724 | 0.476897302 | 0.620194415 | 0.001069746 | 0.015935637 | 0.028068822 |
| Dst | S | 4680 | 0.009807473 | 0.212927551 | 0.139718896 | 0.129911423 | -0.073208655 | 0.934803287 | 0.054966907 | 0.32984005 | 0.358398599 | 0.53458214 |
| Il1rap | S | 556 | 0.212128635 | -0.002953841 | 0.06214412 | -0.149984515 | 0.06509796 | 0.013149422 | 0.97018458 | 0.348305374 | 0.015031623 | 0.330028254 |
| Chd6 | S | 1357 | 0.025694419 | 0.130467673 | 0.198340682 | 0.172646263 | 0.067873009 | 0.715271132 | 0.195254964 | 0.118332508 | 0.162013879 | 0.559557561 |
| Sash1 | S | 163 | -0.184008474 | 0.348112964 | 0.221202888 | 0.405211362 | -0.126910076 | 0.013666542 | 0.013527673 | 0.032707508 | 0.008509284 | 0.229888398 |
| Foxj2 | S | 46 | -0.005363567 | 0.319065198 | 0.023329567 | 0.028693134 | -0.29573563 | 0.955540007 | 0.01547173 | 0.824853892 | 0.694271103 | 0.018873432 |
| Sp100 | T | 209 | -0.030782216 | 0.063910872 | 0.226167353 | 0.25694957 | 0.162256482 | 0.594796968 | 0.313831922 | 0.007027375 | 0.000186985 | 0.009665097 |
| Sap30bp | S | 18 | 0.168220584 | 0.044596363 | 0.046656987 | -0.121563596 | 0.002060624 | 0.019875435 | 0.41458209 | 0.44440311 | 0.103963654 | 0.972938945 |
| Acin1 | T | 253 | 0.108898978 | 0.086634775 | 0.134847361 | 0.025948383 | 0.048212586 | 0.254775813 | 0.387927937 | 0.122442628 | 0.690949652 | 0.531083419 |
| Snip1 | S | 89 | 0.028870282 | 0.036240586 | 0.081494704 | 0.052624422 | 0.045254118 | 0.224217011 | 0.299581786 | 0.000496398 | 0.077253125 | 0.223281125 |
| Ak1 | S | 38 | 0.113624056 | 0.117003319 | 0.030594214 | -0.083029842 | -0.086409105 | 0.136744443 | 0.058068068 | 0.472158112 | 0.244848763 | 0.148076935 |
| Ptpn21 | S | 673 | -0.029920901 | 0.113051743 | 0.09811314 | 0.128034041 | -0.014938603 | 0.342627625 | 0.004267865 | 0.026323743 | 0.013502686 | 0.592403112 |
| Rfc1 | S | 155 | 0.134777196 | 0.054177704 | 0.036156786 | -0.098620409 | -0.018020918 | 0.015310934 | 0.209963526 | 0.482893038 | 0.072739067 | 0.64587776 |
| Gpkow | S | 485 | 0.051433105 | 0.076608195 | 0.059758806 | 0.008325701 | -0.016849389 | 0.133499251 | 0.053246715 | 0.062020416 | 0.793393471 | 0.602450903 |

| | | | | | | | | | | | | |
|---|---|---|---|---|---|---|---|---|---|---|---|---|
| Dbn1 | S | 385 | 0.116111285 | -0.00930516 | 0.060546403 | -0.055564883 | 0.069851563 | 0.003242039 | 0.799856717 | 0.045112046 | 0.001915717 | 0.111754199 |
| Zgpat | S | 275 | 0.185111888 | 0.237178684 | 0.055388484 | -0.129723403 | -0.1817902 | 0.482025299 | 0.193848833 | 0.444557136 | 0.612637374 | 0.285779381 |
| Zgpat | S | 276 | 0.185111888 | 0.237178684 | 0.055388484 | -0.129723403 | -0.1817902 | 0.482025299 | 0.193848833 | 0.444557136 | 0.612637374 | 0.285779381 |
| Zgpat | S | 278 | 0.185111888 | 0.237178684 | 0.055388484 | -0.129723403 | -0.1817902 | 0.482025299 | 0.193848833 | 0.444557136 | 0.612637374 | 0.285779381 |
| Zgpat | T | 280 | 0.185111888 | 0.237178684 | 0.055388484 | -0.129723403 | -0.1817902 | 0.482025299 | 0.193848833 | 0.444557136 | 0.612637374 | 0.285779381 |
| Mtdh | S | 454 | 0.017732679 | 0.081345608 | 0.117903795 | 0.100171116 | 0.036558187 | 0.701737897 | 0.036378097 | 0.044994386 | 0.100208477 | 0.391216843 |
| Pacsin3 | S | 319 | 0.12469969 | 0.0881393 | 0.028655713 | -0.096043977 | -0.059483587 | 0.057494648 | 0.066839901 | 0.616936128 | 0.178142459 | 0.324747973 |
| Mark2 | S | 453 | 0.075843818 | 0.121054766 | 0.080811852 | 0.004968034 | -0.040242914 | 0.160805053 | 0.072191523 | 0.416935714 | 0.956820538 | 0.675995002 |
| Ssh1 | S | 1035 | 0.203880838 | 0.025299022 | 0.109170078 | -0.09471076 | 0.083871056 | 0.172041467 | 0.716289041 | 0.092016909 | 0.457520341 | 0.278709518 |
| Samhd1 | T | 310 | -0.1075349 | 0.123126831 | 0.205723286 | 0.313258186 | 0.082596455 | 0.035647371 | 0.021576547 | 0.002437733 | 0.004398056 | 0.059892103 |
| Eif4g1 | S | 1081 | -0.150028753 | 0.331072172 | 0.090955784 | 0.240984537 | -0.240116387 | 0.012520323 | 0.000103886 | 0.451456352 | 0.121088949 | 0.124309356 |
| Adar | S | 238 | 0.004670575 | 0.033983246 | 0.308212869 | 0.303542294 | 0.274229622 | 0.962622703 | 0.749543731 | 0.021453715 | 0.009991888 | 0.008147957 |
| Ubr5 | S | 2479 | 0.063298183 | 0.118187143 | 0.054771022 | -0.008527161 | -0.063416121 | 0.187933082 | 0.050390017 | 0.237639658 | 0.860281303 | 0.252769775 |
| Map3k3 | S | 237 | 0.091207944 | 0.229948538 | -0.014381433 | -0.105589376 | -0.24432997 | 0.270913885 | 0.029252349 | 0.860607079 | 0.215563429 | 0.033426945 |
| Syne2 | S | 793 | 0.022553692 | -0.02733868 | 0.248395921 | 0.225842229 | 0.275734601 | 0.839910204 | 0.826212612 | 0.002773384 | 0.127856893 | 0.113510842 |
| Egfr | S | 1028 | 0.03065205 | 0.167730235 | 0.219425918 | 0.188773868 | 0.051695682 | 0.790064573 | 0.320836327 | 0.123088054 | 0.160936007 | 0.745763649 |
| Foxp1 | S | 653 | -0.026119518 | 0.080711403 | 0.221340681 | 0.247460199 | 0.140629278 | 0.527028253 | 0.154392138 | 0.018436257 | 0.008621902 | 0.059343148 |
| Mrps36 | S | 60 | 0.135720691 | 0.041151655 | 0.108329337 | -0.027391354 | 0.067177683 | 0.08434066 | 0.484590064 | 0.158585954 | 0.711934266 | 0.344484317 |
| Prpf4b | S | 381 | 0.102996775 | 0.023783328 | 0.162017146 | 0.059020371 | 0.138233817 | 0.439555707 | 0.72977904 | 0.015599968 | 0.633669731 | 0.081642485 |
| Abca1 | S | 1169 | 0.231830859 | -0.04290919 | 0.214487659 | -0.0173432 | 0.257396849 | 0.114461101 | 0.725108309 | 0.179350455 | 0.907372658 | 0.161736101 |
| Atg16l2 | S | 308 | 0.168646208 | 0.311967823 | -0.135265686 | -0.303911894 | -0.44723351 | 0.211615213 | 0.047447247 | 0.540631645 | 0.20971851 | 0.10380615 |
| Srrm2 | S | 1339 | 0.035450174 | 0.032797903 | 0.111136593 | 0.075686419 | 0.078338689 | 0.149824264 | 0.069724104 | 0.024082338 | 0.071951214 | 0.081719589 |
| Mprip | S | 292 | 0.07783236 | 0.026524324 | 0.128384303 | 0.050551943 | 0.101859979 | 0.141685214 | 0.297155273 | 0.045932797 | 0.367866667 | 0.084734066 |
| Tra2a Tra2b | S | 270;265 | 0.011053193 | 0.127368598 | 0.110718005 | 0.099664812 | -0.016650593 | 0.681007212 | 0.043638046 | 0.103260819 | 0.140428287 | 0.781928751 |
| Hdgfl2 | S | 628 | -0.045289473 | 0.131095027 | 0.187162285 | 0.232451758 | 0.056067258 | 0.35335812 | 0.067515873 | 0.027445202 | 0.018933106 | 0.422443294 |
| Wrap53 | S | 27 | 0.045090058 | 0.045839688 | 0.108325074 | 0.063235016 | 0.062485386 | 0.364298574 | 0.062117781 | 0.026598005 | 0.251159288 | 0.12256217 |
| Ncoa5 | S | 377 | 0.053726001 | 0.022902759 | 0.18598582 | 0.132259819 | 0.163083062 | 0.16371727 | 0.349397806 | 0.057062941 | 0.1148372 | 0.087618812 |
| Dctn1 | T | 108 | 0.057653026 | 0.096718141 | 0.05805949 | 0.000406465 | -0.038658651 | 0.129125247 | 0.066695964 | 0.134890727 | 0.984830846 | 0.33675056 |
| Scap | S | 850 | -0.081546796 | 0.157652266 | 0.144568619 | 0.226115415 | -0.013083647 | 0.021999557 | 0.000967573 | 0.031319064 | 0.022955674 | 0.763592262 |
| Cdyl | S | 83 | 0.108245126 | 0.130101084 | -0.004089764 | -0.11233489 | -0.134190849 | 0.143742865 | 0.013200119 | 0.959828918 | 0.269630259 | 0.19156828 |
| Srrm2 | S | 851 | 0.02214775 | 0.054334232 | 0.111167661 | 0.089019911 | 0.056833429 | 0.410370625 | 0.318990825 | 0.00292828 | 0.016807126 | 0.299936956 |
| Lurap1l | S | 69 | 0.196731812 | 0.105004533 | 0.089622881 | -0.107108931 | -0.015381652 | 0.400498007 | 0.259703262 | 0.176769897 | 0.626335885 | 0.861341524 |
| Ubr4 | S | 1655 | -0.020336392 | 0.381231973 | 0.34071002 | 0.361046412 | -0.040521954 | 0.914089227 | 0.261547882 | 0.449572497 | 0.419038909 | 0.930146623 |
| Reps1 | S | 170 | 0.050480819 | 0.064245139 | 0.043252455 | -0.007228363 | -0.020992684 | 0.015172464 | 0.070948 | 0.091511449 | 0.722544066 | 0.479283503 |
| Nfic | S | 305 | 0.017367048 | 0.079763878 | 0.063154691 | 0.045787643 | -0.016609188 | 0.634620118 | 0.02526211 | 0.006635094 | 0.265034155 | 0.491013145 |
| Zfp638 | S | 486 | 0.062005345 | 0.131393622 | 0.128522828 | 0.066517483 | -0.002870794 | 0.602100894 | 0.073494172 | 0.236965355 | 0.62634696 | 0.976506713 |
| Zfp638 | S | 488 | 0.062005345 | 0.131393622 | 0.128522828 | 0.066517483 | -0.002870794 | 0.602100894 | 0.073494172 | 0.236965355 | 0.62634696 | 0.976506713 |
| Rc3h1 | S | 462 | 0.194094018 | 0.054325702 | 0.076834447 | -0.117259571 | 0.022508746 | 0.093219922 | 0.313742169 | 0.405891444 | 0.304290131 | 0.772649898 |
| Tmpo | T | 298 | 0.094746813 | 0.084522414 | 0.034777434 | -0.059969379 | -0.04974498 | 0.199542961 | 0.005616988 | 0.571817244 | 0.453390766 | 0.428775205 |
| Las1l | S | 238 | -0.014637848 | 0.111442454 | 0.180188668 | 0.194826517 | 0.068746215 | 0.823649044 | 0.171905006 | 0.02817403 | 0.010006132 | 0.288901911 |
| Ticrr | T | 1186 | 0.113161034 | 0.049084368 | 0.217794654 | 0.10463362 | 0.168710286 | 0.210364836 | 0.66639808 | 0.16128134 | 0.443986722 | 0.297525141 |
| Scrib | S | 436 | 0.150096131 | 0.21794533 | 0.051154205 | -0.098941927 | -0.166791125 | 0.21183241 | 0.237456317 | 0.650283221 | 0.461432591 | 0.363906116 |
| Pgam1 | S | 14 | 0.073238735 | 0.148156905 | 0.029784344 | -0.043454391 | -0.11837256 | 0.058537009 | 0.07313508 | 0.65499326 | 0.498383798 | 0.180353967 |
| Wrnip1 | S | 153 | 0.066220727 | 0.024856667 | 0.077220591 | 0.010999865 | 0.052363924 | 0.014651048 | 0.252788222 | 0.052647328 | 0.698027908 | 0.14182391 |
| Xrcc1 | S | 460 | 0.069673747 | 0.192802543 | 0.039599752 | -0.030073995 | -0.153202791 | 0.385052287 | 0.042992642 | 0.553948338 | 0.558283559 | 0.076077601 |
| Ppp1r12a | S | 870 | 0.015134506 | 0.05702366 | 0.122580219 | 0.107445713 | 0.065556559 | 0.616199677 | 0.092592806 | 0.012152336 | 0.027016073 | 0.087843948 |
| Iws1 | S | 366 | 0.051410634 | 0.03060315 | 0.080733141 | 0.029322507 | 0.050129991 | 0.104164915 | 0.246672204 | 0.006179169 | 0.302985501 | 0.098268748 |
| Mybl2 | T | 267 | 0.151778087 | -0.00299613 | 0.103759662 | -0.048018425 | 0.106755792 | 0.037095331 | 0.942319591 | 0.094932379 | 0.430853162 | 0.084413852 |
| Cobll1 | S | 1215 | 0.120183029 | 0.163616294 | -0.070585077 | -0.190768106 | -0.23420137 | 0.110942756 | 0.008749789 | 0.545509973 | 0.177751685 | 0.126563562 |
| Rab7 | S | 72 | 0.22533145 | -0.004541316 | 0.040975357 | -0.184356094 | 0.045516673 | 0.007246659 | 0.940785332 | 0.575519876 | 0.037612151 | 0.431226256 |
| Mpp1 | S | 52 | 0.027901279 | 0.193732419 | 0.115582348 | 0.087681069 | -0.07815007 | 0.808215793 | 0.078023535 | 0.237859097 | 0.414446418 | 0.357791991 |
| Ccnl1 | S | 380 | 0.029202432 | 0.056407727 | 0.06604425 | 0.036841818 | 0.009636523 | 0.39076483 | 0.018298285 | 0.010885173 | 0.279387433 | 0.435444836 |
| Rnf168 | S | 262 | -0.060458594 | 0.109497536 | 0.185351732 | 0.245810326 | 0.075854196 | 0.204523782 | 0.004580647 | 0.056975556 | 0.02060981 | 0.270234767 |
| Dmtn | S | 226 | 0.032322365 | 0.042705251 | 0.103690772 | 0.071368407 | 0.06098552 | 0.374979845 | 0.235565313 | 0.003803778 | 0.091946078 | 0.111474272 |
| Dnajc5 | S | 8 | 0.073055834 | 0.078811785 | 0.094603532 | 0.021547698 | 0.015791747 | 0.54998933 | 0.04013607 | 0.092851098 | 0.856755817 | 0.724411147 |
| Cbx3 | S | 95 | -0.012112342 | 0.0479813 | 0.108600116 | 0.120712458 | 0.060618816 | 0.501912417 | 0.001782405 | 0.007729737 | 0.004706111 | 0.04761565 |
| Ttf1 | S | 457 | -0.005512568 | 0.085460166 | 0.190412357 | 0.195924925 | 0.104952191 | 0.866269213 | 0.14760224 | 0.031845023 | 0.017134636 | 0.123504291 |
| Ttf1 | S | 460 | -0.005512568 | 0.085460166 | 0.190412357 | 0.195924925 | 0.104952191 | 0.866269213 | 0.14760224 | 0.031845023 | 0.017134636 | 0.123504291 |
| Arhgap31 | S | 1163 | 0.056536309 | 0.234503017 | -0.044295677 | -0.100831985 | -0.278798694 | 0.294711598 | 0.01018938 | 0.746471431 | 0.47050756 | 0.121712589 |
| Zfp654 | S | 566 | 0.174242267 | 0.064584824 | 0.217751009 | 0.043508742 | 0.153166185 | 0.274444584 | 0.732173005 | 0.225677141 | 0.695098587 | 0.361854998 |
| Zfp654 | S | 570 | 0.174242267 | 0.064584824 | 0.217751009 | 0.043508742 | 0.153166185 | 0.274444584 | 0.732173005 | 0.225677141 | 0.695098587 | 0.361854998 |
| Mki67 | S | 958 | 0.129583115 | 0.342906991 | -0.108908903 | -0.238492018 | -0.451815894 | 0.448954921 | 0.024131074 | 0.521021529 | 0.247304778 | 0.047427181 |
| Mki67 | T | 1956 | 0.176595041 | 0.091742192 | 0.072070873 | -0.104524168 | -0.019671319 | 0.065848749 | 0.3862194 | 0.583023163 | 0.456913663 | 0.892509327 |
| Farp1 | S | 902 | -0.003878612 | 0.059993699 | 0.150178554 | 0.154057166 | 0.090184856 | 0.943215278 | 0.120143991 | 0.008699019 | 0.057870808 | 0.066811343 |
| Taf3 | S | 229 | 0.256711298 | 0.107551344 | -0.008088857 | -0.264800155 | -0.115640201 | 0.212246846 | 0.040350148 | 0.875151397 | 0.19869603 | 0.096443092 |
| Thrap3 | S | 55 | -0.027679219 | 0.062610326 | 0.205903042 | 0.233582261 | 0.143292716 | 0.415223061 | 0.128214234 | 0.009190695 | 0.023066639 | 0.043093835 |
| Mpp6 | S | 110 | 0.035892569 | 0.135535077 | 0.077834199 | 0.04194163 | -0.057700878 | 0.595373847 | 0.114495048 | 0.036900084 | 0.533671457 | 0.405815932 |
| Atp2a2 Atp2a3 | T | 441;441 | -0.000714263 | 0.145244185 | 0.266834758 | 0.267549022 | 0.121590573 | 0.995141941 | 0.2533714 | 0.128207915 | 0.116424011 | 0.368579871 |
| Fbxo38 | S | 722 | 0.147862903 | 0.12320862 | 0.173562408 | 0.025699505 | 0.050353787 | 0.305647321 | 0.360537287 | 0.376471753 | 0.887287075 | 0.7766506 |
| Kif14 | S | 386 | -0.060628161 | 0.166408667 | 0.106968177 | 0.167596338 | -0.059440489 | 0.025873163 | 0.000149481 | 0.157774007 | 0.073211811 | 0.360994308 |
| Slc12a4 Slc12a6 Slc1 | T | 991;926;926 | 0.13711512 | 0.015136242 | 0.160462443 | 0.023347323 | 0.145326201 | 0.089624652 | 0.57163269 | 0.188925677 | 0.82515219 | 0.224692504 |
| Prkd1 Prkd3 | S | 748;734 | 0.024405522 | 0.079402529 | 0.168963308 | 0.144557786 | 0.08956078 | 0.438343479 | 0.176911101 | 0.069387084 | 0.095630777 | 0.253306915 |
| Epb41l2 | T | 192 | 0.01935763 | 0.169913999 | 0.146849455 | 0.127491825 | -0.023064545 | 0.608169304 | 0.101066415 | 0.233926743 | 0.284951333 | 0.844663311 |
| Ino80b | S | 147 | 0.01389289 | 0.082049822 | 0.150993767 | 0.137100877 | 0.068943946 | 0.654498582 | 0.050527617 | 0.083672249 | 0.091037184 | 0.301520764 |
| Tra2a | S | 80 | 0.016842972 | 0.091222265 | 0.131387586 | 0.114544614 | 0.040165321 | 0.808043645 | 0.106567983 | 0.032348728 | 0.13928169 | 0.154644675 |
| Kat7 | S | 59 | -0.046124465 | 0.106948355 | 0.199391972 | 0.245516437 | 0.092443618 | 0.449624709 | 0.031622426 | 0.052189868 | 0.024771809 | 0.196206922 |
| Snx16 | S | 108 | 0.04646024 | 0.103122681 | 0.107129132 | 0.060668891 | 0.004006451 | 0.384391754 | 0.080541503 | 0.135900547 | 0.285884785 | 0.936241583 |
| Phldb1 | S | 553 | 0.353650193 | -0.001308602 | -0.039921062 | -0.393571256 | -0.038612461 | 0.005337452 | 0.992231628 | 0.466796488 | 0.006396232 | 0.77387662 |
| Tut7 | S | 645 | 0.0041201 | 0.07972207 | 0.122975818 | 0.118855718 | 0.043253748 | 0.922366745 | 0.050510948 | 0.024166487 | 0.042169301 | 0.280429192 |
| Mgrn1 | S | 428 | 0.120103741 | 0.062097892 | -0.001434367 | -0.121538108 | -0.063532259 | 0.005943501 | 0.073064901 | 0.981889593 | 0.135355775 | 0.344163883 |
| Eml4 | S | 906 | 0.124975643 | 0.09032864 | 0.027460374 | -0.09751527 | -0.062868266 | 0.158298253 | 0.024725243 | 0.56877425 | 0.248723284 | 0.210054053 |
| Ranbp10 | S | 451 | 0.090654296 | 0.1132587 | 0.135499365 | 0.044845069 | 0.022240666 | 0.461705374 | 0.277195658 | 0.119756846 | 0.701906423 | 0.813794955 |
| Mki67 | S | 2257 | 0.043666716 | 0.129863882 | 0.086108763 | 0.042442047 | -0.043755119 | 0.59586969 | 0.04847847 | 0.149148757 | 0.546689151 | 0.079726339 |
| Map3k3 | S | 355 | 0.08952821 | 0.261078469 | -0.040616304 | -0.130144514 | -0.301694773 | 0.379532022 | 0.018967819 | 0.656893733 | 0.262959426 | 0.027920065 |
| Ctnnbl1 | S | 545 | -0.021274706 | 0.087140026 | 0.104682858 | 0.125957564 | 0.017542832 | 0.845760888 | 0.001213825 | 0.110428837 | 0.321444839 | 0.709847 |
| Rbbp6 | T | 985 | 0.08580297 | 0.057751276 | 0.161898336 | 0.076095365 | 0.104147059 | 0.236406888 | 0.426786106 | 0.102409817 | 0.368346421 | 0.24725455 |
| Rock1 | S | 1105 | -0.004107024 | 0.068867394 | 0.136600948 | 0.140707972 | 0.067733554 | 0.72057113 | 0.015968126 | 0.0329367 | 0.027683448 | 0.118404655 |
| Tpm1 | S | 174 | -0.050266921 | 0.05916291 | 0.216610746 | 0.266877667 | 0.157447836 | 0.240302591 | 0.252819343 | 0.00196094 | 0.003498875 | 0.016687332 |
| Zc3h18 | S | 888 | 0.011805493 | 0.031508044 | 0.121301497 | 0.109496005 | 0.089793453 | 0.816444156 | 0.198938057 | 0.001167533 | 0.114151227 | 0.01150419 |
| Herpud2 | S | 94 | -0.082225957 | 0.207346174 | 0.199282166 | 0.281508123 | -0.008064008 | 0.128161886 | 0.004093979 | 0.253703321 | 0.154171367 | 0.955848489 |
| Rbmx Rbmxl1 | S | 323;326 | 0.064284338 | 0.06361296 | 0.122501433 | 0.058217094 | 0.058888472 | 0.589320551 | 0.106765383 | 0.044440471 | 0.626128872 | 0.232501509 |
| Srrm2 | S | 1685 | 0.051469175 | 0.011327888 | 0.085171654 | 0.033702479 | 0.073843766 | 0.06016076 | 0.657110151 | 0.001922454 | 0.16219845 | 0.040393639 |
| Kif27 | S | 999 | -0.00030496 | 0.094186847 | 0.06171833 | 0.06202329 | -0.032468517 | 0.993483745 | 0.008384107 | 0.015066995 | 0.178033537 | 0.200138694 |
| Srrm2 | T | 2689 | 0.056680914 | 0.017262126 | 0.093911681 | 0.037230768 | 0.076649556 | 0.019971752 | 0.634851945 | 0.016945193 | 0.192650306 | 0.109263986 |

| | | | | | | | | | | | | | |
|---|---|---|---|---|---|---|---|---|---|---|---|---|---|
| Trpm7 | S | | 1403 | -0.07809338 | 0.154116844 | 0.157643026 | 0.235736407 | 0.003526182 | 0.198969952 | 0.024152801 | 0.02195011 | 0.013350776 | 0.940745084 |
| Srrm1 | S | | 946 | 0.015687684 | 0.072835912 | 0.091467902 | 0.075780218 | 0.01863199 | 0.56486649 | 0.032319688 | 0.024307982 | 0.049812523 | 0.442277462 |
| Sipa1l1 | S | | 1181 | -0.031957682 | 0.04472053 | 0.19345941 | 0.225417092 | 0.148738881 | 0.472298908 | 0.054943438 | 0.007567909 | 0.008478644 | 0.032442632 |
| Thrap3 | S | | 681 | -0.044255498 | 0.07132159 | 0.180732008 | 0.224987506 | 0.109410418 | 0.256000757 | 0.027664301 | 0.009548829 | 0.005965025 | 0.055649288 |
| Araf | T | | 181 | 0.186538566 | 0.106453703 | -0.032170724 | -0.218709289 | -0.138624427 | 0.035753603 | 0.119339745 | 0.789622597 | 0.150936418 | 0.305364766 |
| Mavs | S | | 157 | 0.068116882 | 0.018400456 | 0.140956995 | 0.072840113 | 0.122556539 | 0.362355564 | 0.662052669 | 0.007842471 | 0.337506928 | 0.039422887 |
| Ppp6r1 | S | | 662 | 0.217105081 | 0.156442284 | 0.007988561 | -0.20911652 | -0.148453723 | 0.106860957 | 0.251158768 | 0.972120381 | 0.382360919 | 0.521653177 |
| Ppp6r1 | S | | 665 | 0.217105081 | 0.156442284 | 0.007988561 | -0.20911652 | -0.148453723 | 0.106860957 | 0.251158768 | 0.972120381 | 0.382360919 | 0.521653177 |
| Ppp6r1 | S | | 668 | 0.217105081 | 0.156442284 | 0.007988561 | -0.20911652 | -0.148453723 | 0.106860957 | 0.251158768 | 0.972120381 | 0.382360919 | 0.521653177 |
| Ppp6r1 | T | | 666 | 0.217105081 | 0.156442284 | 0.007988561 | -0.20911652 | -0.148453723 | 0.106860957 | 0.251158768 | 0.972120381 | 0.382360919 | 0.521653177 |
| Eva1b | T | | 86 | 0.045507735 | 0.123313936 | 0.113591882 | 0.068084147 | -0.009722054 | 0.635719972 | 0.099782324 | 0.111152902 | 0.51195839 | 0.890306668 |
| Ncor1 | S | | 1274 | 0.034170693 | 0.041889043 | 0.154838858 | 0.120668165 | 0.112949815 | 0.463488131 | 0.629304445 | 0.007145076 | 0.024939172 | 0.243301424 |
| Cavin2 | S | | 363 | 0.006442886 | 0.10303551 | 0.013744031 | 0.007301145 | -0.089291479 | 0.661167732 | 3.69022E-05 | 0.491769726 | 0.712095534 | 0.019820936 |
| Fgd3 | S | | 589 | -0.018660893 | 0.074910595 | 0.25578829 | 0.274449182 | 0.180877694 | 0.575909171 | 0.08809493 | 0.059189567 | 0.049139791 | 0.111772722 |
| Lbr | S | | 90 | 0.080534166 | 0.085235662 | 0.091182693 | 0.010648527 | 0.005947031 | 0.240049208 | 0.166955501 | 0.11411762 | 0.833037195 | 0.880391879 |
| Ralbp1 | T | | 60 | -0.00429844 | 0.212447039 | 0.070465292 | 0.074763732 | -0.141981747 | 0.907190561 | 0.057404185 | 0.086020104 | 0.04575393 | 0.145544327 |
| Per2 | S | | 1231 | 0.089651085 | -0.062220309 | 0.366969297 | 0.277318213 | 0.429189607 | 0.522349427 | 0.564600193 | 0.032465822 | 0.09768516 | 0.029085606 |
| Elac2 | S | | 189 | 0.162534344 | 0.25942058 | -0.141808901 | -0.304343245 | -0.401229481 | 0.19917805 | 0.017032228 | 0.388908794 | 0.124664669 | 0.070800902 |
| Nucks1 | S | | 61 | -0.041274133 | 0.190266147 | 0.089010133 | 0.130284266 | -0.101256015 | 0.41410596 | 0.015538797 | 0.079785036 | 0.029013431 | 0.109884878 |
| Sntb2 | S | | 367 | -0.142735971 | 0.161777202 | 0.282149313 | 0.424885284 | 0.120372111 | 0.080610585 | 0.075882162 | 0.011062016 | 0.004298459 | 0.190251617 |
| Mki67 | T | | 2101 | 0.202435996 | 0.143455437 | -0.010678228 | -0.213114224 | -0.154133665 | 0.193669372 | 0.056680561 | 0.830646005 | 0.174210262 | 0.059551481 |
| Kdm4b | S | | 453 | 0.087067008 | 0.071209692 | 0.388531253 | 0.301464245 | 0.317321561 | 0.28674383 | 0.761844505 | 0.27870727 | 0.373682117 | 0.398693105 |
| Rbm3 | S | | 133 | 0.563129665 | 0.183871991 | -0.43610686 | -0.999236524 | -0.61997885 | 0.029863223 | 0.279443834 | 0.386363155 | 0.116894175 | 0.248379397 |
| Cfap97 | S | | 161 | -0.077244096 | 0.179624763 | 0.204240737 | 0.281484833 | 0.024615974 | 0.076413991 | 0.005580677 | 0.124414389 | 0.070641362 | 0.804480771 |
| Sulf2 | S | | 503 | 0.041644941 | 0.172586948 | 0.12831724 | 0.086672299 | -0.044269708 | 0.716642003 | 0.076478244 | 0.309520953 | 0.544459999 | 0.712202523 |
| Mid2 | S | | 116 | 0.078615324 | 0.039817446 | 0.220869846 | 0.142254522 | 0.1810524 | 0.397825823 | 0.750750027 | 0.056906998 | 0.161954753 | 0.197352858 |
| Cep170 | S | | 963 | 0.246229089 | 0.053237572 | -0.041048756 | -0.287277845 | -0.094286328 | 0.006203622 | 0.487736095 | 0.604583279 | 0.013789595 | 0.249740228 |
| Vim | S | | 430 | 0.143630058 | 0.051235951 | 0.080607216 | -0.063022843 | 0.029371264 | 0.057815407 | 0.496488992 | 0.236024424 | 0.049695668 | 0.559881325 |
| Ddx42 | S | | 96 | 0.095382651 | 0.09265261 | 0.089907457 | -0.005475194 | -0.002745153 | 0.178720309 | 0.19870396 | 0.204642442 | 0.940189679 | 0.970553388 |
| Nedd4l | S | | 516 | 0.06606497 | 0.047511098 | 0.116646191 | 0.050581221 | 0.069135093 | 0.054190097 | 0.279443834 | 0.133804855 | 0.429513469 | 0.322860595 |
| Slain2 | T | | 314 | 0.215762476 | 0.131779066 | 0.017090172 | -0.198672304 | -0.114688894 | 0.164724288 | 0.155247403 | 0.865583175 | 0.190838063 | 0.192572253 |
| Nr2f6 | S | | 41 | 0.059041239 | 0.077938892 | 0.123832596 | 0.064791357 | 0.045893704 | 0.293915287 | 0.253432321 | 0.070696387 | 0.304908207 | 0.505580039 |
| Atg9a | S | | 14 | -0.005692937 | 0.1044874 | 0.120576341 | 0.126269278 | 0.016088941 | 0.944088364 | 0.035072592 | 0.05317387 | 0.187022556 | 0.690375381 |
| Git1 | S | | 419 | 0.003775733 | 0.145904658 | 0.081667486 | 0.077891753 | -0.064237172 | 0.9135681 | 0.017736032 | 0.166590875 | 0.181431058 | 0.279550611 |
| Zfp608 | S | | 962 | 0.195784654 | -0.020359195 | 0.137678745 | -0.05810591 | 0.158037939 | 0.05109088 | 0.785302655 | 0.194159002 | 0.572644583 | 0.168383576 |
| Srrm2 | T | | 2350 | 0.072498877 | 0.065827313 | 0.078087431 | 0.005588554 | 0.012260118 | 0.078392593 | 0.114485355 | 0.190267205 | 0.903152507 | 0.796051926 |
| Hnrnpd | S | | 82 | -0.096859971 | 0.072347495 | 0.189555354 | 0.286415325 | 0.117207859 | 0.132314499 | 0.067948227 | 0.000745448 | 0.010092821 | 0.018184533 |
| Trp53bp1 | S | | 297 | 0.080665317 | 0.265692808 | 0.163793833 | 0.083128516 | -0.101898976 | 0.642814892 | 0.394481396 | 0.245181768 | 0.600203511 | 0.72093907 |
| Trp53bp1 | S | | 298 | 0.080665317 | 0.265692808 | 0.163793833 | 0.083128516 | -0.101898976 | 0.642814892 | 0.394481396 | 0.245181768 | 0.600203511 | 0.72093907 |
| Trp53bp1 | S | | 303 | 0.080665317 | 0.265692808 | 0.163793833 | 0.083128516 | -0.101898976 | 0.642814892 | 0.394481396 | 0.245181768 | 0.600203511 | 0.72093907 |
| Trp53bp1 | S | | 304 | 0.080665317 | 0.265692808 | 0.163793833 | 0.083128516 | -0.101898976 | 0.642814892 | 0.394481396 | 0.245181768 | 0.600203511 | 0.72093907 |
| Trp53bp1 | T | | 305 | 0.080665317 | 0.265692808 | 0.163793833 | 0.083128516 | -0.101898976 | 0.642814892 | 0.394481396 | 0.245181768 | 0.600203511 | 0.72093907 |
| Rai1 | T | | 1062 | 0.175676165 | -0.014860026 | 0.080789967 | -0.094886198 | 0.095649993 | 0.019631571 | 0.875709813 | 0.169122482 | 0.161563866 | 0.3699707 |
| Kdm1a | S | | 132 | 0.01785563 | 0.043194944 | 0.107800713 | 0.089945083 | 0.064605769 | 0.470978074 | 0.06827669 | 0.009436823 | 0.026441829 | 0.045026161 |
| Fcho2 | S | | 507 | -0.054663895 | 0.146905261 | 0.089068522 | 0.143732417 | -0.057836739 | 0.254289477 | 0.000912323 | 0.131847615 | 0.05109853 | 0.274161865 |
| Mki67 | S | | 1187 | 0.089753989 | 0.081858986 | 0.039504632 | -0.050249357 | -0.042354354 | 0.212578973 | 0.025880081 | 0.218411102 | 0.435950316 | 0.18852551 |
| Macf1 | S | | 7311 | 0.093368067 | 0.209321782 | 0.111714517 | 0.018346451 | -0.097607265 | 0.44135174 | 0.118519571 | 0.647416161 | 0.941611997 | 0.697545269 |
| Gigyf1 | T | | 598 | -0.572877146 | 0.335177748 | 0.771849633 | 1.344726779 | 0.436671885 | 0.453439497 | 0.49033954 | 0.094757306 | 0.172612801 | 0.424849662 |
| Gigyf1 | S | | 755 | -0.104206396 | 0.184159177 | 0.192095846 | 0.296302243 | 0.00793667 | 0.056583064 | 0.004305062 | 0.072827639 | 0.032919935 | 0.91481932 |
| Srsf4 | S | | 457 | 0.029820692 | 0.153665832 | 0.140419957 | 0.110599264 | -0.013245875 | 0.71657371 | 0.04423612 | 0.374096442 | 0.477994992 | 0.925749297 |
| Snw1 | S | | 224 | 0.041176439 | 0.031504427 | 0.074572294 | 0.033395855 | 0.043067867 | 0.040581512 | 0.189756252 | 0.01256539 | 0.13749861 | 0.111943901 |
| Hirip3 | S | | 398 | 0.289669746 | 0.030896439 | 0.041634665 | -0.248035081 | 0.010738227 | 0.086628269 | 0.354216548 | 0.617438431 | 0.109560451 | 0.894952995 |
| Pcm1 | S | | 1370 | 0.128319215 | 0.130972036 | 0.103715871 | -0.024603345 | -0.027256165 | 0.240897884 | 0.163127725 | 0.553658277 | 0.887107063 | 0.870563036 |
| Adgra2 | S | | 1113 | -0.039661696 | 0.132539447 | 0.142942105 | 0.1826038 | 0.010402658 | 0.5438868 | 0.056282131 | 0.032416148 | 0.039868019 | 0.853862963 |
| Trp53bp1 | S | | 822 | -0.030460368 | 0.08369334 | 0.228053543 | 0.258513911 | 0.144360203 | 0.649879945 | 0.131333621 | 0.037812793 | 0.030468264 | 0.110607067 |
| Slc23a2 | S | | 81 | -0.026752902 | 0.149180886 | 0.150980397 | 0.177733298 | 0.001799511 | 0.678970882 | 0.154137248 | 0.027939449 | 0.023581163 | 0.982244847 |
| Esyt2 | S | | 660 | 0.020055477 | 0.113752883 | 0.062345485 | 0.042290008 | -0.051407398 | 0.505919622 | 0.017254474 | 0.118209732 | 0.272667308 | 0.220593264 |
| Nab2 | S | | 479 | 0.023542208 | 0.116507247 | 0.142445022 | 0.118902813 | 0.025937775 | 0.697597229 | 0.06861401 | 0.160960869 | 0.218593566 | 0.74987101 |
| Son | Y | | 1815 | 0.052534581 | 0.1844224 | 0.272069941 | 0.219535359 | 0.08764754 | 0.651645253 | 0.204915159 | 0.444005869 | 0.523704971 | 0.792609219 |
| Raly | S | | 270 | 0.085560963 | 0.054189424 | 0.045203311 | -0.040357652 | -0.008986113 | 0.012061115 | 0.172785411 | 0.287729571 | 0.33104506 | 0.837459943 |
| Srrm2 | S | | 1038 | 0.001390017 | 0.030664614 | 0.132594986 | 0.131204969 | 0.101930372 | 0.981412585 | 0.22849615 | 0.001232967 | 0.112481109 | 0.007254171 |
| Evi2a | S | | 203 | 0.18259652 | 0.119928362 | 0.076812147 | -0.105784372 | -0.043116215 | 0.087965468 | 0.46193852 | 0.625879243 | 0.466030201 | 0.810968651 |
| Ubr5 | S | | 2476 | 0.076239291 | 0.157295235 | 0.042279073 | -0.033960219 | -0.115016162 | 0.237337725 | 0.043698312 | 0.681395736 | 0.727036808 | 0.295172945 |
| Adar | S | | 140 | 0.001263526 | 0.083173618 | 0.129014769 | 0.127751243 | 0.045841151 | 0.98952133 | 0.056612128 | 0.031542664 | 0.256071036 | 0.310357627 |
| Pxn | S | | 320 | 0.09342153 | 0.200617007 | 0.011136953 | -0.082284577 | -0.189480055 | 0.369455415 | 0.037433527 | 0.847345306 | 0.409535949 | 0.051361675 |
| Trp53bp1 | S | | 1755 | 0.086483773 | 0.146079373 | -0.010842004 | -0.097325777 | -0.156921377 | 0.503889837 | 0.002175833 | 0.918968358 | 0.527712917 | 0.228408846 |
| Mtdh | S | | 565 | 0.101681035 | 0.161077031 | 0.009291984 | -0.092389052 | -0.151785047 | 0.316385074 | 0.02193879 | 0.891614072 | 0.384421515 | 0.08823554 |
| Smarcad1 | S | | 145 | 0.048436807 | 0.019839333 | 0.090368228 | 0.041931421 | 0.070528896 | 0.155457779 | 0.354509096 | 0.003473277 | 0.19040297 | 0.005999382 |
| Smarcad1 | S | | 34 | 0.142050619 | 0.082807442 | 0.067181153 | -0.074869466 | -0.015626289 | 0.228175396 | 0.088375313 | 0.390746281 | 0.514803128 | 0.834577244 |
| Il16 | S | | 319 | -0.013042551 | 0.313307135 | 0.045756149 | 0.0587987 | -0.267550986 | 0.821948335 | 0.028070853 | 0.616384483 | 0.509474412 | 0.059994233 |
| Srrm1 | S | | 528 | 0.133712885 | 0.032861809 | 0.081289277 | -0.052423608 | 0.048427468 | 0.045916045 | 0.636350083 | 0.150356601 | 0.362917384 | 0.495514695 |
| Kifc3 | S | | 782 | 0.084078743 | 0.136136393 | 0.008462352 | -0.075616391 | -0.127674041 | 0.203569692 | 0.014656068 | 0.885402542 | 0.334028504 | 0.114217335 |
| Ncaph2 | S | | 120 | -0.046701491 | 0.125754233 | 0.29489665 | 0.341598141 | 0.169142417 | 0.663025821 | 0.342764455 | 0.054167806 | 0.040013161 | 0.251255158 |
| Stt3b | S | | 496 | 0.042707036 | 0.025901372 | 0.176943491 | 0.134236455 | 0.15104212 | 0.311975587 | 0.56817661 | 0.024193101 | 0.06262043 | 0.047432251 |
| Snx2 | S | | 185 | 0.01807721 | 0.332249842 | -0.016243888 | -0.034321098 | -0.34849373 | 0.875088661 | 0.019332067 | 0.910421482 | 0.781931371 | 0.052129053 |
| Cdk2 Cdk3 | T | 14;14 | | 0.173565205 | 0.07024648 | 0.04511532 | -0.128449885 | -0.02513116 | 0.053584553 | 0.334050399 | 0.54825503 | 0.064846 | 0.539337939 |
| Bcl9l | S | | 88 | 0.022674055 | 0.076853491 | 0.08287464 | 0.060200585 | 0.006021149 | 0.468906177 | 0.011447345 | 0.099701797 | 0.202663576 | 0.868418793 |
| Phactr4 | T | | 425 | 0.017319706 | 0.184185465 | 0.128355893 | 0.111036187 | -0.055829573 | 0.91334012 | 0.128382544 | 0.143035873 | 0.490240515 | 0.562713598 |
| Vdac1 | S | | 117 | 0.027016074 | 0.021068189 | 0.141226729 | 0.114210654 | 0.12015854 | 0.549607126 | 0.505789845 | 0.00295484 | 0.05268271 | 0.007164556 |
| Tpx2 | S | | 121 | -0.066815717 | 0.114894607 | 0.068695603 | 0.13551132 | -0.046199004 | 0.466863336 | 0.000830641 | 0.036902387 | 0.206161292 | 0.113164038 |
| Mavs | S | | 179 | 0.114558871 | 0.126289453 | 0.109567262 | -0.004991609 | -0.016722191 | 0.377396421 | 0.365627473 | 0.141866057 | 0.966256939 | 0.894786688 |
| Tbc1d10b | S | | 644 | 0.006958894 | 0.097378361 | 0.132453726 | 0.125494832 | 0.035075365 | 0.890775876 | 0.316032224 | 0.009004762 | 0.05525281 | 0.681421539 |
| Gpatch8 | S | | 912 | 0.064910518 | 0.036583056 | 0.193837641 | 0.128927123 | 0.157254585 | 0.339869367 | 0.52010803 | 0.062361146 | 0.177066142 | 0.110072942 |
| Cic | S | | 2147 | 0.016637079 | 0.026626003 | 0.130050634 | 0.113413556 | 0.103424631 | 0.475795782 | 0.368668501 | 0.002666973 | 0.009676978 | 0.025740058 |
| Add1 | S | | 355 | 0.088699185 | 0.080098591 | 0.097735319 | 0.009036135 | 0.017636729 | 0.344204215 | 0.256687115 | 0.066778555 | 0.916460021 | 0.78664014 |
| Arglu1 | S | | 75 | 0.032801153 | 0.047269685 | 0.111514206 | 0.078713053 | 0.064244521 | 0.390963442 | 0.231177431 | 0.010533802 | 0.049223719 | 0.079500092 |
| Rpgrip1l | S | | 985 | 0.029952862 | 0.090821424 | 0.199888902 | 0.169936039 | 0.109067477 | 0.812349805 | 0.24799887 | 0.076730549 | 0.247441311 | 0.261923891 |
| Rpgrip1l | S | | 989 | 0.029952862 | 0.090821424 | 0.199888902 | 0.169936039 | 0.109067477 | 0.812349805 | 0.24799887 | 0.076730549 | 0.247441311 | 0.261923891 |
| Mast2 | S | | 14 | -0.107465912 | 0.14977899 | 0.196494313 | 0.303960225 | 0.046715323 | 0.083779068 | 0.019881987 | 0.016018228 | 0.005639798 | 0.428176884 |
| Bclaf3 | S | | 20 | 0.024603184 | 0.129869267 | 0.135533947 | 0.110930763 | 0.00566468 | 0.517528581 | 0.07155618 | 0.243862999 | 0.318251056 | 0.956254145 |
| Frmd6 | Y | | 607 | 0.000743568 | 0.400248429 | -0.052197917 | -0.052941485 | -0.452446346 | 0.996644533 | 0.0203907 | 0.825440554 | 0.823133478 | 0.12971851 |
| Gtf2i | S | | 823 | 0.108213732 | 0.035749571 | 0.144424762 | 0.036211031 | 0.108675191 | 0.36215299 | 0.735478419 | 0.066201849 | 0.746111734 | 0.107258036 |
| Sntb2 | S | | 75 | -0.009779135 | 0.077875764 | 0.108969888 | 0.118749023 | 0.031094125 | 0.610936745 | 0.015869264 | 0.020628338 | 0.014158413 | 0.315636799 |
| Zfp513 | S | | 85 | 0.032852519 | 0.051496297 | 0.074554553 | 0.041702033 | 0.023058256 | 0.160827942 | 0.197683927 | 0.006425286 | 0.076603579 | 0.494997976 |

| | | | | | | | | | | | | |
|---|---|---|---|---|---|---|---|---|---|---|---|---|
| Sipa1l1 | S | 1151 | 0.203668303 | 0.061468909 | -0.014453319 | -0.218121622 | -0.075922228 | 0.010784089 | 0.430006025 | 0.884551791 | 0.103599008 | 0.463689188 |
| Micall2 | S | 657 | 0.107266579 | -0.067933081 | 0.189886942 | 0.082620363 | 0.257820024 | 0.046602659 | 0.370722051 | 0.026042946 | 0.20610694 | 0.023917502 |
| Kansl3 | S | 539 | 0.082290579 | 0.129629357 | -0.026742886 | -0.109033465 | -0.156372243 | 0.09469359 | 0.00686033 | 0.697074443 | 0.193793007 | 0.099912598 |
| Taf3 | S | 791 | -0.008175884 | 0.046543507 | 0.201610791 | 0.209786675 | 0.155067283 | 0.929229823 | 0.598017301 | 0.006010922 | 0.090327833 | 0.138220703 |
| Hnrnpu | S | 247 | 0.042370022 | 0.032422655 | 0.238642373 | 0.196272351 | 0.206219719 | 0.595811579 | 0.583416667 | 0.04351024 | 0.087422198 | 0.089017133 |
| E4f1 | S | 408 | 0.084517881 | 0.017658966 | 0.287889814 | 0.203371933 | 0.270230848 | 0.544716414 | 0.929394667 | 0.066789438 | 0.188178737 | 0.233188243 |
| Zmym4 | S | 1242 | 0.105502264 | 0.053518596 | 0.210722603 | 0.105220339 | 0.157204008 | 0.429581255 | 0.638840215 | 0.103580748 | 0.288868499 | 0.041554183 |
| Prrc2a | S | 378 | 0.051060005 | 0.077632158 | 0.051661988 | 0.000601983 | -0.02597017 | 0.15937061 | 0.011965708 | 0.292103461 | 0.989566873 | 0.549579936 |
| Micall2 | S | 659 | 0.155659038 | -0.025805756 | 0.082682187 | -0.072976851 | 0.108487943 | 0.009845025 | 0.771686332 | 0.177471954 | 0.17793438 | 0.271783628 |
| Shroom4 | S | 369 | -0.057529635 | 0.04075058 | 0.134409273 | 0.191938908 | 0.093658693 | 0.346200951 | 0.078496026 | 0.000203207 | 0.049876358 | 0.003355589 |
| Cep164 | S | 227 | -0.093594552 | 0.308964049 | 0.034315822 | 0.127910374 | -0.274648227 | 0.452072497 | 0.003210523 | 0.811795165 | 0.447740424 | 0.1356587 |
| Mcm2 | S | 21 | -0.015573375 | 0.079661251 | 0.09397376 | 0.109547135 | 0.014312509 | 0.516583128 | 0.000560619 | 0.092553779 | 0.057357949 | 0.707833753 |
| Arid1a | T | 1600 | -0.146620129 | 0.257880662 | 0.168694552 | 0.315314681 | -0.089186111 | 0.000501081 | 0.010678869 | 0.000612577 | 0.00050414 | 0.151662024 |
| Asap1 | T | 852 | 0.060117509 | 0.165827829 | 0.071613068 | 0.011495558 | -0.094214761 | 0.371326918 | 0.070112625 | 0.461270247 | 0.898730684 | 0.363868629 |
| Sin3a | S | 861 | -0.024494542 | 0.101442804 | 0.113431495 | 0.137926037 | 0.011988691 | 0.56344712 | 0.057973954 | 0.00979679 | 0.024862171 | 0.765299781 |
| Gkap1 | S | 23 | -0.052741056 | 0.111825882 | 0.190431805 | 0.243172861 | 0.078605922 | 0.383780963 | 0.024340623 | 0.064424169 | 0.03510656 | 0.321761727 |
| Foxk1 | S | 209 | -0.046135485 | 0.066356484 | 0.127014112 | 0.173149597 | 0.060657628 | 0.525659844 | 0.148533447 | 0.00073072 | 0.099792378 | 0.184927637 |
| Kdm2b | S | 26 | 0.183209966 | 0.10005629 | 0.037006364 | -0.146203602 | -0.063049926 | 0.115795317 | 0.246159737 | 0.600816414 | 0.188994744 | 0.448248821 |
| Parn | T | 554 | 0.083253613 | 0.020637481 | 0.086240746 | 0.002987133 | 0.065603265 | 0.130583761 | 0.434119032 | 0.015972136 | 0.943963106 | 0.024011045 |
| Kank2 | T | 178 | 0.020753247 | 0.119946063 | 0.091856456 | 0.071103209 | -0.028089607 | 0.759942775 | 0.034542678 | 0.135352043 | 0.306504993 | 0.511962663 |
| Arhgap21 | S | 1847 | 0.189669145 | 0.072614926 | 0.039008553 | -0.150660592 | -0.033606372 | 0.04322899 | 0.357517655 | 0.79816324 | 0.34236179 | 0.80973219 |
| Zwilch | S | 196 | 0.094750459 | 0.14805805 | 0.021641774 | -0.073108685 | -0.126416276 | 0.318915402 | 0.022236508 | 0.85081821 | 0.579869334 | 0.323692286 |
| Ccnl2 | S | 377 | -0.019653524 | 0.17185332 | 0.145990271 | 0.165643795 | -0.025863049 | 0.783744702 | 0.146604248 | 0.063965009 | 0.062118773 | 0.792422751 |
| Ccnl2 | S | 379 | -0.019653524 | 0.17185332 | 0.145990271 | 0.165643795 | -0.025863049 | 0.783744702 | 0.146604248 | 0.063965009 | 0.062118773 | 0.792422751 |
| Rpl6 | T | 153 | 0.59988529 | 0.226288714 | 0.405169136 | -0.194716154 | 0.178880422 | 0.115420996 | NA | NA | NA | NA |
| Cfap97 | S | 160 | -0.060957992 | 0.141162459 | 0.153410892 | 0.214368884 | 0.012248433 | 0.335017665 | 0.029762076 | 0.04422266 | 0.021287314 | 0.82604494 |
| Cfap97 | T | 155 | -0.060957992 | 0.141162459 | 0.153410892 | 0.214368884 | 0.012248433 | 0.335017665 | 0.029762076 | 0.04422266 | 0.021287314 | 0.82604494 |
| Ktn1 | S | 77 | 0.137307318 | -0.009602035 | 0.069777252 | -0.067530066 | 0.079379288 | 0.0010609 | 0.900489413 | 0.564507022 | 0.57431103 | 0.552326433 |
| Retreg3 | S | 274 | 0.138636298 | 0.068628561 | 0.073940275 | -0.064696023 | 0.005311714 | 0.173693723 | 0.224384912 | 0.241121209 | 0.47175278 | 0.914629515 |
| Srrm2 | S | 2648 | 0.028815456 | 0.03673516 | 0.061403514 | 0.032588057 | 0.024668353 | 0.104901581 | 0.048400952 | 0.007487788 | 0.039796824 | 0.08673538 |
| Bclaf3 | S | 15 | 0.032364322 | 0.069507148 | 0.126132508 | 0.093768186 | 0.05662536 | 0.326451434 | 0.091093101 | 0.100901413 | 0.175567925 | 0.367851442 |
| Gpatch8 | S | 451 | 0.015394222 | 0.064474182 | 0.154633229 | 0.139239006 | 0.090159046 | 0.754030812 | 0.226675262 | 0.023713466 | 0.050511328 | 0.149462934 |
| Tra2a | S | 82 | 0.671990061 | -0.007836351 | 0.072173552 | -0.59981651 | 0.080009903 | 0.309613619 | 0.858705732 | 0.331769841 | 0.351536785 | 0.285362818 |
| Retreg3 | T | 310 | 0.059187577 | 0.032927232 | 0.179969938 | 0.120782361 | 0.147042707 | 0.206788126 | 0.464334738 | 0.082124394 | 0.178924375 | 0.123472621 |
| Epb41l1 | S | 868 | 0.074122444 | 0.246964097 | -0.013004902 | -0.087127345 | -0.259968999 | 0.257661738 | 0.049544471 | 0.86404873 | 0.190244242 | 0.049235972 |
| Tut4 | S | 131 | -0.04487994 | 0.101482856 | 0.14207077 | 0.18695071 | 0.040587915 | 0.011274647 | 0.017876278 | 0.001877437 | 0.003644167 | 0.198288627 |
| Kif20a | T | 238 | 0.188252962 | 0.143168719 | 0.099106556 | -0.089146406 | -0.044062163 | 0.279797511 | 0.363159588 | 0.494248996 | 0.447166548 | 0.601886387 |
| Tnk2 | S | 102 | -0.078711479 | 0.047165737 | 0.234671764 | 0.313383242 | 0.187506027 | 0.030757822 | 0.278063304 | 0.000204212 | 0.003308278 | 0.008130346 |
| Ranbp2 | S | 757 | 0.138662728 | 0.145581299 | 0.048708102 | -0.089954626 | -0.096873196 | 0.22835795 | 0.200403391 | 0.448513416 | 0.408050634 | 0.365599206 |
| Mat2b | S | 282 | -0.062772728 | 0.191655418 | 0.072036066 | 0.134808794 | -0.119619352 | 0.339565754 | 0.002204707 | 0.260188952 | 0.115474201 | 0.103727114 |
| Dact3 | S | 165 | 0.055459224 | 0.01101666 | 0.211782853 | 0.15632363 | 0.200766193 | 0.728343585 | 0.858631056 | 0.012880981 | 0.359712928 | 0.00117627 |
| Exoc1 | S | 470 | 0.011631156 | 0.045707875 | 0.195360069 | 0.183728913 | 0.149652194 | 0.853677149 | 0.459096169 | 0.015533454 | 0.013811904 | 0.022470458 |
| Ncor1 | S | 2134 | -0.043523276 | 0.10735448 | 0.156268568 | 0.199791844 | 0.048914088 | 0.53193367 | 0.075646435 | 0.021882758 | 0.030373539 | 0.236939195 |
| Safb2 | S | 645 | 0.060267786 | 0.110821409 | 0.119631827 | 0.059364041 | 0.008810418 | 0.547490593 | 0.215434342 | 0.095430946 | 0.549053031 | 0.91130298 |
| Kmt2d | S | 706 | 0.30935491 | 0.249626147 | 0.104920486 | -0.204434423 | -0.144705661 | 0.287542924 | 0.349607701 | NA | NA | NA |
| Farp1 | S | 892 | -0.02033699 | 0.075877985 | 0.099209686 | 0.119546676 | 0.023331701 | 0.286325471 | 0.001543992 | 0.029493292 | 0.021010253 | 0.426431685 |
| Map4 | S | 1046 | 0.02600468 | 0.114731113 | 0.054190712 | 0.028186031 | -0.060540402 | 0.213341057 | 0.034417679 | 0.13082447 | 0.356651839 | 0.2006409 |
| Afdn | S | 1173 | -0.031171409 | 0.042875881 | 0.148067687 | 0.179239096 | 0.105191806 | 0.226042985 | 0.149463883 | 0.000492362 | 0.000571578 | 0.008786102 |
| Supt6 | S | 1701 | 0.029440064 | 0.077857942 | 0.122581471 | 0.093141406 | 0.044723528 | 0.192250111 | 0.200636853 | 0.078795122 | 0.146512404 | 0.493864296 |
| Ccne2 | S | 339 | 0.029977491 | 0.086156108 | 0.122128814 | 0.092151324 | 0.035972707 | 0.583482862 | 0.202847321 | 0.037257615 | 0.071650254 | 0.498832318 |
| Yap1 | T | 104 | 0.173031381 | 0.031949125 | 0.118946639 | -0.054084742 | 0.086997514 | 0.1669787 | 0.352615168 | 0.27354893 | 0.667096733 | 0.392603426 |
| Ccny | S | 25 | 0.009400568 | 0.109206197 | 0.105131721 | 0.095731153 | -0.004074476 | 0.833746175 | 0.043001265 | 0.089260892 | 0.121690456 | 0.936628948 |
| Rbm12b1 Rbm12b2 | S | 277;277 | 0.004504801 | 0.025992452 | 0.115291049 | 0.110786248 | 0.089298597 | 0.842570145 | 0.208826784 | 0.000862813 | 0.004493254 | 0.005441572 |
| Rbm12b1 Rbm12b2 | S | 279;279 | 0.004504801 | 0.025992452 | 0.115291049 | 0.110786248 | 0.089298597 | 0.842570145 | 0.208826784 | 0.000862813 | 0.004493254 | 0.005441572 |
| Prr12 | S | 648 | -0.046331477 | 0.083298603 | 0.211176152 | 0.257507629 | 0.127877549 | 0.369247953 | 0.185424446 | 0.011537666 | 0.009043782 | 0.094431892 |
| Akap12 | S | 683 | 0.042974467 | 0.320248239 | -0.090465944 | -0.133440411 | -0.410714183 | 0.550135431 | 0.004367847 | 0.257779426 | 0.035348981 | 0.002262579 |
| Ncoa6 | S | 1094 | 0.152870022 | 0.081294087 | 0.062883791 | -0.089986231 | -0.018410296 | 0.127944311 | 0.533412712 | 0.154975275 | 0.303787678 | 0.880881883 |
| Mki67 | T | 1295 | 0.025753141 | 0.177306109 | 0.095716046 | 0.069962905 | -0.081590063 | 0.745141719 | 0.109534767 | 0.166458334 | 0.361087258 | 0.381759165 |
| Srrm2 | T | 456 | -0.011445278 | 0.09201716 | 0.062386631 | 0.073831909 | -0.029630529 | 0.865647593 | 0.005319114 | 0.027382217 | 0.339664247 | 0.192980104 |
| Fxr2 | S | 604 | 0.090549788 | 0.068197727 | 0.056326273 | -0.034223515 | -0.011871454 | 0.140123317 | 0.079057924 | 0.227380724 | 0.549939847 | 0.780837259 |
| Mprip | S | 617 | 0.074507206 | 0.028635485 | 0.114865261 | 0.040358055 | 0.086229776 | 0.163700297 | 0.583794856 | 0.029937344 | 0.287332835 | 0.10170585 |
| Brpf1 | S | 1074 | -0.032915845 | 0.04021383 | 0.214561738 | 0.247477583 | 0.174347908 | 0.612353314 | 0.498059043 | 0.004119791 | 0.028122785 | 0.048229454 |
| Uimc1 | S | 379 | 0.023276271 | 0.018436374 | 0.102572065 | 0.079295794 | 0.084135691 | 0.543890016 | 0.266949771 | 0.000864727 | 0.112773494 | 0.005324224 |
| Srek1ip1 | S | 143 | 0.003760907 | 0.080592037 | 0.159959504 | 0.156198597 | 0.079367467 | 0.932901923 | 0.275703399 | 0.020792224 | 0.032086072 | 0.299358559 |
| Prkd3 | S | 31 | 0.028091871 | 0.117788885 | 0.075733184 | 0.047641312 | -0.042055701 | 0.525535803 | 0.013437644 | 0.359517984 | 0.548190303 | 0.583289202 |
| D5Ertd579e | S | 1295 | -0.146320248 | 0.118231057 | 0.23599525 | 0.382315498 | 0.117764193 | 0.028697998 | 0.111535932 | 0.0005148 | 0.00180453 | 0.117485977 |
| Mcl1 | S | 140 | -0.022527622 | 0.056537243 | 0.141098039 | 0.16362566 | 0.084560796 | 0.737846684 | 0.076464182 | 0.007899688 | 0.084034828 | 0.056131881 |
| Mef2a | S | 485 | -0.00853683 | 0.074000329 | 0.097826601 | 0.106363431 | 0.023826272 | 0.710667667 | 0.072308926 | 0.003895276 | 0.00868909 | 0.466561834 |
| Srek1 | S | 465 | -0.065739296 | 0.06031452 | 0.128712901 | 0.194452197 | 0.068398381 | 0.348445231 | 0.004133554 | 0.005999778 | 0.058112008 | 0.040364836 |
| Tom1l2 | S | 479 | 0.084744504 | 0.055537224 | 0.189445944 | 0.10470144 | 0.13390872 | 0.280912844 | 0.444143759 | 0.166087634 | 0.400354612 | 0.295422386 |
| Mphosph10 | S | 172 | 0.130675912 | -0.003644986 | 0.117528709 | -0.013147202 | 0.121173695 | 0.228314884 | 0.959476754 | 0.016139348 | 0.884077777 | 0.167176562 |
| Ezh1 | S | 416 | 0.046262538 | 0.041878274 | 0.396020894 | 0.349758356 | 0.35414262 | 0.862407051 | 0.840012549 | 0.099837759 | 0.268609141 | 0.184631875 |
| Rbm15b | S | 606 | 0.026717868 | 0.069010453 | 0.119042309 | 0.092324441 | 0.050031857 | 0.392212125 | 0.046832341 | 0.116742236 | 0.185558196 | 0.41478376 |
| Crebbp Ep300 | S | 2325;2363 | 0.026378403 | 0.083077786 | 0.151952233 | 0.12557383 | 0.068874447 | 0.659234018 | 0.515670764 | 0.017521709 | 0.057613703 | 0.5758015 |
| Gnas | S | 1091;352 | -0.008581893 | 0.082986397 | 0.120662208 | 0.129244101 | 0.037675811 | 0.82397267 | 0.051630616 | 0.024726325 | 0.026810419 | 0.353063625 |
| Cenpk | S | 196 | 0.068599425 | 0.270215258 | 0.187131594 | 0.118532169 | -0.083083664 | 0.859843994 | 0.19547924 | 0.507090494 | 0.782669309 | 0.768287608 |
| Srrm2 | S | 1576 | 0.03335455 | 0.081881512 | 0.052702323 | 0.019347773 | -0.029179189 | 0.380101521 | 0.006117635 | 0.188167259 | 0.647633748 | 0.402168863 |
| Cdc26 | S | 42 | -0.012184171 | 0.096838343 | 0.082106049 | 0.09429022 | -0.014732294 | 0.663259285 | 0.057720054 | 0.004429905 | 0.027647525 | 0.675118005 |
| Pcdh18 | S | 877 | 0.122240391 | 0.097898771 | 0.046984419 | -0.075255972 | -0.050914352 | 0.022872129 | 0.429301127 | 0.602896429 | 0.407527901 | 0.699071735 |
| Pcdh18 | Y | 884 | 0.122240391 | 0.097898771 | 0.046984419 | -0.075255972 | -0.050914352 | 0.022872129 | 0.429301127 | 0.602896429 | 0.407527901 | 0.699071735 |
| Kif23 | T | 184 | 0.352076369 | 0.061414552 | -0.126936639 | -0.479013008 | -0.188351192 | 0.004343098 | 0.594562804 | 0.215371219 | 0.013856135 | 0.139800465 |
| Srsf2 | S | 187 | 0.079221018 | 0.098700811 | 0.038890619 | -0.040330399 | -0.059810191 | 0.107655443 | 0.038853996 | 0.627728698 | 0.607623346 | 0.455084441 |
| Srgap2 | S | 427 | -0.073654655 | 0.158237529 | 0.159152456 | 0.232807111 | 0.000914927 | 0.196477958 | 0.069423312 | 0.015087551 | 0.002712741 | 0.988097987 |
| Nin | S | 1794 | 0.147991332 | 0.117742082 | 0.030351595 | -0.117639737 | -0.087390487 | 0.217988391 | 0.110345668 | 0.543277468 | 0.301226172 | 0.183922217 |
| Pkn1 | T | 778 | 0.024056508 | 0.127969064 | 0.146521651 | 0.122465142 | 0.018552587 | 0.838791828 | 0.043291212 | 0.298233592 | 0.448503955 | 0.880448249 |
| Ckap4 | S | 67 | 0.119654924 | 0.104717293 | 0.041015028 | -0.078639895 | -0.063702265 | 0.089163888 | 0.140285351 | 0.669020628 | 0.430108922 | 0.519067164 |
| Pacsin3 | S | 354 | -0.00621121 | 0.19691272 | 0.041058604 | 0.047269814 | -0.155854116 | 0.932897069 | 0.008289534 | 0.500649105 | 0.494784775 | 0.023466512 |
| Taf5l | S | 230 | -0.032478314 | 0.089120402 | 0.267998938 | 0.300477252 | 0.178878536 | 0.701866258 | 0.225453908 | 0.062695863 | 0.051755876 | 0.160529626 |
| Kmt2b | S | 1044 | -0.05622289 | 0.110561981 | 0.161477607 | 0.217700497 | 0.050915626 | 0.304908921 | 0.267962512 | 0.002167844 | 0.019792619 | 0.562434834 |
| Zfp608 | S | 1452 | 0.065574282 | 0.060635705 | 0.076467991 | 0.01089371 | 0.015832286 | 0.118798141 | 0.132678449 | 0.1312761 | 0.792953704 | 0.697790335 |
| Prkar2b | S | 85 | -0.022054847 | 0.085375721 | 0.118118275 | 0.140173122 | 0.032742554 | 0.504311288 | 0.012364536 | 0.035422741 | 0.020690416 | 0.360124425 |
| Raf1 | S | 29 | -0.001260452 | 0.106583318 | 0.08913305 | 0.090393502 | -0.017450268 | 0.98934522 | 0.024937441 | 0.065044469 | 0.37741425 | 0.613595112 |
| Srrm2 | S | 250 | 0.039040445 | 0.052110225 | 0.130626128 | 0.091585683 | 0.078515903 | 0.256523859 | 0.067373491 | 0.135094765 | 0.244987022 | 0.302262728 |
| Etv1 | S | 94 | -0.066633037 | 0.177794031 | 0.108590603 | 0.17522364 | -0.069203428 | 0.643622101 | 0.026994678 | 0.113051148 | 0.275873687 | 0.262691002 |

| | | | | | | | | | | | | |
|---|---|---|---|---|---|---|---|---|---|---|---|---|
| Kdm1a | T | 105 | 0.069114596 | 0.141221885 | -0.02613867 | -0.095253265 | -0.167360555 | 0.258677545 | 0.001850556 | 0.502645825 | 0.155057833 | 0.010374671 |
| Myl12b | T | 135 | -7.02244E-05 | 0.101133998 | 0.086736884 | 0.086807108 | -0.014397115 | 0.998964551 | 0.040439948 | 0.032646744 | 0.182833503 | 0.706996489 |
| Ccne2 | S | 383 | 0.113345797 | 0.084179495 | 0.092798883 | -0.020546914 | 0.008619388 | 0.190939981 | 0.214487157 | 0.329848507 | 0.833250188 | 0.921964065 |
| Srrm1 | S | 177 | 0.121303253 | 0.034233091 | 0.05684591 | -0.064457343 | 0.022612819 | 0.016300521 | 0.285152626 | 0.494613557 | 0.448406009 | 0.774291521 |
| Srrm1 | S | 179 | 0.121303253 | 0.034233091 | 0.05684591 | -0.064457343 | 0.022612819 | 0.016300521 | 0.285152626 | 0.494613557 | 0.448406009 | 0.774291521 |
| Srrm1 | S | 181 | 0.121303253 | 0.034233091 | 0.05684591 | -0.064457343 | 0.022612819 | 0.016300521 | 0.285152626 | 0.494613557 | 0.448406009 | 0.774291521 |
| Elk1 | S | 325 | 0.193817044 | 0.032615252 | 0.037048907 | -0.156768137 | 0.004433655 | 0.028188316 | 0.538213294 | 0.58939886 | 0.087013432 | 0.944376824 |
| Sp100 | S | 314 | -0.078521464 | 0.07320714 | 0.21835653 | 0.296877993 | 0.145149389 | 0.109270434 | 0.13067958 | 0.002476144 | 1.46824E-05 | 0.000285646 |
| Igf2bp2 | S | 164 | -0.003929338 | 0.130389981 | 0.05990027 | 0.063829608 | -0.070489711 | 0.898836602 | 0.004330344 | 0.243107151 | 0.203739261 | 0.173341087 |
| Ncbp3 | S | 495 | -0.001475983 | 0.068953563 | 0.095488609 | 0.096964591 | 0.026535046 | 0.962844085 | 0.028948197 | 0.016464422 | 0.03337238 | 0.354666343 |
| Vcp | S | 705 | 0.051204164 | 0.143546998 | 0.047630211 | -0.003573953 | -0.095916787 | 0.218992777 | 0.060440862 | 0.435668801 | 0.949367324 | 0.212126138 |
| Rap1gap | S | 606 | -0.094095462 | 0.296231229 | 0.138593542 | 0.232689004 | -0.157637687 | 0.556927989 | 0.027354872 | 0.490309728 | 0.301783446 | 0.421413173 |
| Ranbp3 | S | 463 | -0.015326639 | 0.048843554 | 0.078783616 | 0.094110255 | 0.029940062 | 0.519765025 | 0.030480273 | 0.000303574 | 0.033204818 | 0.127368272 |
| Gas2l1 | S | 292 | -0.023399145 | 0.147981944 | 0.1118867 | 0.135285846 | -0.036095244 | 0.585638091 | 0.05557287 | 0.05552353 | 0.032665671 | 0.540681277 |
| Ncoa3 | S | 544 | 0.089364707 | 0.237305338 | -0.05413891 | -0.143503617 | -0.291444249 | 0.465779016 | 0.012213873 | 0.573782436 | 0.276817263 | 0.041535283 |
| Srrm2 | S | 438 | 0.05497044 | 0.05431534 | 0.155288838 | 0.100318398 | 0.100973498 | 0.43093082 | 0.329652213 | 0.066045511 | 0.191832358 | 0.165171795 |
| Srrm2 | S | 1151 | 0.041327068 | 0.025174304 | 0.098653171 | 0.057326103 | 0.073478867 | 0.182554172 | 0.364387753 | 0.008526512 | 0.046088941 | 0.006548295 |
| Golm1 | S | 213 | 0.037988428 | 0.078719078 | 0.063931731 | 0.025943303 | -0.014787347 | 0.179532355 | 0.025864604 | 0.206482095 | 0.565898497 | 0.738793497 |
| Mycbp2 | S | 2970 | -0.099354737 | 0.107324588 | 0.174641828 | 0.273996565 | 0.06731724 | 0.166272039 | 0.049177189 | 0.004947143 | 0.026509261 | 0.108734474 |
| Ets2 | S | 309 | 0.108872144 | 0.116050834 | 0.075941537 | -0.032930607 | -0.040109297 | 0.091404297 | 0.212298353 | 0.668757137 | 0.848504188 | 0.823263511 |
| Scrib | S | 974 | -0.074461652 | 0.248653222 | 0.177981081 | 0.252442733 | -0.07067214 | 0.639559641 | 0.045306341 | 0.381441121 | 0.285231857 | 0.712246925 |
| Setdb2 | S | 119 | 0.04161308 | 0.104200054 | 0.074395542 | 0.032782462 | -0.029804512 | 0.456078349 | 0.06515831 | 0.123496268 | 0.48180655 | 0.445749271 |
| Fam117a | S | 305 | -0.188172061 | 0.394223606 | 0.221242569 | 0.40941463 | -0.172981037 | 0.47700055 | 0.038586389 | 0.576596643 | 0.354853577 | 0.652258381 |
| Fam117a | S | 318 | -0.188172061 | 0.394223606 | 0.221242569 | 0.40941463 | -0.172981037 | 0.47700055 | 0.038586389 | 0.576596643 | 0.354853577 | 0.652258381 |
| Fam117a | S | 326 | -0.188172061 | 0.394223606 | 0.221242569 | 0.40941463 | -0.172981037 | 0.47700055 | 0.038586389 | 0.576596643 | 0.354853577 | 0.652258381 |
| Lasp1 | T | 104 | 0.268294354 | -0.015365477 | 0.013227569 | -0.255066785 | 0.028593046 | 0.010547662 | 0.885781513 | 0.852332266 | 0.01389669 | 0.763588232 |
| Pcyt1a | S | 315 | -0.037662019 | 0.169504393 | 0.020379808 | 0.058041827 | -0.149124585 | 0.52127233 | 0.000387251 | 0.552682783 | 0.346839332 | 0.011252559 |
| Cers2 | S | 341 | 0.043901319 | 0.049169 | 0.103910841 | 0.060009523 | 0.054741842 | 0.244102391 | 0.135434968 | 0.057537421 | 0.220983561 | 0.230641507 |
| Cers2 | S | 348 | 0.043901319 | 0.049169 | 0.103910841 | 0.060009523 | 0.054741842 | 0.244102391 | 0.135434968 | 0.057537421 | 0.220983561 | 0.230641507 |
| Cers2 | T | 346 | 0.043901319 | 0.049169 | 0.103910841 | 0.060009523 | 0.054741842 | 0.244102391 | 0.135434968 | 0.057537421 | 0.220983561 | 0.230641507 |
| Swsap1 | S | 262 | -0.003935296 | 0.07025098 | 0.217099885 | 0.221035181 | 0.146848905 | 0.972013078 | 0.423455143 | 0.030158256 | 0.104754398 | 0.057943524 |
| Svil | S | 960 | 0.004702766 | 0.213616571 | 0.082579231 | 0.077876466 | -0.131037339 | 0.946596494 | 0.050502565 | 0.355655549 | 0.432650946 | 0.227190519 |
| Dennd1b | S | 530 | -0.040979482 | 0.085264334 | 0.204390093 | 0.245369575 | 0.119125759 | 0.416303692 | 0.433632318 | 0.005627535 | 0.003690082 | 0.290109842 |
| Dennd1b | S | 533 | -0.040979482 | 0.085264334 | 0.204390093 | 0.245369575 | 0.119125759 | 0.416303692 | 0.433632318 | 0.005627535 | 0.003690082 | 0.290109842 |
| Trim28 | S | 51 | -0.006479523 | 0.045777569 | 0.195919308 | 0.202398831 | 0.150141739 | 0.8521115 | 0.414975803 | 0.01123533 | 0.00768246 | 0.057284826 |
| Med24 | S | 871 | 0.05290612 | 0.067630129 | 0.050572842 | -0.002333278 | -0.017057287 | 0.025794999 | 0.128531113 | 0.18185374 | 0.938281963 | 0.68591044 |
| Ep300 | S | 1037 | 0.022884278 | 0.102828256 | 0.147858399 | 0.124974121 | 0.045030143 | 0.748233972 | 0.107714562 | 0.133365977 | 0.209442361 | 0.582490242 |
| Arid4b | S | 710 | -0.18222157 | 0.500640264 | 0.379545691 | 0.561767262 | -0.121094572 | 0.637605086 | 0.327677351 | 0.334567084 | 0.203730108 | 0.803496212 |
| Arid4b | S | 712 | -0.18222157 | 0.500640264 | 0.379545691 | 0.561767262 | -0.121094572 | 0.637605086 | 0.327677351 | 0.334567084 | 0.203730108 | 0.803496212 |
| Arid4b | S | 713 | -0.18222157 | 0.500640264 | 0.379545691 | 0.561767262 | -0.121094572 | 0.637605086 | 0.327677351 | 0.334567084 | 0.203730108 | 0.803496212 |
| Arid4b | S | 717 | -0.18222157 | 0.500640264 | 0.379545691 | 0.561767262 | -0.121094572 | 0.637605086 | 0.327677351 | 0.334567084 | 0.203730108 | 0.803496212 |
| Xrcc1 | S | 467 | 0.081073935 | 0.150837658 | 0.031031109 | -0.050042826 | -0.119806549 | 0.148626751 | 0.147511215 | 0.371978246 | 0.296178879 | 0.22670036 |
| Xrcc1 | T | 469 | 0.081073935 | 0.150837658 | 0.031031109 | -0.050042826 | -0.119806549 | 0.148626751 | 0.147511215 | 0.371978246 | 0.296178879 | 0.22670036 |
| Evi5 | S | 102 | -0.020141996 | 0.177180497 | 0.171829864 | 0.19197186 | -0.005350633 | 0.794201049 | 0.057443582 | 0.288535166 | 0.243464706 | 0.967994144 |
| Srrm2 | S | 1577 | 0.033110622 | 0.079268097 | 0.05225601 | 0.019145388 | -0.027012087 | 0.372857151 | 0.00712479 | 0.181599819 | 0.640791405 | 0.419555264 |
| Sf3b1 | T | 326 | 0.048275991 | 0.025632927 | 0.108955126 | 0.060679136 | 0.083322199 | 0.128097207 | 0.37253964 | 0.02581837 | 0.137477878 | 0.066134579 |
| Tns2 | S | 102 | -0.006574515 | 0.399448602 | 0.030607894 | 0.03718241 | -0.368840707 | 0.94503689 | 0.066641369 | 0.792599757 | 0.764404822 | 0.083078072 |
| Fcho2 | S | 487 | 0.004032277 | 0.126849758 | 0.097159608 | 0.093127331 | -0.02969015 | 0.894429328 | 0.062903967 | 0.08316341 | 0.087611184 | 0.563813328 |
| Ppig | S | 395 | 0.019699014 | 0.080828744 | 0.068800606 | 0.049101592 | -0.012028138 | 0.554492703 | 0.003431471 | 0.226913386 | 0.378849568 | 0.799441622 |
| Zfp367 | S | 21 | 0.005952542 | 0.08135231 | 0.214216439 | 0.208263897 | 0.132864129 | 0.936556643 | 0.097390332 | 0.139616091 | 0.149514519 | 0.292377293 |
| Trio | S | 1818 | 0.048900407 | 0.061502195 | 0.265999585 | 0.217099178 | 0.20449739 | 0.397963843 | 0.617235907 | 0.142620776 | 0.204638633 | 0.261879666 |
| Kmt2a | S | 1233 | 0.104585203 | 0.057045465 | 0.017382815 | -0.087202387 | -0.03966265 | 0.002473366 | 0.375297027 | 0.722352989 | 0.164643058 | 0.579079299 |
| Ctr9 | S | 1085 | 0.218438651 | 0.095896314 | -0.040780407 | -0.259219058 | -0.136676722 | 0.031863104 | 0.305777667 | 0.776717966 | 0.137534388 | 0.354367898 |
| Zfp746 | S | 323 | -0.027690923 | 0.154952605 | 0.285807271 | 0.313498194 | 0.130854666 | 0.89043849 | 0.236229339 | 0.191912216 | 0.236051424 | 0.503803285 |
| Pitpnm1 | S | 373 | 0.033519993 | 0.190507554 | 0.005960701 | -0.027559292 | -0.184546853 | 0.690520603 | 0.007442725 | 0.902644475 | 0.709455176 | 0.001047042 |
| Luzp1 | S | 395 | -0.019046156 | 0.071147054 | 0.103218242 | 0.122264398 | 0.032071188 | 0.523090799 | 0.032097646 | 0.006737747 | 0.005183911 | 0.062678556 |
| Stt3b | S | 495 | 0.012324265 | 0.026128511 | 0.172802164 | 0.160477899 | 0.146673653 | 0.750720157 | 0.47283772 | 0.008987409 | 0.018628433 | 0.022350835 |
| Brca1 | S | 1149 | -0.040793151 | 0.087118136 | 0.148613808 | 0.189406959 | 0.061495673 | 0.435761932 | 0.129996825 | 0.006574484 | 0.020087804 | 0.253745419 |
| Nsd3 | S | 1433 | -0.027120524 | 0.131925703 | 0.105004626 | 0.132125149 | -0.026921078 | 0.636344058 | 0.015931789 | 0.11842817 | 0.10906813 | 0.629407666 |
| Hivep2 | S | 2284 | 0.085074654 | 0.120984363 | 0.176046892 | 0.090972237 | 0.055062529 | 0.580435534 | 0.355574255 | 0.199759992 | 0.586968 | 0.700351175 |
| Lrrc1 | S | 428 | 0.192444285 | 0.04106297 | 0.165633229 | -0.026811056 | 0.124570259 | 0.095823731 | 0.828654246 | 0.49503933 | 0.905306234 | 0.643660019 |
| Ncoa2 | S | 29 | 0.050772234 | 0.025484079 | 0.113820107 | 0.063047874 | 0.088336029 | 0.06513322 | 0.298968319 | 0.068177568 | 0.233045581 | 0.138922974 |
| Tmpo | T | 112;112 | 0.231300164 | 0.044233884 | 0.018226437 | -0.213073727 | -0.026007447 | 0.046191364 | 0.438553411 | 0.785645676 | 0.065236807 | 0.583801674 |
| Cic | S | 1049 | -0.102569651 | 0.083478624 | 0.197783033 | 0.300352684 | 0.114304409 | 0.107974597 | 0.076097766 | 0.001868865 | 0.00606532 | 0.021310108 |
| Cic | S | 1053 | -0.102569651 | 0.083478624 | 0.197783033 | 0.300352684 | 0.114304409 | 0.107974597 | 0.076097766 | 0.001868865 | 0.00606532 | 0.021310108 |
| Bcar3 | S | 284 | -0.007931195 | 0.077484242 | 0.143019596 | 0.15095079 | 0.065535354 | 0.845813028 | 0.215309956 | 0.012480697 | 0.02092505 | 0.290915296 |
| Tasor2 | S | 1082 | 0.089504938 | 0.112053137 | 0.093685179 | 0.004180241 | -0.018367958 | 0.343599474 | 0.110700445 | 0.397429181 | 0.972253142 | 0.862875799 |
| Peak1 | S | 723 | 0.101335339 | 0.23443238 | -0.026386816 | -0.127722155 | -0.260819196 | 0.257924986 | 0.06156916 | 0.786127049 | 0.265640214 | 0.071055686 |
| Top2b | S | 1387 | -0.063459703 | 0.104113689 | 0.136800322 | 0.200260024 | 0.032686633 | 0.374435912 | 0.017741926 | 0.027571362 | 0.047458864 | 0.464892854 |
| Sh3bp5l | S | 377 | -0.045802742 | 0.227790895 | 0.09299734 | 0.138800082 | -0.134793554 | 0.54856299 | 0.02103782 | 0.317233019 | 0.140403361 | 0.157405033 |
| Mki67 | T | 2283 | 0.051919109 | 0.151560247 | 0.075929147 | 0.024012038 | -0.0756311 | 0.495501049 | 0.077566606 | 0.342945898 | 0.598854363 | 0.151357677 |
| Filip1l | S | 990 | 0.008219612 | 0.119644148 | 0.217879317 | 0.209659705 | 0.098235169 | 0.911105448 | 0.147874028 | 0.199882934 | 0.218448745 | 0.496746658 |
| Zfp608 | S | 626 | 0.047897796 | 0.014282741 | 0.158205692 | 0.110307896 | 0.143922951 | 0.203491556 | 0.583119562 | 0.033246923 | 0.083728748 | 0.048034774 |
| Fam92a | T | 351 | 0.088564171 | 0.107160741 | 0.035787488 | -0.052776683 | -0.071373253 | 0.535818904 | 0.012391554 | 0.418557151 | 0.703659236 | 0.123170074 |
| Son | S | 1809 | 0.039470118 | 0.166871665 | 0.236336488 | 0.19686637 | 0.069464824 | 0.760030891 | 0.153573523 | 0.450311105 | 0.528423648 | 0.813427849 |
| Zmym4 | S | 1543 | 0.04501391 | 0.046821644 | 0.07064182 | 0.02562791 | 0.023820176 | 0.262701206 | 0.079866796 | 0.026010473 | 0.471800512 | 0.215613095 |
| Sash1 | S | 534 | -0.000781084 | 0.106310653 | 0.174738858 | 0.175519943 | 0.068428205 | 0.990958592 | 0.107441788 | 0.116708951 | 0.117121948 | 0.43978758 |
| Incenp | S | 248 | 0.024627741 | 0.162035023 | 0.397051649 | 0.372423908 | 0.235016625 | 0.622087699 | 0.451306012 | 0.368291373 | 0.392862256 | 0.58619043 |
| Farp2 Frmd7 | S | 344;302 | 0.206416294 | 0.04203965 | -0.006882561 | -0.213298856 | -0.048922211 | 0.016981955 | 0.356466498 | 0.887817152 | 0.020163484 | 0.341417288 |
| Rsrc1 | T | 8 | 0.018767526 | 0.108277113 | 0.077598725 | 0.0588312 | -0.030678388 | 0.737121727 | 0.01802891 | 0.186207568 | 0.360993267 | 0.518568371 |
| Fam219a | T | 85 | 0.030357847 | -0.005816109 | 0.172541323 | 0.142183477 | 0.178357432 | 0.408500945 | 0.890517397 | 0.004945015 | 0.016715742 | 0.011195808 |
| Tnrc6c | S | 1358 | 0.078965344 | -0.010022008 | 0.15980693 | 0.080841586 | 0.169828938 | 0.562243248 | 0.92656966 | 0.005595783 | 0.549170257 | 0.20350033 |
| Mkrn2 | T | 138 | 0.072715351 | 0.080108984 | 0.158073212 | 0.085357861 | 0.077964228 | 0.334944114 | 0.4696465 | 0.130259439 | 0.39198372 | 0.528394553 |
| Pak4 | T | 207 | 0.05113816 | 0.148462562 | 0.040835147 | -0.010303012 | -0.107627415 | 0.254014766 | 0.054457428 | 0.478289872 | 0.856793874 | 0.16208008 |
| Dock7 | S | 905 | -0.075973818 | 0.249453905 | 0.09327873 | 0.169252548 | -0.156175175 | 0.673354111 | 0.025837099 | 0.348766647 | 0.36406683 | 0.070982099 |
| Ipo5 | S | 827 | -0.00034116 | 0.154797862 | 0.134380447 | 0.134721607 | -0.020417415 | 0.993372948 | 0.121684559 | 0.128277899 | 0.13634299 | 0.826885744 |
| Nfic | T | 293 | 0.030608098 | 0.119417558 | 0.160460175 | 0.129852077 | 0.041042617 | 0.706555938 | 0.26446719 | 0.111834512 | 0.222866067 | 0.707069646 |
| Rbm12 | S | 420 | 0.023649641 | 0.044938189 | 0.076775443 | 0.053125801 | 0.031837254 | 0.604593427 | 0.027216205 | 0.011994443 | 0.292797461 | 0.159329292 |
| Top2b | S | 1539 | -0.00334198 | 0.058922913 | 0.144025057 | 0.147367036 | 0.085102144 | 0.938288291 | 0.015632118 | 0.073401458 | 0.06814742 | 0.203997508 |
| Scrib | S | 1566 | 0.066237485 | 0.115799372 | 0.069367149 | 0.003129664 | -0.046432223 | 0.328440172 | 0.045884459 | 0.483221947 | 0.973825789 | 0.608155008 |
| Klf16 | S | 245 | 0.012705653 | 0.15426424 | 0.104844547 | 0.092138894 | -0.049419694 | 0.867086237 | 0.037786902 | 0.33303364 | 0.381573167 | 0.594337166 |
| Gigyf1 | S | 756 | -0.063816773 | 0.146062796 | 0.139924054 | 0.203740827 | -0.006138742 | 0.215870657 | 0.004260073 | 0.153165365 | 0.071587979 | 0.935007953 |
| Dock1 | S | 1826 | 0.132327343 | 0.009896545 | 0.058309882 | -0.074017461 | 0.048413337 | 0.028318057 | 0.894224433 | 0.092631983 | 0.138124455 | 0.52725029 |
| Cep170b | S | 563 | 0.217177529 | 0.10253899 | 0.000493969 | -0.21668356 | -0.102045021 | 0.096070676 | 0.248897518 | 0.99685684 | 0.148527717 | 0.384485355 |

| | | | | | | | | | | | | |
|---|---|---|---|---|---|---|---|---|---|---|---|---|
| Cep170b | T | 567 | 0.217177529 | 0.10253899 | 0.000493969 | -0.21668356 | -0.102045021 | 0.096070676 | 0.248897518 | 0.99685684 | 0.148527717 | 0.384485355 |
| Rubcn | S | 936 | 0.575358456 | 0.052543129 | 0.025541868 | -0.549816589 | -0.027001261 | 0.205026703 | 0.854072182 | 0.91974176 | 0.21692567 | 0.911030239 |
| Pkn2 | T | 957 | -0.027727748 | 0.096730775 | 0.118004631 | 0.145732378 | 0.021273856 | 0.304150523 | 0.015555532 | 0.031378222 | 0.013866976 | 0.551995781 |
| Hnrnpul2 | S | 159 | -0.047735921 | 0.151344593 | 0.239020372 | 0.286756293 | 0.087675779 | 0.038336615 | 0.07970752 | 0.042811802 | 0.030499238 | 0.300977103 |
| Nktr | S | 245 | 0.096932952 | -0.005282724 | 0.095891153 | -0.001041798 | 0.101173877 | 0.279838705 | 0.78073148 | 0.001857466 | 0.988940185 | 0.00096168 |
| Resf1 | S | 1482 | -0.006371659 | 0.14174214 | 0.191413678 | 0.197785337 | 0.049671537 | 0.9583226 | 0.190972085 | 0.131232648 | 0.20821348 | 0.680941788 |
| Nes | S | 1084 | -0.004974306 | 0.11281021 | 0.198569571 | 0.203543877 | 0.085759362 | 0.954382669 | 0.109260366 | 0.15698879 | 0.159073324 | 0.46617147 |
| Cttn | S | 418 | -0.092808534 | 0.197207538 | 0.182836357 | 0.275644891 | -0.014371181 | 0.001882642 | 0.004972769 | 0.023091848 | 0.010707196 | 0.763873245 |
| Atad5 | S | 1201 | 0.150980255 | 0.169058203 | -0.029970059 | -0.180950314 | -0.199028262 | 0.226830037 | 0.056887933 | 0.808220383 | 0.256469694 | 0.184129786 |
| Hp1bp3 | S | 515 | 0.072226441 | 0.131945226 | 0.064747937 | -0.007478504 | -0.067197288 | 0.387772506 | 0.072470548 | 0.415009924 | 0.937325482 | 0.432215193 |
| Rplp2 | S | 16 | 0.264697697 | 0.046541674 | 0.146185182 | -0.118512515 | 0.099643508 | 0.174576231 | 0.825071878 | 0.509301413 | 0.622230632 | 0.707867196 |
| Srrm2 | S | 1556 | 0.053653601 | 0.092751152 | 0.057953115 | 0.004299514 | -0.034798037 | 0.283966234 | 0.036090097 | 0.276233019 | 0.939161379 | 0.495811562 |
| Pelp1 | S | 749 | -0.002650096 | 0.082475391 | 0.065332826 | 0.067982921 | -0.017142566 | 0.916458278 | 0.00233058 | 0.091156684 | 0.094645112 | 0.562916049 |
| Gas2 | T | 282 | -0.113122752 | 0.345749209 | 0.328251154 | 0.441373906 | -0.017498054 | 0.284437258 | 0.1366808 | 0.29356459 | 0.196453781 | 0.953663214 |
| Tspyl1 | S | 51 | 0.055593137 | 0.035312958 | 0.091334948 | 0.035741811 | 0.056021989 | 0.164998814 | 0.30757862 | 0.029946837 | 0.188342007 | 0.004354305 |
| Cntrob | S | 149 | 0.279563961 | 0.137804455 | 0.123645666 | -0.155918295 | -0.014158789 | 0.392772441 | 0.487500712 | 0.577383753 | 0.621785499 | 0.943390005 |
| Luc7l2 | S | 353 | -0.02260523 | 0.090037841 | 0.230610851 | 0.253216081 | 0.14057301 | 0.655423389 | 0.479273135 | 0.024143525 | 0.01940349 | 0.31217702 |
| Luc7l2 | S | 358 | -0.02260523 | 0.090037841 | 0.230610851 | 0.253216081 | 0.14057301 | 0.655423389 | 0.479273135 | 0.024143525 | 0.01940349 | 0.31217702 |
| Plcd1 | T | 457 | 0.033309401 | 0.15471647 | 0.048497138 | 0.015187737 | -0.106219333 | 0.711902872 | 0.012879688 | 0.592987321 | 0.888164848 | 0.274996432 |
| Map7d1 | S | 115 | 0.078934408 | 0.096268392 | 0.018385167 | -0.06054924 | -0.077883225 | 0.136707644 | 0.02249011 | 0.672484382 | 0.241841642 | 0.100827362 |
| Trio | S | 1785 | -0.138059201 | 0.16034114 | 0.238401644 | 0.376460845 | 0.078060503 | 0.05469434 | 0.083572122 | 0.006953242 | 8.09458E-05 | 0.280324592 |
| Gpatch8 | S | 648 | 0.008172201 | 0.069104078 | 0.107858428 | 0.099686226 | 0.03875435 | 0.838984983 | 0.223680652 | 0.007526387 | 0.06705646 | 0.453390645 |
| Nsrp1 | S | 33 | 0.025292948 | 0.05797831 | 0.075422091 | 0.050129143 | 0.017443781 | 0.539012031 | 0.048612871 | 0.020505691 | 0.268912248 | 0.503033007 |
| Snip1 | S | 59 | 0.10942384 | 0.191333279 | 0.014779537 | -0.094644303 | -0.176553742 | 0.191424036 | 0.141484454 | 0.858698385 | 0.295845547 | 0.177279334 |
| Rpl23 LOC100044627 | S | 17 | 0.442592085 | 0.049463573 | -0.130123263 | -0.572715349 | -0.179586837 | 0.020608017 | 0.731132928 | 0.388414912 | 0.002776951 | 0.032173596 |
| Cdk16 | S | 138 | 0.027489184 | 0.235221083 | -0.02178449 | -0.049273674 | -0.257005573 | 0.792234551 | 0.008546969 | 0.872822433 | 0.738705099 | 0.140571024 |
| Zbtb7c | S | 215 | 0.042609048 | 0.07962487 | 0.111830132 | 0.069221084 | 0.032205261 | 0.429390564 | 0.111181022 | 0.13192215 | 0.335278674 | 0.611926843 |
| Snap23 | S | 110 | 0.030945043 | 0.001628782 | 0.101958821 | 0.071013779 | 0.10033004 | 0.055445376 | 0.911614706 | 0.002875442 | 0.033381538 | 0.009126971 |
| Srrm1 | S | 427 | 0.132720382 | 0.04732531 | 0.013292709 | -0.119427672 | -0.0340326 | 0.028345826 | 0.087368875 | 0.787536246 | 0.08580047 | 0.499012242 |
| Ccdc88a | S | 1822 | 0.038337947 | 0.169206839 | 0.056935491 | 0.018597544 | -0.112271348 | 0.588233275 | 0.029689973 | 0.60429737 | 0.864888507 | 0.335499068 |
| Ccdc88a | T | 1821 | 0.038337947 | 0.169206839 | 0.056935491 | 0.018597544 | -0.112271348 | 0.588233275 | 0.029689973 | 0.60429737 | 0.864888507 | 0.335499068 |
| Gpr50 | S | 7 | 0.090948173 | 0.097041483 | -0.034016045 | -0.124964218 | -0.131057528 | 0.015818918 | 0.034570308 | 0.734061881 | 0.281592975 | 0.263823378 |
| Foxj2 | S | 170 | -0.024812505 | 0.136067629 | 0.12247801 | 0.147290515 | -0.013589619 | 0.605263832 | 0.044412747 | 0.090283096 | 0.06706699 | 0.828987538 |
| Cenpc1 | S | 612 | 0.088529569 | 0.105200802 | 0.040193353 | -0.048336216 | -0.065007448 | 0.126875813 | 0.081269671 | 0.61327933 | 0.484787628 | 0.370800493 |
| Cope | S | 10 | -0.01528243 | 0.288996232 | 0.07515275 | 0.09043518 | -0.213843482 | 0.910886082 | 0.07003783 | 0.45710854 | 0.466642518 | 0.13800407 |
| Ubr5 | S | 606 | 0.110954308 | 0.053652076 | 0.057086194 | -0.053868114 | 0.003434117 | 0.15036324 | 0.251065511 | 0.116565334 | 0.412701542 | 0.930118199 |
| Srrm2 | S | 2073 | 0.074596605 | 0.068336483 | 0.044022087 | -0.030574518 | -0.024314396 | 0.137331905 | 0.038295761 | 0.321221484 | 0.535841048 | 0.519444584 |
| Fam133b | S | 244 | 0.061970601 | 0.116417635 | 0.063389571 | 0.00141897 | -0.053028064 | 0.204278612 | 0.111592666 | 0.340873351 | 0.980368209 | 0.467108594 |
| Rbm39 | S | 100 | -0.013207154 | 0.090547811 | 0.102874807 | 0.11608196 | 0.012326995 | 0.811114195 | 0.061772426 | 0.020940231 | 0.115934357 | 0.754809187 |
| Ralbp1 | S | 62 | 0.026764595 | 0.105658944 | 0.002381155 | -0.02438344 | -0.103277789 | 0.196595362 | 0.000946095 | 0.92386126 | 0.284818075 | 0.012047595 |
| Wdhd1 | S | 821 | 0.005226209 | 0.079011004 | 0.12045853 | 0.115232321 | 0.041447526 | 0.88818046 | 0.175902823 | 0.020459611 | 0.045312461 | 0.443720145 |
| Zc3h18 | S | 483 | 0.046645046 | 0.007961747 | 0.195622878 | 0.148977832 | 0.187661131 | 0.488120815 | 0.894379971 | 0.021470691 | 0.088794008 | 0.039098276 |
| Spin1 | S | 199 | -0.100061647 | 0.117680577 | 0.210163652 | 0.310225298 | 0.092483075 | 0.527367638 | 0.087683335 | 0.048654102 | 0.120948465 | 0.269744249 |
| Irf2bp2 | S | 300 | -0.005846977 | 0.10320888 | 0.114719306 | 0.120566283 | 0.011510426 | 0.938868331 | 0.090042212 | 0.043548458 | 0.188609596 | 0.827108819 |
| Mrc2 | S | 1452 | 0.017477536 | 0.214481014 | -0.072975776 | -0.090453312 | -0.28745679 | 0.676804064 | 0.000550337 | 0.358409246 | 0.273385225 | 0.031405044 |
| Macf1 | S | 5551 | -0.011454462 | 0.084857422 | 0.119934154 | 0.131388615 | 0.035076732 | 0.717565427 | 0.070941725 | 0.024927226 | 0.023360581 | 0.416913649 |
| Tmem245 | S | 324 | 0.174032589 | 0.052367304 | 0.019686377 | -0.154346212 | -0.032680928 | 0.021400143 | 0.493951793 | 0.794159469 | 0.073305729 | 0.6586034 |
| Dcaf6 | S | 673 | 0.021203026 | 0.05623732 | 0.145635539 | 0.124432513 | 0.089398219 | 0.551954612 | 0.096437031 | 0.082654728 | 0.119774329 | 0.215984619 |
| Dcaf6 | T | 670 | 0.021203026 | 0.05623732 | 0.145635539 | 0.124432513 | 0.089398219 | 0.551954612 | 0.096437031 | 0.082654728 | 0.119774329 | 0.215984619 |
| Tab2 | S | 580 | -0.010392408 | 0.126865218 | 0.15946362 | 0.169856028 | 0.032598402 | 0.897348374 | 0.043001008 | 0.23733039 | 0.227165931 | 0.773813599 |
| Vim | S | 83 | 0.018221324 | 0.076360129 | 0.107730615 | 0.089509291 | 0.031370486 | 0.674486071 | 0.217380922 | 0.019559443 | 0.077196094 | 0.535892022 |
| Epb41 | S | 556 | 0.022537678 | 0.052633106 | 0.150375447 | 0.127837769 | 0.09774234 | 0.63364956 | 0.472920918 | 0.018016366 | 0.011094523 | 0.190124317 |
| Zfp318 | S | 291 | -0.016112008 | 0.159019188 | 0.058073118 | 0.074185126 | -0.100946071 | 0.777495923 | 0.005981632 | 0.376813984 | 0.298102856 | 0.159133084 |
| Zfp318 | S | 292 | -0.016112008 | 0.159019188 | 0.058073118 | 0.074185126 | -0.100946071 | 0.777495923 | 0.005981632 | 0.376813984 | 0.298102856 | 0.159133084 |
| Zfp318 | S | 293 | -0.016112008 | 0.159019188 | 0.058073118 | 0.074185126 | -0.100946071 | 0.777495923 | 0.005981632 | 0.376813984 | 0.298102856 | 0.159133084 |
| Cdk14 | S | 119 | 0.101856755 | 0.250947142 | -0.044510659 | -0.146367413 | -0.2954578 | 0.407671957 | 0.037564863 | 0.671194556 | 0.279230635 | 0.041784398 |
| Srrm2 | T | 233 | 0.041270432 | 0.058747312 | 0.132845197 | 0.091574766 | 0.074097885 | 0.61338681 | 0.059771883 | 0.123337019 | 0.358064311 | 0.305346137 |
| Dnmt1 | S | 15 | -0.180968383 | 0.243075116 | 0.213804064 | 0.394772447 | -0.029271052 | 0.006232796 | 0.001091253 | 0.073492605 | 0.02159718 | 0.730289941 |
| Mpp6 | S | 106 | -0.013924949 | 0.078281958 | 0.095433075 | 0.109358024 | 0.017151116 | 0.599393982 | 0.00753033 | 0.046252048 | 0.036628293 | 0.604285361 |
| Zeb2 | S | 780 | 0.164726411 | 0.027594044 | 0.05288939 | -0.111837021 | 0.025295345 | 0.153555836 | 0.454782215 | 0.072593753 | 0.277450051 | 0.442924596 |
| Farp1 | S | 427 | 0.087258281 | 0.005264702 | 0.079308634 | -0.007949648 | 0.074043932 | 0.122765275 | 0.809825614 | 0.008601129 | 0.853082409 | 0.008401567 |
| Ncor1 | S | 1011 | -0.025118804 | 0.081490391 | 0.252719962 | 0.277838766 | 0.171229571 | 0.699974089 | 0.336189047 | 0.049398793 | 0.061532693 | 0.139131746 |
| Bag3 | S | 285 | 0.01048899 | 0.134828138 | 0.067250811 | 0.05676182 | -0.067577327 | 0.744500087 | 0.065512099 | 0.079405497 | 0.025538555 | 0.261294163 |
| Dyrk1a | S | 748 | -0.008361702 | 0.105399798 | 0.205079566 | 0.213441268 | 0.099679768 | 0.91057341 | 0.132258648 | 0.135111075 | 0.12446773 | 0.382211553 |
| Whamm | S | 787 | 0.103673604 | 0.054358564 | 0.045499798 | -0.058173806 | -0.008858766 | 0.040198921 | 0.154642218 | 0.49628129 | 0.391813432 | 0.883006485 |
| Camk2d | S | 319 | -0.039007367 | 0.33849001 | -0.004578041 | 0.034429326 | -0.343068051 | 0.704089177 | 0.013929214 | 0.971002059 | 0.762441774 | 0.039763035 |
| Mov10 | S | 266 | -0.004032336 | 0.161001637 | 0.036436239 | 0.040468576 | -0.124565398 | 0.935294791 | 0.003304024 | 0.624190871 | 0.60250288 | 0.164167085 |
| Lbr | S | 69 | -0.033281018 | 0.07800879 | 0.117808911 | 0.151089929 | 0.039800121 | 0.174373796 | 0.013044671 | 0.010495927 | 0.019754285 | 0.214904872 |
| Srrm2 | S | 1747 | 0.132354291 | -0.010357244 | 0.107231752 | -0.02512254 | 0.117588996 | 0.037627248 | 0.90442533 | 0.155714256 | 0.709016591 | 0.269568534 |
| Nf1 | S | 2517 | 0.06364849 | 0.089337041 | 0.136756381 | 0.073107891 | 0.04741934 | 0.385849111 | 0.208592246 | 0.223453187 | 0.501626716 | 0.647122209 |
| Rpl5 | S | 185 | 0.030973881 | 0.032652808 | 0.186977145 | 0.156003264 | 0.154324337 | 0.694725297 | 0.515791884 | 0.027530669 | 0.12287424 | 0.057642094 |
| Arhgap17 | S | 739 | -0.00090102 | 0.12937408 | 0.114074591 | 0.11497561 | -0.015299489 | 0.985395995 | 0.104012484 | 0.081076785 | 0.080582171 | 0.822188557 |
| Chd8 | S | 1980 | -0.071703317 | 0.172278307 | 0.113647822 | 0.185351139 | -0.058630485 | 0.354316442 | 0.036305316 | 0.046895851 | 0.071087911 | 0.316027928 |
| Ephx1 | S | 73 | 0.018192029 | 0.084014547 | 0.066079759 | 0.04788773 | -0.017934789 | 0.777029585 | 0.000804912 | 0.445156616 | 0.621281723 | 0.821877406 |
| Nap1l4 | S | 49 | 0.049302744 | 0.123407463 | 0.055120722 | 0.005817978 | -0.068286742 | 0.294010939 | 0.046619571 | 0.445220075 | 0.920104013 | 0.326386935 |
| Ino80b | T | 211 | 0.111676181 | 0.07184428 | 0.212217282 | 0.100541101 | 0.140373002 | 0.431426079 | 0.306268904 | 0.347688712 | 0.654065664 | 0.50428206 |
| Nfic | S | 302 | 0.038178513 | 0.131112992 | 0.056217168 | 0.018038655 | -0.074895823 | 0.254017591 | 0.163369633 | 0.105989227 | 0.486760817 | 0.360865433 |
| Glcci1 | S | 131 | 0.089230034 | 0.118895873 | 0.078978894 | -0.01025114 | -0.039916978 | 0.657549821 | 0.280812379 | 0.048290452 | 0.95805648 | 0.677482365 |
| Rfc1 | S | 363 | 0.016774985 | 0.065491118 | 0.117583296 | 0.100808312 | 0.052092178 | 0.712616713 | 0.151355973 | 0.029538731 | 0.053465528 | 0.197858457 |
| Rfc1 | S | 357 | 0.016774985 | 0.065491118 | 0.117583296 | 0.100808312 | 0.052092178 | 0.712616713 | 0.151355973 | 0.029538731 | 0.053465528 | 0.197858457 |
| Thrap3 | S | 750 | -0.044972625 | 0.098474949 | 0.239131562 | 0.284104187 | 0.140656613 | 0.412971765 | 0.133306228 | 0.058273903 | 0.031868349 | 0.16824919 |
| Ubxn4 | S | 454 | 0.082894154 | 0.130703165 | -0.024380551 | -0.107274704 | -0.155083715 | 0.12959743 | 0.010449749 | 0.590272455 | 0.06898761 | 0.013342097 |
| Safb Safb2 | S 235;229 | | 0.034799637 | -0.013887576 | 0.151163575 | 0.116363939 | 0.165051151 | 0.401604649 | 0.766031977 | 0.002459975 | 0.027200182 | 0.023585726 |
| Sorbs2 | S | 319 | 0.130653387 | -0.034169899 | 0.075987752 | -0.054665635 | 0.110157651 | 0.026212348 | 0.480835514 | 0.022141926 | 0.180802078 | 0.090722427 |
| Srrm2 | S | 472 | 0.097601296 | 0.004510577 | 0.151539752 | 0.053938455 | 0.147029175 | 0.269059062 | 0.950659018 | 0.040688406 | 0.528808756 | 0.119731636 |
| Wee1 | S | 150 | 0.03317241 | 0.203096241 | -0.011089004 | -0.044261414 | -0.214185245 | 0.573946707 | 0.009216754 | 0.851525934 | 0.273936889 | 0.003457887 |
| Gas2l1 | S | 482 | -0.05473221 | 0.116850081 | 0.207683026 | 0.262415236 | 0.090832944 | 0.02039799 | 0.003619389 | 0.082316347 | 0.05724468 | 0.305859252 |
| Hmgcr | S | 871 | -0.048512852 | 0.054048669 | 0.145573361 | 0.194086213 | 0.091524691 | 0.228098984 | 0.193086785 | 0.000605076 | 0.013342761 | 0.071625919 |
| Akap13 | S | 2692 | 0.06595155 | 0.007487761 | 0.182381811 | 0.116430261 | 0.17489405 | 0.393890904 | 0.923684519 | 0.030854359 | 0.088578777 | 0.052303248 |
| Mrnip | S | 139 | 0.008506663 | 0.231830342 | 0.103495734 | 0.094989071 | -0.128334608 | 0.932334299 | 0.109664651 | 0.353214003 | 0.349102361 | 0.324907968 |
| Git1 | S | 580 | 0.073936191 | 0.03918872 | 0.079524613 | 0.005588421 | 0.040335892 | 0.23159993 | 0.281365137 | 0.034977493 | 0.912009619 | 0.167613949 |
| Zc3h13 | S | 898 | 0.101880089 | 0.065743454 | 0.071369298 | -0.030510791 | 0.005625844 | 0.06513722 | 0.366777074 | 0.323329009 | 0.649935662 | 0.944380358 |
| Ell | S | 310 | 0.171376223 | 0.070413305 | -0.035636462 | -0.207012684 | -0.106049767 | 0.020263037 | 0.169274247 | 0.741009731 | 0.133551087 | 0.358733541 |
| Toe1 | S | 349 | 0.022889247 | 0.10259987 | 0.026803371 | 0.003914124 | -0.075796499 | 0.394690796 | 0.001523102 | 0.660417832 | 0.947782581 | 0.274434266 |

| | | | | | | | | | | | | |
|---|---|---|---|---|---|---|---|---|---|---|---|---|
| Lbr | S | 73 | -0.076159952 | 0.144902556 | 0.15160976 | 0.227769712 | 0.006707204 | 0.459754974 | 0.145274819 | 0.017794094 | 0.103334977 | 0.92874138 |
| Cwc22 | S | 862 | 0.032649378 | 0.086861839 | 0.18335026 | 0.150700881 | 0.096488421 | 0.715968727 | 0.570542991 | 0.049800722 | 0.153261181 | 0.53294743 |
| Pnisr | S | 563 | -0.013774456 | 0.066393196 | 0.162564191 | 0.176338647 | 0.096170995 | 0.745195562 | 0.149489956 | 0.024840537 | 0.033375316 | 0.11894907 |
| Hirip3 | T | 141 | 0.013436798 | 0.056402285 | 0.094627342 | 0.081190544 | 0.038225057 | 0.645150293 | 0.039934526 | 0.035155547 | 0.065055982 | 0.264017292 |
| Nab2 | S | 6 | 0.042008469 | 0.05442606 | 0.030969192 | -0.011039276 | -0.023456868 | 0.00058612 | 0.053175464 | 0.692788303 | 0.88563665 | 0.764888704 |
| Arid5b | S | 712 | -0.009710962 | 0.08699309 | 0.12502329 | 0.134734252 | 0.0380302 | 0.897836769 | 0.093893597 | 0.036328181 | 0.136163058 | 0.353874644 |
| Trim2 | T | 371 | 0.047997061 | 0.120347944 | 0.063635456 | 0.015638394 | -0.056712489 | 0.666200842 | 0.03546466 | 0.263752715 | 0.890983404 | 0.338007399 |
| Scaf1 | S | 240 | 0.036494419 | 0.114229503 | 0.070442803 | 0.033948384 | -0.0437867 | 0.514902725 | 0.082673272 | 0.13521886 | 0.523680884 | 0.4204571 |
| Ttc28 | S | 2102 | 0.030941618 | 0.136517348 | 0.130291465 | 0.099349847 | -0.006225883 | 0.711858554 | 0.164711253 | 0.189073343 | 0.211828928 | 0.937314496 |
| Eif4g3 | S | 1576 | 0.002559366 | 0.153715075 | 0.046115443 | 0.043556077 | -0.107599633 | 0.948346519 | 0.020224288 | 0.186143445 | 0.348477299 | 0.055148049 |
| Rbl1 | S | 640 | 0.054829071 | 0.133769874 | 0.048545545 | -0.006283525 | -0.085224328 | 0.584847536 | 0.020283771 | 0.550654431 | 0.954556246 | 0.312575007 |
| Nav1 | S | 1379 | 0.049919394 | 0.147837405 | 0.043043676 | -0.006875718 | -0.104793729 | 0.374730938 | 0.061069447 | 0.396895521 | 0.867059471 | 0.135676109 |
| Itga4 | S | 1030 | 0.015083104 | 0.163270528 | 0.095995236 | 0.080912132 | -0.067275292 | 0.785090937 | 0.099829728 | 0.205411047 | 0.297131255 | 0.459234674 |
| Srrm2 | S | 1750 | 0.119476718 | -0.012884281 | 0.111248832 | -0.008227885 | 0.124133113 | 0.034871938 | 0.88341678 | 0.137506281 | 0.896761258 | 0.248732734 |
| Klc2 | S | 505 | 0.031231982 | 0.162043362 | 0.117081464 | 0.085849482 | -0.044961898 | 0.765840535 | 0.0675728 | 0.438439039 | 0.578616636 | 0.746417006 |
| Fhod1 | S | 551 | 0.073972692 | 0.10175091 | 0.119344184 | 0.045371492 | 0.017593275 | 0.347678343 | 0.220827043 | 0.277482896 | 0.639278117 | 0.855565602 |
| Map3k3 | S | 175 | 0.010667136 | 0.104534641 | 0.087705759 | 0.077038623 | -0.016828882 | 0.822960894 | 0.049239981 | 0.093487058 | 0.056240367 | 0.478716843 |
| Map3k3 | S | 176 | 0.010667136 | 0.104534641 | 0.087705759 | 0.077038623 | -0.016828882 | 0.822960894 | 0.049239981 | 0.093487058 | 0.056240367 | 0.478716843 |
| Hnrnpc | S | 241 | 0.005050499 | 0.108978693 | 0.124221671 | 0.119171171 | 0.015242977 | 0.9320881 | 0.07655549 | 0.119174474 | 0.138530641 | 0.812335211 |
| Mcm6 | S | 699 | 0.089510491 | 0.062755024 | 0.113945441 | 0.02443495 | 0.051190417 | 0.267124711 | 0.126689551 | 0.324411302 | 0.830164064 | 0.621152424 |
| Bud13 | S | 251 | 0.111564465 | 0.0895733 | 0.041456892 | -0.070107573 | -0.048116408 | 0.130642757 | 0.151317102 | 0.4840921 | 0.267833483 | 0.331538814 |
| Pnpla7 | S | 1280 | 0.00719964 | 0.211939928 | 0.122543103 | 0.115343463 | -0.089396824 | 0.951796046 | 0.110711826 | 0.348843722 | 0.448136121 | 0.530927613 |
| Pnpla7 | T | 1284 | 0.00719964 | 0.211939928 | 0.122543103 | 0.115343463 | -0.089396824 | 0.951796046 | 0.110711826 | 0.348843722 | 0.448136121 | 0.530927613 |
| Spice1 | S | 769 | 0.058942725 | 0.097468991 | 0.065238016 | 0.006295292 | -0.032230975 | 0.287055483 | 0.072888423 | 0.288939986 | 0.907312776 | 0.529052335 |
| Dennd4c | S | 1246 | 0.032060854 | 0.097617839 | 0.093424424 | 0.06136357 | -0.004193415 | 0.725472917 | 0.027603386 | 0.236615598 | 0.561444147 | 0.950540955 |
| Dennd4c | S | 1248 | 0.032060854 | 0.097617839 | 0.093424424 | 0.06136357 | -0.004193415 | 0.725472917 | 0.027603386 | 0.236615598 | 0.561444147 | 0.950540955 |
| Atn1 | S | 168 | 0.128812506 | 0.048646493 | -0.016431859 | -0.145244365 | -0.065078352 | 0.013156677 | 0.059700882 | 0.821494589 | 0.134507735 | 0.410554585 |
| Chaf1b | S | 465 | 0.022367559 | 0.130528576 | 0.20569497 | 0.183327411 | 0.075166394 | 0.877398092 | 0.134565116 | 0.304507924 | 0.401130975 | 0.677333481 |
| Iws1 | S | 250 | 0.038895686 | 0.027406855 | 0.109655007 | 0.070759321 | 0.082248152 | 0.070358793 | 0.153229678 | 0.091577976 | 0.197888256 | 0.155357318 |
| Taf1 | S | 1817 | -0.006292121 | 0.104468493 | 0.159143901 | 0.165436022 | 0.054675408 | 0.914323223 | 0.154307841 | 0.075104807 | 0.072952287 | 0.484784937 |
| Taf1 | Y | 1824 | -0.006292121 | 0.104468493 | 0.159143901 | 0.165436022 | 0.054675408 | 0.914323223 | 0.154307841 | 0.075104807 | 0.072952287 | 0.484784937 |
| Cdk12 | S | 382 | 0.079154351 | -0.02178716 | 0.135854874 | 0.056700523 | 0.157642034 | 0.039547364 | 0.550756821 | 0.037382238 | 0.258352725 | 0.030527469 |
| Cdk12 | S | 384 | 0.079154351 | -0.02178716 | 0.135854874 | 0.056700523 | 0.157642034 | 0.039547364 | 0.550756821 | 0.037382238 | 0.258352725 | 0.030527469 |
| Ino80 | S | 1492 | -0.026831128 | 0.134996639 | 0.172661185 | 0.199492313 | 0.037664546 | 0.836371047 | 0.17429387 | 0.091095288 | 0.159361469 | 0.3477927 |
| Zmynd11 | S | 403 | 0.048573253 | 0.095922301 | 0.094402968 | 0.045829716 | -0.001519333 | 0.295452886 | 0.114956588 | 0.252654374 | 0.539030226 | 0.984092384 |
| Tcf20 | T | 1362 | 0.221259789 | 0.101106358 | -0.095107454 | -0.316367243 | -0.196213812 | 0.02532884 | 0.189466391 | 0.509769701 | 0.091652156 | 0.21947375 |
| Sgpl1 | S | 564 | 0.033130913 | 0.211485002 | 0.009878298 | -0.023252614 | -0.201606704 | 0.554618751 | 0.023544158 | 0.921996192 | 0.822864942 | 0.12848688 |
| Mcm2 | S | 27 | 0.00618474 | 0.069567523 | 0.085041096 | 0.078856356 | 0.015473573 | 0.809591656 | 0.019323061 | 0.054195916 | 0.071185445 | 0.621009512 |
| N4bp1 | S | 549 | 0.091080312 | 0.122888852 | 0.151291674 | 0.060211361 | 0.028402822 | 0.622474163 | 0.170209069 | 0.377993712 | 0.775114562 | 0.846160567 |
| Mecp2 | S | 350 | 0.236643705 | 0.034108261 | 0.105009159 | -0.131634546 | 0.070900898 | 0.153545122 | 0.781077503 | 0.425172836 | 0.332763243 | 0.428620271 |
| Iqce | S | 604 | 0.41703988 | 0.053709806 | -0.105572921 | -0.5226128 | -0.159282727 | 0.064541447 | 0.635806906 | 0.765668572 | 0.219951533 | 0.652822228 |
| Kif16b | S | 662 | -0.007210911 | 0.172211314 | 0.060283488 | 0.067494398 | -0.111927827 | 0.936278512 | 0.036319151 | 0.176846966 | 0.466898528 | 0.120913298 |
| Arhgef40 | S | 1478 | -0.043946262 | 0.0866622 | 0.213652149 | 0.25759841 | 0.126989948 | 0.745070646 | 0.296712539 | 0.035043292 | 0.120899649 | 0.107574569 |
| Srrm1 | S | 524 | 0.266202883 | -0.096243273 | 0.037330062 | -0.228872821 | 0.133573334 | 0.002175724 | 0.231049343 | 0.525701902 | 0.003107982 | 0.105078755 |
| Erh | S | 24 | 0.103439469 | 0.115291304 | -0.040574309 | -0.144013778 | -0.155865614 | 0.025847491 | 0.064600761 | 0.699402647 | 0.243823707 | 0.216023716 |
| Kdm2a | S | 868 | 0.176985022 | 0.05637451 | -0.035656669 | -0.212641691 | -0.092031179 | 0.009219653 | 0.237905869 | 0.58168154 | 0.021675492 | 0.171196944 |
| Kif14 | S | 283 | 0.185763868 | 0.028319988 | -0.016332791 | -0.202096659 | -0.044652779 | 0.004628803 | 0.645667152 | 0.833716445 | 0.06597795 | 0.588087715 |
| Reps1 | T | 543 | 0.063366676 | 0.170221305 | 0.060352997 | -0.003013679 | -0.109868308 | 0.524757988 | 0.122517955 | 0.337352686 | 0.973746099 | 0.265402336 |
| Plk2 | S | 386 | 0.058729293 | 0.107868441 | 0.070263635 | 0.011534342 | -0.037604806 | 0.271864014 | 0.071516269 | 0.425626459 | 0.888396985 | 0.654782359 |
| Arhgap23 | S | 552 | -0.074312635 | 0.21791767 | 0.163898999 | 0.238211634 | -0.054018671 | 0.321657654 | 0.055825364 | 0.155691257 | 0.072879234 | 0.628097874 |
| Bod1l | S | 3010 | 0.033028445 | 0.047858631 | 0.049187295 | 0.01615885 | 0.001328664 | 0.034950294 | 0.037256831 | 0.11761858 | 0.514314388 | 0.959152392 |
| Hmga2 | S | 44 | 0.050811611 | 0.078252 | 0.047151618 | -0.003659993 | -0.031100382 | 0.279422459 | 0.038408952 | 0.158986873 | 0.918687388 | 0.040975722 |
| Anapc1 | S | 46 | 0.004741662 | 0.155038365 | 0.181391426 | 0.176649764 | 0.026353062 | 0.94882391 | 0.201436124 | 0.201378531 | 0.208154766 | 0.845862724 |
| Ripk1 | S | 313 | 0.055980836 | 0.108547763 | 0.031820646 | -0.02416019 | -0.076727117 | 0.204808774 | 0.032001087 | 0.522973523 | 0.623114697 | 0.169101961 |
| Srrm2 | S | 924 | 0.031170886 | 0.054591655 | 0.078325188 | 0.047154302 | 0.023733533 | 0.311277694 | 0.066266974 | 0.054661523 | 0.221465243 | 0.471847996 |
| Thap7 | S | 162 | 0.054437694 | 0.193396148 | 0.028578288 | -0.025859406 | -0.16481786 | 0.696825678 | 0.031218863 | 0.738466155 | 0.852635636 | 0.091567223 |
| Aven | S | 84 | -0.073672623 | 0.21612694 | 0.143766349 | 0.217438971 | -0.072360592 | 0.339714561 | 0.141904868 | 0.038508531 | 0.042920126 | 0.538409699 |
| Aven | T | 86 | -0.073672623 | 0.21612694 | 0.143766349 | 0.217438971 | -0.072360592 | 0.339714561 | 0.141904868 | 0.038508531 | 0.042920126 | 0.538409699 |
| Denn2b | S | 571 | -0.08833244 | 0.122189913 | 0.271513492 | 0.359845932 | 0.149323579 | 0.627038854 | 0.489525608 | 0.033731704 | 0.11770106 | 0.4049575 |
| Pdlim7 | S | 209 | 0.302883762 | -0.010648872 | -0.031337371 | -0.334221133 | -0.0206885 | 0.00894316 | 0.907620452 | 0.67228461 | 0.012323899 | 0.757812899 |
| Rps6ka1 Rps6ka3 Rps | S | 227;221;252 | 0.036222284 | 0.21371896 | -0.001966658 | -0.038188942 | -0.215685618 | 0.65639999 | 0.01685905 | 0.980023731 | 0.668507003 | 0.044530447 |
| Snrnp48 | S | 204 | 0.083207257 | 0.236284659 | -0.074985802 | -0.158193058 | -0.311270461 | 0.544055591 | 0.013776385 | 0.649258009 | 0.409944026 | 0.144777168 |
| Rtn4 | S | 913 | 0.411799817 | 0.141067664 | -0.344780217 | -0.756580034 | -0.48584788 | 0.020415826 | 0.3286852 | 0.338699521 | 0.107991472 | 0.21570615 |
| Thrap3 | S | 669 | -0.02135613 | 0.081320601 | 0.137793088 | 0.159149218 | 0.056472487 | 0.512729702 | 0.035760228 | 0.046658702 | 0.034248033 | 0.287448437 |
| Clcn7 | S | 9 | -0.055985888 | 0.274816132 | 0.087407342 | 0.14339323 | -0.18740879 | 0.459441193 | 0.046531904 | 0.263646829 | 0.045205914 | 0.128279293 |
| Cacnb2 | T | 549 | 0.028545198 | 0.08810873 | 0.142583884 | 0.114038685 | 0.054475153 | 0.724545495 | 0.281775566 | 0.066965668 | 0.14601111 | 0.402077624 |
| Irs2 | T | 1189 | 0.040061828 | 0.093959317 | 0.11050191 | 0.070440083 | 0.016542593 | 0.477634147 | 0.167473215 | 0.13672589 | 0.311014829 | 0.81171177 |
| Arid1a | S | 697 | 0.031542287 | 0.037876745 | 0.135243893 | 0.103701605 | 0.097367147 | 0.44420219 | 0.348872406 | 0.029290831 | 0.065596438 | 0.07908791 |
| Twf2 | S | 349 | 0.117164423 | 0.171916962 | -0.058595516 | -0.175759939 | -0.230512477 | 0.097105659 | 0.029290942 | 0.340667179 | 0.015621942 | 0.008634925 |
| Foxo3 | S | 293 | -0.078864278 | 0.140620631 | 0.167586419 | 0.246450697 | 0.026965788 | 0.366457335 | 0.035863225 | 0.081784295 | 0.047610566 | 0.714378558 |
| Ncapd2 | S | 1358 | -0.064204837 | 0.099447855 | 0.085791253 | 0.149996089 | -0.013656602 | 0.466122047 | 0.005230887 | 0.029839026 | 0.162255535 | 0.638687556 |
| Ncapd2 | S | 1361 | -0.064204837 | 0.099447855 | 0.085791253 | 0.149996089 | -0.013656602 | 0.466122047 | 0.005230887 | 0.029839026 | 0.162255535 | 0.638687556 |
| Ncapd2 | S | 1362 | -0.064204837 | 0.099447855 | 0.085791253 | 0.149996089 | -0.013656602 | 0.466122047 | 0.005230887 | 0.029839026 | 0.162255535 | 0.638687556 |
| Arhgap31 | S | 1055 | 0.047644194 | 0.146631871 | 0.065059964 | 0.01741577 | -0.081571907 | 0.474923744 | 0.083035037 | 0.36023197 | 0.724232436 | 0.232298426 |
| Mtmr2 | S | 46 | 0.223280424 | 0.204997373 | -0.001023391 | -0.224303815 | -0.206020764 | 0.361100613 | 0.199202353 | 0.994635973 | 0.365616977 | 0.229361635 |
| Mtmr2 | S | 49 | 0.223280424 | 0.204997373 | -0.001023391 | -0.224303815 | -0.206020764 | 0.361100613 | 0.199202353 | 0.994635973 | 0.365616977 | 0.229361635 |
| Map2k1 | S | 385 | -0.205563604 | 0.445359363 | 0.1827677 | 0.388331303 | -0.262591663 | 0.497608437 | 0.090281251 | 0.428567775 | 0.178778964 | 0.013958115 |
| Nol8 | S | 1063 | 0.100643802 | 0.122955581 | 0.003878455 | -0.096765347 | -0.119077126 | 0.143531118 | 0.057326252 | 0.946831618 | 0.099623752 | 0.024971371 |
| Nol8 | S | 1064 | 0.100643802 | 0.122955581 | 0.003878455 | -0.096765347 | -0.119077126 | 0.143531118 | 0.057326252 | 0.946831618 | 0.099623752 | 0.024971371 |
| Nol8 | S | 1065 | 0.100643802 | 0.122955581 | 0.003878455 | -0.096765347 | -0.119077126 | 0.143531118 | 0.057326252 | 0.946831618 | 0.099623752 | 0.024971371 |
| Nol8 | S | 1080 | 0.100643802 | 0.122955581 | 0.003878455 | -0.096765347 | -0.119077126 | 0.143531118 | 0.057326252 | 0.946831618 | 0.099623752 | 0.024971371 |
| Nfib | S | 569 | 0.02074831 | 0.037466627 | 0.2991482 | 0.27839989 | 0.261681574 | 0.933967212 | 0.826610481 | 0.056677551 | 0.311455274 | 0.167649686 |
| Fndc3b | S | 261 | 0.138605397 | 0.050719644 | 0.002429037 | -0.13617636 | -0.048290607 | 0.010936221 | 0.296649667 | 0.965704466 | 0.06054842 | 0.381550656 |
| Rbm10 | S | 736 | 0.07834137 | 0.098387337 | -0.012592257 | -0.090933627 | -0.110979595 | 0.056799686 | 0.017904 | 0.78845582 | 0.142439867 | 0.096694692 |
| Rbm10 | S | 738 | 0.07834137 | 0.098387337 | -0.012592257 | -0.090933627 | -0.110979595 | 0.056799686 | 0.017904 | 0.78845582 | 0.142439867 | 0.096694692 |
| Gcfc2 | S | 85 | 0.04311571 | 0.035747193 | 0.13390546 | 0.09078975 | 0.098158267 | 0.282141959 | 0.503649083 | 0.039382553 | 0.117974882 | 0.153531691 |
| Gcfc2 | T | 86 | 0.04311571 | 0.035747193 | 0.13390546 | 0.09078975 | 0.098158267 | 0.282141959 | 0.503649083 | 0.039382553 | 0.117974882 | 0.153531691 |
| Shroom4 | S | 460 | -0.014066457 | 0.054496122 | 0.149632259 | 0.163698715 | 0.095136137 | 0.856194518 | 0.13102414 | 0.020605799 | 0.122625897 | 0.066501079 |
| Nek1 | S | 953 | 0.006942413 | 0.110115218 | 0.102788324 | 0.095845911 | -0.007326894 | 0.92412092 | 0.037079719 | 0.176231038 | 0.274563681 | 0.899645462 |
| U2surp | T | 919 | -0.022655184 | 0.04523329 | 0.131446512 | 0.154101696 | 0.086213221 | 0.544597769 | 0.037145746 | 0.009823646 | 0.016771116 | 0.050529479 |
| Ralgapa1 | T | 753 | -0.004663804 | 0.068998921 | 0.095823493 | 0.100487297 | 0.026824572 | 0.876521037 | 0.090943874 | 0.009768356 | 0.00884851 | 0.390438699 |
| Itpr1 | S | 1755 | 0.133699381 | 0.151479971 | -0.032215453 | -0.165914834 | -0.183695424 | 0.167050542 | 0.061044774 | 0.768110714 | 0.179489486 | 0.147564333 |
| Rreb1 | S | 1230 | 0.016104833 | 0.131493072 | 0.071019121 | 0.054914287 | -0.060473952 | 0.874307168 | 0.073583499 | 0.088015268 | 0.594340645 | 0.311057607 |
| Fam117b | S | 216 | 0.200906427 | 0.027503464 | 0.108049441 | -0.092856986 | 0.080545977 | 0.123304729 | 0.824191131 | 0.373627566 | 0.369187635 | 0.455486917 |

| | | | | | | | | | | | | |
|---|---|---|---|---|---|---|---|---|---|---|---|---|
| Slbp | T | 171 | -0.058489122 | 0.252310498 | 0.115955711 | 0.174444833 | -0.136354786 | 0.615585312 | 0.070491878 | 0.255411579 | 0.113303873 | 0.207070918 |
| Setd2 | S | 131 | 0.050598396 | 0.00665015 | 0.132686596 | 0.0820882 | 0.126036446 | 0.117449415 | 0.798441194 | 0.030842947 | 0.089768135 | 0.028409991 |
| Ccdc88a | S | 1562 | 0.065522741 | 0.134808836 | 0.016114327 | -0.049408414 | -0.118694509 | 0.442872643 | 0.015259813 | 0.76785542 | 0.572138379 | 0.085096681 |
| Aak1 | S | 635 | 0.127908953 | 0.150039385 | -0.02852184 | -0.156430793 | -0.178561225 | 0.313708298 | 0.01800319 | 0.631679377 | 0.23822685 | 0.021282764 |
| Bzw2 | S | 419 | 0.118444928 | 0.086486713 | 0.215045719 | 0.096600791 | 0.128559006 | 0.530667039 | 0.637472007 | 0.21456621 | 0.612124457 | 0.498597629 |
| Arhgef12 | S | 341 | 0.188446775 | 0.164651663 | -0.088139893 | -0.276586667 | -0.252791555 | 0.059287112 | 0.083703227 | 0.307166339 | 0.003474598 | 0.000853918 |
| Purg | S | 342 | 0.050817349 | 0.131773988 | 0.069190033 | 0.018372684 | -0.062583954 | 0.434034154 | 0.073058504 | 0.396831597 | 0.767987534 | 0.357706248 |
| Tnrc18 | S | 1889 | -0.010720201 | 0.085738977 | 0.125010414 | 0.135730615 | 0.039271437 | 0.853164641 | 0.122067099 | 0.029971189 | 0.090273033 | 0.456707377 |
| Nktr | S | 866 | 0.086746102 | 0.04397649 | 0.139184972 | 0.052438869 | 0.095208482 | 0.227924356 | 0.505636289 | 0.149876008 | 0.561427263 | 0.315553798 |
| Sap30l | S | 92 | -0.041324229 | 0.109392041 | 0.159001688 | 0.200325917 | 0.049609647 | 0.391499372 | 0.013426941 | 0.143630799 | 0.09153315 | 0.563199374 |
| Cbarp | S | 312 | 0.011829877 | 0.095920829 | 0.077715668 | 0.065885792 | -0.01820516 | 0.856043685 | 0.087181849 | 0.031410239 | 0.345024626 | 0.658726775 |
| Ank | S | 470 | 0.010280951 | 0.124183844 | 0.108814329 | 0.098533377 | -0.015369515 | 0.934395365 | 0.051272487 | 0.211784584 | 0.468040255 | 0.824841845 |
| Ank | T | 473 | 0.010280951 | 0.124183844 | 0.108814329 | 0.098533377 | -0.015369515 | 0.934395365 | 0.051272487 | 0.211784584 | 0.468040255 | 0.824841845 |
| Rbm15 | S | 670 | 0.072486821 | 0.143567791 | 0.034822498 | -0.037664323 | -0.108745293 | 0.241028297 | 0.060669168 | 0.795501193 | 0.766760139 | 0.433358616 |
| Rbm15 | S | 674 | 0.072486821 | 0.143567791 | 0.034822498 | -0.037664323 | -0.108745293 | 0.241028297 | 0.060669168 | 0.795501193 | 0.766760139 | 0.433358616 |
| Rbm15 | T | 668 | 0.072486821 | 0.143567791 | 0.034822498 | -0.037664323 | -0.108745293 | 0.241028297 | 0.060669168 | 0.795501193 | 0.766760139 | 0.433358616 |
| Psma3 | S | 250 | 0.008652332 | 0.119529461 | 0.032525436 | 0.023873104 | -0.087004024 | 0.69518956 | 0.004203645 | 0.370312398 | 0.49642315 | 0.066327019 |
| Arhgef17 | T | 610 | 0.191744531 | -0.019065892 | 0.133145857 | -0.058598675 | 0.152211748 | 0.122245342 | 0.86155368 | 0.202035055 | 0.446015696 | 0.097732316 |
| Rad18 | T | 190 | 0.063648878 | 0.013124181 | 0.158505205 | 0.094856327 | 0.145381024 | 0.09758075 | 0.895105781 | 0.08874807 | 0.239116948 | 0.238700186 |
| Tnks1bp1 | S | 1241 | 0.055870021 | 0.174683434 | 0.044746577 | -0.011123444 | -0.129936857 | 0.624306665 | 0.061406475 | 0.476583995 | 0.912990016 | 0.116478258 |
| AW554918 | S | 471 | -0.001461711 | 0.028178323 | 0.132164275 | 0.133625986 | 0.103985952 | 0.970153904 | 0.377316362 | 0.00273693 | 0.022963986 | 0.012074289 |
| Specc1l | S | 385 | 0.037635216 | 0.087116784 | 0.03476756 | -0.002867656 | -0.052349224 | 0.200639884 | 0.018423175 | 0.293863916 | 0.906028284 | 0.104952873 |
| Eloa | S | 125 | -0.001393428 | 0.09224797 | 0.101192646 | 0.102586074 | 0.008944676 | 0.9624677 | 0.125892854 | 0.024978394 | 0.035598901 | 0.855969646 |
| Ythdc1 | S | 146 | 0.019327368 | 0.09923961 | 0.100438055 | 0.081110687 | 0.001198446 | 0.759689139 | 0.044088928 | 0.185309683 | 0.32739379 | 0.985018618 |
| Prkar2b | S | 83 | -0.02718892 | 0.092414844 | 0.106714282 | 0.133903201 | 0.014299437 | 0.497003079 | 0.013371527 | 0.060756564 | 0.039478713 | 0.700093556 |
| Mcm4 | T | 19 | -0.013375107 | 0.072672739 | 0.137942793 | 0.1513179 | 0.065270054 | 0.692624924 | 0.046193548 | 0.052700813 | 0.051336262 | 0.252982513 |
| Ddx42 | Y | 183 | -0.0453263 | 0.078039773 | 0.06838404 | 0.11371034 | -0.009655733 | 0.478067907 | 0.005030587 | 0.007323488 | 0.157760436 | 0.472126752 |
| Bcl2l11 | S | 83 | -0.092210636 | 0.227036574 | 0.116346534 | 0.208557171 | -0.11069004 | 0.323178396 | 0.04337574 | 0.099802399 | 0.068595042 | 0.218122738 |
| Pak4 | S | 476 | -0.005752005 | 0.097001518 | 0.049915829 | 0.055667833 | -0.047085689 | 0.882381089 | 0.001334065 | 0.246183132 | 0.28485 | 0.26787614 |
| Dync1i2 | S | 51 | 0.077931709 | 0.075943307 | 0.067470704 | -0.010461005 | -0.008472604 | 0.097437792 | 0.349277151 | 0.198518619 | 0.831384212 | 0.913093856 |
| Zhx1 | S | 47 | 0.06716647 | 0.085860668 | 0.127130256 | 0.059963786 | 0.041269588 | 0.568301847 | 0.221730546 | 0.157957126 | 0.607744583 | 0.548820776 |
| Zhx1 | S | 48 | 0.06716647 | 0.085860668 | 0.127130256 | 0.059963786 | 0.041269588 | 0.568301847 | 0.221730546 | 0.157957126 | 0.607744583 | 0.548820776 |
| Pkmyt1 | S | 481 | 0.172291267 | 0.273247779 | -0.282160675 | -0.454451942 | -0.555408454 | 0.158876358 | 0.04513612 | 0.496764842 | 0.311079088 | 0.240953185 |
| Sphk1 | T | 226 | 0.050688004 | 0.20892525 | 0.029395571 | -0.021292432 | -0.179529678 | 0.676123601 | 0.042836503 | 0.775129835 | 0.869893801 | 0.136953812 |
| Nfic | S | 343 | 0.08115295 | 0.073739624 | 0.0402253 | -0.04092765 | -0.033514324 | 0.083739156 | 0.123624568 | 0.380227321 | 0.184337989 | 0.291730632 |
| Mia3 | S | 1705 | 0.043050277 | 0.097758639 | 0.017653305 | -0.025396972 | -0.080105334 | 0.303456526 | 0.003551653 | 0.763428857 | 0.689298217 | 0.242898235 |
| 4833420G17Rik | S | 547 | 0.049838557 | 0.030141413 | 0.183381516 | 0.133542959 | 0.153240102 | 0.305228599 | 0.572816843 | 0.089904419 | 0.169855439 | 0.132952825 |
| 4833420G17Rik | T | 546 | 0.049838557 | 0.030141413 | 0.183381516 | 0.133542959 | 0.153240102 | 0.305228599 | 0.572816843 | 0.089904419 | 0.169855439 | 0.132952825 |
| Lrrfip2 | S | 92 | 0.043950356 | 0.122814159 | 0.034234268 | -0.009716088 | -0.088579891 | 0.536916199 | 0.025408856 | 0.310501703 | 0.880271256 | 0.077907288 |
| Rreb1 | S | 1667 | -0.032318758 | -0.049274651 | 0.299430171 | 0.331748929 | 0.348704822 | 0.535831432 | 0.430461399 | 0.001692352 | 0.005595259 | 0.002385794 |
| Accs | T | 316 | 0.057039779 | 0.222553017 | -0.081634632 | -0.138674411 | -0.304187648 | 0.412769823 | 0.001855348 | 0.184813948 | 0.098501242 | 0.004437404 |
| Lpin1 | S | 328 | -0.060172786 | 0.132474616 | 0.091702454 | 0.151875241 | -0.040772162 | 0.338326487 | 0.010250998 | 0.05684512 | 0.057469088 | 0.198405707 |
| Smarca4 | T | 609 | 0.044295478 | -0.04048968 | 0.322413547 | 0.278118069 | 0.362903227 | 0.789159524 | 0.728404617 | 0.033382983 | 0.180572098 | 0.045475805 |
| Ndrg3 | S | 352 | 0.192134282 | 0.084739738 | -0.024923992 | -0.217058274 | -0.10966373 | 0.140964961 | 0.031702873 | 0.566273077 | 0.109012245 | 0.056208306 |
| Jam2 | S | 277 | 0.186022881 | -0.009517825 | -0.051392338 | -0.23741522 | -0.041874513 | 8.15042E-05 | 0.773670475 | 0.328830603 | 0.019120672 | 0.430748731 |
| Rbm15b | S | 107 | 0.011340533 | 0.017069342 | 0.141607258 | 0.130266725 | 0.124537916 | 0.663297787 | 0.419230923 | 0.006498471 | 0.012366066 | 0.017305202 |
| Ranbp3 | S | 33 | 0.020069504 | 0.059414566 | 0.049260565 | 0.029191061 | -0.010154001 | 0.132366711 | 0.008177792 | 0.164500857 | 0.34606252 | 0.724067864 |
| Pcm1 | S | 864 | -0.113933991 | 0.120007088 | 0.199491428 | 0.313425419 | 0.07948434 | 0.019510462 | 0.01498484 | 0.007111641 | 0.00475504 | 0.139698368 |
| Ppig | S | 431 | 0.016094432 | 0.075001215 | 0.160635261 | 0.14454083 | 0.085634046 | 0.819207786 | 0.259871309 | 0.067122939 | 0.139609916 | 0.277712838 |
| Dync1i2 | S | 95 | 0.075685962 | 0.098286355 | 0.025283817 | -0.050402145 | -0.073002538 | 0.26477204 | 0.022770519 | 0.639039603 | 0.489943304 | 0.233924555 |
| Arhgap21 | S | 1092 | 0.032289922 | -0.005102614 | 0.141331012 | 0.109041089 | 0.146433625 | 0.595723781 | 0.917411693 | 0.002365802 | 0.145770358 | 0.047599888 |
| Prrc2c | T | 2645 | 0.006807378 | 0.098549013 | 0.136272608 | 0.12946523 | 0.037723595 | 0.951401564 | 0.233516473 | 0.051511258 | 0.28750856 | 0.580929705 |
| Prkd3 | S | 27 | 0.017344419 | 0.122176159 | 0.09203153 | 0.07468711 | -0.03014463 | 0.699580015 | 0.049380438 | 0.268682422 | 0.361298681 | 0.698480899 |
| Wbp11 | S | 353 | 0.076737817 | 0.051556893 | 0.123038374 | 0.046300557 | 0.071481481 | 0.207061978 | 0.291579213 | 0.217885248 | 0.606615449 | 0.42632599 |
| Wbp11 | S | 361 | 0.076737817 | 0.051556893 | 0.123038374 | 0.046300557 | 0.071481481 | 0.207061978 | 0.291579213 | 0.217885248 | 0.606615449 | 0.42632599 |
| Klc4 | S | 590 | -0.010369696 | 0.160063435 | 0.096552321 | 0.106922017 | -0.063511114 | 0.832038986 | 0.039304289 | 0.269063949 | 0.22710864 | 0.459890081 |
| Tlk2 | S | 41 | 0.210756125 | 0.129815745 | 0.024702585 | -0.186053541 | -0.10511316 | 0.244064046 | 0.223606691 | 0.918417825 | 0.487966553 | 0.658669484 |
| Tlk2 | T | 52 | 0.210756125 | 0.129815745 | 0.024702585 | -0.186053541 | -0.10511316 | 0.244064046 | 0.223606691 | 0.918417825 | 0.487966553 | 0.658669484 |
| Srrm2 | T | 1848 | 0.184557639 | -0.015692289 | 0.231842583 | 0.047284944 | 0.247534872 | 0.179382112 | 0.870175756 | 0.318474372 | 0.822946864 | 0.290036834 |
| Lrif1 | T | 717 | -0.0259197 | 0.256460939 | 0.034886542 | 0.060806242 | -0.221574397 | 0.79063857 | 0.021368071 | 0.680642418 | 0.530587389 | 0.047222331 |
| Adam17 | S | 770 | 0.100760456 | 0.147580356 | 0.088376706 | -0.01238375 | -0.05920365 | 0.327231962 | 0.148059873 | 0.73629923 | 0.961545525 | 0.818024854 |
| Adam17 | S | 773 | 0.100760456 | 0.147580356 | 0.088376706 | -0.01238375 | -0.05920365 | 0.327231962 | 0.148059873 | 0.73629923 | 0.961545525 | 0.818024854 |
| Adam17 | T | 771 | 0.100760456 | 0.147580356 | 0.088376706 | -0.01238375 | -0.05920365 | 0.327231962 | 0.148059873 | 0.73629923 | 0.961545525 | 0.818024854 |
| Dync1i2 | T | 89 | 0.161432846 | -0.036843392 | 0.082417155 | -0.079015691 | 0.119260547 | 0.025351146 | 0.406671264 | 0.092719734 | 0.182659672 | 0.047229692 |
| Pcyt1a | S | 362 | 0.060864558 | 0.080329756 | 0.032484632 | -0.028379926 | -0.047845124 | 0.247309788 | 0.01216951 | 0.528328297 | 0.642530044 | 0.373383151 |
| Cdc25b | S | 15 | 0.057423499 | 0.071769647 | 0.081748694 | 0.024325196 | 0.009979048 | 0.300515556 | 0.04185514 | 0.355763844 | 0.779884979 | 0.900048057 |
| Rasa3 | S | 528 | 0.039153653 | 0.234253005 | -0.156516795 | -0.195670448 | -0.3907698 | 0.718000026 | 6.33407E-05 | 0.109730085 | 0.164075323 | 0.021259956 |
| Dzip3 | S | 613 | -0.007108582 | 0.076340833 | 0.107939597 | 0.115048179 | 0.031598764 | 0.857803503 | 0.041054746 | 0.049346132 | 0.054004048 | 0.443823303 |
| Cul4b | T | 15 | 0.001568652 | 0.2127263 | 0.023200779 | 0.021632128 | -0.18952552 | 0.984717132 | 0.016183983 | 0.710822829 | 0.74837744 | 0.020987408 |
| Alyref | S | 205 | 0.282783442 | 0.080690239 | -0.137157464 | -0.419940906 | -0.217847703 | 0.012531437 | 0.323338383 | 0.340203064 | 0.040702996 | 0.168540014 |
| Bclaf1 | S | 296 | 0.10326996 | 0.09733133 | -0.038320885 | -0.141590846 | -0.135652215 | 0.029709983 | 0.057297385 | 0.7701844 | 0.333482819 | 0.349251279 |
| Ints5 | S | 278 | 0.112351989 | 0.111741297 | 0.034182892 | -0.078169097 | -0.077558405 | 0.127703637 | 0.22295767 | 0.565897355 | 0.276080556 | 0.377015808 |
| Mia3 | S | 1915 | 0.077992397 | 0.05068473 | 0.070951464 | -0.007040933 | 0.020266733 | 0.242146337 | 0.160462138 | 0.113989449 | 0.907693618 | 0.61087893 |
| Dennd2a | S | 302 | 0.083531371 | 0.104222431 | -0.001020304 | -0.084551675 | -0.105242735 | 0.04087243 | 0.077071009 | 0.983644241 | 0.157967977 | 0.12357533 |
| Pds5b | S | 1356 | 0.094686388 | 0.019653649 | 0.014008953 | -0.080677435 | -0.005644696 | 0.000748619 | 0.663730294 | 0.395056612 | 0.001036152 | 0.895544157 |
| Borcs6 | T | 201 | 0.103747441 | 0.244985609 | -0.077832847 | -0.181580288 | -0.322818456 | 0.16098125 | 0.040029624 | 0.329463042 | 0.03424637 | 0.021015275 |
| Lpar1 | S | 341 | 0.065111799 | 0.097762659 | 0.102481234 | 0.037369436 | 0.004718575 | 0.366013276 | 0.210342147 | 0.239932801 | 0.498551059 | 0.933105739 |
| Pdlim5 | S | 228 | -0.031349036 | 0.243442872 | 0.042357378 | 0.073706414 | -0.201085494 | 0.652342873 | 0.014110515 | 0.688650119 | 0.430983935 | 0.099025104 |
| Atxn3 | S | 268 | 0.003363814 | 0.15198653 | 0.093729998 | 0.090366185 | -0.058256532 | 0.949621632 | 0.052809325 | 0.277453747 | 0.297706218 | 0.500252599 |
| Pacs1 | S | 409 | 0.033882735 | 0.085402319 | 0.175715566 | 0.141832832 | 0.090313247 | 0.673657659 | 0.19764337 | 0.189966175 | 0.285718835 | 0.448309033 |
| Srrm2 | S | 452 | 0.013838109 | 0.04365023 | 0.089603724 | 0.075765615 | 0.045953495 | 0.490219789 | 0.064715124 | 0.022609243 | 0.042228771 | 0.141498015 |
| E4f1 | S | 49 | 0.051438035 | 0.309211448 | -0.047859683 | -0.099297718 | -0.357071131 | 0.824087646 | 0.02916806 | 0.748926668 | 0.671029797 | 0.05491368 |
| Srsf2 | S | 191 | 0.057499009 | 0.036533934 | 0.104223098 | 0.04672409 | 0.067689165 | 0.129176554 | 0.296743398 | 0.116153037 | 0.391124253 | 0.251412245 |
| Arhgap28 | S | 53 | -0.014990015 | 0.151923852 | 0.103030217 | 0.118020232 | -0.048893635 | 0.839347326 | 0.052736374 | 0.195914546 | 0.116646058 | 0.399945815 |
| Rbm25 | S | 698 | 0.016242778 | 0.112408072 | 0.169603882 | 0.153361103 | 0.05719581 | 0.813637256 | 0.218114462 | 0.162514763 | 0.200569808 | 0.613105049 |
| Srrm2 | S | 2351 | 0.09667001 | 0.097886078 | 0.09360697 | -0.003063039 | -0.004279108 | 0.253705829 | 0.254348936 | 0.390984422 | 0.969184241 | 0.957656688 |
| Vim | S | 420 | 0.006728017 | 0.049832283 | 0.197855597 | 0.19112758 | 0.148023314 | 0.909886051 | 0.387714364 | 0.044733691 | 0.057782033 | 0.110635717 |
| Magi3 | S | 829 | 0.064029351 | 0.073848121 | 0.079019014 | 0.014989663 | 0.005170893 | 0.319230927 | 0.081924644 | 0.260302921 | 0.842813168 | 0.934353956 |
| Zeb2 | S | 353 | 0.064967271 | 0.066758958 | 0.026154964 | -0.038812306 | -0.040603993 | 0.039604209 | 0.076817301 | 0.402695852 | 0.25261358 | 0.279431406 |
| Srrm2 | S | 1438 | 0.028980285 | 0.020453191 | 0.106600305 | 0.07762002 | 0.086147114 | 0.247922605 | 0.227858 | 0.018745531 | 0.054483467 | 0.050149915 |
| Ncoa6 | S | 2022 | 0.020093469 | 0.037732523 | 0.274143471 | 0.254050002 | 0.236410947 | 0.777153743 | 0.200617985 | 0.160382126 | 0.173490982 | 0.20275171 |
| Ncoa6 | T | 2024 | 0.020093469 | 0.037732523 | 0.274143471 | 0.254050002 | 0.236410947 | 0.777153743 | 0.200617985 | 0.160382126 | 0.173490982 | 0.20275171 |
| Bclaf1 | S | 387 | 0.004085762 | 0.086246895 | 0.200061975 | 0.195976213 | 0.11381508 | 0.969186354 | 0.213010108 | 0.121719634 | 0.165619357 | 0.306632348 |
| Cybc1 | S | 176 | -0.050725959 | 0.028243491 | 0.245152279 | 0.295878237 | 0.216908788 | 0.372228929 | 0.566629741 | 0.005863961 | 0.00596131 | 0.015022342 |

| | | | | | | | | | | | | |
|---|---|---|---|---|---|---|---|---|---|---|---|---|
| Tcf20 | S | 603 | 0.038073979 | 0.072024387 | 0.115891026 | 0.077817047 | 0.043866639 | 0.673485859 | 0.082771023 | 0.138674686 | 0.446827584 | 0.494624941 |
| Steap3 | S | 486 | 0.010039061 | 0.077625568 | 0.086090395 | 0.076051334 | 0.008464827 | 0.759559311 | 0.216361236 | 0.010334385 | 0.057996389 | 0.869934703 |
| Srrm2 | S | 1572 | 0.021766904 | 0.092934545 | 0.050658699 | 0.028891795 | -0.042275846 | 0.589240345 | 0.0096785 | 0.261221668 | 0.561110242 | 0.335065339 |
| Btf3l4 | S | 153 | 0.052417295 | 0.198528504 | 0.093563121 | 0.041145825 | -0.104965383 | 0.712025028 | 0.153989099 | 0.428841942 | 0.794290165 | 0.464344737 |
| Magi3 | S | 1232 | 0.023494582 | 0.15486607 | 0.052771777 | 0.029277195 | -0.102094292 | 0.774764846 | 0.033356162 | 0.396042151 | 0.750190184 | 0.177354986 |
| Dennd1a | S | 538 | -0.001583221 | 0.147976295 | 0.069149294 | 0.070732515 | -0.078827001 | 0.98354795 | 0.020263954 | 0.349667862 | 0.408091342 | 0.254526377 |
| Prpf4b | S | 62 | 0.005677181 | 0.075656876 | 0.15656032 | 0.150883139 | 0.080903444 | 0.926604792 | 0.130820826 | 0.092256284 | 0.120112175 | 0.302646349 |
| Bap1 | S | 395 | 0.096580287 | 0.120803918 | 0.039955893 | -0.056624393 | -0.080848024 | 0.080272763 | 0.203203939 | 0.810691708 | 0.730387914 | 0.644919013 |
| Palld | S | 1143 | -0.005323654 | 0.14710817 | 0.044687712 | 0.050011366 | -0.102420458 | 0.882154786 | 0.004632184 | 0.565451389 | 0.508056251 | 0.23654246 |
| Runx1 | T | 207 | 0.006616007 | 0.234896932 | 0.07248665 | 0.065870643 | -0.162410282 | 0.926751119 | 0.082860949 | 0.465330106 | 0.456871397 | 0.211865055 |
| Prpf4b | S | 143 | 0.056367716 | 0.033241603 | 0.101456933 | 0.045089217 | 0.068215329 | 0.32052401 | 0.086450362 | 0.0991143 | 0.466426775 | 0.186610614 |
| Prpf4b | S | 145 | 0.056367716 | 0.033241603 | 0.101456933 | 0.045089217 | 0.068215329 | 0.32052401 | 0.086450362 | 0.0991143 | 0.466426775 | 0.186610614 |
| Tcf12 | S | 386 | -0.07830362 | 0.156687564 | 0.112812216 | 0.191115837 | -0.043875348 | 0.121439064 | 0.024672241 | 0.01929368 | 0.011301921 | 0.383134255 |
| Arhgap31 | S | 1359 | -0.015396717 | 0.119397953 | 0.108733205 | 0.124129922 | -0.010664749 | 0.817223088 | 0.01628962 | 0.270343657 | 0.247543583 | 0.899641382 |
| Abi1 | T | 200 | 0.036685223 | 0.133601934 | 0.037216614 | 0.000531391 | -0.09638532 | 0.382217484 | 0.030578738 | 0.540694919 | 0.992098838 | 0.164212981 |
| Triobp | S | 1101 | 0.121645457 | 0.091172882 | -0.001622685 | -0.123268141 | -0.092795567 | 0.055330838 | 0.125795936 | 0.974844139 | 0.057199128 | 0.116138021 |
| Szt2 | S | 2135 | 0.088083692 | 0.090914161 | -0.000468625 | -0.088552316 | -0.091382786 | 0.054467903 | 0.052738669 | 0.992660949 | 0.125109082 | 0.11814915 |
| Palb2 | S | 559 | 0.019368969 | 0.008032942 | 0.157808097 | 0.138439127 | 0.149775154 | 0.684477988 | 0.863685012 | 0.005838973 | 0.025125749 | 0.018261013 |
| Actn1 Actn4 | S | 368;348 | 0.070439923 | 0.034804273 | 0.048559265 | -0.021880659 | 0.013754992 | 0.038387592 | 0.21376478 | 0.139082387 | 0.493019847 | 0.659863421 |
| Snrpd2 | S | 9 | 0.103404758 | 0.043937299 | 0.063511021 | -0.039893737 | 0.019573721 | 0.073020538 | 0.191467283 | 0.432803334 | 0.624105364 | 0.796957 |
| Prrc2a | Y | 1092 | 0.228758002 | 0.047361173 | -0.014493548 | -0.24325155 | -0.061854721 | 0.031222427 | 0.558027607 | 0.912400619 | 0.122195409 | 0.605730631 |
| Smn1 | S | 25 | 0.039145472 | 0.069049698 | 0.047911627 | 0.008766155 | -0.021138072 | 0.10537774 | 0.09639622 | 0.115654521 | 0.744524717 | 0.55254682 |
| Bcl9l | S | 25 | -0.036966161 | 0.084631303 | 0.147328759 | 0.18429492 | 0.062697456 | 0.295603301 | 0.094730164 | 0.015211263 | 0.007824422 | 0.201228449 |
| Hnrnpf Hnrnph1 | S | 310;310 | -0.017083645 | 0.16931437 | 0.111475335 | 0.128558981 | -0.057839035 | 0.846280904 | 0.035174558 | 0.399821427 | 0.370861565 | 0.64696569 |
| Huwe1 | S | 2917 | 0.020740466 | 0.171321183 | 0.088903786 | 0.06816332 | -0.082417397 | 0.859131931 | 0.138103506 | 0.19645281 | 0.534047193 | 0.385103726 |
| Asap2 | S | 704 | -0.043796236 | 0.088450404 | 0.122786885 | 0.166583121 | 0.03433648 | 0.055702104 | 0.012125055 | 0.010330342 | 0.007667956 | 0.295707176 |
| 4932438A13Rik | S | 3653 | 0.079422128 | -0.00711796 | 0.095343477 | 0.015921349 | 0.102461438 | 0.028970368 | 0.822202738 | 0.055422114 | 0.666074638 | 0.054992095 |
| Synrg | S | 801 | -0.017513742 | 0.174892714 | -0.02346334 | -0.005949597 | -0.198356054 | 0.805032013 | 0.000411662 | 0.717705446 | 0.944416837 | 0.058522487 |
| Klf6 | S | 171 | 0.031431724 | 0.046052895 | 0.091979758 | 0.060548033 | 0.045926862 | 0.55987835 | 0.338490592 | 0.008919614 | 0.296263746 | 0.333007088 |
| Plekhg3 | S | 758 | 0.057050784 | 0.111279129 | 0.03623621 | -0.020814574 | -0.075042919 | 0.382782954 | 0.022716755 | 0.608847543 | 0.804117314 | 0.337324462 |
| Kmt2d | S | 727 | -0.007799906 | 0.138187792 | 0.073207868 | 0.081007774 | -0.064979924 | 0.801866485 | 0.074512698 | 0.056039332 | 0.013425306 | 0.300745035 |
| Rbm15 | S | 293 | -0.019415282 | 0.086197161 | 0.130913996 | 0.150329278 | 0.044716835 | 0.437286814 | 0.218383506 | 0.009484944 | 0.009862445 | 0.480169498 |
| Tra2a | Y | 85 | 0.033817655 | 0.08104872 | 0.097794283 | 0.063976628 | 0.016745563 | 0.643246135 | 0.158313411 | 0.071810646 | 0.353584212 | 0.634339573 |
| Fbxl20 | T | 417 | -0.027028666 | 0.12323747 | 0.138892254 | 0.16592092 | 0.015654784 | 0.455727752 | 0.106382165 | 0.054947547 | 0.052662856 | 0.821365926 |
| Faap20 | S | 119 | 0.013193118 | 0.057104337 | 0.135279791 | 0.122086673 | 0.078175454 | 0.69268307 | 0.340998589 | 0.027569796 | 0.056004776 | 0.230407868 |
| Pms2 | S | 569 | 0.239422347 | 0.083450524 | -0.083587716 | -0.323010063 | -0.167038239 | 0.032483925 | 0.338561132 | 0.643356489 | 0.145871714 | 0.369423961 |
| Ahi1 | S | 974 | -0.045498645 | 0.155799875 | 0.082742861 | 0.128241506 | -0.073057014 | 0.192810485 | 0.032081348 | 0.023957327 | 0.009004625 | 0.201948322 |
| Pcm1 | S | 867 | -0.038636425 | 0.055433198 | 0.151501003 | 0.190137429 | 0.096067805 | 0.476486246 | 0.264674797 | 0.004155894 | 0.025264021 | 0.073641031 |
| Pcm1 | S | 870 | -0.038636425 | 0.055433198 | 0.151501003 | 0.190137429 | 0.096067805 | 0.476486246 | 0.264674797 | 0.004155894 | 0.025264021 | 0.073641031 |
| Wnk1 | S | 2133 | -0.079950703 | 0.141802671 | 0.19602435 | 0.275975054 | 0.05422168 | 0.474982295 | 0.214088653 | 0.041407729 | 0.05615463 | 0.5858797 |
| Srrm2 | S | 911 | 0.009765136 | 0.201383656 | 0.144723389 | 0.134958253 | -0.056660267 | 0.96524829 | 0.217315439 | 0.26252794 | 0.573501518 | 0.72110678 |
| Taok1 | S | 965 | 0.101420492 | 0.128567127 | -0.028725655 | -0.130146147 | -0.157292782 | 0.092667494 | 0.035804763 | 0.53629655 | 0.038036833 | 0.015588453 |
| Dtna | S | 666 | -0.008178941 | 0.132892175 | 0.116183284 | 0.124362225 | -0.01670889 | 0.929824314 | 0.255693695 | 0.040525817 | 0.241844576 | 0.866657101 |
| Cttnbp2nl | S | 481 | 0.098098989 | -0.014706243 | 0.107665437 | 0.009566448 | 0.12237168 | 0.051905787 | 0.650091428 | 0.058777506 | 0.836766299 | 0.05738888 |
| Chaf1b | S | 416 | -0.045787156 | 0.31602306 | -0.075758981 | -0.029971826 | -0.391782042 | 0.447137106 | 0.001971626 | 0.590356426 | 0.816515153 | 0.063175728 |
| Pdlim2 | S | 74 | 0.017549408 | 0.061703304 | 0.084035505 | 0.066486096 | 0.0223322 | 0.621198666 | 0.182029192 | 0.014697462 | 0.109230461 | 0.548128587 |
| Adgra2 | S | 1273 | -0.053301982 | 0.181224966 | 0.112809588 | 0.16611157 | -0.068415378 | 0.453435106 | 0.068306105 | 0.091884336 | 0.039552281 | 0.363088364 |
| Erf | S | 187 | -0.008680868 | 0.085921709 | 0.093333201 | 0.10201407 | 0.007411492 | 0.82444794 | 0.100743444 | 0.019098132 | 0.0394943 | 0.845619322 |
| Trim44 | S | 338 | 0.001919521 | 0.068635693 | 0.097347735 | 0.095428214 | 0.028712042 | 0.967830475 | 0.024000279 | 0.079021018 | 0.138376473 | 0.489709432 |
| Srrm2 | T | 474 | 0.087983714 | 0.014434662 | 0.132228299 | 0.044244585 | 0.117793637 | 0.303398974 | 0.845673845 | 0.046398979 | 0.590055366 | 0.188163045 |
| Mcm3 | S | 611 | 0.248772453 | 0.178686005 | -0.129252159 | -0.378024612 | -0.307938165 | 0.096718297 | 0.034839138 | 0.117103637 | 0.030907269 | 0.012464407 |
| Dennd2a | S | 512 | -0.01210103 | 0.039685279 | 0.146842721 | 0.15894375 | 0.107157441 | 0.822587247 | 0.399021903 | 0.00594829 | 0.042036177 | 0.044970138 |
| Dennd2a | S | 514 | -0.01210103 | 0.039685279 | 0.146842721 | 0.15894375 | 0.107157441 | 0.822587247 | 0.399021903 | 0.00594829 | 0.042036177 | 0.044970138 |
| Dennd2a | S | 516 | -0.01210103 | 0.039685279 | 0.146842721 | 0.15894375 | 0.107157441 | 0.822587247 | 0.399021903 | 0.00594829 | 0.042036177 | 0.044970138 |
| Cers2 | S | 349 | 0.043920444 | 0.046568151 | 0.095650749 | 0.051730305 | 0.049082598 | 0.278822293 | 0.134304215 | 0.102164421 | 0.336955216 | 0.322622322 |
| Ep300 | T | 887 | 0.090138391 | 0.064339898 | 0.052560255 | -0.037578136 | -0.011779643 | 0.024216803 | 0.50387549 | 0.396450558 | 0.532837414 | 0.906125106 |
| Sde2 | S | 308 | -0.029920935 | 0.106382439 | 0.126923246 | 0.156844181 | 0.020540806 | 0.784500824 | 0.084487638 | 0.069459887 | 0.237690031 | 0.709333861 |
| Mki67 | S | 1587 | 0.135107029 | 0.065982152 | 0.016748085 | -0.118358943 | -0.049234067 | 0.052068858 | 0.19073844 | 0.880052315 | 0.343253171 | 0.663597423 |
| Srsf6 | S | 45 | 0.022507461 | 0.110342984 | 0.127991721 | 0.105484261 | 0.017648738 | 0.734818604 | 0.14747904 | 0.183508781 | 0.214045007 | 0.811834584 |
| Dido1 | S | 151 | -0.050825146 | 0.105166823 | 0.150612156 | 0.201437302 | 0.045445333 | 0.054963389 | 0.045041288 | 0.011338428 | 0.013768104 | 0.331548652 |
| Prrx1 | S | 86 | 0.020476767 | 0.083483432 | 0.159766588 | 0.13928982 | 0.076283155 | 0.84854109 | 0.165438038 | 0.137785578 | 0.301796482 | 0.408642013 |
| Foxk1 | S | 235 | 0.208608624 | 0.542570031 | -0.019032412 | -0.227641036 | -0.561602443 | NA | 0.136500545 | 0.950082823 | NA | 0.055351461 |
| Ubr5 | Y | 1740 | -0.173240491 | 0.385613896 | 0.121911039 | 0.29515153 | -0.263702857 | 0.481883748 | 0.162219293 | 0.103241819 | 0.277244895 | 0.288821318 |
| Akap12 | S | 811 | 0.040673383 | 0.066986859 | 0.144124237 | 0.103450854 | 0.077137378 | 0.741767961 | 0.197635034 | 0.106804343 | 0.445131094 | 0.313704622 |
| Rai1 | S | 1105 | 0.057597261 | 0.035253257 | 0.07654964 | 0.01895238 | 0.041296383 | 0.05627351 | 0.185767706 | 0.189772918 | 0.693238681 | 0.419094155 |
| Rtn4 | S | 186 | 0.052861724 | 0.118912094 | 0.112790693 | 0.059928968 | -0.006121402 | 0.446352122 | 0.118873962 | 0.457295598 | 0.662767497 | 0.963444123 |
| Srrm1 | S | 169 | 0.082661883 | 0.009187185 | 0.095592821 | 0.012930938 | 0.086405636 | 0.337815206 | 0.860295068 | 0.009123603 | 0.864054963 | 0.16968665 |
| Srrm1 | S | 171 | 0.082661883 | 0.009187185 | 0.095592821 | 0.012930938 | 0.086405636 | 0.337815206 | 0.860295068 | 0.009123603 | 0.864054963 | 0.16968665 |
| Brd4 | T | 296 | 0.078001024 | 0.295857119 | -0.055160404 | -0.133161428 | -0.351017522 | 0.596889021 | 0.037203481 | 0.553380669 | 0.417554167 | 0.028892465 |
| Dido1 | S | 1726 | -0.079542291 | 0.087493782 | 0.233843787 | 0.313386078 | 0.146350005 | 0.156766114 | 0.287987592 | 0.008176892 | 0.004680393 | 0.126738436 |
| Map3k4 | T | 451 | 0.098470203 | 0.112990864 | 0.073243999 | -0.025226203 | -0.039746865 | 0.38752027 | 0.146352355 | 0.472104278 | 0.842039743 | 0.698136331 |
| Iws1 | S | 345 | 0.015758726 | 0.016948349 | 0.095323727 | 0.079565001 | 0.078375379 | 0.234019981 | 0.248963807 | 0.005773906 | 0.019026245 | 0.016134402 |
| Cenpu | S | 186 | 0.013553971 | 0.086710133 | 0.108756467 | 0.095202496 | 0.022046334 | 0.624937108 | 0.114152526 | 0.108342693 | 0.138264854 | 0.708631491 |
| Tef | S | 138 | 0.032379863 | 0.167482837 | 0.0412101 | 0.008830237 | -0.126272737 | 0.669733623 | 0.063996288 | 0.33365996 | 0.894522075 | 0.1444722 |
| Arid1a | S | 773 | -0.005649524 | 0.087287608 | 0.066649095 | 0.072298619 | -0.020638514 | 0.893103065 | 0.007115167 | 0.123638016 | 0.177357387 | 0.553819875 |
| Arhgap21 | S | 1728 | 0.035159503 | 0.097961894 | 0.098493988 | 0.063334485 | 0.000532094 | 0.689919527 | 0.318593998 | 0.04345725 | 0.476539228 | 0.995029292 |
| Ssr1 | S | 268 | 0.029442989 | 0.094234434 | 0.065144009 | 0.03570102 | -0.029090425 | 0.53795713 | 0.059870782 | 0.145407712 | 0.310399103 | 0.28356075 |
| Scaf1 | S | 676 | 0.002652102 | 0.049787644 | 0.086991364 | 0.084339263 | 0.037203721 | 0.91362524 | 0.037330563 | 0.019692253 | 0.037268284 | 0.12360625 |
| Scaf1 | S | 682 | 0.002652102 | 0.049787644 | 0.086991364 | 0.084339263 | 0.037203721 | 0.91362524 | 0.037330563 | 0.019692253 | 0.037268284 | 0.12360625 |
| Synrg | S | 152 | 0.091655442 | 0.236993748 | -0.093773762 | -0.185429204 | -0.330767511 | 0.383449251 | 0.029154769 | 0.631969667 | 0.37411457 | 0.167158308 |
| Synrg | S | 153 | 0.091655442 | 0.236993748 | -0.093773762 | -0.185429204 | -0.330767511 | 0.383449251 | 0.029154769 | 0.631969667 | 0.37411457 | 0.167158308 |
| Rfx1 | S | 113 | -0.026419856 | 0.087045074 | 0.180625078 | 0.207044934 | 0.093580005 | 0.54714738 | 0.108346476 | 0.07129622 | 0.056704801 | 0.260920999 |
| Parg | S | 297 | -0.026511348 | 0.131520182 | 0.047282992 | 0.073794339 | -0.08423719 | 0.728749462 | 0.002728246 | 0.378523945 | 0.394926511 | 0.160568445 |
| Epn2 | S | 443 | 0.10450581 | 0.016063784 | 0.057847335 | -0.046658475 | 0.041783552 | 0.042976206 | 0.537798488 | 0.152025685 | 0.300870969 | 0.242199332 |
| Baz1b | T | 711 | 0.012927941 | 0.130539639 | 0.130413297 | 0.117485356 | -0.000126341 | 0.905783401 | 0.064785601 | 0.350268816 | 0.439524776 | 0.999163657 |
| Spp1 | S | 234 | 0.189869604 | 0.038264098 | 0.012492096 | -0.177377508 | -0.025772002 | 0.036747092 | 0.49646876 | 0.835841401 | 0.058000824 | 0.673481174 |
| Slk | S | 643 | 0.046157183 | 0.109154862 | 0.044955107 | -0.001202076 | -0.064199754 | 0.30060568 | 0.075112125 | 0.290062969 | 0.970013988 | 0.212743128 |
| Prpf4b | S | 387 | -0.030425214 | 0.161522612 | 0.282858953 | 0.313284167 | 0.121336341 | 0.294910399 | 0.15712171 | 0.225931661 | 0.196976229 | 0.554094239 |
| Tpr | T | 715 | 0.27770947 | -0.063573622 | 0.145122109 | -0.132587361 | 0.208695731 | 0.077001362 | 0.658334235 | 0.490601251 | 0.536898831 | 0.3616785 |
| Sphk1 | S | 225 | 0.026297839 | 0.176620661 | 0.030511855 | 0.004214016 | -0.146108806 | 0.747850426 | 0.025283499 | 0.679572143 | 0.961081639 | 0.096298098 |
| Usp20 | S | 415 | 0.074151982 | 0.060882945 | 0.049711618 | -0.024440365 | -0.011171328 | 0.080921317 | 0.157979164 | 0.329843659 | 0.607721438 | 0.816900481 |
| Rnf217 | S | 27 | 0.087565368 | 0.015164274 | 0.077989475 | -0.009575893 | 0.0628252 | 0.038332986 | 0.75928773 | 0.125817663 | 0.819831546 | 0.279235516 |
| Ctdnep1 | S | 49 | 0.062330608 | -0.00121613 | 0.125875832 | 0.063545225 | 0.127091962 | 0.160634174 | 0.974150114 | 0.028753828 | 0.201263579 | 0.03484586 |
| Larp4b | S | 601 | -0.016041587 | 0.069455203 | 0.093273527 | 0.109315113 | 0.023818323 | 0.568345267 | 0.016892206 | 0.033503268 | 0.026005573 | 0.442771114 |

| | | | | | | | | | | | | |
|---|---|---|---|---|---|---|---|---|---|---|---|---|
| Ppig | S | 288 | 0.01167281 | 0.076193038 | 0.058678116 | 0.047005307 | -0.017514922 | 0.688596601 | 0.018600906 | 0.082432078 | 0.128455154 | 0.433905556 |
| Bard1 | S | 397 | 0.126566917 | 0.02231854 | 0.052275946 | -0.074290971 | 0.029957406 | 0.097651517 | 0.491301508 | 0.122880564 | 0.253917908 | 0.370975192 |
| Zcchc8 | S | 501 | 0.115757831 | 0.288649511 | -0.037097055 | -0.152854886 | -0.325746566 | 0.538254361 | 0.106547036 | 0.868995076 | 0.527460712 | 0.200691777 |
| Chchd2 | T | 13 | 0.144501029 | 0.046443945 | 0.014951113 | -0.129549916 | -0.031492832 | 0.022453036 | 0.36996748 | 0.879357104 | 0.243085042 | 0.736902844 |
| Smarcad1 | S | 144 | 0.024854542 | 0.016914962 | 0.09804398 | 0.073189438 | 0.081129019 | 0.431011342 | 0.460079534 | 0.003785953 | 0.063447484 | 0.001092346 |
| Anks1 | S | 903 | 0.001239846 | 0.146536216 | 0.067844891 | 0.066605045 | -0.078691325 | 0.977033409 | 0.014251379 | 0.542121497 | 0.537836804 | 0.477331935 |
| Srrm2 | S | 810 | 0.001593153 | 0.101737828 | 0.087652848 | 0.086059695 | -0.01408498 | 0.968092537 | 0.094534617 | 0.057644507 | 0.120376166 | 0.76778852 |
| Cav2 | Y | 19 | -0.075959629 | 0.059275178 | 0.222712516 | 0.298672145 | 0.163437338 | 0.066070419 | 0.068366493 | 0.006924193 | 0.003488965 | 0.029725355 |
| Mki67 | T | 1952 | 0.159480049 | 0.280618015 | -0.07285182 | -0.23233187 | -0.353469835 | 0.341591915 | 0.107274083 | 0.549907403 | 0.155335177 | 0.059746418 |
| Sltm | S | 1017 | 0.004167232 | 0.040349581 | 0.171578267 | 0.167411035 | 0.131228686 | 0.94978158 | 0.584838382 | 0.017229325 | 0.033540682 | 0.100318041 |
| Oasl2 | S | 375 | -0.013642117 | 0.044089946 | 0.263656279 | 0.277298396 | 0.219566333 | 0.895691374 | 0.683344844 | 0.044643806 | 0.01136145 | 0.015867801 |
| Sorbs2 | S | 111 | -0.091537475 | -0.069340364 | 0.304297749 | 0.395835224 | 0.373638113 | 0.54671208 | 0.443675902 | 0.000837214 | 0.081472724 | 0.021898959 |
| Sipa1l2 | S | 161 | -0.014505516 | 0.018916154 | 0.219731966 | 0.234237481 | 0.200815812 | 0.91736574 | 0.810191135 | 0.01412473 | 0.178147497 | 0.057675718 |
| Cavin2 | T | 368 | 0.039777994 | 0.054737513 | 0.035164895 | -0.004613099 | -0.019572618 | 0.120552274 | 0.00688012 | 0.385665061 | 0.905065121 | 0.601546258 |
| Acd | S | 313 | 0.090323982 | 0.042754105 | 0.08729756 | -0.003026422 | 0.044543455 | 0.323781568 | 0.54779163 | 0.047735928 | 0.970007663 | 0.522559674 |
| Acd | S | 317 | 0.090323982 | 0.042754105 | 0.08729756 | -0.003026422 | 0.044543455 | 0.323781568 | 0.54779163 | 0.047735928 | 0.970007663 | 0.522559674 |
| Tesk1 | S | 439 | -0.003251401 | 0.050985409 | 0.139138401 | 0.142389802 | 0.088152993 | 0.960419055 | 0.290828302 | 0.016711504 | 0.086440434 | 0.023567566 |
| Tesk1 | S | 442 | -0.003251401 | 0.050985409 | 0.139138401 | 0.142389802 | 0.088152993 | 0.960419055 | 0.290828302 | 0.016711504 | 0.086440434 | 0.023567566 |
| Fmnl1 | S | 184 | 0.044039968 | 0.113757315 | 0.099345352 | 0.055305384 | -0.014411963 | 0.278837203 | 0.145847003 | 0.431943603 | 0.638320267 | 0.906764572 |
| Nes | S | 957 | -0.079589112 | 0.089846601 | 0.225381744 | 0.304970856 | 0.135535143 | 0.20856015 | 0.327347818 | 0.009012503 | 0.006984543 | 0.184794944 |
| Atrx | S | 1975 | -0.045270289 | 0.133936436 | 0.222638094 | 0.267908383 | 0.088701658 | 0.483000091 | 0.135426137 | 0.143223735 | 0.115687779 | 0.494460094 |
| Srpk2 | S | 8 | 0.125413771 | -0.030720867 | 0.16249138 | 0.037077609 | 0.193212247 | 0.32010767 | 0.65489817 | 0.037798867 | 0.748395925 | 0.043824231 |
| Adnp | S | 98 | 0.0245397 | 0.123062908 | 0.059882086 | 0.035342386 | -0.063180822 | 0.653558631 | 0.030687702 | 0.393287228 | 0.651878991 | 0.382244761 |
| Clspn | S | 1265 | 0.094227774 | 0.012028488 | 0.052385251 | -0.041842523 | 0.040356763 | 0.00803097 | 0.799325217 | 0.256017667 | 0.340575126 | 0.473342675 |
| Smim13 | S | 59 | 0.049804143 | 0.055214628 | 0.068326951 | 0.018522808 | 0.013112323 | 0.282086775 | 0.223875444 | 0.048599038 | 0.671577921 | 0.751868174 |
| Smim13 | S | 61 | 0.049804143 | 0.055214628 | 0.068326951 | 0.018522808 | 0.013112323 | 0.282086775 | 0.223875444 | 0.048599038 | 0.671577921 | 0.751868174 |
| Ifi35 | S | 36 | -0.238184762 | 0.285711136 | 0.208234793 | 0.446419555 | -0.077476343 | 0.014015977 | 0.003178345 | 0.083455629 | 0.011352142 | 0.407407534 |
| Supt6 | S | 1708 | 0.189451023 | 0.225950295 | -0.063180293 | -0.252631316 | -0.289130589 | 0.418971887 | 0.087690288 | 0.683529854 | 0.333874944 | 0.142505406 |
| Srrm2 | S | 1225 | 0.077745963 | 0.031956028 | 0.118359731 | 0.040613768 | 0.086403703 | 0.1942324 | 0.677915424 | 0.098248459 | 0.541335056 | 0.342019525 |
| Abi2 | S | 177 | 0.031158991 | 0.146482233 | 0.020156322 | -0.011002669 | -0.126325911 | 0.463416731 | 0.02410976 | 0.615380562 | 0.790144695 | 0.047723519 |
| Kmt2d | S | 2253 | 0.135770372 | 0.012680383 | 0.167569299 | 0.031798927 | 0.154888916 | 0.131422361 | 0.880567406 | 0.33711417 | 0.84206319 | 0.374649002 |
| Gtse1 | T | 406 | 0.020835707 | 0.161613345 | 0.000755093 | -0.020080614 | -0.160858252 | 0.514910273 | 0.009229764 | 0.981747352 | 0.5825962 | 0.013063507 |
| Syne2 | T | 4195 | -0.025604598 | 0.21632689 | 0.115693882 | 0.14129848 | -0.100633007 | 0.73679549 | 0.159394012 | 0.174555664 | 0.175944592 | 0.451323896 |
| Cdan1 | S | 160 | -0.111543817 | 0.13105078 | 0.207902504 | 0.319446321 | 0.076851724 | 0.344563951 | 0.148984927 | 0.034792953 | 0.046941482 | 0.386532929 |
| Pard3 | S | 1172 | 0.013297415 | 0.202151165 | 0.340022767 | 0.326725352 | 0.137871602 | 0.83227821 | 0.436447081 | 0.435701116 | 0.45050501 | 0.756249184 |
| Enah | S | 740 | 0.220444237 | 0.086793101 | -0.015520607 | -0.235964844 | -0.102313708 | 0.156727687 | 0.163202648 | 0.874755988 | 0.152194958 | 0.322837418 |
| Kctd12 | S | 178 | 0.047044992 | 0.027785037 | 0.130212736 | 0.083167744 | 0.1024277 | 0.706878312 | 0.510968404 | 0.017556083 | 0.512480232 | 0.004404724 |
| Srrm1 | T | 745 | 0.034404529 | 0.0669042 | 0.062596233 | 0.028191704 | -0.004307968 | 0.519079645 | 0.005798874 | 0.361781059 | 0.705345544 | 0.943744148 |
| Cenpv | S | 18 | 0.098928011 | -0.018858211 | 0.079510518 | -0.019417494 | 0.098368729 | 0.014151285 | 0.547347715 | 0.081239321 | 0.561790273 | 0.046759844 |
| Ccdc88a | S | 233 | -0.020206194 | 0.170779658 | 0.074044017 | 0.094250211 | -0.096735642 | 0.724067871 | 0.0200461 | 0.421997152 | 0.309184752 | 0.301000766 |
| Atp13a1 | S | 22 | 0.089403302 | 0.090479076 | 0.019776587 | -0.069626715 | -0.070702489 | 0.117718238 | 0.089303584 | 0.705528555 | 0.180722372 | 0.143631313 |
| Itsn1 | S | 335 | -0.023343423 | 0.075848262 | 0.083094892 | 0.106438315 | 0.00724663 | 0.21697844 | 0.141534446 | 0.000530392 | 0.009164076 | 0.843244424 |
| Xpnpep3 | S | 263 | -0.124815625 | 0.156919775 | 0.238800138 | 0.363615762 | 0.081880363 | 0.060615737 | 0.050697953 | 0.034869641 | 0.019685591 | 0.348886949 |
| Xpnpep3 | S | 273 | -0.124815625 | 0.156919775 | 0.238800138 | 0.363615762 | 0.081880363 | 0.060615737 | 0.050697953 | 0.034869641 | 0.019685591 | 0.348886949 |
| Xpnpep3 | T | 262 | -0.124815625 | 0.156919775 | 0.238800138 | 0.363615762 | 0.081880363 | 0.060615737 | 0.050697953 | 0.034869641 | 0.019685591 | 0.348886949 |
| Xpnpep3 | T | 269 | -0.124815625 | 0.156919775 | 0.238800138 | 0.363615762 | 0.081880363 | 0.060615737 | 0.050697953 | 0.034869641 | 0.019685591 | 0.348886949 |
| Ahdc1 | S | 1060 | -0.0221153 | -0.009059084 | 0.185925937 | 0.208041237 | 0.194985021 | 0.486795112 | 0.918313936 | 0.000904283 | 0.002364011 | 0.117921068 |
| Rbl1 | S | 696 | -0.084038401 | 0.145417687 | 0.253857929 | 0.337896331 | 0.108440242 | 0.392181567 | 0.038331626 | 0.266607414 | 0.171380222 | 0.58749486 |
| Actn4 | S | 657 | -0.015773225 | 0.034453754 | 0.193350884 | 0.20912411 | 0.158897131 | 0.789192092 | 0.528811526 | 0.015093844 | 0.02493585 | 0.045635515 |
| Hjurp A730008H23Ril | S | 567 | 0.041888524 | 0.080746161 | 0.04640895 | 0.004520426 | -0.034337211 | 0.108137907 | 0.071515971 | 0.350340991 | 0.914842179 | 0.501000122 |
| Ctnnd1 | S | 125 | -0.01725631 | 0.208429911 | 0.053173104 | 0.070429414 | -0.155256807 | 0.935564726 | 0.021089308 | 0.748993992 | 0.778670036 | 0.388494223 |
| Adgrl3 | S | 1268 | -0.014656806 | 0.018005678 | 0.174637761 | 0.189294566 | 0.156632082 | 0.724228348 | 0.678774433 | 0.004472301 | 0.000614636 | 0.004616278 |
| Tiam2 | S | 1604 | 0.011150425 | 0.052866611 | 0.106946456 | 0.095796031 | 0.054079845 | 0.641841608 | 0.444530206 | 0.009077325 | 0.020518862 | 0.435963506 |
| Carmil1 | S | 1103 | 0.136517303 | 0.071761575 | 0.011583839 | -0.124933463 | -0.060177736 | 0.068809678 | 0.202675671 | 0.854332325 | 0.093030153 | 0.263752039 |
| Radil | S | 235 | 0.114313921 | 0.136812751 | -0.005997029 | -0.12031095 | -0.14280978 | 0.175231746 | 0.101305279 | 0.930687243 | 0.207275332 | 0.135814765 |
| Brf1 | S | 552 | -0.022218088 | 0.108761524 | 0.054243119 | 0.076461207 | -0.054518405 | 0.569766914 | 0.00101521 | 0.375718917 | 0.260452297 | 0.371299129 |
| Limd1 | S | 308 | 0.083104189 | 0.005205608 | 0.140043652 | 0.056939463 | 0.134838044 | 0.240859144 | 0.865269687 | 0.068378489 | 0.454636177 | 0.097157114 |
| Ppp1r12a | S | 861 | 0.014797938 | 0.040120843 | 0.053939181 | 0.039141243 | 0.013818338 | 0.466718442 | 0.030031351 | 0.009225253 | 0.081982239 | 0.187053763 |
| Psen1 | S | 329 | 0.07234333 | 0.08765019 | 0.001426996 | -0.070916334 | -0.086223194 | 0.075299211 | 0.033780024 | 0.971539876 | 0.090716069 | 0.053405604 |
| Huwe1 | S | 3932 | 0.074153934 | 0.263654623 | -0.051108875 | -0.125262808 | -0.314763498 | 0.717818898 | 0.031784302 | 0.730514507 | 0.570854794 | 0.087367117 |
| Zfp462 | S | 1083 | -0.058529045 | 0.100649526 | 0.135298895 | 0.19382794 | 0.034649369 | 0.339935949 | 0.019762945 | 0.066185004 | 0.037759599 | 0.519382693 |
| Cdk13 | S | 401 | 0.059564983 | 0.018671914 | 0.157988788 | 0.098423805 | 0.139316874 | 0.578776379 | 0.673438462 | 0.03867874 | 0.396269825 | 0.062708644 |
| Snip1 | S | 91 | 0.024726873 | 0.027187417 | 0.062680986 | 0.037954113 | 0.035493569 | 0.255603089 | 0.490046137 | 0.000956043 | 0.121443968 | 0.383776627 |
| Gkap1 | S | 25 | -0.010904843 | 0.08806137 | 0.127797491 | 0.138702334 | 0.039736121 | 0.831011719 | 0.06286359 | 0.107254264 | 0.09400461 | 0.518267063 |
| Mastl | S | 560 | 0.181888747 | 0.054578416 | 0.029742558 | -0.152146188 | -0.024835857 | 0.037970467 | 0.660422226 | 0.88474391 | 0.475302881 | 0.90814797 |
| Dennd5a | T | 1079 | -0.042835223 | 0.316192842 | 0.086124793 | 0.128960016 | -0.230068049 | 0.75323616 | 0.131645456 | 0.372840866 | 0.401786255 | 0.227672387 |
| Nup98 | S | 1098 | 0.096330343 | -0.01883845 | 0.170826961 | 0.074496618 | 0.189665411 | 0.116710187 | 0.854432147 | 0.138093677 | 0.439385767 | 0.168965146 |
| Chd4 | T | 510 | 0.038879381 | 0.16065387 | 0.095614017 | 0.056734636 | -0.065039853 | 0.480437714 | 0.107271272 | 0.618651147 | 0.759116095 | 0.735476723 |
| Arhgap35 | S | 975 | 0.025584676 | 0.123694222 | 0.070837665 | 0.045252988 | -0.052856557 | 0.4696138 | 0.050898944 | 0.456762992 | 0.612720318 | 0.576237892 |
| Kif23 | S | 685 | 0.090188823 | 0.098067623 | 0.050791829 | -0.039396994 | -0.047275794 | 0.096501349 | 0.304514435 | 0.52509457 | 0.630405971 | 0.652620644 |
| Prkaa1 Prkaa2 | S | 187;176 | 0.034069062 | 0.169177773 | 0.185756689 | 0.151687627 | 0.016578916 | 0.763913265 | 0.320797277 | 0.34083143 | 0.444973168 | 0.93730572 |
| Otud7b | T | 730 | -0.09082901 | 0.108198141 | 0.197535761 | 0.288364772 | 0.08933762 | 0.164995182 | 0.085384712 | 0.02170498 | 0.006922432 | 0.160342793 |
| Morc2a | S | 859 | 0.120960148 | 0.032969072 | 0.041156548 | -0.0798036 | 0.008187476 | 0.067999207 | 0.190875716 | 0.465993291 | 0.251546101 | 0.873055642 |
| Mphosph10 | S | 164 | 0.212456146 | -0.056816744 | 0.112273814 | -0.100182332 | 0.169090558 | 0.258890867 | 0.494974888 | 0.024563853 | 0.543063399 | 0.112274626 |
| Triobp | S | 1603 | 0.02772206 | 0.087737298 | 0.132729497 | 0.105007437 | 0.044992199 | 0.775639806 | 0.183757057 | 0.139409641 | 0.353177505 | 0.570781393 |
| U2surp | S | 948 | 0.020336676 | 0.043801097 | 0.100154842 | 0.079818166 | 0.056353745 | 0.655936748 | 0.115506172 | 0.034566471 | 0.159738175 | 0.154373257 |
| Aak1 | S | 676 | 0.048762112 | 0.110053114 | -0.006154502 | -0.054916614 | -0.116207616 | 0.162033884 | 0.009460227 | 0.832439405 | 0.089707351 | 0.009428567 |
| Sin3b | S | 115 | 0.06083849 | 0.191006546 | 0.016493379 | -0.04434511 | -0.174513167 | 0.469129306 | 0.068992276 | 0.854147776 | 0.483371218 | 0.066535416 |
| Spag9 | T | 365 | 0.02167162 | 0.291095527 | -0.025453384 | -0.047125004 | -0.316548911 | 0.844464151 | 0.038002073 | 0.881457784 | 0.740737571 | 0.118012969 |
| Cic | T | 2221 | 0.022555074 | 0.039919197 | 0.099831606 | 0.077276532 | 0.059912409 | 0.561781239 | 0.152642668 | 0.03012975 | 0.117307589 | 0.125672937 |
| Pola2 | S | 152 | -0.00502691 | 0.107252151 | 0.041939779 | 0.046966689 | -0.065312372 | 0.806091736 | 0.002249315 | 0.373066154 | 0.330631475 | 0.209518019 |
| Ncoa7 | S | 214 | -0.015948746 | 0.142799781 | 0.082574805 | 0.098523551 | -0.060224976 | 0.87582169 | 0.034382991 | 0.264780749 | 0.36341821 | 0.314323101 |
| Kmt2b | S | 1040 | -0.035295348 | 0.134710751 | 0.07025082 | 0.105546168 | -0.064459931 | 0.377949968 | 0.017014072 | 0.077948904 | 0.028075304 | 0.143656886 |
| Hirip3 | T | 135 | 0.032244439 | 0.066064111 | 0.041606376 | 0.009361937 | -0.024457735 | 0.298329323 | 0.013600004 | 0.226869098 | 0.782295437 | 0.408804354 |
| Armc10 | S | 43 | -0.038913379 | 0.006605041 | 0.204125429 | 0.243038807 | 0.197520387 | 0.129479416 | 0.836023799 | 0.000631289 | 0.003142932 | 0.003057912 |
| Bnc2 | S | 431 | 0.098684855 | -0.037978812 | 0.158540973 | 0.059856118 | 0.196519785 | 0.008531966 | 0.050843521 | 0.082384796 | 0.365841127 | 0.057721509 |
| Nde1 | S | 282 | 0.21956771 | 0.133091512 | -0.156534017 | -0.376101727 | -0.289625529 | 0.003215083 | 0.020557774 | 0.008507686 | 0.002355522 | 0.00116663 |
| Zc3h13 | S | 1256 | 0.063099104 | 0.021579216 | 0.117993062 | 0.054893958 | 0.096413846 | 0.260781088 | 0.709662659 | 0.047261918 | 0.335568545 | 0.156540132 |
| Flnb | S | 1988 | 0.102717931 | 0.046854174 | 0.027461199 | -0.075256732 | -0.019392975 | 0.021919749 | 0.383144567 | 0.495890301 | 0.137590909 | 0.7349332 |
| Mageh1 | T | 55 | -0.10817594 | -0.013958422 | 0.281233293 | 0.389409233 | 0.295191715 | 0.50223235 | 0.876463213 | 0.002636652 | 0.095177801 | 0.046520659 |
| Tjp2 | T | 1045 | 0.087168064 | 0.089948135 | 0.134713087 | 0.047545023 | 0.044764952 | 0.482532008 | 0.42883591 | 0.224808672 | 0.683168975 | 0.670818825 |
| Atf7ip | S | 290 | 0.001066545 | 0.254009245 | 0.040409504 | 0.039342958 | -0.213599741 | 0.995323001 | 0.057587227 | 0.689850675 | 0.824463721 | 0.09282027 |
| Per3 | S | 909 | 0.068386242 | 0.177090247 | 0.048467981 | -0.01991826 | -0.128622266 | 0.739408419 | 0.053665511 | 0.774140141 | 0.933293437 | 0.460330237 |
| Per3 | S | 910 | 0.068386242 | 0.177090247 | 0.048467981 | -0.01991826 | -0.128622266 | 0.739408419 | 0.053665511 | 0.774140141 | 0.933293437 | 0.460330237 |

| | | | | | | | | | | | | | |
|---|---|---|---|---|---|---|---|---|---|---|---|---|---|
| Per3 | S | | 911 | 0.068386242 | 0.177090247 | 0.048467981 | -0.01991826 | -0.128622266 | 0.739408419 | 0.053665511 | 0.774140141 | 0.933293437 | 0.460330237 |
| Tmpo | S | | 183 | -0.141769233 | 0.164896308 | 0.163632765 | 0.305401997 | -0.001263543 | 0.048358256 | 0.004642943 | 0.055146068 | 0.009971789 | 0.983323531 |
| Mtfr1l | S | | 234 | -0.019062759 | 0.176901009 | 0.141306489 | 0.160369248 | -0.035594519 | 0.63549758 | 0.095863815 | 0.281549438 | 0.235925376 | 0.776537432 |
| Nav1 | S | | 315 | -0.007850396 | 0.123899122 | 0.083300554 | 0.091150951 | -0.040598568 | 0.811517081 | 0.032745161 | 0.204366394 | 0.176795125 | 0.514176159 |
| Sh3bp5l | S | | 349 | -0.046074548 | 0.205341609 | 0.063512929 | 0.109587477 | -0.14182868 | 0.452268731 | 0.012106245 | 0.490343462 | 0.266483953 | 0.175675353 |
| Erbin | S | | 1125 | -0.033745325 | 0.097674687 | 0.091290393 | 0.125035719 | -0.006384294 | 0.182563842 | 0.00737963 | 0.052191511 | 0.039671442 | 0.850980145 |
| Xrn2 | S | | 501 | 0.003247603 | 0.045683902 | 0.084486756 | 0.081239153 | 0.038802854 | 0.926442549 | 0.08904107 | 0.010018491 | 0.071514084 | 0.03765956 |
| Ccm2 | S | | 425 | 0.06243897 | 0.150447419 | 0.02205997 | -0.040379001 | -0.128387449 | 0.18524877 | 0.130791591 | 0.670063098 | 0.484462951 | 0.179954637 |
| Tjp2 | S | | 1136 | -0.040716323 | -0.027179641 | 0.201805317 | 0.24252164 | 0.228984958 | 0.078618281 | 0.181027089 | 1.26136E-05 | 0.000110384 | 5.52069E-05 |
| Golga1 | S | | 36 | 0.05583181 | 0.062024317 | 0.152696268 | 0.096864458 | 0.090671951 | 0.56440941 | 0.484721612 | 0.109877605 | 0.324973491 | 0.309501939 |
| Fbxo38 | S | | 607 | -0.007370042 | 0.114313911 | 0.099586775 | 0.106956817 | -0.014727136 | 0.908593919 | 0.041505515 | 0.210415951 | 0.243815877 | 0.833441536 |
| Pura | T | | 182 | 0.061838901 | 0.165980393 | 0.060900231 | -0.00093867 | -0.105080161 | 0.469814351 | 0.178374032 | 0.453693924 | 0.990120619 | 0.348603562 |
| Cherp | S | | 822 | 0.08630947 | 0.013894717 | 0.028884531 | -0.057424939 | 0.014989814 | 0.002139067 | 0.453127019 | 0.521694431 | 0.251018206 | 0.720991208 |
| Rprd2 | S | | 926 | 0.099523776 | 0.371089871 | -0.00387591 | -0.103399685 | -0.374965781 | 0.56699255 | 0.230165355 | 0.991610964 | 0.776749108 | 0.385720456 |
| Mllt3 | S | | 420 | -0.03993226 | 0.004886799 | 0.159032583 | 0.198964842 | 0.154145784 | 0.280590456 | 0.880703024 | 0.000183367 | 0.014498623 | 0.021202989 |
| Bclaf1 | S | | 183 | -0.007201099 | 0.015420249 | 0.152085001 | 0.1592861 | 0.136664753 | 0.864594219 | 0.722633445 | 0.003358274 | 0.012265427 | 0.022393607 |
| Rbm8a Rbm8a2 | S | | 166 | 0.131733122 | 0.025595422 | 0.083885146 | -0.047847976 | 0.058289724 | 0.158512174 | 0.64304531 | 0.196903099 | 0.5740953 | 0.394251321 |
| Rbm8a Rbm8a2 | S | | 168 | 0.131733122 | 0.025595422 | 0.083885146 | -0.047847976 | 0.058289724 | 0.158512174 | 0.64304531 | 0.196903099 | 0.5740953 | 0.394251321 |
| Mybl2 | S | | 396 | 0.21122277 | 0.053623649 | 0.169177135 | -0.042045635 | 0.115553486 | 0.315900483 | 0.657871608 | 0.536160837 | 0.882950547 | 0.653023086 |
| Mybl2 | T | | 408 | 0.21122277 | 0.053623649 | 0.169177135 | -0.042045635 | 0.115553486 | 0.315900483 | 0.657871608 | 0.536160837 | 0.882950547 | 0.653023086 |
| Gemin8 | S | | 120 | 0.011019904 | 0.061511158 | 0.080314328 | 0.069294424 | 0.01880317 | 0.701067863 | 0.366351107 | 0.004678839 | 0.08008699 | 0.760353285 |
| Gemin8 | S | | 122 | 0.011019904 | 0.061511158 | 0.080314328 | 0.069294424 | 0.01880317 | 0.701067863 | 0.366351107 | 0.004678839 | 0.08008699 | 0.760353285 |
| Usp39 | S | | 81 | 0.017754593 | 0.034428845 | 0.089516187 | 0.071761593 | 0.055087342 | 0.555630584 | 0.350456366 | 0.006147867 | 0.074711863 | 0.181028415 |
| Dock1 | S | | 1743 | 0.062505294 | 0.066570228 | 0.051753198 | -0.010752096 | -0.01481703 | 0.040193734 | 0.231812364 | 0.437065893 | 0.862249471 | 0.835672759 |
| Carmil1 | S | | 1255 | 0.088068126 | 0.066290396 | 0.035949221 | -0.052118906 | -0.030341175 | 0.127060424 | 0.140522589 | 0.407387721 | 0.254917733 | 0.180148162 |
| Atrx | S | | 669 | -0.009056192 | 0.145937116 | 0.074295974 | 0.083352166 | -0.071641142 | 0.819317704 | 0.027415018 | 0.337376817 | 0.301372386 | 0.368444301 |
| Bckdk | T | | 32 | 0.006412029 | 0.084400495 | 0.250562091 | 0.244150062 | 0.166161597 | 0.95163953 | 0.391917925 | 0.169764902 | 0.173870304 | 0.306797142 |
| Dlg5 | S | | 132 | 0.065113807 | -0.025191766 | 0.202725047 | 0.13761124 | 0.227916814 | 0.284146694 | 0.557612772 | 0.042401038 | 0.124949309 | 0.054454231 |
| Plec | S | | 1729 | -0.140930608 | 0.178025846 | 0.137685248 | 0.278615857 | -0.040340598 | 0.029172726 | 0.006305818 | 0.022844462 | 0.00246719 | 0.23549488 |
| Wapl | S | | 226 | -0.016252103 | 0.071164883 | 0.078843092 | 0.095095195 | 0.007678209 | 0.728571162 | 0.105293576 | 0.005894525 | 0.128445564 | 0.813317611 |
| Ring1 | S | | 248 | -0.014972935 | 0.126464533 | 0.162693282 | 0.177666217 | 0.036228749 | 0.919339517 | 0.121629548 | 0.237105687 | 0.321874548 | 0.755991968 |
| Ppig | S | | 685 | -0.026348319 | 0.080275694 | 0.1008495 | 0.127197819 | 0.020573806 | 0.46470061 | 0.00344888 | 0.157699485 | 0.098365223 | 0.70950768 |
| Arfgef2 | S | | 621 | 0.113354557 | 0.184401932 | -0.032737573 | -0.146092129 | -0.217139505 | 0.228014587 | 0.104063356 | 0.775117186 | 0.287098039 | 0.150405103 |
| Dido1 | S | | 1016 | -0.034927142 | 0.046565707 | 0.150317317 | 0.185244458 | 0.10375161 | 0.341720814 | 0.050641643 | 0.016594999 | 0.009795029 | 0.0547498 |
| Gpkow | S | | 144 | 0.032362799 | 0.013052159 | 0.120186415 | 0.087823616 | 0.107134256 | 0.667827502 | 0.710189824 | 0.006147149 | 0.29260392 | 0.026276313 |
| Apbb2 | S | | 409 | -0.025278264 | 0.093684861 | 0.159559791 | 0.184838055 | 0.06587493 | 0.324692216 | 0.068999808 | 0.082182878 | 0.068198848 | 0.362796833 |
| Rbmx2 | S | | 227 | 0.048277558 | 0.041954293 | 0.128700972 | 0.080423414 | 0.086746679 | 0.372477066 | 0.583971735 | 0.064164364 | 0.236675071 | 0.324780723 |
| U2surp | S | | 946 | 0.021182481 | 0.04231907 | 0.100723176 | 0.079540695 | 0.058404106 | 0.662905607 | 0.126320884 | 0.038307477 | 0.183487707 | 0.153699193 |
| Akap13 | S | | 1908 | 0.044382604 | 0.151832139 | 0.101419311 | 0.057036706 | -0.050412828 | 0.568610179 | 0.154587589 | 0.483625658 | 0.686092422 | 0.732580053 |
| Tjp1 | S | | 841 | -0.011713174 | 0.096914955 | 0.174913476 | 0.18662665 | 0.077998521 | 0.929857395 | 0.375900347 | 0.069353584 | 0.241756062 | 0.494103295 |
| Bclaf1 | S | | 177 | -0.003678224 | 0.053643385 | 0.092632211 | 0.096310434 | 0.038988826 | 0.900954235 | 0.123941927 | 0.010928453 | 0.011517309 | 0.161063213 |
| 4921524J17Rik | S | | 50 | 0.113731725 | 0.032543482 | 0.019708852 | -0.094022873 | -0.01283463 | 0.023771525 | 0.338933685 | 0.470083295 | 0.058036485 | 0.618467038 |
| Sp110 | S | | 177 | 0.15717415 | 0.105338135 | 0.097567862 | -0.059606288 | -0.007770274 | 0.157624164 | 0.551175452 | 0.73912153 | 0.840036113 | 0.980423774 |
| Mapkbp1 | S | | 1252 | -0.009508522 | 0.106322828 | 0.089123102 | 0.098631624 | -0.017199726 | 0.875661819 | 0.092730408 | 0.064875008 | 0.145414621 | 0.696693572 |
| Usp7 | S | | 964 | 0.188885175 | 0.053713061 | 0.020478437 | -0.168406738 | -0.033234624 | 0.066529555 | 0.554431222 | 0.800205441 | 0.065427031 | 0.633810004 |
| Rbm39 | S | | 97 | -0.016652837 | 0.025276475 | 0.107801923 | 0.12445476 | 0.082525449 | 0.703944616 | 0.015137889 | 0.009074604 | 0.064614236 | 0.031642599 |
| Srsf2 | S | | 189 | 0.055547034 | 0.034374843 | 0.101451808 | 0.045904774 | 0.067076964 | 0.157105037 | 0.346225711 | 0.136865261 | 0.415987091 | 0.270542748 |
| Pdlim2 | S | | 87 | 0.048501044 | 0.055168008 | 0.017535287 | -0.030965757 | -0.037632721 | 0.031049653 | 0.014660568 | 0.769777392 | 0.609469651 | 0.54040907 |
| Myo18a | S | | 2037 | -0.006867024 | 0.061009789 | 0.068103887 | 0.07497091 | 0.007094098 | 0.821671107 | 0.00505648 | 0.066717897 | 0.088030282 | 0.781515606 |
| Strn3 | S | | 257 | -0.014149251 | 0.034071251 | 0.083001165 | 0.097150416 | 0.048929914 | 0.746797729 | 0.114878623 | 0.001073166 | 0.113512163 | 0.0409089 |
| Phactr4 | S | | 174 | 0.025356681 | 0.137458919 | 0.008041886 | -0.017314795 | -0.129417033 | 0.443570217 | 0.013230286 | 0.769351206 | 0.591131077 | 0.021035618 |
| Psmd1 | T | | 311 | 0.011500549 | 0.063003023 | 0.089755693 | 0.078255145 | 0.02675267 | 0.627477018 | 0.134784641 | 0.042815064 | 0.077510406 | 0.512668121 |
| Srrm2 | S | | 1048 | -0.001407292 | 0.017437672 | 0.10742581 | 0.108833102 | 0.089988138 | 0.980039692 | 0.390459168 | 0.001468739 | 0.148765831 | 0.000478531 |
| Anapc2 | T | | 481 | -0.024668677 | 0.172593864 | 0.201677288 | 0.226345965 | 0.029083424 | 0.895414226 | 0.383258661 | 0.401513051 | 0.892977054 | |
| Kmt2a | T | | 504 | -0.015265528 | 0.098168143 | 0.067832782 | 0.08309831 | -0.030335362 | 0.855619743 | 0.014174503 | 0.145528953 | 0.37349171 | 0.454195572 |
| Tmem87b | S | | 470 | 0.085779869 | -0.045806414 | 0.114655889 | 0.02887602 | 0.160462303 | 0.043743135 | 0.431412787 | 0.028985965 | 0.472721835 | 0.047601358 |
| Slc12a4 | S | | 24 | 0.10156252 | 0.05471078 | 0.093230186 | -0.008332334 | 0.038519406 | 0.305840648 | 0.421764816 | 0.187841269 | 0.925391377 | 0.543732825 |
| Cry1 | S | | 183 | 0.198911494 | 0.025830826 | 0.056379171 | -0.142532323 | 0.030548346 | 0.071815746 | 0.79467577 | 0.642441143 | 0.196606118 | 0.747887242 |
| Brd9 | S | | 103 | -0.036916044 | 0.099302131 | 0.10149322 | 0.138409265 | 0.002191089 | 0.36829174 | 0.011656245 | 0.097737146 | 0.04846866 | 0.961346599 |
| Irf2bp2 | S | | 440 | 0.023030545 | 0.040876899 | 0.087422453 | 0.064391908 | 0.046545555 | 0.361539484 | 0.09557205 | 0.06033434 | 0.131419841 | 0.232461386 |
| Srrm2 | S | | 470 | 0.085640855 | 0.0134017 | 0.123635796 | 0.037994941 | 0.110234097 | 0.30054699 | 0.850982254 | 0.059355284 | 0.634260059 | 0.200842042 |
| U2surp | S | | 974 | 0.035440637 | -0.009981886 | 0.112981304 | 0.077540668 | 0.122963191 | 0.570601326 | 0.854935009 | 0.001513056 | 0.267309482 | 0.107767798 |
| Ranbp3 | S | | 472 | -0.00158334 | 0.084057647 | 0.129959655 | 0.131542994 | 0.045902007 | 0.974067224 | 0.128877931 | 0.098022455 | 0.102983577 | 0.472523547 |
| Srek1 | S | | 389 | 0.111255902 | 0.021016265 | 0.138160109 | 0.026904207 | 0.117143844 | 0.343856096 | 0.811658102 | 0.114663441 | 0.814727709 | 0.286281054 |
| Srek1 | S | | 391 | 0.111255902 | 0.021016265 | 0.138160109 | 0.026904207 | 0.117143844 | 0.343856096 | 0.811658102 | 0.114663441 | 0.814727709 | 0.286281054 |
| Srek1 | S | | 393 | 0.111255902 | 0.021016265 | 0.138160109 | 0.026904207 | 0.117143844 | 0.343856096 | 0.811658102 | 0.114663441 | 0.814727709 | 0.286281054 |
| Eif3g | T | | 41 | 0.039668377 | -0.040526578 | 0.136091343 | 0.096422966 | 0.17661792 | 0.208132512 | 0.293391329 | 0.001708964 | 0.018456169 | 0.008279497 |
| Irf2bp1 | S | | 66 | 0.065064374 | 0.19463346 | 0.037558226 | -0.027506149 | -0.157075234 | 0.355751248 | 0.145087817 | 0.823295804 | 0.863767665 | 0.407015475 |
| Itprid2 | S | | 1251 | -0.088172856 | 0.207145931 | 0.073555835 | 0.161728691 | -0.133590096 | 0.030305245 | 0.000589933 | 0.366802242 | 0.120690923 | 0.157473597 |
| Mta1 | S | | 576 | 0.205079268 | 0.037782739 | -0.101988111 | -0.307067379 | -0.13977085 | 0.000524985 | 0.300940764 | 0.086027267 | 0.012585606 | 0.043070141 |
| Usp24 | S | | 1138 | -0.026789393 | 0.053466689 | 0.063623813 | 0.090413206 | 0.010157124 | 0.260204827 | 0.00260605 | 0.004960504 | 0.01780368 | 0.462674095 |
| Ptov1 | S | | 36 | -0.017186623 | 0.166654236 | 0.0036875 | 0.020874123 | -0.162966735 | 0.782317709 | 0.002335403 | 0.941553547 | 0.750858737 | 0.029333601 |
| Arhgap21 | T | | 1846 | 0.150508813 | 0.091723068 | 0.033381002 | -0.117127811 | -0.058342065 | 0.090264577 | 0.427208908 | 0.850015126 | 0.505123026 | 0.74704927 |
| Dlg4 | S | | 295 | 0.028960737 | 0.063057059 | 0.062347217 | 0.03338648 | -0.000709842 | 0.615222344 | 0.067136806 | 0.056565813 | 0.558458808 | 0.978646781 |
| Tti2 | S | | 504 | 0.045390585 | 0.144776365 | 0.015921025 | -0.02946956 | -0.12885534 | 0.61854937 | 0.020791944 | 0.810651852 | 0.75799927 | 0.105725478 |
| Resf1 | S | | 1514 | -0.073911725 | 0.165601457 | 0.110456102 | 0.184367826 | -0.055145356 | 0.046763113 | 0.020383258 | 0.028467582 | 0.008335207 | 0.299230524 |
| Resf1 | S | | 1518 | -0.073911725 | 0.165601457 | 0.110456102 | 0.184367826 | -0.055145356 | 0.046763113 | 0.020383258 | 0.028467582 | 0.008335207 | 0.299230524 |
| Raver1 Gm38431 | S | | 626 | 0.009301622 | 0.049741913 | 0.075046267 | 0.065744645 | 0.025304354 | 0.615122745 | 0.042441552 | 0.040640907 | 0.076717018 | 0.36237989 |
| Actb Actg1 | Y | 218;218 | | 0.269591337 | -0.054032305 | 0.207498917 | -0.06209242 | 0.261531222 | 0.159031536 | 0.682330394 | 0.474532314 | 0.827797751 | 0.376658623 |
| Gys1 | S | | 711 | -0.038566131 | 0.212312967 | 0.119117198 | 0.157683329 | -0.09319577 | 0.650940572 | 0.060643374 | 0.444594586 | 0.328362993 | 0.547714881 |
| Pkp4 | S | | 272 | 0.082828337 | 0.038553923 | 0.061908052 | -0.020920284 | 0.02335413 | 0.093277364 | 0.290101631 | 0.253967296 | 0.669708931 | 0.587422378 |
| Tjp2 | S | | 947 | 0.041492508 | 0.059397608 | 0.140357718 | 0.09886521 | 0.08096011 | 0.369242786 | 0.209187521 | 0.244852056 | 0.376136368 | 0.453469353 |
| Slu7 | S | | 512 | -0.028445041 | 0.241180433 | 0.136447418 | 0.164892459 | -0.104733015 | 0.803002241 | 0.239940646 | 0.234732352 | 0.268362993 | 0.577279908 |
| Tbc1d10b | T | | 140 | 0.081918501 | 0.119959784 | 0.053208721 | -0.028709781 | -0.066751064 | 0.444879053 | 0.090843914 | 0.624701672 | 0.828357177 | 0.555163331 |
| Zbtb11 Rpl24 | S | | 473 | 0.025197651 | 0.078321331 | 0.102213306 | 0.077015655 | 0.023891976 | 0.455792956 | 0.138973868 | 0.180526009 | 0.280187075 | 0.72782572 |
| Zic5 | S | | 537 | 0.449114084 | -0.061020149 | -0.102895642 | -0.552009726 | -0.041875494 | 0.020754196 | 0.750058715 | 0.585456792 | 0.03148279 | 0.836605229 |
| Zic5 | S | | 541 | 0.449114084 | -0.061020149 | -0.102895642 | -0.552009726 | -0.041875494 | 0.020754196 | 0.750058715 | 0.585456792 | 0.03148279 | 0.836605229 |
| Zic5 | S | | 559 | 0.449114084 | -0.061020149 | -0.102895642 | -0.552009726 | -0.041875494 | 0.020754196 | 0.750058715 | 0.585456792 | 0.03148279 | 0.836605229 |
| Srrm2 | S | | 1535 | 0.042173942 | 0.01057427 | 0.059193517 | 0.017019575 | 0.048619247 | 0.01862256 | 0.831290828 | 0.01531486 | 0.297595878 | 0.374188919 |
| Chchd2 | S | | 41 | 0.139542837 | 0.01930394 | 0.102768614 | -0.036774223 | 0.083464673 | 0.140515494 | 0.717205802 | 0.307755432 | 0.726418789 | 0.408981765 |
| Ksr1 | S | | 518 | 0.160803581 | 0.183632148 | 0.011756967 | -0.149046613 | -0.171875181 | 0.316128272 | 0.259710495 | 0.955016608 | 0.532025498 | 0.474340318 |
| Cobll1 | S | | 300 | -0.074224894 | 0.182240045 | 0.080162975 | 0.154387868 | -0.10207707 | 0.208287093 | 0.018181556 | 0.091565439 | 0.0447956 | 0.101431198 |
| Atrx | S | | 956 | -0.024355789 | 0.075910928 | 0.135977744 | 0.160333533 | 0.060066816 | 0.5820448 | 0.199992539 | 0.022963865 | 0.013418877 | 0.278445799 |
| Nhsl1 | S | | 340 | 0.116795056 | 0.126844198 | 0.033191002 | -0.083604054 | -0.093653196 | 0.204191744 | 0.233227941 | 0.782817947 | 0.475664386 | 0.457410004 |

| | | | | | | | | | | | | |
|---|---|---|---|---|---|---|---|---|---|---|---|---|
| Srsf4 | S | 179 | 0.035813828 | 0.045984912 | 0.114910255 | 0.079096427 | 0.068925343 | 0.588286009 | 0.174703887 | 0.087813651 | 0.307583557 | 0.240712824 |
| Hacd3 | S | 114 | 0.042860992 | 0.010931403 | 0.137603311 | 0.094742319 | 0.126671908 | 0.186892116 | 0.756686728 | 0.057913199 | 0.148866391 | 0.078580073 |
| Klf6 | S | 192 | -0.013247175 | 0.151377777 | 0.045948644 | 0.059195819 | -0.105429134 | 0.763160059 | 0.01541527 | 0.368012839 | 0.174211547 | 0.059974771 |
| Tubg1 Tubg2 | S | 284;284 | -0.07236494 | 0.097636465 | 0.188780204 | 0.261145144 | 0.091143739 | 0.32530725 | 0.10505359 | 0.044965109 | 0.026497056 | 0.238289853 |
| Ctnna1 | S | 641 | -0.02471427 | 0.11721777 | 0.071381767 | 0.096096037 | -0.045836003 | 0.562957104 | 0.02034325 | 0.115614333 | 0.092074562 | 0.308351215 |
| Mki67 | S | 2052 | 0.179864965 | 0.088108445 | 0.077774677 | -0.102090287 | -0.010333768 | 0.335098828 | 0.6071545 | 0.324956813 | 0.549687592 | 0.948476142 |
| Ythdc1 | T | 148 | 0.0215988 | 0.088965818 | 0.109982614 | 0.088383815 | 0.021016796 | 0.772273954 | 0.090999577 | 0.213223011 | 0.366394461 | 0.782356361 |
| Prr12 | S | 1372 | -0.000117387 | 0.139094019 | 0.13309496 | 0.133212347 | -0.005999059 | 0.997742649 | 0.08198815 | 0.378995925 | 0.378221427 | 0.966025141 |
| Prr12 | S | 1373 | -0.000117387 | 0.139094019 | 0.13309496 | 0.133212347 | -0.005999059 | 0.997742649 | 0.08198815 | 0.378995925 | 0.378221427 | 0.966025141 |
| Srrm1 | S | 550 | 0.068148651 | 0.049203477 | 0.095149308 | 0.027000656 | 0.04594583 | 0.333399827 | 0.169209069 | 0.220782158 | 0.74290733 | 0.500371584 |
| Gas2l3 | S | 367 | 0.162428108 | 0.043869881 | 0.002620574 | -0.159807534 | -0.041249307 | 0.022529389 | 0.581914122 | 0.96744986 | 0.028610979 | 0.574666169 |
| Cpsf6 | T | 407 | 0.012307731 | 0.058699197 | 0.106408826 | 0.094101094 | 0.047709629 | 0.750657775 | 0.098678715 | 0.078798177 | 0.112244643 | 0.327277108 |
| Scrib | T | 1527 | 0.013197282 | 0.007018517 | 0.340807902 | 0.32761062 | 0.333789385 | 0.892764445 | 0.957349584 | 0.096912996 | 0.124655508 | 0.10905594 |
| Acbd3 | T | 125 | 0.025560591 | 0.159439296 | 0.037364936 | 0.011804345 | -0.12207436 | 0.425967306 | 0.085727835 | 0.443748957 | 0.78372899 | 0.154270203 |
| Trp53bp2 | S | 697 | -0.044428363 | 0.045499593 | 0.176939659 | 0.221368022 | 0.131440065 | 0.275626614 | 0.213073605 | 0.008260459 | 0.006179574 | 0.031555487 |
| Polr2a | S | 1850 | 0.101540878 | 0.055451293 | 0.038654474 | -0.062886404 | -0.016796819 | 0.100981144 | 0.164974956 | 0.565117447 | 0.393506538 | 0.785030627 |
| Arid4a | S | 793 | -0.05259798 | 0.070096386 | 0.22568629 | 0.27828427 | 0.155589904 | 0.699555813 | 0.194222302 | 0.085477327 | 0.128224644 | 0.168890789 |
| Resf1 | S | 1142 | 0.0083227 | 0.105121331 | 0.071541866 | 0.063219166 | -0.033579465 | 0.861715288 | 0.04985235 | 0.180620583 | 0.175877083 | 0.410737173 |
| Supt6 | T | 1697 | -0.011547886 | 0.175267755 | 0.077044922 | 0.088592808 | -0.098222833 | 0.737629473 | 0.05404412 | 0.33398981 | 0.277735615 | 0.286833132 |
| Nav3 | S | 1504 | 0.083249083 | 0.133417857 | -0.006110231 | -0.089359314 | -0.139528088 | 0.575381648 | 0.01552737 | 0.96517567 | 0.635015219 | 0.371515725 |
| Nrip1 | S | 808 | 0.145895121 | 0.059216679 | -0.033311281 | -0.179206401 | -0.09252796 | 0.007571152 | 0.572781219 | 0.822208405 | 0.296197577 | 0.586410657 |
| Nfat5 | S | 559 | 0.107333989 | 0.03493613 | 0.037281899 | -0.070052091 | 0.002345769 | 0.026481355 | 0.383898896 | 0.60123697 | 0.344909478 | 0.971948779 |
| Akt1 | T | 448 | 0.064463572 | 0.109503482 | 0.022074697 | -0.042388874 | -0.087428785 | 0.401465051 | 0.027357445 | 0.779607042 | 0.647702104 | 0.300831757 |
| Api5 | S | 464 | -0.027682579 | 0.092510136 | 0.081511623 | 0.109194202 | -0.010998513 | 0.384348985 | 0.009655976 | 0.081597303 | 0.040957891 | 0.748825539 |
| Trio | S | 1786 | -0.115018507 | 0.169706642 | 0.178421822 | 0.293440329 | 0.008715181 | 0.035348497 | 0.058790923 | 0.014246878 | 0.013225205 | 0.898839583 |
| Ercc5 | S | 382 | 0.203997769 | 0.089203361 | 0.040772772 | -0.163224996 | -0.048430589 | 0.175892527 | 0.521815065 | 0.752889073 | 0.335732451 | 0.770591297 |
| Cdk12 | S | 332 | 0.024705144 | 0.016326851 | 0.171417073 | 0.146711929 | 0.155090222 | 0.762662159 | 0.758829431 | 0.025148715 | 0.11765524 | 0.034990379 |
| Fnbp1l | S | 488 | -0.016453639 | 0.201571262 | -0.01918431 | -0.00273067 | -0.220755571 | 0.760596717 | 0.003629572 | 0.860699736 | 0.979477249 | 0.131793882 |
| Arhgap21 | S | 453 | 0.008964276 | 0.115206071 | 0.052131307 | 0.043167031 | -0.063074764 | 0.865940736 | 0.011791161 | 0.492500908 | 0.58508296 | 0.402505054 |
| Amfr | S | 509 | -0.042786399 | 0.249334897 | 0.099279229 | 0.142065627 | -0.150055668 | 0.643613518 | 0.209057668 | 0.124257628 | 0.195138221 | 0.394403352 |
| Nktr | S | 470 | -0.013583393 | 0.009121026 | 0.162446906 | 0.1760303 | 0.15332588 | 0.843324574 | 0.842475982 | 0.003878695 | 0.07752267 | 0.02022865 |
| Rflnb | S | 197 | 0.128433222 | 0.227207467 | 0.058012397 | -0.070420826 | -0.169195071 | 0.555821684 | 0.247742242 | 0.832425457 | 0.774978167 | 0.478059902 |
| Rprd2 | S | 612 | -0.010715477 | 0.013401494 | 0.2225907 | 0.233306177 | 0.209189206 | 0.838228351 | 0.870481967 | 0.018698965 | 0.017908929 | 0.08073323 |
| Cwc22 | S | 26 | 0.002744293 | 0.067941223 | 0.115333586 | 0.112589294 | 0.047392363 | 0.953347081 | 0.389620294 | 0.019014247 | 0.064959556 | 0.531142955 |
| Cep350 | S | 1822 | -0.004023013 | 0.019888906 | 0.203637718 | 0.207660731 | 0.183748812 | 0.964330655 | 0.65380868 | 0.026382916 | 0.085877983 | 0.06474934 |
| Cep350 | S | 1824 | -0.004023013 | 0.019888906 | 0.203637718 | 0.207660731 | 0.183748812 | 0.964330655 | 0.65380868 | 0.026382916 | 0.085877983 | 0.06474934 |
| Maf1 | S | 68 | -0.039379138 | 0.198076033 | 0.058653252 | 0.09803239 | -0.139422781 | 0.480159986 | 0.014787323 | 0.51752668 | 0.30620601 | 0.181969525 |
| Prpf4b | Y | 849 | 0.026899713 | 0.01848794 | 0.102040311 | 0.075140598 | 0.083552371 | 0.653515731 | 0.454214809 | 0.006918548 | 0.264976535 | 0.018560332 |
| Srrm2 | S | 2019 | 0.023465992 | 0.035275297 | 0.107959895 | 0.084493903 | 0.072684598 | 0.342024289 | 0.276697763 | 0.050069584 | 0.10504242 | 0.140202394 |
| Srrm2 | S | 2023 | 0.023465992 | 0.035275297 | 0.107959895 | 0.084493903 | 0.072684598 | 0.342024289 | 0.276697763 | 0.050069584 | 0.10504242 | 0.140202394 |
| Zbtb44 | S | 194 | -0.022671029 | 0.051636335 | 0.173391985 | 0.196063014 | 0.12175565 | 0.566116027 | 0.462773747 | 0.01487314 | 0.012175717 | 0.153446124 |
| Rgl3 | S | 506 | 0.011112459 | 0.166091773 | 0.077968261 | 0.066855802 | -0.088123512 | 0.862220975 | 0.158597067 | 0.215130139 | 0.263025793 | 0.390736758 |
| Rgl3 | S | 510 | 0.011112459 | 0.166091773 | 0.077968261 | 0.066855802 | -0.088123512 | 0.862220975 | 0.158597067 | 0.215130139 | 0.263025793 | 0.390736758 |
| Rnf214 | S | 500 | -0.072882087 | 0.064557242 | 0.161780502 | 0.234662589 | 0.09722326 | 0.514902314 | 0.219623554 | 0.012025928 | 0.111320265 | 0.041377244 |
| Ints12 | S | 126 | 0.025344082 | 0.048754076 | 0.062071757 | 0.036727675 | 0.01331768 | 0.480776836 | 0.126360905 | 0.022747586 | 0.34028556 | 0.629965886 |
| Kars | S | 588 | -0.025401297 | 0.120312108 | 0.07877578 | 0.104177077 | -0.041536328 | 0.396992964 | 0.009331202 | 0.227565914 | 0.144797469 | 0.483772357 |
| Mpp6 | S | 109 | -0.025338564 | 0.089201179 | 0.155170709 | 0.180509273 | 0.065969531 | 0.380482987 | 0.11361879 | 0.063598113 | 0.050069872 | 0.336989428 |
| Sltm | S | 506 | -0.065143932 | 0.177673584 | 0.101734669 | 0.166878601 | -0.075938916 | 0.445924157 | 0.029182414 | 0.252285837 | 0.139445866 | 0.387330661 |
| Specc1l | S | 948 | 0.023831515 | 0.113968156 | 0.078692319 | 0.054860804 | -0.035275837 | 0.575680779 | 0.09981446 | 0.295350328 | 0.454418424 | 0.648133204 |
| Polr2a | S | 1899 | 0.057625455 | 0.062818947 | 0.05706705 | -0.000558406 | -0.005751897 | 0.210789099 | 0.128324224 | 0.257758762 | 0.991343381 | 0.904731452 |
| Rbm15 | S | 126 | -0.004007913 | 0.084267449 | 0.090996552 | 0.095004465 | 0.006729104 | 0.887045438 | 0.04481341 | 0.111083764 | 0.103974727 | 0.886356003 |
| Actb Actg1 | S | 232;232 | -0.062678848 | -0.030108132 | 0.340998129 | 0.403676977 | 0.371106261 | 0.844362984 | 0.802105281 | 0.025116074 | 0.271572651 | 0.01061272 |
| Mical1 | S | 777 | -0.077072756 | 0.169727816 | 0.147279699 | 0.224352455 | -0.022448116 | 0.027145648 | 0.142115833 | 0.004726194 | 0.001961814 | 0.796605764 |
| Baz1b | S | 158 | 0.016483468 | 0.094013659 | 0.03364753 | 0.017164062 | -0.060366129 | 0.600692081 | 0.025791368 | 0.140889788 | 0.526068122 | 0.112956165 |
| Cactin | S | 82 | 0.003261879 | 0.094859574 | 0.036285734 | 0.033023855 | -0.05857384 | 0.89791465 | 0.011411087 | 0.145079185 | 0.217025118 | 0.068821715 |
| Fam122a | S | 273 | 0.028344467 | 0.095138174 | 0.248780667 | 0.2204362 | 0.153642493 | 0.784743668 | 0.420379599 | 0.277521643 | 0.321976733 | 0.471530454 |
| Smn1 | S | 28 | 0.03848308 | 0.082901782 | 0.02641445 | -0.01206863 | -0.056487332 | 0.103937156 | 0.064406335 | 0.36869525 | 0.648234213 | 0.175098905 |
| Jcad | S | 941 | 0.153139369 | 0.095906795 | 0.052521932 | -0.100617437 | -0.043384863 | 0.280264213 | 0.337100856 | 0.577939383 | 0.421821294 | 0.549754094 |
| Snx29 | S | 268 | -0.073871843 | 0.22767981 | 0.113520142 | 0.187391984 | -0.114159669 | 0.357994008 | 0.072970359 | 0.185419363 | 0.08319768 | 0.304750558 |
| Mki67 | T | 1215 | 0.08531723 | 0.056725871 | 0.019560372 | -0.065756858 | -0.037165499 | 0.072073935 | 0.087748047 | 0.669569421 | 0.233450922 | 0.431004378 |
| Arhgap21 | S | 1501 | -0.001516287 | 0.075759052 | 0.131538417 | 0.133054703 | 0.055779365 | 0.979053499 | 0.116395656 | 0.112772975 | 0.122969091 | 0.410514324 |
| Trp53bp1 | T | 536 | 0.039233013 | 0.036010974 | 0.152916503 | 0.11368349 | 0.116905529 | 0.185327218 | 0.344486353 | 0.220954054 | 0.326859137 | 0.316157115 |
| Prpf40b | S | 831 | -0.030746926 | 0.03439814 | 0.159994857 | 0.190741783 | 0.125596717 | 0.585711432 | 0.57087615 | 0.004891632 | 0.025097153 | 0.090212115 |
| Dtl | S | 511 | 0.026711411 | 0.055226992 | 0.069427097 | 0.042715687 | 0.014200106 | 0.342188252 | 0.069365069 | 0.124948697 | 0.29799815 | 0.703606082 |
| Dtl | T | 515 | 0.026711411 | 0.055226992 | 0.069427097 | 0.042715687 | 0.014200106 | 0.342188252 | 0.069365069 | 0.124948697 | 0.29799815 | 0.703606082 |
| Tomm34 | S | 186 | 0.221876433 | 0.119761709 | -0.163083981 | -0.384960414 | -0.28284569 | 0.027885672 | 0.187915924 | 0.36447417 | 0.095782555 | 0.16642725 |
| Prpf40a | S | 910 | 0.107095556 | -0.031526699 | 0.121223997 | 0.014128441 | 0.152750696 | 0.2003484 | 0.737144918 | 0.043767283 | 0.836647415 | 0.170897825 |
| Prpf40a | S | 914 | 0.107095556 | -0.031526699 | 0.121223997 | 0.014128441 | 0.152750696 | 0.2003484 | 0.737144918 | 0.043767283 | 0.836647415 | 0.170897825 |
| Surf6 | S | 55 | 0.074163047 | -0.021164881 | 0.154632365 | 0.080469317 | 0.175797246 | 0.273885624 | 0.704369439 | 0.038779771 | 0.277530399 | 0.039524435 |
| Fanci | S | 797 | 0.078817652 | 0.104129933 | 0.250256814 | 0.171439163 | 0.146126881 | 0.500485296 | 0.536788567 | 0.454184504 | 0.596102717 | 0.660943402 |
| Fanci | S | 800 | 0.078817652 | 0.104129933 | 0.250256814 | 0.171439163 | 0.146126881 | 0.500485296 | 0.536788567 | 0.454184504 | 0.596102717 | 0.660943402 |
| Fanci | S | 802 | 0.078817652 | 0.104129933 | 0.250256814 | 0.171439163 | 0.146126881 | 0.500485296 | 0.536788567 | 0.454184504 | 0.596102717 | 0.660943402 |
| Fanci | T | 796 | 0.078817652 | 0.104129933 | 0.250256814 | 0.171439163 | 0.146126881 | 0.500485296 | 0.536788567 | 0.454184504 | 0.596102717 | 0.660943402 |
| Usp9x | S | 1593 | -0.065105927 | 0.158665266 | 0.06348867 | 0.128594596 | -0.095176597 | 0.333605449 | 0.016692588 | 0.087014705 | 0.113611997 | 0.084589639 |
| Iws1 | S | 183 | 0.025812802 | 0.029556149 | 0.099652386 | 0.073839584 | 0.070096238 | 0.431730827 | 0.351447694 | 0.024175848 | 0.076244084 | 0.080713763 |
| Srrm2 | S | 1752 | 0.141808533 | 0.006175342 | 0.142959045 | 0.001150512 | 0.136783703 | 0.302041758 | 0.968870235 | 0.167676241 | 0.993166677 | 0.440439352 |
| Ccar2 | S | 458 | 0.020198104 | 0.071818054 | 0.155638214 | 0.13544011 | 0.083820159 | 0.869525086 | 0.392893341 | 0.085372021 | 0.305119226 | 0.301216502 |
| Arpc5l | S | 64 | 0.035236495 | 0.1198079 | 0.046276743 | 0.011040247 | -0.073531157 | 0.43602001 | 0.132914084 | 0.211804808 | 0.818513353 | 0.280253895 |
| Brip1 | T | 1130 | -0.01850169 | 0.076135239 | 0.228408711 | 0.246910401 | 0.152273472 | 0.862832245 | 0.496171931 | 0.084343627 | 0.030965921 | 0.099614517 |
| Zdhhc8 | Y | 576 | -0.020367781 | 0.157918378 | 0.142227426 | 0.162595207 | -0.015690952 | 0.705885532 | 0.144081632 | 0.22499265 | 0.18253923 | 0.893601129 |
| Nktr | S | 1195 | -0.016635502 | 0.042525877 | 0.130088549 | 0.146724051 | 0.087562672 | 0.765559648 | 0.195134378 | 0.016180582 | 0.067821345 | 0.066445413 |
| Rictor | S | 21 | -0.006877666 | 0.048640482 | 0.067646853 | 0.074524519 | 0.019006371 | 0.715161998 | 0.000972879 | 0.098049738 | 0.067186191 | 0.506873611 |
| Dtnbp1 | S | 340 | -0.005229008 | 0.063292048 | 0.087514286 | 0.092743294 | 0.024222238 | 0.910156625 | 0.135185581 | 0.017475605 | 0.12505099 | 0.519445126 |
| Dtnbp1 | S | 343 | -0.005229008 | 0.063292048 | 0.087514286 | 0.092743294 | 0.024222238 | 0.910156625 | 0.135185581 | 0.017475605 | 0.12505099 | 0.519445126 |
| Sik2 | S | 576 | -0.119830963 | 0.147792446 | 0.240649341 | 0.360480304 | 0.092856895 | 0.096349952 | 0.034814552 | 0.094369342 | 0.038821987 | 0.414673198 |
| Scaf1 | S | 821 | -0.015259522 | 0.015532458 | 0.135924755 | 0.151184277 | 0.120392297 | 0.555478528 | 0.647810451 | 0.001482426 | 0.004098856 | 0.027597871 |
| Glud1 | T | 410 | 0.021774511 | 0.030449533 | 0.131155075 | 0.109380565 | 0.100705543 | 0.512862872 | 0.374138673 | 0.046336375 | 0.080221462 | 0.09672178 |
| Dek | S | 306 | -0.032421888 | 0.044190693 | 0.221492722 | 0.25391461 | 0.177302029 | 0.609782646 | 0.360612467 | 0.040600164 | 0.032012863 | 0.087576505 |
| Dek | S | 308 | -0.032421888 | 0.044190693 | 0.221492722 | 0.25391461 | 0.177302029 | 0.609782646 | 0.360612467 | 0.040600164 | 0.032012863 | 0.087576505 |
| Dek | S | 311 | -0.032421888 | 0.044190693 | 0.221492722 | 0.25391461 | 0.177302029 | 0.609782646 | 0.360612467 | 0.040600164 | 0.032012863 | 0.087576505 |
| Dek | S | 312 | -0.032421888 | 0.044190693 | 0.221492722 | 0.25391461 | 0.177302029 | 0.609782646 | 0.360612467 | 0.040600164 | 0.032012863 | 0.087576505 |
| Map4k4 | S | 619 | 0.032779498 | 0.0662556 | 0.035054416 | 0.002274918 | -0.031201184 | 0.267708174 | 0.057930685 | 0.097829545 | 0.929161248 | 0.273783343 |
| Bcl7c | S | 122 | -0.042866964 | 0.095078266 | 0.217770124 | 0.260637088 | 0.122691858 | 0.622940651 | 0.331381138 | 0.077791125 | 0.062774626 | 0.30657846 |
| Srrm2 | S | 724 | 0.176231041 | -0.056209581 | 0.096555002 | -0.079676039 | 0.152764583 | 0.034574951 | 0.612691933 | 0.338910203 | 0.431241497 | 0.268459047 |

| | | | | | | | | | | | | |
|---|---|---|---|---|---|---|---|---|---|---|---|---|
| Faf1 | S | 319 | -0.011894268 | 0.070077825 | 0.106872969 | 0.118767237 | 0.036795143 | 0.750456679 | 0.075048541 | 0.053312955 | 0.047091066 | 0.402185831 |
| Shroom4 | S | 658 | 0.064028521 | 0.105771332 | 0.039001604 | -0.025026918 | -0.066769729 | 0.180222765 | 0.134929554 | 0.62333173 | 0.732782247 | 0.438172746 |
| Mlh1 | S | 422 | 0.029387174 | 0.026390651 | 0.11725044 | 0.087863266 | 0.090859789 | 0.358502089 | 0.462380884 | 0.041462879 | 0.111896069 | 0.087982907 |
| Bclaf1 | S | 314 | 0.097762736 | 0.062233011 | 0.016764691 | -0.080998045 | -0.04546832 | 0.069983643 | 0.13422005 | 0.784859141 | 0.26929655 | 0.494868158 |
| Cdc27 | S | 370 | 0.227607642 | -0.017997963 | 0.220130557 | -0.007477085 | 0.23812852 | 0.466975715 | 0.886839365 | 0.244234439 | 0.980652602 | 0.215671781 |
| Tra2b | S | 102 | -0.012003549 | 0.069132324 | 0.183975946 | 0.195979495 | 0.114843622 | 0.851081885 | 0.498786451 | 0.046449686 | 0.041537145 | 0.294235148 |
| Arhgap21 | S | 1424 | -0.086759588 | 0.08904667 | 0.146062745 | 0.232822333 | 0.057016075 | 0.164535021 | 0.017827725 | 0.02786044 | 0.014609528 | 0.251482123 |
| Rbm12 | S | 422 | 0.040746495 | 0.039636492 | 0.035151131 | -0.005595364 | -0.004485361 | 0.152971239 | 0.00842319 | 0.27081506 | 0.863201764 | 0.869634091 |
| Rbm12 | S | 424 | 0.040746495 | 0.039636492 | 0.035151131 | -0.005595364 | -0.004485361 | 0.152971239 | 0.00842319 | 0.27081506 | 0.863201764 | 0.869634091 |
| Hdgf | S | 165 | -0.030498268 | 0.128289998 | 0.075168152 | 0.10566642 | -0.053121845 | 0.049639791 | 0.053105169 | 0.00657 | 0.005946087 | 0.25991578 |
| Fcho2 | S | 403 | 0.047406133 | 0.089389592 | 0.036362136 | -0.011043998 | -0.053027457 | 0.377600826 | 0.032588337 | 0.507188329 | 0.850782148 | 0.315272088 |
| D830031N03Rik | S | 436 | 0.047002523 | 0.025414887 | 0.074116581 | 0.027114058 | 0.048701695 | 0.324563446 | 0.361400037 | 0.021288577 | 0.520558762 | 0.040403145 |
| Nav1 | T | 1835 | 0.174549356 | -0.05545391 | 0.112390423 | -0.062158933 | 0.167844333 | 0.06281273 | 0.574955 | 0.23231279 | 0.479575103 | 0.150778164 |
| Tln1 | Y | 1116 | 0.105725483 | 0.088446358 | 0.042010926 | -0.063714557 | -0.046435432 | 0.243290926 | 0.206742971 | 0.524697849 | 0.417252789 | 0.383449781 |
| Pakap | S | 799 | 0.151702337 | 0.089788037 | 0.007755884 | -0.143946454 | -0.082032153 | 0.097059875 | 0.347367734 | 0.942595097 | 0.266691061 | 0.523575394 |
| Mki67 | S | 1955 | 0.161415538 | 0.309642689 | -0.093699536 | -0.255115074 | -0.403342225 | 0.387511562 | 0.130973619 | 0.485247336 | 0.166474873 | 0.079769841 |
| Atrx | S | 1309 | 0.006933807 | 0.078787262 | 0.082175508 | 0.075241702 | 0.003388246 | 0.87349341 | 0.039866386 | 0.158357423 | 0.209824998 | 0.941367928 |
| Atrx | S | 1311 | 0.006933807 | 0.078787262 | 0.082175508 | 0.075241702 | 0.003388246 | 0.87349341 | 0.039866386 | 0.158357423 | 0.209824998 | 0.941367928 |
| Atrx | S | 1313 | 0.006933807 | 0.078787262 | 0.082175508 | 0.075241702 | 0.003388246 | 0.87349341 | 0.039866386 | 0.158357423 | 0.209824998 | 0.941367928 |
| Shb | S | 357 | -0.020707129 | 0.11397945 | 0.110088784 | 0.130795913 | -0.003890666 | 0.798891476 | 0.095464232 | 0.129780218 | 0.158271677 | 0.946851575 |
| Shb | S | 359 | -0.020707129 | 0.11397945 | 0.110088784 | 0.130795913 | -0.003890666 | 0.798891476 | 0.095464232 | 0.129780218 | 0.158271677 | 0.946851575 |
| Sorbs2 | S | 379 | 0.021680949 | 0.031974262 | 0.075659125 | 0.053978176 | 0.043684863 | 0.339513985 | 0.214383669 | 0.018881101 | 0.056450074 | 0.11414148 |
| Eif3b | S | 37 | 0.011187768 | 0.068233617 | 0.108421171 | 0.097233403 | 0.040187554 | 0.727225742 | 0.353599996 | 0.037445436 | 0.060639696 | 0.573422986 |
| Kdm5c | S | 897 | 0.009368938 | 0.018480675 | 0.118793348 | 0.109424409 | 0.100312673 | 0.79322757 | 0.724651567 | 0.004664733 | 0.039643218 | 0.135814409 |
| Terf2ip | S | 200 | 0.055031203 | 0.05093604 | 0.042120887 | -0.012910316 | -0.008815153 | 0.217827465 | 0.054459993 | 0.234464533 | 0.76802016 | 0.785688808 |
| Setd1a | S | 521 | 0.058251562 | 0.042625506 | 0.075128282 | 0.01687672 | 0.032502776 | 0.16470361 | 0.288836887 | 0.166394091 | 0.737492127 | 0.530895707 |
| Msl3 | S | 404 | 0.097158325 | 0.056456719 | 0.060909834 | -0.036248491 | 0.004453115 | 0.179395041 | 0.343252931 | 0.310351303 | 0.564426781 | 0.931654154 |
| Mtmr10 | T | 766 | 0.136936103 | 0.224933258 | -0.110123811 | -0.247059914 | -0.335057069 | 0.383143832 | 0.045039531 | 0.499656515 | 0.225599222 | 0.101678212 |
| Usp24 | S | 2074 | 0.084020625 | 0.056705235 | 0.013375046 | -0.070645579 | -0.043330189 | 0.056264839 | 0.109838941 | 0.653166388 | 0.110688205 | 0.262944914 |
| Ythdc1 | S | 425 | -0.0170494 | 0.086014112 | 0.074581347 | 0.091630747 | -0.011432765 | 0.82708238 | 0.017653261 | 0.133180536 | 0.308211742 | 0.776991487 |
| Pde4a | S | 145 | -0.027358584 | 0.221963201 | 0.058769771 | 0.086128354 | -0.16319343 | 0.807045747 | 0.041467719 | 0.660629083 | 0.592401204 | 0.290684305 |
| Srek1ip1 | S | 144 | -0.015202312 | 0.072084254 | 0.18093218 | 0.196134492 | 0.108847926 | 0.823797563 | 0.299947965 | 0.077893081 | 0.076946587 | 0.247959145 |
| Tns3 | S | 752 | -0.014513794 | 0.102147259 | 0.115869995 | 0.130383788 | 0.013722736 | 0.594538974 | 0.064003745 | 0.140874484 | 0.124394321 | 0.836993467 |
| U2af2 | T | 133 | 0.034368983 | 0.043585619 | 0.15325355 | 0.118884568 | 0.109667932 | 0.459581554 | 0.308856272 | 0.160282977 | 0.242066538 | 0.26984925 |
| Larp1 | S | 494 | 0.112211214 | 0.113744346 | -0.008888285 | -0.121099499 | -0.122632632 | 0.29611367 | 0.048383278 | 0.855084628 | 0.263560606 | 0.005624934 |
| Dact3 | S | 303 | 0.006873076 | 0.143723297 | 0.020657423 | 0.013784347 | -0.123065874 | 0.881235711 | 0.00979479 | 0.765948834 | 0.834944759 | 0.138861059 |
| Numa1 | S | 196 | 0.263897094 | -0.113272497 | 0.103365303 | -0.160531791 | 0.216637799 | 0.055478294 | 0.659341049 | 0.513403946 | 0.292396987 | 0.42869658 |
| Numa1 | T | 209 | 0.263897094 | -0.113272497 | 0.103365303 | -0.160531791 | 0.216637799 | 0.055478294 | 0.659341049 | 0.513403946 | 0.292396987 | 0.42869658 |
| Fndc3a | S | 207 | 0.003248576 | -0.038175509 | 0.23226111 | 0.229012535 | 0.270436619 | 0.96642097 | 0.699436471 | 0.010885257 | 0.016709684 | 0.048168633 |
| Kmt2b | T | 1490 | 0.079789754 | 0.125329212 | -0.017195742 | -0.096985495 | -0.142524954 | 0.195098864 | 0.05547391 | 0.886114517 | 0.450650591 | 0.296786492 |
| Tmpo | S | 353 | 0.055685218 | 0.126534967 | 0.082964542 | 0.027279324 | -0.043570425 | 0.70590709 | 0.307048284 | 0.158133671 | 0.845365515 | 0.683228157 |
| Epn1 | T | 459 | 0.047396685 | 0.205161655 | 0.0400527 | -0.007343986 | -0.165108956 | 0.739429616 | 0.101067132 | 0.733007306 | 0.947291081 | 0.05896678 |
| Srrm2 | S | 725 | 0.174895809 | -0.059881998 | 0.096847939 | -0.07804787 | 0.156729937 | 0.032930969 | 0.569600918 | 0.330449184 | 0.429661866 | 0.237236244 |
| Pias1 | S | 488 | 0.018809257 | 0.106210602 | 0.055835512 | 0.037026255 | -0.05037509 | 0.496430748 | 0.049791204 | 0.366884243 | 0.533221824 | 0.43463677 |
| Ehbp1l1 | S | 284 | 0.103108882 | 0.067547447 | -0.007777348 | -0.11088623 | -0.075324795 | 0.096089851 | 0.048965279 | 0.907281962 | 0.18593801 | 0.306761671 |
| Rabgap1 | S | 494 | 0.124235273 | 0.189127883 | -0.029661857 | -0.153897129 | -0.21878974 | 0.433537353 | 0.097358998 | 0.842019384 | 0.422725118 | 0.209531591 |
| Dbn1 | S | 431 | 0.118888596 | 0.159814513 | -0.023472029 | -0.142360625 | -0.183286542 | 0.23241133 | 0.1571277 | 0.914454171 | 0.515817467 | 0.421692366 |
| Suds3 | S | 237 | 0.003131021 | 0.039084803 | 0.182301223 | 0.179170202 | 0.143216419 | 0.948214488 | 0.377281621 | 0.064751367 | 0.074288543 | 0.123929 |
| Snx19 | S | 379 | 0.018307245 | -0.027627885 | 0.238082207 | 0.219774962 | 0.265710092 | 0.782107068 | 0.861822085 | 0.030558162 | 0.046292293 | 0.176016925 |
| Lin54 | T | 280 | 0.177201122 | 0.078865356 | -0.000954671 | -0.178155793 | -0.079820027 | 0.063579898 | 0.54870212 | 0.99444092 | 0.235894304 | 0.615261312 |
| Usp6nl | S | 394 | 0.015644855 | 0.050144183 | 0.064330523 | 0.048685668 | 0.014186339 | 0.585022344 | 0.080478068 | 0.033744799 | 0.171931457 | 0.584474802 |
| Nktr | S | 611 | 0.048731886 | 0.071424042 | 0.164330371 | 0.115598485 | 0.092906329 | 0.489334676 | 0.510459267 | 0.207772967 | 0.350030053 | 0.503631233 |
| Brd4 | S | 1067 | 0.005716911 | 0.059626617 | 0.098268437 | 0.092551526 | 0.03864182 | 0.830146325 | 0.337681878 | 0.018363646 | 0.027710655 | 0.512788898 |
| Slc7a6os | S | 156 | 0.00924687 | 0.326356999 | -0.053835986 | -0.063082857 | -0.380192985 | 0.934406107 | 0.036173642 | 0.650336026 | 0.256512765 | 0.014946893 |
| Smarca4 | S | 1593 | 0.005136594 | 0.051443488 | 0.149726078 | 0.144589484 | 0.09828259 | 0.916866969 | 0.170348665 | 0.097389291 | 0.110366546 | 0.22040444 |
| Smarca4 | S | 1597 | 0.005136594 | 0.051443488 | 0.149726078 | 0.144589484 | 0.09828259 | 0.916866969 | 0.170348665 | 0.097389291 | 0.110366546 | 0.22040444 |
| Smarca4 | S | 1606 | 0.005136594 | 0.051443488 | 0.149726078 | 0.144589484 | 0.09828259 | 0.916866969 | 0.170348665 | 0.097389291 | 0.110366546 | 0.22040444 |
| Smarca4 | S | 1608 | 0.005136594 | 0.051443488 | 0.149726078 | 0.144589484 | 0.09828259 | 0.916866969 | 0.170348665 | 0.097389291 | 0.110366546 | 0.22040444 |
| Smarca4 | S | 1610 | 0.005136594 | 0.051443488 | 0.149726078 | 0.144589484 | 0.09828259 | 0.916866969 | 0.170348665 | 0.097389291 | 0.110366546 | 0.22040444 |
| Vim | S | 55 | 0.101016739 | 0.079658465 | -0.091629427 | -0.192646166 | -0.171287892 | 0.019982545 | 0.006460673 | 0.262538756 | 0.07043825 | 0.098245868 |
| Nop56 | S | 513 | 0.067821651 | 0.032213621 | 0.074319603 | 0.006497953 | 0.042105982 | 0.26389565 | 0.422018677 | 0.069710093 | 0.893046419 | 0.156184975 |
| Nfix | S | 288 | 0.072966112 | 0.035490525 | 0.094937636 | 0.021971524 | 0.059447111 | 0.190719413 | 0.562176771 | 0.153954318 | 0.653614019 | 0.327628064 |
| Iws1 | S | 306 | 0.010687123 | 0.004890506 | 0.14294672 | 0.132259597 | 0.138056214 | 0.651174051 | 0.721888388 | 0.010826566 | 0.012853422 | 0.024863221 |
| Mov10 | T | 253 | -0.04704104 | 0.055629977 | 0.168424054 | 0.215465094 | 0.112794077 | 0.628294108 | 0.365524282 | 0.017924027 | 0.090577498 | 0.062093639 |
| Mov10 | T | 254 | -0.04704104 | 0.055629977 | 0.168424054 | 0.215465094 | 0.112794077 | 0.628294108 | 0.365524282 | 0.017924027 | 0.090577498 | 0.062093639 |
| Med26 | S | 337 | 0.072690089 | 0.110587927 | 0.01453228 | -0.058157809 | -0.096055647 | 0.151412427 | 0.109789053 | 0.859341104 | 0.486820362 | 0.304949186 |
| Smarcc1 | S | 327 | -0.005556889 | 0.017653001 | 0.101101225 | 0.106658114 | 0.083448225 | 0.883343053 | 0.575050608 | 0.000946284 | 0.064472875 | 0.058756688 |
| Mia3 | S | 1697 | -0.070368575 | 0.152649372 | 0.170018362 | 0.240386937 | 0.01736899 | 0.588596413 | 0.304379645 | 0.066788065 | 0.127691949 | 0.886321901 |
| Mia3 | S | 1700 | -0.070368575 | 0.152649372 | 0.170018362 | 0.240386937 | 0.01736899 | 0.588596413 | 0.304379645 | 0.066788065 | 0.127691949 | 0.886321901 |
| Cavin2 | T | 276 | 0.13487134 | 0.061095754 | 0.038080912 | -0.096790428 | -0.023014843 | 0.139099924 | 0.450348114 | 0.448935317 | 0.249429017 | 0.759055009 |
| Trip12 | S | 241 | 0.048434629 | 0.004056814 | 0.078506863 | 0.030072234 | 0.074450049 | 0.073380162 | 0.910880473 | 0.025926898 | 0.281189378 | 0.109116893 |
| Fnbp1l | S | 501 | -0.03943814 | 0.180485318 | 0.053639978 | 0.093078118 | -0.126845339 | 0.37752138 | 0.006364129 | 0.65342047 | 0.44591134 | 0.327128044 |
| Reps1 | S | 118 | 0.005566862 | 0.073558739 | 0.187202247 | 0.181635385 | 0.113643508 | 0.912909785 | 0.275788176 | 0.165845192 | 0.175097699 | 0.3504141 |
| Srrm2 | S | 1996 | 0.106017324 | 0.053762665 | 0.042362204 | -0.06365512 | -0.011400462 | 0.075984654 | 0.30014273 | 0.661205992 | 0.528094887 | 0.906207989 |
| Srrm2 | S | 1998 | 0.106017324 | 0.053762665 | 0.042362204 | -0.06365512 | -0.011400462 | 0.075984654 | 0.30014273 | 0.661205992 | 0.528094887 | 0.906207989 |
| Tjp1 | S | 178 | 0.044709614 | 0.038260366 | 0.09107051 | 0.046360897 | 0.052810145 | 0.243291178 | 0.231761937 | 0.142470153 | 0.393713601 | 0.328414532 |
| Mcm4 | S | 32 | 0.072695294 | 0.122343755 | -0.03448278 | -0.107178073 | -0.156826535 | 0.231131258 | 0.019081314 | 0.689675737 | 0.269413925 | 0.149116333 |
| Pan3 | S | 300 | 0.133506492 | -0.040038378 | 0.043941286 | -0.089565206 | 0.083979665 | 0.003126035 | 0.401522135 | 0.323611561 | 0.083979907 | 0.139273202 |
| Clasp1 | S | 556 | 0.044401012 | 0.238539201 | -0.042062249 | -0.086463261 | -0.280601449 | 0.666251817 | 0.036692441 | 0.734726674 | 0.502286111 | 0.075784894 |
| Tbc1d10b | S | 647 | -0.007810991 | 0.086266382 | 0.097459948 | 0.105270939 | 0.011193566 | 0.821423738 | 0.334796186 | 0.014981163 | 0.014736897 | 0.885913742 |
| Camsap1 | S | 569 | 0.139233229 | 0.039543745 | 0.017279673 | -0.121953556 | -0.022264072 | 0.063208696 | 0.318192361 | 0.729862342 | 0.107479446 | 0.69171421 |
| Hectd1 | S | 1538 | -0.040871921 | 0.153613815 | 0.136665106 | 0.177537027 | -0.016948709 | 0.680066514 | 0.103246159 | 0.241766957 | 0.206204872 | 0.879076094 |
| Gzf1 | S | 193 | -0.012105728 | 0.08497548 | 0.078134129 | 0.090239857 | -0.006841351 | 0.731632323 | 0.061703391 | 0.05235548 | 0.058844833 | 0.856869387 |
| Kmt2a | S | 2934 | -0.018896127 | 0.032871531 | 0.221982107 | 0.240878234 | 0.189110575 | 0.8108387 | 0.75015567 | 0.03298022 | 0.026619992 | 0.116628231 |
| Nek1 | S | 966 | 0.000596442 | 0.091892236 | 0.043622767 | 0.043026325 | -0.04826947 | 0.986342027 | 0.006413708 | 0.328078965 | 0.360156968 | 0.274010035 |
| Brca1 | S | 1297 | -0.032567351 | 0.075390399 | 0.085900566 | 0.118467917 | 0.010510167 | 0.310487953 | 0.108663501 | 0.003878165 | 0.019911793 | 0.770151241 |
| Tnrc6b | S | 1489 | 0.290362907 | 0.118509427 | -0.049645603 | -0.340008509 | -0.16815503 | 0.263280578 | 0.240251549 | 0.71466529 | 0.210978458 | 0.21912345 |
| Prrc2a | S | 1090 | 0.127581308 | 0.083217576 | 0.005420322 | -0.122160986 | -0.077797254 | 0.059252692 | 0.383527458 | 0.918890029 | 0.066598039 | 0.402564278 |
| Pcif1 | S | 19 | 0.00023472 | 0.178849211 | 0.121214568 | 0.120979848 | -0.057634643 | 0.997373041 | 0.154243917 | 0.418161135 | 0.428193835 | 0.71495963 |
| Srrm2 | S | 722 | 0.180946575 | -0.060023222 | 0.100930868 | -0.080015707 | 0.160954089 | 0.054466094 | 0.583341676 | 0.280842105 | 0.416920903 | 0.230103725 |
| Sntb2 | S | 188 | -0.013151759 | 0.116996648 | 0.117709271 | 0.130861029 | 0.000712623 | 0.899030251 | 0.032320194 | 0.448462099 | 0.444554741 | 0.996013773 |
| Champ1 | S | 204 | -0.031737199 | 0.005584108 | 0.213251661 | 0.24498886 | 0.207667553 | 0.560817641 | 0.88625478 | 0.008148124 | 0.010590845 | 0.018689947 |
| Srpr | S | 177 | 0.028638673 | 0.025948434 | 0.132430325 | 0.103791652 | 0.106481891 | 0.656944121 | 0.555667163 | 0.033755699 | 0.179116425 | 0.081536789 |
| Als2 | S | 1456 | -0.005088639 | 0.140986355 | 0.147977404 | 0.153066044 | 0.00699105 | 0.927157056 | 0.234664085 | 0.224658241 | 0.21350825 | 0.956453771 |

| | | | | | | | | | | | | |
|---|---|---|---|---|---|---|---|---|---|---|---|---|
| Ncor2 | S | 2152 | -0.003646881 | -0.056896457 | 0.243832788 | 0.247479669 | 0.300729245 | 0.953257143 | 0.432175816 | 0.005124946 | 0.003215482 | 0.006289701 |
| Slc9a3r1 | S | 275 | 0.01837841 | 0.09303214 | 0.063718851 | 0.045340442 | -0.029313288 | 0.516011196 | 0.038037935 | 0.406165092 | 0.541410372 | 0.689432667 |
| Ncl | T | 307 | 0.127358173 | 0.122934434 | 0.066049605 | -0.061308568 | -0.056884829 | 0.404752054 | 0.344972844 | 0.546034947 | 0.67220261 | 0.642700145 |
| Rbm15b | S | 9 | 0.22966877 | 0.123443883 | -0.023333371 | -0.253002142 | -0.146777255 | 0.302087021 | 0.207678993 | 0.839212579 | 0.267874946 | 0.202441279 |
| Clspn | S | 823 | 0.106195157 | 0.015758571 | 0.054957564 | -0.051237593 | 0.039198993 | 0.035411103 | 0.642794744 | 0.408859816 | 0.448049622 | 0.542955015 |
| Igf2bp2 | S | 162 | 0.025758431 | 0.103345087 | 0.032845024 | 0.007086594 | -0.070500063 | 0.495461739 | 0.026876659 | 0.481717648 | 0.856454299 | 0.142752631 |
| Lemd2 | S | 507 | 0.045451433 | 0.050780651 | 0.061529281 | 0.016077848 | 0.010748629 | 0.268605326 | 0.217492802 | 0.098590659 | 0.594726139 | 0.713389198 |
| Itga4 | S | 1028 | 0.056289786 | 0.040443991 | 0.080609652 | 0.024319867 | 0.040165661 | 0.204660722 | 0.465825628 | 0.105673222 | 0.484559626 | 0.43015687 |
| Phactr4 | S | 285 | 0.124623118 | 0.046708722 | 0.002418411 | -0.122204706 | -0.044290311 | 0.030907676 | 0.317446436 | 0.9655327 | 0.052991034 | 0.338761056 |
| Rbm39 | S | 127 | 0.080086909 | -0.009255512 | 0.201920771 | 0.121833863 | 0.211176283 | 0.31968177 | 0.88080974 | 0.139301727 | 0.320391805 | 0.140914511 |
| Prrc2a | T | 377 | 0.034498537 | 0.056179159 | 0.079204852 | 0.044706315 | 0.023025693 | 0.584285875 | 0.243369972 | 0.053942757 | 0.491264246 | 0.61298639 |
| Gas2l1 | S | 306 | 0.013549036 | 0.111968571 | 0.026166579 | 0.012617544 | -0.085801992 | 0.759146596 | 0.006850967 | 0.730757494 | 0.870664793 | 0.307297992 |
| Nhsl2 | T | 1032 | -0.126721885 | 0.037162507 | 0.233135257 | 0.359857143 | 0.19597275 | 0.259147072 | 0.012554734 | 0.028932078 | 0.028935374 | 0.044027736 |
| Huwe1 | S | 1907 | -0.009130195 | 0.072809587 | 0.048110803 | 0.057240998 | -0.024698784 | 0.805639136 | 0.029864842 | 0.018856425 | 0.203791344 | 0.30985563 |
| Ppig | S | 429 | 0.020270461 | 0.025624044 | 0.163226174 | 0.142955713 | 0.137602131 | 0.830992915 | 0.675346141 | 0.036065988 | 0.219156763 | 0.10525225 |
| Zfp553 | S | 371 | 0.049734837 | 0.084574951 | 0.230858037 | 0.181123201 | 0.146283086 | 0.786040676 | 0.545331963 | 0.254692005 | 0.448576336 | 0.479571633 |
| Mrgbp | S | 191 | -0.029829058 | 0.110625783 | 0.090861248 | 0.120690306 | -0.019764534 | 0.600822194 | 0.127923374 | 0.038531862 | 0.101889032 | 0.73607095 |
| 1700025G04Rik | S | 95 | 0.0299257 | 0.049051781 | 0.083151307 | 0.053225607 | 0.034099525 | 0.430927971 | 0.308745409 | 0.050355318 | 0.049230942 | 0.373747414 |
| Il6st | T | 668 | 0.025105411 | 0.033662347 | 0.147207152 | 0.122101741 | 0.113544805 | 0.756815721 | 0.575668968 | 0.04444943 | 0.146127229 | 0.002854388 |
| Iws1 | S | 343 | 0.009387473 | 0.003594629 | 0.090192815 | 0.080805342 | 0.086598186 | 0.452290072 | 0.775035807 | 0.001172067 | 0.004942412 | 0.003699518 |
| Unkl | S | 15 | 0.096044989 | 0.041396964 | 0.040155202 | -0.055889787 | -0.001241762 | 0.032892321 | 0.54927481 | 0.501844326 | 0.380168746 | 0.987568192 |
| Fam117b | S | 302 | 0.046046129 | 0.15950287 | 0.110376579 | 0.064330451 | -0.04912629 | 0.551897325 | 0.470409725 | 0.262944826 | 0.41406867 | 0.813488465 |
| Hirip3 | S | 396 | 0.025090757 | 0.061537658 | 0.027940051 | 0.002849295 | -0.033597607 | 0.531253645 | 0.007322961 | 0.17645367 | 0.939637199 | 0.078364713 |
| Trp53bp1 | S | 1968 | 0.159105816 | 0.17885742 | -0.111178723 | -0.270284539 | -0.290036143 | 0.132490819 | 0.019687637 | 0.097128812 | 0.036702625 | 0.007118674 |
| Naa10 | S | 205 | -0.017174625 | 0.165541828 | 0.059863867 | 0.077038492 | -0.10567796 | 0.741383969 | 0.050981752 | 0.308161634 | 0.179782548 | 0.16578206 |
| Utp18 | T | 200 | 0.120542548 | 0.011736621 | 0.043445977 | -0.077096571 | 0.031709356 | 0.049893898 | 0.807182921 | 0.254588135 | 0.15901033 | 0.542183313 |
| Bud13 | S | 404 | 0.036536224 | 0.043609445 | 0.077603327 | 0.041067103 | 0.033993882 | 0.47821242 | 0.128720395 | 0.09040679 | 0.452839891 | 0.355713095 |
| Bud13 | S | 405 | 0.036536224 | 0.043609445 | 0.077603327 | 0.041067103 | 0.033993882 | 0.47821242 | 0.128720395 | 0.09040679 | 0.452839891 | 0.355713095 |
| Bud13 | S | 407 | 0.036536224 | 0.043609445 | 0.077603327 | 0.041067103 | 0.033993882 | 0.47821242 | 0.128720395 | 0.09040679 | 0.452839891 | 0.355713095 |
| Bud13 | S | 410 | 0.036536224 | 0.043609445 | 0.077603327 | 0.041067103 | 0.033993882 | 0.47821242 | 0.128720395 | 0.09040679 | 0.452839891 | 0.355713095 |
| Mertk | S | 538 | 0.064330417 | 0.247863656 | 0.060952281 | -0.003378136 | -0.186911375 | 0.757220883 | 0.218264106 | 0.70185473 | 0.988482888 | 0.389845945 |
| Mki67 | S | 2934 | 0.209723077 | 0.113722897 | -0.052811762 | -0.262534838 | -0.166534658 | 0.101328919 | 0.190629261 | 0.35975885 | 0.0576757 | 0.095564953 |
| Rpl37 Rpl37rt | T | 17 | 0.498175136 | 0.015991798 | -0.154734655 | -0.652909791 | -0.170726452 | 0.082977669 | 0.909983462 | 0.650532682 | 0.1310333 | 0.60147409 |
| Wrnip1 | S | 91 | 0.007701178 | 0.043441416 | 0.127235088 | 0.119533909 | 0.083793672 | 0.847040872 | 0.304566281 | 0.043565037 | 0.062831406 | 0.145887117 |
| Srrm1 | S | 429 | 0.11148255 | 0.033178815 | 0.021017082 | -0.090465469 | -0.012161734 | 0.062660932 | 0.176776625 | 0.630560559 | 0.141900566 | 0.771585103 |
| Mis18bp1 | T | 893 | 0.135461439 | 0.203969564 | 0.036384651 | -0.099076788 | -0.167584913 | 0.537791606 | 0.25661962 | 0.848559769 | 0.651876048 | 0.364661583 |
| Dlc1 | S | 406 | 0.157094342 | 0.080433815 | 0.010521096 | -0.146573245 | -0.069912719 | 0.118211703 | 0.370286249 | 0.907344071 | 0.212604977 | 0.523648066 |
| Nrcam | S | 1243 | 0.027668057 | 0.127751799 | 0.04828743 | 0.020619374 | -0.079464369 | 0.680881194 | 0.056941144 | 0.491988737 | 0.806347441 | 0.317547088 |
| Lmnb1 | T | 21 | -0.018355704 | 0.063841106 | 0.078538521 | 0.096894225 | 0.014697415 | 0.656960145 | 0.076629478 | 0.014650612 | 0.080487416 | 0.565502211 |
| Kif1c | S | 1089 | 0.054879991 | 0.206756689 | -0.003051276 | -0.057931267 | -0.209807964 | 0.600875209 | 0.074450095 | 0.982085746 | 0.694774842 | 0.199398118 |
| Samhd1 | T | 634 | 0.011389549 | 0.044765838 | 0.089754802 | 0.078365253 | 0.044988963 | 0.687274429 | 0.191560034 | 0.032319443 | 0.056478943 | 0.162643187 |
| Bicd2 | S | 578 | 0.193294137 | 0.021395762 | -0.092537012 | -0.285831149 | -0.113932773 | 0.002878296 | 0.73356224 | 0.501476294 | 0.119491175 | 0.428352205 |
| Tubgcp3 | S | 513 | 0.137071393 | 0.177746349 | -0.078162125 | -0.215233518 | -0.255908473 | 0.255988086 | 0.099269974 | 0.657811687 | 0.276249911 | 0.207661231 |
| Mtfr1l | S | 231 | -0.065147662 | 0.240225417 | 0.19405035 | 0.259198012 | -0.046175068 | 0.200979989 | 0.151039748 | 0.220731749 | 0.138551474 | 0.783092377 |
| Scap | S | 837 | -0.012295896 | 0.167337618 | 0.072223707 | 0.084519603 | -0.095113911 | 0.837119166 | 0.083595129 | 0.301887291 | 0.217498707 | 0.276986562 |
| Map3k1 | S | 245 | 0.084287664 | 0.098733009 | 0.282814388 | 0.198526724 | 0.184081379 | 0.634735695 | 0.554790768 | 0.481688773 | 0.605349101 | 0.627784902 |
| Map3k1 | S | 247 | 0.084287664 | 0.098733009 | 0.282814388 | 0.198526724 | 0.184081379 | 0.634735695 | 0.554790768 | 0.481688773 | 0.605349101 | 0.627784902 |
| Trp53bp1 | S | 80 | -0.109216095 | 0.133166125 | 0.620295061 | 0.729511155 | 0.487128935 | 0.462100753 | 0.597541575 | 0.255956259 | 0.201042221 | 0.355839422 |
| Trp53bp1 | S | 82 | -0.109216095 | 0.133166125 | 0.620295061 | 0.729511155 | 0.487128935 | 0.462100753 | 0.597541575 | 0.255956259 | 0.201042221 | 0.355839422 |
| Trp53bp1 | S | 83 | -0.109216095 | 0.133166125 | 0.620295061 | 0.729511155 | 0.487128935 | 0.462100753 | 0.597541575 | 0.255956259 | 0.201042221 | 0.355839422 |
| Nr3c1 | T | 159 | 0.065477373 | 0.079179155 | 0.013924443 | -0.05155293 | -0.065254712 | 0.059764765 | 0.152229888 | 0.669872228 | 0.068443997 | 0.210419036 |
| Slc20a2 | S | 433 | 0.140210178 | 0.040922286 | -0.105464794 | -0.245674972 | -0.14638708 | 0.011719078 | 0.015902237 | 0.399271249 | 0.12575257 | 0.2777252 |
| Wrnip1 | S | 92 | 0.007312301 | 0.041816693 | 0.118730624 | 0.111418323 | 0.07691393 | 0.863767218 | 0.262350285 | 0.039269059 | 0.072081491 | 0.135454084 |
| Srrm2 | S | 484 | 0.030572527 | 0.02517777 | 0.100500702 | 0.069928205 | 0.075322962 | 0.71575944 | 0.459547535 | 0.015427877 | 0.431062305 | 0.061409426 |
| Kat7 | S | 52 | 0.100819112 | 0.029493915 | 0.051386009 | -0.049433103 | 0.021892095 | 0.069553584 | 0.434854411 | 0.4124534 | 0.445238822 | 0.702682291 |
| Kat7 | S | 55 | 0.100819112 | 0.029493915 | 0.051386009 | -0.049433103 | 0.021892095 | 0.069553584 | 0.434854411 | 0.4124534 | 0.445238822 | 0.702682291 |
| Nup107 | S | 87 | 0.11429015 | -0.04704482 | 0.116520803 | 0.002230653 | 0.163565623 | 0.082125877 | 0.4647181 | 0.079068576 | 0.939419535 | 0.017687599 |
| Srrm2 | S | 792 | 0.158039044 | -0.051379306 | 0.072884604 | -0.08515444 | 0.124263909 | 0.00975161 | 0.413154794 | 0.436357214 | 0.367345626 | 0.234739473 |
| Araf | S | 255 | 0.306659391 | 0.163006359 | -0.129104201 | -0.435763592 | -0.292110561 | 0.196805929 | 0.19892942 | 0.427822222 | 0.100244495 | 0.10561891 |
| Wdhd1 | S | 377 | -0.10100507 | 0.033561908 | 0.205172399 | 0.306177469 | 0.171610491 | 0.298876604 | 0.573077205 | 0.002733488 | 0.038295808 | 0.037059663 |
| Ofd1 | S | 664 | 0.068931647 | 0.101390109 | 0.013802374 | -0.055129272 | -0.087587735 | 0.306157931 | 0.04883336 | 0.841389256 | 0.47157312 | 0.233372097 |
| Mpp1 | S | 57 | 0.047428435 | 0.083929252 | 0.057274551 | 0.009846115 | -0.026654701 | 0.369077789 | 0.09880174 | 0.39900162 | 0.867026315 | 0.631334525 |
| Clspn | S | 1302 | 0.165542411 | 0.019212244 | 0.045560127 | -0.119982284 | 0.026347883 | 0.067002399 | 0.809325747 | 0.591710632 | 0.169719155 | 0.733638945 |
| Micall1 | S | 480 | 0.181423704 | 0.061955876 | -0.025202631 | -0.206626334 | -0.087158507 | 0.035939574 | 0.485387629 | 0.666009986 | 0.034747767 | 0.375371006 |
| Nppb | S | 52 | 0.043910439 | 0.10541727 | 0.025830197 | -0.018080242 | -0.079587073 | 0.39578938 | 0.04127136 | 0.648582397 | 0.765063181 | 0.207807629 |
| Sipa1l1 | S | 1369 | 0.046758609 | -0.036212719 | 0.217929468 | 0.17117086 | 0.254142187 | 0.323085215 | 0.596109174 | 0.055591218 | 0.112761203 | 0.041317466 |
| Pcbp1 | S | 190 | -0.023909828 | 0.082405794 | 0.066479825 | 0.090389652 | -0.015925969 | 0.397234432 | 0.001255985 | 0.275190828 | 0.172250424 | 0.760196184 |
| Arhgap27 | S | 238 | -0.018308166 | 0.078999462 | 0.21479852 | 0.233106686 | 0.135799059 | 0.941301886 | 0.709560776 | 0.070532152 | 0.387946262 | 0.527686807 |
| Arhgap27 | T | 237 | -0.018308166 | 0.078999462 | 0.21479852 | 0.233106686 | 0.135799059 | 0.941301886 | 0.709560776 | 0.070532152 | 0.387946262 | 0.527686807 |
| Usp42 | S | 1125 | 0.178834592 | -0.080100211 | 0.073515289 | -0.105319303 | 0.1536155 | 0.004445064 | 0.1430419 | 0.302796459 | 0.162687806 | 0.074266956 |
| Macf1 | S | 2823 | 0.15080043 | 0.246831074 | -0.169091093 | -0.319891523 | -0.415922168 | 0.476471507 | 0.023834604 | 0.371917538 | 0.228029219 | 0.090476448 |
| Ccnl1 | S | 508 | 0.031063834 | 0.022904337 | 0.077945341 | 0.046881507 | 0.055041004 | 0.215427501 | 0.336498005 | 0.035851741 | 0.150465474 | 0.104535132 |
| Ccnl1 | S | 510 | 0.031063834 | 0.022904337 | 0.077945341 | 0.046881507 | 0.055041004 | 0.215427501 | 0.336498005 | 0.035851741 | 0.150465474 | 0.104535132 |
| Hp1bp3 | S | 513 | -0.056337773 | 0.057400195 | 0.181782877 | 0.238120651 | 0.124382683 | 0.470394527 | 0.151023576 | 0.040886702 | 0.041359587 | 0.116104364 |
| Prkab2 | S | 107 | 0.017505005 | 0.060040196 | 0.121543902 | 0.104038898 | 0.061503706 | 0.746899407 | 0.530617224 | 0.042716985 | 0.050718787 | 0.493309026 |
| Phtf2 | S | 293 | -0.040495691 | 0.062002962 | 0.199721066 | 0.240216757 | 0.137718104 | 0.723887148 | 0.708562784 | 0.02606848 | 0.096351376 | 0.423459392 |
| Phtf2 | S | 297 | -0.040495691 | 0.062002962 | 0.199721066 | 0.240216757 | 0.137718104 | 0.723887148 | 0.708562784 | 0.02606848 | 0.096351376 | 0.423459392 |
| Phtf2 | S | 298 | -0.040495691 | 0.062002962 | 0.199721066 | 0.240216757 | 0.137718104 | 0.723887148 | 0.708562784 | 0.02606848 | 0.096351376 | 0.423459392 |
| Xrn2 | S | 499 | -0.00032006 | 0.016509837 | 0.101500979 | 0.101821039 | 0.084991142 | 0.99201277 | 0.367285323 | 0.003470971 | 0.035148436 | 0.022161018 |
| Ccnl1 | S | 358 | -0.004997602 | 0.035530507 | 0.127231945 | 0.132229547 | 0.091701438 | 0.882013017 | 0.223355573 | 0.02675887 | 0.030027007 | 0.091747934 |
| Rbm12b2 | S | 555 | 0.041025601 | 0.028094212 | 0.133847371 | 0.092821771 | 0.105753159 | 0.564506435 | 0.453233119 | 0.07965579 | 0.283280185 | 0.133398736 |
| Vim | S | 49 | -0.041502816 | 0.083304157 | 0.124181171 | 0.165683987 | 0.040877013 | 0.371738683 | 0.06133292 | 0.052806813 | 0.027968926 | 0.409598708 |
| Brf1 | S | 449 | 0.061812969 | 0.147198341 | 0.025365526 | -0.036447443 | -0.121832815 | 0.745860499 | 0.069100431 | 0.555711641 | 0.846848209 | 0.118841343 |
| Cep350 | S | 1944 | 0.101335247 | 0.040147441 | 0.070469935 | -0.030865312 | 0.030322495 | 0.122850675 | 0.543222852 | 0.365745626 | 0.656165643 | 0.679254293 |
| Pml | T | 535 | -0.044158346 | 0.093186657 | 0.057292256 | 0.101450601 | -0.035894401 | 0.342366181 | 0.003381678 | 0.088146649 | 0.084079334 | 0.232552898 |
| Bclaf1 | S | 289 | 0.008145361 | 0.021512042 | 0.071032894 | 0.062887533 | 0.049520852 | 0.784382664 | 0.183722252 | 0.00272793 | 0.108369174 | 0.009629345 |
| Prpf40a | S | 879 | 0.00142224 | 0.049862483 | 0.193510399 | 0.192088159 | 0.143647916 | 0.98016408 | 0.37124703 | 0.108908248 | 0.110831518 | 0.191883089 |
| Slc9a3r1 | T | 288 | -0.067304908 | 0.060633356 | 0.131257911 | 0.198562819 | 0.070624556 | 0.025528368 | 0.037728119 | 0.001307658 | 0.000882718 | 0.033660486 |
| Atp5a1 | S | 477 | 0.137593604 | -0.056301483 | 0.19591522 | 0.058321616 | 0.252216703 | 0.16940119 | 0.631067218 | 0.214661106 | 0.684267948 | 0.164663922 |
| Chd8 | T | 2206 | -0.076144298 | 0.09429684 | 0.080857604 | 0.157001902 | -0.013439236 | 0.341358984 | 0.003092165 | 0.101491931 | 0.108525277 | 0.718711651 |
| Rai1 | S | 1332 | 0.019313158 | 0.05608423 | 0.072726948 | 0.053413789 | 0.016642717 | 0.376213186 | 0.099907823 | 0.130366287 | 0.230279471 | 0.687025159 |
| Htra2 | S | 212 | 0.08129575 | 0.141516069 | 0.075320369 | -0.005975382 | -0.066195701 | 0.656745846 | 0.226596343 | 0.472075124 | 0.972289228 | 0.49781281 |
| Heatr6 | S | 646 | -0.066937365 | 0.15484117 | 0.114620868 | 0.181558233 | -0.040220302 | 0.407610283 | 0.132647618 | 0.060687821 | 0.086516954 | 0.621760097 |
| Ap2a1 | T | 650 | 0.007387138 | 0.076438612 | 0.123658447 | 0.11627131 | 0.047219835 | 0.843245489 | 0.189975104 | 0.145132091 | 0.168392188 | 0.535874806 |

| | | | | | | | | | | | | |
|---|---|---|---|---|---|---|---|---|---|---|---|---|
| Spry2 | S | 114 | 0.093090587 | 0.11418447 | -0.075456751 | -0.168547338 | -0.189641221 | 0.024993506 | 0.091609761 | 0.500967297 | 0.203627468 | 0.16202859 |
| Atp5b | S | 529 | 0.069576403 | 0.112386908 | 0.020800713 | -0.04877569 | -0.091586195 | 0.261169323 | 0.102216814 | 0.801664539 | 0.49246504 | 0.251508328 |
| Nol9 | S | 88 | 0.105446266 | 0.117572229 | -0.014624643 | -0.120070909 | -0.132196872 | 0.19468924 | 0.106247888 | 0.84792898 | 0.200082915 | 0.141752127 |
| Vim | S | 72 | 0.126574057 | 0.026645542 | -0.03580563 | -0.162379688 | -0.062451172 | 0.001614495 | 0.311713013 | 0.330394908 | 0.010347297 | 0.115494975 |
| Acbd3 | S | 126 | 0.069657946 | 0.200854644 | -0.025256999 | -0.094914946 | -0.226111643 | 0.517538743 | 0.071261474 | 0.852443916 | 0.444079235 | 0.134726178 |
| 3110082I17Rik | S | 52 | 0.062997302 | 0.063440779 | 0.001706453 | -0.061290849 | -0.061734325 | 0.00722343 | 0.21026998 | 0.97216642 | 0.279046173 | 0.32691993 |
| Rbm15b | T | 628 | 0.026957774 | 0.057099257 | 0.076343106 | 0.049385332 | 0.01924385 | 0.500454225 | 0.169961554 | 0.095242532 | 0.130872816 | 0.478982813 |
| Sfswap | S | 841 | 0.12994886 | 0.052287865 | -0.026640607 | -0.156589467 | -0.078928472 | 0.015269207 | 0.34571841 | 0.713178207 | 0.103402885 | 0.31254718 |
| Sfswap | S | 843 | 0.12994886 | 0.052287865 | -0.026640607 | -0.156589467 | -0.078928472 | 0.015269207 | 0.34571841 | 0.713178207 | 0.103402885 | 0.31254718 |
| Cstf3 | S | 691 | -0.010290018 | 0.019456646 | 0.138265563 | 0.148555581 | 0.118808917 | 0.772762236 | 0.36552901 | 0.011049407 | 0.017870103 | 0.02599525 |
| Itsn1 | S | 334 | -0.035575012 | 0.069021068 | 0.082501573 | 0.118076586 | 0.013480506 | 0.482986635 | 0.027364354 | 0.029118177 | 0.077401991 | 0.599726843 |
| Tlk1 | S | 159 | 0.076782877 | 0.114564593 | 0.002181782 | -0.074601095 | -0.112382811 | 0.290341502 | 0.062920369 | 0.980651841 | 0.449488215 | 0.258559633 |
| Mcm2 | S | 140 | 0.017612741 | 0.060625636 | 0.04663709 | 0.029024349 | -0.013988547 | 0.419483997 | 0.02747706 | 0.171708958 | 0.374423579 | 0.641383007 |
| Bcl9 | S | 19 | -0.035180236 | 0.076886717 | 0.179036133 | 0.214216369 | 0.102149415 | 0.437551114 | 0.188878007 | 0.070467617 | 0.039071731 | 0.216842482 |
| Irf2bp2 | S | 389 | 0.01552433 | 0.039126327 | 0.080692452 | 0.065168121 | 0.041566125 | 0.753631951 | 0.129617504 | 0.031848371 | 0.260305421 | 0.191636525 |
| Ksr1 | T | 411 | 0.024256123 | 0.173015989 | 0.010082929 | -0.014173194 | -0.16293306 | 0.651808329 | 0.040307686 | 0.92179655 | 0.88940513 | 0.19272525 |
| Usp39 | S | 46 | 0.086201234 | 0.067278321 | 0.007359235 | -0.078841999 | -0.059919086 | 0.14685843 | 0.055346522 | 0.902321735 | 0.27892436 | 0.342190017 |
| Nes | S | 1290 | 0.077310822 | -0.016694954 | 0.096764439 | 0.019453617 | 0.113459393 | 0.041074845 | 0.568425786 | 0.097150485 | 0.672717411 | 0.075863737 |
| Pofut2 | S | 318 | 0.039598235 | 0.116989357 | 0.076692688 | 0.037094452 | -0.04029667 | 0.685433336 | 0.259582709 | 0.194165926 | 0.707823112 | 0.668644704 |
| Pofut2 | T | 322 | 0.039598235 | 0.116989357 | 0.076692688 | 0.037094452 | -0.04029667 | 0.685433336 | 0.259582709 | 0.194165926 | 0.707823112 | 0.668644704 |
| Wdr60 | S | 252 | -0.048665874 | 0.210334272 | 0.072121372 | 0.120787246 | -0.1382129 | 0.222682963 | 0.023634209 | 0.401143187 | 0.20601167 | 0.174204871 |
| Mastl | S | 442 | 0.055438732 | 0.099779224 | -0.027814861 | -0.083253593 | -0.127594085 | 0.197282537 | 0.007649242 | 0.576925448 | 0.176926195 | 0.069517712 |
| Mknk1 | S | 396 | 0.065787428 | 0.211940908 | -0.044362491 | -0.110149919 | -0.256303399 | 0.619694951 | 0.048510094 | 0.77848774 | 0.552281335 | 0.181663808 |
| Arfgef2 | T | 623 | 0.117953619 | 0.188323956 | -0.065873917 | -0.183827536 | -0.254197873 | 0.249525624 | 0.104258082 | 0.59372645 | 0.224103252 | 0.119681474 |
| Fhdc1 | S | 501 | -0.01837912 | 0.083764071 | 0.08388965 | 0.102268771 | 0.00012558 | 0.555463906 | 0.028761548 | 0.109726943 | 0.074703351 | 0.997504799 |
| Cavin1 | T | 378 | 0.06476031 | 0.113723587 | 0.044084349 | -0.020675961 | -0.069639238 | 0.34971074 | 0.096925542 | 0.81294533 | 0.910627209 | 0.708395658 |
| BC005624 | S | 24 | 0.098807713 | 0.180809156 | -0.036492386 | -0.135300098 | -0.217301542 | 0.390251108 | 0.093561257 | 0.809377989 | 0.438160369 | 0.229278534 |
| Mfap1a Mfap1b | T | 267;267 | 0.010810902 | 0.019399156 | 0.127963217 | 0.117152315 | 0.108564062 | 0.786213415 | 0.620867245 | 0.016576051 | 0.005977355 | 0.014976755 |
| Smim13 | T | 63 | 0.051377786 | 0.046901823 | 0.058045064 | 0.006667278 | 0.011143241 | 0.22179946 | 0.302046087 | 0.112851857 | 0.869522297 | 0.801569625 |
| Fam122b | S | 130 | 0.004288796 | 0.0578657 | 0.186928037 | 0.18263924 | 0.129062337 | 0.942093365 | 0.534844661 | 0.094452833 | 0.101857765 | 0.276092005 |
| Fam122b | S | 145 | 0.004288796 | 0.0578657 | 0.186928037 | 0.18263924 | 0.129062337 | 0.942093365 | 0.534844661 | 0.094452833 | 0.101857765 | 0.276092005 |
| Zfhx3 | S | 1600 | 0.036181385 | 0.033354625 | 0.080230823 | 0.044049438 | 0.046876198 | 0.089830099 | 0.114994344 | 0.311651949 | 0.54044858 | 0.517667293 |
| Prr12 | S | 1123 | 0.026424094 | 0.093041189 | 0.169462072 | 0.143037978 | 0.076420883 | 0.879590999 | 0.402115986 | 0.202368767 | 0.456859819 | 0.551987023 |
| Rgs12 | S | 661 | -0.038002078 | 0.133359737 | 0.133939394 | 0.171941471 | 0.000579657 | 0.620351779 | 0.028425585 | 0.477152514 | 0.382883098 | 0.997348479 |
| Rfc1 | T | 109 | 0.086949046 | 0.073318742 | 0.171700353 | 0.084751308 | 0.098381611 | 0.340655219 | 0.616442643 | 0.44353189 | 0.690724927 | 0.674189554 |
| Mdm1 | S | 556 | 0.031361261 | -0.017024335 | 0.208475166 | 0.177113905 | 0.225499501 | 0.508196485 | 0.740657167 | 0.048798979 | 0.063702989 | 0.034185766 |
| Ptk2 | Y | 397 | -0.050385287 | 0.127037354 | 0.073379831 | 0.123765118 | -0.053657524 | 0.466931231 | 0.061375558 | 0.050580645 | 0.142254488 | 0.316641153 |
| Kif1b | S | 1454 | -0.152509241 | 0.20005771 | 0.180239171 | 0.332748411 | -0.01981854 | 0.023288292 | 0.027956134 | 0.034787139 | 0.015199067 | 0.781154402 |
| Anapc1 | S | 555 | -0.001735717 | 0.051351386 | 0.103870758 | 0.105606475 | 0.052519373 | 0.933529751 | 0.2083749 | 0.033586149 | 0.037197267 | 0.248698701 |
| Cobl | S | 272 | -0.105436145 | 0.036416399 | 0.230876212 | 0.336312357 | 0.194459813 | 0.37703354 | 0.671286772 | 0.007235335 | 0.059840568 | 0.074876748 |
| Rev1 | S | 291 | 0.06273908 | 0.016165638 | 0.09362168 | 0.0308826 | 0.077456042 | 0.046020863 | 0.627667245 | 0.290614033 | 0.687820659 | 0.364236534 |
| Prpf38a | S | 292 | 0.151741612 | 0.078668224 | 0.087956695 | -0.063784916 | 0.009288472 | 0.498580066 | 0.442802396 | 0.390463425 | 0.76741419 | 0.928455802 |
| Kif23 | S | 687 | -0.006092756 | 0.114244019 | 0.247058851 | 0.253151607 | 0.132814832 | 0.973545428 | 0.535842042 | 0.22012173 | 0.263832359 | 0.527680679 |
| Vim | T | 63 | 0.053852408 | 0.033286865 | 0.171357137 | 0.117504729 | 0.138070273 | 0.714996001 | 0.689180664 | 0.106295648 | 0.420155524 | 0.102223657 |
| Lrrc47 | S | 430 | 0.432735128 | 0.209977882 | -0.512497021 | -0.94523215 | -0.722474903 | 0.037140904 | 0.297954267 | 0.258616134 | 0.0990038 | 0.151739871 |
| Cdc23 | T | 582 | -0.164347616 | 0.249699792 | 0.142744702 | 0.307092318 | -0.10695509 | 0.113355442 | 0.013899902 | 0.273642315 | 0.066007442 | 0.393698084 |
| Fnbp4 | S | 120 | 0.015837736 | 0.029840678 | 0.090001661 | 0.074163925 | 0.060160983 | 0.483977479 | 0.016565848 | 0.157399116 | 0.207597876 | 0.282143528 |
| Cdk14 | T | 125 | -0.032439073 | 0.114905631 | 0.146210776 | 0.17864985 | 0.031305146 | 0.674558887 | 0.105367046 | 0.222243398 | 0.161485958 | 0.757676627 |
| Med13 Med13l | T | 1010;1045 | -0.077528576 | -0.013695934 | 0.407740217 | 0.485268794 | 0.421436151 | 0.654122175 | 0.881160713 | 0.055091873 | 0.058583992 | 0.069788848 |
| Cav1 | Y | 42 | -0.056657173 | 0.086023023 | 0.182751835 | 0.239409007 | 0.096728812 | 0.501163788 | 0.087887868 | 0.135383592 | 0.085231538 | 0.353927617 |
| Gorasp2 | S | 451 | 0.050653247 | 0.045560287 | 0.102965397 | 0.05231215 | 0.05740511 | 0.273569559 | 0.283599323 | 0.268238679 | 0.538299791 | 0.498486656 |
| Huwe1 | S | 3135 | 0.029138746 | 0.047311762 | 0.131756713 | 0.102617967 | 0.084444951 | 0.664446314 | 0.227599348 | 0.162378727 | 0.29192521 | 0.315620104 |
| Myo9a | S | 1300 | 0.045250459 | 0.039774483 | 0.04722157 | 0.001971111 | 0.007447087 | 0.216229438 | 0.15653725 | 0.10236752 | 0.953701747 | 0.790888541 |
| Spen | S | 3446 | 0.035694623 | 0.085525622 | 0.058932352 | 0.02323773 | -0.02659327 | 0.458893512 | 0.20617949 | 0.160012856 | 0.604761044 | 0.649770107 |
| Spen | T | 3447 | 0.035694623 | 0.085525622 | 0.058932352 | 0.02323773 | -0.02659327 | 0.458893512 | 0.20617949 | 0.160012856 | 0.604761044 | 0.649770107 |
| Exoc2 | S | 432 | -0.013270167 | 0.093371739 | 0.065730367 | 0.079000534 | -0.027641371 | 0.745933765 | 0.025304176 | 0.173747555 | 0.152099737 | 0.523493613 |
| Tmem237 | S | 227 | 0.081187035 | 0.068747491 | 0.048126308 | -0.033060727 | -0.020621183 | 0.100910666 | 0.485210291 | 0.406347192 | 0.463275685 | 0.825576552 |
| Srrm2 | S | 2535 | 0.003664069 | 0.048188501 | 0.086531218 | 0.082867149 | 0.038342717 | 0.918086786 | 0.097638277 | 0.049468337 | 0.103286379 | 0.250102728 |
| Polr2a | S | 1966 | 0.013237358 | 0.112514143 | 0.080969412 | 0.067732054 | -0.031544731 | 0.901127683 | 0.181762901 | 0.156788706 | 0.530578002 | 0.655119016 |
| Sec62 | S | 313 | -0.109936026 | 0.080412191 | 0.192223638 | 0.302159664 | 0.111811447 | 0.288519117 | 0.178431394 | 0.021434729 | 0.03682989 | 0.105348274 |
| Nrcam | T | 1173 | 0.14394582 | 0.044567867 | 0.070737213 | -0.073208607 | 0.026169346 | 0.188331523 | 0.570847315 | 0.524894767 | 0.50204153 | 0.761871706 |
| Adam9 | S | 300 | -0.030723348 | 0.048128431 | 0.094133544 | 0.124856892 | 0.046005113 | 0.677185182 | 0.174623365 | 0.007251482 | 0.177772545 | 0.15583508 |
| Map1b | T | 1323 | -0.032085744 | 0.1655034 | 0.087389905 | 0.119475649 | -0.078113495 | 0.718793704 | 0.070312986 | 0.353899081 | 0.30818205 | 0.440441297 |
| Fam219a | S | 44 | 0.031138134 | 0.076717908 | 0.070660369 | 0.039522235 | -0.006057538 | 0.57213901 | 0.219694365 | 0.118478917 | 0.49674684 | 0.915140138 |
| Tnrc18 | S | 348 | 0.107239058 | 0.035472884 | 0.02984672 | -0.077392338 | -0.005626164 | 0.055868053 | 0.469764212 | 0.514787352 | 0.102233193 | 0.883380393 |
| Tcof1 | T | 324 | 0.287836807 | -0.020160421 | -0.033330273 | -0.32116708 | -0.013169852 | 0.028901466 | 0.868145706 | 0.758729338 | 0.006392229 | 0.88195078 |
| Faf1 | T | 268 | -0.10525253 | 0.161037074 | 0.173490938 | 0.278743467 | 0.012453863 | 0.293730664 | 0.242585578 | 0.041819196 | 0.044024481 | 0.917468481 |
| 4932438A13Rik | S | 1305 | 0.111539195 | 0.009328545 | 0.027516337 | -0.084022858 | 0.018187792 | 0.016029194 | 0.759450856 | 0.536787111 | 0.116927383 | 0.663381542 |
| A430005L14Rik | S | 180 | 0.06588103 | 0.064927637 | 0.040703437 | -0.025177594 | -0.0242242 | 0.355126866 | 0.187785848 | 0.167884238 | 0.69048815 | 0.544553146 |
| Kansl2 | T | 131 | 0.071917659 | 0.068099033 | 0.126649049 | 0.054731389 | 0.058550016 | 0.403608569 | 0.485469712 | 0.315438904 | 0.604833489 | 0.607071717 |
| Clasp1 | S | 636 | 0.075801506 | 0.094074885 | 0.004702705 | -0.071098801 | -0.08937218 | 0.213590037 | 0.069999425 | 0.938616465 | 0.305910671 | 0.181046337 |
| Uhrf1bp1l | S | 986 | 0.072041807 | 0.06099412 | 0.085222443 | 0.013180636 | 0.024228323 | 0.238076136 | 0.547993465 | 0.24801475 | 0.785497352 | 0.798326091 |
| Jmy | S | 75 | 0.006066121 | 0.069042699 | 0.091714302 | 0.085648181 | 0.022671603 | 0.878387474 | 0.185706367 | 0.073907392 | 0.0536203 | 0.577035605 |
| Erbb2 | T | 702 | 0.042303731 | 0.068481938 | 0.104900997 | 0.062597266 | 0.036419059 | 0.568969088 | 0.376963547 | 0.151944535 | 0.438772393 | 0.646346612 |
| Cdk17 | S | 105 | 0.075786455 | 0.063505961 | 0.030644928 | -0.045141527 | -0.032861032 | 0.212079513 | 0.080011624 | 0.654082969 | 0.562275665 | 0.629499506 |
| Med1 | S | 1000 | 0.076407041 | -0.000334104 | 0.059828668 | -0.016578373 | 0.060162772 | 0.018971219 | 0.996598559 | 0.20622239 | 0.685666531 | 0.486960927 |
| Srrm2 | S | 1840 | 0.049058916 | 0.023613005 | 0.128315742 | 0.079256826 | 0.104702737 | 0.367099564 | 0.651727825 | 0.108283448 | 0.23510117 | 0.150995134 |
| Carmil1 | S | 1070 | 0.00192127 | 0.309686325 | 0.030322388 | 0.028401117 | -0.279363937 | 0.99144741 | 0.153828423 | 0.763305825 | 0.875011107 | 0.187361108 |
| Tmem131 | S | 1153 | 0.039856152 | 0.133288708 | 0.057379893 | 0.017523741 | -0.075908815 | 0.657380916 | 0.102460445 | 0.584568232 | 0.868827114 | 0.449360552 |
| Pi4ka | S | 262 | 0.106861047 | 0.10233997 | -0.038566975 | -0.145428022 | -0.140906945 | 0.192465788 | 0.030309928 | 0.505708112 | 0.121690489 | 0.076310392 |
| Ybx3 | S | 195 | 0.023704749 | 0.057010701 | 0.034470829 | 0.010766079 | -0.022539872 | 0.480841337 | 0.043450584 | 0.072151343 | 0.725754198 | 0.295130455 |
| Nop2 | S | 179 | 0.060875384 | 0.043686179 | 0.033194524 | -0.027680859 | -0.010491654 | 0.088794157 | 0.235904718 | 0.231452574 | 0.366424093 | 0.746450863 |
| Mast1 Mast2 | S | 1345;1268 | 0.039938006 | 0.119714771 | 0.039197299 | -0.000740706 | -0.080517471 | 0.474504842 | 0.119474423 | 0.443243413 | 0.986079978 | 0.232993183 |
| Pde4a | S | 333 | -0.092576152 | 0.220462549 | 0.114751032 | 0.207327184 | -0.105711518 | 0.035383283 | 0.016977853 | 0.153654983 | 0.061136821 | 0.213917668 |
| Steap3 | S | 20 | 0.052702307 | 0.079415365 | 0.01224005 | -0.040462257 | -0.067175315 | 0.350866065 | 0.021355125 | 0.687788185 | 0.447291126 | 0.033619037 |
| Fam126b | S | 521 | 0.137219371 | 0.054773993 | 0.042704812 | -0.094514558 | -0.01206918 | 0.13981701 | 0.49922471 | 0.629274441 | 0.351174823 | 0.897365257 |
| AW554918 | S | 558 | 0.070361406 | -0.00367948 | 0.067325556 | -0.00303585 | 0.071005037 | 0.035129442 | 0.888567035 | 0.103728629 | 0.924061162 | 0.091441124 |
| Ahctf1 | T | 1954 | 0.059150771 | 0.094086864 | 0.055778046 | -0.003372725 | -0.038308818 | 0.262983565 | 0.248141491 | 0.448192487 | 0.963517418 | 0.667432642 |
| Depdc5 | S | 503 | 0.046794716 | 0.024072182 | 0.14923081 | 0.102436094 | 0.125158628 | 0.625750846 | 0.668584659 | 0.083844842 | 0.352576363 | 0.144083713 |
| Cramp1l | S | 1245 | 0.325676176 | 0.260776004 | -0.133790054 | -0.45946623 | -0.394566058 | 0.335623705 | 0.481198438 | 0.788328203 | 0.415041264 | 0.490797642 |
| Arid4b | S | 675 | -0.015960542 | 0.000408658 | 0.228777974 | 0.244738515 | 0.228369315 | 0.914325517 | 0.99481985 | 0.031215935 | 0.190592435 | 0.037924018 |
| Rbm15b | S | 111 | 0.09219969 | 0.011693034 | 0.09408749 | 0.0018878 | 0.082394456 | 0.286256942 | 0.859047975 | 0.093333598 | 0.980965112 | 0.277482743 |
| Lmna | S | 614 | 0.036173916 | 0.034086208 | 0.094511878 | 0.058337962 | 0.06042567 | 0.2686571 | 0.468228816 | 0.09203887 | 0.243373104 | 0.298843091 |
| Arhgef12 | T | 189 | 0.093558394 | 0.047806173 | 0.100154266 | 0.006595872 | 0.052348094 | 0.269451574 | 0.590622349 | 0.305581351 | 0.948027406 | 0.632879603 |

| | | | | | | | | | | | | | |
|---|---|---|---|---|---|---|---|---|---|---|---|---|---|
| Nup133 | S | | 593 | -0.057726504 | 0.139069045 | 0.157263769 | 0.214990273 | 0.018194724 | 0.460077811 | 0.286932703 | 0.069214081 | 0.021094778 | 0.871671892 |
| Trp53bp1 | S | | 1094 | -0.004777396 | 0.052195682 | 0.085989493 | 0.090766889 | 0.033793811 | 0.931233598 | 0.106163476 | 0.038841918 | 0.179470829 | 0.320203239 |
| Tcea1 | S | | 96 | -0.182653367 | 0.284323187 | 0.17716847 | 0.359821837 | -0.107154717 | 0.015518888 | 0.022062019 | 0.071914367 | 0.017563559 | 0.290644885 |
| Add1 | T | | 429 | 0.119866415 | 0.090909145 | -0.035409402 | -0.155275817 | -0.126318547 | 0.031135073 | 0.199911882 | 0.41853983 | 0.007002379 | 0.108589585 |
| Phf8-ps Phf8 | S | | 615 | 0.051168436 | 0.056443137 | 0.164051423 | 0.112882987 | 0.107608286 | 0.60368849 | 0.406968804 | 0.267304484 | 0.444788869 | 0.430184989 |
| Slc25a31 Slc25a4 Slc | T | 84;97;84 | | -0.080446529 | 0.120848845 | 0.134423836 | 0.214870365 | 0.013574991 | 0.617240748 | 0.163086889 | 0.082061353 | 0.240188852 | 0.811334112 |
| Prpf38b | S | | 290 | 0.108029696 | -0.069361952 | 0.162099714 | 0.054070018 | 0.231461666 | 0.300414951 | 0.260957854 | 0.020224499 | 0.559772175 | 0.005532621 |
| Kank2 | S | | 532 | 0.023559502 | 0.124123616 | 0.006561255 | -0.016998247 | -0.117562361 | 0.355732624 | 0.029610025 | 0.846431062 | 0.576950205 | 0.042886515 |
| Tbc1d2 | S | | 914 | 0.029193733 | 0.029757611 | 0.107146994 | 0.077953261 | 0.077389383 | 0.37587581 | 0.408659097 | 0.081870378 | 0.173930235 | 0.166263616 |
| Ttc7 | S | | 52 | -0.110804205 | 0.171877202 | 0.122600002 | 0.233404208 | -0.0492772 | 0.234870872 | 0.011863583 | 0.312422361 | 0.116228947 | 0.654786851 |
| Tjp1 | S | | 175 | 0.081340916 | 0.052912298 | 0.042704851 | -0.038636065 | -0.010207446 | 0.137144686 | 0.316329499 | 0.387401775 | 0.479573091 | 0.85117723 |
| Farp1 | S | | 408 | 0.100999817 | -0.064741926 | 0.143242149 | 0.042242332 | 0.207984075 | 0.118207536 | 0.277788875 | 0.04249875 | 0.35520125 | 0.006257976 |
| Trip12 | S | | 85 | 0.115654952 | 0.092505594 | 0.017897698 | -0.097757254 | -0.074607895 | 0.226575753 | 0.215455908 | 0.829617938 | 0.333304523 | 0.379634816 |
| Mki67 | S | | 1371 | -0.007318534 | 0.124482116 | 0.051135873 | 0.058454407 | -0.073346243 | 0.896424647 | 0.042899853 | 0.291403218 | 0.379085666 | 0.211909398 |
| Aff4 | S | | 205 | -0.073632131 | 0.09295368 | 0.135658242 | 0.209290373 | 0.042704563 | 0.444650777 | 0.221441689 | 0.022247361 | 0.095738464 | 0.534621431 |
| Thoc5 | T | | 328 | 0.004489857 | -0.031910505 | 0.227035064 | 0.222545207 | 0.25894557 | 0.954084341 | 0.825315276 | 0.025594823 | 0.057153298 | 0.162428822 |
| Tjp2 | S | | 265 | -0.038343246 | -0.073334405 | 0.270795535 | 0.30913878 | 0.344129939 | 0.536384499 | 0.191688194 | 0.001181959 | 0.004580013 | 0.00101802 |
| Inpp5f | S | | 943 | -0.071688114 | 0.129220321 | 0.026848145 | 0.098536259 | -0.102372176 | 0.31405409 | 0.000272002 | 0.656947727 | 0.255460525 | 0.177517552 |
| Iqsec1 | S | | 922 | 0.004897322 | 0.163727631 | 0.053528565 | 0.048631243 | -0.110199067 | 0.945285518 | 0.088207815 | 0.472676439 | 0.333395239 | 0.164990877 |
| Chaf1b | S | | 436 | 0.011045739 | 0.093017158 | 0.093205854 | 0.082160114 | 0.000188696 | 0.815710518 | 0.091427852 | 0.315649247 | 0.373555417 | 0.998228493 |
| Cwc22 | S | | 25 | 0.028201619 | 0.052953136 | 0.097520878 | 0.069319259 | 0.044567743 | 0.554152313 | 0.289582911 | 0.112920728 | 0.275508067 | 0.454661391 |
| Hnrnpk | S | | 379 | 0.052778326 | 0.099996579 | 0.066143329 | 0.013365003 | -0.03385325 | 0.337058271 | 0.336349968 | 0.315518765 | 0.845660713 | 0.740111069 |
| Nsrp1 | S | | 31 | 0.023032054 | 0.050952592 | 0.052805122 | 0.029773068 | 0.00185253 | 0.606222434 | 0.098465498 | 0.052275951 | 0.510209079 | 0.942604897 |
| Prpf38a | S | | 294 | 0.113646605 | 0.072233067 | 0.081483399 | -0.032163206 | 0.009250332 | 0.516776565 | 0.357430372 | 0.333607534 | 0.851613508 | 0.917291754 |
| Prpf38a | S | | 296 | 0.113646605 | 0.072233067 | 0.081483399 | -0.032163206 | 0.009250332 | 0.516776565 | 0.357430372 | 0.333607534 | 0.851613508 | 0.917291754 |
| Cdk10 | T | | 196 | -0.020880544 | 0.01110016 | 0.125915844 | 0.146796389 | 0.114815685 | 0.601542998 | 0.710427976 | 0.001944217 | 0.031378661 | 0.023417799 |
| Plcd1 | S | | 460 | -0.007743909 | 0.121946952 | 0.053524534 | 0.061268443 | -0.068422418 | 0.888182548 | 0.023837387 | 0.472099819 | 0.41615235 | 0.343123778 |
| Cenpc1 | S | | 409 | 0.149952077 | 0.120460984 | -0.02617767 | -0.176129748 | -0.146638655 | 0.235063494 | 0.156711132 | 0.716229695 | 0.185632678 | 0.125091145 |
| Cenpc1 | S | | 412 | 0.149952077 | 0.120460984 | -0.02617767 | -0.176129748 | -0.146638655 | 0.235063494 | 0.156711132 | 0.716229695 | 0.185632678 | 0.125091145 |
| Atp7a | T | | 327 | 0.017263341 | 0.085350196 | 0.083632858 | 0.066369517 | -0.001717338 | 0.855491806 | 0.134601025 | 0.179176335 | 0.522235094 | 0.977428306 |
| Zcchc8 | S | | 503 | 0.037367216 | 0.004579888 | 0.092342102 | 0.054974886 | 0.087762214 | 0.075719887 | 0.82685301 | 0.063817005 | 0.190489936 | 0.075930173 |
| Rab11fip5 | S | | 254 | 0.040297185 | 0.242228068 | -0.075706603 | -0.116003788 | -0.31793467 | 0.720109255 | 0.015654207 | 0.36847005 | 0.301415983 | 0.005466355 |
| Chd8 | S | | 2202 | -0.083909983 | 0.163165081 | 0.108902458 | 0.192812442 | -0.054262622 | 0.6105197 | 0.05776756 | 0.295395504 | 0.302277693 | 0.584651066 |
| Yeats2 | S | | 446 | -0.011324359 | 0.030127408 | 0.134566785 | 0.145891144 | 0.104439377 | 0.690053931 | 0.429838601 | 0.015757637 | 0.015354578 | 0.057763336 |
| Cep135 | S | | 439 | 0.046653217 | 0.020238326 | 0.095842295 | 0.049189078 | 0.075603969 | 0.34240147 | 0.575193827 | 0.064628342 | 0.36574956 | 0.138412345 |
| Tmem63a | S | | 100 | 0.136037755 | 0.281603856 | -0.296857463 | -0.432895218 | -0.578461319 | 0.357670054 | 0.066845806 | 0.574956638 | 0.432972445 | 0.320915489 |
| Cep68 | S | | 459 | -0.007355432 | 0.143341231 | 0.155850361 | 0.163205793 | 0.012509131 | 0.972535922 | 0.263764367 | 0.325026746 | 0.507611694 | 0.93868066 |
| Stag2 | S | | 1058 | 0.022829898 | 0.12900155 | 0.072994501 | 0.050164603 | -0.056007049 | 0.830895447 | 0.103542147 | 0.468982054 | 0.680498368 | 0.561761449 |
| Erc1 Erc2 | S | 318;314 | | -0.002735374 | 0.034241919 | 0.067299847 | 0.070035221 | 0.033057928 | 0.954222158 | 0.20784265 | 0.0033039 | 0.227879905 | 0.218775211 |
| Limk2 | S | | 287 | 0.068993216 | 0.085196107 | 0.021061709 | -0.047931506 | -0.064134398 | 0.204165731 | 0.087083115 | 0.874384148 | 0.726666385 | 0.641036844 |
| Ccdc9b | T | | 514 | 0.056463848 | 0.008920931 | 0.168658139 | 0.112174291 | 0.159737208 | 0.490250649 | 0.940388155 | 0.105440344 | 0.288757915 | 0.269394182 |
| Iws1 | S | | 309 | 0.012490073 | -0.00478235 | 0.133350981 | 0.120860908 | 0.138133331 | 0.59246585 | 0.736337746 | 0.008497733 | 0.012673774 | 0.017138003 |
| Exoc1 | S | | 473 | -0.017796529 | 0.033109337 | 0.163989784 | 0.181786313 | 0.130880447 | 0.764003846 | 0.598985414 | 0.024744714 | 0.011331292 | 0.045319438 |
| Slf2 | S | | 708 | 0.04987909 | 0.134155352 | 0.064803601 | 0.014924511 | -0.069351751 | 0.762343678 | 0.195448097 | 0.366039813 | 0.924605327 | 0.42732003 |
| Pura | S | | 181 | 0.054492684 | 0.158092504 | 0.040042113 | -0.014450571 | -0.118050391 | 0.528239487 | 0.189880552 | 0.611814771 | 0.850160189 | 0.292824751 |
| Aebp2 | S | | 131 | 0.121183803 | -0.013199659 | 0.01575656 | -0.105427244 | 0.028956219 | 0.002359333 | 0.648108509 | 0.804012374 | 0.180125731 | 0.648642519 |
| Lbr | S | | 71 | 0.019846422 | 0.047462468 | 0.08650875 | 0.066662329 | 0.039046282 | 0.57594584 | 0.188722687 | 0.093125134 | 0.157459791 | 0.349177916 |
| Senp1 | S | | 157 | 0.039935103 | -0.122701167 | 0.201415055 | 0.161479952 | 0.324116223 | 0.401863334 | 0.340292753 | 0.006196282 | 0.018364244 | 0.069391461 |
| Ralgapa2 | S | | 373 | 0.107270164 | 0.031739664 | 0.032226514 | -0.07504365 | 0.00048685 | 0.090421871 | 0.161059308 | 0.775627702 | 0.535551909 | 0.996509968 |
| Ralgapa2 | S | | 376 | 0.107270164 | 0.031739664 | 0.032226514 | -0.07504365 | 0.00048685 | 0.090421871 | 0.161059308 | 0.775627702 | 0.535551909 | 0.996509968 |
| Agfg1 | Y | | 327 | -0.115840633 | 0.045792946 | 0.25754895 | 0.373389583 | 0.211756004 | 0.308558433 | 0.458039268 | 0.017812173 | 0.027821858 | 0.041107581 |
| Hps6 | T | | 765 | 0.082708616 | 0.083696783 | 0.085071267 | 0.002362652 | 0.001374485 | 0.475349399 | 0.16804782 | 0.61302958 | 0.98940471 | 0.993028352 |
| Macroh2a1 | S | | 170 | 0.021413218 | 0.109477828 | 0.065301826 | 0.043888608 | -0.044176002 | 0.839253003 | 0.066731234 | 0.436412656 | 0.702244542 | 0.557038717 |
| Atrx | S | | 315 | 0.001975634 | 0.03263869 | 0.094278836 | 0.092303202 | 0.061640146 | 0.938049387 | 0.073849452 | 0.048416582 | 0.052163954 | 0.128335041 |
| Rlim | S | | 163 | 0.098514487 | 0.138222207 | -0.093994329 | -0.192508816 | -0.232216536 | 0.115910937 | 0.010263121 | 0.154133816 | 0.029612943 | 0.020410525 |
| Mast2 | S | | 1367 | -0.115789473 | 0.264059039 | 0.097170972 | 0.212960445 | -0.166888068 | 0.072552923 | 0.014435052 | 0.325280057 | 0.093483958 | 0.139295982 |
| Coil | T | | 122 | 0.092558002 | 0.020346648 | 0.111452728 | 0.018894726 | 0.09110608 | 0.227063253 | 0.624995364 | 0.29541902 | 0.855609965 | 0.373034138 |
| Spout1 | S | | 374 | 0.044860775 | 0.060190108 | 0.016520746 | -0.028340029 | -0.043669362 | 0.313844847 | 0.016168742 | 0.478794194 | 0.507702679 | 0.112762731 |
| Neo1 | S | | 1330 | -0.007745 | 0.027212481 | 0.256524419 | 0.264269419 | 0.229311938 | 0.884454472 | 0.444592814 | 0.133435626 | 0.120029469 | 0.162718686 |
| Fam193b | S | | 764 | 0.01738729 | 0.151402369 | 0.044414735 | 0.027027445 | -0.106987635 | 0.824290077 | 0.071454587 | 0.61796272 | 0.720497 | 0.193299715 |
| Mtmr3 | S | | 288 | 0.007555341 | 0.132144807 | -0.003983718 | -0.011539059 | -0.136128525 | 0.84261823 | 0.006555289 | 0.944874672 | 0.848692406 | 0.08783323 |
| Sipa1l3 | S | | 198 | 0.030800291 | 0.011242469 | 0.171151112 | 0.140350821 | 0.159908642 | 0.71914407 | 0.882584872 | 0.058241915 | 0.074551263 | 0.018673296 |
| Larp7 | S | | 256 | -0.00192502 | 0.049521471 | 0.060162919 | 0.062087939 | 0.010641448 | 0.940248545 | 0.066383325 | 0.021722986 | 0.081057388 | 0.643430572 |
| Aptx | S | | 127 | 0.106815777 | 0.025746225 | 0.039827893 | -0.066987884 | 0.014081668 | 0.061610963 | 0.603128058 | 0.462389094 | 0.163172775 | 0.730786025 |
| Bud13 | S | | 238 | 0.025021862 | 0.02556627 | 0.097030923 | 0.072009061 | 0.071464653 | 0.393083999 | 0.304088368 | 0.07043609 | 0.144314921 | 0.155087034 |
| Rapgef2 | S | | 960 | -0.03073375 | 0.067247863 | 0.086063997 | 0.116797746 | 0.018816134 | 0.49873649 | 0.011678226 | 0.113209974 | 0.080708492 | 0.649661242 |
| Champ1 | S | | 237 | -0.040348693 | 0.051698363 | 0.228400229 | 0.268748923 | 0.176701866 | 0.600591318 | 0.473018957 | 0.069490834 | 0.051696526 | 0.13554094 |
| Champ1 | S | | 243 | -0.040348693 | 0.051698363 | 0.228400229 | 0.268748923 | 0.176701866 | 0.600591318 | 0.473018957 | 0.069490834 | 0.051696526 | 0.13554094 |
| Champ1 | T | | 242 | -0.040348693 | 0.051698363 | 0.228400229 | 0.268748923 | 0.176701866 | 0.600591318 | 0.473018957 | 0.069490834 | 0.051696526 | 0.13554094 |
| Nek4 | S | | 429 | -0.042722853 | 0.17155511 | 0.056860485 | 0.099583338 | -0.114694624 | 0.390056913 | 0.020817464 | 0.430584181 | 0.183674804 | 0.153208497 |
| Iws1 | S | | 307 | 0.014320016 | -0.006680899 | 0.129452687 | 0.115132672 | 0.136133586 | 0.52006513 | 0.633282653 | 0.007979046 | 0.012359889 | 0.013326137 |
| Srrm2 | S | | 476 | 0.074940305 | 0.057810485 | 0.094278804 | 0.019338499 | 0.03646832 | 0.358716515 | 0.552482553 | 0.234370633 | 0.828265012 | 0.727662292 |
| Peak1 | T | | 692 | 0.051985846 | 0.016749517 | 0.064719675 | 0.012733829 | 0.047970158 | 0.135309808 | 0.709946479 | 0.054706727 | 0.696281592 | 0.338736689 |
| Farp1 | S | | 512 | 0.086399955 | 0.043887769 | 0.002700398 | -0.083699557 | -0.041187371 | 0.010999647 | 0.475567824 | 0.76471069 | 0.011952433 | 0.49975504 |
| Zc3h13 | S | | 380 | 0.069955515 | 0.027722886 | 0.051065722 | -0.018889793 | 0.023342836 | 0.027532323 | 0.524466893 | 0.471669537 | 0.774448729 | 0.74511106 |
| Rab3gap1 | S | | 540 | 0.130456538 | 0.120003879 | 0.053195977 | -0.077260561 | -0.066807902 | 0.400660351 | 0.507854471 | 0.470191485 | 0.594540086 | 0.698147404 |
| Cdk12 | S | | 317 | 0.056210895 | 0.041641622 | 0.061575051 | 0.005364156 | 0.019933428 | 0.255672273 | 0.331426445 | 0.15416952 | 0.906931554 | 0.63349876 |
| Lrch3 | S | | 415 | 0.140873082 | 0.201891342 | -0.149587605 | -0.290460687 | -0.351478947 | 0.088763849 | 0.013774049 | 0.043838151 | 0.009232188 | 0.00101074 |
| Efhd2 | S | | 74 | 0.216450339 | 0.050725621 | -0.144723921 | -0.36117426 | -0.195449542 | 0.008763948 | 0.29025619 | 0.27365621 | 0.050873197 | 0.179386736 |
| Dnttip2 | S | | 80 | -0.776050805 | 0.506475975 | -0.274503201 | 0.501547604 | -0.780979176 | NA | 0.25687983 | NA | NA | NA |
| Dnttip2 | T | | 78 | -0.776050805 | 0.506475975 | -0.274503201 | 0.501547604 | -0.780979176 | NA | 0.25687983 | NA | NA | NA |
| Dnttip2 | T | | 82 | -0.776050805 | 0.506475975 | -0.274503201 | 0.501547604 | -0.780979176 | NA | 0.25687983 | NA | NA | NA |
| Hnrnpa2b1 | S | | 341 | 0.081748153 | 0.055882787 | 0.048445127 | -0.033303027 | -0.00743766 | 0.152540336 | 0.339543428 | 0.468098294 | 0.51320172 | 0.888360631 |
| Lemd2 | S | | 150 | -0.093665115 | 0.061454469 | 0.236846149 | 0.330511264 | 0.175391681 | 0.188991489 | 0.290100165 | 0.022356396 | 0.009364031 | 0.056745029 |
| Phldb1 | T | | 524 | 0.165421945 | 0.000279925 | 0.007528217 | -0.157893729 | 0.007248291 | 0.02069818 | 0.994166683 | 0.905752476 | 0.075755627 | 0.908366282 |
| Smarca5 | S | | 65 | 0.026688748 | 0.043601752 | 0.059789161 | 0.033100414 | 0.016187409 | 0.368991805 | 0.080336908 | 0.153941745 | 0.389925642 | 0.623889926 |
| Kdm1b | S | | 253 | -0.014177466 | 0.12492176 | 0.072103146 | 0.086280612 | -0.052818614 | 0.599765421 | 0.177560387 | 0.060773782 | 0.037141556 | 0.499129167 |
| Cul4b | S | | 17 | -0.076759482 | 0.200192342 | 0.063832008 | 0.14059149 | -0.136360334 | 0.512183658 | 0.035956507 | 0.407088966 | 0.23134826 | 0.030807767 |
| Canx | S | | 582 | -0.018348518 | 0.061703619 | 0.079927141 | 0.098275659 | 0.018223522 | 0.502685798 | 0.010279078 | 0.12306245 | 0.080288414 | 0.64456832 |
| Uba1 | S | | 13 | -0.067990008 | 0.146199568 | 0.110687009 | 0.178677017 | -0.035512559 | 0.035526764 | 0.048061797 | 0.025261697 | 0.006378656 | 0.495378487 |
| Specc1l | T | | 52 | 0.032873856 | 0.093986921 | 0.049188268 | 0.016314412 | -0.044798653 | 0.369059486 | 0.202782909 | 0.254215142 | 0.632162752 | 0.496203867 |
| Rplp1 | T | | 84 | 0.060400373 | -0.025240487 | 0.077544983 | 0.01714461 | 0.102785469 | 0.064980132 | 0.595065518 | 0.02469934 | 0.517538604 | 0.095011922 |
| Hirip3 | S | | 238 | 0.006734498 | 0.018369649 | 0.093860015 | 0.087125518 | 0.075490366 | 0.741572659 | 0.637598247 | 0.006116492 | 0.013887955 | 0.125941697 |
| Mtfr1l | S | | 235 | 0.011314825 | 0.003317043 | 0.124482188 | 0.113167363 | 0.121165145 | 0.752808467 | 0.955407285 | 0.009862372 | 0.020739409 | 0.111573693 |

| | | | | | | | | | | | | |
|---|---|---|---|---|---|---|---|---|---|---|---|---|
| Gse1 | S | 116 | 0.161243372 | 0.046609116 | -0.019185399 | -0.180428771 | -0.065794514 | 0.055570798 | 0.260245203 | 0.556737198 | 0.036842041 | 0.170289733 |
| Gse1 | S | 119 | 0.161243372 | 0.046609116 | -0.019185399 | -0.180428771 | -0.065794514 | 0.055570798 | 0.260245203 | 0.556737198 | 0.036842041 | 0.170289733 |
| Gse1 | T | 120 | 0.161243372 | 0.046609116 | -0.019185399 | -0.180428771 | -0.065794514 | 0.055570798 | 0.260245203 | 0.556737198 | 0.036842041 | 0.170289733 |
| Atad5 | T | 616 | 0.01619188 | 0.400350052 | -0.040154472 | -0.056346352 | -0.440504524 | 0.883836589 | 0.116418632 | 0.58853869 | 0.602320082 | 0.102148911 |
| Srrm1 | S | 207 | 0.037474473 | 0.031945303 | 0.094969591 | 0.057495119 | 0.063024289 | 0.204782183 | 0.356460067 | 0.183651048 | 0.366131971 | 0.33531396 |
| Ercc3 | S | 34 | 0.034744943 | 0.051345789 | 0.079942749 | 0.045197806 | 0.02859696 | 0.468990809 | 0.283147288 | 0.123151996 | 0.393591936 | 0.567545602 |
| Cep170 | T | 1333 | 0.088535528 | 0.051005835 | 0.093214281 | 0.004678753 | 0.042208446 | 0.297359969 | 0.561215488 | 0.319197619 | 0.950213288 | 0.606222213 |
| Papola | S | 537 | 0.034101479 | 0.062863122 | 0.02633665 | -0.007764829 | -0.036526472 | 0.231661241 | 0.06174802 | 0.278120187 | 0.684235346 | 0.173765874 |
| Rtn4 | S | 953 | 0.105975202 | 0.013595306 | 0.124772751 | 0.018797549 | 0.111177445 | 0.30583625 | 0.770843094 | 0.2578364 | 0.874577455 | 0.300174396 |
| Kmt2a | T | 1847 | 0.04435546 | 0.118850663 | 0.045660203 | 0.001304743 | -0.07319046 | 0.524933512 | 0.238506596 | 0.274389593 | 0.983534788 | 0.423436182 |
| Rer1 | S | 95 | -0.079148915 | -0.011370865 | 0.209087261 | 0.288236176 | 0.220458126 | 0.479780753 | 0.849367987 | 0.002797704 | 0.075794094 | 0.027203987 |
| Slc26a6 | S | 604 | 0.21620859 | 0.042931842 | 0.077150666 | -0.139057924 | 0.034218824 | 0.318526113 | 0.618877652 | 0.646534194 | 0.555286077 | 0.83753845 |
| Slc26a6 | S | 607 | 0.21620859 | 0.042931842 | 0.077150666 | -0.139057924 | 0.034218824 | 0.318526113 | 0.618877652 | 0.646534194 | 0.555286077 | 0.83753845 |
| Slc26a6 | S | 608 | 0.21620859 | 0.042931842 | 0.077150666 | -0.139057924 | 0.034218824 | 0.318526113 | 0.618877652 | 0.646534194 | 0.555286077 | 0.83753845 |
| Slc26a6 | S | 610 | 0.21620859 | 0.042931842 | 0.077150666 | -0.139057924 | 0.034218824 | 0.318526113 | 0.618877652 | 0.646534194 | 0.555286077 | 0.83753845 |
| Slc26a6 | T | 615 | 0.21620859 | 0.042931842 | 0.077150666 | -0.139057924 | 0.034218824 | 0.318526113 | 0.618877652 | 0.646534194 | 0.555286077 | 0.83753845 |
| Nup107 | S | 735 | 0.071057225 | 0.05075054 | 0.056336065 | -0.01472116 | 0.005585524 | 0.180850776 | 0.242793458 | 0.465352277 | 0.838828245 | 0.934540321 |
| Specc1l | S | 171 | 0.025612481 | 0.140314939 | 0.071784642 | 0.04617216 | -0.068530297 | 0.778001135 | 0.181574114 | 0.405108486 | 0.505592194 | 0.400718569 |
| Arhgef12 | S | 279 | 0.017852725 | 0.087885988 | 0.082927338 | 0.065074613 | -0.00495865 | 0.84480593 | 0.266473502 | 0.110050798 | 0.482017671 | 0.941192477 |
| Atrx | S | 1287 | 0.026401285 | 0.057141282 | 0.092540619 | 0.066139334 | 0.035399336 | 0.593201481 | 0.253465467 | 0.140794237 | 0.208268794 | 0.449312761 |
| Suz12 | S | 548 | 0.025479197 | 0.038808439 | 0.063435718 | 0.037956521 | 0.024627278 | 0.106810685 | 0.313684978 | 0.112369266 | 0.275739616 | 0.559845785 |
| Cep170 | S | 628 | 0.232531824 | 0.030470696 | -0.082449656 | -0.31498148 | -0.112920352 | 0.014393463 | 0.783370951 | 0.451179936 | 0.040358686 | 0.392896407 |
| Hirip3 | S | 364 | 0.123455003 | 0.116011262 | -0.066730328 | -0.190185331 | -0.182741589 | 0.057108602 | 0.282902661 | 0.700004728 | 0.321967892 | 0.348917994 |
| Pard3 | S | 958 | -0.00121254 | 0.125396426 | 0.045655752 | 0.046868292 | -0.079740675 | 0.981699027 | 0.04710355 | 0.38701331 | 0.103808148 | 0.028013006 |
| Atat1 | S | 276 | -0.014502903 | 0.214878484 | -0.008835021 | 0.005667882 | -0.223713506 | 0.835730256 | 0.022646758 | 0.907691837 | 0.926724141 | 0.025129711 |
| Dnm1 | S | 778 | -0.068109267 | 0.150175321 | 0.193609763 | 0.26171903 | 0.043434442 | 0.150211597 | 0.068153065 | 0.215069823 | 0.142037057 | 0.745927163 |
| Fam83h | S | 639 | -0.041356298 | 0.017560576 | 0.266912225 | 0.308268523 | 0.249351649 | 0.672925457 | 0.553870494 | 0.068123111 | 0.056031126 | 0.090831211 |
| Arhgef7 | S | 497 | -0.027703993 | 0.107741033 | 0.095740802 | 0.123444795 | -0.012000232 | 0.582430215 | 0.129927119 | 0.095737944 | 0.028841153 | 0.830272166 |
| Scaf1 | S | 695 | 0.100523841 | -0.011417569 | 0.056364375 | -0.044159466 | 0.067781944 | 0.048514026 | 0.808723934 | 0.166309708 | 0.321707654 | 0.212524999 |
| Smarcad1 | S | 151 | 0.196230457 | -0.030363175 | 0.052077711 | -0.144152746 | 0.082440886 | 0.040806777 | 0.58880462 | 0.632699029 | 0.25206388 | 0.459002989 |
| Ttc28 | S | 2219 | -0.097260694 | 0.121063894 | 0.129591327 | 0.226852022 | 0.008527434 | 0.352510062 | 0.259424211 | 0.015645985 | 0.104978964 | 0.92349581 |
| Srrm1 | S | 389 | 0.050446973 | -0.003896537 | 0.09260053 | 0.042153557 | 0.096497066 | 0.150921208 | 0.87731385 | 0.046415948 | 0.299375307 | 0.046393687 |
| Ascc3 | S | 2197 | 0.029024132 | -0.06792005 | 0.285433026 | 0.256408895 | 0.353353076 | 0.885594925 | 0.763809588 | 0.060365473 | 0.267538246 | 0.193690068 |
| Ercc5 | S | 406 | 0.09969886 | 0.004711031 | 0.180910193 | 0.081211333 | 0.176199162 | 0.356966987 | 0.947657361 | 0.276088675 | 0.610297013 | 0.284605771 |
| Ercc5 | S | 407 | 0.09969886 | 0.004711031 | 0.180910193 | 0.081211333 | 0.176199162 | 0.356966987 | 0.947657361 | 0.276088675 | 0.610297013 | 0.284605771 |
| Sltm | S | 552 | 0.015156423 | 0.030347201 | 0.055157166 | 0.040000743 | 0.024809965 | 0.474466807 | 0.62093397 | 0.00173241 | 0.133320541 | 0.680734623 |
| Cenpc1 | S | 79 | 0.138860211 | 0.0165946 | 0.044705471 | -0.09415474 | 0.02811087 | 0.095616358 | 0.828464482 | 0.43907805 | 0.180392156 | 0.648014722 |
| Chaf1a | S | 293 | -0.057522225 | 0.074082571 | 0.109672836 | 0.167195061 | 0.035590265 | 0.07618856 | 0.031610461 | 0.011841364 | 0.004988159 | 0.206137971 |
| Scaf1 | S | 691 | 0.098293681 | -0.010236473 | 0.056079994 | -0.042213687 | 0.066316467 | 0.052020695 | 0.815703557 | 0.152214549 | 0.333978024 | 0.193764606 |
| Sfr1 | S | 71 | 0.033921171 | 0.08594513 | 0.004042241 | -0.02987893 | -0.081902889 | 0.219307283 | 0.022449234 | 0.905056489 | 0.326186759 | 0.051674754 |
| Sipa1l2 | S | 1549 | -0.008336263 | 0.137923622 | 0.050379191 | 0.058715454 | -0.087544431 | 0.865450444 | 0.030774303 | 0.601813202 | 0.545661535 | 0.389639293 |
| Fam83h | S | 637 | -0.009845553 | 0.087915035 | 0.087211054 | 0.097056607 | -0.000703981 | 0.905109229 | 0.111322019 | 0.14599522 | 0.286954184 | 0.988554098 |
| Iqsec1 | S | 513 | -0.001075927 | 0.131160329 | -0.01455614 | -0.013480212 | -0.145716469 | 0.986927155 | 0.003078354 | 0.844619883 | 0.879104628 | 0.138441746 |
| Ints6 | S | 150 | 0.090060002 | 0.130758851 | -0.005576243 | -0.095636244 | -0.136335094 | 0.507300394 | 0.069366989 | 0.927329335 | 0.481821802 | 0.070949345 |
| Nup93 | T | 766 | 0.091962351 | 0.049499133 | 0.065901349 | -0.026061001 | 0.016402216 | 0.410687258 | 0.411784359 | 0.20826638 | 0.790823383 | 0.678659896 |
| Dhx8 | S | 484 | 0.077604733 | 0.022828941 | 0.042059389 | -0.035545344 | 0.019230448 | 0.103579189 | 0.460248312 | 0.189471001 | 0.369785356 | 0.446695434 |
| Itgav | S | 1042 | 0.029162456 | 0.084745438 | 0.052135722 | 0.022973266 | -0.032609716 | 0.512624404 | 0.097319885 | 0.353345707 | 0.566739931 | 0.444691644 |
| Bcorl1 | S | 593 | 0.153240436 | 0.032733259 | 0.06050293 | -0.092737507 | 0.027769671 | 0.171734258 | 0.756358847 | 0.512199197 | 0.246448327 | 0.700299912 |
| Kif18a | S | 674 | 0.11622623 | 0.043072957 | -0.027850683 | -0.144076912 | -0.070923639 | 0.003545135 | 0.473111309 | 0.341487949 | 0.008105217 | 0.269732626 |
| Nfatc2ip | S | 79 | 0.006252717 | 0.075226546 | 0.041736948 | 0.035484231 | -0.033489599 | 0.894234897 | 0.021043052 | 0.221575655 | 0.464272845 | 0.239849806 |
| Nfatc2ip | S | 81 | 0.006252717 | 0.075226546 | 0.041736948 | 0.035484231 | -0.033489599 | 0.894234897 | 0.021043052 | 0.221575655 | 0.464272845 | 0.239849806 |
| Nfatc2ip | S | 83 | 0.006252717 | 0.075226546 | 0.041736948 | 0.035484231 | -0.033489599 | 0.894234897 | 0.021043052 | 0.221575655 | 0.464272845 | 0.239849806 |
| Cenpc1 | S | 767 | -0.096385452 | 0.175673775 | 0.114049529 | 0.210434981 | -0.061624246 | 0.167629688 | 0.034425342 | 0.154581785 | 0.040124295 | 0.398353504 |
| Serinc1 | S | 351 | 0.029965827 | 0.062628475 | 0.112226951 | 0.082261125 | 0.049598477 | 0.605142968 | 0.120762826 | 0.380684659 | 0.513143845 | 0.673318498 |
| Champ1 | S | 254 | -0.032714111 | 0.025652215 | 0.181974624 | 0.214688735 | 0.156322409 | 0.625162954 | 0.633564116 | 0.022547355 | 0.019493979 | 0.038820621 |
| Taf9 | T | 161 | 0.126965217 | 0.01987001 | 0.091125418 | -0.035839798 | 0.071255408 | 0.200554797 | 0.783679218 | 0.3885102 | 0.755626006 | 0.516494043 |
| Eif3b | T | 45 | 0.125102393 | 0.054508924 | 0.155068774 | 0.029966381 | 0.10055985 | 0.473428247 | 0.768740806 | 0.376854244 | 0.738319089 | 0.417342529 |
| Phkg2 | T | 397 | -0.098332084 | 0.196728953 | 0.10693997 | 0.205272054 | -0.089788983 | 0.599470372 | 0.158794188 | 0.167864395 | 0.321967494 | 0.462328773 |
| Chaf1b | S | 422 | 0.055065344 | 0.128678387 | -0.016295647 | -0.071360991 | -0.144974033 | 0.457936988 | 0.026943935 | 0.705091392 | 0.332216241 | 0.017024796 |
| Srrm2 | S | 797 | 0.105003446 | -0.023409147 | 0.066535586 | -0.038467861 | 0.089944733 | 0.014766033 | 0.54839559 | 0.342131176 | 0.555949809 | 0.23051777 |
| Clspn | S | 787 | 0.139930514 | -0.058356221 | 0.084566952 | -0.055363563 | 0.142923173 | 0.04516374 | 0.26369092 | 0.107502487 | 0.266624324 | 0.010278317 |
| 4833439L19Rik | T | 44 | 0.088084039 | 0.07518363 | 0.059095046 | -0.028988994 | -0.016088585 | 0.353236932 | 0.205825513 | 0.627567977 | 0.830967772 | 0.895164077 |
| Samd4 | S | 420 | 0.000569551 | 0.081526994 | 0.094625981 | 0.09405643 | 0.013098988 | 0.987738878 | 0.19739404 | 0.121434756 | 0.123129822 | 0.830019287 |
| Gorasp2 | S | 443 | 0.073637583 | 0.045750282 | 0.04644675 | -0.027190832 | 0.000696468 | 0.133693427 | 0.349579304 | 0.392079216 | 0.502838935 | 0.986706876 |
| Bcl9l | S | 121 | -0.043182706 | 0.092950819 | 0.114174506 | 0.157357212 | 0.021223687 | 0.086607818 | 0.006925555 | 0.159370721 | 0.093537965 | 0.737824108 |
| Lrp1 | S | 4521 | -0.075037322 | 0.136964148 | 0.114263197 | 0.189300519 | -0.022700951 | 0.395208514 | 0.13528987 | 0.068660375 | 0.090209368 | 0.769201183 |
| Polr2a | S | 1864 | 0.040283421 | 0.011792905 | 0.074440351 | 0.034156931 | 0.062647447 | 0.561331671 | 0.853134211 | 0.006025422 | 0.617248796 | 0.374541923 |
| Zdhhc5 | S | 296 | 0.036252635 | 0.025189599 | 0.104526306 | 0.068273671 | 0.079336707 | 0.294274258 | 0.418640829 | 0.139521224 | 0.272391602 | 0.21442701 |
| Atg2a | S | 1290 | 0.205803631 | 0.253013337 | -0.095977898 | -0.301781529 | -0.348991236 | 0.395930215 | 0.228456776 | 0.560858179 | 0.263320243 | 0.151188104 |
| Trim28 | S | 601 | -0.057292363 | 0.089234686 | 0.100659664 | 0.157952027 | 0.011424979 | 0.335069686 | 0.011985657 | 0.15397654 | 0.073188896 | 0.833624603 |
| Cdk11b | S | 113 | 0.187227941 | -0.10406065 | 0.129472143 | -0.057755798 | 0.233532793 | 0.073040264 | 0.255325685 | 0.15799296 | 0.544093471 | 0.062970998 |
| Zc3h13 | S | 1086 | 0.00652796 | 0.034466079 | 0.110291549 | 0.103763589 | 0.07582547 | 0.882171864 | 0.40951945 | 0.035218093 | 0.054933407 | 0.09944941 |
| Plcb3 | S | 474 | 0.041058164 | 0.11563004 | 0.081022453 | 0.039964288 | -0.034607587 | 0.612644487 | 0.183048392 | 0.531575796 | 0.767316438 | 0.79430044 |
| Arhgef17 | S | 1984 | -0.021719504 | -0.011221465 | 0.210263514 | 0.231983017 | 0.221484979 | 0.843073755 | 0.939992115 | 0.021594368 | 0.103342187 | 0.215639907 |
| Prkd1 | S | 203 | 0.056284539 | 0.083187952 | 0.049521331 | -0.006763208 | -0.033666621 | 0.547668281 | 0.107771384 | 0.460771503 | 0.943840212 | 0.585349245 |
| Mtmr3 | S | 613 | 0.021106505 | 0.034649557 | 0.071776119 | 0.050669614 | 0.037126562 | 0.652103069 | 0.090050097 | 0.079247263 | 0.343722427 | 0.275418468 |
| Atad2 | S | 406 | 0.040316764 | 0.006174995 | 0.096095018 | 0.055778254 | 0.089920023 | 0.455908465 | 0.892876481 | 0.021912785 | 0.301729827 | 0.097642626 |
| Fgfr1op | S | 158 | -0.071357983 | 0.10161895 | 0.146235342 | 0.217593325 | 0.044616392 | 0.620143387 | 0.135384885 | 0.13219711 | 0.197758045 | 0.554267477 |
| Fgfr1op | S | 160 | -0.071357983 | 0.10161895 | 0.146235342 | 0.217593325 | 0.044616392 | 0.620143387 | 0.135384885 | 0.13219711 | 0.197758045 | 0.554267477 |
| Prkab2 | T | 39 | -0.208691645 | 0.14088501 | 0.306919895 | 0.51561154 | 0.166034885 | 0.207874297 | 0.389266064 | 0.034227329 | 0.027358092 | 0.341911605 |
| Sf3b1 | T | 328 | 0.047909303 | -0.014877669 | 0.10397072 | 0.056061417 | 0.118848389 | 0.1691425 | 0.682926356 | 0.033875694 | 0.183518098 | 0.034728663 |
| Tlk2 | S | 46 | 0.141237262 | 0.087423605 | -0.002124294 | -0.143361556 | -0.0895479 | 0.184339749 | 0.296520608 | 0.987217914 | 0.348816828 | 0.523626433 |
| Wdr7 | T | 926 | 0.223746933 | 0.026862854 | -0.057033491 | -0.280780423 | -0.083896345 | 0.016312484 | 0.664023725 | 0.319027839 | 0.010809799 | 0.225284599 |
| Rufy1 | S | 14 | -0.109267957 | 0.007949447 | 0.250889042 | 0.360157 | 0.242939595 | 0.373723941 | 0.946717282 | 0.006979562 | 0.045296101 | 0.112289055 |
| Prrc2a | S | 1002 | 0.145617498 | 0.001942469 | 0.005583147 | -0.140034351 | 0.003640678 | 0.015367113 | 0.969241894 | 0.92104838 | 0.04050686 | 0.938138585 |
| Setd2 | S | 675 | 0.080230344 | 0.049663307 | 0.04887943 | -0.031350915 | -0.000783877 | 0.229644631 | 0.316062383 | 0.351657383 | 0.55212794 | 0.977787933 |
| Plcl1 | T | 557 | -0.065600819 | 0.198177101 | 0.040328576 | 0.105929395 | -0.157848525 | 0.416076935 | 0.014901495 | 0.638128527 | 0.288626806 | 0.120172671 |
| L3mbtl1 | S | 234 | 0.168852639 | 0.036230935 | -0.058189468 | -0.227042107 | -0.094420403 | 0.005960517 | 0.475461143 | 0.243719688 | 0.003712964 | 0.087868384 |
| L3mbtl1 | S | 236 | 0.168852639 | 0.036230935 | -0.058189468 | -0.227042107 | -0.094420403 | 0.005960517 | 0.475461143 | 0.243719688 | 0.003712964 | 0.087868384 |
| L3mbtl1 | S | 239 | 0.168852639 | 0.036230935 | -0.058189468 | -0.227042107 | -0.094420403 | 0.005960517 | 0.475461143 | 0.243719688 | 0.003712964 | 0.087868384 |
| L3mbtl1 | Y | 232 | 0.168852639 | 0.036230935 | -0.058189468 | -0.227042107 | -0.094420403 | 0.005960517 | 0.475461143 | 0.243719688 | 0.003712964 | 0.087868384 |
| Sema6d | S | 711 | 0.055562968 | 0.014683062 | 0.128786274 | 0.073223306 | 0.114103212 | 0.531782343 | 0.903446634 | 0.072597413 | 0.443573069 | 0.396994809 |
| Mki67 | S | 774 | 0.092602277 | 0.170521754 | -0.136098239 | -0.228700516 | -0.306619993 | 0.06292559 | 0.001345836 | 0.014213993 | 0.004741057 | 0.007773586 |
| Hdac4 | S | 562 | -0.026589835 | 0.095851917 | 0.034076105 | 0.06066594 | -0.061775812 | 0.138347111 | 0.003252247 | 0.077545182 | 0.008629894 | 0.032629366 |

| | | | | | | | | | | | | |
|---|---|---|---|---|---|---|---|---|---|---|---|---|
| Ehbp1 | S | 171 | 0.009326874 | 0.103464209 | 0.054973467 | 0.045646593 | -0.048490742 | 0.852634649 | 0.036454006 | 0.544282323 | 0.606053393 | 0.572114511 |
| Ehbp1 | S | 174 | 0.009326874 | 0.103464209 | 0.054973467 | 0.045646593 | -0.048490742 | 0.852634649 | 0.036454006 | 0.544282323 | 0.606053393 | 0.572114511 |
| Ier3 | S | 31 | 0.05667563 | 0.177854666 | -0.072312513 | -0.128988142 | -0.250167179 | 0.373825307 | 0.01620852 | 0.321903021 | 0.090313752 | 0.014855926 |
| Sbf1 | T | 1819 | 0.067548437 | 0.192187683 | 0.128241409 | 0.060692972 | -0.063946274 | 0.690142986 | 0.338204218 | 0.752232687 | 0.878601111 | 0.87446751 |
| Erbin | S | 712 | 0.043494777 | 0.083182707 | 0.074544322 | 0.031049546 | -0.008638385 | 0.538513774 | 0.124411464 | 0.509431767 | 0.792286611 | 0.936912514 |
| Scrib | S | 1529 | -0.156868169 | 0.217419838 | 0.143476146 | 0.300344314 | -0.073943693 | 0.499620471 | 0.191454529 | 0.136119012 | 0.247741034 | 0.581637532 |
| Sipa1l2 | S | 163 | 0.137264651 | -0.006344506 | 0.123717571 | -0.01354708 | 0.130062077 | 0.151990967 | 0.9575768 | 0.454170151 | 0.927025991 | 0.449789224 |
| Zcchc7 | T | 115 | 0.29859284 | 0.00015623 | -0.118664184 | -0.417257024 | -0.118820414 | 0.007404676 | 0.999068534 | 0.248658912 | 0.013779747 | 0.446411085 |
| Cramp1l | S | 795 | 0.20152186 | 0.107471035 | -0.128389733 | -0.329911593 | -0.235860768 | 0.058981763 | 0.125524694 | 0.247173087 | 0.032766871 | 0.078630218 |
| Papolg | S | 649 | 0.148137091 | 0.048851337 | -0.021547257 | -0.169684348 | -0.070398594 | 0.058548165 | 0.458877853 | 0.928659508 | 0.507883146 | 0.77286628 |
| Mff | S | 129 | 0.03418114 | 0.113521903 | 0.003226406 | -0.030954734 | -0.110295497 | 0.413118206 | 0.036435518 | 0.919679298 | 0.471977048 | 0.047072089 |
| Armcx3 | S | 61 | 0.016162404 | -0.02294903 | 0.067581675 | 0.051419271 | 0.090530705 | 0.549407935 | 0.551298701 | 4.78823E-05 | 0.143761939 | 0.103915503 |
| Tnks1bp1 | S | 1463 | 0.056523386 | 0.133631688 | 0.028556224 | -0.027967162 | -0.105075464 | 0.479756841 | 0.153049713 | 0.724183213 | 0.514561615 | 0.127688747 |
| Dact3 | S | 496 | -0.073620238 | 0.16256334 | 0.228574751 | 0.302194989 | 0.066011411 | 0.404444501 | 0.144040576 | 0.339566028 | 0.236922947 | 0.763421759 |
| Ccdc61 | S | 330 | -0.003124588 | 0.021307879 | 0.1170962 | 0.120220788 | 0.09578832 | 0.930448578 | 0.702071431 | 0.010854558 | 0.038045378 | 0.172363138 |
| Wnk1 | S | 185 | 0.116909938 | 0.172448529 | -0.017891319 | -0.134801257 | -0.190339848 | 0.465741308 | 0.166203893 | 0.84039737 | 0.374435144 | 0.114709457 |
| Synm | S | 780 | 0.057128949 | 0.183284341 | 0.020549398 | -0.036579551 | -0.162734942 | 0.792819991 | 0.100464613 | 0.94228755 | 0.911971876 | 0.583294508 |
| Arhgap21 | S | 947 | 0.016979265 | 0.007034165 | 0.140996891 | 0.124017626 | 0.133962726 | 0.873823997 | 0.883628141 | 0.024888447 | 0.301895832 | 0.035113753 |
| Zeb1 | S | 302 | 0.064105927 | 0.06271406 | 0.034190093 | -0.029915834 | -0.028523967 | 0.326369189 | 0.166884279 | 0.327675248 | 0.598278013 | 0.394319574 |
| Prpf4b | S | 240 | -0.00378824 | 0.009699174 | 0.117558506 | 0.121346745 | 0.107859331 | 0.878676974 | 0.711543726 | 0.007078238 | 0.01060902 | 0.017200007 |
| Prpf4b | S | 242 | -0.00378824 | 0.009699174 | 0.117558506 | 0.121346745 | 0.107859331 | 0.878676974 | 0.711543726 | 0.007078238 | 0.01060902 | 0.017200007 |
| Nfib | S | 284 | -0.017888563 | 0.059543868 | 0.085711263 | 0.103599826 | 0.026167395 | 0.499783334 | 0.09558036 | 0.035190409 | 0.024517 | 0.454067538 |
| Ptprg | S | 1180 | 0.077455924 | 0.033560865 | 0.388367953 | 0.31091203 | 0.354807088 | 0.833159721 | 0.902233205 | 0.37127594 | 0.480903528 | 0.370462208 |
| Plekhg3 | S | 1146 | 0.032470744 | 0.013821842 | 0.098035844 | 0.0655651 | 0.084214002 | 0.352928122 | 0.652858447 | 0.05296393 | 0.165191446 | 0.088362695 |
| Ahctf1 | T | 2067 | 0.163839972 | 0.004736208 | 0.060262627 | -0.103577345 | 0.055526419 | 0.117512123 | 0.967546373 | 0.54155757 | 0.255065266 | 0.611788223 |
| Numa1 | S | 795 | -0.00155056 | 0.062025581 | 0.147079639 | 0.148630199 | 0.085054059 | 0.979323667 | 0.357901304 | 0.127637544 | 0.13541894 | 0.350728199 |
| Nes | S | 1861 | 0.010519721 | 0.01568421 | 0.096910186 | 0.086390465 | 0.081225976 | 0.753499344 | 0.628320484 | 0.011691688 | 0.036025449 | 0.034985413 |
| Mrps36 | T | 59 | 0.011298488 | 0.050447616 | 0.134018014 | 0.122719526 | 0.083570398 | 0.787323265 | 0.407261959 | 0.121415869 | 0.14150024 | 0.31373669 |
| Supt6 | T | 1523 | 0.040652339 | 0.032027745 | 0.067642954 | 0.026990614 | 0.035615208 | 0.258417735 | 0.292432096 | 0.14224032 | 0.501155615 | 0.354388837 |
| Zhx3 | S | 941 | -0.054042512 | 0.051072778 | 0.130813607 | 0.184856119 | 0.079740829 | 0.330378737 | 0.256189714 | 0.00989825 | 0.026680551 | 0.061007133 |
| Dclk1 | T | 336 | 0.015672321 | 0.062222666 | 0.028589753 | 0.012917432 | -0.033632913 | 0.309396453 | 0.02133057 | 0.360380823 | 0.651374026 | 0.306408144 |
| Csnk1a1 | T | 321 | 0.391253877 | -0.048413224 | -0.124890391 | -0.516144269 | -0.076477167 | 0.013185538 | 0.756606395 | 0.296458638 | 0.000110711 | 0.543241015 |
| Incenp | T | 270 | 0.13884792 | 0.03160701 | 0.087112134 | -0.051735786 | 0.055505125 | 0.32153993 | 0.838573973 | 0.302119816 | 0.656235464 | 0.698212551 |
| Rcor3 | Y | 229 | -0.004613636 | 0.037555365 | 0.187533934 | 0.19214757 | 0.149978568 | 0.945406559 | 0.426479197 | 0.119282989 | 0.119100135 | 0.188860193 |
| Akap13 | S | 579 | 0.101871775 | 0.104886242 | 0.045703836 | -0.056167939 | -0.059182406 | 0.570545757 | 0.175459607 | 0.699711007 | 0.770149172 | 0.624501998 |
| Tpm1 | S | 87 | 0.047881911 | 0.008448451 | 0.096467931 | 0.04858602 | 0.088019479 | 0.390210822 | 0.725946509 | 0.054733306 | 0.411261167 | 0.081290837 |
| Srrm2 | T | 2077 | 0.053438073 | 0.015916071 | 0.058356604 | 0.004918531 | 0.042440533 | 0.048859315 | 0.709604325 | 0.176919276 | 0.884117862 | 0.39656235 |
| Prkd1 | S | 206 | 0.086518682 | 0.047455603 | 0.02688578 | -0.059632902 | -0.020569823 | 0.297945219 | 0.052042535 | 0.714050651 | 0.536445626 | 0.773925258 |
| Ino80d | S | 354 | 0.025269447 | 0.023522057 | 0.140240946 | 0.114971499 | 0.116718889 | 0.649231151 | 0.65438817 | 0.084335062 | 0.158726374 | 0.14579688 |
| Ubr5 | S | 2022 | 0.053643631 | 0.044555591 | 0.041419367 | -0.012224264 | -0.003136223 | 0.103830575 | 0.258159165 | 0.386321646 | 0.768561626 | 0.94585479 |
| Camk2g | S | 319 | 0.058653016 | 0.366201558 | -0.294170818 | -0.352823834 | -0.660372376 | 0.665522729 | 0.023946997 | 0.382400305 | 0.307300082 | 0.122072502 |
| Gtpbp1 | S | 24 | -0.005119547 | 0.069852277 | 0.030529159 | 0.035648706 | -0.039323118 | 0.85547527 | 0.005610929 | 0.211230442 | 0.258510029 | 0.110546731 |
| Vrk3 | T | 58 | -0.072767068 | 0.090531893 | 0.129368474 | 0.202135542 | 0.03883658 | 0.067033022 | 0.055986011 | 0.017432414 | 0.005598534 | 0.379068785 |
| Dock7 | S | 30 | 0.226479257 | -0.030204357 | -0.028875456 | -0.255354713 | 0.001328901 | 0.008381689 | 0.667373127 | 0.51172759 | 0.010692389 | 0.983599126 |
| Chchd2 | S | 45 | 0.071679234 | 0.067156822 | -0.034780778 | -0.106460011 | -0.1019376 | 0.015559432 | 0.081394887 | 0.453962994 | 0.094275269 | 0.086081254 |
| Zfhx3 | S | 426 | -0.023562266 | -0.007346166 | 0.147619706 | 0.171181972 | 0.154965872 | 0.614057794 | 0.854002977 | 0.002775865 | 0.023713084 | 0.012348476 |
| 1110059E24Rik | S | 136 | 0.036157093 | 0.082995026 | 0.007504917 | -0.028652176 | -0.075490109 | 0.242377692 | 0.03686642 | 0.799478508 | 0.138816358 | 0.036406964 |
| Nes | S | 1860 | 0.01025814 | 0.01541099 | 0.09731274 | 0.0870546 | 0.08190175 | 0.759616155 | 0.637776948 | 0.012288674 | 0.034441046 | 0.035253011 |
| Cdc5l | S | 410 | -0.026692271 | 0.051306972 | 0.259453325 | 0.286145596 | 0.208146354 | 0.78799287 | 0.472733475 | 0.173571035 | 0.148462314 | 0.248825535 |
| Cdc5l | S | 417 | -0.026692271 | 0.051306972 | 0.259453325 | 0.286145596 | 0.208146354 | 0.78799287 | 0.472733475 | 0.173571035 | 0.148462314 | 0.248825535 |
| Top2a | S | 1448 | 0.268765272 | 0.27982367 | -0.239342423 | -0.508107695 | -0.519166093 | 0.183540092 | 0.307746124 | 0.414480736 | 0.146950915 | 0.168850354 |
| Kif2a | S | 431 | 0.404028572 | 0.105525903 | -0.098830433 | -0.502859005 | -0.204356335 | 0.321045171 | 0.496721952 | 0.773195065 | 0.302096856 | 0.563127604 |
| Nav1 | S | 21 | 0.059248547 | 0.000427927 | 0.071260284 | 0.012011737 | 0.070832358 | 0.225789184 | 0.992081311 | 0.033933356 | 0.756400111 | 0.159693645 |
| Arid1b | S | 1563 | 0.006880765 | 0.037709366 | 0.09859481 | 0.091714046 | 0.060885444 | 0.801315067 | 0.28538148 | 0.055930439 | 0.086491441 | 0.180598893 |
| Tmem63a | S | 734 | 0.100488707 | 0.043518638 | 0.025082767 | -0.07540594 | -0.018435871 | 0.062446376 | 0.477593596 | 0.763936855 | 0.401873094 | 0.838292839 |
| Tmem63a | S | 738 | 0.100488707 | 0.043518638 | 0.025082767 | -0.07540594 | -0.018435871 | 0.062446376 | 0.477593596 | 0.763936855 | 0.401873094 | 0.838292839 |
| Stk38 | S | 281 | -0.002247765 | 0.123942896 | 0.04246922 | 0.044716985 | -0.081473676 | 0.96410717 | 0.037304143 | 0.547956319 | 0.551137005 | 0.295946493 |
| Pik3c3 | S | 243 | 0.159618712 | 0.049560345 | 0.027074882 | -0.13254383 | -0.022485463 | 0.129497761 | 0.597728034 | 0.88252782 | 0.48406785 | 0.899614957 |
| Pkp4 | S | 336 | 0.022285998 | 0.012000956 | 0.145520002 | 0.123234004 | 0.133519045 | 0.481811753 | 0.547967025 | 0.101911632 | 0.132180896 | 0.122857392 |
| Srrm1 Srsf1 Tra2a Tr | S | 280;304;223;273 | 0.160160606 | -0.055275187 | 0.185404213 | 0.025243606 | 0.2406794 | 0.245753869 | 0.685012464 | 0.320377578 | 0.876166265 | 0.205176706 |
| Rbl1 | S | 649 | 0.008093512 | 0.144035878 | 0.03189811 | 0.023804597 | -0.112137769 | 0.862588228 | 0.089660312 | 0.42691861 | 0.459484312 | 0.168411902 |
| Specc1l | S | 833 | 0.063435958 | 0.170057985 | -0.052989712 | -0.11642567 | -0.223047697 | 0.526328733 | 0.039697347 | 0.639750616 | 0.352697561 | 0.113678371 |
| Ints12 | S | 127 | 0.047210477 | 0.059014262 | 0.059397213 | 0.012186736 | 0.000382951 | 0.24112345 | 0.339286286 | 0.275993124 | 0.818456858 | 0.995382551 |
| Ncam1 | S | 1005 | -0.012157665 | 0.051820779 | 0.097890547 | 0.110048212 | 0.046069768 | 0.687772746 | 0.060534816 | 0.10064807 | 0.081228948 | 0.34703625 |
| Cobll1 | T | 183 | 0.041111284 | 0.036123749 | 0.052285018 | 0.011173735 | 0.01616127 | 0.222288686 | 0.452198611 | 0.073127583 | 0.685901122 | 0.714083221 |
| Srrm2 | S | 622 | -0.016532305 | 0.126309711 | 0.084359539 | 0.100891844 | -0.041950173 | 0.803088246 | 0.072621442 | 0.414997288 | 0.371222435 | 0.680824849 |
| Srrm2 | S | 626 | -0.016532305 | 0.126309711 | 0.084359539 | 0.100891844 | -0.041950173 | 0.803088246 | 0.072621442 | 0.414997288 | 0.371222435 | 0.680824849 |
| Srrm2 | S | 624 | -0.016532305 | 0.126309711 | 0.084359539 | 0.100891844 | -0.041950173 | 0.803088246 | 0.072621442 | 0.414997288 | 0.371222435 | 0.680824849 |
| Cds2 | T | 30 | -0.050838941 | 0.153638473 | 0.051035525 | 0.101874466 | -0.102602948 | 0.619045169 | 0.023078508 | 0.51920122 | 0.362959986 | 0.200600265 |
| Eml4 | S | 908 | 0.021981042 | 0.050310957 | 0.069386612 | 0.04740557 | 0.019075655 | 0.628941894 | 0.186383153 | 0.104816168 | 0.372853699 | 0.630775839 |
| Mex3d | S | 484 | 0.000376561 | 0.182106773 | -0.019977301 | -0.020353862 | -0.202084074 | 0.994975599 | 0.017150973 | 0.756118165 | 0.727366046 | 0.021582925 |
| Epb41l1 | S | 782 | 0.062955372 | 0.095268446 | 0.05305364 | -0.009901732 | -0.042214806 | 0.436349897 | 0.317717677 | 0.37569942 | 0.887998137 | 0.617067996 |
| Srsf1 | S | 242 | -0.006356983 | 0.015509603 | 0.13028692 | 0.136643904 | 0.114777318 | 0.90958094 | 0.722075827 | 0.01501722 | 0.056575307 | 0.014141834 |
| Fam122a | S | 32 | -0.010221917 | 0.077099948 | 0.048973899 | 0.059195816 | -0.028126049 | 0.886521623 | 0.038186638 | 0.124625477 | 0.437532407 | 0.342749399 |
| Hnrnpd | S | 210 | -0.067080954 | 0.017305505 | 0.170625377 | 0.23770633 | 0.153319872 | 0.363821069 | 0.841665778 | 0.003138303 | 0.037685361 | 0.159366065 |
| Gcfc2 | S | 118 | -0.026755409 | 0.054129357 | 0.131056578 | 0.157811987 | 0.076927221 | 0.720993331 | 0.413163132 | 0.035815677 | 0.096818488 | 0.260446545 |
| Akap12 | S | 350 | -0.005532531 | 0.161775983 | 0.034836422 | 0.040368953 | -0.126939561 | 0.92834825 | 0.07441431 | 0.489739629 | 0.452550035 | 0.13532921 |
| Necap2 | T | 180 | -0.155105094 | 0.328428567 | 0.022430405 | 0.177535499 | -0.305998162 | 0.131217097 | 0.006755759 | 0.828667216 | 0.107625643 | 0.0238876 |
| Map3k20 | S | 706 | -0.018560732 | 0.228030433 | 0.054607066 | 0.073167798 | -0.173423367 | 0.844802918 | 0.135812766 | 0.606212953 | 0.505325515 | 0.24805751 |
| Xrcc1 | S | 228 | 0.136330021 | 0.032832673 | 0.037326535 | -0.099003486 | 0.004493862 | 0.093636823 | 0.841444395 | 0.589021415 | 0.128508643 | 0.977288847 |
| Trim24 | T | 1044 | -0.048926633 | 0.052090668 | 0.229114403 | 0.278041036 | 0.177023734 | 0.079035759 | 0.169045834 | 0.048715912 | 0.034166587 | 0.076396956 |
| Zc3h18 | S | 847 | -0.007425199 | 0.013550966 | 0.084748374 | 0.092173573 | 0.071197408 | 0.681289963 | 0.690297015 | 0.000983764 | 0.005405362 | 0.116320492 |
| Champ1 | S | 231 | -0.143848263 | 0.099685483 | 0.394304945 | 0.538153208 | 0.294619462 | 0.425057386 | 0.456211019 | 0.157082785 | 0.085538124 | 0.252604663 |
| Eaf1 | S | 165 | 0.02341648 | 0.070513241 | 0.050212324 | 0.026795844 | -0.020300917 | 0.559573518 | 0.063125526 | 0.364511634 | 0.600986852 | 0.661618703 |
| Faxc | S | 395 | -0.052003183 | 0.165740857 | 0.066506916 | 0.1185101 | -0.099233941 | 0.559286657 | 0.058147105 | 0.333974508 | 0.268920608 | 0.237089501 |
| Tcp1 | S | 551 | -0.008186342 | 0.090289016 | 0.084616827 | 0.092803168 | -0.00567219 | 0.933161874 | 0.158004404 | 0.153220396 | 0.351611179 | 0.879642044 |
| Brf1 | S | 358 | -0.08653017 | 0.081709375 | 0.219633854 | 0.306164023 | 0.137924479 | 0.485169137 | 0.319122257 | 0.090051752 | 0.071344367 | 0.232008034 |
| Caskin2 | S | 797 | 0.009146498 | 0.111822212 | 0.06528017 | 0.056133672 | -0.046542042 | 0.858744719 | 0.17025159 | 0.25155584 | 0.246115635 | 0.508036212 |
| Arhgap39 | S | 115 | 0.008239953 | 0.081247722 | 0.139592322 | 0.131352369 | 0.0583446 | 0.920759166 | 0.297610068 | 0.246028433 | 0.302505232 | 0.608960407 |
| Zc3h13 | S | 951 | 0.078175461 | 0.025047362 | 0.011704514 | -0.066470947 | -0.013342848 | 0.013663175 | 0.521716084 | 0.616108102 | 0.013036797 | 0.697377368 |
| Zc3h13 | S | 955 | 0.078175461 | 0.025047362 | 0.011704514 | -0.066470947 | -0.013342848 | 0.013663175 | 0.521716084 | 0.616108102 | 0.013036797 | 0.697377368 |
| Mtmr10 | S | 763 | 0.107394751 | 0.21107321 | -0.077473622 | -0.184868372 | -0.288546832 | 0.548465162 | 0.080956309 | 0.595929878 | 0.356968445 | 0.098044855 |
| Tfpt | S | 180 | 0.014249752 | 0.022095476 | 0.071376578 | 0.057126827 | 0.049281103 | 0.752758258 | 0.311768998 | 0.012743834 | 0.258983178 | 0.002388782 |
| Kmt2a | S | 2668 | -0.053491881 | 0.216403751 | 0.058906136 | 0.112398018 | -0.157497615 | 0.736097096 | 0.096295184 | 0.516950544 | 0.519356192 | 0.218941854 |

| | | | | | | | | | | | | | |
|---|---|---|---|---|---|---|---|---|---|---|---|---|---|
| Kmt2a | S | | 2670 | -0.053491881 | 0.216403751 | 0.058906136 | 0.112398018 | -0.157497615 | 0.736097096 | 0.096295184 | 0.516950544 | 0.519356192 | 0.218941854 |
| Kmt2a | T | | 2669 | -0.053491881 | 0.216403751 | 0.058906136 | 0.112398018 | -0.157497615 | 0.736097096 | 0.096295184 | 0.516950544 | 0.519356192 | 0.218941854 |
| Slc9a3r1 | S | | 286 | -0.06177586 | 0.110572962 | 0.13212177 | 0.19389763 | 0.021548808 | 0.009255986 | 0.024274453 | 0.031158502 | 0.013105717 | 0.599028021 |
| Fam122a | S | | 140 | 0.067120616 | 0.133454778 | -0.063759844 | -0.13088046 | -0.197214622 | 0.124917328 | 0.039042907 | 0.37018148 | 0.119145129 | 0.043816072 |
| Fam122a | S | | 144 | 0.067120616 | 0.133454778 | -0.063759844 | -0.13088046 | -0.197214622 | 0.124917328 | 0.039042907 | 0.37018148 | 0.119145129 | 0.043816072 |
| Srrm1 | S | | 779 | 0.019367023 | 0.010366932 | 0.066090565 | 0.046723542 | 0.055723633 | 0.441685383 | 0.541309444 | 0.00688145 | 0.099163759 | 0.008804286 |
| Mettl15 | S | | 358 | 0.051481861 | 0.071259291 | 0.032713052 | -0.018768809 | -0.038546239 | 0.405306938 | 0.065286359 | 0.632113004 | 0.805465367 | 0.552340016 |
| Mprip | S | | 1015 | -0.132240502 | 0.258658462 | 0.137875249 | 0.270115752 | -0.120783213 | 0.021587635 | 0.045452931 | 0.075395955 | 0.012836527 | 0.227436892 |
| Srrt | S | | 67 | -0.045722912 | 0.036028855 | 0.120843132 | 0.166566045 | 0.084814277 | 0.236906478 | 0.261357857 | 0.004064565 | 0.007605657 | 0.020434597 |
| Aco1 | S | | 462 | 0.606172547 | -0.239332196 | 0.188826216 | -0.417346332 | 0.428158411 | 0.324745107 | 0.701703397 | 0.729263034 | 0.449906543 | 0.466467498 |
| Mki67 | T | | 783 | 0.246383661 | 0.123255485 | -0.201611944 | -0.447995605 | -0.324867429 | 0.00453729 | 0.066460966 | 0.011077308 | 3.82971E-05 | 0.000685778 |
| Dbn1 | S | | 142 | 0.095960888 | 0.02045414 | 0.015781361 | -0.080179527 | -0.00467278 | 0.037266818 | 0.487042019 | 0.565390884 | 0.079256583 | 0.742991652 |
| Ybx3 | S | | 52 | -0.047438083 | 0.089987385 | 0.066712322 | 0.114150405 | -0.023275063 | 0.308522142 | 0.012183128 | 0.110207291 | 0.060066267 | 0.497592664 |
| Ctnna1 Ctnna2 | S | 323;321 | | 0.066193536 | 0.04692911 | 0.170939737 | 0.1047462 | 0.124010626 | 0.758141665 | 0.661756195 | 0.228596854 | 0.629946707 | 0.30118826 |
| Mta1 | T | | 564 | 0.259054322 | -0.023980009 | -0.047862686 | -0.306917007 | -0.023882676 | 0.021428182 | 0.813458633 | 0.622533922 | 0.011875802 | 0.765231982 |
| Spp1 | S | | 231 | 0.122909597 | -0.006768241 | -0.051629969 | -0.174539566 | -0.044861727 | 0.00034468 | 0.763854439 | 0.569518709 | 0.146653668 | 0.616631025 |
| Bod1l | S | | 2841 | 0.033819611 | 0.025811537 | 0.114306074 | 0.080486462 | 0.088494536 | 0.4357498 | 0.566124679 | 0.124875644 | 0.238769326 | 0.208247096 |
| Usp9x | S | | 2443 | 0.048382657 | 0.065283203 | 0.032819944 | -0.015562713 | -0.032463259 | 0.281911691 | 0.10234618 | 0.535806082 | 0.790654259 | 0.561065359 |
| Arhgef17 | T | | 1985 | -0.02772522 | -0.003847417 | 0.209344374 | 0.237069593 | 0.213191791 | 0.816358625 | 0.981305539 | 0.029311842 | 0.124154272 | 0.267749545 |
| Nhsl2 | S | | 1028 | -0.14903554 | 0.036054007 | 0.234676826 | 0.383712366 | 0.198622819 | 0.232728265 | 0.052249688 | 0.028077263 | 0.030980738 | 0.046946149 |
| Tubgcp5 | T | | 161 | 0.087024987 | -0.099373484 | 0.203761277 | 0.11673629 | 0.303134761 | 0.417915166 | 0.412904735 | 0.044431284 | 0.268156586 | 0.053562832 |
| Phf10 | S | | 53 | 0.012731343 | 0.059131164 | 0.133819047 | 0.121087704 | 0.074687883 | 0.866227726 | 0.478781256 | 0.128440713 | 0.075081723 | 0.26829715 |
| Prpf4b | S | | 636 | 0.057748652 | 0.025660853 | 0.109943288 | 0.052194636 | 0.084282435 | 0.385396509 | 0.707218309 | 0.163773585 | 0.483261277 | 0.297631281 |
| Slc12a4 | S | | 47 | 0.156373576 | 0.04159156 | -0.049325955 | -0.205699531 | -0.090917515 | 0.008543178 | 0.611277697 | 0.336895444 | 0.006711111 | 0.291931012 |
| Tbc1d4 | S | | 276 | 0.148892468 | -0.055128276 | 0.063554847 | -0.085337621 | 0.118683124 | 0.013326339 | 0.415024717 | 0.557696073 | 0.442122414 | 0.324805669 |
| Srrm2 | S | | 1849 | 0.169178677 | -0.069256655 | 0.210361871 | 0.041183194 | 0.279618526 | 0.148299239 | 0.47654233 | 0.411560313 | 0.859937403 | 0.297080597 |
| Atp7a | S | | 1423 | 0.009401739 | 0.01023588 | 0.094625949 | 0.08522421 | 0.084390069 | 0.813761377 | 0.62736082 | 0.009923763 | 0.099148806 | 0.02485276 |
| Atp7a | S | | 1426 | 0.009401739 | 0.01023588 | 0.094625949 | 0.08522421 | 0.084390069 | 0.813761377 | 0.62736082 | 0.009923763 | 0.099148806 | 0.02485276 |
| Nes | S | | 312 | 0.07853093 | 0.022699179 | 0.048307543 | -0.030223387 | 0.025608364 | 0.196266172 | 0.529254827 | 0.163458259 | 0.566456592 | 0.455246301 |
| Wdr37 | T | | 28 | -0.002434569 | 0.192831454 | 0.094692409 | 0.097126978 | -0.098139045 | 0.983107889 | 0.18183309 | 0.639290173 | 0.619234991 | 0.629255863 |
| Rab1a Rab1b | S | 78;75 | | 0.05552689 | 0.021053733 | 0.080234747 | 0.024707857 | 0.059181014 | 0.267758734 | 0.562616569 | 0.147382188 | 0.659052904 | 0.278585323 |
| Brca2 | S | | 3207 | 0.240495191 | -0.118817072 | 0.125439533 | -0.115055658 | 0.244256604 | 0.092847549 | 0.687269927 | 0.661941721 | 0.675047173 | 0.508816953 |
| Chd4 | S | | 303 | -0.075683783 | 0.077917117 | 0.147363543 | 0.223047326 | 0.069446426 | 0.194105463 | 0.165459335 | 0.021945908 | 0.012317078 | 0.229420098 |
| Tssc4 | S | | 124 | 0.000421892 | 0.012929307 | 0.104982942 | 0.10456105 | 0.092053635 | 0.985752301 | 0.721931669 | 0.00882924 | 0.015136174 | 0.071553277 |
| Tgoln1 Tgoln2 | S | 230;240 | | -0.153578492 | 0.04587759 | 0.313424887 | 0.467003379 | 0.267547297 | 0.0820947 | 0.531185968 | 0.008622528 | 0.003820562 | 0.023510855 |
| Nup133 | T | | 490 | -0.003299268 | 0.073730892 | 0.148388609 | 0.151687877 | 0.074657717 | 0.934890201 | 0.153685286 | 0.314312545 | 0.306225885 | 0.579316441 |
| Srrm2 | S | | 796 | 0.108799923 | -0.037000651 | 0.068733677 | -0.040066246 | 0.105734328 | 0.014637452 | 0.384932502 | 0.3171591 | 0.523218427 | 0.163641136 |
| Kdm5c | S | | 893 | -0.123054276 | 0.32825476 | 0.047252622 | 0.170306898 | -0.281002138 | 0.352244997 | 0.050562877 | 0.664546307 | 0.172510165 | 0.079067472 |
| Lmna | S | | 66 | -0.032972439 | 0.010518674 | 0.189875285 | 0.222847723 | 0.17935661 | 0.665987504 | 0.895734803 | 0.02183955 | 0.01996025 | 0.052700775 |
| Arhgap35 | S | | 1142 | 0.05170659 | 0.226528264 | -0.007089409 | -0.058795998 | -0.233617672 | 0.662895488 | 0.1586813 | 0.938329481 | 0.672316511 | 0.156015109 |
| Zdhhc5 | S | | 380 | 0.017223183 | 0.030162469 | 0.054796233 | 0.03757305 | 0.024633764 | 0.366059728 | 0.238965366 | 0.033724533 | 0.122342429 | 0.348280811 |
| Hdgfl2 | S | | 635 | -0.083283029 | 0.098416322 | 0.105867485 | 0.189150514 | 0.007451163 | 0.018731128 | 0.024318777 | 0.005594749 | 0.001619364 | 0.810755239 |
| Dcaf1 | S | | 202 | 0.109807758 | -0.08692839 | 0.182379033 | 0.072571274 | 0.269307422 | 0.299265992 | 0.262831538 | 0.046952202 | 0.439725976 | 0.008309032 |
| Casc3 | S | | 263 | 0.040840713 | 0.052732978 | 0.039463127 | -0.001377587 | -0.013269852 | 0.372800933 | 0.119974418 | 0.247736746 | 0.970728534 | 0.474478376 |
| Zfp639 | S | | 60 | 0.039999291 | -0.021220584 | 0.091148737 | 0.051149446 | 0.11236932 | 0.206249999 | 0.448937326 | 0.009954945 | 0.085964236 | 0.007060377 |
| Glcci1 | S | | 396 | -0.031501939 | 0.092213818 | 0.090750804 | 0.122252744 | -0.001463013 | 0.608972016 | 0.04835032 | 0.230254959 | 0.158388288 | 0.98114878 |
| Afap1 | T | | 676 | -0.230175322 | 0.642623284 | 0.446267679 | 0.676443001 | -0.196355605 | NA | 0.398840847 | NA | NA | NA |
| Ldlrap1 | S | | 201 | -0.021853162 | 0.129517089 | 0.014734447 | 0.036587609 | -0.114782642 | 0.567531714 | 0.005720032 | 0.771537022 | 0.458282971 | 0.08027598 |
| Supt7l | S | | 106 | 0.100579221 | 0.007807802 | 0.085798516 | -0.014780705 | 0.077990714 | 0.330681418 | 0.889750773 | 0.148495178 | 0.878016088 | 0.26425833 |
| Rai1 | S | | 666 | -0.019613893 | 0.067425743 | 0.099787679 | 0.119401572 | 0.032361936 | 0.535110523 | 0.075688039 | 0.11817554 | 0.097332404 | 0.523761952 |
| Huwe1 | S | | 2391 | 0.041385545 | 0.060065954 | 0.066032994 | 0.024647449 | 0.00596704 | 0.400555837 | 0.12859334 | 0.472606406 | 0.784588772 | 0.944539502 |
| Uba1 | S | | 21 | 0.011663852 | 0.117604586 | 0.157710907 | 0.146047056 | 0.040106321 | 0.951593916 | 0.447522142 | 0.298842809 | 0.477378321 | 0.800116619 |
| Rpa1 | S | | 393 | -0.002953716 | 0.206060384 | -0.027231945 | -0.02427823 | -0.233292329 | 0.933351084 | 0.021536245 | 0.656347323 | 0.692817799 | 0.025092663 |
| Parva | S | | 19 | -0.086759904 | 0.181230272 | 0.088922381 | 0.175682284 | -0.092307891 | 0.099455149 | 0.015778184 | 0.255896713 | 0.071078023 | 0.240050193 |
| Dpf2 | T | | 176 | 0.075652322 | 0.040897865 | 0.028541676 | -0.047110646 | -0.012356189 | 0.088254107 | 0.354211962 | 0.537955848 | 0.203568965 | 0.728134213 |
| Osbpl3 | S | | 250 | 0.072156591 | 0.018213352 | 0.085874075 | 0.013717484 | 0.067660723 | 0.34050723 | 0.737893622 | 0.135633972 | 0.847411723 | 0.262600234 |
| Sf3b1 | S | | 308 | 0.030687044 | 0.100999078 | 0.048003738 | 0.017316695 | -0.052995339 | 0.561328468 | 0.114227101 | 0.564074553 | 0.803980511 | 0.489328862 |
| Mcm3 | S | | 708 | 0.008416763 | 0.04861831 | 0.052041228 | 0.043624464 | 0.003422918 | 0.873898282 | 0.078242325 | 0.06115176 | 0.446471483 | 0.891105796 |
| Amer1 | S | | 534 | -0.128993744 | 0.202084403 | 0.161520624 | 0.290514369 | -0.040563779 | 0.128465751 | 0.182581832 | 0.050378864 | 0.008328729 | 0.742853608 |
| Amer1 | S | | 538 | -0.128993744 | 0.202084403 | 0.161520624 | 0.290514369 | -0.040563779 | 0.128465751 | 0.182581832 | 0.050378864 | 0.008328729 | 0.742853608 |
| Srrm1 | S | | 522 | 0.210539583 | -0.054479144 | -0.025968527 | -0.23650811 | 0.028510618 | 0.006117865 | 0.528448512 | 0.765607797 | 0.062209498 | 0.770769504 |
| Tpm3 | S | | 88 | -0.053239455 | 0.056645748 | 0.148996024 | 0.202235479 | 0.092350276 | 0.603375893 | 0.485130972 | 0.028711046 | 0.126720751 | 0.299291292 |
| Arhgap24 | S | | 380 | -0.009753901 | 0.037582413 | 0.173992482 | 0.183746383 | 0.136410069 | 0.872762921 | 0.5829248 | 0.082019277 | 0.076135648 | 0.149769227 |
| Taf6l | S | | 494 | 0.122297602 | 0.030917493 | 0.043312179 | -0.078985424 | 0.012394686 | 0.114282675 | 0.617154611 | 0.683381017 | 0.490375053 | 0.908662217 |
| Srrm2 | S | | 1749 | 0.130959317 | -0.049017953 | 0.105459179 | -0.025500137 | 0.154477132 | 0.068478641 | 0.597123911 | 0.351596489 | 0.810554655 | 0.252442001 |
| Virma | S | | 138 | -0.007643112 | 0.095585635 | 0.025639435 | 0.033282547 | -0.0699462 | 0.840893568 | 0.010319322 | 0.462145663 | 0.417407045 | 0.078011146 |
| Rbbp6 | S | | 1079 | 0.079498147 | -0.032665243 | 0.089836049 | 0.010337902 | 0.122501292 | 0.106626234 | 0.400613727 | 0.051739954 | 0.725708335 | 0.001760427 |
| Trim2 | S | | 375 | -0.050746268 | 0.143566889 | 0.090442997 | 0.141189266 | -0.053123892 | 0.589709789 | 0.074874067 | 0.327198881 | 0.235419514 | 0.563758182 |
| Rtn4 | S | | 155 | 0.004247454 | 0.080066628 | 0.124743674 | 0.12049622 | 0.044677046 | 0.941024587 | 0.320566824 | 0.203348958 | 0.227106802 | 0.651989783 |
| Pak4 | S | | 266 | 0.023087166 | 0.085273456 | 0.037744764 | 0.014657598 | -0.047528692 | 0.48770663 | 0.07814647 | 0.471896894 | 0.786519994 | 0.413106024 |
| R3hdm2 | S | | 393 | -0.034847093 | 0.127472043 | 0.08247966 | 0.117326753 | -0.044992383 | 0.524926493 | 0.080517252 | 0.23440397 | 0.131518639 | 0.531621599 |
| Eif4g3 | S | | 472 | -0.08538114 | 0.164835977 | 0.083902012 | 0.169283153 | -0.080933965 | 0.233930334 | 0.018966782 | 0.327751072 | 0.089255209 | 0.30713577 |
| Pkp4 | T | | 1016 | 0.007051785 | 0.036910756 | 0.108125316 | 0.101073531 | 0.07121456 | 0.925734954 | 0.228238296 | 0.094343727 | 0.267342532 | 0.215730983 |
| Canx | S | | 56 | -0.047197942 | 0.014988232 | 0.136437397 | 0.183635339 | 0.121449166 | 0.484165886 | 0.602605632 | 0.00443231 | 0.060875385 | 0.021109497 |
| Parp2 | S | | 32 | -0.00930904 | -0.003688077 | 0.100277782 | 0.109586822 | 0.103965859 | 0.541900614 | 0.806678263 | 0.000569083 | 0.002716411 | 0.003524041 |
| Acin1 | S | | 593 | -0.04774019 | 0.107907539 | 0.1196103 | 0.16735049 | 0.011702762 | 0.692172713 | 0.149691207 | 0.197018524 | 0.23882929 | 0.888595451 |
| Cic | T | | 2300 | -0.08917504 | 0.086161608 | 0.199048504 | 0.288223544 | 0.112886896 | 0.00073936 | 0.021279131 | 0.007516977 | 0.002064418 | 0.012614366 |
| Kif1b | S | | 1659 | -0.012063637 | 0.113589592 | 0.055300228 | 0.067363864 | -0.058289364 | 0.814002202 | 0.098260772 | 0.202415032 | 0.173239803 | 0.304181614 |
| Fcho2 | S | | 394 | -0.015894987 | 0.02148456 | 0.138733302 | 0.154628289 | 0.117248742 | 0.595797562 | 0.466428101 | 0.0230475 | 0.020427906 | 0.044313907 |
| 1110059E24Rik | S | | 144 | 0.034719558 | 0.078361873 | 0.008199282 | -0.026520276 | -0.070162591 | 0.251521648 | 0.043630937 | 0.770980117 | 0.189189727 | 0.049429959 |
| 1110059E24Rik | S | | 146 | 0.034719558 | 0.078361873 | 0.008199282 | -0.026520276 | -0.070162591 | 0.251521648 | 0.043630937 | 0.770980117 | 0.189189727 | 0.049429959 |
| Samd1 | S | | 406 | 0.114240776 | 0.007316003 | 0.10977067 | -0.004470106 | 0.102454667 | 0.09192809 | 0.961153416 | 0.640796412 | 0.984386209 | 0.693110292 |
| Ncoa5 | T | | 274 | 0.143847738 | 0.076486341 | -0.07986623 | -0.223713968 | -0.156352571 | 0.003471352 | 0.203143203 | 0.048314138 | 0.000444057 | 0.049010337 |
| Akap13 | S | | 1569 | -0.016380799 | 0.107366797 | 0.072384387 | 0.088765186 | -0.03498241 | 0.863318157 | 0.092019408 | 0.282400649 | 0.39087653 | 0.574211396 |
| Ccnk | S | | 341 | 0.022925654 | 0.011763138 | 0.105153001 | 0.082227347 | 0.093389863 | 0.678345841 | 0.578500373 | 0.038629073 | 0.21697845 | 0.067193472 |
| Ankrd17 | S | | 205 | 0.124264807 | 0.116257162 | 0.090340169 | -0.033924638 | -0.025916994 | 0.464623954 | 0.575222569 | 0.650079703 | 0.863663124 | 0.909226405 |
| Baz1b | S | | 160 | 0.018719608 | 0.138957288 | 0.05515805 | 0.036438442 | -0.083799238 | 0.860153557 | 0.143878646 | 0.532668817 | 0.653158852 | 0.117955805 |
| Pnisr | S | | 561 | 0.02987559 | 0.001547316 | 0.203256128 | 0.173380537 | 0.201708812 | 0.503855063 | 0.97732227 | 0.161057342 | 0.217995658 | 0.166668086 |
| Arhgef17 | S | | 79 | -0.005937035 | -0.006322034 | 0.186332446 | 0.19226948 | 0.192654479 | 0.934604867 | 0.922148017 | 0.032326122 | 0.068827445 | 0.052984338 |
| Ing5 | S | | 118 | 0.049664874 | -0.016902041 | 0.075812484 | 0.02614761 | 0.092714525 | 0.076408783 | 0.557060066 | 0.03759846 | 0.298355816 | 0.021060133 |
| Kat6a | S | | 1242 | 0.104123647 | -0.064730368 | 0.061119173 | -0.043004474 | 0.125849541 | 0.004233926 | 0.262272559 | 0.237704469 | 0.375699281 | 0.091212007 |
| Kat6a | T | | 1240 | 0.104123647 | -0.064730368 | 0.061119173 | -0.043004474 | 0.125849541 | 0.004233926 | 0.262272559 | 0.237704469 | 0.375699281 | 0.091212007 |
| Kat6a | T | | 1241 | 0.104123647 | -0.064730368 | 0.061119173 | -0.043004474 | 0.125849541 | 0.004233926 | 0.262272559 | 0.237704469 | 0.375699281 | 0.091212007 |
| Phrf1 | S | | 867 | 0.104762788 | -0.023431324 | 0.069893191 | -0.034869597 | 0.093324514 | 0.039564782 | 0.543406341 | 0.291378054 | 0.556621494 | 0.178074875 |

| | | | | | | | | | | | | |
|---|---|---|---|---|---|---|---|---|---|---|---|---|
| Cic | S | 778 | -0.011323598 | 0.138798597 | 0.140098936 | 0.151422534 | 0.001300339 | 0.878306976 | 0.240798429 | 0.418893714 | 0.391732111 | 0.994050604 |
| Sympk | S | 1260 | -0.052874577 | 0.007735535 | 0.173126104 | 0.22600068 | 0.165390569 | 0.245841292 | 0.831286371 | 0.003496579 | 0.002701668 | 0.008313989 |
| Ticrr | S | 1195 | 0.213291648 | 0.023003014 | -0.01122537 | -0.224517018 | -0.034228384 | 0.091181567 | 0.801160275 | 0.916869251 | 0.132189521 | 0.779752411 |
| Zfp532 | S | 349 | -0.124468745 | 0.119605453 | 0.134015519 | 0.258484264 | 0.014410066 | 0.168356815 | 0.024009238 | 0.098332064 | 0.033464614 | 0.819744821 |
| Bicral | S | 594 | 0.016710913 | -0.051234875 | 0.200367772 | 0.18365686 | 0.251602647 | 0.816712163 | 0.729260735 | 0.033899855 | 0.031013432 | 0.161039099 |
| Gnptab | S | 1229 | 0.063697454 | 0.026034649 | 0.033818165 | -0.029879289 | 0.007783516 | 0.119419683 | 0.375780587 | 0.218388067 | 0.411641687 | 0.808954149 |
| Lbr | S | 70 | -0.095953783 | 0.096712311 | 0.108084906 | 0.204038689 | 0.011372595 | 0.009431794 | 0.009204822 | 0.005498752 | 0.000729133 | 0.63084525 |
| Nktr | S | 255 | -0.045093708 | -0.027376158 | 0.193745458 | 0.238839166 | 0.221121617 | 0.5429288 | 0.541973369 | 0.003393829 | 0.035787045 | 0.002956664 |
| Nktr | T | 253 | -0.045093708 | -0.027376158 | 0.193745458 | 0.238839166 | 0.221121617 | 0.5429288 | 0.541973369 | 0.003393829 | 0.035787045 | 0.002956664 |
| Prpf38b | S | 487 | 0.004968859 | 0.0930246 | 0.064060673 | 0.059091813 | -0.028963928 | 0.934219597 | 0.128715413 | 0.24817217 | 0.404887039 | 0.634202908 |
| Maged1 | S | 319 | 0.193059392 | 0.092534732 | -0.059401103 | -0.252460495 | -0.151935835 | 0.213571065 | 0.204471813 | 0.589850424 | 0.143555183 | 0.198568486 |
| Baz2a | S | 1383 | 0.095316543 | 0.051887884 | -0.063998534 | -0.159315077 | -0.115886419 | 0.007495356 | 0.176861109 | 0.434316562 | 0.134205877 | 0.21042892 |
| Zc3h13 | S | 198 | 0.01298275 | 0.035147794 | 0.094176473 | 0.081193723 | 0.059028678 | 0.866055324 | 0.180177061 | 0.102626887 | 0.361552466 | 0.240904286 |
| Atrx | S | 992 | -0.011855444 | 0.080447136 | 0.055425298 | 0.067280741 | -0.025021838 | 0.797647893 | 0.069019789 | 0.124978549 | 0.204568542 | 0.504583339 |
| Rere | S | 642 | 0.014372399 | 0.035538512 | 0.047416683 | 0.033044284 | 0.011878171 | 0.738794853 | 0.032395124 | 0.091973691 | 0.482538507 | 0.584297334 |
| Hltf | S | 21 | 0.086240229 | 0.021186687 | 0.067203598 | -0.019036631 | 0.046016911 | 0.129161108 | 0.621259649 | 0.453116351 | 0.82783142 | 0.597600883 |
| AI597479 | S | 182 | -0.104059108 | 0.044312672 | 0.256468686 | 0.360527794 | 0.212156014 | 0.054898492 | 0.30802913 | 0.011360935 | 0.004461361 | 0.024462628 |
| Ssh1 | S | 597 | 0.046758113 | -0.02691562 | 0.124425286 | 0.077667173 | 0.151340906 | 0.192533265 | 0.642335502 | 0.065924612 | 0.189348073 | 0.067069697 |
| Jph2 | S | 462 | 0.024792327 | -0.022889727 | 0.176716342 | 0.151924015 | 0.199606069 | 0.739228388 | 0.479965276 | 0.028446849 | 0.126337705 | 0.017316345 |
| Tcf7l2 | S | 120 | -0.365165537 | 0.182934647 | 0.62192643 | 0.987091967 | 0.438991784 | 0.499692036 | 0.66757978 | 0.212478621 | 0.164378256 | 0.402033471 |
| Sf3b1 | T | 313 | 0.069162012 | 0.0637077 | 0.009952974 | -0.059209038 | -0.053754726 | 0.094478525 | 0.248173396 | 0.797760499 | 0.171028626 | 0.326097466 |
| Htatsf1 | S | 679 | -0.004597306 | 0.053743655 | 0.140391581 | 0.144988887 | 0.086647925 | 0.937743211 | 0.373961129 | 0.128623577 | 0.133480284 | 0.316621484 |
| Rbm17 | S | 222 | 0.007354915 | 0.040633343 | 0.05686271 | 0.049507795 | 0.016229367 | 0.716289034 | 0.071606649 | 0.088062594 | 0.121390895 | 0.513523071 |
| Pcnx3 | S | 496 | -0.031454205 | 0.120347253 | 0.104435211 | 0.135889416 | -0.015912042 | 0.696578374 | 0.185578944 | 0.208715225 | 0.199502668 | 0.862588739 |
| Rtn4 | S | 105 | -0.018151936 | 0.057745501 | 0.062868214 | 0.081020151 | 0.005122714 | 0.624191014 | 0.057845639 | 0.049972302 | 0.074003856 | 0.806139427 |
| Mprip | T | 540 | -0.081818625 | 0.331023969 | 0.017057268 | 0.098875894 | -0.313966701 | 0.444790113 | 0.063063591 | 0.873901163 | 0.48231877 | 0.074360551 |
| Pias1 | S | 485 | -0.027516077 | 0.092772959 | 0.048225011 | 0.075741088 | -0.044547948 | 0.486751383 | 0.016595029 | 0.218431823 | 0.147291183 | 0.268081794 |
| Cdk13 | S | 494 | 0.053048147 | 0.012342511 | 0.126253759 | 0.073205611 | 0.113911247 | 0.608379829 | 0.741348169 | 0.115077127 | 0.517446341 | 0.13730106 |
| Sfswap | S | 865 | 0.123281368 | -0.026693948 | 0.048540795 | -0.074740573 | 0.075234743 | 0.042310264 | 0.682896018 | 0.363597998 | 0.212642329 | 0.307050343 |
| Ufl1 | S | 458 | -0.07194306 | 0.035276348 | 0.122956346 | 0.194899406 | 0.087679999 | 0.299570721 | 0.457201531 | 0.002020064 | 0.056922016 | 0.123874849 |
| Xrcc6 | S | 2 | -0.03424409 | 0.09392664 | 0.058211597 | 0.092455687 | -0.035715043 | 0.133277762 | 0.061746831 | 0.014569394 | 0.009557029 | 0.33043981 |
| Rad9a | S | 385 | 0.007913579 | 0.088172547 | 0.058992565 | 0.051078987 | -0.029179982 | 0.289829701 | 0.181619172 | 0.323609273 | 0.378157518 | 0.669123422 |
| Rnf6 | S | 182 | 0.043393185 | 0.090838716 | -0.003046198 | -0.046439384 | -0.093884915 | 0.518489805 | 0.023458245 | 0.976243813 | 0.685204659 | 0.405180157 |
| Srrm2 | T | 251 | 0.060539247 | 0.014618394 | 0.059587805 | -0.000951443 | 0.04496941 | 0.178353204 | 0.662744352 | 0.138188476 | 0.980570177 | 0.200149961 |
| Mcm9 | S | 742 | -0.026243943 | 0.094038693 | 0.138341006 | 0.164584949 | 0.044302313 | 0.744083679 | 0.495312531 | 0.091817056 | 0.07746713 | 0.733669002 |
| Mcm9 | S | 743 | -0.026243943 | 0.094038693 | 0.138341006 | 0.164584949 | 0.044302313 | 0.744083679 | 0.495312531 | 0.091817056 | 0.07746713 | 0.733669002 |
| Shisa8 | S | 214 | 0.123274181 | -0.021349369 | 0.083642021 | -0.03963216 | 0.10499139 | 0.11160339 | 0.546672118 | 0.257459208 | 0.615624406 | 0.184083224 |
| Shisa8 | S | 218 | 0.123274181 | -0.021349369 | 0.083642021 | -0.03963216 | 0.10499139 | 0.11160339 | 0.546672118 | 0.257459208 | 0.615624406 | 0.184083224 |
| Iqsec1 | S | 415 | 0.029403784 | 0.021429831 | 0.070820908 | 0.041417124 | 0.049391077 | 0.44481887 | 0.643690462 | 0.033080667 | 0.248787585 | 0.295089734 |
| Spen | T | 2458 | 0.084310875 | 0.042943558 | 0.08144128 | -0.002869595 | 0.038497722 | 0.215977145 | 0.641123627 | 0.478900786 | 0.97990082 | 0.766294229 |
| Hdgfl2 | S | 603 | 0.07385374 | 0.027380452 | 0.040868107 | -0.032985633 | 0.013487655 | 0.11303348 | 0.43141909 | 0.415609182 | 0.4886771 | 0.737234601 |
| Add1 | S | 613 | -0.032873054 | 0.188367854 | 0.021133203 | 0.054006257 | -0.167234651 | 0.809485521 | 0.044685092 | 0.841524288 | 0.731316732 | 0.195465989 |
| Srrm2 | S | 349 | 0.024088456 | 0.020727944 | 0.063555053 | 0.039466597 | 0.042827109 | 0.34203621 | 0.36945383 | 0.049224221 | 0.176561445 | 0.13424109 |
| Pcm1 | S | 1259 | 0.034453035 | 0.07881543 | 0.035810663 | 0.001357628 | -0.043004767 | 0.360490521 | 0.14853506 | 0.460578393 | 0.978581376 | 0.462267937 |
| Mex3d | S | 483 | 0.021254621 | 0.201557583 | 0.042080537 | 0.020825916 | -0.159477046 | 0.813775653 | 0.19815249 | 0.775698813 | 0.871929579 | 0.35894993 |
| Zc3hc1 | S | 340 | 0.113266649 | 0.025322145 | 0.021909796 | -0.091356853 | -0.003412349 | 0.049807747 | 0.718567939 | 0.822548322 | 0.395894384 | 0.975125886 |
| Rubcn | S | 198 | 0.119887459 | 0.009250939 | -0.015899742 | -0.135787201 | -0.025150681 | 0.008291188 | 0.877675127 | 0.837342583 | 0.166704882 | 0.770586982 |
| Gpkow | S | 348 | -0.02098625 | 0.030971203 | 0.069066139 | 0.090052389 | 0.038094937 | 0.660306583 | 0.153719996 | 0.006257275 | 0.145412085 | 0.091211865 |
| Hnrnpk | S | 216 | 0.006751933 | 0.108678826 | 0.032023197 | 0.025271263 | -0.07665563 | 0.91983013 | 0.053904824 | 0.484531747 | 0.722000152 | 0.178011927 |
| Syap1 | T | 282 | -0.025070471 | 0.193877938 | 0.170961412 | 0.196031882 | -0.022916526 | 0.793733603 | 0.400762665 | 0.403375336 | 0.357532339 | 0.929788022 |
| Zc3h13 | S | 207 | 0.040889926 | 0.046967374 | 0.058506006 | 0.01761608 | 0.011538631 | 0.245405314 | 0.123215343 | 0.554846752 | 0.852065304 | 0.90114622 |
| Mtdh | S | 110 | -0.005504412 | 0.116878215 | 0.02172526 | 0.027229673 | -0.095152954 | 0.829702206 | 0.028567166 | 0.480040819 | 0.462639746 | 0.047810456 |
| Mycbp2 | S | 2707 | 0.00342933 | 0.060971455 | 0.125261004 | 0.121831674 | 0.064289549 | 0.972881093 | 0.449930199 | 0.127013003 | 0.307178723 | 0.4819638 |
| B230219D22Rik | S | 121 | 0.000671391 | 0.005969399 | 0.166354773 | 0.165683382 | 0.160385374 | 0.995600627 | 0.93084333 | 0.046180992 | 0.220717353 | 0.014220624 |
| Polk | S | 748 | 0.003431256 | 0.116429979 | 0.031192452 | 0.027761196 | -0.085237527 | 0.924525334 | 0.061450184 | 0.48159909 | 0.552967104 | 0.162544646 |
| Aak1 | S | 21 | 0.082555181 | 0.085189059 | 0.027285475 | -0.055269706 | -0.057903584 | 0.246517216 | 0.366303618 | 0.671483385 | 0.486951336 | 0.554467936 |
| Apbb2 | S | 412 | -0.061452153 | 0.106396445 | 0.135944705 | 0.197396858 | 0.02954826 | 0.044317051 | 0.039307181 | 0.078996222 | 0.041283393 | 0.62216453 |
| Ncam1 | S | 887 | 0.015031053 | 0.071768653 | 0.036493192 | 0.02146214 | -0.035275461 | 0.564206102 | 0.071701038 | 0.302541364 | 0.528748339 | 0.37238304 |
| Bmp2k | T | 995 | 0.115211259 | -0.011056816 | 0.02236273 | -0.092848529 | 0.033419546 | 0.020898452 | 0.781149058 | 0.520835013 | 0.044284461 | 0.352072573 |
| Ncoa2 | S | 1072 | 0.120982355 | 0.031654524 | 0.239387991 | 0.118405637 | 0.207733468 | 0.540005062 | 0.919268238 | 0.493213469 | 0.724455677 | 0.609910143 |
| Ppp4r3a | S | 127 | -0.033891247 | 0.059890168 | 0.153232419 | 0.187123666 | 0.093342251 | 0.499314237 | 0.239730824 | 0.097800969 | 0.062470942 | 0.252677639 |
| Eef1d | S | 119 | -0.093686776 | 0.190039163 | 0.010770599 | 0.104457375 | -0.179268563 | 0.134126978 | 0.001308076 | 0.818611736 | 0.13815808 | 0.024471355 |
| Tjp2 | Y | 914 | 0.103439964 | 0.102511697 | 0.094079781 | -0.009360183 | -0.008431916 | 0.502571575 | 0.422094442 | 0.730982787 | 0.97155789 | 0.973222872 |
| Hirip3 | S | 292 | 0.070400304 | -0.052231728 | 0.19136512 | 0.120964816 | 0.243596848 | 0.334155927 | 0.262386873 | 0.065431641 | 0.206584295 | 0.032027162 |
| Hirip3 | S | 295 | 0.070400304 | -0.052231728 | 0.19136512 | 0.120964816 | 0.243596848 | 0.334155927 | 0.262386873 | 0.065431641 | 0.206584295 | 0.032027162 |
| Hirip3 | S | 296 | 0.070400304 | -0.052231728 | 0.19136512 | 0.120964816 | 0.243596848 | 0.334155927 | 0.262386873 | 0.065431641 | 0.206584295 | 0.032027162 |
| Chd8 | S | 1422 | 0.038787145 | 0.0388921 | 0.07768035 | 0.038893205 | 0.03878825 | 0.238109677 | 0.316412471 | 0.343429537 | 0.598668072 | 0.607468006 |
| Chd8 | S | 1426 | 0.038787145 | 0.0388921 | 0.07768035 | 0.038893205 | 0.03878825 | 0.238109677 | 0.316412471 | 0.343429537 | 0.598668072 | 0.607468006 |
| Sp100 | S | 319 | -0.030541086 | 0.037847834 | 0.17483406 | 0.205375146 | 0.136986226 | 0.742348133 | 0.621099949 | 0.062261288 | 0.101432233 | 0.169272687 |
| Srsf7 | S | 192 | -0.079644023 | -0.000132171 | 0.251520264 | 0.331164286 | 0.251652435 | 0.306717747 | 0.998945766 | 0.013266762 | 0.008832566 | 0.06298001 |
| Srsf7 | S | 194 | -0.079644023 | -0.000132171 | 0.251520264 | 0.331164286 | 0.251652435 | 0.306717747 | 0.998945766 | 0.013266762 | 0.008832566 | 0.06298001 |
| Pard3 | S | 378 | 0.085079848 | 0.037037419 | 0.000849455 | -0.084230393 | -0.036187964 | 0.046562208 | 0.269600023 | 0.980639236 | 0.03119539 | 0.149024031 |
| Rb1 | S | 800 | -0.128768014 | 0.115518599 | 0.158584475 | 0.287352488 | 0.043065876 | 0.01090053 | 0.063685026 | 0.004082153 | 6.59612E-05 | 0.359440041 |
| Prpf4b | S | 94 | -0.013848945 | 0.009020607 | 0.109990877 | 0.123839821 | 0.10097027 | 0.709012848 | 0.602193736 | 0.007268381 | 0.033398155 | 0.025381987 |
| Tbc1d10b | S | 129 | -0.021345617 | 0.020469943 | 0.196945945 | 0.218291563 | 0.176476002 | 0.695349286 | 0.757513578 | 0.058739603 | 0.048513876 | 0.09248555 |
| Gli3 | S | 664 | 0.00798389 | 0.032993089 | 0.098119512 | 0.090135622 | 0.065126424 | 0.805991316 | 0.352165553 | 0.072647141 | 0.097377753 | 0.1948013 |
| Map4 | T | 847 | 0.177925936 | 0.071212979 | -0.126777189 | -0.304703125 | -0.197990168 | 0.011669931 | 0.196622848 | 0.14344244 | 0.020112833 | 0.062278642 |
| Tjp2 | S | 270 | -0.024306024 | -0.121397243 | 0.348000389 | 0.372306413 | 0.469397632 | 0.764983318 | 0.110708593 | 0.006128392 | 0.010633671 | 0.003773596 |
| Srrm1 | S | 624 | -0.004954924 | -0.008930301 | 0.073256804 | 0.078211728 | 0.082187105 | 0.843687244 | 0.565697873 | 7.49193E-05 | 0.061569326 | 0.012052785 |
| Ncoa3 | S | 1304 | 0.015188923 | 0.052182951 | 0.061675906 | 0.046486983 | 0.009492955 | 0.458603031 | 0.197911833 | 0.163917035 | 0.260712784 | 0.829490497 |
| Cul4b | T | 106 | 0.103522779 | 0.08558371 | -0.026499082 | -0.130021861 | -0.112082792 | 0.051055077 | 0.137663919 | 0.127899274 | 0.033561961 | 0.08610389 |
| Med1 | S | 996 | 0.095861154 | 0.036761948 | 0.038498687 | -0.057362467 | 0.001736739 | 0.142292887 | 0.628908293 | 0.469804706 | 0.394291652 | 0.982946389 |
| Ostm1 | S | 329 | -0.018497902 | 0.0526032 | 0.123279775 | 0.141777677 | 0.070676576 | 0.557096274 | 0.055705814 | 0.197017014 | 0.154171579 | 0.397075069 |
| Polr2a | S | 1913 | 0.023041117 | 0.038227123 | 0.051439217 | 0.0283981 | 0.013212094 | 0.429455193 | 0.162322852 | 0.12080914 | 0.387321986 | 0.648722305 |
| Nelfa | S | 363 | -0.000296831 | 0.045165324 | 0.089402141 | 0.089698972 | 0.044236818 | 0.995464826 | 0.369387505 | 0.04752467 | 0.125224331 | 0.294304384 |
| Pa2g4 | S | 375 | 0.013728376 | 0.115807641 | 0.028470044 | 0.014741668 | -0.087337597 | 0.822734455 | 0.051172151 | 0.732883234 | 0.854816835 | 0.303189689 |
| Ube2j1 | S | 251 | 0.074996752 | 0.057122248 | 0.026865323 | -0.048131429 | -0.030256924 | 0.175528275 | 0.282744378 | 0.681834315 | 0.359723481 | 0.536422798 |
| Esyt2 | S | 663 | -0.029266703 | 0.004204778 | 0.147541413 | 0.176808116 | 0.143336636 | 0.603758526 | 0.877417308 | 0.009464909 | 0.039540026 | 0.01755328 |
| Phldb2 | S | 73 | 0.142561599 | 0.015025153 | 0.029716291 | -0.112845308 | 0.014691139 | 0.077883944 | 0.888386732 | 0.781074909 | 0.358457964 | 0.917100334 |
| Map7d1 | S | 256 | 0.052572071 | 0.167058654 | -0.032918106 | -0.085490177 | -0.199976761 | 0.454026477 | 0.094840285 | 0.738008645 | 0.434509202 | 0.128541704 |
| Srrm2 | T | 1428 | 0.047643565 | 0.070537123 | -0.000171326 | -0.047814891 | -0.070708449 | 0.256904718 | 0.039767689 | 0.995673503 | 0.244705112 | 0.045893369 |
| Usp34 | S | 2540 | 0.003912343 | -0.029725821 | 0.285807246 | 0.281894903 | 0.315533067 | 0.985794534 | 0.846180168 | 0.11064392 | 0.206130848 | 0.025784681 |
| Ncor1 | S | 2116 | -0.018822943 | 0.103873563 | 0.058282107 | 0.07710505 | -0.045591457 | 0.840988929 | 0.071151616 | 0.304979313 | 0.414186511 | 0.225398178 |
| Sntb2 | S | 213 | -0.104664463 | 0.059876704 | 0.161439779 | 0.266104242 | 0.101563075 | 0.030248234 | 0.087169592 | 0.003611626 | 0.001788644 | 0.031747155 |

| | | | | | | | | | | | | |
|---|---|---|---|---|---|---|---|---|---|---|---|---|
| Ppp1r18 | S | 126 | 0.102534081 | 0.029520122 | 0.033159636 | -0.069374446 | 0.003639513 | 0.115616403 | 0.462328497 | 0.70137401 | 0.463384472 | 0.965529431 |
| Ddx59 | S | 150 | 0.011311984 | 0.0478912 | 0.050833355 | 0.039521371 | 0.002942155 | 0.718672351 | 0.054435984 | 0.181010949 | 0.329381547 | 0.925736694 |
| H2-K1 | T | 350;309;349;350 | 0.012465609 | 0.1249841 | -0.000178159 | -0.012643768 | -0.125162259 | 0.879797141 | 0.022268303 | 0.997723809 | 0.883441477 | 0.10966541 |
| Cep170 | S | 1157 | 0.088427701 | 0.16183059 | -0.014211924 | -0.102639625 | -0.176042514 | 0.470143043 | 0.209982489 | 0.928546749 | 0.560209307 | 0.337359406 |
| Srbd1 | S | 28 | 0.140622376 | 0.033576285 | 0.116994848 | -0.023627528 | 0.083418563 | 0.538001521 | 0.776747087 | 0.370278377 | 0.908562124 | 0.296475563 |
| Srbd1 | S | 29 | 0.140622376 | 0.033576285 | 0.116994848 | -0.023627528 | 0.083418563 | 0.538001521 | 0.776747087 | 0.370278377 | 0.908562124 | 0.296475563 |
| Srbd1 | S | 31 | 0.140622376 | 0.033576285 | 0.116994848 | -0.023627528 | 0.083418563 | 0.538001521 | 0.776747087 | 0.370278377 | 0.908562124 | 0.296475563 |
| Srbd1 | S | 39 | 0.140622376 | 0.033576285 | 0.116994848 | -0.023627528 | 0.083418563 | 0.538001521 | 0.776747087 | 0.370278377 | 0.908562124 | 0.296475563 |
| Osbpl3 | T | 482 | 0.043654758 | 0.176613742 | -0.021541921 | -0.065196678 | -0.198155663 | 0.378903378 | 0.13494779 | 0.782169868 | 0.410882851 | 0.120892491 |
| Srrm1 | S | 209 | 0.04159336 | 0.019588293 | 0.078681973 | 0.037088613 | 0.05909368 | 0.144786268 | 0.547572898 | 0.23912092 | 0.525622033 | 0.350522367 |
| Srrm1 | S | 211 | 0.04159336 | 0.019588293 | 0.078681973 | 0.037088613 | 0.05909368 | 0.144786268 | 0.547572898 | 0.23912092 | 0.525622033 | 0.350522367 |
| Sptbn1 | S | 2127 | 0.086727184 | 0.000987725 | 0.032635877 | -0.054091308 | 0.031648151 | 0.017535605 | 0.983033204 | 0.593591551 | 0.38421696 | 0.631430038 |
| Cep170b | T | 1289 | 0.001326255 | 0.076497446 | 0.054788301 | 0.053462046 | -0.021709146 | 0.965878139 | 0.38107663 | 0.025535697 | 0.147317337 | 0.781569071 |
| Srrm2 | S | 1813 | 0.040245463 | 0.010115663 | 0.132118492 | 0.091873029 | 0.122002829 | 0.32944024 | 0.765169926 | 0.197306195 | 0.329173013 | 0.230259113 |
| Bclaf1 | S | 267 | 0.004215333 | 0.10024966 | 0.032334745 | 0.028119413 | -0.067914915 | 0.948975118 | 0.063302567 | 0.345273752 | 0.66214135 | 0.155029794 |
| Nes | S | 169 | 0.019373927 | 0.033859456 | 0.040979621 | 0.021605694 | 0.007120165 | 0.213826635 | 0.17156003 | 0.09727438 | 0.289897657 | 0.766073934 |
| Polr2a | S | 1920 | 0.061659936 | 0.05595299 | -0.009755191 | -0.071415127 | -0.065708181 | 0.134693794 | 0.04294885 | 0.809818564 | 0.162199087 | 0.167688787 |
| Ncapd2 | T | 1329 | 0.095652888 | 0.029145845 | 0.041682198 | -0.05397069 | 0.012536353 | 0.11992789 | 0.578453552 | 0.590774894 | 0.44487592 | 0.844649096 |
| Trak1 | S | 777 | 0.013946154 | 0.204967724 | 0.02717632 | 0.013230166 | -0.177791404 | 0.884744742 | 0.186478013 | 0.708954022 | 0.876824151 | 0.235452758 |
| Trak1 | S | 782 | 0.013946154 | 0.204967724 | 0.02717632 | 0.013230166 | -0.177791404 | 0.884744742 | 0.186478013 | 0.708954022 | 0.876824151 | 0.235452758 |
| Trak1 | T | 778 | 0.013946154 | 0.204967724 | 0.02717632 | 0.013230166 | -0.177791404 | 0.884744742 | 0.186478013 | 0.708954022 | 0.876824151 | 0.235452758 |
| Sipa1l1 | S | 1567 | 0.02843512 | 0.042604675 | 0.056013305 | 0.027578185 | 0.01340863 | 0.422715793 | 0.072783392 | 0.376180378 | 0.656610088 | 0.814257653 |
| Cnot4 | S | 324 | 0.073522196 | 0.051792024 | -0.001530956 | -0.075053152 | -0.05332298 | 0.07602216 | 0.188467615 | 0.986004659 | 0.413134741 | 0.543770113 |
| Pkmyt1 | S | 486 | 0.118184066 | 0.056021297 | 0.010713055 | -0.107471012 | -0.045308242 | 0.127671759 | 0.473495845 | 0.938310523 | 0.478064725 | 0.760173536 |
| Nek1 | S | 1071 | -0.041056952 | 0.080360798 | 0.114667581 | 0.155724533 | 0.034306783 | 0.355321749 | 0.290364202 | 0.037118584 | 0.006653416 | 0.601180955 |
| Ptpn1 | S | 118 | 0.075406834 | 0.062170332 | -0.030030321 | -0.105437155 | -0.092200653 | 0.084568116 | 0.054671349 | 0.554552913 | 0.098119507 | 0.136806956 |
| Clasrp | S | 285 | 0.038129264 | -0.016508959 | 0.073678097 | 0.035548834 | 0.090187056 | 0.390836484 | 0.447454942 | 0.003334972 | 0.419311926 | 0.020064684 |
| Trp53bp1 | S | 631 | -0.056181324 | 0.203018048 | 0.034717524 | 0.090898849 | -0.168300524 | 0.593852552 | 0.047012848 | 0.725550861 | 0.464251144 | 0.161693289 |
| Trp53bp1 | S | 632 | -0.056181324 | 0.203018048 | 0.034717524 | 0.090898849 | -0.168300524 | 0.593852552 | 0.047012848 | 0.725550861 | 0.464251144 | 0.161693289 |
| Nup133 | S | 479 | 0.12858155 | 0.106969219 | -0.027347173 | -0.155928722 | -0.134316391 | 0.132024683 | 0.402937966 | 0.694768806 | 0.064921049 | 0.299015558 |
| Coil | S | 171 | -0.008503158 | 0.017365263 | 0.118687046 | 0.127190203 | 0.101321782 | 0.87301667 | 0.706424515 | 0.020708067 | 0.060770097 | 0.042297542 |
| AW554918 | S | 549 | 0.048705695 | -0.001390704 | 0.087331717 | 0.038626022 | 0.088722421 | 0.052888134 | 0.978914694 | 0.247660137 | 0.5554596 | 0.279630899 |
| Epb41l1 | T | 550 | 0.008545484 | 0.050318718 | 0.063204926 | 0.054659442 | 0.012886207 | 0.800182739 | 0.068659422 | 0.2165028 | 0.2848294 | 0.758255185 |
| Yap1 | S | 351 | 0.099498612 | 0.073701803 | -0.000841073 | -0.100339684 | -0.074542875 | 0.418005423 | 0.079026393 | 0.965780005 | 0.414486052 | 0.086322612 |
| Spen | S | 1237 | 0.055182317 | -0.018590086 | 0.090316773 | 0.035134456 | 0.108906858 | 0.157851475 | 0.61636954 | 0.065174473 | 0.36016019 | 0.037387116 |
| Dido1 | S | 1250 | -0.012662367 | -0.016204957 | 0.160644518 | 0.173306885 | 0.176849475 | 0.870700075 | 0.713128434 | 0.012946169 | 0.087110773 | 0.011410193 |
| Farp1 | S | 394 | 0.012303061 | 0.147359975 | -0.032347742 | -0.044650803 | -0.179707718 | 0.89540311 | 0.010924003 | 0.573144936 | 0.647260096 | 0.02761064 |
| Zc3h3 | S | 851 | -0.014973296 | 0.085800015 | 0.044180984 | 0.05915428 | -0.04161903 | 0.690927603 | 0.026471625 | 0.288535222 | 0.180677476 | 0.269685588 |
| Rnf2 | S | 41 | 0.140652536 | 0.009456238 | -0.01031204 | -0.150964576 | -0.019768277 | 0.021297281 | 0.910616012 | 0.803385435 | 0.030021159 | 0.824497517 |
| Cic | S | 2304 | -0.065164546 | 0.031757891 | 0.128574828 | 0.193739374 | 0.096816936 | 0.040513424 | 0.023059803 | 0.004774359 | 0.001225111 | 0.017010699 |
| Setd2 | S | 459 | -0.052973858 | 0.03313051 | 0.155238356 | 0.208212213 | 0.122107846 | 0.393754204 | 0.361096839 | 0.025200209 | 0.028077835 | 0.058963408 |
| Setd2 | S | 460 | -0.052973858 | 0.03313051 | 0.155238356 | 0.208212213 | 0.122107846 | 0.393754204 | 0.361096839 | 0.025200209 | 0.028077835 | 0.058963408 |
| Setd2 | S | 462 | -0.052973858 | 0.03313051 | 0.155238356 | 0.208212213 | 0.122107846 | 0.393754204 | 0.361096839 | 0.025200209 | 0.028077835 | 0.058963408 |
| Ift46 | T | 283 | -0.086257357 | 0.161269019 | 0.109532098 | 0.195789455 | -0.051736921 | 0.133157495 | 0.095834025 | 0.085498129 | 0.015630808 | 0.515940469 |
| Cc2d1b | S | 455 | -0.125992752 | 0.361246798 | -0.023459029 | 0.102533724 | -0.384705827 | 0.183849267 | 0.014736581 | 0.876982268 | 0.499851972 | 0.066949513 |
| Zeb1 | S | 998 | -0.108419632 | 0.105249965 | 0.213128703 | 0.321548336 | 0.107878738 | 0.025149275 | 0.056105616 | 0.050349312 | 0.015413476 | 0.182378813 |
| Map4k4 | S | 689 | 0.07936297 | 0.218526998 | -0.105617713 | -0.184980683 | -0.324144711 | 0.549500029 | 0.096844901 | 0.644237478 | 0.423661107 | 0.211310592 |
| Sec62 | T | 375 | 0.042892751 | 0.010366126 | 0.101565309 | 0.058672557 | 0.091199183 | 0.335712418 | 0.710970704 | 0.147719197 | 0.36721081 | 0.186476901 |
| Afap1l2 | S | 165 | 0.139787166 | 0.188884159 | -0.103435134 | -0.243222299 | -0.292319293 | 0.538753432 | 0.102306673 | 0.589674339 | 0.385934297 | 0.205256391 |
| Rere | S | 1258 | 0.088550741 | 0.067676955 | -0.059260425 | -0.147811166 | -0.12693738 | 0.066177119 | 0.126086534 | 0.690189527 | 0.362466329 | 0.421203766 |
| Clasp1 | S | 820 | -0.048639894 | 0.004330026 | 0.199504469 | 0.248144363 | 0.195174443 | 0.518980332 | 0.914903965 | 0.021498323 | 0.033392342 | 0.027282964 |
| Tasor | S | 628 | 0.056894677 | 0.003467334 | 0.152168761 | 0.095274084 | 0.148701426 | 0.282972767 | 0.977627403 | 0.28126668 | 0.45973084 | 0.370015265 |
| Tasor | S | 631 | 0.056894677 | 0.003467334 | 0.152168761 | 0.095274084 | 0.148701426 | 0.282972767 | 0.977627403 | 0.28126668 | 0.45973084 | 0.370015265 |
| Etaa1 | S | 75 | 0.164536478 | 0.036955071 | 0.038247919 | -0.126288558 | 0.001292848 | 0.422755699 | 0.420794204 | 0.509926774 | 0.526693826 | 0.982136141 |
| Tcf20 | T | 1792 | -0.126673658 | 0.14721029 | 0.131341776 | 0.258015434 | -0.015868514 | 0.051325709 | 0.007967924 | 0.166118781 | 0.037286095 | 0.83206332 |
| Brca1 | S | 831 | -0.002248271 | 0.028276649 | 0.097325913 | 0.099574184 | 0.069049264 | 0.967047263 | 0.532135731 | 0.027717198 | 0.132607162 | 0.135571394 |
| Srsf4 | S | 461 | 0.010200124 | 0.133883347 | 0.101741871 | 0.091541748 | -0.032141475 | 0.926791104 | 0.178370478 | 0.655604903 | 0.683068327 | 0.88067154 |
| Ccnl1 | S | 451 | 0.01895846 | 0.037337944 | 0.120739379 | 0.101780919 | 0.083401436 | 0.934223119 | 0.461524 | 0.116565143 | 0.669150015 | 0.258479613 |
| Sik3 | S | 721 | -0.001594433 | 0.070693516 | 0.010758672 | 0.012353115 | -0.059934844 | 0.9622651 | 0.002488306 | 0.617650532 | 0.728174516 | 0.043277503 |
| Ccnt2 | S | 477 | -0.070918169 | 0.105349268 | 0.166079565 | 0.236997734 | 0.060730297 | 0.620111317 | 0.312803463 | 0.162811244 | 0.154772634 | 0.556288781 |
| Son | S | 1956 | 0.10415856 | 0.024445783 | 0.063459895 | -0.040698664 | 0.039014112 | 0.188603141 | 0.710260428 | 0.510683474 | 0.68851941 | 0.69280029 |
| Son | T | 1958 | 0.10415856 | 0.024445783 | 0.063459895 | -0.040698664 | 0.039014112 | 0.188603141 | 0.710260428 | 0.510683474 | 0.68851941 | 0.69280029 |
| Srrm2 | S | 248 | 0.040214518 | 0.022526469 | 0.063329534 | 0.023115016 | 0.040803065 | 0.269140667 | 0.463360379 | 0.14000471 | 0.53817074 | 0.267401768 |
| Sipa1l2 | S | 195 | -3.28202E-05 | 0.030496999 | 0.074803198 | 0.074836018 | 0.044306199 | 0.999560991 | 0.355515107 | 0.021362624 | 0.271332429 | 0.157107612 |
| Ilk | S | 232 | 0.025179578 | 0.230205653 | -0.005019203 | -0.030198782 | -0.235224856 | 0.857065671 | 0.145502111 | 0.955465837 | 0.836928853 | 0.146810385 |
| Ep400 | S | 1009 | 0.016430761 | 0.032373682 | 0.106788544 | 0.090357783 | 0.074414862 | 0.883215464 | 0.607992068 | 0.064567761 | 0.432120531 | 0.194217031 |
| Ep400 | S | 1010 | 0.016430761 | 0.032373682 | 0.106788544 | 0.090357783 | 0.074414862 | 0.883215464 | 0.607992068 | 0.064567761 | 0.432120531 | 0.194217031 |
| Stard8 | S | 733 | -0.071045657 | 0.104746622 | 0.102141317 | 0.173186974 | -0.002605305 | 0.324403415 | 0.223068973 | 0.029842243 | 0.067853722 | 0.971384164 |
| Psen1 | S | 371 | 0.124519666 | 0.066296858 | -0.014227908 | -0.138747574 | -0.080524766 | 0.22003859 | 0.182998773 | 0.793860215 | 0.190607908 | 0.214568316 |
| Topbp1 | S | 657 | -0.053781994 | 0.28976096 | 0.009721622 | 0.063503616 | -0.280039338 | 0.838735204 | 0.114271416 | 0.971801321 | 0.838710067 | 0.31756554 |
| Actn1 | S | 362 | -0.042546633 | 0.072984343 | 0.092197664 | 0.134744297 | 0.019213322 | 0.643962249 | 0.196497784 | 0.063488145 | 0.207126286 | 0.645652201 |
| Srrm1 | S | 526 | 0.189870623 | -0.062914319 | -0.016999375 | -0.206869997 | 0.045914945 | 0.004402513 | 0.390482691 | 0.808839625 | 0.043944942 | 0.581433906 |
| Srrm1 | S | 530 | 0.189870623 | -0.062914319 | -0.016999375 | -0.206869997 | 0.045914945 | 0.004402513 | 0.390482691 | 0.808839625 | 0.043944942 | 0.581433906 |
| Wac | S | 523 | -0.062590116 | -0.037238338 | 0.212826995 | 0.27541711 | 0.250065332 | 0.406523263 | 0.629518179 | 0.004167743 | 0.021564212 | 0.035146868 |
| Scaf1 | S | 442 | -0.066173818 | 0.208384782 | 0.116600157 | 0.182773975 | -0.091784625 | 0.439461367 | 0.106411563 | 0.556739304 | 0.370653507 | 0.64215491 |
| Map4k5 | T | 400 | 0.087472486 | 0.220946845 | -0.151131932 | -0.238604418 | -0.372078777 | 0.216263199 | 0.011585442 | 0.062617662 | 0.007165627 | 0.001720997 |
| Mecp2 | S | 216 | 0.100045547 | 0.013986072 | 0.032708873 | -0.067336674 | 0.018722801 | 0.080007893 | 0.763369677 | 0.517409591 | 0.077030661 | 0.434690987 |
| Camk2g | T | 287 | 0.015926041 | 0.09752802 | 0.002527811 | -0.013398229 | -0.095000208 | 0.491820776 | 0.028410587 | 0.911678831 | 0.622185733 | 0.03155619 |
| Rbm8a Rbm8a2 | S | 46 | 0.073446206 | 0.078087695 | 0.040095323 | -0.033350883 | -0.037992372 | 0.529467706 | 0.200919 | 0.61000981 | 0.780602505 | 0.612838397 |
| Rps6ka4 | S | 682 | 0.026333579 | 0.119788554 | -0.032210531 | -0.05854411 | -0.151999085 | 0.586025121 | 0.018938616 | 0.745492933 | 0.573908445 | 0.207505974 |
| Pwwp3a | S | 345 | 0.139254625 | -0.011784782 | 0.094305735 | -0.04494889 | 0.106090517 | 0.239094195 | 0.897615987 | 0.385359681 | 0.640477127 | 0.200183178 |
| Stard13 | S | 414 | 0.046703346 | 0.125109184 | 0.048482395 | 0.001779048 | -0.076626789 | 0.732164475 | 0.217965179 | 0.61756692 | 0.989223338 | 0.416601444 |
| Brpf1 | S | 459 | -0.03098634 | -0.017103298 | 0.185061864 | 0.216048204 | 0.202165162 | 0.460534797 | 0.625198685 | 0.007266008 | 0.007179769 | 0.016947394 |
| Brpf1 | S | 461 | -0.03098634 | -0.017103298 | 0.185061864 | 0.216048204 | 0.202165162 | 0.460534797 | 0.625198685 | 0.007266008 | 0.007179769 | 0.016947394 |
| Rhbdf2 | S | 147 | -0.006834124 | 0.125984865 | 0.015288272 | 0.022122395 | -0.110696593 | 0.90001258 | 0.02657804 | 0.787106899 | 0.722051447 | 0.104796607 |
| Sde2 | T | 307 | -0.037848433 | 0.082452464 | 0.118869232 | 0.156717665 | 0.036416767 | 0.788254612 | 0.188075878 | 0.205955768 | 0.326593189 | 0.645214688 |
| Uckl1 | T | 546 | -0.032385682 | 0.166312661 | 0.018417973 | 0.050803655 | -0.147894688 | 0.785114967 | 0.040365962 | 0.782409159 | 0.691989362 | 0.109070223 |
| Rgl1 | S | 747 | 0.557239231 | 0.587796668 | -0.883017101 | -1.440256332 | -1.47081377 | 0.188651145 | 0.243183242 | 0.214104178 | 0.08455172 | 0.079907601 |
| Plin3 | T | 220 | 0.034030558 | 0.087855345 | 0.107913364 | 0.073882806 | 0.020058019 | 0.882549576 | 0.579443287 | 0.188297502 | 0.746883029 | 0.891916573 |
| Dyrk1a Dyrk1b | Y | 321;273 | -0.052768387 | 0.094958671 | 0.090175663 | 0.142944049 | -0.004783009 | 0.090112844 | 0.023012952 | 0.089726476 | 0.04340368 | 0.909228198 |
| Afdn | S | 1172 | 0.01465517 | 0.078193226 | 0.053999954 | 0.039344783 | -0.024193272 | 0.75540497 | 0.153622485 | 0.283364681 | 0.07940398 | 0.397335819 |
| Pcnt | S | 2838 | -0.006548695 | 0.268491049 | -0.05494865 | -0.048400258 | -0.323439698 | 0.950889155 | 0.046534712 | 0.548701964 | 0.50449273 | 0.036393724 |
| Sgpp1 | S | 101 | 0.060961156 | 0.014312363 | 0.053514864 | -0.007446292 | 0.039202501 | 0.099974727 | 0.656390035 | 0.296801948 | 0.857286801 | 0.382384477 |
| Cic | S | 2148 | -0.06098432 | 0.034050656 | 0.176991867 | 0.237976187 | 0.142941211 | 0.038758297 | 0.183411112 | 0.00910179 | 0.013208841 | 0.042587473 |

| | | | | | | | | | | | | |
|---|---|---|---|---|---|---|---|---|---|---|---|---|
| Polr2a | S | 1833 | 0.007802131 | 0.049366266 | 0.125796805 | 0.117994675 | 0.07643054 | 0.874962108 | 0.432932375 | 0.17105944 | 0.195078743 | 0.386562854 |
| Clasp1 | S | 688 | 0.040173641 | 0.148440629 | -0.043082307 | -0.083255948 | -0.191522936 | 0.462160984 | 0.053116553 | 0.677615058 | 0.433725085 | 0.137134993 |
| Rbm8a Rbm8a2 | S | 42 | 0.026462167 | 0.068066794 | 0.001250198 | -0.025211969 | -0.066816596 | 0.371657816 | 0.01440108 | 0.954912411 | 0.442535009 | 0.047495352 |
| Dsn1 | S | 344 | 0.028660094 | 0.09681617 | 0.014646051 | -0.014014043 | -0.082170118 | 0.486699147 | 0.081751223 | 0.711321732 | 0.452937377 | 0.09969901 |
| Mapk11 | S | 243 | -0.124940962 | 0.093553056 | 0.100524726 | 0.225465687 | 0.006971669 | 0.168307608 | 0.035189894 | 0.022166528 | 0.06575798 | 0.747729781 |
| Mapk11 | T | 241 | -0.124940962 | 0.093553056 | 0.100524726 | 0.225465687 | 0.006971669 | 0.168307608 | 0.035189894 | 0.022166528 | 0.06575798 | 0.747729781 |
| Dmwd | S | 393 | 0.064790781 | 0.093706645 | 0.106441119 | 0.041650338 | 0.012734475 | 0.684276485 | 0.663106565 | 0.310302804 | 0.80229549 | 0.953026275 |
| Tmpo | T | 334 | 0.070124465 | 0.01403163 | 0.08347434 | 0.013349875 | 0.06944271 | 0.32521256 | 0.812770964 | 0.191900498 | 0.760827213 | 0.020513535 |
| Ranbp3 | S | 40 | 0.014333561 | 0.041897613 | 0.024423295 | 0.010089734 | -0.017474317 | 0.221162747 | 0.016700622 | 0.425276752 | 0.722864887 | 0.556371481 |
| 1700017B05Rik | T | 470 | 0.042488131 | 0.0474841 | 0.071680507 | 0.029192375 | 0.024196407 | 0.232902574 | 0.380879428 | 0.44095864 | 0.731237634 | 0.789739028 |
| Phf8-ps Phf8 | S | 790 | 0.022917755 | 0.131087628 | -0.005361869 | -0.028279624 | -0.136449498 | 0.742089488 | 0.045692153 | 0.965846303 | 0.820311421 | 0.319189739 |
| Wdhd1 | S | 400 | 0.010045872 | -0.008594233 | 0.137814916 | 0.127769044 | 0.146409149 | 0.816741886 | 0.846785932 | 0.02931888 | 0.048500738 | 0.034037739 |
| Lmnb1 | S | 17 | 0.031398496 | 0.05581992 | 0.022181894 | -0.009216602 | -0.033638026 | 0.497447667 | 0.03294603 | 0.521540458 | 0.849179098 | 0.332396805 |
| Cdk13 | S | 438 | 0.02209032 | -0.005072027 | 0.116343635 | 0.094253314 | 0.121415662 | 0.458996276 | 0.905924051 | 0.047369837 | 0.086950606 | 0.064383036 |
| Cdk13 | S | 440 | 0.02209032 | -0.005072027 | 0.116343635 | 0.094253314 | 0.121415662 | 0.458996276 | 0.905924051 | 0.047369837 | 0.086950606 | 0.064383036 |
| Kif18b | S | 677 | 0.172308553 | -0.00440254 | -0.00210427 | -0.174412823 | 0.00229827 | 0.040159464 | 0.829615006 | 0.937240771 | 0.036016664 | 0.926033599 |
| Zfp609 | S | 491 | 0.017290236 | 0.056774904 | 0.044593495 | 0.027303259 | -0.012181409 | 0.585636238 | 0.103777094 | 0.24871907 | 0.354285236 | 0.658268029 |
| Kif1c | S | 674 | -0.022422201 | 0.085626982 | 0.057923472 | 0.080345673 | -0.02770351 | 0.521233357 | 0.079896725 | 0.107653143 | 0.027092233 | 0.452330145 |
| Setd2 | S | 2058 | 0.030802002 | 0.080629197 | 0.090180299 | 0.059378297 | 0.009551101 | 0.704963275 | 0.213241059 | 0.509898486 | 0.666470014 | 0.940319859 |
| Pdlim4 | S | 119 | 0.05562021 | 0.191458995 | -0.076546397 | -0.132166608 | -0.268005393 | 0.408480544 | 0.046844629 | 0.343922019 | 0.093807011 | 0.019736256 |
| Rab3gap2 | S | 812 | 0.026899045 | 0.114954228 | 0.087351384 | 0.060452339 | -0.027602845 | 0.726050482 | 0.347122685 | 0.457857869 | 0.593321903 | 0.839198432 |
| Camk1 | T | 177 | 0.040697495 | 0.224610548 | -0.074170137 | -0.114867632 | -0.298780686 | 0.55086759 | 0.071307804 | 0.510197398 | 0.333613727 | 0.064056314 |
| Vps26b | S | 302 | 0.110155524 | 0.037063396 | -0.046031375 | -0.156186899 | -0.083094771 | 0.012092652 | 0.35612793 | 0.454278471 | 0.066758912 | 0.213854773 |
| Erbin | S | 869 | -0.065712206 | 0.120534428 | 0.064953484 | 0.13066569 | -0.055580944 | 0.138843996 | 0.024228324 | 0.092937056 | 0.022976188 | 0.19859144 |
| Ripk2 | T | 483 | 0.047455331 | 0.073116567 | 0.103070101 | 0.05561477 | 0.029953533 | 0.745851994 | 0.610890311 | 0.221121057 | 0.675582977 | 0.811510075 |
| Mcm3 | S | 672 | -0.045747101 | 0.067953752 | 0.065916125 | 0.111663226 | -0.002037627 | 0.224808678 | 0.101160933 | 0.007392522 | 0.042067687 | 0.947131169 |
| Zfp687 | S | 433 | 0.006735596 | 0.005917481 | 0.109455919 | 0.102720322 | 0.103538437 | 0.819177063 | 0.798187441 | 0.027020423 | 0.040361132 | 0.05597624 |
| Map3k12 | S | 533 | 0.039554829 | 0.051286018 | 0.046566124 | 0.007011295 | -0.004719893 | 0.304220169 | 0.193092741 | 0.44412607 | 0.888572664 | 0.924217055 |
| Mki67 | S | 561 | 0.18317576 | 0.226888304 | -0.289315979 | -0.472491739 | -0.516204283 | 0.076833342 | 0.042763815 | 0.092559091 | 0.028003939 | 0.021678356 |
| Eif3g | S | 42 | -0.000967097 | -0.020601185 | 0.092751481 | 0.093718578 | 0.113352667 | 0.958764197 | 0.091232702 | 8.92365E-05 | 0.023013509 | 0.000949451 |
| Uckl1 | S | 539 | -0.001475547 | 0.091397401 | 0.033133566 | 0.034609113 | -0.058263835 | 0.976007086 | 0.046315477 | 0.387176935 | 0.394624921 | 0.02709045 |
| Pdlim2 | T | 276 | -0.005030314 | 0.034029501 | 0.109330796 | 0.11436111 | 0.075301295 | 0.789198323 | 0.146408837 | 0.12195437 | 0.109692966 | 0.221350895 |
| Fam83h | S | 904 | -0.113301069 | 0.178656652 | 0.061882703 | 0.175183772 | -0.116773949 | 0.185287988 | 0.046618037 | 0.077581625 | 0.096052624 | 0.142619055 |
| Eif4g2 | S | 394 | 0.09773988 | 0.00516748 | 0.018262436 | -0.079477444 | 0.013094955 | 0.026989667 | 0.880446431 | 0.701259594 | 0.132569528 | 0.729513178 |
| Akap12 | S | 1292 | 0.037119635 | 0.301813361 | -0.164654466 | -0.201774101 | -0.466467827 | 0.622961449 | 0.006789501 | 0.071063297 | 0.016446816 | 0.000349708 |
| Faf1 | S | 269 | -0.098776728 | 0.15920288 | 0.058736292 | 0.15751302 | -0.100466588 | 0.026719742 | 0.002912298 | 0.149693009 | 0.003258695 | 0.016600876 |
| Tcof1 | T | 860 | 0.024494083 | 0.056583509 | 0.082063797 | 0.057569714 | 0.025480288 | 0.691179289 | 0.216884582 | 0.308314301 | 0.499644543 | 0.728562246 |
| Reps1 | T | 459 | 0.034116794 | 0.145624428 | 0.020957421 | -0.013159373 | -0.124667007 | 0.638420842 | 0.145019744 | 0.904305133 | 0.935886228 | 0.489093067 |
| Epb41l1 | S | 677 | -0.052612774 | 0.091920284 | 0.132127534 | 0.184740308 | 0.04020725 | 0.270030055 | 0.095483907 | 0.153484358 | 0.08124685 | 0.606677312 |
| Irf2bp1 | S | 436 | 0.02218126 | 0.027106835 | 0.077273301 | 0.05509204 | 0.050166466 | 0.568403527 | 0.491381615 | 0.07379491 | 0.098043823 | 0.126403488 |
| Lmna | S | 392 | 0.051904839 | 0.020978011 | 0.021362666 | -0.030542173 | 0.000384655 | 0.030463926 | 0.348562222 | 0.485027228 | 0.299301956 | 0.988652948 |
| Nfrkb | S | 884 | 0.100730539 | 0.064189387 | -0.021625417 | -0.122355956 | -0.085814804 | 0.076153305 | 0.350970644 | 0.796817206 | 0.227099132 | 0.395745577 |
| Usp34 | T | 3401 | 0.101808371 | -0.050094197 | 0.053090426 | -0.048717945 | 0.103184624 | 0.005786947 | 0.149630726 | 0.222375104 | 0.247321426 | 0.055969915 |
| Nes | S | 485 | 0.090415387 | -0.039780606 | 0.099820521 | 0.009405134 | 0.139601127 | 0.170212441 | 0.389364904 | 0.088843094 | 0.875935875 | 0.038326459 |
| Nfrkb | T | 1285 | -0.005573007 | 0.073326691 | 0.1086883 | 0.114261307 | 0.035361609 | 0.930198676 | 0.432480299 | 0.130844547 | 0.1562923 | 0.704462955 |
| Epb41l3 | S | 804 | 0.035641149 | -0.031856029 | 0.119073338 | 0.083432189 | 0.150929367 | 0.608611831 | 0.550609565 | 0.01583497 | 0.287809281 | 0.0530828 |
| Spen | S | 2491 | 0.116163612 | 0.079086914 | -0.0077949 | -0.123958512 | -0.086881815 | 0.259117867 | 0.224859048 | 0.79850898 | 0.24009108 | 0.195315868 |
| Prpf6 | T | 266 | 0.202733658 | 0.016407362 | 0.004597976 | -0.198135682 | -0.011809387 | 0.14725754 | 0.789667705 | 0.967887799 | 0.215060373 | 0.920229099 |
| Uhrf1 | S | 289 | 0.048347592 | 0.093849893 | -0.049096126 | -0.097443718 | -0.142946019 | 0.202572213 | 0.020515676 | 0.501025225 | 0.229798357 | 0.124408551 |
| Dab2ip | S | 717 | 0.044073541 | 0.082415011 | 0.075010887 | 0.030937345 | -0.007404125 | 0.732757606 | 0.241114862 | 0.470645341 | 0.822369657 | 0.933615421 |
| Mvb12a | S | 200 | -0.02650681 | 0.080208406 | 0.078674622 | 0.105181432 | -0.001533784 | 0.385557446 | 0.016200802 | 0.368997673 | 0.259476121 | 0.984127108 |
| Srrm1 | S | 789 | 0.053001733 | 0.01586713 | 0.022183445 | -0.030818288 | 0.006316315 | 0.01818041 | 0.525796615 | 0.44503611 | 0.281767262 | 0.829686033 |
| Mef2d | S | 437 | 0.08299072 | -0.010962969 | 0.037527477 | -0.045463243 | 0.048490446 | 0.022421549 | 0.767486044 | 0.390237891 | 0.323867409 | 0.348311528 |
| Ticrr | S | 1406 | 0.058404826 | 0.053349197 | 0.049430644 | -0.008974182 | -0.003918553 | 0.365662134 | 0.260832283 | 0.499857845 | 0.911700281 | 0.95649534 |
| Srrm2 | S | 402 | 0.146722908 | 0.042671804 | -0.06854571 | -0.215268618 | -0.111217514 | 0.018986699 | 0.600723384 | 0.509794009 | 0.113709051 | 0.34209108 |
| Srrm2 | S | 406 | 0.146722908 | 0.042671804 | -0.06854571 | -0.215268618 | -0.111217514 | 0.018986699 | 0.600723384 | 0.509794009 | 0.113709051 | 0.34209108 |
| Srrm2 | S | 422 | 0.146722908 | 0.042671804 | -0.06854571 | -0.215268618 | -0.111217514 | 0.018986699 | 0.600723384 | 0.509794009 | 0.113709051 | 0.34209108 |
| Hps4 | S | 478 | -0.015207295 | 0.067390189 | 0.058918768 | 0.074126063 | -0.008471421 | 0.588473659 | 0.103122363 | 0.072587641 | 0.0219187 | 0.790443239 |
| Dcaf1 | S | 999 | 0.033257664 | -0.01209374 | 0.105318278 | 0.072060614 | 0.117412018 | 0.458790312 | 0.751355582 | 0.035737474 | 0.182257318 | 0.044110263 |
| Ints3 | S | 1040 | 0.112835924 | -0.018061152 | 0.027005792 | -0.085830132 | 0.045066945 | 0.023052923 | 0.644498455 | 0.529768207 | 0.090276135 | 0.303194182 |
| Eif1ad | S | 131 | 0.160796379 | 0.084836288 | 0.048154696 | -0.112641683 | -0.036681592 | 0.338214971 | 0.624161241 | 0.886464871 | 0.735594081 | 0.912276461 |
| Eif1ad | S | 135 | 0.160796379 | 0.084836288 | 0.048154696 | -0.112641683 | -0.036681592 | 0.338214971 | 0.624161241 | 0.886464871 | 0.735594081 | 0.912276461 |
| Eif1ad | S | 136 | 0.160796379 | 0.084836288 | 0.048154696 | -0.112641683 | -0.036681592 | 0.338214971 | 0.624161241 | 0.886464871 | 0.735594081 | 0.912276461 |
| Eif1ad | S | 137 | 0.160796379 | 0.084836288 | 0.048154696 | -0.112641683 | -0.036681592 | 0.338214971 | 0.624161241 | 0.886464871 | 0.735594081 | 0.912276461 |
| Hnrnpf | S | 187 | 0.056773654 | -0.033039983 | 0.141399868 | 0.084626214 | 0.174439851 | 0.656537057 | 0.65837651 | 0.047597929 | 0.514700479 | 0.065887886 |
| Nomo1 | S | 1197 | -0.041594292 | 0.136956635 | 0.262526533 | 0.304120826 | 0.125569898 | 0.346943186 | 0.305548279 | 0.385963355 | 0.330323343 | 0.664462108 |
| Mllt3 | S | 430 | -0.029297922 | 0.022531198 | 0.125745921 | 0.155043843 | 0.103214723 | 0.470707056 | 0.649285512 | 0.012401806 | 0.008077876 | 0.079324475 |
| Pi4kb | S | 511 | -0.04419025 | 0.269605134 | 0.000519747 | 0.044709997 | -0.269085387 | 0.641406357 | 0.07681313 | 0.99684482 | 0.692264834 | 0.108598408 |
| Sptbn1 | T | 2194 | 0.047488786 | 0.164386889 | 0.048786017 | 0.001297231 | -0.115600873 | 0.809205889 | 0.239904277 | 0.787545554 | 0.995435267 | 0.535204475 |
| Mastl | S | 362 | 0.038510613 | 0.043460246 | 0.055501733 | 0.01699112 | 0.012041487 | 0.249747661 | 0.295617417 | 0.382129247 | 0.763160417 | 0.840152038 |
| Nup98 | S | 932 | 0.132865273 | 0.019206186 | -0.013793418 | -0.14665869 | -0.032999603 | 0.033516122 | 0.731993245 | 0.790947919 | 0.03792146 | 0.565255318 |
| Racgap1 | T | 585 | 0.223638194 | -0.028674158 | 0.033471292 | -0.190166903 | 0.062145449 | 0.163392873 | 0.821976659 | 0.600604542 | 0.212490081 | 0.640623963 |
| Atf7ip | S | 707 | 0.114612327 | 0.004073485 | 0.012622422 | -0.101989905 | 0.008548937 | 0.036551781 | 0.930655022 | 0.758083424 | 0.040383688 | 0.811531032 |
| Ehbp1l1 | S | 1443 | 0.188616871 | 0.062040534 | -0.176764329 | -0.3653812 | -0.238804863 | 0.006004781 | 0.331734337 | 0.178526386 | 0.045799447 | 0.10175395 |
| Pxn | S | 322 | 0.05378291 | 0.110322249 | 0.004318621 | -0.049464289 | -0.106003628 | 0.395190022 | 0.156291394 | 0.942975118 | 0.415296672 | 0.170121483 |
| Mavs | S | 152 | 0.037659804 | 0.00546289 | 0.069608919 | 0.031949115 | 0.064146029 | 0.481930749 | 0.829659214 | 0.030076076 | 0.52825973 | 0.012727761 |
| Nin | S | 1921 | -0.101851406 | 0.042695012 | 0.169429552 | 0.271280958 | 0.12673454 | 0.075843185 | 0.479955149 | 0.001982594 | 0.005217267 | 0.094861751 |
| Larp1 | S | 138 | -0.069668628 | 0.297646247 | -0.080378275 | -0.010709647 | -0.378024522 | 0.452531936 | 0.009290967 | 0.44881603 | 0.900587491 | 0.015666354 |
| Brpf1 | S | 120 | -0.057309703 | 0.037698497 | 0.309268262 | 0.366577964 | 0.271569765 | 0.295362418 | 0.760072904 | 0.094794019 | 0.079553716 | 0.150165824 |
| Srrm2 | S | 982 | 0.046505776 | 0.02835527 | 0.093295061 | 0.046789285 | 0.064939791 | 0.333417275 | 0.509906992 | 0.319330765 | 0.5918495 | 0.463547607 |
| Gpalpp1 | S | 146 | -0.034454817 | 0.059807993 | 0.107023265 | 0.141478081 | 0.047215272 | 0.295834544 | 0.043835689 | 0.135464011 | 0.072300078 | 0.419263152 |
| Acd | S | 298 | 0.066442433 | 0.005010594 | 0.09653638 | 0.030093947 | 0.091525786 | 0.207188138 | 0.956909495 | 0.277340604 | 0.715943645 | 0.430190195 |
| Afap1 | S | 284 | 0.052268758 | 0.05933093 | 0.02428181 | -0.027986948 | -0.03504912 | 0.284909521 | 0.184178246 | 0.569655936 | 0.475457984 | 0.294936107 |
| Srrm2 | S | 505 | 0.049937847 | -0.043826074 | 0.239130798 | 0.189192951 | 0.282956872 | 0.585828208 | 0.621918022 | 0.152467295 | 0.225325327 | 0.116902395 |
| Srrm2 | S | 507 | 0.049937847 | -0.043826074 | 0.239130798 | 0.189192951 | 0.282956872 | 0.585828208 | 0.621918022 | 0.152467295 | 0.225325327 | 0.116902395 |
| Dtl | S | 654 | 0.041786509 | 0.061145174 | 0.003995764 | -0.037790745 | -0.05714941 | 0.145543289 | 0.084207421 | 0.932831278 | 0.422459465 | 0.269100714 |
| Ncoa2 | S | 487 | 0.089650481 | 0.076203497 | 0.044863097 | -0.044787384 | -0.031340401 | 0.299489769 | 0.647296108 | 0.49133926 | 0.528974975 | 0.843156553 |
| Adgre5 | S | 799 | -0.016700614 | 0.120226411 | -0.007700422 | 0.009000192 | -0.127926833 | 0.828459135 | 0.006764674 | 0.891037846 | 0.914794569 | 0.10074892 |
| Srrm2 | T | 860 | 0.080725129 | 0.007666005 | 0.051372652 | -0.029352477 | 0.043706647 | 0.091424738 | 0.838650417 | 0.415397684 | 0.61331686 | 0.448554957 |
| Map4 | T | 503 | -0.014592302 | 0.102797517 | 0.056267784 | 0.070860086 | -0.046529733 | 0.616038342 | 0.093738066 | 0.249961454 | 0.164782465 | 0.414078291 |
| Nwd2 | S | 923 | -0.074828604 | 0.046408941 | 0.178836297 | 0.2536649 | 0.132427355 | 0.342332992 | 0.23883634 | 0.057895281 | 0.035120348 | 0.132601263 |
| Nwd2 | S | 924 | -0.074828604 | 0.046408941 | 0.178836297 | 0.2536649 | 0.132427355 | 0.342332992 | 0.23883634 | 0.057895281 | 0.035120348 | 0.132601263 |
| Med1 | S | 1158 | 0.123893227 | 0.066527856 | -0.038619874 | -0.162513101 | -0.10514773 | 0.102417807 | 0.341838446 | 0.728930491 | 0.191493205 | 0.336318988 |
| Rnps1 | T | 161 | 0.052210118 | -0.007823603 | 0.171235782 | 0.119025664 | 0.179059385 | 0.503261336 | 0.921118173 | 0.201205541 | 0.326947272 | 0.18147121 |

| | | | | | | | | | | | | |
|---|---|---|---|---|---|---|---|---|---|---|---|---|
| Rsrc1 | S | 116 | -0.309587305 | 0.304522331 | 0.230858488 | 0.540445793 | -0.073663844 | 0.734825769 | 0.237739564 | 0.615286561 | 0.576093221 | 0.85555652 |
| Rsrc1 | S | 118 | -0.309587305 | 0.304522331 | 0.230858488 | 0.540445793 | -0.073663844 | 0.734825769 | 0.237739564 | 0.615286561 | 0.576093221 | 0.85555652 |
| Rsrc1 | S | 120 | -0.309587305 | 0.304522331 | 0.230858488 | 0.540445793 | -0.073663844 | 0.734825769 | 0.237739564 | 0.615286561 | 0.576093221 | 0.85555652 |
| Aimp2 | S | 36 | -0.138947974 | 0.100736273 | 0.12208565 | 0.261033624 | 0.021349377 | 0.332316681 | 0.04927146 | 0.102301623 | 0.125908852 | 0.701625386 |
| Mdc1 | S | 1443 | -0.028990901 | 0.140534171 | 0.046191999 | 0.0751829 | -0.094342172 | 0.796110766 | 0.124595817 | 0.327183311 | 0.517566484 | 0.25417054 |
| Rsrc1 | S | 6 | -0.040470046 | 0.062199438 | 0.064749882 | 0.105219928 | 0.002550444 | 0.26085338 | 0.026952414 | 0.046441468 | 0.030871645 | 0.910068001 |
| Secisbp2l | S | 897 | -0.000923878 | 0.095719869 | 0.003744521 | 0.004668399 | -0.091975348 | 0.968808115 | 0.008797871 | 0.908240884 | 0.890834803 | 0.052402252 |
| Arfgap3 | S | 331 | 0.027417193 | 0.077630243 | 0.060180556 | 0.032763363 | -0.017449687 | 0.597510852 | 0.23131177 | 0.412948838 | 0.660593666 | 0.82308021 |
| Trappc12 | S | 10 | 0.037521718 | 0.063672983 | 0.133885207 | 0.096363489 | 0.070212224 | 0.838173934 | 0.570956008 | 0.274689044 | 0.614410042 | 0.564584187 |
| Trappc12 | S | 15 | 0.037521718 | 0.063672983 | 0.133885207 | 0.096363489 | 0.070212224 | 0.838173934 | 0.570956008 | 0.274689044 | 0.614410042 | 0.564584187 |
| Ubtd2 | T | 230 | 0.20265769 | 0.155788516 | -0.177620325 | -0.380278016 | -0.333408842 | 0.262770585 | 0.210028889 | 0.546586285 | 0.257849531 | 0.300099073 |
| Hivep2 | S | 767 | 0.054641401 | 0.021376775 | 0.153822779 | 0.099181378 | 0.132446004 | 0.542937454 | 0.713106684 | 0.320870057 | 0.519296305 | 0.380984401 |
| Srrm1 | S | 814 | 0.036656986 | 0.035600091 | 0.049719463 | 0.013062477 | 0.014119373 | 0.305162616 | 0.411345301 | 0.165082416 | 0.716537934 | 0.742261368 |
| Srrm2 | S | 926 | 0.029699429 | 0.014293841 | 0.035043576 | 0.005344147 | 0.020749735 | 0.075939492 | 0.432852414 | 0.077986825 | 0.713274734 | 0.271104041 |
| C2cd3 | S | 2028 | -0.08522183 | 0.113777462 | 0.125114613 | 0.210336443 | 0.011337151 | 0.574445753 | 0.322057533 | 0.094615594 | 0.225388565 | 0.910548574 |
| Epc2 | S | 119 | 0.00697234 | 0.083951104 | 0.014090948 | 0.007118608 | -0.069860156 | 0.917157494 | 0.013887245 | 0.767579097 | 0.923822729 | 0.207245543 |
| Gtpbp1 | S | 12 | -0.016963832 | 0.081309605 | 0.035051185 | 0.052015018 | -0.046258419 | 0.607572414 | 0.028839403 | 0.194940495 | 0.13890661 | 0.128488156 |
| Spag9 | S | 1188 | -0.014486159 | 0.246563171 | -0.16912534 | -0.154639181 | -0.415688511 | 0.851776189 | 0.000546415 | 0.096741924 | 0.159292547 | 0.019266527 |
| Thoc1 | S | 560 | -0.038903118 | 0.109343988 | 0.121285226 | 0.160188344 | 0.011941238 | 0.690782845 | 0.286993604 | 0.211647267 | 0.194691506 | 0.911193791 |
| Nfkb2 | T | 425 | 0.020953103 | 0.118967832 | 0.035409037 | 0.014455934 | -0.083558794 | 0.761304418 | 0.110517367 | 0.771009803 | 0.909632629 | 0.52075837 |
| Hipk3 | Y | 359 | -0.018434653 | 0.105482615 | 0.057340199 | 0.075774852 | -0.048142416 | 0.708180963 | 0.159977285 | 0.161267387 | 0.156124311 | 0.453828002 |
| Aak1 | S | 729 | -0.149354316 | 0.403141981 | 0.022762249 | 0.172116565 | -0.380379732 | 0.613224705 | 0.125137216 | 0.937936288 | 0.646715908 | 0.284299696 |
| Nup54 | S | 292 | 0.070041667 | -0.041729648 | 0.055132798 | -0.014908868 | 0.096862446 | 0.000988261 | 0.008853996 | 0.034330933 | 0.409990377 | 0.01873547 |
| Rbm5 | S | 18 | 0.086714955 | 0.077883856 | -0.029854224 | -0.116569179 | -0.10773808 | 0.135752985 | 0.193003084 | 0.757452782 | 0.279259359 | 0.311667176 |
| Spen | S | 1293 | 0.161330388 | -0.018671796 | 0.056431615 | -0.104898772 | 0.075103411 | 0.279937789 | 0.541849044 | 0.134623746 | 0.443512182 | 0.093988387 |
| Aff4 | S | 485 | 0.010833971 | 0.09997681 | 0.058417723 | 0.047583753 | -0.041559087 | 0.885144981 | 0.167399979 | 0.426007065 | 0.451980161 | 0.429598919 |
| Aff4 | S | 486 | 0.010833971 | 0.09997681 | 0.058417723 | 0.047583753 | -0.041559087 | 0.885144981 | 0.167399979 | 0.426007065 | 0.451980161 | 0.429598919 |
| Dnttip2 | S | 255 | 0.091210732 | 0.003860782 | 0.004890132 | -0.0863206 | 0.00102935 | 0.012146839 | 0.891604439 | 0.867105754 | 0.012727789 | 0.961791383 |
| Irf2bp1 | S | 421 | -0.018197224 | 0.053179343 | 0.121449006 | 0.13964623 | 0.068269663 | 0.880014589 | 0.333547218 | 0.14126345 | 0.301340793 | 0.31329847 |
| Srrm2 | S | 482 | 0.033134143 | 0.015262963 | 0.076071945 | 0.042937803 | 0.060808982 | 0.704067619 | 0.646980986 | 0.041908203 | 0.619669036 | 0.058264638 |
| Dclk1 | S | 32 | -0.011052517 | 0.1232207 | -0.022521791 | -0.011469273 | -0.145742491 | 0.819227632 | 0.005769306 | 0.806996082 | 0.905909878 | 0.203881586 |
| Arid4a | S | 564 | 0.017432103 | 0.042720859 | 0.064687933 | 0.04725583 | 0.021967074 | 0.529362159 | 0.240354229 | 0.188678079 | 0.290508459 | 0.617037441 |
| Rnmt | T | 34 | 0.024249382 | 0.07751327 | -0.005006091 | -0.029255473 | -0.082519361 | 0.636416104 | 0.010615509 | 0.937764616 | 0.693854247 | 0.268716111 |
| Camk2d | T | 337 | -0.051502341 | 0.125564197 | 0.030106139 | 0.081608481 | -0.095458058 | 0.408189478 | 0.010052032 | 0.545316927 | 0.254308711 | 0.110490356 |
| Srrm1 | S | 791 | 0.054121342 | 0.018568131 | 0.029362117 | -0.024759225 | 0.010793987 | 0.036569936 | 0.620189785 | 0.404941799 | 0.434760475 | 0.792597503 |
| Epb41 | S | 87 | -0.009168329 | 0.238648427 | 0.03836339 | 0.047531719 | -0.200285037 | 0.826820407 | 0.194320326 | 0.784099049 | 0.736266841 | 0.315848083 |
| Flna | S | 968 | 0.069835707 | 0.088814493 | -0.050653957 | -0.120489665 | -0.139468451 | 0.161820076 | 0.055737516 | 0.515315037 | 0.177605627 | 0.139597812 |
| Otud7b | T | 698 | 0.056010499 | 0.142728787 | -0.025976683 | -0.081987181 | -0.16870547 | 0.234621399 | 0.193821266 | 0.73583497 | 0.30711861 | 0.163598919 |
| Bag3 | S | 280 | -0.007586691 | 0.105718785 | 0.043364715 | 0.050951406 | -0.062354071 | 0.877383443 | 0.082926089 | 0.405917589 | 0.050435056 | 0.102059764 |
| Thoc2 | S | 1364 | -0.002627773 | 0.00565557 | 0.130545408 | 0.133173181 | 0.124889838 | 0.97011788 | 0.828180425 | 0.037270868 | 0.148234749 | 0.041112118 |
| Pbx2 | S | 330 | -0.045131642 | 0.077775062 | 0.1179721 | 0.163103742 | 0.040197038 | 0.582392657 | 0.25571569 | 0.110775824 | 0.068565118 | 0.331784287 |
| Canx | T | 561 | 0.046280987 | -0.025667769 | 0.21646713 | 0.170186142 | 0.242134899 | 0.550973514 | 0.655003673 | 0.17248842 | 0.258133931 | 0.148861235 |
| Vps8 | T | 435 | -0.136173635 | 0.123102566 | 0.197911854 | 0.334085489 | 0.074809288 | 0.09934138 | 0.191029485 | 0.043436118 | 0.006755072 | 0.358548678 |
| Fnbp1l | T | 293 | -0.057636228 | 0.088589907 | 0.044929086 | 0.102565314 | -0.043660821 | 0.446638679 | 0.006802313 | 0.312724563 | 0.232956982 | 0.321837258 |
| Nes | S | 484 | 0.080829248 | -0.039000835 | 0.09411096 | 0.013281712 | 0.133111795 | 0.162246878 | 0.384373882 | 0.080068028 | 0.797425605 | 0.031515473 |
| Lmna | S | 18 | 0.045775392 | 0.084924663 | 0.028105013 | -0.017670379 | -0.05681965 | 0.722399173 | 0.108130912 | 0.593841071 | 0.886406488 | 0.146819087 |
| Cdyl2 | S | 111 | 0.018877823 | 0.063859496 | 0.129973706 | 0.111095882 | 0.06611421 | 0.857400227 | 0.533477791 | 0.251902138 | 0.107173938 | 0.243168997 |
| Cpsf7 | S | 194 | 0.095340735 | -0.026550155 | 0.087078744 | -0.008261991 | 0.113628899 | 0.212018071 | 0.638683604 | 0.157041139 | 0.874770413 | 0.004141875 |
| Rtn4 | S | 185 | 0.343252353 | 0.132842493 | -0.108144492 | -0.451396845 | -0.240986985 | 0.2464668 | 0.862778632 | 0.795567059 | 0.341018614 | 0.762562958 |
| Macf1 | S | 7346 | 0.125273078 | 0.011616331 | -0.023994546 | -0.149267623 | -0.035610876 | 0.008785481 | 0.894945963 | 0.492756286 | 0.000685661 | 0.679556824 |
| Srrm2 | S | 882 | 0.014613541 | 0.023657544 | 0.066873677 | 0.052260137 | 0.043216133 | 0.639979496 | 0.455585321 | 0.043261317 | 0.138826434 | 0.19984436 |
| Hecw2 | S | 1048 | 0.194162181 | -0.056911155 | 0.032436542 | -0.161725639 | 0.089347696 | 0.040942308 | 0.484836836 | 0.776626732 | 0.185837157 | 0.392545572 |
| Usp24 | S | 2601 | -0.101860198 | 0.059492312 | 0.129037695 | 0.230897893 | 0.069545383 | 0.065386671 | 0.177310891 | 0.002430206 | 0.007976082 | 0.134759349 |
| Ercc5 | T | 444 | 0.0391535 | 0.06301142 | 0.032678096 | -0.006475405 | -0.030333324 | 0.325581536 | 0.20068566 | 0.514664179 | 0.904183057 | 0.600415602 |
| Klhl22 | S | 621 | 0.17414895 | 0.144663658 | -0.022225059 | -0.19637401 | -0.166888717 | 0.375667646 | 0.520186948 | 0.909046234 | 0.401064653 | 0.51262193 |
| Arhgef17 | S | 561 | -0.007806938 | 0.121212327 | 0.053642785 | 0.061449722 | -0.067569543 | 0.955992316 | 0.114392049 | 0.545579989 | 0.692880554 | 0.485578488 |
| Card14 | S | 940 | 0.034102354 | 0.01527679 | 0.079527215 | 0.045424861 | 0.064250425 | 0.27139816 | 0.814624526 | 0.124614241 | 0.314843151 | 0.382809748 |
| Card14 | S | 941 | 0.034102354 | 0.01527679 | 0.079527215 | 0.045424861 | 0.064250425 | 0.27139816 | 0.814624526 | 0.124614241 | 0.314843151 | 0.382809748 |
| Sp3 | S | 565 | -0.049000279 | 0.077194138 | 0.263145525 | 0.312145804 | 0.185951386 | 0.31141254 | 0.576045101 | 0.150029423 | 0.103353432 | 0.320794497 |
| Rptor | S | 863 | 0.077206364 | 0.047242202 | 0.003603118 | -0.073603245 | -0.043639084 | 0.219390278 | 0.130335108 | 0.883354168 | 0.236851942 | 0.16972921 |
| Spag9 | S | 493 | -0.031176743 | 0.066167379 | 0.093442883 | 0.124619626 | 0.027275504 | 0.383685362 | 0.085090034 | 0.123476127 | 0.071707155 | 0.561735248 |
| Msl1 | S | 127 | 0.117687376 | -0.012346323 | 0.04076917 | -0.076918206 | 0.053115493 | 0.090067246 | 0.725924186 | 0.361532879 | 0.216289048 | 0.167812418 |
| Nav1 | S | 314 | -0.002740997 | 0.06153953 | 0.039319098 | 0.042060095 | -0.022220432 | 0.908920866 | 0.045859164 | 0.196260128 | 0.193582828 | 0.4601732 |
| Mrtfb | S | 207 | 0.039083278 | 0.030459566 | 0.126873068 | 0.08778979 | 0.096413502 | 0.643147195 | 0.616967657 | 0.249583098 | 0.455666384 | 0.372017297 |
| Tor1aip1 | S | 151 | -0.086985316 | 0.002646765 | 0.332615267 | 0.419600583 | 0.329968502 | 0.583080344 | 0.95919977 | 0.098680116 | 0.079142748 | 0.09994141 |
| Peak1 | S | 1193 | 0.265854393 | 0.139567813 | -0.08713896 | -0.352993353 | -0.226706773 | 0.384701939 | 0.402743946 | 0.663743307 | 0.26495479 | 0.204803216 |
| Rif1 | S | 1538 | 0.057523805 | 0.052843799 | 0.014857801 | -0.042666004 | -0.037985998 | 0.218477075 | 0.119940622 | 0.894693485 | 0.712763523 | 0.737703765 |
| Mdc1 | T | 472 | -0.06460346 | 0.09471593 | 0.19714299 | 0.26174645 | 0.10242706 | 0.336033824 | 0.330763943 | 0.149511165 | 0.087588751 | 0.430914646 |
| Lmtk2 | S | 611 | 0.118233922 | -0.082026907 | 0.101956313 | -0.01627761 | 0.183983219 | 0.121321834 | 0.345803191 | 0.119509435 | 0.81549504 | 0.093142533 |
| Lmtk2 | S | 615 | 0.118233922 | -0.082026907 | 0.101956313 | -0.01627761 | 0.183983219 | 0.121321834 | 0.345803191 | 0.119509435 | 0.81549504 | 0.093142533 |
| Hivep2 | S | 764 | -0.083135501 | 0.056228667 | 0.136918426 | 0.220053927 | 0.080689758 | 0.222349975 | 0.413470909 | 0.009334984 | 0.030318446 | 0.251627253 |
| Med12 | S | 636 | -0.034904898 | 0.02196801 | 0.117790066 | 0.152694964 | 0.095822056 | 0.296076105 | 0.446223661 | 0.009310646 | 0.004422107 | 0.033280512 |
| Akap13 | S | 2365 | 0.027945063 | 0.021924406 | 0.063198107 | 0.035253044 | 0.041273701 | 0.669640416 | 0.458898999 | 0.051745776 | 0.578740439 | 0.06922632 |
| Pard3b | S | 368 | -0.104258088 | 0.02835641 | 0.280577839 | 0.384835927 | 0.252221429 | 0.075837988 | 0.452944603 | 0.021528957 | 0.007764745 | 0.034398661 |
| Kif18b | S | 484 | 0.04085816 | 0.039198981 | 0.038132394 | -0.002725766 | -0.001066587 | 0.27338227 | 0.20279153 | 0.39471043 | 0.946988294 | 0.976356484 |
| Dck | S | 13 | 0.014597284 | 0.116611607 | -0.010823782 | -0.025421066 | -0.127435389 | 0.739886235 | 0.031772698 | 0.909127417 | 0.788679582 | 0.249628519 |
| Sik3 | T | 163 | 0.104655101 | 0.09746623 | 0.016915491 | -0.08773961 | -0.080550739 | 0.351118277 | 0.441319722 | 0.837050717 | 0.472803478 | 0.550649085 |
| Luc7l3 | S | 333 | 0.098223792 | 0.06812721 | -0.017946622 | -0.116170413 | -0.086073832 | 0.065545627 | 0.550245546 | 0.864832081 | 0.314179747 | 0.534508385 |
| Luc7l3 | S | 335 | 0.098223792 | 0.06812721 | -0.017946622 | -0.116170413 | -0.086073832 | 0.065545627 | 0.550245546 | 0.864832081 | 0.314179747 | 0.534508385 |
| Rgs22 | S | 273 | -0.171970509 | 0.209764546 | 0.098551102 | 0.27052161 | -0.111213444 | 0.085528616 | 0.040909249 | 0.07956021 | 0.030558218 | 0.183365542 |
| Rgs22 | S | 275 | -0.171970509 | 0.209764546 | 0.098551102 | 0.27052161 | -0.111213444 | 0.085528616 | 0.040909249 | 0.07956021 | 0.030558218 | 0.183365542 |
| Calu | T | 303 | -0.047124841 | 0.383703315 | -0.085772793 | -0.038647952 | -0.469476108 | 0.692140128 | 0.093398541 | 0.757705579 | 0.891904542 | 0.183101621 |
| Irf2bp2 | S | 343 | 0.02167768 | -0.020199239 | 0.059133042 | 0.037455362 | 0.079332281 | 0.222381352 | 0.067498915 | 0.000157404 | 0.098166894 | 0.006499652 |
| Aff4 | S | 615 | 0.057184292 | 0.048282149 | 0.029312625 | -0.027871667 | -0.018969524 | 0.293161119 | 0.334368893 | 0.365166925 | 0.580370692 | 0.68430787 |
| Bmp2k | T | 981 | -0.065203675 | 0.146570268 | 0.128451009 | 0.193654685 | -0.018119258 | 0.803381448 | 0.422270274 | 0.236712671 | 0.475561735 | 0.911003052 |
| Srrm2 | S | 1305 | 0.025891639 | 0.042701513 | 0.122200799 | 0.09630916 | 0.079499287 | 0.795314737 | 0.5567304 | 0.218602377 | 0.386985342 | 0.366903669 |
| Med26 | S | 314 | -0.022531527 | 0.059099484 | 0.264642945 | 0.287174472 | 0.205543461 | 0.785757073 | 0.583654147 | 0.304051661 | 0.273438114 | 0.410360046 |
| Septin5 | S | 327 | 0.106402468 | 0.090109461 | -0.019677241 | -0.126079709 | -0.109786703 | 0.256096342 | 0.193649022 | 0.604975679 | 0.203894364 | 0.140071189 |
| Gorasp1 | S | 400 | -0.089714077 | 0.291407429 | 0.134559628 | 0.224273705 | -0.156847801 | 0.451282224 | 0.351435317 | 0.340336445 | 0.090563442 | 0.588017048 |
| Cbfa2t2 | S | 255 | 0.046997978 | 0.024155242 | 0.056542954 | 0.009544976 | 0.032387711 | 0.374631659 | 0.460675502 | 0.174948167 | 0.848374108 | 0.349847426 |
| Myl6 | S | 135 | -0.014592183 | 0.109869244 | 0.026673279 | 0.041265462 | -0.083195965 | 0.635451705 | 0.019294197 | 0.746560056 | 0.612011444 | 0.352762773 |
| Incenp | S | 273 | 0.104382784 | 0.029058878 | -0.01801276 | -0.122395544 | -0.047071638 | 0.014810626 | 0.424430257 | 0.259041689 | 0.017873935 | 0.236540638 |
| Senp6 | S | 211 | 0.160077277 | -0.017552348 | 0.093995696 | -0.066081582 | 0.111548043 | 0.318675142 | 0.804079155 | 0.387053154 | 0.682545652 | 0.347820126 |
| Fcho2 | S | 509 | -0.006919446 | 0.105183482 | 0.081597831 | 0.088517277 | -0.023585651 | 0.934711056 | 0.276385627 | 0.288648367 | 0.276526479 | 0.773844405 |

| | | | | | | | | | | | | |
|---|---|---|---|---|---|---|---|---|---|---|---|---|
| Shroom4 | S | 722 | -0.064173478 | 0.048298741 | 0.233082411 | 0.297255889 | 0.18478367 | 0.301052713 | 0.520791274 | 0.0819397 | 0.053330053 | 0.144186834 |
| Tenm3 | S | 207 | 0.156135023 | -0.049302414 | 0.044449863 | -0.11168516 | 0.093752277 | 0.039907303 | 0.444348743 | 0.585834371 | 0.15825348 | 0.208332787 |
| Pum2 | S | 181 | -0.041164854 | 0.064642107 | 0.052180075 | 0.09334493 | -0.012462032 | 0.10386834 | 0.058211938 | 0.003748765 | 0.013889463 | 0.561875967 |
| Rb1 | T | 350 | 0.010308814 | 0.008303588 | 0.103789336 | 0.093480522 | 0.095485748 | 0.84075185 | 0.819128711 | 0.039383054 | 0.114417226 | 0.042937583 |
| Rnf214 | S | 497 | 0.057447891 | 0.065683507 | 0.076167437 | 0.018719546 | 0.01048393 | 0.513632834 | 0.394367973 | 0.531726403 | 0.879960271 | 0.929075654 |
| Zfp668 | T | 45 | 0.049139575 | 0.047330205 | 0.047460125 | -0.00167945 | 0.00012992 | 0.215824584 | 0.38716555 | 0.521983589 | 0.981669968 | 0.998698211 |
| Sf3b1 | T | 248 | 0.054617023 | 0.054493118 | 0.004945287 | -0.049671736 | -0.049547831 | 0.108903793 | 0.218743292 | 0.872315142 | 0.020001132 | 0.211980605 |
| Ctr9 | S | 1074 | 0.178136136 | -0.148585959 | 0.085635852 | -0.092500284 | 0.234221811 | 0.032094979 | 0.277681312 | 0.313209334 | 0.27475482 | 0.132253766 |
| Fam117a | S | 304 | -0.207188829 | 0.211544458 | 0.33619825 | 0.543387079 | 0.124653791 | 0.284496502 | 0.277409721 | 0.283584698 | 0.123656532 | 0.655747473 |
| Bmp2k | S | 927 | 0.138823579 | -0.044354447 | 0.096006152 | -0.042817426 | 0.140360599 | 0.062175463 | 0.320437502 | 0.481279411 | 0.746629542 | 0.331883001 |
| Hnrnpa3 | S | 359 | 0.002662937 | 0.060693386 | 0.054241488 | 0.051578552 | -0.006451898 | 0.966034281 | 0.14156543 | 0.175511103 | 0.403953046 | 0.785981929 |
| Prr12 | S | 330 | -0.029908919 | 0.115606966 | 0.048349995 | 0.078258914 | -0.067256971 | 0.616031024 | 0.075590282 | 0.318832833 | 0.237367391 | 0.257408738 |
| Fmr1 | S | 349 | -0.061670028 | 0.124740732 | 0.047797088 | 0.109467115 | -0.076943644 | 0.264717661 | 0.01838264 | 0.296723187 | 0.095632457 | 0.133172366 |
| Utp14b | S | 443 | 0.117470942 | 0.122471733 | -0.031120807 | -0.148591749 | -0.15359254 | 0.292316713 | 0.360749453 | 0.842061359 | 0.317173193 | 0.334028512 |
| Utp14b | T | 440 | 0.117470942 | 0.122471733 | -0.031120807 | -0.148591749 | -0.15359254 | 0.292316713 | 0.360749453 | 0.842061359 | 0.317173193 | 0.334028512 |
| Senp3 | S | 71 | 0.043373032 | 0.049977184 | 0.101557058 | 0.058184026 | 0.051579874 | 0.457712186 | 0.704919885 | 0.311831305 | 0.550116788 | 0.725600572 |
| Senp3 | S | 73 | 0.043373032 | 0.049977184 | 0.101557058 | 0.058184026 | 0.051579874 | 0.457712186 | 0.704919885 | 0.311831305 | 0.550116788 | 0.725600572 |
| Fubp1 | S | 626 | 0.197501901 | 0.02565982 | 0.040836785 | -0.156665117 | 0.015176965 | 0.460911289 | 0.665481147 | 0.507251806 | 0.547755445 | 0.789207568 |
| Mxra7 | S | 57 | 0.031145333 | 0.102865571 | -0.040339698 | -0.071485031 | -0.143205268 | 0.63525149 | 0.005694766 | 0.415571963 | 0.352579398 | 0.048707718 |
| Zc3h13 | S | 406 | 0.048881353 | 0.044087394 | 0.059154348 | 0.010272996 | 0.015066954 | 0.278987247 | 0.467466399 | 0.419411145 | 0.88727845 | 0.852519737 |
| Fam83d | S | 454 | -0.000336319 | 0.048207958 | 0.089383305 | 0.089719624 | 0.041175347 | 0.997297217 | 0.26311085 | 0.157041158 | 0.423579692 | 0.482058988 |
| Phf12 | T | 670 | 0.020639526 | 0.087104974 | 0.078886649 | 0.058247123 | -0.008218325 | 0.738030729 | 0.274829518 | 0.467471639 | 0.550336148 | 0.93535129 |
| Virma | S | 173 | 0.052053549 | 0.014366297 | 0.045530647 | -0.006522902 | 0.031164349 | 0.10591997 | 0.611140952 | 0.302197027 | 0.851774507 | 0.408607909 |
| Calu | S | 300 | -0.042768865 | 0.493351331 | -0.135262229 | -0.092493365 | -0.62861356 | 0.595901975 | 0.090346205 | 0.557873456 | 0.678231587 | 0.071784681 |
| Wac | S | 525 | -0.058401072 | -0.047463799 | 0.219348188 | 0.277749259 | 0.266811987 | 0.49722421 | 0.545682591 | 0.007901019 | 0.033425047 | 0.027720459 |
| Ddx42 | S | 109 | 0.035058749 | -0.014417893 | 0.082256068 | 0.047197319 | 0.096673961 | 0.377033882 | 0.609773624 | 0.022612871 | 0.271441304 | 0.028562394 |
| Ddx42 | S | 111 | 0.035058749 | -0.014417893 | 0.082256068 | 0.047197319 | 0.096673961 | 0.377033882 | 0.609773624 | 0.022612871 | 0.271441304 | 0.028562394 |
| Snx16 | S | 70 | 0.079580794 | 0.016311689 | -0.007598062 | -0.087178856 | -0.02390975 | 0.014566272 | 0.516208372 | 0.918231473 | 0.301396772 | 0.74307718 |
| Eva1c | S | 431 | 0.135886076 | 0.077544288 | -0.052240623 | -0.188126698 | -0.129784911 | 0.154429164 | 0.352575666 | 0.723163367 | 0.236539214 | 0.374060977 |
| Flna | S | 1010 | 0.091204115 | 0.037042758 | 0.007139661 | -0.084064454 | -0.029903098 | 0.118223142 | 0.403414754 | 0.856066669 | 0.125028961 | 0.294611477 |
| Gigyf2 | S | 237 | 0.04448641 | 0.033794569 | -0.006874564 | -0.051360974 | -0.040669133 | 0.004305419 | 0.300277931 | 0.827453119 | 0.192670388 | 0.327913102 |
| Git1 | S | 570 | 0.052402182 | 0.042749066 | 0.13023679 | 0.077834608 | 0.087487724 | 0.64061918 | 0.661651376 | 0.346025527 | 0.593493628 | 0.526885199 |
| Ano8 | S | 1009 | 0.08959258 | -0.030919128 | 0.074338136 | -0.015254444 | 0.105257264 | 0.088664779 | 0.195817358 | 0.097511949 | 0.743439643 | 0.062336003 |
| Fgd1 | S | 815 | -0.156014724 | 0.146662421 | 0.194138159 | 0.350152883 | 0.047475739 | 0.06475863 | 0.090442061 | 0.07074325 | 0.015000865 | 0.603612674 |
| Arfgef2 | S | 1535 | 0.039301233 | -0.011444904 | 0.078064128 | 0.038762895 | 0.089509032 | 0.30273915 | 0.28280118 | 0.017825799 | 0.319251392 | 0.024030743 |
| Podxl | S | 482 | 0.271607108 | 0.103733609 | -0.002171524 | -0.273778632 | -0.105905133 | 0.40540973 | 0.767829696 | 0.994059798 | 0.212516872 | 0.673761649 |
| Prickle2 | S | 732 | 0.039951643 | 0.136131261 | -0.073106228 | -0.11305787 | -0.209237489 | 0.411549374 | 0.033051891 | 0.539189821 | 0.359303012 | 0.152329354 |
| Samd1 | T | 107 | 0.021814393 | 0.112845413 | -0.030883099 | -0.052697492 | -0.143728512 | 0.706032594 | 0.008328373 | 0.412737946 | 0.389528927 | 0.011073458 |
| Dlgap1 | S | 534 | 0.014604061 | 0.255130829 | -0.081675183 | -0.096279244 | -0.336806013 | 0.823660082 | 0.038664983 | 0.31347344 | 0.211555857 | 0.020691527 |
| Dlgap1 | S | 535 | 0.014604061 | 0.255130829 | -0.081675183 | -0.096279244 | -0.336806013 | 0.823660082 | 0.038664983 | 0.31347344 | 0.211555857 | 0.020691527 |
| Dlgap1 | T | 536 | 0.014604061 | 0.255130829 | -0.081675183 | -0.096279244 | -0.336806013 | 0.823660082 | 0.038664983 | 0.31347344 | 0.211555857 | 0.020691527 |
| Alyref Alyref2 | S | 237;200 | 0.072813051 | 0.008393333 | 0.057776929 | -0.015036122 | 0.049383596 | 0.264041686 | 0.814532717 | 0.203147745 | 0.804295325 | 0.263880218 |
| Smarcad1 | S | 96 | -0.024577566 | 0.032517597 | 0.090820923 | 0.115398489 | 0.058303326 | 0.200364443 | 0.122495128 | 0.020624356 | 0.021587078 | 0.086848326 |
| Mpdz | S | 1803 | 0.029658969 | 0.190361745 | -0.021365372 | -0.051024341 | -0.211727118 | 0.53685493 | 0.162523631 | 0.80297789 | 0.530036317 | 0.145012705 |
| Cwc22 | S | 27 | 0.041187449 | 0.04557971 | 0.042933699 | 0.00174625 | -0.002646011 | 0.492123817 | 0.089342494 | 0.582399381 | 0.984171702 | 0.972115914 |
| Srrm2 | T | 877 | 0.010572674 | 0.012874079 | 0.073330058 | 0.062757384 | 0.060455979 | 0.743095582 | 0.691995927 | 0.021718477 | 0.107886253 | 0.118434438 |
| Kif4 | S | 952 | -0.069075806 | 0.097569471 | 0.034799431 | 0.103875237 | -0.06277004 | 0.424003221 | 0.007652602 | 0.286280386 | 0.266013732 | 0.060946333 |
| Caml | S | 88 | -0.135528813 | 0.183525976 | 0.108243069 | 0.243771882 | -0.075282907 | 0.064029834 | 0.059486413 | 0.064267744 | 0.01217626 | 0.326389522 |
| Myo9b | S | 1175 | -0.038605049 | 0.071478077 | 0.057503219 | 0.096108268 | -0.013974858 | 0.294731634 | 0.023896812 | 0.1125212 | 0.041959839 | 0.58389508 |
| Sf3b1 | T | 299 | 0.069284837 | 0.023720584 | 0.06471046 | -0.004574376 | 0.040989876 | 0.242604506 | 0.698551598 | 0.385631451 | 0.930643215 | 0.495932 |
| Tiam2 | S | 1007 | -0.04780997 | 0.088609788 | 0.14685004 | 0.19466001 | 0.058240252 | 0.617436515 | 0.41075725 | 0.153321083 | 0.131198032 | 0.619566179 |
| Nfic | S | 322 | 0.101653759 | -0.002237997 | 0.039054357 | -0.062599402 | 0.041292353 | 0.092002109 | 0.965102626 | 0.476591677 | 0.217098081 | 0.356738619 |
| Cherp | S | 826 | 0.057297012 | 0.014606297 | 0.022515165 | -0.034781846 | 0.007908868 | 0.040192871 | 0.42439836 | 0.58046439 | 0.408505824 | 0.83198938 |
| Cherp | T | 828 | 0.057297012 | 0.014606297 | 0.022515165 | -0.034781846 | 0.007908868 | 0.040192871 | 0.42439836 | 0.58046439 | 0.408505824 | 0.83198938 |
| Ash1l | S | 729 | 0.067965684 | -0.019711976 | 0.073366416 | 0.005400732 | 0.093078393 | 0.138579301 | 0.587227648 | 0.129587061 | 0.903126779 | 0.073996606 |
| Ptpn12 | S | 612 | -0.448919429 | 0.089206712 | 0.385855792 | 0.834775221 | 0.29664908 | 0.467769044 | 0.587707571 | 0.068452586 | 0.235590027 | 0.178182222 |
| Ptpn12 | S | 613 | -0.448919429 | 0.089206712 | 0.385855792 | 0.834775221 | 0.29664908 | 0.467769044 | 0.587707571 | 0.068452586 | 0.235590027 | 0.178182222 |
| Polr2a | T | 1919 | 0.006798565 | 0.06436116 | 0.054848859 | 0.048050294 | -0.009512301 | 0.805688881 | 0.20037229 | 0.214539227 | 0.239137867 | 0.843956828 |
| Sorbs2 | S | 99 | 0.008580367 | 0.005413494 | 0.148404696 | 0.139824329 | 0.142991202 | 0.708758751 | 0.881544816 | 0.111980605 | 0.119668877 | 0.108444834 |
| Frmd6 | S | 544 | -0.062191203 | 0.128601393 | 0.075583936 | 0.137775139 | -0.053017458 | 0.023927373 | 0.049845102 | 0.031527685 | 0.007834346 | 0.28973833 |
| Strn4 | S | 331 | 0.203684346 | 0.011939871 | 0.005982198 | -0.197702148 | -0.005957673 | 0.153260937 | 0.954128252 | 0.988368485 | 0.637019454 | 0.988881631 |
| Zc3h13 | S | 77 | 0.040802527 | -0.017214757 | 0.089940777 | 0.04913825 | 0.107155534 | 0.412613943 | 0.681932723 | 0.038599595 | 0.356048811 | 0.069784621 |
| Srrm1 | S | 915 | 0.027057862 | 0.006676557 | 0.059084523 | 0.032026661 | 0.052407966 | 0.329936144 | 0.771423963 | 0.040915375 | 0.143227295 | 0.01156537 |
| Dido1 | S | 2092 | 0.066515888 | 0.029307366 | 0.046667725 | -0.019848162 | 0.017360359 | 0.319597434 | 0.571736803 | 0.273860821 | 0.75752384 | 0.755160005 |
| Zc3h13 | T | 375 | 0.024763084 | 0.051620601 | 0.05197042 | 0.027207336 | 0.000349819 | 0.406713327 | 0.288038618 | 0.312019683 | 0.582314018 | 0.995055918 |
| Trim28 | S | 596 | -0.026281691 | 0.140332949 | 0.077076829 | 0.103358521 | -0.06325612 | 0.638558887 | 0.17401042 | 0.411868308 | 0.305615991 | 0.569588325 |
| Tnks1bp1 | S | 887 | 0.028199138 | 0.069909887 | 0.038142646 | 0.009943508 | -0.031767241 | 0.552598786 | 0.205756734 | 0.40498942 | 0.701932185 | 0.402398664 |
| Camk2d | S | 334 | -0.186551365 | 0.314749805 | 0.058934668 | 0.245486033 | -0.255815137 | 0.083226337 | 0.015012626 | 0.582624785 | 0.033926846 | 0.034056039 |
| Eif2ak2 | S | 110 | 0.029788822 | 0.186832779 | -0.060297484 | -0.090086306 | -0.247130263 | 0.61840046 | 0.08802792 | 0.698212483 | 0.565926919 | 0.191892059 |
| Triobp | S | 1614 | 0.037335895 | 0.017484129 | 0.123035562 | 0.085699668 | 0.105551433 | 0.546941237 | 0.785079609 | 0.221607532 | 0.327543199 | 0.25048584 |
| Hsp90aa1 | S | 477 | 0.04521845 | 0.101277594 | 0.002104291 | -0.043114159 | -0.099173303 | 0.482016675 | 0.127459188 | 0.976388532 | 0.4417048 | 0.126066124 |
| Vim | S | 419 | 0.005923064 | -0.01749525 | 0.151268623 | 0.145345559 | 0.168763873 | 0.937589263 | 0.795021401 | 0.043433434 | 0.072166658 | 0.019227408 |
| Rnf10 | S | 41 | 0.01893575 | 0.074168969 | 0.064392827 | 0.045457077 | -0.009776142 | 0.806062643 | 0.265958082 | 0.329153655 | 0.612371698 | 0.895533691 |
| Lrrfip2 | S | 91 | 0.022817031 | 0.077279094 | 0.016882997 | -0.005934034 | -0.060396097 | 0.535594503 | 0.068832988 | 0.744857806 | 0.886650489 | 0.234838313 |
| Zcchc8 | S | 601 | -0.01100766 | -0.04566379 | 0.267826564 | 0.278834224 | 0.31349035 | 0.89410983 | 0.331308572 | 0.047288051 | 0.050302493 | 0.049153512 |
| Pja1 | S | 137 | 0.068224604 | 0.085842278 | 0.019324854 | -0.04889975 | -0.066517424 | 0.377550406 | 0.271308587 | 0.869923121 | 0.63492175 | 0.527095973 |
| Wee1 | T | 190 | 0.034423403 | 0.085994436 | 0.025670548 | -0.008752855 | -0.060323888 | 0.592767813 | 0.108918685 | 0.812209196 | 0.93567784 | 0.57268717 |
| Eif3j1 Eif3j2 | S | 130;132 | 0.047073629 | 0.14676888 | -0.015453844 | -0.062527473 | -0.162222724 | 0.374366747 | 0.213459891 | 0.863520366 | 0.507623753 | 0.223251639 |
| Phf14 | S | 287 | 0.114402228 | -0.050686302 | 0.086164796 | -0.028237433 | 0.136851098 | 0.220929419 | 0.480213039 | 0.128284011 | 0.734492141 | 0.1230386 |
| Tjp2 | S | 267 | -0.064174138 | -0.078447834 | 0.265413278 | 0.329587416 | 0.343861112 | 0.322424534 | 0.188710211 | 0.002198699 | 0.004105712 | 0.002006946 |
| Nfib | S | 194 | 0.089171192 | 0.080730408 | 0.003247888 | -0.085923305 | -0.07748252 | 0.204034373 | 0.43729468 | 0.97879395 | 0.480510991 | 0.569448929 |
| Chd8 | S | 551 | -0.001174727 | 0.010093316 | 0.15650217 | 0.157676898 | 0.146408855 | 0.989865668 | 0.782592699 | 0.102603781 | 0.191360123 | 0.129126617 |
| Chd8 | S | 555 | -0.001174727 | 0.010093316 | 0.15650217 | 0.157676898 | 0.146408855 | 0.989865668 | 0.782592699 | 0.102603781 | 0.191360123 | 0.129126617 |
| Chd8 | S | 564 | -0.001174727 | 0.010093316 | 0.15650217 | 0.157676898 | 0.146408855 | 0.989865668 | 0.782592699 | 0.102603781 | 0.191360123 | 0.129126617 |
| Zfp148 | T | 305 | 0.02808118 | 0.01253647 | 0.074929522 | 0.046848342 | 0.062393052 | 0.561310388 | 0.813878202 | 0.055881275 | 0.375431896 | 0.307478206 |
| Srrm2 | S | 794 | 0.127215025 | -0.053812786 | 0.032018966 | -0.095196059 | 0.085831752 | 0.007529867 | 0.282452422 | 0.660657384 | 0.250946339 | 0.301799366 |
| Gtf3c2 | S | 218 | 0.264473033 | 0.148295048 | -0.202016979 | -0.466490012 | -0.350312027 | 0.109935191 | 0.365984687 | 0.279436668 | 0.059605438 | 0.136482731 |
| Nes | S | 841 | -0.085508868 | 0.044819965 | 0.218521599 | 0.304030467 | 0.173701634 | 0.103437112 | 0.289849567 | 0.037367859 | 0.020746034 | 0.093282857 |
| Slf2 | S | 815 | 0.03695015 | 0.00590818 | 0.115302931 | 0.078352781 | 0.109394751 | 0.414508638 | 0.757707441 | 0.213986107 | 0.365777243 | 0.232161159 |
| Utp18 | S | 115 | 0.06270409 | 0.014554747 | 0.025536236 | -0.037167854 | 0.010981489 | 0.044458053 | 0.63258735 | 0.578465157 | 0.405798048 | 0.799757804 |
| Taf9 | T | 159 | 0.143678199 | -0.115921083 | 0.080173122 | -0.063505078 | 0.196094204 | 0.03388431 | 0.31225817 | 0.29787829 | 0.403381234 | 0.141611074 |
| Ralgapb | T | 724 | 0.118498369 | 0.079523497 | -0.033323036 | -0.151821404 | -0.112846533 | 0.414744912 | 0.031336733 | 0.279347681 | 0.31982177 | 0.002353554 |
| Farp1 | S | 403 | 0.049719769 | -0.024198879 | 0.069912446 | 0.020192677 | 0.094111325 | 0.024276696 | 0.359375965 | 0.143126891 | 0.593806112 | 0.07814385 |
| Sik3 | T | 527 | 0.050623662 | 0.109002529 | -0.022811124 | -0.073434786 | -0.131813653 | 0.573066633 | 0.072582103 | 0.846693248 | 0.579877394 | 0.312497394 |

| | | | | | | | | | | | | |
|---|---|---|---|---|---|---|---|---|---|---|---|---|
| Mcm9 | S | 885 | 0.035330635 | -0.00493082 | 0.113307202 | 0.077976567 | 0.118238022 | 0.282048366 | 0.725595149 | 0.143216831 | 0.25525557 | 0.137052819 |
| Epb41l1 | S | 378 | 0.092356625 | 0.008940431 | 0.04923939 | -0.043117235 | 0.040298958 | 0.178216626 | 0.697285944 | 0.498990539 | 0.601230964 | 0.569225508 |
| Rfc1 | S | 516 | -0.024326292 | 0.059209398 | 0.039569579 | 0.063895871 | -0.019639819 | 0.619696296 | 0.026855928 | 0.126635147 | 0.244063128 | 0.255818975 |
| Clasp1 | T | 793 | -0.027246425 | 0.444892385 | -0.221753634 | -0.194507208 | -0.666646019 | 0.971234515 | 0.184054403 | 0.83603147 | 0.870949277 | 0.569741828 |
| 4933417A18Rik | S | 83 | -0.006002961 | 0.025178641 | 0.089766643 | 0.095769604 | 0.064588002 | 0.72167937 | 0.332275829 | 0.050512369 | 0.056279581 | 0.119818978 |
| Ppp4r2 | S | 159 | 0.099636356 | 0.00596832 | 0.006277914 | -0.093358442 | 0.000309595 | 0.031314077 | 0.887496927 | 0.900582117 | 0.133272961 | 0.995476663 |
| Wwox | S | 14 | 0.012372129 | 0.038684291 | 0.047995761 | 0.035623632 | 0.00931147 | 0.364239593 | 0.319954481 | 0.121884155 | 0.210945931 | 0.8090518 |
| Wwox | T | 12 | 0.012372129 | 0.038684291 | 0.047995761 | 0.035623632 | 0.00931147 | 0.364239593 | 0.319954481 | 0.121884155 | 0.210945931 | 0.8090518 |
| Cfap65 | S | 746 | 0.188061222 | 0.084857634 | 0.084022916 | -0.104038306 | -0.000834718 | 0.634328328 | 0.648370022 | 0.751323616 | 0.802556687 | 0.997377 |
| Cfap65 | T | 749 | 0.188061222 | 0.084857634 | 0.084022916 | -0.104038306 | -0.000834718 | 0.634328328 | 0.648370022 | 0.751323616 | 0.802556687 | 0.997377 |
| Eif3f | S | 262 | 0.16504378 | 0.078005089 | -0.20349044 | -0.368534221 | -0.28149553 | 0.014930269 | 0.130262703 | 0.177833543 | 0.059937263 | 0.106343709 |
| Myh9 | S | 1714 | -0.106982402 | 0.160243125 | 0.150940417 | 0.257922819 | -0.009302709 | 0.00542158 | 0.059626128 | 0.034235901 | 0.009481927 | 0.884234326 |
| Dsn1 | S | 330 | 0.014176533 | 0.067758993 | -0.007681915 | -0.021858449 | -0.075440908 | 0.784348925 | 0.003632992 | 0.836582982 | 0.711098218 | 0.129009004 |
| Champ1 | S | 643 | -0.01065616 | 0.015274574 | 0.109163829 | 0.119819989 | 0.093889256 | 0.679599541 | 0.384206019 | 0.045219859 | 0.032039506 | 0.067968145 |
| Srrm2 | S | 333 | -0.079227913 | 0.050787903 | 0.112755677 | 0.19198359 | 0.061967774 | 0.292624942 | 0.194829225 | 0.020545229 | 0.059559732 | 0.078440391 |
| Kansl3 | T | 538 | 0.175197642 | -0.073945776 | 0.022849676 | -0.152347967 | 0.096795451 | 0.024849446 | 0.533834711 | 0.900627869 | 0.443284911 | 0.63724909 |
| Fbxo38 | S | 723 | -0.206075389 | -0.042164298 | 0.382582743 | 0.588658133 | 0.424747042 | 0.423429845 | 0.814349619 | 0.031180056 | 0.080873396 | 0.071364081 |
| Plekho2 | S | 239 | 0.017587013 | 0.156604284 | -0.032899992 | -0.050487004 | -0.189504276 | 0.847171775 | 0.049741904 | 0.730236644 | 0.610181108 | 0.099533907 |
| Srrm2 | S | 435 | 0.064956377 | -0.000457703 | 0.068659937 | 0.00370356 | 0.06911764 | 0.166356479 | 0.990905585 | 0.274016845 | 0.939768495 | 0.236681016 |
| Fam117b | S | 340 | 0.03951612 | 0.003802207 | 0.05638824 | 0.01687212 | 0.052586033 | 0.272283398 | 0.860119046 | 0.070520914 | 0.633958345 | 0.101361248 |
| Sh3pxd2a | S | 608 | 0.064093732 | -0.051437753 | 0.215193324 | 0.151099592 | 0.266631077 | 0.625017329 | 0.774186822 | 0.211220561 | 0.359013848 | 0.219243922 |
| Prrx1 | S | 63 | -0.0768815 | 0.208681489 | 0.005063525 | 0.081945025 | -0.203617964 | 0.096326476 | 0.005529966 | 0.926015044 | 0.145994028 | 0.015580763 |
| Rcan1 Rcan3 | S | 152;165 | -0.005525617 | 0.065088882 | 0.084332155 | 0.089857772 | 0.019243273 | 0.940459881 | 0.148820339 | 0.336149394 | 0.36324035 | 0.797598577 |
| Mical2 | S | 515 | -0.132123274 | 0.073951956 | 0.153535092 | 0.285658365 | 0.079583135 | 0.050795965 | 0.123187385 | 0.007162892 | 0.005156669 | 0.124572968 |
| Kat7 | S | 166 | -0.029662042 | 0.007066336 | 0.13573497 | 0.165397012 | 0.128668634 | 0.707767209 | 0.918470016 | 0.023549102 | 0.117107433 | 0.148645181 |
| Nfu1 | S | 254 | 0.254280744 | 0.048637182 | 0.043896285 | -0.21038446 | -0.004740897 | 0.406001561 | 0.860802707 | 0.887363364 | 0.572278377 | 0.989365199 |
| Nfu1 | S | 255 | 0.254280744 | 0.048637182 | 0.043896285 | -0.21038446 | -0.004740897 | 0.406001561 | 0.860802707 | 0.887363364 | 0.572278377 | 0.989365199 |
| Bud13 | S | 135 | 0.010521823 | 0.035206858 | 0.074325038 | 0.063803214 | 0.03911818 | 0.559070634 | 0.283850671 | 0.196619904 | 0.252998761 | 0.459089125 |
| Srrm2 | S | 1844 | 0.058246771 | 0.017999054 | 0.100493513 | 0.042246741 | 0.082494459 | 0.651596452 | 0.844113998 | 0.168525736 | 0.744463244 | 0.41020625 |
| Numa1 | T | 1982 | -0.177099635 | -0.038367556 | 0.365520738 | 0.542620373 | 0.403888293 | 0.213470742 | 0.724735741 | 0.012111545 | 0.009522315 | 0.004641371 |
| Fam117b | S | 102 | -0.012502229 | 0.097661887 | 0.018783009 | 0.031285238 | -0.078878877 | 0.787464799 | 0.027002554 | 0.609196318 | 0.430309156 | 0.014961382 |
| Arhgef17 | T | 329 | -0.111597768 | 0.198593022 | 0.08233979 | 0.193937558 | -0.116253232 | 0.546103967 | 0.118655429 | 0.482829936 | 0.337879156 | 0.380099259 |
| Fam177a Fam177a2 | Y | 64 | 0.013770676 | 0.026631883 | 0.092415057 | 0.078644381 | 0.065783174 | 0.811361529 | 0.619642349 | 0.092807789 | 0.132091282 | 0.110902627 |
| Nktr | S | 243 | 0.043693063 | -0.016465514 | 0.071808342 | 0.028115279 | 0.088273856 | 0.478106148 | 0.41570046 | 0.009877489 | 0.634199892 | 0.006668437 |
| Pcnx3 | S | 1955 | 0.00079444 | 0.053167245 | 0.120346253 | 0.119551812 | 0.067179008 | 0.988442945 | 0.490555811 | 0.205842938 | 0.199101413 | 0.468469811 |
| Mef2d | S | 251 | 0.079519801 | 0.00547991 | 0.035039969 | -0.044479832 | 0.029560059 | 0.077582037 | 0.873721096 | 0.501328089 | 0.431620662 | 0.588011599 |
| Rai1 | S | 1116 | 0.025872124 | -0.007817137 | 0.08885691 | 0.062984785 | 0.096674046 | 0.370317253 | 0.831066068 | 0.052794226 | 0.133566759 | 0.067776894 |
| Blm | T | 1311 | 0.11680223 | 0.019509155 | -0.021411803 | -0.138214033 | -0.040920958 | 0.029865463 | 0.70671202 | 0.689322287 | 0.037647917 | 0.443228331 |
| H2-K1 | S | 348;307;347;348 | 0.003122144 | 0.100632491 | -0.007963515 | -0.011085658 | -0.108596005 | 0.953762205 | 0.013406223 | 0.838053954 | 0.836412966 | 0.032553734 |
| Nckipsd | S | 260 | 0.190403715 | -0.025510546 | 0.000519375 | -0.189884339 | 0.026029921 | 0.072026506 | 0.789980803 | 0.994602148 | 0.099804049 | 0.820292172 |
| Gorasp1 | S | 376 | 0.002633657 | 0.034636177 | 0.08078598 | 0.078152323 | 0.046149803 | 0.977451319 | 0.476572261 | 0.06989544 | 0.433508801 | 0.342940325 |
| Jmy | S | 701 | -0.035260094 | 0.080889577 | 0.100731567 | 0.135991661 | 0.019841991 | 0.668750878 | 0.207572628 | 0.214207886 | 0.135506397 | 0.716720974 |
| Jmy | T | 702 | -0.035260094 | 0.080889577 | 0.100731567 | 0.135991661 | 0.019841991 | 0.668750878 | 0.207572628 | 0.214207886 | 0.135506397 | 0.716720974 |
| Myo18a | S | 2032 | 0.071152025 | 0.067506514 | -0.024722993 | -0.095875018 | -0.092229507 | 0.120514068 | 0.162785381 | 0.630019559 | 0.084999472 | 0.077506866 |
| Gpatch8 | S | 437 | -0.028104865 | 0.041361809 | 0.132000675 | 0.16010554 | 0.090638865 | 0.632689527 | 0.180163276 | 0.17764372 | 0.130778787 | 0.307705094 |
| Epb41l2 | S | 543 | -0.061043138 | 0.09277511 | 0.10229524 | 0.163338378 | 0.00952013 | 0.547849819 | 0.310368452 | 0.093679369 | 0.178542098 | 0.910978762 |
| Taf6 | S | 532 | 0.056687038 | 0.038478171 | 0.045304578 | -0.01138246 | 0.006826408 | 0.418923482 | 0.460825914 | 0.289948992 | 0.864325591 | 0.895650726 |
| Zc3h18 | S | 64 | 0.016713382 | -0.006304315 | 0.07650369 | 0.059790308 | 0.082808005 | 0.713411645 | 0.751250748 | 0.009491372 | 0.243800391 | 0.011032539 |
| Trmt6 | S | 289 | 0.053225407 | 0.070212298 | 0.008867336 | -0.04435807 | -0.061344962 | 0.303235662 | 0.168562413 | 0.937385444 | 0.704456768 | 0.603217137 |
| Brip1 | S | 197 | -0.00378034 | 0.023269875 | 0.059827011 | 0.063607351 | 0.036557136 | 0.940899459 | 0.563814313 | 0.009271796 | 0.289057174 | 0.393013432 |
| Rif1 | S | 1828 | 0.10886561 | 0.092998712 | 0.018717395 | -0.090148215 | -0.074281317 | 0.436102162 | 0.438048363 | 0.868503372 | 0.394936245 | 0.279048335 |
| Nsfl1c | S | 272 | -0.023578448 | 0.089434077 | 0.036831173 | 0.060409621 | -0.052602904 | 0.66923903 | 0.058895632 | 0.260411791 | 0.288907604 | 0.174561694 |
| Sp100 | S | 318 | -0.049435246 | -0.004061626 | 0.169721068 | 0.219156314 | 0.173782694 | 0.304827404 | 0.941696722 | 0.009778769 | 0.003653768 | 0.019213421 |
| Lbr | T | 154 | 0.120601401 | -0.029978244 | 0.162184889 | 0.041583487 | 0.192163133 | 0.478394418 | 0.840343133 | 0.369469871 | 0.809802061 | 0.264753022 |
| Zc3h4 | S | 917 | 0.006362566 | 0.042000543 | 0.064228703 | 0.057866137 | 0.022228159 | 0.892656559 | 0.377706836 | 0.085147915 | 0.225651194 | 0.574658131 |
| Tfam | S | 241 | -0.064674548 | 0.065787719 | 0.111531591 | 0.17620614 | 0.045743873 | 0.239622175 | 0.108527695 | 0.067376985 | 0.026705149 | 0.337903018 |
| Capn15 | S | 348 | 0.064963903 | 0.089542084 | -0.019707693 | -0.084671596 | -0.109249777 | 0.203055831 | 0.183161861 | 0.692738038 | 0.106364272 | 0.119734956 |
| Wnk1 | T | 1801 | 0.070765917 | 0.136697794 | -0.085474349 | -0.156240265 | -0.222172143 | 0.333277992 | 0.038957022 | 0.304764724 | 0.100714021 | 0.057257405 |
| Zbtb41 | S | 191 | 0.053356514 | 0.138185573 | 0.022876705 | -0.030479808 | -0.115308868 | 0.659157107 | 0.305966232 | 0.795331945 | 0.804302931 | 0.391747619 |
| Zbtb41 | S | 192 | 0.053356514 | 0.138185573 | 0.022876705 | -0.030479808 | -0.115308868 | 0.659157107 | 0.305966232 | 0.795331945 | 0.804302931 | 0.391747619 |
| Sh3bp5 | S | 354 | -0.139513988 | 0.42884155 | -0.27174377 | -0.132229782 | -0.70058532 | 0.611958817 | 0.15918582 | NA | NA | NA |
| Zc3h4 | S | 92 | -0.148662072 | -0.007170396 | 0.2507049 | 0.399366972 | 0.257875296 | 0.206040349 | 0.918345445 | 0.005854081 | 0.028421165 | 0.00603462 |
| Zc3h4 | S | 94 | -0.148662072 | -0.007170396 | 0.2507049 | 0.399366972 | 0.257875296 | 0.206040349 | 0.918345445 | 0.005854081 | 0.028421165 | 0.00603462 |
| Polr2a | S | 1906 | 0.031528816 | 0.04735997 | 0.018218839 | -0.013309977 | -0.029141131 | 0.170087065 | 0.175298315 | 0.563014198 | 0.659871752 | 0.438254217 |
| Msantd2 | S | 83 | 0.053163844 | 0.059233092 | 0.021048847 | -0.032114997 | -0.038184245 | 0.362147094 | 0.207719577 | 0.692700009 | 0.571158838 | 0.412906569 |
| Macroh2a1 | S | 173 | 0.001128934 | 0.067809075 | 0.064052907 | 0.062923973 | -0.003756168 | 0.988970055 | 0.12989819 | 0.37799092 | 0.521057667 | 0.955828361 |
| Srrm2 | S | 1846 | 0.115748926 | -0.053845331 | 0.16107718 | 0.045328254 | 0.214922512 | 0.274886172 | 0.494354105 | 0.311885827 | 0.762518635 | 0.204517472 |
| Asph | S | 17 | 0.131899794 | -0.00714849 | 0.160005611 | 0.028105818 | 0.167154102 | 0.583540508 | 0.978038914 | 0.422841934 | 0.913105496 | 0.560823671 |
| Peak1 | S | 726 | -0.032247535 | 0.014056818 | 0.152915268 | 0.185162803 | 0.13885845 | 0.742911608 | 0.776718353 | 0.056283751 | 0.128429368 | 0.080310408 |
| Esyt2 | S | 682 | -0.034278689 | 0.021252889 | 0.08060072 | 0.114879409 | 0.059347831 | 0.328983164 | 0.214226148 | 0.006291791 | 0.030088937 | 0.046389453 |
| Srrm1 | S | 781 | 0.013595906 | 0.008457149 | 0.059491066 | 0.04589516 | 0.051033917 | 0.526290738 | 0.671456949 | 0.023263614 | 0.040837435 | 0.023792389 |
| Eif5a Eif5a2 | S | 75;75 | -0.029058344 | -0.004759804 | 0.214154603 | 0.243212947 | 0.218914406 | 0.393996713 | 0.965439697 | 0.049673012 | 0.03423969 | 0.137955451 |
| Eif5a Eif5a2 | T | 76;76 | -0.029058344 | -0.004759804 | 0.214154603 | 0.243212947 | 0.218914406 | 0.393996713 | 0.965439697 | 0.049673012 | 0.03423969 | 0.137955451 |
| Dzip1l | S | 425 | 0.010562875 | 0.02200083 | 0.072138793 | 0.061575918 | 0.050137963 | 0.811280368 | 0.590350721 | 0.047180491 | 0.211859529 | 0.245305941 |
| Dzip1l | S | 426 | 0.010562875 | 0.02200083 | 0.072138793 | 0.061575918 | 0.050137963 | 0.811280368 | 0.590350721 | 0.047180491 | 0.211859529 | 0.245305941 |
| Ankrd11 | S | 408 | 0.016958704 | 0.001361336 | 0.078011173 | 0.061052469 | 0.076649837 | 0.507561407 | 0.942247682 | 0.036710041 | 0.095657593 | 0.045378573 |
| Ankrd11 | S | 411 | 0.016958704 | 0.001361336 | 0.078011173 | 0.061052469 | 0.076649837 | 0.507561407 | 0.942247682 | 0.036710041 | 0.095657593 | 0.045378573 |
| Ankrd11 | T | 410 | 0.016958704 | 0.001361336 | 0.078011173 | 0.061052469 | 0.076649837 | 0.507561407 | 0.942247682 | 0.036710041 | 0.095657593 | 0.045378573 |
| Tomm40 | S | 142 | 0.298006034 | 0.104104388 | -0.101140486 | -0.39914652 | -0.205244874 | 0.346054092 | 0.577657007 | 0.732472578 | 0.304550281 | 0.504358117 |
| Atp6v0a2 | S | 695 | 0.081426921 | -0.007159457 | 0.007622421 | -0.0738045 | 0.014781878 | 0.008206776 | 0.731983002 | 0.768988477 | 0.021685638 | 0.443081176 |
| Proser3 | S | 591 | 0.007645857 | 0.392338918 | NA | NA | NA | NA | 0.372246452 | NA | NA | NA |
| Fbxo46 | S | 273 | 0.16429647 | -0.046348695 | 0.167382093 | 0.003085622 | 0.213730788 | 0.403988611 | 0.697557741 | 0.449717879 | 0.990233517 | 0.367858921 |
| Ccdc61 | S | 332 | -0.00788348 | -0.005571551 | 0.106411786 | 0.114295266 | 0.111983336 | 0.825679734 | 0.923825702 | 0.012592319 | 0.042242321 | 0.147066448 |
| Gabbr1 | S | 438 | 0.061242408 | 0.094194245 | 0.016863702 | -0.044378705 | -0.077330543 | 0.622014054 | 0.180693765 | 0.862163231 | 0.745532667 | 0.440054375 |
| Gabbr1 | S | 442 | 0.061242408 | 0.094194245 | 0.016863702 | -0.044378705 | -0.077330543 | 0.622014054 | 0.180693765 | 0.862163231 | 0.745532667 | 0.440054375 |
| Gabbr1 | T | 441 | 0.061242408 | 0.094194245 | 0.016863702 | -0.044378705 | -0.077330543 | 0.622014054 | 0.180693765 | 0.862163231 | 0.745532667 | 0.440054375 |
| Ticrr | T | 1251 | 0.121615733 | -0.061206588 | 0.087457847 | -0.034157886 | 0.148664435 | 0.049316578 | 0.233510119 | 0.320754528 | 0.680235537 | 0.140650185 |
| Chd4 | S | 508 | -0.055660115 | 0.075717276 | 0.165288808 | 0.220948923 | 0.089571532 | 0.36533159 | 0.233344983 | 0.186662955 | 0.119932567 | 0.409022062 |
| Mcm10 | T | 85 | 0.16053 | 0.014526123 | 0.011814366 | -0.148715635 | -0.002711757 | 0.155017644 | 0.873780717 | 0.923995514 | 0.249599483 | 0.97967932 |
| Ticrr | S | 1180 | 0.00224906 | -0.001376984 | 0.291968581 | 0.28971952 | 0.293345565 | 0.963929688 | 0.98701858 | 0.271913862 | 0.274312282 | 0.268751213 |
| 5730455P16Rik | S | 18 | -0.045230327 | 0.09390154 | 0.043795279 | 0.089025606 | -0.050106262 | 0.291063796 | 0.036619183 | 0.127357987 | 0.086688603 | 0.183914504 |
| Yap1 | T | 95 | -0.036981656 | 0.183161512 | 0.051253163 | 0.088234819 | -0.131908349 | 0.693409913 | 0.13190116 | 0.724211053 | 0.558246087 | 0.412706358 |
| Msl1 | S | 452 | 0.008533609 | 0.02034304 | 0.124018274 | 0.115484665 | 0.103675235 | 0.825175384 | 0.580515464 | 0.16148538 | 0.185150926 | 0.220158405 |
| Dnm1 | S | 777 | -0.078708383 | 0.12134041 | 0.140040104 | 0.218748487 | 0.018699693 | 0.191597962 | 0.06441388 | 0.310188719 | 0.165237182 | 0.873940848 |

| | | | | | | | | | | | | |
|---|---|---|---|---|---|---|---|---|---|---|---|---|
| Ptbp1 | S | 16 | 0.01611129 | 0.051954932 | 0.051710267 | 0.035598977 | -0.000244665 | 0.58783518 | 0.190390297 | 0.345694651 | 0.501665811 | 0.996352851 |
| Zfp91 Gm44505 | S | 106 | -0.100849278 | 0.07663548 | 0.121705961 | 0.222555239 | 0.045070481 | 0.241471114 | 0.192635592 | 0.033259368 | 0.055685517 | 0.318475536 |
| Bmpr2 | S | 863 | -0.032529868 | 0.090708066 | 0.068483381 | 0.101013249 | -0.022224685 | 0.659784649 | 0.117070425 | 0.282169755 | 0.24132444 | 0.710956969 |
| Dennd4c | S | 1096 | -0.008462999 | 0.189932434 | -0.029822314 | -0.021359315 | -0.219754748 | 0.910669301 | 0.037758888 | 0.509400371 | 0.76570418 | 0.033975767 |
| Kat6b | S | 1105 | -0.1251248 | 0.07397655 | 0.249253026 | 0.374377826 | 0.175276476 | 0.238538016 | 0.449265944 | 0.07749963 | 0.024565526 | 0.146961652 |
| Mex3d | T | 497 | 0.005384274 | 0.15210298 | -0.058994292 | -0.064378567 | -0.211097272 | 0.92077022 | 0.012292063 | 0.4364147 | 0.410697668 | 0.048327941 |
| Pde4a | S | 51 | -0.048770328 | 0.254524029 | -0.007950964 | 0.040819364 | -0.262474993 | 0.569090225 | 0.078665501 | 0.947171957 | 0.767486148 | 0.122855468 |
| Nfib | S | 328 | -0.055368931 | 0.042588301 | 0.073951219 | 0.129320149 | 0.031362918 | 0.021355196 | 0.037281683 | 0.0012081 | 0.002953666 | 0.076900735 |
| Nrcam | S | 1223 | -0.000783849 | 0.059602487 | 0.068105897 | 0.068889745 | 0.008503409 | 0.987747231 | 0.261082594 | 0.187375529 | 0.146857765 | 0.831547098 |
| Dst | S | 4378 | 0.157108688 | -0.009188706 | -0.029047519 | -0.186156208 | -0.019858814 | 0.04367498 | 0.944287943 | 0.880042457 | 0.374229733 | 0.924855058 |
| Zcchc8 | S | 599 | 0.123449756 | -0.024313684 | 0.124924379 | 0.001474623 | 0.149238063 | 0.420111031 | 0.88058832 | 0.383800721 | 0.986257377 | 0.238465784 |
| Ate1 | S | 118 | 0.252521386 | 0.174685511 | -0.133053897 | -0.385575283 | -0.307739409 | 0.253262942 | 0.456417229 | 0.415720286 | 0.102025555 | 0.205626822 |
| Ate1 | T | 119 | 0.252521386 | 0.174685511 | -0.133053897 | -0.385575283 | -0.307739409 | 0.253262942 | 0.456417229 | 0.415720286 | 0.102025555 | 0.205626822 |
| Spred2 | S | 168 | 0.117069712 | 0.031150459 | 0.02670183 | -0.090367882 | -0.00444863 | 0.237421664 | 0.63789942 | 0.74598146 | 0.36058506 | 0.948910462 |
| Map1b | S | 561 | 0.037256482 | 0.166627313 | -0.160281632 | -0.197538115 | -0.326908946 | 0.582277688 | 0.007573469 | 0.320326265 | 0.24429672 | 0.116514666 |
| Taf2 | S | 1099 | 0.032512291 | 0.039418036 | 0.059865171 | 0.027352881 | 0.020447135 | 0.41880751 | 0.467782125 | 0.281385296 | 0.61462172 | 0.744770394 |
| Taf2 | S | 1101 | 0.032512291 | 0.039418036 | 0.059865171 | 0.027352881 | 0.020447135 | 0.41880751 | 0.467782125 | 0.281385296 | 0.61462172 | 0.744770394 |
| Taf2 | S | 1103 | 0.032512291 | 0.039418036 | 0.059865171 | 0.027352881 | 0.020447135 | 0.41880751 | 0.467782125 | 0.281385296 | 0.61462172 | 0.744770394 |
| Bcl9l | S | 1417 | 0.098118156 | 0.081614584 | -0.001570967 | -0.099689123 | -0.083185551 | 0.302358381 | 0.38898508 | 0.990488169 | 0.43768943 | 0.511206142 |
| Cep170 | S | 962 | 0.17826036 | -0.007785388 | 0.005298015 | -0.172962346 | 0.013083403 | 0.111250403 | 0.924581855 | 0.957961252 | 0.104976031 | 0.844455872 |
| Zfp106 | S | 1291 | 0.10523349 | 0.072570679 | -0.04533848 | -0.15057197 | -0.11790916 | 0.104868438 | 0.278770609 | 0.505088032 | 0.06810288 | 0.138916438 |
| Gatad2b | S | 135 | 0.015578477 | 0.061738994 | 0.058291461 | 0.042712984 | -0.003447533 | 0.864505897 | 0.27975165 | 0.242119442 | 0.657577077 | 0.953026456 |
| Arhgap31 | S | 1116 | 0.156199558 | -0.09092941 | 0.0680167 | -0.088182858 | 0.15894611 | 0.02664138 | 0.116713247 | 0.254986087 | 0.185905965 | 0.041793025 |
| Zdhhc5 | S | 529 | 0.005503683 | 0.037289846 | 0.074017414 | 0.068513731 | 0.036727567 | 0.79901006 | 0.406392149 | 0.119011084 | 0.148163685 | 0.47243109 |
| Gpam | S | 687 | 0.004397583 | 0.004758143 | 0.057757929 | 0.053360346 | 0.052999786 | 0.83605972 | 0.753751286 | 0.00526137 | 0.060698532 | 0.01126075 |
| Trim11 | S | 85 | 0.139700674 | -0.005024551 | 0.007615643 | -0.132085031 | 0.012640193 | 0.072666049 | 0.925636929 | 0.891189654 | 0.109484877 | 0.84423205 |
| Ppp1r10 | S | 545 | -0.07684717 | 0.027612519 | 0.141239885 | 0.218087055 | 0.113627366 | 0.294542964 | 0.642034184 | 0.009498174 | 0.05179947 | 0.115626584 |
| Med1 | S | 675 | 0.07585524 | -0.019672829 | 0.092380851 | 0.016525612 | 0.11205368 | 0.323148381 | 0.857858458 | 0.215988313 | 0.686916279 | 0.303211652 |
| Prune2 | S | 2326 | -0.116575738 | 0.093029303 | 0.324369709 | 0.440945447 | 0.231340406 | 0.873501089 | 0.639846529 | 0.41281684 | 0.584666714 | 0.549884638 |
| Egfr | S | 1166 | -0.027079943 | 0.125592965 | 0.033827131 | 0.060907074 | -0.091765834 | 0.546251754 | 0.035469766 | 0.768768787 | 0.590310521 | 0.441636805 |
| Kifc3 | S | 815 | -0.009387965 | 0.035451127 | 0.075943394 | 0.085331359 | 0.040492268 | 0.74977952 | 0.42670953 | 0.041415344 | 0.010681428 | 0.312949926 |
| Cep131 | S | 14 | -0.036612209 | 0.134830896 | 0.05820503 | 0.09481724 | -0.076625866 | 0.570498396 | 0.246538659 | 0.168244259 | 0.182418826 | 0.462085366 |
| Cep131 | T | 10 | -0.036612209 | 0.134830896 | 0.05820503 | 0.09481724 | -0.076625866 | 0.570498396 | 0.246538659 | 0.168244259 | 0.182418826 | 0.462085366 |
| Crtc2 | T | 459 | 0.028670807 | 0.061405482 | 0.031489425 | 0.002818617 | -0.029916057 | 0.594226053 | 0.134713765 | 0.539776487 | 0.958778784 | 0.48547611 |
| Tcerg1 | S | 640 | 0.096178038 | 0.03114003 | -0.020520354 | -0.116698392 | -0.051660384 | 0.084389423 | 0.305452818 | 0.799136985 | 0.228616059 | 0.534777244 |
| Pkp4 | S | 280 | -0.003907428 | 0.034881845 | 0.06588539 | 0.069792818 | 0.031003545 | 0.924583595 | 0.101511815 | 0.161757463 | 0.209212914 | 0.435547653 |
| Aplf | S | 149 | -0.018153545 | 0.05133265 | 0.0611447 | 0.079298246 | 0.00981205 | 0.595291348 | 0.160805333 | 0.071308928 | 0.075036635 | 0.770894644 |
| Scrib | S | 1543 | -0.037907737 | 0.077937649 | 0.162974953 | 0.20088269 | 0.085037304 | 0.34165506 | 0.323743596 | 0.194930548 | 0.148760186 | 0.471386591 |
| Palm | S | 157 | 0.094314452 | 0.030844974 | -0.058296104 | -0.152610555 | -0.089141077 | 0.011167312 | 0.280681615 | 0.377318657 | 0.086480143 | 0.217884569 |
| Champ1 | T | 593 | 0.214920888 | -0.035898191 | -0.04490536 | -0.259826248 | -0.009007169 | 0.029464181 | 0.63691278 | 0.676021291 | 0.068860887 | 0.912950095 |
| Itprid2 | T | 632 | -0.001788323 | 0.122730786 | 0.072888957 | 0.07467728 | -0.049841829 | 0.989836715 | 0.283957883 | 0.510967664 | 0.541798643 | 0.512651587 |
| Pde4a Pde4c | S | 140;121 | -0.041936333 | 0.157106771 | 0.047497583 | 0.089433916 | -0.109609188 | 0.542120462 | 0.095977212 | 0.569738472 | 0.313087333 | 0.268789151 |
| Atrx | S | 1335 | -0.070614854 | 0.082817884 | 0.086773134 | 0.157387988 | 0.00395525 | 0.071460491 | 0.103579135 | 0.014054258 | 0.007448436 | 0.920182164 |
| Zfp219 | S | 710 | -0.017111958 | 0.032295157 | 0.113633032 | 0.13074499 | 0.081337874 | 0.669859337 | 0.524918961 | 0.070930231 | 0.057356526 | 0.214238196 |
| Map1s | S | 462 | -0.005229309 | 0.050194258 | 0.029295922 | 0.034525231 | -0.020898336 | 0.683345193 | 0.034686118 | 0.131983 | 0.091172533 | 0.315101721 |
| Ei24 | S | 326 | 0.064742552 | 0.059634453 | -0.023851866 | -0.088594419 | -0.083486319 | 0.124759731 | 0.120390421 | 0.587176058 | 0.116252184 | 0.124950334 |
| Ei24 | S | 330 | 0.064742552 | 0.059634453 | -0.023851866 | -0.088594419 | -0.083486319 | 0.124759731 | 0.120390421 | 0.587176058 | 0.116252184 | 0.124950334 |
| Cwc22 | S | 91 | -0.051225473 | 0.010493883 | 0.173035516 | 0.224260989 | 0.162541634 | 0.466427124 | 0.8788792 | 0.032434733 | 0.030204106 | 0.076972547 |
| Cwc22 | S | 93 | -0.051225473 | 0.010493883 | 0.173035516 | 0.224260989 | 0.162541634 | 0.466427124 | 0.8788792 | 0.032434733 | 0.030204106 | 0.076972547 |
| Srrm2 | S | 2679 | 0.035420909 | -0.000414033 | 0.073795774 | 0.038374865 | 0.074209808 | 0.540910512 | 0.993890437 | 0.054832838 | 0.507854773 | 0.23221327 |
| Nfrkb | S | 228 | 0.003351946 | -0.02931284 | 0.147167914 | 0.143815968 | 0.176480754 | 0.959251963 | 0.625306595 | 0.031506243 | 0.048365856 | 0.01148834 |
| Srrm2 | S | 867 | 0.064749931 | 0.016225674 | 0.04303961 | -0.021710321 | 0.026813936 | 0.129521949 | 0.698346775 | 0.520729795 | 0.709967456 | 0.657341312 |
| Smarcc1 | S | 329 | 0.00145325 | 0.00226037 | 0.073080586 | 0.071627335 | 0.070820215 | 0.969330036 | 0.944405732 | 0.00859396 | 0.133639552 | 0.091233749 |
| Phlpp1 | S | 1503 | -0.000772391 | 0.1294752 | 0.160469762 | 0.161242153 | 0.030994562 | 0.995969827 | 0.624141673 | 0.44113579 | 0.38147585 | 0.907421634 |
| Bclaf1 | S | 688 | 0.118937292 | 0.028946169 | -0.024395681 | -0.143332972 | -0.053341849 | 0.07560064 | 0.600537474 | 0.826876229 | 0.252825809 | 0.622883795 |
| Zfp687 | S | 1211 | 0.081528957 | 0.01396868 | 0.023677034 | -0.057851923 | 0.009708354 | 0.103466138 | 0.777734977 | 0.555159303 | 0.215855863 | 0.838517606 |
| Irf2bp1 | S | 384 | 0.044796946 | 0.069603792 | -0.007742177 | -0.052539124 | -0.077345969 | 0.308761372 | 0.100025968 | 0.936785767 | 0.595539981 | 0.449082783 |
| Map3k5 | S | 1040 | -0.007887994 | 0.016457997 | 0.115855459 | 0.123743453 | 0.099397462 | 0.862879184 | 0.687267138 | 0.065036732 | 0.062303432 | 0.097728402 |
| Rb1 | S | 31 | -0.02925055 | 0.120997705 | 0.033284237 | 0.062534788 | -0.087713468 | 0.419622076 | 0.052163139 | 0.425594976 | 0.126484046 | 0.128458537 |
| Filip1l | S | 1080 | -0.069740973 | 0.119350828 | 0.116749548 | 0.186490521 | -0.00260128 | 0.433926032 | 0.167149323 | 0.27949872 | 0.095276764 | 0.974578977 |
| Srrm2 | T | 831 | 0.018009159 | 0.002427153 | 0.187081875 | 0.169072716 | 0.184654722 | 0.855717273 | 0.979943585 | 0.202702829 | 0.259595128 | 0.222780905 |
| Kmt2a | S | 2100 | -0.064754383 | 0.044323905 | 0.171636681 | 0.236391064 | 0.127312776 | 0.351896404 | 0.410319046 | 0.074481293 | 0.037049606 | 0.150057961 |
| Nfic | S | 194 | 0.013923649 | 0.089273888 | 0.014696385 | 0.000772737 | -0.074577503 | 0.772509479 | 0.092163012 | 0.613008389 | 0.98544059 | 0.14372177 |
| Rb1 | T | 814 | -0.028517275 | 0.108136446 | 0.049425688 | 0.077942964 | -0.058710757 | 0.528103075 | 0.055484877 | 0.512576621 | 0.347227356 | 0.460953463 |
| Rb1 | T | 819 | -0.028517275 | 0.108136446 | 0.049425688 | 0.077942964 | -0.058710757 | 0.528103075 | 0.055484877 | 0.512576621 | 0.347227356 | 0.460953463 |
| Usp20 | S | 369 | 0.251435488 | -0.005670367 | -0.020976062 | -0.27241155 | -0.015305695 | 0.130396051 | 0.914503808 | 0.688800576 | 0.117879399 | 0.738443544 |
| Atrip | S | 224 | 0.058221032 | 0.005943188 | 0.079077472 | 0.020856441 | 0.073134284 | 0.318485311 | 0.932112621 | 0.26059036 | 0.665301189 | 0.284516862 |
| Lpin1 | S | 106 | -0.018919309 | 0.100815951 | 0.136448766 | 0.155368075 | 0.035632815 | 0.833670814 | 0.466598878 | 0.335446109 | 0.270157735 | 0.81895276 |
| Eif3h | S | 191 | 0.135887277 | 0.087401425 | -0.100217487 | -0.236104764 | -0.187618912 | 0.066287325 | 0.188862581 | 0.226812701 | 0.026332575 | 0.059900254 |
| Lnpk | T | 392 | 0.1892226 | -0.038038115 | 0.076444708 | -0.112777891 | 0.114482824 | 0.309271731 | 0.651475175 | 0.414124606 | 0.502562232 | 0.090436714 |
| Rock2 | S | 425 | 0.150583136 | 0.030363501 | 0.012432 | -0.138151136 | -0.0179315 | 0.310668439 | 0.500145508 | 0.889128929 | 0.369448694 | 0.831482014 |
| G3bp2 | S | 225 | 0.021163888 | 0.021902538 | 0.087881543 | 0.066717655 | 0.065979006 | 0.489341492 | 0.506104957 | 0.233882095 | 0.338634621 | 0.346019053 |
| Kdm6a | S | 829 | 0.105844557 | 0.063917221 | -0.029632471 | -0.135477028 | -0.093549692 | 0.172263221 | 0.257116264 | 0.594493304 | 0.113752338 | 0.155792134 |
| Fxr2 | T | 599 | 0.051434831 | 0.053266865 | 0.032665009 | -0.018769821 | -0.020601856 | 0.420983369 | 0.228516728 | 0.688938194 | 0.830790988 | 0.793448592 |
| Ing5 | S | 148 | -0.014247394 | 0.014452045 | 0.095577436 | 0.109824831 | 0.081125391 | 0.683257566 | 0.641042227 | 0.024587542 | 0.035497771 | 0.062713389 |
| Vrk1 | S | 376 | 0.037367948 | 0.051119056 | 0.017180436 | -0.020187512 | -0.03393862 | 0.501030129 | 0.107404279 | 0.53941969 | 0.688498424 | 0.091194417 |
| Fyb | S | 561 | -0.052449313 | -0.016950303 | 0.242601586 | 0.295050899 | 0.259551889 | 0.383071992 | 0.798886466 | 0.038923084 | 0.03457963 | 0.038633783 |
| H3f3a H3f3b Gm742( | S | 32;32 | 0.122809123 | 0.110427464 | -0.159948686 | -0.282757809 | -0.27037615 | 0.015479186 | 0.256749384 | 0.207202714 | 0.078426989 | 0.080190678 |
| Tmem132c | S | 3 | 0.156145223 | -0.027955335 | 0.007815174 | -0.148330049 | 0.035770509 | 0.054058386 | 0.714675095 | 0.934427676 | 0.154289965 | 0.692385232 |
| Snip1 | S | 33 | 0.023915552 | 0.017982159 | 0.045123208 | 0.021207656 | 0.027141049 | 0.5387376 | 0.381216719 | 0.069302068 | 0.559444376 | 0.096791212 |
| Zfp106 | S | 1041 | 0.10546868 | -0.070305282 | 0.101969532 | -0.003499148 | 0.172274814 | 0.158175791 | 0.049394505 | 0.023314309 | 0.953258983 | 0.005389177 |
| Dock7 | T | 909 | 0.14659484 | 0.028500173 | -0.012524574 | -0.159119414 | -0.041024747 | 0.159930871 | 0.616922328 | 0.903155404 | 0.209507144 | 0.66196193 |
| Lyn | Y | 397 | 0.21416478 | 0.230868625 | -0.141565991 | -0.355730771 | -0.372434616 | 0.134417135 | 0.167350336 | 0.031901482 | 0.056773935 | 0.076710288 |
| Top2b | T | 1390 | 0.035533342 | 0.00974951 | 0.089055512 | 0.05352217 | 0.079306002 | 0.651023967 | 0.871886677 | 0.114304415 | 0.448894705 | 0.145287255 |
| Itsn2 | S | 211 | 0.017464581 | -0.015211782 | 0.111147121 | 0.093682539 | 0.126358903 | 0.732205736 | 0.772944121 | 0.044617256 | 0.133047255 | 0.071429899 |
| Eif3b | S | 123 | 0.013597146 | 0.063357677 | 0.028051294 | 0.014454148 | -0.035306383 | 0.539515789 | 0.223252693 | 0.243891196 | 0.258688414 | 0.443220773 |
| Abcc1 | S | 290 | 0.016128756 | 0.029790639 | 0.064406073 | 0.048277317 | 0.034615434 | 0.670921747 | 0.197380671 | 0.250329477 | 0.406465881 | 0.493682235 |
| Bod1l | S | 2898 | 0.013550211 | 0.01281539 | 0.096700276 | 0.083150065 | 0.083884886 | 0.728813592 | 0.82549377 | 0.089096844 | 0.124709011 | 0.218152123 |
| Mdc1 | S | 750 | 0.031592687 | -0.017242579 | 0.086446292 | 0.054853605 | 0.10368887 | 0.569825473 | 0.655468636 | 0.030058404 | 0.330134092 | 0.031450906 |
| Setd5 | S | 81 | -0.136049209 | 0.098974327 | 0.218627735 | 0.354676944 | 0.119653409 | 0.528329595 | 0.641181133 | 0.11828673 | 0.155576011 | 0.560178916 |
| Setd5 | S | 84 | -0.136049209 | 0.098974327 | 0.218627735 | 0.354676944 | 0.119653409 | 0.528329595 | 0.641181133 | 0.11828673 | 0.155576011 | 0.560178916 |
| Zgrf1 | S | 445 | 0.026166404 | 0.049349739 | 0.089592263 | 0.063425859 | 0.040242524 | 0.495628067 | 0.520128252 | 0.408858134 | 0.548162713 | 0.728164055 |
| Kri1 | S | 142 | 0.096390215 | 0.038945181 | -0.036501222 | -0.132891437 | -0.075446403 | 0.061705375 | 0.272585879 | 0.561574428 | 0.09609066 | 0.261170788 |
| Gpatch8 | S | 1037 | 0.020402941 | 0.021189277 | 0.095381834 | 0.074978892 | 0.074192556 | 0.593224111 | 0.733920224 | 0.166155901 | 0.246608745 | 0.336194814 |

| | | | | | | | | | | | | |
|---|---|---|---|---|---|---|---|---|---|---|---|---|
| Secisbp2l | S | 1081 | 0.041879766 | -0.081178129 | 0.245874657 | 0.20399489 | 0.327052785 | 0.729850446 | 0.32363171 | 0.081392466 | 0.190054344 | 0.044166087 |
| Srrm2 | S | 928 | 0.02558631 | 0.01890511 | 0.036807743 | 0.011221434 | 0.017902633 | 0.305302613 | 0.242823343 | 0.161579138 | 0.678291204 | 0.428216338 |
| Bsdc1 | S | 216 | 0.072899125 | 0.130824702 | 0.045335976 | -0.027563149 | -0.085488726 | 0.777813533 | 0.397593644 | 0.75044779 | 0.911195644 | 0.522460175 |
| Cdk13 | S | 349 | -0.043143608 | 0.063479701 | 0.065057536 | 0.108201144 | 0.001577835 | 0.195434624 | 0.055596958 | 0.055401202 | 0.007178762 | 0.914743907 |
| Cers4 | S | 342 | -0.102050148 | 0.063557783 | 0.164697577 | 0.266747725 | 0.101139794 | 0.290422894 | 0.409367227 | 0.04541864 | 0.03535914 | 0.172787535 |
| Cers4 | S | 349 | -0.102050148 | 0.063557783 | 0.164697577 | 0.266747725 | 0.101139794 | 0.290422894 | 0.409367227 | 0.04541864 | 0.03535914 | 0.172787535 |
| Cers4 | S | 350 | -0.102050148 | 0.063557783 | 0.164697577 | 0.266747725 | 0.101139794 | 0.290422894 | 0.409367227 | 0.04541864 | 0.03535914 | 0.172787535 |
| Sipa1l1 | S | 1600 | 0.080432761 | -0.070082 | 0.153457878 | 0.073025117 | 0.223539878 | 0.545477111 | 0.516067369 | 0.10260385 | 0.599502339 | 0.112537634 |
| Kitl | S | 261 | 0.1858594 | 0.066502212 | -0.051732866 | -0.237592266 | -0.118235078 | 0.205297026 | 0.42519097 | 0.518239154 | 0.133014255 | 0.148144931 |
| Fosl2 | S | 317 | -0.051360965 | -0.062100128 | 0.274233911 | 0.325594876 | 0.336334038 | 0.554614816 | 0.553851172 | 0.034553832 | 0.02490569 | 0.023045111 |
| Zbed6 | S | 32 | 0.092325217 | 0.174083891 | 0.004206849 | -0.088118368 | -0.169877042 | 0.784977006 | 0.335311349 | 0.980404653 | 0.802689864 | 0.425838403 |
| Zbed6 | S | 35 | 0.092325217 | 0.174083891 | 0.004206849 | -0.088118368 | -0.169877042 | 0.784977006 | 0.335311349 | 0.980404653 | 0.802689864 | 0.425838403 |
| Zbed6 | T | 37 | 0.092325217 | 0.174083891 | 0.004206849 | -0.088118368 | -0.169877042 | 0.784977006 | 0.335311349 | 0.980404653 | 0.802689864 | 0.425838403 |
| Rb1cc1 | S | 646 | 0.062400225 | 0.091322686 | -0.054947399 | -0.117347625 | -0.146270085 | 0.328008808 | 0.077354476 | 0.652372074 | 0.370926746 | 0.283547992 |
| Tbc1d22a | T | 51 | 0.261521075 | 0.036251059 | -0.277951034 | -0.539472109 | -0.314202094 | 0.000892808 | 0.766907983 | 0.07019607 | 0.019027616 | 0.08374614 |
| Kdm3b | S | 599 | 0.019263086 | 0.090888677 | 0.002307505 | -0.016955582 | -0.088581172 | 0.766266269 | 0.056041646 | 0.954038496 | 0.771591238 | 0.025205856 |
| Virma | T | 184 | -0.002323177 | 0.104744008 | 0.042685824 | 0.045009001 | -0.062058184 | 0.979242902 | 0.089272765 | 0.801258796 | 0.798551741 | 0.712250494 |
| Use1 | S | 165 | 0.051550454 | 0.043432812 | 0.029636252 | -0.021914201 | -0.01379656 | 0.25269964 | 0.4394567 | 0.52596506 | 0.626538735 | 0.803136448 |
| Msl2 | S | 211 | 0.030048281 | 0.03747866 | 0.061637356 | 0.031589075 | 0.024158696 | 0.342922467 | 0.215861278 | 0.611294822 | 0.7874562 | 0.835429287 |
| Nipbl | S | 306 | 0.013698635 | 0.02938586 | 0.082635552 | 0.068936917 | 0.053249691 | 0.77907827 | 0.712660635 | 0.101370139 | 0.217351314 | 0.519760354 |
| Tmcc3 | S | 242 | 0.215644295 | -0.092864166 | -0.015863983 | -0.231508279 | 0.077000182 | 0.010235793 | 0.318934384 | 0.700697163 | 0.010320748 | 0.396731858 |
| Hexim1 | S | 103 | 0.005073388 | 0.057803802 | 0.104612978 | 0.099539589 | 0.046809175 | 0.947817297 | 0.451614132 | 0.28581469 | 0.247213862 | 0.541838814 |
| Tubgcp5 | S | 165 | 0.068651363 | -0.132302711 | 0.180919331 | 0.112267968 | 0.313222042 | 0.462420017 | 0.279524574 | 0.036669393 | 0.216499561 | 0.053125776 |
| Tubgcp5 | S | 168 | 0.068651363 | -0.132302711 | 0.180919331 | 0.112267968 | 0.313222042 | 0.462420017 | 0.279524574 | 0.036669393 | 0.216499561 | 0.053125776 |
| Zgpat | S | 70 | 0.088499075 | 0.030125954 | 0.054264682 | -0.034234393 | 0.024138728 | 0.420297784 | 0.617809487 | 0.495435007 | 0.759466006 | 0.751208068 |
| Mprip | S | 371 | -0.159897137 | 0.169102542 | 0.181787573 | 0.34168471 | 0.012685032 | 0.012621572 | 0.100159985 | 0.028306406 | 0.008683261 | 0.881265197 |
| Myo18a | S | 1064 | 0.020879453 | 0.153200367 | -0.042049279 | -0.062928731 | -0.195249645 | 0.686019625 | 0.049410574 | 0.483023678 | 0.386431274 | 0.038802961 |
| Pdzd8 | S | 968 | -0.054803487 | -0.128617024 | 0.347929054 | 0.402732541 | 0.476546079 | 0.798514277 | 0.355557656 | 0.037811732 | 0.121873573 | 0.000505346 |
| Septin2 | S | 218 | -0.015565592 | 0.066801178 | 0.039313826 | 0.054879419 | -0.027487352 | 0.692720848 | 0.082124572 | 0.182599807 | 0.221688043 | 0.414625447 |
| Dtl | S | 598 | 0.0787021 | -0.014113108 | 0.046917138 | -0.031784962 | 0.061030246 | 0.082540398 | 0.723656032 | 0.370886521 | 0.558661348 | 0.312073577 |
| Col12a1 | S | 1320 | 0.014985099 | 0.000419156 | 0.080883966 | 0.065898867 | 0.08046481 | 0.771046421 | 0.990735848 | 0.032449042 | 0.248222281 | 0.066997012 |
| Pcyt1a | S | 319 | -0.056293227 | 0.122180027 | -0.007337051 | 0.048956176 | -0.129517078 | 0.257491273 | 0.001130413 | 0.774603144 | 0.30698147 | 0.003457689 |
| Rbm8a Rbm8a2 | S | 56 | 0.04657246 | 0.01848748 | 0.055784921 | 0.00921246 | 0.037297441 | 0.263091968 | 0.747095679 | 0.27599274 | 0.831715054 | 0.537494563 |
| Pias2 | S | 499 | 0.077107008 | 0.041148965 | -0.037916598 | -0.115023606 | -0.079065563 | 0.153435661 | 0.056895283 | 0.626304971 | 0.221206644 | 0.353938989 |
| Sash1 | S | 831 | -0.081627464 | 0.139624946 | 0.078556991 | 0.160184455 | -0.061067955 | 0.278140474 | 0.105510296 | 0.204299956 | 0.090594302 | 0.42450364 |
| Atg2a | S | 875 | -0.01437062 | 0.071053694 | 0.070557826 | 0.084928446 | -0.000495868 | 0.837299176 | 0.194767046 | 0.27967544 | 0.23958379 | 0.991028729 |
| Atf7ip | S | 533 | -0.022159352 | 0.020087991 | 0.089167117 | 0.111326469 | 0.069079126 | 0.629236727 | 0.549736972 | 0.022490228 | 0.075877079 | 0.103272986 |
| Nhsl1 | T | 1547 | -0.021719969 | 0.104184709 | 0.089731248 | 0.111451218 | -0.01445346 | 0.780848565 | 0.295472671 | 0.350450864 | 0.329022231 | 0.897494218 |
| Myh9 | S | 1660 | -0.025208967 | 0.081693289 | 0.067853404 | 0.093062371 | -0.013839885 | 0.337096523 | 0.055313749 | 0.344673837 | 0.231350599 | 0.833751544 |
| Larp1 | T | 1048 | 0.384020063 | 0.018382164 | -0.375873153 | -0.759893216 | -0.394255317 | 0.011290273 | 0.864190704 | 0.202349237 | 0.064899664 | 0.197099021 |
| Zfp462 | S | 1736 | 0.045754355 | -0.026017142 | 0.095261516 | 0.049507162 | 0.121278658 | 0.335059792 | 0.614603505 | 0.090970002 | 0.372003546 | 0.095844394 |
| Psip1 | S | 106 | -0.09181109 | 0.030522116 | 0.192360813 | 0.284171903 | 0.161838697 | 0.233076086 | 0.465060906 | 0.035495845 | 0.018989437 | 0.062243995 |
| Slc20a2 | S | 385 | -0.00318652 | 0.054971136 | 0.06595569 | 0.06914221 | 0.010984554 | 0.947968681 | 0.568138724 | 0.06454863 | 0.203511241 | 0.904072285 |
| Pbrm1 | S | 353 | -0.019697445 | 0.038857024 | 0.076252892 | 0.095950337 | 0.037395868 | 0.144594392 | 0.03362109 | 0.098607625 | 0.074703782 | 0.316727917 |
| Pbrm1 | S | 355 | -0.019697445 | 0.038857024 | 0.076252892 | 0.095950337 | 0.037395868 | 0.144594392 | 0.03362109 | 0.098607625 | 0.074703782 | 0.316727917 |
| Prrc2c | S | 1472 | 0.120577107 | 0.022347248 | 0.031583469 | -0.088993638 | 0.00923622 | 0.243227193 | 0.783434386 | 0.722296163 | 0.294655881 | 0.864272419 |
| Prrc2c | S | 1475 | 0.120577107 | 0.022347248 | 0.031583469 | -0.088993638 | 0.00923622 | 0.243227193 | 0.783434386 | 0.722296163 | 0.294655881 | 0.864272419 |
| Prrc2c | T | 1470 | 0.120577107 | 0.022347248 | 0.031583469 | -0.088993638 | 0.00923622 | 0.243227193 | 0.783434386 | 0.722296163 | 0.294655881 | 0.864272419 |
| Zc3h13 | S | 929 | -0.036178015 | 0.019767507 | 0.08676199 | 0.122940005 | 0.066994482 | 0.677735801 | 0.682609141 | 0.012128734 | 0.234684242 | 0.22032465 |
| Gas2 | S | 184 | -0.024574652 | 0.11349184 | 0.040655656 | 0.065230308 | -0.072836184 | 0.891489519 | 0.227738954 | 0.272441732 | 0.721868778 | 0.397031632 |
| Golim4 | S | 328 | 0.190192746 | -0.024887583 | 0.044309077 | -0.145883668 | 0.069196661 | 0.272468282 | 0.323358026 | 0.225543413 | 0.370923973 | 0.096584485 |
| Sun2 | S | 83 | -0.02871018 | 0.004987857 | 0.128219085 | 0.156929265 | 0.123231228 | 0.620894848 | 0.90666614 | 0.026754127 | 0.033445787 | 0.022245169 |
| Riok3 | S | 512 | -0.032863691 | 0.138879915 | 0.027314817 | 0.060178508 | -0.111565099 | 0.562176515 | 0.057606009 | 0.721183742 | 0.405339453 | 0.182978031 |
| Mga | S | 769 | 0.201398361 | 0.010578196 | -0.022249145 | -0.223647506 | -0.032827342 | 0.167127581 | 0.913131297 | 0.913131297 | 0.347382849 | 0.876747803 |
| Ripk2 | T | 481 | 0.048364847 | 0.051983119 | 0.107504696 | 0.059139849 | 0.055521577 | 0.772013201 | 0.722705387 | 0.331872827 | 0.693332897 | 0.657158039 |
| Sec16a | S | 2053 | 0.024928041 | 0.130081828 | -0.045293434 | -0.070221475 | -0.175375262 | 0.774147864 | 0.018440962 | 0.357942311 | 0.429517831 | 0.007359361 |
| Eif4ebp1 | S | 100 | -0.17451145 | 0.177071482 | 0.183850344 | 0.358361794 | 0.006778862 | 0.05290849 | 0.169931483 | 0.050549255 | 0.008877132 | 0.951670479 |
| Frmd6 | S | 372 | -0.00703882 | 0.081911175 | 0.047071832 | 0.054110652 | -0.034839343 | 0.789757757 | 0.195337004 | 0.238493179 | 0.182691003 | 0.548032873 |
| Kmt2c | S | 695 | 0.065379734 | 0.001084768 | 0.0562292 | -0.009150534 | 0.055144432 | 0.21716499 | 0.986234021 | 0.287434563 | 0.794099528 | 0.337431326 |
| Svil | S | 1278 | -0.017121246 | 0.001329911 | 0.097185567 | 0.114306812 | 0.095855656 | 0.70084824 | 0.978432804 | 0.011800794 | 0.073514525 | 0.138479354 |
| Dvl1 | S | 194 | 0.021560858 | 0.096897701 | -0.000603241 | -0.0221641 | -0.097500943 | 0.657052594 | 0.082759495 | 0.988872365 | 0.48701013 | 0.051065666 |
| Map7d1 | S | 114 | 0.022172286 | 0.148946125 | -0.043720812 | -0.065893098 | -0.192666937 | 0.75706971 | 0.057872565 | 0.612618498 | 0.392714944 | 0.058021792 |
| Myo18a | S | 1938 | -0.009906478 | 0.024369098 | 0.073158778 | 0.083065256 | 0.04878968 | 0.785116337 | 0.474680492 | 0.033801676 | 0.057096027 | 0.120567184 |
| Tns2 | S | 811 | 0.140865817 | 0.014919045 | 0.027818662 | -0.113047156 | 0.012899617 | 0.220073529 | 0.916358421 | 0.768231919 | 0.362730743 | 0.932538009 |
| Acd | T | 297 | 0.091490459 | 0.067898983 | 0.029043015 | -0.062447445 | -0.038855969 | 0.487364358 | 0.423639669 | 0.778655968 | 0.670634173 | 0.734206706 |
| Specc1l | S | 54 | 0.036220191 | 0.038433249 | 0.016326055 | -0.019894136 | -0.022107195 | 0.171926475 | 0.26531398 | 0.510898269 | 0.209781855 | 0.434895709 |
| Prpf4b | S | 21 | -0.011030562 | 0.042701884 | 0.085860149 | 0.096890711 | 0.043158265 | 0.852395532 | 0.135582642 | 0.275751134 | 0.273919892 | 0.536588853 |
| Prpf4b | S | 24 | -0.011030562 | 0.042701884 | 0.085860149 | 0.096890711 | 0.043158265 | 0.852395532 | 0.135582642 | 0.275751134 | 0.273919892 | 0.536588853 |
| Tra2a | S | 101 | -0.060214748 | 0.05320574 | 0.17146734 | 0.231682088 | 0.118261601 | 0.570748393 | 0.45350707 | 0.159227939 | 0.112510627 | 0.284676271 |
| Unkl | S | 346 | 0.056189399 | 0.020476222 | 0.117868912 | 0.061679513 | 0.09739269 | 0.469328094 | 0.77783473 | 0.454980846 | 0.665891524 | 0.508165657 |
| Nisch | S | 669 | 0.07834422 | 0.046638747 | -0.053531156 | -0.131875376 | -0.100169903 | 0.01487436 | 0.23217611 | 0.30321967 | 0.063745731 | 0.107799283 |
| Xkr5 | S | 458 | 0.047257151 | 0.020758582 | 0.089617036 | 0.042359885 | 0.068858454 | 0.459732091 | 0.741600912 | 0.338138215 | 0.655661229 | 0.482380671 |
| Sipa1l1 | T | 1552 | -0.009475251 | 0.041699696 | 0.089130855 | 0.098606106 | 0.047431159 | 0.74754014 | 0.173918618 | 0.225415966 | 0.195328724 | 0.4666239 |
| Tmem131 | S | 1601 | 0.099845541 | 0.042181318 | 0.009231597 | -0.090613944 | -0.032949721 | 0.261081852 | 0.445409323 | 0.905205271 | 0.37387274 | 0.684153318 |
| Nuf2 | S | 247 | 0.007887504 | 0.12855172 | -0.070115089 | -0.078002594 | -0.198666809 | 0.828956633 | 0.007289205 | 0.401141582 | 0.356446217 | 0.081794943 |
| Myh9 | T | 1657 | 0.017167693 | 0.079171219 | 0.030311814 | 0.013144121 | -0.048859405 | 0.719997715 | 0.173190497 | 0.557610299 | 0.67755993 | 0.274273011 |
| Ubap2l | S | 625 | 0.116621007 | 0.038548322 | -0.064543493 | -0.1811645 | -0.103091815 | 0.010653323 | 0.307220836 | 0.133512861 | 0.005342762 | 0.026010915 |
| Ncor2 | S | 2371 | -0.071813328 | 0.074526327 | 0.135991389 | 0.207804717 | 0.061465062 | 0.249151159 | 0.115107025 | 0.165848977 | 0.069576085 | 0.457270301 |
| Ralgapb | S | 414 | 0.018403761 | 0.075746164 | 0.047313623 | 0.028909862 | -0.028432541 | 0.621230676 | 0.23059407 | 0.610989896 | 0.74483223 | 0.767182248 |
| Chaf1b | S | 435 | 0.067407333 | 0.006049799 | 0.034062935 | -0.033344398 | 0.028013136 | 0.136498121 | 0.922766161 | 0.343564381 | 0.390191758 | 0.64920903 |
| Chaf1b | T | 440 | 0.067407333 | 0.006049799 | 0.034062935 | -0.033344398 | 0.028013136 | 0.136498121 | 0.922766161 | 0.343564381 | 0.390191758 | 0.64920903 |
| Msl1 | S | 403 | 0.158411383 | -0.036986709 | 0.010591146 | -0.147820237 | 0.047577855 | 0.0675835 | 0.74537973 | 0.921519883 | 0.189864624 | 0.695300274 |
| Msl1 | S | 394 | 0.158411383 | -0.036986709 | 0.010591146 | -0.147820237 | 0.047577855 | 0.0675835 | 0.74537973 | 0.921519883 | 0.189864624 | 0.695300274 |
| Hirip3 | S | 575 | -0.029440662 | 0.072725713 | 0.03910119 | 0.068541852 | -0.033624523 | 0.125971792 | 0.025226987 | 0.088583789 | 0.021383088 | 0.206910803 |
| Nsd2 | T | 110 | 0.079915584 | 0.046755942 | 0.006958803 | -0.072956781 | -0.039797139 | 0.250285604 | 0.343905609 | 0.935817951 | 0.466206388 | 0.661884783 |
| Mast1 Mast2 Mast4 | S | 1337;1436;1260 | 0.273806403 | -0.061416774 | -0.04797685 | -0.321783253 | 0.013439924 | 0.024008635 | 0.462347253 | 0.380608112 | 0.031971092 | 0.84549139 |
| Atad5 | S | 1209 | 0.128032364 | -0.012263536 | -0.000817538 | -0.128849901 | 0.011445999 | 0.04609028 | 0.834156189 | 0.988919544 | 0.088919526 | 0.86221954 |
| Mphosph8 | S | 261 | 0.075886507 | 0.116938832 | -0.022513241 | -0.098399748 | -0.139452073 | 0.429876914 | 0.297287654 | 0.884702781 | 0.567239486 | 0.438221539 |
| Champ1 | S | 642 | -0.023261664 | 0.02018237 | 0.127527967 | 0.150789631 | 0.107345597 | 0.599256739 | 0.433124675 | 0.088320226 | 0.062304369 | 0.134428219 |
| Xrn2 | S | 485 | 0.062752476 | 0.039344804 | -0.012108692 | -0.074861169 | -0.051453496 | 0.090042787 | 0.225546279 | 0.787808042 | 0.132201721 | 0.24899976 |
| Ehbp1l1 | S | 1093 | 0.197824439 | 0.175769139 | -0.173890481 | -0.37171492 | -0.34965962 | 0.067711667 | 0.445501708 | 0.199042468 | 0.037688595 | 0.191473238 |
| Dennd2a | S | 400 | 0.140703311 | 0.058937506 | -0.087173309 | -0.22787662 | -0.146110815 | 0.025479442 | 0.162979128 | 0.080431723 | 0.006374834 | 0.012596807 |
| Lpin1 | T | 105 | -0.119545548 | 0.326216349 | 0.048789356 | 0.168334904 | -0.277426992 | 0.551808118 | 0.20316736 | 0.735625522 | 0.43134866 | 0.270415234 |
| Bclaf3 | S | 205 | 0.045469457 | 0.012275945 | 0.063777254 | 0.018307797 | 0.051501309 | 0.335422443 | 0.59863913 | 0.306914668 | 0.774323969 | 0.39385462 |

| | | | | | | | | | | | | |
|---|---|---|---|---|---|---|---|---|---|---|---|---|
| Pacs1 | S | 527 | 0.09931083 | 0.038603666 | 0.014194656 | -0.085116175 | -0.02440901 | 0.225959455 | 0.605966484 | 0.818660145 | 0.219899377 | 0.661185374 |
| Pacs1 | S | 532 | 0.09931083 | 0.038603666 | 0.014194656 | -0.085116175 | -0.02440901 | 0.225959455 | 0.605966484 | 0.818660145 | 0.219899377 | 0.661185374 |
| Camkk2 | S | 133 | 0.018768689 | 0.077875368 | 0.00152746 | -0.017241229 | -0.076347909 | 0.808927516 | 0.037071317 | 0.968126381 | 0.822444829 | 0.070978032 |
| Camkk2 | S | 137 | 0.018768689 | 0.077875368 | 0.00152746 | -0.017241229 | -0.076347909 | 0.808927516 | 0.037071317 | 0.968126381 | 0.822444829 | 0.070978032 |
| Adgrl2 | S | 1437 | -0.045843811 | 0.007107437 | 0.147875485 | 0.193719295 | 0.140768048 | 0.582963647 | 0.854076169 | 0.036326035 | 0.085807326 | 0.048717667 |
| Taf3 | T | 797 | -0.02236281 | 0.04809025 | 0.130801496 | 0.153164306 | 0.082711246 | 0.748572525 | 0.453881484 | 0.195703213 | 0.154916013 | 0.375862795 |
| Wasl | S | 480 | -0.06373704 | 0.089710616 | 0.071663427 | 0.135400467 | -0.018047189 | 0.299084673 | 0.058898968 | 0.181634175 | 0.081540602 | 0.71060542 |
| Wasl | S | 481 | -0.06373704 | 0.089710616 | 0.071663427 | 0.135400467 | -0.018047189 | 0.299084673 | 0.058898968 | 0.181634175 | 0.081540602 | 0.71060542 |
| Utp18 | S | 118 | 0.060129113 | 0.013426943 | 0.016442052 | -0.043687061 | 0.003015109 | 0.049328661 | 0.675548711 | 0.693082391 | 0.303009486 | 0.941328908 |
| Nedd4l | S | 512 | 0.086998982 | 0.09111473 | -0.051285992 | -0.138284974 | -0.142400722 | 0.109642575 | 0.261111388 | 0.413609466 | 0.083998859 | 0.137671424 |
| Cpeb2 | S | 21 | 0.092547246 | -0.032032024 | 0.020304547 | -0.072242699 | 0.052336571 | 0.004572569 | 0.191111058 | 0.614627084 | 0.136911051 | 0.230348646 |
| Actn4 | S | 382 | 0.001098105 | 0.045365534 | 0.075293013 | 0.074194908 | 0.029927479 | 0.985674966 | 0.307957302 | 0.215939907 | 0.275845814 | 0.530862445 |
| Thoc2 | S | 1516 | -0.056781522 | 0.031503041 | 0.143915914 | 0.200697435 | 0.112412873 | 0.290931688 | 0.573983157 | 0.030965886 | 0.014786426 | 0.098570251 |
| Epn2 | S | 172 | 0.080233556 | 0.013029596 | 0.045955572 | -0.034277984 | 0.032925976 | 0.189168098 | 0.823360476 | 0.578361482 | 0.687026277 | 0.703901568 |
| Kdm4c | S | 395 | -0.027018398 | 0.011421206 | 0.131868053 | 0.158886451 | 0.120446847 | 0.417322642 | 0.77287026 | 0.032017099 | 0.019902916 | 0.056356441 |
| Vim | T | 446 | 0.035440308 | 0.039421491 | 0.069293217 | 0.033852909 | 0.009871726 | 0.639578438 | 0.56474718 | 0.276983984 | 0.552978678 | 0.501341367 |
| Mast2 | S | 1462 | 0.002450506 | 0.058784406 | 0.060097061 | 0.057646555 | 0.001312655 | 0.920592945 | 0.29577005 | 0.242172998 | 0.257276698 | 0.982456234 |
| Mgrn1 | S | 449 | -0.053443526 | 0.025810158 | 0.180494874 | 0.2339384 | 0.154684715 | 0.236374638 | 0.640350049 | 0.046554091 | 0.019349297 | 0.07828677 |
| Mgrn1 | S | 452 | -0.053443526 | 0.025810158 | 0.180494874 | 0.2339384 | 0.154684715 | 0.236374638 | 0.640350049 | 0.046554091 | 0.019349297 | 0.07828677 |
| Mtss2 | S | 575 | -0.130541241 | 0.116538574 | 0.172655403 | 0.303196644 | 0.056116829 | 0.172347086 | 0.234861423 | 0.0923405 | 0.026423593 | 0.572850571 |
| Ino80b | S | 209 | -0.028829495 | 0.133237748 | 0.08891608 | 0.117745575 | -0.044321669 | 0.823847406 | 0.294333724 | 0.511855533 | 0.487246907 | 0.774531036 |
| Ptpn14 | S | 710 | 0.082682311 | 0.056106757 | 0.053483903 | -0.029198408 | -0.002622854 | 0.422819861 | 0.613074995 | 0.692366921 | 0.841249158 | 0.986171938 |
| Ptpn14 | S | 711 | 0.082682311 | 0.056106757 | 0.053483903 | -0.029198408 | -0.002622854 | 0.422819861 | 0.613074995 | 0.692366921 | 0.841249158 | 0.986171938 |
| Ptpn14 | S | 713 | 0.082682311 | 0.056106757 | 0.053483903 | -0.029198408 | -0.002622854 | 0.422819861 | 0.613074995 | 0.692366921 | 0.841249158 | 0.986171938 |
| Sec62 | S | 345 | 0.111833781 | 0.136859226 | -0.053656028 | -0.165489809 | -0.190515254 | 0.509513973 | 0.322751662 | 0.84349969 | 0.55143474 | 0.482576302 |
| Cbx3 | S | 97 | -0.016652388 | 0.012337034 | 0.080126011 | 0.096778399 | 0.067788978 | 0.482759439 | 0.418058796 | 0.015326173 | 0.01543973 | 0.049889719 |
| Tpd52l2 | S | 206 | -0.011915871 | 0.067138327 | 0.02888732 | 0.040803191 | -0.038251007 | 0.713590013 | 0.022319031 | 0.557800299 | 0.429159575 | 0.432110826 |
| Cbx8 | S | 229 | -0.07616213 | 0.107586416 | 0.178470423 | 0.254632553 | 0.070884006 | 0.228429363 | 0.159195221 | 0.30179221 | 0.185843796 | 0.647334676 |
| Zgrf1 | S | 701 | -0.083321723 | 0.029763475 | 0.183190134 | 0.266511857 | 0.153426659 | 0.226007234 | 0.141664252 | 0.078909045 | 0.026174514 | 0.116087628 |
| Inpp5f | S | 830 | 0.041346579 | 0.035728196 | 0.022948621 | -0.018397958 | -0.012779575 | 0.259090726 | 0.239499214 | 0.597768223 | 0.66672682 | 0.740688471 |
| Phip | S | 911 | 0.010999382 | 0.022711554 | 0.094021719 | 0.083022336 | 0.071310164 | 0.801523003 | 0.49662571 | 0.189393286 | 0.25069326 | 0.289851084 |
| Gatad2b | T | 489 | 0.116275659 | 0.039808394 | 0.048934462 | -0.067341197 | 0.009126068 | 0.385658819 | 0.765023752 | 0.690015687 | 0.419251187 | 0.91338269 |
| Rbm12b2 | S | 562 | 0.094879787 | -0.039665836 | 0.113289483 | 0.018409697 | 0.152955319 | 0.338516054 | 0.332429326 | 0.156322876 | 0.854192677 | 0.086815096 |
| Dst | S | 3894 | 0.002897915 | 0.048729102 | 0.041564692 | 0.038666777 | -0.00716441 | 0.936661189 | 0.216987805 | 0.145062582 | 0.297373803 | 0.827664976 |
| Mef2a | S | 444 | -0.055127449 | 0.026574315 | 0.21823594 | 0.27336339 | 0.191661626 | 0.302562644 | 0.567153128 | 0.103447691 | 0.063966629 | 0.135259201 |
| Lyn | T | 504 | -0.299623345 | 0.372305945 | 0.058992209 | 0.358615553 | -0.313313737 | 0.295038385 | 0.048217397 | 0.809537042 | 0.260048954 | 0.224432021 |
| Arhgap21 | S | 1703 | -0.067522176 | 0.094790126 | 0.098064826 | 0.165587002 | 0.003274701 | 0.284087363 | 0.271035444 | 0.071477824 | 0.035393703 | 0.963121443 |
| Stx17 | S | 288 | 0.072913029 | 0.017016761 | 0.027187234 | -0.045725794 | 0.010170474 | 0.134871341 | 0.744650646 | 0.610905205 | 0.405644237 | 0.862099545 |
| Kmt2b | T | 953 | 0.275399097 | -0.094798376 | 0.049121562 | -0.226277534 | 0.143919938 | 0.199981287 | 0.205965896 | 0.181365686 | 0.266279566 | 0.110694742 |
| Srsf4 | S | 327 | -0.018490075 | 0.039412855 | 0.067350699 | 0.085840775 | 0.027937845 | 0.860863745 | 0.562467715 | 0.037549726 | 0.453736223 | 0.678989298 |
| Srsf4 | S | 331 | -0.018490075 | 0.039412855 | 0.067350699 | 0.085840775 | 0.027937845 | 0.860863745 | 0.562467715 | 0.037549726 | 0.453736223 | 0.678989298 |
| Dcaf11 | S | 73 | -0.121723426 | 0.137141934 | 0.087155 | 0.208878427 | -0.049986934 | 0.21742922 | 0.021313492 | 0.427018207 | 0.133644106 | 0.618754017 |
| Dcaf11 | S | 75 | -0.121723426 | 0.137141934 | 0.087155 | 0.208878427 | -0.049986934 | 0.21742922 | 0.021313492 | 0.427018207 | 0.133644106 | 0.618754017 |
| Acta1 Acta2 Actc1 Ac | T | 68;68;67;68 | 0.056027179 | 0.051402724 | -0.011655028 | -0.067682207 | -0.063057752 | 0.166079658 | 0.202313218 | 0.891526773 | 0.444658385 | 0.472470234 |
| Tex30 | S | 110 | -0.024686923 | 0.003833397 | 0.104971498 | 0.129658422 | 0.101138102 | 0.655304568 | 0.932305461 | 0.017276091 | 0.079722134 | 0.079671864 |
| Tex30 | T | 115 | -0.024686923 | 0.003833397 | 0.104971498 | 0.129658422 | 0.101138102 | 0.655304568 | 0.932305461 | 0.017276091 | 0.079722134 | 0.079671864 |
| Ipmk | S | 19 | 0.000964189 | 0.027555927 | 0.065931807 | 0.064967618 | 0.03837588 | 0.96473449 | 0.209399289 | 0.150594755 | 0.158739018 | 0.341360024 |
| Serbp1 | S | 390 | 0.008249432 | 0.094135948 | 0.040046927 | 0.031797495 | -0.054089021 | 0.928162305 | 0.17737571 | 0.620477095 | 0.726912955 | 0.427263871 |
| Cdk13 | S | 1228 | -0.056074943 | 0.141410819 | 0.109233486 | 0.16530843 | -0.032177333 | 0.638019662 | 0.259147215 | 0.467440632 | 0.199118836 | 0.755188583 |
| Cdk13 | T | 1225 | -0.056074943 | 0.141410819 | 0.109233486 | 0.16530843 | -0.032177333 | 0.638019662 | 0.259147215 | 0.467440632 | 0.199118836 | 0.755188583 |
| Phip | S | 1315 | 0.26670368 | -0.077663463 | -0.032417623 | -0.299121303 | 0.045245841 | 0.008005301 | 0.070689666 | 0.666478013 | 0.019192452 | 0.535358454 |
| Dsn1 | S | 331 | 0.037499017 | 0.045254903 | -0.041920657 | -0.079419674 | -0.08717556 | 0.149733261 | 0.000580716 | 0.151355127 | 0.038273096 | 0.042439212 |
| Fnbp1 | S | 358 | 0.000329373 | 0.072979527 | 0.027177331 | 0.026847958 | -0.045802196 | 0.99674154 | 0.113280777 | 0.361346156 | 0.732952448 | 0.241633597 |
| Irs2 | T | 517 | 0.007502394 | 0.015970921 | 0.085707724 | 0.07820533 | 0.069736804 | 0.880641641 | 0.729310533 | 0.081082957 | 0.198863922 | 0.212633917 |
| Plekhg3 | S | 1199 | 0.021378559 | 0.097910044 | 0.004798728 | -0.016579831 | -0.093111316 | 0.658129695 | 0.123558152 | 0.932719327 | 0.779945179 | 0.195976148 |
| Srrm2 | S | 2681 | 0.028010936 | 0.00507859 | 0.069537838 | 0.041526902 | 0.064459248 | 0.633620312 | 0.922587469 | 0.068307271 | 0.494242062 | 0.277013191 |
| Kat7 | T | 90 | 0.071903931 | 0.013250311 | -0.001208897 | -0.073112828 | -0.014459208 | 0.038833158 | 0.549767134 | 0.959437258 | 0.047258785 | 0.532840464 |
| Map1b | S | 1317 | -0.039808959 | 0.108751197 | 0.027797327 | 0.067606286 | -0.08095387 | 0.167701031 | 0.008872071 | 0.609066417 | 0.265447181 | 0.203365036 |
| Pcm1 | S | 1434 | -0.016457816 | -0.022248425 | 0.203107227 | 0.219565043 | 0.225355652 | 0.790343221 | 0.650762119 | 0.075679894 | 0.067401709 | 0.074438912 |
| Nono | S | 149 | 0.013403291 | 0.058546328 | 0.052152337 | 0.038749046 | -0.006393991 | 0.695246659 | 0.314883841 | 0.35006215 | 0.448743697 | 0.918183524 |
| Ints1 | S | 1320 | 0.059327001 | 0.057785964 | 0.012349729 | -0.046977272 | -0.045436235 | 0.330263674 | 0.316795573 | 0.822327656 | 0.492184603 | 0.491536523 |
| Ints1 | S | 1328 | 0.059327001 | 0.057785964 | 0.012349729 | -0.046977272 | -0.045436235 | 0.330263674 | 0.316795573 | 0.822327656 | 0.492184603 | 0.491536523 |
| Setd2 | S | 624 | 0.007573224 | 0.048844564 | 0.06331796 | 0.055744736 | 0.014473396 | 0.76160785 | 0.463314306 | 0.190752743 | 0.238364303 | 0.832736484 |
| Peak1 | S | 727 | 0.059614297 | 0.103626988 | -0.022883729 | -0.082498026 | -0.126510717 | 0.387218104 | 0.168217228 | 0.619868032 | 0.254653088 | 0.116267302 |
| Ppp4r1 | S | 945 | -0.02112101 | 0.049290585 | 0.065703528 | 0.086824538 | 0.016412943 | 0.625876 | 0.205029073 | 0.107516292 | 0.056566161 | 0.395238435 |
| C77080 | S | 419 | -0.117704969 | 0.636451816 | -0.186814496 | -0.069109527 | -0.823266312 | 0.669074725 | 0.089192473 | 0.321870745 | 0.758503757 | 0.071681852 |
| Gigyf2 | S | 30 | 0.267569569 | -0.05591103 | -0.095269865 | -0.362839434 | -0.039358836 | 0.005031242 | 0.612894303 | 0.106284633 | 0.003577712 | 0.709316192 |
| Vim | S | 65 | 0.096896337 | 0.124707382 | -0.137944866 | -0.234841202 | -0.262652248 | 0.040145293 | 0.160391459 | 0.196115163 | 0.075236552 | 0.054380706 |
| Rbmxl1 | S | 244 | 0.085126403 | 0.033930421 | 0.040443166 | -0.044683237 | 0.006512745 | 0.277879706 | 0.728396953 | 0.667337519 | 0.630625009 | 0.952272795 |
| Rbmxl1 | S | 247 | 0.085126403 | 0.033930421 | 0.040443166 | -0.044683237 | 0.006512745 | 0.277879706 | 0.728396953 | 0.667337519 | 0.630625009 | 0.952272795 |
| Mcrs1 | S | 282 | 0.015795811 | -0.043174167 | 0.0990526 | 0.083256789 | 0.142226767 | 0.649009774 | 0.374376337 | 0.008754962 | 0.048655946 | 0.050052275 |
| Hivep2 | S | 1077 | -0.069634969 | -0.010219017 | 0.136865908 | 0.206500877 | 0.147084924 | 0.423702371 | 0.824673604 | 0.005068271 | 0.086804624 | 0.027644303 |
| Atg2a | S | 877 | -0.023756875 | 0.07723269 | 0.064110212 | 0.087867087 | -0.013122478 | 0.731135366 | 0.16480572 | 0.308669129 | 0.207941964 | 0.747123956 |
| Prpf38b | S | 319 | 0.045806036 | 0.042880553 | 0.029445251 | -0.016360786 | -0.013435302 | 0.336904413 | 0.354108311 | 0.577511487 | 0.767958691 | 0.805427541 |
| Prpf38b | S | 321 | 0.045806036 | 0.042880553 | 0.029445251 | -0.016360786 | -0.013435302 | 0.336904413 | 0.354108311 | 0.577511487 | 0.767958691 | 0.805427541 |
| Arhgap21 | S | 41 | -0.04455934 | 0.034656665 | 0.131219798 | 0.175779139 | 0.096563133 | 0.335615302 | 0.353584833 | 0.069164452 | 0.035387959 | 0.150434783 |
| Vcp | S | 775 | 0.149081211 | -0.011941953 | 0.027508694 | -0.121572518 | 0.039450647 | 0.154195365 | 0.900488274 | 0.842261195 | 0.442006834 | 0.800952796 |
| Ralbp1 | S | 48 | 0.04081021 | 0.097722227 | -0.061429555 | -0.102239765 | -0.159151782 | 0.443474237 | 0.003219034 | 0.107771791 | 0.119166574 | 0.019176538 |
| Sfr1 | S | 115 | 0.042667285 | 0.100024356 | -0.093708524 | -0.136375809 | -0.19373288 | 0.148105563 | 0.009062588 | 0.165555567 | 0.085469265 | 0.038985725 |
| Smn1 | T | 22 | 0.000542802 | 0.132137585 | 0.020690957 | 0.020148155 | -0.111446628 | 0.9895528 | 0.143572461 | 0.877544347 | 0.881216555 | 0.459244138 |
| Cdk12 | S | 681 | 0.095279544 | -0.029484686 | 0.020980238 | -0.074299306 | 0.050464924 | 0.040480403 | 0.587426631 | 0.429286489 | 0.087302825 | 0.372113431 |
| Vim | T | 32 | 0.02386785 | 0.030633932 | 0.096886287 | 0.072998437 | 0.066232354 | 0.827932174 | 0.655462686 | 0.225644814 | 0.479409111 | 0.19847244 |
| Tmem209 | S | 147 | -0.009697359 | 0.095746531 | 0.059980933 | 0.069678292 | -0.035765597 | 0.861923289 | 0.249425439 | 0.402619254 | 0.291268844 | 0.654960126 |
| Vim | S | 293 | 0.066201728 | 0.04716941 | 0.049301902 | -0.016899827 | 0.002132492 | 0.492404812 | 0.625443509 | 0.463104876 | 0.8686886 | 0.983723535 |
| Zfp148 | S | 306 | 0.014993943 | 0.042507067 | 0.034743761 | 0.019749818 | -0.007763307 | 0.538996578 | 0.223523129 | 0.276084933 | 0.406011404 | 0.801182175 |
| Atrx | T | 996 | 0.012282529 | 0.048635573 | 0.038381909 | 0.026099379 | -0.010253664 | 0.869099875 | 0.232796687 | 0.191539644 | 0.719599344 | 0.738640995 |
| Fbxw8 | S | 36 | 0.235594717 | 0.107946195 | -0.226912441 | -0.462507158 | -0.334858636 | 0.02328855 | 0.295462688 | 0.086938093 | 0.025923554 | 0.031623711 |
| Palld | T | 784 | 0.033150355 | 0.163457353 | 0.052016438 | 0.018866083 | -0.111440915 | 0.90925262 | 0.340868295 | 0.873789415 | 0.959093692 | 0.712058575 |
| Wrnip1 | T | 85 | 0.023597095 | 0.050658729 | 0.037126733 | 0.013529638 | -0.013531996 | 0.756341348 | 0.303129169 | 0.234518016 | 0.853619112 | 0.746703813 |
| Sgo2a | S | 185 | -0.039709873 | 0.07407828 | 0.061803137 | 0.10151301 | -0.012275143 | 0.683854528 | 0.175080341 | 0.186765303 | 0.333869402 | 0.777619748 |
| Zfp318 | S | 1908 | 0.164917428 | 0.020448655 | -0.034904305 | -0.199821733 | -0.05535296 | 0.106856191 | 0.728525174 | 0.617023382 | 0.069298294 | 0.314888458 |
| Ice1 | S | 1841 | -0.049543317 | -0.045791117 | 0.135877102 | 0.185420419 | 0.181668219 | 0.616986924 | 0.695221097 | 0.005463462 | 0.149913671 | 0.208191263 |
| Phf14 | T | 785 | 0.043526047 | 0.12170885 | -0.02054501 | -0.064071057 | -0.14225386 | 0.566913527 | 0.182638772 | 0.86423765 | 0.597301174 | 0.292033599 |
| Ncor2 | S | 2004 | 0.026630362 | -0.017293809 | 0.131323408 | 0.104693046 | 0.148617217 | 0.810943481 | 0.797237148 | 0.103999102 | 0.39077087 | 0.098688076 |

| | | | | | | | | | | | | |
|---|---|---|---|---|---|---|---|---|---|---|---|---|
| Ncor2 | S | 2012 | 0.026630362 | -0.017293809 | 0.131323408 | 0.104693046 | 0.148617217 | 0.810943481 | 0.797237148 | 0.103999102 | 0.39077087 | 0.098688076 |
| Ncor2 | S | 2016 | 0.026630362 | -0.017293809 | 0.131323408 | 0.104693046 | 0.148617217 | 0.810943481 | 0.797237148 | 0.103999102 | 0.39077087 | 0.098688076 |
| Fbxl20 | S | 421 | -0.130977003 | 0.131927341 | 0.208247881 | 0.339224883 | 0.07632054 | 0.098926448 | 0.20156638 | 0.133212066 | 0.047803537 | 0.547373171 |
| Gtpbp1 | S | 6 | -0.063391845 | 0.102823215 | 0.073195117 | 0.136586962 | -0.029628098 | 0.056302815 | 0.081615226 | 0.044938165 | 0.005260348 | 0.522388123 |
| Setd3 | S | 571 | -0.028879553 | 0.067716076 | 0.102224346 | 0.131103899 | 0.03450827 | 0.298535483 | 0.251289241 | 0.153181814 | 0.092264351 | 0.617291324 |
| Hirip3 | S | 564 | 0.030547783 | 0.066926361 | 0.011448874 | -0.019098909 | -0.055477487 | 0.624274411 | 0.100063234 | 0.843274149 | 0.791949731 | 0.360266879 |
| Hdac5 | S | 600 | -0.026803837 | 0.041668501 | 0.157832481 | 0.184636318 | 0.11616398 | 0.681433613 | 0.600999456 | 0.190041362 | 0.14441824 | 0.32015244 |
| Clspn | S | 1133 | 0.137421625 | 0.007351781 | -0.003703103 | -0.141124727 | -0.011054883 | 0.11878296 | 0.880370839 | 0.932982368 | 0.119118756 | 0.786670068 |
| Son | S | 1723 | -0.006782174 | -0.029360412 | 0.113916844 | 0.120699018 | 0.143277255 | 0.827220371 | 0.276352604 | 0.004424974 | 0.017902967 | 5.14643E-05 |
| Fkbp15 | S | 1159 | -0.038922641 | 0.039283028 | 0.107893176 | 0.146815817 | 0.068610147 | 0.550882521 | 0.467265362 | 0.070418323 | 0.053550797 | 0.065231758 |
| Pcnx3 | S | 1953 | -0.064002553 | 0.057108196 | 0.159863153 | 0.223865706 | 0.102754957 | 0.709775628 | 0.431210383 | 0.230270867 | 0.27640557 | 0.405564389 |
| Ifnar2 | S | 448 | -0.112758061 | 0.230017469 | 0.052063408 | 0.164821469 | -0.177954061 | 0.511006636 | 0.129214641 | 0.669577624 | 0.399135737 | 0.275786916 |
| Ccdc84 | S | 176 | -0.040219556 | 0.062914535 | 0.079114595 | 0.119334151 | 0.016200059 | 0.63262221 | 0.181889388 | 0.188080507 | 0.22279494 | 0.755540637 |
| Slk | S | 344 | 0.035610006 | 0.116543589 | -0.050261607 | -0.085871613 | -0.166805196 | 0.504584019 | 0.027753173 | 0.295137738 | 0.186257486 | 0.021153743 |
| Dst | S | 3896 | 0.038166187 | 0.100643633 | -0.003567265 | -0.041733452 | -0.104210898 | 0.459394334 | 0.212276148 | 0.956029606 | 0.566258648 | 0.255508763 |
| Wwc1 | S | 939 | -0.112653981 | 0.141646591 | 0.155530128 | 0.268184109 | 0.013883537 | 0.005045312 | 0.119561788 | 0.020413683 | 0.003732469 | 0.841367499 |
| Gatad2a | S | 105 | 0.017967634 | 0.038679926 | 0.0704048 | 0.052437165 | 0.031724874 | 0.586177658 | 0.237818243 | 0.500201461 | 0.601453601 | 0.74510405 |
| Cbx4 | S | 291 | 0.02370765 | 0.02613775 | 0.046198407 | 0.022490757 | 0.020060658 | 0.59763871 | 0.417446817 | 0.156096272 | 0.568172017 | 0.340328626 |
| Pak1 | T | 225 | 0.040456431 | 0.074891034 | 0.005523547 | -0.034932884 | -0.069367487 | 0.673187699 | 0.104122503 | 0.955209964 | 0.774109036 | 0.489521295 |
| Dennd2c | S | 83 | 0.091528092 | 0.10339646 | -0.050937657 | -0.142465748 | -0.154334117 | 0.358572336 | 0.279502765 | 0.769682141 | 0.413385109 | 0.379978179 |
| Med1 | S | 771 | -0.046204195 | 0.059130797 | 0.099553834 | 0.145758029 | 0.040423036 | 0.431001266 | 0.197718141 | 0.141897576 | 0.080823064 | 0.496465722 |
| Map7d1 | S | 276 | 0.008366775 | 0.197314916 | 0.00898441 | 0.000617635 | -0.188330506 | 0.944899867 | 0.27884132 | 0.908721704 | 0.995938042 | 0.297798873 |
| Champ1 | S | 795 | 0.118080823 | -0.002566267 | 0.031360749 | -0.086720074 | 0.033927016 | 0.174990387 | 0.9872826 | 0.704083702 | 0.356237702 | 0.837502914 |
| Erf | S | 20 | 0.066601339 | 0.058577028 | -0.029156431 | -0.09575777 | -0.087733459 | 0.192527781 | 0.158231922 | 0.672872096 | 0.214917784 | 0.241155445 |
| Srrm2 | S | 1790 | 0.143109236 | 0.056787972 | -0.084697723 | -0.227806958 | -0.141485695 | 0.077802784 | 0.413202718 | 0.403707012 | 0.064912007 | 0.172717158 |
| Tenm4 | S | 41 | 0.010502609 | 0.029808 | 0.049171775 | 0.038669166 | 0.019363775 | 0.742354046 | 0.330011573 | 0.135761614 | 0.102929966 | 0.194113362 |
| Prrc2c | S | 1219 | 0.015971298 | -0.013543815 | 0.041501887 | 0.025530589 | 0.055045703 | 0.420974054 | 0.549016944 | 0.001351211 | 0.237970599 | 0.090679056 |
| Wrnip1 | S | 465 | 0.015380646 | 0.031400881 | 0.085183652 | 0.069803006 | 0.053782771 | 0.825242933 | 0.610325745 | 0.196899307 | 0.210086188 | 0.150142737 |
| Relch | S | 54 | 0.039802205 | 0.057181234 | 0.089298924 | 0.049496718 | 0.03211769 | 0.702346309 | 0.589963771 | 0.474447602 | 0.578680047 | 0.71796548 |
| Unc119 | S | 41 | -0.027479312 | 0.136345363 | -0.013998741 | 0.013480571 | -0.150344104 | 0.602724205 | 0.021803873 | 0.810425984 | 0.795322268 | 0.038480713 |
| Rps6ka5 | S | 211 | 0.11744671 | -0.091317048 | 0.159303599 | 0.041856888 | 0.250620646 | 0.341257624 | 0.565335613 | 0.325411718 | 0.782796407 | 0.209471834 |
| Maml1 | S | 1019 | 0.106709656 | 0.045139501 | 0.000143252 | -0.106566404 | -0.044996249 | 0.207746973 | 0.647582817 | 0.998496304 | 0.131538733 | 0.599534428 |
| Scarf2 | S | 672 | 0.225099283 | -0.020483145 | 0.048972025 | -0.176127259 | 0.069455169 | 0.32125215 | 0.89930084 | 0.844390759 | 0.503931491 | 0.752299259 |
| Ino80b | S | 144 | -0.054226117 | 0.011665989 | 0.216146377 | 0.270372494 | 0.204480388 | 0.371078972 | 0.81373226 | 0.083411838 | 0.049718224 | 0.099035876 |
| Kdm5b | S | 1405 | -0.085764348 | -0.011321419 | 0.225684914 | 0.311449262 | 0.237006333 | 0.122708183 | 0.807176383 | 0.009765524 | 0.00462138 | 0.011491586 |
| Git2 | S | 505 | -0.01448546 | 0.273100013 | -0.071548597 | -0.057063136 | -0.34464861 | 0.945235062 | 0.156767196 | 0.777468361 | 0.843780746 | 0.247518298 |
| Cwc22 | S | 38 | 0.043065275 | -0.042443434 | 0.059059822 | 0.015994547 | 0.101503256 | 0.425017375 | 0.119503655 | 0.000672549 | 0.747323975 | 0.024587856 |
| Clstn1 | S | 978 | 0.144724901 | -0.024993607 | 0.014217863 | -0.130507038 | 0.03921147 | 0.08271678 | 0.738208568 | 0.918499542 | 0.360767117 | 0.761147155 |
| Snrnp70 | S | 268 | 0.046098694 | -0.032864286 | 0.176693186 | 0.130594492 | 0.209557473 | 0.492410817 | 0.579642835 | 0.241106215 | 0.357746416 | 0.188108857 |
| Srrm2 | S | 876 | 0.025519868 | 0.015907836 | 0.08616855 | 0.060648682 | 0.070260714 | 0.330160819 | 0.656617681 | 0.356368493 | 0.492532991 | 0.441235828 |
| Lmnb2 | S | 427 | 0.009529379 | 0.060035073 | 0.032234047 | 0.022704668 | -0.027801026 | 0.781486688 | 0.13067113 | 0.497329653 | 0.660923208 | 0.586304293 |
| Lsm14b | S | 233 | 0.088670656 | -0.055060528 | 0.104431799 | 0.015761143 | 0.159492327 | 0.272985171 | 0.410095459 | 0.169975085 | 0.799549884 | 0.025748177 |
| Mllt3 | S | 365 | 0.068413541 | -0.23897484 | 0.287510769 | 0.219097228 | 0.526485609 | 0.701554661 | 0.05655297 | 0.005633953 | 0.275531023 | 0.007306152 |
| Vps4a | S | 95 | -0.007681485 | -0.030859147 | 0.096025477 | 0.103706962 | 0.126884624 | 0.721967455 | 0.535304465 | 0.003544514 | 0.007862926 | 0.076793585 |
| Kank2 | S | 248 | -0.123692281 | 0.187236777 | 0.065436096 | 0.189128377 | -0.121800681 | 0.037836533 | 0.021634204 | 0.1775799 | 0.013486093 | 0.086126492 |
| Rap1gap2 | S | 590 | -0.114479602 | 0.13269982 | 0.06781634 | 0.182295943 | -0.064883479 | 0.125285435 | 0.017407402 | 0.262641993 | 0.043754206 | 0.234529256 |
| Rap1gap2 | S | 594 | -0.114479602 | 0.13269982 | 0.06781634 | 0.182295943 | -0.064883479 | 0.125285435 | 0.017407402 | 0.262641993 | 0.043754206 | 0.234529256 |
| Smarca4 | S | 1349 | 0.144065877 | -0.003621809 | -0.019388094 | -0.163453971 | -0.015766286 | 0.084541784 | 0.958684322 | 0.929680064 | 0.478944722 | 0.941280697 |
| Plec | T | 4635 | -0.128875221 | 0.196236431 | 0.067349754 | 0.196224975 | -0.128886676 | 0.070707829 | 0.018023497 | 0.417407512 | 0.053059279 | 0.140258421 |
| Atrx | S | 691 | 0.051758072 | -0.12697496 | 0.230201285 | 0.178443213 | 0.357176246 | 0.500222386 | 0.252780777 | 0.073901193 | 0.140690351 | 0.032617046 |
| Dennd2a | S | 303 | -0.029157925 | 0.071671072 | 0.072251008 | 0.101408934 | 0.000579936 | 0.430471157 | 0.093948328 | 0.310652212 | 0.192101452 | 0.992916839 |
| Zfpm2 | S | 476 | -0.125667393 | 0.114089406 | 0.113923535 | 0.239590928 | -0.000165871 | 0.01458021 | 0.029928053 | 0.027097179 | 0.002594594 | 0.996034697 |
| Rhbdf1 | S | 10 | -0.0782606 | 0.094888222 | 0.132769016 | 0.211029616 | 0.037880793 | 0.166459436 | 0.098032257 | 0.218103158 | 0.091340879 | 0.685737966 |
| Numa1 | S | 198 | -0.026900554 | 0.118758198 | 0.072399965 | 0.099300519 | -0.046358232 | 0.74879721 | 0.268866488 | 0.458682702 | 0.394687196 | 0.697533858 |
| Hmx3 | S | 218 | 0.11828704 | 0.147204512 | -0.099063827 | -0.217350867 | -0.246268338 | 0.335860811 | 0.150042814 | 0.304151543 | 0.094946502 | 0.000796394 |
| Myh9 | S | 1114 | 0.030485485 | 0.058433677 | 0.080921582 | 0.050436097 | 0.022487904 | 0.68325192 | 0.410917808 | 0.584971877 | 0.730861056 | 0.875316288 |
| Piezo1 | S | 1385 | 0.06488787 | 0.058498598 | -0.028628706 | -0.093516576 | -0.087127305 | 0.131460192 | 0.17398097 | 0.509599185 | 0.032855199 | 0.029893378 |
| Cdr2 | T | 405 | 0.263246834 | -0.179350429 | 0.144054091 | -0.119192744 | 0.323404519 | 0.114840194 | 0.219570352 | 0.513610441 | 0.548622581 | 0.176943083 |
| Fyn | S | 21 | 0.016437911 | 0.082800157 | 0.000175492 | -0.016262418 | -0.082624664 | 0.537883984 | 0.095413078 | 0.997474872 | 0.754793151 | 0.215512107 |
| Ap3d1 | S | 784 | -0.022712515 | 0.045416591 | 0.032245915 | 0.05495843 | -0.013170675 | 0.561294923 | 0.022118854 | 0.165673846 | 0.226857397 | 0.528916502 |
| Nes | S | 816 | 0.020944538 | 0.016549896 | 0.034989748 | 0.014045209 | 0.018439852 | 0.263126946 | 0.676018096 | 0.0837608 | 0.154671878 | 0.626365073 |
| Kmt5b | S | 670 | 0.040303148 | -0.019428568 | 0.093372092 | 0.053068944 | 0.11280066 | 0.561532975 | 0.685932727 | 0.088959817 | 0.403610657 | 0.017524657 |
| Ifi205 Ifi211 Mndal | T | 439;353;332 | -0.082758039 | 0.168014315 | 0.096114021 | 0.17887206 | -0.071900294 | 0.021782011 | 0.037223883 | 0.257008443 | 0.095438652 | 0.402284221 |
| Sf3b1 | T | 244 | 0.050052335 | 0.034861636 | -0.004322851 | -0.054375186 | -0.039184487 | 0.050869559 | 0.379821305 | 0.838131808 | 0.003373517 | 0.311849785 |
| Slc23a2 | S | 78 | -0.00367034 | 0.200243688 | 0.007257845 | 0.010928185 | -0.192985843 | 0.974686922 | 0.24492765 | 0.942178729 | 0.901862192 | 0.244750773 |
| Dbn1 | S | 389 | 0.169019069 | -0.060587306 | 0.00502899 | -0.163990079 | 0.065616297 | 0.041120205 | 0.479190516 | 0.936315564 | 0.055496465 | 0.440883749 |
| Dbn1 | S | 390 | 0.169019069 | -0.060587306 | 0.00502899 | -0.163990079 | 0.065616297 | 0.041120205 | 0.479190516 | 0.936315564 | 0.055496465 | 0.440883749 |
| Dtx3l | S | 558 | -0.160930354 | 0.103475259 | 0.198750241 | 0.359680594 | 0.095274982 | 0.082780607 | 0.266011286 | 0.038530035 | 0.003768331 | 0.17309046 |
| Mycbp2 | S | 2911 | -0.044641631 | 0.055815903 | 0.065259593 | 0.109901224 | 0.00944369 | 0.488626665 | 0.141210529 | 0.097821988 | 0.148063007 | 0.542723355 |
| Plekhg5 | T | 137 | -0.155886905 | 0.100760272 | 0.213437742 | 0.369324646 | 0.11267747 | 0.045802362 | 0.143429122 | 0.052915184 | 0.010213743 | 0.220057541 |
| Pelp1 | S | 51 | 0.021408987 | -0.013599368 | 0.091853749 | 0.070444762 | 0.105453117 | 0.684579998 | 0.766031263 | 0.058215371 | 0.161392775 | 0.027846831 |
| Sptbn1 | T | 2158 | -0.035572341 | 0.020106597 | 0.124357578 | 0.159929919 | 0.104250982 | 0.518304721 | 0.56108652 | 0.063330773 | 0.047859432 | 0.113227188 |
| Mtfr1l | S | 100 | -0.002134256 | 0.087112154 | 0.023750285 | 0.02588454 | -0.063361869 | 0.961187344 | 0.095079861 | 0.672866529 | 0.602147525 | 0.260256299 |
| Insr | S | 1340 | -0.121176317 | 0.162131279 | 0.143079205 | 0.264255522 | -0.019052074 | 0.047901735 | 0.132914043 | 0.098930308 | 0.019117222 | 0.837955527 |
| Reps1 | S | 273 | 0.009999931 | 0.185909599 | -0.085280793 | -0.095280724 | -0.271190393 | 0.865786787 | 0.0262072 | 0.299932712 | 0.225486462 | 0.020160372 |
| Paxx | S | 150 | 0.057718506 | 0.063307915 | -0.011802892 | -0.069521397 | -0.075110807 | 0.150656664 | 0.343115485 | 0.803531562 | 0.189988971 | 0.297241525 |
| BC003965 | S | 34 | 0.051245747 | -0.031225179 | 0.041707911 | -0.009537835 | 0.07293309 | 0.022387602 | 0.543851667 | 0.262001491 | 0.765431794 | 0.230941103 |
| Nav1 | S | 477 | -0.073850358 | 0.081227584 | 0.052872669 | 0.126723027 | -0.028354916 | 0.121600484 | 0.0111536 | 0.128769583 | 0.03412571 | 0.37048738 |
| Dock7 | S | 929 | -0.074545102 | 0.156498601 | 0.067795969 | 0.142341071 | -0.088702631 | 0.114386221 | 0.137879678 | 0.113336079 | 0.010785744 | 0.331770879 |
| Slk | S | 340 | 0.015887376 | 0.098730087 | -0.008928748 | -0.024816124 | -0.107658835 | 0.85562041 | 0.059805496 | 0.845855552 | 0.760798761 | 0.009544744 |
| Lmnb1 | S | 406 | -0.038876153 | 0.225561297 | -0.080312529 | -0.041436376 | -0.305873826 | 0.686258862 | 0.043296753 | 0.660259359 | 0.805018926 | 0.168855235 |
| Slk | S | 816 | 0.048551781 | 0.128098409 | -0.043054635 | -0.091606416 | -0.171153044 | 0.603595326 | 0.104068706 | 0.564120195 | 0.386839214 | 0.08571735 |
| Srrm2 | S | 434 | 0.06521302 | -0.008343561 | 0.055076961 | -0.010136059 | 0.063420522 | 0.170778124 | 0.838590471 | 0.362322002 | 0.831059781 | 0.251179575 |
| Gtf2i | S | 694 | 0.103423843 | 0.029175596 | -0.041169172 | -0.144593015 | -0.070344768 | 0.083953082 | 0.387184367 | 0.640952648 | 0.177817162 | 0.439778459 |
| Ubr4 | S | 1646 | -0.080914144 | 0.117291535 | 0.06808639 | 0.149000534 | -0.049205145 | 0.017180988 | 0.011484689 | 0.111433121 | 0.027381998 | 0.232337727 |
| Larp4b | S | 739 | -0.037397417 | 0.042113114 | 0.020015776 | 0.057413193 | -0.022097338 | 0.284244822 | 0.000909416 | 0.109857272 | 0.148996302 | 0.089543925 |
| Afdn | S | 1201 | 0.010726154 | 0.045537373 | 0.033910104 | 0.02318395 | -0.011627269 | 0.837012555 | 0.101227542 | 0.427601163 | 0.693182002 | 0.767009053 |
| Nfat5 | S | 617 | 0.00948566 | 0.124763857 | -0.015857289 | -0.025342949 | -0.140621146 | 0.887466139 | 0.074327963 | 0.753475159 | 0.643384504 | 0.04709585 |
| Kif20b | S | 1694 | 0.178343868 | -0.014596669 | -0.064584242 | -0.24292811 | -0.049987573 | 0.00774378 | 0.752093337 | 0.183795662 | 0.00427769 | 0.069801701 |
| Srrm2 | S | 1179 | 0.005956095 | 0.018985592 | 0.074337474 | 0.068381379 | 0.055351882 | 0.857850362 | 0.539997431 | 0.119498014 | 0.145412289 | 0.204931897 |
| Washc2 | S | 782 | 0.105307793 | 0.049552153 | -0.058845721 | -0.164153514 | -0.108397875 | 0.075992445 | 0.511887841 | 0.621257259 | 0.23018298 | 0.40298373 |
| Aebp2 | S | 204 | -0.032744485 | 0.025308409 | 0.099198853 | 0.131943338 | 0.073890444 | 0.697199427 | 0.551726446 | 0.064629823 | 0.187400111 | 0.134065634 |
| Srrt | S | 74 | -0.04403677 | 0.014788926 | 0.13944722 | 0.18348399 | 0.124658293 | 0.247109376 | 0.680177696 | 0.027518663 | 0.016660702 | 0.047720889 |
| Srrm2 | S | 1343 | 0.007568361 | -0.014728037 | 0.086028801 | 0.07846044 | 0.100756837 | 0.746918061 | 0.427855622 | 0.015250195 | 0.029266092 | 0.02849056 |

| | | | | | | | | | | | | |
|---|---|---|---|---|---|---|---|---|---|---|---|---|
| Pan3 | T | 299 | 0.184841606 | 0.010635705 | -0.027129149 | -0.211970755 | -0.037764854 | 0.12245668 | 0.885307618 | 0.567202644 | 0.095924295 | 0.613271965 |
| Axl | S | 856 | 0.193992604 | -0.047769962 | 0.025279808 | -0.168712796 | 0.07304977 | 0.199467746 | 0.801432429 | 0.783362596 | 0.266966144 | 0.711330051 |
| Jmy | S | 134 | -0.067656215 | 0.119790942 | 0.083951589 | 0.151607803 | -0.035839353 | 0.251275797 | 0.110778727 | 0.27990133 | 0.104590734 | 0.657984224 |
| Kmt2b | S | 1932 | 0.055650367 | 0.023417961 | -0.008857404 | -0.064507771 | -0.032275365 | 0.039881628 | 0.419081987 | 0.893361939 | 0.37096467 | 0.63212042 |
| Ldlr | S | 445 | -0.067770226 | 0.021745608 | 0.175505323 | 0.243275549 | 0.153759715 | 0.322421663 | 0.670643931 | 0.052877365 | 0.0284866 | 0.081643813 |
| Caap1 | S | 307 | -0.00997549 | 0.058187415 | 0.049100434 | 0.059075924 | -0.009086981 | 0.794961733 | 0.159098654 | 0.265291921 | 0.26544171 | 0.842688229 |
| Cenpa | S | 16 | 0.005934388 | -0.014220427 | 0.122716301 | 0.116781913 | 0.136936728 | 0.901264258 | 0.845906397 | 0.078897623 | 0.092446462 | 0.134437433 |
| Mcrip1 | S | 17 | 0.120255474 | -0.021820084 | -0.067313097 | -0.187568571 | -0.045493013 | 0.005864982 | 0.78890125 | 0.618065585 | 0.241341538 | 0.755478316 |
| Nfrkb | S | 226 | 0.04376031 | 0.010983712 | 0.114333444 | 0.070573134 | 0.103349732 | 0.654862875 | 0.870407919 | 0.324188068 | 0.582809373 | 0.377060787 |
| Zc3hc1 | S | 24 | 0.082740427 | 0.053491721 | -0.062379694 | -0.145120121 | -0.115871415 | 0.059003293 | 0.063138733 | 0.164385117 | 0.021926065 | 0.054694095 |
| Vrk3 | S | 115 | -0.118205611 | 0.114376607 | 0.190448871 | 0.308654482 | 0.076072264 | 0.056175281 | 0.199644986 | 0.084079754 | 0.02106833 | 0.436144738 |
| Wdr76 | S | 146 | 0.185721834 | -0.007718772 | 0.039319681 | -0.146402154 | 0.047038453 | 0.357990374 | 0.831674949 | 0.63690948 | 0.46103449 | 0.570260891 |
| Mphosph8 | T | 84 | 0.093709874 | -0.00667592 | 0.043024625 | -0.050685249 | 0.049700545 | 0.188700668 | 0.921412668 | 0.551658514 | 0.550398823 | 0.586302468 |
| Nelfe | S | 179 | -0.007845159 | 0.004596979 | 0.08122539 | 0.08907055 | 0.076628412 | 0.74289091 | 0.833005738 | 0.023767532 | 0.027352598 | 0.037482965 |
| Gpsm2 | S | 408 | -0.084155579 | 0.174973213 | 0.063640868 | 0.147796447 | -0.111332345 | 0.083738067 | 0.107963393 | 0.124009286 | 0.015630374 | 0.248057411 |
| Casp8ap2 | S | 1532 | 0.035104932 | 0.007110301 | 0.070320986 | 0.035216053 | 0.063210685 | 0.549336477 | 0.745556786 | 0.191572954 | 0.59420635 | 0.232020548 |
| Herc2 | S | 1943 | 0.053977566 | 0.052768945 | -0.006026346 | -0.060003912 | -0.058795291 | 0.168633341 | 0.315524566 | 0.899647014 | 0.250624856 | 0.327712309 |
| Zbtb20 | T | 304 | 0.105905926 | 0.074106886 | 0.028723014 | -0.077182912 | -0.045383872 | 0.507742398 | 0.697737859 | 0.70890118 | 0.624850423 | 0.811039083 |
| Srrm2 | S | 1557 | 0.026127416 | 0.034623881 | 0.024249626 | -0.00187779 | -0.010374255 | 0.364105373 | 0.223856269 | 0.444111547 | 0.956765509 | 0.759429878 |
| Nsrp1 | S | 254 | -0.039435299 | 0.029754184 | 0.067219966 | 0.106655264 | 0.037465782 | 0.303231262 | 0.191810767 | 0.016399612 | 0.045962337 | 0.069847478 |
| Bmp2k | S | 702 | 0.192473479 | 0.044327368 | -0.102462912 | -0.294936391 | -0.14679028 | 0.050713837 | 0.456741194 | 0.187112239 | 0.015889733 | 0.068514116 |
| Tcf20 | S | 1162 | 0.075001003 | -0.045954306 | 0.152471814 | 0.077470811 | 0.19842612 | 0.522332951 | 0.53315638 | 0.222531522 | 0.562189183 | 0.136332186 |
| Samd4 | S | 578 | 0.115099975 | -0.146229924 | 0.137882867 | 0.022782891 | 0.284112791 | 0.099000121 | 0.089433996 | 0.068756938 | 0.586559369 | 0.010811552 |
| Daxx | T | 523 | 0.024905639 | 0.042057389 | 0.011411745 | -0.013493893 | -0.030645644 | 0.397279619 | 0.091102288 | 0.72858287 | 0.690192466 | 0.340985218 |
| Bcl9l | S | 1449 | 0.05931858 | 0.123356523 | -0.022791596 | -0.082110175 | -0.146148118 | 0.376123247 | 0.376014033 | 0.858144647 | 0.538562777 | 0.393148193 |
| Wipf2 | S | 235 | 0.164476884 | 0.001751229 | -0.021081625 | -0.185558509 | -0.022832853 | 0.115650197 | 0.977371299 | 0.776190642 | 0.108164259 | 0.775139999 |
| Sh3kbp1 | S | 630 | -0.008085482 | 0.198569255 | -0.056057032 | -0.04797155 | -0.254626287 | 0.959276658 | 0.079917917 | 0.62876382 | 0.786623221 | 0.099522144 |
| Phf20 | S | 900 | 0.047588631 | 0.025168208 | 0.055502598 | 0.007913967 | 0.03033439 | 0.587849522 | 0.647036505 | 0.282695237 | 0.924911804 | 0.567251283 |
| Cherp | S | 824 | 0.057829648 | 0.007444983 | 0.031443875 | -0.026385773 | 0.023998892 | 0.151542587 | 0.78476778 | 0.463109163 | 0.568816315 | 0.564849573 |
| Htatsf1 | T | 109 | 0.132584358 | -0.072919033 | 0.097381519 | -0.035202839 | 0.170300552 | 0.288113679 | 0.582806407 | 0.349344576 | 0.661184681 | 0.176710482 |
| Six5 | S | 65 | -0.048743169 | 0.060497169 | 0.110960099 | 0.159703267 | 0.05046293 | 0.733146277 | 0.451673081 | 0.186977092 | 0.296107157 | 0.417080733 |
| Hgsnat | S | 238 | 0.104207273 | 0.014135111 | -0.009697141 | -0.113904414 | -0.023832253 | 0.095553563 | 0.742162935 | 0.898937612 | 0.216001161 | 0.756764897 |
| Slc39a6 | S | 191 | 0.071454131 | 0.012078122 | 0.044919283 | -0.026534848 | 0.03284116 | 0.378746916 | 0.77723492 | 0.380665573 | 0.713509669 | 0.333519282 |
| Aktip | S | 30 | 0.054705455 | 0.09049797 | -0.020681832 | -0.075387286 | -0.111179802 | 0.419409269 | 0.200112391 | 0.773815693 | 0.200519068 | 0.089712256 |
| Sipa1l3 | S | 1358 | 0.022638539 | 0.007496159 | 0.054118981 | 0.031480443 | 0.046622822 | 0.503265228 | 0.803769059 | 0.089477365 | 0.323855582 | 0.130294708 |
| Parg | S | 311 | 0.098914758 | 0.106038053 | -0.088692799 | -0.187607556 | -0.194730852 | 0.114324491 | 0.086971897 | 0.106653957 | 0.024583669 | 0.019649183 |
| Ncor1 | S | 1207 | -0.033713382 | 0.03481571 | 0.071510926 | 0.105224308 | 0.036695217 | 0.517583302 | 0.248463296 | 0.055074816 | 0.101479454 | 0.157912302 |
| Cyfip1 | T | 1234 | -0.01025002 | 0.125035693 | 0.057418988 | 0.067669008 | -0.067616705 | 0.854589291 | 0.390277934 | 0.419813579 | 0.375879297 | 0.631793375 |
| Zc3h18 | S | 852 | 0.020685824 | 0.011171319 | 0.06521675 | 0.044530926 | 0.054045431 | 0.705380564 | 0.799331655 | 0.095826917 | 0.442287326 | 0.274548351 |
| Atxn2l | S | 32 | 0.162070953 | 0.077636292 | -0.121121397 | -0.28319235 | -0.198757689 | 0.084023699 | 0.432889156 | 0.337705411 | 0.073094548 | 0.168309352 |
| Scyl1 | S | 17 | 0.059283747 | 0.035619288 | 0.024345834 | -0.034937914 | -0.011273455 | 0.19198311 | 0.75029379 | 0.598191933 | 0.439003857 | 0.918919917 |
| Prkab1 | T | 106 | -0.031242358 | 0.224427483 | -0.116492203 | -0.085249844 | -0.340919685 | 0.781459135 | 0.035171689 | 0.576580497 | 0.680845675 | 0.175725364 |
| Zc3h13 | S | 913 | 0.048118862 | 0.02596312 | 0.035995237 | -0.012123625 | 0.010032117 | 0.32545555 | 0.411206687 | 0.574368533 | 0.861452863 | 0.874252438 |
| Setx | S | 870 | 0.164240411 | -0.030721083 | 0.025610762 | -0.138629648 | 0.056331846 | 0.172481579 | 0.724496324 | 0.740483334 | 0.25702408 | 0.595869918 |
| Setx | S | 871 | 0.164240411 | -0.030721083 | 0.025610762 | -0.138629648 | 0.056331846 | 0.172481579 | 0.724496324 | 0.740483334 | 0.25702408 | 0.595869918 |
| Setx | S | 872 | 0.164240411 | -0.030721083 | 0.025610762 | -0.138629648 | 0.056331846 | 0.172481579 | 0.724496324 | 0.740483334 | 0.25702408 | 0.595869918 |
| Tex2 | S | 733 | 0.036253967 | 0.00292221 | 0.11136385 | 0.075109883 | 0.10844164 | 0.767820231 | 0.976303803 | 0.206443644 | 0.551802078 | 0.311678741 |
| Lyst | S | 2207 | 0.099364575 | 0.313009872 | -0.129458441 | -0.228823016 | -0.442468312 | 0.681567565 | 0.174520934 | 0.274125407 | 0.364266908 | 0.097637687 |
| Slx4 | S | 417 | 0.009015029 | 0.040563806 | 0.084030464 | 0.075015436 | 0.043466659 | 0.872504822 | 0.365542659 | 0.373182642 | 0.442242263 | 0.627418279 |
| Ncaph | S | 189 | 0.001789521 | 0.049335603 | 0.049026186 | 0.047236664 | -0.000309417 | 0.972312838 | 0.395048723 | 0.152422407 | 0.35223627 | 0.994960508 |
| Vim | Y | 61 | 0.075484596 | -0.021136575 | 0.048000207 | -0.02748439 | 0.069136781 | 0.111887324 | 0.519841225 | 0.315701266 | 0.556880088 | 0.154540685 |
| Ssh1 | S | 928 | 0.005120114 | 0.083076329 | 0.070760338 | 0.065640225 | -0.012315991 | 0.920915809 | 0.571450064 | 0.309528107 | 0.363224375 | 0.932523133 |
| Clk3 | S | 173 | 0.135947701 | -0.002248659 | 0.012661635 | -0.123286066 | 0.014910293 | 0.166566392 | 0.979020975 | 0.896293728 | 0.317246295 | 0.900572178 |
| Clk3 | S | 175 | 0.135947701 | -0.002248659 | 0.012661635 | -0.123286066 | 0.014910293 | 0.166566392 | 0.979020975 | 0.896293728 | 0.317246295 | 0.900572178 |
| Ppp4r1 | S | 948 | -0.016619672 | 0.049352138 | 0.062165371 | 0.078785043 | 0.012813233 | 0.716495242 | 0.259242668 | 0.156395397 | 0.067112567 | 0.585433147 |
| Srrm1 | S | 645 | 0.009810273 | 0.001433335 | 0.078774213 | 0.06896394 | 0.077340878 | 0.753005948 | 0.959923537 | 0.065879079 | 0.12601807 | 0.086686217 |
| Clk2 | S | 141 | 0.016958734 | 0.010755434 | 0.078947066 | 0.061988331 | 0.068191632 | 0.560615306 | 0.700590096 | 0.189608039 | 0.276854181 | 0.241093898 |
| Shb | T | 210 | 0.026190694 | 0.055685772 | 0.005856317 | -0.020334377 | -0.049829455 | 0.534288174 | 0.084642323 | 0.909709523 | 0.710052166 | 0.346765126 |
| Flna | S | 2284 | -0.041331646 | 0.020856765 | 0.072568672 | 0.113900318 | 0.051711907 | 0.03766724 | 0.313740369 | 0.000591054 | 0.003089496 | 0.059720168 |
| Srrm2 | T | 1926 | 0.074541689 | -0.018908682 | 0.107336635 | 0.032794946 | 0.126245317 | 0.344104009 | 0.794556423 | 0.41887201 | 0.790164545 | 0.344265369 |
| Sipa1 | S | 51 | 0.097366619 | -0.016184689 | -0.023370927 | -0.120737547 | -0.007186238 | 0.003037605 | 0.517557758 | 0.504753808 | 0.030831717 | 0.820710467 |
| Ncor1 | T | 568 | 0.102771079 | -0.009017814 | 0.058111809 | -0.04465927 | 0.067129624 | 0.297772237 | 0.913590105 | 0.51610368 | 0.639082422 | 0.442925486 |
| Hmgn3 | S | 78 | -0.027493894 | 0.04679639 | 0.111449158 | 0.138943053 | 0.064652768 | 0.490401008 | 0.267925315 | 0.218395422 | 0.150530331 | 0.4327659 |
| Cactin | S | 148 | 0.191959066 | 0.015672181 | -0.014500778 | -0.206459844 | -0.030172959 | 0.233195749 | 0.861116145 | 0.796650155 | 0.208487968 | 0.747402579 |
| Srrm1 | S | 702 | 0.046278715 | -0.021449327 | 0.061361194 | 0.015082479 | 0.082810521 | 0.137672945 | 0.470921199 | 0.145682805 | 0.66245983 | 0.072855401 |
| Nedd4l | S | 508 | 0.076250585 | 0.020384466 | 0.000936602 | -0.075313984 | -0.019447864 | 0.08690988 | 0.703143658 | 0.989976518 | 0.369437566 | 0.817478239 |
| Gpr149 | S | 544 | 0.149457472 | -0.171046176 | 0.145358644 | -0.004098828 | 0.31640482 | 0.066131034 | 0.040800126 | 0.070100875 | 0.905381144 | 0.000388608 |
| Matr3 | S | 630 | 0.021240977 | -0.014783642 | 0.159596625 | 0.138355649 | 0.174380267 | 0.82501207 | 0.866716321 | 0.20756709 | 0.284833209 | 0.183750529 |
| Brms1 | S | 237 | 0.040535811 | 0.018092698 | 0.045410942 | 0.004875132 | 0.027318244 | 0.262932214 | 0.600882428 | 0.451874071 | 0.92407009 | 0.606323341 |
| Rpp25l | S | 16 | 0.014206695 | 0.051593324 | 0.077091183 | 0.062884487 | 0.025497858 | 0.946106159 | 0.473221493 | 0.326506171 | 0.768606081 | 0.74197248 |
| Exoc1 | S | 472 | 0.042268973 | 0.024526435 | 0.079186364 | 0.036917391 | 0.054659929 | 0.401746289 | 0.753221834 | 0.487380817 | 0.724901384 | 0.642962636 |
| Knl1 | S | 1270 | 0.020806317 | 0.053536515 | 0.047280922 | 0.026474606 | -0.006255593 | 0.790028942 | 0.40029954 | 0.338268975 | 0.725171307 | 0.911867099 |
| Wdhd1 | S | 333 | -0.080407621 | 0.142574763 | 0.186806505 | 0.267214126 | 0.044231742 | 0.672447434 | 0.727252093 | 0.286591442 | 0.176857525 | 0.911386663 |
| Wdhd1 | S | 334 | -0.080407621 | 0.142574763 | 0.186806505 | 0.267214126 | 0.044231742 | 0.672447434 | 0.727252093 | 0.286591442 | 0.176857525 | 0.911386663 |
| Wapl | S | 81 | 0.015342643 | 0.017084703 | 0.11413993 | 0.098797287 | 0.097055227 | 0.84758417 | 0.792579871 | 0.239014337 | 0.344454247 | 0.311522592 |
| Tmem245 | S | 12 | 0.082278808 | -0.032968775 | 0.051174024 | -0.031104784 | 0.084142798 | 0.161580883 | 0.385749586 | 0.157951543 | 0.510863802 | 0.029443549 |
| Pard3 | S | 187 | 0.047254155 | 0.012937438 | 0.056529661 | 0.009275506 | 0.043592223 | 0.384571707 | 0.833323347 | 0.31298717 | 0.876800434 | 0.531329723 |
| Wac | S | 62 | -0.006706791 | 0.072230111 | 0.0299865 | 0.03669329 | -0.042243611 | 0.919644634 | 0.158011794 | 0.356291576 | 0.588030911 | 0.358425732 |
| Atf7ip | S | 285 | -0.042852191 | 0.158965193 | 0.04331222 | 0.086164411 | -0.115652973 | 0.64785862 | 0.212353712 | 0.565395502 | 0.435018003 | 0.354202767 |
| Pds5b | S | 1166 | 0.099039222 | 0.012384173 | -0.046674521 | -0.145713743 | -0.059058694 | 0.015177209 | 0.654666409 | 0.42510321 | 0.069333815 | 0.329325812 |
| Npm1 | T | 277 | 0.004843353 | -0.002212154 | 0.211603851 | 0.206760499 | 0.213816005 | 0.98710681 | 0.992272714 | 0.289240621 | 0.491493106 | 0.341490458 |
| Mcm2 | S | 53 | 0.140066473 | -0.068661891 | 0.16384356 | 0.023777087 | 0.232505451 | 0.558558891 | 0.754840144 | 0.435538683 | 0.921427374 | 0.33590704 |
| Gtf2f1 | T | 437 | 0.067764894 | -0.023274247 | 0.06533188 | -0.002433014 | 0.088606127 | 0.477832712 | 0.898607924 | 0.167169895 | 0.978326032 | 0.636938901 |
| Ep400 | T | 3065 | 0.108982042 | -0.053357323 | 0.0249359 | -0.084046141 | 0.078293224 | 0.003034185 | 0.059936887 | 0.624947102 | 0.170903543 | 0.191033845 |
| Mief1 | S | 59 | -0.048886027 | -0.004541583 | 0.083686822 | 0.132572849 | 0.088228405 | 0.333438725 | 0.902971899 | 0.000862431 | 0.069942662 | 0.098996069 |
| Gmip | S | 479 | 0.078491973 | -0.019035751 | 0.12120879 | 0.042716818 | 0.140244542 | 0.625846178 | 0.823039151 | 0.295797447 | 0.8047968 | 0.27204663 |
| Extl3 | S | 361 | -0.07331948 | 0.059568594 | 0.21257946 | 0.28589894 | 0.153010866 | 0.012179268 | 0.336918652 | 0.039466837 | 0.02609974 | 0.093480799 |
| Stag1 | S | 45 | 0.764426438 | 0.047929647 | -0.231758571 | -0.996185009 | -0.279688218 | 0.319055348 | 0.673234423 | 0.261754995 | 0.224709885 | 0.197539948 |
| Agap1 | S | 521 | -0.022437614 | 0.056196882 | 0.043756145 | 0.066193759 | -0.012440737 | 0.31934217 | 0.127039215 | 0.078264486 | 0.025727931 | 0.682475413 |
| Fam53c | S | 122 | -0.044656077 | 0.14247679 | 0.020419417 | 0.065075494 | -0.122057373 | 0.580234764 | 0.085397509 | 0.741462652 | 0.441539632 | 0.142881982 |
| Aak1 | T | 618 | -0.019271929 | 0.082175147 | 0.031798322 | 0.051070251 | -0.050376825 | 0.423218168 | 0.067496849 | 0.393389365 | 0.209326608 | 0.267066637 |
| Mex3d | T | 490 | 0.030223115 | 0.129219212 | -0.060355519 | -0.090578633 | -0.189574731 | 0.677419623 | 0.052652002 | 0.477222014 | 0.356821304 | 0.083304844 |
| Mex3d | T | 491 | 0.030223115 | 0.129219212 | -0.060355519 | -0.090578633 | -0.189574731 | 0.677419623 | 0.052652002 | 0.477222014 | 0.356821304 | 0.083304844 |
| Ccdc6 | S | 412 | 0.147092806 | 0.144282137 | -0.291127835 | -0.438220641 | -0.435409971 | 0.086801352 | 0.145752273 | 0.275259743 | 0.152769994 | 0.149902218 |

| | | | | | | | | | | | | |
|---|---|---|---|---|---|---|---|---|---|---|---|---|
| Tmcc3 | S | 174 | 0.154524826 | -0.019138449 | -0.022818534 | -0.17734336 | -0.003680085 | 0.047945652 | 0.824599352 | 0.579585544 | 0.05492628 | 0.963003622 |
| Pum1 | S | 212 | 0.073020869 | -0.052971535 | 0.084578077 | 0.011557208 | 0.137549613 | 0.438906674 | 0.528920009 | 0.09021818 | 0.888925592 | 0.164721781 |
| Ncaph2 | T | 305 | 0.088401205 | 0.051468206 | 0.049487332 | -0.038913874 | -0.001980874 | 0.482261534 | 0.697241595 | 0.738950302 | 0.78035827 | 0.989126964 |
| Sos1 | S | 1082 | 0.017210092 | 0.033593032 | 0.119667244 | 0.102457152 | 0.086074212 | 0.816418342 | 0.66984428 | 0.387029195 | 0.446863423 | 0.520344238 |
| Rgl1 | S | 520 | -0.196683707 | 0.299396639 | 0.093331864 | 0.29001557 | -0.206064775 | 0.110279926 | 0.066751224 | 0.457322533 | 0.074492873 | 0.201482667 |
| Irf2bp2 | S | 439 | 0.051885397 | 0.056738466 | 0.002550413 | -0.049334984 | -0.054188053 | 0.348436926 | 0.280121526 | 0.966742411 | 0.424456546 | 0.367872956 |
| Arhgef7 | S | 776 | 0.02211267 | 0.096019563 | -0.057315549 | -0.079428218 | -0.153335112 | 0.554969708 | 0.028141723 | 0.539716011 | 0.407004019 | 0.175944804 |
| Clip1 | S | 1317 | -0.022711809 | 0.074872153 | 0.033391321 | 0.05610313 | -0.041480832 | 0.548241058 | 0.039499716 | 0.57512172 | 0.365286884 | 0.472693147 |
| Phf2 | S | 899 | 0.021825647 | 0.049629749 | 0.007496062 | -0.014329586 | -0.042133687 | 0.541827215 | 0.072202612 | 0.856206982 | 0.754786107 | 0.335694532 |
| Ehmt1 | T | 78 | -0.042505567 | 0.048743376 | 0.104449604 | 0.146955171 | 0.055706228 | 0.415890789 | 0.482336608 | 0.078581497 | 0.00895156 | 0.365010849 |
| Pdcl | S | 20 | 0.007895292 | 0.067149621 | -0.009874093 | -0.017769385 | -0.077023713 | 0.822023545 | 0.019400128 | 0.826942326 | 0.715421843 | 0.164632563 |
| Pdcl | S | 25 | 0.007895292 | 0.067149621 | -0.009874093 | -0.017769385 | -0.077023713 | 0.822023545 | 0.019400128 | 0.826942326 | 0.715421843 | 0.164632563 |
| Ranbp10 | S | 355 | 0.128595988 | 0.082330491 | -0.10204012 | -0.230636107 | -0.184370611 | 0.247929015 | 0.291031842 | 0.590443846 | 0.273180863 | 0.354079448 |
| Rad21 | S | 175 | -0.003794618 | 0.00914132 | 0.128028805 | 0.131823422 | 0.118887485 | 0.898965259 | 0.8278493 | 0.154547646 | 0.161223876 | 0.179230436 |
| Ppig | S | 534 | 0.105403797 | -0.152392225 | 0.18018832 | 0.074784523 | 0.332580546 | 0.327496845 | 0.371697771 | 0.180189364 | 0.503034197 | 0.102521284 |
| Nol9 | T | 91 | 0.120400888 | 0.062143299 | -0.035616694 | -0.156017582 | -0.097759992 | 0.185481269 | 0.148370613 | 0.340534594 | 0.119590374 | 0.180512998 |
| Gja1 | S | 369 | 0.234780639 | -0.001807645 | 0.004786948 | -0.229993692 | 0.006594593 | 0.339921267 | 0.929761759 | 0.931609196 | 0.347578826 | 0.906057497 |
| Aff4 | S | 701 | 0.044975724 | -0.019500929 | 0.04908649 | 0.004110766 | 0.068587419 | 0.090027978 | 0.420965087 | 0.139240498 | 0.860642597 | 0.056728352 |
| Olfr143 | T | 5 | 0.030086236 | 0.082448338 | -0.017755334 | -0.04784157 | -0.100203672 | 0.623911667 | 0.083355719 | 0.807665185 | 0.542270907 | 0.215557889 |
| Syde1 | S | 235 | 0.036155856 | 0.063688285 | -0.01193363 | -0.048089486 | -0.075621915 | 0.411897867 | 0.09198802 | 0.748716188 | 0.269157235 | 0.05823842 |
| Rsl1d1 | S | 6 | 0.033391163 | 0.013570154 | 0.088301547 | 0.054910385 | 0.074731393 | 0.440200065 | 0.807647026 | 0.385923755 | 0.5722242 | 0.470503423 |
| Lin9 | S | 76 | 0.060304004 | 0.029370395 | 0.02804961 | -0.032254394 | -0.001320785 | 0.532026543 | 0.512970509 | 0.339077681 | 0.724130749 | 0.972135479 |
| Mtmr1 | T | 659 | 0.082946782 | 0.166554949 | -0.105053387 | -0.188000168 | -0.271608335 | 0.537634818 | 0.125633496 | 0.410274732 | 0.224258245 | 0.069313511 |
| Mycbp2 | S | 2968 | -0.043221303 | 0.01656748 | 0.096678162 | 0.139899465 | 0.080110682 | 0.268149955 | 0.617717062 | 0.011388878 | 0.016630983 | 0.031675783 |
| Mycbp2 | T | 2969 | -0.043221303 | 0.01656748 | 0.096678162 | 0.139899465 | 0.080110682 | 0.268149955 | 0.617717062 | 0.011388878 | 0.016630983 | 0.031675783 |
| Rbm26 | S | 523 | 0.207234781 | -0.06610226 | -0.020341936 | -0.227576717 | 0.045760324 | 0.035059947 | 0.37888208 | 0.776300358 | 0.025010089 | 0.141545757 |
| Phf12 | S | 661 | -5.3567E-05 | 0.0807896 | 0.039579878 | 0.039633445 | -0.041209721 | 0.998969894 | 0.260283359 | 0.447411713 | 0.471484027 | 0.566153391 |
| Glcci1 | S | 395 | -0.060050181 | 0.084527305 | 0.074518169 | 0.13456835 | -0.010009136 | 0.293621531 | 0.122408464 | 0.179543741 | 0.076620903 | 0.856977644 |
| Cttnbp2nl | S | 556 | 0.052783912 | -0.058855915 | 0.038178752 | -0.01460516 | 0.097034667 | 0.0184667 | 0.142768235 | 0.019708124 | 0.362699265 | 0.053220344 |
| Farp2 | S | 884 | 0.11558104 | -0.031887026 | -0.003072229 | -0.118653269 | 0.028814797 | 0.018695104 | 0.437163709 | 0.97588038 | 0.303144481 | 0.774968123 |
| C77080 | S | 886 | -0.011024843 | 0.049082145 | 0.043661512 | 0.054686356 | -0.005420633 | 0.726780429 | 0.187352846 | 0.167312809 | 0.051270595 | 0.811922868 |
| Rlim | S | 229 | 0.031227269 | 0.010661444 | 0.03538766 | 0.00416039 | 0.024726216 | 0.313868551 | 0.795802719 | 0.133088065 | 0.868542529 | 0.538973022 |
| Bcl9l | S | 102 | 0.096316985 | 0.025885089 | 0.017074909 | -0.079242077 | -0.008810181 | 0.277716028 | 0.761488154 | 0.743772597 | 0.347975121 | 0.913438693 |
| Eif3b | S | 40 | 0.039797824 | 0.021973173 | 0.034298638 | -0.005499186 | 0.012325465 | 0.136009297 | 0.694219736 | 0.588372226 | 0.927006844 | 0.868489909 |
| Map1a | T | 1633 | 0.018678951 | 0.04445706 | 0.047108258 | 0.028429307 | 0.002651198 | 0.700427556 | 0.391763813 | 0.356216609 | 0.631307697 | 0.964043273 |
| Spen | S | 1022 | -0.004602166 | -0.009346388 | 0.126708321 | 0.131310486 | 0.136054708 | 0.918438179 | 0.77774709 | 0.07424161 | 0.075272077 | 0.05148666 |
| Afdn | S | 1182 | -0.024806151 | 0.022288637 | 0.05900395 | 0.083810101 | 0.036715313 | 0.09658205 | 0.290846442 | 0.003303193 | 0.000800706 | 0.11238578 |
| Tra2b | S | 16 | 0.27044191 | -0.038879108 | -0.111031969 | -0.381473879 | -0.072152861 | 0.038928697 | 0.710136529 | 0.406895065 | 0.046292627 | 0.620055097 |
| Tra2b | S | 18 | 0.27044191 | -0.038879108 | -0.111031969 | -0.381473879 | -0.072152861 | 0.038928697 | 0.710136529 | 0.406895065 | 0.046292627 | 0.620055097 |
| Tjp1 | S | 179 | 0.001810308 | -0.001089168 | 0.121980847 | 0.120170539 | 0.123070015 | 0.961405921 | 0.975122379 | 0.127179848 | 0.13961343 | 0.13950945 |
| Pik3c3 | S | 244 | 0.089665836 | 0.005573821 | 0.035188658 | -0.054477177 | 0.029614838 | 0.185751962 | 0.933539114 | 0.767778766 | 0.656860414 | 0.809634261 |
| Iqgap1 | T | 582 | 0.147599931 | -0.070054137 | 0.061847892 | -0.085752039 | 0.131902029 | 0.22150608 | 0.616813845 | 0.469447518 | 0.4805172 | 0.401569027 |
| Srsf4 | S | 270 | 0.0404926 | 0.106722154 | -0.010504396 | -0.050996996 | -0.11722655 | 0.652620793 | 0.212781808 | 0.895201685 | 0.59697616 | 0.218859244 |
| Kat7 | S | 164 | -0.016548195 | 0.064109085 | 0.091540779 | 0.108088973 | 0.027431694 | 0.571662288 | 0.297273869 | 0.324485492 | 0.263569933 | 0.76421632 |
| Arhgap21 | S | 1425 | 0.068565005 | 0.040118853 | -0.008737827 | -0.077302832 | -0.04885668 | 0.178019978 | 0.384524149 | 0.858527327 | 0.078766837 | 0.179634536 |
| Thoc2 | S | 1514 | -0.151573566 | 0.184706495 | 0.105127521 | 0.256701088 | -0.079578973 | 0.246745784 | 0.084212041 | 0.455091294 | 0.160485767 | 0.580802445 |
| Tprn | S | 184 | 0.074571641 | -0.056929256 | 0.055760283 | -0.018811358 | 0.112689539 | 0.046028773 | 0.074710623 | 0.050542954 | 0.510362324 | 0.009864247 |
| Atf7ip | S | 885 | 0.14748896 | 0.048603179 | -0.09268574 | -0.2401747 | -0.141288919 | 0.107707272 | 0.530773704 | 0.520596635 | 0.169298756 | 0.366890843 |
| Trip12 | S | 75 | -0.031189518 | 0.077167585 | 0.060935295 | 0.092124813 | -0.01623229 | 0.376431762 | 0.40391513 | 0.061052962 | 0.034466683 | 0.8449971 |
| Pbrm1 | Y | 351 | -0.109673691 | 0.074388627 | 0.141075647 | 0.250749338 | 0.06668702 | 0.520388902 | 0.582490999 | 0.100546775 | 0.201719229 | 0.60571504 |
| Rtn4 | S | 81 | 0.019843352 | 0.054962055 | 0.04654698 | 0.026703628 | -0.008415075 | 0.717343664 | 0.327887158 | 0.524600793 | 0.64279251 | 0.879800474 |
| Recql5 | S | 773 | 0.075907349 | 0.030141516 | 0.02064911 | -0.055258239 | -0.009492407 | 0.449533371 | 0.530943778 | 0.514244024 | 0.566817484 | 0.829986961 |
| Hmga1 Hmga1b | T | 53 | 0.051350365 | -0.003635256 | 0.035313735 | -0.01603663 | 0.038948992 | 0.135373953 | 0.905346792 | 0.322817376 | 0.449032685 | 0.135724883 |
| Sptbn1 | S | 2137 | 0.004390018 | 0.024399439 | 0.038730129 | 0.034340111 | 0.01433069 | 0.835266591 | 0.597214935 | 0.034917423 | 0.144210417 | 0.743956141 |
| Zbtb7a | S | 537 | 0.015504783 | 0.019107636 | 0.058961709 | 0.043456926 | 0.039854073 | 0.710002824 | 0.61450018 | 0.15602701 | 0.2947726 | 0.280734182 |
| Slirp | S | 105 | 0.09538571 | 0.016338629 | -0.059230888 | -0.154616597 | -0.075569516 | 0.021466727 | 0.606567723 | 0.52475085 | 0.174119573 | 0.423342404 |
| Prune2 | S | 594 | 0.07508299 | 0.077378957 | -0.055041541 | -0.130124531 | -0.132420498 | 0.213463324 | 0.18156169 | 0.470186217 | 0.124670692 | 0.11935004 |
| Krcc1 | S | 180 | 0.015386419 | -0.015514051 | 0.101190242 | 0.085803822 | 0.116704293 | 0.612753367 | 0.46994892 | 0.06805717 | 0.103336297 | 0.049627084 |
| Dnm2 | S | 764 | -0.010892392 | 0.154479039 | -0.029683387 | -0.018790995 | -0.184162426 | 0.850187629 | 0.071030607 | 0.727891975 | 0.795646251 | 0.082564847 |
| Emsy | T | 171 | 0.025567689 | -0.082144201 | 0.130285029 | 0.104717341 | 0.212429231 | 0.345533542 | 0.023294523 | 0.002260622 | 0.011260489 | 0.00040289 |
| Hirip3 | S | 134 | 0.014102494 | 0.051585418 | 0.006754734 | -0.007347761 | -0.044830684 | 0.68417497 | 0.047699545 | 0.875961 | 0.874389123 | 0.327106674 |
| Phip | T | 910 | -0.020627482 | 0.010970788 | 0.086873634 | 0.107501116 | 0.075902846 | 0.781712745 | 0.827428857 | 0.042096497 | 0.228571229 | 0.210622887 |
| Cdkn1b | S | 178 | -0.020218519 | 0.092743659 | 0.060605347 | 0.080823866 | -0.032138312 | 0.632174443 | 0.297317523 | 0.328613435 | 0.242200337 | 0.716588082 |
| Ociad1 | T | 146 | 0.012813038 | 0.058312876 | 0.02006701 | 0.007253972 | -0.038245865 | 0.693383804 | 0.149903984 | 0.622086626 | 0.875660813 | 0.430470704 |
| Rexo1 | S | 905 | 0.147487364 | 0.032432913 | -0.072232074 | -0.219719437 | -0.104664987 | 0.072338544 | 0.767039614 | 0.52229562 | 0.114888994 | 0.412635278 |
| Abl1 | S | 569 | 0.056910541 | 0.032927236 | 0.053292727 | -0.003617814 | 0.020365492 | 0.506587211 | 0.715037849 | 0.48470313 | 0.970410912 | 0.843921606 |
| Mettl16 | S | 297 | -0.042308093 | 0.068434066 | 0.039744334 | 0.082052427 | -0.028689732 | 0.562089934 | 0.140944781 | 0.123945675 | 0.306629799 | 0.466336857 |
| Nup133 | S | 500 | 0.096033514 | -0.040930862 | 0.077411564 | -0.018621951 | 0.118342426 | 0.110515059 | 0.270393406 | 0.391395896 | 0.82995377 | 0.230562078 |
| Srrm2 | S | 1439 | 0.038127752 | -0.005974671 | 0.058741922 | 0.02061417 | 0.064716593 | 0.305520657 | 0.745008235 | 0.161491662 | 0.629986987 | 0.143922537 |
| Bag3 | S | 281 | -0.049734009 | 0.133675397 | 0.011359006 | 0.061093016 | -0.122316391 | 0.194056494 | 0.020028426 | 0.727271698 | 0.0651942 | 0.03995983 |
| Filip1l | S | 972 | 0.107166699 | -0.023332306 | 0.09695453 | -0.010212169 | 0.120286836 | 0.421836015 | 0.767334933 | 0.466134304 | 0.946456865 | 0.362664919 |
| Ppp6r3 | S | 822 | 0.085819339 | -0.04553696 | 0.085098044 | -0.000721296 | 0.130635004 | 0.253336118 | 0.673692149 | 0.376478269 | 0.992854841 | 0.282558604 |
| Ehbp1l1 | S | 1040 | 0.082294887 | 0.045994988 | -0.039102819 | -0.121397705 | -0.085097806 | 0.086375802 | 0.149076694 | 0.208585168 | 0.034937753 | 0.024556798 |
| Chd8 | S | 2213 | -0.056180594 | 0.098041381 | 0.030635113 | 0.086815707 | -0.067406268 | 0.027042953 | 0.005286303 | 0.176084373 | 0.014081679 | 0.03386142 |
| Sox4 | S | 312 | -0.191438637 | 0.00857138 | 0.3294847 | 0.520923337 | 0.32091332 | 0.398177045 | 0.95765673 | 0.086624612 | 0.080676273 | 0.147171738 |
| Sox4 | S | 316 | -0.191438637 | 0.00857138 | 0.3294847 | 0.520923337 | 0.32091332 | 0.398177045 | 0.95765673 | 0.086624612 | 0.080676273 | 0.147171738 |
| Sox4 | S | 320 | -0.191438637 | 0.00857138 | 0.3294847 | 0.520923337 | 0.32091332 | 0.398177045 | 0.95765673 | 0.086624612 | 0.080676273 | 0.147171738 |
| Nup88 | S | 47 | 0.066218804 | -0.009888113 | -0.002405067 | -0.068623871 | 0.007483046 | 0.007037467 | 0.651969073 | 0.951621997 | 0.166825585 | 0.8497147 |
| Nav3 | S | 1859 | 0.262870847 | 0.05721928 | -0.226530701 | -0.489401549 | -0.283749981 | 0.074704258 | 0.870667107 | 0.4624774 | 0.176969421 | 0.504218644 |
| Nes | S | 862 | 0.122609641 | 0.048532325 | -0.068749318 | -0.191358959 | -0.117281643 | 0.04639301 | 0.383331917 | 0.199816615 | 0.006021658 | 0.037519325 |
| Bcl9l | S | 1014 | -0.05559888 | 0.071757623 | 0.110650255 | 0.166249135 | 0.038892632 | 0.211656637 | 0.218682576 | 0.148291598 | 0.063694035 | 0.585664584 |
| Dact3 | S | 268 | 0.087875018 | 0.076191193 | 0.004570383 | -0.083304634 | -0.07162081 | 0.563101786 | 0.422221761 | 0.953277629 | 0.575498577 | 0.423877362 |
| Baz1b | S | 161 | 0.017591593 | 0.088496341 | -0.004814864 | -0.022406457 | -0.093311205 | 0.692079147 | 0.114442331 | 0.897687809 | 0.456495971 | 0.098508277 |
| Utp3 | S | 37 | 0.027863754 | 0.013588449 | 0.011407859 | -0.016455895 | -0.002180591 | 0.212047031 | 0.048289585 | 0.532281473 | 0.49124205 | 0.898960049 |
| Terf1 | S | 23 | 0.021780903 | 0.085147595 | 0.028467737 | 0.006686835 | -0.056679858 | 0.883030934 | 0.253963293 | 0.695289192 | 0.963256502 | 0.402790084 |
| Aff4 | S | 482 | -0.024853826 | 0.050595279 | 0.062825956 | 0.087679782 | 0.012230677 | 0.478696603 | 0.08551791 | 0.294127634 | 0.179974585 | 0.814078313 |
| Myo9a | S | 1832 | -0.04617055 | 0.078729744 | 0.055792427 | 0.101962977 | -0.022937317 | 0.286547036 | 0.081504307 | 0.220097693 | 0.023606579 | 0.4263686 |
| Pdcl | T | 19 | 0.005730103 | 0.068676685 | -0.007359921 | -0.013090024 | -0.076036605 | 0.873035175 | 0.026777602 | 0.869159091 | 0.785106612 | 0.157376985 |
| Prpf4b | S | 341 | -0.101291726 | 0.062439499 | 0.146927631 | 0.248219356 | 0.084488132 | 0.121111702 | 0.142606304 | 0.081126328 | 0.022077628 | 0.240248976 |
| Peak1 | S | 791 | -0.005824723 | 0.056595934 | 0.022034499 | 0.027859222 | -0.034561435 | 0.692193509 | 0.133919676 | 0.155979312 | 0.027873349 | 0.298767937 |
| Plec | S | 4649 | -0.035315077 | 0.040072692 | 0.096704623 | 0.1320197 | 0.05663193 | 0.333118935 | 0.271931962 | 0.110422348 | 0.064365781 | 0.272245799 |
| Sos1 | S | 1164 | -0.018754786 | 0.015456969 | 0.08059808 | 0.099352866 | 0.065141111 | 0.577789619 | 0.446217589 | 0.055537943 | 0.04974823 | 0.107708739 |
| Mcm6 | S | 704 | 0.049106631 | 0.03842737 | 0.001970656 | -0.047135975 | -0.036456715 | 0.157637744 | 0.427153514 | 0.941244426 | 0.163316567 | 0.440753168 |
| Rbm5 | S | 59 | -0.00884253 | -0.013913375 | 0.140838976 | 0.149681506 | 0.15475235 | 0.828627283 | 0.594843072 | 0.066081136 | 0.058766862 | 0.067055557 |

| | | | | | | | | | | | | |
|---|---|---|---|---|---|---|---|---|---|---|---|---|
| Cav2 | S | 23 | -0.126444814 | -0.016919671 | 0.322206833 | 0.448651648 | 0.339126504 | 0.065671074 | 0.77748297 | 0.017143969 | 0.011004426 | 0.023242736 |
| Mars1 | S | 827 | -0.077678729 | 0.082001401 | 0.083707801 | 0.161386531 | 0.0017064 | 0.17072024 | 0.036981794 | 0.245152329 | 0.072368034 | 0.977497813 |
| Pard3 | S | 383 | 0.096877727 | 0.015642212 | -0.035428212 | -0.132305938 | -0.051070423 | 0.026482915 | 0.624122775 | 0.372681549 | 0.012819618 | 0.147743763 |
| Pak1 | S | 220 | 0.079430089 | 0.032189205 | -0.001220172 | -0.080650261 | -0.033409377 | 0.146397603 | 0.712159773 | 0.986388992 | 0.266342157 | 0.7208998 |
| Chd5 | S | 185 | 0.005593997 | 0.069371871 | 0.048498552 | 0.042904555 | -0.020873319 | 0.899072631 | 0.306645189 | 0.513580339 | 0.578690052 | 0.804907039 |
| Chd5 | T | 189 | 0.005593997 | 0.069371871 | 0.048498552 | 0.042904555 | -0.020873319 | 0.899072631 | 0.306645189 | 0.513580339 | 0.578690052 | 0.804907039 |
| Rassf5 | S | 177 | 0.011962846 | 0.090637939 | 0.000469192 | -0.011493654 | -0.090168746 | 0.826409129 | 0.117506914 | 0.991471518 | 0.777531267 | 0.08396084 |
| Acin1 | S | 1003 | 0.010911056 | 0.01252115 | 0.061545948 | 0.050634892 | 0.049024799 | 0.688430034 | 0.614322702 | 0.139250039 | 0.1945092 | 0.202721188 |
| Arhgap31 | S | 235 | 0.185827331 | 0.321755698 | -0.294043911 | -0.479871241 | -0.615799608 | 0.64062993 | 0.238646128 | 0.461784842 | 0.308866855 | 0.155390683 |
| Scrib | S | 1561 | -0.034101787 | 0.066873345 | 0.062188243 | 0.09629003 | -0.004685101 | 0.815136962 | 0.254438073 | 0.30744874 | 0.514952082 | 0.861769557 |
| Atad5 | S | 87 | -0.028820047 | 0.069062964 | 0.067956729 | 0.096776776 | -0.001106234 | 0.461160789 | 0.09587525 | 0.439513952 | 0.296740101 | 0.989035018 |
| Git2 | S | 517 | 0.033336504 | 0.001696831 | 0.104628146 | 0.071291642 | 0.102931315 | 0.752462354 | 0.987804441 | 0.238953949 | 0.450910182 | 0.332246948 |
| Itgav | T | 1044 | 0.040807895 | 0.128703955 | 0.001157031 | -0.039650865 | -0.127546924 | 0.655048289 | 0.376246068 | 0.993485559 | 0.771603614 | 0.459083886 |
| Mast2 | S | 32 | -0.071700273 | 0.098672776 | 0.087973486 | 0.159673759 | -0.01069929 | 0.137580458 | 0.122285731 | 0.14713428 | 0.033550266 | 0.858621385 |
| Atf7ip | S | 57 | -0.023883004 | 0.017966054 | 0.10365482 | 0.127537825 | 0.085688766 | 0.624643254 | 0.732167388 | 0.073666759 | 0.021370029 | 0.092558957 |
| Ptk2b | Y | 579 | 0.034770903 | 0.373031489 | -0.145721462 | -0.180492365 | -0.518752951 | 0.82032651 | 0.277435676 | 0.570664912 | 0.499745889 | 0.192954319 |
| Ptk2b | Y | 580 | 0.034770903 | 0.373031489 | -0.145721462 | -0.180492365 | -0.518752951 | 0.82032651 | 0.277435676 | 0.570664912 | 0.499745889 | 0.192954319 |
| Osbpl5 | S | 311 | -0.096199788 | 0.006055406 | 0.213010749 | 0.309210538 | 0.206955343 | 0.503664264 | 0.91789857 | 0.08610614 | 0.10264417 | 0.093730832 |
| Senp1 | S | 159 | 0.034681904 | -0.027541726 | 0.08779948 | 0.053117577 | 0.115341207 | 0.582684228 | 0.652286268 | 0.090923687 | 0.35696465 | 0.093105497 |
| Itpr1 | S | 1588 | 0.002529251 | 0.129179018 | -0.031680694 | -0.034209945 | -0.160859712 | 0.948286154 | 0.062387336 | 0.619014211 | 0.609003142 | 0.076287187 |
| Srrm1 | T | 633 | 0.002286204 | -0.00817486 | 0.062656148 | 0.060369944 | 0.070831008 | 0.911196209 | 0.720252 | 0.012475057 | 0.014362547 | 0.025238685 |
| Zfhx3 | S | 2933 | 0.055912127 | 0.055948709 | 0.090877954 | 0.034965827 | 0.034929245 | 0.726606376 | 0.777627372 | 0.546304027 | 0.856581321 | 0.876041625 |
| Tut4 | S | 102 | 0.061800024 | 0.036428111 | -0.026184811 | -0.087984835 | -0.062612922 | 0.128387154 | 0.215493405 | 0.651187534 | 0.209285614 | 0.33113092 |
| Sh3pxd2a | T | 256 | 0.106040118 | 0.336215921 | -0.108331939 | -0.214372057 | -0.44454786 | 0.635044509 | 0.224856787 | 0.16571726 | 0.368287832 | 0.153471334 |
| Nfic | S | 192 | 0.02412734 | 0.057562702 | 0.058260712 | 0.034133371 | 0.00069801 | 0.774028509 | 0.503804834 | 0.472104919 | 0.55533665 | 0.990372973 |
| Dock6 | S | 878 | -0.151422165 | 0.293407448 | 0.061115734 | 0.212537899 | -0.232291714 | 0.36690707 | 0.190233892 | 0.597819028 | 0.236998635 | 0.277592797 |
| Ranbp10 | S | 365 | 0.066928798 | -0.002773472 | 0.079610161 | 0.012681363 | 0.082383633 | 0.419196749 | 0.979603449 | 0.450964626 | 0.881651185 | 0.471875945 |
| Afap1l2 | S | 144 | -0.094258433 | 0.15737457 | 0.0721845 | 0.166442933 | -0.08519007 | 0.204187335 | 0.074012298 | 0.473449411 | 0.131360523 | 0.381334475 |
| Cdv3 | S | 82 | 0.229979388 | -0.151911642 | 0.003763642 | -0.226215746 | 0.155675283 | 0.07290861 | 0.554819072 | 0.980900386 | 0.195509208 | 0.562517612 |
| Abca1 | S | 1141 | 0.075827151 | -0.009396095 | 0.060895763 | -0.014931389 | 0.070291857 | 0.288910574 | 0.871247658 | 0.470426037 | 0.870025907 | 0.441972048 |
| Usp24 | S | 2044 | -0.093761482 | 0.129189763 | 0.059618135 | 0.153379617 | -0.069571628 | 0.081905543 | 0.041647704 | 0.163479125 | 0.018786791 | 0.179669712 |
| Ttc21a | S | 141 | 0.090909794 | 0.094072967 | -0.077100426 | -0.168010221 | -0.171173393 | 0.032551706 | 0.086020705 | 0.018723997 | 0.006791595 | 0.025232137 |
| Ttc21a | Y | 147 | 0.090909794 | 0.094072967 | -0.077100426 | -0.168010221 | -0.171173393 | 0.032551706 | 0.086020705 | 0.018723997 | 0.006791595 | 0.025232137 |
| Zfp592 | S | 142 | 0.015025186 | -0.021212609 | 0.19730673 | 0.182281545 | 0.218519339 | 0.851555141 | 0.816187831 | 0.28057198 | 0.311503527 | 0.246303738 |
| Zfp592 | S | 145 | 0.015025186 | -0.021212609 | 0.19730673 | 0.182281545 | 0.218519339 | 0.851555141 | 0.816187831 | 0.28057198 | 0.311503527 | 0.246303738 |
| Zfp592 | S | 146 | 0.015025186 | -0.021212609 | 0.19730673 | 0.182281545 | 0.218519339 | 0.851555141 | 0.816187831 | 0.28057198 | 0.311503527 | 0.246303738 |
| Hoxc10 | T | 237 | -0.02636988 | 0.036187402 | 0.093905595 | 0.120275475 | 0.057718192 | 0.629606395 | 0.397566951 | 0.164400679 | 0.120318637 | 0.345374193 |
| Fcho2 | S | 681 | 0.066115462 | 0.015995022 | 0.027704438 | -0.038411024 | 0.011709416 | 0.201207644 | 0.756807216 | 0.738692746 | 0.6069054 | 0.874508248 |
| Frmd8 | S | 447 | 0.090669366 | 0.064288815 | 0.006958863 | -0.083710503 | -0.057329953 | 0.581172406 | 0.388391445 | 0.966830841 | 0.687378151 | 0.725931233 |
| Tcf20 | S | 1363 | 0.100177541 | 0.091558542 | -0.080113902 | -0.180291443 | -0.171672444 | 0.195056631 | 0.208736642 | 0.325788451 | 0.02949229 | 0.047747871 |
| Srrm2 | S | 1834 | 0.019935194 | 0.024591657 | 0.066368111 | 0.046432918 | 0.041776454 | 0.576161307 | 0.515617141 | 0.387723257 | 0.509249924 | 0.550848 |
| Poldip3 | S | 368 | 0.105133651 | 0.016454226 | -0.052455185 | -0.157588836 | -0.06890941 | 0.017787163 | 0.650122409 | 0.298502089 | 0.038776872 | 0.165817162 |
| Scrib | S | 948 | 0.037898582 | 0.034115708 | 0.039204026 | 0.001305444 | 0.005088317 | 0.564760209 | 0.372731171 | 0.553251688 | 0.986078408 | 0.928184823 |
| Trp53bp1 | T | 1644 | 0.153655478 | -0.040300654 | 0.033798909 | -0.119856569 | 0.074099564 | 0.171919277 | 0.380770018 | 0.46977441 | 0.257064868 | 0.088884979 |
| Fam117b | S | 190 | -0.085593945 | 0.167232813 | 0.050647876 | 0.136241821 | -0.116584938 | 0.093493076 | 0.049521267 | 0.371680293 | 0.042464391 | 0.131305432 |
| Sh2b3 | S | 129 | 0.06583418 | 0.038825625 | -0.003005627 | -0.068839807 | -0.041831253 | 0.208299557 | 0.472018404 | 0.969227138 | 0.366705886 | 0.569465477 |
| Arhgap23 | S | 607 | -0.055764468 | 0.130239921 | 0.042771039 | 0.098535507 | -0.087468882 | 0.298611742 | 0.090638088 | 0.459232553 | 0.062493723 | 0.193897037 |
| Gpatch8 | T | 650 | 0.137117136 | 0.00472032 | 0.013130595 | -0.123986541 | 0.008410276 | 0.269165264 | 0.925466783 | 0.892765795 | 0.364544123 | 0.927265222 |
| Macf1 | S | 4483 | 0.005799733 | 0.001579047 | 0.091202588 | 0.085402855 | 0.089623541 | 0.922580375 | 0.97914613 | 0.104742865 | 0.134931546 | 0.12916429 |
| Sulf2 | S | 843 | -0.022258512 | 0.094955712 | 0.021091798 | 0.04335031 | -0.073863914 | 0.467158281 | 0.053976111 | 0.712912302 | 0.475253091 | 0.271854941 |
| Rere | S | 594 | -0.020557147 | 0.071274142 | 0.054474775 | 0.075031922 | -0.016799367 | 0.458272785 | 0.10387358 | 0.459919225 | 0.339121465 | 0.815305369 |
| Tmem209 | T | 115 | 0.017879021 | 0.05344384 | 0.027478918 | 0.009599897 | -0.025964921 | 0.640424412 | 0.196994332 | 0.710777529 | 0.892126588 | 0.718525599 |
| Nipbl | S | 1083 | 0.029080831 | 0.051439945 | 0.006890247 | -0.022190583 | -0.044549698 | 0.543709807 | 0.124462974 | 0.942551033 | 0.823380115 | 0.647055699 |
| Nipbl | S | 1084 | 0.029080831 | 0.051439945 | 0.006890247 | -0.022190583 | -0.044549698 | 0.543709807 | 0.124462974 | 0.942551033 | 0.823380115 | 0.647055699 |
| Nipbl | S | 1090 | 0.029080831 | 0.051439945 | 0.006890247 | -0.022190583 | -0.044549698 | 0.543709807 | 0.124462974 | 0.942551033 | 0.823380115 | 0.647055699 |
| Psma1 | S | 177 | 0.01386669 | -0.021412952 | 0.120031374 | 0.106164684 | 0.141444326 | 0.793804209 | 0.862207625 | 0.146838767 | 0.174889299 | 0.311366649 |
| Arhgap21 | S | 686 | 0.070102935 | 0.068092753 | 0.056662699 | -0.013440236 | -0.011430054 | 0.839255407 | 0.562562876 | 0.596801365 | 0.96911015 | 0.929226763 |
| Arhgap21 | T | 685 | 0.070102935 | 0.068092753 | 0.056662699 | -0.013440236 | -0.011430054 | 0.839255407 | 0.562562876 | 0.596801365 | 0.96911015 | 0.929226763 |
| Jmjd1c | S | 992 | -0.074805128 | 0.04350806 | 0.126972738 | 0.201777866 | 0.083464678 | 0.439637478 | 0.629566896 | 0.067069204 | 0.100601201 | 0.393969754 |
| Ccdc6 | S | 360 | -0.001604999 | 0.025040513 | 0.094356365 | 0.095961364 | 0.069315852 | 0.990805293 | 0.628280465 | 0.224152596 | 0.531689057 | 0.381288168 |
| Pcm1 | S | 65 | 0.073040603 | -0.011397971 | 0.041210498 | -0.031830105 | 0.052608469 | 0.460302822 | 0.391370491 | 0.055750172 | 0.731822423 | 0.048556817 |
| Tasor2 | S | 1538 | 0.071523944 | 0.039303393 | -0.016471284 | -0.087995228 | -0.055774677 | 0.103560145 | 0.597623325 | 0.769657409 | 0.189349662 | 0.509167204 |
| Sp1 | S | 61 | -0.035110626 | 0.06141632 | 0.099035493 | 0.13414612 | 0.037619173 | 0.45032299 | 0.398999435 | 0.184846112 | 0.100211464 | 0.635897 |
| Ppp6r1 | S | 636 | 0.02255933 | 0.011074244 | 0.060354358 | 0.037795027 | 0.049280113 | 0.685967338 | 0.619855882 | 0.195540902 | 0.548217359 | 0.268013348 |
| Brpf1 | S | 49 | 0.053415081 | 0.10843845 | -0.030639166 | -0.084054247 | -0.139077615 | 0.417321757 | 0.330992048 | 0.795741793 | 0.517079599 | 0.35751738 |
| Wrnip1 | T | 87 | 0.019639946 | 0.057322734 | 0.050857437 | 0.03121749 | -0.006465297 | 0.859671303 | 0.41323262 | 0.459264136 | 0.754940256 | 0.869276574 |
| Pgrmc2 | T | 205 | -0.019265109 | -0.021757256 | 0.16895274 | 0.188217849 | 0.190709996 | 0.733472356 | 0.406977853 | 0.057314418 | 0.051012855 | 0.062043667 |
| Lemd3 | S | 261 | 0.011721214 | 0.033839945 | 0.06311566 | 0.051394647 | 0.029275915 | 0.832469564 | 0.356116266 | 0.388415549 | 0.498329078 | 0.649048236 |
| Lrch3 | S | 324 | -0.053625755 | 0.02456007 | 0.136757565 | 0.19038332 | 0.112197494 | 0.299033401 | 0.630402866 | 0.057214381 | 0.026690792 | 0.114306196 |
| Srrm2 | T | 1153 | -0.00787838 | 0.01351473 | 0.062844536 | 0.070722916 | 0.049329806 | 0.894366093 | 0.832193581 | 0.035879568 | 0.289697771 | 0.457577181 |
| Tubgcp3 | S | 510 | -0.018436599 | 0.172797303 | -0.03981414 | -0.021377542 | -0.212611444 | 0.869223362 | 0.089779531 | 0.708526503 | 0.872894621 | 0.122283057 |
| Acin1 | S | 647 | 0.14169265 | 0.109042764 | -0.092187482 | -0.233880132 | -0.201230246 | 0.139044537 | 0.313584773 | 0.1658488 | 0.0432937 | 0.117650407 |
| Macroh2a2 | S | 129 | -0.041846231 | 0.076460556 | 0.068155273 | 0.110001504 | -0.008305283 | 0.149229438 | 0.204157623 | 0.083982401 | 0.024011094 | 0.874687802 |
| Pbxip1 | S | 164 | 0.073358005 | 0.159331597 | -0.104057015 | -0.177415019 | -0.263388611 | 0.306440753 | 0.143010758 | 0.260849214 | 0.113495348 | 0.055884183 |
| Saal1 | S | 6 | 0.030534238 | 0.034631907 | 0.048257533 | 0.017723295 | 0.013625626 | 0.76134219 | 0.441878908 | 0.375401963 | 0.863955175 | 0.802996039 |
| Saal1 | S | 14 | 0.030534238 | 0.034631907 | 0.048257533 | 0.017723295 | 0.013625626 | 0.76134219 | 0.441878908 | 0.375401963 | 0.863955175 | 0.802996039 |
| Smarca4 | S | 657 | 0.046542902 | 0.094225615 | 0.002189799 | -0.044353104 | -0.092035817 | 0.515426651 | 0.406688022 | 0.955445498 | 0.518706999 | 0.413156894 |
| Smarca4 | S | 660 | 0.046542902 | 0.094225615 | 0.002189799 | -0.044353104 | -0.092035817 | 0.515426651 | 0.406688022 | 0.955445498 | 0.518706999 | 0.413156894 |
| Smarca4 | S | 662 | 0.046542902 | 0.094225615 | 0.002189799 | -0.044353104 | -0.092035817 | 0.515426651 | 0.406688022 | 0.955445498 | 0.518706999 | 0.413156894 |
| Cav2 | S | 20 | -0.126997395 | -0.018749365 | 0.326572236 | 0.453569631 | 0.3453216 | 0.063432115 | 0.763757526 | 0.018767588 | 0.011994386 | 0.022391009 |
| Phf14 | S | 291 | 0.198324359 | 0.05158565 | -0.068029908 | -0.266354267 | -0.119615558 | 0.249856164 | 0.769277181 | 0.672692073 | 0.184493896 | 0.540592801 |
| Snap23 | S | 20 | 0.118455228 | -0.054947371 | -0.03717015 | -0.155625378 | 0.017777222 | 0.000740316 | 0.28572225 | 0.396295022 | 0.036251922 | 0.744034953 |
| Iws1 | S | 185 | 0.016749521 | -0.018924225 | 0.063933242 | 0.04718372 | 0.082857466 | 0.527252673 | 0.417280095 | 0.019933756 | 0.10073521 | 0.009842499 |
| Aak1 | T | 671 | -0.096276691 | 0.12919995 | 0.083672125 | 0.179948816 | -0.045527825 | 0.254646429 | 0.112093714 | 0.325748926 | 0.043806826 | 0.441981012 |
| Cwc22 | S | 903 | -0.054457495 | 0.07908003 | 0.057814555 | 0.112272049 | -0.021265475 | 0.198435881 | 0.068187287 | 0.187882011 | 0.057442665 | 0.625414178 |
| Ifi203 Mndal | S | 109;109 | -0.037067833 | 0.011020919 | 0.090209738 | 0.127277571 | 0.079188819 | 0.676760017 | 0.777675799 | 0.03833089 | 0.223872177 | 0.04612487 |
| Oat | S | 188 | 0.026374117 | 0.070142047 | 0.040276342 | 0.013902225 | -0.029865704 | 0.701810528 | 0.4488623 | 0.610322543 | 0.785816765 | 0.706209031 |
| Acaca | S | 49 | 0.090686304 | 0.139161054 | -0.090905735 | -0.181592039 | -0.230066789 | 0.679867991 | 0.113248995 | 0.467114127 | 0.441530508 | 0.115171056 |
| Arhgap12 | S | 444 | 0.01934925 | 0.046128971 | 0.021673886 | 0.002324636 | -0.024455085 | 0.758701046 | 0.227795973 | 0.404895673 | 0.97119643 | 0.515902419 |
| Map1s | T | 659 | 0.059679256 | 0.028589319 | 0.003102235 | -0.056577021 | -0.025487084 | 0.148115266 | 0.583398413 | 0.950658798 | 0.230496922 | 0.629746721 |
| Iqsec2 | S | 383 | -0.09998037 | 0.118334887 | 0.084287396 | 0.184267766 | -0.034047491 | 0.02578263 | 0.024135926 | 0.129814375 | 0.022262234 | 0.474917374 |
| Kif15 | S | 568 | 0.049839528 | 0.037655457 | 0.037666949 | -0.012172579 | 1.14921E-05 | 0.486763769 | 0.566434386 | 0.602330151 | 0.866737172 | 0.99986431 |
| Sltm | S | 421 | -0.012717032 | 0.141183027 | -0.040789514 | -0.028072482 | -0.181972541 | 0.89577421 | 0.070279912 | 0.776506064 | 0.849868359 | 0.25872099 |
| Fzr1 | S | 70 | 0.050392828 | 0.065602508 | -0.057803131 | -0.108195959 | -0.123405639 | 0.139729142 | 0.060707901 | 0.282191495 | 0.092458221 | 0.080673233 |

| | | | | | | | | | | | | |
|---|---|---|---|---|---|---|---|---|---|---|---|---|
| Agap1 | S | 422 | 0.177244345 | -0.046170801 | -0.061942797 | -0.239187142 | -0.015771996 | 0.010625766 | 0.311957099 | 0.549852039 | 0.0879166 | 0.866504598 |
| Arhgap21 | S | 1655 | 0.010008904 | 0.018527781 | 0.049338589 | 0.039329685 | 0.030810808 | 0.759054554 | 0.573422704 | 0.127464645 | 0.216708406 | 0.313320338 |
| Rbms1 | T | 208 | -0.014226668 | 0.062867661 | 0.059742052 | 0.07396872 | -0.003125609 | 0.820110943 | 0.319209867 | 0.328661852 | 0.140690182 | 0.936939002 |
| Ncbp1 | S | 674 | -0.012515179 | 0.035277791 | 0.094441041 | 0.10695622 | 0.05916325 | 0.768330067 | 0.293531118 | 0.333433979 | 0.287112857 | 0.513023404 |
| Samd4b | S | 271 | 0.102024996 | 0.049679899 | -0.025527243 | -0.127552239 | -0.075207141 | 0.285747019 | 0.181323741 | 0.295950069 | 0.212424974 | 0.075627035 |
| Fkbpl | S | 38 | 0.185761682 | 0.131176128 | -0.269251569 | -0.455013251 | -0.400427697 | 0.103197964 | 0.265817745 | 0.269201123 | 0.116549672 | 0.143167527 |
| Eif3b | S | 120 | -0.000506104 | 0.061613277 | 0.021116735 | 0.021622839 | -0.040496542 | 0.982436446 | 0.180041883 | 0.344900614 | 0.18132913 | 0.327685574 |
| Csnk1d | T | 347 | 0.044899924 | 0.01701217 | 0.127058872 | 0.082158948 | 0.110046702 | 0.4755451 | 0.839883793 | 0.64817235 | 0.765185255 | 0.695165652 |
| Kirrel | S | 592 | 0.006600841 | 0.087720095 | 0.035503537 | 0.028902696 | -0.052216558 | 0.861280461 | 0.276358027 | 0.734960711 | 0.782386929 | 0.656672361 |
| Kif1c | S | 676 | -0.035054722 | 0.088484696 | 0.034073802 | 0.069128524 | -0.054410894 | 0.342449151 | 0.077606515 | 0.347250855 | 0.063893793 | 0.207229019 |
| Ybx3 | S | 193 | 0.01159743 | 0.048682671 | 0.009590484 | -0.002006945 | -0.039092187 | 0.782300527 | 0.067438087 | 0.721966434 | 0.960034801 | 0.116013756 |
| Ybx3 | S | 196 | 0.01159743 | 0.048682671 | 0.009590484 | -0.002006945 | -0.039092187 | 0.782300527 | 0.067438087 | 0.721966434 | 0.960034801 | 0.116013756 |
| Srrm1 | S | 378 | 0.026162671 | 0.009351641 | 0.071826413 | 0.045663741 | 0.062474772 | 0.563329997 | 0.818668258 | 0.27215444 | 0.429055148 | 0.294543457 |
| Snrnp200 | S | 225 | -0.0131009 | 0.031947386 | 0.084449004 | 0.097549904 | 0.052501617 | 0.69230464 | 0.341464554 | 0.224571261 | 0.181062415 | 0.39801987 |
| Chd8 | S | 2220 | 0.024493857 | 0.052896976 | 0.079360097 | 0.05486624 | 0.026463121 | 0.764158509 | 0.626044468 | 0.506620664 | 0.633103034 | 0.838959271 |
| Ehbp1l1 | T | 278 | 0.084504893 | 0.00649928 | 0.017428692 | -0.067076201 | 0.010929413 | 0.274800615 | 0.861502742 | 0.508382206 | 0.36197245 | 0.728097128 |
| Xkr5 | S | 517 | 0.024361588 | 0.059677227 | 0.002782211 | -0.021579378 | -0.056895017 | 0.726669079 | 0.09827279 | 0.966072355 | 0.799408097 | 0.410291919 |
| Ppp1r18 | S | 212 | 0.041080755 | -0.01432743 | 0.067229141 | 0.026148386 | 0.081556571 | 0.432548366 | 0.586190197 | 0.127981392 | 0.622550673 | 0.092649203 |
| Supt5 | T | 1028 | 0.071869012 | 0.005919946 | 0.027981627 | -0.043887386 | 0.022061681 | 0.313802396 | 0.840215332 | 0.452954793 | 0.519708409 | 0.536686665 |
| Phf20l1 | S | 584 | 0.02392003 | 0.058741595 | 0.063716604 | 0.039796574 | 0.004975009 | 0.755832198 | 0.538326634 | 0.545306874 | 0.71872356 | 0.966802056 |
| Nop58 | T | 34 | -0.070096484 | 0.060748011 | 0.137981836 | 0.20807832 | 0.077233825 | 0.729395932 | 0.427271859 | 0.332194294 | 0.367389338 | 0.544011646 |
| Ppp6r3 | S | 823 | 0.155774392 | -0.021676109 | -0.040680935 | -0.196455327 | -0.019004826 | 0.026453385 | 0.704661723 | 0.386383735 | 0.01645052 | 0.727873967 |
| Nucks1 | S | 58 | -0.090752423 | 0.141131281 | 0.046119899 | 0.136872322 | -0.095011382 | 0.143030538 | 0.047127918 | 0.312436716 | 0.05411178 | 0.134719658 |
| Tmcc3 | T | 176 | 0.094886441 | 0.023835545 | 0.001595697 | -0.093290744 | -0.022239848 | 0.182036912 | 0.85943853 | 0.982254111 | 0.148702013 | 0.864411082 |
| Kirrel | S | 715 | -0.073722519 | 0.076430844 | 0.139723163 | 0.213445682 | 0.063292318 | 0.310021864 | 0.571793908 | 0.110730991 | 0.038747379 | 0.639840292 |
| Mndal | S | 177 | -0.047191946 | 0.02025054 | 0.137163662 | 0.184355609 | 0.116913123 | 0.319208686 | 0.640141803 | 0.069852159 | 0.037989859 | 0.111730449 |
| Atp2b1 | S | 1177 | -0.128125834 | 0.059233966 | 0.16022836 | 0.288354194 | 0.100994394 | 0.367038245 | 0.469242759 | 0.095348735 | 0.099513045 | 0.275944641 |
| Pcdh18 | S | 778 | -0.096492647 | 0.04485466 | 0.183297173 | 0.27978982 | 0.138442512 | 0.008924303 | 0.167889587 | 0.017875505 | 0.01491578 | 0.051019031 |
| Dck | S | 16 | -0.084859044 | 0.045159892 | 0.10179308 | 0.186652124 | 0.056633188 | 0.202639858 | 0.197155576 | 0.044067184 | 0.036630823 | 0.202815824 |
| Tenm3 | S | 116 | 0.202150733 | -0.022895148 | -0.110712542 | -0.312863275 | -0.087817394 | 0.049929405 | 0.827653318 | 0.578386303 | 0.187673631 | 0.661697825 |
| Snrpa1 | S | 236 | 0.122430894 | 0.038272571 | -0.101921269 | -0.224352163 | -0.14019384 | 0.028333331 | 0.482346834 | 0.277936007 | 0.069656206 | 0.167806403 |
| Fam126b | S | 519 | 0.096709863 | -0.008728977 | 0.048715723 | -0.04799414 | 0.0574447 | 0.28393132 | 0.940409248 | 0.668603454 | 0.600302332 | 0.630109101 |
| Dnttip2 | S | 248 | 0.02027602 | 0.018418517 | 0.037062963 | 0.016786943 | 0.018644446 | 0.316451522 | 0.300998641 | 0.483413822 | 0.736627851 | 0.706717948 |
| Osbpl3 | S | 483 | 0.025045543 | 0.157763866 | -0.085269643 | -0.110315186 | -0.243033509 | 0.72620106 | 0.043248956 | 0.281499137 | 0.1350023 | 0.015534632 |
| Wdr11 | S | 959 | -0.053066954 | -0.000282098 | 0.172669063 | 0.225736017 | 0.172951161 | 0.461746021 | 0.996069841 | 0.075473031 | 0.049482305 | 0.082743209 |
| Tjp2 | S | 263 | -0.080054386 | -0.114539382 | 0.340465056 | 0.420519442 | 0.455004438 | 0.208558542 | 0.15493098 | 0.008090953 | 0.007531966 | 0.004714042 |
| Zfp646 | S | 1412 | -0.135300332 | 0.053146414 | 0.203509631 | 0.338809963 | 0.150363217 | 0.0589756 | 0.203337271 | 0.046015291 | 0.010780718 | 0.106139348 |
| Szt2 | T | 2433 | 0.075290911 | -0.002730414 | 0.04449692 | -0.030793991 | 0.047227334 | 0.285543623 | 0.950277265 | 0.529410667 | 0.711610012 | 0.524301777 |
| Rbmxl1 | S | 121 | 0.170463051 | -0.068631743 | 0.060767104 | -0.109695947 | 0.129398847 | 0.211118669 | 0.353431801 | 0.464674909 | 0.379131989 | 0.095902886 |
| AW554918 | S | 521 | 0.025434963 | 0.007048198 | 0.069041569 | 0.043606606 | 0.061993371 | 0.723703906 | 0.890023547 | 0.180814359 | 0.532238666 | 0.211521891 |
| AW554918 | S | 524 | 0.025434963 | 0.007048198 | 0.069041569 | 0.043606606 | 0.061993371 | 0.723703906 | 0.890023547 | 0.180814359 | 0.532238666 | 0.211521891 |
| Tanc1 | S | 267 | 0.065228681 | -0.007200805 | 0.072193953 | 0.006965272 | 0.079394758 | 0.415281196 | 0.906276795 | 0.420545732 | 0.932545042 | 0.315924807 |
| Cenpk | T | 197 | 0.115623909 | -0.081489458 | 0.082026055 | -0.033597855 | 0.163515513 | 0.298653334 | 0.487067211 | 0.262802937 | 0.723707787 | 0.198515299 |
| Aak1 | S | 622 | -0.022784746 | 0.079369566 | 0.031326277 | 0.054111023 | -0.048043289 | 0.323425976 | 0.071256549 | 0.418320207 | 0.206298191 | 0.297017637 |
| Rab11fip5 | T | 305 | 0.060591929 | -0.023822654 | 0.082482781 | 0.021890851 | 0.106305435 | 0.174567554 | 0.325402771 | 0.334343727 | 0.782859761 | 0.242061978 |
| Nktr | S | 609 | 0.223561428 | 0.316372143 | -0.164181553 | -0.387742981 | -0.480553695 | 0.246559725 | 0.24997731 | 0.029781299 | 0.102077694 | 0.133705505 |
| Farp1 | S | 899 | 0.011437431 | 0.051860345 | 0.041818603 | 0.030381172 | -0.010041742 | 0.916976201 | 0.436610275 | 0.303252266 | 0.776884275 | 0.859822862 |
| Ccm2 | S | 452 | 0.024050769 | 0.088916248 | 0.049580394 | 0.025529624 | -0.039335854 | 0.871946631 | 0.544898571 | 0.550007373 | 0.861960677 | 0.779482857 |
| Sf3b1 | T | 350 | -0.029854346 | 0.012827574 | 0.146294764 | 0.17614911 | 0.133467191 | 0.422749498 | 0.780365995 | 0.112414563 | 0.071305182 | 0.133067441 |
| Brd4 | S | 1086 | -0.043168545 | 0.055155056 | 0.132511507 | 0.175680052 | 0.077356451 | 0.34773786 | 0.412761809 | 0.211341944 | 0.132911654 | 0.446788532 |
| Socs6 | S | 159 | -0.055583605 | 0.123988957 | 0.035006019 | 0.090589624 | -0.088982938 | 0.307029116 | 0.050868506 | 0.709358981 | 0.369628921 | 0.377652189 |
| Zfp638 | S | 1099 | 0.006824189 | -0.005299227 | 0.08861689 | 0.081792701 | 0.093916118 | 0.870114874 | 0.861067123 | 0.093622388 | 0.130425879 | 0.096741788 |
| Rbm17 | S | 62 | 0.025515776 | 0.189464952 | -0.086644579 | -0.112160355 | -0.27610953 | 0.881949185 | 0.17789376 | 0.722564881 | 0.651534121 | 0.287705019 |
| Rev1 | S | 1106 | 0.019627315 | 0.050447225 | 0.095115177 | 0.075487861 | 0.044667951 | 0.834604985 | 0.691829982 | 0.47859246 | 0.589705188 | 0.777499931 |
| Ubxn7 | S | 266 | -0.035694825 | 0.075024581 | 0.053274474 | 0.0889693 | -0.021750106 | 0.557807641 | 0.105705178 | 0.49335682 | 0.298507502 | 0.753087471 |
| Mcm4 | S | 119 | 0.086860189 | -0.005257439 | 0.038293616 | -0.048566573 | 0.043551055 | 0.279773846 | 0.903111977 | 0.552447341 | 0.567739203 | 0.507415706 |
| Irs2 | T | 401 | -0.105462595 | 0.315915903 | 0.024597398 | 0.130059993 | -0.291318505 | 0.455120852 | 0.23439208 | 0.814661196 | 0.379616314 | 0.264314892 |
| Ccnl2 | S | 345 | 0.028703368 | -0.018783337 | 0.039138293 | 0.010434925 | 0.05792163 | 0.090148486 | 0.229941754 | 0.034712148 | 0.042613769 | 5.57436E-05 |
| Fnbp1l | T | 496 | 0.024623337 | 0.274194481 | -0.271142727 | -0.295766065 | -0.545337209 | 0.902280959 | 0.027023395 | 0.281538355 | 0.306647485 | 0.087816834 |
| Dhx9 | S | 323 | 0.121645566 | -0.052733437 | 0.048536895 | -0.07310867 | 0.101270333 | 0.122836443 | 0.538045381 | 0.689202997 | 0.535192738 | 0.423676289 |
| Tnrc6b | S | 422 | 0.138530977 | 0.08067257 | -0.093235804 | -0.23176678 | -0.173908374 | 0.155285366 | 0.223131638 | 0.183295386 | 0.047586474 | 0.029318136 |
| Tdrd3 | S | 349 | -0.084441618 | 0.095504635 | 0.092975478 | 0.177417096 | -0.002529157 | 0.250995475 | 0.177084235 | 0.189883304 | 0.068724653 | 0.972274327 |
| Pacc1 | S | 9 | 0.108562074 | -0.071942267 | 0.070854223 | -0.037707851 | 0.14279649 | 0.115289985 | 0.035907574 | 0.048690132 | 0.497570006 | 0.005157286 |
| Ash2l | S | 96 | 0.163860969 | 0.025378461 | 0.042918552 | -0.120942418 | 0.01754009 | 0.653290915 | 0.811688891 | 0.711918245 | 0.739369915 | 0.887634257 |
| Zmym4 | S | 306 | -0.052213382 | 0.012680084 | 0.208101525 | 0.260314908 | 0.195421442 | 0.694216663 | 0.935443767 | 0.214167851 | 0.142169975 | 0.289770668 |
| Atrx | S | 1135 | -0.053305161 | 0.022555083 | 0.11882856 | 0.172133721 | 0.096273478 | 0.294480927 | 0.594819983 | 0.046297807 | 0.023822782 | 0.095122652 |
| Sos1 | T | 1076 | -0.069378441 | 0.090302281 | 0.148472131 | 0.217850572 | 0.05816985 | 0.168219366 | 0.314197484 | 0.204565273 | 0.100404477 | 0.613733376 |
| Hcfc1 | S | 1516 | 0.020523308 | 0.028262592 | 0.020052217 | -0.000471091 | -0.008210376 | 0.517074102 | 0.228601673 | 0.396288991 | 0.987977509 | 0.723956453 |
| Jade2 | T | 662 | -0.006561569 | 0.042452251 | 0.085533902 | 0.092095471 | 0.043081651 | 0.939447727 | 0.59183456 | 0.284665876 | 0.177154266 | 0.322206229 |
| Akap12 | S | 234 | 0.012574948 | 0.127846823 | -0.065125316 | -0.077700264 | -0.192972139 | 0.73723835 | 0.011313058 | 0.133428716 | 0.060758762 | 0.003563371 |
| Zdhhc8 | S | 534 | -0.003343182 | 0.033488734 | 0.060312729 | 0.063655911 | 0.026823995 | 0.892568514 | 0.418130566 | 0.203410915 | 0.189394326 | 0.602810138 |
| Sgip1 | S | 300 | 0.271322005 | 0.190810894 | -0.2576876 | -0.529009605 | -0.448498494 | 0.323735909 | 0.347357732 | 0.314771373 | 0.117796834 | 0.116154245 |
| Hbs1l | T | 210 | 0.060495692 | 0.141714473 | -0.120022116 | -0.180517808 | -0.261736589 | 0.532105853 | 0.044079135 | 0.256805673 | 0.157137629 | 0.061255053 |
| Slc12a7 | S | 30 | 0.032123236 | -0.013269482 | 0.074744326 | 0.042621091 | 0.088013808 | 0.348840062 | 0.670278104 | 0.20255159 | 0.425781891 | 0.153425088 |
| Cdc5l | T | 404 | -0.044430639 | 0.069139812 | 0.13377612 | 0.178206759 | 0.064636309 | 0.484616498 | 0.443903278 | 0.305023234 | 0.200980082 | 0.604480885 |
| Cdc5l | T | 411 | -0.044430639 | 0.069139812 | 0.13377612 | 0.178206759 | 0.064636309 | 0.484616498 | 0.443903278 | 0.305023234 | 0.200980082 | 0.604480885 |
| Cdc5l | T | 415 | -0.044430639 | 0.069139812 | 0.13377612 | 0.178206759 | 0.064636309 | 0.484616498 | 0.443903278 | 0.305023234 | 0.200980082 | 0.604480885 |
| Map1a | S | 2591 | 0.021371114 | 0.059983171 | -0.010554993 | -0.031926107 | -0.070538164 | 0.405721551 | 0.123470747 | 0.877688385 | 0.639699302 | 0.353226603 |
| Ercc6l2 | S | 1387 | 0.051172493 | 0.077637967 | -0.000719835 | -0.051892328 | -0.078357802 | 0.637135538 | 0.300184055 | 0.995991715 | 0.732239946 | 0.571420197 |
| Rpusd1 | S | 271 | -0.011611094 | 0.125221798 | -0.001327498 | 0.010283596 | -0.126549296 | 0.873428942 | 0.135587981 | 0.981277641 | 0.89508373 | 0.149931028 |
| Papolg | S | 651 | 0.034317609 | 0.03400461 | 0.010069904 | -0.024247705 | -0.023934706 | 0.257597952 | 0.326447672 | 0.822554287 | 0.611667255 | 0.629989157 |
| Ints3 | S | 1038 | 0.109679266 | -0.033082998 | -0.004471788 | -0.114151055 | 0.028611209 | 0.026521071 | 0.473885458 | 0.925340783 | 0.046775711 | 0.535119583 |
| Srrm1 | S | 723 | 0.068130822 | -0.012280461 | 0.08012788 | 0.011997057 | 0.092408341 | 0.606393364 | 0.70389541 | 0.244548456 | 0.928030293 | 0.198408522 |
| Srrm1 | S | 725 | 0.06930002 | -0.013979815 | 0.08077111 | 0.01147109 | 0.094750925 | 0.626515729 | 0.678775239 | 0.222573438 | 0.935478948 | 0.172601464 |
| Srrm2 | S | 433 | 0.061458904 | -0.011125585 | 0.045912985 | -0.015545919 | 0.05703857 | 0.197008828 | 0.786190597 | 0.453985786 | 0.757026498 | 0.304019208 |
| Zfp280c | S | 142 | -0.114907952 | 0.075495798 | 0.117522293 | 0.232430245 | 0.042026495 | 0.15891191 | 0.200552786 | 0.065664219 | 0.028755326 | 0.400888454 |
| Dock7 | T | 186 | 0.052522804 | 0.008279611 | 0.075004899 | 0.022482095 | 0.066725289 | 0.739882206 | 0.936384324 | 0.289514081 | 0.886707929 | 0.543322841 |
| Sorbs2 | S | 28 | -0.04499257 | 0.046272234 | 0.106480046 | 0.151472617 | 0.060207812 | 0.292391407 | 0.303532367 | 0.144638722 | 0.08242983 | 0.332860855 |
| Cherp | S | 831 | 0.018695032 | 0.066936219 | 0.039461693 | 0.020766661 | -0.027474526 | 0.881850926 | 0.437457426 | 0.482362374 | 0.870123907 | 0.745006834 |
| Irf2bp1 | S | 186 | 0.129660048 | -0.072728877 | 0.054269763 | -0.075390285 | 0.12699864 | 0.086253451 | 0.238693629 | 0.43642267 | 0.349296444 | 0.139447994 |
| Aff4 | T | 523 | 0.004090743 | 0.052587008 | 0.065932829 | 0.061842086 | 0.013345821 | 0.954899147 | 0.53339222 | 0.366113108 | 0.287025695 | 0.845341135 |
| Bcl9 | S | 291 | 0.015904915 | 0.025016562 | 0.150606261 | 0.134701346 | 0.125589699 | 0.915528069 | 0.858440716 | 0.436334474 | 0.428396164 | 0.441761569 |
| Usp2 | S | 260 | 0.031407916 | 0.02770641 | 0.024389942 | -0.007017973 | -0.003316467 | 0.303298456 | 0.463611489 | 0.593687221 | 0.864702705 | 0.942435306 |
| Polr2a | T | 1877 | 0.123296721 | 0.091722772 | -0.020040418 | -0.143337139 | -0.111763189 | 0.479230236 | 0.615478239 | 0.835739417 | 0.42151302 | 0.547788972 |

| | | | | | | | | | | | | |
|---|---|---|---|---|---|---|---|---|---|---|---|---|
| Tbx3 | S | 407 | 0.121860136 | -0.034113372 | 0.052522454 | -0.069337682 | 0.086635825 | 0.182054179 | 0.58949293 | 0.722032611 | 0.65996245 | 0.579353588 |
| Srrm2 | S | 396 | 0.056879011 | 0.002892539 | 0.042713229 | -0.014165782 | 0.03982069 | 0.309777735 | 0.905365786 | 0.480367807 | 0.836757706 | 0.503954946 |
| Son | S | 1794 | 0.124725555 | -0.026383172 | 0.024693548 | -0.100032007 | 0.05107672 | 0.183511533 | 0.827794808 | 0.853314011 | 0.497081187 | 0.755266261 |
| Nmt1 | S | 31 | 0.020261591 | 0.061258334 | 0.016457701 | -0.00380389 | -0.044800632 | 0.576968516 | 0.421814934 | 0.442375853 | 0.908047997 | 0.539055302 |
| Lemd3 | S | 112 | 0.043225873 | 0.096169549 | -0.002901956 | -0.046127829 | -0.099071505 | 0.79026602 | 0.264910428 | 0.981132319 | 0.798093877 | 0.43915315 |
| Yju2 | S | 220 | 0.024801851 | 0.047709101 | 0.008420908 | -0.016380943 | -0.039288193 | 0.568663425 | 0.175596901 | 0.794468032 | 0.687420431 | 0.224712549 |
| Ncaph2 | S | 317 | 0.104356259 | 0.126876733 | -0.030269906 | -0.134626165 | -0.157146639 | 0.643341885 | 0.441750419 | 0.781170115 | 0.572543443 | 0.388998892 |
| E4f1 | S | 397 | -0.124487912 | 0.127390306 | 0.119020057 | 0.243507969 | -0.008370248 | 0.576745869 | 0.204463407 | 0.561516664 | 0.371960795 | 0.963720069 |
| E4f1 | S | 399 | -0.124487912 | 0.127390306 | 0.119020057 | 0.243507969 | -0.008370248 | 0.576745869 | 0.204463407 | 0.561516664 | 0.371960795 | 0.963720069 |
| E4f1 | S | 400 | -0.124487912 | 0.127390306 | 0.119020057 | 0.243507969 | -0.008370248 | 0.576745869 | 0.204463407 | 0.561516664 | 0.371960795 | 0.963720069 |
| Pum1 | S | 229 | 0.08608868 | -0.020640396 | 0.045491693 | -0.040596987 | 0.066132089 | 0.241697219 | 0.636138675 | 0.426262507 | 0.593119626 | 0.318166635 |
| Ncor2 | T | 1406 | -0.001700185 | -0.06589835 | 0.262939018 | 0.264639203 | 0.328837368 | 0.985658127 | 0.308430185 | 0.129544552 | 0.133110647 | 0.097225903 |
| Dock7 | S | 910 | 0.065442137 | 0.133070267 | -0.079762586 | -0.145204724 | -0.212832853 | 0.489269658 | 0.095321522 | 0.263651811 | 0.19633236 | 0.041277742 |
| Arhgap35 | S | 1134 | -0.00733362 | 0.09895174 | 0.029614658 | 0.036948278 | -0.069337082 | 0.906497535 | 0.225087714 | 0.751376497 | 0.723235543 | 0.526656109 |
| Zfp668 | S | 10 | 0.025934191 | -0.002030161 | 0.059144808 | 0.033210617 | 0.061174969 | 0.345573119 | 0.911281012 | 0.225886774 | 0.461149102 | 0.222140437 |
| Kansl1 | S | 759 | -0.142322364 | 0.090778779 | 0.189665903 | 0.331988267 | 0.098887125 | 0.177262314 | 0.155178121 | 0.226405363 | 0.078767976 | 0.475486836 |
| Phf10 | S | 49 | -0.01016143 | -0.026868893 | 0.127612304 | 0.137773734 | 0.154481197 | 0.865278881 | 0.672202858 | 0.065221057 | 0.00609026 | 0.015749427 |
| Trerf1 | S | 961 | 0.089209116 | -0.049766772 | 0.113261911 | 0.024052795 | 0.163028683 | 0.651992351 | 0.767237218 | 0.302382484 | 0.902593879 | 0.374241286 |
| Trerf1 | T | 960 | 0.089209116 | -0.049766772 | 0.113261911 | 0.024052795 | 0.163028683 | 0.651992351 | 0.767237218 | 0.302382484 | 0.902593879 | 0.374241286 |
| Tpd52l2 | T | 187 | 0.200673914 | 0.055426818 | -0.192379127 | -0.393053041 | -0.247805945 | 0.067869752 | 0.579343426 | 0.332664869 | 0.107469818 | 0.232457499 |
| Vps13c | S | 873 | 0.053117828 | 0.056610099 | 0.005992985 | -0.047124843 | -0.050617114 | 0.619627771 | 0.30238158 | 0.872874017 | 0.658425332 | 0.358837202 |
| Ppp1r12a | T | 858 | 0.040645426 | 0.014710755 | 0.054582616 | 0.01393719 | 0.039871861 | 0.467759353 | 0.86481971 | 0.392184866 | 0.830888558 | 0.66938879 |
| Myo18a | S | 1970 | -0.018082255 | 0.011681854 | 0.096616537 | 0.114698792 | 0.084934683 | 0.650005146 | 0.743745217 | 0.094496693 | 0.066657291 | 0.127530104 |
| Sf3a3 | T | 475 | 0.133275255 | 0.014497563 | -0.024422689 | -0.157697945 | -0.038920252 | 0.170960857 | 0.760098059 | 0.662414876 | 0.12504327 | 0.358419472 |
| Cdk13 | S | 384 | 0.007529668 | -0.013544189 | 0.103147051 | 0.095617383 | 0.116691239 | 0.880511919 | 0.757592271 | 0.109283875 | 0.128899874 | 0.076897999 |
| Sltm | T | 345 | -0.111147994 | 0.121295259 | 0.139678888 | 0.250826881 | 0.018383629 | 0.120329781 | 0.216623874 | 0.164762192 | 0.044524989 | 0.859183656 |
| Pcm1 | S | 1301 | 0.057415582 | 0.026935462 | 0.03338971 | -0.024025872 | 0.006454248 | 0.448536934 | 0.678458209 | 0.603269918 | 0.689889438 | 0.888796059 |
| Puf60 | T | 319 | 0.144623815 | 0.066704204 | -0.10419815 | -0.248821965 | -0.170902354 | 0.270996643 | 0.574962942 | 0.730919732 | 0.445310361 | 0.586333618 |
| Sptan1 | S | 982 | 0.062787628 | 0.060926996 | -0.03493957 | -0.097727198 | -0.095866566 | 0.406502214 | 0.173651081 | 0.667516643 | 0.334247471 | 0.28743801 |
| Vps26a | S | 315 | 0.117738264 | 0.019914374 | 0.010339076 | -0.107399188 | -0.009575298 | 0.355202501 | 0.781382308 | 0.935482895 | 0.511610577 | 0.944485426 |
| Rai1 | S | 1343 | 0.037973853 | 0.007836365 | 0.067643774 | 0.029669921 | 0.059807409 | 0.659006057 | 0.863899402 | 0.299920344 | 0.751529469 | 0.376342022 |
| Klf6 | S | 146 | -0.297748948 | 0.195605223 | 0.138949731 | 0.436698679 | -0.056655493 | 0.382539796 | 0.067890884 | 0.628554913 | 0.292557246 | 0.837951224 |
| Gigyf2 | T | 25 | 0.263427417 | -0.108385193 | -0.063838612 | -0.327266029 | 0.044546581 | 0.023190409 | 0.353299912 | 0.483111151 | 0.016621873 | 0.683823021 |
| Arhgap12 | T | 102 | 0.107626174 | 0.068014013 | -0.010537562 | -0.118163737 | -0.078551576 | 0.49128874 | 0.574100482 | 0.933510807 | 0.373563373 | 0.316509783 |
| Msantd2 | S | 436 | -0.012491825 | 0.026214684 | 0.081547036 | 0.094038861 | 0.055332352 | 0.843552203 | 0.536182351 | 0.200702571 | 0.231607567 | 0.349354946 |
| Farp2 | T | 28 | 0.14239817 | -0.109156908 | -0.003230278 | -0.145628448 | 0.10592663 | 0.004013648 | 0.186276805 | 0.93660603 | 0.014180114 | 0.198163988 |
| Rai1 | S | 996 | 0.037764408 | 0.038719124 | 0.020368401 | -0.017396007 | -0.018350723 | 0.385312328 | 0.433844017 | 0.77001798 | 0.785880486 | 0.785176959 |
| Soga1 | S | 1015 | -0.061150244 | 0.056461529 | 0.06818054 | 0.129330784 | 0.011719011 | 0.004220322 | 0.005907957 | 0.03306409 | 0.014818954 | 0.581738076 |
| Efr3a | S | 367 | 0.187292228 | -0.00217958 | -0.14726263 | -0.334554857 | -0.14508305 | 0.033131024 | 0.976423968 | 0.43622611 | 0.147725009 | 0.44125159 |
| Srrm2 | S | 949 | -0.034307893 | 0.046257461 | 0.089337232 | 0.123645126 | 0.043079771 | 0.571151241 | 0.296913557 | 0.26648671 | 0.178956774 | 0.557628245 |
| Acin1 | S | 391 | -0.045662118 | 0.031559596 | 0.112948349 | 0.158610466 | 0.081388753 | 0.082395729 | 0.241562872 | 0.057822268 | 0.026983831 | 0.120830284 |
| Micall1 | S | 649 | 0.111860788 | -0.008119685 | 0.033700641 | -0.078160147 | 0.041820326 | 0.368866712 | 0.948098136 | 0.613568153 | 0.484989123 | 0.716014237 |
| Pdcl | S | 296 | 0.025683115 | 0.086275159 | -0.019101372 | -0.044784487 | -0.105376531 | 0.776449703 | 0.090239431 | 0.624206862 | 0.617238019 | 0.045104932 |
| Msantd2 | S | 48 | 0.008345544 | 0.02340632 | 0.076386472 | 0.068040928 | 0.052980152 | 0.813441475 | 0.701449256 | 0.287337294 | 0.337103148 | 0.510981382 |
| Nop56 | S | 461 | 0.071807453 | -0.018008183 | 0.058936189 | -0.012871263 | 0.076944372 | 0.242731086 | 0.852567342 | 0.592804652 | 0.906540346 | 0.56396939 |
| Nop56 | S | 462 | 0.071807453 | -0.018008183 | 0.058936189 | -0.012871263 | 0.076944372 | 0.242731086 | 0.852567342 | 0.592804652 | 0.906540346 | 0.56396939 |
| Irf2bp2 | S | 443 | 0.078664169 | -0.00702703 | -0.001323298 | -0.079987466 | 0.005703732 | 0.053825153 | 0.854897605 | 0.975188596 | 0.092284954 | 0.87138255 |
| Ubl7 | S | 230 | -0.01236174 | 0.048817181 | 0.046482081 | 0.058843821 | -0.0023351 | 0.824967861 | 0.217146485 | 0.368746235 | 0.382438315 | 0.962587418 |
| Ttf2 | S | 119 | 0.093278803 | 0.036394986 | -0.058384771 | -0.151663574 | -0.094779757 | 0.033740465 | 0.525819267 | 0.272168498 | 0.035758163 | 0.181561619 |
| Mis18bp1 | S | 997 | 0.018950154 | -0.024102373 | 0.064617967 | 0.045667813 | 0.088720339 | 0.636676361 | 0.82995207 | 0.070738256 | 0.311024047 | 0.462553218 |
| Nf1 | S | 2819 | 0.047157349 | 0.072331333 | -0.014581979 | -0.061739328 | -0.086913312 | 0.419382107 | 0.231551464 | 0.582886746 | 0.310327362 | 0.174159674 |
| Prdx5 | T | 93 | -0.015611131 | 0.115176943 | 0.0835381 | 0.099149231 | -0.031638843 | 0.887680742 | 0.467717349 | 0.660443648 | 0.581682879 | 0.874001743 |
| Ncbp3 | S | 25 | -0.003240637 | 0.060151082 | 0.042241649 | 0.045482287 | -0.017909433 | 0.910779914 | 0.199229773 | 0.608907112 | 0.584593363 | 0.830980911 |
| Zc3h13 | S | 924 | -0.033067481 | 0.019980278 | 0.068396978 | 0.101464459 | 0.0484167 | 0.736912619 | 0.621786592 | 0.057679193 | 0.353581922 | 0.28111734 |
| Prpf4b | S | 88 | -0.022149111 | -0.00771923 | 0.080572615 | 0.102721726 | 0.088291845 | 0.512971271 | 0.650316112 | 0.008773476 | 0.04026407 | 0.013347291 |
| Nup98 | S | 934 | 0.066703545 | -0.000123354 | 0.00987472 | -0.056828824 | 0.009998074 | 0.111639668 | 0.99756829 | 0.723123974 | 0.178721683 | 0.818741119 |
| Arhgef12 | S | 309 | 0.002368786 | 0.024943142 | 0.04751187 | 0.045143085 | 0.022568728 | 0.931982997 | 0.442937051 | 0.185387241 | 0.229130779 | 0.545459048 |
| Fanci | T | 948 | 0.016397589 | -0.025809705 | 0.104338812 | 0.087941223 | 0.130148517 | 0.865117269 | 0.817971573 | 0.129728676 | 0.342247847 | 0.27473985 |
| Lmnb1 | S | 409 | -0.047721495 | 0.239027923 | -0.118909983 | -0.071188489 | -0.357937906 | 0.673388 | 0.058042319 | 0.593182575 | 0.729788079 | 0.179356621 |
| Gtf3c2 | S | 219 | 0.015766347 | -0.000502961 | 0.063126861 | 0.047360514 | 0.063629821 | 0.745191076 | 0.987428781 | 0.110958708 | 0.31031944 | 0.055806471 |
| Rptor | S | 859 | -0.065722744 | 0.062355654 | 0.076243552 | 0.141966295 | 0.013887898 | 0.389519462 | 0.188081208 | 0.175561818 | 0.107267258 | 0.697230606 |
| Lig3 | S | 243 | 0.195517891 | -0.00121872 | -0.12072212 | -0.316240012 | -0.1195034 | 0.015886315 | 0.983788547 | 0.176479003 | 0.020997064 | 0.156093892 |
| Anln | S | 974 | -0.052881109 | 0.005900995 | 0.181794267 | 0.234675377 | 0.175893272 | 0.458547126 | 0.85067572 | 0.129278303 | 0.074997947 | 0.144711229 |
| Anln | S | 979 | -0.052881109 | 0.005900995 | 0.181794267 | 0.234675377 | 0.175893272 | 0.458547126 | 0.85067572 | 0.129278303 | 0.074997947 | 0.144711229 |
| Ubxn8 | S | 141 | 0.168948325 | -0.048315272 | -0.007194979 | -0.176143303 | 0.041120293 | 0.076815169 | 0.456834728 | 0.908586102 | 0.069547586 | 0.310403241 |
| Zeb1 | S | 657 | 0.064482965 | 0.084882595 | -0.072595029 | -0.137077994 | -0.157477624 | 0.589974253 | 0.092359985 | 0.586199248 | 0.405599648 | 0.287746834 |
| Cwc22 | T | 905 | -0.041359322 | 0.069340134 | 0.094349736 | 0.135709058 | 0.025009602 | 0.554383422 | 0.494389441 | 0.230602926 | 0.110268795 | 0.800545622 |
| Ep400 | S | 735 | 0.006955933 | -0.009798751 | 0.08359222 | 0.076636288 | 0.093390971 | 0.833292294 | 0.778736409 | 0.088020753 | 0.109409365 | 0.071062041 |
| Prpf4b | S | 328 | -0.020748116 | -0.061575479 | 0.253706096 | 0.274454211 | 0.315281574 | 0.755442203 | 0.227931069 | 0.073186359 | 0.063585361 | 0.063966099 |
| Rb1cc1 | S | 649 | 0.066783593 | 0.059698904 | 0.008333166 | -0.058450427 | -0.051365737 | 0.573290349 | 0.466973268 | 0.895318715 | 0.619790682 | 0.530627731 |
| Cit | S | 1348 | 0.072029548 | 0.030800082 | -0.011048093 | -0.083077641 | -0.041848175 | 0.238331022 | 0.462631579 | 0.869928069 | 0.272066401 | 0.501931485 |
| Tcof1 | S | 1191 | -0.017499692 | 0.005256258 | 0.064786404 | 0.082286096 | 0.059530146 | 0.621021732 | 0.811475047 | 0.023134656 | 0.07407209 | 0.020140615 |
| Cenpu | T | 174 | 0.080631553 | -0.039113176 | 0.015669011 | -0.064962543 | 0.054782186 | 0.037455208 | 0.567491427 | 0.731287628 | 0.180545283 | 0.450253818 |
| Sfr1 | S | 67 | 0.023279866 | 0.057154742 | -0.023657527 | -0.046937393 | -0.080812269 | 0.298593512 | 0.040097496 | 0.335464305 | 0.043757157 | 0.010145549 |
| Paxbp1 | S | 264 | -0.007426276 | -0.015495026 | 0.097994932 | 0.105421208 | 0.113489957 | 0.839928603 | 0.552649848 | 0.037802845 | 0.047933755 | 0.037412729 |
| Qars | S | 70 | -0.053902785 | 0.2298551 | -0.112869255 | -0.058966469 | -0.342724355 | 0.485196721 | 0.023687036 | 0.461456592 | 0.666159273 | 0.088719509 |
| Apc | S | 1040 | 0.007083194 | 0.056733957 | 0.044970371 | 0.037887177 | -0.011763586 | 0.931233836 | 0.356613263 | 0.524704554 | 0.63275304 | 0.827709459 |
| Trpm4 | S | 1207 | -0.357653362 | 0.061189747 | 0.21427915 | 0.571932513 | 0.153089403 | 0.342618233 | 0.727123578 | 0.02028141 | 0.18679593 | 0.402325085 |
| 2310033P09Rik | S | 232 | 0.048352123 | 0.015142516 | 0.007465905 | -0.040886218 | -0.007676611 | 0.156970082 | 0.604521747 | 0.733681021 | 0.208570073 | 0.775312158 |
| Arhgef40 | S | 1485 | -0.034713599 | 0.091829547 | 0.022945858 | 0.057659457 | -0.068883689 | 0.472725542 | 0.099700265 | 0.448529115 | 0.28207731 | 0.18315797 |
| Pdha1 | S | 300 | 0.020468494 | 0.065369719 | -0.011565999 | -0.032034493 | -0.076935719 | 0.604588941 | 0.103516818 | 0.788435319 | 0.369265013 | 0.089732479 |
| Yap1 | S | 90 | -0.000436768 | 0.170483282 | -0.091184686 | -0.090747918 | -0.261667968 | 0.994323495 | 0.07134764 | 0.544913877 | 0.549289744 | 0.157530747 |
| Mbd2 | S | 44 | -0.080808691 | 0.156382537 | 0.025096957 | 0.105905648 | -0.13128558 | 0.219182564 | 0.044527378 | 0.741551925 | 0.149285103 | 0.100208798 |
| Jph2 | S | 479 | 0.098142816 | -0.075990013 | 0.056855735 | -0.041287082 | 0.132845747 | 0.060503952 | 0.104422188 | 0.159120955 | 0.353620337 | 0.024946308 |
| Jph2 | T | 483 | 0.098142816 | -0.075990013 | 0.056855735 | -0.041287082 | 0.132845747 | 0.060503952 | 0.104422188 | 0.159120955 | 0.353620337 | 0.024946308 |
| Mta2 | S | 435 | 0.012039289 | -0.009845418 | 0.068168051 | 0.056128762 | 0.07801347 | 0.492930134 | 0.604146069 | 0.083143373 | 0.139399521 | 0.0691184 |
| Dusp6 Dusp7 | S | 331;372 | -0.146129629 | 0.224331765 | 0.099212075 | 0.245341704 | -0.12511969 | 0.099925988 | 0.106095808 | 0.393397459 | 0.088211863 | 0.37272407 |
| Tsc2 | S | 1365 | -0.006784561 | 0.179311881 | -0.115604322 | -0.108819761 | -0.294916203 | 0.906420998 | 0.030381466 | 0.383986917 | 0.405190773 | 0.086931436 |
| Zc3h13 | S | 209 | 0.031858132 | 0.017296418 | 0.011219077 | -0.020639055 | -0.006077341 | 0.116166814 | 0.390811427 | 0.844244473 | 0.721214843 | 0.915459615 |
| Kdm2a | S | 444 | 0.03017565 | 0.064384624 | -0.02541683 | -0.05559248 | -0.089801454 | 0.222020112 | 0.180672478 | 0.609960787 | 0.298705107 | 0.162206899 |
| Afdn | S | 561 | -0.109350415 | 0.113195563 | 0.084136856 | 0.193487271 | -0.029058707 | 0.035923254 | 0.031556888 | 0.135059323 | 0.014510092 | 0.453867863 |
| Frmd8 | S | 409 | 0.062493823 | 0.118692253 | -0.096505737 | -0.158999561 | -0.21519799 | 0.442392312 | 0.122280302 | 0.411754858 | 0.22265109 | 0.124349176 |
| Kif1c | S | 1036 | 0.01127828 | 0.063363591 | 0.044596688 | 0.033318408 | -0.018766903 | 0.871315953 | 0.419965579 | 0.568840918 | 0.558298897 | 0.771050273 |
| Spty2d1 | S | 483 | 0.182088064 | 0.041968837 | -0.069331672 | -0.251419736 | -0.111300509 | 0.272322804 | 0.605922192 | 0.563164127 | 0.174455214 | 0.336045111 |

| | | | | | | | | | | | | |
|---|---|---|---|---|---|---|---|---|---|---|---|---|
| Mrtfa | S | 548 | 0.014280045 | -0.027600911 | 0.089231064 | 0.074951019 | 0.116831976 | 0.710363495 | 0.517127064 | 0.062754223 | 0.113999229 | 0.043879712 |
| Ccar1 | S | 829 | -0.083277739 | 0.015603809 | 0.214475288 | 0.297753026 | 0.198871479 | 0.303211456 | 0.709564541 | 0.120562601 | 0.057360751 | 0.150483041 |
| Arid4a | S | 1022 | -0.122443188 | 0.246668447 | 0.08870762 | 0.211150807 | -0.157960827 | 0.195996719 | 0.187197931 | 0.536432756 | 0.185622927 | 0.407530043 |
| Arid4a | S | 1025 | -0.122443188 | 0.246668447 | 0.08870762 | 0.211150807 | -0.157960827 | 0.195996719 | 0.187197931 | 0.536432756 | 0.185622927 | 0.407530043 |
| Rb1cc1 | T | 663 | 0.087583674 | 0.110028561 | -0.031494787 | -0.119078461 | -0.141523348 | 0.524412379 | 0.325911408 | 0.451235572 | 0.407744051 | 0.234379568 |
| Dnttip2 | S | 108 | 0.158289226 | -0.011288258 | -0.037800744 | -0.19608997 | -0.026512486 | 0.074456126 | 0.827456475 | 0.488248586 | 0.047923552 | 0.511303475 |
| Phactr4 | S | 620 | 0.026241001 | 0.007821701 | 0.051866969 | 0.025625968 | 0.044045267 | 0.355018669 | 0.906360633 | 0.336441657 | 0.600395449 | 0.563072974 |
| Prpf31 | T | 455 | 0.006296703 | 0.01537031 | 0.087487355 | 0.081190652 | 0.072117045 | 0.906903579 | 0.803528688 | 0.251805212 | 0.242571583 | 0.316279807 |
| Fancm | S | 1379 | 0.064704266 | -0.002012604 | 0.058587374 | -0.006116892 | 0.060599978 | 0.631748504 | 0.972504043 | 0.301136966 | 0.963191689 | 0.363230377 |
| Fancm | S | 1381 | 0.064704266 | -0.002012604 | 0.058587374 | -0.006116892 | 0.060599978 | 0.631748504 | 0.972504043 | 0.301136966 | 0.963191689 | 0.363230377 |
| Fancm | T | 1378 | 0.064704266 | -0.002012604 | 0.058587374 | -0.006116892 | 0.060599978 | 0.631748504 | 0.972504043 | 0.301136966 | 0.963191689 | 0.363230377 |
| Gtf3c1 | T | 1196 | 0.096922305 | -0.102635555 | 0.082708793 | -0.014213513 | 0.185344348 | 0.222673293 | 0.495133903 | 0.296793016 | 0.856075177 | 0.263327447 |
| Tbc1d10b | S | 142 | 0.116160605 | 0.021636308 | -0.087554103 | -0.203714708 | -0.109190412 | 0.026572687 | 0.680818708 | 0.331669476 | 0.082462653 | 0.250851075 |
| Rgs12 | S | 850 | 0.09890721 | 0.033832382 | -0.055413447 | -0.154320657 | -0.089245828 | 0.068274238 | 0.462787073 | 0.331837417 | 0.02476781 | 0.102040811 |
| Wbp4 | S | 230 | 0.099689041 | -0.080150518 | 0.055520964 | -0.044168077 | 0.135671482 | 0.016167985 | 0.05081369 | 0.253189444 | 0.30346951 | 0.032251972 |
| Mdc1 | T | 942 | -0.149876475 | 0.099107656 | 0.163735605 | 0.313612081 | 0.064627949 | 0.045854688 | 0.101037058 | 0.087414179 | 0.014244492 | 0.410735367 |
| Ralgapa1 | S | 733 | -0.05513004 | 0.206655929 | -0.008670758 | 0.046459282 | -0.215326687 | 0.612378775 | 0.16131488 | 0.9491535 | 0.697241555 | 0.178289827 |
| Palm | S | 87 | 0.033965208 | 0.029660299 | -0.001690075 | -0.035655283 | -0.031350374 | 0.285281331 | 0.166677882 | 0.95637579 | 0.320527009 | 0.303403101 |
| Pcyt1a | S | 347 | 0.115286084 | 0.085743288 | -0.167886039 | -0.283172124 | -0.253629327 | 0.167189366 | 0.015365819 | 0.097856287 | 0.028473127 | 0.053966439 |
| Tns2 | S | 825 | 0.075651916 | 0.036289273 | -0.00406022 | -0.079712136 | -0.040349493 | 0.258839028 | 0.664817239 | 0.954595585 | 0.339585994 | 0.67129722 |
| Rpl8 | S | 14 | 0.336371848 | 0.079685799 | -0.173405962 | -0.50977781 | -0.253091761 | 0.127322966 | 0.465263301 | 0.086174835 | 0.058574135 | 0.054069934 |
| Mon2 | S | 1213 | 0.04251539 | 0.152522007 | -0.102263899 | -0.144779289 | -0.254785906 | 0.405900765 | 0.056902933 | 0.164722215 | 0.072351301 | 0.017642733 |
| Bsdc1 | S | 415 | 0.103034461 | -0.087448175 | 0.045521913 | -0.057512548 | 0.132970088 | 0.115902885 | 0.669160679 | 0.635427698 | 0.549827306 | 0.539124547 |
| Zkscan1 | S | 354 | -0.013490228 | 0.089901577 | 0.038357885 | 0.051848113 | -0.051543692 | 0.857683733 | 0.225897683 | 0.741015219 | 0.683126302 | 0.675653513 |
| Rai14 | S | 300 | 0.118621419 | 0.033841664 | -0.0920085 | -0.21062992 | -0.125850164 | 0.047743293 | 0.533960119 | 0.330615989 | 0.091006541 | 0.202211866 |
| Tasor | S | 836 | -0.021698647 | -0.023652532 | 0.111914096 | 0.133612742 | 0.135566628 | 0.5563053 | 0.501515845 | 0.018705232 | 0.00547621 | 0.00777552 |
| Brpf1 | S | 84 | 0.099064422 | -0.015852957 | 0.006537536 | -0.092526886 | 0.022390492 | 0.121265281 | 0.83348547 | 0.92038921 | 0.231685866 | 0.792391165 |
| Eif3b | S | 84 | -0.004555348 | 0.047133333 | 0.070601621 | 0.075156969 | 0.023468288 | 0.950533965 | 0.36736643 | 0.489604032 | 0.509800971 | 0.814398538 |
| Iqsec1 | S | 923 | 0.043560354 | 0.062307759 | -0.018299103 | -0.061859457 | -0.080606862 | 0.245034144 | 0.315002478 | 0.647313551 | 0.084022214 | 0.199314257 |
| Hirip3 | T | 391 | 0.038455457 | 0.022793878 | 0.021297915 | -0.017157542 | -0.001495964 | 0.397521565 | 0.608429014 | 0.447010788 | 0.692954823 | 0.972675827 |
| Lats2 | S | 262 | -0.098585477 | 0.185035036 | 0.044690295 | 0.143275772 | -0.140344741 | 0.370285996 | 0.141845949 | 0.655753346 | 0.259440462 | 0.278125829 |
| Zc3h13 | Y | 373 | 0.108387505 | -0.036501044 | 0.008037744 | -0.100349762 | 0.044538788 | 0.088147329 | 0.706578685 | 0.92611713 | 0.321446033 | 0.711089453 |
| Srsf4 Srsf5 Srsf6 | S | 117;119;113 | -0.069620989 | -0.02639695 | 0.231874184 | 0.301495173 | 0.258271135 | 0.636559265 | 0.633154116 | 0.11203985 | 0.125680006 | 0.106192418 |
| Zeb2 | S | 142 | 0.014926124 | 0.117712824 | 0.022017021 | 0.007090898 | -0.095695802 | 0.857671956 | 0.436160998 | 0.828328486 | 0.924477357 | 0.510658005 |
| Arhgap42 | S | 680 | 0.086334702 | 0.236495245 | -0.122982339 | -0.209317041 | -0.359477584 | 0.653220085 | 0.341765582 | 0.550393794 | 0.161163027 | 0.133143977 |
| 1110004F10Rik | S | 17 | -0.003700327 | 0.104157007 | -0.014979459 | -0.011279131 | -0.119136465 | 0.937500112 | 0.095894937 | 0.821613212 | 0.87735027 | 0.165868613 |
| Nbeal2 | S | 2424 | -0.074709283 | 0.101700354 | 0.110535801 | 0.185245084 | 0.008835447 | 0.271876419 | 0.491815314 | 0.138504954 | 0.057951043 | 0.950982477 |
| Nbeal2 | S | 2431 | -0.074709283 | 0.101700354 | 0.110535801 | 0.185245084 | 0.008835447 | 0.271876419 | 0.491815314 | 0.138504954 | 0.057951043 | 0.950982477 |
| Nbeal2 | Y | 2420 | -0.074709283 | 0.101700354 | 0.110535801 | 0.185245084 | 0.008835447 | 0.271876419 | 0.491815314 | 0.138504954 | 0.057951043 | 0.950982477 |
| Abl2 | S | 621 | 0.154102213 | 0.045420558 | -0.116296358 | -0.270398571 | -0.161716917 | 0.0119008 | 0.524075305 | 0.058997222 | 0.004531348 | 0.07862984 |
| Gigyf2 | S | 385 | 0.005154318 | 0.000689713 | 0.071452077 | 0.06629776 | 0.070762364 | 0.954834763 | 0.989877772 | 0.099991407 | 0.481457965 | 0.214895122 |
| Sin3a | T | 436 | 0.014438594 | 0.017579379 | 0.036726311 | 0.022287717 | 0.019146932 | 0.505456479 | 0.565791428 | 0.25589473 | 0.442373009 | 0.580400255 |
| Fam120a | S | 485 | -0.028948259 | 0.113427437 | 0.015594517 | 0.044542776 | -0.09783292 | 0.686926062 | 0.138116693 | 0.833199729 | 0.607391531 | 0.271757647 |
| Spp1 | T | 41 | -0.150275058 | 0.005639856 | 0.242800708 | 0.393075766 | 0.237160852 | 0.695195613 | 0.982985898 | 0.178498368 | 0.353809435 | 0.418258755 |
| Spen | S | 1302 | -0.033668528 | 0.031625765 | 0.079868872 | 0.1135374 | 0.048243108 | 0.489037801 | 0.565453779 | 0.084989852 | 0.064985676 | 0.394450391 |
| Sf3b1 | T | 227 | -0.002969297 | 0.011773748 | 0.057341937 | 0.060311234 | 0.045568189 | 0.92667807 | 0.684845662 | 0.090673095 | 0.048694535 | 0.044522971 |
| Bag2 | S | 26 | -0.237665108 | 0.083151035 | 0.149560393 | 0.387225501 | 0.066409357 | 0.066215371 | 0.094166309 | 0.016630781 | 0.025152841 | 0.129991364 |
| Nedd4l | S | 504 | 0.089037784 | 0.089840085 | -0.066281946 | -0.15531973 | -0.156122031 | 0.147149078 | 0.349003576 | 0.261574548 | 0.058515697 | 0.158987841 |
| Scyl2 | S | 677 | -0.097973158 | 0.245940566 | -0.0030472 | 0.094925957 | -0.248987767 | 0.128042466 | 0.042555888 | 0.966789565 | 0.212340659 | 0.048723622 |
| Pdcl | S | 18 | 0.006516833 | 0.065150652 | -0.017016857 | -0.02353369 | -0.082167509 | 0.84783772 | 0.033298204 | 0.677756895 | 0.595923312 | 0.108387972 |
| Synj1 | S | 1350 | -0.009825126 | 0.075245673 | -0.008579936 | 0.00124519 | -0.083825608 | 0.82252517 | 0.028276098 | 0.729715127 | 0.974255022 | 0.019546004 |
| Arhgap21 | S | 1426 | -0.03557272 | 0.056163804 | 0.094488057 | 0.130060777 | 0.038324253 | 0.002886204 | 0.086890182 | 0.065319085 | 0.039077795 | 0.321140519 |
| Myc | S | 164 | 0.107886019 | -0.029984509 | 0.030971076 | -0.076914943 | 0.060955585 | 0.234405728 | 0.674110831 | 0.555995563 | 0.376127861 | 0.423736598 |
| Strn3 | S | 229 | 0.100861914 | 0.039938432 | -0.070955364 | -0.171817277 | -0.110893796 | 0.062718205 | 0.275235022 | 0.207564424 | 0.026004558 | 0.084966545 |
| Hirip3 | S | 595 | 0.045974923 | 0.027653207 | -0.006503061 | -0.052477985 | -0.034156268 | 0.107089006 | 0.535240542 | 0.866356602 | 0.200597811 | 0.481543483 |
| Hirip3 | S | 596 | 0.045974923 | 0.027653207 | -0.006503061 | -0.052477985 | -0.034156268 | 0.107089006 | 0.535240542 | 0.866356602 | 0.200597811 | 0.481543483 |
| Hirip3 | S | 600 | 0.045974923 | 0.027653207 | -0.006503061 | -0.052477985 | -0.034156268 | 0.107089006 | 0.535240542 | 0.866356602 | 0.200597811 | 0.481543483 |
| Plekhg3 | S | 1158 | 0.025317053 | 0.012933748 | 0.028959749 | 0.003642696 | 0.016026001 | 0.432104535 | 0.477023403 | 0.338118529 | 0.921299706 | 0.592239022 |
| Slk | S | 341 | 0.019142931 | 0.111153102 | -0.045863391 | -0.065006322 | -0.157016493 | 0.776156309 | 0.063235026 | 0.407633682 | 0.40628323 | 0.047198365 |
| Ttc3 | S | 1060 | 0.016272942 | 0.070132558 | -0.014970809 | -0.031243751 | -0.085103367 | 0.684665079 | 0.098069219 | 0.777261664 | 0.50530911 | 0.139034166 |
| Pcm1 | S | 68 | 0.027611623 | 0.014406979 | 0.041218459 | 0.013606836 | 0.02681148 | 0.563158589 | 0.62412151 | 0.337776834 | 0.777206605 | 0.442632837 |
| Polq | S | 1596 | -0.03237962 | 0.049439593 | 0.210245766 | 0.242625387 | 0.160806174 | 0.811007233 | 0.832503276 | 0.454883424 | 0.398527885 | 0.616475639 |
| Wdr81 | S | 682 | 0.062479024 | 0.108972893 | -0.043572825 | -0.106051849 | -0.152545718 | 0.554163841 | 0.242569505 | 0.533561511 | 0.367102175 | 0.155940147 |
| Nap1l4 | S | 54 | 0.026533465 | 0.116354562 | -0.027506639 | -0.054040104 | -0.143861201 | 0.729895651 | 0.143119363 | 0.452946268 | 0.485625646 | 0.102259904 |
| Leo1 | S | 280 | 0.058182795 | 0.044639925 | -0.033866644 | -0.092049439 | -0.078506569 | 0.043279159 | 0.342023591 | 0.218613917 | 0.010294159 | 0.142009506 |
| Nes | S | 1216 | -0.020299699 | 0.015713978 | 0.07678043 | 0.09708013 | 0.061066452 | 0.497070597 | 0.644796415 | 0.067377323 | 0.045823809 | 0.174140742 |
| Tmcc1 | S | 378 | -0.036892197 | -0.014465673 | 0.161024792 | 0.197916989 | 0.175490465 | 0.641070546 | 0.824861832 | 0.107670615 | 0.088925149 | 0.098277078 |
| Apc | S | 2533 | 0.294218118 | 0.040934609 | -0.054765849 | -0.348983967 | -0.095700458 | 0.594494504 | 0.684240571 | 0.839912055 | 0.553564831 | 0.730332788 |
| Arhgap31 | S | 1413 | -0.017119906 | 0.140606489 | 0.001485633 | 0.018605539 | -0.139120855 | 0.892817652 | 0.225330571 | 0.987421157 | 0.8532651 | 0.138429561 |
| Mis18bp1 | S | 478 | 0.072927135 | 0.037315695 | -0.015322525 | -0.08824966 | -0.052638221 | 0.271144016 | 0.51557014 | 0.859837916 | 0.311333214 | 0.504351031 |
| Jade2 | T | 738 | 0.079384672 | 0.048565449 | 0.038676196 | -0.040708476 | -0.009889253 | 0.581864045 | 0.718086938 | 0.835196696 | 0.810085489 | 0.95142945 |
| Atg2a | T | 560 | 0.013102045 | 0.083663847 | 0.084253106 | 0.071151061 | 0.000589259 | 0.87776282 | 0.668494301 | 0.643510157 | 0.694350649 | 0.998016943 |
| Atg2a | T | 562 | 0.013102045 | 0.083663847 | 0.084253106 | 0.071151061 | 0.000589259 | 0.87776282 | 0.668494301 | 0.643510157 | 0.694350649 | 0.998016943 |
| Zfp318 | S | 2140 | 0.012266559 | -0.051966051 | 0.124620245 | 0.112353686 | 0.176586296 | 0.882786762 | 0.164858601 | 0.023382328 | 0.244950741 | 0.011226419 |
| Reps1 | S | 539 | -0.084969577 | 0.118295459 | 0.010796289 | 0.095765866 | -0.10749917 | 0.567818661 | 0.050170029 | 0.853190801 | 0.520011448 | 0.067684545 |
| Ccar1 | S | 336 | 0.054053772 | -0.055466612 | 0.09264011 | 0.038586338 | 0.148106721 | 0.219038044 | 0.354529993 | 0.227915641 | 0.575538296 | 0.107144274 |
| Ccar1 | S | 340 | 0.054053772 | -0.055466612 | 0.09264011 | 0.038586338 | 0.148106721 | 0.219038044 | 0.354529993 | 0.227915641 | 0.575538296 | 0.107144274 |
| Pcif1 | S | 143 | -0.057367088 | 0.050353643 | 0.138432016 | 0.195799104 | 0.088078373 | 0.756531595 | 0.671213851 | 0.312120792 | 0.333302403 | 0.493921167 |
| Cobl | S | 1128 | -0.111243645 | 0.022367403 | 0.108022889 | 0.219266533 | 0.085655485 | 0.214510416 | 0.557002734 | 0.008702512 | 0.063223202 | 0.080779835 |
| Pde2a | S | 204 | 0.013013869 | 0.247914786 | -0.195845599 | -0.208859468 | -0.443760385 | 0.891758814 | 0.01961494 | 0.150004152 | 0.141858223 | 0.021366709 |
| Reep3 | S | 152 | -0.009851364 | 0.136639319 | -0.016173583 | -0.006322219 | -0.152812902 | 0.894183282 | 0.158256459 | 0.843021011 | 0.939434301 | 0.155030881 |
| Phf23 | S | 330 | -0.010439626 | 0.077522008 | 0.03613413 | 0.046573757 | -0.041387877 | 0.92208456 | 0.301757297 | 0.582032052 | 0.673039409 | 0.583861076 |
| Mical3 | S | 1179 | 0.132280796 | -0.043959107 | 0.012991929 | -0.119288867 | 0.056951036 | 0.09187984 | 0.367920936 | 0.736386139 | 0.118855416 | 0.269512779 |
| Macf1 | S | 7296 | 0.004801615 | 0.049881854 | 0.038396268 | 0.033594652 | -0.011485586 | 0.867743808 | 0.383258772 | 0.469016099 | 0.510508953 | 0.857793165 |
| Usf1 Usf2 | S | 222;186 | 0.045889425 | 0.044246251 | -0.002974446 | -0.04886387 | -0.047220697 | 0.507718355 | 0.248508681 | 0.971143287 | 0.616807192 | 0.580703621 |
| Yaf2 | S | 172 | 0.067985263 | 0.001232235 | 0.047084566 | -0.020900697 | 0.045852331 | 0.475120426 | 0.977091197 | 0.484779404 | 0.836486315 | 0.517797437 |
| Map7d1 | S | 275 | 0.085858536 | 0.137289477 | -0.072122241 | -0.157980777 | -0.209411718 | 0.431573556 | 0.333487653 | 0.44366259 | 0.236745572 | 0.195351411 |
| Hspa5 | S | 650 | -0.017774112 | 0.024275929 | 0.043284735 | 0.061058846 | 0.019008805 | 0.756864795 | 0.310559225 | 0.099704269 | 0.329827763 | 0.253850613 |
| Mycbp2 | S | 2916 | -0.042838559 | 0.102020608 | -0.023903908 | 0.01893465 | -0.125924516 | 0.596252201 | 0.014053778 | 0.675306684 | 0.825534955 | 0.103983605 |
| Aebp2 | S | 199 | -0.020596855 | 0.04247098 | 0.026871962 | 0.047468817 | -0.015599018 | 0.684939331 | 0.09652991 | 0.273355697 | 0.373724522 | 0.392085996 |
| Zfp148 | S | 665 | 0.062820446 | 0.031925901 | 0.010894135 | -0.05192631 | -0.021031765 | 0.457811233 | 0.485085363 | 0.867074321 | 0.547249949 | 0.692744808 |
| Prrc2a | T | 798 | 0.245831019 | -0.042428289 | -0.110165846 | -0.355996865 | -0.067737557 | 0.019164873 | 0.594968035 | 0.194546354 | 0.002236655 | 0.070192483 |
| Clasrp | S | 639 | 0.054273958 | -0.081901782 | 0.122811729 | 0.068537771 | 0.204713511 | 0.629584873 | 0.349105568 | 0.086729366 | 0.533031475 | 0.056644153 |
| Mis18bp1 | S | 985 | 0.027913039 | 0.003533174 | 0.068840347 | 0.040927307 | 0.065307172 | 0.751901335 | 0.931994041 | 0.253761346 | 0.669696889 | 0.303494866 |

| | | | | | | | | | | | | |
|---|---|---|---|---|---|---|---|---|---|---|---|---|
| Brat1 | S | 735 | 0.068607524 | 0.071321835 | -0.069074802 | -0.137682326 | -0.140396637 | 0.099822593 | 0.062004291 | 0.068165401 | 0.020481993 | 0.000116489 |
| Foxm1 | T | 658 | 0.09185421 | 0.052818944 | 0.000447586 | -0.091406624 | -0.052371358 | 0.408209333 | 0.746347213 | 0.998440436 | 0.688969617 | 0.833126275 |
| Acbd5 | S | 187 | -0.077544937 | -0.010913121 | 0.175362366 | 0.252907303 | 0.186275487 | 0.331889176 | 0.823562334 | 0.040349026 | 0.031661122 | 0.039912781 |
| Acbd5 | S | 191 | -0.077544937 | -0.010913121 | 0.175362366 | 0.252907303 | 0.186275487 | 0.331889176 | 0.823562334 | 0.040349026 | 0.031661122 | 0.039912781 |
| Nelfa | T | 157 | 0.067041768 | 0.047809009 | 0.015172228 | -0.05186954 | -0.032636781 | 0.630033111 | 0.500653349 | 0.825731949 | 0.692073509 | 0.496806386 |
| Zfp768 | S | 65 | -0.037270574 | -0.033043745 | 0.180635111 | 0.217905685 | 0.213678856 | 0.551475565 | 0.728289104 | 0.088232776 | 0.075692508 | 0.084647301 |
| Zfp768 | S | 72 | -0.037270574 | -0.033043745 | 0.180635111 | 0.217905685 | 0.213678856 | 0.551475565 | 0.728289104 | 0.088232776 | 0.075692508 | 0.084647301 |
| Zfp768 | S | 79 | -0.037270574 | -0.033043745 | 0.180635111 | 0.217905685 | 0.213678856 | 0.551475565 | 0.728289104 | 0.088232776 | 0.075692508 | 0.084647301 |
| Xrcc4 | S | 320 | 0.043746511 | 0.043710163 | 0.015039421 | -0.028707089 | -0.028670742 | 0.642990443 | 0.318958624 | 0.85617855 | 0.797171184 | 0.728450326 |
| Syde1 | S | 244 | 0.038240975 | 0.055778812 | -0.018785405 | -0.057026381 | -0.074564218 | 0.445484623 | 0.153613166 | 0.657754857 | 0.262162407 | 0.084267184 |
| Srrm1 | S | 647 | 0.001923523 | -0.00894886 | 0.075377991 | 0.073454468 | 0.084326851 | 0.949175661 | 0.759864837 | 0.068227346 | 0.101820184 | 0.068098323 |
| Evl | S | 367 | -0.202075842 | 0.13040214 | 0.176808244 | 0.378884086 | 0.046406104 | 0.000594569 | 0.077215015 | 0.001792671 | 0.000192985 | 0.424393388 |
| Chchd3 | S | 113 | 0.168460767 | -0.109717868 | -0.045057394 | -0.213518161 | 0.064660474 | 0.015847457 | 0.425250123 | 0.729872929 | 0.182399717 | 0.695094556 |
| Thoc5 | S | 312 | 0.033589336 | -0.028789726 | 0.075391269 | 0.041801932 | 0.104180994 | 0.343189159 | 0.337373019 | 0.123256434 | 0.343002855 | 0.063169643 |
| Thoc5 | S | 314 | 0.033589336 | -0.028789726 | 0.075391269 | 0.041801932 | 0.104180994 | 0.343189159 | 0.337373019 | 0.123256434 | 0.343002855 | 0.063169643 |
| Cav2 | Y | 27 | -0.114839333 | -0.043911224 | 0.185732008 | 0.300571342 | 0.229643232 | 0.13923917 | 0.536075027 | 0.002620538 | 0.018480645 | 0.039530534 |
| Amfr | T | 548 | -0.244344523 | 0.121370706 | 0.165933503 | 0.410278026 | 0.044562797 | 0.592800498 | 0.46629722 | 0.275558029 | 0.391462929 | 0.672372689 |
| Rock2 | S | 1134 | -0.060785446 | 0.103335428 | 0.052933599 | 0.113719045 | -0.050401828 | 0.12332034 | 0.067160089 | 0.344173861 | 0.088988287 | 0.395064258 |
| Zc3h13 | T | 263 | -0.030211712 | -0.042174554 | 0.151832196 | 0.182043908 | 0.19400675 | 0.551895619 | 0.40889398 | 0.027869223 | 0.016100091 | 0.013158461 |
| Piezo1 | T | 1917 | 0.132388684 | -0.019051683 | -0.06043765 | -0.192826334 | -0.041385967 | 0.01351062 | 0.611454557 | 0.357715378 | 0.039187738 | 0.494911719 |
| Tiam2 | T | 1606 | -0.022873278 | 0.045115252 | 0.06897778 | 0.091851058 | 0.023862528 | 0.349705308 | 0.235797961 | 0.251974747 | 0.163317221 | 0.671336607 |
| Arid1b | S | 454 | 0.077692991 | -0.002768238 | -0.002178234 | -0.079871224 | 0.000590004 | 0.093189135 | 0.933228192 | 0.976400247 | 0.324095884 | 0.993254242 |
| Mki67 | T | 1766 | -0.003265791 | 0.048371525 | 0.037997813 | 0.041263603 | -0.010373712 | 0.934523245 | 0.789495711 | 0.625640851 | 0.612255292 | 0.892802191 |
| AW554918 | S | 64 | 0.051255706 | -0.025089393 | 0.058691112 | 0.007435406 | 0.083780505 | 0.467152243 | 0.652605835 | 0.187078291 | 0.90757193 | 0.164646184 |
| AW554918 | S | 67 | 0.051255706 | -0.025089393 | 0.058691112 | 0.007435406 | 0.083780505 | 0.467152243 | 0.652605835 | 0.187078291 | 0.90757193 | 0.164646184 |
| Slmap | S | 235 | 0.056383211 | 0.017860793 | 0.045847754 | -0.010535457 | 0.027986961 | 0.523784817 | 0.773614559 | 0.616566231 | 0.915143505 | 0.732533086 |
| Kmt2a | S | 2282 | -0.045413872 | 0.12033705 | -0.006270171 | 0.039143701 | -0.126607221 | 0.267798635 | 0.0210852 | 0.868731224 | 0.029732415 | 0.018846931 |
| Gpbp1 | S | 49 | 0.02648784 | 0.028696412 | 0.03827726 | 0.01178942 | 0.009580848 | 0.758330768 | 0.361230314 | 0.553500611 | 0.904025574 | 0.879615486 |
| Snrpa1 | S | 239 | 0.08461388 | 0.036594621 | -0.071257152 | -0.155871032 | -0.107851773 | 0.02340923 | 0.452620592 | 0.222844739 | 0.04656572 | 0.132201267 |
| Nes | S | 1114 | 0.073087631 | 0.012286297 | 0.026640047 | -0.046447584 | 0.01435375 | 0.386354491 | 0.789495711 | 0.745236454 | 0.647271223 | 0.860696584 |
| Gon4l | S | 1412 | 0.071589554 | -0.039826504 | 0.048741968 | -0.022847586 | 0.088568472 | 0.149415209 | 0.478983923 | 0.41187346 | 0.660543843 | 0.185034661 |
| Tmem131l | S | 1267 | 0.026868549 | 0.080162872 | -0.041107758 | -0.067976306 | -0.121270629 | 0.570503152 | 0.06952562 | 0.455868123 | 0.284131811 | 0.082096094 |
| Retreg2 | T | 325 | -0.013056454 | -0.02028383 | 0.114732426 | 0.12778888 | 0.135016256 | 0.752759915 | 0.5897069 | 0.059231363 | 0.044705942 | 0.056034219 |
| Ncbp3 | S | 407 | 0.127249712 | -0.022724852 | -0.002109572 | -0.129359284 | 0.02061528 | 0.074477487 | 0.331117922 | 0.938735699 | 0.079090831 | 0.325618848 |
| Smc6 | S | 1095 | -0.016839603 | -0.016787631 | 0.102891406 | 0.119731008 | 0.119679037 | 0.575008967 | 0.686899169 | 0.037935666 | 0.03914659 | 0.038209352 |
| Tor1aip1 | S | 140 | 0.13433265 | 0.007957718 | -0.013293481 | -0.147626131 | -0.021251199 | 0.28702725 | 0.770679052 | 0.820928021 | 0.257740392 | 0.725544026 |
| Kdm2a | S | 28 | 0.006939696 | 0.022302474 | 0.017583101 | 0.010643405 | -0.004719373 | 0.802979353 | 0.079103533 | 0.403997589 | 0.735350947 | 0.81348303 |
| Klhl42 | S | 159 | 0.021334602 | -0.033182184 | 0.186484697 | 0.165150095 | 0.219666881 | 0.925247051 | 0.82907819 | 0.352755679 | 0.466839764 | 0.204901162 |
| Kat7 | S | 101 | 0.00612745 | 0.055971243 | -0.005636516 | -0.011763966 | -0.061607759 | 0.825725811 | 0.062130212 | 0.886684352 | 0.75770623 | 0.171521613 |
| Kat7 | S | 102 | 0.00612745 | 0.055971243 | -0.005636516 | -0.011763966 | -0.061607759 | 0.825725811 | 0.062130212 | 0.886684352 | 0.75770623 | 0.171521613 |
| Kat7 | S | 104 | 0.00612745 | 0.055971243 | -0.005636516 | -0.011763966 | -0.061607759 | 0.825725811 | 0.062130212 | 0.886684352 | 0.75770623 | 0.171521613 |
| Rad50 | T | 690 | -0.043991595 | 0.060311318 | 0.095448069 | 0.139439664 | 0.03513675 | 0.70901615 | 0.447069519 | 0.378868609 | 0.285701979 | 0.684641578 |
| 5730455P16Rik | S | 19 | -0.138216592 | 0.291340025 | -0.030237278 | 0.107979314 | -0.321577304 | 0.144596897 | 0.030697757 | 0.768623538 | 0.210668283 | 0.026369119 |
| Xpo4 | S | 464 | 0.165206603 | -0.046961399 | 0.051743668 | -0.113462935 | 0.098705067 | 0.404846559 | 0.660188417 | 0.630994775 | 0.563288655 | 0.43061725 |
| Glcci1 | S | 163 | -0.279549981 | 0.115271949 | 0.165664053 | 0.445214034 | 0.050392104 | 0.293328122 | 0.147492153 | 0.200091874 | 0.14177147 | 0.63666208 |
| Cnst | S | 34 | 0.015627675 | 0.011463559 | 0.08280252 | 0.067174845 | 0.071338961 | 0.872112282 | 0.866922263 | 0.305028551 | 0.527308755 | 0.388461874 |
| Cds2 | S | 20 | -0.131563132 | 0.078310497 | 0.105535044 | 0.237098175 | 0.027224547 | 0.057184867 | 0.044311552 | 0.07813584 | 0.012766591 | 0.556020268 |
| Sorbs2 | S | 251 | 0.033726708 | -0.044885382 | 0.068027226 | 0.034300517 | 0.112912608 | 0.373986127 | 0.230391713 | 0.037159087 | 0.354059807 | 0.026555508 |
| Srsf5 | S | 226 | 0.035836543 | 0.037673619 | 0.034419354 | -0.001417189 | -0.003254265 | 0.59338105 | 0.517519363 | 0.731123183 | 0.989571211 | 0.975146904 |
| Srsf5 | S | 228 | 0.035836543 | 0.037673619 | 0.034419354 | -0.001417189 | -0.003254265 | 0.59338105 | 0.517519363 | 0.731123183 | 0.989571211 | 0.975146904 |
| Ccdc61 | T | 334 | -0.007949913 | 0.006390602 | 0.084499135 | 0.092449048 | 0.078108533 | 0.92680365 | 0.882345148 | 0.158956524 | 0.359935935 | 0.224752228 |
| Tmem63a | S | 101 | 0.035603599 | 0.063435457 | 0.037185141 | 0.001581542 | -0.026250315 | 0.659854013 | 0.573458812 | 0.812610706 | 0.991136882 | 0.868283218 |
| Purb | S | 112 | 0.265256041 | -0.009839583 | -0.180373485 | -0.445629526 | -0.170533902 | 0.028025685 | 0.946478976 | 0.183274851 | 0.019587016 | 0.316794225 |
| Mdc1 | S | 461 | -0.012512011 | -0.00308147 | 0.091710412 | 0.104222423 | 0.094791883 | 0.609820236 | 0.915942355 | 0.071122535 | 0.053111436 | 0.070709623 |
| Akap8 | S | 325 | 0.044615603 | -0.042769908 | 0.073418972 | 0.028803369 | 0.11618888 | 0.183592874 | 0.197659385 | 0.132664854 | 0.444660058 | 0.054588674 |
| Bclaf3 | T | 409 | 0.016221108 | 0.00201625 | 0.060802219 | 0.04458111 | 0.058785968 | 0.69088155 | 0.952007194 | 0.207293081 | 0.350327964 | 0.200707082 |
| Dbndd2 | S | 142 | -0.05661059 | 0.083771376 | 0.038728703 | 0.095339293 | -0.045042673 | 0.624661119 | 0.109756684 | 0.64814014 | 0.465989385 | 0.584943664 |
| Tmf1 | S | 334 | 0.050346598 | -0.043703978 | 0.074767105 | 0.024420507 | 0.118471083 | 0.693557513 | 0.669844715 | 0.126667042 | 0.840599505 | 0.290980067 |
| Srrm2 | Y | 2647 | 0.00700824 | 0.007998521 | 0.076121998 | 0.069113758 | 0.068123477 | 0.904158102 | 0.866017364 | 0.227655745 | 0.350744108 | 0.307976399 |
| Utp18 | S | 114 | 0.054704569 | -0.000507567 | 0.003240311 | -0.051464259 | 0.003747878 | 0.067381671 | 0.986157792 | 0.936292208 | 0.245274598 | 0.925735597 |
| Ptpn14 | S | 593 | 0.015168194 | 0.049148233 | -0.007952733 | -0.023120927 | -0.057100966 | 0.628256178 | 0.083842451 | 0.841768429 | 0.567427657 | 0.193990683 |
| Srrm2 | S | 1832 | 0.014439065 | 0.035804275 | 0.033880383 | 0.019441318 | -0.001923892 | 0.670921818 | 0.311783614 | 0.673282685 | 0.796063124 | 0.979402808 |
| Cdr2 | T | 312 | 0.030998057 | 0.116120545 | -0.064509916 | -0.095507973 | -0.180630462 | 0.767660117 | 0.168135646 | 0.703671156 | 0.587043342 | 0.32263642 |
| Uimc1 | S | 44 | -0.039816694 | 0.044510396 | 0.094922407 | 0.134739101 | 0.050412011 | 0.454084475 | 0.552033325 | 0.170912236 | 0.101271899 | 0.555218701 |
| Uimc1 | S | 46 | -0.039816694 | 0.044510396 | 0.094922407 | 0.134739101 | 0.050412011 | 0.454084475 | 0.552033325 | 0.170912236 | 0.101271899 | 0.555218701 |
| Rai1 | S | 453 | 0.134791547 | 0.011358758 | 0.033533123 | -0.101258424 | 0.022174365 | 0.512525412 | 0.948147931 | 0.865988943 | 0.646197646 | 0.909550784 |
| Disp2 | S | 1317 | 0.013248358 | -0.04440208 | 0.237178772 | 0.223930414 | 0.281580852 | 0.879888955 | 0.610135098 | 0.347034641 | 0.367596916 | 0.281634146 |
| Wwp2 | S | 211 | 0.143065105 | -0.055701575 | -0.030879579 | -0.173944684 | 0.024821996 | 0.011334505 | 0.223926242 | 0.635800692 | 0.066801111 | 0.669268254 |
| Dock11 | S | 1237 | -0.166450926 | -0.023822217 | 0.276674008 | 0.443124934 | 0.300496225 | 0.130459405 | 0.784852669 | 0.018379372 | 0.00521788 | 0.001846028 |
| Prkd3 | T | 535 | -0.098729206 | 0.036728359 | 0.113664413 | 0.212393619 | 0.076936054 | 0.567028227 | 0.717310481 | 0.093145205 | 0.271733542 | 0.449478801 |
| Edc4 | S | 733 | 0.0025183 | 0.011172528 | 0.050174038 | 0.047655737 | 0.039001509 | 0.915435205 | 0.607905308 | 0.159338665 | 0.18903531 | 0.251382796 |
| Kmt2d | S | 2299 | 0.041693634 | -0.015394799 | 0.047573616 | 0.005879982 | 0.062968415 | 0.373168426 | 0.692067802 | 0.226979035 | 0.846739899 | 0.04249132 |
| Kif1c | T | 1080 | -0.016019453 | 0.035844184 | 0.028354085 | 0.044373538 | -0.0074901 | 0.663423776 | 0.168930024 | 0.177901527 | 0.271573025 | 0.729541369 |
| Kmt2b | S | 1037 | -0.026761002 | 0.072899718 | 0.023249891 | 0.050010893 | -0.049649827 | 0.498965758 | 0.140602341 | 0.377239584 | 0.241814607 | 0.272187317 |
| Sec23ip | S | 738 | 0.085317126 | 0.111057167 | -0.095428452 | -0.180745578 | -0.206485619 | 0.410463223 | 0.033235403 | 0.074046481 | 0.144722716 | 0.004906829 |
| Zfp280d | S | 557 | -0.030155893 | 0.0296449 | 0.112744909 | 0.142900802 | 0.083100008 | 0.842970439 | 0.700541793 | 0.315158934 | 0.39392966 | 0.403508384 |
| Rbm25 | S | 578 | -0.037720764 | -0.007350752 | 0.148388664 | 0.186109428 | 0.155739416 | 0.2829477 | 0.855982682 | 0.061478216 | 0.050237153 | 0.060366178 |
| Akap8 | S | 320 | 0.045787636 | -0.042913523 | 0.071711335 | 0.0259237 | 0.114624858 | 0.17163448 | 0.19399604 | 0.144890047 | 0.493621104 | 0.059898022 |
| Pomgnt1 | S | 248 | 0.058402865 | 0.086904026 | -0.066298113 | -0.124700979 | -0.153202139 | 0.222412521 | 0.340919767 | 0.509992396 | 0.268639803 | 0.232783819 |
| Git2 | S | 563 | 0.012212645 | 0.066283919 | 0.059658729 | 0.047446084 | -0.00662519 | 0.864101204 | 0.432997734 | 0.808868771 | 0.847329652 | 0.97862097 |
| Dnajc21 | T | 377 | 0.060064263 | 0.041412477 | 0.021252619 | -0.038811644 | -0.020159858 | 0.485015292 | 0.668446266 | 0.879793032 | 0.795919421 | 0.896509798 |
| Osbpl3 | T | 248 | 0.028879088 | -0.036906211 | 0.060364065 | 0.031484977 | 0.097270276 | 0.384205631 | 0.646436685 | 0.130911747 | 0.38382664 | 0.283031741 |
| Espl1 | S | 238 | 0.054356526 | 0.056233506 | -0.003105078 | -0.057461603 | -0.059338583 | 0.281622866 | 0.702700727 | 0.94170448 | 0.206403553 | 0.684655801 |
| Slc12a4 | S | 88 | 0.063948408 | -0.005782172 | 0.103744481 | 0.039796073 | 0.109526654 | 0.483867341 | 0.966880185 | 0.711274906 | 0.884613428 | 0.707128044 |
| Srrm2 | T | 1390 | 0.134894806 | -0.068961976 | -0.018204376 | -0.153099182 | 0.0507576 | 0.007818578 | 0.217680362 | 0.603887464 | 0.008645713 | 0.334757033 |
| Git1 | S | 423 | 0.005219898 | 0.086404381 | -0.024448428 | -0.029668326 | -0.110852809 | 0.888181628 | 0.090710852 | 0.706318921 | 0.657013491 | 0.167890085 |
| Rbm5 | S | 621 | 0.012940369 | 0.008550093 | 0.055561653 | 0.042621284 | 0.047011561 | 0.672654502 | 0.804405035 | 0.260978729 | 0.391135309 | 0.367564783 |
| Tnks1bp1 | S | 1705 | -0.052957273 | 0.068182095 | 0.07116421 | 0.124121482 | 0.002982114 | 0.620592103 | 0.184986519 | 0.545608935 | 0.395763057 | 0.978659783 |
| Nmt2 | S | 66 | 0.047605611 | 0.037323379 | -0.001863069 | -0.049468681 | -0.039186448 | 0.316629543 | 0.477722332 | 0.92088617 | 0.304199046 | 0.462093096 |
| Ahctf1 | S | 1191 | -0.259133678 | 0.326683805 | 0.124492578 | 0.383626257 | -0.202191227 | 0.621370881 | 0.333374959 | 0.854852153 | 0.624346054 | 0.769869053 |
| Ahctf1 | S | 1192 | -0.259133678 | 0.326683805 | 0.124492578 | 0.383626257 | -0.202191227 | 0.621370881 | 0.333374959 | 0.854852153 | 0.624346054 | 0.769869053 |
| Arhgef18 | S | 973 | -0.079857903 | -0.018766417 | 0.162308256 | 0.242166159 | 0.181074673 | 0.517220197 | 0.785616312 | 0.056002157 | 0.121760699 | 0.06968839 |
| Rab11fip5 | S | 251 | -0.013043771 | 0.13766698 | 0.006443046 | 0.019486818 | -0.131223934 | 0.914725715 | 0.296117083 | 0.978576541 | 0.928715857 | 0.567442011 |
| Nup214 | S | 922 | 0.066483622 | 0.006504952 | -0.003600798 | -0.07008442 | -0.010105749 | 0.12567892 | 0.851981757 | 0.916380308 | 0.11112825 | 0.754564242 |

| | | | | | | | | | | | | |
|---|---|---|---|---|---|---|---|---|---|---|---|---|
| Dbr1 | S | 520 | 0.040068978 | 0.073582696 | -0.040913778 | -0.080982756 | -0.114496474 | 0.581382797 | 0.061232253 | 0.323370638 | 0.298599649 | 0.023675509 |
| Limk2 | S | 289 | -0.015937073 | 0.175557983 | -0.085405641 | -0.069468568 | -0.260963624 | 0.83444832 | 0.08238574 | 0.558882232 | 0.610804544 | 0.135934238 |
| Gatad2a | S | 110 | 0.052501283 | -0.032180809 | 0.045167132 | -0.007334152 | 0.077347941 | 0.154883181 | 0.290473836 | 0.195228185 | 0.790930712 | 0.029235514 |
| Uba1 | S | 810 | -0.056035429 | 0.165743448 | -0.017911241 | 0.038124188 | -0.183654689 | 0.383786261 | 0.061612683 | 0.800873889 | 0.607831809 | 0.070368402 |
| Sqstm1 | S | 28 | 0.064242106 | -0.016838657 | 0.024453173 | -0.039788933 | 0.041291829 | 0.161091061 | 0.78919382 | 0.634178842 | 0.379723929 | 0.520428133 |
| Arid5b | S | 566 | 0.002852738 | -0.056897663 | 0.103290699 | 0.100437961 | 0.160188363 | 0.946745689 | 0.559880866 | 0.051263328 | 0.078703774 | 0.174694864 |
| Ncaph | S | 662 | -0.144458262 | 0.154232742 | 0.080111093 | 0.224569354 | -0.074121649 | 0.03100009 | 0.011134335 | 0.430308342 | 0.087879359 | 0.448562785 |
| Srrm1 | S | 635 | 0.033116614 | -0.002254496 | 0.022901497 | -0.010215117 | 0.025155993 | 0.160471237 | 0.907842516 | 0.359936828 | 0.681044863 | 0.315078965 |
| Osbpl8 | S | 91 | 0.043425366 | -0.05701204 | 0.076243358 | 0.032817992 | 0.133255399 | 0.436534055 | 0.455965656 | 0.115096495 | 0.441802181 | 0.139272132 |
| Bud13 | S | 373 | -0.031501286 | 0.009777788 | 0.068383017 | 0.099884304 | 0.05860523 | 0.135558353 | 0.620885218 | 0.009178053 | 0.00318153 | 0.021564167 |
| Mapt | S | 696 | 1.141257657 | -0.152958058 | 0.018628468 | -1.122629188 | 0.171586526 | 0.322545694 | 0.012580622 | 0.795130183 | 0.328508111 | 0.077603652 |
| Mphosph8 | S | 51 | 0.056923184 | 0.00655975 | 0.004038583 | -0.0528846 | -0.002521167 | 0.139943147 | 0.880268748 | 0.905610292 | 0.075040218 | 0.943025343 |
| Nectin1 | S | 506 | 0.115421635 | 0.091180954 | -0.205271092 | -0.320692728 | -0.296452047 | 0.067958827 | 0.054438173 | 0.098991729 | 0.034686564 | 0.050473745 |
| Dpf2 | S | 142 | -0.159580882 | 0.020571369 | 0.21939082 | 0.378971702 | 0.198819451 | 0.053036973 | 0.700485287 | 0.012310958 | 0.003649454 | 0.02771844 |
| Ssrp1 | S | 673 | 0.011767266 | 0.008162334 | 0.067821068 | 0.056053802 | 0.059658734 | 0.768370364 | 0.715144175 | 0.328313859 | 0.42955036 | 0.378680533 |
| Arhgef17 | S | 458 | -0.049917046 | -0.168943143 | 0.222762054 | 0.272679099 | 0.391705196 | 0.358901235 | 0.098819355 | 0.001074494 | 0.006350555 | 0.015244821 |
| Kif4 | S | 811 | 0.054578511 | 0.012098506 | 0.002227673 | -0.052350838 | -0.009870833 | 0.161711344 | 0.761070517 | 0.961982233 | 0.29226267 | 0.837179974 |
| Cdca3 | S | 31 | 0.081646977 | -0.001647214 | 0.002553681 | -0.079093297 | 0.004200895 | 0.172838854 | 0.982207258 | 0.97049513 | 0.286275216 | 0.959319525 |
| Ptpn14 | S | 578 | 0.076704615 | 0.072338732 | -0.06360331 | -0.140307924 | -0.135942042 | 0.300704888 | 0.150172308 | 0.261488837 | 0.113757478 | 0.057617079 |
| Cdv3 | T | 182 | 0.026238068 | 0.092144444 | -0.041812159 | -0.068050227 | -0.133956603 | 0.558951209 | 0.139813162 | 0.511034038 | 0.250891009 | 0.076289845 |
| Cltc | S | 412 | -0.112477694 | 0.006312456 | 0.286050224 | 0.398527918 | 0.279737768 | 0.123554722 | 0.8890868 | 0.08512951 | 0.031826451 | 0.07866775 |
| Iws1 | S | 248 | 0.016105312 | -0.015648996 | 0.071938427 | 0.055833115 | 0.087587423 | 0.383040656 | 0.426235121 | 0.146328469 | 0.223652266 | 0.100255327 |
| Rab12 | S | 105 | 0.017110073 | 0.121364352 | -0.059300838 | -0.076410912 | -0.180665191 | 0.757155617 | 0.056520046 | 0.269409015 | 0.073604938 | 0.009366112 |
| Kmt2a | S | 151 | -0.005550211 | 0.017872052 | 0.056609087 | 0.062159298 | 0.038737035 | 0.864358906 | 0.694650225 | 0.149451281 | 0.111318084 | 0.411356477 |
| Tshz3 | S | 463 | 0.033496601 | 0.019150084 | 0.034713084 | 0.001216483 | 0.015563 | 0.695461868 | 0.538655678 | 0.475804131 | 0.988897676 | 0.719992632 |
| Hivep3 | S | 898 | 0.05498803 | 0.056253314 | -0.005823287 | -0.060811316 | -0.0620766 | 0.488979171 | 0.502250914 | 0.959060573 | 0.64158714 | 0.639818363 |
| Syde1 | S | 60 | 0.104045962 | 0.011261935 | -0.042481523 | -0.146527485 | -0.053743458 | 0.117855688 | 0.840967692 | 0.645237008 | 0.175431002 | 0.558697678 |
| Iws1 | T | 299 | 0.045228948 | -0.01197783 | 0.073485295 | 0.028256347 | 0.085463125 | 0.563181038 | 0.859549271 | 0.396585291 | 0.746674689 | 0.320809018 |
| Luzp1 | S | 660 | 0.011753646 | -0.009115562 | 0.058467742 | 0.046714096 | 0.067583304 | 0.813177512 | 0.779663138 | 0.08828597 | 0.379328042 | 0.089245651 |
| Atrx | S | 768 | -0.013430519 | 0.000260469 | 0.085412586 | 0.098843106 | 0.085152118 | 0.783035916 | 0.995281789 | 0.109970912 | 0.08116739 | 0.091258703 |
| Glis2 | S | 54 | -0.006810477 | -0.005452337 | 0.083301759 | 0.090112236 | 0.088754096 | 0.835932621 | 0.861150549 | 0.096664807 | 0.079979207 | 0.085586678 |
| Smg6 | S | 869 | -0.024805709 | 0.070500116 | 0.04487484 | 0.069680549 | -0.025625276 | 0.515316772 | 0.178215157 | 0.578626403 | 0.389151897 | 0.742214124 |
| Smg6 | S | 873 | -0.024805709 | 0.070500116 | 0.04487484 | 0.069680549 | -0.025625276 | 0.515316772 | 0.178215157 | 0.578626403 | 0.389151897 | 0.742214124 |
| Fgd1 | S | 364 | -0.050260375 | 0.111403612 | 0.051683399 | 0.101943774 | -0.059720213 | 0.437695308 | 0.163042742 | 0.715947921 | 0.473475327 | 0.667740588 |
| Hnrnpk | S | 430 | -0.213227502 | 0.264060529 | 0.164269146 | 0.377496648 | -0.099791382 | 0.080871121 | 0.256258617 | 0.205603894 | 0.033533708 | 0.639639945 |
| Acin1 | S | 413 | -0.115386836 | 0.062820324 | 0.16926897 | 0.284655806 | 0.106448646 | 0.014191966 | 0.049153681 | 0.087182718 | 0.027545373 | 0.209262805 |
| Bcorl1 | S | 1448 | 0.004662427 | -0.024510737 | 0.074039864 | 0.069377438 | 0.098550601 | 0.913536482 | 0.704007465 | 0.065682283 | 0.140892543 | 0.190950856 |
| Atrx | S | 1512 | -0.026034085 | 0.001663042 | 0.067514296 | 0.093548381 | 0.065851255 | 0.572957714 | 0.95030309 | 0.029493665 | 0.109810217 | 0.026637178 |
| Lmna | S | 390 | -0.007998739 | 0.009443589 | 0.048342977 | 0.056341716 | 0.038899388 | 0.648644417 | 0.592546065 | 0.063570921 | 0.047471959 | 0.118791085 |
| Cdca7l | S | 21 | 0.02528219 | 0.021175431 | 0.055630254 | 0.030348065 | 0.034454823 | 0.659232311 | 0.564863292 | 0.619612787 | 0.788772229 | 0.749653763 |
| Atad5 | S | 1188 | 0.038894506 | 0.055656801 | -0.028725906 | -0.067620412 | -0.084382707 | 0.518369189 | 0.095315642 | 0.476989382 | 0.293827869 | 0.079852656 |
| Fgd1 | T | 710 | -0.018279349 | 0.055641583 | 0.007452809 | 0.025732158 | -0.048188773 | 0.296850243 | 0.014754829 | 0.805590794 | 0.390764725 | 0.177383715 |
| Erbin | S | 1076 | 0.020336441 | 0.056378928 | -0.009140241 | -0.029476682 | -0.065519169 | 0.645894662 | 0.162585461 | 0.846906646 | 0.509336914 | 0.156936915 |
| Tnk2 | S | 772 | 0.02648174 | 0.03068958 | 0.036026196 | 0.009544456 | 0.005336616 | 0.721513731 | 0.584058407 | 0.519457289 | 0.904496668 | 0.933342299 |
| Tmx1 | S | 245 | -0.004167286 | 0.01695245 | 0.047435213 | 0.051602499 | 0.030482763 | 0.882372595 | 0.414026711 | 0.202555107 | 0.196298245 | 0.361498895 |
| Arhgef40 | T | 249 | 0.048717534 | 0.087856544 | 0.00539344 | -0.043324094 | -0.082463104 | 0.873689503 | 0.397232929 | 0.986834617 | 0.917051776 | 0.803077809 |
| Gpatch8 | S | 1018 | -0.054387327 | 0.009042904 | 0.120988469 | 0.175375795 | 0.111945564 | 0.184990909 | 0.695127375 | 0.041894494 | 0.014715061 | 0.050645834 |
| Serpini2 | T | 52 | 0.00991378 | 0.075985146 | -0.012638899 | -0.022552679 | -0.088624045 | 0.793514648 | 0.143125769 | 0.763114357 | 0.629810604 | 0.136027272 |
| Serpini2 | T | 53 | 0.00991378 | 0.075985146 | -0.012638899 | -0.022552679 | -0.088624045 | 0.793514648 | 0.143125769 | 0.763114357 | 0.629810604 | 0.136027272 |
| Amotl2 | S | 664 | -0.109119036 | 0.100353777 | 0.177193542 | 0.286312578 | 0.076839765 | 0.387658673 | 0.576179008 | 0.303221853 | 0.153732553 | 0.714279192 |
| Mtres1 | S | 111 | 0.041476835 | 0.002872204 | 0.038326569 | -0.003150266 | 0.035454365 | 0.551229736 | 0.973487104 | 0.325345582 | 0.960139053 | 0.676770045 |
| Mepce | S | 307 | 0.08934028 | 0.014563839 | -0.048340221 | -0.137680501 | -0.06290406 | 0.052805096 | 0.830739438 | 0.467782962 | 0.096423455 | 0.440623886 |
| Tnks1bp1 | S | 1657 | -0.074657497 | 0.081011271 | 0.049018188 | 0.123675685 | -0.031993082 | 0.051095277 | 0.180295529 | 0.010733876 | 0.019769698 | 0.520105925 |
| Kif18a | S | 744 | 0.03416004 | 0.031611173 | 0.01202228 | -0.02213776 | -0.019588892 | 0.450445722 | 0.466820093 | 0.79696499 | 0.46056093 | 0.472920319 |
| Nrcam | S | 1242 | 0.046302072 | 0.048043261 | -0.015079 | -0.061381072 | -0.063122261 | 0.402794209 | 0.335913677 | 0.774559932 | 0.384687817 | 0.348183484 |
| Rapgef2 | S | 498 | -0.139977145 | 0.317860807 | -0.094099357 | 0.045877789 | -0.411960163 | 0.361580108 | 0.0598807 | 0.662703055 | 0.829587108 | 0.12069586 |
| Dync1i2 | S | 98 | -0.058771191 | 0.085490437 | 0.055409539 | 0.11418073 | -0.030080898 | 0.504367343 | 0.254610362 | 0.384998919 | 0.246257095 | 0.684063127 |
| Rad21 | S | 549 | -0.153153713 | -0.081223321 | 0.290515519 | 0.443669232 | 0.37173884 | 0.210070906 | 0.261112192 | 0.007674103 | 0.01794378 | 0.005637328 |
| Arid4b | S | 1186 | -0.079307821 | 0.053316573 | 0.129167569 | 0.20847539 | 0.075850996 | 0.397128157 | 0.597794417 | 0.174797379 | 0.04941822 | 0.420111806 |
| Pard3b | S | 801 | 0.037645672 | 0.04033787 | 0.046159592 | 0.00851392 | 0.005821721 | 0.799976453 | 0.756894044 | 0.527757318 | 0.951649399 | 0.961486883 |
| Foxm1 | S | 727 | 0.05843848 | -0.001542283 | 0.054741699 | -0.003696781 | 0.056283982 | 0.536186097 | 0.982352019 | 0.539628078 | 0.97084562 | 0.512404816 |
| Adgrl3 | S | 1508 | -0.0629033 | 0.046668779 | 0.09000895 | 0.15291225 | 0.043340171 | 0.421922823 | 0.244860544 | 0.264409624 | 0.141816927 | 0.53845819 |
| Dtl | T | 428 | 0.02151764 | 0.027555148 | 0.064169065 | 0.042651425 | 0.036613917 | 0.713775571 | 0.72138433 | 0.573692008 | 0.715950547 | 0.768482762 |
| Gapvd1 | T | 762 | -0.026571228 | 0.067383862 | 0.020926989 | 0.047498217 | -0.046456873 | 0.220552433 | 0.038964809 | 0.611852369 | 0.285882672 | 0.304008484 |
| Gapvd1 | S | 761 | -0.039425418 | 0.03270273 | 0.07529355 | 0.114718968 | 0.04259082 | 0.747146993 | 0.721550011 | 0.158491052 | 0.394952528 | 0.658407161 |
| Gtf2i | S | 831 | 0.020952322 | -0.01255868 | 0.064282146 | 0.043329824 | 0.076840827 | 0.768777869 | 0.690816823 | 0.129797126 | 0.545748388 | 0.070365279 |
| Polr3e | S | 161 | 0.038562579 | -0.015602451 | 0.026836875 | -0.011725704 | 0.042439326 | 0.136668362 | 0.370867853 | 0.178390401 | 0.604997864 | 0.055468604 |
| Prpf4b | S | 258 | -0.009575778 | -0.012512355 | 0.094691258 | 0.104267036 | 0.107203613 | 0.832965922 | 0.703309577 | 0.091518077 | 0.110046011 | 0.073325207 |
| Mki67 | S | 276 | 0.042697517 | 0.094943818 | -0.080651552 | -0.123349069 | -0.17559537 | 0.296270215 | 0.016197621 | 0.053484967 | 0.022942432 | 0.014109363 |
| Huwe1 | S | 1395 | -0.012982806 | 0.0713039 | 0.002189103 | 0.015171909 | -0.069114797 | 0.609339823 | 0.069027741 | 0.946513026 | 0.589315163 | 0.098087452 |
| Trp53bp1 | S | 983 | -0.074317806 | 0.095974586 | 0.078490143 | 0.152807949 | -0.017484443 | 0.439450931 | 0.133873891 | 0.344438925 | 0.194649749 | 0.866167794 |
| Kmt2b | T | 485 | 0.209428265 | -0.051050642 | -0.053369994 | -0.262798259 | -0.002319352 | 0.068411675 | 0.641489183 | 0.365733748 | 0.06027935 | 0.980914756 |
| Ncor1 | S | 1598 | -0.033471197 | 0.069818452 | 0.089477346 | 0.122948542 | 0.019658893 | 0.48007151 | 0.376820768 | 0.462241765 | 0.329850153 | 0.870763872 |
| Arhgef12 | S | 304 | 0.05031596 | 0.039420752 | -0.008528686 | -0.058844646 | -0.047949438 | 0.401284393 | 0.418387819 | 0.879852628 | 0.437994515 | 0.488714309 |
| Kmt2b | S | 849 | 0.095095082 | -0.009106625 | -0.032246664 | -0.127341747 | -0.023140039 | 0.022272855 | 0.798486289 | 0.325956445 | 0.002819092 | 0.319058659 |
| Mier1 | S | 482 | -0.008550332 | 0.028861848 | 0.070948593 | 0.079498925 | 0.042086745 | 0.809903934 | 0.530237774 | 0.349370297 | 0.308322132 | 0.577783497 |
| Kmt2b | S | 2346 | 0.109492894 | -0.032271671 | -0.030858667 | -0.140351561 | 0.001413005 | 0.012379273 | 0.389792686 | 0.466171225 | 0.022554822 | 0.972446934 |
| Cep170 | S | 378 | -0.087718451 | 0.113838745 | 0.15916173 | 0.246880182 | 0.045322985 | 0.008429747 | 0.097405057 | 0.211337729 | 0.107698534 | 0.679327318 |
| Plec | S | 2866 | 0.100263411 | -0.022682109 | 0.111699821 | 0.01143641 | 0.134381931 | 0.719521167 | 0.850985101 | 0.587258355 | 0.971133786 | 0.543462416 |
| Sgo1 | S | 120 | 0.025145536 | 0.044477701 | -0.008814661 | -0.033960197 | -0.053292362 | 0.597542485 | 0.123530457 | 0.71855359 | 0.461361941 | 0.03246275 |
| Lrch4 | S | 281 | 0.035546342 | -0.047320625 | 0.097813523 | 0.062267181 | 0.145134147 | 0.38948298 | 0.341234357 | 0.183775717 | 0.341659351 | 0.082239444 |
| Chd2 | S | 1727 | 0.110328182 | -0.019993075 | -0.038079889 | -0.148408072 | -0.018086815 | 0.04867756 | 0.671715831 | 0.66886833 | 0.162127238 | 0.823951567 |
| Vps51 | S | 640 | -0.036621864 | 0.088144376 | 0.049129401 | 0.085751265 | -0.039014975 | 0.687589612 | 0.338880674 | 0.544558095 | 0.326918921 | 0.627668972 |
| Wbp11 | S | 237 | 0.039119759 | 0.012676426 | 0.003465499 | -0.03565426 | -0.009210927 | 0.18534389 | 0.571948504 | 0.85259314 | 0.212504939 | 0.65076694 |
| Anks1 | S | 640 | -0.077193107 | 0.065270551 | 0.139612259 | 0.216805366 | 0.074341709 | 0.020829335 | 0.469295369 | 0.033495302 | 0.017990488 | 0.428971121 |
| Telo2 | S | 457 | -0.024450405 | 0.024052989 | 0.070743696 | 0.095194101 | 0.046690708 | 0.360988748 | 0.52081476 | 0.108252823 | 0.056852943 | 0.316951607 |
| Nectin1 | S | 421 | 0.038835688 | 0.024279349 | 0.014945522 | -0.023890165 | -0.009333827 | 0.441883128 | 0.506085104 | 0.804001263 | 0.723575856 | 0.877083378 |
| Ociad1 | S | 122 | -0.15276526 | 0.055246742 | 0.149167205 | 0.301932465 | 0.093920464 | 0.054911048 | 0.340024347 | 0.016031237 | 0.008110942 | 0.156318394 |
| Phf14 | S | 283 | 0.002670046 | 0.049898414 | 0.002933593 | 0.000263546 | -0.046964822 | 0.945783291 | 0.100969802 | 0.944496577 | 0.995837381 | 0.31934559 |
| Srpk1 | S | 39 | 0.13223073 | -0.06244882 | 0.031279769 | -0.100950961 | 0.093728589 | 0.078963267 | 0.151699925 | 0.597272293 | 0.17942153 | 0.151187132 |
| Zc3h18 | S | 80 | 0.037264319 | 0.055229902 | 0.021653163 | -0.015611156 | -0.033576739 | 0.747876087 | 0.524167351 | 0.796531634 | 0.882418033 | 0.636518224 |
| Trpm7 | S | 1300 | -0.100483729 | 0.059151069 | 0.232050125 | 0.332533854 | 0.172899056 | 0.217267379 | 0.667196427 | 0.212068557 | 0.124759484 | 0.366828408 |
| Afap1 | S | 669 | 0.078194245 | -0.016276416 | -0.024321347 | -0.102515592 | -0.008044931 | 0.002974282 | 0.377190008 | 0.242680793 | 0.006929905 | 0.60095271 |
| Spag9 | S | 728 | -0.032079364 | 0.141293493 | -0.019567094 | 0.01251227 | -0.160860587 | 0.819238962 | 0.186090056 | 0.90183843 | 0.942335362 | 0.326897647 |

| | | | | | | | | | | | | |
|---|---|---|---|---|---|---|---|---|---|---|---|---|
| Polr2a | S | 1836 | 0.000709388 | 0.013257265 | 0.043917481 | 0.043208093 | 0.030660216 | 0.93758576 | 0.489739171 | 0.21927886 | 0.226600268 | 0.369750203 |
| Dcaf10 | S | 98 | -0.069954471 | 0.060531594 | 0.091963216 | 0.161917687 | 0.031431622 | 0.645825421 | 0.536993269 | 0.278165215 | 0.338714161 | 0.765976732 |
| Amfr | S | 552 | -0.287598632 | 0.119115979 | 0.147979624 | 0.435578256 | 0.028863645 | 0.537308865 | 0.267533766 | 0.361029131 | 0.37903846 | 0.839853875 |
| Sfswap | S | 283 | -0.030849723 | 0.008589043 | 0.118431419 | 0.149281142 | 0.109842375 | 0.551326872 | 0.876988222 | 0.163027428 | 0.104738086 | 0.174348385 |
| Psip1 | T | 122 | -0.113626546 | -0.023468317 | 0.234779291 | 0.348405837 | 0.258247608 | 0.339042682 | 0.774246255 | 0.070349979 | 0.043385875 | 0.063122737 |
| Map1a | S | 995 | -0.004497743 | 0.029477342 | 0.043551484 | 0.048049227 | 0.014074142 | 0.869112464 | 0.390370909 | 0.329673586 | 0.271659533 | 0.739005914 |
| Sf3b1 | S | 332 | 0.016209552 | -0.03970708 | 0.076430356 | 0.060220803 | 0.116137436 | 0.675360302 | 0.275384773 | 0.034286531 | 0.15852421 | 0.019583906 |
| Map4 | S | 449 | 0.092129209 | 0.105379127 | -0.107798557 | -0.199927765 | -0.213177684 | 0.174380004 | 0.248875937 | 0.189299689 | 0.035268731 | 0.054402195 |
| Rpl34 Rpl34-ps1 | S | 12 | 0.237957764 | 0.101646504 | -0.231756544 | -0.469714308 | -0.333403048 | 0.088048447 | 0.417722332 | 0.122161135 | 0.009864984 | 0.02547745 |
| Mpdz | S | 908 | -0.071567681 | 0.105536535 | 0.000682477 | 0.072250157 | -0.104854059 | 0.555005774 | 0.049325001 | 0.994437997 | 0.613238099 | 0.343048706 |
| Nup98 | S | 623 | 0.011041243 | 0.027091127 | 0.028629717 | 0.017588474 | 0.00153859 | 0.752784055 | 0.432477861 | 0.43979023 | 0.628233812 | 0.964475346 |
| Ubap2l | S | 490 | 0.055798691 | 0.006635028 | -0.001831868 | -0.057630559 | -0.008466896 | 0.136605043 | 0.826352654 | 0.956377677 | 0.196384493 | 0.829103985 |
| Map3k5 | S | 1310 | 0.051717249 | 0.097604322 | -0.019691047 | -0.071408295 | -0.117295368 | 0.745211121 | 0.408702488 | 0.83536871 | 0.682824396 | 0.398171393 |
| Dffa | S | 314 | 0.003332428 | 0.06634567 | 0.00239863 | -0.000933798 | -0.063947041 | 0.945842947 | 0.182310289 | 0.963772103 | 0.987863321 | 0.307544677 |
| Psmd7 | T | 149 | 0.074134717 | -0.025731581 | 0.016188024 | -0.057946692 | 0.041919605 | 0.074133818 | 0.597740111 | 0.929218451 | 0.753907749 | 0.820974335 |
| Ski | S | 477 | 0.008577035 | 0.046204447 | 0.011975826 | 0.003398791 | -0.034228621 | 0.774357289 | 0.230335771 | 0.803344073 | 0.948699054 | 0.537405279 |
| Serinc1 | S | 364 | 0.032606422 | 0.063762327 | -0.042421703 | -0.075028126 | -0.10618403 | 0.262550909 | 0.153654751 | 0.404601243 | 0.19636659 | 0.107378511 |
| Rtn4 | S | 857 | 0.052686343 | -0.030342323 | 0.066219546 | 0.013533203 | 0.096561869 | 0.209098844 | 0.299845961 | 0.354556808 | 0.842142355 | 0.216286553 |
| Zfp106 | S | 1381 | 0.002467918 | 0.015707139 | 0.062059756 | 0.059591838 | 0.046352617 | 0.943328957 | 0.447491352 | 0.41531953 | 0.44011987 | 0.524576409 |
| Cdk13 | S | 590 | 0.029808185 | 0.024224464 | 0.012736968 | -0.017071216 | -0.011487496 | 0.475677312 | 0.4613104 | 0.667205026 | 0.661044818 | 0.697050421 |
| Atad5 | S | 739 | -0.001119363 | 0.070185162 | 0.008681584 | 0.009800947 | -0.061503578 | 0.992956934 | 0.211825815 | 0.900982279 | 0.942236754 | 0.433050696 |
| Anapc1 | S | 1349 | 0.051099301 | -0.0306701 | 0.034995395 | -0.016103906 | 0.065665495 | 0.307436178 | 0.471173906 | 0.137853306 | 0.708929806 | 0.178875063 |
| Vcl | S | 290 | -0.092393988 | 0.087612431 | 0.091150973 | 0.183544961 | 0.003538541 | 0.16025655 | 0.153284036 | 0.237265642 | 0.050399242 | 0.956879254 |
| Arfgef2 | S | 276 | 0.014794037 | 0.041863418 | -0.00178296 | -0.016576997 | -0.043646379 | 0.646123845 | 0.144983221 | 0.958562526 | 0.653230509 | 0.231970874 |
| Suz12 | S | 585 | 0.036121521 | -0.042397354 | 0.064334746 | 0.028213225 | 0.106732099 | 0.440354714 | 0.25734622 | 0.061376193 | 0.489305973 | 0.02924306 |
| H2-K1 | S | 354;313;353;354 | 0.063609698 | 0.081930933 | -0.062538856 | -0.126148554 | -0.144469789 | 0.228664341 | 0.091151566 | 0.084100778 | 0.058279659 | 0.021876962 |
| Son | S | 1973 | 0.038400842 | 0.000733994 | 0.060883963 | 0.022483121 | 0.060149969 | 0.56260511 | 0.987226608 | 0.504717289 | 0.809877141 | 0.496343503 |
| Camkk2 | S | 129 | -0.00655447 | 0.062404097 | -0.010510239 | -0.003955769 | -0.072914336 | 0.935470966 | 0.054838625 | 0.794117616 | 0.961558687 | 0.120723531 |
| Taf6l | S | 500 | 0.07911665 | -0.02515287 | 0.030823606 | -0.048293044 | 0.055976476 | 0.112057682 | 0.309478023 | 0.671353674 | 0.538270367 | 0.467564086 |
| Rai1 | T | 455 | 0.100483452 | -0.01978369 | -0.004862564 | -0.105346016 | 0.014921126 | 0.153154142 | 0.814845205 | 0.95209192 | 0.165895724 | 0.852102769 |
| Myo9a | S | 2296 | -0.026290059 | 0.078834957 | 0.031506656 | 0.057796716 | -0.047328301 | 0.603246162 | 0.250259522 | 0.537319248 | 0.359222579 | 0.5002221 |
| Stk10 | S | 437 | -0.012946298 | 0.037386582 | 0.037676653 | 0.050622951 | 0.000290071 | 0.848871847 | 0.311677818 | 0.375523587 | 0.458876856 | 0.991401343 |
| Tbc1d15 | T | 669 | 0.096901279 | -0.04612774 | 0.082527023 | -0.014374256 | 0.128654763 | 0.593473415 | 0.623455797 | 0.393337891 | 0.936785839 | 0.265824175 |
| Wnk1 | S | 2006 | 0.024454287 | 0.102963068 | -0.046654045 | -0.071108332 | -0.149617112 | 0.662460978 | 0.206463272 | 0.621880948 | 0.425708157 | 0.17136488 |
| Rpl37 Rpl37rt | S | 95 | 0.195148675 | 0.029988601 | -0.0730678 | -0.268216475 | -0.103056401 | 0.208199337 | 0.674272302 | 0.273820495 | 0.114781744 | 0.256478745 |
| Srrm2 | S | 829 | 0.018473882 | -0.007779893 | 0.042247085 | 0.023773202 | 0.050026978 | 0.360793034 | 0.761494698 | 0.194371449 | 0.428055129 | 0.174450758 |
| Srrm1 | S | 602 | 0.055352067 | 0.018403731 | 0.009961293 | -0.045390773 | -0.008442438 | 0.365872818 | 0.706559771 | 0.864895465 | 0.419956158 | 0.841674061 |
| Srrm1 | T | 600 | 0.055352067 | 0.018403731 | 0.009961293 | -0.045390773 | -0.008442438 | 0.365872818 | 0.706559771 | 0.864895465 | 0.419956158 | 0.841674061 |
| Abhd16a | T | 32 | 0.01047185 | -0.023964672 | 0.157610768 | 0.147138918 | 0.18157544 | 0.918035087 | 0.838046369 | 0.383103799 | 0.434729203 | 0.360281087 |
| Epb41l2 | S | 15 | 0.070752731 | 0.053586396 | -0.025861498 | -0.096614229 | -0.079447893 | 0.404278715 | 0.516840708 | 0.743210035 | 0.062758653 | 0.071694643 |
| Yeats2 | S | 1244 | -0.004922077 | 0.00365406 | 0.156028462 | 0.160950539 | 0.152374402 | 0.971356233 | 0.981708485 | 0.441874485 | 0.363145403 | 0.41037812 |
| Fam114a1 | T | 197 | 0.066700092 | 0.081375837 | -0.056903325 | -0.123603416 | -0.138279161 | 0.226366572 | 0.194524296 | 0.153146946 | 0.060784593 | 0.06679166 |
| H4c1 H4c11 H4c12 H4 | S | 48 | 0.017454737 | 0.074850106 | -0.026645214 | -0.044099951 | -0.10149532 | 0.758304109 | 0.103039273 | 0.535451804 | 0.379403613 | 0.014618816 |
| Dtl | S | 446 | 0.096370091 | 0.048085296 | -0.072715778 | -0.169085869 | -0.120801074 | 0.214438762 | 0.651222472 | 0.651988698 | 0.325863928 | 0.482201811 |
| Arhgap24 | S | 368 | -0.203245156 | 0.070470154 | 0.227108972 | 0.430354128 | 0.156638818 | 0.00297911 | 0.226268247 | 0.004166766 | 0.000842686 | 0.034608424 |
| Macf1 | S | 3541 | 0.007051646 | 0.089758989 | -0.000253465 | -0.007305111 | -0.090012454 | 0.902052967 | 0.30694278 | 0.998239814 | 0.949469328 | 0.490245603 |
| Macf1 | T | 3549 | 0.007051646 | 0.089758989 | -0.000253465 | -0.007305111 | -0.090012454 | 0.902052967 | 0.30694278 | 0.998239814 | 0.949469328 | 0.490245603 |
| Cdc42ep3 | S | 89 | -0.026705242 | 0.058207091 | 0.075396304 | 0.102101545 | 0.017189213 | 0.682537912 | 0.554839364 | 0.386712005 | 0.197177479 | 0.857102739 |
| Mcm3 | S | 732 | -0.035027717 | 0.033730651 | 0.053122334 | 0.088150051 | 0.019391683 | 0.437679018 | 0.396502077 | 0.107171493 | 0.108128306 | 0.614549043 |
| Farp2 | S | 897 | 0.041888159 | -0.042493182 | 0.084433291 | 0.042545133 | 0.126926474 | 0.460770732 | 0.539325581 | 0.269397532 | 0.498396603 | 0.130321719 |
| Mki67 | T | 1924 | 0.121064794 | 0.073585278 | -0.069980973 | -0.191045767 | -0.14356625 | 0.076882096 | 0.241427035 | 0.017988088 | 0.036241983 | 0.080655849 |
| Map4k5 | S | 434 | 0.069520553 | -0.051451633 | 0.037696293 | -0.031824261 | 0.089147926 | 0.200528058 | 0.411385839 | 0.246846407 | 0.488846131 | 0.197844322 |
| Eva1b | T | 95 | -0.082759141 | 0.234912818 | 0.001337388 | 0.084096529 | -0.233575429 | 0.399510717 | 0.19333753 | 0.986095461 | 0.419752173 | 0.195945892 |
| Cep170b | S | 360 | 0.015675188 | 0.015214659 | 0.017496771 | 0.001821584 | 0.002282112 | 0.433150038 | 0.314421736 | 0.547681563 | 0.948293164 | 0.928852114 |
| Srrm2 | T | 2021 | 0.003176755 | 0.001033516 | 0.065871703 | 0.062694949 | 0.064838188 | 0.905896969 | 0.977978725 | 0.197185063 | 0.210875258 | 0.219331326 |
| Sec31a | S | 531 | -0.003944402 | 0.024421587 | 0.029988202 | 0.033932603 | 0.005566615 | 0.926106517 | 0.484479219 | 0.161293096 | 0.454816943 | 0.867764247 |
| Pdzd8 | S | 973 | -0.111291877 | -0.085096034 | 0.26141165 | 0.372703527 | 0.346507684 | 0.518451826 | 0.481231039 | 0.055599523 | 0.083774373 | 0.00434443 |
| Ranbp2 | S | 2505 | 0.120463569 | -0.05946192 | -0.001819086 | -0.122282655 | 0.057642833 | 0.021279094 | 0.170281121 | 0.968986124 | 0.029494524 | 0.149538972 |
| Sec62 | S | 335 | 0.005489737 | -0.063900338 | 0.215535701 | 0.210045964 | 0.279436039 | 0.957828316 | 0.541594487 | 0.270483587 | 0.291733418 | 0.185949328 |
| Septin10 | S | 414 | -0.054896737 | 0.17885002 | -0.075632343 | -0.020735607 | -0.254482363 | 0.657057965 | 0.057329715 | 0.542869073 | 0.889209365 | 0.104766778 |
| Rad18 | S | 99 | -0.057748251 | 0.036688366 | 0.042143225 | 0.099891476 | 0.005454859 | 0.442022829 | 0.100633344 | 0.166033668 | 0.234172492 | 0.816392594 |
| Nav1 | S | 1251 | 0.027534353 | 0.030585092 | -0.01396353 | -0.041497883 | -0.044548621 | 0.518667406 | 0.097825606 | 0.817559184 | 0.547267852 | 0.482318054 |
| Rab12 | T | 107 | 0.034152939 | 0.113634521 | -0.066819231 | -0.10097217 | -0.180453752 | 0.58593225 | 0.115414359 | 0.302769142 | 0.045978204 | 0.013418566 |
| Sec62 | T | 343 | 0.069629582 | -0.056417917 | 0.116635764 | 0.047006182 | 0.173053681 | 0.537338965 | 0.363095387 | 0.292202311 | 0.723091167 | 0.159901593 |
| Mndal | S | 170 | -0.027243849 | -0.015488207 | 0.120475344 | 0.147719193 | 0.135963551 | 0.671362367 | 0.750847056 | 0.107387621 | 0.0708987 | 0.085708911 |
| Fam189b | S | 494 | 0.05163018 | 0.006982496 | 0.041104408 | -0.010525772 | 0.034121913 | 0.476361513 | 0.956567743 | 0.646213136 | 0.912125555 | 0.809027501 |
| Mbd1 | S | 264 | 0.073394268 | -0.006341818 | 0.012431859 | -0.060962409 | 0.018773677 | 0.268278792 | 0.937529732 | 0.775915261 | 0.354671401 | 0.819356193 |
| Zfp592 | S | 461 | 0.02052807 | 0.025440111 | 0.033499107 | 0.012971036 | 0.008058996 | 0.735912763 | 0.595236799 | 0.500114714 | 0.805149457 | 0.819328032 |
| Srsf10 | S | 213 | 0.01108214 | 0.024311448 | 0.055609811 | 0.044527671 | 0.031298363 | 0.870399088 | 0.593682373 | 0.506280803 | 0.640941995 | 0.707019882 |
| Tenm3 | S | 202 | 0.138354063 | -0.010901559 | -0.060784156 | -0.199138219 | -0.049882597 | 0.071216844 | 0.899492158 | 0.436863338 | 0.058560596 | 0.614066942 |
| Srrm2 | S | 1825 | -0.048982066 | 0.017744424 | 0.140943817 | 0.189925882 | 0.123199393 | 0.408882083 | 0.754784129 | 0.188794804 | 0.109255983 | 0.241532627 |
| Foxp4 | S | 269 | 0.104564051 | -0.021015979 | 0.006069033 | -0.098495018 | 0.027085011 | 0.216192683 | 0.796853453 | 0.964092577 | 0.484410727 | 0.841696433 |
| Srrm1 | S | 704 | 0.035117857 | -0.036838841 | 0.06228896 | 0.027171103 | 0.099127801 | 0.240081089 | 0.227713919 | 0.131815526 | 0.445020824 | 0.04348624 |
| Srsf6 | S | 316 | 0.034803313 | -0.047949151 | 0.099249852 | 0.064446539 | 0.147199004 | 0.452078199 | 0.219939725 | 0.13131089 | 0.280264942 | 0.070938215 |
| Rabl6 | S | 594 | -0.043827494 | 0.032999748 | 0.061913904 | 0.105741398 | 0.028914156 | 0.274503823 | 0.224577318 | 0.121268117 | 0.055873279 | 0.422137012 |
| Rabl6 | T | 597 | -0.043827494 | 0.032999748 | 0.061913904 | 0.105741398 | 0.028914156 | 0.274503823 | 0.224577318 | 0.121268117 | 0.055873279 | 0.422137012 |
| Clspn | S | 424 | 0.114282153 | -0.037536016 | -0.023490824 | -0.137772977 | 0.014045192 | 0.009615057 | 0.180612133 | 0.488019202 | 0.010224211 | 0.599689451 |
| Ccnl1 | S | 378 | -0.035947787 | 0.0599242 | 0.032442595 | 0.068390382 | -0.027481605 | 0.217763942 | 0.09900072 | 0.26701756 | 0.034311702 | 0.362363363 |
| Nes | S | 1287 | 0.056244174 | -0.042986959 | 0.043339893 | -0.012904281 | 0.086326852 | 0.07342599 | 0.11754146 | 0.197547328 | 0.690401052 | 0.044815059 |
| Vcl | T | 719 | -0.055309445 | 0.172148551 | 0.016103107 | 0.071412553 | -0.156045444 | 0.426352513 | 0.191159063 | 0.951348888 | 0.784961012 | 0.577293649 |
| Ubxn7 | S | 258 | -0.09128786 | 0.133316896 | 0.016592215 | 0.107880075 | -0.116724681 | 0.345996026 | 0.068337057 | 0.816957033 | 0.248976575 | 0.078168828 |
| Zc3h18 | S | 45 | 0.027701859 | -0.02342008 | 0.095174833 | 0.067472974 | 0.118594913 | 0.441490239 | 0.261651015 | 0.235342141 | 0.370929346 | 0.16921474 |
| Tmem209 | S | 260 | 0.070526326 | 0.009559952 | 0.032112602 | -0.038413725 | 0.022552649 | 0.447824145 | 0.946802435 | 0.806836441 | 0.731930156 | 0.883223587 |
| Ppp4r3a | S | 117 | 0.056436607 | -0.001232152 | -0.002139453 | -0.05857606 | -0.000907301 | 0.093389034 | 0.986293243 | 0.860840397 | 0.100556418 | 0.989844388 |
| Kansl2 | S | 168 | 0.039211327 | 0.001631756 | 0.046207808 | 0.006996482 | 0.044576052 | 0.586050484 | 0.98601728 | 0.442460464 | 0.92336704 | 0.643651731 |
| Kansl2 | S | 172 | 0.039211327 | 0.001631756 | 0.046207808 | 0.006996482 | 0.044576052 | 0.586050484 | 0.98601728 | 0.442460464 | 0.92336704 | 0.643651731 |
| Kansl2 | S | 175 | 0.039211327 | 0.001631756 | 0.046207808 | 0.006996482 | 0.044576052 | 0.586050484 | 0.98601728 | 0.442460464 | 0.92336704 | 0.643651731 |
| Mycbp2 | S | 181 | -0.044810596 | 0.098594381 | 0.00373949 | 0.048550086 | -0.094854891 | 0.493806989 | 0.071921005 | 0.962800431 | 0.560158069 | 0.262766399 |
| Mycbp2 | S | 183 | -0.044810596 | 0.098594381 | 0.00373949 | 0.048550086 | -0.094854891 | 0.493806989 | 0.071921005 | 0.962800431 | 0.560158069 | 0.262766399 |
| Mybl2 | S | 455 | 0.033403506 | -0.107919512 | 0.210916977 | 0.17751347 | 0.318836489 | 0.686448198 | 0.151956082 | 0.110209251 | 0.158977232 | 0.060341254 |
| Arid1a | S | 384 | -0.022187231 | 0.03711644 | 0.047943154 | 0.070130385 | 0.010826714 | 0.594987691 | 0.248282207 | 0.354352026 | 0.206973569 | 0.796581512 |
| Itpr2 | S | 1160 | -0.038547717 | 0.088515705 | 0.024009038 | 0.062556755 | -0.064506667 | 0.649291121 | 0.19067206 | 0.722070799 | 0.535172418 | 0.435064849 |
| Vcp | S | 787 | 0.075650669 | -0.002177164 | 0.000627318 | -0.075023352 | 0.002804481 | 0.19684179 | 0.939603895 | 0.982605288 | 0.202437631 | 0.77034232 |
| Kdm1a | S | 850 | 0.039455417 | 0.031458378 | -0.014444625 | -0.053900041 | -0.045903003 | 0.251334151 | 0.440398011 | 0.828894539 | 0.444639565 | 0.520014907 |

| | | | | | | | | | | | | | |
|---|---|---|---|---|---|---|---|---|---|---|---|---|---|
| Wdr76 | S | | 141 | 0.01514786 | 0.028467584 | 0.038752571 | 0.023604712 | 0.010284987 | 0.91581201 | 0.635398124 | 0.386895695 | 0.869658597 | 0.864204418 |
| Wdr76 | S | | 142 | 0.01514786 | 0.028467584 | 0.038752571 | 0.023604712 | 0.010284987 | 0.91581201 | 0.635398124 | 0.386895695 | 0.869658597 | 0.864204418 |
| Srsf1 | S | | 201 | -0.019394358 | -0.042034425 | 0.132878681 | 0.152273039 | 0.174913105 | 0.739470033 | 0.343354823 | 0.050278231 | 0.049780768 | 0.032689796 |
| Atf7ip | S | | 288 | -0.109497792 | 0.074158572 | 0.09848748 | 0.207985271 | 0.024328908 | 0.506368687 | 0.310386662 | 0.353138223 | 0.254780182 | 0.772294456 |
| Tra2b | T | | 203 | 0.099827281 | -0.088707095 | 0.135871928 | 0.036044647 | 0.224579022 | 0.100287751 | 0.096463647 | 0.475843462 | 0.839953788 | 0.283145038 |
| Spen | S | | 190 | -0.05824982 | -0.015332861 | 0.152491562 | 0.210741382 | 0.167824423 | 0.398428154 | 0.759505685 | 0.069501177 | 0.041757861 | 0.059945197 |
| Apc | S | | 2260 | -0.025613568 | -0.028318382 | 0.095009677 | 0.120623245 | 0.123328059 | 0.694605006 | 0.587582498 | 0.03685194 | 0.140163646 | 0.074351296 |
| Phrf1 | S | | 1201 | 0.006674241 | -0.03146361 | 0.055281019 | 0.048606778 | 0.086744629 | 0.810256767 | 0.429618424 | 0.019688679 | 0.139547012 | 0.095496389 |
| Phactr4 | S | | 175 | -0.001469601 | 0.091472915 | -0.048802435 | -0.047332834 | -0.14027535 | 0.969591851 | 0.058496991 | 0.581115447 | 0.578868312 | 0.180571913 |
| Zfand5 | S | | 48 | 0.046676357 | -0.014068283 | 0.024712814 | -0.021963543 | 0.038781097 | 0.235354211 | 0.655498671 | 0.367393235 | 0.50263046 | 0.180917589 |
| Rbm12b1 Rbm12b2 | S | 641;643 | | 0.007563992 | 0.007057872 | 0.059955637 | 0.052391645 | 0.052897765 | 0.922296152 | 0.840511861 | 0.270166088 | 0.53159846 | 0.296399408 |
| Cbfb | S | | 173 | -0.01421989 | 0.065392778 | 0.02168217 | 0.035902059 | -0.043710608 | 0.608423658 | 0.170496881 | 0.727544987 | 0.563280087 | 0.51922243 |
| Sugp1 | S | | 483 | -0.066418687 | 0.07693361 | 0.057119655 | 0.123538342 | -0.019813955 | 0.006213292 | 0.013189264 | 0.12754728 | 0.02290238 | 0.5375178 |
| Nes | S | | 1291 | 0.019156256 | -0.066896321 | 0.084426971 | 0.065270715 | 0.151323292 | 0.638991775 | 0.29453661 | 0.034883949 | 0.189741869 | 0.069043327 |
| Hnrnpc | S | | 306 | 0.030423392 | -0.045628687 | 0.118077975 | 0.087654583 | 0.163706662 | 0.583612154 | 0.534531464 | 0.294895777 | 0.39694486 | 0.17608123 |
| Nectin3 | S | | 465 | -0.022858358 | -0.014094345 | 0.114851456 | 0.137709814 | 0.128945801 | 0.573484147 | 0.758475396 | 0.100653521 | 0.076643035 | 0.077734205 |
| Cep170 | S | | 870 | -0.070972971 | 0.057777314 | 0.063676705 | 0.134649676 | 0.005899392 | 0.184956427 | 0.147412114 | 0.143306878 | 0.043587175 | 0.871439127 |
| Slc12a7 | S | | 33 | 0.01133974 | 0.005978026 | 0.06159714 | 0.0502574 | 0.055619114 | 0.760211531 | 0.832867558 | 0.364726543 | 0.459456127 | 0.404167447 |
| Fbxo6 | S | | 276 | -0.060117358 | 0.142981629 | -0.021749946 | 0.038367412 | -0.164731575 | 0.183890525 | 0.022056841 | 0.7371094 | 0.538423148 | 0.060015155 |
| Fggy | S | | 398 | 0.176280762 | 0.168493744 | -0.068789008 | -0.24506977 | -0.237282752 | 0.707245531 | 0.754182622 | 0.789838526 | 0.627035574 | 0.676385862 |
| Hivep1 | S | | 1739 | -0.14932305 | 0.067716423 | 0.19770676 | 0.34702981 | 0.129990336 | 0.159118971 | 0.464695478 | 0.131156175 | 0.026821871 | 0.231242072 |
| Wdr26 | S | | 638 | 0.106120423 | -0.018457583 | 0.02012658 | -0.085993843 | 0.038584163 | 0.332293414 | 0.737440746 | 0.782247797 | 0.450398178 | 0.615407867 |
| Fndc3b | S | | 208 | 0.009310619 | -0.008163063 | 0.055307406 | 0.045996787 | 0.063470469 | 0.750386014 | 0.757774597 | 0.134485747 | 0.235352462 | 0.113587412 |
| Tob1 | S | | 205 | -0.146253931 | 0.137454527 | 0.049834984 | 0.196088916 | -0.087619542 | 0.163866805 | 0.024063626 | 0.691957707 | 0.206614804 | 0.493998125 |
| Tob1 | T | | 204 | -0.146253931 | 0.137454527 | 0.049834984 | 0.196088916 | -0.087619542 | 0.163866805 | 0.024063626 | 0.691957707 | 0.206614804 | 0.493998125 |
| Ubxn4 | S | | 127 | 0.002141695 | -0.027323172 | 0.125879138 | 0.123737442 | 0.15320231 | 0.989913713 | 0.818346563 | 0.269763028 | 0.488220042 | 0.250459313 |
| Pitpnm1 | T | | 369 | 0.450223079 | -0.151766078 | -0.043798655 | -0.494021734 | 0.107967423 | 0.206842519 | 0.308212735 | 0.854670867 | 0.202826359 | 0.669571324 |
| Nedd4l | T | | 502 | 0.056087774 | 0.086587173 | -0.063400765 | -0.119488539 | -0.149987938 | 0.527299274 | 0.301593881 | 0.617097839 | 0.398129642 | 0.294423122 |
| Dlg5 | S | | 295 | 0.046698787 | 0.019997403 | -0.008579899 | -0.055278685 | -0.028577302 | 0.232546528 | 0.566715173 | 0.894217777 | 0.397954742 | 0.636891149 |
| Tjp2 | S | | 948 | -0.04248754 | -0.107658019 | 0.198457011 | 0.240944552 | 0.30611503 | 0.462965827 | 0.025292081 | 0.001326558 | 0.014938563 | 0.000533184 |
| Asxl2 | S | | 524 | 0.122849783 | -0.033797337 | -5.33245E-06 | -0.122855115 | 0.033792005 | 0.152862643 | 0.541571572 | 0.999913128 | 0.153024069 | 0.47481383 |
| Ccar2 | S | | 686 | 0.058436444 | 0.02675476 | 0.06040429 | 0.001967846 | 0.033649531 | 0.81910131 | 0.860365395 | 0.637506128 | 0.993849223 | 0.828683388 |
| Zc3h3 | S | | 855 | -0.001285961 | 0.107041627 | -0.029365678 | -0.028079717 | -0.136407305 | 0.981689871 | 0.173343605 | 0.749920426 | 0.76192196 | 0.220848505 |
| Zfp646 | S | | 1411 | -0.183118835 | 0.098221836 | 0.201513208 | 0.384632044 | 0.103291372 | 0.024885542 | 0.079939729 | 0.114991641 | 0.022992025 | 0.327291065 |
| Hirip3 | S | | 231 | -0.111040812 | 0.015353661 | 0.110359813 | 0.221400625 | 0.095006152 | 0.036671299 | 0.717927605 | 0.001046538 | 0.009536328 | 0.099997448 |
| Hirip3 | S | | 234 | -0.111040812 | 0.015353661 | 0.110359813 | 0.221400625 | 0.095006152 | 0.036671299 | 0.717927605 | 0.001046538 | 0.009536328 | 0.099997448 |
| Rbm12b1 Rbm12b2 | S | 376;376 | | -0.015248186 | -0.034416865 | 0.16654362 | 0.181791805 | 0.200960485 | 0.693539033 | 0.425163044 | 0.130873265 | 0.114395148 | 0.091519501 |
| Safb | S | | 506 | -0.000227777 | 0.014814093 | 0.063905055 | 0.064132832 | 0.049090962 | 0.996178188 | 0.701392168 | 0.336846229 | 0.334343535 | 0.418133801 |
| Pdlim7 | T | | 96 | -0.020255968 | 0.021387796 | 0.057432104 | 0.077688071 | 0.036044308 | 0.153227363 | 0.146380711 | 0.154632681 | 0.099926365 | 0.314180667 |
| Rpp25l | S | | 100 | 0.156791053 | -0.075850935 | -0.00618452 | -0.162975573 | 0.069666415 | 0.121363414 | 0.514139776 | 0.959615376 | 0.211656124 | 0.597706005 |
| Akap12 | S | | 1190 | -0.037038814 | 0.320658997 | -0.094931832 | -0.057893018 | -0.415590829 | 0.669581874 | 0.174113097 | 0.405062884 | 0.535728859 | 0.108365162 |
| Setd2 | S | | 633 | 0.019935477 | -0.021888105 | 0.070186498 | 0.050251021 | 0.092074603 | 0.248005162 | 0.427690308 | 0.269338537 | 0.396007734 | 0.178215073 |
| Incenp | T | | 195 | 0.030316755 | 0.029785724 | 0.003229781 | -0.027086974 | -0.026555943 | 0.450303167 | 0.479412141 | 0.753496686 | 0.491529301 | 0.52062916 |
| Phip | S | | 1476 | 0.027478467 | -0.013916566 | 0.047064476 | 0.019586009 | 0.060981042 | 0.503610082 | 0.676986354 | 0.213194364 | 0.631850554 | 0.126148582 |
| Cdc23 | S | | 578 | -0.137102928 | 0.15276162 | 0.116092063 | 0.253194991 | -0.036669557 | 0.021790709 | 0.049085535 | 0.290166057 | 0.076133067 | 0.719999938 |
| Map4 | S | | 506 | -0.016878159 | 0.062138249 | 0.019826171 | 0.03670433 | -0.042312078 | 0.658505401 | 0.170626692 | 0.69246651 | 0.513911022 | 0.461343328 |
| Amot | T | | 1084 | -0.131744398 | 0.34375751 | -0.079377046 | 0.052367352 | -0.423134556 | 0.167111921 | 0.022450084 | 0.266148861 | 0.458845831 | 0.028520909 |
| Nup98 | S | | 653 | 0.084786927 | -0.043868475 | 0.030138206 | -0.054648721 | 0.074006681 | 0.154687363 | 0.437591931 | 0.594599363 | 0.092308418 | 0.04813419 |
| Cdca7l | T | | 153 | 0.086929191 | -0.008933654 | 0.010494266 | -0.076434925 | 0.01942792 | 0.309902014 | 0.843714666 | 0.769878842 | 0.353310927 | 0.565500614 |
| Ppp1r18 | S | | 59 | 0.028910725 | 0.019884872 | 0.030852106 | 0.001941381 | 0.010967234 | 0.512381768 | 0.73589179 | 0.654114459 | 0.976454336 | 0.883936082 |
| Tom1l1 | S | | 170 | -0.013697975 | 0.130703076 | -0.071922189 | -0.058224214 | -0.202625265 | 0.78301142 | 0.035646549 | 0.360756936 | 0.400807981 | 0.053219021 |
| Zc3h13 | S | | 110 | 0.058708373 | -0.011176077 | 0.032229607 | -0.026478765 | 0.043405684 | 0.411947684 | 0.801840202 | 0.486662227 | 0.679164456 | 0.245088887 |
| C77080 | S | | 560 | -0.034776791 | 0.143347245 | 0.01249653 | 0.047273321 | -0.130850715 | 0.726417322 | 0.318993862 | 0.924459007 | 0.707106226 | 0.410732659 |
| Scrib | S | | 1287 | 0.080110393 | -0.000199621 | -0.01511816 | -0.095228553 | -0.014918539 | 0.154779618 | 0.995752803 | 0.817803525 | 0.223574521 | 0.813780422 |
| Ncbp3 | T | | 408 | 0.141003692 | -0.041042002 | -0.012468485 | -0.153472177 | 0.028573517 | 0.019481863 | 0.067991603 | 0.578380599 | 0.022672553 | 0.130971186 |
| Leo1 | S | | 274 | 0.026218757 | -0.021723415 | 0.033657828 | 0.007439071 | 0.055381242 | 0.48999176 | 0.393766132 | 0.035370521 | 0.835456259 | 0.090456649 |
| Tada3 | S | | 338 | 0.02518686 | -0.056876821 | 0.09632854 | 0.07114168 | 0.153205361 | 0.700792739 | 0.434712445 | 0.119287189 | 0.333484729 | 0.094351787 |
| Bcl9l | S | | 994 | -0.064810005 | -0.027788179 | 0.140032558 | 0.204842564 | 0.167820737 | 0.241248352 | 0.483054227 | 0.013747037 | 0.012667144 | 0.008711322 |
| Mta3 | S | | 547 | -0.011796453 | 0.057402352 | 0.006917496 | 0.018713949 | -0.050484856 | 0.863201582 | 0.140607321 | 0.836789518 | 0.781564919 | 0.171833468 |
| Rbmxl1 | S | | 170 | 0.095005047 | -0.110922863 | 0.117975956 | 0.022970909 | 0.228898819 | 0.483616649 | 0.34977763 | 0.244534725 | 0.872366108 | 0.126680189 |
| Akap13 | S | | 2462 | -0.080159032 | 0.255027627 | 0.014105648 | 0.09426468 | -0.240921979 | 0.431542597 | 0.320810767 | 0.838302416 | 0.37054033 | 0.342681561 |
| Fam120a | S | | 1038 | -0.168541053 | 0.078952505 | 0.193175645 | 0.361716699 | 0.11422314 | 0.022727436 | 0.154131026 | 0.05746342 | 0.011126258 | 0.189092729 |
| Fam120a | S | | 1039 | -0.168541053 | 0.078952505 | 0.193175645 | 0.361716699 | 0.11422314 | 0.022727436 | 0.154131026 | 0.05746342 | 0.011126258 | 0.189092729 |
| Fam120a | S | | 1042 | -0.168541053 | 0.078952505 | 0.193175645 | 0.361716699 | 0.11422314 | 0.022727436 | 0.154131026 | 0.05746342 | 0.011126258 | 0.189092729 |
| Mki67 | S | | 1100 | 0.034880635 | -0.043577141 | 0.062290735 | 0.0274101 | 0.105867876 | 0.403971345 | 0.581819597 | 0.226185367 | 0.528076343 | 0.233505553 |
| Atrx | S | | 1339 | -0.058829594 | 0.115533564 | 0.025181331 | 0.084010925 | -0.090352233 | 0.152433706 | 0.0859708 | 0.518093115 | 0.025942308 | 0.149427574 |
| Nudc | T | | 99 | 0.138884055 | -0.020332466 | -0.011799519 | -0.150683574 | 0.008532947 | 0.263146831 | 0.815733726 | 0.884807442 | 0.245239448 | 0.927457102 |
| Trim33 | S | | 877 | 0.068099224 | -0.009972146 | -0.004680848 | -0.072780071 | 0.005291298 | 0.09634144 | 0.76806523 | 0.918788025 | 0.12255371 | 0.877487731 |
| H1f2 | S | | 113 | 0.113507998 | 0.053899615 | -0.053071427 | -0.166579425 | -0.106971042 | 0.367281261 | 0.659777285 | 0.620991494 | 0.26274379 | 0.458245772 |
| Vta1 | Y | | 280 | -0.05449914 | 0.174046203 | 0.017535755 | 0.072034895 | -0.156510448 | 0.443735215 | 0.279203015 | 0.774298329 | 0.301931688 | 0.319550414 |
| Foxc1 | S | | 320 | -0.029502884 | 0.049149159 | 0.064342231 | 0.093845115 | 0.015193072 | 0.651943881 | 0.518893332 | 0.349540616 | 0.038283824 | 0.770435325 |
| Ticrr | T | | 1814 | 0.008366317 | 0.075573733 | 0.050421017 | 0.0420547 | -0.025152716 | 0.911923129 | 0.577641481 | 0.833724817 | 0.858286673 | 0.920417137 |
| Srrm2 | T | | 1836 | 0.014937122 | 0.022406311 | 0.038753763 | 0.023816641 | 0.016347452 | 0.718102858 | 0.567026892 | 0.585325057 | 0.714483603 | 0.796444376 |
| Fam110b | S | | 255 | 0.018885078 | 0.055890004 | 0.015761671 | -0.003123407 | -0.040128333 | 0.779313002 | 0.466566483 | 0.842698894 | 0.969934709 | 0.653551309 |
| Msantd4 | S | | 152 | 0.033715536 | -0.014621314 | 0.074484776 | 0.04076924 | 0.08910609 | 0.790532221 | 0.851636833 | 0.333016226 | 0.760227559 | 0.352460074 |
| Hsp90b1 | T | | 44 | -0.027018062 | 0.046266778 | 0.06559974 | 0.092617802 | 0.019332962 | 0.661902163 | 0.411080514 | 0.453272673 | 0.284615025 | 0.793431625 |
| Mzt2 | S | | 140 | 0.041647578 | 0.144534261 | -0.090898652 | -0.132546231 | -0.235432913 | 0.692838714 | 0.192489389 | 0.404701417 | 0.218809224 | 0.058666095 |
| Pimreg | S | | 187 | -0.039705211 | 0.095902761 | -0.032433518 | 0.007271693 | -0.128336279 | 0.499228406 | 0.011659626 | 0.439963743 | 0.901636056 | 0.048085698 |
| Arhgap21 | S | | 437 | 0.02933067 | 0.044538404 | -0.007064943 | -0.036395613 | -0.051603347 | 0.6941965 | 0.253372583 | 0.902390428 | 0.665784067 | 0.400870698 |
| Eif4g3 | S | | 267 | -0.104590005 | 0.097549818 | 0.106159888 | 0.210749893 | 0.00861007 | 0.026996846 | 0.067455184 | 0.182167993 | 0.047493795 | 0.899587509 |
| Cdk13 | S | | 662 | -0.038961956 | 0.068820062 | 0.100334944 | 0.139296901 | 0.031514882 | 0.494350037 | 0.288099258 | 0.68229866 | 0.577772633 | 0.895858114 |
| Thumpd1 | S | | 86 | -0.000169881 | 0.018243592 | 0.030910432 | 0.031080313 | 0.01266684 | 0.994445928 | 0.468421789 | 0.253286514 | 0.311647851 | 0.660604412 |
| Thumpd1 | S | | 88 | -0.000169881 | 0.018243592 | 0.030910432 | 0.031080313 | 0.01266684 | 0.994445928 | 0.468421789 | 0.253286514 | 0.311647851 | 0.660604412 |
| Mast4 | S | | 1395 | -0.076838252 | 0.056190709 | 0.076399359 | 0.153237612 | 0.020208651 | 0.262082511 | 0.309641881 | 0.141365058 | 0.074221432 | 0.728294603 |
| Clk4 | S | | 136 | 0.086990487 | -0.00759412 | -0.010351327 | -0.097341815 | -0.002757207 | 0.154278134 | 0.815688689 | 0.845755971 | 0.170321899 | 0.954883634 |
| Clk4 | S | | 138 | 0.086990487 | -0.00759412 | -0.010351327 | -0.097341815 | -0.002757207 | 0.154278134 | 0.815688689 | 0.845755971 | 0.170321899 | 0.954883634 |
| Poll | S | | 167 | 0.066087823 | -0.055197715 | 0.03561708 | -0.030470743 | 0.090814795 | 0.080239613 | 0.303691748 | 0.442912635 | 0.524181263 | 0.169495685 |
| Myh9 | S | | 876 | 0.008971309 | 0.058614599 | 0.048487834 | 0.039516524 | -0.010126765 | 0.934476027 | 0.593887595 | 0.734461129 | 0.812788429 | 0.950925852 |
| Srrm2 | S | | 778 | 0.008978582 | 0.007804803 | 0.019451958 | 0.010473375 | 0.011647154 | 0.687864575 | 0.567372124 | 0.135768671 | 0.635682768 | 0.383201967 |
| Sltm | S | | 311 | -0.022304791 | -0.019619799 | 0.081309554 | 0.103614345 | 0.100929353 | 0.488745452 | 0.616280091 | 0.025818665 | 0.019849494 | 0.05191424 |
| Dcp1a | S | | 372 | -0.099345137 | 0.04863525 | 0.120669477 | 0.220014614 | 0.072034227 | 0.019063273 | 0.271683559 | 0.017880408 | 0.003221861 | 0.158681617 |
| Ran | S | | 135 | 0.101379889 | 0.035844088 | -0.002819789 | -0.104199678 | -0.038663877 | 0.537769074 | 0.837503117 | 0.969805439 | 0.540642713 | 0.831232522 |
| Huwe1 | T | | 3930 | 0.021502317 | 0.006669745 | 0.016359343 | -0.005142974 | 0.009689598 | 0.395326424 | 0.714124602 | 0.342123281 | 0.813678711 | 0.508650268 |
| Kat6a | S | | 954 | 0.023073696 | 0.002014042 | 0.057165325 | 0.034091629 | 0.055151283 | 0.713540048 | 0.971968194 | 0.411720481 | 0.581741882 | 0.354998669 |

| | | | | | | | | | | | | |
|---|---|---|---|---|---|---|---|---|---|---|---|---|
| Atxn7l3 | S | 281 | -0.046140227 | 0.022873767 | 0.088410207 | 0.134550434 | 0.065536439 | 0.167732193 | 0.384239133 | 0.091383218 | 0.033321619 | 0.173222044 |
| Hivep2 | S | 169 | -0.028233489 | 0.032318385 | 0.073637541 | 0.101871029 | 0.041319156 | 0.576023212 | 0.455783575 | 0.328908549 | 0.204511147 | 0.543784895 |
| Cdk13 | S | 664 | 0.025833454 | -3.03467E-05 | 0.067461116 | 0.041627662 | 0.067491463 | 0.796117866 | 0.999555767 | 0.414172949 | 0.706410534 | 0.401374502 |
| Mpzl1 | S | 221 | 0.058737238 | -0.054800887 | 0.055365361 | -0.003371877 | 0.110166248 | 0.313024484 | 0.438043273 | 0.262955388 | 0.955551686 | 0.187211027 |
| Zfp592 | S | 346 | -0.023274946 | 0.016621623 | 0.084794651 | 0.108069598 | 0.068173028 | 0.547000259 | 0.638965924 | 0.230893612 | 0.152242815 | 0.303292648 |
| Plekho2 | S | 403 | 0.033021559 | -0.008276768 | 0.069579342 | 0.036557783 | 0.07785611 | 0.67189515 | 0.935684405 | 0.471149575 | 0.734821811 | 0.543529231 |
| Srrm2 | S | 351 | 0.019690883 | 0.000747583 | 0.026465134 | 0.006774251 | 0.025717551 | 0.3703816 | 0.970756685 | 0.312774467 | 0.780918096 | 0.314231663 |
| Pcnt | S | 1959 | 0.091089576 | 0.019356066 | -0.002530848 | -0.093620424 | -0.021886914 | 0.412639698 | 0.915462538 | 0.988241365 | 0.586452217 | 0.919323553 |
| Tsc2 | S | 671 | 0.034274306 | -0.005829455 | 0.034795043 | 0.000520738 | 0.040624498 | 0.407232598 | 0.886638632 | 0.432547359 | 0.991942073 | 0.465690417 |
| Med1 | S | 1484 | 0.053774707 | -0.002619485 | 0.025809009 | -0.027965698 | 0.028428494 | 0.39597205 | 0.960444588 | 0.642703233 | 0.687729135 | 0.658651669 |
| Map11 | S | 517 | 0.041753707 | 0.07175514 | -0.026623585 | -0.068377292 | -0.098378724 | 0.590876893 | 0.425372949 | 0.803337547 | 0.523330105 | 0.397866772 |
| Lrch1 | S | 518 | -0.034028526 | 0.009265751 | 0.08711593 | 0.121144456 | 0.077850179 | 0.486650556 | 0.89581022 | 0.1040181 | 0.043470431 | 0.303593807 |
| Lrch1 | S | 522 | -0.034028526 | 0.009265751 | 0.08711593 | 0.121144456 | 0.077850179 | 0.486650556 | 0.89581022 | 0.1040181 | 0.043470431 | 0.303593807 |
| Ercc6l | S | 997 | -0.058917417 | 0.137294902 | 0.017455365 | 0.076372781 | -0.119839537 | 0.360746813 | 0.147187474 | 0.85892856 | 0.435350028 | 0.293065042 |
| Ppp4r1 | S | 949 | -0.024491321 | 0.044053871 | 0.039202589 | 0.06369391 | -0.004851282 | 0.523262591 | 0.270257362 | 0.341872865 | 0.062561484 | 0.854165103 |
| Wnk1 | S | 2055 | -0.165647861 | 0.084569853 | 0.140677268 | 0.306325129 | 0.056107415 | 0.359402701 | 0.384657049 | 0.242735 | 0.141109091 | 0.584797534 |
| Wnk1 | S | 2056 | -0.165647861 | 0.084569853 | 0.140677268 | 0.306325129 | 0.056107415 | 0.359402701 | 0.384657049 | 0.242735 | 0.141109091 | 0.584797534 |
| Rad50 | S | 635 | 0.000438775 | 0.001022846 | 0.085361841 | 0.084923067 | 0.084338995 | 0.985572574 | 0.971049544 | 0.322687861 | 0.3248941 | 0.327494053 |
| Zeb2 | T | 141 | -0.050565806 | 0.120770545 | 0.05232534 | 0.102891146 | -0.068445205 | 0.520734094 | 0.293084385 | 0.775604925 | 0.57374101 | 0.717278868 |
| Snrpa1 | T | 250 | 0.064930502 | 0.025112842 | -0.003982933 | -0.068913434 | -0.029095774 | 0.312266017 | 0.860764596 | 0.924166193 | 0.283247187 | 0.838430446 |
| Gen1 | S | 636 | 0.043254853 | 0.064318372 | -0.008213743 | -0.051468595 | -0.072532114 | 0.806243091 | 0.386567336 | 0.925443092 | 0.783233784 | 0.485076888 |
| Ankrd17 | S | 2037 | 0.009785917 | 0.058878034 | -0.046587719 | -0.056373635 | -0.105465753 | 0.586517702 | 0.033468535 | 0.419693279 | 0.342462219 | 0.13754854 |
| Ttc14 | S | 709 | -0.002632863 | -0.059428322 | 0.122141637 | 0.1247745 | 0.181569959 | 0.969339185 | 0.185269985 | 0.03376868 | 0.157414727 | 0.014303596 |
| Dido1 | S | 1307 | -0.143142114 | -0.078459338 | 0.26180027 | 0.404942383 | 0.340259608 | 0.590787857 | 0.582120581 | 0.087859469 | 0.199735778 | 0.077633027 |
| Inpp5f | S | 941 | -0.071554039 | 0.120284746 | -0.012143639 | 0.0594104 | -0.132428385 | 0.072227572 | 0.003502408 | 0.636039629 | 0.121336592 | 0.005497459 |
| Prpf40a | S | 899 | -0.069984842 | 0.008872453 | 0.131529515 | 0.201514357 | 0.122657062 | 0.254572315 | 0.854479501 | 0.071548817 | 0.031334473 | 0.10434303 |
| Peak1 | S | 565 | -0.024537592 | 0.04259252 | 0.06395017 | 0.088487762 | 0.021357651 | 0.439379887 | 0.354178327 | 0.380611131 | 0.259435993 | 0.769812041 |
| Nf2 | S | 518 | -0.100817702 | -0.040011168 | 0.240904127 | 0.341721829 | 0.280915295 | 0.21581871 | 0.623947206 | 0.050865522 | 0.022957669 | 0.036767582 |
| Iqgap1 | S | 648 | 0.047408873 | 0.048090433 | -0.051401864 | -0.098810737 | -0.099492297 | 0.271734966 | 0.268456224 | 0.519205918 | 0.249494104 | 0.247009956 |
| Sf3b1 | T | 207 | 0.07640421 | 0.010950937 | -0.016922695 | -0.093326905 | -0.027873633 | 0.191748576 | 0.722299067 | 0.563094576 | 0.136505848 | 0.331148417 |
| Safb2 | S | 95 | 0.067498955 | 0.012060761 | -0.020225795 | -0.087724749 | -0.032286556 | 0.155599257 | 0.49266039 | 0.430444147 | 0.09304134 | 0.215864061 |
| Enah | S | 144 | 0.062955568 | 0.018860386 | 0.032300491 | -0.030655077 | 0.013440105 | 0.68371376 | 0.829550568 | 0.72422132 | 0.848007915 | 0.896645712 |
| Etv6 | S | 215 | -0.012414464 | 0.010620321 | 0.068360334 | 0.080774798 | 0.057740012 | 0.649299926 | 0.719234766 | 0.20298506 | 0.153742643 | 0.263173451 |
| Mllt3 | S | 413 | -0.066963399 | -0.012021958 | 0.109550351 | 0.17651375 | 0.121572309 | 0.067310753 | 0.767095338 | 0.00303699 | 0.010827272 | 0.057763952 |
| Luzp1 | S | 746 | 0.14649254 | -0.08754336 | 0.020391519 | -0.126101021 | 0.107934879 | 0.109750207 | 0.32892679 | 0.846479388 | 0.199547703 | 0.250457037 |
| Cep55 | S | 23 | -0.10432079 | 0.200851332 | -0.034408773 | 0.069912017 | -0.235260104 | 0.133551522 | 0.020897115 | 0.712097036 | 0.423491124 | 0.054532584 |
| Hdac6 | S | 59 | -0.041398472 | 0.10207419 | -0.02711636 | 0.014282112 | -0.12919055 | 0.511264155 | 0.055605454 | 0.870140163 | 0.93241877 | 0.467299195 |
| Tfe3 | S | 553 | 0.031709169 | 0.043006093 | -0.005566627 | -0.037275796 | -0.04857272 | 0.543546798 | 0.384943905 | 0.878749782 | 0.481223123 | 0.337219874 |
| Psen1 | S | 367 | 0.014064833 | 0.01780657 | 0.023893281 | 0.009828447 | 0.006086711 | 0.738118701 | 0.446862022 | 0.494586998 | 0.840290146 | 0.865397858 |
| Usp34 | S | 2525 | -0.028910337 | 0.010706548 | 0.086245645 | 0.115155982 | 0.075539098 | 0.491923265 | 0.72527733 | 0.157594722 | 0.097386515 | 0.202946073 |
| Arhgef12 | S | 1327 | 0.080666383 | -0.001995643 | -0.000552139 | -0.081218522 | 0.001443504 | 0.273805889 | 0.97621594 | 0.994083845 | 0.146209114 | 0.972838491 |
| Cep350 | S | 874 | 0.052367694 | -0.056984388 | 0.074250487 | 0.021882793 | 0.131234874 | 0.561538134 | 0.375598365 | 0.149454208 | 0.801053354 | 0.088176623 |
| Snx29 | S | 642 | 0.092326144 | 0.032183515 | -0.037864807 | -0.130190952 | -0.070048322 | 0.360188745 | 0.631536105 | 0.702813459 | 0.264611198 | 0.439170708 |
| Snx29 | S | 645 | 0.092326144 | 0.032183515 | -0.037864807 | -0.130190952 | -0.070048322 | 0.360188745 | 0.631536105 | 0.702813459 | 0.264611198 | 0.439170708 |
| Snx29 | S | 649 | 0.092326144 | 0.032183515 | -0.037864807 | -0.130190952 | -0.070048322 | 0.360188745 | 0.631536105 | 0.702813459 | 0.264611198 | 0.439170708 |
| Ptpn13 | S | 1621 | -0.163824639 | 0.208386192 | -0.025987252 | 0.137837387 | -0.234373444 | 0.295821346 | 0.033511698 | 0.771576642 | 0.354200127 | 0.020967597 |
| Jmjd1c | S | 1039 | -0.031856023 | 0.064162138 | 0.071902608 | 0.103758631 | 0.00774047 | 0.056889638 | 0.336503311 | 0.181552406 | 0.104809781 | 0.90897946 |
| Phldb1 | T | 518 | 0.065883561 | 0.062614736 | -0.063988272 | -0.129871833 | -0.126603008 | 0.237058219 | 0.353404283 | 0.373607362 | 0.126944499 | 0.159572428 |
| Atad2 | S | 806 | -0.070264068 | 0.06820277 | 0.106928928 | 0.177192995 | 0.038726157 | 0.417671972 | 0.427411879 | 0.387075108 | 0.159811215 | 0.693001962 |
| Selenon | S | 13 | -0.056873338 | 0.04210523 | 0.064009432 | 0.12088277 | 0.021904202 | 0.33178966 | 0.0719232 | 0.440720742 | 0.216273479 | 0.774162348 |
| Mcrip1 | T | 23 | 0.163679891 | 0.075680883 | -0.145508647 | -0.309188538 | -0.22118953 | 0.169422904 | 0.355684933 | 0.170361586 | 0.044217143 | 0.066560363 |
| Anp32b | S | 129 | 0.016939615 | 0.027472757 | 0.025666194 | 0.00872658 | -0.001806563 | 0.633648901 | 0.674183443 | 0.57597772 | 0.812193541 | 0.977667989 |
| Srrm1 | S | 943 | 0.007383272 | 0.014884788 | 0.029288332 | 0.02190506 | 0.014403545 | 0.725186231 | 0.627731644 | 0.335748003 | 0.437714621 | 0.673821047 |
| Jmy | S | 704 | -0.054549673 | 0.039723504 | 0.108156494 | 0.162706168 | 0.068432991 | 0.27862217 | 0.382708655 | 0.265308707 | 0.136026515 | 0.443654201 |
| Eif4ebp2 | Y | 34 | 0.058716538 | 0.070654431 | -0.07509233 | -0.133808868 | -0.145746761 | 0.651423451 | 0.137234318 | 0.514021832 | 0.407003066 | 0.256576798 |
| Birc2 | S | 153 | 0.164272022 | -0.060504903 | -0.068041709 | -0.23231373 | -0.007536806 | 0.045687674 | 0.676808505 | 0.402402604 | 0.036711562 | 0.958879286 |
| Snrnp70 | S | 226 | 0.054326384 | 0.005924498 | -0.003877304 | -0.058203688 | -0.009801802 | 0.215480985 | 0.799315439 | 0.936772341 | 0.318743121 | 0.833331075 |
| Fip1l1 | S | 479 | -0.012687028 | 0.020054147 | 0.022271742 | 0.034958769 | 0.002217595 | 0.510887015 | 0.051509542 | 0.327559682 | 0.208518513 | 0.912357167 |
| Phf8-ps Phf8 | S | 768 | 0.177649701 | -0.033084439 | -0.041019908 | -0.218669609 | -0.007935469 | 0.086275143 | 0.576954851 | 0.320761905 | 0.063936165 | 0.881258608 |
| Eif2s1 | S | 158 | 0.050097833 | 0.000819119 | 0.025525559 | -0.024572275 | 0.02470644 | 0.411921774 | 0.994557558 | 0.698552991 | 0.680690146 | 0.836933462 |
| Srrm2 | S | 2368 | 0.109368616 | -0.031550326 | 0.004231839 | -0.105136778 | 0.035782164 | 0.193889182 | 0.584026201 | 0.947015802 | 0.230394429 | 0.579111811 |
| 1110059G10Rik | S | 150 | -0.011726696 | 0.148972417 | -0.055774841 | -0.044048146 | -0.204747258 | 0.853903293 | 0.098230585 | 0.366184202 | 0.521735989 | 0.049408028 |
| Limd1 | S | 413 | 0.102755538 | 0.007520183 | -0.01510738 | -0.117862919 | -0.022627563 | 0.241734535 | 0.795991502 | 0.414122192 | 0.202737294 | 0.412406145 |
| Sh3pxd2a | Y | 619 | 0.004880253 | 0.071429279 | 0.012561251 | 0.007680998 | -0.058868028 | 0.93718167 | 0.403430693 | 0.905654648 | 0.934033994 | 0.585419388 |
| Mdc1 | S | 1111 | -0.038218368 | 0.123723794 | -0.043674154 | -0.005455786 | -0.167397948 | 0.728489841 | 0.077104175 | 0.616339702 | 0.963039921 | 0.106403327 |
| Rmi1 | T | 278 | 0.183724209 | 0.036200464 | -0.132501418 | -0.316225627 | -0.168701882 | 0.117156907 | 0.628634014 | 0.217054593 | 0.042367246 | 0.152746859 |
| Hnrnpa1 | S | 6 | -0.053067663 | 0.020105431 | 0.09520231 | 0.148269972 | 0.075096878 | 0.206667182 | 0.538751496 | 0.084895334 | 0.02619679 | 0.149460393 |
| Cwc22 | S | 874 | 0.08911822 | 0.031504282 | -0.05772656 | -0.14684478 | -0.089230842 | 0.204984757 | 0.627092398 | 0.534114478 | 0.143877154 | 0.312549235 |
| Jarid2 | S | 331 | 0.105725793 | -0.000272804 | -0.043861129 | -0.149586922 | -0.043588325 | 0.140625324 | 0.996438085 | 0.605140153 | 0.110044199 | 0.548824165 |
| Arhgap12 | S | 584 | 0.042182712 | 0.01881346 | 0.003609543 | -0.038573169 | -0.015203917 | 0.342252441 | 0.690582691 | 0.973676044 | 0.726006164 | 0.889839819 |
| Rock1 | S | 1102 | -0.093434417 | 0.086241811 | 0.104420153 | 0.197854571 | 0.018178342 | 0.365338001 | 0.256830123 | 0.497016045 | 0.240877243 | 0.893234989 |
| Mki67 | T | 2045 | 0.082106129 | 0.050333605 | -0.049914714 | -0.132020843 | -0.100248319 | 0.241830544 | 0.427047041 | 0.306818531 | 0.099561685 | 0.159075784 |
| Akap13 | T | 2431 | 0.081674569 | 0.027536114 | -0.036012325 | -0.117686894 | -0.063548439 | 0.307668971 | 0.660777566 | 0.705571589 | 0.249344066 | 0.467459592 |
| Mon1a | S | 91 | -0.153418648 | 0.161155184 | 0.077169547 | 0.230588195 | -0.083985637 | 0.118572329 | 0.083157019 | 0.462934636 | 0.056199994 | 0.349892194 |
| Macf1 | S | 6904 | 0.0183218 | 0.009856801 | 0.028862904 | 0.010541104 | 0.019006103 | 0.60750963 | 0.831662382 | 0.38245055 | 0.765410374 | 0.68429594 |
| Tor1aip1 | T | 180 | 0.006093005 | 0.026186462 | 0.051364011 | 0.045271006 | 0.025177549 | 0.910871342 | 0.683418159 | 0.52930025 | 0.596983551 | 0.777820508 |
| Srsf10 | S | 131 | -0.00698792 | -0.079040252 | 0.228439875 | 0.235427795 | 0.307480127 | 0.943664542 | 0.393300826 | 0.211633781 | 0.19777337 | 0.136650839 |
| Atp8b2 | S | 1187 | -0.050168175 | 0.10173021 | -0.003182326 | 0.04698585 | -0.104912536 | 0.301693036 | 0.042622469 | 0.903630918 | 0.319808443 | 0.047862785 |
| Bub3 | S | 211 | -0.099144055 | 0.108979055 | 0.079759092 | 0.178903147 | -0.029219964 | 0.034869083 | 0.037894699 | 0.366664533 | 0.107128529 | 0.722413183 |
| Dis3 | S | 215 | 0.036808796 | 0.023714556 | 0.016238622 | -0.020570174 | -0.007475934 | 0.477538058 | 0.7209203 | 0.856048919 | 0.797800574 | 0.932536355 |
| Pelp1 | T | 49 | -0.116911449 | 0.043962359 | 0.118787046 | 0.235698495 | 0.074824686 | 0.286455716 | 0.40440382 | 0.137193734 | 0.08317043 | 0.32522797 |
| Akap1 | S | 55 | 0.060618997 | 0.007397135 | -0.02959685 | -0.090215847 | -0.036993985 | 0.031995723 | 0.677023088 | 0.232661363 | 0.015609977 | 0.13562901 |
| Mier3 | S | 156 | 0.123090915 | -0.033294052 | -0.039927832 | -0.163018747 | -0.00663378 | 0.051856279 | 0.526523961 | 0.549336141 | 0.040873913 | 0.893865052 |
| Zc3h18 | S | 863 | 0.072783486 | 0.009701432 | 0.001651828 | -0.071131658 | -0.008049605 | 0.411568627 | 0.837741352 | 0.970783866 | 0.435829486 | 0.887428771 |
| Pabpc1 | S | 120 | 0.592022264 | -0.218546785 | -0.110189645 | -0.702211909 | 0.10835714 | 0.117042735 | 0.175083888 | 0.677081264 | 0.095670246 | 0.661908427 |
| Dnmt3a | S | 102 | -0.094228694 | 0.004540831 | 0.146884592 | 0.241113286 | 0.142343761 | 0.263101735 | 0.951600685 | 0.060374888 | 0.033350063 | 0.061383471 |
| Fzr1 | S | 72 | 0.044077379 | -0.019576769 | 0.029631758 | -0.014445621 | 0.049208527 | 0.185843812 | 0.567511273 | 0.539608842 | 0.771007035 | 0.368959642 |
| Ctnnd1 | S | 288 | -0.011956395 | 0.081413434 | -0.008385616 | 0.003570779 | -0.08979905 | 0.795526965 | 0.15872117 | 0.880643373 | 0.942055724 | 0.158427454 |
| Rock1 | S | 1098 | 0.019330634 | 0.024971361 | 0.024820669 | 0.005490035 | -0.000150692 | 0.743013995 | 0.475321597 | 0.721424211 | 0.947006641 | 0.998304807 |
| Ftsj3 | S | 446 | 0.218156124 | -0.209771542 | 0.087686128 | -0.130469995 | 0.297457671 | 0.239433233 | 0.422229692 | 0.722471091 | 0.537111142 | 0.301444387 |
| Lmnb2 | S | 432 | -0.152814592 | 0.166464855 | 0.008918355 | 0.161732948 | -0.157546499 | 0.634437713 | 0.170792333 | 0.941765535 | 0.61402226 | 0.190338138 |
| 1700017B05Rik | S | 471 | 0.029692458 | -0.012935367 | 0.042559521 | 0.012867063 | 0.055494887 | 0.389553933 | 0.742904128 | 0.40359518 | 0.791026284 | 0.320650426 |
| Osbpl8 | S | 314 | 0.144250992 | -0.061140116 | -0.035226854 | -0.179477847 | 0.025913262 | 0.012490527 | 0.30433346 | 0.238771608 | 0.013924633 | 0.620592835 |
| Dtl | S | 678 | -0.09938551 | -0.078357945 | 0.216733347 | 0.316118856 | 0.295091292 | 0.204098817 | 0.284885765 | 0.008922156 | 0.007479157 | 0.006289459 |

| | | | | | | | | | | | | |
|---|---|---|---|---|---|---|---|---|---|---|---|---|
| Erbb2 | S | 999 | 0.013270017 | 0.067774277 | 0.005994777 | -0.00727524 | -0.0617795 | 0.90124162 | 0.412720768 | 0.9474556 | 0.94986999 | 0.520393979 |
| Srrm2 | S | 1419 | 0.034605657 | 0.099576381 | -0.079806679 | -0.114412336 | -0.17938306 | 0.475033322 | 0.032969958 | 0.084968492 | 0.042952016 | 0.003267188 |
| Mtdh | T | 96 | 0.161316383 | -0.012243209 | -0.034161658 | -0.195478042 | -0.021918449 | 0.224224719 | 0.701696783 | 0.630190134 | 0.166983513 | 0.739338864 |
| Prrc2c | S | 1221 | 0.034804561 | -0.011512093 | 0.014425141 | -0.020379421 | 0.025937234 | 0.131199724 | 0.526980812 | 0.371572151 | 0.328464145 | 0.186108474 |
| Brat1 | S | 736 | 0.057939582 | -0.011265325 | -0.00427156 | -0.062211142 | 0.006993764 | 0.111995874 | 0.880698177 | 0.918155227 | 0.21648223 | 0.930117131 |
| Ampd3 | S | 85 | -0.08347971 | 0.189227498 | -0.051135099 | 0.03234461 | -0.240362597 | 0.285999351 | 0.039039983 | 0.600617015 | 0.739478113 | 0.063905469 |
| Srrm1 | S | 626 | -0.018974826 | -0.024021583 | 0.056080617 | 0.075055443 | 0.0801022 | 0.461057346 | 0.162611087 | 0.00088442 | 0.058879466 | 0.009383102 |
| Mki67 | T | 2162 | 0.097399114 | 0.001221866 | -0.050197538 | -0.147596652 | -0.051419404 | 0.055016057 | 0.975149317 | 0.317878029 | 0.025974914 | 0.260484492 |
| Tmem115 | S | 339 | 0.28021839 | -0.244059223 | 0.207641326 | -0.072577065 | 0.451700548 | 0.505075743 | 0.46654717 | 0.665225329 | 0.893396015 | 0.411538578 |
| Ddx41 | S | 21 | -0.020284677 | -0.024650177 | 0.119887988 | 0.140172666 | 0.144538165 | 0.657116859 | 0.586526649 | 0.120193054 | 0.088931613 | 0.083071973 |
| Cenpc1 | S | 154 | 0.038876526 | 0.003504029 | 0.013986431 | -0.024890095 | 0.010482402 | 0.276465678 | 0.918241522 | 0.798292995 | 0.640067165 | 0.841033201 |
| Rbbp6 | T | 1275 | -0.098162817 | 0.026336578 | 0.113061737 | 0.211224555 | 0.08672516 | 0.393770273 | 0.565610011 | 0.133961702 | 0.130483632 | 0.230026869 |
| Ranbp2 | S | 2641 | -0.062423662 | 0.028734396 | 0.104421897 | 0.166845559 | 0.075687501 | 0.318086335 | 0.719847352 | 0.124071793 | 0.007890893 | 0.299655499 |
| Scamp1 | S | 43 | 0.093366884 | -0.038580073 | 0.012885535 | -0.080481349 | 0.051465607 | 0.215129168 | 0.663714495 | 0.86442914 | 0.271862072 | 0.55401117 |
| Actn1 | Y | 193 | 0.00547865 | -0.042401648 | 0.103367545 | 0.097888895 | 0.145769193 | 0.865556963 | 0.344330473 | 0.107520945 | 0.127900159 | 0.051535618 |
| Setd6 | S | 22 | 0.073621308 | 0.129257537 | -0.177788104 | -0.251409413 | -0.307045641 | 0.124887766 | 0.56566873 | 0.467797651 | 0.334453932 | 0.326590712 |
| Rbbp6 | S | 361 | 0.071306792 | 0.042844754 | -0.057393828 | -0.12870062 | -0.100238581 | 0.143082121 | 0.547447726 | 0.373239533 | 0.084463869 | 0.219436652 |
| Tjp2 | S | 404 | -0.051304012 | -0.099604954 | 0.178125437 | 0.229429449 | 0.277730391 | 0.216767266 | 0.070583356 | 0.001286592 | 0.002065278 | 0.005870145 |
| Hmgn1 | S | 84 | 0.064088378 | -0.005146188 | -0.012052733 | -0.076141112 | -0.006906545 | 0.110093746 | 0.901531907 | 0.755849248 | 0.102686314 | 0.874253452 |
| Ric1 | T | 991 | -0.00408057 | 0.090262522 | -0.045876556 | -0.041795986 | -0.136139078 | 0.951332838 | 0.08967384 | 0.581618575 | 0.653823467 | 0.169465725 |
| Trak1 | S | 905 | 0.083682397 | 0.02001602 | -0.04224814 | -0.125930537 | -0.06226416 | 0.106205342 | 0.84680939 | 0.401278019 | 0.053886169 | 0.565053273 |
| Prkd3 | S | 415 | 0.033698801 | 0.00670167 | 0.018512092 | -0.015186709 | 0.011810422 | 0.370401796 | 0.865280806 | 0.706458309 | 0.707599524 | 0.78186811 |
| Hnrnpc | S | 113 | 0.018508344 | 0.114799999 | -0.044122115 | -0.062630459 | -0.158922114 | 0.840950236 | 0.39747579 | 0.839610143 | 0.780242248 | 0.513824152 |
| Fip1l1 | S | 487 | -0.021418694 | 0.02816031 | 0.033128506 | 0.0545472 | 0.004968196 | 0.474411192 | 0.073603756 | 0.509458706 | 0.330189959 | 0.915954821 |
| Alms1 Alms1-ps2 | S | 73 | -0.118723753 | -0.030559774 | 0.155297928 | 0.274021681 | 0.185857702 | 0.328808407 | 0.550054616 | 0.017801605 | 0.086530908 | 0.006838399 |
| Alms1 Alms1-ps2 | S | 74 | -0.118723753 | -0.030559774 | 0.155297928 | 0.274021681 | 0.185857702 | 0.328808407 | 0.550054616 | 0.017801605 | 0.086530908 | 0.006838399 |
| Ncl | S | 425 | 0.110301754 | -0.01211864 | 0.021919702 | -0.088382051 | 0.034038342 | 0.477978444 | 0.667929576 | 0.764568501 | 0.577016262 | 0.638314866 |
| Actr10 | S | 414 | 0.110593222 | 0.101906807 | -0.158633176 | -0.269226398 | -0.260539983 | 0.349292433 | 0.298102087 | 0.36347363 | 0.167473494 | 0.175287975 |
| Kri1 | S | 95 | 0.064070684 | -0.031415002 | 0.037847439 | -0.026223245 | 0.069262441 | 0.282090982 | 0.602447811 | 0.627632313 | 0.713494086 | 0.372006873 |
| Kri1 | S | 96 | 0.064070684 | -0.031415002 | 0.037847439 | -0.026223245 | 0.069262441 | 0.282090982 | 0.602447811 | 0.627632313 | 0.713494086 | 0.372006873 |
| Kri1 | S | 97 | 0.064070684 | -0.031415002 | 0.037847439 | -0.026223245 | 0.069262441 | 0.282090982 | 0.602447811 | 0.627632313 | 0.713494086 | 0.372006873 |
| Kri1 | S | 99 | 0.064070684 | -0.031415002 | 0.037847439 | -0.026223245 | 0.069262441 | 0.282090982 | 0.602447811 | 0.627632313 | 0.713494086 | 0.372006873 |
| Apc | S | 2242 | 0.160417946 | -0.023263539 | -0.044275604 | -0.20469355 | -0.021012065 | 0.141425795 | 0.690682116 | 0.496258095 | 0.086659802 | 0.611588636 |
| Apc | S | 2244 | 0.160417946 | -0.023263539 | -0.044275604 | -0.20469355 | -0.021012065 | 0.141425795 | 0.690682116 | 0.496258095 | 0.086659802 | 0.611588636 |
| Dag1 | S | 886 | 0.10836383 | 0.007190031 | -0.051176243 | -0.159540073 | -0.058366274 | 0.166764918 | 0.817027886 | 0.501804592 | 0.121642354 | 0.439686088 |
| Hnrnpc | S | 119 | 0.112984309 | 0.109381728 | -0.260686465 | -0.373670774 | -0.370068192 | 0.271080443 | 0.2385724 | 0.399333467 | 0.262709322 | 0.266977858 |
| Prkd1 | S | 351 | -0.032215418 | 0.099099461 | -0.012249215 | 0.019966203 | -0.111348677 | 0.468335998 | 0.095662049 | 0.901464511 | 0.830007925 | 0.305538365 |
| Srrm2 | S | 777 | 0.02714329 | 0.008602642 | 0.005140003 | -0.022003286 | -0.003462639 | 0.33593223 | 0.514497923 | 0.70555799 | 0.407383465 | 0.620491563 |
| Recql5 | S | 489 | 0.043742229 | -0.032435139 | 0.054404896 | 0.010662667 | 0.086840035 | 0.427925919 | 0.343571976 | 0.229724774 | 0.856048075 | 0.108621388 |
| Recql5 | S | 492 | 0.043742229 | -0.032435139 | 0.054404896 | 0.010662667 | 0.086840035 | 0.427925919 | 0.343571976 | 0.229724774 | 0.856048075 | 0.108621388 |
| Brd9 | S | 565 | -0.002510138 | 0.002650854 | 0.041553492 | 0.04406363 | 0.038902638 | 0.945052288 | 0.920563443 | 0.140015114 | 0.225286924 | 0.051591672 |
| Leo1 | S | 659 | 0.018606451 | -0.017888846 | 0.042597595 | 0.023991143 | 0.060486441 | 0.333781905 | 0.32397427 | 0.143146341 | 0.365584602 | 0.068619935 |
| Med19 | S | 194 | 0.000812499 | 0.079991121 | -0.025957991 | -0.02677049 | -0.105949111 | 0.991455279 | 0.207977336 | 0.833723456 | 0.842421377 | 0.435024485 |
| Ubxn2b | S | 56 | 0.142923435 | -0.033138398 | -0.056606161 | -0.199529596 | -0.023467764 | 0.044415785 | 0.635486673 | 0.313357433 | 0.024641897 | 0.750661469 |
| Ubn1 | S | 834 | 0.141837393 | -0.092582134 | 0.023424219 | -0.118413175 | 0.116006352 | 0.132686472 | 0.319493894 | 0.773767351 | 0.045296788 | 0.090476519 |
| Rab3gap2 | S | 566 | 0.027842481 | 0.116728357 | -0.031616971 | -0.059459452 | -0.148345328 | 0.739484412 | 0.376685108 | 0.52974574 | 0.510791072 | 0.286147393 |
| Cir1 | S | 427 | -0.082421921 | 0.072810093 | 0.07654304 | 0.158964961 | 0.003732946 | 0.071159986 | 0.193446394 | 0.113261321 | 0.018743343 | 0.94387681 |
| Cope | S | 13 | -0.000774468 | 0.092170902 | -0.013046814 | -0.012272346 | -0.105217716 | 0.986160357 | 0.258076158 | 0.735523042 | 0.762643109 | 0.211591099 |
| Cast | S | 442 | 0.07068301 | -0.013182966 | 0.013445414 | -0.057237597 | 0.026628379 | 0.30535844 | 0.924119045 | 0.938444782 | 0.734015652 | 0.89281647 |
| Tpr | T | 2135 | 0.280947232 | -0.093835157 | -0.075332223 | -0.356279455 | 0.018502934 | 0.070304593 | 0.364324704 | 0.381024975 | 0.050202721 | 0.809904367 |
| Phf6 | S | 145 | 0.030536871 | 0.031701318 | -0.007129775 | -0.037666646 | -0.038831092 | 0.686704468 | 0.218551815 | 0.793313984 | 0.620260016 | 0.140238213 |
| Dmap1 | S | 460 | 0.064173082 | -0.044669496 | 0.03829149 | -0.025881592 | 0.082960986 | 0.096977726 | 0.300685692 | 0.641296198 | 0.753260672 | 0.362890334 |
| Spire1 | S | 229 | -0.063114389 | 0.059070989 | 0.125509889 | 0.188624278 | 0.0664389 | 0.314984175 | 0.454476035 | 0.400920605 | 0.247624875 | 0.653053331 |
| Abhd2 | S | 415 | -0.020036679 | 0.037611935 | 0.039274075 | 0.059310755 | 0.00166214 | 0.795339571 | 0.264036207 | 0.657728242 | 0.581604476 | 0.984469819 |
| Nbn | S | 58 | 0.022701089 | 0.034385081 | 0.01172726 | -0.010973829 | -0.022657821 | 0.736764075 | 0.514586308 | 0.8109805 | 0.83971444 | 0.465773773 |
| Ptpn14 | S | 762 | -0.051458206 | 0.094542671 | -0.007431341 | 0.044026866 | -0.101974012 | 0.151469488 | 0.013954352 | 0.805670421 | 0.235293987 | 0.023903182 |
| Sorbs1 | S | 471 | 0.002961185 | -0.028525583 | 0.07868614 | 0.075724955 | 0.107211723 | 0.963759246 | 0.739565386 | 0.182259273 | 0.324763355 | 0.280692923 |
| Snip1 | S | 120 | -0.01167118 | -0.008380991 | 0.101169892 | 0.112841071 | 0.109550882 | 0.925593716 | 0.929362305 | 0.322787607 | 0.387171789 | 0.274213055 |
| Nup98 | S | 1085 | 0.025941463 | -0.339028459 | 0.362220612 | 0.336279149 | 0.701249071 | 0.927970721 | 0.419552106 | 0.184874268 | 0.173357784 | 0.139282872 |
| Akirin2 | S | 31 | 0.040788151 | 0.103801691 | -0.065290232 | -0.106078384 | -0.169091923 | 0.817092357 | 0.244928515 | 0.576259056 | 0.567965517 | 0.166081002 |
| Cep170b | T | 1345 | 0.087247213 | 0.021872079 | -0.027913209 | -0.115160422 | -0.049785288 | 0.32238557 | 0.895591179 | 0.794023433 | 0.328306024 | 0.780029536 |
| Vcpip1 | S | 746 | -0.020734592 | 0.058537028 | 0.012015684 | 0.032750275 | -0.046521344 | 0.405629023 | 0.133248099 | 0.615624844 | 0.058737584 | 0.193355165 |
| Setx | S | 507 | -0.024649654 | -0.016607456 | 0.091074945 | 0.115724599 | 0.107682401 | 0.588814088 | 0.84048099 | 0.107243016 | 0.067419132 | 0.257901946 |
| Eif6 | S | 239 | -0.063227081 | 0.08987476 | 0.020291452 | 0.083518533 | -0.069583308 | 0.134613882 | 0.033024731 | 0.699493348 | 0.184401828 | 0.241950973 |
| Dicer1 | S | 413 | 0.099371751 | -0.021572199 | -0.031936103 | -0.131307854 | -0.010363904 | 0.07597169 | 0.712209022 | 0.549360243 | 0.024016971 | 0.834975432 |
| Dicer1 | S | 415 | 0.099371751 | -0.021572199 | -0.031936103 | -0.131307854 | -0.010363904 | 0.07597169 | 0.712209022 | 0.549360243 | 0.024016971 | 0.834975432 |
| Srrm1 | S | 664 | -0.004990028 | 0.014143908 | 0.040887219 | 0.045877247 | 0.026743311 | 0.869246815 | 0.514270106 | 0.337432346 | 0.321029074 | 0.505851082 |
| C77080 | S | 909 | -0.05965239 | 0.021375767 | 0.081468037 | 0.141120427 | 0.060092269 | 0.113510812 | 0.646362372 | 0.026309005 | 0.009541621 | 0.219737982 |
| Prrc2a | S | 1145 | 0.090125461 | -0.061809178 | 0.011004341 | -0.07912112 | 0.072813519 | 0.132356847 | 0.531327669 | 0.786350634 | 0.149809157 | 0.453371346 |
| Arhgef17 | S | 142 | -0.043794557 | 0.057324033 | 0.042965836 | 0.086760394 | -0.014358197 | 0.165834673 | 0.078267692 | 0.474770347 | 0.20174383 | 0.797081623 |
| Npm1 | S | 143 | 0.274180926 | -0.120895711 | -0.104585704 | -0.37876663 | 0.016310007 | 0.001412137 | 0.14675901 | 0.04038934 | 0.000955691 | 0.811425076 |
| Jund | S | 253 | 0.051482762 | -0.050474471 | 0.062610935 | 0.011128172 | 0.113085406 | 0.05080546 | 0.049857291 | 0.291617232 | 0.826766964 | 0.12322354 |
| Ndc1 | S | 438 | 0.019352154 | -0.154170683 | 0.174598533 | 0.155246379 | 0.328769216 | 0.914521649 | 0.35798146 | 0.098269702 | 0.394267478 | 0.112433257 |
| Ndc1 | S | 444 | 0.019352154 | -0.154170683 | 0.174598533 | 0.155246379 | 0.328769216 | 0.914521649 | 0.35798146 | 0.098269702 | 0.394267478 | 0.112433257 |
| Spindoc | T | 50 | -0.058656016 | 0.040532756 | 0.105361203 | 0.164017218 | 0.064828447 | 0.217468612 | 0.484613219 | 0.206646902 | 0.092133716 | 0.412544395 |
| Zc3h13 | S | 265 | -0.021147009 | -0.047885783 | 0.121654872 | 0.142801882 | 0.169540655 | 0.627657814 | 0.292736999 | 0.04179568 | 0.024493775 | 0.014514453 |
| Ap2m1 | S | 45 | 0.10787874 | 0.08385636 | -0.093320526 | -0.201199265 | -0.177176885 | 0.199955717 | 0.505472369 | 0.219232789 | 0.036191103 | 0.203518241 |
| Samd1 | S | 150 | -0.007472651 | 0.055601044 | -0.010741599 | -0.003268948 | -0.066342643 | 0.675345569 | 0.066833713 | 0.758524978 | 0.919113753 | 0.129632333 |
| Mki67 | S | 1773 | 0.052723992 | -0.025792549 | 0.060113263 | 0.00738927 | 0.085905812 | 0.445928188 | 0.598923544 | 0.555995216 | 0.943870031 | 0.413718548 |
| Virma | S | 1578 | 0.029513329 | -0.014111922 | 0.041976353 | 0.012463024 | 0.056088275 | 0.380463557 | 0.435016819 | 0.320070793 | 0.772658688 | 0.217156533 |
| Ints1 | S | 1365 | 0.016744269 | 0.05569034 | 0.020333708 | 0.00358944 | -0.035356632 | 0.88305863 | 0.595522979 | 0.819751453 | 0.975066149 | 0.737429901 |
| Tbc1d9b | S | 1090 | -0.033019879 | 0.078468741 | 0.019286293 | 0.052306173 | -0.059182448 | 0.448854054 | 0.175334479 | 0.763684297 | 0.379667862 | 0.367613045 |
| Itprid2 | S | 1059 | 0.015624818 | -0.035193846 | 0.103102986 | 0.087478168 | 0.138296831 | 0.718438804 | 0.389271028 | 0.230884181 | 0.294188215 | 0.142795705 |
| Hirip3 | S | 328 | -0.072074198 | 0.050086157 | 0.090266409 | 0.162340607 | 0.040180252 | 0.062370721 | 0.431781062 | 0.049869901 | 0.01046642 | 0.531141493 |
| Hirip3 | S | 331 | -0.072074198 | 0.050086157 | 0.090266409 | 0.162340607 | 0.040180252 | 0.062370721 | 0.431781062 | 0.049869901 | 0.01046642 | 0.531141493 |
| Hirip3 | S | 332 | -0.072074198 | 0.050086157 | 0.090266409 | 0.162340607 | 0.040180252 | 0.062370721 | 0.431781062 | 0.049869901 | 0.01046642 | 0.531141493 |
| Map1a | S | 1205 | 0.023808795 | 0.064378413 | -0.028880524 | -0.052689319 | -0.093258937 | 0.668247534 | 0.242002947 | 0.596345198 | 0.458098705 | 0.195228024 |
| Med13 | T | 326 | -0.087367096 | 0.041471003 | 0.051692807 | 0.139059903 | 0.010221804 | 0.278682586 | 0.194642566 | 0.092531188 | 0.138427758 | 0.699151091 |
| Srsf6 | S | 314 | 0.040974086 | -0.058830135 | 0.095370389 | 0.054396302 | 0.154200524 | 0.44163776 | 0.184687553 | 0.172506146 | 0.392305922 | 0.076167413 |
| Pcyt1a | S | 343 | 0.076853315 | 0.082117425 | -0.150425193 | -0.227278509 | -0.232542618 | 0.078643052 | 0.070755042 | 0.070512364 | 0.020935999 | 0.019417947 |
| Ep300 | S | 1036 | -0.039597694 | 0.051897181 | 0.071536119 | 0.111133813 | 0.019638938 | 0.624760801 | 0.368095831 | 0.619097815 | 0.474809411 | 0.888860827 |
| AW554918 | T | 548 | 0.03412169 | 0.001032935 | 0.058110101 | 0.023988412 | 0.057077166 | 0.630777554 | 0.987246976 | 0.689128323 | 0.872484296 | 0.703284221 |
| Rps14 | S | 139 | 0.20407373 | 0.119886014 | -0.307700582 | -0.511774312 | -0.427586596 | 0.005907996 | 0.375068824 | 0.021017677 | 0.009197904 | 0.033715825 |
| Usp20 | S | 132 | -0.01965925 | 0.000543117 | 0.065245291 | 0.084904541 | 0.064702174 | 0.305611507 | 0.983829674 | 0.062396194 | 0.031254932 | 0.099740009 |

| | | | | | | | | | | | | |
|---|---|---|---|---|---|---|---|---|---|---|---|---|
| Usp20 | S | 134 | -0.01965925 | 0.000543117 | 0.065245291 | 0.084904541 | 0.064702174 | 0.305611507 | 0.983829674 | 0.062396194 | 0.031254932 | 0.099740009 |
| Tenm3 | T | 201 | 0.103648836 | 0.010243924 | -0.080952606 | -0.184601443 | -0.09119653 | 0.041868427 | 0.904579212 | 0.260416198 | 0.052428313 | 0.378638688 |
| Gys1 | S | 728 | 0.083882847 | -0.013432767 | 0.040264124 | -0.043618723 | 0.053696891 | 0.565439765 | 0.906767097 | 0.807053211 | 0.822368532 | 0.764888444 |
| Pask | S | 692 | 0.05246168 | 0.099466459 | -0.04049824 | -0.092959921 | -0.139964699 | 0.75613167 | 0.486557949 | 0.727148458 | 0.517802008 | 0.224057953 |
| Aurkb | T | 237 | 0.073742412 | 0.105114367 | -0.116841732 | -0.190584143 | -0.221956098 | 0.331637998 | 0.229350106 | 0.247877707 | 0.104447064 | 0.078797949 |
| Cpsf1 | S | 763 | 0.070543171 | -0.025675506 | 0.002905275 | -0.067637896 | 0.028580781 | 0.131946383 | 0.596054653 | 0.951671539 | 0.217997197 | 0.601188048 |
| Cpsf1 | S | 764 | 0.070543171 | -0.025675506 | 0.002905275 | -0.067637896 | 0.028580781 | 0.131946383 | 0.596054653 | 0.951671539 | 0.217997197 | 0.601188048 |
| Pdlim2 | S | 96 | 0.241003678 | 0.016017274 | -0.048559903 | -0.289563581 | -0.064577177 | 0.599515595 | 0.929097399 | 0.776602222 | 0.524276124 | 0.580497196 |
| Inpp5f | S | 946 | 0.020571584 | 0.142805239 | -0.059798957 | -0.080370541 | -0.202604196 | 0.463048466 | 0.098323714 | 0.047530148 | 0.014790772 | 0.061456828 |
| Srrm2 | S | 888 | 0.020135342 | 0.009185046 | 0.032641143 | 0.0125058 | 0.023456097 | 0.6280019 | 0.86252135 | 0.545291176 | 0.767173481 | 0.661738899 |
| Bclaf1 | S | 883 | 0.199694003 | 0.04435711 | -0.090133753 | -0.289827756 | -0.134490863 | 0.549044351 | 0.760167918 | 0.692461704 | 0.428465457 | 0.544352326 |
| Srrm1 | S | 424 | 0.017598873 | -0.03789513 | 0.095645376 | 0.078046503 | 0.133540506 | 0.787877782 | 0.082146732 | 0.060974107 | 0.30486927 | 0.043354006 |
| Incenp | S | 239 | 0.123116804 | 0.207134438 | -0.183569502 | -0.306686306 | -0.39070394 | 0.166412261 | 0.253892837 | 0.055448479 | 0.010041648 | 0.084539837 |
| Srrm1 | S | 797 | 0.075743916 | -0.006171366 | -0.009601573 | -0.085345488 | -0.003430207 | 0.129623286 | 0.601455764 | 0.656818131 | 0.097839334 | 0.861038743 |
| Rem1 | S | 288 | -0.07465593 | 0.213136452 | -0.086583478 | -0.011927548 | -0.29971993 | 0.528694131 | 0.075718626 | 0.486317833 | 0.932479268 | 0.072122935 |
| Ticrr | S | 599 | 0.025152137 | -0.031270707 | 0.049314477 | 0.02416234 | 0.080585184 | 0.390967713 | 0.398060495 | 0.182355175 | 0.40048571 | 0.0676792 |
| Thoc2 | S | 1450 | 0.01069197 | 0.036848143 | 0.008906833 | -0.001785137 | -0.02794131 | 0.774265299 | 0.397114546 | 0.858500787 | 0.974543405 | 0.635016296 |
| Ppip5k2 | S | 44 | -0.146971096 | 0.128436528 | 0.103132966 | 0.250104062 | -0.025303562 | 0.007970579 | 0.028961235 | 0.150582932 | 0.024571591 | 0.676955184 |
| Ccdc61 | S | 379 | -0.009236838 | -0.051465429 | 0.180546057 | 0.189782895 | 0.232011486 | 0.908696804 | 0.562510646 | 0.286024482 | 0.263711357 | 0.195195372 |
| Usp34 | S | 3396 | 0.017787159 | 0.006474334 | 0.028247055 | 0.010459896 | 0.021772721 | 0.529773285 | 0.860074174 | 0.520173861 | 0.786983581 | 0.628934689 |
| Glud1 | S | 128 | -0.029178248 | 0.040982507 | 0.045555701 | 0.074733949 | 0.004573194 | 0.256838203 | 0.236362648 | 0.315101275 | 0.144327436 | 0.917758189 |
| Trps1 | S | 90 | 0.013329609 | -0.135694799 | 0.228893025 | 0.215563416 | 0.364587825 | 0.845752923 | 0.102266872 | 0.069002354 | 0.088574693 | 0.02247851 |
| Vim | S | 144 | 0.048409561 | 0.041242004 | -0.068644682 | -0.117054243 | -0.109886686 | 0.138098902 | 0.233794641 | 0.31574115 | 0.140133922 | 0.156232955 |
| Cwc22 | S | 829 | 0.059981851 | 0.019760859 | -0.063276853 | -0.123258704 | -0.083037712 | 0.053942218 | 0.20097803 | 0.190552397 | 0.046674957 | 0.13118534 |
| Atrx | S | 1223 | -0.003140443 | -0.056031983 | 0.157034867 | 0.160175311 | 0.21306685 | 0.933765515 | 0.154810248 | 0.093055741 | 0.096294703 | 0.054884316 |
| Usp10 | S | 364 | 0.062296124 | -0.003316408 | -0.005203083 | -0.067499207 | -0.001886676 | 0.191248827 | 0.890009607 | 0.843673015 | 0.171108169 | 0.942968207 |
| Kank2 | S | 89 | -0.015477286 | -0.017923116 | 0.104822713 | 0.120299999 | 0.122745829 | 0.784051247 | 0.797988506 | 0.223079415 | 0.169637638 | 0.190442913 |
| Champ1 | T | 447 | 0.050614156 | 0.022250693 | -0.037517648 | -0.088131804 | -0.059768341 | 0.022166629 | 0.647161943 | 0.151750706 | 0.017588016 | 0.277541149 |
| Epb41l1 | T | 685 | -0.08131617 | 0.060696618 | 0.139500304 | 0.220816474 | 0.078803686 | 0.088677224 | 0.284825676 | 0.251382717 | 0.120666896 | 0.481124363 |
| Praf2 | S | 178 | -0.789879041 | -0.766607148 | 0.092622226 | 0.882501267 | 0.859229374 | NA | NA | 0.452689319 | NA | NA |
| Rps6ka5 | S | 862 | -0.078131001 | 0.080422651 | 0.074855721 | 0.152986721 | -0.005566931 | 0.474566239 | 0.329853862 | 0.545378744 | 0.250285322 | 0.955560457 |
| Iqgap1 | S | 360 | -0.001514836 | 0.01229928 | 0.054148901 | 0.055663737 | 0.041849622 | 0.979652216 | 0.830020455 | 0.384211109 | 0.401165579 | 0.506828838 |
| Psen1 | T | 370 | 0.001681147 | 0.020708048 | 0.023848006 | 0.022166859 | 0.003139958 | 0.955794949 | 0.41150861 | 0.516886455 | 0.596697489 | 0.933223096 |
| Zcchc24 | S | 93 | 0.02417362 | -0.029956821 | 0.174724503 | 0.150550883 | 0.204681324 | 0.8841292 | 0.908745572 | 0.661501315 | 0.713890132 | 0.647559886 |
| Zcchc24 | S | 97 | 0.02417362 | -0.029956821 | 0.174724503 | 0.150550883 | 0.204681324 | 0.8841292 | 0.908745572 | 0.661501315 | 0.713890132 | 0.647559886 |
| Dck | S | 74 | -0.008999215 | 0.031022083 | 0.02989039 | 0.038889605 | -0.001131693 | 0.715245431 | 0.321636896 | 0.533608448 | 0.442729046 | 0.981749307 |
| Pdzd8 | S | 490 | -0.056038883 | -0.011483432 | 0.157504851 | 0.213543734 | 0.168988283 | 0.530756372 | 0.870382142 | 0.207364421 | 0.126669837 | 0.184110863 |
| Arhgap31 | S | 935 | 0.015146371 | 0.082235772 | -0.029556034 | -0.044702405 | -0.111791806 | 0.788009602 | 0.298398243 | 0.68035928 | 0.442240261 | 0.179693949 |
| Tmem74 | S | 127 | 0.042104292 | -0.071006561 | 0.162182957 | 0.120078665 | 0.233189517 | 0.587585787 | 0.731187113 | 0.643767558 | 0.726553018 | 0.54414104 |
| Arhgef1 | S | 775 | 0.037274755 | -0.075029821 | 0.106683442 | 0.069408687 | 0.181713263 | 0.401333397 | 0.132434564 | 0.123162516 | 0.242794135 | 0.035114397 |
| Plpp3 | S | 19 | 0.154839266 | 0.129283159 | -0.247173525 | -0.402012791 | -0.376456684 | 0.10216586 | 0.156064624 | 0.066982494 | 0.025546719 | 0.031555209 |
| Neurl4 | S | 504 | 0.261837672 | -0.046330094 | -0.245231818 | -0.50706949 | -0.198901724 | 0.006723287 | 0.57043878 | 0.123402204 | 0.026649149 | 0.188073485 |
| Trpm7 | S | 1360 | -0.018973556 | -0.013237923 | 0.107413894 | 0.126387451 | 0.120651817 | 0.733904691 | 0.662584016 | 0.197856278 | 0.166555285 | 0.168551477 |
| Nes | S | 481 | 0.075342274 | -0.050393606 | 0.040131036 | -0.035211238 | 0.090524642 | 0.261262501 | 0.339747346 | 0.438532857 | 0.593621876 | 0.155779436 |
| Eif2b5 | S | 221 | 0.059537953 | 0.282546995 | -0.25920529 | -0.318743243 | -0.541752285 | 0.792662946 | 0.115258507 | 0.213504685 | 0.187339948 | 0.032821252 |
| Hnrnpa2b1 | S | 259 | 0.015258696 | 0.004104023 | 0.035756533 | 0.020497837 | 0.031652509 | 0.706754798 | 0.930415758 | 0.419379115 | 0.573488898 | 0.478137268 |
| Ddx1 | S | 481 | -0.008790866 | 0.021689089 | 0.016394607 | 0.025185472 | -0.005294483 | 0.846003125 | 0.240129212 | 0.364927749 | 0.575620047 | 0.305380838 |
| Clasp1 | S | 590 | -0.127267385 | 0.218691697 | 0.056602583 | 0.183869968 | -0.162089114 | 0.170619853 | 0.247440761 | 0.536386597 | 0.083024653 | 0.365325922 |
| Clasp2 | S | 509 | -0.014095726 | 0.115060296 | -0.044822595 | -0.030726869 | -0.159882892 | 0.765047622 | 0.118348084 | 0.531689672 | 0.638204791 | 0.088190475 |
| Clasp2 | S | 513 | -0.014095726 | 0.115060296 | -0.044822595 | -0.030726869 | -0.159882892 | 0.765047622 | 0.118348084 | 0.531689672 | 0.638204791 | 0.088190475 |
| Gapvd1 | S | 758 | -0.030432451 | 0.066669956 | 0.014280346 | 0.044712798 | -0.052389609 | 0.161302436 | 0.057437841 | 0.74399821 | 0.336883952 | 0.288494537 |
| Tnpo2 | S | 354 | 0.077202349 | 0.097482529 | -0.107768912 | -0.184971262 | -0.205251441 | 0.647687583 | 0.321260923 | 0.559799924 | 0.426259578 | 0.316626811 |
| Cdyl | S | 196 | 0.082210983 | 0.028616927 | -0.058494001 | -0.140704984 | -0.087110928 | 0.367900932 | 0.353508442 | 0.526573566 | 0.245812677 | 0.367480005 |
| Tsc2 | S | 1388 | -0.231412565 | 0.129150854 | 0.144754496 | 0.376167061 | 0.015603642 | 0.125406847 | 0.259160513 | 0.132543215 | 0.038360337 | 0.889866656 |
| Srek1 | S | 371 | 0.090009592 | -0.119450729 | 0.161017201 | 0.071007608 | 0.28046793 | 0.363331575 | 0.35736361 | 0.542267088 | 0.771994294 | 0.321199274 |
| Srek1 | S | 374 | 0.090009592 | -0.119450729 | 0.161017201 | 0.071007608 | 0.28046793 | 0.363331575 | 0.35736361 | 0.542267088 | 0.771994294 | 0.321199274 |
| Srek1 | S | 375 | 0.090009592 | -0.119450729 | 0.161017201 | 0.071007608 | 0.28046793 | 0.363331575 | 0.35736361 | 0.542267088 | 0.771994294 | 0.321199274 |
| Srek1 | S | 377 | 0.090009592 | -0.119450729 | 0.161017201 | 0.071007608 | 0.28046793 | 0.363331575 | 0.35736361 | 0.542267088 | 0.771994294 | 0.321199274 |
| Pard3b | S | 1182 | -0.13561645 | 0.054049035 | 0.159631631 | 0.295248081 | 0.105582596 | 0.029162055 | 0.085338242 | 0.110767924 | 0.023707302 | 0.225521362 |
| Srrm2 | T | 1370 | 0.043050609 | -0.005106796 | 0.004111913 | -0.038938695 | 0.00921871 | 0.149419201 | 0.730255371 | 0.917489449 | 0.393998195 | 0.816511289 |
| Etl4 | T | 1840 | -0.096569461 | 0.282991345 | -0.077248094 | 0.019321367 | -0.360239439 | 0.208839022 | 0.078372308 | 0.610467153 | 0.897531761 | 0.090170333 |
| Zc3hc1 | S | 393 | 0.063357408 | -0.014769316 | 0.014080372 | -0.049277036 | 0.028849688 | 0.329055508 | 0.78434151 | 0.78115014 | 0.309189489 | 0.274533902 |
| Myo9a | S | 1736 | -0.097671579 | 0.049869828 | 0.118438431 | 0.21611001 | 0.068568603 | 0.155178635 | 0.406856607 | 0.143839975 | 0.02538222 | 0.289213019 |
| Kmt2a | S | 163 | -0.00257148 | 0.067234048 | -0.013866166 | -0.011294686 | -0.081100214 | 0.955768192 | 0.168109425 | 0.694579338 | 0.796952021 | 0.110611807 |
| Kmt2a | S | 159 | -0.00257148 | 0.067234048 | -0.013866166 | -0.011294686 | -0.081100214 | 0.955768192 | 0.168109425 | 0.694579338 | 0.796952021 | 0.110611807 |
| Nbea | S | 2128 | -0.019422841 | 0.019156245 | 0.056640426 | 0.076063266 | 0.037484181 | 0.827876229 | 0.655425774 | 0.378071248 | 0.433265962 | 0.494860532 |
| Prpf4b | S | 278 | -0.063109248 | 0.003329125 | 0.110123275 | 0.173232522 | 0.106794149 | 0.23870343 | 0.910616789 | 0.066390753 | 0.029580287 | 0.071272773 |
| Stard13 | S | 411 | -0.055831725 | 0.113682177 | -0.024831313 | 0.031000412 | -0.138513491 | 0.50510298 | 0.071729082 | 0.728823819 | 0.703896031 | 0.096195104 |
| Polr2a | S | 1843 | -0.008207696 | 0.040205337 | -0.000535485 | 0.007672211 | -0.040740822 | 0.896342017 | 0.113885632 | 0.988843169 | 0.907613595 | 0.299764637 |
| Polr2a | S | 1857 | -0.008207696 | 0.040205337 | -0.000535485 | 0.007672211 | -0.040740822 | 0.896342017 | 0.113885632 | 0.988843169 | 0.907613595 | 0.299764637 |
| Rnmt | S | 11 | -0.040712656 | 0.024151114 | 0.053939063 | 0.094651719 | 0.029787949 | 0.645368511 | 0.764657308 | 0.157119431 | 0.332006324 | 0.715488209 |
| Mcm2 | S | 26 | 0.026969143 | 0.043243304 | -0.030180884 | -0.057150027 | -0.073424188 | 0.289737974 | 0.12592761 | 0.202925963 | 0.028051809 | 0.027100446 |
| Zfp692 | S | 243 | 0.074098849 | -0.004605776 | -0.003495427 | -0.077594276 | 0.001110349 | 0.336812382 | 0.971392325 | 0.973460599 | 0.429596143 | 0.99350645 |
| Pold3 | S | 148 | -0.061584084 | 0.098059004 | 0.008633885 | 0.070217969 | -0.089425119 | 0.078537609 | 0.020027187 | 0.837769088 | 0.12977948 | 0.071100524 |
| Slx4 | S | 447 | -0.022325905 | -0.017804502 | 0.072888928 | 0.095214832 | 0.09069343 | 0.827484936 | 0.791873705 | 0.121920405 | 0.397150794 | 0.24831181 |
| Tra2a | S | 20 | -0.01311361 | 0.007983858 | 0.063663868 | 0.076777478 | 0.055680009 | 0.835451054 | 0.881123759 | 0.295092011 | 0.148215434 | 0.143902413 |
| Casp8 | S | 188 | -0.07199977 | 0.216432878 | -0.105315889 | -0.033316118 | -0.321748766 | 0.529953814 | 0.088560411 | 0.556080043 | 0.8334399 | 0.124798925 |
| Srrm1 | T | 834 | 0.024129791 | -0.019244126 | 0.043095502 | 0.018965711 | 0.062339627 | 0.361609049 | 0.155235627 | 0.133572721 | 0.531604916 | 0.067527523 |
| Atrx | S | 212 | 0.013931673 | -0.022000422 | 0.072134101 | 0.058202428 | 0.094134523 | 0.775871361 | 0.770918505 | 0.343547496 | 0.388104852 | 0.289244255 |
| Ehmt2 | S | 403 | 0.225610606 | -0.177551265 | -0.031887173 | -0.257497779 | 0.145664092 | 0.02304511 | 0.173838342 | 0.712562592 | 0.018037946 | 0.238548902 |
| Bud13 | S | 389 | -0.009169795 | 0.081594165 | 0.011425374 | 0.020595169 | -0.070168791 | 0.820082112 | 0.446238356 | 0.825275968 | 0.728453008 | 0.522365343 |
| Bud13 | S | 391 | -0.009169795 | 0.081594165 | 0.011425374 | 0.020595169 | -0.070168791 | 0.820082112 | 0.446238356 | 0.825275968 | 0.728453008 | 0.522365343 |
| Bud13 | S | 394 | -0.009169795 | 0.081594165 | 0.011425374 | 0.020595169 | -0.070168791 | 0.820082112 | 0.446238356 | 0.825275968 | 0.728453008 | 0.522365343 |
| Slc30a1 | S | 463 | 0.052758171 | -0.085474565 | 0.043250984 | -0.009507187 | 0.128725549 | 0.39861139 | 0.60116266 | 0.238707905 | 0.859679394 | 0.448756 |
| Bcl9l | S | 984 | -0.056164371 | -0.01881732 | 0.121009662 | 0.177174033 | 0.139826983 | 0.294479595 | 0.602738034 | 0.041738469 | 0.028018373 | 0.031677361 |
| Kmt5b | S | 630 | 0.047703574 | -1.65816E-05 | 0.02924019 | -0.018463384 | 0.029256771 | 0.551579823 | 0.999847951 | 0.752419588 | 0.846419724 | 0.774220635 |
| Scrib | S | 432 | 0.124320156 | 0.080896155 | -0.203167014 | -0.32748717 | -0.284063169 | 0.126776357 | 0.170720338 | 0.145940233 | 0.052325363 | 0.084222912 |
| Piezo1 | S | 1390 | 0.064344077 | 0.053057538 | -0.077819315 | -0.142163393 | -0.130876853 | 0.12774875 | 0.198001644 | 0.130776991 | 0.030901829 | 0.032423057 |
| Bcl9 | T | 315 | 0.039881086 | -0.056955468 | 0.085047487 | 0.045166401 | 0.142002955 | 0.381610888 | 0.00561249 | 0.013256028 | 0.338321413 | 0.007852371 |
| Zcchc8 | T | 495 | -0.076675824 | 0.152325612 | -0.034194772 | 0.042481052 | -0.186520384 | 0.439243466 | 0.10092231 | 0.826008094 | 0.775121435 | 0.26877913 |
| Zcchc8 | T | 499 | -0.076675824 | 0.152325612 | -0.034194772 | 0.042481052 | -0.186520384 | 0.439243466 | 0.10092231 | 0.826008094 | 0.775121435 | 0.26877913 |
| Polr2a | S | 1868 | 0.056799654 | -0.057819313 | 0.035560923 | -0.02123873 | 0.093380237 | 0.219897313 | 0.170050834 | 0.101387439 | 0.596536499 | 0.066949877 |
| Myh9 | S | 1304 | 0.058408101 | 0.046714358 | -0.013690249 | -0.07209835 | -0.060404607 | 0.664407346 | 0.596517196 | 0.823030728 | 0.575803768 | 0.42672477 |
| Nup98 | S | 888 | -0.002700477 | -0.060196732 | 0.123127251 | 0.125827728 | 0.183323982 | 0.947460318 | 0.114774089 | 0.039338154 | 0.043362994 | 0.020595197 |

| | | | | | | | | | | | | |
|---|---|---|---|---|---|---|---|---|---|---|---|---|
| Flna | S | 1459 | -0.000110422 | 0.060515898 | -0.013265836 | -0.013155414 | -0.073781734 | 0.997903284 | 0.207454374 | 0.828024838 | 0.807065054 | 0.243929633 |
| Tusc2 | S | 50 | 0.022431503 | 0.048829255 | -0.020744285 | -0.043175788 | -0.06957354 | 0.592737037 | 0.229529128 | 0.508563879 | 0.27493523 | 0.089539509 |
| Cyth2 | T | 58 | 0.158467976 | -0.035343217 | -0.048872345 | -0.20734032 | -0.013529127 | 0.08113232 | 0.710912339 | 0.242228297 | 0.055119935 | 0.881451422 |
| Med13 | S | 1028 | 0.09220461 | 0.014112494 | -0.045388851 | -0.137593461 | -0.059501345 | 0.289293553 | 0.764884372 | 0.640993216 | 0.240494503 | 0.52313462 |
| Nab1 | S | 327 | 0.031144507 | 0.073820876 | -0.07195672 | -0.103101227 | -0.145777597 | 0.448537575 | 0.109898826 | 0.276076644 | 0.147714049 | 0.078445718 |
| Hdlbp | S | 944 | -0.015733372 | 0.081569082 | -0.012806401 | 0.002926971 | -0.094375484 | 0.810465445 | 0.215926389 | 0.889588153 | 0.97347822 | 0.321273869 |
| Washc2 | S | 873 | -0.001164502 | 0.129634813 | -0.04463129 | -0.043466788 | -0.174266103 | 0.986492438 | 0.17677206 | 0.362891426 | 0.490783351 | 0.107398802 |
| Phip | S | 879 | 0.019003057 | -0.062149141 | 0.094643475 | 0.075640417 | 0.156792616 | 0.538447462 | 0.028646023 | 0.017543234 | 0.067637115 | 0.00498962 |
| Phip | S | 880 | 0.019003057 | -0.062149141 | 0.094643475 | 0.075640417 | 0.156792616 | 0.538447462 | 0.028646023 | 0.017543234 | 0.067637115 | 0.00498962 |
| Phip | S | 881 | 0.019003057 | -0.062149141 | 0.094643475 | 0.075640417 | 0.156792616 | 0.538447462 | 0.028646023 | 0.017543234 | 0.067637115 | 0.00498962 |
| Atp2b1 Atp2b2 | S | 1127;1105 | -0.078003011 | 0.118218064 | 0.04358571 | 0.121588721 | -0.074632354 | 0.252555448 | 0.188197446 | 0.636395018 | 0.192694229 | 0.416754739 |
| Sptbn1 | S | 2340 | 0.086441716 | -0.059626797 | -0.002208278 | -0.088649994 | 0.057418519 | 0.014088243 | 0.143787747 | 0.955699921 | 0.079345661 | 0.215555453 |
| Nck1 | S | 85 | 0.022455794 | 0.101898007 | -0.090011194 | -0.112466988 | -0.191909201 | 0.830977936 | 0.039463237 | 0.198156343 | 0.352276799 | 0.03914326 |
| Otud7b | S | 471 | 0.014053009 | 0.013862792 | 0.072130455 | 0.058077446 | 0.058267663 | 0.870924864 | 0.907948879 | 0.651330568 | 0.721369735 | 0.743884697 |
| Atad5 | S | 727 | 0.025742365 | 0.030811913 | 0.002086242 | -0.023656123 | -0.028725671 | 0.723585774 | 0.46965367 | 0.926688043 | 0.741537477 | 0.483815368 |
| Esyt2 | S | 669 | 0.020687116 | 0.051174607 | 0.048540564 | 0.027853448 | -0.002634043 | 0.927598502 | 0.796011304 | 0.785066922 | 0.911031755 | 0.990573508 |
| Akap12 | S | 584 | 0.00666209 | 0.147678908 | -0.098902889 | -0.105564978 | -0.246581797 | 0.909590773 | 0.040013794 | 0.118996012 | 0.022438783 | 0.003116301 |
| Setd1a | T | 523 | -0.009570292 | 0.021083465 | 0.048804127 | 0.058374418 | 0.027720662 | 0.87448339 | 0.699152055 | 0.425118574 | 0.427680468 | 0.67572772 |
| Sf3b1 | T | 426 | 0.015879981 | -0.049966448 | 0.079889586 | 0.064009605 | 0.129856034 | 0.726378946 | 0.228836759 | 0.074474047 | 0.114979106 | 0.001719107 |
| Gatad2a | S | 96 | 0.044197199 | -0.034788433 | 0.036363286 | -0.007833913 | 0.071151719 | 0.205474256 | 0.233782559 | 0.278464899 | 0.788218629 | 0.043199279 |
| Akap1 | S | 856 | 0.120741724 | -0.049285489 | 0.070277905 | -0.050463819 | 0.119563394 | 0.6600161 | 0.866190311 | 0.828050877 | 0.836445054 | 0.65619932 |
| Ptpn21 | S | 589 | -0.021836371 | 0.036568315 | 0.041996624 | 0.063832996 | 0.00542831 | 0.678058902 | 0.414805365 | 0.512093017 | 0.382534096 | 0.932279241 |
| Cdk13 | S | 226 | 0.061365485 | -0.021798818 | 0.018615104 | -0.042750381 | 0.040413922 | 0.310512679 | 0.583526594 | 0.614843086 | 0.434015899 | 0.221203973 |
| Wee1 | S | 165 | 0.000677427 | -0.00552546 | 0.042681112 | 0.042003685 | 0.048206572 | 0.981644155 | 0.822577833 | 0.151351266 | 0.113776929 | 0.057704859 |
| Rbm5 | S | 624 | -0.065156159 | -0.025662741 | 0.115008673 | 0.180164832 | 0.140671414 | 0.15786707 | 0.385629232 | 0.006810256 | 0.011769771 | 0.003817107 |
| Phf8-ps Phf8 | S | 817 | 0.044173241 | -0.019183136 | 0.030297178 | -0.013876063 | 0.049480314 | 0.188416274 | 0.461007306 | 0.696901392 | 0.859505477 | 0.541525966 |
| Rtn4 | S | 80 | 0.0896321 | 0.051508194 | -0.08147179 | -0.171103889 | -0.132979983 | 0.244668689 | 0.352575489 | 0.256800844 | 0.08995818 | 0.121912473 |
| Hnrnpc | S | 162 | 0.176861806 | 0.035591689 | -0.211592636 | -0.388454442 | -0.247184324 | 0.133011992 | 0.663303988 | 0.361037217 | 0.151582623 | 0.302891185 |
| Tnrc6a | S | 724 | 0.007485809 | 0.065147652 | -0.021125309 | -0.028611117 | -0.08627296 | 0.91871433 | 0.185445381 | 0.597942806 | 0.710357333 | 0.131351054 |
| Ecm1 | S | 84 | 0.113754182 | 0.04017541 | -0.067417152 | -0.181171334 | -0.107592562 | 0.451219198 | 0.682984434 | 0.6431233 | 0.329574168 | 0.472677723 |
| Ecm1 | S | 85 | 0.113754182 | 0.04017541 | -0.067417152 | -0.181171334 | -0.107592562 | 0.451219198 | 0.682984434 | 0.6431233 | 0.329574168 | 0.472677723 |
| Sec31a | S | 526 | -0.024528427 | 0.029340253 | 0.027807361 | 0.052335788 | -0.001532892 | 0.545980479 | 0.392985402 | 0.210669964 | 0.255565097 | 0.963318669 |
| Tmem209 | S | 98 | -0.007561107 | 0.023463734 | 0.042708558 | 0.050269665 | 0.019244824 | 0.909178845 | 0.707304693 | 0.437207002 | 0.379355602 | 0.68872486 |
| Pagr1a | S | 91 | 0.006485668 | 0.009293708 | 0.052816853 | 0.046331185 | 0.043523145 | 0.816283742 | 0.850921033 | 0.555349319 | 0.598559129 | 0.640397237 |
| Stxbp5 | S | 783 | -0.007286904 | 0.084611065 | -0.061714503 | -0.054427599 | -0.146325568 | 0.828606929 | 0.049940591 | 0.471661945 | 0.521604195 | 0.158790871 |
| Dclk1 | S | 334 | -0.024538334 | 0.03267275 | 0.032612081 | 0.057150414 | -6.06691E-05 | 0.83946647 | 0.363682713 | 0.572502696 | 0.649920063 | 0.998989971 |
| Srrm2 | S | 1718 | -0.003837295 | -0.025567177 | 0.104681655 | 0.10851895 | 0.130248833 | 0.933254733 | 0.394000337 | 0.184790409 | 0.183933434 | 0.131676396 |
| Srrm2 | S | 1720 | -0.003837295 | -0.025567177 | 0.104681655 | 0.10851895 | 0.130248833 | 0.933254733 | 0.394000337 | 0.184790409 | 0.183933434 | 0.131676396 |
| Cwc22 | S | 106 | 0.001780746 | -0.03988664 | 0.072052567 | 0.070271822 | 0.111939207 | 0.936449171 | 0.060365811 | 0.010476905 | 0.01978532 | 0.006032744 |
| Ccser2 | S | 222 | -0.088704279 | 0.075366558 | 0.056014755 | 0.144719034 | -0.019351803 | 0.364027655 | 0.221725217 | 0.449146432 | 0.166430721 | 0.706562441 |
| Zfp516 | S | 912 | 0.060016973 | -0.014784262 | 0.005565348 | -0.054451625 | 0.02034961 | 0.187066123 | 0.613554071 | 0.949286203 | 0.562989851 | 0.818894526 |
| Phf14 | S | 295 | 0.033163937 | 0.055558239 | -0.03802976 | -0.071193697 | -0.093587999 | 0.800321411 | 0.285879824 | 0.76431979 | 0.669646824 | 0.478839835 |
| Rai1 | S | 875 | 0.029315866 | -0.015611115 | 0.034073878 | 0.004758012 | 0.049684992 | 0.550039849 | 0.746538225 | 0.371126444 | 0.908043571 | 0.280222549 |
| Crispld2 | S | 75 | 0.046082232 | -0.039083981 | 0.075745292 | 0.02966306 | 0.114829273 | 0.601839015 | 0.425669258 | 0.36130479 | 0.774740212 | 0.209379313 |
| Crispld2 | T | 81 | 0.046082232 | -0.039083981 | 0.075745292 | 0.02966306 | 0.114829273 | 0.601839015 | 0.425669258 | 0.36130479 | 0.774740212 | 0.209379313 |
| Crispld2 | Y | 71 | 0.046082232 | -0.039083981 | 0.075745292 | 0.02966306 | 0.114829273 | 0.601839015 | 0.425669258 | 0.36130479 | 0.774740212 | 0.209379313 |
| Chd2 | S | 1213 | 0.05180718 | 0.028568697 | -0.022611309 | -0.074418489 | -0.051180006 | 0.368380083 | 0.597905208 | 0.638295562 | 0.251968817 | 0.401549053 |
| Gorab | S | 335 | -0.021997121 | 0.027729481 | 0.017640056 | 0.039637176 | -0.010089426 | 0.696357913 | 0.16469024 | 0.482747845 | 0.507293613 | 0.686399542 |
| M6pr | S | 268 | 0.032536246 | 0.053783666 | -0.08870071 | -0.121236957 | -0.142484376 | 0.301996932 | 0.114435103 | 0.384357295 | 0.266865637 | 0.212679854 |
| Tnks1bp1 | S | 796 | 0.016772159 | 0.072384198 | -0.035713994 | -0.052486152 | -0.108098191 | 0.801251045 | 0.147953196 | 0.381851958 | 0.43182271 | 0.045770302 |
| Fcho2 | S | 402 | 0.007569941 | -0.018366807 | 0.059309999 | 0.051740059 | 0.077676806 | 0.822673493 | 0.727919669 | 0.253636704 | 0.284510021 | 0.224003993 |
| Cbx3 | S | 93 | -0.021981935 | 0.012834243 | 0.046749888 | 0.068731823 | 0.033915645 | 0.392336552 | 0.477198254 | 0.159012368 | 0.076849755 | 0.264006134 |
| Trip10 | S | 296 | -0.000358991 | 0.020107786 | 0.03567784 | 0.036036831 | 0.015570054 | 0.997029149 | 0.795825659 | 0.401762934 | 0.719244805 | 0.846638287 |
| Prkd3 | S | 213 | -0.078254611 | 0.015520424 | 0.108816232 | 0.187070843 | 0.093295807 | 0.760210187 | 0.930125388 | 0.304143824 | 0.495313821 | 0.625459952 |
| Prkd3 | S | 216 | -0.078254611 | 0.015520424 | 0.108816232 | 0.187070843 | 0.093295807 | 0.760210187 | 0.930125388 | 0.304143824 | 0.495313821 | 0.625459952 |
| Zfp646 | S | 523 | 0.039225926 | -0.006296234 | 0.02135882 | -0.017867106 | 0.027655054 | 0.349085193 | 0.879435834 | 0.784162961 | 0.800583862 | 0.701454662 |
| Cad | S | 64 | 0.280955633 | 0.162786098 | -0.422035455 | -0.702991088 | -0.584821553 | 0.208470256 | 0.740187264 | 0.380870827 | 0.194731923 | 0.353496392 |
| Akap13 | S | 1839 | -0.031199564 | 0.055457229 | 0.02534578 | 0.056545344 | -0.030111449 | 0.526484894 | 0.292066483 | 0.530491096 | 0.232566212 | 0.505998937 |
| Samd4 | T | 625 | 0.069721835 | -0.013526378 | 0.002409581 | -0.067312254 | 0.015935958 | 0.246813704 | 0.605285963 | 0.8968421 | 0.262900653 | 0.476412621 |
| Mark2 | S | 390 | 0.032674896 | -0.047621111 | 0.078187633 | 0.045512737 | 0.125808744 | 0.600772271 | 0.451725229 | 0.288856565 | 0.533233072 | 0.134098935 |
| Ppp1r18 | S | 234 | 0.00370791 | -0.01832961 | 0.07404511 | 0.070337199 | 0.09237472 | 0.904088537 | 0.09667543 | 0.078986926 | 0.122140743 | 0.056891333 |
| Med13 | S | 330 | -0.046041246 | 0.024343211 | 0.053380145 | 0.099421391 | 0.029036934 | 0.582581998 | 0.78390839 | 0.135539689 | 0.288771072 | 0.746357051 |
| Zfp638 | S | 1399 | 0.043854168 | -0.043008824 | 0.046755605 | 0.002901437 | 0.089764429 | 0.316334139 | 0.333227759 | 0.295994272 | 0.950304316 | 0.112846316 |
| Tbc1d2b | S | 959 | -0.03137682 | 0.09372278 | -0.011047062 | 0.020329759 | -0.104769841 | 0.609623477 | 0.171749089 | 0.889929955 | 0.756332641 | 0.187018007 |
| Gatad2a | S | 103 | 0.053655576 | -0.035656715 | 0.029885388 | -0.023770188 | 0.065542103 | 0.204994357 | 0.277391459 | 0.389831776 | 0.498587789 | 0.046212325 |
| Nsd1 | S | 2369 | -0.051621033 | 0.021649048 | 0.067119687 | 0.11874072 | 0.045470638 | 0.594044295 | 0.706481995 | 0.211896211 | 0.280666836 | 0.494458369 |
| Ppfibp1 | T | 581 | 0.002697762 | 0.403772724 | -0.281661066 | -0.284358828 | -0.68543379 | 0.988457221 | 0.196023553 | 0.31390364 | 0.330134868 | 0.086396263 |
| Usp47 | S | 832 | 0.022356087 | 0.09312317 | -0.077330247 | -0.099686333 | -0.170453417 | 0.702609895 | 0.154790881 | 0.415374017 | 0.287705103 | 0.119218849 |
| Scaf8 | S | 276 | -0.060728742 | 0.022884903 | 0.140907861 | 0.201636603 | 0.118022958 | 0.13056948 | 0.787175767 | 0.144168649 | 0.07672702 | 0.286183787 |
| Map3k4 | S | 59 | -0.028667262 | 0.062662824 | -0.017475188 | 0.011192074 | -0.080138013 | 0.662262207 | 0.071456277 | 0.849850324 | 0.913617252 | 0.420634907 |
| Axin1 | S | 492 | -0.011788755 | 0.066172159 | 0.016002064 | 0.027790819 | -0.050170095 | 0.701360938 | 0.354108434 | 0.87032453 | 0.773623974 | 0.646888547 |
| Nmnat1 | S | 117 | 0.041497604 | 0.065204258 | -0.070407979 | -0.111905583 | -0.135612237 | 0.509544211 | 0.210763347 | 0.436989645 | 0.240808126 | 0.18216995 |
| Set | S | 28 | 0.090606912 | -0.053745139 | 0.030574936 | -0.060031976 | 0.084320075 | 0.335290783 | 0.324897651 | 0.366805314 | 0.499023969 | 0.167974935 |
| Thoc2 | S | 1486 | 0.341620041 | -0.115664836 | -0.202657974 | -0.544278014 | -0.086993138 | 0.041909795 | 0.438269066 | 0.308173024 | 0.047185797 | 0.674211847 |
| Frmd6 | S | 605 | -0.130742711 | 0.27788634 | -0.040152868 | 0.090589842 | -0.318039208 | 0.140491519 | 0.057695734 | 0.612992322 | 0.342476893 | 0.041518471 |
| Srrm2 | S | 1372 | 0.112262956 | -0.021580417 | -0.051894788 | -0.164157744 | -0.030314371 | 0.012159672 | 0.162906472 | 0.266034105 | 0.022358151 | 0.471977419 |
| Peak1 | S | 786 | 0.038654682 | -0.012393738 | 0.02267107 | -0.015983613 | 0.035064808 | 0.365504456 | 0.599063831 | 0.498372754 | 0.729135683 | 0.355116554 |
| Sfr1 | T | 129 | 0.171736289 | -0.009688966 | -0.113998013 | -0.285734302 | -0.104309046 | 0.019371994 | 0.891511749 | 0.07221098 | 0.002270479 | 0.150053702 |
| Map2 | S | 1013 | -0.167444026 | 0.238942753 | 0.016395072 | 0.183839099 | -0.222547681 | 0.007435889 | 0.007113232 | 0.801463756 | 0.04633409 | 0.024358327 |
| Atrx | S | 1140 | -0.033878595 | -0.018660496 | 0.111442955 | 0.14532155 | 0.130103452 | 0.333098298 | 0.744531971 | 0.098741364 | 0.051732728 | 0.112642049 |
| Acin1 | S | 468 | -0.020750013 | -0.037670861 | 0.11947163 | 0.140221643 | 0.157142491 | 0.690545589 | 0.385398919 | 0.106206381 | 0.086514715 | 0.059362192 |
| Glyr1 | S | 151 | 0.024370737 | -0.001772786 | 0.044266081 | 0.019895944 | 0.046039467 | 0.744653125 | 0.951383906 | 0.541580487 | 0.831776545 | 0.530997747 |
| Brap | S | 116 | 0.004390644 | 0.051467421 | -0.019535749 | -0.023926393 | -0.07100317 | 0.903226661 | 0.204640777 | 0.795858372 | 0.732105883 | 0.363075032 |
| Stk10 | S | 191 | 0.021561578 | 0.040051046 | -0.007159058 | -0.028720636 | -0.047210104 | 0.630182504 | 0.467245804 | 0.845937766 | 0.544035649 | 0.410435123 |
| Nfix | S | 250 | 0.138446116 | -0.14951256 | 0.130608038 | -0.007838077 | 0.280120598 | 0.626130347 | 0.421755982 | 0.403929886 | 0.976838368 | 0.152659462 |
| Srrm1 | S | 616 | 0.057545909 | -0.03233298 | -0.003325495 | -0.060871404 | 0.029007486 | 0.008480622 | 0.113127855 | 0.893785048 | 0.069103526 | 0.278270025 |
| Ktn1 | S | 124 | 0.016618144 | -0.048170408 | 0.098657172 | 0.082039028 | 0.14682758 | 0.748683256 | 0.25677248 | 0.159374558 | 0.237068809 | 0.073096024 |
| Sirt1 | S | 46 | -0.053436116 | 0.043360808 | 0.09177606 | 0.145212176 | 0.048415251 | 0.199659007 | 0.317267186 | 0.352954897 | 0.189556573 | 0.59601249 |
| Ddi2 | S | 121 | 0.033031826 | 0.032770676 | -0.02211814 | -0.055149966 | -0.054888815 | 0.56654026 | 0.265422718 | 0.597700322 | 0.382662785 | 0.210373941 |
| Inpp5f | S | 827 | 0.078584506 | -0.046966717 | 0.021216618 | -0.057367888 | 0.068183334 | 0.202916356 | 0.484559348 | 0.798726512 | 0.433522051 | 0.387959219 |
| Peak1 | S | 282 | -0.01150434 | 0.024573876 | 0.028134745 | 0.039639085 | 0.003560868 | 0.728839907 | 0.482804025 | 0.381003166 | 0.350145556 | 0.929302356 |
| Tmpo | S | 264 | 0.062116216 | -0.026392221 | 0.044229894 | -0.017886322 | 0.070622114 | 0.536383093 | 0.539774601 | 0.524563679 | 0.866143739 | 0.333722365 |
| Snip1 | S | 18 | 0.014487386 | -0.003134382 | 0.029227881 | 0.014740495 | 0.032362263 | 0.464845181 | 0.806627797 | 0.423040734 | 0.681471099 | 0.3805974 |
| Pikfyve | S | 329 | 0.031136942 | -0.050752605 | 0.062319281 | 0.031182339 | 0.113071886 | 0.48482437 | 0.661664132 | 0.420011075 | 0.681108913 | 0.393925299 |

| | | | | | | | | | | | | |
|---|---|---|---|---|---|---|---|---|---|---|---|---|
| Huwe1 | S | 3663 | 0.17197958 | 0.033417584 | -0.173397648 | -0.345377228 | -0.206815232 | 0.115815409 | 0.547304504 | 0.219185059 | 0.054640389 | 0.165615958 |
| Rbm10 | S | 733 | 0.008843147 | -0.028974012 | 0.056670571 | 0.047827424 | 0.085644583 | 0.770193315 | 0.471859276 | 0.134397655 | 0.249504086 | 0.109399473 |
| Rbm15b | T | 529 | 0.021460381 | -0.061614027 | 0.100732042 | 0.079271661 | 0.162346069 | 0.617995948 | 0.33212798 | 0.209579601 | 0.307924793 | 0.09639962 |
| Myc | S | 277 | 0.12419272 | 0.058773624 | -0.159641266 | -0.283833986 | -0.21841489 | 0.357284536 | 0.744271016 | 0.671872124 | 0.466405377 | 0.574636714 |
| Srrm2 | T | 2553 | 0.31363305 | -0.050205196 | -0.104204881 | -0.417837931 | -0.053999685 | 0.104590923 | 0.590352142 | 0.153459273 | 0.066272389 | 0.523553652 |
| Smchd1 | S | 1975 | 0.011598513 | 0.026227764 | 0.000208002 | -0.01139051 | -0.026019761 | 0.74363094 | 0.304002114 | 0.995148268 | 0.757697549 | 0.388046472 |
| Mpdz | S | 1066 | 0.012022416 | 0.093580926 | -0.054039924 | -0.066062341 | -0.14762085 | 0.860683211 | 0.152412948 | 0.388046338 | 0.227024572 | 0.00838601 |
| Snd1 | S | 426 | -0.06256789 | 0.034111521 | 0.057013366 | 0.119581257 | 0.022901845 | 0.149637452 | 0.234197273 | 0.105806786 | 0.034205508 | 0.492831509 |
| Tbx3 | S | 373 | 0.048734565 | -0.070903664 | 0.057798852 | 0.009064287 | 0.128702517 | 0.193628908 | 0.051769341 | 0.053835071 | 0.751365285 | 0.009981874 |
| Nes | S | 479 | 0.074921879 | -0.06051484 | 0.036554508 | -0.038367371 | 0.097069349 | 0.37274747 | 0.378149667 | 0.312704668 | 0.611074622 | 0.197223634 |
| Unc119 | S | 39 | -0.058781763 | 0.062702081 | 0.038025396 | 0.096807158 | -0.024676685 | 0.387958511 | 0.203918676 | 0.529291663 | 0.204447099 | 0.637403004 |
| Cdk11b | T | 740 | -0.013323713 | -0.01521313 | 0.073187533 | 0.086511246 | 0.088400663 | 0.764962202 | 0.72304364 | 0.161970931 | 0.083682942 | 0.07454263 |
| Myl12b Myl9 | S | 29;29 | 0.049600206 | -0.08377579 | 0.079994697 | 0.030394492 | 0.163770487 | 0.552980102 | 0.376235612 | 0.216840742 | 0.712837957 | 0.135312339 |
| Pard3 | S | 871 | 0.11625729 | -0.020044818 | -0.059317172 | -0.175574462 | -0.039272354 | 0.027011082 | 0.637962947 | 0.180813422 | 0.002618757 | 0.165415858 |
| Sf3b1 | T | 436 | 0.02740959 | -0.020582158 | 0.035722744 | 0.008313155 | 0.056304903 | 0.471030429 | 0.443286627 | 0.257369529 | 0.826882264 | 0.105254534 |
| Mospd3 | S | 24 | -0.023139514 | -0.055211684 | 0.113227252 | 0.136366766 | 0.168438936 | 0.804580809 | 0.362259632 | 0.075007277 | 0.204777039 | 0.028597625 |
| Rexo1 | S | 353 | 0.005872328 | 0.014623659 | 0.04043761 | 0.034565282 | 0.02581395 | 0.882160099 | 0.605077405 | 0.696512745 | 0.741114976 | 0.79957662 |
| Srrm2 | S | 1269 | -0.017399332 | -0.03861361 | 0.085407957 | 0.102807289 | 0.124021567 | 0.618589391 | 0.247938499 | 0.026444096 | 0.033348739 | 0.012585993 |
| Kif23 | S | 904 | 0.022285443 | 0.011780871 | -0.015685585 | -0.037971028 | -0.027466456 | 0.208385198 | 0.242526394 | 0.645458536 | 0.320982826 | 0.442159531 |
| Hdac5 | S | 488 | -0.171411959 | 0.052089555 | 0.205098605 | 0.376510564 | 0.15300905 | 0.126182986 | 0.493085293 | 0.138310645 | 0.032901831 | 0.228701831 |
| Ahdc1 | S | 593 | -0.047777651 | 0.045362901 | 0.051820044 | 0.099597694 | 0.006457143 | 0.120990019 | 0.306215928 | 0.140424051 | 0.022684569 | 0.865882167 |
| Cby1 | S | 20 | -0.024340235 | 0.089864617 | -0.054597536 | -0.030257302 | -0.144462153 | 0.540913364 | 0.017396516 | 0.323240578 | 0.570588369 | 0.06551998 |
| Herc2 | T | 1945 | 0.02711066 | 0.065487139 | -0.044234459 | -0.071345119 | -0.109721598 | 0.574047159 | 0.229575814 | 0.403193487 | 0.184897222 | 0.082422493 |
| Ehmt1 | S | 81 | -0.001124212 | 0.006364527 | 0.04192798 | 0.043052193 | 0.035563454 | 0.97961744 | 0.883607427 | 0.351511572 | 0.386912311 | 0.463260576 |
| Tmem209 | S | 194 | 0.100895824 | 0.042603856 | -0.064428621 | -0.165324445 | -0.107032476 | 0.282692379 | 0.842480004 | 0.502871769 | 0.037710095 | 0.610584933 |
| Tmem209 | S | 198 | 0.100895824 | 0.042603856 | -0.064428621 | -0.165324445 | -0.107032476 | 0.282692379 | 0.842480004 | 0.502871769 | 0.037710095 | 0.610584933 |
| Srrm1 | S | 793 | 0.059390677 | -0.011691174 | -0.015356321 | -0.074746998 | -0.003665147 | 0.062731728 | 0.491869925 | 0.638989712 | 0.088074554 | 0.903439661 |
| Satb2 | S | 7 | 0.000837762 | 0.011206466 | 0.043754443 | 0.042916681 | 0.032547977 | 0.991807992 | 0.838534502 | 0.460095603 | 0.56101834 | 0.371811597 |
| H2bc1 H2bc11 H2bc1 | S | 88;88;88;89;88;88 | 0.20881412 | -0.071793809 | -0.13123671 | -0.34005083 | -0.059442901 | 0.007498199 | 0.248201973 | 0.194955248 | 0.038898061 | 0.494390746 |
| Relb | S | 96 | 0.033561857 | 0.002993124 | 0.042213799 | 0.008651943 | 0.039220676 | 0.69508843 | 0.958210778 | 0.752840813 | 0.95037413 | 0.762068749 |
| Pds5a | T | 1207 | -0.00207926 | 0.054157466 | 0.007506277 | 0.009585536 | -0.046651189 | 0.946178362 | 0.412624718 | 0.932105789 | 0.915468111 | 0.647848023 |
| Dbn1 | S | 658 | 0.076402427 | -0.024759484 | -0.006013655 | -0.082416082 | 0.01874583 | 0.139947491 | 0.475891895 | 0.929887288 | 0.285466301 | 0.771691283 |
| Fam126a | S | 433 | -0.180428638 | 0.251538083 | 0.005338128 | 0.185766766 | -0.246199955 | 0.248289284 | 0.128234541 | 0.965769246 | 0.195979489 | 0.11603857 |
| Fbxl3 | S | 8 | 0.104700331 | -0.034300988 | 0.05168423 | -0.053016101 | 0.085985218 | 0.683781154 | 0.562471078 | 0.564205959 | 0.837821825 | 0.385847951 |
| Fbxl3 | S | 12 | 0.104700331 | -0.034300988 | 0.05168423 | -0.053016101 | 0.085985218 | 0.683781154 | 0.562471078 | 0.564205959 | 0.837821825 | 0.385847951 |
| Son | S | 1808 | 0.055001601 | -0.066273869 | 0.054970615 | -3.09861E-05 | 0.121244483 | 0.262497672 | 0.236670105 | 0.269303596 | 0.999524226 | 0.084103535 |
| Lpin3 | S | 408 | -0.047815513 | 0.249305913 | -0.159597267 | -0.111781754 | -0.408903179 | 0.612828061 | 0.03832795 | 0.185858557 | 0.223543801 | 0.009316797 |
| Eif3a | S | 1149 | 0.022175547 | 0.033701181 | -0.007076023 | -0.02925157 | -0.040777204 | 0.585086436 | 0.510719655 | 0.886757414 | 0.61188786 | 0.526412453 |
| Stub1 | S | 20 | -0.012999722 | 0.071263485 | -0.011013363 | 0.001986359 | -0.082276848 | 0.478912263 | 0.121713441 | 0.726616956 | 0.952462169 | 0.10357124 |
| Tsc22d4 | S | 189 | 0.026059704 | -0.038293928 | 0.040309676 | 0.014249972 | 0.078603604 | 0.229528647 | 0.347589261 | 0.252544843 | 0.637502943 | 0.120247481 |
| Pask | S | 1039 | -0.163210262 | 0.177835138 | 0.050738342 | 0.213948604 | -0.127096796 | 0.06014238 | 0.036395353 | 0.681376182 | 0.149081323 | 0.328020844 |
| Abl1 | S | 909 | 0.010879254 | 0.027298637 | 0.026478766 | 0.015599512 | -0.000819871 | 0.880213968 | 0.675252931 | 0.743828479 | 0.871550582 | 0.992842876 |
| Akap13 | S | 1613 | -0.083840627 | 0.137583303 | 0.007875851 | 0.091716479 | -0.129707451 | 0.188323091 | 0.094270313 | 0.867832291 | 0.135713194 | 0.112061093 |
| Cux1 | S | 1054 | 0.267597238 | -0.070205961 | 0.061146443 | -0.206450795 | 0.131352404 | 0.485791539 | 0.116119581 | 0.593787604 | 0.585755944 | 0.29798704 |
| Phlpp1 | S | 304 | 0.020870765 | 0.037160432 | -0.021330326 | -0.042201091 | -0.058490758 | 0.464474511 | 0.302562279 | 0.578099812 | 0.256946914 | 0.17527792 |
| Cep170b | S | 843 | 0.01283868 | 0.003640242 | 0.026232981 | 0.013394301 | 0.022592739 | 0.610978782 | 0.915165228 | 0.516470602 | 0.720531897 | 0.603975758 |
| Med17 | S | 55 | 0.211117484 | 0.015087398 | -0.147861084 | -0.358978569 | -0.162948483 | 0.331597152 | 0.876510257 | 0.54788513 | 0.242133607 | 0.507061286 |
| Bag6 | S | 995 | -0.062492863 | 0.067861083 | 0.033227113 | 0.095719976 | -0.03463397 | 0.070400878 | 0.0351193 | 0.464961099 | 0.1044917 | 0.460013561 |
| Fnip1 | S | 259 | -0.062415193 | 0.073662891 | 0.065014531 | 0.127429724 | -0.00864836 | 0.799790879 | 0.497668276 | 0.801174453 | 0.688371618 | 0.971653285 |
| Cdk12 | S | 302 | -0.043476095 | 0.038944326 | 0.014845957 | 0.058322052 | -0.024098369 | 0.369646875 | 0.035489738 | 0.677805678 | 0.293250615 | 0.502325279 |
| Usp36 | S | 916 | 0.233572237 | -0.150020277 | -0.055360219 | -0.288932456 | 0.094660058 | 0.022065716 | 0.226334015 | 0.343556061 | 0.020839454 | 0.396903463 |
| Eapp | S | 109 | -0.007364613 | 0.001105393 | 0.040463062 | 0.047827675 | 0.03935767 | 0.880249084 | 0.956684239 | 0.208984727 | 0.378716135 | 0.204021557 |
| Eapp | S | 111 | -0.007364613 | 0.001105393 | 0.040463062 | 0.047827675 | 0.03935767 | 0.880249084 | 0.956684239 | 0.208984727 | 0.378716135 | 0.204021557 |
| Sf3b1 | T | 273 | 0.017710118 | 0.041204198 | 0.027169104 | 0.009458986 | -0.014035094 | 0.90727923 | 0.598754572 | 0.940087287 | 0.979867164 | 0.968896518 |
| Fam83d | S | 17 | 0.072095963 | 0.119992019 | -0.105141503 | -0.177237465 | -0.225133522 | 0.548906032 | 0.282900456 | 0.273195822 | 0.139157816 | 0.04951666 |
| Slk | S | 666 | -0.025928002 | 0.085152462 | -0.012388264 | 0.013539738 | -0.097540726 | 0.514559659 | 0.152698696 | 0.869450478 | 0.849979469 | 0.261990914 |
| Aup1 | T | 367 | 0.063473677 | -0.140521638 | 0.127086644 | 0.063612968 | 0.267608283 | 0.299210865 | 0.387439291 | 0.457660752 | 0.695897017 | 0.228354706 |
| Tex2 | S | 795 | -0.020621376 | 0.031324532 | 0.067629948 | 0.088251324 | 0.036305417 | 0.815215178 | 0.759085673 | 0.557883582 | 0.443354667 | 0.763099697 |
| Luc7l2 | S | 384 | -0.127509083 | 0.028584014 | 0.088788395 | 0.216297478 | 0.060204381 | 0.425167102 | 0.63245228 | 0.110243651 | 0.228547369 | 0.375745738 |
| Abi2 | S | 210 | 0.132514311 | -0.059701973 | -0.01431843 | -0.146832741 | 0.045383543 | 0.079347876 | 0.210682904 | 0.851909596 | 0.150004695 | 0.585598864 |
| Baz1b | S | 424 | 0.024492748 | 0.034606125 | 0.014403061 | -0.010089687 | -0.020203064 | 0.81357932 | 0.679828217 | 0.895673108 | 0.931306579 | 0.842555742 |
| Sh3pxd2a | S | 1034 | -0.036766035 | 0.023193172 | 0.144614565 | 0.1813806 | 0.121421393 | 0.207534515 | 0.633530706 | 0.40013047 | 0.314251519 | 0.470933694 |
| Elp2 | S | 485 | 0.033021235 | 0.047952445 | -0.010472915 | -0.04349415 | -0.05842536 | 0.803547962 | 0.54684977 | 0.923179324 | 0.763604583 | 0.575248319 |
| Aff4 | S | 689 | 0.054718616 | 0.060884628 | -0.06073182 | -0.115450436 | -0.121616448 | 0.608335265 | 0.275187745 | 0.453901311 | 0.339664294 | 0.168788853 |
| Myadm | S | 15 | 0.04469548 | 0.027524385 | -0.027012051 | -0.071707531 | -0.054536436 | 0.259392214 | 0.579644788 | 0.474119389 | 0.059123121 | 0.263009767 |
| Marcksl1 | S | 22 | 0.741742179 | -0.158018422 | -0.022226421 | -0.7639686 | 0.135792002 | 0.353588787 | 0.124424999 | 0.757963073 | 0.342519306 | 0.174362472 |
| Xrn2 | S | 487 | 0.074025399 | 0.001724033 | -0.041138923 | -0.115164322 | -0.042862956 | 0.113653346 | 0.954326146 | 0.436722649 | 0.080431346 | 0.384914073 |
| Champ1 | S | 622 | 0.081247698 | -0.076653727 | 0.049651785 | -0.031595913 | 0.126305512 | 0.165012479 | 0.226783071 | 0.520820457 | 0.649298039 | 0.143599783 |
| Tmem87a | S | 540 | 0.011582732 | -0.022986731 | 0.041717084 | 0.030134352 | 0.064703815 | 0.748987959 | 0.69695132 | 0.202074439 | 0.470426645 | 0.331048861 |
| Map3k1 | S | 518 | 0.050864841 | 0.032649781 | -0.043165017 | -0.094029858 | -0.075814798 | 0.371362409 | 0.282852021 | 0.386812919 | 0.166698597 | 0.167679967 |
| Dtl | S | 675 | 0.007130493 | -0.004653823 | 0.063119412 | 0.055988919 | 0.067773235 | 0.886332744 | 0.940570545 | 0.489971823 | 0.509401916 | 0.457867614 |
| Srrm1 | S | 684 | 0.030740905 | -0.016962914 | 0.0555396 | 0.024798694 | 0.072502513 | 0.615861139 | 0.806669345 | 0.620051534 | 0.801741696 | 0.500198171 |
| Cic | T | 1078 | -0.089204064 | 0.073943187 | 0.086967775 | 0.176171839 | 0.013024588 | 0.175983519 | 0.281364666 | 0.326992479 | 0.099802724 | 0.881302312 |
| Mia2 | S | 1190 | -0.057900866 | 0.009378392 | 0.060190156 | 0.118091022 | 0.050811764 | 0.1100688 | 0.61993382 | 0.012333855 | 0.021898465 | 0.03352543 |
| Cenpc1 | S | 363 | 0.03645193 | -0.040076312 | 0.039371892 | 0.002919962 | 0.079448204 | 0.440091378 | 0.623070046 | 0.406667038 | 0.957423167 | 0.379494539 |
| Ccne2 | S | 21 | -0.015890048 | 0.016179789 | 0.040912342 | 0.056802389 | 0.024732553 | 0.511749348 | 0.567631768 | 0.271875011 | 0.154186185 | 0.499013411 |
| Cep350 | S | 1641 | -0.25772355 | -0.034885408 | 0.201318592 | 0.459042142 | 0.236204 | 0.065804522 | 0.784679953 | 0.001365761 | 0.027938572 | 0.162830864 |
| Zc3h13 | S | 907 | 0.038283695 | 0.010098499 | -0.010271572 | -0.048555267 | -0.02037007 | 0.347372572 | 0.670346178 | 0.843451618 | 0.417021492 | 0.691692887 |
| Hdac2 | S | 422 | 0.009814523 | 0.001836769 | 0.013382492 | 0.003567969 | 0.011545722 | 0.567495082 | 0.867100766 | 0.271128217 | 0.810958667 | 0.148971251 |
| Srrm1 | S | 539 | -0.018414502 | 0.021194319 | 0.023262848 | 0.04167735 | 0.002068529 | 0.632042132 | 0.463672606 | 0.222128995 | 0.309983127 | 0.933600204 |
| Atn1 | S | 112 | 0.050141942 | -0.028353499 | 0.041674339 | -0.008467603 | 0.070027837 | 0.451037058 | 0.542159987 | 0.565069588 | 0.913345388 | 0.334631455 |
| Kmt2a | S | 1860 | 0.064124472 | -0.003806794 | -0.037579342 | -0.101703814 | -0.033772548 | 0.051284212 | 0.907582059 | 0.362520903 | 0.058437615 | 0.43681863 |
| Pak1 | T | 146 | -0.08523532 | 0.258670998 | -0.056012121 | 0.029223199 | -0.314683118 | 0.260587671 | 0.113926047 | 0.527927609 | 0.709350426 | 0.076739311 |
| Phax | S | 56 | 0.015284961 | 0.084878385 | -0.029043585 | -0.044328546 | -0.11392197 | 0.910837971 | 0.412002153 | 0.754657298 | 0.729148574 | 0.229965887 |
| Phax | S | 57 | 0.015284961 | 0.084878385 | -0.029043585 | -0.044328546 | -0.11392197 | 0.910837971 | 0.412002153 | 0.754657298 | 0.729148574 | 0.229965887 |
| Phax | S | 60 | 0.015284961 | 0.084878385 | -0.029043585 | -0.044328546 | -0.11392197 | 0.910837971 | 0.412002153 | 0.754657298 | 0.729148574 | 0.229965887 |
| Phax | S | 63 | 0.015284961 | 0.084878385 | -0.029043585 | -0.044328546 | -0.11392197 | 0.910837971 | 0.412002153 | 0.754657298 | 0.729148574 | 0.229965887 |
| Ncoa3 | S | 850 | -0.00971346 | 0.068354884 | 0.046741757 | 0.056455217 | -0.021613128 | 0.910558018 | 0.749837495 | 0.767582547 | 0.720255727 | 0.928167271 |
| Cd34 | S | 343 | -0.099991361 | 0.094143412 | 0.038360725 | 0.138352086 | -0.055782687 | 0.148588893 | 0.079439124 | 0.473185472 | 0.061057142 | 0.167627299 |
| Misp | S | 416 | 0.215850193 | -0.046974844 | -0.164089653 | -0.379939845 | -0.117114809 | 0.100754873 | 0.747189933 | 0.442525983 | 0.136663602 | 0.5931975 |
| Misp | S | 419 | 0.215850193 | -0.046974844 | -0.164089653 | -0.379939845 | -0.117114809 | 0.100754873 | 0.747189933 | 0.442525983 | 0.136663602 | 0.5931975 |
| Misp | T | 420 | 0.215850193 | -0.046974844 | -0.164089653 | -0.379939845 | -0.117114809 | 0.100754873 | 0.747189933 | 0.442525983 | 0.136663602 | 0.5931975 |
| Misp | T | 423 | 0.215850193 | -0.046974844 | -0.164089653 | -0.379939845 | -0.117114809 | 0.100754873 | 0.747189933 | 0.442525983 | 0.136663602 | 0.5931975 |
| Nxf1 | S | 187 | 0.045346385 | 0.024451892 | -0.023347971 | -0.068694355 | -0.047799863 | 0.258759905 | 0.553563433 | 0.396733954 | 0.112189548 | 0.256477146 |

| | | | | | | | | | | | | |
|---|---|---|---|---|---|---|---|---|---|---|---|---|
| Elob | S | 94 | -0.001343728 | 0.096800157 | 0.031150623 | 0.032494351 | -0.065649534 | 0.992863769 | 0.787087603 | 0.847511701 | 0.814588364 | 0.852247983 |
| Nfib | T | 286 | -0.109668399 | 0.146024467 | 0.014860014 | 0.124528413 | -0.131164453 | 0.126480065 | 0.056995213 | 0.843609691 | 0.137493474 | 0.124380018 |
| Mical1 | S | 949 | -0.032343802 | 0.035258035 | 0.060478632 | 0.092822433 | 0.025220597 | 0.732050388 | 0.722084085 | 0.468493579 | 0.216499083 | 0.728100827 |
| Hsf3 | S | 371 | 0.008553682 | -0.02330361 | 0.083307251 | 0.074753569 | 0.10661086 | 0.921506751 | 0.596447316 | 0.329555192 | 0.474332777 | 0.228580038 |
| Hsf3 | Y | 369 | 0.008553682 | -0.02330361 | 0.083307251 | 0.074753569 | 0.10661086 | 0.921506751 | 0.596447316 | 0.329555192 | 0.474332777 | 0.228580038 |
| Chd4 | T | 1546 | 0.002128984 | -0.034779193 | 0.062524618 | 0.060395634 | 0.097303811 | 0.934912276 | 0.605311958 | 0.155979738 | 0.163153457 | 0.217313215 |
| Sec16a | S | 2058 | -0.029910953 | 0.111779235 | -0.060373093 | -0.030462139 | -0.172152328 | 0.607661524 | 0.034648337 | 0.297329333 | 0.643474078 | 0.031893114 |
| Psmd2 | T | 20 | 0.065937544 | 0.207465317 | -0.211754765 | -0.277692309 | -0.419220081 | 0.426920858 | 0.026027635 | 0.024370532 | 0.014307315 | 5.00988E-05 |
| Dync1li2 | T | 202 | -0.223905481 | 0.1739683 | 0.091401522 | 0.315307003 | -0.082566777 | 0.251276637 | 0.197117469 | 0.520870112 | 0.15022186 | 0.570109448 |
| Eif3b | S | 68 | 0.016068035 | -0.017957413 | 0.030169413 | 0.014101378 | 0.048126826 | 0.465254543 | 0.63709819 | 0.271087464 | 0.486445556 | 0.242749868 |
| Phrf1 | S | 982 | -0.003626415 | 0.037596606 | 0.016020918 | 0.019647332 | -0.021575688 | 0.836676496 | 0.415092194 | 0.78485848 | 0.740227045 | 0.749643393 |
| Utp18 | S | 201 | 0.054098748 | -0.035447695 | -0.001439844 | -0.055538591 | 0.034007851 | 0.046921303 | 0.399527299 | 0.958125978 | 0.085742465 | 0.425954256 |
| Prpf40a | S | 881 | -0.036743023 | -0.015516776 | 0.131093717 | 0.16783674 | 0.146610493 | 0.474106558 | 0.767904439 | 0.217351112 | 0.143563877 | 0.183602519 |
| Kank2 | S | 550 | 0.044890692 | -0.043813429 | 0.083145848 | 0.038255156 | 0.126959277 | 0.528317327 | 0.583155227 | 0.613491218 | 0.808344735 | 0.455294402 |
| Champ1 | S | 448 | 0.055329316 | -0.092555747 | 0.038661737 | -0.016667579 | 0.131217484 | 0.025371139 | 0.147828188 | 0.405956099 | 0.689999685 | 0.080059775 |
| Nkap | S | 9 | -0.008810959 | -0.003112071 | 0.042097248 | 0.050908207 | 0.045209318 | 0.854872335 | 0.855045086 | 0.163690042 | 0.347514825 | 0.144096364 |
| Gnl3 | S | 525 | 0.123428116 | -0.016568324 | -0.095249758 | -0.218677874 | -0.078681434 | 0.194478875 | 0.885581226 | 0.689389158 | 0.389864265 | 0.743778439 |
| Ralgapa1 | S | 739 | 0.050027316 | -0.022061658 | 0.009520744 | -0.040506572 | 0.031582402 | 0.154326226 | 0.44536645 | 0.816049578 | 0.311050769 | 0.390823439 |
| Zmym2 | T | 1375 | 0.063319951 | -0.021940723 | 0.022382044 | -0.040937907 | 0.044322767 | 0.431989534 | 0.60323168 | 0.691667991 | 0.634308952 | 0.479396708 |
| Map3k11 | S | 525 | 0.102824287 | -0.017552839 | 0.007438003 | -0.095386284 | 0.024990843 | 0.565237921 | 0.899731789 | 0.963632599 | 0.634699149 | 0.883358973 |
| Pak4 | S | 104 | -0.056649562 | 0.061870084 | 0.046921742 | 0.103571304 | -0.014948342 | 0.061442767 | 0.101677318 | 0.30166562 | 0.071359119 | 0.73359378 |
| Ank3 | S | 907 | 0.168262115 | 0.147937599 | -0.327604165 | -0.49586628 | -0.475541764 | 0.407707231 | 0.483933628 | 0.464975344 | 0.295522731 | 0.311765706 |
| Dclre1c | S | 385 | -0.033709135 | 0.084627078 | 0.007801638 | 0.041510773 | -0.07682544 | 0.43728461 | 0.234722408 | 0.877317643 | 0.450329905 | 0.303265384 |
| H3f3a H3f3b Gm742( | S | 29;29 | 0.060937274 | 0.047131087 | -0.132095511 | -0.193032785 | -0.179226598 | 0.003556758 | 0.459448575 | 0.090436376 | 0.045932748 | 0.059981478 |
| Traf5 | S | 19 | -0.0430102 | 0.100674344 | 0.004354636 | 0.047364836 | -0.096319708 | 0.796384182 | 0.399660741 | 0.981030917 | 0.816979991 | 0.587301305 |
| Arglu1 | S | 74 | 0.005916884 | -0.022020932 | 0.085979811 | 0.080062927 | 0.108000743 | 0.947585694 | 0.610122283 | 0.350559353 | 0.491226385 | 0.268018414 |
| Srrm2 | S | 875 | 0.008036037 | -0.049615441 | 0.090049036 | 0.082012999 | 0.139664477 | 0.757909379 | 0.528441216 | 0.268158525 | 0.303145904 | 0.189071176 |
| Fam160a2 | S | 900 | 0.06444161 | -0.053516098 | 0.100489683 | 0.036048074 | 0.154005782 | 0.5463932 | 0.301037877 | 0.485072499 | 0.817086327 | 0.316482411 |
| Trappc12 | S | 83 | 0.032629615 | -0.013137039 | 0.041823149 | 0.009193534 | 0.054960188 | 0.695938923 | 0.82821157 | 0.565356445 | 0.926775237 | 0.530505195 |
| Trappc12 | S | 87 | 0.032629615 | -0.013137039 | 0.041823149 | 0.009193534 | 0.054960188 | 0.695938923 | 0.82821157 | 0.565356445 | 0.926775237 | 0.530505195 |
| Map1a | S | 1634 | -0.040891247 | 0.030121678 | 0.03823879 | 0.079130037 | 0.008117112 | 0.279384841 | 0.357543199 | 0.157936702 | 0.082647993 | 0.795234643 |
| Fam117b | S | 452 | -0.055299402 | 0.088372597 | 0.019500471 | 0.074799873 | -0.068872127 | 0.255412077 | 0.141231587 | 0.766311861 | 0.309424154 | 0.362473924 |
| Dennd2a | S | 397 | 0.108056206 | 0.033522997 | -0.095762675 | -0.203818881 | -0.129285672 | 0.135292682 | 0.416647622 | 0.152974703 | 0.03309663 | 0.088511635 |
| Pgls | S | 49 | 0.171554595 | -0.003287233 | -0.054312396 | -0.225866991 | -0.051025163 | 0.537478355 | 0.981725637 | 0.811136639 | 0.486946167 | 0.827105744 |
| Dcaf5 | S | 528 | 0.182365037 | 0.003735181 | -0.144669266 | -0.327034302 | -0.148404447 | 0.300689901 | 0.983848621 | 0.617635014 | 0.301930422 | 0.61949088 |
| Cdk11b | S | 741 | 0.014146394 | -0.034539455 | 0.069784077 | 0.055637683 | 0.104323532 | 0.817031444 | 0.534535045 | 0.262682206 | 0.269441175 | 0.048330763 |
| Numb | S | 241 | -0.009156717 | 0.056655878 | 0.048606932 | 0.057763649 | -0.008048946 | 0.91879399 | 0.75286816 | 0.772631285 | 0.722052598 | 0.970021509 |
| Srsf5 | S | 230 | 0.036663781 | -0.004585897 | 0.039849997 | 0.003186216 | 0.044435894 | 0.67778055 | 0.954720212 | 0.756414432 | 0.982160598 | 0.752608694 |
| Ric1 | T | 995 | 0.031308554 | 0.081047669 | -0.072136147 | -0.103444701 | -0.153183815 | 0.585973777 | 0.145315023 | 0.253082086 | 0.161287186 | 0.05361441 |
| Tomm22 | S | 45 | 0.025621648 | -0.10560059 | 0.088148678 | 0.06252703 | 0.193749268 | 0.440088695 | 0.104787561 | 0.031246708 | 0.041477384 | 0.03019739 |
| Rfc1 | S | 266 | 0.04450912 | 0.019976997 | -0.023420621 | -0.06792974 | -0.043397618 | 0.351815928 | 0.169637299 | 0.198232841 | 0.200754815 | 0.041405505 |
| Frmd8 | S | 448 | -0.059467958 | 0.067313096 | 0.027025436 | 0.086493394 | -0.04028766 | 0.510908739 | 0.214713315 | 0.754763095 | 0.432367604 | 0.634776349 |
| Tcof1 | S | 386 | 0.011654346 | 0.041805449 | -0.023680016 | -0.035334362 | -0.065485465 | 0.77628905 | 0.319998234 | 0.843160013 | 0.765591889 | 0.590821961 |
| Smpd4 | S | 130 | 0.014306026 | -0.013601376 | 0.055277944 | 0.040971918 | 0.06887932 | 0.651476285 | 0.337354349 | 0.316937768 | 0.461397323 | 0.240940053 |
| Mrpl1 | S | 85 | 0.009587672 | -0.001477845 | 0.032166009 | 0.022578337 | 0.033643854 | 0.825395106 | 0.964638891 | 0.395745711 | 0.5432659 | 0.196202597 |
| Kdm6a | S | 818 | 0.095687386 | 0.013246324 | -0.069610639 | -0.165298026 | -0.082856963 | 0.194593696 | 0.807611711 | 0.434688046 | 0.127336983 | 0.357102705 |
| Fam53c | S | 236 | -0.02057177 | 0.010010211 | 0.06359205 | 0.08416382 | 0.053581839 | 0.854231537 | 0.899580903 | 0.428343274 | 0.480966062 | 0.534847725 |
| Mfap1a Mfap1b | S | 132;132 | -0.023781353 | -0.020664231 | 0.092764074 | 0.116545427 | 0.113428304 | 0.591916694 | 0.684778487 | 0.167194649 | 0.103693769 | 0.095076792 |
| Mfap1a Mfap1b | S | 133;133 | -0.023781353 | -0.020664231 | 0.092764074 | 0.116545427 | 0.113428304 | 0.591916694 | 0.684778487 | 0.167194649 | 0.103693769 | 0.095076792 |
| Ticrr | S | 1002 | -0.030107998 | 0.009357784 | 0.053934939 | 0.084042937 | 0.044577155 | 0.635962007 | 0.83038956 | 0.22840592 | 0.211995882 | 0.222103508 |
| Usp34 | T | 3387 | 0.074384873 | -0.05601585 | 0.026177592 | -0.048207281 | 0.082193442 | 0.199820203 | 0.341557064 | 0.651023985 | 0.183800245 | 0.073450138 |
| Sipa1l3 | S | 394 | -0.011441989 | 0.081988736 | -0.0494648 | -0.038022812 | -0.131453536 | 0.893618931 | 0.093638646 | 0.451480874 | 0.680970295 | 0.093764329 |
| Hnrnpa0 | S | 68 | -0.273754903 | -0.034586763 | 0.167240784 | 0.440995688 | 0.201827548 | 0.366399345 | 0.529716437 | 0.009888723 | 0.202475932 | 0.008132797 |
| Pkp4 | S | 326 | 0.047398992 | -0.072350425 | 0.082471252 | 0.03507226 | 0.154821676 | 0.086743829 | 0.101929286 | 0.407658664 | 0.70042714 | 0.180099767 |
| Cad | S | 1407 | -0.035383465 | 0.293431836 | -0.217031759 | -0.181648293 | -0.510463594 | 0.784271787 | 0.088469893 | 0.254158211 | 0.270835681 | 0.031052172 |
| Fnbp1l | S | 525 | 0.037201765 | 0.006959245 | 0.000452776 | -0.036748988 | -0.006506468 | 0.458036491 | 0.856984365 | 0.992041364 | 0.486170296 | 0.877570921 |
| Bcr | S | 237 | -0.100907881 | 0.047232679 | 0.327358259 | 0.428266139 | 0.28012558 | 0.375077823 | 0.884675665 | 0.548445612 | 0.444834205 | 0.631464308 |
| Epb41l3 | S | 585 | 0.099509129 | 0.008920331 | -0.074067694 | -0.173576822 | -0.082988025 | 0.183458121 | 0.915491114 | 0.477676427 | 0.166384458 | 0.490325083 |
| Nfic | S | 273 | 0.088776344 | -0.012674114 | -0.023665057 | -0.112441401 | -0.010990943 | 0.304443776 | 0.87823108 | 0.763601486 | 0.047417812 | 0.754615078 |
| Atn1 | S | 119 | -0.049376797 | -0.009536376 | 0.100453302 | 0.149830099 | 0.109989678 | 0.560552323 | 0.801914888 | 0.173730552 | 0.153893035 | 0.156315859 |
| Arhgap32 | S | 1797 | 0.009809233 | 0.047665489 | -0.002065475 | -0.011874709 | -0.049730964 | 0.932142443 | 0.489182023 | 0.978899298 | 0.923503236 | 0.562652218 |
| Rictor | S | 1283 | -0.023562169 | 0.050575969 | 0.026926696 | 0.050488865 | -0.023649273 | 0.673159286 | 0.472589337 | 0.658430915 | 0.440325019 | 0.749020407 |
| Ppp1r18 | S | 412 | 0.05757236 | -0.024925308 | 0.002658054 | -0.054914305 | 0.027583363 | 0.107636869 | 0.313816295 | 0.915934018 | 0.149236313 | 0.389880142 |
| Rbm27 | S | 120 | 0.035453069 | -0.028484403 | 0.030414749 | -0.00503832 | 0.058899152 | 0.188673761 | 0.142120842 | 0.283634146 | 0.855038259 | 0.088092343 |
| Arhgdia | S | 47 | 0.075932115 | 0.037360003 | -0.057895935 | -0.13382805 | -0.095255938 | 0.237223211 | 0.444517931 | 0.208249992 | 0.071851766 | 0.045690193 |
| Carmil1 | S | 1289 | 0.090245869 | 0.109798354 | -0.1494096 | -0.239655468 | -0.259207953 | 0.175313199 | 0.048175679 | 0.018812725 | 0.01195416 | 0.002092581 |
| Sh3pxd2a | S | 812 | -0.016636627 | 0.191199904 | -0.128286628 | -0.111650001 | -0.319486532 | 0.885768343 | 0.140994793 | 0.355611177 | 0.2494586 | 0.039440941 |
| Zfp292 | S | 654 | 0.042702228 | -0.023312542 | 0.034583308 | -0.00811892 | 0.05789585 | 0.473275771 | 0.744640098 | 0.718126248 | 0.928195346 | 0.562516235 |
| Hdac2 | S | 424 | 0.008718078 | -0.000302727 | 0.014976963 | 0.006258885 | 0.015279691 | 0.611840009 | 0.976916212 | 0.270422798 | 0.703813929 | 0.187822919 |
| Zbtb20 | S | 431 | -0.027487319 | -0.086516676 | 0.162469203 | 0.189956522 | 0.24898588 | 0.828981837 | 0.390949001 | 0.145671586 | 0.176633972 | 0.042171022 |
| Eif4b | S | 488 | -0.058462016 | 0.066532443 | 0.033717416 | 0.092179433 | -0.032815027 | 0.561466024 | 0.338788194 | 0.66325962 | 0.34806799 | 0.554528023 |
| Trim24 | S | 37 | -0.141699089 | -0.001470779 | 0.205867522 | 0.347566611 | 0.207338301 | 0.231410363 | 0.979017668 | 0.118131873 | 0.044421489 | 0.13098055 |
| Coro1b | S | 419 | 0.001917554 | 0.06054241 | -0.014454471 | -0.016372025 | -0.074996881 | 0.974543711 | 0.342705314 | 0.782362517 | 0.764346837 | 0.227406917 |
| Ncor2 | T | 549 | -0.022035019 | 0.015967157 | 0.048239497 | 0.070274516 | 0.03227234 | 0.53479728 | 0.645825111 | 0.315362701 | 0.145763035 | 0.431795016 |
| Abca1 | S | 1296 | 0.069881749 | -0.007916686 | -0.008116219 | -0.077997968 | -0.000199533 | 0.301712005 | 0.705253447 | 0.861740183 | 0.295836758 | 0.996494421 |
| Maoa | S | 388 | -0.027756704 | 0.036476616 | 0.01979055 | 0.047547254 | -0.016686066 | 0.7097633 | 0.326127104 | 0.619551717 | 0.515947546 | 0.475342542 |
| Acsl4 | T | 679 | -0.013568552 | -0.04381317 | 0.073910094 | 0.087478646 | 0.117723265 | 0.871897623 | 0.584131718 | 0.116648262 | 0.347920484 | 0.205950177 |
| Srrm1 | S | 591 | 0.007114474 | 0.00790686 | 0.02141125 | 0.014296776 | 0.013504391 | 0.701750646 | 0.842602519 | 0.542002659 | 0.665496246 | 0.770631025 |
| Srrm1 | T | 593 | 0.007114474 | 0.00790686 | 0.02141125 | 0.014296776 | 0.013504391 | 0.701750646 | 0.842602519 | 0.542002659 | 0.665496246 | 0.770631025 |
| Tomm40 | S | 169 | -0.041989049 | 0.029143603 | 0.058686653 | 0.100675703 | 0.02954305 | 0.528213006 | 0.582476318 | 0.376311256 | 0.193273285 | 0.613886933 |
| Trim2 | S | 456 | -0.025057851 | 0.070216404 | 0.007651456 | 0.032709307 | -0.062564948 | 0.600116789 | 0.314795062 | 0.894603588 | 0.438929604 | 0.338251089 |
| Top2a | S | 1521 | 0.156526189 | 0.041379266 | -0.1329226 | -0.289448789 | -0.174301867 | 0.146456156 | 0.303980496 | 0.087211277 | 0.034240854 | 0.048385752 |
| Tmcc3 | S | 4 | 0.076555374 | -0.00323448 | -0.01618279 | -0.092738164 | -0.01294831 | 0.355696818 | 0.949199038 | 0.753687946 | 0.306251704 | 0.842716803 |
| Rps6kc1 | S | 634 | 0.047826894 | 0.001093858 | -0.005615195 | -0.053442089 | -0.006709053 | 0.38177023 | 0.97512188 | 0.932718254 | 0.504805687 | 0.924955156 |
| Git1 | S | 422 | -0.031785564 | 0.094603975 | -0.062501981 | -0.030716417 | -0.157105956 | 0.38531963 | 0.021368925 | 0.432074053 | 0.690290715 | 0.120779552 |
| Gpsm1 | S | 653 | 0.042684884 | 0.051064727 | -0.062098356 | -0.10478324 | -0.113163084 | 0.570800218 | 0.175876739 | 0.35696628 | 0.252064195 | 0.146338766 |
| Stmn1 | S | 16 | -0.00739425 | 0.106605926 | -0.034318427 | -0.026924177 | -0.140924353 | 0.976853751 | 0.415619976 | 0.77614611 | 0.918607282 | 0.360661784 |
| Pds5b | T | 1368 | 0.093353033 | -0.029313318 | -0.042174355 | -0.135527388 | -0.012861037 | 0.009815932 | 0.563391243 | 0.077566479 | 0.006527092 | 0.785731308 |
| Nup93 | T | 236 | 0.06682447 | -0.001582502 | 0.017816477 | -0.049007994 | 0.019398978 | 0.716206919 | 0.986335824 | 0.848918244 | 0.79078189 | 0.844711376 |
| Polr2a | T | 1912 | 0.09743358 | 0.066716881 | -0.100962215 | -0.198395795 | -0.167679095 | 0.1568009 | 0.229165362 | 0.055521947 | 0.028671577 | 0.020089869 |
| Sf3b1 | S | 400 | 0.084316683 | -0.035369388 | 0.008295587 | -0.076021096 | 0.043664975 | 0.182485099 | 0.285752024 | 0.898589591 | 0.340785481 | 0.514761875 |
| Ppig | S | 693 | 0.057513963 | 0.035069901 | -0.032677323 | -0.090191286 | -0.067747224 | 0.576192205 | 0.352727166 | 0.406604397 | 0.39762647 | 0.037570106 |
| Ttc7b | S | 677 | 0.042846127 | 0.048569472 | -0.053174555 | -0.096020681 | -0.101744026 | 0.779634081 | 0.37674674 | 0.748760641 | 0.641537366 | 0.54472172 |
| Srsf10 | S | 133 | -0.003314085 | -0.064091977 | 0.155924679 | 0.159238763 | 0.220016656 | 0.965698143 | 0.381155538 | 0.263667316 | 0.24570258 | 0.154989655 |

| | | | | | | | | | | | | |
|---|---|---|---|---|---|---|---|---|---|---|---|---|
| Tra2a | S | 94 | -0.014787168 | -0.042834924 | 0.111107667 | 0.125894834 | 0.153942591 | 0.752720886 | 0.330814416 | 0.126902268 | 0.106606783 | 0.063629592 |
| Itgb1bp1 | S | 41 | -0.013758704 | 0.043656083 | -0.008158164 | 0.00560054 | -0.051814248 | 0.643086547 | 0.106919501 | 0.832446304 | 0.882886203 | 0.21586884 |
| Rbm17 | S | 155 | 0.012658921 | -0.017009794 | 0.029729802 | 0.017070881 | 0.046739597 | 0.810972557 | 0.521030168 | 0.123200416 | 0.742172611 | 0.12247941 |
| Bcl7b | S | 120 | 0.189253142 | -0.020964185 | -0.108704801 | -0.297957943 | -0.087740616 | 0.016984594 | 0.742625057 | 0.034688138 | 0.005286655 | 0.225041641 |
| R3hcc1l | S | 337 | -0.036372297 | 0.071647086 | 0.010960215 | 0.047332512 | -0.060686871 | 0.818780099 | 0.412400379 | 0.931446549 | 0.794956256 | 0.647586564 |
| R3hcc1l | S | 339 | -0.036372297 | 0.071647086 | 0.010960215 | 0.047332512 | -0.060686871 | 0.818780099 | 0.412400379 | 0.931446549 | 0.794956256 | 0.647586564 |
| R3hcc1l | S | 340 | -0.036372297 | 0.071647086 | 0.010960215 | 0.047332512 | -0.060686871 | 0.818780099 | 0.412400379 | 0.931446549 | 0.794956256 | 0.647586564 |
| Tmf1 | S | 340 | -0.026980222 | 0.007805286 | 0.07659117 | 0.103571392 | 0.068785884 | 0.075789881 | 0.718102639 | 0.11493206 | 0.07369948 | 0.140441225 |
| Phf10 | S | 55 | -0.068010181 | 0.010743874 | 0.097188826 | 0.165199007 | 0.086444953 | 0.239242845 | 0.86095683 | 0.107397028 | 0.023552911 | 0.183119965 |
| Shroom4 | S | 1243 | -0.000163669 | 0.059310204 | -0.027984718 | -0.027821049 | -0.087294922 | 0.998499427 | 0.313291065 | 0.847717705 | 0.859525985 | 0.566442644 |
| Serpinh1 | S | 90 | 0.075933258 | 0.051101818 | -0.065919057 | -0.141852316 | -0.117020875 | 0.48785632 | 0.590595556 | 0.532618381 | 0.23752651 | 0.268337474 |
| Brca1 | S | 1127 | 0.008325763 | 0.053894836 | -5.71545E-05 | -0.008382917 | -0.053951991 | 0.895914558 | 0.608923896 | 0.999436207 | 0.893797072 | 0.605170652 |
| Cxxc1 | S | 138 | 0.007271059 | -0.037000338 | 0.109749641 | 0.102478582 | 0.14674998 | 0.937945451 | 0.724641934 | 0.488824971 | 0.51431339 | 0.380136438 |
| Utp14a | T | 565 | 0.193602312 | -0.058061955 | -0.009123336 | -0.202725648 | 0.048938619 | 0.494769442 | 0.657622909 | 0.955258314 | 0.467584662 | 0.664681897 |
| G3bp2 | S | 141 | -0.033331902 | 0.020117359 | 0.058748475 | 0.092080377 | 0.038631116 | 0.540781572 | 0.670809082 | 0.357141353 | 0.159443181 | 0.467134906 |
| Arhgap23 | S | 617 | -0.053003819 | 0.108726049 | -0.022873109 | 0.03013071 | -0.131599158 | 0.395229768 | 0.111644188 | 0.786232877 | 0.6636129 | 0.142342712 |
| Ccm2 | S | 181 | -0.03417427 | 0.044794105 | 0.049395596 | 0.083569866 | 0.004601492 | 0.679593351 | 0.533885157 | 0.674507952 | 0.530519736 | 0.970311492 |
| Nes | S | 1418 | -0.047265765 | 0.003627771 | 0.072763758 | 0.120029522 | 0.069135986 | 0.46571673 | 0.969135657 | 0.150630243 | 0.103510879 | 0.472028381 |
| Tepsin | S | 400 | -0.014212782 | 0.117173461 | -0.060981955 | -0.046769173 | -0.178155416 | 0.866026164 | 0.107027031 | 0.28996077 | 0.61162774 | 0.047370546 |
| Nsun2 | S | 723 | -0.104507681 | 0.03415772 | 0.084786946 | 0.189294628 | 0.050629226 | 0.090148612 | 0.28114771 | 0.062414298 | 0.019566587 | 0.210215495 |
| Eif4g1 | S | 1147 | -0.081880959 | 0.047996715 | 0.049043256 | 0.130924215 | 0.001046541 | 0.129799929 | 0.174121967 | 0.139466246 | 0.046255733 | 0.973571294 |
| Synpo | S | 258 | -0.190837218 | 0.238455471 | 0.085553207 | 0.276390425 | -0.152902264 | 0.059265263 | 0.120195263 | 0.582153771 | 0.14124661 | 0.391620621 |
| Zhx1 | S | 26 | 0.032691952 | -0.050286354 | 0.099681519 | 0.066989567 | 0.149967873 | 0.814697471 | 0.685152442 | 0.540881807 | 0.726046941 | 0.425893211 |
| Zfhx3 | S | 2237 | -0.034761241 | -0.068028914 | 0.176252123 | 0.211013364 | 0.244281037 | 0.685344175 | 0.133153891 | 0.074694018 | 0.083733462 | 0.051679517 |
| Sipa1l3 | S | 166 | 0.043053037 | 0.010902031 | -0.022044652 | -0.06509769 | -0.032946683 | 0.268573746 | 0.737117903 | 0.608674009 | 0.206250667 | 0.464645684 |
| Cttn | T | 401 | -0.210287154 | 0.080757031 | 0.16701309 | 0.377300244 | 0.086256059 | 0.005372404 | 0.061045443 | 0.030611546 | 0.001813605 | 0.142306262 |
| Mybl2 | T | 443 | 0.03324014 | 0.018661657 | 0.014155871 | -0.019084269 | -0.004505787 | 0.787484764 | 0.83661171 | 0.877696431 | 0.88319029 | 0.964520186 |
| Srrm2 | S | 865 | 0.067978945 | -0.019004996 | -0.034984127 | -0.102963072 | -0.015979131 | 0.085173176 | 0.677794895 | 0.601144306 | 0.186023912 | 0.806860357 |
| Top1 | S | 114 | 0.080111856 | -0.020654347 | -0.034693142 | -0.114804998 | -0.014038795 | 0.02118644 | 0.599976912 | 0.134434184 | 0.009026371 | 0.713717379 |
| Fam53a | S | 265 | 0.005913943 | 0.028546988 | -0.008280278 | -0.014194221 | -0.036827266 | 0.874418339 | 0.179565281 | 0.642783224 | 0.688752646 | 0.049491991 |
| Cdc27 | T | 369 | -0.121789805 | 0.144156323 | 0.051388172 | 0.173177977 | -0.092768151 | 0.341032458 | 0.339817766 | 0.620590871 | 0.237221246 | 0.54571906 |
| Tmem135 | S | 221 | 0.006705647 | -0.028832866 | 0.075404917 | 0.06869927 | 0.104237783 | 0.928570037 | 0.744009465 | 0.411334941 | 0.314055542 | 0.217610615 |
| Tmem135 | T | 216 | 0.006705647 | -0.028832866 | 0.075404917 | 0.06869927 | 0.104237783 | 0.928570037 | 0.744009465 | 0.411334941 | 0.314055542 | 0.217610615 |
| Tmem94 | S | 221 | -0.096460872 | -0.003511662 | 0.139628713 | 0.236089585 | 0.143140376 | 0.624369087 | 0.932944434 | 0.287530229 | 0.303805275 | 0.279748804 |
| Tmem94 | S | 225 | -0.096460872 | -0.003511662 | 0.139628713 | 0.236089585 | 0.143140376 | 0.624369087 | 0.932944434 | 0.287530229 | 0.303805275 | 0.279748804 |
| Pias1 | T | 508 | 0.085471166 | 0.062479442 | -0.15512923 | -0.240600396 | -0.217608672 | 0.270912823 | 0.627610183 | 0.523676382 | 0.355548197 | 0.410317298 |
| Srek1 | S | 486 | -0.025006579 | 0.006106146 | 0.063862642 | 0.088869221 | 0.057756496 | 0.429145522 | 0.872449961 | 0.234215305 | 0.135924394 | 0.311554115 |
| Hnrnpc | S | 115 | -0.084100826 | 0.03477965 | 0.119646128 | 0.203746954 | 0.084866478 | 0.50129768 | 0.820476183 | 0.368732451 | 0.235503393 | 0.64397672 |
| Map1a | T | 1717 | 0.053854291 | 0.014447638 | -0.042388488 | -0.096242779 | -0.056836127 | 0.172873531 | 0.843951977 | 0.506144729 | 0.201203621 | 0.531095567 |
| Stxbp5l | S | 1162 | -0.017546748 | -0.086237025 | 0.116900254 | 0.134447002 | 0.203137279 | 0.82309023 | 0.483240208 | 0.122607918 | 0.157851563 | 0.162979197 |
| Stxbp5l | S | 1167 | -0.017546748 | -0.086237025 | 0.116900254 | 0.134447002 | 0.203137279 | 0.82309023 | 0.483240208 | 0.122607918 | 0.157851563 | 0.162979197 |
| Stxbp5l | T | 1156 | -0.017546748 | -0.086237025 | 0.116900254 | 0.134447002 | 0.203137279 | 0.82309023 | 0.483240208 | 0.122607918 | 0.157851563 | 0.162979197 |
| Stxbp5l | T | 1161 | -0.017546748 | -0.086237025 | 0.116900254 | 0.134447002 | 0.203137279 | 0.82309023 | 0.483240208 | 0.122607918 | 0.157851563 | 0.162979197 |
| Ehbp1l1 | S | 191 | 0.067070486 | -0.00216501 | -0.015255982 | -0.082326468 | -0.013090972 | 0.339937324 | 0.96159906 | 0.699291412 | 0.274380867 | 0.803879439 |
| Med13l | S | 392 | -0.104039915 | 0.122809368 | -0.013007837 | 0.091032077 | -0.135817205 | 0.059088515 | 0.00798837 | 0.873444078 | 0.320282368 | 0.180788185 |
| Sipa1l1 | S | 1654 | 0.055204438 | -0.054478084 | 0.065187404 | 0.009982966 | 0.119665488 | 0.596084186 | 0.357092324 | 0.374941525 | 0.918841468 | 0.104504748 |
| Zc3h13 | S | 909 | -0.011632027 | 0.011689651 | 0.028437382 | 0.040069409 | 0.016747729 | 0.732549499 | 0.592825644 | 0.376059904 | 0.341967416 | 0.598808404 |
| Prrc2a | T | 608 | 0.105070832 | 0.025562849 | -0.090008054 | -0.195078886 | -0.115570903 | 0.343688093 | 0.740304436 | 0.53250042 | 0.229996907 | 0.410957008 |
| Safb | S | 626 | -0.105441618 | 0.093369489 | 0.08541251 | 0.190854128 | -0.00795698 | 0.080425988 | 0.411245322 | 0.137230733 | 0.001251215 | 0.936097913 |
| Srrm2 | T | 2056 | -0.068184149 | 0.030207002 | 0.051989209 | 0.120173357 | 0.021782206 | 0.622926051 | 0.604802769 | 0.330274524 | 0.415301057 | 0.727188095 |
| Lysmd3 | S | 55 | -0.022418025 | -0.024361826 | 0.081098594 | 0.103516619 | 0.10546042 | 0.796530471 | 0.482733359 | 0.156237035 | 0.302430925 | 0.092645205 |
| Zc3h14 | S | 671 | 0.059844517 | 0.020847512 | -0.051945796 | -0.111790314 | -0.072793309 | 0.100601445 | 0.812820942 | 0.291407065 | 0.070035323 | 0.452938541 |
| Slc4a8 | S | 1052 | 0.075119502 | 0.005396704 | -0.020559185 | -0.095678687 | -0.025955889 | 0.518640734 | 0.922319539 | 0.840181485 | 0.493612636 | 0.800089843 |
| Sfswap | S | 831 | 0.008970104 | -0.002051999 | 0.036627276 | 0.027657172 | 0.038679275 | 0.807997368 | 0.923221543 | 0.531706846 | 0.65105381 | 0.503640483 |
| Epb41l2 | S | 395 | -0.030503255 | 0.022790048 | 0.05693006 | 0.087433314 | 0.034140011 | 0.594229565 | 0.60456014 | 0.48947748 | 0.323427862 | 0.656270409 |
| Zfp638 | S | 1635 | 0.024639615 | 0.084087835 | -0.052217197 | -0.076856811 | -0.136305032 | 0.791586278 | 0.487338562 | 0.690582428 | 0.576606786 | 0.386404743 |
| Kif4 | S | 942 | -0.000922045 | 0.012904232 | 0.044802933 | 0.045724978 | 0.031898702 | 0.996199472 | 0.908348192 | 0.608509186 | 0.815816204 | 0.783577949 |
| Larp4b | S | 453 | 0.174022073 | -0.021744451 | -0.133063907 | -0.30708598 | -0.111319456 | 0.005651548 | 0.710818093 | 0.030575205 | 0.003231606 | 0.092448673 |
| Zfp609 | T | 618 | 0.059746151 | 0.101800582 | -0.111065291 | -0.170811442 | -0.212865873 | 0.723343993 | 0.34390474 | 0.475459661 | 0.415209298 | 0.231368801 |
| Arhgap28 | S | 640 | -0.12898263 | 0.161868113 | 0.04444689 | 0.17342952 | -0.117421222 | 0.097651216 | 0.104632752 | 0.705097492 | 0.20869352 | 0.376363814 |
| Arhgap35 | Y | 1105 | -0.003034721 | 0.062766822 | -0.022164974 | -0.019130252 | -0.084931796 | 0.960583023 | 0.321618631 | 0.776342546 | 0.787319118 | 0.275967878 |
| Rin1 | S | 340 | -0.052701639 | 0.137738533 | 0.001643223 | 0.054344862 | -0.13609531 | 0.269218664 | 0.24772868 | 0.983882552 | 0.510029466 | 0.285323911 |
| Trim24 | S | 772 | -0.034314309 | -0.004565919 | 0.122100857 | 0.156415166 | 0.126666776 | 0.723686654 | 0.975427786 | 0.486747746 | 0.411736343 | 0.549794901 |
| Yeats2 | S | 124 | -0.002466426 | -0.014936819 | 0.079972679 | 0.082439105 | 0.094909498 | 0.970910528 | 0.710880452 | 0.428602403 | 0.441923726 | 0.353533413 |
| Kdm2a | S | 718 | 0.01030474 | 0.012217001 | 0.00400331 | -0.00630143 | -0.00821369 | 0.646559049 | 0.523835001 | 0.862737559 | 0.807508811 | 0.725919464 |
| Ppig | S | 714 | -0.069499278 | 0.023157665 | 0.089275717 | 0.158774995 | 0.066118052 | 0.183171968 | 0.453578018 | 0.185568674 | 0.055022135 | 0.290267146 |
| Cep170 | S | 922 | 0.039687789 | -0.013206784 | -0.01017532 | -0.049863109 | 0.003031464 | 0.122107995 | 0.688082765 | 0.859132516 | 0.408550599 | 0.958364829 |
| Gen1 | S | 794 | 0.099629744 | 0.03062451 | -0.096066903 | -0.195696646 | -0.126691413 | 0.043493309 | 0.403860277 | 0.044742532 | 0.003317354 | 0.021225373 |
| Gen1 | S | 795 | 0.099629744 | 0.03062451 | -0.096066903 | -0.195696646 | -0.126691413 | 0.043493309 | 0.403860277 | 0.044742532 | 0.003317354 | 0.021225373 |
| Fmnl3 | T | 189 | -0.066116336 | 0.118368618 | -0.010231875 | 0.05588446 | -0.128600493 | 0.465171673 | 0.219062761 | 0.961316185 | 0.787676174 | 0.551345356 |
| Elf2 | S | 191 | -0.031556847 | 0.053816587 | -0.013412743 | 0.018144104 | -0.06722933 | 0.347241507 | 0.03518257 | 0.739161671 | 0.684636232 | 0.168706894 |
| Gpm6b | S | 320 | -0.020881605 | 0.300846984 | -0.077089467 | -0.056207861 | -0.377936451 | 0.66475669 | 0.184582986 | 0.07878239 | 0.320532315 | 0.126039065 |
| Plekhg3 | S | 1155 | 0.031375682 | -0.008945363 | 0.010548252 | -0.020827429 | 0.019493615 | 0.34871166 | 0.636428032 | 0.712759355 | 0.587475942 | 0.537283092 |
| Fam177a Fam177a2 | S | 65 | -0.024843781 | -0.019637875 | 0.092512454 | 0.117356235 | 0.112150329 | 0.560071588 | 0.565229166 | 0.171881289 | 0.108167575 | 0.129743936 |
| Fam177a Fam177a2 | T | 66 | -0.024843781 | -0.019637875 | 0.092512454 | 0.117356235 | 0.112150329 | 0.560071588 | 0.565229166 | 0.171881289 | 0.108167575 | 0.129743936 |
| Cdk12 | S | 264 | -0.049993104 | -0.030884104 | 0.109457382 | 0.159450486 | 0.140341486 | 0.366323493 | 0.667793837 | 0.090096683 | 0.052020515 | 0.134542305 |
| Kif20a | S | 542 | 0.033084908 | -0.039710355 | 0.030298372 | -0.002786536 | 0.070008727 | 0.181687697 | 0.417912414 | 0.592500343 | 0.957174369 | 0.305141361 |
| Dnttip2 | S | 261 | 0.097548039 | -0.025809809 | -0.025696367 | -0.123244407 | 0.000113442 | 0.06443397 | 0.322867905 | 0.328136301 | 0.039785012 | 0.996224803 |
| Aspm | S | 1071 | -0.018013979 | 0.060440329 | 0.004931487 | 0.022945466 | -0.055508843 | 0.834966286 | 0.472479751 | 0.91745932 | 0.777295696 | 0.482849977 |
| Sbno1 | S | 690 | 0.035520873 | 0.118010753 | -0.105838419 | -0.141359292 | -0.223849172 | 0.910421844 | 0.331180712 | 0.573977498 | 0.669181398 | 0.242732577 |
| Sbno1 | S | 691 | 0.035520873 | 0.118010753 | -0.105838419 | -0.141359292 | -0.223849172 | 0.910421844 | 0.331180712 | 0.573977498 | 0.669181398 | 0.242732577 |
| Micall2 | S | 150 | 0.023897578 | 0.017761486 | -0.018233077 | -0.042130655 | -0.035994563 | 0.310774067 | 0.495126236 | 0.561792256 | 0.225172838 | 0.305370924 |
| Arhgef10 | S | 1291 | -0.027530043 | 0.148085936 | -0.087132029 | -0.059601987 | -0.235217966 | 0.752606507 | 0.081255944 | 0.25505382 | 0.517853251 | 0.029844682 |
| Lrrfip1 | S | 346 | -0.114803832 | 0.108114025 | 0.123799842 | 0.238603674 | 0.015685818 | 0.000219577 | 0.041798201 | 0.070235322 | 0.022035425 | 0.762092749 |
| Atf7ip | S | 593 | -0.030117543 | 0.011989881 | 0.029287715 | 0.059405258 | 0.017297834 | 0.595297109 | 0.743374045 | 0.137288056 | 0.337120894 | 0.640170351 |
| Scnm1 | S | 218 | -0.046106816 | 0.064112085 | -0.01094899 | 0.035157826 | -0.075061076 | 0.62453943 | 0.135680774 | 0.89329595 | 0.747271408 | 0.377341058 |
| Huwe1 | T | 3927 | 0.016459674 | 0.001768614 | 0.005755659 | -0.010704015 | 0.003987045 | 0.413422582 | 0.923399259 | 0.730033971 | 0.546342409 | 0.803259935 |
| Cic | S | 1061 | -0.208237291 | -0.02156034 | 0.29444041 | 0.502677701 | 0.31600075 | 0.310594095 | 0.87435014 | 0.170659972 | 0.077714021 | 0.167043222 |
| Unc119 | S | 37 | -0.070035574 | 0.071601622 | 0.024036802 | 0.094072376 | -0.04756482 | 0.272215206 | 0.1320463 | 0.686421888 | 0.198686867 | 0.396619702 |
| Sun2 | T | 9 | -0.090060368 | 0.176496614 | -0.055626754 | 0.034433613 | -0.232123368 | 0.248352347 | 0.064520499 | 0.603437967 | 0.740597313 | 0.089819512 |
| Atrx | S | 940 | -0.056604111 | 0.011854157 | 0.07691312 | 0.133517232 | 0.065058964 | 0.052243911 | 0.580175845 | 0.037765509 | 0.008392964 | 0.068500872 |
| Rftn2 | S | 228 | 0.047172611 | -0.015200955 | -0.004527077 | -0.051699687 | 0.010673878 | 0.233746563 | 0.72631183 | 0.925393551 | 0.242840966 | 0.805730596 |
| Akt3 | T | 305 | 0.159206363 | -0.157670758 | 0.01898743 | -0.140218933 | 0.176658187 | 0.03548759 | 0.082159114 | 0.76357942 | 0.032603115 | 0.05888013 |
| Zfp106 | S | 953 | -0.024755961 | 0.023830881 | 0.046731552 | 0.071487512 | 0.02290067 | 0.80189805 | 0.855773169 | 0.477825033 | 0.465022883 | 0.857263509 |

| | | | | | | | | | | | | |
|---|---|---|---|---|---|---|---|---|---|---|---|---|
| Frs2 | S | 503 | 0.001380871 | 0.091990903 | -0.039649782 | -0.041030653 | -0.131640685 | 0.962741432 | 0.076964228 | 0.074489051 | 0.237698559 | 0.03265846 |
| Trp53i11 | S | 14 | 1.087003713 | 0.100666812 | -0.301302769 | -1.388306482 | -0.401969581 | 0.359419153 | 0.159851714 | 0.003290789 | 0.270784668 | 0.001715148 |
| Epb41l2 | S | 492 | -0.040422091 | 0.024140313 | 0.050821453 | 0.091243544 | 0.02668114 | 0.111507258 | 0.338630948 | 0.154951196 | 0.040897253 | 0.389254819 |
| Map1a | S | 1694 | -0.033714207 | 0.096306669 | -0.02544211 | 0.008272097 | -0.121748779 | 0.543011007 | 0.143564246 | 0.581098925 | 0.825276119 | 0.079210446 |
| Srrm2 | S | 2052 | -0.03766689 | 0.035992265 | 0.027130705 | 0.064797595 | -0.00886156 | 0.789333327 | 0.502472842 | 0.650566769 | 0.655935754 | 0.885685469 |
| Ighmbp2 | S | 800 | 0.00355546 | 0.015052745 | 0.043554213 | 0.039998754 | 0.028501468 | 0.967800286 | 0.843564318 | 0.759931106 | 0.774686175 | 0.831272027 |
| Apc | S | 1662 | 0.13105424 | -0.069954233 | 0.056321479 | -0.074732761 | 0.126275712 | 0.469473816 | 0.248058763 | 0.605837106 | 0.686040147 | 0.274665205 |
| Tfip11 | S | 96 | -0.011133487 | -0.025171691 | 0.065583604 | 0.076717091 | 0.090755295 | 0.810233105 | 0.496937787 | 0.152032786 | 0.127643077 | 0.047290535 |
| Tfip11 | S | 99 | -0.011133487 | -0.025171691 | 0.065583604 | 0.076717091 | 0.090755295 | 0.810233105 | 0.496937787 | 0.152032786 | 0.127643077 | 0.047290535 |
| Wdr12 | T | 221 | 0.019998971 | 0.046454509 | -0.0346619 | -0.054660871 | -0.081116409 | 0.522916515 | 0.196915656 | 0.253040162 | 0.106634552 | 0.051217043 |
| Prkd3 | S | 391 | -0.06560687 | 0.059122897 | 0.071923055 | 0.137529925 | 0.012800159 | 0.096439419 | 0.249718458 | 0.348551516 | 0.133626829 | 0.860100836 |
| Ncoa6 | S | 1485 | 0.070369519 | -0.011520293 | -0.034826744 | -0.105196263 | -0.023306451 | 0.199332678 | 0.815417391 | 0.668551542 | 0.230708733 | 0.754519318 |
| Eif2ak2 | S | 223 | 0.017229898 | -0.004930234 | 0.02271083 | 0.005480932 | 0.027641064 | 0.588298444 | 0.881068249 | 0.648163117 | 0.89429413 | 0.531674126 |
| Srrm1 Srsf1 Tra2a Tr | S | 282;306;225;275 | 0.043063578 | -0.062224019 | 0.063279426 | 0.020215849 | 0.125503445 | 0.486781558 | 0.410628152 | 0.436966886 | 0.81543296 | 0.225374381 |
| Srrm1 Srsf1 Tra2a Tr | S | 284;308;227;277 | 0.043063578 | -0.062224019 | 0.063279426 | 0.020215849 | 0.125503445 | 0.486781558 | 0.410628152 | 0.436966886 | 0.81543296 | 0.225374381 |
| Rtn4 | T | 156 | 0.06044273 | -0.061350762 | 0.038854431 | -0.021588299 | 0.100205193 | 0.293634167 | 0.355432226 | 0.563177195 | 0.632679125 | 0.126136438 |
| Cdca3 | T | 37 | 0.040785649 | -0.042081931 | 0.039227798 | -0.00155785 | 0.081309729 | 0.432785541 | 0.485782361 | 0.524823751 | 0.975768319 | 0.217615693 |
| Med1 | S | 770 | -0.04914163 | 0.031618451 | 0.041276703 | 0.090418334 | 0.009658253 | 0.481399719 | 0.387749083 | 0.432789364 | 0.2456537 | 0.821989286 |
| Lrrfip1 | S | 84 | -0.125876478 | 0.179024456 | -0.037649146 | 0.088227332 | -0.216673602 | 0.177894841 | 0.036731063 | 0.622128739 | 0.329441641 | 0.032679759 |
| Zeb2 | T | 188 | -0.118029069 | 0.165764692 | 0.055810247 | 0.173839316 | -0.109954446 | 0.430675066 | 0.386714455 | 0.871970313 | 0.632560255 | 0.765282584 |
| Prrc2a | S | 1068 | 0.07045239 | 0.021952522 | -0.040507297 | -0.110959688 | -0.06245982 | 0.352899489 | 0.785341751 | 0.469637598 | 0.19546425 | 0.475820761 |
| Hnrnpc | S | 268 | 0.01277027 | -0.053202409 | 0.091174715 | 0.078404445 | 0.144377124 | 0.768357266 | 0.274593128 | 0.217488675 | 0.253751206 | 0.086021608 |
| Map1s | S | 785 | 0.04874978 | 0.088746999 | -0.069531473 | -0.118281253 | -0.158278471 | 0.32146684 | 0.412956123 | 0.176877995 | 0.015262792 | 0.195174416 |
| Fam122b | S | 115 | 0.005823596 | 0.096388346 | -0.091860187 | -0.097683784 | -0.188248534 | 0.90257224 | 0.043006374 | 0.164638794 | 0.152296504 | 0.033429638 |
| Fam122b | S | 119 | 0.005823596 | 0.096388346 | -0.091860187 | -0.097683784 | -0.188248534 | 0.90257224 | 0.043006374 | 0.164638794 | 0.152296504 | 0.033429638 |
| Zbtb20 | S | 353 | -0.102765916 | 0.09540385 | 0.043529514 | 0.14629543 | -0.051874335 | 0.043761826 | 0.036996105 | 0.498267863 | 0.073409956 | 0.394358823 |
| Acd | S | 315 | 0.032656503 | -0.014182432 | 0.013745717 | -0.018910785 | 0.027928149 | 0.553340578 | 0.926208569 | 0.757576563 | 0.679539484 | 0.853634034 |
| Mark3 | S | 540 | -0.016509569 | 0.128931034 | -0.112203795 | -0.095694226 | -0.241134829 | 0.848043122 | 0.155798497 | 0.522855995 | 0.562058167 | 0.219904194 |
| Gab2 | S | 319 | -0.000200464 | -0.000606861 | 0.090472053 | 0.090672517 | 0.091078914 | 0.998943012 | 0.996719432 | 0.727872978 | 0.7133834 | 0.710396858 |
| Gab2 | T | 316 | -0.000200464 | -0.000606861 | 0.090472053 | 0.090672517 | 0.091078914 | 0.998943012 | 0.996719432 | 0.727872978 | 0.7133834 | 0.710396858 |
| Lig1 | S | 188 | -0.167510303 | 0.098986612 | 0.107689296 | 0.2751996 | 0.008702685 | 0.007691563 | 0.08757824 | 0.044240354 | 0.00010803 | 0.804608303 |
| Taf1 | S | 328 | -0.082222984 | 0.040506045 | 0.135353778 | 0.217576762 | 0.094847733 | 0.095825332 | 0.702456256 | 0.173760963 | 0.070187547 | 0.449423817 |
| Srsf2 | S | 221 | -0.006347108 | -0.039445375 | 0.092246752 | 0.09859386 | 0.131692127 | 0.915661729 | 0.643366305 | 0.325018017 | 0.300949667 | 0.242620719 |
| Mdc1 | T | 1113 | -0.009065083 | 0.076850338 | -0.029125544 | -0.020060461 | -0.105975882 | 0.702166016 | 0.210825974 | 0.594530216 | 0.716627443 | 0.167868104 |
| Rcan1 Rcan3 | S | 148;161 | -0.223712495 | 0.169607656 | 0.065987702 | 0.289700197 | -0.103619954 | 0.42224425 | 0.26832871 | 0.764809579 | 0.369200996 | 0.652877807 |
| Rab11fip5 | S | 296 | 0.196089328 | -0.101595752 | -0.085511149 | -0.281600476 | 0.016084603 | 0.032786825 | 0.317368561 | 0.322164447 | 0.01330855 | 0.859172577 |
| Washc2 | S | 343 | 0.062703503 | -0.012320121 | -0.024308688 | -0.087012191 | -0.011988567 | 0.271595288 | 0.849430558 | 0.788994652 | 0.383062598 | 0.902974878 |
| Lyst | S | 2203 | -0.023039472 | 0.106319988 | -0.043680462 | -0.02064099 | -0.150000451 | 0.738754114 | 0.127207561 | 0.330514449 | 0.748016147 | 0.059562605 |
| Map1a | T | 1472 | -0.042948076 | 0.101695833 | -0.03729242 | 0.005655656 | -0.138988253 | 0.636665364 | 0.177799516 | 0.70084592 | 0.95726489 | 0.193935191 |
| Raly | S | 248 | 0.024947134 | 0.034501278 | 0.000446753 | -0.024500381 | -0.034054526 | 0.86885408 | 0.729013062 | 0.997622646 | 0.887598142 | 0.803775762 |
| Zc3h14 | S | 409 | 0.028412963 | -0.012009197 | 0.032796748 | 0.004383785 | 0.044805944 | 0.563608222 | 0.619791317 | 0.682288416 | 0.95926609 | 0.584315489 |
| C77080 | S | 883 | -0.019259466 | 0.018596398 | 0.029064974 | 0.04832444 | 0.010468576 | 0.56986433 | 0.619910398 | 0.355225318 | 0.078335124 | 0.685533507 |
| Tcof1 | S | 1303 | -0.015279241 | 0.01957695 | 0.015964448 | 0.031243689 | -0.003612502 | 0.734593276 | 0.392164931 | 0.555801514 | 0.511812815 | 0.887175401 |
| Atxn2l | S | 27 | 0.047428225 | 0.00926058 | -0.0369261 | -0.084354325 | -0.04618668 | 0.104117421 | 0.688825075 | 0.260605591 | 0.042843452 | 0.163801085 |
| Sorbs2 | S | 82 | 0.020847548 | -0.039162076 | 0.042166311 | 0.021318763 | 0.081328387 | 0.260683387 | 0.481898365 | 0.523774497 | 0.73465218 | 0.316126848 |
| Camsap2 | S | 992 | 0.100185187 | -0.003448439 | -0.057870046 | -0.158055233 | -0.054421607 | 0.032553826 | 0.904916165 | 0.05963237 | 0.011083396 | 0.087416174 |
| Pcm1 | S | 780 | 0.038180535 | 0.008899844 | -0.042742192 | -0.080922727 | -0.051642035 | 0.089559131 | 0.879496731 | 0.491944205 | 0.248419794 | 0.512068906 |
| Dennd2a | S | 327 | -0.000319006 | 0.084439778 | -0.05160135 | -0.051282345 | -0.136041128 | 0.996987823 | 0.214240563 | 0.471741096 | 0.553503417 | 0.088966263 |
| Ncor2 | S | 1749 | 0.061492121 | 0.023996612 | -0.09707367 | -0.158565791 | -0.121070283 | 0.058416422 | 0.291400417 | 0.15268368 | 0.056296796 | 0.106849584 |
| Mtcl1 | S | 255 | 0.038594176 | 0.011756368 | -0.018959703 | -0.05755388 | -0.030716072 | 0.303037486 | 0.873723439 | 0.641410462 | 0.106877827 | 0.671422626 |
| Ncor2 | S | 1220 | 0.079670623 | -0.029766801 | -0.059184638 | -0.138855261 | -0.029417838 | 0.008904273 | 0.584777362 | 0.168542729 | 0.026760375 | 0.615815563 |
| Sp100 | S | 248 | -0.011892798 | 0.011106057 | 0.05521403 | 0.067106828 | 0.044107973 | 0.896941028 | 0.901565049 | 0.616869896 | 0.430328764 | 0.583901015 |
| Zfp638 | S | 868 | -0.003630069 | 0.007626298 | 0.035552112 | 0.039182181 | 0.027925814 | 0.963164576 | 0.896489313 | 0.558367958 | 0.644802147 | 0.676351318 |
| Prpf38a | S | 193 | -0.015719253 | -0.069830789 | 0.142571453 | 0.158290706 | 0.212402242 | 0.796764126 | 0.265301414 | 0.144496995 | 0.114660838 | 0.061571901 |
| Prpf38a | S | 194 | -0.015719253 | -0.069830789 | 0.142571453 | 0.158290706 | 0.212402242 | 0.796764126 | 0.265301414 | 0.144496995 | 0.114660838 | 0.061571901 |
| Cenpc1 | S | 258 | 0.052156866 | 0.020609623 | -0.035727737 | -0.087884603 | -0.05633736 | 0.405298352 | 0.770485953 | 0.623446048 | 0.322797749 | 0.542483256 |
| Hdac4 | S | 465 | -0.056074518 | 0.105418654 | 0.005819843 | 0.061894361 | -0.099598812 | 0.344546242 | 0.256000269 | 0.891640829 | 0.322307518 | 0.280429472 |
| Pi4k2a | S | 47 | 0.010871131 | 0.02767284 | -0.015558791 | -0.026429921 | -0.04323163 | 0.718181084 | 0.315028253 | 0.673228762 | 0.437782551 | 0.225420185 |
| Enox2 | S | 360 | -0.113712757 | 0.077098869 | 0.067612363 | 0.18132512 | -0.009486506 | 0.374694527 | 0.42239701 | 0.440296666 | 0.2067107 | 0.92533346 |
| Rbmx2 | S | 297 | 0.122283961 | -0.185316956 | 0.173613027 | 0.051329066 | 0.358929983 | 0.232358362 | 0.084280734 | 0.374029899 | 0.77717428 | 0.126840321 |
| Rbmx2 | S | 299 | 0.122283961 | -0.185316956 | 0.173613027 | 0.051329066 | 0.358929983 | 0.232358362 | 0.084280734 | 0.374029899 | 0.77717428 | 0.126840321 |
| Pds5a | S | 1194 | 0.100526217 | 0.135940655 | -0.269279553 | -0.369805769 | -0.405220208 | 0.35504529 | 0.128284025 | 0.168574304 | 0.090917382 | 0.079781331 |
| Anapc2 | S | 489 | -0.184126269 | -0.001423821 | 0.213839801 | 0.39796607 | 0.215263621 | 0.179625277 | 0.994152797 | 0.086382339 | 0.030384786 | 0.323163024 |
| Abi2 | T | 297 | 0.042399457 | 0.099000564 | -0.068536433 | -0.11093589 | -0.167536997 | 0.476355225 | 0.227462425 | 0.094874076 | 0.115557222 | 0.095073799 |
| Tmf1 | S | 138 | 0.085207154 | 0.014618995 | -0.113768674 | -0.198975828 | -0.128387669 | 0.062904268 | 0.696368209 | 0.243299654 | 0.101640393 | 0.210201065 |
| BC005624 | S | 261 | -0.061692957 | 0.059253432 | 0.015285315 | 0.076978272 | -0.043968117 | 0.244705487 | 0.116058538 | 0.556949648 | 0.171718899 | 0.235931638 |
| Ctdp1 | S | 459 | 0.064958756 | 0.019366744 | -0.107852693 | -0.172811449 | -0.127219437 | 0.041177577 | 0.540750359 | 0.214913415 | 0.10598567 | 0.167770672 |
| Cic | T | 672 | -0.024990515 | 0.050629336 | 0.004531762 | 0.029522277 | -0.046097575 | 0.695413771 | 0.338735356 | 0.929115146 | 0.632684849 | 0.361379052 |
| Pcyt1a | S | 350 | 0.051338952 | 0.069793669 | -0.07563182 | -0.126970772 | -0.145425489 | 0.204444324 | 0.304927538 | 0.100325938 | 0.033668028 | 0.085318083 |
| Mrpl55 | S | 84 | -0.02147531 | -0.101407662 | 0.14158114 | 0.163056451 | 0.242988802 | 0.845433152 | 0.515313024 | 0.20223651 | 0.081387433 | 0.151018169 |
| Acin1 | S | 389 | -0.079239442 | 0.034083122 | 0.122071818 | 0.201311261 | 0.087988696 | 0.018219888 | 0.347168546 | 0.100329647 | 0.041153574 | 0.192303708 |
| Rcor1 | S | 254 | -0.098262793 | -0.025529864 | 0.150381949 | 0.248644742 | 0.175911813 | 0.086580978 | 0.649994288 | 0.023393416 | 0.003549047 | 0.009241804 |
| Abi2 | S | 221 | 0.070063475 | 0.014848653 | -0.046024843 | -0.116088317 | -0.060873495 | 0.135429405 | 0.601269667 | 0.132139806 | 0.046437785 | 0.029057473 |
| Ccar2 | S | 612 | -0.051430656 | 0.036882065 | 0.078383437 | 0.129814093 | 0.041501373 | 0.070349882 | 0.252653559 | 0.317502308 | 0.152766676 | 0.568810853 |
| Cdc42bpa | S | 1708 | -0.04414605 | 0.058469867 | 0.038830861 | 0.082976911 | -0.019639006 | 0.576462921 | 0.584606759 | 0.629533564 | 0.395734558 | 0.864188235 |
| Micall2 | S | 832 | 0.144314756 | -0.006470432 | -0.084158218 | -0.228472974 | -0.077687786 | 0.150140496 | 0.860470615 | 0.259931396 | 0.057932093 | 0.296861397 |
| Brca1 | S | 686 | 0.0683358 | -1.53626E-05 | -0.013405132 | -0.081740931 | -0.013389769 | 0.443873448 | 0.999724084 | 0.617544601 | 0.375174256 | 0.77580137 |
| Arhgap10 | S | 589 | -0.041385337 | -0.018812789 | 0.079211451 | 0.120596788 | 0.09802424 | 0.59230786 | 0.8494033 | 0.198696573 | 0.135707697 | 0.3267212 |
| Zc2hc1a | T | 242 | 0.051399925 | 0.056749295 | -0.061009413 | -0.112409338 | -0.117758708 | 0.429953824 | 0.267799663 | 0.173465344 | 0.130308522 | 0.046233183 |
| Srrm2 | S | 1360 | -0.002351325 | -0.022081259 | 0.070653159 | 0.073004484 | 0.092734417 | 0.945045055 | 0.50109196 | 0.305358218 | 0.284800388 | 0.20623412 |
| Lyst | S | 113 | 0.077352008 | 0.009291611 | -0.030490752 | -0.10784276 | -0.039782363 | 0.552550465 | 0.91106983 | 0.767703147 | 0.46036128 | 0.712628531 |
| Parp14 | S | 1407 | -0.122460347 | 0.027505852 | 0.139784549 | 0.262244896 | 0.112278696 | 0.208404119 | 0.784023436 | 0.155298201 | 0.002897729 | 0.143292511 |
| Ranbp10 | S | 490 | 0.031614825 | -0.167768673 | 0.178527215 | 0.14691239 | 0.346295888 | 0.856169009 | 0.14339315 | 0.090963956 | 0.399476375 | 0.018513044 |
| Ptms | S | 100 | -0.07356951 | 0.185509158 | -0.100813543 | -0.027244033 | -0.286322701 | 0.679100029 | 0.093938097 | 0.317112197 | 0.878070004 | 0.040173156 |
| Shmt2 | S | 50 | 0.019262765 | -0.070224515 | 0.110979675 | 0.091716909 | 0.181204189 | 0.783723853 | 0.406959911 | 0.362589379 | 0.466331048 | 0.201377682 |
| Stmn1 | S | 63 | -0.160895972 | 0.21769647 | -0.043023113 | 0.117872859 | -0.260719583 | 0.200248212 | 0.048658352 | 0.593646623 | 0.334149787 | 0.039293297 |
| Cdc42ep1 | S | 207 | -0.118704265 | 0.074255166 | 0.074422414 | 0.19312668 | 0.000167249 | 0.021004827 | 0.018470391 | 0.335201258 | 0.06475913 | 0.998036346 |
| Mki67 | T | 1335 | 0.077242344 | -0.019291718 | -0.042232827 | -0.119475171 | -0.022941109 | 0.082398428 | 0.715139753 | 0.365352925 | 0.027182549 | 0.642312273 |
| Srrm2 | S | 850 | -0.080641444 | 0.040471584 | 0.089882652 | 0.170524096 | 0.049411068 | 0.000137721 | 0.126122903 | 0.00371729 | 0.001986966 | 0.08890256 |
| Prrc2b | T | 823 | 0.25172968 | -0.139129808 | -0.117032813 | -0.368762493 | 0.022096995 | 0.075388657 | 0.518845058 | 0.345253874 | 0.00619421 | 0.906526891 |
| Srrm2 | S | 886 | 0.005133207 | -0.037597773 | 0.0545421 | 0.049408893 | 0.092139873 | 0.883207246 | 0.272154115 | 0.109268757 | 0.195489597 | 0.034418862 |
| Sf3b1 | T | 211 | 0.022444903 | 0.001493906 | -0.000104366 | -0.022549268 | -0.001598272 | 0.39108294 | 0.969682984 | 0.99640371 | 0.426936074 | 0.968486472 |
| Srrm2 | T | 841 | 0.087665301 | -0.086798927 | 0.033162725 | -0.054502576 | 0.119961651 | 0.138314995 | 0.17122169 | 0.619449323 | 0.367556624 | 0.10551144 |
| Tgoln1 Tgoln2 | T | 208;218 | 0.060712421 | -0.031182212 | 0.006329229 | -0.054383192 | 0.03751144 | 0.384565085 | 0.675628793 | 0.93721991 | 0.467740925 | 0.632931461 |

| | | | | | | | | | | | | |
|---|---|---|---|---|---|---|---|---|---|---|---|---|
| Nfrkb | S | 178 | -0.205873758 | 0.198782542 | -0.013260418 | 0.19261334 | -0.212042959 | 0.149679642 | 0.034031608 | 0.864239313 | 0.17526536 | 0.048919893 |
| H1f0 | T | 153 | 0.217747629 | 0.023938652 | -0.208675255 | -0.426422884 | -0.232613907 | 0.002521811 | 0.562473074 | 0.002766732 | 0.000480903 | 0.004070467 |
| Zzz3 | S | 138 | -0.189249594 | -0.01810368 | 0.234122418 | 0.423372012 | 0.252226098 | 0.05352904 | 0.649646326 | 0.016161015 | 0.004709703 | 0.016961294 |
| Mecom | S | 64 | -0.095312045 | 0.044684386 | 0.09014386 | 0.185455905 | 0.045459474 | 0.17764202 | 0.50258981 | 0.200998443 | 0.028692365 | 0.456530059 |
| Dnajc8 | S | 35 | -0.005015931 | 0.078148701 | -0.042039035 | -0.037023104 | -0.120187736 | 0.919497154 | 0.22024947 | 0.488388405 | 0.566128729 | 0.134632983 |
| Tmpo | S | 306 | 0.005907445 | 0.06127564 | -0.03465006 | -0.040557505 | -0.095925699 | 0.963432116 | 0.393023556 | 0.72654649 | 0.779122744 | 0.369539091 |
| Hmgn1 | S | 7 | -0.009848095 | 0.027729646 | 0.008309823 | 0.018157918 | -0.019419823 | 0.791558961 | 0.413597123 | 0.823891708 | 0.610431207 | 0.539230165 |
| Pank1 | S | 11 | 0.032663476 | 0.128047931 | -0.13639922 | -0.169062696 | -0.264447151 | 0.721430698 | 0.20915811 | 0.305050075 | 0.184204712 | 0.076839577 |
| Ptov1 | S | 53 | -0.079791776 | 0.107989937 | 0.0114866 | 0.091278376 | -0.096503337 | 0.199891494 | 0.129569614 | 0.889792351 | 0.345770325 | 0.328477046 |
| Ankrd17 | S | 15 | 0.032743506 | 0.020798071 | -0.020222239 | -0.052965745 | -0.041020309 | 0.504689943 | 0.58088101 | 0.561604343 | 0.320240939 | 0.332612514 |
| Mef2d | S | 464 | -0.015380472 | -0.02501992 | 0.0866944 | 0.102074872 | 0.11171432 | 0.832444857 | 0.6360123 | 0.336498271 | 0.334132789 | 0.254612043 |
| Mef2d | S | 465 | -0.015380472 | -0.02501992 | 0.0866944 | 0.102074872 | 0.11171432 | 0.832444857 | 0.6360123 | 0.336498271 | 0.334132789 | 0.254612043 |
| Srrm2 | S | 1798 | -0.009242323 | 0.015516705 | 0.013233556 | 0.022475879 | -0.00228315 | 0.703308974 | 0.439448746 | 0.530442263 | 0.296685602 | 0.862801564 |
| Rars | S | 370 | -0.02711762 | 0.156552765 | -0.051591057 | -0.024473437 | -0.208143822 | 0.901429645 | 0.557370834 | 0.81677585 | 0.92337157 | 0.485046381 |
| Kdm2a | T | 713 | 0.022022483 | 0.041069811 | -0.040692786 | -0.062715268 | -0.081762596 | 0.430349114 | 0.152594308 | 0.173424943 | 0.033183065 | 0.012647894 |
| Dzip1l | S | 731 | 0.122910887 | -0.075089147 | -0.073227743 | -0.196138631 | 0.001861403 | 0.128621851 | 0.616163596 | 0.691564045 | 0.327419302 | 0.992829869 |
| Lats2 | S | 156 | -0.032681098 | 0.058415549 | -0.013704835 | 0.018976263 | -0.072120384 | 0.634859121 | 0.178087443 | 0.839847761 | 0.827952781 | 0.35021128 |
| Hirip3 | S | 385 | -0.046111026 | 0.019694005 | 0.045081358 | 0.091192384 | 0.025387353 | 0.372027282 | 0.72158158 | 0.176355385 | 0.130669847 | 0.647764901 |
| Git1 | S | 426 | -0.043489574 | 0.090234126 | -0.033124726 | 0.010364848 | -0.123358853 | 0.412511991 | 0.081985072 | 0.585471676 | 0.868565339 | 0.097403681 |
| Pcm1 | S | 957 | 0.024690515 | -0.031357171 | 0.033301795 | 0.00861128 | 0.064658966 | 0.449290995 | 0.340162151 | 0.34209207 | 0.613740906 | 0.03124874 |
| Igf2bp2 | S | 178 | 0.159584736 | -0.108172878 | 0.052735366 | -0.10684937 | 0.160908245 | 0.263612238 | 0.199182804 | 0.82129607 | 0.672814511 | 0.517053568 |
| Srrt | S | 319 | -0.122784285 | 0.03248042 | 0.182614275 | 0.30539856 | 0.150133855 | 0.007985734 | 0.420986996 | 0.046791193 | 0.017239097 | 0.080083593 |
| Parva | S | 14 | -0.072847959 | 0.105712698 | 0.000661475 | 0.073509434 | -0.105051223 | 0.169054747 | 0.091712537 | 0.991762358 | 0.259708962 | 0.152638243 |
| Dock7 | S | 180 | 0.000230906 | -0.00021508 | 0.022335799 | 0.022104893 | 0.022550879 | 0.995389481 | 0.991225292 | 0.388760811 | 0.608569395 | 0.394696717 |
| Mcm9 | S | 1079 | 0.005148539 | 0.091905629 | -0.039737952 | -0.044886491 | -0.131643581 | 0.975095449 | 0.33276748 | 0.424407077 | 0.786860488 | 0.199782824 |
| Sh3bp5l | S | 30 | -0.076549776 | 0.056919882 | 0.047893105 | 0.124442881 | -0.009026777 | 0.267421677 | 0.440831703 | 0.281161845 | 0.108811817 | 0.891361214 |
| Prkd1 | S | 407 | -0.062527706 | 0.032638678 | 0.070160653 | 0.132688359 | 0.037521976 | 0.223200804 | 0.700881897 | 0.186865489 | 0.020832168 | 0.644125618 |
| Prkd3 | S | 395 | -0.062527706 | 0.032638678 | 0.070160653 | 0.132688359 | 0.037521976 | 0.223200804 | 0.700881897 | 0.186865489 | 0.020832168 | 0.644125618 |
| Faf2 | S | 355 | -0.000468341 | -0.014057158 | 0.029453684 | 0.029922024 | 0.043510842 | 0.995043162 | 0.813542772 | 0.257975092 | 0.699453039 | 0.493233776 |
| Ptpn14 | T | 579 | -0.027606524 | 0.093471079 | -0.050714877 | -0.023108353 | -0.144185956 | 0.414530178 | 0.120560585 | 0.661869114 | 0.836735731 | 0.277449956 |
| Yeats2 | S | 625 | -0.006268791 | -0.05007029 | 0.095958151 | 0.102226942 | 0.146028441 | 0.888010786 | 0.281803358 | 0.155071314 | 0.13976301 | 0.062153078 |
| Huwe1 | S | 3818 | -0.005754379 | 0.021076575 | 0.027727577 | 0.033481957 | 0.006651002 | 0.932949462 | 0.852883463 | 0.668215956 | 0.688065306 | 0.956095129 |
| Numbl | S | 224 | -0.139493388 | 0.117487885 | 0.066168761 | 0.205662149 | -0.051319124 | 0.078732073 | 0.121944596 | 0.413084486 | 0.024714999 | 0.367242622 |
| Numbl | S | 228 | -0.139493388 | 0.117487885 | 0.066168761 | 0.205662149 | -0.051319124 | 0.078732073 | 0.121944596 | 0.413084486 | 0.024714999 | 0.367242622 |
| Zc3h4 | S | 1114 | 0.04673268 | -0.073968496 | 0.078616007 | 0.031883328 | 0.152584504 | 0.428226633 | 0.023597771 | 0.060003618 | 0.584177197 | 0.010614667 |
| Tap1 | S | 464 | -0.054561256 | 0.000440879 | 0.082032299 | 0.136593555 | 0.08159142 | 0.321125652 | 0.985747525 | 0.157588681 | 0.074441023 | 0.168152209 |
| Klf16 | S | 237 | 0.100745664 | -0.074631969 | 0.023641742 | -0.077103922 | 0.098273711 | 0.409372378 | 0.528646549 | 0.854220834 | 0.630851286 | 0.545508023 |
| Kif4 | S | 395 | 0.095166268 | -0.024679519 | -0.06611948 | -0.161285748 | -0.041439961 | 0.035510292 | 0.47072525 | 0.29307133 | 0.052697254 | 0.458301053 |
| Ap3d1 | S | 685 | -0.030515705 | -0.069939852 | 0.13320205 | 0.163717755 | 0.203141903 | 0.538048156 | 0.245337189 | 0.071212319 | 0.043769397 | 0.025765058 |
| Usp20 | S | 342 | 0.025572429 | 0.081390198 | -0.052776522 | -0.078348951 | -0.13416672 | 0.496754128 | 0.382058616 | 0.24730708 | 0.089848984 | 0.196237792 |
| Brd4 | S | 1073 | -0.058329757 | 0.03564285 | 0.054240912 | 0.112570669 | 0.018598062 | 0.404095388 | 0.614421398 | 0.34521301 | 0.189497222 | 0.81434223 |
| Phf10 | S | 52 | -0.018396503 | -0.05740417 | 0.089427277 | 0.107823781 | 0.146831447 | 0.668865335 | 0.333292026 | 0.077421306 | 0.002176999 | 0.049166943 |
| Pkd2 | S | 827 | 0.084343467 | 0.070409935 | -0.091058323 | -0.17540179 | -0.161468258 | 0.076917883 | 0.385240848 | 0.023860106 | 0.019679721 | 0.122334679 |
| Tra2a | S | 98 | -0.057212244 | -0.0233205 | 0.140919432 | 0.198131676 | 0.164239932 | 0.240882167 | 0.588022065 | 0.128581537 | 0.06079168 | 0.094281827 |
| Srrm1 | S | 561 | -0.019693543 | -0.022977077 | 0.07675511 | 0.096448653 | 0.099732187 | 0.518999946 | 0.387569916 | 0.122129143 | 0.077695282 | 0.072914923 |
| Srrm1 | S | 563 | -0.019693543 | -0.022977077 | 0.07675511 | 0.096448653 | 0.099732187 | 0.518999946 | 0.387569916 | 0.122129143 | 0.077695282 | 0.072914923 |
| Cavin2 | S | 359 | -0.024172067 | 0.069658707 | -0.005980596 | 0.018191471 | -0.075639304 | 0.073400307 | 0.044113819 | 0.395183055 | 0.145212827 | 0.042698126 |
| Adgra2 | T | 974 | -0.040495069 | 0.079012088 | 0.042467725 | 0.082962794 | -0.036544363 | 0.306076939 | 0.430521418 | 0.785170617 | 0.601058628 | 0.828851199 |
| Dmpk | S | 379 | -0.049188398 | 0.052288334 | 0.044565744 | 0.093754141 | -0.007722591 | 0.474149093 | 0.50663888 | 0.630200629 | 0.27954831 | 0.926829071 |
| Trip6 | S | 92 | -0.145542865 | 0.160698393 | 0.064092601 | 0.209635466 | -0.096605792 | 0.059503366 | 0.096938574 | 0.64138182 | 0.193003877 | 0.493384857 |
| Ccny | S | 73 | -0.042221448 | 0.184635966 | -0.096723182 | -0.054501733 | -0.281359147 | 0.46976685 | 0.056578265 | 0.176158467 | 0.258684077 | 0.017892123 |
| Cic | S | 1364 | 0.062167537 | 0.019593104 | -0.072670019 | -0.134837556 | -0.092263123 | 0.400384178 | 0.540900065 | 0.549461186 | 0.323601192 | 0.4558531 |
| Reps1 | S | 534 | -0.186803467 | 0.132860617 | 0.122771815 | 0.309575283 | -0.010088801 | 0.23958205 | 0.501180786 | 0.361843469 | 0.061248582 | 0.953549274 |
| Afap1 | T | 547 | 0.130196608 | -0.008392593 | -0.097739761 | -0.227936368 | -0.089347168 | 0.064613038 | 0.880115847 | 0.157394709 | 0.007418102 | 0.104875009 |
| Epb41l1 | S | 546 | 0.028693891 | -0.005331126 | 0.001672939 | -0.027020952 | 0.007004065 | 0.416918118 | 0.878440245 | 0.971015628 | 0.504036412 | 0.857527987 |
| Zc3h13 | S | 703 | 0.003226911 | 0.029139311 | -0.000697236 | -0.003924148 | -0.029836547 | 0.951588574 | 0.565414077 | 0.990254872 | 0.932817971 | 0.504163558 |
| Nfrkb | S | 176 | -0.158960706 | 0.08154435 | 0.021485661 | 0.180446366 | -0.060058689 | 0.342603734 | 0.098183195 | 0.534773193 | 0.295249633 | 0.214341404 |
| Trio | S | 1809 | 0.06417732 | 0.060986274 | -0.105756181 | -0.169934131 | -0.166743085 | 0.10926881 | 0.129780615 | 0.036503608 | 0.00850754 | 0.008871457 |
| Herc2 | S | 2929 | 0.027769924 | 0.03385978 | -0.055767331 | -0.083537254 | -0.08962711 | 0.644744408 | 0.281719152 | 0.560177421 | 0.424950818 | 0.372625759 |
| Mcm6 | S | 762 | -0.029463677 | 0.017325219 | 0.041372119 | 0.070835796 | 0.0240469 | 0.268423492 | 0.567721662 | 0.261770086 | 0.097813578 | 0.525010524 |
| Ralgapa2 | S | 819 | 0.095314949 | 0.009572175 | -0.053782853 | -0.149097802 | -0.063355028 | 0.370139465 | 0.917424739 | 0.477876422 | 0.217155271 | 0.544579938 |
| Ralgapa2 | S | 820 | 0.095314949 | 0.009572175 | -0.053782853 | -0.149097802 | -0.063355028 | 0.370139465 | 0.917424739 | 0.477876422 | 0.217155271 | 0.544579938 |
| Slc9a3r1 | S | 285 | -0.125780832 | 0.136097369 | 0.047112952 | 0.172893785 | -0.088984417 | 0.016980444 | 0.045161501 | 0.415790913 | 0.041181434 | 0.207328504 |
| Tram1 | S | 365 | 0.090718104 | -0.031896613 | -0.042694912 | -0.133413016 | -0.010798299 | 0.107175859 | 0.424795427 | 0.596996068 | 0.158810928 | 0.880696533 |
| Bod1l | S | 1794 | -0.077335687 | 0.070536439 | 0.032830575 | 0.110166263 | -0.037705864 | 0.107074438 | 0.099033724 | 0.511901281 | 0.076152509 | 0.43013531 |
| Sf3a1 | S | 329 | -0.09967074 | 0.063441745 | 0.06235241 | 0.16202315 | -0.001089335 | 0.168387767 | 0.354070372 | 0.234532872 | 0.063218549 | 0.987217615 |
| Chd4 | S | 79 | -0.04550457 | -0.008181209 | 0.086495417 | 0.131999986 | 0.094676625 | 0.638313007 | 0.827784097 | 0.310466638 | 0.274677264 | 0.27678977 |
| Cbl | S | 692 | 0.018942776 | 0.024907015 | -0.013004344 | -0.031947119 | -0.037911359 | 0.698841301 | 0.586614299 | 0.745040154 | 0.386640887 | 0.241778321 |
| Srsf6 | S | 310 | 0.011337998 | -0.003585319 | 0.029958269 | 0.018620271 | 0.033543588 | 0.859130261 | 0.923648798 | 0.662171002 | 0.816457089 | 0.613444807 |
| Ranbp2 | T | 2476 | 0.096438297 | -0.047622417 | -0.016023272 | -0.112461569 | 0.031599146 | 0.492916325 | 0.839092276 | 0.929832326 | 0.600033564 | 0.909354929 |
| Ap3d1 | T | 758 | 0.138726285 | -0.053855481 | 0.021626875 | -0.11709941 | 0.075482357 | 0.522065615 | 0.206666832 | 0.476414382 | 0.582574514 | 0.104853687 |
| Hspa12b | S | 46 | 0.021760603 | 0.02292246 | -0.02699447 | -0.048755073 | -0.04991693 | 0.634872212 | 0.393687413 | 0.635994252 | 0.44778122 | 0.389340306 |
| Hspa12b | T | 42 | 0.021760603 | 0.02292246 | -0.02699447 | -0.048755073 | -0.04991693 | 0.634872212 | 0.393687413 | 0.635994252 | 0.44778122 | 0.389340306 |
| Apc | S | 2671 | 0.046595108 | -0.039024641 | 0.019957565 | -0.026637543 | 0.058982205 | 0.437141035 | 0.573527435 | 0.691120945 | 0.488925098 | 0.32470469 |
| Hmbs | S | 28 | 0.084013274 | 0.090730007 | -0.099313456 | -0.183326729 | -0.190043462 | 0.294529879 | 0.362528322 | 0.089759668 | 0.070563297 | 0.116552399 |
| Hmbs | T | 35 | 0.084013274 | 0.090730007 | -0.099313456 | -0.183326729 | -0.190043462 | 0.294529879 | 0.362528322 | 0.089759668 | 0.070563297 | 0.116552399 |
| Hmbs | T | 37 | 0.084013274 | 0.090730007 | -0.099313456 | -0.183326729 | -0.190043462 | 0.294529879 | 0.362528322 | 0.089759668 | 0.070563297 | 0.116552399 |
| Smarcad1 | S | 97 | -0.007809739 | -0.005256191 | 0.029643978 | 0.037453717 | 0.034900169 | 0.626523561 | 0.717299686 | 0.151123788 | 0.075375161 | 0.092345247 |
| Ash2l | S | 618 | -0.03308419 | -0.070723378 | 0.130268845 | 0.163353035 | 0.200992223 | 0.707267413 | 0.330478747 | 0.124141519 | 0.093095323 | 0.023486726 |
| Nucks1 | S | 79 | -0.161891472 | 0.067403159 | 0.065556268 | 0.22744774 | -0.001846891 | 0.095512474 | 0.32090576 | 0.032395863 | 0.047543036 | 0.975500821 |
| Tjp1 | S | 1614 | 0.032975523 | 0.022013023 | -0.019729687 | -0.05270521 | -0.04174271 | 0.315436847 | 0.538608134 | 0.202279515 | 0.157058173 | 0.28618705 |
| Pcm1 | S | 1092 | 0.041216054 | -0.048179697 | 0.04990993 | 0.008693876 | 0.098089627 | 0.567257096 | 0.457023046 | 0.540137488 | 0.898979548 | 0.188265208 |
| Peak1 | T | 571 | -0.074750479 | 0.080258597 | 0.019501308 | 0.094251787 | -0.060757289 | 0.569097878 | 0.375120519 | 0.83868149 | 0.46423886 | 0.459850032 |
| Trim65 | S | 89 | -0.031614871 | 0.190527291 | -0.130609208 | -0.098994337 | -0.3211365 | 0.785217966 | 0.230903707 | 0.484776158 | 0.559673413 | 0.141888952 |
| Irs2 | S | 362 | -0.146368722 | 0.059967227 | 0.100152952 | 0.246521674 | 0.040185725 | 0.030475598 | 0.293590438 | 0.03599805 | 0.010210554 | 0.414721728 |
| Dnttip2 | T | 252 | 0.114770795 | -0.08200304 | 0.028103276 | -0.086667519 | 0.110106316 | 0.342923631 | 0.367646062 | 0.843127088 | 0.603135818 | 0.485643997 |
| Nf1 | S | 866 | 0.009562705 | 0.005919817 | 0.014427113 | 0.004864408 | 0.008507296 | 0.874809635 | 0.892216754 | 0.701578909 | 0.929596816 | 0.804992047 |
| Fmnl2 | S | 171 | -0.059793114 | 0.030312113 | 0.042562745 | 0.102355859 | 0.012250632 | 0.069579374 | 0.034010596 | 0.347839325 | 0.080023932 | 0.760586966 |
| Epn1 | T | 493 | -0.038706008 | 0.080790775 | -0.015163123 | 0.023542885 | -0.095953898 | 0.730058886 | 0.341594305 | 0.840632505 | 0.850094937 | 0.353455691 |
| Srrm1 | S | 795 | 0.084877857 | -0.011510797 | -0.02484892 | -0.109726777 | -0.013338123 | 0.146677784 | 0.359557428 | 0.346092548 | 0.082817309 | 0.57529102 |
| Akap13 | S | 867 | 0.080877203 | 0.089401487 | -0.093305624 | -0.174182827 | -0.182707111 | 0.636867976 | 0.715606232 | 0.563087571 | 0.221082204 | 0.436621814 |
| Rbm12b1 Rbm12b2 | S | 254;254 | -0.012925219 | -0.011712635 | 0.030120711 | 0.04304593 | 0.041833346 | 0.716525112 | 0.640154776 | 0.089129113 | 0.28282494 | 0.156474816 |
| Arid4b | T | 876 | 0.029161341 | -0.042820997 | 0.043360828 | 0.014199487 | 0.086181825 | 0.484877808 | 0.218383453 | 0.259458774 | 0.678448849 | 0.030898511 |
| Nup88 | T | 526 | -0.018358803 | 0.009657035 | 0.031084778 | 0.049443582 | 0.021427743 | 0.598598662 | 0.727306435 | 0.380043315 | 0.233359128 | 0.516581756 |

| | | | | | | | | | | | | | |
|---|---|---|---|---|---|---|---|---|---|---|---|---|---|
| Cog3 | S | | 204 | 0.084819929 | 0.030819238 | -0.063690527 | -0.148510455 | -0.094509765 | 0.390939224 | 0.609216969 | 0.306858354 | 0.190400072 | 0.224452626 |
| Rrbp1 | S | | 1472 | 0.0349779 | -0.021888513 | -0.006957081 | -0.041934981 | 0.014931433 | 0.1354535 | 0.68275006 | 0.777546524 | 0.08145499 | 0.772345997 |
| Hipk1 Hipk2 | Y | 352;361 | | -0.081009627 | 0.080124109 | 0.04946877 | 0.130478398 | -0.030655339 | 0.057278008 | 0.138605029 | 0.394569623 | 0.082505381 | 0.593505108 |
| Prpf4b | S | | 266 | 0.021608677 | 0.07682965 | -0.095642743 | -0.11725142 | -0.172472393 | 0.826889372 | 0.333103473 | 0.592339031 | 0.513296725 | 0.352561928 |
| Srsf6 | S | | 81 | 0.056402946 | 0.02565836 | -0.07807852 | -0.134481467 | -0.103736881 | 0.410571186 | 0.570558122 | 0.558213692 | 0.35627791 | 0.450944788 |
| Tango6 | S | | 561 | -0.007001053 | 0.020746242 | 0.010331436 | 0.017332488 | -0.010414806 | 0.802108679 | 0.497000577 | 0.790251141 | 0.578148137 | 0.743746874 |
| Snrpa1 | T | | 180 | 0.105673863 | -0.006422208 | -0.097184045 | -0.202857908 | -0.090761836 | 0.031435691 | 0.912468169 | 0.124478022 | 0.019994041 | 0.231237373 |
| Pard3 | S | | 25 | 0.062127874 | -0.03472928 | 0.009593542 | -0.052534331 | 0.044322822 | 0.400638404 | 0.627493037 | 0.962148098 | 0.791829696 | 0.823088671 |
| Vgll4 | S | | 58 | 0.09760328 | -0.045742477 | -0.02112606 | -0.11872934 | 0.024616417 | 0.15239107 | 0.359408219 | 0.723988923 | 0.097020406 | 0.584968679 |
| Srgap2 | S | | 795 | -0.036774739 | 0.054802901 | -0.037584226 | -0.000809487 | -0.092387127 | 0.550796163 | 0.064199192 | 0.722604606 | 0.994242635 | 0.416385514 |
| Srrm2 | S | | 1278 | -0.003883225 | -0.004404348 | 0.022064196 | 0.02594742 | 0.026468544 | 0.793100627 | 0.83797044 | 0.204335389 | 0.061276035 | 0.218429813 |
| Brwd1 | S | | 1810 | 0.072730486 | 0.032216475 | -0.047211621 | -0.119942107 | -0.079428096 | 0.275648733 | 0.481334364 | 0.099002333 | 0.133567976 | 0.143971311 |
| Brwd1 | S | | 1812 | 0.072730486 | 0.032216475 | -0.047211621 | -0.119942107 | -0.079428096 | 0.275648733 | 0.481334364 | 0.099002333 | 0.133567976 | 0.143971311 |
| Scarf2 | S | | 683 | 0.036954505 | 0.027747937 | -0.022675661 | -0.059630166 | -0.050423597 | 0.664942819 | 0.525623224 | 0.48956256 | 0.490206871 | 0.227803379 |
| Npat | S | | 1249 | 0.050767481 | 0.002824122 | -0.012252064 | -0.063019546 | -0.015076186 | 0.509253385 | 0.937094245 | 0.673030215 | 0.418448011 | 0.620600034 |
| Rock1 | T | | 1101 | -0.075529311 | 0.000155329 | 0.108121469 | 0.183650781 | 0.10796614 | 0.142898172 | 0.996820811 | 0.088868413 | 0.024892998 | 0.099986968 |
| Mcm2 | S | | 40 | -0.093942527 | 0.003091632 | 0.125263488 | 0.219206014 | 0.122171856 | 0.454351108 | 0.975653071 | 0.292746295 | 0.107016752 | 0.220788984 |
| Osbpl5 | S | | 746 | -0.040882135 | -0.005940074 | 0.102908981 | 0.143791116 | 0.108849056 | 0.300312148 | 0.902167923 | 0.254840476 | 0.152149538 | 0.2392717 |
| Nop14 | S | | 146 | -0.015954597 | -0.049921674 | 0.08700845 | 0.102963047 | 0.136930123 | 0.93181712 | 0.594478451 | 0.301130724 | 0.604578262 | 0.244708536 |
| Nop14 | S | | 148 | -0.015954597 | -0.049921674 | 0.08700845 | 0.102963047 | 0.136930123 | 0.93181712 | 0.594478451 | 0.301130724 | 0.604578262 | 0.244708536 |
| Tln1 | S | | 1225 | -0.026584352 | 0.065987259 | -0.022212635 | 0.004371717 | -0.088199893 | 0.659739219 | 0.194559931 | 0.664721132 | 0.949972247 | 0.185492582 |
| Son | S | | 1910 | 0.017261964 | -0.01465381 | 0.034159215 | 0.016897251 | 0.048813025 | 0.675782122 | 0.528044496 | 0.551086811 | 0.790233613 | 0.41968162 |
| Son | S | | 1912 | 0.017261964 | -0.01465381 | 0.034159215 | 0.016897251 | 0.048813025 | 0.675782122 | 0.528044496 | 0.551086811 | 0.790233613 | 0.41968162 |
| Champ1 | S | | 464 | 0.025733302 | -0.090458405 | 0.047413749 | 0.021680446 | 0.137872153 | 0.337581197 | 0.117692871 | 0.033708532 | 0.353448443 | 0.062316382 |
| Adgra2 | S | | 1085 | 0.042956879 | -0.044871572 | 0.024784941 | -0.018171939 | 0.069656512 | 0.540596405 | 0.655784671 | 0.651468419 | 0.722656404 | 0.467465206 |
| Prpf6 | T | | 227 | 0.037940015 | 0.049310519 | -0.059559302 | -0.097499317 | -0.108869821 | 0.934678681 | 0.886155818 | NA | NA | NA |
| Aak1 | T | | 604 | -0.000584692 | 0.037538452 | -0.016082292 | -0.0154976 | -0.053620744 | 0.989684897 | 0.367639413 | 0.777229926 | 0.781233178 | 0.345750098 |
| Arhgap35 | S | | 1179 | -0.052436704 | 0.143017968 | -0.047349592 | 0.005087111 | -0.19036756 | 0.313085822 | 0.056631446 | 0.223014467 | 0.912209438 | 0.031830603 |
| Setd2 | S | | 707 | 0.126883528 | -0.106284953 | 0.031113653 | -0.095769875 | 0.137398606 | 0.216909593 | 0.179649364 | 0.657700329 | 0.353116146 | 0.141872806 |
| Gas2l1 | S | | 316 | -0.136346944 | 0.201732617 | -0.037139756 | 0.099207188 | -0.238872373 | 0.175890944 | 0.084118157 | 0.793450971 | 0.443155754 | 0.134048068 |
| Tbc1d15 | S | | 32 | -0.043243965 | 0.052886219 | 0.030377058 | 0.073621023 | -0.022509161 | 0.604049711 | 0.523453667 | 0.767019667 | 0.41561672 | 0.790738346 |
| Ncor2 | S | | 550 | -0.023491069 | 0.024903964 | 0.029942426 | 0.053433495 | 0.005038462 | 0.421065492 | 0.394066564 | 0.631584416 | 0.399294633 | 0.929793853 |
| Ncor2 | T | | 546 | -0.023491069 | 0.024903964 | 0.029942426 | 0.053433495 | 0.005038462 | 0.421065492 | 0.394066564 | 0.631584416 | 0.399294633 | 0.929793853 |
| 4933427D14Rik | S | | 663 | -0.029551235 | -0.056837165 | 0.142687151 | 0.172238386 | 0.199524317 | 0.633947471 | 0.318148027 | 0.17582907 | 0.129029117 | 0.092156122 |
| Setd1a | S | | 1255 | -0.071969494 | 0.031881856 | 0.082438624 | 0.154408117 | 0.050556768 | 0.446117571 | 0.52560085 | 0.542435547 | 0.318606658 | 0.696729373 |
| Afap1 | S | | 549 | 0.085051262 | 0.020395801 | -0.093121865 | -0.178173127 | -0.113517666 | 0.068889713 | 0.69810963 | 0.129964237 | 0.036499336 | 0.089286593 |
| Lmnb1 | S | | 29 | -0.031790963 | -0.037040307 | 0.112103315 | 0.143894279 | 0.149143622 | 0.649685092 | 0.667234943 | 0.301474062 | 0.19273957 | 0.201801882 |
| Mta2 | T | | 534 | 0.217388863 | -0.130197452 | -0.08123489 | -0.298623752 | 0.048962562 | 0.038839792 | 0.228688985 | 0.392697992 | 0.015204096 | 0.543411011 |
| Wnk1 | S | | 2027 | -0.037369089 | 0.082098404 | -0.038285996 | -0.000916906 | -0.120384399 | 0.599636009 | 0.106791076 | 0.495637866 | 0.990385101 | 0.089385968 |
| Sh3bp1 | S | | 632 | -0.034492743 | -0.009772903 | 0.070383191 | 0.104875935 | 0.080156094 | 0.419084953 | 0.79148751 | 0.180247386 | 0.096006261 | 0.157829464 |
| Tnks1bp1 | S | | 1022 | 0.136853494 | 0.000180695 | -0.102477216 | -0.23933071 | -0.102657911 | 0.188318902 | 0.995429771 | 0.29688654 | 0.081184099 | 0.297473227 |
| Soga1 | S | | 1195 | 0.159763122 | -0.076129088 | -0.067998465 | -0.227761587 | 0.008130623 | 0.018599714 | 0.185363782 | 0.235055049 | 0.004679466 | 0.804521886 |
| Pum2 | S | | 177 | -0.203125561 | 0.298308451 | 0.028085976 | 0.231211536 | -0.270222476 | 0.032412328 | 0.077186038 | 0.768191739 | 0.05386498 | 0.104278167 |
| Dck | T | | 14 | -0.10214678 | 0.070810783 | 0.047353621 | 0.149500401 | -0.023457162 | 0.069526468 | 0.12406779 | 0.231980775 | 0.034967704 | 0.392145076 |
| Camkk1 | S | | 458 | -0.031800142 | 0.225685472 | -0.172171963 | -0.140371821 | -0.397857435 | 0.784063946 | 0.105681333 | 0.235800295 | 0.201466466 | 0.016320267 |
| Cep170 | S | | 1102 | -0.030006834 | 0.046332241 | -0.012604635 | 0.017402198 | -0.058936876 | 0.551085275 | 0.121597327 | 0.79671799 | 0.774179365 | 0.277455276 |
| Srrm1 | S | | 572 | 0.007830744 | -0.019460945 | 0.030118649 | 0.022287905 | 0.049579595 | 0.791321859 | 0.423921881 | 0.226232962 | 0.459474672 | 0.082667171 |
| Srrm1 | S | | 574 | 0.007830744 | -0.019460945 | 0.030118649 | 0.022287905 | 0.049579595 | 0.791321859 | 0.423921881 | 0.226232962 | 0.459474672 | 0.082667171 |
| Ppig | S | | 532 | 0.025137799 | -0.032921566 | 0.039483212 | 0.014345413 | 0.072404777 | 0.773930373 | 0.735754641 | 0.585855677 | 0.855411255 | 0.443270442 |
| Trp53bp1 | S | | 716 | 0.033750421 | -0.02122958 | 0.009558696 | -0.024191725 | 0.030788276 | 0.389612722 | 0.713770014 | 0.904126444 | 0.743969004 | 0.711799885 |
| Raly | S | | 235 | -0.051270044 | -0.099772535 | 0.136859463 | 0.188129507 | 0.236631998 | 0.655503825 | 0.549589699 | 0.142605718 | 0.134459784 | 0.207874866 |
| Nab1 | S | | 400 | 0.007295215 | 0.064352339 | -0.060285738 | -0.067580953 | -0.124638077 | 0.877650511 | 0.20158798 | 0.424649673 | 0.371059474 | 0.148433789 |
| Srrm1 | S | | 836 | 0.037182293 | -0.036841977 | 0.044613289 | 0.007430997 | 0.081455266 | 0.444231269 | 0.254331055 | 0.501354598 | 0.91319772 | 0.257704323 |
| Prpf31 | S | | 450 | 0.001889609 | 0.007001643 | 0.023444564 | 0.021554955 | 0.016442921 | 0.963910737 | 0.900002679 | 0.714928357 | 0.72786015 | 0.815016644 |
| Cherp | S | | 714 | 0.04892697 | 0.023306382 | -0.005960237 | -0.054887207 | -0.029266619 | 0.792240751 | 0.953771921 | 0.957932912 | 0.730238204 | 0.940438022 |
| Gorasp1 | S | | 422 | 0.204545038 | -0.118878306 | -0.077023695 | -0.281568734 | 0.041854611 | 0.232774847 | 0.559985048 | 0.717931784 | 0.268633524 | 0.871082793 |
| Sirt2 | T | | 27 | -0.073507933 | 0.086161414 | 0.01469745 | 0.088205382 | -0.071463964 | 0.081157107 | 0.054218317 | 0.831625368 | 0.256387343 | 0.337590625 |
| Nhej1 | S | | 264 | -0.063264064 | 0.057024136 | 0.077547688 | 0.140811752 | 0.020523552 | 0.118590783 | 0.336718428 | 0.471250795 | 0.243572234 | 0.844448515 |
| Asf1b | S | | 198 | -0.04523644 | -0.042240783 | 0.114628572 | 0.159865012 | 0.156869355 | 0.397506245 | 0.616446162 | 0.146231659 | 0.077511854 | 0.152119254 |
| Cdk5rap2 | S | | 1243 | 0.027984791 | -0.020085893 | 0.012265929 | -0.015718862 | 0.032351822 | 0.207590947 | 0.266210435 | 0.629073074 | 0.581472443 | 0.270044171 |
| Thoc5 | S | | 307 | -0.033485415 | -0.009729018 | 0.077894277 | 0.111379692 | 0.087623295 | 0.469962572 | 0.807830836 | 0.254565838 | 0.126268477 | 0.199277098 |
| Prpf40a | S | | 934 | -0.017112912 | -0.042590973 | 0.086967297 | 0.104080208 | 0.12955827 | 0.589766646 | 0.178243984 | 0.064921679 | 0.042006218 | 0.039820533 |
| Trim44 | S | | 341 | -0.040937285 | 0.007710361 | 0.046666407 | 0.087603692 | 0.038956045 | 0.18138763 | 0.74503193 | 0.086645809 | 0.030739443 | 0.162800204 |
| Dclk1 | S | | 307 | -0.073386033 | 0.128888541 | -0.027262747 | 0.046123286 | -0.156151288 | 0.224811638 | 0.100793025 | 0.731969884 | 0.519686237 | 0.103210089 |
| Dnajc5 | S | | 12 | -0.022799684 | 0.027338337 | 0.027659459 | 0.050459143 | 0.000321123 | 0.584470874 | 0.528189977 | 0.70323287 | 0.459497119 | 0.995973341 |
| Washc2 | S | | 1173 | 0.015298674 | 0.024352669 | -0.021915249 | -0.037213922 | -0.046267917 | 0.590902227 | 0.468967677 | 0.600464949 | 0.409669222 | 0.338378865 |
| Ube4b | T | | 234 | -0.19024782 | 0.108630587 | 0.102109541 | 0.292357361 | -0.006521046 | 0.02028884 | 0.06879486 | 0.148144308 | 0.008821419 | 0.908319704 |
| Ranbp2 | T | | 2342 | 0.094751174 | -0.002194851 | -0.066349013 | -0.161100187 | -0.064154161 | 0.056879056 | 0.954975416 | 0.099088544 | 0.013718944 | 0.150194023 |
| Prrc2c | S | | 1516 | -0.003798159 | 0.016978458 | 0.016425596 | 0.020223755 | -0.000552862 | 0.934281883 | 0.755019405 | 0.76606768 | 0.578359618 | 0.990177016 |
| Trappc9 | S | | 953 | -0.021558069 | -0.007267306 | 0.038468347 | 0.060026416 | 0.045735653 | 0.417675189 | 0.723225424 | 0.090033227 | 0.06021153 | 0.041197232 |
| Lrba | S | | 994 | 0.097773755 | 0.002443618 | -0.053713944 | -0.1514877 | -0.056157562 | 0.381409368 | 0.957599248 | 0.477186987 | 0.230846209 | 0.450461666 |
| Trip4 | S | | 404 | 0.084070664 | 0.054761006 | -0.067235269 | -0.151305933 | -0.121996276 | 0.467742363 | 0.562761112 | 0.233864566 | 0.233313492 | 0.23142357 |
| Dzip1l | S | | 726 | 0.132562642 | -0.077251061 | -0.075191987 | -0.207754629 | 0.002059074 | 0.107148195 | 0.409627896 | 0.69839584 | 0.337168336 | 0.991691289 |
| Pbrm1 | S | | 509 | 0.084421568 | -0.001785592 | -0.040737941 | -0.125159509 | -0.038952349 | 0.297013022 | 0.977800302 | 0.38605002 | 0.165824588 | 0.579840667 |
| Ubr4 | S | | 1651 | -0.087039808 | 0.106691324 | 0.024067717 | 0.111107525 | -0.082623607 | 0.028138361 | 0.049287062 | 0.618225433 | 0.072343617 | 0.156864713 |
| Carmil1 | S | | 1323 | 0.100757705 | 0.024931049 | -0.099196488 | -0.199954193 | -0.124127536 | 0.208398214 | 0.613534001 | 0.213724737 | 0.067198296 | 0.153158324 |
| Asph | S | | 19 | -0.164940264 | 0.003281626 | 0.188365117 | 0.353305381 | 0.185083491 | 0.441913358 | 0.983111791 | 0.300227117 | 0.12749605 | 0.213944704 |
| Dock1 | S | | 1823 | 0.052545932 | -0.057798011 | 0.034387746 | -0.018158186 | 0.092185757 | 0.698812272 | 0.69234695 | 0.634926349 | 0.890832823 | 0.53482427 |
| Phldb1 | S | | 541 | 0.164734701 | -0.031790041 | -0.139436241 | -0.304170942 | -0.107646199 | 0.001499785 | 0.66112192 | 0.009709727 | 0.001124611 | 0.202590591 |
| Zfp513 | S | | 540 | -0.030027588 | -0.035579443 | 0.102982681 | 0.133010269 | 0.138562123 | 0.559806036 | 0.249660978 | 0.116684101 | 0.084919598 | 0.071655173 |
| Tra2b | S | | 20 | 0.202534546 | -0.141736121 | -0.024388093 | -0.226922639 | 0.117348028 | 0.139459425 | 0.283763881 | 0.701341202 | 0.11469601 | 0.358836938 |
| Il17ra | S | | 603 | -0.101556359 | 0.139786239 | -0.029128344 | 0.072428015 | -0.168914583 | 0.502462001 | 0.268535859 | 0.922607693 | 0.81840998 | 0.59281569 |
| Aspscr1 | S | | 196 | 0.000595023 | 0.061259741 | -0.021478445 | -0.022073469 | -0.082738186 | 0.995258932 | 0.574529013 | 0.794925672 | 0.843970618 | 0.493967122 |
| Erf | S | | 190 | -0.045647845 | 0.045456683 | 0.020705998 | 0.066353843 | -0.024750685 | 0.253633622 | 0.253223785 | 0.52344027 | 0.082343434 | 0.356566915 |
| Azi2 | S | | 314 | 0.157769222 | -0.074757866 | -0.064711041 | -0.222480264 | 0.010046825 | 0.133011305 | 0.4885605 | 0.481550748 | 0.085706256 | 0.931243253 |
| Aff4 | S | | 698 | 0.025855058 | -0.0403011 | 0.019399439 | -0.00645562 | 0.059700538 | 0.083502658 | 0.087218034 | 0.206123457 | 0.574162778 | 0.034861391 |
| Gins2 | S | | 182 | -0.004443898 | 0.274740907 | -0.128709612 | -0.124265714 | -0.403450519 | 0.966994617 | 0.30209798 | 0.212513957 | 0.134529211 | 0.180144529 |
| Srrm1 | T | | 404 | 0.005560932 | -0.016910813 | 0.02357467 | 0.018013738 | 0.040485483 | 0.839553494 | 0.578072851 | 0.253528614 | 0.461627426 | 0.204133696 |
| Emg1 | T | | 25 | 0.054190641 | -0.070210126 | 0.053044566 | -0.001146075 | 0.123254692 | 0.25422699 | 0.063510085 | 0.222978938 | 0.978732183 | 0.03008667 |
| Asap1 | T | | 1066 | -0.018109005 | 0.047645058 | 0.010806038 | 0.028915042 | -0.03683902 | 0.845791465 | 0.661059802 | 0.923976456 | 0.810129685 | 0.77807659 |
| Anks1 | S | | 663 | -0.01488894 | 0.088212928 | -0.037746355 | -0.022857415 | -0.125959283 | 0.606239145 | 0.275422659 | 0.610541077 | 0.753529136 | 0.217751033 |
| Gorasp1 | S | | 367 | -0.084173816 | 0.029544441 | 0.045356202 | 0.129530018 | 0.015811761 | 0.554085352 | 0.581316487 | 0.376121604 | 0.379979765 | 0.526024716 |
| Irf2bpl | S | | 636 | 0.084574565 | 0.054046835 | -0.105910045 | -0.19048461 | -0.15995688 | 0.380289749 | 0.248020506 | 0.152536654 | 0.108913281 | 0.063975151 |
| Eif3b | T | | 72 | 0.02966481 | -0.00522644 | -0.012537574 | -0.042202384 | -0.007311134 | 0.213976348 | 0.833474911 | 0.68542995 | 0.216982027 | 0.816778624 |

| | | | | | | | | | | | | |
|---|---|---|---|---|---|---|---|---|---|---|---|---|
| Eif5b | S | 187 | 0.019041213 | 0.027732076 | -0.050443744 | -0.069484957 | -0.07817582 | 0.251562542 | 0.134420264 | 0.201828928 | 0.118446393 | 0.092157363 |
| H2-K1 | S | 351;310;350;351 | 0.054266233 | 0.057579324 | -0.070324211 | -0.124590444 | -0.127903535 | 0.17779422 | 0.283557991 | 0.047750261 | 0.031537098 | 0.077260969 |
| Fnip1 | S | 181 | -0.002075219 | 0.107894255 | -0.069424756 | -0.067349537 | -0.177319011 | 0.985746909 | 0.06083293 | 0.068419554 | 0.572343919 | 0.025697852 |
| Prkd1 | S | 403 | -0.069422276 | 0.055333328 | 0.068898517 | 0.138320793 | 0.013565189 | 0.080697319 | 0.265860139 | 0.375407639 | 0.138551918 | 0.853491634 |
| Cenpt | S | 233 | 0.048661249 | 0.170531681 | -0.140109173 | -0.188770422 | -0.310640854 | 0.902955224 | 0.743377803 | 0.786602683 | 0.755435324 | 0.649005346 |
| Lrrfip1 | S | 3 | -0.098154224 | 0.091010457 | 0.012129978 | 0.110284202 | -0.078880478 | 0.066921302 | 0.030937961 | 0.709054646 | 0.054817583 | 0.046530481 |
| Nfia | S | 303 | -0.010098333 | 0.002223256 | 0.027119677 | 0.03721801 | 0.024896421 | 0.876365536 | 0.955785127 | 0.494866274 | 0.549299548 | 0.410100595 |
| Osbpl3 | S | 166 | 0.14684475 | -0.008631192 | -0.180808075 | -0.327652825 | -0.172176883 | 0.04084504 | 0.899199825 | 0.194999587 | 0.064187208 | 0.214379958 |
| Srsf1 | S | 205 | -0.003541466 | -0.050821256 | 0.123069061 | 0.126610527 | 0.173890317 | 0.956093945 | 0.42276078 | 0.405987666 | 0.384236823 | 0.267085009 |
| 1700030K09Rik | S | 380 | 0.008694294 | -0.025321287 | 0.069082802 | 0.060388508 | 0.094404088 | 0.950945835 | 0.653040354 | 0.580097654 | 0.722612862 | 0.457387633 |
| 1700030K09Rik | T | 379 | 0.008694294 | -0.025321287 | 0.069082802 | 0.060388508 | 0.094404088 | 0.950945835 | 0.653040354 | 0.580097654 | 0.722612862 | 0.457387633 |
| Flot1 | S | 19 | -0.088644861 | -0.054896763 | 0.220431102 | 0.309075963 | 0.275327865 | 0.158311088 | 0.534388954 | 0.107852873 | 0.053044896 | 0.07103807 |
| Plekhg3 | S | 423 | 0.161560683 | -0.013934005 | -0.027623123 | -0.189183805 | -0.013689117 | 0.433204924 | 0.730645103 | 0.267901547 | 0.372394441 | 0.733490892 |
| Tra2a | S | 96 | -0.059685275 | -0.026197341 | 0.13968873 | 0.199374005 | 0.165886071 | 0.244793006 | 0.532324836 | 0.134943249 | 0.061726853 | 0.095734839 |
| Rabl6 | S | 578 | -0.080196302 | 0.013040481 | 0.15901918 | 0.239215482 | 0.145978698 | 0.915115644 | 0.974027996 | 0.822631444 | 0.786332377 | 0.830477541 |
| Rabl6 | T | 580 | -0.080196302 | 0.013040481 | 0.15901918 | 0.239215482 | 0.145978698 | 0.915115644 | 0.974027996 | 0.822631444 | 0.786332377 | 0.830477541 |
| Tcf12 | S | 67 | 0.046017915 | -0.032117252 | -0.013294546 | -0.059312462 | 0.018822706 | 0.155264235 | 0.644990698 | 0.736843838 | 0.143699546 | 0.784375827 |
| Washc2 | T | 322 | 0.069648306 | -0.066134972 | 0.01445715 | -0.055191156 | 0.080592122 | 0.056737703 | 0.072664753 | 0.648367683 | 0.018141797 | 0.009633722 |
| Thop1 | S | 16 | -0.049303332 | 0.045977458 | 0.034467115 | 0.083770447 | -0.011510343 | 0.356938588 | 0.417426392 | 0.591243411 | 0.183000433 | 0.839578871 |
| Med7 | S | 195 | -0.170701098 | 0.026887235 | 0.162981982 | 0.33368308 | 0.136094747 | 0.062556475 | 0.611984967 | 0.057567789 | 0.009928863 | 0.104887794 |
| Zzz3 | S | 137 | -0.112885433 | -0.019600331 | 0.082769121 | 0.195654554 | 0.102369452 | 0.299442126 | 0.500349316 | 0.013983705 | 0.133535007 | 0.021878197 |
| Zzz3 | S | 142 | -0.112885433 | -0.019600331 | 0.082769121 | 0.195654554 | 0.102369452 | 0.299442126 | 0.500349316 | 0.013983705 | 0.133535007 | 0.021878197 |
| Asap1 | S | 854 | -0.009672702 | 0.04482219 | -0.026927613 | -0.017254911 | -0.071749803 | 0.81912706 | 0.109396538 | 0.377389958 | 0.71389021 | 0.077080171 |
| Nipbl | T | 599 | -0.036709533 | -0.049948096 | 0.101299084 | 0.138008617 | 0.15124718 | 0.50609444 | 0.530495566 | 0.142005114 | 0.051794838 | 0.089981884 |
| Wee1 | S | 66 | -0.088218618 | 0.059077944 | 0.044971441 | 0.133190059 | -0.014106503 | 0.081380154 | 0.304211204 | 0.112526638 | 0.036948566 | 0.778110051 |
| Washc2 | S | 538 | 0.066198986 | 0.101533275 | -0.095416626 | -0.161615612 | -0.1969499 | 0.526717848 | 0.432055791 | 0.202817459 | 0.175826576 | 0.180414185 |
| Hexim1 | Y | 288 | -0.055930633 | 0.072835483 | -0.001331695 | 0.054598939 | -0.074167177 | 0.036407323 | 0.016912879 | 0.962910881 | 0.127826987 | 0.063495074 |
| Ahctf1 | T | 1824 | 0.128951266 | -0.09050457 | -0.115193436 | -0.244144702 | -0.024688866 | 0.036051751 | 0.413209088 | 0.545110022 | 0.259843008 | 0.899181605 |
| Nhp2 | S | 11 | 0.209568762 | -0.084804028 | -0.154622484 | -0.364191245 | -0.069818456 | 0.037748369 | 0.334623275 | 0.324578892 | 0.073098713 | 0.623564804 |
| Kif18b | S | 725 | 0.036250833 | -0.000853132 | -0.006737172 | -0.042988005 | -0.00588404 | 0.616206936 | 0.990846833 | 0.925375543 | 0.083917379 | 0.847551464 |
| Arhgef18 | S | 1381 | 0.107885475 | -0.058026558 | -0.118563238 | -0.226448713 | -0.06053668 | 0.071805319 | 0.70753898 | 0.503974494 | 0.257463921 | 0.77380412 |
| Kmt2a | T | 2170 | -0.006541623 | 0.000727538 | 0.056415349 | 0.062957172 | 0.055687811 | 0.945345451 | 0.993914987 | 0.764265767 | 0.74374119 | 0.771926607 |
| Syde1 | S | 231 | 0.035480941 | 0.033116257 | -0.046719483 | -0.082200424 | -0.07983574 | 0.521227038 | 0.330722967 | 0.298923533 | 0.194275069 | 0.098216767 |
| Srsf10 | S | 106 | 0.057772795 | 0.001225721 | -0.0333556 | -0.091128395 | -0.03458132 | 0.228669015 | 0.971365511 | 0.33101367 | 0.0856748 | 0.240795613 |
| Supt16 | S | 1013 | 0.044516296 | 0.044715502 | -0.073120361 | -0.117636657 | -0.117835863 | 0.559448888 | 0.471891229 | 0.459798342 | 0.24888152 | 0.236749191 |
| Atrx | S | 828 | -0.106838685 | -0.034897647 | 0.211510442 | 0.318349127 | 0.246408089 | 0.290766237 | 0.557869875 | 0.170570168 | 0.076163488 | 0.133923461 |
| Atrx | S | 829 | -0.106838685 | -0.034897647 | 0.211510442 | 0.318349127 | 0.246408089 | 0.290766237 | 0.557869875 | 0.170570168 | 0.076163488 | 0.133923461 |
| Ythdc2 | S | 1294 | 0.056899141 | -0.032272315 | -0.010356249 | -0.067255391 | 0.021916065 | 0.280071687 | 0.5870135 | 0.90323915 | 0.444224667 | 0.801663015 |
| Frmd4a | S | 353 | -0.034676668 | 0.065750123 | -0.017813106 | 0.016863562 | -0.083563229 | 0.783283312 | 0.325988577 | 0.734285728 | 0.890423296 | 0.19976849 |
| Plekhg3 | S | 766 | -0.013162889 | -0.101189094 | 0.141493443 | 0.154656331 | 0.242682537 | 0.820468321 | 0.159484507 | 0.088130534 | 0.08771552 | 0.02700339 |
| Ppfibp1 | S | 444 | -0.036301257 | -0.082778969 | 0.100488093 | 0.13678935 | 0.183267062 | 0.534770325 | 0.28489477 | 0.036357407 | 0.072198903 | 0.077206025 |
| Arhgap22 | S | 387 | 0.062546801 | 0.015007707 | -0.038626908 | -0.101173709 | -0.053634614 | 0.278747252 | 0.723765432 | 0.202232525 | 0.130048181 | 0.220894576 |
| Ncor2 | S | 1449 | 0.013924424 | 0.04036292 | -0.035149033 | -0.049073458 | -0.075511954 | 0.772915264 | 0.523027719 | 0.673760082 | 0.544913235 | 0.402801266 |
| Abcc1 | S | 878 | 0.068305364 | 0.070596168 | -0.100985078 | -0.169290442 | -0.171581246 | 0.514064497 | 0.524877306 | 0.356365577 | 0.206107088 | 0.215901303 |
| Abcc1 | S | 882 | 0.068305364 | 0.070596168 | -0.100985078 | -0.169290442 | -0.171581246 | 0.514064497 | 0.524877306 | 0.356365577 | 0.206107088 | 0.215901303 |
| Arfgef2 | S | 218 | 0.076397316 | -0.002739458 | -0.035705248 | -0.112102564 | -0.03296579 | 0.385524504 | 0.931218303 | 0.522558766 | 0.255042023 | 0.552478047 |
| Ctbp2 | S | 365 | -0.078348359 | -0.038346515 | 0.110641438 | 0.188989797 | 0.148987953 | 0.569835334 | 0.761527465 | 0.210098996 | 0.217040376 | 0.274827291 |
| Usp42 | S | 1227 | 0.023854651 | 0.004622017 | -0.000940632 | -0.024795283 | -0.005562648 | 0.727542627 | 0.91581686 | 0.983626086 | 0.687913987 | 0.844716645 |
| Usp42 | S | 1230 | 0.023854651 | 0.004622017 | -0.000940632 | -0.024795283 | -0.005562648 | 0.727542627 | 0.91581686 | 0.983626086 | 0.687913987 | 0.844716645 |
| Usp42 | S | 1234 | 0.023854651 | 0.004622017 | -0.000940632 | -0.024795283 | -0.005562648 | 0.727542627 | 0.91581686 | 0.983626086 | 0.687913987 | 0.844716645 |
| Irs1 | S | 325 | 0.149583026 | 0.013318517 | -0.066582987 | -0.216166013 | -0.079901504 | 0.498531594 | 0.804033603 | 0.325163438 | 0.356531623 | 0.268847171 |
| Slk | T | 812 | -0.036674847 | 0.028015654 | 0.055047133 | 0.09172198 | 0.027031479 | 0.311215948 | 0.686084939 | 0.459008387 | 0.250658407 | 0.752871844 |
| Pbrm1 | S | 648 | -0.035906208 | 0.011497577 | 0.022147012 | 0.058053219 | 0.010649434 | 0.514112372 | 0.537741057 | 0.260164332 | 0.322387173 | 0.189304744 |
| Pola2 | S | 141 | -0.092745011 | 0.074814508 | 0.030955316 | 0.123700327 | -0.043859192 | 0.330889479 | 0.30179167 | 0.611260454 | 0.226962879 | 0.547434271 |
| Bag6 | S | 1139 | 0.06533919 | 0.004302758 | -0.054749317 | -0.120088507 | -0.059052075 | 0.142640485 | 0.918931699 | 0.255658206 | 0.060734472 | 0.296428704 |
| 2810004N23Rik | S | 67 | 0.028522528 | -0.017299928 | 0.005849014 | -0.022673514 | 0.023148941 | 0.327481944 | 0.515558226 | 0.848737134 | 0.429889003 | 0.394864849 |
| Fras1 | T | 1234 | 0.031054827 | -0.005201475 | -0.007366108 | -0.038420935 | -0.002164633 | 0.346541433 | 0.682084351 | 0.849072094 | 0.415552066 | 0.955046589 |
| Fras1 | T | 1236 | 0.031054827 | -0.005201475 | -0.007366108 | -0.038420935 | -0.002164633 | 0.346541433 | 0.682084351 | 0.849072094 | 0.415552066 | 0.955046589 |
| Fras1 | T | 1237 | 0.031054827 | -0.005201475 | -0.007366108 | -0.038420935 | -0.002164633 | 0.346541433 | 0.682084351 | 0.849072094 | 0.415552066 | 0.955046589 |
| Nefm | S | 605 | 0.077433308 | -0.007098042 | -0.050041763 | -0.127475071 | -0.042943721 | 0.152948174 | 0.915498552 | 0.274270452 | 0.030127963 | 0.475968999 |
| 1110059G10Rik | S | 50 | -0.11581566 | 0.088478921 | 0.049744706 | 0.165560366 | -0.038734215 | 0.161594963 | 0.203305506 | 0.499581378 | 0.113159706 | 0.644068162 |
| Nsfl1c | S | 114 | -0.035288618 | 0.056031699 | 0.011340446 | 0.046629064 | -0.044691253 | 0.603860777 | 0.528333852 | 0.829790237 | 0.498631021 | 0.609665328 |
| Inf2 | S | 23 | 0.125430543 | -0.045757805 | -0.00091452 | -0.126345063 | 0.044843285 | 0.417721696 | 0.3278465 | 0.986373337 | 0.416401072 | 0.404552882 |
| Morc2a | S | 614 | 0.058495552 | -0.058754637 | 0.01519537 | -0.043300182 | 0.073950008 | 0.261816233 | 0.383282777 | 0.798639704 | 0.53232763 | 0.366847829 |
| Cbx3 | S | 176 | 0.001347651 | -0.005039319 | 0.024619747 | 0.023272097 | 0.029659067 | 0.971314066 | 0.919016159 | 0.584804104 | 0.518144523 | 0.541501301 |
| Edc4 | S | 712 | 0.104950977 | 0.006117086 | -0.085116042 | -0.190067019 | -0.091233128 | 0.176632049 | 0.911375022 | 0.237203497 | 0.041507974 | 0.154392708 |
| Triobp | S | 1604 | 0.050478932 | -0.030229655 | -0.009979489 | -0.060458421 | 0.020250166 | 0.126558826 | 0.392664591 | 0.741571116 | 0.058867054 | 0.508394382 |
| Rnf6 | S | 183 | 0.039984501 | 0.015896878 | -0.031722358 | -0.071706859 | -0.047619236 | 0.566327477 | 0.69897432 | 0.605759779 | 0.37121215 | 0.421812898 |
| Phldb1 | S | 520 | 0.094328048 | 0.000459647 | -0.062538121 | -0.156866169 | -0.062997768 | 0.009982466 | 0.984462342 | 0.006426405 | 0.004514828 | 0.050445759 |
| Gas2 | S | 289 | 0.164172943 | -0.011878954 | -0.172752901 | -0.336925844 | -0.160873947 | 0.107136233 | 0.858218948 | 0.289259947 | 0.089247703 | 0.313061852 |
| Srrm2 | S | 2054 | -0.096025027 | 0.030592101 | 0.055937417 | 0.151962445 | 0.025345317 | 0.535144225 | 0.665563542 | 0.377617329 | 0.355062513 | 0.719466115 |
| Atrx | S | 1979 | -0.011412473 | -0.004853945 | 0.051374968 | 0.062787441 | 0.056228913 | 0.854944569 | 0.948678099 | 0.616590016 | 0.504128205 | 0.571694713 |
| Cep170b | S | 1122 | -0.010114127 | 0.073375063 | -0.042844148 | -0.032730021 | -0.116219211 | 0.828667871 | 0.284728877 | 0.546842367 | 0.656690123 | 0.203773252 |
| Tjp1 | S | 899 | 0.217520592 | -0.001524147 | -0.200969847 | -0.418490439 | -0.1994457 | 0.195890159 | 0.986345502 | 0.299677032 | 0.09054658 | 0.300346069 |
| Atad2 | T | 801 | 0.055093986 | -0.050115404 | 0.021963755 | -0.033130231 | 0.072079159 | 0.466950987 | 0.529913548 | 0.789470031 | 0.564649698 | 0.283750861 |
| Coil | S | 146 | 0.031364671 | -0.015076803 | 0.01029798 | -0.021066691 | 0.025374783 | 0.579617621 | 0.660818675 | 0.809469005 | 0.730633217 | 0.575978777 |
| Fam204a | S | 155 | -0.063223384 | 0.003824112 | 0.101605361 | 0.164828745 | 0.09778125 | 0.226607179 | 0.936663146 | 0.225726368 | 0.08968641 | 0.228818591 |
| Cep170 | S | 1229 | -0.007383503 | 0.01515613 | 0.012462482 | 0.019845985 | -0.002693648 | 0.883026357 | 0.72293351 | 0.761430952 | 0.649361559 | 0.935119807 |
| Prrc2a | T | 1110 | -0.024208853 | -0.11047719 | 0.130372047 | 0.154580901 | 0.240849237 | 0.654614773 | 0.189995887 | 0.048713739 | 0.021855582 | 0.031039667 |
| Sash1 | S | 435 | -0.138300312 | 0.136996677 | 0.039906259 | 0.178206571 | -0.097090418 | 0.126636436 | 0.158011418 | 0.655933222 | 0.073093558 | 0.288797389 |
| Fto | S | 352 | 0.166527023 | -0.054957076 | -0.081154579 | -0.247681602 | -0.026197504 | 0.106063984 | 0.402406872 | 0.354873466 | 0.045883489 | 0.710027952 |
| Lmna | S | 395 | -0.000634844 | 0.004463426 | 0.009691609 | 0.010326453 | 0.005228183 | 0.969160053 | 0.673868606 | 0.691522886 | 0.686382626 | 0.819051398 |
| Nhsl1 | T | 139 | 0.076268256 | -0.064901432 | 0.016487679 | -0.059780577 | 0.081389111 | 0.426524752 | 0.491538872 | 0.821227058 | 0.462985373 | 0.330745285 |
| Sfr1 | T | 70 | 0.043594106 | 0.043139879 | -0.062500884 | -0.10609499 | -0.105640764 | 0.248169367 | 0.344145295 | 0.11989106 | 0.005189229 | 0.042679711 |
| Tubgcp6 | S | 1345 | -0.039032823 | 0.031033505 | 0.020042988 | 0.059075811 | -0.010990517 | 0.455934213 | 0.283607082 | 0.763282153 | 0.445139123 | 0.865077251 |
| Sox4 | S | 315 | -0.189132044 | -0.138554516 | 0.273894916 | 0.46302696 | 0.412449433 | 0.50113154 | 0.543493168 | 0.129634249 | 0.16826423 | 0.143322549 |
| Prrc2a | T | 825 | 0.101010451 | -0.114285853 | -0.00599564 | -0.107006091 | 0.108290213 | 0.055224617 | 0.208310497 | 0.925258694 | 0.150106835 | 0.255962979 |
| Soga1 | S | 1300 | -0.118426783 | -0.019383313 | 0.096636468 | 0.21506325 | 0.116019781 | 0.176664866 | 0.53373208 | 0.011135912 | 0.05847324 | 0.015988426 |
| Rala | S | 183 | -0.064459094 | 0.004434642 | 0.061916902 | 0.126375995 | 0.057482259 | 0.290704254 | 0.890254228 | 0.130176942 | 0.090605111 | 0.203332529 |
| Set | S | 132 | -0.04009392 | 0.037061182 | 0.018377745 | 0.058471664 | -0.018683437 | 0.453802614 | 0.453201263 | 0.565167466 | 0.273735048 | 0.656250622 |
| Pxn | Y | 88 | 0.028485362 | 0.087255681 | -0.098237 | -0.126722362 | -0.185492681 | 0.775419339 | 0.305920857 | 0.370035315 | 0.28312847 | 0.117477773 |
| Nktr | S | 1139 | -0.129740213 | 0.159305215 | -0.067241957 | 0.062498256 | -0.226547172 | 0.183201523 | 0.016569985 | 0.409109014 | 0.519004117 | 0.048457936 |
| Arhgef5 | S | 720 | -0.049832027 | 0.018008102 | 0.049817835 | 0.099649861 | 0.031809733 | 0.161250172 | 0.579774971 | 0.172464175 | 0.018670373 | 0.273306984 |
| Pop5 | S | 157 | -0.023736036 | 0.013794646 | 0.031953197 | 0.055689233 | 0.018158551 | 0.319416842 | 0.707081454 | 0.330820008 | 0.116477747 | 0.643207922 |

| | | | | | | | | | | | | |
|---|---|---|---|---|---|---|---|---|---|---|---|---|
| Secisbp2l | S | 251 | -0.065288136 | 0.098837844 | -0.021586613 | 0.043701523 | -0.120424457 | 0.559944938 | 0.294133358 | 0.841415176 | 0.722494127 | 0.296442762 |
| Cntrob | S | 782 | 0.034615538 | 0.066520532 | -0.065377925 | -0.099993463 | -0.131898457 | 0.805368293 | 0.648721769 | 0.661713363 | 0.553946965 | 0.450723903 |
| Prrc2c | S | 1222 | 0.039777392 | -0.032267054 | 0.007665882 | -0.03211151 | 0.039932936 | 0.091317568 | 0.102723184 | 0.681103401 | 0.183524177 | 0.092785394 |
| Rpa1 | S | 39 | -0.011928629 | 0.007055718 | 0.025005249 | 0.036933878 | 0.017949531 | 0.792107009 | 0.940760751 | 0.542760907 | 0.385942549 | 0.847741375 |
| Cdca3 | S | 43 | 0.053419555 | -0.019742624 | -0.027707679 | -0.081127233 | -0.007965055 | 0.236614713 | 0.760020294 | 0.634685877 | 0.222456723 | 0.914706671 |
| Usp38 | S | 680 | 0.022912172 | 0.043387974 | -0.062083373 | -0.084995545 | -0.105471347 | 0.53476685 | 0.613007636 | 0.613718038 | 0.498438988 | 0.452048417 |
| Zeb2 | S | 705 | 0.02126315 | 0.036746474 | -0.052626549 | -0.0738897 | -0.089373024 | 0.545139591 | 0.190119355 | 0.251275127 | 0.139118721 | 0.103147235 |
| Klf16 | T | 239 | -0.076475335 | 0.252488812 | -0.09170167 | -0.015226334 | -0.344190482 | 0.230160559 | 0.170686444 | 0.526514853 | 0.910669248 | 0.115878575 |
| Tmem209 | T | 150 | -0.041691883 | 0.013336865 | 0.054522269 | 0.096214152 | 0.041185404 | 0.540525614 | 0.798797857 | 0.496765984 | 0.244305286 | 0.53777397 |
| Brd9 | S | 567 | -0.013283039 | -0.002612687 | 0.035341771 | 0.04862481 | 0.037954458 | 0.796408811 | 0.95552452 | 0.505146889 | 0.287979411 | 0.336866972 |
| Prune2 | S | 2710 | -0.009955164 | 0.022349504 | 0.002410546 | 0.01236571 | -0.019938959 | 0.753338168 | 0.467483035 | 0.940146287 | 0.61618686 | 0.396079794 |
| Coro1b | S | 413 | 0.06791584 | 0.013792643 | -0.039256399 | -0.10717224 | -0.053049042 | 0.583310272 | 0.913373055 | 0.61626835 | 0.39247482 | 0.670455852 |
| Wee1 | S | 262 | 0.071264833 | -0.012469289 | -0.070666794 | -0.141931627 | -0.058197506 | 0.025046678 | 0.593299269 | 0.18601128 | 0.045709491 | 0.253691467 |
| Wee1 | S | 269 | 0.071264833 | -0.012469289 | -0.070666794 | -0.141931627 | -0.058197506 | 0.025046678 | 0.593299269 | 0.18601128 | 0.045709491 | 0.253691467 |
| Nfatc2ip | S | 49 | -0.011134614 | 0.017699676 | 0.017187369 | 0.028321983 | -0.000512307 | 0.904862182 | 0.81988433 | 0.804529082 | 0.796509964 | 0.995793067 |
| Nfatc2ip | S | 51 | -0.011134614 | 0.017699676 | 0.017187369 | 0.028321983 | -0.000512307 | 0.904862182 | 0.81988433 | 0.804529082 | 0.796509964 | 0.995793067 |
| Sema6d | S | 973 | 0.003271851 | -0.016817952 | 0.03191002 | 0.028638169 | 0.048727971 | 0.905110456 | 0.531131554 | 0.395188596 | 0.387719517 | 0.185826479 |
| Srrm1 | S | 196 | -0.233801898 | 0.257536198 | -0.117061548 | 0.11674035 | -0.374597746 | 0.693845283 | 0.276716439 | 0.719415993 | 0.847929781 | 0.273814335 |
| Srrm1 | S | 198 | -0.233801898 | 0.257536198 | -0.117061548 | 0.11674035 | -0.374597746 | 0.693845283 | 0.276716439 | 0.719415993 | 0.847929781 | 0.273814335 |
| Srrm1 | S | 200 | -0.233801898 | 0.257536198 | -0.117061548 | 0.11674035 | -0.374597746 | 0.693845283 | 0.276716439 | 0.719415993 | 0.847929781 | 0.273814335 |
| Hirip3 | S | 80 | -0.015142976 | -0.007120296 | 0.026698498 | 0.041841474 | 0.033818794 | 0.470648266 | 0.534009094 | 0.079255526 | 0.112406313 | 0.051667559 |
| Zbtb44 | S | 191 | -0.082395747 | -0.037652747 | 0.137182734 | 0.219578481 | 0.174835481 | 0.417152116 | 0.702295591 | 0.19950616 | 0.115985352 | 0.191874146 |
| Nab2 | S | 171 | 0.015022887 | 0.043923765 | -0.044246926 | -0.059269813 | -0.088170691 | 0.719626276 | 0.177539999 | 0.2176957 | 0.240989152 | 0.057673701 |
| Lmbrd1 | S | 528 | 0.001781401 | 0.034946651 | -0.019330672 | -0.021112073 | -0.054277323 | 0.927754848 | 0.353647563 | 0.497651167 | 0.497865835 | 0.221492572 |
| Arhgap21 | S | 1905 | 0.006924181 | 0.049274119 | -0.064990573 | -0.071914754 | -0.114264692 | 0.923974445 | 0.261840773 | 0.557716513 | 0.545961342 | 0.333642515 |
| Arhgap21 | T | 1901 | 0.006924181 | 0.049274119 | -0.064990573 | -0.071914754 | -0.114264692 | 0.923974445 | 0.261840773 | 0.557716513 | 0.545961342 | 0.333642515 |
| Cep350 | S | 2108 | 0.013575189 | -0.074786241 | 0.051505818 | 0.037930629 | 0.126292059 | 0.735494826 | 0.297826733 | 0.133849178 | 0.4040249 | 0.130369084 |
| Larp1 | T | 626 | -0.065126058 | 0.000881187 | 0.081497753 | 0.146623811 | 0.080616566 | 0.138455453 | 0.978821814 | 0.093400899 | 0.016867527 | 0.087553694 |
| Trip12 | S | 238 | 0.06828405 | -0.065044342 | -0.049511521 | -0.117795571 | 0.015532822 | 0.004192023 | 0.226766735 | 0.390721981 | 0.11959259 | 0.807292514 |
| Bicral | S | 673 | -0.071297441 | 0.102352559 | -0.046068056 | 0.025229385 | -0.148420615 | 0.53849636 | 0.135064771 | 0.582364025 | 0.838129947 | 0.144021113 |
| Pogz | S | 1334 | -0.084361378 | 0.021387983 | 0.072855437 | 0.157216815 | 0.051467454 | 0.335345882 | 0.681531714 | 0.296022844 | 0.127467736 | 0.418499061 |
| Nup133 | S | 58 | -0.030972785 | 0.076268167 | -0.013198843 | 0.017773942 | -0.08946701 | 0.750099614 | 0.529284449 | 0.879356323 | 0.854044725 | 0.465872392 |
| Nup133 | S | 59 | -0.030972785 | 0.076268167 | -0.013198843 | 0.017773942 | -0.08946701 | 0.750099614 | 0.529284449 | 0.879356323 | 0.854044725 | 0.465872392 |
| Nup133 | T | 62 | -0.030972785 | 0.076268167 | -0.013198843 | 0.017773942 | -0.08946701 | 0.750099614 | 0.529284449 | 0.879356323 | 0.854044725 | 0.465872392 |
| Fam171b | T | 584 | -0.163138238 | 0.070300287 | 0.119403511 | 0.282541748 | 0.049103223 | 0.041795638 | 0.376084657 | 0.085461248 | 0.009508249 | 0.52505797 |
| Myo18a | S | 1066 | -0.004063984 | -0.048914498 | 0.073782822 | 0.077846806 | 0.12269732 | 0.968560128 | 0.752999754 | 0.5396207 | 0.588896136 | 0.505296657 |
| Prpf40a | S | 929 | -0.020595581 | -0.043212838 | 0.086110976 | 0.106706557 | 0.129323813 | 0.524945545 | 0.182053037 | 0.076960119 | 0.046887834 | 0.044260421 |
| Prpf40a | S | 931 | -0.020595581 | -0.043212838 | 0.086110976 | 0.106706557 | 0.129323813 | 0.524945545 | 0.182053037 | 0.076960119 | 0.046887834 | 0.044260421 |
| Cpsf3 | S | 659 | -0.128292461 | 0.078571986 | 0.034021181 | 0.162313642 | -0.044550805 | 0.248945817 | 0.213007956 | 0.483845454 | 0.174313384 | 0.414990449 |
| Fto | S | 249 | -0.014213707 | 0.10677644 | -0.117097586 | -0.102883878 | -0.223874025 | 0.927556039 | 0.169974603 | 0.419332803 | 0.595005208 | 0.181806901 |
| Fto | S | 253 | -0.014213707 | 0.10677644 | -0.117097586 | -0.102883878 | -0.223874025 | 0.927556039 | 0.169974603 | 0.419332803 | 0.595005208 | 0.181806901 |
| Vcl | T | 754 | -0.12572386 | 0.083063651 | 0.083139885 | 0.208863745 | 7.62337E-05 | 0.014460311 | 0.04324071 | 0.33361332 | 0.071016703 | 0.99919429 |
| Il10rb | S | 334 | 0.031761127 | -0.011182172 | -0.001901314 | -0.033662441 | 0.009280857 | 0.528130509 | 0.788902681 | 0.964856407 | 0.4065891 | 0.723262366 |
| Fyb | S | 28 | -0.060944192 | -0.111406048 | 0.217890033 | 0.278834224 | 0.329296081 | 0.337034135 | 0.121195192 | 0.044019822 | 0.030456946 | 0.016648207 |
| Aco2 | S | 559 | 0.001615183 | -0.022817307 | 0.031371241 | 0.029756058 | 0.054188548 | 0.984135949 | 0.796426725 | 0.490292974 | 0.720352975 | 0.555594656 |
| Src | S | 17 | -0.083543402 | 0.02521831 | 0.085616996 | 0.169160399 | 0.060398686 | 0.228663015 | 0.542402076 | 0.327360372 | 0.117575349 | 0.47435367 |
| Stxbp5 | S | 786 | -0.023765643 | 0.07443496 | -0.05903222 | -0.035266577 | -0.13346718 | 0.280506044 | 0.038463178 | 0.387817729 | 0.581553361 | 0.117524396 |
| Stk3 | S | 316 | 0.025764111 | -0.008726048 | -0.006734156 | -0.032498267 | 0.001991892 | 0.314439343 | 0.727855177 | 0.808374945 | 0.261542054 | 0.940499744 |
| Rbm12b1 Rbm12b2 | S | 681;683 | 0.052541154 | -0.027132789 | -0.00347378 | -0.056014934 | 0.023659009 | 0.304375374 | 0.418998327 | 0.942183309 | 0.339807882 | 0.603845672 |
| Casc3 | S | 262 | -0.064401248 | 0.065036121 | 0.022163173 | 0.086564421 | -0.042872949 | 0.222824034 | 0.245070901 | 0.691430977 | 0.222948129 | 0.524576842 |
| Ctnnd1 | S | 268 | -0.065072795 | 0.119226687 | -0.040529236 | 0.024543559 | -0.159755923 | 0.355722576 | 0.14742423 | 0.631378034 | 0.729120005 | 0.09394507 |
| Fam193a | S | 293 | -0.017490636 | -0.007545879 | 0.026504681 | 0.043995317 | 0.03405056 | 0.611135008 | 0.815956199 | 0.178024435 | 0.247124162 | 0.325830558 |
| Etv6 | S | 240 | -0.029475305 | -0.004539032 | 0.052527969 | 0.082003274 | 0.057067001 | 0.28426241 | 0.918459389 | 0.19288022 | 0.07473842 | 0.276007324 |
| Cep170 | T | 376 | 0.01800352 | -0.010789206 | 0.017674366 | -0.000329155 | 0.028463572 | 0.817233079 | 0.871468817 | 0.819218718 | 0.996674838 | 0.682288087 |
| Haus7 | S | 169 | 0.029492506 | -0.093442902 | 0.059284633 | 0.029792127 | 0.152727535 | 0.827483296 | 0.753370169 | 0.61749957 | 0.704237041 | 0.598337508 |
| Ulk1 | S | 757 | 0.01918252 | 0.093086559 | -0.127068523 | -0.146251043 | -0.220155082 | 0.742700124 | 0.123522672 | 0.224509571 | 0.179909282 | 0.080572684 |
| Phf20 | S | 522 | -0.00799255 | 0.004914888 | 0.021910354 | 0.029902905 | 0.016995467 | 0.756419611 | 0.86272038 | 0.62647865 | 0.534431459 | 0.724201731 |
| Ssrp1 | S | 671 | -0.006546324 | -0.033539869 | 0.061861627 | 0.068407951 | 0.095401496 | 0.747697135 | 0.18254924 | 0.11828585 | 0.09984161 | 0.044335253 |
| Asap1 | S | 732 | -0.23587595 | 0.115008031 | 0.095529939 | 0.331405889 | -0.019478092 | 0.02825744 | 0.073156264 | 0.121410943 | 0.006543243 | 0.738695486 |
| Telo2 | S | 450 | -0.051685398 | 0.143467527 | -0.104460594 | -0.052775196 | -0.247928121 | 0.724618473 | 0.202142826 | 0.497918262 | 0.777357347 | 0.179616053 |
| Telo2 | S | 451 | -0.051685398 | 0.143467527 | -0.104460594 | -0.052775196 | -0.247928121 | 0.724618473 | 0.202142826 | 0.497918262 | 0.777357347 | 0.179616053 |
| Telo2 | S | 453 | -0.051685398 | 0.143467527 | -0.104460594 | -0.052775196 | -0.247928121 | 0.724618473 | 0.202142826 | 0.497918262 | 0.777357347 | 0.179616053 |
| Arhgef7 | S | 655 | -0.020608875 | 0.025830652 | 0.01893139 | 0.039540265 | -0.006899262 | 0.476564122 | 0.640052096 | 0.614123625 | 0.236580409 | 0.897319591 |
| Strn3 | S | 202 | 0.059760309 | 0.010640495 | -0.063779159 | -0.123539468 | -0.074419654 | 0.314712997 | 0.849899087 | 0.452431211 | 0.205674991 | 0.409475065 |
| Rrp12 | S | 1081 | 0.091891479 | -0.015873272 | -0.03466353 | -0.126555009 | -0.018790258 | 0.217096196 | 0.444868793 | 0.385510732 | 0.123644176 | 0.606784706 |
| Pum2 | T | 174 | 0.031657525 | 0.01590674 | -0.020567234 | -0.052224758 | -0.036473974 | 0.494698225 | 0.844620341 | 0.487041238 | 0.287511644 | 0.658461411 |
| Pcif1 | S | 8 | 0.083905987 | -0.074133923 | -0.012932941 | -0.096838928 | 0.061200982 | 0.115372614 | 0.291081592 | 0.861090479 | 0.236426625 | 0.470486505 |
| Ccdc9b | S | 190 | 0.028020151 | 0.017670138 | -0.02232541 | -0.050345561 | -0.039995548 | 0.492622187 | 0.383673314 | 0.192043993 | 0.261393972 | 0.069960852 |
| Baz2a | S | 135 | -0.057247134 | 0.084634659 | -0.00982662 | 0.047420514 | -0.094461278 | 0.394449084 | 0.247196524 | 0.886480377 | 0.512166539 | 0.24073159 |
| Mink1 | S | 746 | -0.031133026 | 0.017747159 | 0.028933214 | 0.06006624 | 0.011186055 | 0.214741676 | 0.228782187 | 0.529015769 | 0.251189768 | 0.797867453 |
| Cd2bp2 | T | 244 | -0.013139252 | 0.089207605 | -0.057076771 | -0.043937519 | -0.146284376 | 0.827607112 | 0.303976222 | 0.50424703 | 0.60759873 | 0.184280667 |
| Osbpl6 | S | 190 | 0.080689302 | 0.046067496 | -0.105788268 | -0.18647757 | -0.151855764 | 0.09387259 | 0.262149288 | 0.030706184 | 0.00655175 | 0.008743845 |
| Rras2 | S | 186 | -0.031985957 | 0.015478071 | 0.022798927 | 0.054784884 | 0.007320856 | 0.216128201 | 0.411796653 | 0.232613521 | 0.063704152 | 0.516268001 |
| Dlgap5 | S | 780 | 0.04098477 | 0.016328037 | -0.020136816 | -0.061121586 | -0.036464853 | 0.732429662 | 0.721870446 | 0.561449518 | 0.617555299 | 0.45705329 |
| Atxn2l | T | 31 | 0.036377789 | 8.66497E-05 | -0.016905847 | -0.053283636 | -0.016992497 | 0.544061361 | 0.997640553 | 0.740756038 | 0.453465136 | 0.737994116 |
| Tcof1 | S | 1216 | 0.015591884 | -0.010882067 | 0.007966686 | -0.007625197 | 0.018848753 | 0.418580405 | 0.567072499 | 0.819825951 | 0.831810047 | 0.608079228 |
| Map3k4 | S | 77 | -0.048768427 | 0.061141399 | -0.029929994 | 0.018838433 | -0.091071393 | 0.558555405 | 0.149351481 | 0.74110715 | 0.863852439 | 0.360178573 |
| Kmt2b | S | 1927 | 0.049802322 | -0.03039995 | -0.0052041 | -0.055006421 | 0.025195851 | 0.088175571 | 0.123852252 | 0.936717374 | 0.440455533 | 0.703345784 |
| Dnmt1 | S | 717 | 0.016076484 | 0.021016481 | -0.028797912 | -0.044874396 | -0.049814393 | 0.57259269 | 0.528535796 | 0.540015996 | 0.376701039 | 0.347905419 |
| Slx4 | S | 1046 | 0.051824138 | 0.080411963 | -0.083484053 | -0.135308191 | -0.163896016 | 0.822143768 | 0.73513772 | 0.675375019 | 0.593665046 | 0.530305771 |
| Topors | S | 553 | -0.007787609 | 0.015164733 | 0.019276537 | 0.027064146 | 0.004111804 | 0.526515498 | 0.760404215 | 0.700183374 | 0.591603136 | 0.949018543 |
| Srrm2 | T | 787 | -0.074408262 | -0.041435492 | 0.124044896 | 0.198453159 | 0.165480388 | 0.223975955 | 0.267276109 | 0.035410865 | 0.025156569 | 0.01787336 |
| Smcr8 | S | 515 | 0.10095118 | -0.030302669 | -0.077929767 | -0.178880947 | -0.047627099 | 0.257567709 | 0.805330753 | 0.559105702 | 0.222331234 | 0.753240979 |
| Smcr8 | T | 510 | 0.10095118 | -0.030302669 | -0.077929767 | -0.178880947 | -0.047627099 | 0.257567709 | 0.805330753 | 0.559105702 | 0.222331234 | 0.753240979 |
| Wdr47 | S | 296 | 0.23918505 | -0.027585382 | -0.228253509 | -0.467438559 | -0.200668127 | 0.272628986 | 0.858457641 | 0.467549674 | 0.200970672 | 0.517919616 |
| Atxn2 | S | 435 | -0.015120483 | 0.018168644 | 0.009197799 | 0.024318282 | -0.008970844 | 0.775055732 | 0.599207253 | 0.827912288 | 0.681233065 | 0.839535336 |
| Plec | S | 4633 | -0.06391953 | 0.067852872 | -0.007534556 | 0.056384973 | -0.075387429 | 0.123638812 | 0.034160977 | 0.833251158 | 0.189210476 | 0.081624632 |
| Smap2 | S | 239 | 0.001933351 | 0.029228283 | -0.009372006 | -0.011305357 | -0.038600289 | 0.987919383 | 0.720796957 | 0.925353612 | 0.928541693 | 0.636751474 |
| Scaf4 | S | 154 | -0.018660326 | -0.037210548 | 0.054041006 | 0.072701332 | 0.091251554 | 0.585213191 | 0.40429443 | 0.091826667 | 0.081372242 | 0.095157575 |
| Ulk2 | S | 799 | 0.095125395 | -0.057578824 | 0.671015532 | 0.575890137 | 0.728594357 | 0.778580814 | 0.897814643 | NA | NA | NA |
| Adnp | S | 708 | 0.009126731 | 0.007281884 | -0.00057579 | -0.009702521 | -0.007857675 | 0.793716395 | 0.867984289 | 0.988527862 | 0.808035956 | 0.868305361 |
| Srrm2 | S | 1497 | 0.069356842 | -0.100752134 | 0.037021519 | -0.032335323 | 0.137773653 | 0.092454277 | 0.064683119 | 0.360685845 | 0.3477222 | 0.028623126 |
| Srrm1 | S | 816 | 0.012387111 | -0.023548682 | 0.022424857 | 0.010037746 | 0.045973539 | 0.787488474 | 0.207944799 | 0.131650342 | 0.823826658 | 0.040903091 |
| Fam174c | S | 96 | 0.034207584 | 0.006169146 | -0.039318247 | -0.073525831 | -0.045487393 | 0.17635623 | 0.885243839 | 0.371631954 | 0.150190139 | 0.384752171 |

| | | | | | | | | | | | | |
|---|---|---|---|---|---|---|---|---|---|---|---|---|
| Peak1 | S | 569 | 0.003966508 | -0.020559062 | 0.039927562 | 0.035961053 | 0.060486624 | 0.604914914 | 0.378544837 | 0.483304792 | 0.522232629 | 0.326066686 |
| Akap13 | S | 2361 | 0.001989954 | -0.002095708 | 0.010762584 | 0.00877263 | 0.012858292 | 0.976995625 | 0.921570832 | 0.554404912 | 0.896977327 | 0.395394699 |
| Synrg | S | 974 | 0.081536825 | 0.035931724 | -0.093642161 | -0.175178986 | -0.129573886 | 0.071020028 | 0.363972942 | 0.026938498 | 0.00664132 | 0.011247515 |
| Rbmx Rbmxl1 | S | 216;219 | -0.037160744 | -0.018451639 | 0.045653813 | 0.082814557 | 0.064105452 | 0.580433052 | 0.601117655 | 0.11841623 | 0.268604924 | 0.103567523 |
| Acin1 | T | 417 | -0.216924124 | 0.071959321 | 0.099454369 | 0.316378493 | 0.027495048 | 0.379060464 | 0.224385421 | 0.65868473 | 0.309358855 | 0.899332491 |
| Smarcad1 | S | 37 | 0.126105705 | -0.052249815 | -0.0965843 | -0.222690005 | -0.044334485 | 0.062277957 | 0.444123261 | 0.363136835 | 0.095500527 | 0.675596775 |
| Matr3 | S | 598 | -0.052583517 | -0.018484742 | 0.100046508 | 0.152630025 | 0.11853125 | 0.209182957 | 0.595395512 | 0.123733455 | 0.048195159 | 0.092864016 |
| Etv1 | S | 191 | -0.05312508 | 0.12722891 | -0.015859013 | 0.037266067 | -0.143087922 | 0.298473819 | 0.303181376 | 0.773692998 | 0.395283228 | 0.258220561 |
| Dst | S | 4987 | 0.04251781 | -0.090777698 | 0.050715918 | 0.008198108 | 0.141493616 | 0.782030905 | 0.603244264 | 0.512134616 | 0.958049455 | 0.441904656 |
| Iqsec1 | S | 510 | -0.093011818 | 0.054446895 | 0.024217282 | 0.1172291 | -0.030229613 | 0.236926022 | 0.133503267 | 0.623307463 | 0.174441603 | 0.556533611 |
| Epb41l3 | S | 63 | -0.068337229 | 0.054516408 | 0.028434493 | 0.096771722 | -0.026081915 | 0.272123812 | 0.269049375 | 0.690090925 | 0.270111378 | 0.728812072 |
| Nktr | S | 1060 | 0.051297057 | -0.055303637 | -0.014852911 | -0.066149968 | 0.040450726 | 0.098643725 | 0.484480694 | 0.69819401 | 0.139728574 | 0.609018812 |
| Cep170b | S | 819 | 0.033019357 | -0.040015778 | 0.035631719 | 0.002612362 | 0.075647497 | 0.686569176 | 0.740243279 | 0.87614805 | 0.990962409 | 0.75664935 |
| Srrm2 | S | 1400 | 0.043891671 | -0.02849709 | 0.01060287 | -0.033288801 | 0.039099961 | 0.268576339 | 0.177769033 | 0.776064708 | 0.472896478 | 0.328643562 |
| Mis18bp1 | S | 305 | -0.014680746 | 0.005647229 | 0.03309179 | 0.047772536 | 0.027444561 | 0.73301845 | 0.8926437 | 0.683231072 | 0.544692319 | 0.718586201 |
| Sipa1l3 | S | 1550 | 0.140136905 | -0.056921999 | -0.116785361 | -0.256922265 | -0.059863362 | 0.08647372 | 0.576161556 | 0.374822469 | 0.110867158 | 0.669601471 |
| Tmem74 | S | 66 | -0.077541637 | 0.022194897 | 0.029630946 | 0.107172583 | 0.007436048 | 0.783606522 | 0.672894252 | 0.646552312 | 0.705805471 | 0.862210236 |
| Eif5b | S | 215 | 0.006605079 | 0.011278421 | -0.009572613 | -0.016177692 | -0.020851034 | 0.835592275 | 0.51582015 | 0.760810479 | 0.689240906 | 0.527220041 |
| Srsf4 | S | 297 | -0.05141738 | -0.061069926 | 0.130545345 | 0.181962725 | 0.191615271 | 0.522129941 | 0.441449347 | 0.176444493 | 0.05346686 | 0.047584677 |
| Irf2bp2 | S | 438 | 0.044315167 | -0.041901882 | -0.009696948 | -0.054012115 | 0.032204934 | 0.107998858 | 0.436930455 | 0.702682069 | 0.121418155 | 0.552850465 |
| Map4 | S | 512 | 0.011306539 | 0.019712905 | -0.014823194 | -0.026129734 | -0.034536099 | 0.873411346 | 0.677490472 | 0.798122651 | 0.694408385 | 0.41668434 |
| Prpf4b | S | 231 | 0.109971815 | -0.14025452 | 0.048125426 | -0.061846389 | 0.188379946 | 0.386688069 | 0.245133251 | 0.47358698 | 0.599314042 | 0.148270425 |
| Nes | T | 316 | 0.073755347 | -0.024513811 | -0.023381129 | -0.097136476 | 0.001132682 | 0.131613058 | 0.220585638 | 0.509847381 | 0.084622071 | 0.970831898 |
| Zc3h18 | T | 62 | 0.123579332 | -0.073803305 | 0.033756691 | -0.089822641 | 0.107559996 | 0.28649342 | 0.05606482 | 0.574209229 | 0.428561065 | 0.142679904 |
| Srrm2 | S | 2062 | -0.22736195 | 0.027786918 | 0.127822665 | 0.355184616 | 0.100035748 | 0.309223956 | 0.79152703 | 0.13635937 | 0.163548744 | 0.336996456 |
| Map1b | S | 2207 | 0.068530748 | -0.009109516 | -0.026050655 | -0.094581403 | -0.016941139 | 0.399640617 | 0.837566724 | 0.461169291 | 0.272229146 | 0.68812951 |
| Scaf1 | S | 718 | 0.009117455 | -0.060011373 | 0.079725873 | 0.070608419 | 0.139737246 | 0.900601293 | 0.370196181 | 0.369691096 | 0.471359842 | 0.17387087 |
| Ralgapa1 | S | 860 | -0.117865207 | 0.096288523 | 0.081412484 | 0.199277692 | -0.014876039 | 0.047231316 | 0.182009837 | 0.427849392 | 0.121129905 | 0.879385782 |
| Gm13889 | S | 217 | 0.010379495 | 0.10201198 | -0.092769589 | -0.103149085 | -0.194781569 | 0.857007893 | 0.251066773 | 0.272155393 | 0.228384883 | 0.088420688 |
| Nes | S | 324 | -0.000123123 | -0.013241584 | 0.023076662 | 0.023199785 | 0.036318246 | 0.998332015 | 0.814460933 | 0.532725951 | 0.676961403 | 0.503071326 |
| Mllt10 | S | 436 | -0.082030208 | 0.100526709 | 0.00388385 | 0.085914058 | -0.096642859 | 0.155944066 | 0.143514573 | 0.932955356 | 0.136974955 | 0.155485873 |
| Hnrnpk | S | 284 | 0.037810254 | -0.016145127 | -0.017221769 | -0.055032023 | -0.001076642 | 0.022493412 | 0.335301307 | 0.157663863 | 0.008195753 | 0.937566265 |
| Sltm | S | 589 | -0.086670103 | -0.020852434 | 0.107831046 | 0.194501148 | 0.12868348 | 0.350431463 | 0.458907768 | 0.101727979 | 0.094696737 | 0.084417763 |
| Pnisr | S | 313 | 0.084831709 | -0.063029997 | -0.009826764 | -0.094658473 | 0.053203232 | 0.268964891 | 0.405497377 | 0.924432866 | 0.425715323 | 0.645664003 |
| H1f8 | S | 20 | 0.016048966 | 0.085279789 | -0.078717408 | -0.094766374 | -0.163997196 | 0.702277556 | 0.108182372 | 0.082098562 | 0.020682813 | 0.015860271 |
| Sf3b1 | T | 223 | -0.028466846 | 0.029523959 | 0.002471498 | 0.030938344 | -0.027052461 | 0.40249691 | 0.204330893 | 0.949311807 | 0.479536502 | 0.480352163 |
| Socs5 | S | 271 | 0.019044097 | -0.027707946 | 0.019095931 | 5.18337E-05 | 0.046803877 | 0.631357976 | 0.594703063 | 0.634023093 | 0.999033159 | 0.405910584 |
| Dgcr8 | S | 377 | 0.010012363 | -0.02188993 | 0.023523344 | 0.013510981 | 0.045413273 | 0.57153164 | 0.314554726 | 0.379889592 | 0.551241865 | 0.123837747 |
| Rfc1 | S | 281 | 0.038953742 | -0.002106902 | -0.013882238 | -0.05283598 | -0.011775336 | 0.522195284 | 0.914883164 | 0.638279772 | 0.407538829 | 0.649477777 |
| Stxbp5 | T | 1040 | 0.258204284 | -0.113973632 | -0.169847623 | -0.428051907 | -0.055873991 | 0.095315163 | 0.535473184 | 0.309978829 | 0.058508263 | 0.780755043 |
| Prrc2a | T | 1085 | 0.1365669 | 0.162503877 | -0.13042167 | -0.26698857 | -0.292925547 | 0.179275051 | 0.170654711 | 0.000926409 | 0.057734894 | 0.063669044 |
| Atr | S | 431 | 0.018536311 | -0.026814394 | 0.018815015 | 0.000278703 | 0.045629409 | 0.483128201 | 0.480091132 | 0.64947915 | 0.994730082 | 0.37143237 |
| Cdc42ep1 | S | 377 | -0.127122326 | -0.039573976 | 0.300843136 | 0.427965463 | 0.340417112 | 0.11350743 | 0.468686375 | 0.143387798 | 0.0696369 | 0.11693458 |
| Upf1 | S | 1122 | 0.065765769 | -0.007068885 | -0.045169179 | -0.110934948 | -0.038100294 | 0.317475438 | 0.910548819 | 0.473721941 | 0.10300671 | 0.50711127 |
| Psmd1 | S | 315 | 0.008395467 | -0.019897817 | 0.024200271 | 0.015804804 | 0.044098088 | 0.727002657 | 0.388948891 | 0.392195309 | 0.555100915 | 0.139987703 |
| Cdk14 | S | 134 | -0.054878958 | 0.088863367 | -0.022720501 | 0.032158457 | -0.111583868 | 0.581938193 | 0.364281729 | 0.843612971 | 0.783115565 | 0.356845723 |
| Golga4 | S | 1440 | -0.023729163 | 0.059799258 | -0.009393097 | 0.014336066 | -0.069192354 | 0.647632497 | 0.49654638 | 0.856583061 | 0.582427027 | 0.402365096 |
| Golga4 | S | 1445 | -0.023729163 | 0.059799258 | -0.009393097 | 0.014336066 | -0.069192354 | 0.647632497 | 0.49654638 | 0.856583061 | 0.582427027 | 0.402365096 |
| Dnaja4 | S | 77 | 0.392045527 | -0.005548718 | -0.494587015 | -0.886632542 | -0.489038296 | 0.245049638 | 0.986208065 | 0.404746512 | 0.193747011 | 0.433610166 |
| Zc3h14 | S | 620 | 0.022336524 | -0.031344677 | 0.013531416 | -0.008805108 | 0.044876093 | 0.496044005 | 0.545849685 | 0.6427314 | 0.638014359 | 0.370375186 |
| Cdk13 | S | 326 | -0.027105753 | 0.040504158 | -0.00154535 | 0.025560403 | -0.042049508 | 0.305258671 | 0.20944173 | 0.96316794 | 0.417730274 | 0.251921446 |
| Zranb2 | T | 303 | -0.17020312 | 0.015077712 | 0.11289642 | 0.283099539 | 0.097818708 | 0.399657555 | 0.741705123 | 0.255616652 | 0.212161993 | 0.306668456 |
| Cdk13 | S | 316 | -0.028032582 | 0.04503379 | -0.008166692 | 0.01986589 | -0.053200482 | 0.354144213 | 0.191415871 | 0.83909719 | 0.594557615 | 0.220773159 |
| Cdk13 | S | 318 | -0.028032582 | 0.04503379 | -0.008166692 | 0.01986589 | -0.053200482 | 0.354144213 | 0.191415871 | 0.83909719 | 0.594557615 | 0.220773159 |
| Specc1l | S | 832 | 0.038874361 | 0.067227193 | -0.08792115 | -0.126795511 | -0.155148343 | 0.596898353 | 0.338859114 | 0.252968595 | 0.062593384 | 0.028054128 |
| Zc3hav1 | S | 592 | 0.105103171 | -0.014468705 | -0.132748319 | -0.237851489 | -0.118279613 | 0.098137072 | 0.675837067 | 0.386735021 | 0.179477093 | 0.431880982 |
| Knl1 | S | 794 | 0.022603917 | 0.026146842 | -0.024077168 | -0.046681084 | -0.05022401 | 0.757347646 | 0.806217374 | 0.707673346 | 0.385557009 | 0.606458228 |
| Tle3 | T | 285 | -0.024848337 | 0.058957095 | -0.010906339 | 0.013941998 | -0.069863434 | 0.687851744 | 0.481628048 | 0.808700661 | 0.836481056 | 0.430044156 |
| Cep170 | S | 829 | 0.032835011 | -0.013404656 | -0.013915045 | -0.046750056 | -0.000510389 | 0.212652977 | 0.614956645 | 0.61519729 | 0.143259572 | 0.985813706 |
| Vcp | T | 791 | 0.029278186 | 0.022588968 | -0.020902721 | -0.050180907 | -0.043491689 | 0.699712891 | 0.758292579 | 0.507690227 | 0.527654513 | 0.570797617 |
| Wtip | S | 29 | -0.140005591 | 0.121550773 | 0.051749027 | 0.191754618 | -0.069801747 | 0.007766779 | 0.017004348 | 0.522770359 | 0.089461561 | 0.411059539 |
| Magi3 | S | 236 | -0.098986878 | -0.029119542 | 0.220206837 | 0.319193715 | 0.249326379 | 0.170613218 | 0.742557517 | 0.224455532 | 0.124700355 | 0.18601589 |
| Snapc4 | S | 1301 | 0.08938471 | -0.055776682 | -0.002763588 | -0.092148298 | 0.053013094 | 0.637173377 | 0.73874637 | 0.971768198 | 0.618130025 | 0.74101 |
| Snapc4 | S | 1309 | 0.08938471 | -0.055776682 | -0.002763588 | -0.092148298 | 0.053013094 | 0.637173377 | 0.73874637 | 0.971768198 | 0.618130025 | 0.74101 |
| Srrm2 | S | 1631 | 0.022733489 | 0.002821721 | -0.019332548 | -0.042066037 | -0.022154269 | 0.370945613 | 0.931657444 | 0.580296884 | 0.25770509 | 0.577202006 |
| Ncor1 | S | 1004 | -0.101203509 | 0.076805477 | 0.027658943 | 0.128862452 | -0.049146534 | 0.479404705 | 0.593521513 | 0.521489165 | 0.382808872 | 0.724288133 |
| Rbmx Rbmxl1 | S | 325;328 | -0.007797703 | -0.048996253 | 0.084315255 | 0.092112958 | 0.133311508 | 0.954988122 | 0.440300611 | 0.39209599 | 0.559529665 | 0.228401213 |
| Aff4 | S | 156 | -0.007797703 | -0.048996253 | 0.084315255 | 0.092112958 | 0.133311508 | 0.954988122 | 0.440300611 | 0.39209599 | 0.559529665 | 0.228401213 |
| Des Krt7 Krt8 Prph Vi | S | 409;409;413;395; | 0.211691596 | -0.091512171 | 0.021190657 | -0.190500939 | 0.112702828 | 0.45833179 | 0.163301062 | 0.833210677 | 0.503953933 | 0.291895589 |
| Ndc1 | S | 443 | 0.177778664 | -0.141129872 | 0.039969357 | -0.137809307 | 0.181099229 | 0.382249335 | 0.286206499 | 0.89110551 | 0.659797722 | 0.539476251 |
| Srrm2 | S | 1924 | 0.009824073 | -0.039378776 | 0.070240417 | 0.060416344 | 0.109619194 | 0.882120765 | 0.363816092 | 0.485476162 | 0.566830694 | 0.30436619 |
| Zc3h13 | S | 1027 | 0.046292461 | 0.023383627 | -0.071774052 | -0.118066513 | -0.095157679 | 0.705185784 | 0.824679093 | 0.779588668 | 0.663679761 | 0.720039858 |
| Reps1 | S | 174 | -0.008672268 | 0.010494504 | 0.016819933 | 0.025492201 | 0.006325429 | 0.889700979 | 0.849142617 | 0.856782259 | 0.806386859 | 0.949679846 |
| Bak1 | S | 19 | 0.091933664 | -0.070652603 | 0.007338422 | -0.084595242 | 0.077991025 | 0.157393677 | 0.121061023 | 0.852740135 | 0.174029373 | 0.019737464 |
| Dock5 | S | 1801 | -0.039440592 | 0.007383308 | 0.055775898 | 0.09521649 | 0.04839259 | 0.249194248 | 0.745500755 | 0.334769815 | 0.194358232 | 0.384912972 |
| Htatsf1 | S | 446 | -0.165478167 | -0.006756085 | 0.161138197 | 0.326616364 | 0.167894282 | 0.151899282 | 0.919843956 | 0.077718533 | 0.0312433 | 0.094017705 |
| Sh3pxd2a | S | 607 | -0.079193233 | 0.020987243 | 0.083194103 | 0.162387336 | 0.06220686 | 0.278080318 | 0.783495247 | 0.32441642 | 0.060325304 | 0.391326119 |
| Lmna | T | 10 | -0.029608006 | 0.050608085 | -0.023185247 | 0.006422759 | -0.073793333 | 0.71475264 | 0.364669025 | 0.822667257 | 0.953474935 | 0.47352054 |
| Ubap2l | S | 487 | 0.034830578 | 0.008019106 | -0.026799813 | -0.061630391 | -0.034818919 | 0.1861108 | 0.695220049 | 0.122872991 | 0.05631469 | 0.132329813 |
| Phrf1 | S | 97 | 0.083942425 | 0.049250425 | -0.086702023 | -0.170644448 | -0.135952448 | 0.526815662 | 0.769873732 | 0.38746076 | 0.259642958 | 0.456094929 |
| Per1 | S | 592 | -0.113763457 | 0.029124571 | 0.084729759 | 0.198493216 | 0.055605188 | 0.249926705 | 0.682516051 | 0.23481779 | 0.081544055 | 0.400818348 |
| Rab6a Rab6b | S | 77;77 | 0.088398399 | -0.009362596 | -0.099934057 | -0.188332456 | -0.09057146 | 0.301582732 | 0.913541686 | 0.566193142 | 0.315845409 | 0.601274359 |
| Pla2g4a | S | 434 | -0.120944249 | 0.138037174 | -0.036906991 | 0.084037257 | -0.174944166 | 0.432781734 | 0.15230788 | 0.616582848 | 0.574030801 | 0.099003448 |
| Map1b | S | 1788 | -0.077190795 | 0.06704033 | 0.030760348 | 0.107951143 | -0.036279982 | 0.058228519 | 0.10250109 | 0.551616265 | 0.091376841 | 0.478330646 |
| Akap12 | S | 631 | 0.068226765 | -0.005727509 | -0.073761216 | -0.14198798 | -0.068033707 | 0.19401763 | 0.953917855 | 0.398358857 | 0.151073362 | 0.551889466 |
| Phrf1 | S | 1190 | 0.022813965 | -0.05507391 | 0.035967165 | 0.0131532 | 0.091041075 | 0.576436626 | 0.302663896 | 0.32191879 | 0.757194052 | 0.136453189 |
| Gprc5a | S | 344 | 0.152670845 | -0.113537317 | -0.028531456 | -0.181202302 | 0.085005861 | 0.014948544 | 0.04861271 | 0.417968269 | 0.008717808 | 0.104901799 |
| Gja5 | S | 281 | -0.061135573 | 0.035802816 | 0.037774012 | 0.098909585 | 0.001971196 | 0.096321582 | 0.2413865 | 0.312070145 | 0.033663394 | 0.945955945 |
| Pwp1 | S | 58 | 0.04775233 | -0.026153533 | -0.002702029 | -0.050454359 | 0.023451504 | 0.228674802 | 0.219118341 | 0.964328141 | 0.455788397 | 0.69954104 |
| Cpsf7 | S | 197 | 0.022945709 | -0.009960503 | 0.010534056 | -0.012411653 | 0.020494559 | 0.873123122 | 0.92613795 | 0.929061293 | 0.942443539 | 0.888765554 |
| Srrm2 | S | 2660 | 0.043141387 | -0.047564583 | 0.017516552 | -0.025624835 | 0.065081135 | 0.068407521 | 0.047311662 | 0.58793808 | 0.397593688 | 0.107906489 |
| Sec61b | S | 17 | 0.025386308 | 0.008602824 | -0.021186696 | -0.046573004 | -0.02978952 | 0.717245946 | 0.899133375 | 0.810949243 | 0.577580983 | 0.713007227 |
| Pacs1 | T | 44 | 0.095277947 | 0.033917265 | -0.096538613 | -0.19181656 | -0.130455878 | 0.241504907 | 0.816668758 | 0.19804708 | 0.027700963 | 0.392027296 |
| Ptprg | S | 1432 | 0.099139308 | -0.129079891 | 0.045519549 | -0.05361976 | 0.174599439 | 0.475328763 | 0.439106427 | 0.768072502 | 0.750874435 | 0.374891322 |

| | | | | | | | | | | | | |
|---|---|---|---|---|---|---|---|---|---|---|---|---|
| Mapkbp1 | S | 756 | -0.016062021 | 0.039232176 | -0.017248908 | -0.001186887 | -0.056481083 | 0.688880113 | 0.236647792 | 0.526214392 | 0.974583706 | 0.1005414 |
| Fubp1 | T | 149 | -0.112002884 | 0.071634936 | 0.027807497 | 0.139810382 | -0.043827439 | 0.271771023 | 0.24377692 | 0.600181799 | 0.191497809 | 0.393950739 |
| Atxn2 | S | 593 | 0.05410703 | -0.036546766 | -0.007831238 | -0.061938268 | 0.028715528 | 0.247553996 | 0.434568274 | 0.770466558 | 0.191706615 | 0.489232688 |
| Arl2 | S | 72 | -0.057882371 | 0.125221758 | -0.060273389 | -0.002391017 | -0.185495147 | 0.367160284 | 0.200701307 | 0.679423337 | 0.986131557 | 0.26551243 |
| Sipa1l1 | S | 288 | -0.015899681 | 0.254025632 | -0.179345105 | -0.163445424 | -0.433370737 | 0.919472612 | 0.068483433 | 0.049547104 | 0.348957077 | 0.017513087 |
| Snx2 | S | 93 | 0.02017103 | 0.079451726 | -0.078188155 | -0.098359184 | -0.157639881 | 0.857369432 | 0.511074652 | 0.501269615 | 0.219133369 | 0.115081898 |
| Slain2 | S | 316 | 0.001419792 | 0.007298388 | 0.001391257 | -2.85343E-05 | -0.00590713 | 0.970570733 | 0.804864424 | 0.956261404 | 0.999282548 | 0.749468887 |
| Mphosph8 | S | 271 | 0.009819785 | -0.04084443 | 0.063574224 | 0.05375444 | 0.104418655 | 0.88032763 | 0.601767254 | 0.65933191 | 0.700204833 | 0.48386663 |
| Mcm2 | T | 158 | 0.432905609 | -0.034336288 | -0.104432512 | -0.537338121 | -0.070096225 | 0.458452977 | 0.622661109 | 0.216866046 | 0.375106792 | 0.370432241 |
| Itprid2 | S | 738 | -0.052673097 | 0.002266632 | 0.062475336 | 0.115148433 | 0.060208704 | 0.24925685 | 0.956799898 | 0.205896173 | 0.015918662 | 0.094265317 |
| Supt20 | S | 504 | -0.039123474 | 0.078984016 | -0.019243356 | 0.019880119 | -0.098227371 | 0.453662926 | 0.32755659 | 0.789350842 | 0.751125771 | 0.272373105 |
| Ppp1r35 | S | 53 | -0.017320175 | 0.041022043 | -0.02524763 | -0.007927455 | -0.066269673 | 0.817163782 | 0.321131772 | 0.653642325 | 0.924866658 | 0.299964811 |
| Anapc1 | S | 569 | -0.016057063 | 0.038815438 | -0.050813524 | -0.03475646 | -0.089628962 | 0.898627975 | 0.316833377 | 0.764889953 | 0.859494661 | 0.605454581 |
| Yju2 | S | 309 | 0.049745129 | 0.038297824 | -0.106772254 | -0.156517383 | -0.145070078 | 0.250847923 | 0.524446281 | 0.284055671 | 0.158608022 | 0.187324002 |
| Yju2 | S | 312 | 0.049745129 | 0.038297824 | -0.106772254 | -0.156517383 | -0.145070078 | 0.250847923 | 0.524446281 | 0.284055671 | 0.158608022 | 0.187324002 |
| Vars | S | 1225 | -0.067390237 | -0.001098429 | 0.081319252 | 0.14870949 | 0.082417681 | 0.241752393 | 0.977372619 | 0.190472586 | 0.067554468 | 0.195951245 |
| Abcc1 | S | 884 | -0.057869236 | 0.045732702 | 0.039071893 | 0.096941129 | -0.006660809 | 0.066692038 | 0.198951656 | 0.448648293 | 0.124558136 | 0.889747949 |
| Wiz | S | 1368 | 0.04770861 | 0.004949861 | -0.037526583 | -0.085235193 | -0.042476445 | 0.557058718 | 0.953817215 | 0.639387853 | 0.356659872 | 0.649648364 |
| Zcchc8 | T | 474 | 0.129713873 | 0.074198614 | -0.169980938 | -0.29969481 | -0.244179552 | 0.042161168 | 0.594839439 | 0.026756611 | 0.005057714 | 0.164126075 |
| Zcchc8 | T | 482 | 0.129713873 | 0.074198614 | -0.169980938 | -0.29969481 | -0.244179552 | 0.042161168 | 0.594839439 | 0.026756611 | 0.005057714 | 0.164126075 |
| Zcchc8 | T | 488 | 0.129713873 | 0.074198614 | -0.169980938 | -0.29969481 | -0.244179552 | 0.042161168 | 0.594839439 | 0.026756611 | 0.005057714 | 0.164126075 |
| Akap12 | S | 1645 | -0.100780681 | 0.155642942 | -0.054196539 | 0.046584142 | -0.209839481 | 0.364095806 | 0.184795463 | 0.692713999 | 0.637612878 | 0.124006717 |
| Ets1 | S | 251 | 0.033182848 | -0.005549149 | -0.003773537 | -0.036956385 | 0.001775613 | 0.764058163 | 0.87628731 | 0.94105416 | 0.740895018 | 0.966004464 |
| Fancm | S | 1174 | -0.029055293 | -0.008550219 | 0.062557195 | 0.091612487 | 0.071107414 | 0.568175832 | 0.8755609 | 0.480530725 | 0.33994476 | 0.452949934 |
| Ampd3 | S | 87 | -0.113353937 | 0.187082392 | -0.005332409 | 0.108021528 | -0.192414801 | 0.066027964 | 0.122224758 | 0.928804926 | 0.057902036 | 0.121709761 |
| Slirp | T | 104 | -0.000850345 | 0.006421472 | 0.00526576 | 0.006116105 | -0.001155711 | 0.971391073 | 0.847303968 | 0.893202704 | 0.880279611 | 0.97981278 |
| Map4 | T | 96 | -0.009369637 | 0.094134144 | -0.072447138 | -0.063077501 | -0.166581282 | 0.791767764 | 0.205285378 | 0.319560906 | 0.36311237 | 0.09047241 |
| Srrm2 | T | 2268 | 0.028031693 | 0.000593768 | -0.006384551 | -0.034416244 | -0.006978319 | 0.770592894 | 0.980456783 | 0.806170236 | 0.720543245 | 0.746046127 |
| Wdr62 | S | 966 | 0.088146121 | -0.04399835 | -0.049711305 | -0.137857426 | -0.005712955 | 0.164047151 | 0.305024184 | 0.752532822 | 0.417896027 | 0.970222671 |
| Wdr62 | S | 972 | 0.088146121 | -0.04399835 | -0.049711305 | -0.137857426 | -0.005712955 | 0.164047151 | 0.305024184 | 0.752532822 | 0.417896027 | 0.970222671 |
| Aff1 | S | 197 | -0.042693083 | 0.039011156 | 0.03199559 | 0.074688673 | -0.007015567 | 0.835908406 | 0.809565681 | 0.929476941 | 0.85065528 | 0.985288402 |
| Syap1 | S | 298 | 0.040432225 | -0.044511567 | 0.031578213 | -0.008854012 | 0.07608978 | 0.801034251 | 0.683125419 | 0.786242545 | 0.953215621 | 0.436620179 |
| Lbr | S | 88 | 0.061642092 | -0.037149987 | -0.009392213 | -0.071034304 | 0.027757775 | 0.454483613 | 0.591210764 | 0.913520854 | 0.397090832 | 0.688617714 |
| Exosc9 | S | 393 | -0.060352843 | -0.084444527 | 0.213655547 | 0.274008389 | 0.298100074 | 0.416891798 | 0.24059359 | 0.157813941 | 0.105923428 | 0.099174203 |
| Exosc9 | S | 395 | -0.060352843 | -0.084444527 | 0.213655547 | 0.274008389 | 0.298100074 | 0.416891798 | 0.24059359 | 0.157813941 | 0.105923428 | 0.099174203 |
| Cep350 | S | 468 | -0.068505815 | -0.00510369 | 0.104347602 | 0.172853417 | 0.109451292 | 0.30283555 | 0.918856119 | 0.292597394 | 0.13361362 | 0.278752483 |
| Slc12a7 | Y | 63 | 0.004749256 | -0.037029352 | 0.047250743 | 0.042501487 | 0.084280095 | 0.959204475 | 0.583181086 | 0.509533331 | 0.5816421 | 0.066600067 |
| Npepps | S | 919 | -0.041368135 | 0.048491484 | 0.0150689 | 0.056437035 | -0.033422585 | 0.276137035 | 0.357119373 | 0.796872443 | 0.33026982 | 0.588932683 |
| Nop56 | S | 529 | 0.036659046 | 0.000232042 | -0.025309617 | -0.061968663 | -0.025541659 | 0.310894872 | 0.994092858 | 0.377985307 | 0.145542313 | 0.476684006 |
| Prrc2c | S | 2644 | -0.015339633 | 0.032043205 | -0.006080703 | 0.00925893 | -0.038123908 | 0.714562099 | 0.488031331 | 0.870621712 | 0.801279481 | 0.377957124 |
| Epha2 | S | 900 | 0.249011487 | -0.189402371 | -0.074362019 | -0.323373506 | 0.115040352 | 0.086998716 | 0.253952886 | 0.52249638 | 0.063880046 | 0.490426112 |
| Inpp5k | S | 301 | 0.032151756 | 0.014681007 | -0.031951841 | -0.064103597 | -0.046632848 | 0.611978474 | 0.799525248 | 0.574477665 | 0.324094963 | 0.421656234 |
| Smn1 | S | 10 | 0.018098282 | -0.04076471 | 0.041876828 | 0.023778546 | 0.082641538 | 0.638877512 | 0.406895834 | 0.579508881 | 0.748355756 | 0.325667502 |
| Mdc1 | S | 1169 | -0.074502339 | 0.036875017 | 0.067020683 | 0.141523022 | 0.030145666 | 0.459947066 | 0.67629119 | 0.649182195 | 0.402899906 | 0.847224001 |
| Tcf20 | S | 900 | -0.004023153 | 0.069029534 | -0.056942845 | -0.052919693 | -0.125972379 | 0.951393821 | 0.087410334 | 0.121648271 | 0.42902395 | 0.007041735 |
| Nes | S | 819 | 0.018786967 | 0.021896035 | -0.029436539 | -0.048223506 | -0.051332574 | 0.701189954 | 0.5638938 | 0.492452129 | 0.273670436 | 0.120039412 |
| Ctdspl2 | S | 104 | 0.050550754 | 0.032740774 | -0.069918537 | -0.120469291 | -0.102659311 | 0.320114841 | 0.603797583 | 0.23257523 | 0.048491879 | 0.136845251 |
| Sipa1l1 | T | 209 | -0.15920028 | 0.113348441 | 0.129321344 | 0.288521624 | 0.015972903 | 0.00061983 | 0.03240611 | 0.133989326 | 0.026750677 | 0.807708904 |
| Rai14 | T | 295 | 0.054890184 | -0.040493009 | 0.006902996 | -0.047987188 | 0.047396005 | 0.647326029 | 0.662988381 | 0.929176804 | 0.652644058 | 0.517786914 |
| Hivep3 | S | 489 | -0.082709958 | 0.071350418 | 0.053099185 | 0.135809142 | -0.018251233 | 0.0008865 | 0.03143774 | 0.157311931 | 0.031714729 | 0.593194315 |
| Nktr | S | 563 | -0.101445735 | -0.038900984 | 0.140369974 | 0.241815709 | 0.179270958 | 0.081601711 | 0.394174817 | 0.019768394 | 0.005601297 | 0.009541125 |
| Nktr | S | 565 | -0.101445735 | -0.038900984 | 0.140369974 | 0.241815709 | 0.179270958 | 0.081601711 | 0.394174817 | 0.019768394 | 0.005601297 | 0.009541125 |
| Wdr44 | S | 472 | -0.013052517 | 0.03274175 | -0.030656385 | -0.017603868 | -0.063398135 | 0.769037291 | 0.174344371 | 0.560923133 | 0.766803327 | 0.271830926 |
| Dock7 | S | 896 | 0.027839497 | 0.052889201 | -0.078921322 | -0.106760829 | -0.131810533 | 0.211889409 | 0.120218336 | 0.044014685 | 0.028203503 | 0.013293822 |
| Mink1 | S | 644 | -0.04885216 | 0.051049555 | 0.036761598 | 0.085613757 | -0.014287958 | 0.223102606 | 0.445535457 | 0.705194135 | 0.410797797 | 0.891959585 |
| Lca5 | S | 7 | 0.003811301 | 0.045807375 | -0.03289147 | -0.036702771 | -0.078698845 | 0.905549588 | 0.35175442 | 0.323883017 | 0.017785623 | 0.139638135 |
| Acin1 | S | 898 | -0.044190814 | -0.015390241 | 0.055196059 | 0.099386873 | 0.0705863 | 0.254410191 | 0.533146303 | 0.063610352 | 0.045420937 | 0.018795151 |
| Mdc1 | T | 1480 | 0.080427748 | -0.047695917 | -0.022701534 | -0.103129282 | 0.024994383 | 0.459935676 | 0.67923502 | 0.812485772 | 0.165390297 | 0.7341506 |
| Dcp1a | S | 542 | 0.027184523 | -0.01264806 | 0.001764931 | -0.025419592 | 0.014412991 | 0.730074645 | 0.782846638 | 0.961173178 | 0.728451362 | 0.657818691 |
| Mdfic | S | 141 | -0.001189336 | 0.021497163 | -0.003792383 | -0.002603047 | -0.025289546 | 0.977696198 | 0.81674404 | 0.937557772 | 0.944309079 | 0.781126509 |
| Med16 | S | 314 | 0.15127717 | -0.004365055 | -0.1249552 | -0.27623237 | -0.120590145 | 0.183688337 | 0.925006715 | 0.207437663 | 0.059698052 | 0.220429671 |
| Yap1 | S | 94 | 0.035342291 | 0.015315812 | -0.051937053 | -0.087279344 | -0.067252865 | 0.364725284 | 0.723747969 | 0.411783422 | 0.217565901 | 0.332671432 |
| Vdac2 | S | 116 | 0.029158465 | -0.037593512 | 0.02053492 | -0.008623545 | 0.058128432 | 0.478990397 | 0.373309112 | 0.671881938 | 0.839485411 | 0.228389558 |
| Hnrnpul1 | S | 513 | 0.05919775 | 0.013323223 | -0.041090693 | -0.100288443 | -0.054413915 | 0.711054549 | 0.853333921 | 0.60483073 | 0.5307558 | 0.366018571 |
| Srrm2 | T | 1800 | -0.010806358 | 0.00858972 | 0.007869375 | 0.018675733 | -0.000720345 | 0.67153471 | 0.679936112 | 0.710340796 | 0.386562028 | 0.956454353 |
| Srrm1 | S | 694 | -0.034474377 | -0.001642569 | 0.042505356 | 0.076979733 | 0.044147925 | 0.20380062 | 0.932690084 | 0.141231452 | 0.046990907 | 0.150142043 |
| Kmt2b | S | 866 | 0.084305303 | -0.068120058 | -0.035953557 | -0.12025886 | 0.032166501 | 0.004049052 | 0.06720437 | 0.296253175 | 0.024719418 | 0.39299654 |
| Wwtr1 | S | 89 | -0.10183233 | 0.13028345 | -0.040126867 | 0.061705463 | -0.170410317 | 0.387358412 | 0.170522822 | 0.673497161 | 0.521193458 | 0.040809978 |
| Prkaa1 Prkaa2 | T | 183;172 | -0.02984582 | 0.049854952 | -0.003008385 | 0.026837435 | -0.052863337 | 0.55054387 | 0.481108771 | 0.935090092 | 0.630247655 | 0.476716894 |
| Ncor2 | S | 1445 | 0.002596414 | 0.031659453 | -0.030402462 | -0.032998876 | -0.062061915 | 0.963110381 | 0.56792405 | 0.7168124 | 0.659845882 | 0.427744791 |
| Pacsin3 | S | 358 | -0.018199381 | 0.173417999 | -0.178816908 | -0.160617528 | -0.352234908 | 0.887401096 | 0.104557301 | 0.200487655 | 0.273051765 | 0.042897566 |
| Trp53bp1 | T | 1645 | 0.110291762 | 0.000597221 | -0.068718095 | -0.179009857 | -0.069315316 | 0.16923543 | 0.972598812 | 0.093114296 | 0.060471886 | 0.099535268 |
| Ilf2 | S | 385 | -0.088499254 | 0.142419068 | -0.0316381 | 0.056861154 | -0.174057168 | 0.238633976 | 0.150024154 | 0.62807109 | 0.305778593 | 0.093021 |
| Phrf1 | S | 1379 | 0.087586401 | -0.116654915 | 0.038277324 | -0.049309077 | 0.154932239 | 0.329900555 | 0.261634354 | 0.66150963 | 0.493394205 | 0.127108476 |
| Cdc5l | T | 424 | 0.035959922 | -0.016365341 | -0.01196538 | -0.047925302 | 0.004399961 | 0.438648469 | 0.790053945 | 0.677895897 | 0.347255912 | 0.944918304 |
| Nipbl | S | 350 | -0.033321998 | -0.024268964 | 0.062814136 | 0.096136134 | 0.0870831 | 0.615186323 | 0.611566573 | 0.292828341 | 0.173890002 | 0.112805543 |
| Gpsm1 | S | 538 | -0.310205995 | 0.104237282 | 0.19701195 | 0.507217945 | 0.092774668 | 0.340750098 | 0.732867074 | 0.375707708 | 0.155063872 | 0.752936678 |
| Lin9 | T | 96 | 0.051322955 | -0.04668306 | -0.002550576 | -0.053873531 | 0.044132483 | 0.203424863 | 0.328174332 | 0.897032072 | 0.190879981 | 0.354211518 |
| Csnk1g3 | S | 367 | 0.058035157 | -0.048958797 | 0.005194464 | -0.052840693 | 0.054153261 | 0.53585392 | 0.571271263 | 0.950151946 | 0.632235949 | 0.609030397 |
| Tmem245 | S | 329 | 0.116955544 | -0.073838609 | -0.02132661 | -0.138282154 | 0.052511999 | 0.135722716 | 0.183665844 | 0.691400921 | 0.106876345 | 0.384846261 |
| Chordc1 | T | 199 | -0.067519037 | 0.076048738 | 0.004864638 | 0.072383675 | -0.071184101 | 0.244623143 | 0.235066964 | 0.941105589 | 0.31444504 | 0.341625563 |
| Dst | S | 7348 | 0.129527989 | 0.027180795 | -0.155160662 | -0.284688651 | -0.182341457 | 0.283648294 | 0.828111288 | 0.29550293 | 0.115942505 | 0.289691979 |
| Gak | S | 824 | -0.101491483 | 0.097935196 | 0.003259383 | 0.104750866 | -0.094675813 | 0.112839975 | 0.079422564 | 0.948506501 | 0.150198925 | 0.160931815 |
| Slc23a2 | T | 79 | -0.087404718 | -0.04828667 | 0.117561448 | 0.204966165 | 0.165848118 | 0.42740884 | 0.728160034 | 0.197685047 | 0.099430009 | 0.269446788 |
| Sart1 | S | 492 | -0.019953572 | -0.058273562 | 0.120033516 | 0.139987088 | 0.178307078 | 0.643361665 | 0.18819765 | 0.191052327 | 0.15125462 | 0.101340481 |
| Spout1 | S | 378 | 0.069153069 | 0.006131023 | -0.053212127 | -0.122365196 | -0.059343149 | 0.047504691 | 0.773800206 | 0.019116525 | 0.00981809 | 0.051229485 |
| Hp1bp3 | S | 431 | -0.003648171 | 0.031608645 | -0.023269969 | -0.019621798 | -0.054878615 | 0.979468678 | 0.691699471 | 0.857599402 | 0.907106603 | 0.671916482 |
| Hp1bp3 | S | 443 | -0.003648171 | 0.031608645 | -0.023269969 | -0.019621798 | -0.054878615 | 0.979468678 | 0.691699471 | 0.857599402 | 0.907106603 | 0.671916482 |
| Hp1bp3 | S | 444 | -0.003648171 | 0.031608645 | -0.023269969 | -0.019621798 | -0.054878615 | 0.979468678 | 0.691699471 | 0.857599402 | 0.907106603 | 0.671916482 |
| Hp1bp3 | S | 447 | -0.003648171 | 0.031608645 | -0.023269969 | -0.019621798 | -0.054878615 | 0.979468678 | 0.691699471 | 0.857599402 | 0.907106603 | 0.671916482 |
| Ticrr | S | 1655 | 0.01323934 | -0.119223899 | 0.107192417 | 0.093953077 | 0.226416316 | 0.903267447 | 0.48419384 | 0.440209765 | 0.52251531 | 0.261255929 |
| Ticrr | T | 1653 | 0.01323934 | -0.119223899 | 0.107192417 | 0.093953077 | 0.226416316 | 0.903267447 | 0.48419384 | 0.440209765 | 0.52251531 | 0.261255929 |
| Tcf4 | S | 514 | 0.025399599 | -0.024063005 | -0.023408068 | -0.048807667 | 0.000654938 | 0.165414985 | 0.725159054 | 0.724615583 | 0.486311912 | 0.994036829 |
| Pnisr | S | 650 | 0.000797666 | -0.021210215 | 0.041711296 | 0.04091363 | 0.06292151 | 0.973356468 | 0.553196637 | 0.598680137 | 0.608793459 | 0.457811148 |

| | | | | | | | | | | | | |
|---|---|---|---|---|---|---|---|---|---|---|---|---|
| Mki67 | S | 1130 | 0.012761412 | 0.075635601 | -0.080531137 | -0.093292549 | -0.156166738 | 0.843408 | 0.101031442 | 0.106444528 | 0.182333682 | 0.011834164 |
| S100a11 | S | 11 | 0.011146598 | -0.063052098 | 0.07444664 | 0.063300042 | 0.137498738 | 0.96591236 | 0.745353312 | 0.725150768 | 0.787805889 | 0.3659486 |
| Slc20a2 | S | 435 | 0.219853091 | -0.061272251 | -0.088375685 | -0.308228776 | -0.027103434 | 0.11020246 | 0.092034559 | 0.422288566 | 0.065064623 | 0.786673945 |
| Zfp91 Gm44505 | S | 86 | -0.135176497 | 0.016353359 | 0.084929164 | 0.220105661 | 0.068575806 | 0.16832827 | 0.767072412 | 0.080600592 | 0.079445375 | 0.190847855 |
| Cic | S | 1511 | -0.057832358 | -0.037537517 | 0.097310888 | 0.155143246 | 0.134848405 | 0.616065373 | 0.770063762 | 0.387603794 | 0.183099781 | 0.293972752 |
| Zzef1 | S | 1515 | 0.012905125 | 0.044813216 | -0.039020496 | -0.051925621 | -0.083833712 | 0.821966253 | 0.374021828 | 0.278148296 | 0.38648505 | 0.136347651 |
| Eif5b | S | 184 | 0.028595229 | 0.014511471 | -0.055846853 | -0.084442082 | -0.070358323 | 0.130976044 | 0.420505335 | 0.166461762 | 0.080337622 | 0.11356064 |
| Nomo1 | S | 1196 | 0.013264978 | -0.028227833 | 0.002179219 | -0.011085759 | 0.030407052 | 0.180253739 | 0.588786288 | 0.971117288 | 0.854547597 | 0.683148743 |
| Ino80c | S | 26 | -0.050954917 | 0.041911501 | 0.020892153 | 0.07184707 | -0.021019349 | 0.243694081 | 0.391930082 | 0.583750344 | 0.160654189 | 0.677287634 |
| Gpatch8 | S | 753 | -0.028486705 | -0.023584572 | 0.045045936 | 0.073532641 | 0.068630508 | 0.439688401 | 0.285786261 | 0.055815143 | 0.107731171 | 0.008428553 |
| Eif3h | S | 17 | 0.097372183 | 0.011680911 | -0.061488706 | -0.158860889 | -0.073169617 | 0.220421766 | 0.869044478 | 0.085694634 | 0.098295844 | 0.341265195 |
| Pdcl | S | 293 | 0.001737476 | 0.073221843 | -0.065239474 | -0.06697695 | -0.138461317 | 0.982106697 | 0.169083881 | 0.18627794 | 0.409136791 | 0.03203568 |
| Nes | S | 478 | 0.070507521 | -0.061862498 | 0.009747685 | -0.060759836 | 0.071610183 | 0.306985532 | 0.25386976 | 0.863574855 | 0.425939011 | 0.295004014 |
| Il6st | S | 778 | 0.198038902 | -0.173687443 | 0.017026966 | -0.181011936 | 0.190714409 | 0.402984711 | 0.327037222 | 0.858004686 | 0.441141226 | 0.295184423 |
| Bud13 | S | 121 | 0.034492818 | -0.027692265 | 0.010818051 | -0.023674766 | 0.038510316 | 0.738379412 | 0.696437906 | 0.907093684 | 0.839151634 | 0.681194602 |
| Arhgef5 | S | 1110 | -0.039206845 | 0.02918508 | 0.007953463 | 0.047160308 | -0.021231616 | 0.551417848 | 0.463180484 | 0.839590939 | 0.451048632 | 0.43084709 |
| Top2b | S | 1400 | -0.035670032 | 0.012625453 | 0.026941923 | 0.062611955 | 0.01431647 | 0.484088375 | 0.738948517 | 0.516345234 | 0.221317829 | 0.650335225 |
| Tjp2 | S | 239 | -0.061867203 | -0.053933214 | 0.215459721 | 0.277326924 | 0.269392935 | 0.022800878 | 0.020006656 | 0.011875899 | 0.006600548 | 0.009397041 |
| Dkc1 | S | 453 | 0.05082343 | -0.011525481 | -0.013502184 | -0.064325614 | -0.001976703 | 0.495528017 | 0.642519819 | 0.598745917 | 0.398575858 | 0.900973298 |
| Ski | S | 340 | 0.049265558 | 0.034155619 | -0.064667628 | -0.113933186 | -0.098823247 | 0.585790916 | 0.641907052 | 0.376297163 | 0.226727228 | 0.183977011 |
| Dlg5 | S | 264 | 0.039905356 | -0.077443907 | 0.05582481 | 0.015919454 | 0.133268717 | 0.434781756 | 0.075860444 | 0.202580263 | 0.701852882 | 0.013117624 |
| Bcl9l | S | 988 | -0.073896537 | -0.043201328 | 0.124127405 | 0.198023941 | 0.167328733 | 0.339232621 | 0.308544719 | 0.093883245 | 0.060836226 | 0.043622929 |
| Stk4 | S | 320 | -0.015173503 | 0.056015865 | -0.053767145 | -0.038543962 | -0.10973333 | 0.767792912 | 0.059270639 | 0.185162031 | 0.507405011 | 0.038707445 |
| Cep250 | T | 1226 | -0.010093102 | 0.129947264 | -0.139636386 | -0.129543284 | -0.26958365 | 0.971230041 | 0.553793148 | 0.692676516 | 0.728437322 | 0.446112684 |
| 2810004N23Rik | S | 106 | 0.027111543 | -0.056289184 | 0.027285138 | 0.000173595 | 0.083574321 | 0.077816192 | 0.013366484 | 0.079885623 | 0.97393541 | 0.00888898 |
| Eif4g3 | S | 266 | -0.014364263 | 0.054907338 | -0.029350577 | -0.014986315 | -0.084257915 | 0.839234436 | 0.481612919 | 0.625086996 | 0.851718576 | 0.343500339 |
| Eif4g3 | T | 265 | -0.014364263 | 0.054907338 | -0.029350577 | -0.014986315 | -0.084257915 | 0.839234436 | 0.481612919 | 0.625086996 | 0.851718576 | 0.343500339 |
| Ptpn1 | S | 392 | 0.022627757 | 0.055397095 | -0.080709619 | -0.103337376 | -0.136106714 | 0.488824124 | 0.5662508 | 0.398979435 | 0.301596995 | 0.281873416 |
| Dennd1b | S | 517 | -0.16583883 | 0.12250489 | 0.077506439 | 0.243345269 | -0.044998452 | 0.012924622 | 0.112377502 | 0.180763403 | 0.004319766 | 0.461682236 |
| Usp8 | S | 535 | 0.033668869 | 0.017080037 | -0.034469977 | -0.068138846 | -0.051550014 | 0.674263884 | 0.842684385 | 0.580274203 | 0.303691749 | 0.469614312 |
| Skap2 | Y | 237 | -0.033384826 | 0.071278392 | -0.057414007 | -0.024029181 | -0.128692398 | 0.848948842 | 0.467753148 | 0.756693979 | 0.912469801 | 0.484574567 |
| Arhgef18 | S | 1336 | -0.054392399 | 0.067052821 | -0.001811083 | 0.052581316 | -0.068863904 | 0.319909036 | 0.296218843 | 0.988245354 | 0.660186749 | 0.578893511 |
| Wac | S | 12 | 0.112198903 | -0.128970628 | 0.018518289 | -0.093680614 | 0.147488917 | 0.132708794 | 0.133674063 | 0.794737801 | 0.223147404 | 0.112627558 |
| Cnot2 | S | 391 | 0.078866292 | -0.19547485 | 0.151651573 | 0.072785282 | 0.347126424 | 0.557883394 | 0.157843174 | 0.316321721 | 0.668811321 | 0.081300701 |
| Slk | S | 377 | 0.06173825 | -0.01345746 | -0.076163901 | -0.137902151 | -0.062706442 | 0.123378984 | 0.794539117 | 0.376782973 | 0.166824679 | 0.468927961 |
| Cep170b | S | 653 | 0.082264935 | -0.027924669 | -0.03692513 | -0.119190065 | -0.009000461 | 0.32340689 | 0.680621587 | 0.455349742 | 0.187483214 | 0.892779039 |
| Pds5b | S | 1360 | 0.010426585 | -0.013716726 | 0.013785585 | 0.003358999 | 0.027502311 | 0.871947916 | 0.783338902 | 0.831136782 | 0.967062469 | 0.704221668 |
| Evl | S | 352 | -0.133242759 | 0.098070465 | 0.091693875 | 0.224936633 | -0.006376591 | 0.08181243 | 0.439662622 | 0.318226103 | 0.047298585 | 0.959087554 |
| Dcp2 | S | 246 | 0.006485207 | -0.024945222 | 0.023828896 | 0.017343688 | 0.048774118 | 0.865772061 | 0.598639218 | 0.585621346 | 0.738505666 | 0.412851655 |
| Dcp2 | S | 247 | 0.006485207 | -0.024945222 | 0.023828896 | 0.017343688 | 0.048774118 | 0.865772061 | 0.598639218 | 0.585621346 | 0.738505666 | 0.412851655 |
| Dcp2 | S | 249 | 0.006485207 | -0.024945222 | 0.023828896 | 0.017343688 | 0.048774118 | 0.865772061 | 0.598639218 | 0.585621346 | 0.738505666 | 0.412851655 |
| Mark3 | S | 489 | 0.046404103 | -0.003065019 | -0.039870408 | -0.086274511 | -0.036805389 | 0.437736997 | 0.921224727 | 0.570286812 | 0.305375703 | 0.587553662 |
| Anks1 | Y | 472 | -0.091909578 | 0.065883107 | 0.044407447 | 0.136317025 | -0.02147566 | 0.147823632 | 0.263793876 | 0.591592347 | 0.171231317 | 0.799209944 |
| Zc3h13 | S | 921 | -0.049062915 | -0.001800153 | 0.034329857 | 0.083392771 | 0.03613001 | 0.455977024 | 0.968462832 | 0.205010901 | 0.248992881 | 0.422013099 |
| Bcar1 | Y | 253 | -0.042330059 | 0.017511896 | 0.023488498 | 0.065818557 | 0.005976602 | 0.564815977 | 0.649543563 | 0.665918612 | 0.397309818 | 0.890704684 |
| Adrb2 | S | 246 | -0.13926915 | -0.010470589 | 0.093226714 | 0.232495864 | 0.103697303 | 0.213649916 | 0.867259283 | 0.048427269 | 0.092989939 | 0.151347381 |
| Smarcad1 | S | 33 | -0.022537164 | 7.23232E-05 | 0.033944474 | 0.056481638 | 0.033872151 | 0.557677687 | 0.999241869 | 0.546265364 | 0.359561743 | 0.694712183 |
| Map1b | S | 989 | 0.035962466 | 0.052851228 | -0.081766641 | -0.117729108 | -0.134617869 | 0.689183485 | 0.583810386 | 0.447759428 | 0.297887132 | 0.25895361 |
| Mcm2 | S | 108 | -0.047475754 | 0.028468038 | 0.030339404 | 0.077815158 | 0.001871366 | 0.41011453 | 0.719922784 | 0.541783693 | 0.108244878 | 0.977414572 |
| Khdc4 | S | 570 | 0.12719209 | -0.077377536 | -0.087579838 | -0.214771928 | -0.010202302 | 0.009195741 | 0.350448675 | 0.089117228 | 0.014385574 | 0.893069416 |
| Pde4a | S | 204 | -0.07671688 | 0.043299197 | 0.052285714 | 0.129002594 | 0.008986518 | 0.069862664 | 0.144386991 | 0.503214004 | 0.16697089 | 0.900034284 |
| Khdc4 | S | 569 | 0.123775408 | -0.033584734 | -0.099609598 | -0.223385006 | -0.066024865 | 0.101407463 | 0.588011723 | 0.249804186 | 0.05183984 | 0.458649463 |
| Mcm10 | T | 861 | 0.050627911 | 0.053698751 | -0.08378643 | -0.134414341 | -0.137485181 | 0.561347408 | 0.384858503 | 0.192478191 | 0.150209856 | 0.004025497 |
| Rb1cc1 | S | 652 | 0.02977264 | 0.065829649 | -0.088640451 | -0.118413091 | -0.154470101 | 0.586608899 | 0.312210094 | 0.199950501 | 0.088178213 | 0.055941163 |
| Chtop | T | 243 | 0.043612396 | 0.010621611 | -0.056691875 | -0.100304271 | -0.067313486 | 0.297791796 | 0.847485372 | 0.341515958 | 0.119935456 | 0.29448073 |
| Nemf | S | 417 | 0.050895753 | 0.0348321 | -0.071812664 | -0.122708417 | -0.106644764 | 0.42225933 | 0.286324159 | 0.115988225 | 0.110883979 | 0.043077598 |
| Sart1 | S | 480 | -0.020997608 | -0.058833665 | 0.118454675 | 0.139452283 | 0.17728834 | 0.624161488 | 0.179641874 | 0.191989912 | 0.149209702 | 0.100476216 |
| Tcof1 | T | 1114 | 0.115278368 | -0.009755812 | -0.086792739 | -0.202071107 | -0.077036927 | 0.167938684 | 0.820721462 | 0.181286523 | 0.049565466 | 0.195110063 |
| Telo2 | S | 465 | 0.045188757 | -0.033690702 | 0.004311549 | -0.040877208 | 0.038002251 | 0.62552461 | 0.578554945 | 0.951393704 | 0.627983076 | 0.428762981 |
| Nes | S | 1562 | 0.069628207 | -0.038967196 | -0.052427853 | -0.122056061 | -0.013460657 | 0.017187389 | 0.18695934 | 0.297339695 | 0.080115046 | 0.765257496 |
| Arhgap32 | S | 830 | 0.024600476 | -0.009992192 | -0.011123398 | -0.035723874 | -0.001131206 | 0.415729517 | 0.624016867 | 0.76695698 | 0.394937451 | 0.974198329 |
| Rem1 | S | 290 | -0.108007955 | 0.201115718 | -0.071632684 | 0.036375271 | -0.272748402 | 0.345805515 | 0.166685842 | 0.454503585 | 0.770788435 | 0.09661791 |
| Med13 | T | 329 | -0.128692947 | 0.058598794 | 0.050005468 | 0.178698415 | -0.008593327 | 0.197558296 | 0.350715083 | 0.338084607 | 0.109394245 | 0.881058338 |
| Usp8 | S | 681 | -0.051085422 | 0.073090291 | -0.008670528 | 0.042414894 | -0.081760819 | 0.410663329 | 0.374072433 | 0.893742315 | 0.567553107 | 0.370495752 |
| Akap17b | S | 753 | -0.051946503 | -0.002551801 | 0.056715441 | 0.108661944 | 0.059267241 | 0.137418122 | 0.968318155 | 0.110832655 | 0.001209503 | 0.373756645 |
| Akap17b | S | 754 | -0.051946503 | -0.002551801 | 0.056715441 | 0.108661944 | 0.059267241 | 0.137418122 | 0.968318155 | 0.110832655 | 0.001209503 | 0.373756645 |
| G3bp1 | S | 231 | 0.024190735 | 0.038200434 | -0.055313127 | -0.079503862 | -0.09351356 | 0.542020645 | 0.243491159 | 0.137885047 | 0.080406217 | 0.02294521 |
| Wnk1 | S | 2060 | -0.036179356 | 0.073150905 | -0.126703155 | -0.090523799 | -0.19985406 | 0.333350128 | 0.001748968 | 0.189158068 | 0.302583082 | 0.090829307 |
| Tubgcp3 | T | 214 | 0.031993415 | 0.010219704 | -0.023718251 | -0.055711666 | -0.033937956 | 0.328397684 | 0.711259222 | 0.117230348 | 0.144837725 | 0.268726914 |
| Birc2 | S | 152 | 0.153900375 | -0.123663548 | -0.026251944 | -0.180152319 | 0.097411604 | 0.051079696 | 0.07586326 | 0.735893111 | 0.094519384 | 0.290560084 |
| Cobll1 | S | 384 | 0.035789299 | 0.125265014 | -0.135061209 | -0.170850508 | -0.260326222 | 0.645204963 | 0.147687152 | 0.074221544 | 0.045596282 | 0.017932665 |
| Prkar1a | S | 77 | 0.088151878 | 0.099068409 | -0.178629317 | -0.266781195 | -0.277697725 | 0.28764648 | 0.347680074 | 0.106079546 | 0.042867154 | 0.056568747 |
| Irs1 | S | 414 | -0.10068781 | 0.001746939 | 0.107431977 | 0.208119787 | 0.105685039 | 0.452267218 | 0.965783021 | 0.440827327 | 0.253031375 | 0.448562522 |
| Tmcc3 | S | 15 | 0.004829177 | 0.038782005 | -0.023831587 | -0.028660764 | -0.062613592 | 0.921335917 | 0.588447972 | 0.429607778 | 0.557478835 | 0.401843271 |
| Mia3 | S | 1754 | 0.137105054 | -0.010647789 | -0.054113967 | -0.191219021 | -0.043466179 | 0.54020375 | 0.816037855 | 0.399391596 | 0.414569553 | 0.489419036 |
| Mphosph10 | S | 346 | -0.036324708 | -0.061879875 | 0.101364456 | 0.137689164 | 0.163244331 | 0.547337489 | 0.157632672 | 0.07916037 | 0.067204559 | 0.024099064 |
| Pawr | S | 243 | -0.037913912 | -0.052570203 | 0.11128452 | 0.149198432 | 0.163854723 | 0.817382344 | 0.539260247 | 0.473438277 | 0.469477351 | 0.324066514 |
| Sh3pxd2a | S | 419 | 0.120176185 | 0.063673904 | -0.159573135 | -0.27974932 | -0.223247039 | 0.456632949 | 0.670814618 | 0.287751048 | 0.10303723 | 0.141255858 |
| Tmem209 | T | 114 | 0.057290788 | -0.171194141 | 0.130259491 | 0.072968703 | 0.301453631 | 0.766586528 | 0.217576809 | 0.2786817 | 0.688246406 | 0.044754082 |
| Zfp318 | S | 2110 | 0.039625989 | -0.062875134 | 0.035659475 | -0.003966514 | 0.098534609 | 0.657299494 | 0.478505053 | 0.727776921 | 0.973759813 | 0.427545499 |
| Fam193b | S | 776 | 0.002857875 | -0.005085511 | 0.006756537 | 0.003898661 | 0.011842047 | 0.968453312 | 0.886737779 | 0.832109988 | 0.953951746 | 0.573181772 |
| Wdr62 | S | 32 | 0.006204436 | 0.068806993 | -0.073374612 | -0.079579048 | -0.142181605 | 0.926785056 | 0.135849059 | 0.150994724 | 0.313308604 | 0.036678195 |
| Srrm1 | S | 401 | 0.01355733 | -0.011971113 | 0.002102708 | -0.011454622 | 0.014073821 | 0.619097491 | 0.636067576 | 0.926216804 | 0.599171254 | 0.466884951 |
| Dop1a | S | 1140 | 0.014008034 | -0.040899493 | 0.038573499 | 0.024565464 | 0.079472991 | 0.823299566 | 0.409430464 | 0.52172226 | 0.744567541 | 0.260542091 |
| Ubap2l | S | 513 | 0.015029802 | 0.137736087 | -0.138282593 | -0.153312395 | -0.276018679 | 0.811979694 | 0.184061078 | 0.141235832 | 0.10465845 | 0.037012319 |
| Cds2 | S | 32 | -0.08159309 | 0.010932617 | 0.060662583 | 0.142255674 | 0.049729966 | 0.331334923 | 0.692979653 | 0.34186706 | 0.155687291 | 0.40960589 |
| Kifc3 | S | 86 | -0.061082084 | 0.035660114 | 0.02995957 | 0.091041653 | -0.005700544 | 0.11986844 | 0.264884267 | 0.4251696 | 0.063545711 | 0.868951666 |
| Itsn2 | S | 841 | 0.037428846 | -0.018713426 | -0.032848405 | -0.070277251 | -0.014134979 | 0.241432535 | 0.757387007 | 0.54696397 | 0.229701028 | 0.836120252 |
| Fbxo38 | S | 730 | 0.087835734 | -0.033812305 | -0.046075092 | -0.133910826 | -0.012262787 | 0.686329845 | 0.833343664 | 0.85250116 | 0.636854191 | 0.960313275 |
| Cic | T | 1051 | -0.114766476 | -0.02833206 | 0.135525737 | 0.250292213 | 0.163857797 | 0.123539691 | 0.625040056 | 0.057244696 | 0.018530981 | 0.055427749 |
| Akap13 | T | 1910 | -0.080726143 | 0.117871525 | -0.043825435 | 0.036900708 | -0.161696959 | 0.111493395 | 0.096492128 | 0.84174989 | 0.866255779 | 0.491074425 |
| Cwc22 | S | 97 | -0.089734749 | -0.043601753 | 0.128904271 | 0.21863902 | 0.172506024 | 0.199489882 | 0.54369907 | 0.087678609 | 0.028122483 | 0.070890188 |
| Arhgap17 | S | 570 | 0.05485351 | 0.008751647 | -0.077661431 | -0.132514941 | -0.086413078 | 0.552483202 | 0.914851684 | 0.623375541 | 0.426506062 | 0.584136634 |
| Wbp4 | S | 228 | 0.08550198 | -0.075673381 | 0.005567332 | -0.079934648 | 0.081240712 | 0.018365111 | 0.010186998 | 0.922079132 | 0.239481208 | 0.234624154 |

| | | | | | | | | | | | | | |
|---|---|---|---|---|---|---|---|---|---|---|---|---|---|
| Mia3 | S | | 772 | -0.180080921 | -0.014865658 | 0.152379182 | 0.332460103 | 0.167244841 | 0.064401081 | 0.767729796 | 0.018101586 | 0.024769228 | 0.002238729 |
| Fam122a | S | | 34 | -0.04176973 | 0.028844147 | 0.016456979 | 0.058226708 | -0.012387169 | 0.43441622 | 0.544324968 | 0.743132816 | 0.28755423 | 0.786093698 |
| Fam122a | S | | 45 | -0.04176973 | 0.028844147 | 0.016456979 | 0.058226708 | -0.012387169 | 0.43441622 | 0.544324968 | 0.743132816 | 0.28755423 | 0.786093698 |
| Ccnl1 | S | | 361 | -0.094054249 | 0.034672654 | 0.117801849 | 0.211856098 | 0.083129195 | 0.020276144 | 0.386970351 | 0.193940103 | 0.069433393 | 0.321884214 |
| Pop5 | S | | 159 | 0.009385212 | 0.011094672 | -0.015778568 | -0.025163781 | -0.02687324 | 0.662068661 | 0.583465334 | 0.376837403 | 0.226061711 | 0.167242471 |
| Tmem119 | S | | 164 | -0.0225841 | 0.015482712 | 0.009272245 | 0.031856345 | -0.006210467 | 0.754457266 | 0.769119687 | 0.898968585 | 0.671994298 | 0.914187368 |
| Tmem119 | T | | 175 | -0.0225841 | 0.015482712 | 0.009272245 | 0.031856345 | -0.006210467 | 0.754457266 | 0.769119687 | 0.898968585 | 0.671994298 | 0.914187368 |
| Pnn | S | | 380 | 0.039020124 | -0.03771826 | -0.001221815 | -0.040241939 | 0.036496445 | 0.330720309 | 0.393642716 | 0.980471156 | 0.343626072 | 0.413606027 |
| Arhgef2 | S | | 951 | -0.104674178 | -0.017375378 | 0.146611663 | 0.251285841 | 0.163987041 | 0.415128633 | 0.820617447 | 0.379015056 | 0.205355916 | 0.33872578 |
| Arhgef2 | S | | 952 | -0.104674178 | -0.017375378 | 0.146611663 | 0.251285841 | 0.163987041 | 0.415128633 | 0.820617447 | 0.379015056 | 0.205355916 | 0.33872578 |
| Arhgef17 | S | | 953 | -0.02302701 | 0.147877927 | -0.24736745 | -0.22434044 | -0.395245377 | 0.794695376 | 0.042490372 | 0.237992823 | 0.274348398 | 0.10965465 |
| Rnf216 | S | | 407 | -0.088162471 | 0.047303937 | 0.020059429 | 0.1082219 | -0.027244508 | 0.468754469 | 0.503315259 | 0.649894783 | 0.379859323 | 0.643413183 |
| Sema6d | T | | 773 | 0.174890604 | -0.077292793 | -0.126803801 | -0.301694406 | -0.049511009 | 0.089740403 | 0.348899187 | 0.381870057 | 0.093469831 | 0.711498837 |
| Dhx29 | S | | 522 | 0.222766194 | -0.094239515 | -0.156149216 | -0.378915411 | -0.061909701 | 0.468598566 | 0.769551757 | 0.69304423 | 0.415823148 | 0.89260974 |
| Khdrbs1 | S | | 38 | -0.065801286 | -0.047049345 | 0.201818438 | 0.267619724 | 0.248867782 | 0.210115964 | 0.545901409 | 0.284616033 | 0.196261984 | 0.211582291 |
| Ncor2 | S | | 938 | 0.061791758 | -0.011526189 | -0.035547039 | -0.097338798 | -0.02402085 | 0.213280427 | 0.583753864 | 0.269517218 | 0.091714826 | 0.398210037 |
| Ap2m1 | T | | 156 | -0.013889188 | 0.030919185 | -0.017316667 | -0.003427479 | -0.048235851 | 0.798162964 | 0.410273788 | 0.587830366 | 0.946089816 | 0.173986848 |
| Apc | S | | 2674 | 0.077888358 | -0.058815008 | -0.032910857 | -0.110799215 | 0.025904151 | 0.104307488 | 0.367353981 | 0.446752629 | 0.027868597 | 0.655478863 |
| Mrtfa | S | | 605 | 0.098861768 | -0.008917738 | -0.073322954 | -0.172184723 | -0.064405217 | 0.306211953 | 0.757461784 | 0.346483983 | 0.145057641 | 0.399197031 |
| Nav1 | S | | 756 | 0.110997567 | -0.036664957 | -0.044382205 | -0.155379772 | -0.007717248 | 0.459106725 | 0.564965966 | 0.658940908 | 0.336155835 | 0.926760143 |
| Rock2 | S | | 1137 | -0.097863762 | 0.084444452 | 0.028549277 | 0.126413039 | -0.055895175 | 0.072901291 | 0.131290904 | 0.678095541 | 0.134110561 | 0.447343469 |
| Chd7 | S | | 2171 | 0.065575617 | -0.09468042 | 0.072064027 | 0.00648841 | 0.166744447 | 0.116485799 | 0.002795836 | 0.129643279 | 0.883243419 | 0.024395349 |
| Tjap1 | S | | 330 | 0.147807517 | -0.010036952 | -0.179659213 | -0.32746673 | -0.169622261 | 0.051019377 | 0.855588876 | 0.125448277 | 0.03411127 | 0.156809561 |
| Uba2 | S | | 207 | -0.090286668 | 0.006911487 | 0.11828476 | 0.208571428 | 0.111373273 | 0.074080836 | 0.861918393 | 0.160124832 | 0.049695244 | 0.182748601 |
| Mast4 | S | | 1417 | -0.074690406 | 0.006725538 | 0.084824382 | 0.159514788 | 0.078098843 | 0.030326924 | 0.876580565 | 0.056348394 | 0.010485762 | 0.144602738 |
| Inpp5f | S | | 879 | -0.027543678 | 0.076570933 | -0.037676457 | -0.010132779 | -0.11424739 | 0.19758899 | 0.164921469 | 0.560875326 | 0.866947012 | 0.162940844 |
| Vcpip1 | S | | 997 | -0.023606228 | 0.023776221 | 0.012559755 | 0.036165982 | -0.011216466 | 0.330089259 | 0.534030755 | 0.805593171 | 0.483013551 | 0.837685731 |
| Vcl | S | | 346 | -0.122037611 | 0.09228824 | 0.069881627 | 0.191919238 | -0.022406612 | 0.083003775 | 0.240066424 | 0.562278584 | 0.173965997 | 0.848702472 |
| Srrm2 | S | | 2656 | -0.005313673 | -0.039146242 | 0.064132261 | 0.069445935 | 0.103278503 | 0.804369149 | 0.077520478 | 0.12875459 | 0.114538165 | 0.054470299 |
| Rad54l2 | S | | 1168 | -0.014215305 | -0.016706431 | 0.050070857 | 0.064286162 | 0.066777288 | 0.712268734 | 0.686150825 | 0.597153827 | 0.510005198 | 0.497617372 |
| Nipbl | S | | 1144 | -0.052707399 | 0.032402436 | 0.006865762 | 0.059573161 | -0.025536674 | 0.495073864 | 0.35762188 | 0.938106858 | 0.581564892 | 0.776740252 |
| Nipbl | S | | 1146 | -0.052707399 | 0.032402436 | 0.006865762 | 0.059573161 | -0.025536674 | 0.495073864 | 0.35762188 | 0.938106858 | 0.581564892 | 0.776740252 |
| Nipbl | S | | 1148 | -0.052707399 | 0.032402436 | 0.006865762 | 0.059573161 | -0.025536674 | 0.495073864 | 0.35762188 | 0.938106858 | 0.581564892 | 0.776740252 |
| Nipbl | S | | 1154 | -0.052707399 | 0.032402436 | 0.006865762 | 0.059573161 | -0.025536674 | 0.495073864 | 0.35762188 | 0.938106858 | 0.581564892 | 0.776740252 |
| Irf2bp1 | S | | 453 | 0.013055558 | -0.009232262 | -0.00590394 | -0.018959497 | 0.003328322 | 0.811447219 | 0.935923201 | 0.954126436 | 0.841456923 | 0.980064901 |
| Prpf40a | T | | 928 | -0.022783928 | -0.052261559 | 0.087864804 | 0.110648732 | 0.140126363 | 0.515465126 | 0.155395194 | 0.104368745 | 0.064952686 | 0.04688223 |
| Seh1l | S | | 190 | 0.051683702 | 0.020377536 | -0.100482805 | -0.152166507 | -0.120860341 | 0.422924629 | 0.548526394 | 0.382347897 | 0.231658759 | 0.310755461 |
| Nsd2 | S | | 376 | -0.099687054 | -0.008815935 | 0.114175435 | 0.213862488 | 0.122991369 | 0.437078758 | 0.9535714 | 0.41188673 | 0.17658663 | 0.464830987 |
| Nsd2 | S | | 377 | -0.099687054 | -0.008815935 | 0.114175435 | 0.213862488 | 0.122991369 | 0.437078758 | 0.9535714 | 0.41188673 | 0.17658663 | 0.464830987 |
| Nsd2 | S | | 380 | -0.099687054 | -0.008815935 | 0.114175435 | 0.213862488 | 0.122991369 | 0.437078758 | 0.9535714 | 0.41188673 | 0.17658663 | 0.464830987 |
| Ndc1 | S | | 499 | 0.154026429 | -0.071220975 | -0.122061883 | -0.276088312 | -0.050840908 | 0.175845474 | 0.628746739 | 0.419442681 | 0.114822776 | 0.76286554 |
| Ap1b1 | S | | 783 | -0.184491688 | 0.409448488 | -0.36918829 | -0.184696602 | -0.778636778 | 0.442745944 | 0.205661671 | 0.638107607 | 0.8110634 | 0.365365975 |
| Eif5b | S | | 183 | 0.020420865 | 0.018948153 | -0.056534545 | -0.07695541 | -0.075482698 | 0.232294007 | 0.298752712 | 0.174186077 | 0.103276589 | 0.104218137 |
| E2f6 | S | | 254 | 0.043914737 | 0.021520515 | -0.060117652 | -0.104032389 | -0.081638167 | 0.280388029 | 0.492179184 | 0.129998467 | 0.036291137 | 0.044022476 |
| Dennd2a | S | | 401 | 0.0754091 | 0.008108487 | -0.104352041 | -0.179761141 | -0.112460528 | 0.389784414 | 0.870142313 | 0.445334587 | 0.243456549 | 0.409898225 |
| Pkp4 | S | | 509 | 0.033302719 | -0.002335492 | -0.041078644 | -0.074381363 | -0.038743152 | 0.556932393 | 0.969143982 | 0.697564784 | 0.479486415 | 0.705351523 |
| Pkp4 | S | | 511 | 0.033302719 | -0.002335492 | -0.041078644 | -0.074381363 | -0.038743152 | 0.556932393 | 0.969143982 | 0.697564784 | 0.479486415 | 0.705351523 |
| Pkp4 | S | | 514 | 0.033302719 | -0.002335492 | -0.041078644 | -0.074381363 | -0.038743152 | 0.556932393 | 0.969143982 | 0.697564784 | 0.479486415 | 0.705351523 |
| Mars1 | T | | 895 | 0.110384392 | -0.039468814 | -0.097880788 | -0.208265179 | -0.058411973 | 0.095086613 | 0.592940901 | 0.272645368 | 0.063995349 | 0.482688549 |
| D1Ertd622e | S | | 42 | 0.346112123 | -0.119077385 | -0.352722287 | -0.69883441 | -0.233644902 | 0.031485109 | 0.315447792 | 0.235512302 | 0.069688373 | 0.389077494 |
| Washc2 | S | | 1333 | 0.037165165 | -0.070697738 | 0.040507832 | 0.003342667 | 0.11120557 | 0.189570841 | 0.043128693 | 0.264793297 | 0.899142226 | 0.019608115 |
| Pds5b | S | | 1415 | -0.168276183 | 0.153073586 | 0.04080308 | 0.209079262 | -0.112270506 | 0.072241488 | 0.108932238 | 0.81446507 | 0.279674412 | 0.522155579 |
| Tenm3 | S | | 199 | 0.099058888 | -0.01500672 | -0.133125856 | -0.232184744 | -0.118119136 | 0.027632973 | 0.873187644 | 0.110015953 | 0.029144054 | 0.301181566 |
| Setd2 | T | | 1827 | -0.020515711 | 0.011367171 | 0.020239405 | 0.040755116 | 0.008872234 | 0.423183912 | 0.813616377 | 0.651786229 | 0.376077739 | 0.875881953 |
| Mki67 | S | | 1825 | 0.041562165 | 0.017053089 | -0.038182808 | -0.079744973 | -0.055235897 | 0.388103127 | 0.662468884 | 0.141021293 | 0.154318707 | 0.214789269 |
| Rai14 | T | | 249 | 0.117424797 | -0.016147675 | -0.088130482 | -0.205555279 | -0.071982807 | 0.066742382 | 0.552576192 | 0.093151688 | 0.015920425 | 0.149358849 |
| Itprid2 | S | | 270 | -0.121177068 | 0.183106844 | -0.071622821 | 0.049554247 | -0.254729665 | 0.106139246 | 0.026451446 | 0.320032141 | 0.508432928 | 0.019415704 |
| Nav1 | S | | 1012 | -0.015186107 | -0.016544902 | 0.037563859 | 0.052750776 | 0.05410876 | 0.812037902 | 0.863450399 | 0.708945797 | 0.582305136 | 0.641369773 |
| Chd4 | S | | 421 | -0.00813394 | -0.020698778 | 0.039010835 | 0.047144775 | 0.059709613 | 0.794154664 | 0.23087815 | 0.271529003 | 0.263946326 | 0.138587065 |
| Pask | S | | 488 | 0.046468912 | 0.008844683 | -0.04196004 | -0.088428952 | -0.050804723 | 0.739123251 | 0.945916796 | 0.656333612 | 0.502523059 | 0.668053999 |
| Strada | S | | 313 | -0.040666842 | 0.108894629 | -0.092278604 | -0.051611762 | -0.201173233 | 0.809250155 | 0.223010325 | 0.367897983 | 0.769125661 | 0.103714695 |
| Ranbp3 | S | | 490 | 0.019068337 | -0.029143613 | 0.013883599 | -0.005184738 | 0.043027213 | 0.300778082 | 0.247967149 | 0.785910867 | 0.917043627 | 0.434337978 |
| Ctu1 | S | | 378 | -0.076517904 | 0.103605997 | 0.005003043 | 0.081520947 | -0.098602954 | 0.209234838 | 0.331458951 | 0.954426285 | 0.359672369 | 0.399634728 |
| Bcorl1 | S | | 1474 | 0.078184156 | -0.047488336 | -0.035448856 | -0.113633012 | 0.01203948 | 0.149610146 | 0.338987454 | 0.521243158 | 0.081445519 | 0.811277552 |
| Mcm2 | S | | 41 | -0.028082848 | -0.064690934 | 0.089112539 | 0.117195387 | 0.153803474 | 0.679867405 | 0.195414978 | 0.118803572 | 0.148143734 | 0.028035257 |
| 2610507B11Rik | T | | 1269 | 4.35402E-05 | -0.058771323 | 0.073876188 | 0.073832647 | 0.132647511 | 0.999673095 | 0.517954773 | 0.572168849 | 0.634300237 | 0.380030311 |
| Atrip | S | | 97 | 0.034593673 | -0.001422613 | -0.032469005 | -0.067062678 | -0.031046392 | 0.215347572 | 0.958417534 | 0.235835782 | 0.063385032 | 0.326683722 |
| Xrcc4 | S | | 319 | 0.002214375 | 0.009391962 | -0.013079374 | -0.015293748 | -0.022471336 | 0.976685008 | 0.812813481 | 0.847577571 | 0.857913342 | 0.711990177 |
| Asxl2 | S | | 373 | 0.045945737 | -0.070086413 | 0.037434228 | -0.008511509 | 0.10752064 | 0.348400981 | 0.060972624 | 0.2539676 | 0.825168947 | 0.002610543 |
| Ptpn21 | S | | 538 | -0.010478615 | 0.019854042 | -0.001548161 | 0.008930454 | -0.021402203 | 0.580168543 | 0.681628177 | 0.96020476 | 0.770246148 | 0.682393875 |
| Mia2 | S | | 1178 | 0.020658967 | -0.045919437 | 0.044106517 | 0.02344755 | 0.090025954 | 0.870076884 | 0.339741336 | 0.510053192 | 0.854505786 | 0.183470269 |
| Ash1l | S | | 1678 | 0.068019026 | -0.049789321 | 0.003942426 | -0.0640766 | 0.053731747 | 0.320090851 | 0.175209753 | 0.908825591 | 0.328929268 | 0.019180744 |
| Taf1 | S | | 1173 | 0.001988636 | 0.006985899 | -0.007772766 | -0.009761402 | -0.014758664 | 0.982933153 | 0.924345444 | 0.91491571 | 0.906485147 | 0.80430066 |
| Taf1 | T | | 1175 | 0.001988636 | 0.006985899 | -0.007772766 | -0.009761402 | -0.014758664 | 0.982933153 | 0.924345444 | 0.91491571 | 0.906485147 | 0.80430066 |
| Nup98 | S | | 612 | 0.007326988 | -0.035924705 | 0.036294866 | 0.028967877 | 0.07221957 | 0.911020331 | 0.589587444 | 0.699745844 | 0.751464546 | 0.450066018 |
| Ncam1 | S | | 770 | 0.085613359 | -0.036256786 | -0.04956981 | -0.135183169 | -0.013313024 | 0.028036676 | 0.231497168 | 0.125492206 | 0.004445503 | 0.383268162 |
| Dido1 | T | | 1305 | -0.020966958 | -0.067485103 | 0.099523083 | 0.120490041 | 0.167008185 | 0.904630481 | 0.488781984 | 0.488399916 | 0.561680818 | 0.299953062 |
| Rtn4 | S | | 165 | 0.009760208 | -0.00443075 | -0.007312053 | -0.017072261 | -0.002881303 | 0.73525761 | 0.856066741 | 0.901124323 | 0.769842802 | 0.959198802 |
| Gphn | S | | 280 | 0.096447312 | -0.006422325 | -0.179190656 | -0.275637968 | -0.172768331 | 0.101489485 | 0.869409799 | 0.340247429 | 0.191470441 | 0.35438071 |
| --- | S | --- | | -0.044693847 | -0.782386299 | -0.693880728 | -0.649186881 | 0.088505572 | 0.549829033 | 4.70601E-09 | 0.005834032 | 0.00204778 | 0.312488474 |
| --- | S | --- | | -0.96221903 | -0.71153625 | -1.77604281 | -0.81382378 | -1.06450656 | 0.037225468 | 0.209078669 | 0.012101809 | 0.134555323 | 0.112202468 |
| --- | S | --- | | -0.647723583 | -0.629496556 | -1.431926919 | -0.784203336 | -0.802430363 | 0.030902353 | 0.163362632 | 0.009025936 | 0.070554884 | 0.116624815 |
| --- | S | --- | | -0.229441137 | -0.20687115 | -0.513286206 | -0.283845069 | -0.306415056 | 0.00379262 | 0.004310788 | 1.67778E-05 | 0.002849295 | 0.002707836 |
| --- | S | --- | | -0.609558802 | -0.523614638 | -0.92466346 | -0.315104658 | -0.401048822 | 0.032021293 | 0.090553679 | 0.019612146 | 0.271945149 | 0.213544439 |
| --- | S | --- | | -0.75302385 | -0.514131985 | -0.740424042 | 0.012599808 | -0.226292056 | 0.015496396 | 0.103662614 | 0.03088477 | 0.949154715 | 0.398303835 |
| --- | T | --- | | -0.170438061 | -0.32036004 | -0.28062972 | -0.11019166 | 0.03973032 | 0.009141367 | 2.75837E-05 | 6.80467E-05 | 0.056777104 | 0.102725465 |
| --- | S | --- | | -0.09653924 | -0.538322748 | -0.818620112 | -0.722080871 | -0.280297363 | 0.252043321 | 0.016471452 | 0.001800134 | 0.001159448 | 0.060399812 |
| --- | S | --- | | -0.594934892 | -0.688738866 | -0.453647493 | 0.1412874 | 0.235091374 | 0.01331665 | 0.03109586 | 0.185108002 | 0.624796515 | 0.472134001 |
| --- | T | --- | | -1.183508672 | -0.675370545 | -0.399046574 | 0.784462098 | 0.276323971 | 0.033957756 | 0.098644903 | 0.479631826 | 0.194869202 | 0.564570547 |
| --- | S | --- | | -0.192551775 | -0.324846038 | -0.747331343 | -0.554779568 | -0.422485306 | 0.167928295 | 0.113728314 | 0.000815061 | 0.002206265 | 0.061097748 |
| --- | S | --- | | 0.026563886 | -0.350156547 | -0.353264844 | -0.37982873 | -0.003108297 | 0.768418831 | 0.000572269 | 8.00817E-06 | 0.033855045 | 0.937118258 |
| --- | S | --- | | 0.026563886 | -0.350156547 | -0.353264844 | -0.37982873 | -0.003108297 | 0.768418831 | 0.000572269 | 8.00817E-06 | 0.033855045 | 0.937118258 |
| --- | S | --- | | 0.026563886 | -0.350156547 | -0.353264844 | -0.37982873 | -0.003108297 | 0.768418831 | 0.000572269 | 8.00817E-06 | 0.033855045 | 0.937118258 |
| --- | S | | 138 | -0.681398264 | -0.518819717 | -0.60192848 | 0.079469784 | -0.083108763 | 0.035578361 | 0.117458384 | 0.224686971 | 0.848599977 | 0.846973283 |

| | | | | | | | | | | | | | |
|---|---|---|---|---|---|---|---|---|---|---|---|---|---|
| --- | S | --- | | -0.583052648 | -0.227094841 | -0.60464831 | -0.021595663 | -0.377553469 | 0.004972707 | 0.358026845 | 0.175364598 | 0.95113637 | 0.360780723 |
| --- | S | --- | | -0.264783435 | -0.188543078 | -0.33709704 | -0.072313605 | -0.148553962 | 0.014310916 | 0.048335482 | 0.003892798 | 0.375792735 | 0.105712128 |
| --- | S | | 323 | -0.552119527 | -0.394150225 | -0.326121545 | 0.225997982 | 0.068028681 | 0.021058223 | 0.083799848 | 0.334512376 | 0.479854937 | 0.826173305 |
| --- | S | --- | | -0.235061553 | -0.218094003 | -0.54568913 | -0.310627577 | -0.327595127 | 0.153585075 | 0.186313099 | 0.005375137 | 0.04602057 | 0.045174377 |
| --- | S | --- | | -0.069279185 | -0.25684571 | -0.35049146 | -0.281212275 | -0.09364575 | 0.122285736 | 0.005085199 | 0.00467787 | 0.014820215 | 0.212761338 |
| --- | S | --- | | -0.119318229 | -0.285564577 | -0.516814412 | -0.397496183 | -0.231249835 | 0.151093617 | 0.07638696 | 0.008019594 | 0.030550143 | 0.156372285 |
| --- | S | --- | | 0.011876362 | -0.264634905 | -0.425921663 | -0.437798025 | -0.161266758 | 0.82391612 | 0.024789774 | 0.000230969 | 0.001020138 | 0.104632802 |
| --- | S | | 325 | -0.493151937 | -0.33224297 | -0.259367256 | 0.233784681 | 0.072875714 | 0.02168791 | 0.097639377 | 0.361356167 | 0.400488554 | 0.783147907 |
| --- | S | | 320 | -0.492735235 | -0.32970737 | -0.260465553 | 0.232269682 | 0.069241816 | 0.022792027 | 0.107516267 | 0.373528633 | 0.416074064 | 0.800279273 |
| --- | S | --- | | -0.124582518 | -0.163777043 | -0.202715778 | -0.07813326 | -0.038938734 | 0.001322703 | 0.004176088 | 0.027133583 | 0.203651294 | 0.484222644 |
| --- | S | --- | | -0.099874568 | -0.139812853 | -0.194505766 | -0.094631198 | -0.054692914 | 0.034919186 | 0.054726612 | 2.64666E-05 | 0.057434193 | 0.303781345 |
| --- | S | --- | | -0.101696892 | -0.137427381 | -0.189584832 | -0.087887939 | -0.05215745 | 0.016334026 | 0.004646115 | 0.001053893 | 0.027907951 | 0.113332489 |
| --- | S | --- | | -0.101696892 | -0.137427381 | -0.189584832 | -0.087887939 | -0.05215745 | 0.016334026 | 0.004646115 | 0.001053893 | 0.027907951 | 0.113332489 |
| --- | S | --- | | -0.054545768 | -0.166204747 | -0.225422183 | -0.170876415 | -0.059217435 | 0.221715654 | 0.004298088 | 0.000354021 | 0.022381045 | 0.138935696 |
| --- | S | --- | | -0.054545768 | -0.166204747 | -0.225422183 | -0.170876415 | -0.059217435 | 0.221715654 | 0.004298088 | 0.000354021 | 0.022381045 | 0.138935696 |
| --- | S | --- | | 0.011639013 | -0.221817455 | -0.223845222 | -0.235484235 | -0.002027767 | 0.554544802 | 9.4471E-05 | 0.008467609 | 0.008517864 | 0.956722041 |
| --- | S | --- | | 0.011639013 | -0.221817455 | -0.223845222 | -0.235484235 | -0.002027767 | 0.554544802 | 9.4471E-05 | 0.008467609 | 0.008517864 | 0.956722041 |
| --- | T | --- | | -0.031146795 | -0.201154225 | -0.140878265 | -0.109731469 | 0.06027596 | 0.468666252 | 0.00015808 | 0.003069952 | 0.050990535 | 0.071312608 |
| --- | S | --- | | -0.175577975 | -0.157887533 | -0.333339627 | -0.157761653 | -0.175452095 | 0.063867173 | 0.211952131 | 0.021346866 | 0.160827238 | 0.195089419 |
| --- | S | --- | | -0.171929741 | -0.167349964 | -0.33504195 | -0.163112209 | -0.167691986 | 0.079508249 | 0.203978154 | 0.019841442 | 0.144040807 | 0.216277218 |
| --- | S | --- | | -0.274086043 | -0.144375974 | -0.073464026 | 0.200622017 | 0.070911949 | 0.001023922 | 0.018650884 | 0.660309046 | 0.290552491 | 0.665002092 |
| --- | S | --- | | -0.274086043 | -0.144375974 | -0.073464026 | 0.200622017 | 0.070911949 | 0.001023922 | 0.018650884 | 0.660309046 | 0.290552491 | 0.665002092 |
| --- | S | --- | | -0.20948273 | -0.064691758 | -0.145131509 | 0.064351222 | -0.080439751 | 0.000127589 | 0.367075942 | 0.048969597 | 0.234151726 | 0.310082861 |
| --- | S | --- | | -0.331962655 | -0.057610427 | -0.338596987 | -0.006634332 | -0.28098656 | 0.018632901 | 0.732498469 | 0.066748166 | 0.960201974 | 0.1702311 |
| --- | S | --- | | -0.331962655 | -0.057610427 | -0.338596987 | -0.006634332 | -0.28098656 | 0.018632901 | 0.732498469 | 0.066748166 | 0.960201974 | 0.1702311 |
| --- | S | --- | | -0.331962655 | -0.057610427 | -0.338596987 | -0.006634332 | -0.28098656 | 0.018632901 | 0.732498469 | 0.066748166 | 0.960201974 | 0.1702311 |
| --- | S | --- | | -0.331962655 | -0.057610427 | -0.338596987 | -0.006634332 | -0.28098656 | 0.018632901 | 0.732498469 | 0.066748166 | 0.960201974 | 0.1702311 |
| --- | S | | 60 | -0.272083377 | -0.174312808 | -0.182564181 | 0.089519196 | -0.008251373 | 0.015501272 | 0.074034678 | 0.325733011 | 0.588893653 | 0.958395961 |
| --- | S | --- | | -0.140572878 | -0.123785528 | -0.381954151 | -0.241381273 | -0.258168623 | 0.264147241 | 0.23744256 | 0.011662883 | 0.08201926 | 0.045359189 |
| --- | S | --- | | 0.02758768 | -0.29561717 | -0.174036386 | -0.201624065 | 0.121580784 | 0.785699654 | 0.000362402 | 0.062084711 | 0.125652958 | 0.151740344 |
| --- | S | --- | | 0.02758768 | -0.29561717 | -0.174036386 | -0.201624065 | 0.121580784 | 0.785699654 | 0.000362402 | 0.062084711 | 0.125652958 | 0.151740344 |
| --- | S | | 481 | -0.058492079 | -0.21241233 | -0.305317403 | -0.246825323 | -0.092905072 | 0.674629687 | 0.063453595 | 0.008254145 | 0.134470462 | 0.281579725 |
| --- | S | --- | | -0.064309514 | -0.253792037 | -0.242414249 | -0.178104735 | 0.011377787 | 0.658108534 | 0.021702117 | 0.019604166 | 0.251588048 | 0.852101099 |
| --- | S | --- | | -0.17627726 | -0.103654088 | -0.335520496 | -0.159243236 | -0.231866409 | 0.23398485 | 0.337925547 | 0.010930532 | 0.249418731 | 0.038485403 |
| --- | S | --- | | 0.198381287 | -0.498062602 | -0.388861311 | -0.587242598 | 0.109201291 | 0.001601689 | 0.002990946 | 0.00374331 | 0.006282111 | 0.231062967 |
| --- | S | --- | | -0.031793184 | -0.218952579 | -0.125271593 | -0.093478409 | 0.093680986 | 0.459562947 | 0.001430176 | 0.022276084 | 0.030793527 | 0.033579123 |
| --- | S | --- | | -0.132225996 | -0.156701056 | -0.166818891 | -0.034592896 | -0.010117836 | 0.073325717 | 0.031133052 | 0.071636403 | 0.654192977 | 0.889911924 |
| --- | S | --- | | -0.052615924 | -0.095818794 | -0.15498293 | -0.102367005 | -0.059164135 | 0.121704595 | 0.015171629 | 0.002203274 | 0.014689464 | 0.068465322 |
| --- | S | --- | | -0.127782952 | -0.193690675 | -0.066418993 | 0.061363958 | 0.127271682 | 0.123636767 | 0.00221139 | 0.195595156 | 0.388869728 | 0.049532335 |
| --- | S | --- | | -0.12340745 | -0.054390751 | -0.105098145 | 0.018309305 | -0.050707394 | 0.00013602 | 0.108145942 | 0.096335509 | 0.682772313 | 0.328963756 |
| --- | S | --- | | -0.075134408 | -0.126780784 | -0.149326076 | -0.074191668 | -0.022545292 | 0.07525744 | 0.006457233 | 0.15599964 | 0.406863457 | 0.782240987 |
| --- | S | --- | | 0.015510604 | -0.222393665 | -0.189179442 | -0.204690045 | 0.033214223 | 0.703208695 | 0.005342335 | 0.066700064 | 0.061026264 | 0.665766354 |
| --- | S | --- | | -0.137276946 | -0.02765146 | -0.180949855 | -0.043672909 | -0.153298395 | 0.018685078 | 0.732267544 | 0.020835122 | 0.42795881 | 0.128689289 |
| --- | S | | 696 | -0.119862695 | -0.198868013 | 0.053729115 | 0.173591811 | 0.252597128 | 0.049912017 | 0.002473133 | 0.714074774 | 0.299330865 | 0.17731338 |
| --- | S | --- | | -0.05011819 | -0.218873835 | -0.049338071 | 0.000780119 | 0.169535764 | 0.49711565 | 0.005206515 | 0.413614969 | 0.992116599 | 0.044647539 |
| --- | S | | 324 | -0.319425156 | -0.081002567 | -0.037955957 | 0.281469199 | 0.04304661 | 0.020299518 | 0.57692676 | 0.829029282 | 0.164366251 | 0.813240166 |
| --- | T | --- | | -0.256019686 | 0.162561934 | -0.443951729 | -0.187932043 | -0.606513662 | 0.108589809 | 0.531127208 | 0.122550021 | 0.432769878 | 0.094319435 |
| --- | S | --- | | -0.229791779 | -0.015699635 | -0.197242814 | 0.032548965 | -0.181543179 | 0.094084256 | 0.901023599 | 0.148956509 | 0.786731091 | 0.196004759 |
| --- | T | --- | | -0.03083911 | -0.05733052 | -0.244406281 | -0.213567171 | -0.187075761 | 0.474679884 | 0.372558197 | 0.021657843 | 0.039295432 | 0.059043224 |
| --- | S | --- | | -0.113796641 | -0.296982216 | 0.135581307 | 0.249377948 | 0.432563522 | 0.125493231 | 0.024920526 | 0.631760259 | 0.409477231 | 0.208266339 |
| --- | T | --- | | 0.043696797 | -0.144435338 | -0.127768338 | -0.171465135 | 0.01666705 | 0.311305489 | 0.011845218 | 0.019588305 | 0.000905974 | 0.389989955 |
| --- | T | --- | | -0.202399199 | -0.019447829 | -0.107650101 | 0.094749097 | -0.088202273 | 0.041589177 | 0.807980117 | 0.343355724 | 0.338162245 | 0.360436117 |
| --- | S | --- | | -0.10358715 | -0.101648029 | -0.205062812 | -0.101475661 | -0.103414783 | 0.346627384 | 0.403668693 | 0.266517326 | 0.563401623 | 0.567927784 |
| --- | S | --- | | -0.10358715 | -0.101648029 | -0.205062812 | -0.101475661 | -0.103414783 | 0.346627384 | 0.403668693 | 0.266517326 | 0.563401623 | 0.567927784 |
| --- | S | --- | | -0.10358715 | -0.101648029 | -0.205062812 | -0.101475661 | -0.103414783 | 0.346627384 | 0.403668693 | 0.266517326 | 0.563401623 | 0.567927784 |
| --- | S | --- | | -0.118201424 | -0.091894779 | -0.087094749 | 0.031106675 | 0.00480003 | 0.108154482 | 0.255108561 | 0.410936899 | 0.72316141 | 0.959331451 |
| --- | S | --- | | -0.04121709 | -0.064730422 | -0.145681763 | -0.104464673 | -0.08095134 | 0.547584687 | 0.271735432 | 0.044427872 | 0.157185032 | 0.172690294 |
| --- | S | --- | | -0.04121709 | -0.064730422 | -0.145681763 | -0.104464673 | -0.08095134 | 0.547584687 | 0.271735432 | 0.044427872 | 0.157185032 | 0.172690294 |
| --- | S | | 7 | -0.265078197 | 0.069510012 | -0.137296467 | 0.12778173 | -0.206806479 | 0.017991532 | 0.416754765 | 0.472921643 | 0.503302907 | 0.312094571 |
| --- | S | | 8 | -0.265078197 | 0.069510012 | -0.137296467 | 0.12778173 | -0.206806479 | 0.017991532 | 0.416754765 | 0.472921643 | 0.503302907 | 0.312094571 |
| --- | S | | 10 | -0.265078197 | 0.069510012 | -0.137296467 | 0.12778173 | -0.206806479 | 0.017991532 | 0.416754765 | 0.472921643 | 0.503302907 | 0.312094571 |
| --- | S | --- | | 0.116026729 | -0.217993169 | -0.425872601 | -0.54189933 | -0.207879432 | 0.785873528 | 0.689395256 | 0.131056712 | 0.258423474 | 0.701081483 |
| --- | T | --- | | -0.175191626 | -0.129636124 | 0.018720739 | 0.193912365 | 0.148356863 | 0.089016402 | 0.234541486 | 0.876272434 | 0.180119852 | 0.297499353 |
| --- | S | --- | | 0.003219711 | -0.044501259 | -0.254368104 | -0.257587815 | -0.209866845 | 0.920600065 | 0.452713306 | 0.068718501 | 0.070154174 | 0.09347582 |
| --- | S | --- | | 0.003219711 | -0.044501259 | -0.254368104 | -0.257587815 | -0.209866845 | 0.920600065 | 0.452713306 | 0.068718501 | 0.070154174 | 0.09347582 |
| --- | S | --- | | -0.083697766 | 0.005545665 | -0.111891988 | -0.028194222 | -0.117437652 | 0.151055352 | 0.90240046 | 0.087382895 | 0.637828896 | 0.086120955 |
| --- | T | --- | | -0.094258626 | 0.027785282 | -0.18449297 | -0.090234344 | -0.212278251 | 0.250383247 | 0.771212604 | 0.134657674 | 0.414150455 | 0.126049604 |
| --- | S | --- | | -0.123594355 | -0.054300668 | -0.062976602 | 0.060617753 | -0.008675934 | 0.18920721 | 0.535744561 | 0.382405371 | 0.437966633 | 0.906299683 |
| --- | S | --- | | -0.038744743 | 0.039211706 | -0.160117763 | -0.12137302 | -0.199329469 | 0.432531267 | 0.693404969 | 0.015786123 | 0.053905085 | 0.122970712 |
| --- | T | --- | | 0.06439352 | -0.196608029 | -0.132108343 | -0.196501864 | 0.064499686 | 0.389857161 | 0.10038069 | 0.229040193 | 0.100948218 | 0.574401117 |
| --- | T | --- | | -0.464210995 | -0.197600865 | 0.278563038 | 0.742774033 | 0.476163903 | 0.156169642 | 0.39146654 | 0.414254612 | 0.07315625 | 0.177179343 |
| --- | S | --- | | -0.053647656 | -0.019865536 | -0.098574716 | -0.04492706 | -0.07870918 | 0.578128425 | 0.631077007 | 0.072899351 | 0.635152642 | 0.104703933 |
| --- | S | --- | | -0.053647656 | -0.019865536 | -0.098574716 | -0.04492706 | -0.07870918 | 0.578128425 | 0.631077007 | 0.072899351 | 0.635152642 | 0.104703933 |
| --- | T | --- | | -0.053647656 | -0.019865536 | -0.098574716 | -0.04492706 | -0.07870918 | 0.578128425 | 0.631077007 | 0.072899351 | 0.635152642 | 0.104703933 |
| --- | S | --- | | -0.089665487 | -0.011583652 | -0.072119948 | 0.017545539 | -0.060536296 | 0.150231486 | 0.818555385 | 0.283408646 | 0.787802055 | 0.351477842 |
| --- | S | | 55 | -0.15614192 | 0.009454236 | -0.014220116 | 0.141921804 | -0.023674352 | 0.024104372 | 0.857065371 | 0.940372353 | 0.481097391 | 0.899360966 |
| --- | S | --- | | -0.177073648 | 0.021186476 | -0.057458918 | 0.11961473 | -0.078645394 | 0.149079275 | 0.797808743 | 0.423741805 | 0.273269715 | 0.260257291 |
| --- | S | --- | | -0.007388161 | -0.041306475 | -0.076711256 | -0.069323095 | -0.035404782 | 0.795271917 | 0.276582336 | 0.093171004 | 0.123969678 | 0.416142044 |
| --- | S | --- | | 0.085565307 | -0.180711551 | -0.048336005 | -0.133901312 | 0.132375546 | 0.126162803 | 0.04817105 | 0.096962168 | 0.049599941 | 0.096572691 |
| --- | S | --- | | 0.168683209 | 0.046050101 | -0.279676594 | -0.448359803 | -0.325726695 | 0.279640614 | 0.331333571 | 0.001836267 | 0.050340649 | 0.004887943 |
| --- | S | --- | | -0.260720142 | -0.000362515 | -0.021970848 | 0.238749294 | -0.021608333 | 0.258291679 | 0.998550331 | 0.947382203 | 0.478334802 | 0.94448265 |
| --- | S | --- | | -0.089358457 | 0.046414065 | -0.099644699 | -0.010286242 | -0.146058764 | 0.27075312 | 0.637403157 | 0.158179327 | 0.898456701 | 0.211298423 |
| --- | S | --- | | -0.089358457 | 0.046414065 | -0.099644699 | -0.010286242 | -0.146058764 | 0.27075312 | 0.637403157 | 0.158179327 | 0.898456701 | 0.211298423 |
| --- | S | --- | | 0.091961018 | -0.13615403 | -0.084061449 | -0.176022467 | 0.052092581 | 0.060057468 | 0.067024641 | 0.079115275 | 0.00187756 | 0.365162331 |
| --- | S | --- | | -0.112343438 | -0.082501185 | 0.049685557 | 0.162028995 | 0.132186741 | 0.118221296 | 0.536985908 | 0.572048362 | 0.148962775 | 0.387090431 |
| --- | S | --- | | -0.112343438 | -0.082501185 | 0.049685557 | 0.162028995 | 0.132186741 | 0.118221296 | 0.536985908 | 0.572048362 | 0.148962775 | 0.387090431 |
| --- | S | --- | | 0.06464991 | -0.098541318 | -0.046421626 | -0.111071536 | 0.052119692 | 0.560586631 | 0.035181268 | 0.63674373 | 0.413719383 | 0.586614837 |
| --- | S | --- | | -0.046666185 | -0.102931664 | 0.038765299 | 0.085431485 | 0.141696963 | 0.539278496 | 0.079439379 | 0.332128419 | 0.282832772 | 0.045993849 |
| --- | S | --- | | -0.048688107 | 0.041192362 | -0.120945911 | -0.072257804 | -0.162138273 | 0.715108959 | 0.741204517 | 0.225666788 | 0.596018845 | 0.246587486 |
| --- | S | --- | | -0.152823231 | 0.092726726 | 0.01863056 | 0.171453791 | -0.074096166 | 0.007649411 | 0.291888597 | 0.682488399 | 0.007279344 | 0.376026249 |
| --- | S | --- | | 0.000847712 | -0.057074639 | -0.022298536 | -0.023146248 | 0.034776103 | 0.987512365 | 0.175438678 | 0.6574719 | 0.666969499 | 0.400113108 |
| --- | S | --- | | 0.000847712 | -0.057074639 | -0.022298536 | -0.023146248 | 0.034776103 | 0.987512365 | 0.175438678 | 0.6574719 | 0.666969499 | 0.400113108 |
| --- | S | --- | | -0.102829759 | 0.050055785 | -0.022571188 | 0.080258571 | -0.072626973 | 0.006080274 | 0.01186334 | 0.425309216 | 0.045772226 | 0.064001864 |
| --- | S | --- | | -0.102829759 | 0.050055785 | -0.022571188 | 0.080258571 | -0.072626973 | 0.006080274 | 0.01186334 | 0.425309216 | 0.045772226 | 0.064001864 |

| | | | | | | | | | | | | |
|---|---|---|---|---|---|---|---|---|---|---|---|---|
| --- | S | --- | -0.102829759 | 0.050055785 | -0.022571188 | 0.080258571 | -0.072626973 | 0.006080274 | 0.01186334 | 0.425309216 | 0.045772226 | 0.064001864 |
| --- | S | 432 | -0.037937824 | 0.102767807 | -0.182874198 | -0.144936374 | -0.285642004 | 0.761548068 | 0.173082698 | 0.131393743 | 0.34011743 | 0.047839839 |
| --- | S | 518 | -0.191246203 | 0.281210301 | -0.219108341 | -0.027862138 | -0.500318642 | 0.530558705 | 0.335554715 | 0.35497535 | 0.898763286 | 0.091766963 |
| --- | S | --- | -0.013986504 | 0.008610536 | -0.066382476 | -0.052395972 | -0.074993012 | 0.777159383 | 0.895030481 | 0.446746628 | 0.504407162 | 0.396297565 |
| --- | T | --- | -0.013986504 | 0.008610536 | -0.066382476 | -0.052395972 | -0.074993012 | 0.777159383 | 0.895030481 | 0.446746628 | 0.504407162 | 0.396297565 |
| --- | S | --- | -0.032109741 | -0.129458659 | 0.063002867 | 0.095112608 | 0.192461527 | 0.863538787 | 0.57010332 | 0.805600969 | 0.679036418 | 0.467711167 |
| --- | T | --- | -0.032109741 | -0.129458659 | 0.063002867 | 0.095112608 | 0.192461527 | 0.863538787 | 0.57010332 | 0.805600969 | 0.679036418 | 0.467711167 |
| --- | T | --- | -0.084317378 | 0.014136981 | 0.01697249 | 0.101289868 | 0.002835509 | 0.149767981 | 0.666187451 | 0.466374099 | 0.123765475 | 0.91078553 |
| --- | S | 517 | -0.154192224 | 0.299387621 | -0.222827072 | -0.068634849 | -0.522214693 | 0.547011115 | 0.281359642 | 0.314166055 | 0.689577782 | 0.075352538 |
| --- | S | --- | -0.049125048 | -0.03221549 | 0.04657074 | 0.095695788 | 0.07878623 | 0.397363892 | 0.503525373 | 0.439367912 | 0.124997492 | 0.152688596 |
| --- | S | --- | -0.049125048 | -0.03221549 | 0.04657074 | 0.095695788 | 0.07878623 | 0.397363892 | 0.503525373 | 0.439367912 | 0.124997492 | 0.152688596 |
| --- | S | --- | -0.071031453 | 0.021180566 | 0.032713933 | 0.103745386 | 0.011533367 | 0.093564638 | 0.681025616 | 0.458440308 | 0.068377494 | 0.83590978 |
| --- | S | --- | -0.068292198 | 0.037594742 | 0.030512449 | 0.098804647 | -0.007082293 | 0.272997324 | 0.582350685 | 0.695428217 | 0.271020566 | 0.935157592 |
| --- | S | --- | -0.151549978 | 0.166455183 | -0.033813485 | 0.117736494 | -0.200268668 | 0.117975973 | 0.100480187 | 0.842644542 | 0.491900048 | 0.284440723 |
| --- | T | --- | -0.151549978 | 0.166455183 | -0.033813485 | 0.117736494 | -0.200268668 | 0.117975973 | 0.100480187 | 0.842644542 | 0.491900048 | 0.284440723 |
| --- | S | --- | -0.027856766 | 0.000236243 | 0.01610062 | 0.043957386 | 0.015864377 | 0.663012311 | 0.992561427 | 0.650304975 | 0.513872241 | 0.628569572 |
| --- | S | --- | -0.027856766 | 0.000236243 | 0.01610062 | 0.043957386 | 0.015864377 | 0.663012311 | 0.992561427 | 0.650304975 | 0.513872241 | 0.628569572 |
| --- | T | --- | -0.027856766 | 0.000236243 | 0.01610062 | 0.043957386 | 0.015864377 | 0.663012311 | 0.992561427 | 0.650304975 | 0.513872241 | 0.628569572 |
| --- | S | --- | -0.113725144 | 0.859820139 | 1.429776129 | 1.543501273 | 0.56995599 | 0.545485904 | 0.000661408 | 7.70606E-05 | 0.004247265 | 0.011640559 |
| --- | S | --- | 0.168587784 | 0.497100818 | 0.652440157 | 0.483852373 | 0.155339339 | 0.30848275 | 0.004450796 | 0.007011908 | 0.035623577 | 0.30268337 |
| --- | S | --- | 0.051381374 | 0.287468059 | 0.315137575 | 0.263756201 | 0.027669516 | 0.299550806 | 0.000136209 | 0.000341594 | 0.005735859 | 0.460142353 |
| --- | S | --- | 0.261933108 | 0.376037572 | 0.202568353 | -0.059364755 | -0.173469219 | 0.069553607 | 0.000163547 | 0.220656774 | 0.70333333 | 0.274984788 |
| --- | S | --- | 0.259993074 | 0.186885599 | 0.244946143 | -0.01504693 | 0.058060544 | 0.001808407 | 0.002811206 | 0.088063306 | 0.88280343 | 0.574859556 |
| --- | S | --- | 0.062395766 | 0.310149019 | 0.308161117 | 0.245765351 | -0.001987902 | 0.284315298 | 0.000106157 | 0.01835393 | 0.036629301 | 0.977708633 |
| --- | T | --- | 0.062395766 | 0.310149019 | 0.308161117 | 0.245765351 | -0.001987902 | 0.284315298 | 0.000106157 | 0.01835393 | 0.036629301 | 0.977708633 |
| --- | S | --- | 0.047944175 | 0.190708227 | 0.362382776 | 0.314438601 | 0.171674549 | 0.539785759 | 0.005621564 | 0.00022119 | 0.017259504 | 0.011441538 |
| --- | S | --- | 0.160906507 | 0.114587137 | 0.405583241 | 0.244676734 | 0.290996104 | 0.167276463 | 0.179407123 | 0.000354488 | 0.085679863 | 0.025600573 |
| --- | S | --- | 0.028478274 | 0.132172805 | 0.292229098 | 0.263750824 | 0.160056293 | 0.573630639 | 0.005648821 | 4.9173E-05 | 0.009080128 | 0.004463472 |
| --- | S | --- | 0.028478274 | 0.132172805 | 0.292229098 | 0.263750824 | 0.160056293 | 0.573630639 | 0.005648821 | 4.9173E-05 | 0.009080128 | 0.004463472 |
| --- | S | --- | 0.046794193 | 0.297982007 | 0.321233827 | 0.274439634 | 0.023251819 | 0.361804398 | 0.042551153 | 0.001853297 | 0.007587062 | 0.810570325 |
| --- | S | --- | -0.023654977 | 0.222871531 | 0.350515861 | 0.374170837 | 0.127644329 | 0.740782222 | 0.008870003 | 0.000626243 | 0.007110138 | 0.046782875 |
| --- | S | --- | 0.023127162 | 0.936884581 | 1.404395536 | 1.381268374 | 0.467510955 | 0.947161968 | 0.338858147 | 0.234063005 | 0.242397404 | 0.701374223 |
| --- | S | --- | 0.050173097 | 0.145657055 | 0.237885568 | 0.187712471 | 0.092228513 | 0.132327936 | 0.027349116 | 0.000220469 | 0.001289556 | 0.119484401 |
| --- | S | --- | 0.155617663 | 0.238861208 | 0.583107801 | 0.427490138 | 0.344246593 | 0.308781187 | 0.241656962 | 0.031944491 | 0.094999153 | 0.154179016 |
| --- | S | --- | -0.044450011 | 0.270264847 | 0.347434199 | 0.39188421 | 0.077169352 | 0.34049208 | 0.000992607 | 0.01238926 | 0.01467311 | 0.348949377 |
| --- | S | --- | -0.044450011 | 0.270264847 | 0.347434199 | 0.39188421 | 0.077169352 | 0.34049208 | 0.000992607 | 0.01238926 | 0.01467311 | 0.348949377 |
| --- | S | --- | 0.160010989 | 0.149265822 | 0.161565643 | 0.001554654 | 0.012299821 | 0.028967696 | 0.020529555 | 0.015195325 | 0.971986528 | 0.627584032 |
| --- | T | --- | 0.160010989 | 0.149265822 | 0.161565643 | 0.001554654 | 0.012299821 | 0.028967696 | 0.020529555 | 0.015195325 | 0.971986528 | 0.627584032 |
| --- | S | --- | 0.099487966 | 0.332332048 | 0.044206898 | -0.055281068 | -0.28812515 | 0.255455586 | 0.000173043 | 0.75006327 | 0.707211047 | 0.13422204 |
| --- | S | --- | 0.014695277 | 0.220237066 | 0.262471576 | 0.2477763 | 0.042234511 | 0.782692146 | 0.005639925 | 0.028014364 | 0.044178769 | 0.59967938 |
| --- | S | --- | -0.032632293 | 0.59239124 | 1.233019153 | 1.265651445 | 0.640627912 | 0.904169138 | 0.408257984 | 0.238179165 | 0.230859006 | 0.535832949 |
| --- | S | --- | -0.048708764 | 0.693564415 | 1.136182419 | 1.184891183 | 0.442618004 | 0.859941422 | 0.401729025 | 0.295769126 | 0.281840198 | 0.694218436 |
| --- | S | --- | -0.122033037 | 0.142123144 | 0.291052434 | 0.41308547 | 0.14892929 | 0.068401633 | 0.008009188 | 0.000147881 | 0.002336725 | 0.011605577 |
| --- | S | --- | -0.001995629 | 0.121617079 | 0.196649888 | 0.198645517 | 0.075032809 | 0.947477856 | 0.004836232 | 0.003836375 | 0.004395039 | 0.067995076 |
| --- | S | 273 | 0.053853046 | 0.105341417 | 0.201750955 | 0.147897908 | 0.096409537 | 0.354975861 | 0.160034354 | 0.001410624 | 0.063153087 | 0.194259256 |
| --- | T | 263 | 0.053853046 | 0.105341417 | 0.201750955 | 0.147897908 | 0.096409537 | 0.354975861 | 0.160034354 | 0.001410624 | 0.063153087 | 0.194259256 |
| --- | S | --- | 0.144897267 | 0.188803779 | 0.200290368 | 0.055393102 | 0.01148659 | 0.25511733 | 0.077165197 | 0.14450395 | 0.689268631 | 0.923316555 |
| --- | S | --- | 0.363892466 | 0.180026638 | -0.032378041 | -0.396270507 | -0.212404679 | 0.015774798 | 0.140305152 | 0.855475106 | 0.082702105 | 0.253909121 |
| --- | S | --- | 0.159650137 | 0.312849974 | 0.100213933 | -0.059436204 | -0.212636041 | 0.157572139 | 0.076201383 | 0.424699352 | 0.632014948 | 0.208393404 |
| --- | S | --- | 0.159650137 | 0.312849974 | 0.100213933 | -0.059436204 | -0.212636041 | 0.157572139 | 0.076201383 | 0.424699352 | 0.632014948 | 0.208393404 |
| --- | S | --- | 0.123419307 | 0.140037383 | 0.134276261 | 0.010856955 | -0.005761121 | 0.07960953 | 0.043579987 | 0.166209194 | 0.894637841 | 0.942343372 |
| --- | S | --- | -0.006693934 | 0.150424137 | 0.200508945 | 0.207202878 | 0.050084808 | 0.910014691 | 0.014317644 | 0.027523139 | 0.032212453 | 0.420378762 |
| --- | T | --- | -0.006693934 | 0.150424137 | 0.200508945 | 0.207202878 | 0.050084808 | 0.910014691 | 0.014317644 | 0.027523139 | 0.032212453 | 0.420378762 |
| --- | S | --- | 0.036161947 | 0.130637269 | 0.164889662 | 0.128727714 | 0.034252393 | 0.66154327 | 0.012067828 | 0.043765219 | 0.205131568 | 0.553077585 |
| --- | T | --- | -0.035660327 | 0.126665551 | 0.121239498 | 0.156899825 | -0.005426053 | 0.474281166 | 0.001930319 | 0.003220652 | 0.039796544 | 0.775781352 |
| --- | S | --- | 0.143309406 | 0.176561575 | 0.094411973 | -0.048897433 | -0.082149602 | 0.074675221 | 0.076731633 | 0.522753377 | 0.725871613 | 0.57805724 |
| --- | S | --- | 0.001747363 | 0.109113939 | 0.188095628 | 0.186348264 | 0.078981689 | 0.973011154 | 0.060182561 | 0.008573139 | 0.006222449 | 0.076750781 |
| --- | S | --- | 0.070121396 | 0.086703516 | 0.21011807 | 0.139996674 | 0.123414554 | 0.110716451 | 0.157158196 | 0.00956542 | 0.206825103 | 0.253956713 |
| --- | T | --- | 0.024160483 | 0.177996054 | 0.034066811 | 0.009906329 | -0.143929243 | 0.737879832 | 0.000162665 | 0.537244861 | 0.902512431 | 0.078807763 |
| --- | S | --- | -0.12243478 | 0.207783413 | 0.170029147 | 0.292463927 | -0.037754266 | 0.416078202 | 0.039589491 | 0.005432043 | 0.129559929 | 0.556843501 |
| --- | S | --- | -0.12243478 | 0.207783413 | 0.170029147 | 0.292463927 | -0.037754266 | 0.416078202 | 0.039589491 | 0.005432043 | 0.129559929 | 0.556843501 |
| --- | T | --- | -0.017274811 | 0.161890142 | 0.148548356 | 0.165823167 | -0.013341786 | 0.790924515 | 0.032973434 | 0.031154287 | 0.038233335 | 0.800710278 |
| --- | S | --- | 0.098807513 | 0.14690858 | 0.174211614 | 0.075404101 | 0.027303034 | 0.466458009 | 0.161280774 | 0.149863738 | 0.536145586 | 0.71604509 |
| --- | T | --- | 0.024162685 | 0.258726469 | 0.160484325 | 0.13632164 | -0.098242143 | 0.618704105 | 0.229367942 | 0.194827451 | 0.252048414 | 0.614048553 |
| --- | S | --- | 0.040029661 | 0.098829419 | 0.146799254 | 0.106769593 | 0.047969835 | 0.33163943 | 0.105334653 | 0.125750059 | 0.213958952 | 0.545666826 |
| --- | S | --- | 0.040029661 | 0.098829419 | 0.146799254 | 0.106769593 | 0.047969835 | 0.33163943 | 0.105334653 | 0.125750059 | 0.213958952 | 0.545666826 |
| --- | S | --- | -0.044960105 | 0.172829982 | 0.208732057 | 0.253692162 | 0.035902075 | 0.769279456 | 0.143109812 | 0.061969565 | 0.152096093 | 0.67545648 |
| --- | T | --- | -0.044960105 | 0.172829982 | 0.208732057 | 0.253692162 | 0.035902075 | 0.769279456 | 0.143109812 | 0.061969565 | 0.152096093 | 0.67545648 |
| --- | S | --- | -0.077149838 | 0.190697096 | 0.071756178 | 0.148906015 | -0.118940918 | 0.216121323 | 0.000325212 | 0.247325291 | 0.076437819 | 0.10702102 |
| --- | S | --- | 0.011809175 | 0.179080168 | 0.086524021 | 0.074714846 | -0.092556147 | 0.818391269 | 0.062111407 | 0.110763689 | 0.234506287 | 0.240023745 |
| --- | S | --- | 0.011809175 | 0.179080168 | 0.086524021 | 0.074714846 | -0.092556147 | 0.818391269 | 0.062111407 | 0.110763689 | 0.234506287 | 0.240023745 |
| --- | S | --- | 0.06113751 | 0.095220569 | 0.093892815 | 0.032755305 | -0.001327755 | 0.32177329 | 0.027772661 | 0.337105796 | 0.730472562 | 0.987657123 |
| --- | S | --- | 0.061608538 | 0.064704899 | 0.112458504 | 0.050849966 | 0.047753605 | 0.333637573 | 0.345957626 | 0.051905226 | 0.431954799 | 0.485777649 |
| --- | S | --- | 0.009149982 | 0.189614033 | 0.084338606 | 0.075188623 | -0.105275427 | 0.838451435 | 0.07670036 | 0.252406865 | 0.302896444 | 0.276119254 |
| --- | T | --- | 0.003869417 | 0.026198945 | 0.13206 | 0.128190583 | 0.105861056 | 0.913089218 | 0.465398068 | 0.0053401 | 0.029038001 | 0.043646534 |
| --- | S | --- | 0.019097378 | 0.17871086 | 0.062998734 | 0.043901356 | -0.115712127 | 0.889868895 | 0.055785537 | 0.552250865 | 0.755879689 | 0.247606804 |
| --- | T | --- | 0.019097378 | 0.17871086 | 0.062998734 | 0.043901356 | -0.115712127 | 0.889868895 | 0.055785537 | 0.552250865 | 0.755879689 | 0.247606804 |
| --- | S | --- | -0.084747062 | 0.128600831 | 0.214871873 | 0.299618935 | 0.086271042 | 0.319869392 | 0.0932868 | 0.097408395 | 0.045341563 | 0.401548358 |
| --- | T | --- | -0.084747062 | 0.128600831 | 0.214871873 | 0.299618935 | 0.086271042 | 0.319869392 | 0.0932868 | 0.097408395 | 0.045341563 | 0.401548358 |
| --- | S | --- | -0.012767911 | 0.179181984 | 0.032000446 | 0.044768357 | -0.147181538 | 0.890348935 | 0.023696678 | 0.460616008 | 0.64435634 | 0.047121513 |
| --- | S | --- | -0.10295407 | 0.182935134 | 0.125465963 | 0.228420033 | -0.05746917 | 0.086727937 | 0.007982647 | 0.139193432 | 0.033797501 | 0.433462126 |
| --- | S | --- | 0.037986455 | 0.14207915 | 0.044224509 | 0.006238053 | -0.097854641 | 0.460817015 | 0.131674547 | 0.535884473 | 0.911615905 | 0.28216782 |
| --- | T | --- | 0.037986455 | 0.14207915 | 0.044224509 | 0.006238053 | -0.097854641 | 0.460817015 | 0.131674547 | 0.535884473 | 0.911615905 | 0.28216782 |
| --- | S | --- | 0.289430723 | 0.013985693 | 0.004153419 | -0.285277304 | -0.009832274 | 0.125025132 | 0.864400977 | 0.967842135 | 0.134216235 | 0.91956027 |
| --- | S | --- | 0.16940724 | -0.093263087 | 0.04803885 | -0.12136839 | 0.141301937 | 0.00593343 | 0.332365792 | 0.373735572 | 0.057056099 | 0.179814148 |
| --- | S | --- | 0.16940724 | -0.093263087 | 0.04803885 | -0.12136839 | 0.141301937 | 0.00593343 | 0.332365792 | 0.373735572 | 0.057056099 | 0.179814148 |
| --- | S | --- | 0.16940724 | -0.093263087 | 0.04803885 | -0.12136839 | 0.141301937 | 0.00593343 | 0.332365792 | 0.373735572 | 0.057056099 | 0.179814148 |
| --- | S | --- | 0.071263986 | 0.064656332 | 0.095014766 | 0.02375078 | 0.030358433 | 0.38924608 | 0.430319193 | 0.320790201 | 0.705597923 | 0.630029028 |
| --- | S | --- | -0.070798731 | 0.144385315 | 0.074909553 | 0.145708285 | -0.069475761 | 0.10997318 | 0.003481669 | 0.36965087 | 0.139805384 | 0.396849308 |
| --- | S | --- | 0.10928975 | 0.085264191 | 0.096877204 | -0.012412546 | 0.011613013 | 0.49184415 | 0.484250733 | 0.611262934 | 0.948537937 | 0.945314106 |
| --- | S | --- | 0.012261846 | 0.027716897 | 0.056707826 | 0.04444598 | 0.028990929 | 0.3385494 | 0.069167441 | 0.107596442 | 0.173264754 | 0.321129329 |
| --- | S | --- | -0.156218862 | 0.059530517 | 0.186456625 | 0.342675487 | 0.126926108 | 0.186566669 | 0.185414922 | 0.009562072 | 0.037962603 | 0.045645149 |
| --- | T | --- | 0.147409428 | 0.025351256 | 0.057662001 | -0.089747427 | 0.032310745 | 0.205014288 | 0.837796913 | 0.583833033 | 0.25866863 | 0.734896238 |
| --- | T | --- | 0.147409428 | 0.025351256 | 0.057662001 | -0.089747427 | 0.032310745 | 0.205014288 | 0.837796913 | 0.583833033 | 0.25866863 | 0.734896238 |

| | | | | | | | | | | | | |
|---|---|---|---|---|---|---|---|---|---|---|---|---|
| --- | S | --- | 0.07153546 | -0.083201654 | 0.148352241 | 0.076816782 | 0.231553895 | 0.551471194 | 0.443762635 | 0.017341829 | 0.520843855 | 0.102029216 |
| --- | S | --- | 0.01603125 | 0.059449198 | 0.033602286 | 0.017571036 | -0.025846913 | 0.653578881 | 0.033358127 | 0.711743527 | 0.84844028 | 0.774059813 |
| --- | S | --- | 0.093673466 | 0.075343172 | 0.002122639 | -0.091550827 | -0.073220533 | 0.371408271 | 0.269733627 | 0.972032194 | 0.349017423 | 0.110965126 |
| --- | S | --- | -0.052467966 | -0.001676938 | 0.145407454 | 0.19787542 | 0.147084392 | 0.160532404 | 0.988310435 | 0.004166583 | 0.00109323 | 0.273456925 |
| --- | S | --- | 0.010256369 | 0.049456896 | 0.04652984 | 0.036273471 | -0.002927056 | 0.778003262 | 0.291819176 | 0.183710337 | 0.272376499 | 0.940836839 |
| --- | S | --- | 0.095469555 | -0.033558721 | 0.178921912 | 0.083452357 | 0.212480633 | 0.587592167 | 0.909722314 | 0.437103814 | 0.714086629 | 0.523265077 |
| --- | S | --- | -0.057546096 | 0.030411751 | 0.198701521 | 0.256247617 | 0.168289771 | 0.25472982 | 0.603918694 | 0.090125219 | 0.061304989 | 0.13336209 |
| --- | S | --- | -0.057546096 | 0.030411751 | 0.198701521 | 0.256247617 | 0.168289771 | 0.25472982 | 0.603918694 | 0.090125219 | 0.061304989 | 0.13336209 |
| --- | S | --- | -0.057546096 | 0.030411751 | 0.198701521 | 0.256247617 | 0.168289771 | 0.25472982 | 0.603918694 | 0.090125219 | 0.061304989 | 0.13336209 |
| --- | S | --- | -0.058528489 | 0.105875543 | 0.036786683 | 0.095315173 | -0.06908886 | 0.019692773 | 0.001620822 | 0.429567951 | 0.111234254 | 0.191506723 |
| --- | S | --- | -0.05665584 | 0.003485848 | 0.108061971 | 0.164717811 | 0.104576123 | 0.181463845 | 0.878542936 | 0.002288104 | 0.016487381 | 0.004787559 |
| --- | S | --- | -0.354006901 | 0.360561782 | 0.232222493 | 0.586229394 | -0.128339289 | 0.010525263 | 0.083054597 | 0.052305927 | 0.000581062 | 0.426716897 |
| --- | T | --- | 0.050281779 | -0.015650486 | 0.083201887 | 0.032920107 | 0.098852372 | 0.408172511 | 0.776444526 | 0.271312102 | 0.603443377 | 0.167681288 |
| --- | S | --- | 0.0587819 | -0.043816457 | 0.092638274 | 0.033856374 | 0.136454731 | 0.244873263 | 0.313141176 | 0.154130596 | 0.579885869 | 0.069630053 |
| --- | S | --- | 0.144636457 | 0.029189717 | -0.063625993 | -0.20826245 | -0.092815711 | 0.092873084 | 0.716032542 | 0.479688829 | 0.048159747 | 0.199549419 |
| --- | S | --- | -0.132591514 | 0.142693501 | 0.090667183 | 0.223258697 | -0.052026317 | 0.222749793 | 0.273182793 | 0.128303769 | 0.083532119 | 0.65508161 |
| --- | S | --- | -0.132591514 | 0.142693501 | 0.090667183 | 0.223258697 | -0.052026317 | 0.222749793 | 0.273182793 | 0.128303769 | 0.083532119 | 0.65508161 |
| --- | T | --- | -0.132591514 | 0.142693501 | 0.090667183 | 0.223258697 | -0.052026317 | 0.222749793 | 0.273182793 | 0.128303769 | 0.083532119 | 0.65508161 |
| --- | T | --- | 0.033172974 | -0.0453523 | 0.089674419 | 0.056501444 | 0.135026719 | 0.443888923 | 0.437334885 | 0.129348869 | 0.268306633 | 0.070366212 |
| --- | S | --- | 0.364986391 | 0.003030153 | -0.165737494 | -0.530723885 | -0.168767647 | 0.032176786 | 0.968255296 | 0.032316987 | 0.013070266 | 0.083669705 |
| --- | S | 400;416 | 0.322790256 | -0.096299583 | 0.081336986 | -0.24145327 | 0.177636568 | 0.44341658 | 0.244480565 | 0.561293295 | 0.557432105 | 0.235378222 |
| --- | S | --- | -0.100507976 | 0.173967273 | 0.042758322 | 0.143266298 | -0.131208951 | 0.161572759 | 0.156450491 | 0.467983625 | 0.071513588 | 0.252019609 |
| --- | S | --- | 0.352554863 | -0.003346768 | -0.175013181 | -0.527568044 | -0.171666413 | 0.030609735 | 0.963914017 | 0.021821069 | 0.011915705 | 0.07317617 |
| --- | S | --- | 0.012699198 | 0.04326867 | -0.014583556 | -0.027282754 | -0.057852226 | 0.890372021 | 0.41560105 | 0.876522477 | 0.818988032 | 0.561178809 |
| --- | S | --- | -0.048029773 | 0.055949168 | 0.060558115 | 0.108587888 | 0.004608947 | 0.405916347 | 0.59176716 | 0.586043143 | 0.376959131 | 0.97315036 |
| --- | S | --- | -0.048029773 | 0.055949168 | 0.060558115 | 0.108587888 | 0.004608947 | 0.405916347 | 0.59176716 | 0.586043143 | 0.376959131 | 0.97315036 |
| --- | S | --- | -0.078048441 | 0.044535587 | 0.034981842 | 0.113030282 | -0.009553745 | 0.471558731 | 0.359416003 | 0.527025515 | 0.335874875 | 0.875538457 |
| --- | S | --- | 0.918826436 | -0.269517005 | -0.945280698 | -1.864107135 | -0.675763693 | 0.012949741 | 0.368725966 | 0.122509599 | 0.033938051 | 0.227754591 |
| --- | S | --- | 0.918826436 | -0.269517005 | -0.945280698 | -1.864107135 | -0.675763693 | 0.012949741 | 0.368725966 | 0.122509599 | 0.033938051 | 0.227754591 |
| --- | T | --- | -0.033106353 | 0.053857947 | -0.006268071 | 0.026838282 | -0.060126018 | 0.568368391 | 0.484785974 | 0.929319674 | 0.603425374 | 0.411671955 |
| --- | S | --- | -0.25787609 | 0.249328094 | 0.123913326 | 0.381789417 | -0.125414767 | 0.008744279 | 0.097696563 | 0.319598321 | 0.035768734 | 0.397150597 |
| --- | S | --- | -0.25787609 | 0.249328094 | 0.123913326 | 0.381789417 | -0.125414767 | 0.008744279 | 0.097696563 | 0.319598321 | 0.035768734 | 0.397150597 |
| --- | S | --- | -0.035071138 | -0.000832592 | 0.040545619 | 0.075616758 | 0.041378211 | 0.328236222 | 0.985637577 | 0.292817924 | 0.070712819 | 0.385381853 |
| --- | S | --- | -0.035071138 | -0.000832592 | 0.040545619 | 0.075616758 | 0.041378211 | 0.328236222 | 0.985637577 | 0.292817924 | 0.070712819 | 0.385381853 |
| --- | S | --- | 0.012911654 | -0.030407424 | 0.016299456 | 0.003387802 | 0.04670688 | 0.60353112 | 0.424248629 | 0.532849018 | 0.894545118 | 0.254441014 |
| --- | S | --- | 0.012911654 | -0.030407424 | 0.016299456 | 0.003387802 | 0.04670688 | 0.60353112 | 0.424248629 | 0.532849018 | 0.894545118 | 0.254441014 |
| --- | T | --- | 0.012911654 | -0.030407424 | 0.016299456 | 0.003387802 | 0.04670688 | 0.60353112 | 0.424248629 | 0.532849018 | 0.894545118 | 0.254441014 |
| --- | T | --- | 0.012911654 | -0.030407424 | 0.016299456 | 0.003387802 | 0.04670688 | 0.60353112 | 0.424248629 | 0.532849018 | 0.894545118 | 0.254441014 |
