## Supplementary Table 2 (pY) for "The lipid phosphatase activity of PTEN dampens FRA1 expression via AKT/mTOR signaling to suppress melanoma"

| | | | *Log2 ratio vs. C124S+GFP* | | | WT vs. other | | *T-tests* | | | | |
|---|---|---|---|---|---|---|---|---|---|---|---|---|
| | | | G129E | Y138L | WT | WT | WT | G129E | Y138L | WT | WT | WT |
| | | | vs | vs | vs | vs | vs | vs | vs | vs | vs | vs |
| | | | C124S+GFP | C124S+GFP | C124S+GFP | G129E | Y138L | C124S+GFP | C124S+GFP | C124S+GFP | G129E | Y138L |
| **Symbol** | **Amino Ac** | **Positions Within Proteins** | **G129EvsC124S+GFP _log2ratio** | **Y138LvsC124S+GFP _log2ratio** | **WTvsC124S+GFP _log2ratio** | **WTvsG129E _log2ratio** | **WTvsY138L _log2ratio** | **G129EvsC124S+GFP _pvalue** | **Y138LvsC124S+GFP _pvalue** | **WTvsC124S+GFP _pvalue** | **WTvsG129E _pvalue** | **WTvsY138L _pvalue** |
| Prpf18 | Y | 42 | -0.702208371 | -0.944194707 | -1.690543069 | -0.988334697 | -0.746348361 | 0.013055133 | 0.003313089 | 0.000229093 | 0.005267556 | 0.013727808 |
| Chd1 Chd2 | Y | 1066;1068 | -0.677134837 | -1.032691698 | -1.672958901 | -0.995824064 | -0.640267203 | 0.02893225 | 0.005684634 | 0.000126762 | 0.010036678 | 0.038660351 |
| Zc3h18 | Y | 381 | -0.934253234 | -1.123145818 | -1.576822847 | -0.642569613 | -0.45367703 | 0.007395558 | 0.020040428 | 0.000640268 | 0.020033837 | 0.20321029 |
| Epha2 | Y | 595 | -0.596689203 | -0.918991118 | -1.237604849 | -0.640915646 | -0.318613731 | 0.023364705 | 0.007353952 | 0.002717808 | 0.025630181 | 0.164305009 |
| Stam2 | Y | 371 | -0.607509257 | -0.688073814 | -1.291705762 | -0.684196505 | -0.603631948 | 0.046962516 | 0.036534254 | 0.000754201 | 0.036971142 | 0.055839673 |
| Map1b | S | 1391 | -0.946885695 | -1.151927141 | -1.190474385 | -0.24358869 | -0.038547244 | 0.002309573 | 0.004736156 | 0.002672244 | 0.262986676 | 0.878811435 |
| Syap1 | Y | 76 | -0.730136292 | -0.796402045 | -0.972704445 | -0.242568153 | -0.1763024 | 0.002104684 | 0.001112514 | 0.000423506 | 0.080516501 | 0.124183332 |
| Plin2 | Y | 215 | -0.669666012 | -0.921231055 | -1.143374989 | -0.473708977 | -0.222143935 | 0.010519959 | 0.001373261 | 0.000581091 | 0.03867661 | 0.15568685 |
| Uba1 | Y | 55 | -1.141955637 | -1.031249133 | -1.259579495 | -0.117623857 | -0.228330362 | 0.007984961 | 0.017781283 | 0.007930193 | 0.664662636 | 0.462766436 |
| Cdv3 | Y | 213 | -0.808580424 | -0.886240825 | -1.442248161 | -0.633667738 | -0.556007336 | 0.008129781 | 0.023479126 | 0.00195943 | 0.051711311 | 0.112049966 |
| Map1b | S | 1395 | -0.892681293 | -0.981425973 | -1.021115452 | -0.128434159 | -0.03968948 | 0.002947537 | 0.003109333 | 0.008538887 | 0.560108691 | 0.883974297 |
| Stac2 | Y | 205 | -0.589396469 | -0.794473119 | -1.15449949 | -0.565103021 | -0.360026371 | 0.003247343 | 0.006816807 | 0.00138872 | 0.029810556 | 0.121012911 |
| Lrp1 | T | 4473 | -0.662790085 | -0.708431071 | -1.123099173 | -0.460309088 | -0.414668102 | 0.007911742 | 0.025119244 | 0.000439536 | 0.031333542 | 0.120429938 |
| 2310011J03Rik | Y | 63 | -0.681515345 | -0.81051437 | -1.153193721 | -0.471678376 | -0.342679352 | 0.014839325 | 0.004252587 | 0.003821235 | 0.093917908 | 0.169733091 |
| Cdv3 | S | 214 | -0.762423764 | -0.869197577 | -1.44674024 | -0.684316476 | -0.577542663 | 0.016465722 | 0.048755796 | 0.003565847 | 0.059280413 | 0.154908208 |
| Map1b | T | 1905 | -0.622836456 | -0.880104811 | -0.913360791 | -0.290524336 | -0.033255981 | 0.009909818 | 0.000603646 | 0.016734617 | 0.259442699 | 0.877688069 |
| Tuba1a Tuba1b Tuba1c Tu | Y | 103;103;103;103; | -0.795677335 | -0.876100062 | -1.195920673 | -0.400243339 | -0.319820612 | 0.020901921 | 0.014077627 | 0.006804966 | 0.185110501 | 0.269164637 |
| Rbmx Rbmxl1 | Y | 332;335 | -0.70079697 | -0.824815938 | -1.168722306 | -0.467925336 | -0.343906369 | 0.018817112 | 0.031566363 | 0.001430164 | 0.049489793 | 0.236540552 |
| Arpin | Y | 5 | -0.827038867 | -0.854630904 | -1.160952235 | -0.333913368 | -0.306321331 | 0.020785966 | 0.01762658 | 0.008359516 | 0.22014986 | 0.263389648 |
| Ppp1r11 | Y | 69 | -0.841488362 | -0.768240492 | -1.279957818 | -0.438469456 | -0.511717326 | 0.015870832 | 0.042069614 | 0.005682903 | 0.146361783 | 0.140893917 |
| Tagln2 Tagln3 | Y | 8;8 | -0.637027917 | -0.753689875 | -1.167602703 | -0.530574786 | -0.413912828 | 0.017842905 | 0.034898551 | 0.001323032 | 0.026921236 | 0.157833074 |
| Usp47 | Y | 174 | -0.723616548 | -0.721214762 | -1.097452043 | -0.373835495 | -0.376237281 | 0.014686967 | 0.019088819 | 0.003660943 | 0.131453404 | 0.145544215 |
| Nosip | Y | 14 | -0.73565117 | -0.828523117 | -1.171672514 | -0.436021343 | -0.343149396 | 0.032065049 | 0.011766147 | 0.007174363 | 0.179100083 | 0.23904001 |
| Cdv3 | Y | 280 | -0.768581548 | -0.907076357 | -0.63546787 | 0.133113678 | 0.271608487 | 0.001967885 | 0.031631064 | 0.00372814 | 0.160152373 | 0.339449747 |
| Inppl1 | Y | 672 | -0.642559795 | -0.758547906 | -0.835590337 | -0.193030542 | -0.077042431 | 0.00876242 | 0.011610967 | 0.003714513 | 0.234979386 | 0.691120424 |
| Svbp | Y | 40 | -0.86017554 | -0.826347787 | -1.083924608 | -0.223749068 | -0.257576821 | 0.013075968 | 0.043424726 | 0.0124901 | 0.407137822 | 0.445732994 |
| Map1b | Y | 1902 | -0.641423449 | -0.84738717 | -0.672562204 | -0.031138755 | 0.174824966 | 0.00982979 | 0.002655846 | 0.010868999 | 0.815320416 | 0.285217585 |
| Cnn2 | Y | 146 | -0.766840059 | -0.725638101 | -0.968475592 | -0.201635532 | -0.242837491 | 0.017058547 | 0.020447602 | 0.006467976 | 0.082687312 | 0.144152694 |
| Tex2 | Y | 298 | -0.952552699 | -0.948107214 | -0.948577932 | 0.003974767 | -0.000470718 | 0.041946607 | 0.027449189 | 0.017355185 | 0.991001858 | 0.998834261 |
| Nisch | Y | 1382 | -0.630909681 | -0.63003033 | -0.884412483 | -0.253502802 | -0.254382153 | 0.003122006 | 0.005402852 | 0.033706222 | 0.351762032 | 0.354124456 |
| Map1b | Y | 1405 | -0.656250465 | -0.724062337 | -0.786221686 | -0.129971221 | -0.062159349 | 0.017547592 | 0.009446199 | 0.008571584 | 0.53685142 | 0.754849044 |
| Uba1 Uba1y | Y | 60;59 | -0.651207703 | -0.728399451 | -1.029302124 | -0.37809442 | -0.300902673 | 0.046406 | 0.036412672 | 0.00506278 | 0.161317251 | 0.262114149 |
| Pebp1 | Y | 181 | -0.645999106 | -0.681965004 | -0.890081067 | -0.244081962 | -0.208116063 | 0.036499749 | 0.029745702 | 0.005992477 | 0.251628546 | 0.310437263 |
| Pacsin2 | Y | 388 | -0.623991383 | -0.638603841 | -0.70484814 | -0.080856756 | -0.066244298 | 0.011529694 | 0.014106856 | 0.009408894 | 0.56198202 | 0.701253304 |
| Cttn | T | 401 | -0.596995828 | -0.624532108 | -0.648785296 | -0.051789468 | -0.024253187 | 0.008570627 | 0.035416484 | 0.009023903 | 0.686829689 | 0.908937015 |
| Peak1 | Y | 632 | 0.73191117 | 0.676737148 | 1.143141173 | 0.411230003 | 0.466404025 | 0.042101798 | 0.047577888 | 0.018758153 | 0.254602863 | 0.201993303 |
| Atg3 | Y | 18 | -1.060270934 | -1.217140491 | -1.732682382 | -0.672411448 | -0.515541892 | 0.0021362 | 0.022375792 | 0.122910688 | 0.434142797 | 0.543462098 |
| Zfp638 | S | 727 | -0.646398737 | -1.066439763 | -1.202292961 | -0.555894224 | -0.135853198 | 0.006469117 | 0.009371342 | 0.073016137 | 0.268472426 | 0.762709189 |
| Zfp638 | T | 726 | -0.646398737 | -1.066439763 | -1.202292961 | -0.555894224 | -0.135853198 | 0.006469117 | 0.009371342 | 0.073016137 | 0.268472426 | 0.762709189 |
| Zfp638 | T | 734 | -0.646398737 | -1.066439763 | -1.202292961 | -0.555894224 | -0.135853198 | 0.006469117 | 0.009371342 | 0.073016137 | 0.268472426 | 0.762709189 |
| Zfp638 | Y | 731 | -0.646398737 | -1.066439763 | -1.202292961 | -0.555894224 | -0.135853198 | 0.006469117 | 0.009371342 | 0.073016137 | 0.268472426 | 0.762709189 |
| Met | S | 1234 | -0.979636288 | -0.979348905 | -0.668042348 | 0.31159394 | 0.311306556 | 0.014856927 | 0.025271569 | 0.061145315 | 0.152557763 | 0.300309156 |
| Met | Y | 1232 | -0.979636288 | -0.979348905 | -0.668042348 | 0.31159394 | 0.311306556 | 0.014856927 | 0.025271569 | 0.061145315 | 0.152557763 | 0.300309156 |
| Met | Y | 1233 | -0.979636288 | -0.979348905 | -0.668042348 | 0.31159394 | 0.311306556 | 0.014856927 | 0.025271569 | 0.061145315 | 0.152557763 | 0.300309156 |
| Fam120a | Y | 429 | -0.864017209 | -0.640778086 | -0.723739476 | 0.140277733 | -0.08296139 | 0.015557386 | 0.046298811 | 0.082759239 | 0.636720783 | 0.780144598 |
| Papss1 | Y | 30 | -0.670209223 | -0.663343747 | -1.409397007 | -0.739187784 | -0.74605326 | 0.033343986 | 0.067238805 | 0.00191713 | 0.032171301 | 0.053402348 |
| Rbm15 | Y | 335 | -0.757587049 | -0.847617042 | -1.428240329 | -0.67065328 | -0.580623286 | 0.022273455 | 0.057233316 | 0.007391455 | 0.089746915 | 0.177752088 |
| Tab3 | Y | 617 | -0.604838606 | -0.511693178 | -0.939336701 | -0.334498095 | -0.427643522 | 0.00707851 | 0.011654152 | 0.003848558 | 0.120141371 | 0.078582559 |
| Rtn4 | Y | 696 | -0.681570847 | -0.520626326 | -1.116092201 | -0.434521354 | -0.595465874 | 0.010072486 | 0.060265757 | 0.009707695 | 0.150996063 | 0.082509813 |
| Arfgap2 | Y | 444 | -0.625269747 | -0.576187381 | -0.819559946 | -0.194290199 | -0.243372564 | 0.011859101 | 0.021025435 | 0.004709986 | 0.322334052 | 0.245186229 |
| Anxa1 | Y | 21 | -0.753435062 | -0.514337984 | -1.198874609 | -0.445439547 | -0.684536625 | 0.006828705 | 0.164332466 | 0.01424354 | 0.18448635 | 0.115650277 |
| Arhgef10l | Y | 152 | -0.762621539 | -0.469807376 | -0.659970711 | 0.102650828 | -0.190163335 | 0.003810089 | 0.119527355 | 0.024375853 | 0.575512384 | 0.488382198 |
| Hnrnpk | Y | 280 | -0.601779184 | -0.57425896 | -0.728279322 | -0.126500138 | -0.154020363 | 0.025722313 | 0.03424177 | 0.019035911 | 0.472811261 | 0.446607732 |
| Cttn | S | 418 | -0.650872196 | -0.515780725 | -0.604995453 | 0.045876743 | -0.089214727 | 0.004243288 | 0.059075561 | 0.034477483 | 0.80073677 | 0.711822786 |
| Epb41l2 | T | 597 | -0.651392134 | -0.434051705 | -0.750778378 | -0.099386245 | -0.316726674 | 0.011601481 | 0.069884772 | 0.034431496 | 0.683780195 | 0.256358195 |
| Slc25a31 Slc25a4 Slc25a5 | Y | 81;81;94 | 0.644400055 | 0.458153312 | 0.666899278 | 0.022499224 | 0.208745966 | 0.008450559 | 0.084286457 | 0.005099686 | 0.861417577 | 0.325281379 |
| H4c1 H4c11 H4c12 H4c14 | Y | 89 | 0.706406918 | 0.515674483 | 0.630092961 | -0.076313957 | 0.114418478 | 0.014249423 | 0.031567897 | 0.021047144 | 0.705809983 | 0.569419481 |
| Rps26 | Y | 62 | 0.802942744 | 0.537176103 | 0.597962317 | -0.204980427 | 0.060786213 | 0.014681182 | 0.124811235 | 0.029914184 | 0.366626375 | 0.827728989 |
| Cdk16 Cdk17 | T | 175;202 | 0.613398028 | 0.402240028 | 0.613607718 | 0.00020969 | 0.211367691 | 0.032551089 | 0.083028171 | 0.015103491 | 0.999136826 | 0.307609285 |
| Tom1l2 | Y | 192 | 1.080845559 | 0.432451707 | 0.756250881 | -0.324594678 | 0.323799174 | 0.049046411 | 0.439532737 | 0.027261179 | 0.42171122 | 0.55572458 |
| Shb | Y | 113 | -0.652429937 | 0.035831174 | 0.130745791 | 0.783175728 | 0.094914617 | 0.003323104 | 0.835978369 | 0.411117948 | 0.002898604 | 0.554602998 |
| Aaas | S | 495 | 0.626318785 | 0.624316658 | -0.087207792 | -0.713526577 | -0.71152445 | 0.035246489 | 0.168301481 | 0.702342884 | 0.014729387 | 0.136047066 |
| Cttn | Y | 421 | -0.697734582 | -0.376625883 | -0.556394848 | 0.141339735 | -0.179768965 | 0.004628945 | 0.141200728 | 0.07668672 | 0.540522981 | 0.5168143 |
| Slc20a2 | Y | 377 | -0.671135722 | -0.549016012 | -0.41154892 | 0.259586802 | 0.137467092 | 0.021638133 | 0.040553644 | 0.093353731 | 0.086519613 | 0.340944438 |
| Cttn | S | 417 | -0.65698801 | -0.367812864 | -0.516842498 | 0.140145512 | -0.149029634 | 0.004780121 | 0.132309814 | 0.073130353 | 0.511980281 | 0.557887204 |
| Fam102b | Y | 332 | -0.822221177 | -0.291553626 | -0.639524535 | 0.182696642 | -0.347970909 | 0.010053288 | 0.28914192 | 0.104074339 | 0.56353777 | 0.323185019 |
| Tjp2 | Y | 1156 | -0.593000218 | -0.544310379 | -0.120383275 | 0.472616944 | 0.423927104 | 0.005243648 | 0.011641906 | 0.40541776 | 0.006142394 | 0.025218115 |
| Fgfr1 | Y | 653 | -0.809146726 | -0.412921254 | -0.568640921 | 0.240505804 | -0.155719667 | 0.03026368 | 0.207263658 | 0.102903119 | 0.434873789 | 0.61361699 |
| Stxbp3 | Y | 521 | -0.670214495 | -0.500755915 | -0.348115696 | 0.3220988 | 0.15264022 | 0.014995642 | 0.055322369 | 0.241325151 | 0.246804861 | 0.561636236 |
| Shb | Y | 262 | -0.759115754 | -0.332320353 | -0.266914478 | 0.492201276 | 0.065405874 | 0.004228124 | 0.133842806 | 0.142758859 | 0.016465237 | 0.680816237 |
| Arhgap39 | Y | 444 | -0.769760131 | -0.590773587 | -0.345440407 | 0.424319724 | 0.24533318 | 0.043713101 | 0.11218714 | 0.245888675 | 0.229244909 | 0.482548316 |
| Epb41l1 | Y | 68 | -0.717897254 | -0.218881862 | -0.451116242 | 0.266781012 | -0.23223438 | 0.040156757 | 0.451011792 | 0.035410915 | 0.309760269 | 0.408649006 |
| Txnl4b | Y | 146 | -1.081683876 | -0.376993553 | -0.435363942 | 0.646319934 | -0.058370389 | 0.003897601 | NA | NA | NA | NA |
| Rpl36a Rpl36al | S | 32 | 1.191738052 | 0.682078695 | 0.508391293 | -0.683346759 | -0.173687401 | 0.006783721 | 0.050809776 | 0.168760359 | 0.065727773 | 0.509149924 |
| Rpl36a Rpl36al | Y | 34 | 1.090522583 | 0.524199305 | 0.475767341 | -0.614755242 | -0.048431964 | 0.005621446 | 0.095308067 | 0.243450781 | 0.143195447 | 0.886896531 |
| Bcar1 | S | 359 | 0.817563967 | 0.422403639 | 0.496005172 | -0.321558794 | 0.073601534 | 0.020518641 | 0.064721314 | 0.13534048 | 0.318464108 | 0.791922072 |
| Rpl31 | Y | 108 | 0.723060796 | 0.552893548 | 0.6985092 | -0.024551597 | 0.145615652 | 0.040342701 | 0.141157411 | 0.108033083 | 0.934674658 | 0.688949394 |
| Safb | Y | 802 | 0.593180402 | 0.162202555 | 0.455450907 | -0.137729495 | 0.293248352 | 0.002449084 | 0.628843244 | 0.018734273 | 0.320100022 | 0.40894053 |
| Fyb | Y | 816 | 0.66740441 | 0.364219894 | 0.120478771 | -0.546925639 | -0.243741123 | 0.019920826 | 0.021713875 | 0.446256381 | 0.042662008 | 0.138285803 |
| Pabpc1 | Y | 116 | 0.707050651 | 0.39031876 | 0.166781563 | -0.540269088 | -0.223537197 | 0.020590882 | 0.151267273 | 0.568115382 | 0.090117776 | 0.393513802 |

| | | | | | | | | | | | | |
|---|---|---|---|---|---|---|---|---|---|---|---|---|
| Actn1 Actn4 | S | 264;244 | 0.820530035 | 0.174182037 | 0.367520108 | -0.453009927 | 0.193338072 | 0.03267785 | 0.465005544 | 0.278705796 | 0.233425432 | 0.548500866 |
| Rps9 | Y | 165 | 0.610487503 | 0.028639468 | 0.129403936 | -0.481083567 | 0.100764467 | 0.004589337 | 0.921599413 | 0.267837751 | 0.015663732 | 0.735601885 |
| Ubap2l | Y | 130 | 0.619410966 | 0.014326336 | 0.05682271 | -0.562588257 | 0.042496374 | 0.027384525 | 0.954166933 | 0.504769421 | 0.045789986 | 0.862602914 |
| Hnrnpf Hnrnph1 | Y | 306;306 | 0.779724511 | -0.265404941 | 0.057670778 | -0.722053733 | 0.323075719 | 0.004300897 | 0.250271363 | 0.844068794 | 0.077531185 | 0.308826553 |
| Eef2 | Y | 443 | 0.063312942 | -0.797028356 | -0.888691689 | -0.952004631 | -0.091663333 | 0.629771493 | 2.2831E-07 | 3.63614E-07 | 0.010899782 | 0.034936937 |
| Fosl1 | Y | 18 | -0.439527626 | -1.258581548 | -1.931333692 | -1.491806066 | -0.672752143 | 0.021182058 | 0.008579602 | 0.005556647 | 0.011372775 | 0.083456698 |
| Rps21 | Y | 53 | -0.35414073 | -1.006858796 | -1.350342134 | -0.996201403 | -0.343483338 | 0.096759182 | 0.002498154 | 0.00073592 | 0.00578654 | 0.144157838 |
| Snw1 | Y | 113 | -0.516957946 | -0.875163953 | -1.350823349 | -0.833865403 | -0.475659397 | 0.03648016 | 0.011909693 | 0.000907343 | 0.000197452 | 0.093011235 |
| Tacc3 | Y | 349 | -0.581571831 | -0.787223453 | -1.209020964 | -0.627449133 | -0.421797511 | 0.020703697 | 0.013437007 | 0.001059979 | 0.001530892 | 0.092851053 |
| Gpatch1 | Y | 322 | -0.503030081 | -0.703867352 | -1.112599943 | -0.609569861 | -0.408732591 | 0.022913012 | 0.005948442 | 0.001899705 | 0.029180091 | 0.077663649 |
| Mllt11 | Y | 9 | -0.381820135 | -0.864732482 | -1.143620331 | -0.761800196 | -0.27888785 | 0.108236077 | 0.005357959 | 0.001080335 | 0.011631319 | 0.179927207 |
| Sdc2 | S | 188 | -0.195866034 | -0.799774891 | -1.00841141 | -0.812545376 | -0.208636519 | 0.123465371 | 0.000936574 | 0.000386137 | 0.006312101 | 0.139669337 |
| Grasp | Y | 236 | 0.160353524 | -0.948031696 | -0.988407758 | -1.148761282 | -0.040376063 | 0.115026631 | 5.9221E-05 | 7.44222E-05 | 0.001568822 | 0.640503303 |
| Cbl | S | 673 | -0.348830782 | -0.660018557 | -0.941107195 | -0.592276412 | -0.281088638 | 0.055070323 | 0.004141447 | 0.001061314 | 0.009855739 | 0.083995079 |
| Spag7 | Y | 189 | -0.279882153 | -0.587415207 | -0.994060448 | -0.714178295 | -0.406645241 | 0.130705333 | 0.01264314 | 0.005419668 | 0.025881835 | 0.104861615 |
| Ccdc88a | Y | 1767 | -0.201642308 | -0.59534344 | -0.789170153 | -0.587527845 | -0.193826713 | 0.097741501 | 0.000489643 | 0.016020374 | 0.027057777 | 0.276934729 |
| Arap2 | Y | 440 | -0.176796476 | -0.648970025 | -0.91606696 | -0.739270485 | -0.267096935 | 0.11431002 | 0.009561846 | 0.008997625 | 0.014518061 | 0.18748036 |
| Arap2 | Y | 378 | 0.019765007 | -0.676250438 | -0.733738507 | -0.753503514 | -0.057488069 | 0.857106806 | 4.58404E-05 | 0.00814062 | 0.004796283 | 0.622858068 |
| Prrc2c | Y | 17 | -0.098336652 | -0.715592549 | -0.695752434 | -0.597415781 | 0.019840116 | 0.46966426 | 0.003496486 | 0.002414396 | 0.009594082 | 0.863621211 |
| Rpl37 Rpl37rt | Y | 39 | 0.020430899 | -0.883837898 | -0.655863541 | -0.67629444 | 0.227974356 | 0.877060538 | 0.009394056 | 0.038010557 | 0.034746857 | 0.356692298 |
| Irs2 | S | 732 | 0.418190863 | 2.675224611 | 3.070651198 | 2.652460335 | 0.395426587 | 0.159895931 | 0.000118262 | 1.22683E-05 | 0.001204532 | 0.038730561 |
| Pdgfra | Y | 754 | 0.278872881 | 1.26784603 | 1.103430675 | 0.824557794 | -0.164415355 | 0.120995033 | 1.14783E-05 | 2.59011E-06 | 0.014688589 | 0.07802249 |
| Stat5a | Y | 694 | 0.035470923 | 1.249738506 | 1.603239726 | 1.567768803 | 0.35350122 | 0.829348811 | 4.4829E-05 | 9.35174E-06 | 0.001264573 | 0.016546196 |
| Crim1 | Y | 1020 | 0.354874108 | 1.243953621 | 1.701518306 | 1.346644198 | 0.457564685 | 0.027449681 | 0.003376607 | 0.000423752 | 0.000612867 | 0.048307698 |
| Pdgfra | Y | 762 | 0.358165074 | 1.053450104 | 1.139187883 | 0.781022809 | 0.085737779 | 0.048686835 | 0.00036351 | 7.64982E-05 | 0.004956732 | 0.453904416 |
| Nedd9 | Y | 630 | 0.394943783 | 0.833236239 | 0.985391238 | 0.590447455 | 0.152154999 | 0.031603432 | 9.76727E-05 | 0.000118644 | 0.025102132 | 0.096045477 |
| Numa1 | Y | 1603 | 0.367610941 | 0.820942733 | 1.157558491 | 0.789947551 | 0.336615758 | 0.024831981 | 0.001354697 | 0.000297855 | 0.004042337 | 0.069600788 |
| Tns2 | Y | 705 | 0.183731748 | 1.027342632 | 1.03606021 | 0.852328461 | 0.008717578 | 0.203975867 | 0.000235613 | 0.000209924 | 0.001176382 | 0.938894401 |
| Aplp2 | Y | 694 | 0.352764345 | 0.992750949 | 1.313237836 | 0.96047349 | 0.320486887 | 0.054006223 | 0.003556459 | 0.001037599 | 0.006411611 | 0.175569813 |
| Tns2 | Y | 483 | 0.150380646 | 0.705978721 | 0.78816054 | 0.637779894 | 0.082181818 | 0.010380931 | 0.000340121 | 2.88738E-06 | 0.000134595 | 0.245855714 |
| Stat5b | Y | 699 | 0.012938557 | 0.900182408 | 1.412966171 | 1.400027613 | 0.512783762 | 0.939514282 | 0.000671008 | 4.20988E-05 | 0.00139584 | 0.006576917 |
| Irs1 | Y | 628 | 0.158682975 | 1.075146898 | 0.924417973 | 0.765734998 | -0.150728925 | 0.259126186 | 0.000256709 | 0.000548349 | 0.001800664 | 0.179634719 |
| Nedd9 | Y | 260 | 0.262977664 | 0.714998943 | 0.911660577 | 0.648682913 | 0.196661634 | 0.022162671 | 0.001278532 | 0.001110785 | 0.009374927 | 0.188901848 |
| Aplp2 | Y | 699 | 0.287038703 | 0.766979643 | 1.100024853 | 0.81298615 | 0.33304521 | 0.076740521 | 0.002734013 | 0.002407241 | 0.00619901 | 0.090912303 |
| Ephb3 | Y | 595 | 0.215062401 | 0.676744545 | 0.848860185 | 0.633797784 | 0.17211564 | 0.296626499 | 0.010809735 | 0.000402627 | 0.033540215 | 0.313493764 |
| Nup35 | Y | 299 | -0.555938814 | -0.854406607 | -1.034411616 | -0.478472803 | -0.180005009 | 0.006191953 | 0.001497512 | 0.000366136 | 0.012988941 | 0.238374126 |
| Tra2b | Y | 260 | -0.681793687 | -0.756371362 | -1.48173083 | -0.799937143 | -0.725359468 | 0.05906208 | 0.039327272 | 0.000107338 | 0.050044752 | 0.056109604 |
| Abl2 | Y | 719 | -0.519246096 | -1.087650084 | -0.852561001 | -0.333314906 | 0.235089083 | 0.01472811 | 0.000507207 | 0.011711543 | 0.164595006 | 0.281241585 |
| Hnrnpab | Y | 269 | -0.573223556 | -0.746183892 | -1.241514601 | -0.668291045 | -0.495330709 | 0.019758862 | 0.006889365 | 0.002594032 | 0.05140064 | 0.085658257 |
| Map1b | Y | 1758 | -0.573577235 | -0.738370703 | -1.136454667 | -0.562877432 | -0.398083964 | 0.019505872 | 0.012644991 | 0.001046371 | 0.023923647 | 0.087452836 |
| Pdlim5 | Y | 251 | -0.519963781 | -0.737801006 | -0.923763012 | -0.403799231 | -0.185962006 | 0.008663146 | 0.005069211 | 0.001640393 | 0.048302022 | 0.308371269 |
| Mapk6 | Y | 467 | -0.245630197 | -0.785888716 | -0.60209496 | -0.356464763 | 0.183793756 | 0.00887413 | 9.86855E-06 | 0.003641575 | 0.024371822 | 0.122151565 |
| Ubap2 | Y | 854 | -0.562840351 | -0.69400582 | -0.630347142 | -0.067506791 | 0.063658678 | 0.056969888 | 0.000387026 | 0.000500601 | 0.728041263 | 0.298584505 |
| Tdp1 | Y | 46 | -0.56774098 | -0.747642612 | -1.1203777 | -0.55263672 | -0.372735087 | 0.049178954 | 0.013874266 | 0.002469522 | 0.035797595 | 0.065401158 |
| Ptpn14 | Y | 485 | -0.387558557 | -0.646217411 | -0.959980376 | -0.572421819 | -0.313762965 | 0.02403497 | 0.008905043 | 0.000423822 | 0.001236786 | 0.093291781 |
| Snap29 | Y | 160 | -0.573018321 | -0.746226749 | -1.091682839 | -0.518664518 | -0.34545609 | 0.013377117 | 0.005913298 | 0.015031068 | 0.126213334 | 0.248161978 |
| Sec31a | Y | 803 | -0.579793811 | -0.700424948 | -0.863767873 | -0.283974062 | -0.163342925 | 0.013451719 | 0.010367601 | 0.002198648 | 0.072432724 | 0.347488643 |
| Tpx2 | Y | 518 | -0.475892569 | -0.622154716 | -1.026518199 | -0.55062563 | -0.404363483 | 0.02817406 | 0.008994089 | 0.001207385 | 0.020633657 | 0.050733344 |
| Cnn3 | T | 180 | -0.509522471 | -0.800873795 | -0.881314846 | -0.371792375 | -0.080441051 | 0.014174183 | 0.006132511 | 0.011175232 | 0.134372976 | 0.718318055 |
| Dapp1 | Y | 139 | -0.482689695 | -0.820456712 | -0.990578618 | -0.507888923 | -0.170121906 | 0.010182002 | 0.008886185 | 0.017997888 | 0.08613021 | 0.488543725 |
| Map1b | Y | 1059 | -0.430674013 | -0.585495796 | -0.666975466 | -0.236301453 | -0.08147967 | 0.008285627 | 0.002239719 | 0.002658543 | 0.098618661 | 0.496938131 |
| Epha2 | T | 588 | -0.51122058 | -0.987033175 | -0.676073221 | -0.164852641 | 0.310959953 | 0.040250731 | 0.004760106 | 0.014868415 | 0.204809881 | 0.066372803 |
| Pik3r1 | Y | 607 | -0.401328878 | -0.871934384 | -0.960881766 | -0.559552848 | -0.088947342 | 0.026492337 | 0.006868003 | 0.01889127 | 0.044345483 | 0.651195181 |
| Szrd1 | Y | 95 | -0.25826983 | -0.683686377 | -0.651121636 | -0.392851806 | 0.032564742 | 0.203665573 | 0.000497666 | 0.000270742 | 0.098571315 | 0.57661528 |
| Cnn2 Cnn3 | Y | 12;10 | -0.559481692 | -0.638114313 | -1.005064264 | -0.445582573 | -0.366949952 | 0.058388978 | 0.036557307 | 0.002105267 | 0.114325815 | 0.153069365 |
| Jcad | Y | 972 | -0.543929672 | -0.60955547 | -0.931935983 | -0.38800631 | -0.322380513 | 0.027201625 | 0.030793624 | 0.00284426 | 0.033908632 | 0.13600422 |
| Sugt1 | Y | 288 | -0.47612084 | -0.682949387 | -0.909657872 | -0.433537031 | -0.226708485 | 0.038763071 | 0.02186511 | 0.002659961 | 0.029548105 | 0.270353513 |
| Cbl | Y | 672 | -0.286263118 | -0.598852685 | -0.8279331 | -0.541669982 | -0.229080416 | 0.05095543 | 0.004164649 | 0.000743516 | 0.013848735 | 0.113308698 |
| Yipf5 | Y | 39 | -0.47840985 | -0.896983508 | -1.103284345 | -0.624874494 | -0.206300837 | 0.065018447 | 0.021519997 | 0.015908957 | 0.090692716 | 0.533704079 |
| Eif4g1 | Y | 600 | -0.469421132 | -0.626630153 | -0.666536871 | -0.197115739 | -0.039906719 | 0.018551881 | 0.002055441 | 0.013075199 | 0.298071509 | 0.793128467 |
| Polr2a | Y | 1832 | -0.562203283 | -0.66407159 | -0.93026114 | -0.368057857 | -0.26618955 | 0.030305397 | 0.014743327 | 0.023082195 | 0.222806424 | 0.338755156 |
| BC005624 | Y | 147 | -0.478363924 | -0.672995748 | -1.091509006 | -0.613145082 | -0.418513258 | 0.074005489 | 0.045366077 | 0.00708085 | 0.054773851 | 0.171485142 |
| U2surp | Y | 117 | -0.554561764 | -0.585487968 | -1.165427692 | -0.610865928 | -0.579939724 | 0.112294245 | 0.04323382 | 0.006704866 | 0.100933877 | 0.074217452 |
| Lasp1 | Y | 152 | -0.495472827 | -0.640174662 | -0.817425436 | -0.321952609 | -0.177250774 | 0.023635071 | 0.024857698 | 0.024570588 | 0.232922752 | 0.506044387 |
| Rbm14 | Y | 645 | -0.47016071 | -0.63103687 | -0.888130228 | -0.417969518 | -0.257093358 | 0.17638776 | 0.006137458 | 0.018505499 | 0.244576558 | 0.305925057 |
| Fmnl3 | Y | 203 | -0.31087039 | -0.71254412 | -0.804458264 | -0.493587874 | -0.091914144 | 0.141112097 | 0.015333752 | 0.007226463 | 0.035216574 | 0.612805114 |
| Rbm3 | Y | 147 | -0.400952355 | -0.590957763 | -0.74134613 | -0.340393775 | -0.150388367 | 0.055703563 | 0.015246168 | 0.012068958 | 0.129795662 | 0.388694651 |
| Fmnl3 | S | 204 | -0.441388392 | -0.749217841 | -0.708229701 | -0.266841309 | 0.04098814 | 0.106371368 | 0.012119856 | 0.022145996 | 0.259508945 | 0.808895285 |
| Bag3 | Y | 246 | -0.399622387 | -0.639677147 | -0.706624873 | -0.307002486 | -0.066947726 | 0.027362113 | 0.020815455 | 0.030913756 | 0.211319541 | 0.779249742 |
| Pard3 | S | 1254 | -0.267927275 | -0.620500942 | -0.673355248 | -0.405427973 | -0.052854306 | 0.13389601 | 0.003949033 | 0.0134651 | 0.076711775 | 0.74822433 |
| Ppp1r12a | S | 909 | -0.380845079 | -0.585026292 | -0.675912296 | -0.295067217 | -0.090886003 | 0.065631552 | 0.028992286 | 0.01006314 | 0.079458905 | 0.616663438 |
| Rin1 | S | 280 | -0.314008587 | -0.600992007 | -0.620408382 | -0.306399795 | -0.019416375 | 0.132039723 | 0.040691905 | 0.010282043 | 0.130994727 | 0.926423767 |
| Nedd9 | Y | 628 | 0.449159895 | 0.853902236 | 0.993304374 | 0.544144479 | 0.139402138 | 0.018148091 | 0.000173272 | 0.000204722 | 0.025114854 | 0.088204276 |
| Irs2 | Y | 649 | 0.337511025 | 0.937397886 | 0.894141443 | 0.556630418 | -0.043256443 | 0.042121738 | 4.7712E-05 | 0.00010343 | 0.01101117 | 0.600184554 |
| Pdgfra | Y | 768 | 0.684462869 | 1.245657545 | 1.164684054 | 0.480221184 | -0.080973491 | 0.124748437 | 0.010651439 | 0.000707056 | 0.2138629 | 0.677098364 |
| Afap1l2 | Y | 359 | 0.401289438 | 0.791404195 | 0.869112326 | 0.467822888 | 0.077708131 | 0.129239797 | 0.000266658 | 0.000104754 | 0.101225525 | 0.423263568 |
| Arhgap42 | Y | 759 | 0.243541864 | 0.72087379 | 0.757068355 | 0.513526491 | 0.036194565 | 0.106851913 | 0.00011644 | 2.98432E-05 | 0.020146545 | 0.61203977 |
| Nedd9 | Y | 344 | 0.296752572 | 0.833340465 | 0.767961292 | 0.47120872 | -0.065379173 | 0.058210795 | 0.000592371 | 0.001030108 | 0.006780234 | 0.43169472 |
| F2r | Y | 425 | 0.45610205 | 0.604430198 | 0.706152365 | 0.250050315 | 0.101722167 | 0.015252629 | 0.010171417 | 0.000246204 | 0.097145882 | 0.458766555 |
| Fam228a | S | 173 | 0.157350726 | 0.679179501 | 0.740445703 | 0.583094977 | 0.061266203 | 0.064573226 | 0.000543922 | 0.000167027 | 0.000682821 | 0.444138854 |
| Mxra8 | Y | 400 | 0.21358726 | 0.624165275 | 0.676016087 | 0.462428826 | 0.051850812 | 0.040888757 | 0.000457709 | 0.000178286 | 0.004204461 | 0.52267562 |
| Ephb3 | Y | 919 | 0.573853786 | 0.623028143 | 0.792256275 | 0.218402489 | 0.169228132 | 0.17993319 | 0.000767847 | 0.001402671 | 0.537135831 | 0.196671537 |
| Laptm4a | Y | 215 | 0.659109732 | 0.724346592 | 1.095616205 | 0.436506473 | 0.371269613 | 0.184701156 | 0.007912969 | 0.005228515 | 0.339000528 | 0.154702647 |
| Pdgfra | Y | 1018 | 0.454359686 | 0.90383817 | 0.726481601 | 0.272121915 | -0.177356569 | 0.245462899 | 0.000254142 | 0.014492877 | 0.453055962 | 0.324022816 |
| Hgs | Y | 132 | 0.285187887 | 0.591807138 | 0.657864809 | 0.372676922 | 0.06605767 | 0.034558119 | 0.006084394 | 5.51761E-05 | 0.030509319 | 0.515563008 |
| Irs2 | Y | 814 | 0.131345011 | 0.642241242 | 0.644163187 | 0.512818176 | 0.001921945 | 0.112108103 | 0.00020231 | 0.00026295 | 0.0072965 | 0.979553643 |
| 9530068E07Rik | Y | 257 | 0.506714404 | 0.813096126 | 0.967568483 | 0.460854079 | 0.154472357 | 0.013578963 | 0.019719853 | 0.025463831 | 0.116227032 | 0.556206194 |
| Ptpra | Y | 825 | 0.267193009 | 0.635866113 | 0.698152728 | 0.43095972 | 0.062286615 | 0.145489352 | 0.000789067 | 0.000686275 | 0.049874418 | 0.476891949 |
| Laptm4a | Y | 230 | 0.395975821 | 0.650486352 | 0.846155528 | 0.450179707 | 0.195669176 | 0.00573982 | 0.021836007 | 0.007564806 | 0.047647539 | 0.340931642 |
| App | T | 761 | 0.466702431 | 0.623973562 | 0.690182174 | 0.223479744 | 0.066208612 | 0.47554298 | 0.001789522 | 0.000405682 | 0.717023532 | 0.512959334 |
| Prkaca Prkacb | T | 196;196 | 0.635636784 | 0.687052956 | 0.761043052 | 0.125406268 | 0.073990096 | 0.08242874 | 0.016043634 | 0.015136335 | 0.672208407 | 0.743192088 |

| | | | | | | | | | | | | | |
|---|---|---|---|---|---|---|---|---|---|---|---|---|---|
| Magoh Magohb | Y | 40;40 | | 0.160010807 | 0.600977049 | 0.716858688 | 0.556847881 | 0.11588164 | 0.034207232 | 0.003111825 | 0.001773613 | 0.0027343 | 0.253219013 |
| 2610507B11Rik | S | | 110 | 0.200487503 | 0.708472934 | 0.684822112 | 0.484334609 | -0.023650822 | 0.16495902 | 0.001979283 | 0.004442941 | 0.022008507 | 0.842515328 |
| Atp1a1 | Y | | 55 | -0.952373924 | -0.972634759 | -0.98625859 | -0.033884666 | -0.013623831 | 0.202873607 | 0.023461732 | 0.054683445 | 0.95822374 | 0.967696697 |
| Usp15 | Y | | 274 | -0.629581669 | -0.972946728 | -0.676135456 | -0.046553787 | 0.296811273 | 0.05872935 | 0.013372231 | 0.120755145 | 0.887306792 | 0.388364191 |
| Eps8 | Y | | 612 | -0.499234726 | -0.618135697 | -0.371234692 | 0.128000034 | 0.246901005 | 0.009530903 | 0.005101525 | 0.048227542 | 0.411455842 | 0.161602694 |
| Gpr149 | Y | | 230 | -0.215234441 | -0.791926416 | -0.465726966 | -0.250492525 | 0.32619945 | 0.252251164 | 0.001960786 | 0.018301683 | 0.160690717 | 0.038528129 |
| Ybx1 | Y | | 160 | -0.495838146 | -0.727039284 | -0.55069727 | -0.054859124 | 0.176342013 | 0.081967181 | 0.037926103 | 0.057702618 | 0.684608389 | 0.425156763 |
| Myo10 | Y | | 1878 | 0.032293605 | -0.595181451 | -0.448885115 | -0.48117872 | 0.146296336 | 0.538179566 | 0.005273532 | 0.00024107 | 0.000838404 | 0.194487346 |
| Baiap2 | S | | 353 | -0.381217657 | -0.679892596 | -0.211702974 | 0.169514683 | 0.468189622 | 0.042272631 | 0.005961393 | 0.216080473 | 0.186883131 | 0.0257983 |
| Tjp2 | Y | | 1095 | -0.369221592 | -0.637389671 | -0.105208445 | 0.264013147 | 0.532181226 | 0.021788407 | 0.004059957 | 0.41393744 | 0.023043857 | 0.011720158 |
| Fip1l1 | Y | | 497 | 0.052553109 | -0.704334672 | -0.42513171 | -0.477684819 | 0.279202962 | 0.524252115 | 0.002005599 | 0.022783762 | 0.020208686 | 0.090106161 |
| Ncoa5 | Y | | 107 | -0.15697958 | -0.728515265 | -0.054131353 | 0.102848226 | 0.674383912 | 0.492313975 | 0.015340892 | 0.886467059 | 0.804907241 | 0.171897062 |
| Srsf7 | Y | | 146 | 0.341437267 | -0.635380783 | -0.054922176 | -0.396359443 | 0.580458607 | 0.058998348 | 0.036213856 | 0.873295284 | 0.310863125 | 0.180352441 |
| Asap2 | T | | 765 | 0.542127136 | 0.987419136 | 0.66222228 | 0.120095144 | -0.325196856 | 0.136666274 | 0.009003794 | 0.069949417 | 0.712730421 | 0.252852947 |
| Tln2 | Y | | 1666 | 0.217975663 | 0.589637623 | 0.376645744 | 0.158670081 | -0.212991879 | 0.013295764 | 0.005900157 | 0.001167022 | 0.006263587 | 0.123007788 |
| Tln2 | S | | 1667 | 0.36323703 | 0.883012573 | 0.296812052 | -0.066424978 | -0.58620052 | 0.03510274 | 0.001338074 | 0.275541885 | 0.76904664 | 0.077890377 |
| Dcbld2 | Y | | 727 | 0.147854746 | 0.677843725 | 0.543781081 | 0.395926335 | -0.134062645 | 0.148245853 | 0.003225491 | 0.009352055 | 0.050834424 | 0.378919828 |
| Yes1 | S | | 137 | 0.334154054 | 0.699955184 | 0.771830646 | 0.437676592 | 0.071875462 | 0.474157998 | 0.015313965 | 0.075353003 | 0.39502253 | 0.805819916 |
| Nsfl1c | S | | 176 | 0.409902239 | 0.651392626 | 0.272160152 | -0.137742088 | -0.379232474 | 0.017387139 | 0.041203895 | 0.184825456 | 0.417677151 | 0.168758241 |
| Ptk2b | Y | | 580 | 0.131015469 | 0.594216962 | 0.182160593 | 0.051145125 | -0.412056368 | 0.250199058 | 0.028273496 | 0.241500184 | 0.661010433 | 0.085412587 |
| Ptk2b | Y | | 579 | 0.121518575 | 0.601486622 | 0.146218187 | 0.024699612 | -0.455268435 | 0.315901032 | 0.02893743 | 0.383929693 | 0.8519187 | 0.074436206 |
| Sirpa | Y | | 507 | 0.140241839 | 0.667700514 | 0.030658476 | -0.109583363 | -0.637042038 | 0.524479298 | 0.01894911 | 0.950863079 | 0.819885185 | 0.263917641 |
| Ddx24 | S | | 270 | -0.110742772 | 0.61625881 | 0.217354196 | 0.328096968 | -0.398904615 | 0.444477703 | 0.01173386 | 0.498241912 | 0.33264807 | 0.261119202 |
| Ddx24 | S | | 288 | -0.110742772 | 0.61625881 | 0.217354196 | 0.328096968 | -0.398904615 | 0.444477703 | 0.01173386 | 0.498241912 | 0.33264807 | 0.261119202 |
| Ddx24 | T | | 271 | -0.110742772 | 0.61625881 | 0.217354196 | 0.328096968 | -0.398904615 | 0.444477703 | 0.01173386 | 0.498241912 | 0.33264807 | 0.261119202 |
| Eps8 | Y | | 539 | 0.338535527 | 0.555343956 | 1.225705456 | 0.887169929 | 0.670361499 | 0.034762563 | 0.015660123 | 0.000509401 | 0.004084408 | 0.013691156 |
| Ceacam1 Ceacam2 | Y | 514;515 | | 0.038732573 | -0.182634418 | 0.793183814 | 0.754451241 | 0.975818232 | 0.841795955 | 0.352065411 | 0.007365507 | 0.002703438 | 0.004040738 |
| Stam | Y | | 384 | -0.353992102 | -0.52023971 | -1.093347841 | -0.739355739 | -0.573108131 | 0.074782626 | 0.060149307 | 0.000316765 | 0.006683849 | 0.052594521 |
| Tmem63a | T | | 773 | -0.293930131 | -0.549881398 | -1.042472102 | -0.748541971 | -0.492590704 | 0.149370045 | 0.06536765 | 0.00015542 | 0.014843877 | 0.091512204 |
| Arap2 | Y | | 77 | -0.087646778 | -0.518770656 | -0.673418698 | -0.58577192 | -0.154648042 | 0.169026371 | 0.005762553 | 1.5886E-05 | 0.000487449 | 0.133542026 |
| Tbc1d15 | Y | | 215 | -0.411345582 | -0.565434199 | -1.014003844 | -0.602658262 | -0.448569645 | 0.05270843 | 0.042033247 | 0.006130817 | 0.046999323 | 0.116013154 |
| Tbc1d15 | S | | 205 | -0.37283806 | -0.591383505 | -1.032973967 | -0.660135907 | -0.441590462 | 0.151463215 | 0.051077958 | 0.005806079 | 0.013556998 | 0.041748735 |
| Prc1 | Y | | 464 | -0.234168689 | -0.458608674 | -0.918146233 | -0.683977545 | -0.459537559 | 0.196212821 | 0.029836777 | 0.00084126 | 0.013323369 | 0.034429287 |
| Ddr2 | Y | | 471 | -0.061177085 | -0.460306532 | -0.6882294 | -0.627052315 | -0.227922868 | 0.552448922 | 0.000969353 | 0.000371221 | 0.006056421 | 0.026748519 |
| Otud4 | S | | 1042 | -0.073194396 | -0.46881076 | -0.700524062 | -0.627329666 | -0.231713302 | 0.585004861 | 0.001902155 | 0.001633025 | 0.009519055 | 0.088852877 |
| Epha2 | T | | 775 | -0.096050714 | -0.286650141 | -0.778533096 | -0.682482382 | -0.491882955 | 0.544485722 | 0.027141867 | 0.000507015 | 0.02659163 | 0.006007862 |
| Pard3 | S | | 1172 | 0.004899098 | -0.434380613 | -0.598233844 | -0.603132942 | -0.163853231 | 0.958109107 | 0.049942197 | 9.73489E-05 | 0.004352577 | 0.302002352 |
| Pard3 | Y | | 1171 | 0.004899098 | -0.434380613 | -0.598233844 | -0.603132942 | -0.163853231 | 0.958109107 | 0.049942197 | 9.73489E-05 | 0.004352577 | 0.302002352 |
| Khdrbs1 Khdrbs2 | Y | 440;346 | | 0.125484008 | -0.32277973 | -0.847793319 | -0.973277327 | -0.525013589 | 0.413074935 | 0.131901913 | 0.000152713 | 0.006028021 | 0.05427995 |
| Ddr2 | S | | 461 | 0.127022411 | -0.534915293 | -0.722534838 | -0.849557249 | -0.187619545 | 0.405663911 | 0.059754566 | 0.00181119 | 0.00250277 | 0.390568064 |
| Hspa8 | Y | | 115 | -0.010483932 | -0.328086039 | -0.625768198 | -0.615284266 | -0.297682159 | 0.888822274 | 0.001004202 | 0.035018023 | 0.034167723 | 0.163048158 |
| Epha2 | Y | | 773 | 0.016942284 | -0.214829651 | -0.599915671 | -0.616857955 | -0.38508602 | 0.890858351 | 0.215247621 | 6.89801E-05 | 0.017116682 | 0.077580122 |
| Khsrp | Y | | 318 | -0.074334756 | -0.272582023 | -0.698242474 | -0.623907718 | -0.425660451 | 0.582947541 | 0.126329972 | 0.019469202 | 0.031137281 | 0.087937116 |
| Magi3 | Y | | 356 | 0.383242219 | 0.579933765 | 0.99276001 | 0.609517792 | 0.412826245 | 0.012159357 | 0.005078351 | 0.00077279 | 0.009548152 | 0.036690167 |
| Pdgfrb | Y | | 856 | 0.224357205 | 0.437241868 | 0.83845818 | 0.614100975 | 0.401216312 | 0.112603497 | 0.014460501 | 7.31762E-05 | 0.007749291 | 0.02534285 |
| Nedd9 | Y | | 165 | 0.034281249 | 0.447183238 | 0.70254554 | 0.668264291 | 0.255362302 | 0.783767468 | 0.004158867 | 0.000496663 | 0.008805095 | 0.04626039 |
| Ephb3 | Y | | 588 | 0.0897671 | 0.530034081 | 0.821639032 | 0.731871932 | 0.291604951 | 0.352468308 | 0.001712921 | 0.022846963 | 0.045423804 | 0.224662497 |
| Jak2 | Y | | 570 | 0.10157819 | 0.45118693 | 0.762365338 | 0.660787148 | 0.311178408 | 0.478233622 | 0.001458877 | 0.008624821 | 0.015090633 | 0.105037399 |
| Tjp2 | Y | | 218 | -0.116018064 | 0.385496345 | 0.809695128 | 0.925713192 | 0.424198783 | 0.400689127 | 0.036992936 | 0.000680331 | 0.000998637 | 0.029542327 |
| Cav1 | Y | | 25 | 0.187771112 | 0.22708851 | 0.773863536 | 0.586092423 | 0.546775026 | 0.121492696 | 0.158358815 | 0.022140432 | 0.043574494 | 0.049529699 |
| Fyb | Y | | 791 | 0.140916824 | 0.178274588 | 0.770618655 | 0.62970183 | 0.592344066 | 0.305111826 | 0.485098172 | 0.003556514 | 0.011689727 | 0.081129089 |
| Agfg1 | Y | | 327 | -0.044207869 | 0.166360589 | 0.659572808 | 0.703780677 | 0.493212219 | 0.779767157 | 0.295681484 | 0.048093378 | 0.043381082 | 0.10143448 |
| Ubap2 | Y | | 853 | -0.555919028 | -0.18340425 | -0.877385763 | -0.321466735 | -0.693981513 | 0.197522279 | 0.370185476 | 0.00825017 | 0.40969165 | 0.022848923 |
| Cwc22 | Y | | 47 | -0.422843541 | -0.598120925 | -0.879354209 | -0.456510668 | -0.281233284 | 0.000566525 | 0.053156408 | 1.1021E-05 | 0.000303534 | 0.220458441 |
| Sf3b1 | Y | | 44 | -0.504134526 | -0.835932198 | -1.038813658 | -0.534679132 | -0.20288146 | 0.034793697 | 0.084623382 | 0.000112513 | 0.040306187 | 0.550115407 |
| Map3k3 | Y | | 155 | -0.537440559 | -0.559043813 | -0.907868727 | -0.370428168 | -0.348824914 | 0.021203019 | 0.015459511 | 0.002681519 | 0.089272126 | 0.096607782 |
| Igbp1 | Y | | 278 | -0.394337546 | -0.569331033 | -0.844316868 | -0.449979322 | -0.274985834 | 0.025126891 | 0.006739323 | 0.001604516 | 0.025048638 | 0.099142296 |
| Cnn2 | Y | | 231 | -0.538047883 | -0.528601274 | -0.954630734 | -0.41658285 | -0.42602946 | 0.044556551 | 0.053265661 | 0.002185088 | 0.076880296 | 0.083544543 |
| Eif4h | Y | | 12 | -0.556088801 | -0.667556437 | -0.78127071 | -0.225181909 | -0.113714273 | 0.015877058 | 0.070615575 | 0.003580733 | 0.095306958 | 0.666130615 |
| Plec | Y | | 4619 | -0.324527252 | -0.527994929 | -0.596476528 | -0.271949276 | -0.068481598 | 0.00981848 | 0.001302672 | 0.003798425 | 0.07372141 | 0.54496502 |
| Tagln2 Tagln3 | Y | 192;192 | | -0.475705981 | -0.549020044 | -0.87277179 | -0.397065808 | -0.323751746 | 0.02922022 | 0.033205122 | 0.003705477 | 0.060458168 | 0.139786447 |
| Pkp4 | S | | 1136 | -0.555965936 | -0.55304577 | -0.631483518 | -0.075517582 | -0.078437747 | 0.012512076 | 0.008580632 | 0.011228357 | 0.660202544 | 0.630885346 |
| Polr2a | Y | | 1916 | -0.500976563 | -0.379363489 | -0.62259298 | -0.121616417 | -0.243229491 | 0.006258893 | 0.01572401 | 0.002122513 | 0.227350671 | 0.025143985 |
| Strap | Y | | 342 | -0.584850199 | -0.655935797 | -1.065199928 | -0.480349729 | -0.409264131 | 0.039846107 | 0.098033938 | 0.01305373 | 0.12819934 | 0.263091722 |
| Vcp | Y | | 805 | -0.462071755 | -0.494706891 | -0.668192092 | -0.206120337 | -0.1734852 | 0.010610504 | 0.013046564 | 0.006947452 | 0.210101514 | 0.301451149 |
| Dbnl | Y | | 140 | -0.544260023 | -0.487851434 | -0.89279687 | -0.348536847 | -0.404945436 | 0.052592476 | 0.050403472 | 0.003262263 | 0.133648648 | 0.053906441 |
| Tjap1 | Y | | 337 | -0.382681687 | -0.568008817 | -0.706549689 | -0.323868002 | -0.138540872 | 0.034341855 | 0.008552824 | 0.004733816 | 0.067762518 | 0.345044113 |
| Rasa3 | Y | | 822 | -0.291610472 | -0.565721665 | -0.590426866 | -0.298816394 | -0.024705201 | 0.031049412 | 0.000691367 | 0.001514645 | 0.011575834 | 0.830727818 |
| Cdv3 | Y | | 120 | -0.55585396 | -0.507810003 | -0.734323032 | -0.178469071 | -0.226513029 | 0.010501821 | 0.080390916 | 0.008417226 | 0.304506417 | 0.363823683 |
| Skint9 | S | | 129 | -0.521450538 | -0.535342419 | -0.712588755 | -0.191138217 | -0.177246335 | 0.009812895 | 0.098015907 | 0.005832006 | 0.238102258 | 0.500403415 |
| Skint9 | S | | 132 | -0.521450538 | -0.535342419 | -0.712588755 | -0.191138217 | -0.177246335 | 0.009812895 | 0.098015907 | 0.005832006 | 0.238102258 | 0.500403415 |
| Ppp1r12b | Y | | 712 | -0.316568526 | -0.565243645 | -0.786899694 | -0.470331168 | -0.221656049 | 0.050533608 | 0.005901785 | 0.009971367 | 0.076053525 | 0.257993813 |
| Fip1l1 | Y | | 128 | -0.283876851 | -0.418139904 | -0.850568232 | -0.566691381 | -0.432428329 | 0.077748146 | 0.025432535 | 0.001027459 | 0.004666461 | 0.014392179 |
| Erbin | Y | | 1097 | -0.518324409 | -0.492917514 | -0.661823052 | -0.143498643 | -0.168905537 | 0.018133796 | 0.064159628 | 0.004364991 | 0.316116375 | 0.406917327 |
| Pkn2 | Y | | 623 | -0.572995583 | -0.425525189 | -0.698607445 | -0.125611862 | -0.273082256 | 0.010031457 | 0.115368378 | 0.005221053 | 0.355477149 | 0.253719587 |
| Mpzl1 | S | | 261 | -0.264805295 | -0.564696828 | -0.698708179 | -0.433902884 | -0.134011351 | 0.054007967 | 0.02145513 | 0.001373506 | 0.013747963 | 0.428305895 |
| Ercc3 | Y | | 56 | -0.63266058 | -0.469508169 | -0.974981584 | -0.342321004 | -0.505473415 | 0.051519837 | 0.16394562 | 0.007295035 | 0.1769627 | 0.119936074 |
| Cnn2 | Y | | 184 | -0.382835989 | -0.449774 | -0.78479042 | -0.401954432 | -0.335016421 | 0.053837058 | 0.034385551 | 0.002544717 | 0.027919296 | 0.057883467 |
| Ppp1r12a | Y | | 760 | -0.549456702 | -0.553605144 | -0.756597276 | -0.207140574 | -0.202992132 | 0.033190108 | 0.062171149 | 0.012069567 | 0.227037992 | 0.372353061 |
| Ppp2r5a | Y | | 474 | -0.500467897 | -0.441456629 | -0.726986919 | -0.226519022 | -0.28553029 | 0.021057004 | 0.063014786 | 0.005882718 | 0.185007035 | 0.17426985 |
| Palld | Y | | 747 | -0.42373586 | -0.45785003 | -0.700022487 | -0.276286627 | -0.242172457 | 0.020152055 | 0.022427591 | 0.009012727 | 0.15518682 | 0.202279154 |
| Cnn3 | Y | | 261 | -0.365397602 | -0.441125972 | -0.851518022 | -0.48612042 | -0.41039205 | 0.038292242 | 0.082565595 | 0.002421756 | 0.028710371 | 0.106918547 |
| Ppp1r12a | Y | | 910 | -0.421771544 | -0.513264731 | -0.654765585 | -0.232994041 | -0.141500854 | 0.029844249 | 0.024683509 | 0.006840145 | 0.009928149 | 0.325762241 |
| Iqsec2 | Y | | 123 | -0.358656437 | -0.493317803 | -0.842856513 | -0.484200075 | -0.349538709 | 0.048838727 | 0.031175279 | 0.00717915 | 0.051289553 | 0.132395705 |
| Rtn4 | S | | 694 | -0.577077105 | -0.400879623 | -0.886514184 | -0.309437079 | -0.485634561 | 0.020641786 | 0.108855482 | 0.015607667 | 0.237088177 | 0.111153261 |
| Lars | Y | | 475 | -0.720667983 | -0.560058898 | -1.096706373 | -0.37603839 | -0.536647475 | 0.10682914 | 0.166901431 | 0.019972806 | 0.372388928 | 0.214303056 |
| Tom1l2 | T | | 403 | -0.477438522 | -0.337482625 | -0.816885204 | -0.339446683 | -0.47940258 | 0.041169754 | 0.127244915 | 0.002013365 | 0.089815693 | 0.046764813 |
| Otud4 | Y | | 1040 | -0.122371246 | -0.490325746 | -0.688681979 | -0.566310732 | -0.198356233 | 0.386235036 | 0.000172118 | 0.00610418 | 0.017465853 | 0.140189296 |
| Fer | Y | | 402 | -0.51389055 | -0.451287903 | -0.68150171 | -0.16761116 | -0.230213807 | 0.004544973 | 0.079258353 | 0.03558942 | 0.432270075 | 0.369611403 |
| Inppl1 | Y | | 1136 | -0.235909302 | -0.471434133 | -0.594740555 | -0.358831253 | -0.123306422 | 0.101294972 | 0.00100545 | 0.009054646 | 0.043205007 | 0.288076901 |
| Dnm3 | Y | | 622 | -0.463591613 | -0.44889921 | -0.757242884 | -0.293651271 | -0.308343674 | 0.037577168 | 0.078115964 | 0.009826765 | 0.141388293 | 0.182559131 |

| | | | | | | | | | | | | |
|---|---|---|---|---|---|---|---|---|---|---|---|---|
| Zc3hav1 | Y | 508 | -0.44021199 | -0.440875742 | -0.8831389 | -0.44292691 | -0.442263158 | 0.027008374 | 0.047945139 | 0.031315528 | 0.164910641 | 0.166512656 |
| Cnn3 | Y | 182 | -0.419714348 | -0.445941423 | -0.785785328 | -0.36607098 | -0.339843906 | 0.098279284 | 0.049090938 | 0.006317953 | 0.113003002 | 0.034875661 |
| Ubap2l | Y | 855 | -0.334640329 | -0.463964638 | -0.795501077 | -0.460860747 | -0.331536439 | 0.217542554 | 0.030881272 | 0.004073905 | 0.111001556 | 0.071879372 |
| Ppp1r12a | Y | 764 | -0.196964299 | -0.540817406 | -0.618301838 | -0.421337539 | -0.077484432 | 0.047418585 | 0.001576752 | 0.037948696 | 0.101596399 | 0.674938293 |
| Arhgap5 | Y | 1109 | -0.466480963 | -0.412973486 | -0.755876076 | -0.289395113 | -0.34290259 | 0.026896626 | 0.076196486 | 0.020368268 | 0.211285433 | 0.177980788 |
| Son | Y | 2210 | -0.321273682 | -0.423411545 | -0.612199867 | -0.290926185 | -0.188788323 | 0.047835296 | 0.001763434 | 0.041267891 | 0.198205743 | 0.347292689 |
| Lnpep | Y | 70 | -0.468404247 | -0.497759516 | -0.611308082 | -0.142903835 | -0.113548566 | 0.064314982 | 0.017503896 | 0.029653156 | 0.539434556 | 0.567523867 |
| Sgsm3 | Y | 608 | -0.374864989 | -0.323960989 | -0.609331311 | -0.234466322 | -0.285370323 | 0.038877462 | 0.061973671 | 0.001824071 | 0.128855 | 0.08313591 |
| Plec | Y | 3040 | -0.349096845 | -0.288334016 | -0.764975894 | -0.415879049 | -0.476641878 | 0.054279969 | 0.107192426 | 0.002640407 | 0.019449709 | 0.013110048 |
| Atn1 Rere | Y | 874;1251 | -0.297614443 | -0.515331551 | -0.671923906 | -0.374309463 | -0.156592355 | 0.162781482 | 0.03795264 | 0.009428077 | 0.05048184 | 0.32270887 |
| Crip2 | Y | 198 | -0.361228325 | -0.28059453 | -0.673195189 | -0.311966864 | -0.392600658 | 0.017455075 | 0.053789279 | 0.015434854 | 0.12638027 | 0.07659167 |
| Otud4 | Y | 173 | -0.505224817 | -0.476855463 | -0.619512862 | -0.114288045 | -0.142657399 | 0.063794221 | 0.099253869 | 0.027107712 | 0.630785288 | 0.578876479 |
| Stam2 | Y | 374 | -0.301021931 | -0.337367755 | -0.855033234 | -0.554011303 | -0.517665479 | 0.105082586 | 0.234466731 | 0.00364258 | 0.02488144 | 0.105509645 |
| Tom1l2 | Y | 404 | -0.338878206 | -0.305656485 | -0.67267391 | -0.333795704 | -0.367017425 | 0.047140472 | 0.152650913 | 0.003834572 | 0.037764921 | 0.09410948 |
| Pla2g4a | Y | 534 | -0.225306153 | -0.313542093 | -0.663235965 | -0.437929812 | -0.349693872 | 0.203182198 | 0.108721454 | 0.000410006 | 0.055128396 | 0.093300771 |
| Faf2 | Y | 79 | -0.299234861 | -0.405976435 | -0.651461305 | -0.352226445 | -0.245484871 | 0.099441004 | 0.059494256 | 0.00871696 | 0.051800653 | 0.172492425 |
| Crip2 | Y | 77 | -0.330561233 | -0.3677846 | -0.74053566 | -0.409974427 | -0.37275106 | 0.224317407 | 0.0531855 | 0.008182085 | 0.159462762 | 0.085079117 |
| Yipf5 | S | 45 | -0.185283403 | -0.77926742 | -0.797147317 | -0.611863914 | -0.017879897 | 0.193774496 | 0.068915247 | 0.038773253 | 0.073875585 | 0.957191674 |
| Tmem63a | Y | 774 | -0.174819406 | -0.413066865 | -0.713209501 | -0.538390095 | -0.300142636 | 0.194511659 | 0.050223929 | 0.004650861 | 0.014107352 | 0.10848478 |
| Spp1 | S | 33 | -0.226096835 | -0.468530531 | -0.813068758 | -0.586971923 | -0.344538227 | 0.123492546 | 0.068388528 | 0.020787992 | 0.068642253 | 0.215862501 |
| Cr1l | Y | 441 | -0.104328502 | -0.387726969 | -0.641224913 | -0.536896411 | -0.253497944 | 0.4524325 | 0.023988275 | 0.001300976 | 0.016921376 | 0.082590446 |
| Trim28 | Y | 517 | -0.312517356 | -0.380814277 | -0.602356506 | -0.28983915 | -0.221542229 | 0.06036097 | 0.043218529 | 0.023070573 | 0.160847032 | 0.258347515 |
| Epb41l1 | Y | 862 | -0.368119622 | -0.525734426 | -0.793227326 | -0.425107704 | -0.2674929 | 0.108967978 | 0.181543292 | 0.029333056 | 0.148095328 | 0.468619368 |
| Ddr2 | S | 469 | -0.095505197 | -0.457539325 | -0.63967774 | -0.544172543 | -0.182138414 | 0.506021406 | 0.000794749 | 0.036172614 | 0.048827125 | 0.350187523 |
| Fbl | Y | 124 | -0.152348928 | -0.292760046 | -0.695621734 | -0.543272806 | -0.402861687 | 0.344322187 | 0.043611234 | 0.001573423 | 0.016778376 | 0.018953396 |
| Epb41l2 | Y | 606 | -0.357938793 | -0.287412479 | -0.606374132 | -0.24843534 | -0.318961653 | 0.015955168 | 0.093501691 | 0.033597162 | 0.235375211 | 0.165520253 |
| Fyn | S | 21 | -0.252352907 | -0.30424352 | -0.671598223 | -0.419245316 | -0.367354702 | 0.070328956 | 0.080884077 | 0.011116316 | 0.052590966 | 0.085660114 |
| Iqsec2 | Y | 89 | -0.236681958 | -0.409367676 | -0.811379498 | -0.57469754 | -0.402011822 | 0.157843237 | 0.103924567 | 0.022509326 | 0.039235156 | 0.130047593 |
| Anxa1 | S | 27 | -0.327486306 | -0.264500442 | -0.676322459 | -0.348836153 | -0.411822018 | 0.019871227 | 0.127416691 | 0.042168429 | 0.176184578 | 0.131865908 |
| Atp1a1 | Y | 260 | -0.415817916 | -0.371358845 | -0.585002911 | -0.169184995 | -0.213644066 | 0.385931293 | 0.084560115 | 0.021749477 | 0.698130927 | 0.150244022 |
| Anln | S | 653 | -0.30601074 | -0.260589108 | -0.802942294 | -0.496931554 | -0.542353185 | 0.32148306 | 0.162073477 | 0.020642202 | 0.1659604 | 0.072024182 |
| Plec | Y | 3369 | -0.169866728 | -0.229209774 | -0.646821245 | -0.476954517 | -0.417611471 | 0.114377006 | 0.117360991 | 0.012497067 | 0.045866268 | 0.054639485 |
| Plin3 | Y | 39 | -0.123723787 | -0.253020189 | -0.685253797 | -0.561530009 | -0.432233608 | 0.640532346 | 0.115998766 | 0.002661892 | 0.105562656 | 0.026470454 |
| Phtf1 | S | 569 | -0.192161985 | -0.251096689 | -0.661536627 | -0.469374641 | -0.410439937 | 0.383892438 | 0.30778089 | 0.010287605 | 0.087307241 | 0.142383309 |
| Phtf1 | T | 568 | -0.192161985 | -0.251096689 | -0.661536627 | -0.469374641 | -0.410439937 | 0.383892438 | 0.30778089 | 0.010287605 | 0.087307241 | 0.142383309 |
| Acap2 | Y | 742 | -0.099848463 | -0.287230065 | -0.631568256 | -0.531719793 | -0.344338191 | 0.179399865 | 0.086538172 | 0.035129673 | 0.060908649 | 0.133768575 |
| Paics | Y | 22 | -0.308403422 | 0.018385778 | -0.604534177 | -0.296130755 | -0.622919954 | 0.057439159 | 0.944054728 | 0.013981163 | 0.12324284 | 0.08522755 |
| Serpinh1 | Y | 134 | 0.158680674 | -0.323145546 | -0.688572585 | -0.847253259 | -0.365427039 | 0.705397414 | 0.107623749 | 0.00784282 | 0.128024215 | 0.030984649 |
| Usp9x | Y | 2380 | 0.328410571 | 0.535465262 | 0.625117855 | 0.296707285 | 0.089652594 | 0.014447756 | 0.006633584 | 0.000132147 | 0.026068902 | 0.412179401 |
| Magi1 | Y | 373 | 0.259773666 | 0.45048954 | 0.756116018 | 0.496342352 | 0.305626478 | 0.087796714 | 0.004856838 | 7.46092E-05 | 0.018367921 | 0.025987436 |
| Flnb | Y | 1530 | 0.807279644 | 0.485491638 | 0.642583172 | -0.164696471 | 0.157091535 | 0.175466931 | 0.033734934 | 3.89417E-05 | 0.717205592 | 0.29953659 |
| Dyrk1a | Y | 145 | 0.240100841 | 0.444777714 | 0.641440227 | 0.401339386 | 0.196662513 | 0.018922759 | 0.005828403 | 0.000563649 | 0.017876302 | 0.107273791 |
| Tns3 | S | 332 | 0.373819108 | 0.511433581 | 0.682189769 | 0.308370661 | 0.170756188 | 0.322045887 | 0.000553901 | 0.002615949 | 0.397328547 | 0.168493735 |
| Tns3 | Y | 354 | 0.373819108 | 0.511433581 | 0.682189769 | 0.308370661 | 0.170756188 | 0.322045887 | 0.000553901 | 0.002615949 | 0.397328547 | 0.168493735 |
| Egfr | Y | 1197 | 0.106992847 | 0.500193073 | 0.649309225 | 0.542316378 | 0.149116152 | 0.209823101 | 0.000845933 | 0.000243842 | 0.001483527 | 0.084034502 |
| Axl | Y | 773 | 0.517811218 | 0.497460535 | 0.671138875 | 0.153327656 | 0.17367834 | 0.148648056 | 0.013240084 | 0.002117587 | 0.594791491 | 0.242687345 |
| Laptm4a | T | 214 | 0.51015673 | 0.525871272 | 0.774030239 | 0.26387351 | 0.248158968 | 0.186972448 | 0.01839873 | 0.003475234 | 0.435897452 | 0.190263699 |
| Sorbs2 | Y | 971 | 0.421425901 | 0.359964833 | 0.653096314 | 0.231670413 | 0.293131481 | 0.021653806 | 0.008376239 | 0.010280138 | 0.181526322 | 0.095238601 |
| Mxra8 | T | 402 | 0.201095319 | 0.547537141 | 0.659501188 | 0.458405869 | 0.111964048 | 0.010267061 | 0.003888358 | 0.0202046 | 0.061165855 | 0.472827751 |
| Laptm4a | Y | 210 | 0.503853901 | 0.561380102 | 0.771992772 | 0.268138871 | 0.21061267 | 0.132679212 | 0.045689809 | 0.006455289 | 0.357261949 | 0.348223116 |
| Hipk3 | S | 362 | 0.356851201 | 0.429760954 | 0.59122852 | 0.23437732 | 0.161467567 | 0.00953743 | 0.008975774 | 0.020881894 | 0.165361888 | 0.307803168 |
| Vta1 | Y | 280 | 0.38711587 | 0.539898534 | 0.609128469 | 0.222012599 | 0.069229935 | 0.177875723 | 0.009853635 | 0.004860274 | 0.37700354 | 0.594149898 |
| Actn1 Actn2 Actn3 Actn4 | Y | 339;326;332;319 | 0.262767197 | 0.343739442 | 0.754375753 | 0.491608556 | 0.410636311 | 0.068691967 | 0.033922587 | 0.001137605 | 0.014325514 | 0.027573058 |
| Itga3 | Y | 1030 | 0.215057704 | 0.45562066 | 0.778329998 | 0.563269294 | 0.322706338 | 0.174075752 | 0.002170916 | 0.010990522 | 0.028796066 | 0.109934941 |
| Pard3 | Y | 489 | 0.314183465 | 0.392602277 | 0.604207395 | 0.29002393 | 0.211605118 | 0.011843552 | 0.004940961 | 0.025544926 | 0.154964222 | 0.25629832 |
| Peak1 | S | 640 | 0.738122104 | 0.630177965 | 0.672809945 | -0.065312159 | 0.04263198 | 0.207852652 | 0.057947113 | 0.018914241 | 0.892317853 | 0.870178428 |
| Rnh1 | Y | 146 | 0.727563506 | 0.346819261 | 0.769313473 | 0.041749967 | 0.422494213 | 0.145588022 | 0.438506614 | 0.001510621 | 0.910534277 | 0.359303617 |
| Pik3r2 | Y | 447 | 0.078448752 | 0.579468377 | 0.648195888 | 0.569747136 | 0.068727512 | 0.471445586 | 0.001592157 | 0.009952046 | 0.027119277 | 0.628915843 |
| Sh3d19 | Y | 738 | 0.440019244 | 0.533862805 | 0.644813734 | 0.20479449 | 0.110950928 | 0.180432657 | 0.022348704 | 0.018844099 | 0.485854812 | 0.542986845 |
| Cilk1 | T | 157 | 0.326593475 | 0.367199866 | 0.709102728 | 0.382509253 | 0.341902862 | 0.011428173 | 0.069077556 | 0.021952088 | 0.116923127 | 0.152124974 |
| Cilk1 | Y | 159 | 0.326593475 | 0.367199866 | 0.709102728 | 0.382509253 | 0.341902862 | 0.011428173 | 0.069077556 | 0.021952088 | 0.116923127 | 0.152124974 |
| Ctnna1 | Y | 619 | 0.339959309 | 0.391867565 | 0.710549502 | 0.370590193 | 0.318681937 | 0.054422487 | 0.131113819 | 0.005612888 | 0.068196692 | 0.205181952 |
| Dlg3 | Y | 705 | 0.284635853 | 0.362855347 | 0.616651812 | 0.332015959 | 0.253796465 | 0.090268955 | 0.010939929 | 0.01212685 | 0.097569994 | 0.140885694 |
| Peak1 | Y | 528 | 0.260046974 | 0.47338525 | 0.720672678 | 0.460625704 | 0.247287428 | 0.024902027 | 0.087565194 | 0.021488864 | 0.049606228 | 0.291626589 |
| Snrnp70 | Y | 126 | 0.438208956 | 0.221995812 | 0.587747971 | 0.149539014 | 0.365752158 | 0.008666797 | 0.039298295 | 0.031904168 | 0.413076993 | 0.105758819 |
| Stat3 | Y | 705 | 0.10084719 | 0.463722951 | 0.595784944 | 0.494937755 | 0.132061993 | 0.478380652 | 0.001477798 | 0.015169546 | 0.034658138 | 0.366486199 |
| Dab2 | Y | 170 | 0.301921581 | 0.210412335 | 0.671364666 | 0.369443085 | 0.460952331 | 0.026942549 | 0.407005602 | 0.001358904 | 0.024207402 | 0.131947782 |
| Ctnna1 Ctnna2 | Y | 419;417 | 0.340721079 | 0.340145536 | 0.83820227 | 0.497481191 | 0.498056734 | 0.07642826 | 0.07097121 | 0.035719905 | 0.131634855 | 0.131916877 |
| Lyn | Y | 508 | 0.227628846 | 0.4022197 | 0.662088621 | 0.434459776 | 0.259868921 | 0.41128608 | 0.026859617 | 0.005208289 | 0.169085129 | 0.126662297 |
| Ncl | Y | 526 | 0.301511622 | 0.464630176 | 0.687680404 | 0.386168782 | 0.223050228 | 0.120639151 | 0.061706132 | 0.026887606 | 0.134829924 | 0.367268381 |
| Kank2 | Y | 362 | 0.295858591 | 0.317783876 | 0.703724233 | 0.407865642 | 0.385940356 | 0.133215262 | 0.249406448 | 0.00296308 | 0.078619072 | 0.191327098 |
| Tjp1 | Y | 1360 | 0.205770235 | 0.242590961 | 0.708926541 | 0.503156306 | 0.46633558 | 0.055273255 | 0.124121994 | 0.006880978 | 0.026776127 | 0.035431863 |
| Lpp | Y | 45 | 0.066062448 | 0.371590547 | 0.640316857 | 0.57425441 | 0.268726311 | 0.622711352 | 0.002858431 | 0.01086801 | 0.018320978 | 0.095664415 |
| Pitpnm2 | Y | 1083 | 0.221462582 | 0.19631804 | 0.592504071 | 0.371041489 | 0.396186031 | 0.200121195 | 0.05711937 | 0.049616026 | 0.148514095 | 0.134652605 |
| Ctnna1 | Y | 177 | 0.155069433 | 0.197602672 | 0.602568855 | 0.447499421 | 0.404966183 | 0.031932894 | 0.303083309 | 0.029282246 | 0.06164766 | 0.103189512 |
| Dlg1 | Y | 399 | 0.118854542 | 0.29966272 | 0.789485763 | 0.670631221 | 0.489823042 | 0.571086015 | 0.131083332 | 0.043096007 | 0.073371764 | 0.156268723 |
| Trim16 | Y | 523 | 0.04315253 | 0.294560969 | 0.786775673 | 0.743623143 | 0.492214704 | 0.901948084 | 0.370874124 | 0.006914983 | 0.179760715 | 0.216199101 |
| Cct2 | Y | 297 | 0.454319719 | 0.130680331 | 0.667203065 | 0.212883347 | 0.536522734 | 0.357069237 | 0.838669856 | 0.039563659 | 0.621189865 | 0.424579253 |
| Dapk3 | Y | 314 | 0.11312971 | 0.043969947 | 0.586656469 | 0.473526759 | 0.542686522 | 0.491785635 | 0.860150265 | 0.010531548 | 0.039971982 | 0.101101462 |
| Ptprj | Y | 1212 | 0.308508897 | 0.188602142 | -0.534316453 | -0.84282535 | -0.722918595 | 0.324302954 | 0.154424818 | 0.062148426 | 0.049927527 | 0.043619868 |
| Ybx1 | S | 207 | 0.116452537 | -0.36913699 | -0.507421843 | -0.623874381 | -0.138284853 | 0.248726857 | 0.009419371 | 0.002490948 | 0.002843255 | 0.127997393 |
| Ereg | Y | 149 | 0.127288569 | -0.388969793 | -0.570704481 | -0.69799305 | -0.181734688 | 0.402967713 | 0.058816668 | 0.055864213 | 0.031506301 | 0.431726402 |
| Ctnnd1 | Y | 257 | 0.055116105 | -0.108463202 | -0.582260973 | -0.637377078 | -0.473797771 | 0.698018538 | 0.341267099 | 0.030840249 | 0.020903142 | 0.041388432 |
| Baz1a | Y | 1295 | 0.242891076 | -0.515888096 | -0.39542639 | -0.638317467 | 0.120461706 | 0.037482859 | 0.043727108 | 0.020095947 | 0.005181356 | 0.500718898 |
| Pkp4 | Y | 1141 | 0.105218416 | -0.091268685 | -0.483816887 | -0.589035303 | -0.392548202 | 0.394567614 | 0.473605756 | 0.015036958 | 0.010907594 | 0.042344021 |
| Bclaf1 | Y | 80 | 0.250732358 | -0.409580407 | -0.408493122 | -0.659225481 | 0.001087284 | 0.012940408 | 0.073036467 | 0.020390396 | 0.011593857 | 0.994994549 |
| Dyrk3 | Y | 368 | 0.389937712 | -0.223494152 | -0.208597741 | -0.598535453 | 0.014896411 | 0.036112568 | 0.169913831 | 0.084898189 | 0.009426304 | 0.917612715 |
| Ptpn11 | Y | 542 | -0.033838161 | 0.433768939 | 0.566542468 | 0.600380629 | 0.132773529 | 0.73781537 | 4.34216E-05 | 7.57999E-05 | 0.010253216 | 0.043237037 |
| Gab1 | Y | 628 | -0.185093894 | 0.343198829 | 0.502458439 | 0.687552333 | 0.159259609 | 0.017124222 | 0.05970681 | 0.000143364 | 9.84015E-05 | 0.260742005 |
| Ephb4 | Y | 574 | -0.186500427 | 0.16095189 | 0.469713673 | 0.6562141 | 0.308761783 | 0.121047133 | 0.173753915 | 0.019970087 | 0.010909327 | 0.076711584 |
| Gab1 | Y | 660 | -0.338113827 | 0.18139242 | 0.341488659 | 0.679602486 | 0.160096239 | 0.029131743 | 0.15472436 | 0.061564688 | 0.017935809 | 0.254480002 |

| | | | | | | | | | | | | |
|---|---|---|---|---|---|---|---|---|---|---|---|---|
| Shb | Y | 240 | -0.259233734 | 0.072854236 | 0.354964082 | 0.614197817 | 0.282109847 | 0.030224118 | 0.421029384 | 0.030751878 | 0.014758894 | 0.081656549 |
| Zc3h13 | Y | 669 | 0.407954397 | -0.229768809 | -0.295575074 | -0.703529471 | -0.065806264 | 0.016124303 | 0.301724411 | 0.158586916 | 0.034752099 | 0.772318513 |
| Gab1 | Y | 407 | -0.291335096 | -0.366536051 | -0.562620886 | -0.271285789 | -0.196084834 | 0.003259337 | 0.001231084 | 0.000113292 | 0.005372682 | 0.019928681 |
| Srrt | Y | 23 | -0.543443973 | -1.022332543 | -0.455996745 | 0.087447228 | 0.566335798 | 0.002739006 | 0.074667154 | 0.006504517 | 0.147567506 | 0.217181623 |
| Eif3l | Y | 36 | -0.454835302 | -0.551538756 | -0.505613382 | -0.05077808 | 0.045925375 | 0.013385115 | 0.005944645 | 0.008811482 | 0.485572232 | 0.558944901 |
| Cuedc2 | Y | 272 | -0.426696258 | -0.521713516 | -0.521413516 | -0.094717258 | 0.0003 | 0.005219922 | 0.001588194 | 0.046761885 | 0.594433844 | 0.99854322 |
| Map1b | Y | 2038 | -0.549226572 | -0.557861049 | -0.546000552 | 0.00322602 | 0.011860498 | 0.004363822 | 0.045387913 | 0.013344666 | 0.979456058 | 0.953756708 |
| Bag3 | Y | 253 | -0.314557818 | -0.44546101 | -0.560223007 | -0.245665188 | -0.114761997 | 0.007644261 | 0.002078155 | 0.011819091 | 0.127481145 | 0.380394089 |
| Gprc5a | S | 344 | -0.121261694 | -0.509932336 | -0.564295365 | -0.443033671 | -0.05436303 | 0.202667427 | 0.001045715 | 0.000419769 | 0.003140481 | 0.476808591 |
| Mxra7 | Y | 117 | -0.367165648 | -0.451934639 | -0.410018748 | -0.0428531 | 0.041915891 | 0.008925494 | 0.00766023 | 0.001967932 | 0.602006446 | 0.660257182 |
| Cnn3 | T | 184 | -0.603005248 | -0.822213974 | -0.878625546 | -0.275620298 | -0.056411572 | 0.079949679 | 0.050584941 | 0.065381444 | 0.480311191 | 0.887284468 |
| Dennd2a | Y | 346 | -0.392297928 | -0.506002122 | -0.494960792 | -0.102662864 | 0.01104133 | 0.013151744 | 0.011064116 | 0.013073404 | 0.41087576 | 0.93782596 |
| Iqschfp Schip1 | Y | 276;351 | -0.544252668 | -0.364289954 | -0.709005828 | -0.16475316 | -0.344715874 | 0.042594018 | 0.003091414 | 0.054443875 | 0.527006966 | 0.212235239 |
| Son | S | 2208 | -0.224092241 | -0.418072003 | -0.526143035 | -0.302050794 | -0.108071032 | 0.022623726 | 0.027341084 | 0.000557053 | 0.010350642 | 0.400090805 |
| Rbm3 | Y | 151 | -0.473739552 | -0.395650448 | -0.420203636 | 0.053535916 | -0.024553188 | 0.007214934 | 0.013223366 | 0.021122153 | 0.662484482 | 0.831583583 |
| Efnb2 | Y | 307 | -0.189567314 | -0.494285289 | -0.518002691 | -0.328435377 | -0.023717402 | 0.104474743 | 0.002471233 | 0.003809012 | 0.024943768 | 0.792944611 |
| Snd1 | Y | 908 | -0.250132277 | -0.437467448 | -0.414853332 | -0.164721055 | 0.022614116 | 0.022617038 | 0.003030448 | 0.004394912 | 0.091523529 | 0.793590613 |
| Rin1 | Y | 35 | -0.1993901 | -0.383477563 | -0.562090683 | -0.362700583 | -0.17861312 | 0.069171211 | 0.000794377 | 0.008970328 | 0.035064413 | 0.177361504 |
| Ppp1r12a | T | 759 | -0.543815629 | -0.653807802 | -0.677288429 | -0.1334728 | -0.023480627 | 0.08829387 | 0.051195697 | 0.050955119 | 0.506540233 | 0.904344294 |
| Inppl1 | Y | 887 | -0.270368119 | -0.32933075 | -0.438596692 | -0.168228573 | -0.109265942 | 0.02269709 | 0.0139512 | 0.001608155 | 0.078346699 | 0.22353986 |
| Eps8 | Y | 45 | -0.392744801 | -0.345042269 | -0.569526821 | -0.17678202 | -0.224484552 | 0.004784011 | 0.100047112 | 0.016473945 | 0.262218445 | 0.272294284 |
| Lrp1 | Y | 4474 | -0.326189784 | -0.371055025 | -0.508660607 | -0.182470823 | -0.137605582 | 0.070332629 | 0.019605558 | 0.003225614 | 0.229138891 | 0.226634579 |
| Epha2 | T | 594 | -0.288194518 | -0.569670993 | -0.316628636 | -0.028434118 | 0.253042357 | 0.038853437 | 0.002557465 | 0.034095305 | 0.712126007 | 0.035546771 |
| Abi1 | Y | 213 | -0.291817817 | -0.341376759 | -0.489848797 | -0.19803098 | -0.148472038 | 0.027391564 | 0.018678304 | 0.00379784 | 0.117660603 | 0.220406141 |
| Ubap2 | Y | 1124 | -0.227322256 | -0.289903508 | -0.551214344 | -0.323892088 | -0.261310835 | 0.042083881 | 0.028488889 | 0.001020692 | 0.005424594 | 0.0341836 |
| Lpp | Y | 297 | -0.583798079 | -0.360811682 | -0.475499749 | 0.108298331 | -0.114688066 | 0.011938565 | 0.044927951 | 0.066097415 | 0.591294186 | 0.533391688 |
| Lpp | Y | 298 | -0.583798079 | -0.360811682 | -0.475499749 | 0.108298331 | -0.114688066 | 0.011938565 | 0.044927951 | 0.066097415 | 0.591294186 | 0.533391688 |
| Map1b | Y | 1970 | -0.392759503 | -0.394805604 | -0.380202506 | 0.012556997 | 0.014603098 | 0.00750785 | 0.008845911 | 0.078914806 | 0.934690355 | 0.922745573 |
| Eps8 | Y | 524 | -0.389297654 | -0.572878305 | -0.477879319 | -0.088581665 | 0.094998986 | 0.004077843 | 0.070459465 | 0.11654055 | 0.692736804 | 0.737882165 |
| Abi1 | Y | 198 | -0.351494447 | -0.353816761 | -0.317005752 | 0.034488695 | 0.036811009 | 0.007296828 | 0.008548658 | 0.032943822 | 0.722851114 | 0.690621949 |
| Abi1 | T | 215 | -0.307111959 | -0.332720462 | -0.517707419 | -0.21059546 | -0.184986957 | 0.077079704 | 0.03784876 | 0.003252701 | 0.184898842 | 0.184640189 |
| Syk | Y | 317 | -0.50754764 | -0.420232488 | -0.823220655 | -0.315673015 | -0.402988167 | 0.047725554 | 0.146076487 | 0.06930884 | 0.36661082 | 0.290245303 |
| Rbmx Rbmxl1 | Y | 310;313 | -0.279950399 | -0.361797948 | -0.507336406 | -0.227386008 | -0.145538458 | 0.058548656 | 0.025080132 | 0.007809656 | 0.020824979 | 0.081567405 |
| Pard3 | Y | 1123 | -0.275807542 | -0.340659292 | -0.375915304 | -0.100107762 | -0.035256011 | 0.075619863 | 0.000210322 | 0.047661708 | 0.482978457 | 0.751151642 |
| Rbm3 | Y | 124 | -0.191679887 | -0.416163986 | -0.417118409 | -0.225438523 | -0.000954423 | 0.042496983 | 0.001856622 | 0.029168572 | 0.146680455 | 0.993673459 |
| Cttn | Y | 154 | -0.53827166 | -0.328747892 | -0.44251862 | 0.09575304 | -0.113770728 | 0.026267138 | 0.078142908 | 0.026669559 | 0.532550329 | 0.318502986 |
| Rps27a | Y | 105 | -0.21386708 | -0.416106379 | -0.504561903 | -0.290694823 | -0.088455525 | 0.339202534 | 0.008174575 | 0.003653259 | 0.221438399 | 0.149645056 |
| Rps27a | Y | 106 | -0.21386708 | -0.416106379 | -0.504561903 | -0.290694823 | -0.088455525 | 0.339202534 | 0.008174575 | 0.003653259 | 0.221438399 | 0.149645056 |
| Tuba1a Tuba1b Tuba1c Tu | Y | 272;272;272;272; | -0.260792901 | -0.422361831 | -0.564023161 | -0.30323026 | -0.14166133 | 0.160597451 | 0.0254262 | 0.008885438 | 0.094333359 | 0.167269636 |
| Cttn | Y | 191 | -0.448146093 | -0.360523888 | -0.249495072 | 0.198651021 | 0.111028816 | 0.005895013 | 0.01329984 | 0.100366287 | 0.155107408 | 0.367537331 |
| Plec | Y | 4622 | -0.045861599 | -0.498988179 | -0.279239439 | -0.23337784 | 0.219748741 | 0.439686121 | 8.04202E-05 | 0.008650779 | 0.02203627 | 0.028625362 |
| Plcg1 | Y | 771 | -0.386818499 | -0.363169454 | -0.504136878 | -0.117318379 | -0.140967424 | 0.015825187 | 0.051411018 | 0.058261919 | 0.543950955 | 0.497947989 |
| Lrp6 | Y | 1541 | -0.358128769 | -0.282150665 | -0.470215425 | -0.112086656 | -0.188064759 | 0.010669788 | 0.033548991 | 0.037565502 | 0.465187761 | 0.271136191 |
| Cdk5rap2 | Y | 560 | -0.205512713 | -0.421279784 | -0.492906677 | -0.287393964 | -0.071626893 | 0.137674788 | 0.0156953 | 0.008607693 | 0.026145768 | 0.414369589 |
| Abi1 | T | 200 | -0.356196353 | -0.381833091 | -0.223552219 | 0.132644133 | 0.158280871 | 0.006483624 | 0.00489456 | 0.097889705 | 0.242368958 | 0.186560964 |
| Ptpn11 | Y | 580 | -0.416465632 | -0.262302451 | -0.467238408 | -0.050772776 | -0.204935957 | 0.014641105 | 0.104791904 | 0.016990799 | 0.667620661 | 0.180657972 |
| Mink1 | Y | 509 | -0.497993632 | -0.366195192 | -0.530767034 | -0.032773402 | -0.164571841 | 0.04419037 | 0.028359183 | 0.109515577 | 0.900913813 | 0.519106895 |
| Lbr | Y | 176 | -0.139074096 | -0.303743912 | -0.300378788 | -0.161304691 | 0.003365124 | 0.027647175 | 0.000213179 | 0.012307347 | 0.052976013 | 0.952266669 |
| Cttn | Y | 162 | -0.404085317 | -0.36538028 | -0.509622535 | -0.105537218 | -0.144242255 | 0.043343407 | 0.062246003 | 0.03396271 | 0.483832577 | 0.368328295 |
| Myo1e | Y | 988 | -0.25567468 | -0.425250996 | -0.320779413 | -0.065104733 | 0.104471582 | 0.036402433 | 0.005937779 | 0.034374594 | 0.501138703 | 0.312973475 |
| Ddr2 | Y | 740 | -0.259547623 | -0.380002185 | -0.479604865 | -0.220057242 | -0.09960268 | 0.100628394 | 0.016341157 | 0.018327119 | 0.176661411 | 0.436090124 |
| Map1b | Y | 2019 | -0.305438715 | -0.491119333 | -0.364616229 | -0.059177515 | 0.126503104 | 0.098697174 | 0.00460613 | 0.113805354 | 0.768495213 | 0.487368265 |
| Baiap2 | Y | 506 | -0.233683763 | -0.303180801 | -0.513881622 | -0.280197859 | -0.210700821 | 0.019535953 | 0.033066475 | 0.027059863 | 0.121946922 | 0.213389205 |
| Fer | Y | 715 | -0.372272044 | -0.231307442 | -0.437469171 | -0.065197127 | -0.206161729 | 0.005090026 | 0.274099093 | 0.012528488 | 0.547970859 | 0.327106433 |
| Tank | Y | 108 | -0.254399033 | -0.485333505 | -0.484601377 | -0.230202344 | 0.000732128 | 0.127399396 | 0.016744478 | 0.051914448 | 0.25113523 | 0.996763178 |
| Mia2 | Y | 1321 | -0.186907115 | -0.271921678 | -0.392808093 | -0.205900977 | -0.120886415 | 0.044485713 | 0.012439235 | 0.004129605 | 0.046845649 | 0.16687891 |
| Fyn | Y | 28 | -0.171015123 | -0.254064819 | -0.497848262 | -0.326833139 | -0.243783444 | 0.105703006 | 0.086343431 | 0.000715662 | 0.017025507 | 0.099645087 |
| Spred1 | S | 291 | -0.443152599 | -0.46238191 | -0.151541155 | 0.291611445 | 0.310840755 | 0.023811962 | 0.005475677 | 0.27231158 | 0.096411034 | 0.047406117 |
| Ybx1 | Y | 143 | -0.061503278 | -0.450047767 | -0.558235024 | -0.496731746 | -0.108187257 | 0.556526302 | 0.027675676 | 0.002912602 | 0.007551578 | 0.465225591 |
| Myo9b | Y | 1886 | -0.378671888 | -0.418434386 | -0.313327717 | 0.065344171 | 0.105106668 | 0.031105686 | 0.030850732 | 0.071255114 | 0.515771096 | 0.446031522 |
| Tnks1bp1 | Y | 202 | -0.22714222 | -0.531295293 | -0.4148295 | -0.18768728 | 0.116465793 | 0.086141267 | 0.005985968 | 0.170166682 | 0.464891021 | 0.642062674 |
| Nhsl1 | Y | 339 | -0.161107574 | -0.32561115 | -0.491221264 | -0.33011369 | -0.165610114 | 0.119289017 | 0.074488904 | 0.002233576 | 0.018329931 | 0.272069413 |
| Hnrnpk | S | 379 | -0.110216321 | -0.284185253 | -0.545041076 | -0.434824755 | -0.260855822 | 0.307223596 | 0.020413215 | 0.002247755 | 0.010024926 | 0.043372253 |
| Ica1 | Y | 261 | -0.333531881 | -0.1725689 | -0.485676537 | -0.152144657 | -0.313107637 | 0.012717512 | 0.201093434 | 0.010910351 | 0.244737096 | 0.070573734 |
| Nrp1 | S | 921 | -0.188575206 | -0.301746736 | -0.539561884 | -0.350986677 | -0.237815147 | 0.082702634 | 0.082990706 | 0.006468016 | 0.047061713 | 0.163448874 |
| Rassf8 | Y | 418 | -0.363438056 | -0.788656847 | -0.624019528 | -0.260581472 | 0.164637319 | 0.335688026 | 0.100647554 | 0.090331447 | 0.51057006 | 0.662875997 |
| Ctnnd1 | Y | 904 | -0.385636876 | -0.268508923 | -0.462305123 | -0.076668247 | -0.1937962 | 0.035175379 | 0.188191511 | 0.016435386 | 0.548139095 | 0.292520576 |
| Baiap2 | Y | 354 | -0.201652854 | -0.404197399 | -0.269972891 | -0.068320037 | 0.134224508 | 0.036131637 | 0.009061825 | 0.027738004 | 0.423402034 | 0.236750932 |
| Lpp | Y | 333 | -0.311346363 | -0.325447076 | -0.336010678 | -0.024664316 | -0.010563603 | 0.032452318 | 0.030254266 | 0.031736117 | 0.764391523 | 0.89220842 |
| Denn2b | Y | 485 | -0.35142224 | -0.384394849 | -0.274937548 | 0.076484692 | 0.109457301 | 0.088068306 | 0.014688937 | 0.032833495 | 0.612461317 | 0.259667259 |
| Pi4ka | S | 2090 | -0.150745691 | -0.308772696 | -0.473075246 | -0.322329555 | -0.164302551 | 0.504507747 | 0.015833548 | 0.002287963 | 0.212365038 | 0.037411193 |
| Cttn | S | 407 | -0.497386474 | -0.265255156 | -0.420286735 | 0.077099739 | -0.155031579 | 0.009750749 | 0.176200844 | 0.125658078 | 0.712944247 | 0.519945759 |
| Dctn2 | Y | 6 | -0.343496972 | -0.436773483 | -0.363927184 | -0.020430212 | 0.0728463 | 0.056213261 | 0.028097705 | 0.122085031 | 0.903566935 | 0.686049461 |
| Gdi2 | Y | 442 | -0.318387113 | -0.233297654 | -0.467259149 | -0.148872036 | -0.233961496 | 0.031593567 | 0.220642726 | 0.010009825 | 0.233369102 | 0.21852826 |
| 1110004F10Rik | Y | 84 | -0.226472014 | -0.26870026 | -0.441311247 | -0.214839232 | -0.172610987 | 0.056783209 | 0.119005626 | 0.004645673 | 0.012534205 | 0.251129225 |
| Cttn | Y | 104 | -0.182351124 | -0.431145407 | -0.419290855 | -0.236939731 | 0.011854552 | 0.099871547 | 0.041504006 | 0.019908241 | 0.108475789 | 0.939148093 |
| Acap2 | T | 741 | -0.121551628 | -0.243613864 | -0.498995117 | -0.377443489 | -0.255381253 | 0.251347994 | 0.066739047 | 0.000884801 | 0.031650151 | 0.082081353 |
| Anks1 | Y | 850 | -0.245496186 | -0.361349617 | -0.32685069 | -0.081354504 | 0.034498927 | 0.061236452 | 0.002905453 | 0.1340924 | 0.649729346 | 0.832025438 |
| Cnn3 | Y | 322 | -0.239366531 | -0.202637801 | -0.335608033 | -0.096241502 | -0.132970232 | 0.017349066 | 0.032121888 | 0.008536392 | 0.232911368 | 0.129337988 |
| Tjp2 | S | 213 | -0.420907629 | -0.370247416 | -0.087613002 | 0.333294627 | 0.282634413 | 0.002574866 | 0.016169388 | 0.457096777 | 0.035150655 | 0.065287107 |
| Dok1 | Y | 450 | -0.307811284 | -0.256485674 | -0.44637864 | -0.138567356 | -0.189892966 | 0.01957345 | 0.131124885 | 0.029521443 | 0.300758105 | 0.260248268 |
| Rbm3 | Y | 139 | -0.052495642 | -0.511850319 | -0.316120978 | -0.263625336 | 0.195729341 | 0.582984695 | 0.001009063 | 0.037616882 | 0.067815002 | 0.132226662 |
| Mapk14 | T | 180 | -0.39237185 | -0.246850184 | -0.318195716 | 0.074176133 | -0.071345533 | 0.012167475 | 0.073430629 | 0.064451923 | 0.551343734 | 0.577162663 |
| Spred1 | Y | 293 | -0.477901142 | -0.243288393 | -0.386587885 | 0.091313256 | -0.143299493 | 0.014917239 | 0.315764828 | 0.043456057 | 0.564048895 | 0.545470219 |
| Pard3 | Y | 1076 | -0.368975696 | -0.449972965 | -0.221150847 | 0.147824849 | 0.228822118 | 0.032624253 | 0.043676957 | 0.094155933 | 0.260361849 | 0.205509496 |
| Arhgef12 | Y | 66 | -0.248494647 | -0.300898161 | -0.53536085 | -0.286866203 | -0.234462689 | 0.160940706 | 0.129321958 | 0.00998225 | 0.074571972 | 0.171160095 |
| Rin1 | Y | 283 | -0.190143547 | -0.546396328 | -0.290087474 | -0.099943927 | 0.256308854 | 0.183616749 | 0.006827568 | 0.104972762 | 0.4230713 | 0.103232478 |
| Pard3b | Y | 1080 | -0.282747251 | -0.368844915 | -0.268069522 | 0.014677729 | 0.100775393 | 0.045871653 | 0.013024633 | 0.079289288 | 0.900822625 | 0.364537449 |
| Mapk14 | Y | 182 | -0.196925316 | -0.220649718 | -0.312860237 | -0.115934921 | -0.092210519 | 0.014132189 | 0.010826555 | 0.020356781 | 0.228508089 | 0.31801601 |
| Dcbld2 | Y | 560 | -0.143511826 | -0.198398034 | -0.563663135 | -0.420151308 | -0.365265101 | 0.293137239 | 0.17035864 | 0.000814993 | 0.034441009 | 0.048460799 |
| Ptpn6 | Y | 536 | -0.427546615 | -0.238198002 | -0.391970988 | 0.035575627 | -0.153772986 | 0.019837746 | 0.227899413 | 0.040236261 | 0.814786571 | 0.437285302 |

| | | | | | | | | | | | | |
|---|---|---|---|---|---|---|---|---|---|---|---|---|
| Lurap1l | Y | 218 | -0.214864511 | -0.397715216 | -0.510178731 | -0.29531422 | -0.112463515 | 0.013824496 | 0.098837274 | 0.100265102 | 0.25103675 | 0.659378537 |
| Snx9 | Y | 177 | -0.362728829 | -0.448690866 | -0.403264696 | -0.040535867 | 0.04542617 | 0.053614593 | 0.101718713 | 0.104170964 | 0.814139538 | 0.852139566 |
| Xkr5 | Y | 472 | -0.226470068 | -0.267633499 | -0.509660625 | -0.283190558 | -0.242027126 | 0.140156712 | 0.195559084 | 0.004891548 | 0.094784653 | 0.235337844 |
| Mpp5 | Y | 243 | -0.079928321 | -0.327581068 | -0.337426397 | -0.257498076 | -0.009845329 | 0.213424706 | 0.05233647 | 0.000376733 | 0.013411935 | 0.92492437 |
| Carhsp1 | S | 53 | -0.24345807 | -0.171771935 | -0.497008903 | -0.253550832 | -0.325236968 | 0.095560594 | 0.128190632 | 0.005310683 | 0.086060602 | 0.031456142 |
| Gab1 | Y | 259 | -0.094572917 | -0.300161589 | -0.463861465 | -0.369288548 | -0.163699876 | 0.046857688 | 0.004742585 | 0.057287772 | 0.087084555 | 0.309259319 |
| Ctnnd1 | S | 230 | -0.255046444 | -0.228882744 | -0.434073651 | -0.179027208 | -0.205190908 | 0.092927821 | 0.07136036 | 0.011509831 | 0.228029393 | 0.138901585 |
| Slc37a2 | Y | 262 | -0.37461635 | -0.547685807 | -0.680287655 | -0.305671305 | -0.132601848 | 0.607857522 | 0.050683798 | 0.105623788 | 0.682809623 | 0.680873149 |
| Dok1 | S | 415 | -0.226629323 | -0.437174122 | -0.233670085 | -0.007040762 | 0.203504037 | 0.083989554 | 0.009181567 | 0.082815184 | 0.914408605 | 0.034072136 |
| Rbm3 | Y | 132 | -0.165772782 | -0.351347761 | -0.336308406 | -0.170535624 | 0.015039355 | 0.061659274 | 0.008704847 | 0.066028031 | 0.25112565 | 0.91107492 |
| Ddr2 | Y | 735 | -0.132027508 | -0.306631369 | -0.520023049 | -0.387995542 | -0.213391681 | 0.331253902 | 0.036192462 | 0.011393201 | 0.035845679 | 0.145573478 |
| Iqsec1 | Y | 909 | -0.32601271 | -0.211681919 | -0.433340082 | -0.107327371 | -0.221658162 | 0.067585517 | 0.093605591 | 0.025585394 | 0.498778654 | 0.156912306 |
| Riok1 | Y | 465 | -0.149632588 | -0.376961824 | -0.566003839 | -0.41637125 | -0.189042014 | 0.515273247 | 0.074418939 | 0.010847112 | 0.123584861 | 0.299913755 |
| Sh3pxd2b | Y | 661 | -0.173236924 | -0.390566604 | -0.783466078 | -0.610229154 | -0.392899474 | 0.187873621 | 0.116401518 | 0.068412329 | 0.096686449 | 0.229152619 |
| Cttn | Y | 265 | -0.187606033 | -0.274990255 | -0.275993073 | -0.08838704 | -0.001002818 | 0.032016834 | 0.042995503 | 0.007155754 | 0.083615274 | 0.991185746 |
| Atp2b1 Atp2b2 | Y | 1107;1129 | -0.379644449 | -0.252137247 | -0.351745305 | 0.027899144 | -0.099608058 | 0.091513473 | 0.094105836 | 0.022105615 | 0.870574294 | 0.420594038 |
| Cnn2 | Y | 295 | -0.0739836 | -0.220267001 | -0.418371713 | -0.344388114 | -0.198104713 | 0.724262946 | 0.04604856 | 0.000182417 | 0.193295857 | 0.067040513 |
| Tjp1 | Y | 1164 | -0.453506066 | -0.452240059 | -0.215477714 | 0.238028352 | 0.236762344 | 0.021971976 | 0.090707847 | 0.299374852 | 0.251861805 | 0.345490596 |
| Axl | Y | 860 | -0.316424424 | -0.277188501 | -0.181378579 | 0.135045845 | 0.095809922 | 0.01156428 | 0.021895691 | 0.099731054 | 0.114783274 | 0.211900031 |
| Mtmr10 | Y | 704 | -0.072690389 | -0.195326751 | -0.414905095 | -0.342214706 | -0.219578344 | 0.539046947 | 0.024322119 | 0.000429445 | 0.056698232 | 0.015311175 |
| Cttn | Y | 228 | -0.39654932 | -0.227480499 | -0.210324162 | 0.186225158 | 0.017156337 | 0.009478543 | 0.068685846 | 0.086397631 | 0.029722956 | 0.7866701 |
| Ociad1 | Y | 201 | -0.018075025 | -0.398992208 | -0.301460215 | -0.28338519 | 0.097531993 | 0.788087579 | 0.002566002 | 0.006468418 | 0.000776457 | 0.020338719 |
| Srsf2 | Y | 115 | -0.032537666 | -0.368496599 | -0.228406611 | -0.195868945 | 0.140089987 | 0.795885731 | 0.000537892 | 0.00574618 | 0.200329181 | 0.008405232 |
| Efnb1 | Y | 312 | -0.271773712 | -0.291195064 | -0.3990764 | -0.127302688 | -0.107881336 | 0.213398384 | 0.093585988 | 0.012029581 | 0.495251189 | 0.412322619 |
| Lurap1l | Y | 217 | -0.023788959 | -0.325443662 | -0.556949043 | -0.533160083 | -0.231505381 | 0.7878099 | 0.010033277 | 0.018250481 | 0.018620921 | 0.14753565 |
| Bet1 | Y | 18 | -0.271990391 | -0.175590383 | -0.347333106 | -0.075342715 | -0.171742723 | 0.06773193 | 0.169441589 | 0.005392184 | 0.508586084 | 0.167933375 |
| Klc1 | Y | 448 | 0.016185374 | -0.343476968 | -0.547352725 | -0.5635381 | -0.203875757 | 0.911806164 | 0.040838103 | 0.004370899 | 0.023943674 | 0.168467741 |
| Cttn | Y | 334 | -0.320754144 | -0.238427515 | -0.256338148 | 0.064415996 | -0.017910633 | 0.021432756 | 0.078737952 | 0.062889251 | 0.502766816 | 0.863740445 |
| Sipa1l1 | S | 1564 | -0.247823991 | -0.227771278 | -0.339615818 | -0.091791827 | -0.11184454 | 0.019263078 | 0.122861931 | 0.040060345 | 0.440985784 | 0.448162449 |
| Sipa1l1 | Y | 1569 | -0.247823991 | -0.227771278 | -0.339615818 | -0.091791827 | -0.11184454 | 0.019263078 | 0.122861931 | 0.040060345 | 0.440985784 | 0.448162449 |
| Cdc42ep1 | Y | 349 | -0.513512041 | -0.081707799 | -1.016013431 | -0.50250139 | -0.934305632 | 0.056254977 | 0.799210706 | 0.16635202 | 0.419173213 | 0.191958875 |
| Slc6a8 | Y | 11 | -0.068655068 | -0.292845795 | -0.364043053 | -0.295387985 | -0.071197257 | 0.439379594 | 0.002239101 | 0.030967053 | 0.057913158 | 0.503290997 |
| Hnrnpul1 | Y | 111 | -0.326967512 | -0.360901484 | -0.317367421 | 0.009600091 | 0.043534066 | 0.104128468 | 0.04658608 | 0.143151935 | 0.958514152 | 0.77534306 |
| Abi2 | Y | 207 | -0.141951131 | -0.310517972 | -0.310522192 | -0.168571061 | -4.22024E-06 | 0.232285698 | 0.041931604 | 0.008753131 | 0.155833972 | 0.999968105 |
| Wee1 | Y | 19 | -0.447019706 | -0.24154567 | -0.132624901 | 0.314394806 | 0.108920769 | 0.008741734 | 0.032977079 | 0.513864061 | 0.187978202 | 0.58605148 |
| Itsn2 | Y | 922 | -0.175618482 | -0.237097809 | -0.33533623 | -0.159717748 | -0.098238421 | 0.034215507 | 0.085044351 | 0.02425618 | 0.148978989 | 0.424813788 |
| Ctnnd1 | Y | 280 | -0.152836363 | -0.149836055 | -0.529491222 | -0.376654859 | -0.379655167 | 0.171552594 | 0.278115079 | 0.004993608 | 0.024667491 | 0.041088271 |
| D1Pas1 Ddx3x Ddx3y | Y | 69;70;67 | -0.123976666 | -0.420881964 | -0.386380186 | -0.26240352 | 0.034501778 | 0.284949088 | 0.048186901 | 0.044744335 | 0.125456755 | 0.841820189 |
| Gprc5a | Y | 346 | -0.140427249 | -0.231493855 | -0.5487194 | -0.408292151 | -0.317225545 | 0.351027957 | 0.173660545 | 0.009519572 | 0.046251266 | 0.102416922 |
| Cdc20 | Y | 152 | -0.003957132 | -0.285950998 | -0.474049357 | -0.470092224 | -0.188098359 | 0.96458862 | 0.020444497 | 0.006276119 | 0.011072589 | 0.126980179 |
| Baiap2 | Y | 338 | -0.049063917 | -0.379681524 | -0.556349893 | -0.507285975 | -0.176668369 | 0.554966609 | 0.075940592 | 0.022383283 | 0.015425653 | 0.306951337 |
| Myl6 | Y | 86 | -0.261557849 | -0.173711687 | -0.315392647 | -0.053834798 | -0.14168096 | 0.048252858 | 0.158621071 | 0.015264248 | 0.514879727 | 0.173322946 |
| Eif4enif1 | Y | 781 | 0.062594992 | -0.492745936 | -0.423378461 | -0.485973454 | 0.069367475 | 0.762135749 | 0.013613165 | 0.021777116 | 0.08450971 | 0.481971497 |
| Mapk14 | T | 185 | -0.158749025 | -0.197104844 | -0.817112289 | -0.658363264 | -0.620007444 | 0.134212304 | 0.050703345 | 0.142379341 | 0.199809553 | 0.219036965 |
| Dlg5 | Y | 1197 | -0.133426349 | -0.495456974 | -0.305771085 | -0.172344736 | 0.189685888 | 0.155900334 | 0.044915328 | 0.099973605 | 0.278839206 | 0.346581031 |
| Emd | Y | 100 | -0.090914768 | -0.234680522 | -0.563816811 | -0.472902043 | -0.329136289 | 0.266995176 | 0.085235748 | 0.022094604 | 0.053423331 | 0.096156352 |
| Tuba1a Tuba1b Tuba1c Tu | Y | 224;224;224;224; | -0.112485762 | -0.381923599 | -0.369875168 | -0.257389406 | 0.012048431 | 0.661905551 | 0.014244531 | 0.043179215 | 0.353970698 | 0.917220361 |
| Pkp4 | Y | 477 | -0.192799344 | -0.291318989 | -0.184468221 | 0.008331123 | 0.106850768 | 0.045608405 | 0.011532465 | 0.098524286 | 0.919698307 | 0.264904855 |
| Hnrnpa1 | Y | 289 | 0.00942614 | -0.3556684 | -0.439554196 | -0.448980336 | -0.083885797 | 0.960485131 | 0.021318504 | 0.010582974 | 0.075792675 | 0.35315697 |
| Ddr2 | Y | 481 | -0.104992634 | -0.261679757 | -0.190991031 | -0.085998398 | 0.070688726 | 0.108716017 | 0.011092738 | 0.006830273 | 0.135292969 | 0.28408523 |
| Memo1 | Y | 210 | -0.109669044 | -0.252942216 | -0.270422369 | -0.160753325 | -0.017480153 | 0.042412795 | 0.027390362 | 0.026070508 | 0.109396714 | 0.848599512 |
| Map1b | Y | 2021 | -0.350468227 | -0.181781956 | -0.205029909 | 0.145438318 | -0.023247953 | 0.064236224 | 0.054429678 | 0.037245915 | 0.323386905 | 0.739856623 |
| Tmem106b | Y | 51 | -0.221463829 | -0.310574826 | -0.196671494 | 0.024792335 | 0.113903332 | 0.054781596 | 0.014133976 | 0.187829158 | 0.838834563 | 0.370515557 |
| Chchd3 | Y | 53 | 0.054412792 | -0.357822406 | -0.367380587 | -0.42179338 | -0.009558181 | 0.552260534 | 0.006927833 | 0.00684848 | 0.009304742 | 0.916433104 |
| Cttn | Y | 302 | -0.363931139 | -0.193692329 | -0.301600399 | 0.06233074 | -0.10790807 | 0.036603584 | 0.189909233 | 0.094938404 | 0.696439947 | 0.510054485 |
| Igf1r Insr | Y | 1167;1179 | -0.417701576 | -0.229560273 | -0.208289157 | 0.209412419 | 0.021271116 | 0.013176438 | 0.195752555 | 0.233663578 | 0.197469471 | 0.900682041 |
| Ptprk | Y | 870 | -0.207174988 | -0.179907156 | -0.440644648 | -0.23346966 | -0.260737492 | 0.121802166 | 0.278561866 | 0.01495212 | 0.090385138 | 0.136639748 |
| Mpp5 | S | 245 | -0.132781818 | -0.332513987 | -0.678059648 | -0.54527783 | -0.345545661 | 0.434800773 | 0.106177742 | 0.084316639 | 0.120527843 | 0.269372515 |
| Nectin3 | Y | 511 | -0.088107149 | -0.234403862 | -0.296143541 | -0.208036392 | -0.061739679 | 0.278057555 | 0.051703636 | 0.002819744 | 0.039374935 | 0.490098923 |
| Pard3b | Y | 1170 | -0.161911447 | -0.233945187 | -0.270674064 | -0.108762617 | -0.036728878 | 0.045584802 | 0.074449714 | 0.025473453 | 0.237962722 | 0.738029044 |
| Ctnnd1 | Y | 228 | -0.154677362 | -0.112352428 | -0.347677397 | -0.193000035 | -0.235324969 | 0.059176525 | 0.054116143 | 0.011845562 | 0.07349321 | 0.045684469 |
| Samd4 | Y | 171 | -0.066780109 | -0.343718057 | -0.394486714 | -0.327706605 | -0.050768657 | 0.476635304 | 0.020283573 | 0.05235074 | 0.078189245 | 0.725161182 |
| Herc2 | Y | 4798 | -0.178402654 | -0.335794464 | -0.248117381 | -0.069714726 | 0.087677083 | 0.143834628 | 0.009294804 | 0.215790771 | 0.6924537 | 0.600806726 |
| Baiap2 | Y | 472 | -0.183221162 | -0.14220049 | -0.43431664 | -0.251095477 | -0.29211615 | 0.184626664 | 0.126811696 | 0.014380842 | 0.109099341 | 0.039146515 |
| Pttg1ip | Y | 171 | -0.190510972 | -0.184362932 | -0.482759725 | -0.292248752 | -0.298396793 | 0.258155291 | 0.233188387 | 0.015655636 | 0.055279612 | 0.018739273 |
| Fes | Y | 713 | -0.156811688 | -0.185742826 | -0.371311886 | -0.214500198 | -0.18556906 | 0.042057523 | 0.157719093 | 0.02975053 | 0.125447607 | 0.213747558 |
| Baiap2 | S | 493 | -0.203774804 | -0.222694159 | -0.229055529 | -0.025280725 | -0.00636137 | 0.089390206 | 0.058341523 | 0.023961615 | 0.795660342 | 0.944737802 |
| Rbmx Rbmxl1 | Y | 211;214 | -0.176866814 | -0.270709675 | -0.280261164 | -0.10339435 | -0.00955149 | 0.143118955 | 0.053362596 | 0.04416535 | 0.023506792 | 0.881391643 |
| Anln | Y | 666 | 0.037231282 | -0.328703431 | -0.438807807 | -0.476039089 | -0.110104376 | 0.872302369 | 0.014901435 | 0.020733574 | 0.125715357 | 0.395712983 |
| Ybx1 | Y | 206 | 0.021023229 | -0.479789537 | -0.312394225 | -0.333417453 | 0.167395313 | 0.873991397 | 0.034739156 | 0.011175449 | 0.063980912 | 0.301516782 |
| Tkt | Y | 275 | -0.132610791 | -0.203323252 | -0.421247819 | -0.288637028 | -0.217924567 | 0.575245547 | 0.127798814 | 0.005685657 | 0.27861494 | 0.122877514 |
| Ttc1 | Y | 280 | -0.230658104 | -0.142035458 | -0.345150554 | -0.114492449 | -0.203115095 | 0.014858957 | 0.061057262 | 0.178566212 | 0.583148498 | 0.365219531 |
| Mertk Tyro3 | Y | 748;675 | -0.246102977 | -0.217502924 | -0.172293408 | 0.073809569 | 0.045209516 | 0.027769822 | 0.045223243 | 0.103123408 | 0.27881072 | 0.533618538 |
| Xaf1 | Y | 235 | -0.418539114 | -0.19579007 | -0.147950869 | 0.270588245 | 0.047839201 | 0.009053613 | 0.217374788 | 0.280229827 | 0.082722273 | 0.740873762 |
| Rbm3 | S | 133 | -0.105835985 | -0.366199919 | -0.28942143 | -0.183585445 | 0.076778489 | 0.24237653 | 0.02084635 | 0.1169246 | 0.260731798 | 0.622728899 |
| Timp3 | Y | 70 | -0.195110276 | -0.177107258 | -0.323560782 | -0.128450507 | -0.146453525 | 0.158059805 | 0.224405488 | 0.008143092 | 0.301856679 | 0.288939576 |
| Tagln | Y | 193 | -0.223552849 | -0.307339265 | -0.384968764 | -0.161415915 | -0.0776295 | 0.22272062 | 0.117329856 | 0.089480329 | 0.283310544 | 0.569167324 |
| Macf1 | Y | 2930 | -0.32332116 | -0.233539987 | -0.12986587 | 0.19345529 | 0.103674117 | 0.021585371 | 0.05366403 | 0.274800922 | 0.091977075 | 0.247087035 |
| Map4 | Y | 34 | -0.430952273 | -0.61574232 | -0.20066095 | 0.230291324 | 0.41508137 | 0.349260676 | 0.060463924 | 0.539528886 | 0.629286518 | 0.273057516 |
| Lima1 | Y | 190 | -0.162087929 | -0.454711883 | -0.056307196 | 0.105780732 | 0.398404687 | 0.11459835 | 0.00449482 | 0.584499542 | 0.195877579 | 0.018571792 |
| Hdgf | Y | 100 | -0.225629328 | -0.318363732 | -0.658437282 | -0.432807953 | -0.34007355 | 0.403827609 | 0.277902855 | 0.100175208 | 0.250948744 | 0.355519035 |
| Cttn | Y | 199 | -0.28410273 | -0.304837362 | -0.165701466 | 0.118401263 | 0.139135895 | 0.061242979 | 0.055273269 | 0.25259056 | 0.224250904 | 0.211823037 |
| Syk | Y | 520 | -0.244618028 | -0.23321492 | -0.162417547 | 0.082200481 | 0.070797374 | 0.025258172 | 0.048039164 | 0.210638121 | 0.463158677 | 0.547471188 |
| Ptk2 | T | 406 | -0.342928891 | -0.124906079 | -0.298827201 | 0.04410169 | -0.173921122 | 0.007782141 | 0.692629385 | 0.128562956 | 0.772738773 | 0.604653807 |
| Afap1l2 | Y | 124 | -0.547906868 | -0.098045452 | -0.173793915 | 0.374112953 | -0.075748463 | 0.013211458 | 0.46683341 | 0.282888096 | 0.074813625 | 0.632488767 |
| Spred1 | Y | 295 | -0.280847748 | -0.135472227 | -0.217025438 | 0.063822311 | -0.081553211 | 0.005158569 | 0.33951638 | 0.127533207 | 0.578899436 | 0.598006224 |
| Dennd2a | Y | 242 | -0.120209256 | -0.196671598 | -0.372306754 | -0.252097498 | -0.175635157 | 0.274364097 | 0.018426355 | 0.074530858 | 0.159080461 | 0.26760721 |
| Igf1r Insr | Y | 1163;1175 | -0.356561759 | -0.188560997 | -0.182206092 | 0.174355667 | 0.006354904 | 0.020520966 | 0.178036285 | 0.256518496 | 0.231141688 | 0.962963463 |
| Pkn2 | S | 619 | -0.29920988 | -0.231483468 | -0.29743437 | 0.001775511 | -0.065950902 | 0.199493943 | 0.166644728 | 0.06880684 | 0.992256594 | 0.554706865 |
| Pkn2 | S | 621 | -0.29920988 | -0.231483468 | -0.29743437 | 0.001775511 | -0.065950902 | 0.199493943 | 0.166644728 | 0.06880684 | 0.992256594 | 0.554706865 |
| Ptbp3 | Y | 98 | -0.16656321 | -0.187492833 | -0.473585026 | -0.307021816 | -0.286092193 | 0.507146224 | 0.115600736 | 0.037031533 | 0.281032609 | 0.127784211 |

| | | | | | | | | | | | | |
|---|---|---|---|---|---|---|---|---|---|---|---|---|
| Dok1 | Y | 361 | -0.285660812 | -0.170817925 | -0.104611281 | 0.181049531 | 0.066206644 | 0.006373571 | 0.045752884 | 0.438502498 | 0.215526399 | 0.591756235 |
| Slc38a2 | Y | 41 | -0.143541824 | -0.182765291 | -0.198706393 | -0.055164569 | -0.015941103 | 0.076373484 | 0.12418478 | 0.007337684 | 0.407350894 | 0.862388434 |
| Rps3a1 | Y | 256 | -0.250719293 | -0.117734186 | -0.185962823 | 0.06475647 | -0.068228637 | 0.015251069 | 0.224179532 | 0.042364317 | 0.063114365 | 0.359111269 |
| Rcn2 | Y | 314 | -0.22673146 | -0.170738728 | -0.481801016 | -0.255069556 | -0.311062287 | 0.383636093 | 0.34262866 | 0.027541983 | 0.32241949 | 0.098489467 |
| Slc20a2 | Y | 344 | -0.190566707 | -0.382719601 | -0.095972252 | 0.094594456 | 0.286747349 | 0.099742696 | 0.010316652 | 0.606610428 | 0.581668799 | 0.181635353 |
| Mertk | T | 945 | -0.147198351 | -0.362064961 | -0.493085011 | -0.34588666 | -0.13102005 | 0.70285011 | 0.117591454 | 0.092635863 | 0.423710616 | 0.603027688 |
| Clasp2 | Y | 1014 | -0.115592952 | -0.28230657 | -0.3682567 | -0.252663748 | -0.08595013 | 0.366494721 | 0.075298143 | 0.06762887 | 0.158345837 | 0.594189204 |
| Ptpn14 | T | 497 | -0.167009974 | -0.438838124 | -0.354024794 | -0.18701482 | 0.084813329 | 0.468544321 | 0.137614669 | 0.10606111 | 0.46102362 | 0.743433451 |
| Shb | Y | 330 | -0.27735757 | -0.193676552 | -0.163602186 | 0.113755384 | 0.030074366 | 0.020905916 | 0.183302613 | 0.141906767 | 0.24352056 | 0.811292146 |
| Bcar3 | Y | 260 | -0.117089039 | -0.351904165 | -0.243649028 | -0.126559989 | 0.108255137 | 0.208735473 | 0.012588872 | 0.375838032 | 0.618558781 | 0.671052113 |
| Baiap2 | Y | 492 | -0.205578442 | -0.177570862 | -0.219319126 | -0.013740684 | -0.041748264 | 0.015119317 | 0.122114791 | 0.179102599 | 0.916642265 | 0.773331754 |
| Dok1 | Y | 340 | -0.139372044 | -0.222774758 | -0.346590898 | -0.207218853 | -0.12381614 | 0.192012912 | 0.175564732 | 0.04431457 | 0.146128158 | 0.43235752 |
| Tec | Y | 518 | 0.028007444 | -0.245019566 | -0.414235408 | -0.442242852 | -0.169215842 | 0.801532375 | 0.030223292 | 0.01788258 | 0.022941497 | 0.183784598 |
| Slc12a4 | Y | 17 | -0.198309858 | -0.263511008 | -0.125914014 | 0.072395844 | 0.137596994 | 0.108557962 | 0.024656614 | 0.218206973 | 0.452474239 | 0.096088661 |
| Sdc1 | Y | 310 | -0.118475283 | -0.076232431 | -0.450511272 | -0.332035989 | -0.374278841 | 0.228552181 | 0.535230304 | 0.010015602 | 0.032132739 | 0.042798485 |
| Dennd2a | Y | 363 | 0.06808581 | -0.24457741 | -0.400768467 | -0.468854276 | -0.156191056 | 0.867276224 | 0.058345752 | 0.011938343 | 0.317944366 | 0.219068575 |
| Ttyh2 | Y | 424 | -0.002121196 | -0.419850102 | -0.285130811 | -0.283009615 | 0.134719291 | 0.985132369 | 0.057560154 | 0.037702522 | 0.0556313 | 0.425346969 |
| Ppp1cb | Y | 306 | -0.118487708 | -0.187989163 | -0.462804052 | -0.344316345 | -0.27481489 | 0.503922555 | 0.274463105 | 0.028202494 | 0.07096283 | 0.099121774 |
| Rpl8 | Y | 133 | -0.159263033 | -0.227105242 | -0.199279288 | -0.040016254 | 0.027825954 | 0.156871684 | 0.054620683 | 0.082524038 | 0.558580492 | 0.619456829 |
| Ints7 | Y | 935 | -0.199751301 | -0.16337435 | -0.164391509 | 0.035359792 | -0.001017159 | 0.00896977 | 0.123353016 | 0.221694627 | 0.749681125 | 0.993584867 |
| Rbmx Rbmxl1 | Y | 203;206 | 0.071292946 | -0.279680964 | -0.33342061 | -0.404713556 | -0.053739646 | 0.471647745 | 0.007290341 | 0.026739598 | 0.019391076 | 0.587703725 |
| Mink1 | Y | 913 | -0.401118184 | -0.097229874 | -0.381252846 | 0.019865339 | -0.284022972 | 0.026136817 | 0.592443055 | 0.427679069 | 0.963973879 | 0.544619767 |
| Iap | T | 494 | -0.068243085 | -0.456319843 | -0.070329575 | -0.00208649 | 0.385990268 | 0.619607231 | 0.00234606 | 0.726460765 | 0.992047234 | 0.136655545 |
| Tjp2 | Y | 222 | -0.411957469 | -0.222309628 | 0.118858217 | 0.530815686 | 0.341167845 | 0.003088405 | 0.042563218 | 0.279016214 | 0.005079425 | 0.017257543 |
| Dok1 | Y | 314 | -0.094942833 | -0.159498229 | -0.111416822 | -0.01647399 | 0.048081407 | 0.048856744 | 0.024067668 | 0.015355722 | 0.626266317 | 0.329429581 |
| Ptpn13 | T | 323 | -0.208889666 | -0.185181524 | -0.276698735 | -0.067809069 | -0.091517212 | 0.344678653 | 0.233240933 | 0.027575792 | 0.738342594 | 0.526689587 |
| Pdlim1 | Y | 149 | -0.219520138 | -0.166675357 | -0.383620312 | -0.164100174 | -0.216944954 | 0.433674767 | 0.300471694 | 0.04231048 | 0.539061528 | 0.160002747 |
| Axl | Y | 696 | -0.257538684 | -0.202519486 | -0.110419243 | 0.14711944 | 0.092100243 | 0.044545644 | 0.079617391 | 0.297388438 | 0.096068038 | 0.1548376 |
| Actr3 | Y | 233 | -0.168629243 | -0.347455702 | -0.213618124 | -0.044988881 | 0.133837577 | 0.148926584 | 0.192655203 | 0.141649913 | 0.726554562 | 0.566781737 |
| Eif4b | Y | 316 | -0.220790405 | -0.344537162 | -0.100273802 | 0.120516603 | 0.244263359 | 0.253856612 | 0.019453642 | 0.566119783 | 0.555880358 | 0.194632991 |
| Pi4ka | T | 2092 | -0.175964596 | -0.261985248 | -0.454590167 | -0.278625571 | -0.192604919 | 0.556637064 | 0.272226308 | 0.08926593 | 0.404999567 | 0.467763761 |
| Crtc1 | Y | 133 | -0.13603674 | -0.188765851 | -0.20693622 | -0.07089948 | -0.018170369 | 0.31909844 | 0.02691569 | 0.07615772 | 0.599710912 | 0.821608777 |
| Nav3 | S | 995 | 0.086381553 | -0.200607071 | -0.423724656 | -0.510106209 | -0.223117585 | 0.710694844 | 0.048437357 | 0.015969955 | 0.111689048 | 0.08954292 |
| Nav3 | T | 1000 | 0.086381553 | -0.200607071 | -0.423724656 | -0.510106209 | -0.223117585 | 0.710694844 | 0.048437357 | 0.015969955 | 0.111689048 | 0.08954292 |
| Tjp2 | Y | 1127 | -0.265618023 | -0.24804741 | -0.103456971 | 0.162161052 | 0.14459044 | 0.019882945 | 0.252524926 | 0.380293368 | 0.181866724 | 0.476298087 |
| Rtn4 | Y | 367 | 0.204855625 | -0.157532416 | -0.529699147 | -0.734554771 | -0.372166731 | 0.703667725 | 0.289294552 | 0.003185689 | 0.2522895 | 0.039810525 |
| Dnaja1 | Y | 376 | -0.22617087 | -0.32149735 | -0.229430163 | -0.003259293 | 0.092067187 | 0.233384004 | 0.146509448 | 0.271594059 | 0.981725057 | 0.599952653 |
| Nrp1 | Y | 920 | 0.05730045 | -0.142330407 | -0.276858177 | -0.334158627 | -0.13452777 | 0.247830781 | 0.013693496 | 0.000781056 | 0.003004489 | 0.013184429 |
| Nectin3 | Y | 510 | -0.105913335 | -0.169613223 | -0.390539425 | -0.28462609 | -0.220926202 | 0.275735495 | 0.198780724 | 0.073606829 | 0.148536139 | 0.243062493 |
| Aph1a | Y | 256 | 0.032714432 | -0.137361152 | -0.377269302 | -0.409983734 | -0.23990815 | 0.717129382 | 0.200470844 | 0.002073466 | 0.009671092 | 0.058693852 |
| Ctnnd1 | Y | 96 | -0.058211998 | -0.107334281 | -0.38687868 | -0.328666682 | -0.279544398 | 0.596305234 | 0.05273747 | 0.032790539 | 0.056945877 | 0.06215272 |
| Ctnnd1 | Y | 248 | -0.137574926 | -0.080659252 | -0.142769585 | -0.005194659 | -0.062110333 | 0.135782202 | 0.272471277 | 0.000613166 | 0.93727741 | 0.369891176 |
| Nfya | Y | 265 | 0.002807724 | -0.231215086 | -0.36899087 | -0.371798593 | -0.137775784 | 0.991847905 | 0.138635236 | 0.019695416 | 0.248331481 | 0.325945826 |
| Kirrel | S | 606 | -0.200090609 | -0.152772871 | -0.250955446 | -0.050864837 | -0.098182575 | 0.126376167 | 0.405004708 | 0.066388857 | 0.630107273 | 0.567111321 |
| Cherp | Y | 906 | -0.074037914 | -0.250518115 | -0.312889997 | -0.238852083 | -0.062371882 | 0.595700993 | 0.046224667 | 0.155193224 | 0.263830988 | 0.72616972 |
| Mapk11 | Y | 182 | -0.145096019 | -0.125456047 | -0.110699209 | 0.03439681 | 0.014756838 | 0.041458573 | 0.025192042 | 0.138481164 | 0.614188761 | 0.812493987 |
| Dennd2a | Y | 376 | -0.042919181 | -0.204577762 | -0.27658915 | -0.23366997 | -0.072011388 | 0.890522174 | 0.089917222 | 0.02173354 | 0.480083784 | 0.459030624 |
| Sdc1 | T | 304 | -0.138249322 | -0.077736762 | -0.258516651 | -0.12026733 | -0.180779889 | 0.178587218 | 0.465674096 | 0.01205332 | 0.193657677 | 0.132160593 |
| Vti1b | Y | 112 | -0.088698423 | -0.239273699 | -0.244190255 | -0.155491831 | -0.004916556 | 0.572538721 | 0.087973795 | 0.067452813 | 0.317736754 | 0.962499603 |
| Cav1 | T | 15 | -0.158423997 | -0.293151091 | 0.004697278 | 0.163121275 | 0.297848369 | 0.077905412 | 0.009364334 | 0.966384878 | 0.180652479 | 0.05149726 |
| Tnks1bp1 | T | 201 | -0.139557652 | -0.396875312 | -0.25512898 | -0.115571328 | 0.141746332 | 0.257607521 | 0.189951263 | 0.321283762 | 0.616741352 | 0.646318194 |
| Gemin5 | Y | 1038 | -0.064480947 | -0.527748006 | -0.178851173 | -0.114370225 | 0.348896833 | 0.735321355 | 0.124530568 | 0.141407301 | 0.570751398 | 0.240190263 |
| Acap2 | Y | 734 | 0.029257276 | -0.219321052 | -0.314753811 | -0.344011087 | -0.095432759 | 0.942592704 | 0.078896668 | 0.026920666 | 0.432780325 | 0.189936942 |
| Calu | Y | 275 | -0.279360518 | -0.084521137 | -0.506280343 | -0.226919825 | -0.421759206 | 0.318888453 | 0.770137045 | 0.110358193 | 0.270189735 | 0.110633421 |
| Cbl | Y | 698 | -0.148710197 | -0.106855703 | -0.183672547 | -0.034962349 | -0.076816844 | 0.055536741 | 0.176517125 | 0.078545766 | 0.642660985 | 0.370272042 |
| Ddr2 | T | 470 | 0.036052763 | -0.109728872 | -0.514161952 | -0.550214715 | -0.40443308 | 0.771541205 | 0.191171909 | 0.017981049 | 0.018397131 | 0.062095136 |
| Cmtm4 | Y | 187 | -0.275239524 | -0.105028682 | -0.102735565 | 0.172503959 | 0.002293117 | 0.007728245 | 0.498648985 | 0.34036614 | 0.143871908 | 0.988387861 |
| Micall1 | Y | 692 | 0.117918357 | -0.128178231 | -0.442658374 | -0.560576731 | -0.314480144 | 0.432228123 | 0.224162086 | 0.002614769 | 0.036473926 | 0.031227306 |
| Adgrl3 | Y | 1482 | -0.123226324 | -0.204194549 | -0.257032186 | -0.133805861 | -0.052837637 | 0.259983484 | 0.316903631 | 0.066107014 | 0.294209479 | 0.782313796 |
| Fus | Y | 232 | 0.046521241 | -0.249415214 | -0.194288123 | -0.240809364 | 0.05512709 | 0.535367405 | 0.010397568 | 0.029178496 | 0.015680842 | 0.29912535 |
| Mapk12 | Y | 185 | -0.327732471 | -0.190903881 | 0.017460067 | 0.345192538 | 0.208363948 | 0.012442611 | 0.199247314 | 0.904450916 | 0.088039837 | 0.23736538 |
| Pik3r1 | S | 505 | 0.029640903 | -0.227053932 | -0.201046194 | -0.230687097 | 0.026007738 | 0.808291313 | 0.113991337 | 0.003032509 | 0.15187922 | 0.803546328 |
| Ptpn6 | Y | 564 | -0.437036661 | -0.128475065 | -0.368382149 | 0.068654513 | -0.239907084 | 0.209271108 | 0.70170943 | 0.356222522 | 0.792524677 | 0.419964841 |
| Eno1 Eno1b Eno2 Eno3 | Y | 44;44;44 | -0.016624071 | -0.143235757 | -0.29741824 | -0.28079417 | -0.154182483 | 0.903564653 | 0.028121748 | 0.06847944 | 0.138726874 | 0.214972003 |
| Cep170 | Y | 955 | 0.087912782 | -0.217425113 | -0.305228842 | -0.393141624 | -0.087803729 | 0.287935766 | 0.073647441 | 0.007386926 | 0.007498187 | 0.388513294 |
| Afdn | Y | 1230 | -0.221700593 | -0.230711014 | -0.061629039 | 0.160071554 | 0.169081975 | 0.069522896 | 0.108175962 | 0.675262731 | 0.26217104 | 0.273923287 |
| Dok1 | Y | 376 | -0.185136556 | -0.179423645 | -0.121221087 | 0.06391547 | 0.058202558 | 0.082017683 | 0.132085705 | 0.359365855 | 0.561882675 | 0.63077409 |
| Acsl5 | Y | 562 | -0.196507704 | -0.140253888 | -0.216399449 | -0.019891745 | -0.076145561 | 0.244673862 | 0.399889382 | 0.074102766 | 0.891955873 | 0.624087982 |
| Tmem63a | S | 767 | 0.069002091 | -0.266347611 | -0.338374958 | -0.407377049 | -0.072027347 | 0.449594815 | 0.074354768 | 0.041469191 | 0.024179915 | 0.594907181 |
| Specc1l | Y | 909 | 0.016886062 | -0.116309821 | -0.266040854 | -0.282926916 | -0.149731033 | 0.799074256 | 0.286387608 | 0.001525749 | 0.018049983 | 0.199166152 |
| Cfl1 | Y | 68 | -0.090253297 | -0.215996015 | -0.336864407 | -0.24661111 | -0.120868392 | 0.716724089 | 0.216324937 | 0.099450538 | 0.372118915 | 0.520000935 |
| Iqsec2 | Y | 1119 | -0.060634035 | -0.077998193 | -0.5296547 | -0.469020665 | -0.451656507 | 0.696307624 | 0.64515946 | 0.043814892 | 0.062807684 | 0.078750817 |
| Akap12 | S | 350 | -0.289508778 | 0.095091758 | -0.445861845 | -0.156353068 | -0.540953604 | 0.462254422 | 0.472864865 | 0.011412374 | 0.672750131 | 0.003685354 |
| Kif20a | Y | 867 | 0.060032237 | -0.422194883 | -0.310247649 | -0.370279886 | 0.111947234 | 0.371592716 | 0.087441085 | 0.101190096 | 0.08383507 | 0.586972418 |
| Kirrel | S | 605 | -0.162039552 | -0.172032379 | -0.231145652 | -0.0691061 | -0.059113273 | 0.212524848 | 0.249246915 | 0.185864478 | 0.648361372 | 0.718142266 |
| Epb41l1 | Y | 343 | -0.029947413 | -0.169745834 | -0.270228769 | -0.240281356 | -0.100482935 | 0.768968859 | 0.117257522 | 0.045642789 | 0.067675915 | 0.336647084 |
| Sorbs2 | Y | 312 | 0.113709244 | -0.21686266 | -0.258734933 | -0.372444178 | -0.041872274 | 0.436839481 | 0.014036245 | 0.030905124 | 0.061841223 | 0.59538394 |
| Lpp | Y | 245 | -0.03731439 | -0.156879278 | -0.575567625 | -0.538253235 | -0.418688346 | 0.826843933 | 0.261618118 | 0.143820473 | 0.154257543 | 0.231415125 |
| Isoc1 | Y | 159 | -0.406138302 | -0.09526981 | -0.446150881 | -0.04001258 | -0.350881072 | 0.321943061 | 0.674962947 | 0.31944232 | 0.934458783 | 0.418134521 |
| Ubap2l | Y | 878 | -0.00324066 | -0.202820446 | -0.225871733 | -0.222631073 | -0.023051287 | 0.961681081 | 0.065410922 | 0.053220274 | 0.075156194 | 0.80947247 |
| Ephb3 Ephb4 | Y | 609;596 | -0.043441631 | -0.190496119 | -0.244006855 | -0.200565224 | -0.053510736 | 0.686011032 | 0.120255887 | 0.070796633 | 0.155403901 | 0.660577753 |
| Arhgap39 | Y | 423 | -0.178010977 | -0.084047677 | -0.129458501 | 0.048552476 | -0.045410824 | 0.123352646 | 0.036077002 | 0.246161842 | 0.680770412 | 0.633250162 |
| Kirrel | Y | 604 | -0.028886714 | -0.113936732 | -0.236186469 | -0.207299755 | -0.122249737 | 0.664603591 | 0.280161826 | 0.00764744 | 0.031321934 | 0.256274194 |
| Papss2 | Y | 21 | -0.188394918 | -0.30944891 | -0.334273539 | -0.145878621 | -0.024824629 | 0.349494858 | 0.366971743 | 0.446930419 | 0.729692374 | 0.957964407 |
| Adgrl2 | Y | 1434 | -0.185046857 | -0.174996491 | -0.114045195 | 0.071001662 | 0.060951297 | 0.088196628 | 0.227477359 | 0.421425088 | 0.613017957 | 0.705939215 |
| Pkp4 | Y | 493 | -0.195770601 | -0.109784245 | -0.06001089 | 0.135759711 | 0.049773355 | 0.018215855 | 0.280426559 | 0.408817304 | 0.070778046 | 0.587054086 |
| Dok1 | Y | 408 | -0.168055897 | -0.124274013 | -0.125972602 | 0.042083294 | -0.00169859 | 0.114562666 | 0.16590986 | 0.25926779 | 0.674865201 | 0.984582021 |
| Slc4a7 | Y | 49 | -0.05391512 | -0.156458572 | -0.162708929 | -0.108793808 | -0.006250357 | 0.604208155 | 0.082264878 | 0.050464076 | 0.286891838 | 0.908446943 |
| Gab2 | T | 262 | -0.193081004 | -0.115127995 | -0.079221468 | 0.113859535 | 0.035906527 | 0.013633508 | 0.385661893 | 0.5125394 | 0.367800413 | 0.81338576 |
| Rbm3 | S | 143 | -0.233421527 | -0.20519989 | -0.135448758 | 0.097972769 | 0.069751132 | 0.28547498 | 0.193247526 | 0.436485719 | 0.663978732 | 0.702582817 |
| Peg3 | S | 160 | -0.174017696 | -0.207199012 | -0.134641359 | 0.039376336 | 0.072557652 | 0.304011192 | 0.126465222 | 0.429148377 | 0.834997305 | 0.661889318 |

| | | | | | | | | | | | | | |
|---|---|---|---|---|---|---|---|---|---|---|---|---|---|
| Peg3 | T | | 156 | -0.174017696 | -0.207199012 | -0.134641359 | 0.039376336 | 0.072557652 | 0.304011192 | 0.126465222 | 0.429148377 | 0.834997305 | 0.661889318 |
| Peg3 | T | | 161 | -0.174017696 | -0.207199012 | -0.134641359 | 0.039376336 | 0.072557652 | 0.304011192 | 0.126465222 | 0.429148377 | 0.834997305 | 0.661889318 |
| Peg3 | Y | | 158 | -0.174017696 | -0.207199012 | -0.134641359 | 0.039376336 | 0.072557652 | 0.304011192 | 0.126465222 | 0.429148377 | 0.834997305 | 0.661889318 |
| Epb41l3 | Y | | 479 | -0.071818879 | -0.147970762 | -0.219946243 | -0.148127365 | -0.071975481 | 0.54761612 | 0.475974268 | 0.028439831 | 0.227987423 | 0.710134293 |
| Stip1 | Y | | 354 | 0.085383163 | -0.272215182 | -0.199576464 | -0.284959627 | 0.072638718 | 0.539623521 | 0.013168928 | 0.202650549 | 0.135951099 | 0.58408413 |
| Dennd2a | S | | 358 | 0.071560853 | -0.152936766 | -0.271630651 | -0.343191505 | -0.118693885 | 0.852358122 | 0.177875802 | 0.02367102 | 0.413984979 | 0.26063066 |
| Snap23 | Y | | 138 | -0.288142699 | -0.02263342 | -0.050709352 | 0.237433347 | -0.028075932 | 0.008340533 | 0.750353162 | 0.636736806 | 0.073765985 | 0.75527971 |
| Supt5 | Y | | 759 | -0.028734784 | -0.024489946 | -0.278273605 | -0.249538821 | -0.253783659 | 0.722263541 | 0.779018139 | 0.004802521 | 0.02994429 | 0.039936999 |
| Ncaph | Y | | 625 | 0.141879458 | -0.334451365 | -0.280532031 | -0.42241149 | 0.053919334 | 0.105441038 | 0.016366474 | 0.11264262 | 0.046245891 | 0.707613866 |
| Pkp4 | Y | | 1166 | -0.301205804 | -0.173707986 | 0.051547059 | 0.352752863 | 0.225255045 | 0.04140238 | 0.175868515 | 0.647914402 | 0.004619988 | 0.013711295 |
| Dctn2 | Y | | 314 | -0.151450646 | -0.340420775 | -0.076685162 | 0.074765483 | 0.263735613 | 0.558501188 | 0.101347548 | 0.595976237 | 0.750997254 | 0.150039069 |
| Fermt2 | Y | | 185 | 0.036155131 | -0.205600993 | -0.294105586 | -0.330260717 | -0.088504593 | 0.826475415 | 0.092643344 | 0.139843077 | 0.149085546 | 0.589184825 |
| Prag1 | Y | | 391 | 0.048928795 | -0.347662893 | -0.185886828 | -0.234815624 | 0.161776064 | 0.679906334 | 0.049660257 | 0.275380978 | 0.222441722 | 0.356079244 |
| Ctnnd1 | Y | | 208 | -0.144162885 | -0.055292739 | -0.188457986 | -0.044295101 | -0.133165247 | 0.173370946 | 0.604788517 | 0.07297034 | 0.477583589 | 0.172547888 |
| Pag1 | Y | | 224 | -0.297820663 | 0.196937025 | -0.204708319 | 0.093112344 | -0.401645344 | 0.011027469 | 0.385278713 | 0.13628295 | 0.439862165 | 0.138536218 |
| Dnaja1 | Y | | 381 | -0.114746815 | -0.346131412 | -0.146034538 | -0.031287723 | 0.200096875 | 0.564839746 | 0.179690425 | 0.461943238 | 0.785261057 | 0.319056603 |
| Iqgap1 | Y | | 172 | -0.040407656 | -0.117967193 | -0.315041071 | -0.274633415 | -0.197073878 | 0.873102445 | 0.444736084 | 0.047638239 | 0.311063235 | 0.14599615 |
| Pard3b | S | | 990 | -0.136826371 | -0.07045462 | -0.121961531 | 0.01486484 | -0.051506911 | 0.079683458 | 0.511797651 | 0.099430828 | 0.821853324 | 0.624370675 |
| Rbm3 | Y | | 126 | 0.005959454 | -0.180583786 | -0.542227603 | -0.548187057 | -0.361643817 | 0.969056416 | 0.262073567 | 0.260548024 | 0.257176335 | 0.415850015 |
| Tjp2 | Y | | 183 | -0.314548136 | -0.181741578 | 0.084783099 | 0.399331235 | 0.266524677 | 0.013888421 | 0.375690592 | 0.474404182 | 0.022468804 | 0.22681639 |
| Itgb1 | Y | | 783 | 0.028417906 | -0.035856617 | -0.254381825 | -0.282799731 | -0.218525209 | 0.706721655 | 0.477057351 | 0.001840249 | 0.024230948 | 0.006421768 |
| Myo10 | S | | 1135 | -0.121586986 | -0.246083829 | -0.094284099 | 0.027302887 | 0.15179973 | 0.25363854 | 0.188800462 | 0.453950318 | 0.842318812 | 0.405167207 |
| Myo10 | Y | | 1132 | -0.121586986 | -0.246083829 | -0.094284099 | 0.027302887 | 0.15179973 | 0.25363854 | 0.188800462 | 0.453950318 | 0.842318812 | 0.405167207 |
| Pkp4 | Y | | 371 | 0.014377534 | -0.066011469 | -0.196594473 | -0.210972007 | -0.130583003 | 0.855767475 | 0.00661364 | 0.041591965 | 0.072962642 | 0.101400206 |
| Ctnnd1 | T | | 304 | -0.026518665 | -0.066943399 | -0.333422539 | -0.306903874 | -0.26647914 | 0.829621718 | 0.511174327 | 0.041673013 | 0.087088343 | 0.088838359 |
| Eif3c | Y | | 911 | 0.111791077 | -0.268997948 | -0.228035678 | -0.339826755 | 0.040962271 | 0.493326664 | 0.036299112 | 0.161427418 | 0.100751881 | 0.74002971 |
| Hnrnpk | Y | | 323 | 0.177035031 | -0.191610751 | -0.397348966 | -0.574383996 | -0.205738214 | 0.304599301 | 0.126911029 | 0.042046757 | 0.024866468 | 0.194013932 |
| Abi2 | Y | | 304 | -0.088280768 | -0.096433872 | -0.222089232 | -0.133808464 | -0.12565536 | 0.376974307 | 0.414466756 | 0.111365995 | 0.239207964 | 0.311510364 |
| Rack1 | Y | | 52 | 0.098450671 | -0.329975297 | -0.201588573 | -0.300039244 | 0.128386724 | 0.35976274 | 0.06677305 | 0.123497145 | 0.063867197 | 0.387424246 |
| Eps8 | T | | 602 | -0.096875884 | -0.257701649 | 0.096204853 | 0.193080737 | 0.353906502 | 0.12288427 | 0.004386514 | 0.259928863 | 0.067468132 | 0.01163769 |
| Sh3pxd2a | S | | 566 | 0.036332376 | -0.339761011 | -0.161408302 | -0.197740678 | 0.178352709 | 0.848612425 | 0.110215042 | 0.242471112 | 0.360539665 | 0.350255575 |
| Rpl35a | Y | | 34 | 0.085229918 | -0.344860279 | -0.189502565 | -0.274732483 | 0.155357713 | 0.509635776 | 0.038933345 | 0.354539024 | 0.190529629 | 0.390749602 |
| Flt1 | Y | | 1053 | -0.095774265 | -0.166646302 | -0.011550913 | 0.084223352 | 0.15509539 | 0.135138351 | 0.024718232 | 0.909207571 | 0.407866817 | 0.187981684 |
| Met | Y | | 1001 | -0.103303127 | -0.041519965 | -0.220091314 | -0.116788187 | -0.178571349 | 0.178909105 | 0.613852822 | 0.117581477 | 0.334090488 | 0.185179977 |
| Slc12a4 | Y | | 61 | -0.273288136 | -0.28165538 | 0.147592507 | 0.420880643 | 0.429247887 | 0.09797298 | 0.07542533 | 0.330465815 | 0.015124741 | 0.015827325 |
| Ddx5 | Y | | 514 | -0.040738033 | -0.134930041 | -0.37517869 | -0.334440658 | -0.240248649 | 0.769334293 | 0.189353103 | 0.305580832 | 0.353425062 | 0.478475658 |
| Ahcy Gm4737 | Y | | 193 | 0.172104394 | -0.299275293 | -0.159336257 | -0.331440651 | 0.139939036 | 0.032275003 | 0.00036825 | 0.248271653 | 0.06344342 | 0.295406761 |
| Abl1 Abl2 | Y | 439;393 | | -0.010046816 | -0.090703418 | -0.126801112 | -0.116754296 | -0.036097695 | 0.824105107 | 0.066376544 | 0.02414055 | 0.039134572 | 0.154780007 |
| Pdlim1 | S | | 144 | 0.081465616 | -0.353205967 | -0.099690173 | -0.18115579 | 0.253515793 | 0.73114067 | 0.028253506 | 0.505452435 | 0.457276589 | 0.095160569 |
| Pdlim1 | T | | 146 | 0.081465616 | -0.353205967 | -0.099690173 | -0.18115579 | 0.253515793 | 0.73114067 | 0.028253506 | 0.505452435 | 0.457276589 | 0.095160569 |
| Sirpa | Y | | 464 | -0.079777341 | -0.053672038 | -0.157798534 | -0.078021193 | -0.104126497 | 0.164722668 | 0.741810922 | 0.038453059 | 0.2418634 | 0.541587118 |
| Vav2 | Y | | 142 | 0.104800605 | -0.177464269 | -0.36852927 | -0.473329876 | -0.191065002 | 0.315500577 | 0.137009253 | 0.077031735 | 0.05065512 | 0.273287629 |
| Arhgap12 | Y | | 388 | -0.10519211 | -0.004234822 | -0.186428973 | -0.081236863 | -0.18219415 | 0.08151334 | 0.955168529 | 0.086549416 | 0.358798982 | 0.11274289 |
| Ptk2 | S | | 843 | -0.05974678 | -0.228074873 | -0.005145778 | 0.054601002 | 0.222929096 | 0.392209274 | 0.021496791 | 0.943958466 | 0.333641918 | 0.021916885 |
| Phax | Y | | 48 | -0.098910976 | -0.104961889 | -0.184109698 | -0.085198722 | -0.079147809 | 0.42501647 | 0.337290314 | 0.19377892 | 0.563174494 | 0.562579808 |
| Pkp4 | Y | | 1093 | -0.20212382 | -0.097818475 | -0.028060826 | 0.174062994 | 0.069757649 | 0.085933369 | 0.250292359 | 0.699084365 | 0.119139917 | 0.316749495 |
| Flnb | Y | | 2502 | -0.100133191 | -0.155241044 | -0.028133902 | 0.071999288 | 0.127107142 | 0.203277708 | 0.068322931 | 0.755394039 | 0.383586157 | 0.166256577 |
| Ubap2 | S | | 1001 | 0.149562724 | -0.592060825 | -0.290525909 | -0.440088632 | 0.301534917 | 0.78633676 | 0.23908581 | 0.588831268 | 0.34915826 | 0.389097226 |
| Ubap2 | S | | 1002 | 0.149562724 | -0.592060825 | -0.290525909 | -0.440088632 | 0.301534917 | 0.78633676 | 0.23908581 | 0.588831268 | 0.34915826 | 0.389097226 |
| Yipf5 | Y | | 44 | -0.216000533 | 0.004978329 | -0.246462878 | -0.030462344 | -0.251441206 | 0.373404536 | 0.978571852 | 0.163980714 | 0.895750532 | 0.241632034 |
| Rbm3 | Y | | 115 | -0.039642209 | -0.184074895 | -0.187988483 | -0.148346273 | -0.003913587 | 0.781648934 | 0.20681779 | 0.294964602 | 0.370241827 | 0.979117975 |
| Fermt2 | Y | | 179 | -0.073528314 | -0.001216279 | -0.262575856 | -0.189047541 | -0.261359577 | 0.551817126 | 0.98347538 | 0.047117395 | 0.20407275 | 0.068884525 |
| Acsl5 | Y | | 70 | -0.051205483 | -0.462734036 | -0.209921925 | -0.158716442 | 0.252812112 | 0.923417403 | 0.295526528 | 0.666786623 | 0.788611091 | 0.607976001 |
| Frs2 | Y | | 349 | -0.098734146 | -0.123526596 | -0.184506557 | -0.085772411 | -0.060979961 | 0.715168043 | 0.291782311 | 0.212984935 | 0.751520269 | 0.613711696 |
| Cbl | S | | 692 | -0.122472699 | -0.026816594 | -0.181780286 | -0.059307587 | -0.154963692 | 0.092909944 | 0.764804918 | 0.285036227 | 0.68476942 | 0.354252455 |
| Tchp | Y | | 9 | 0.040157667 | -0.305840678 | -0.161175155 | -0.201332822 | 0.144665523 | 0.549036347 | 0.065733902 | 0.50196043 | 0.414514429 | 0.557999759 |
| Rbm8a Rbm8a2 | Y | | 54 | -0.063488814 | -0.225684464 | -0.129472245 | -0.065983431 | 0.096212219 | 0.660863084 | 0.204161952 | 0.445010174 | 0.671180049 | 0.581235169 |
| Lpp | Y | | 302 | -0.197615999 | 0.015111804 | -0.07203496 | 0.12558104 | -0.087146764 | 0.01078515 | 0.84575569 | 0.630771672 | 0.419009399 | 0.571519437 |
| Bcar1 | Y | | 238 | -0.058433544 | -0.048816386 | -0.206016147 | -0.147582603 | -0.157199762 | 0.429465202 | 0.375601356 | 0.157442996 | 0.273778904 | 0.251732726 |
| Mertk | Y | | 924 | 0.008307265 | -0.07930386 | -0.204800702 | -0.213107968 | -0.125496842 | 0.976778507 | 0.191413983 | 0.086114328 | 0.489156324 | 0.225917468 |
| Cttnbp2nl | Y | | 343 | -0.16186664 | -0.049041874 | -0.131660811 | 0.030205829 | -0.082618936 | 0.272331918 | 0.579192636 | 0.247038009 | 0.831199818 | 0.439549257 |
| Rtn4 | Y | | 26 | 0.071921772 | -0.105227625 | -0.391915217 | -0.46383699 | -0.286687592 | 0.862113713 | 0.683640886 | 0.102871377 | 0.358025113 | 0.28369288 |
| Gab2 | Y | | 263 | -0.127401476 | -0.199314599 | 0.021722712 | 0.149124189 | 0.221037311 | 0.250449622 | 0.115725186 | 0.875240844 | 0.322824404 | 0.18079373 |
| Prkaa1 | Y | | 500 | -0.226894257 | 0.015601222 | -0.104676988 | 0.12221727 | -0.12027821 | 0.092514855 | 0.883548356 | 0.386152964 | 0.332123101 | 0.305508588 |
| Ubap2 | Y | | 998 | 0.157242655 | -0.519334062 | -0.192732363 | -0.349975017 | 0.326601699 | 0.702363644 | 0.185731284 | 0.615999239 | 0.295005313 | 0.202425762 |
| Sh3pxd2a | Y | | 619 | 0.292119341 | -0.02196141 | -0.242477145 | -0.534596486 | -0.220515735 | 0.499559871 | 0.838098048 | 0.000415698 | 0.272873982 | 0.133827788 |
| Gprc5a | Y | | 349 | -0.045432164 | -0.022209093 | -0.173091498 | -0.127659334 | -0.150882404 | 0.481884926 | 0.738609426 | 0.051874763 | 0.101406015 | 0.072049358 |
| Zdhhc8 | Y | | 601 | -0.047101016 | -0.058941505 | -0.269490561 | -0.222389545 | -0.210549056 | 0.757244996 | 0.448321304 | 0.220694085 | 0.335997225 | 0.308831438 |
| Frmd4a | Y | | 843 | -0.12248933 | -0.108198712 | -0.080872765 | 0.041616565 | 0.027325947 | 0.197424578 | 0.408430958 | 0.612130352 | 0.799399738 | 0.880054996 |
| Snx12 | Y | | 23 | 0.090334432 | -0.007468479 | -0.341511034 | -0.431845465 | -0.334042555 | 0.758665222 | 0.935018297 | 0.03059118 | 0.225153196 | 0.038620291 |
| Sdc2 | Y | | 192 | -0.134622208 | -0.030283481 | -0.102931764 | 0.031690444 | -0.072648284 | 0.148758549 | 0.700245737 | 0.317860266 | 0.699312098 | 0.373828738 |
| Asap1 | S | | 781 | -0.179059398 | -0.138655032 | -0.394936572 | -0.215877174 | -0.256281539 | 0.884933472 | 0.61989409 | 0.477883676 | 0.863953912 | 0.621189974 |
| Arhgef2 | Y | | 893 | 0.066625503 | -0.094099839 | -0.211206797 | -0.2778323 | -0.117106958 | 0.522302057 | 0.270420879 | 0.049128455 | 0.030779296 | 0.110000333 |
| Mapk1 | Y | | 185 | 0.122103852 | -0.135635188 | -0.259058865 | -0.381162717 | -0.123423678 | 0.294856638 | 0.221115282 | 0.053097464 | 0.024994944 | 0.297801468 |
| Zdhhc5 Zdhhc8 | Y | 758;711 | | -0.059675713 | -0.071310919 | -0.112034015 | -0.052358302 | -0.040723095 | 0.575244376 | 0.162393086 | 0.248677088 | 0.66123203 | 0.621126595 |
| Cttn | Y | | 84 | -0.153001349 | -0.245664002 | 0.044461083 | 0.197462432 | 0.290125085 | 0.321898126 | 0.146700634 | 0.5018517 | 0.230192715 | 0.098010112 |
| Gja1 | S | | 306 | -0.063087533 | -0.033537403 | -0.155280964 | -0.092193431 | -0.121743561 | 0.609672106 | 0.641647351 | 0.085071243 | 0.444446923 | 0.082571181 |
| Caprin1 | Y | | 668 | 0.150533499 | -0.233917475 | -0.228821849 | -0.379355348 | 0.005095626 | 0.059399213 | 0.164321269 | 0.026852027 | 0.01187843 | 0.970345961 |
| Spred2 | Y | | 246 | -0.141406309 | 0.020049466 | -0.147660484 | -0.006254175 | -0.16770995 | 0.194955102 | 0.863321406 | 0.24554104 | 0.960767795 | 0.277820823 |
| Bcar1 | Y | | 228 | -0.038813086 | -0.050504055 | -0.141572203 | -0.102759117 | -0.091068148 | 0.529200089 | 0.485286935 | 0.139450428 | 0.228398676 | 0.299160774 |
| Ddr2 | Y | | 739 | -0.129170132 | -0.068404956 | -0.030438595 | 0.098731537 | 0.037966361 | 0.130976908 | 0.436080688 | 0.626480558 | 0.164607731 | 0.590927684 |
| Kif23 | Y | | 583 | 0.034564734 | -0.166614224 | -0.139671801 | -0.174236535 | 0.026942422 | 0.745383296 | 0.168386881 | 0.285979193 | 0.195666183 | 0.82533032 |
| Rbm14 | Y | | 665 | -0.071093995 | -0.054929972 | -0.271960376 | -0.200866381 | -0.217030404 | 0.621325788 | 0.793426296 | 0.309572411 | 0.447270682 | 0.465622112 |
| Cfl1 | Y | | 140 | -0.056478845 | -0.026261486 | -0.088737107 | -0.032258263 | -0.062475621 | 0.234333183 | 0.523628952 | 0.111958624 | 0.47576564 | 0.182747976 |
| Rab34 | Y | | 247 | -0.058891308 | 0.001468818 | -0.168570369 | -0.109679061 | -0.170039187 | 0.481149227 | 0.988708388 | 0.082144237 | 0.029961624 | 0.122786877 |
| Dok1 | Y | | 336 | 0.030367476 | -0.186968143 | -0.11572697 | -0.146094446 | 0.071241173 | 0.907142185 | 0.204283689 | 0.332478806 | 0.598204278 | 0.631379596 |
| Grasp | Y | | 55 | 0.475609674 | -0.397673373 | -0.257410052 | -0.733019726 | 0.140263321 | 0.256116759 | 0.155826981 | 0.092356106 | 0.127698257 | 0.551712879 |
| Cttn | Y | | 141 | -0.174806225 | -0.095517653 | 0.02434899 | 0.199155215 | 0.119866642 | 0.142951347 | 0.361855163 | 0.861714052 | 0.164192854 | 0.340117256 |
| Tor1aip1 | Y | | 67 | 0.041240238 | 0.119628519 | -0.402679895 | -0.443920133 | -0.522308413 | 0.749949916 | 0.343202366 | 0.015152996 | 0.021675174 | 0.009941551 |
| Gab1 | Y | | 285 | -0.513686862 | 0.012255927 | -0.06116439 | 0.452522472 | -0.073420317 | 0.370323505 | 0.947407517 | 0.728852071 | 0.422374366 | 0.738459547 |
| Gsto1 | Y | | 228 | 0.088281813 | -0.42123689 | -0.027024112 | -0.115305924 | 0.394212778 | 0.812571397 | 0.145377132 | 0.90044012 | 0.750772882 | 0.147622886 |

| | | | | | | | | | | | | |
|---|---|---|---|---|---|---|---|---|---|---|---|---|
| Arhgef40 | Y | 242 | 0.461390165 | -0.214486641 | -0.268081095 | -0.72947126 | -0.053594455 | 0.531405009 | 0.130863073 | 0.161770794 | 0.356444538 | 0.753089976 |
| Kifbp | S | 229 | -0.12288003 | -0.112174418 | 0.012819261 | 0.135699291 | 0.12499368 | 0.18790762 | 0.234704456 | 0.887597394 | 0.068601363 | 0.096222695 |
| Kifbp | T | 230 | -0.12288003 | -0.112174418 | 0.012819261 | 0.135699291 | 0.12499368 | 0.18790762 | 0.234704456 | 0.887597394 | 0.068601363 | 0.096222695 |
| Kifbp | Y | 226 | -0.12288003 | -0.112174418 | 0.012819261 | 0.135699291 | 0.12499368 | 0.18790762 | 0.234704456 | 0.887597394 | 0.068601363 | 0.096222695 |
| Erbb2 | Y | 878 | 0.068366195 | 0.144379519 | -0.36511138 | -0.433477575 | -0.509490899 | 0.665760243 | 0.558776905 | 0.011919556 | 0.052246939 | 0.119083938 |
| Ephb3 Ephb4 | T | 608;595 | 0.081362543 | -0.020732463 | -0.160800843 | -0.242163386 | -0.14006838 | 0.707776445 | 0.636930518 | 0.007760529 | 0.324087492 | 0.017122016 |
| Pitpna | Y | 141 | -0.075800859 | -0.037458561 | -0.115481243 | -0.039680384 | -0.078022682 | 0.402405465 | 0.66507511 | 0.225966866 | 0.701299988 | 0.462705411 |
| Rin1 | Y | 670 | 0.252798254 | -0.225611789 | -0.246981187 | -0.499779442 | -0.021369399 | 0.120282864 | 0.128558787 | 0.057082674 | 0.021865333 | 0.86874367 |
| Flrt2 | Y | 588 | -0.323948992 | -0.095225377 | 0.152152098 | 0.47610109 | 0.247377475 | 0.067136968 | 0.588085268 | 0.39680739 | 0.038532474 | 0.166064925 |
| C77080 | Y | 239 | -0.139694111 | 0.03220588 | -0.167639051 | -0.02794494 | -0.199844931 | 0.326076789 | 0.838257596 | 0.280500988 | 0.867240284 | 0.319706177 |
| Spred2 | Y | 263 | -0.313690273 | 0.156970258 | -0.133941482 | 0.179748791 | -0.29091174 | 0.139107244 | 0.132399234 | 0.440133037 | 0.328836567 | 0.149596684 |
| Sh3pxd2a | Y | 557 | 0.214573414 | -0.213866916 | -0.129075713 | -0.343649127 | 0.084791202 | 0.565090526 | 0.104389246 | 0.238673309 | 0.384330895 | 0.439619012 |
| Epha7 | Y | 791 | 0.151302464 | -0.059731821 | -0.252418642 | -0.403721106 | -0.192686821 | 0.227925857 | 0.507105378 | 0.014532523 | 0.029618579 | 0.050871133 |
| Gja1 | S | 279 | 0.221143457 | -0.257272511 | -0.246323206 | -0.467466663 | 0.010949306 | 0.472794593 | 0.146519063 | 0.423273878 | 0.231564474 | 0.967220454 |
| Ppp1ca | Y | 306 | -0.023892751 | -0.087926479 | -0.045616557 | -0.021723806 | 0.042309922 | 0.531723317 | 0.071860352 | 0.605815439 | 0.791378177 | 0.620939509 |
| Actr3 | Y | 231 | 0.049097586 | -0.252302902 | -0.080013485 | -0.129111071 | 0.172289417 | 0.636910772 | 0.159615477 | 0.541158963 | 0.353848415 | 0.335422121 |
| Bcr | Y | 178 | -0.150726279 | -0.008501302 | -0.003714259 | 0.14701202 | 0.004787043 | 0.040521964 | 0.904571522 | 0.951697238 | 0.03544301 | 0.939230067 |
| Ptk2b | T | 850 | -0.230167736 | 0.25825179 | -0.031452399 | 0.198715336 | -0.28970419 | 0.002424098 | 0.229526555 | 0.422496843 | 0.013995184 | 0.198204427 |
| Esyt1 | Y | 111 | 0.029997421 | -0.077631591 | -0.269508291 | -0.299505712 | -0.1918767 | 0.733657023 | 0.54428347 | 0.273424475 | 0.239906513 | 0.414026665 |
| Vti1b | Y | 115 | -0.080828024 | -0.118597468 | -0.011239416 | 0.069588608 | 0.107358052 | 0.380794858 | 0.280999147 | 0.887446226 | 0.523366418 | 0.372406419 |
| Vim | Y | 53 | -0.054535475 | 0.021396024 | -0.245732018 | -0.191196543 | -0.267128042 | 0.572362861 | 0.855808654 | 0.247679333 | 0.349000497 | 0.230304219 |
| Dok1 | S | 290 | 0.082700388 | -0.148302233 | -0.161771236 | -0.244471623 | -0.013469002 | 0.804445648 | 0.384720128 | 0.275285452 | 0.500035779 | 0.941941579 |
| Wasl | Y | 172 | 0.052388465 | -0.092889883 | -0.126438657 | -0.178827122 | -0.033548774 | 0.584072781 | 0.317608902 | 0.102545618 | 0.11586429 | 0.690909796 |
| Otud4 | Y | 169 | -0.344509967 | -0.183334551 | 0.074355911 | 0.418865878 | 0.257690463 | 0.498422153 | 0.38471473 | 0.539046781 | 0.420579118 | 0.232956793 |
| Wipf2 | Y | 74 | -0.124741453 | -0.063775305 | 0.037798635 | 0.162540088 | 0.10157394 | 0.050858096 | 0.457126637 | 0.57516063 | 0.05761217 | 0.26391105 |
| Pik3r1 Pik3r3 | Y | 467;199 | 0.164543195 | -0.200332673 | -0.167300421 | -0.331843616 | 0.033032252 | 0.065536551 | 0.051564797 | 0.130856557 | 0.037967244 | 0.714615965 |
| Abi1 | S | 267 | 0.106512755 | -0.079896752 | -0.218497197 | -0.325009952 | -0.138600445 | 0.660516345 | 0.511665578 | 0.14156157 | 0.250076615 | 0.357295698 |
| Dok1 | Y | 397 | -0.288677675 | 0.004408661 | 0.064376465 | 0.353054141 | 0.059967805 | 0.077653701 | 0.97887726 | 0.608293262 | 0.033783033 | 0.682639825 |
| Slc35b3 | Y | 11 | -0.009050458 | -0.087139793 | -0.105324344 | -0.096273886 | -0.018184551 | 0.92251804 | 0.387722015 | 0.333462117 | 0.382649411 | 0.866006625 |
| Mpzl1 | Y | 242 | 0.415922927 | -0.103950548 | -0.388647599 | -0.804570526 | -0.284697052 | 0.295382766 | 0.245195379 | 0.082856727 | 0.097110733 | 0.15899846 |
| Trp53bp2 | Y | 609 | 0.062718289 | -0.060506262 | -0.214508466 | -0.277226754 | -0.154002203 | 0.442659093 | 0.584376488 | 0.102939224 | 0.058291447 | 0.251944244 |
| Eps8 | Y | 601 | -0.098517744 | -0.225214198 | 0.103042841 | 0.201560585 | 0.328257039 | 0.190106071 | 0.080331648 | 0.080638624 | 0.037167343 | 0.045123956 |
| Hars | Y | 115 | 0.166570615 | -0.178015239 | -0.230140519 | -0.396711134 | -0.05212528 | 0.105356954 | 0.088101327 | 0.222532437 | 0.091535222 | 0.733827019 |
| Asap2 | Y | 727 | -0.037213995 | -0.023459498 | -0.148521214 | -0.111307219 | -0.125061716 | 0.746482969 | 0.832860723 | 0.245065402 | 0.458411936 | 0.403022896 |
| Mapk1 | T | 183 | 0.199267855 | -0.211887186 | -0.225334834 | -0.424602689 | -0.013447647 | 0.239848456 | 0.243675 | 0.194991919 | 0.047740661 | 0.937032733 |
| Cct4 | Y | 24 | 0.260918404 | -0.20215972 | -0.244853682 | -0.505772085 | -0.042693962 | 0.064640198 | 0.250673302 | 0.031661914 | 0.012493091 | 0.771626266 |
| Eps8 | Y | 674 | -0.269668959 | -0.206955756 | 0.287318249 | 0.556987208 | 0.494274005 | 0.066922289 | 0.078528519 | 0.035432023 | 0.007311188 | 0.004998726 |
| Arhgap42 | Y | 342 | 0.011917429 | 0.027887486 | -0.158527539 | -0.170444968 | -0.186415026 | 0.905598119 | 0.560232815 | 0.028278332 | 0.183107493 | 0.024067189 |
| Dst | Y | 555 | -0.242219511 | -0.376883699 | 0.290726974 | 0.532946485 | 0.667610673 | 0.564269378 | 0.276584406 | 0.450796875 | 0.259672841 | 0.120260047 |
| Plcg1 | Y | 1253 | -0.360381796 | 0.120493925 | 0.056978706 | 0.417360502 | -0.063515218 | 0.05739895 | 0.445915897 | 0.762352597 | 0.062057421 | 0.651531491 |
| Tes | Y | 236 | 0.138702127 | -0.218003843 | -0.036760263 | -0.17546239 | 0.18124358 | 0.035500094 | 0.004247416 | 0.519637472 | 0.020895184 | 0.01710441 |
| Hnrnpa3 | Y | 361 | -0.091037347 | -0.016950008 | -0.101887856 | -0.010850509 | -0.084937848 | 0.566355874 | 0.909901898 | 0.453615945 | 0.932609026 | 0.493649528 |
| Asap2 | S | 773 | 0.372519104 | -0.072531714 | -0.452428416 | -0.824947521 | -0.379896702 | 0.24858233 | 0.563190864 | 0.084945794 | 0.051408156 | 0.130572384 |
| Epha2 | T | 774 | 0.050973033 | -0.028303788 | -0.13785215 | -0.188825183 | -0.109548362 | 0.576016516 | 0.723764693 | 0.076722658 | 0.079824578 | 0.146867193 |
| Bcar3 | Y | 694 | 0.075155754 | -0.336067542 | -0.026353483 | -0.101509237 | 0.309714058 | 0.690311949 | 0.33815213 | 0.804937832 | 0.558646279 | 0.368195213 |
| Axl | Y | 475 | -0.023321652 | -0.095725542 | -0.013310325 | 0.010011327 | 0.082415217 | 0.730161175 | 0.215236962 | 0.875814086 | 0.907298055 | 0.379267003 |
| Caprin1 | Y | 660 | -0.026822792 | -0.128565574 | -0.073222475 | -0.046399683 | 0.055343099 | 0.730113493 | 0.629159442 | 0.673237924 | 0.781816801 | 0.847131026 |
| Emd | Y | 161 | 0.002175395 | 0.047436409 | -0.139530619 | -0.141706014 | -0.186967028 | 0.981645593 | 0.652799657 | 0.056137357 | 0.222206886 | 0.159686282 |
| Hnrnpa2b1 | Y | 336 | -0.009806226 | -0.061183274 | -0.052096469 | -0.042290244 | 0.009086805 | 0.917790649 | 0.296034943 | 0.546303404 | 0.687578109 | 0.901657367 |
| Epha2 | T | 772 | 0.123552869 | -0.027948101 | -0.199090885 | -0.322643754 | -0.171142784 | 0.214559724 | 0.736647773 | 0.036540696 | 0.028111149 | 0.054292002 |
| Etl4 | Y | 393 | -0.1383024 | 0.060942373 | -0.047701432 | 0.090600968 | -0.108643805 | 0.16678738 | 0.510747926 | 0.651643883 | 0.301296614 | 0.230011863 |
| Rpl7a | Y | 181 | 0.160507581 | -0.075090554 | -0.198440068 | -0.358947649 | -0.123349514 | 0.286351761 | 0.595279043 | 0.099975498 | 0.053224594 | 0.32979995 |
| Map2k3 Map2k6 | Y | 203;214 | -0.208388498 | -0.100198774 | 0.045971747 | 0.254360245 | 0.146170521 | 0.609305693 | 0.672242464 | 0.780984691 | 0.542149536 | 0.558612461 |
| Tnk2 | Y | 284 | -0.084107102 | -0.06812533 | 0.04070759 | 0.124814692 | 0.10883292 | 0.337156159 | 0.400645643 | 0.624257073 | 0.227490799 | 0.265051717 |
| Lrig2 | Y | 911 | 0.047419482 | -0.010595453 | -0.129462168 | -0.17688165 | -0.118866715 | 0.525011474 | 0.884213716 | 0.102933321 | 0.079752373 | 0.19187935 |
| Pard3b | Y | 998 | -0.091990273 | 0.075354282 | -0.061738333 | 0.03025194 | -0.137092615 | 0.231089047 | 0.573417847 | 0.400027637 | 0.702937488 | 0.345834171 |
| Gja1 | S | 282 | 0.226620004 | -0.138205335 | -0.175580844 | -0.402200847 | -0.037375508 | 0.338626338 | 0.392056947 | 0.199125598 | 0.142833573 | 0.818886531 |
| Larp1 | Y | 336 | 0.431663311 | -0.323059275 | -0.1154245 | -0.547087811 | 0.207634774 | 0.236940984 | 0.16086668 | 0.223761624 | 0.164249586 | 0.310253596 |
| Nphp4 | S | 1199 | -0.193271786 | 0.073403839 | 0.020001503 | 0.213273289 | -0.053402337 | 0.081937566 | 0.440799258 | 0.842198285 | 0.114980788 | 0.645025208 |
| Nphp4 | S | 1202 | -0.193271786 | 0.073403839 | 0.020001503 | 0.213273289 | -0.053402337 | 0.081937566 | 0.440799258 | 0.842198285 | 0.114980788 | 0.645025208 |
| Mark3 | Y | 508 | 0.097824406 | -0.023179637 | -0.224508597 | -0.322333003 | -0.201328959 | 0.582691578 | 0.884527989 | 0.245100461 | 0.074441781 | 0.170876462 |
| Asap1 | Y | 308 | 0.056570712 | -0.071265974 | -0.151491345 | -0.208062057 | -0.080225371 | 0.395680093 | 0.461955207 | 0.402182245 | 0.278325495 | 0.64835403 |
| Psmd11 | Y | 415 | 0.053185872 | -0.087100652 | -0.075754366 | -0.128940237 | 0.011346287 | 0.147277922 | 0.169311287 | 0.309678252 | 0.143849425 | 0.879318339 |
| Abl2 | Y | 568 | 0.293230209 | -0.105996837 | -0.143756751 | -0.43698696 | -0.037759914 | 0.343457273 | 0.435322115 | 0.066943627 | 0.205605667 | 0.766643363 |
| Zyx | Y | 308 | -0.045406644 | 0.031584947 | -0.120205723 | -0.074799079 | -0.15179067 | 0.475941479 | 0.644762031 | 0.487270523 | 0.648723674 | 0.392084822 |
| Antxr1 | Y | 423 | -0.03414679 | -0.042537117 | -0.015033201 | 0.019113589 | 0.027503916 | 0.648651177 | 0.503957748 | 0.746398439 | 0.794214591 | 0.655851519 |
| Pkm | Y | 390 | 0.152633742 | -0.159578558 | -0.070370304 | -0.223004046 | 0.089208255 | 0.232910071 | 0.176331981 | 0.278405466 | 0.127761799 | 0.380589787 |
| Megf10 | Y | 1061 | -0.10113315 | -0.027227881 | 0.060197308 | 0.161330458 | 0.087425189 | 0.146905666 | 0.609838557 | 0.454400682 | 0.118595822 | 0.327482927 |
| Fam171b | Y | 521 | 0.126900862 | 0.189962487 | -0.282019076 | -0.408919938 | -0.471981563 | 0.394140403 | 0.126532231 | 0.002505334 | 0.062312697 | 0.015097541 |
| Crcp | Y | 47 | 0.027106029 | -0.119469869 | -0.013653191 | -0.04075922 | 0.105816678 | 0.584571958 | 0.286095315 | 0.810802381 | 0.434145091 | 0.335573724 |
| Gpi1 | Y | 92 | 0.144501075 | -0.215982642 | -0.040438656 | -0.184939731 | 0.175543986 | 0.193844033 | 0.131709194 | 0.625251642 | 0.131839508 | 0.20291682 |
| G6pdx | Y | 401 | -0.011792458 | -0.030185926 | -0.044628976 | -0.032836518 | -0.01444305 | 0.809966193 | 0.511396191 | 0.725134649 | 0.798079834 | 0.909139941 |
| Iqgap1 | Y | 801 | -0.04537442 | 0.020417014 | -0.056218853 | -0.010844433 | -0.076635867 | 0.583696199 | 0.811236465 | 0.506338684 | 0.896915218 | 0.402236461 |
| Fermt2 | Y | 378 | -0.217323439 | 0.330298859 | -0.121942205 | 0.095381234 | -0.452241064 | 0.691261118 | NA | NA | NA | NA |
| Hnrnpa3 | Y | 365 | 0.100061901 | -0.184298106 | -0.034155767 | -0.134217668 | 0.150142339 | 0.35019216 | 0.253481712 | 0.759812603 | 0.331306579 | 0.373873648 |
| Pakap | Y | 439 | 0.114804503 | -0.129121441 | -0.069841391 | -0.184645894 | 0.05928005 | 0.08328767 | 0.044196355 | 0.647842678 | 0.285590055 | 0.694642274 |
| AA986860 | S | 130 | 0.08867573 | -0.221182334 | 0.066195083 | -0.022480647 | 0.287377417 | 0.409646599 | 0.092777998 | 0.634273118 | 0.8683714 | 0.100559192 |
| Pkp4 | Y | 1130 | -0.15571786 | -0.08334141 | 0.142807626 | 0.298525485 | 0.226149036 | 0.18812407 | 0.598383953 | 0.241840344 | 0.035461099 | 0.194921506 |
| Bcar1 | T | 389 | -0.092050877 | 0.022772582 | -0.009786151 | 0.082264726 | -0.032558732 | 0.349250406 | 0.807963448 | 0.969370587 | 0.746494514 | 0.896886736 |
| Zdhhc5 | Y | 456 | 0.143091681 | -0.35019273 | 0.088080628 | -0.055011053 | 0.438273358 | 0.661718746 | 0.236991708 | 0.610809071 | 0.853596864 | 0.156220387 |
| Abi1 | Y | 306 | -0.205170603 | -0.39906347 | 1.057754262 | 1.262924865 | 1.456817732 | 0.727844928 | NA | NA | NA | NA |
| Tes | Y | 239 | 0.029179033 | -0.083306517 | -0.018342199 | -0.047521232 | 0.064964318 | 0.79779316 | 0.405618735 | 0.844391409 | 0.556254286 | 0.173606249 |
| Rin1 | Y | 690 | 0.182593722 | -0.131837308 | -0.084972936 | -0.267566658 | 0.046864372 | 0.23970457 | 0.211763603 | 0.33858749 | 0.124365427 | 0.64439243 |
| Dhx8 | T | 740 | 0.061207486 | 0.074072278 | -0.263650396 | -0.324857882 | -0.337722675 | 0.580515019 | 0.601281305 | 0.2341123 | 0.167104723 | 0.155851018 |
| Dhx8 | T | 743 | 0.061207486 | 0.074072278 | -0.263650396 | -0.324857882 | -0.337722675 | 0.580515019 | 0.601281305 | 0.2341123 | 0.167104723 | 0.155851018 |
| Mcm7 | Y | 600 | 0.273882008 | -0.255965766 | -0.160339182 | -0.43422119 | 0.095626584 | 0.12862844 | 0.191437506 | 0.51909332 | 0.141795757 | 0.70507868 |
| Myo10 | Y | 1138 | -0.015511914 | -0.104023808 | 0.060436917 | 0.075948831 | 0.164460725 | 0.851577941 | 0.348079479 | 0.630777066 | 0.508651666 | 0.238424201 |
| Ghr | Y | 639 | -0.092627389 | 0.008305016 | 0.094123455 | 0.186750845 | 0.085818439 | 0.050464672 | 0.897363641 | 0.377924354 | 0.145211156 | 0.445010005 |
| Map4k3 | Y | 509 | -0.052378318 | -0.002893792 | -0.027445211 | 0.024933107 | -0.024551419 | 0.76384655 | 0.992698072 | 0.837801822 | 0.893830184 | 0.939331574 |
| Pik3r1 | Y | 556 | 0.182263743 | -0.174275953 | -0.102495757 | -0.284759501 | 0.071780196 | 0.072032658 | 0.150514152 | 0.398992124 | 0.057905666 | 0.555376668 |
| Abl1 Abl2 | Y | 218;172 | 0.241071737 | -0.156532781 | -0.117072957 | -0.358144694 | 0.039459824 | 0.128549887 | 0.340998338 | 0.137747992 | 0.054121276 | 0.793047924 |

| | | | | | | | | | | | | |
|---|---|---|---|---|---|---|---|---|---|---|---|---|
| Erbin | Y | 1019 | -0.076383615 | -0.001466867 | 0.03028934 | 0.106672955 | 0.031756207 | 0.393232597 | 0.993058495 | 0.79993642 | 0.372178516 | 0.859367021 |
| Arhgdia | T | 132 | -0.035616617 | 0.108161846 | -0.213746349 | -0.178129732 | -0.321908195 | 0.667923393 | 0.220073352 | 0.355396794 | 0.425904606 | 0.207988428 |
| Arhgef10 | Y | 628 | -0.013275877 | -0.098583702 | 0.077679068 | 0.090954945 | 0.176262771 | 0.886193657 | 0.331623352 | 0.627883667 | 0.533883837 | 0.281179001 |
| Dcbld1 | Y | 413 | 0.110018265 | -0.254520578 | 0.043093683 | -0.066924582 | 0.297614261 | 0.558973101 | 0.347984568 | 0.84051864 | 0.788630544 | 0.340785477 |
| Pfn1 | Y | 129;129 | -0.01881147 | -0.240312131 | 0.213020759 | 0.231832229 | 0.45333289 | 0.851558016 | 0.062262359 | 0.097989957 | 0.094053439 | 0.01426532 |
| Ldlr | Y | 830 | 0.059610877 | -0.130051045 | 0.04779457 | -0.011816307 | 0.177845616 | 0.496151412 | 0.159248911 | 0.738288498 | 0.931328064 | 0.257887977 |
| Mgrn1 | Y | 389 | -0.138664737 | -0.332585432 | 0.023988882 | 0.162653619 | 0.356574314 | 0.826667804 | NA | NA | NA | NA |
| Rbm39 | S | 97 | 0.188711448 | -0.118612948 | -0.113168097 | -0.301879545 | 0.005444851 | 0.15234384 | 0.376225412 | 0.217990902 | 0.051839165 | 0.961671912 |
| Rbm39 | Y | 99 | 0.188711448 | -0.118612948 | -0.113168097 | -0.301879545 | 0.005444851 | 0.15234384 | 0.376225412 | 0.217990902 | 0.051839165 | 0.961671912 |
| Abl2 | Y | 231 | 0.12164261 | -0.253239788 | 0.051494807 | -0.070147803 | 0.304734595 | 0.284061088 | 0.175844599 | 0.779214981 | 0.70971966 | 0.204309581 |
| Pik3r2 | Y | 458 | 0.078632953 | -0.099910938 | -0.026394288 | -0.10502724 | 0.07351665 | 0.14740724 | 0.14553039 | 0.65219364 | 0.084039981 | 0.236854048 |
| Pdcd5 | Y | 80 | 0.042742616 | -0.020991461 | -0.068146493 | -0.110889108 | -0.047155031 | 0.63655079 | 0.824613139 | 0.469444586 | 0.299340576 | 0.652953599 |
| Eif4b | Y | 237 | 0.25651773 | -0.323044751 | -0.013107876 | -0.269625606 | 0.309936875 | 0.042124194 | 0.042603853 | 0.906776953 | 0.045262208 | 0.05484148 |
| Ddb1 | Y | 718 | 0.082897145 | -0.107083665 | -0.069885959 | -0.152783104 | 0.037197705 | 0.053487076 | 0.209512715 | 0.482060702 | 0.192483866 | 0.730467902 |
| Epha2 | Y | 589 | 0.01456895 | -0.030186842 | -0.022796123 | -0.037365073 | 0.007390719 | 0.771393916 | 0.654221225 | 0.723089953 | 0.450561646 | 0.906981793 |
| Kirrel3 | Y | 689 | -0.101326383 | 0.036159642 | 0.047376743 | 0.148703127 | 0.011217102 | 0.2484679 | 0.650173409 | 0.690082677 | 0.236595391 | 0.916554598 |
| Myo10 | T | 1131 | 0.036426193 | 0.000758668 | -0.082556969 | -0.118983162 | -0.083315636 | 0.709305096 | 0.994881824 | 0.526575567 | 0.41933453 | 0.595348827 |
| Vasp | Y | 39 | 0.15450536 | -0.087307443 | -0.126532098 | -0.281037458 | -0.039224655 | 0.024373863 | 0.437641628 | 0.058220365 | 0.004485646 | 0.711941897 |
| Vim | S | 420 | 0.164090595 | -0.143889912 | -0.134680748 | -0.298771343 | 0.009209164 | 0.700868677 | 0.804187655 | 0.804142207 | 0.53308282 | 0.987776369 |
| Anxa6 | Y | 201 | NA | -0.08384143 | -0.240697334 | NA | -0.156855904 | NA | 0.881783097 | NA | NA | NA |
| Nhsl1 | S | 444 | 0.079853268 | -0.050134569 | -0.094632732 | -0.174486 | -0.044498163 | 0.497059647 | 0.670597493 | 0.679919188 | 0.447022727 | 0.835692726 |
| Eps15 | Y | 850 | 0.189526756 | 0.014016355 | -0.179491438 | -0.369018194 | -0.193507794 | 0.482208298 | 0.930353324 | 0.249932421 | 0.213203656 | 0.209396021 |
| Pxdn | S | 868 | -0.136209813 | 0.052720355 | 0.004191104 | 0.140400917 | -0.04852925 | 0.574494689 | 0.571033365 | 0.952267794 | 0.564241925 | 0.607949112 |
| Wasl | Y | 253 | -0.15659447 | 0.177243386 | -0.067328126 | 0.089266344 | -0.244571511 | 0.147338814 | 0.107923029 | 0.668482618 | 0.55290633 | 0.171894065 |
| Kirrel | Y | 654 | 0.037638068 | -0.032520477 | -0.028698617 | -0.066336685 | 0.003821861 | 0.499141585 | 0.577599728 | 0.659067415 | 0.227399334 | 0.940248143 |
| Fam171b | Y | 569 | 0.210705893 | 0.08756401 | -0.246033622 | -0.456739514 | -0.333597632 | 0.137671542 | 0.595746816 | 0.028113812 | 0.034682959 | 0.122667384 |
| Myh9 | Y | 11 | 0.077024329 | -0.234249606 | 0.247468806 | 0.170444476 | 0.481718411 | 0.70161664 | 0.120316687 | 0.423120907 | 0.583549571 | 0.1801032 |
| Rack1 | T | 50 | 0.324929616 | -0.280578055 | 0.038437301 | -0.286492315 | 0.319015356 | 0.314203662 | 0.101018432 | 0.616704591 | 0.3617085 | 0.067495774 |
| Pkp4 | Y | 1154 | -0.04454604 | 0.049297729 | -0.046550415 | -0.002004375 | -0.095848144 | 0.691818335 | 0.510059259 | 0.788239358 | 0.991280513 | 0.584524558 |
| Afdn | Y | 584 | -0.138008496 | -0.066665736 | 0.184168596 | 0.322177092 | 0.250834332 | 0.206309252 | 0.656534682 | 0.144519267 | 0.028700939 | 0.129326289 |
| Mpzl1 | Y | 264 | 0.099352567 | -0.080430126 | -0.030836409 | -0.130188976 | 0.049593717 | 0.058559188 | 0.115715363 | 0.477110675 | 0.016487442 | 0.218945707 |
| Hnrnpf | Y | 210 | 0.17420877 | -0.092025018 | -0.07544284 | -0.24965161 | 0.016582179 | 0.219305879 | 0.532721115 | 0.283594199 | 0.114960991 | 0.908879952 |
| Arap2 | Y | 332 | 0.383254554 | -0.080196195 | -0.11539802 | -0.498652575 | -0.035201826 | 0.297086252 | 0.532348515 | 0.126684794 | 0.210671426 | 0.765382269 |
| Txnrd1 | Y | 245 | 0.059716237 | -0.045901623 | -0.053424201 | -0.113140438 | -0.007522578 | 0.022303651 | 0.320686 | 0.236506476 | 0.068611183 | 0.881755621 |
| Jcad | Y | 113 | 0.103519132 | 0.014760702 | -0.100309507 | -0.203828639 | -0.115070209 | 0.426898426 | 0.785291175 | 0.245505081 | 0.185010855 | 0.216381793 |
| Hsp90aa1 Hsp90ab1 | Y | 285;276 | 0.362430771 | -0.167577809 | -0.103628813 | -0.466059584 | 0.063948996 | 0.300655673 | 0.315338616 | 0.586244838 | 0.217869922 | 0.76260531 |
| Epn2 | Y | 186 | -0.072246427 | -0.005930971 | 0.077002014 | 0.149248441 | 0.082932985 | 0.110777413 | 0.901939456 | 0.111460107 | 0.008781142 | 0.106210449 |
| Myh9 | Y | 9 | 0.051038555 | -0.195858215 | 0.259607495 | 0.20856894 | 0.45546571 | 0.799943583 | 0.170757765 | 0.421560243 | 0.524275583 | 0.211541282 |
| Vim | S | 419 | 1.245096021 | 1.035359435 | NA | NA | NA | NA | NA | NA | NA | NA |
| Flnb | S | 1528 | NA | NA | 0.293835474 | NA | NA | NA | NA | NA | NA | NA |
| Pik3r2 | Y | 74 | NA | NA | NA | NA | NA | NA | NA | NA | NA | NA |
| Lyn | Y | 316 | 0.101000984 | NA | 0.226996628 | 0.125995644 | NA | NA | NA | NA | NA | NA |
| Jak1 | Y | 412 | NA | NA | NA | NA | NA | NA | NA | NA | NA | NA |
| Rars | Y | 536 | NA | NA | NA | NA | NA | NA | NA | NA | NA | NA |
| Rexo2 | Y | 122 | 0.172015501 | 0.428981292 | NA | NA | NA | NA | NA | NA | NA | NA |
| Vps35 | Y | 791 | NA | NA | NA | NA | NA | NA | NA | NA | NA | NA |
| Ptpra | S | 823 | 0.538697551 | 0.581317581 | 0.542407731 | 0.00371018 | -0.03890985 | 0.008597173 | 0.004510431 | 0.020807891 | 0.981062983 | 0.786094434 |
| Emg1 | Y | 222 | 0.536626676 | 0.353505354 | 0.528647913 | -0.007978762 | 0.17514256 | 0.039890325 | 0.008177122 | 0.000463342 | 0.959617272 | 0.090655848 |
| Rras2 | Y | 105 | 0.327324724 | 0.39558661 | 0.570897338 | 0.243572614 | 0.175310728 | 0.007351732 | 0.006936801 | 0.002563948 | 0.068320381 | 0.153937454 |
| Flna | Y | 373 | 0.298434549 | 0.546676408 | 0.468961466 | 0.170526917 | -0.077714942 | 0.019978422 | 0.002113483 | 0.004306876 | 0.108694781 | 0.356925359 |
| Nck2 | S | 90 | 0.351228052 | 0.525885286 | 0.483103811 | 0.131875758 | -0.042781476 | 0.085731221 | 0.002312116 | 0.001428474 | 0.412374398 | 0.64334733 |
| Clip3 | Y | 280 | 1.017432069 | 0.91833086 | 0.56394582 | -0.453486249 | -0.35438504 | 0.128827623 | 0.064127523 | 0.03726623 | 0.389712476 | 0.318361362 |
| Anxa2 | Y | 333 | 0.352069431 | 0.516641295 | 0.563444575 | 0.211375144 | 0.04680328 | 0.017486854 | 0.047399719 | 0.001988284 | 0.033904396 | 0.787176189 |
| Nedd9 | Y | 213 | 0.305755888 | 0.532202523 | 0.524269708 | 0.21851382 | -0.007932815 | 0.216778229 | 0.002424526 | 0.001619132 | 0.34129922 | 0.936772982 |
| Nedd9 | Y | 222 | 0.305755888 | 0.532202523 | 0.524269708 | 0.21851382 | -0.007932815 | 0.216778229 | 0.002424526 | 0.001619132 | 0.34129922 | 0.936772982 |
| Asap1 | Y | 782 | 0.820510855 | 0.814479237 | 0.95972636 | 0.139215505 | 0.145247123 | 0.051089607 | 0.234647778 | 0.060315636 | 0.687600789 | 0.787833491 |
| Nedd9 | T | 218 | 0.324424436 | 0.48914308 | 0.441187657 | 0.116763221 | -0.047955423 | 0.33384765 | 0.002682671 | 0.000161735 | 0.695138544 | 0.529245607 |
| Tln1 | Y | 1116 | 0.400494506 | 0.38224149 | 0.447342032 | 0.046847526 | 0.065100542 | 0.005958562 | 0.016067702 | 0.004957572 | 0.614841038 | 0.556980781 |
| Kank2 | Y | 111 | 0.242731765 | 0.321197552 | 0.384127862 | 0.141396096 | 0.06293031 | 0.001197696 | 0.037631952 | 0.000101532 | 0.01176552 | 0.514007786 |
| Pcdhga4 | Y | 894 | 0.181984888 | 0.444647156 | 0.525743605 | 0.343758717 | 0.081096449 | 0.018632698 | 0.050201727 | 0.000260778 | 0.004176587 | 0.577681284 |
| Shc1 | Y | 423 | 0.296134955 | 0.40277957 | 0.455455586 | 0.15932063 | 0.052676015 | 0.226641392 | 0.029134777 | 2.24643E-05 | 0.455156512 | 0.603354506 |
| Tns2 | S | 481 | 0.001219537 | 0.564052389 | 0.534704203 | 0.533484667 | -0.029348185 | 0.990143433 | 0.00036794 | 0.000850499 | 0.004023152 | 0.69037723 |
| Anxa2 | Y | 317 | 0.346771611 | 0.380536277 | 0.345510779 | -0.001260831 | -0.035025498 | 0.004550405 | 0.00383978 | 0.013464898 | 0.988354784 | 0.70200847 |
| Sdc2 | Y | 201 | 0.074884963 | 0.464869052 | 0.400451352 | 0.325566389 | -0.0644177 | 0.314587726 | 8.07922E-05 | 0.001143865 | 0.009429557 | 0.283553458 |
| Dnaja2 | Y | 69 | 0.322203515 | 0.321564764 | 0.264127149 | -0.058076366 | -0.057437615 | 0.002535698 | 0.004493509 | 0.001402654 | 0.3190146 | 0.365235277 |
| Prpf4b | Y | 849 | 0.528590655 | 0.355582599 | 0.447921122 | -0.080669533 | 0.092338524 | 0.022999203 | 0.0294137 | 0.010045953 | 0.60990286 | 0.48143445 |
| Elmo2 | Y | 48 | 0.336520082 | 0.471636648 | 0.557919675 | 0.221399593 | 0.086283027 | 0.042606903 | 0.008425052 | 0.031341951 | 0.238254116 | 0.601685307 |
| Peak1 | S | 612 | 0.755775619 | 0.49595268 | 0.421322684 | -0.334452935 | -0.074629997 | 0.353181543 | 0.012533618 | 0.005555505 | 0.649230966 | 0.541201318 |
| Esyt1 | Y | 997 | 0.559989424 | 0.425754622 | 0.485845311 | -0.074144113 | 0.060090689 | 0.192638807 | 0.098097424 | 0.000710003 | 0.824152945 | 0.738297505 |
| Lyn | Y | 397 | 0.472837888 | 0.567937164 | 0.123499176 | -0.349338712 | -0.444437988 | 0.00537107 | 0.002342993 | 0.311102263 | 0.009898799 | 0.004321788 |
| H2-K1 | Y | 301;341;342;342 | 0.093613536 | 0.487526837 | 0.516406335 | 0.422792799 | 0.028879498 | 0.235022685 | 0.002720148 | 0.00294122 | 0.012341901 | 0.778097675 |
| Afap1l2 | S | 414 | 0.467163894 | 0.567414576 | 0.43175007 | -0.035413824 | -0.135664506 | 0.045556023 | 0.010805025 | 0.10074819 | 0.874767203 | 0.530771134 |
| Cep55 | Y | 377 | 0.22303621 | 0.393064355 | 0.484660035 | 0.261623825 | 0.091595681 | 0.037494805 | 0.01856078 | 0.002882159 | 0.033385391 | 0.391715173 |
| Dlg5 | S | 430 | 0.309066718 | 0.34271447 | 0.524579085 | 0.215512366 | 0.181864614 | 0.006760381 | 0.047495562 | 0.013535542 | 0.150081865 | 0.263491113 |
| Stat6 | Y | 641 | 0.042121802 | 0.310398475 | 0.454062187 | 0.411940386 | 0.143663713 | 0.572678589 | 0.00056929 | 6.622E-05 | 0.008439889 | 0.020213101 |
| H2-D1 H2-L | Y | 154;344;344 | 0.123317951 | 0.377707388 | 0.381455315 | 0.258137365 | 0.003747927 | 0.084606013 | 0.001262702 | 0.000992158 | 0.009740315 | 0.940672944 |
| Hipk3 | Y | 359 | 0.277110228 | 0.380907937 | 0.489473735 | 0.212363506 | 0.108565798 | 0.010192198 | 0.010254706 | 0.030235543 | 0.168200484 | 0.435335825 |
| Layn | Y | 380 | 0.24067151 | 0.293747999 | 0.314625172 | 0.073953662 | 0.020877173 | 0.007254296 | 0.001012975 | 0.008130633 | 0.321037079 | 0.74882805 |
| Anks1 | S | 462 | 0.448626556 | 0.364294636 | 0.483597105 | 0.034970549 | 0.119302469 | 0.035032929 | 0.169098042 | 0.002360869 | 0.795681601 | 0.578654952 |
| Sorbs2 | Y | 62 | 0.284448515 | 0.316085579 | 0.515170571 | 0.230722056 | 0.199084992 | 0.049335043 | 0.093882857 | 0.000997231 | 0.087042137 | 0.229037456 |
| Anxa2 | Y | 318 | 0.269205528 | 0.263890466 | 0.35028054 | 0.081075012 | 0.086390074 | 0.016668803 | 0.005670562 | 0.002611908 | 0.327060216 | 0.211551429 |
| Cdk1 | T | 14 | 0.482850059 | 0.406152517 | 0.375884239 | -0.10696582 | -0.030268278 | 0.060247158 | 0.035273519 | 0.009710332 | 0.55713205 | 0.813504964 |
| Git2 | Y | 541 | 0.462603679 | 0.452894677 | 0.554639263 | 0.092035584 | 0.101744586 | 0.036674443 | 0.145510583 | 0.02108604 | 0.340416461 | 0.667046755 |
| Sgms2 | Y | 59 | 0.143065607 | 0.437517036 | 0.464067673 | 0.321002066 | 0.026550637 | 0.213246316 | 0.002470229 | 0.005922806 | 0.034488684 | 0.776787627 |
| Peak1 | S | 1203 | 0.226326983 | 0.410395976 | 0.425112818 | 0.198785835 | 0.014716843 | 0.193647791 | 0.054315823 | 0.000171379 | 0.237778181 | 0.91034871 |
| Ptk2 | S | 574 | 0.152734495 | 0.472209894 | 0.456705612 | 0.303971117 | -0.015504282 | 0.141161305 | 0.018544874 | 0.00178261 | 0.026166632 | 0.896716678 |
| Kirrel | Y | 679 | 0.174358904 | 0.247339993 | 0.476100808 | 0.301741904 | 0.228760815 | 0.008230563 | 0.019633451 | 0.002836996 | 0.029172228 | 0.044253214 |
| Vta1 | S | 283 | 0.390428425 | 0.370633839 | 0.449191857 | 0.058763432 | 0.078558019 | 0.363261517 | 0.02312815 | 0.001017196 | 0.877417405 | 0.47019764 |
| Peak1 | Y | 1204 | 0.237282366 | 0.42671223 | 0.488811631 | 0.251529265 | 0.062099401 | 0.213361473 | 0.023345608 | 0.002896091 | 0.190800324 | 0.605856217 |
| Irs2 | Y | 758 | 0.843596754 | 0.44260084 | 0.58101826 | -0.262578494 | 0.13841742 | 0.13198286 | 0.151205827 | 0.046210563 | 0.547716107 | 0.623398265 |
| Tns3 | Y | 849 | 0.438567015 | 0.388388138 | 0.413461295 | -0.025105721 | 0.025073156 | 0.322410201 | 0.023841329 | 0.001588714 | 0.947448377 | 0.802065453 |

| | | | | | | | | | | | | | |
|---|---|---|---|---|---|---|---|---|---|---|---|---|---|
| Nck1 | Y | | 105 | 0.411810727 | 0.430609805 | 0.385713899 | -0.026096828 | -0.044895906 | 0.197659811 | 0.011270715 | 0.009947443 | 0.919576441 | 0.69287234 |
| Hipk1 Hipk2 | Y | 361;352 | | 0.253390633 | 0.349313516 | 0.44996994 | 0.196579308 | 0.100656425 | 0.057311463 | 0.018182621 | 0.006857143 | 0.138730312 | 0.384936869 |
| Prpf4b | S | | 839 | 0.595429887 | 0.500454411 | 0.510251947 | -0.08517794 | 0.009797536 | 0.2769613 | 0.0710937 | 0.012479116 | 0.855227749 | 0.962475224 |
| Naxd | Y | | 81 | 0.355593735 | 0.459789484 | 0.459321177 | 0.103727442 | -0.000468307 | 0.233011355 | 0.016692155 | 0.012723013 | 0.683686587 | 0.997006531 |
| Sdcbp | Y | | 47 | 0.362730955 | 0.333527107 | 0.583275661 | 0.220544706 | 0.249748555 | 0.20889223 | 0.005737102 | 0.028546075 | 0.423115513 | 0.196195347 |
| Cavin2 | Y | | 103 | 0.197606198 | 0.297963208 | 0.584255402 | 0.386649203 | 0.286292194 | 0.124037638 | 0.04214551 | 0.00237598 | 0.014952948 | 0.045085431 |
| Cyfip1 Cyfip2 | Y | 836;835 | | 0.670936495 | 0.449388901 | 0.583410535 | -0.08752596 | 0.134021633 | 0.107572546 | 0.193555887 | 0.035052155 | 0.790627751 | 0.658524388 |
| Caskin2 | Y | | 253 | 0.505439009 | 0.394047694 | 0.411217327 | -0.094221682 | 0.017169633 | 0.331939467 | 0.031644971 | 0.003076835 | 0.835501225 | 0.882133963 |
| Hgs | Y | | 216 | 0.124733497 | 0.301831178 | 0.491574089 | 0.366840591 | 0.18974291 | 0.213452399 | 0.053285611 | 0.000156847 | 0.019899402 | 0.139049708 |
| Tln1 | Y | | 1737 | 0.458787097 | 0.358974723 | 0.565977651 | 0.107190554 | 0.207002928 | 0.201885442 | 0.05767137 | 0.011838211 | 0.723181104 | 0.249717632 |
| Esyt1 | Y | | 809 | 0.267548422 | 0.335694749 | 0.520187479 | 0.252639056 | 0.18449273 | 0.154381583 | 0.072610914 | 0.002120794 | 0.173983109 | 0.242273089 |
| Src | Y | | 192 | 0.464724395 | 0.268435922 | 0.453244682 | -0.011479713 | 0.18480876 | 0.007436935 | 0.145812039 | 0.045939271 | 0.941028225 | 0.350277246 |
| Myh9 | Y | | 1408 | 0.278177752 | 0.323559663 | 0.414117915 | 0.135940163 | 0.090558252 | 0.018531074 | 0.023453613 | 0.026459787 | 0.290942479 | 0.482957114 |
| Ptk2 | Y | | 576 | 0.130789001 | 0.289255147 | 0.254191169 | 0.123402168 | -0.035063978 | 0.029283476 | 0.000939585 | 0.001631054 | 0.022597901 | 0.312524214 |
| Unc5b | Y | | 457 | 0.467272277 | 0.23637843 | 0.558152794 | 0.090880517 | 0.321774364 | 0.097935404 | 0.121535452 | 0.009379876 | 0.683578731 | 0.061387419 |
| Myh9 | Y | | 754 | 0.269661022 | 0.230932825 | 0.486875519 | 0.217214497 | 0.255942693 | 0.050658097 | 0.016951584 | 0.010570299 | 0.139131239 | 0.080361365 |
| Bcar1 | Y | | 271 | 0.879526484 | 0.482177196 | 0.305893406 | -0.573633079 | -0.17628379 | 0.239186286 | 0.035284165 | 0.04879605 | 0.396219092 | 0.29212129 |
| Sh2b3 | Y | | 536 | 0.162862403 | 0.199684716 | 0.481205337 | 0.318342934 | 0.28152062 | 0.049742937 | 0.066305647 | 0.000485534 | 0.001637956 | 0.028653616 |
| Dnajc10 | Y | | 751 | 0.381511474 | 0.36559898 | 0.344575768 | -0.036935706 | -0.021023212 | 0.133015188 | 0.002515058 | 0.058198153 | 0.860572748 | 0.866770098 |
| Gcnt4 | S | | 289 | 0.400921452 | 0.383757689 | 0.323238299 | -0.077683154 | -0.060519391 | 0.043254433 | 0.005087705 | 0.140689842 | 0.695744577 | 0.721120762 |
| Gcnt4 | S | | 303 | 0.400921452 | 0.383757689 | 0.323238299 | -0.077683154 | -0.060519391 | 0.043254433 | 0.005087705 | 0.140689842 | 0.695744577 | 0.721120762 |
| Gcnt4 | T | | 286 | 0.400921452 | 0.383757689 | 0.323238299 | -0.077683154 | -0.060519391 | 0.043254433 | 0.005087705 | 0.140689842 | 0.695744577 | 0.721120762 |
| Gcnt4 | Y | | 305 | 0.400921452 | 0.383757689 | 0.323238299 | -0.077683154 | -0.060519391 | 0.043254433 | 0.005087705 | 0.140689842 | 0.695744577 | 0.721120762 |
| Unk | Y | | 237 | 0.468355517 | 0.235261594 | 0.41693686 | -0.051418657 | 0.181675267 | 0.0131332 | 0.316129226 | 0.010152817 | 0.644255084 | 0.407872173 |
| Atp5a1 | Y | | 337 | 0.343644035 | 0.467473258 | 0.749836649 | 0.406192614 | 0.282363391 | 0.025936515 | 0.064197021 | 0.215605809 | 0.440761462 | 0.583724256 |
| Micall2 | Y | | 59 | 0.336760358 | 0.243802745 | 0.550876727 | 0.214116369 | 0.307073982 | 0.059045531 | 0.027484741 | 0.029077465 | 0.27247416 | 0.135959396 |
| Hgs | Y | | 286 | 0.119706946 | 0.358353797 | 0.474839869 | 0.355132922 | 0.116486072 | 0.325343874 | 0.060199572 | 0.000387447 | 0.058327305 | 0.390695883 |
| Bcar1 | Y | | 331 | 0.472458586 | 0.299991865 | 0.320905935 | -0.15155265 | 0.02091407 | 0.0210652 | 0.053543633 | 0.042522143 | 0.269089307 | 0.743109651 |
| Git1 | S | | 397 | 0.1371783 | 0.420273281 | 0.422584856 | 0.285406556 | 0.002311576 | 0.345587691 | 0.010298995 | 0.004338023 | 0.089797062 | 0.98092436 |
| Git1 | Y | | 392 | 0.1371783 | 0.420273281 | 0.422584856 | 0.285406556 | 0.002311576 | 0.345587691 | 0.010298995 | 0.004338023 | 0.089797062 | 0.98092436 |
| Myh10 Myh14 Myh9 | Y | 400;407;430 | | 0.240705911 | 0.284642169 | 0.466739643 | 0.226033732 | 0.182097474 | 0.05363587 | 0.011638867 | 0.025642177 | 0.163949586 | 0.225661416 |
| Tmod3 | Y | | 86 | 0.344959544 | 0.240458536 | 0.409841824 | 0.06488228 | 0.169383288 | 0.023471085 | 0.025999004 | 0.030269203 | 0.642678598 | 0.233688029 |
| Cdk2 Cdk3 | T | 14;14 | | 0.493637436 | 0.338306232 | 0.413833006 | -0.079804431 | 0.075526773 | 0.081800695 | 0.037925003 | 0.053253404 | 0.728598114 | 0.646366843 |
| Actn1 | Y | | 193 | 0.287818515 | 0.207593742 | 0.579381376 | 0.291562861 | 0.371787634 | 0.007738948 | 0.198235797 | 0.021120837 | 0.118699177 | 0.084932088 |
| Unk | Y | | 236 | 0.348137616 | 0.215337829 | 0.370051506 | 0.02191389 | 0.154713676 | 0.009554808 | 0.308890098 | 0.002650384 | 0.766870849 | 0.434939713 |
| Afap1l2 | Y | | 413 | 0.509345438 | 0.482277685 | 0.37382773 | -0.135517708 | -0.108449955 | 0.15715319 | 0.010539963 | 0.19723506 | 0.691060227 | 0.652078896 |
| Pxn | Y | | 76 | 0.065040563 | 0.321175042 | 0.479658492 | 0.414617929 | 0.15848345 | 0.500437187 | 0.022478857 | 0.000491292 | 0.021075981 | 0.101328155 |
| Nck2 | Y | | 110 | 0.273445025 | 0.372480374 | 0.328987424 | 0.055542399 | -0.043492951 | 0.299175097 | 0.008936272 | 0.004834597 | 0.807795354 | 0.598995059 |
| Crkl | Y | | 251 | 0.025604461 | 0.233887624 | 0.342945182 | 0.317340721 | 0.109057559 | 0.768995567 | 0.000693629 | 5.49903E-05 | 0.041663351 | 0.035859399 |
| Prpf31 | Y | | 205 | 0.437557366 | 0.240712516 | 0.361384918 | -0.076172447 | 0.120672402 | 0.043478916 | 0.005905667 | 0.088293481 | 0.67643517 | 0.44260047 |
| Dyrk1a Dyrk1b | Y | 321;273 | | 0.239371174 | 0.328335491 | 0.40631579 | 0.166944616 | 0.077980299 | 0.007089352 | 0.106873356 | 0.024984648 | 0.184216409 | 0.637897295 |
| Tjp1 | Y | | 1145 | 0.06642119 | 0.208047434 | 0.354888066 | 0.288466875 | 0.146840631 | 0.505708419 | 0.00153369 | 0.000114269 | 0.059776 | 0.009554171 |
| Mtcp1 | Y | | 25 | 0.19476839 | 0.291716162 | 0.436207526 | 0.241439136 | 0.144491364 | 0.050808117 | 0.064711783 | 0.005869081 | 0.039984059 | 0.24813312 |
| Cdc42ep1 | S | | 121 | 0.506205115 | 0.471078916 | 0.063476313 | -0.442728802 | -0.407602603 | 0.017600055 | 0.007023964 | 0.570156396 | 0.041858755 | 0.010323971 |
| Stam | Y | | 219 | 0.527362859 | 0.387063271 | 0.487238228 | -0.04012463 | 0.100174957 | 0.126614756 | 0.14598994 | 0.043979641 | 0.885991877 | 0.678265654 |
| Sirpa | Y | | 440 | 0.447890663 | 0.428232053 | 0.181016645 | -0.266874018 | -0.247215408 | 0.078812333 | 0.003252306 | 0.250275132 | 0.237360334 | 0.142119496 |
| Blk Lyn | Y | 194;182 | | 0.188889172 | 0.283750994 | 0.431845876 | 0.242956704 | 0.148094882 | 0.068608915 | 0.094381443 | 0.003002364 | 0.045557199 | 0.296930435 |
| Cpd | Y | | 1341 | 0.301792732 | 0.439025744 | 0.417918356 | 0.116125624 | -0.021107387 | 0.137051957 | 0.138662862 | 0.009558853 | 0.499671246 | 0.926632979 |
| Vim | Y | | 319 | 0.464667192 | 0.287452474 | 0.555805181 | 0.091137989 | 0.268352707 | 0.087600135 | 0.163216697 | 0.044968761 | 0.707367133 | 0.264964278 |
| Ptk2 | Y | | 720 | 0.141017391 | 0.30852719 | 0.534711157 | 0.393693766 | 0.226183967 | 0.351484883 | 0.037892815 | 0.004310151 | 0.034937729 | 0.024569694 |
| Anxa2 | Y | | 238 | 0.21930687 | 0.292834825 | 0.36207153 | 0.142764661 | 0.069236705 | 0.093565706 | 0.089528196 | 0.001510611 | 0.218692438 | 0.584817035 |
| Ndc80 | Y | | 458 | 0.395147134 | 0.274228658 | 0.327760672 | -0.067386462 | 0.053532014 | 0.025327061 | 0.045417889 | 0.072817894 | 0.663973902 | 0.713439111 |
| Erbin | Y | | 917 | 0.93196216 | 0.576889064 | 0.617958086 | -0.314004074 | 0.041069022 | 0.089262175 | 0.470149915 | 0.283089895 | 0.5930934 | 0.95713636 |
| Notch2 | Y | | 2342 | 0.476940187 | 0.34802831 | 0.437578589 | -0.039361597 | 0.089550279 | 0.386878472 | 0.064908482 | 0.011995961 | 0.936417442 | 0.52813385 |
| Peak1 | Y | | 638 | 0.482601184 | 0.240689168 | 0.682814463 | 0.20021328 | 0.442125295 | 0.105381712 | 0.179128414 | 0.069452759 | 0.525434865 | 0.168922752 |
| Anxa2 | Y | | 30 | 0.256632351 | 0.237790265 | 0.409754501 | 0.153122149 | 0.171964236 | 0.021048864 | 0.008583854 | 0.097904503 | 0.418834976 | 0.368324835 |
| Anxa2 | Y | | 316 | 0.170411793 | 0.295738953 | 0.357488183 | 0.18707639 | 0.06174923 | 0.08632986 | 0.095951807 | 0.001155148 | 0.068609524 | 0.637673151 |
| Ephb3 | Y | | 787 | 0.434091347 | 0.291054899 | 0.455906788 | 0.021815441 | 0.164851889 | 0.306179217 | 0.200989598 | 0.002880056 | 0.951923527 | 0.411307421 |
| Fyb | Y | | 793 | 0.112471341 | 0.167742897 | 0.535207405 | 0.422736064 | 0.367464507 | 0.235165924 | 0.309082662 | 0.000137898 | 0.012410829 | 0.0893939 |
| Gsk3a Gsk3b | S | 219;282 | | 0.243452193 | 0.36937933 | 0.217195545 | -0.026256648 | -0.152183785 | 0.029737679 | 0.01811279 | 0.027674016 | 0.687664031 | 0.188064504 |
| Lasp1 Nebl | Y | 57;57 | | 0.177272946 | 0.274832888 | 0.252764537 | 0.075491591 | -0.022068351 | 0.033328142 | 0.007065556 | 0.010187763 | 0.056192527 | 0.627120522 |
| Actb Actg1 | Y | 218;218 | | 0.388101501 | 0.347321577 | 0.37122612 | -0.016875381 | 0.023904543 | 0.120194769 | 0.078767466 | 0.025199765 | 0.931850558 | 0.877713469 |
| Vim | Y | | 117 | 0.376810786 | 0.202704003 | 0.647404323 | 0.270593537 | 0.444700321 | 0.049975151 | 0.16321073 | 0.06102482 | 0.317294638 | 0.146470423 |
| Hdlbp | Y | | 437 | 0.302008438 | 0.264731289 | 0.492632037 | 0.190623599 | 0.227900747 | 0.035196703 | 0.149920565 | 0.03493623 | 0.275968058 | 0.263906879 |
| Ighm | S | | 172 | 0.206871668 | 1.160891873 | 1.567337531 | 1.360465863 | 0.406445658 | 0.679609114 | 0.29536813 | 0.228147473 | 0.279646139 | 0.760607952 |
| Ighm | T | | 173 | 0.206871668 | 1.160891873 | 1.567337531 | 1.360465863 | 0.406445658 | 0.679609114 | 0.29536813 | 0.228147473 | 0.279646139 | 0.760607952 |
| Ighm | T | | 176 | 0.206871668 | 1.160891873 | 1.567337531 | 1.360465863 | 0.406445658 | 0.679609114 | 0.29536813 | 0.228147473 | 0.279646139 | 0.760607952 |
| Cdkl5 | Y | | 171 | 0.104993182 | 0.262787526 | 0.427837433 | 0.322844251 | 0.165049907 | 0.099976179 | 0.017878091 | 0.006223013 | 0.034806208 | 0.134428915 |
| Qars | Y | | 248 | 0.533728343 | 0.237144284 | 0.534165149 | 0.000436806 | 0.297020865 | 0.080227571 | 0.422882739 | 0.028559198 | 0.998437122 | 0.302561162 |
| Cltc | Y | | 899 | 0.302445823 | 0.260633543 | 0.649931434 | 0.347485612 | 0.389297892 | 0.104511574 | 0.059800979 | 0.074987111 | 0.247958214 | 0.20719815 |
| Dnah2 | T | | 633 | 0.244378796 | 0.299777295 | 0.436247873 | 0.191869077 | 0.136470578 | 0.19638277 | 0.066073797 | 0.008638412 | 0.289451063 | 0.321915991 |
| Dnah2 | T | | 638 | 0.244378796 | 0.299777295 | 0.436247873 | 0.191869077 | 0.136470578 | 0.19638277 | 0.066073797 | 0.008638412 | 0.289451063 | 0.321915991 |
| Dnah2 | T | | 640 | 0.244378796 | 0.299777295 | 0.436247873 | 0.191869077 | 0.136470578 | 0.19638277 | 0.066073797 | 0.008638412 | 0.289451063 | 0.321915991 |
| Nck1 | S | | 85 | 0.444987681 | 0.501107611 | 0.19565614 | -0.249331541 | -0.305451471 | 0.078520309 | 0.020449476 | 0.270498967 | 0.244719084 | 0.079312529 |
| Cavin1 | Y | | 158 | 0.223526529 | 0.197748779 | 0.556624986 | 0.333098457 | 0.358876207 | 0.128504722 | 0.1152939 | 0.008689007 | 0.068792922 | 0.046062054 |
| Fgd6 | Y | | 717 | 0.165395821 | 0.233552778 | 0.557742435 | 0.392346614 | 0.324189657 | 0.255329994 | 0.091395534 | 0.004971862 | 0.022076546 | 0.045945387 |
| Dlg5 | Y | | 429 | 0.365386669 | 0.197272079 | 0.44199906 | 0.07661239 | 0.244726981 | 0.022112875 | 0.123124525 | 0.05869818 | 0.658224404 | 0.210235384 |
| Ephb3 | T | | 786 | 0.241210711 | 0.461235623 | 0.294901417 | 0.053690706 | -0.166334206 | 0.366921894 | 0.003404411 | 0.097228206 | 0.833021891 | 0.250072536 |
| Actn4 | Y | | 213 | 0.323107481 | 0.138076204 | 0.511166496 | 0.188059015 | 0.373090292 | 0.013676051 | 0.393391695 | 0.022124862 | 0.237801128 | 0.087443292 |
| Egfr | Y | | 1092 | 0.276382673 | 0.307407361 | 0.486554288 | 0.210171615 | 0.179146928 | 0.223394777 | 0.093371601 | 0.01473212 | 0.336223278 | 0.291417053 |
| Ncl | Y | | 464 | 0.437361356 | 0.095256062 | 0.323637656 | -0.1137237 | 0.228381594 | 0.007406888 | 0.415894356 | 0.014176093 | 0.304728024 | 0.092292942 |
| Bmpr2 | Y | | 708 | 0.478628661 | 0.567553658 | 0.415078898 | -0.063549763 | -0.15247476 | 0.467569344 | 0.018324387 | 0.207783426 | 0.920969316 | 0.601915017 |
| Cuedc1 | Y | | 268 | 0.173384927 | 0.21083628 | 0.472236454 | 0.298851527 | 0.261400173 | 0.021780231 | 0.088626457 | 0.01698979 | 0.053876842 | 0.083881319 |
| Plec | Y | | 3185 | 0.347690608 | 0.24205347 | 0.500536947 | 0.152846339 | 0.258483477 | 0.057530665 | 0.138694033 | 0.050127265 | 0.429222091 | 0.213689581 |
| Ptk2 | Y | | 397 | 0.152223277 | 0.220809881 | 0.276629329 | 0.124406052 | 0.055819449 | 0.078712125 | 0.008473651 | 0.003140868 | 0.114526093 | 0.179450572 |
| Matr3 | S | | 598 | 0.250810957 | 0.367019121 | 0.41886092 | 0.168049963 | 0.051841799 | 0.136683749 | 0.073569465 | 0.028155855 | 0.11978499 | 0.689630532 |
| Fyn Yes1 | Y | 214;221 | | 0.362382804 | 0.17928065 | 0.453758909 | 0.091376106 | 0.27447826 | 0.029649102 | 0.225867914 | 0.032769719 | 0.564922917 | 0.148690007 |
| Septin7 | Y | | 29 | 0.394798851 | 0.302660766 | 0.429994144 | 0.035195293 | 0.127333378 | 0.031447145 | 0.201073009 | 0.089997698 | 0.85621936 | 0.603121984 |
| Cdk1 | Y | | 15 | 0.316287396 | 0.287024742 | 0.188135892 | -0.128151504 | -0.09888885 | 0.042697076 | 0.029291267 | 0.019537918 | 0.275523054 | 0.295760252 |
| Brk1 | Y | | 63 | 0.238891459 | 0.235880087 | 0.303807691 | 0.064916232 | 0.067927605 | 0.085082623 | 0.114009466 | 0.001964575 | 0.538089479 | 0.560450976 |
| Ptk2 | T | | 575 | 0.183882482 | 0.257758728 | 0.280252553 | 0.096370071 | 0.022493824 | 0.059614891 | 0.010174428 | 0.020094049 | 0.293005584 | 0.753751476 |

| | | | | | | | | | | | | |
|---|---|---|---|---|---|---|---|---|---|---|---|---|
| Sipa1l3 | T | 1162 | 0.104495109 | 0.295281645 | 0.341502815 | 0.237007706 | 0.04622117 | 0.218623283 | 0.017771115 | 0.004326699 | 0.036423282 | 0.598667548 |
| Sipa1l1 | Y | 1598 | 0.682167455 | 0.212846464 | 0.38857948 | -0.293587975 | 0.175733016 | 0.245899961 | 0.096423985 | 0.030922406 | 0.563232755 | 0.213764415 |
| Bcar1 | Y | 266 | 0.779956595 | 0.380640419 | 0.384554999 | -0.395401595 | 0.00391458 | 0.404308111 | 0.068881882 | 0.076748494 | 0.649715225 | 0.98306983 |
| Coro1c | Y | 301 | 0.332196511 | 0.178496777 | 0.346957726 | 0.014761215 | 0.168460949 | 0.040874665 | 0.19948159 | 0.009535531 | 0.900635529 | 0.227700957 |
| Egfr | Y | 1110 | 0.100596936 | 0.278376596 | 0.335642059 | 0.235045123 | 0.057265463 | 0.429652657 | 0.017461542 | 0.001462525 | 0.130310191 | 0.468129112 |
| H2bc11 H2bc13 H2bc15 H | Y | 43;43;43;43;43;43 | 0.53313998 | 0.20176178 | 0.338022288 | -0.195117693 | 0.136260508 | 0.171846466 | 0.118609253 | 0.013854013 | 0.53427311 | 0.27596724 |
| Ppfibp1 | Y | 711 | 0.207420536 | 0.2495942 | 0.432989923 | 0.225569387 | 0.183395723 | 0.231332677 | 0.247555453 | 0.001454431 | 0.203419257 | 0.364154311 |
| Arhgap35 | Y | 1087 | 0.414658084 | 0.30036455 | 0.249299636 | -0.165538448 | -0.051064915 | 0.261581086 | 0.049550481 | 0.006521927 | 0.602883308 | 0.626570977 |
| Cdk2 Cdk3 | Y | 15;15 | 0.307554726 | 0.24307067 | 0.208647455 | -0.098907271 | -0.034423215 | 0.07632922 | 0.030501291 | 0.009113635 | 0.438602262 | 0.652064947 |
| Bcar1 | Y | 253 | 0.311374097 | 0.254487473 | 0.212812747 | -0.098561349 | -0.041674726 | 0.014923952 | 0.04890586 | 0.061152325 | 0.165437959 | 0.624965892 |
| Actb Actg1 | Y | 294;294 | 0.286018008 | 0.216868438 | 0.241493878 | -0.044524129 | 0.02462544 | 0.022709429 | 0.06411983 | 0.019501481 | 0.61362205 | 0.784783977 |
| H2bc11 H2bc13 H2bc15 H | Y | 41;41;41;41;41;41 | 0.65714212 | 0.201464858 | 0.356278311 | -0.300863809 | 0.154813453 | 0.180936268 | 0.201813245 | 0.023278358 | 0.464096577 | 0.300510913 |
| Hspe1 | Y | 76 | 0.375737225 | 0.23018601 | 0.549247657 | 0.173510432 | 0.319061647 | 0.079572167 | 0.353150824 | 0.044103905 | 0.347338349 | 0.209732239 |
| Rpl27a | Y | 52 | 0.432516382 | 0.206800192 | 0.183953536 | -0.248562846 | -0.022846656 | 0.00547002 | 0.091439087 | 0.177557479 | 0.08963752 | 0.846081818 |
| Crkl | Y | 132 | 0.229440628 | 0.272097227 | 0.268577081 | 0.039136453 | -0.003520146 | 0.343070142 | 0.012323103 | 0.005885593 | 0.853966267 | 0.95532973 |
| Rbck1 | S | 326 | 0.487188849 | 0.292073184 | 0.351360224 | -0.135828625 | 0.059287039 | 0.288866924 | 0.179108756 | 0.010108518 | 0.731323796 | 0.739977955 |
| Rbck1 | T | 327 | 0.487188849 | 0.292073184 | 0.351360224 | -0.135828625 | 0.059287039 | 0.288866924 | 0.179108756 | 0.010108518 | 0.731323796 | 0.739977955 |
| Tln1 | Y | 1777 | 0.149048654 | 0.416185462 | 0.28798036 | 0.138931706 | -0.128205102 | 0.563533799 | 0.063633084 | 0.001217201 | 0.587190576 | 0.403396146 |
| Bckdha | Y | 288 | 0.224668683 | 0.214720812 | 0.580633544 | 0.355964861 | 0.365912733 | 0.114977008 | 0.228794748 | 0.022230931 | 0.094549258 | 0.110158877 |
| Anks1 | Y | 472 | 0.285155179 | 0.225052815 | 0.346793777 | 0.061638597 | 0.121740961 | 0.088404966 | 0.213131947 | 0.006722548 | 0.616353989 | 0.440887955 |
| Vdac1 | Y | 208 | 0.21902138 | 0.154801339 | 0.52330596 | 0.30428458 | 0.368504621 | 0.152855332 | 0.47696544 | 0.002735578 | 0.068601043 | 0.156625609 |
| Abi1 | Y | 428 | 0.210007289 | 0.268708609 | 0.370148658 | 0.160141369 | 0.101440049 | 0.134351244 | 0.051032785 | 0.022177433 | 0.229113363 | 0.379979942 |
| Unk | S | 240 | 0.25231373 | 0.184179191 | 0.405605508 | 0.153291778 | 0.221426317 | 0.090494472 | 0.336312452 | 0.003826066 | 0.24689261 | 0.265150101 |
| Gab2 | Y | 321 | 0.158777005 | 0.411555707 | 0.346825118 | 0.188048114 | -0.064730588 | 0.660572734 | 0.046982213 | 0.005250508 | 0.605616208 | 0.645459775 |
| Vim | Y | 358 | 0.540303581 | 0.285110203 | 0.284242577 | -0.256061004 | -0.000867626 | 0.187946666 | 0.114021555 | 0.033461388 | 0.457686287 | 0.995294729 |
| Bcar1 | Y | 376 | 0.495418344 | 0.259575769 | 0.227880639 | -0.267537705 | -0.03169513 | 0.163219455 | 0.017905644 | 0.085610567 | 0.378498185 | 0.733097454 |
| Asap2 | Y | 766 | 0.326223427 | 0.31455483 | 0.218252319 | -0.107971108 | -0.096302512 | 0.230376875 | 0.013340494 | 0.033770583 | 0.639724129 | 0.293669731 |
| Rbck1 | Y | 328 | 0.271591911 | 0.278908293 | 0.40726222 | 0.135670309 | 0.128353927 | 0.164492682 | 0.167629202 | 0.015113365 | 0.425316409 | 0.460923859 |
| Nck2 | T | 92 | 0.208562234 | 0.364088651 | 0.284844463 | 0.076282229 | -0.079244189 | 0.212306198 | 0.036583243 | 0.02189149 | 0.608066624 | 0.511364889 |
| Pard3b | S | 1082 | 0.071184059 | 0.221723653 | 0.416098146 | 0.344914088 | 0.194374493 | 0.463174965 | 0.020148966 | 0.003413429 | 0.017844462 | 0.058907971 |
| Eif5a | Y | 98 | 0.356044942 | 0.369257679 | 0.313778166 | -0.042266776 | -0.055479513 | 0.364800755 | 0.079681955 | 0.017043763 | 0.903540186 | 0.720527251 |
| Fgfr1 | Y | 585 | 0.054254006 | 0.401200555 | 0.261658334 | 0.207404328 | -0.139542221 | 0.333269149 | 0.001617036 | 0.06717702 | 0.104749501 | 0.219498554 |
| Kirrel | Y | 685 | 0.148783905 | 0.13071391 | 0.307191483 | 0.158407578 | 0.176477573 | 0.037275045 | 0.071287124 | 0.001688611 | 0.015496719 | 0.01895927 |
| Ptk2 | Y | 577 | 0.170707982 | 0.193910379 | 0.229985546 | 0.059277564 | 0.036075166 | 0.034610547 | 0.016587555 | 0.009431978 | 0.292758508 | 0.409832351 |
| Psmb4 | Y | 102 | 0.471686175 | 0.208631006 | 0.334008753 | -0.137677422 | 0.125377747 | 0.195191174 | 0.104616063 | 0.024211011 | 0.642562782 | 0.324789365 |
| Snd1 | Y | 109 | 0.29958609 | 0.174899878 | 0.410478891 | 0.110892801 | 0.235579013 | 0.087338301 | 0.193936485 | 0.019894214 | 0.478341414 | 0.137274873 |
| Pxn | Y | 118 | 0.078686158 | 0.380816251 | 0.300037428 | 0.221351269 | -0.080778824 | 0.30802442 | 0.002032399 | 0.111336399 | 0.198504183 | 0.570507475 |
| Set | Y | 145 | 0.357308119 | 0.095185411 | 0.454781045 | 0.097472926 | 0.359595634 | 0.141524391 | 0.383712924 | 0.006692818 | 0.62387521 | 0.023320628 |
| Rps10 | Y | 12 | 0.368387097 | 0.164426456 | 0.358747536 | -0.009639561 | 0.19432108 | 0.073310821 | 0.244248259 | 0.020147239 | 0.950842049 | 0.175571738 |
| Bst2 | Y | 8 | 0.441249089 | 0.162470324 | 0.491796172 | 0.050547082 | 0.329325847 | 0.289591423 | 0.436832392 | 0.008289313 | 0.888175232 | 0.17303117 |
| Pcdh18 | Y | 743 | 0.237092786 | 0.205050217 | 0.416641628 | 0.179548843 | 0.211591411 | 0.106239798 | 0.228894952 | 0.01187227 | 0.176007839 | 0.206077696 |
| Crkl | T | 130 | 0.055565485 | 0.374960289 | 0.446157627 | 0.390592142 | 0.071197338 | 0.686644026 | 0.019828879 | 0.025731168 | 0.042834652 | 0.568840474 |
| Fes | Y | 456 | 0.648246871 | 0.247726142 | 0.246219876 | -0.402026995 | -0.001506266 | 0.165518495 | 0.104634326 | 0.074320092 | 0.328955225 | 0.986595384 |
| Ccnl1 | S | 380 | 0.285756069 | 0.159105436 | 0.567101663 | 0.281345594 | 0.407996227 | 0.256719937 | 0.493871981 | 0.007878686 | 0.25659351 | 0.131662971 |
| Ccnl1 | S | 382 | 0.285756069 | 0.159105436 | 0.567101663 | 0.281345594 | 0.407996227 | 0.256719937 | 0.493871981 | 0.007878686 | 0.25659351 | 0.131662971 |
| Fgr Fyn Yes1 | Y | 185;168;192 | 0.135426664 | 0.302042823 | 0.204776631 | 0.069349967 | -0.097266192 | 0.087035163 | 0.001032835 | 0.172893157 | 0.585918299 | 0.446821159 |
| Plec | S | 4649 | 0.06396975 | 0.41066575 | 0.396055162 | 0.332085411 | -0.014610588 | 0.650290732 | 0.050638846 | 0.011615119 | 0.060149935 | 0.92278707 |
| Rsrp1 | Y | 157 | 0.198330055 | 0.22318654 | 0.529974447 | 0.331644392 | 0.306787907 | 0.171084522 | 0.194413263 | 0.024326848 | 0.100303186 | 0.143559186 |
| Sirpa | Y | 505 | 0.065609404 | 0.365796716 | 0.191922633 | 0.126313229 | -0.173874083 | 0.3412092 | 0.002089745 | 0.028105387 | 0.043810579 | 0.033749906 |
| Pak1 Pak2 | Y | 453;474 | 0.467544751 | 0.211877667 | 0.272368087 | -0.195176665 | 0.06049042 | 0.10195149 | 0.119183114 | 0.05488604 | 0.397234271 | 0.600335112 |
| Etl4 | Y | 397 | 0.143719473 | 0.269254492 | 0.398539023 | 0.25481955 | 0.12928453 | 0.165010138 | 0.016263285 | 0.069347865 | 0.144341665 | 0.381034657 |
| Vcl | Y | 100 | 0.380638615 | 0.298091447 | 0.394467521 | 0.013828906 | 0.096376074 | 0.129132492 | 0.19267935 | 0.079547777 | 0.937226861 | 0.550018519 |
| Cpne3 | S | 260 | 0.332141454 | 0.152687895 | 0.301344282 | -0.030797172 | 0.148656387 | 0.039662073 | 0.194093769 | 0.026845582 | 0.783784828 | 0.143372279 |
| Cpne3 | S | 264 | 0.332141454 | 0.152687895 | 0.301344282 | -0.030797172 | 0.148656387 | 0.039662073 | 0.194093769 | 0.026845582 | 0.783784828 | 0.143372279 |
| Cpne3 | Y | 261 | 0.332141454 | 0.152687895 | 0.301344282 | -0.030797172 | 0.148656387 | 0.039662073 | 0.194093769 | 0.026845582 | 0.783784828 | 0.143372279 |
| Crim1 | Y | 1034 | -0.068309711 | 0.344025478 | 0.476743814 | 0.545053525 | 0.132718337 | 0.49805449 | 0.073639165 | 0.000810233 | 0.016319102 | 0.364500384 |
| Ankrd13b | S | 622 | -0.043938144 | 0.369295275 | 0.419321015 | 0.463259159 | 0.05002574 | 0.776526966 | 0.001834975 | 0.043487359 | 0.054529455 | 0.699869085 |
| Ankrd13b | T | 624 | -0.043938144 | 0.369295275 | 0.419321015 | 0.463259159 | 0.05002574 | 0.776526966 | 0.001834975 | 0.043487359 | 0.054529455 | 0.699869085 |
| Cit | Y | 426 | 0.385415129 | 0.181524037 | 0.275047256 | -0.110367873 | 0.093523219 | 0.015434617 | 0.214443556 | 0.119688049 | 0.476439743 | 0.571431819 |
| Crk | Y | 251 | 0.08931741 | 0.314527553 | 0.24048478 | 0.15116737 | -0.074042773 | 0.489286242 | 0.0019611 | 0.032719657 | 0.285880275 | 0.353584699 |
| Il1rap | S | 563 | 0.322825869 | 0.275464998 | 0.113726885 | -0.209098984 | -0.161738113 | 0.012243498 | 0.023726018 | 0.344464616 | 0.10280221 | 0.162386606 |
| Peak1 | S | 791 | 0.21892361 | 0.282929242 | 0.232206893 | 0.013283282 | -0.05072235 | 0.439773583 | 0.050362181 | 0.001814917 | 0.958959368 | 0.584852025 |
| Vim | Y | 11 | 0.375199631 | 0.175142888 | 0.387151916 | 0.011952284 | 0.212009028 | 0.196013885 | 0.247054891 | 0.016173698 | 0.959906024 | 0.19776886 |
| Slc30a4 | Y | 356 | 0.521259062 | 0.532071177 | 0.301488422 | -0.21977064 | -0.230582756 | 0.459211914 | 0.08595273 | 0.148315665 | 0.73632649 | 0.336371689 |
| Flii | Y | 737 | 0.390098803 | 0.423022964 | 0.54582186 | 0.155723057 | 0.122798896 | 0.325789176 | 0.213757954 | 0.126053861 | 0.581377596 | 0.390641233 |
| Tnip2 | Y | 231 | 0.132040003 | 0.233274653 | 0.340944219 | 0.208904216 | 0.107669565 | 0.011589188 | 0.05543269 | 0.078170353 | 0.192405206 | 0.457091756 |
| Pard3b | Y | 953 | 0.211510504 | 0.278295933 | 0.208067047 | -0.003443458 | -0.070228887 | 0.17410075 | 0.008670014 | 0.065885998 | 0.979501968 | 0.398519838 |
| Tln1 | Y | 70 | 0.238433121 | 0.118470073 | 0.297937995 | 0.059504874 | 0.179467923 | 0.010515346 | 0.176971888 | 0.032649648 | 0.523806591 | 0.132530404 |
| Dpysl2 Dpysl3 | Y | 32;32 | 0.351420381 | 0.313476662 | 0.263452106 | -0.087968274 | -0.050024556 | 0.288746718 | 0.047100199 | 0.059725236 | 0.762552597 | 0.703300231 |
| Bcar1 | Y | 414 | 0.520143437 | 0.380475313 | 0.179850082 | -0.340293355 | -0.200625231 | 0.316462109 | 0.016350201 | 0.290577949 | 0.483219487 | 0.212807753 |
| Epha2 | Y | 961 | 0.38015499 | 0.132501544 | 0.602976331 | 0.22282134 | 0.470474786 | 0.104267083 | 0.33678214 | 0.098617597 | 0.464108128 | 0.164909497 |
| Plcg1 | Y | 783 | 0.093025424 | 0.306288645 | 0.51317023 | 0.420144806 | 0.206881585 | 0.509540881 | 0.033104408 | 0.05591914 | 0.078871003 | 0.267372902 |
| Ehbp1l1 | Y | 1555 | 0.460256778 | 0.390312584 | 0.427598179 | -0.032658598 | 0.037285595 | 0.21699389 | 0.267363368 | 0.132029505 | 0.910707236 | 0.892435266 |
| Gsk3a Gsk3b | Y | 216;279 | 0.175107394 | 0.218430035 | 0.141278935 | -0.033828459 | -0.0771511 | 0.010677637 | 0.032795559 | 0.023276603 | 0.33679097 | 0.300186058 |
| H2bc1 H2bc11 H2bc13 H2 | Y | 84;84;84;84;84;84 | 0.388304173 | 0.222501893 | 0.327063933 | -0.06124024 | 0.10456204 | 0.087782899 | 0.058637209 | 0.207467403 | 0.805156973 | 0.629141042 |
| Ackr3 | Y | 354 | 0.343506305 | 0.070704084 | 0.181960613 | -0.161545691 | 0.111256529 | 0.000972306 | 0.19326034 | 0.144587281 | 0.182108118 | 0.311835259 |
| Rps17 | S | 113 | 0.497570514 | 0.432892302 | 0.283238495 | -0.214332019 | -0.149653807 | 0.199944715 | 0.15270728 | 0.203566654 | 0.524195512 | 0.553592459 |
| Myh9 | Y | 1805 | 0.197905807 | 0.262575664 | 0.439469019 | 0.241563212 | 0.176893355 | 0.084466279 | 0.177942605 | 0.067685575 | 0.2119806 | 0.405727687 |
| Psma2 | Y | 57 | 0.203866324 | 0.133912141 | 0.336188355 | 0.132322031 | 0.202276214 | 0.022550791 | 0.223763643 | 0.02261119 | 0.219987559 | 0.131780126 |
| Plec | Y | 3516 | 0.313936162 | 0.100344595 | 0.522433407 | 0.208497246 | 0.422088812 | 0.076400279 | 0.499305194 | 0.041784755 | 0.293288526 | 0.080386377 |
| Stam2 | Y | 211 | 0.291424469 | 0.347932552 | 0.344209752 | 0.052785283 | -0.003722799 | 0.162169384 | 0.282999857 | 0.038593473 | 0.751566603 | 0.989285167 |
| Eif3b | Y | 328 | 0.243304353 | 0.01305642 | 0.140869935 | -0.102434418 | 0.127813515 | 7.96504E-05 | 0.907962816 | 0.003351723 | 0.015962352 | 0.323381311 |
| Fyn Yes1 | Y | 213;220 | 0.239027323 | 0.191425244 | 0.350969901 | 0.111942578 | 0.159544657 | 0.091422775 | 0.187674667 | 0.027401933 | 0.411868187 | 0.291687957 |
| Bcar1 | Y | 366 | 0.457442144 | 0.238802351 | 0.193388836 | -0.264053308 | -0.045413515 | 0.197602464 | 0.029922998 | 0.100259934 | 0.397687149 | 0.564988739 |
| Tln1 | Y | 127 | 0.237291966 | 0.122377767 | 0.330926959 | 0.093634994 | 0.208549192 | 0.0521955 | 0.426629609 | 0.006971969 | 0.290702934 | 0.210131102 |
| Epb41l2 | S | 569 | 0.475171331 | 0.322928922 | 0.205659476 | -0.269511855 | -0.117269446 | 0.406045571 | 0.035915307 | 0.067379593 | 0.614118789 | 0.304641816 |
| Iqgap1 | Y | 831 | 0.318393491 | 0.256737379 | 0.161241631 | -0.15715186 | -0.095495748 | 0.004797999 | 0.111758722 | 0.412250484 | 0.419432904 | 0.639142037 |
| Ccdc22 | Y | 474 | 0.263526265 | 0.218154958 | 0.44777718 | 0.184250914 | 0.229622221 | 0.062114429 | 0.10072818 | 0.188053005 | 0.518956061 | 0.434023215 |
| Axl | Y | 753 | 0.382179366 | 0.219318891 | 0.294691001 | -0.087488364 | 0.07537211 | 0.114161843 | 0.234533097 | 0.045788933 | 0.636179051 | 0.611501438 |
| Nolc1 | S | 701 | 0.494530067 | 0.255297264 | 0.136442733 | -0.358087334 | -0.118854531 | 0.014027043 | 0.248508807 | 0.350162105 | 0.050562179 | 0.55205475 |
| Plec | Y | 4162 | 0.248917038 | 0.207664903 | 0.328530726 | 0.079613688 | 0.120865823 | 0.061142596 | 0.085441431 | 0.0970251 | 0.605999238 | 0.437703161 |

| | | | | | | | | | | | | |
|---|---|---|---|---|---|---|---|---|---|---|---|---|
| Msn | Y | 116 | 0.406258188 | 0.223312701 | 0.405350069 | -0.000908119 | 0.182037368 | 0.231382934 | 0.362069688 | 0.034418276 | 0.997378546 | 0.408618185 |
| Plec | Y | 3784 | 0.204322888 | 0.249982362 | 0.348227877 | 0.143904989 | 0.098245514 | 0.075842139 | 0.115432365 | 0.074616184 | 0.359884537 | 0.561571289 |
| Itgb1 | Y | 795 | 0.06741675 | 0.373917458 | 0.194685022 | 0.127268272 | -0.179232436 | 0.367449956 | 0.003415062 | 0.089873753 | 0.193371555 | 0.102675616 |
| Shmt2 | Y | 100 | 0.264524631 | 0.108250137 | 0.444473211 | 0.17994858 | 0.336223074 | 0.017864243 | 0.498945484 | 0.067498663 | 0.327187669 | 0.149970766 |
| Pcbp1 | S | 173 | 0.538533702 | 0.132642215 | 0.29896803 | -0.239565672 | 0.166325815 | 0.035154807 | 0.618836767 | 0.120471565 | 0.191232378 | 0.488860439 |
| Set | Y | 139 | 0.501039248 | 0.492526864 | 0.433243704 | -0.067795544 | -0.059283161 | 0.16917937 | 0.28580949 | 0.365496473 | 0.874896359 | 0.905630202 |
| Unc93b1 | S | 194 | 0.200990722 | 0.443142118 | 0.552161812 | 0.351171089 | 0.109019694 | 0.487951136 | 0.170511004 | 0.10312641 | 0.013887332 | 0.326475878 |
| Bcar3 | Y | 424 | 0.31382616 | 0.211713281 | 0.190228021 | -0.123598139 | -0.021485261 | 0.309370882 | 0.014528157 | 0.02825821 | 0.651441203 | 0.744988355 |
| H4c1 H4c11 H4c12 H4c14 | Y | 52 | 0.597507434 | 0.213009377 | 0.248745943 | -0.348761492 | 0.035736565 | 0.12631303 | 0.27841764 | 0.10908469 | 0.282805572 | 0.850220439 |
| Rps2 | Y | 133 | 0.375774339 | 0.270568348 | 0.263735541 | -0.112038798 | -0.006832807 | 0.08509921 | 0.163724161 | 0.150767158 | 0.457503458 | 0.956618662 |
| Kirrel | Y | 638 | 0.249407806 | 0.053389132 | 0.373785497 | 0.124377691 | 0.320396365 | 0.071805796 | 0.432967079 | 0.008671708 | 0.31160549 | 0.028242351 |
| Actn1 | Y | 681 | 0.093842809 | 0.316278794 | 0.418491083 | 0.324648274 | 0.10221229 | 0.16018647 | 0.046716638 | 0.114001395 | 0.183843575 | 0.621200773 |
| Flna | Y | 1604 | 0.148053936 | 0.34377868 | 0.294372936 | 0.146319001 | -0.049405744 | 0.184916778 | 0.102185521 | 0.045345244 | 0.17177425 | 0.764620623 |
| H3c13 H3c14 H3c15 H3c2 | Y | 42;42;42;42 | 1.217490399 | 0.103028495 | 0.354485622 | -0.863004777 | 0.251457127 | 0.247228529 | 0.3948725 | 0.229234904 | 0.372273827 | 0.359270415 |
| Anks1 | S | 461 | 0.292569719 | 0.144893208 | 0.402240297 | 0.109670579 | 0.257347089 | 0.18940119 | 0.467243027 | 0.014510048 | 0.568112601 | 0.228655503 |
| Prrc2a | Y | 635 | 0.199908418 | 0.289908713 | 0.241815361 | 0.041906943 | -0.048093352 | 0.103187809 | 0.148093635 | 0.031634019 | 0.690893207 | 0.765693338 |
| Atic | Y | 290 | 0.439159858 | 0.328430958 | 0.322780956 | -0.116378902 | -0.005650001 | 0.542510988 | 0.025242241 | 0.212026905 | 0.867767268 | 0.978316611 |
| Vcl | Y | 692 | 0.077141222 | 0.218126209 | 0.194231849 | 0.117090627 | -0.02389436 | 0.056999864 | 0.086002785 | 0.001512661 | 0.00597917 | 0.78178201 |
| Eif1a Eif1ax | Y | 106;106 | 0.438094715 | 0.059580911 | 0.346444375 | -0.09165034 | 0.286863464 | 0.039465072 | 0.696749556 | 0.062718242 | 0.599735614 | 0.136083139 |
| Iqgap1 | Y | 654 | 0.356395423 | 0.276178729 | 0.263915432 | -0.092479991 | -0.012263297 | 0.264848328 | 0.093798807 | 0.078517733 | 0.741546436 | 0.933326727 |
| Tpm1 | Y | 162 | 0.13081887 | 0.179843797 | 0.327573004 | 0.196754134 | 0.147729207 | 0.34956022 | 0.228158744 | 0.002019708 | 0.197613248 | 0.299096289 |
| Bcar3 | Y | 206 | 0.234759185 | 0.093045068 | 0.355176237 | 0.120417051 | 0.262131169 | 0.117716794 | 0.259621978 | 0.013000052 | 0.355590422 | 0.038461806 |
| Rps24 | Y | 81 | 0.37122606 | 0.133556796 | 0.285154721 | -0.08607134 | 0.151597924 | 0.010940746 | 0.513445355 | 0.169294801 | 0.624599654 | 0.525056983 |
| Arpc3 | Y | 47 | -0.047574445 | 0.406114272 | 0.354928703 | 0.402503148 | -0.051185569 | 0.819628181 | 0.09534744 | 0.003292813 | 0.149652689 | 0.765133415 |
| Caskin2 | T | 256 | 0.204890928 | 0.444283716 | 0.378990622 | 0.174099694 | -0.065293094 | 0.329185006 | 0.254790478 | 0.055912732 | 0.331474601 | 0.8425199 |
| Sfpq | S | 375 | 0.716996168 | 0.055799984 | 0.568358386 | -0.148637782 | 0.512558402 | 0.318604568 | 0.934061999 | 0.043954922 | 0.812420577 | 0.47004618 |
| Eps15l1 | Y | 562 | 0.295154243 | 0.000183582 | 0.427846419 | 0.132692176 | 0.427662837 | 0.40377529 | 0.998446649 | 0.000686872 | 0.68456522 | 0.013737904 |
| Calm1 Calm2 Calm3 | Y | 100;100;100 | 0.267698873 | 0.132889704 | 0.194928206 | -0.072770667 | 0.062038502 | 0.016133666 | 0.203574168 | 0.048674712 | 0.156829112 | 0.414561658 |
| Cav1 | Y | 42 | 0.197928318 | 0.142754978 | 0.49172043 | 0.293792112 | 0.348965452 | 0.17771257 | 0.300868141 | 0.035830753 | 0.13418177 | 0.091157888 |
| Acta1 Actb Actc1 Actg1 | Y | 91;91;93;93 | 0.408333772 | 0.087050022 | 0.310453744 | -0.097880028 | 0.223403721 | 0.13731377 | 0.32396939 | 0.026103931 | 0.650423508 | 0.077408463 |
| Ubap2l | Y | 854 | 0.768151129 | 0.541140409 | 0.007103757 | -0.761047372 | -0.534036652 | 0.116841026 | 0.170279648 | 0.983800578 | 0.093514115 | 0.017261777 |
| Pacsin2 | Y | 76 | 0.221356204 | 0.189428977 | 0.294981832 | 0.073625628 | 0.105552855 | 0.177474148 | 0.074884654 | 0.046529202 | 0.631286433 | 0.376554585 |
| Ap2b1 | Y | 737 | 0.19118551 | 0.141728263 | 0.297274425 | 0.106088914 | 0.155546162 | 0.163700989 | 0.181515616 | 0.009067514 | 0.390585915 | 0.1567057 |
| Src | Y | 92 | 0.157895488 | 0.315452971 | 0.294611568 | 0.13671608 | -0.020841403 | 0.680292548 | 0.020332702 | 0.066102504 | 0.723152343 | 0.869557402 |
| Add2 | Y | 489 | 0.520271897 | 0.153254865 | 0.247898649 | -0.272373248 | 0.094643784 | 0.24505987 | 0.216318565 | 0.038144277 | 0.487276817 | 0.441719754 |
| Skap2 | Y | 260 | 0.291836363 | 0.099028507 | 0.270251673 | -0.02158469 | 0.171223166 | 0.02678567 | 0.521833852 | 0.034924852 | 0.710355912 | 0.262322483 |
| Cdc42ep1 | Y | 130 | 0.407269813 | 0.273752278 | 0.032485086 | -0.374784728 | -0.241267192 | 0.113176754 | 0.007026254 | 0.619754699 | 0.143131225 | 0.007199924 |
| Rps3 | Y | 120 | 0.292060743 | 0.102735836 | 0.193967267 | -0.098093477 | 0.091231431 | 0.007000391 | 0.396297555 | 0.073158466 | 0.292698652 | 0.465985016 |
| Crk | Y | 136 | 0.092410942 | 0.233454684 | 0.316838319 | 0.224427377 | 0.083383636 | 0.18937937 | 0.045537042 | 0.051819187 | 0.136292857 | 0.510397571 |
| Pik3r2 | Y | 443 | 0.110744528 | 0.142952479 | 0.295012108 | 0.18426758 | 0.152059629 | 0.15323964 | 0.295374143 | 0.002698003 | 0.031399596 | 0.26888018 |
| Tmem59 | Y | 277 | 0.400253311 | 0.085840126 | 0.422118818 | 0.021865507 | 0.336278692 | 0.068310842 | 0.647678767 | 0.118092869 | 0.920124659 | 0.18250751 |
| Bcar1 | S | 407 | 0.62525919 | 0.177714084 | 0.125609374 | -0.499649817 | -0.05210471 | 0.306305211 | 0.034728776 | 0.097770937 | 0.390486334 | 0.46447201 |
| Kirrel | Y | 657 | 0.12977001 | 0.125336791 | 0.290442365 | 0.160672355 | 0.165105574 | 0.142705375 | 0.118748942 | 0.010658654 | 0.108504543 | 0.08471516 |
| Flnb | Y | 155 | 0.293084282 | 0.124953839 | 0.508493663 | 0.21540938 | 0.383539824 | 0.312767666 | 0.441481434 | 0.043334159 | 0.461897611 | 0.107728862 |
| Flnb | Y | 511 | 0.385759016 | 0.220318838 | 0.236500891 | -0.149258125 | 0.016182052 | 0.247150411 | 0.140491038 | 0.080947203 | 0.607853756 | 0.905784016 |
| Plec | Y | 407 | 0.410161312 | 0.148202204 | 0.317383718 | -0.092777595 | 0.169181514 | 0.091572482 | 0.489248442 | 0.109894288 | 0.557605256 | 0.316005497 |
| Cltc | Y | 634 | 0.420528424 | 0.060603032 | 0.544539689 | 0.124011265 | 0.483936656 | 0.100135262 | 0.832274369 | 0.125549957 | 0.64513708 | 0.176754679 |
| Ldlr | Y | 847 | 0.175460998 | 0.177888338 | 0.351753035 | 0.176292037 | 0.173864697 | 0.093139353 | 0.166878455 | 0.080388717 | 0.269507497 | 0.298924828 |
| Rplp0 | Y | 24 | 0.530818583 | 0.135339087 | 0.187337129 | -0.343481454 | 0.051998042 | 0.122422503 | 0.309158039 | 0.077908838 | 0.246235702 | 0.677808232 |
| Il6st | Y | 914 | 0.079726969 | 0.14952813 | 0.321653681 | 0.241926711 | 0.172125551 | 0.331532691 | 0.098582176 | 0.007158795 | 0.003544291 | 0.010966471 |
| Pcdhga4 | Y | 877 | 0.107962017 | 0.247210053 | 0.20885919 | 0.100897173 | -0.038350863 | 0.066477421 | 0.099006221 | 0.036281802 | 0.201827091 | 0.74363114 |
| Cep170 | Y | 176 | 0.366688886 | 0.282711272 | 0.255279552 | -0.111409334 | -0.02743172 | 0.460669545 | 0.1509361 | 0.054856739 | 0.810281793 | 0.859587414 |
| Socs6 | Y | 356 | 0.217506945 | 0.159903639 | 0.239923718 | 0.022416773 | 0.080020079 | 0.477282933 | 0.126620352 | 0.004103568 | 0.936947616 | 0.366931737 |
| Eif3l | Y | 415 | 0.387990818 | 0.094920807 | 0.332118559 | -0.055872259 | 0.237197752 | 0.222495566 | 0.537930757 | 0.023798505 | 0.832686685 | 0.191396428 |
| Golim4 | Y | 633 | 0.176003342 | 0.047993535 | 0.553513242 | 0.377509901 | 0.505519707 | 0.259731028 | 0.819043594 | 0.014353732 | 0.041786007 | 0.0574754 |
| Hipk1 Hipk2 | S | 364;355 | 0.212466033 | 0.192073103 | 0.340902373 | 0.12843634 | 0.14882927 | 0.193516678 | 0.169988429 | 0.073499633 | 0.470882967 | 0.374551706 |
| Gja1 | Y | 286 | 0.318078597 | 0.30199527 | 0.179096672 | -0.138981925 | -0.122898598 | 0.439847425 | 0.031053218 | 0.15073175 | 0.718890207 | 0.307852129 |
| Plec | Y | 2833 | 0.362986653 | 0.235782642 | 0.142945586 | -0.220041068 | -0.092837057 | 0.025983051 | 0.347225849 | 0.227581809 | 0.105594927 | 0.682403054 |
| Ptk2b | Y | 418 | 0.13860264 | 0.283324055 | 0.480988219 | 0.342385579 | 0.197664164 | 0.234053618 | 0.275074996 | 0.098619148 | 0.201912709 | 0.495967665 |
| Cavin1 | Y | 310 | 0.217367231 | 0.077448474 | 0.274106553 | 0.056739323 | 0.196658079 | 0.05894174 | 0.260352585 | 0.025938102 | 0.533324648 | 0.059426039 |
| Sipa1l3 | Y | 1163 | 0.136574465 | 0.176753374 | 0.298803662 | 0.162229197 | 0.122050288 | 0.085498896 | 0.14705155 | 0.05010384 | 0.188915755 | 0.358014051 |
| Sik3 | S | 493 | 0.291505093 | 0.544040764 | 0.042068728 | -0.249436365 | -0.501972037 | 0.0630332 | 0.108784316 | 0.74870712 | 0.112372293 | 0.129257484 |
| Dcbld1 | Y | 332 | 0.6529211 | -0.029546208 | 0.353401005 | -0.299520095 | 0.382947213 | 0.249601302 | 0.922734082 | 0.016656851 | 0.542492295 | 0.277346636 |
| Sorbs2 | Y | 231 | 0.231208197 | 0.111217789 | 0.492328423 | 0.261120226 | 0.381110634 | 0.152916916 | 0.503929605 | 0.062341653 | 0.223378659 | 0.124590332 |
| Myh9 | Y | 933 | 0.146101682 | 0.18214849 | 0.267832718 | 0.121731036 | 0.085684228 | 0.044163735 | 0.23522522 | 0.04393054 | 0.228877854 | 0.555066633 |
| Npdc1 | Y | 258 | 0.26950613 | 0.217545488 | 0.448180064 | 0.178673934 | 0.230634576 | 0.181724659 | 0.111367823 | 0.305920829 | 0.655862793 | 0.561189405 |
| Zdhhc8 | Y | 576 | 0.26181162 | 0.214316434 | 0.207023231 | -0.054788388 | -0.007293202 | 0.353309418 | 0.050664623 | 0.059085675 | 0.829146418 | 0.938051084 |
| Hsp90b1 | Y | 652 | 0.459501669 | 0.250435628 | 0.08581579 | -0.373685879 | -0.164619838 | 0.412885636 | 0.007856865 | 0.251399129 | 0.492241471 | 0.054770049 |
| Sdcbp | Y | 22 | 0.401042946 | 0.276480196 | 0.27410918 | -0.126933766 | -0.002371017 | 0.218036402 | 0.281794361 | 0.159156221 | 0.664159419 | 0.992344648 |
| Flnb | Y | 904 | 0.197203802 | 0.143675213 | 0.375650366 | 0.178446565 | 0.231975153 | 0.106796638 | 0.216485385 | 0.095335848 | 0.339602654 | 0.238744914 |
| Rnf213 | Y | 4666 | 0.36673605 | 0.000872294 | 0.393763237 | 0.027027187 | 0.392890943 | 0.173619269 | 0.996219956 | 0.016920393 | 0.90053235 | 0.090463604 |
| Timm23 | Y | 52 | 0.352236447 | 0.095941742 | 0.325741788 | -0.02649466 | 0.229800046 | 0.196579748 | 0.669735111 | 0.021716207 | 0.905533416 | 0.340747631 |
| Hdlbp | Y | 582 | 0.545707077 | 0.186063594 | 0.262508715 | -0.283198362 | 0.07644512 | 0.163319702 | 0.512398511 | 0.138819729 | 0.396807411 | 0.789290916 |
| Ptbp1 | Y | 126 | 0.346408057 | 0.099283334 | 0.135558206 | -0.21084985 | 0.036274872 | 0.076623227 | 0.061573504 | 0.074939149 | 0.168624011 | 0.532596017 |
| Spry2 | Y | 55 | 0.09851933 | 0.134437485 | 0.368833751 | 0.270314421 | 0.234396266 | 0.331783596 | 0.321525964 | 0.007922064 | 0.052582208 | 0.137391535 |
| Dcbld2 | S | 719 | 0.080708278 | 0.329585197 | 0.202367886 | 0.121659608 | -0.127217311 | 0.143168544 | 0.014713803 | 0.324095252 | 0.519758404 | 0.512731932 |
| Traf4 | Y | 204 | 0.197706558 | 0.03192908 | 0.235623445 | 0.037916886 | 0.203694365 | 0.018348887 | 0.496831731 | 0.013295344 | 0.577851476 | 0.036786358 |
| Mapk9 | Y | 185 | 0.221769185 | 0.230915965 | 0.212737935 | -0.00903125 | -0.018178029 | 0.148512321 | 0.116467469 | 0.106353455 | 0.943251114 | 0.878712029 |
| Rpsa | Y | 139 | 0.409749134 | 0.184591227 | 0.188576024 | -0.22117311 | 0.003984797 | 0.328526639 | 0.195123184 | 0.028385829 | 0.562127215 | 0.973340984 |
| Taok1 | Y | 610 | 0.199183755 | 0.084072169 | 0.348412832 | 0.149229077 | 0.264340663 | 0.040764719 | 0.472370988 | 0.058649444 | 0.274894606 | 0.125012233 |
| Denn2b | Y | 498 | 0.203719313 | 0.106139569 | 0.340365839 | 0.136646526 | 0.23422627 | 0.635272831 | 0.519259745 | 0.001695028 | 0.745721492 | 0.21693568 |
| Dcbld2 | Y | 744 | 0.35321152 | 0.209881831 | 0.089033238 | -0.264178282 | -0.120848593 | 0.084367724 | 0.045371659 | 0.362920131 | 0.149848429 | 0.269030526 |
| Sema4c | Y | 790 | 0.113896976 | 0.146378841 | 0.702844618 | 0.588947642 | 0.556465777 | 0.436779773 | 0.309119131 | 0.085521331 | 0.099629139 | 0.112625014 |
| Pacsin2 | T | 78 | 0.4086356 | 0.035952251 | 0.30345704 | -0.105178559 | 0.267504789 | 0.020247251 | 0.736268387 | 0.230473192 | 0.635642308 | 0.276217814 |
| Phb2 | Y | 248 | 0.171676147 | 0.222621717 | 0.20453618 | 0.032860033 | -0.018085537 | 0.096239424 | 0.166356009 | 0.057265817 | 0.538648857 | 0.884696676 |
| Tns3 | Y | 584 | -0.035296389 | 0.36379861 | 0.216163476 | 0.251459864 | -0.147635135 | 0.691387208 | 0.001519824 | 0.182733274 | 0.141466341 | 0.318179402 |
| Ctnnd1 | Y | 335 | 0.648424417 | 0.388286362 | 0.122238087 | -0.526186329 | -0.266048275 | 0.3100776 | 0.1430459 | 0.507490038 | 0.391222251 | 0.308394333 |
| Hspa9 | S | 162 | 0.53344553 | 0.231285688 | 0.167800868 | -0.365644661 | -0.06348482 | 0.049957123 | 0.273678191 | 0.769699282 | 0.533149105 | 0.908142495 |
| Ptk2 | Y | 407 | 0.170596982 | 0.217343015 | 0.172541732 | 0.001944749 | -0.044801283 | 0.163584893 | 0.032748857 | 0.12159165 | 0.985643458 | 0.584016365 |
| Arhgef2 | S | 885 | 0.630837916 | 0.375011018 | 0.207209446 | -0.42362847 | -0.167801572 | 0.101790172 | 0.50136448 | 0.632663226 | 0.362794059 | 0.778057686 |
| Kirrel | T | 764 | 0.274423864 | 0.18583586 | 0.18339854 | -0.091025324 | -0.002437321 | 0.053990385 | 0.208049239 | 0.16635878 | 0.493991923 | 0.98654201 |

| | | | | | | | | | | | | |
|---|---|---|---|---|---|---|---|---|---|---|---|---|
| Afdn | Y | 1655 | 0.257855678 | 0.094430708 | 0.311185126 | 0.053329448 | 0.216754418 | 0.064239901 | 0.262379434 | 0.12226418 | 0.750486749 | 0.234417306 |
| Gab2 | Y | 290 | 0.019733768 | 0.380803357 | 0.250419744 | 0.230685976 | -0.130383613 | 0.849959471 | 0.091768279 | 0.019200985 | 0.072642792 | 0.451320408 |
| Frk | Y | 53 | 0.313069279 | -0.005841763 | 0.569375365 | 0.256306086 | 0.575217128 | 0.2252227 | 0.978534185 | 0.044938387 | 0.351819549 | 0.074572842 |
| Pkm | Y | 370 | 0.373088537 | 0.23015011 | 0.213501989 | -0.159586548 | -0.016648121 | 0.224292598 | 0.138474889 | 0.215232297 | 0.552544766 | 0.875078966 |
| Tmem200a | Y | 185 | 0.276710959 | 0.129693317 | 0.376395584 | 0.099684625 | 0.246702267 | 0.13250578 | 0.536475262 | 0.09067339 | 0.597309698 | 0.298663966 |
| Irs2 | Y | 181 | 0.468632265 | 0.111250539 | 0.188325661 | -0.280306604 | 0.077075122 | 0.192182472 | 0.287923483 | 0.059470482 | 0.372770703 | 0.452012881 |
| Tra2b | Y | 221 | 0.345908284 | 0.125701727 | 0.284202866 | -0.061705417 | 0.15850114 | 0.090065763 | 0.463085073 | 0.134691555 | 0.7136954 | 0.341832237 |
| Tra2b | Y | 222 | 0.345908284 | 0.125701727 | 0.284202866 | -0.061705417 | 0.15850114 | 0.090065763 | 0.463085073 | 0.134691555 | 0.7136954 | 0.341832237 |
| Ctnnb1 | Y | 489 | 0.132383968 | 0.154757144 | 0.32718042 | 0.194796453 | 0.172423277 | 0.251741869 | 0.187123818 | 0.036671891 | 0.170674625 | 0.211548391 |
| Dock1 | Y | 1811 | 0.331341488 | 0.171831151 | 0.323760627 | -0.007580861 | 0.151929476 | 0.23101176 | 0.362129419 | 0.103064422 | 0.976118191 | 0.471638495 |
| Nectin1 | Y | 466 | 0.256449195 | 0.193359232 | 0.084746323 | -0.171702872 | -0.108612909 | 0.0204181 | 0.115045738 | 0.288393517 | 3.6521E-05 | 0.282072834 |
| Nhlrc2 | Y | 403 | 0.000917077 | 0.185215816 | 0.218766491 | 0.217849413 | 0.033550675 | 0.970183708 | 0.000498029 | 0.06388368 | 0.073357695 | 0.657826446 |
| Vim | Y | 30 | 0.219637008 | 0.125113651 | 0.525969385 | 0.306332377 | 0.400855734 | 0.280915696 | 0.395274003 | 0.102978902 | 0.289459758 | 0.170522906 |
| Chmp5 | Y | 94 | 0.38350588 | 0.281933376 | 0.168693331 | -0.214812549 | -0.113240045 | 0.473251319 | 0.09221064 | 0.114227298 | 0.672827511 | 0.417427539 |
| Ppp3ca Ppp3cb Ppp3cc | Y | 284;288;297 | 0.287972116 | 0.236238154 | 0.502348617 | 0.214376501 | 0.266110463 | 0.431843296 | 0.391245493 | 0.139298207 | 0.515834524 | 0.267902563 |
| Mapk9 | T | 183 | 0.383105695 | 0.282892243 | 0.068666393 | -0.314439303 | -0.214225851 | 0.142454115 | 0.054402477 | 0.607644292 | 0.204424933 | 0.077975902 |
| Rps13 | Y | 38 | 0.274191194 | 0.15582055 | 0.049650715 | -0.224540478 | -0.106169835 | 0.001748146 | 0.298628292 | 0.441499599 | 0.019537977 | 0.451798094 |
| Arhgap35 | Y | 1105 | 0.189361067 | 0.228777107 | 0.182473756 | -0.006887311 | -0.046303351 | 0.11548855 | 0.097047024 | 0.165378585 | 0.956032272 | 0.72417652 |
| Tln1 | Y | 26 | 0.276529166 | 0.052212941 | 0.138040883 | -0.138488283 | 0.085827942 | 0.005065313 | 0.661193978 | 0.083255324 | 0.058044886 | 0.470402973 |
| Unk | Y | 232 | 0.143507875 | 0.154105994 | 0.437526923 | 0.294019048 | 0.283420929 | 0.413339526 | 0.374874339 | 0.038918645 | 0.175588208 | 0.183094007 |
| Peak1 | Y | 796 | 0.067331308 | 0.266022181 | 0.207128625 | 0.139797316 | -0.058893557 | 0.613376404 | 0.060722022 | 0.02152898 | 0.341768893 | 0.554211771 |
| Eif3l | Y | 89 | 0.269236417 | 0.151389707 | 0.478103513 | 0.208867096 | 0.326713806 | 0.141937176 | 0.5025578 | 0.191945193 | 0.513796688 | 0.357926172 |
| Eif3b | Y | 438 | 0.529760743 | 0.140839277 | 0.147670591 | -0.382090152 | 0.006831314 | 0.137576259 | 0.127058669 | 0.393852373 | 0.239017223 | 0.964910424 |
| Arhgap32 | Y | 1523 | 0.19683077 | 0.133508571 | 0.187376935 | -0.009453835 | 0.053868365 | 0.052920461 | 0.242987292 | 0.05444812 | 0.847662747 | 0.541693057 |
| Ctnnd1 | Y | 174 | 0.261330851 | 0.217202707 | 0.118844696 | -0.142486155 | -0.098358012 | 0.505848812 | 0.011575303 | 0.09659312 | 0.704896779 | 0.180589902 |
| Pear1 | Y | 923 | 0.431164163 | 0.243303149 | 0.28454677 | -0.146617392 | 0.041243621 | 0.308798319 | 0.331409165 | 0.184952455 | 0.703140051 | 0.862603302 |
| Anxa2 | Y | 199 | 0.316346208 | 0.083080429 | 0.227916711 | -0.088429497 | 0.144836282 | 0.200200721 | 0.355915676 | 0.02554052 | 0.664155836 | 0.103852187 |
| Bst2 | Y | 6 | 0.518149261 | 0.151518789 | 0.233404614 | -0.284744647 | 0.081885825 | 0.292274267 | 0.355957698 | 0.108383762 | 0.521783321 | 0.553261017 |
| Nectin1 | T | 468 | 0.380225116 | 0.184824827 | -0.03108217 | -0.411307286 | -0.215906997 | 0.008237234 | 0.113831684 | 0.832938324 | 0.047802993 | 0.181768277 |
| Sfpq | Y | 373 | 0.786104547 | 0.284608142 | 0.127793772 | -0.658310775 | -0.15681437 | 0.171143146 | 0.367752258 | 0.737251552 | 0.239309741 | 0.624560545 |
| Itga4 | S | 1028 | 0.154476395 | 0.261662265 | 0.262637708 | 0.108161312 | 0.000975443 | 0.318102507 | 0.075189524 | 0.185624711 | 0.549153899 | 0.995168083 |
| Itga4 | Y | 1031 | 0.154476395 | 0.261662265 | 0.262637708 | 0.108161312 | 0.000975443 | 0.318102507 | 0.075189524 | 0.185624711 | 0.549153899 | 0.995168083 |
| Hspa4 | Y | 336 | 0.208978272 | 0.176977197 | 0.10560688 | -0.103371392 | -0.071370317 | 0.063943869 | 0.057454847 | 0.172691911 | 0.269350435 | 0.337201665 |
| Srrm1 | T | 376 | 0.170868782 | 0.096232426 | 0.407848025 | 0.236979242 | 0.311615599 | 0.393277804 | 0.641788393 | 0.017040309 | 0.237059106 | 0.177841896 |
| Kirrel | Y | 777 | 0.302796469 | 0.009916342 | 0.181499976 | -0.121296492 | 0.171583634 | 0.014667697 | 0.907058751 | 0.052281489 | 0.194962705 | 0.08313359 |
| Eif4a1 Eif4a2 Eif4a3 Eif4a | Y | 50;51;56 | 0.343716215 | 0.137519382 | 0.095657087 | -0.248059128 | -0.041862295 | 0.014060966 | 0.385766782 | 0.34783952 | 0.005951212 | 0.753287504 |
| AW554918 | Y | 268 | 0.128912193 | 0.104018393 | 0.335382145 | 0.206469952 | 0.231363752 | 0.108011083 | 0.462330575 | 0.032701907 | 0.122923447 | 0.176007417 |
| Flna | Y | 791 | 0.075065182 | 0.136211896 | 0.398627884 | 0.323562702 | 0.262415988 | 0.4518922 | 0.423000539 | 0.01462747 | 0.041169052 | 0.160997202 |
| Eif3c | Y | 882 | 0.259106647 | 0.071219231 | 0.283339484 | 0.024232837 | 0.212120253 | 0.081116463 | 0.607057264 | 0.062220636 | 0.857564742 | 0.218440595 |
| Rin1 | Y | 621 | 0.155511093 | 0.136760675 | 0.109202369 | -0.046308724 | -0.027558306 | 0.040010787 | 0.048059631 | 0.079198718 | 0.337609748 | 0.465155744 |
| Itsn2 | Y | 554 | 0.196493333 | 0.048354886 | 0.425241239 | 0.228747906 | 0.376886353 | 0.136353921 | 0.58985369 | 0.061127976 | 0.183227636 | 0.058018141 |
| Ifitm3 | Y | 27 | 0.217017687 | 0.216635695 | 0.175651613 | -0.041366075 | -0.040984082 | 0.186333592 | 0.090097038 | 0.180187539 | 0.75487462 | 0.614289336 |
| Npdc1 | Y | 256 | 0.148936974 | 0.145884629 | 0.42310322 | 0.274166246 | 0.277218591 | 0.101697668 | 0.512199189 | 0.107663369 | 0.230020292 | 0.319999353 |
| Cobll1 | Y | 961 | 0.31955373 | 0.133531295 | 0.185902543 | -0.133651186 | 0.052371248 | 0.109349507 | 0.243284608 | 0.162391102 | 0.44227182 | 0.674129264 |
| Septin10 | Y | 329 | 0.14551765 | 0.017519593 | 0.484328555 | 0.338810905 | 0.466808962 | 0.301619579 | 0.91030709 | 0.017702736 | 0.074752207 | 0.044598621 |
| Afap1 | Y | 554 | 0.221127261 | 0.142238183 | 0.224675475 | 0.003548214 | 0.082437292 | 0.053647065 | 0.287607042 | 0.172254753 | 0.978332958 | 0.592937941 |
| Flt1 | Y | 1048 | 0.120354809 | 0.116546061 | 0.398648188 | 0.278293379 | 0.282102127 | 0.285028369 | 0.342549718 | 0.045884095 | 0.117771758 | 0.121703913 |
| Mapk10 Mapk8 | T | 183;221 | 0.500003164 | 0.192806389 | 0.128244183 | -0.371758981 | -0.064562206 | 0.178373749 | 0.249006979 | 0.281163177 | 0.279748375 | 0.641179053 |
| Triobp | Y | 1974 | 0.265349812 | 0.079402201 | 0.126796183 | -0.138553629 | 0.047393981 | 0.013246356 | 0.466589449 | 0.118185542 | 0.110310508 | 0.645815839 |
| Cdh2 | Y | 820 | 0.085489859 | 0.24287105 | 0.201862394 | 0.116372535 | -0.041008656 | 0.305815729 | 0.017174335 | 0.257677626 | 0.470304834 | 0.78292933 |
| Nectin3 | Y | 438 | 0.089408393 | 0.157659631 | 0.283865687 | 0.194457294 | 0.126206055 | 0.438997512 | 0.204545798 | 0.019697156 | 0.099983217 | 0.23898427 |
| Tubb1 Tubb2a Tubb2b Tul | Y | 159;159;159;159; | 0.89385458 | 0.002895317 | 0.270962152 | -0.622892428 | 0.268066834 | 0.361158378 | 0.973688684 | 0.063493091 | 0.49977057 | 0.072441487 |
| Plxna1 | Y | 1606 | 0.116474177 | 0.184655492 | 0.15954871 | 0.043074532 | -0.025106782 | 0.110430189 | 0.028275451 | 0.197262689 | 0.666030441 | 0.79848495 |
| Rps25 | Y | 55 | 0.295805568 | 0.102244235 | 0.261881973 | -0.033923595 | 0.159637738 | 0.063645877 | 0.412980213 | 0.227551652 | 0.859703235 | 0.419120192 |
| Tln1 | Y | 1893 | 0.14618768 | 0.085804629 | 0.25367026 | 0.107482581 | 0.167865631 | 0.116772934 | 0.392151706 | 0.02300071 | 0.079285206 | 0.079580345 |
| Glg1 | Y | 379 | 0.183827949 | 0.173671351 | 0.368069809 | 0.18424186 | 0.194398458 | 0.172723407 | 0.291019854 | 0.183015489 | 0.445317706 | 0.442492145 |
| Gjc1 | Y | 324 | 0.07716667 | 0.172862312 | 0.25332813 | 0.176161459 | 0.080465818 | 0.455481912 | 0.221792869 | 0.011967033 | 0.120474199 | 0.511232758 |
| Fnbp1l | Y | 291 | 0.298629416 | 0.141657285 | 0.190476618 | -0.108152798 | 0.048819333 | 0.358220678 | 0.29487426 | 0.023825036 | 0.711138124 | 0.684646394 |
| Pard3 | T | 1170 | 0.147472176 | 0.141265298 | 0.20824085 | 0.060768674 | 0.066975553 | 0.036481285 | 0.224236821 | 0.127904344 | 0.571707 | 0.611414752 |
| Eda2r | Y | 289 | 0.630057263 | 0.283485295 | 0.128393996 | -0.501663268 | -0.155091299 | 0.371347106 | 0.198269853 | 0.398470288 | 0.459398403 | 0.419213237 |
| Ephb3 Ephb4 | Y | 603;590 | 0.08818474 | 0.139291767 | 0.365554999 | 0.277370259 | 0.226263232 | 0.631922842 | 0.327494372 | 0.01807637 | 0.198677978 | 0.159693677 |
| Inppl1 | Y | 987 | 0.279960743 | 0.111408049 | 0.351966866 | 0.072006123 | 0.240558817 | 0.345257526 | 0.425541248 | 0.077576975 | 0.797859677 | 0.208691899 |
| Sema4b | Y | 773 | 0.36649203 | 0.081354603 | 0.106594151 | -0.259897879 | 0.025239548 | 0.019272856 | 0.516889054 | 0.289433107 | 0.065201432 | 0.784730687 |
| Cdk20 | Y | 160 | 0.23590397 | 0.131968631 | 0.419311237 | 0.183407267 | 0.287342607 | 0.237839979 | 0.518836063 | 0.132756017 | 0.408932452 | 0.242618803 |
| Psmc1 | Y | 25 | 0.138958115 | 0.149536632 | 0.18715004 | 0.048191924 | 0.037613408 | 0.055151681 | 0.137566892 | 0.138221407 | 0.637865849 | 0.740179292 |
| Vars | Y | 468 | 0.239045744 | -0.037951876 | 0.231343842 | -0.007701902 | 0.269295718 | 0.01709819 | 0.815276669 | 0.033224578 | 0.915627849 | 0.173774291 |
| Afap1 | S | 549 | 0.203799426 | 0.142715763 | 0.214375103 | 0.010575677 | 0.07165934 | 0.085288693 | 0.242320869 | 0.162806432 | 0.924825681 | 0.573675084 |
| Spry1 | T | 54 | -0.106226044 | 0.211512547 | 0.441756727 | 0.547982771 | 0.23024418 | 0.438526364 | 0.291147716 | 0.003932855 | 0.01664528 | 0.253699193 |
| Cbl Cblb | Y | 369;363 | 0.309692765 | 0.015546553 | 0.307831741 | -0.001861024 | 0.292285189 | 0.284558086 | 0.779173401 | 0.021527527 | 0.994060477 | 0.042139036 |
| Gars | Y | 443 | 0.416285861 | 0.178421254 | 0.207778606 | -0.208507255 | 0.029357352 | 0.313481438 | 0.296305271 | 0.166043642 | 0.58044186 | 0.860906831 |
| Cav1 | Y | 14 | 0.137346425 | 0.233725173 | 0.306126918 | 0.168780493 | 0.072401746 | 0.306180535 | 0.125283319 | 0.231507672 | 0.45314577 | 0.738030286 |
| Flt1 | Y | 795 | 0.580850497 | 0.177201956 | 0.252269245 | -0.328581252 | 0.075067289 | 0.345300395 | 0.450574743 | 0.235201659 | 0.567209315 | 0.767306436 |
| Pard3b | Y | 933 | 0.460745714 | 0.125884298 | 0.210577788 | -0.250167927 | 0.08469349 | 0.455594308 | 0.152329994 | 0.115224792 | 0.669049576 | 0.431585539 |
| Hsp90aa1 Hsp90ab1 | Y | 61;56 | 0.217836656 | 0.079678463 | 0.34962075 | 0.131784094 | 0.269942288 | 0.116069648 | 0.356166777 | 0.169093927 | 0.546160505 | 0.256383053 |
| Vim | S | 56 | 0.145942737 | 0.117304688 | 0.240000752 | 0.094058016 | 0.122696065 | 0.107274385 | 0.289126527 | 0.067706038 | 0.387665535 | 0.337121015 |
| Elmo2 | Y | 729 | 0.129527661 | 0.07491835 | 0.309899584 | 0.180371923 | 0.234981234 | 0.218088449 | 0.360729186 | 0.031601939 | 0.159353446 | 0.07098724 |
| Megf10 | Y | 1048 | -0.138352157 | 0.178070209 | 0.415079702 | 0.553431859 | 0.237009493 | 0.103712009 | 0.062775809 | 0.00194434 | 0.001135734 | 0.023884801 |
| Eif4a1 Eif4a2 | Y | 70;71 | 0.430199244 | 0.068638598 | 0.176114065 | -0.254085179 | 0.107475467 | 0.086039765 | 0.505064173 | 0.235091471 | 0.248428913 | 0.378301805 |
| Ddx3x | Y | 301 | 0.115403057 | 0.121292878 | 0.223912704 | 0.108509647 | 0.102619826 | 0.101154602 | 0.304896854 | 0.040240748 | 0.221540039 | 0.398697022 |
| Tpm3 | Y | 163 | 0.093139126 | 0.173846971 | 0.213702113 | 0.120562987 | 0.039855142 | 0.256804658 | 0.119304994 | 0.061352624 | 0.232983345 | 0.689188206 |
| Tln2 | Y | 1780 | 0.056104293 | 0.305154079 | 0.404483921 | 0.348379628 | 0.099329842 | 0.857598483 | 0.133336001 | 0.161712201 | 0.334906596 | 0.675044766 |
| Psma2 | Y | 101 | 0.533794501 | 0.257793766 | 0.135296099 | -0.398498403 | -0.122497667 | 0.421295335 | 0.235304352 | 0.234863166 | 0.531407058 | 0.51026605 |
| Vim | S | 29 | 0.184549159 | 0.17501627 | 0.368428166 | 0.183879008 | 0.193411896 | 0.291068481 | 0.343555561 | 0.160737964 | 0.442002879 | 0.433709424 |
| Ptpn11 | Y | 62 | 0.087493869 | 0.162253875 | 0.145733098 | 0.05823923 | -0.016520777 | 0.070301201 | 0.083999717 | 0.077964719 | 0.367948745 | 0.840574191 |
| Gapdh | Y | 328 | 0.24930519 | 0.294226884 | 0.097215494 | -0.152089696 | -0.19701139 | 0.446886804 | 0.062154885 | 0.240341297 | 0.626633778 | 0.132291558 |
| Nck2 | S | 104 | 0.295104062 | 0.27976809 | 0.131079814 | -0.164024248 | -0.148688276 | 0.392261897 | 0.226770395 | 0.098852284 | 0.610535598 | 0.4655773 |
| Cnot9 | Y | 23 | 0.944092945 | 0.600197962 | 0.289368682 | -0.654724263 | -0.31082928 | NA | 0.087234453 | 0.406462799 | NA | 0.402365597 |
| Dyrk2 Dyrk4 | Y | 380;379 | 0.289401928 | 0.07755549 | 0.130845986 | -0.158555942 | 0.053290496 | 0.026622479 | 0.46441941 | 0.209905631 | 0.14939572 | 0.588919534 |
| Zdhhc8 | S | 534 | 0.134547275 | 0.089435792 | 0.244533941 | 0.109986666 | 0.155098149 | 0.055963061 | 0.251790891 | 0.105524476 | 0.371069511 | 0.246062608 |
| Snw1 | Y | 430 | 0.518501315 | 0.16416596 | 0.041031743 | -0.477469572 | -0.123134217 | 0.364793998 | 0.018836154 | 0.462774452 | 0.396371422 | 0.048225113 |
| Vcl | Y | 822 | 0.060495356 | 0.157738431 | 0.204963661 | 0.144468305 | 0.04722523 | 0.262768242 | 0.16188043 | 0.021616731 | 0.07871993 | 0.628066364 |

| | | | | | | | | | | | | |
|---|---|---|---|---|---|---|---|---|---|---|---|---|
| Cyfip1 Cyfip2 | Y | 108;108 | 0.20671257 | 0.022315593 | 0.408095832 | 0.201383261 | 0.385780238 | 0.103195161 | 0.872477644 | 0.102137951 | 0.333398416 | 0.121044557 |
| Ran | Y | 147 | 0.264814376 | -0.009748596 | 0.236670225 | -0.028144151 | 0.246418822 | 0.020452805 | 0.926456168 | 0.114031828 | 0.80983466 | 0.130521968 |
| Psma2 | Y | 76 | 0.155152603 | 0.084523969 | 0.311125654 | 0.155973052 | 0.226601685 | 0.060125242 | 0.193498284 | 0.235270483 | 0.495916783 | 0.349569141 |
| Acta1 Acta2 Actb Actc1 A | Y | 53;53;55;55;54;5 | 0.41433259 | 0.112503021 | 0.2689018 | -0.14543079 | 0.156398778 | 0.342683571 | 0.349057867 | 0.17050848 | 0.718250651 | 0.393440022 |
| Rbbp6 | Y | 884 | 0.552197549 | 0.216253908 | 0.254787035 | -0.297410514 | 0.038533127 | 0.203520017 | 0.56427734 | 0.520445376 | 0.506879822 | 0.925147371 |
| Sema4c | S | 364 | 0.37846376 | 0.149944278 | 0.150719269 | -0.227744491 | 0.000774991 | 0.084754426 | 0.411419005 | 0.429588605 | 0.257224729 | 0.996397215 |
| Sema4c | T | 359 | 0.37846376 | 0.149944278 | 0.150719269 | -0.227744491 | 0.000774991 | 0.084754426 | 0.411419005 | 0.429588605 | 0.257224729 | 0.996397215 |
| Sema4c | Y | 358 | 0.37846376 | 0.149944278 | 0.150719269 | -0.227744491 | 0.000774991 | 0.084754426 | 0.411419005 | 0.429588605 | 0.257224729 | 0.996397215 |
| Irs2 | Y | 734 | -0.174320174 | 0.365916636 | 0.264224009 | 0.438544183 | -0.101692627 | 0.172830309 | 0.007944796 | 0.044044484 | 0.016234582 | 0.334153552 |
| Ghr | Y | 545 | 0.047284181 | 0.119489313 | 0.299765493 | 0.252481312 | 0.18027618 | 0.581502034 | 0.327410744 | 0.015403762 | 0.01564811 | 0.073586694 |
| Zdhhc20 | Y | 327 | 0.185519765 | 0.087551605 | 0.159970075 | -0.02554969 | 0.07241847 | 0.076900595 | 0.327009993 | 0.05954645 | 0.726726083 | 0.314541789 |
| Abcf1 | Y | 242 | 0.259244864 | -0.027835085 | 0.163516664 | -0.095728201 | 0.191351749 | 0.002627298 | 0.69333099 | 0.176063997 | 0.369704504 | 0.134837955 |
| Psmc3 | T | 134 | 0.430121918 | -0.069236634 | 0.125848467 | -0.304273451 | 0.195085101 | 0.002860451 | 0.742677575 | 0.659589466 | 0.3326049 | 0.550902755 |
| Anxa2 | Y | 188 | 0.15938938 | 0.186170387 | 0.191520836 | 0.032131455 | 0.005350449 | 0.239558021 | 0.151033676 | 0.158958571 | 0.81441831 | 0.966854632 |
| Hsp90aa1 | Y | 605 | 0.378197278 | 0.223655833 | 0.158668824 | -0.219528454 | -0.064987008 | 0.382051771 | 0.305421546 | 0.155647442 | 0.587936297 | 0.744582158 |
| Psmb5 | Y | 220 | 0.080489935 | 0.278359873 | 0.25254049 | 0.172050555 | -0.025819383 | 0.749825384 | 0.144300923 | 0.096252346 | 0.485959006 | 0.849615505 |
| Rpl27 LOC108167922 | Y | 49 | 0.395619473 | 0.050206559 | 0.044549236 | -0.351070237 | -0.005657323 | 0.006393948 | 0.77831718 | 0.737380406 | 0.065120713 | 0.977537079 |
| Adam10 | Y | 742 | 0.220434369 | 0.128975685 | 0.229406768 | 0.008972398 | 0.100431083 | 0.105693294 | 0.365918137 | 0.235188323 | 0.957444931 | 0.582268256 |
| Septin9 | Y | 276 | 0.204425922 | 0.078310023 | 0.336007488 | 0.131581566 | 0.257697464 | 0.452367617 | 0.617166461 | 0.039502759 | 0.620652773 | 0.186242157 |
| Nck2 | S | 94 | 0.078968363 | 0.204356344 | 0.246189551 | 0.167221188 | 0.041833207 | 0.680518629 | 0.182998704 | 0.045515548 | 0.41069755 | 0.756943819 |
| Psmd14 | Y | 32 | 0.628137077 | 0.025083162 | 0.146014813 | -0.482122264 | 0.120931651 | 0.0704427 | 0.899769882 | 0.564327696 | 0.169041821 | 0.659628767 |
| Myof | Y | 969 | 0.585896355 | 0.316489014 | 0.271555501 | -0.314340855 | -0.044933514 | 0.256826726 | 0.741389666 | 0.505729972 | 0.454821838 | 0.960846773 |
| Tspan3 | Y | 252 | 0.164072556 | 0.385030112 | 0.220455353 | 0.056382797 | -0.164574759 | 0.756199193 | 0.082587429 | 0.430143635 | 0.916771438 | 0.530305116 |
| Septin2 | Y | 17 | 0.148344854 | 0.182561631 | 0.125882878 | -0.022461976 | -0.056678753 | 0.193265168 | 0.120160585 | 0.138904544 | 0.791583209 | 0.516424199 |
| Pik3r2 | T | 576 | 0.323843705 | -0.061057661 | 0.216031026 | -0.107812679 | 0.277088687 | 0.034142282 | 0.788624864 | 0.111887667 | 0.131367241 | 0.265463841 |
| Pik3r2 | Y | 571 | 0.323843705 | -0.061057661 | 0.216031026 | -0.107812679 | 0.277088687 | 0.034142282 | 0.788624864 | 0.111887667 | 0.131367241 | 0.265463841 |
| Pxn | S | 83 | 0.103961572 | 0.119362508 | 0.246891075 | 0.142929503 | 0.127528567 | 0.237191242 | 0.32836004 | 0.050719775 | 0.195958149 | 0.334704333 |
| Kirrel | Y | 693 | -0.08169489 | 0.209714716 | 0.179105432 | 0.260800322 | -0.030609284 | 0.576445977 | 0.021253867 | 0.008752244 | 0.160906613 | 0.59364169 |
| Nectin1 | Y | 395 | 0.154424111 | 0.250214152 | 0.186509709 | 0.032085598 | -0.063704442 | 0.252349324 | 0.305502913 | 0.147113673 | 0.681306918 | 0.758123974 |
| Src | Y | 237 | 0.224624687 | 0.02056501 | 0.525522539 | 0.300897852 | 0.504957528 | 0.397243609 | 0.945755456 | 0.095165985 | 0.274366827 | 0.147387608 |
| Smc6 | Y | 467 | 0.234578141 | 0.144213041 | 0.308750339 | 0.074172198 | 0.164537298 | 0.388899467 | 0.366558845 | 0.169732734 | 0.78702626 | 0.411942203 |
| Arhgap35 | S | 1070 | 0.415435279 | 0.195558448 | 0.063251907 | -0.352183372 | -0.132306541 | 0.188105503 | 0.258091321 | 0.393195187 | 0.244704311 | 0.4041028 |
| Mapk1 | Y | 34 | 0.35215585 | 0.011466386 | 0.489538521 | 0.137382671 | 0.478072135 | 0.362055633 | 0.964371404 | 0.147055475 | 0.730523766 | 0.199918774 |
| H4c1 H4c11 H4c12 H4c14 | Y | 73 | 1.459788908 | 0.020354061 | 0.076843378 | -1.38294553 | 0.056489317 | 0.278237447 | 0.904298307 | 0.693810205 | 0.296417225 | 0.804147823 |
| Kirrel | S | 715 | -0.024382058 | 0.11118323 | 0.367750545 | 0.392132603 | 0.256567315 | 0.880372109 | 0.337507753 | 0.011780266 | 0.071938853 | 0.049011888 |
| Hgs | Y | 334 | 0.040289596 | 0.29185847 | 0.105500667 | 0.065211071 | -0.186357803 | 0.683805844 | 0.006736459 | 0.585000905 | 0.733624407 | 0.360652747 |
| Rnf213 | Y | 1717 | 0.352270062 | 0.025305674 | 0.241793293 | -0.110476769 | 0.21648762 | 0.09094749 | 0.869500704 | 0.243467734 | 0.553776304 | 0.230621732 |
| Adgrl3 | Y | 1447 | 0.408216821 | 0.256845428 | 0.091103983 | -0.317112838 | -0.165741445 | 0.363354618 | 0.166118515 | 0.531766941 | 0.463442314 | 0.353065802 |
| Plec | Y | 1103 | 0.095272925 | 0.100477471 | 0.075903485 | -0.01936944 | -0.024573986 | 0.019191146 | 0.038920157 | 0.155523816 | 0.641782406 | 0.607181964 |
| Rbm22 | Y | 33 | 0.393214872 | 0.230533682 | 0.301702373 | -0.091512498 | 0.071168691 | 0.371880999 | 0.625367763 | 0.298602815 | 0.812918677 | 0.870745396 |
| Ywhag | Y | 117 | 0.139985283 | 0.230559685 | 0.3012907 | 0.161305417 | 0.070731014 | 0.719702071 | 0.309933 | 0.11548274 | 0.673427216 | 0.713570025 |
| Bcar1 | Y | 291 | 0.192631119 | 0.177978368 | 0.13777193 | -0.054859189 | -0.040206438 | 0.113288166 | 0.33810859 | 0.225396387 | 0.114279081 | 0.792875701 |
| Rhot1 | Y | 478 | 0.158000501 | 0.154157185 | 0.204481472 | 0.046480971 | 0.050324288 | 0.330367695 | 0.342131947 | 0.082082359 | 0.732771225 | 0.713163973 |
| Frk | Y | 504 | 0.06583425 | 0.051080286 | 0.319608236 | 0.253773986 | 0.26852795 | 0.699294186 | 0.706146134 | 0.007485921 | 0.210019573 | 0.125548574 |
| Snx13 | T | 691 | 0.190706854 | 0.111580837 | 0.223629383 | 0.032922529 | 0.112048546 | 0.114882253 | 0.234857532 | 0.342987188 | 0.876227172 | 0.596912766 |
| Skap2 | Y | 197 | 0.132041849 | -0.016874218 | 0.506104224 | 0.374062376 | 0.522978442 | 0.40899428 | 0.923620868 | 0.051890653 | 0.114071424 | 0.054822051 |
| Pgk1 | Y | 196 | 0.195170903 | 0.057773729 | 0.120430025 | -0.074740878 | 0.062656296 | 0.03057275 | 0.181646667 | 0.286725354 | 0.494653348 | 0.53527663 |
| Spred1 | Y | 212 | -0.001551487 | 0.230873073 | 0.248204237 | 0.249755724 | 0.017331164 | 0.989245993 | 0.12311968 | 0.06037005 | 0.117695316 | 0.89470883 |
| Zdhhc8 | Y | 538 | 0.098195889 | 0.083280223 | 0.215673021 | 0.117477132 | 0.132392799 | 0.105695087 | 0.256771556 | 0.100408499 | 0.289370934 | 0.255049629 |
| Rgs17 | Y | 137 | 0.337073066 | 0.106616178 | 0.308835772 | -0.028237294 | 0.202219594 | 0.151736705 | 0.625709411 | 0.471261824 | 0.942093582 | 0.614791595 |
| Bcar3 | T | 410 | 0.34680445 | 0.191554999 | 0.257935482 | -0.088868968 | 0.066380483 | 0.427428774 | 0.535101933 | 0.234708366 | 0.830273311 | 0.830512831 |
| Ptk2 | Y | 570 | 0.025261803 | 0.118859168 | 0.417093627 | 0.391831824 | 0.298234459 | 0.708824242 | 0.457510945 | 0.05522019 | 0.061571509 | 0.160925045 |
| Itprid2 | Y | 731 | 0.329289538 | 0.001401658 | 0.243419738 | -0.0858698 | 0.24201808 | 0.097435752 | 0.995746292 | 0.207469805 | 0.60153959 | 0.377938063 |
| Larp1 | S | 525 | 0.429229679 | 0.367310421 | 0.022495989 | -0.40673369 | -0.344814432 | 0.282854306 | 0.252730979 | 0.914825346 | 0.297610996 | 0.262020526 |
| Hnrnpa1 | Y | 167 | 0.324427072 | -0.018889782 | 0.25613343 | -0.068293642 | 0.275023212 | 0.065725584 | 0.927167656 | 0.257195691 | 0.732906949 | 0.288637949 |
| Cep89 | Y | 154 | 0.160259045 | 0.063261431 | 0.154547397 | -0.005711648 | 0.091285966 | 0.186863696 | 0.489818944 | 0.024484218 | 0.952343687 | 0.31774956 |
| Arhgef10 | Y | 93 | 0.182265003 | 0.133565084 | 0.283518852 | 0.101253849 | 0.149953768 | 0.653983103 | 0.531558434 | 0.051078006 | 0.795860475 | 0.45693877 |
| Fyb | Y | 375 | 0.322108808 | 0.087005951 | 0.062417952 | -0.259690856 | -0.024587999 | 0.033710209 | 0.461733021 | 0.602417389 | 0.066762525 | 0.812982269 |
| Efnb1 Efnb2 | Y | 342;333 | 0.505624655 | 0.22613139 | 0.036535554 | -0.469089101 | -0.189595836 | 0.36585928 | 0.12954716 | 0.812641613 | 0.396061506 | 0.274238793 |
| Stat3 | Y | 45 | 0.368023498 | 0.171494922 | 0.07186148 | -0.296162018 | -0.099633442 | 0.155919728 | 0.233636983 | 0.745034009 | 0.310110305 | 0.678070368 |
| Cdc40 | Y | 85 | 0.234563388 | 0.267671713 | 0.21616293 | -0.018400459 | -0.051508784 | 0.744658468 | 0.20273015 | 0.255121547 | 0.978970992 | 0.660403658 |
| Plcg1 | Y | 775 | 0.120255546 | 0.247339567 | 0.269669391 | 0.149413845 | 0.022329824 | 0.722136132 | 0.231617324 | 0.201243497 | 0.679028687 | 0.920339148 |
| Anapc5 | Y | 706 | 0.297260063 | -0.101357182 | 0.289380422 | -0.007879641 | 0.390737605 | 0.022059122 | 0.291889888 | 0.148076992 | 0.960103216 | 0.095466392 |
| Hnrnpk | Y | 449 | 0.432048463 | 0.053919412 | 0.100521188 | -0.331527274 | 0.046601776 | 0.038708956 | 0.829480938 | 0.757333065 | 0.343655374 | 0.895697349 |
| Slc38a2 | Y | 20 | 0.189414994 | 0.167957115 | 0.169846694 | -0.0195683 | 0.001889579 | 0.299546353 | 0.218471154 | 0.285972441 | 0.919407611 | 0.990782437 |
| Uba1 | Y | 542 | 0.450405287 | 0.192502402 | -0.255796887 | -0.706202174 | -0.448299289 | 0.012473125 | 0.158916853 | 0.296309226 | 0.055346204 | 0.1212874 |
| Pik3r1 | Y | 580 | 0.381197719 | 0.087617782 | 0.106388862 | -0.274808857 | 0.01877108 | 0.239819562 | 0.448676166 | 0.167130185 | 0.360415494 | 0.844057626 |
| Ddx3x | Y | 104 | 0.296739929 | -0.009116074 | 0.12020041 | -0.176539518 | 0.129316485 | 0.031550791 | 0.95111771 | 0.191008426 | 0.141976111 | 0.425308091 |
| Shc1 | S | 422 | 0.097397913 | 0.429918904 | 0.101603506 | 0.004205593 | -0.328315398 | 0.576037263 | 0.074183507 | 0.805827727 | 0.991534071 | 0.450828256 |
| Pcnx3 | Y | 279 | 0.009625967 | 0.154251527 | 0.300078033 | 0.290452065 | 0.145826505 | 0.910489882 | 0.139718784 | 0.090438572 | 0.08451955 | 0.298144054 |
| Sorbs2 | Y | 922 | 0.064656601 | 0.045495072 | 0.307591167 | 0.242934566 | 0.262096095 | 0.413049881 | 0.698553409 | 0.025468429 | 0.05580033 | 0.10229338 |
| Kirrel | Y | 719 | 0.166298113 | 0.111508975 | 0.197735845 | 0.031437732 | 0.086226871 | 0.219530344 | 0.413893993 | 0.16625436 | 0.559408437 | 0.285866522 |
| Bcar1 | Y | 556 | 0.179819744 | 0.175437005 | 0.185100483 | 0.005280738 | 0.009663478 | 0.296334082 | 0.349812706 | 0.242988898 | 0.976504943 | 0.959999641 |
| Mapk7 | Y | 221 | 0.247457565 | 0.091134234 | 0.053693499 | -0.193764066 | -0.037440735 | 0.084320381 | 0.124787894 | 0.520478241 | 0.153002447 | 0.616087598 |
| Abi1 | S | 282 | 0.334635043 | 0.277780451 | 0.063195879 | -0.271439164 | -0.214584572 | 0.391832422 | 0.207059066 | 0.59460919 | 0.476198275 | 0.313414305 |
| Dcbld2 | Y | 616 | 0.287908173 | 0.0613295 | 0.095400341 | -0.192507832 | 0.034070841 | 0.235810768 | 0.445815896 | 0.05313042 | 0.3826874 | 0.637996052 |
| Micu1 | S | 124 | 0.197085693 | 0.057917383 | 0.091134345 | -0.105951347 | 0.033216963 | 0.022026031 | 0.518277327 | 0.308667361 | 0.211336943 | 0.718444628 |
| Micu1 | T | 125 | 0.197085693 | 0.057917383 | 0.091134345 | -0.105951347 | 0.033216963 | 0.022026031 | 0.518277327 | 0.308667361 | 0.211336943 | 0.718444628 |
| Micu1 | Y | 123 | 0.197085693 | 0.057917383 | 0.091134345 | -0.105951347 | 0.033216963 | 0.022026031 | 0.518277327 | 0.308667361 | 0.211336943 | 0.718444628 |
| Arhgap35 | S | 1106 | 0.209143202 | 0.468567468 | -0.227031351 | -0.436174553 | -0.695598819 | 0.216327665 | 0.068536488 | 0.647350579 | 0.408249677 | 0.233304703 |
| Naa10 Naa11 | Y | 137;137 | 0.188755828 | 0.06966344 | 0.080807857 | -0.107947971 | 0.011144417 | 0.050101942 | 0.370924445 | 0.214918819 | 0.171426426 | 0.834032547 |
| Eif3m | Y | 351 | 0.603041204 | -0.015474987 | 0.052016148 | -0.551025056 | 0.067491135 | 0.064524051 | 0.955638161 | 0.803150245 | 0.095730032 | 0.83100815 |
| Eno1 Eno1b | Y | 287 | 0.579615723 | 0.051492668 | 0.215829589 | -0.363786134 | 0.164336921 | 0.347714552 | 0.829526352 | 0.336181016 | 0.530711795 | 0.514843721 |
| Acot7 | Y | 195 | 0.027084031 | 0.142690229 | 0.197511078 | 0.170427047 | 0.054820849 | 0.692455841 | 0.059898691 | 0.135160422 | 0.182422604 | 0.612670083 |
| Rplp2 | S | 17 | 0.256278358 | 0.274305215 | 0.339683719 | 0.083405362 | 0.065378505 | 0.583868514 | 0.554309966 | 0.378645133 | 0.808565671 | 0.845695404 |
| Gja1 | Y | 137 | 0.34498106 | -0.036856578 | 0.175668608 | -0.169312452 | 0.212525186 | 0.052854407 | 0.807813443 | 0.372570083 | 0.415213638 | 0.340136225 |
| Igf1r | Y | 1352 | -0.000425413 | 0.147024163 | 0.274733294 | 0.275158707 | 0.127709131 | 0.996974185 | 0.244473433 | 0.059543034 | 0.086733013 | 0.359349035 |
| Tfrc | Y | 20 | 0.243858024 | 0.052096152 | 0.195178456 | -0.048679567 | 0.143082305 | 0.316995018 | 0.667570578 | 0.108313637 | 0.824215309 | 0.284556275 |
| Rbmxl1 | Y | 222 | 0.165011206 | 0.151102651 | 0.245490712 | 0.080479506 | 0.094388061 | 0.292729442 | 0.577933939 | 0.242671457 | 0.685604222 | 0.7444278 |
| Erbb2 | Y | 1197 | 0.203744447 | 0.264021053 | -0.132501704 | -0.336246151 | -0.396522757 | 0.398507054 | 0.002579743 | 0.370926082 | 0.220763219 | 0.06744493 |
| Gja1 | Y | 267 | 0.479048876 | 0.125569383 | 0.068690408 | -0.410358468 | -0.056878976 | 0.369852392 | 0.394215153 | 0.333529712 | 0.429428572 | 0.671653385 |

| | | | | | | | | | | | | |
|---|---|---|---|---|---|---|---|---|---|---|---|---|
| Efnb1 Efnb2 | Y | 328;319 | 0.802124312 | 0.182235991 | -0.050771319 | -0.852895631 | -0.23300731 | 0.344334868 | 0.30937473 | 0.879993852 | 0.324678687 | 0.493261521 |
| H1f2 H1f5 | S | 173;170 | 0.168067718 | 0.341415032 | 0.110158466 | -0.057909252 | -0.231256565 | 0.540625568 | 0.200788149 | 0.586216383 | 0.782498489 | 0.261256327 |
| Cdk1 | Y | 19 | -0.021372011 | 0.277412947 | 0.059616967 | 0.080988978 | -0.21779598 | 0.772477779 | 0.004075998 | 0.718075864 | 0.627209752 | 0.253773389 |
| Lyn | Y | 32 | 0.091303721 | 0.263203885 | -0.057043516 | -0.148347237 | -0.320247401 | 0.343828124 | 0.002134189 | 0.212838662 | 0.177498106 | 2.36116E-05 |
| Mapk10 Mapk8 | Y | 185;223 | 0.250174161 | 0.127480852 | 0.084866199 | -0.165307962 | -0.042614653 | 0.169400688 | 0.391436799 | 0.418334046 | 0.320434583 | 0.755820651 |
| Arhgap35 | S | 1085 | 0.346601637 | 0.151764309 | 0.010905278 | -0.33569636 | -0.140859032 | 0.116712198 | 0.373403257 | 0.902615744 | 0.142579061 | 0.378194881 |
| Cap1 | Y | 353 | 0.157212705 | -0.028032202 | 0.145363173 | -0.011849532 | 0.173395375 | 0.035415162 | 0.727611559 | 0.055268791 | 0.823236716 | 0.077590715 |
| Mvb12a | Y | 208 | 0.291725049 | 0.172915979 | 0.088850242 | -0.202874807 | -0.084065737 | 0.417530272 | 0.302863467 | 0.321492615 | 0.552977487 | 0.56685055 |
| Psmb6 | S | 57 | 0.25820974 | 0.000854279 | 0.214400428 | -0.043809312 | 0.213546149 | 0.145785804 | 0.997037179 | 0.228492304 | 0.810033941 | 0.413780807 |
| Skap2 | S | 247 | 0.412998561 | -0.092752023 | 0.159139681 | -0.25385888 | 0.251891704 | 0.027724574 | 0.330028309 | 0.44361999 | 0.266490684 | 0.258257784 |
| Nedd9 | Y | 91 | -0.108476076 | 0.244427933 | 0.372005492 | 0.480481569 | 0.127577559 | 0.33631277 | 0.226682159 | 0.068979252 | 0.037970614 | 0.538344514 |
| Anxa5 | Y | 92 | 0.027944575 | 0.218881781 | 0.105568847 | 0.077624272 | -0.113312934 | 0.734575502 | 0.055698377 | 0.283411294 | 0.386183641 | 0.275566847 |
| Plec | Y | 491 | 0.165185314 | 0.059423594 | 0.152703001 | -0.012482313 | 0.093279407 | 0.101491683 | 0.658764562 | 0.222109409 | 0.910064459 | 0.533177865 |
| Eif4b | Y | 285 | 0.322664509 | -0.109185377 | 0.237455166 | -0.085209343 | 0.346640542 | 0.169301931 | 0.605411636 | 0.10692672 | 0.662538568 | 0.157452602 |
| Eif3c | Y | 879 | 0.252429516 | -0.008721662 | 0.116891646 | -0.13553787 | 0.125613307 | 0.016345977 | 0.932889669 | 0.56857582 | 0.517738715 | 0.561362724 |
| Ncoa5 | Y | 16 | 0.333757103 | 0.046926651 | 0.082055901 | -0.251701202 | 0.03512925 | 0.135763211 | 0.709888321 | 0.317194602 | 0.224963403 | 0.760317107 |
| Eif4g2 | Y | 29 | 0.06096893 | -0.009973694 | 0.167266276 | 0.106297346 | 0.17723997 | 0.219450694 | 0.917774354 | 0.003594382 | 0.075141098 | 0.16279599 |
| Cdk5 | Y | 15 | -0.052654543 | 0.147077942 | 0.161283855 | 0.213938398 | 0.014205914 | 0.342353122 | 0.033163153 | 0.024750886 | 0.002761729 | 0.649583642 |
| Afap1l2 | Y | 383 | 0.119238184 | 0.047219288 | 0.383007896 | 0.263769712 | 0.335788608 | 0.69196903 | 0.902771357 | 0.089550926 | 0.415173701 | 0.424170716 |
| Actb Actg1 | Y | 198;198 | 0.449985661 | 0.027478248 | 0.060715078 | -0.389270583 | 0.03323683 | 0.305857937 | 0.665664781 | 0.072223338 | 0.359638957 | 0.593982624 |
| Stam2 | Y | 185 | 0.139203843 | -0.018410496 | 0.072517212 | -0.066686631 | 0.090927708 | 0.000934621 | 0.787729759 | 0.254147668 | 0.284794902 | 0.296179963 |
| Lpp | Y | 301 | -0.046695556 | 0.141069958 | 0.195808421 | 0.242503978 | 0.054738464 | 0.475825148 | 0.067991381 | 0.036725352 | 0.027307499 | 0.460654515 |
| Copa | Y | 458 | 0.376513647 | -0.012578317 | 0.21838098 | -0.158132667 | 0.230959297 | 0.332711393 | 0.978383787 | 0.190688239 | 0.65234228 | 0.625787202 |
| Pkm | Y | 148 | 0.112256868 | 0.251539611 | 0.052419 | -0.059837868 | -0.199120611 | 0.521637856 | 0.080803688 | 0.62826855 | 0.707872442 | 0.116969565 |
| Hnrnpa2b1 | Y | 174 | 0.280715918 | -0.087168893 | 0.343861737 | 0.063145819 | 0.43103063 | 0.197483403 | 0.683402575 | 0.214461582 | 0.745315259 | 0.105389758 |
| Cavin2 | Y | 392 | 0.107379156 | 0.118870681 | 0.124020726 | 0.01664157 | 0.005150045 | 0.32825279 | 0.222388031 | 0.198504533 | 0.858102654 | 0.946444394 |
| Tln1 | Y | 436 | -0.084573171 | 0.066758914 | 0.232939188 | 0.317512359 | 0.166180274 | 0.376982808 | 0.083274808 | 0.006052053 | 0.033079299 | 0.030791612 |
| Gja1 | Y | 313 | 0.031634666 | 0.109855262 | 0.151088163 | 0.119453498 | 0.041232901 | 0.558633846 | 0.31377016 | 0.034486322 | 0.041339009 | 0.670870083 |
| Pard3 | Y | 199 | -0.003104152 | 0.016639174 | 0.366217948 | 0.3693221 | 0.349578774 | 0.984570999 | 0.909529437 | 0.023987311 | 0.060091918 | 0.043228632 |
| Rpl10a | Y | 11 | 0.28640943 | -0.079272456 | -0.030247892 | -0.316657322 | 0.049024564 | 0.000485378 | 0.618480088 | 0.800890264 | 0.082833955 | 0.787021382 |
| Prrc2c | Y | 1191 | 0.294704869 | 0.181426973 | 0.014570273 | -0.280134596 | -0.1668567 | 0.221854537 | 0.258552639 | 0.923007901 | 0.248505225 | 0.323083167 |
| Jak1 | Y | 567 | 0.279960968 | 0.139124798 | 0.024241485 | -0.255719482 | -0.114883313 | 0.096835575 | 0.456806735 | 0.888014337 | 0.218943953 | 0.595738611 |
| Nedd9 | Y | 163 | 0.162586633 | 0.020210088 | 0.214922282 | 0.05233565 | 0.194712195 | 0.13450994 | 0.843438966 | 0.229920501 | 0.730942004 | 0.26261153 |
| Hsp90ab1 | Y | 596 | 0.464251054 | 0.110014104 | 0.049981032 | -0.414270021 | -0.060033072 | 0.43801752 | 0.314060381 | 0.396213706 | 0.481598819 | 0.551945629 |
| Kirrel | Y | 756 | 0.091081776 | 0.021358988 | 0.111155665 | 0.020073889 | 0.089796677 | 0.080217564 | 0.624421981 | 0.035232795 | 0.512327747 | 0.024303494 |
| Pikfyve | Y | 1085 | 0.350799239 | 0.168379613 | 0.173547723 | -0.177251516 | 0.00516811 | 0.434642444 | 0.626327263 | 0.464402499 | 0.691822113 | 0.988880665 |
| Pabpc1 | Y | 364 | 0.233321637 | 0.008842377 | 0.060803813 | -0.172517824 | 0.051961436 | 0.024442664 | 0.922161925 | 0.443276884 | 0.075241778 | 0.565348317 |
| Ngef | Y | 177 | 0.458679499 | -0.644621227 | 0.297233016 | -0.161446482 | 0.941854243 | 0.261750531 | NA | 0.596896323 | 0.731913192 | NA |
| Vim | Y | 276 | 0.095512309 | 0.054409574 | 0.218342562 | 0.122830253 | 0.163932988 | 0.633590315 | 0.462718917 | 0.070628114 | 0.551143042 | 0.1344559 |
| Fyb | Y | 559 | 0.014965185 | 0.020774912 | 0.351070242 | 0.336105057 | 0.330295331 | 0.873933642 | 0.78860716 | 0.041730175 | 0.052000515 | 0.059087594 |
| Ptk2 | Y | 861 | 0.118057606 | 0.090819204 | 0.175702588 | 0.057644982 | 0.084883384 | 0.297443649 | 0.436977991 | 0.223282102 | 0.565873807 | 0.438866274 |
| Pard3 | Y | 1094 | 0.435883845 | 0.155044941 | 0.026168354 | -0.409715491 | -0.128876587 | 0.339575302 | 0.340508974 | 0.841995776 | 0.363896299 | 0.436961737 |
| Lpp | Y | 403 | 0.151976987 | 0.029282273 | 0.183782586 | 0.031805599 | 0.154500313 | 0.145519041 | 0.794270707 | 0.223574362 | 0.813145249 | 0.323222708 |
| Ppp1ca | T | 320 | 0.196754706 | 0.0303421 | 0.08646393 | -0.110290775 | 0.056121831 | 0.040415211 | 0.857124015 | 0.395030594 | 0.237908619 | 0.742013933 |
| Nxf1 | Y | 74 | -0.082865551 | 0.186584396 | 0.227322847 | 0.310188397 | 0.040738451 | 0.409520908 | 0.104649391 | 0.075331299 | 0.027164051 | 0.680777773 |
| Skap2 | Y | 237 | 0.441017028 | 0.045883505 | 0.075686725 | -0.365330303 | 0.02980322 | 0.258188836 | 0.595121009 | 0.557764211 | 0.328326731 | 0.779066094 |
| Shank1 | T | 158 | 0.190991565 | 0.017344851 | 0.110772271 | -0.080219293 | 0.09342742 | 0.039731786 | 0.800333553 | 0.483978215 | 0.588724095 | 0.530390183 |
| Shank1 | Y | 156 | 0.190991565 | 0.017344851 | 0.110772271 | -0.080219293 | 0.09342742 | 0.039731786 | 0.800333553 | 0.483978215 | 0.588724095 | 0.530390183 |
| Dok1 | Y | 295 | 0.084763626 | 0.024926073 | 0.204982242 | 0.120218616 | 0.180056169 | 0.715713447 | 0.76157286 | 0.024178248 | 0.610725269 | 0.077272033 |
| Adam19 | Y | 799 | 0.257469744 | -0.036539684 | 0.094722746 | -0.162746998 | 0.13126243 | 0.026032035 | 0.740612341 | 0.506886537 | 0.295961755 | 0.416451532 |
| Pard3 | S | 1112 | 0.454758014 | 0.178017135 | -0.039161701 | -0.493919715 | -0.217178836 | 0.287196604 | 0.302519437 | 0.778198552 | 0.257840816 | 0.227946955 |
| N4bp3 | Y | 79 | 0.180554726 | 0.014882827 | 0.104646221 | -0.075908505 | 0.089763394 | 0.021498577 | 0.902549642 | 0.568838877 | 0.675560864 | 0.659326954 |
| Gab2 | S | 319 | 0.33380767 | 0.131982124 | 0.190651156 | -0.143156515 | 0.058669032 | 0.551550735 | 0.573752504 | 0.452694409 | 0.786797704 | 0.743403172 |
| Lamtor1 | Y | 138 | 0.411888944 | -0.274543655 | 0.251392355 | -0.160496589 | 0.52593601 | 0.058649403 | 0.339320057 | 0.295921404 | 0.50840498 | 0.141835453 |
| Errfi1 | Y | 209 | 0.407613863 | 0.150791662 | -0.038187093 | -0.445800956 | -0.188978755 | 0.325059687 | 0.125474586 | 0.526838652 | 0.293379092 | 0.08339302 |
| Actb Actbl2 Actg1 | Y | 166;166;167 | 0.222475596 | 0.138100104 | -0.005006193 | -0.227481789 | -0.143106296 | 0.207028821 | 0.155712671 | 0.925765632 | 0.196555145 | 0.151644781 |
| Eml4 | Y | 237 | 0.007852218 | -0.019106986 | 0.40313668 | 0.395284461 | 0.422243666 | 0.945662356 | 0.914284134 | 0.051413063 | 0.074032506 | 0.073199934 |
| Des Ina Nefm Vim | Y | 383;387;379;382 | 0.101084504 | 0.009327939 | 0.198206372 | 0.097121868 | 0.188878434 | 0.366416972 | 0.917057615 | 0.070730383 | 0.287760976 | 0.021134403 |
| Sdc3 | Y | 441 | 0.018734683 | 0.29359128 | -0.032665304 | -0.051399987 | -0.326256583 | 0.825928409 | 0.013919934 | 0.680700321 | 0.46367994 | 0.010536026 |
| Arhgap12 | S | 238 | 0.167688527 | 0.044318597 | 0.071807415 | -0.095881112 | 0.027488818 | 0.125388702 | 0.417779263 | 0.339090156 | 0.340107841 | 0.653012223 |
| Snx3 | Y | 22 | 0.703002318 | 0.154390015 | -0.073886779 | -0.776889097 | -0.228276794 | 0.410245845 | 0.279489573 | 0.664632396 | 0.371428553 | 0.157426669 |
| Arhgef10 | Y | 215 | 0.498176447 | 0.272423276 | -0.129424605 | -0.627601052 | -0.401847881 | 0.365951974 | 0.306247633 | 0.672613974 | 0.280288448 | 0.218852375 |
| Jcad | Y | 296 | 0.150269226 | 0.158389098 | 0.115300097 | -0.034969129 | -0.043089001 | 0.349627782 | 0.505001199 | 0.430403509 | 0.701939336 | 0.8308486 |
| Gja1 | Y | 265 | 0.294878073 | 0.040751475 | 0.09083948 | -0.204038593 | 0.050088005 | 0.338916078 | 0.748708601 | 0.24372889 | 0.482337702 | 0.705133184 |
| Peak1 | Y | 662 | -0.00070331 | 0.169071947 | 0.125847564 | 0.126550875 | -0.043224383 | 0.990132844 | 0.152144037 | 0.174487192 | 0.181680671 | 0.693254458 |
| Arhgap12 | T | 229 | 0.162676162 | 0.049856728 | 0.061077373 | -0.101598789 | 0.011220645 | 0.139807931 | 0.357594701 | 0.397291075 | 0.325239058 | 0.859472384 |
| Hnrnpa3 | Y | 374 | 0.200223239 | 0.154190574 | -0.060006359 | -0.260229598 | -0.214196933 | 0.033666014 | 0.504420465 | 0.785187 | 0.302669657 | 0.469556187 |
| Acly | Y | 672 | 0.193206664 | 0.039642822 | 0.032399255 | -0.160807409 | -0.007243567 | 0.057833156 | 0.381188359 | 0.811835728 | 0.311438033 | 0.957807596 |
| Tgfbr1 | Y | 182 | 0.079137261 | 0.097519971 | 0.396656594 | 0.317519334 | 0.299136623 | 0.886372035 | 0.751771339 | 0.2496155 | 0.6002823 | 0.451653077 |
| Maco1 | T | 503 | 0.062377185 | -0.014237726 | 0.169876523 | 0.107499338 | 0.184114249 | 0.480764319 | 0.864393214 | 0.017167405 | 0.264323024 | 0.097598982 |
| Maco1 | T | 505 | 0.062377185 | -0.014237726 | 0.169876523 | 0.107499338 | 0.184114249 | 0.480764319 | 0.864393214 | 0.017167405 | 0.264323024 | 0.097598982 |
| Ass1 | Y | 322 | 0.290329507 | 0.075379084 | 0.083398444 | -0.206931062 | 0.00801936 | 0.406890667 | 0.571811525 | 0.383874624 | 0.532948993 | 0.938428356 |
| Adam9 | Y | 841 | 0.173483886 | 0.098866359 | 0.06054607 | -0.112937816 | -0.038320289 | 0.524348616 | 0.197820135 | 0.340121994 | 0.66712524 | 0.534010327 |
| Ddx17 Ddx5 | Y | 202;200 | 0.257165827 | -0.096766413 | 0.056733624 | -0.200432204 | 0.153500037 | 0.008793019 | 0.501555322 | 0.694873851 | 0.226639401 | 0.40233079 |
| Arhgef7 | Y | 811 | 0.230808352 | -0.084498175 | 0.138914085 | -0.091894267 | 0.22341226 | 0.077736237 | 0.372953105 | 0.168174493 | 0.336834624 | 0.010702805 |
| Tubb2a Tubb2b Tubb3 Tul | Y | 340;340;340;340; | 0.317117577 | -0.027295718 | 0.068306157 | -0.24881142 | 0.095601875 | 0.104484085 | 0.739039977 | 0.488761842 | 0.156229164 | 0.423394611 |
| Ptprk Ptprm | Y | 928;929 | -0.063214042 | 0.081836157 | 0.274012949 | 0.337226991 | 0.192176792 | 0.626499973 | 0.554159675 | 0.043709508 | 0.035852128 | 0.150993878 |
| Lmna | S | 22 | 0.101583964 | 0.071937613 | 0.177608501 | 0.076024537 | 0.105670887 | 0.442393191 | 0.66364496 | 0.228044994 | 0.57962925 | 0.538637677 |
| Megf10 | S | 1059 | -0.015569347 | 0.042934594 | 0.164877922 | 0.180447269 | 0.121943328 | 0.810623407 | 0.325517675 | 0.019985587 | 0.050754097 | 0.056970848 |
| Irs1 | Y | 935 | -0.176944541 | 0.221793965 | 0.251809867 | 0.428754408 | 0.030015902 | 0.208443562 | 0.076477724 | 0.120022655 | 0.031622714 | 0.803916565 |
| Wdr6 | Y | 332 | 0.2877555 | -0.036615566 | 0.064059864 | -0.223695636 | 0.10067543 | 0.078685058 | 0.737143543 | 0.524885487 | 0.151915469 | 0.427807676 |
| Ppp1r12a | Y | 496 | 0.158260592 | -0.023976641 | 0.125941571 | -0.032319021 | 0.149918212 | 0.222135951 | 0.826483442 | 0.106597018 | 0.777956334 | 0.247659752 |
| Pik3r1 Pik3r3 | Y | 470;202 | 0.166870791 | -0.03596555 | 0.048361253 | -0.118509537 | 0.084326804 | 0.009682815 | 0.677644457 | 0.47712276 | 0.128057546 | 0.381592513 |
| Bcar1 | Y | 391 | 0.073261428 | 0.054219304 | 0.11566979 | 0.042408362 | 0.061450486 | 0.315265768 | 0.388140715 | 0.205916006 | 0.59370931 | 0.418141565 |
| Fyb | Y | 807 | 0.041741392 | 0.44706679 | -0.18039361 | -0.222135002 | -0.6274604 | 0.912614776 | 0.036424965 | 0.514058149 | 0.598049687 | 0.078826797 |
| Kirrel | Y | 637 | 0.032747226 | -0.002542618 | 0.179132163 | 0.146384938 | 0.181674781 | 0.641926154 | 0.954416079 | 0.023212605 | 0.092550949 | 0.037450771 |
| Atxn2l | Y | 347 | 0.093601199 | 0.054267962 | 0.096516101 | 0.002914902 | 0.042248138 | 0.213814894 | 0.542539055 | 0.256341961 | 0.972264515 | 0.670433947 |
| Hnrnpul1 | Y | 511 | -0.020906428 | -0.023309545 | 0.262054338 | 0.282960765 | 0.285363883 | 0.864595038 | 0.81571845 | 0.015232574 | 0.095673458 | 0.054318179 |
| Pxn | Y | 88 | -0.025279275 | 0.124929992 | 0.180652801 | 0.205932076 | 0.055722809 | 0.640297366 | 0.213905749 | 0.146387411 | 0.104702935 | 0.627519663 |
| Srsf1 | Y | 189 | 0.21628015 | -0.122336709 | 0.276176253 | 0.059896103 | 0.398512962 | 0.236257998 | 0.746948755 | 0.341631368 | 0.820280503 | 0.365903434 |
| Flt1 | Y | 1169 | 0.034538358 | -0.079922285 | 0.529262683 | 0.494724325 | 0.609184968 | 0.911246582 | 0.794811202 | 0.156360069 | 0.247039775 | 0.167682813 |

| | | | | | | | | | | | | |
|---|---|---|---|---|---|---|---|---|---|---|---|---|
| H1f4 | S | 187 | 0.215681766 | 0.276052446 | -0.017187862 | -0.232869628 | -0.293240308 | 0.508070864 | 0.322575239 | 0.932179311 | 0.428902464 | 0.230593336 |
| Spata5 | Y | 701 | 0.133869229 | -0.063632209 | 0.257889921 | 0.124020692 | 0.32152213 | 0.388248575 | 0.778916565 | 0.17807657 | 0.416018943 | 0.204403454 |
| Ptpre | Y | 695 | 0.311781335 | 0.009667721 | 0.060626231 | -0.251155104 | 0.05095851 | 0.215972765 | 0.932144343 | 0.603720495 | 0.291038328 | 0.167526139 |
| Wipf1 | Y | 72 | 0.29135273 | 0.185542086 | -0.051096326 | -0.342449055 | -0.236638412 | 0.385974152 | 0.238700345 | 0.635209954 | 0.3244521 | 0.174130923 |
| Eef1a2 Eef1a1 Eef1a1-ps | Y | 141;141 | 0.125941573 | 0.067526352 | -0.01587816 | -0.141819733 | -0.083404513 | 0.046315348 | 0.196012716 | 0.761315995 | 0.028407046 | 0.093625095 |
| Tnk2 | Y | 874 | 0.082737752 | -0.013120001 | 0.196797004 | 0.114059252 | 0.209917006 | 0.222254815 | 0.879526453 | 0.18724832 | 0.391724445 | 0.177900751 |
| Cdk16 Cdk17 | Y | 176;203 | 0.087116258 | 0.126324943 | -0.074808799 | -0.161925057 | -0.201133742 | 0.101898078 | 0.00207775 | 0.071323628 | 0.021270523 | 0.005224627 |
| Ephb4 | Y | 774 | -0.093200494 | 0.17506953 | 0.149208663 | 0.242409157 | -0.025860866 | 0.329332713 | 0.067426284 | 0.167369974 | 0.042395214 | 0.727248846 |
| Rps27l Rps27 Rps27rt | Y | 31;31 | 0.399472616 | -0.051666025 | -0.096904449 | -0.496377066 | -0.045238424 | 0.010066485 | 0.706966325 | 0.303848467 | 0.009336708 | 0.707134721 |
| Pik3r1 | Y | 452 | 0.06394481 | 0.046179658 | 0.101062108 | 0.037117298 | 0.05488245 | 0.272602399 | 0.478765847 | 0.202413177 | 0.606804627 | 0.488533313 |
| Ctnnd1 | Y | 302 | 0.231054126 | 0.102641632 | -0.041906751 | -0.272960877 | -0.144548383 | 0.282175283 | 0.084265278 | 0.500027889 | 0.22320336 | 0.077649869 |
| Nars | Y | 550 | 0.164868374 | -0.026022722 | 0.115466669 | -0.049401705 | 0.141489392 | 0.112920579 | 0.761030973 | 0.332225618 | 0.681293325 | 0.273461975 |
| Map1a | Y | 1942 | 0.261145799 | 0.188039043 | -0.084117247 | -0.345263047 | -0.272156291 | 0.329403317 | 0.106062765 | 0.304336452 | 0.228751312 | 0.047946129 |
| Mapk3 | T | 203 | 0.228281247 | 0.019914808 | 0.055619047 | -0.1726622 | 0.035704238 | 0.110815616 | 0.914137625 | 0.773740362 | 0.388894736 | 0.873703343 |
| Iqgap1 | Y | 1510 | 0.084138282 | -0.041613179 | 0.186048429 | 0.101910147 | 0.227661608 | 0.425034601 | 0.666899587 | 0.058676842 | 0.262339823 | 0.042836694 |
| Cdk14 | Y | 146 | 0.043569823 | 0.042399706 | 0.162329762 | 0.118759939 | 0.119930056 | 0.697080783 | 0.537907378 | 0.133030495 | 0.352089735 | 0.223200335 |
| Aldoa Aldoc | Y | 204;204 | 0.137541687 | -0.066184874 | 0.202765433 | 0.065223746 | 0.268950307 | 0.147209645 | 0.550920762 | 0.25031413 | 0.675844885 | 0.168808958 |
| Ctps | Y | 473 | 0.200127852 | 0.013522209 | 0.017259696 | -0.182868156 | 0.003737487 | 0.066027195 | 0.857351923 | 0.813826636 | 0.074590822 | 0.934111157 |
| Plpbp | Y | 69 | -0.048218888 | 0.251671652 | 0.299303492 | 0.347522381 | 0.047631841 | 0.900003397 | 0.554978956 | 0.394910671 | 0.292380319 | 0.89148093 |
| Nck1 | S | 96 | 0.517001886 | 0.13872625 | -0.123086466 | -0.640088351 | -0.261812716 | 0.26795536 | 0.215593248 | 0.196671343 | 0.202594131 | 0.041118419 |
| Dennd2a | T | 356 | 0.39039705 | 0.01892264 | 0.104956279 | -0.285440771 | 0.08603364 | 0.489815379 | 0.903659165 | 0.508512794 | 0.604978257 | 0.632887373 |
| Pitrm1 | Y | 771 | 0.059895554 | 0.015369486 | 0.20935961 | 0.149464056 | 0.193990123 | 0.442568586 | 0.888583318 | 0.206756479 | 0.333772057 | 0.261065843 |
| Eif4b | Y | 211 | 0.166988675 | 0.062228273 | 0.101453483 | -0.065535192 | 0.03922521 | 0.399719111 | 0.695351158 | 0.446227278 | 0.748124606 | 0.829601233 |
| Vim | Y | 61 | 0.032164029 | -0.005744902 | 0.203210328 | 0.171046299 | 0.20895523 | 0.583519431 | 0.948811552 | 0.082849153 | 0.131076044 | 0.102575446 |
| Eef1a1 Eef1a1-ps1 | Y | 177 | 0.193731088 | 0.071566538 | 0.011233412 | -0.182497676 | -0.060333126 | 0.386778316 | 0.190556079 | 0.91365797 | 0.421782502 | 0.550495518 |
| Git1 | Y | 554 | 0.325296566 | 0.02241469 | -0.006504269 | -0.331800835 | -0.028918959 | 0.176827105 | 0.892382563 | 0.977066468 | 0.246478471 | 0.900499509 |
| Inppl1 | Y | 1161 | 0.266996039 | 0.128993721 | -0.017205652 | -0.284201691 | -0.146199373 | 0.487400277 | 0.312717883 | 0.915362645 | 0.476030997 | 0.445974049 |
| Srsf9 | Y | 71 | 0.222107802 | -0.085068327 | 0.045496429 | -0.176611374 | 0.130564756 | 0.024885053 | 0.40312953 | 0.655978721 | 0.122057298 | 0.265688841 |
| Nes | Y | 662 | 0.223459588 | 0.101538314 | 0.101141921 | -0.122317667 | -0.000396393 | 0.568680937 | 0.72954609 | 0.589799405 | 0.756861847 | 0.998983443 |
| Coro1b | Y | 420 | 0.020294021 | 0.007247686 | 0.196654616 | 0.176360595 | 0.18940693 | 0.787747885 | 0.952249093 | 0.110620351 | 0.151752603 | 0.225973519 |
| Anxa1 | Y | 207 | -0.07365017 | 0.1203407 | 0.147309472 | 0.220959642 | 0.026968772 | 0.407882473 | 0.146381403 | 0.16492408 | 0.079472172 | 0.770755842 |
| Eef1d | Y | 182 | 0.130364595 | 0.027406111 | 0.052832975 | -0.07753162 | 0.025426864 | 0.204124071 | 0.637319888 | 0.568735499 | 0.495517288 | 0.782308134 |
| Tnk2 | Y | 842 | -0.021787352 | -0.049303273 | 0.249205203 | 0.270992556 | 0.298508476 | 0.699406819 | 0.588308719 | 0.020302778 | 0.025683407 | 0.029348755 |
| Tln1 | Y | 1445 | 0.435965159 | -0.242945281 | 0.309271511 | -0.126693648 | 0.552216792 | 0.54508884 | 0.529546131 | 0.303909944 | 0.847908716 | 0.154390006 |
| Arhgap12 | Y | 241 | 0.116683929 | 0.035332173 | 0.027998664 | -0.088685265 | -0.007333509 | 0.152554225 | 0.495049095 | 0.69234199 | 0.303463456 | 0.909751181 |
| Ctnnd1 | Y | 296 | 0.2366391 | 0.12526675 | -0.140151099 | -0.376790199 | -0.265417848 | 0.146062077 | 0.05514292 | 0.114172442 | 0.05324142 | 0.024466061 |
| Dnajb1 Dnajb4 | Y | 172;176 | 0.113688584 | -0.063997723 | 0.123386521 | 0.009697937 | 0.187384244 | 0.256633061 | 0.501505408 | 0.095412777 | 0.910270359 | 0.10014987 |
| Sh2b3 | Y | 245 | -0.085639331 | -0.329377239 | 0.544372886 | 0.630012217 | 0.873750126 | 0.81442269 | 0.389695629 | 0.028358745 | 0.173574609 | 0.084759214 |
| Git2 | S | 535 | 0.047284901 | 0.200953065 | 0.027360743 | -0.019924158 | -0.173592322 | 0.697193586 | 0.284276431 | 0.84646411 | 0.870320935 | 0.349829205 |
| Il6ra | Y | 449 | 0.284385937 | -0.001832819 | -0.260096844 | -0.544482781 | -0.258264025 | 0.006410784 | 0.991733671 | 0.373568709 | 0.135549444 | 0.407589486 |
| Eif2s1 | Y | 150 | 0.217066326 | -0.1331773 | 0.069321024 | -0.147745302 | 0.202498324 | 0.011155657 | 0.186592692 | 0.665039656 | 0.384784467 | 0.270321043 |
| Rpl38 | Y | 43 | 0.33238022 | -0.027949597 | -0.145140457 | -0.477520677 | -0.117190861 | 0.011071256 | 0.80669136 | 0.215712475 | 0.009491993 | 0.365631451 |
| Hnrnpa1 | Y | 314 | -0.059537233 | 0.118683604 | 0.166974958 | 0.226512191 | 0.048291354 | 0.7159108 | 0.499083548 | 0.188815868 | 0.249933052 | 0.790200096 |
| Kirrel | T | 751 | 0.066344486 | 0.011972757 | 0.067106758 | 0.000762272 | 0.055134001 | 0.254137974 | 0.875322772 | 0.197914916 | 0.987689099 | 0.473846998 |
| Ptk2 | T | 394 | 0.002367182 | 0.582601171 | -0.216124286 | -0.218491468 | -0.798725457 | 0.994526318 | 0.232429184 | 0.73149479 | 0.720711562 | 0.270026451 |
| Tjp1 | Y | 1177 | -0.089032084 | -0.041614596 | 0.32556059 | 0.414592674 | 0.367175187 | 0.423744006 | 0.271492526 | 0.014564576 | 0.022732375 | 0.017842403 |
| Rbmxl1 | Y | 263 | 0.279902128 | -0.138321076 | 0.037357259 | -0.242544869 | 0.175678335 | 0.056572345 | 0.385591405 | 0.63590325 | 0.083436793 | 0.298038076 |
| Pdlim1 | Y | 142 | 0.137260035 | 0.113245104 | 0.022374762 | -0.114885274 | -0.090870342 | 0.63408228 | 0.36896913 | 0.787922648 | 0.689253191 | 0.477272666 |
| Pfas | Y | 967 | 0.104748385 | -0.020353382 | 0.094498794 | -0.010249591 | 0.114852176 | 0.210235261 | 0.874616563 | 0.37014911 | 0.919576336 | 0.439496344 |
| Kirrel | Y | 762 | -0.007124601 | 0.177062077 | 0.010228397 | 0.017352998 | -0.166833681 | 0.931552319 | 0.086846948 | 0.957413197 | 0.929367452 | 0.427941859 |
| Fchsd2 | Y | 708 | -0.086765554 | -0.013529908 | 0.27268454 | 0.359450094 | 0.286214448 | 0.171271425 | 0.809610208 | 0.012413931 | 0.019840257 | 0.028981231 |
| Il6st | S | 903 | 0.002264686 | -0.026942622 | 0.303899329 | 0.301634643 | 0.330841952 | 0.986683139 | 0.837088786 | 0.192757658 | 0.191334453 | 0.163336067 |
| Kirrel | Y | 628 | 0.056168025 | 0.037003458 | 0.100585704 | 0.044417679 | 0.063582246 | 0.581960737 | 0.65946891 | 0.309531282 | 0.654764 | 0.450211756 |
| Psma5 | Y | 26 | -0.067407988 | 0.526002082 | 0.668853136 | 0.736261124 | 0.142851054 | NA | 0.460922492 | NA | NA | NA |
| Scarf2 | Y | 615 | -0.122858055 | 0.239027016 | -0.047665819 | 0.075192235 | -0.286692836 | 0.343188307 | 0.003343755 | 0.533646943 | 0.563549826 | 0.025907808 |
| Tmprss13 | Y | 214;229;520 | 1.069640043 | -0.060821731 | -0.121729968 | -1.19137001 | -0.060908236 | 0.334953252 | 0.594949663 | 0.53016266 | 0.294434281 | 0.762748144 |
| Slc12a4 | Y | 62 | -0.049139291 | -0.023364469 | 0.224239969 | 0.27337926 | 0.247604438 | 0.683111317 | 0.877021078 | 0.057453452 | 0.055720091 | 0.155054399 |
| Rpl11 | Y | 92 | 0.291756445 | -0.101513756 | 0.118182587 | -0.173573858 | 0.219696342 | 0.426669638 | 0.636037361 | 0.41457698 | 0.612357467 | 0.302710242 |
| Prrc2a | Y | 1013 | 0.209705622 | -0.26307859 | 0.194768763 | -0.014936859 | 0.457847353 | 0.237788862 | 0.543289632 | 0.343834568 | 0.935917067 | 0.327784956 |
| Tuba1a Tuba1b Tuba1c Tu | Y | 357;357;357;357; | 0.158223952 | 0.007034524 | 0.008700127 | -0.149523825 | 0.001665602 | 0.14987196 | 0.924003018 | 0.902665662 | 0.15624913 | 0.976493045 |
| Nans | Y | 71 | -0.029451753 | 0.058385661 | 0.071157299 | 0.100609051 | 0.012771637 | 0.427023547 | 0.05418133 | 0.187678434 | 0.103751277 | 0.777658252 |
| Afap1l2 | Y | 54 | 0.208015717 | -0.152501001 | 0.068249634 | -0.139766083 | 0.220750634 | 0.022809612 | 0.298550556 | 0.778388362 | 0.571087287 | 0.415797701 |
| Axl | Y | 697 | -0.07164553 | 0.042816901 | 0.164254329 | 0.235899859 | 0.121437428 | 0.416230888 | 0.537945946 | 0.062303437 | 0.025813043 | 0.051112256 |
| Eif4b | Y | 270 | 0.184525751 | -0.063538931 | 0.077932283 | -0.106593467 | 0.141471214 | 0.265648122 | 0.587057379 | 0.415919149 | 0.495808193 | 0.280776466 |
| Calm1 Calm2 Calm3 | Y | 139;139;139 | -0.132864751 | 0.162644208 | 0.13805489 | 0.270919642 | -0.024589318 | 0.813345121 | 0.548416152 | 0.271732411 | 0.635246908 | 0.920557463 |
| Mvb12a | S | 200 | 0.277952205 | 0.040908202 | 0.001830204 | -0.276122002 | -0.039077998 | 0.456890265 | 0.799475716 | 0.966178821 | 0.459751113 | 0.810561462 |
| Clock | Y | 331 | 0.229614131 | -0.200682618 | 0.178115973 | -0.051498158 | 0.378798591 | 0.231961853 | 0.412454052 | 0.39395035 | 0.821185883 | 0.210008234 |
| Map3k4 | Y | 1545 | 0.038415243 | 0.072911986 | 0.099092109 | 0.060676866 | 0.026180123 | 0.777196427 | 0.57803764 | 0.51175894 | 0.551536032 | 0.773100382 |
| Cnn2 | Y | 298 | 0.208045495 | 0.159839687 | -0.036932703 | -0.244978199 | -0.19677239 | 0.656401797 | 0.459730776 | 0.790151455 | 0.60746205 | 0.399184325 |
| Slc41a2 | Y | 113 | 0.194034711 | 0.010308316 | -0.132373641 | -0.326408352 | -0.142681957 | 0.020969167 | 0.921522709 | 0.408707115 | 0.112284959 | 0.404692558 |
| Rpl3 | Y | 307 | 0.181667799 | -0.192819953 | 0.057859604 | -0.123808194 | 0.250679557 | 0.001777791 | 0.113992199 | 0.779581922 | 0.564505202 | 0.29962776 |
| Stk4 | S | 40 | 0.040010513 | 0.190884504 | -0.129472388 | -0.169482901 | -0.320356892 | 0.713027535 | 0.080449884 | 0.579549589 | 0.48187747 | 0.23357348 |
| Stk4 | S | 43 | 0.040010513 | 0.190884504 | -0.129472388 | -0.169482901 | -0.320356892 | 0.713027535 | 0.080449884 | 0.579549589 | 0.48187747 | 0.23357348 |
| Vim | T | 37 | 0.378456001 | 0.011255248 | -0.051662824 | -0.430118824 | -0.062918072 | 0.372961291 | 0.965307707 | 0.842705625 | 0.305896631 | 0.768638389 |
| Psmb6 | Y | 58 | 0.106454889 | -0.151107703 | 0.254409041 | 0.147954152 | 0.405516745 | 0.544718404 | 0.305402992 | 0.143462666 | 0.439681203 | 0.051479824 |
| Ctnnd1 | S | 288 | 0.307736555 | 0.169159915 | -0.201042092 | -0.508778647 | -0.370202007 | 0.311768539 | 0.147704002 | 0.130304601 | 0.146001693 | 0.03480304 |
| Prrc2c | Y | 1166 | 0.103880065 | -0.119291448 | 0.16973548 | 0.065855415 | 0.289026929 | 0.419073432 | 0.677753496 | 0.318009686 | 0.660081857 | 0.357991381 |
| Tjp1 | Y | 1521 | 0.017442964 | -0.033062254 | 0.116661693 | 0.099218729 | 0.149723947 | 0.794402388 | 0.651583049 | 0.056217948 | 0.164988215 | 0.092010057 |
| Trim25 | Y | 277 | 0.410468962 | -0.985962496 | 0.516478353 | 0.106009391 | 1.502440848 | 0.197193297 | 0.196449466 | 0.161594675 | 0.678055664 | 0.092564033 |
| Abi1 | Y | 284 | 0.167291065 | -0.506103635 | 0.168173322 | 0.000882257 | 0.674276957 | 0.777444814 | NA | 0.560626406 | 0.998763345 | NA |
| Layn | Y | 292 | 0.264170394 | -0.010968843 | -0.090441476 | -0.35461187 | -0.079472632 | 0.100093325 | 0.914153271 | 0.502858233 | 0.073941157 | 0.60471931 |
| Nutm1 | S | 817 | 0.053767862 | 0.084767778 | -0.023450307 | -0.077218169 | -0.108218085 | 0.558532255 | 0.146260786 | 0.664425293 | 0.387221821 | 0.01955853 |
| Nutm1 | S | 824 | 0.053767862 | 0.084767778 | -0.023450307 | -0.077218169 | -0.108218085 | 0.558532255 | 0.146260786 | 0.664425293 | 0.387221821 | 0.01955853 |
| Ptk2b | Y | 849 | -0.147691829 | 0.25989904 | 0.066302787 | 0.213994617 | -0.193596253 | 0.195216553 | 0.170572067 | 0.296801728 | 0.097653809 | 0.268371161 |
| Eno1 Eno1b Eno2 | Y | 25;25 | 0.0673881 | 0.064083385 | 0.000965881 | -0.066422219 | -0.063117504 | 0.532114173 | 0.360963062 | 0.988393325 | 0.484771155 | 0.009407244 |
| Idh1 | Y | 135 | -0.126455166 | 0.195413169 | 0.123196613 | 0.249651779 | -0.072216557 | 0.776896793 | 0.361454937 | 0.734006002 | 0.619944637 | 0.825836902 |
| Pgam1 Pgam2 | Y | 92;92 | 0.034086756 | 0.056462625 | 0.015714869 | -0.018371887 | -0.040747756 | 0.427906742 | 0.413750004 | 0.764215098 | 0.749309852 | 0.588964892 |
| Rasa3 | Y | 66 | 0.286495404 | 0.071070773 | -0.125861446 | -0.41235685 | -0.196932218 | 0.188511119 | 0.683213967 | 0.257230139 | 0.096263403 | 0.297977693 |
| Tjp1 | Y | 830 | 0.123333042 | -0.069055 | 0.07560267 | -0.047730372 | 0.14465767 | 0.194898386 | 0.468305918 | 0.528583478 | 0.7086633 | 0.307736555 |
| Pard3 | Y | 719 | 0.094806006 | 0.030955205 | -0.045104591 | -0.139910597 | -0.076059796 | 0.081818706 | 0.579614755 | 0.483621896 | 0.071012749 | 0.243649671 |
| Pkp4 | Y | 464 | -0.019214728 | -0.083868545 | 0.233848374 | 0.253063102 | 0.317716918 | 0.784475854 | 0.38947329 | 0.072423641 | 0.07778834 | 0.035269989 |

| | | | | | | | | | | | | | |
|---|---|---|---|---|---|---|---|---|---|---|---|---|---|
| Arhgef5 | Y | | 1081 | 0.209797818 | 0.153432621 | -0.179383111 | -0.38918093 | -0.332815733 | 0.560096952 | 0.269584355 | 0.478928846 | 0.354114293 | 0.246365428 |
| Ubr5 | Y | | 1740 | 0.207885679 | 0.028925931 | -0.009025782 | -0.216911462 | -0.037951713 | 0.520174188 | 0.785990771 | 0.937593924 | 0.513304709 | 0.795418606 |
| Rpl15 | T | | 80 | 0.459470579 | -0.234378159 | -0.046518814 | -0.505989394 | 0.187859345 | 0.117100759 | 0.417205148 | 0.745423491 | 0.092695987 | 0.518771763 |
| Rps10 | T | | 126 | 0.320493295 | -0.142115302 | -0.104627104 | -0.425120399 | 0.037488198 | 0.006090718 | 0.099790943 | 0.462603278 | 0.04609689 | 0.767249422 |
| Ptk2 | S | | 390 | 0.227116458 | -0.037459898 | 0.025501801 | -0.201614657 | 0.062961699 | 0.335730877 | 0.550505165 | 0.822980397 | 0.397622084 | 0.603936481 |
| Sema6d | Y | | 738 | 0.246787066 | -0.150469887 | -0.068435131 | -0.315222197 | 0.082034756 | 0.004676816 | 0.167585565 | 0.505790675 | 0.043102531 | 0.493950198 |
| Prrc2c | Y | | 1167 | 0.109032291 | -0.118870716 | 0.14363318 | 0.034600888 | 0.262503896 | 0.36837621 | 0.392808708 | 0.31753468 | 0.817495023 | 0.157477499 |
| Bcar1 | Y | | 132 | 0.075503277 | 0.035624295 | -0.007046008 | -0.082549285 | -0.042670304 | 0.318577438 | 0.600545334 | 0.886020717 | 0.273629225 | 0.51706309 |
| Sptan1 | Y | | 1176 | 0.182373394 | 0.107818295 | -0.115421506 | -0.2977949 | -0.223239802 | 0.40626653 | 0.291885464 | 0.301704357 | 0.215000704 | 0.041809089 |
| Dcbld2 | Y | | 710 | 0.017761459 | 0.178060931 | -0.140929923 | -0.158691383 | -0.318990855 | 0.901734776 | 0.070411581 | 0.438426417 | 0.431531036 | 0.150077869 |
| Anxa2 | Y | | 24 | 0.184966942 | -0.033794008 | -0.093001944 | -0.277968886 | -0.059207936 | 0.04457437 | 0.702428333 | 0.474311376 | 0.09979202 | 0.667987728 |
| Hspa8 | Y | | 107 | 0.222734279 | -0.093523898 | -0.04544593 | -0.268180209 | 0.048077968 | 0.049836 | 0.437483954 | 0.563654314 | 0.029182406 | 0.636231379 |
| Plcg1 | T | | 766 | 0.345819814 | 0.084532764 | -0.195917657 | -0.541737471 | -0.280450421 | 0.545073037 | 0.504582173 | 0.590240347 | 0.400561314 | 0.455589871 |
| Ccdc88a | Y | | 1801 | 0.021767549 | 0.018065541 | 0.060955465 | 0.039187917 | 0.042889918 | 0.7715036 | 0.759466973 | 0.417328524 | 0.610350081 | 0.497814014 |
| Ube3a | Y | | 506 | -0.077422192 | -0.111736855 | 0.339430045 | 0.416852236 | 0.4511669 | 0.885861504 | 0.840068854 | 0.343535321 | 0.494302936 | 0.470613085 |
| Sdc4 | Y | | 197 | 0.049661223 | 0.011514891 | 0.035521511 | -0.014139712 | 0.02400662 | 0.484302568 | 0.857656612 | 0.614086254 | 0.814254307 | 0.658420099 |
| Antxr1 Antxr2 | Y | 381;381 | | 0.155761492 | -0.001595604 | -0.093136059 | -0.248897551 | -0.091540455 | 0.010665421 | 0.977562549 | 0.087937969 | 0.003302243 | 0.152932447 |
| Sipa1l1 | Y | | 1133 | 0.127043681 | -0.025834551 | 0.042941925 | -0.084101756 | 0.068776476 | 0.479794157 | 0.841657864 | 0.724514816 | 0.622052604 | 0.574244379 |
| Acaca | Y | | 1369 | 0.197124142 | -0.133986459 | 0.051577422 | -0.14554672 | 0.185563881 | 0.09789893 | 0.231469056 | 0.756235472 | 0.365801072 | 0.274451692 |
| Nhsl1 | S | | 512 | 0.270656732 | -0.151448822 | -0.094555136 | -0.365211869 | 0.056893686 | 0.004183839 | 0.064626475 | 0.597302537 | 0.127347704 | 0.743477948 |
| Nhsl1 | Y | | 517 | 0.270656732 | -0.151448822 | -0.094555136 | -0.365211869 | 0.056893686 | 0.004183839 | 0.064626475 | 0.597302537 | 0.127347704 | 0.743477948 |
| Zdhhc5 | S | | 529 | 0.070318626 | -0.031652622 | 0.029758874 | -0.040559752 | 0.061411496 | 0.154016247 | 0.503885995 | 0.532457521 | 0.345076085 | 0.194439974 |
| Sh2b3 | Y | | 519 | -0.084585043 | -0.003885714 | 0.280679485 | 0.365264528 | 0.284565199 | 0.41432039 | 0.983459492 | 0.241291839 | 0.158538381 | 0.292840824 |
| Zc3hav1 | Y | | 184 | 0.113637587 | -0.038857694 | 0.006520678 | -0.107116909 | 0.045378372 | 0.178249252 | 0.709438717 | 0.940590144 | 0.232969661 | 0.669637506 |
| Pakap | S | | 440 | 0.219099819 | -0.050816182 | -0.130077199 | -0.349177018 | -0.079261017 | 0.082931983 | 0.736834018 | 0.57613625 | 0.196542724 | 0.736670472 |
| Cdk16 Cdk17 | T | 178;205 | | 0.048039491 | 0.041055411 | -0.079111238 | -0.127150728 | -0.120166648 | 0.11937739 | 0.042262921 | 0.241310789 | 0.106022027 | 0.131903956 |
| Gapdh | Y | | 316 | 0.370944744 | -0.065468773 | -0.029603235 | -0.400547979 | 0.035865538 | 0.283953266 | 0.474130957 | 0.560405404 | 0.259110543 | 0.6547624 |
| Adk | Y | | 341 | 0.206487399 | -0.035161794 | -0.081664378 | -0.288151778 | -0.046502584 | 0.12682631 | 0.755296736 | 0.539337264 | 0.112180219 | 0.766187999 |
| Orai1 | Y | | 303 | 0.042184753 | 0.014772691 | 0.029795451 | -0.012389301 | 0.01502276 | 0.526548262 | 0.899663262 | 0.620563973 | 0.709181418 | 0.887389657 |
| Pdha1 | S | | 293 | -0.0090669 | 0.04613429 | 0.11703615 | 0.12610305 | 0.07090186 | 0.959701115 | 0.680074343 | 0.690775316 | 0.694989481 | 0.810631631 |
| Pdha1 | S | | 301 | -0.0090669 | 0.04613429 | 0.11703615 | 0.12610305 | 0.07090186 | 0.959701115 | 0.680074343 | 0.690775316 | 0.694989481 | 0.810631631 |
| Rpl15 | Y | | 59 | 0.268321557 | -0.155591701 | -0.023429632 | -0.291751189 | 0.13216207 | 0.142541093 | 0.445403656 | 0.871224032 | 0.164978545 | 0.556606999 |
| Ass1 | Y | | 113 | 0.056313982 | 0.271456168 | -0.241531209 | -0.297845191 | -0.512987377 | 0.535907503 | 0.029602014 | 0.048704812 | 0.020464194 | 0.003676347 |
| Jcad | S | | 115 | 0.116838812 | 0.087320255 | -0.085030773 | -0.201869585 | -0.172351028 | 0.40858864 | 0.544642559 | 0.472083518 | 0.132878514 | 0.201014469 |
| Lemd2 | Y | | 113 | 0.344805558 | 0.097371093 | -0.147868034 | -0.492673592 | -0.245239127 | 0.522207606 | 0.629442979 | 0.352512943 | 0.382169278 | 0.230390033 |
| Mapk3 | Y | | 205 | 0.057206631 | 0.067880311 | -0.052211115 | -0.109417746 | -0.120091425 | 0.51815781 | 0.422906032 | 0.653908223 | 0.38433248 | 0.33726131 |
| Sh3pxd2a | S | | 607 | 0.559733982 | 0.030474833 | -0.125855078 | -0.68558906 | -0.156329911 | 0.566897948 | 0.89186867 | 0.381729386 | 0.492496306 | 0.503272107 |
| Shb | S | | 304 | -0.189719986 | -0.03826939 | 0.200275511 | 0.389995496 | 0.238544901 | 0.469014208 | 0.659154866 | 0.048105417 | 0.20525737 | 0.017987606 |
| Shb | S | | 291 | -0.189719986 | -0.03826939 | 0.200275511 | 0.389995496 | 0.238544901 | 0.469014208 | 0.659154866 | 0.048105417 | 0.20525737 | 0.017987606 |
| Kank2 | Y | | 100 | -0.105761002 | 0.119703969 | 0.066106522 | 0.171867524 | -0.053597447 | 0.306178975 | 0.207906752 | 0.565374054 | 0.152678415 | 0.575270525 |
| Eprs | Y | | 754 | 0.109522002 | 0.002438417 | -0.01795153 | -0.127473532 | -0.020389947 | 0.397019897 | 0.986090379 | 0.863462826 | 0.282285757 | 0.871246324 |
| Eef1b2 | Y | | 126 | 0.109807459 | -0.029283921 | -0.008826493 | -0.118633952 | 0.020457428 | 0.207065349 | 0.612936984 | 0.933703081 | 0.310359272 | 0.829741545 |
| Ttc21a | S | | 140 | -0.055431725 | -0.000893883 | 0.172651805 | 0.228083529 | 0.173545688 | 0.622513125 | 0.989412827 | 0.364529512 | 0.264577225 | 0.358000835 |
| Ttc21a | S | | 141 | -0.055431725 | -0.000893883 | 0.172651805 | 0.228083529 | 0.173545688 | 0.622513125 | 0.989412827 | 0.364529512 | 0.264577225 | 0.358000835 |
| Ttc21a | S | | 143 | -0.055431725 | -0.000893883 | 0.172651805 | 0.228083529 | 0.173545688 | 0.622513125 | 0.989412827 | 0.364529512 | 0.264577225 | 0.358000835 |
| Fyb | S | | 561 | -0.043732111 | -0.093055245 | 0.262928172 | 0.306660284 | 0.355983418 | 0.727193627 | 0.403408623 | 0.203483165 | 0.150982897 | 0.123869122 |
| Tut7 | Y | | 77 | 0.181110862 | -0.129593816 | 0.016247469 | -0.164863393 | 0.145841285 | 0.082546529 | 0.161973408 | 0.848207144 | 0.060009466 | 0.049191362 |
| Csmd1 | S | | 1094 | 0.131580838 | 0.037844864 | -0.086496792 | -0.21807763 | -0.124341657 | 0.327614237 | 0.557208145 | 0.355983525 | 0.156152584 | 0.192515971 |
| Csmd1 | T | | 1084 | 0.131580838 | 0.037844864 | -0.086496792 | -0.21807763 | -0.124341657 | 0.327614237 | 0.557208145 | 0.355983525 | 0.156152584 | 0.192515971 |
| Zdhhc5 | Y | | 533 | 0.04944465 | -0.026895083 | 0.022640407 | -0.026804243 | 0.04953549 | 0.278739389 | 0.562145656 | 0.610623388 | 0.488958849 | 0.261313984 |
| Adgrl3 | Y | | 1483 | 0.150516371 | -0.124583968 | 0.017560721 | -0.13295565 | 0.142144689 | 0.10684425 | 0.252783946 | 0.817444047 | 0.075242686 | 0.170850592 |
| Rpl27 LOC108167922 | S | | 39 | 0.340453227 | -0.151996481 | -0.115298631 | -0.455751858 | 0.036697851 | 0.093236366 | 0.399719532 | 0.407699386 | 0.051119535 | 0.841250232 |
| Mvb12a | Y | | 202 | 0.277230177 | -0.048877755 | -0.043229236 | -0.320489413 | 0.00561852 | 0.476759374 | 0.811377377 | 0.590940917 | 0.419602739 | 0.977319794 |
| Zdhhc8 | S | | 523 | 0.093559673 | 0.040542637 | -0.021309102 | -0.114868775 | -0.061851739 | 0.745664234 | 0.903773717 | 0.939508941 | 0.525914786 | 0.809047355 |
| H2ax | Y | | 143 | -0.035518025 | 0.158953818 | -0.021655261 | 0.013862764 | -0.18060908 | 0.682231742 | 0.486954922 | 0.887253383 | 0.927554827 | 0.472387679 |
| Drg2 | Y | | 37 | 0.03285738 | -0.071187679 | 0.111083068 | 0.078225688 | 0.182270746 | 0.772350712 | 0.576308355 | 0.504748402 | 0.659326039 | 0.345784646 |
| Caskin2 | Y | | 384 | 0.182178204 | -0.145484404 | 0.035445406 | -0.146732799 | 0.18092981 | 0.261470195 | 0.305200694 | 0.804186902 | 0.39714293 | 0.27439656 |
| Vim | Y | | 38 | 0.036421319 | -0.234814834 | 0.301521854 | 0.265100534 | 0.536336688 | 0.633245363 | 0.249696355 | 0.298083901 | 0.346775316 | 0.120622385 |
| Rpl7 | Y | | 161 | 0.230718208 | -0.135151346 | -0.21386046 | -0.444578668 | -0.078709113 | 0.124035073 | 0.701616177 | 0.627068174 | 0.356929061 | 0.878359644 |
| Arhgdia | Y | | 133 | 0.023100509 | 0.101641123 | -0.075262674 | -0.098363183 | -0.176903797 | 0.735187735 | 0.18244381 | 0.182393291 | 0.156269622 | 0.051755595 |
| Ndufa8 | Y | | 142 | 0.352047188 | -0.186240433 | -0.1506533 | -0.502700488 | 0.035587134 | 0.027802807 | 0.227314356 | 0.250819291 | 0.011961847 | 0.799693273 |
| Lamtor1 | Y | | 140 | 0.081540439 | -0.09721842 | 0.122440883 | 0.040900444 | 0.219659303 | 0.797476539 | NA | NA | NA | NA |
| Pgam1 Pgam2 | T | 96;96 | | -0.004371736 | 0.235733479 | -0.373141927 | -0.36877019 | -0.608875406 | 0.972226296 | 0.1574322 | 0.483131274 | 0.485668436 | 0.293958046 |
| Zc3h13 | Y | | 663 | 0.30703239 | -0.252694821 | 0.018969555 | -0.288062834 | 0.271664377 | 0.117248259 | 0.111180938 | 0.862805343 | 0.137314204 | 0.116006811 |
| Tars | Y | | 297 | 0.052446761 | -0.03089148 | -0.001371694 | -0.053818455 | 0.029519786 | 0.433707216 | 0.780939537 | 0.991024823 | 0.65871118 | 0.838061145 |
| Phldb2 | Y | | 147 | 0.262146842 | -0.23662878 | -0.014771437 | -0.276918279 | 0.221857343 | 0.09323276 | 0.176643413 | 0.925814717 | 0.149836012 | 0.259388311 |
| Cttnbp2nl | Y | | 447 | 0.057057965 | -0.017956017 | -0.016144216 | -0.07320218 | 0.001811802 | 0.355114682 | 0.782541136 | 0.756193731 | 0.166151288 | 0.972299442 |
| Eef1a2 Eef1a1 Eef1a1-ps: | Y | 29;29 | | 0.328964934 | -0.141083162 | -0.146748499 | -0.475713433 | -0.005665337 | 0.079574209 | 0.144929585 | 0.545752549 | 0.123902733 | 0.980037963 |
| Nme7 | Y | | 16 | 0.002845243 | -0.027027925 | 0.054770579 | 0.051925336 | 0.081798504 | 0.969108478 | 0.60063107 | 0.504214505 | 0.574994239 | 0.32977659 |
| Iqgap1 | T | | 178 | 0.058848648 | 0.065389135 | -0.07452897 | -0.133377618 | -0.139918105 | 0.840464854 | 0.649762722 | 0.634501443 | 0.660507408 | 0.40727216 |
| Slc38a2 | Y | | 28 | -0.001649854 | 0.000704869 | 0.017779258 | 0.019429112 | 0.017074388 | 0.99292003 | 0.992119315 | 0.759874848 | 0.917602443 | 0.82504245 |
| Plcb1 | Y | | 803 | -0.094863813 | 0.283034102 | -0.196738559 | -0.101874746 | -0.479772661 | 0.681314407 | 0.342245342 | 0.641103348 | 0.818044226 | 0.33257765 |
| Rps27a | Y | | 148 | 0.117903846 | -0.123707727 | 0.045059854 | -0.072843992 | 0.168767581 | 0.335564462 | 0.224189658 | 0.731820587 | 0.615197883 | 0.244012031 |
| Cavin1 | Y | | 318 | -0.046902098 | -0.030605572 | 0.099093442 | 0.14599554 | 0.129699014 | 0.713887415 | 0.814804169 | 0.537076344 | 0.279441582 | 0.331225354 |
| Rps10 | Y | | 127 | 0.365264783 | -0.364568803 | -0.114154875 | -0.479419658 | 0.250413928 | 0.03373992 | 0.179840152 | 0.492570001 | 0.033386534 | 0.331271809 |
| Sema5a | Y | | 1074 | 0.171058255 | -0.097553812 | -0.12460221 | -0.295660464 | -0.027048397 | 0.281496469 | 0.535474371 | 0.757055289 | 0.481539096 | 0.945341545 |
| Pdlim1 | Y | | 319 | -0.025371152 | 0.028342507 | 0.011297211 | 0.036668363 | -0.017045296 | 0.824511231 | 0.855869192 | 0.912047579 | 0.567910765 | 0.894156728 |
| Mfap3 | Y | | 277 | 0.114177273 | -0.068480316 | -0.042363368 | -0.156540641 | 0.026116949 | 0.135700187 | 0.258290312 | 0.596502223 | 0.092520287 | 0.690040021 |
| Ttyh2 | Y | | 494 | 0.139390843 | -0.106934534 | -0.077700851 | -0.217091695 | 0.029233683 | 0.106120654 | 0.393086674 | 0.541161696 | 0.162377027 | 0.848462018 |
| Dbi | Y | | 29 | 0.075967921 | 0.068768578 | -0.135641259 | -0.21160918 | -0.204409837 | 0.474558841 | 0.553589584 | 0.280518593 | 0.095291894 | 0.122447732 |
| Adam9 | Y | | 804 | -0.072879888 | 0.09101706 | -0.019106622 | 0.053773265 | -0.110123682 | 0.369323956 | 0.247500858 | 0.801664634 | 0.366913499 | 0.075380945 |
| Kirrel | Y | | 753 | -0.041562083 | 0.180771126 | -0.212235257 | -0.170673174 | -0.393006383 | 0.849632742 | 0.377926713 | 0.606374796 | 0.659854648 | 0.351204281 |
| Lamp1 | Y | | 403 | -0.004645752 | -0.112856438 | 0.117906201 | 0.12255953 | 0.230762639 | 0.965999022 | 0.33382636 | 0.310535337 | 0.035058464 | 0.006902468 |
| Sdc1 | Y | | 300 | 0.048019989 | 0.01406117 | -0.052163991 | -0.10018398 | -0.066225161 | 0.617642821 | 0.893385 | 0.507466127 | 0.298571953 | 0.519173468 |
| Prag1 | S | | 394 | 0.068497602 | -0.040001814 | -0.052291196 | -0.120788798 | -0.012289382 | 0.445664763 | 0.787841005 | 0.708189784 | 0.409173225 | 0.944444968 |
| Megf10 | Y | | 1111 | -0.133771471 | 0.020748386 | 0.175870602 | 0.309642073 | 0.155122216 | 0.11663466 | 0.802506243 | 0.28535785 | 0.112048949 | 0.344898559 |
| --- | Y | --- | | -0.437466579 | -0.703529041 | -1.08963564 | -0.652169061 | -0.386106599 | 0.066203956 | 0.052524688 | 0.000357764 | 0.023084611 | 0.197226388 |
| --- | Y | --- | | -0.437466579 | -0.703529041 | -1.08963564 | -0.652169061 | -0.386106599 | 0.066203956 | 0.052524688 | 0.000357764 | 0.023084611 | 0.197226388 |
| --- | Y | --- | | -0.633331758 | -0.600132797 | -0.624372694 | 0.008959064 | -0.024239896 | 0.010214671 | 0.043139695 | 0.010612771 | 0.923518516 | 0.901417403 |
| --- | Y | | 106 | -0.987968293 | -0.94584164 | -1.023376327 | -0.035408033 | -0.077534687 | 0.159064516 | 0.173300743 | 0.147916884 | 0.771017875 | 0.495736435 |
| --- | Y | --- | | -0.444759887 | -0.387965547 | -0.379036755 | 0.065723132 | 0.008928793 | 0.012705437 | 0.162783683 | 0.030899754 | 0.510169739 | 0.968226803 |

| | | | | | | | | | | | | |
|---|---|---|---|---|---|---|---|---|---|---|---|---|
| --- | Y | --- | 0.043345444 | -0.164219752 | -0.681053156 | -0.724398599 | -0.516833403 | 0.793341307 | 0.306666577 | 0.002393967 | 0.014751313 | 0.027239572 |
| --- | Y | --- | -0.041340239 | -0.164763136 | -0.274212586 | -0.232872347 | -0.109449449 | 0.777610714 | 0.045538027 | 0.031521999 | 0.186379197 | 0.249870436 |
| --- | Y | --- | 0.184762286 | -0.215992996 | -0.301608859 | -0.486371146 | -0.085615864 | 0.612538241 | 0.055128954 | 0.047579936 | 0.253603174 | 0.463981557 |
| --- | S | --- | -0.000971426 | -0.64353601 | 0.219102209 | 0.220073635 | 0.862638219 | 0.997087431 | 0.116052333 | 0.509797518 | 0.411879423 | 0.057663373 |
| --- | T | --- | -0.000971426 | -0.64353601 | 0.219102209 | 0.220073635 | 0.862638219 | 0.997087431 | 0.116052333 | 0.509797518 | 0.411879423 | 0.057663373 |
| --- | Y | --- | -0.388582188 | 0.109839685 | 0.168442673 | 0.557024861 | 0.058602988 | 0.004024268 | 0.159317403 | 0.188779885 | 0.008989523 | 0.580194392 |
| --- | Y | --- | 0.195973651 | -0.176636319 | -0.140405999 | -0.33637965 | 0.03623032 | 0.223974034 | 0.446727255 | 0.479798394 | 0.117153076 | 0.876705261 |
| --- | Y | --- | 0.195973651 | -0.176636319 | -0.140405999 | -0.33637965 | 0.03623032 | 0.223974034 | 0.446727255 | 0.479798394 | 0.117153076 | 0.876705261 |
| --- | Y | --- | 0.282509112 | -0.30380116 | -0.110905804 | -0.393414916 | 0.192895356 | 0.006636901 | 0.093605077 | 0.271364886 | 0.015426912 | 0.231092558 |
| --- | S | --- | 0.475038575 | 0.305182169 | 0.458139546 | -0.016899029 | 0.152957377 | 0.006574384 | 0.048534817 | 0.008080014 | 0.853893851 | 0.190270798 |
| --- | S | --- | 0.475038575 | 0.305182169 | 0.458139546 | -0.016899029 | 0.152957377 | 0.006574384 | 0.048534817 | 0.008080014 | 0.853893851 | 0.190270798 |
| --- | S | --- | 0.475038575 | 0.305182169 | 0.458139546 | -0.016899029 | 0.152957377 | 0.006574384 | 0.048534817 | 0.008080014 | 0.853893851 | 0.190270798 |
| --- | T | --- | 0.475038575 | 0.305182169 | 0.458139546 | -0.016899029 | 0.152957377 | 0.006574384 | 0.048534817 | 0.008080014 | 0.853893851 | 0.190270798 |
| --- | Y | --- | 0.243751563 | 0.456192107 | 0.394317659 | 0.150566096 | -0.061874448 | 0.139200917 | 0.004022084 | 0.010975622 | 0.303197399 | 0.47703826 |
| --- | Y | --- | 0.22558535 | 0.328553146 | 0.34927137 | 0.12368602 | 0.020718224 | 0.017898926 | 0.00377395 | 0.040661495 | 0.321450834 | 0.853330388 |
| --- | Y | --- | 0.142365843 | 0.327117432 | 0.458987462 | 0.316621619 | 0.13187003 | 0.068053096 | 0.033214328 | 0.003560578 | 0.015706151 | 0.25959376 |
| --- | Y | --- | 0.115907218 | 0.304219121 | 0.313935653 | 0.198028435 | 0.009716532 | 0.335246061 | 0.008769465 | 0.038605693 | 0.181152091 | 0.927012416 |
| --- | Y | --- | 0.115907218 | 0.304219121 | 0.313935653 | 0.198028435 | 0.009716532 | 0.335246061 | 0.008769465 | 0.038605693 | 0.181152091 | 0.927012416 |
| --- | S | --- | 0.209324217 | 0.184631644 | 0.251043029 | 0.041718813 | 0.066411386 | 0.047960489 | 0.118292143 | 0.024359375 | 0.646795161 | 0.531481348 |
| --- | T | --- | 0.289016773 | 0.182386404 | 0.245936876 | -0.043079898 | 0.063550472 | 0.105258884 | 0.216070188 | 0.075457594 | 0.759981606 | 0.650598781 |
| --- | Y | --- | 0.205600398 | 0.120341725 | 0.105024713 | -0.100575685 | -0.015317013 | 0.190802452 | 0.097785505 | 0.181548744 | 0.469131702 | 0.831654869 |
| --- | T | --- | 0.659223242 | 0.062137719 | -0.162746741 | -0.821969983 | -0.224884459 | 0.059310111 | 0.81403973 | 0.817454578 | 0.297304298 | 0.740406225 |
| --- | T | --- | 0.659223242 | 0.062137719 | -0.162746741 | -0.821969983 | -0.224884459 | 0.059310111 | 0.81403973 | 0.817454578 | 0.297304298 | 0.740406225 |
| --- | Y | --- | 0.178692502 | 0.421668005 | 0.057358546 | -0.121333956 | -0.364309459 | 0.625583189 | 0.207355395 | 0.809485916 | 0.68751776 | 0.204088579 |
| --- | S | --- | 0.022526531 | 0.147127882 | -0.006785161 | -0.029311693 | -0.153913044 | 0.507852087 | 0.000708253 | 0.948779511 | 0.785889493 | 0.236719686 |
| --- | S | --- | 0.022526531 | 0.147127882 | -0.006785161 | -0.029311693 | -0.153913044 | 0.507852087 | 0.000708253 | 0.948779511 | 0.785889493 | 0.236719686 |
| --- | S | --- | 0.022526531 | 0.147127882 | -0.006785161 | -0.029311693 | -0.153913044 | 0.507852087 | 0.000708253 | 0.948779511 | 0.785889493 | 0.236719686 |
| --- | Y | --- | 0.427628916 | 0.121887917 | -0.116150534 | -0.543779451 | -0.238038452 | 0.108938127 | 0.604902502 | 0.835309743 | 0.363196458 | 0.661820586 |
| --- | Y | --- | 0.163634108 | 0.001843257 | -0.168434315 | -0.332068423 | -0.170277572 | 0.155733654 | 0.987647372 | 0.664315354 | 0.418297191 | 0.658285813 |
| --- | Y | --- | 0.048693811 | -0.015291172 | 0.061040822 | 0.012347012 | 0.076331994 | 0.824509212 | 0.954637709 | 0.750937858 | 0.935578057 | 0.735814219 |
